# microRNA profiling in the Weddell Seal suggests novel regulatory mechanisms contributing to diving adaptation

**DOI:** 10.1101/851394

**Authors:** Luca Penso-Dolfin, Wilfried Haerty, Allyson Hindle, Federica Di Palma

## Abstract

The Weddell Seal (*Leptonychotes weddelli*) represents a remarkable example of adaptation to diving among marine mammals. This species is capable of diving >900 m deep and remaining underwater for more than 60 minutes. A number of key physiological specializations have been identified, including the low levels of aerobic, lipid-based metabolism under hypoxia, significant increase in oxygen storage in blood and muscle; high blood volume and extreme cardiovascular control. These adaptations have been linked to increased abundance of key proteins, suggesting an important, yet still understudied role for gene reprogramming.

In this study, we investigate the possibility that post-transcriptional gene regulation by microRNAs (miRNAs) has contributed to the adaptive evolution of diving capacities in the Weddell Seal.

Using small RNA data across 4 tissues (cortex, heart, muscle and plasma), in 3 biological replicates, we generate the first miRNA annotation in this species, consisting of 559 high confidence, manually curated miRNA loci. Evolutionary analyses of miRNA gain and loss highlight a high number of Weddell seal specific miRNAs.

416 miRNAs were differentially expressed (DE) among tissues, whereas 83 miRNAs were differentially expressed (DE) across all tissues between pups and adults and 221 miRNAs demonstrated developmental changes in specific tissues only. mRNA targets of these altered miRNAs identify possible protective mechanisms in individual tissues, particularly relevant to hypoxia tolerance, anti-apoptotic pathways, and nitric oxide signal transduction. Novel, lineage-specific miRNAs associated with developmental changes target genes with roles in angiogenesis and vasoregulatory signaling.

Altogether, we provide an overview of miRNA composition and evolution in the Weddell seal, and the first insights into their possible role in the specialization to diving.

## Background

The Antarctic Weddell Seal (*Leptonychotes weddelli*) is a deep diving marine mammal, capable of pursuing prey to depths >900 m and remaining underwater for more than 60 minutes [1, 2]. Due to their exceptional diving ability and accessibility on the fast ice during their breeding season, the Weddell seal is one of the best-studied divers in the world. The pinniped lineage recolonized the marine environment ∼25 mya [3] and over this evolutionary time have become specialized to their aquatic habitat. These specializations encompass morphology and physiology; in particular, the extreme cardiovascular physiology of diving mammals is central to their capacity for long-duration diving. The well-developed dive response of marine mammals, including Weddell seals, is characterized by cardiovascular adjustments to lower heart rate and reduce peripheral blood flow during submergence. These adjustments depress tissue oxygen use by restricting its availability to peripheral vascular beds and conserving it for critical central tissues such as the brain and heart. Previous work has also highlighted several complementary traits that support breath-hold hunting in seals, for example: the preference for aerobic, lipid-based metabolism under hypoxia [4–6]; and extremely high oxygen stores in blood and muscle via enhanced haemoglobin and myoglobin [7–9].

Pinnipeds provide a fascinating model system in which to study the development of diving ability and hypoxia tolerance in mammals. Only adult seals are elite divers – unlike cetacean calves, pinniped pups are born on land. Development of the adult diving phenotype has been linked to changes in key proteins (e.g. respiratory pigments), tissue iron content and metabolic enzyme levels [5]. However, the details of the extent of tissue-specific maturation to refine local blood flow [10], metabolic control, and to combat negative effects of hypoxia exposure are still to be elucidated. Interestingly, pup physiology develops during weaning and throughout a post-weaning fast, including cardiac ontogeny to develop the fine-scale control of bradycardia observed in adults. This maturation can begin before pups first enter the water, which suggests an important role for gene reprogramming. The contribution of post-transcriptional gene regulation in the development of hypoxia tolerance and dive capacity has not been investigated.

MicroRNAs (miRNAs) are considered one of the key gene regulators in animals, conferring temporo-spatial precision in the regulation of gene expression. These short (∼22 nt) non-coding RNAs are involved in fundamental processes such as embryonic development and tissue differentiation [11–14] and likely play important roles in seasonal and development transitions involving gene reprogramming. MiRNAs reduce translation by binding to the 3’ region of complementary RNA, resulting in direct translational repression or mRNA degradation [15–17]. MiRNAs appear critical to the development of tissue-specific phenotypes and evolutionary adjustments in gene expression [18, 19]. For example, differential miNAR profiles in the highland yak compared to the lowland cow are enriched for hypoxia signaling pathways in the respiratory and cardiovascular systems – a key component of altitude adaptation [20]. These small non-coding RNAs also regulate seasonal phenotypic shifts, including metabolic depression that accompanies hibernation, anoxia tolerance and estivation in several vertebrates, although the exact miRNAs that may regulate these transitions appear species-specific [21, 22]. MiRNAs are also implicated in protection against environmental stresses such as hypoxia, extreme temperature and nutrient limitation in animals and plants [23–25].

In this study, we investigate a potential role for miRNAs in Weddell seal maturation, by post-transcriptional regulation of genes involved in development of the dive response and hypoxia tolerance. We provide the first comprehensive dataset of high quality, manually curated miRNA loci for this species, and use this dataset to investigate: 1) the patterns of differential expression of miRNAs across four tissues in pups and adults with a range of hypoxia sensitivity; 2) the putative mRNA targets for each miRNA; and 3) pathway analyses for the targets of differentially expressed miRNAs, focusing primarily of significant expression differences between pups and adults that could explain development of diving capacity.

## Results

### Sequencing, alignment, and annotation

As an initial quality check, we mapped all adapter-trimmed reads against the *LepWed1.0* genome assembly, with no gaps or mismatches allowed. Approximately 70% of reads obtained from tissues aligned perfectly to the Weddell seal genome and only 43-55% of reads from plasma samples were perfectly aligned (Fig. S1). These results illuminated a large portion of the read data that was unmapped to the Weddell seal genome (Fig. S2), ranging from 291,923 unmapped reads in a cortex sample to 2,076,932 in a plasma sample. Plasma, in particular, was over-represented in the unmapped data, with 5 of 6 plasma samples containing >1,000,000 unmapped reads. As several unaligned, putative miRNAs were highly expressed, we further identified reads that did not map to the Weddell seal genome, but were perfectly aligned to a miRNA hairpin sequence annotated in miRBase [26]. For example, the sequence TGAGATGAAGCACTGTAGCT was abundant in our dataset, with 532,097-1,014,236 reads in each sample. This sequence did not map to the Weddell seal genome but was identified as miR-143-3p. Thus, assembly incompleteness appears to reduce the percentage of small RNA reads that could be mapped to the Weddell seal genome, with the greatest impact on plasma samples.

In order to annotate high confidence miRNA loci, we manually curated the set of predictions provided by *miRCat2* [27] and *miRDeep2* [28]. This led to a final set of 559 loci (union of high confidence predictions for each tool; Table S1; Fig. S3-S5). Among these, 329 corresponded to a *miRBase* annotated hairpin sequence, while the remaining 230 represent novel miRNAs. Tissue specific expression plots for all novel and miRBase-annotated miRNAs were also generated (Fig. S5, Tables S2-S4).

We examined variation in the relative abundance of miRNA-5p and miRNA-3p across all samples (Fig. S6) and did not identify any clear examples of arm switching (i.e. changes of the most abundant miRNA strand). We also used our annotation to identify adenylated/uridylated miRNA isoforms and to investigate their representation in each sample. Plasma displayed the highest rates of 3’ end modification, with up to 12% of all reads corresponding to a modified isoform (Fig. S1).

### Differential miRNA expression among tissues

416 miRNAs were differentially expressed (DE) in a single sample type compared to all others. Among these, 74 were DE in two tissues (Fig. S7). 50% of tissue-specific DE occurred in the brain, with 31% of significant results in plasma. A smaller set of loci were uniquely elevated or depressed in heart (8%) and muscle (11%; Table 1). These two contractile tissues were also the most similar to each other when expression data was viewed as a heatmap (Fig. 1), although all 4 tissue types were segregated by hierarchical clustering. Principal Component Analysis (PCA) clearly separated samples by tissue along PC1 and PC2 (Fig. 2A). As with the heatmap, PCA points to greater similarity between the contractile tissues, heart and muscle, relative to plasma and brain. Moreover, brain is clearly separated from the other tissues along the PC1 axis, with limited inter-individual variability, especially for PC2 (Tables S6, S7). Random forests analyses identified 6 miRNA inputs that were able to separate the four tissue types from each other with zero error, including one novel, unmapped Weddell seal miRNA novel-4-3p, whose expression is significantly upregulated in plasma and downregulated in muscle (Table 2; Fig 3A). Four of these classifiers were upregulated in heart (miR-490-3p, miR-499-5p, miR-30e-5p and miR-30d-5p) and four were downregulated in plasma (miR-95-3p, miR-499-5p, miR-30a-5p and miR-30d-5p; Table S8).

**Fig.1.**
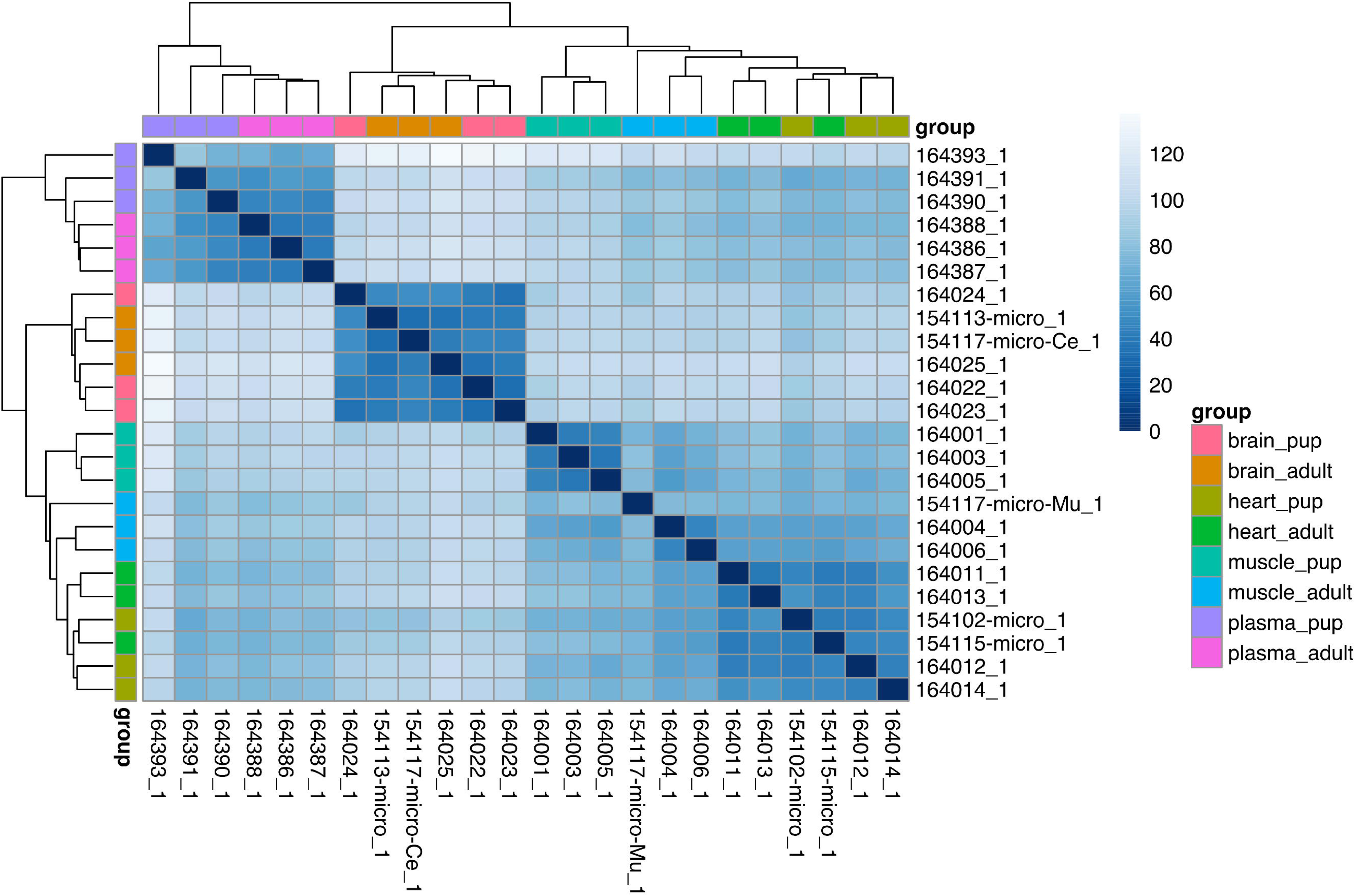
Heatmap of sample distances based on microRNA expression across 4 tissues. Gradient of blue corresponds to distance, while samples are color coded based on the tissue origin and developmental stage of the individual.

**Fig.2.**
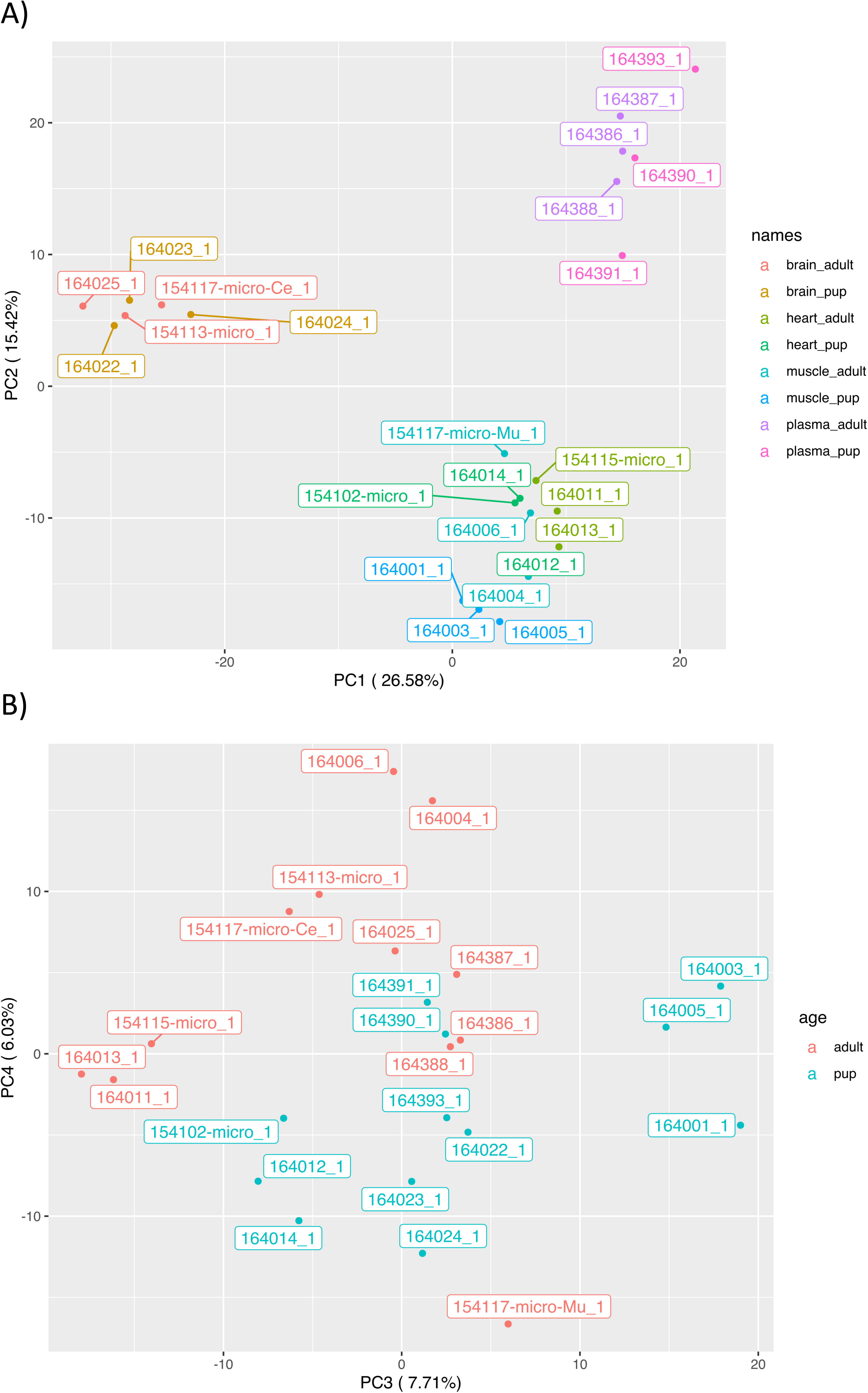
PCA plot based on microRNA expression across all 24 samples. A) PCA plot of principal components 1 and 2. Colour labels are based on both the tissue origin and developmental stage of the individual. B) PCA plot of principal components 3 and 4. Colour labels separate the two developmental stages (pup and adult).

**Fig.3.**
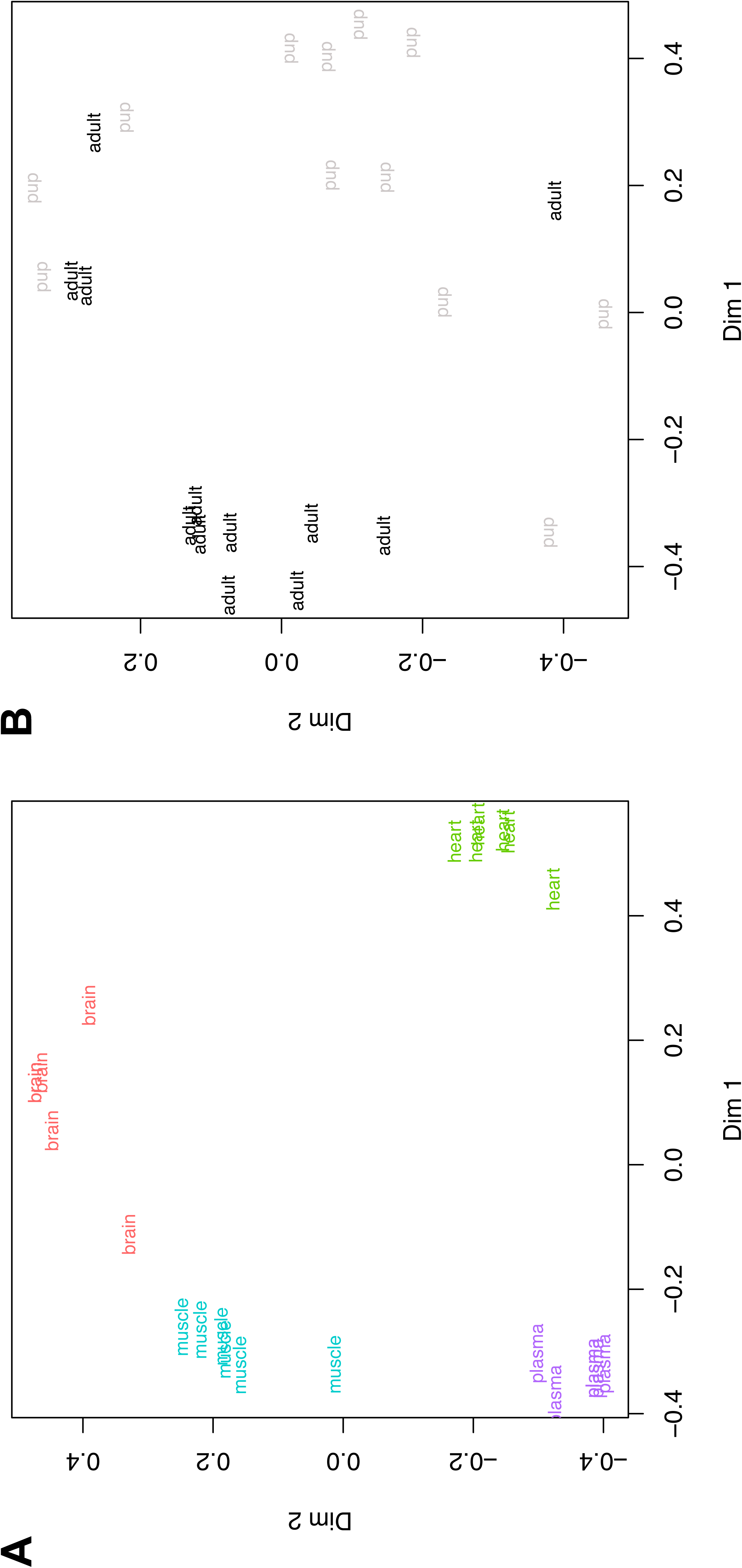
Random Forests plots classifying Weddell seal sample types based on a minimal set of microRNA abundance inputs. Panel A demonstrates clustering among tissue types (n=6 samples per tissue) using 10 microRNA inputs. Panel B demonstrates clustering between developmental stages (n=12 pup versus adult samples).

**Table 1.**
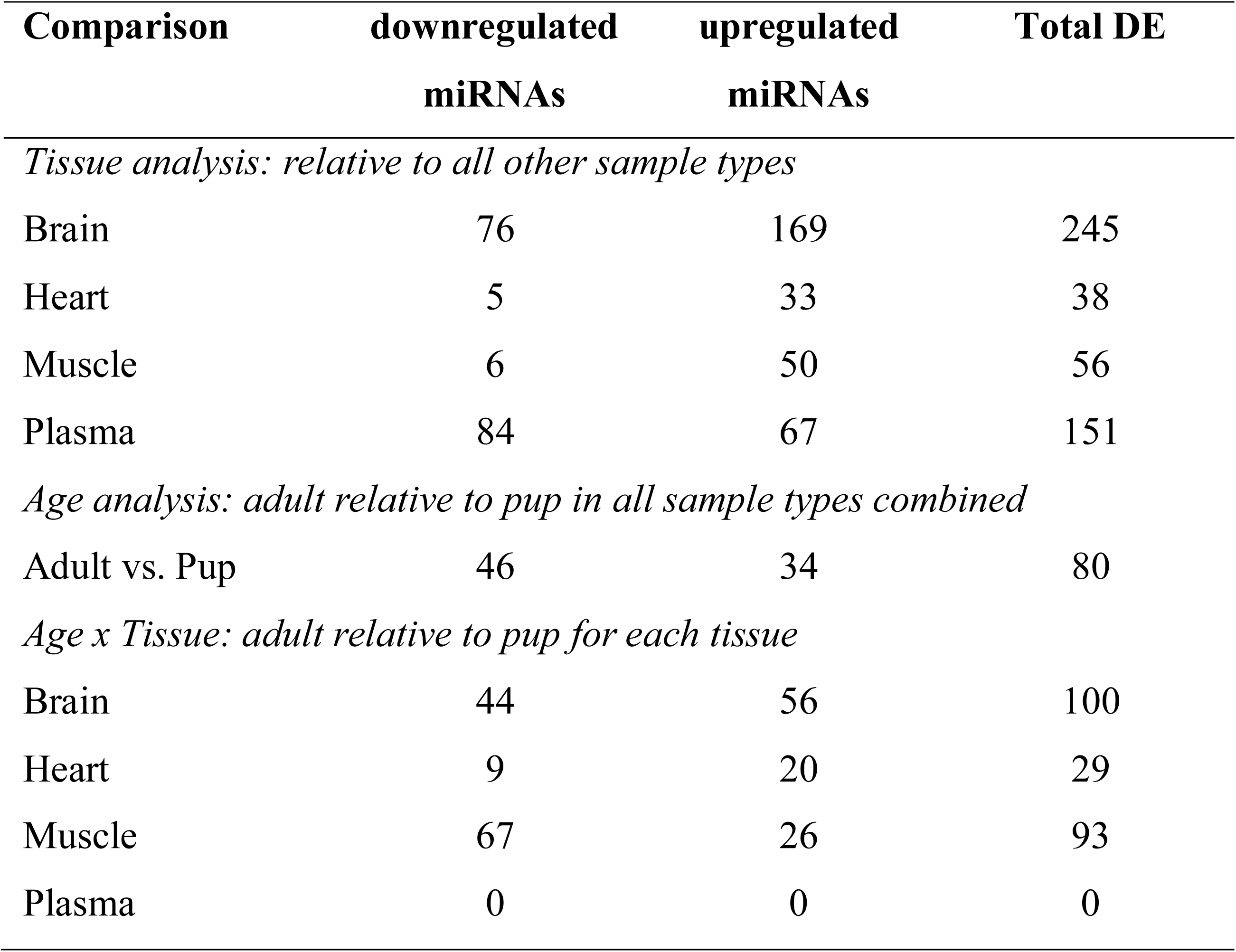
Differential expression (DE) of miRNAs across tissues, with developmental stage, and in tissue-specific developmental comparisons.

| Comparison | downregulated<br>miRNAs | upregulated<br>miRNAs | Total DE |
| --- | --- | --- | --- |
| *Tissue analysis: relative to all other sample types* |  |  |  |
| Brain | 76 | 169 | 245 |
| Heart | 5 | 33 | 38 |
| Muscle | 6 | 50 | 56 |
| Plasma | 84 | 67 | 151 |
| *Age analysis: adult relative to pup in all sample types combined* |  |  |  |
| Adult vs. Pup | 46 | 34 | 80 |
| *Age x Tissue: adult relative to pup for each tissue* |  |  |  |
| Brain | 44 | 56 | 100 |
| Heart | 9 | 20 | 29 |
| Muscle | 67 | 26 | 93 |
| Plasma | 0 | 0 | 0 |

**Table 2.** MiRNAs that best discriminate Weddell seal tissue sampling locations or developmental stages by Random Forests analysis.

| **MicroRNA Inputs** |
| --- |
| *Tissue Selection* |
| lwe-miR-499_loc1-5p |
| novel_4-3p |
| lwe-miR-490_loc1-3p |
| lwe-miR-95_loc1-3p |
| lwe-miR-30e_miRDeep2_loc1-5p |
| lwe-miR-30d_miRDeep2_loc1-5p |
| *Age Selection* |
| lwe-miR-29a_loc1-3p |
| lwe-miR-542_loc1-5p |

Note: Abundance (raw read counts) of the listed miRNA inputs were identified as those that best clustered the four tissues (brain, heart, skeletal muscle and plasma) or the two developmental groups (adult versus pup). Supervised classification in Random Forests analyses of the four tissue types had zero error, clustering of samples by developmental stage had $0.125 \pm 0.068$ out of bag error.

To shed light on the biological role of miRNAs of interest, we mapped mRNA targets of DE miRNAs to significantly enriched gene pathways. MiRNA downregulation within a tissue could infer higher local expression of target mRNAs. Therefore, the activity of pathways associated with upregulated miRNAs are potentially lowered, while pathways associated with downregulated miRNAs and their targets are potentially enhanced.

MiR-499-5p was the most DE locus in the dataset, a Random Forests classifier, and has the highest expression in heart (Table S8). mRNA targets of miR-499-5p, predicted to have lower expression in heart than other seal tissues include *Jade1*, a pro-apoptotic factor. miR-490-3p was also highly expressed in the heart, was a Random Forests classifier, and was among the five most significant DE changes in the dataset (Table S8). miR-490-3p targets ion channels and transporters, including members of the KCN and SLC gene families (*Kcng3*, *Kcnj15*, *Kcne2*, *Kcne4*, *Kcnmb1*, *Kcnip3*, *Kcnk12*, *Slc1A3*, *Slc9a1*, *Slc9a2*), associated with pathway enrichments in potassium ion transmembrane transport (GO:0071804, 0071805). High relative expressions of both miRNAs are consistent with prior miRNA biomarker studies in heart [29, 30], and do not differ between pups and adults. Although KEGG disease pathways were excluded from the presented data, it is noteworthy that pathway analysis identified enrichment of three cardiac disease pathways (hsa05410: Hypertrophic cardiomyopathy, hsa05414: Dilated cardiomyopathy, hsa05412: Arrhythmogenic right ventricular cardiomyopathy) associated with mRNA targets of miRNAs that were significantly upregulated in hearts of Weddell seals.

The brain had more DE miRNAs than any other tissue, impacting the largest number of mRNA targets. In general, targets of DE miRNAs in the seal brain were enriched for neuronal and developmental processes (Tables S8-S13). The largest pairwise fold change occurred in miR-488-3p (brain > plasma), which targets steroid hormone receptors including *Oxtr*, *Ghrhr*, and *Pgrmc2*. High relative expression in brain may downregulate the steroid hormone response pathway (GO:0048545).

MiR-296-3p was among 6 downregulated miRNAs in seal swimming muscle. Enhanced pathways associated with targets of this miRNA include those expected for skeletal muscle, related to ryanodine receptor redox state regulation, calcium handling and the sarcoplasmic reticulum (GO:0060314, 0033017, 0016529, 1901019, 0050848), as well as response to extracellular acidic pH (*Rab11fip5*, *Rab11b*, *Impact*, *Asic1*, *Asic2*, GO:0010447). We also identified elevated miR-206-3p in muscle, which [31] has been identified as a *MyomiR*, with expression restricted to this tissue. In contrast to heart, miR-490-3p was lowest in muscle, pointing to an enrichment of target ion channels and potassium transport. Both strands of miR-10 are upregulated in muscle. Although the 5p strand is most abundant (average 5p read counts: 2,233,312 in pups and 1,634,627 in adults; average 3p read counts: 413.6 in pups, 212 in adults), 3p has large tissue-specific fold changes (23-62X) compared to other tissues, downregulating predicted targets related to the regulation of blood circulation (GO:1903522), including *Bves*, *Casq2*, and *Nos1*.

Of the Random Forest classifiers DE in plasma (Table 2), miR-95-3p had the largest sample-specific downregulation (20-53X lower than heart, muscle, and brain; Table S8). miR-95-3p targets LDL receptor related protein (*Lrp1*) as well as the beta-subunit of guanylyl cyclase (*Gucy1b1*). miR-339-3p, also downregulated in plasma compared to other tissues, targets several key genes in hypoxia sensing (GO:0036293, GO:0070482, GO:0001666) including *Epas1*, *Vhl*, *Hif1an*, and *Cygb*, but does not change with age (Fig. S8). Several miRNAs elevated in Weddell seal plasma versus other tissues (miR-18a-5p, miR-221-3p, and miR-34-5p) target pathways regulating blood vessel and vascular remodeling (e.g. GO:0001974), with miR-34-5p specifically targeting *Epas1* and angiotensin (*Agt*).

### Differential miRNA expression with development

83 miRNAs were DE between adults and pups across all tissues (Table 1). Within each tissue, heatmap visualization also separates pups from adults (Fig. 1). An exception is the miRNA profile of adult skeletal muscle, which clusters more closely with heart samples than with muscle from pups. PC3 and PC4 provide the best developmental clustering (Fig. 2B) but explain only 13.7% of variation (Table S7). Supervised clustering by Random Forests (Fig.3B) identifies 2 miRNA inputs (miR-29a-3p, miR-542-5p) that separate the samples by developmental stage for all tissues combined (Table 3; Fig. 3B). MiR-29a-3p is significantly upregulated in heart and muscle of adults, whereas miR-542-5p is downregulated in adult brain and muscle (Table S10). Brain and skeletal muscle miRNAs had large developmental differences among the four sample types investigated (Table 1). Only 13% of significant adult-pup differences occurred in the heart and there were no developmental differences in plasma miRNAs (Table 1).

**Table 3.** The number of unique mRNA targets of differentially expressed miRNAs in four target tissues, with developmental stage, and in tissue-specific developmental comparisons.

| **Targets of DE**<br>**miRNAs** | **Targets of downregulated**<br>**miRNAs** | **Targets of upregulated**<br>**miRNAs** |
| --- | --- | --- |
| *Tissue analysis: relative to all other sample types* |  |  |
| **Brain** | 7450 | 10172 |
| **Heart** | 580 | 3962 |
| **Muscle** | 1643 | 4668 |
| **Plasma** | 7426 | 7249 |
| *Age analysis: adult relative to pup in all sample types combined* |  |  |
| **Adult vs. Pup** | 4125 | 4572 |
| *Age x Tissue: adult relative to pup for each tissue* |  |  |
| **Brain** | 4560 | 9111 |
| **Heart** | 920 | 2830 |
| **Muscle** | 5900 | 4144 |
| **Plasma** | 0 | 0 |

Predicted targets of miR-29a-3p that would be reduced in the transition between terrestrial pup and diving adult include components of the mitochondrial electron transport system (*Cox4i1*, *Cox10*, *Ndufs7*, *Ndor1*, *Sdhaf2*), genes related to iron regulation (*Tfrc*, *Ireb2*) and negative regulators of hypoxia signaling (*Hif3a*, *Vhl*). Conversely, higher miR-542-5p in pups is predicted to enhance the abundance of mRNAs that are likely relevant in the adult phenotype. Genes of interest include the mitochondrial citrate transporter *Slc25a1*, as well as genes known to regulate hematopoiesis (*Cd44*), anaerobic glycolysis (*Ldha*), and obesity/lipolysis (*Plin1*).

### High number of lineage specific miRNA orthogroups

Recent studies have highlighted the high rate of novel miRNA gains in mammals [32]. We identified 874 loci clusters in a dataset that included Weddell seal, cow, dog, horse, pig, rabbit, mouse, and human. These were considered miRNA orthogroups and were evaluated to infer the evolutionary patterns of gain and loss across the phylogenetic tree (Fig. 4). High net gain rates in both the dog and the Weddell Seal lineages suggest dynamic miRNA evolution. For the seal, we also observe a high number of lineage specific losses. This may reflect purifying selection on young, selectively neutral miRNAs, however this result might be biased by differences in assembly completeness between the *CanFam3.1* dog reference [33] and the *LepWed1.0* genome.

**Fig. 4.**
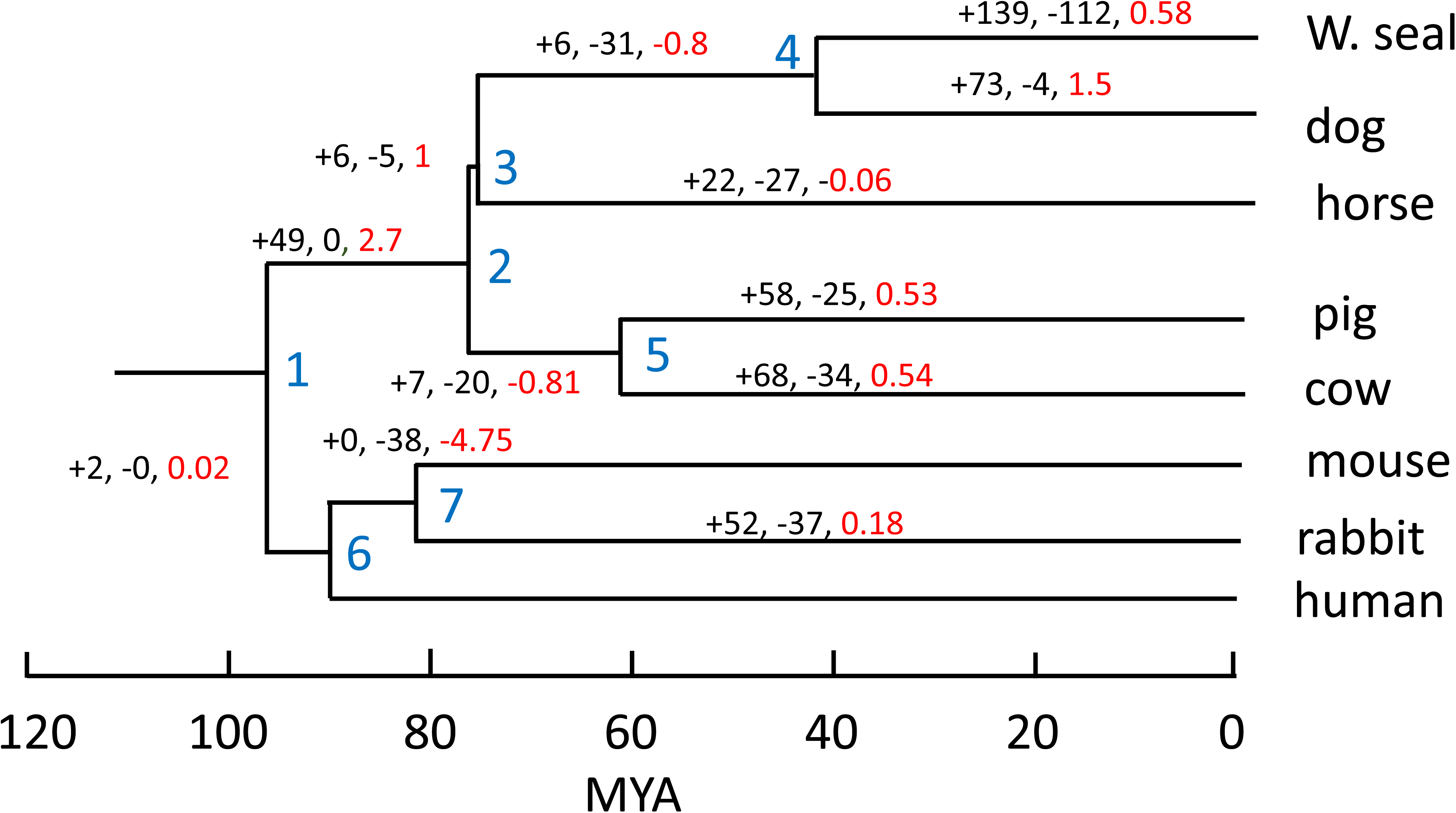
The gain and loss of mammalian miRNA orthogroups in 8 mammals, inferred by Dollo parsimony and synteny analyses. Number of gained (+) and lost (-) orthogroups are listed for each branch of the tree (black). Red numbers represent the branch-specific net gain rate of orthogroups per million year.

Novel Weddell seal miRNAs are more likely to be tissue-specific, as we observed a significantly higher proportion of tissue-specific miRNA families in the novel set compared to the miRBase set (z test, cortex: p=0.0001; heart: p<10^-4^; muscle: p<10^-4^; plasma: p<10^-4^; Fig. 5). This aligns with previous findings, which suggest that young miRNAs are initially expressed in a single or few tissues, then become more broadly expressed later in their evolutionary history [34].

**Fig. 5.**
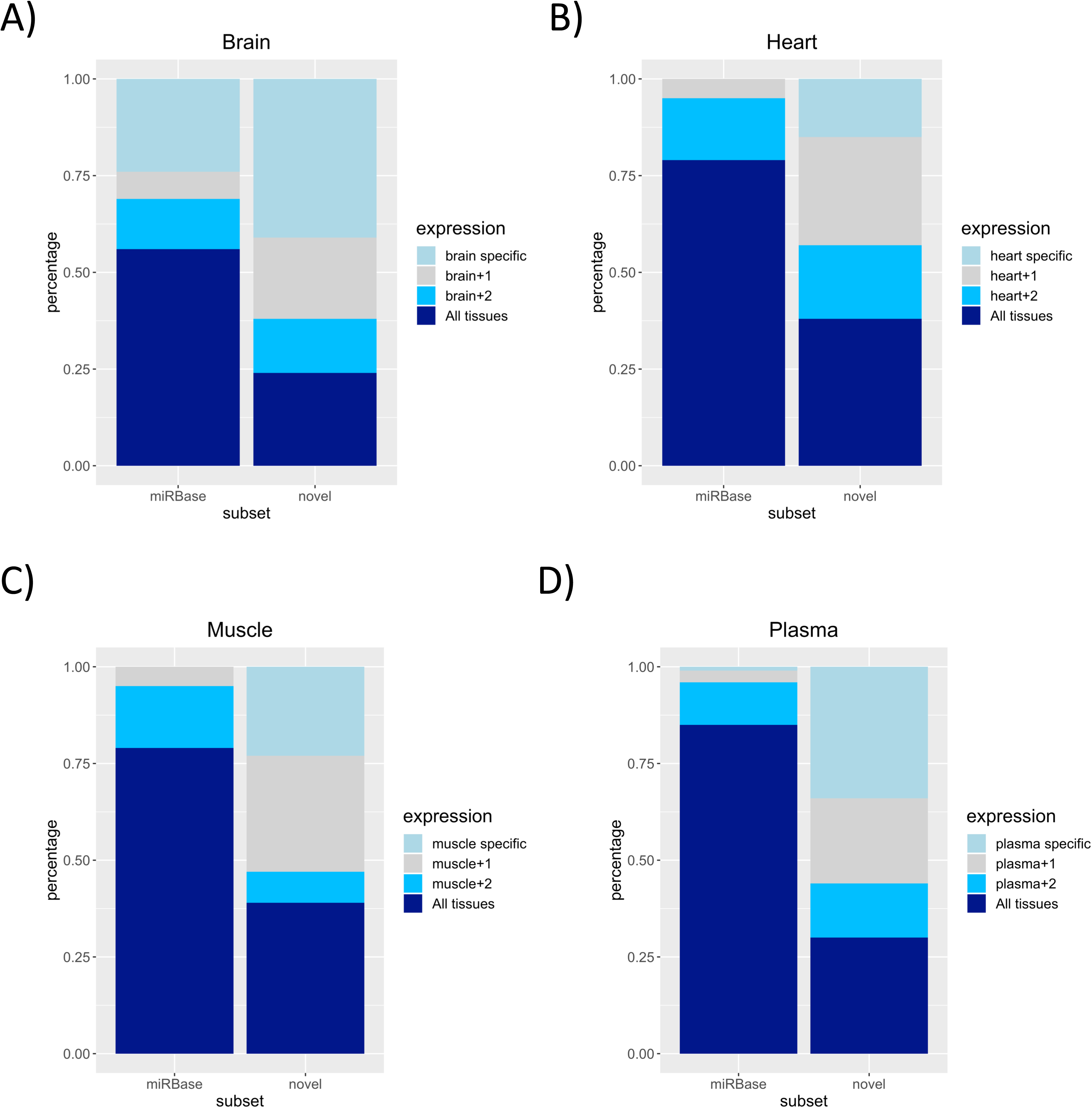
Expression patterns of novel and miRBase orthogroups in 4 sample types. The “miRBase” category contains orthogroups that contain annotated miRNAs, whereas orthogroups with unannotated members are classified as “Novel”. Orthogroups with mixed composition (containing both novel and miRBase genes) were classified as “miRBase”. Colouring denotes the percent of miRNA families expressed in each tissue that are tissue-specific, expressed in the tissue of interest and one (+1) or two (+2) other sample types, or are expressed in all tissues.

Predicted mRNA targets of novel Weddell seal miRNAs have a functional overlap with targets of annotated miRNAs in this dataset, as evidenced by similarities in pathway enrichments for tissue-specific up-and down-regulated miRNA targets, regardless of whether miRNAs were previously annotated or novel in Weddell seals (Tables S11-13). For example, VEGF signaling was targeted by both novel and annotated miRNAs in the seal heart, highlighting the importance of vascular development in cardiac tissue. We specifically investigated pathways identified from novel Weddell seal miRNAs, to evaluate the capability for species-specific gene regulation and manifestation of phenotype.

We compared pathways targeted by DE novel versus miRBase-annotated miRNAs, identifying unique pathway targets of novel miRNAs in each tissue. 37 pathways were enriched (p<0.05, see Methods) for the targets of miRNAs that were highest in the brain and 31 pathways were enriched for targets of muscle-elevated miRNAs, which are primarily signaling pathways (Tables S12, S13). Conversely, only two pathways in each tissue were associated with tissue-specifically downregulated miRNAs. Only a limited selection of pathways was enriched by novel miRNAs that were not similarly identified in pathway enrichments derived from annotated miRNAs. Several of these pathways are associated with lipid metabolism (Peroxisome in brain, ABC transporters in heart) and inflammatory signaling (Cytokine-cytokine receptor and Jak-stat signaling in plasma, CAMs in skeletal muscle), both important elements of the Weddell seal seal phenotype (Fig. S9).

Specific novel miRNAs also indicate potential to control key biochemical and physiological features of the Weddell seal. Novel-15-3p, highly expressed in the Weddell seal heart, is associated with cardiomyopathy through its target dystrophin. Novel-10-3p is upregulated in plasma and is predicted to target 610 mRNAs, including those with vasoactive (*Nos1*, *Nos1ap*), iron modulating (*Hmox1*, *Hfe*), and lipid metabolic functions (*Lep*).

*Targets of differentially expressed miRNAs control physiological changes associated with elite diving* Within each tissue, age related differences in gene expression, driven by miRNA regulation detected in this dataset likely support established developmental patterns of mammals generally [35, 36]. Beyond this, we examined the 221 miRNAs that are DE between adult and pup in either the brain, heart, or muscle specifically, which may regulate the development of the diving phenotype (Table S11). No age-specific differences were identified in plasma miRNAs (Table 1).

Several pathway enrichments and specific predicted target mRNAs related to hypoxia signaling were detected in this dataset. The main developmental signal linked to hypoxia responses was the expression of miR-424-3p, which is significantly reduced in the heart overall, and additionally downregulated in the adult seal compared to pup. *Egln3* is a prolyl hydroxylase involved in HIF1α degradation under normoxic conditions and is predicted to be targeted by miR-424-3p. *Egln3* also hydroxylates pyruvate kinase (PKM) in hypoxia, restricting glycolysis.

Each tissue type revealed DE miRNAs that target nitric oxide (NO) signaling (Fig. 6). Differentially upregulated miRNAs target the beta-1 subunit of guanylyl cyclase (*Gucy1b1*), which would decrease local expression of the heterodimeric enzyme in the brain, heart and muscle overall, whereas miRNAs targeting *Gucy1b1* were downregulated in plasma (Fig. 6). Within the brain, an additional upregulation of novel-25-5p and novel-119-5p was detected in adult seals, which also target *Gucy1b1* (Table S8). Guanylyl cyclase is a major target of NO and has previously been shown to be reduced in seal tissues seal tissues relative to terrestrial mammals [10]. The alpha-1 subunit (*Gucy1a1*) is also targeted by several DE miRNAs, including novel-128-5p (Fig. 6). Novel-128-5p is highly expressed in plasma and is also upregulated overall with age (Tables S8, S9). In addition to miRNAs that may regulate endothelial NO signal transduction via guanylyl cyclase, a number of miRNAs in this dataset target the NO synthase (*Nos1*) directly (Fig. 6), with age-related DE pointing to increased expression in adult muscle (miRNAs 377-5p, 214-5p, cfa-543-3p) but decreased expression in adult brain (miRNAs 34-5p, 1306-5p).

**Fig. 6.**
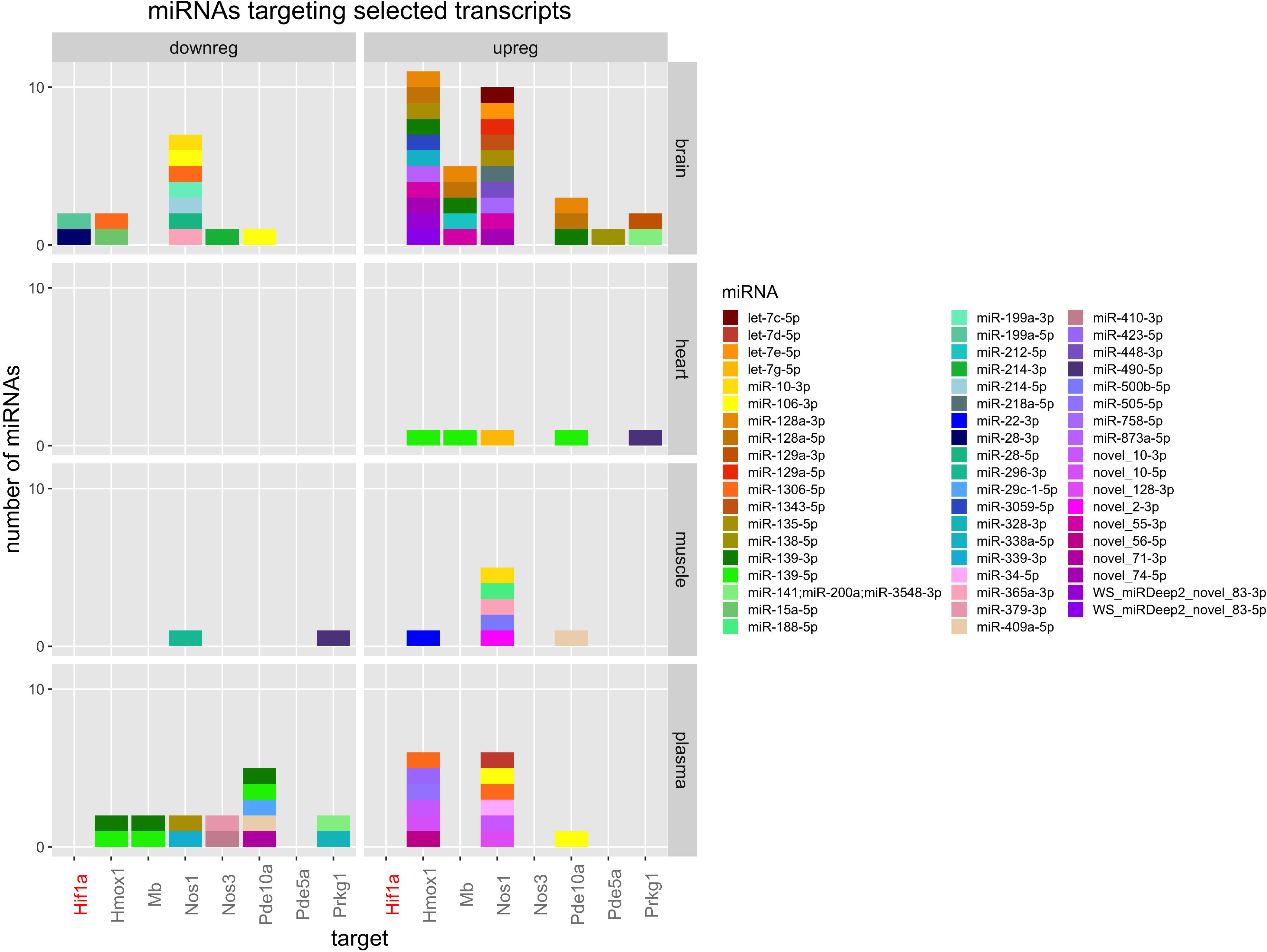
Selected targets of miRNAs DE between tissues. Rows correspond to tissues, columns to the direction of the expression change. Groups of miRNAs targeting the same gene in the same DE group are represented by stacked columns, with a height proportional to the number of different miRNAs. Each miRNA is assigned a specific color label (see legend).

Contrary to central tissues such as the heart and brain, muscles experience reduced and potentially episodic blood flow during diving [37, 38]. Targets of age-increased miR-218a-3p are enriched for ubiquitin/proteasome pathways, suggesting a less robust response to protein damage with development, as well as reduced apoptosis signaling (miR-29c-3p, miR-129a-3p).

Of 137 novel miRNA loci (133 orthogroups) identified in gain and loss evolution analyses as specific to the Weddell Seal lineage, 9 mature miRNAs are DE between adults and pups in the brain, 4 are DE with age in the heart, and 6 are DE with age in muscle (Table S10). Four of the novel miRNAs upregulated in the adult brain and 2 that are upregulated in the adult heart are predicted to target *Vash1*, an angiogenesis inhibitor selective to endothelial cells. Three miRNAs are upregulated in both the heart and skeletal muscle of adult seals and the predicted mRNA targets of these three miRNAs include *Ednra*, the receptor for endothelin 1 (a potent vasoconstrictor), as well as several members of the lipid transporter ABCA gene family (*Abca8*, *Abca10*, *Abca12*).

## Discussion

In this study, we provide the first annotation of the Weddell seal miRNAome, by manually curating the computational predictions of *MiRCat2* and *MiRDeep2*. We identified 559 high confidence miRNA loci; 146 of these represent novel miRNAs absent from *miRBase*. Evolutionary and comparative analyses highlight high lineage-specific miRNA gains in the Weddell Seal, compared to other mammals. Expression of these novel miRNAs tends to be tissue-specific, relative to *miRbase*-annotated miRNAs that are more widely expressed. mRNA targets were assigned to DE miRNAs using a highly conservative approach. As a result, differential miRNA expression by tissue identifies known miRNA biomarkers that have been linked to tissue-specific mRNAs and function, as well as previously undocumented mRNA signals of known and novel miRNAs that are likely associated with seal-specific physiology.

### Protective mechanisms relevant to diving in each tissue

Differential expression of cardiac miRNAs are the most limited in the dataset but represent some of the largest magnitude changes, which highlight miRNAs related to ischemic stress. In cardiomyocytes, miR-499-5p is protective against myocardial infarction due to anti-apoptotic action [39]. In particular, the pro-apoptotic *Jade1* target is predicted to be reduced in the seal heart due to high miR-499-5p, suggesting a protective mechanism against cardiac ischemia [40]. Circulating miR-499 is considered a biomarker of myocardial infarction [30] and this miRNA was not elevated in plasma nor affected by age, suggesting no underlying heart damage in seals despite the intense cardiac regulation involved in the dive response, and the reported potential for cardiac dysregulation in the conflict between diving and for exercise [41]. miR-499 is also important in ischemic postconditioning following cardiac challenge by ischemia-reperfusion [42]. MiR-30d-5p oxer-expression limits ischemic injury and has demonstrated experimental ischemic protection in other tissues [43]. MiR-490-3p is anti-atherosclerotic in human coronary artery smooth muscle cells [44].

Contrary to a central tissue such as the heart, skeletal muscles draw down local oxygen stores during diving [37, 38]. Exercise requirements of swimming muscle such as the *Longissimus dorsi* may be high as seals hunt below the ice, increasing reliance on anaerobic ATP production. *Asic3*, an acid channel and sensor for lactic acidosis [45] is predicted to be elevated in Weddell seal muscles via downregulation of miR-206-3p. Activation of *Asic3* contributes to neurally controlled local vasoconstriction, which would facilitate high peripheral vascular resistance and maintenance of the dive response despite hypoxia. Protective elements such as intrinsically high antioxidants have previously been identified in seal muscles [46, 47], and muscle miRNAs that are upregulated with age suggest dampening of other pathways that promote cell stress (ubiquitin/proteasome pathways, apoptosis signaling).

Brain and plasma have the most DE among miRNAs, likely to due with the high heterogeneity of tissue types found in the brain, and the contribution of many tissues releasing miRNAs into the venous circulation.

Deep diving Antarctic phocids, including Weddell seals, demonstrate hypercortisolemia, however extremely high cortisol binding capabilities produce low free circulating cortisol [48]. These physiological findings are speculated to be a component of protection for the deep diving brain, possibility preventing high pressure nervous syndrome. It is potentially relevant that the largest DE miRNA in the brain (miR-488-3p) also targets steroid hormone receptors.

### Protective mechanisms induced with age and development

Seal pups are novice divers, limited biochemically and physiologically from elite diving [49]. One important component of seal pup maturation is the development of body oxygen stores. Adults maintain aerobic metabolism during routine diving by large oxygen storage abilities of the blood and muscle [50–52]. Rapid synthesis of respiratory pigments (hemoglobin and myoglobin) requires iron, and seal pups do not appear to meet iron needs through intake during the nursing period, requiring mobilization of internal irons reserves [53, 54]. Both *Tfrc* and *Ireb2* respond to low cellular iron levels [55, 56], and their expression could be diminished by miRNAs during development. This may be a regulating factor that minimizes the consequences of high iron turnover and depletion through respiratory pigment biosynthesis.

Increased skeletal muscle LDH activities to exploit anaerobic metabolism in extended diving have been shown in developing seals [57], however the ability to detect statistical differences between pups and adults appears to depend on the species-specific maturity of pups at birth, or pup age at the time of sampling [58]. No significant developmental difference in LDH enzyme activity was detected in Weddell seal *Longissimus dorsi* when 3-5 week old pups were compared to adults [59], however it is possible that this difference exists for younger pups (∼1 week old in this study). Downregulated elements of the skeletal muscle electron transport system with age, predicted from this study, are however consistent with significantly reduced mitochondrial volume densities in in Weddell seal adults compared to pups from the same study [59].

### Regulation of perfusion

Tissue-specific control of perfusion is also important to cardiovascular regulation during diving [38, 60–63]. In terrestrial animals, the vasodilator nitric oxide (NO) improves perfusion and oxygen delivery in hypoxia. In diving seals, a NO response could be detrimental, as high peripheral vascular resistance is required to maintain central arterial pressure with submergence bradycardia. Indeed, downregulation of the nitric oxide-guanylyl cyclase-cGMP signal transduction pathway is limited in Weddell seals [10], which may facilitate central nervous system control of the dive response, but also reduce production of harmful peroxynitrites in the presence of reactive oxygen species. The number of DE miRNAs targeting this pathway, in particular novel miRNAs strongly suggest that lineage-specific miRNA regulation contributes to control of local vasoregulators in the Weddell seal, and perhaps the evolution of diving capabilities. The most significant mRNA target of the pathway in this dataset is NO synthase 1 (*Nos1*). Although it is described as the neuronal form, *Nos1* has wide expression across tissues and in additional to functions in the brain, plays a role in systemic vascular smooth muscle [64] and myocardial function. Furthermore, *Nos1* is linked to calcium handling, as a regulator [65] and in its activation [66], and calcium signaling is enriched for mRNA targets of DE miRNAs in all tissues.

### Regulation of hypoxia tolerance

Despite extensive physiological adaptations to increase body oxygen stores and behavioral selection for aerobic diving, Weddell seals can experience dramatic, global hypoxemia during submergence [63]. This dataset detected a suite of miRNAs that target elements of HIF1α signaling, but only few miRNAs targeted HIF1α itself. For example, miR-424-3p, which regulates HIF1α through interactions with proteins in the ubiquitin-ligase system, as well as promotes angiogenesis *in vitro* as well as in mice, is significantly reduced in heart compared to other tissues and further downregulated in the adult heart compared to the pup. Although low at baseline in Weddell seals, this miRNA is hypoxia-responsive in human endothelial cells [67]. Further, miR-29a-3p expression is increased in heart and muscle of adults versus pups, which is predicted to decrease expression of *Hif3a* and *Vhl*, both negative regulators of hypoxia signal transduction by HIF1α. Similar miRNA targeting of hypoxia signaling pathways was highlighted as an adaptation to high altitude in the yak, however a distinct set of miRNAs of interest were identified [20].

### Novel microRNAs as evolutionary avenues to regulate lineage-specific function

Our results confirmed previous observations of highly dynamic miRNA evolution, and additionally highlighted high net gain rates in both the dog and the Weddell Seal lineages. Novel, Weddell seal-specific miRNAs are evolutionarily young, and may provide key insights into the evolution of the diving phenotype. Similar pathway targets were identified in the hypoxia tolerant diver compared to the high-altitude, hypoxia-tolerant yak; however, they appear to be under the control of different sets of DE miRNAs in the 2 species. This strongly suggests that lineage-specific regulation is crucial to developing phenotypic adaptation in individual species. Additionally, novel miRNAs highlight mRNA targets that were previously overlooked. ∼50% of novel miRNAs which are increased in the adult brain and heart target *Vash1*, an angiogenesis inhibitor selective to endothelial cells. These data could be interpreted to underscore the importance of maintaining vascularization and perfusion in two critical tissues. Three miRNAs are upregulated in both the heart and skeletal muscle of adult seals, implicating them as potential targets for future studies related to contractile tissue function, particularly as they share the potential to regulate the expression of the receptor for endothelin 1.

### Limitations and considerations

This analysis relied on developmental differences between pup and adult Weddell seals to highlight biological significance. Key differences among the tissues are informative towards the biology, but are difficult to interpret in the context of species-specific adaptations without comparison to background(s) of tissue-specific miRNAomes in other species. As with any genome study, this analysis is limited by the quality and completeness of available sequences. miRNA sequences and expression levels were generated specifically for this study, and subject to strict quality controls. The detection of miRNA targets relied on the 3’ UTR multiple alignments available in the *TargetScan7* database [68]. While our results were not experimentally confirmed, our approach was highly conservative, requiring miRNA-mRNA interactions to be independently predicted by three different algorithms. Moreover, miRNAs are capable of propagating regulatory signals by affecting the expression of other miRNAs [69], which were not considered. Functional confirmation of these interactions and their biological roles in the Weddell seal would be important future work towards understanding the molecular basis of the diving phenotype.

## Conclusions

In this study, we take the first steps towards answering the question of whether post-transcriptional regulation by miRNAs has played a role in the evolution of diving capacities in the Weddell Seal. To address this, we have performed differential expression analyses, and identified significant changes among tissues and between developmental stages. Targets of differentially expressed miRNAs are enriched for GO categories relevant to Weddell seal physiology, suggesting that miRNAs have indeed contributed to the evolution of extreme diving capabilities, and providing a new set of mechanistic hypotheses for future follow-ups. We also present a thorough miRNA annotation of the Weddell seal genome, representing a valuable resource for future biomedical and evolutionary studies. Elite diving seals have long been considered a potential model system in which to address human therapies associated with hypoxia and hyperlipidemia, however these links have been tentative due to the lack of available genomic resources.

Recent studies in vertebrates have highlighted expression divergence between vertebrate lineages, and the crucial role of gene regulation in the evolution of adaptive phenotypes [70–76]. The short sequence of a miRNA can quickly be gained or lost in an evolutionary lineage, possibly leading to the rapid acquisition of novel regulatory patterns. This study provides further indication of their evolutionary potential. It is yet another step towards a wider description of miRNA evolutionary dynamics in mammals, which can further clarify the contribution of gene regulatory mechanisms in the evolution of adaptive traits.

## Supporting information

Supplementary Table S1 small RNA reads alignments

## Methods

### Animals and sample collection

Samples were collected from Weddell seal adults and pups near McMurdo Station, Antarctica as part of a larger study [10]. Tissue samples (heart right ventricle, *longissimus dorsi* swimming muscle, cerebral cortex) were collected at necropsy from animals found dead of natural causes on the sea ice (pups were generally ∼1 week of age). Tissues were rinsed in cold PBS, blotted dry, then either snap frozen in liquid nitrogen or preserved in an RNA stabilization solution (RNAlater, ThermoScientific AM7021). Venous blood samples were collected into EDTA-vacutainers from healthy adults and weaned pups during a brief handling period as previously described [10]. Plasma was isolated from whole blood by centrifugation (3000*g* for 15min) within an hour of collection in Antarctica. All animal handling and sample export was conducted under appropriate scientific authorizations (National Marine Fisheries Service #19439, #18662, ACA #2106-005) and approved by the Massachusetts General Hospital IACUC.

### Library preparation and sequencing

We extracted RNA from a total of 24 samples (n=3 adult and n=3 pup samples from each of the 3 tissues plus plasma). Total tissue RNA was isolated from tissues using the *miRNeasy Mini* kit (Qiagen #74104), then enriched for small RNAs using the *RNeasy MinElute Cleanup* kit (Qiagen #74204). miRNAs were extracted from plasma (n=3 adults, n=3 weaned pups) using a miRNeasy Serum/Plasma kit (Qiagen #217184). Enriched fractions were sent to *Macrogen* (South Korea), for small RNA library preparation and sequencing. Libraries were constructed for 24 high quality samples (minimum 100 ng RNA). All 24 libraries were barcoded, then pooled for sequencing using *Illumina HiSeq 2500*, with 10% *PhiX* sequencing control (Illumina FC-110-1003) included to increase sequence diversity. Resulting read data from small RNAs were validated using FASTQC software [77].

### microRNA annotation

MiRNAs were annotated from the combined set of 24 libraries using *miRCat2* [27] and *miRDeep2* [28], generating two independent sets of putative miRNA loci. The genomic coordinates predicted by *miRCat2* and *MiRDeep2* were then merged using *Bedtools merge* to generate a non-overlapping set of loci. We then aligned small RNA reads of each library to our predicted hairpins. This generated a set of high confidence miRNA loci, following the methodology described in [78]. We annotated miRNA loci within the Weddell seal genome based on a combined set of small RNA libraries across all tissues and developmental stages using both *miRCat2* [27] and *miRDeep2* [28] (Fig. S3). Predictions were then filtered based on the following criteria: evidence of both miR-3p and miR-5p expression in small RNA reads; alignments against the predicted loci with no gaps or mismatches allowed (Tables S1-S4); miR-like hairpin secondary structure (as predicted using the *Vienna-RNA* package) [79]; and a minimum of 10 reads mapping perfectly to the predicted locus.

### Clustering analyses

Dendrograms based on hierarchical clustering were generated in *RStudio* version 1.2.1335 and plotted as a heatmap using the *R* package *DeSeq2* [80]. Strand specific miRNA expression data were used as input, after transformation with the *varianceStabilizingTransformation* function. Principal Components (PC) Analyses was performed in *RStudio* version 1.2.1335 using the *prcomp* function. Similar to the clustering analysis, transformed strand specific miRNA expression data was used as input. Supervised clustering was conducted using Random Forests [81] in variable selection mode to identify the minimum miRNA inputs required to best separate the samples by either tissue type (combining pup and adult data) or developmental stage (combining data from all tissues). Analysis was implemented using the *varSelRF* package [82] to eliminate the least important input variables based on 100,000 trees in the first forest and 50,000 trees in all subsequent iterations. 10 replicates of each analysis were performed, and miRNA inputs appearing in ≥3 outputs are reported. MDS plots were generated to visualize group clustering based on minimum input variable sets identified by Random Forests.

### Gain and loss evolution

To investigate miRNA evolution, we used our new Weddell seal annotation, the miRNA annotations of five domestic mammals presented in [78, 83], as well as the available miRBase annotations for human and mouse. All hairpin sequences were clustered using *CD-hit* [83] with 80% minimum identity. Following the method described in [78], we identified syntenic regions between pairs of genomes, thus recovering conserved microRNA loci which had been missed in one or more species (likely due to a lack of sequencing depth). In order to do so, all miRNA loci annotated in each species were aligned, using BLASTN, against the latest genome assemblies of the remaining 7 species. BLAST hits to each assembly were filtered for an e-valueD≤D10^−6^ and an alignment length of at least 40 nucleotides. These hits were treated as putative homologous miRNA loci. We then identified the closest protein coding gene upstream and downstream of the selected hits, as well as the gene containing the hit for all intragenic hits. Genes surrounding or containing the query (for example, a Weddell seal miRNA) and the subject sequences (for example, a BLAST hit in the canine genome) were compared, checking for the presence of at least one homologous pair of genes with conserved synteny structure (i.e. same upstream or downstream gene, both on the same or on the opposite strand with respect to the miRNA/BLAST hit). Any BLAST hit supported by synteny conservation of at least one protein coding gene was then considered an orthologous miRNA locus. The presence absence matrix of orthogroups across all 8 species was then updated, considering these synteny-supported orthologous loci as miRNA genes. We then used *Dollo* parsimony to infer the evolutionary patterns of gain and loss for each orthogroup across the phylogenetic tree. The patterns of gain and loss of miRNA clusters across the phylogeny was estimated using *dollop* from the package *phylip-3.696*.

### Differential expression analyses

We used *DeSeq2* [80] to identify miRNAs that were differentially expressed (DE) among tissues or between developmental stages (pup versus adult). Developmental stage was input as a fixed factor for the tissue comparison analysis. We adopted a conservative approach to identify tissue-specific DE miRNAs, identifying only the most highly (or lowly) expressed miRNAs by requiring that a given miRNA be similarly up-or down-regulated in all pairwise comparisons with the other tissues (i.e. a miRNA was considered up-regulated in brain if it was higher in brain compared to each of heart, muscle, and plasma). Moreover, for the DE miRNA to be considered in downstream analyses, we required the total number of mapped reads across biological replicates to be 30 or more in at least one triplet of biological replicates (e.g., a total of 30 reads across the 3 pup brain samples). This requirement filtered 11 upregulated miRNA strands (3 in brain, 1 in muscle and 7 in plasma) from further analysis. DE miRNA are presented as adult state relative to pup (*i.e.* an upregulated miRNA is higher in adults), and were evaluated first a single pairwise comparison, with tissue included as a fixed factor, and then in each tissue alone. Among the list of DE miRNAs between pup and adult in each tissue, one miRNA strand (downregulated in brain) was filtered based on the minimum read count criterion. *DeSeq2* results were corrected for false discovery rate using the method of Benjamini-Hochberg [84], and miRNAs were considered differentially expressed if FDR-corrected q value ≤ 0.05.

### Target predictions, gene ontology, and pathway analyses

We predicted miRNA target sites by comparing miRNA sequences (for 3p and 5p forms) from our new annotation against the UTR sequence alignments from 72 vertebrates available from *TargetScan7* [68] (7mer and 8mer interactions with a *weighted context++ score* ≤-0.1), *miRanda* [85] (score > 140), and *Pita* [86] (7mer and 8mer interactions with no mismatches and a score ≤-10). Only miRNA:target interactions (*i.e.* a specific miRNA targeting a particular 3’UTR site) predicted by all 3 tools were retained for gene enrichment analysis.

Gene ontology (GO) analyses were performed using the R package *gProfileR* [87] or with DAVID Functional Annotation [88, 89]. We performed GO analysis on each set of mRNA targets for DE miRNAs, using FDR p-value correction, compared to a custom background, represented by the complete (non-redundant) list of genes used for target prediction analysis. This was obtained by mapping the list of human Ensembl transcript ids (ENST) used for the target prediction analyses to the corresponding gene id (ENSG) using gene tables available from *TargetScan7*. To evaluate enriched pathways on all targets of DE miRNAs in a single category (e.g. upregulated in brain) we used DAVID (default settings, p<0.05) and applied the default human background. Significantly enriched pathways were identified by either KEGG classifiers with human disease categories removed or GO Biological Processes.

## Abbreviations

miRNA: microRNA
FDR: False Discovery Rate
ENST: Ensembl transcript ID
ENSG: Ensembl gene ID
DE: Differential expression/Differentially expressed

## Declarations

### Ethics approval and consent to participate

Handling of *L. weddellii* individuals and sample export was conducted under appropriate scientific authorizations (National Marine Fisheries Service #19439, #18662, ACA #2016-005) and approved by the Massachusetts General Hospital IACUC. Consent to participate is not applicable, since our study does not involve human subjects.

### Consent for publication

Not applicable.

### Availability of data and material

Sequencing data used in this study have been submitted to the SRA archive, under accession PRJNA542051.

### Authors’ contributions

LPD: data curation, investigation, formal analysis, methodology, visualisation, writing (original draft). WH: investigation, methodology, supervision, writing (review). AH: sample collection, data curation, conceptualization, investigation, methodology, analysis, writing (review). FDP: conceptualisation, investigation, methodology, supervision, funding acquisition, writing (review). All named authors have read and approved the final manuscript.

## Acknowledgements

Logistical support for tissue acquisition in Antarctica was provided by the U.S. National Science Foundation through the U.S. Antarctic Program. We gratefully acknowledge field team members for assistance. Thanks also to Valerio Joe Utzeri (Universita’ di Bologna, Italy) and Tarang Mehta (Earlham institute, UK) for the useful discussion on total RNA quality control and purification of miRNA enriched fraction. LPD, WH and FDP were supported by the BBSRC, Institute Strategic Programme Grant (BB/J004669/1); WH and FDP are supported by the BBSRC Core Strategic Programme Grant (BB/P016774/1). The Data Infrastructure group at EI are funded in part by EI’s BBSRC Core Strategic Programme (BBS/E/T/000PR9817). AGH was supported by NSF #1443554/1921491.

## Supplementary Material

**Supplementary Figure S1.** Read statistics considering different stages of the bioinformatic analysis: number of raw adapter trimmed reads, number of trimmed reads with a perfect match to the genome.

**Supplementary Figure S2.** Percentage of genome matching reads (no gap or mismatch) compared to the percentage of unmapped reads having a perfect match to a miRBase annotated miRNA.

**Supplementary Figure S3.** Venn diagram showing the intersection between miRCat2 and miRDeep2 predictions

**Supplementary Figure S4.** Secondary structure plots for all 559 annotated miRNA loci, as predicted using the *ViennaRNA* package. Green and purple colors highlight the miRNA-5p and miRNA-3p sequences, respectively.

**Supplementary Figure S5.** Abundance plots for all 559 annotated miRNA loci. Abundance (y axis) along the hairpin sequence (x axis) is defined as reads per million genome matching. Colors separate tissues, while fill and dotted lines denote pup and adult individuals, respectively.

**Supplementary Figure S6.** Comparison of miRNA-5p and miRNA-3p abundance (reads per million genome matching) for all 559 annotated miRNA loci. Colors separate tissues, while different shapes separate pup and adult individuals.

**Supplementary Figure S7.** Venn diagrams showing the overlap between sets of DE miRNAs for each comparison: “Brain”, “Heart”, “Muscle”, “Plasma” and “Adult vs. Pup”.

**Supplementary Figure S8.** Selected GO accessions enriched in the targets of miR-339-3p. The color gradient reflects the calculated q-value; different GO accessions are listed along the x-axis, while the fold change enrichment is shown on the y-axis.

**Supplementary Figure S9.** Venn diagrams demonstrate overlap between enriched pathways of mRNA targets predicted to vary across 4 sample types in Weddell seals. Enriched KEGG pathways were identified based on miRNAs significantly up (+) or down (-) regulated in each tissue, compared to all others. They were then categorized as pathways related to previously annotated, miRBase miRNAs versus novel miRNAs identified in the Weddell seal (novel +, novel -). For each direction of regulation, the identities of enriched pathways targeted by novel miRNAs that were not detected from annotated miRNAs with the same direction of differential expression are listed for each tissue.

**Supplementary Table S1:** small RNA reads aligned against the final set of 559 high confidence miRNA loci. For each locus, the full hairpin sequence is shown, followed by the set of reads (one per line) perfectly matching the locus (with the corresponding abundance) and the predictive secondary structure of the miRNA hairpin.

**Supplementary Table S2:** Number of reads mapped to each miRNA strand across all loci and in all samples

**Supplementary Table S3:** number of reads mapped to each miRNA locus across all samples

**Supplementary Table S4:** Abundance (: number of mapped reads) at each nucleotide position for each miRNA locus across all samples

**Supplementary Table S5.** Naming scheme for 24 samples from Weddell seals

**Supplementary Table S6.** rotation matrix from the PCA analysis of miRNA expression data.

**Supplementary Table S7.** Basic statistics for the PCA analysis on miRNA expression data.

**Supplementary Table S8.** Tissue-specific differential expression statistics considering all pairwise tissue comparisons (brain_heart, brain_muscle, brain_plasma, heart_muscle, heart_plasma, muscle_plasma). Fold change refers to changes from tissue_1 to tissue_2, as defined by column H

**Supplementary Table S9.** Age-specific differential expression statistics considering all tissues together. Fold change refers to changes from pup to adult (i.e. positive sign indicates upregulation in adult)

**Supplementary Table S10.** Tissue-specific differential expression statistics for developmental stage (adult versus pup)

**Supplementary Table S11.** Significant pathway enrichments in brain, heart, and muscle for mRNA targets of all microRNAs differentially expressed in Weddell seal maturation

**Supplementary Table S12.** Enriched KEGG pathways associated with mRNA targets of novel miRNAs upregulated (+) or downregulated (-) in four Weddell seal sample types

**Supplementary Table S13.** Number of significant pathway enrichments annotated to mRNA targets of novel Weddell seal microRNAs that were elevated (+) or decreased (-) in four sample types.

## References

1. Heerah K, Andrews-Goff V, Williams G, Sultan E, Hindell M, Patterson T, Charrassin J-B: Ecology of Weddell seals during winter: Influence of environmental parameters on their foraging behaviour. Deep Sea Research Part II: Topical Studies in Oceanography 2013, 88:23–33.

2. Zapol WM: Diving adaptations of the Weddell seal. Scientific American 1987, 256(6):100–107.

3. Uhen MD: Evolution of marine mammals: back to the sea after 300 million years. The Anatomical Record: Advances in Integrative Anatomy and Evolutionary Biology: Advances in Integrative Anatomy and Evolutionary Biology 2007, 290(6):514–522.

4. De Miranda MA, Schlater AE, Green TL, Kanatous SB: In the face of hypoxia: myoglobin increases in response to hypoxic conditions and lipid supplementation in cultured Weddell seal skeletal muscle cells. Journal of Experimental Biology 2012, 215(5):806–813.

5. Kanatous S, Davis R, Watson R, Polasek L, Williams T, Mathieu-Costello O: Aerobic capacities in the skeletal muscles of Weddell seals: key to longer dive durations? J Exp Biol 2002, 205(23):3601–3608.

6. Trumble SJ, Noren SR, Cornick LA, Hawke TJ, Kanatous SB: Age-related differences in skeletal muscle lipid profiles of Weddell seals: clues to developmental changes. Journal of Experimental Biology 2010, 213(10):1676–1684.

7. Mirceta S, Signore AV, Burns JM, Cossins AR, Campbell KL, Berenbrink M: Evolution of mammalian diving capacity traced by myoglobin net surface charge. Science 2013, 340(6138):1234192.

8. Ponganis PJ, Meir JU, Williams CL: In pursuit of Irving and Scholander: a review of oxygen store management in seals and penguins. J Exp Biol 2011, 214(20):3325–3339.

9. Scholander PF: The master switch of life. Scientific American 1963, 209(6):92–107.

10. Hindle AG, Allen KN, Batten AJ, Hückstädt LA, Turner-Maier J, Schulberg SA, Johnson J, Karlsson E, Lindblad-Toh K, Costa DP: Low guanylyl cyclase activity in Weddell seals: implications for peripheral vasoconstriction and perfusion of the brain during diving. American Journal of Physiology-Regulatory, Integrative and Comparative Physiology 2019, 316(6):R704–R715.

11. Brennecke J, Hipfner DR, Stark A, Russell RB, Cohen SM: Bantam encodes a developmentally regulated microRNA that controls cell proliferation and regulates the proapoptotic gene hid in Drosophila. Cell 2003, 113(1):25–36.

12. Bushati N, Stark A, Brennecke J, Cohen SM: Temporal reciprocity of miRNAs and their targets during the maternal-to-zygotic transition in Drosophila. Current Biology 2008, 18(7):501–506.

13. Lee RC, Feinbaum RL, Ambros V: The C. elegans heterochronic gene lin-4 encodes small RNAs with antisense complementarity to lin-14. cell 1993, 75(5):843–854.

14. Xu P, Vernooy SY, Guo M, Hay BA: The Drosophila microRNA Mir-14 suppresses cell death and is required for normal fat metabolism. Current Biology 2003, 13(9):790–795.

15. Bartel DP: MicroRNAs: genomics, biogenesis, mechanism, and function. cell 2004, 116(2):281–297.

16. Carthew RW, Sontheimer EJ: Origins and mechanisms of miRNAs and siRNAs. Cell 2009, 136(4):642–655.

17. Niwa R, Slack FJ: The evolution of animal microRNA function. Current opinion in genetics & development 2007, 17(2):145–150.

18. Farh KK-H, Grimson A, Jan C, Lewis BP, Johnston WK, Lim LP, Burge CB, Bartel DP: The widespread impact of mammalian MicroRNAs on mRNA repression and evolution. Science 2005, 310(5755):1817–1821.

19. Sevignani C, Calin GA, Siracusa LD, Croce CM: Mammalian microRNAs: a small world for fine-tuning gene expression. Mammalian genome 2006, 17(3):189–202.

20. Guan J, Long K, Ma J, Zhang J, He D, Jin L, Tang Q, Jiang A, Wang X, Hu Y: Comparative analysis of the microRNA transcriptome between yak and cattle provides insight into high-altitude adaptation. PeerJ 2017, 5:e3959.

21. Biggar KK, Storey KB: Functional impact of microRNA regulation in models of extreme stress adaptation. Journal of molecular cell biology 2018, 10(2):93–101.

22. Riggs CL, Summers A, Warren DE, Nilsson GE, Lefevre S, Dowd W, Milton S, Podrabsky JE: Small non-coding RNA expression and vertebrate anoxia tolerance. Frontiers in genetics 2018, 9:230.

23. Hadj-Moussa H, Logan SM, Seibel BA, Storey KB: Potential role for microRNA in regulating hypoxia-induced metabolic suppression in jumbo squids. Biochimica et Biophysica Acta (BBA)-Gene Regulatory Mechanisms 2018, 1861(6):586–593.

24. Lyons PJ, Lang-Ouellette D: CryomiRs: towards the identification of a cold-associated family of microRNAs. Comparative Biochemistry and Physiology Part D: Genomics and Proteomics 2013, 8(4):358–364.

25. Nehammer C, Podolska A, Mackowiak SD, Kagias K, Pocock R: Specific microRNAs regulate heat stress responses in Caenorhabditis elegans. Scientific reports 2015, 5:8866.

26. Castellini MA, Davis RW, Davis M, Horning M: Antarctic marine life under the McMurdo Ice Shelf at White Island: A link between nutrient influx and seal population. Polar Biol 1984, 2(4):229–231.

27. Paicu C, Mohorianu I, Stocks M, Xu P, Coince A, Billmeier M, Dalmay T, Moulton V, Moxon S: miRCat2: accurate prediction of plant and animal microRNAs from next-generation sequencing datasets. Bioinformatics 2017, 33(16):2446–2454.

28. Friedländer MR, Mackowiak SD, Li N, Chen W, Rajewsky N: miRDeep2 accurately identifies known and hundreds of novel microRNA genes in seven animal clades. Nucleic acids research 2011, 40(1):37–52.

29. Ji X, Takahashi R, Hiura Y, Hirokawa G, Fukushima Y, Iwai N: Plasma miR-208 as a biomarker of myocardial injury. Clinical chemistry 2009, 55(11):1944–1949.

30. Xin Y, Yang C, Han Z: Circulating miR-499 as a potential biomarker for acute myocardial infarction. Annals of translational medicine 2016, 4(7).

31. Zhang S, Chen N: Regulatory role of MicroRNAs in muscle atrophy during exercise intervention. International journal of molecular sciences 2018, 19(2):405.

32. Meunier J, Lemoine F, Soumillon M, Liechti A, Weier M, Guschanski K, Hu H, Khaitovich P, Kaessmann H: Birth and expression evolution of mammalian microRNA genes. Genome Res 2013, 23(1):34–45.

33. Hoeppner MP, Lundquist A, Pirun M, Meadows JR, Zamani N, Johnson J, Sundström G, Cook A, FitzGerald MG, Swofford R: An improved canine genome and a comprehensive catalogue of coding genes and non-coding transcripts. PloS one 2014, 9(3):e91172.

34. Meunier J, Lemoine F, Soumillon M, Liechti A, Weier M, Guschanski K, Hu H, Khaitovich P, Kaessmann H: Birth and expression evolution of mammalian microRNA genes. Genome research 2013, 23(1):34–45.

35. Lee CT, Risom T, Strauss WM: MicroRNAs in mammalian development. Birth Defects Research Part C: Embryo Today: Reviews 2006, 78(2):129–139.

36. Song L, Tuan RS: MicroRNAs and cell differentiation in mammalian development. Birth Defects Research Part C: Embryo Today: Reviews 2006, 78(2):140–149.

37. Davis RW, Kanatous SB: Convective oxygen transport and tissue oxygen consumption in Weddell seals during aerobic dives. Journal of Experimental Biology 1999, 202(9):1091–1113.

38. Guyton GP, Stanek KS, Schneider RC, Hochachka PW, Hurford WE, Zapol DG, Liggins GC, Zapol WM: Myoglobin saturation in free-diving Weddell seals. Journal of Applied Physiology 1995, 79(4):1148–1155.

39. Li Y, Lu J, Bao X, Wang X, Wu J, Li X, Hong W: MiR-499-5p protects cardiomyocytes against ischaemic injury via anti-apoptosis by targeting PDCD4. Oncotarget 2016, 7(24):35607.

40. Tang J, Zhuang S: Histone acetylation and DNA methylation in ischemia/reperfusion injury. Clinical Science 2019, 133(4):597–609.

41. Williams TM, Fuiman LA, Kendall T, Berry P, Richter B, Noren SR, Thometz N, Shattock MJ, Farrell E, Stamper AM: Exercise at depth alters bradycardia and incidence of cardiac anomalies in deep-diving marine mammals. Nature communications 2015, 6:6055.

42. Zhu J, Yao K, Wang Q, Guo J, Shi H, Ma L, Liu H, Gao W, Zou Y, Ge J: Ischemic postconditioning-regulated miR-499 protects the rat heart against ischemia/reperfusion injury by inhibiting apoptosis through PDCD4. Cellular Physiology and Biochemistry 2016, 39(6):2364–2380.

43. Jiang M, Wang H, Jin M, Yang X, Ji H, Jiang Y, Zhang H, Wu F, Wu G, Lai X: Exosomes from MiR-30d-5p-ADSCs reverse acute ischemic stroke-induced, autophagy-mediated brain injury by promoting M2 microglial/macrophage polarization. Cellular Physiology and Biochemistry 2018, 47(2):864–878.

44. Sun Y, Chen D, Cao L, Zhang R, Zhou J, Chen H, Li Y, Li M, Cao J, Wang Z: MiR-490-3p modulates the proliferation of vascular smooth muscle cells induced by ox-LDL through targeting PAPP-A. Cardiovascular research 2013, 100(2):272–279.

45. Birdsong WT, Fierro L, Williams FG, Spelta V, Naves LA, Knowles M, Marsh-Haffner J, Adelman JP, Almers W, Elde RP: Sensing muscle ischemia: coincident detection of acid and ATP via interplay of two ion channels. Neuron 2010, 68(4):739–749.

46. Allen K, Hindle A, Vázquez-Medina JP, Lawler JM, Mellish J-AE, Horning M: Age-and Muscle-Specific Oxidative Stress Management Strategies in a Long-Lived Diver, the Weddell Seal. The FASEB Journal 2018, 32(1_supplement):861.865–861.865.

47. Vázquez-Medina JP, Zenteno-Savín T, Elsner R: Antioxidant enzymes in ringed seal tissues: potential protection against dive-associated ischemia/reperfusion. Comparative Biochemistry and Physiology Part C: Toxicology & Pharmacology 2006, 142(3-4):198–204.

48. Liggins GC, France JT, Schneider RC, Knox BS, Zapol WM: Concentrations, metabolic clearance rates, production rates and plasma binding of cortisol in Antarctic phocid seals. European Journal of Endocrinology 1993, 129(4):356–359.

49. Burns JM: The development of diving behavior in juvenile Weddell seals: pushing physiological limits in order to survive. Canadian Journal of Zoology 1999, 77(5):737–747.

50. Kooyman G, Castellini M, Davis R, Maue R: Aerobic diving limits of immature Weddell seals. Journal of Comparative Physiology B: Biochemical, Systemic, and Environmental Physiology 1983, 151(2):171–174.

51. Kooyman G, Wahrenbrock E, Castellini M, Davis R, Sinnett E: Aerobic and anaerobic metabolism during voluntary diving in Weddell seals: Evidence of preferred pathways from blood chemsitry and behavior. Journal of Comparative Physiology B: Biochemical, Systemic, and Environmental Physiology 1980, 138(4):335–346.

52. Ponganis PJ, Kooyman GL, Castellini MA: Determinants of the aerobic dive limit of Weddell seals: analysis of diving metabolic rates, postdive end tidal PO2’s, and blood and muscle oxygen stores. Physiological Zoology 1993, 66(5):732–749.

53. Burns J, Hammill M: Does iron availability limit oxygen store development in seal pups. In: Proceedings of the 4th CPB Meeting in Africa: MARA: 2008. 417–428.

54. Geiseler SJ, Blix AS, Burns JM, Folkow LP: Rapid postnatal development of myoglobin from large liver iron stores in hooded seals. Journal of Experimental Biology 2013, 216(10):1793–1798.

55. Camaschella C, Roetto A, De Gobbi M: Genetic haemochromatosis: genes and mutations associated with iron loading. Best Practice & Research Clinical Haematology 2002, 15(2):261–276.

56. LaVaute T, Smith S, Cooperman S, Iwai K, Land W, Meyron-Holtz E, Drake SK, Miller G, Abu-Asab M, Tsokos M: Targeted deletion of the gene encoding iron regulatory protein-2 causes misregulation of iron metabolism and neurodegenerative disease in mice. Nature genetics 2001, 27(2):209.

57. Prewitt J, Freistroffer D, Schreer J, Hammill M, Burns J: Postnatal development of muscle biochemistry in nursing harbor seal (Phoca vitulina) pups: limitations to diving behavior? Journal of Comparative Physiology B 2010, 180(5):757–766.

58. Burns J, Skomp N, Bishop N, Lestyk K, Hammill M: Development of aerobic and anaerobic metabolism in cardiac and skeletal muscles from harp and hooded seals. Journal of Experimental Biology 2010, 213(5):740–748.

59. Kanatous S, Hawke T, Trumble S, Pearson L, Watson R, Garry D, Williams T, Davis R: The ontogeny of aerobic and diving capacity in the skeletal muscles of Weddell seals. Journal of Experimental Biology 2008, 211(16):2559–2565.

60. Davis R, Castellini M, Kooyman G, Maue R: Renal glomerular filtration rate and hepatic blood flow during voluntary diving in Weddell seals. American Journal of Physiology-Regulatory, Integrative and Comparative Physiology 1983, 245(5):R743–R748.

61. Liggins G, Qvist J, Hochachka P, Murphy B, Creasy R, Schneider R, Snider M, Zapol W: Fetal cardiovascular and metabolic responses to simulated diving in the Weddell seal. Journal of Applied Physiology 1980, 49(3):424–430.

62. McKnight JC, Bennett KA, Bronkhorst M, Russell DJ, Balfour S, Milne R, Bivins M, Moss SE, Colier W, Hall AJ: Shining new light on mammalian diving physiology using wearable near-infrared spectroscopy. PLoS biology 2019, 17(6):e3000306.

63. Zapol WM, Liggins G, Schneider RC, Qvist J, Snider MT, Creasy RK, Hochachka PW: Regional blood flow during simulated diving in the conscious Weddell seal. Journal of Applied Physiology 1979, 47(5):968–973.

64. Brophy CM, Knoepp L, Xin J, Pollock JS: Functional expression of NOS 1 in vascular smooth muscle. American Journal of Physiology-Heart and Circulatory Physiology 2000, 278(3):H991–H997.

65. Khan SA, Skaf MW, Harrison RW, Lee K, Minhas KM, Kumar A, Fradley M, Shoukas AA, Berkowitz DE, Hare JM: Nitric oxide regulation of myocardial contractility and calcium cycling: independent impact of neuronal and endothelial nitric oxide synthases. Circulation research 2003, 92(12):1322–1329.

66. Weissman BA, Jones CL, Liu Q, Gross SS: Activation and inactivation of neuronal nitric oxide synthase: characterization of Ca2+-dependent [125I] Calmodulin binding. European journal of pharmacology 2002, 435(1):9–18.

67. Ghosh G, Subramanian IV, Adhikari N, Zhang X, Joshi HP, Basi D, Chandrashekhar Y, Hall JL, Roy S, Zeng Y: Hypoxia-induced microRNA-424 expression in human endothelial cells regulates HIF-α isoforms and promotes angiogenesis. The Journal of clinical investigation 2010, 120(11):4141–4154.

68. Agarwal V, Bell GW, Nam J-W, Bartel DP: Predicting effective microRNA target sites in mammalian mRNAs elife 2015, 4:e05005.

69. Matkovich SJ, Hu Y, Dorn GW: Regulation of cardiac microRNAs by cardiac microRNAs. Circulation research 2013, 113(1):62–71.

70. Carneiro M, Rubin CJ, Di Palma F, Albert FW, Alfoldi J, Barrio AM, Pielberg G, Rafati N, Sayyab S, Turner-Maier J et al: Rabbit genome analysis reveals a polygenic basis for phenotypic change during domestication. Science 2014, 345(6200):1074–1079.

71. Brawand D, Wagner CE, Li YI, Malinsky M, Keller I, Fan S, Simakov O, Ng AY, Lim ZW, Bezault E et al: The genomic substrate for adaptive radiation in African cichlid fish. Nature 2014, 513(7518):375–381.

72. Mehta TK, Koch C, Nash W, Knaack SA, Sudhakar P, Olbei M, Bastkowski S, Penso-Dolfin L, Korcsmaros T, Haerty W et al: Evolution of regulatory networks controlling adaptive traits in cichlids. bioRxiv 2018.

73. Babbitt CC, Fedrigo O, Pfefferle AD, Boyle AP, Horvath JE, Furey TS, Wray GA: Both noncoding and protein-coding RNAs contribute to gene expression evolution in the primate brain. Genome Biol Evol 2010, 2:67–79.

74. Khaitovich P, Enard W, Lachmann M, Paabo S: Evolution of primate gene expression. Nat Rev Genet 2006, 7(9):693–702.

75. Hu HY, Guo S, Xi J, Yan Z, Fu N, Zhang X, Menzel C, Liang H, Yang H, Zhao M et al: MicroRNA expression and regulation in human, chimpanzee, and macaque brains. PLoS Genet 2011, 7(10):e1002327.

76. Jordan IK, Marino-Ramirez L, Koonin EV: Evolutionary significance of gene expression divergence. Gene 2005, 345(1):119–126.

77. Davis RW, Hagey W, Horning M: Monitoring the behavior and multi-dimensional movements of Weddell seals using an animal-borne video and data recorder. Memoirs of National Institute of Polar Research Special issue 2004, 58:148–154.

78. Penso-Dolfin L, Moxon S, Haerty W, Di Palma F: The evolutionary dynamics of microRNAs in domestic mammals. Scientific reports 2018, 8(1):17050.

79. Hofacker IL: R NA Secondary Structure Analysis Using the Vienna RNA Package. Current protocols in bioinformatics 2009, 26(1):12.12.11–12.12.16.

80. Love MI, Huber W, Anders S: Moderated estimation of fold change and dispersion for RNA-seq data with DESeq2. Genome Biol 2014, 15(12):550.

81. Breiman L: Random forests. Machine learning 2001, 45(1):5–32.

82. Diaz-Uriarte R: GeneSrF and varSelRF: a web-based tool and R package for gene selection and classification using random forest. BMC bioinformatics 2007, 8(1):328.

83. Fu L, Niu B, Zhu Z, Wu S, Li W: CD-HIT: accelerated for clustering the next-generation sequencing data. Bioinformatics 2012, 28(23):3150–3152.

84. Benjamini Y, Hochberg Y: Controlling the false discovery rate: a practical and powerful approach to multiple testing. Journal of the Royal statistical society: series B (Methodological) 1995, 57(1):289–300.

85. Enright AJ, John B, Gaul U, Tuschl T, Sander C, Marks DS: MicroRNA targets in Drosophila. Genome biology 2003, 5(1):R1.

86. Kertesz M, Iovino N, Unnerstall U, Gaul U, Segal E: The role of site accessibility in microRNA target recognition. Nature genetics 2007, 39(10):1278.

87. Raudvere U, Kolberg L, Kuzmin I, Arak T, Adler P, Peterson H, Vilo J: g: Profiler: a web server for functional enrichment analysis and conversions of gene lists (2019 update). Nucleic acids research 2019.

88. Huang DW, Sherman BT, Lempicki RA: Bioinformatics enrichment tools: paths toward the comprehensive functional analysis of large gene lists. Nucleic acids research 2008, 37(1):1–13.

89. Sherman BT, Lempicki RA: Systematic and integrative analysis of large gene lists using DAVID bioinformatics resources. Nature protocols 2009, 4(1):44–57.

