## Supplementary Figures for "microRNA profiling in the Weddell Seal suggests novel regulatory mechanisms contributing to diving adaptation"

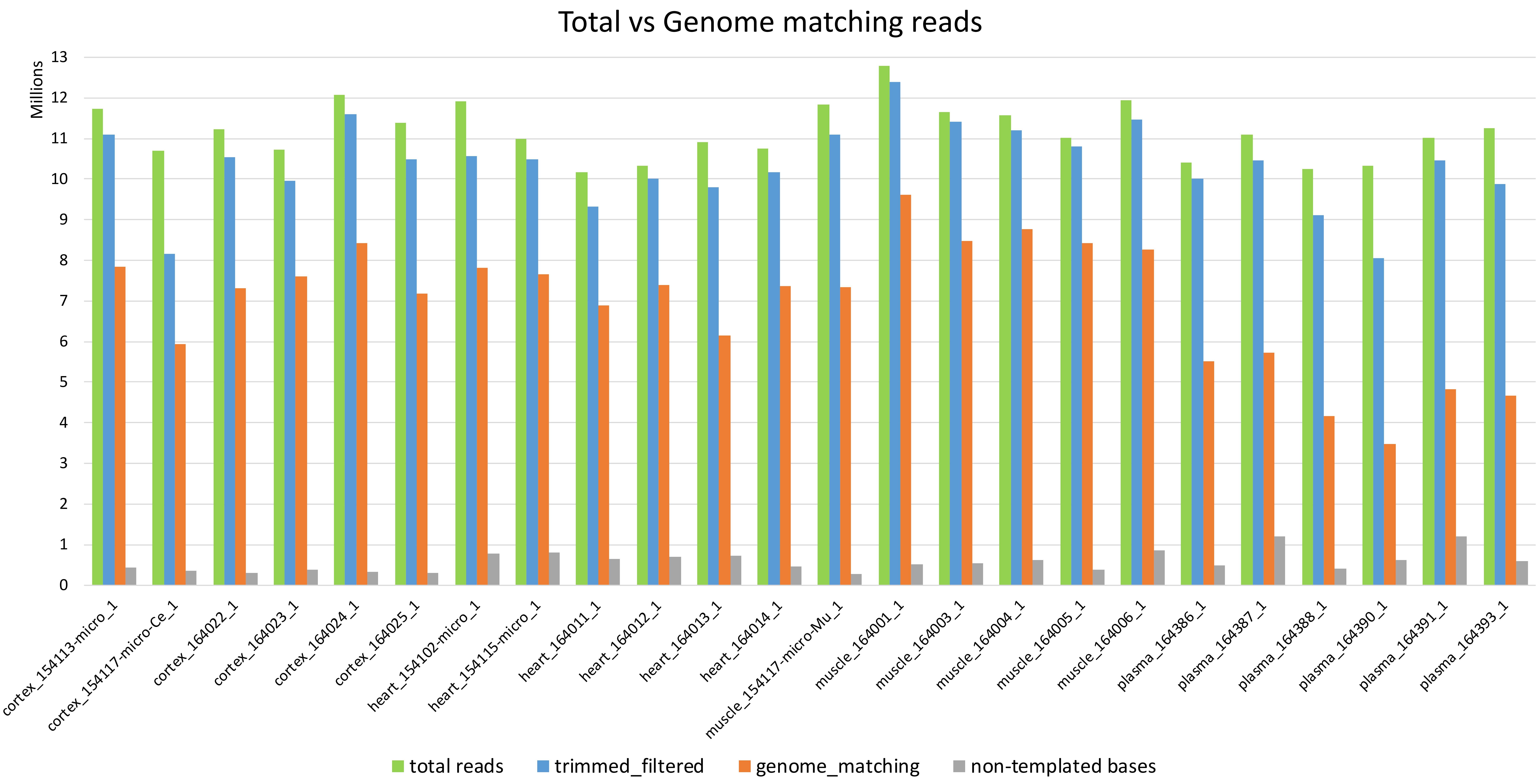


Supplementary Figure S1. Read statistics considering different stages of the bioinformatic analysis: number of raw adapter trimmed reads, number of trimmed reads with a perfect match to the genome.

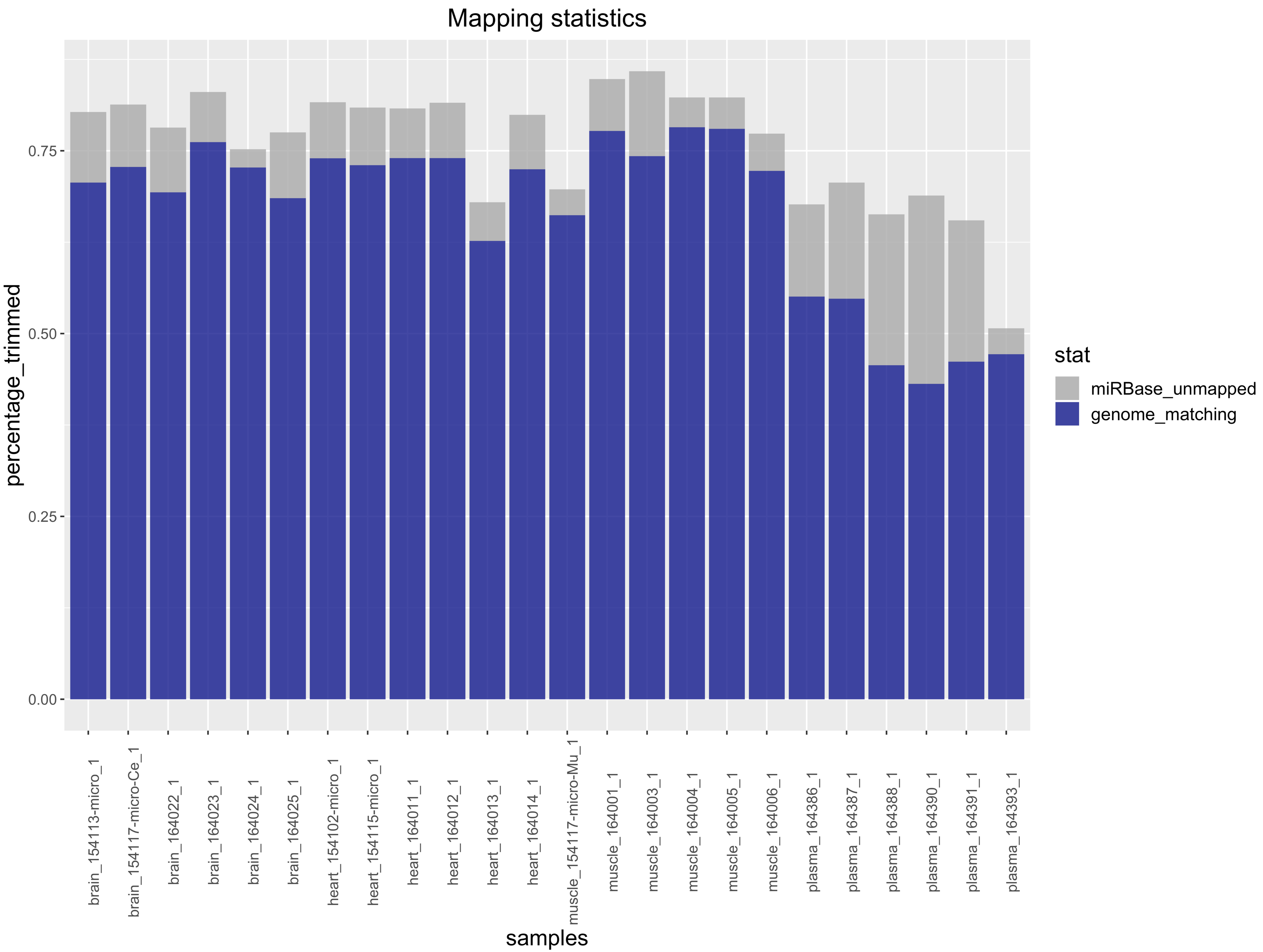


Supplementary Figure S2. Percentage of genome matching reads (no gap or mismatch) compared to the percentage of unmapped reads having a perfect match to a miRBase annotated miRNA.

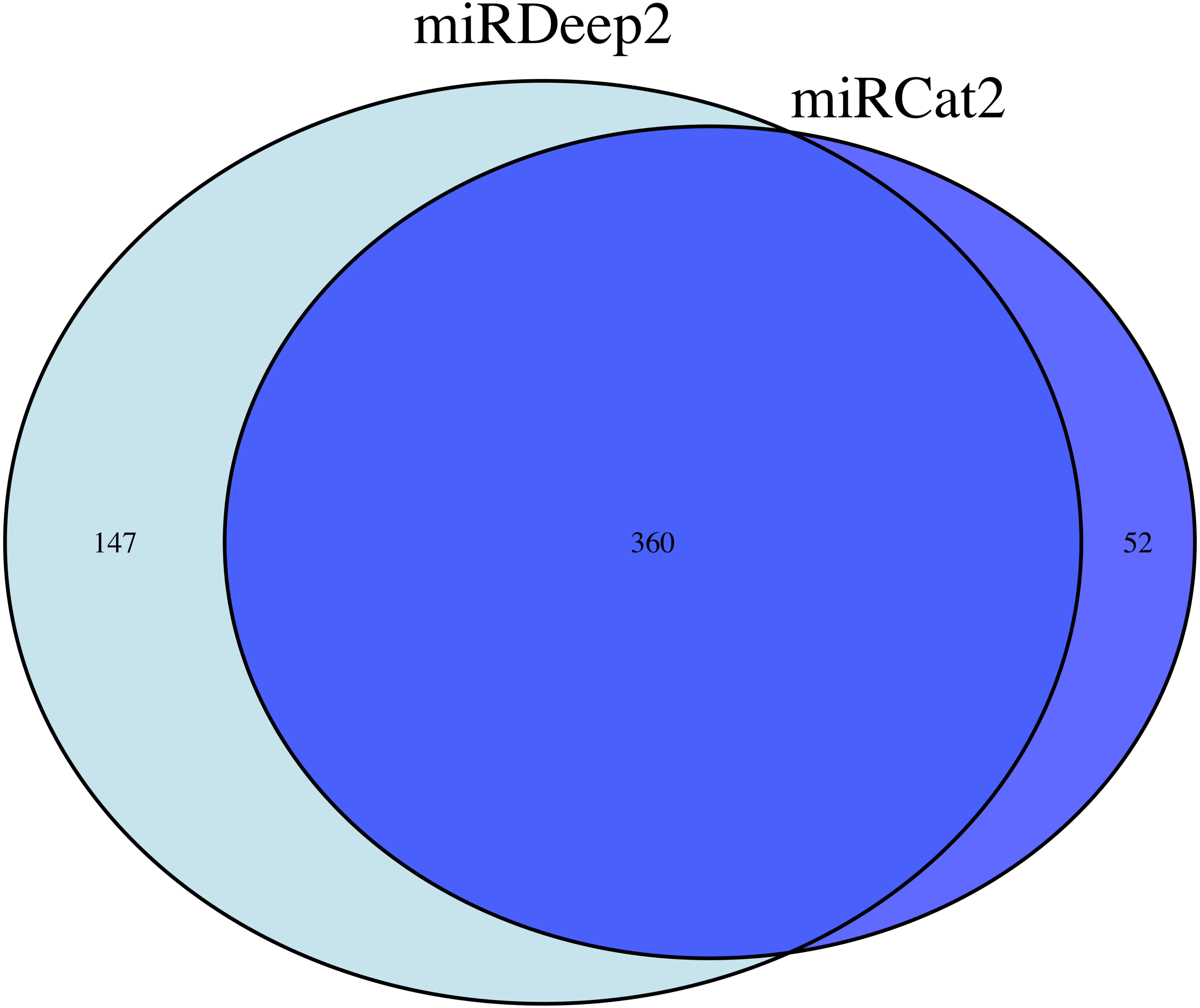


Supplementary Figure S3. Venn diagram showing the intersection between miRCat2 and miRDeep2 predictions

Supplementary Figure S4. Secondary structure plots for all 559 annotated miRNA loci, as predicted using the *ViennaRNA* package. Green and purple colors highlight the miRNA-5p and miRNA-3p sequences, respectively.

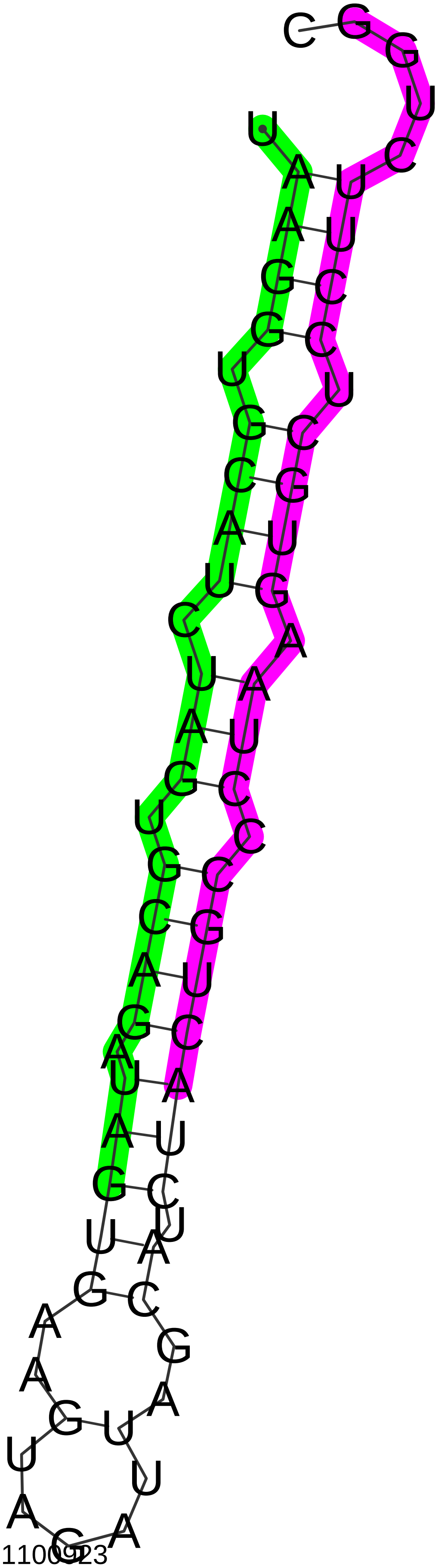

NW_006383042.1_1100859-1100923

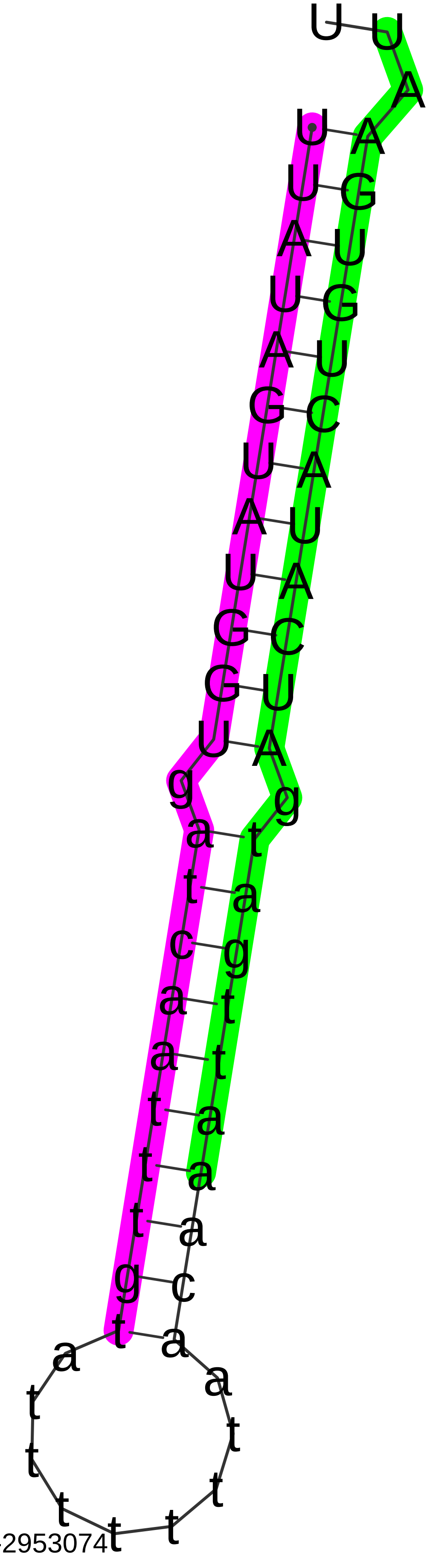

NW_006383067.1_2953017-2953074

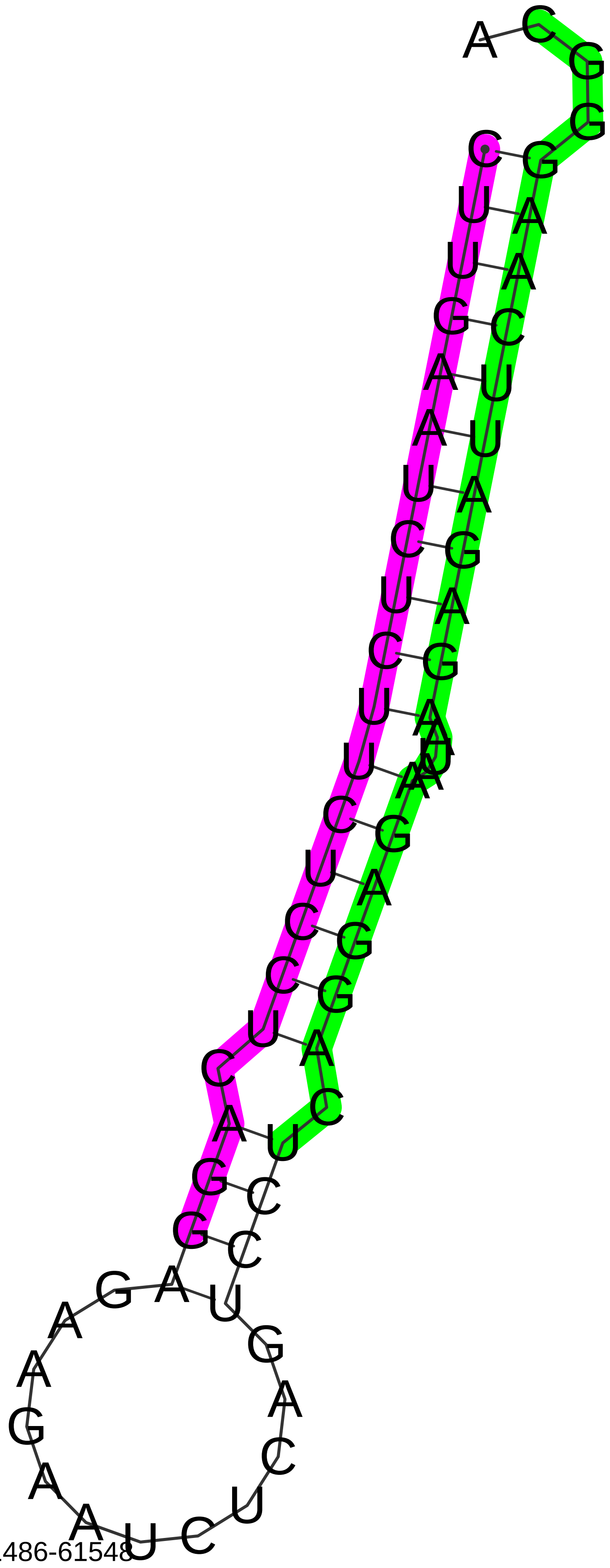

NW_006383070.1_61486-61548

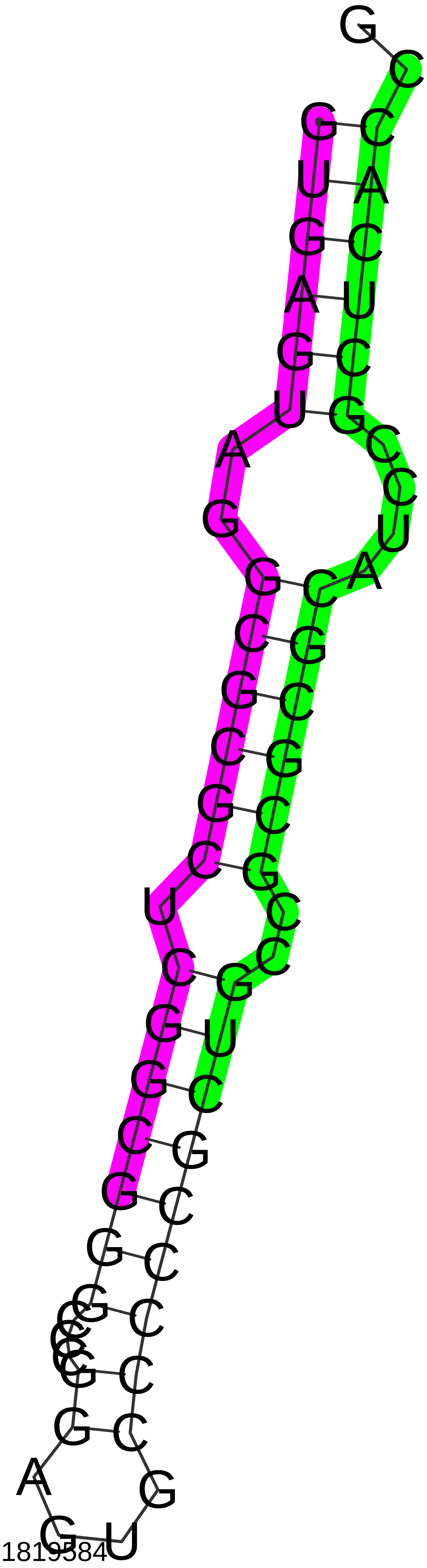

NW_006383074.1_1819525-1819584

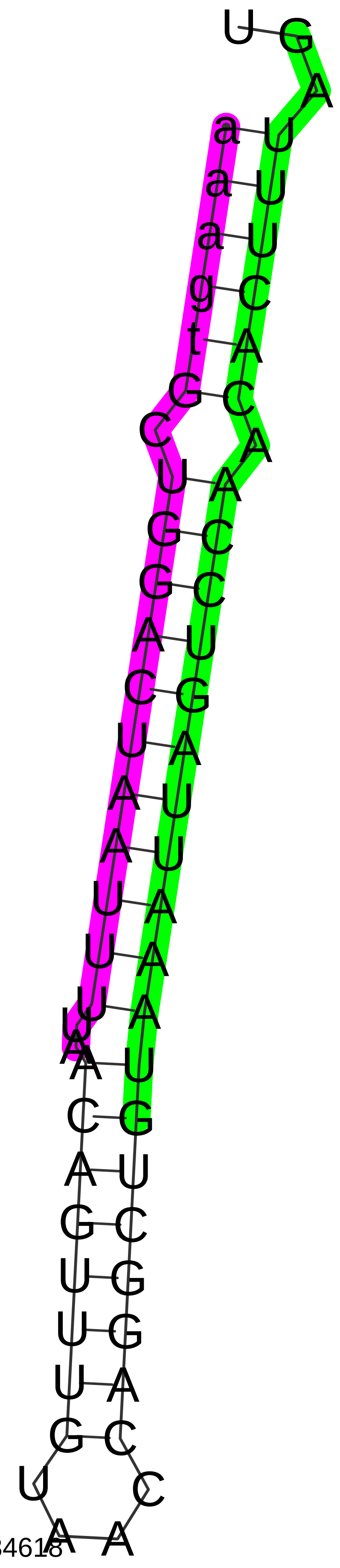

NW_006383089.1_1434558-1434618

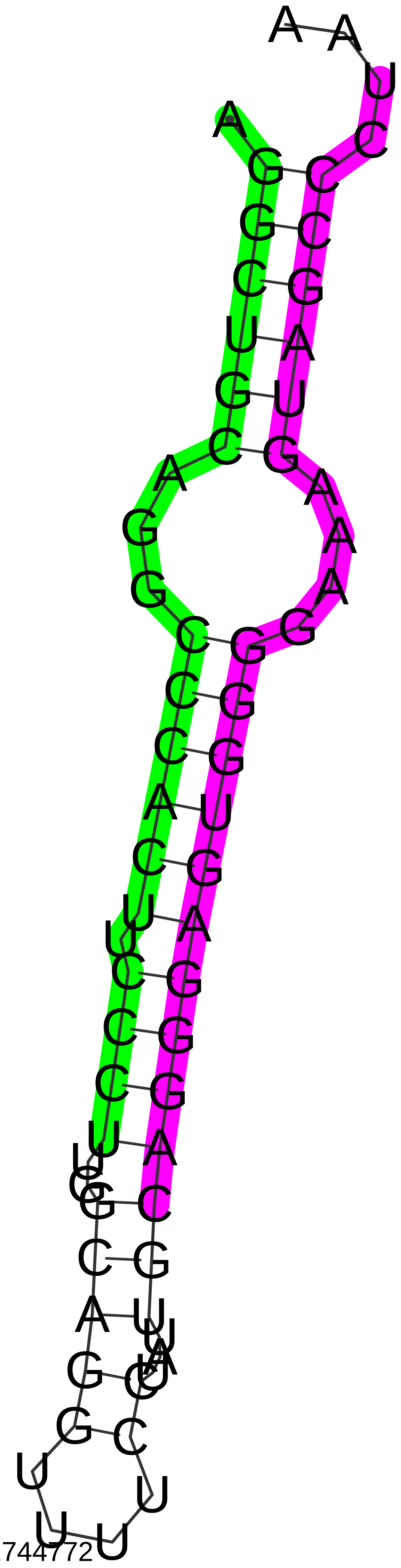

NW_006383124.1_2744709-2744772

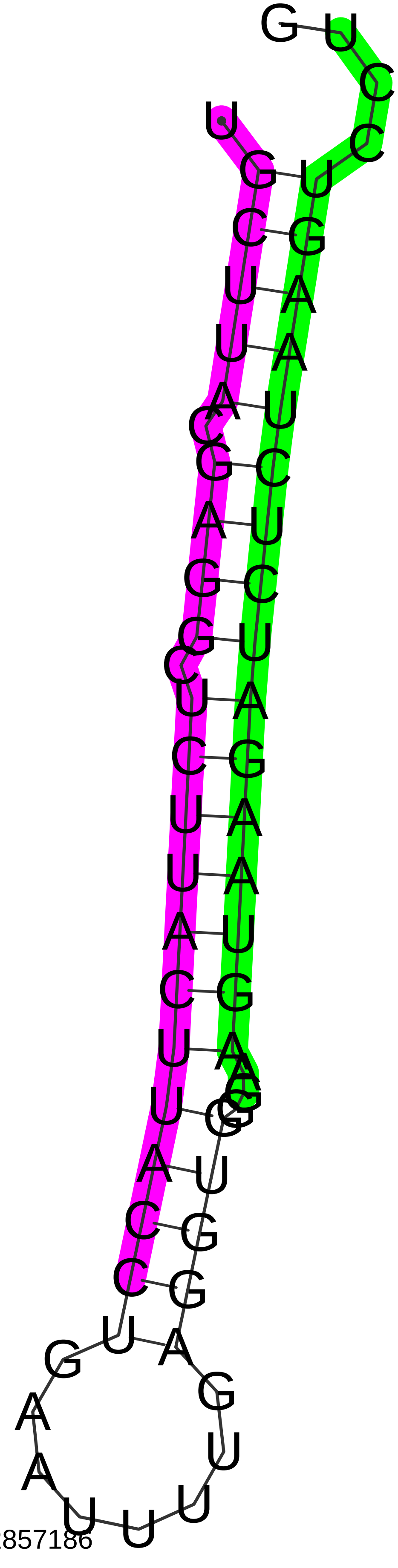

NW_006383133.1_2857127-2857186

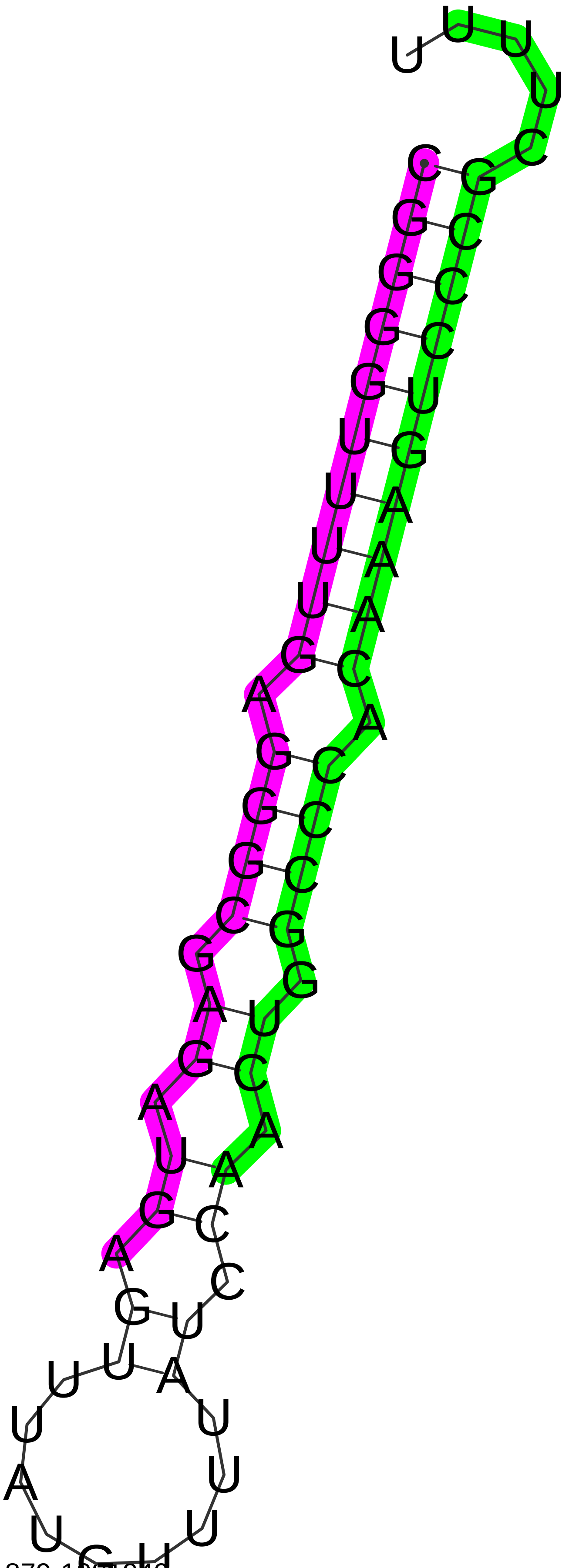

NW_006383147.1_1081879-1081940

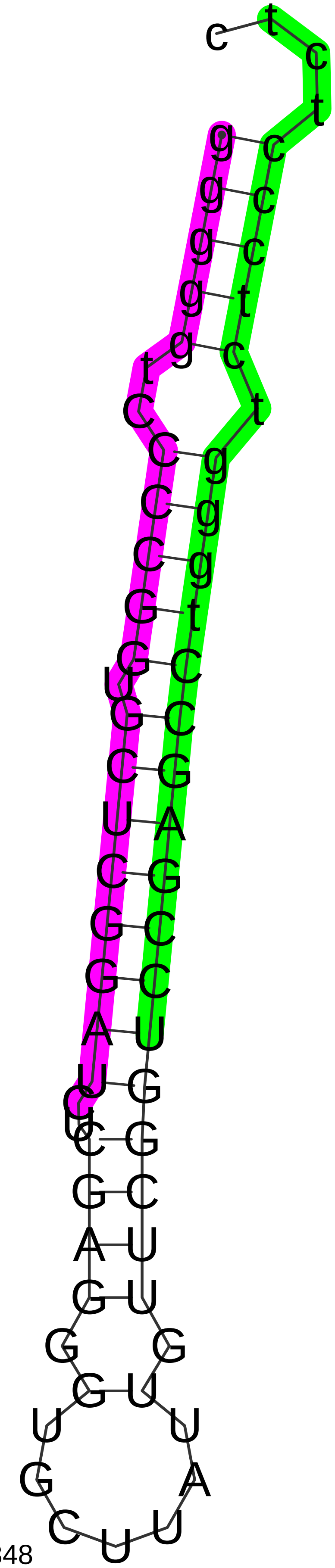

NW_006383149.1_599284-599348

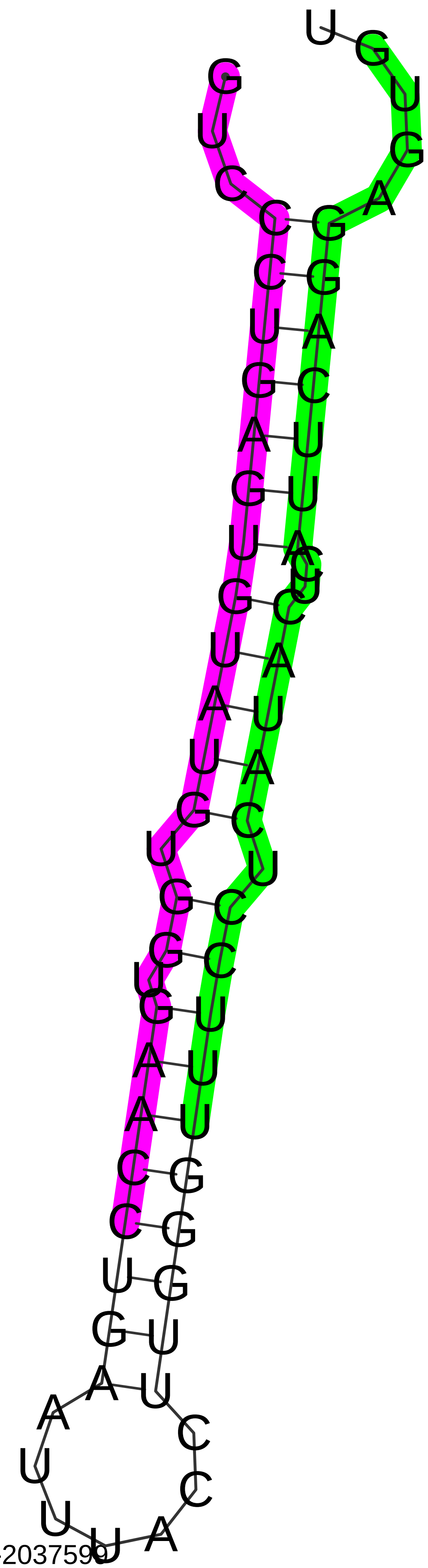

NW_006383150.1_2037536-2037599

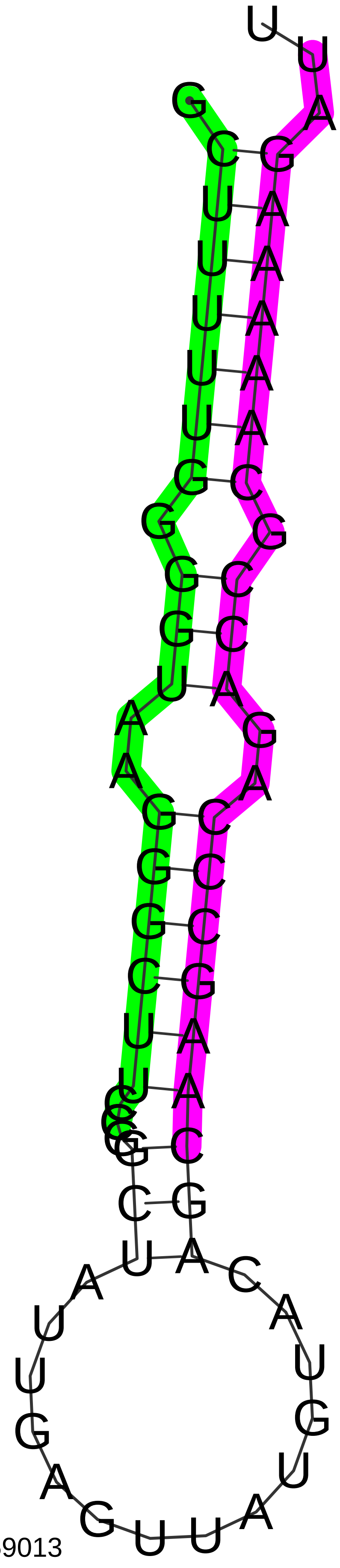

NW_006383150.1_2058949-2059013

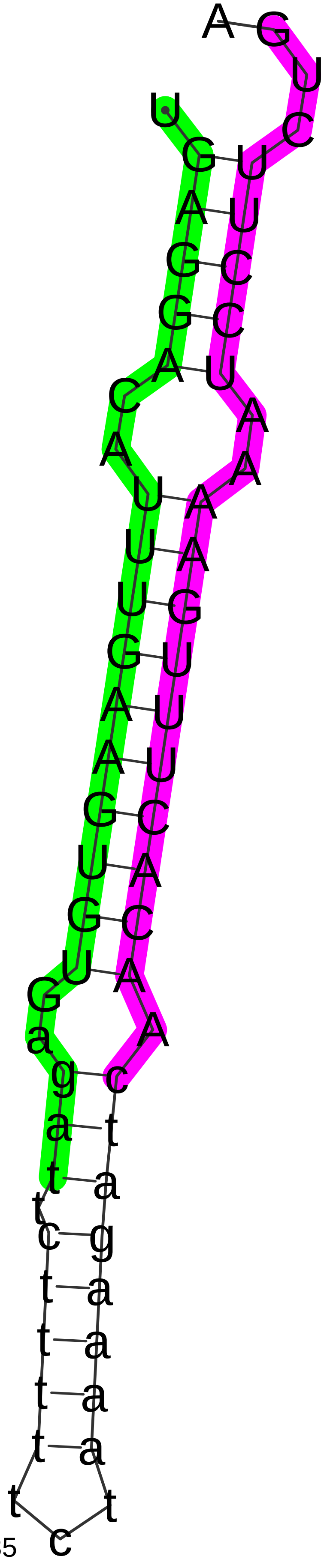

NW_006383167.1_228324-228385

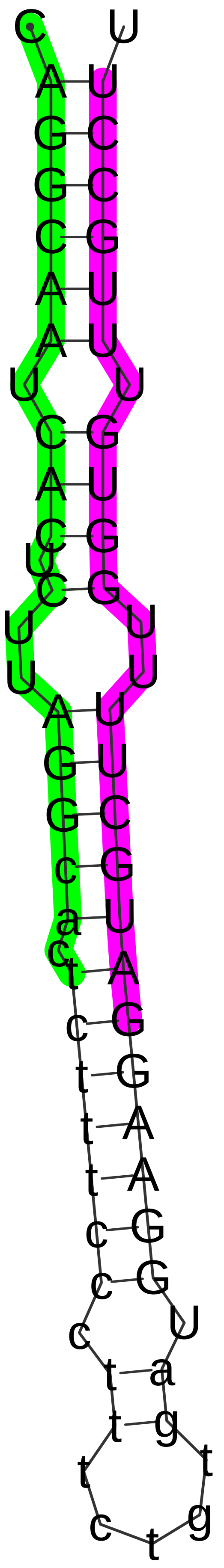

NW_006383171.1_2125908-2125972

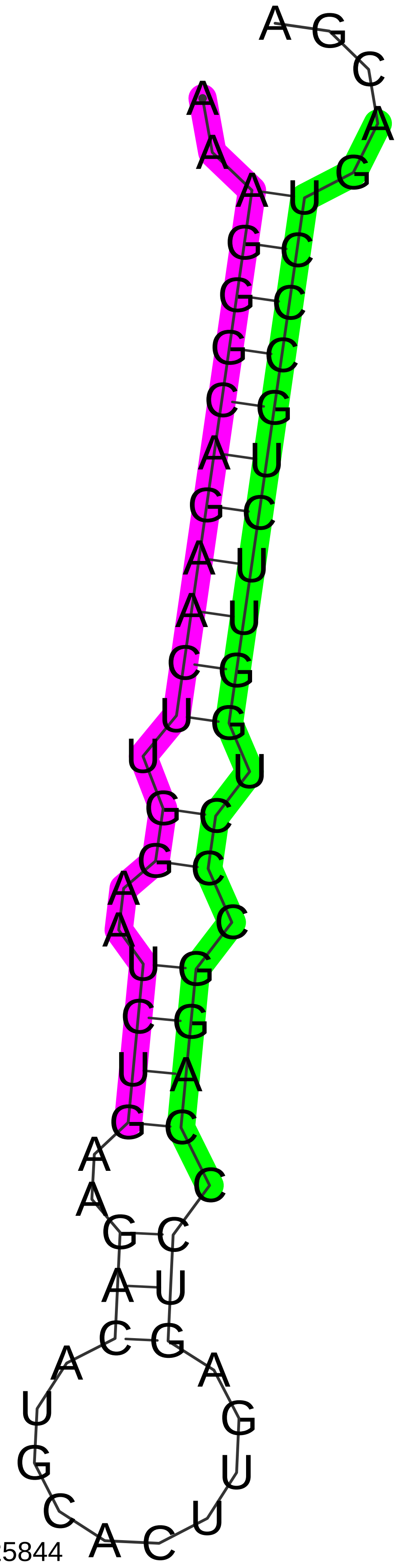

NW_006383177.1_425780-425844

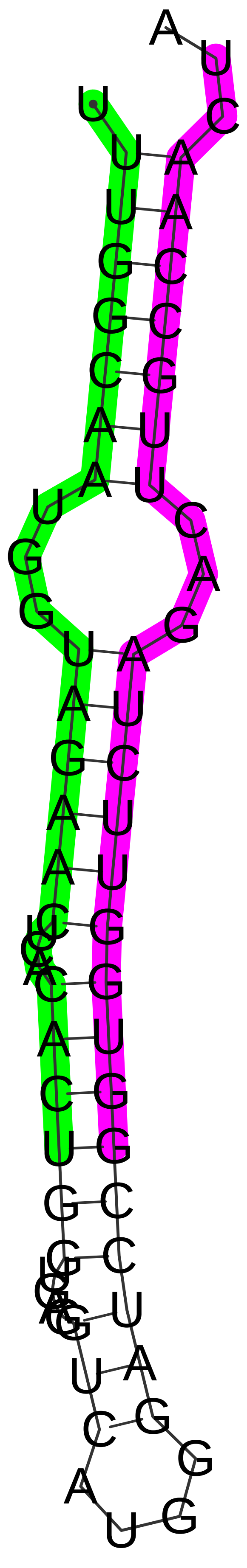

NW_006383179.1_648325-648389

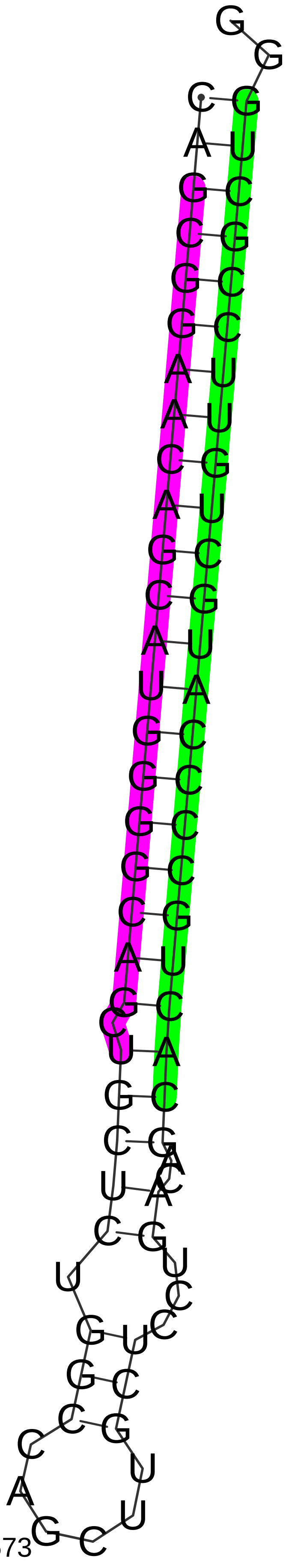

NW_006383225.1_1800601-1800673

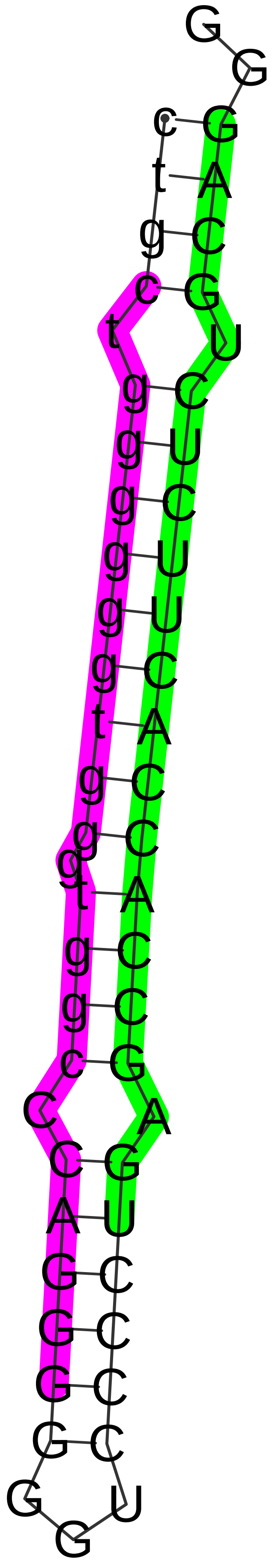

NW_006383245.1_173778-173833

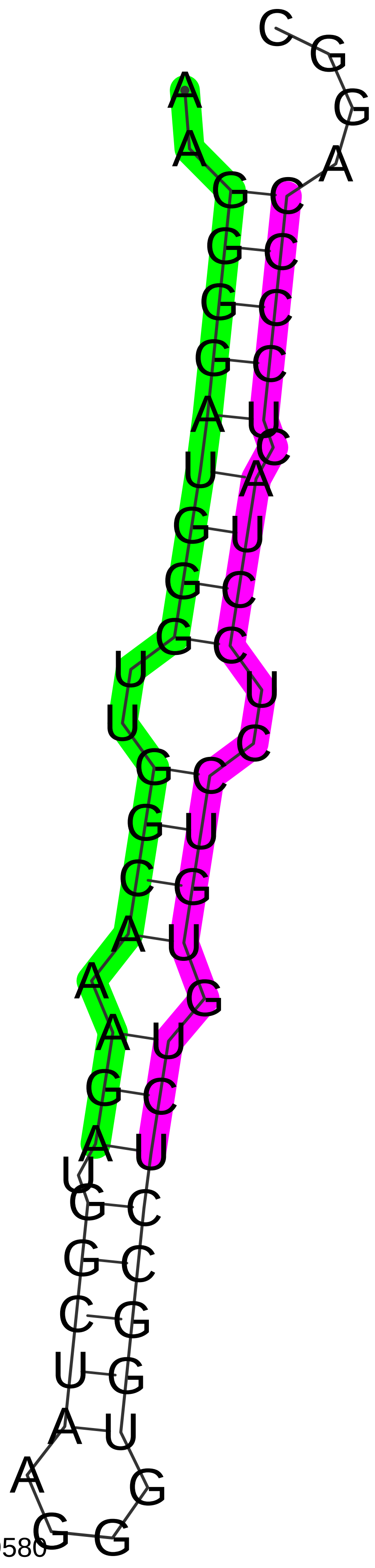

NW_006383287.1_119521-119580

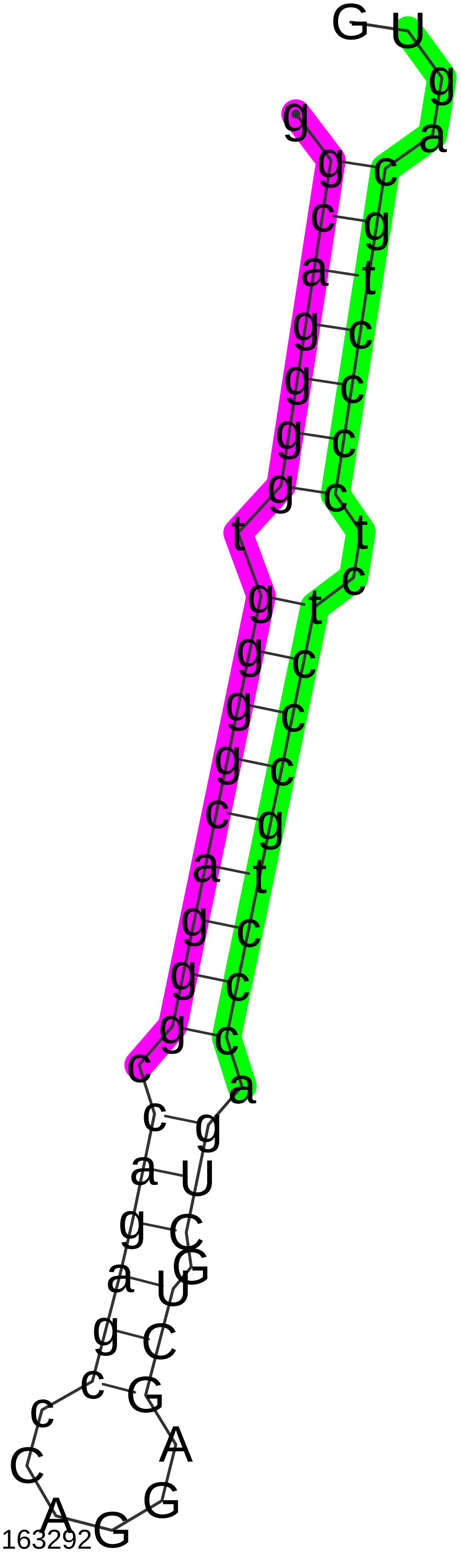

NW_006383292.1_163232-163292

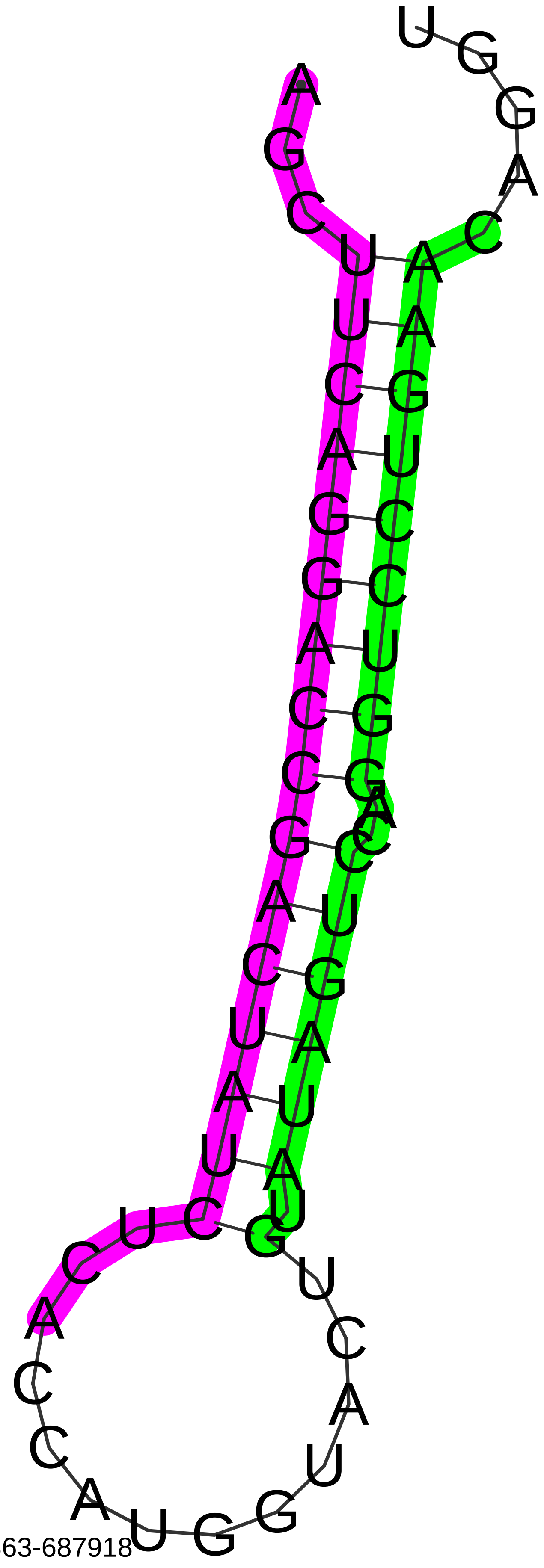

NW_006383299.1_687863-687918

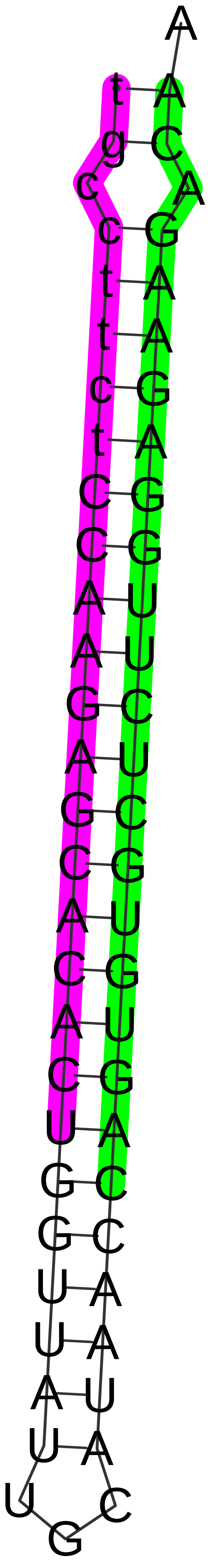

NW_006383307.1_1237775-1237832

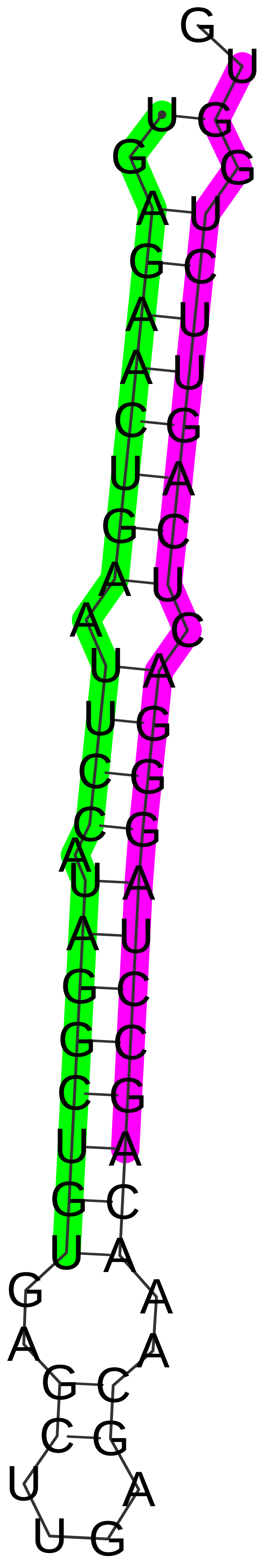

NW_006383308.1_343980-344040

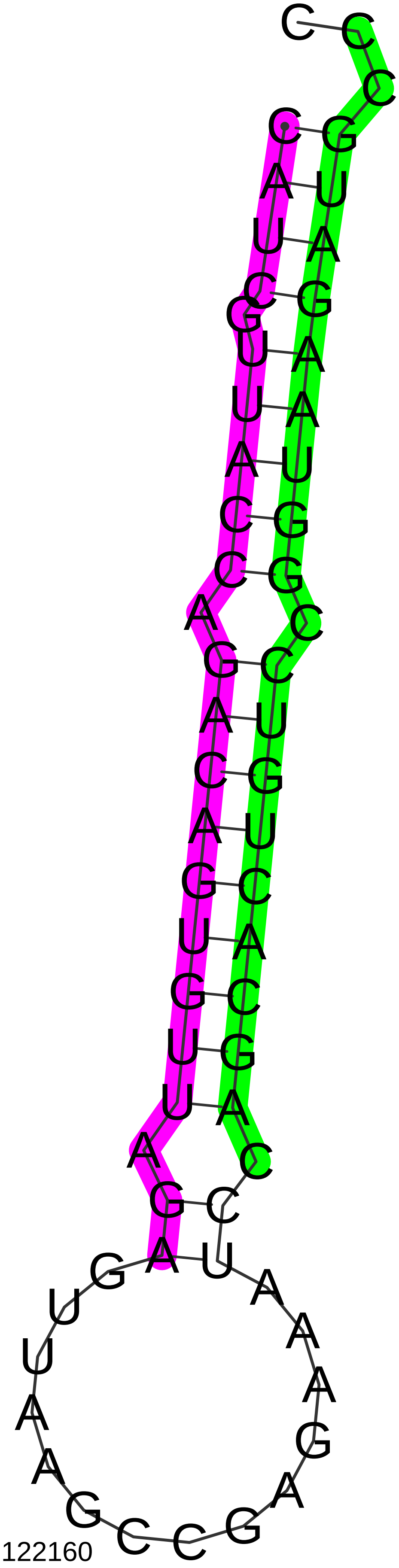

NW_006383401.1_1122099-1122160

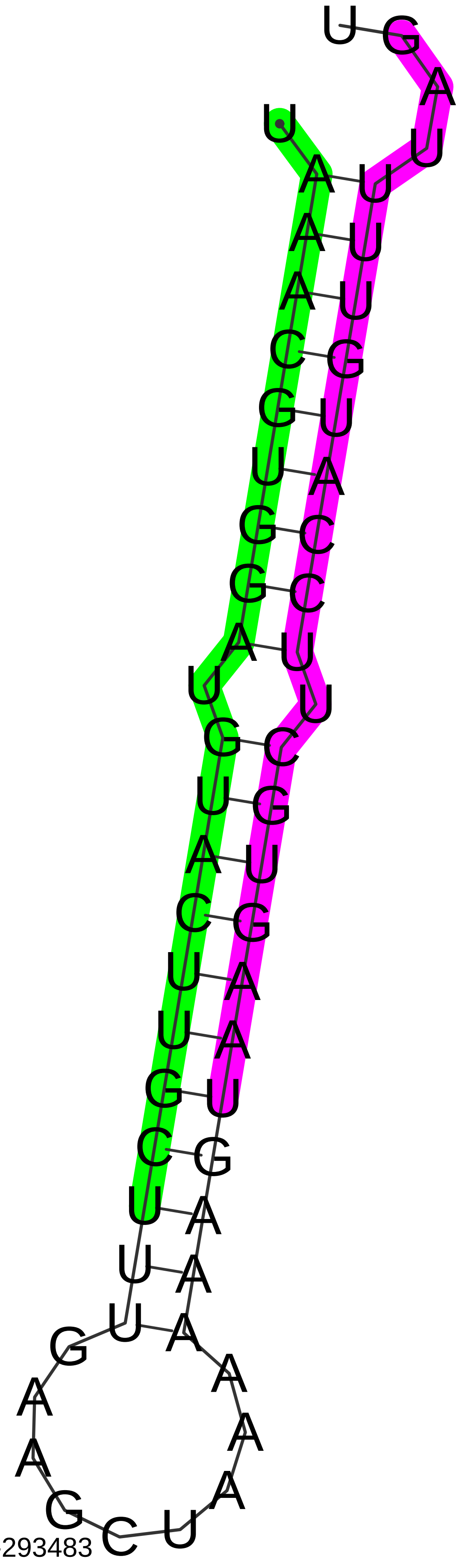

NW_006383431.1_293428-293483

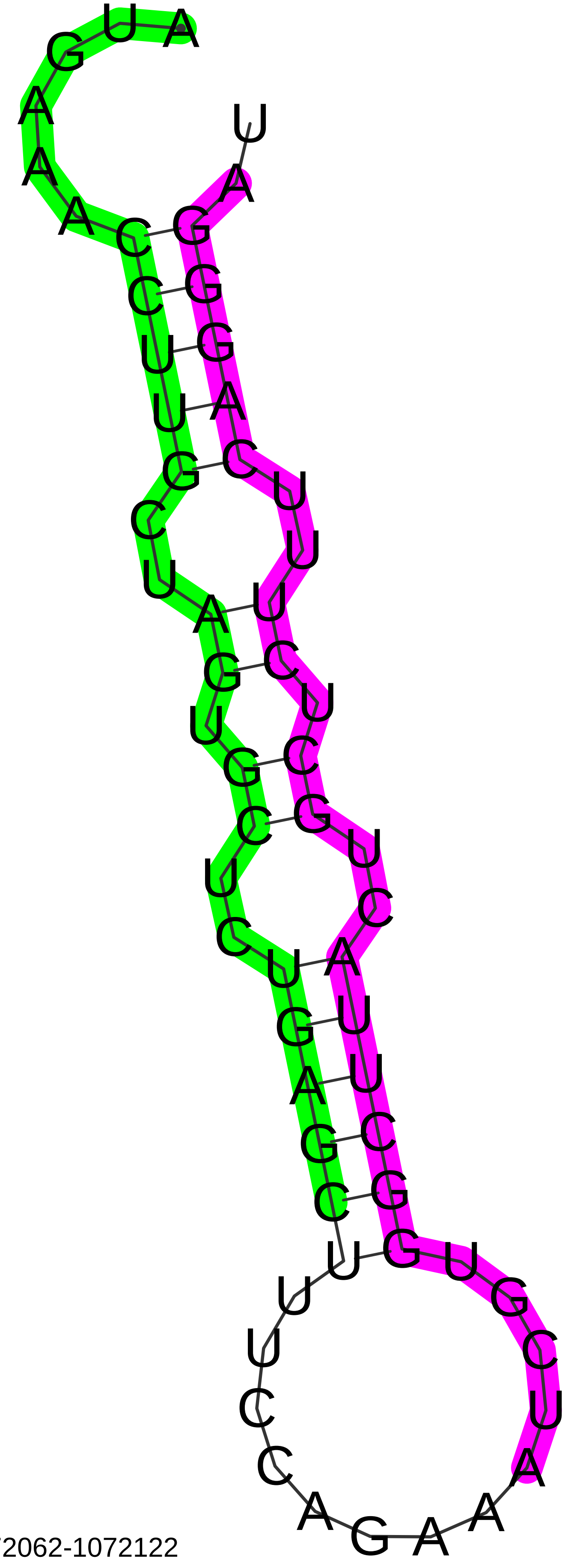

NW_006383432.1_1072062-1072122

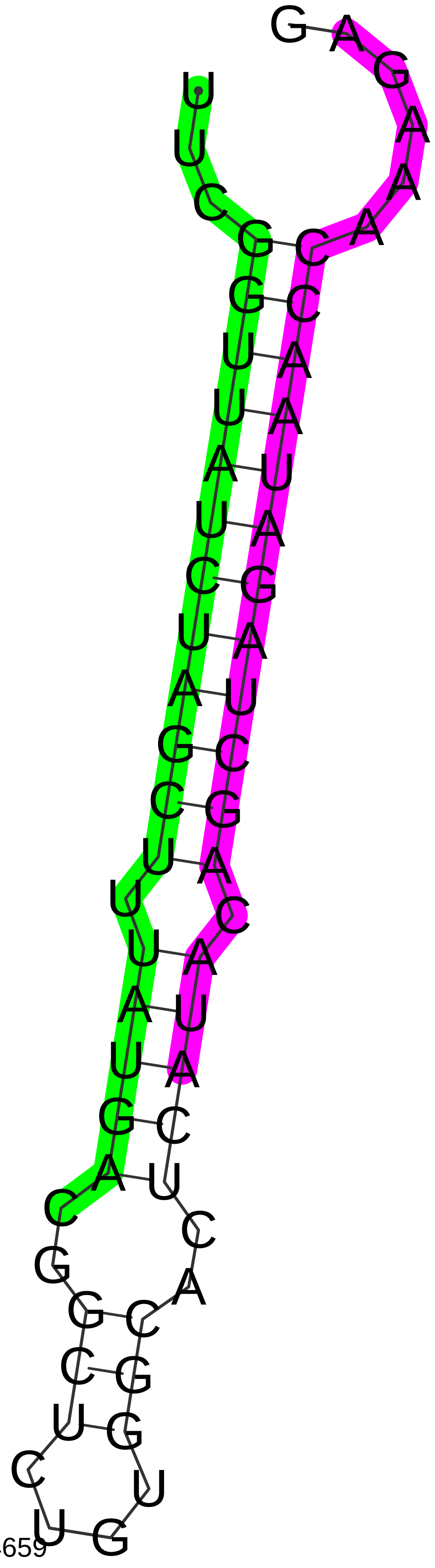

NW_006383462.1_54601-54659

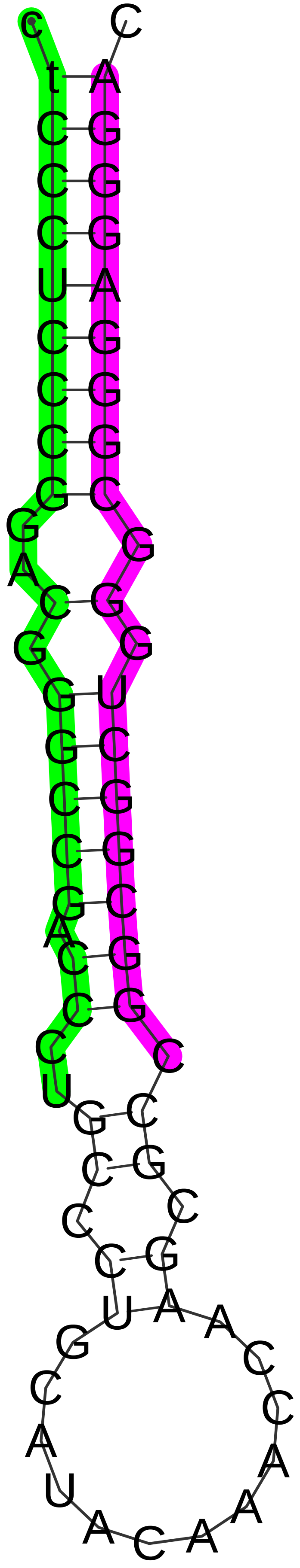

NW_006383462.1_53808-53874

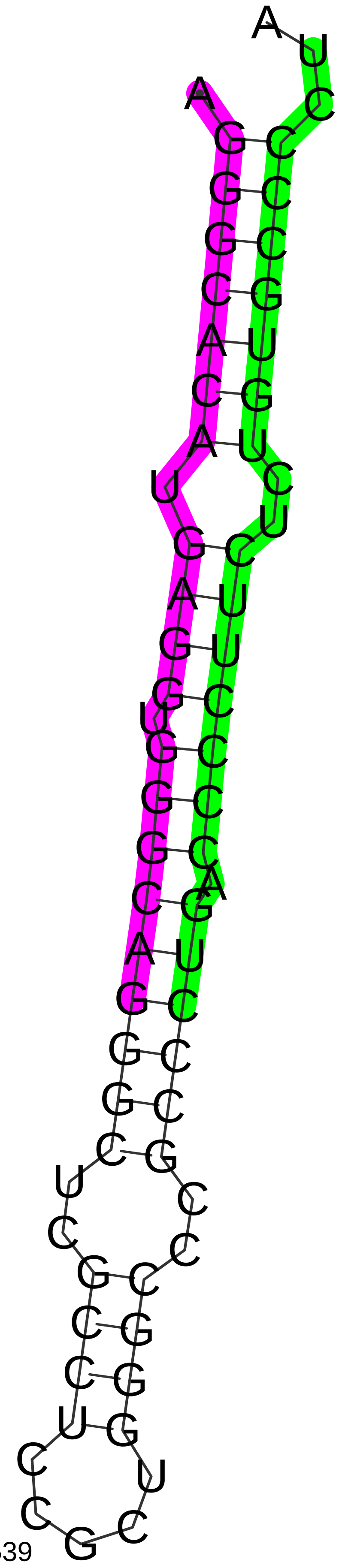

NW_006383464.1_555474-555539

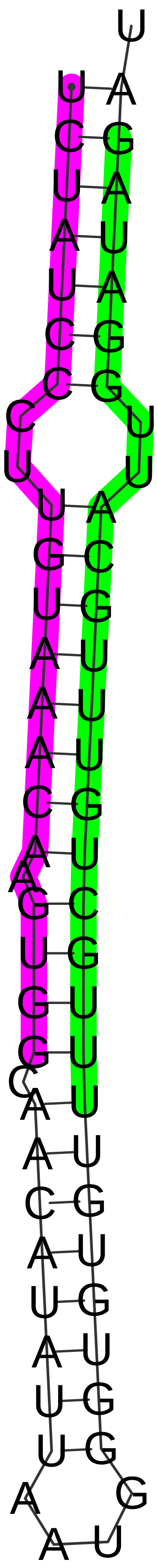

NW_006383473.1_304795-304859

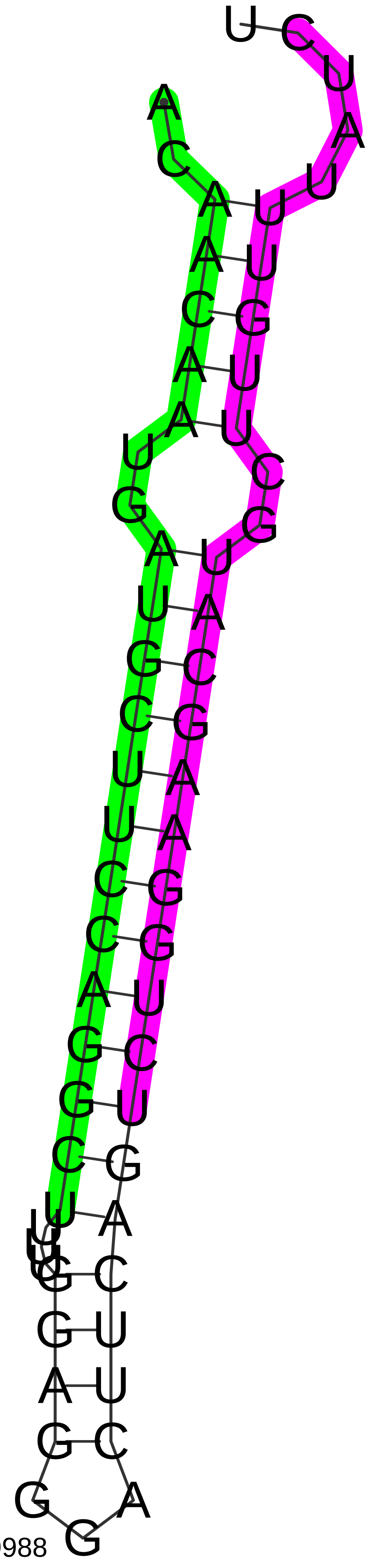

NW_006383507.1_669928-669988

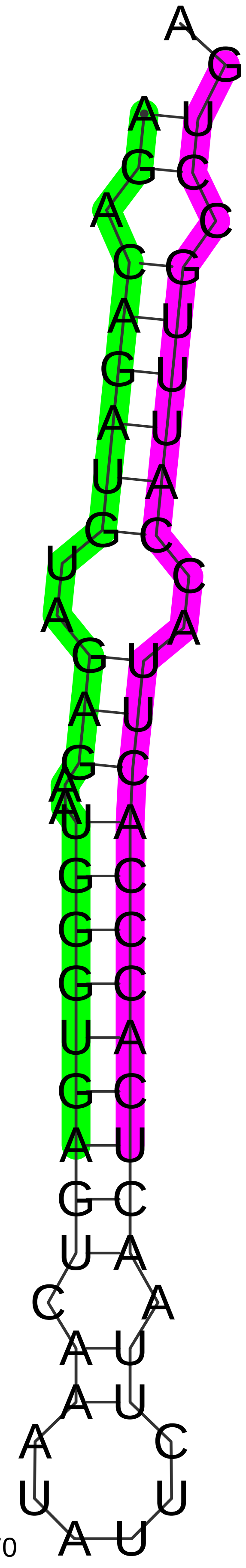

NW_006383509.1_1010609-1010670

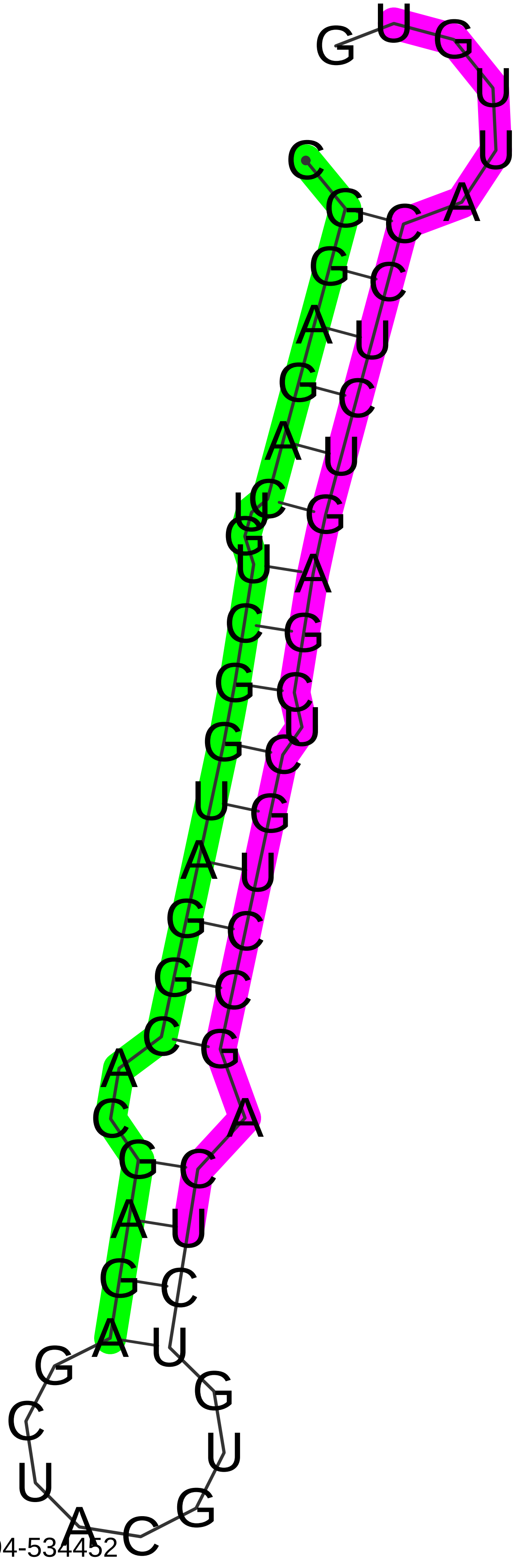

NW_006383515.1_534394-534452

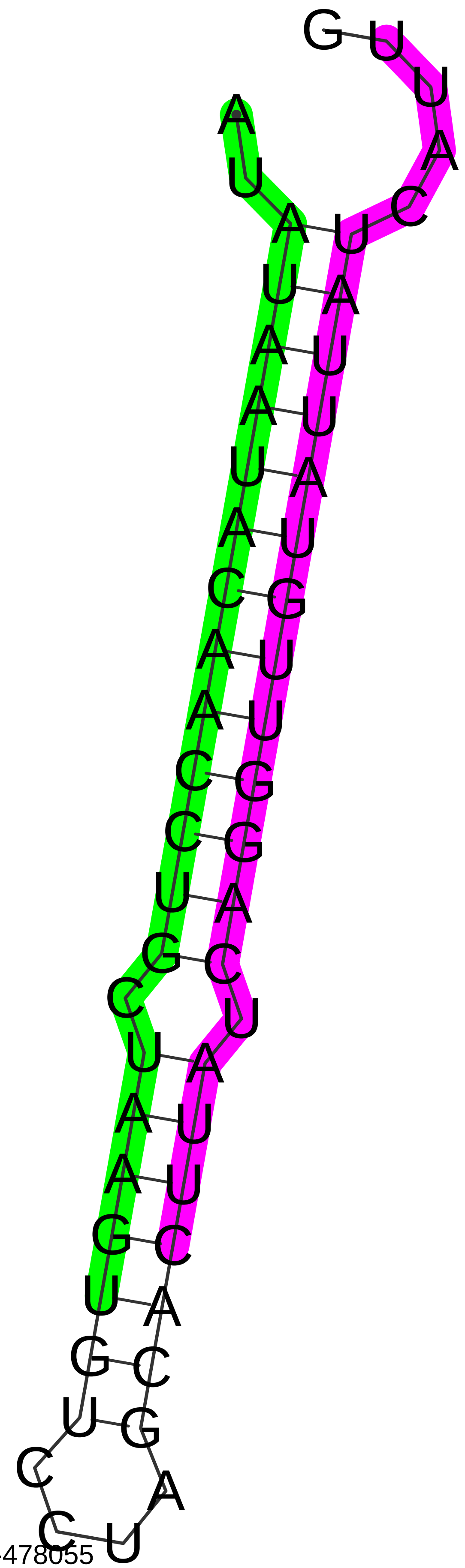


NW_006383521.1_478003-478055

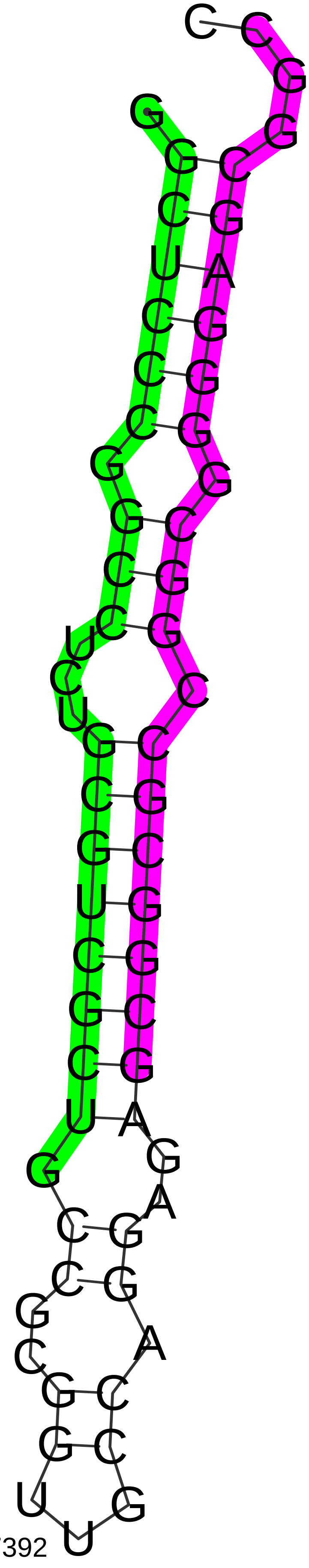

NW_006383530.1_1147331-1147392

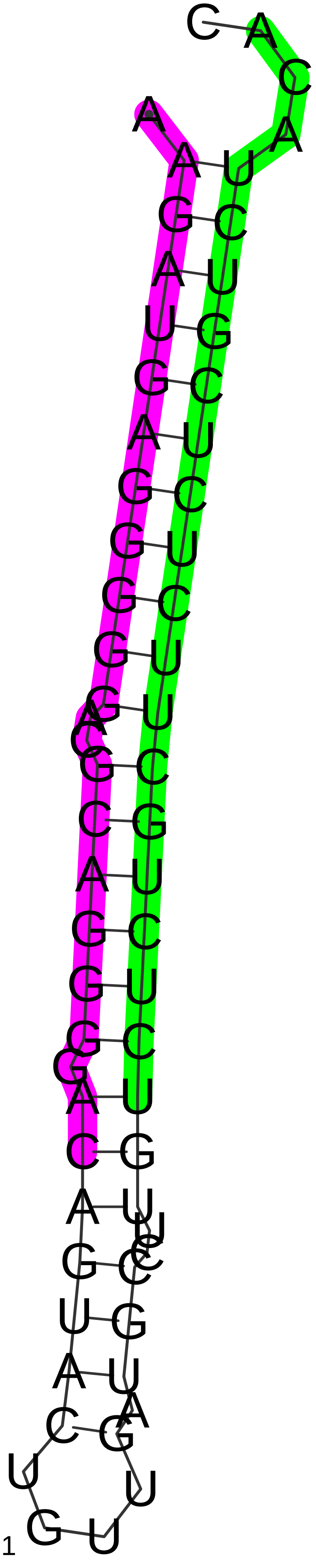

NW_006383552.1_184369-184431

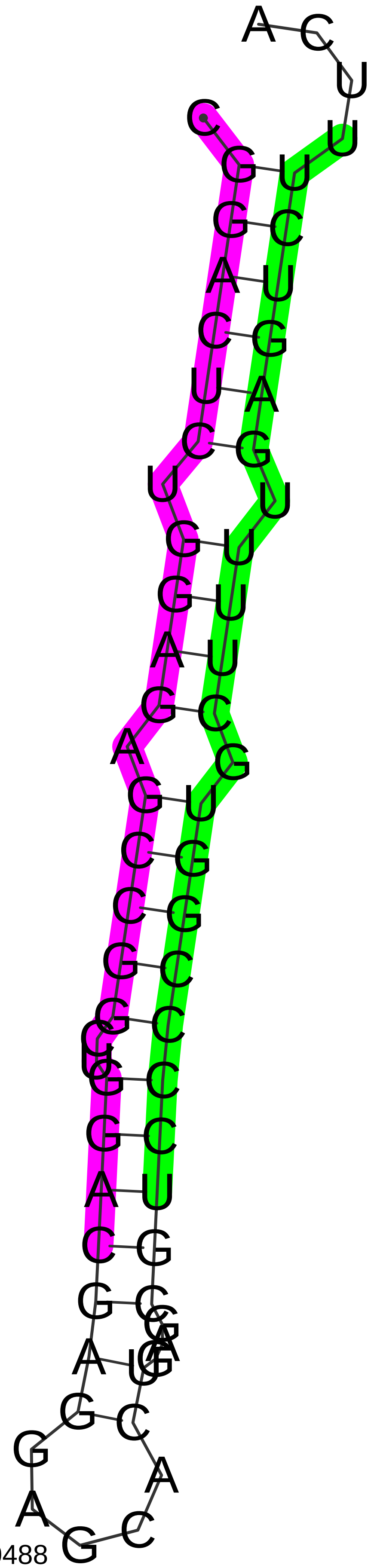

NW_006383573.1_740426-740488

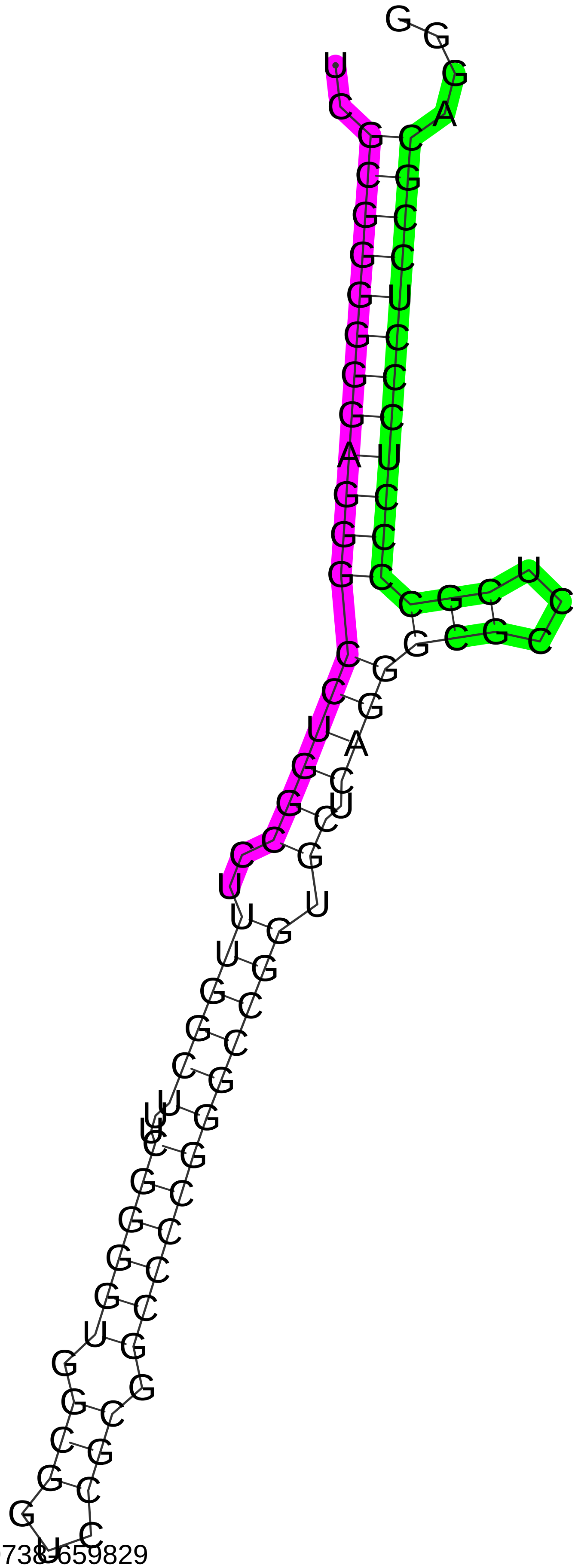

NW_006383590.1_659738-659829

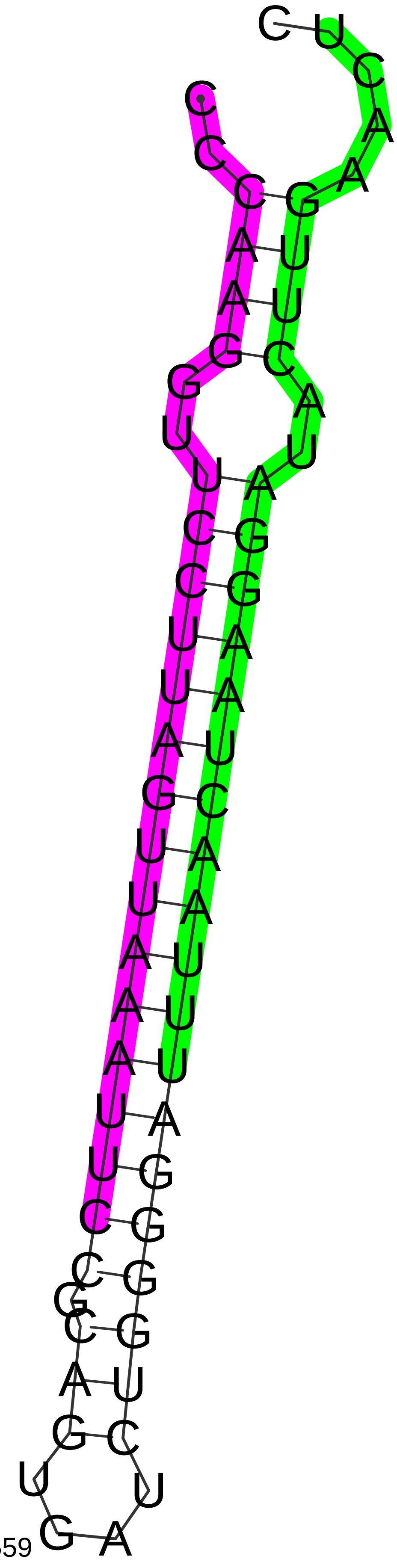

NW_006383612.1_13498-13559

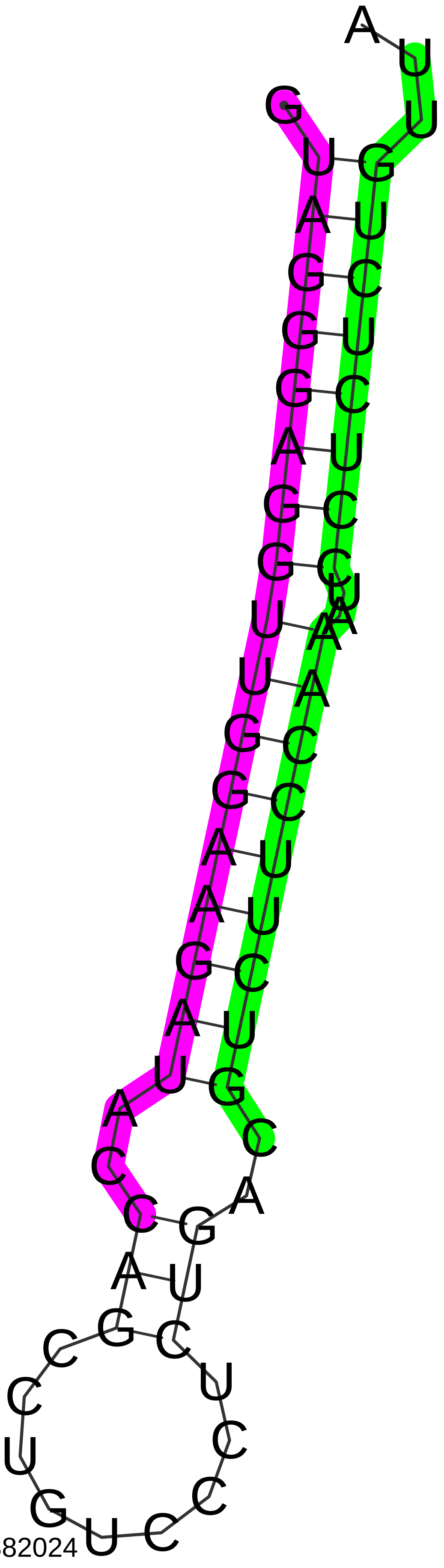

NW_006383625.1_381966-382024

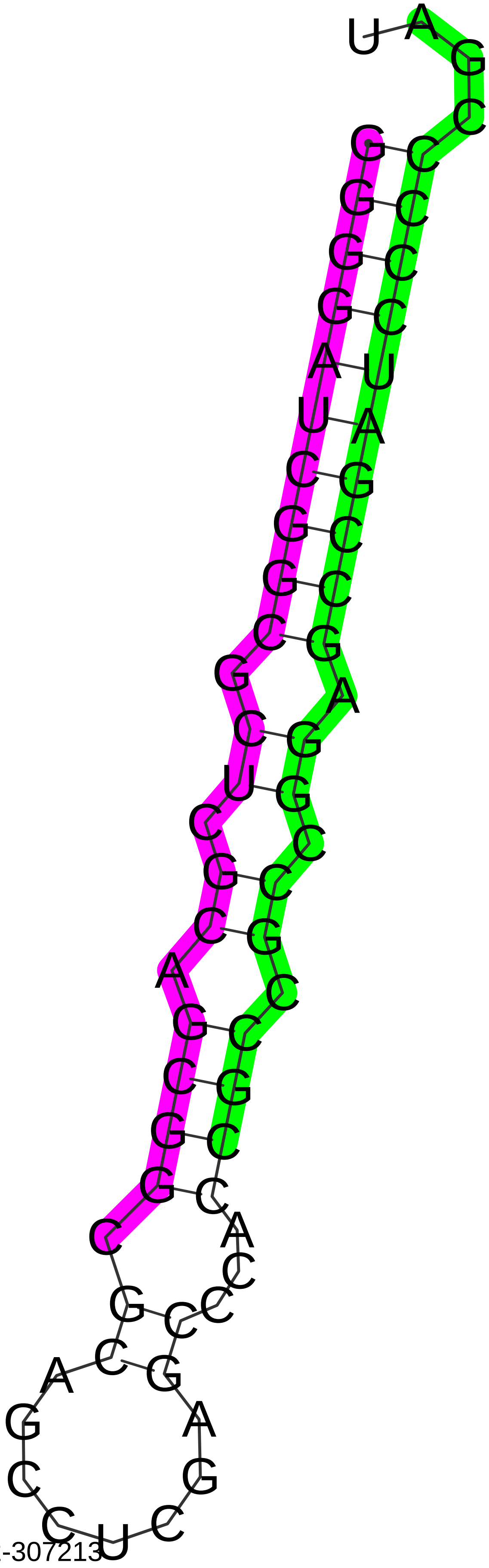

NW_006383681.1_307152-307213

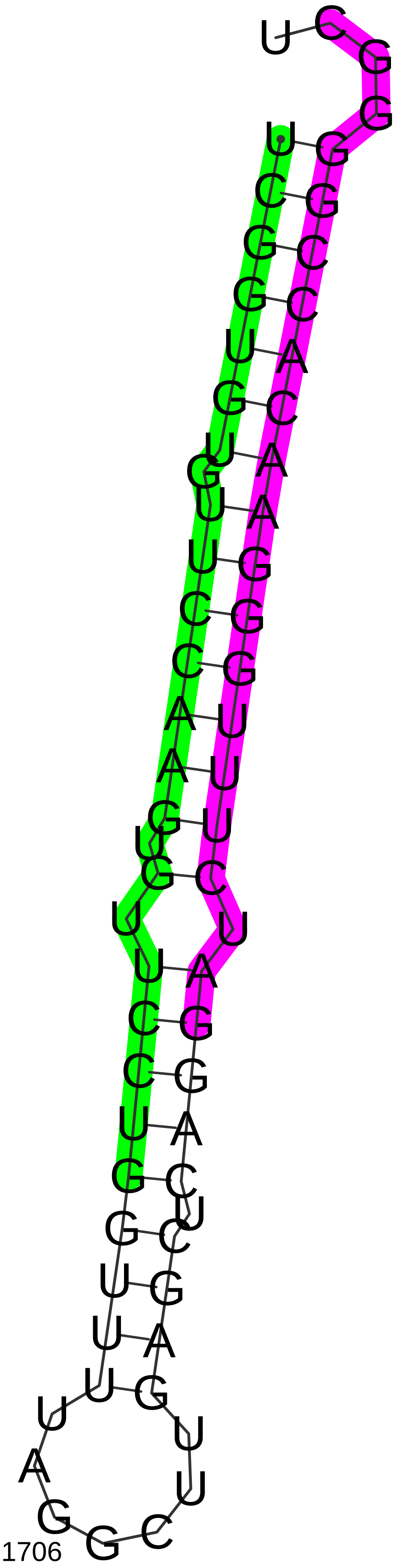

NW_006383712.1_611643-611706

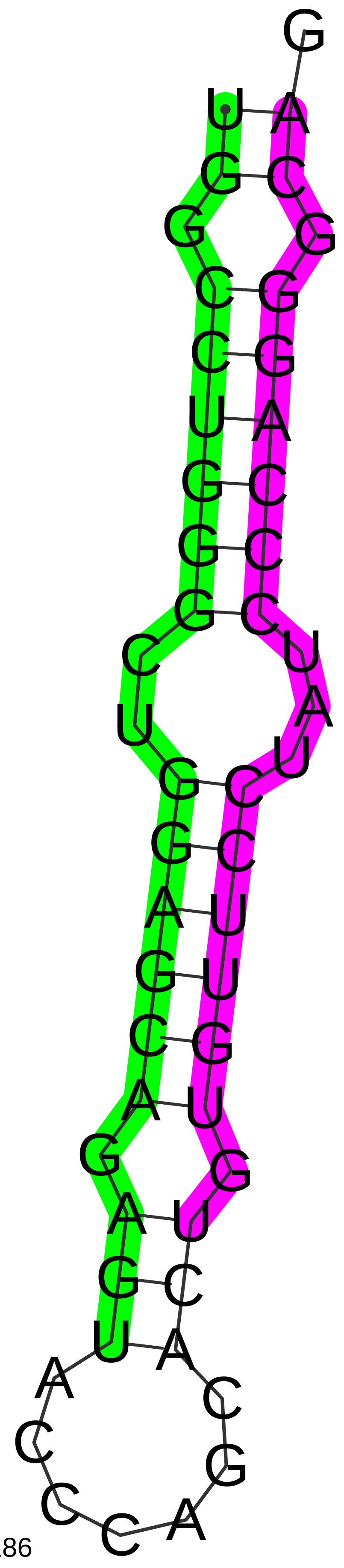

NW_006383719.1_219136-219186

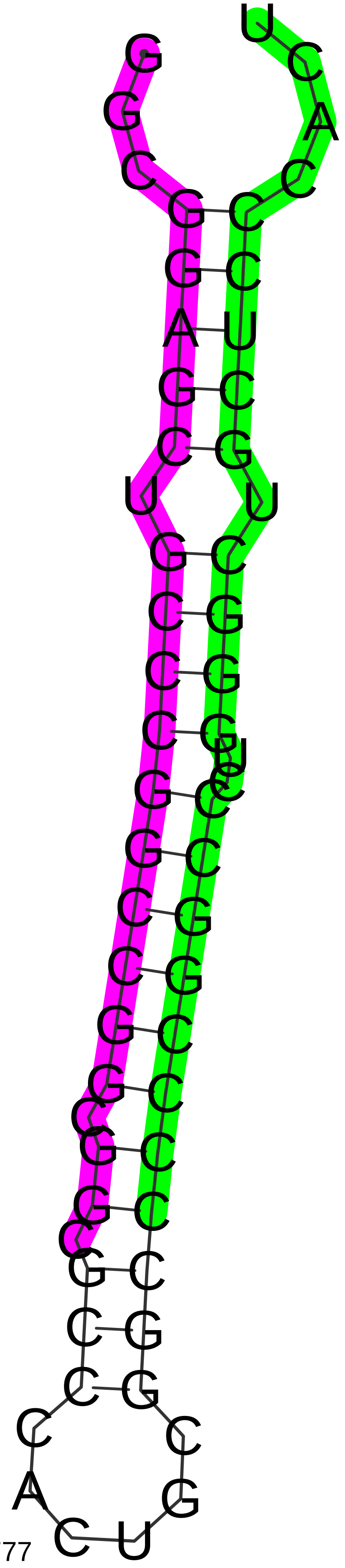

NW_006383721.1_804719-804777

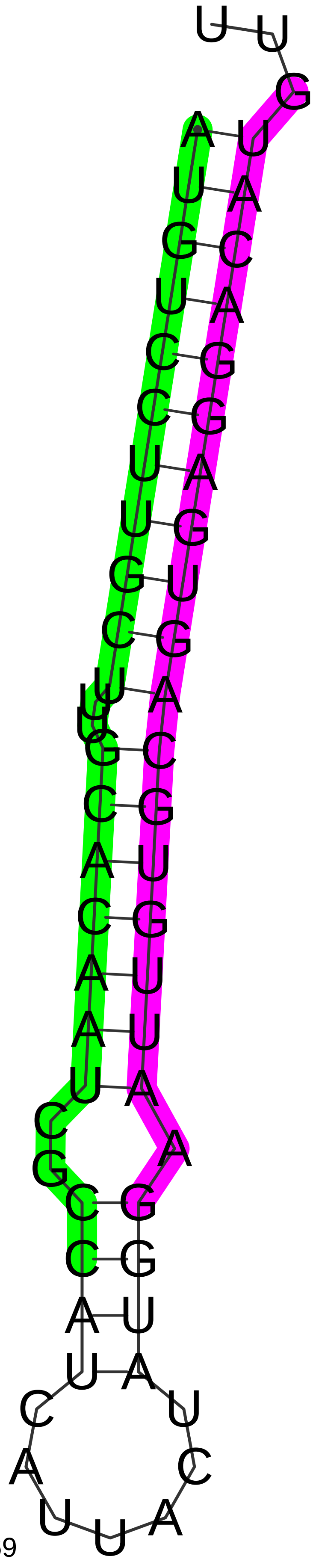

NW_006383731.1_533811-533869

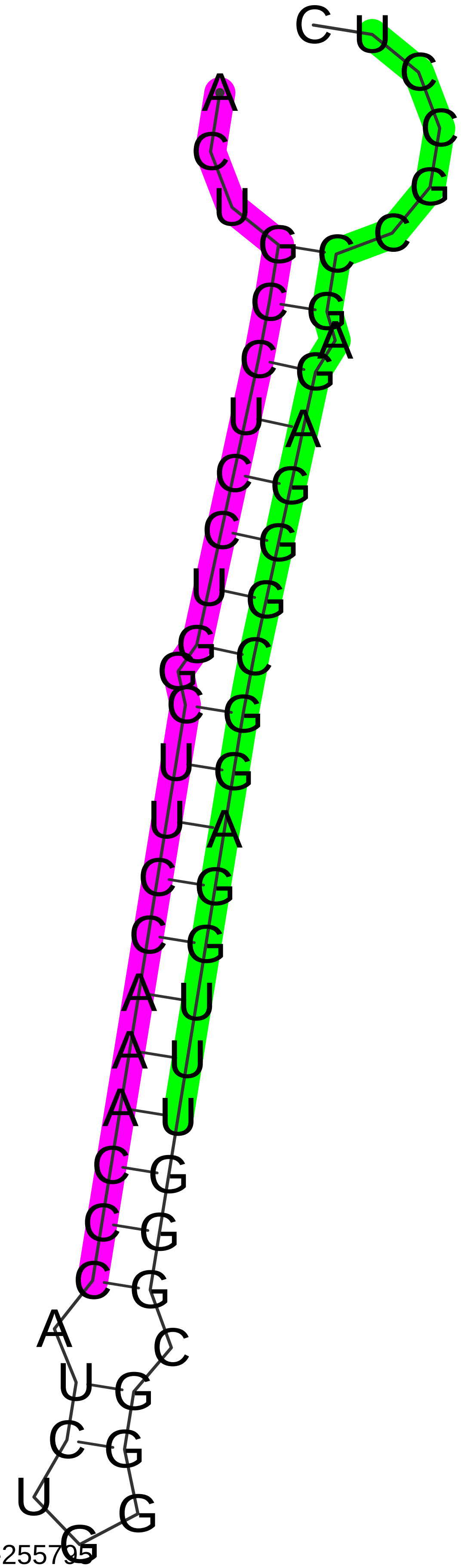

NW_006383764.1_255738-255795

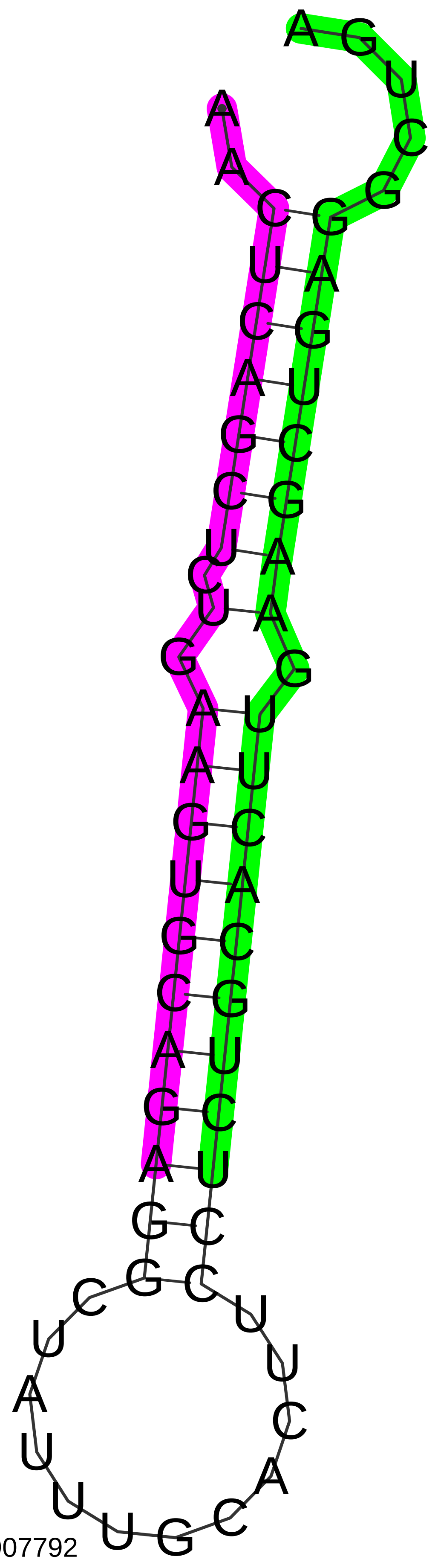

NW_006383800.1_907733-907792

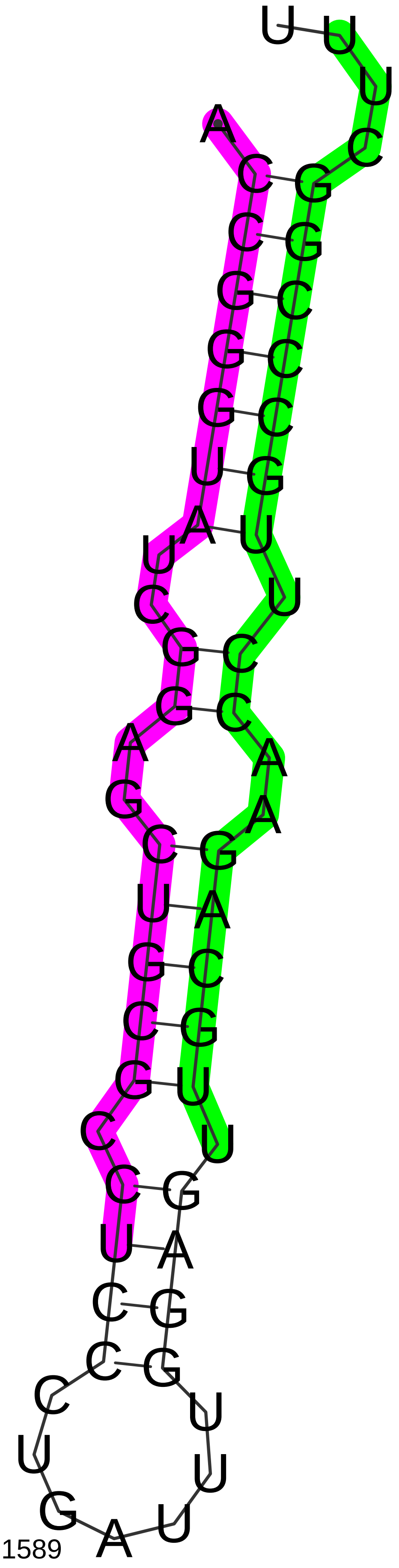

NW_006383813.1_111533-111589

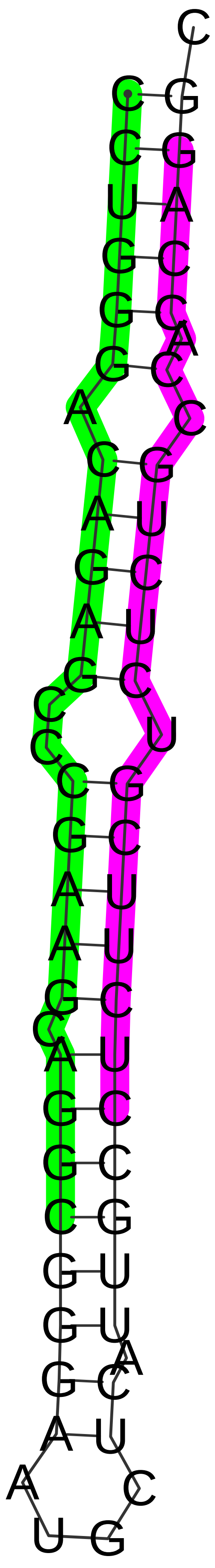

NW_006383816.1_59793-59853

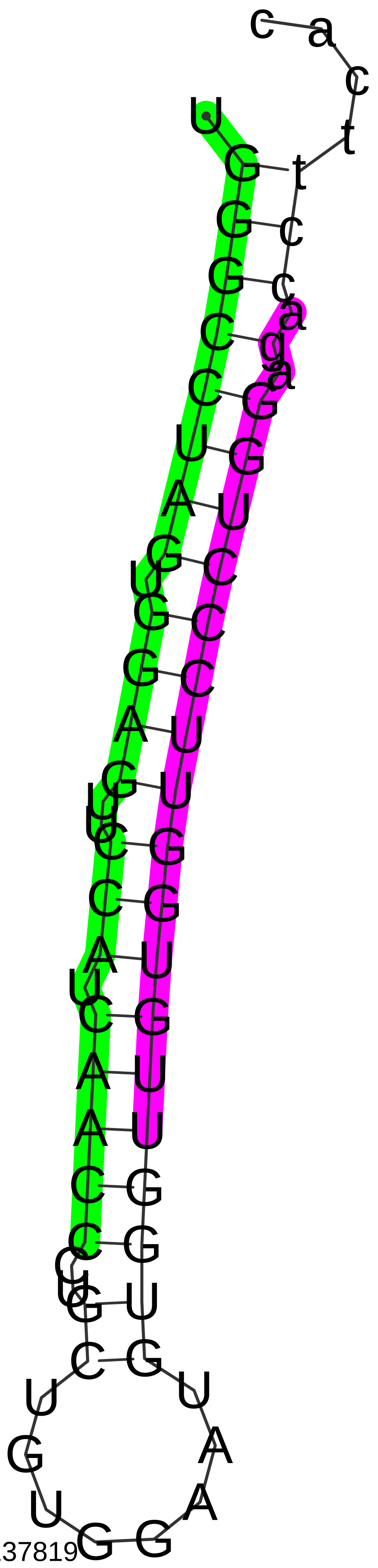

NW_006383846.1_1137755-1137819

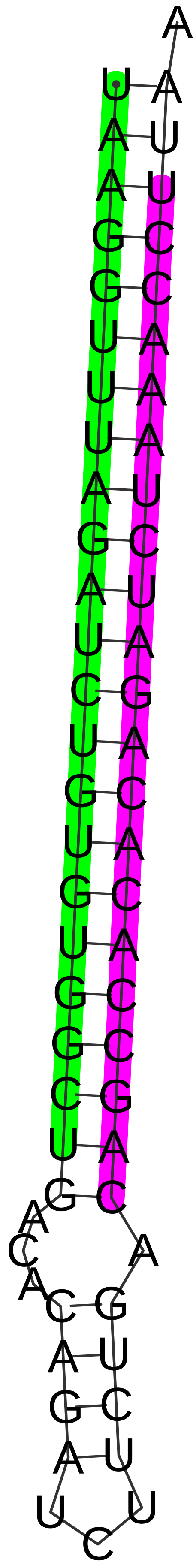

NW_006383860.1_273279-273340

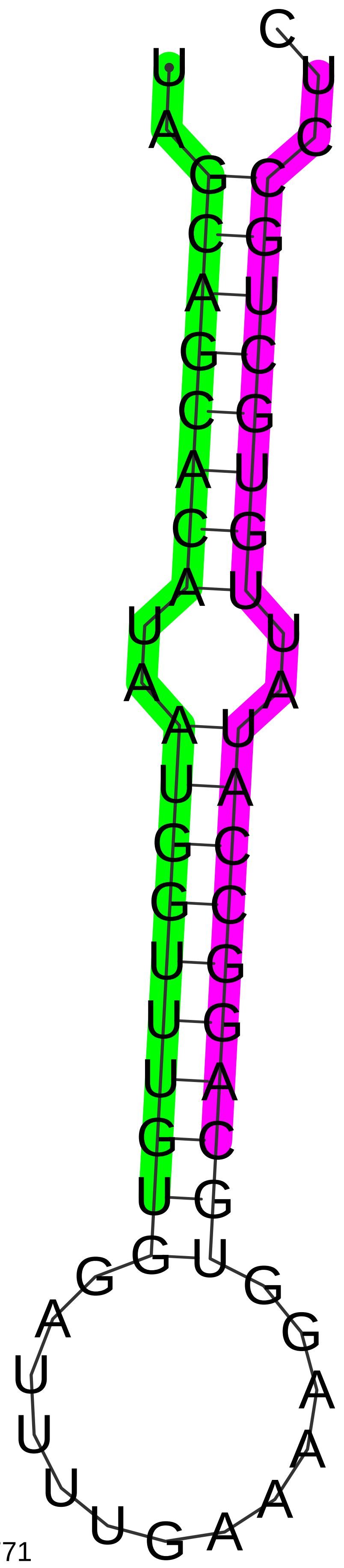

NW_006383872.1_162714-162771

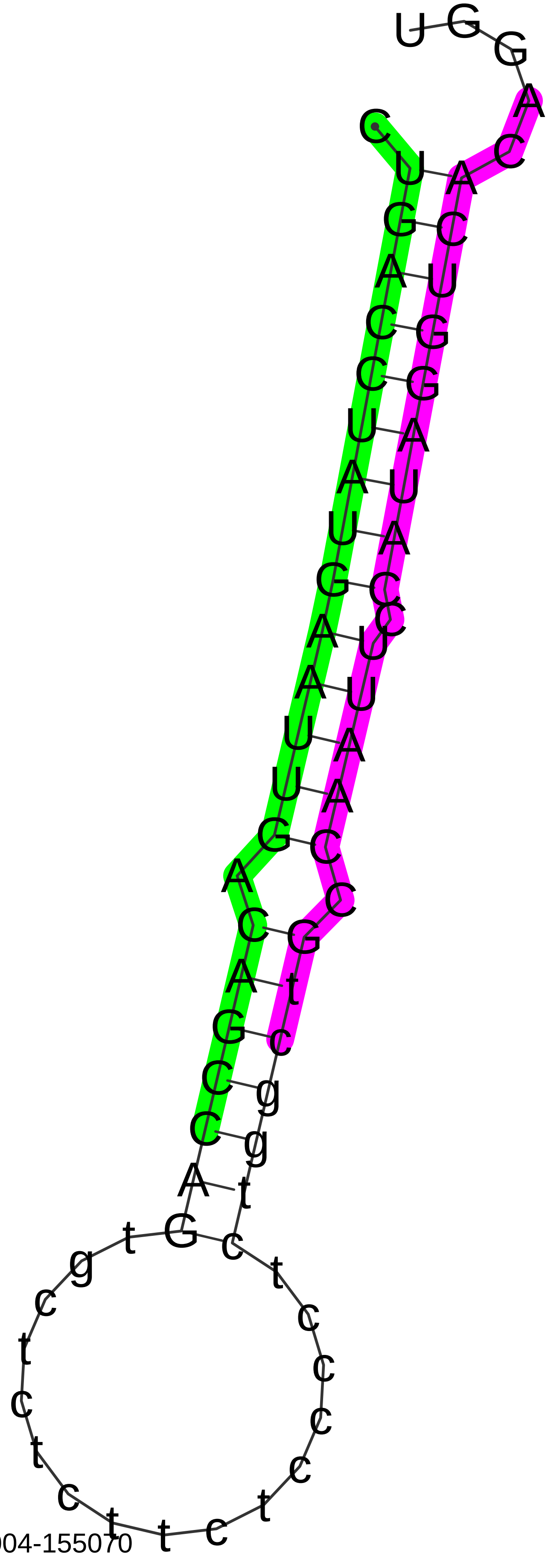

NW_006383874.1_155004-155070

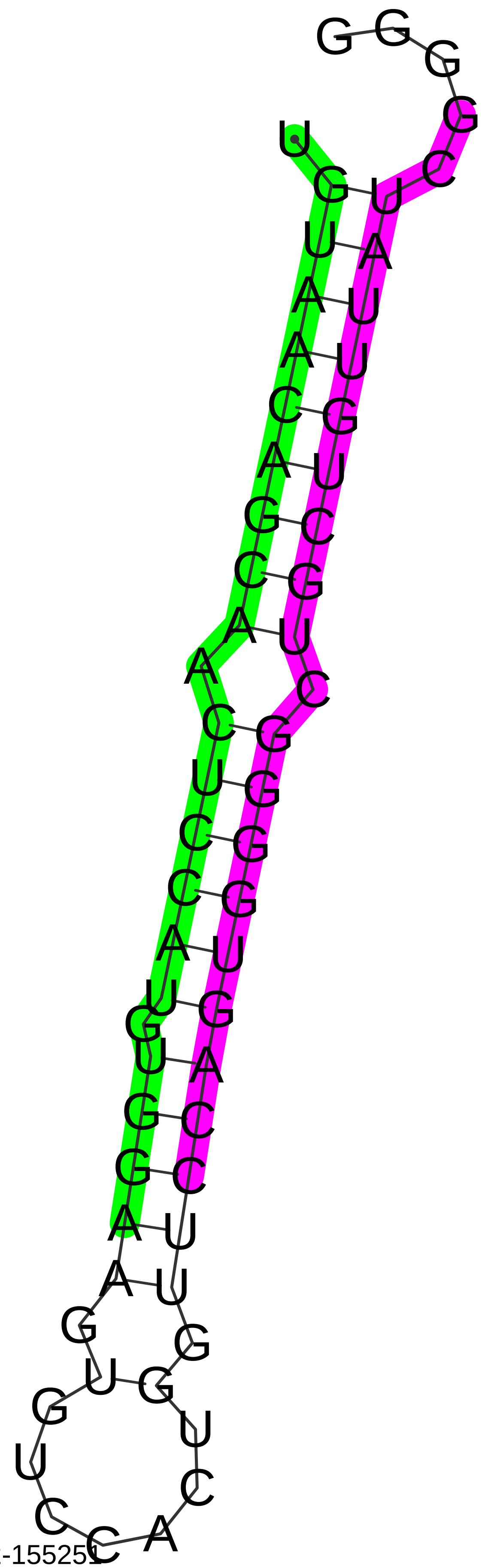

NW_006383874.1_155192-155251

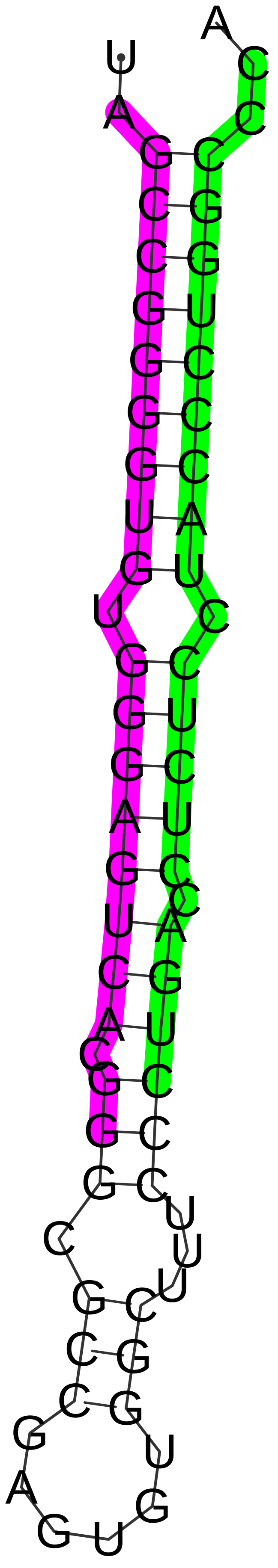

NW_006383874.1_173398-173462

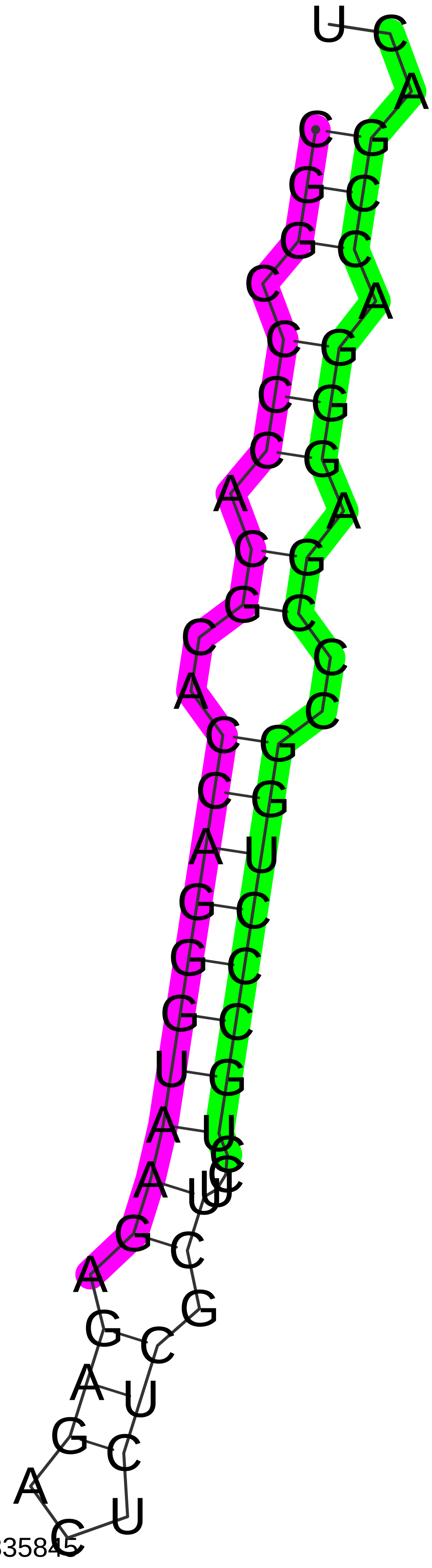

NW_006383886.1_835785-835845

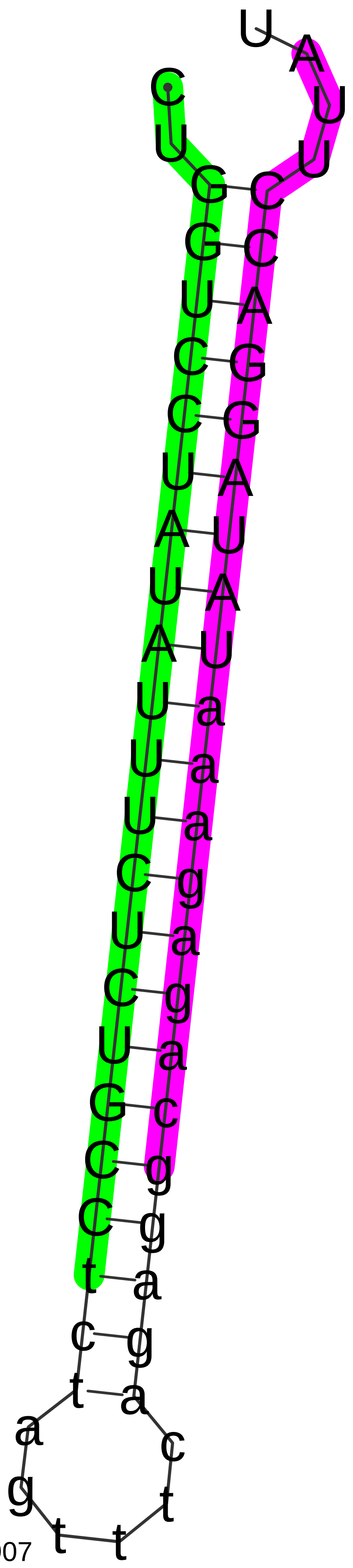


NW_006383902.1_322852-322907

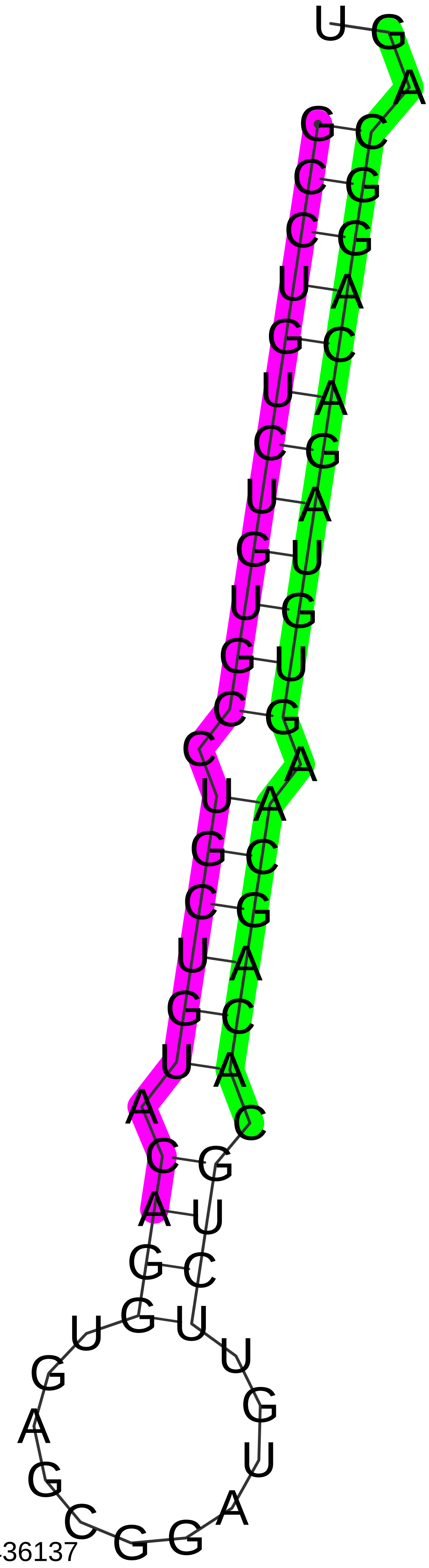

NW_006383916.1_436076-436137

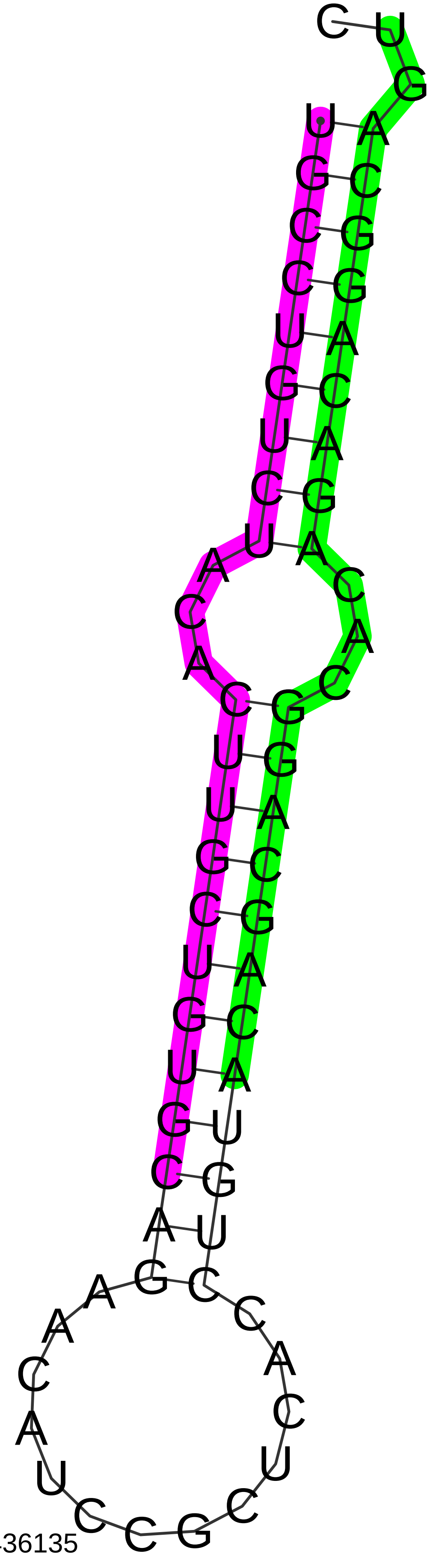

NW_006383916.1_436072-436135

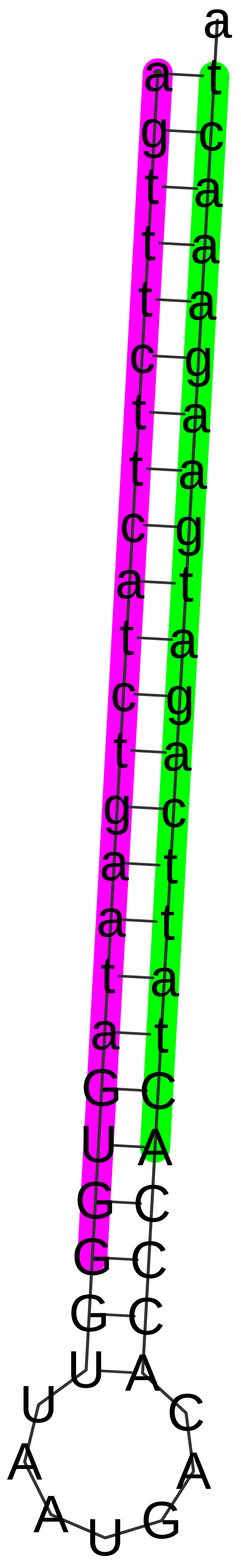

NW_006383927.1_824975-825030

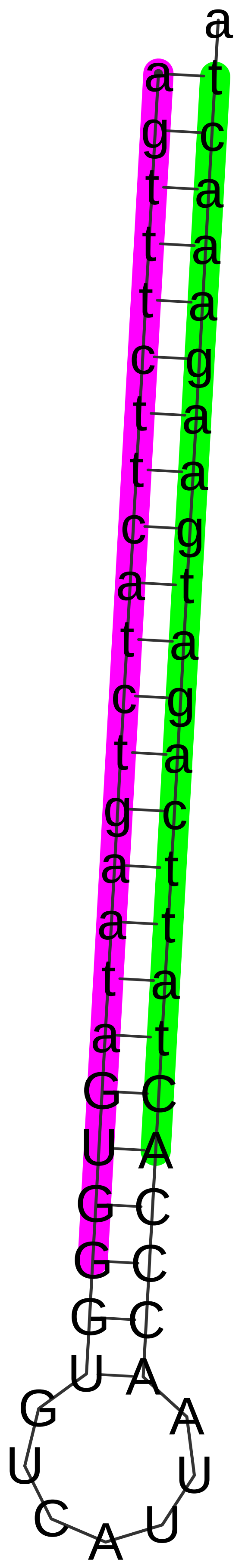


NW_006383927.1_824974-825029

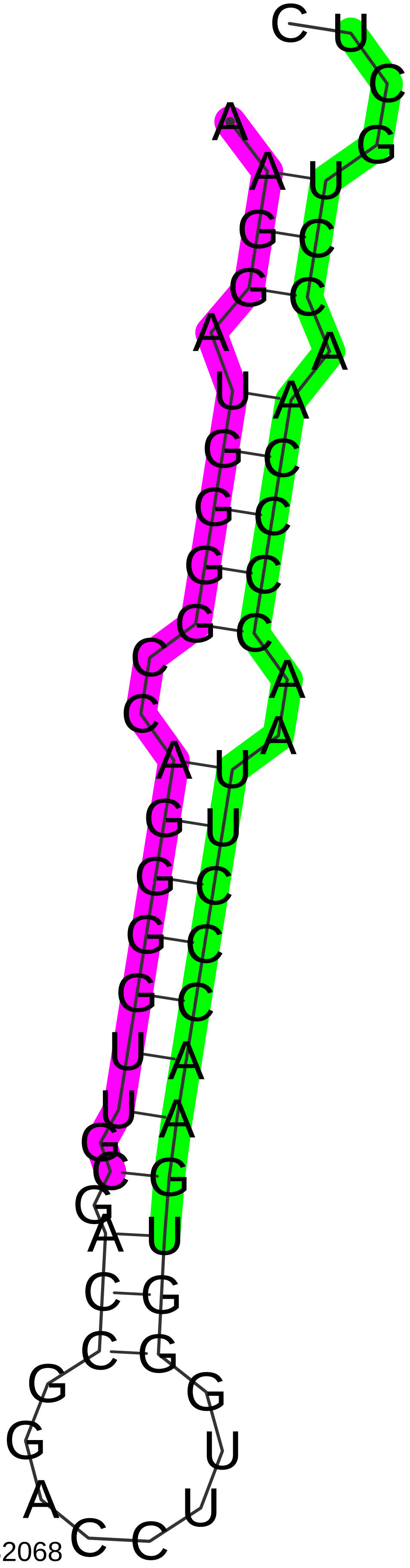

NW_006383948.1_632010-632068

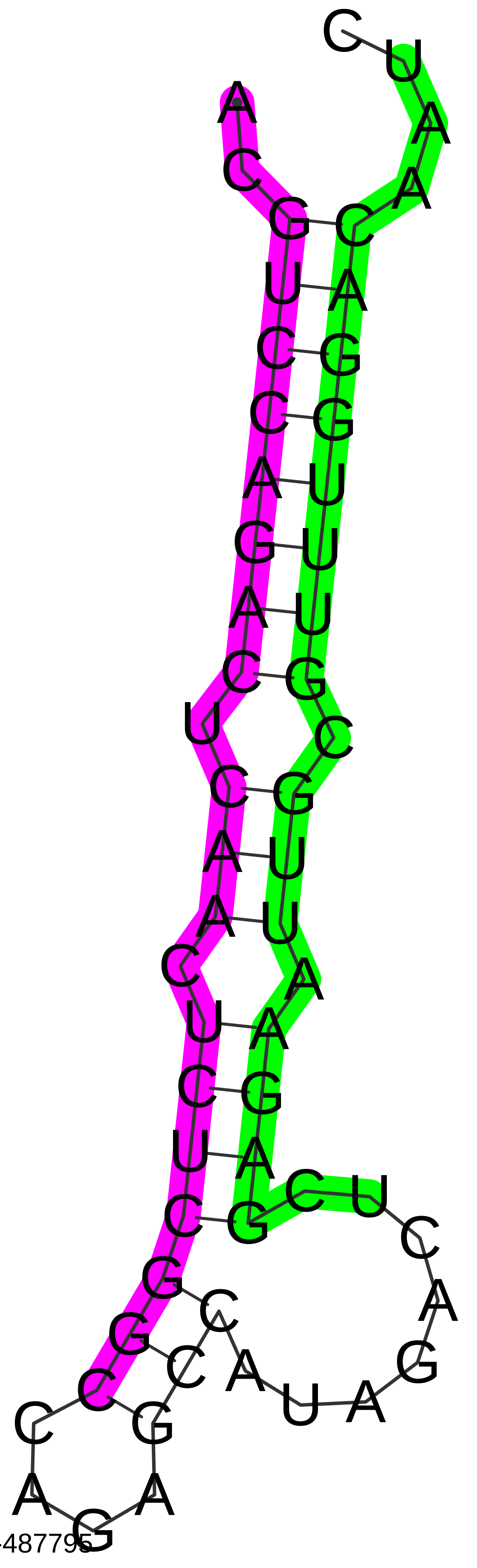

NW_006383968.1_487738-487795

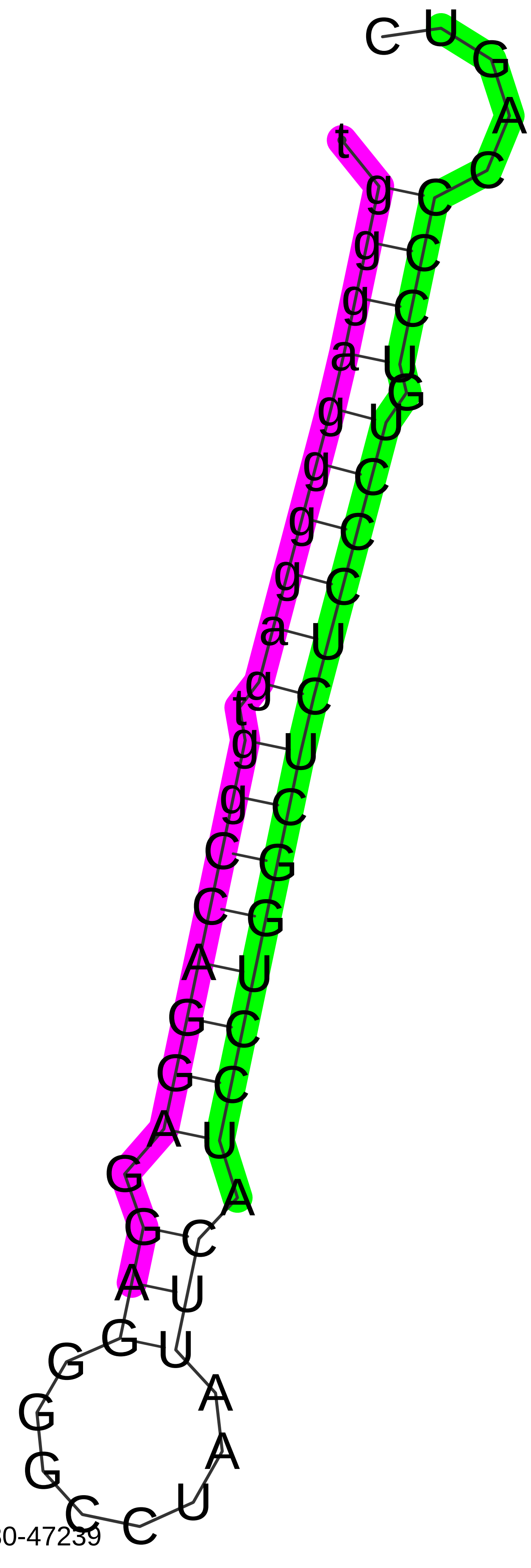

NW_006383971.1_47180-47239

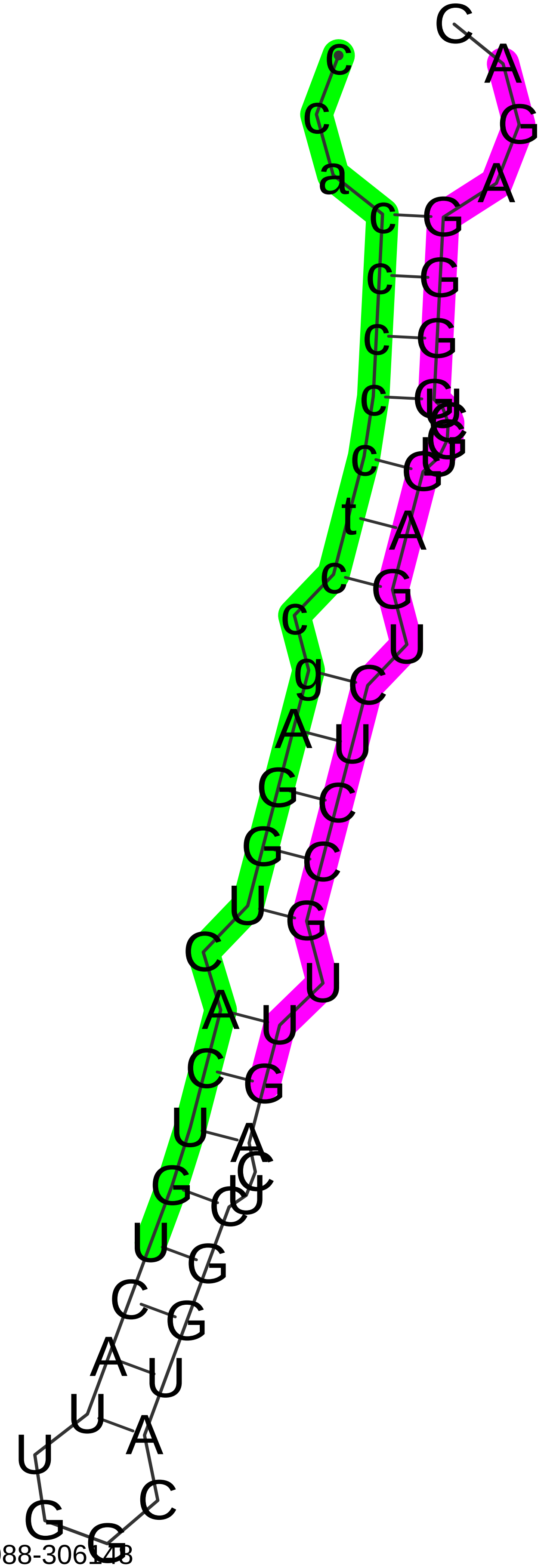

NW_006383984.1_306088-306148

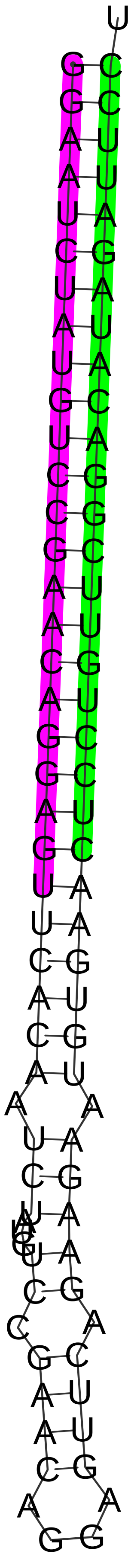

NW_006384008.1_338747-338832

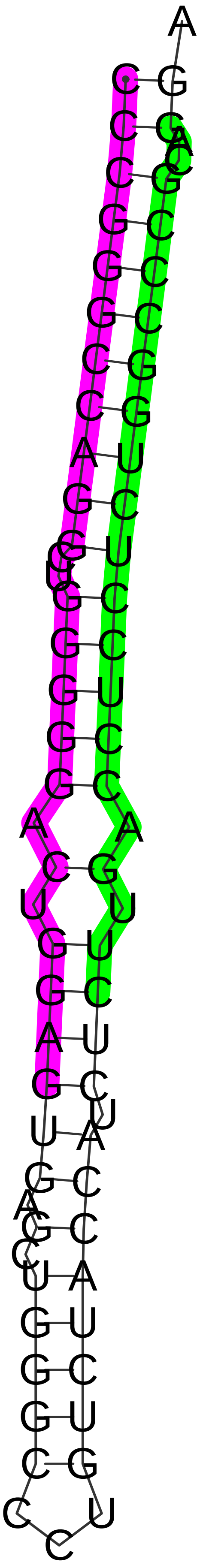

NW_006384039.1_487896-487968

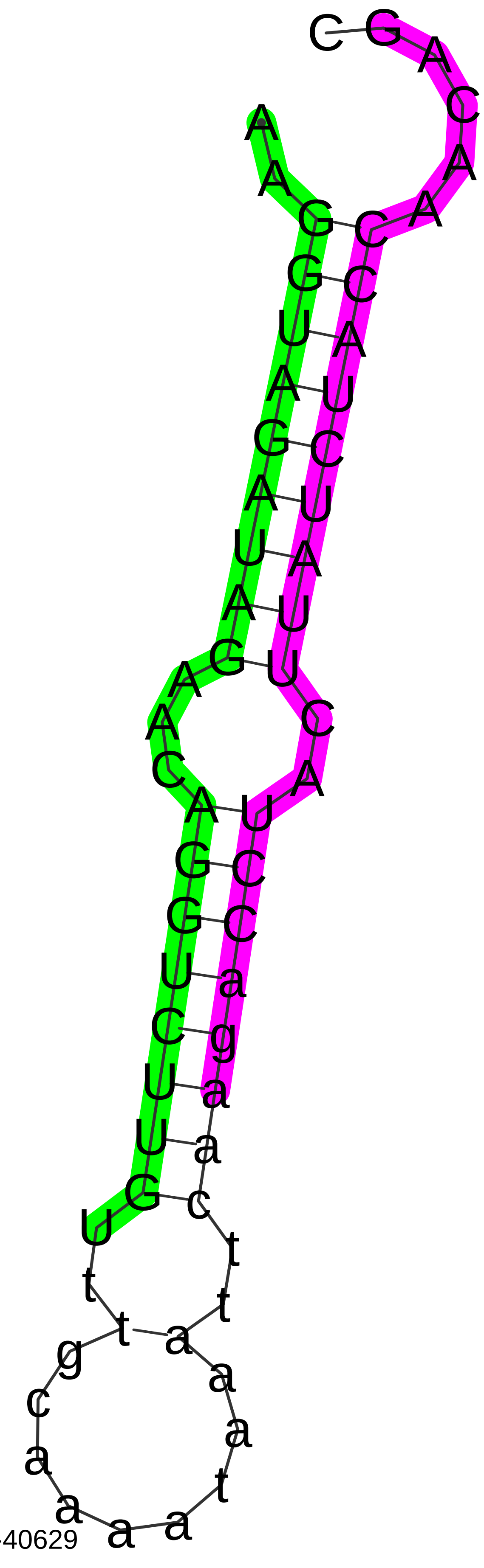

NW_006384074.1_40568-40629

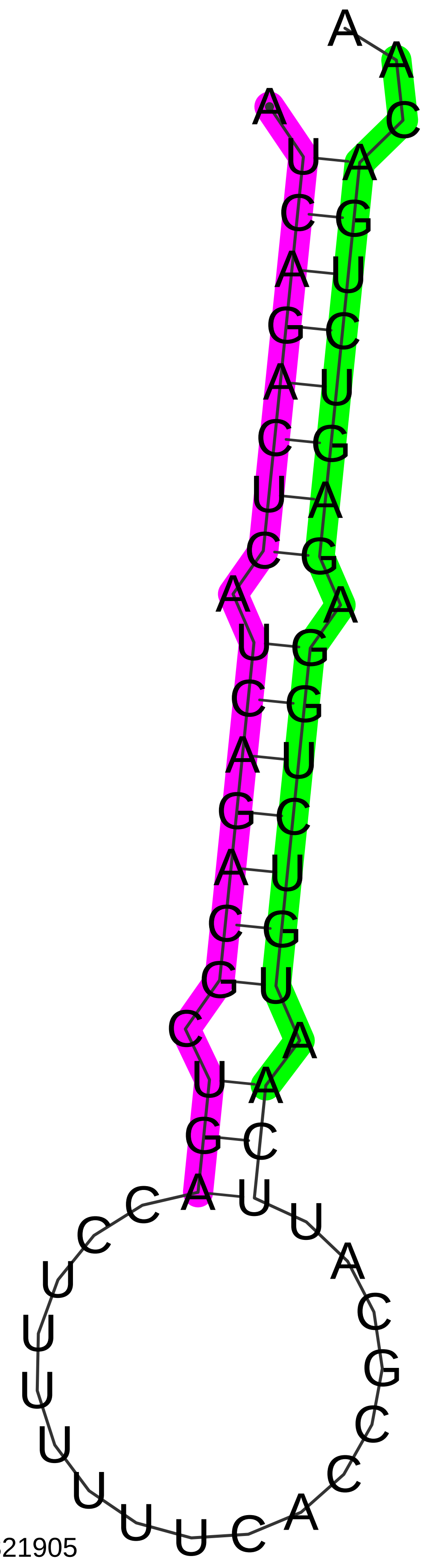

NW_006384099.1_821845-821905

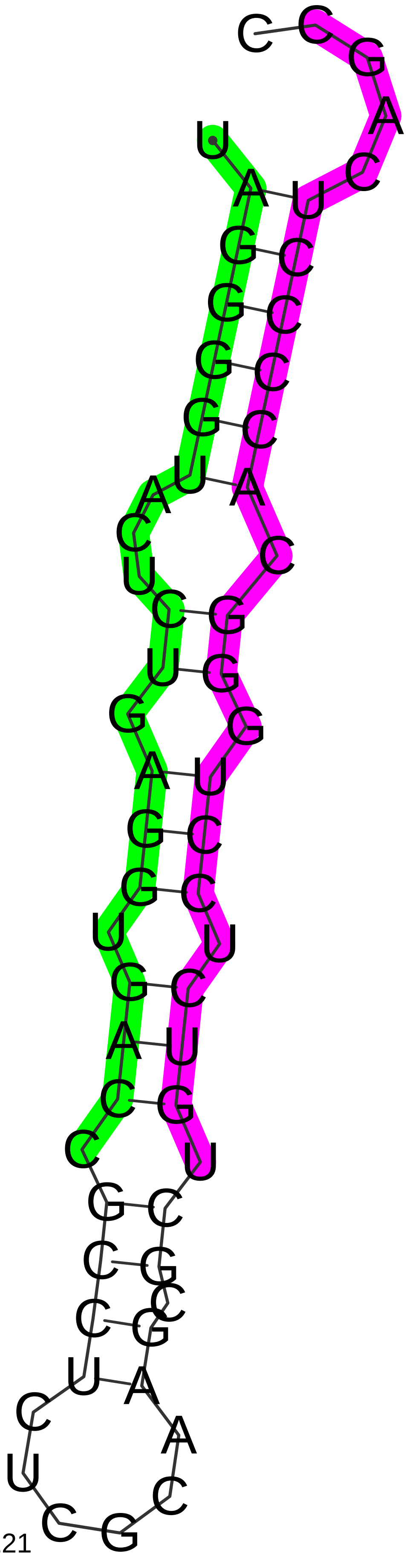

NW_006384131.1_90163-90221

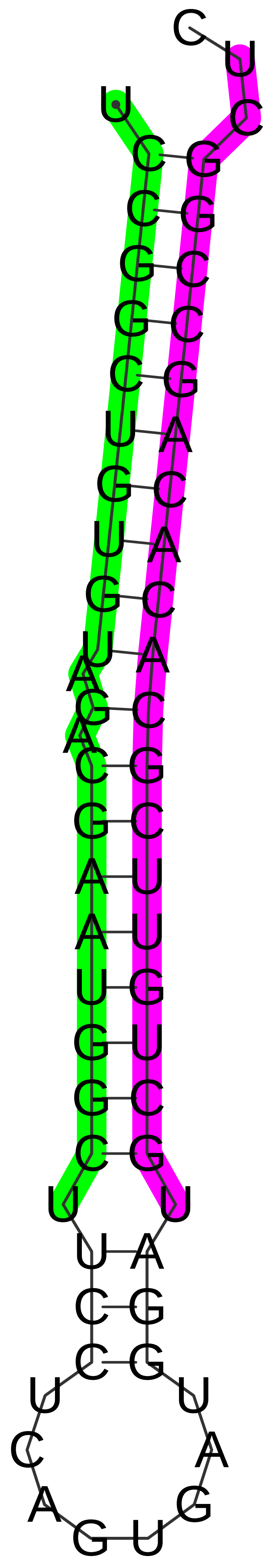

NW_006384141.1_598043-598102

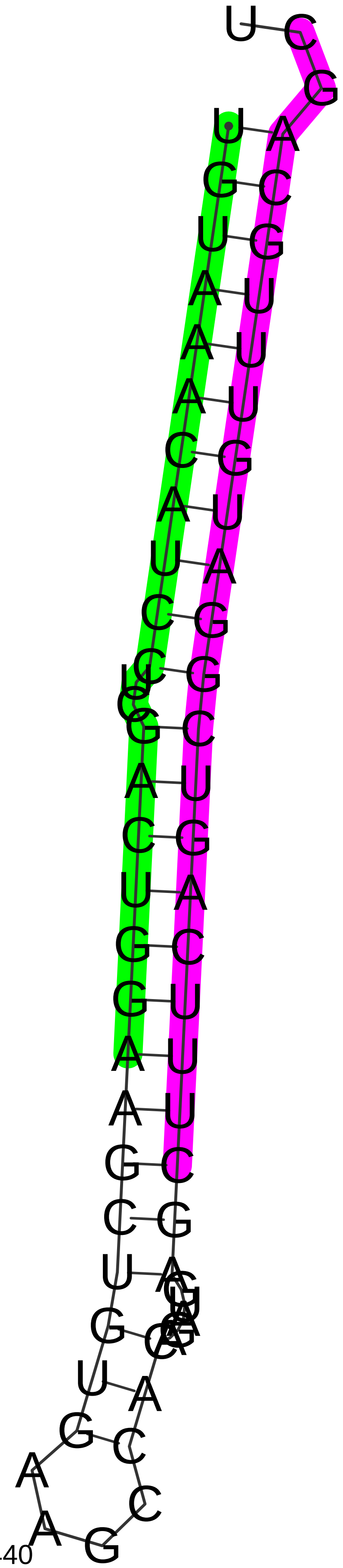

NW_006384161.1_766377-766440

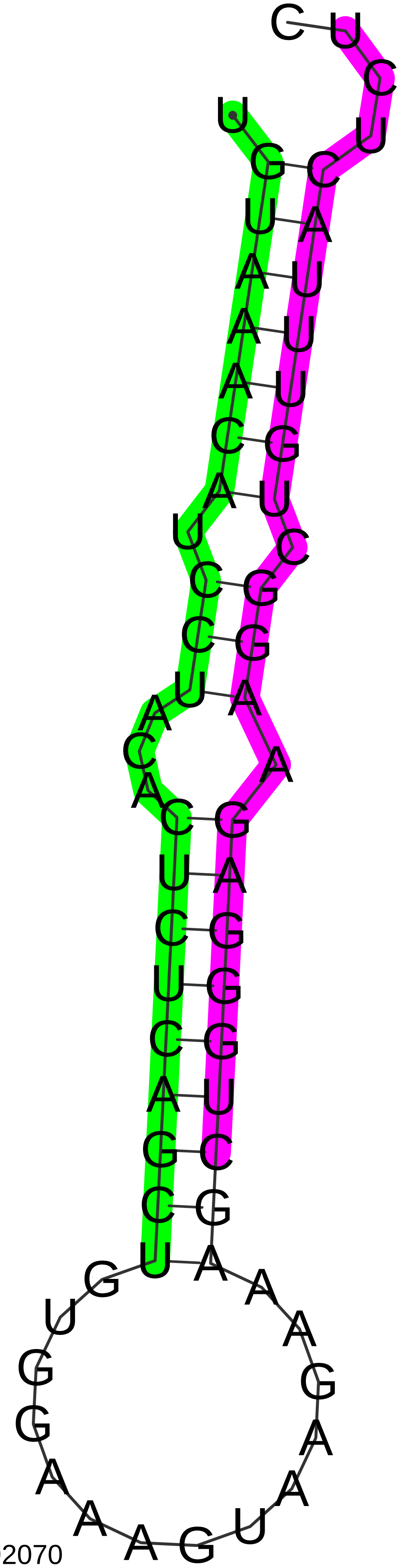


NW_006384161.1_792008-792070

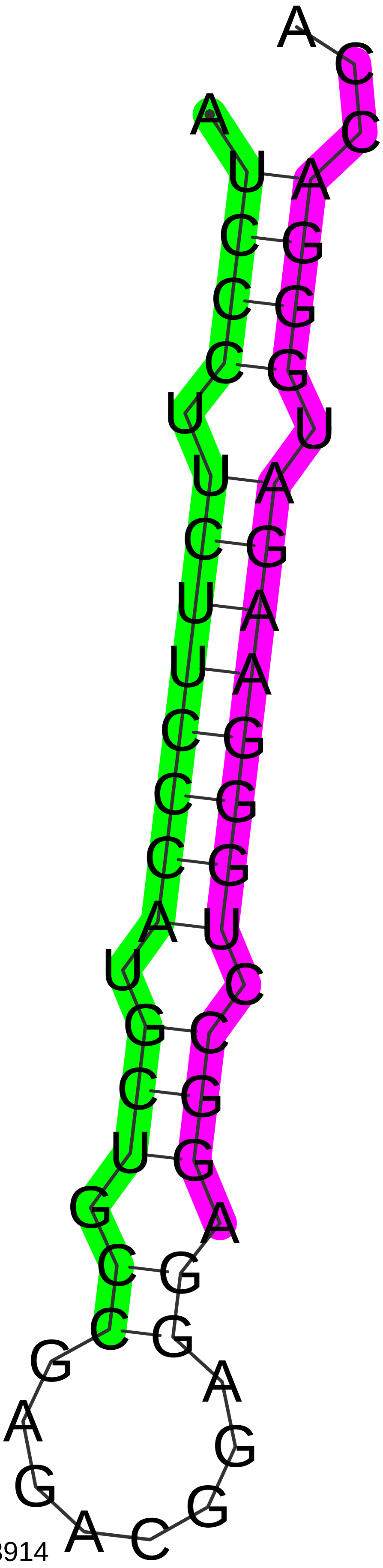

NW_006384171.1_793863-793914

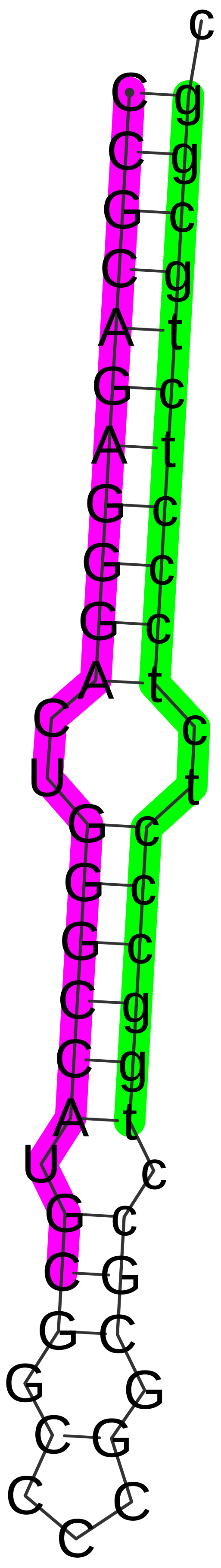

NW_006384171.1_596299-596352

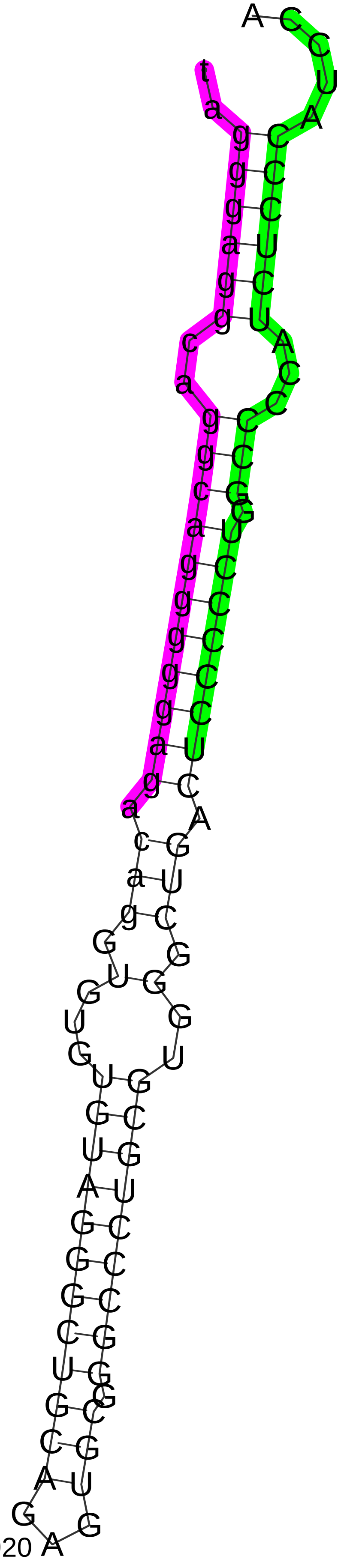

NW_006384180.1_158829-158920

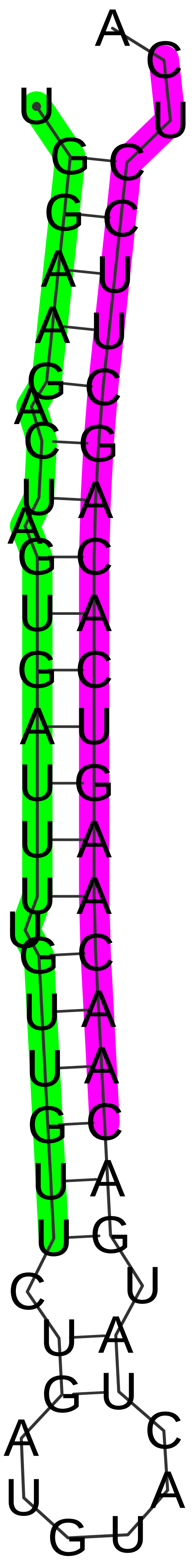

NW_006384182.1_60146-60204

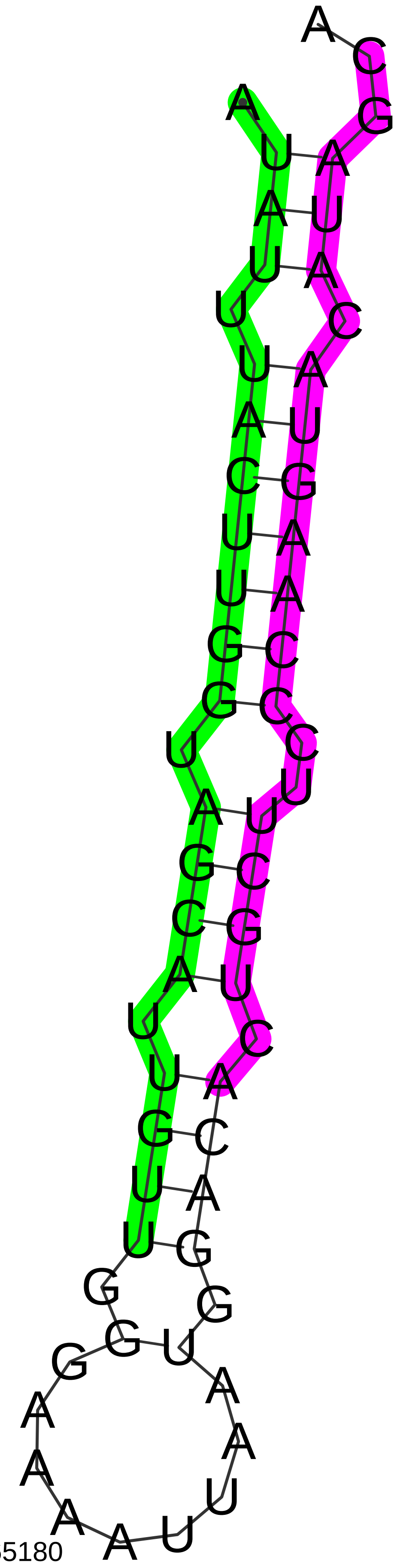

NW_006384198.1_865121-865180

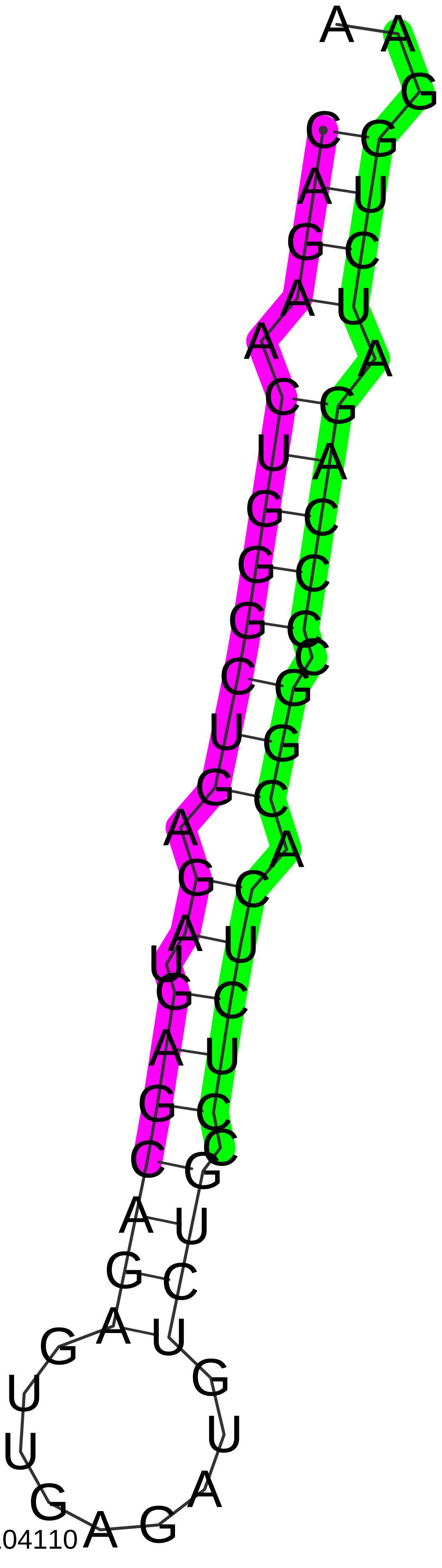

NW_006384260.1_104050-104110

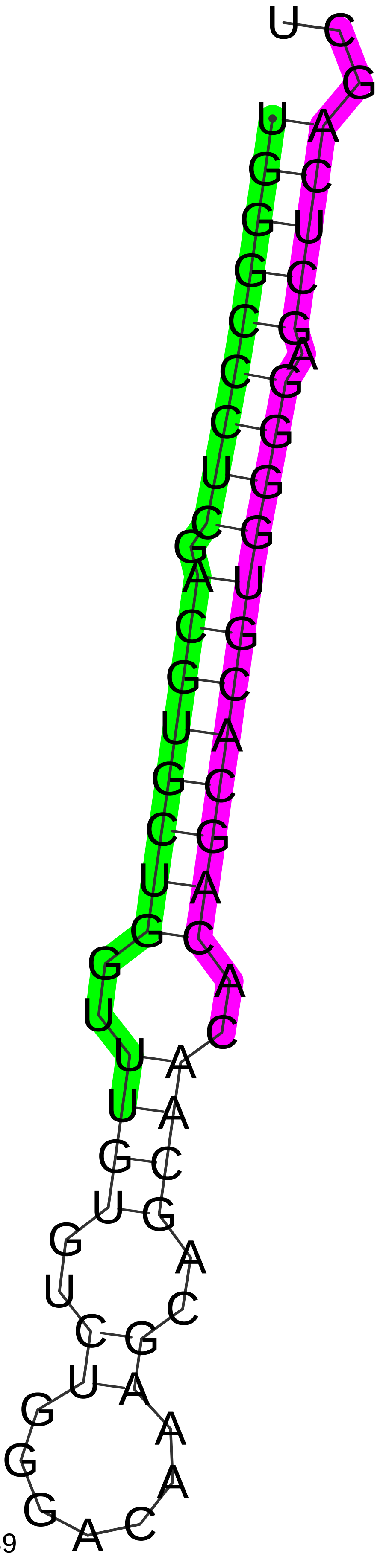

NW_006384301.1_74574-74639

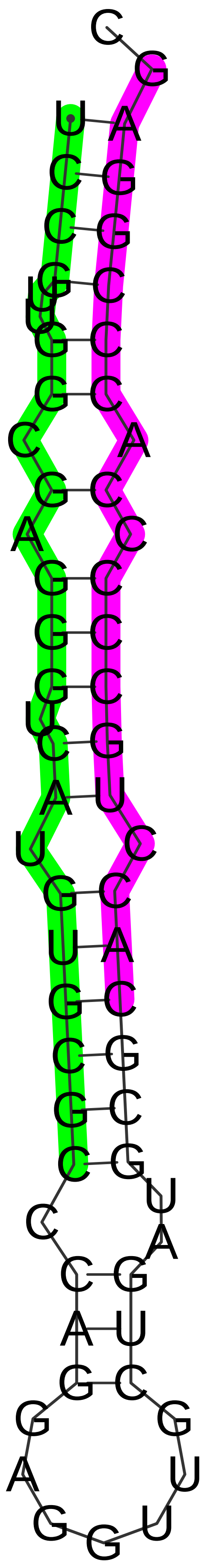

NW_006384367.1_558162-558224

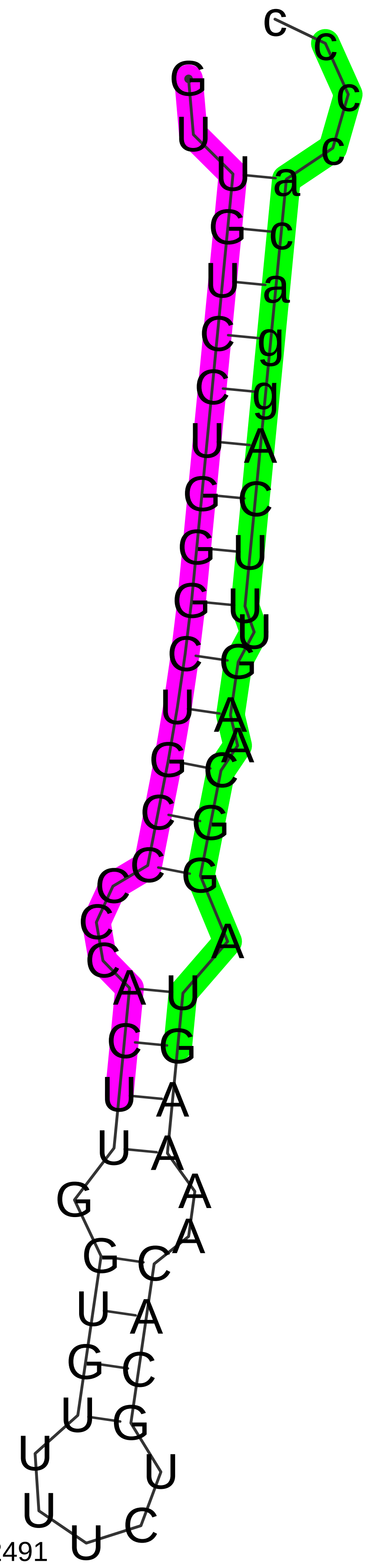

NW_006384417.1_462428-462491

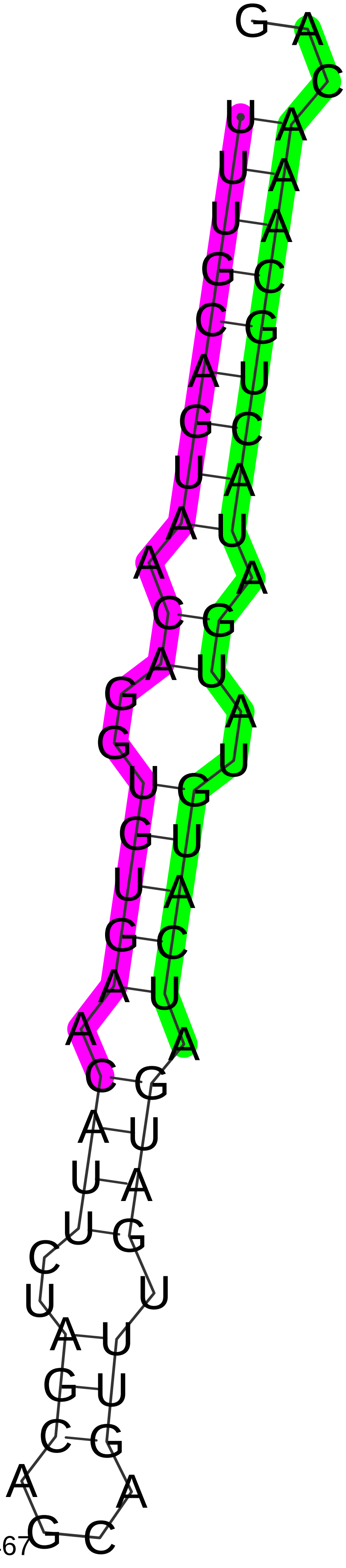

NW_006384417.1_459404-459467

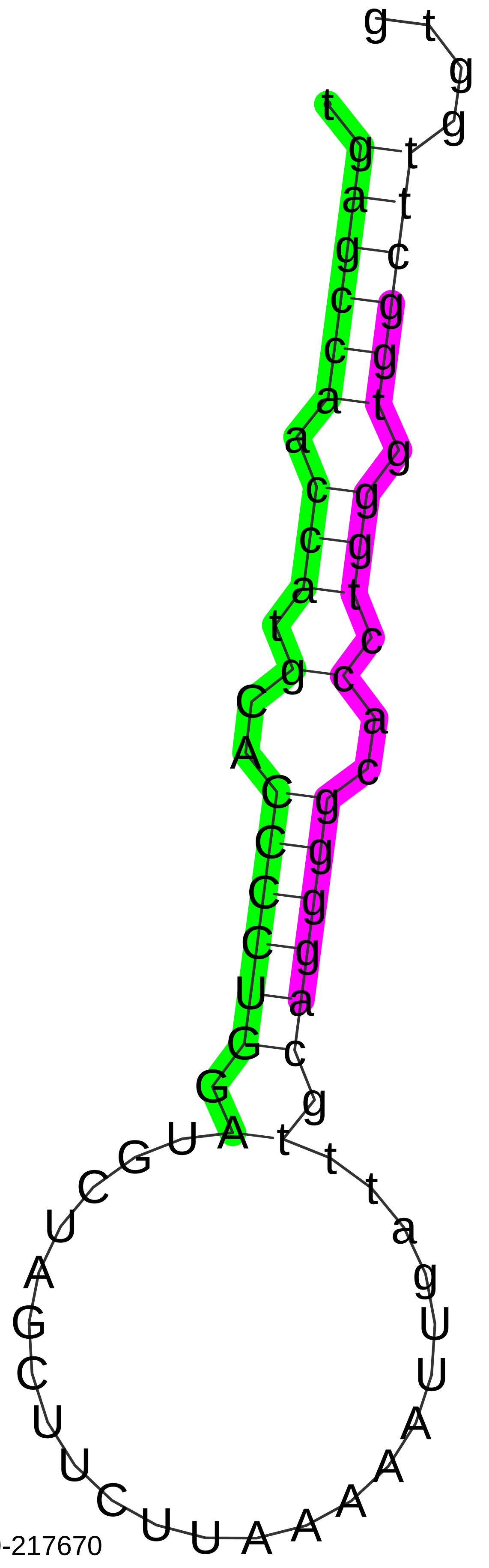

NW_006384440.1_217599-217670

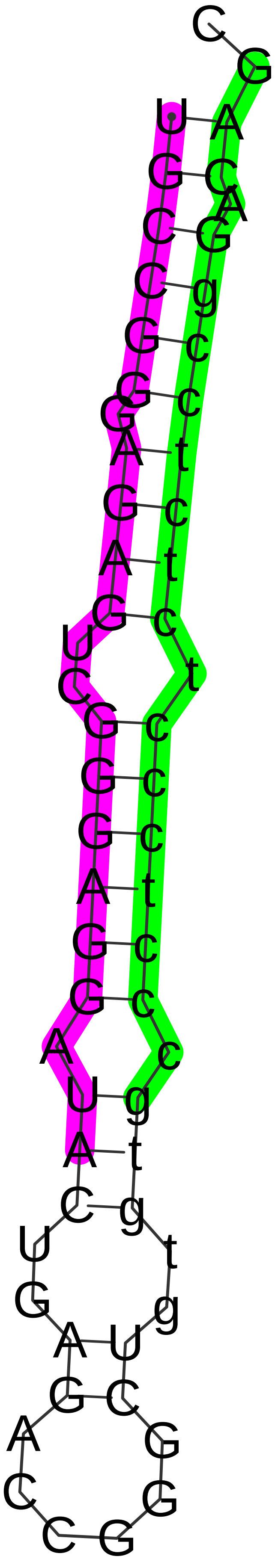

NW_006384491.1_13121-13181

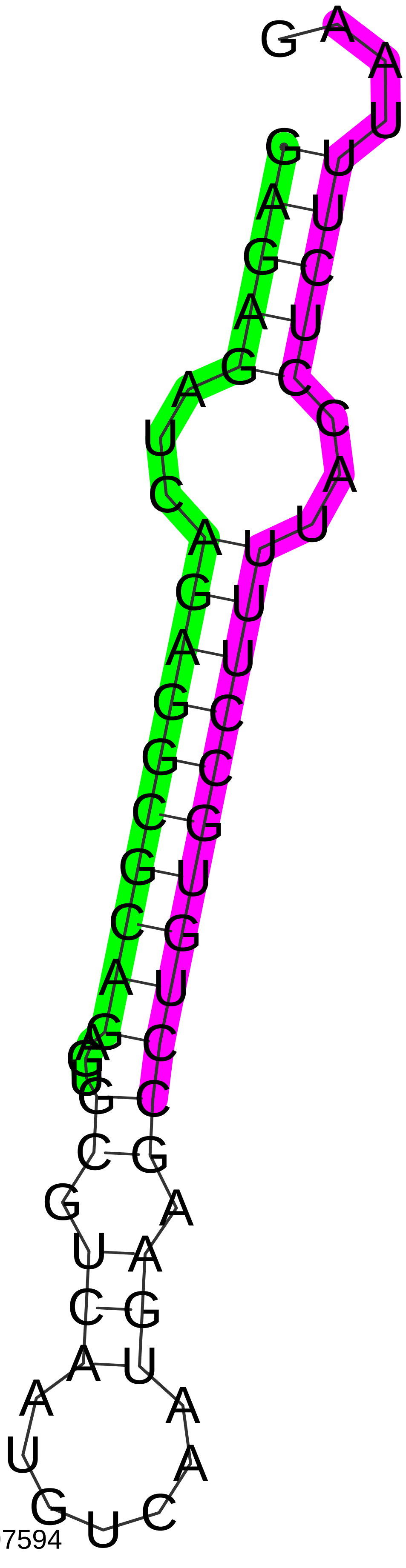

NW_006384494.1_297533-297594

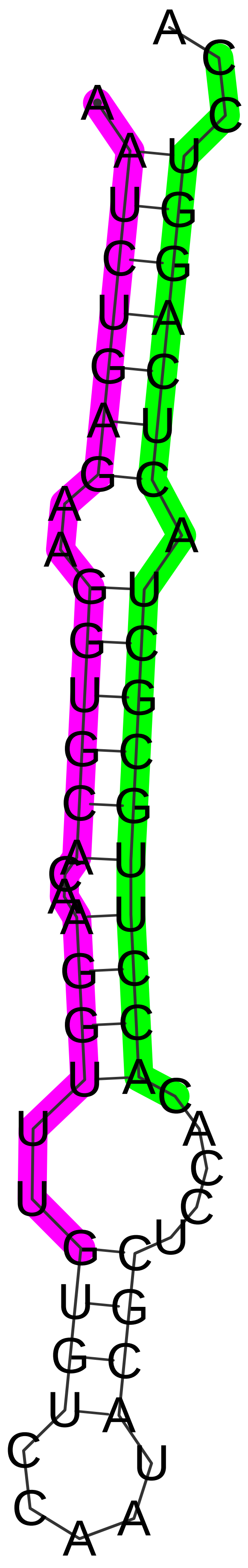

NW_006384502.1_95000-95062

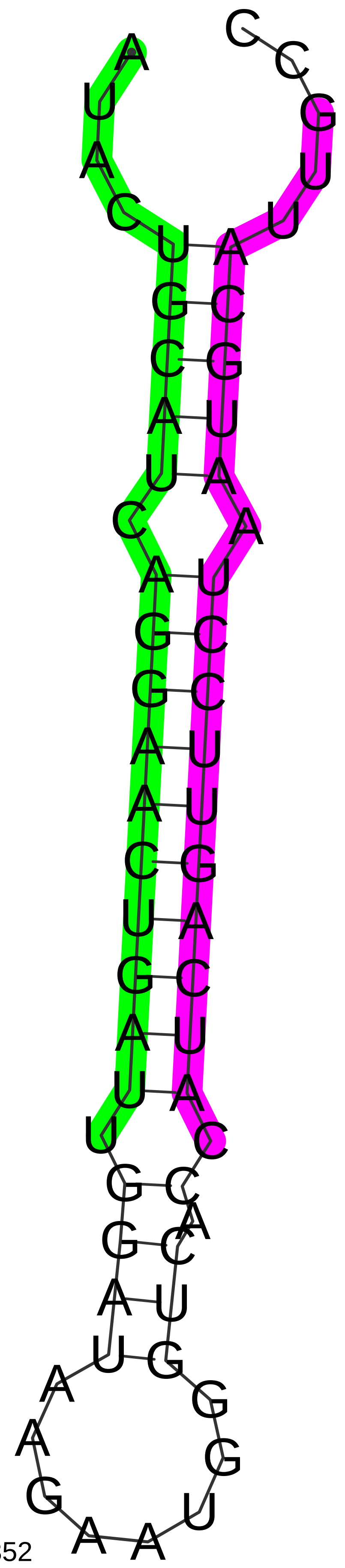
NW_006384532.1_523293-523352

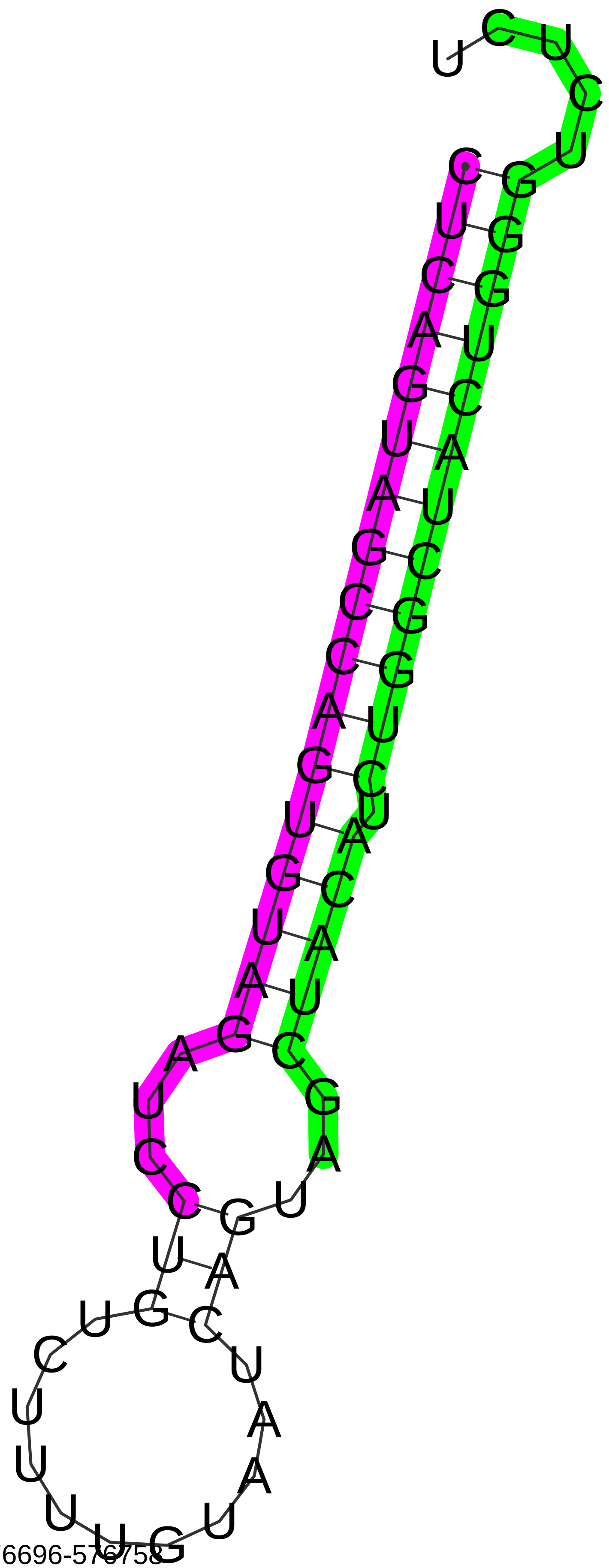

NW_006384555.1_576696-576758

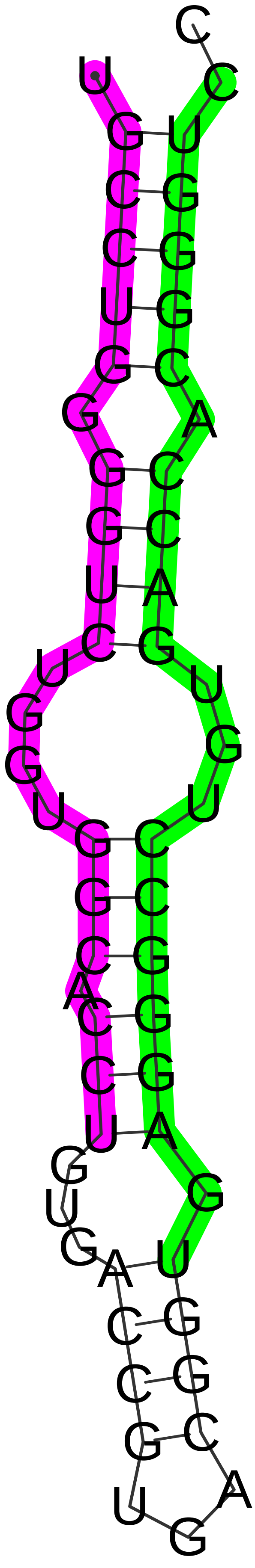

NW_006384577.1_596391-596448

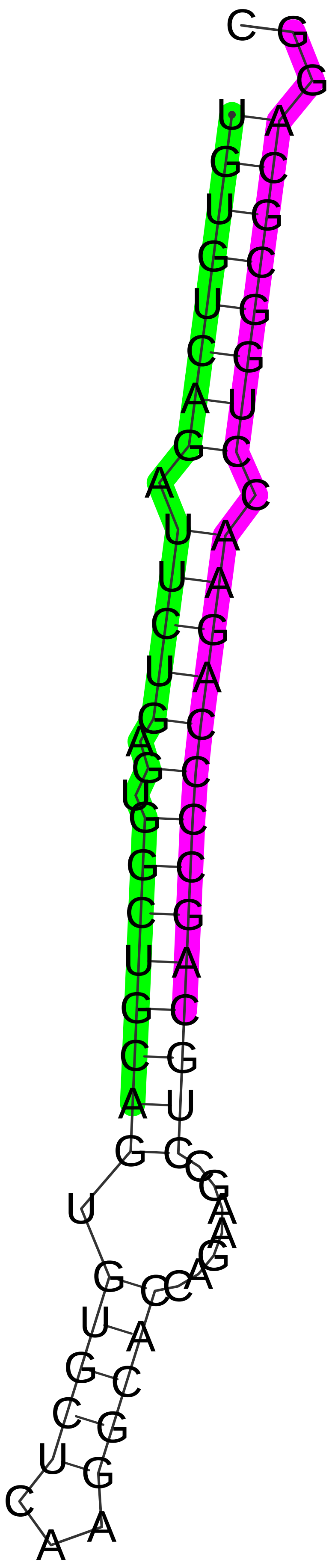

NW_006384581.1_73257-73328

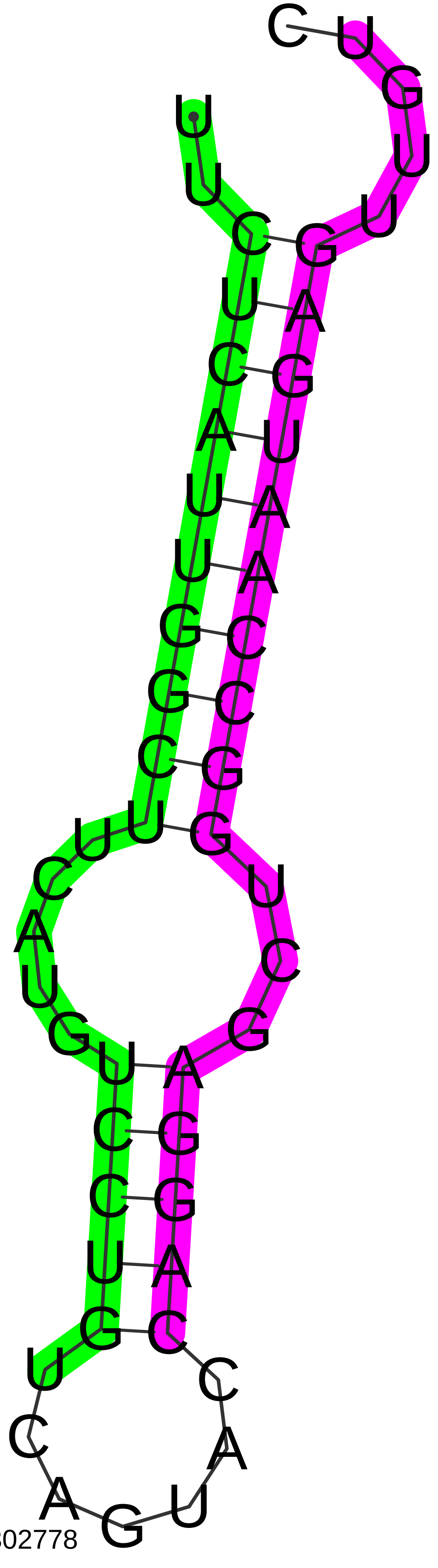

NW_006384586.1_302727-302778

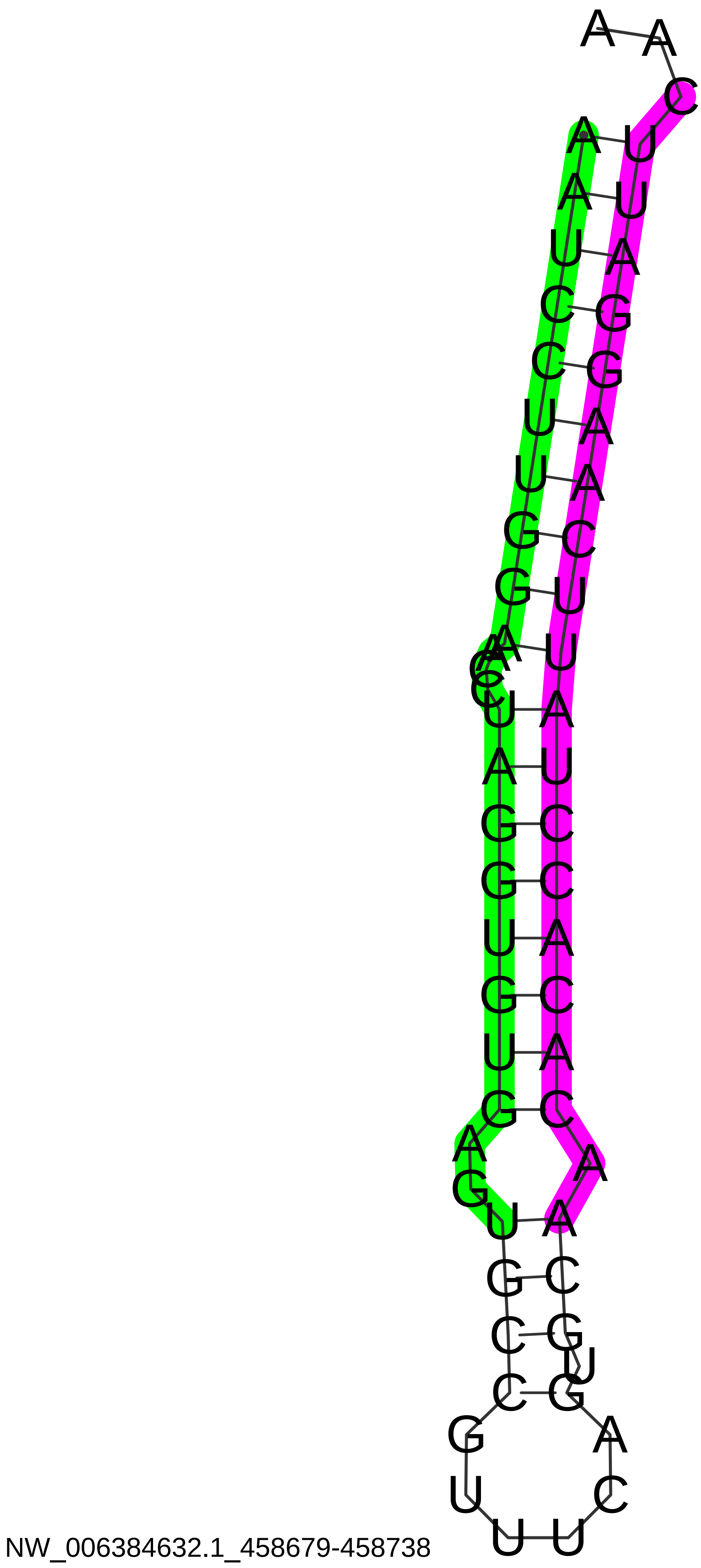

NW_006384632.1_458679-458738

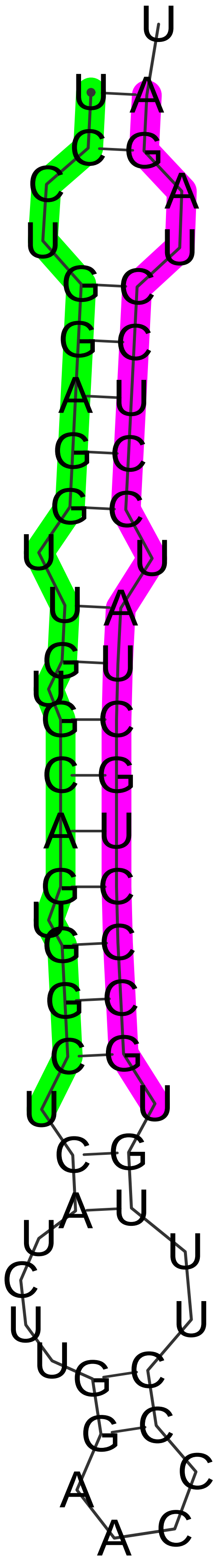

NW_006384689.1_99494-99554

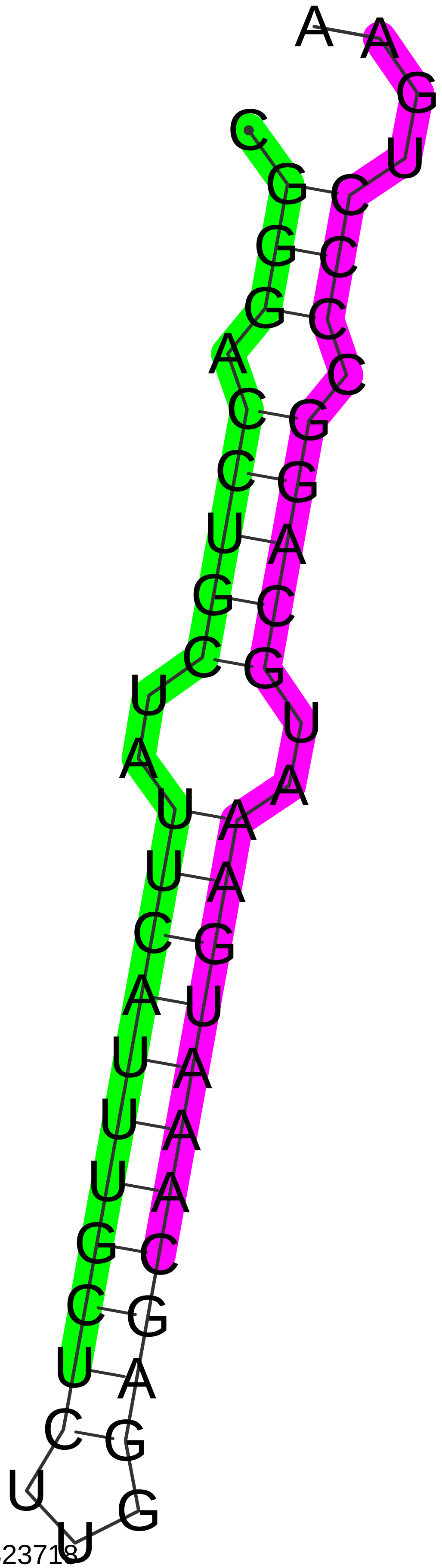

NW_006384749.1_323667-323718

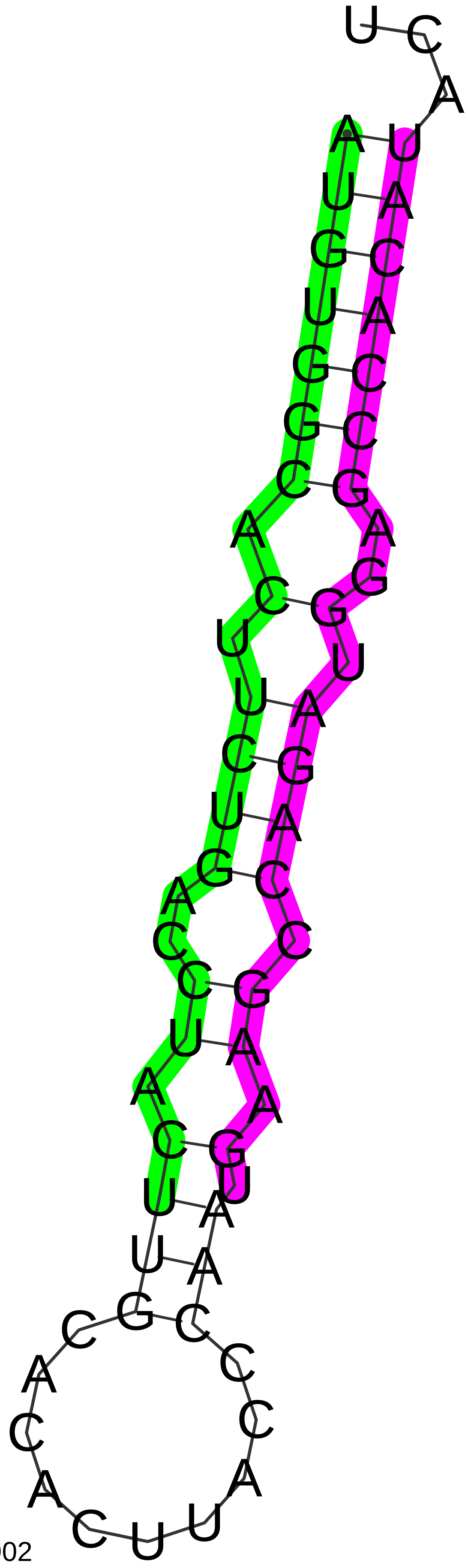

NW_006384770.1_9843-9902

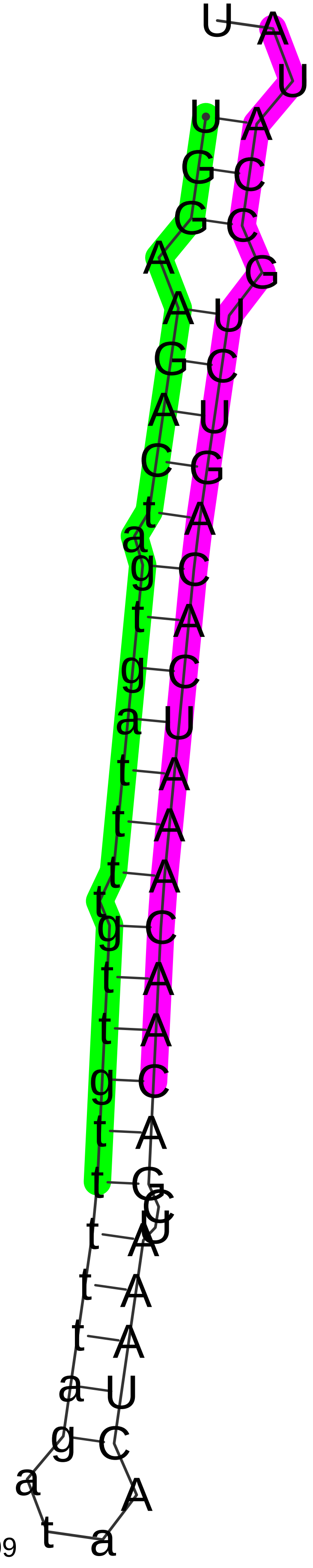

NW_006384820.1_394145-394209

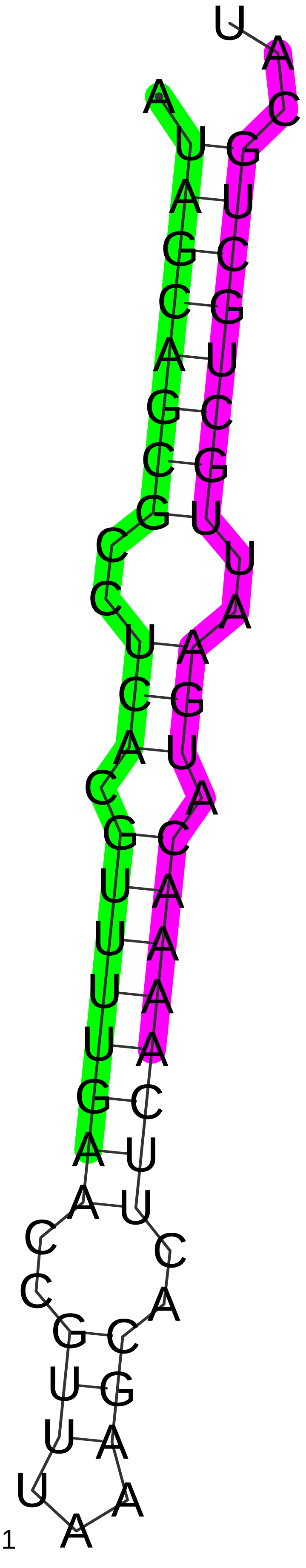

NW_006384929.1_165941-166001

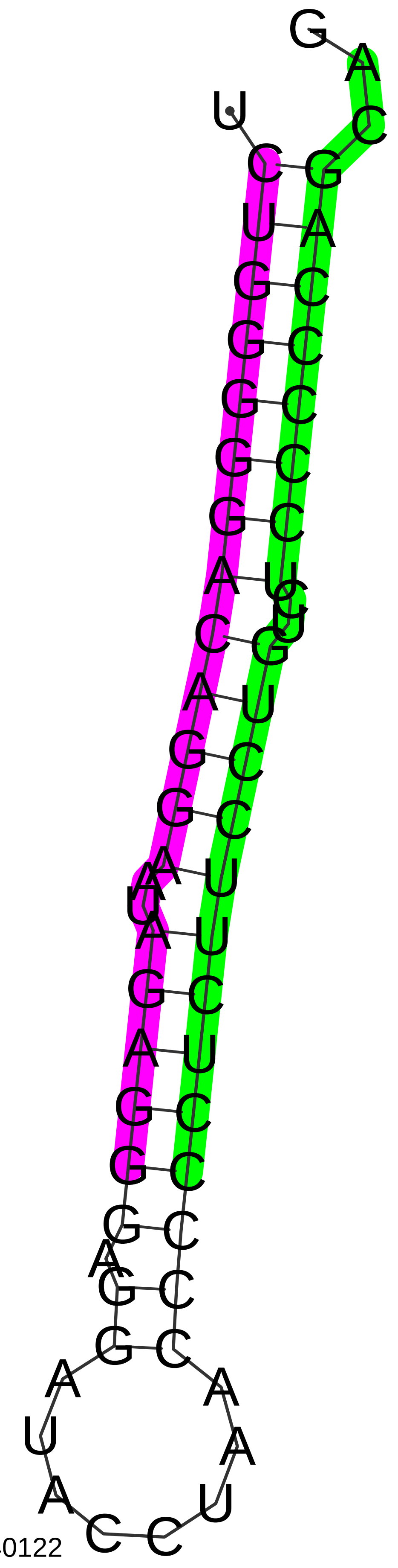

NW_006384946.1_440064-440122

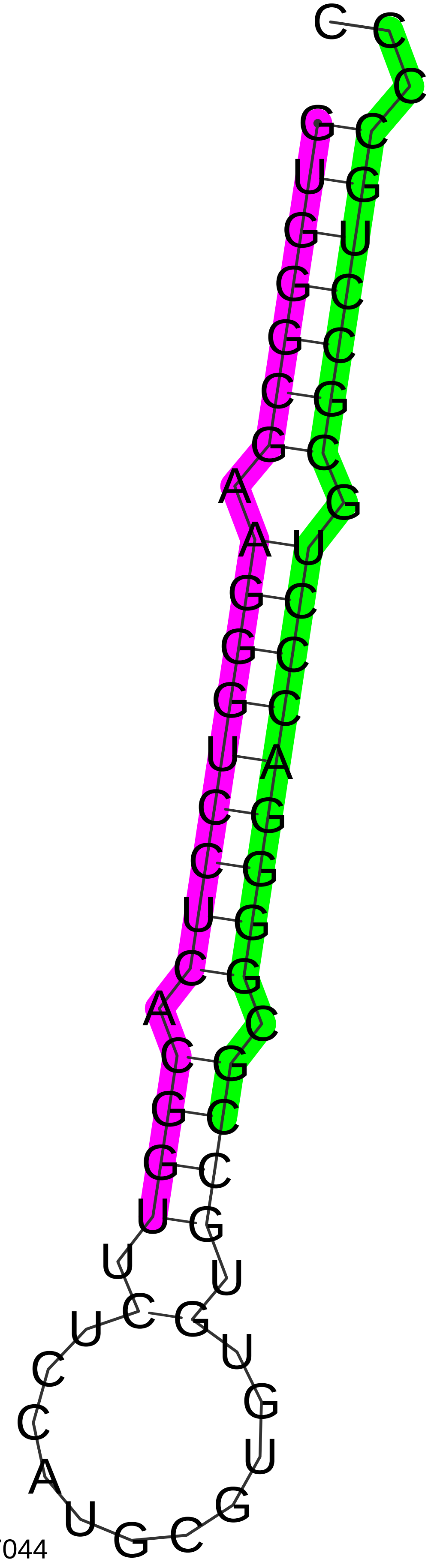

NW_006384996.1_56983-57044

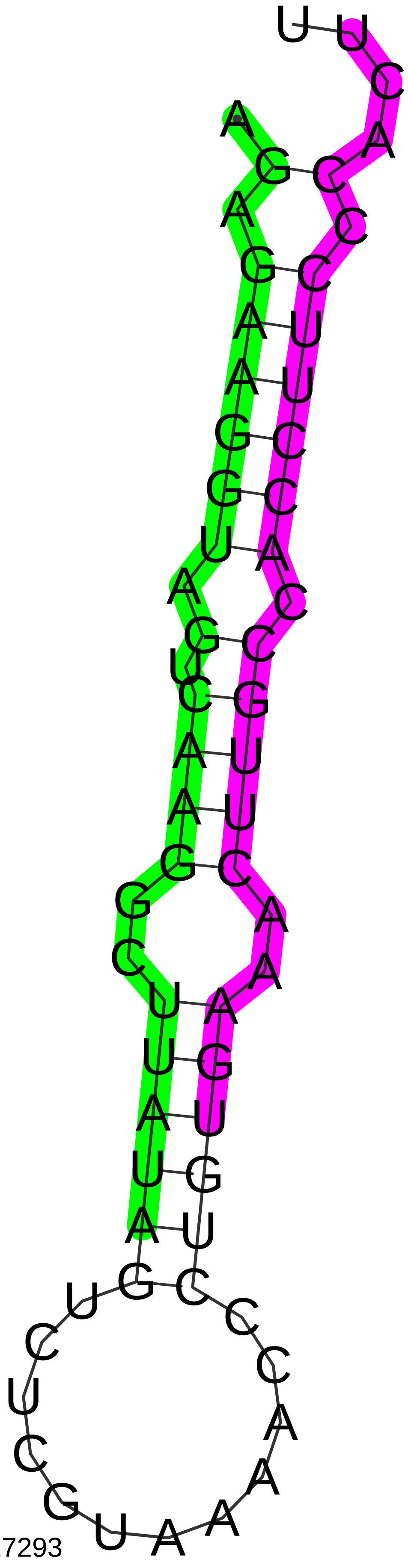

NW_006384999.1_217232-217293

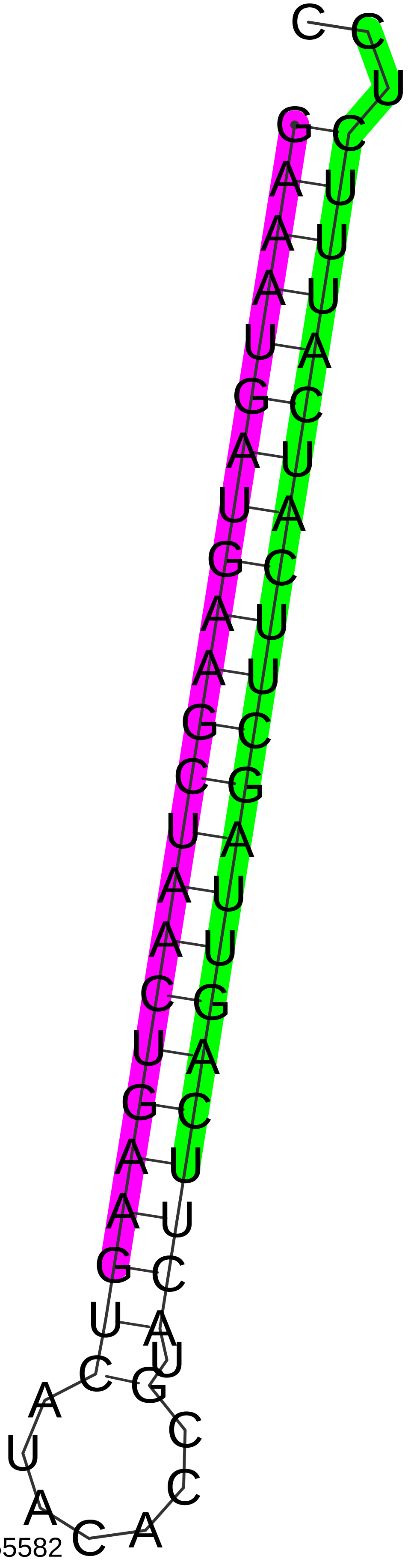

NW_006385011.1_455524-455582

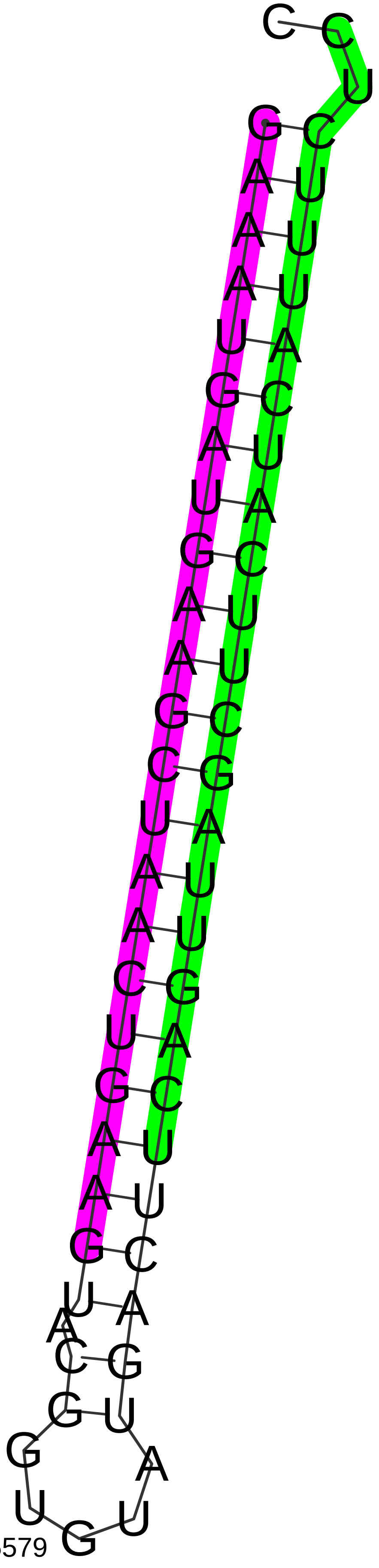

NW_006385011.1_455521-455579

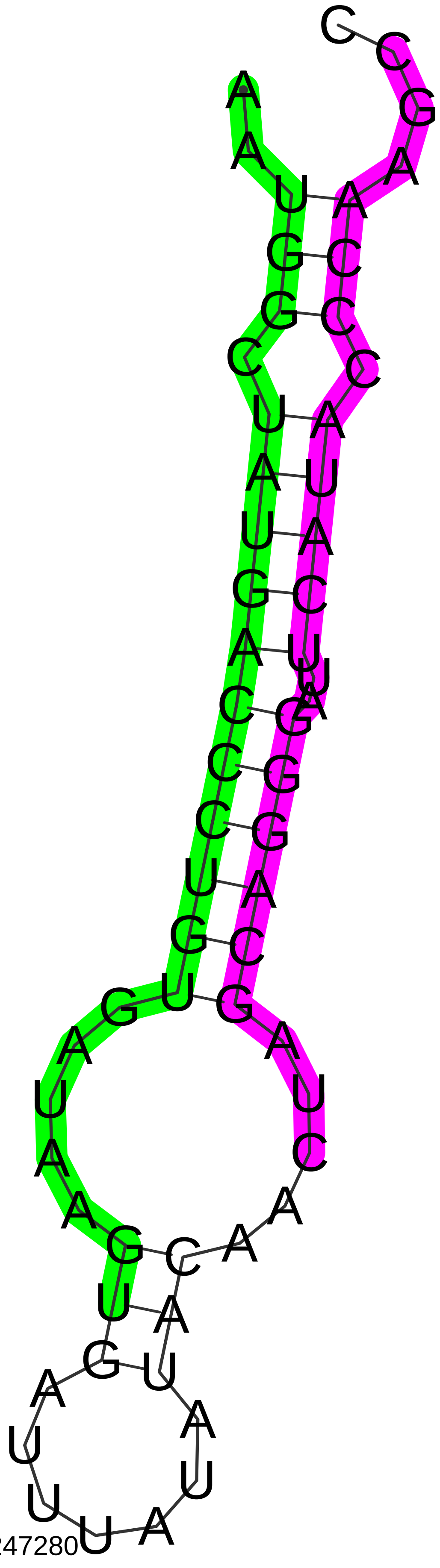

NW_006385054.1_247220-247280

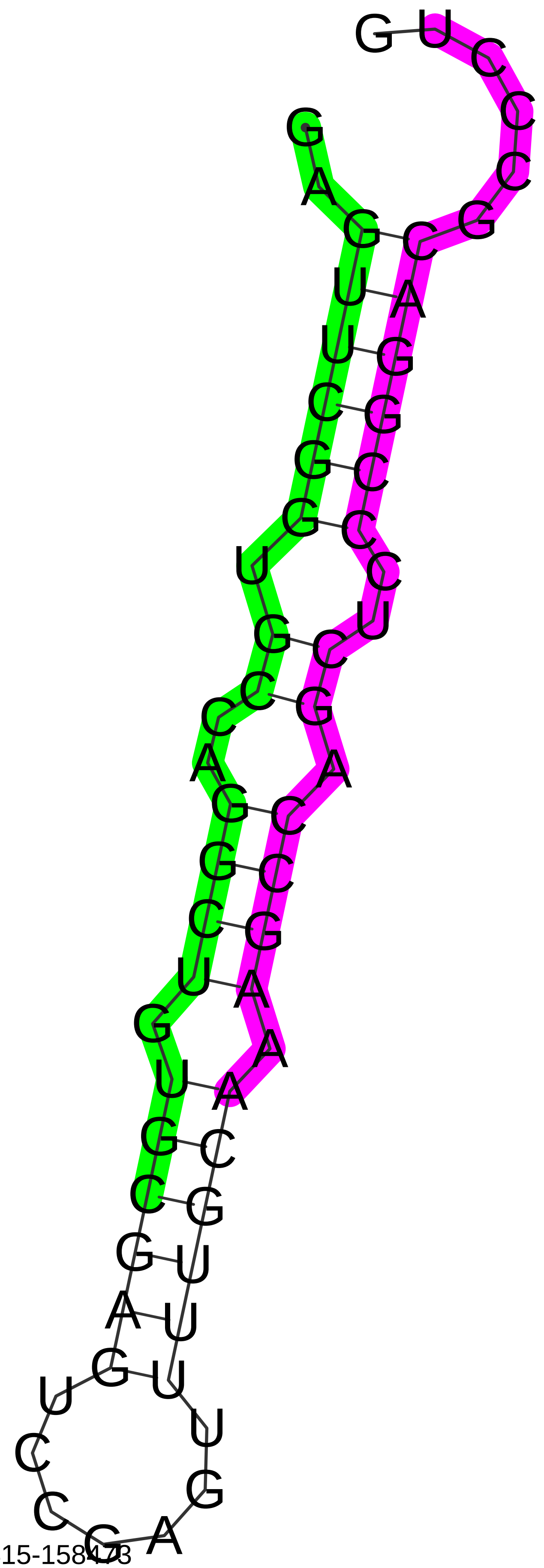

NW_006385069.1_158415-158473

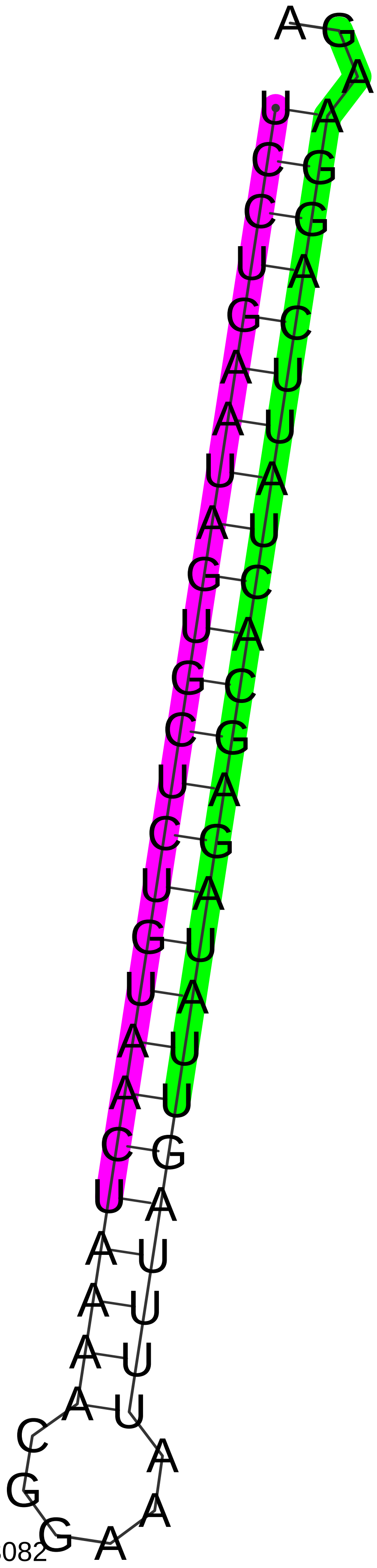

NW_006385135.1_263022-263082

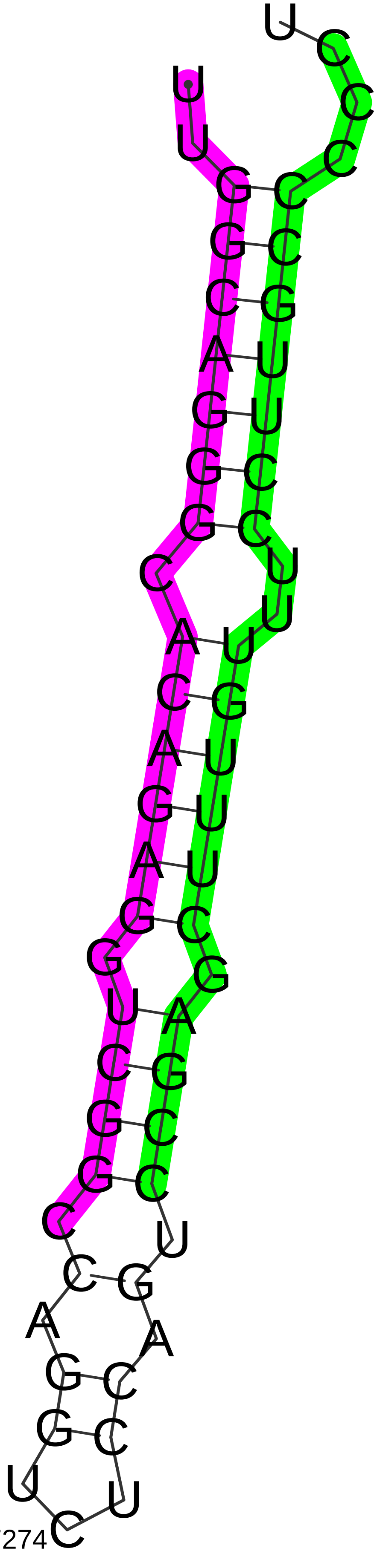

NW_006385183.1_317217-317274

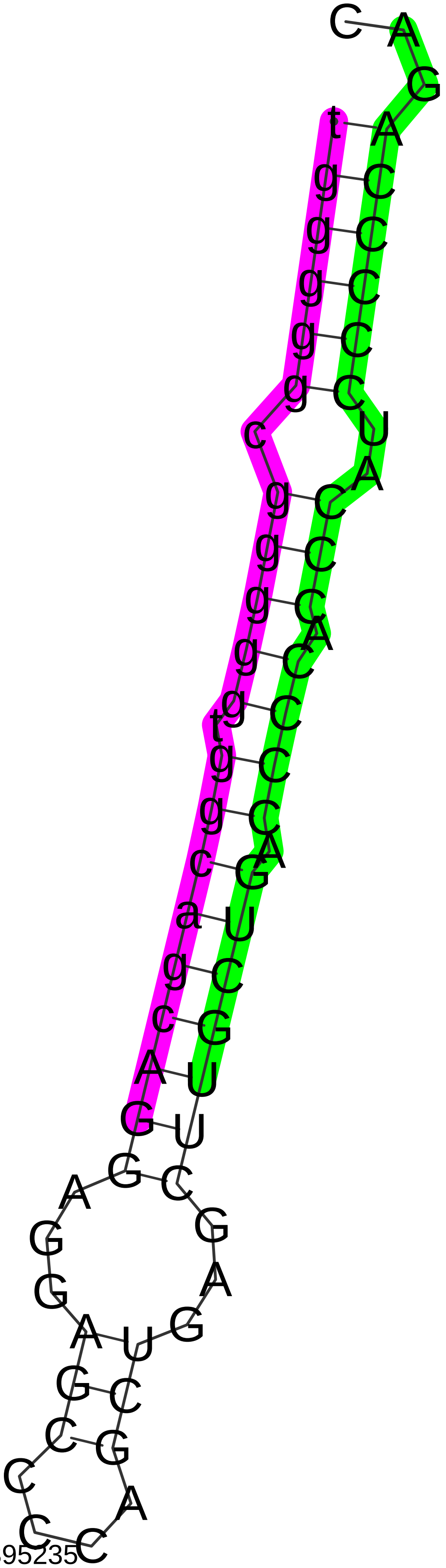

NW_006385255.1_395171-395235

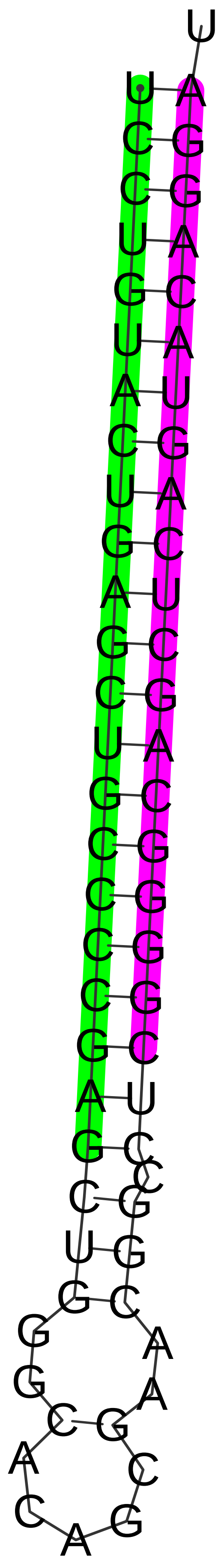

NW_006385323.1_423228-423290

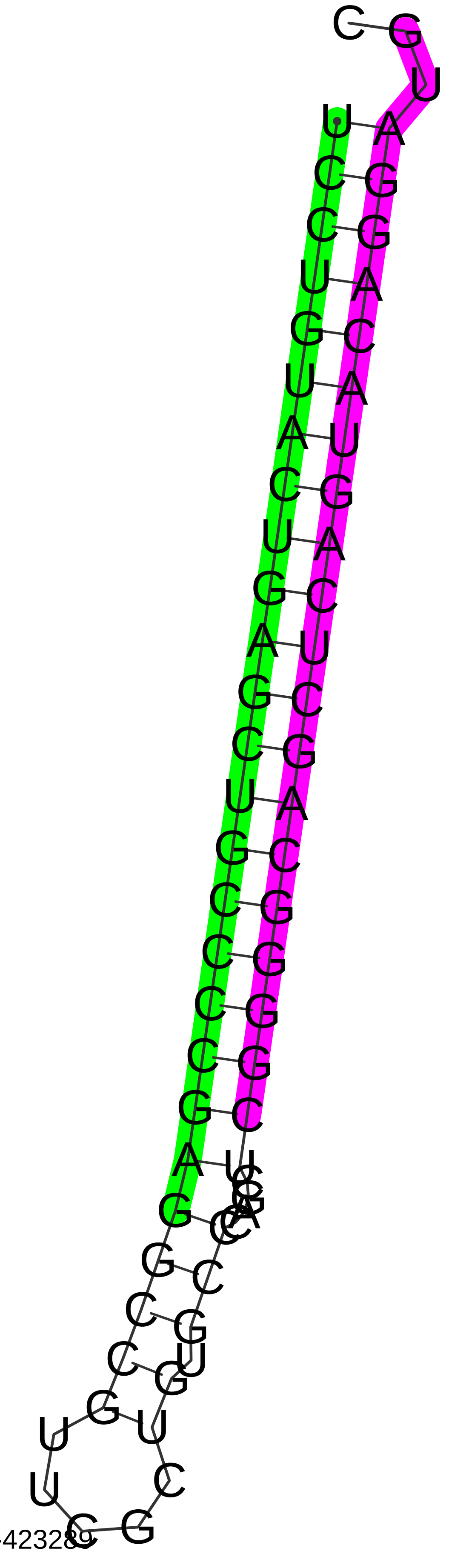

NW_006385323.1_423225-423289

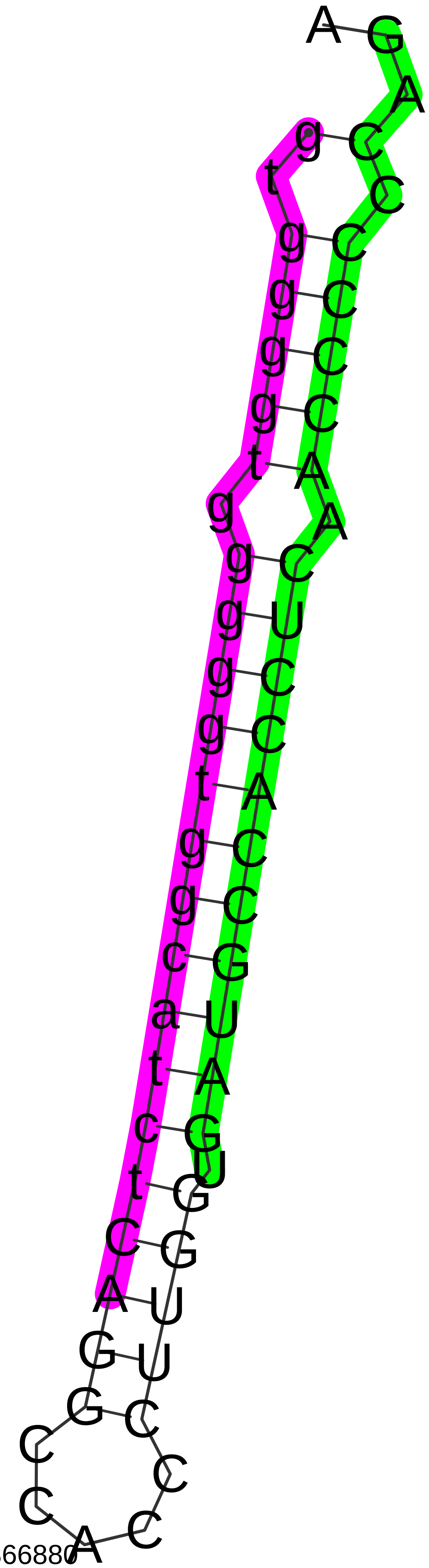

NW_006385327.1_366824-366880

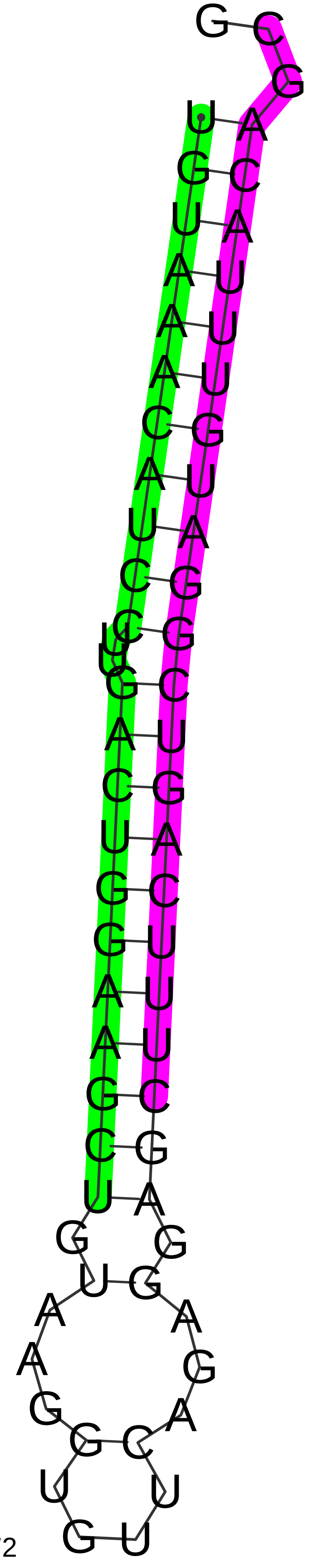

NW_006385528.1_207608-207672

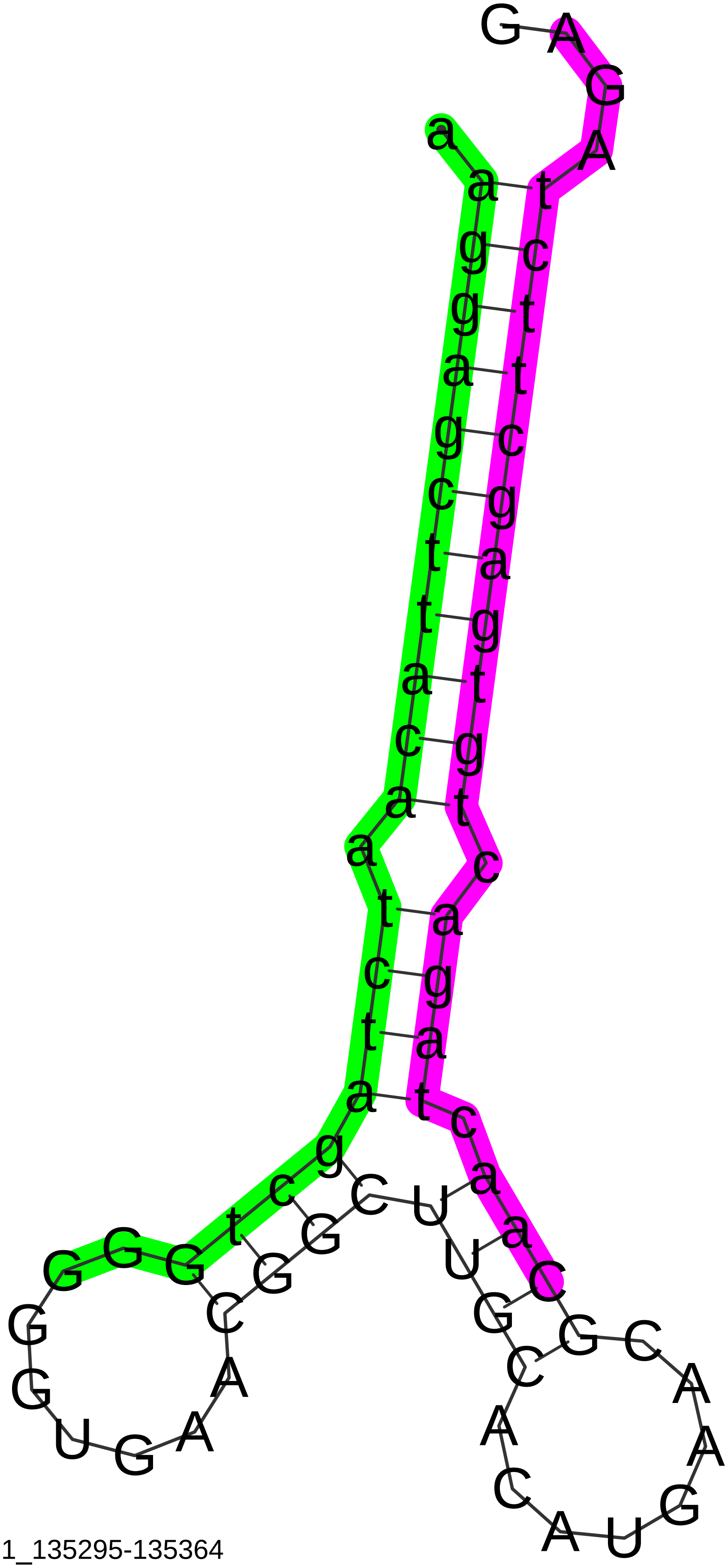

NW_006385537.1_135295-135364

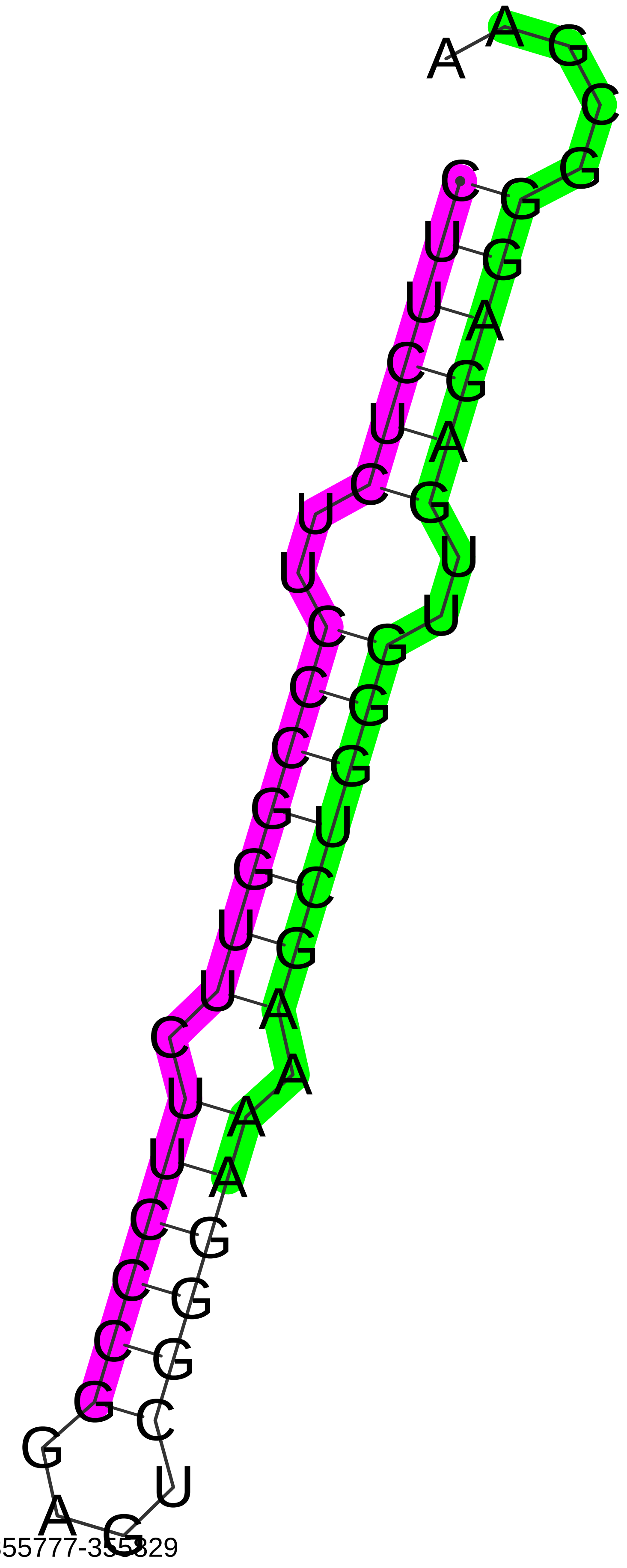

NW_006385625.1_355777-355829

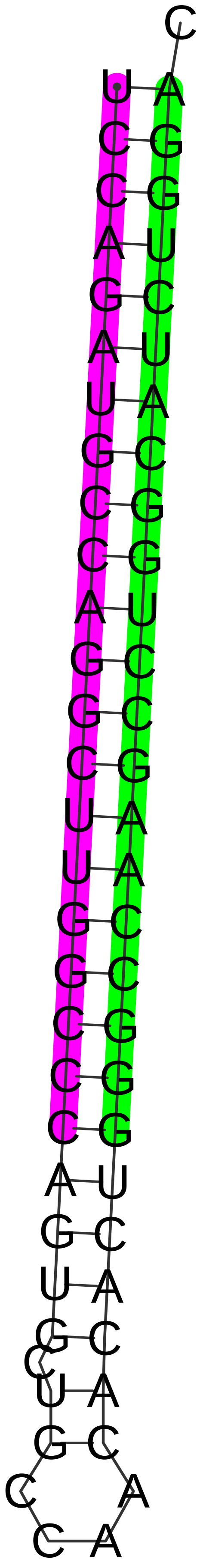

NW_006385655.1_136438-136497

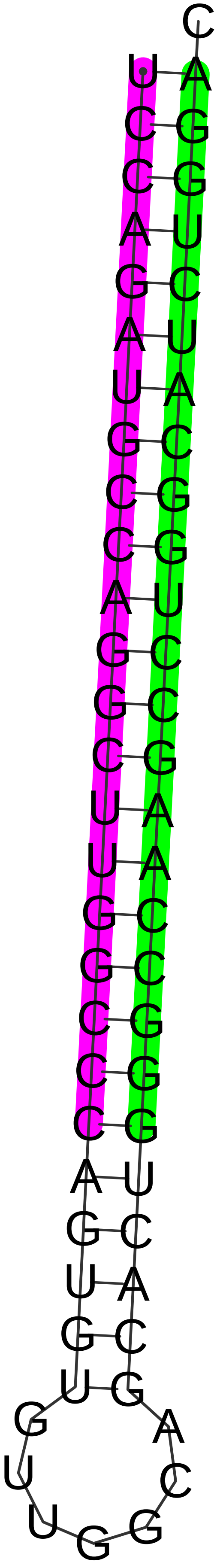

NW_006385655.1_136437-136496

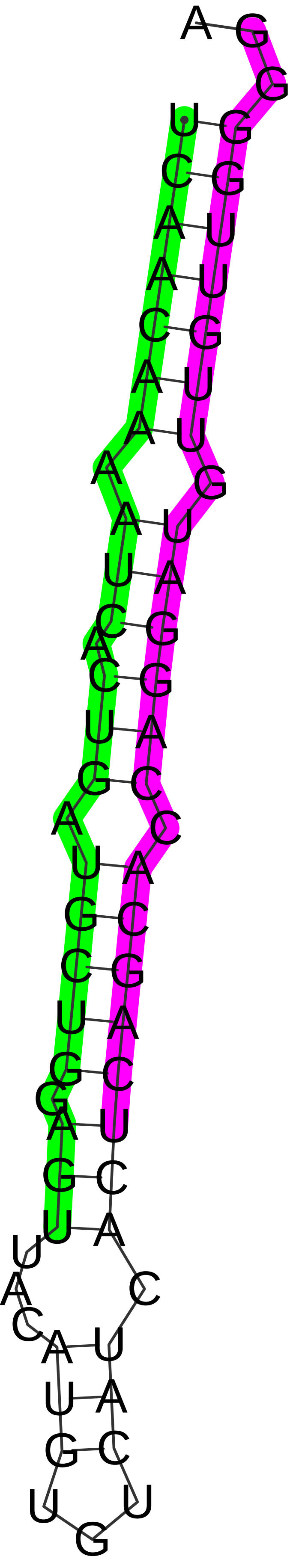

NW_006385655.1_333054-333117

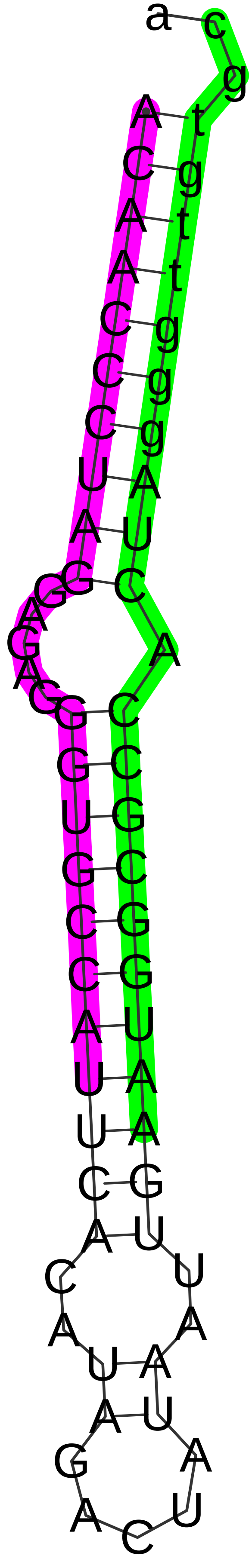

NW_006385734.1_152833-152896

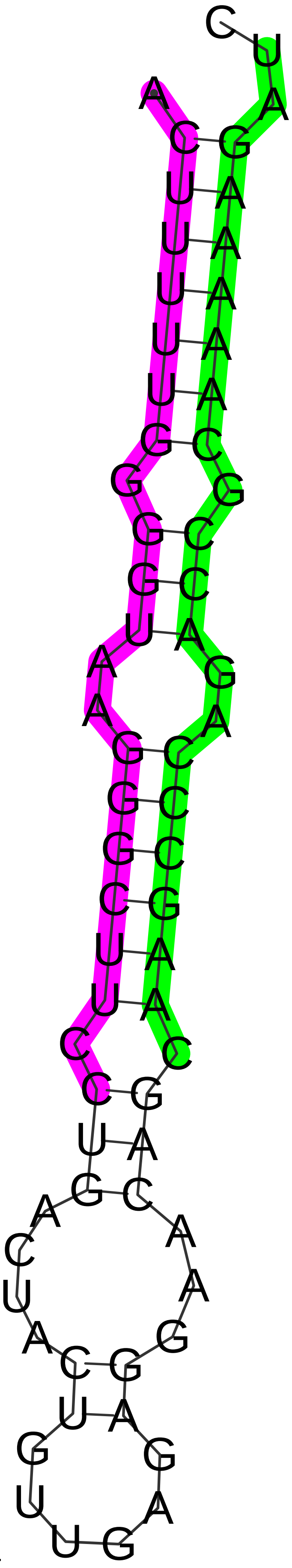

NW_006385739.1_154825-154891

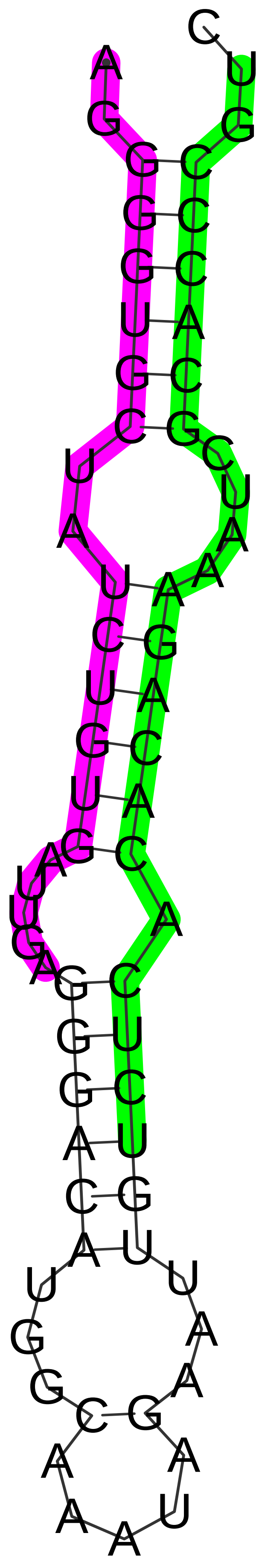

NW_006385810.1_286621-286686

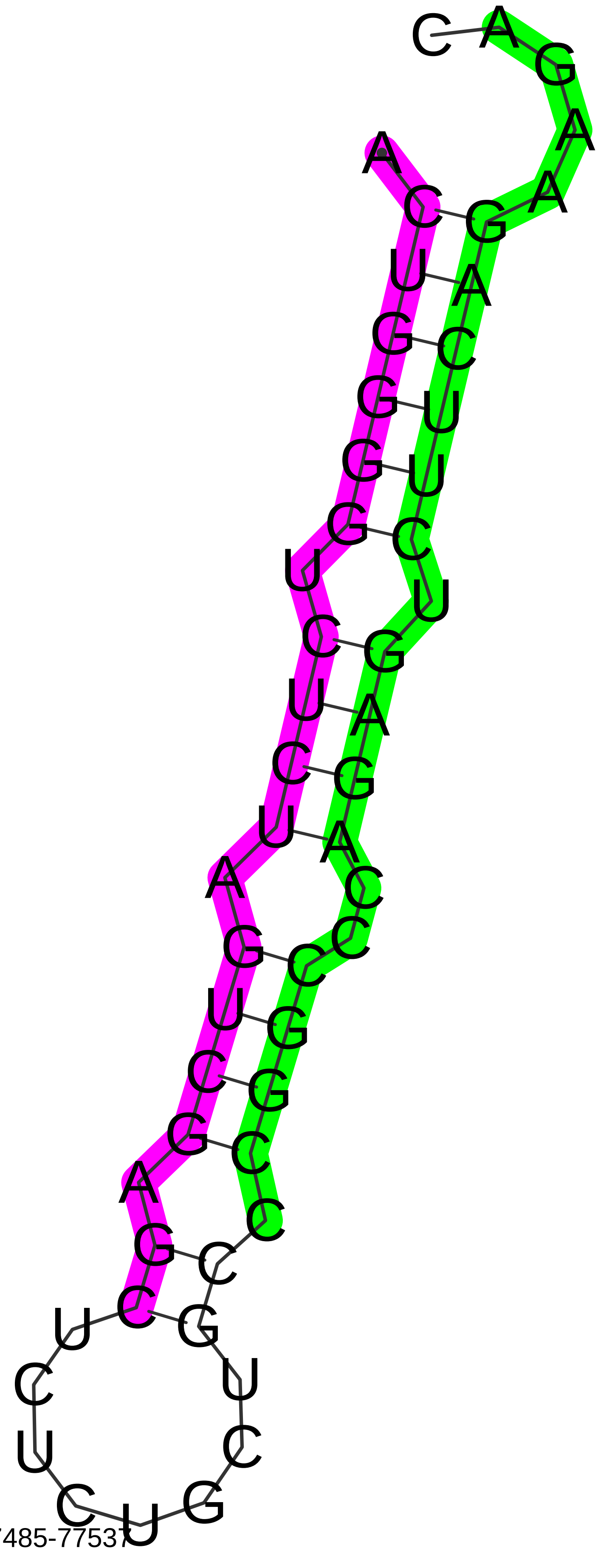

NW_006385875.1_77485-77537

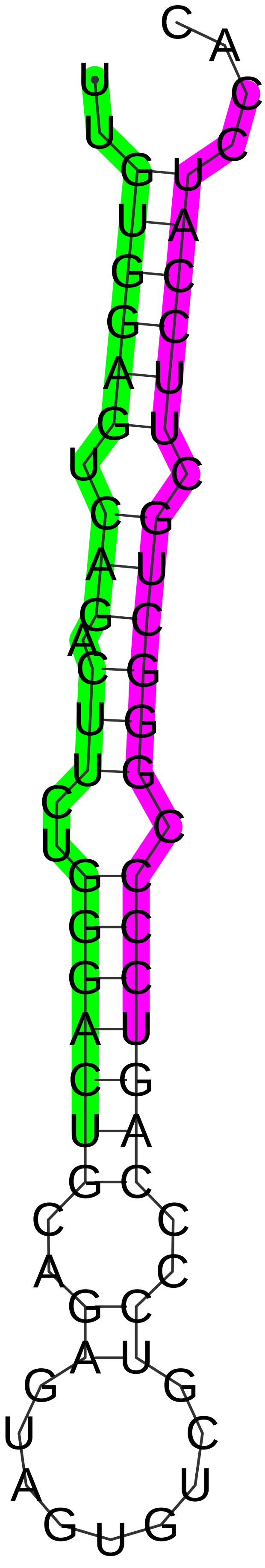

NW_006385878.1_280194-280260

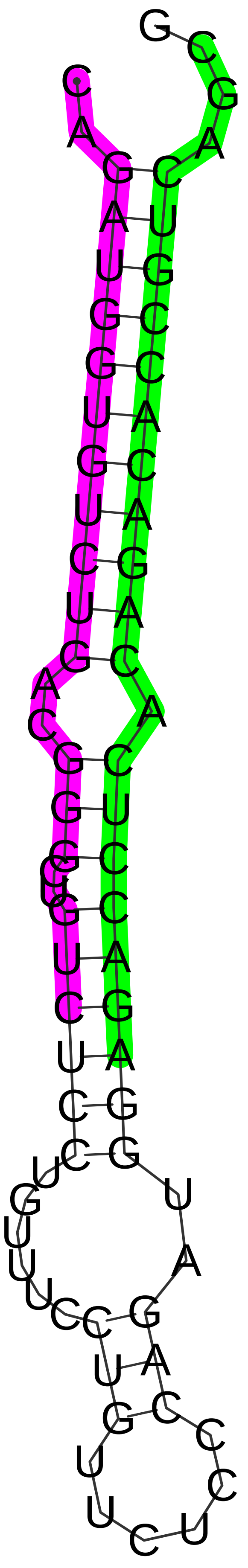

NW_006386136.1_173724-173794

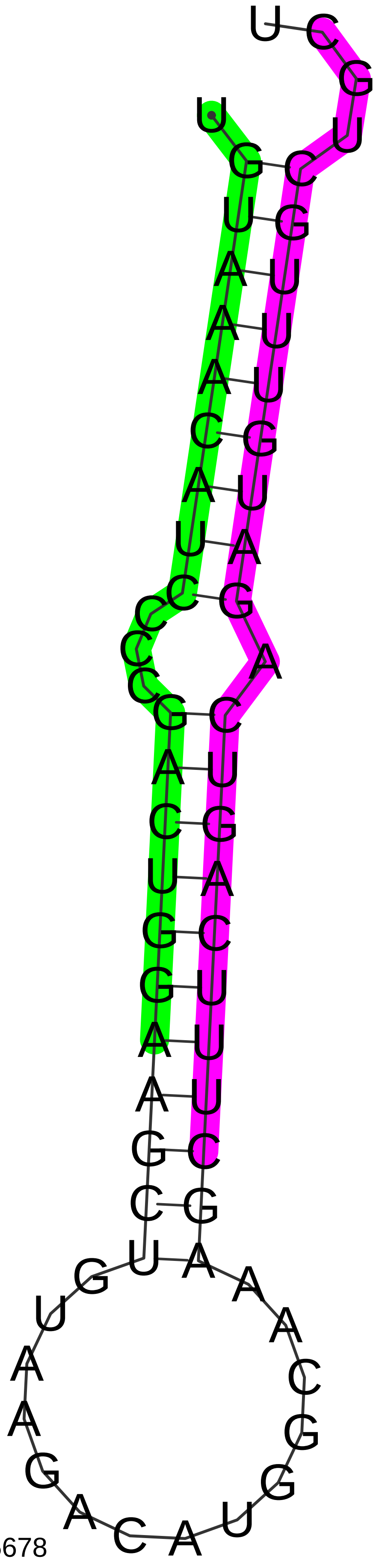

NW_006386213.1_115616-115678

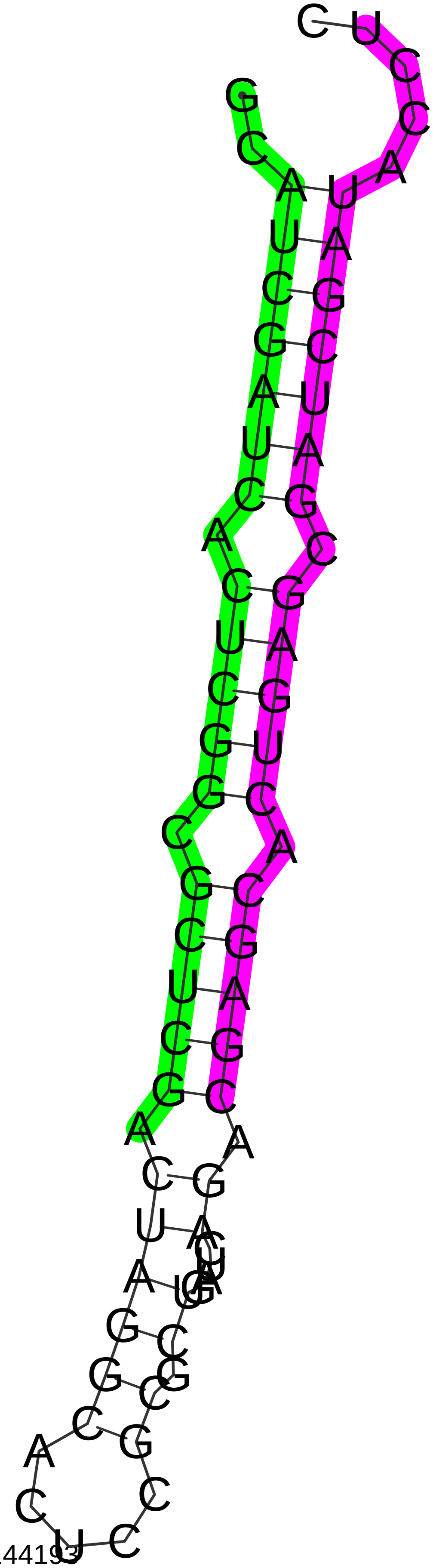

NW_006386216.1_144125-144193

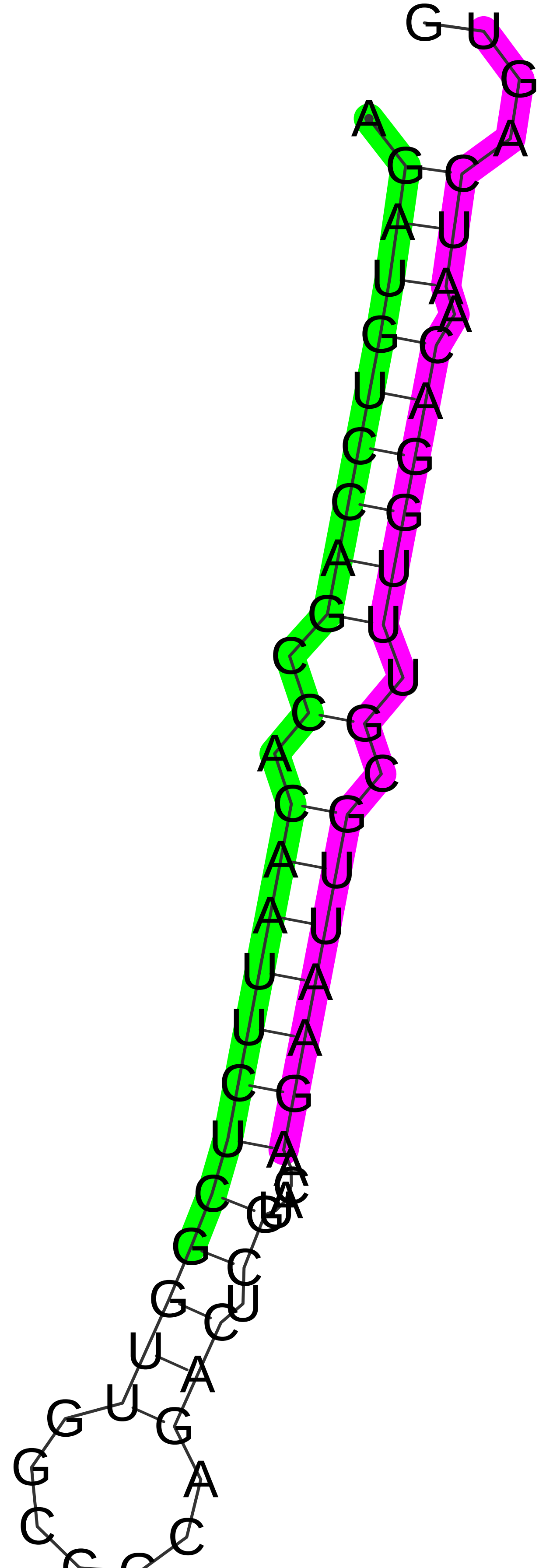

NW_006386229.1_83509-83574

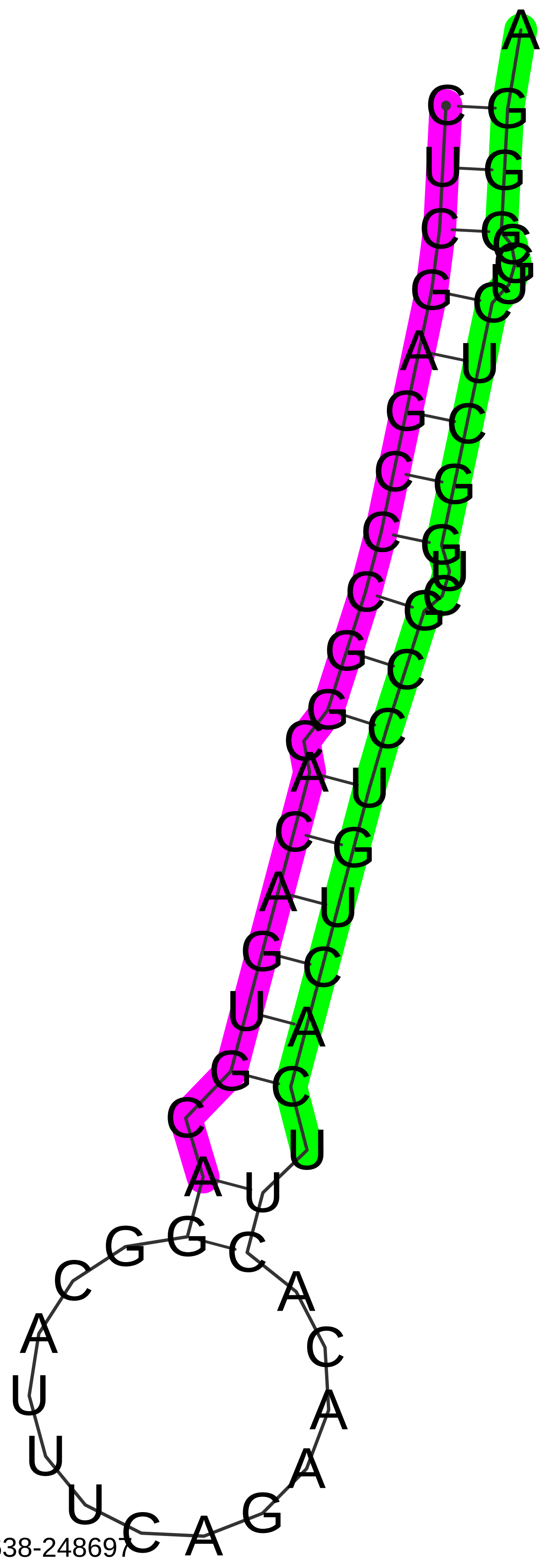

NW_006386284.1_248638-248697

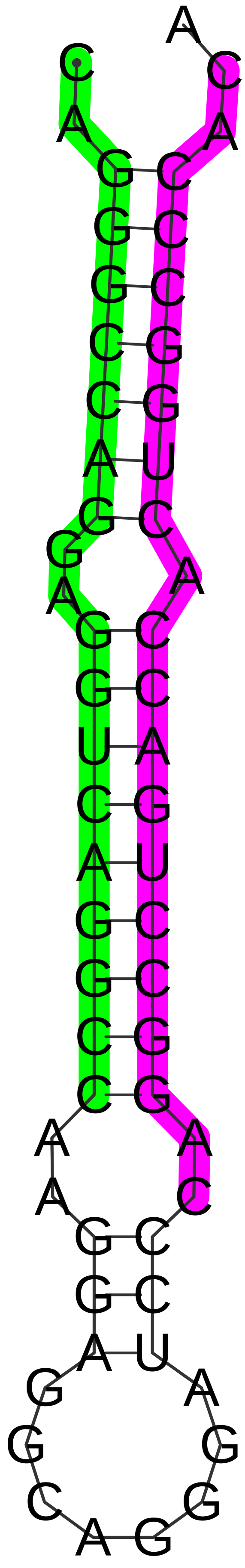

NW_006386296.1_93032-93089

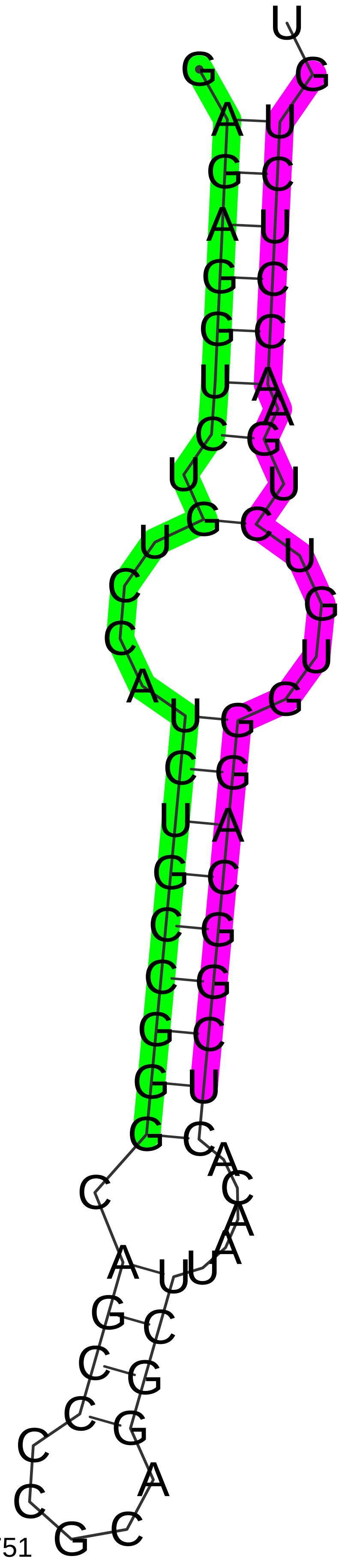

NW_006386338.1_136685-136751

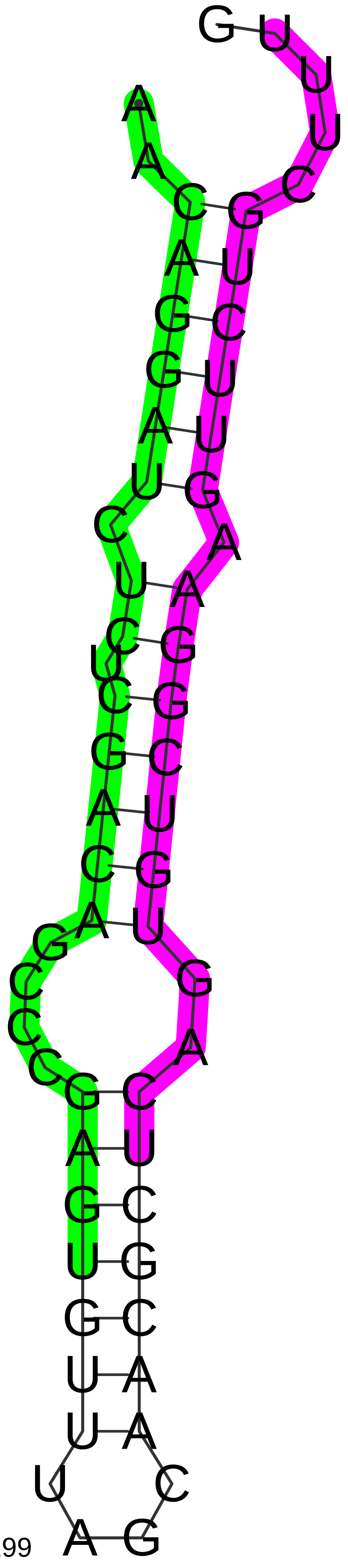

NW_006386347.1_236240-236299

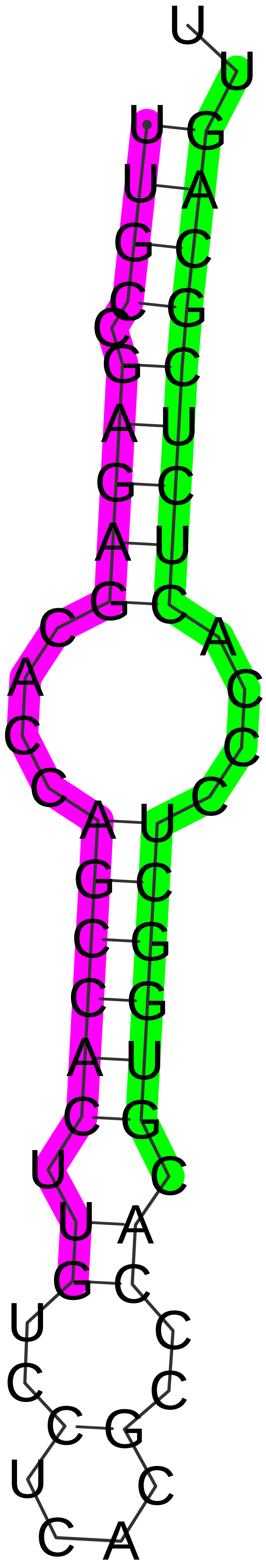

NW_006386363.1_63737-63793

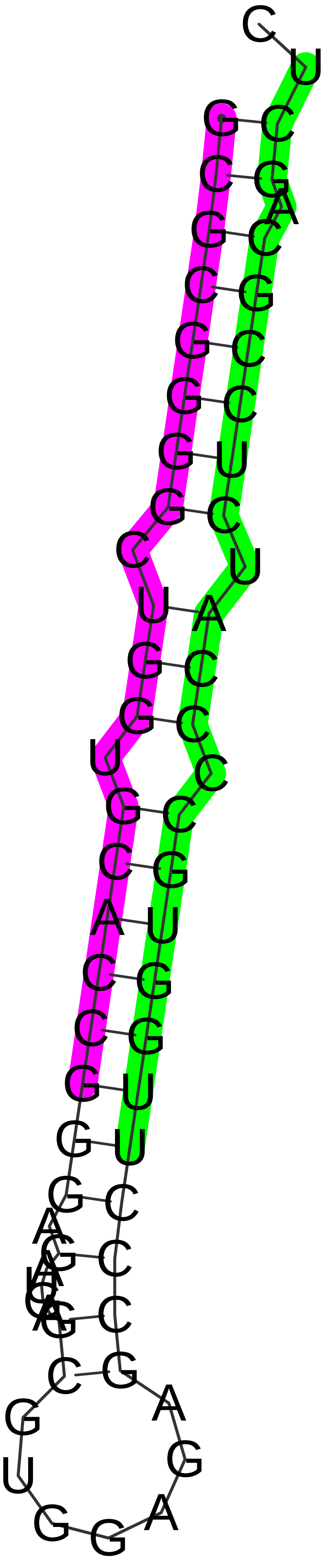

NW_006386592.1_43159-43221

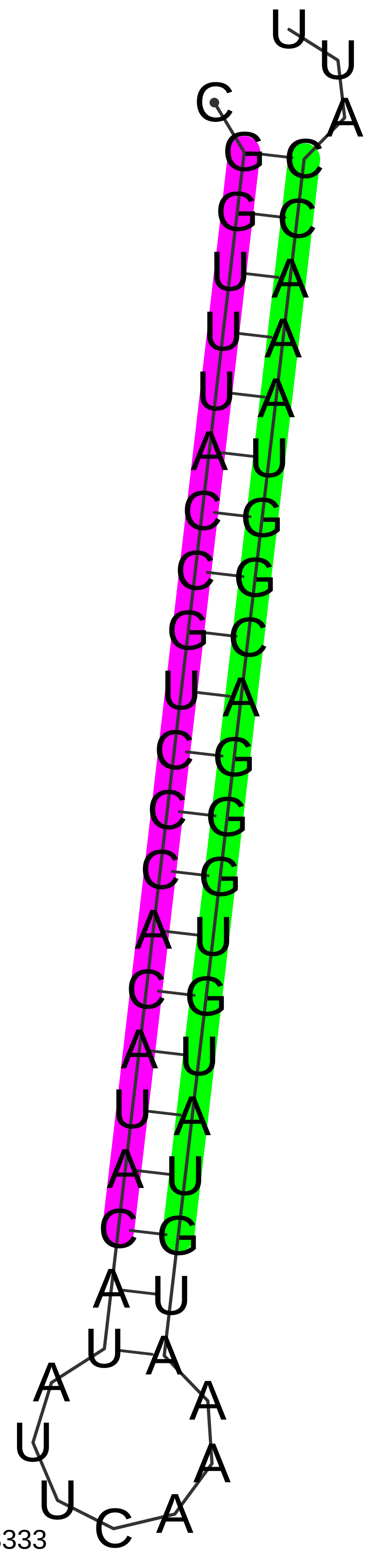


NW_006386594.1_113281-113333

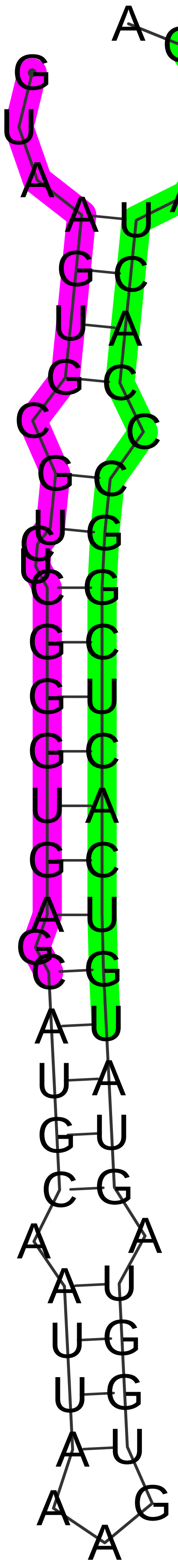

NW_006386594.1_84818-84879

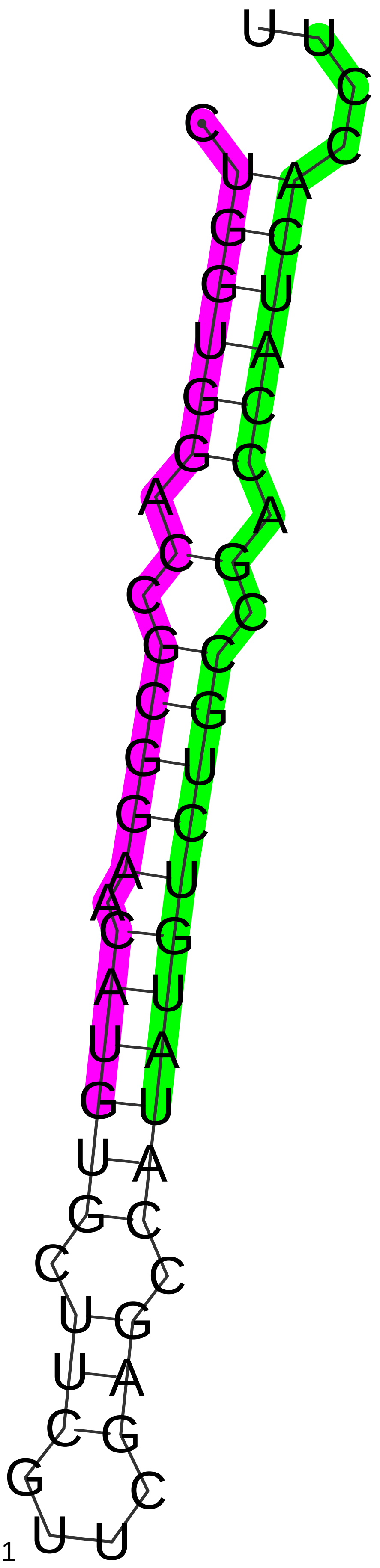

NW_006386594.1_98034-98091

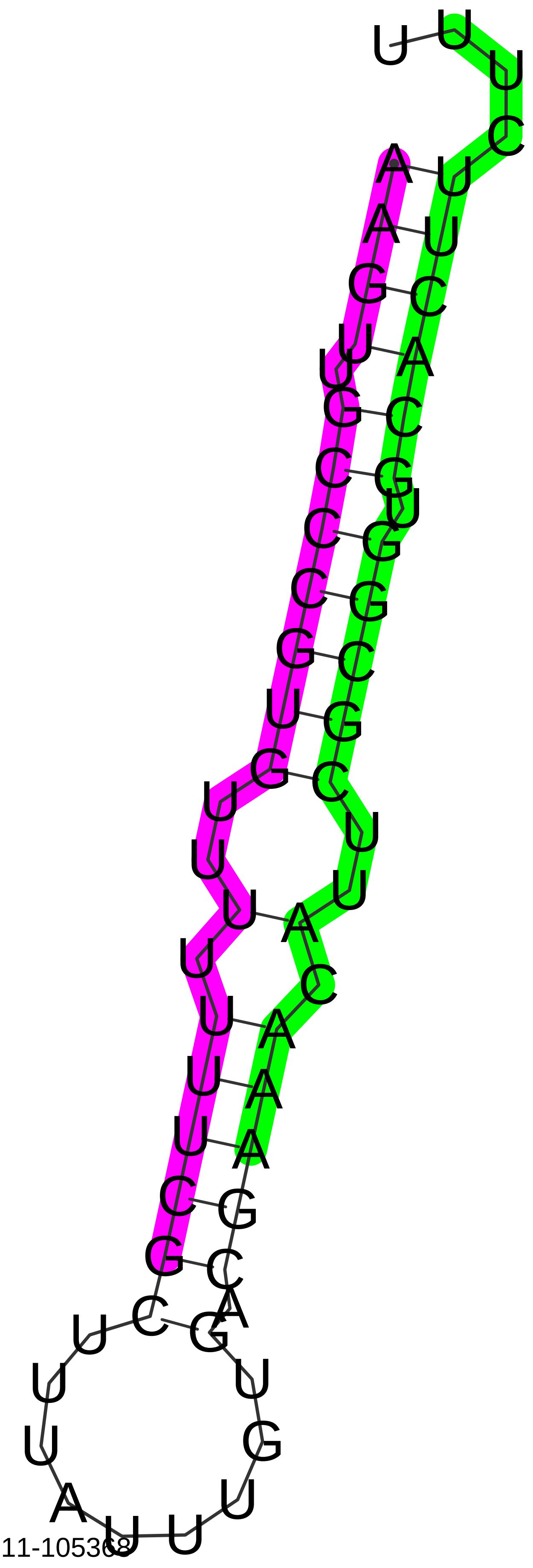

NW_006386594.1_105311-105368

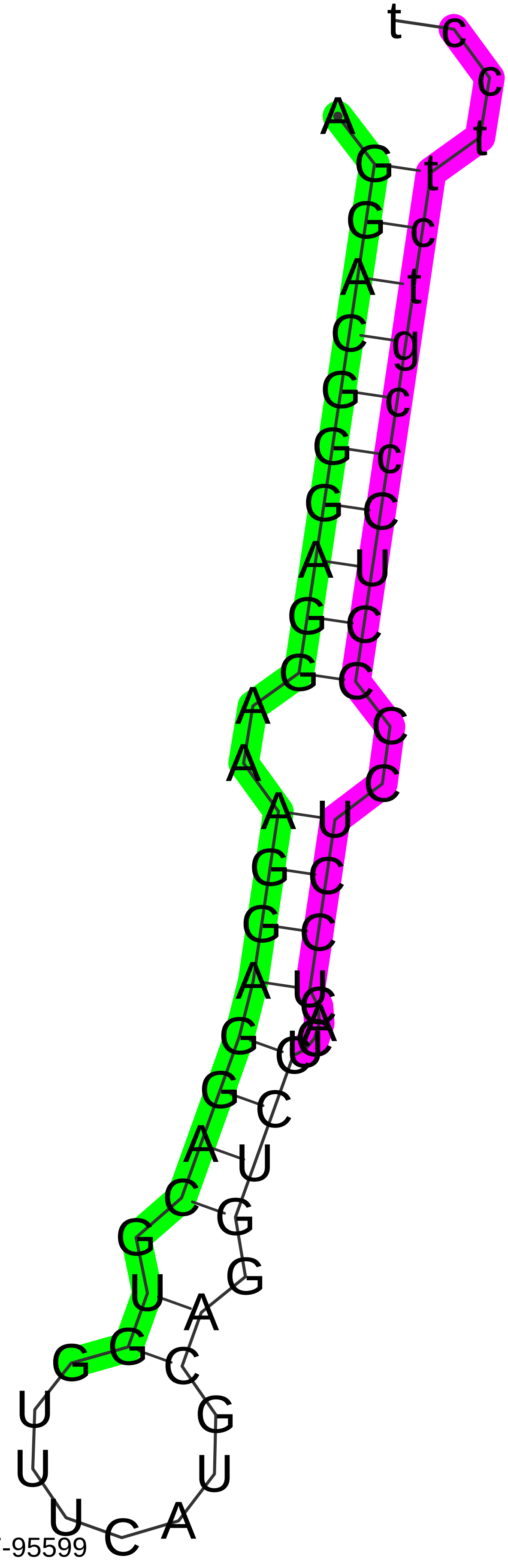

NW_006386681.1_95537-95599

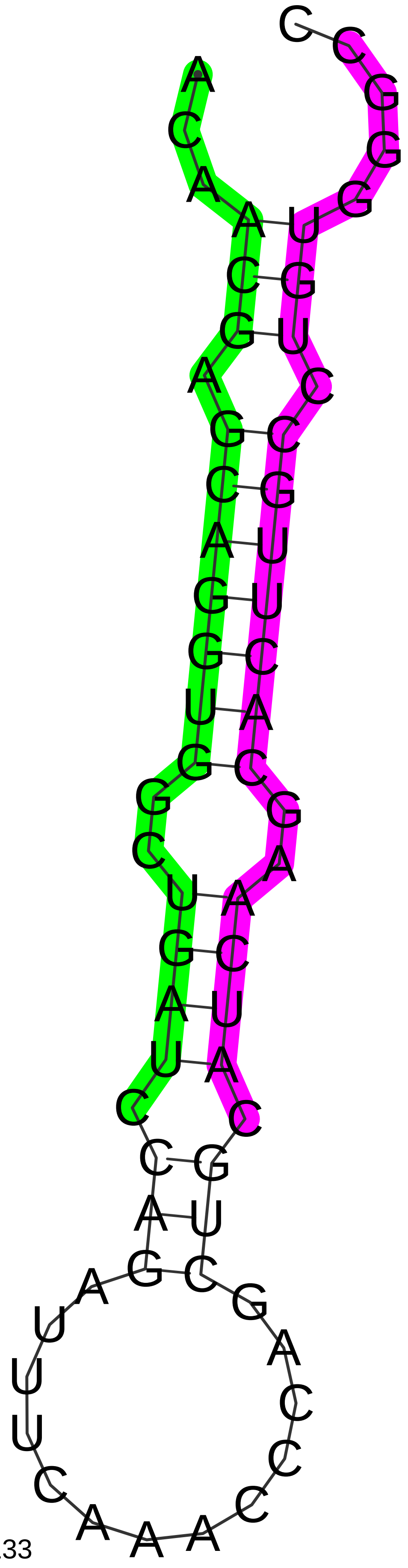

NW_006386689.1_71071-71133

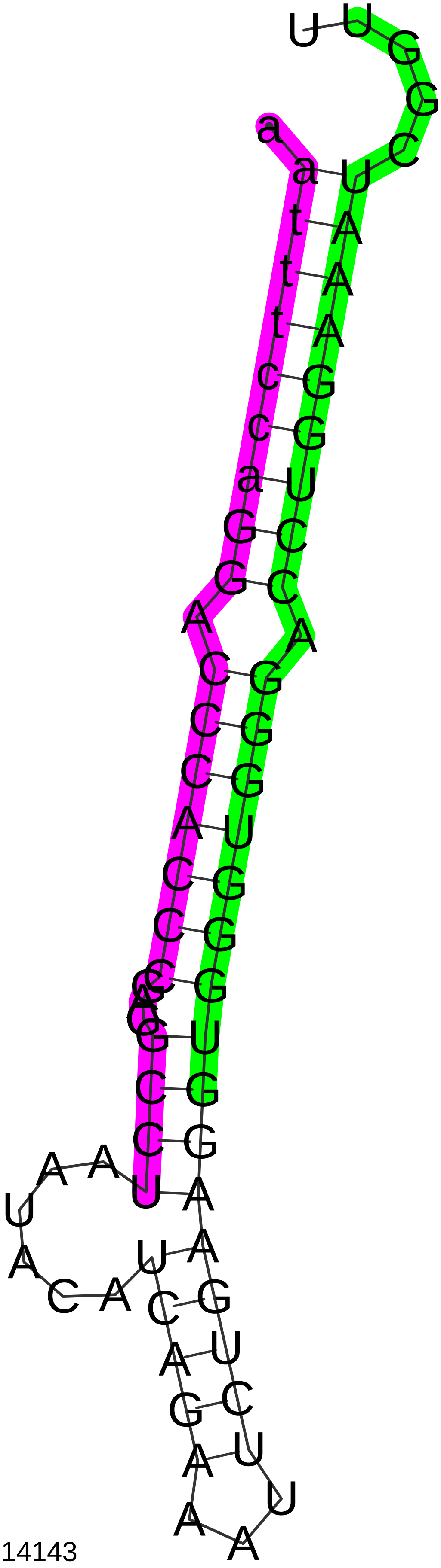

NW_006386974.1_114074-114143

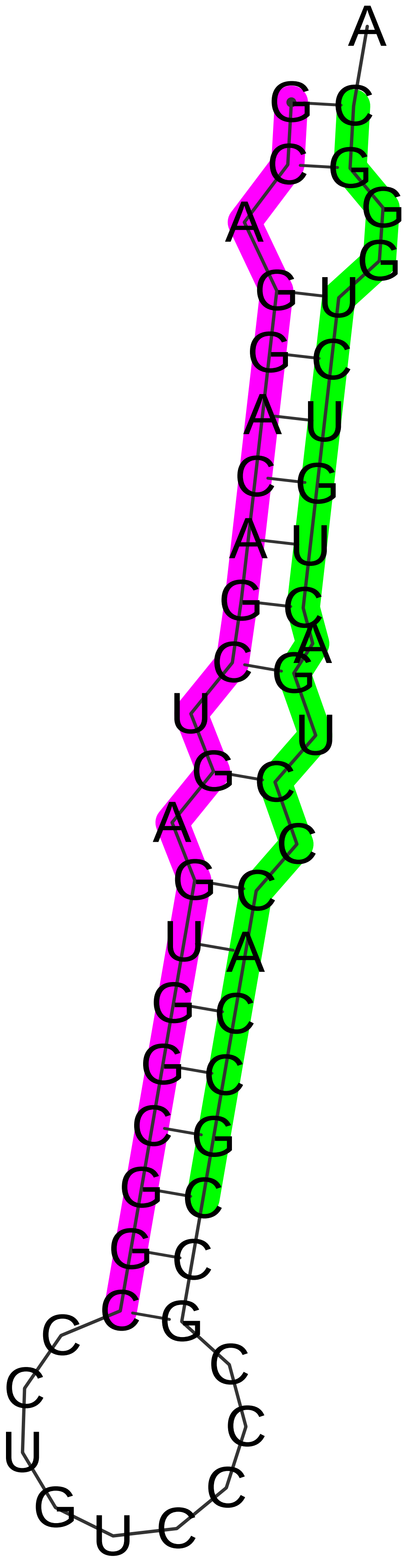

NW_006387221.1_2223-2276

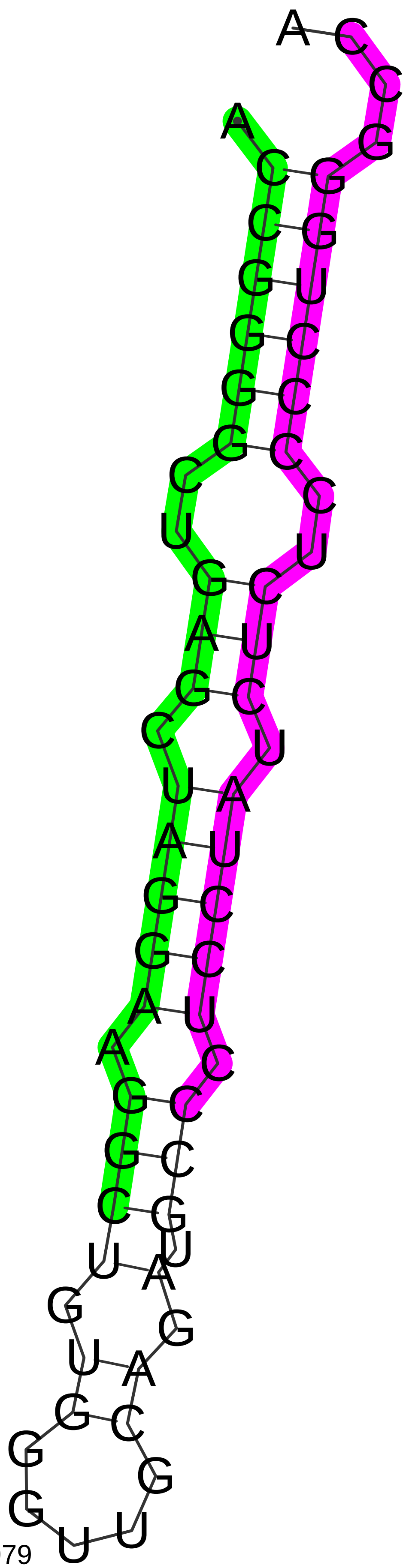


NW_006387535.1_49919-49979

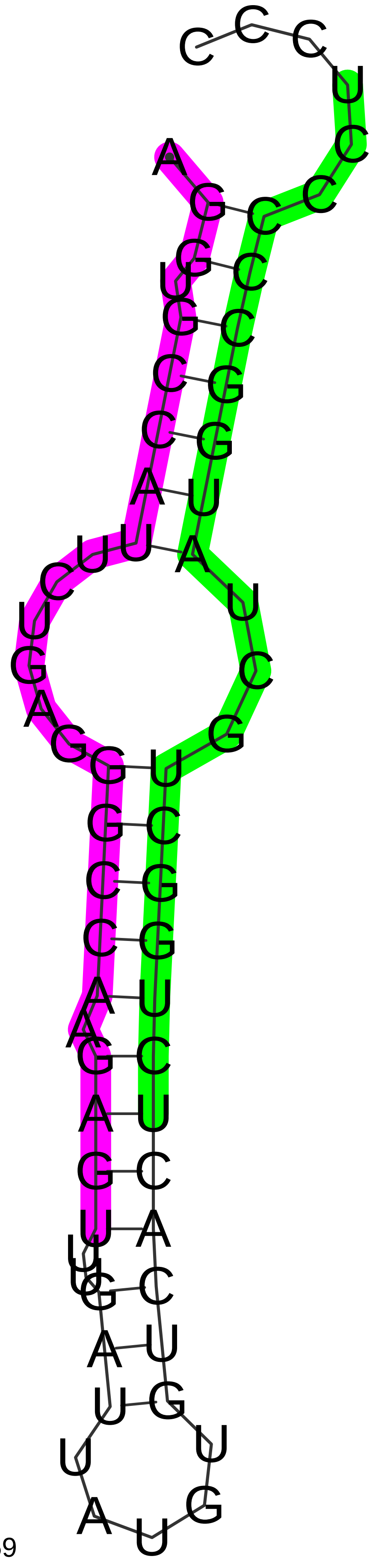

NW_006387597.1_60797-60859

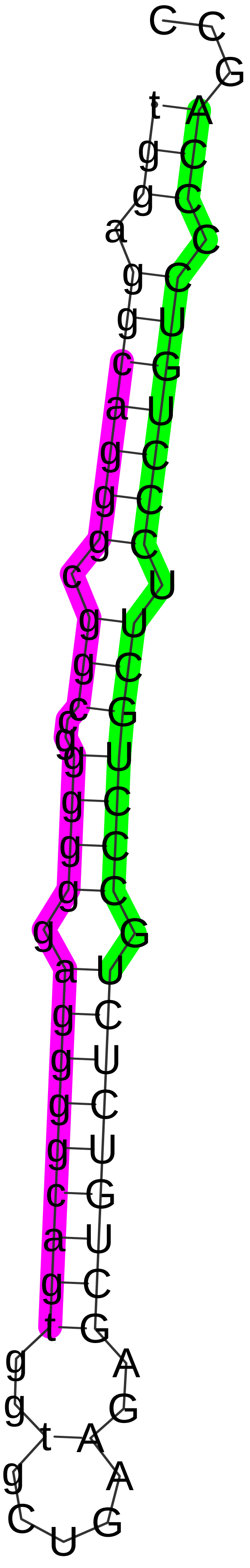

NW_006387694.1_51935-52008

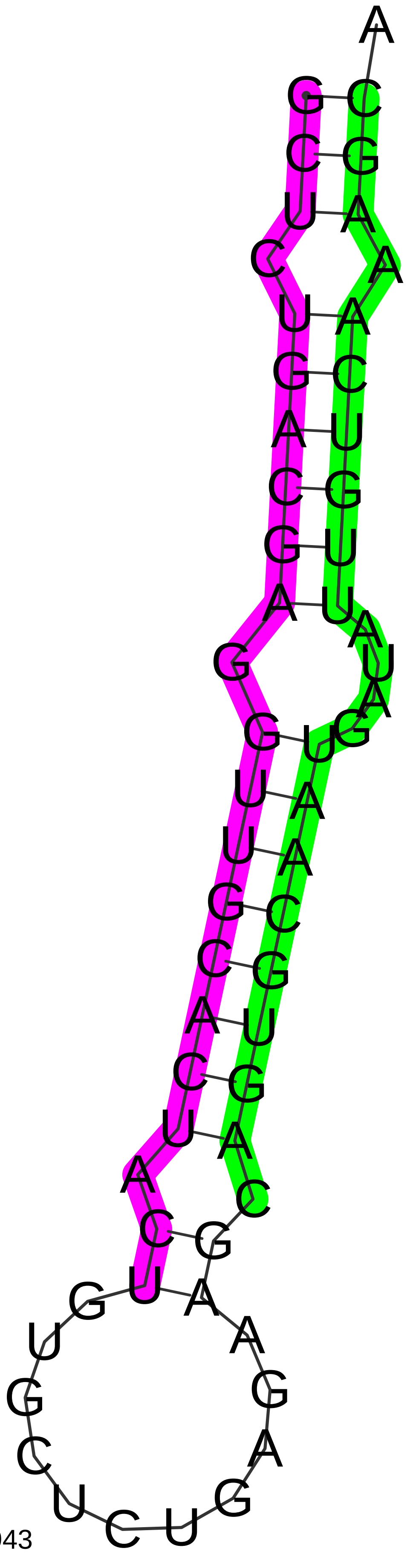

NW_006387848.1_42885-42943

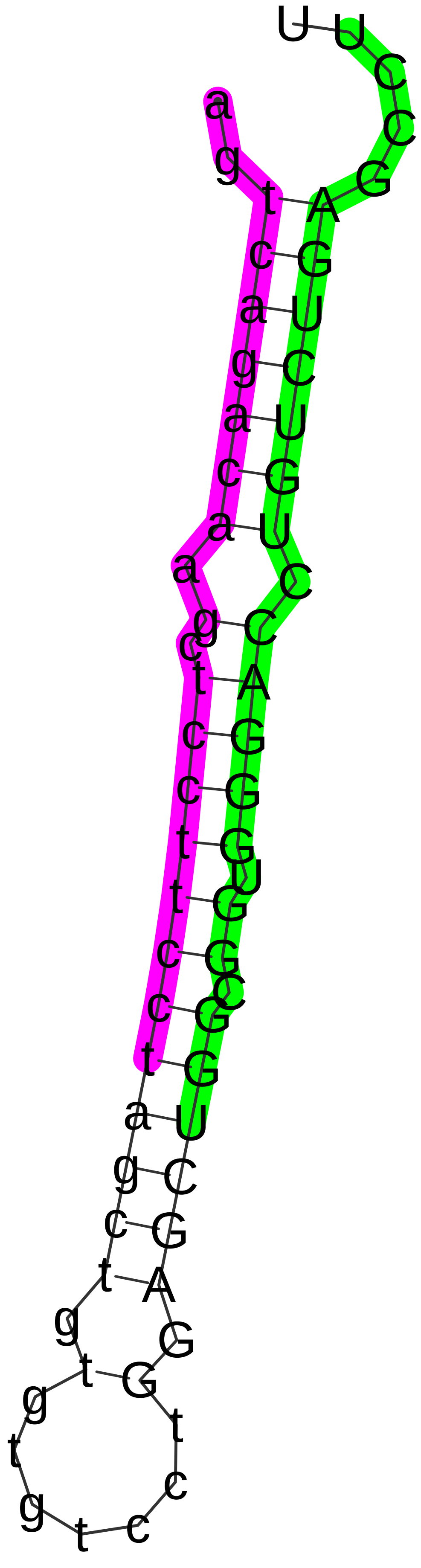

NW_006389115.1_541-603

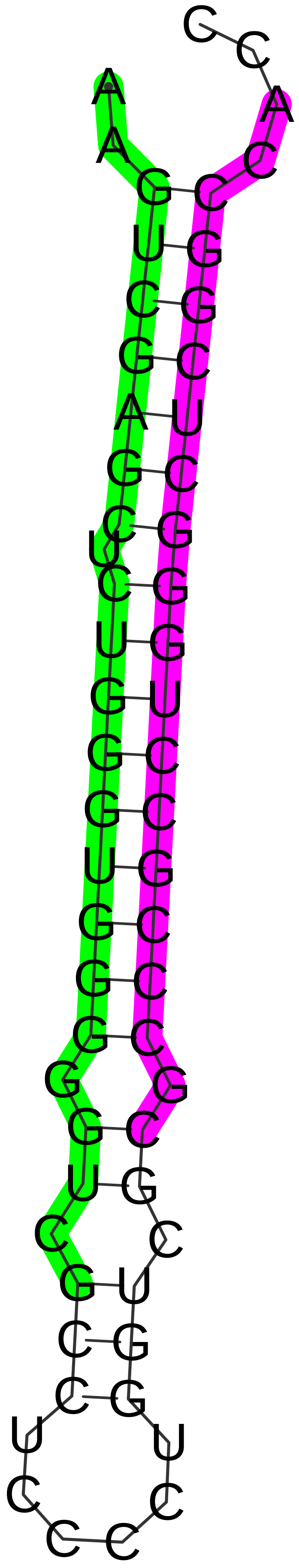

NW_006390488.1_12459-12517

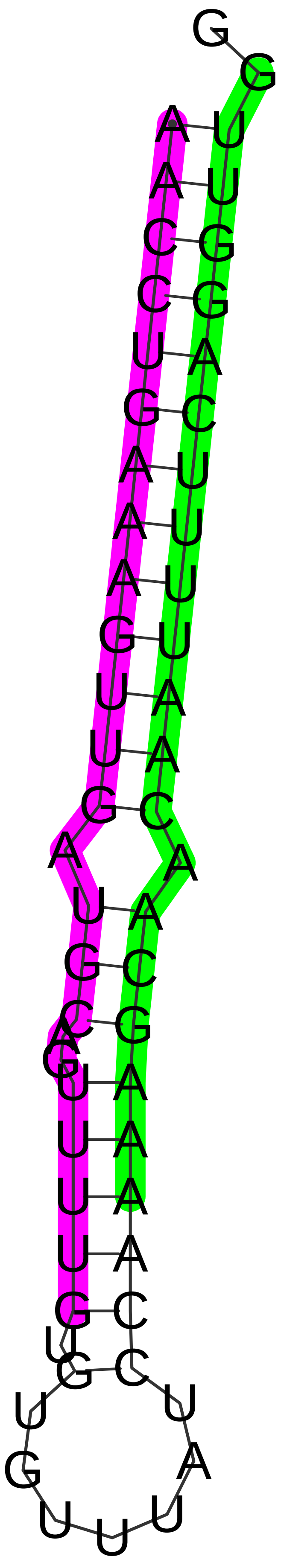

NW_006390522.1_8996-9053

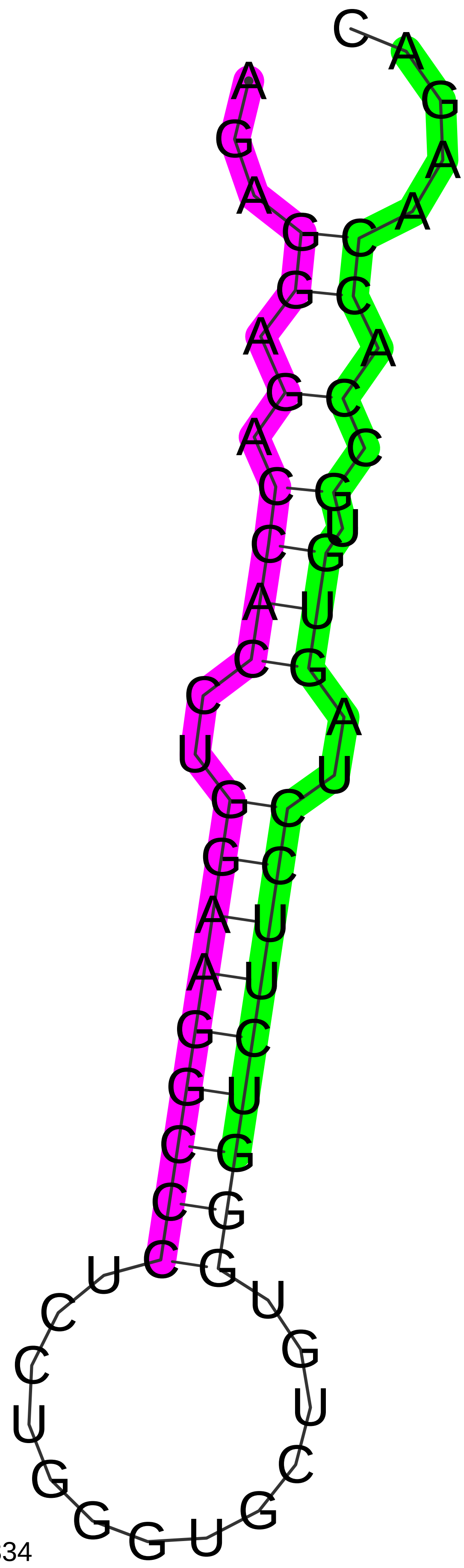

NW_006390703.1_5773-5834

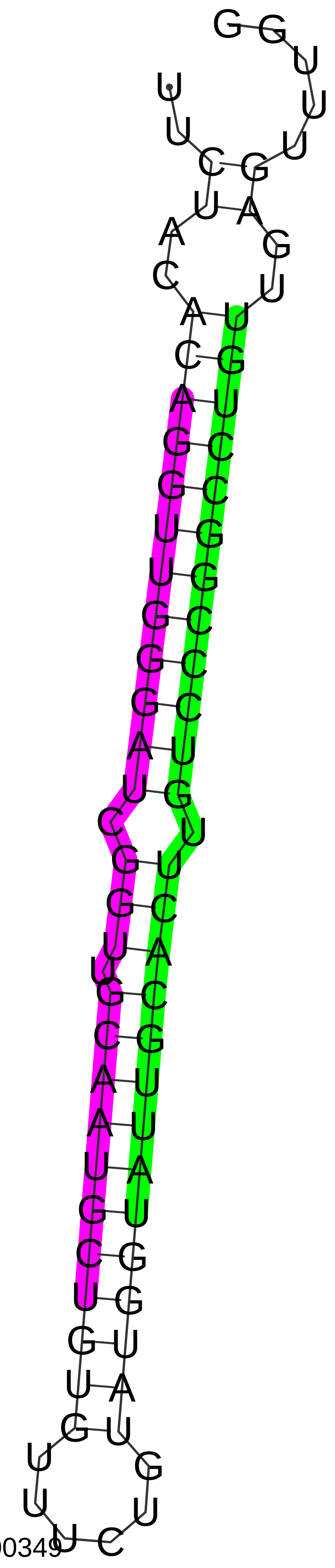

NW_006383042.1_1100274-1100349

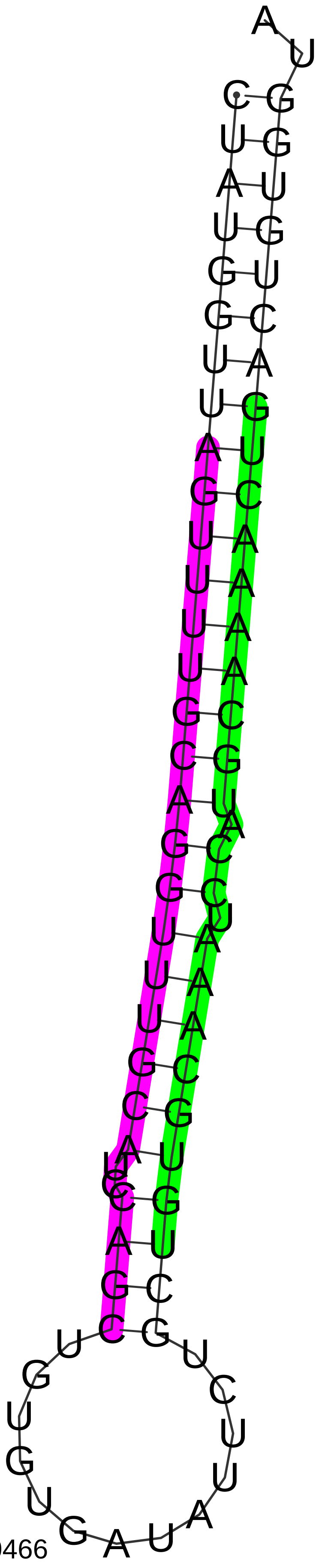

NW_006383042.1_1100390-1100466

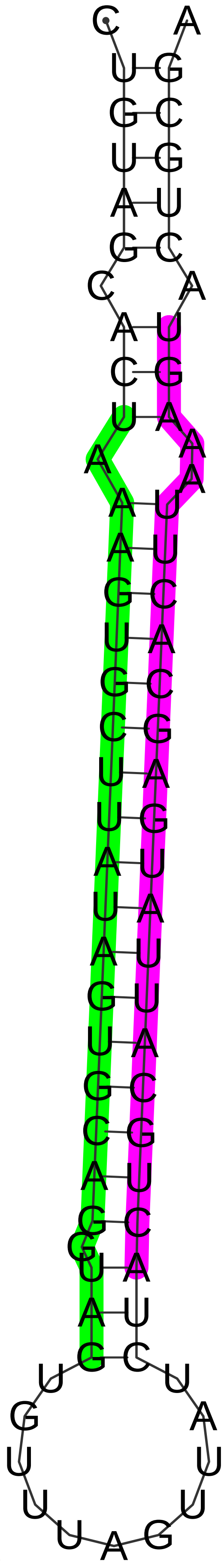

NW_006383042.1_1100528-1100602

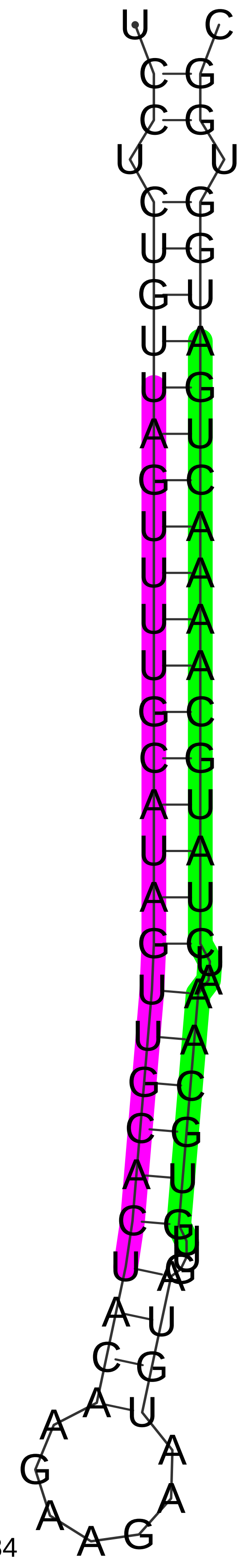

NW_006383042.1_1100711-1100784

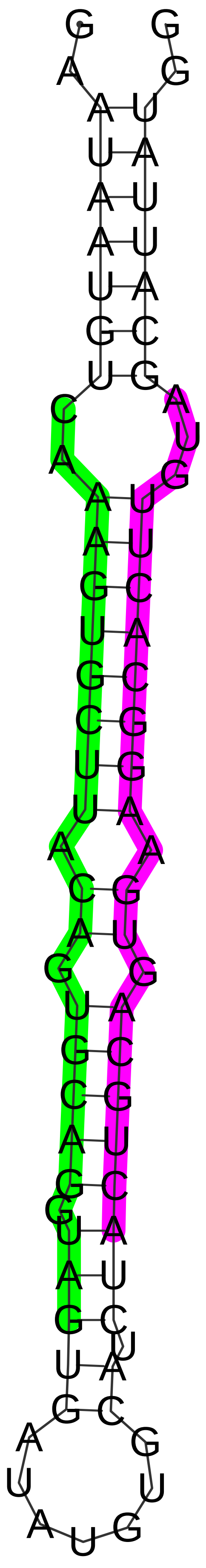

NW_006383042.1_1100995-1101070

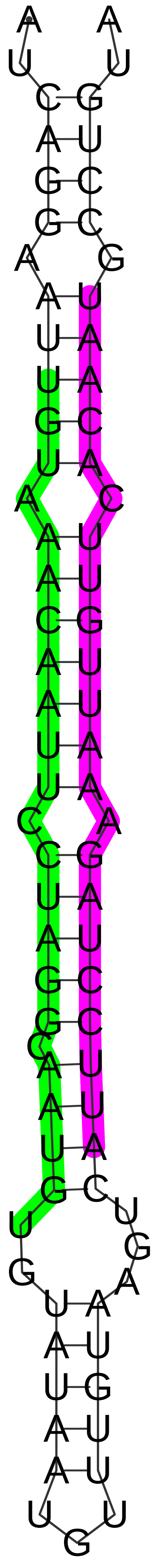

NW_006383043.1_1492335-1492413

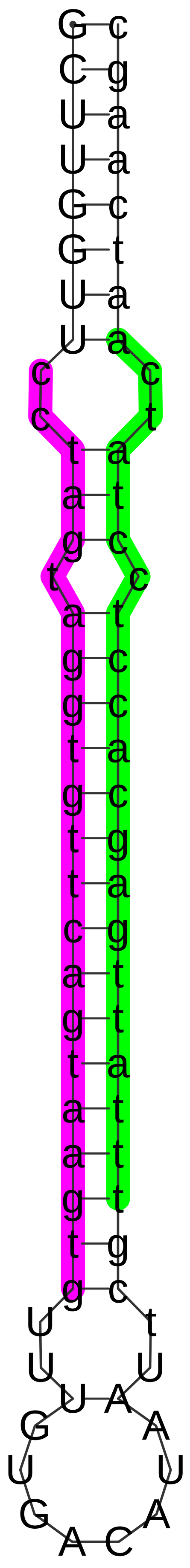

NW_006383043.1_1552314-1552387

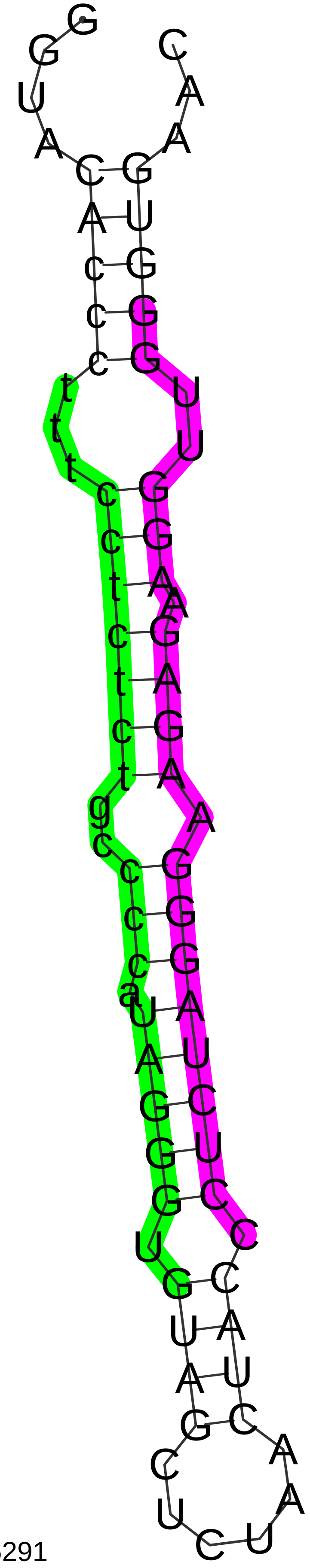

NW_006383047.1_5256219-5256291

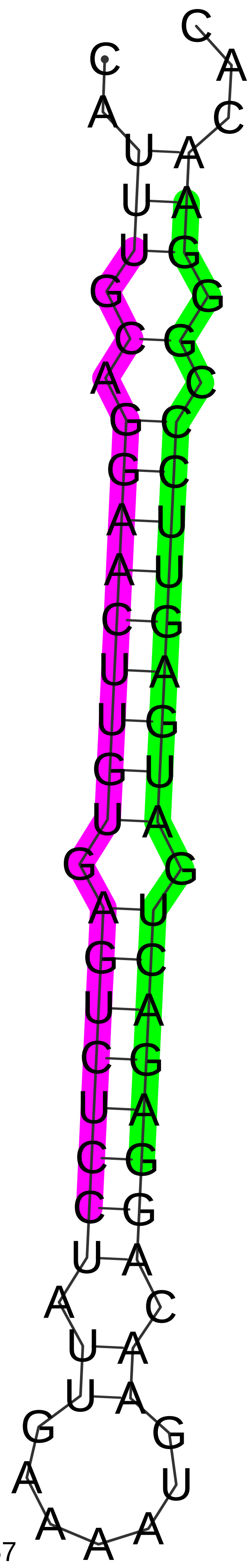

NW_006383062.1_3365692-3365757

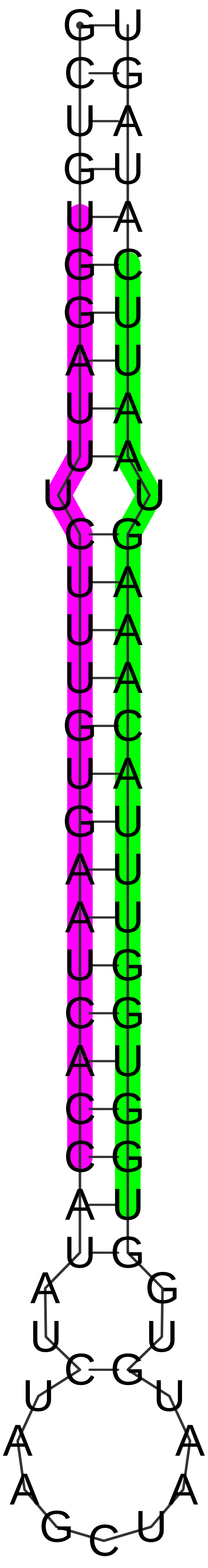

NW_006383062.1_3392514-3392582

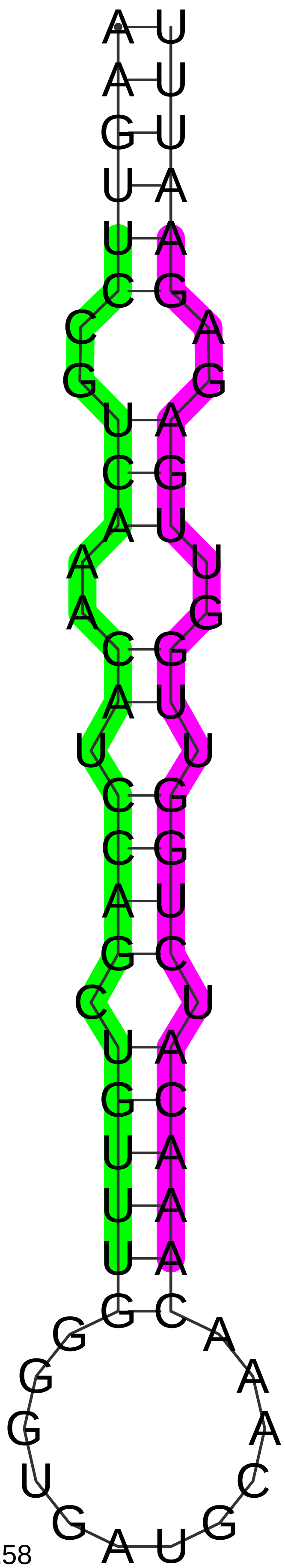


NW_006383063.1_5092193-5092258

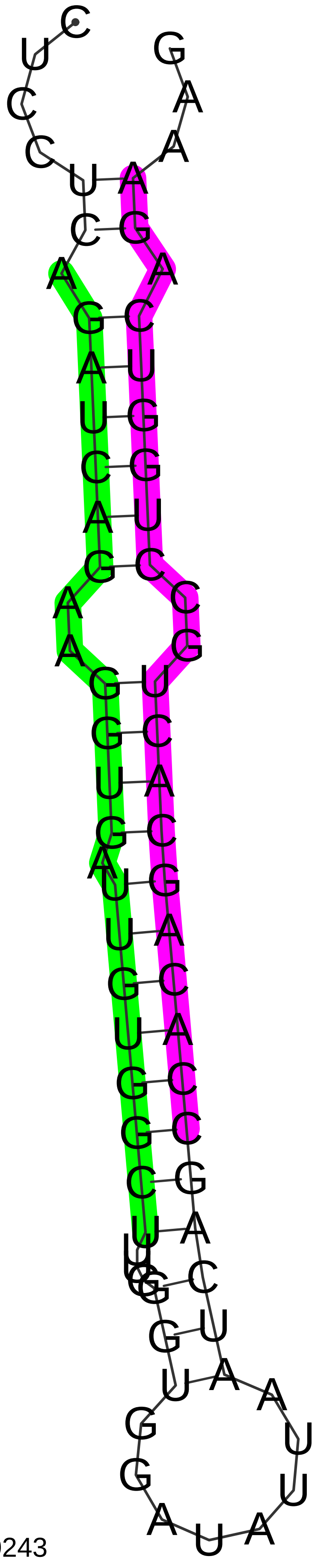

NW_006383069.1_2570173-2570243

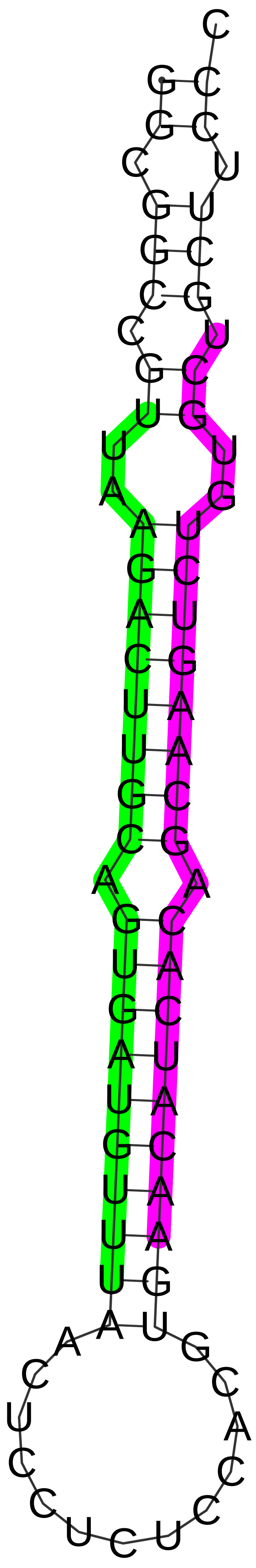

NW_006383074.1_10055-10128

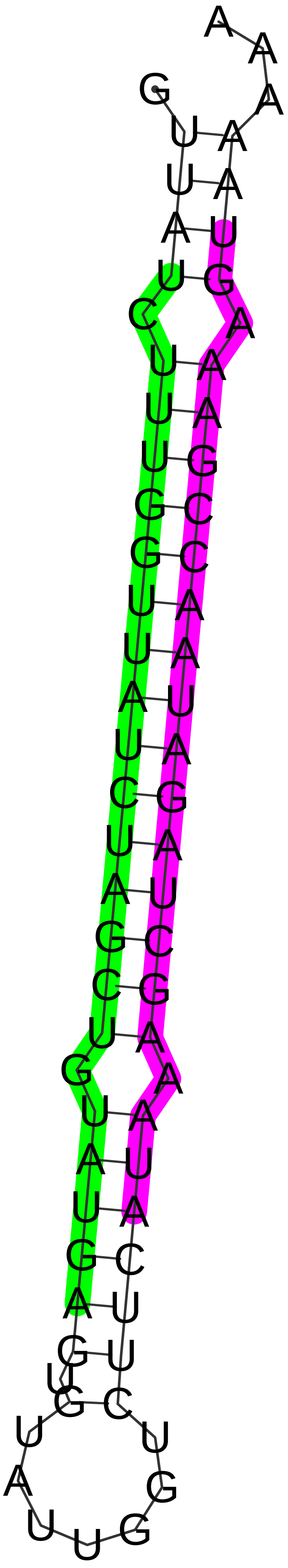

NW_006383098.1_16059-16126

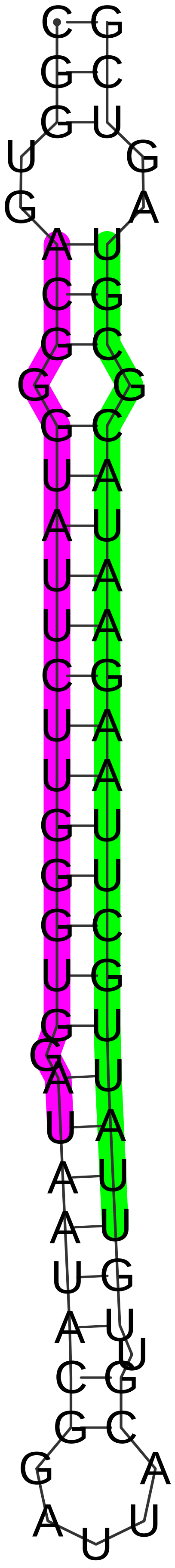

NW_006383103.1_346373-346439

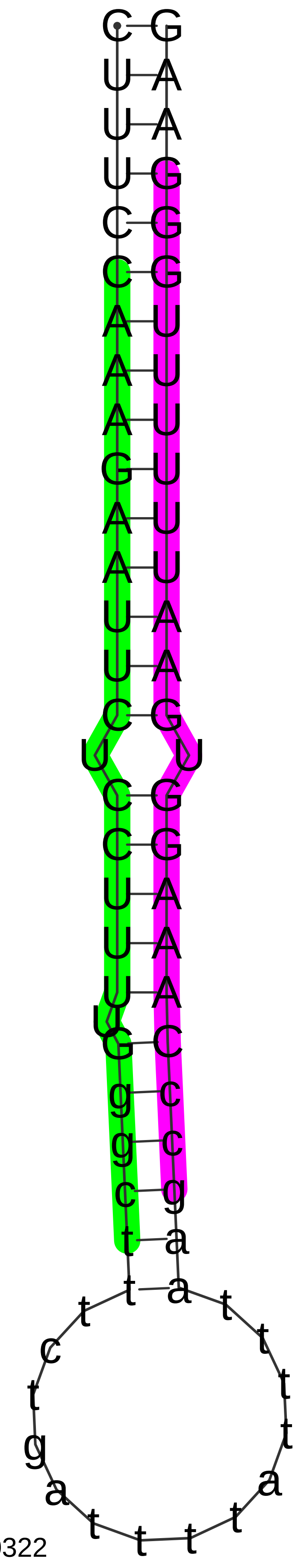


NW_006383105.1_3390254-3390322

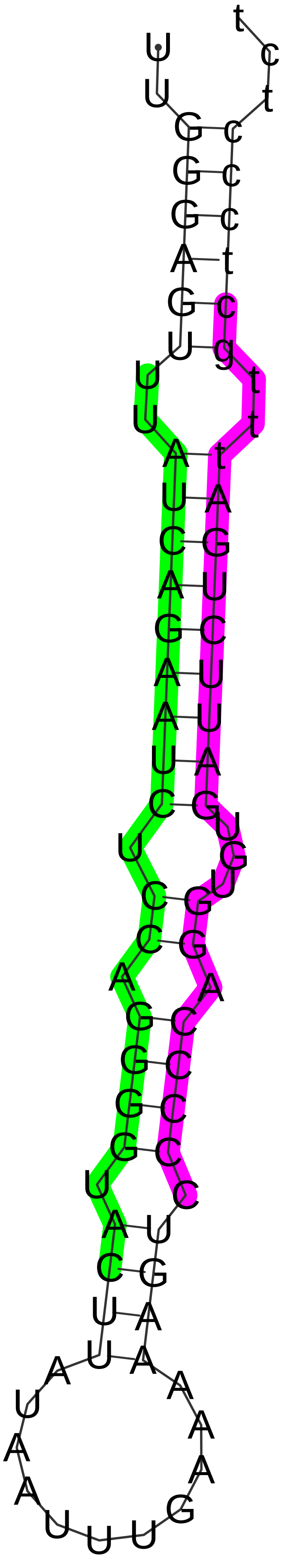

NW_006383110.1_824129-824206

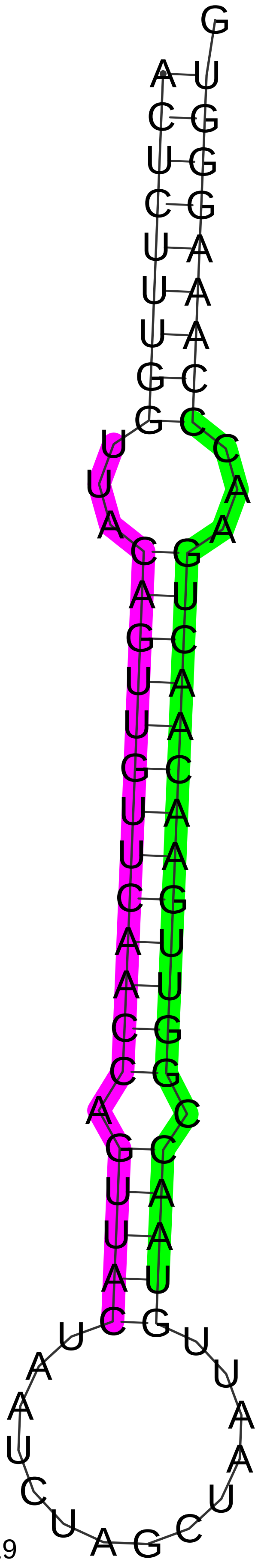

NW_006383112.1_1440143-1440219

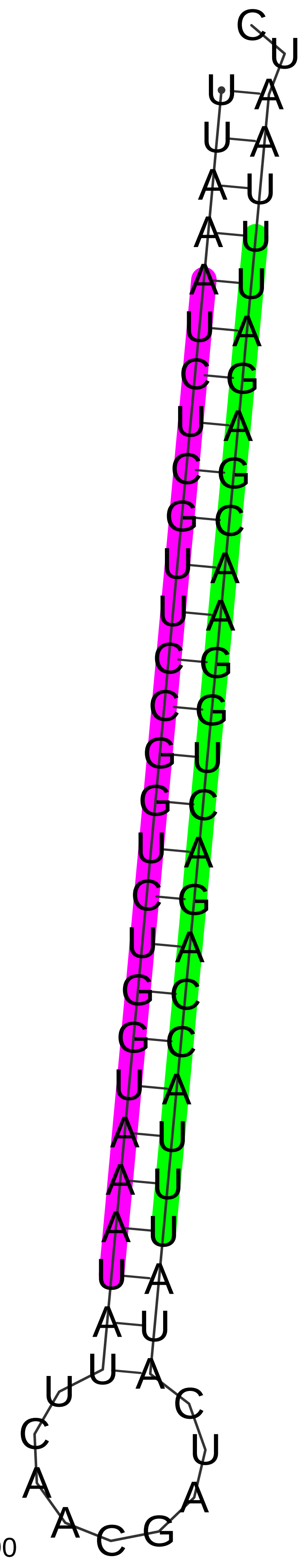

NW_006383112.1_603324-603390

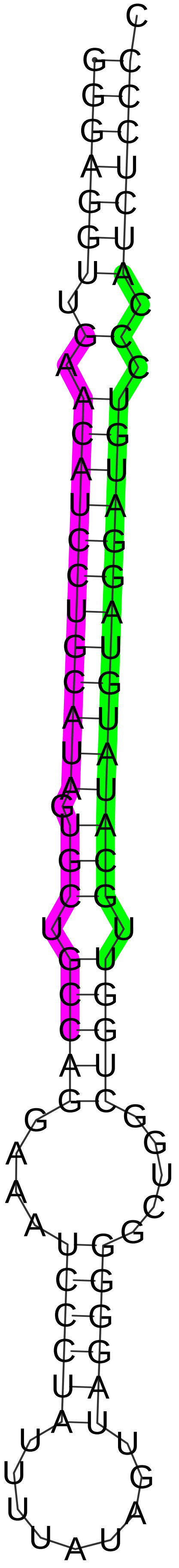

NW_006383124.1_2558105-2558198

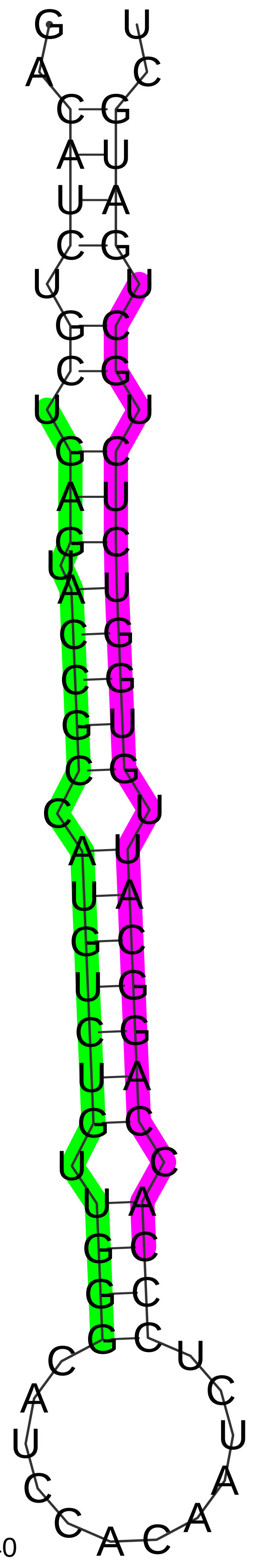

NW_006383124.1_2626066-2626140

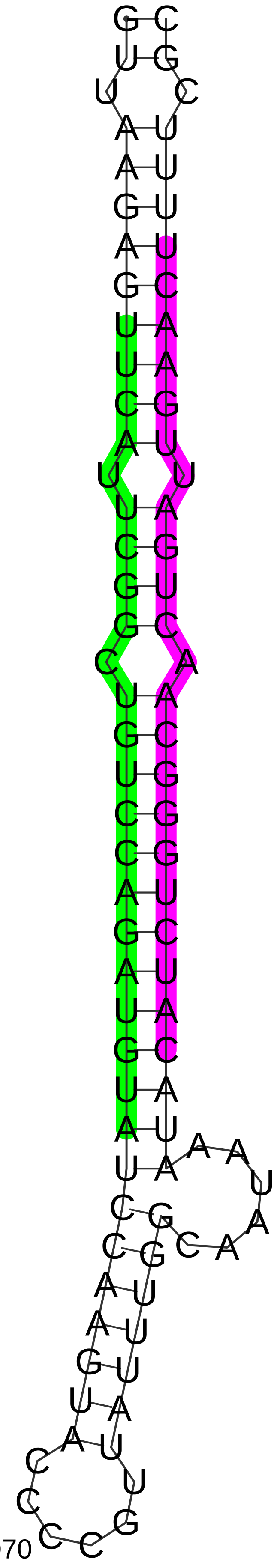

NW_006383124.1_2662983-2663070

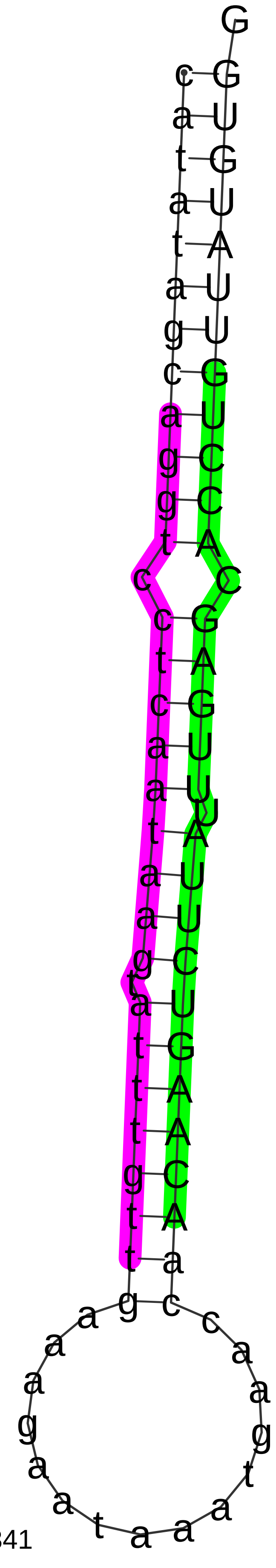

NW_006383124.1_2731264-2731341

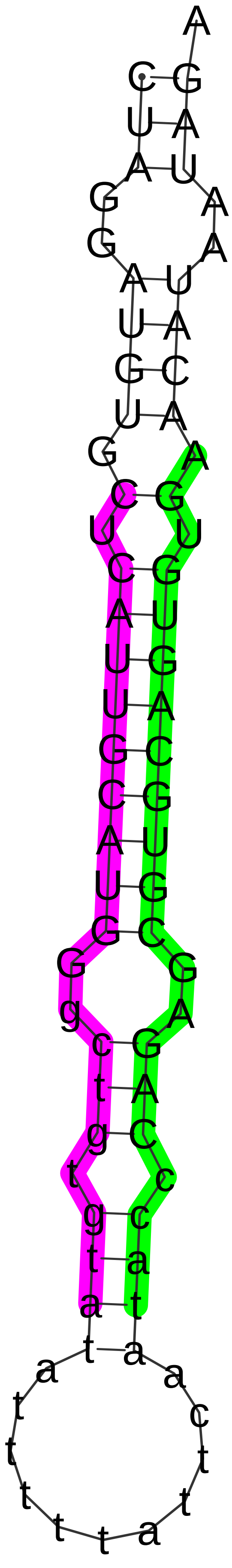

NW_006383124.1_2732441-2732514

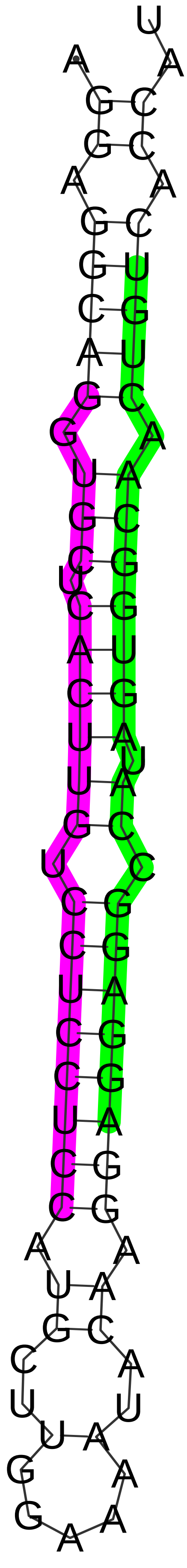

NW_006383124.1_2741972-2742047

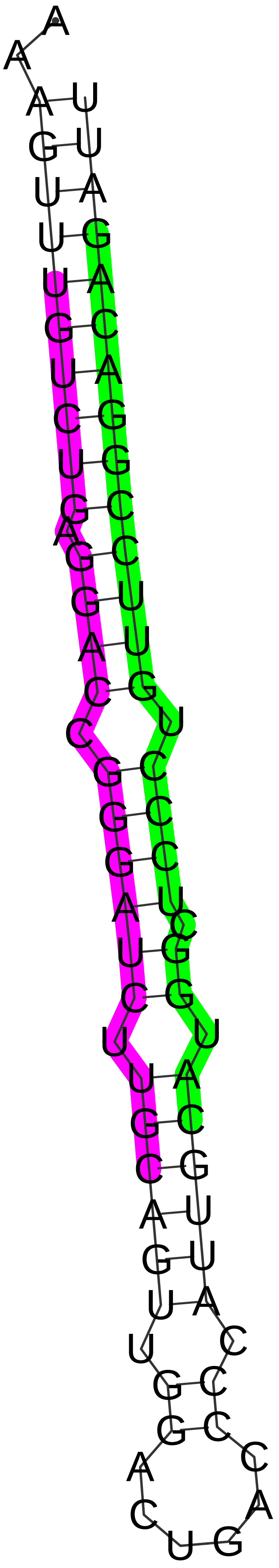

NW_006383138.1_492012-492083

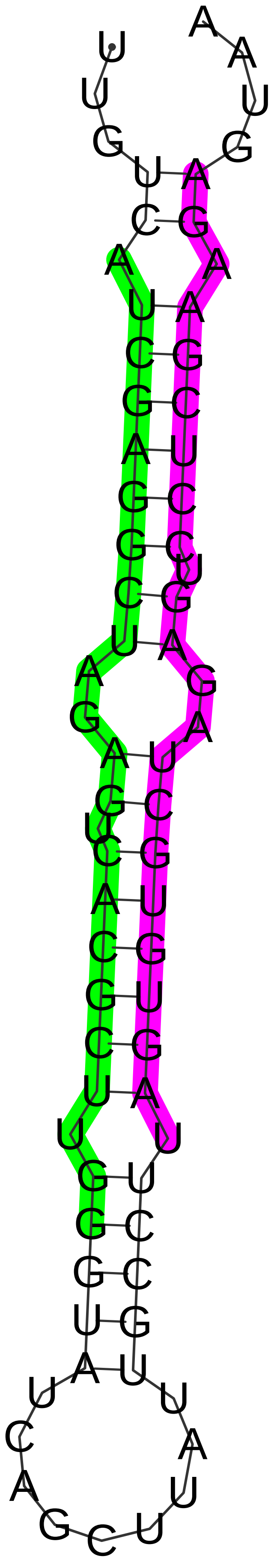

NW_006383139.1_339159-339230

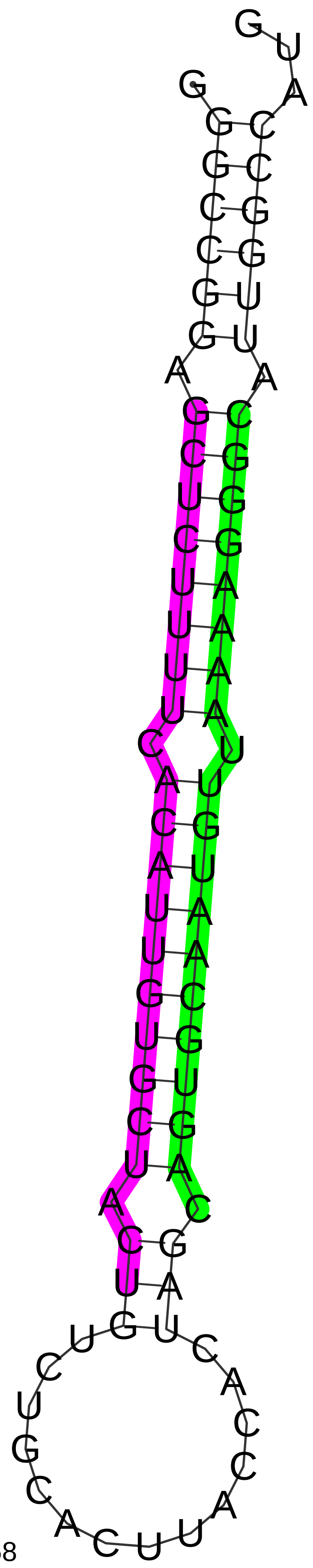

NW_006383141.1_813891-813968

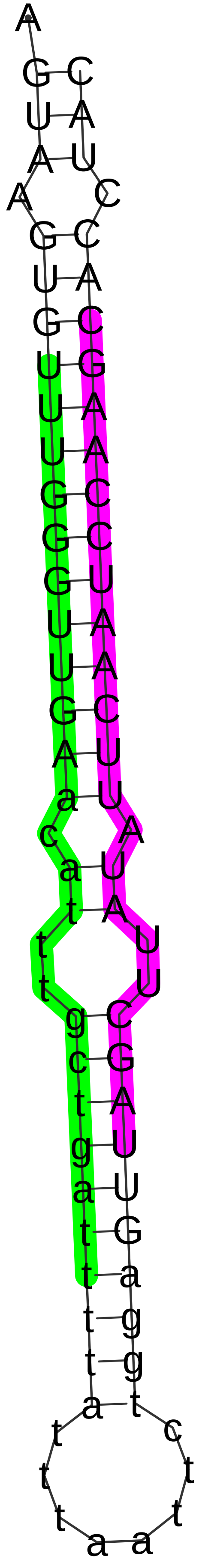

NW_006383145.1_735556-735630

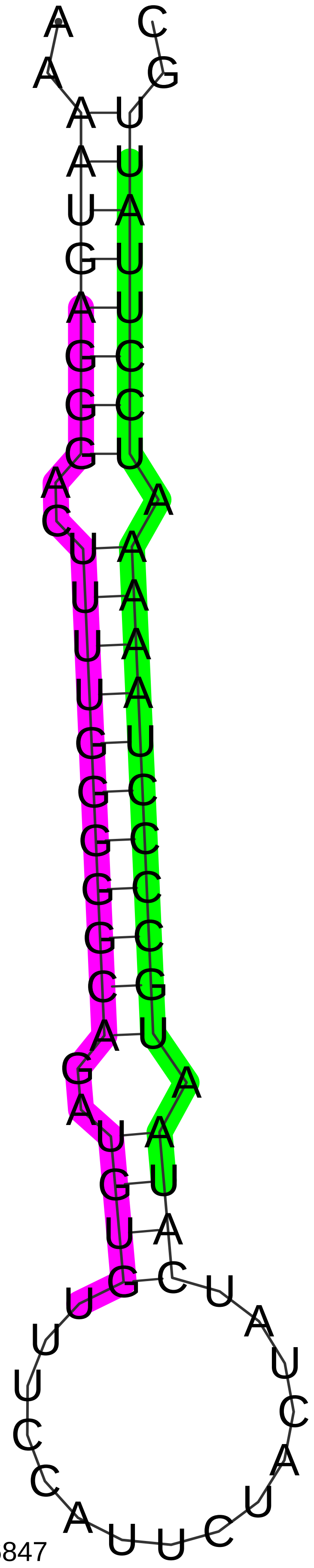

NW_006383147.1_1086777-1086847

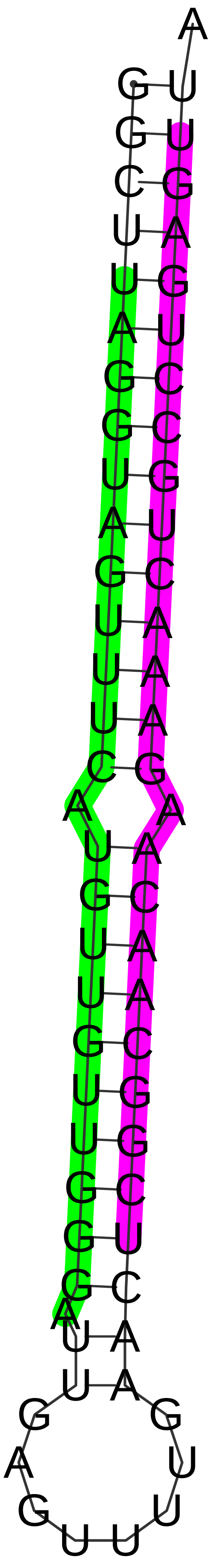

NW_006383149.1_557159-557224

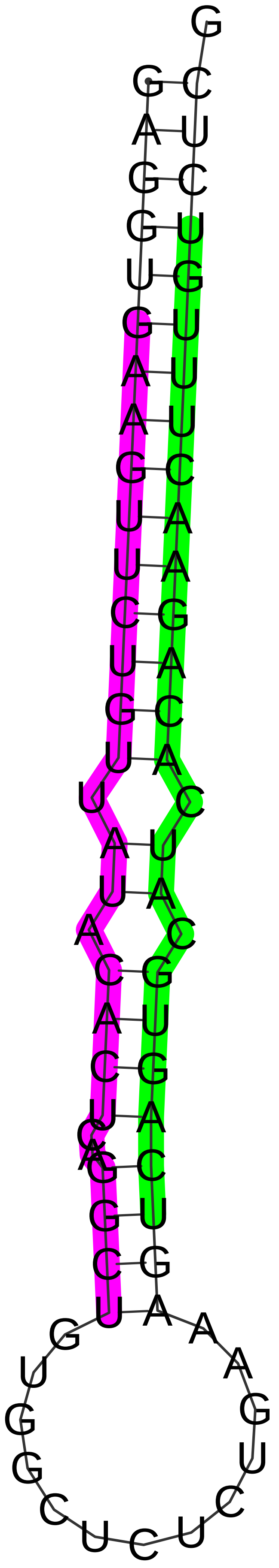

NW_006383149.1_866533-866602

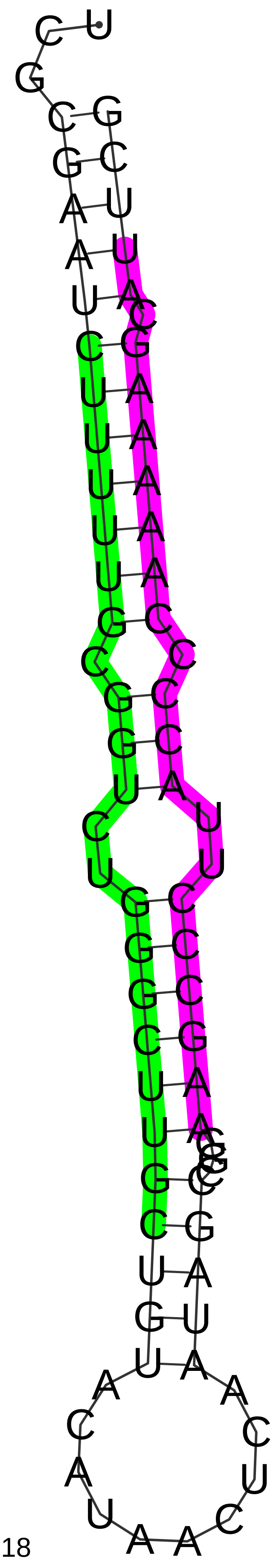

NW_006383150.1_2058944-2059018

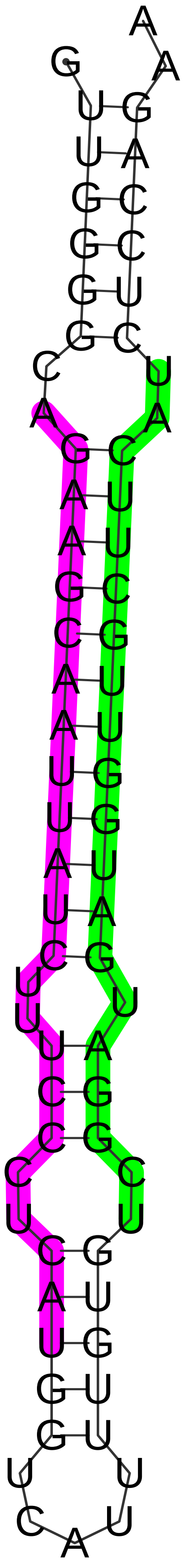

NW_006383153.1_1972479-1972549

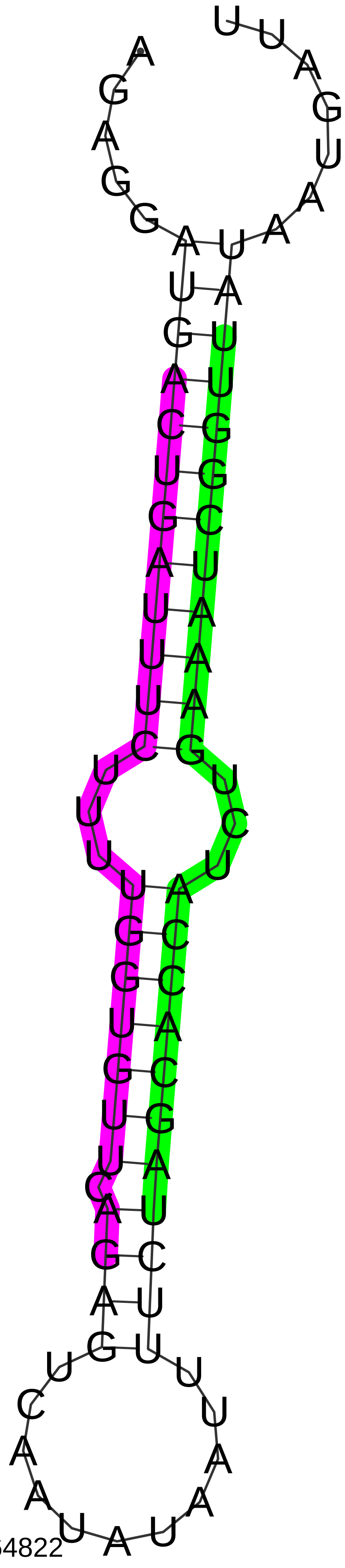

NW_006383179.1_1564747-1564822

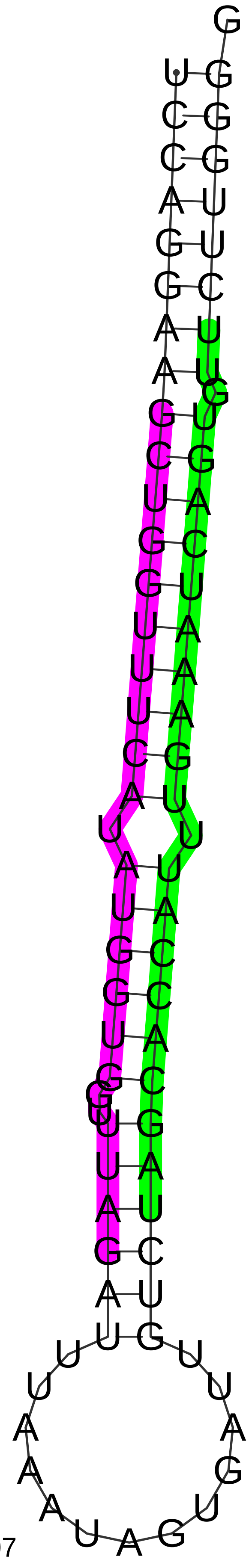

NW_006383179.1_1565129-1565207

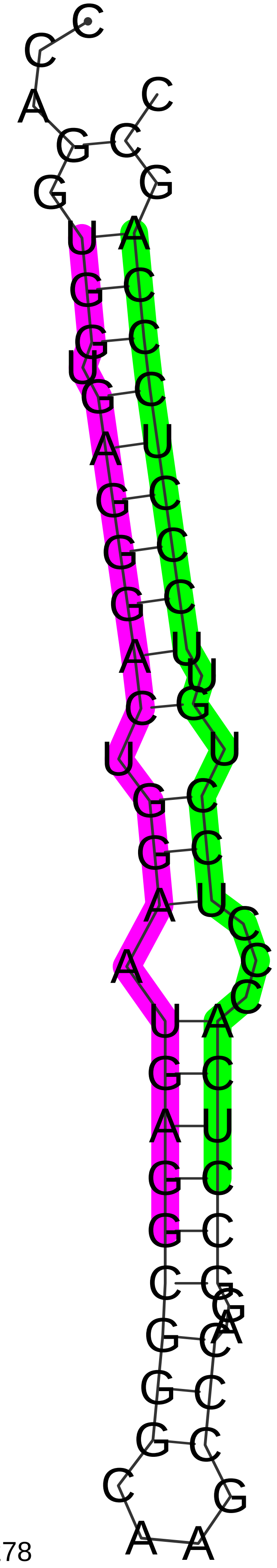

NW_006383180.1_1211213-1211278

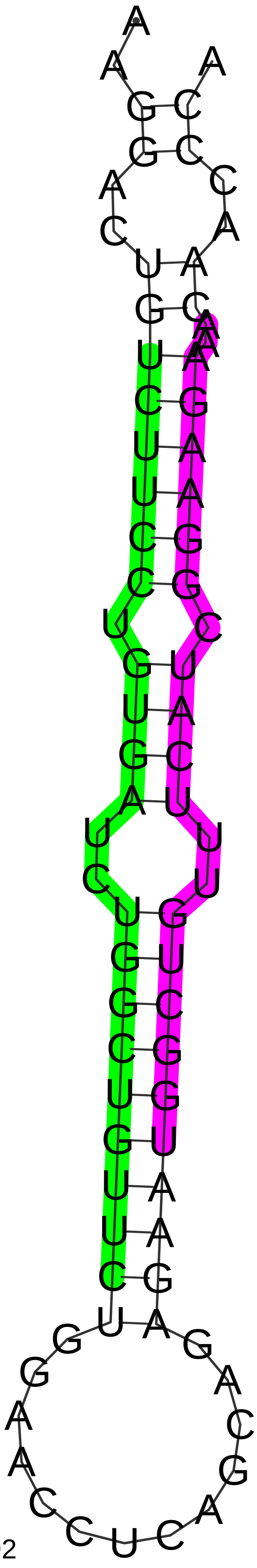

NW_006383180.1_1328017-1328092

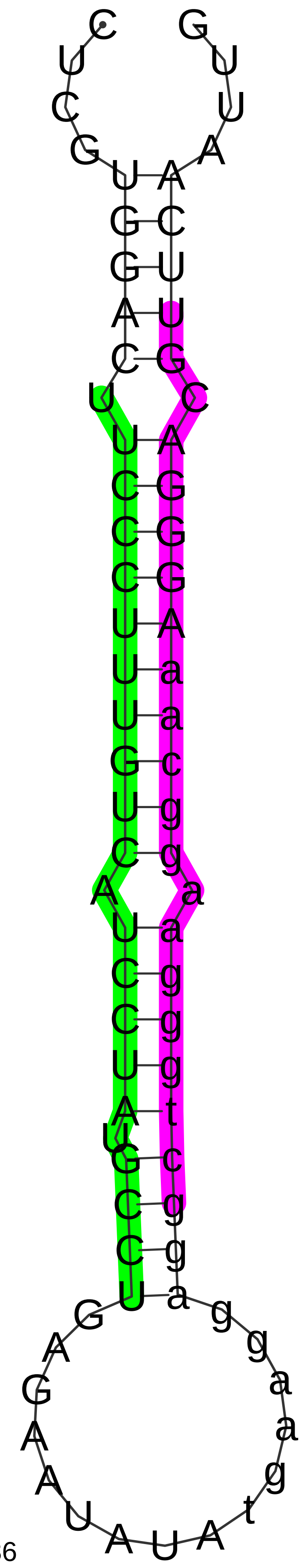


NW_006383196.1_747911-747986

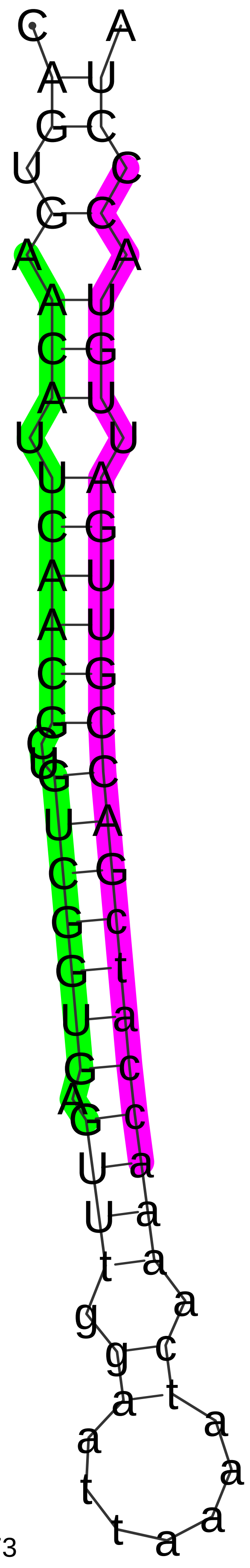

NW_006383202.1_1492504-1492573

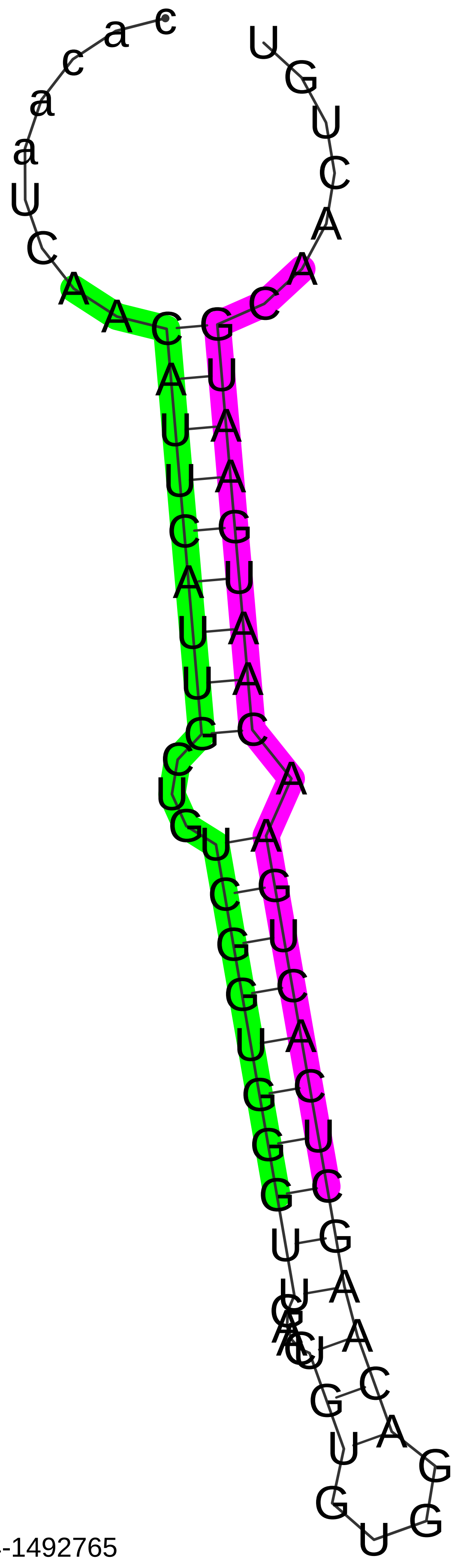

NW_006383202.1_1492694-1492765

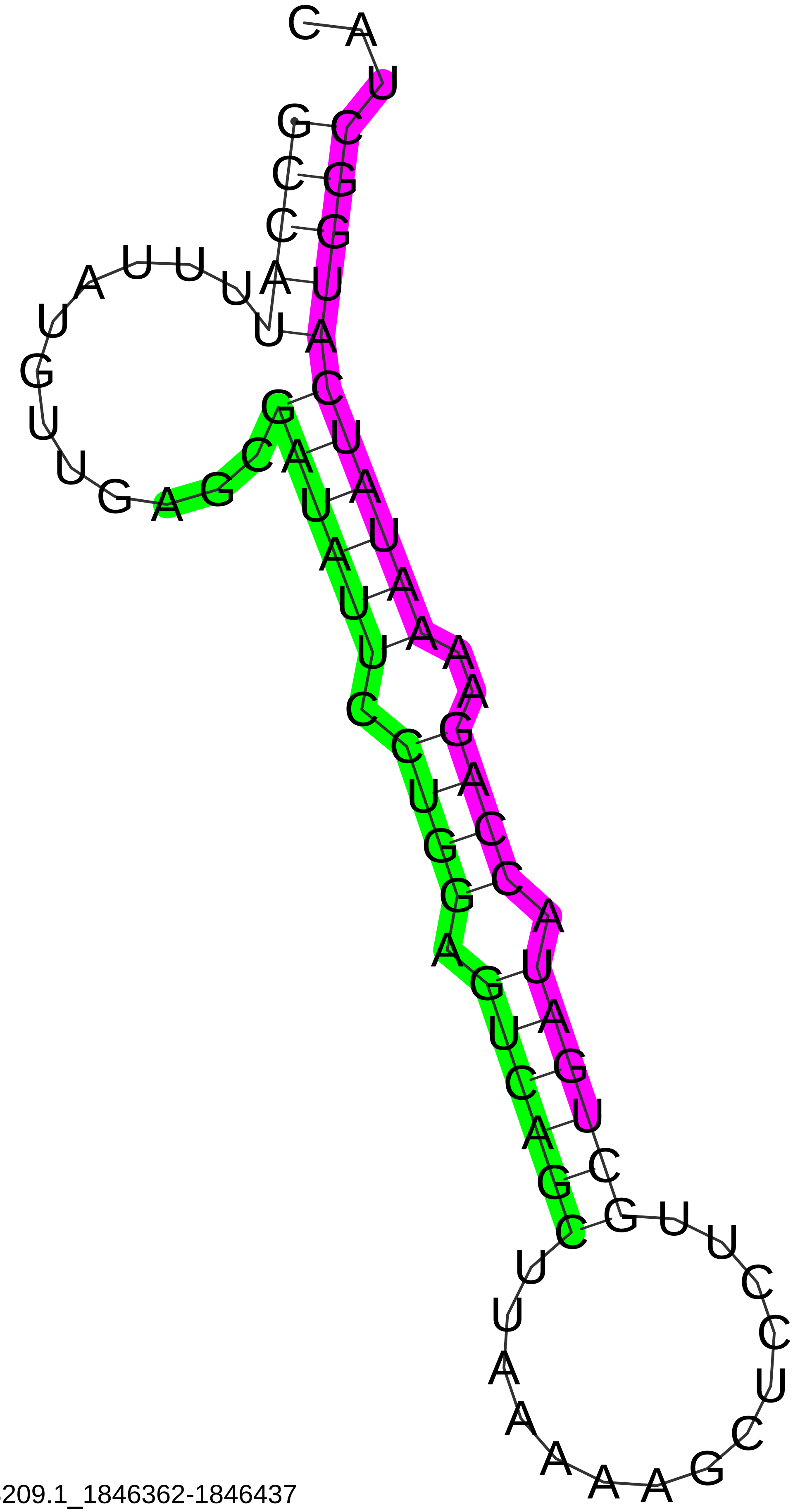

NW_006383209.1_1846362-1846437

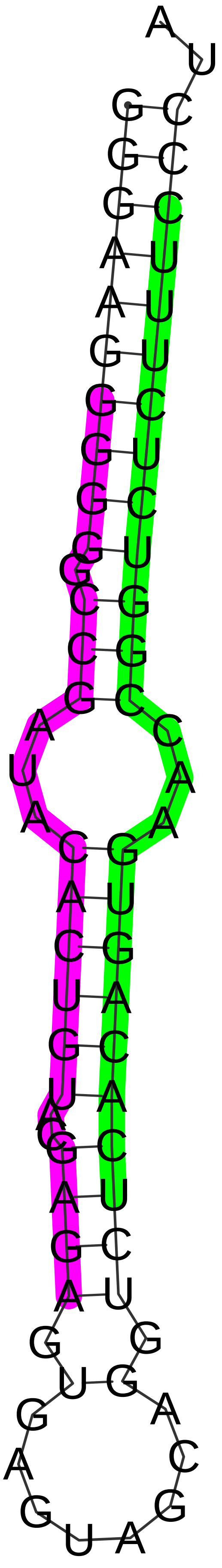

NW_006383213.1_1411071-1411139

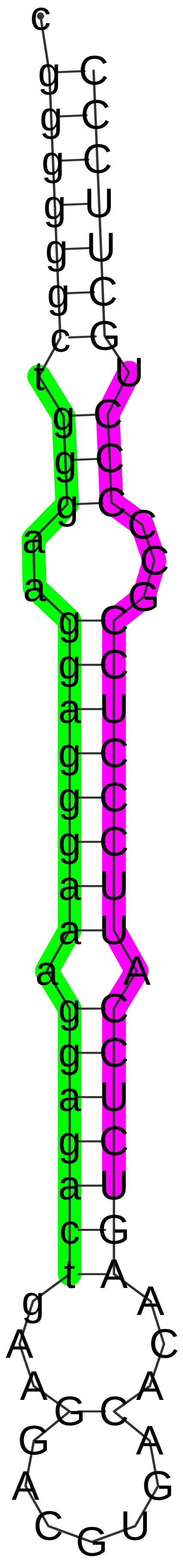


NW_006383218.1_839394-839468

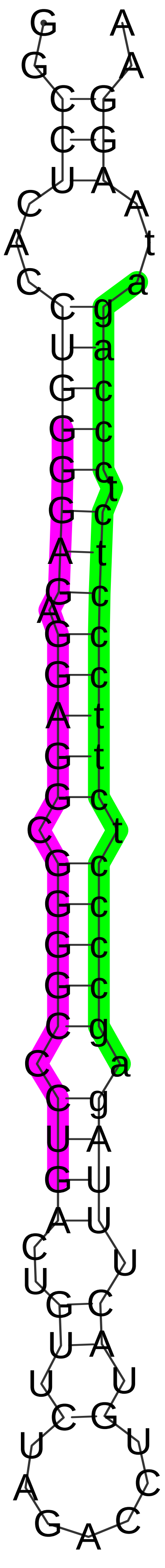

NW_006383229.1_164758-164841

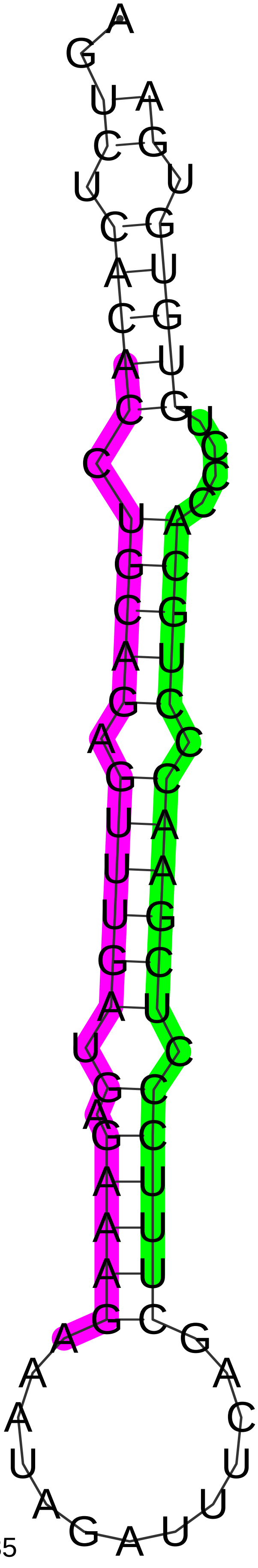

NW_006383232.1_1171461-1171535

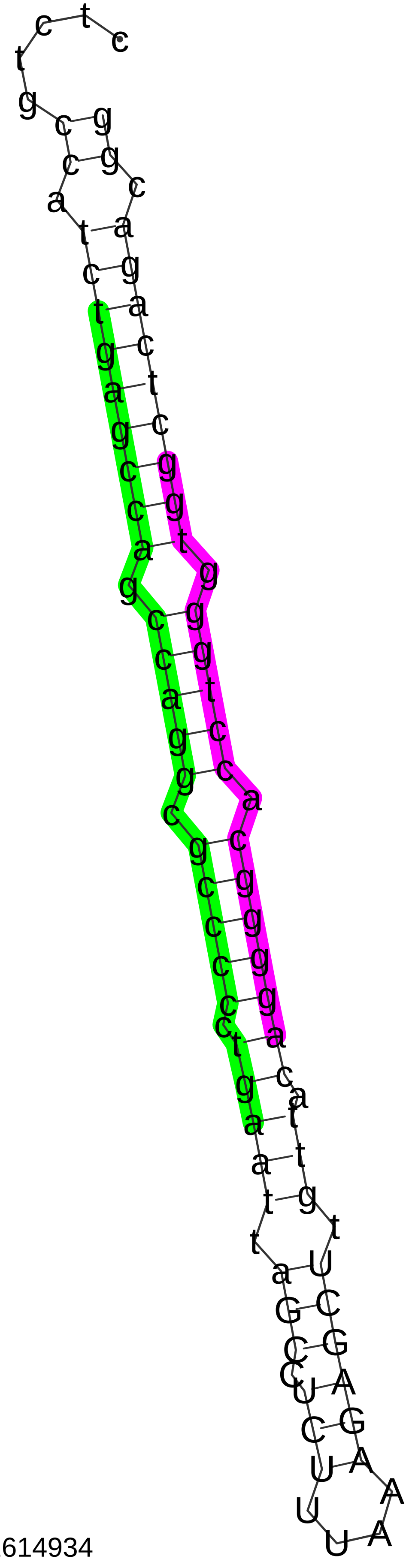

NW_006383238.1_1614851-1614934

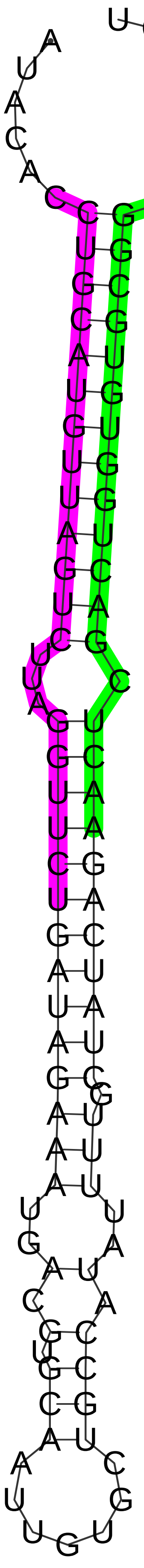

NW_006383243.1_112161-112257

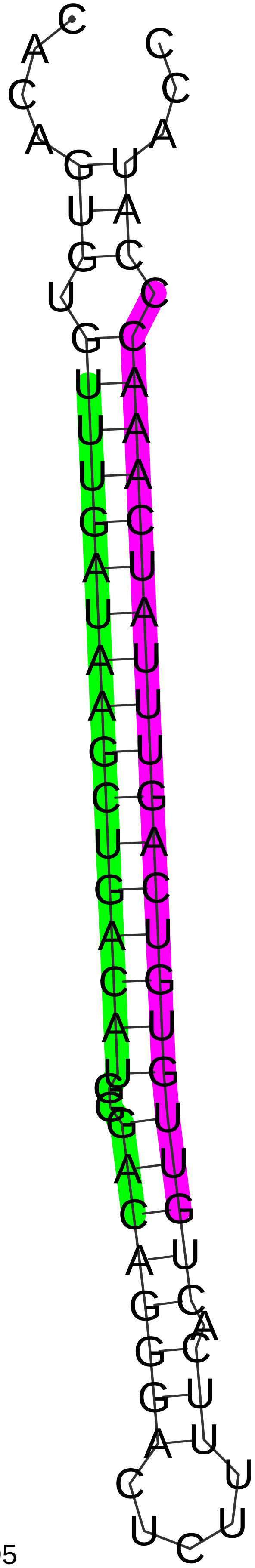

NW_006383249.1_1517123-1517195

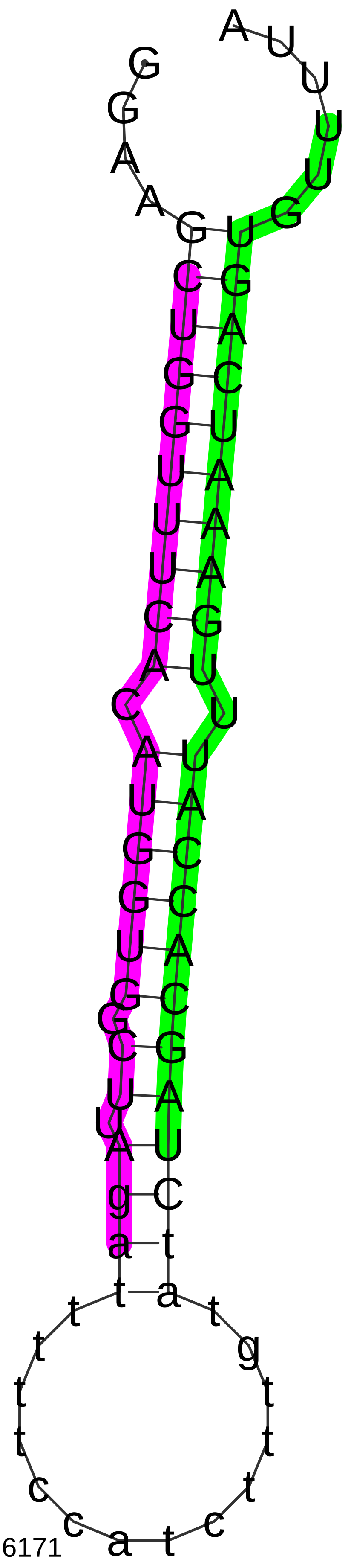

NW_006383259.1_1126100-1126171

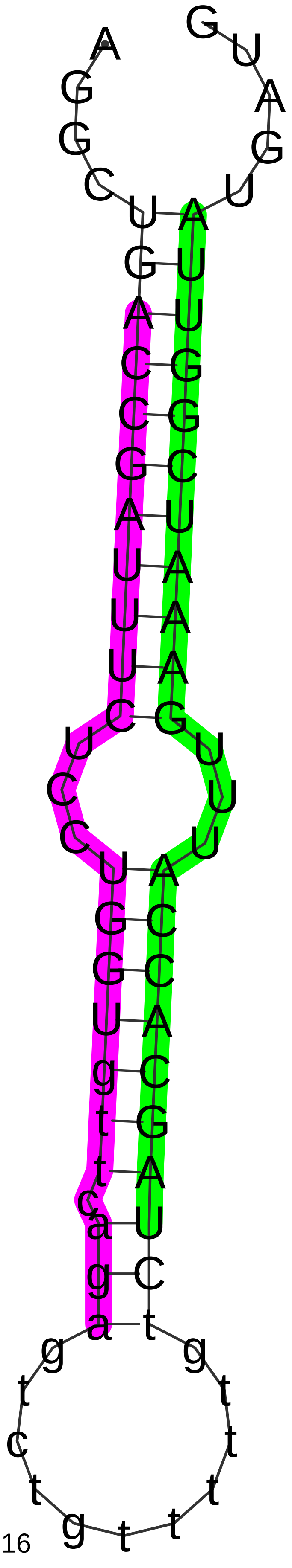

NW_006383259.1_1126648-1126716

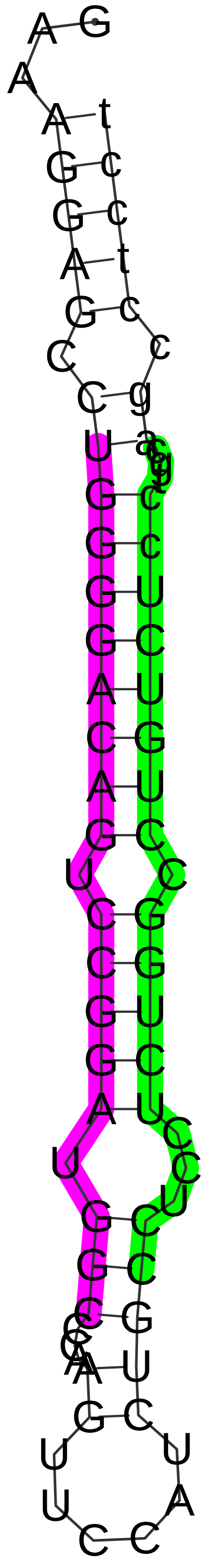

NW_006383262.1_1731879-1731951

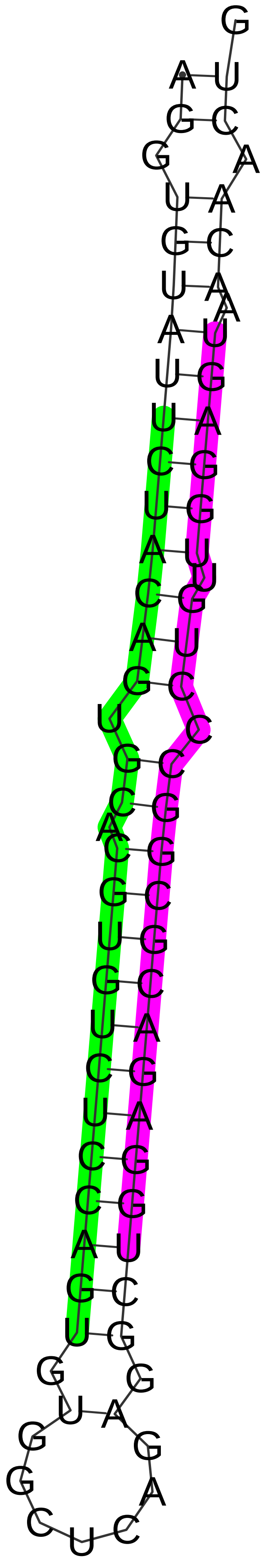

NW_006383262.1_239764-239838

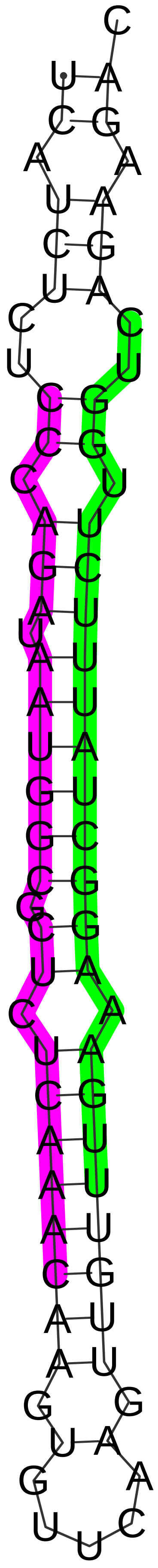

NW_006383264.1_1297503-1297576

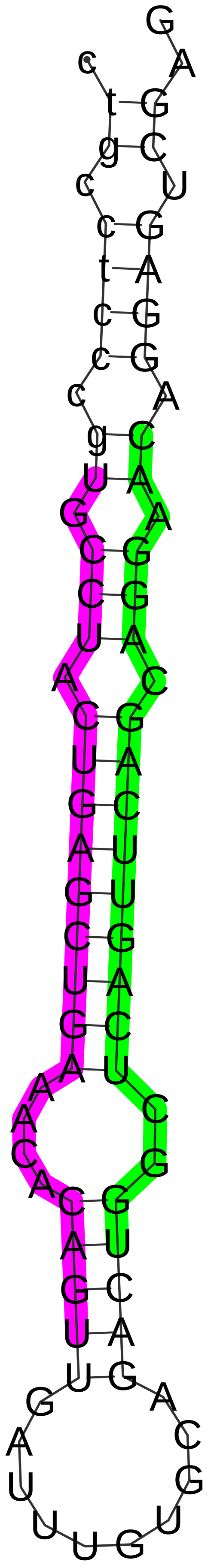

NW_006383268.1_1271580-1271656

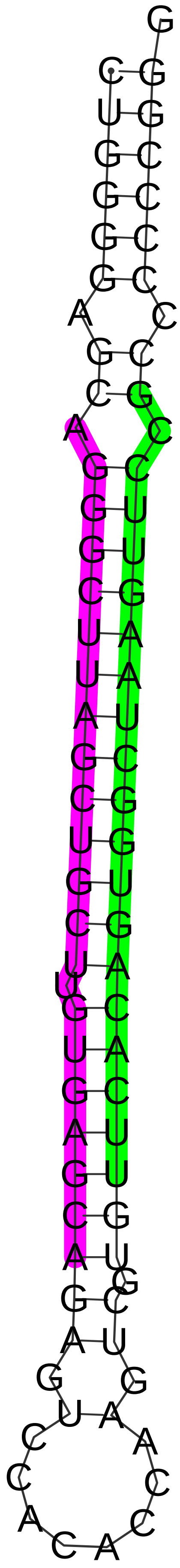

NW_006383268.1_1271720-1271798

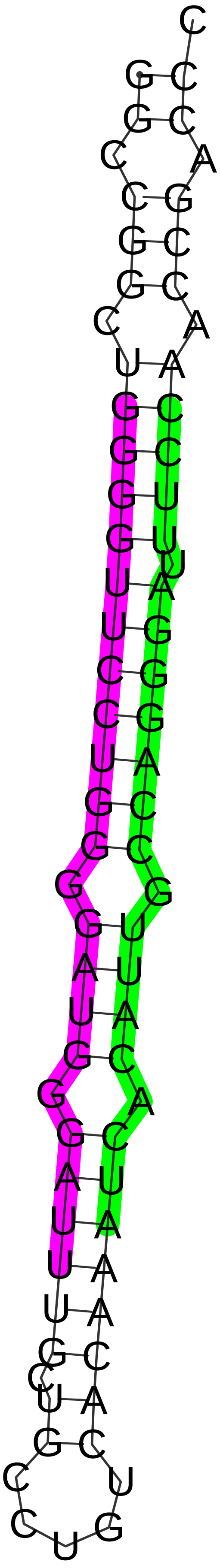

NW_006383268.1_1271906-1271979

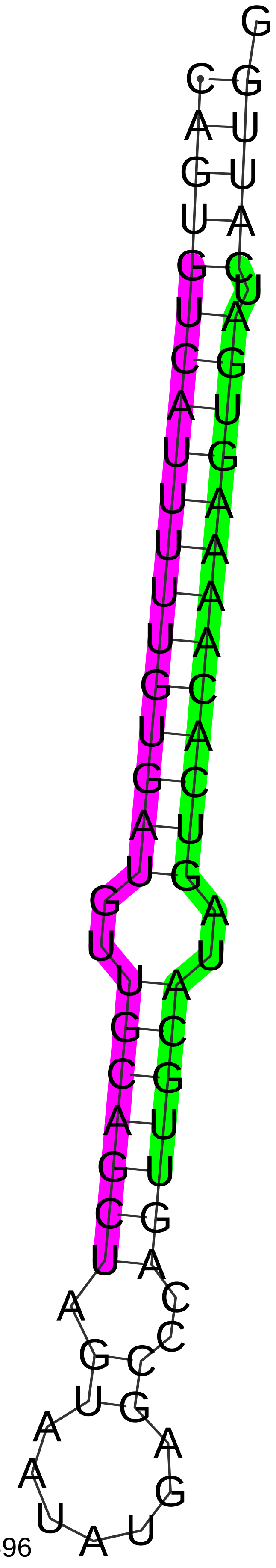

NW_006383283.1_1282527-1282596

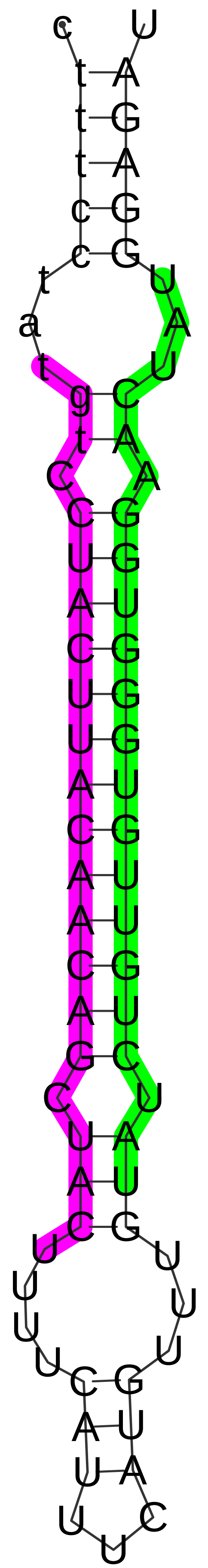


NW_006383297.1_1283494-1283567

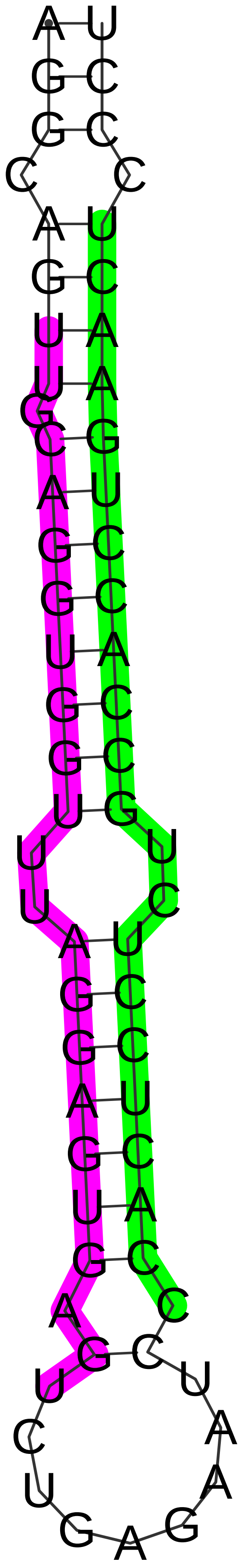

NW_006383297.1_487454-487517

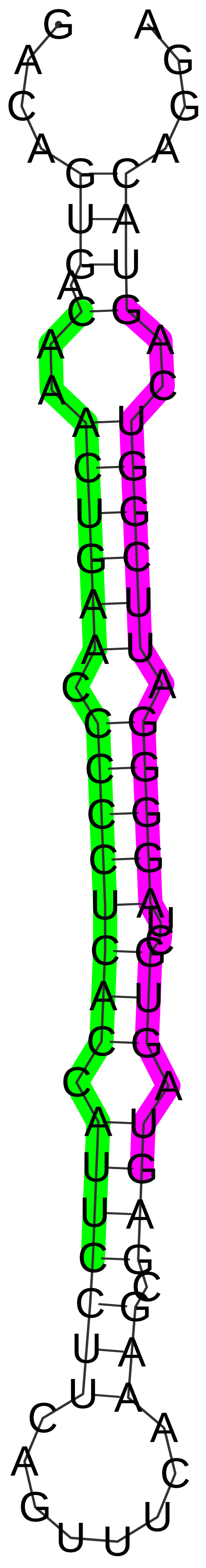

NW_006383303.1_684351-684428

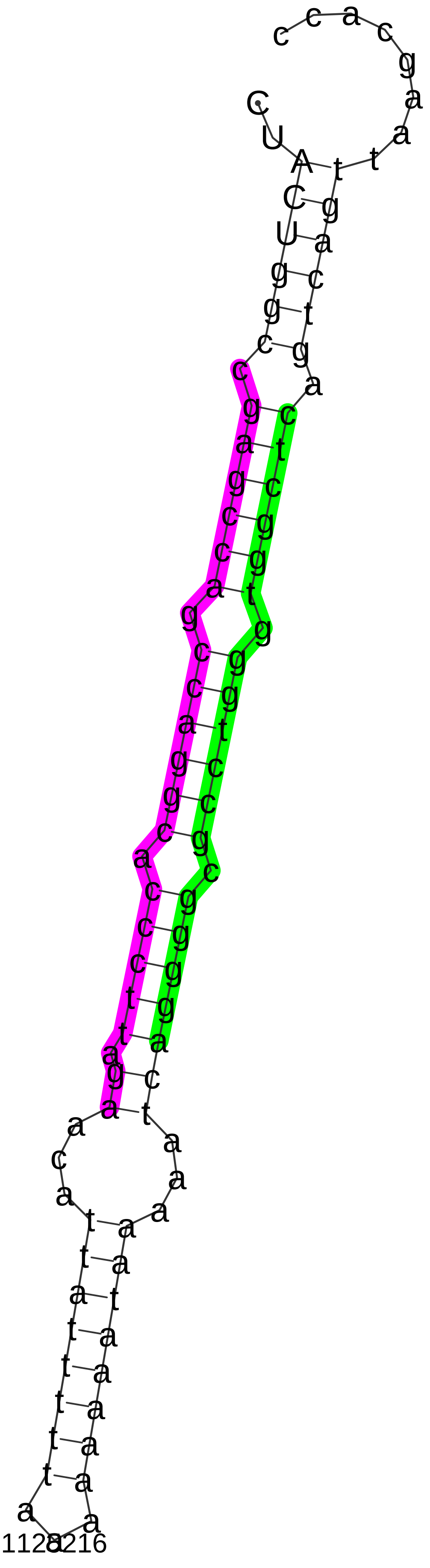

NW_006383309.1_1125125-112[illegible]216

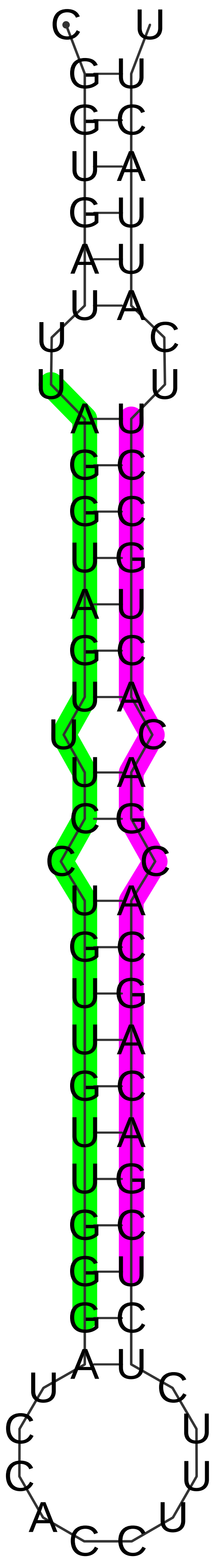

NW_006383334.1_1485765-1485836

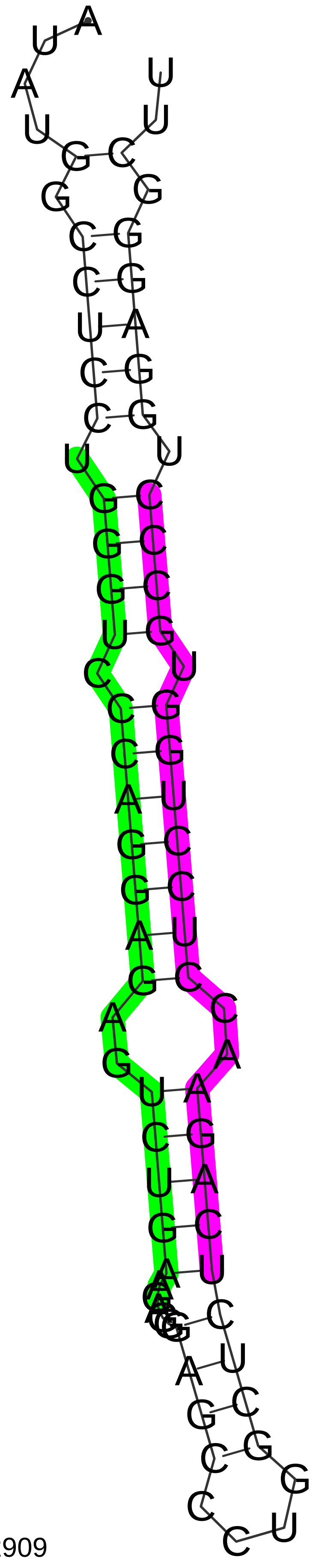

NW_006383336.1_1272833-1272909

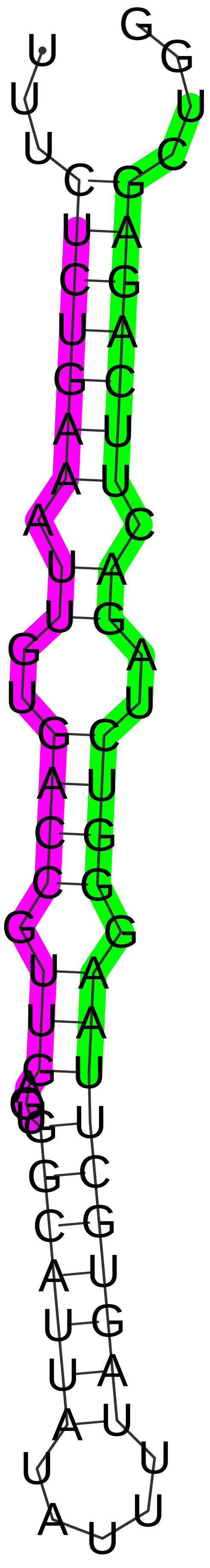

NW_006383340.1_1344314-1344382

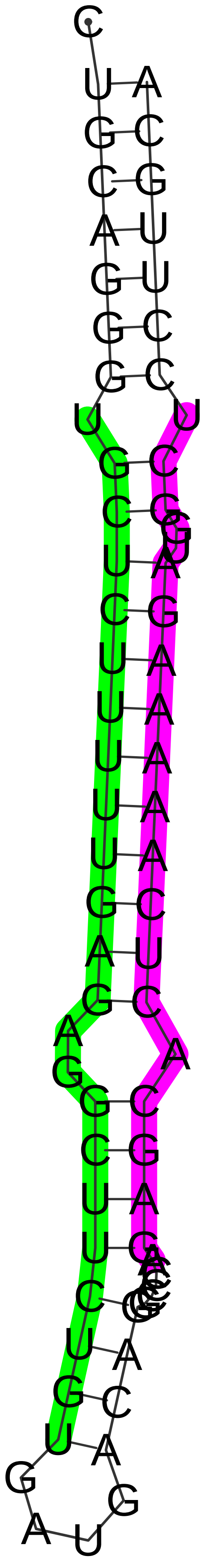

NW_006383353.1_1241921-1241991

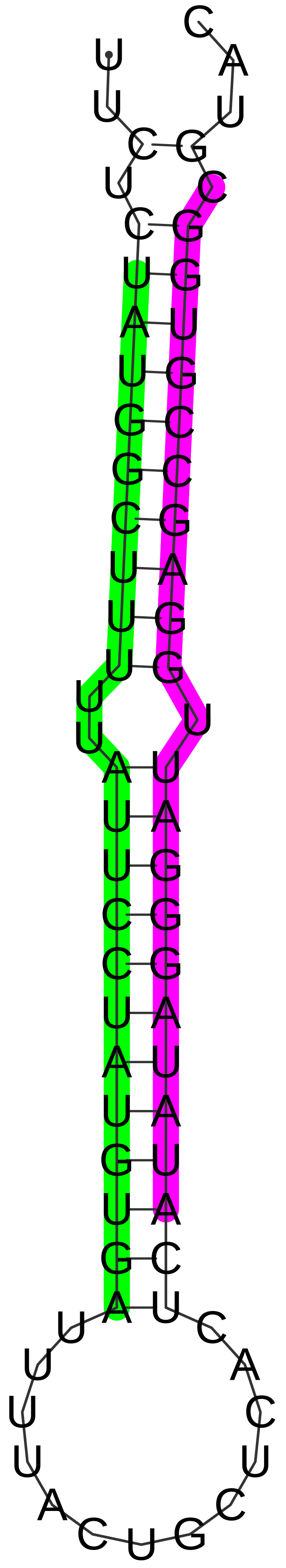

NW_006383353.1_193550-193618

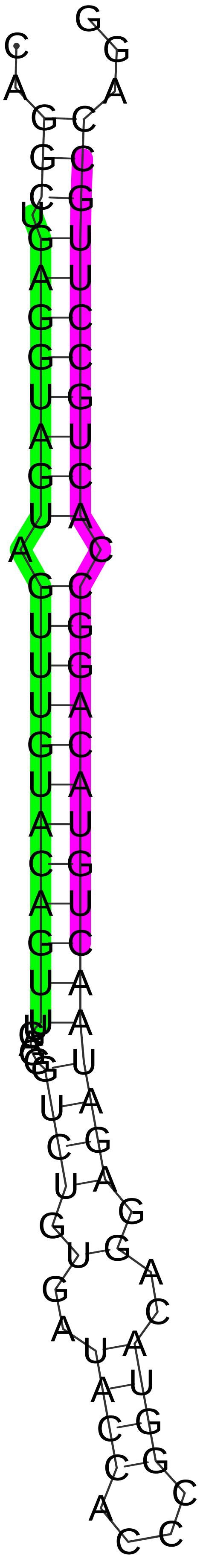

NW_006383353.1_233261-233347

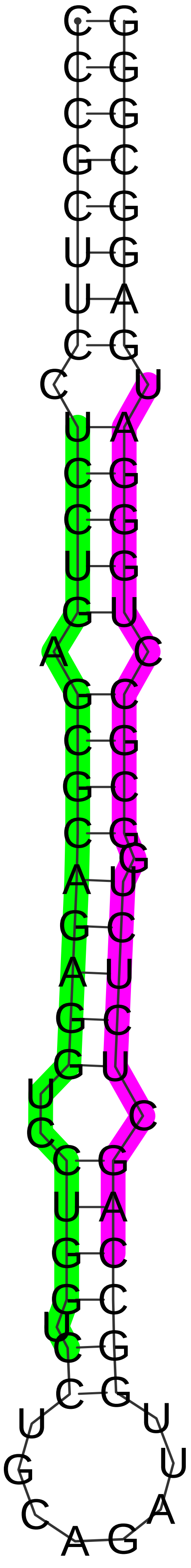

NW_006383358.1_1185095-1185167

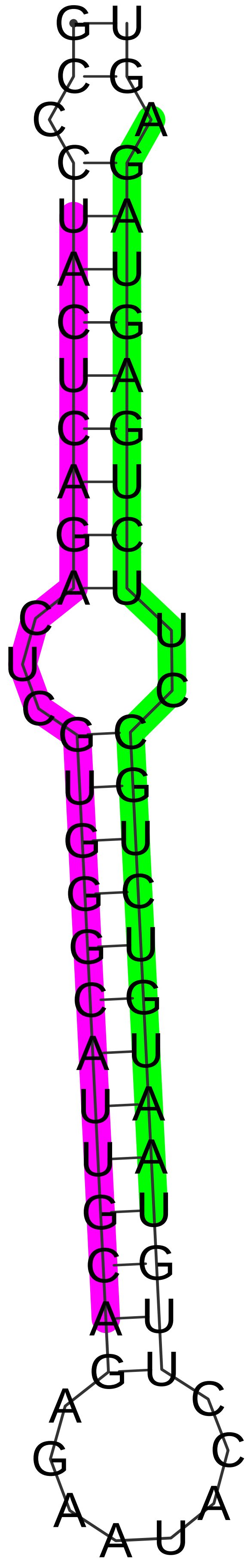

NW_006383360.1_418160-418222

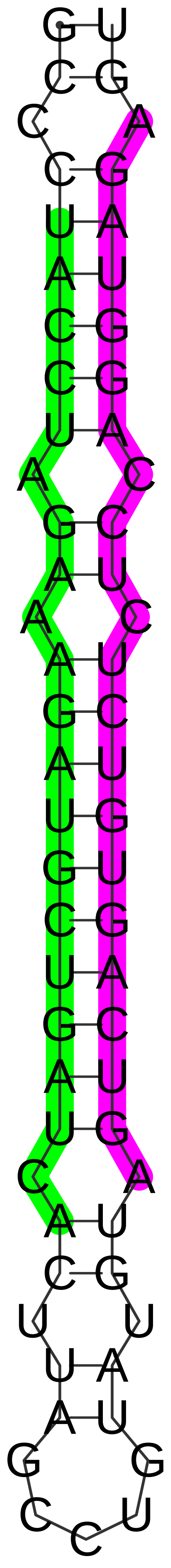

NW_006383360.1_431089-431151

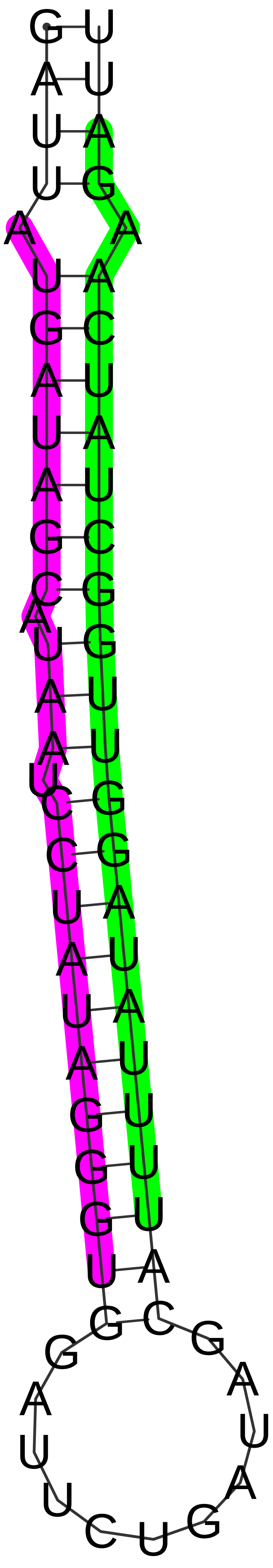

NW_006383364.1_653631-653695

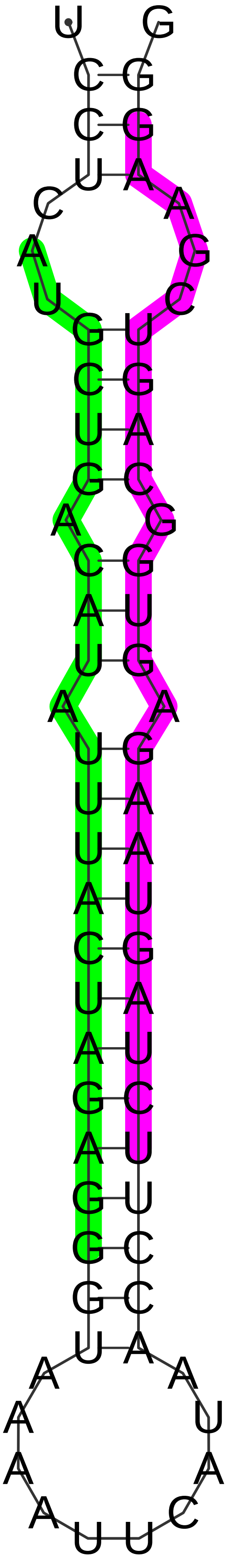

NW_006383375.1_756226-756293

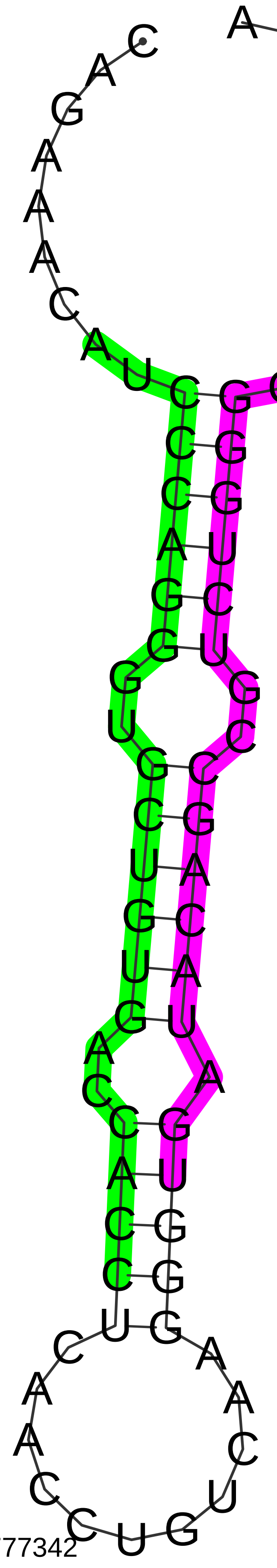

NW_006383383.1_777271-777342

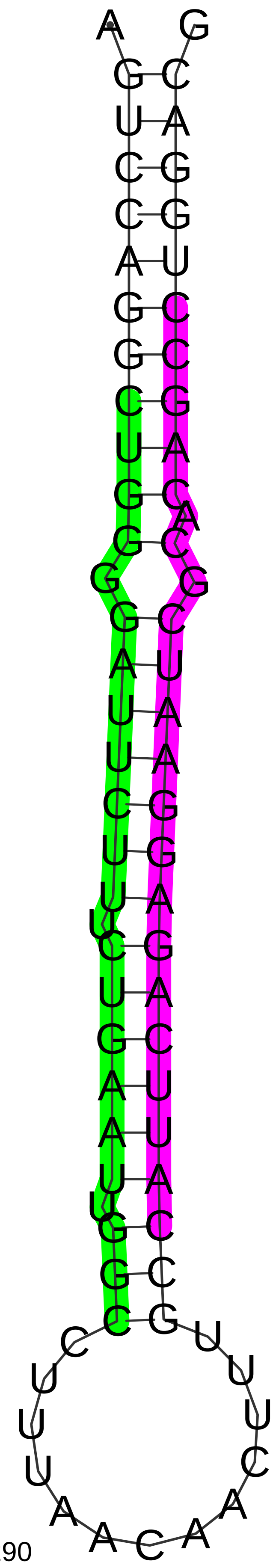

NW_006383384.1_1246217-1246290

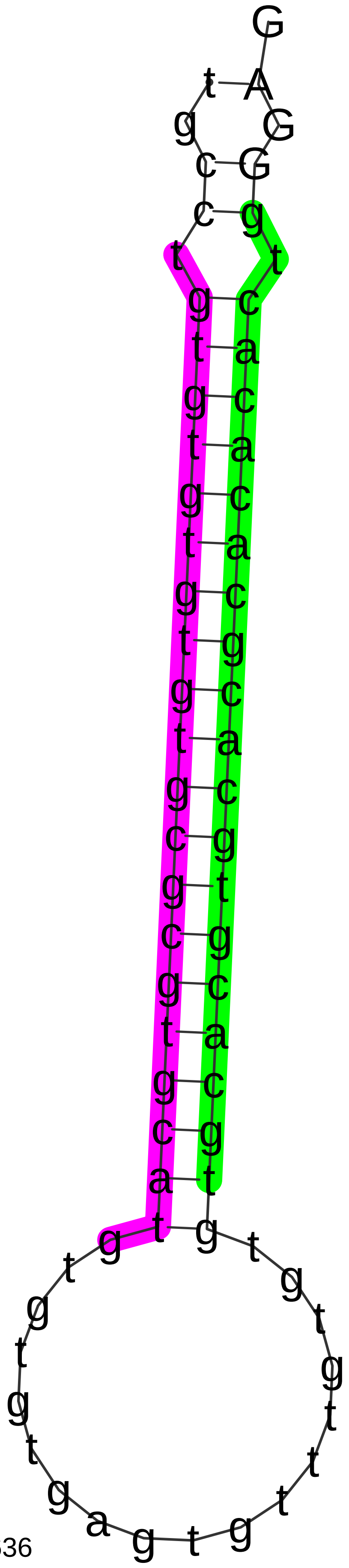

NW_006383399.1_356568-356636

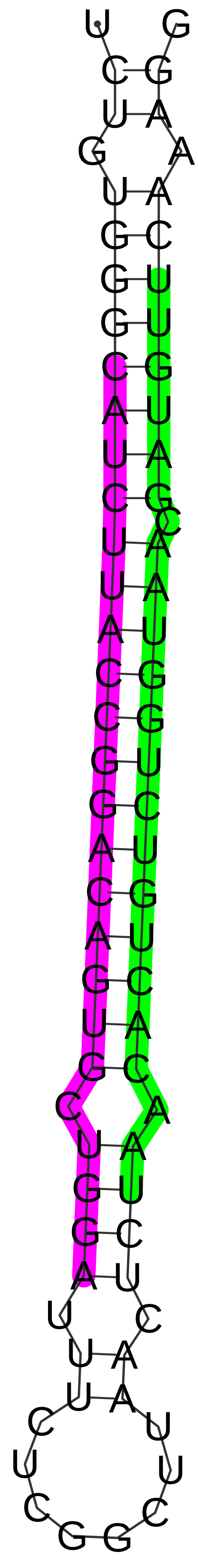

NW_006383401.1_1122091-1122165

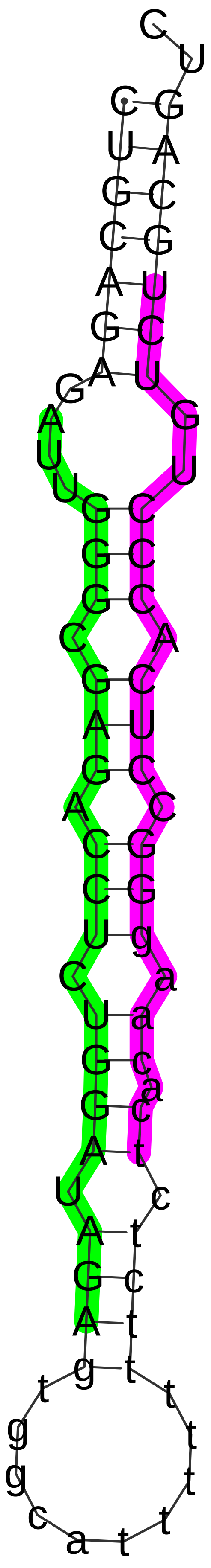

NW_006383413.1_468784-468858

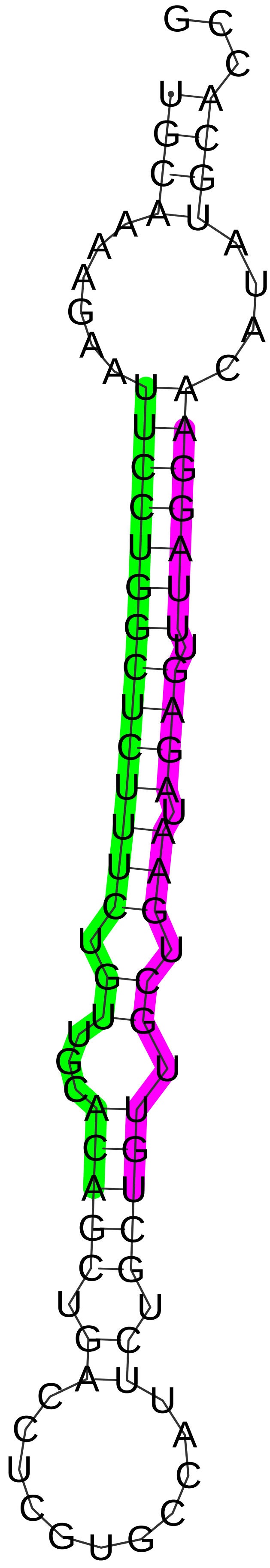

NW_006383427.1_447273-447360

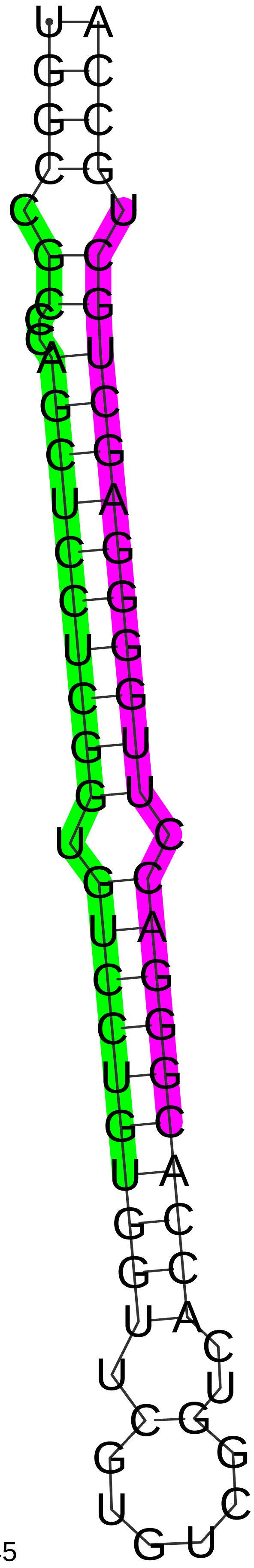

NW_006383436.1_1320377-1320445

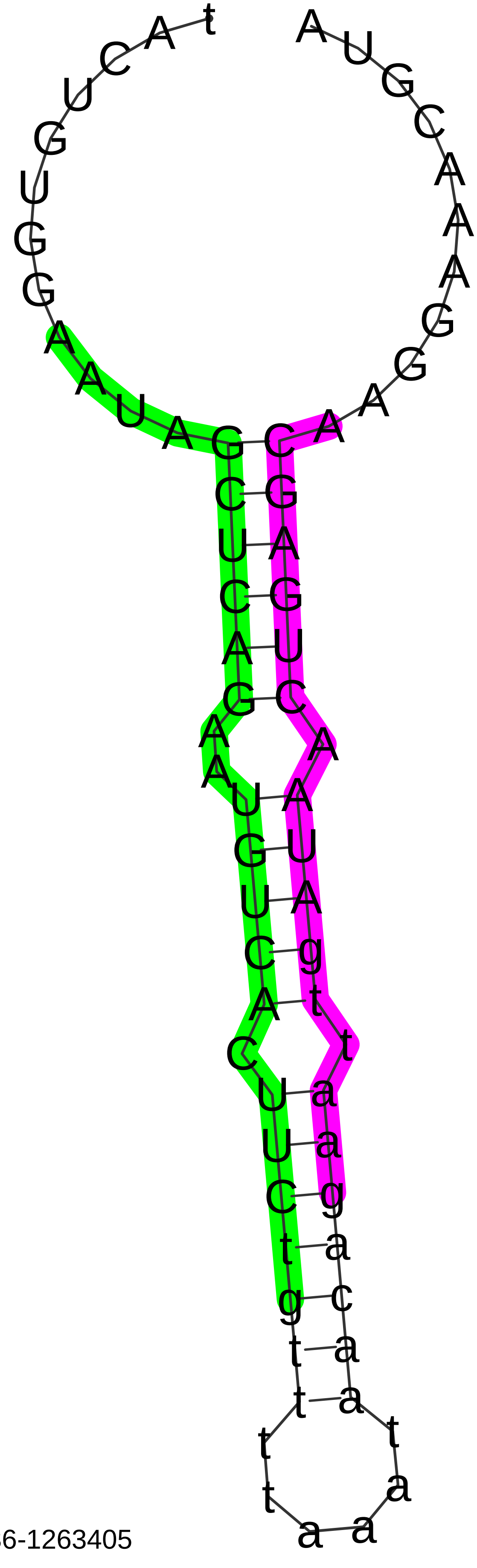

NW_006383438.1_1263336-1263405

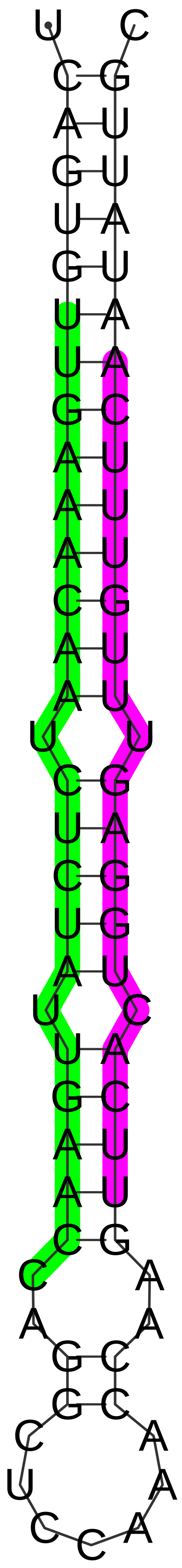

NW_006383440.1_899488-899556

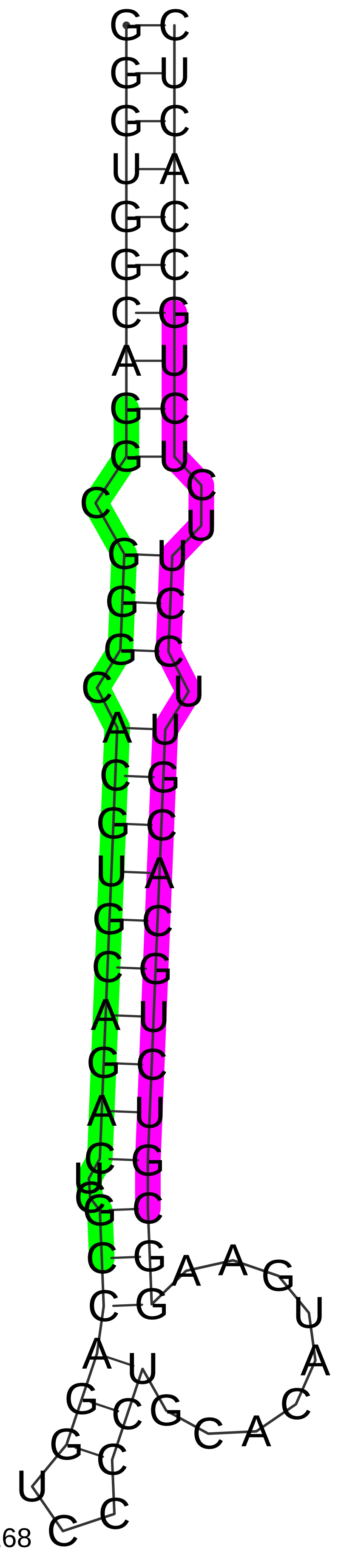

NW_006383442.1_180092-180168

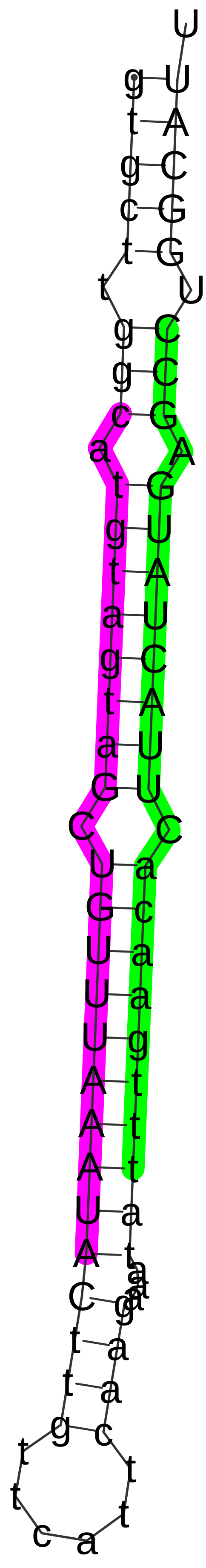

NW_006383446.1_672619-672694

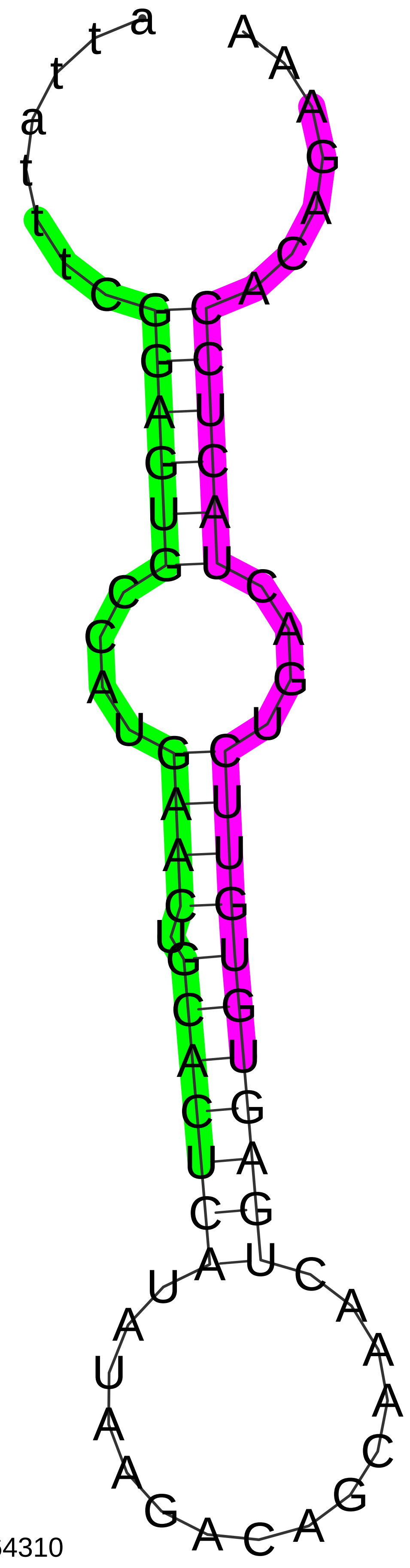

NW_006383446.1_864238-864310

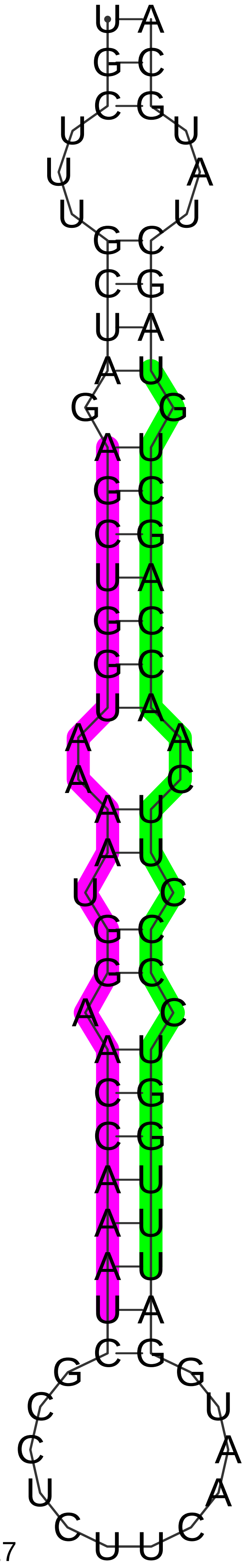

NW_006383454.1_1129248-1129327

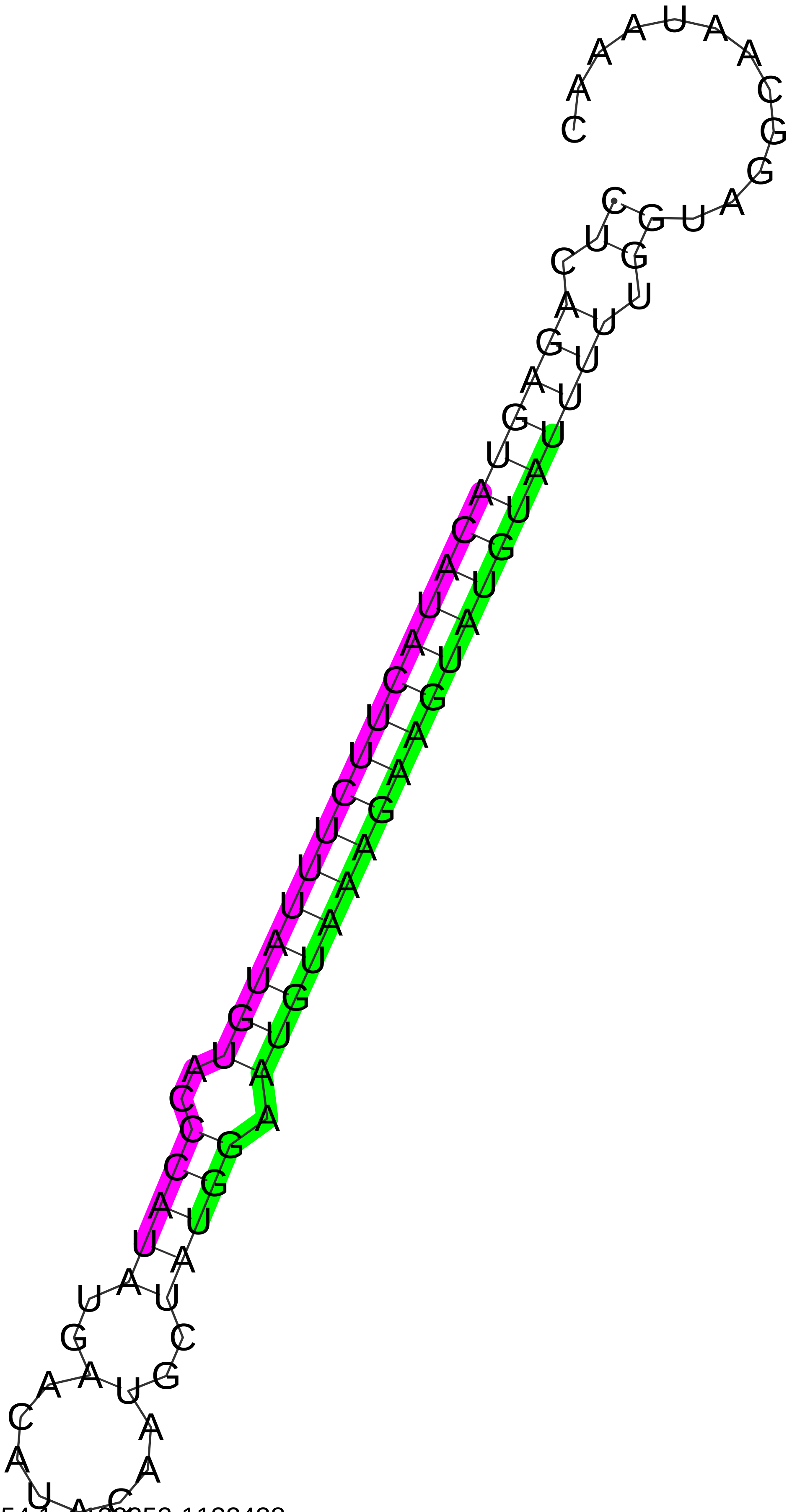

NW_006383454.1_1132352-1132438

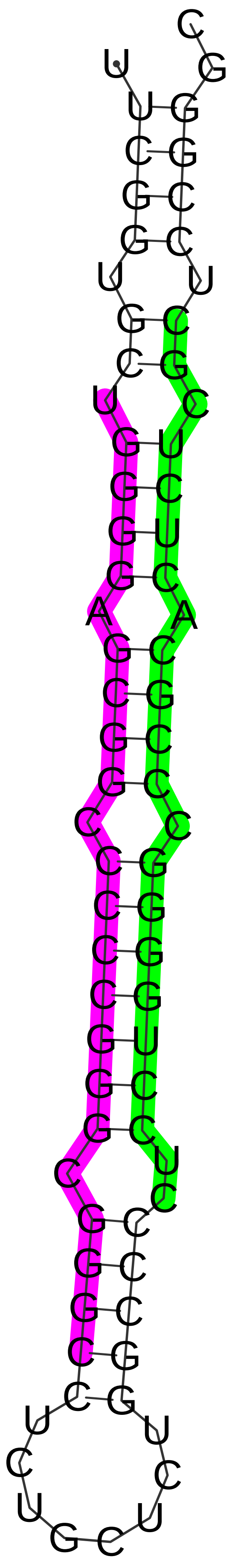

NW_006383455.1_533628-533701

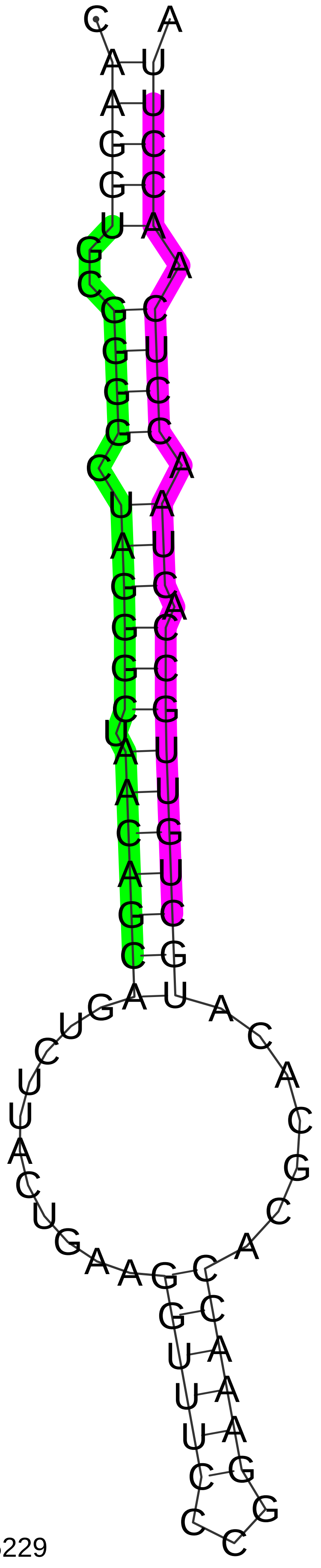

NW_006383458.1_1325144-1325229

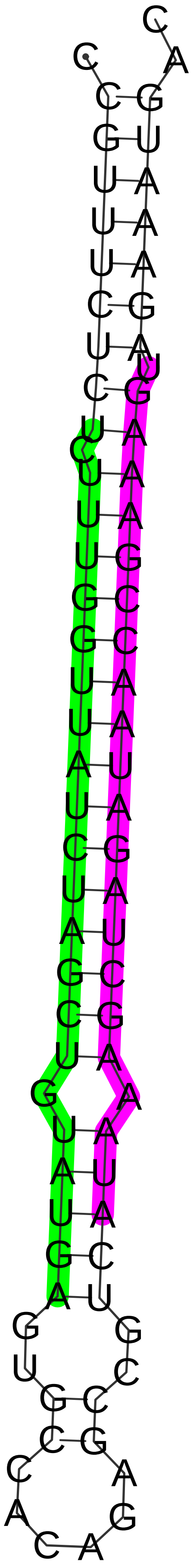

NW_006383462.1_54589-54667

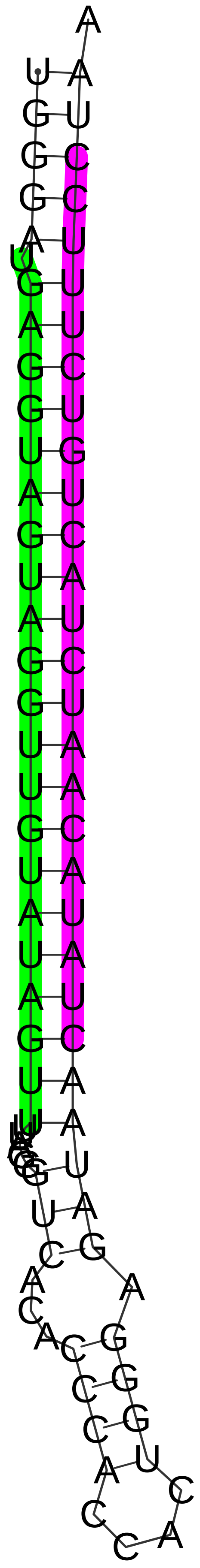

NW_006383491.1_313875-313955

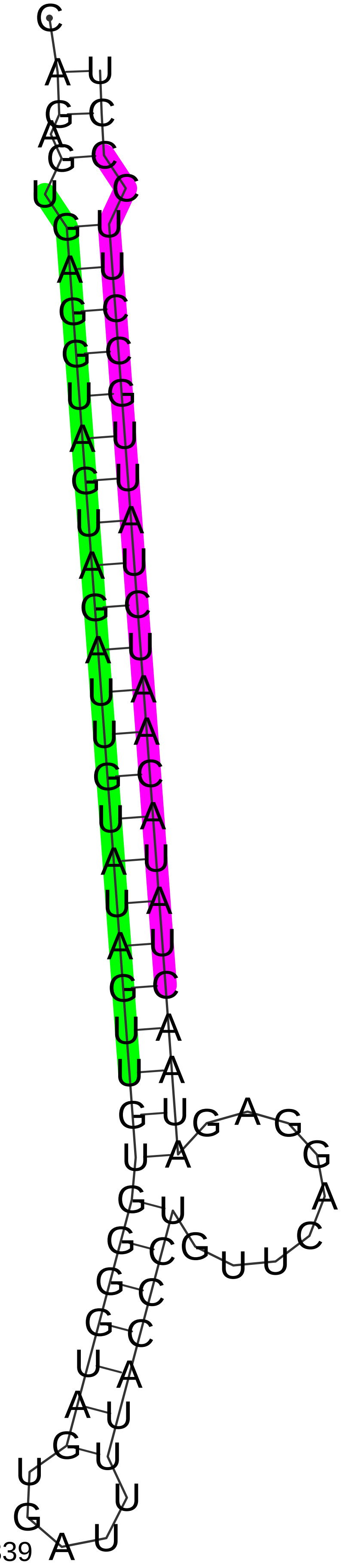

NW_006383491.1_314256-314339

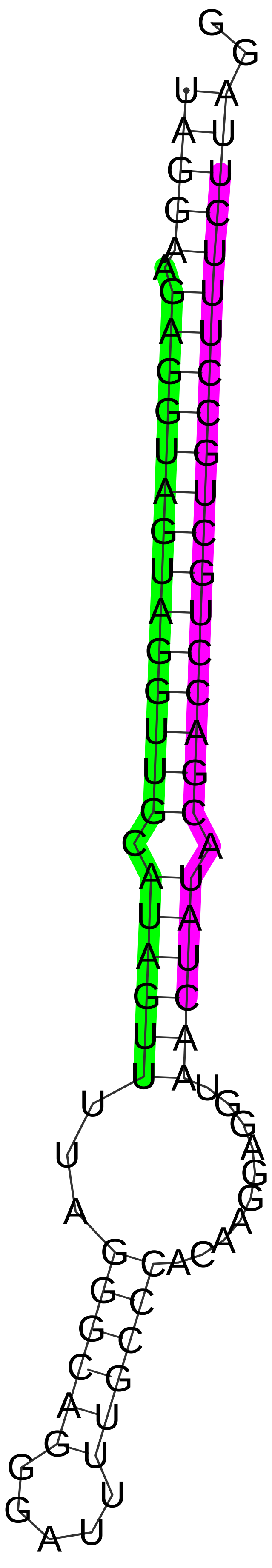

NW_006383491.1_316216-316300

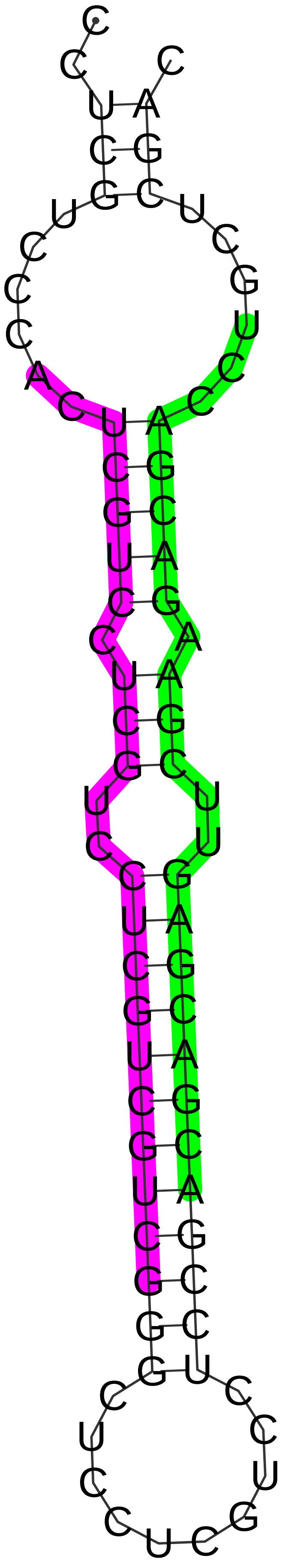

NW_006383504.1_409940-410016

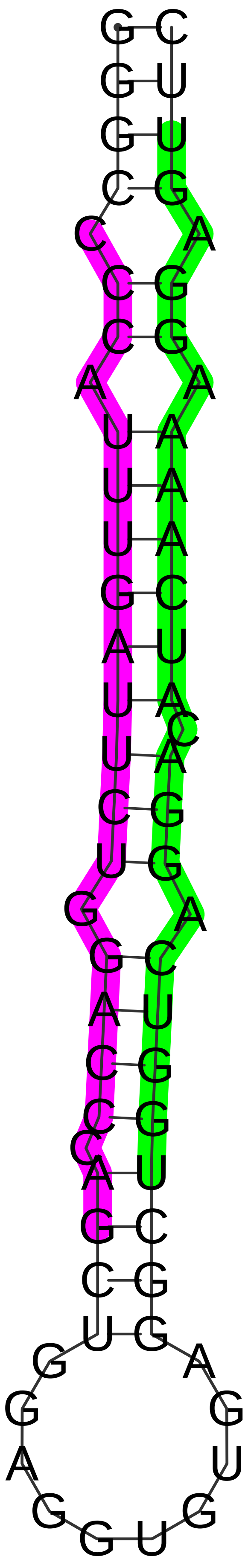

NW_006383520.1_228618-228681

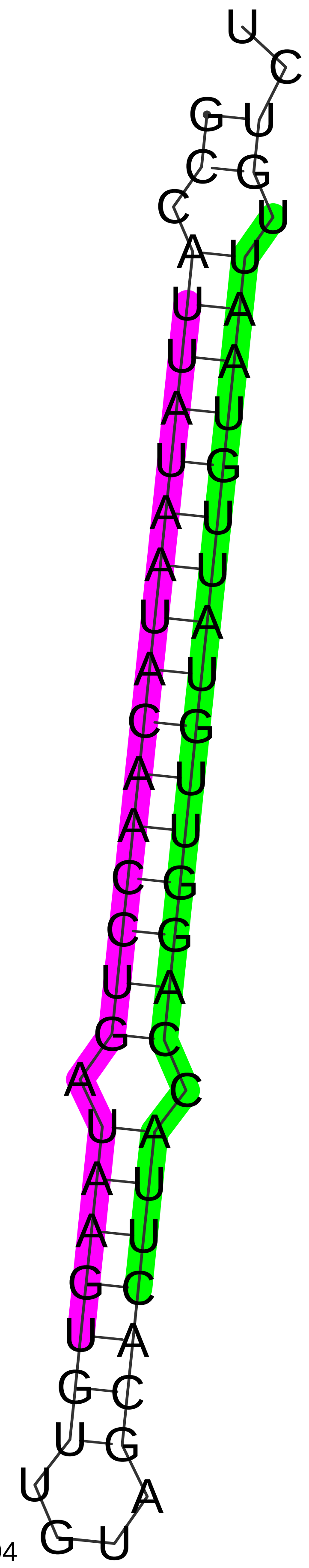

NW_006383521.1_425335-425394

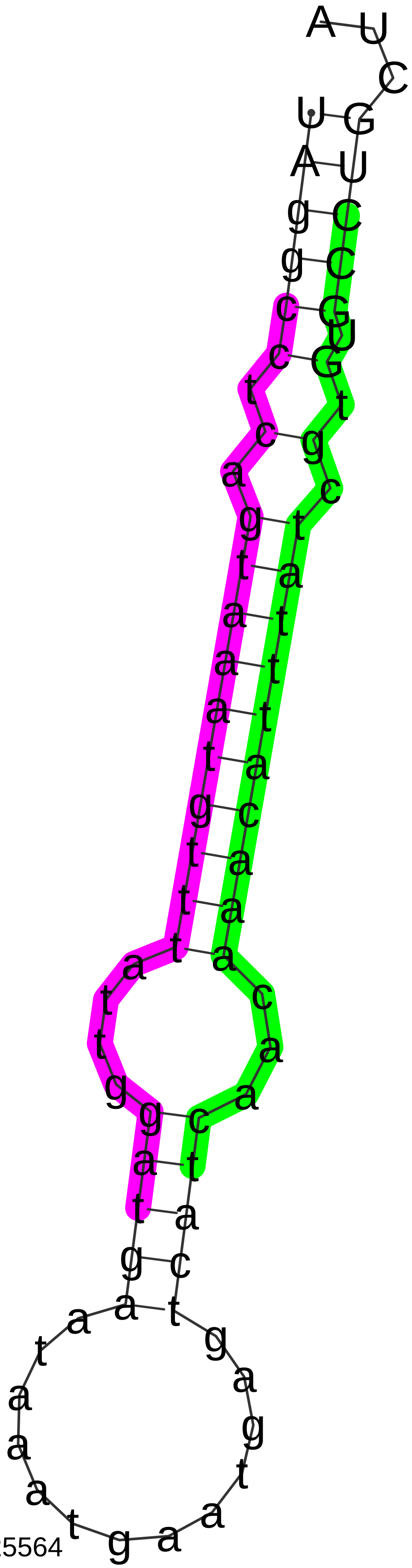

NW_006383521.1_425493-425564

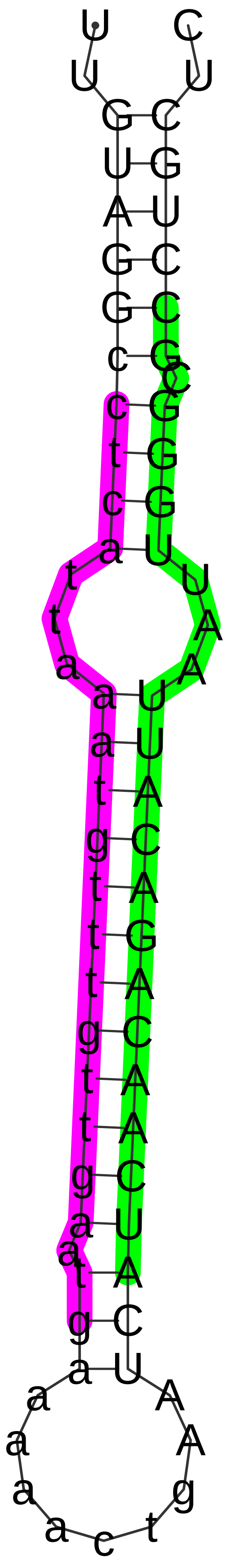

NW_006383521.1_478153-478223

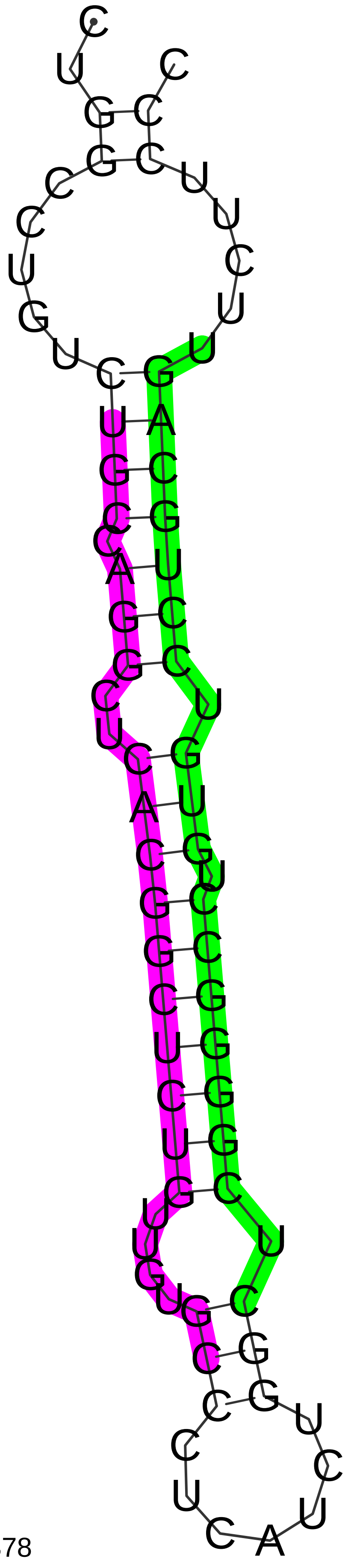

NW_006383524.1_616305-616378

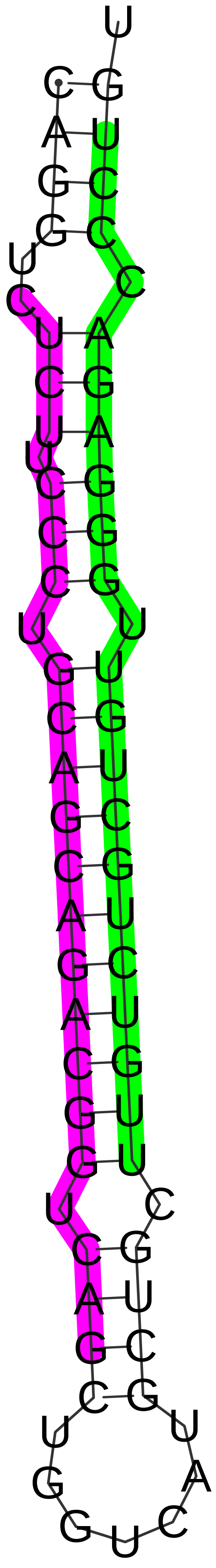

NW_006383532.1_1332945-1333010

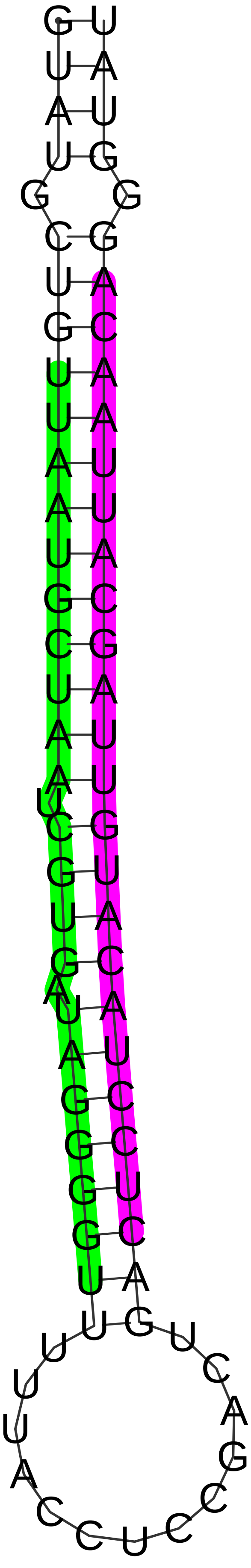


NW_006383537.1_17968-18042

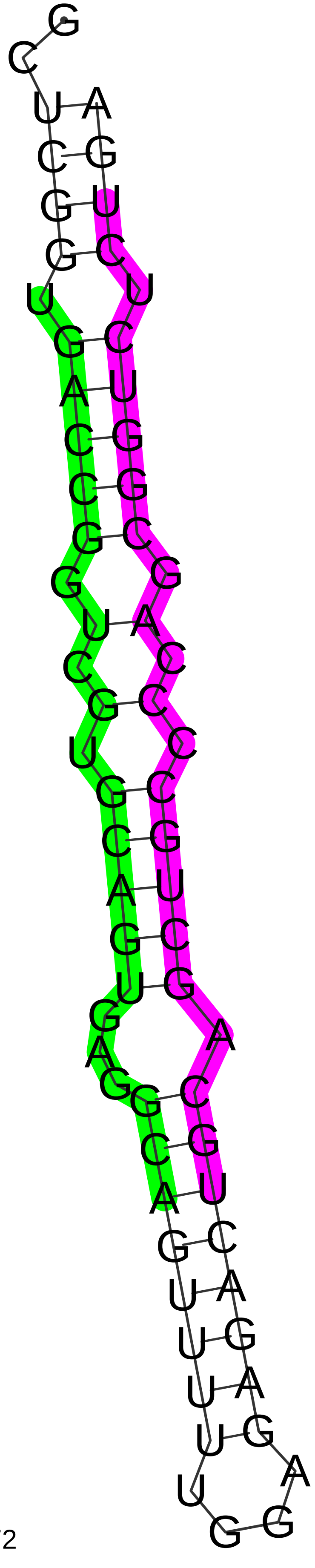

NW_006383554.1_263507-263572

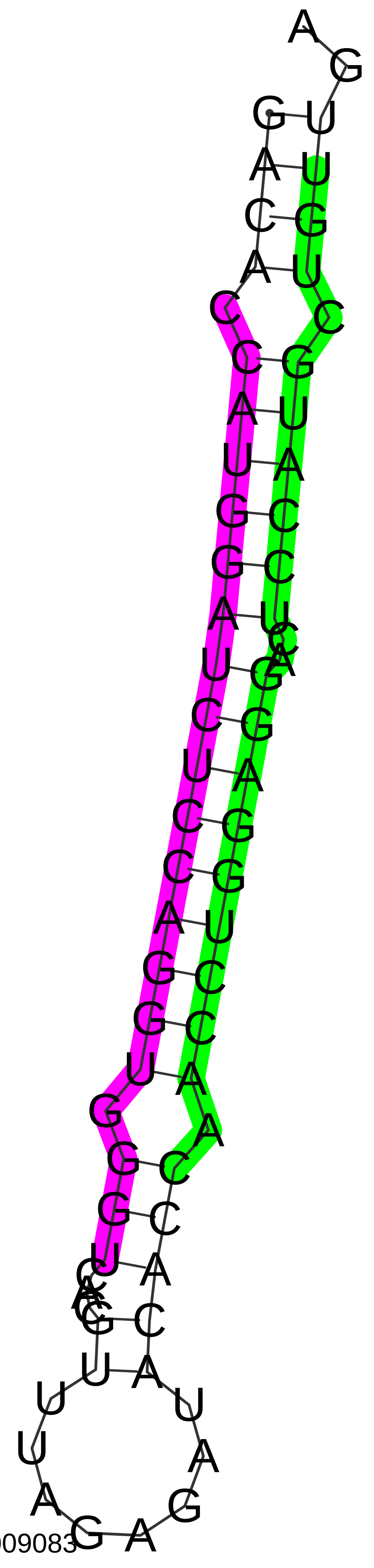

NW_006383587.1_1009017-1009083

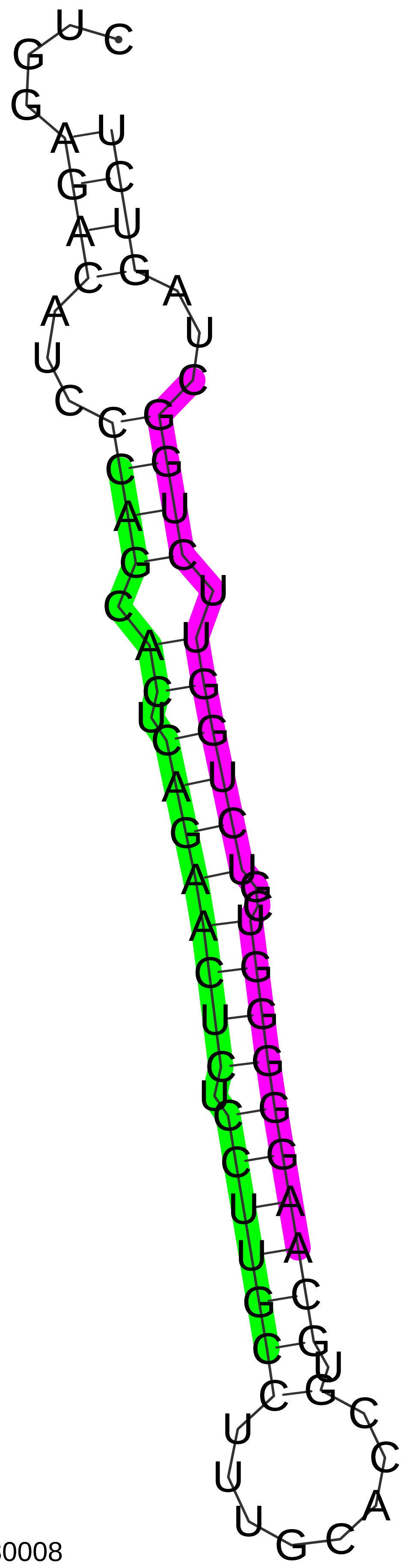

NW_006383608.1_279934-280008

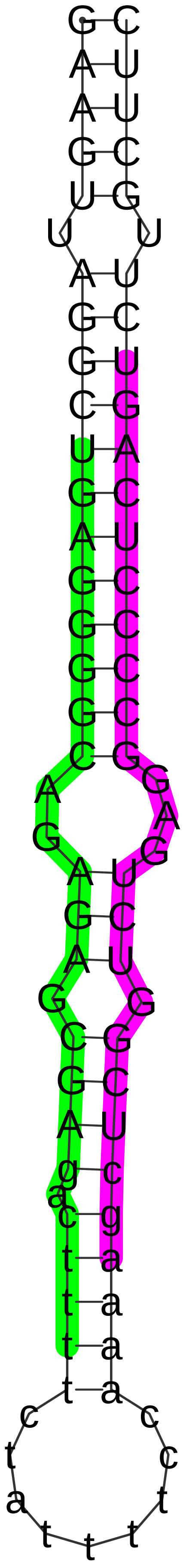

NW_006383610.1_750877-750953

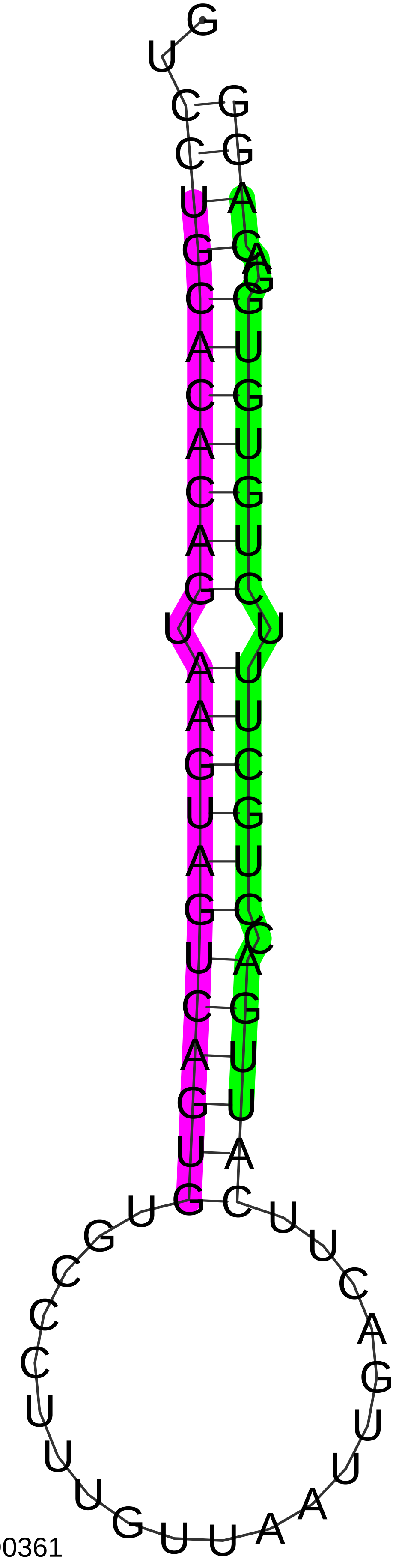

NW_006383615.1_390289-390361

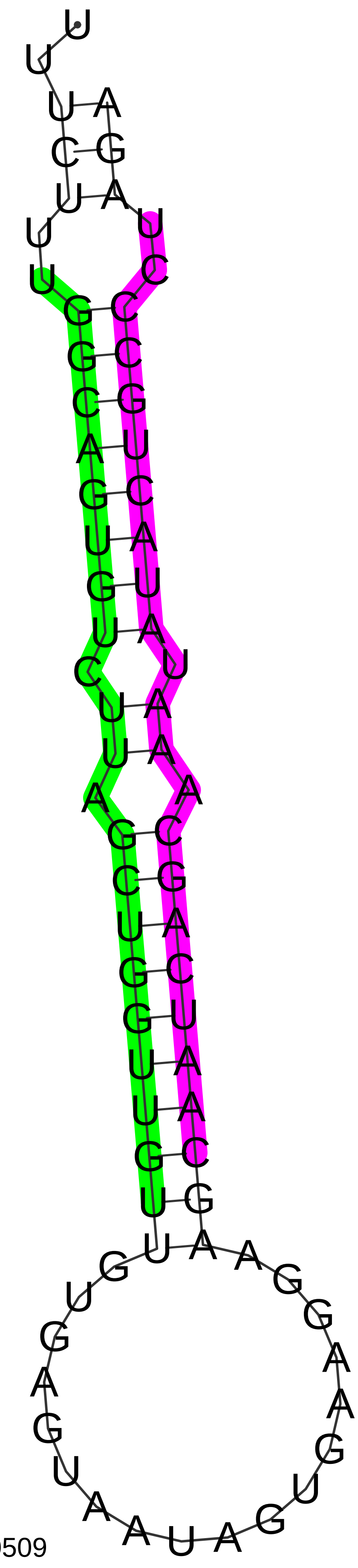

NW_006383621.1_230436-230509

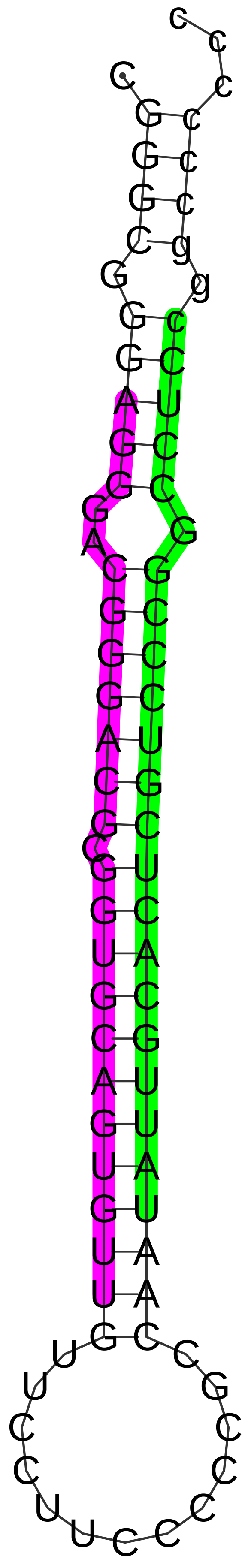

NW_006383630.1_502422-502500

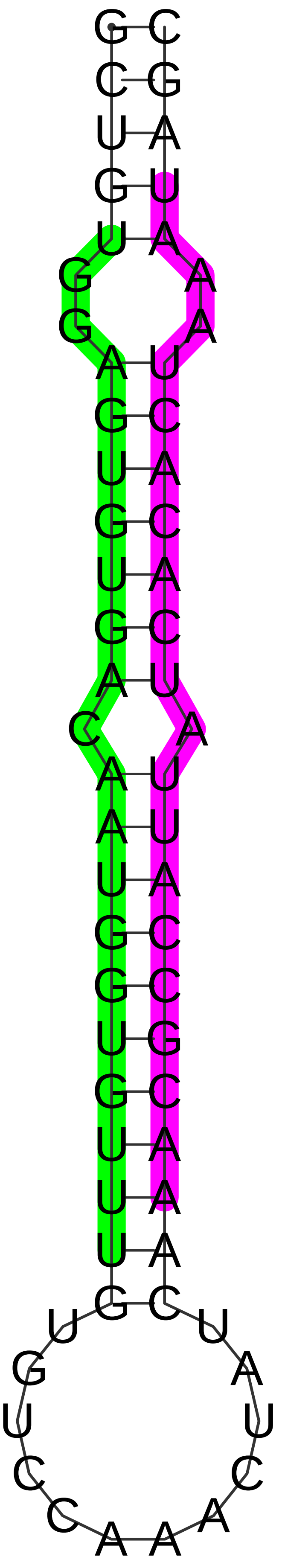


NW_006383633.1_394069-394132

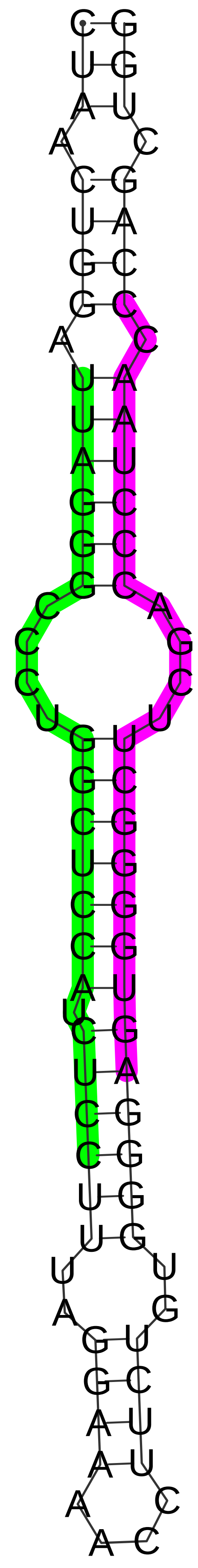

NW_006383643.1_684915-684995

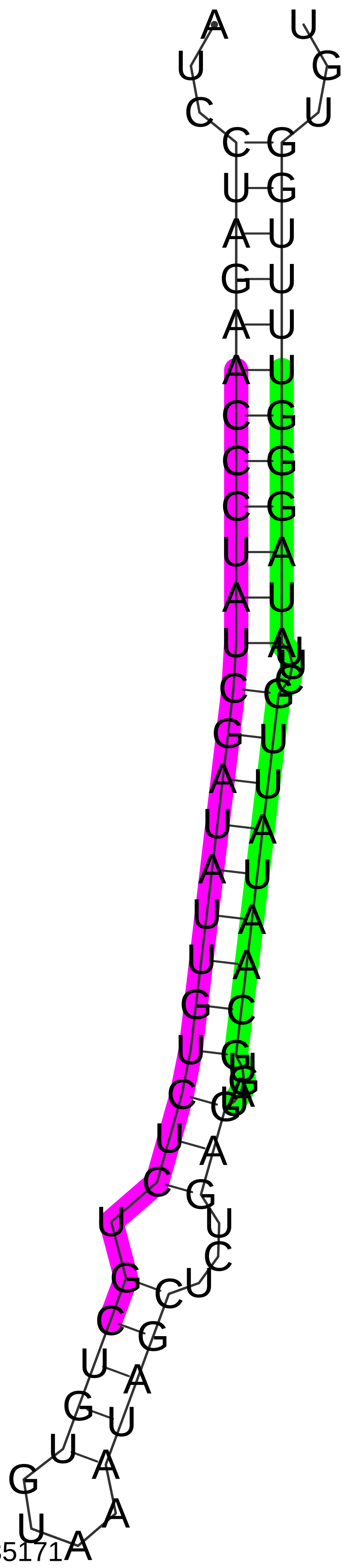

NW_006383655.1_1035093-1035171

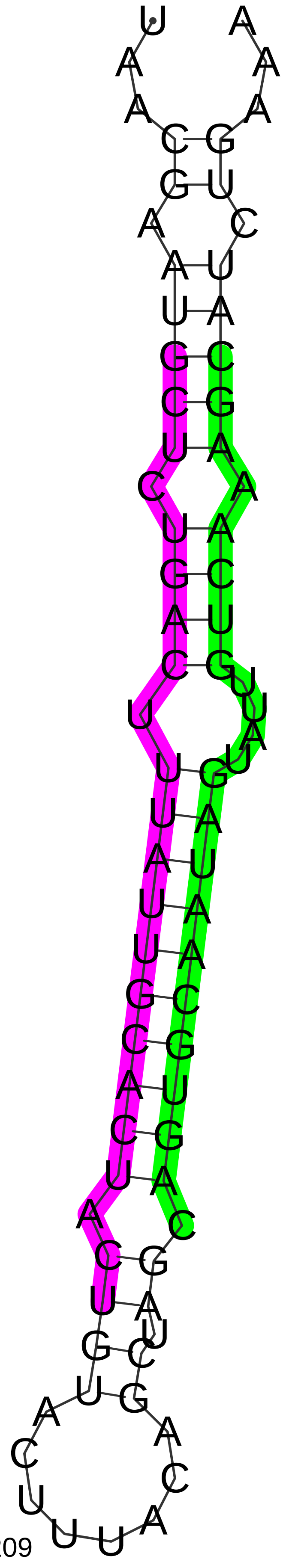

NW_006383655.1_1044134-1044209

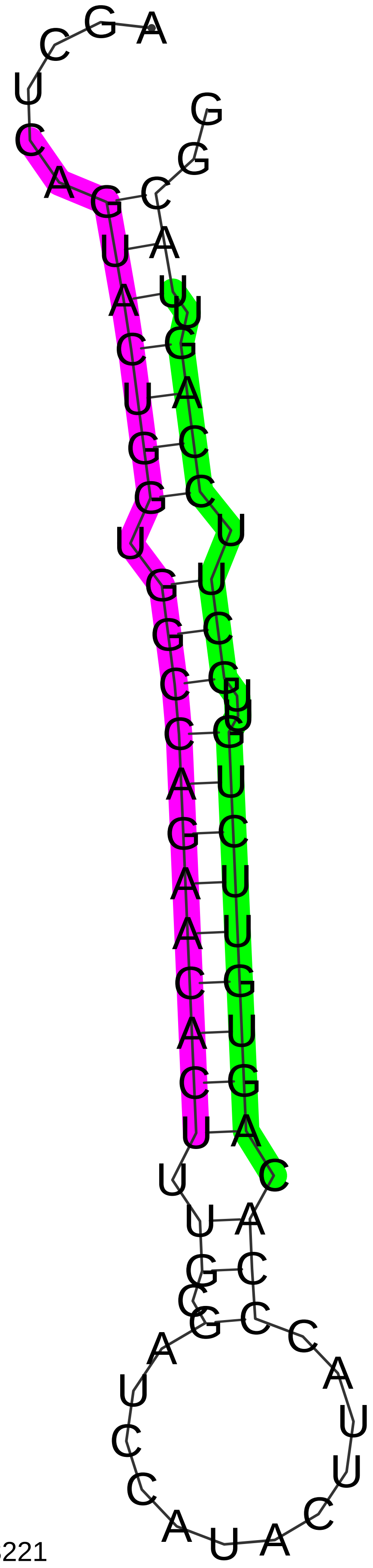

NW_006383666.1_403150-403221

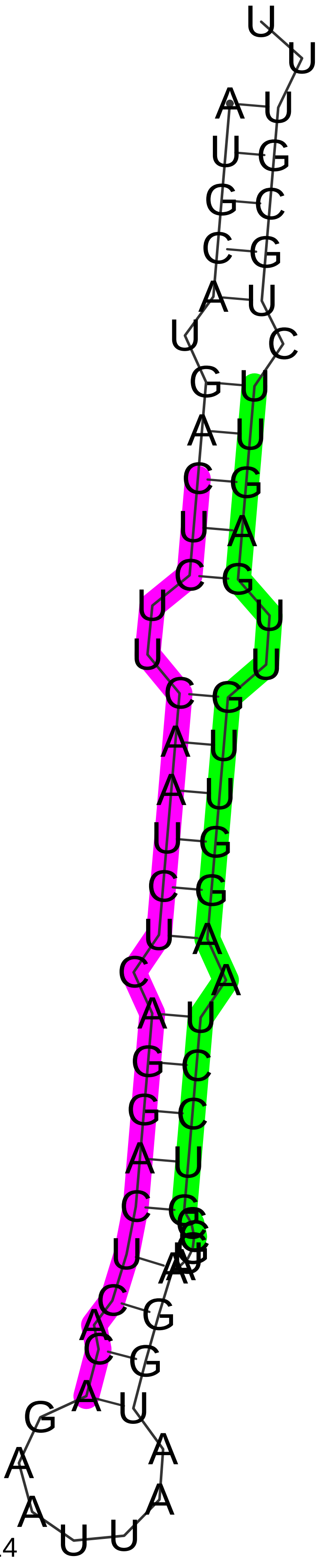

NW_006383667.1_122342-122414

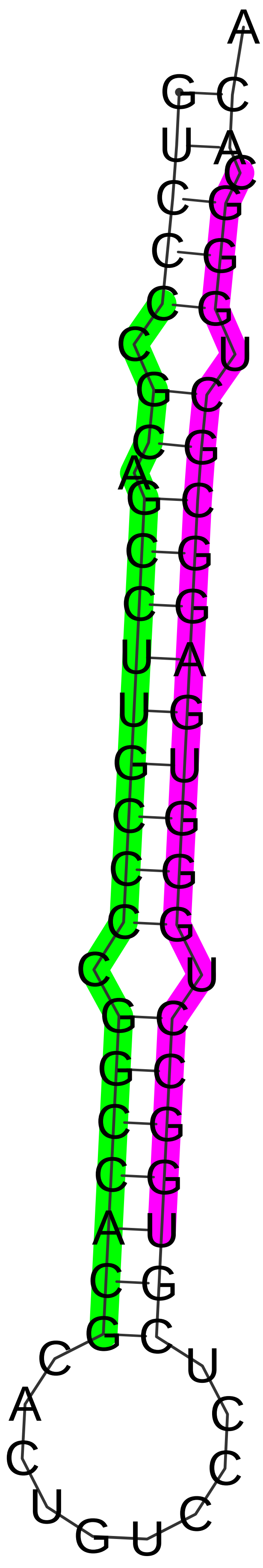

NW_006383707.1_391890-391952

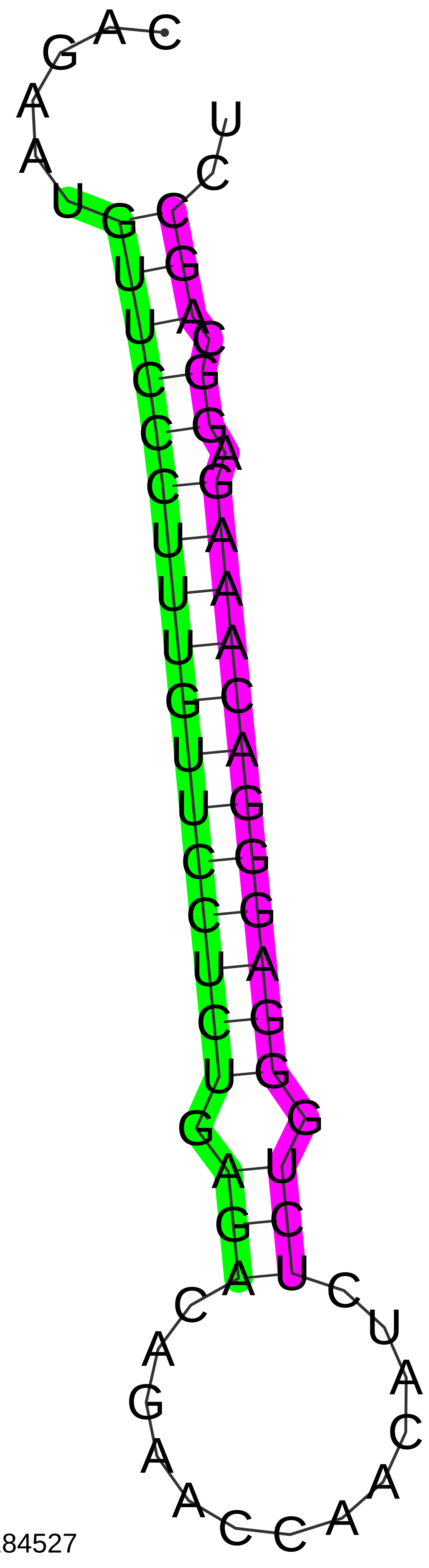

NW_006383723.1_284463-284527

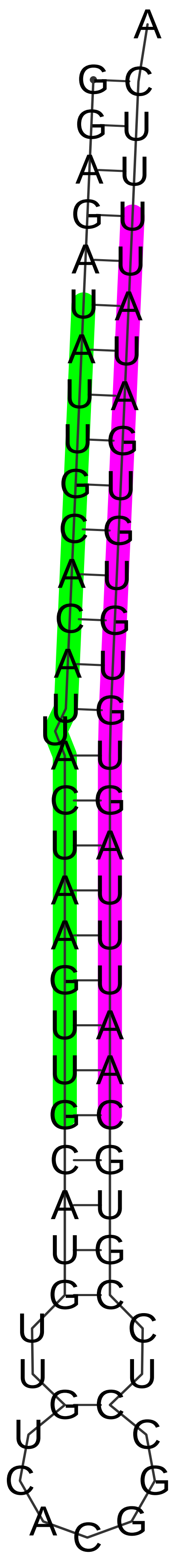

NW_006383740.1_673024-673094

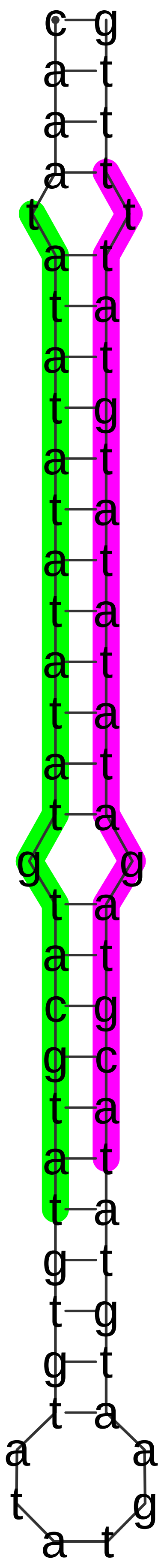

NW_006383751.1_584359-584422

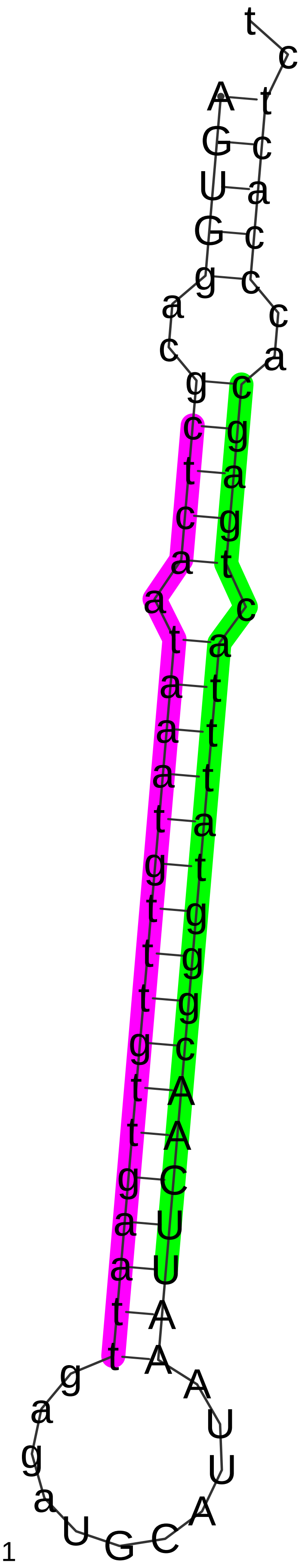

NW_006383760.1_891829-891901

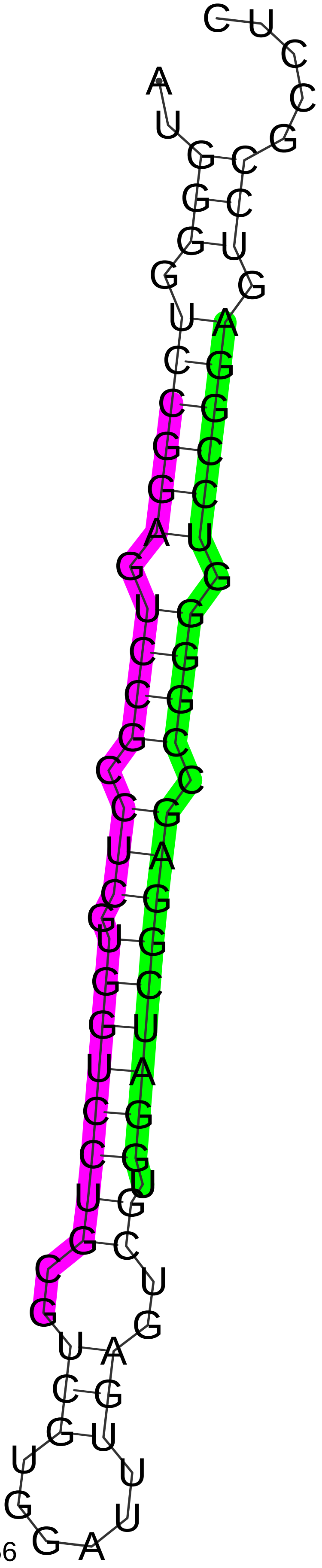

NW_006383782.1_339578-339656

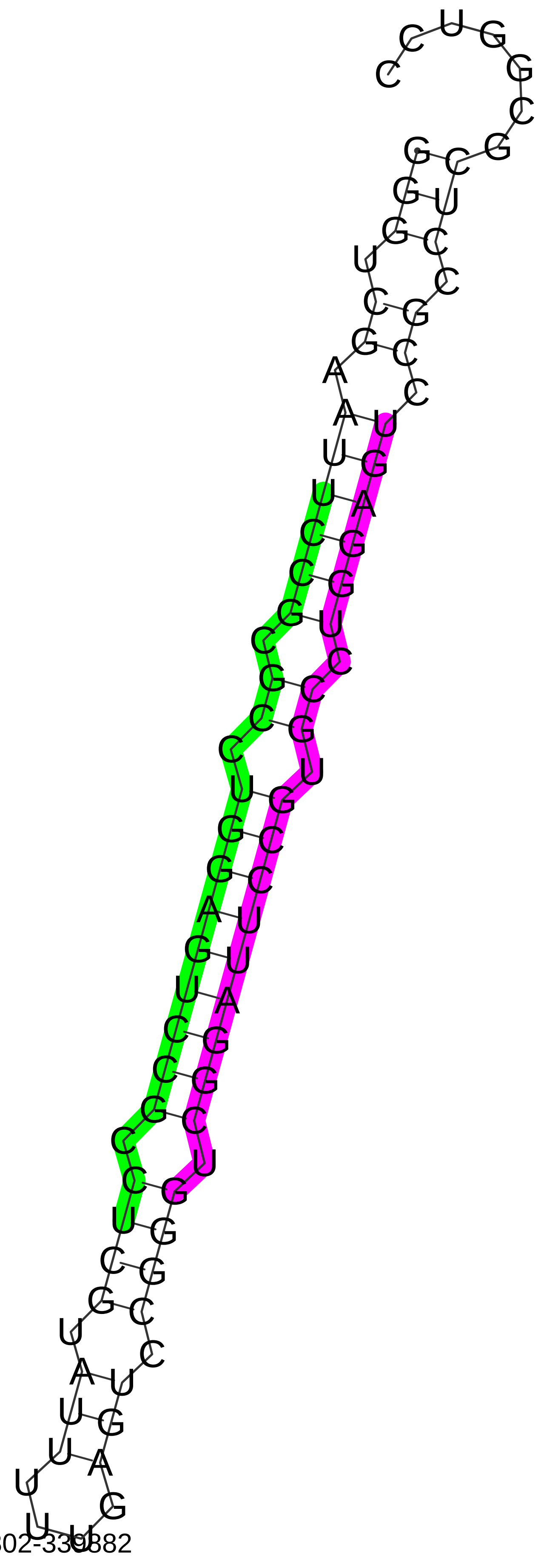

NW_006383782.1_339802-339882

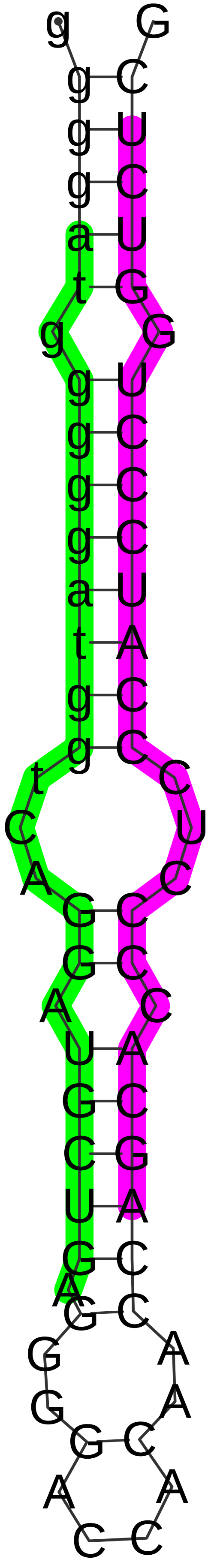

NW_006383788.1_911387-911451

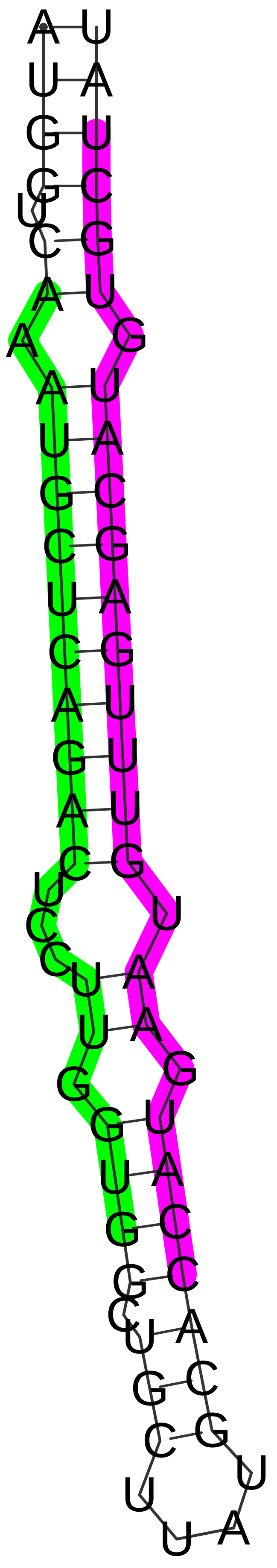

NW_006383799.1_237716-237779

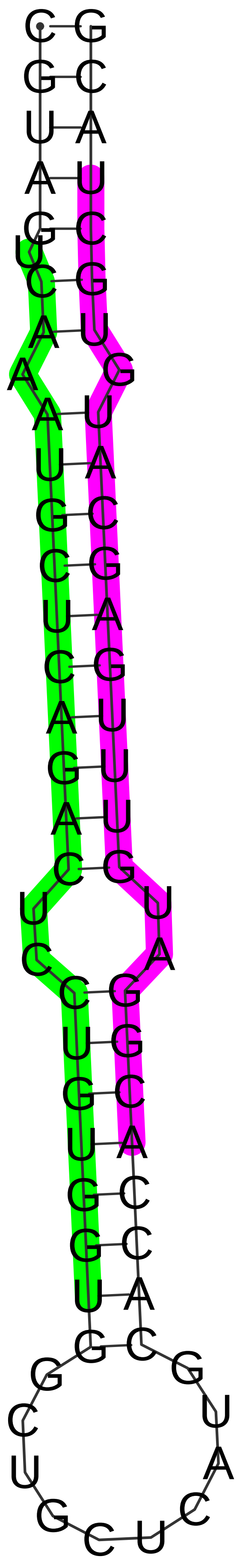

NW_006383799.1_240107-240173

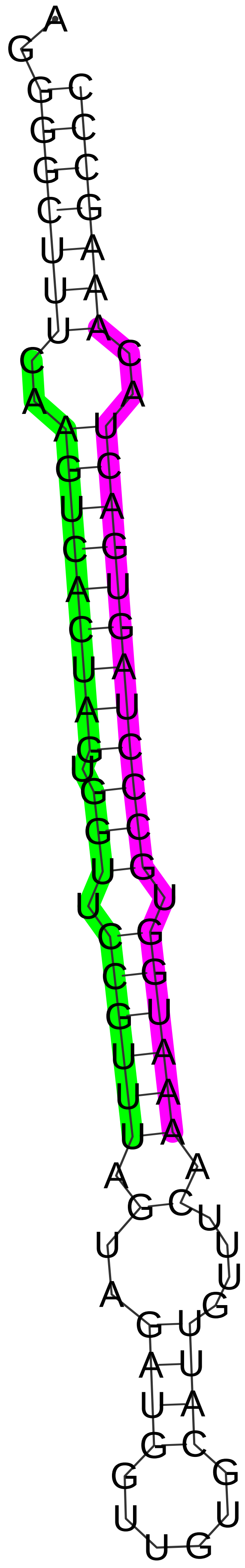

NW_006383799.1_718674-718756

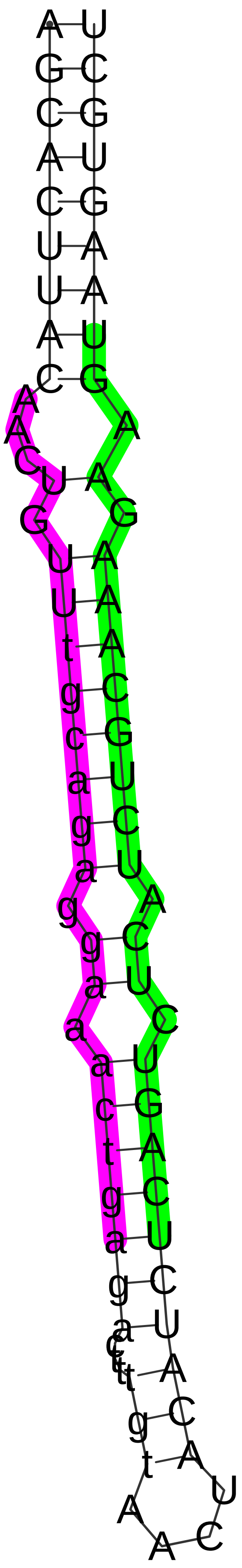

NW_006383799.1_719705-719781

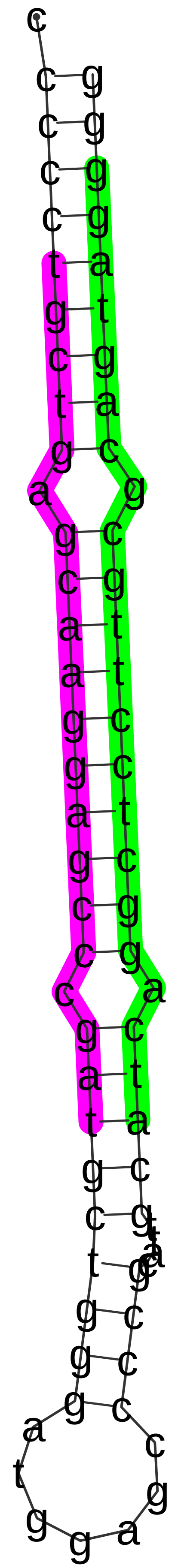

NW_006383817.1_229841-229912

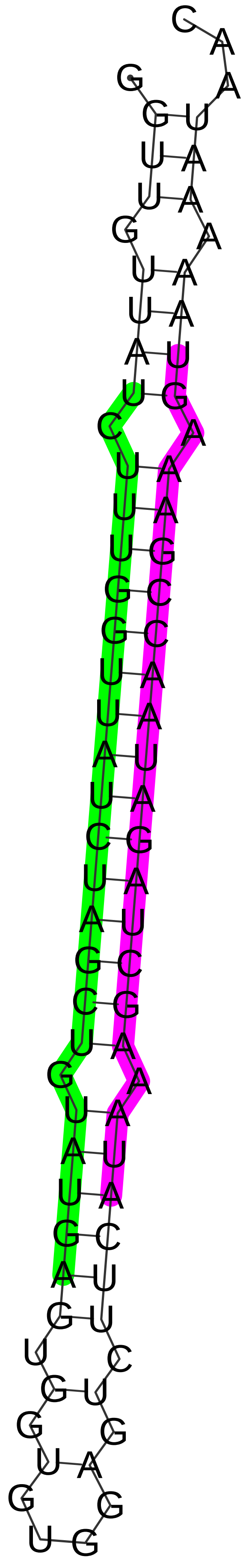

NW_006383833.1_517449-517526

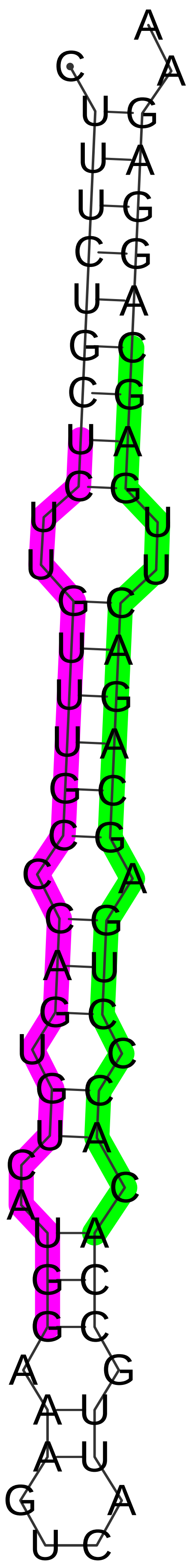

NW_006383853.1_117077-117146

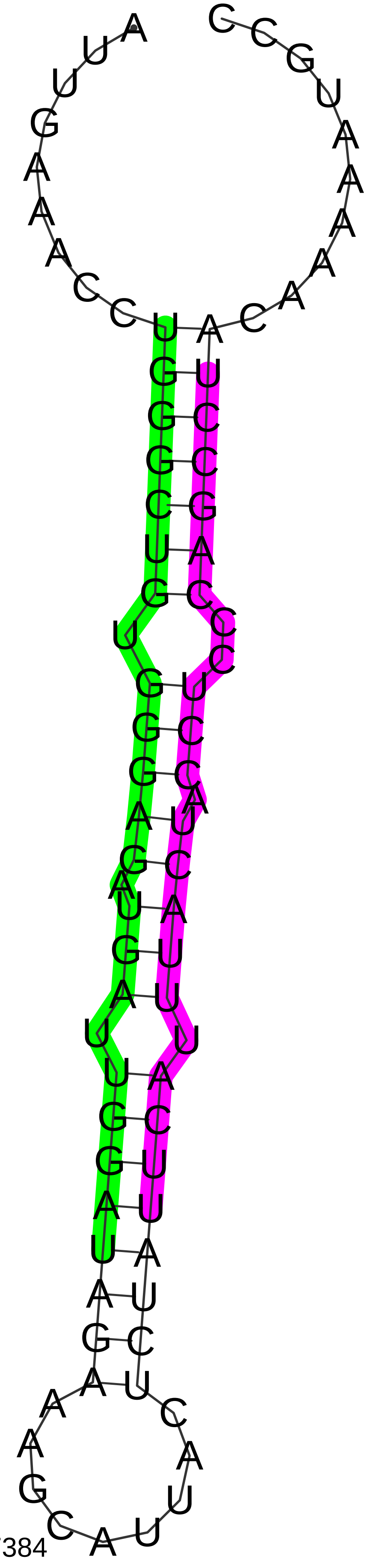


NW_006383854.1_957304-957384

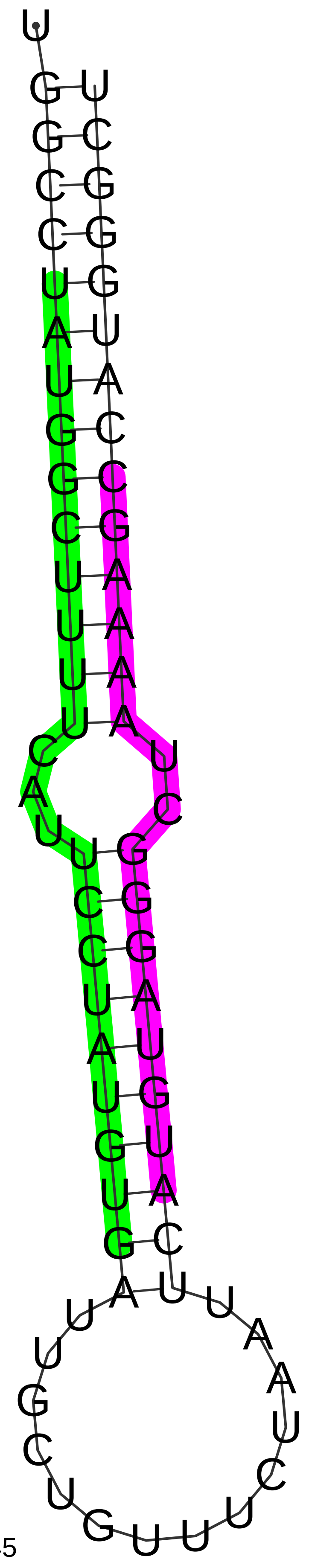


NW_006383867.1_425878-425945

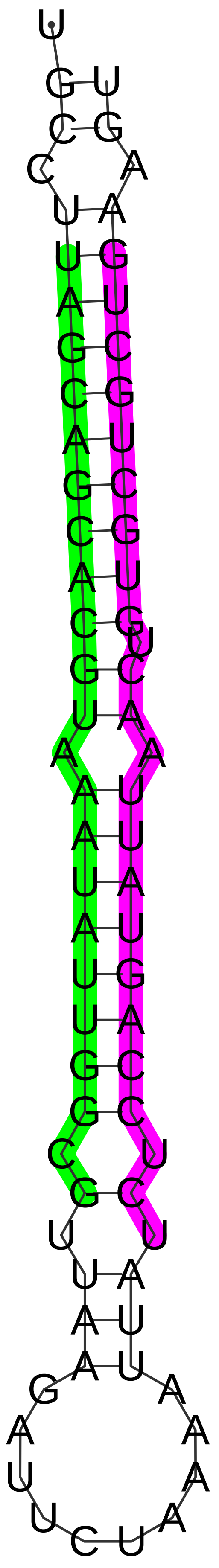

NW_006383872.1_162849-162920

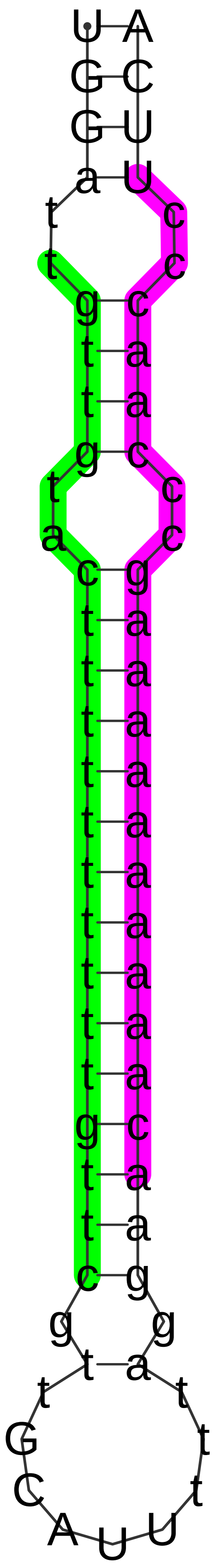

NW_006383872.1_210236-210302

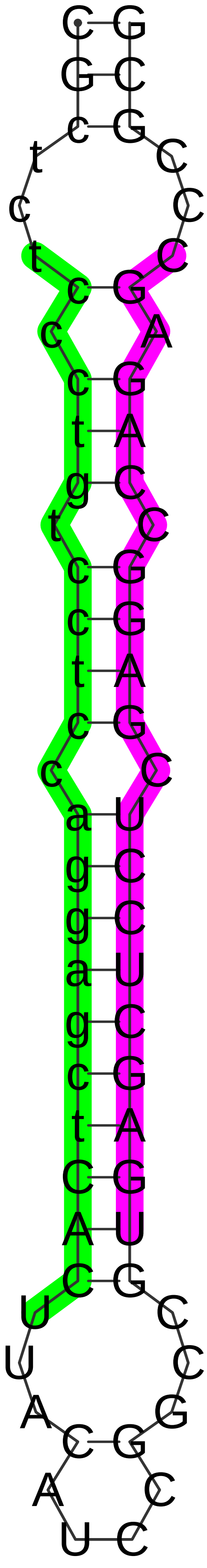


NW_006383882.1_526906-526971

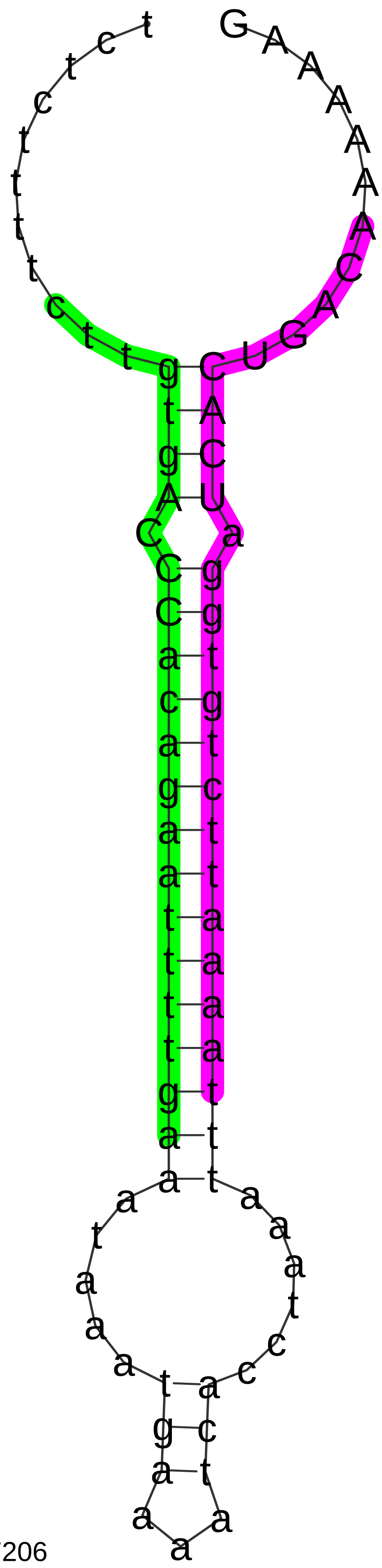

NW_006383888.1_797125-797206

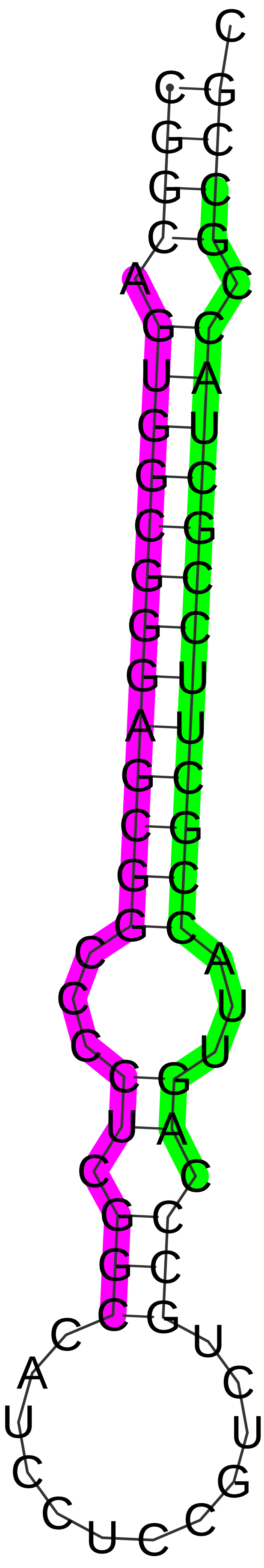

NW_006383891.1_780145-780211

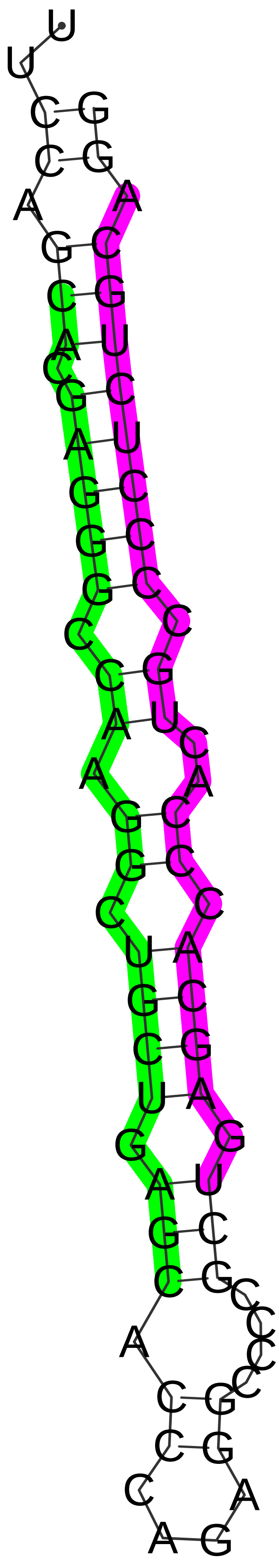

NW_006383894.1_85818-85886

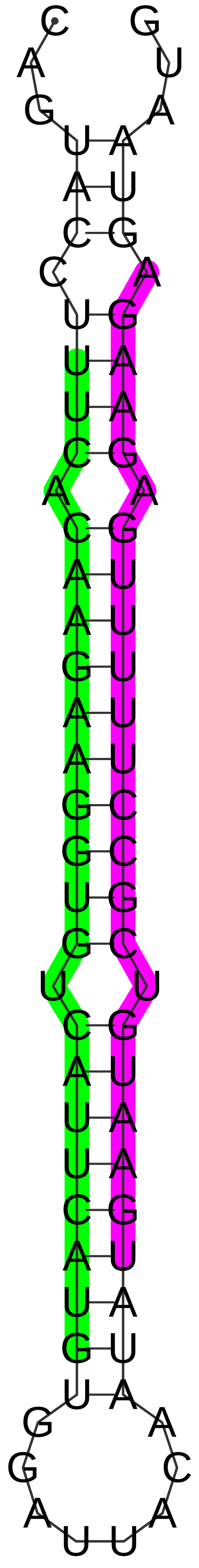

NW_006383898.1_696677-696748

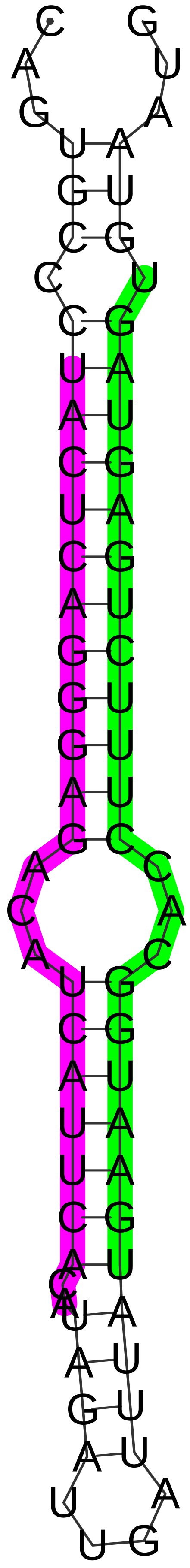

NW_006383898.1_724411-724482

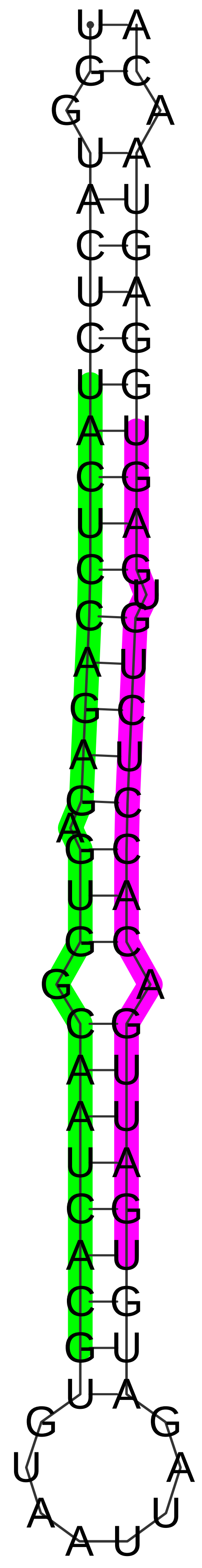

NW_006383898.1_767391-767462

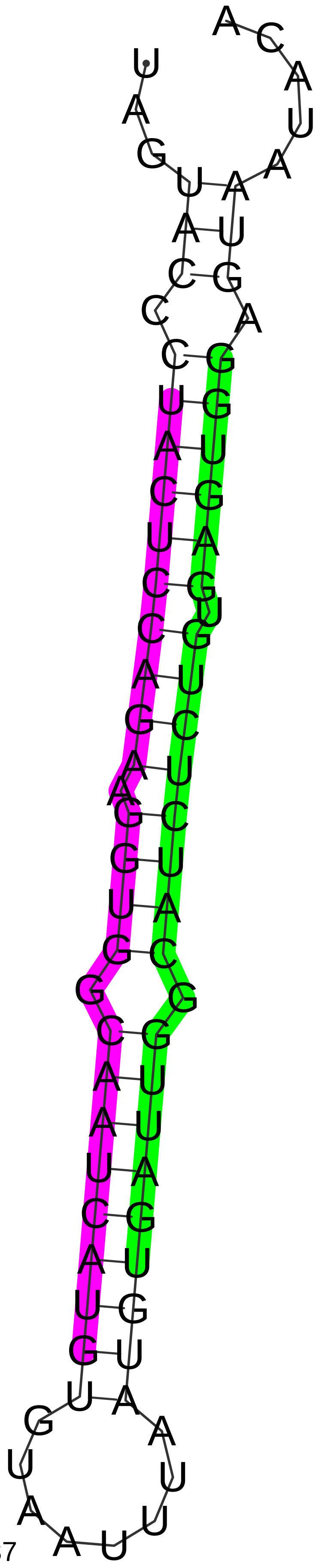

NW_006383898.1_776964-777037

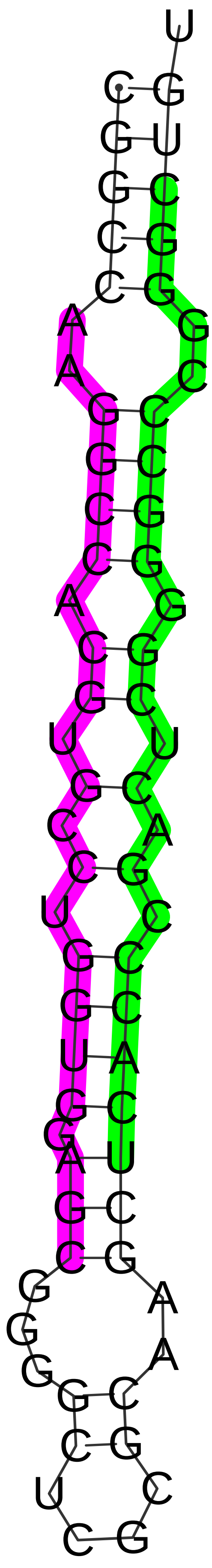

NW_006383901.1_154165-154231

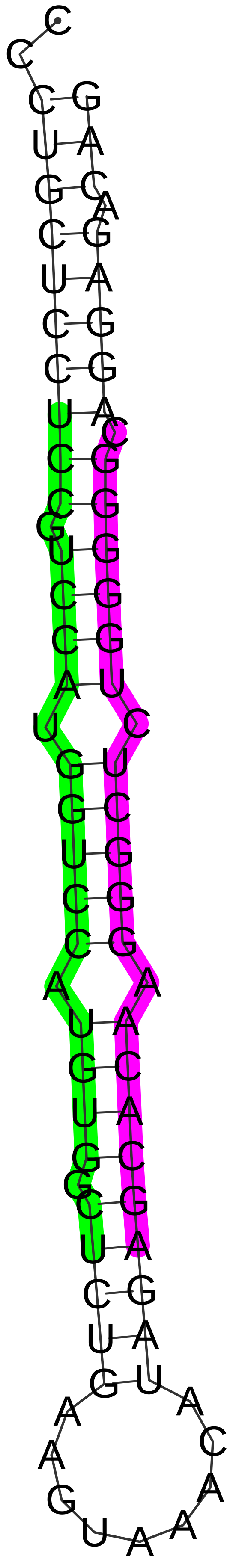


NW_006383907.1_499909-499983

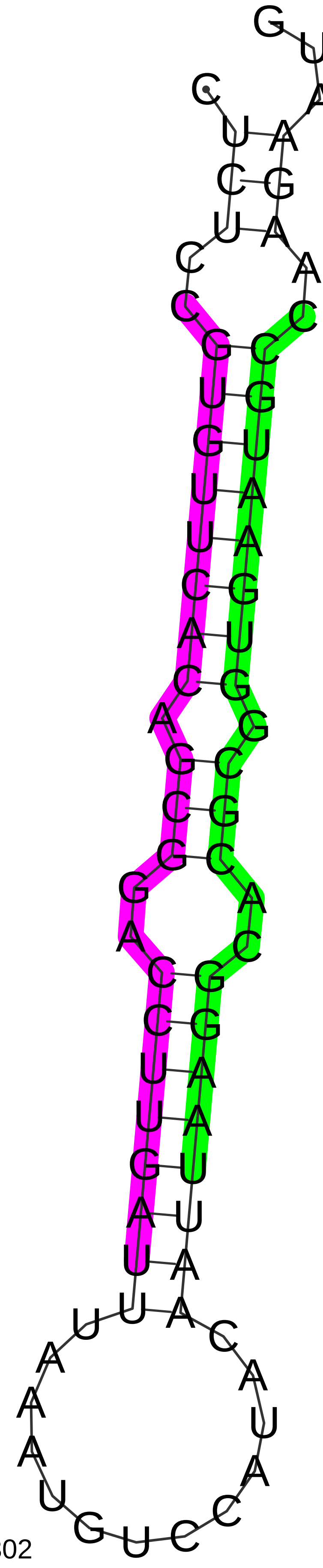

NW_006383915.1_118232-118302

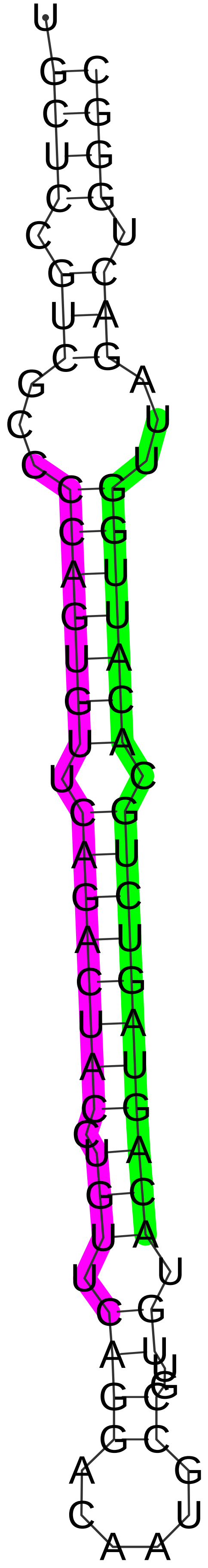

NW_006383916.1_441854-441933

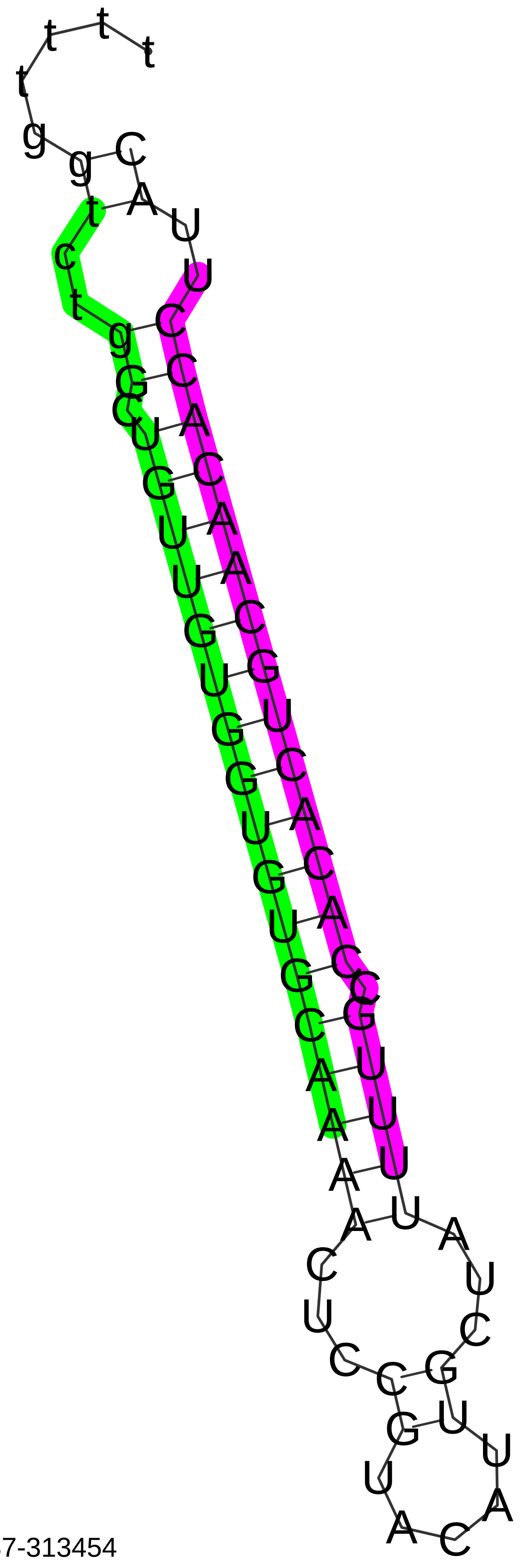

NW_006383930.1_313387-313454

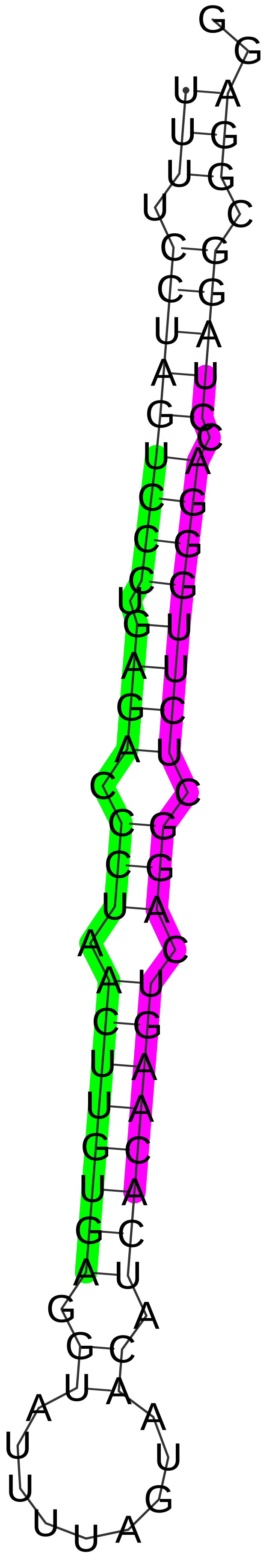

NW_006383936.1_716297-716375

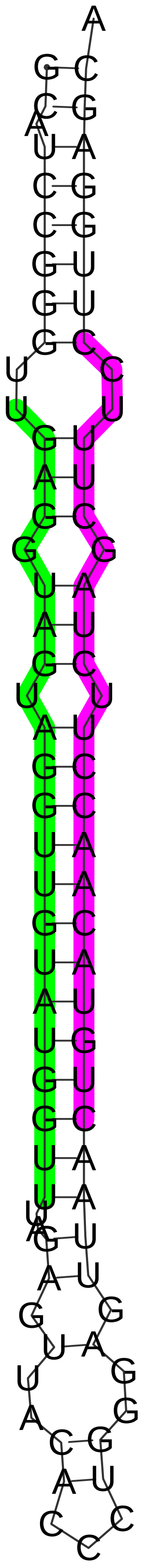

NW_006383936.1_760910-760994

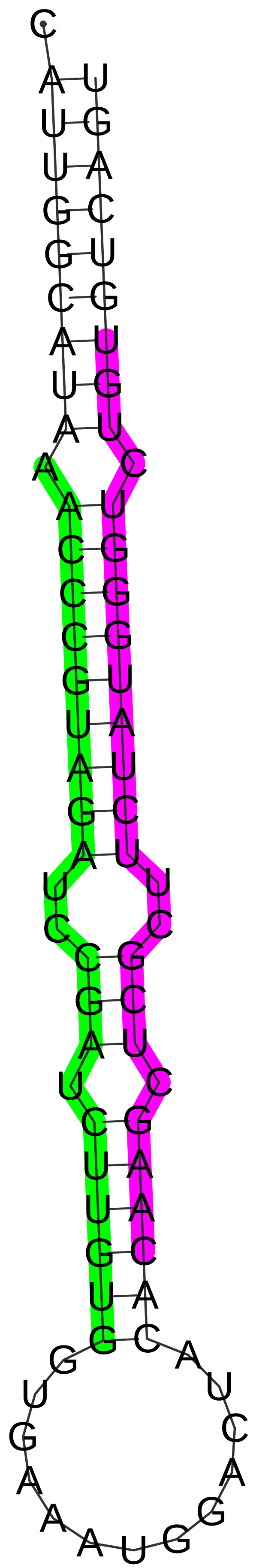

NW_006383936.1_761636-761711

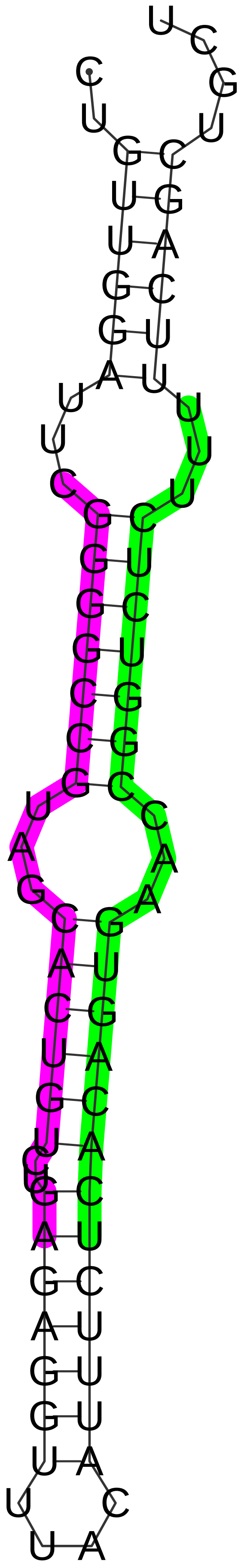

NW_006383941.1_227958-228033

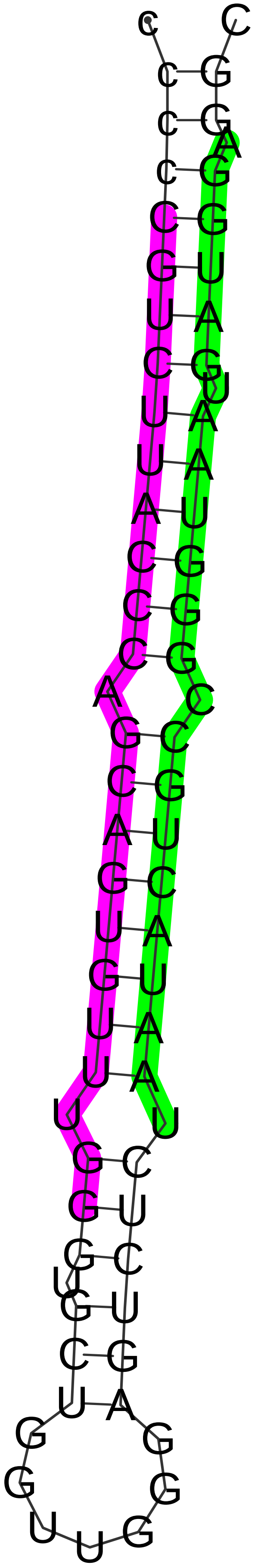


NW_006383948.1_628662-628731

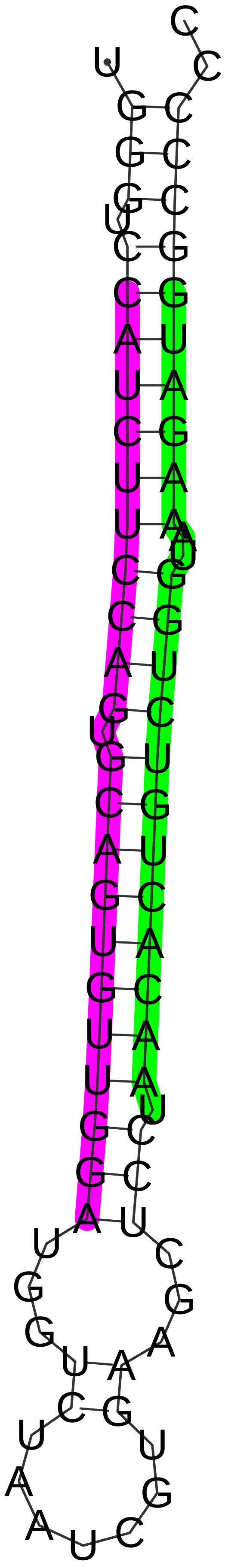

NW_006383948.1_629082-629156

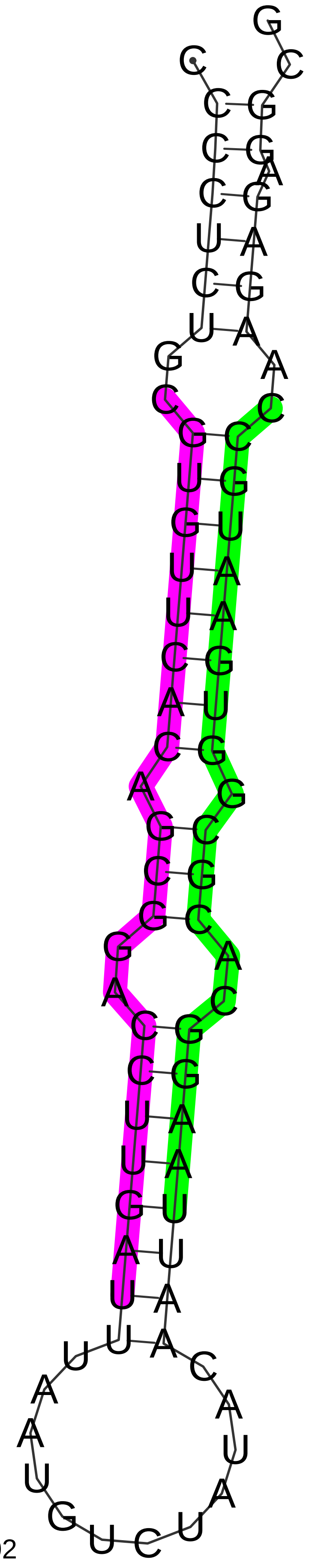

NW_006383949.1_401427-401502

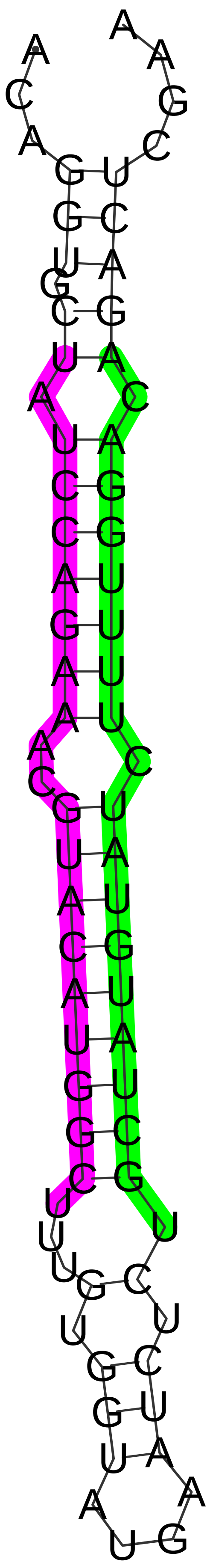

NW_006383954.1_601063-601135

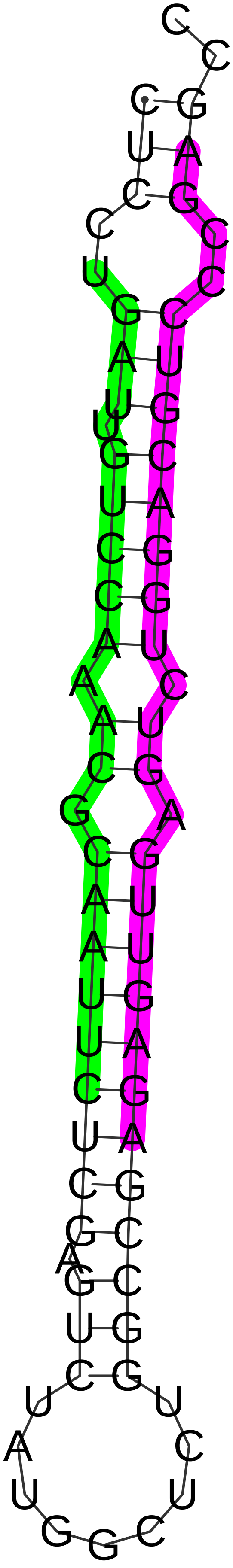

NW_006383968.1_487733-487803

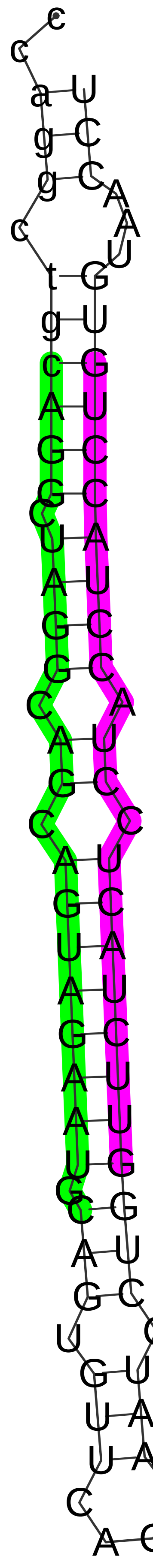

NW_006383974.1_403081-403156

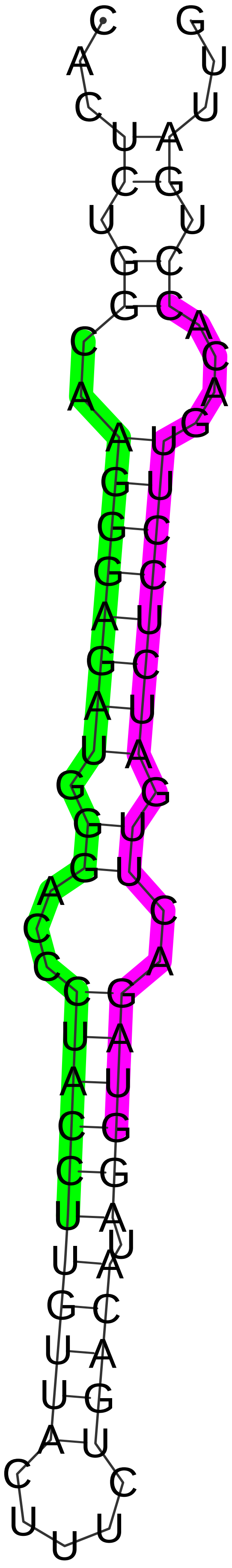

NW_006383985.1_464352-464428

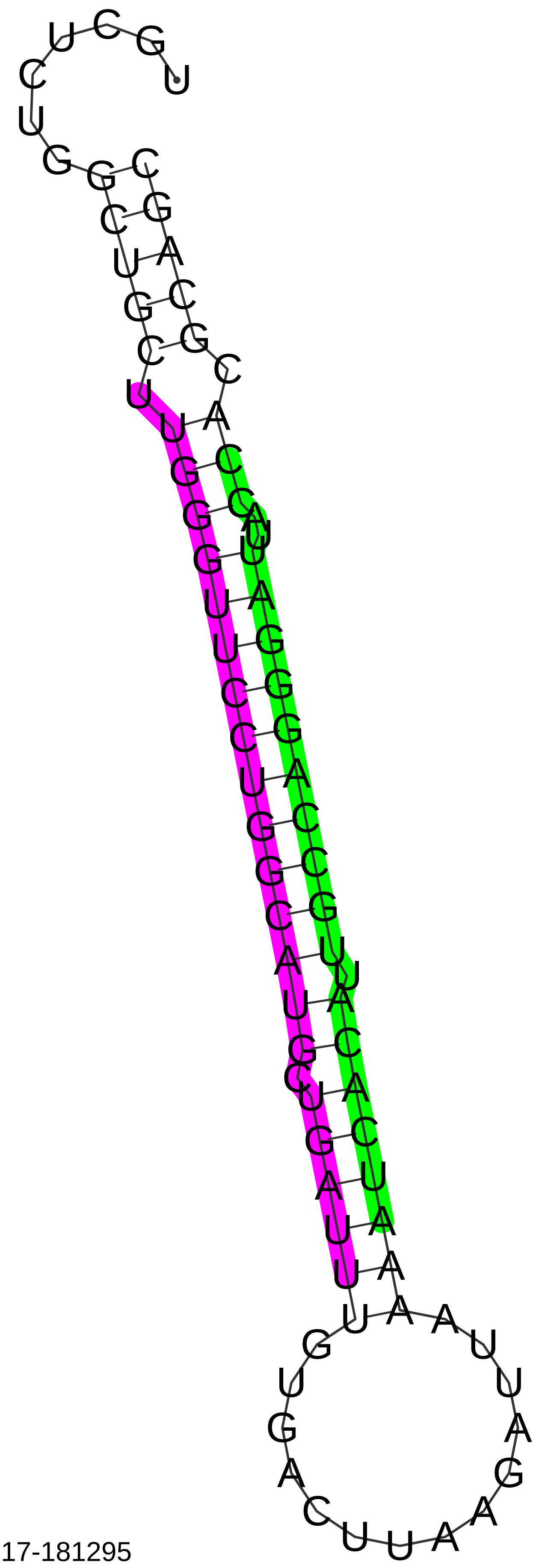

NW_006383994.1_181217-181295

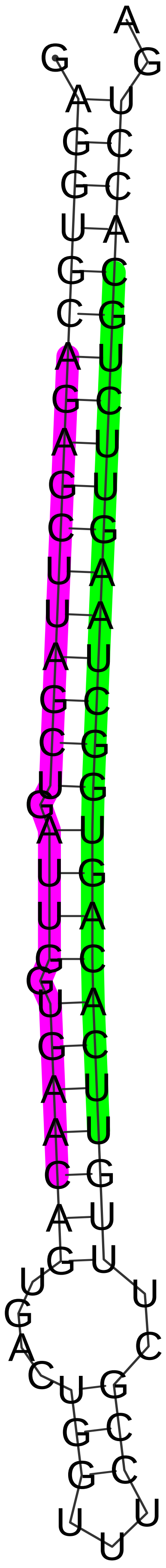

NW_006383994.1_181453-181528

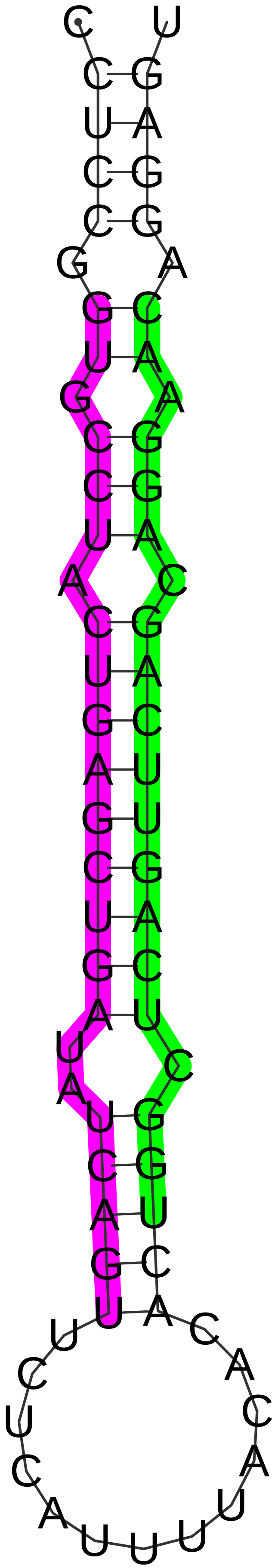


NW_006383994.1_181965-182034

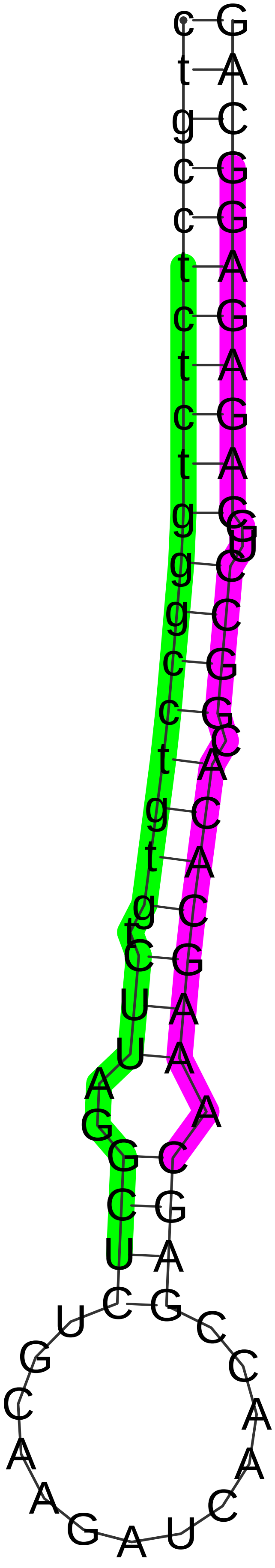

NW_006384004.1_195739-195810

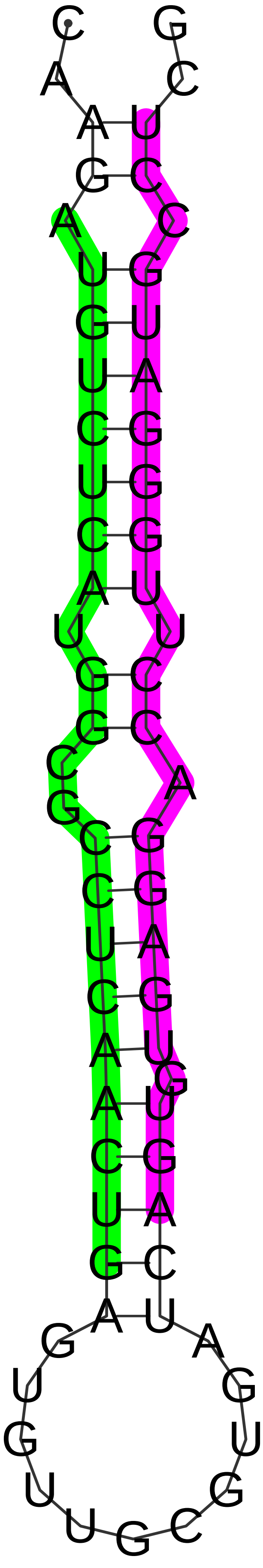

NW_006384005.1_250883-250947

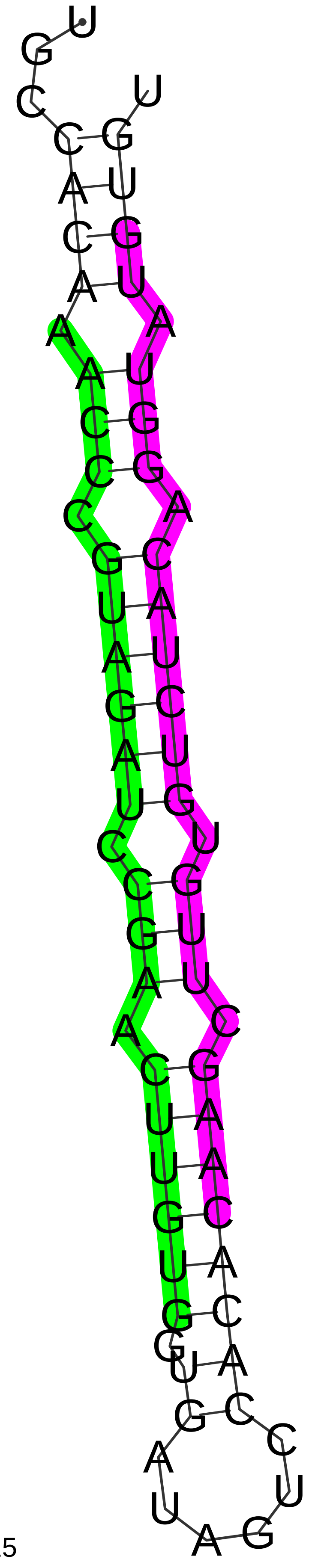

NW_006384005.1_283849-283915

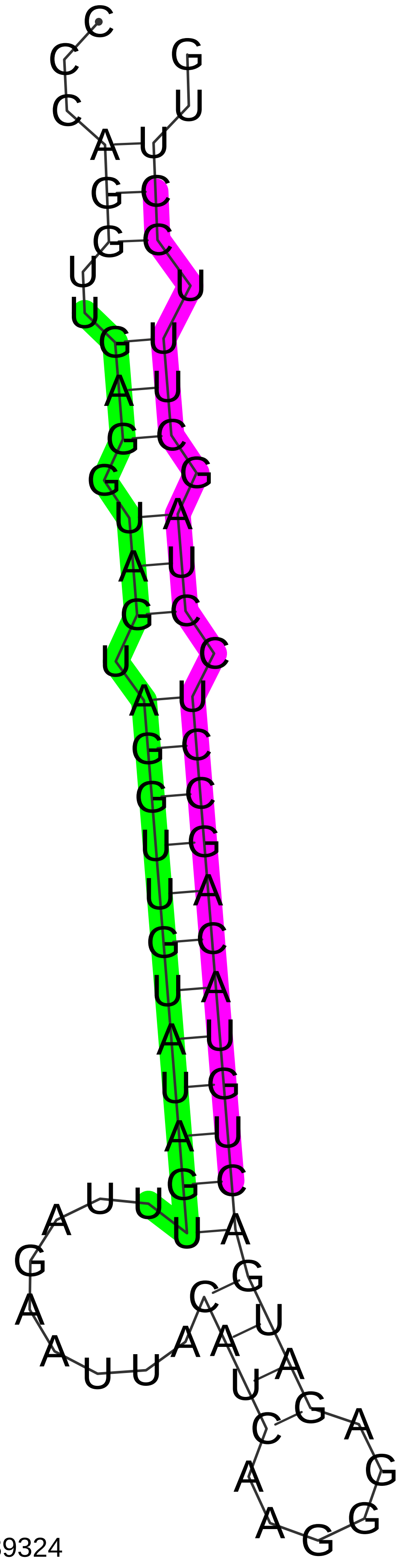

NW_006384005.1_289248-289324

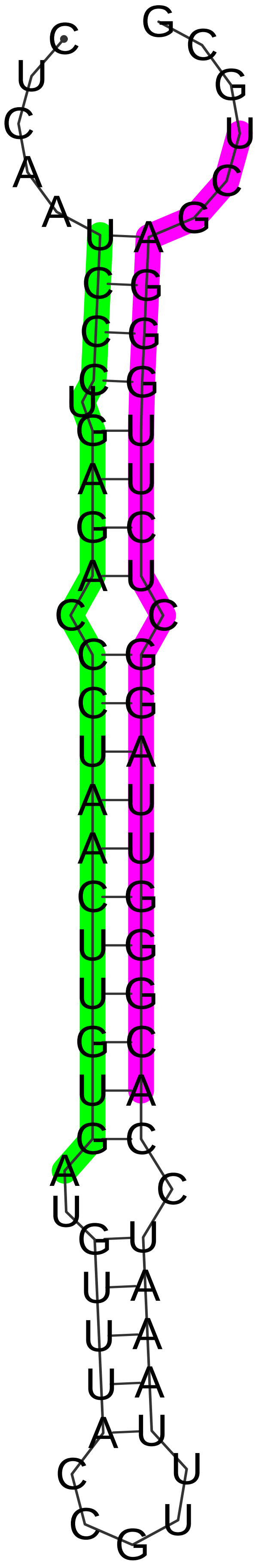

NW_006384005.1_332132-332201

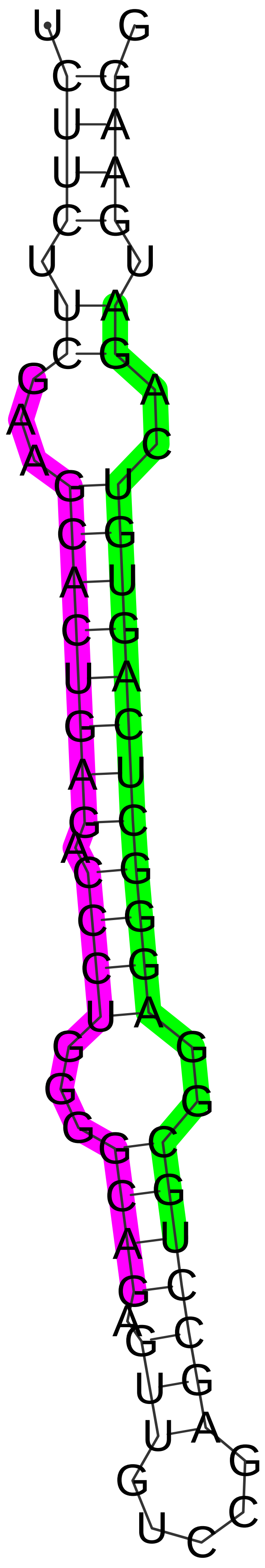

NW_006384018.1_659309-659379

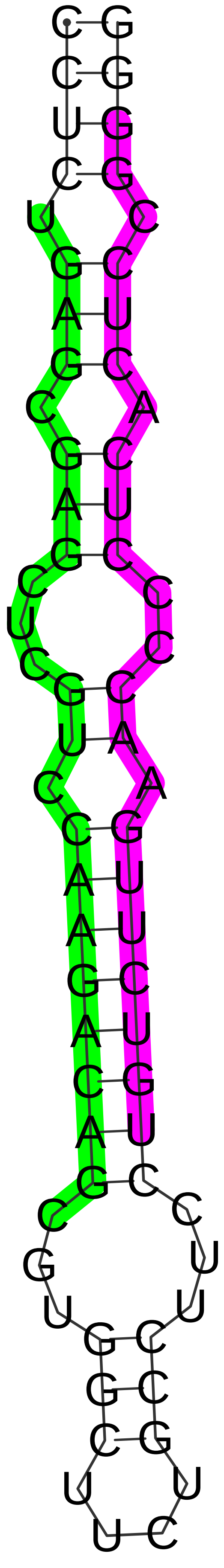

NW_006384039.1_594640-594706

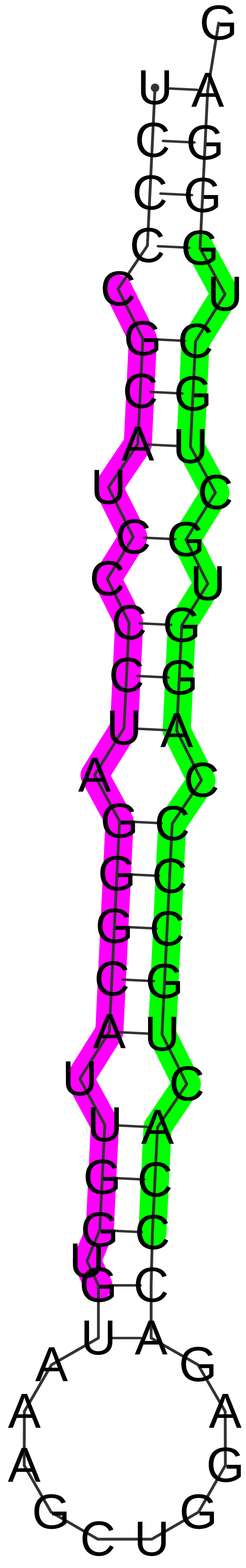

NW_006384039.1_729093-729156

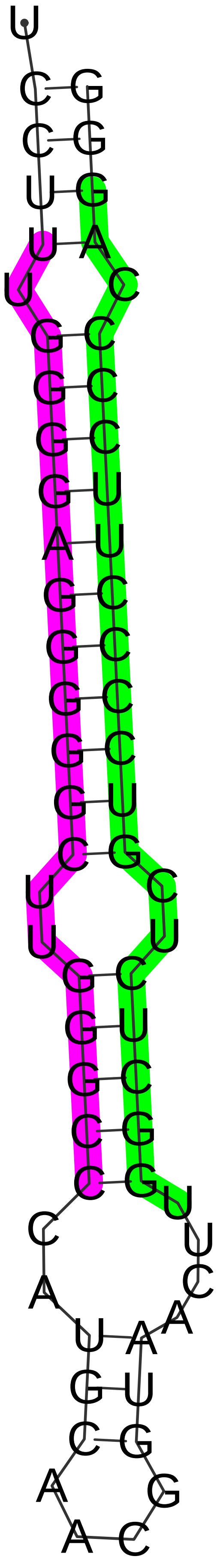

NW_006384046.1_38674-38736

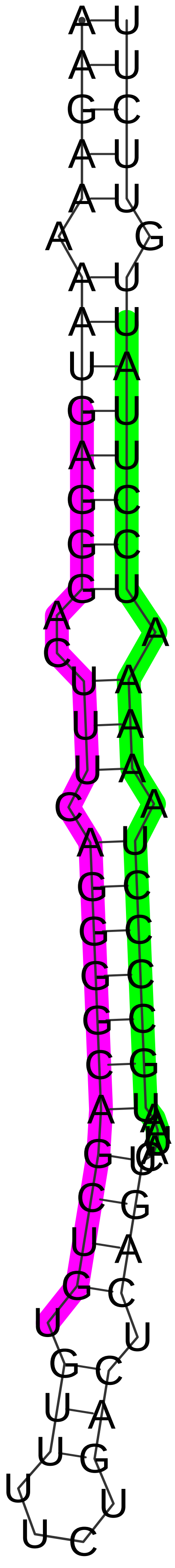

NW_006384048.1_635685-635762

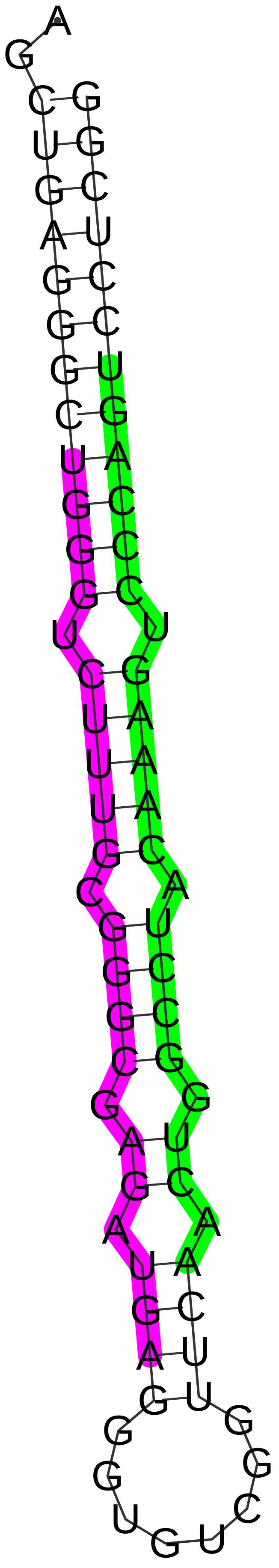

NW_006384048.1_648023-648094

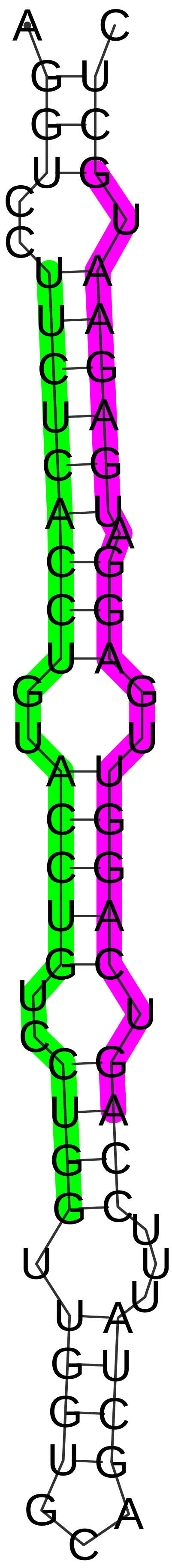

NW_006384051.1_757355-757424

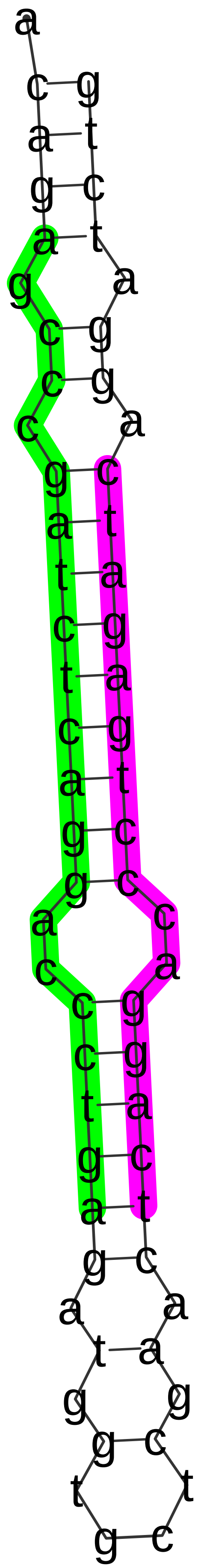

NW_006384057.1_332068-332130

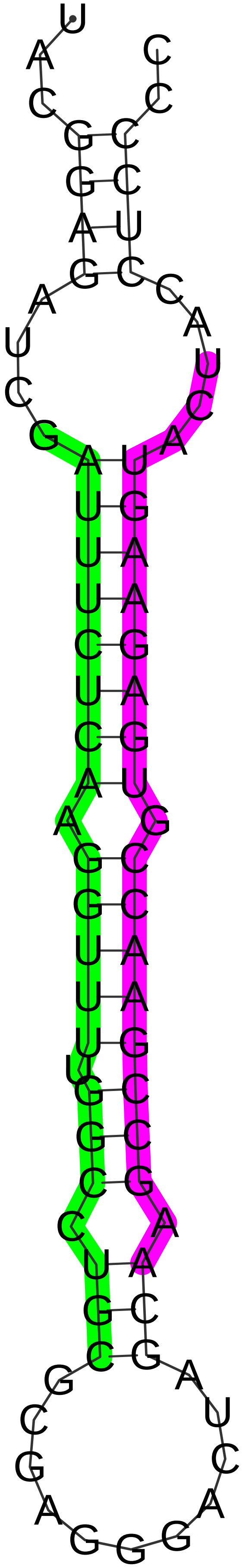


NW_006384057.1_635557-635632

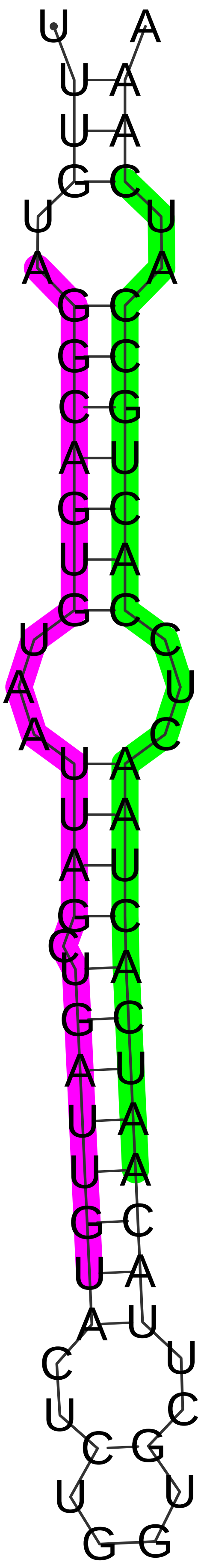

NW_006384065.1_419969-420035

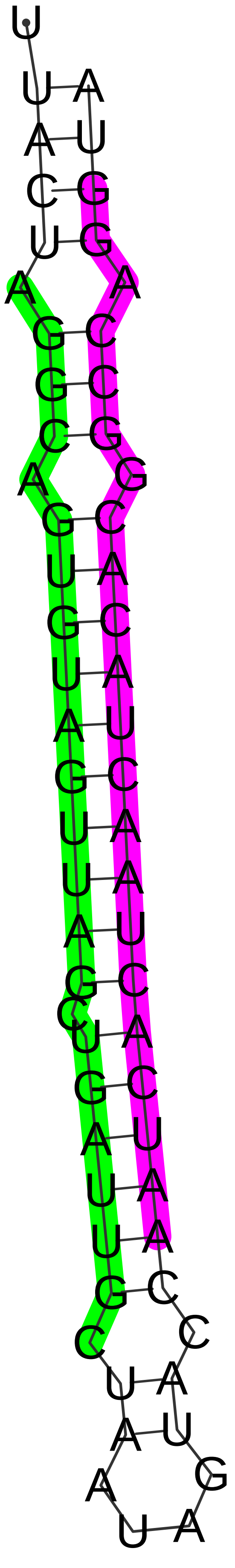

NW_006384065.1_420573-420634

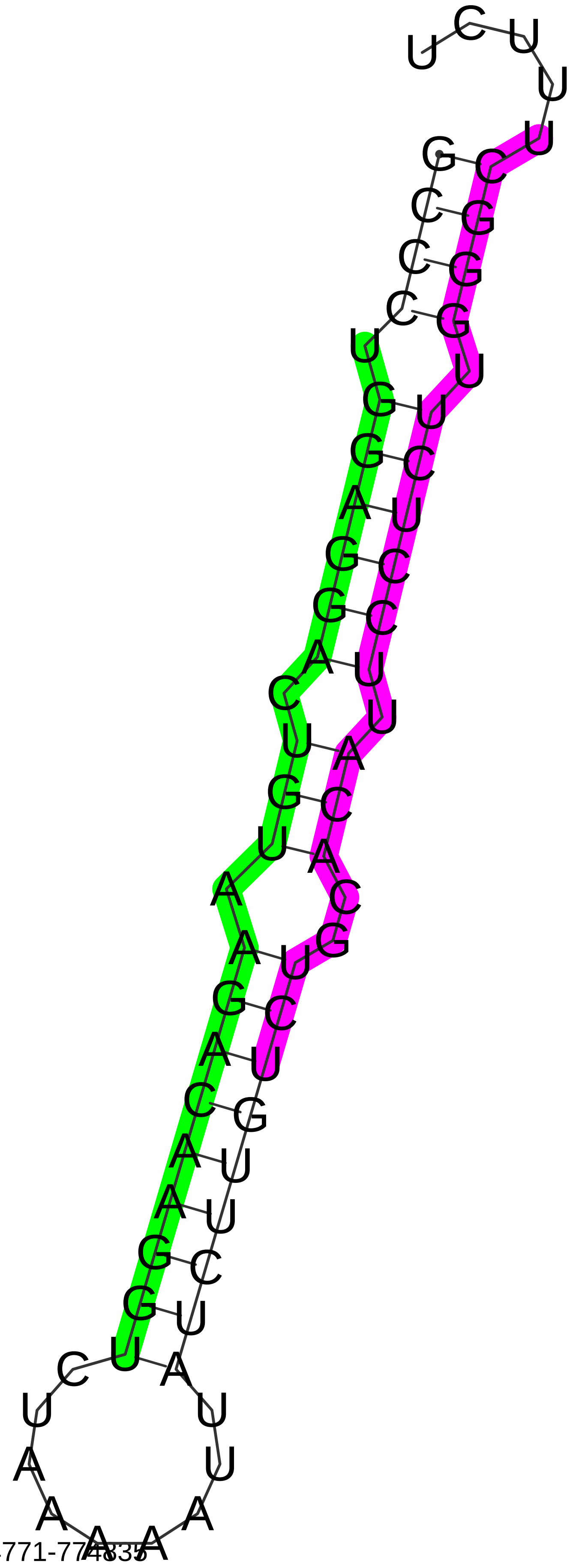

NW_006384078.1_774771-774835

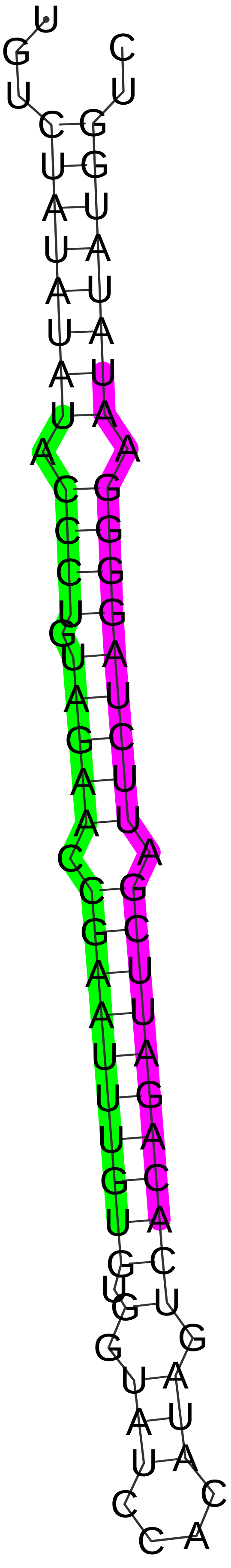

NW_006384104.1_325411-325489

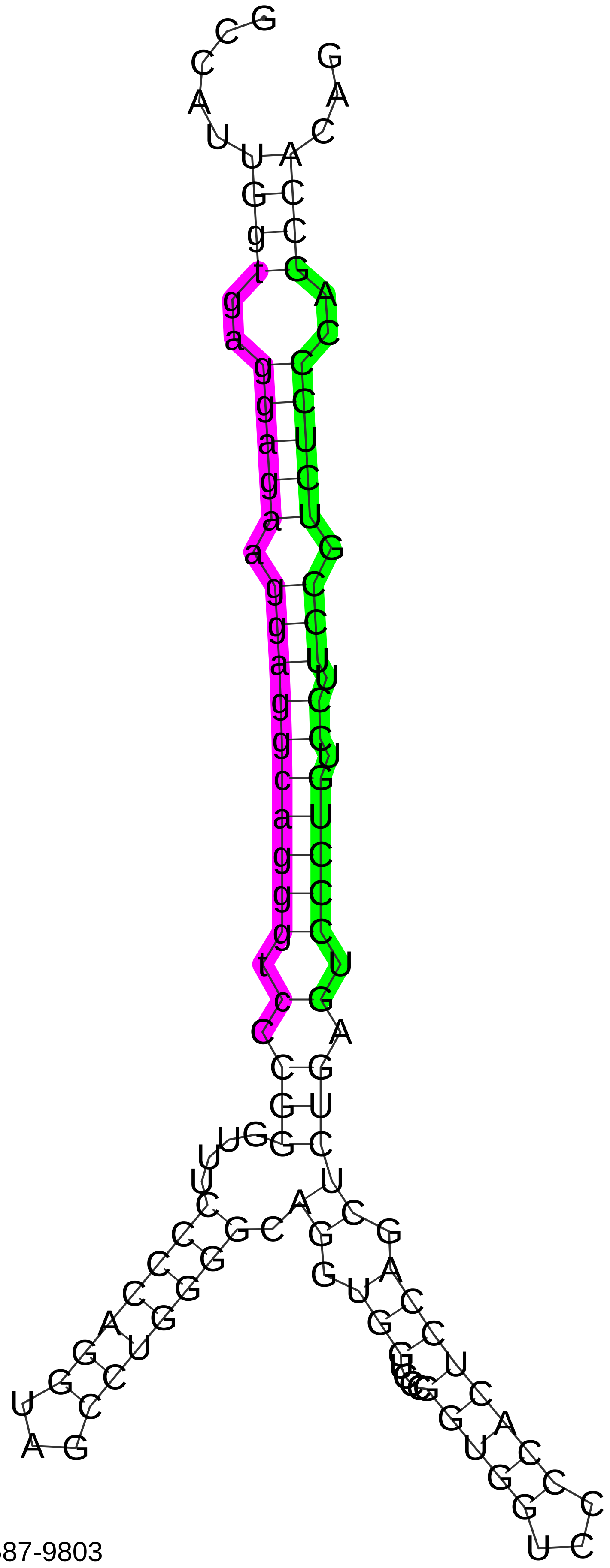

NW_006384107.1_9687-9803

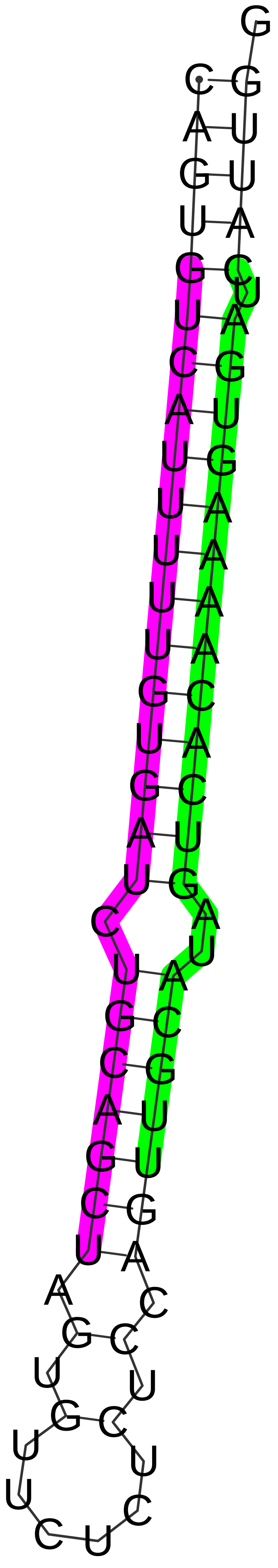

NW_006384119.1_530096-530164

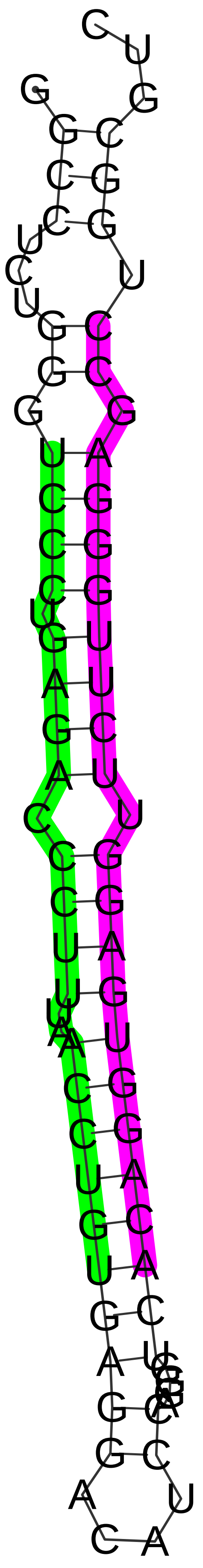

NW_006384123.1_81494-81570

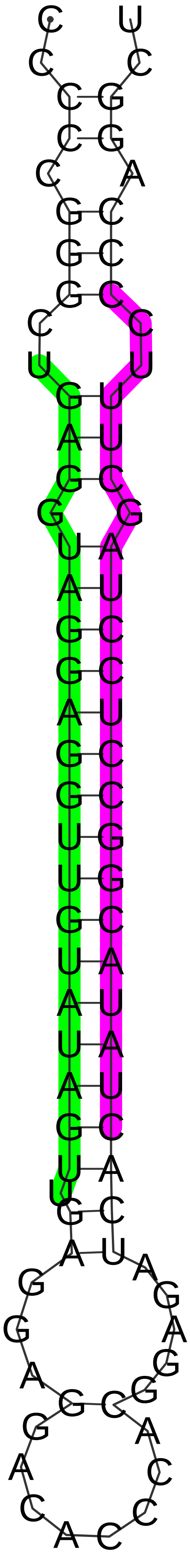

NW_006384123.1_81957-82039

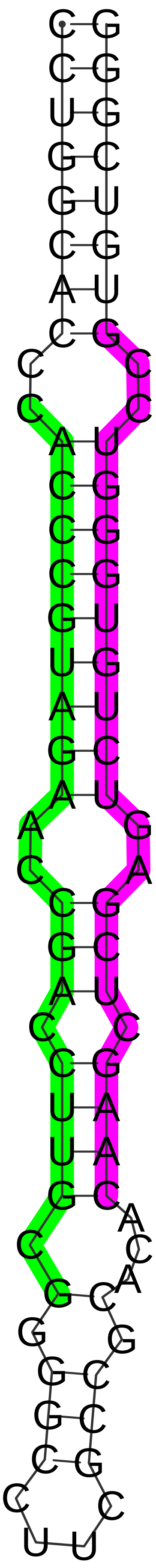

NW_006384123.1_82142-82217

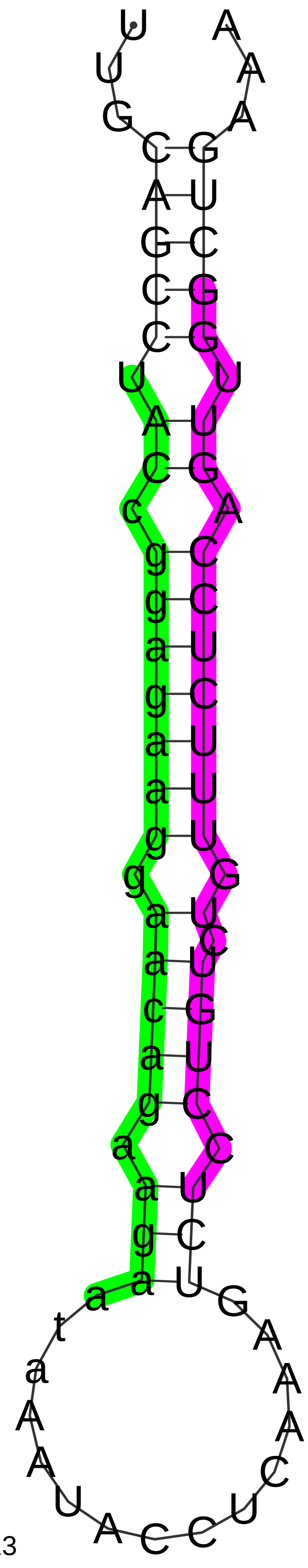

NW_006384133.1_164640-164713

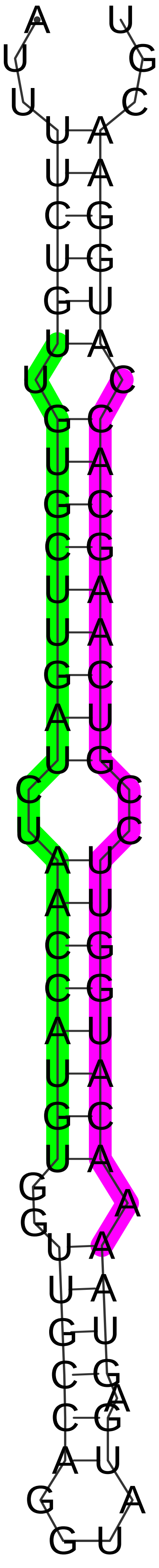

NW_006384138.1_48616-48693

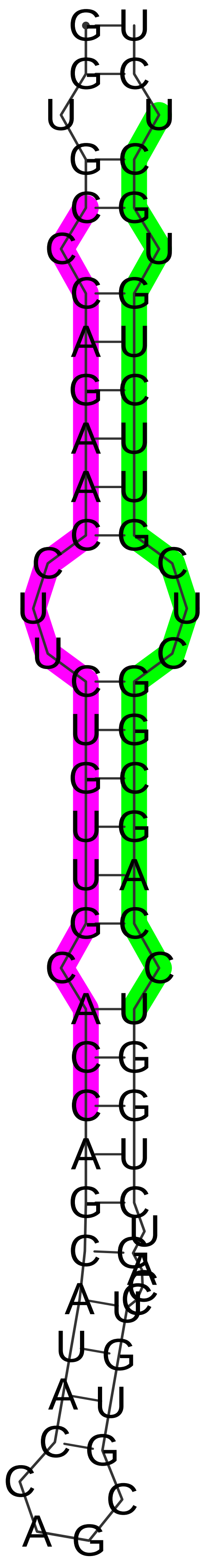

NW_006384142.1_18872-18943

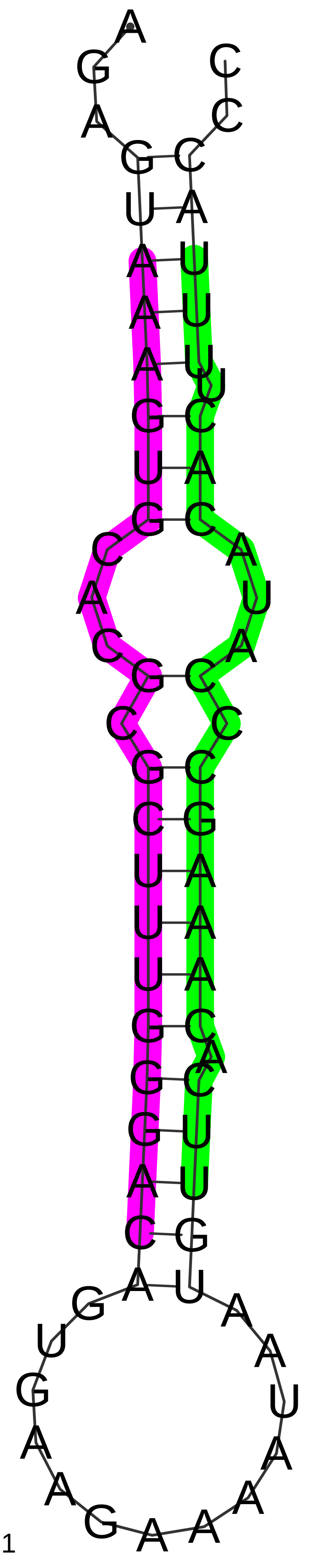

NW_006384158.1_542924-542991

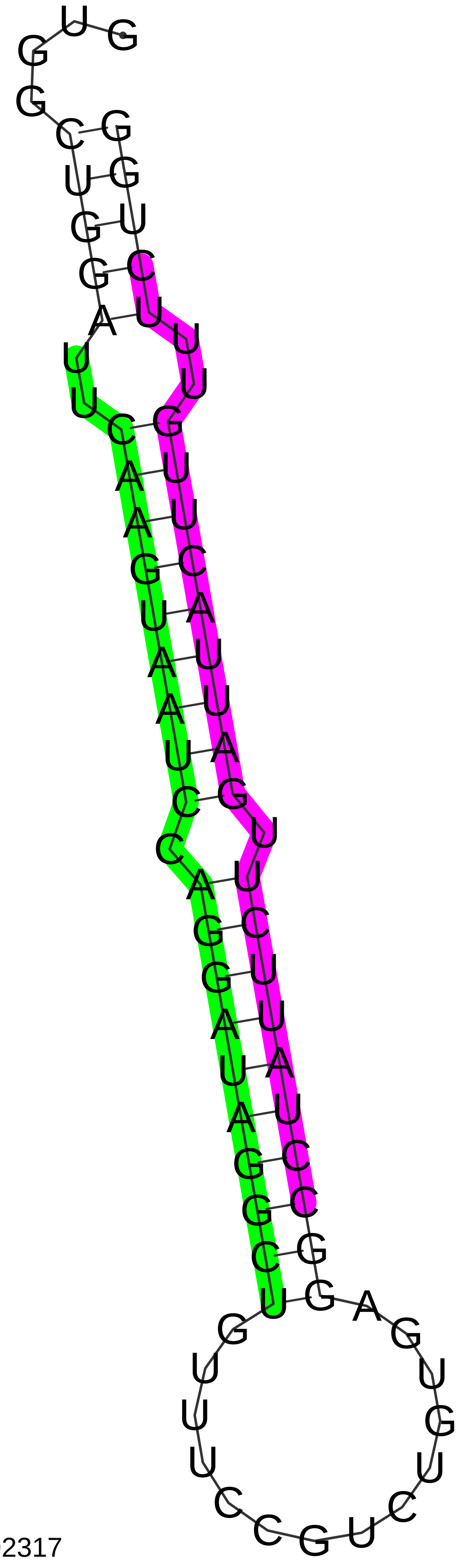

NW_006384165.1_92246-92317

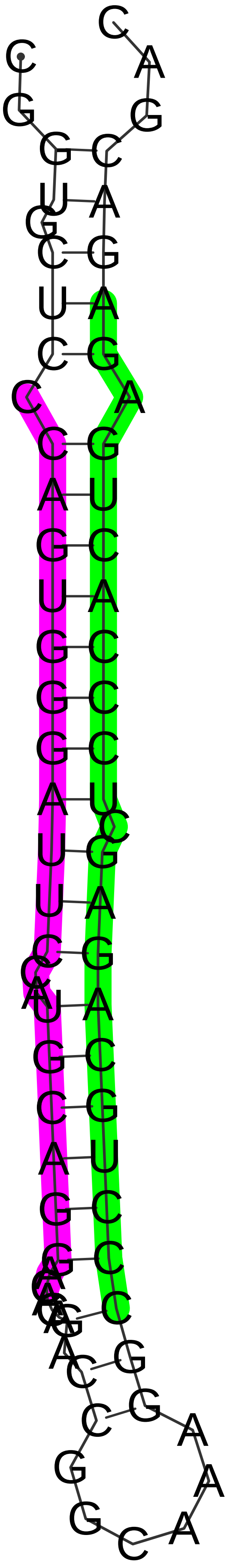

NW_006384171.1_518199-518271

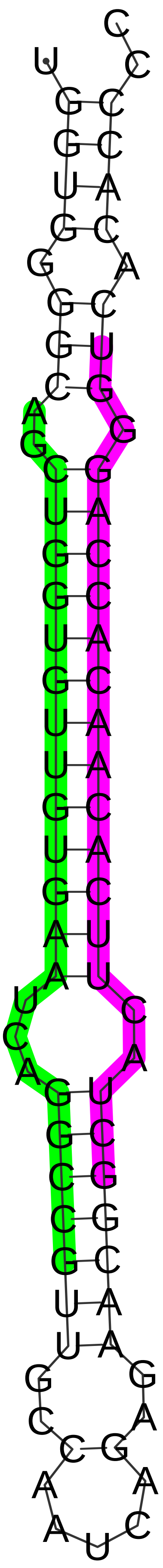

NW_006384196.1_552204-552281

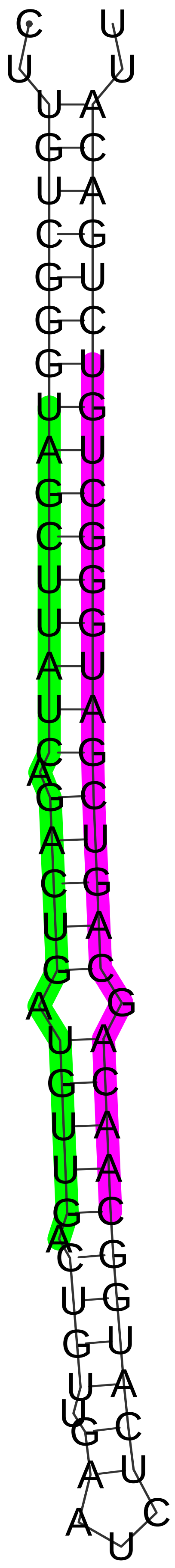

NW_006384200.1_480859-480934

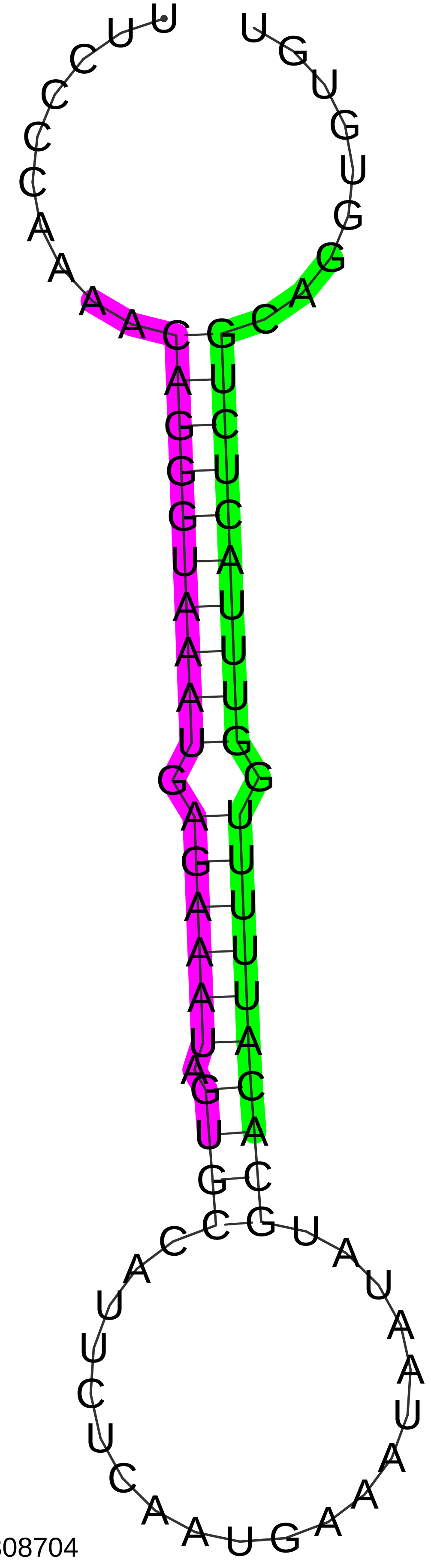

NW_006384206.1_308623-308704

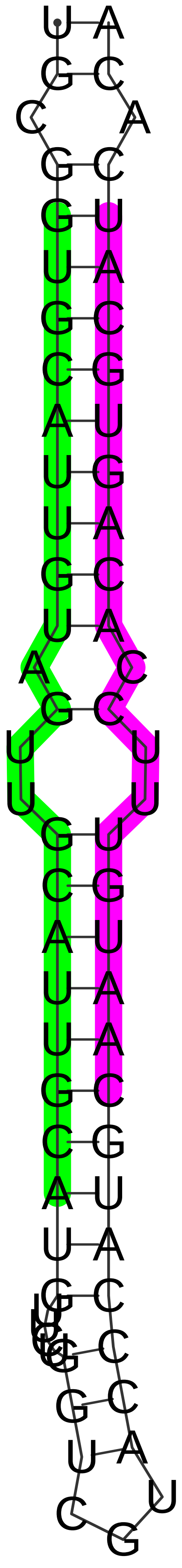

NW_006384209.1_547819-547885

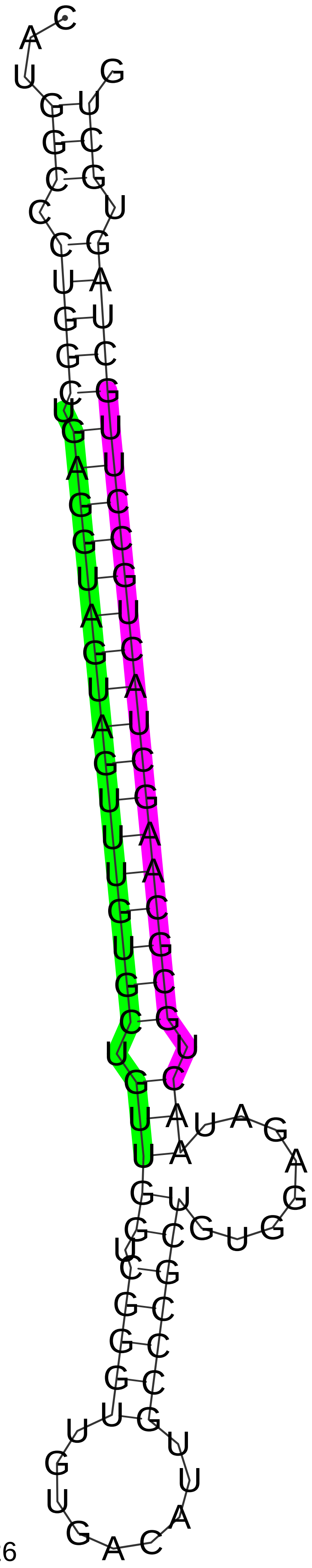

NW_006384263.1_164130-164226

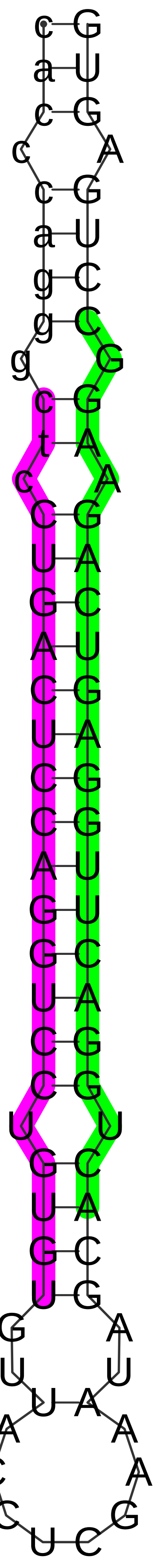
NW_006384269.1_415800-415875

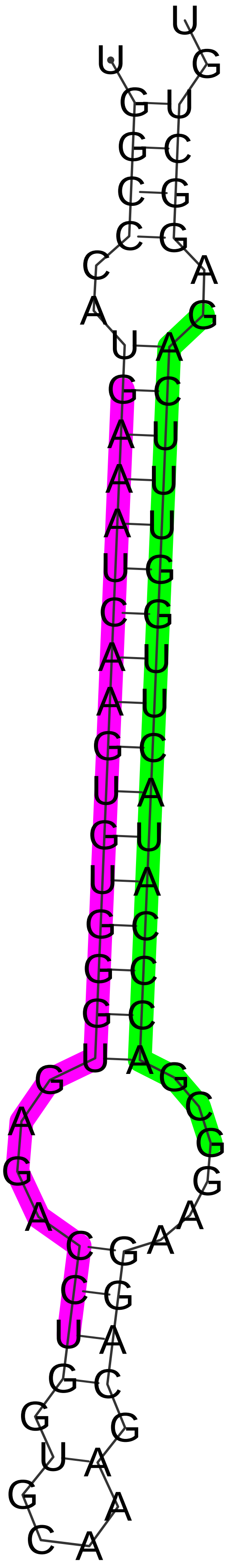

NW_006384280.1_538532-538606

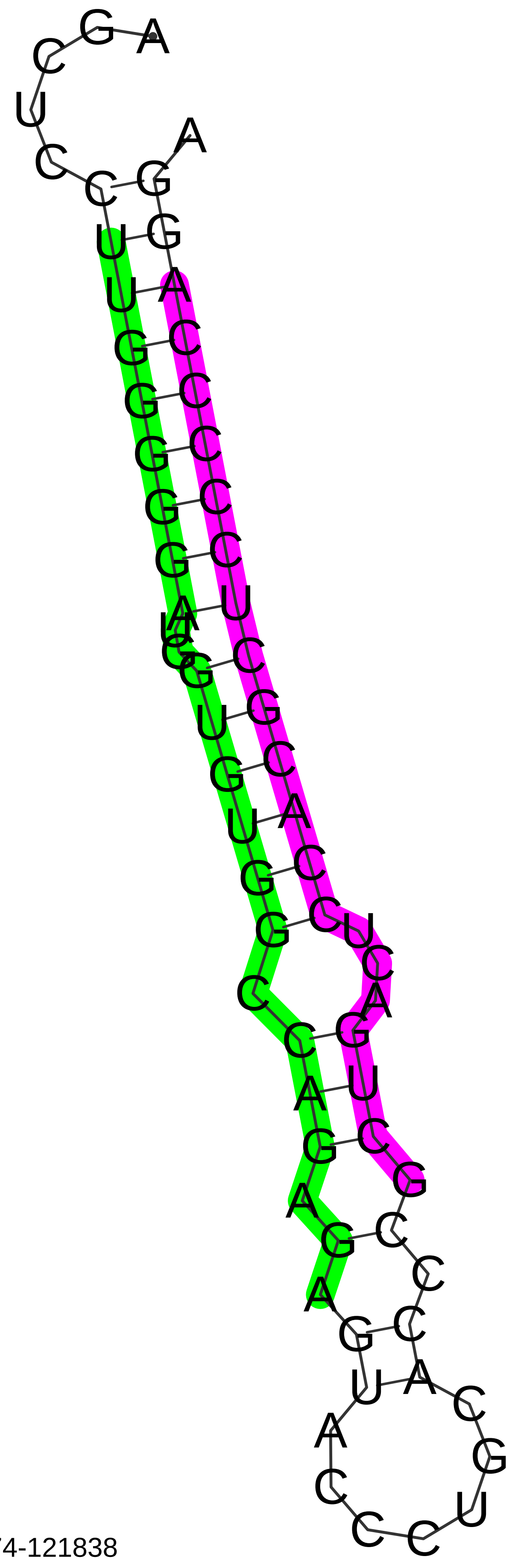

NW_006384281.1_121774-121838

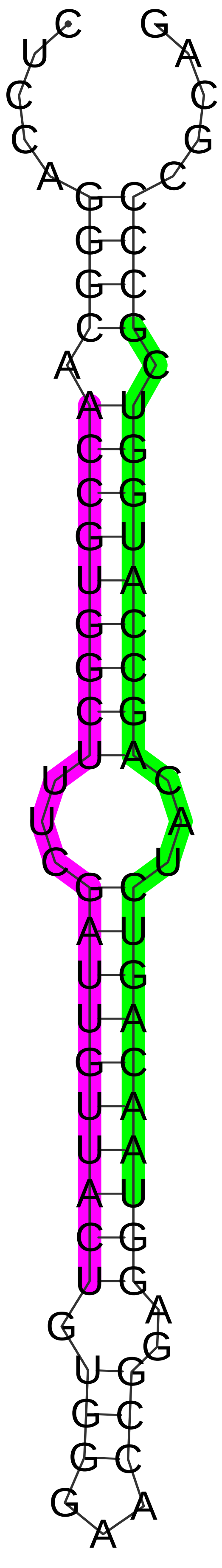


NW_006384289.1_46176-46251

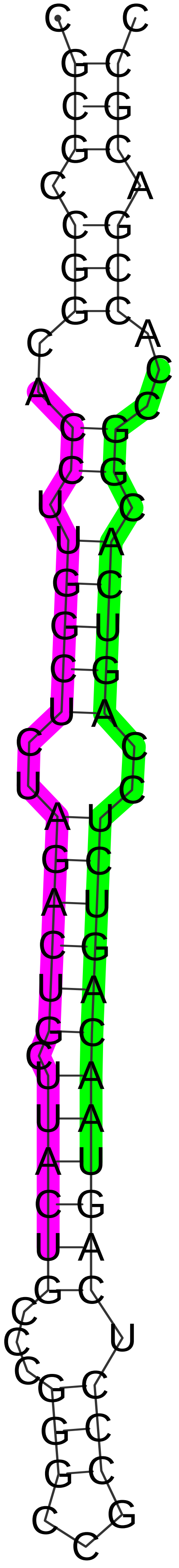

NW_006384289.1_46578-46656

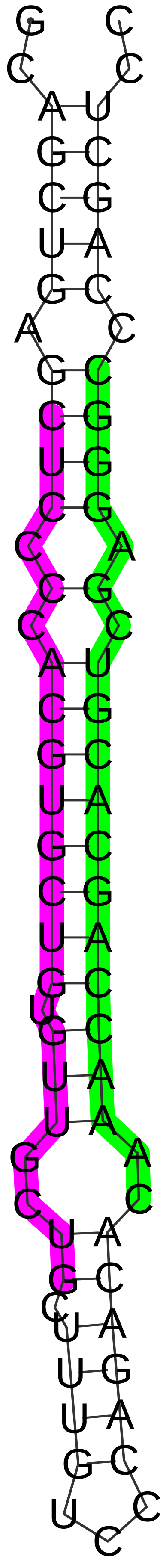

NW_006384301.1_74569-74641

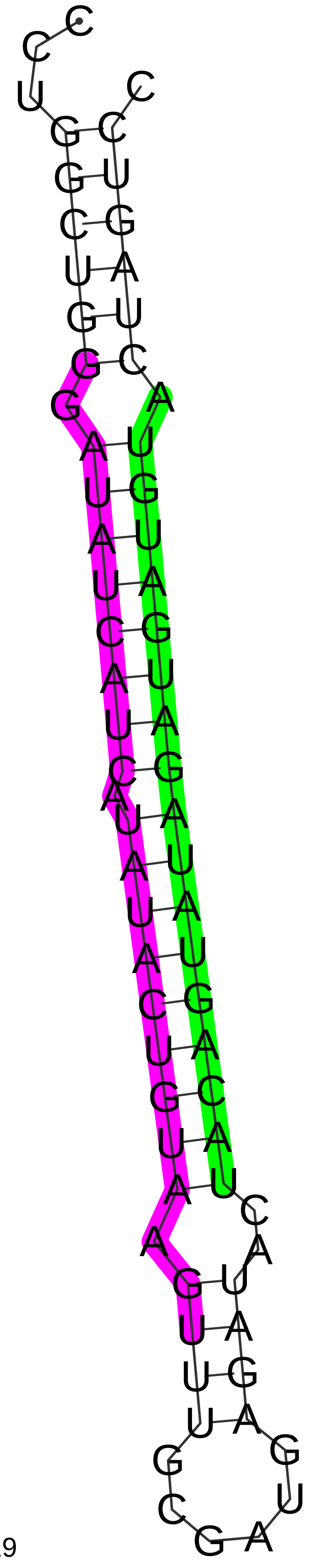

NW_006384301.1_778250-778319

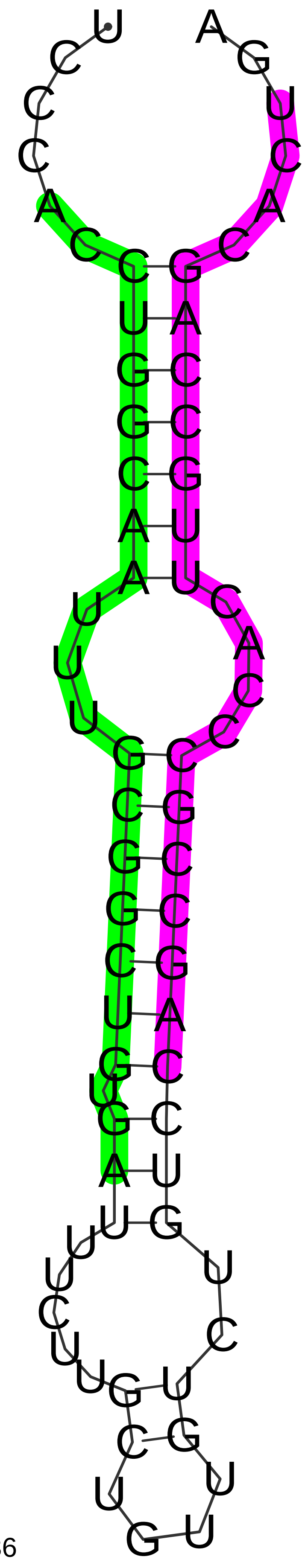

NW_006384303.1_481368-481436

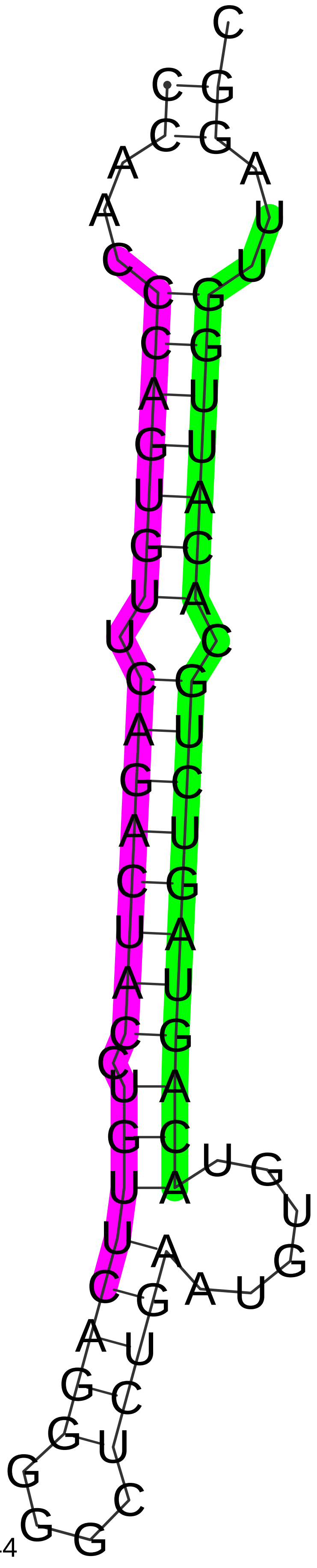

NW_006384317.1_166775-166844

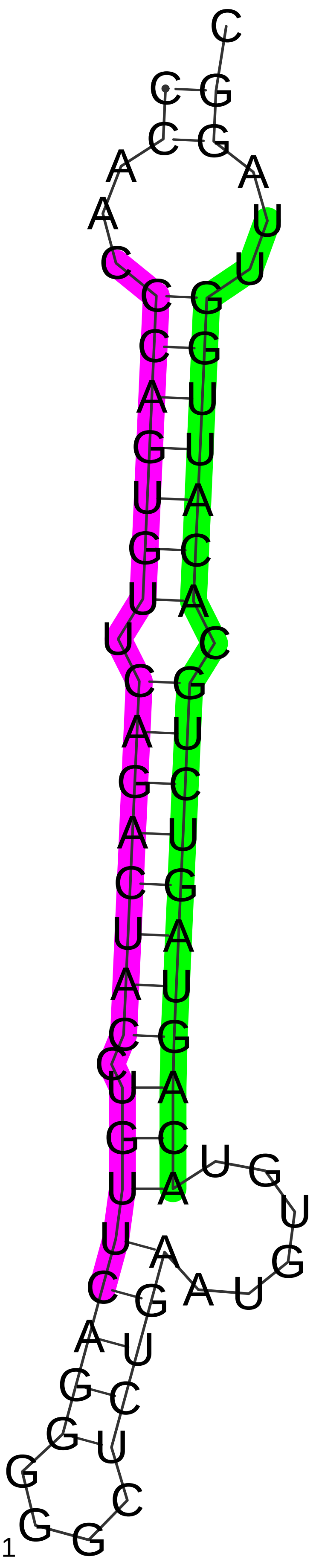

NW_006384317.1_200562-200631

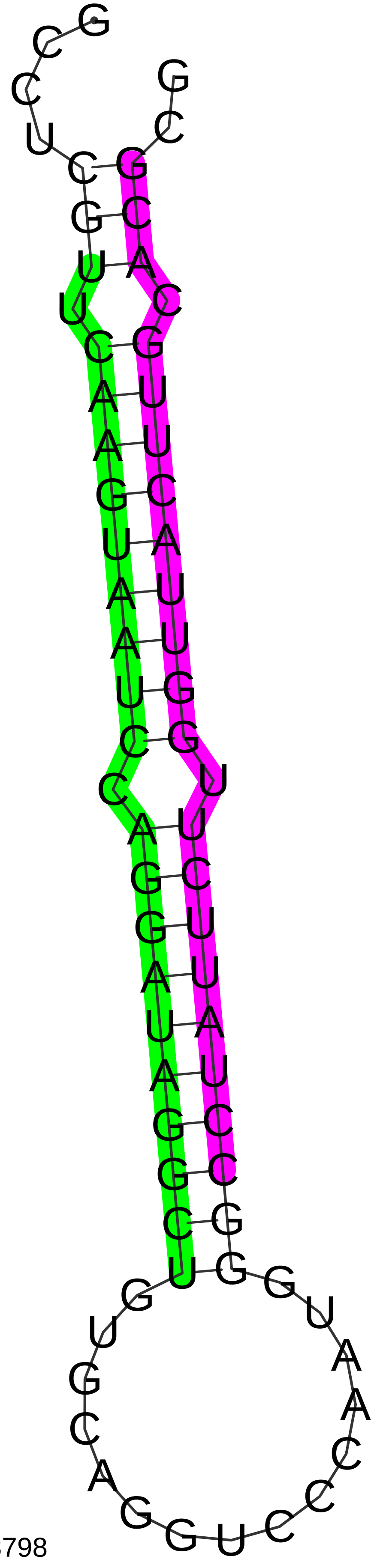

NW_006384324.1_498730-498798

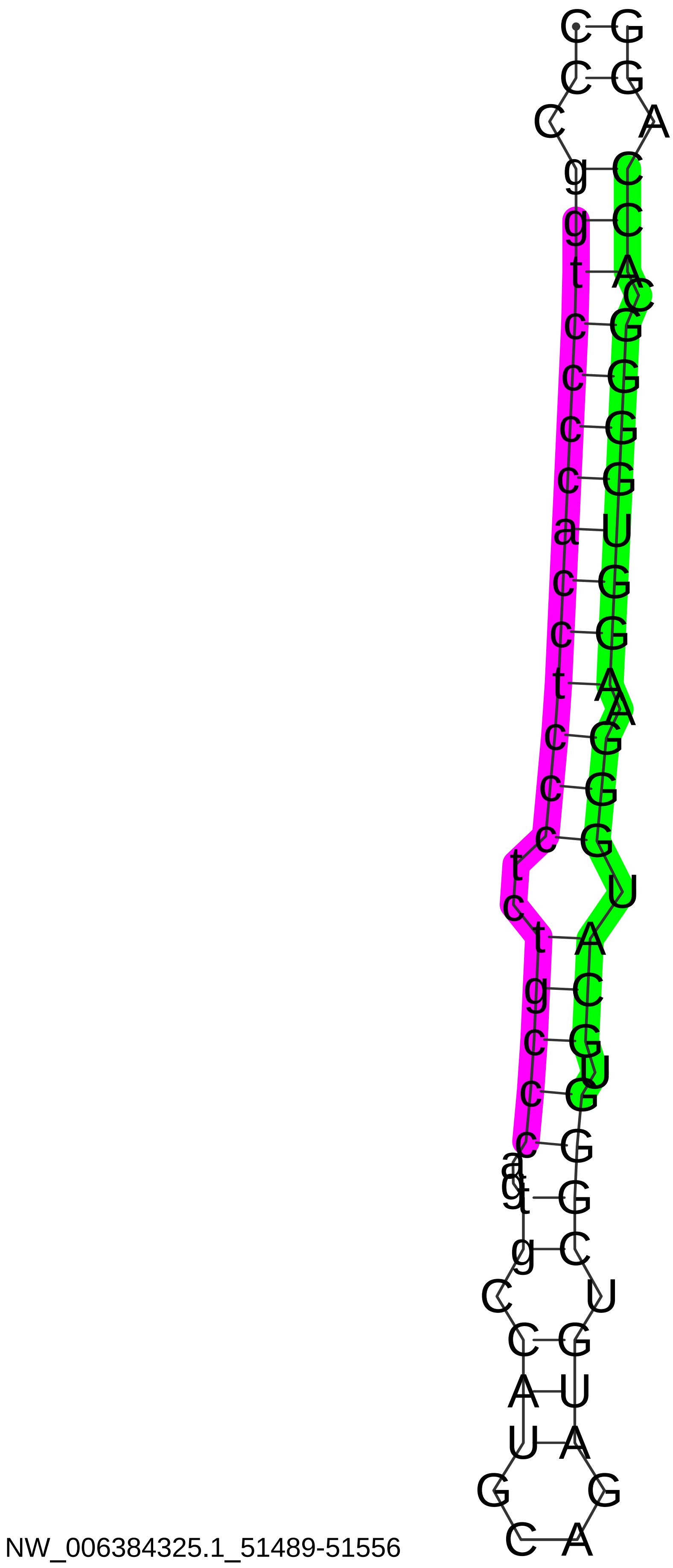
NW_006384325.1_51489-51556

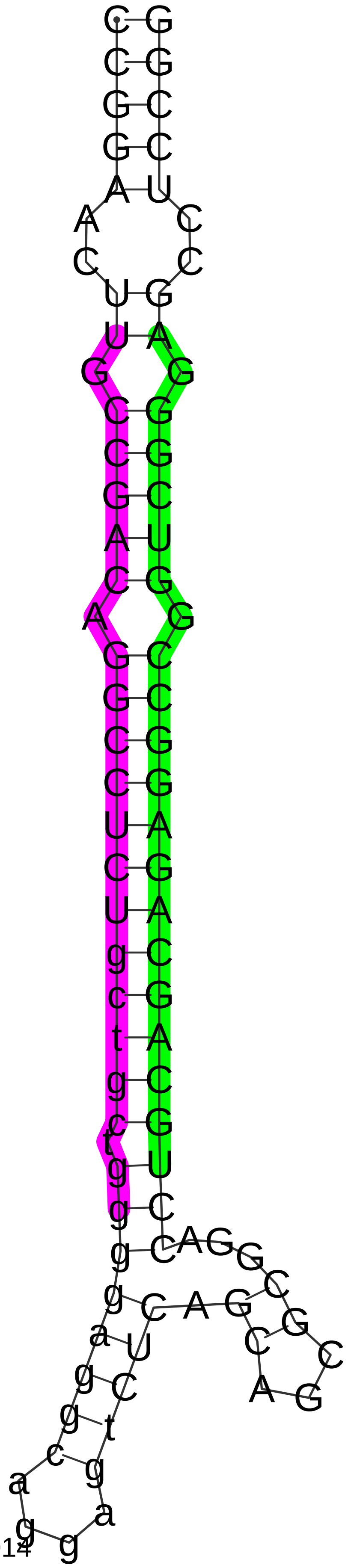

NW_006384333.1_515927-516014

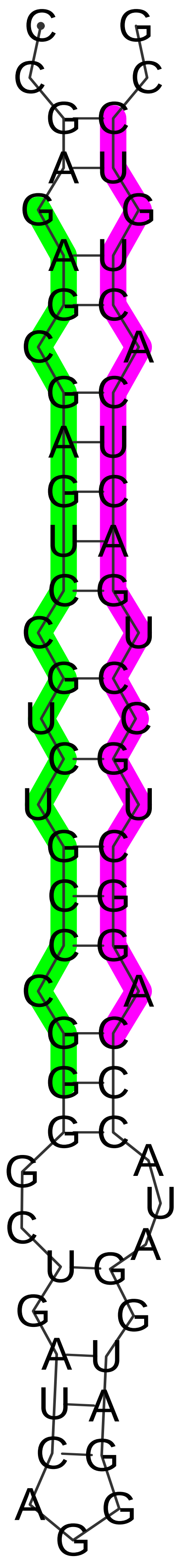

NW_006384335.1_648224-648291

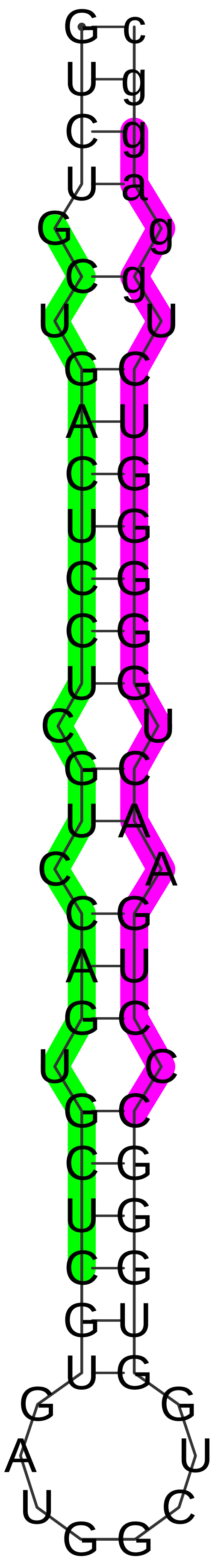
NW_006384355.1_196334-196397

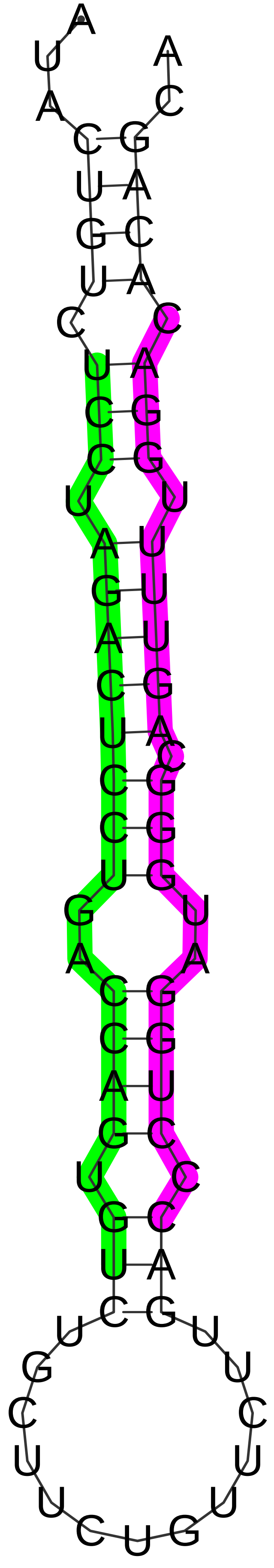

NW_006384369.1_574852-574924

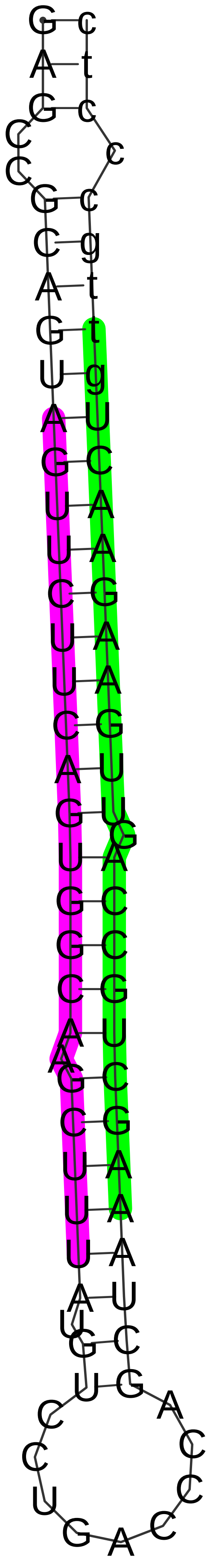

NW_006384375.1_374328-374404

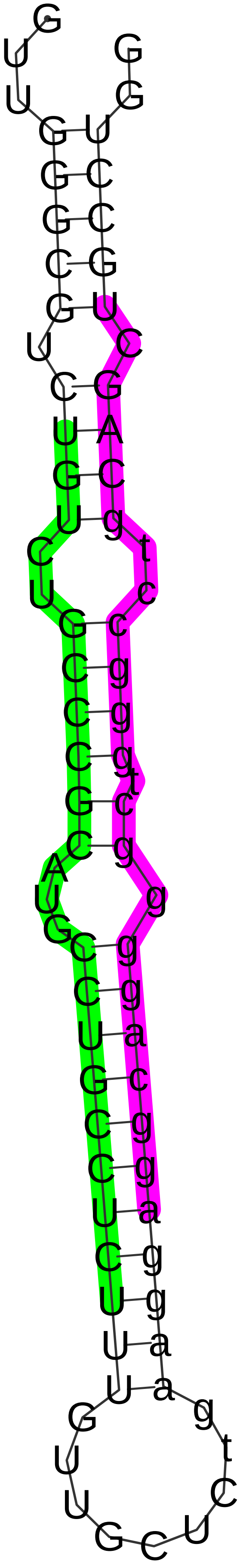

NW_006384383.1_261802-261878

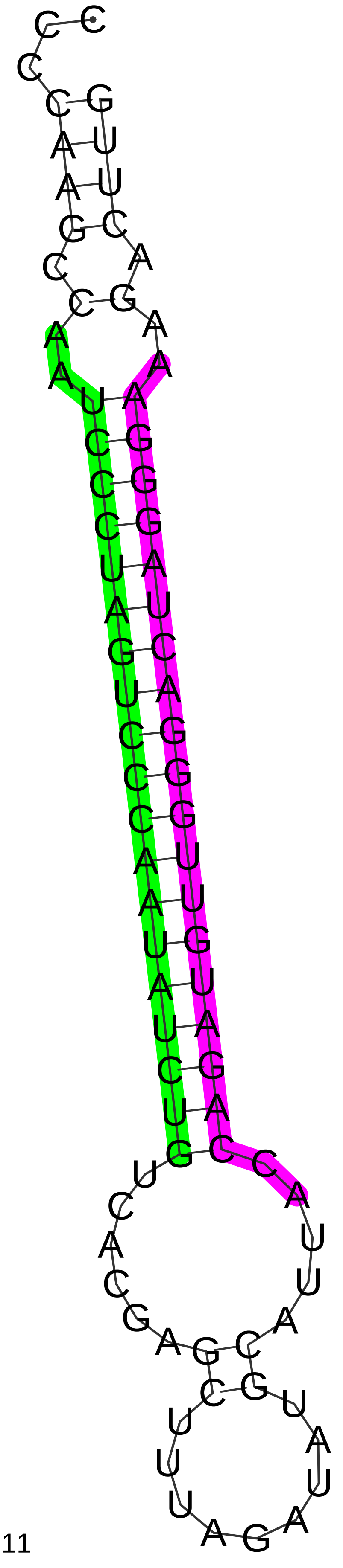

NW_006384385.1_697731-697811

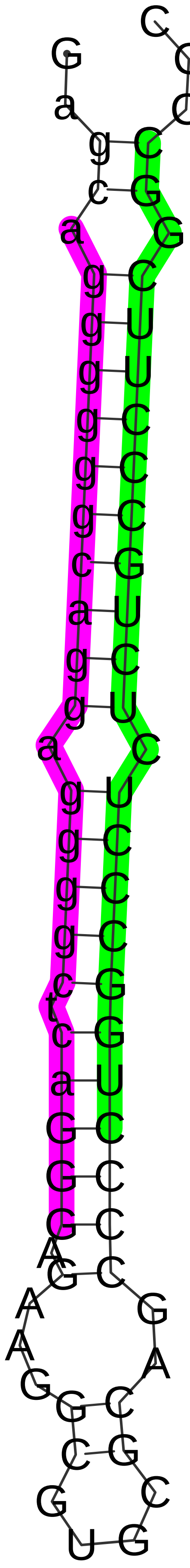

NW_006384393.1_346898-346967

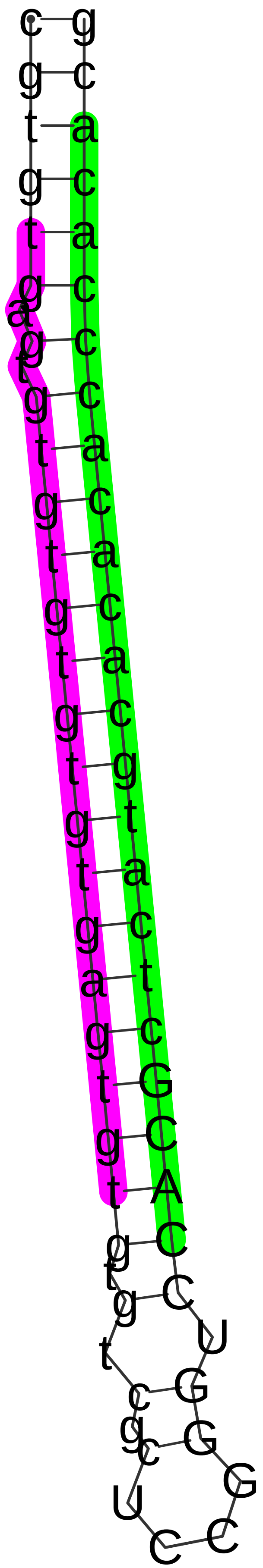

NW_006384413.1_571878-571941

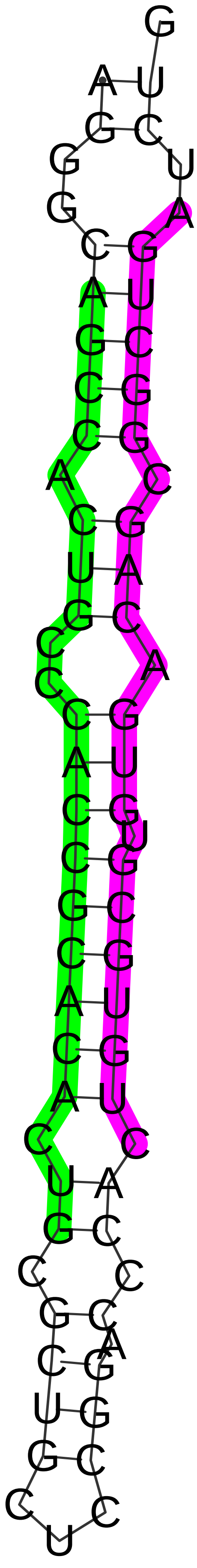

NW_006384434.1_258304-258372

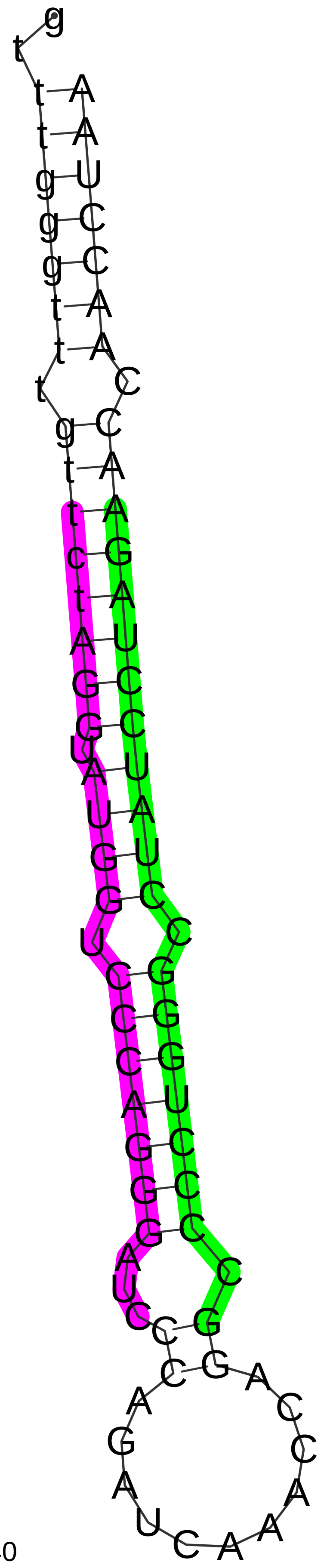

NW_006384440.1_148863-148940

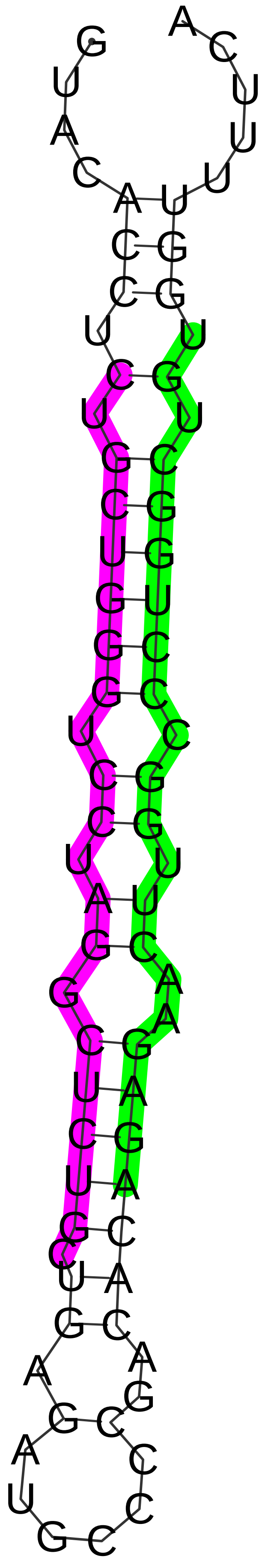

NW_006384482.1_167929-168002

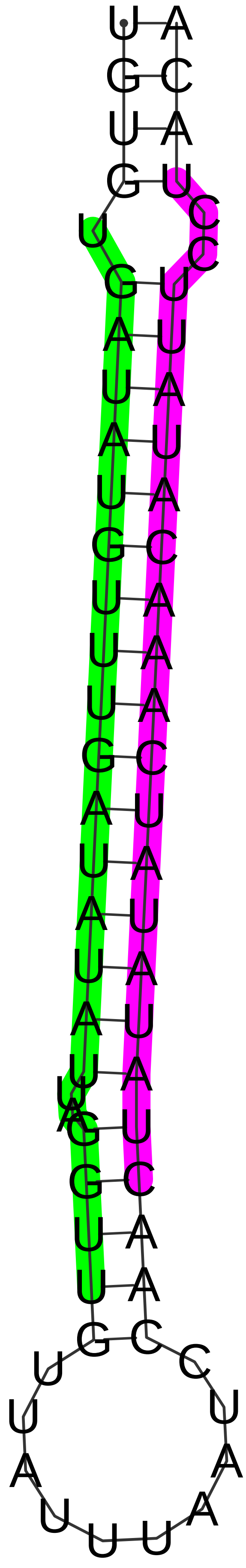

NW_006384487.1_387439-387503

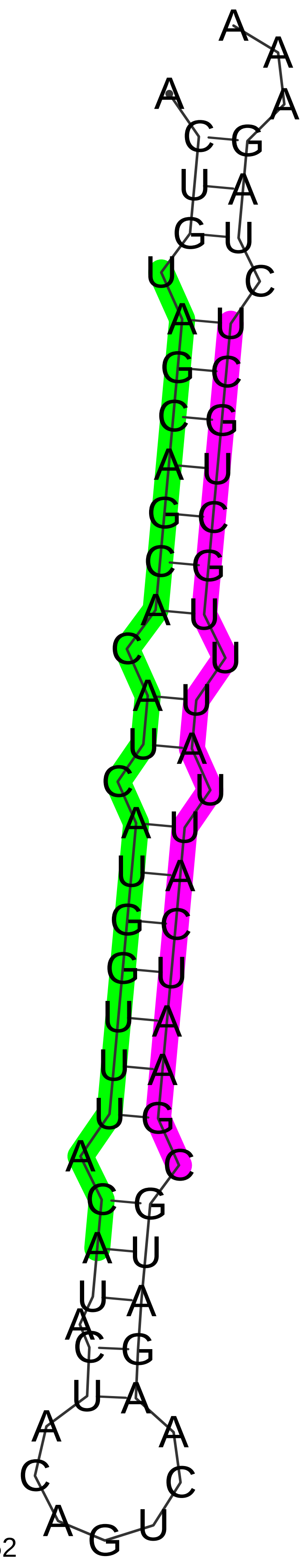

NW_006384492.1_457185-457252

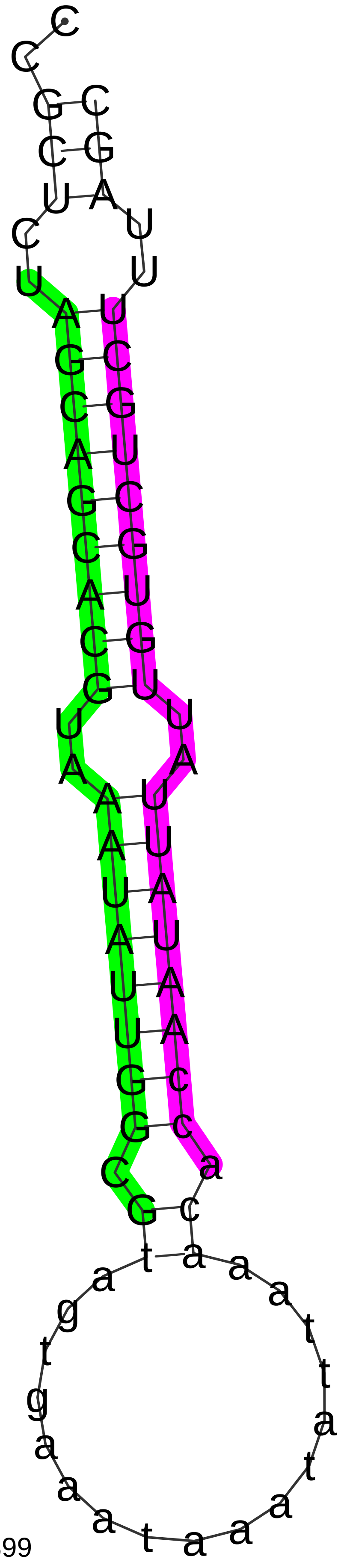

NW_006384492.1_457327-457399

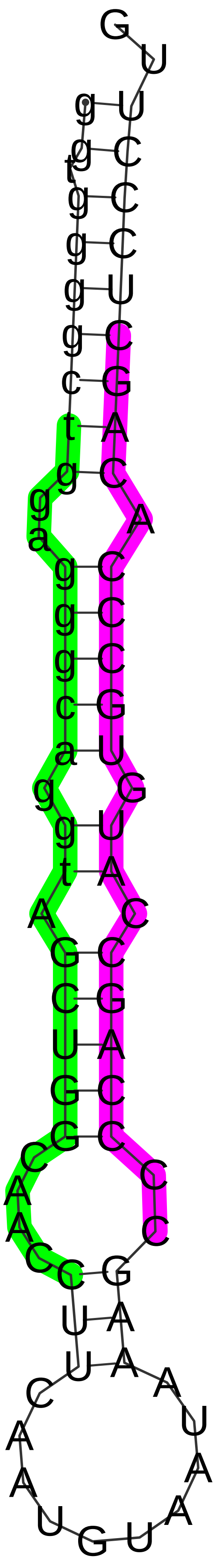

NW_006384502.1_370562-370635

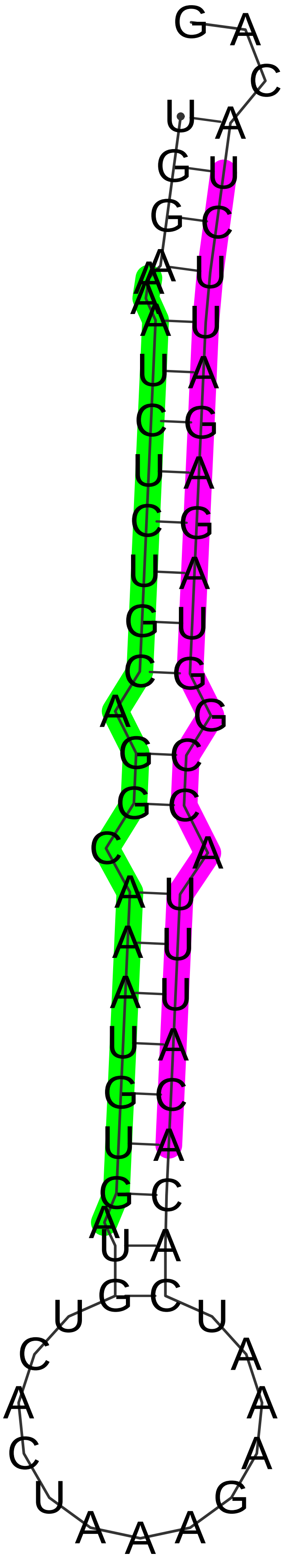

NW_006384532.1_505635-505703

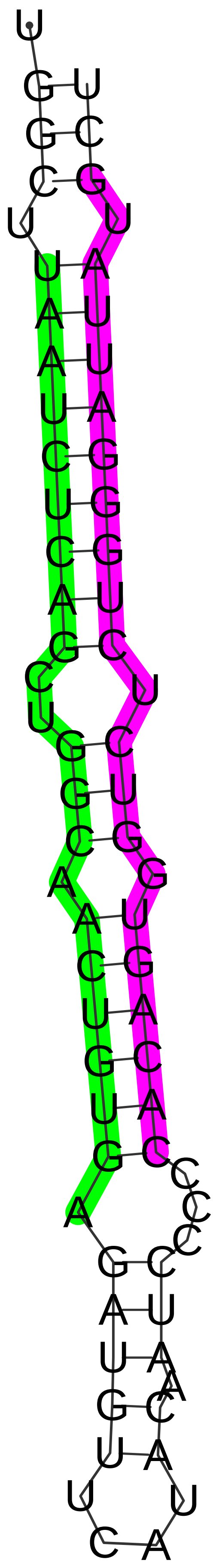

NW_006384532.1_516589-516657

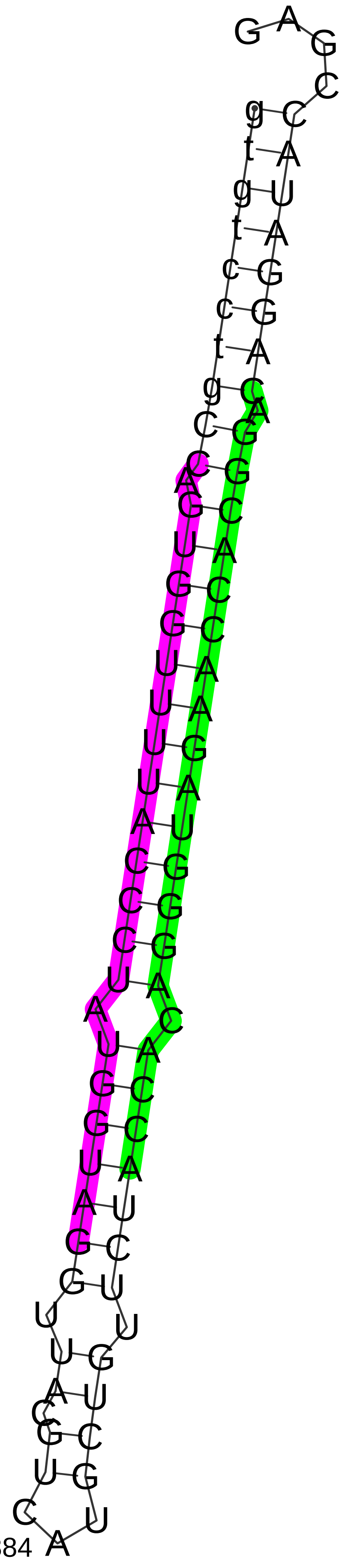

NW_006384546.1_587803-587884

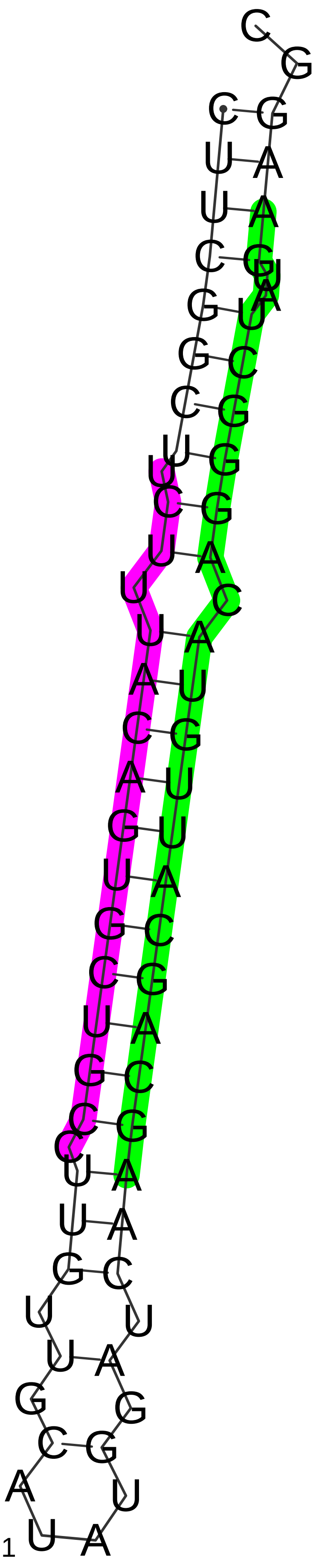

NW_006384553.1_422334-422401

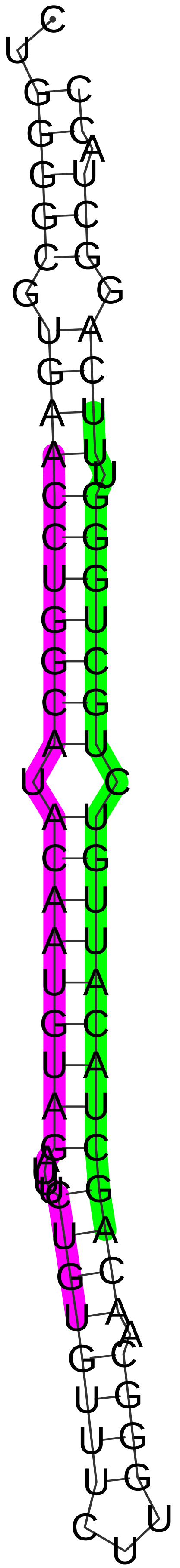

NW_006384555.1_576055-576136

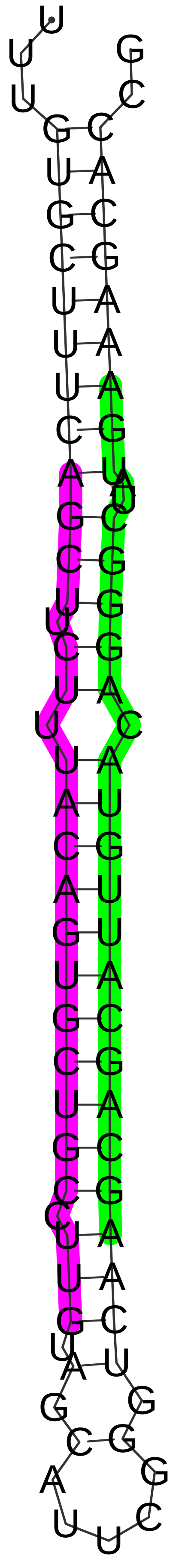

NW_006384559.1_366258-366336

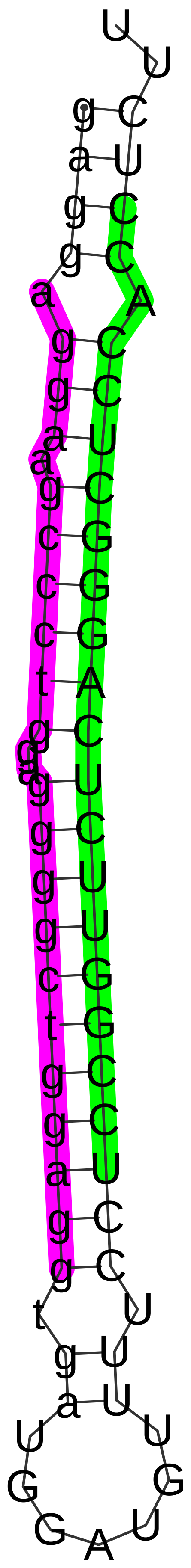

NW_006384571.1_125445-125512

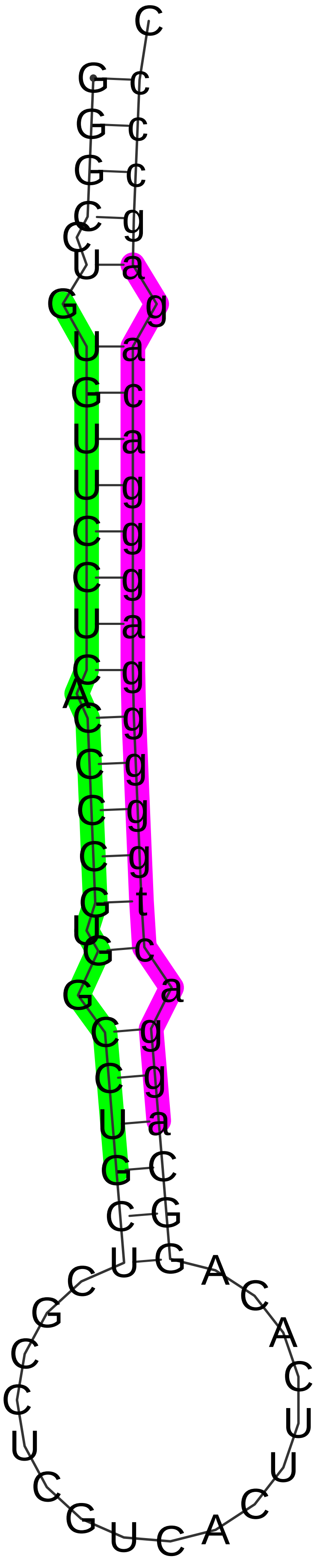

NW_006384578.1_88409-88483

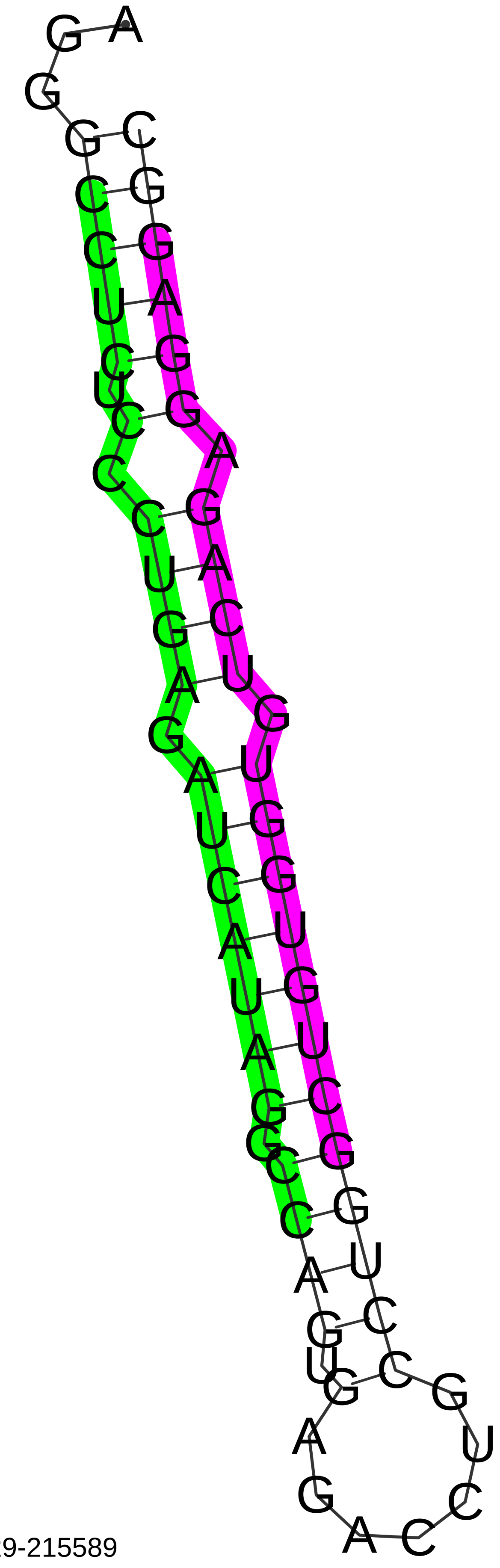

NW_006384592.1_215529-215589

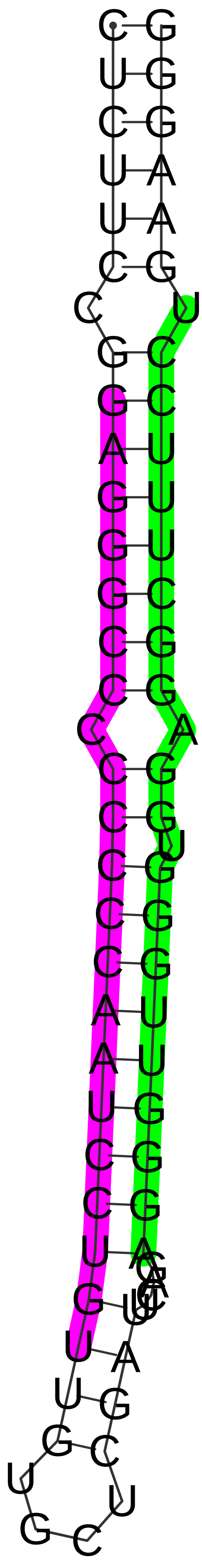

NW_006384598.1_334489-334559

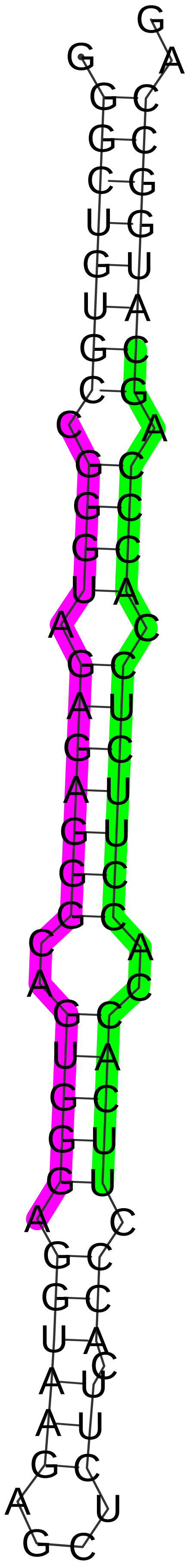

NW_006384601.1_267152-267229

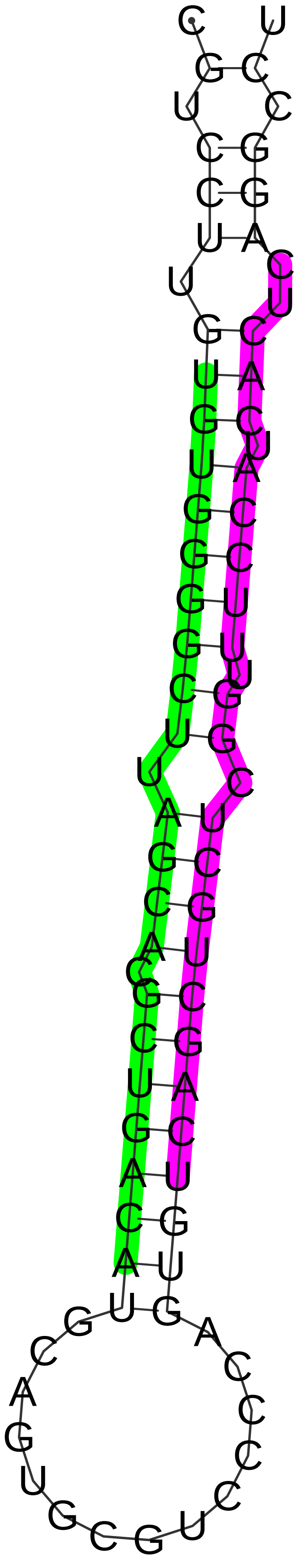

NW_006384619.1_46052-46129

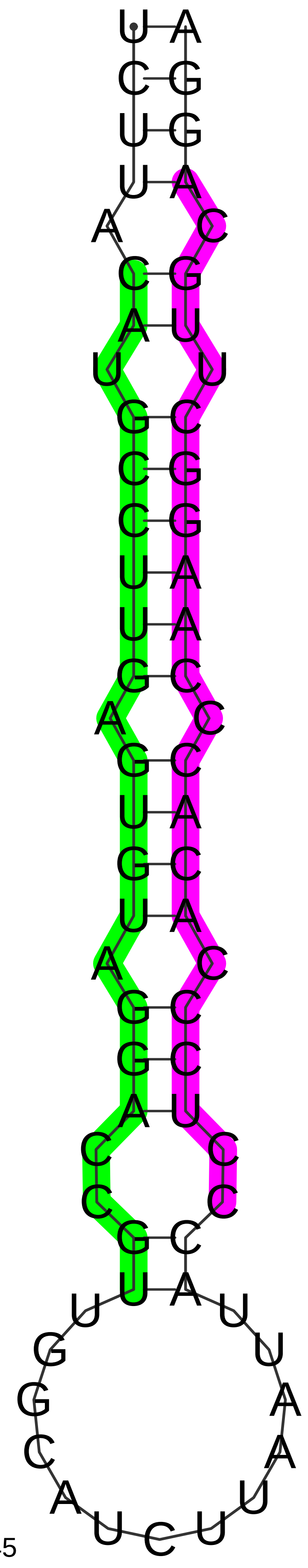

NW_006384632.1_453179-453245

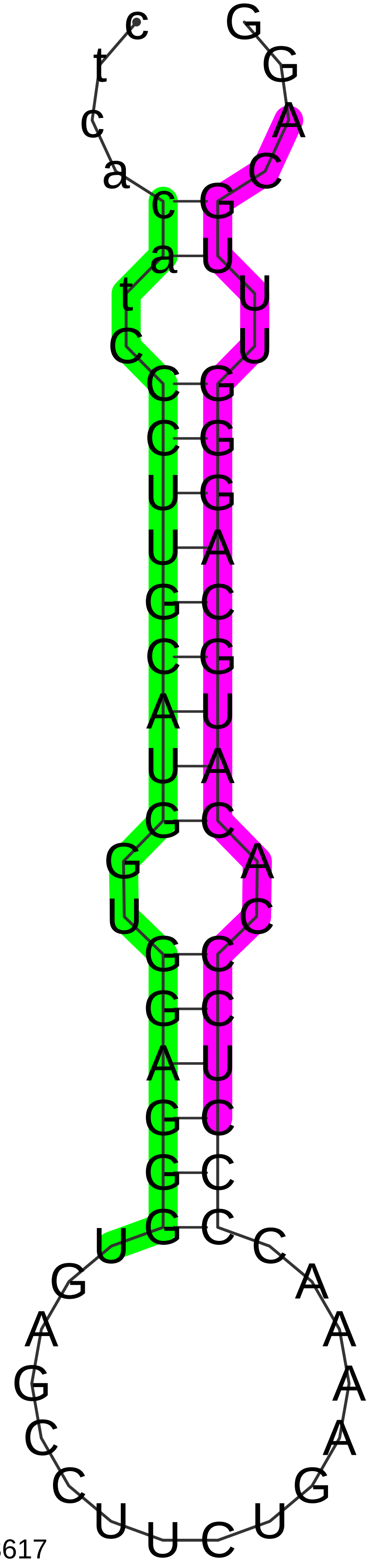

NW_006384632.1_453552-453617

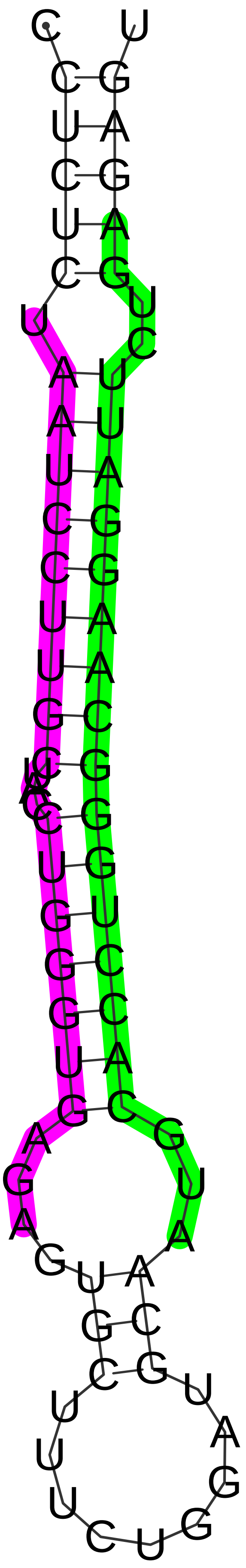

NW_006384632.1_458198-458269

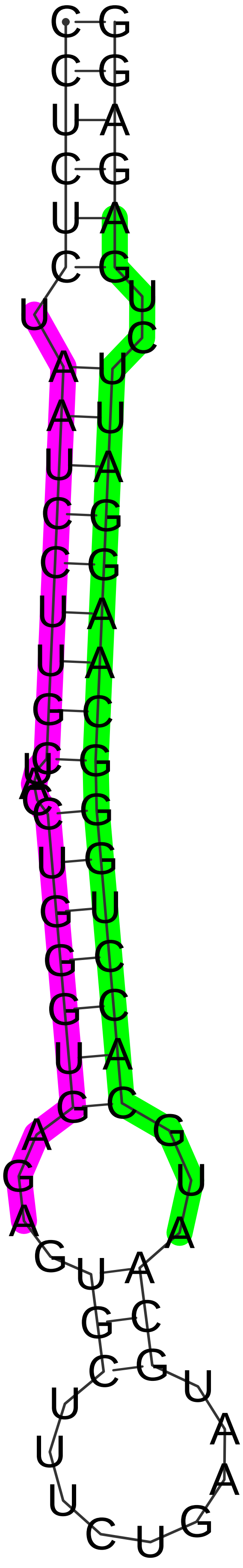

NW_006384632.1_459998-460069

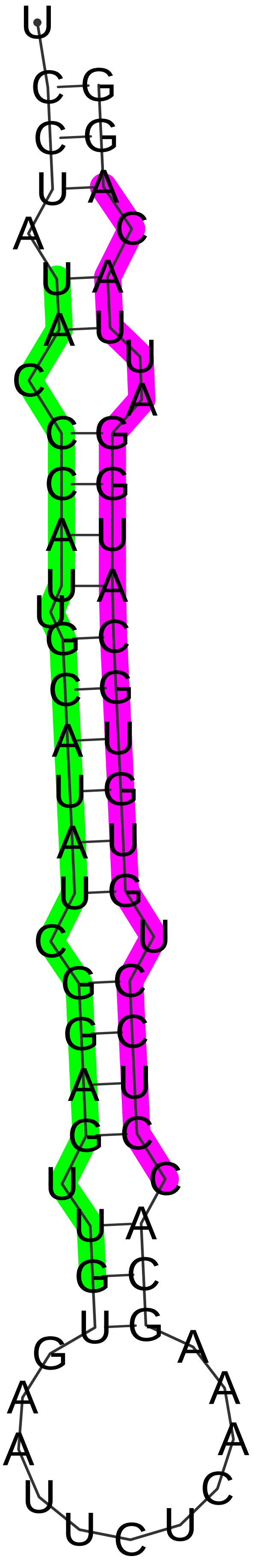

NW_006384632.1_461421-461486

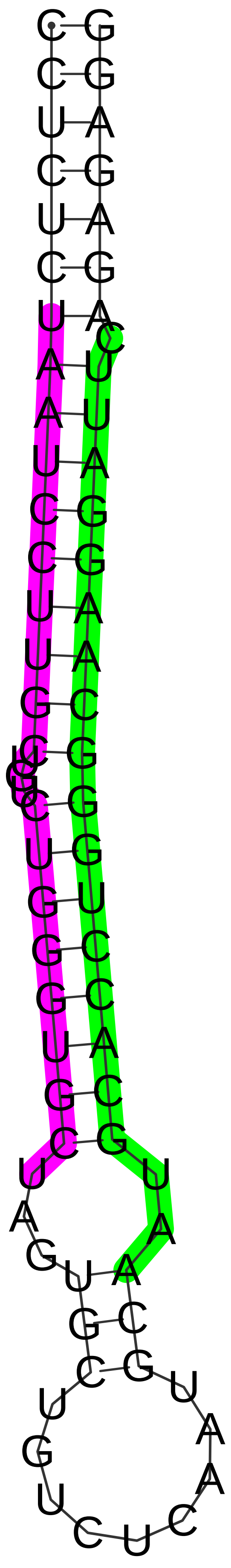

NW_006384632.1_462737-462808

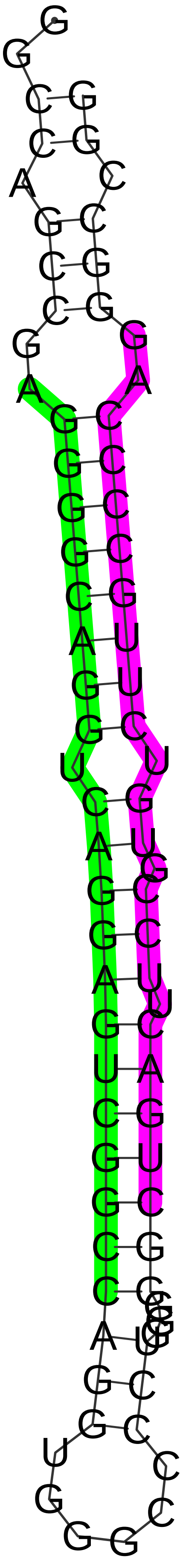

NW_006384645.1_470478-470554

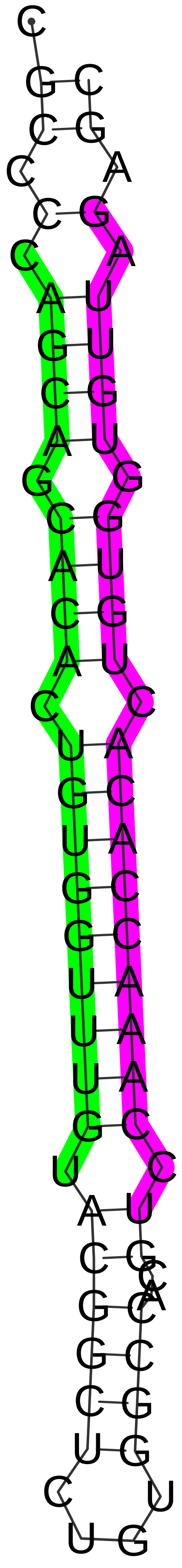

NW_006384645.1_97966-98034

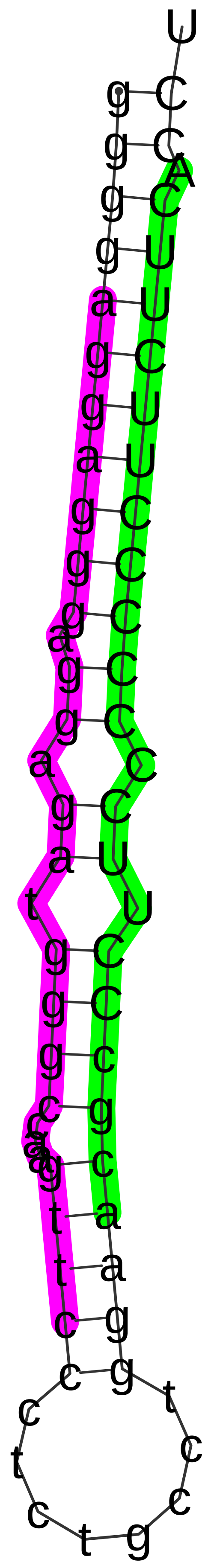

NW_006384648.1_127122-127187

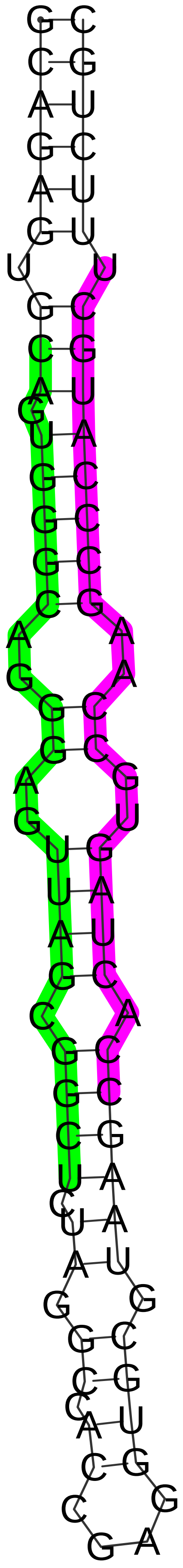

NW_006384754.1_145121-145201

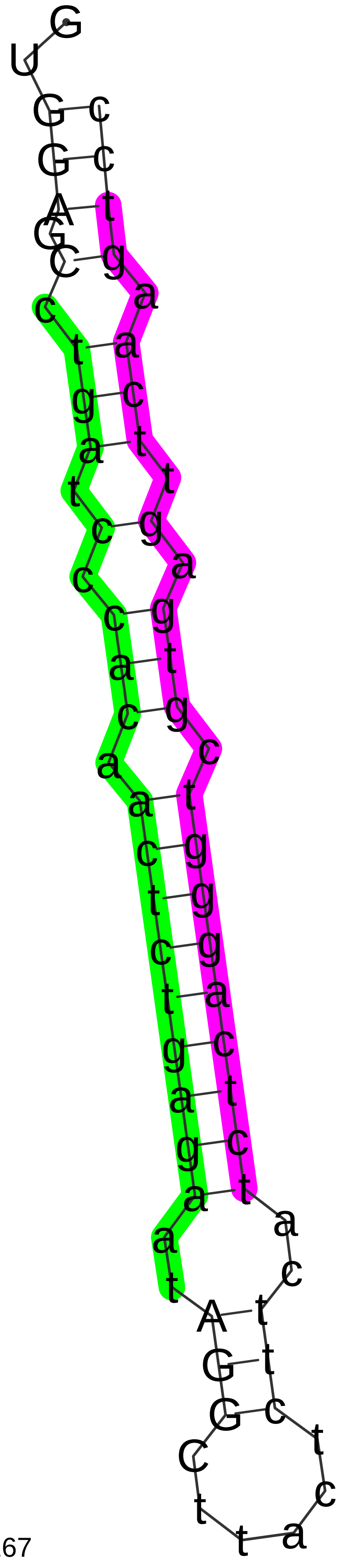

NW_006384761.1_258201-258267

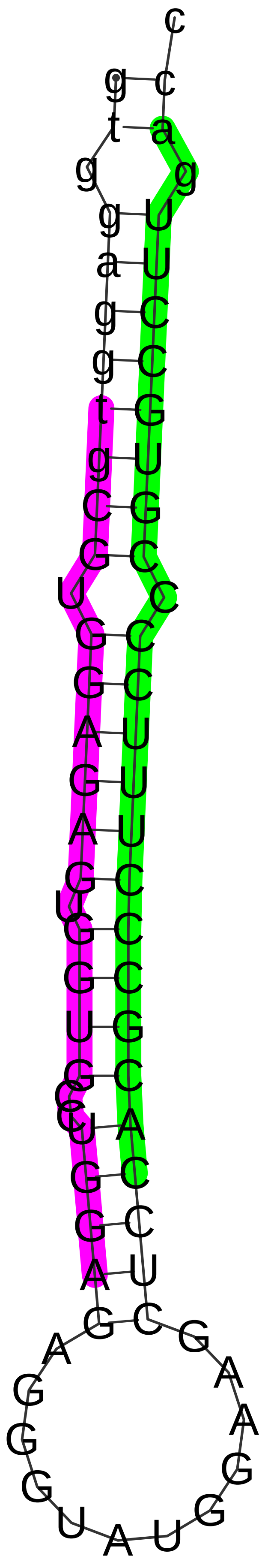

NW_006384796.1_428646-428715

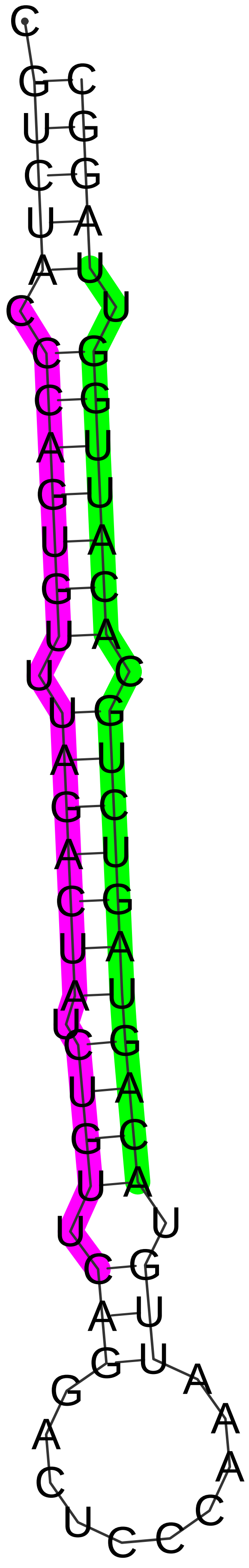

NW_006384809.1_148404-148473

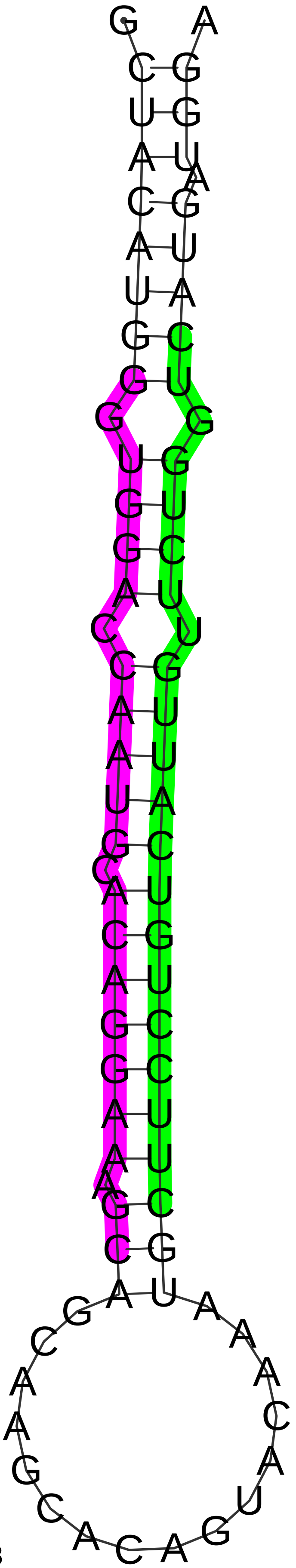

NW_006384838.1_368180-368258

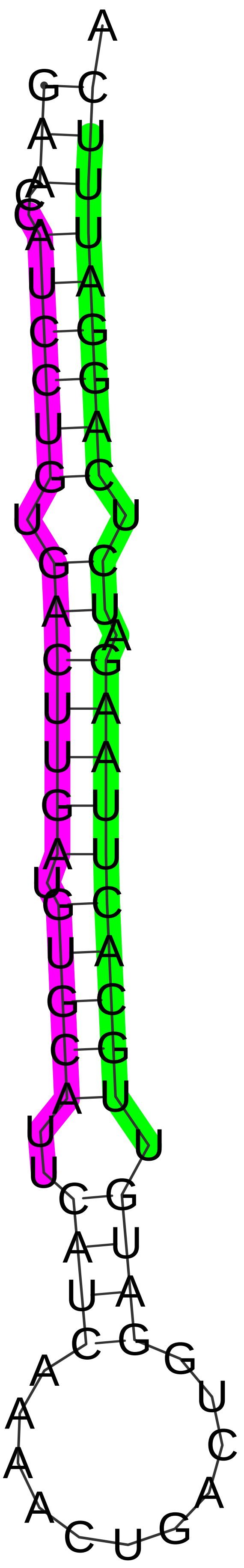

NW_006384862.1_332026-332096

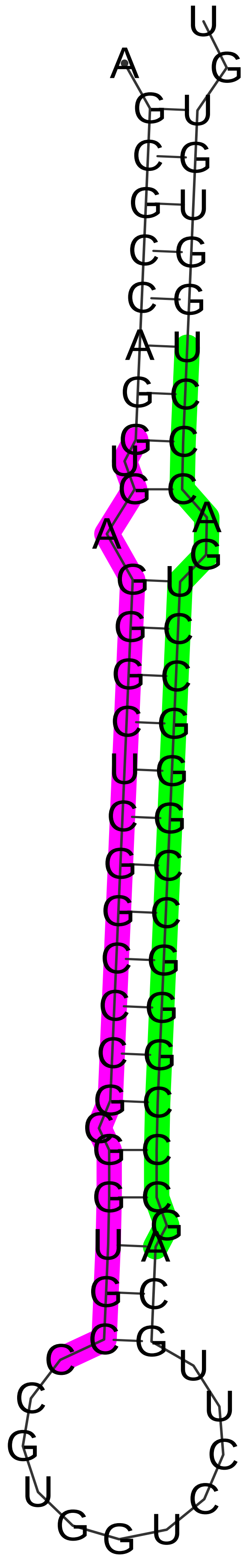

NW_006384870.1_82609-82680

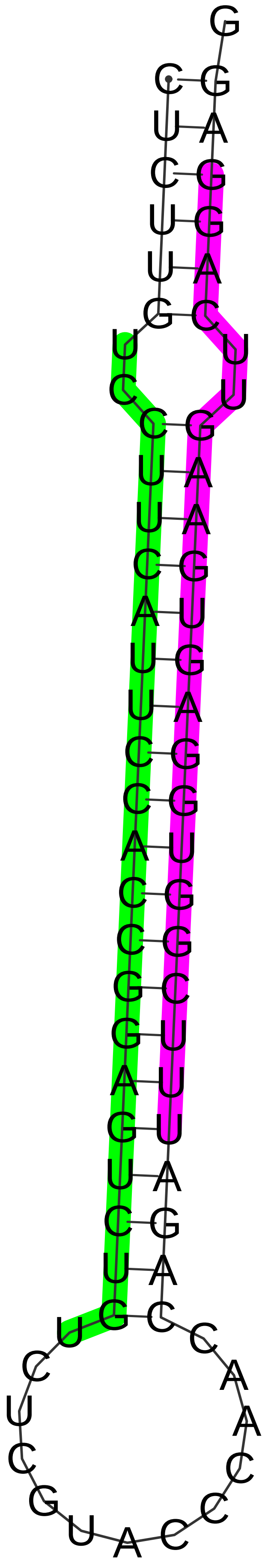

NW_006384878.1_481156-481225

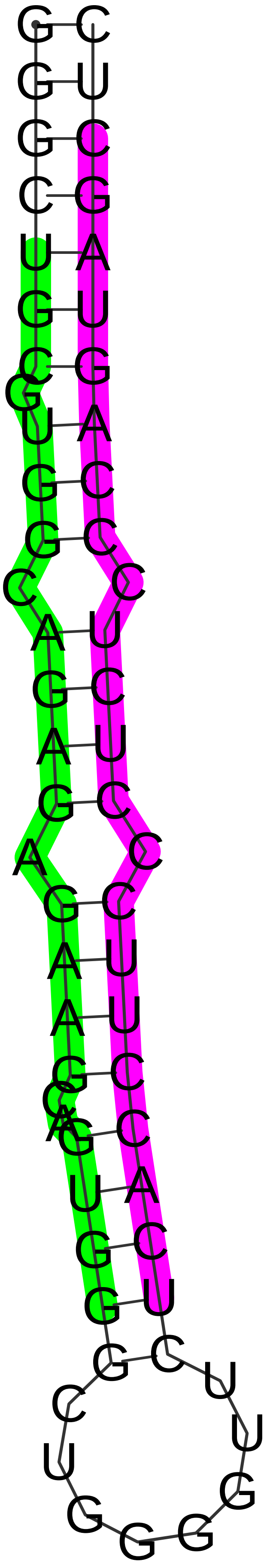

NW_006384908.1_167600-167660

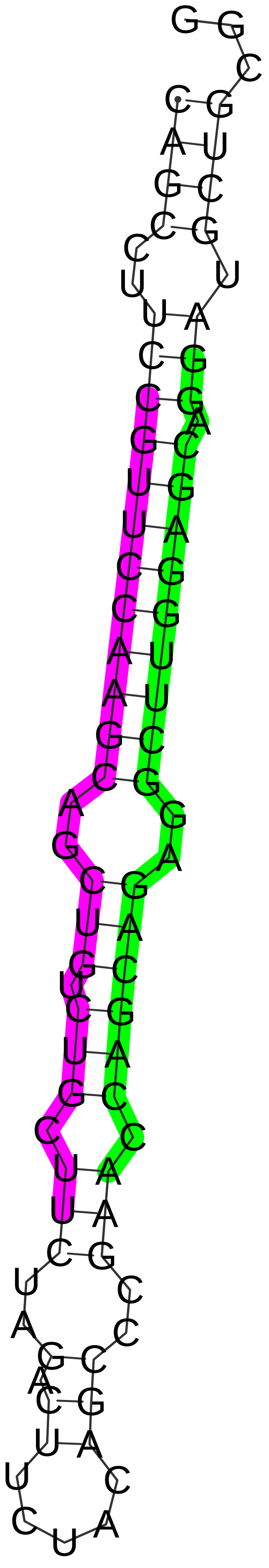

NW_006384928.1_506090-506169

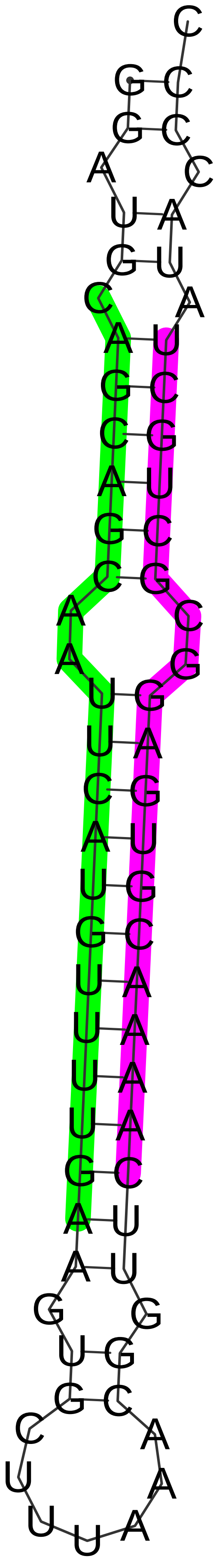

NW_006384929.1_165939-166006

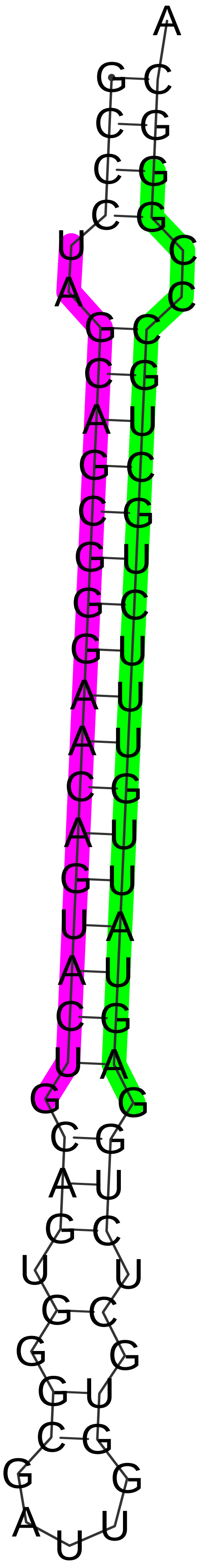

NW_006384929.1_166266-166335

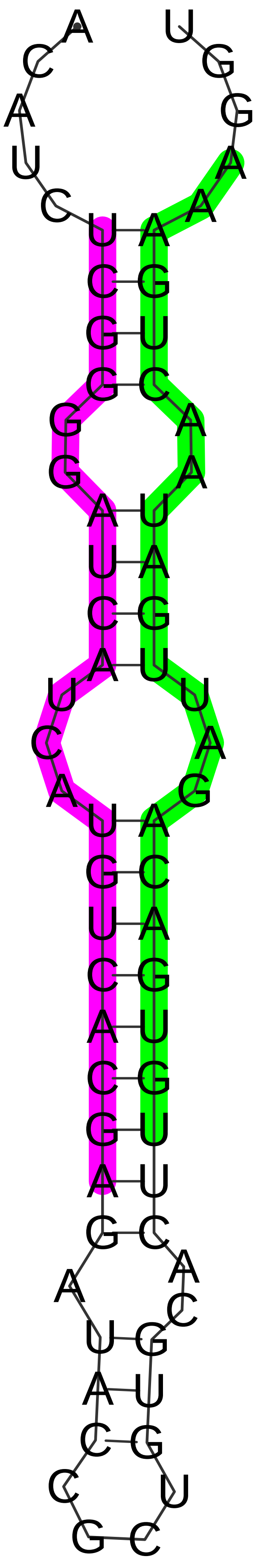

NW_006384929.1_171170-171236

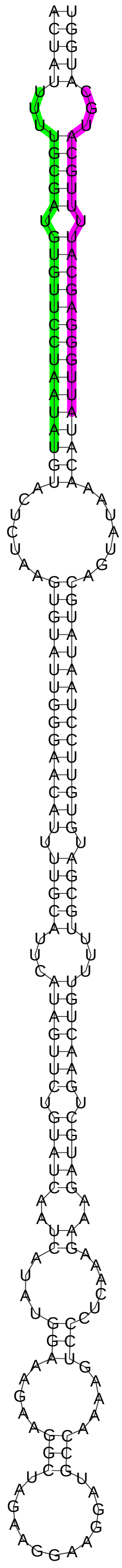

NW_006384929.1_171952-172150

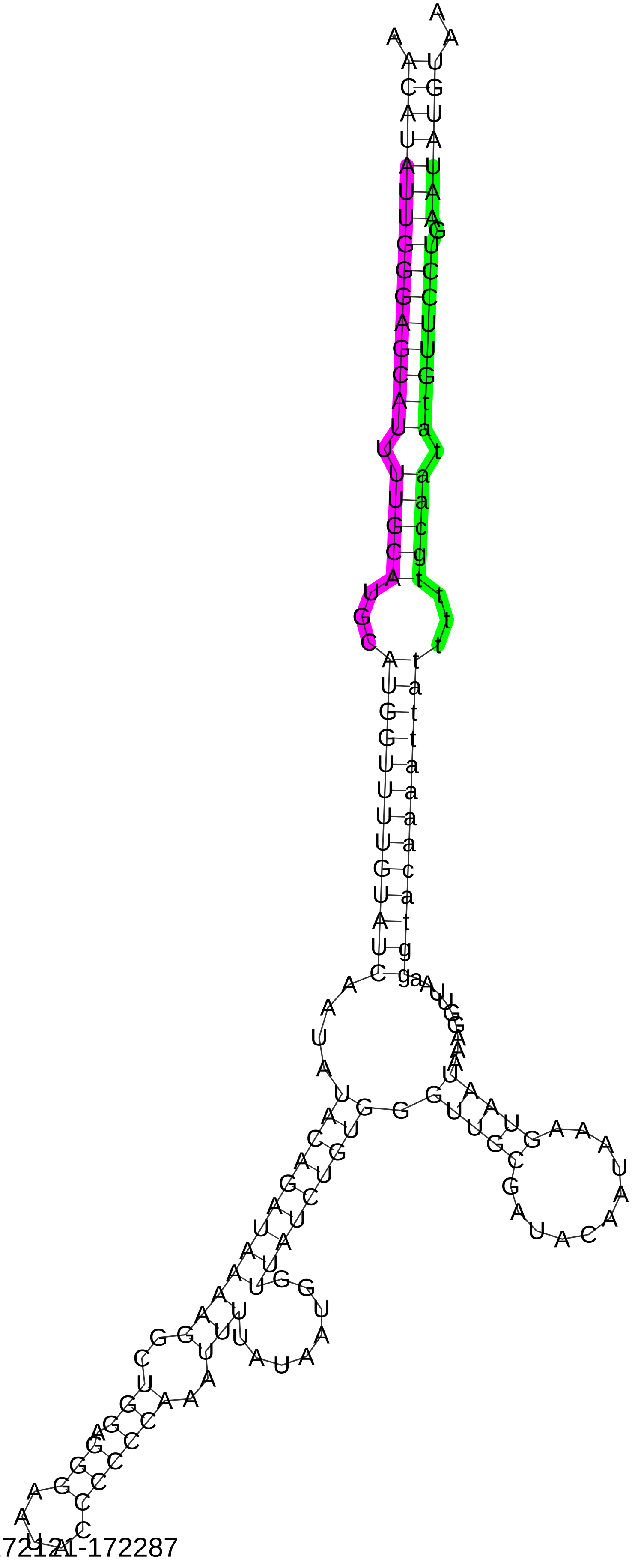

NW_006384929.1_172121-172287

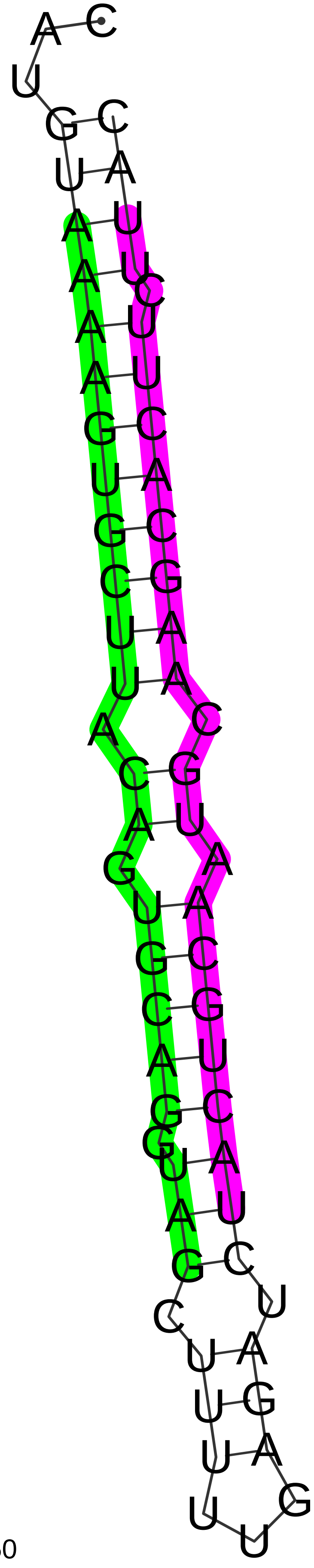

NW_006384929.1_443287-443350

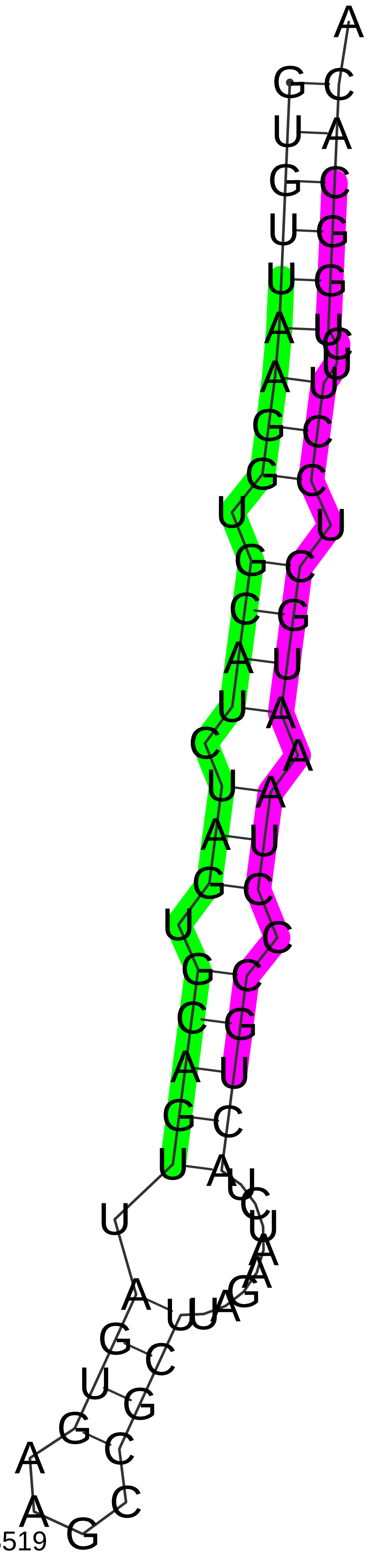

NW_006384929.1_443448-443519

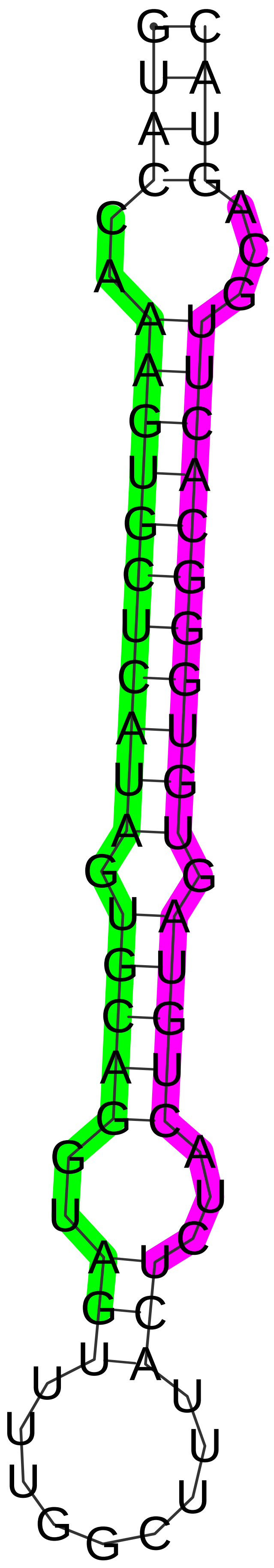

NW_006384929.1_443684-443750

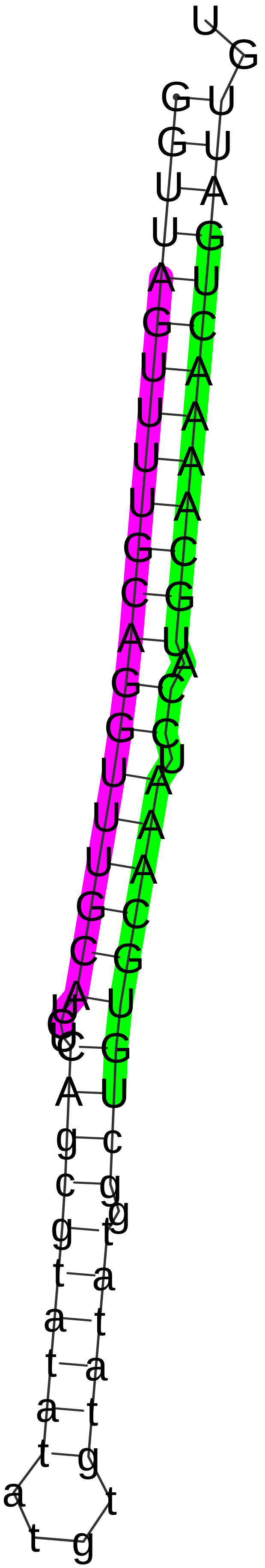

NW_006384929.1_443808-443881

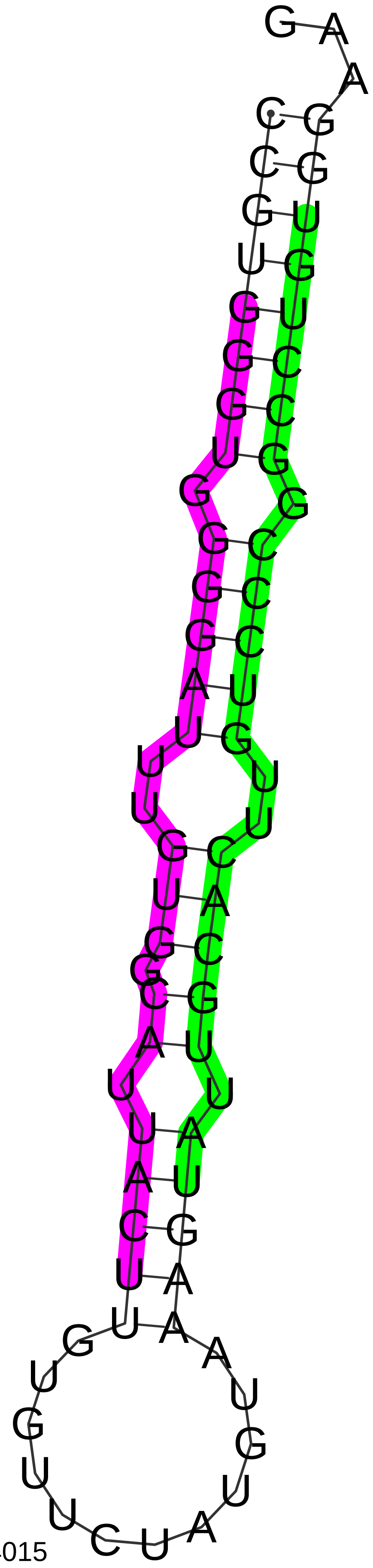

NW_006384929.1_443946-444015

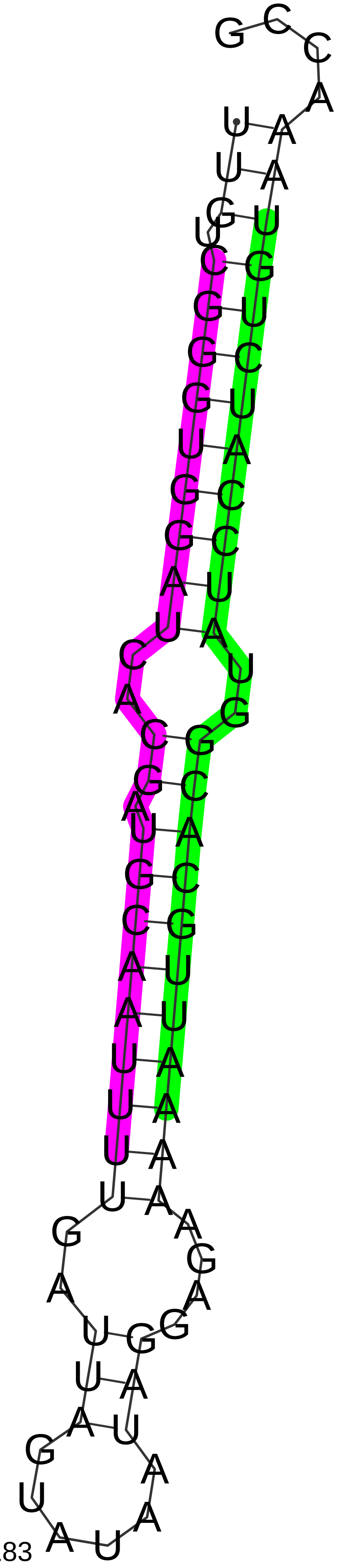

NW_006384929.1_444110-444183

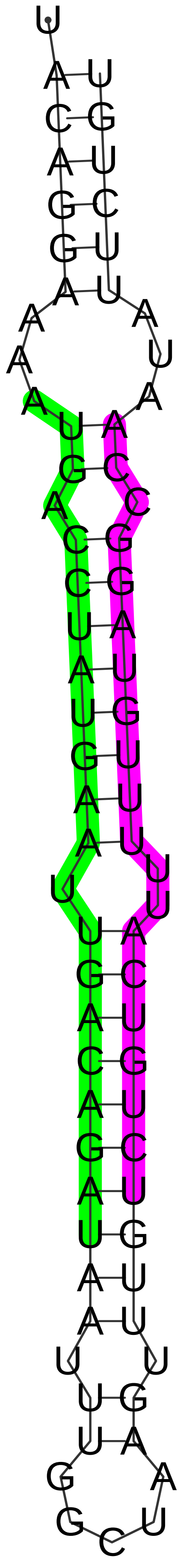

NW_006384939.1_216597-216671

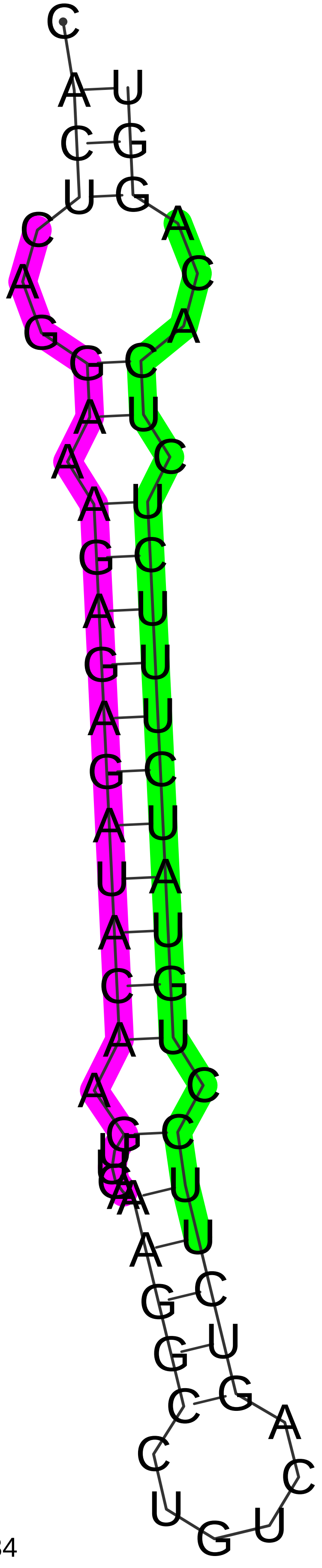

NW_006384939.1_245870-245934

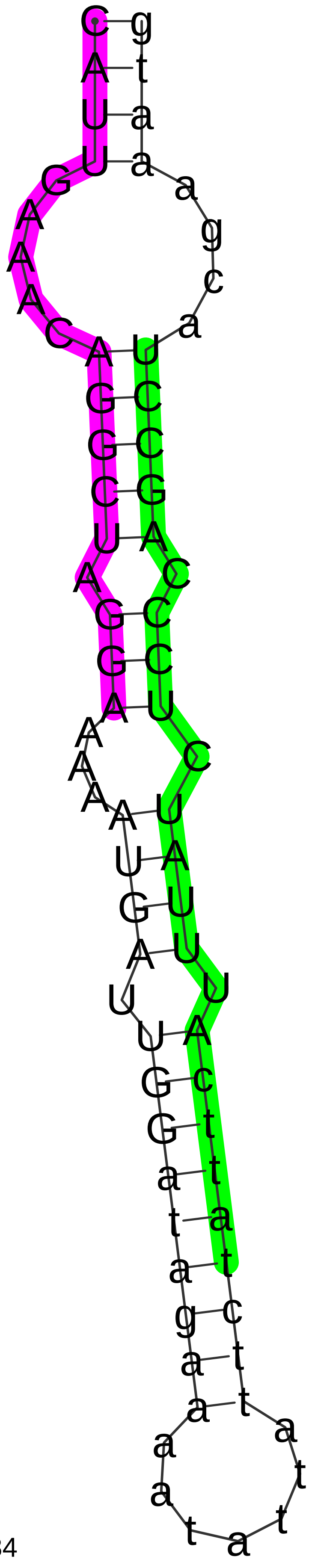

NW_006384939.1_295911-295984

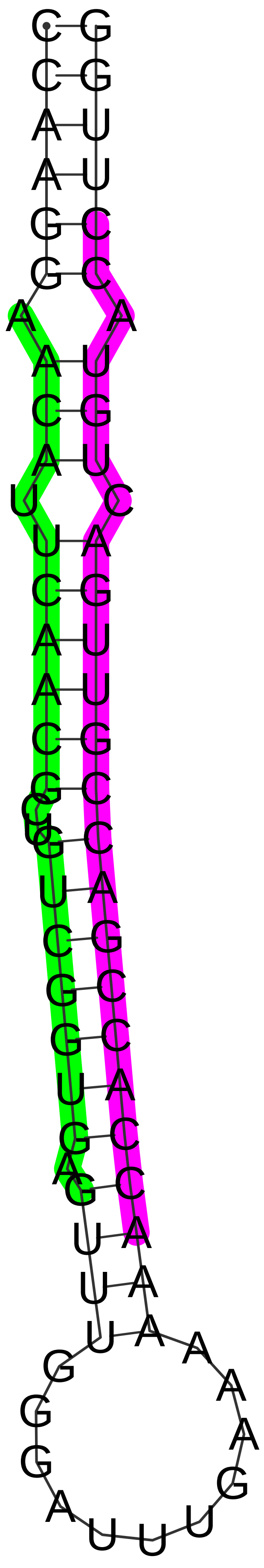

NW_006384957.1_375517-375586

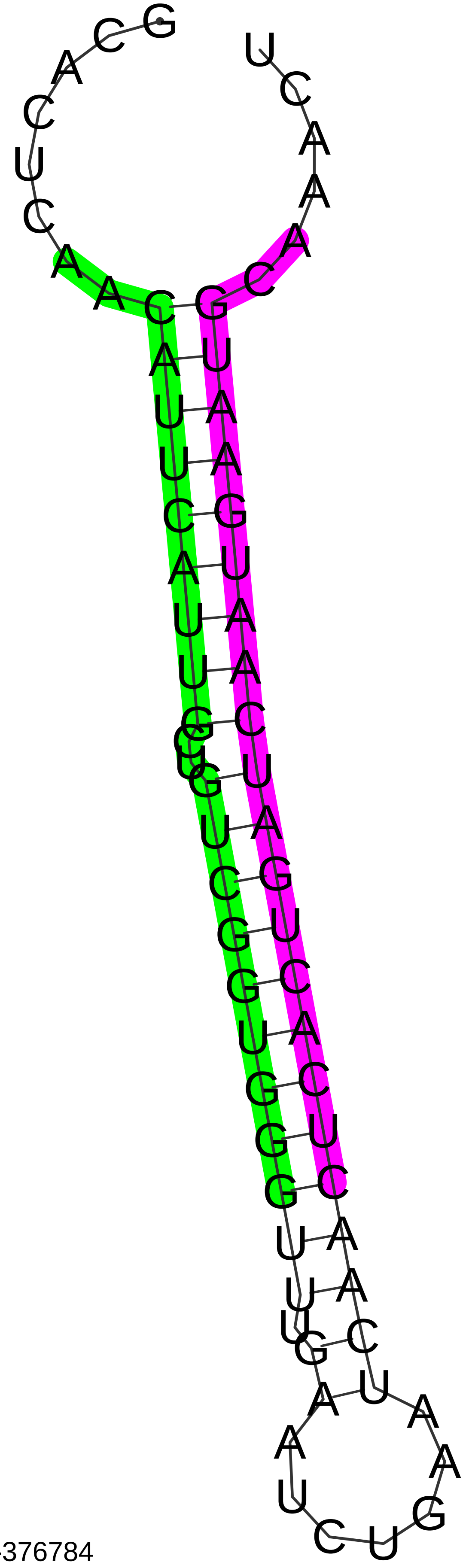

NW_006384957.1_376717-376784

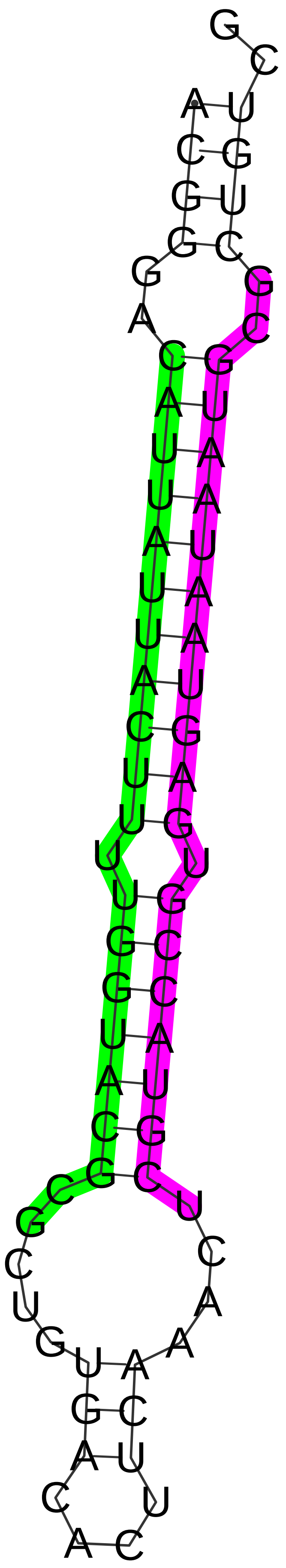

NW_006384974.1_327197-327267

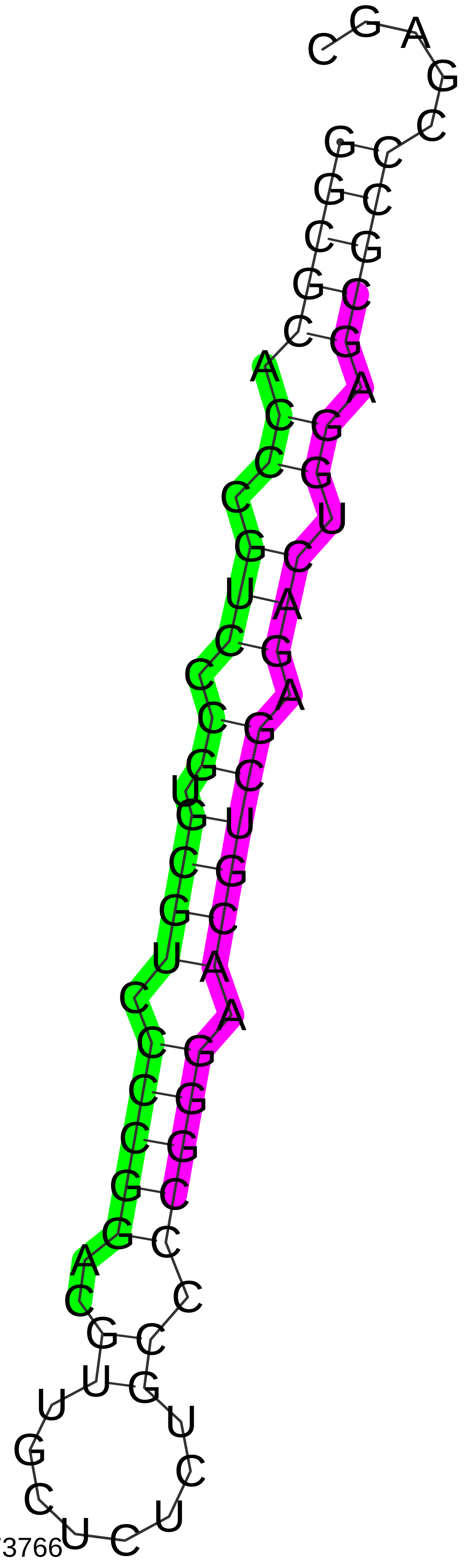

NW_006384993.1_73696-73766

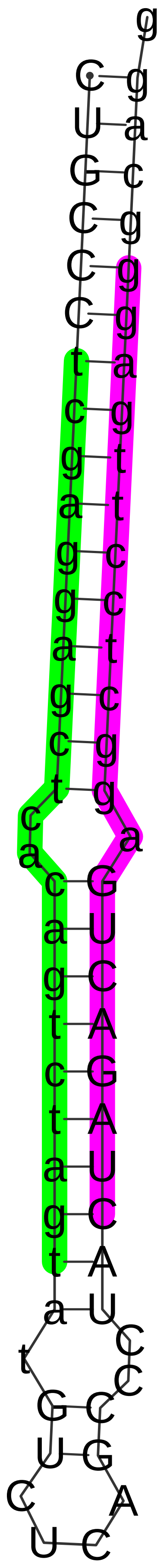

NW_006385004.1_411363-411429

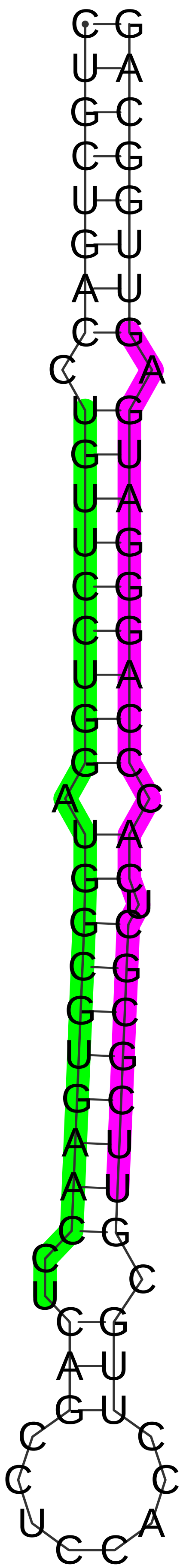

NW_006385011.1_171096-171171

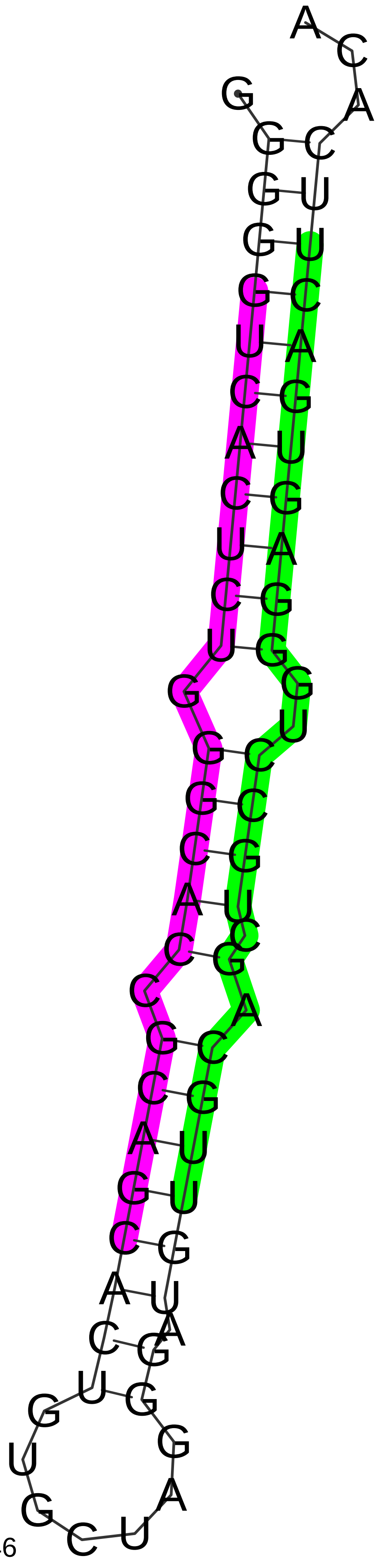

NW_006385057.1_80781-80846

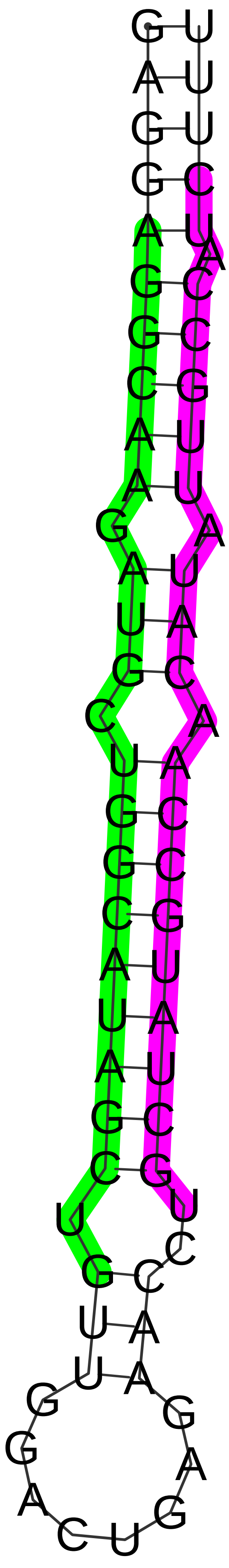

NW_006385061.1_474608-474673

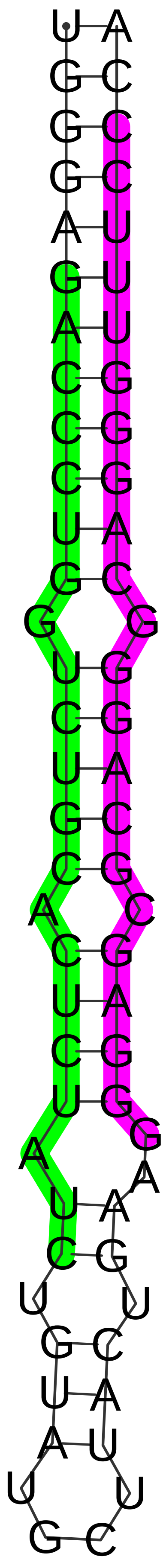

NW_006385065.1_472727-472791

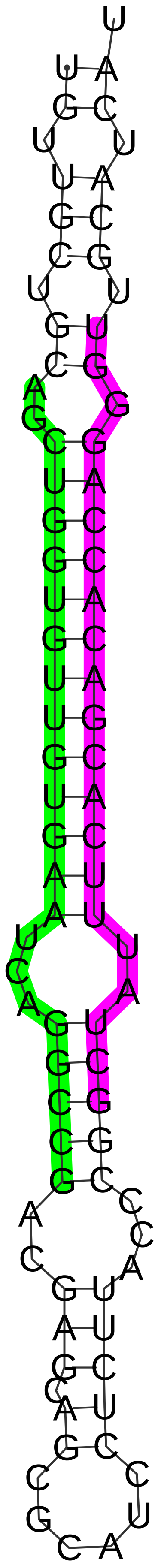

NW_006385073.1_372659-372743

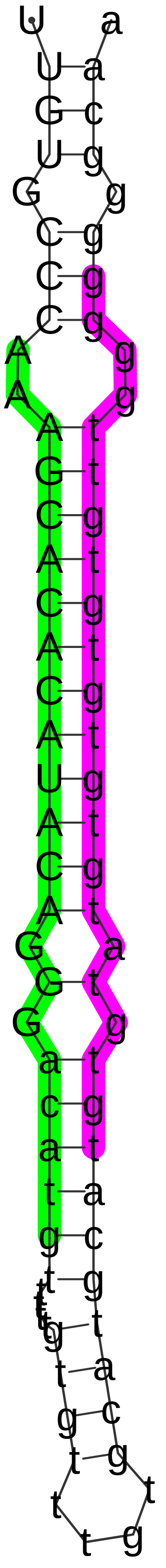

NW_006385100.1_325141-325218

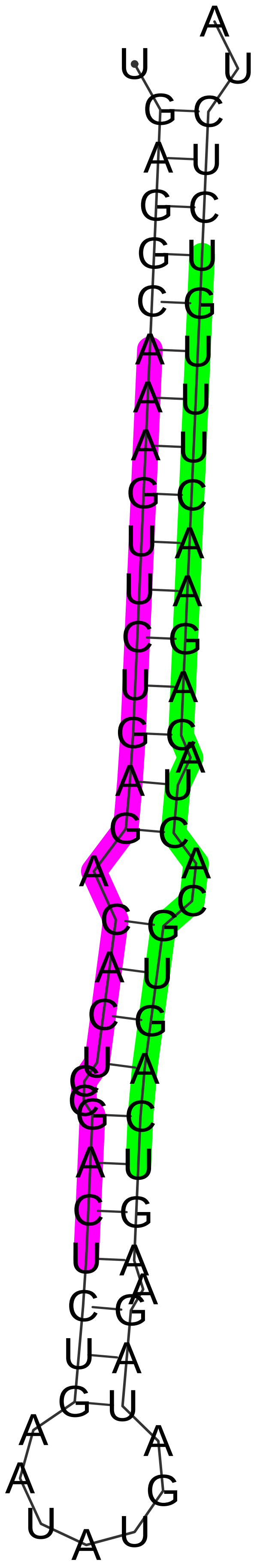

NW_006385126.1_177695-177765

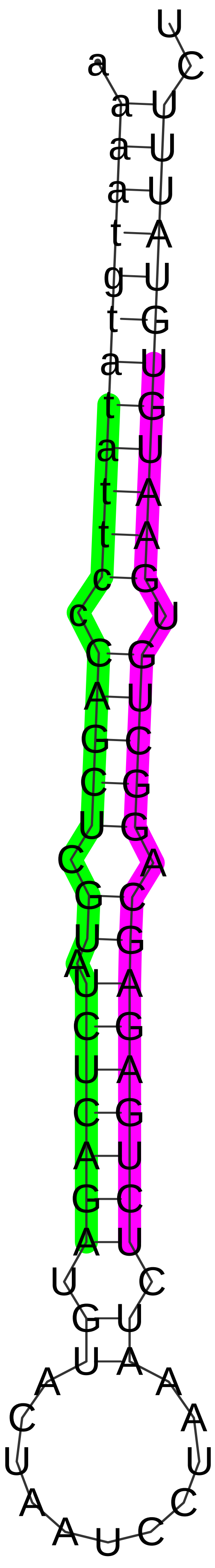

NW_006385169.1_124454-124530

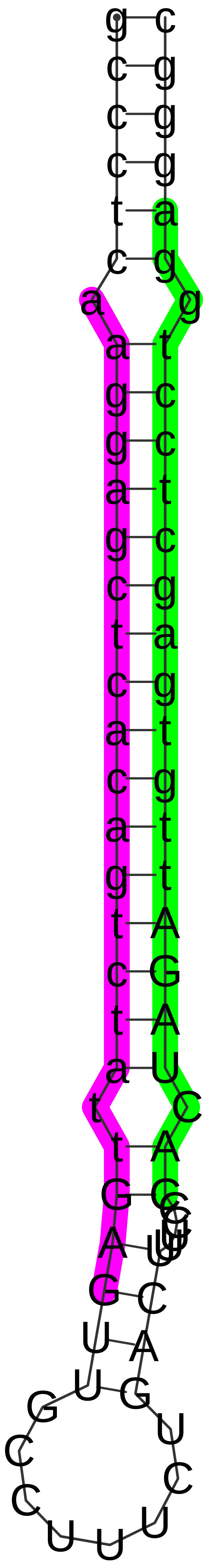

NW_006385178.1_312557-312628

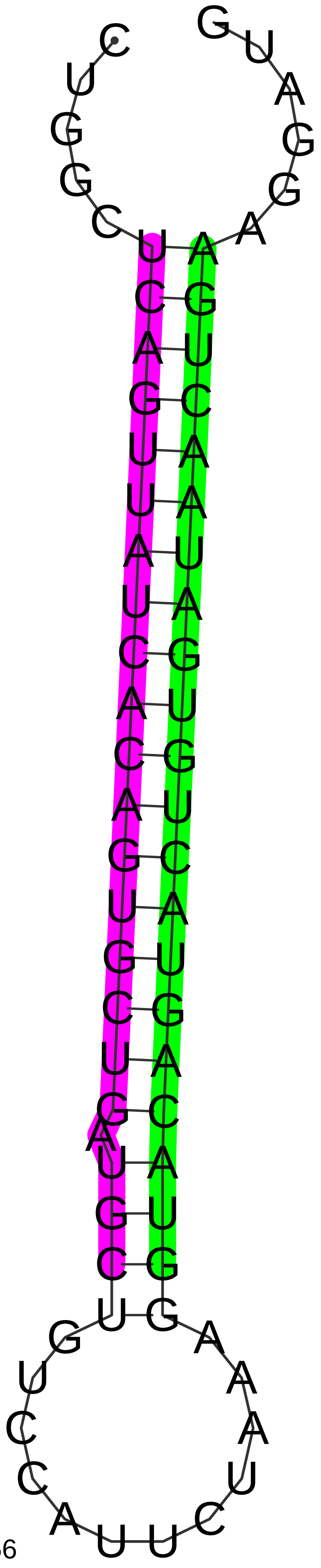

NW_006385197.1_293489-293556

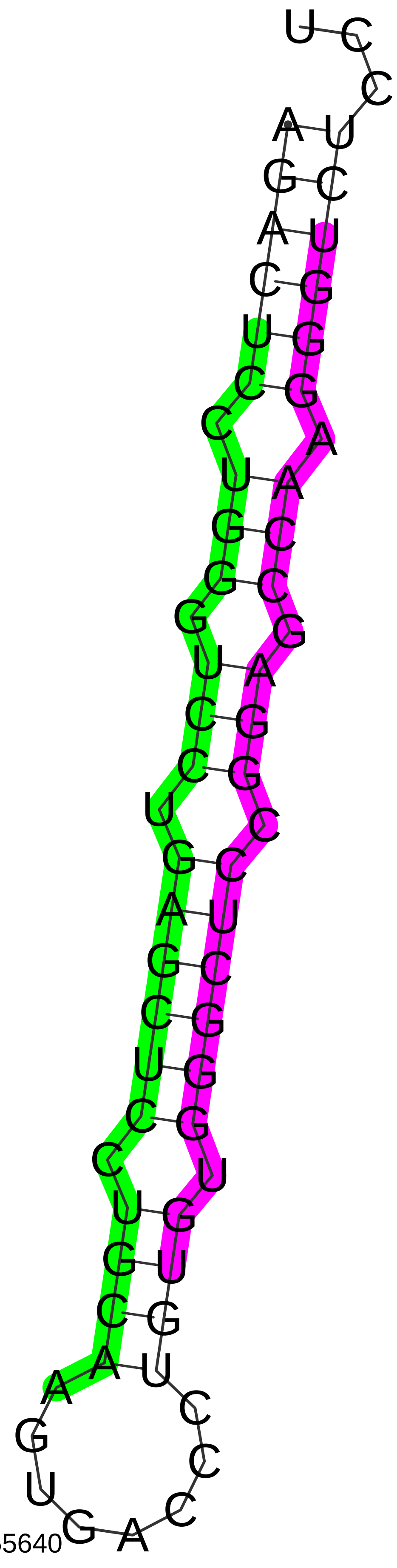

NW_006385210.1_155578-155640

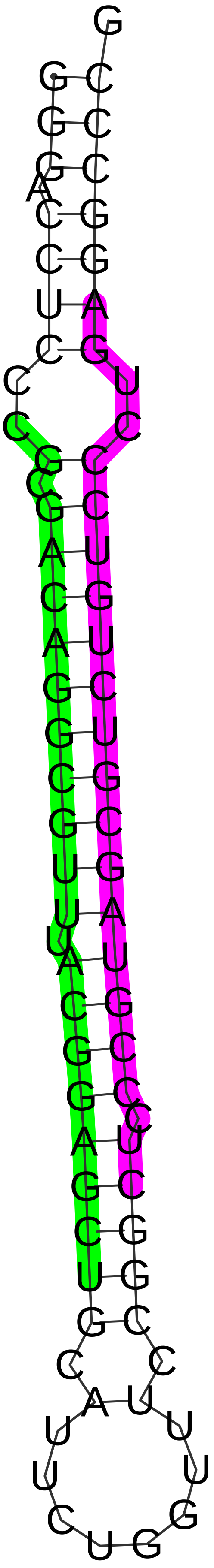

NW_006385222.1_379221-379295

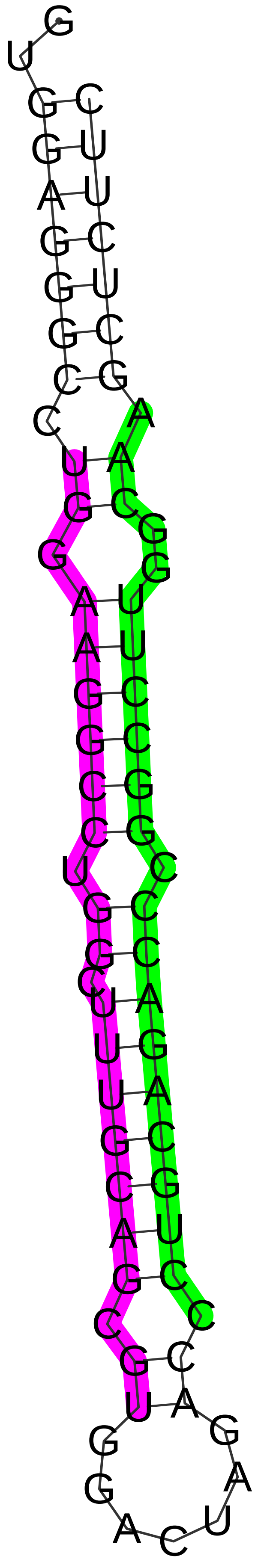

NW_006385227.1_19608-19678

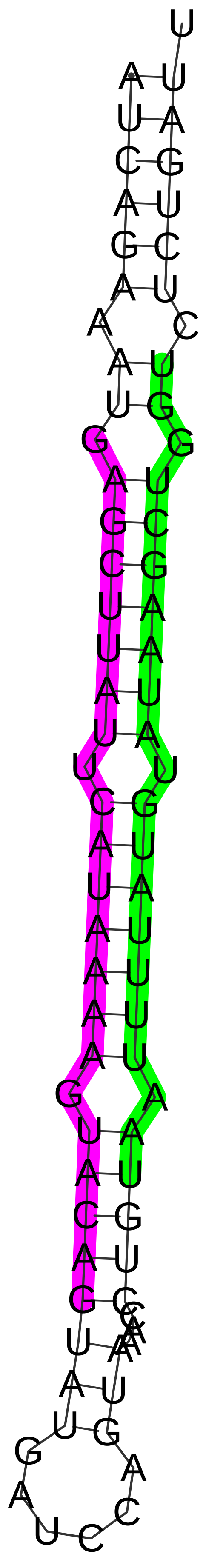

NW_006385227.1_299489-299566

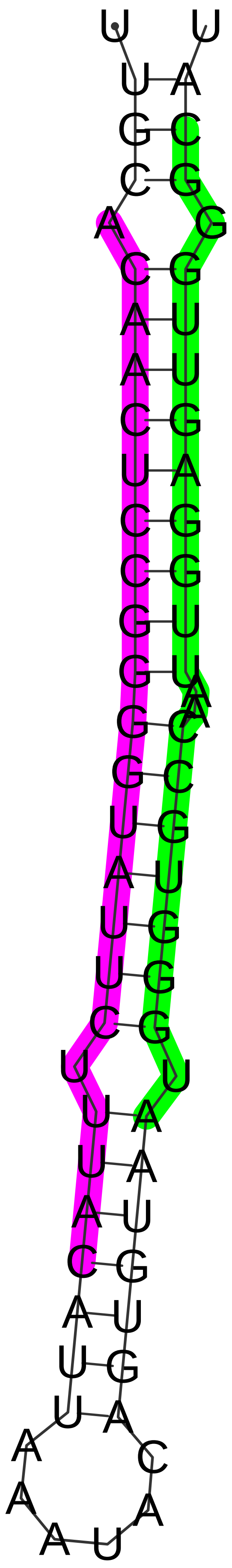

NW_006385228.1_176422-176487

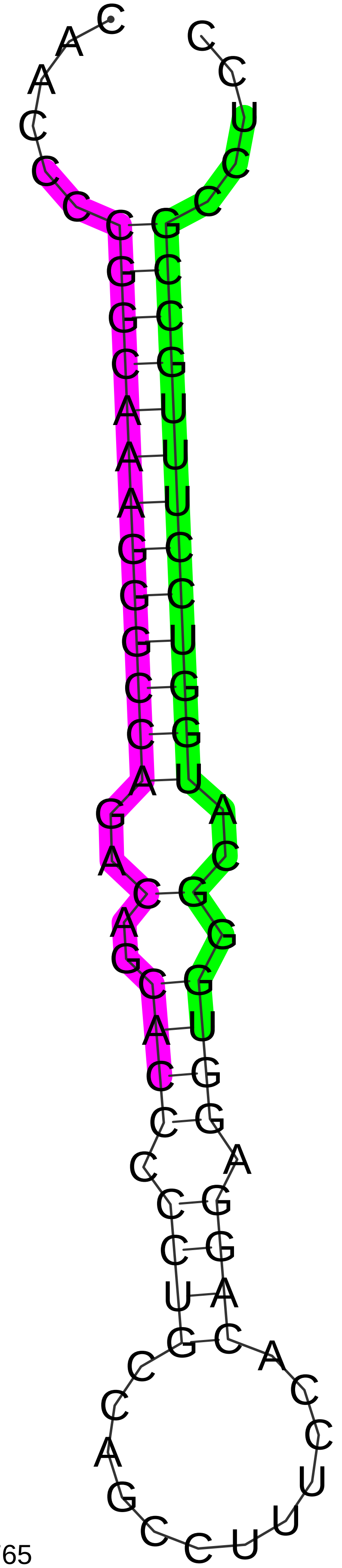


NW_006385241.1_347690-347765

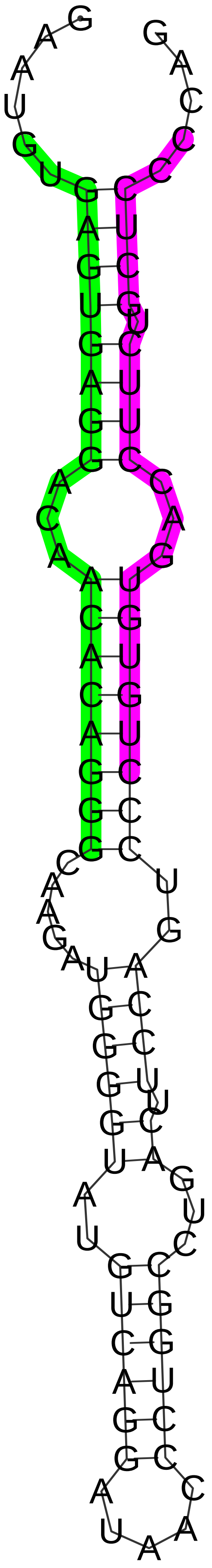

NW_006385327.1_316490-316581

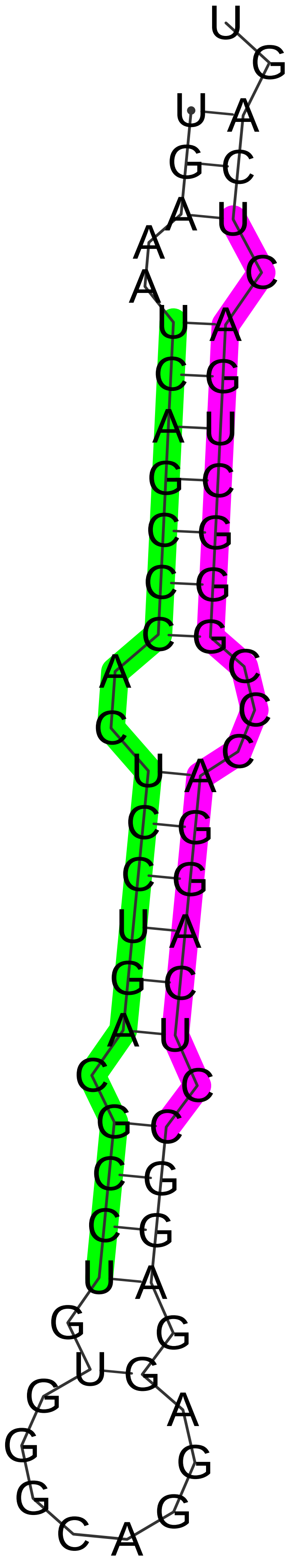

NW_006385330.1_54603-54666

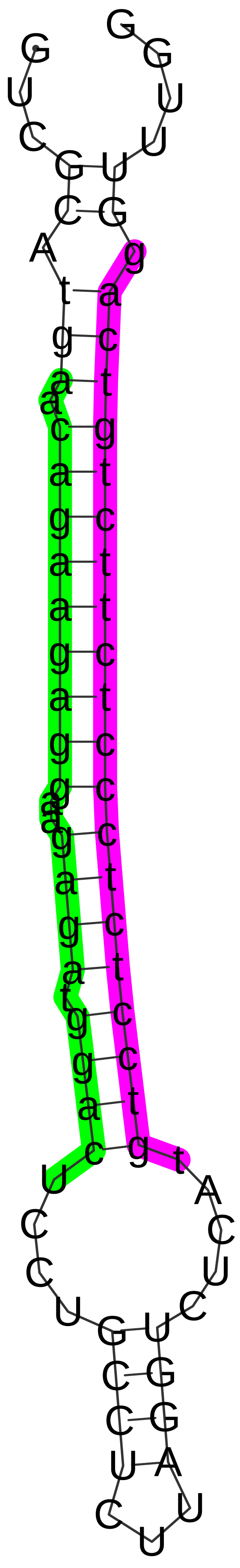

NW_006385363.1_164587-164663

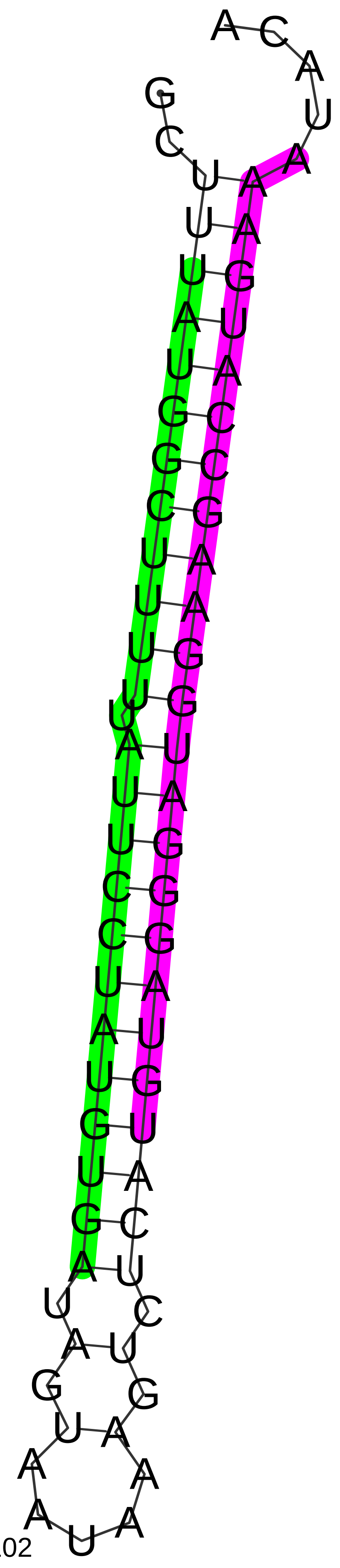

NW_006385438.1_273034-273102

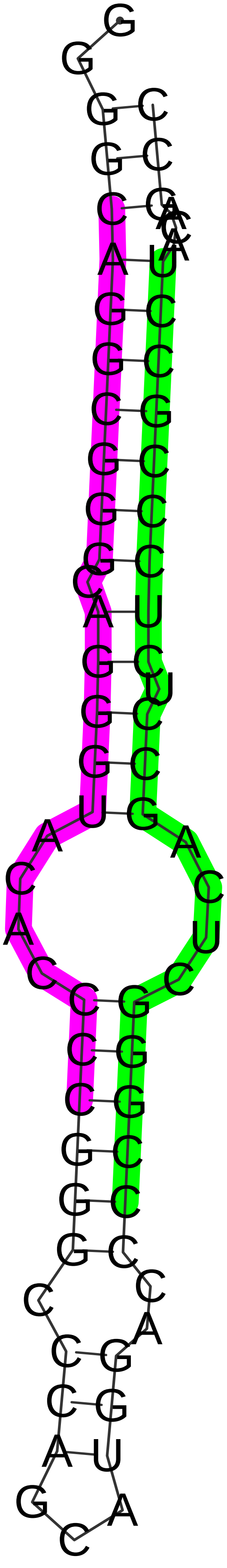

NW_006385452.1_313272-313340

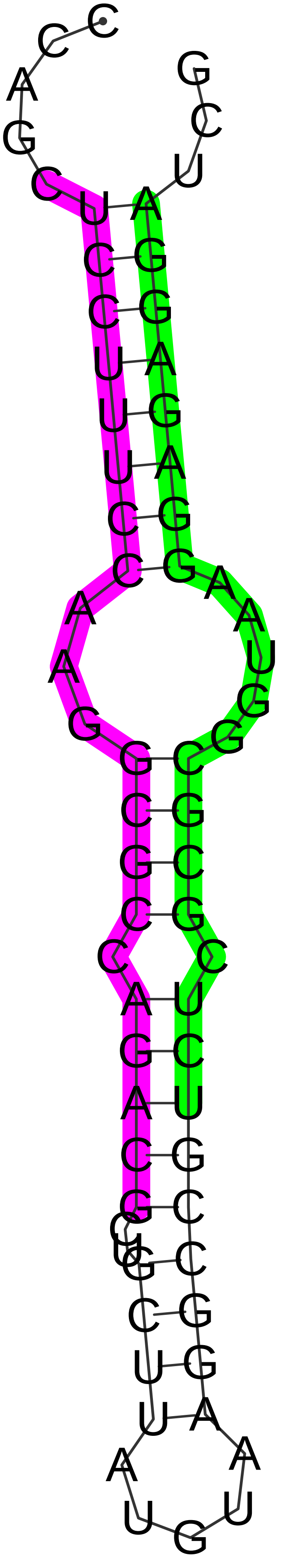

NW_006385454.1_217799-217865

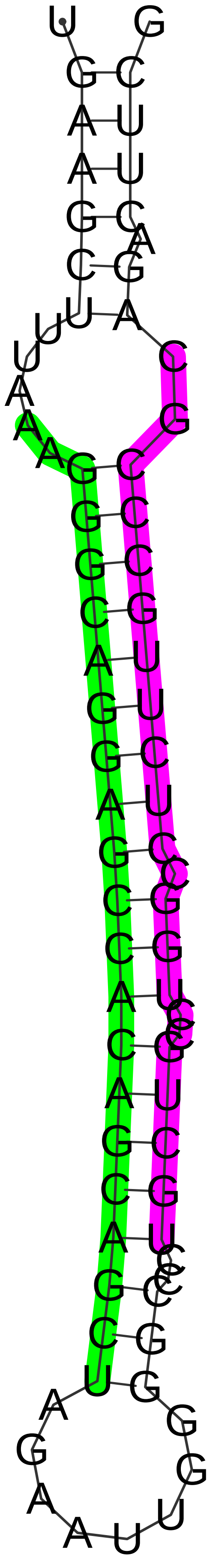

NW_006385495.1_54469-54543

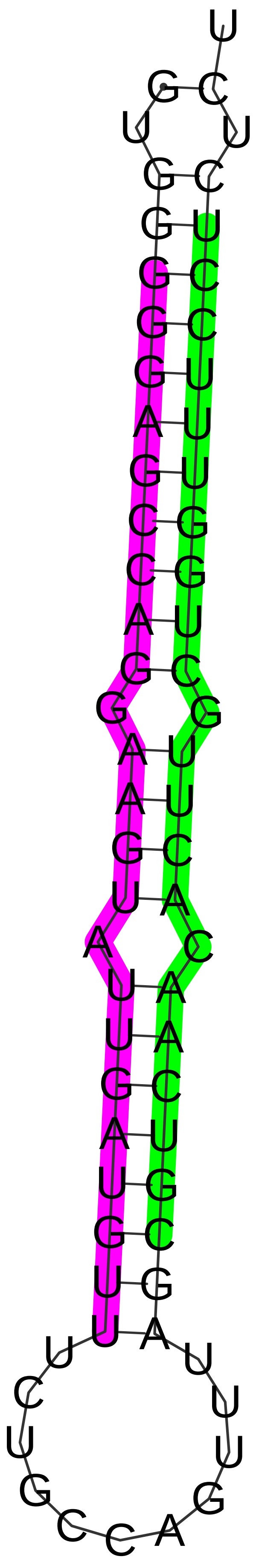

NW_006385515.1_282318-282383

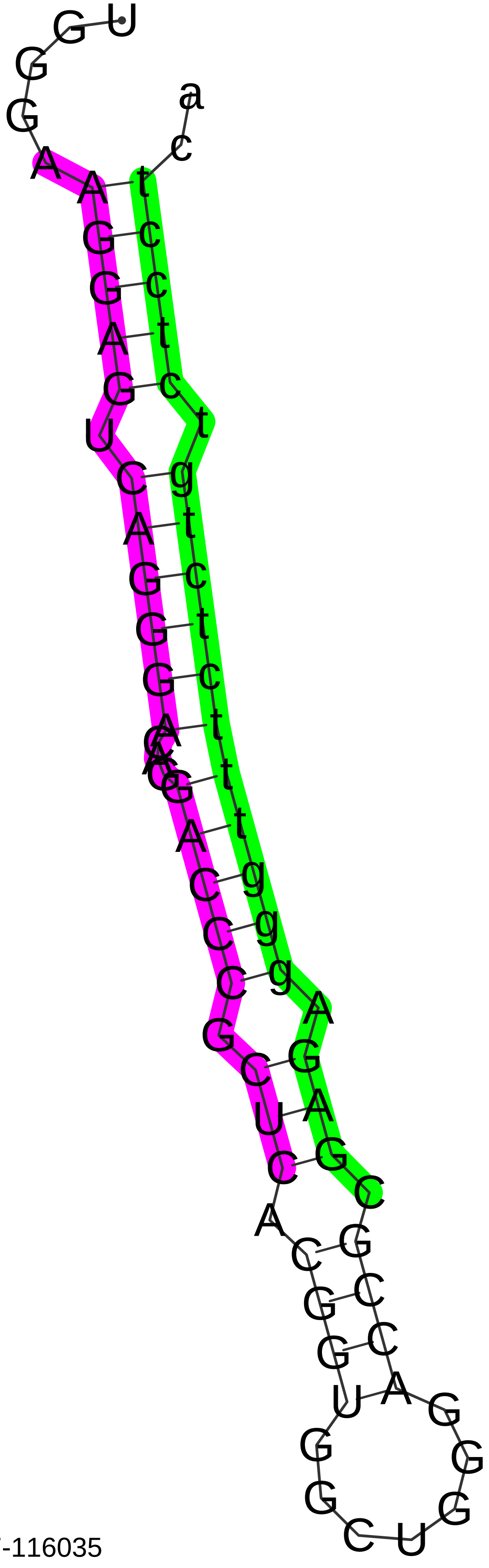

NW_006385565.1_115967-116035

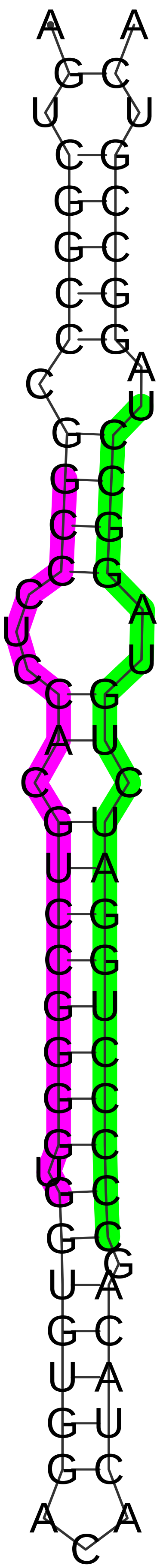

NW_006385575.1_371090-371162

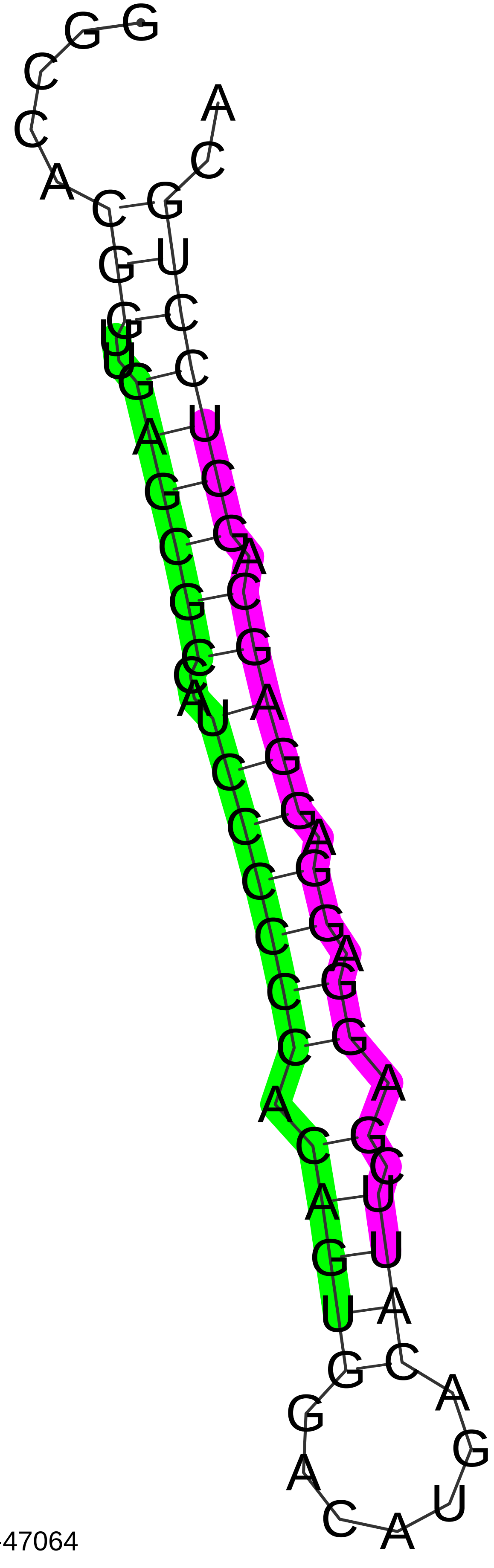

NW_006385575.1_46999-47064

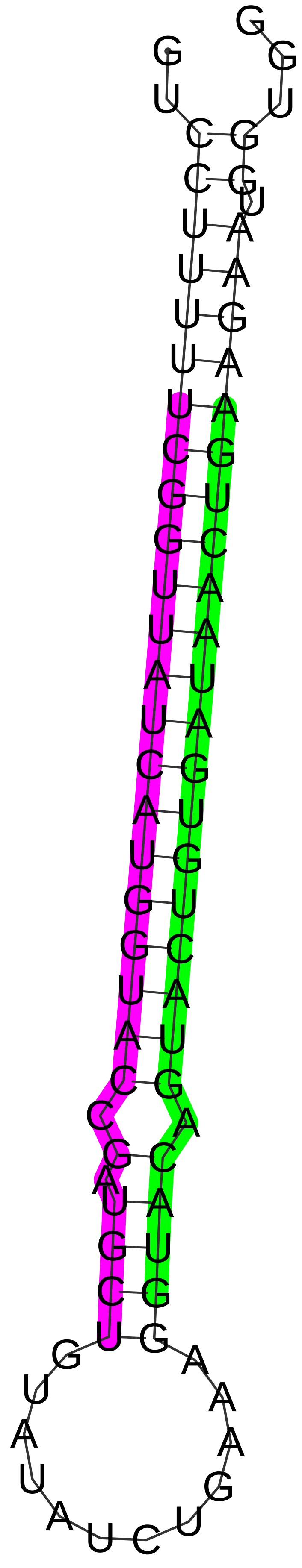

NW_006385576.1_15772-15846

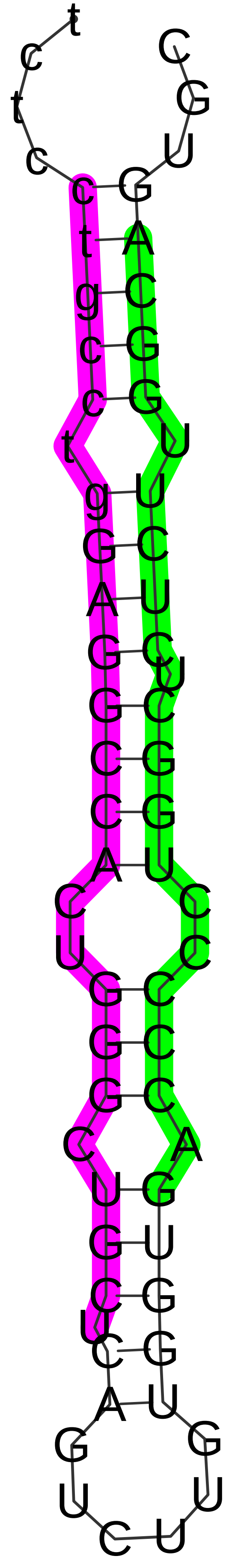

NW_006385581.1_255663-255727

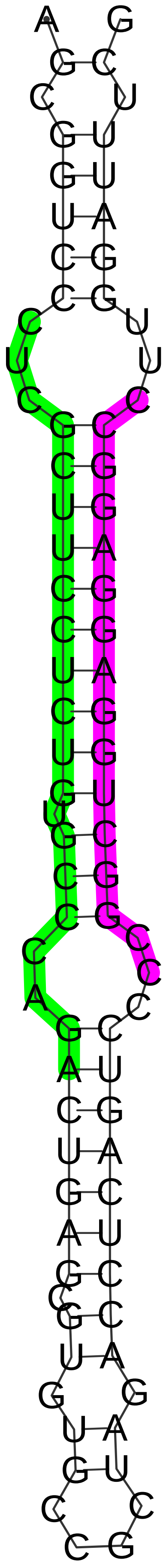

NW_006385624.1_202914-202996

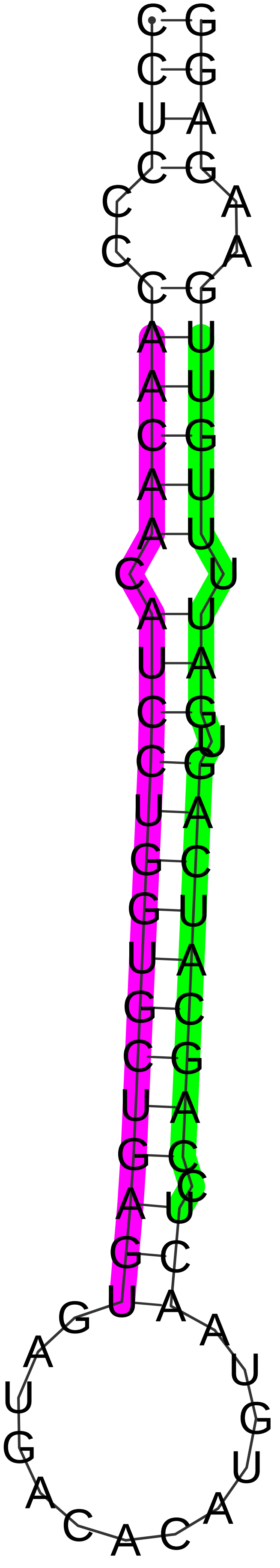

NW_006385655.1_333052-333122

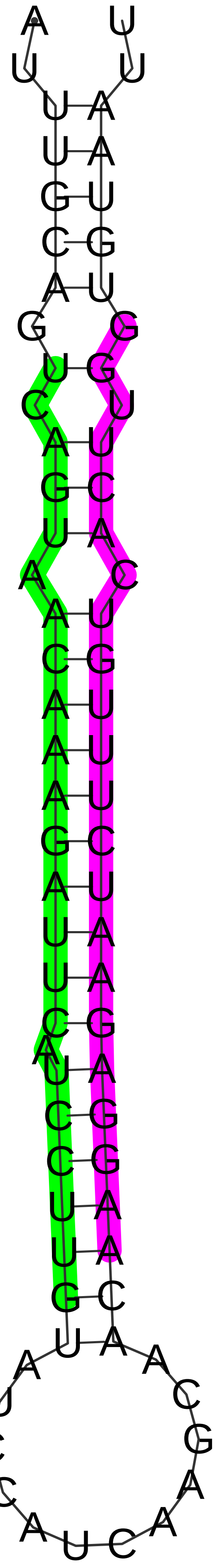

NW_006385724.1_55587-55661

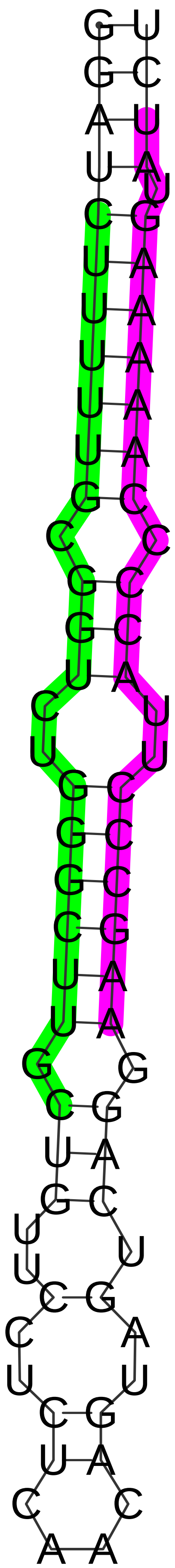

NW_006385739.1_154824-154895

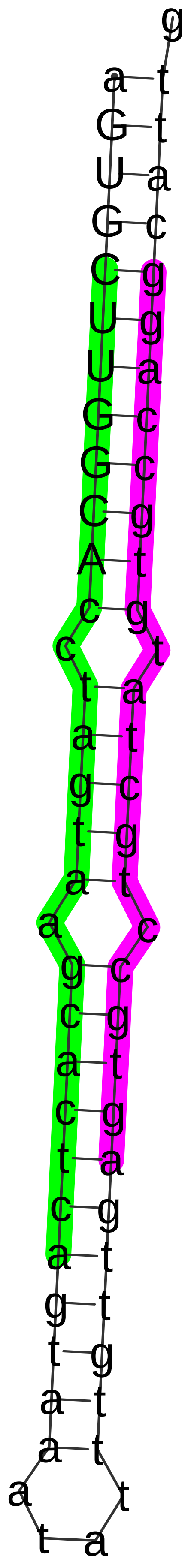

NW_006385755.1_37533-37597

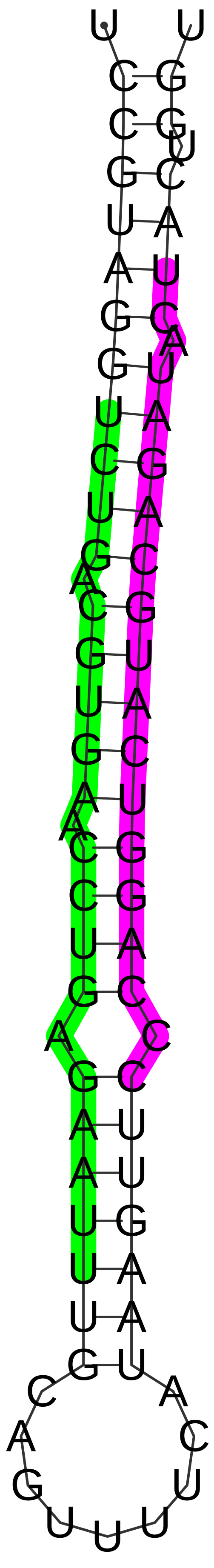

NW_006385787.1_180720-180790

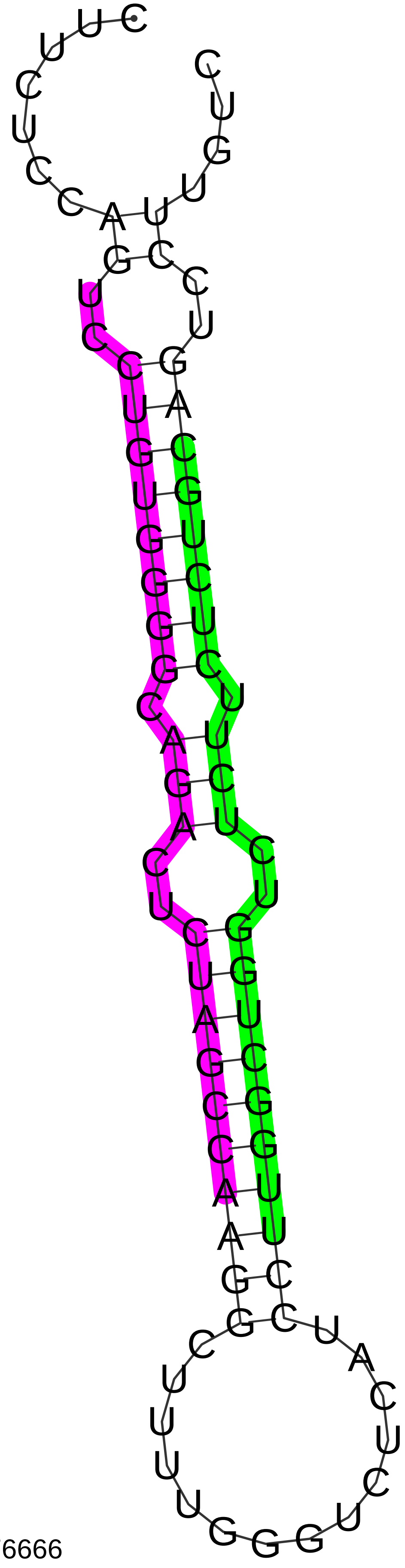

NW_006385832.1_176586-176666

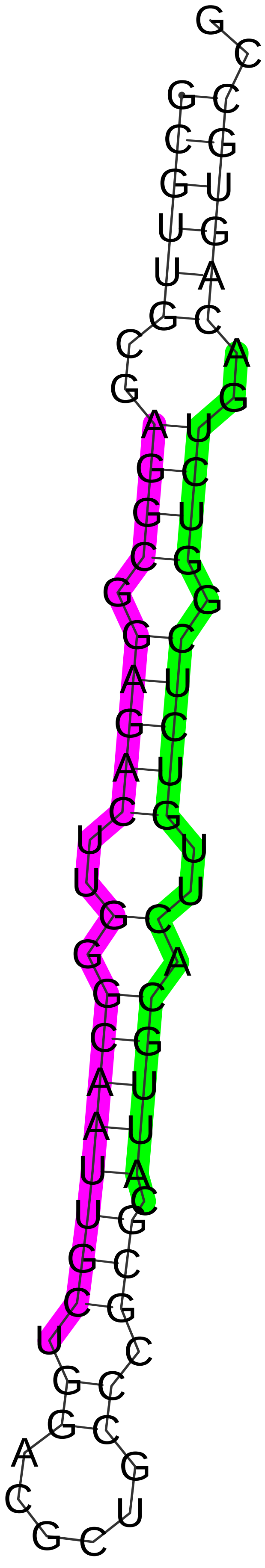

NW_006385874.1_351581-351655

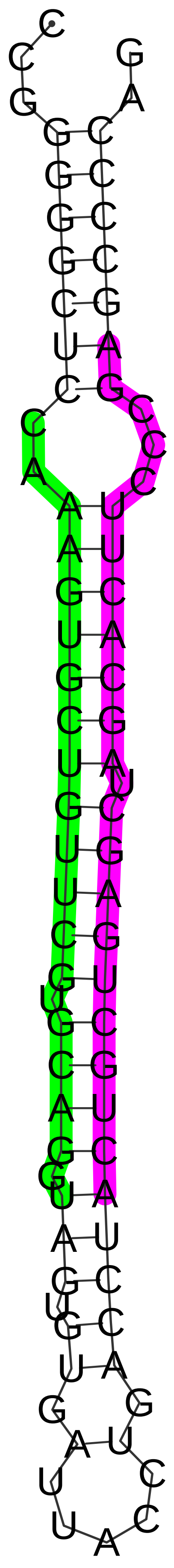

NW_006385874.1_351786-351864

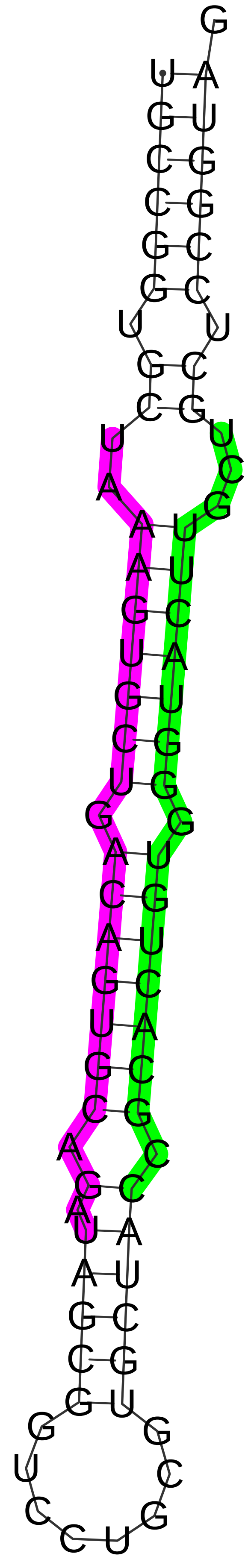

NW_006385874.1_352009-352085

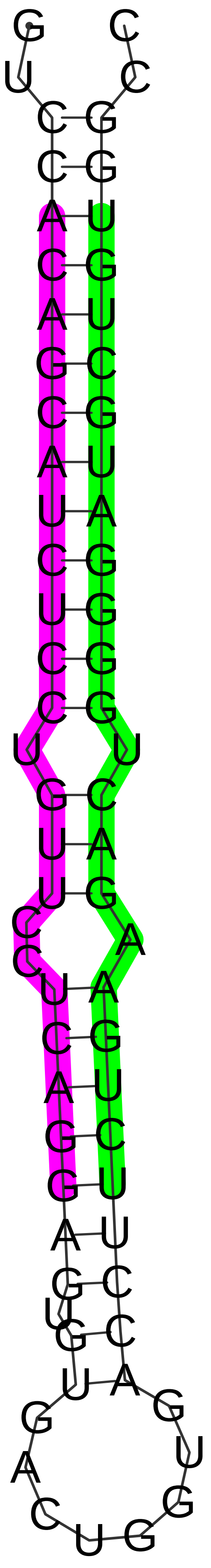

NW_006385933.1_71806-71873

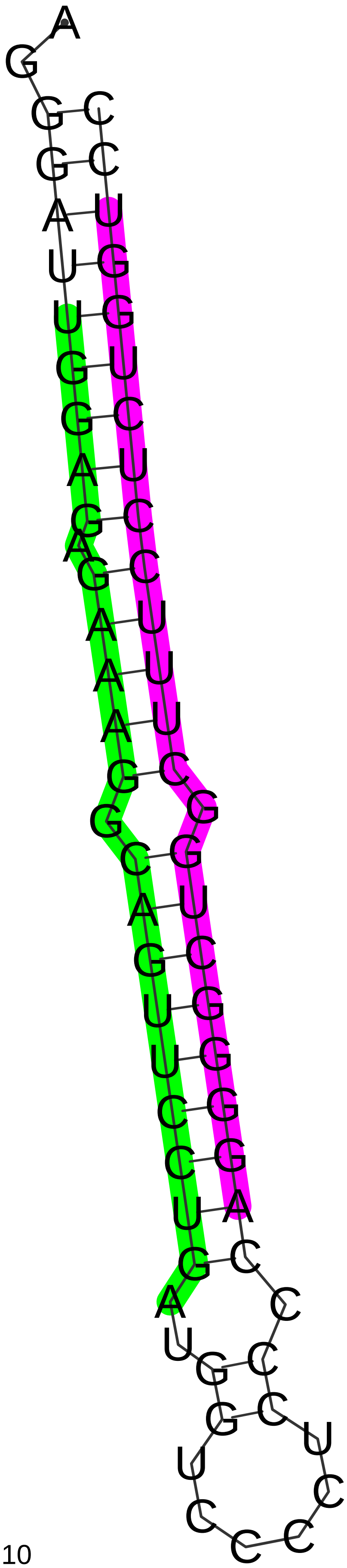

NW_006386020.1_203147-203210

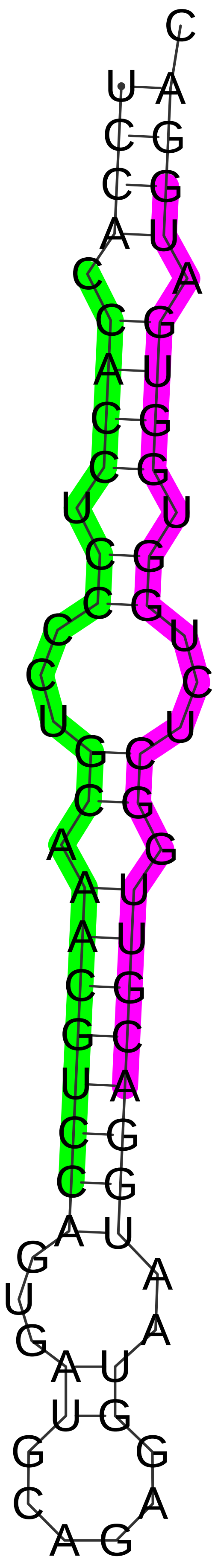

NW_006386020.1_243597-243664

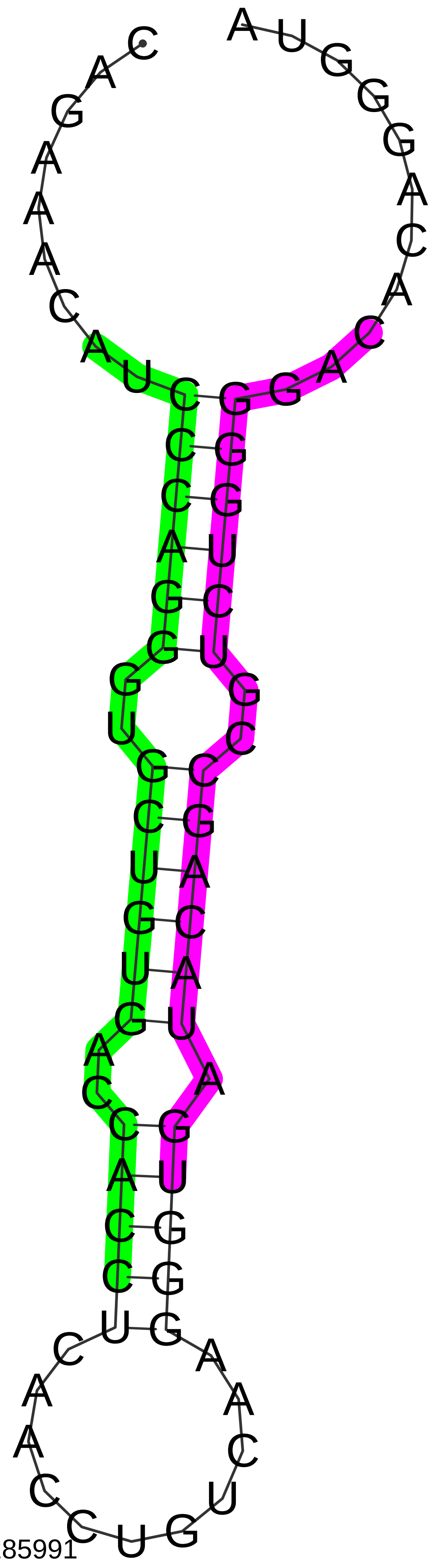

NW_006386020.1_285920-285991

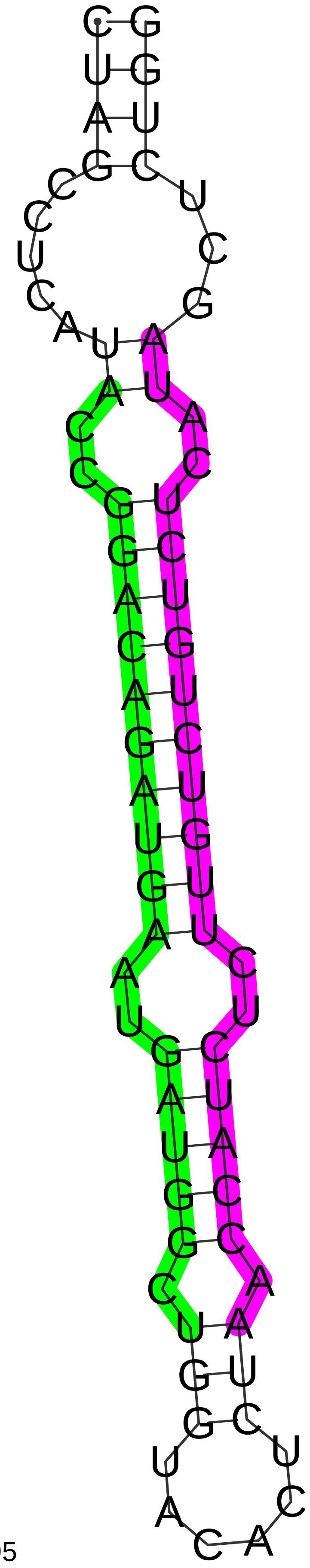

NW_006386061.1_185324-185395

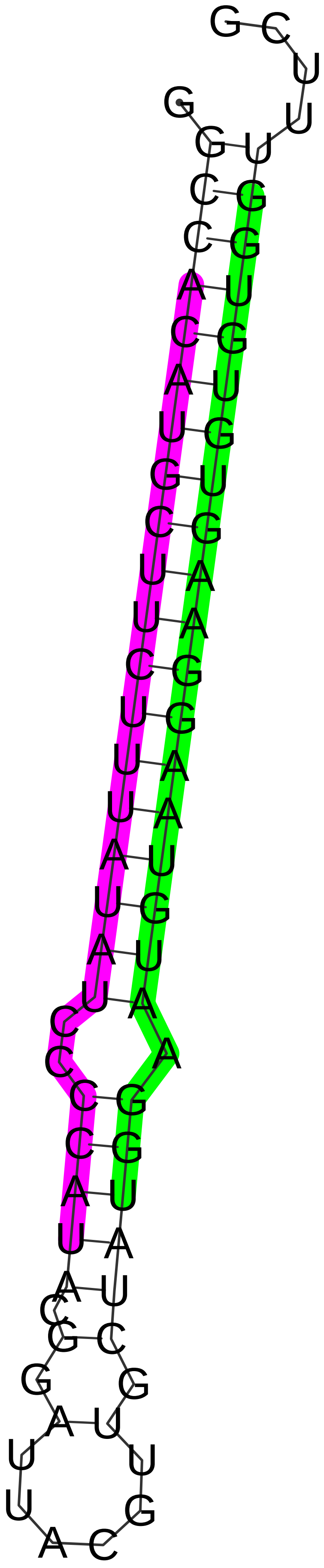

NW_006386069.1_77507-77575

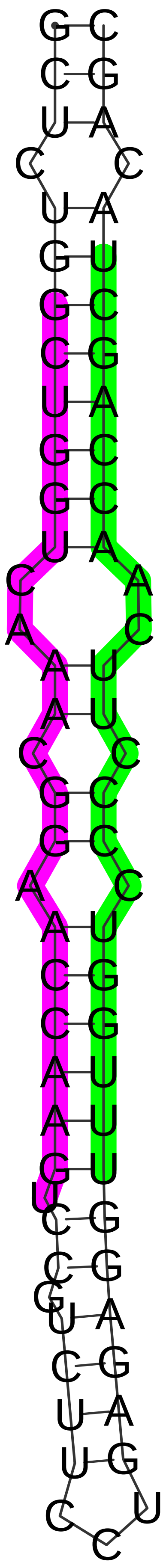

NW_006386069.1_81929-81997

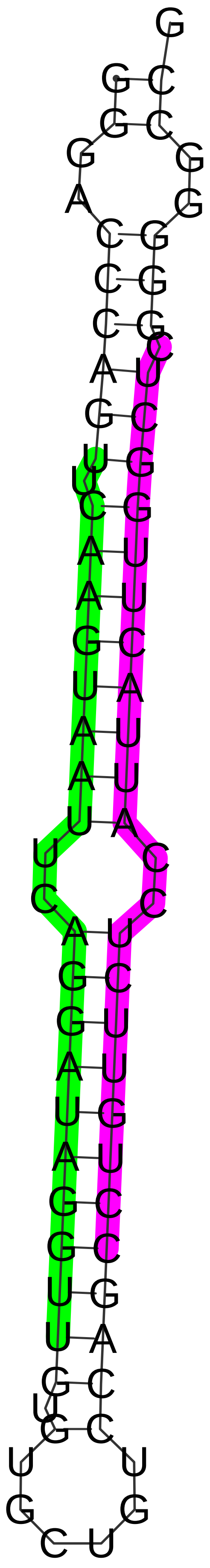

NW_006386073.1_248918-248991

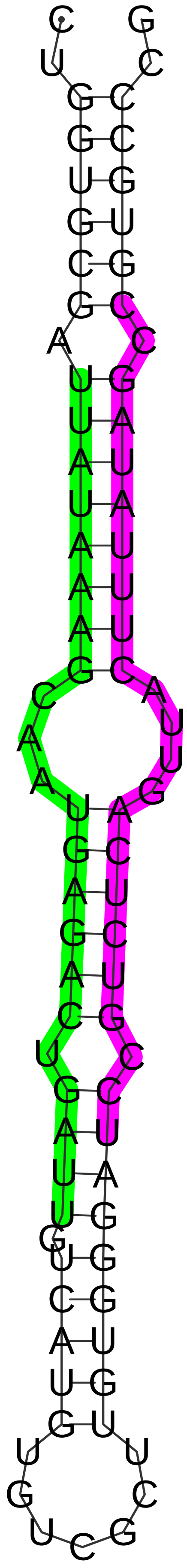


NW_006386076.1_4132-4212

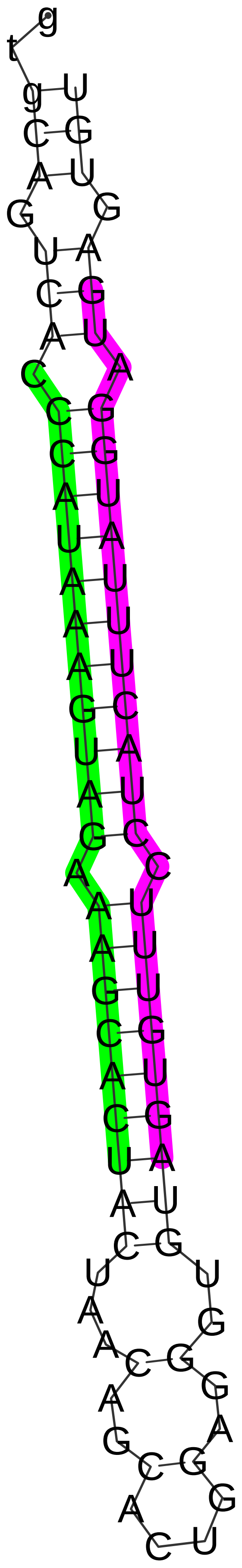

NW_006386120.1_52198-52274

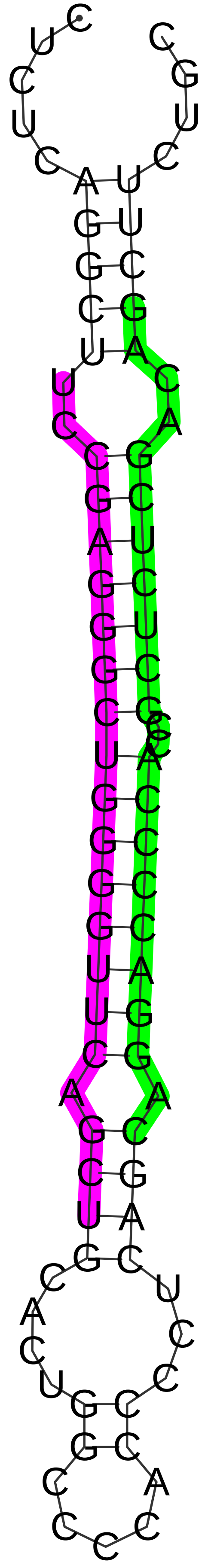

NW_006386192.1_95548-95628

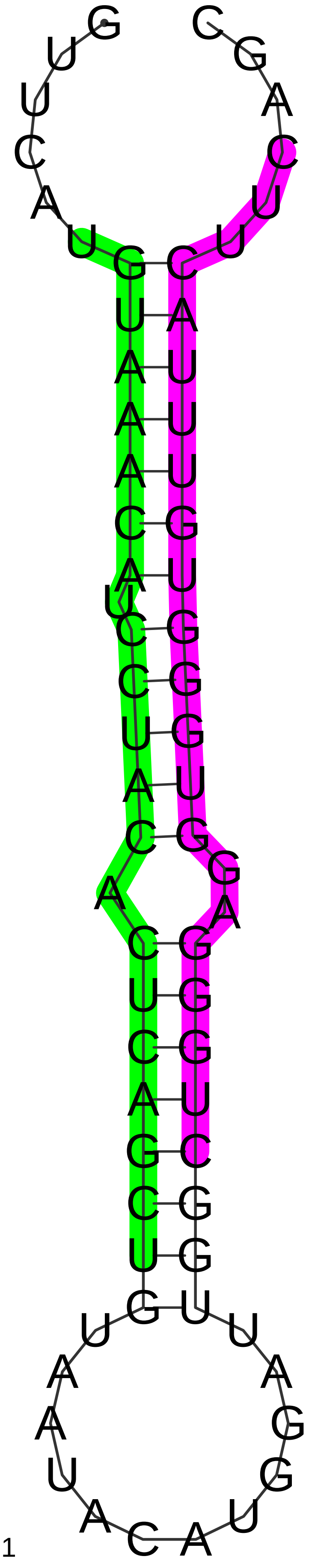

NW_006386213.1_119894-119961

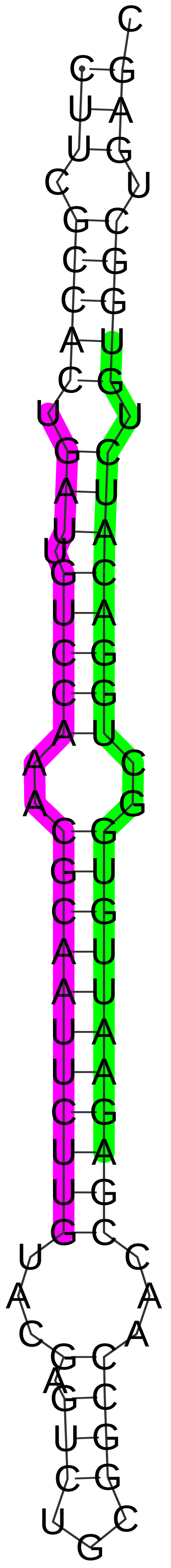

NW_006386229.1_83499-83580

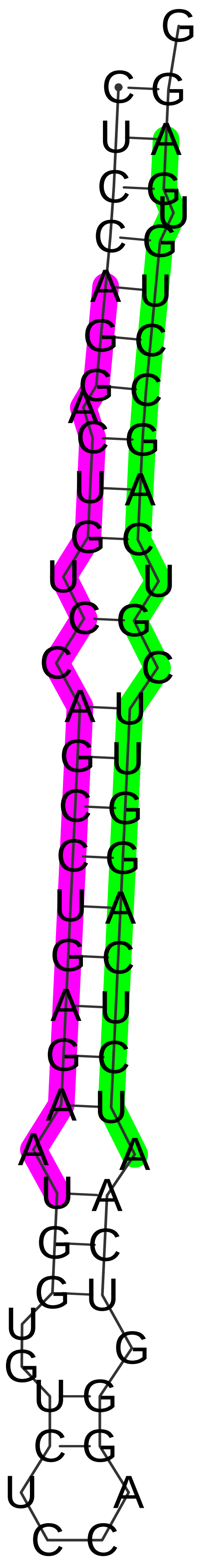

NW_006386233.1_199512-199577

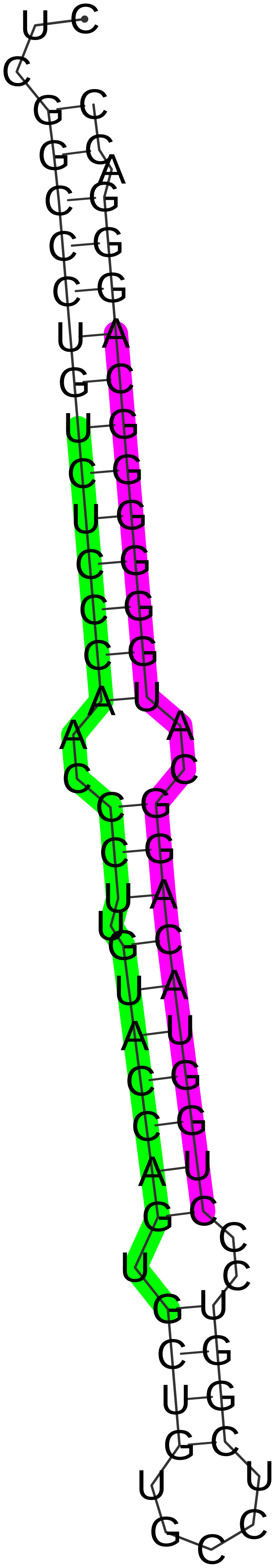

NW_006386261.1_121099-121171

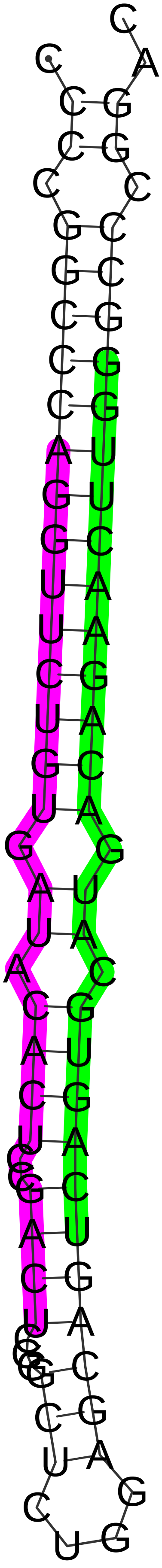

NW_006386302.1_60526-60601

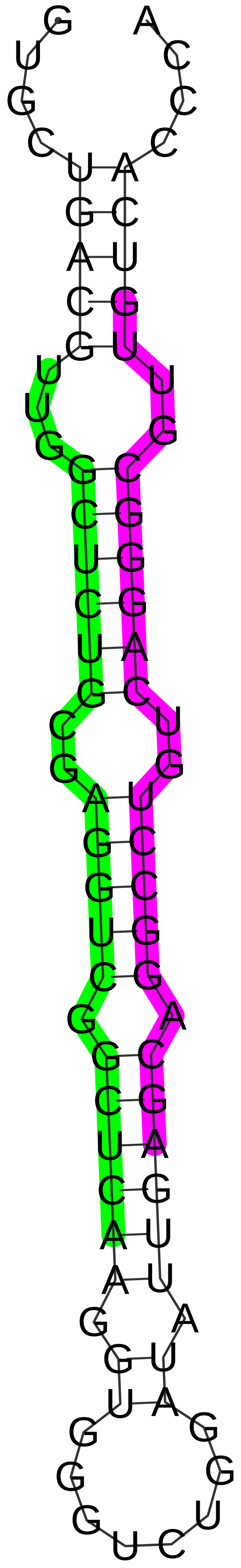

NW_006386311.1_82706-82782

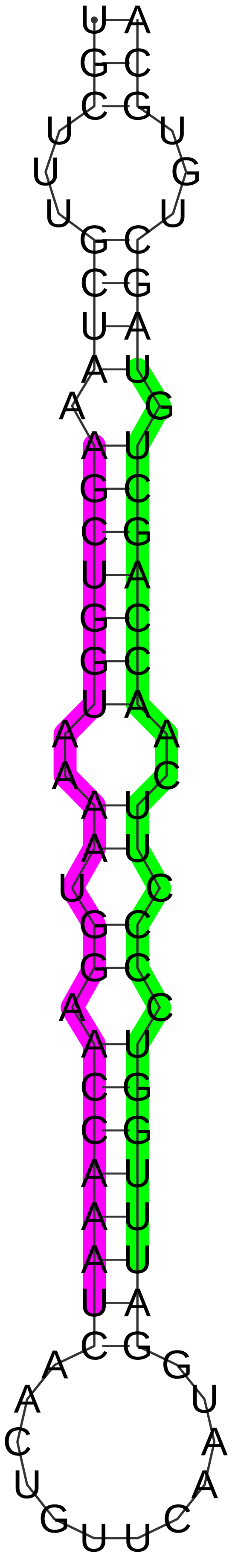

NW_006386338.1_147416-147495

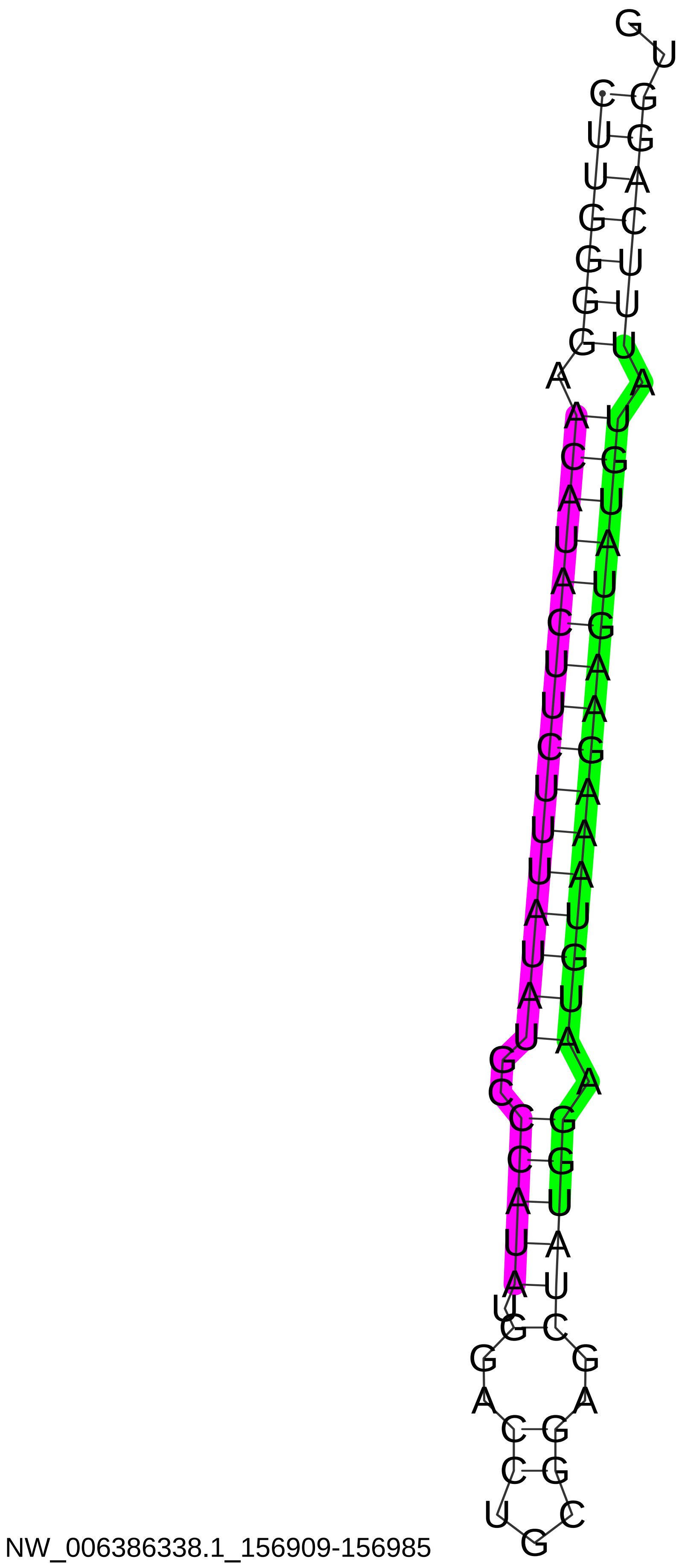

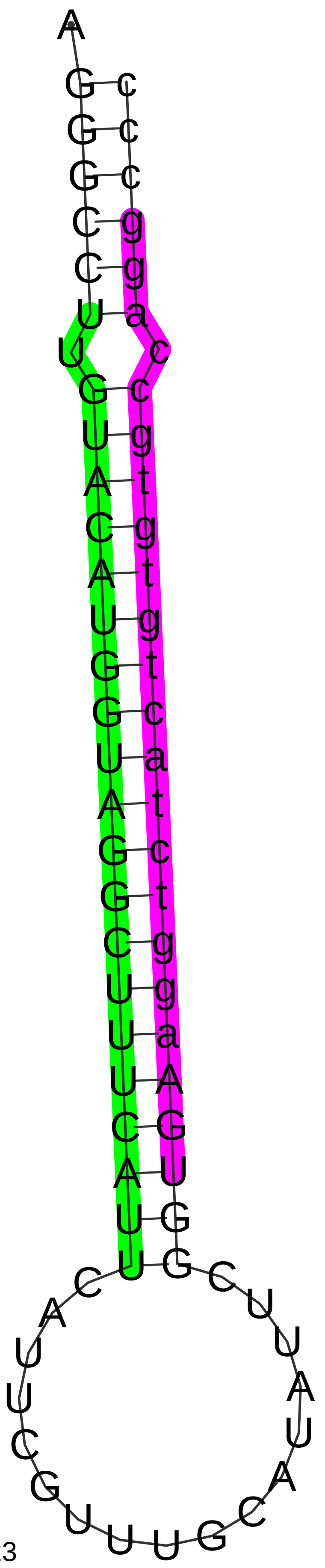

NW_006386394.1_141452-141523

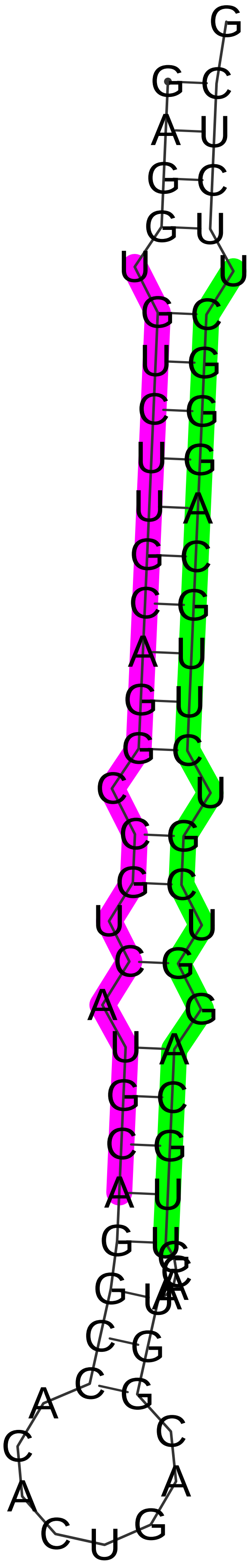

NW_006386394.1_151337-151407

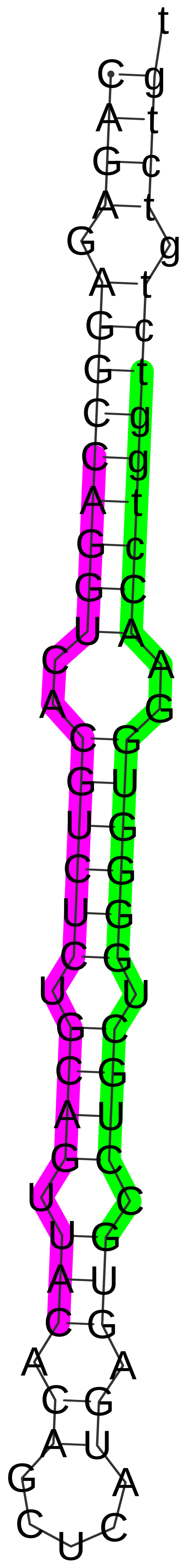

NW_006386394.1_181200-181273

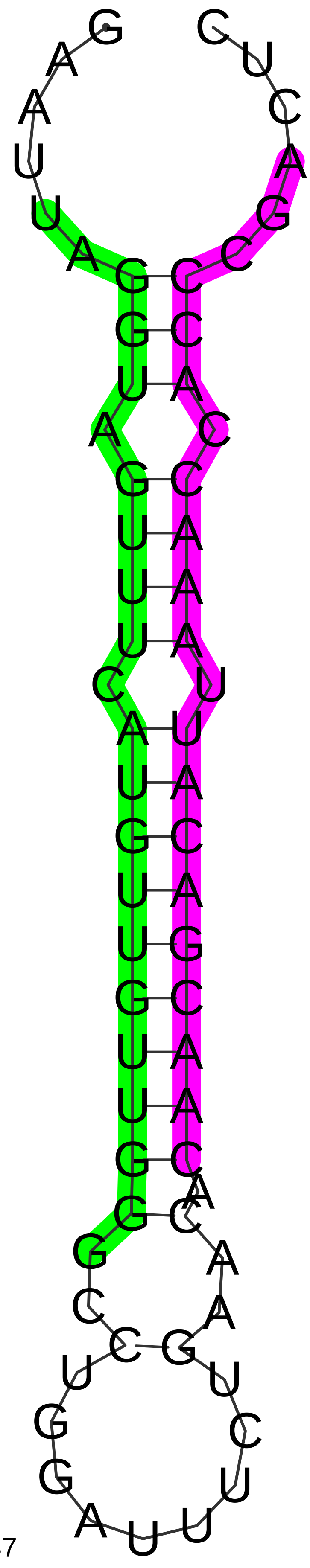

NW_006386434.1_105972-106037

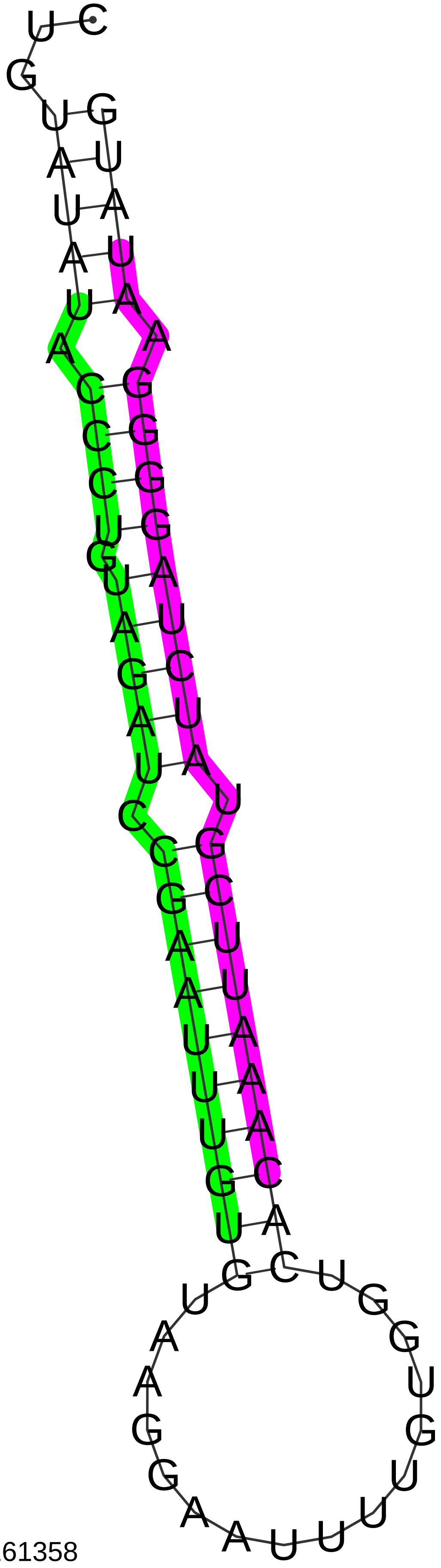


NW_006386434.1_161287-161358

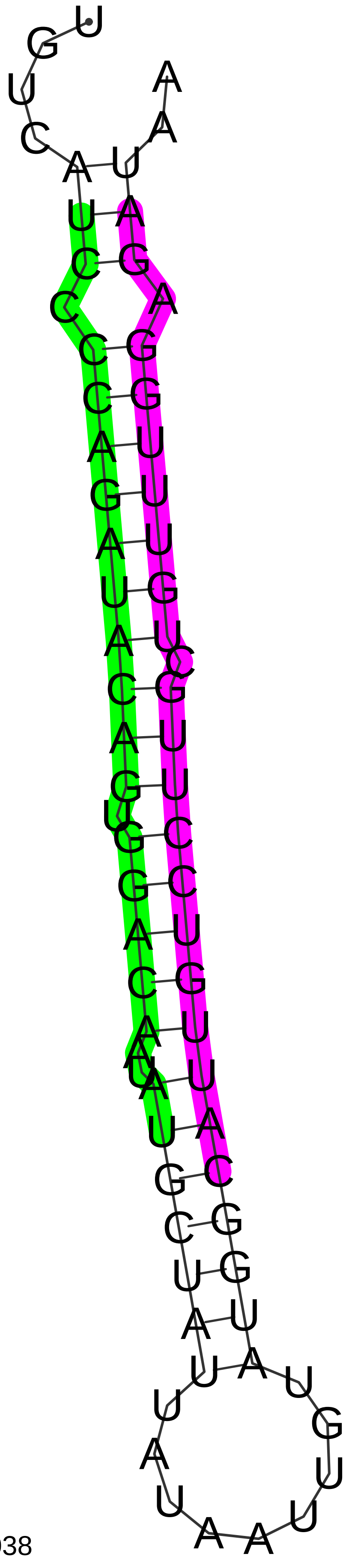

NW_006386495.1_204968-205038

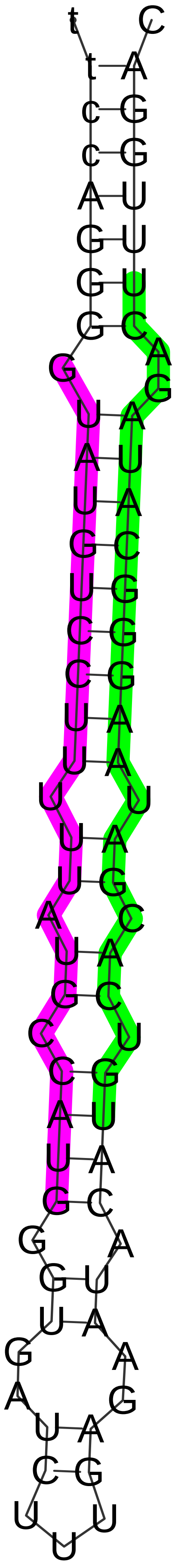

NW_006386502.1_148722-148797

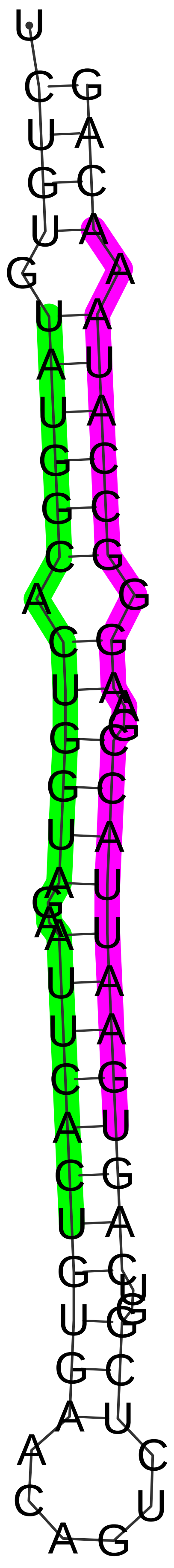

NW_006386524.1_144692-144762

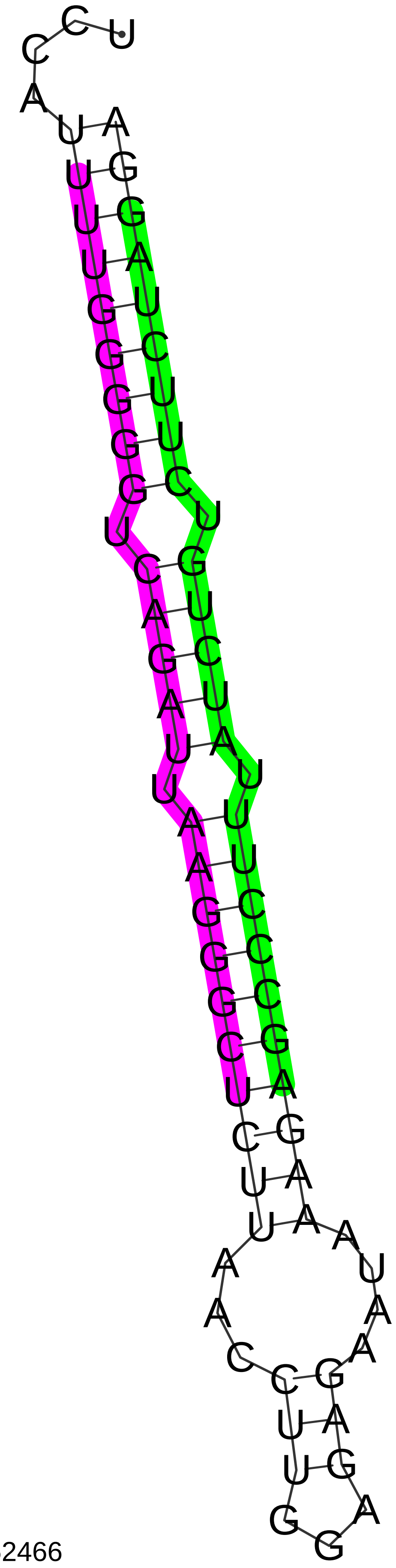

NW_006386541.1_162395-162466

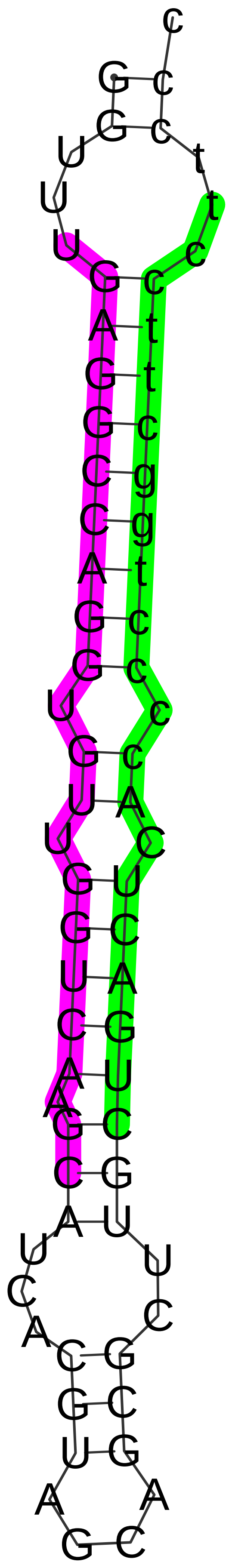

NW_006386567.1_171234-171302

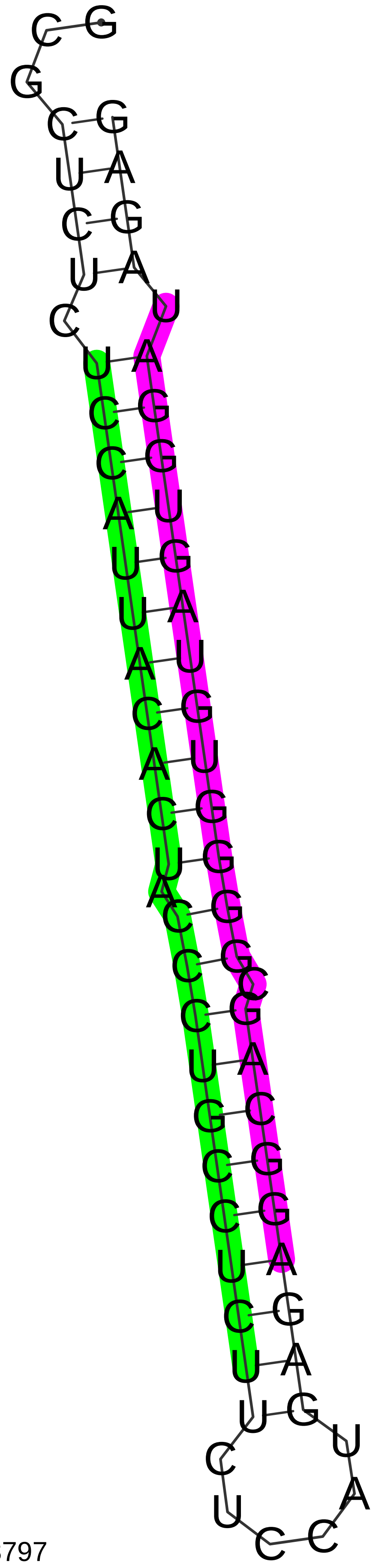

NW_006386574.1_208733-208797

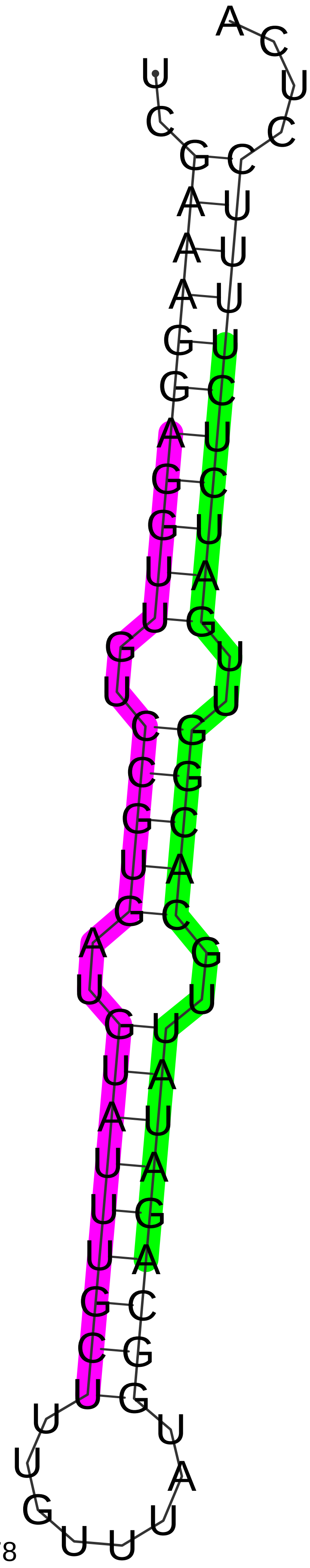


NW_006386594.1_101507-101578

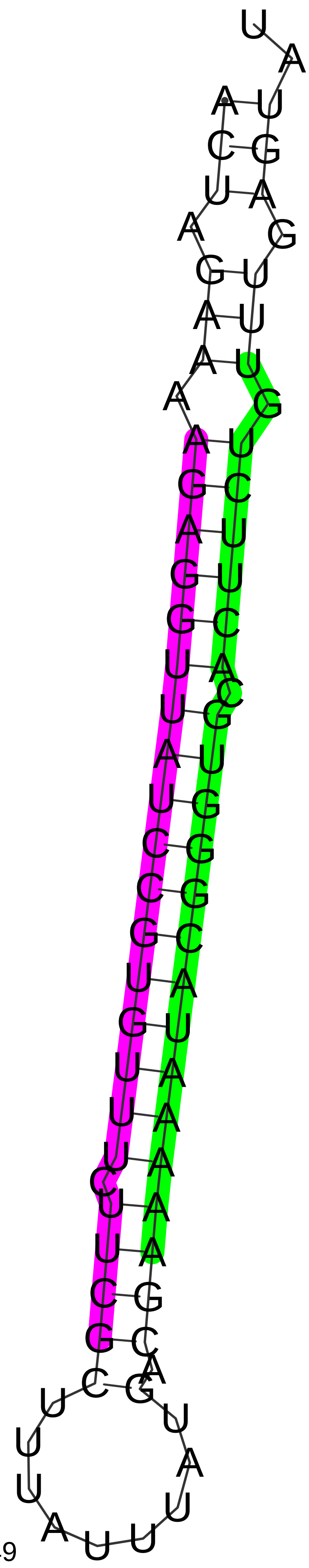

NW_006386594.1_102376-102449

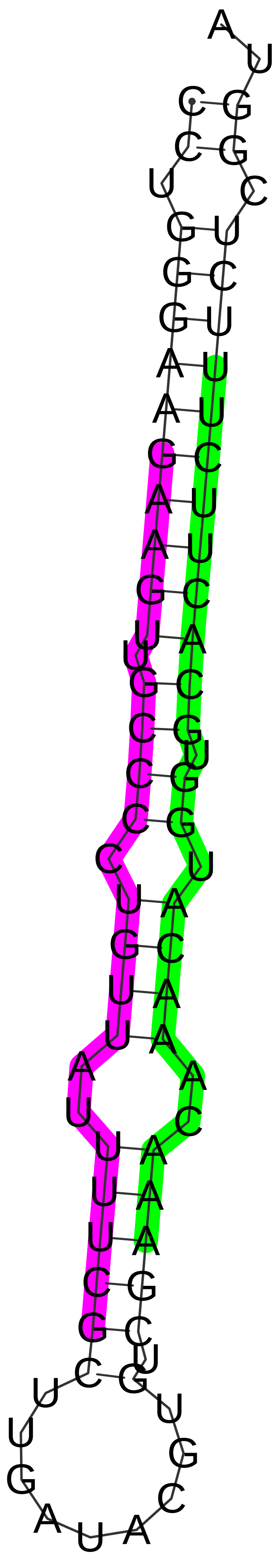

NW_006386594.1_103532-103605

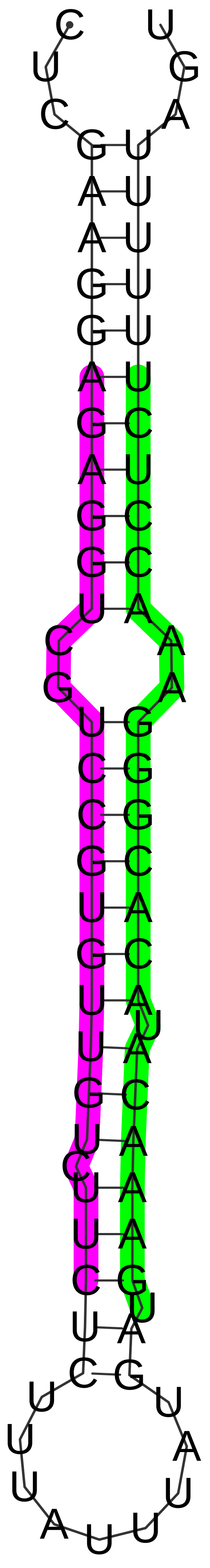

NW_006386594.1_107567-107640

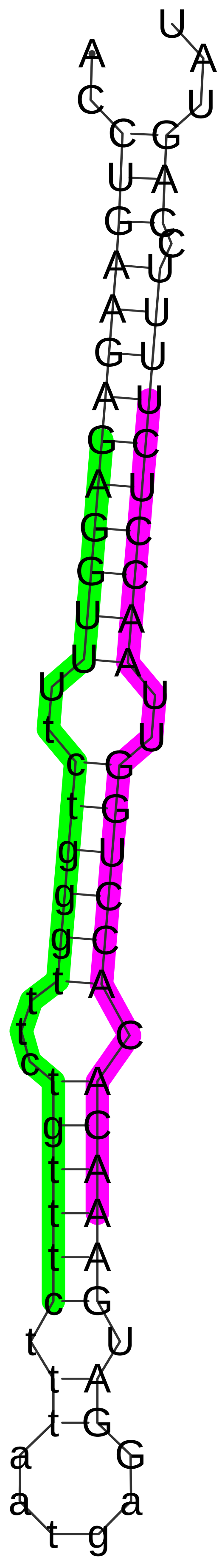

NW_006386594.1_110308-110383

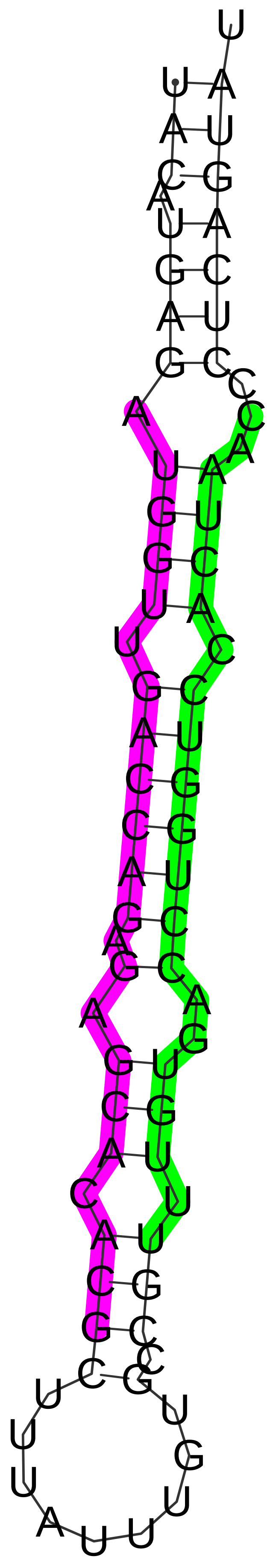

NW_006386594.1_111060-111133

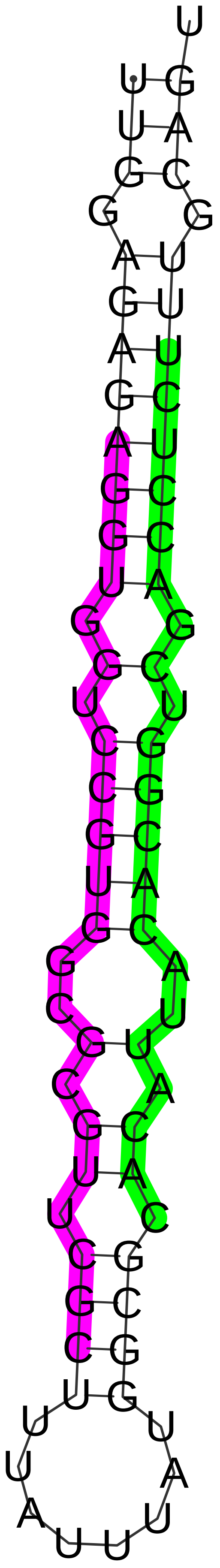

NW_006386594.1_111364-111434

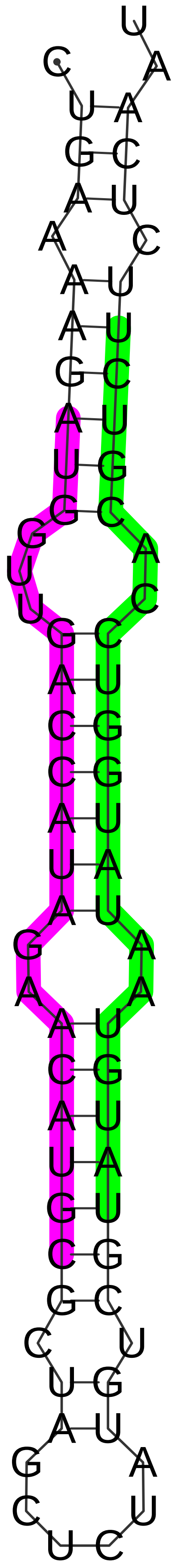

NW_006386594.1_112069-112140

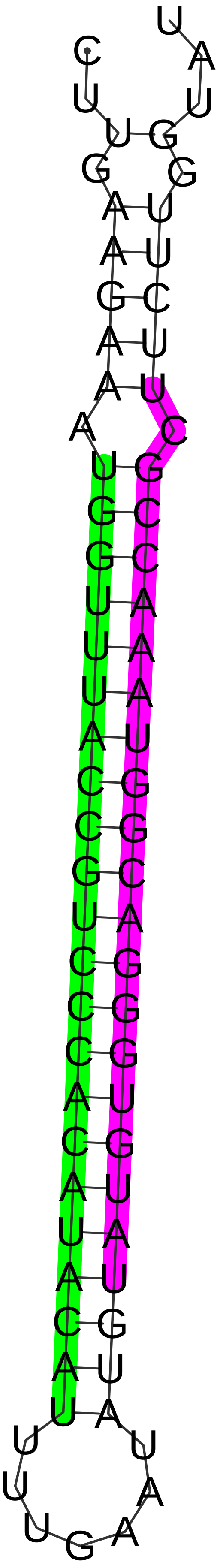

NW_006386594.1_113270-113341

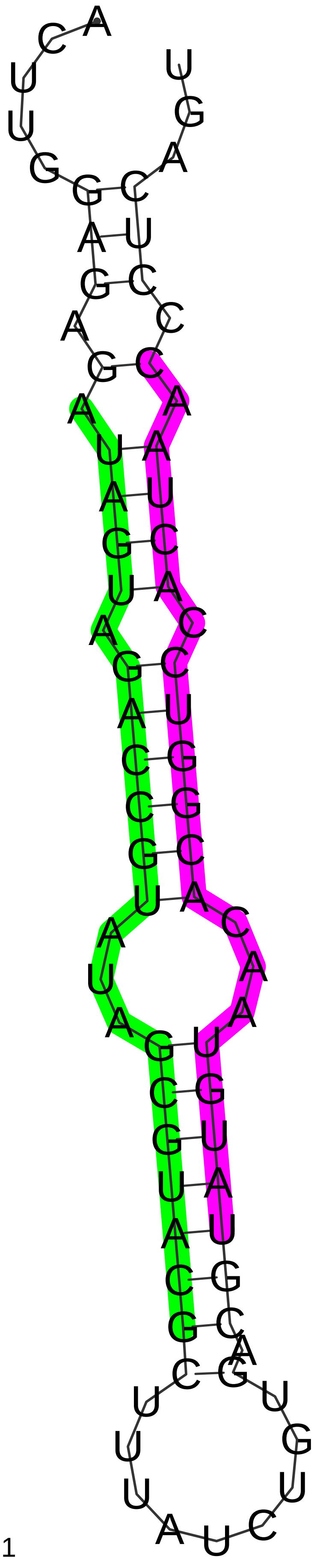

NW_006386594.1_113728-113801

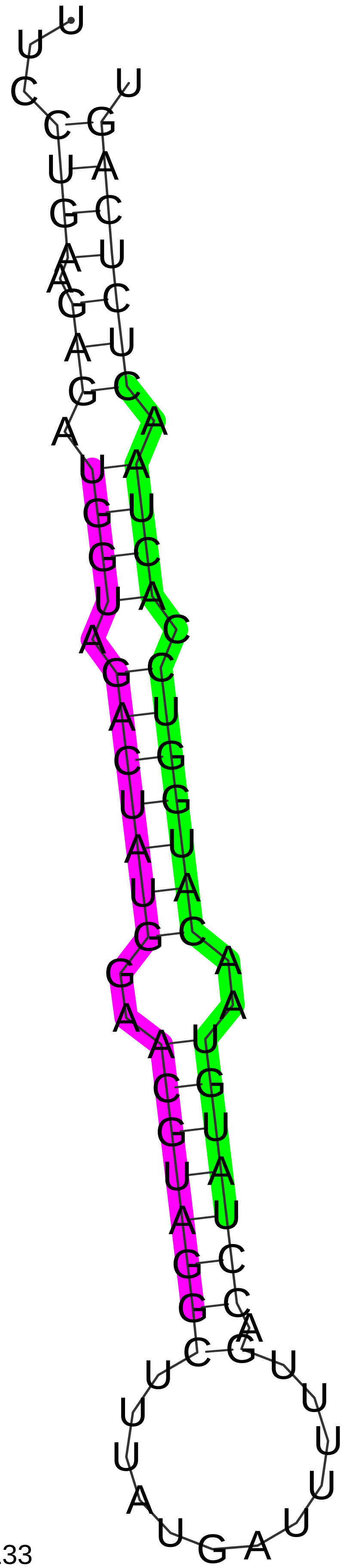

NW_006386594.1_115056-115133

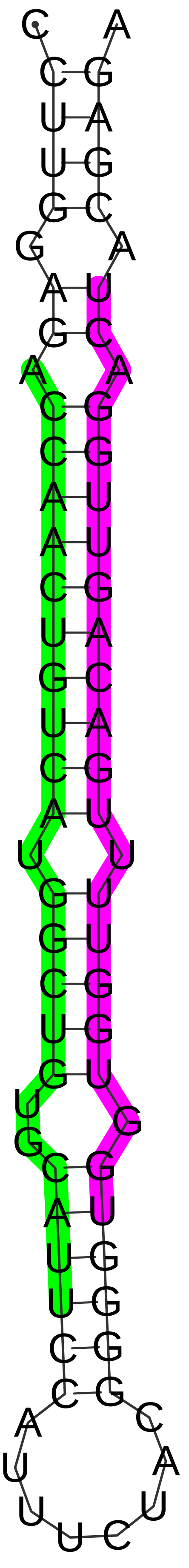

NW_006386594.1_115647-115719

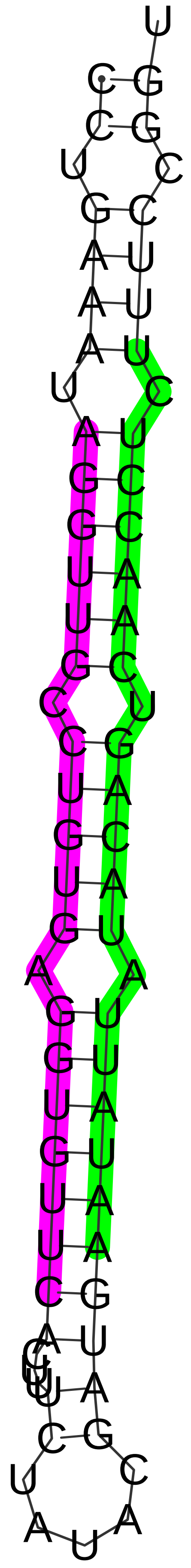

NW_006386594.1_73799-73869

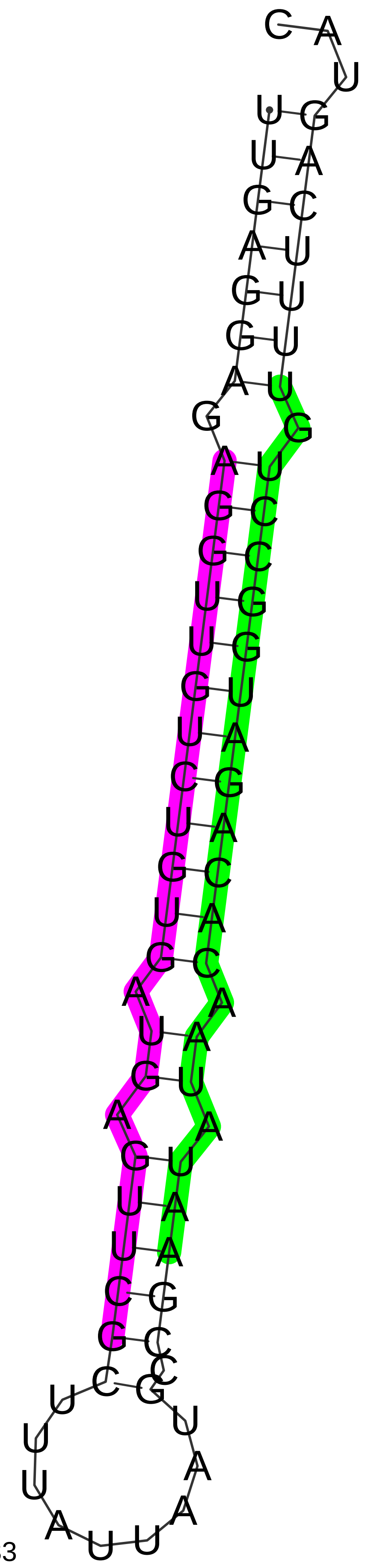

NW_006386594.1_74561-74633

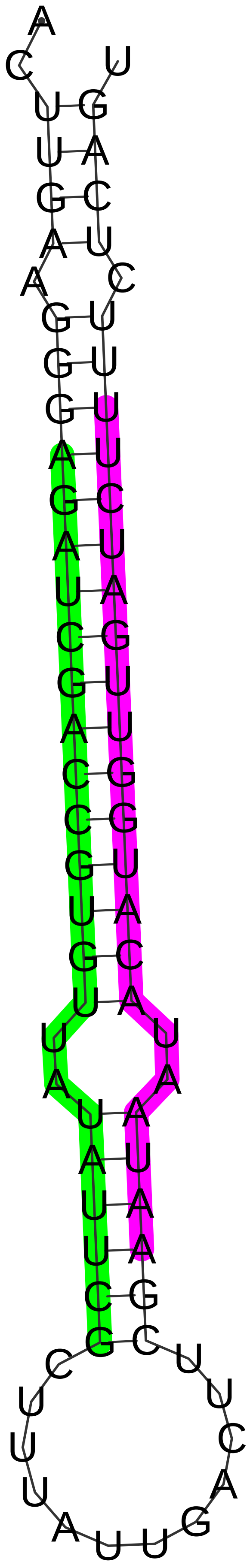

NW_006386594.1_74915-74987

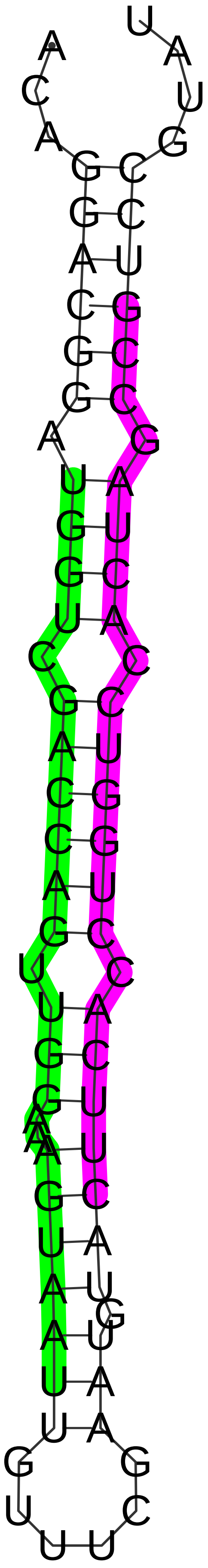

NW_006386594.1_75053-75126

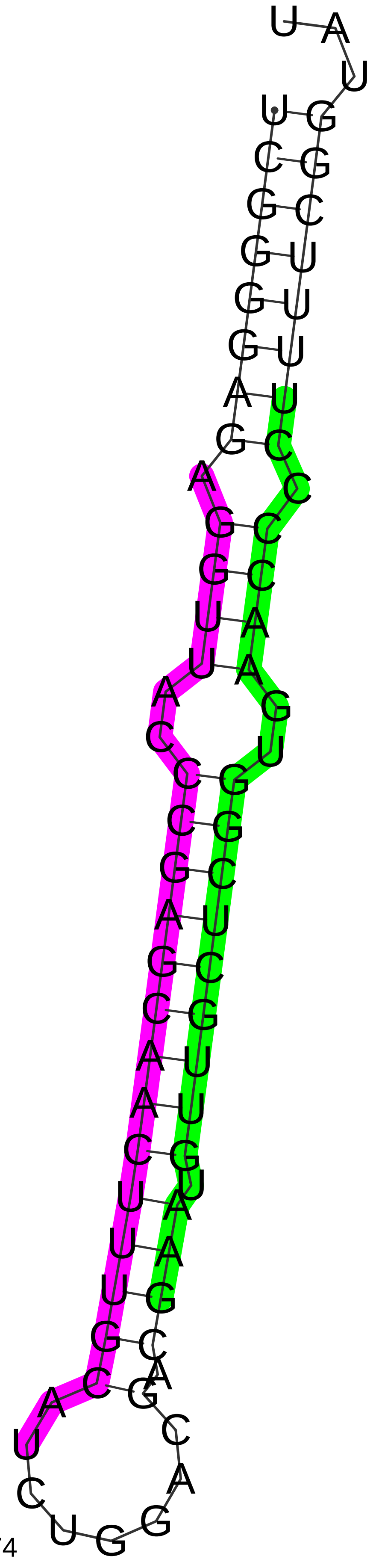

NW_006386594.1_75204-75274

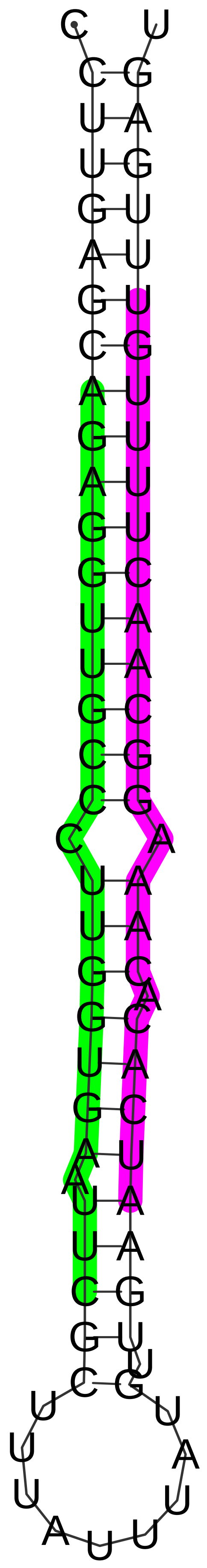

NW_006386594.1_78068-78141

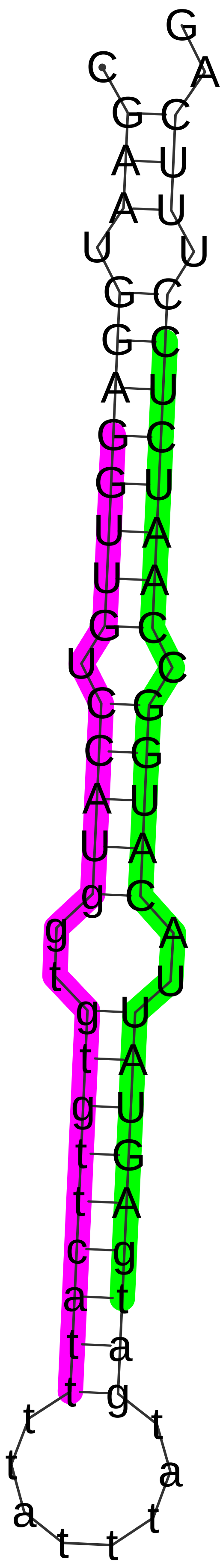

NW_006386594.1_79632-79700

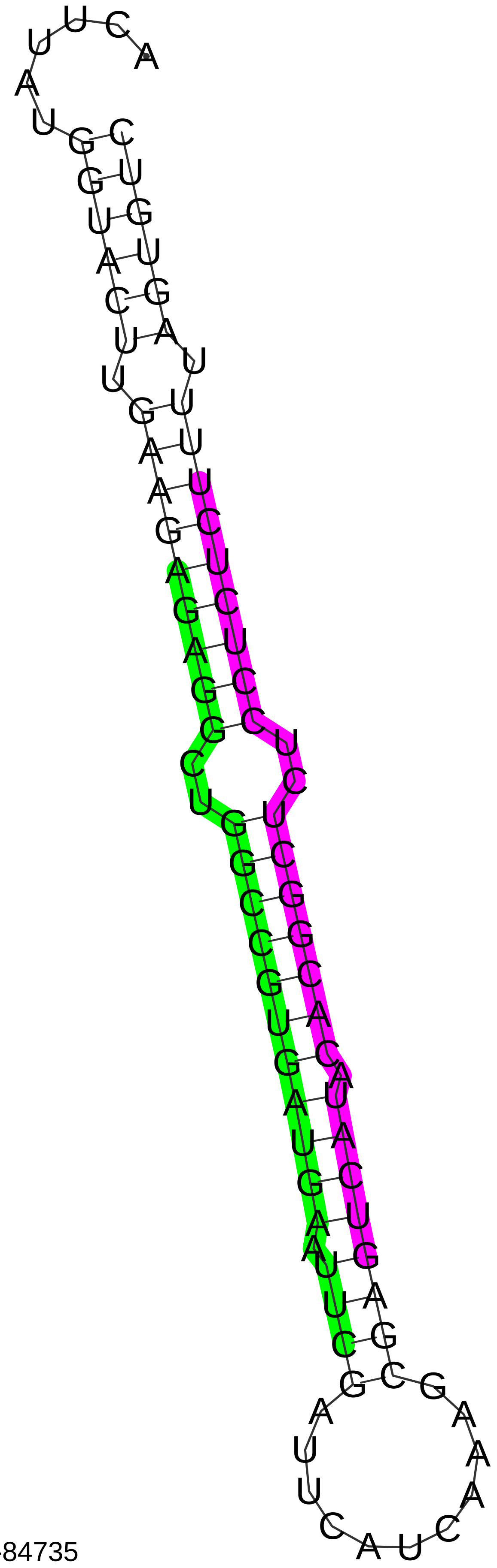

NW_006386594.1_84651-84735

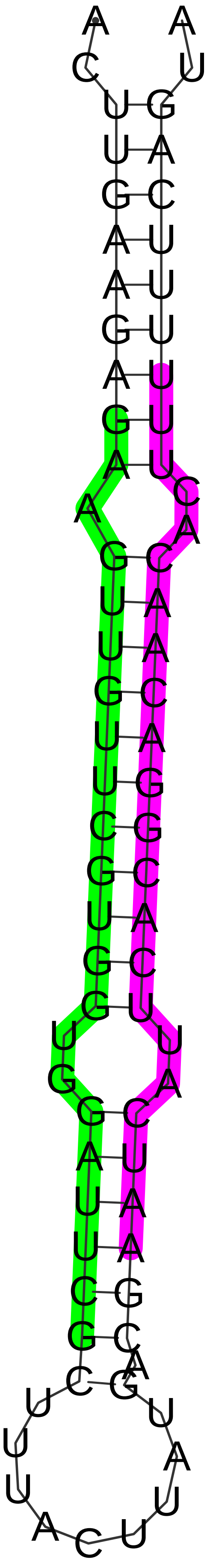

NW_006386594.1_85756-85830

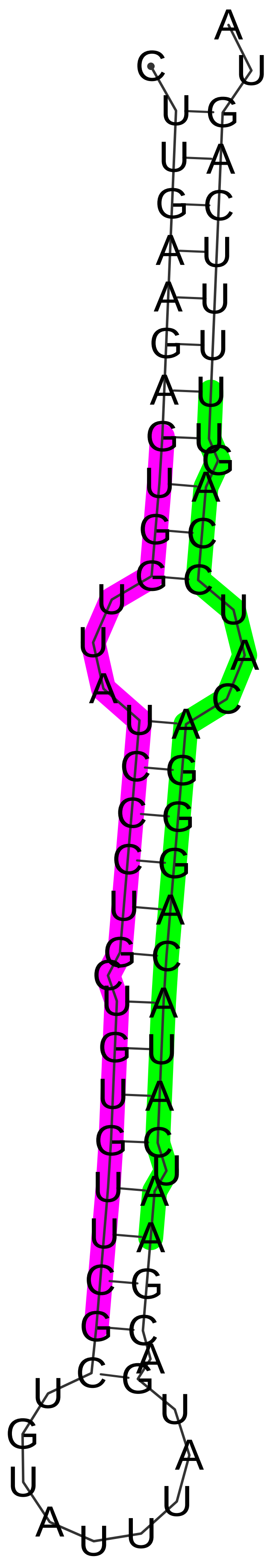

NW_006386594.1_87812-87885

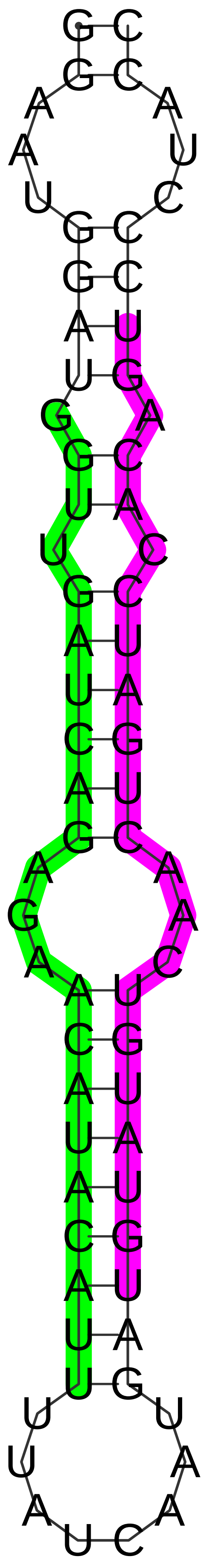

NW_006386594.1_88239-88308

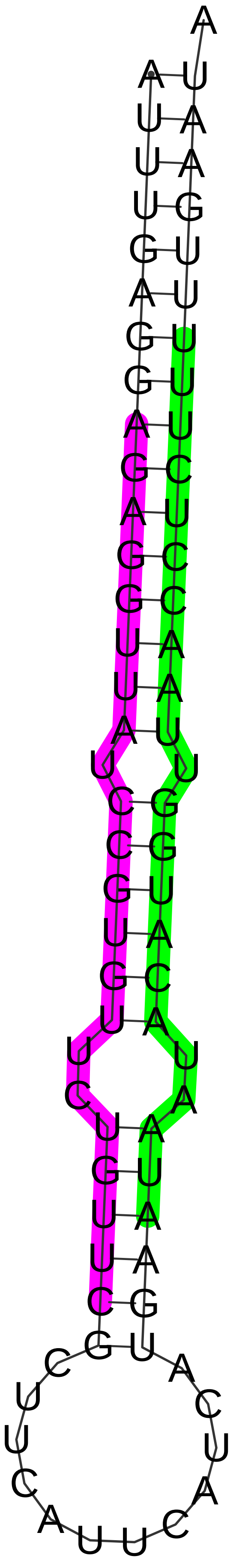

NW_006386594.1_90355-90429

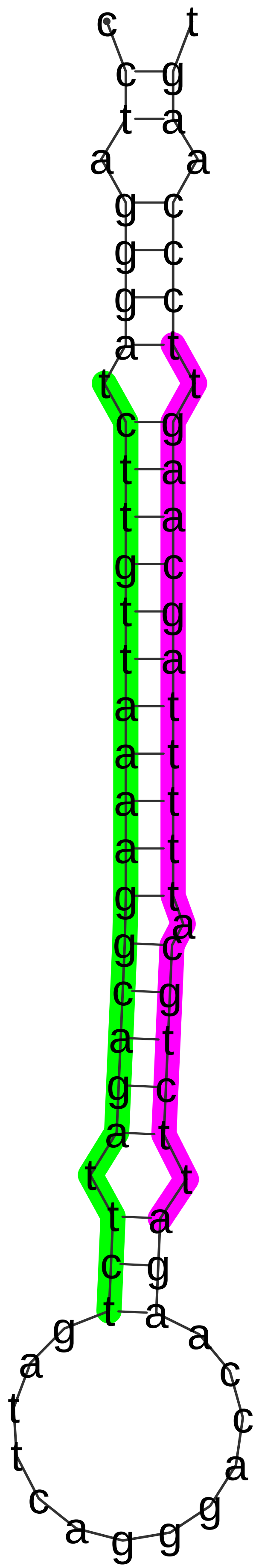

NW_006386594.1_91242-91313

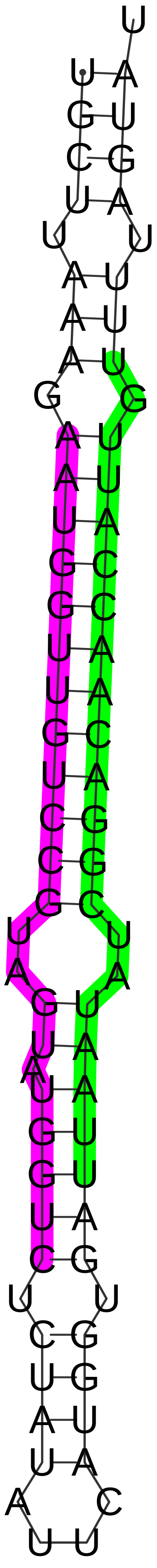

NW_006386594.1_91985-92060

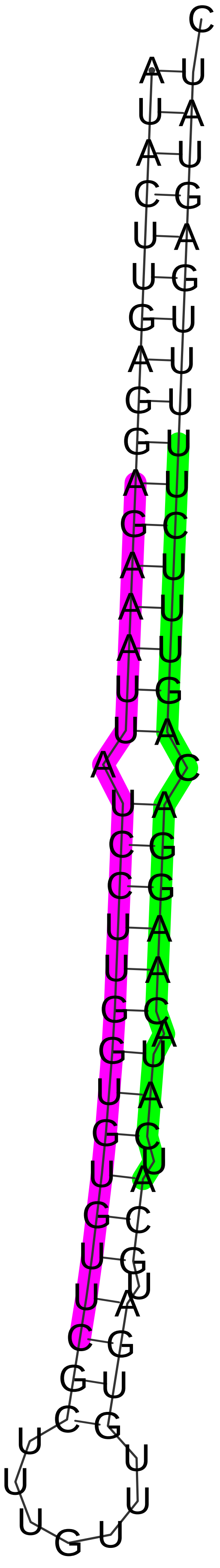

NW_006386594.1_92511-92589

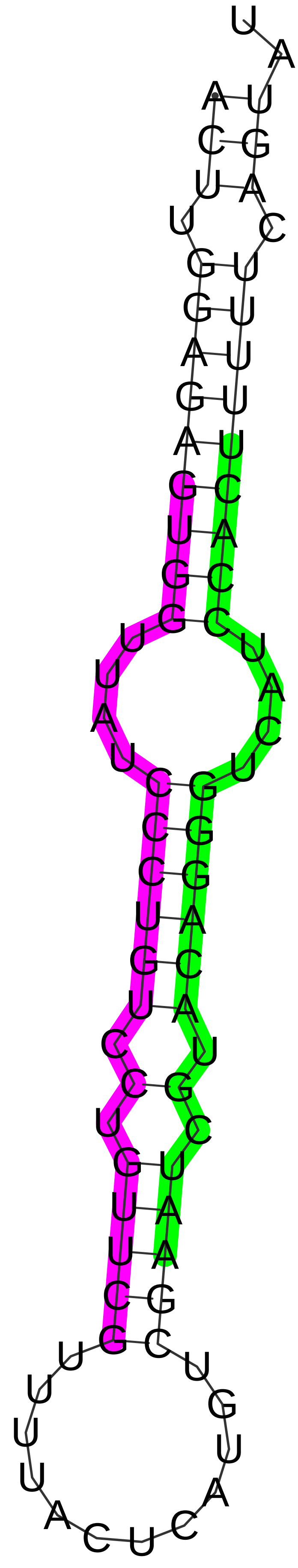

NW_006386594.1_93331-93406

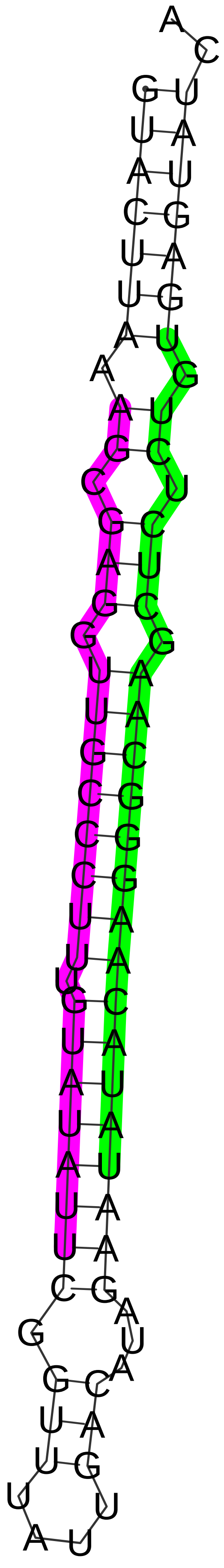

NW_006386594.1_93849-93927

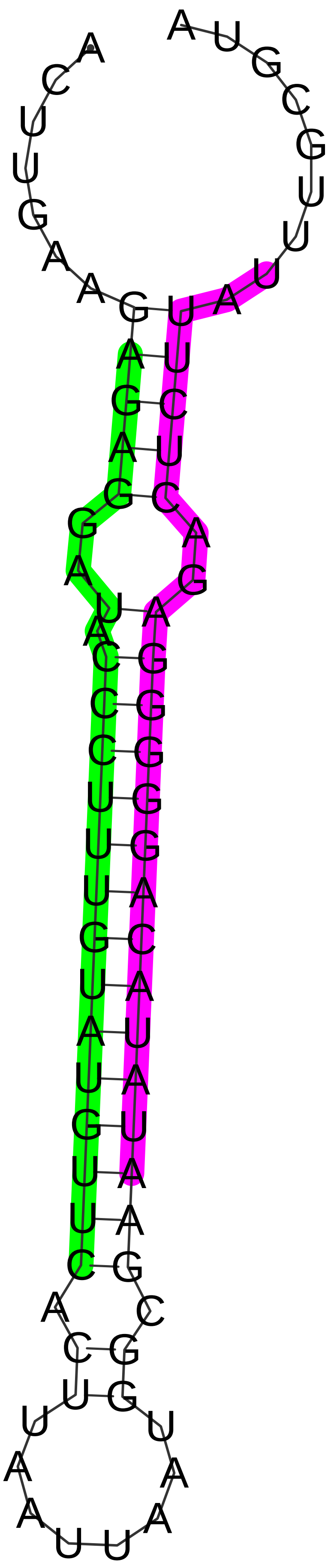

NW_006386594.1_95601-95675

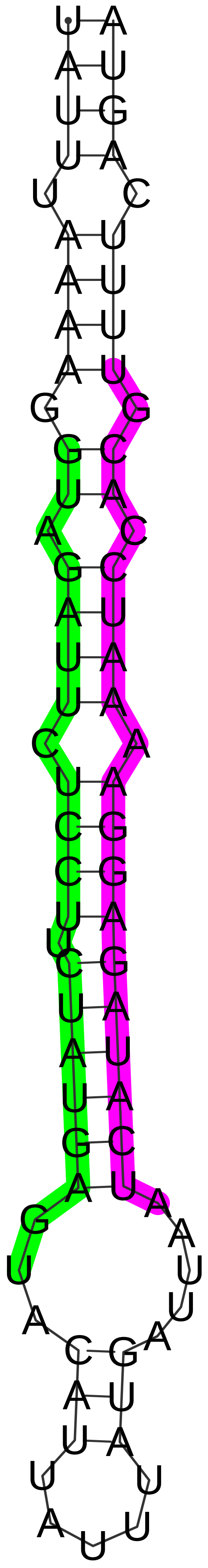

NW_006386594.1_97410-97485

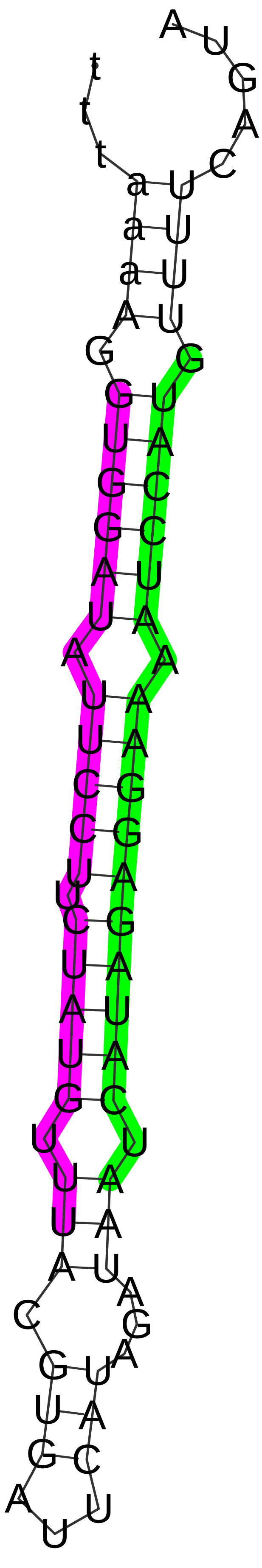

NW_006386594.1_97803-97876

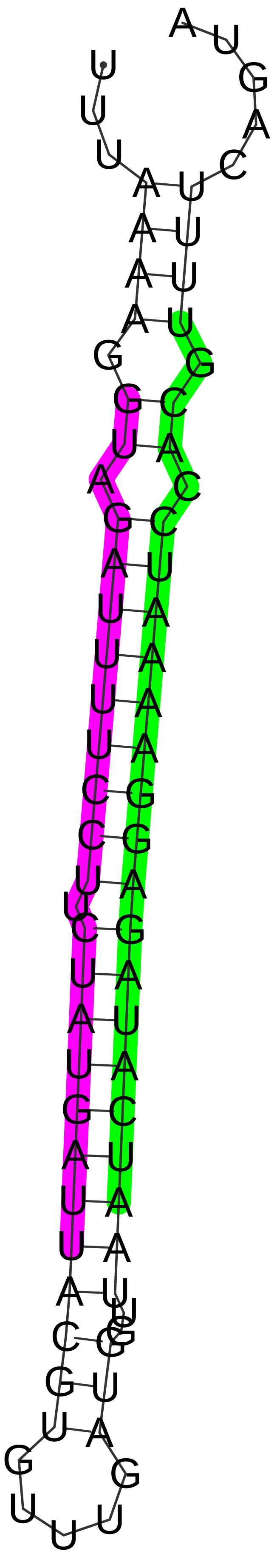

NW_006386594.1_98181-98254

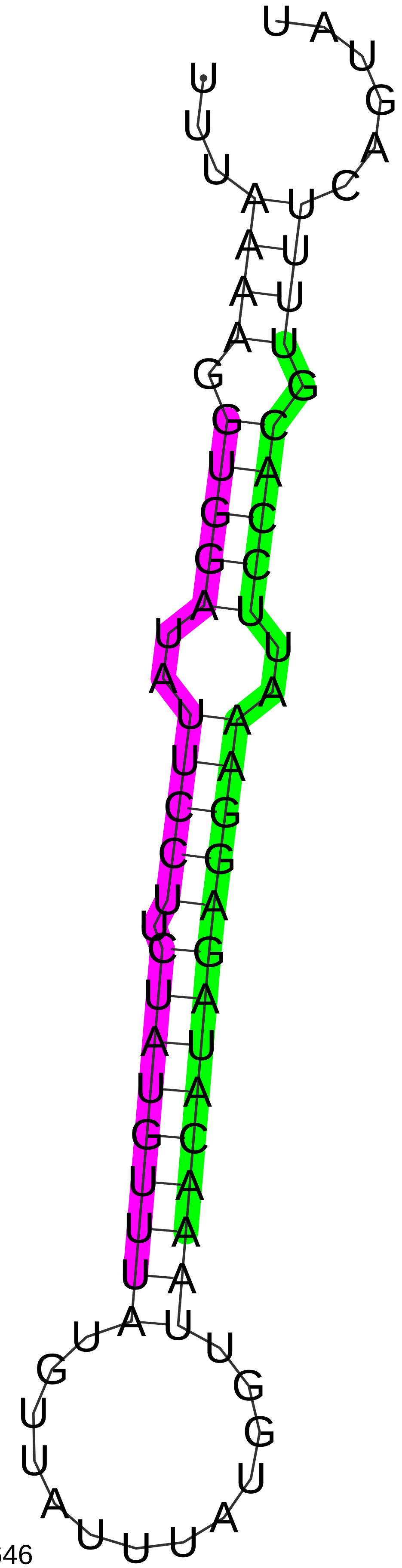

NW_006386594.1_98572-98646

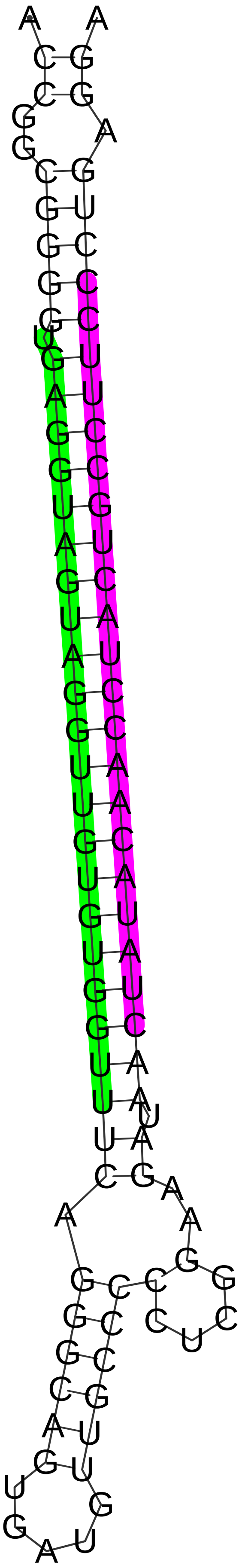

NW_006386630.1_100517-100609

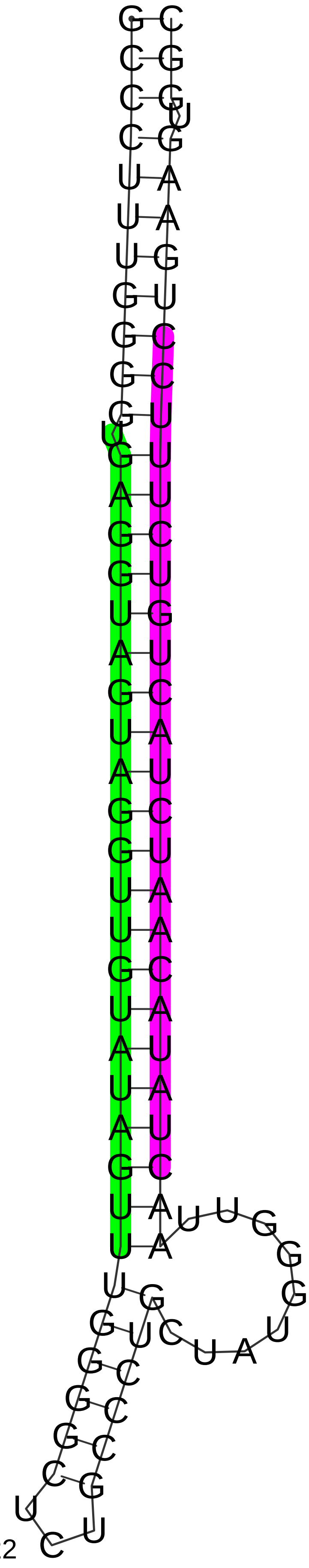

NW_006386630.1_101433-101522

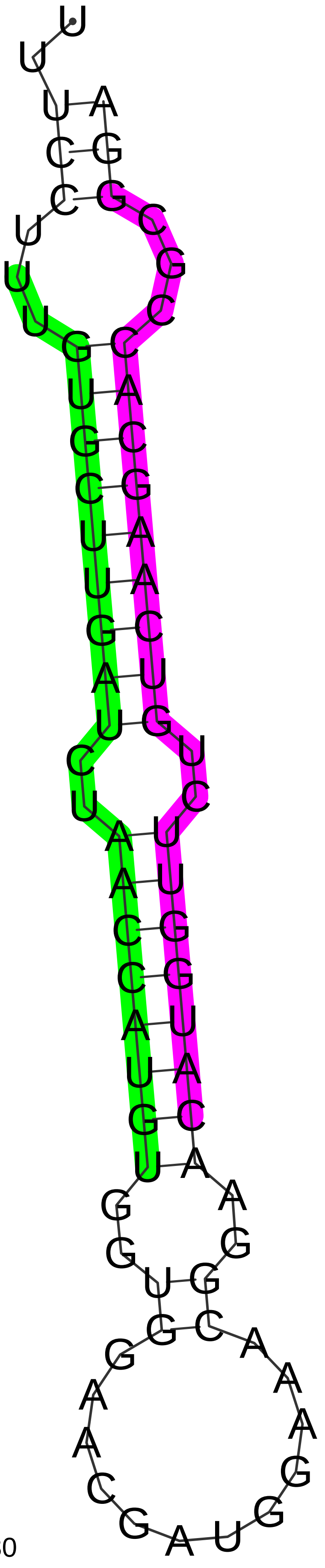

NW_006386638.1_191509-191580

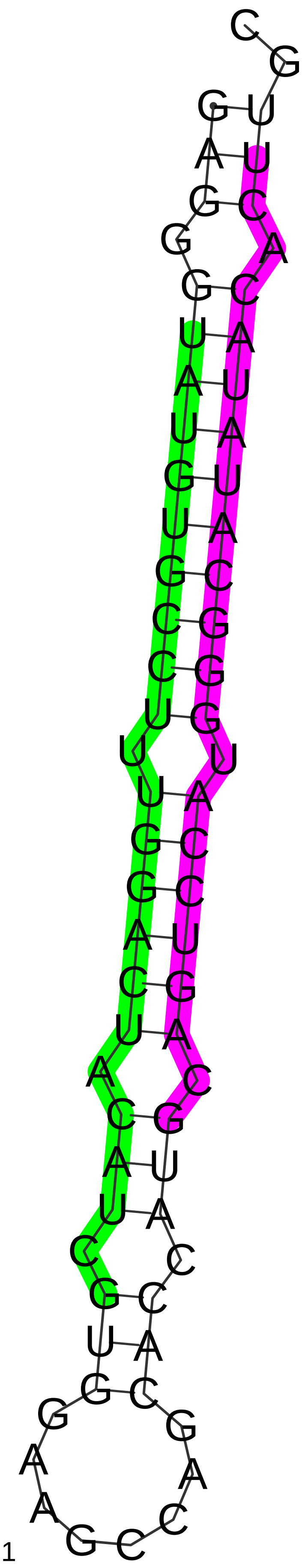

NW_006386641.1_145534-145601

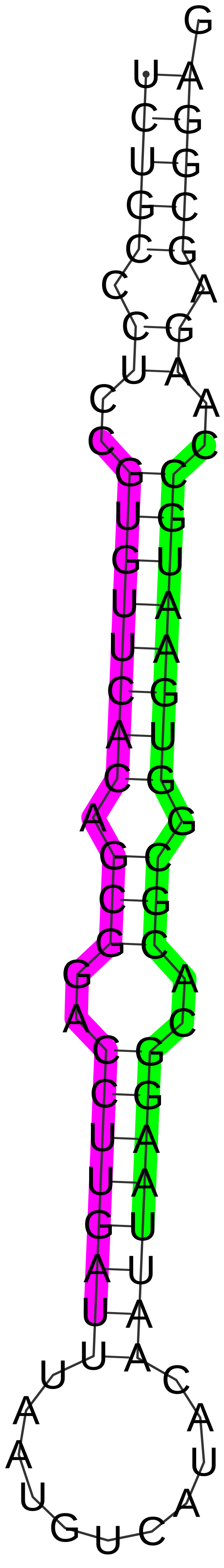

NW_006386794.1_162811-162886

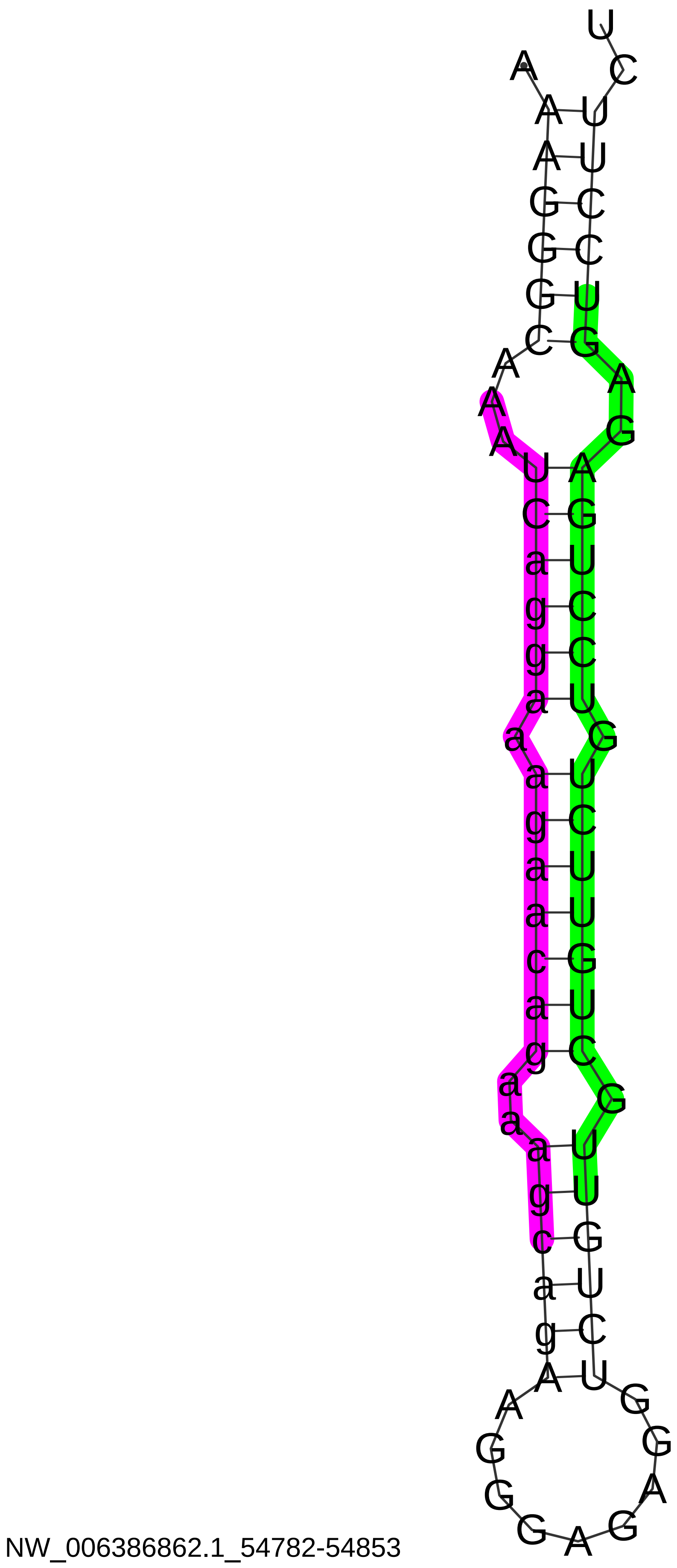

NW_006386862.1_54782-54853

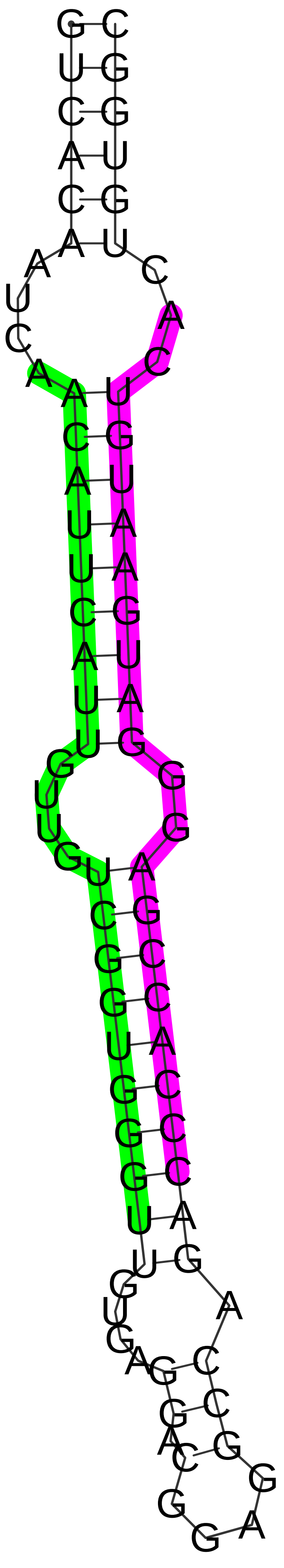

NW_006386921.1_105806-105884

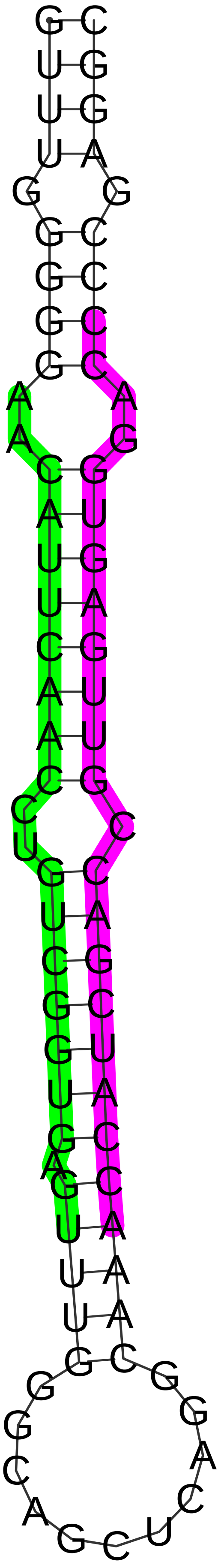

NW_006386921.1_105988-106064

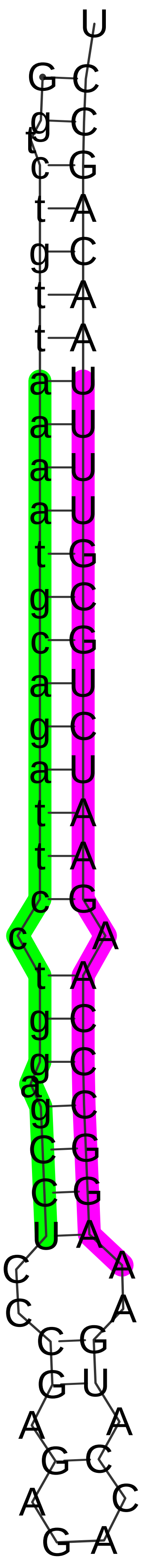

NW_006386957.1_107549-107623

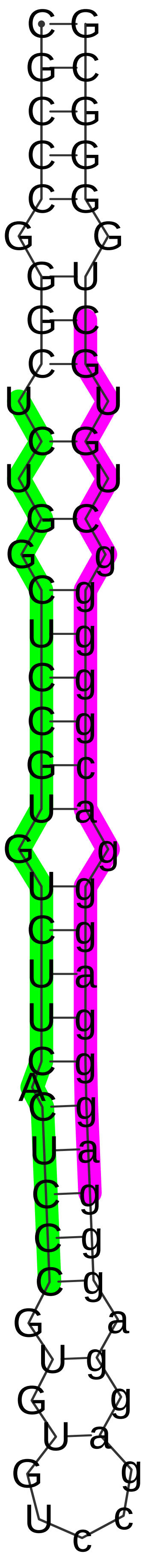

NW_006386972.1_58510-58585

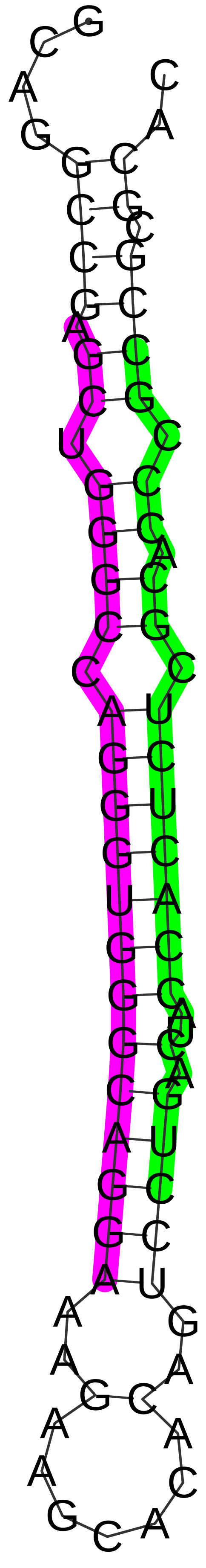

NW_006386972.1_58650-58724

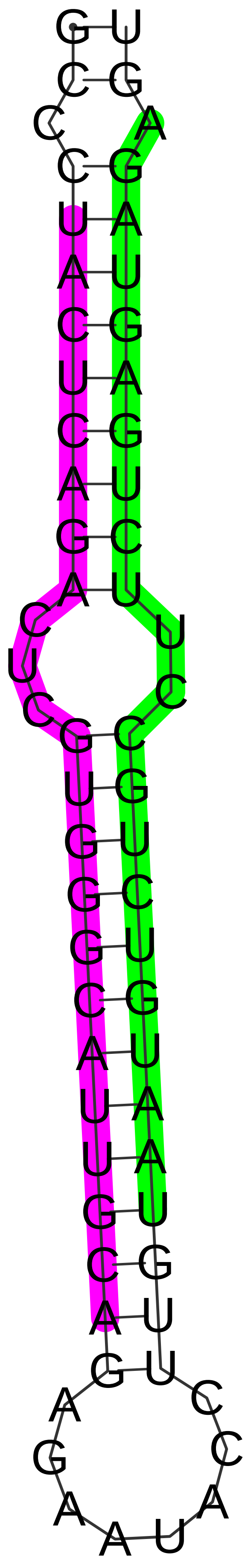

NW_006387102.1_111227-111289

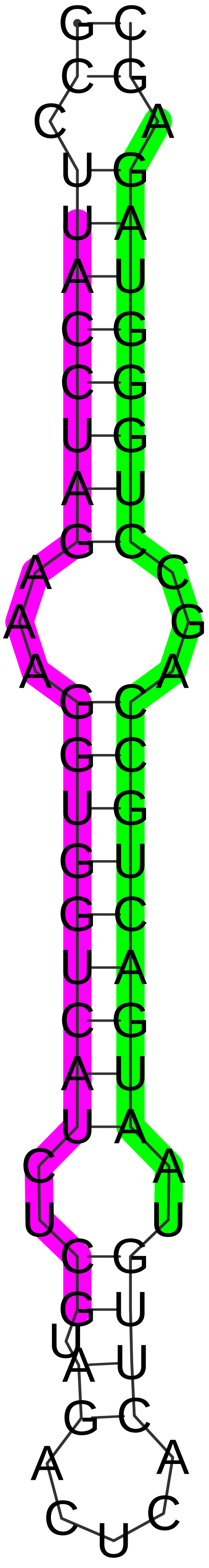

NW_006387102.1_115767-115830

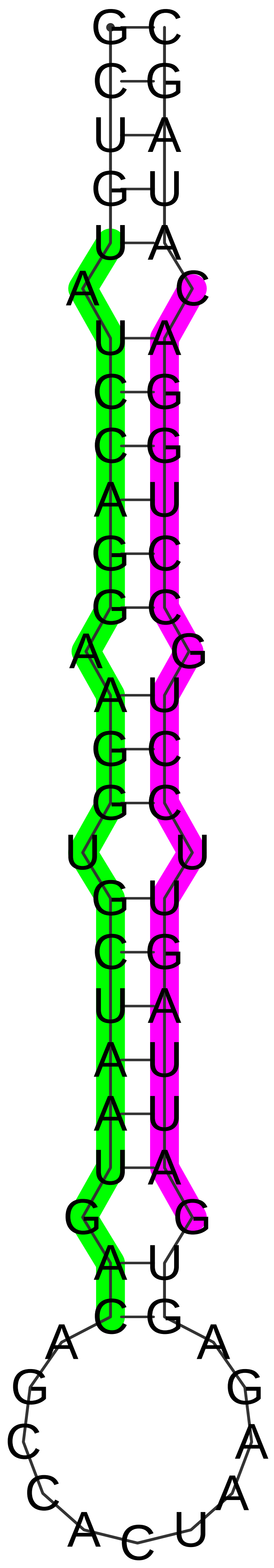

NW_006387102.1_116275-116337

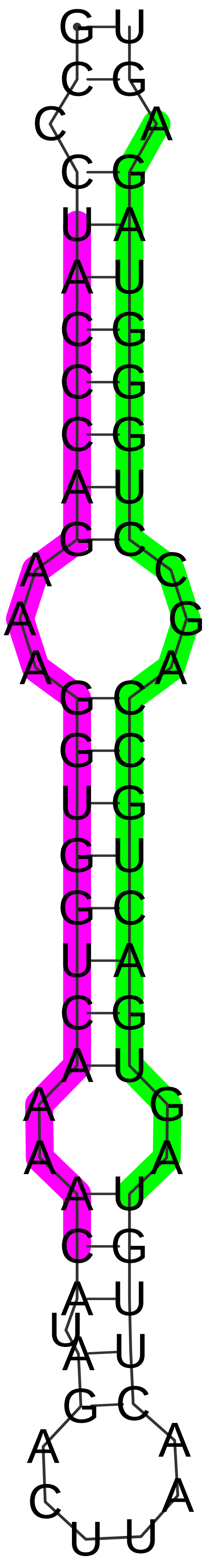

NW_006387102.1_72691-72755

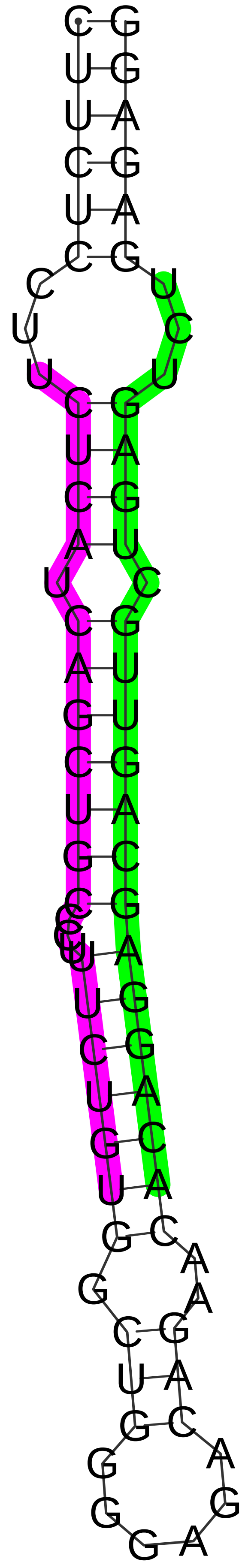

NW_006387121.1_73553-73626

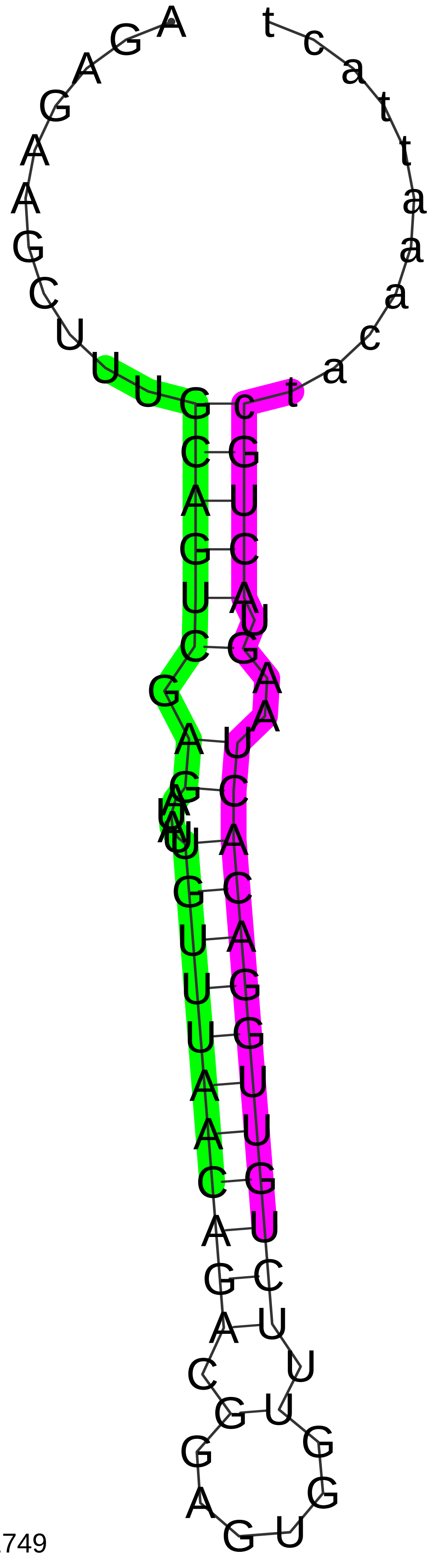

NW_006387130.1_51672-51749

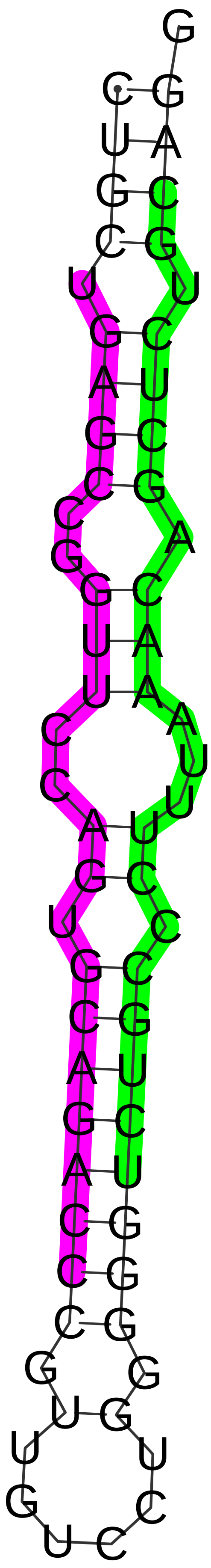

NW_006387384.1_15221-15285

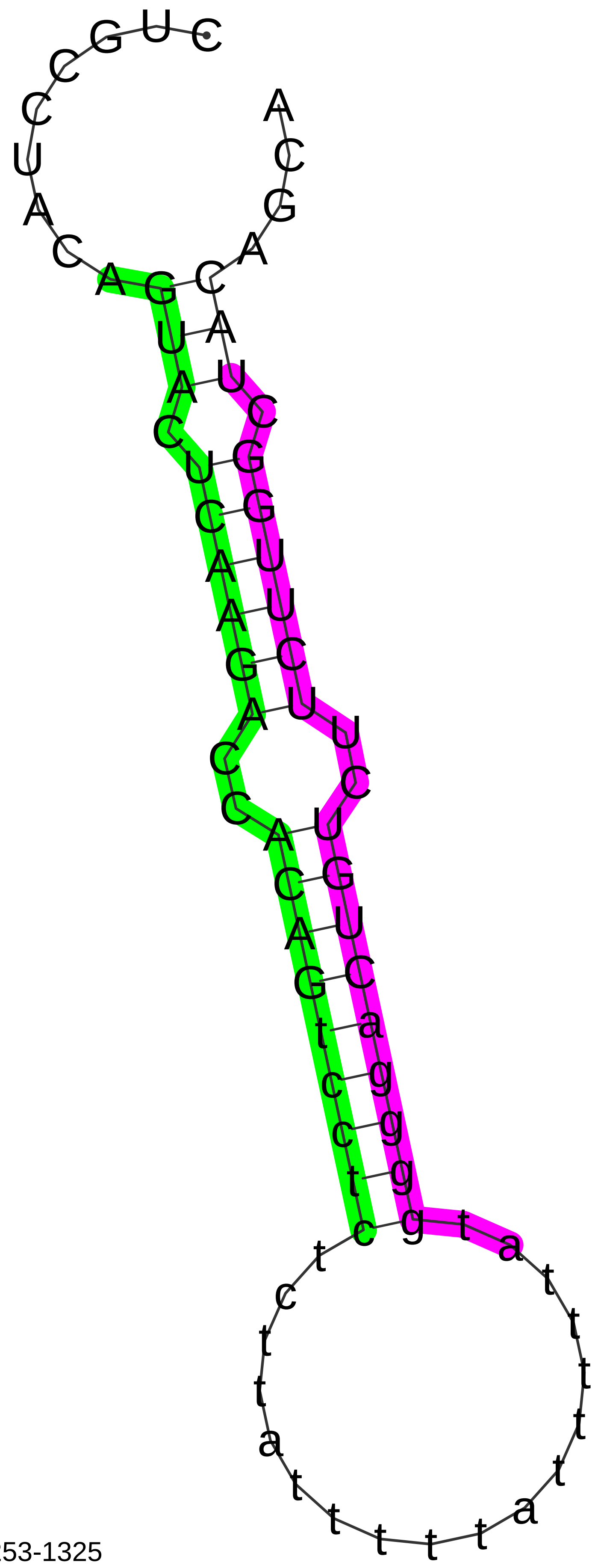

NW_006387434.1_1253-1325

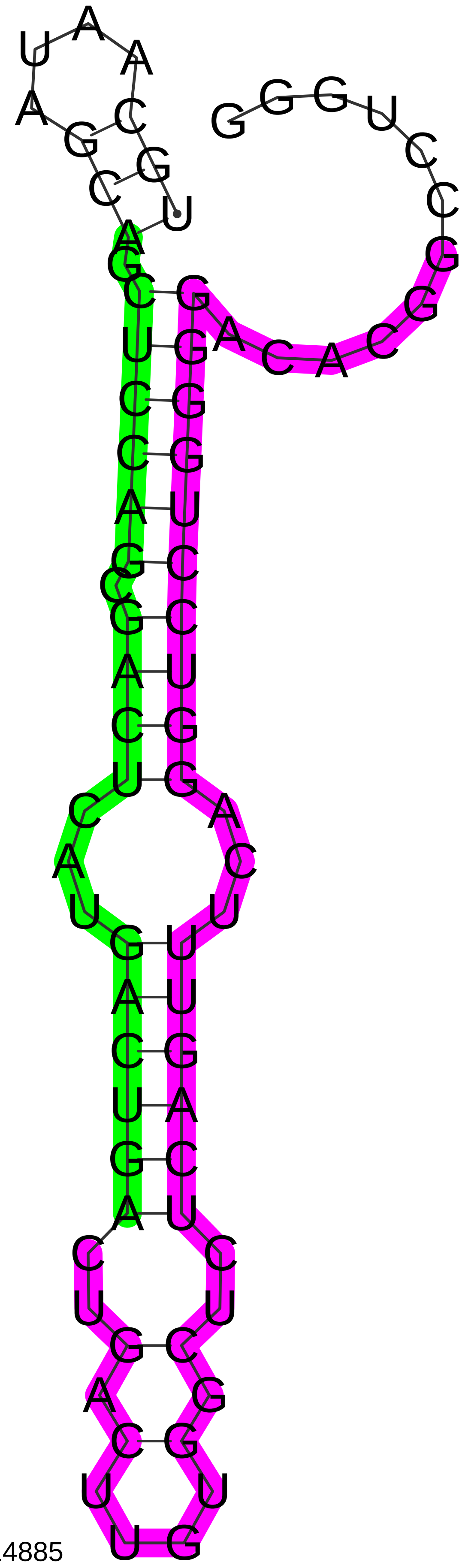

NW_006387494.1_14810-14885

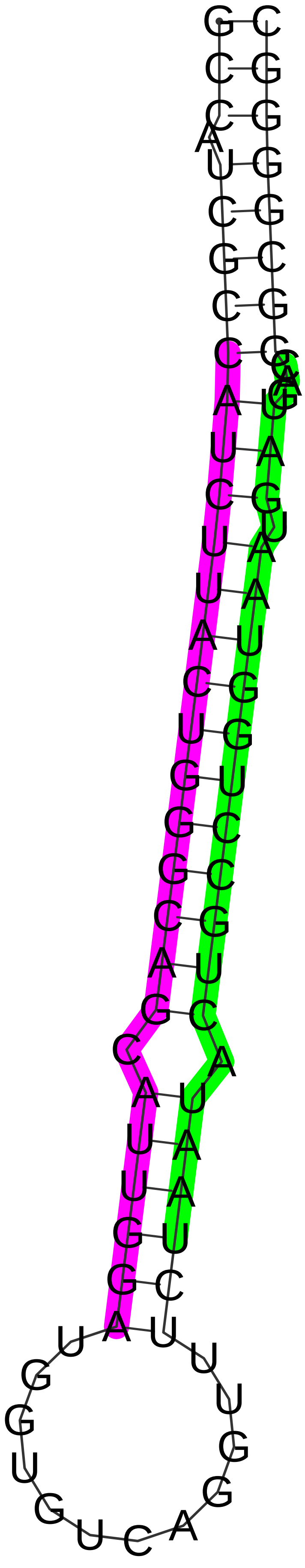

NW_006387595.1_61723-61797

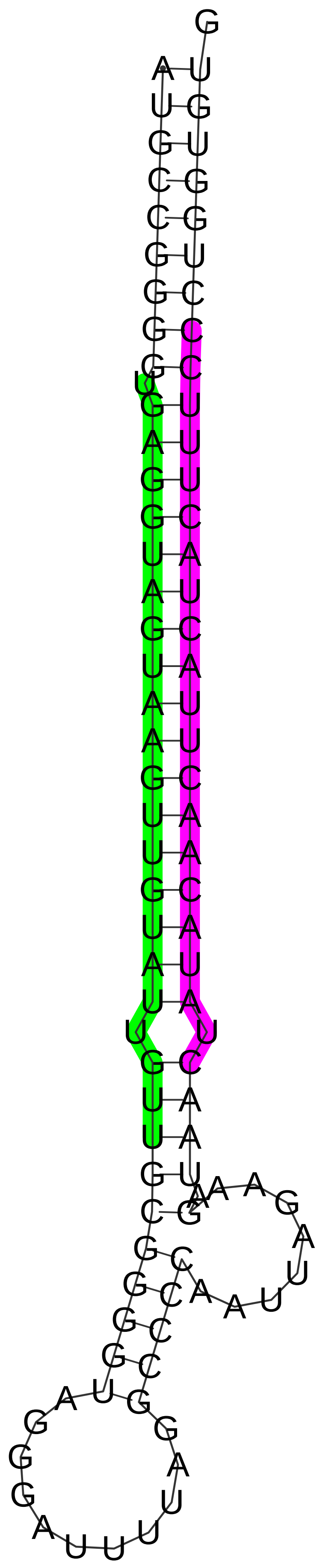

NW_006387656.1_89677-89772

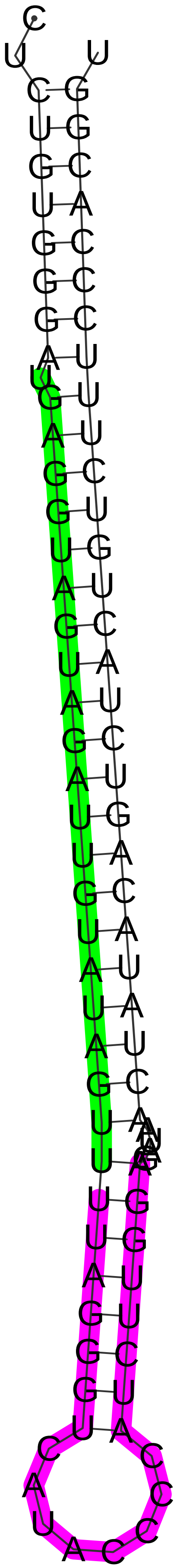

NW_006387656.1_90539-90626

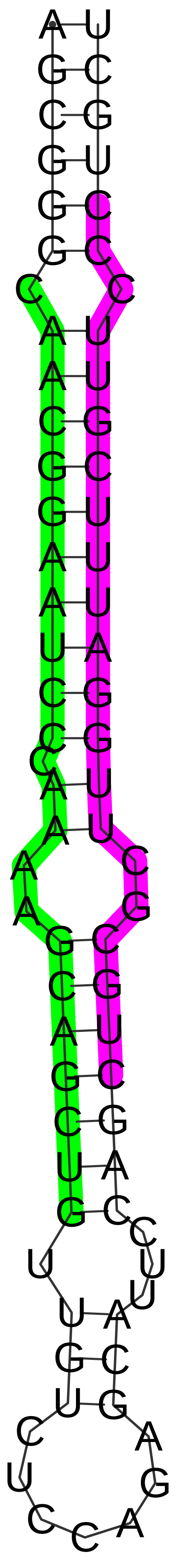

NW_006387742.1_18692-18765

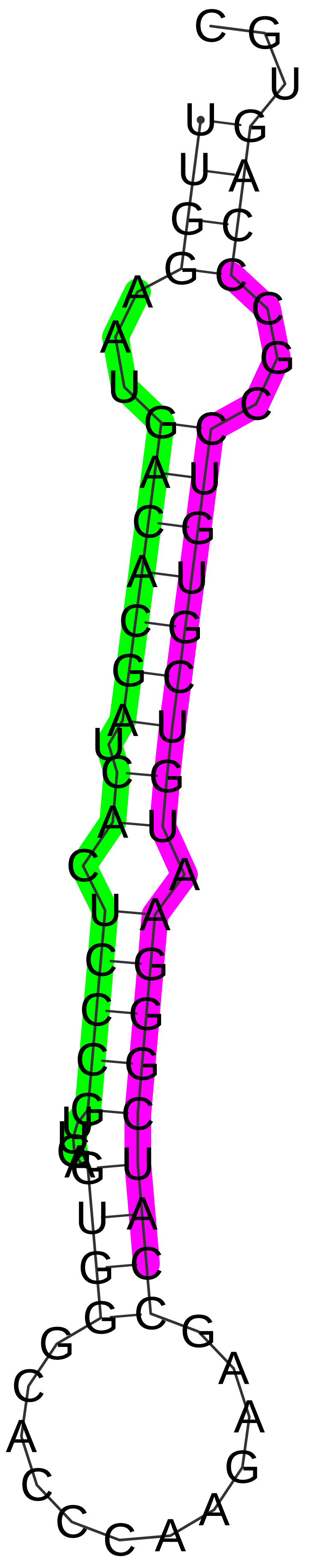

NW_006387742.1_19167-19238

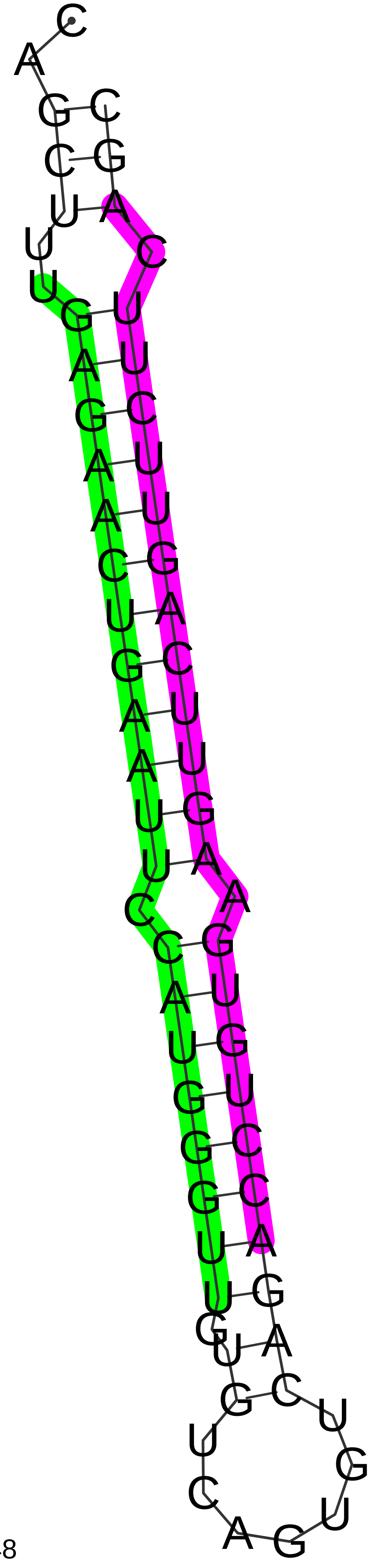

NW_006387754.1_58984-59048

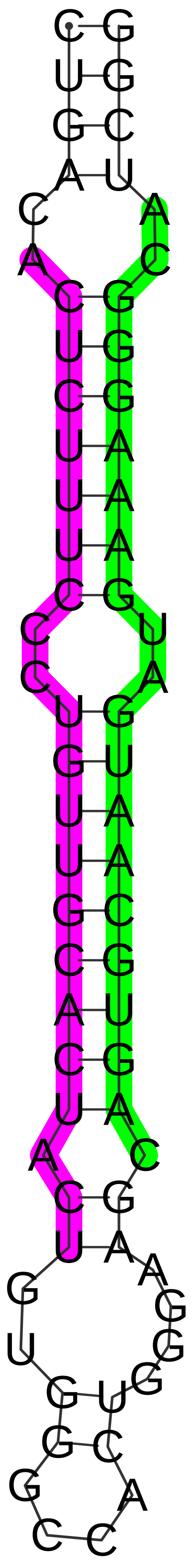

NW_006387848.1_43221-43288

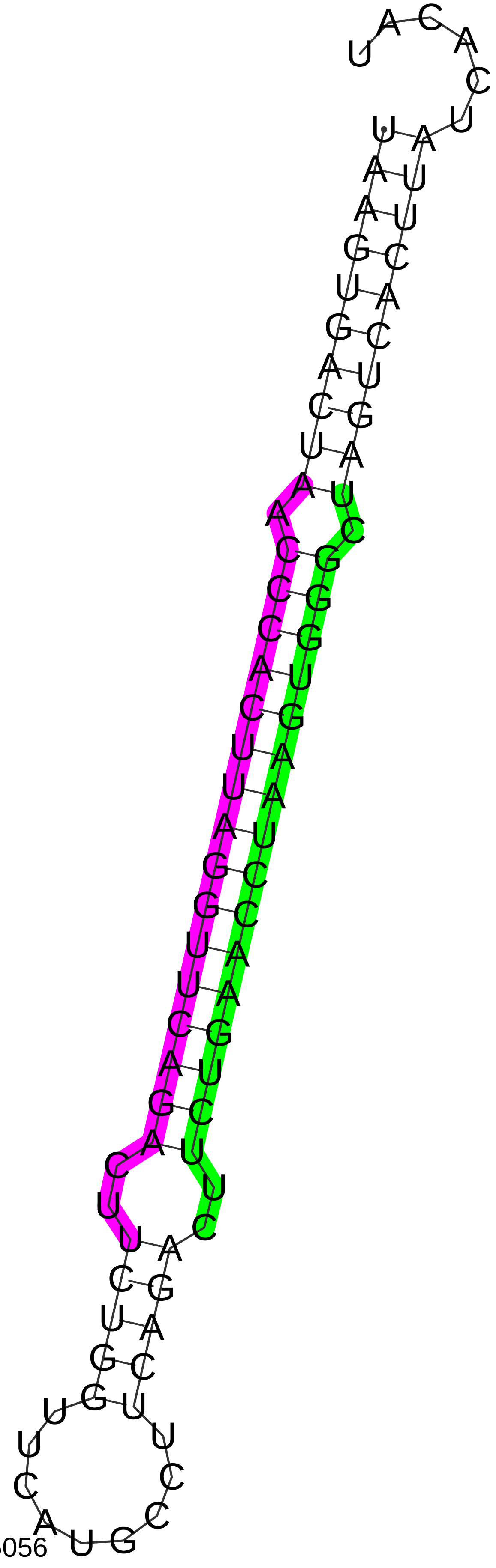

NW_006387925.1_5974-6056

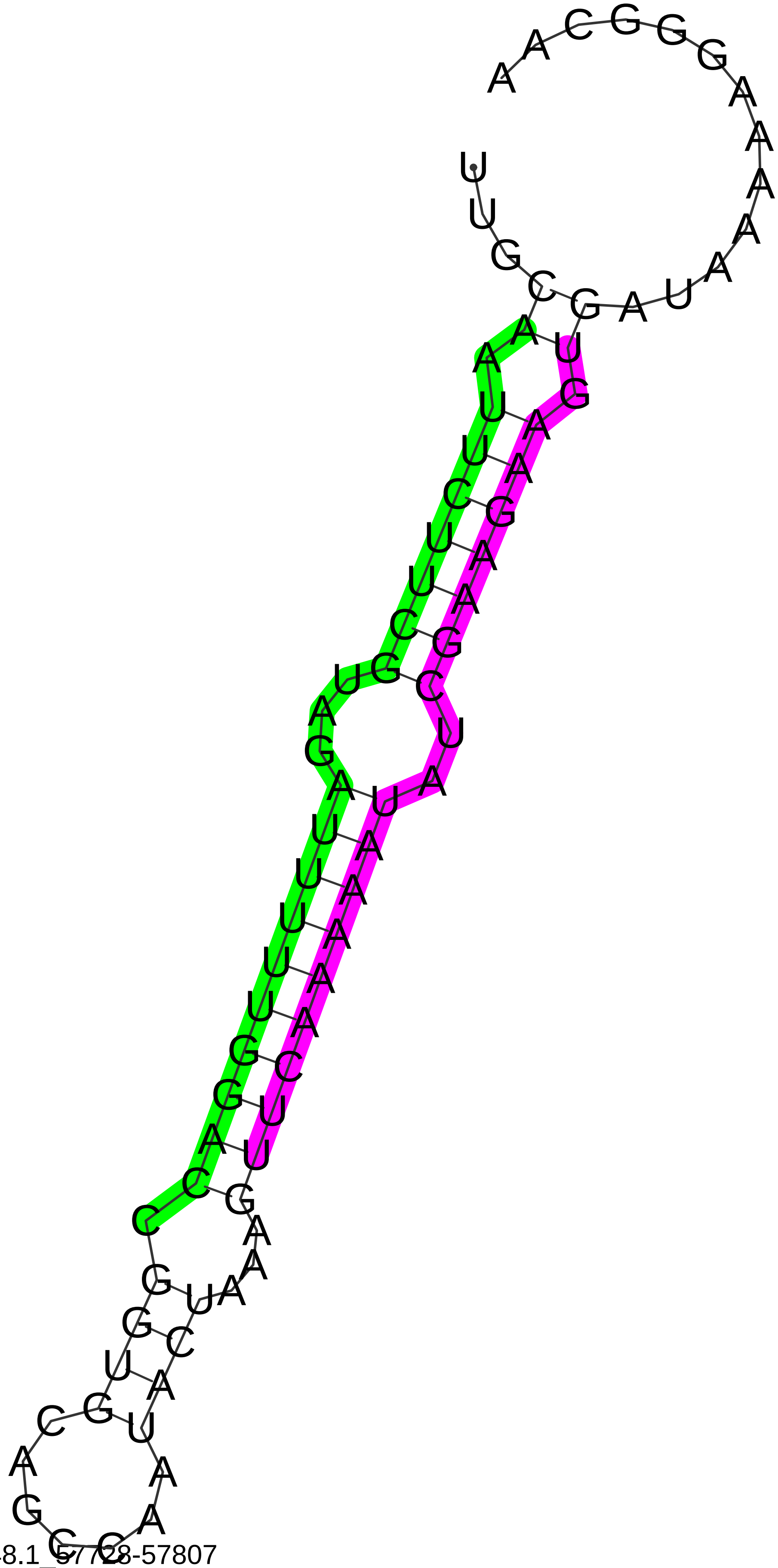

NW_006388248.1_57728-57807

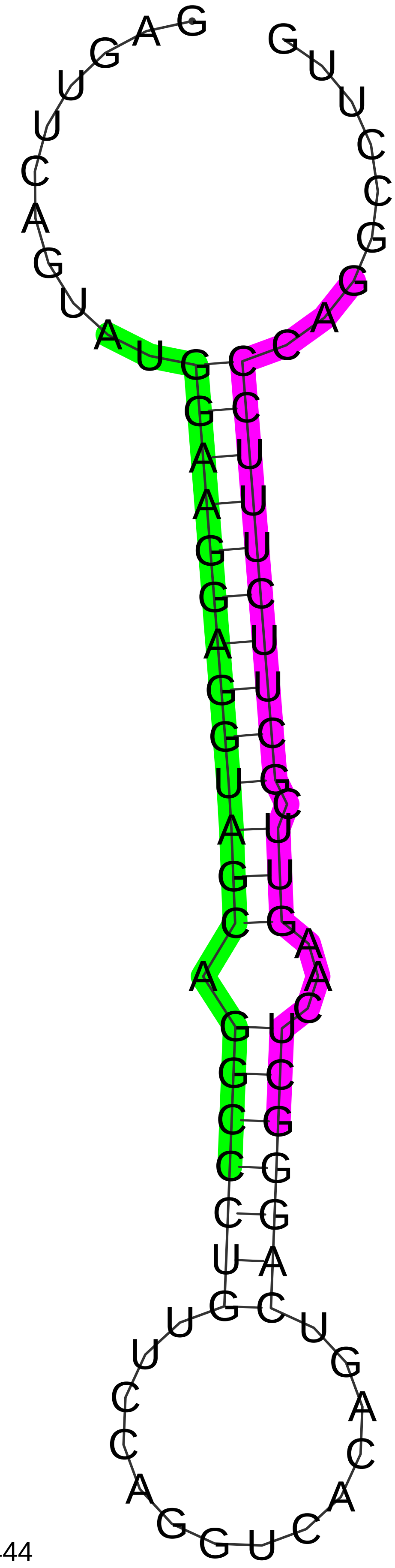

NW_006388286.1_42366-42444

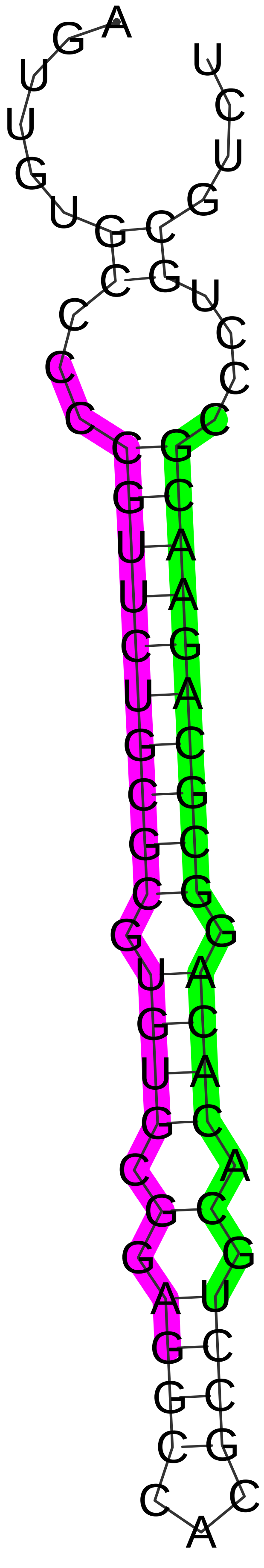

NW_006388351.1_8795-8862

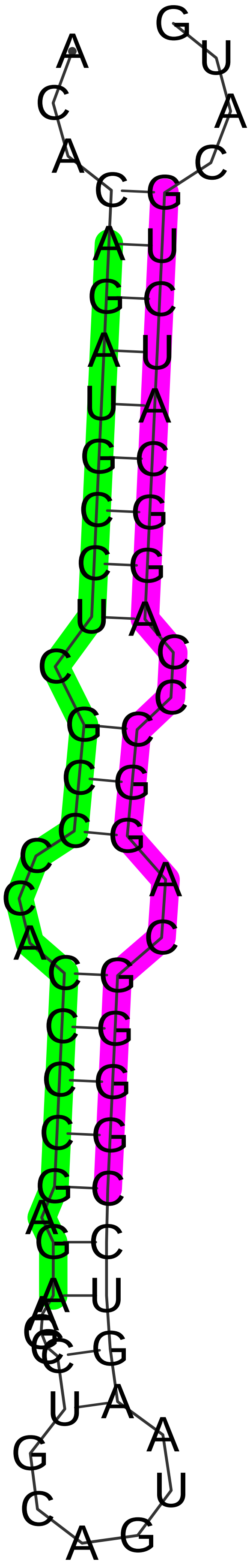

NW_006388531.1_47237-47303

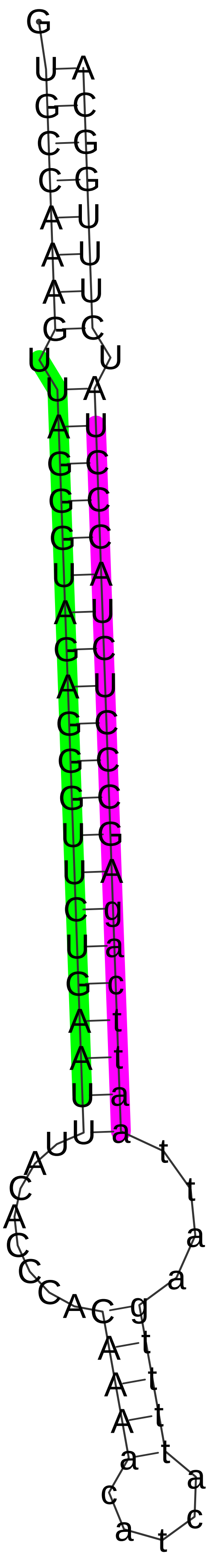

NW_006388751.1_25747-25834

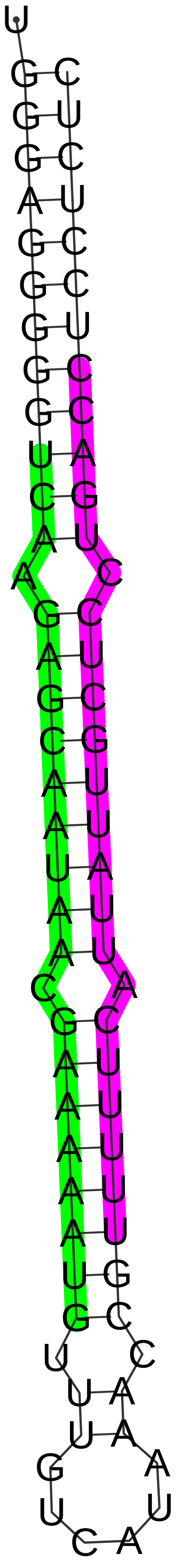

NW_006389291.1_40222-40296

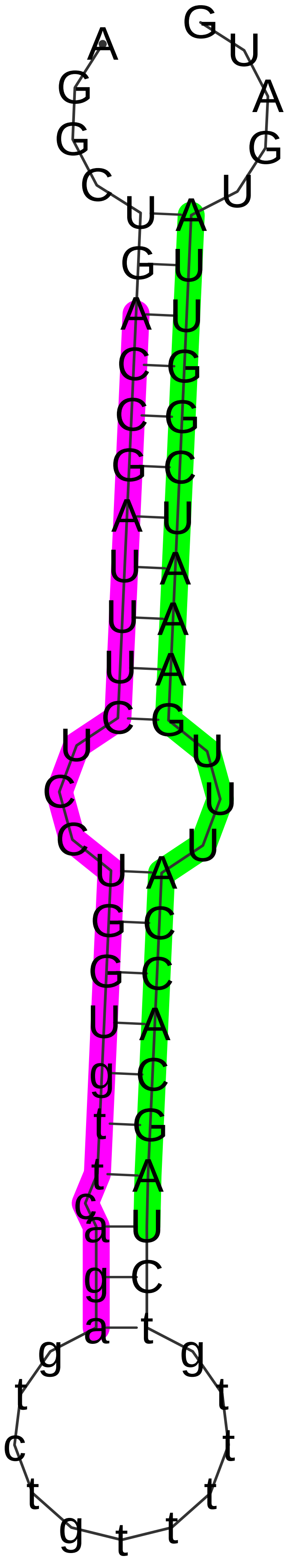

NW_006389358.1_86178-86246

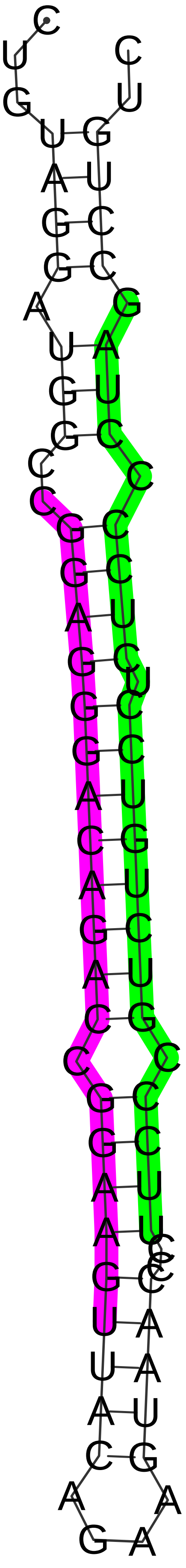

NW_006390155.1_1137-1211

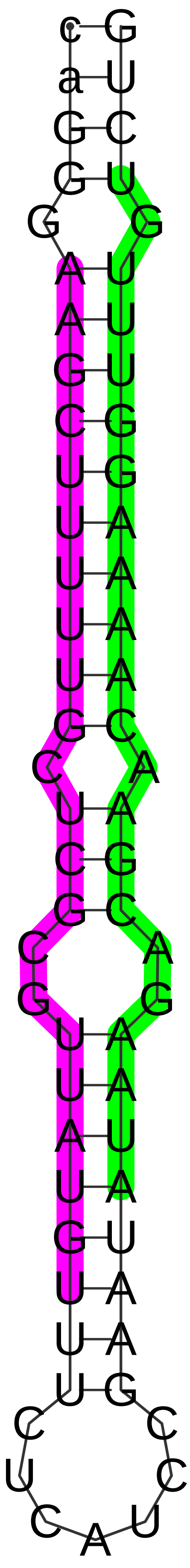


NW_006390703.1_5599-5663

Supplementary Figure S5. Abundance plots for all 559 annotated miRNA loci. Abundance (y axis) along the hairpin sequence (x axis) is defined as reads per million genome matching. Colors separate tissues, while fill and dotted lines denote pup and adult individuals, respectively.

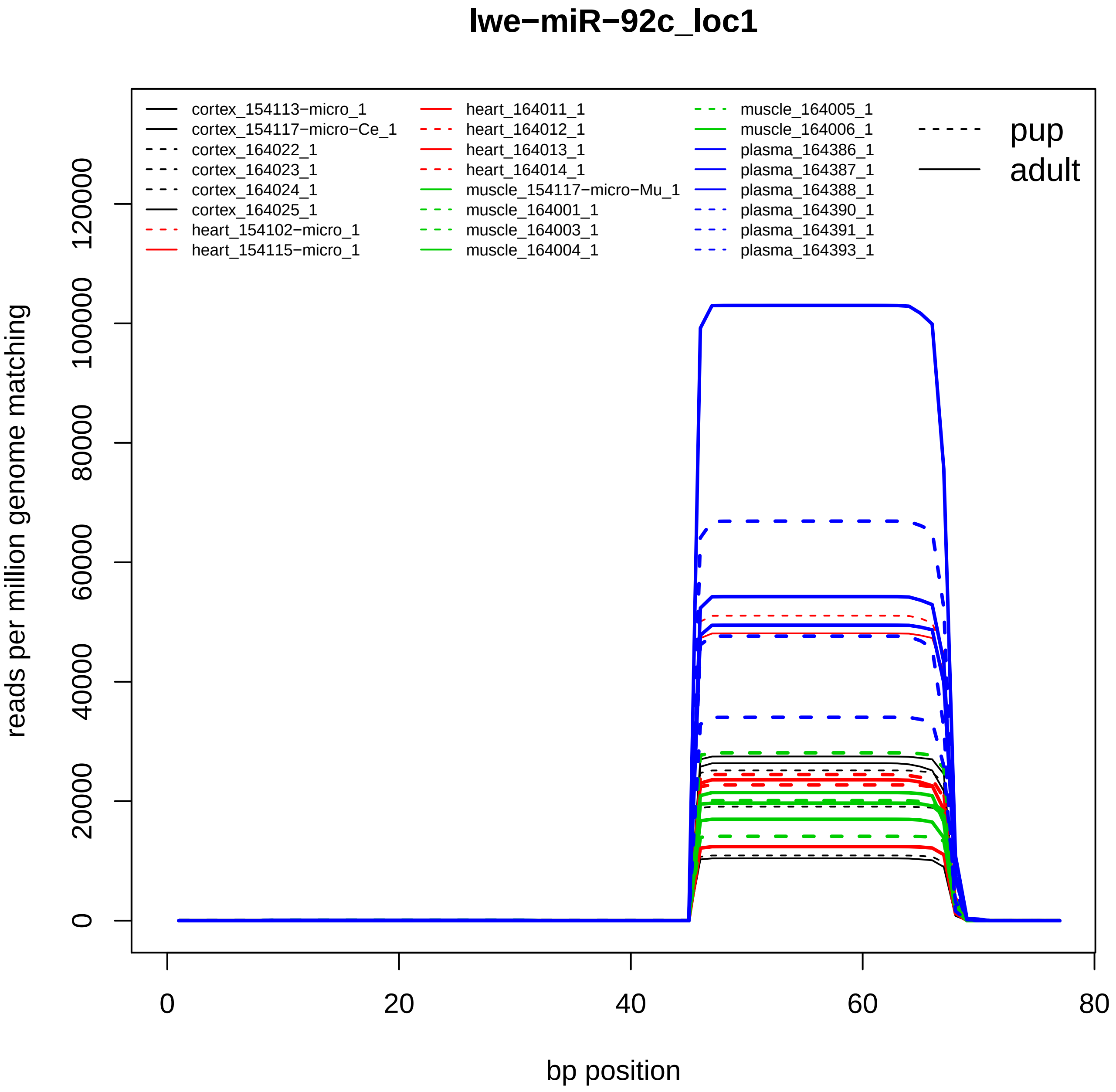
lwe−miR−92c_loc1
cortex_154113−micro_1
cortex_154117−micro−Ce_1
cortex_164022_1
cortex_164023_1
cortex_164024_1
cortex_164025_1
heart_154102−micro_1
heart_154115−micro_1
heart_164011_1
heart_164012_1
heart_164013_1
heart_164014_1
muscle_154117−micro−Mu_1
muscle_164001_1
muscle_164003_1
muscle_164004_1
muscle_164005_1
muscle_164006_1
plasma_164386_1
plasma_164387_1
plasma_164388_1
plasma_164390_1
plasma_164391_1
plasma_164393_1
pup
adult
reads per million genome matching
0
20000
40000
60000
80000
100000
120000
0
20
40
60
80
bp position

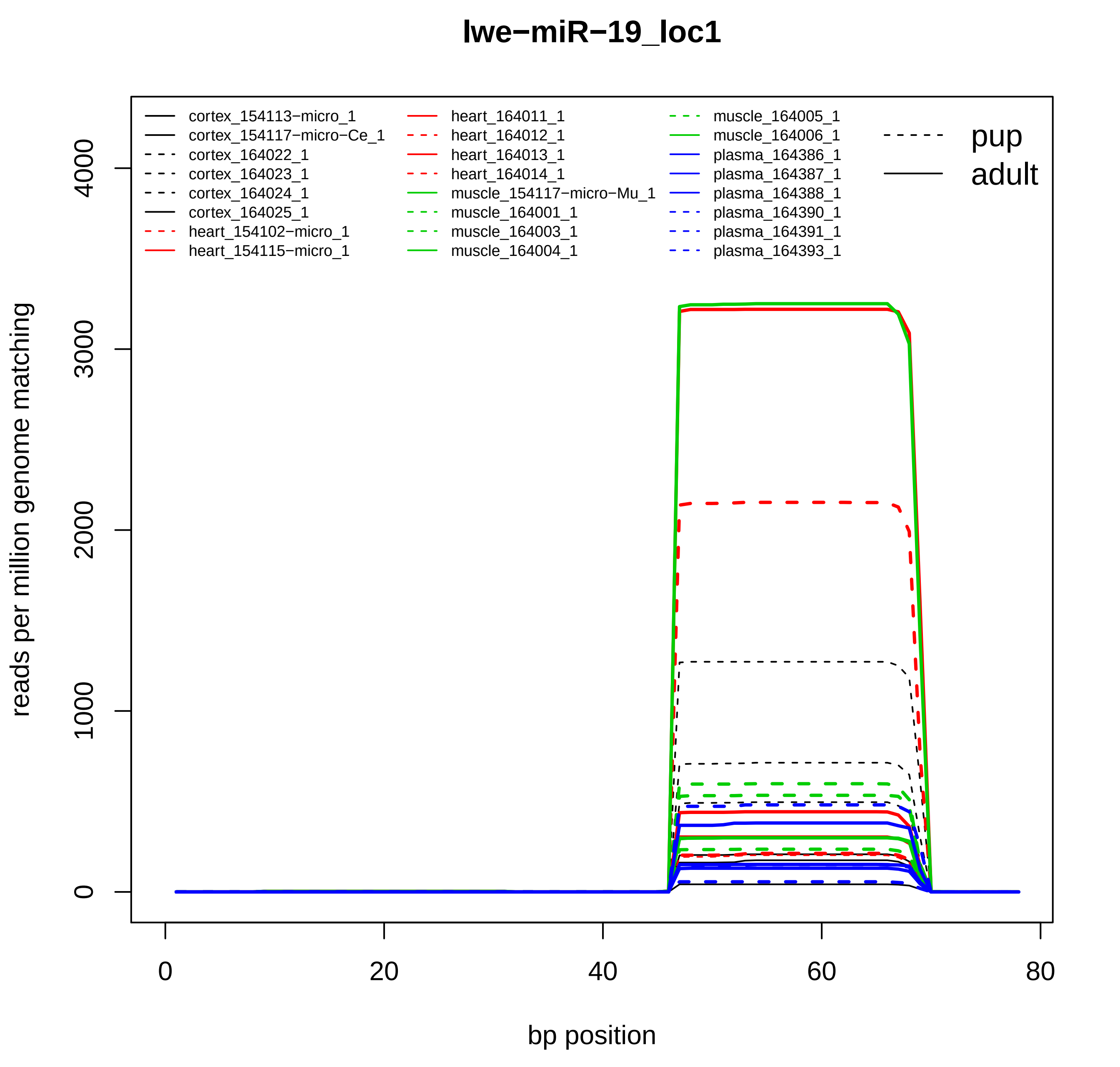
lwe−miR−19_loc1
cortex_154113−micro_1
cortex_154117−micro−Ce_1
cortex_164022_1
cortex_164023_1
cortex_164024_1
cortex_164025_1
heart_154102−micro_1
heart_154115−micro_1
heart_164011_1
heart_164012_1
heart_164013_1
heart_164014_1
muscle_154117−micro−Mu_1
muscle_164001_1
muscle_164003_1
muscle_164004_1
muscle_164005_1
muscle_164006_1
plasma_164386_1
plasma_164387_1
plasma_164388_1
plasma_164390_1
plasma_164391_1
plasma_164393_1
pup
adult
reads per million genome matching
0
1000
2000
3000
4000
0
20
40
60
80
bp position

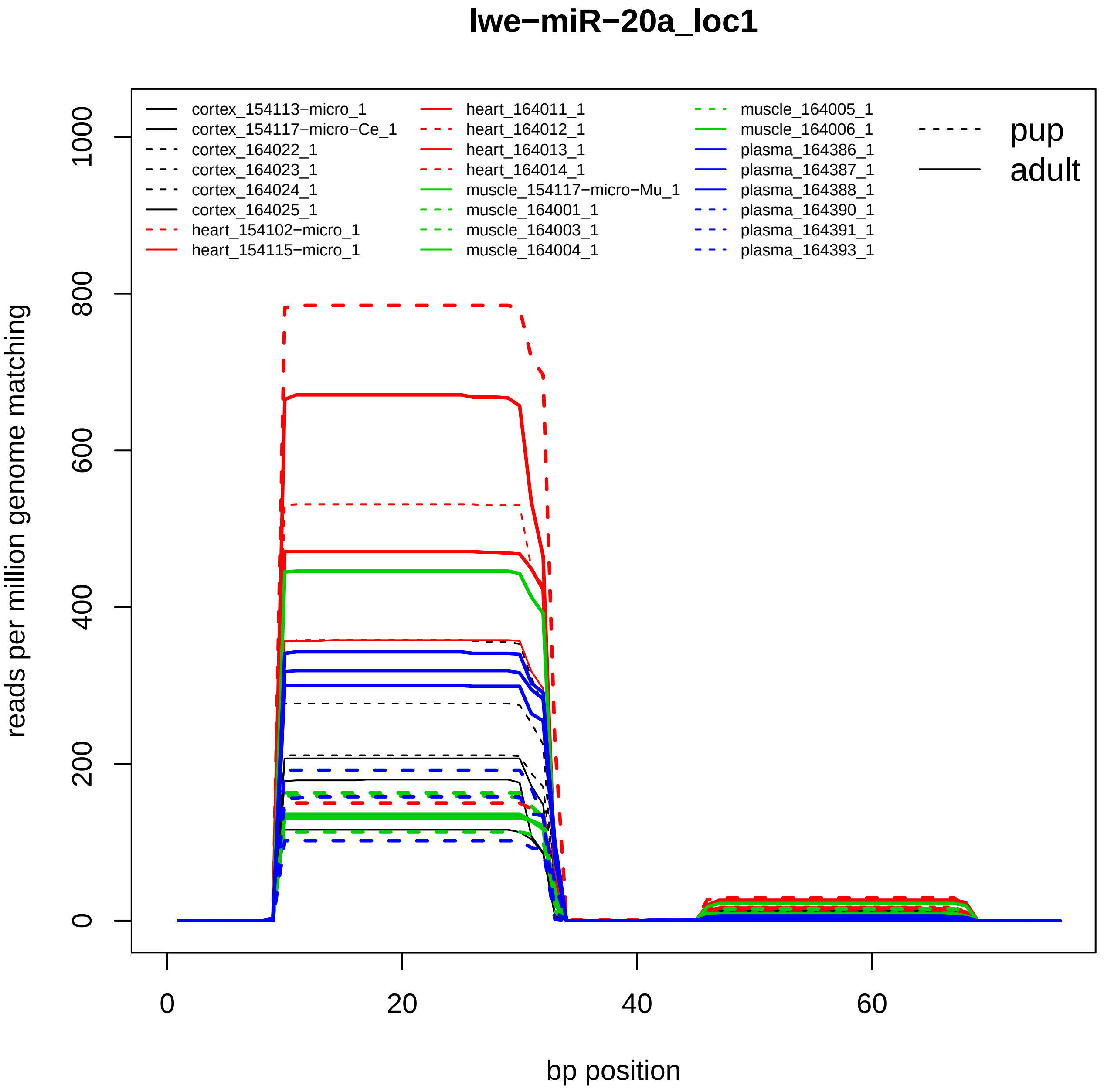

lwe−miR−20a_loc1
cortex_154113−micro_1
cortex_154117−micro−Ce_1
cortex_164022_1
cortex_164023_1
cortex_164024_1
cortex_164025_1
heart_154102−micro_1
heart_154115−micro_1
heart_164011_1
heart_164012_1
heart_164013_1
heart_164014_1
muscle_154117−micro−Mu_1
muscle_164001_1
muscle_164003_1
muscle_164004_1
muscle_164005_1
muscle_164006_1
plasma_164386_1
plasma_164387_1
plasma_164388_1
plasma_164390_1
plasma_164391_1
plasma_164393_1
pup
adult
reads per million genome matching
bp position
0
200
400
600
800
1000
0
20
40
60

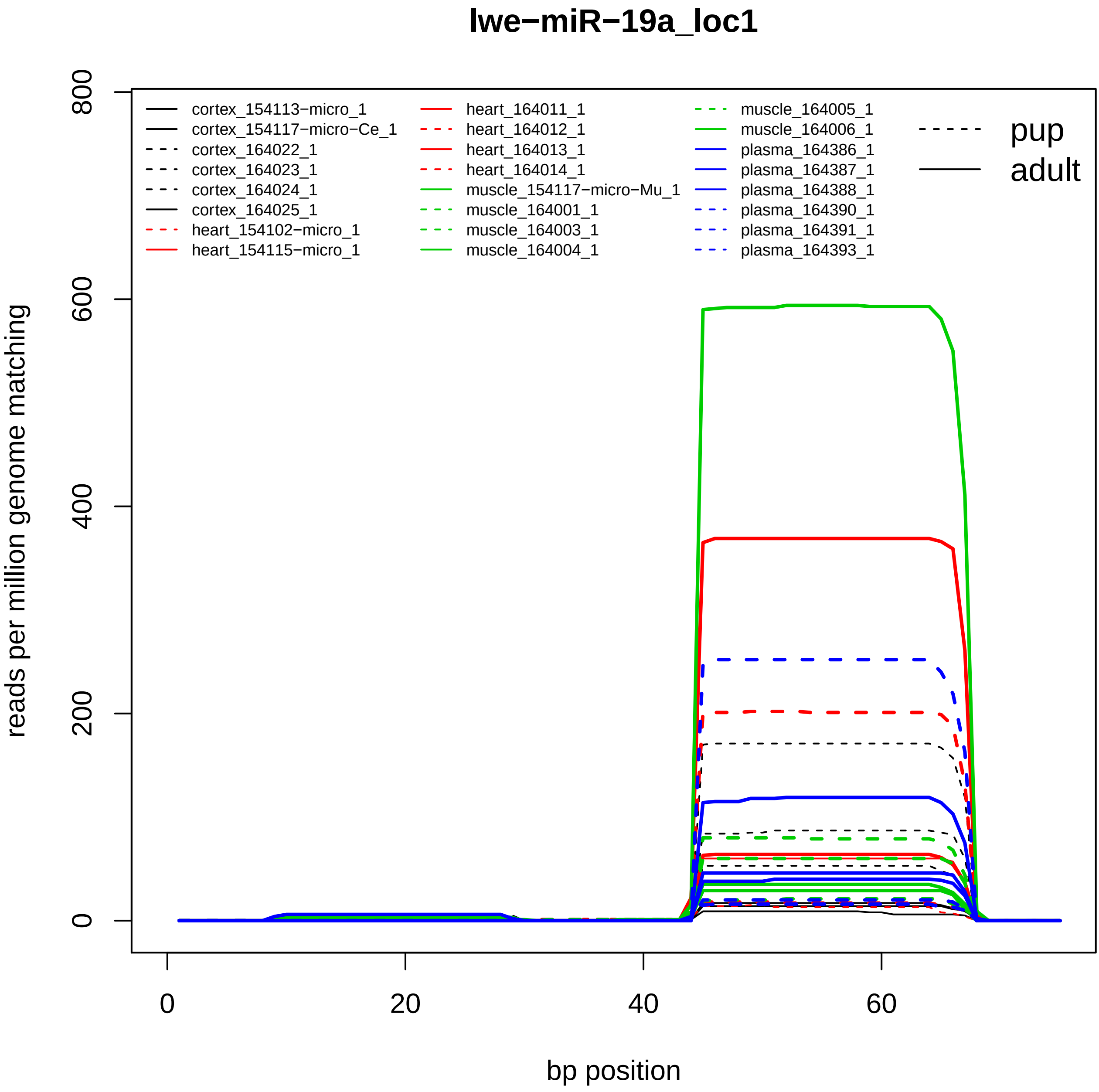
lwe−miR−19a_loc1
reads per million genome matching
bp position
0
200
400
600
800
0
20
40
60
cortex_154113−micro_1
cortex_154117−micro−Ce_1
cortex_164022_1
cortex_164023_1
cortex_164024_1
cortex_164025_1
heart_154102−micro_1
heart_154115−micro_1
heart_164011_1
heart_164012_1
heart_164013_1
heart_164014_1
muscle_154117−micro−Mu_1
muscle_164001_1
muscle_164003_1
muscle_164004_1
muscle_164005_1
muscle_164006_1
plasma_164386_1
plasma_164387_1
plasma_164388_1
plasma_164390_1
plasma_164391_1
plasma_164393_1
pup
adult

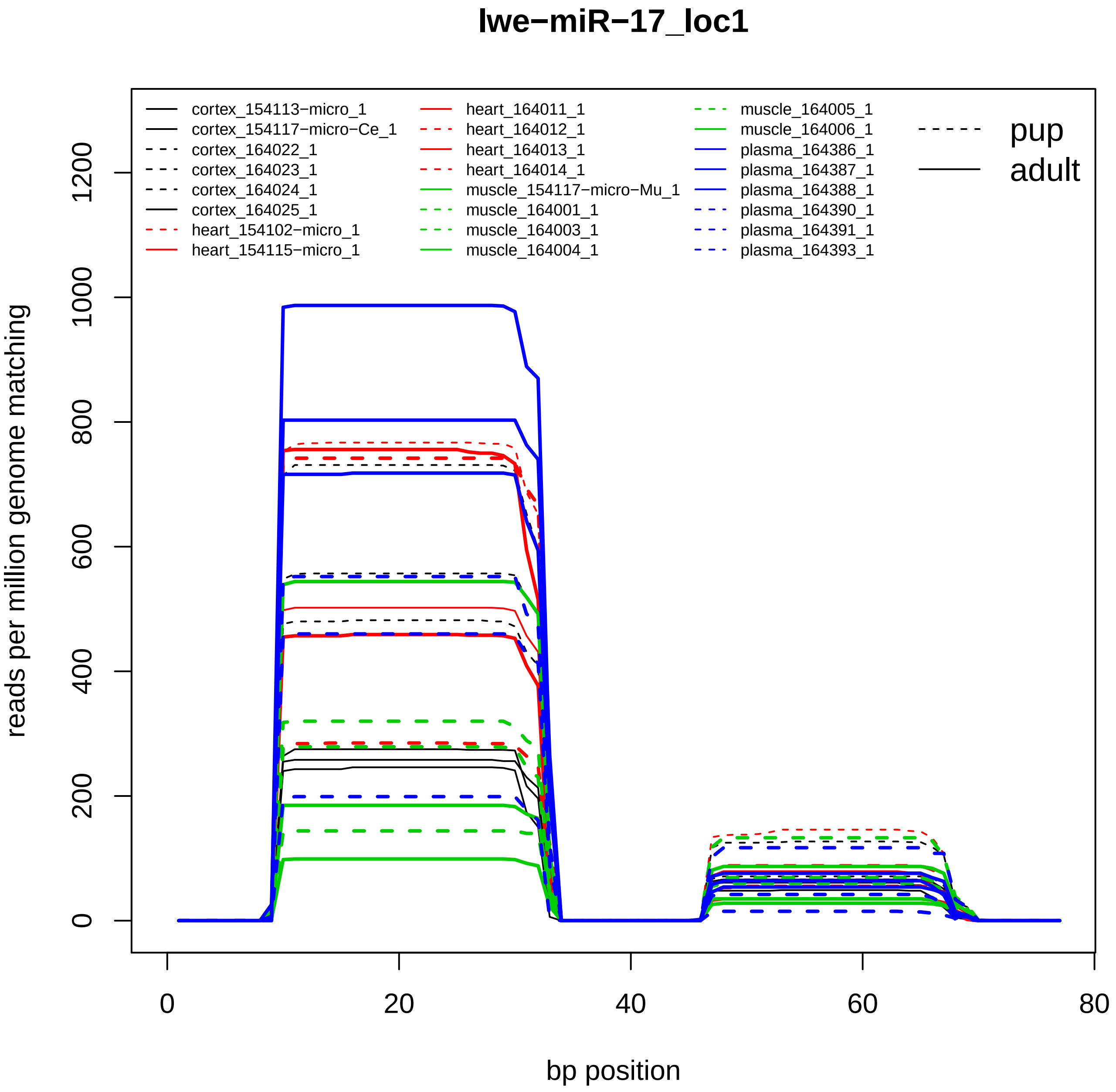
lwe−miR−17_loc1
reads per million genome matching
bp position
0
20
40
60
80
0
200
400
600
800
1000
1200
cortex_154113−micro_1
cortex_154117−micro−Ce_1
cortex_164022_1
cortex_164023_1
cortex_164024_1
cortex_164025_1
heart_154102−micro_1
heart_154115−micro_1
heart_164011_1
heart_164012_1
heart_164013_1
heart_164014_1
muscle_154117−micro−Mu_1
muscle_164001_1
muscle_164003_1
muscle_164004_1
muscle_164005_1
muscle_164006_1
plasma_164386_1
plasma_164387_1
plasma_164388_1
plasma_164390_1
plasma_164391_1
plasma_164393_1
pup
adult

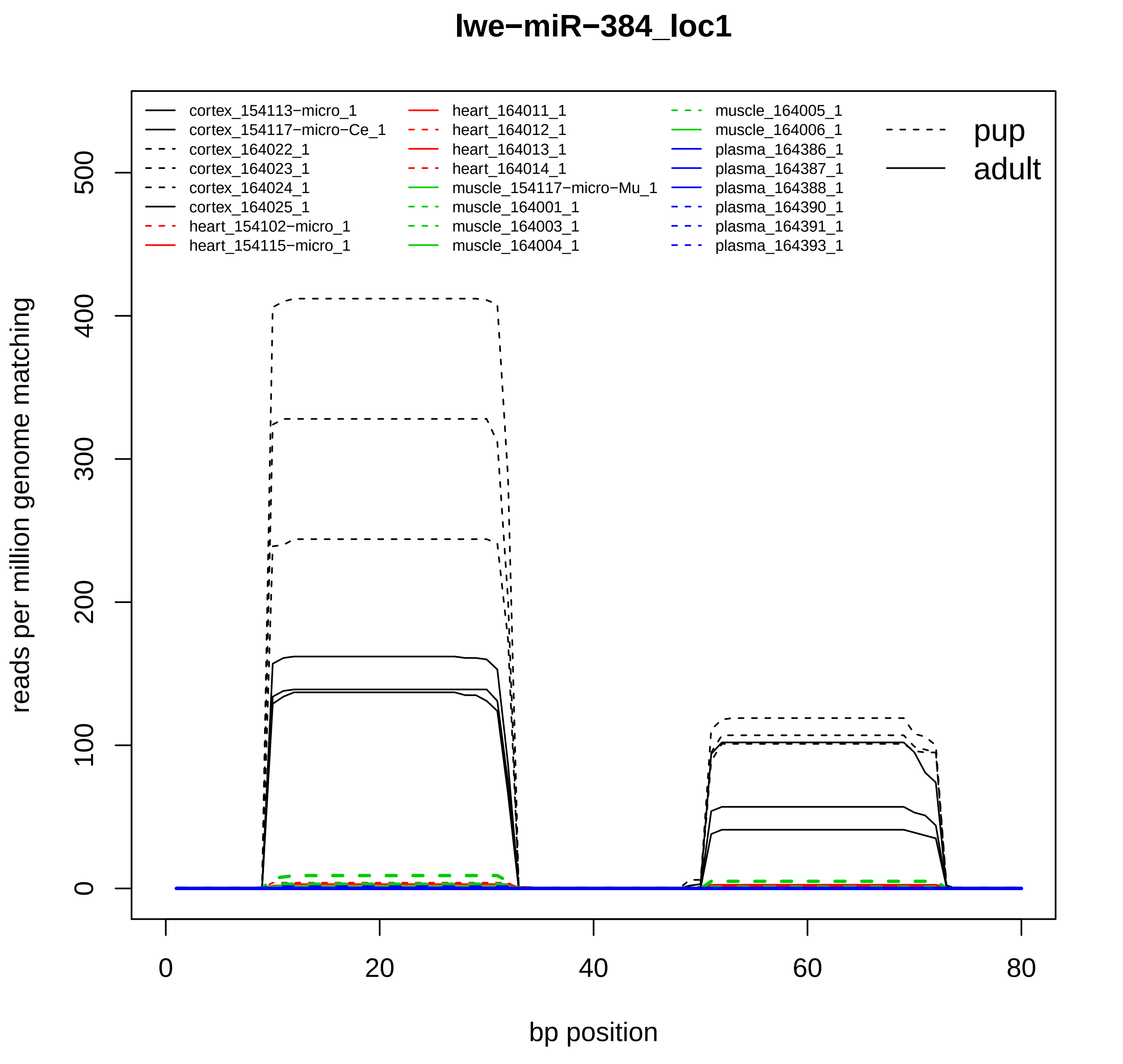

lwe−miR−384_loc1
cortex_154113−micro_1
cortex_154117−micro−Ce_1
cortex_164022_1
cortex_164023_1
cortex_164024_1
cortex_164025_1
heart_154102−micro_1
heart_154115−micro_1
heart_164011_1
heart_164012_1
heart_164013_1
heart_164014_1
muscle_154117−micro−Mu_1
muscle_164001_1
muscle_164003_1
muscle_164004_1
muscle_164005_1
muscle_164006_1
plasma_164386_1
plasma_164387_1
plasma_164388_1
plasma_164390_1
plasma_164391_1
plasma_164393_1
pup
adult
reads per million genome matching
0
100
200
300
400
500
bp position
0
20
40
60
80

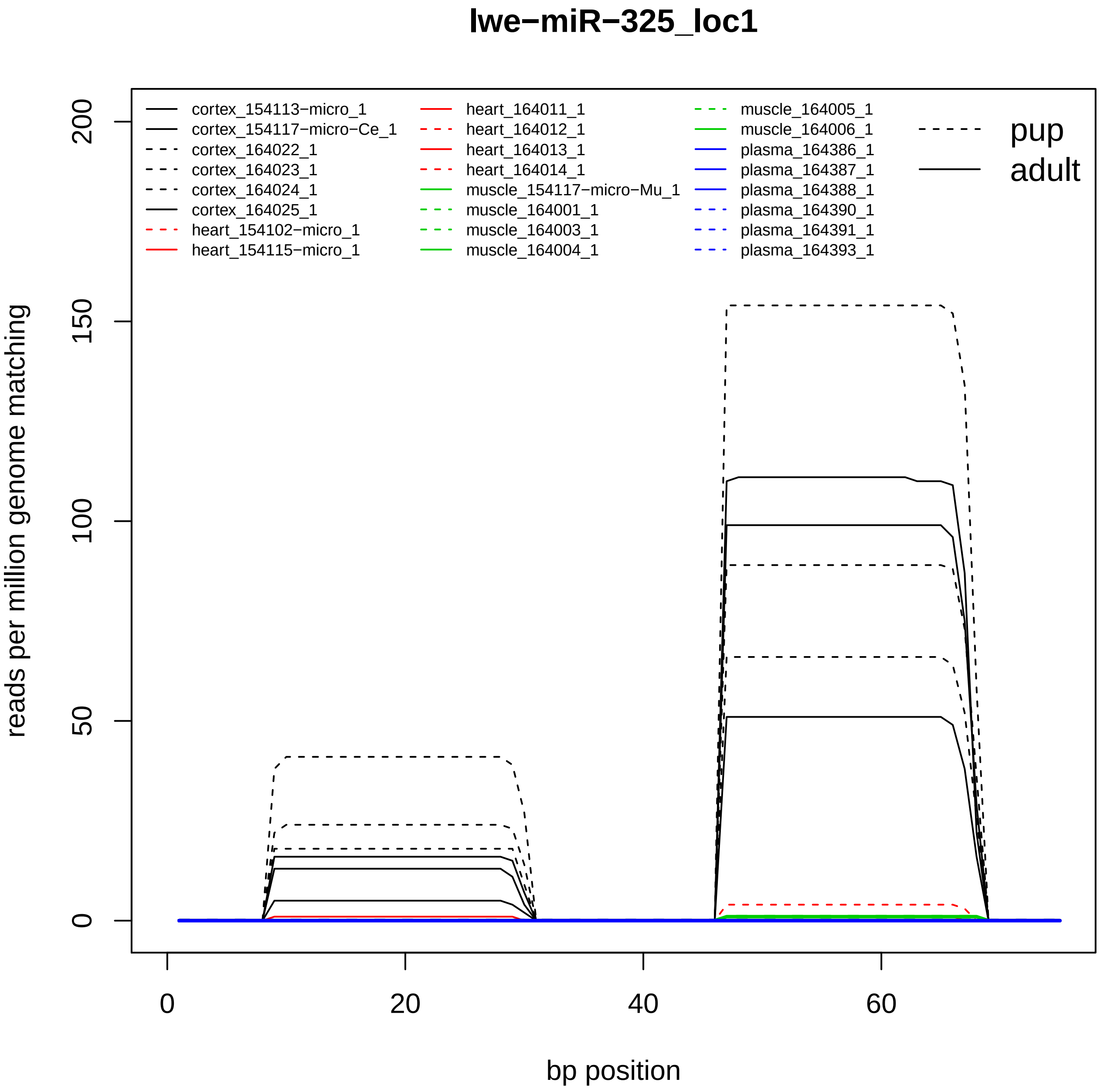
lwe−miR−325_loc1
cortex_154113−micro_1
cortex_154117−micro−Ce_1
cortex_164022_1
cortex_164023_1
cortex_164024_1
cortex_164025_1
heart_154102−micro_1
heart_154115−micro_1
heart_164011_1
heart_164012_1
heart_164013_1
heart_164014_1
muscle_154117−micro−Mu_1
muscle_164001_1
muscle_164003_1
muscle_164004_1
muscle_164005_1
muscle_164006_1
plasma_164386_1
plasma_164387_1
plasma_164388_1
plasma_164390_1
plasma_164391_1
plasma_164393_1
pup
adult
reads per million genome matching
0
50
100
150
200
bp position
0
20
40
60

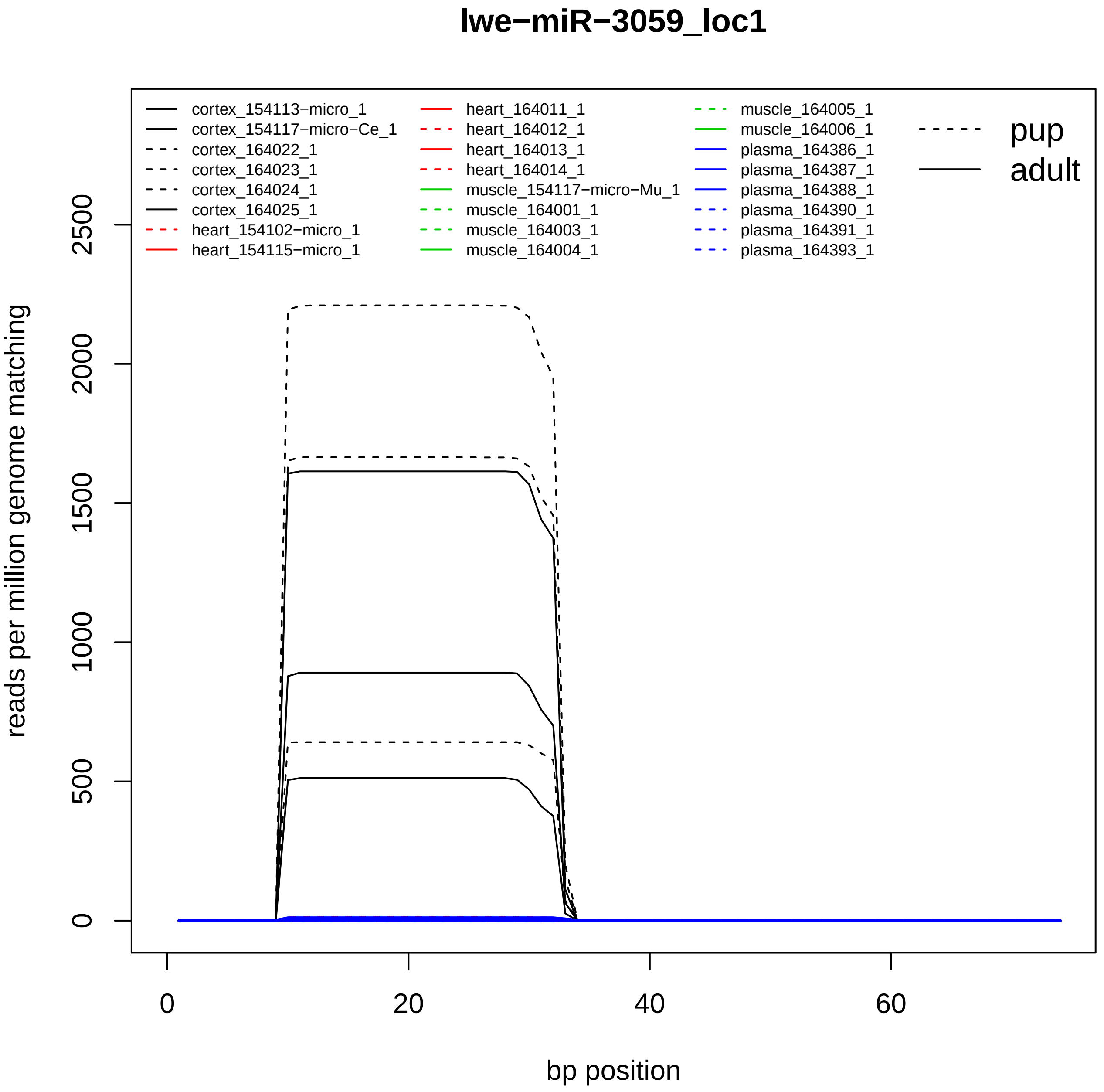
lwe−miR−3059_loc1
cortex_154113−micro_1
cortex_154117−micro−Ce_1
cortex_164022_1
cortex_164023_1
cortex_164024_1
cortex_164025_1
heart_154102−micro_1
heart_154115−micro_1
heart_164011_1
heart_164012_1
heart_164013_1
heart_164014_1
muscle_154117−micro−Mu_1
muscle_164001_1
muscle_164003_1
muscle_164004_1
muscle_164005_1
muscle_164006_1
plasma_164386_1
plasma_164387_1
plasma_164388_1
plasma_164390_1
plasma_164391_1
plasma_164393_1
pup
adult
reads per million genome matching
0
500
1000
1500
2000
2500
0
20
40
60
bp position

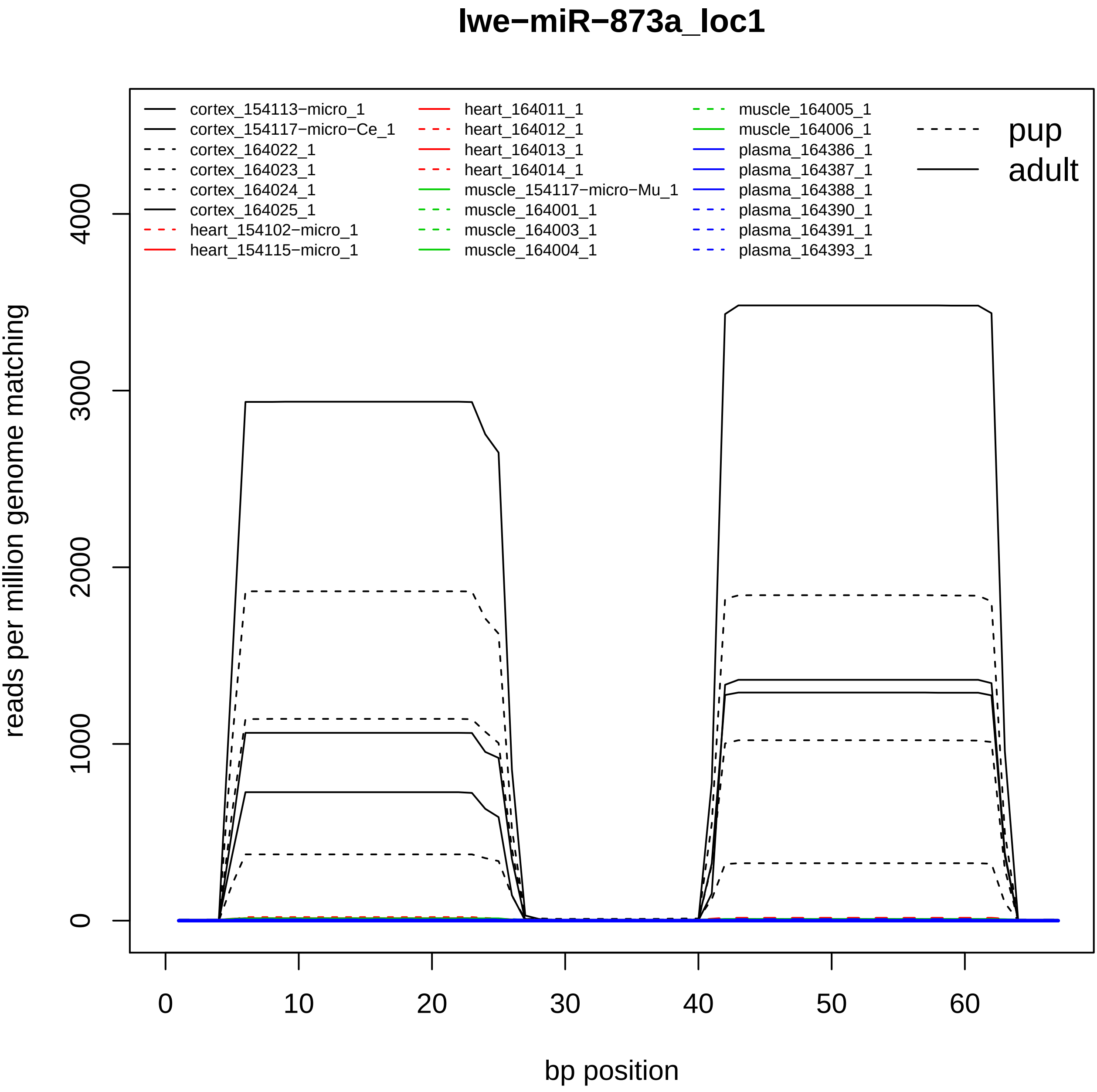
lwe−miR−873a_loc1
cortex_154113−micro_1
cortex_154117−micro−Ce_1
cortex_164022_1
cortex_164023_1
cortex_164024_1
cortex_164025_1
heart_154102−micro_1
heart_154115−micro_1
heart_164011_1
heart_164012_1
heart_164013_1
heart_164014_1
muscle_154117−micro−Mu_1
muscle_164001_1
muscle_164003_1
muscle_164004_1
muscle_164005_1
muscle_164006_1
plasma_164386_1
plasma_164387_1
plasma_164388_1
plasma_164390_1
plasma_164391_1
plasma_164393_1
pup
adult
reads per million genome matching
0
1000
2000
3000
4000
bp position
0
10
20
30
40
50
60

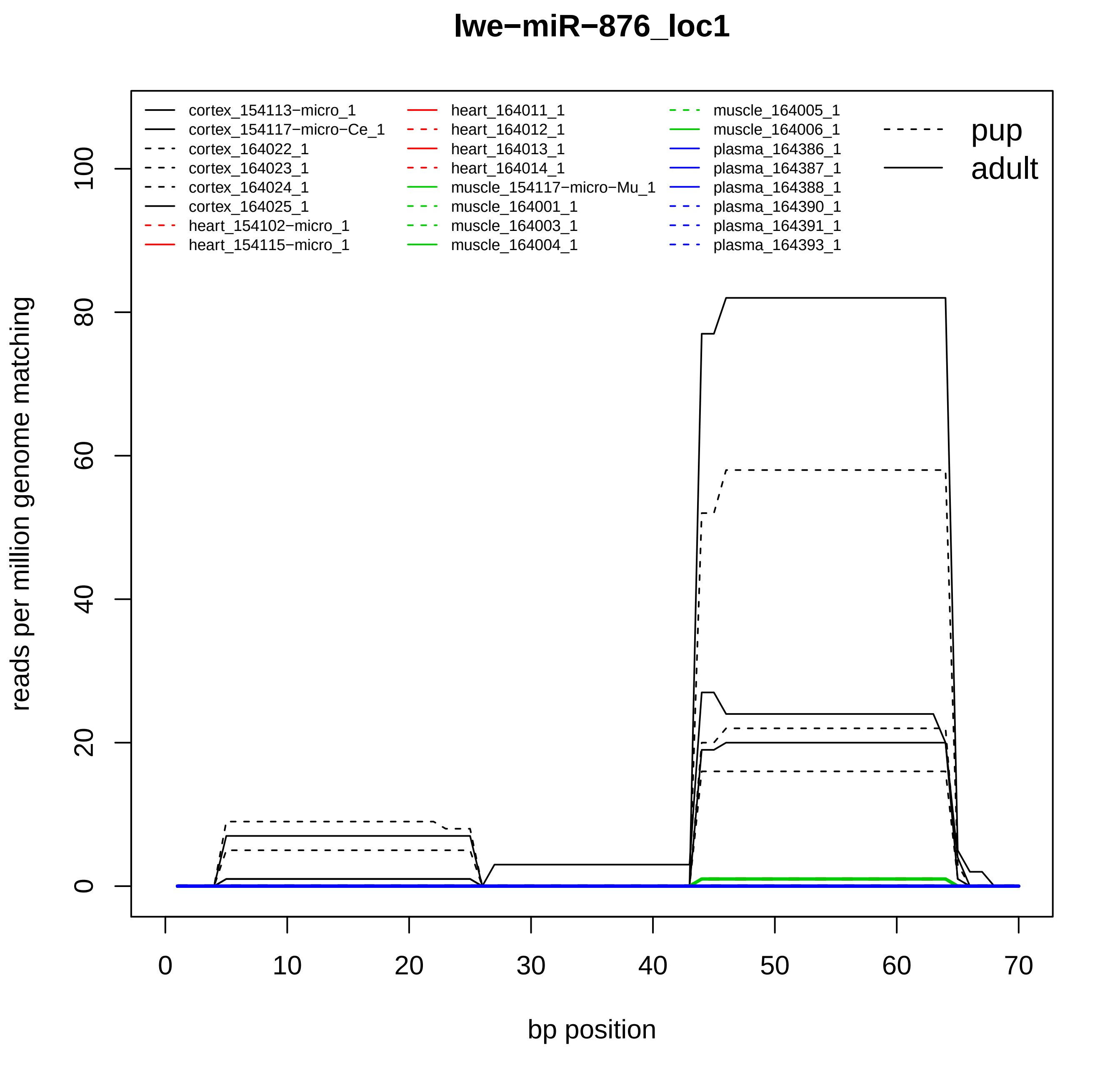
lwe−miR−876_loc1
cortex_154113−micro_1
cortex_154117−micro−Ce_1
cortex_164022_1
cortex_164023_1
cortex_164024_1
cortex_164025_1
heart_154102−micro_1
heart_154115−micro_1
heart_164011_1
heart_164012_1
heart_164013_1
heart_164014_1
muscle_154117−micro−Mu_1
muscle_164001_1
muscle_164003_1
muscle_164004_1
muscle_164005_1
muscle_164006_1
plasma_164386_1
plasma_164387_1
plasma_164388_1
plasma_164390_1
plasma_164391_1
plasma_164393_1
pup
adult
reads per million genome matching
0
20
40
60
80
100
0
10
20
30
40
50
60
70
bp position

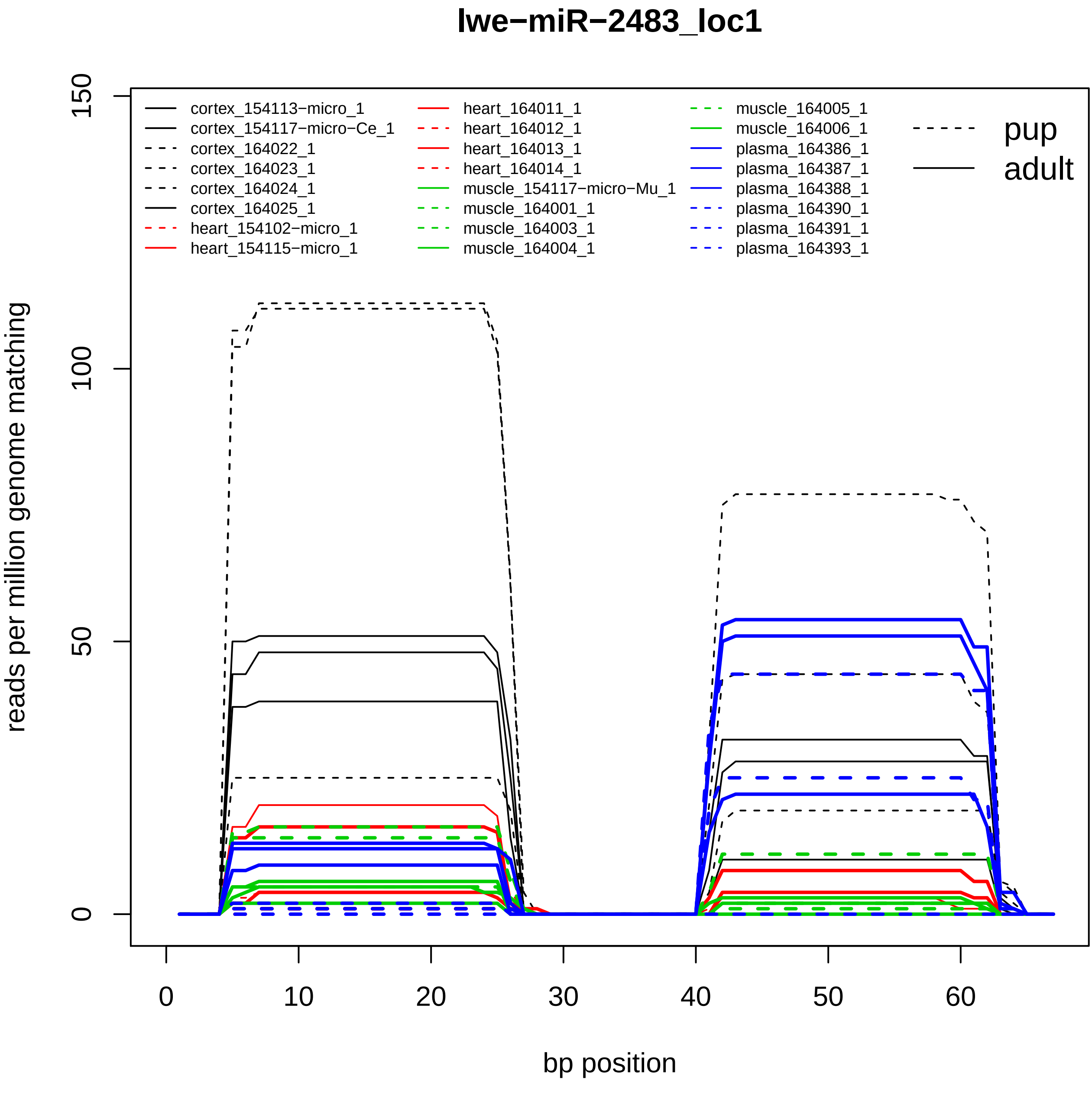
lwe−miR−2483_loc1
reads per million genome matching
bp position
0
50
100
150
0
10
20
30
40
50
60
cortex_154113−micro_1
cortex_154117−micro−Ce_1
cortex_164022_1
cortex_164023_1
cortex_164024_1
cortex_164025_1
heart_154102−micro_1
heart_154115−micro_1
heart_164011_1
heart_164012_1
heart_164013_1
heart_164014_1
muscle_154117−micro−Mu_1
muscle_164001_1
muscle_164003_1
muscle_164004_1
muscle_164005_1
muscle_164006_1
plasma_164386_1
plasma_164387_1
plasma_164388_1
plasma_164390_1
plasma_164391_1
plasma_164393_1
pup
adult

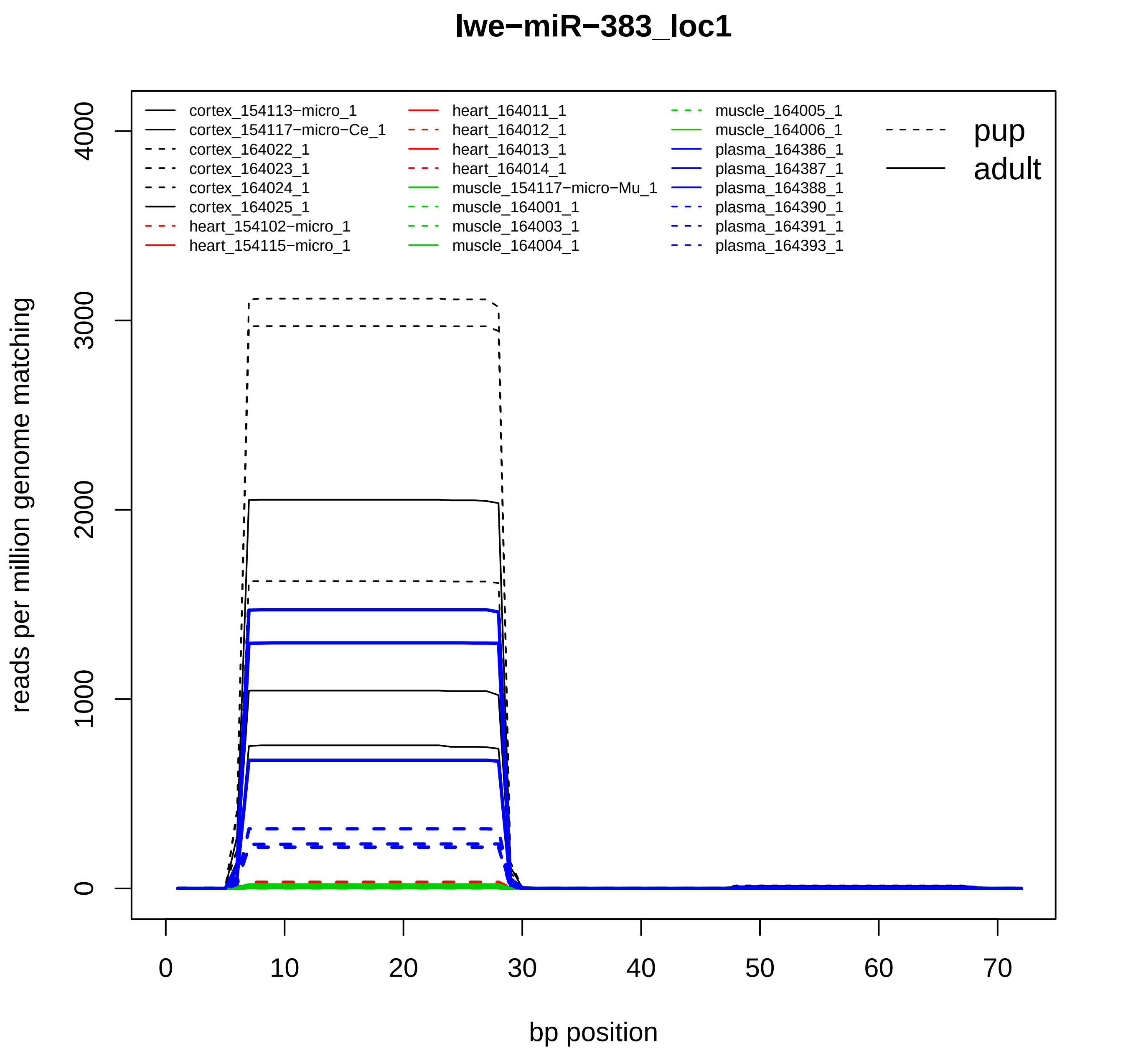

lwe−miR−383_loc1
cortex_154113−micro_1
cortex_154117−micro−Ce_1
cortex_164022_1
cortex_164023_1
cortex_164024_1
cortex_164025_1
heart_154102−micro_1
heart_154115−micro_1
heart_164011_1
heart_164012_1
heart_164013_1
heart_164014_1
muscle_154117−micro−Mu_1
muscle_164001_1
muscle_164003_1
muscle_164004_1
muscle_164005_1
muscle_164006_1
plasma_164386_1
plasma_164387_1
plasma_164388_1
plasma_164390_1
plasma_164391_1
plasma_164393_1
pup
adult
reads per million genome matching
0
1000
2000
3000
4000
0
10
20
30
40
50
60
70
bp position

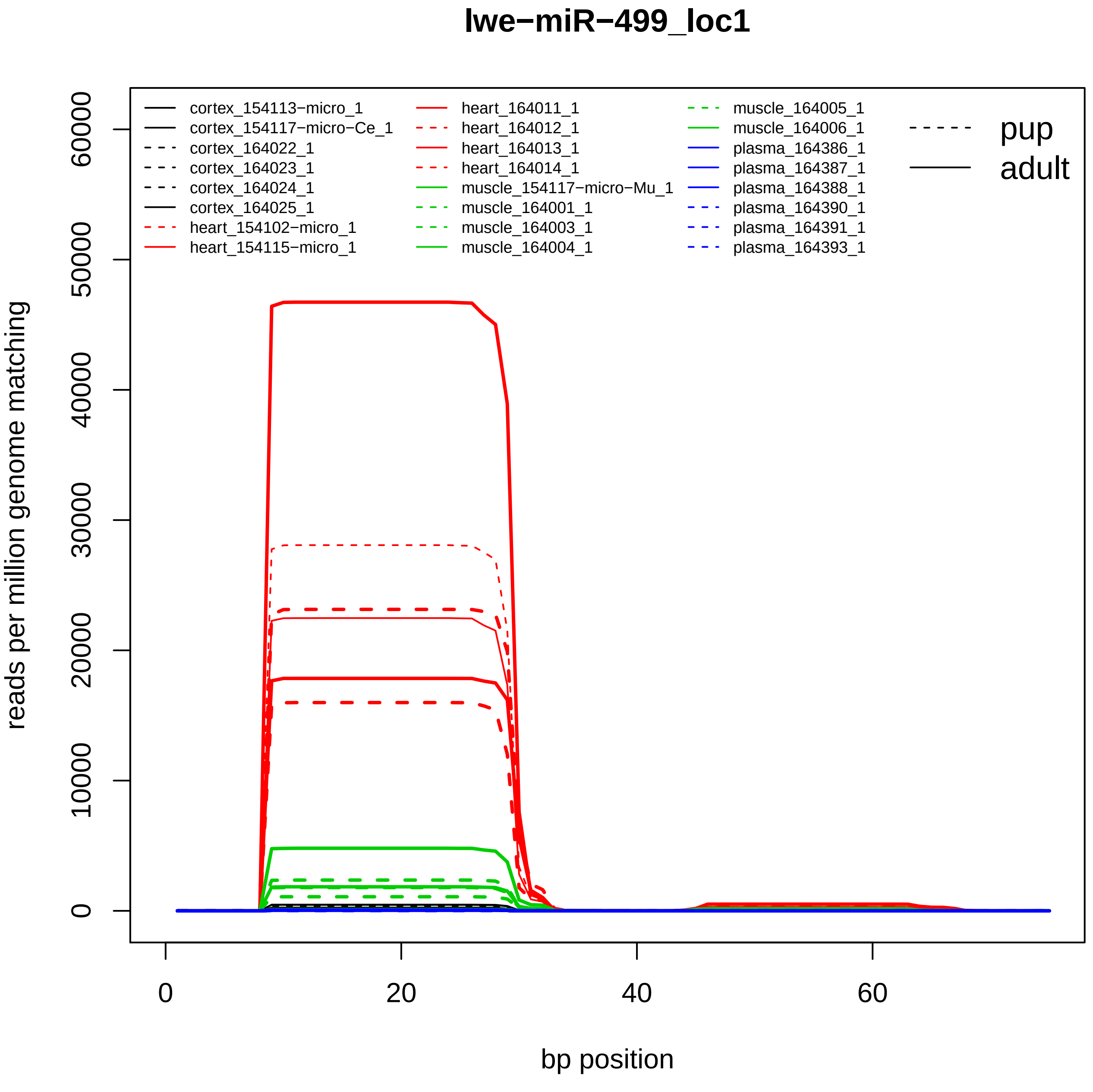

lwe−miR−499_loc1
cortex_154113−micro_1
cortex_154117−micro−Ce_1
cortex_164022_1
cortex_164023_1
cortex_164024_1
cortex_164025_1
heart_154102−micro_1
heart_154115−micro_1
heart_164011_1
heart_164012_1
heart_164013_1
heart_164014_1
muscle_154117−micro−Mu_1
muscle_164001_1
muscle_164003_1
muscle_164004_1
muscle_164005_1
muscle_164006_1
plasma_164386_1
plasma_164387_1
plasma_164388_1
plasma_164390_1
plasma_164391_1
plasma_164393_1
pup
adult
reads per million genome matching
0
10000
20000
30000
40000
50000
60000
0
20
40
60
bp position

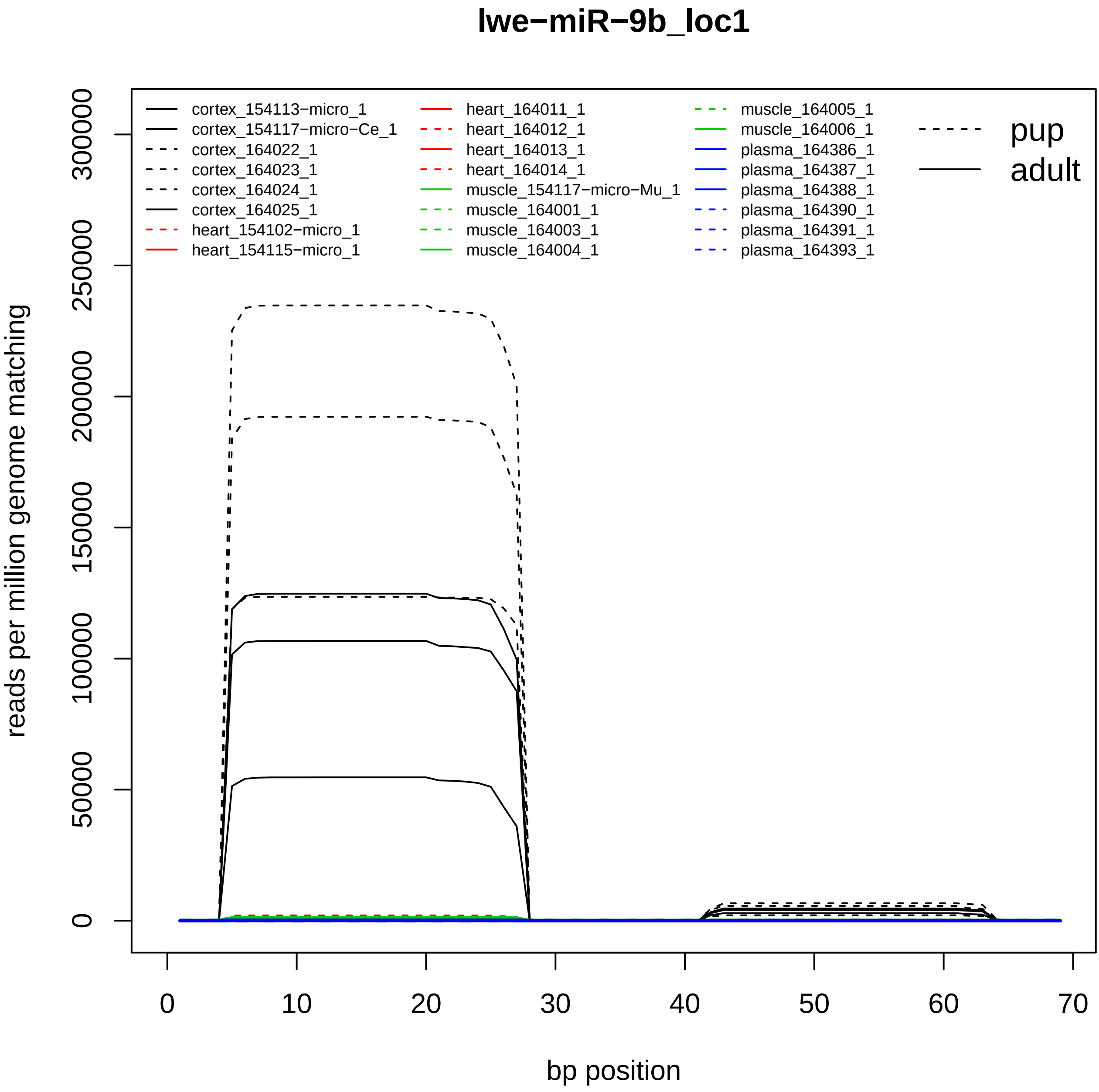

lwe−miR−9b_loc1
cortex_154113−micro_1
cortex_154117−micro−Ce_1
cortex_164022_1
cortex_164023_1
cortex_164024_1
cortex_164025_1
heart_154102−micro_1
heart_154115−micro_1
heart_164011_1
heart_164012_1
heart_164013_1
heart_164014_1
muscle_154117−micro−Mu_1
muscle_164001_1
muscle_164003_1
muscle_164004_1
muscle_164005_1
muscle_164006_1
plasma_164386_1
plasma_164387_1
plasma_164388_1
plasma_164390_1
plasma_164391_1
plasma_164393_1
pup
adult
reads per million genome matching
0
50000
100000
150000
200000
250000
300000
bp position
0
10
20
30
40
50
60
70

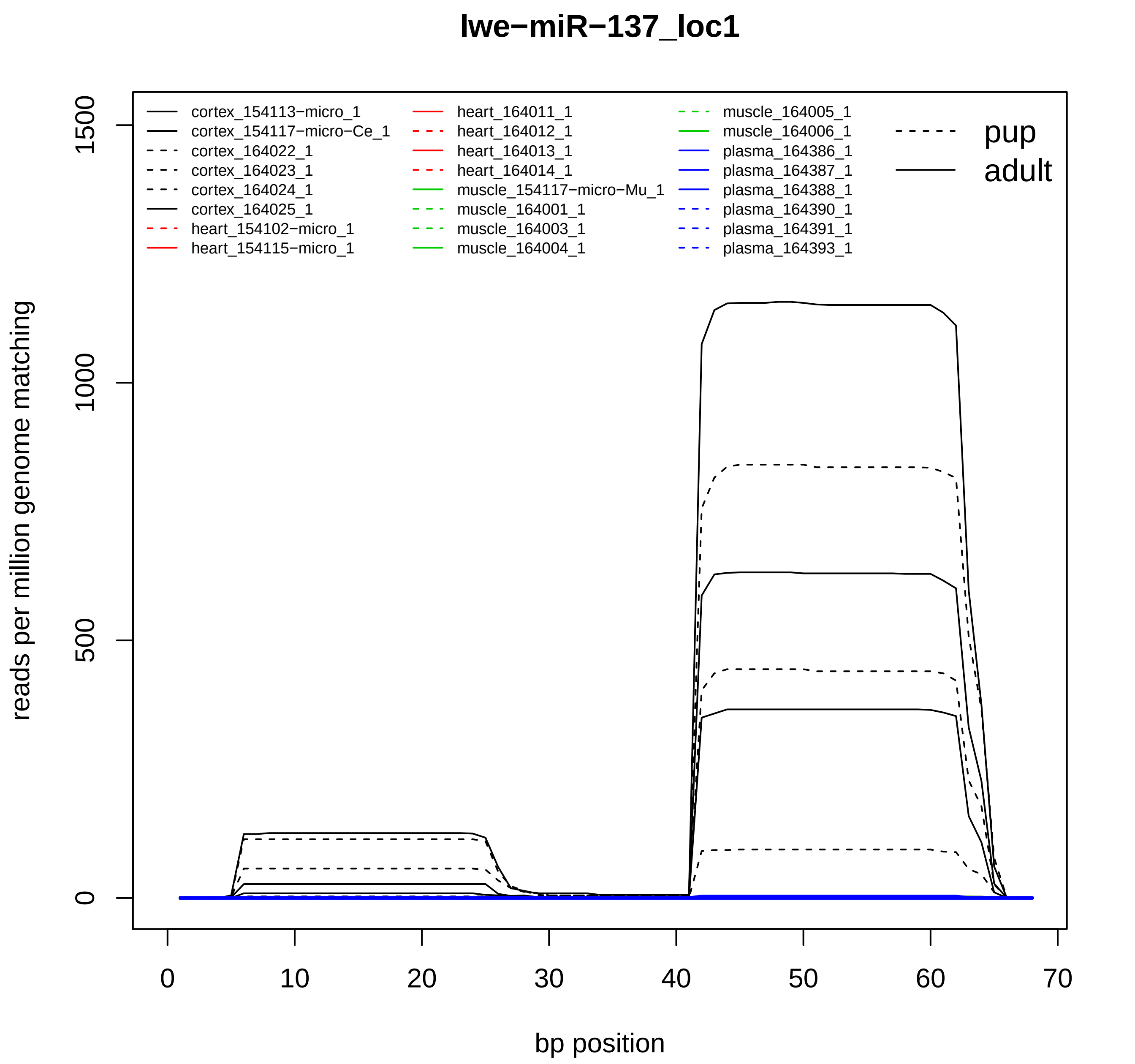
lwe−miR−137_loc1
cortex_154113−micro_1
cortex_154117−micro−Ce_1
cortex_164022_1
cortex_164023_1
cortex_164024_1
cortex_164025_1
heart_154102−micro_1
heart_154115−micro_1
heart_164011_1
heart_164012_1
heart_164013_1
heart_164014_1
muscle_154117−micro−Mu_1
muscle_164001_1
muscle_164003_1
muscle_164004_1
muscle_164005_1
muscle_164006_1
plasma_164386_1
plasma_164387_1
plasma_164388_1
plasma_164390_1
plasma_164391_1
plasma_164393_1
pup
adult
reads per million genome matching
0
500
1000
1500
0
10
20
30
40
50
60
70
bp position

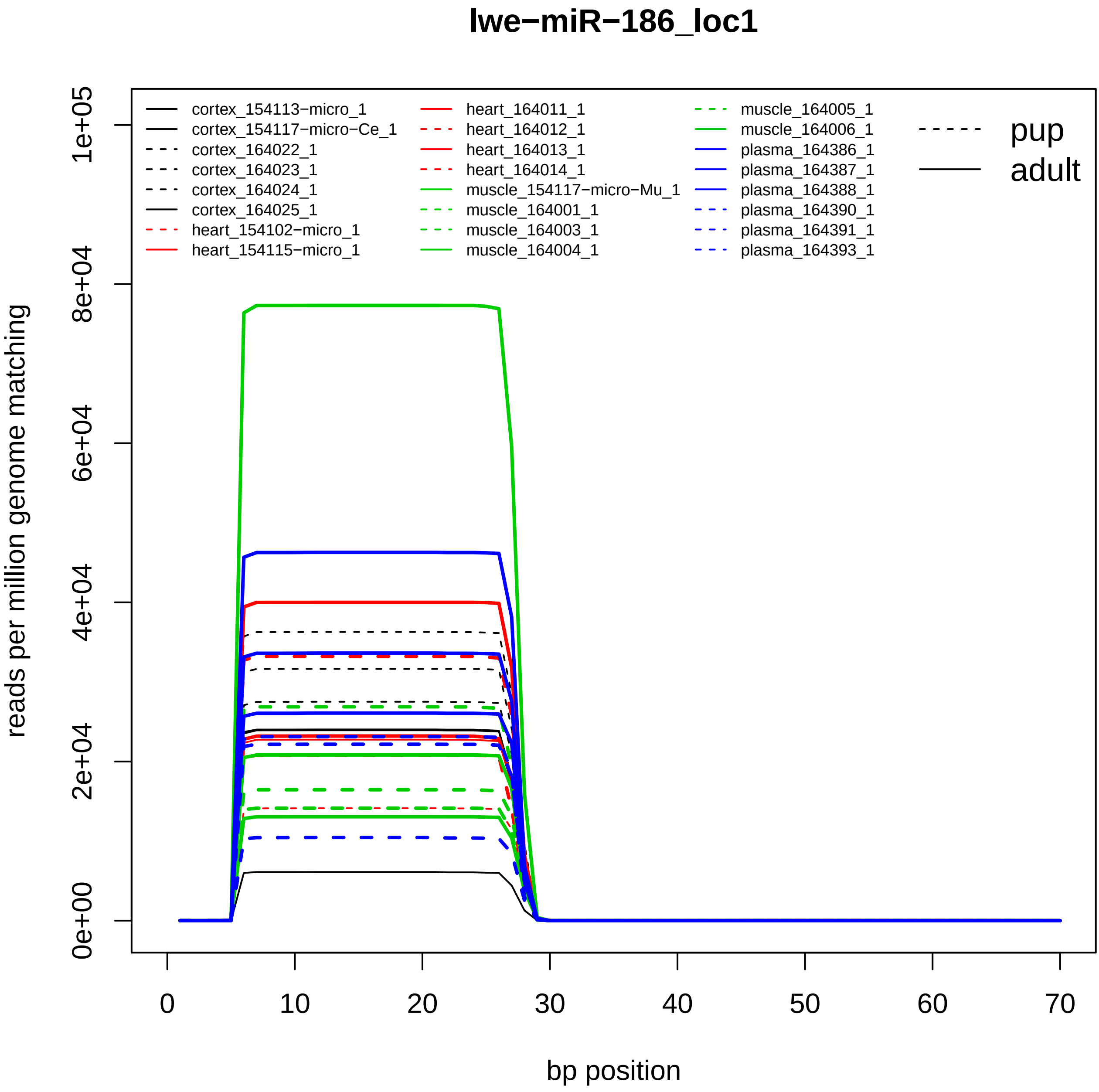
lwe−miR−186_loc1
cortex_154113−micro_1
cortex_154117−micro−Ce_1
cortex_164022_1
cortex_164023_1
cortex_164024_1
cortex_164025_1
heart_154102−micro_1
heart_154115−micro_1
heart_164011_1
heart_164012_1
heart_164013_1
heart_164014_1
muscle_154117−micro−Mu_1
muscle_164001_1
muscle_164003_1
muscle_164004_1
muscle_164005_1
muscle_164006_1
plasma_164386_1
plasma_164387_1
plasma_164388_1
plasma_164390_1
plasma_164391_1
plasma_164393_1
pup
adult
reads per million genome matching
0e+00
2e+04
4e+04
6e+04
8e+04
1e+05
bp position
0
10
20
30
40
50
60
70

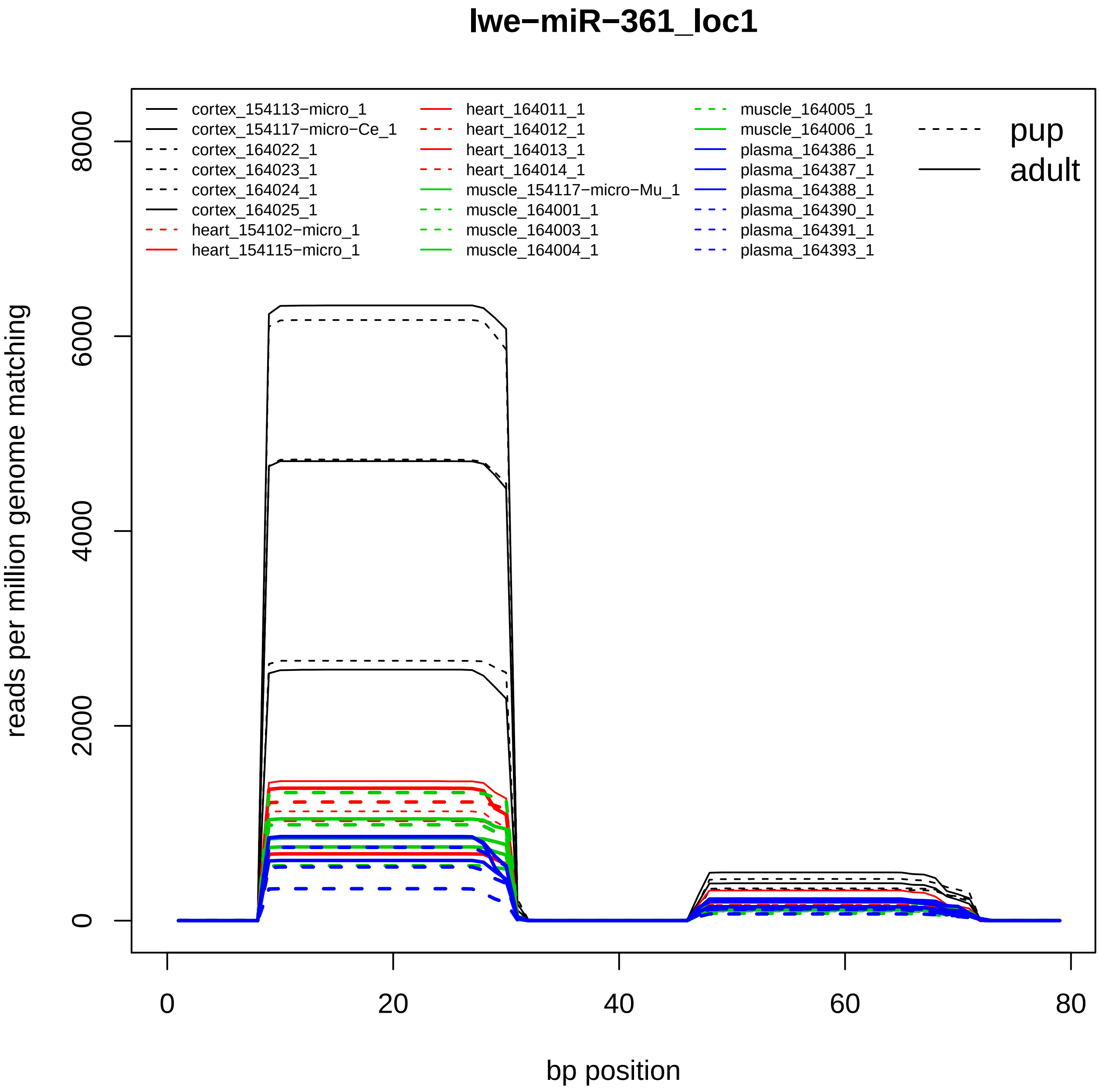
lwe−miR−361_loc1
cortex_154113−micro_1
cortex_154117−micro−Ce_1
cortex_164022_1
cortex_164023_1
cortex_164024_1
cortex_164025_1
heart_154102−micro_1
heart_154115−micro_1
heart_164011_1
heart_164012_1
heart_164013_1
heart_164014_1
muscle_154117−micro−Mu_1
muscle_164001_1
muscle_164003_1
muscle_164004_1
muscle_164005_1
muscle_164006_1
plasma_164386_1
plasma_164387_1
plasma_164388_1
plasma_164390_1
plasma_164391_1
plasma_164393_1
pup
adult
reads per million genome matching
0
2000
4000
6000
8000
0
20
40
60
80
bp position

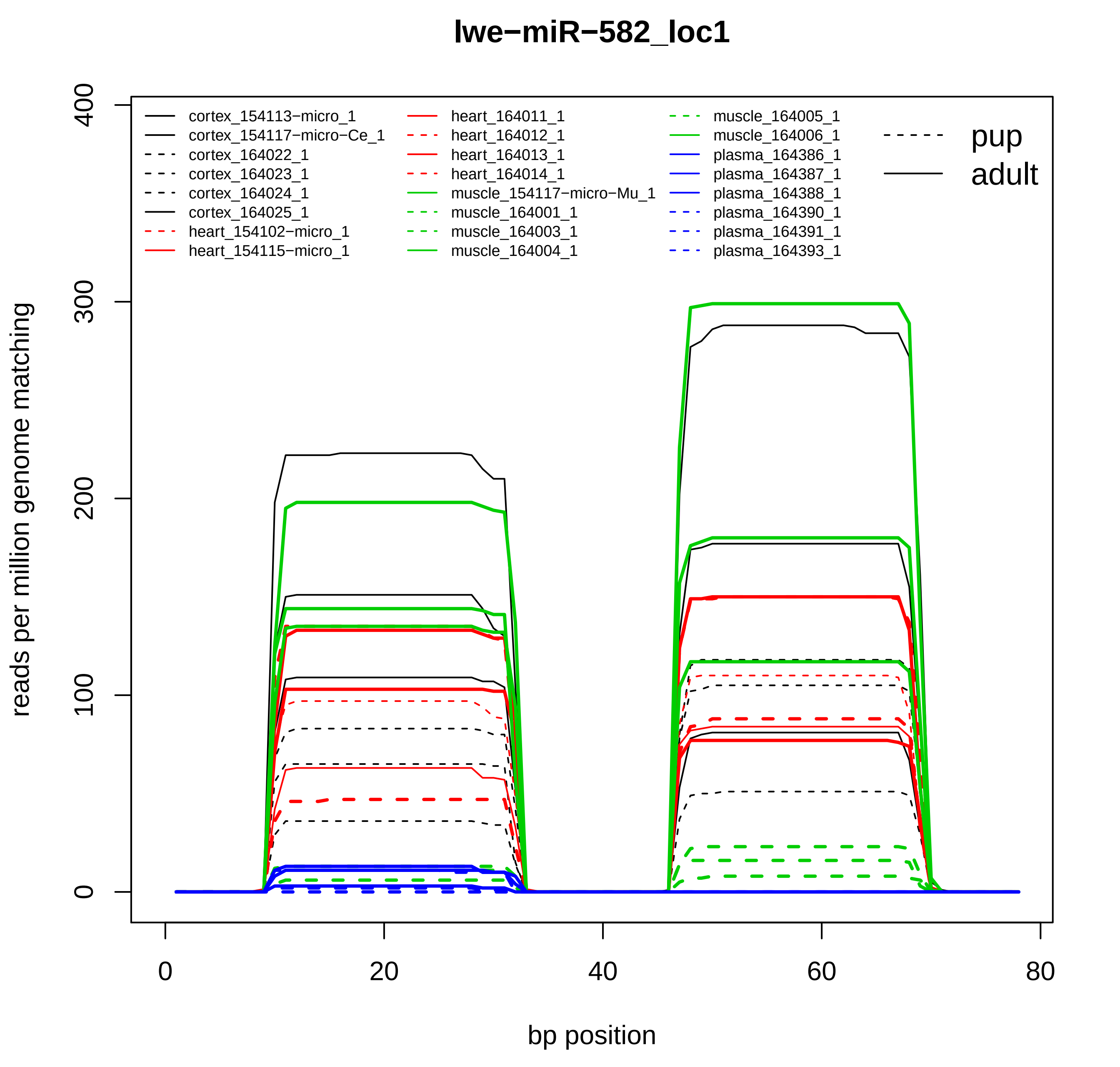
lwe−miR−582_loc1
reads per million genome matching
bp position
0
100
200
300
400
0
20
40
60
80
cortex_154113−micro_1
cortex_154117−micro−Ce_1
cortex_164022_1
cortex_164023_1
cortex_164024_1
cortex_164025_1
heart_154102−micro_1
heart_154115−micro_1
heart_164011_1
heart_164012_1
heart_164013_1
heart_164014_1
muscle_154117−micro−Mu_1
muscle_164001_1
muscle_164003_1
muscle_164004_1
muscle_164005_1
muscle_164006_1
plasma_164386_1
plasma_164387_1
plasma_164388_1
plasma_164390_1
plasma_164391_1
plasma_164393_1
pup
adult

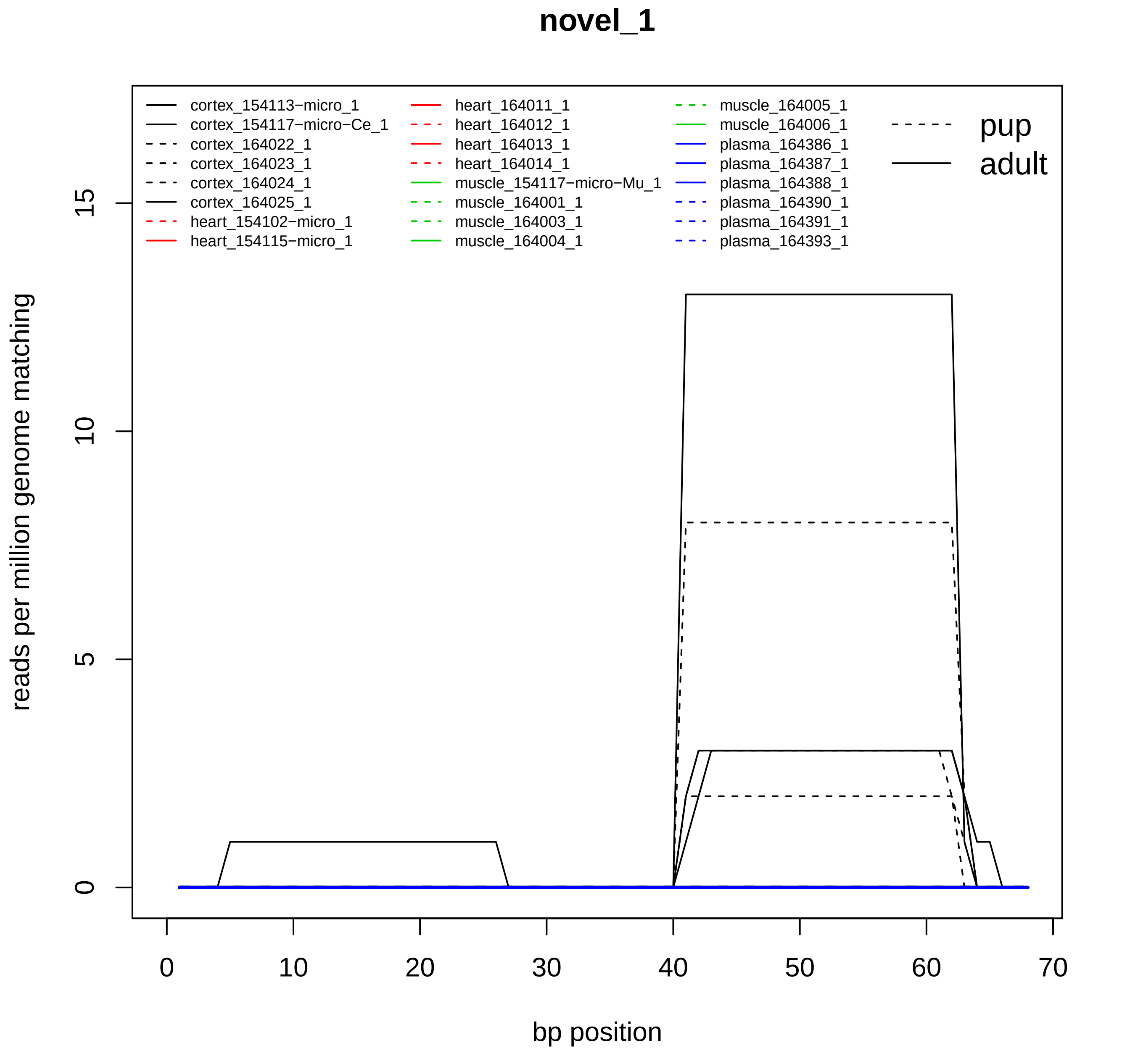
novel_1
cortex_154113−micro_1
cortex_154117−micro−Ce_1
cortex_164022_1
cortex_164023_1
cortex_164024_1
cortex_164025_1
heart_154102−micro_1
heart_154115−micro_1
heart_164011_1
heart_164012_1
heart_164013_1
heart_164014_1
muscle_154117−micro−Mu_1
muscle_164001_1
muscle_164003_1
muscle_164004_1
muscle_164005_1
muscle_164006_1
plasma_164386_1
plasma_164387_1
plasma_164388_1
plasma_164390_1
plasma_164391_1
plasma_164393_1
pup
adult
reads per million genome matching
0
5
10
15
0
10
20
30
40
50
60
70
bp position

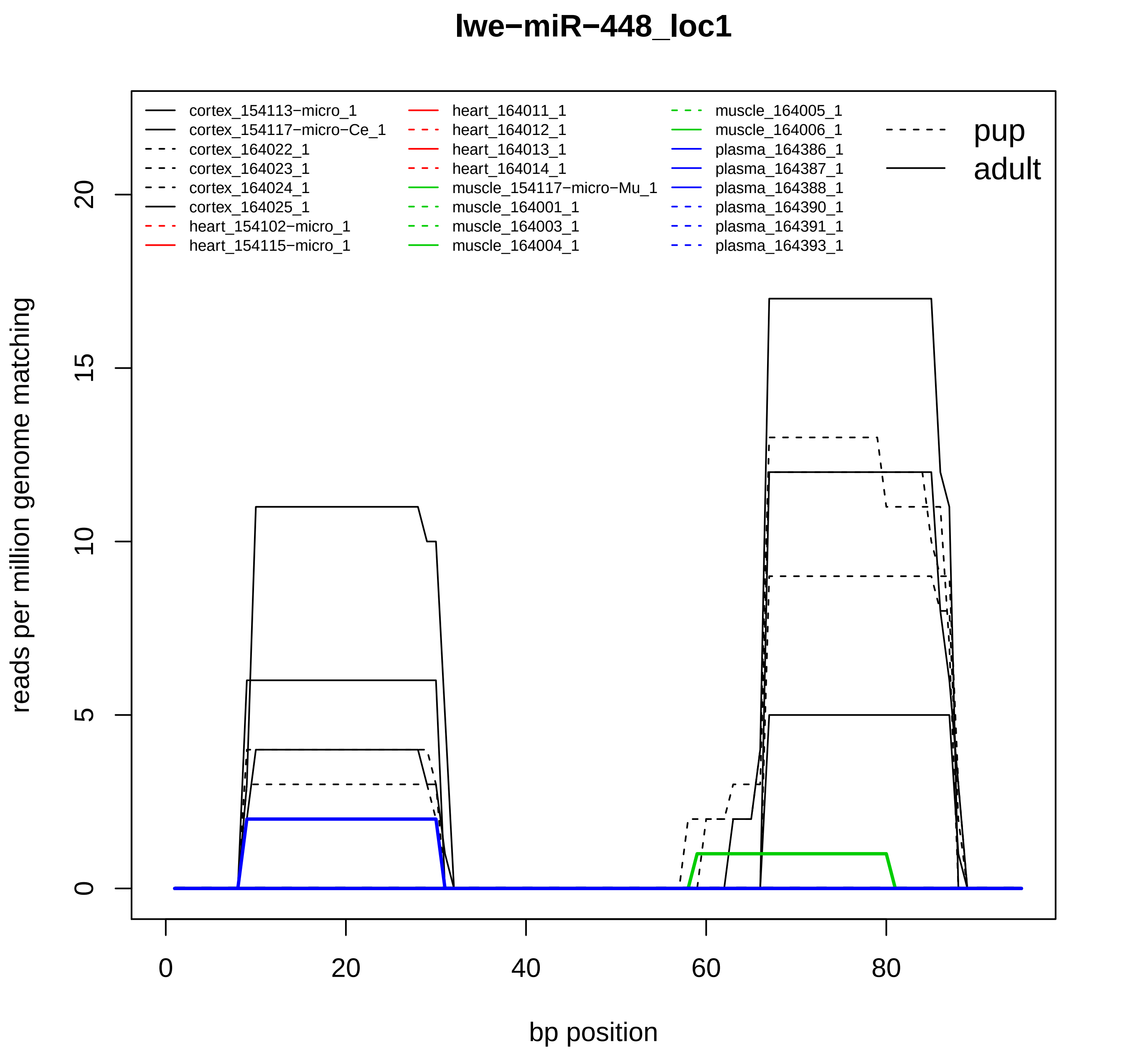

lwe−miR−448_loc1
cortex_154113−micro_1
cortex_154117−micro−Ce_1
cortex_164022_1
cortex_164023_1
cortex_164024_1
cortex_164025_1
heart_154102−micro_1
heart_154115−micro_1
heart_164011_1
heart_164012_1
heart_164013_1
heart_164014_1
muscle_154117−micro−Mu_1
muscle_164001_1
muscle_164003_1
muscle_164004_1
muscle_164005_1
muscle_164006_1
plasma_164386_1
plasma_164387_1
plasma_164388_1
plasma_164390_1
plasma_164391_1
plasma_164393_1
pup
adult
reads per million genome matching
0
5
10
15
20
0
20
40
60
80
bp position

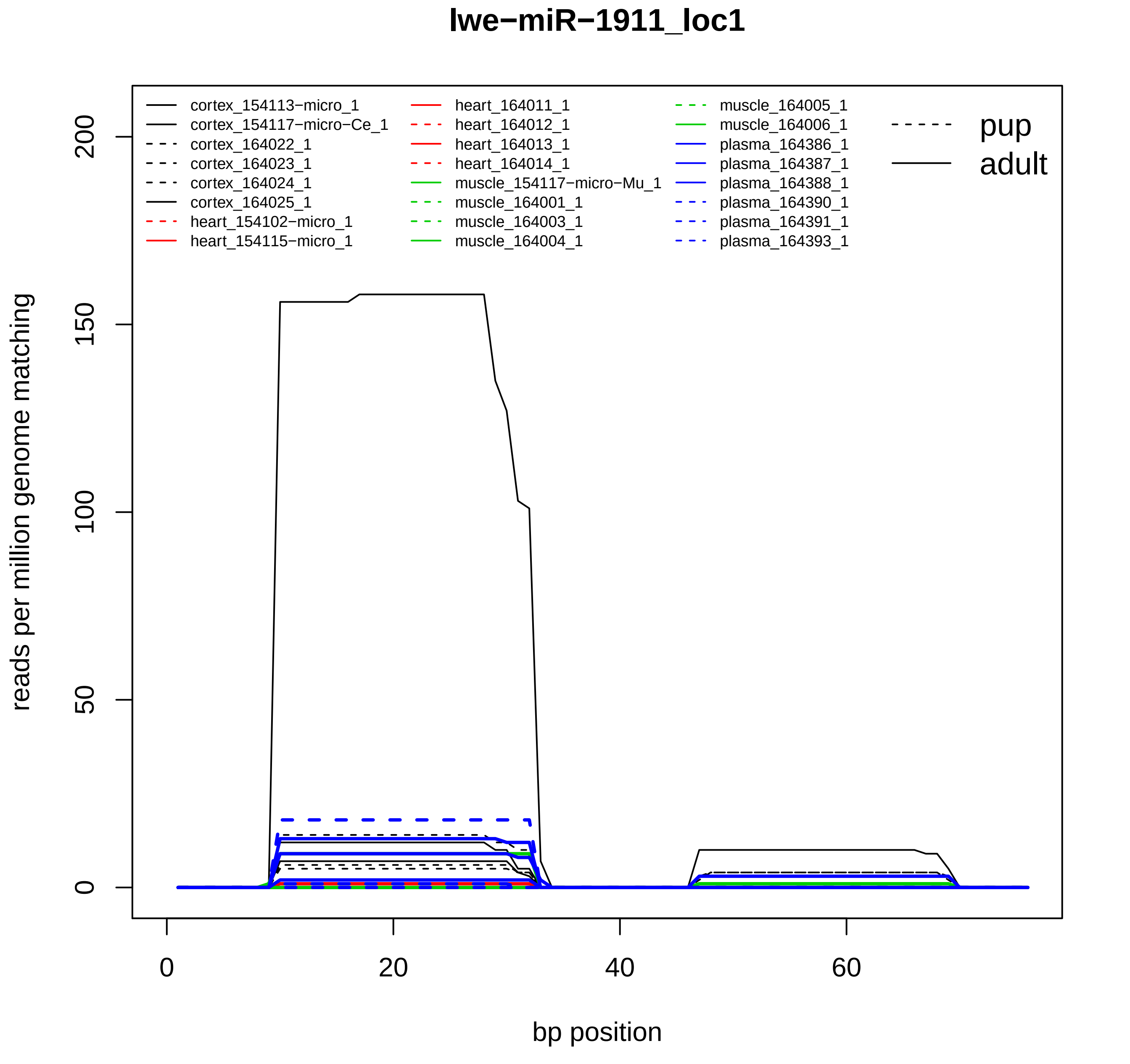
lwe−miR−1911_loc1
cortex_154113−micro_1
cortex_154117−micro−Ce_1
cortex_164022_1
cortex_164023_1
cortex_164024_1
cortex_164025_1
heart_154102−micro_1
heart_154115−micro_1
heart_164011_1
heart_164012_1
heart_164013_1
heart_164014_1
muscle_154117−micro−Mu_1
muscle_164001_1
muscle_164003_1
muscle_164004_1
muscle_164005_1
muscle_164006_1
plasma_164386_1
plasma_164387_1
plasma_164388_1
plasma_164390_1
plasma_164391_1
plasma_164393_1
pup
adult
reads per million genome matching
0
50
100
150
200
0
20
40
60
bp position

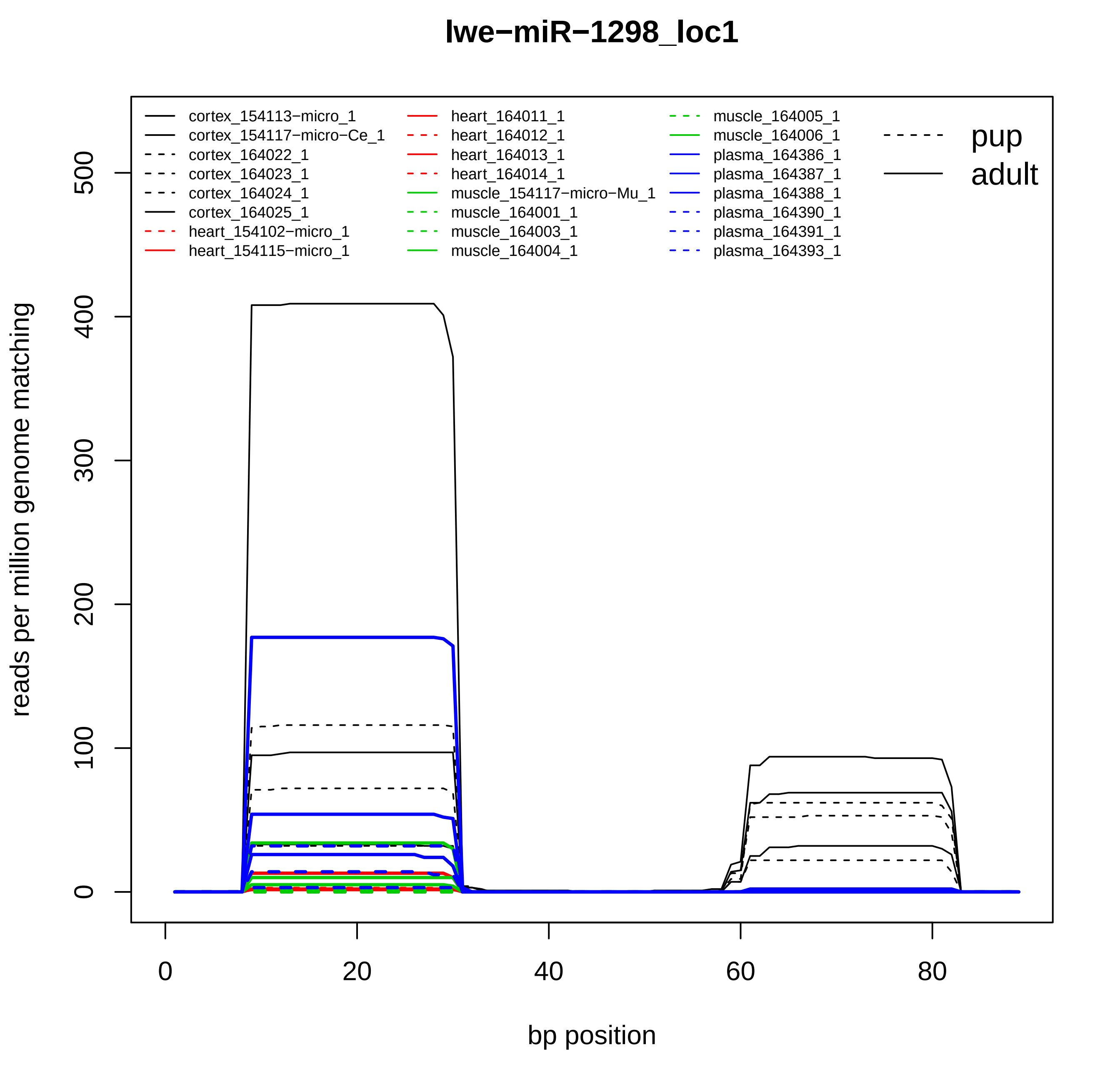

lwe−miR−1298_loc1
cortex_154113−micro_1
cortex_154117−micro−Ce_1
cortex_164022_1
cortex_164023_1
cortex_164024_1
cortex_164025_1
heart_154102−micro_1
heart_154115−micro_1
heart_164011_1
heart_164012_1
heart_164013_1
heart_164014_1
muscle_154117−micro−Mu_1
muscle_164001_1
muscle_164003_1
muscle_164004_1
muscle_164005_1
muscle_164006_1
plasma_164386_1
plasma_164387_1
plasma_164388_1
plasma_164390_1
plasma_164391_1
plasma_164393_1
pup
adult
reads per million genome matching
0
100
200
300
400
500
bp position
0
20
40
60
80

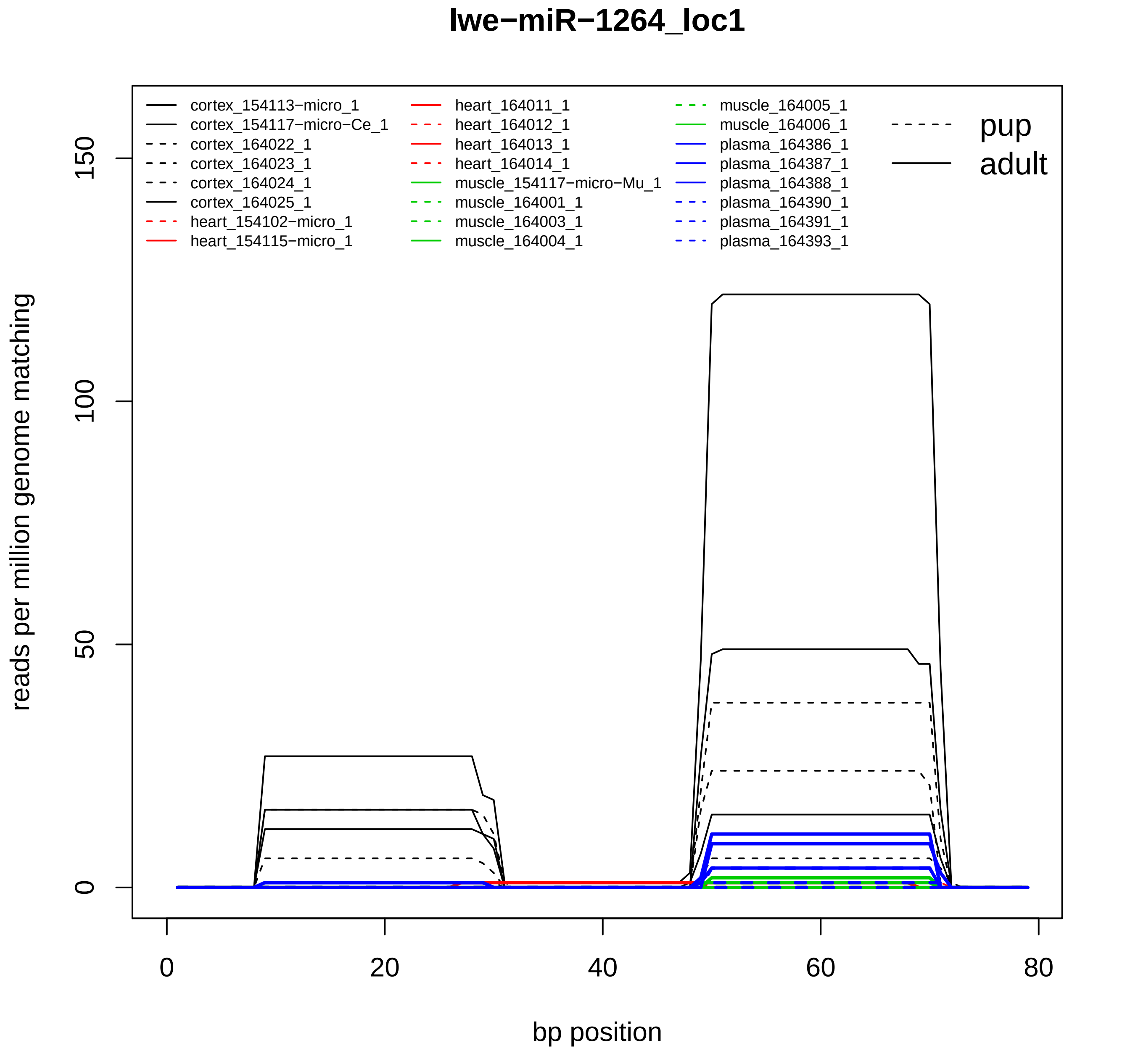
lwe−miR−1264_loc1
cortex_154113−micro_1
cortex_154117−micro−Ce_1
cortex_164022_1
cortex_164023_1
cortex_164024_1
cortex_164025_1
heart_154102−micro_1
heart_154115−micro_1
heart_164011_1
heart_164012_1
heart_164013_1
heart_164014_1
muscle_154117−micro−Mu_1
muscle_164001_1
muscle_164003_1
muscle_164004_1
muscle_164005_1
muscle_164006_1
plasma_164386_1
plasma_164387_1
plasma_164388_1
plasma_164390_1
plasma_164391_1
plasma_164393_1
pup
adult
reads per million genome matching
0
50
100
150
bp position
0
20
40
60
80

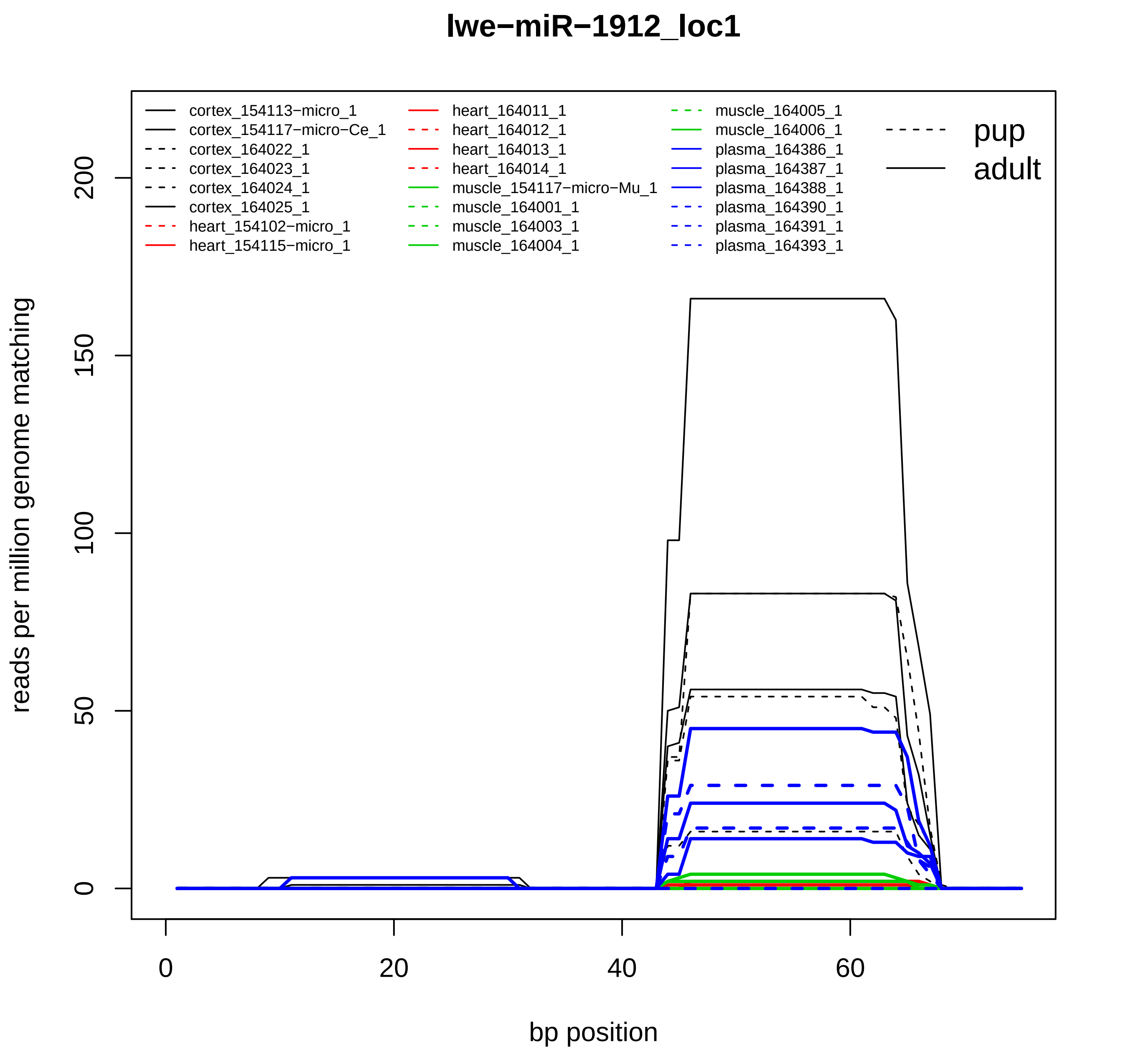

lwe−miR−1912_loc1
cortex_154113−micro_1
cortex_154117−micro−Ce_1
cortex_164022_1
cortex_164023_1
cortex_164024_1
cortex_164025_1
heart_154102−micro_1
heart_154115−micro_1
heart_164011_1
heart_164012_1
heart_164013_1
heart_164014_1
muscle_154117−micro−Mu_1
muscle_164001_1
muscle_164003_1
muscle_164004_1
muscle_164005_1
muscle_164006_1
plasma_164386_1
plasma_164387_1
plasma_164388_1
plasma_164390_1
plasma_164391_1
plasma_164393_1
pup
adult
reads per million genome matching
0
50
100
150
200
0
20
40
60
bp position

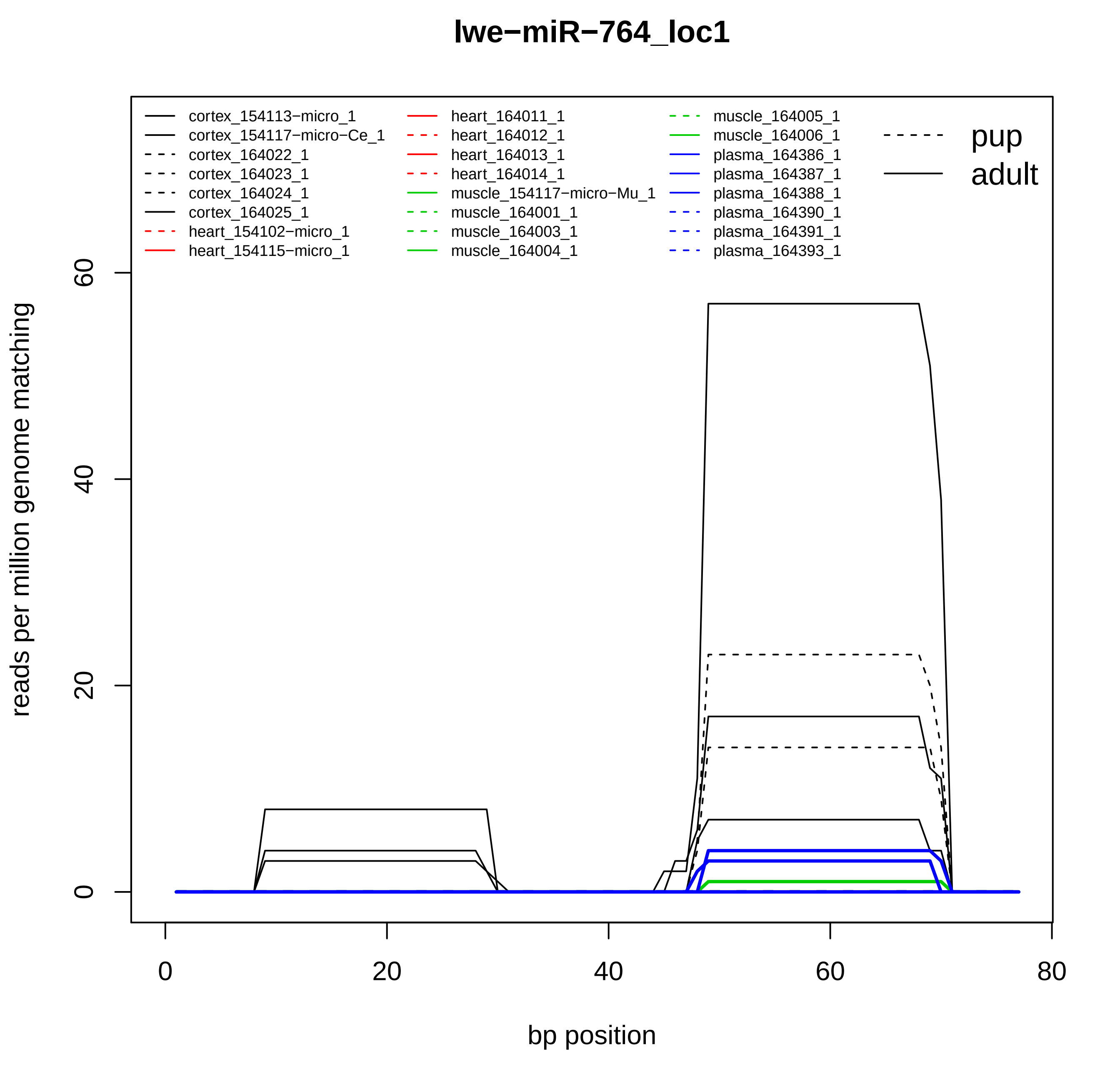

lwe−miR−764_loc1
cortex_154113−micro_1
cortex_154117−micro−Ce_1
cortex_164022_1
cortex_164023_1
cortex_164024_1
cortex_164025_1
heart_154102−micro_1
heart_154115−micro_1
heart_164011_1
heart_164012_1
heart_164013_1
heart_164014_1
muscle_154117−micro−Mu_1
muscle_164001_1
muscle_164003_1
muscle_164004_1
muscle_164005_1
muscle_164006_1
plasma_164386_1
plasma_164387_1
plasma_164388_1
plasma_164390_1
plasma_164391_1
plasma_164393_1
pup
adult
reads per million genome matching
0
20
40
60
0
20
40
60
80
bp position

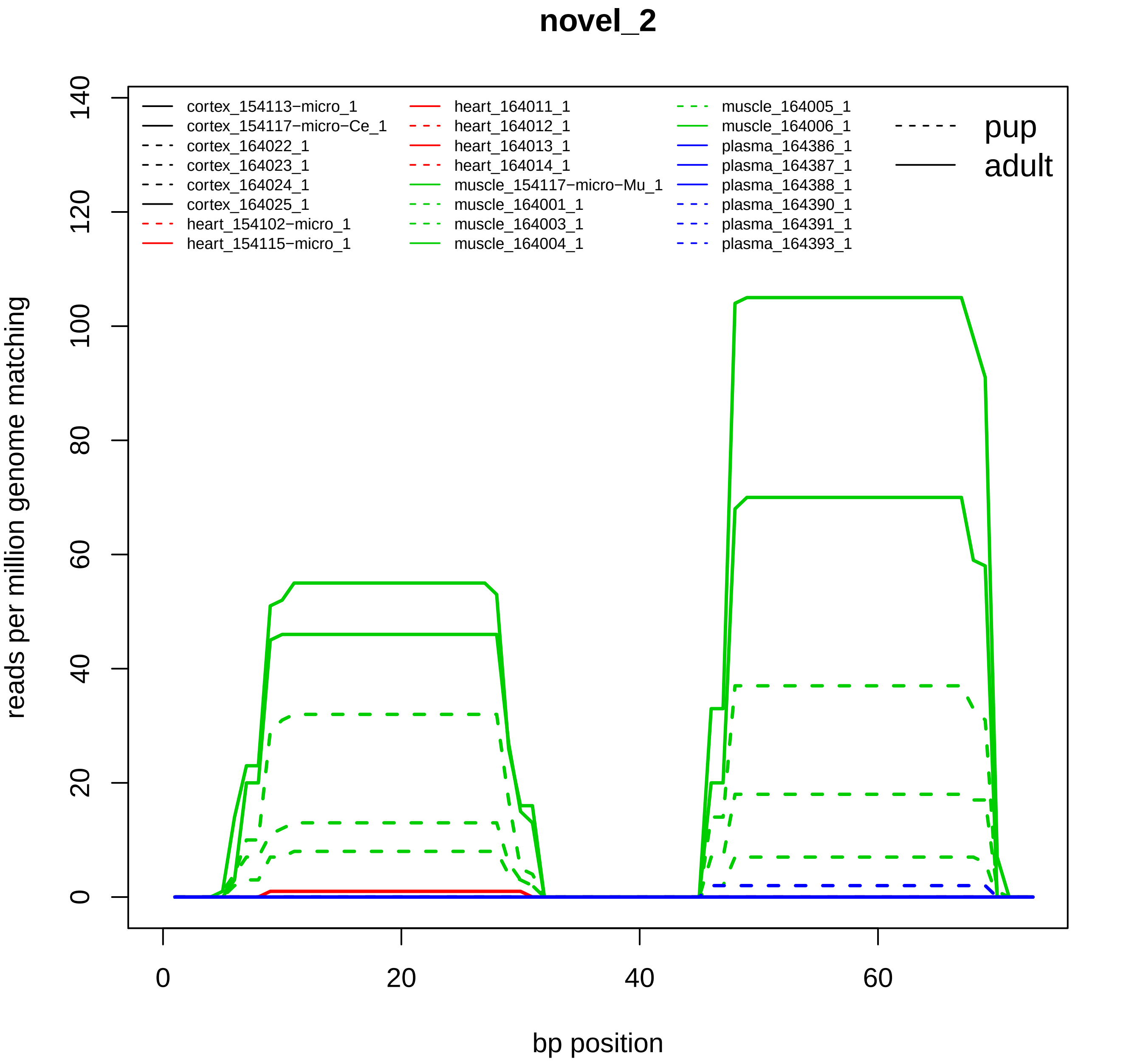

novel_2
cortex_154113−micro_1
cortex_154117−micro−Ce_1
cortex_164022_1
cortex_164023_1
cortex_164024_1
cortex_164025_1
heart_154102−micro_1
heart_154115−micro_1
heart_164011_1
heart_164012_1
heart_164013_1
heart_164014_1
muscle_154117−micro−Mu_1
muscle_164001_1
muscle_164003_1
muscle_164004_1
muscle_164005_1
muscle_164006_1
plasma_164386_1
plasma_164387_1
plasma_164388_1
plasma_164390_1
plasma_164391_1
plasma_164393_1
pup
adult
reads per million genome matching
0
20
40
60
80
100
120
140
0
20
40
60
bp position

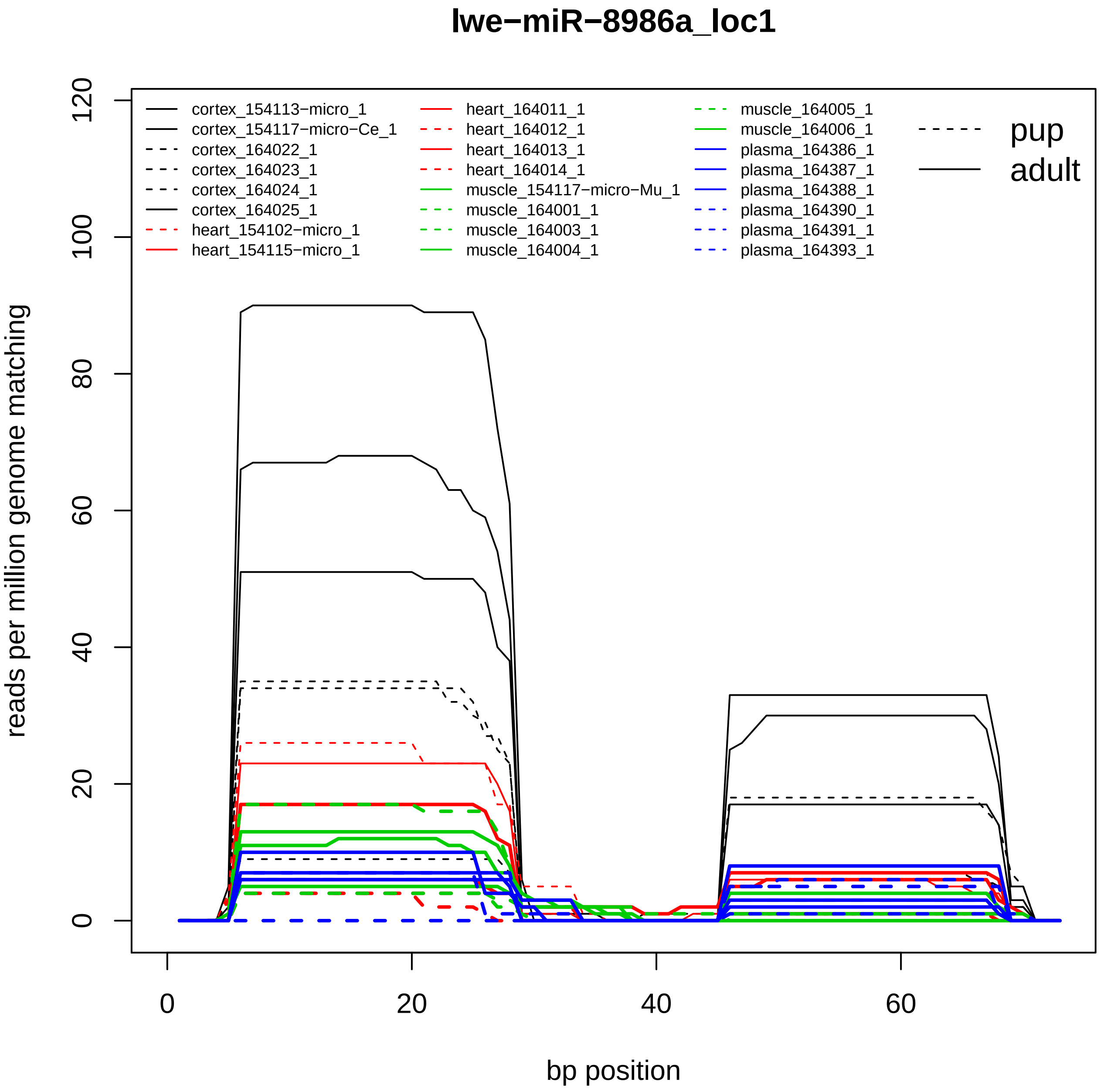

lwe−miR−8986a_loc1
reads per million genome matching
bp position
0
20
40
60
80
100
120
0
20
40
60
cortex_154113−micro_1
cortex_154117−micro−Ce_1
cortex_164022_1
cortex_164023_1
cortex_164024_1
cortex_164025_1
heart_154102−micro_1
heart_154115−micro_1
heart_164011_1
heart_164012_1
heart_164013_1
heart_164014_1
muscle_154117−micro−Mu_1
muscle_164001_1
muscle_164003_1
muscle_164004_1
muscle_164005_1
muscle_164006_1
plasma_164386_1
plasma_164387_1
plasma_164388_1
plasma_164390_1
plasma_164391_1
plasma_164393_1
pup
adult

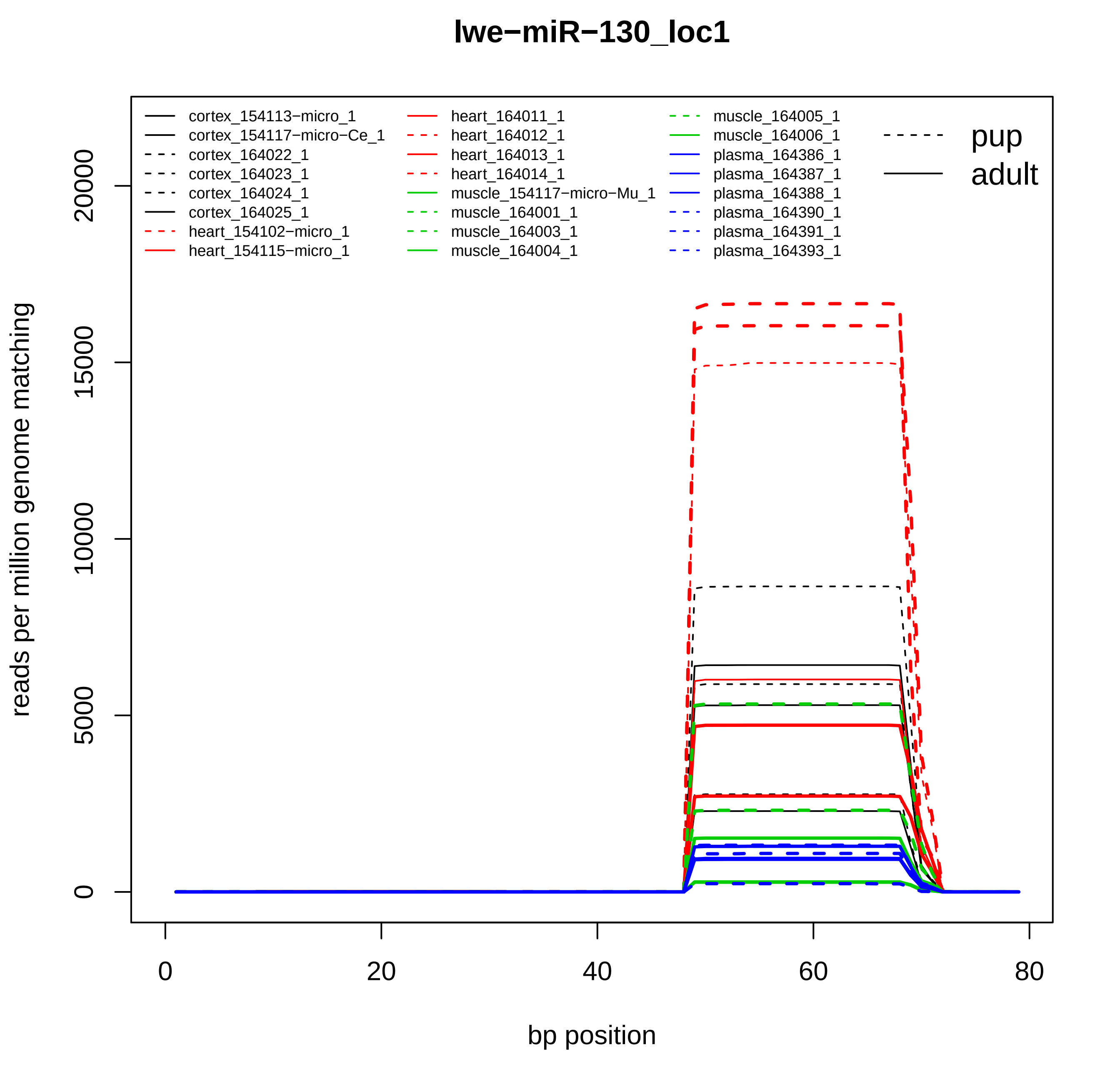

lwe−miR−130_loc1
cortex_154113−micro_1
cortex_154117−micro−Ce_1
cortex_164022_1
cortex_164023_1
cortex_164024_1
cortex_164025_1
heart_154102−micro_1
heart_154115−micro_1
heart_164011_1
heart_164012_1
heart_164013_1
heart_164014_1
muscle_154117−micro−Mu_1
muscle_164001_1
muscle_164003_1
muscle_164004_1
muscle_164005_1
muscle_164006_1
plasma_164386_1
plasma_164387_1
plasma_164388_1
plasma_164390_1
plasma_164391_1
plasma_164393_1
pup
adult
reads per million genome matching
0
5000
10000
15000
20000
bp position
0
20
40
60
80

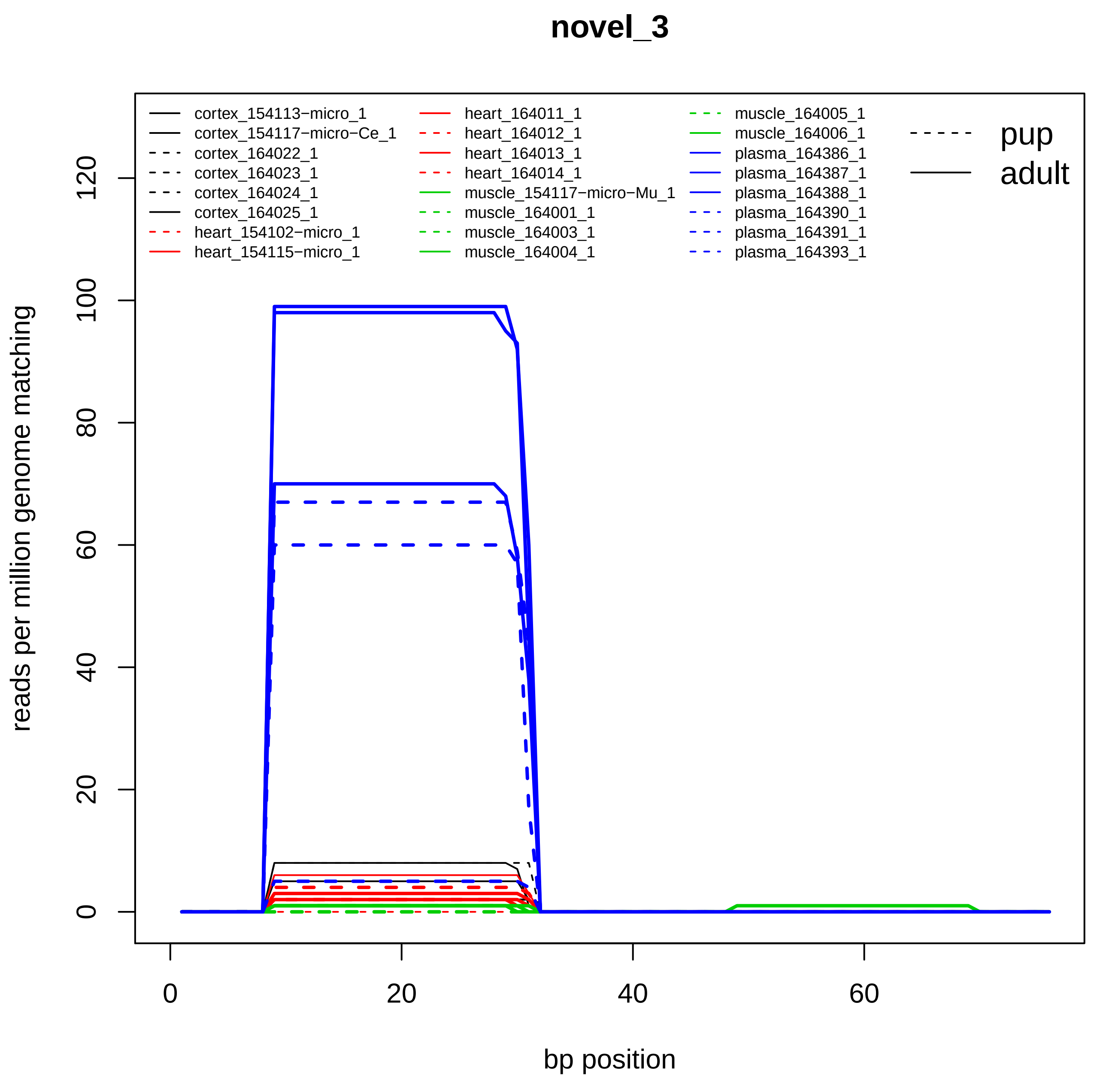
novel_3
cortex_154113−micro_1
cortex_154117−micro−Ce_1
cortex_164022_1
cortex_164023_1
cortex_164024_1
cortex_164025_1
heart_154102−micro_1
heart_154115−micro_1
heart_164011_1
heart_164012_1
heart_164013_1
heart_164014_1
muscle_154117−micro−Mu_1
muscle_164001_1
muscle_164003_1
muscle_164004_1
muscle_164005_1
muscle_164006_1
plasma_164386_1
plasma_164387_1
plasma_164388_1
plasma_164390_1
plasma_164391_1
plasma_164393_1
pup
adult
reads per million genome matching
0
20
40
60
80
100
120
0
20
40
60
bp position

### lwe−miR−365a_loc1

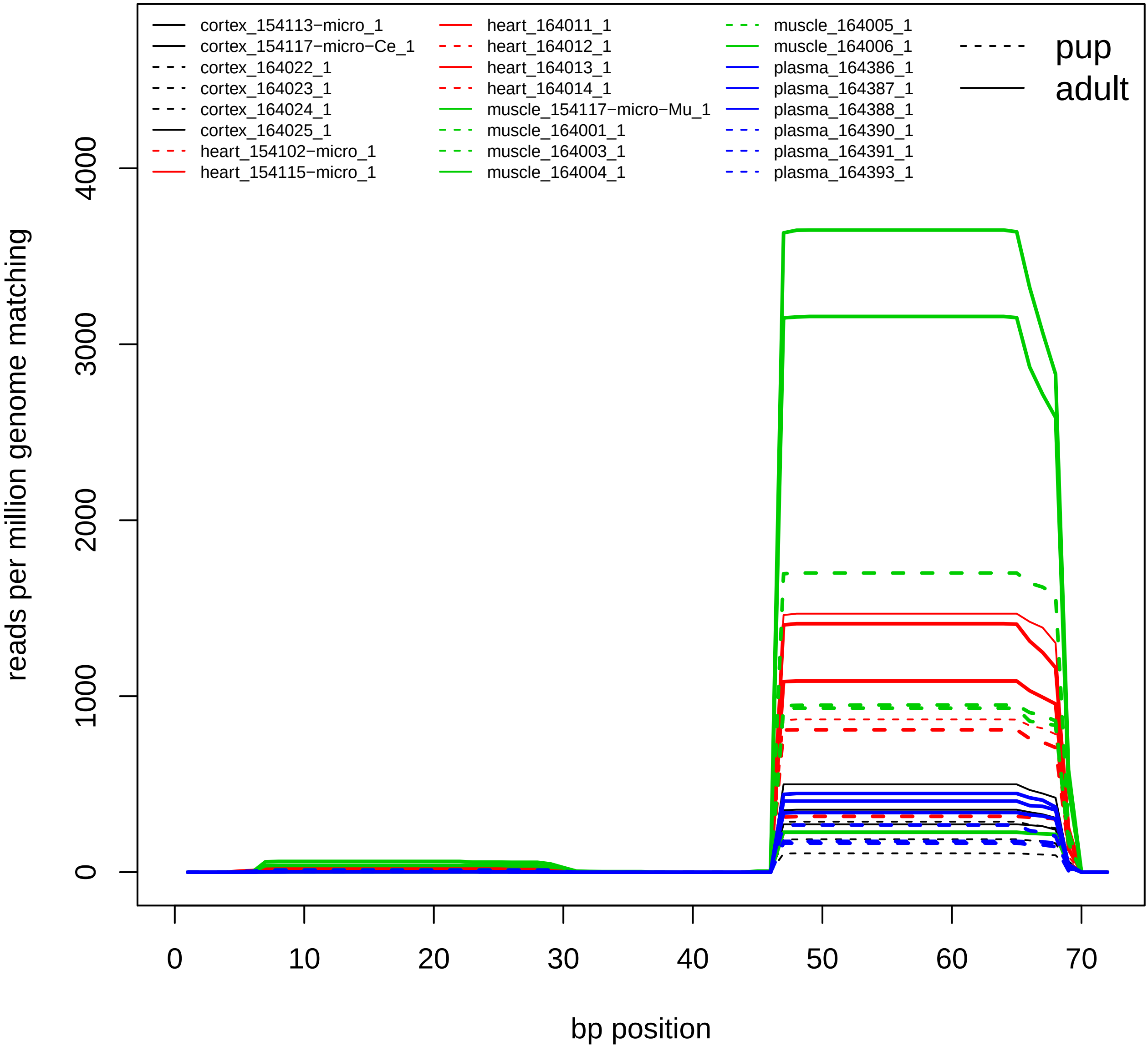

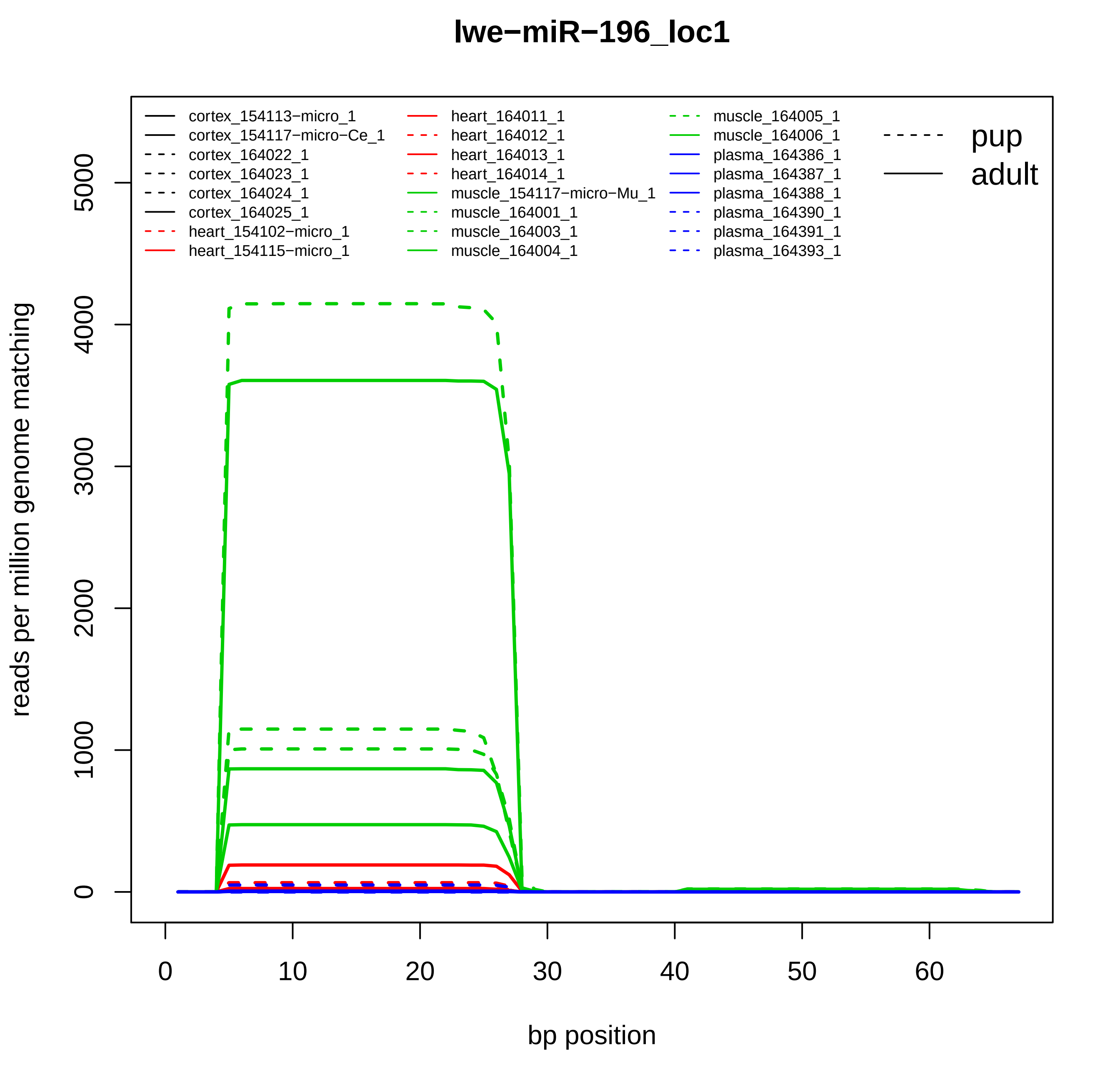
lwe−miR−196_loc1
cortex_154113−micro_1
cortex_154117−micro−Ce_1
cortex_164022_1
cortex_164023_1
cortex_164024_1
cortex_164025_1
heart_154102−micro_1
heart_154115−micro_1
heart_164011_1
heart_164012_1
heart_164013_1
heart_164014_1
muscle_154117−micro−Mu_1
muscle_164001_1
muscle_164003_1
muscle_164004_1
muscle_164005_1
muscle_164006_1
plasma_164386_1
plasma_164387_1
plasma_164388_1
plasma_164390_1
plasma_164391_1
plasma_164393_1
pup
adult
reads per million genome matching
0
1000
2000
3000
4000
5000
0
10
20
30
40
50
60
bp position

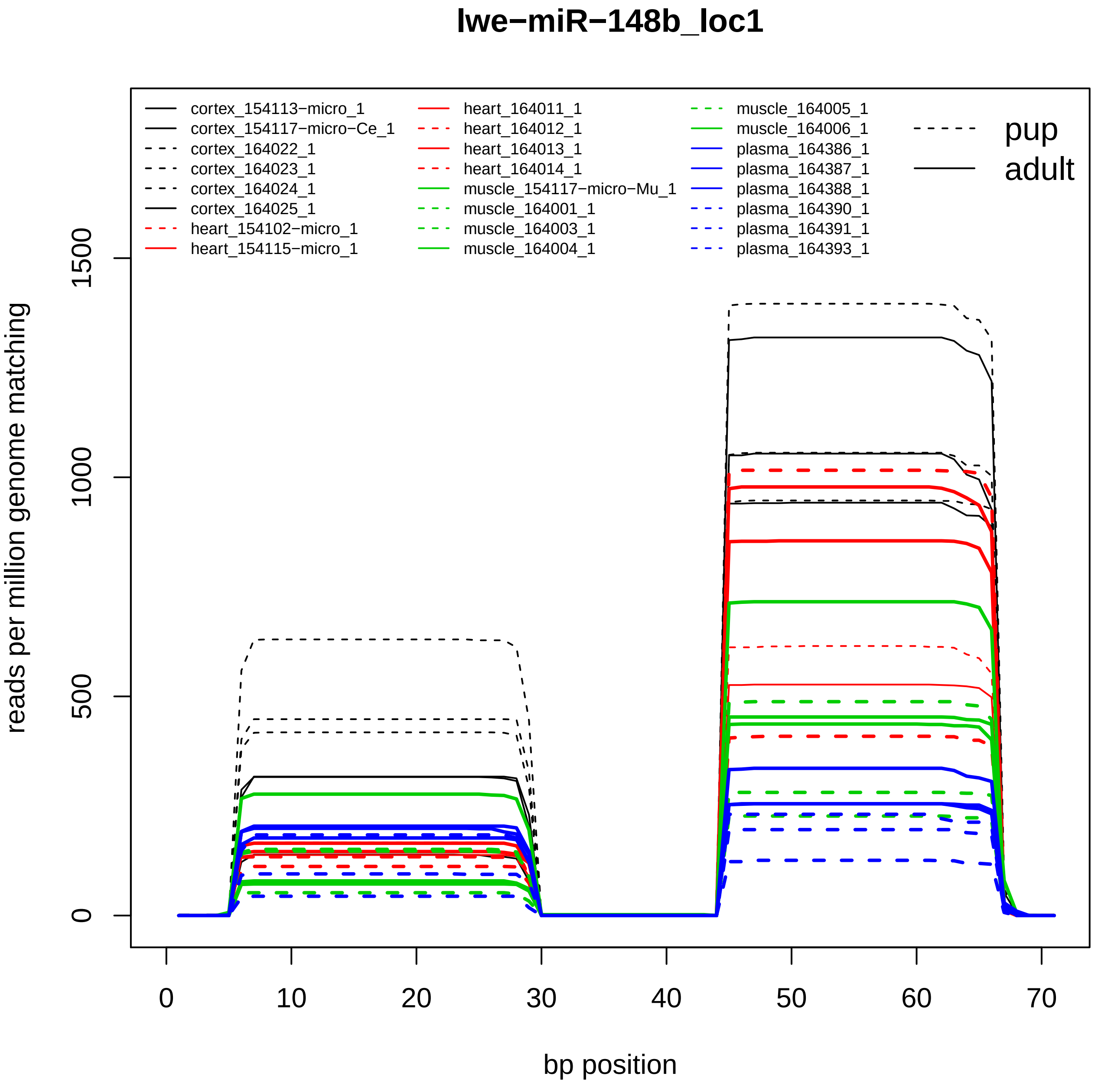
lwe−miR−148b_loc1
cortex_154113−micro_1
cortex_154117−micro−Ce_1
cortex_164022_1
cortex_164023_1
cortex_164024_1
cortex_164025_1
heart_154102−micro_1
heart_154115−micro_1
heart_164011_1
heart_164012_1
heart_164013_1
heart_164014_1
muscle_154117−micro−Mu_1
muscle_164001_1
muscle_164003_1
muscle_164004_1
muscle_164005_1
muscle_164006_1
plasma_164386_1
plasma_164387_1
plasma_164388_1
plasma_164390_1
plasma_164391_1
plasma_164393_1
pup
adult
reads per million genome matching
0
500
1000
1500
bp position
0
10
20
30
40
50
60
70

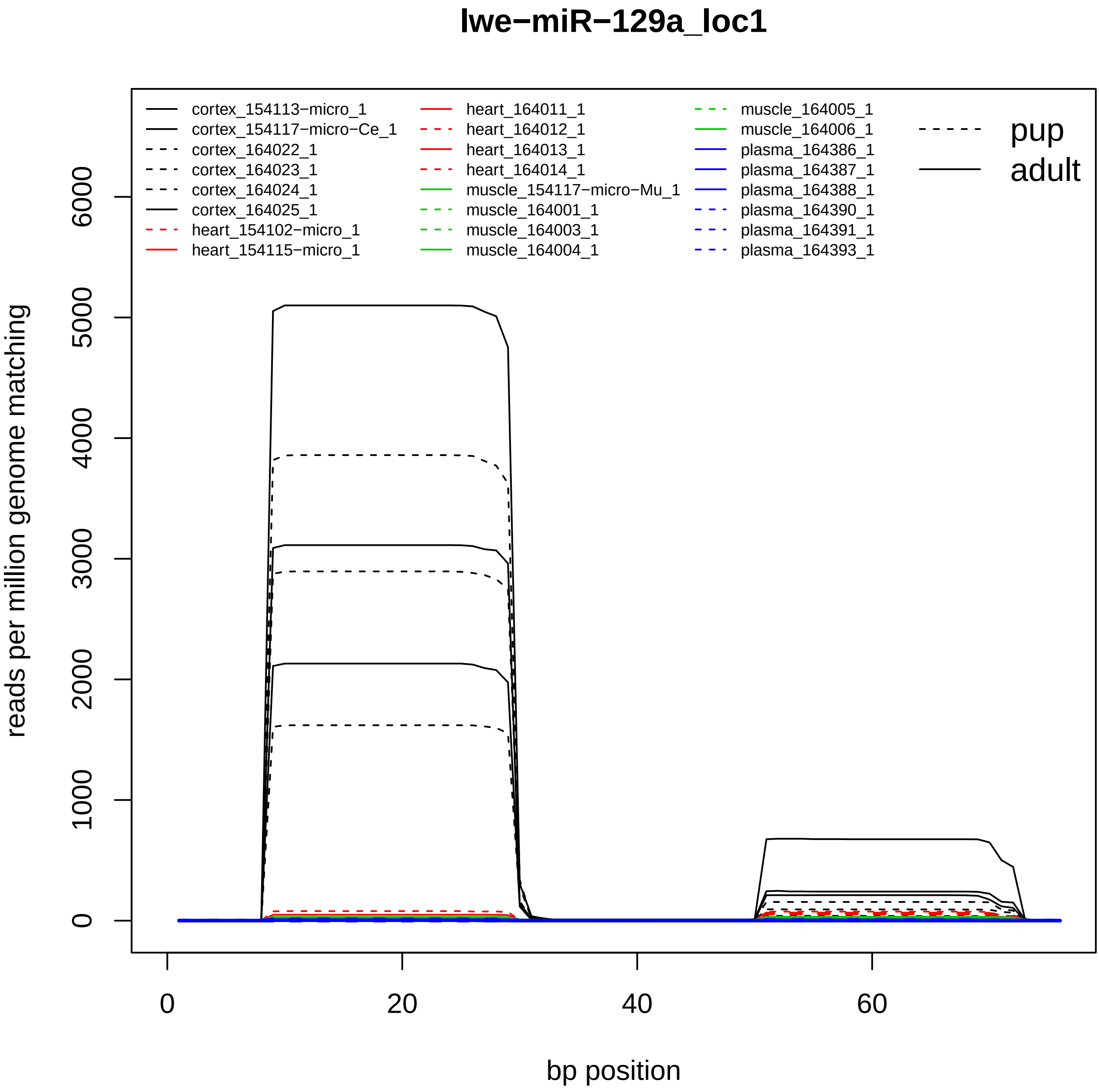
lwe−miR−129a_loc1
cortex_154113−micro_1
cortex_154117−micro−Ce_1
cortex_164022_1
cortex_164023_1
cortex_164024_1
cortex_164025_1
heart_154102−micro_1
heart_154115−micro_1
heart_164011_1
heart_164012_1
heart_164013_1
heart_164014_1
muscle_154117−micro−Mu_1
muscle_164001_1
muscle_164003_1
muscle_164004_1
muscle_164005_1
muscle_164006_1
plasma_164386_1
plasma_164387_1
plasma_164388_1
plasma_164390_1
plasma_164391_1
plasma_164393_1
pup
adult
reads per million genome matching
0
1000
2000
3000
4000
5000
6000
0
20
40
60
bp position

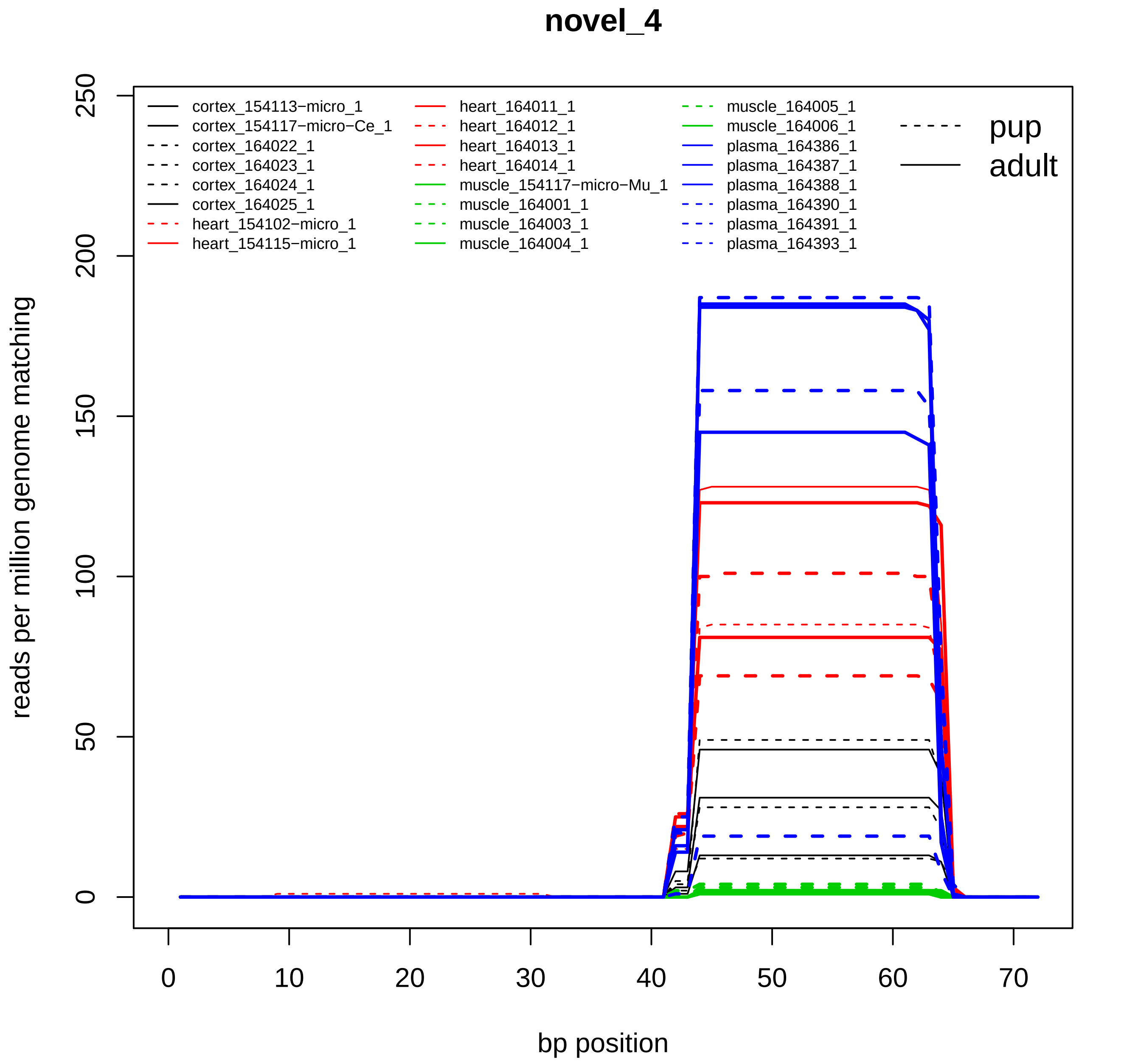
novel_4
cortex_154113−micro_1
cortex_154117−micro−Ce_1
cortex_164022_1
cortex_164023_1
cortex_164024_1
cortex_164025_1
heart_154102−micro_1
heart_154115−micro_1
heart_164011_1
heart_164012_1
heart_164013_1
heart_164014_1
muscle_154117−micro−Mu_1
muscle_164001_1
muscle_164003_1
muscle_164004_1
muscle_164005_1
muscle_164006_1
plasma_164386_1
plasma_164387_1
plasma_164388_1
plasma_164390_1
plasma_164391_1
plasma_164393_1
pup
adult
reads per million genome matching
bp position
0
50
100
150
200
250
0
10
20
30
40
50
60
70

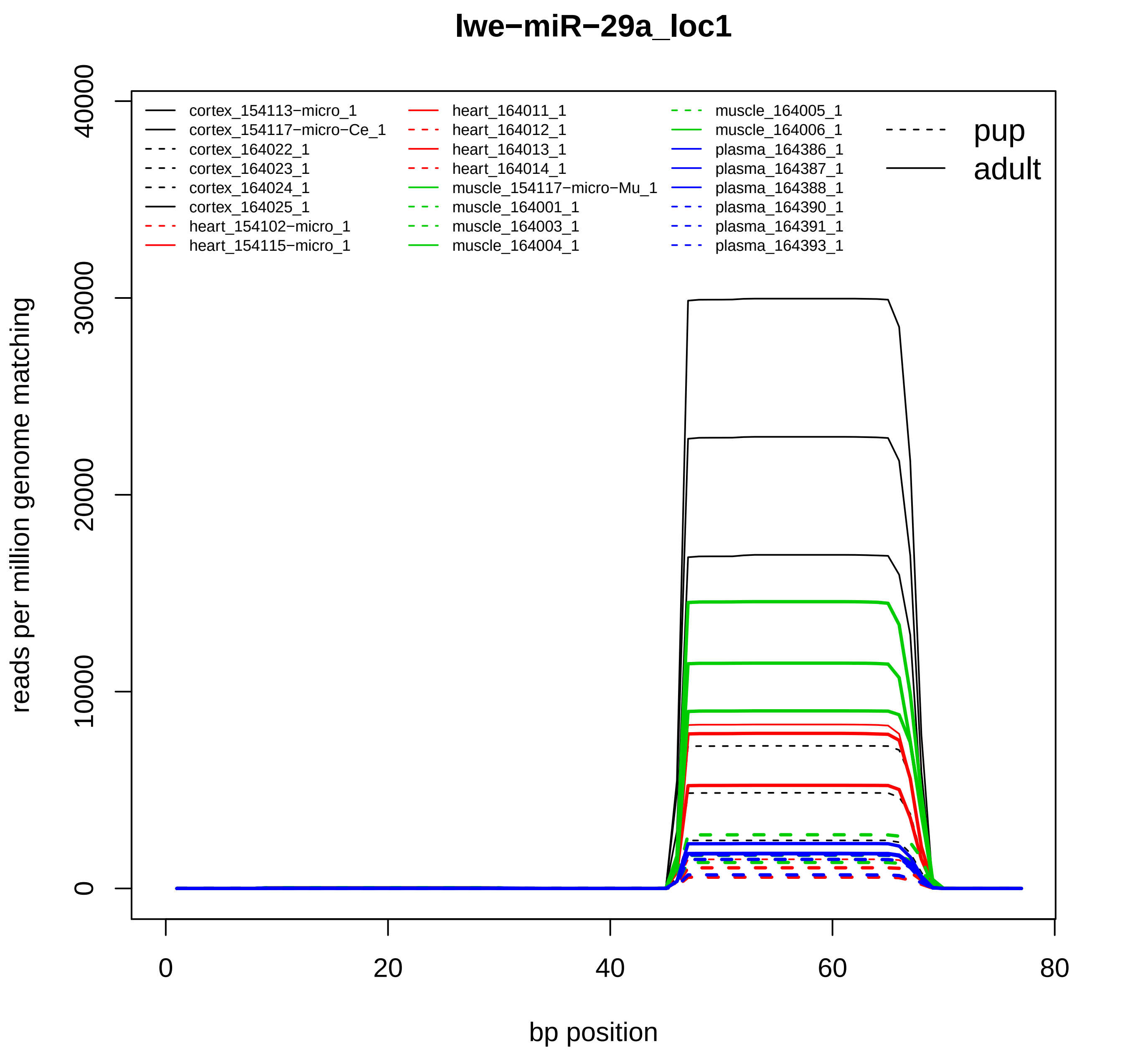
lwe−miR−29a_loc1
cortex_154113−micro_1
cortex_154117−micro−Ce_1
cortex_164022_1
cortex_164023_1
cortex_164024_1
cortex_164025_1
heart_154102−micro_1
heart_154115−micro_1
heart_164011_1
heart_164012_1
heart_164013_1
heart_164014_1
muscle_154117−micro−Mu_1
muscle_164001_1
muscle_164003_1
muscle_164004_1
muscle_164005_1
muscle_164006_1
plasma_164386_1
plasma_164387_1
plasma_164388_1
plasma_164390_1
plasma_164391_1
plasma_164393_1
pup
adult
reads per million genome matching
0
10000
20000
30000
40000
bp position
0
20
40
60
80

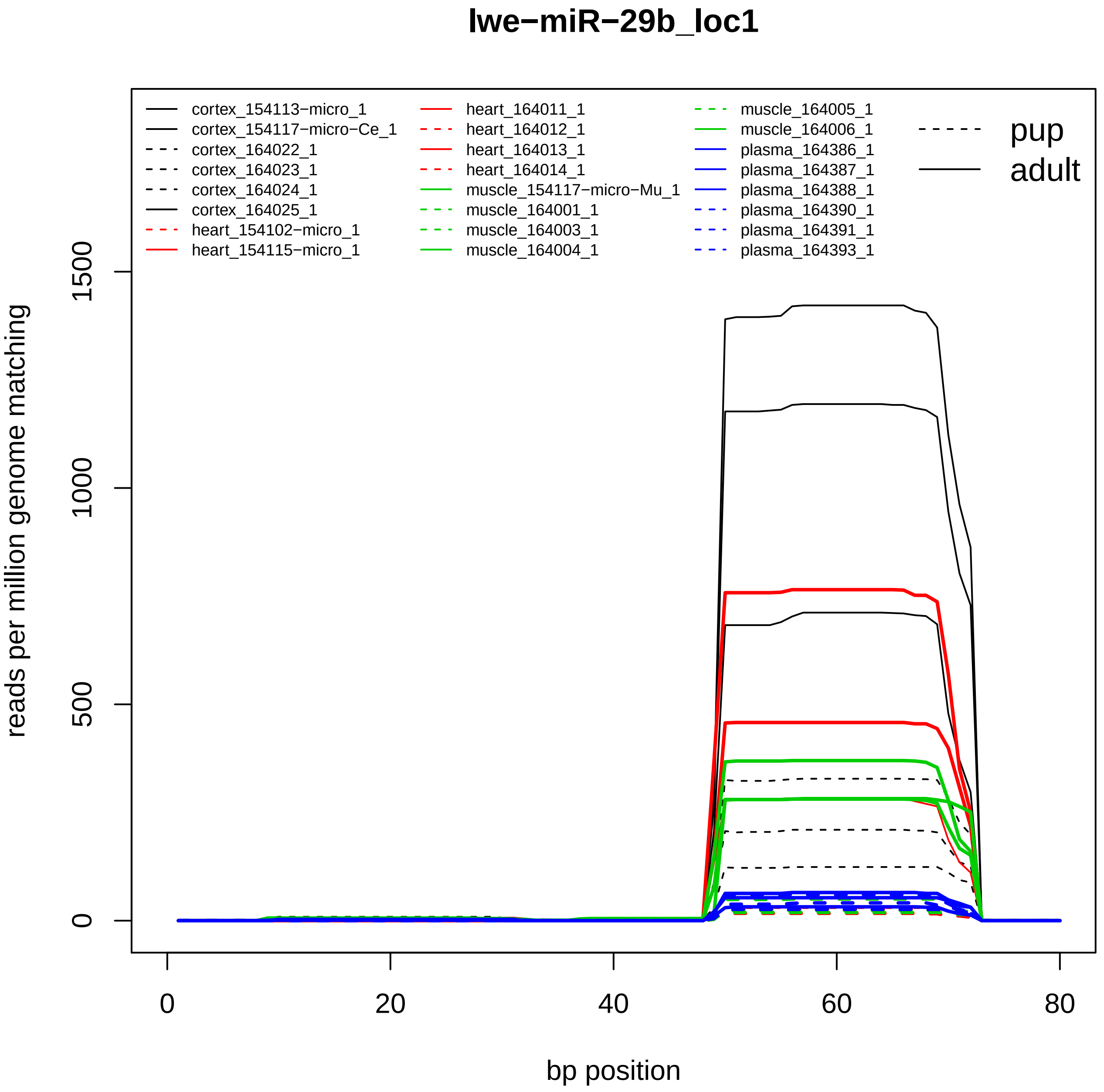
lwe−miR−29b_loc1
cortex_154113−micro_1
cortex_154117−micro−Ce_1
cortex_164022_1
cortex_164023_1
cortex_164024_1
cortex_164025_1
heart_154102−micro_1
heart_154115−micro_1
heart_164011_1
heart_164012_1
heart_164013_1
heart_164014_1
muscle_154117−micro−Mu_1
muscle_164001_1
muscle_164003_1
muscle_164004_1
muscle_164005_1
muscle_164006_1
plasma_164386_1
plasma_164387_1
plasma_164388_1
plasma_164390_1
plasma_164391_1
plasma_164393_1
pup
adult
reads per million genome matching
0
500
1000
1500
0
20
40
60
80
bp position

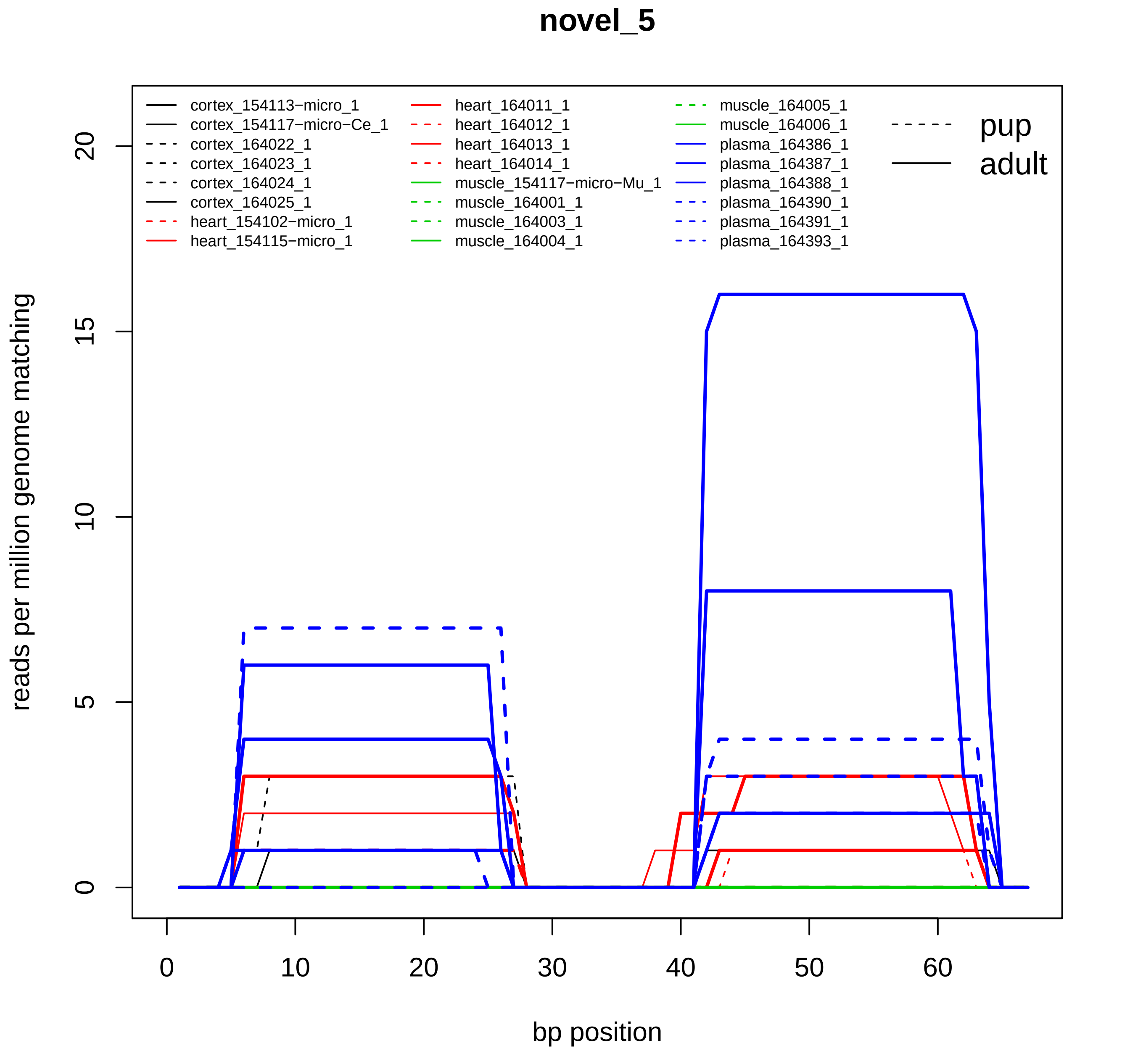
novel_5
cortex_154113−micro_1
cortex_154117−micro−Ce_1
cortex_164022_1
cortex_164023_1
cortex_164024_1
cortex_164025_1
heart_154102−micro_1
heart_154115−micro_1
heart_164011_1
heart_164012_1
heart_164013_1
heart_164014_1
muscle_154117−micro−Mu_1
muscle_164001_1
muscle_164003_1
muscle_164004_1
muscle_164005_1
muscle_164006_1
plasma_164386_1
plasma_164387_1
plasma_164388_1
plasma_164390_1
plasma_164391_1
plasma_164393_1
pup
adult
reads per million genome matching
0
5
10
15
20
0
10
20
30
40
50
60
bp position

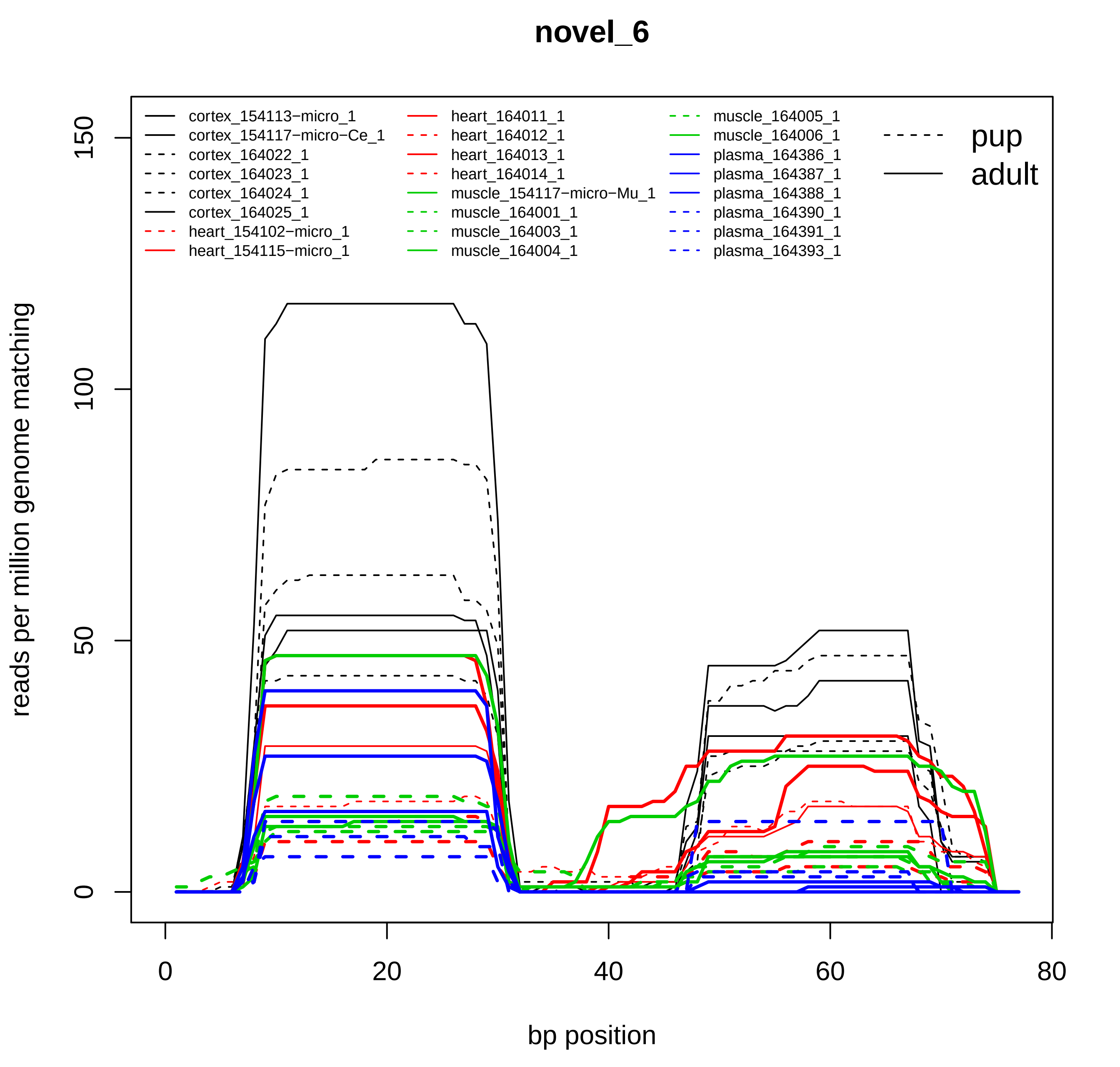

novel_6
reads per million genome matching
bp position
0
20
40
60
80
0
50
100
150
cortex_154113−micro_1
cortex_154117−micro−Ce_1
cortex_164022_1
cortex_164023_1
cortex_164024_1
cortex_164025_1
heart_154102−micro_1
heart_154115−micro_1
heart_164011_1
heart_164012_1
heart_164013_1
heart_164014_1
muscle_154117−micro−Mu_1
muscle_164001_1
muscle_164003_1
muscle_164004_1
muscle_164005_1
muscle_164006_1
plasma_164386_1
plasma_164387_1
plasma_164388_1
plasma_164390_1
plasma_164391_1
plasma_164393_1
pup
adult

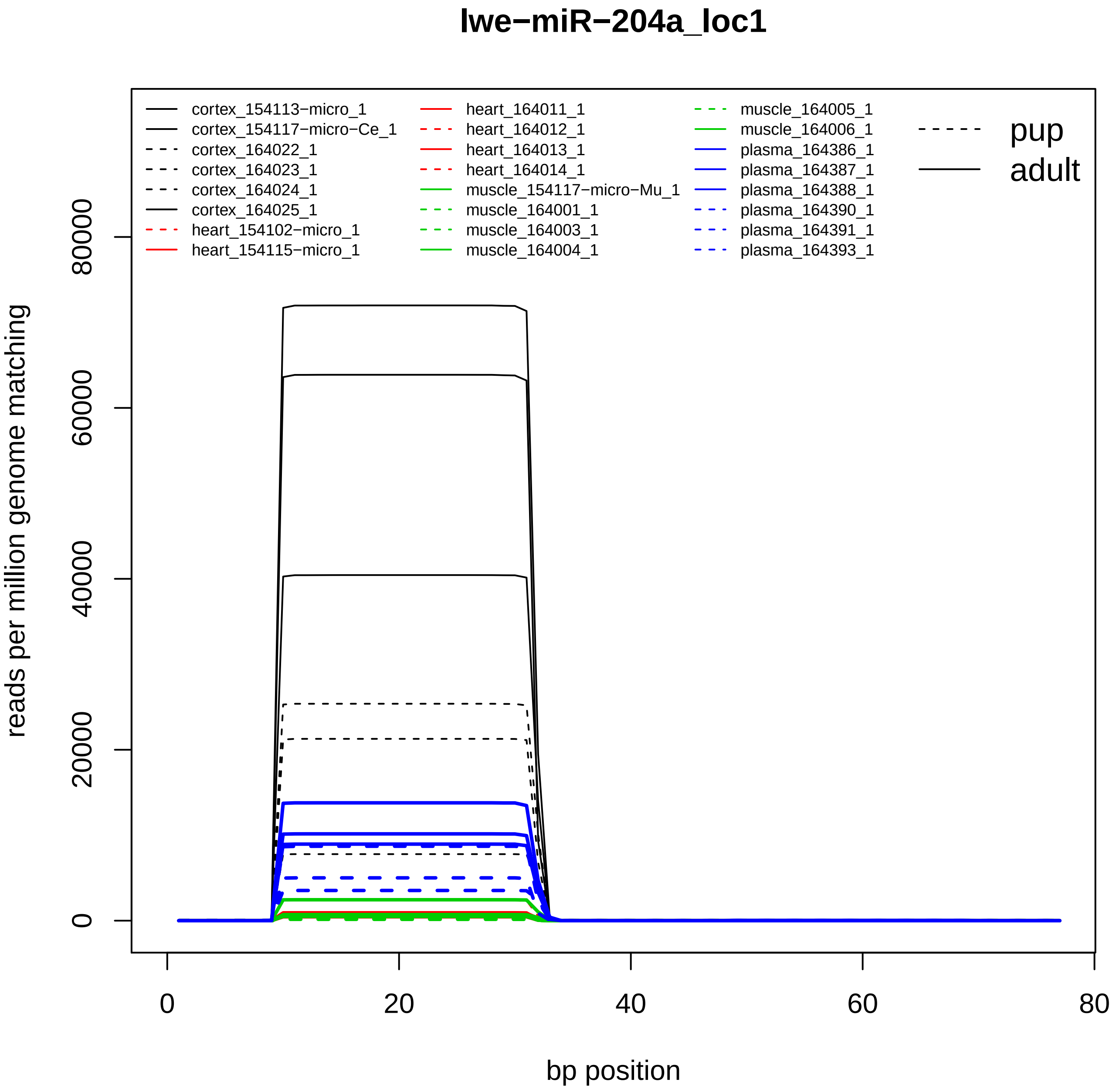

lwe−miR−204a_loc1
cortex_154113−micro_1
cortex_154117−micro−Ce_1
cortex_164022_1
cortex_164023_1
cortex_164024_1
cortex_164025_1
heart_154102−micro_1
heart_154115−micro_1
heart_164011_1
heart_164012_1
heart_164013_1
heart_164014_1
muscle_154117−micro−Mu_1
muscle_164001_1
muscle_164003_1
muscle_164004_1
muscle_164005_1
muscle_164006_1
plasma_164386_1
plasma_164387_1
plasma_164388_1
plasma_164390_1
plasma_164391_1
plasma_164393_1
pup
adult
reads per million genome matching
0
20000
40000
60000
80000
bp position
0
20
40
60
80

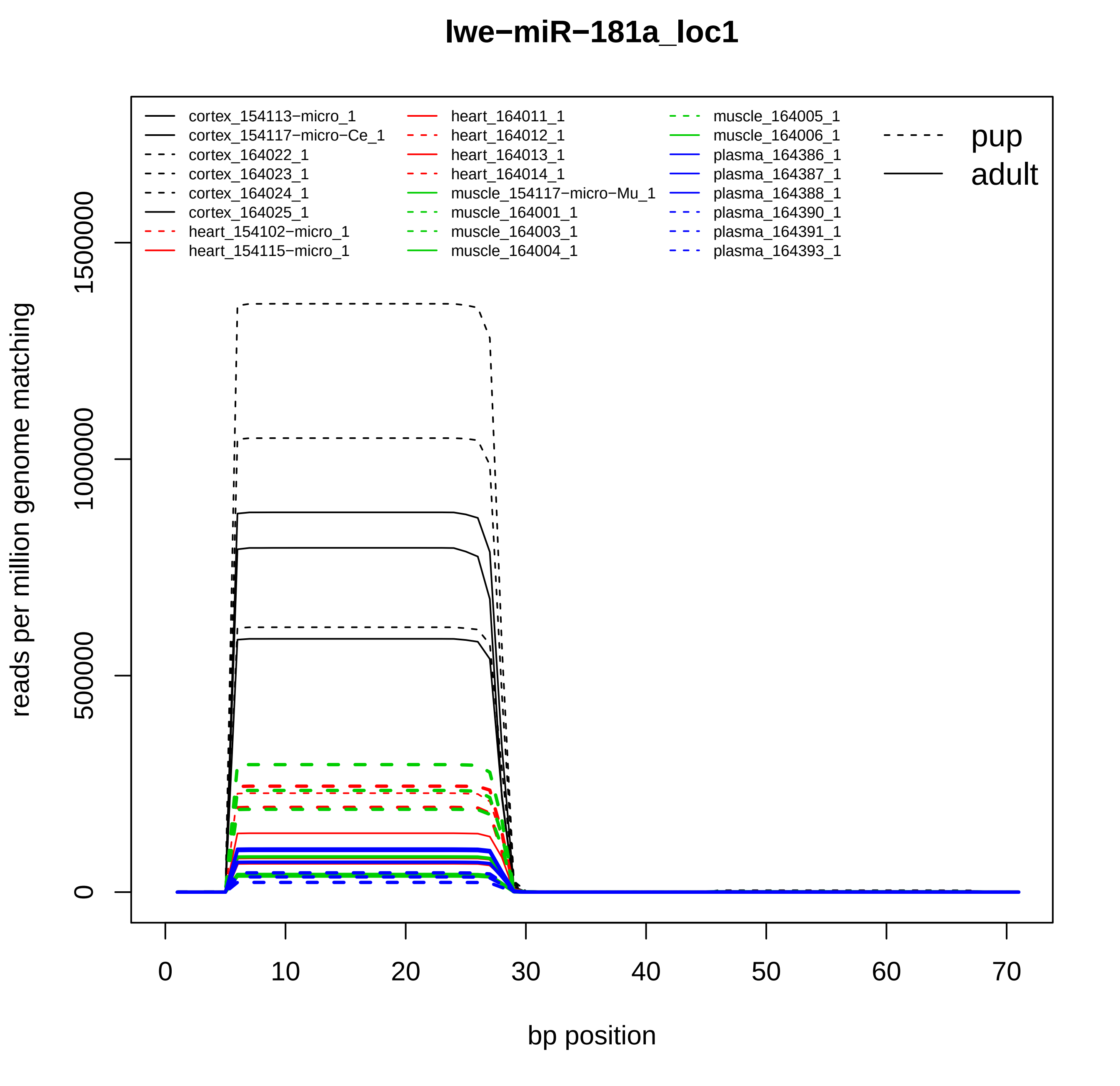
lwe−miR−181a_loc1
reads per million genome matching
bp position
0
500000
1000000
1500000
0
10
20
30
40
50
60
70
cortex_154113−micro_1
cortex_154117−micro−Ce_1
cortex_164022_1
cortex_164023_1
cortex_164024_1
cortex_164025_1
heart_154102−micro_1
heart_154115−micro_1
heart_164011_1
heart_164012_1
heart_164013_1
heart_164014_1
muscle_154117−micro−Mu_1
muscle_164001_1
muscle_164003_1
muscle_164004_1
muscle_164005_1
muscle_164006_1
plasma_164386_1
plasma_164387_1
plasma_164388_1
plasma_164390_1
plasma_164391_1
plasma_164393_1
pup
adult

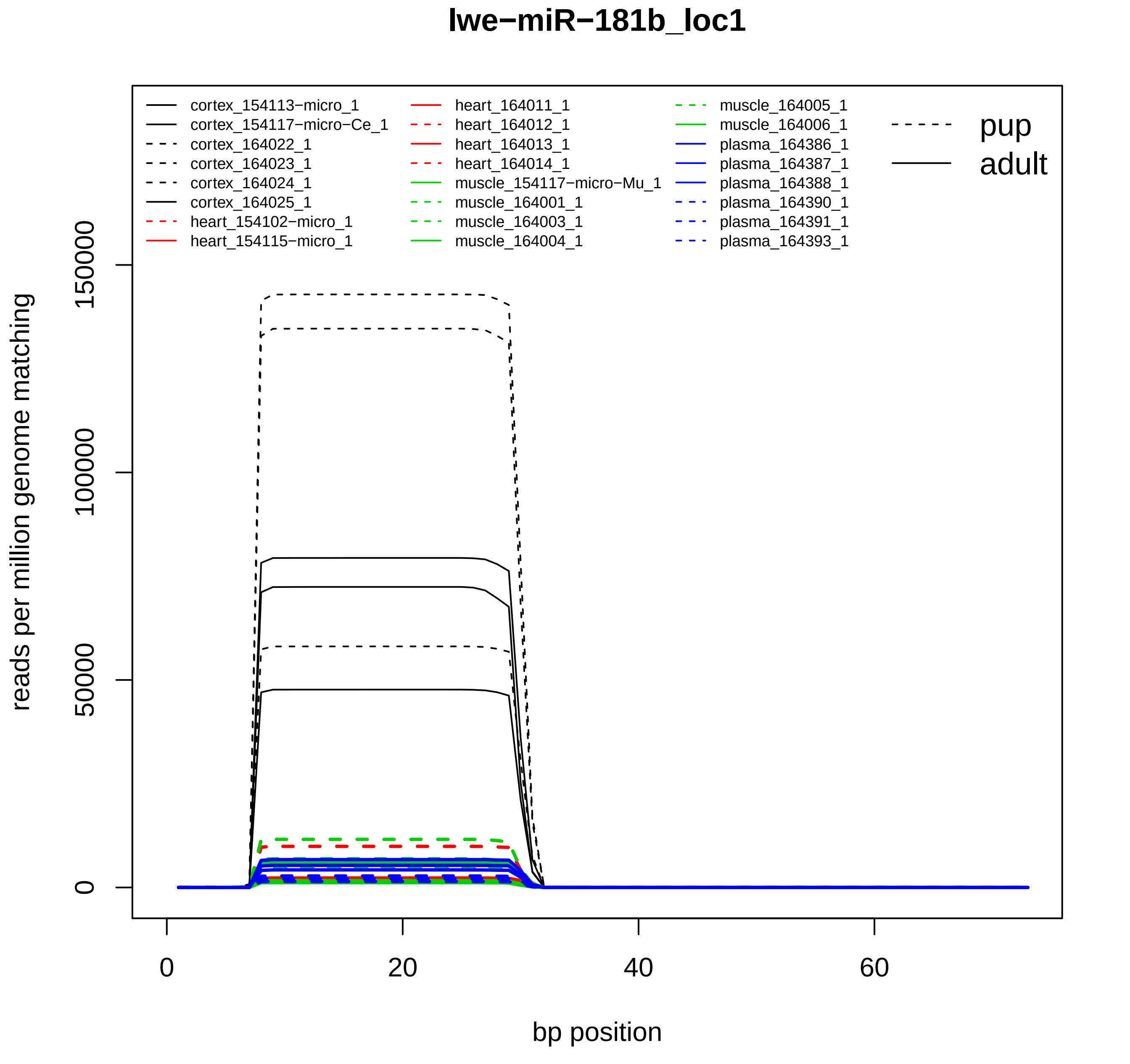
lwe−miR−181b_loc1
cortex_154113−micro_1
cortex_154117−micro−Ce_1
cortex_164022_1
cortex_164023_1
cortex_164024_1
cortex_164025_1
heart_154102−micro_1
heart_154115−micro_1
heart_164011_1
heart_164012_1
heart_164013_1
heart_164014_1
muscle_154117−micro−Mu_1
muscle_164001_1
muscle_164003_1
muscle_164004_1
muscle_164005_1
muscle_164006_1
plasma_164386_1
plasma_164387_1
plasma_164388_1
plasma_164390_1
plasma_164391_1
plasma_164393_1
pup
adult
reads per million genome matching
0
50000
100000
150000
bp position
0
20
40
60

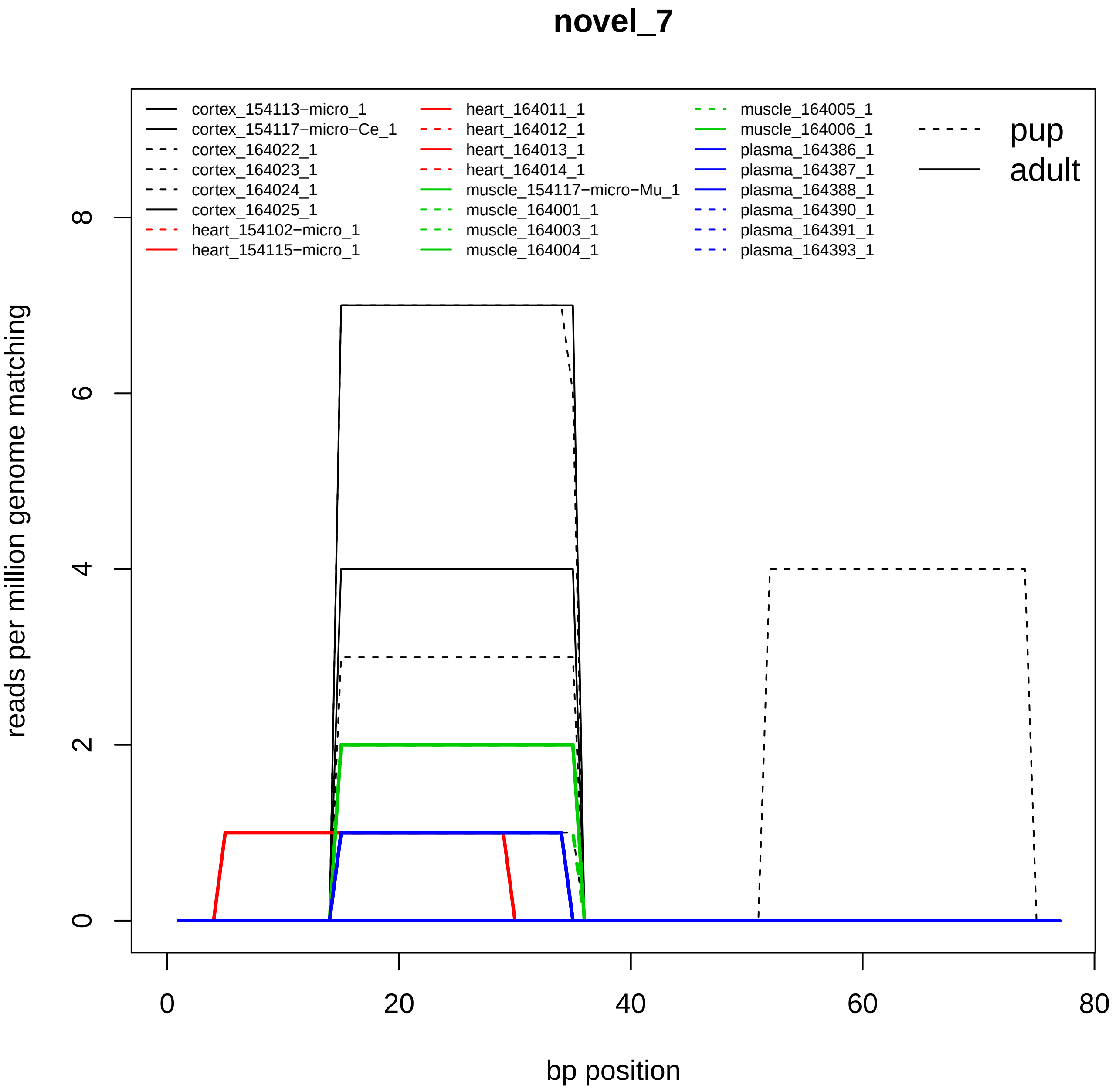
novel_7
cortex_154113−micro_1
cortex_154117−micro−Ce_1
cortex_164022_1
cortex_164023_1
cortex_164024_1
cortex_164025_1
heart_154102−micro_1
heart_154115−micro_1
heart_164011_1
heart_164012_1
heart_164013_1
heart_164014_1
muscle_154117−micro−Mu_1
muscle_164001_1
muscle_164003_1
muscle_164004_1
muscle_164005_1
muscle_164006_1
plasma_164386_1
plasma_164387_1
plasma_164388_1
plasma_164390_1
plasma_164391_1
plasma_164393_1
pup
adult
reads per million genome matching
0
2
4
6
8
0
20
40
60
80
bp position

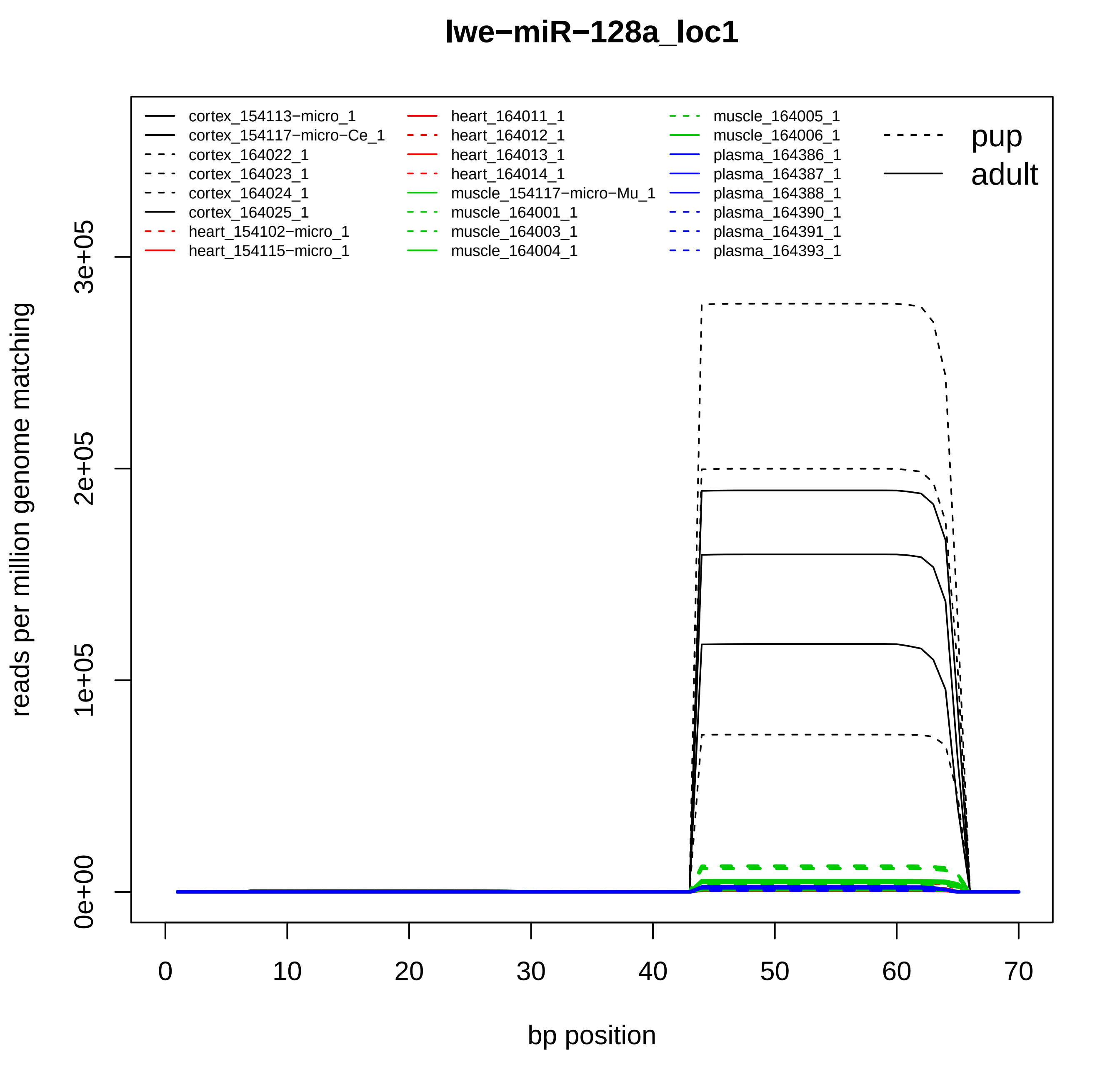
lwe−miR−128a_loc1
cortex_154113−micro_1
cortex_154117−micro−Ce_1
cortex_164022_1
cortex_164023_1
cortex_164024_1
cortex_164025_1
heart_154102−micro_1
heart_154115−micro_1
heart_164011_1
heart_164012_1
heart_164013_1
heart_164014_1
muscle_154117−micro−Mu_1
muscle_164001_1
muscle_164003_1
muscle_164004_1
muscle_164005_1
muscle_164006_1
plasma_164386_1
plasma_164387_1
plasma_164388_1
plasma_164390_1
plasma_164391_1
plasma_164393_1
pup
adult
reads per million genome matching
0e+00
1e+05
2e+05
3e+05
0
10
20
30
40
50
60
70
bp position

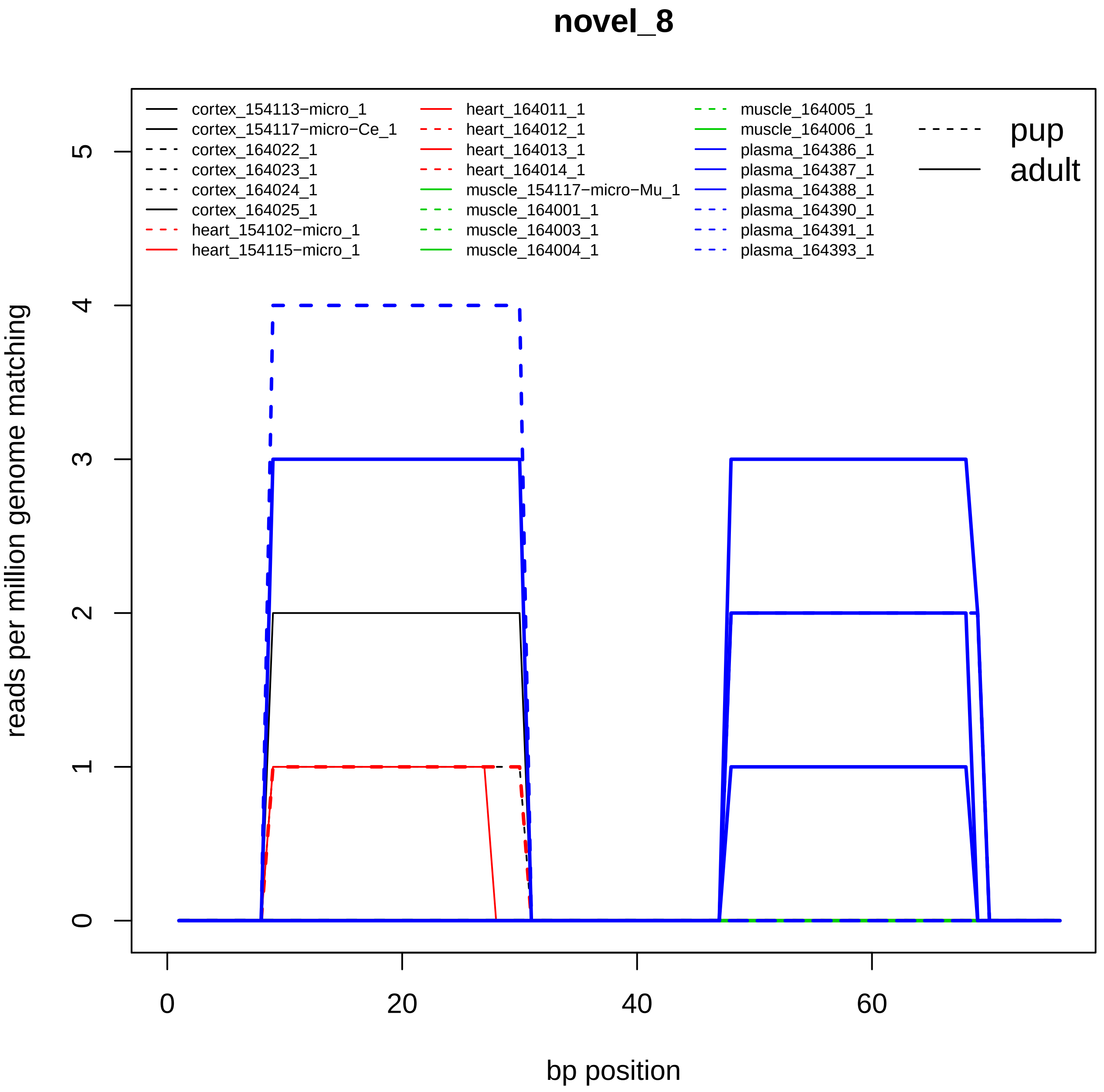

novel_8
cortex_154113−micro_1
cortex_154117−micro−Ce_1
cortex_164022_1
cortex_164023_1
cortex_164024_1
cortex_164025_1
heart_154102−micro_1
heart_154115−micro_1
heart_164011_1
heart_164012_1
heart_164013_1
heart_164014_1
muscle_154117−micro−Mu_1
muscle_164001_1
muscle_164003_1
muscle_164004_1
muscle_164005_1
muscle_164006_1
plasma_164386_1
plasma_164387_1
plasma_164388_1
plasma_164390_1
plasma_164391_1
plasma_164393_1
pup
adult
reads per million genome matching
0
1
2
3
4
5
0
20
40
60
bp position

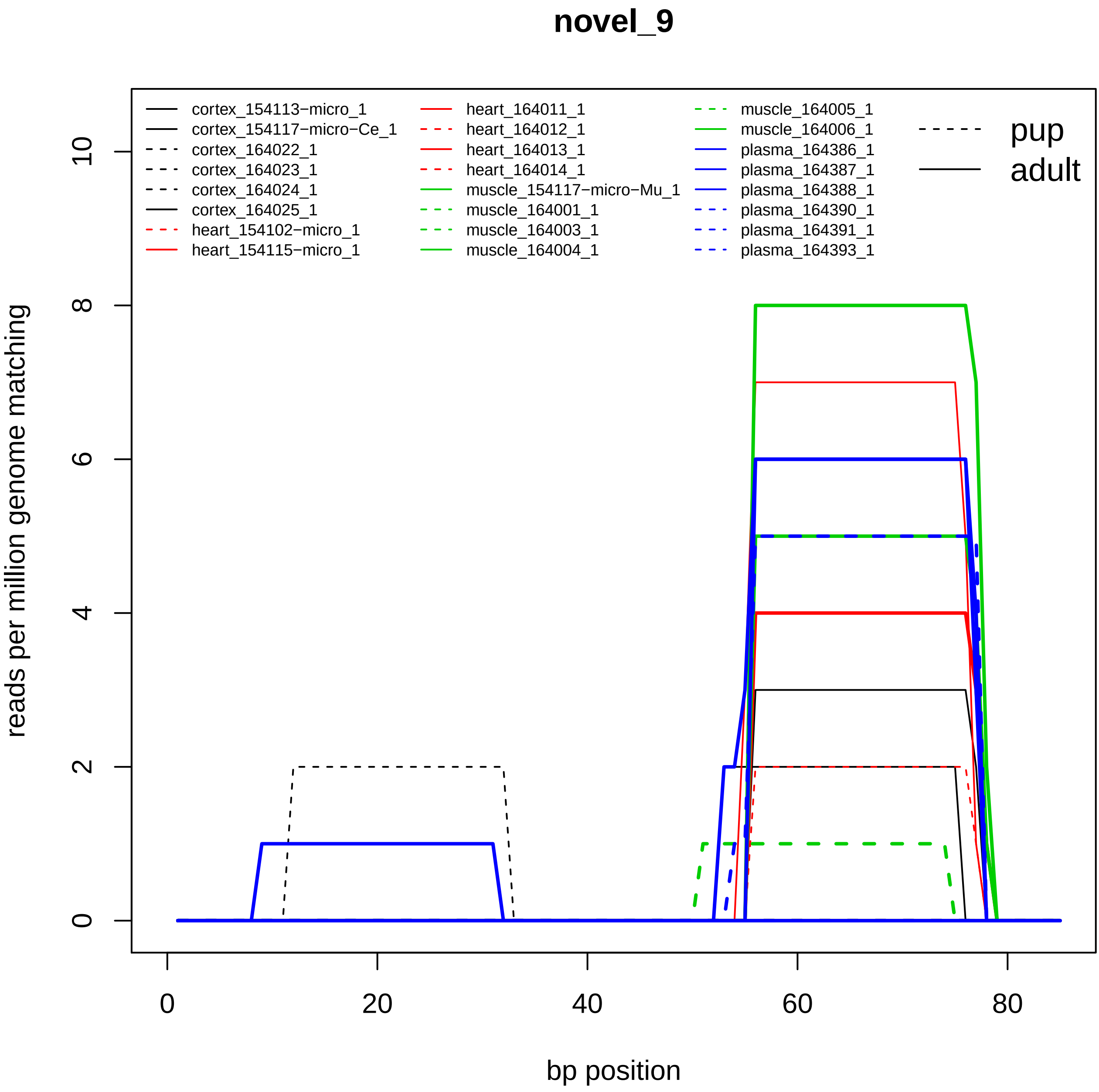

novel_9
cortex_154113−micro_1
cortex_154117−micro−Ce_1
cortex_164022_1
cortex_164023_1
cortex_164024_1
cortex_164025_1
heart_154102−micro_1
heart_154115−micro_1
heart_164011_1
heart_164012_1
heart_164013_1
heart_164014_1
muscle_154117−micro−Mu_1
muscle_164001_1
muscle_164003_1
muscle_164004_1
muscle_164005_1
muscle_164006_1
plasma_164386_1
plasma_164387_1
plasma_164388_1
plasma_164390_1
plasma_164391_1
plasma_164393_1
pup
adult
reads per million genome matching
0
2
4
6
8
10
0
20
40
60
80
bp position

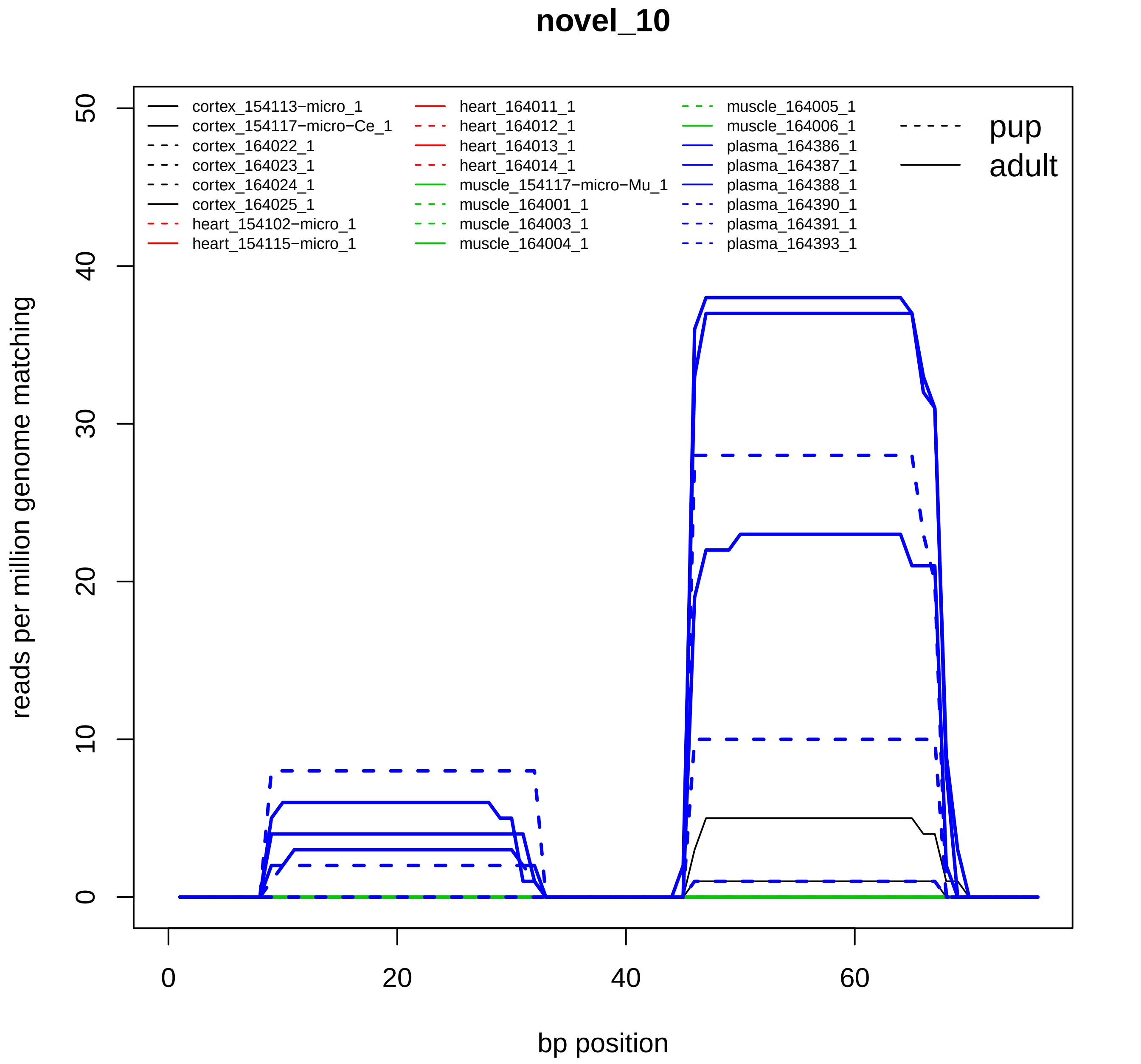
novel_10
cortex_154113−micro_1
cortex_154117−micro−Ce_1
cortex_164022_1
cortex_164023_1
cortex_164024_1
cortex_164025_1
heart_154102−micro_1
heart_154115−micro_1
heart_164011_1
heart_164012_1
heart_164013_1
heart_164014_1
muscle_154117−micro−Mu_1
muscle_164001_1
muscle_164003_1
muscle_164004_1
muscle_164005_1
muscle_164006_1
plasma_164386_1
plasma_164387_1
plasma_164388_1
plasma_164390_1
plasma_164391_1
plasma_164393_1
pup
adult
reads per million genome matching
bp position
0
10
20
30
40
50
60

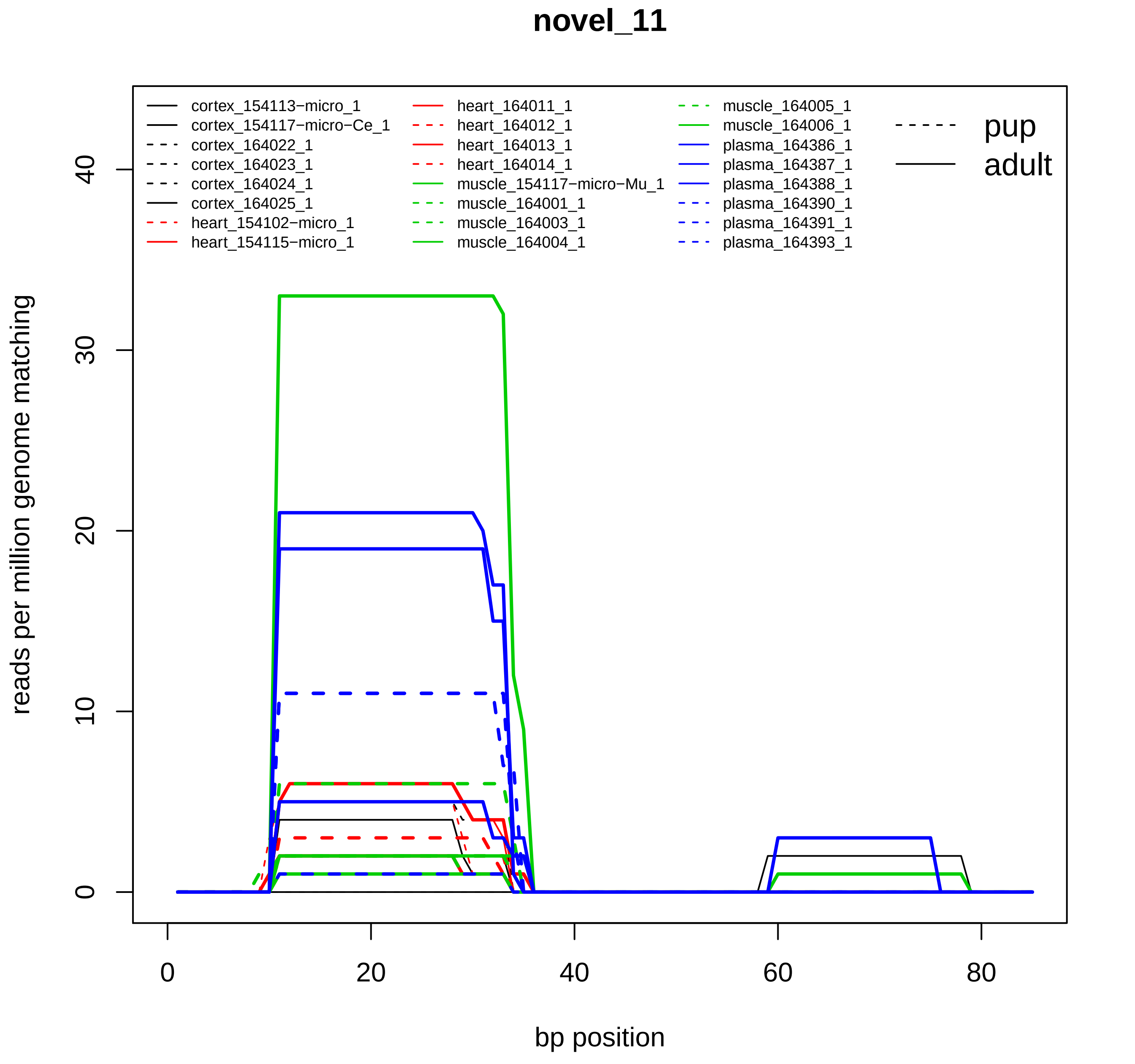

novel_11
cortex_154113−micro_1
cortex_154117−micro−Ce_1
cortex_164022_1
cortex_164023_1
cortex_164024_1
cortex_164025_1
heart_154102−micro_1
heart_154115−micro_1
heart_164011_1
heart_164012_1
heart_164013_1
heart_164014_1
muscle_154117−micro−Mu_1
muscle_164001_1
muscle_164003_1
muscle_164004_1
muscle_164005_1
muscle_164006_1
plasma_164386_1
plasma_164387_1
plasma_164388_1
plasma_164390_1
plasma_164391_1
plasma_164393_1
pup
adult
reads per million genome matching
0
10
20
30
40
0
20
40
60
80
bp position

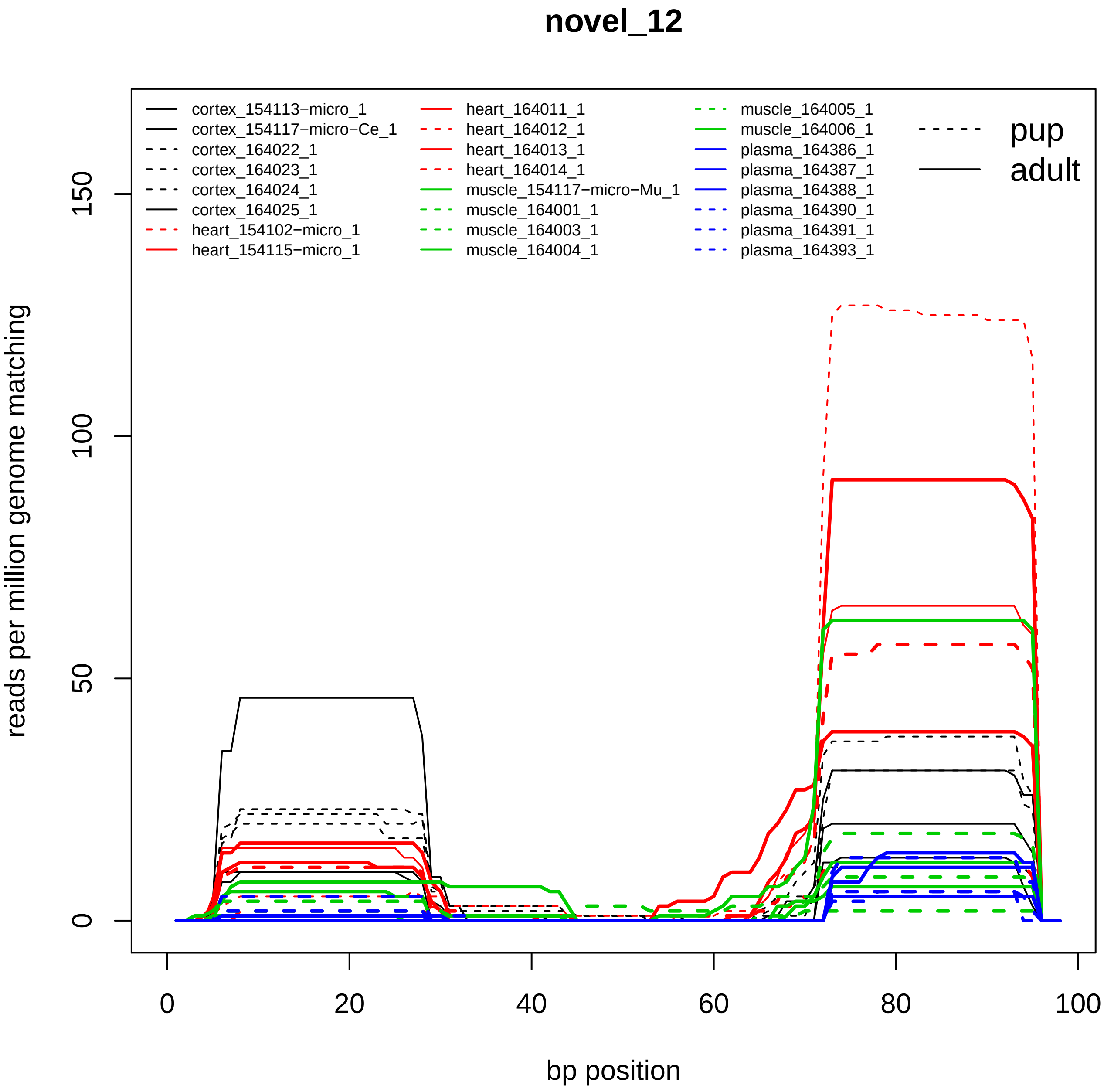

novel_12
cortex_154113−micro_1
cortex_154117−micro−Ce_1
cortex_164022_1
cortex_164023_1
cortex_164024_1
cortex_164025_1
heart_154102−micro_1
heart_154115−micro_1
heart_164011_1
heart_164012_1
heart_164013_1
heart_164014_1
muscle_154117−micro−Mu_1
muscle_164001_1
muscle_164003_1
muscle_164004_1
muscle_164005_1
muscle_164006_1
plasma_164386_1
plasma_164387_1
plasma_164388_1
plasma_164390_1
plasma_164391_1
plasma_164393_1
pup
adult
reads per million genome matching
0
50
100
150
bp position
0
20
40
60
80
100

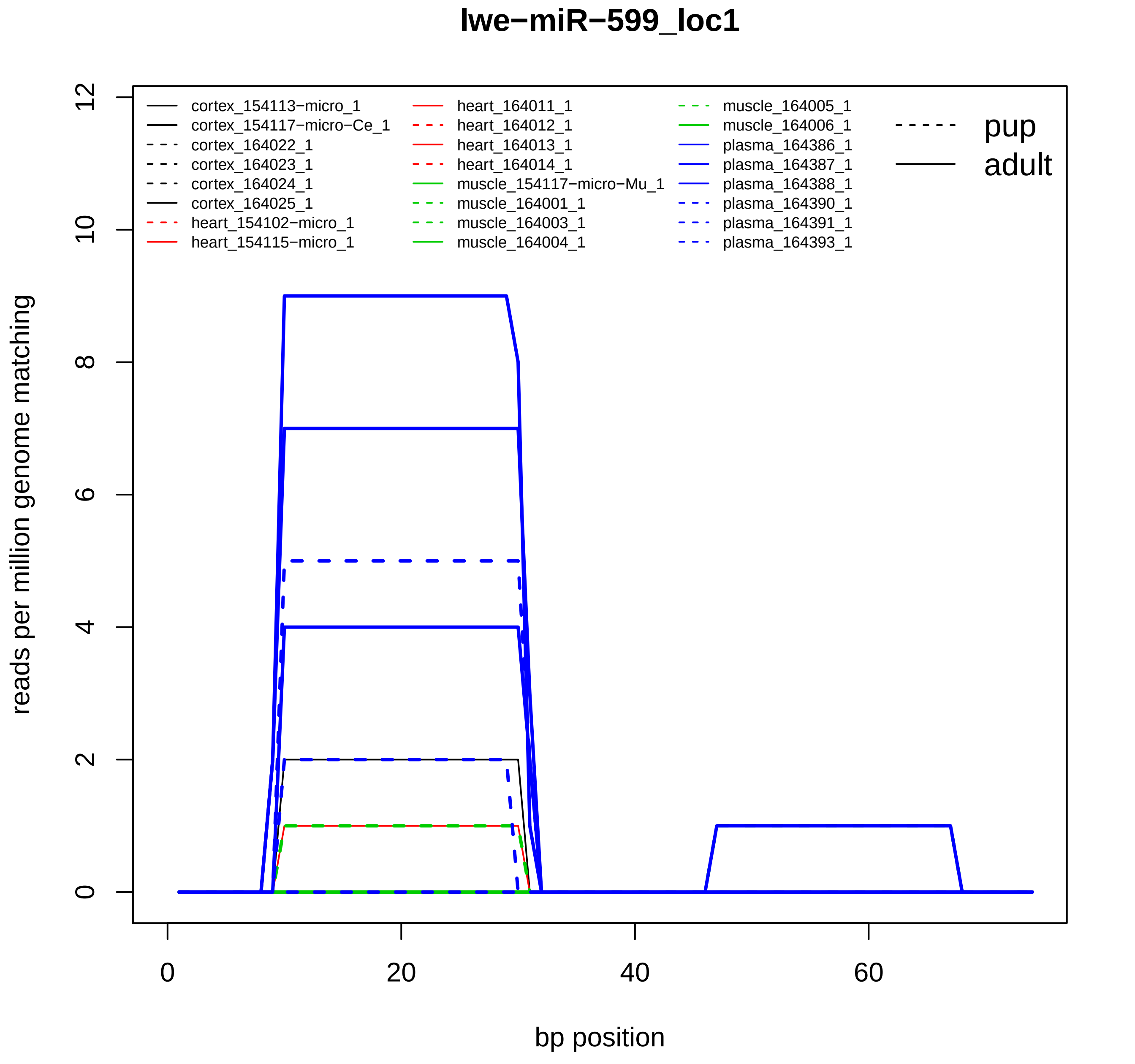
lwe−miR−599_loc1
cortex_154113−micro_1
cortex_154117−micro−Ce_1
cortex_164022_1
cortex_164023_1
cortex_164024_1
cortex_164025_1
heart_154102−micro_1
heart_154115−micro_1
heart_164011_1
heart_164012_1
heart_164013_1
heart_164014_1
muscle_154117−micro−Mu_1
muscle_164001_1
muscle_164003_1
muscle_164004_1
muscle_164005_1
muscle_164006_1
plasma_164386_1
plasma_164387_1
plasma_164388_1
plasma_164390_1
plasma_164391_1
plasma_164393_1
pup
adult
reads per million genome matching
bp position
0
2
4
6
8
10
12
0
20
40
60

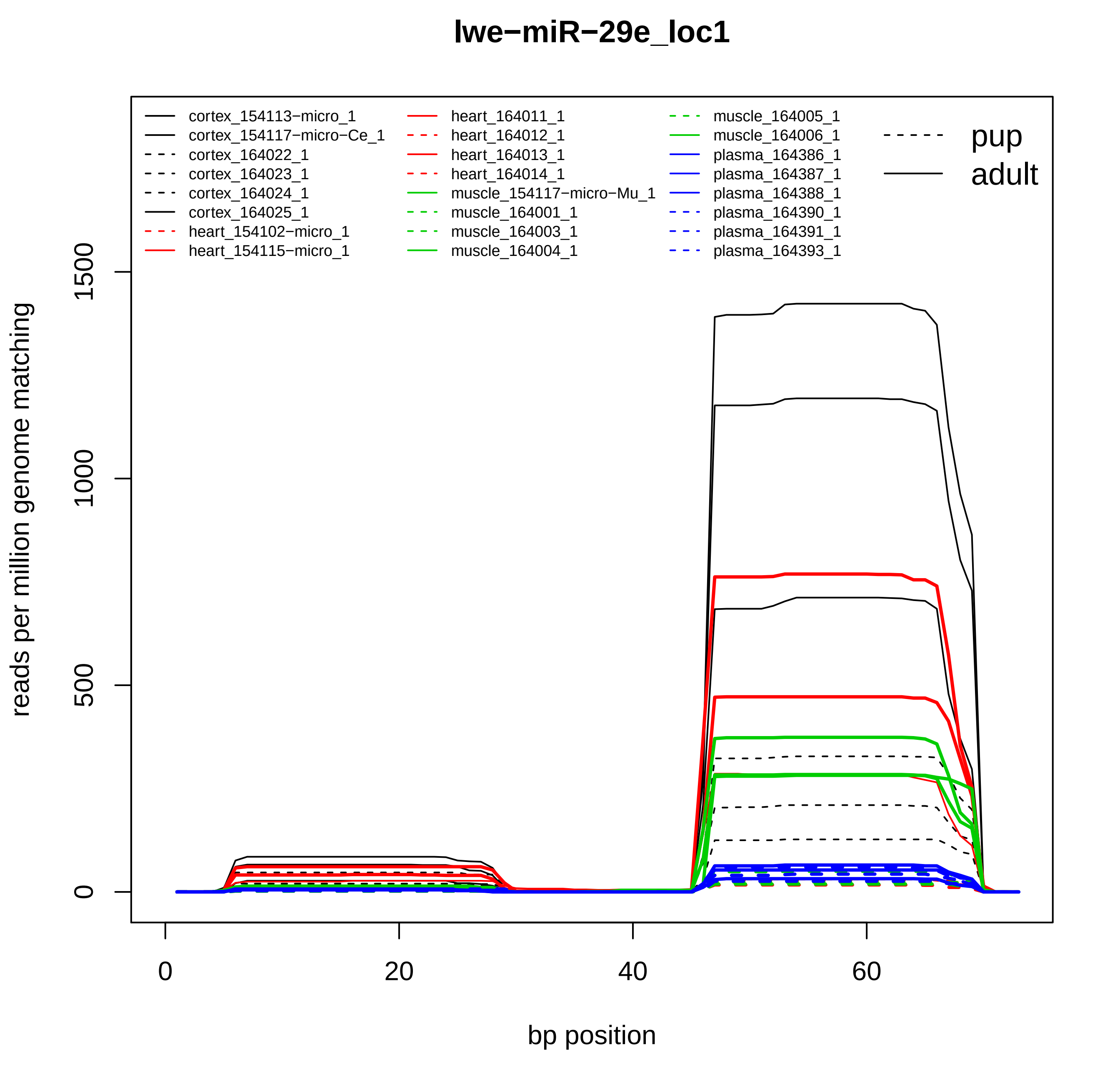
lwe−miR−29e_loc1
cortex_154113−micro_1
cortex_154117−micro−Ce_1
cortex_164022_1
cortex_164023_1
cortex_164024_1
cortex_164025_1
heart_154102−micro_1
heart_154115−micro_1
heart_164011_1
heart_164012_1
heart_164013_1
heart_164014_1
muscle_154117−micro−Mu_1
muscle_164001_1
muscle_164003_1
muscle_164004_1
muscle_164005_1
muscle_164006_1
plasma_164386_1
plasma_164387_1
plasma_164388_1
plasma_164390_1
plasma_164391_1
plasma_164393_1
pup
adult
reads per million genome matching
0
500
1000
1500
0
20
40
60
bp position

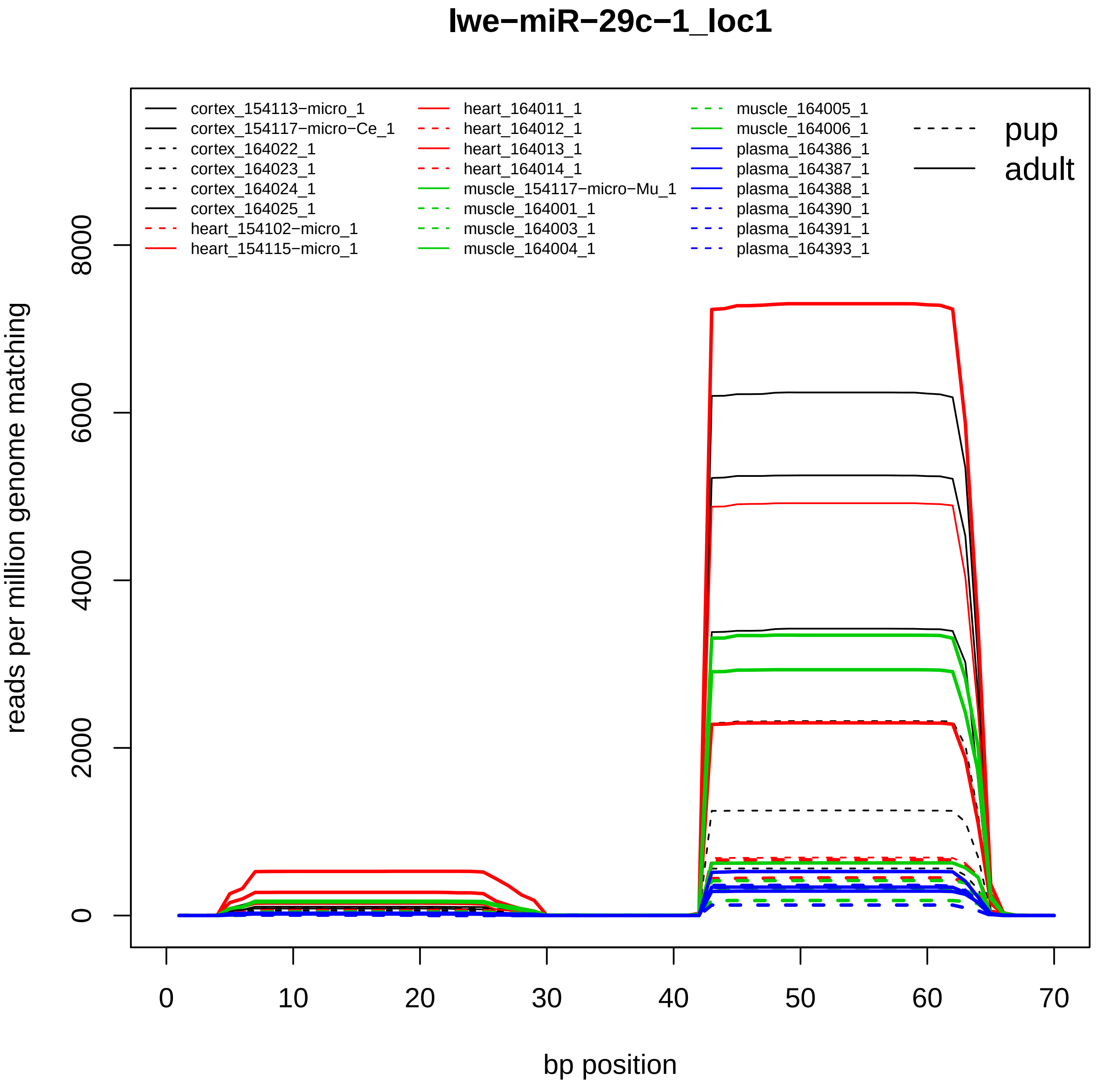

lwe−miR−29c−1_loc1
cortex_154113−micro_1
cortex_154117−micro−Ce_1
cortex_164022_1
cortex_164023_1
cortex_164024_1
cortex_164025_1
heart_154102−micro_1
heart_154115−micro_1
heart_164011_1
heart_164012_1
heart_164013_1
heart_164014_1
muscle_154117−micro−Mu_1
muscle_164001_1
muscle_164003_1
muscle_164004_1
muscle_164005_1
muscle_164006_1
plasma_164386_1
plasma_164387_1
plasma_164388_1
plasma_164390_1
plasma_164391_1
plasma_164393_1
pup
adult
reads per million genome matching
0
2000
4000
6000
8000
0
10
20
30
40
50
60
70
bp position

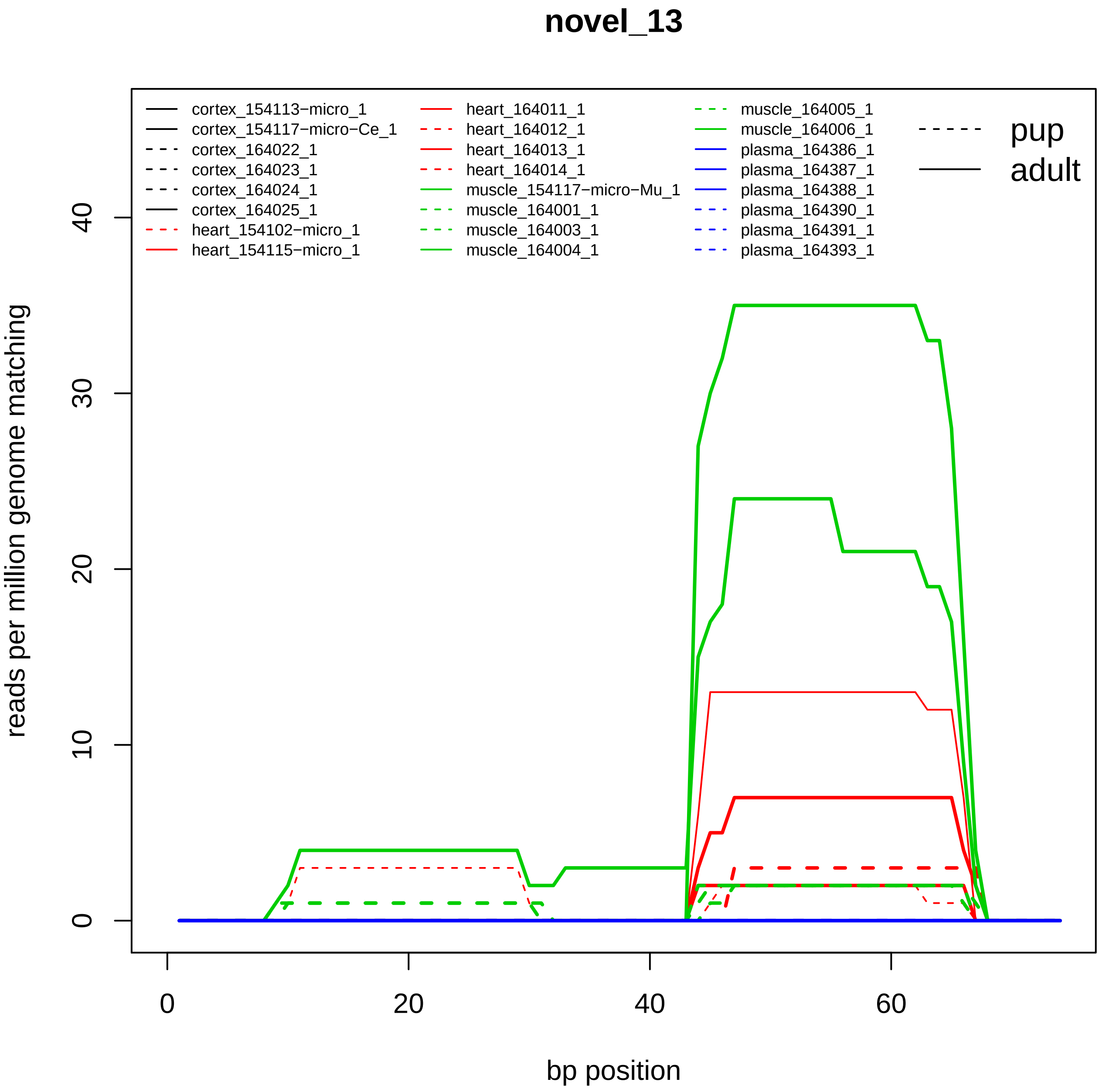

novel_13
cortex_154113−micro_1
cortex_154117−micro−Ce_1
cortex_164022_1
cortex_164023_1
cortex_164024_1
cortex_164025_1
heart_154102−micro_1
heart_154115−micro_1
heart_164011_1
heart_164012_1
heart_164013_1
heart_164014_1
muscle_154117−micro−Mu_1
muscle_164001_1
muscle_164003_1
muscle_164004_1
muscle_164005_1
muscle_164006_1
plasma_164386_1
plasma_164387_1
plasma_164388_1
plasma_164390_1
plasma_164391_1
plasma_164393_1
pup
adult
reads per million genome matching
0
10
20
30
40
0
20
40
60
bp position

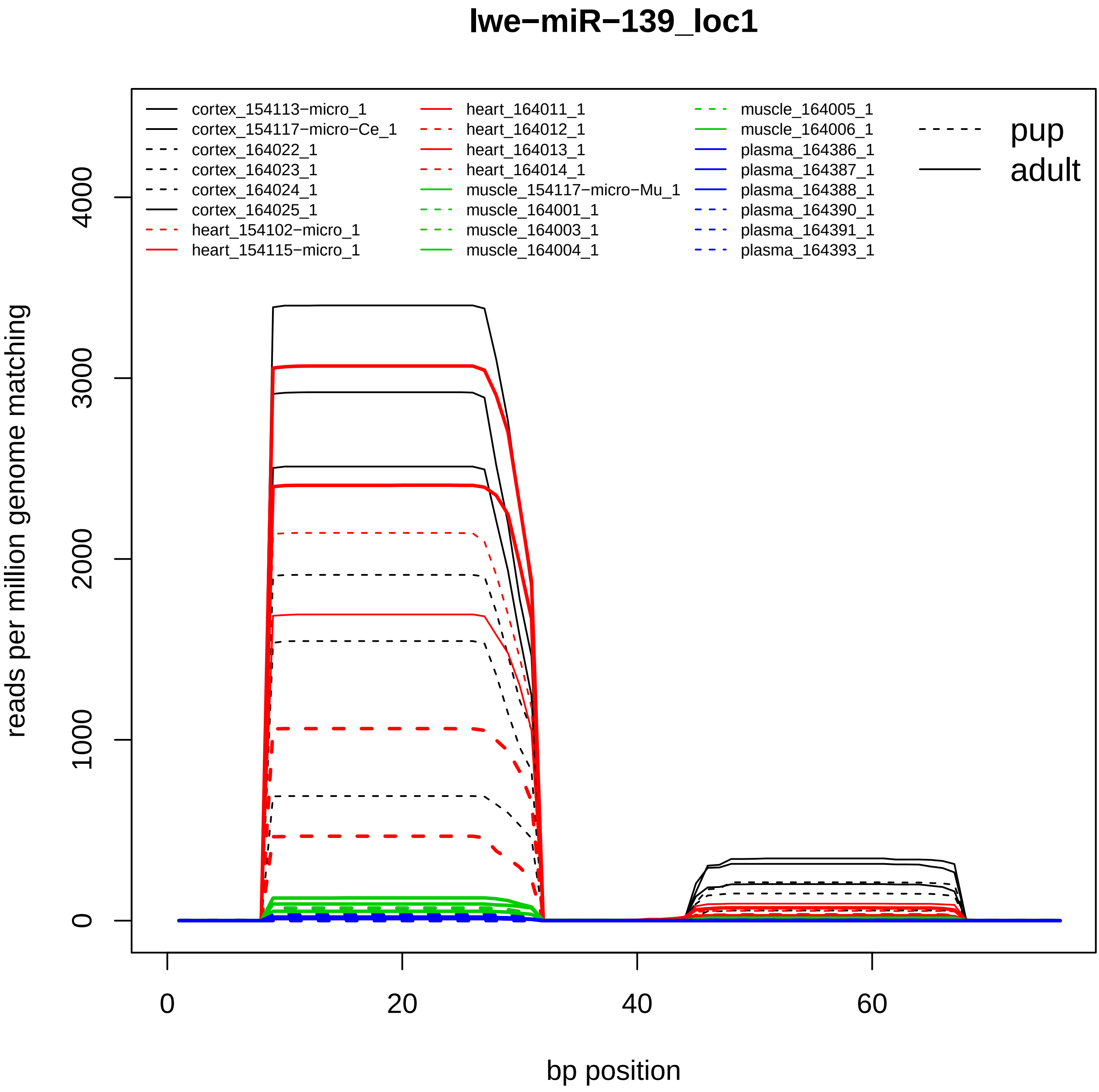
lwe−miR−139_loc1
cortex_154113−micro_1
cortex_154117−micro−Ce_1
cortex_164022_1
cortex_164023_1
cortex_164024_1
cortex_164025_1
heart_154102−micro_1
heart_154115−micro_1
heart_164011_1
heart_164012_1
heart_164013_1
heart_164014_1
muscle_154117−micro−Mu_1
muscle_164001_1
muscle_164003_1
muscle_164004_1
muscle_164005_1
muscle_164006_1
plasma_164386_1
plasma_164387_1
plasma_164388_1
plasma_164390_1
plasma_164391_1
plasma_164393_1
pup
adult
reads per million genome matching
0
1000
2000
3000
4000
0
20
40
60
bp position

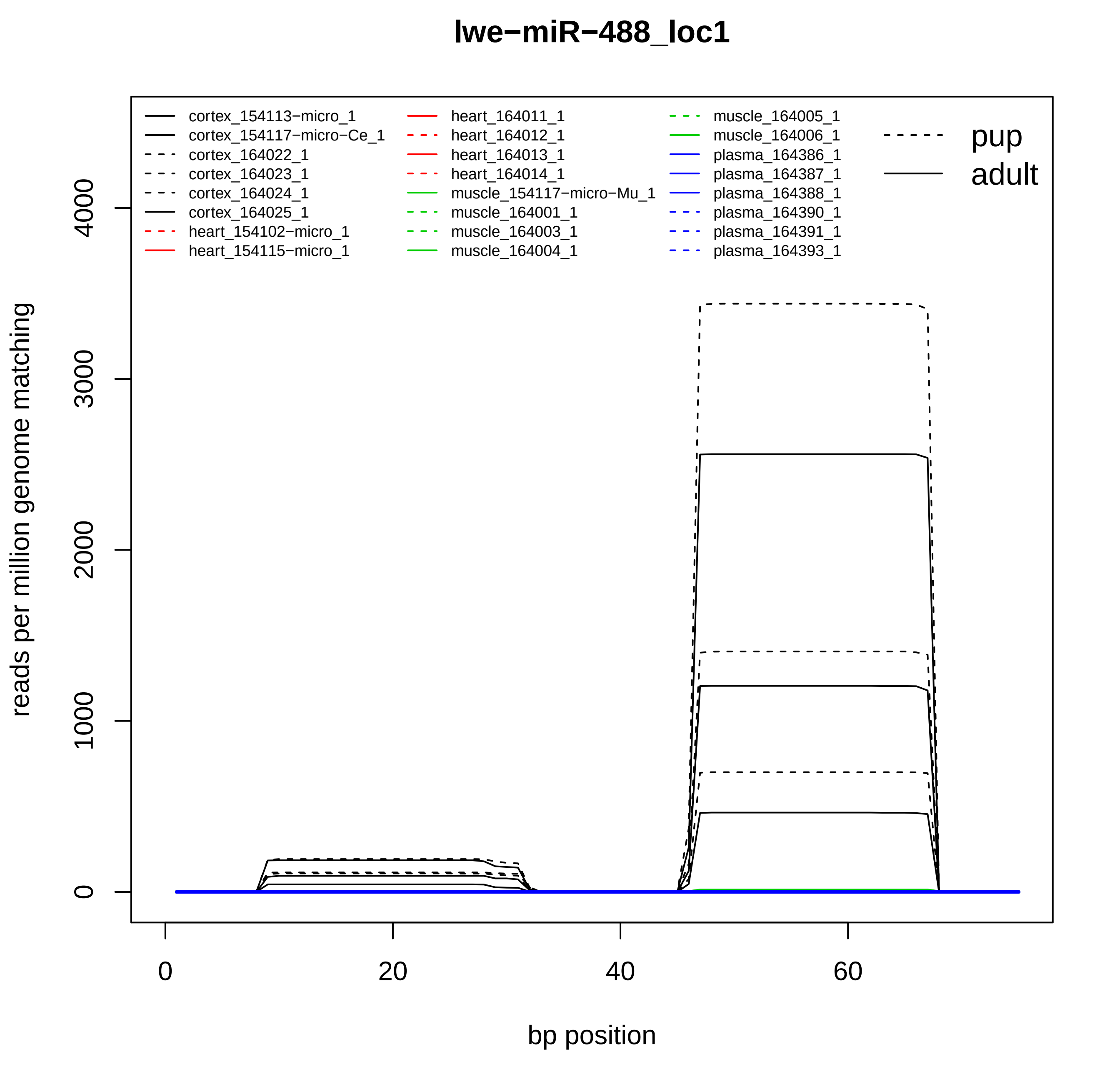
lwe−miR−488_loc1
cortex_154113−micro_1
cortex_154117−micro−Ce_1
cortex_164022_1
cortex_164023_1
cortex_164024_1
cortex_164025_1
heart_154102−micro_1
heart_154115−micro_1
heart_164011_1
heart_164012_1
heart_164013_1
heart_164014_1
muscle_154117−micro−Mu_1
muscle_164001_1
muscle_164003_1
muscle_164004_1
muscle_164005_1
muscle_164006_1
plasma_164386_1
plasma_164387_1
plasma_164388_1
plasma_164390_1
plasma_164391_1
plasma_164393_1
pup
adult
reads per million genome matching
0
1000
2000
3000
4000
0
20
40
60
bp position

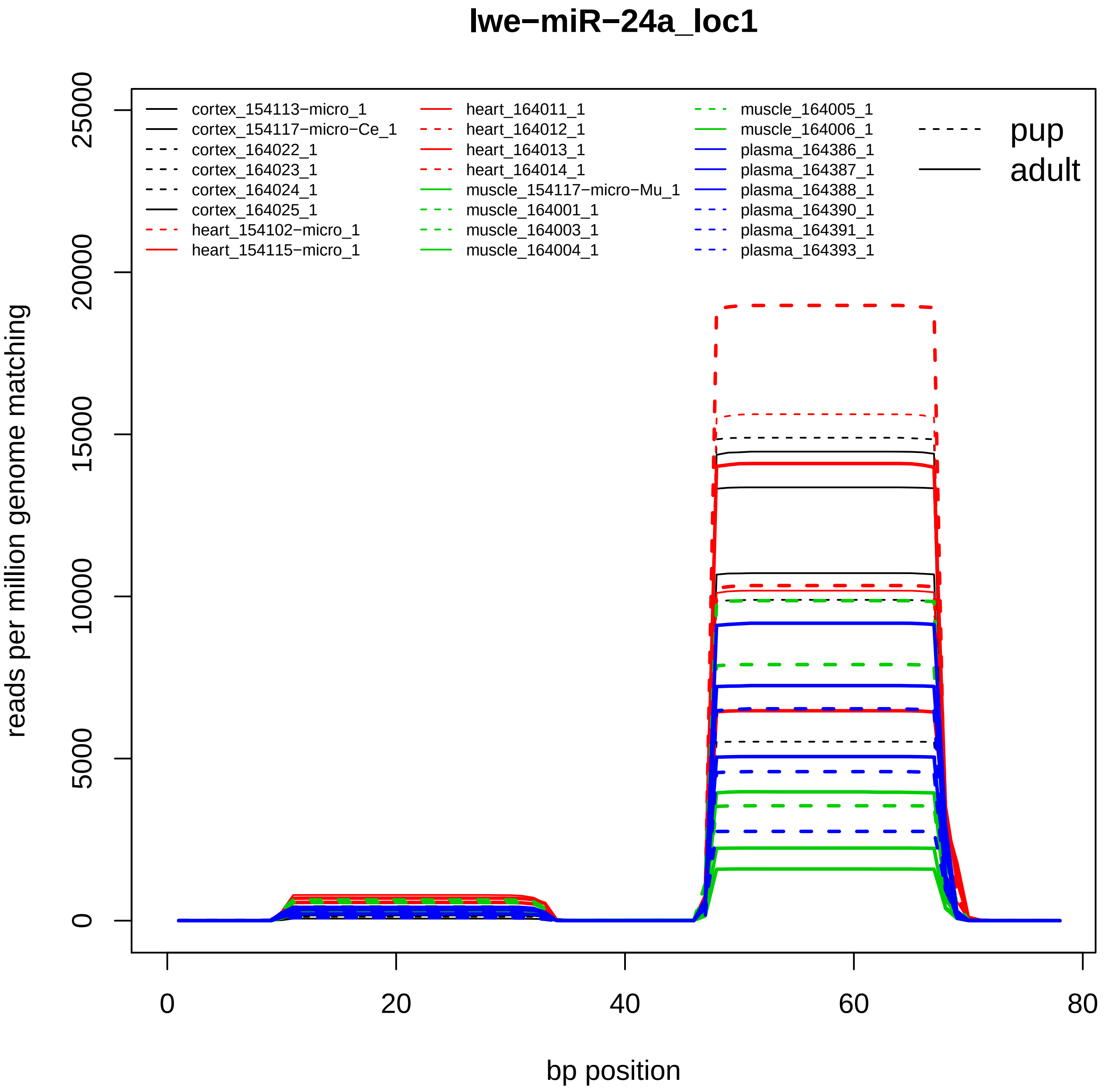

lwe−miR−24a_loc1
cortex_154113−micro_1
cortex_154117−micro−Ce_1
cortex_164022_1
cortex_164023_1
cortex_164024_1
cortex_164025_1
heart_154102−micro_1
heart_154115−micro_1
heart_164011_1
heart_164012_1
heart_164013_1
heart_164014_1
muscle_154117−micro−Mu_1
muscle_164001_1
muscle_164003_1
muscle_164004_1
muscle_164005_1
muscle_164006_1
plasma_164386_1
plasma_164387_1
plasma_164388_1
plasma_164390_1
plasma_164391_1
plasma_164393_1
pup
adult
reads per million genome matching
0
5000
10000
15000
20000
25000
0
20
40
60
80
bp position

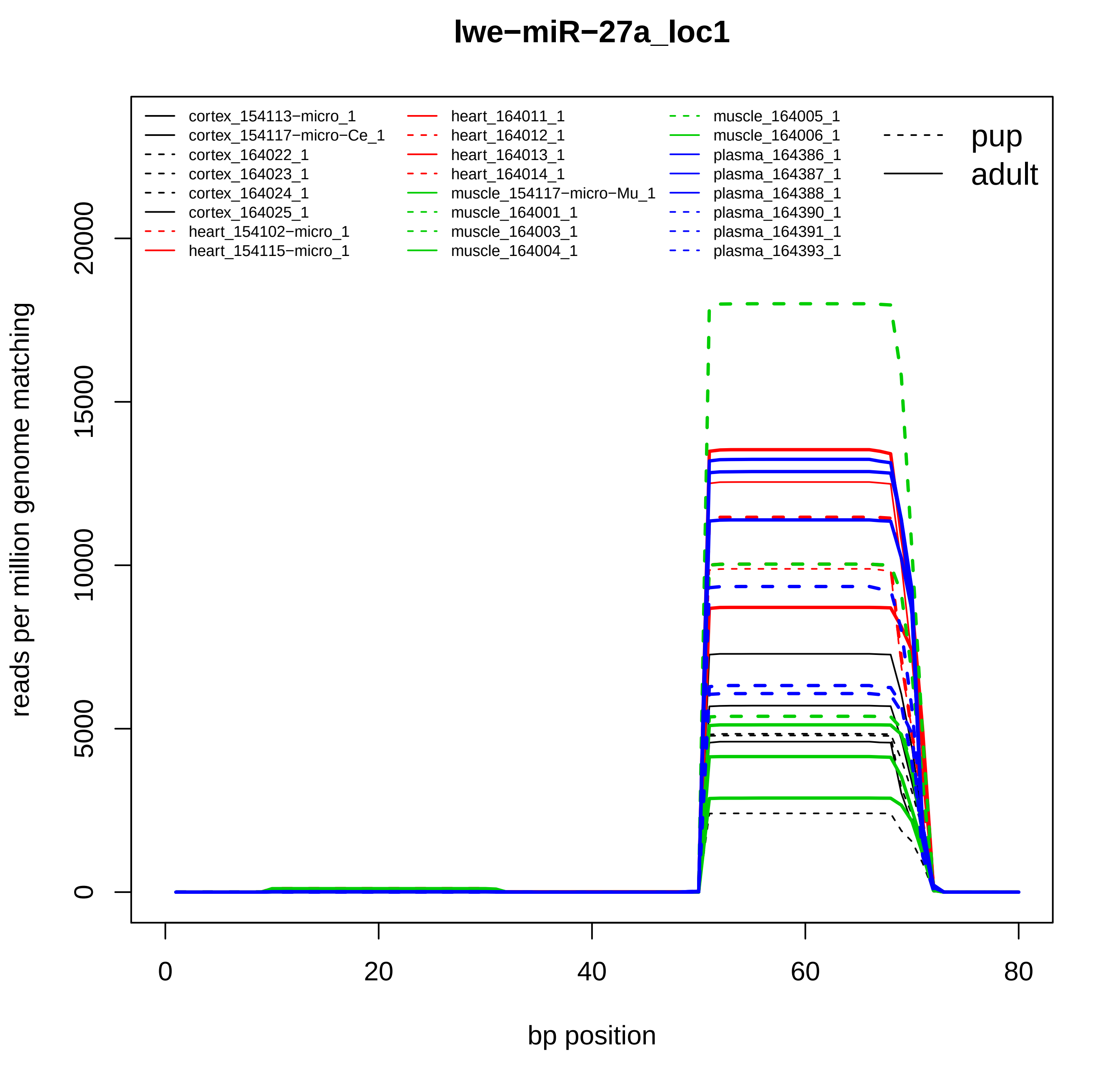
lwe−miR−27a_loc1
cortex_154113−micro_1
cortex_154117−micro−Ce_1
cortex_164022_1
cortex_164023_1
cortex_164024_1
cortex_164025_1
heart_154102−micro_1
heart_154115−micro_1
heart_164011_1
heart_164012_1
heart_164013_1
heart_164014_1
muscle_154117−micro−Mu_1
muscle_164001_1
muscle_164003_1
muscle_164004_1
muscle_164005_1
muscle_164006_1
plasma_164386_1
plasma_164387_1
plasma_164388_1
plasma_164390_1
plasma_164391_1
plasma_164393_1
pup
adult
reads per million genome matching
0
5000
10000
15000
20000
0
20
40
60
80
bp position

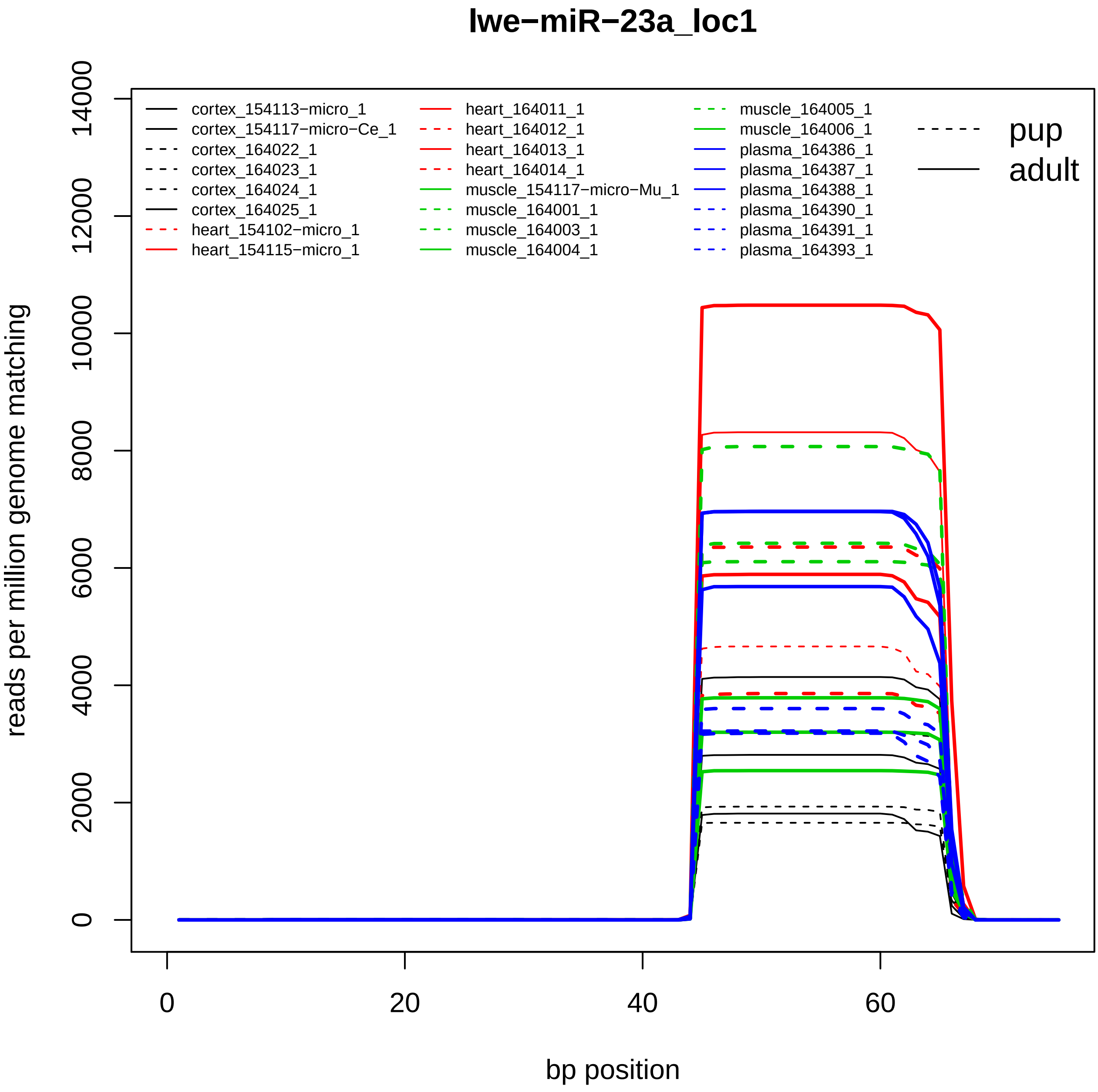
lwe−miR−23a_loc1
reads per million genome matching
bp position
0
2000
4000
6000
8000
10000
12000
14000
0
20
40
60
cortex_154113−micro_1
cortex_154117−micro−Ce_1
cortex_164022_1
cortex_164023_1
cortex_164024_1
cortex_164025_1
heart_154102−micro_1
heart_154115−micro_1
heart_164011_1
heart_164012_1
heart_164013_1
heart_164014_1
muscle_154117−micro−Mu_1
muscle_164001_1
muscle_164003_1
muscle_164004_1
muscle_164005_1
muscle_164006_1
plasma_164386_1
plasma_164387_1
plasma_164388_1
plasma_164390_1
plasma_164391_1
plasma_164393_1
pup
adult

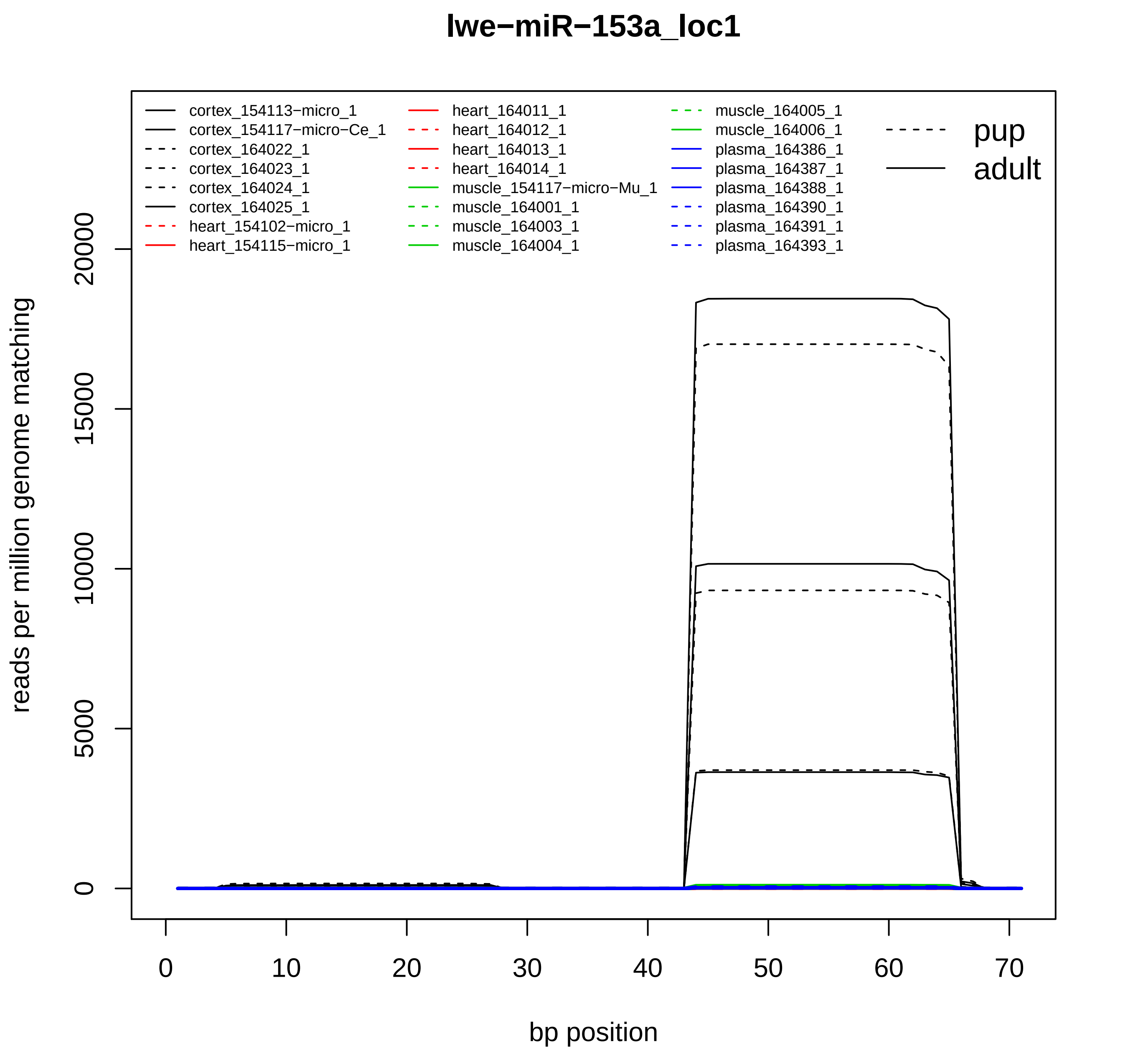
lwe−miR−153a_loc1
cortex_154113−micro_1
cortex_154117−micro−Ce_1
cortex_164022_1
cortex_164023_1
cortex_164024_1
cortex_164025_1
heart_154102−micro_1
heart_154115−micro_1
heart_164011_1
heart_164012_1
heart_164013_1
heart_164014_1
muscle_154117−micro−Mu_1
muscle_164001_1
muscle_164003_1
muscle_164004_1
muscle_164005_1
muscle_164006_1
plasma_164386_1
plasma_164387_1
plasma_164388_1
plasma_164390_1
plasma_164391_1
plasma_164393_1
pup
adult
reads per million genome matching
0
5000
10000
15000
20000
bp position
0
10
20
30
40
50
60
70

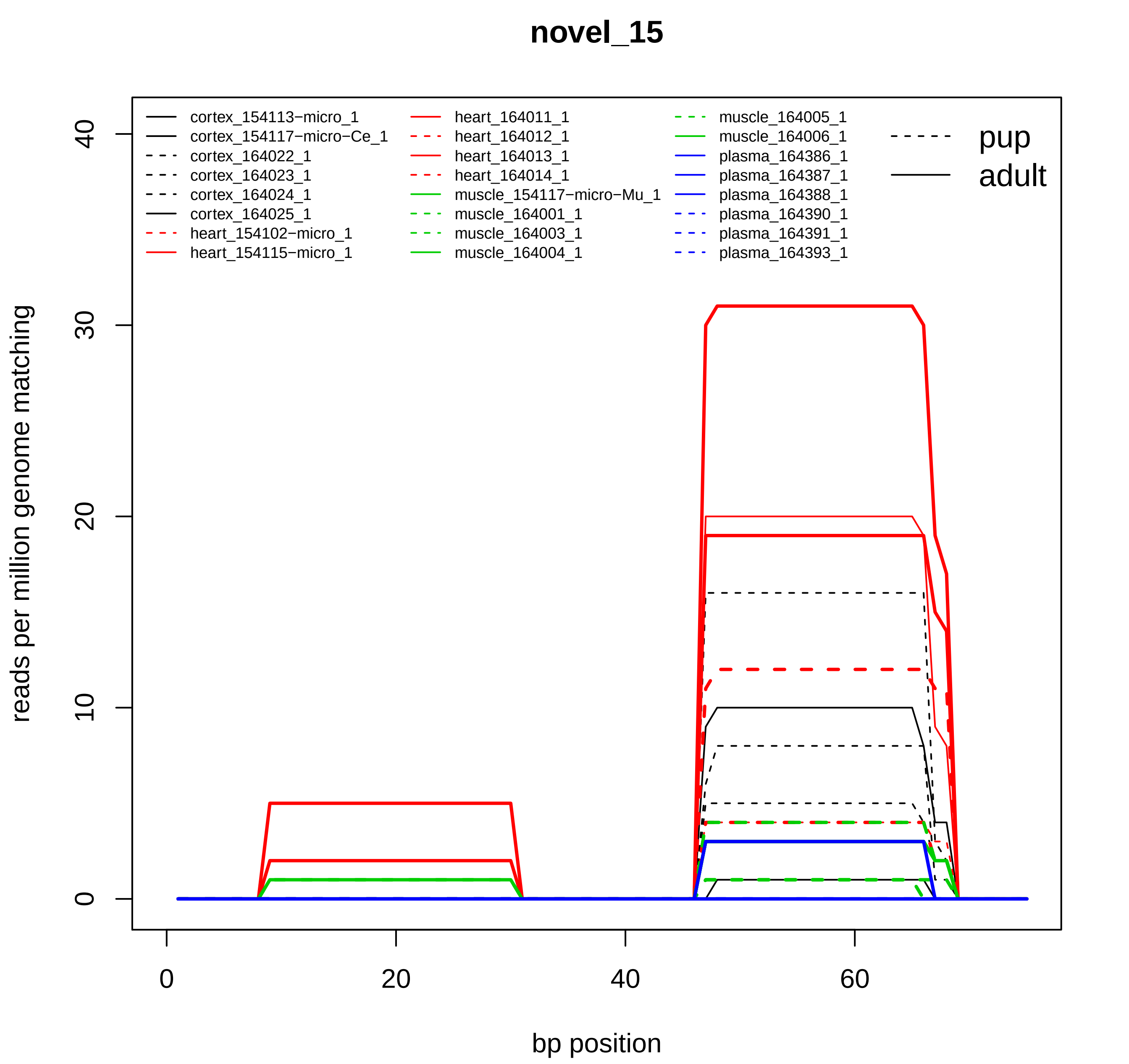
novel_15
cortex_154113−micro_1
cortex_154117−micro−Ce_1
cortex_164022_1
cortex_164023_1
cortex_164024_1
cortex_164025_1
heart_154102−micro_1
heart_154115−micro_1
heart_164011_1
heart_164012_1
heart_164013_1
heart_164014_1
muscle_154117−micro−Mu_1
muscle_164001_1
muscle_164003_1
muscle_164004_1
muscle_164005_1
muscle_164006_1
plasma_164386_1
plasma_164387_1
plasma_164388_1
plasma_164390_1
plasma_164391_1
plasma_164393_1
pup
adult
reads per million genome matching
0
10
20
30
40
0
20
40
60
bp position

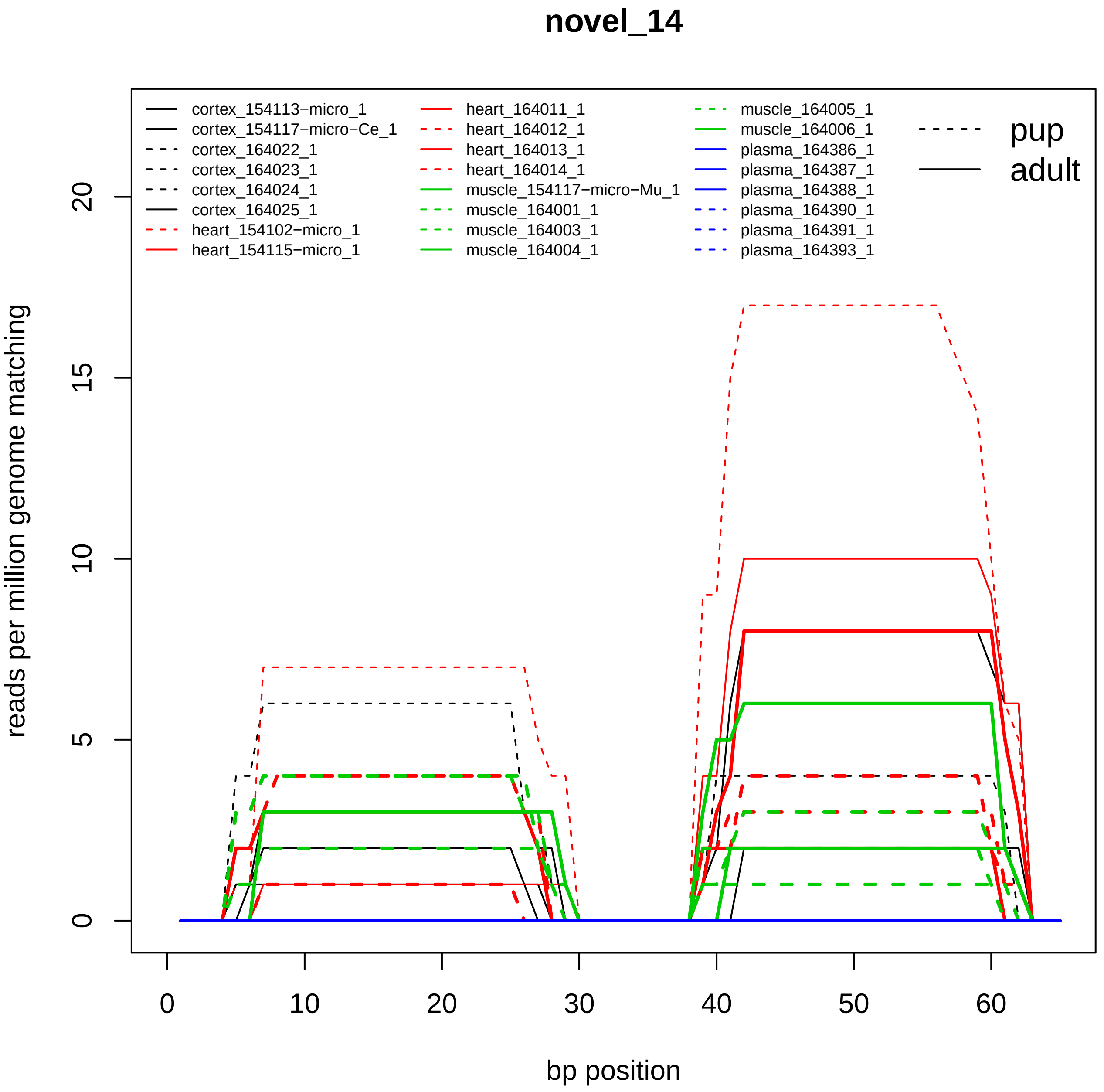
novel_14
cortex_154113−micro_1
cortex_154117−micro−Ce_1
cortex_164022_1
cortex_164023_1
cortex_164024_1
cortex_164025_1
heart_154102−micro_1
heart_154115−micro_1
heart_164011_1
heart_164012_1
heart_164013_1
heart_164014_1
muscle_154117−micro−Mu_1
muscle_164001_1
muscle_164003_1
muscle_164004_1
muscle_164005_1
muscle_164006_1
plasma_164386_1
plasma_164387_1
plasma_164388_1
plasma_164390_1
plasma_164391_1
plasma_164393_1
pup
adult
reads per million genome matching
0
5
10
15
20
0
10
20
30
40
50
60
bp position

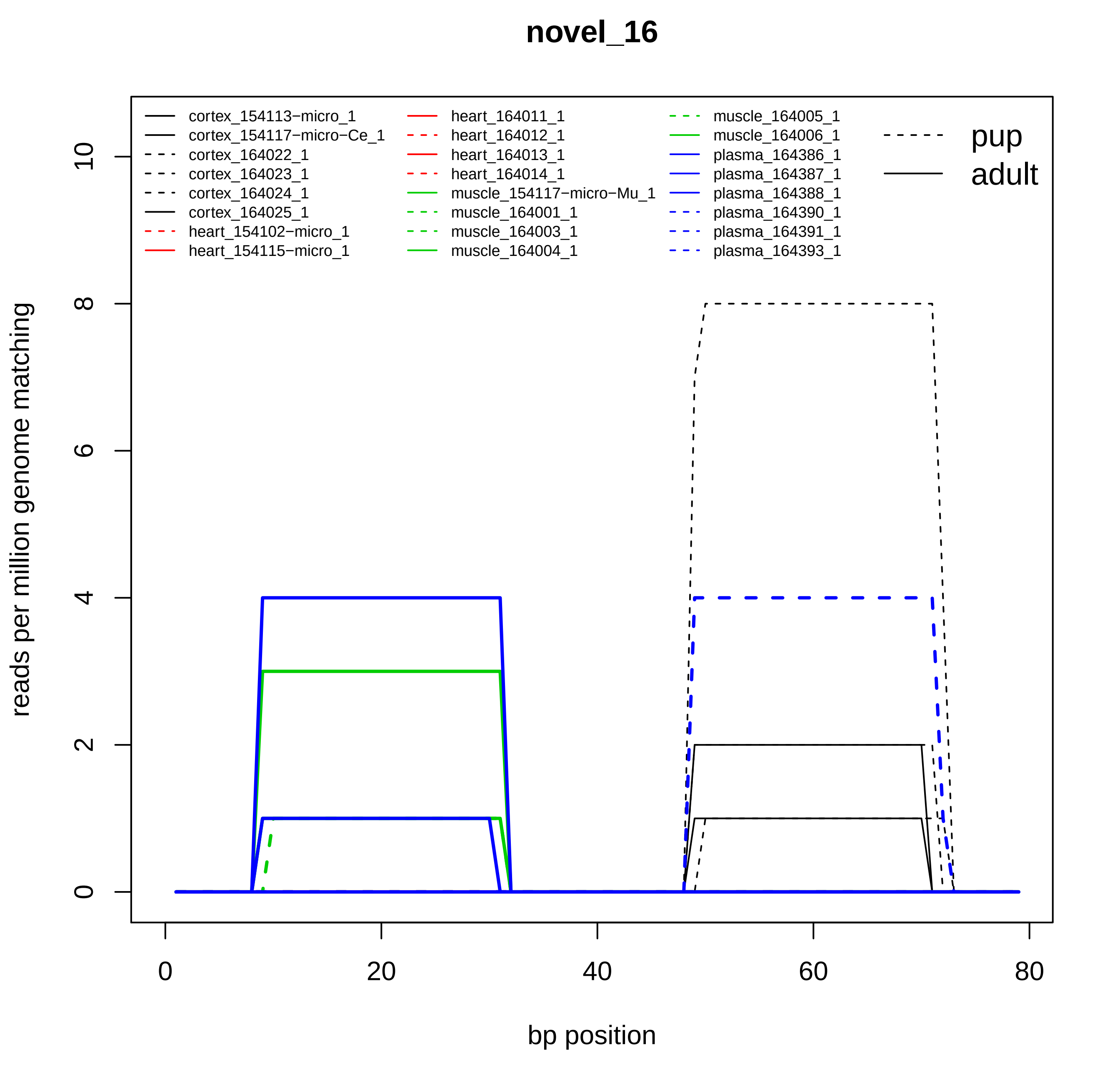
novel_16
cortex_154113−micro_1
cortex_154117−micro−Ce_1
cortex_164022_1
cortex_164023_1
cortex_164024_1
cortex_164025_1
heart_154102−micro_1
heart_154115−micro_1
heart_164011_1
heart_164012_1
heart_164013_1
heart_164014_1
muscle_154117−micro−Mu_1
muscle_164001_1
muscle_164003_1
muscle_164004_1
muscle_164005_1
muscle_164006_1
plasma_164386_1
plasma_164387_1
plasma_164388_1
plasma_164390_1
plasma_164391_1
plasma_164393_1
pup
adult
reads per million genome matching
bp position
0
2
4
6
8
10
0
20
40
60
80

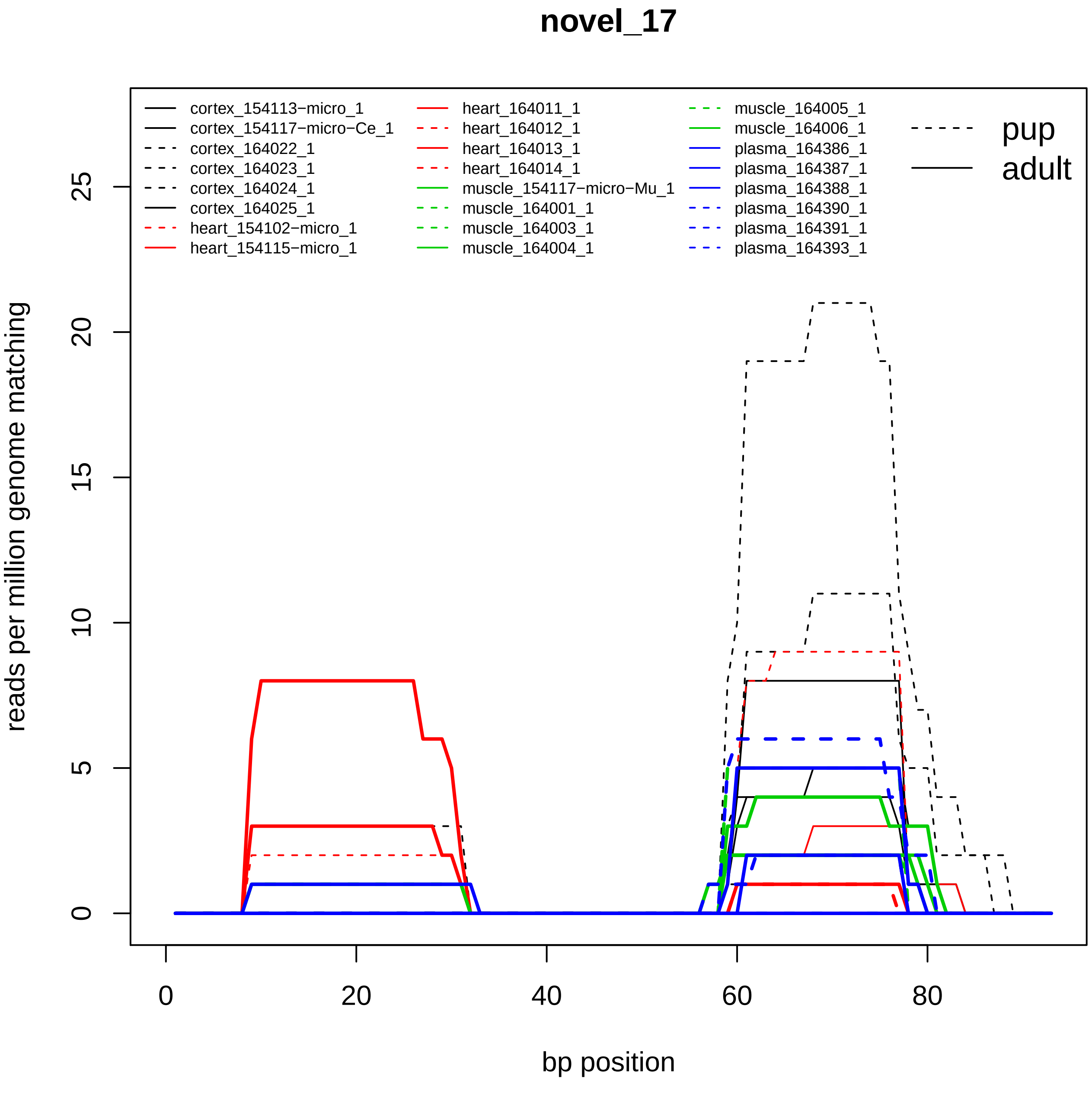

novel_17
cortex_154113−micro_1
cortex_154117−micro−Ce_1
cortex_164022_1
cortex_164023_1
cortex_164024_1
cortex_164025_1
heart_154102−micro_1
heart_154115−micro_1
heart_164011_1
heart_164012_1
heart_164013_1
heart_164014_1
muscle_154117−micro−Mu_1
muscle_164001_1
muscle_164003_1
muscle_164004_1
muscle_164005_1
muscle_164006_1
plasma_164386_1
plasma_164387_1
plasma_164388_1
plasma_164390_1
plasma_164391_1
plasma_164393_1
pup
adult
reads per million genome matching
0
5
10
15
20
25
0
20
40
60
80
bp position

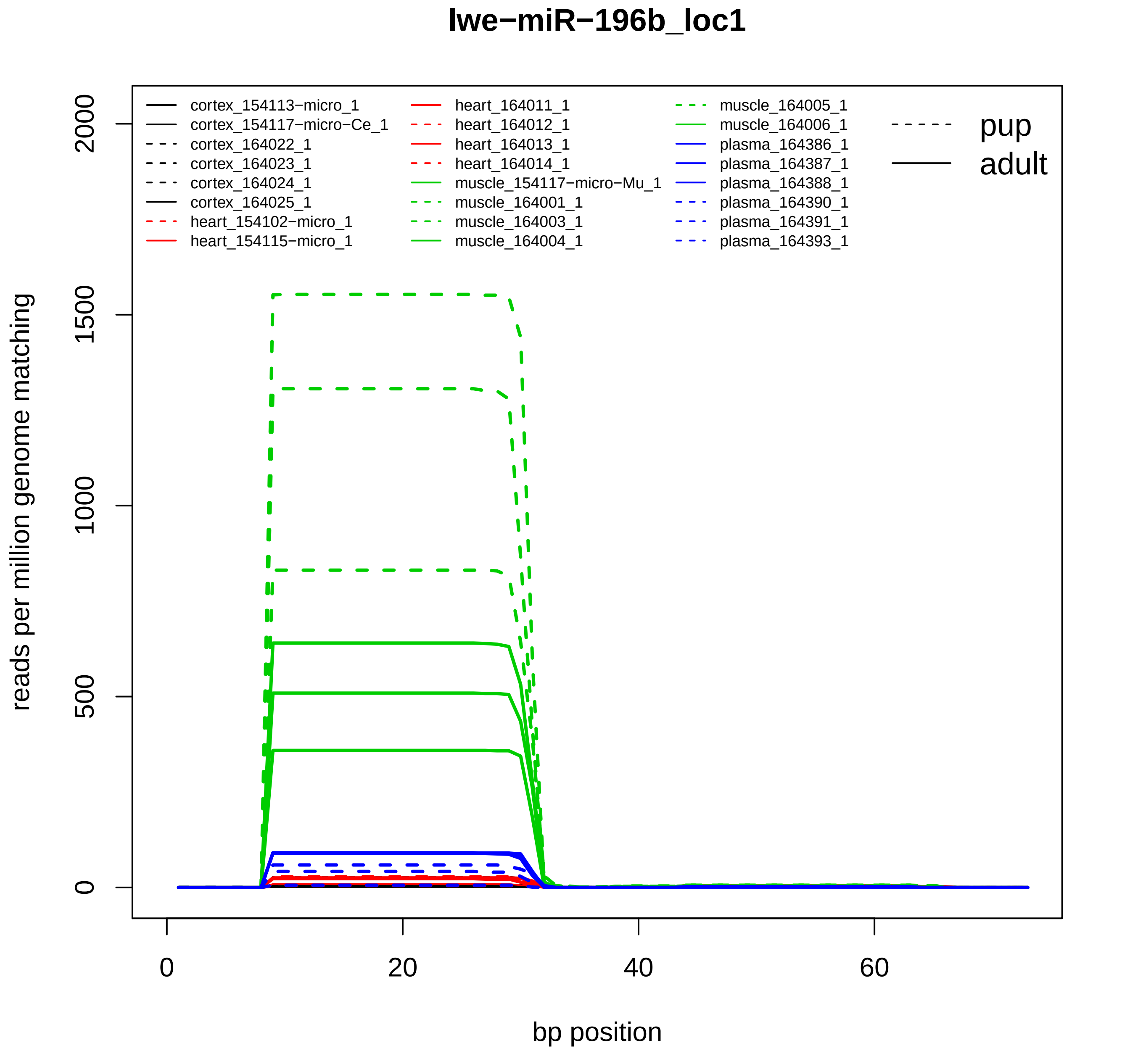
lwe−miR−196b_loc1
reads per million genome matching
bp position
0
500
1000
1500
2000
0
20
40
60
cortex_154113−micro_1
cortex_154117−micro−Ce_1
cortex_164022_1
cortex_164023_1
cortex_164024_1
cortex_164025_1
heart_154102−micro_1
heart_154115−micro_1
heart_164011_1
heart_164012_1
heart_164013_1
heart_164014_1
muscle_154117−micro−Mu_1
muscle_164001_1
muscle_164003_1
muscle_164004_1
muscle_164005_1
muscle_164006_1
plasma_164386_1
plasma_164387_1
plasma_164388_1
plasma_164390_1
plasma_164391_1
plasma_164393_1
pup
adult

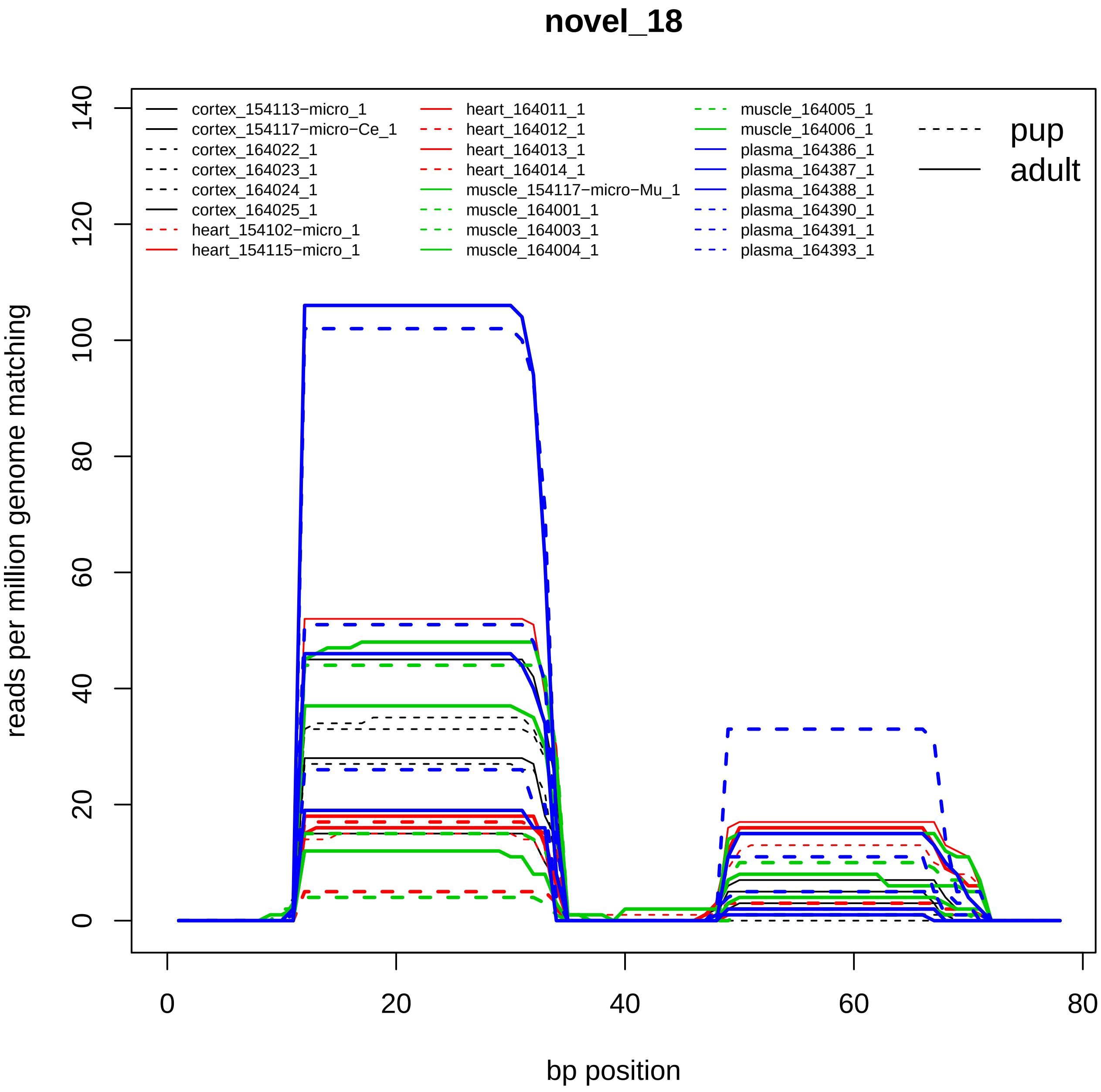

novel_18
cortex_154113−micro_1
cortex_154117−micro−Ce_1
cortex_164022_1
cortex_164023_1
cortex_164024_1
cortex_164025_1
heart_154102−micro_1
heart_154115−micro_1
heart_164011_1
heart_164012_1
heart_164013_1
heart_164014_1
muscle_154117−micro−Mu_1
muscle_164001_1
muscle_164003_1
muscle_164004_1
muscle_164005_1
muscle_164006_1
plasma_164386_1
plasma_164387_1
plasma_164388_1
plasma_164390_1
plasma_164391_1
plasma_164393_1
pup
adult
reads per million genome matching
bp position
0
20
40
60
80
100
120
140

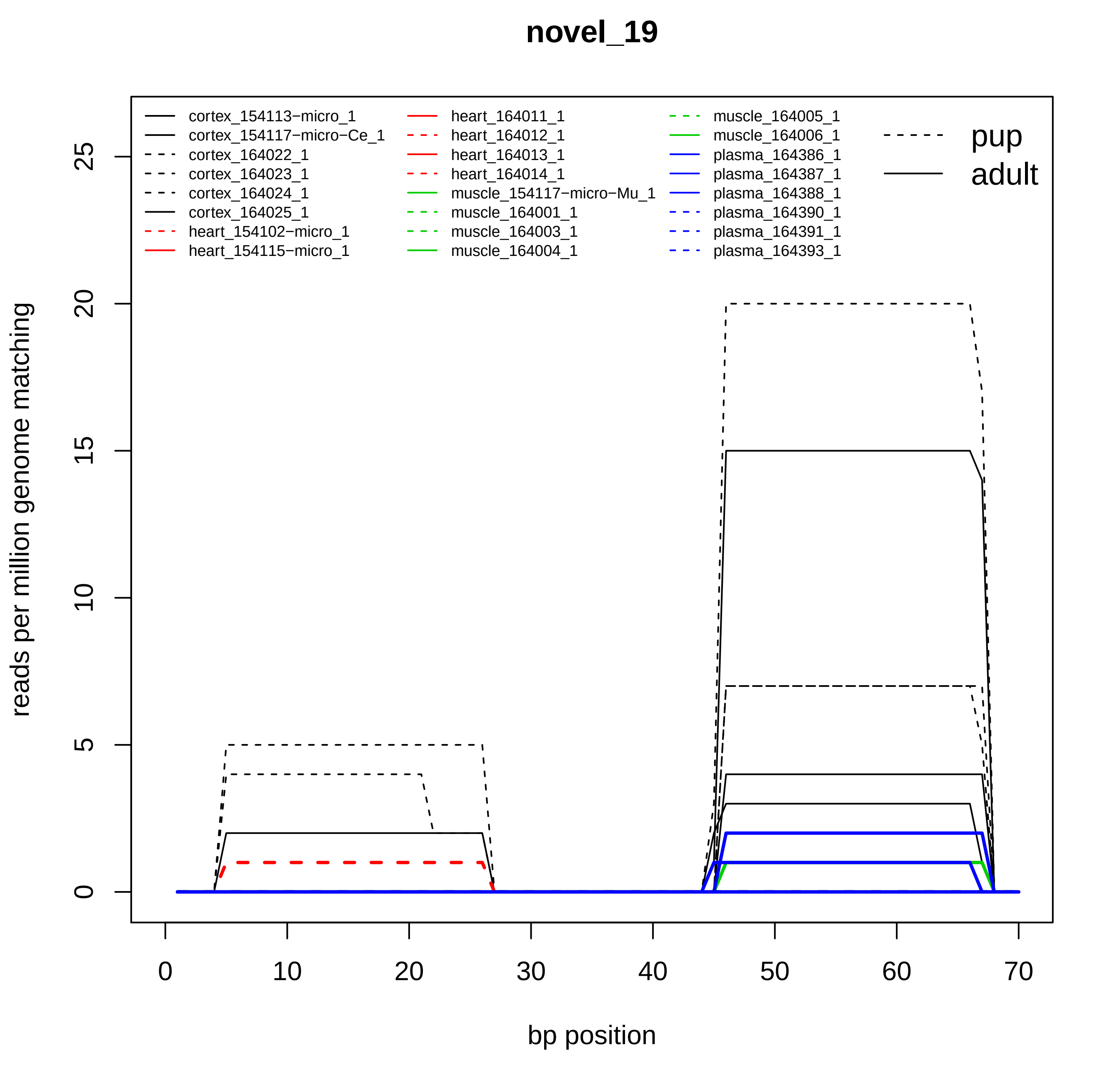
novel_19
cortex_154113−micro_1
cortex_154117−micro−Ce_1
cortex_164022_1
cortex_164023_1
cortex_164024_1
cortex_164025_1
heart_154102−micro_1
heart_154115−micro_1
heart_164011_1
heart_164012_1
heart_164013_1
heart_164014_1
muscle_154117−micro−Mu_1
muscle_164001_1
muscle_164003_1
muscle_164004_1
muscle_164005_1
muscle_164006_1
plasma_164386_1
plasma_164387_1
plasma_164388_1
plasma_164390_1
plasma_164391_1
plasma_164393_1
pup
adult
reads per million genome matching
0
5
10
15
20
25
0
10
20
30
40
50
60
70
bp position

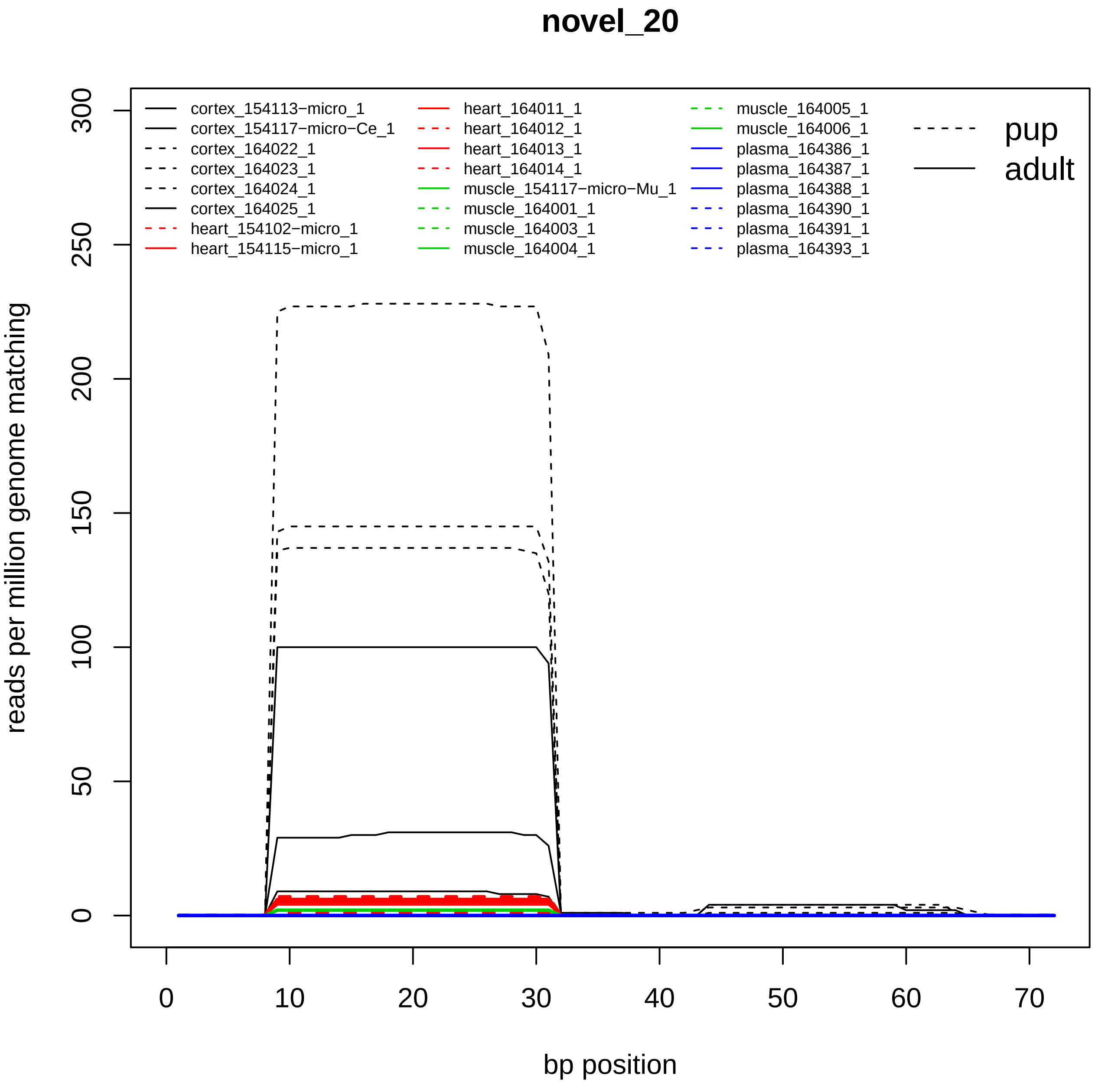

novel_20
cortex_154113−micro_1
cortex_154117−micro−Ce_1
cortex_164022_1
cortex_164023_1
cortex_164024_1
cortex_164025_1
heart_154102−micro_1
heart_154115−micro_1
heart_164011_1
heart_164012_1
heart_164013_1
heart_164014_1
muscle_154117−micro−Mu_1
muscle_164001_1
muscle_164003_1
muscle_164004_1
muscle_164005_1
muscle_164006_1
plasma_164386_1
plasma_164387_1
plasma_164388_1
plasma_164390_1
plasma_164391_1
plasma_164393_1
pup
adult
reads per million genome matching
0
50
100
150
200
250
300
0
10
20
30
40
50
60
70
bp position

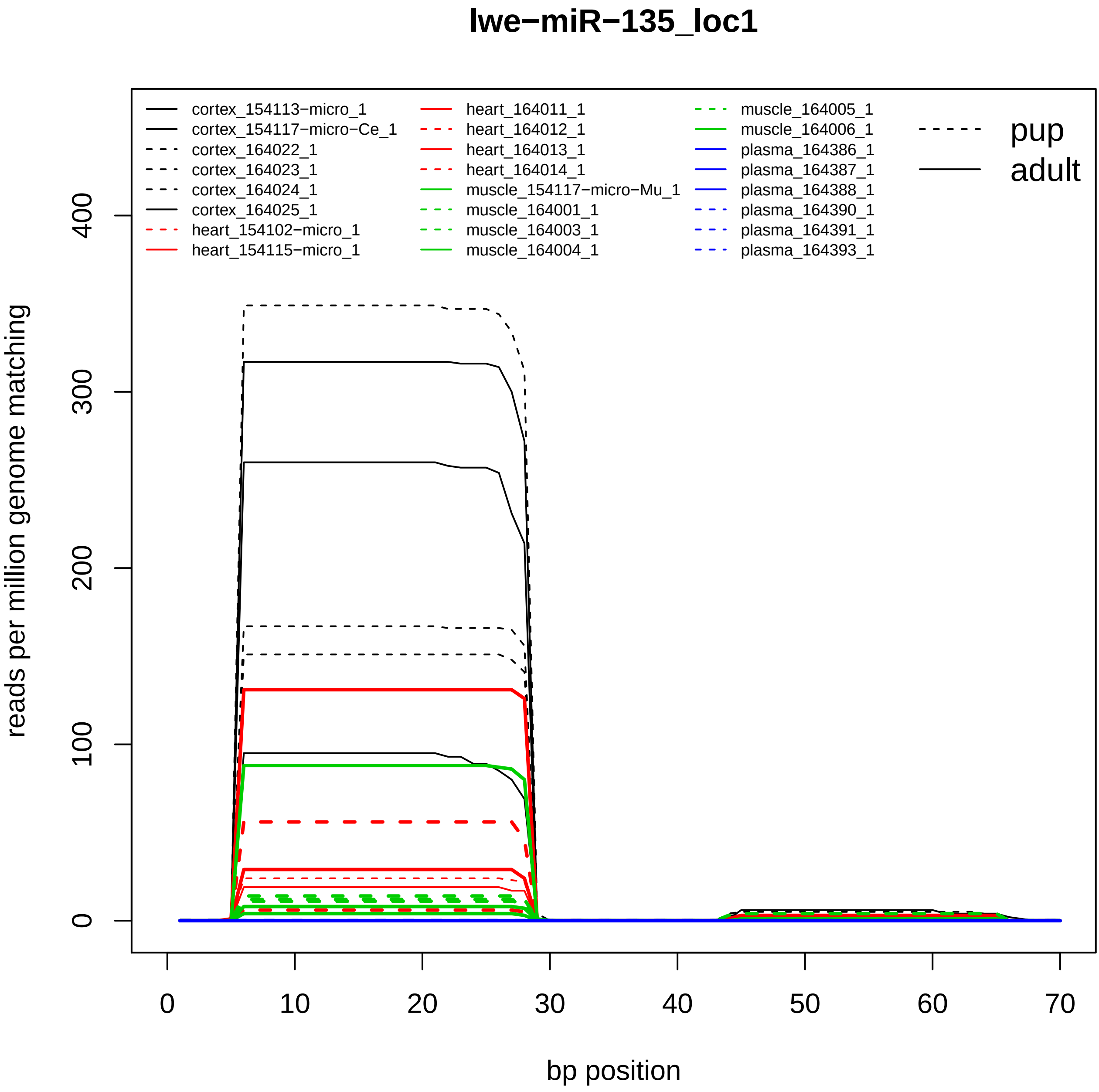

lwe−miR−135_loc1
cortex_154113−micro_1
cortex_154117−micro−Ce_1
cortex_164022_1
cortex_164023_1
cortex_164024_1
cortex_164025_1
heart_154102−micro_1
heart_154115−micro_1
heart_164011_1
heart_164012_1
heart_164013_1
heart_164014_1
muscle_154117−micro−Mu_1
muscle_164001_1
muscle_164003_1
muscle_164004_1
muscle_164005_1
muscle_164006_1
plasma_164386_1
plasma_164387_1
plasma_164388_1
plasma_164390_1
plasma_164391_1
plasma_164393_1
pup
adult
reads per million genome matching
0
100
200
300
400
bp position
0
10
20
30
40
50
60
70

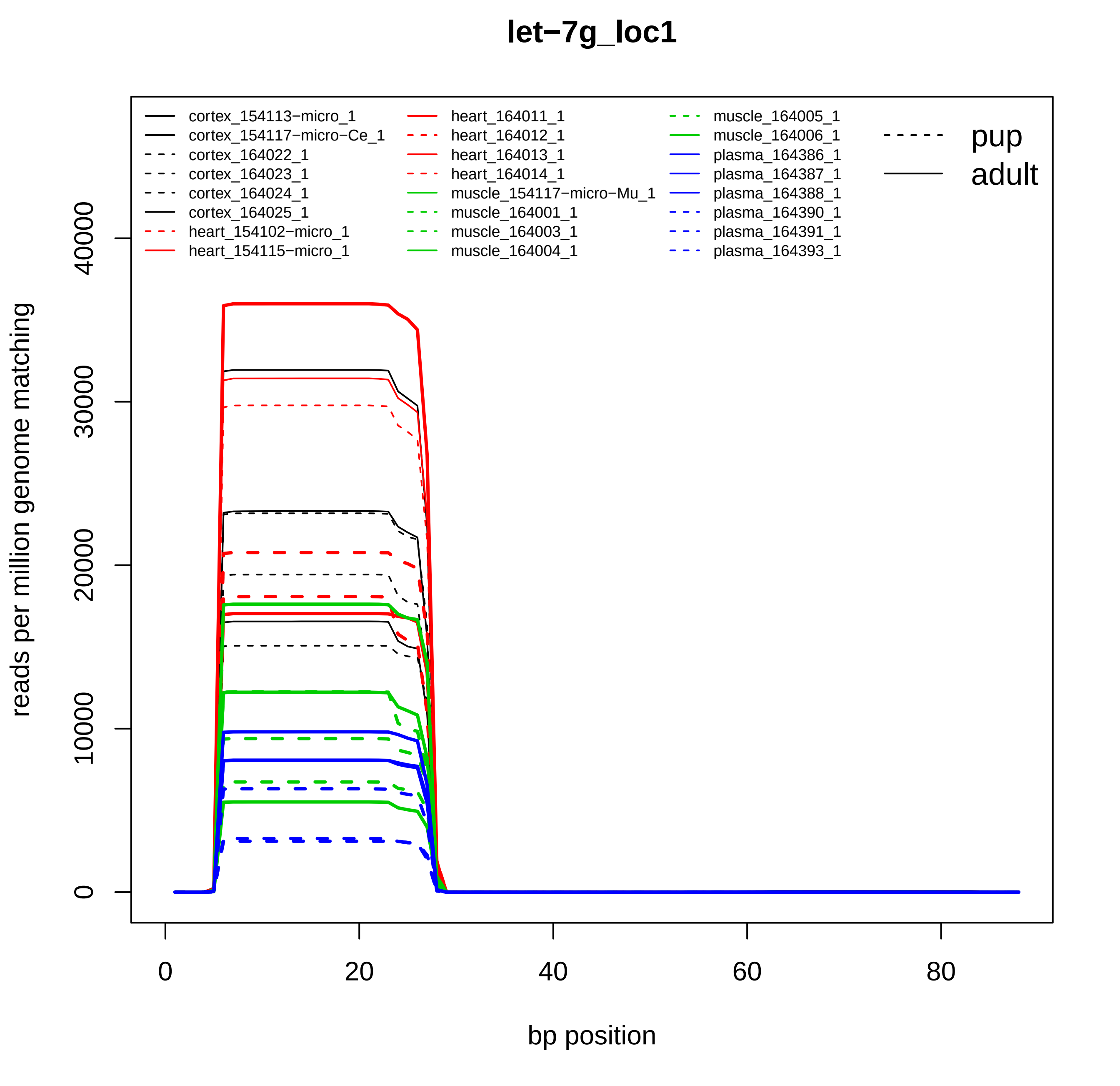

let−7g_loc1
cortex_154113−micro_1
cortex_154117−micro−Ce_1
cortex_164022_1
cortex_164023_1
cortex_164024_1
cortex_164025_1
heart_154102−micro_1
heart_154115−micro_1
heart_164011_1
heart_164012_1
heart_164013_1
heart_164014_1
muscle_154117−micro−Mu_1
muscle_164001_1
muscle_164003_1
muscle_164004_1
muscle_164005_1
muscle_164006_1
plasma_164386_1
plasma_164387_1
plasma_164388_1
plasma_164390_1
plasma_164391_1
plasma_164393_1
pup
adult
reads per million genome matching
0
10000
20000
30000
40000
0
20
40
60
80
bp position

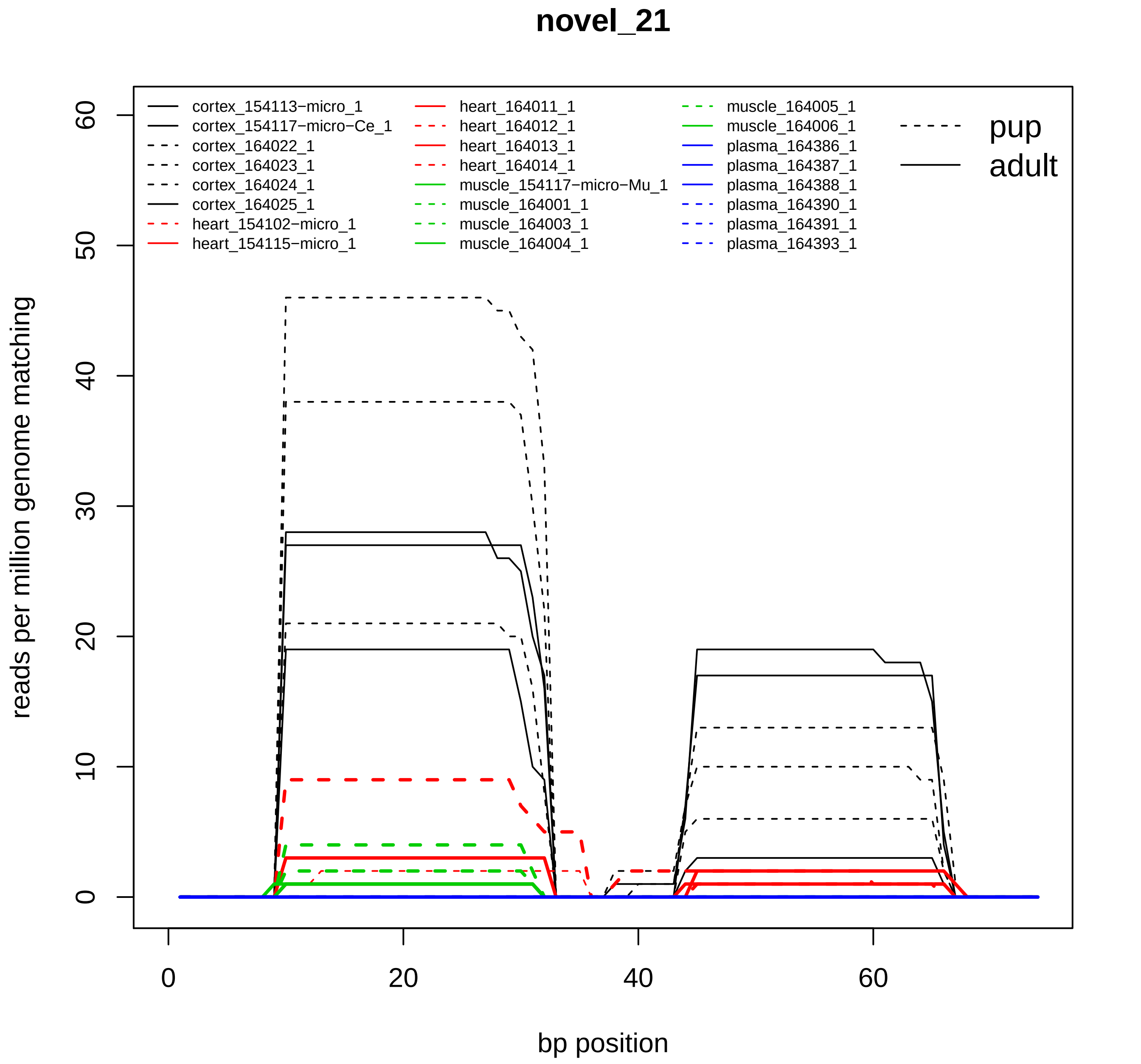
novel_21
reads per million genome matching
bp position
0
10
20
30
40
50
60
0
20
40
60
cortex_154113−micro_1
cortex_154117−micro−Ce_1
cortex_164022_1
cortex_164023_1
cortex_164024_1
cortex_164025_1
heart_154102−micro_1
heart_154115−micro_1
heart_164011_1
heart_164012_1
heart_164013_1
heart_164014_1
muscle_154117−micro−Mu_1
muscle_164001_1
muscle_164003_1
muscle_164004_1
muscle_164005_1
muscle_164006_1
plasma_164386_1
plasma_164387_1
plasma_164388_1
plasma_164390_1
plasma_164391_1
plasma_164393_1
pup
adult

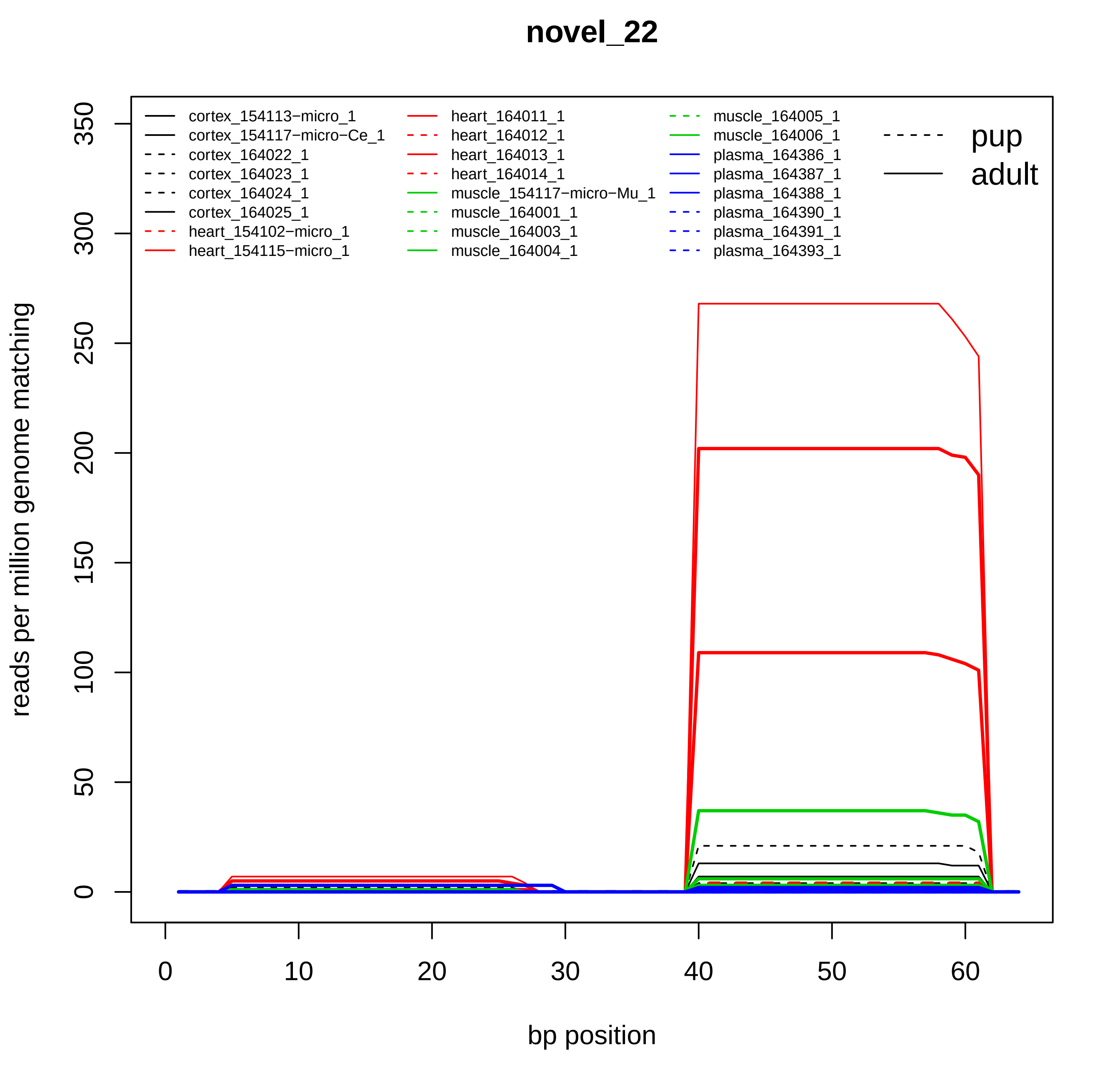

novel_22
cortex_154113−micro_1
cortex_154117−micro−Ce_1
cortex_164022_1
cortex_164023_1
cortex_164024_1
cortex_164025_1
heart_154102−micro_1
heart_154115−micro_1
heart_164011_1
heart_164012_1
heart_164013_1
heart_164014_1
muscle_154117−micro−Mu_1
muscle_164001_1
muscle_164003_1
muscle_164004_1
muscle_164005_1
muscle_164006_1
plasma_164386_1
plasma_164387_1
plasma_164388_1
plasma_164390_1
plasma_164391_1
plasma_164393_1
pup
adult
reads per million genome matching
0
50
100
150
200
250
300
350
0
10
20
30
40
50
60
bp position

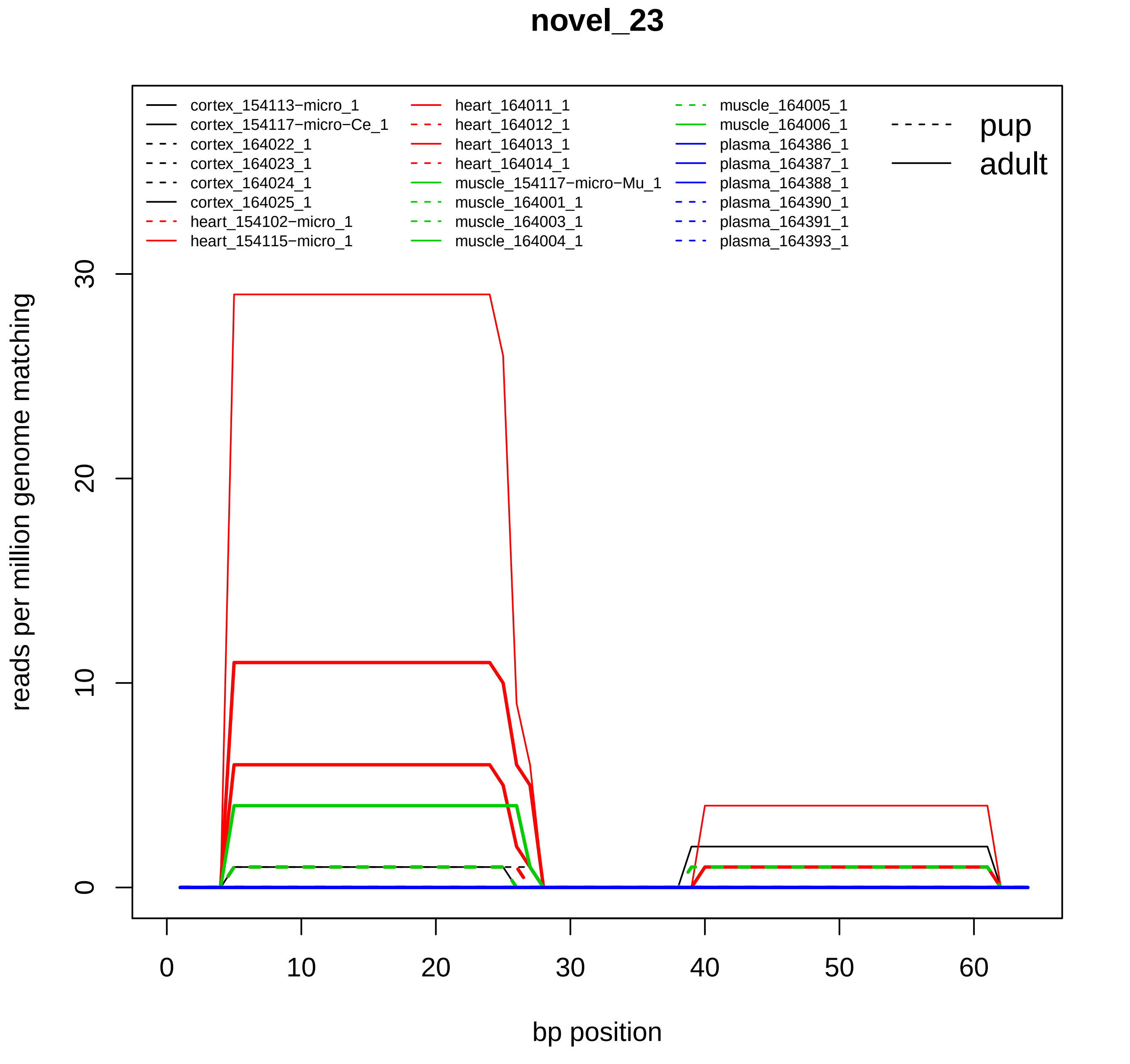
novel_23
cortex_154113−micro_1
cortex_154117−micro−Ce_1
cortex_164022_1
cortex_164023_1
cortex_164024_1
cortex_164025_1
heart_154102−micro_1
heart_154115−micro_1
heart_164011_1
heart_164012_1
heart_164013_1
heart_164014_1
muscle_154117−micro−Mu_1
muscle_164001_1
muscle_164003_1
muscle_164004_1
muscle_164005_1
muscle_164006_1
plasma_164386_1
plasma_164387_1
plasma_164388_1
plasma_164390_1
plasma_164391_1
plasma_164393_1
pup
adult
reads per million genome matching
0
10
20
30
bp position
0
10
20
30
40
50
60

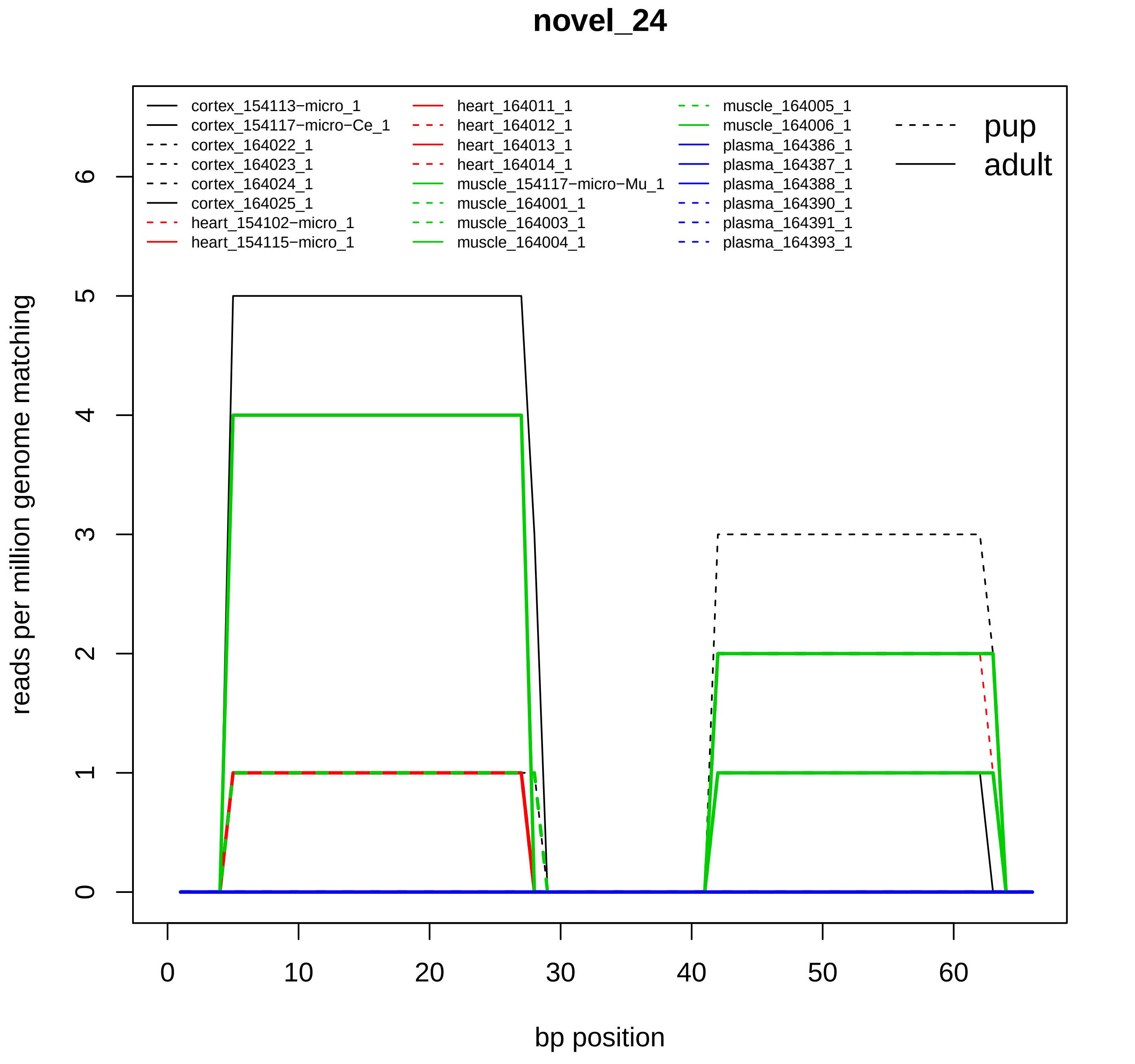
novel_24
cortex_154113−micro_1
cortex_154117−micro−Ce_1
cortex_164022_1
cortex_164023_1
cortex_164024_1
cortex_164025_1
heart_154102−micro_1
heart_154115−micro_1
heart_164011_1
heart_164012_1
heart_164013_1
heart_164014_1
muscle_154117−micro−Mu_1
muscle_164001_1
muscle_164003_1
muscle_164004_1
muscle_164005_1
muscle_164006_1
plasma_164386_1
plasma_164387_1
plasma_164388_1
plasma_164390_1
plasma_164391_1
plasma_164393_1
pup
adult
reads per million genome matching
bp position
0
1
2
3
4
5
6
0
10
20
30
40
50
60

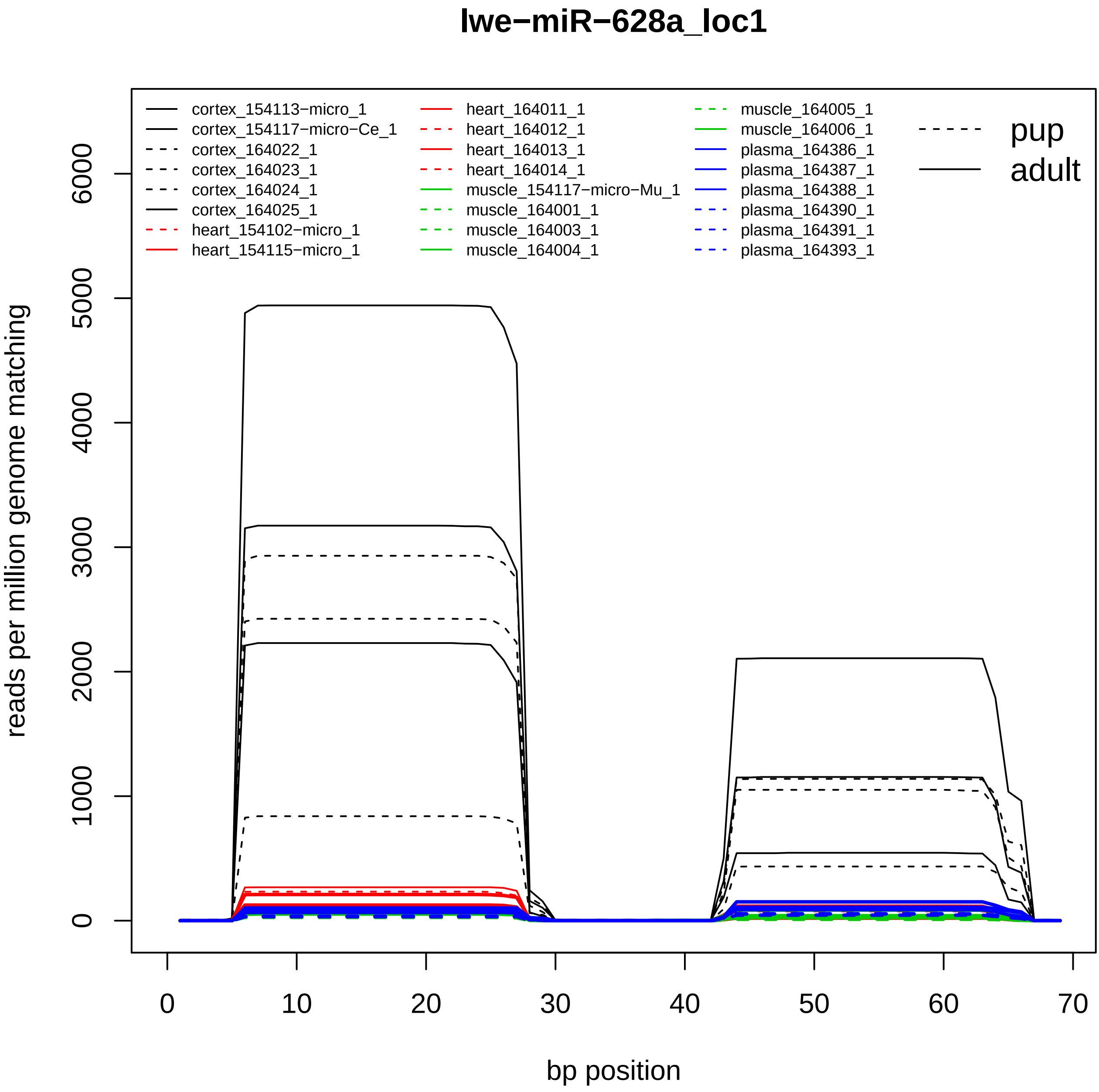
lwe−miR−628a_loc1
cortex_154113−micro_1
cortex_154117−micro−Ce_1
cortex_164022_1
cortex_164023_1
cortex_164024_1
cortex_164025_1
heart_154102−micro_1
heart_154115−micro_1
heart_164011_1
heart_164012_1
heart_164013_1
heart_164014_1
muscle_154117−micro−Mu_1
muscle_164001_1
muscle_164003_1
muscle_164004_1
muscle_164005_1
muscle_164006_1
plasma_164386_1
plasma_164387_1
plasma_164388_1
plasma_164390_1
plasma_164391_1
plasma_164393_1
pup
adult
reads per million genome matching
0
1000
2000
3000
4000
5000
6000
bp position
0
10
20
30
40
50
60
70

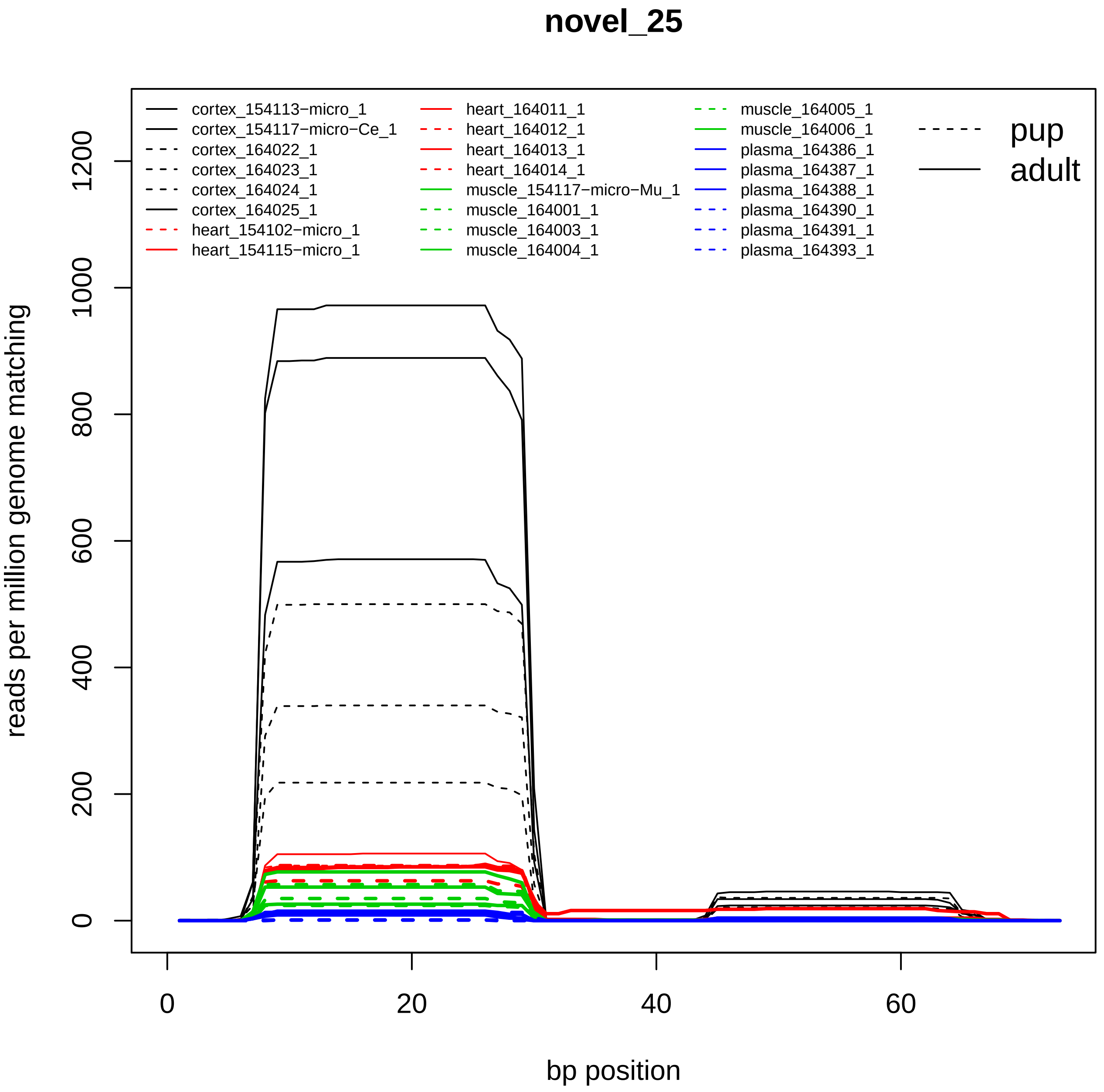

novel_25
cortex_154113−micro_1
cortex_154117−micro−Ce_1
cortex_164022_1
cortex_164023_1
cortex_164024_1
cortex_164025_1
heart_154102−micro_1
heart_154115−micro_1
heart_164011_1
heart_164012_1
heart_164013_1
heart_164014_1
muscle_154117−micro−Mu_1
muscle_164001_1
muscle_164003_1
muscle_164004_1
muscle_164005_1
muscle_164006_1
plasma_164386_1
plasma_164387_1
plasma_164388_1
plasma_164390_1
plasma_164391_1
plasma_164393_1
pup
adult
reads per million genome matching
0
200
400
600
800
1000
1200
bp position
0
20
40
60

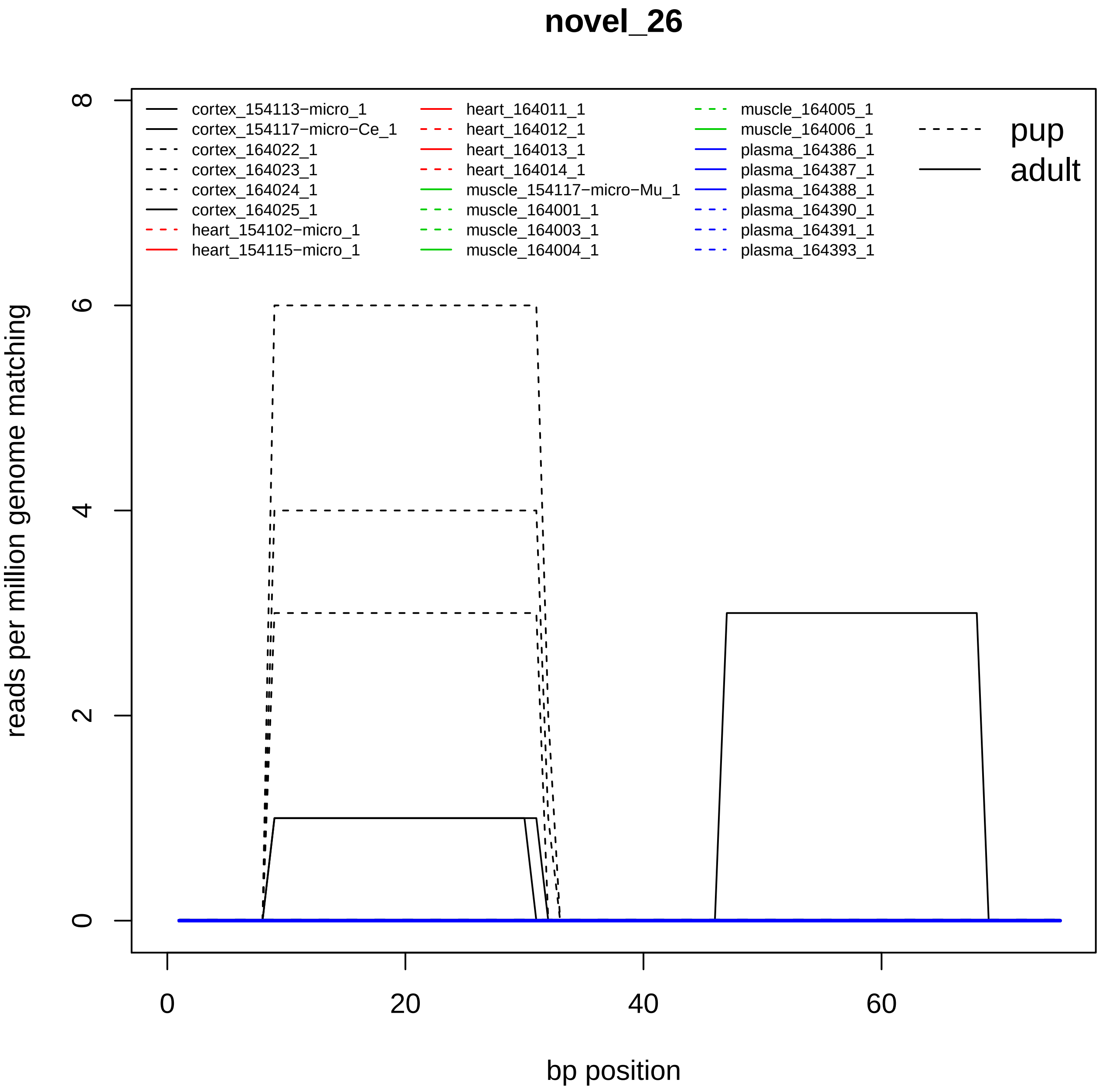
novel_26
cortex_154113−micro_1
cortex_154117−micro−Ce_1
cortex_164022_1
cortex_164023_1
cortex_164024_1
cortex_164025_1
heart_154102−micro_1
heart_154115−micro_1
heart_164011_1
heart_164012_1
heart_164013_1
heart_164014_1
muscle_154117−micro−Mu_1
muscle_164001_1
muscle_164003_1
muscle_164004_1
muscle_164005_1
muscle_164006_1
plasma_164386_1
plasma_164387_1
plasma_164388_1
plasma_164390_1
plasma_164391_1
plasma_164393_1
pup
adult
reads per million genome matching
0
2
4
6
8
0
20
40
60
bp position

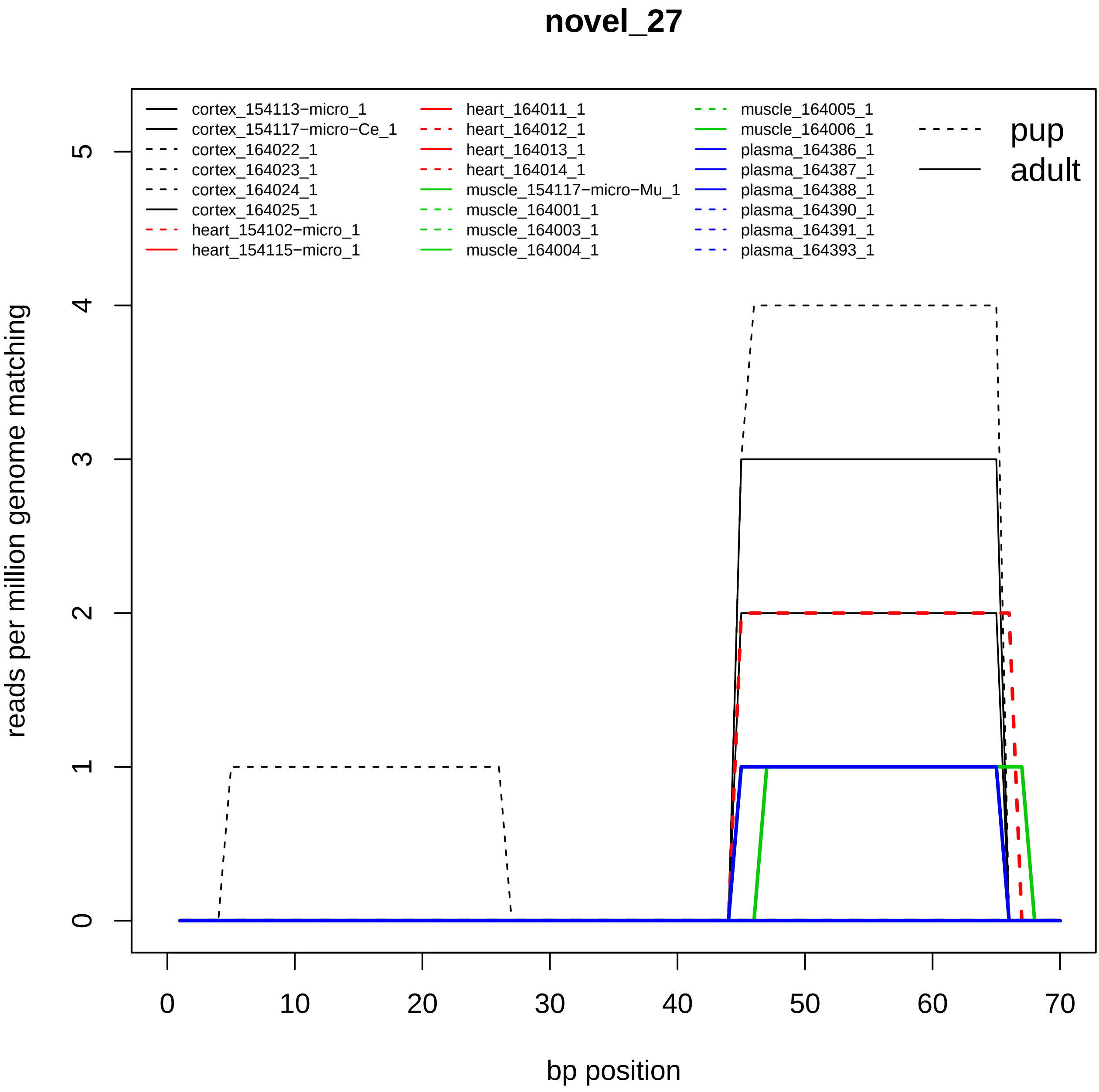
novel_27
cortex_154113−micro_1
cortex_154117−micro−Ce_1
cortex_164022_1
cortex_164023_1
cortex_164024_1
cortex_164025_1
heart_154102−micro_1
heart_154115−micro_1
heart_164011_1
heart_164012_1
heart_164013_1
heart_164014_1
muscle_154117−micro−Mu_1
muscle_164001_1
muscle_164003_1
muscle_164004_1
muscle_164005_1
muscle_164006_1
plasma_164386_1
plasma_164387_1
plasma_164388_1
plasma_164390_1
plasma_164391_1
plasma_164393_1
pup
adult
reads per million genome matching
bp position
0
1
2
3
4
5
0
10
20
30
40
50
60
70

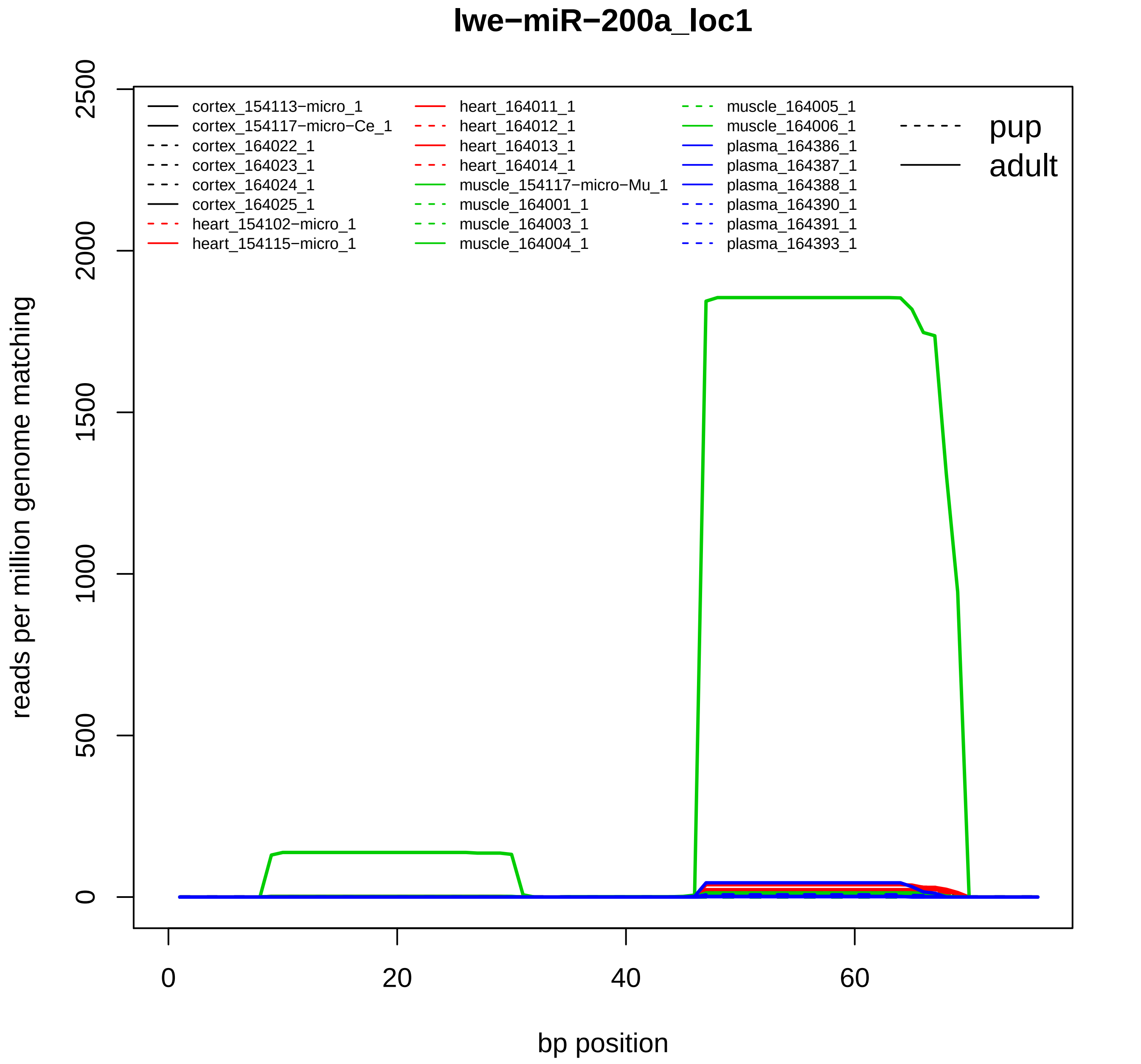

lwe−miR−200a_loc1
cortex_154113−micro_1
cortex_154117−micro−Ce_1
cortex_164022_1
cortex_164023_1
cortex_164024_1
cortex_164025_1
heart_154102−micro_1
heart_154115−micro_1
heart_164011_1
heart_164012_1
heart_164013_1
heart_164014_1
muscle_154117−micro−Mu_1
muscle_164001_1
muscle_164003_1
muscle_164004_1
muscle_164005_1
muscle_164006_1
plasma_164386_1
plasma_164387_1
plasma_164388_1
plasma_164390_1
plasma_164391_1
plasma_164393_1
pup
adult
reads per million genome matching
0
500
1000
1500
2000
2500
0
20
40
60
bp position

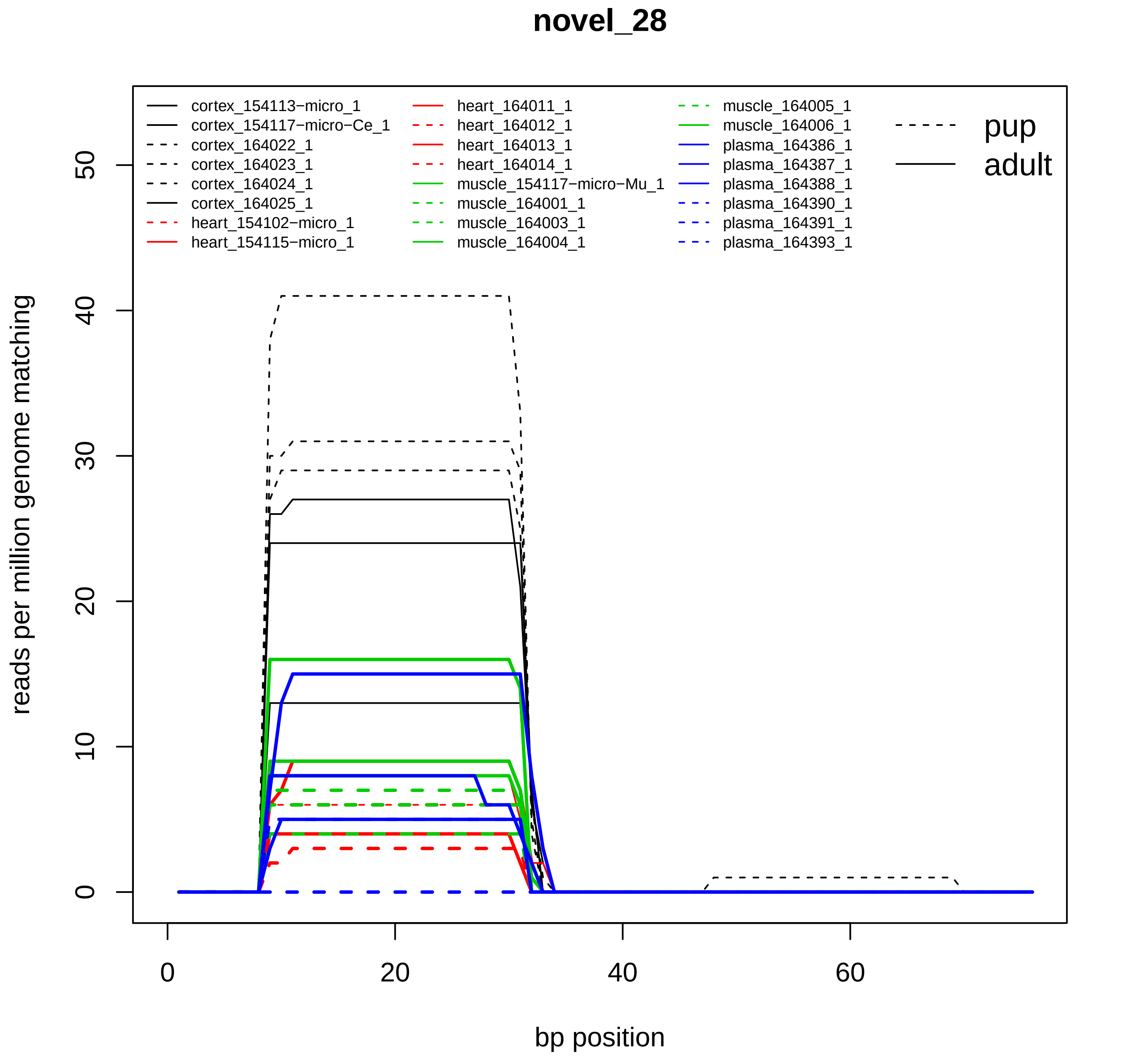

novel_28
cortex_154113−micro_1
cortex_154117−micro−Ce_1
cortex_164022_1
cortex_164023_1
cortex_164024_1
cortex_164025_1
heart_154102−micro_1
heart_154115−micro_1
heart_164011_1
heart_164012_1
heart_164013_1
heart_164014_1
muscle_154117−micro−Mu_1
muscle_164001_1
muscle_164003_1
muscle_164004_1
muscle_164005_1
muscle_164006_1
plasma_164386_1
plasma_164387_1
plasma_164388_1
plasma_164390_1
plasma_164391_1
plasma_164393_1
pup
adult
reads per million genome matching
0
10
20
30
40
50
0
20
40
60
bp position

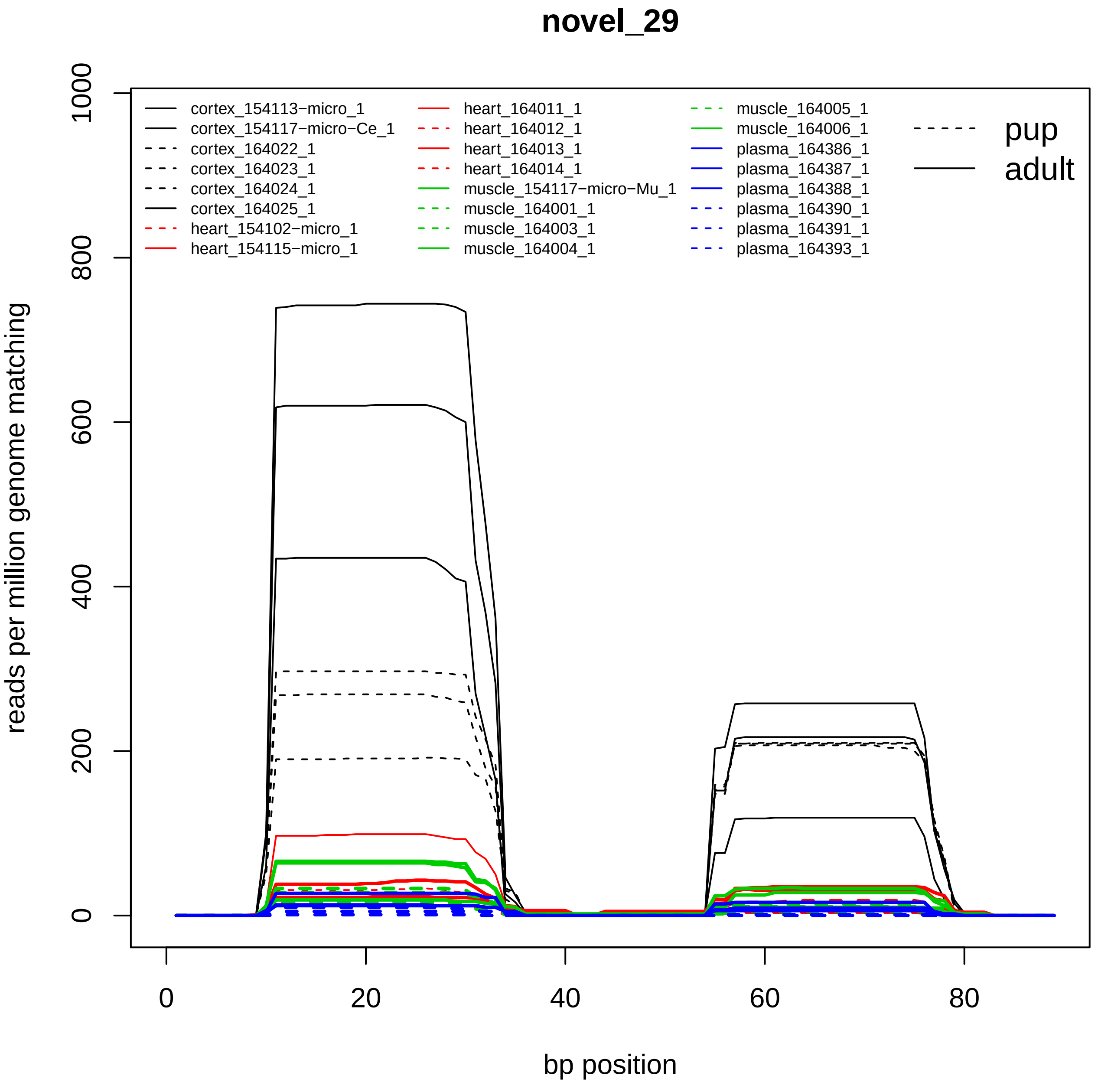

novel_29
reads per million genome matching
bp position
0
200
400
600
800
1000
0
20
40
60
80
cortex_154113−micro_1
cortex_154117−micro−Ce_1
cortex_164022_1
cortex_164023_1
cortex_164024_1
cortex_164025_1
heart_154102−micro_1
heart_154115−micro_1
heart_164011_1
heart_164012_1
heart_164013_1
heart_164014_1
muscle_154117−micro−Mu_1
muscle_164001_1
muscle_164003_1
muscle_164004_1
muscle_164005_1
muscle_164006_1
plasma_164386_1
plasma_164387_1
plasma_164388_1
plasma_164390_1
plasma_164391_1
plasma_164393_1
pup
adult

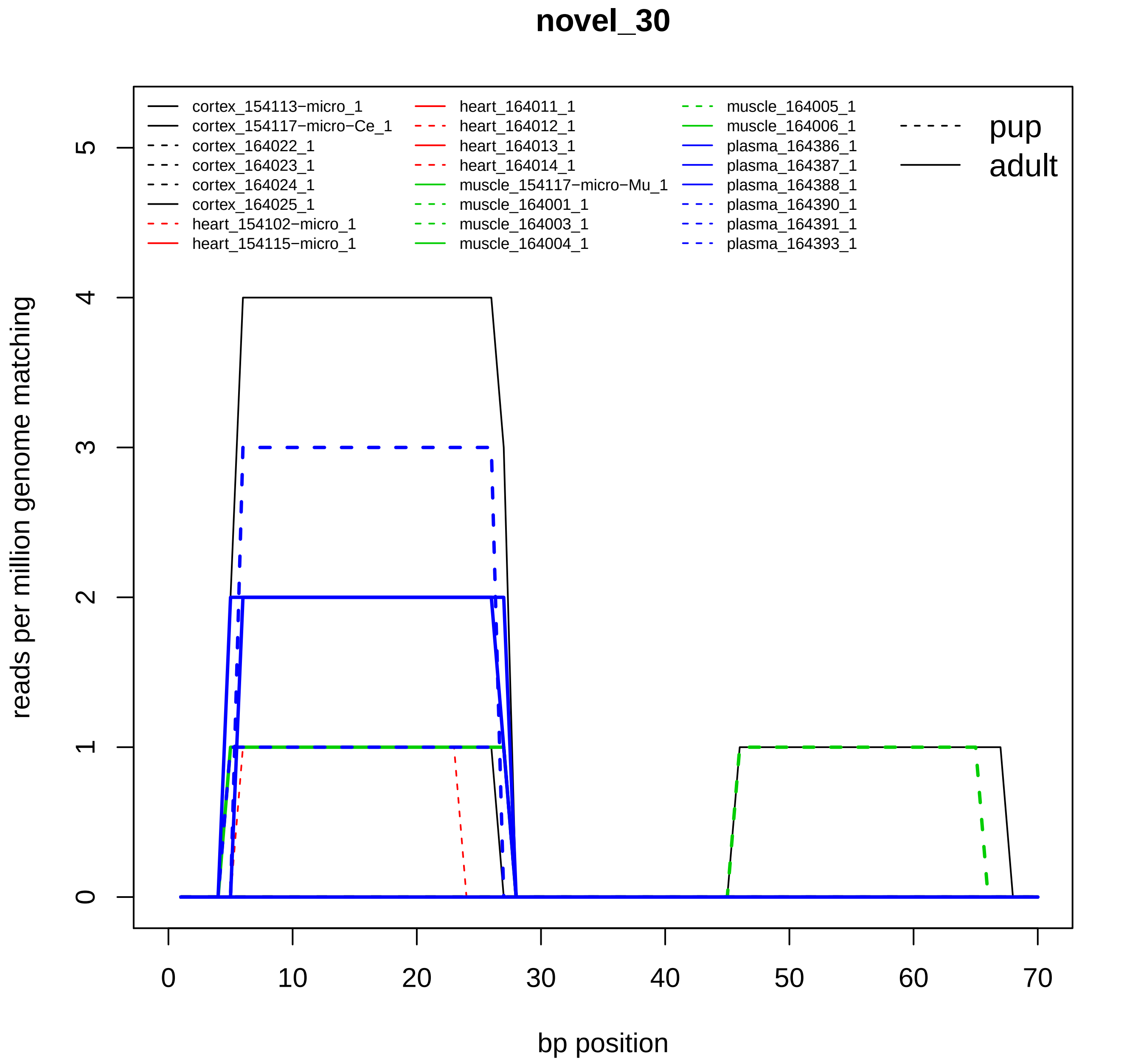

novel_30
cortex_154113−micro_1
cortex_154117−micro−Ce_1
cortex_164022_1
cortex_164023_1
cortex_164024_1
cortex_164025_1
heart_154102−micro_1
heart_154115−micro_1
heart_164011_1
heart_164012_1
heart_164013_1
heart_164014_1
muscle_154117−micro−Mu_1
muscle_164001_1
muscle_164003_1
muscle_164004_1
muscle_164005_1
muscle_164006_1
plasma_164386_1
plasma_164387_1
plasma_164388_1
plasma_164390_1
plasma_164391_1
plasma_164393_1
pup
adult
reads per million genome matching
0
1
2
3
4
5
bp position
0
10
20
30
40
50
60
70

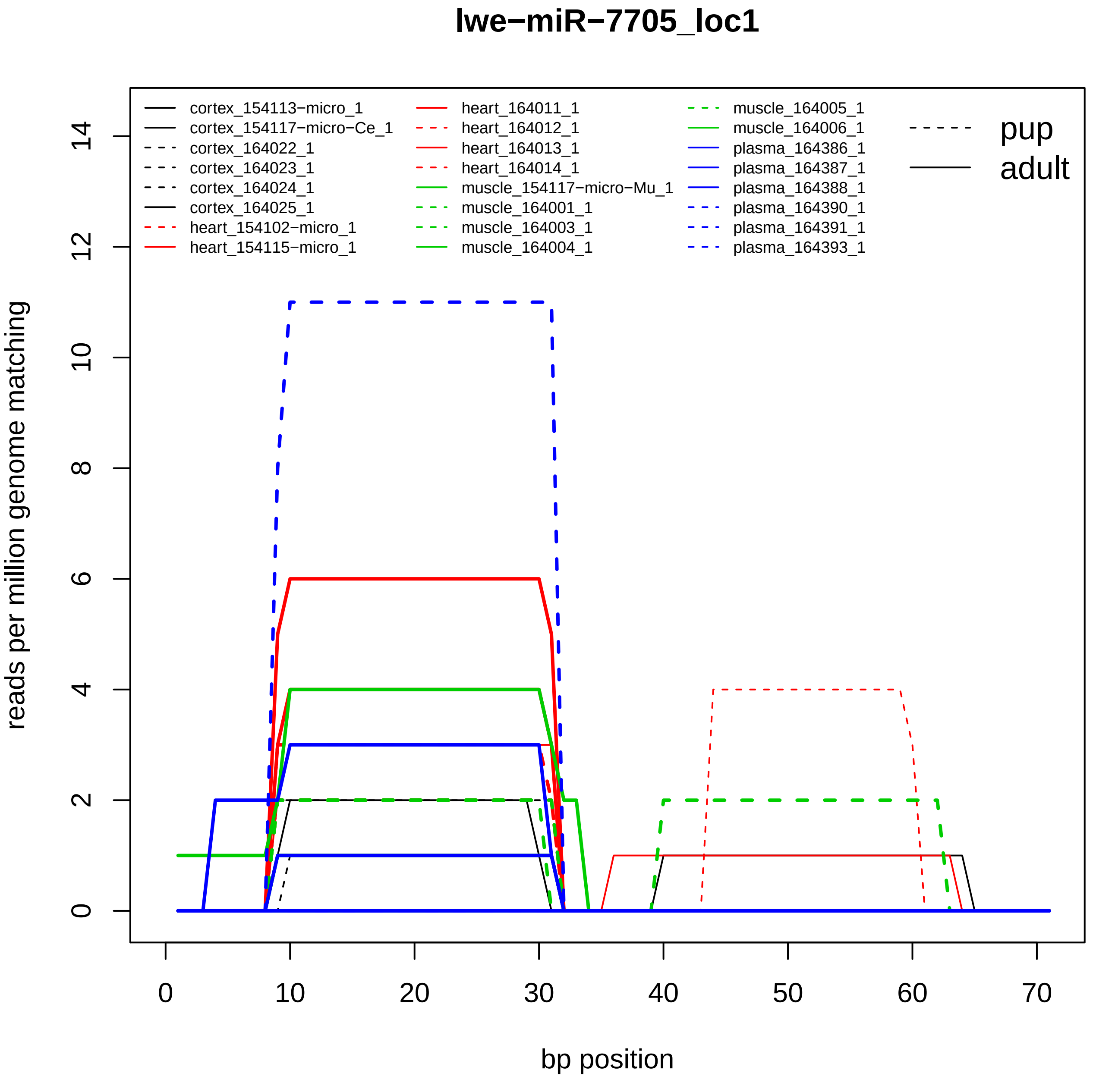
lwe−miR−7705_loc1
reads per million genome matching
bp position
0
2
4
6
8
10
12
14
0
10
20
30
40
50
60
70
cortex_154113−micro_1
cortex_154117−micro−Ce_1
cortex_164022_1
cortex_164023_1
cortex_164024_1
cortex_164025_1
heart_154102−micro_1
heart_154115−micro_1
heart_164011_1
heart_164012_1
heart_164013_1
heart_164014_1
muscle_154117−micro−Mu_1
muscle_164001_1
muscle_164003_1
muscle_164004_1
muscle_164005_1
muscle_164006_1
plasma_164386_1
plasma_164387_1
plasma_164388_1
plasma_164390_1
plasma_164391_1
plasma_164393_1
pup
adult

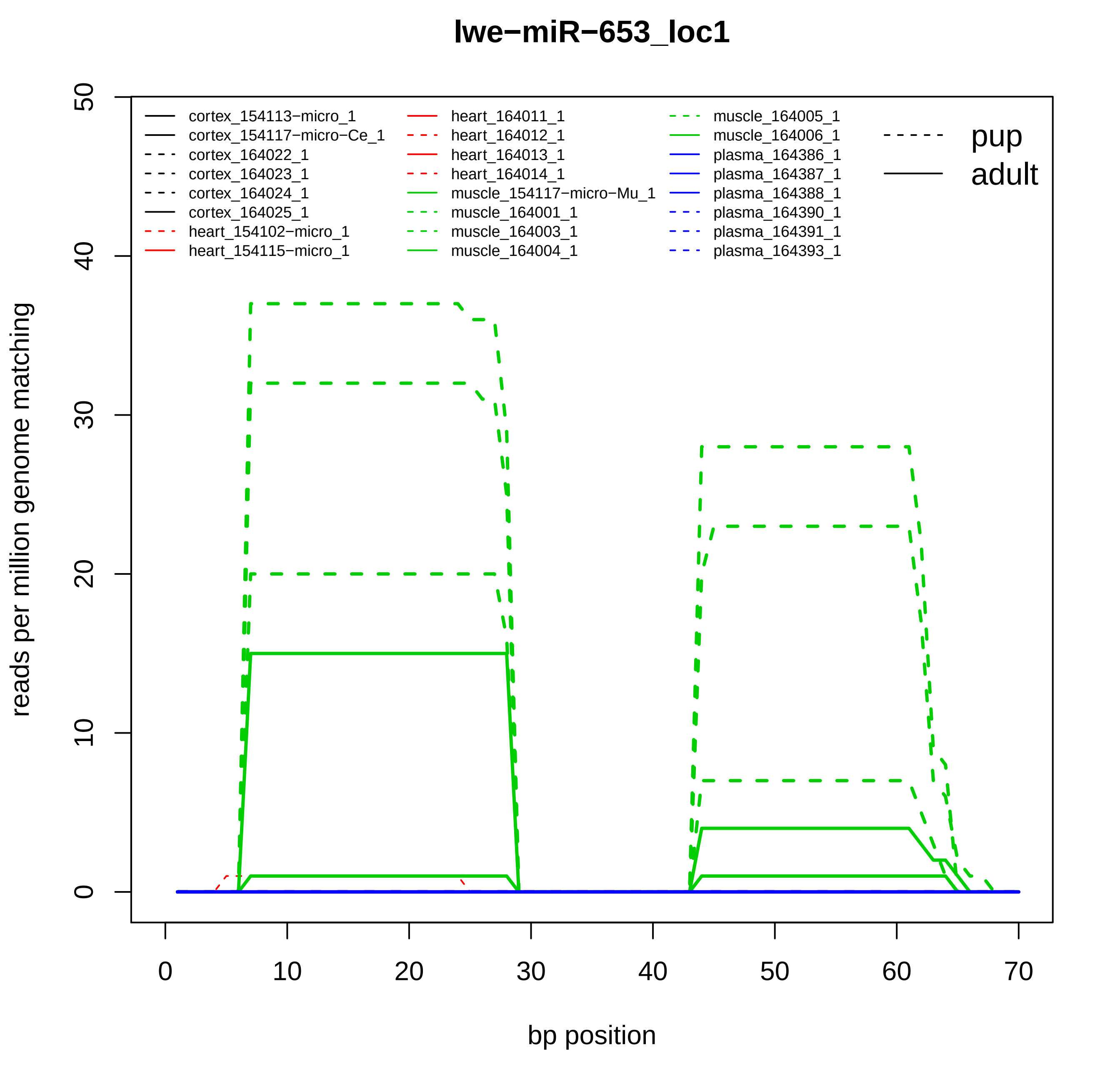
lwe−miR−653_loc1
reads per million genome matching
bp position
0
10
20
30
40
50
60
70
cortex_154113−micro_1
cortex_154117−micro−Ce_1
cortex_164022_1
cortex_164023_1
cortex_164024_1
cortex_164025_1
heart_154102−micro_1
heart_154115−micro_1
heart_164011_1
heart_164012_1
heart_164013_1
heart_164014_1
muscle_154117−micro−Mu_1
muscle_164001_1
muscle_164003_1
muscle_164004_1
muscle_164005_1
muscle_164006_1
plasma_164386_1
plasma_164387_1
plasma_164388_1
plasma_164390_1
plasma_164391_1
plasma_164393_1
pup
adult

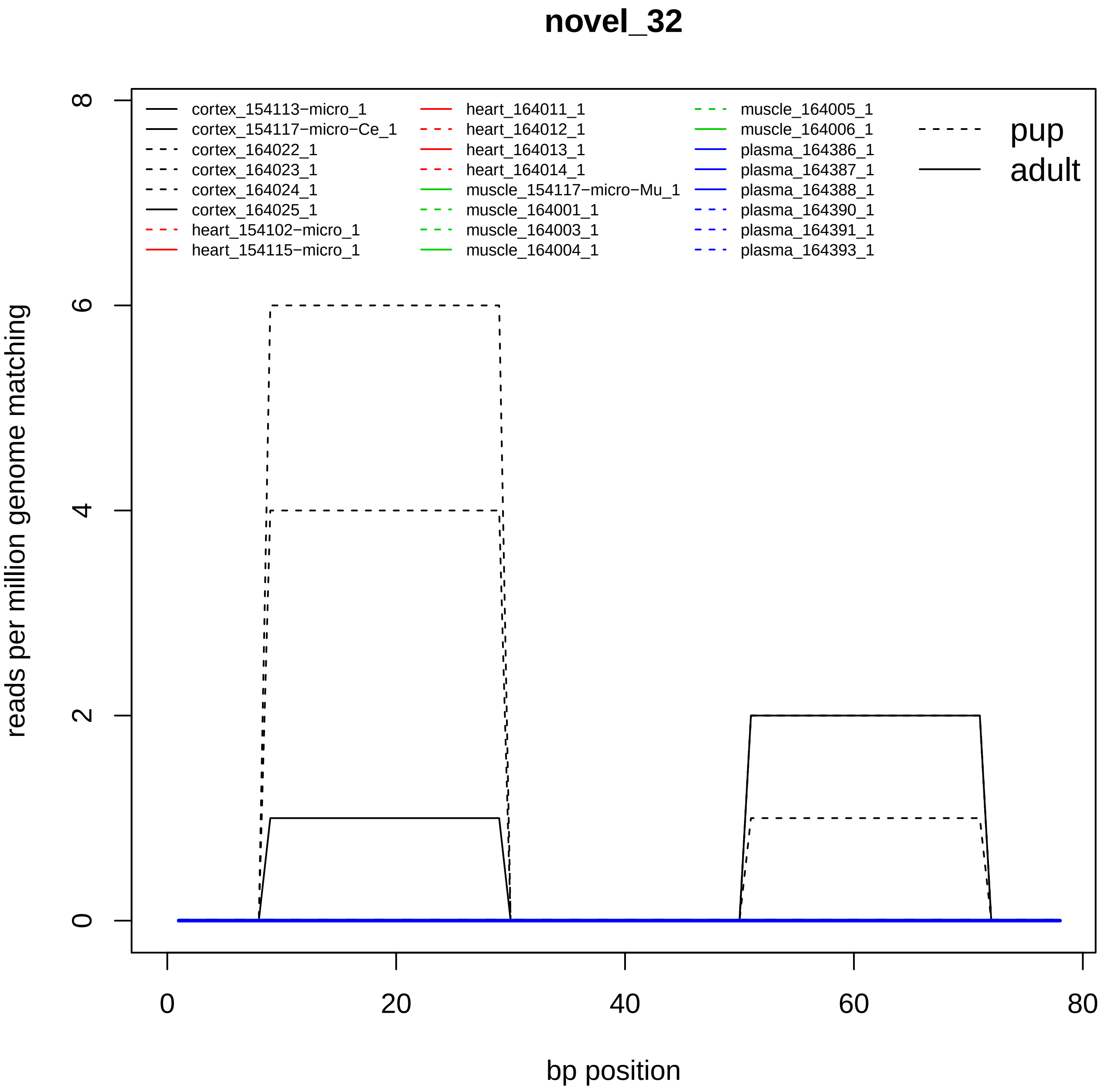
novel_32
cortex_154113−micro_1
cortex_154117−micro−Ce_1
cortex_164022_1
cortex_164023_1
cortex_164024_1
cortex_164025_1
heart_154102−micro_1
heart_154115−micro_1
heart_164011_1
heart_164012_1
heart_164013_1
heart_164014_1
muscle_154117−micro−Mu_1
muscle_164001_1
muscle_164003_1
muscle_164004_1
muscle_164005_1
muscle_164006_1
plasma_164386_1
plasma_164387_1
plasma_164388_1
plasma_164390_1
plasma_164391_1
plasma_164393_1
pup
adult
reads per million genome matching
bp position
0
2
4
6
8
0
20
40
60
80

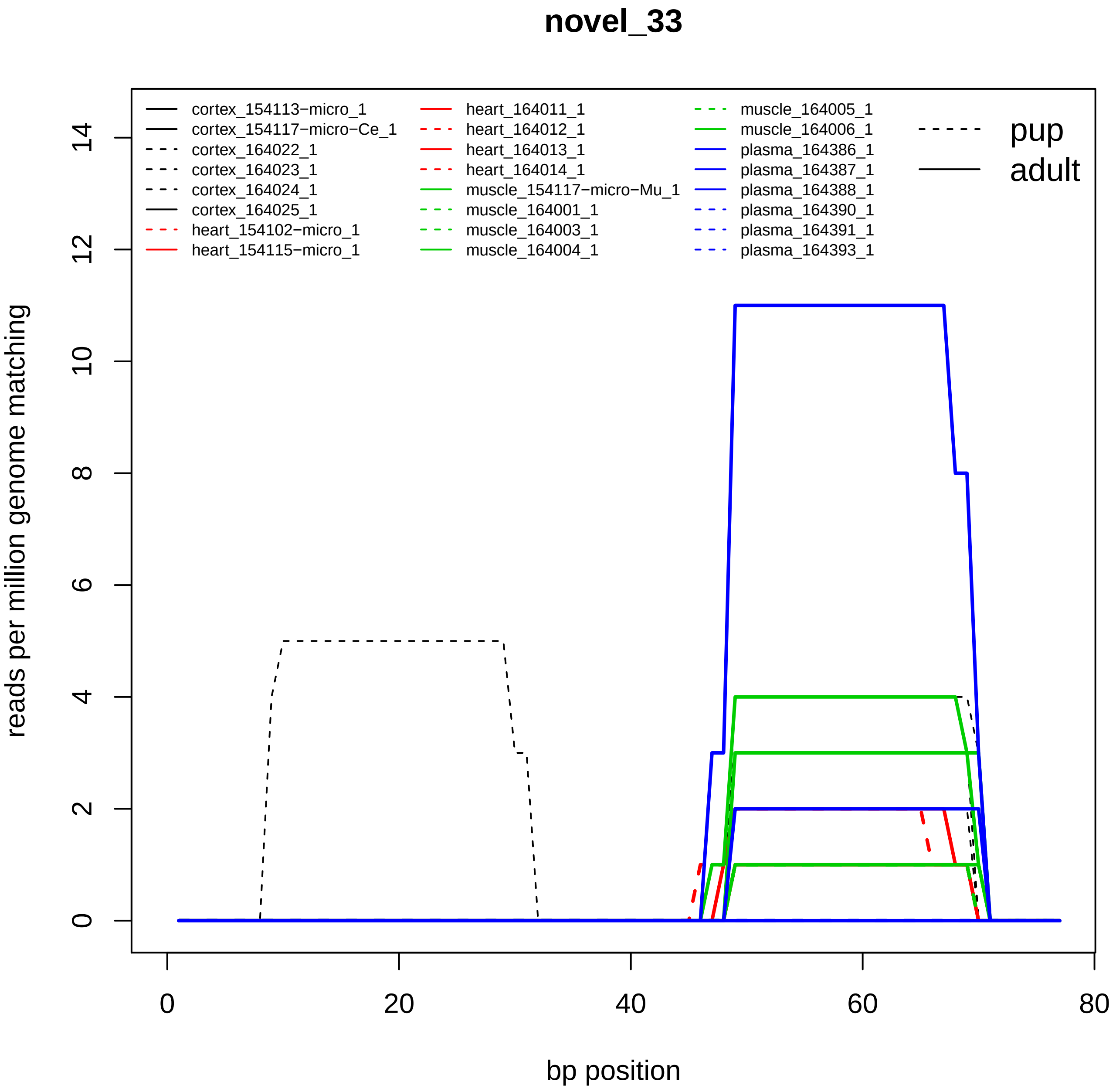

novel_33
cortex_154113−micro_1
cortex_154117−micro−Ce_1
cortex_164022_1
cortex_164023_1
cortex_164024_1
cortex_164025_1
heart_154102−micro_1
heart_154115−micro_1
heart_164011_1
heart_164012_1
heart_164013_1
heart_164014_1
muscle_154117−micro−Mu_1
muscle_164001_1
muscle_164003_1
muscle_164004_1
muscle_164005_1
muscle_164006_1
plasma_164386_1
plasma_164387_1
plasma_164388_1
plasma_164390_1
plasma_164391_1
plasma_164393_1
pup
adult
reads per million genome matching
0
2
4
6
8
10
12
14
bp position
0
20
40
60
80

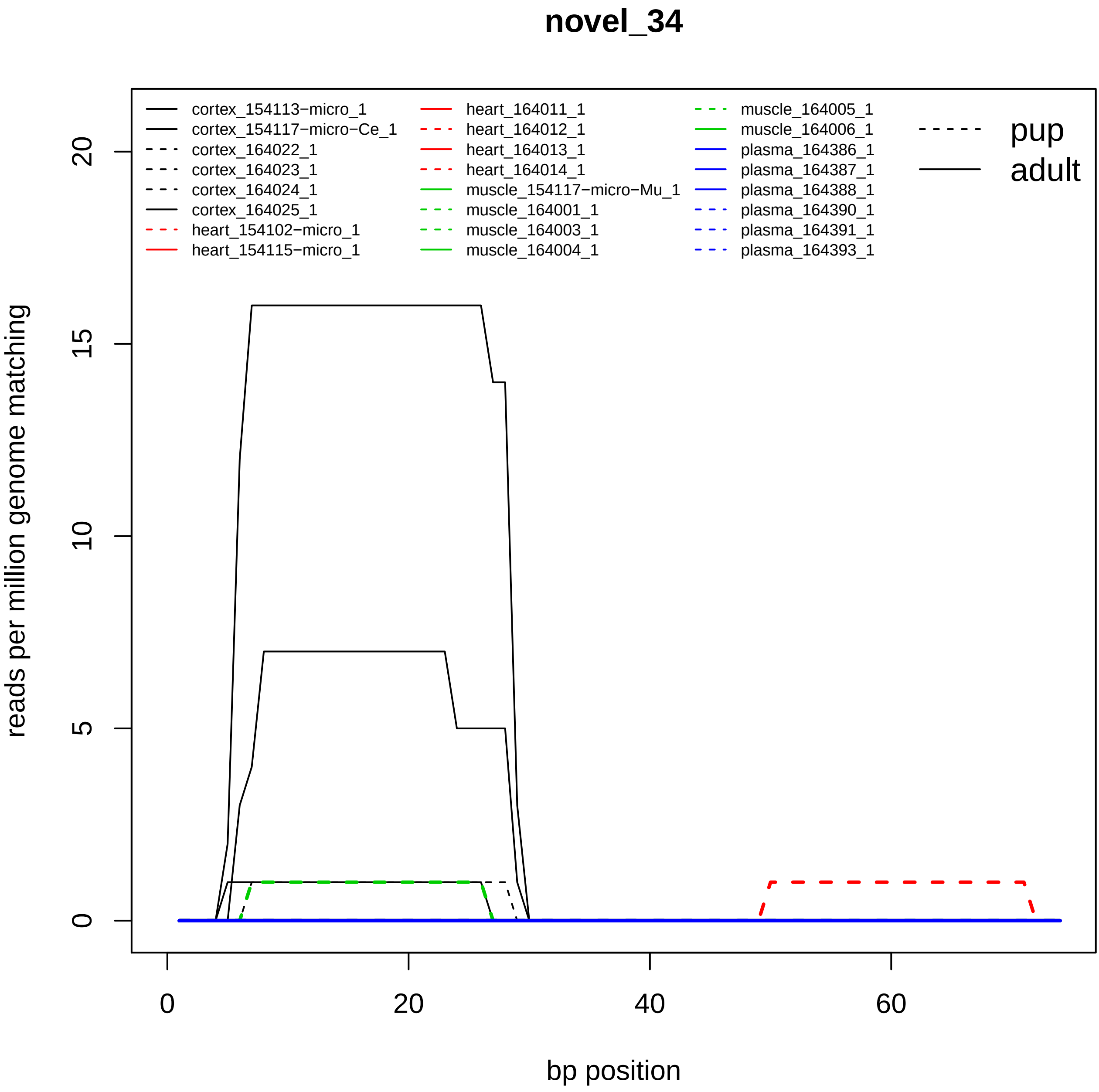

novel_34
cortex_154113−micro_1
cortex_154117−micro−Ce_1
cortex_164022_1
cortex_164023_1
cortex_164024_1
cortex_164025_1
heart_154102−micro_1
heart_154115−micro_1
heart_164011_1
heart_164012_1
heart_164013_1
heart_164014_1
muscle_154117−micro−Mu_1
muscle_164001_1
muscle_164003_1
muscle_164004_1
muscle_164005_1
muscle_164006_1
plasma_164386_1
plasma_164387_1
plasma_164388_1
plasma_164390_1
plasma_164391_1
plasma_164393_1
pup
adult
reads per million genome matching
0
5
10
15
20
0
20
40
60
bp position

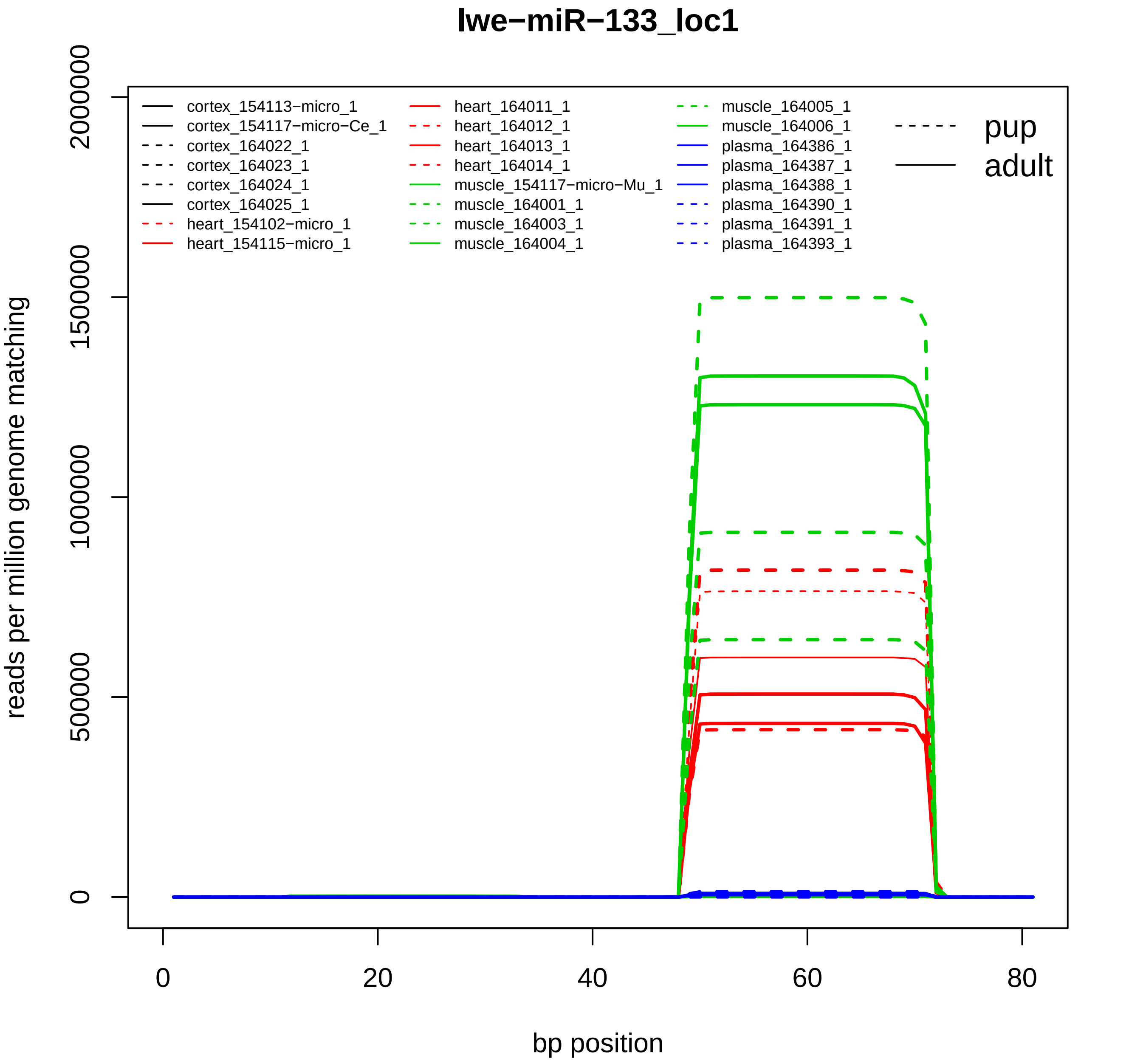

lwe−miR−133_loc1
cortex_154113−micro_1
cortex_154117−micro−Ce_1
cortex_164022_1
cortex_164023_1
cortex_164024_1
cortex_164025_1
heart_154102−micro_1
heart_154115−micro_1
heart_164011_1
heart_164012_1
heart_164013_1
heart_164014_1
muscle_154117−micro−Mu_1
muscle_164001_1
muscle_164003_1
muscle_164004_1
muscle_164005_1
muscle_164006_1
plasma_164386_1
plasma_164387_1
plasma_164388_1
plasma_164390_1
plasma_164391_1
plasma_164393_1
pup
adult
reads per million genome matching
0
500000
1000000
1500000
2000000
bp position
0
20
40
60
80

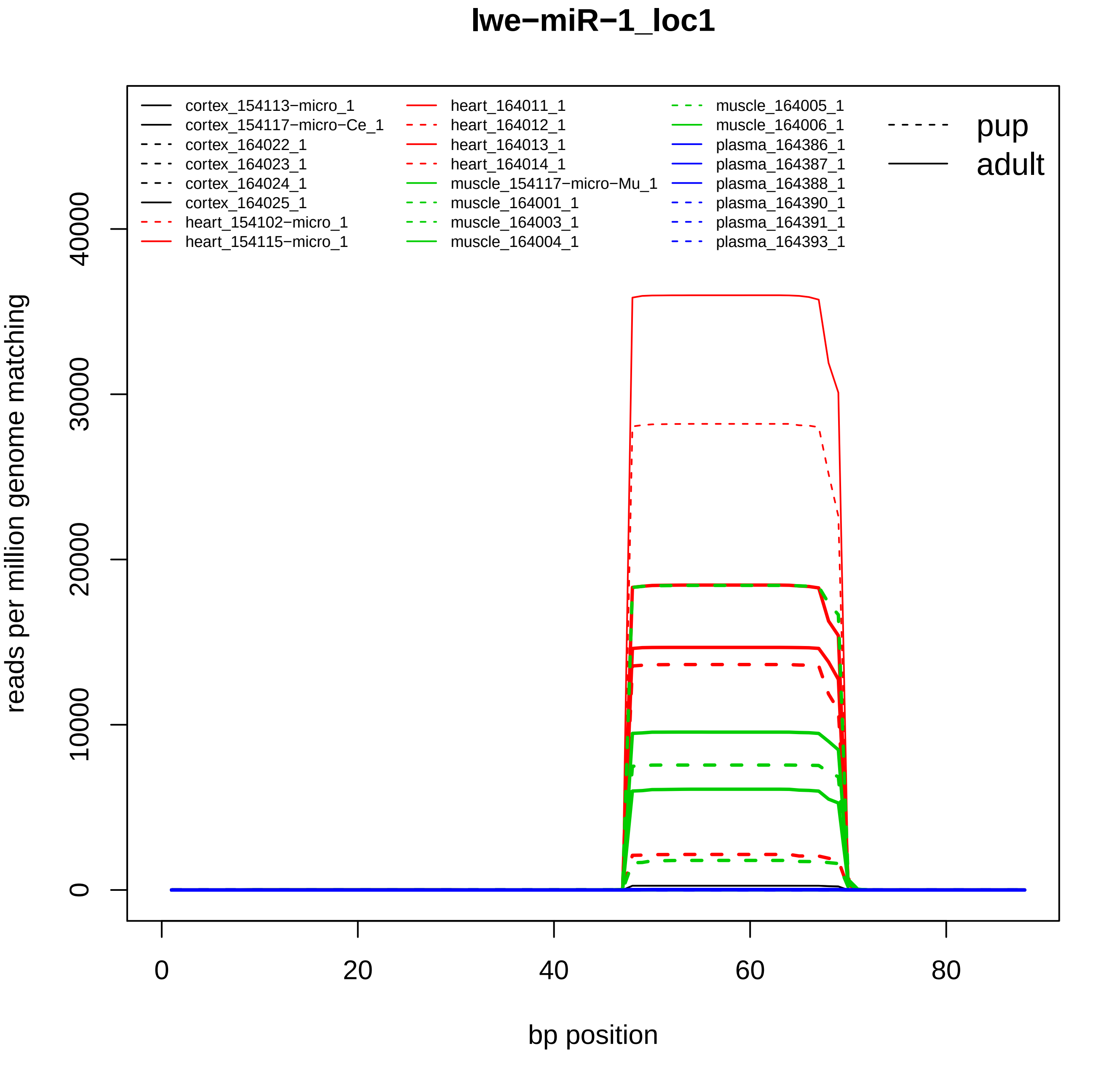

lwe−miR−1_loc1
cortex_154113−micro_1
cortex_154117−micro−Ce_1
cortex_164022_1
cortex_164023_1
cortex_164024_1
cortex_164025_1
heart_154102−micro_1
heart_154115−micro_1
heart_164011_1
heart_164012_1
heart_164013_1
heart_164014_1
muscle_154117−micro−Mu_1
muscle_164001_1
muscle_164003_1
muscle_164004_1
muscle_164005_1
muscle_164006_1
plasma_164386_1
plasma_164387_1
plasma_164388_1
plasma_164390_1
plasma_164391_1
plasma_164393_1
pup
adult
reads per million genome matching
0
10000
20000
30000
40000
0
20
40
60
80
bp position

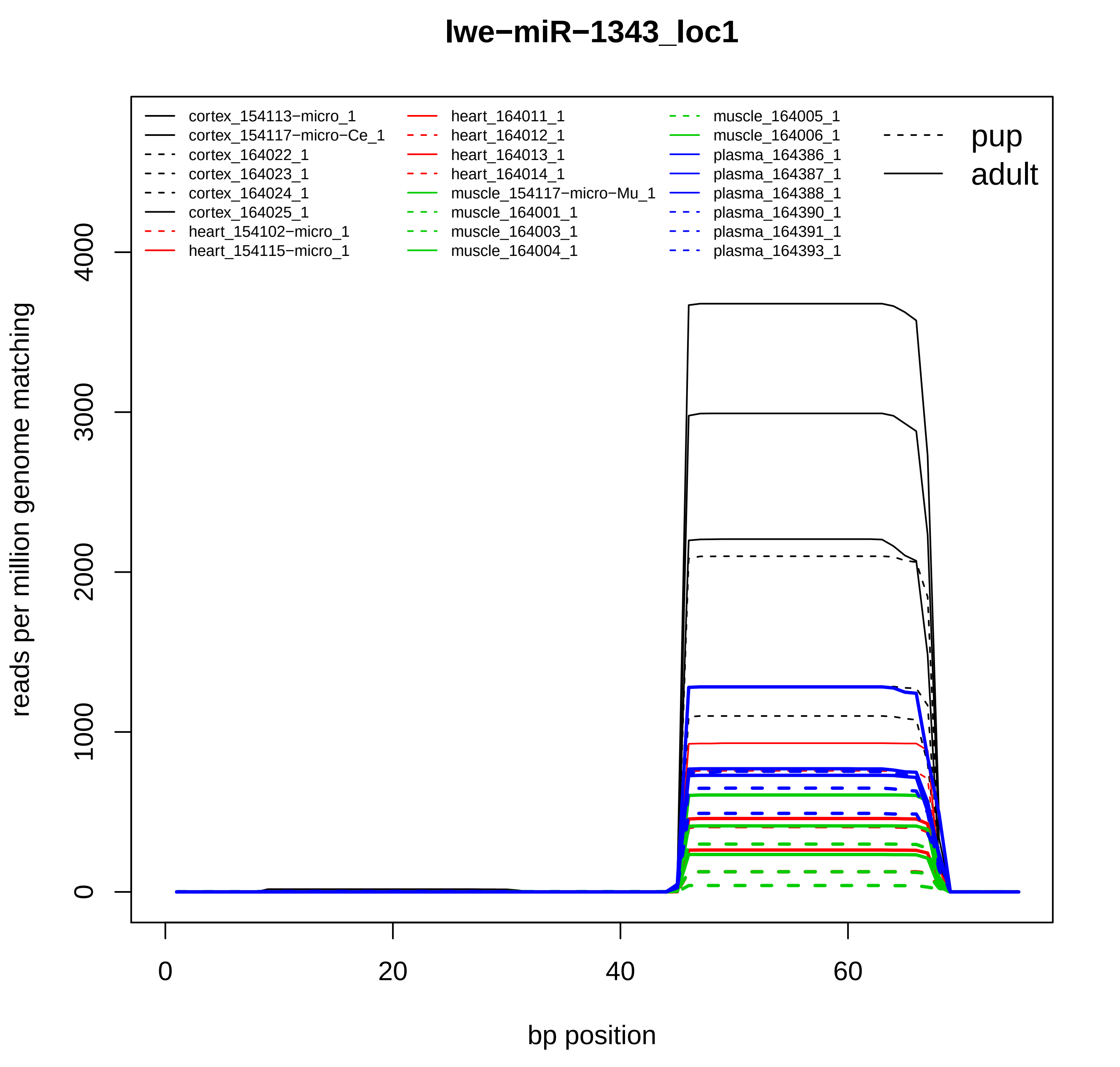
lwe−miR−1343_loc1
cortex_154113−micro_1
cortex_154117−micro−Ce_1
cortex_164022_1
cortex_164023_1
cortex_164024_1
cortex_164025_1
heart_154102−micro_1
heart_154115−micro_1
heart_164011_1
heart_164012_1
heart_164013_1
heart_164014_1
muscle_154117−micro−Mu_1
muscle_164001_1
muscle_164003_1
muscle_164004_1
muscle_164005_1
muscle_164006_1
plasma_164386_1
plasma_164387_1
plasma_164388_1
plasma_164390_1
plasma_164391_1
plasma_164393_1
pup
adult
reads per million genome matching
0
1000
2000
3000
4000
bp position
0
20
40
60

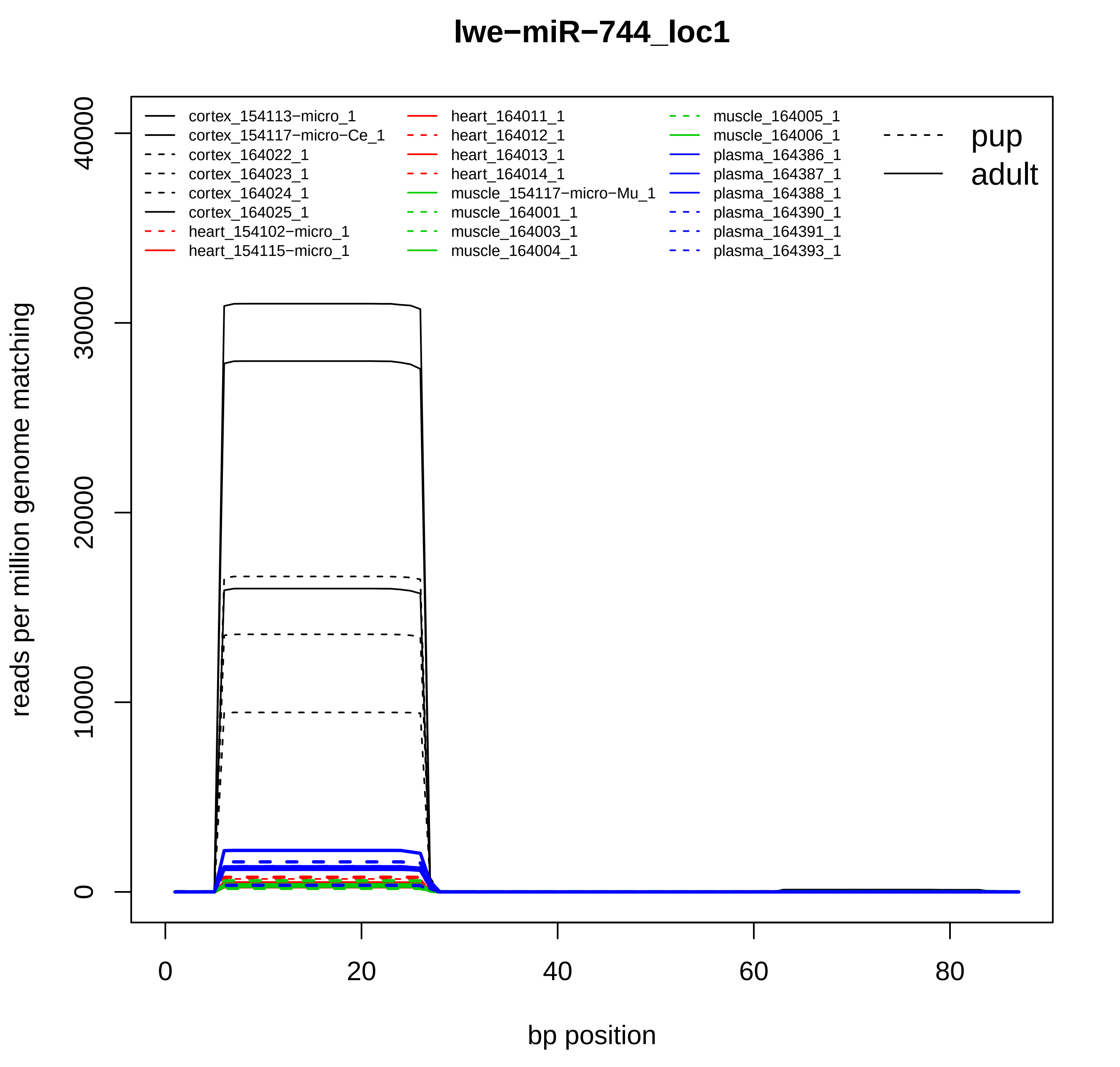

lwe−miR−744_loc1
reads per million genome matching
bp position
0
10000
20000
30000
40000
0
20
40
60
80
cortex_154113−micro_1
cortex_154117−micro−Ce_1
cortex_164022_1
cortex_164023_1
cortex_164024_1
cortex_164025_1
heart_154102−micro_1
heart_154115−micro_1
heart_164011_1
heart_164012_1
heart_164013_1
heart_164014_1
muscle_154117−micro−Mu_1
muscle_164001_1
muscle_164003_1
muscle_164004_1
muscle_164005_1
muscle_164006_1
plasma_164386_1
plasma_164387_1
plasma_164388_1
plasma_164390_1
plasma_164391_1
plasma_164393_1
pup
adult

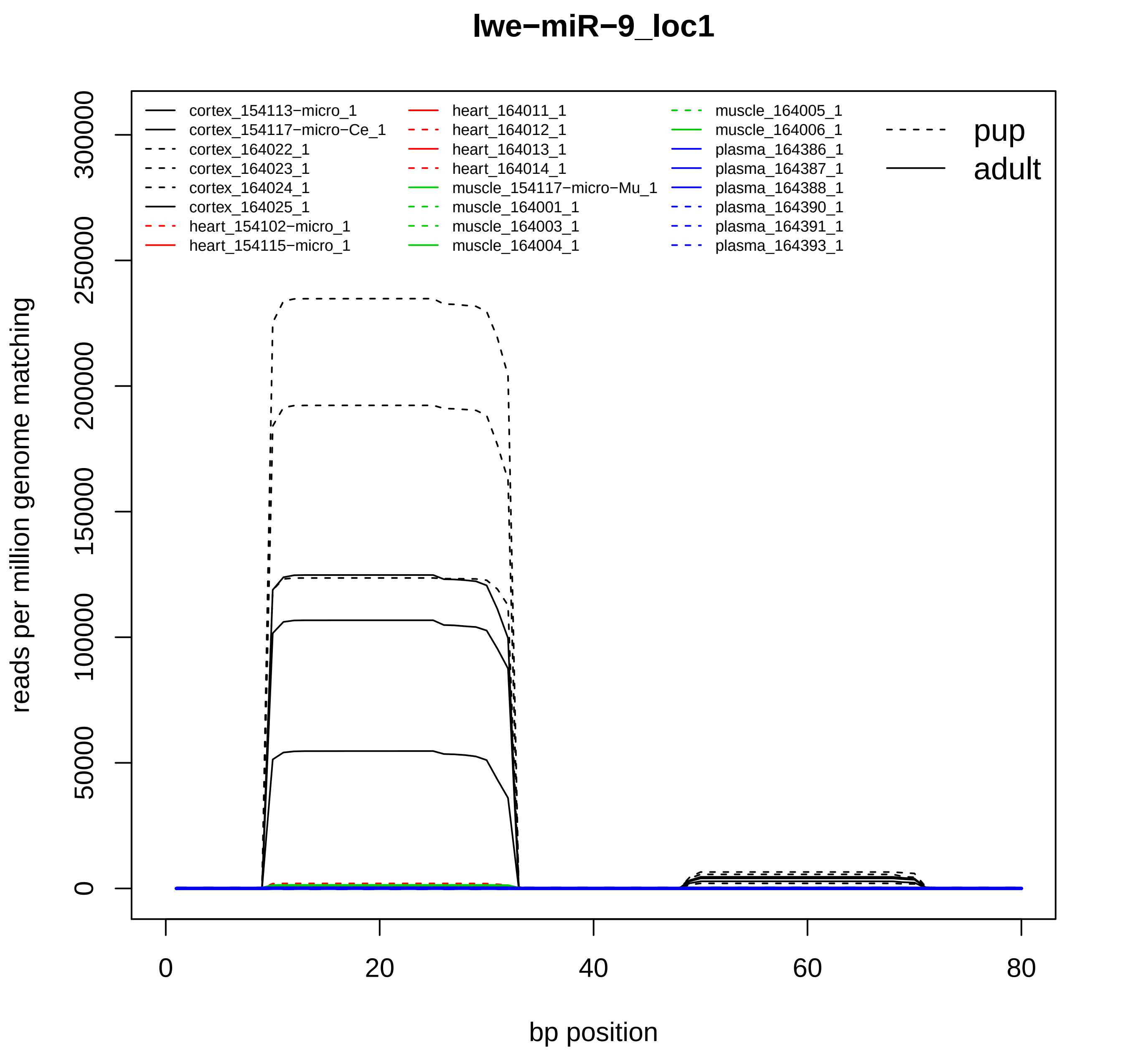
lwe−miR−9_loc1
cortex_154113−micro_1
cortex_154117−micro−Ce_1
cortex_164022_1
cortex_164023_1
cortex_164024_1
cortex_164025_1
heart_154102−micro_1
heart_154115−micro_1
heart_164011_1
heart_164012_1
heart_164013_1
heart_164014_1
muscle_154117−micro−Mu_1
muscle_164001_1
muscle_164003_1
muscle_164004_1
muscle_164005_1
muscle_164006_1
plasma_164386_1
plasma_164387_1
plasma_164388_1
plasma_164390_1
plasma_164391_1
plasma_164393_1
pup
adult
reads per million genome matching
0
50000
100000
150000
200000
250000
300000
0
20
40
60
80
bp position

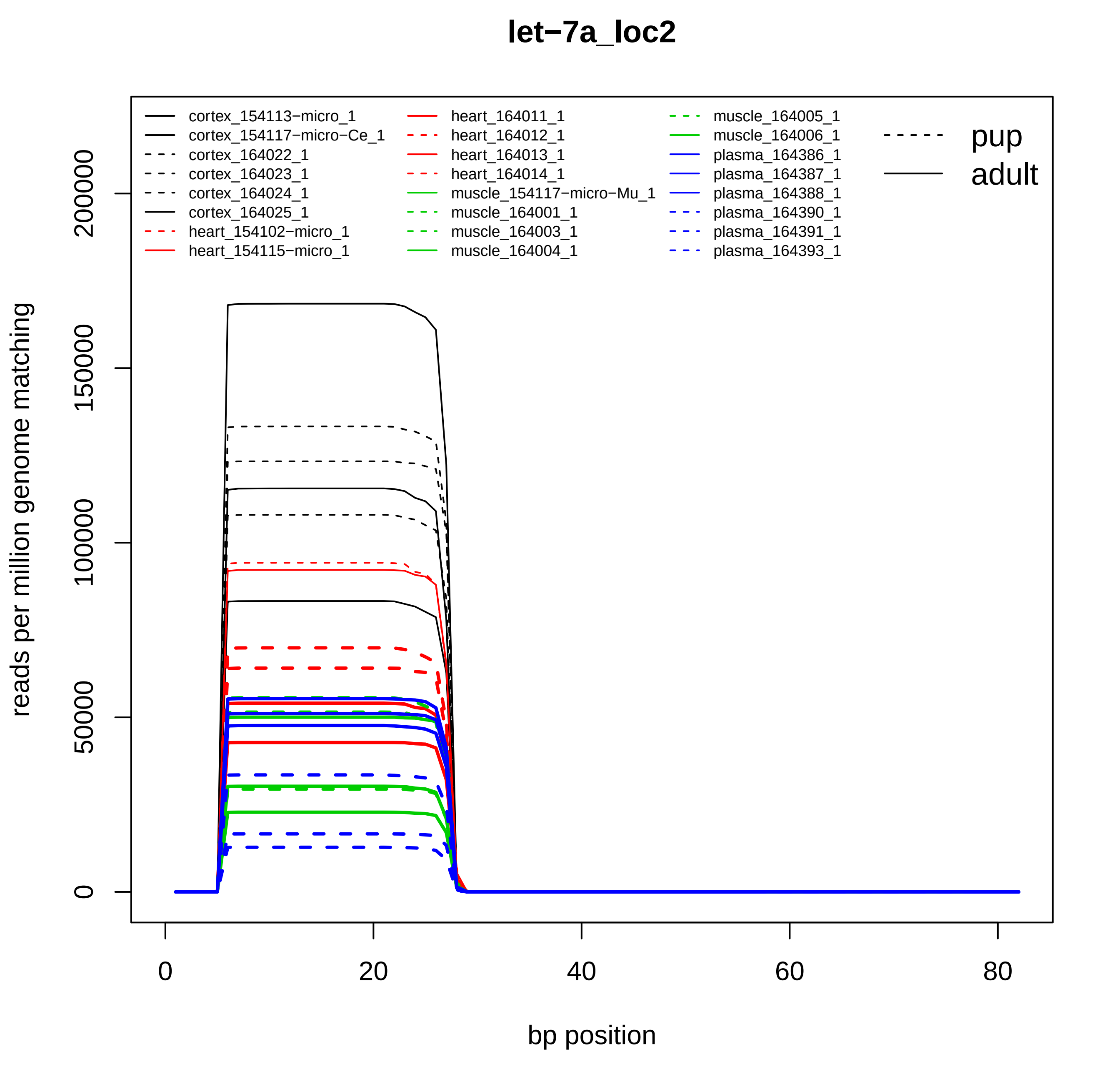
let−7a_loc2
cortex_154113−micro_1
cortex_154117−micro−Ce_1
cortex_164022_1
cortex_164023_1
cortex_164024_1
cortex_164025_1
heart_154102−micro_1
heart_154115−micro_1
heart_164011_1
heart_164012_1
heart_164013_1
heart_164014_1
muscle_154117−micro−Mu_1
muscle_164001_1
muscle_164003_1
muscle_164004_1
muscle_164005_1
muscle_164006_1
plasma_164386_1
plasma_164387_1
plasma_164388_1
plasma_164390_1
plasma_164391_1
plasma_164393_1
pup
adult
reads per million genome matching
0
50000
100000
150000
200000
bp position
0
20
40
60
80

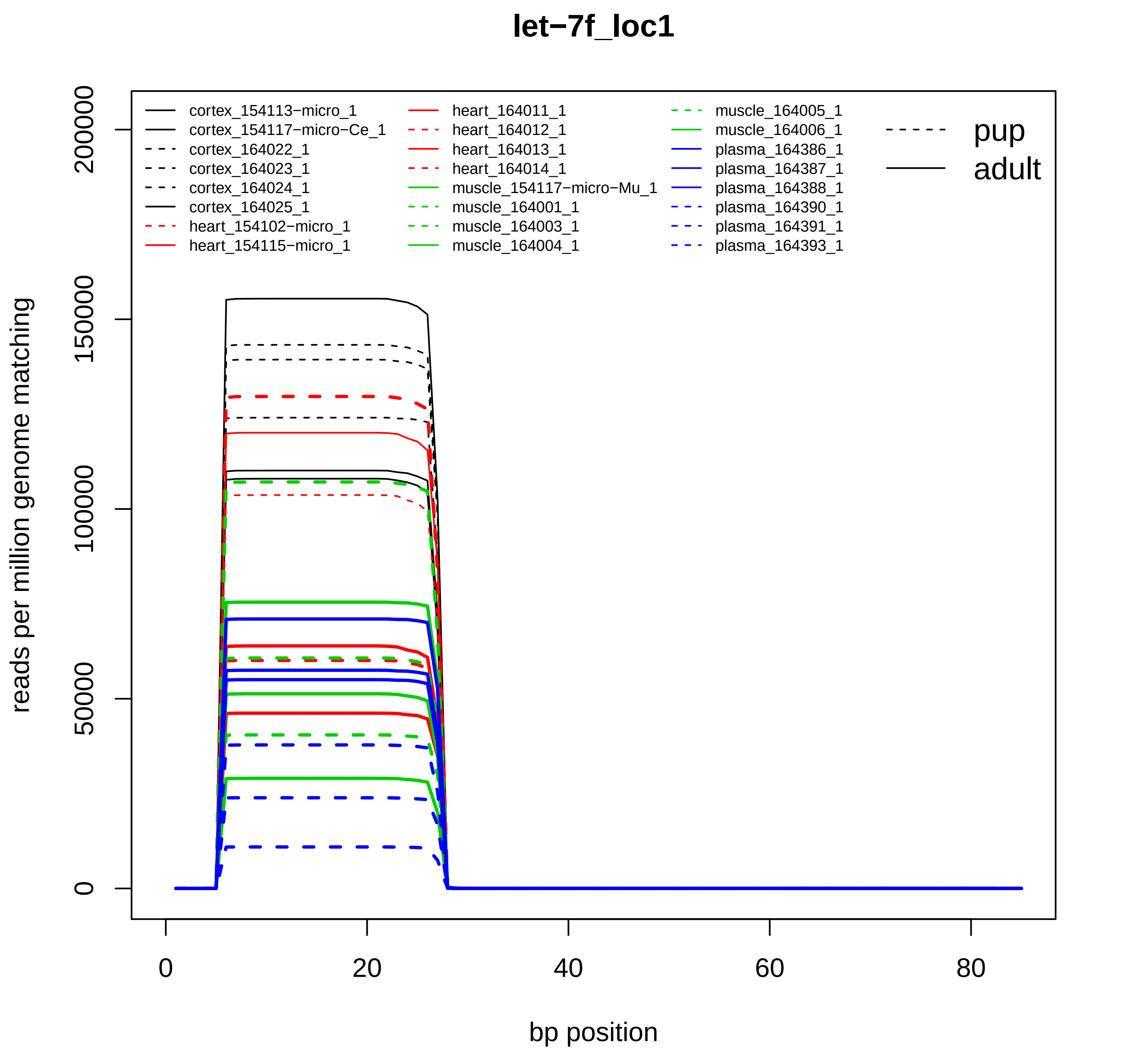

let−7f_loc1
cortex_154113−micro_1
cortex_154117−micro−Ce_1
cortex_164022_1
cortex_164023_1
cortex_164024_1
cortex_164025_1
heart_154102−micro_1
heart_154115−micro_1
heart_164011_1
heart_164012_1
heart_164013_1
heart_164014_1
muscle_154117−micro−Mu_1
muscle_164001_1
muscle_164003_1
muscle_164004_1
muscle_164005_1
muscle_164006_1
plasma_164386_1
plasma_164387_1
plasma_164388_1
plasma_164390_1
plasma_164391_1
plasma_164393_1
pup
adult
reads per million genome matching
0
50000
100000
150000
200000
bp position
0
20
40
60
80

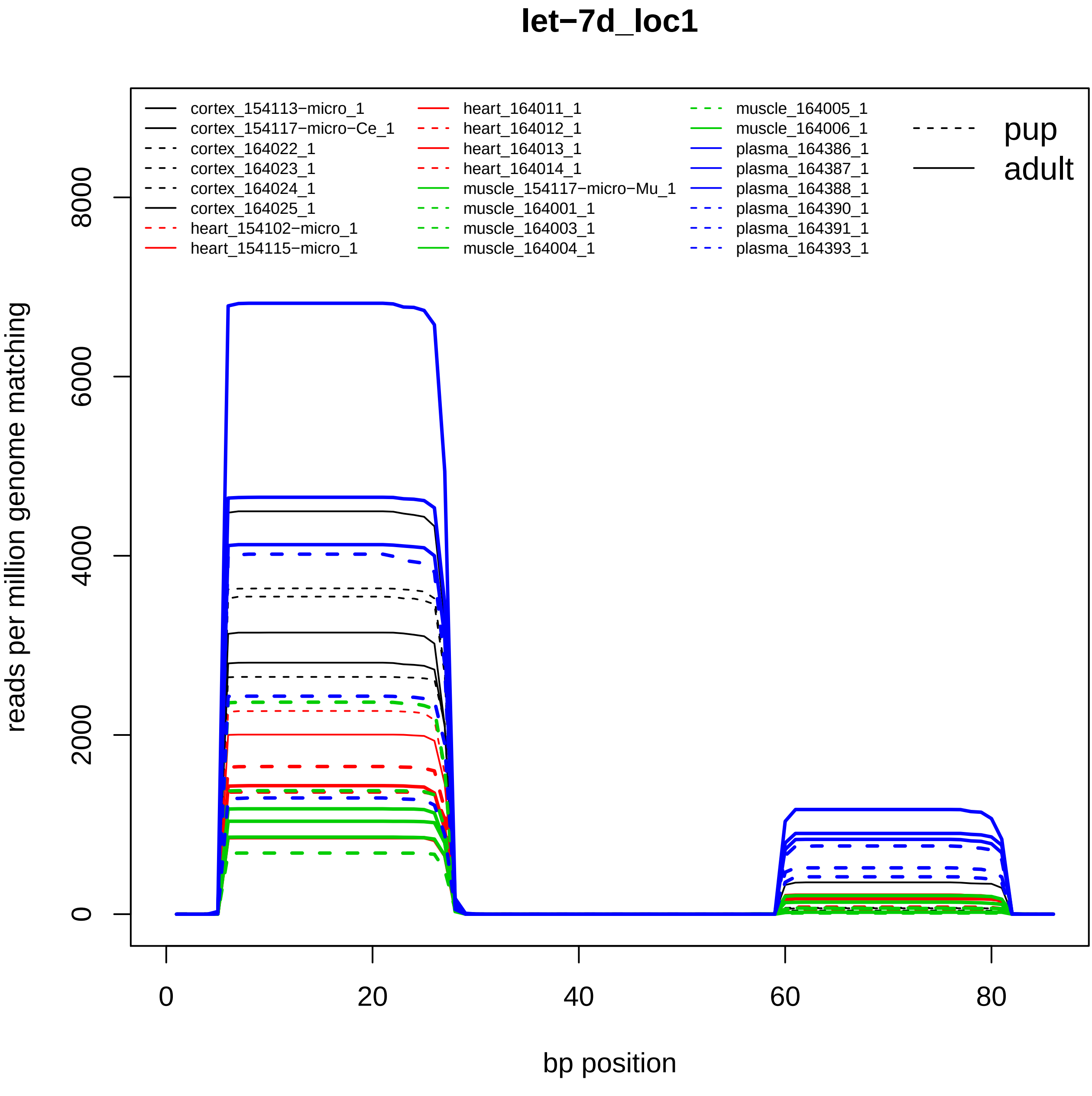
let−7d_loc1
cortex_154113−micro_1
cortex_154117−micro−Ce_1
cortex_164022_1
cortex_164023_1
cortex_164024_1
cortex_164025_1
heart_154102−micro_1
heart_154115−micro_1
heart_164011_1
heart_164012_1
heart_164013_1
heart_164014_1
muscle_154117−micro−Mu_1
muscle_164001_1
muscle_164003_1
muscle_164004_1
muscle_164005_1
muscle_164006_1
plasma_164386_1
plasma_164387_1
plasma_164388_1
plasma_164390_1
plasma_164391_1
plasma_164393_1
pup
adult
reads per million genome matching
0
2000
4000
6000
8000
0
20
40
60
80
bp position

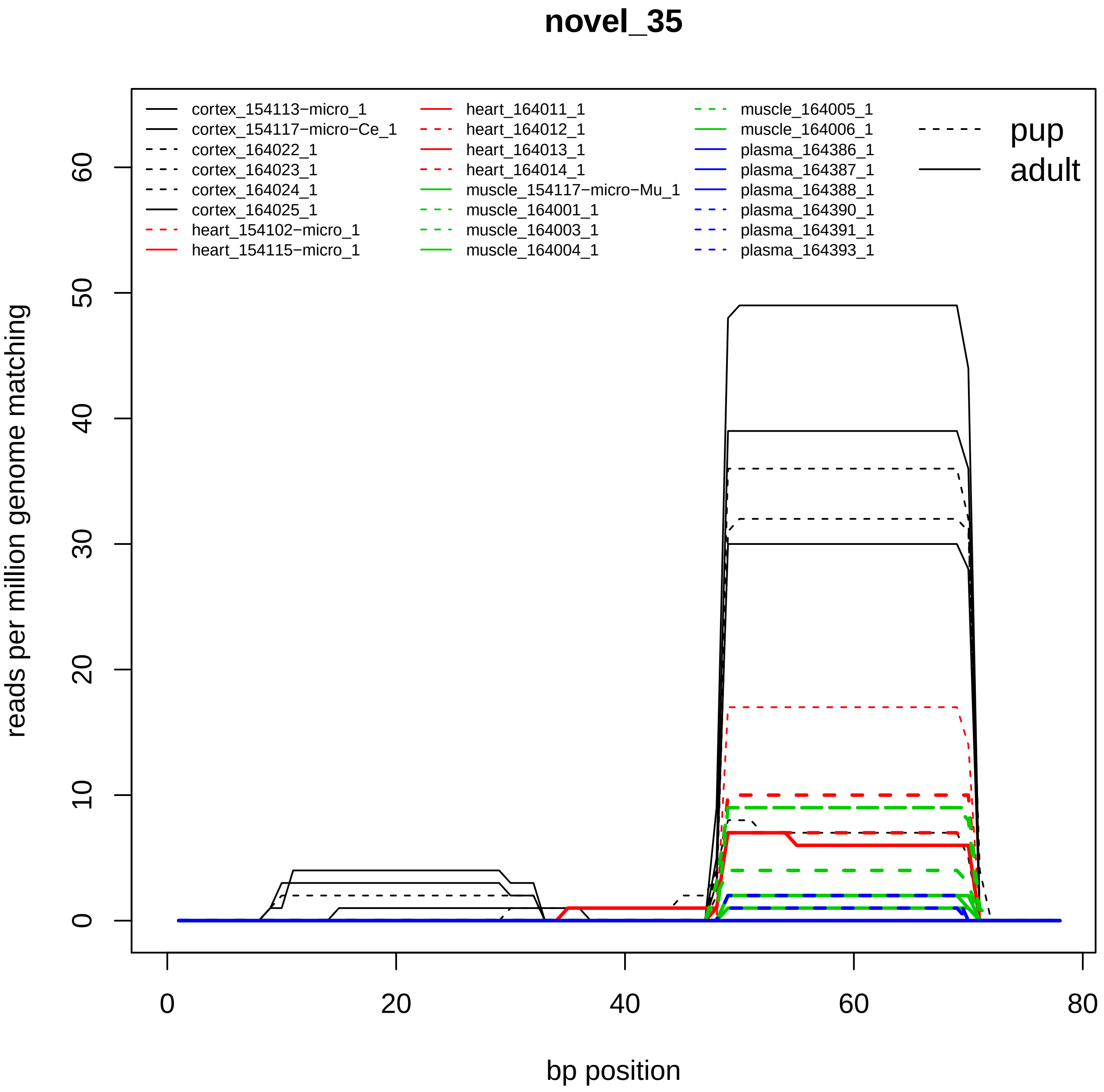
novel_35
reads per million genome matching
bp position
0
10
20
30
40
50
60
0
20
40
60
80
cortex_154113−micro_1
cortex_154117−micro−Ce_1
cortex_164022_1
cortex_164023_1
cortex_164024_1
cortex_164025_1
heart_154102−micro_1
heart_154115−micro_1
heart_164011_1
heart_164012_1
heart_164013_1
heart_164014_1
muscle_154117−micro−Mu_1
muscle_164001_1
muscle_164003_1
muscle_164004_1
muscle_164005_1
muscle_164006_1
plasma_164386_1
plasma_164387_1
plasma_164388_1
plasma_164390_1
plasma_164391_1
plasma_164393_1
pup
adult

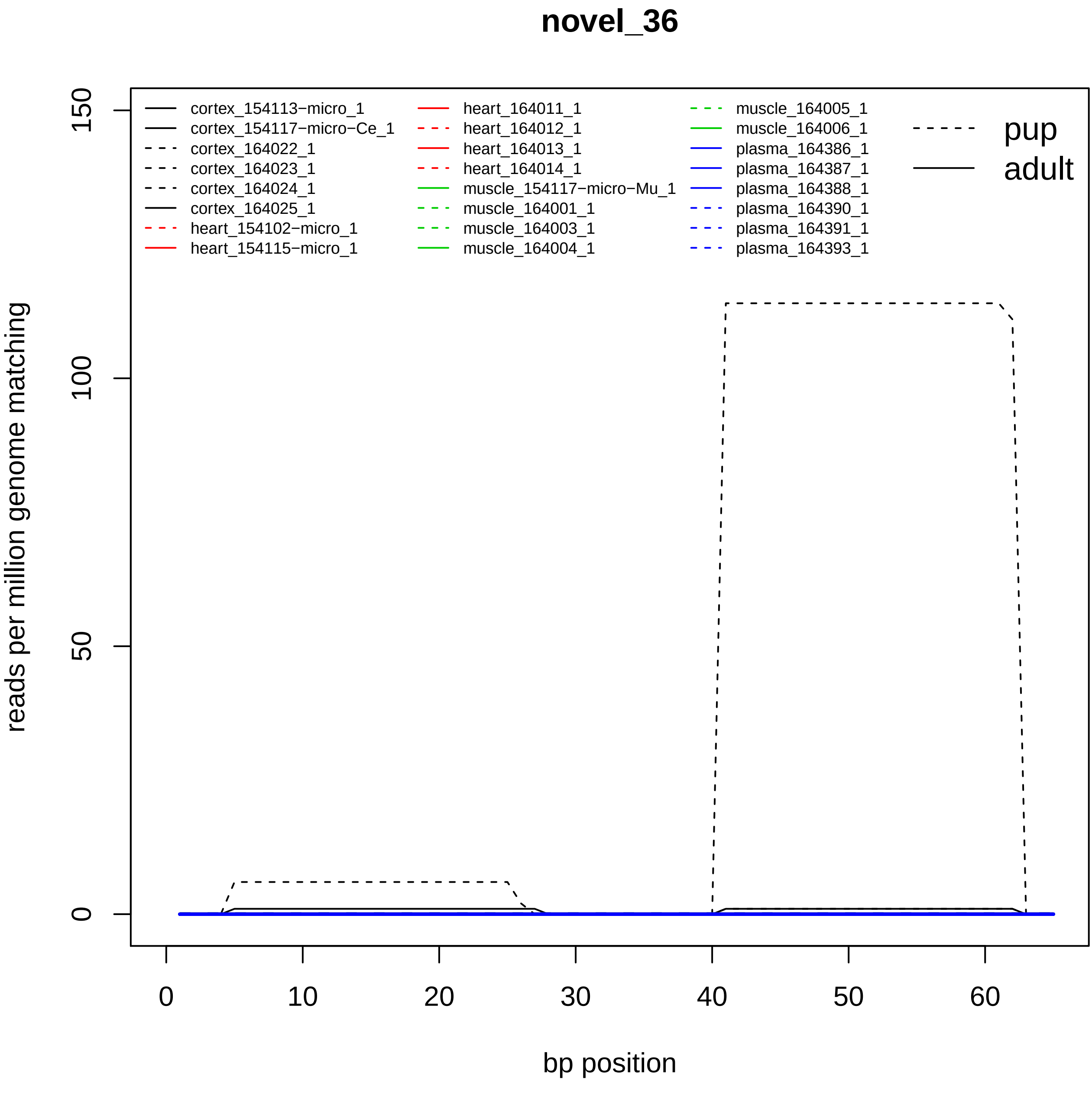

novel_36
cortex_154113−micro_1
cortex_154117−micro−Ce_1
cortex_164022_1
cortex_164023_1
cortex_164024_1
cortex_164025_1
heart_154102−micro_1
heart_154115−micro_1
heart_164011_1
heart_164012_1
heart_164013_1
heart_164014_1
muscle_154117−micro−Mu_1
muscle_164001_1
muscle_164003_1
muscle_164004_1
muscle_164005_1
muscle_164006_1
plasma_164386_1
plasma_164387_1
plasma_164388_1
plasma_164390_1
plasma_164391_1
plasma_164393_1
pup
adult
reads per million genome matching
bp position
0
50
100
150
0
10
20
30
40
50
60

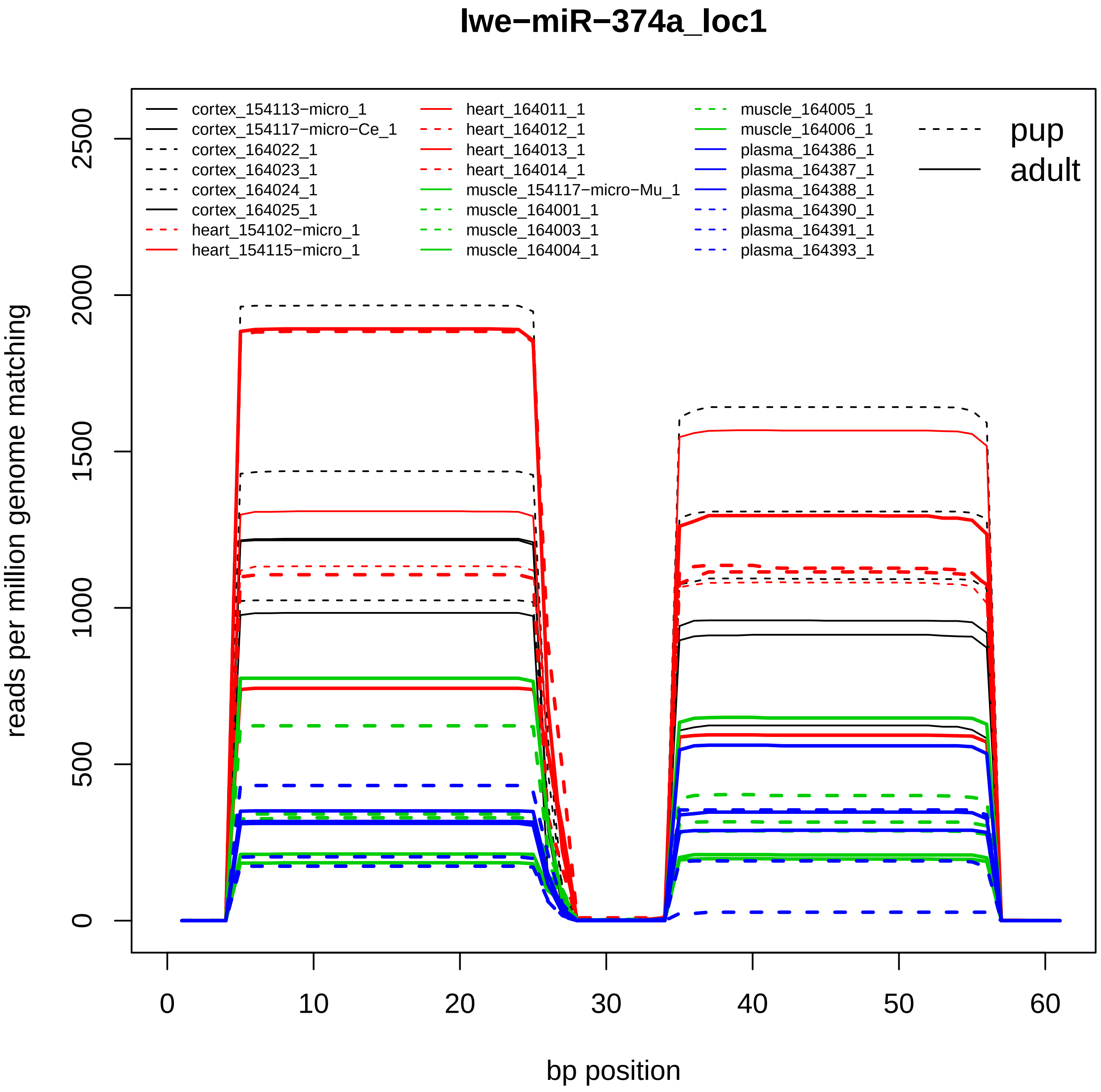
lwe−miR−374a_loc1
cortex_154113−micro_1
cortex_154117−micro−Ce_1
cortex_164022_1
cortex_164023_1
cortex_164024_1
cortex_164025_1
heart_154102−micro_1
heart_154115−micro_1
heart_164011_1
heart_164012_1
heart_164013_1
heart_164014_1
muscle_154117−micro−Mu_1
muscle_164001_1
muscle_164003_1
muscle_164004_1
muscle_164005_1
muscle_164006_1
plasma_164386_1
plasma_164387_1
plasma_164388_1
plasma_164390_1
plasma_164391_1
plasma_164393_1
pup
adult
reads per million genome matching
bp position
0
500
1000
1500
2000
2500
0
10
20
30
40
50
60

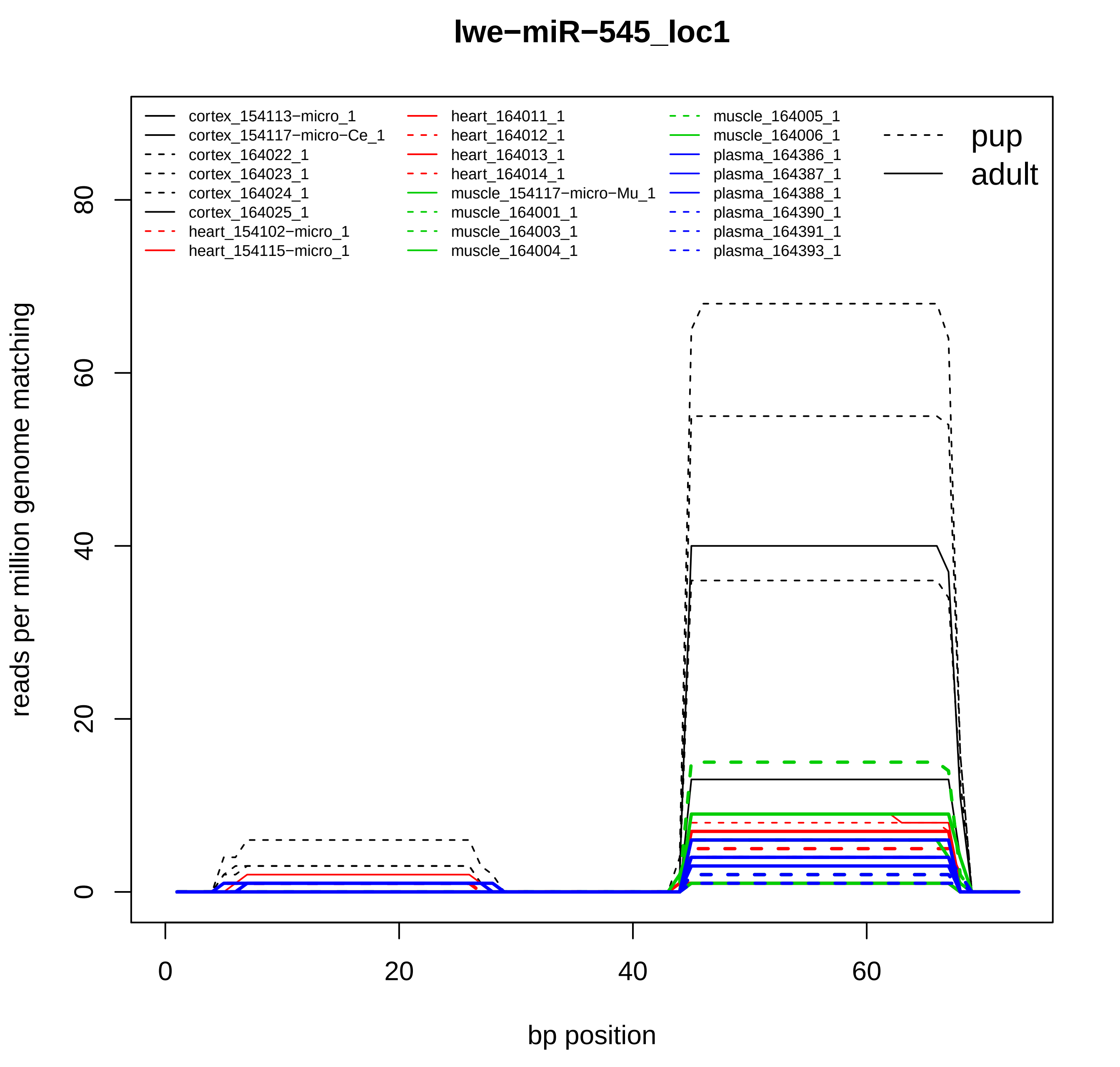

lwe−miR−545_loc1
cortex_154113−micro_1
cortex_154117−micro−Ce_1
cortex_164022_1
cortex_164023_1
cortex_164024_1
cortex_164025_1
heart_154102−micro_1
heart_154115−micro_1
heart_164011_1
heart_164012_1
heart_164013_1
heart_164014_1
muscle_154117−micro−Mu_1
muscle_164001_1
muscle_164003_1
muscle_164004_1
muscle_164005_1
muscle_164006_1
plasma_164386_1
plasma_164387_1
plasma_164388_1
plasma_164390_1
plasma_164391_1
plasma_164393_1
pup
adult
reads per million genome matching
bp position
0
20
40
60
80

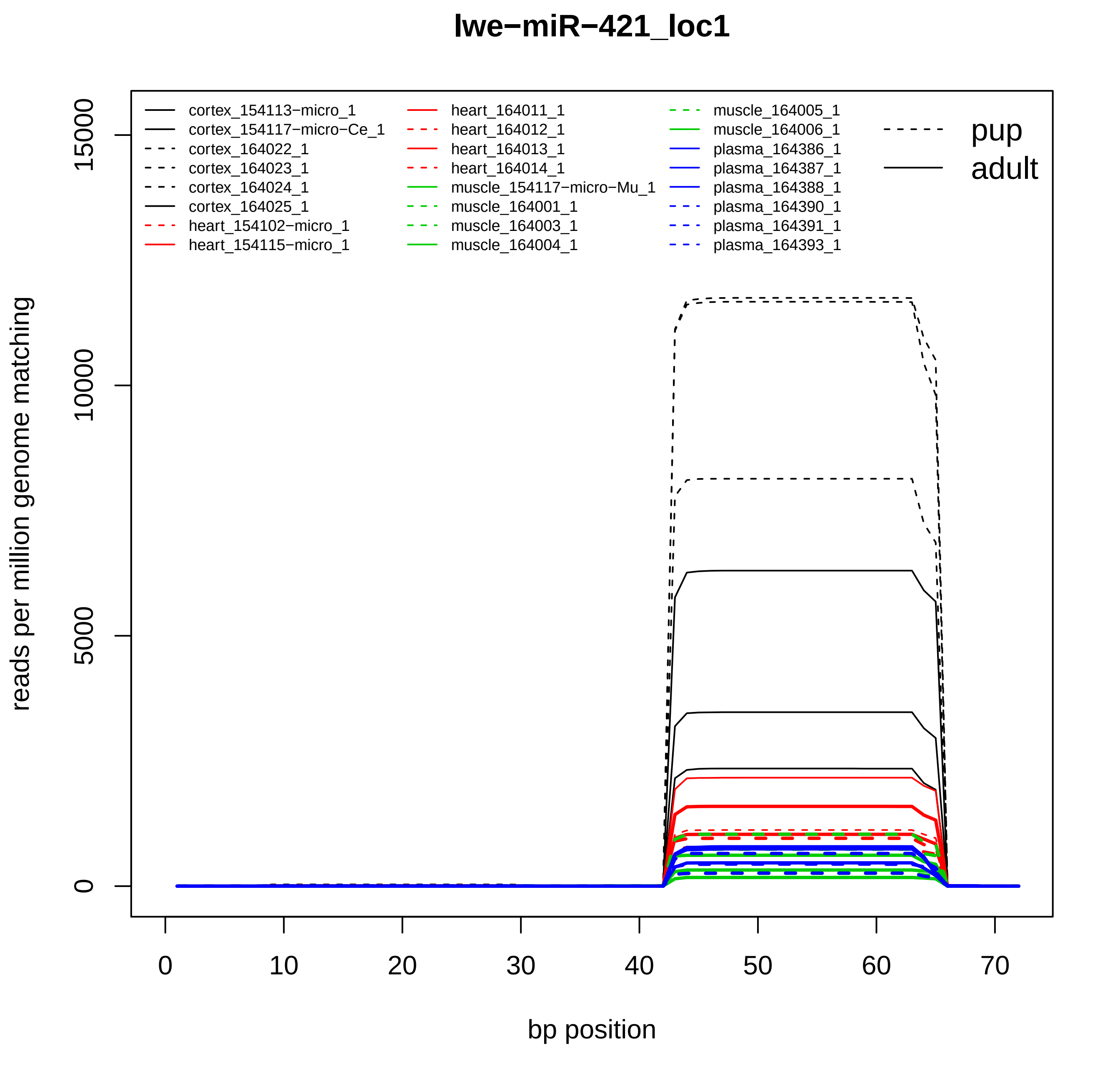

lwe−miR−421_loc1
cortex_154113−micro_1
cortex_154117−micro−Ce_1
cortex_164022_1
cortex_164023_1
cortex_164024_1
cortex_164025_1
heart_154102−micro_1
heart_154115−micro_1
heart_164011_1
heart_164012_1
heart_164013_1
heart_164014_1
muscle_154117−micro−Mu_1
muscle_164001_1
muscle_164003_1
muscle_164004_1
muscle_164005_1
muscle_164006_1
plasma_164386_1
plasma_164387_1
plasma_164388_1
plasma_164390_1
plasma_164391_1
plasma_164393_1
pup
adult
reads per million genome matching
0
5000
10000
15000
0
10
20
30
40
50
60
70
bp position

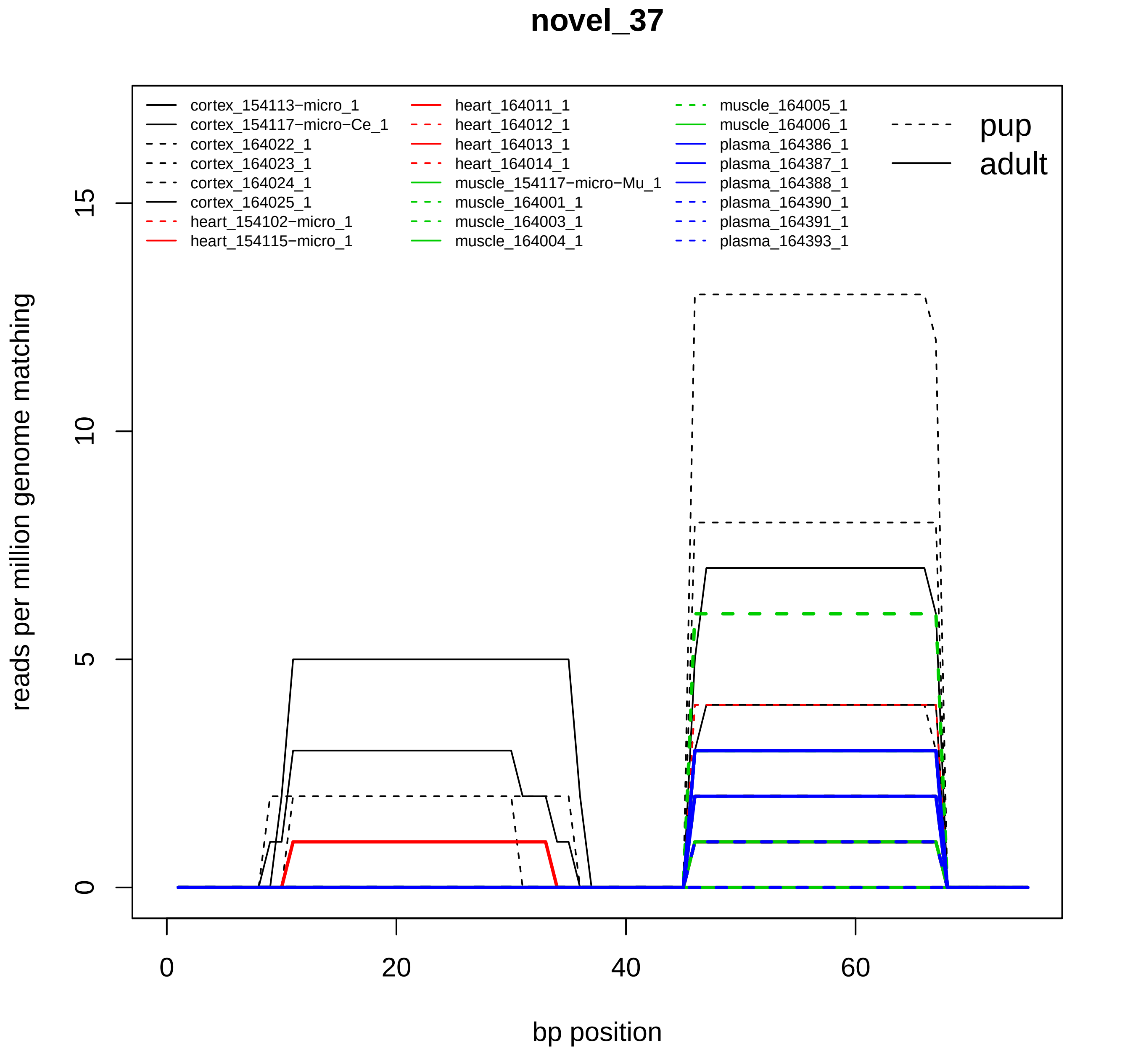
novel_37
cortex_154113−micro_1
cortex_154117−micro−Ce_1
cortex_164022_1
cortex_164023_1
cortex_164024_1
cortex_164025_1
heart_154102−micro_1
heart_154115−micro_1
heart_164011_1
heart_164012_1
heart_164013_1
heart_164014_1
muscle_154117−micro−Mu_1
muscle_164001_1
muscle_164003_1
muscle_164004_1
muscle_164005_1
muscle_164006_1
plasma_164386_1
plasma_164387_1
plasma_164388_1
plasma_164390_1
plasma_164391_1
plasma_164393_1
pup
adult
reads per million genome matching
0
5
10
15
0
20
40
60
bp position

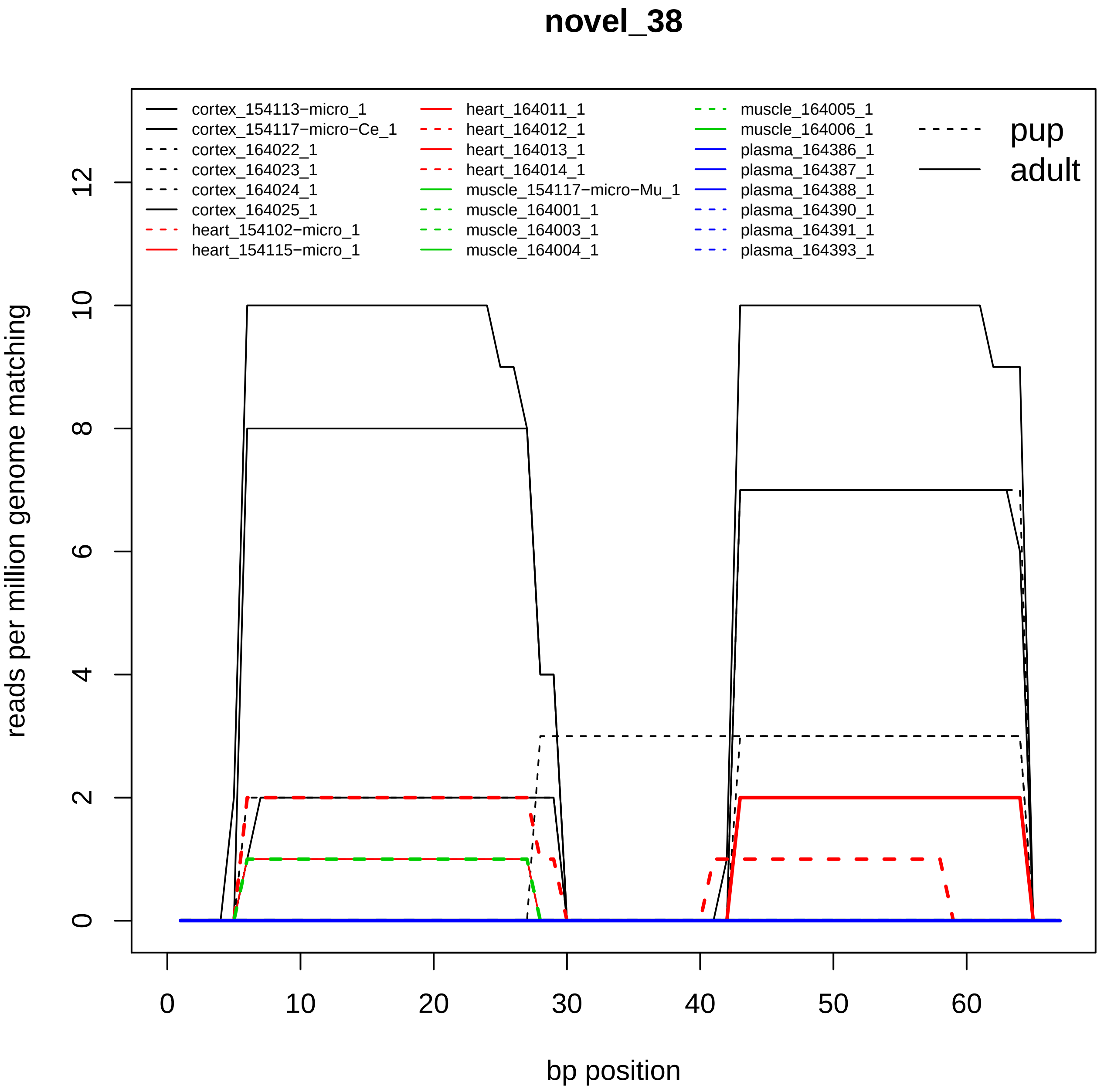

novel_38
cortex_154113−micro_1
cortex_154117−micro−Ce_1
cortex_164022_1
cortex_164023_1
cortex_164024_1
cortex_164025_1
heart_154102−micro_1
heart_154115−micro_1
heart_164011_1
heart_164012_1
heart_164013_1
heart_164014_1
muscle_154117−micro−Mu_1
muscle_164001_1
muscle_164003_1
muscle_164004_1
muscle_164005_1
muscle_164006_1
plasma_164386_1
plasma_164387_1
plasma_164388_1
plasma_164390_1
plasma_164391_1
plasma_164393_1
pup
adult
reads per million genome matching
0
2
4
6
8
10
12
0
10
20
30
40
50
60
bp position

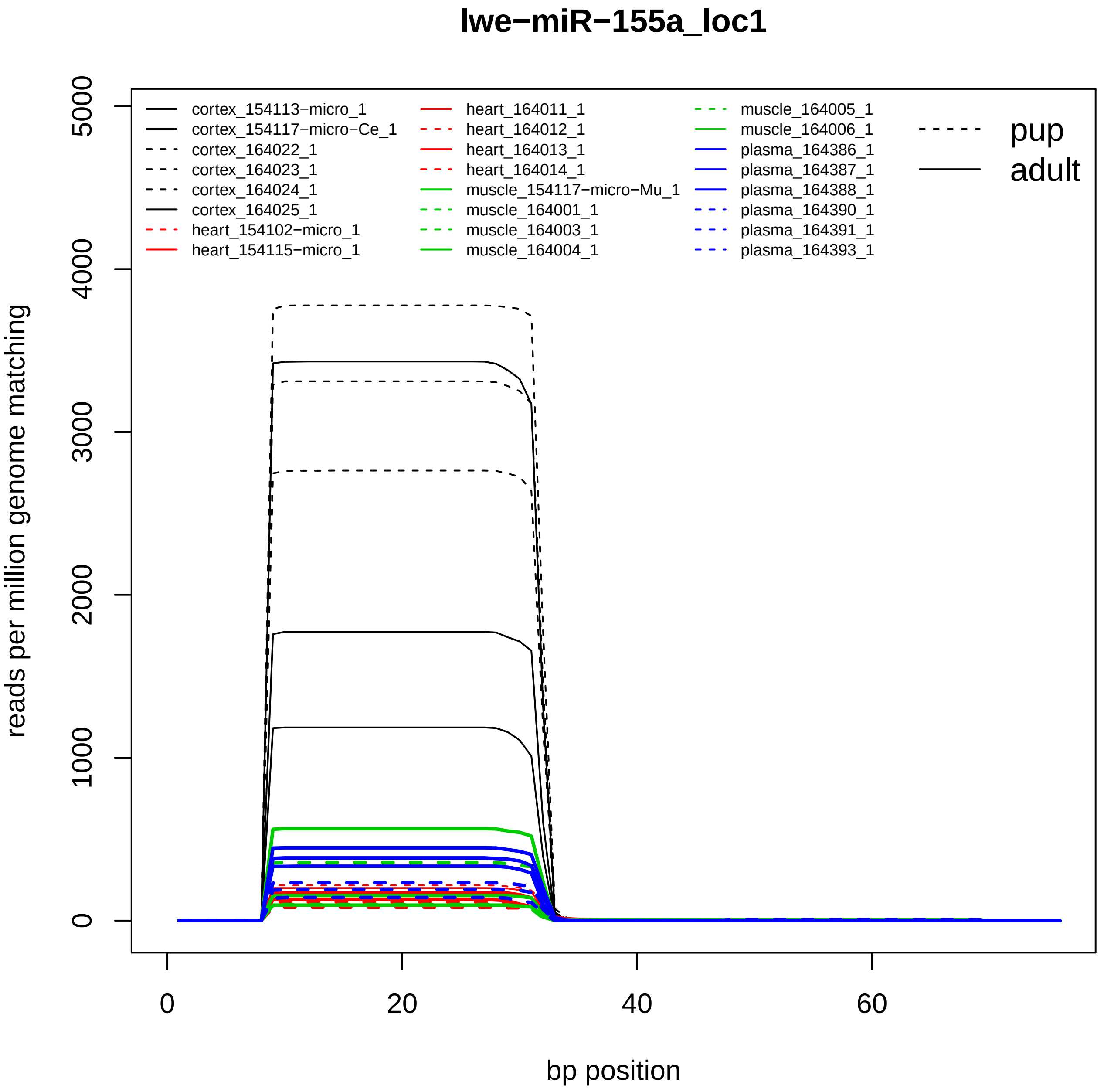
lwe−miR−155a_loc1
cortex_154113−micro_1
cortex_154117−micro−Ce_1
cortex_164022_1
cortex_164023_1
cortex_164024_1
cortex_164025_1
heart_154102−micro_1
heart_154115−micro_1
heart_164011_1
heart_164012_1
heart_164013_1
heart_164014_1
muscle_154117−micro−Mu_1
muscle_164001_1
muscle_164003_1
muscle_164004_1
muscle_164005_1
muscle_164006_1
plasma_164386_1
plasma_164387_1
plasma_164388_1
plasma_164390_1
plasma_164391_1
plasma_164393_1
pup
adult
reads per million genome matching
0
1000
2000
3000
4000
5000
bp position
0
20
40
60

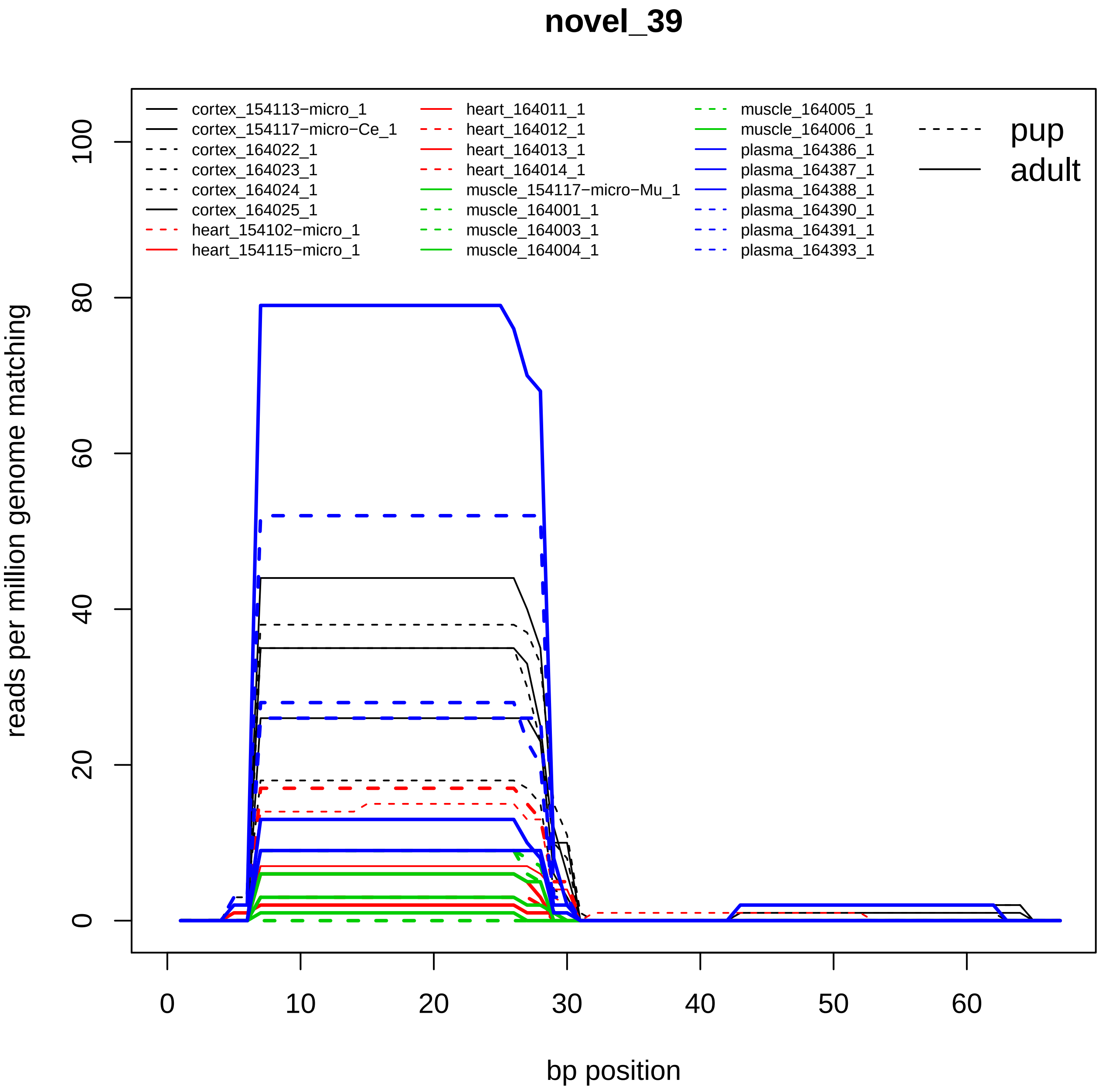
novel_39
cortex_154113−micro_1
cortex_154117−micro−Ce_1
cortex_164022_1
cortex_164023_1
cortex_164024_1
cortex_164025_1
heart_154102−micro_1
heart_154115−micro_1
heart_164011_1
heart_164012_1
heart_164013_1
heart_164014_1
muscle_154117−micro−Mu_1
muscle_164001_1
muscle_164003_1
muscle_164004_1
muscle_164005_1
muscle_164006_1
plasma_164386_1
plasma_164387_1
plasma_164388_1
plasma_164390_1
plasma_164391_1
plasma_164393_1
pup
adult
reads per million genome matching
bp position
0
20
40
60
80
100
0
10
20
30
40
50
60

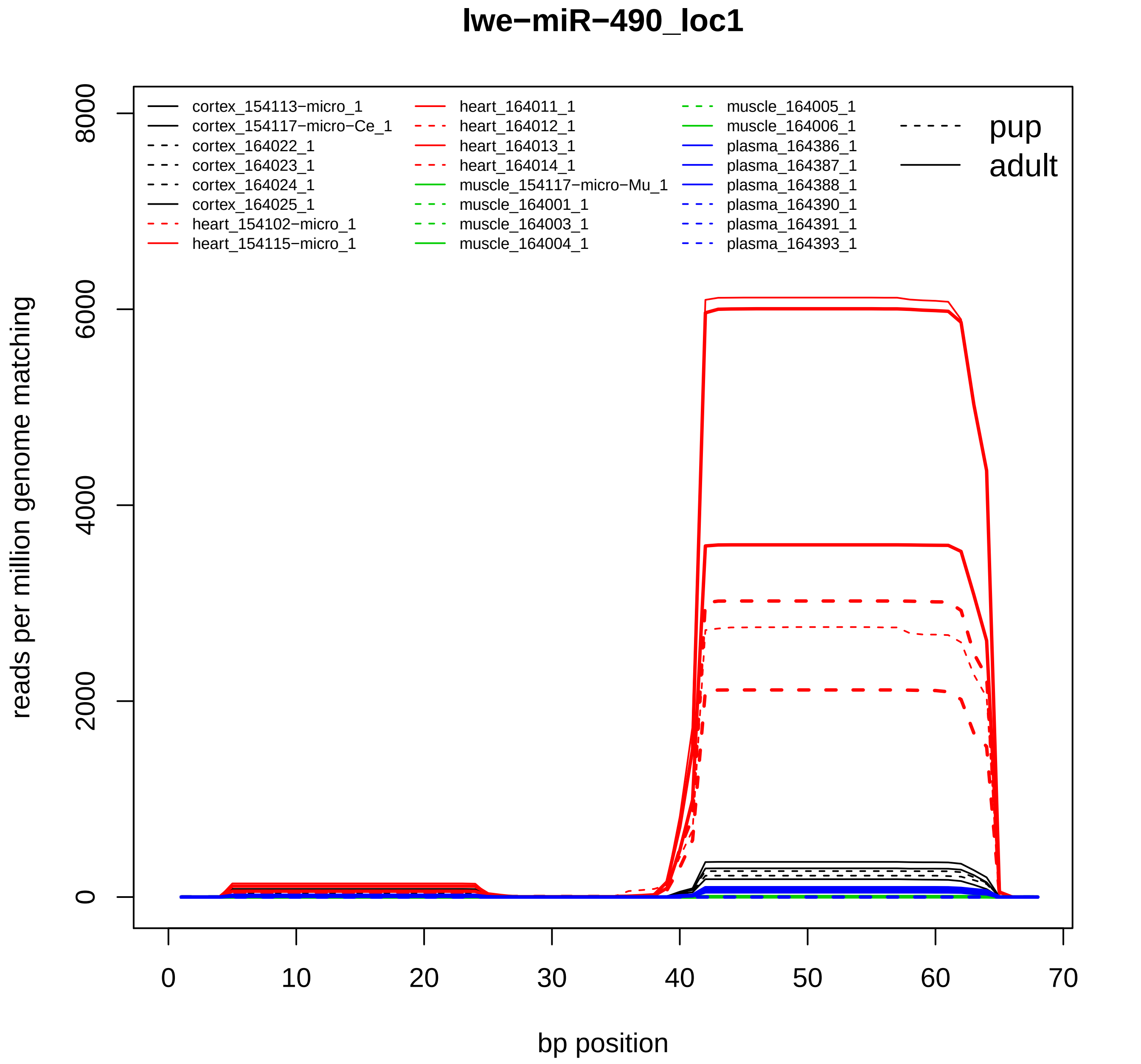
lwe−miR−490_loc1
cortex_154113−micro_1
cortex_154117−micro−Ce_1
cortex_164022_1
cortex_164023_1
cortex_164024_1
cortex_164025_1
heart_154102−micro_1
heart_154115−micro_1
heart_164011_1
heart_164012_1
heart_164013_1
heart_164014_1
muscle_154117−micro−Mu_1
muscle_164001_1
muscle_164003_1
muscle_164004_1
muscle_164005_1
muscle_164006_1
plasma_164386_1
plasma_164387_1
plasma_164388_1
plasma_164390_1
plasma_164391_1
plasma_164393_1
pup
adult
reads per million genome matching
0
2000
4000
6000
8000
bp position
0
10
20
30
40
50
60
70

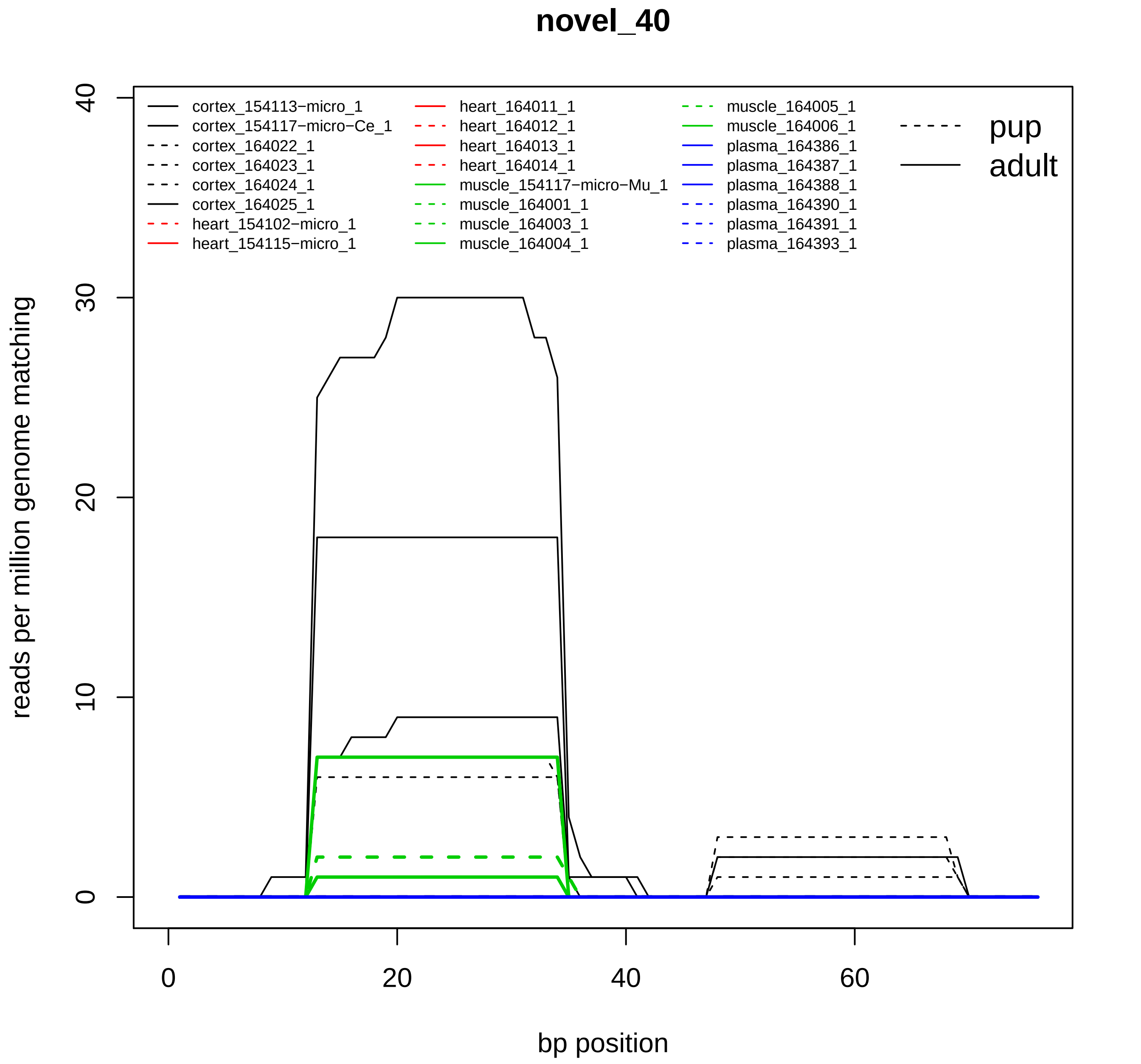
novel_40
reads per million genome matching
bp position
0
10
20
30
40
0
20
40
60
cortex_154113−micro_1
cortex_154117−micro−Ce_1
cortex_164022_1
cortex_164023_1
cortex_164024_1
cortex_164025_1
heart_154102−micro_1
heart_154115−micro_1
heart_164011_1
heart_164012_1
heart_164013_1
heart_164014_1
muscle_154117−micro−Mu_1
muscle_164001_1
muscle_164003_1
muscle_164004_1
muscle_164005_1
muscle_164006_1
plasma_164386_1
plasma_164387_1
plasma_164388_1
plasma_164390_1
plasma_164391_1
plasma_164393_1
pup
adult

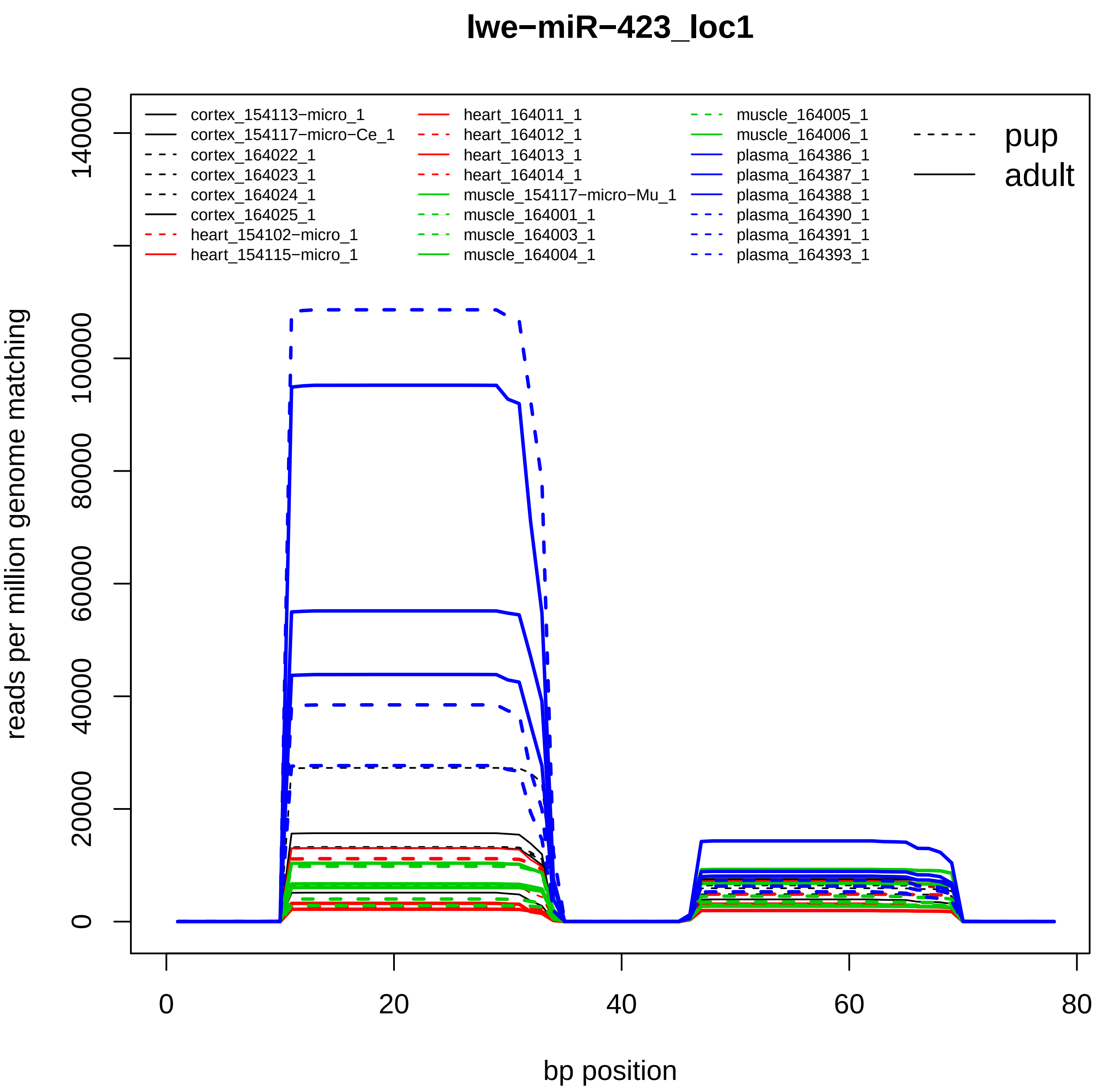
lwe−miR−423_loc1
reads per million genome matching
bp position
0
20000
40000
60000
80000
100000
140000
0
20
40
60
80
cortex_154113−micro_1
cortex_154117−micro−Ce_1
cortex_164022_1
cortex_164023_1
cortex_164024_1
cortex_164025_1
heart_154102−micro_1
heart_154115−micro_1
heart_164011_1
heart_164012_1
heart_164013_1
heart_164014_1
muscle_154117−micro−Mu_1
muscle_164001_1
muscle_164003_1
muscle_164004_1
muscle_164005_1
muscle_164006_1
plasma_164386_1
plasma_164387_1
plasma_164388_1
plasma_164390_1
plasma_164391_1
plasma_164393_1
pup
adult

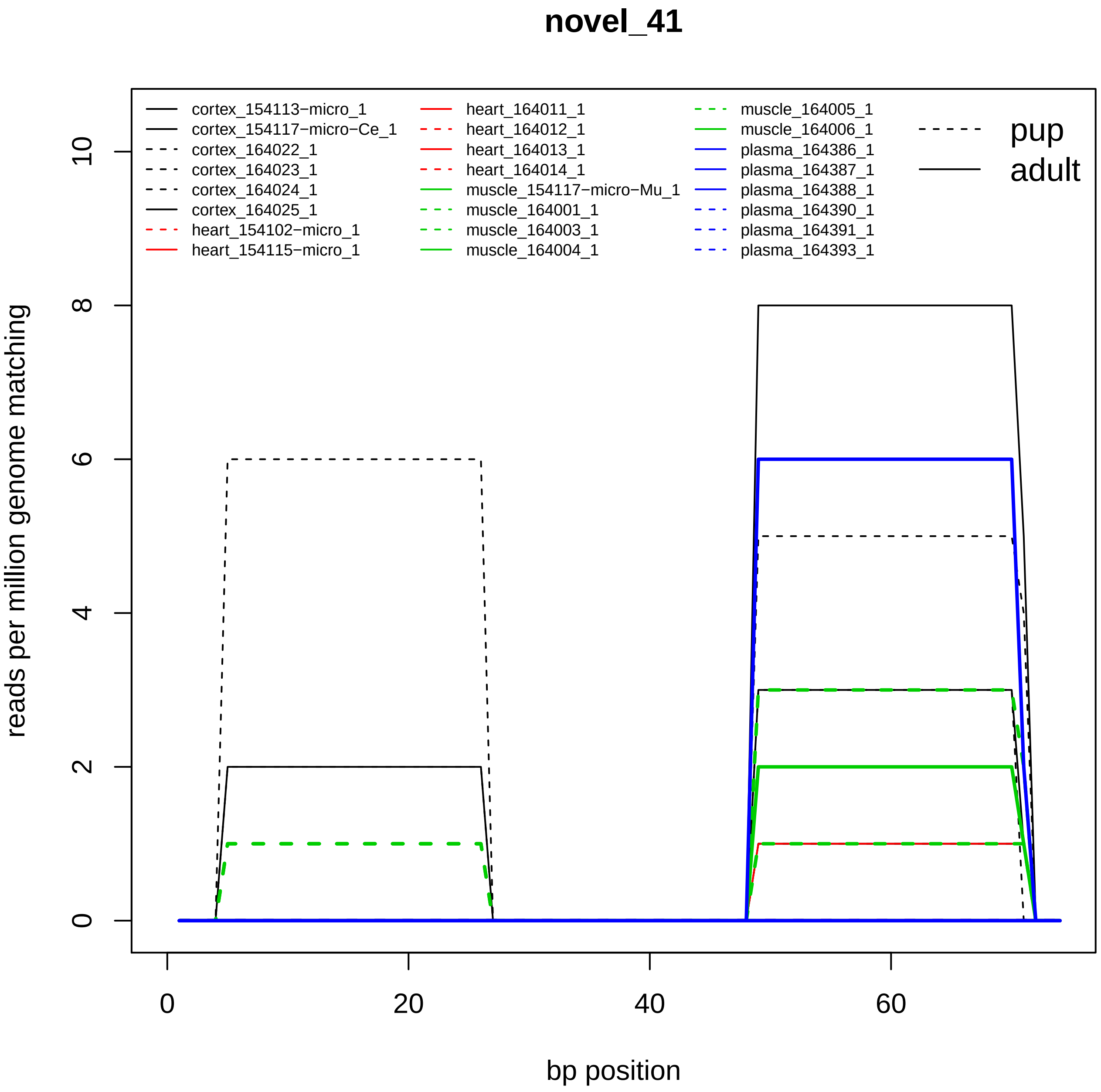
novel_41
cortex_154113−micro_1
cortex_154117−micro−Ce_1
cortex_164022_1
cortex_164023_1
cortex_164024_1
cortex_164025_1
heart_154102−micro_1
heart_154115−micro_1
heart_164011_1
heart_164012_1
heart_164013_1
heart_164014_1
muscle_154117−micro−Mu_1
muscle_164001_1
muscle_164003_1
muscle_164004_1
muscle_164005_1
muscle_164006_1
plasma_164386_1
plasma_164387_1
plasma_164388_1
plasma_164390_1
plasma_164391_1
plasma_164393_1
pup
adult
reads per million genome matching
bp position
0
2
4
6
8
10
0
20
40
60

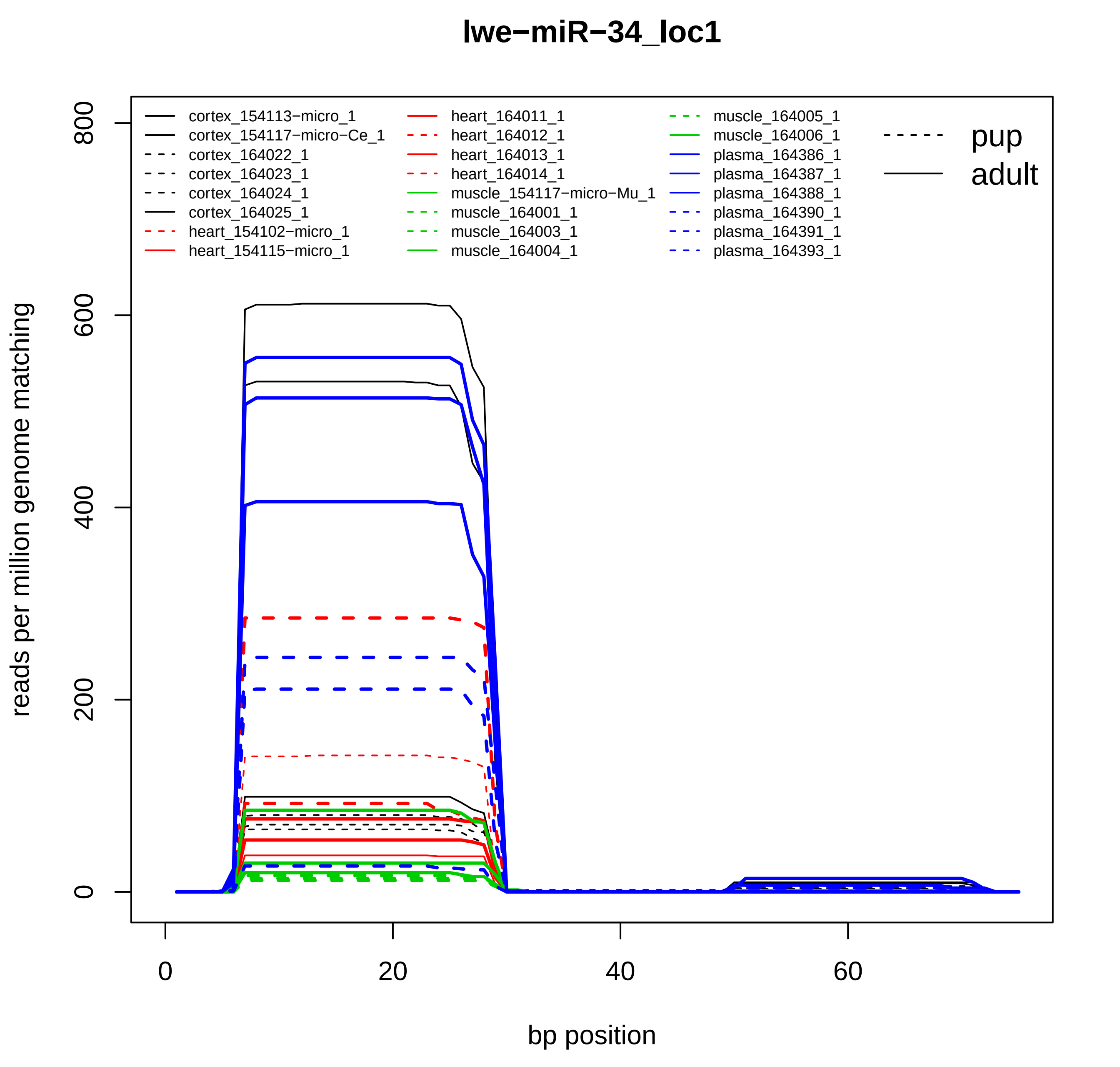
lwe−miR−34_loc1
cortex_154113−micro_1
cortex_154117−micro−Ce_1
cortex_164022_1
cortex_164023_1
cortex_164024_1
cortex_164025_1
heart_154102−micro_1
heart_154115−micro_1
heart_164011_1
heart_164012_1
heart_164013_1
heart_164014_1
muscle_154117−micro−Mu_1
muscle_164001_1
muscle_164003_1
muscle_164004_1
muscle_164005_1
muscle_164006_1
plasma_164386_1
plasma_164387_1
plasma_164388_1
plasma_164390_1
plasma_164391_1
plasma_164393_1
pup
adult
reads per million genome matching
bp position
0
200
400
600
800
0
20
40
60

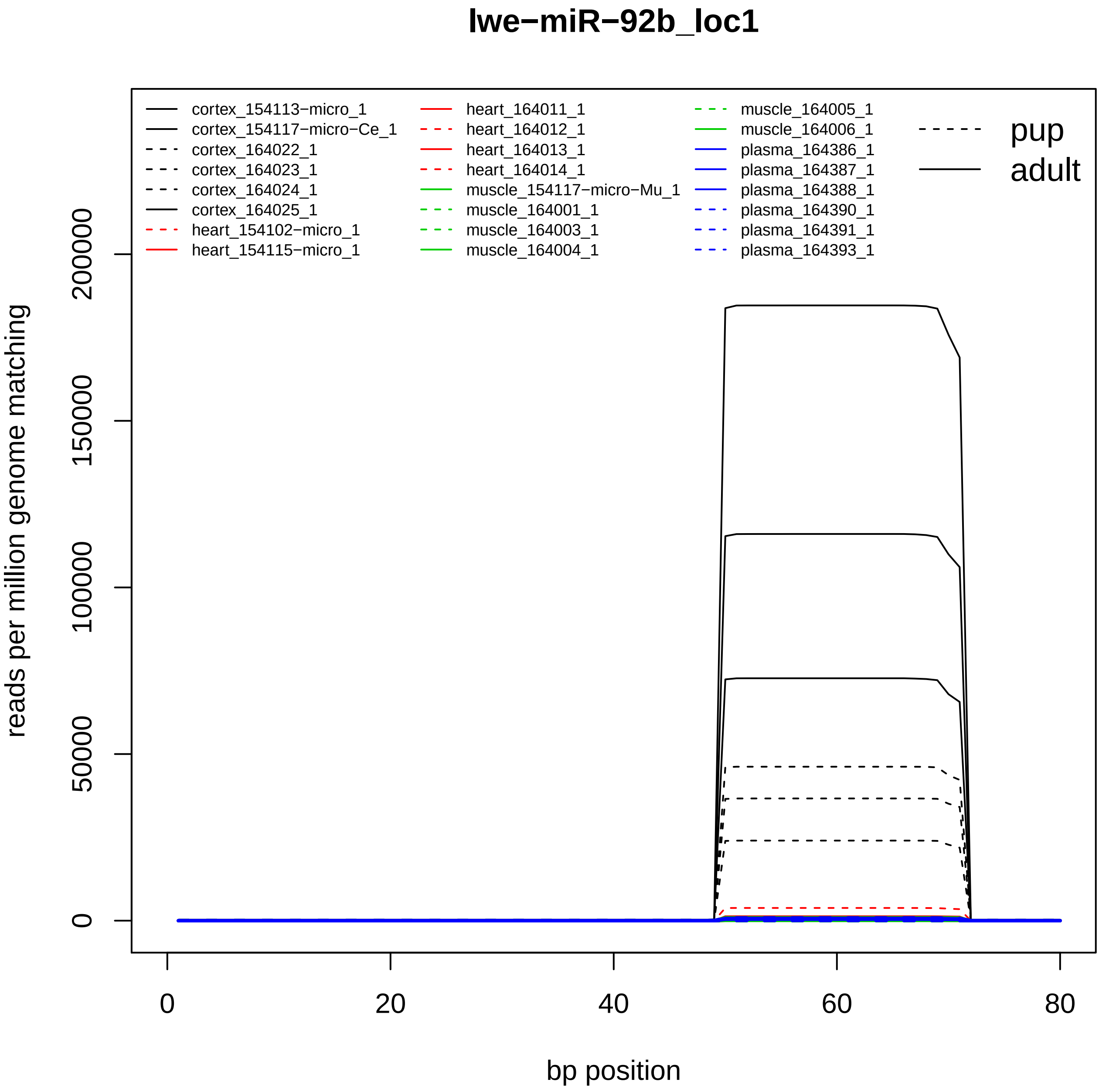
lwe−miR−92b_loc1
cortex_154113−micro_1
cortex_154117−micro−Ce_1
cortex_164022_1
cortex_164023_1
cortex_164024_1
cortex_164025_1
heart_154102−micro_1
heart_154115−micro_1
heart_164011_1
heart_164012_1
heart_164013_1
heart_164014_1
muscle_154117−micro−Mu_1
muscle_164001_1
muscle_164003_1
muscle_164004_1
muscle_164005_1
muscle_164006_1
plasma_164386_1
plasma_164387_1
plasma_164388_1
plasma_164390_1
plasma_164391_1
plasma_164393_1
pup
adult
reads per million genome matching
0
50000
100000
150000
200000
bp position
0
20
40
60
80

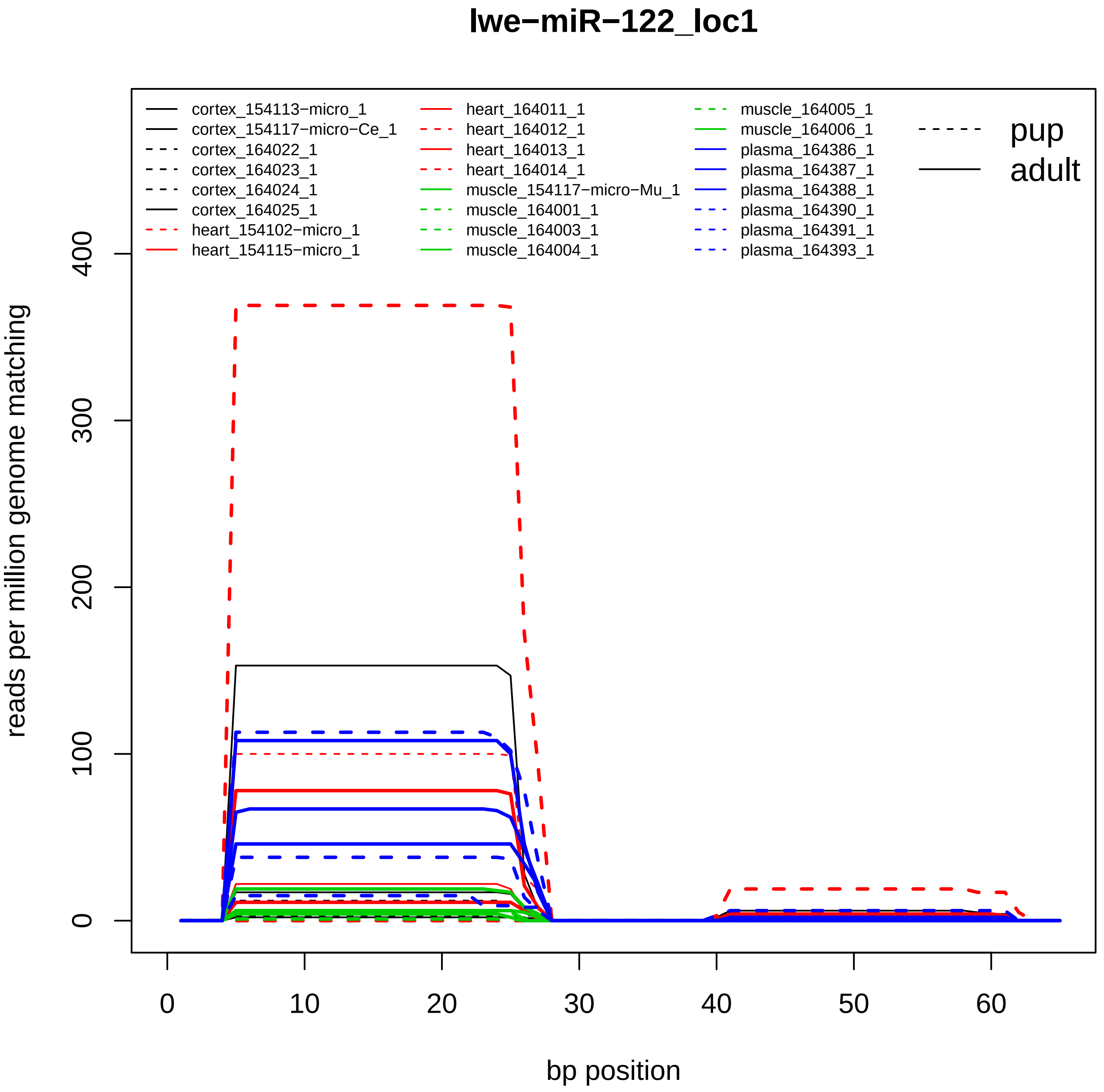

lwe−miR−122_loc1
cortex_154113−micro_1
cortex_154117−micro−Ce_1
cortex_164022_1
cortex_164023_1
cortex_164024_1
cortex_164025_1
heart_154102−micro_1
heart_154115−micro_1
heart_164011_1
heart_164012_1
heart_164013_1
heart_164014_1
muscle_154117−micro−Mu_1
muscle_164001_1
muscle_164003_1
muscle_164004_1
muscle_164005_1
muscle_164006_1
plasma_164386_1
plasma_164387_1
plasma_164388_1
plasma_164390_1
plasma_164391_1
plasma_164393_1
pup
adult
reads per million genome matching
0
100
200
300
400
0
10
20
30
40
50
60
bp position

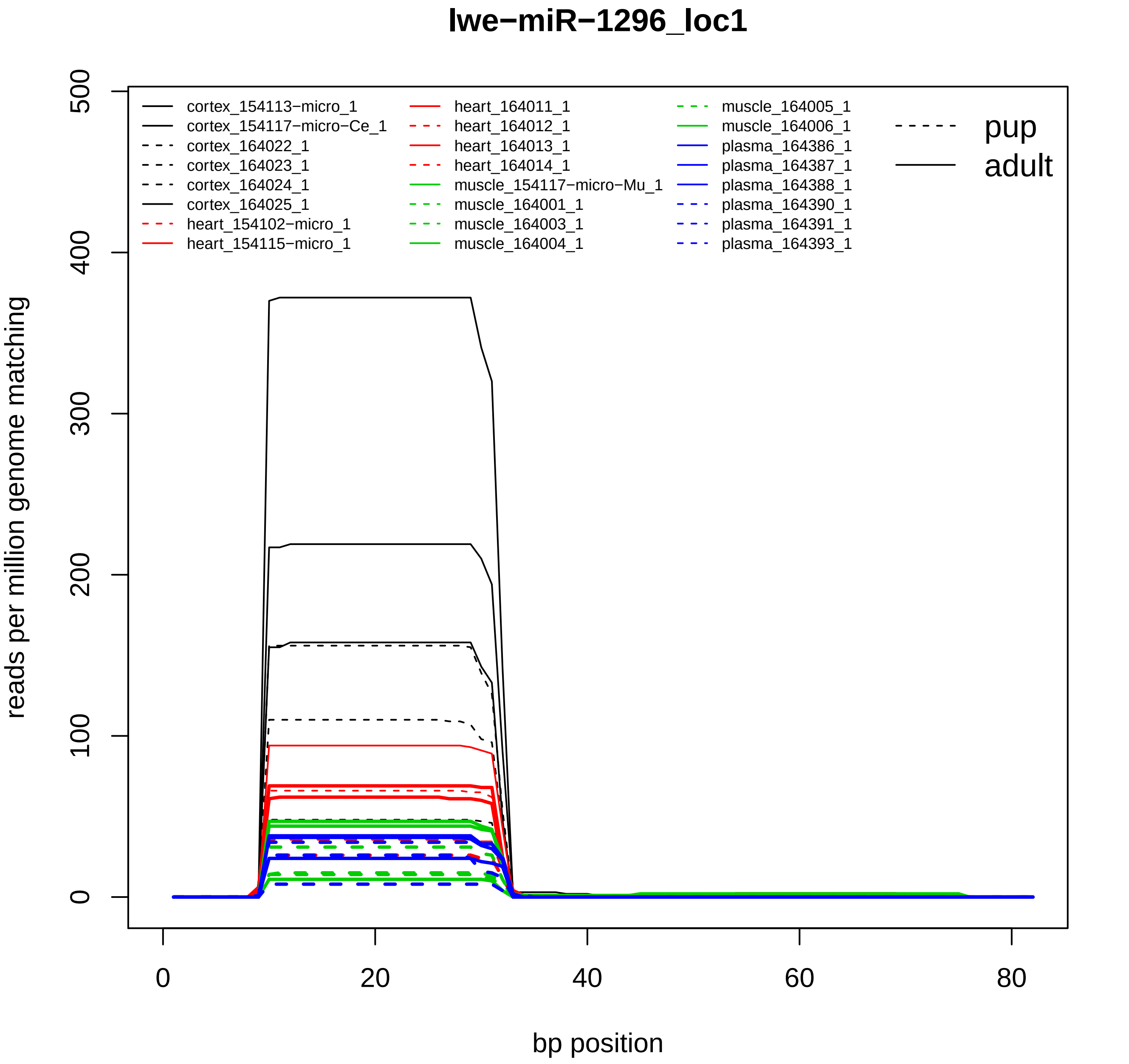

lwe−miR−1296_loc1
reads per million genome matching
bp position
0
100
200
300
400
500
0
20
40
60
80
cortex_154113−micro_1
cortex_154117−micro−Ce_1
cortex_164022_1
cortex_164023_1
cortex_164024_1
cortex_164025_1
heart_154102−micro_1
heart_154115−micro_1
heart_164011_1
heart_164012_1
heart_164013_1
heart_164014_1
muscle_154117−micro−Mu_1
muscle_164001_1
muscle_164003_1
muscle_164004_1
muscle_164005_1
muscle_164006_1
plasma_164386_1
plasma_164387_1
plasma_164388_1
plasma_164390_1
plasma_164391_1
plasma_164393_1
pup
adult

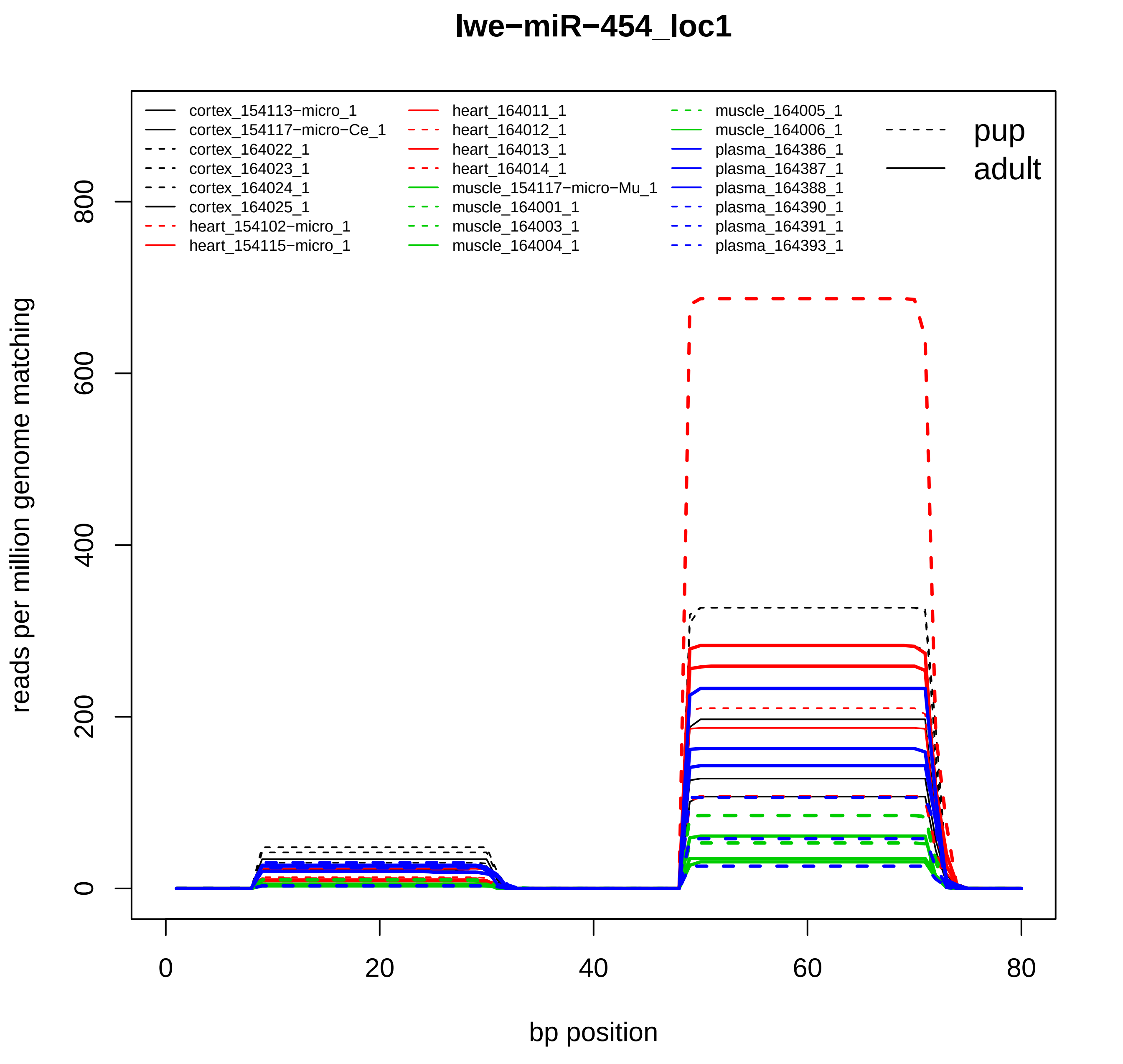

lwe−miR−454_loc1
cortex_154113−micro_1
cortex_154117−micro−Ce_1
cortex_164022_1
cortex_164023_1
cortex_164024_1
cortex_164025_1
heart_154102−micro_1
heart_154115−micro_1
heart_164011_1
heart_164012_1
heart_164013_1
heart_164014_1
muscle_154117−micro−Mu_1
muscle_164001_1
muscle_164003_1
muscle_164004_1
muscle_164005_1
muscle_164006_1
plasma_164386_1
plasma_164387_1
plasma_164388_1
plasma_164390_1
plasma_164391_1
plasma_164393_1
pup
adult
reads per million genome matching
0
200
400
600
800
0
20
40
60
80
bp position

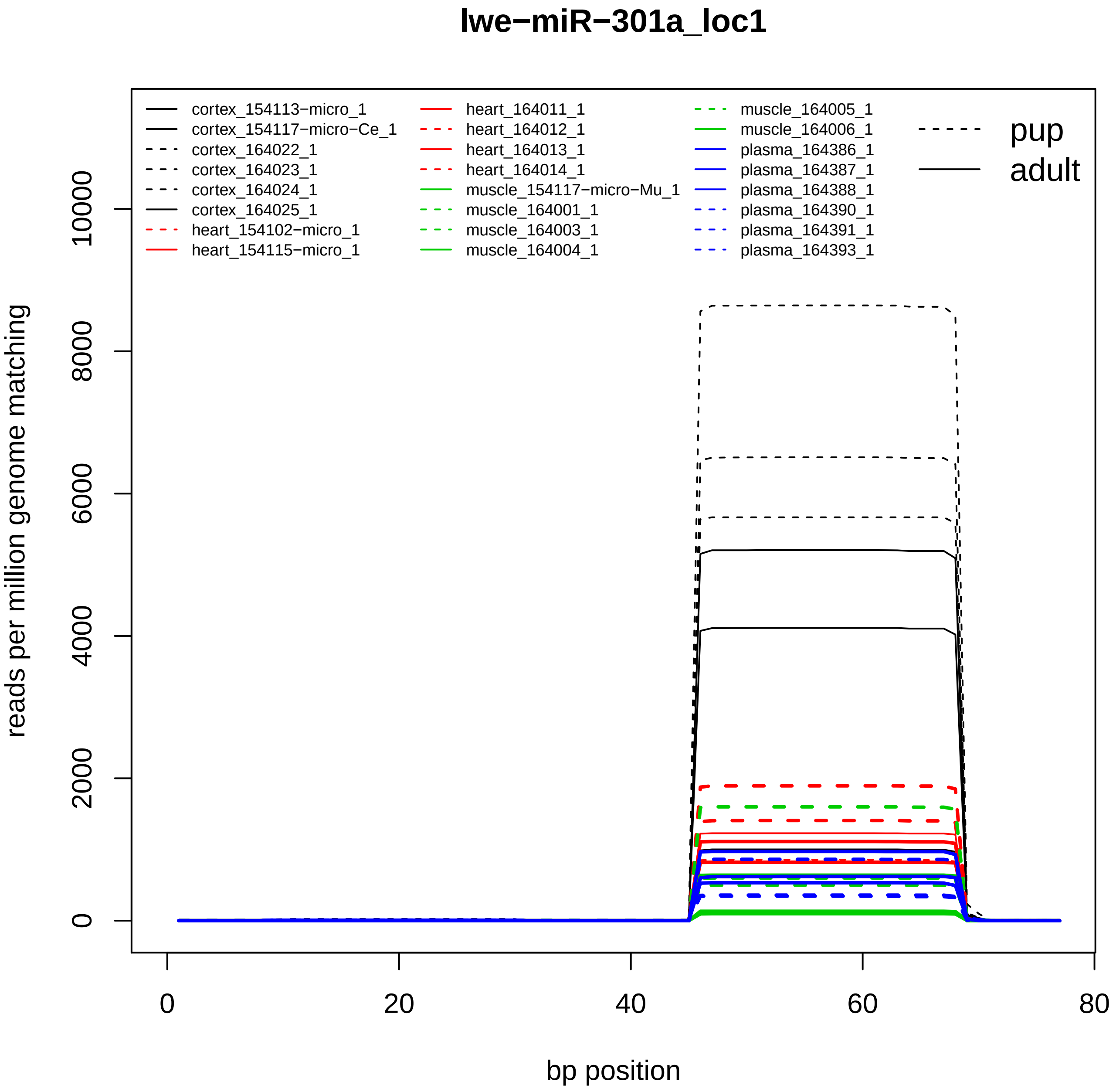

lwe−miR−301a_loc1
cortex_154113−micro_1
cortex_154117−micro−Ce_1
cortex_164022_1
cortex_164023_1
cortex_164024_1
cortex_164025_1
heart_154102−micro_1
heart_154115−micro_1
heart_164011_1
heart_164012_1
heart_164013_1
heart_164014_1
muscle_154117−micro−Mu_1
muscle_164001_1
muscle_164003_1
muscle_164004_1
muscle_164005_1
muscle_164006_1
plasma_164386_1
plasma_164387_1
plasma_164388_1
plasma_164390_1
plasma_164391_1
plasma_164393_1
pup
adult
reads per million genome matching
0
2000
4000
6000
8000
10000
bp position
0
20
40
60
80

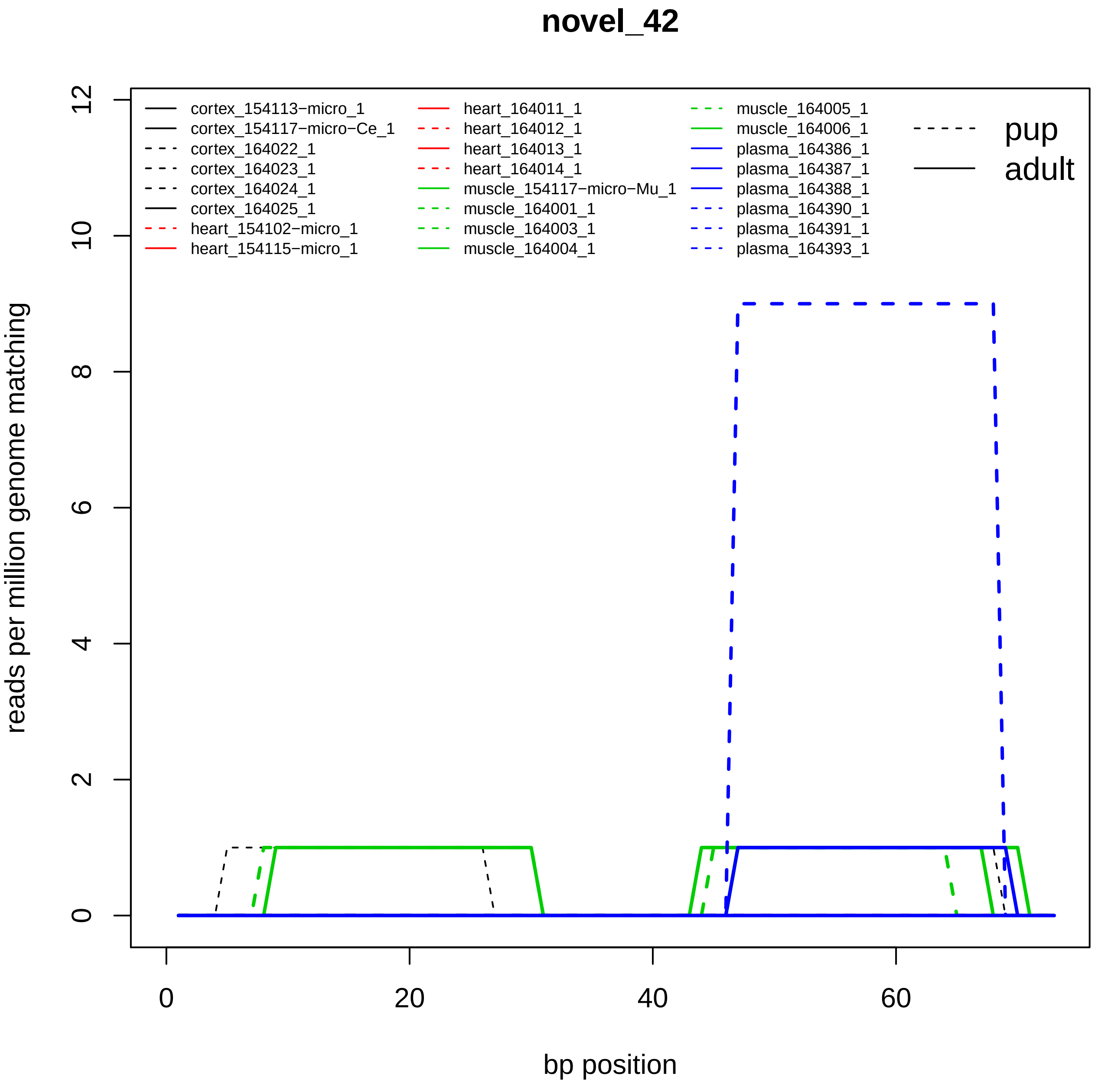
novel_42
cortex_154113−micro_1
cortex_154117−micro−Ce_1
cortex_164022_1
cortex_164023_1
cortex_164024_1
cortex_164025_1
heart_154102−micro_1
heart_154115−micro_1
heart_164011_1
heart_164012_1
heart_164013_1
heart_164014_1
muscle_154117−micro−Mu_1
muscle_164001_1
muscle_164003_1
muscle_164004_1
muscle_164005_1
muscle_164006_1
plasma_164386_1
plasma_164387_1
plasma_164388_1
plasma_164390_1
plasma_164391_1
plasma_164393_1
pup
adult
reads per million genome matching
0
2
4
6
8
10
12
bp position
0
20
40
60

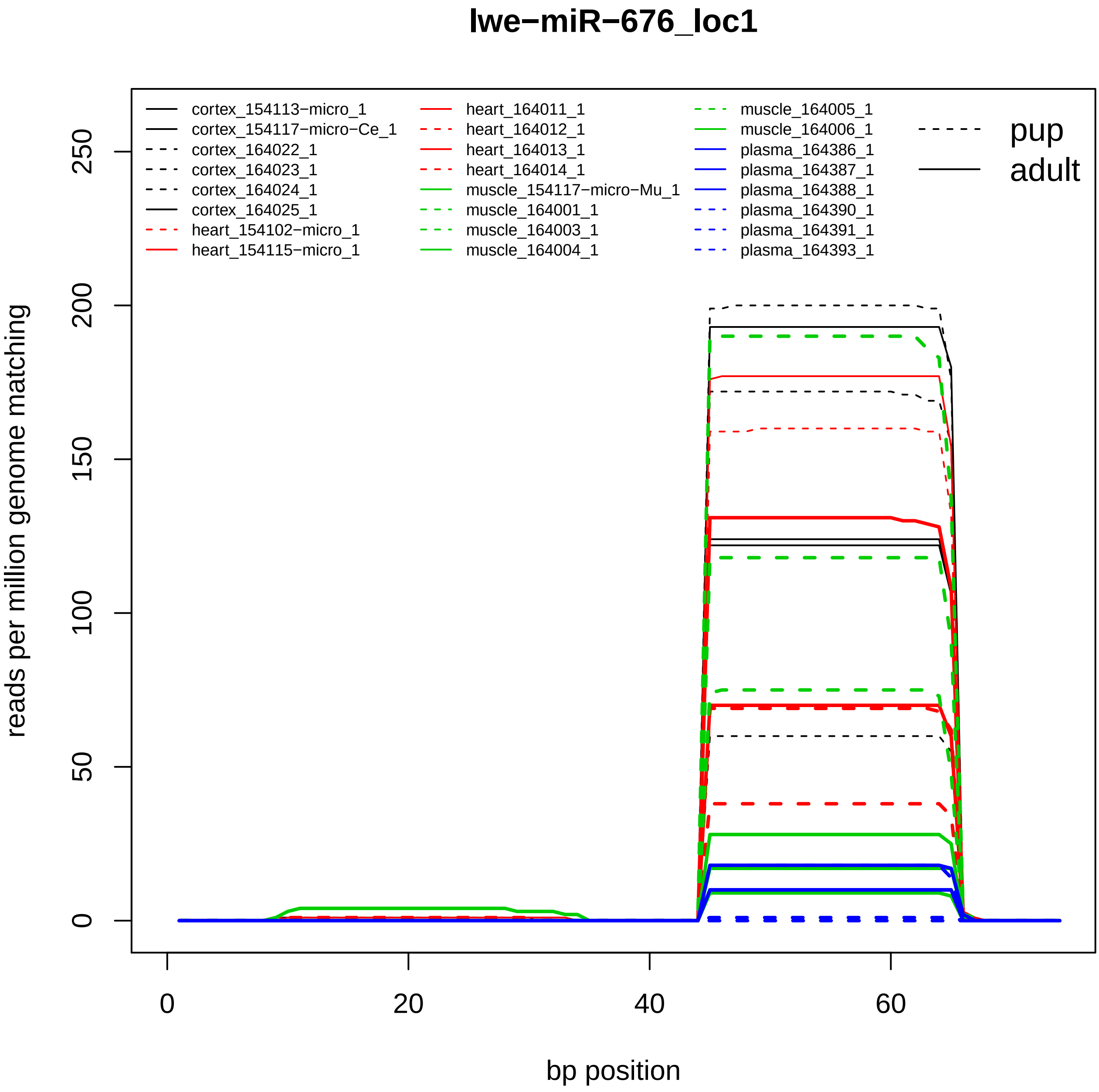

lwe−miR−676_loc1
cortex_154113−micro_1
cortex_154117−micro−Ce_1
cortex_164022_1
cortex_164023_1
cortex_164024_1
cortex_164025_1
heart_154102−micro_1
heart_154115−micro_1
heart_164011_1
heart_164012_1
heart_164013_1
heart_164014_1
muscle_154117−micro−Mu_1
muscle_164001_1
muscle_164003_1
muscle_164004_1
muscle_164005_1
muscle_164006_1
plasma_164386_1
plasma_164387_1
plasma_164388_1
plasma_164390_1
plasma_164391_1
plasma_164393_1
pup
adult
reads per million genome matching
0
50
100
150
200
250
0
20
40
60
bp position

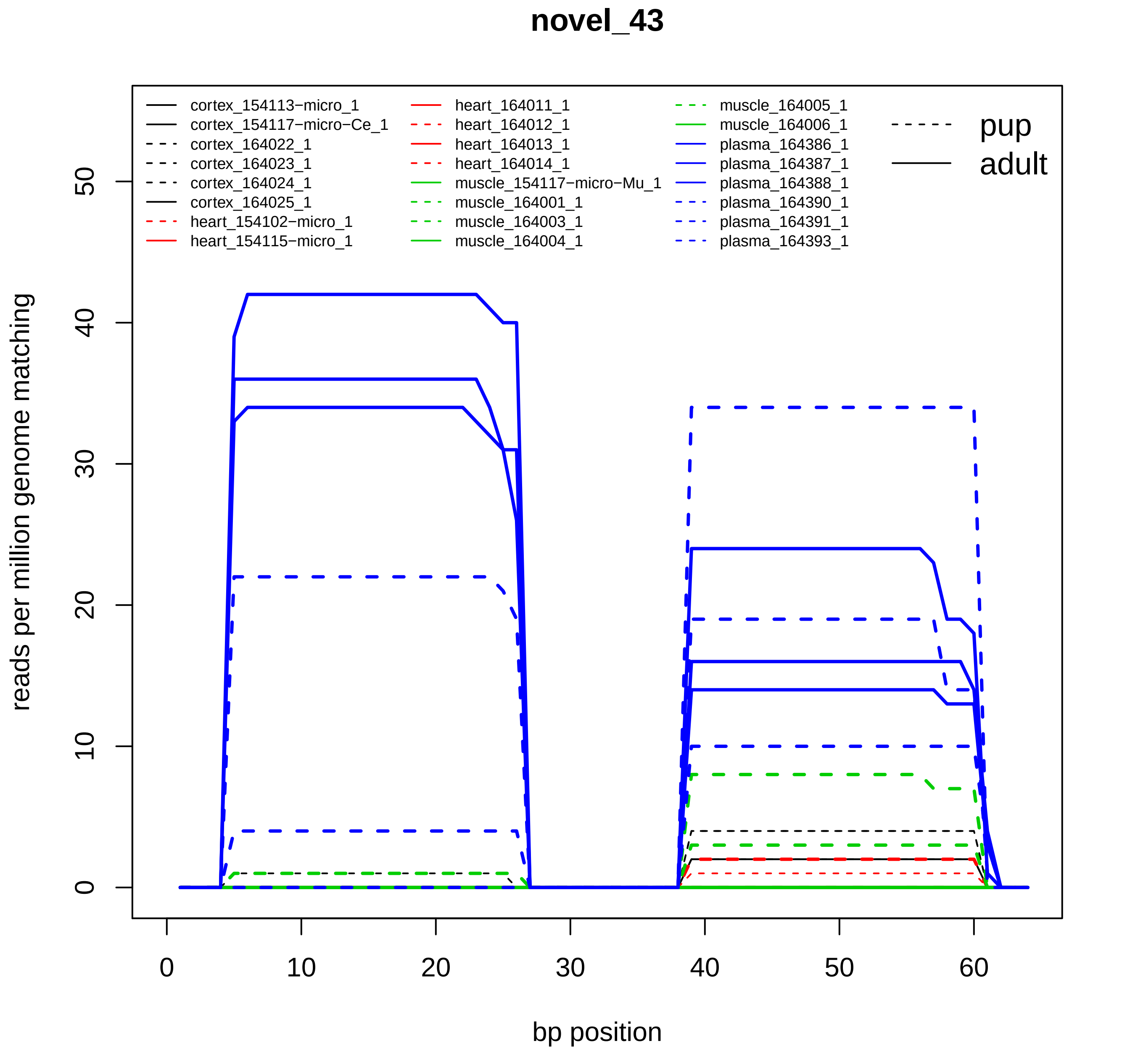

novel_43
cortex_154113−micro_1
cortex_154117−micro−Ce_1
cortex_164022_1
cortex_164023_1
cortex_164024_1
cortex_164025_1
heart_154102−micro_1
heart_154115−micro_1
heart_164011_1
heart_164012_1
heart_164013_1
heart_164014_1
muscle_154117−micro−Mu_1
muscle_164001_1
muscle_164003_1
muscle_164004_1
muscle_164005_1
muscle_164006_1
plasma_164386_1
plasma_164387_1
plasma_164388_1
plasma_164390_1
plasma_164391_1
plasma_164393_1
pup
adult
reads per million genome matching
0
10
20
30
40
50
0
10
20
30
40
50
60
bp position

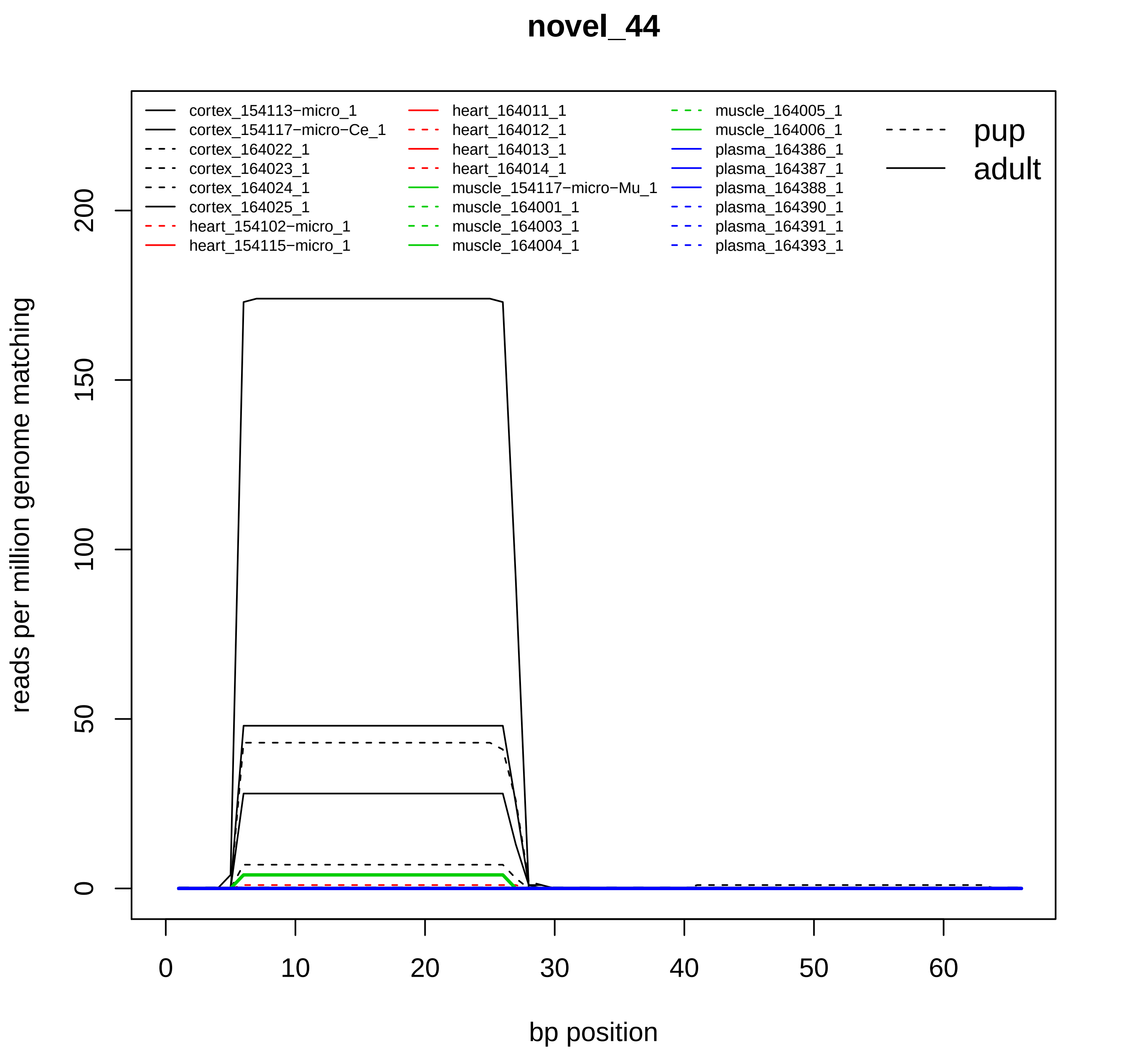
novel_44
cortex_154113−micro_1
cortex_154117−micro−Ce_1
cortex_164022_1
cortex_164023_1
cortex_164024_1
cortex_164025_1
heart_154102−micro_1
heart_154115−micro_1
heart_164011_1
heart_164012_1
heart_164013_1
heart_164014_1
muscle_154117−micro−Mu_1
muscle_164001_1
muscle_164003_1
muscle_164004_1
muscle_164005_1
muscle_164006_1
plasma_164386_1
plasma_164387_1
plasma_164388_1
plasma_164390_1
plasma_164391_1
plasma_164393_1
pup
adult
reads per million genome matching
0
50
100
150
200
0
10
20
30
40
50
60
bp position

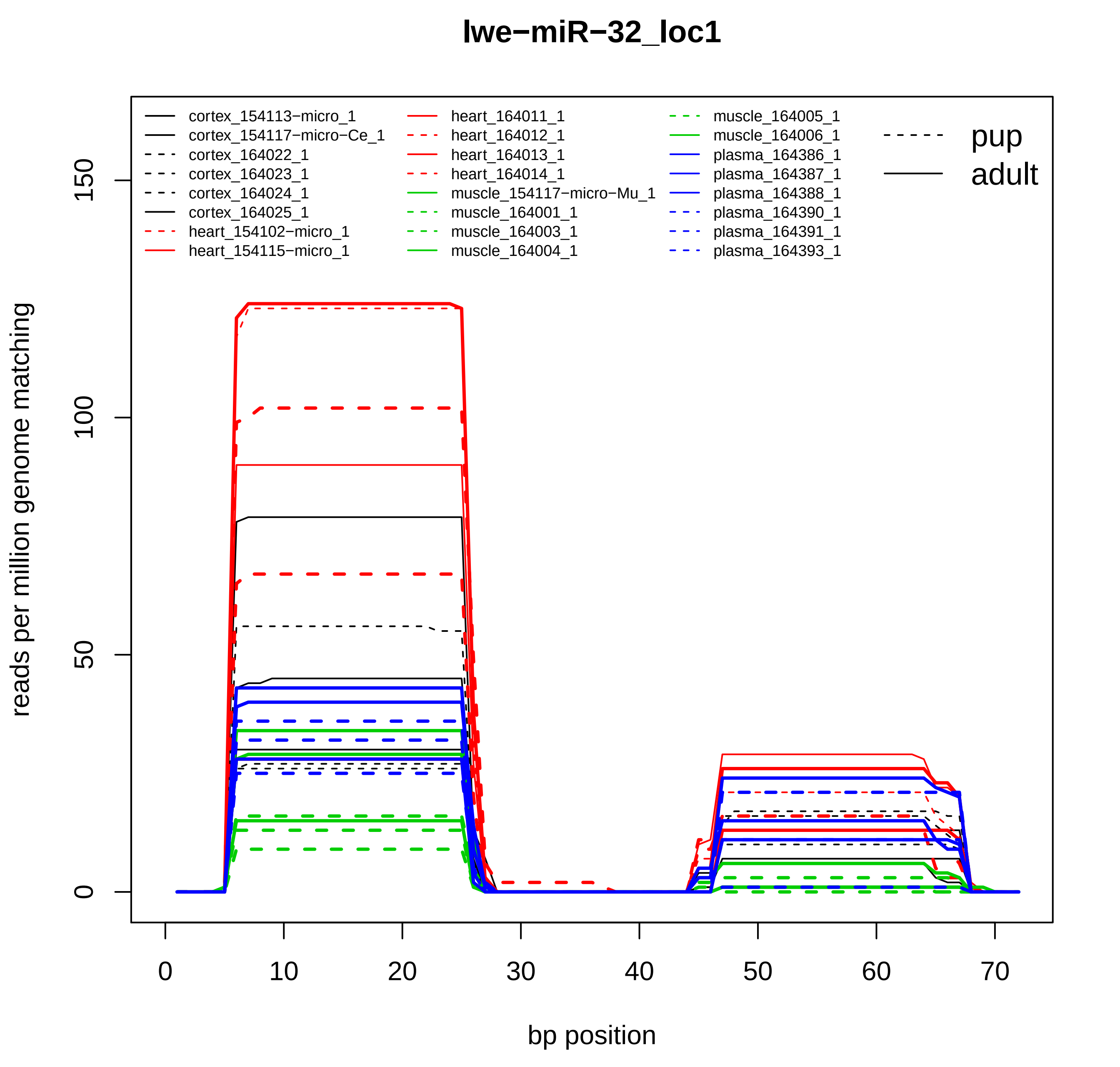
lwe−miR−32_loc1
cortex_154113−micro_1
cortex_154117−micro−Ce_1
cortex_164022_1
cortex_164023_1
cortex_164024_1
cortex_164025_1
heart_154102−micro_1
heart_154115−micro_1
heart_164011_1
heart_164012_1
heart_164013_1
heart_164014_1
muscle_154117−micro−Mu_1
muscle_164001_1
muscle_164003_1
muscle_164004_1
muscle_164005_1
muscle_164006_1
plasma_164386_1
plasma_164387_1
plasma_164388_1
plasma_164390_1
plasma_164391_1
plasma_164393_1
pup
adult
reads per million genome matching
0
50
100
150
bp position
0
10
20
30
40
50
60
70

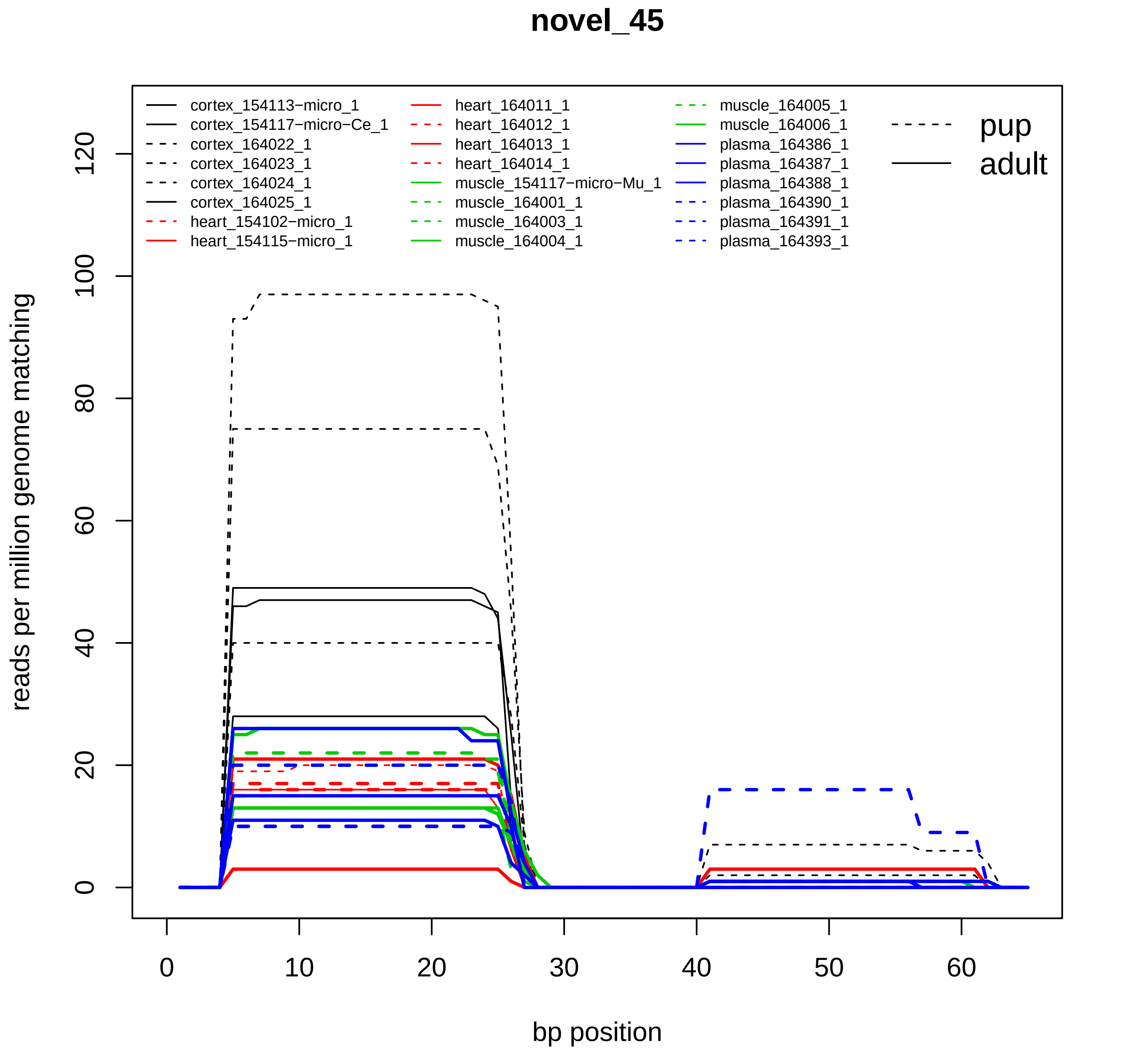
novel_45
cortex_154113−micro_1
cortex_154117−micro−Ce_1
cortex_164022_1
cortex_164023_1
cortex_164024_1
cortex_164025_1
heart_154102−micro_1
heart_154115−micro_1
heart_164011_1
heart_164012_1
heart_164013_1
heart_164014_1
muscle_154117−micro−Mu_1
muscle_164001_1
muscle_164003_1
muscle_164004_1
muscle_164005_1
muscle_164006_1
plasma_164386_1
plasma_164387_1
plasma_164388_1
plasma_164390_1
plasma_164391_1
plasma_164393_1
pup
adult
reads per million genome matching
bp position
0
20
40
60
80
100
120
0
10
20
30
40
50
60

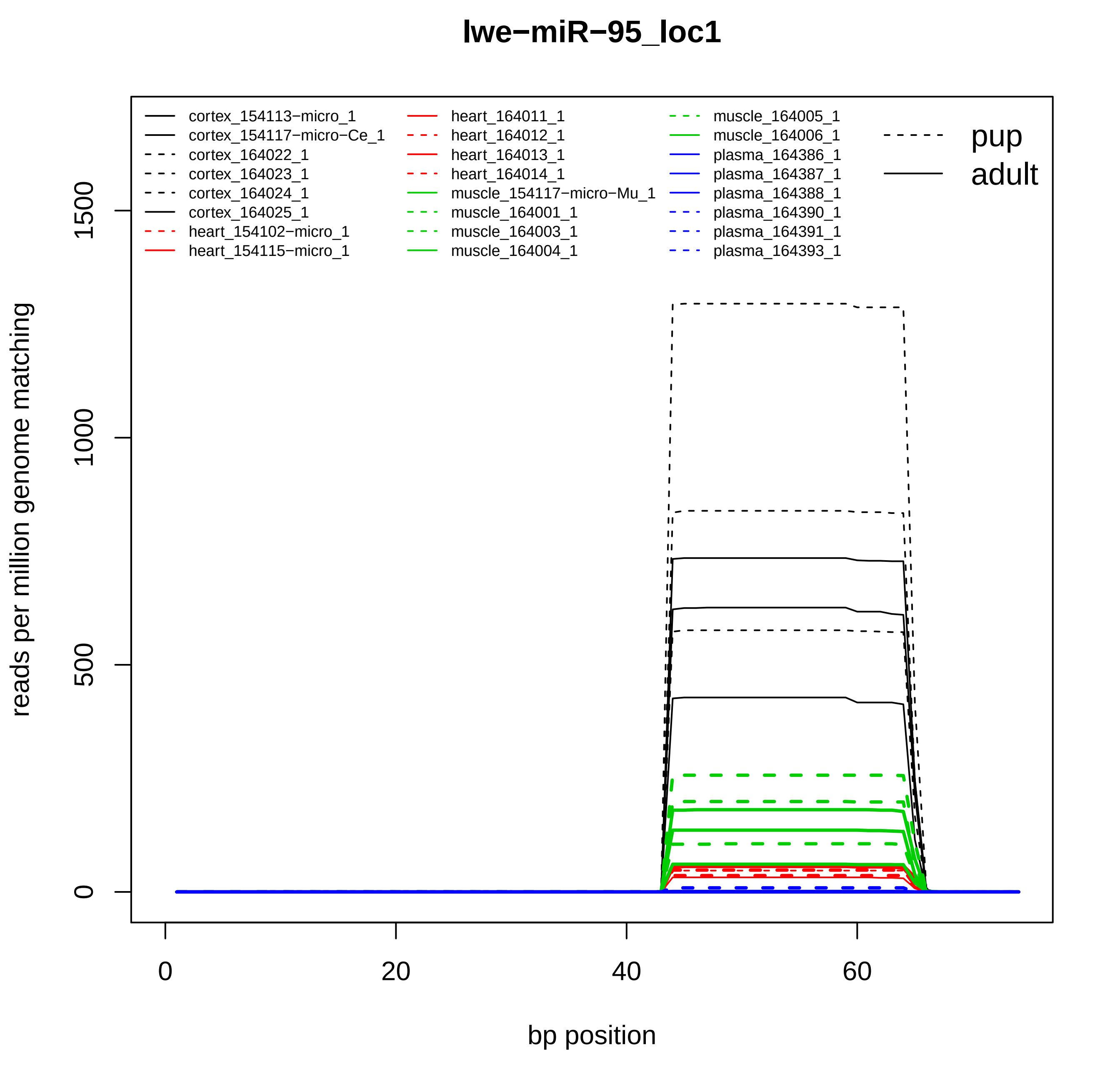
lwe−miR−95_loc1
cortex_154113−micro_1
cortex_154117−micro−Ce_1
cortex_164022_1
cortex_164023_1
cortex_164024_1
cortex_164025_1
heart_154102−micro_1
heart_154115−micro_1
heart_164011_1
heart_164012_1
heart_164013_1
heart_164014_1
muscle_154117−micro−Mu_1
muscle_164001_1
muscle_164003_1
muscle_164004_1
muscle_164005_1
muscle_164006_1
plasma_164386_1
plasma_164387_1
plasma_164388_1
plasma_164390_1
plasma_164391_1
plasma_164393_1
pup
adult
reads per million genome matching
0
500
1000
1500
0
20
40
60
bp position

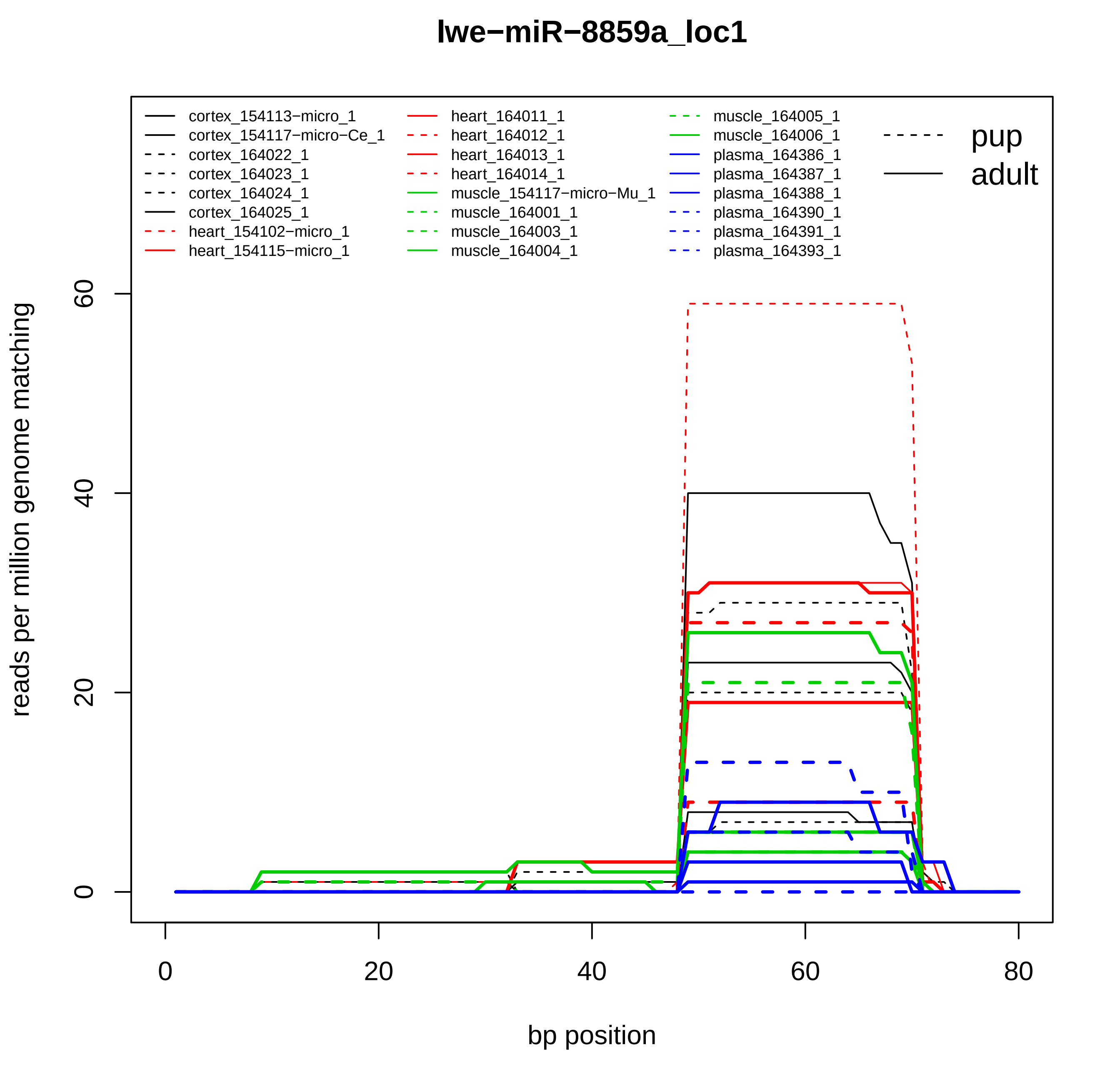

lwe−miR−8859a_loc1
cortex_154113−micro_1
cortex_154117−micro−Ce_1
cortex_164022_1
cortex_164023_1
cortex_164024_1
cortex_164025_1
heart_154102−micro_1
heart_154115−micro_1
heart_164011_1
heart_164012_1
heart_164013_1
heart_164014_1
muscle_154117−micro−Mu_1
muscle_164001_1
muscle_164003_1
muscle_164004_1
muscle_164005_1
muscle_164006_1
plasma_164386_1
plasma_164387_1
plasma_164388_1
plasma_164390_1
plasma_164391_1
plasma_164393_1
pup
adult
reads per million genome matching
0
20
40
60
0
20
40
60
80
bp position

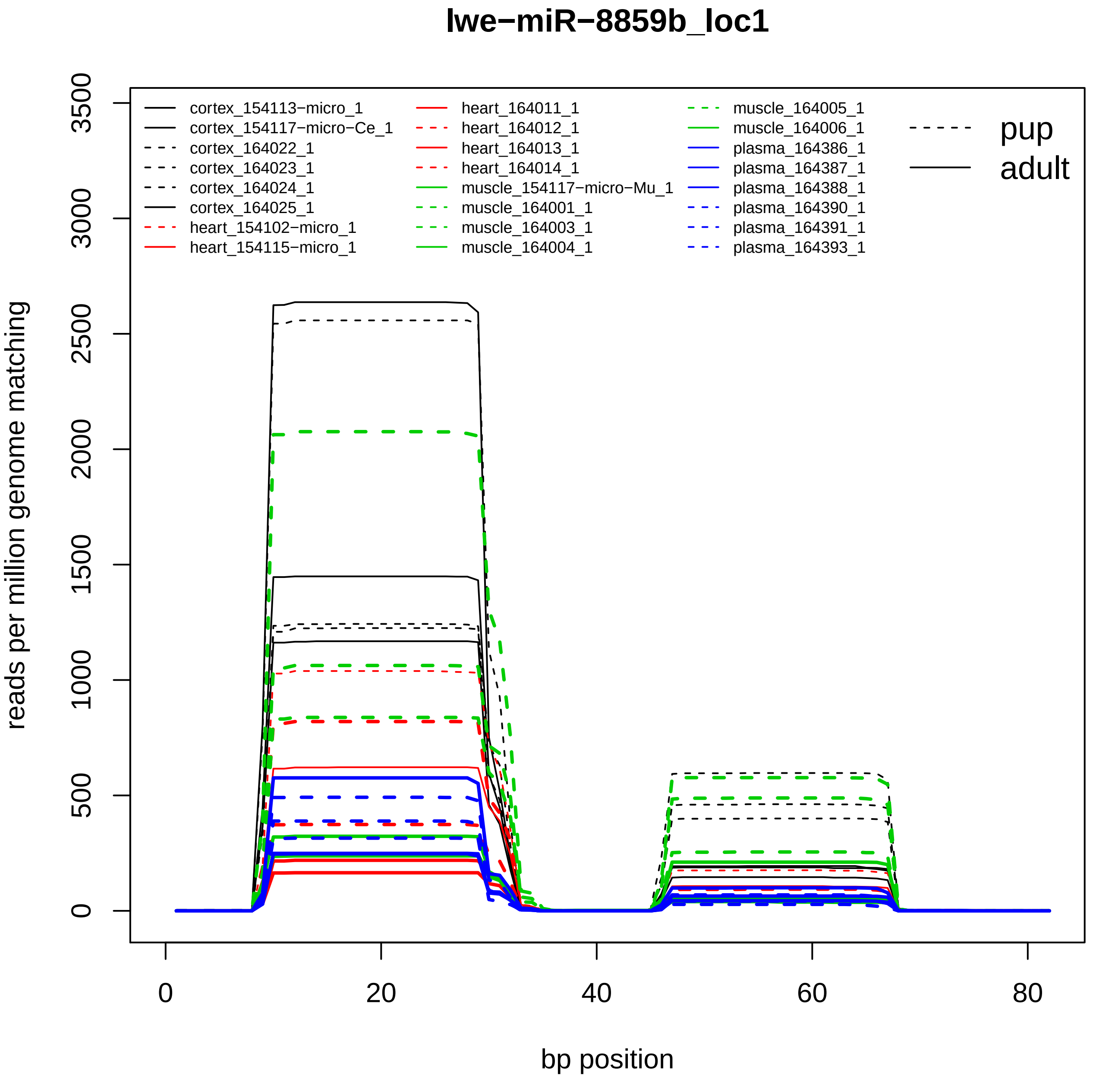

lwe−miR−8859b_loc1
cortex_154113−micro_1
cortex_154117−micro−Ce_1
cortex_164022_1
cortex_164023_1
cortex_164024_1
cortex_164025_1
heart_154102−micro_1
heart_154115−micro_1
heart_164011_1
heart_164012_1
heart_164013_1
heart_164014_1
muscle_154117−micro−Mu_1
muscle_164001_1
muscle_164003_1
muscle_164004_1
muscle_164005_1
muscle_164006_1
plasma_164386_1
plasma_164387_1
plasma_164388_1
plasma_164390_1
plasma_164391_1
plasma_164393_1
pup
adult
reads per million genome matching
0
500
1000
1500
2000
2500
3000
3500
bp position
0
20
40
60
80

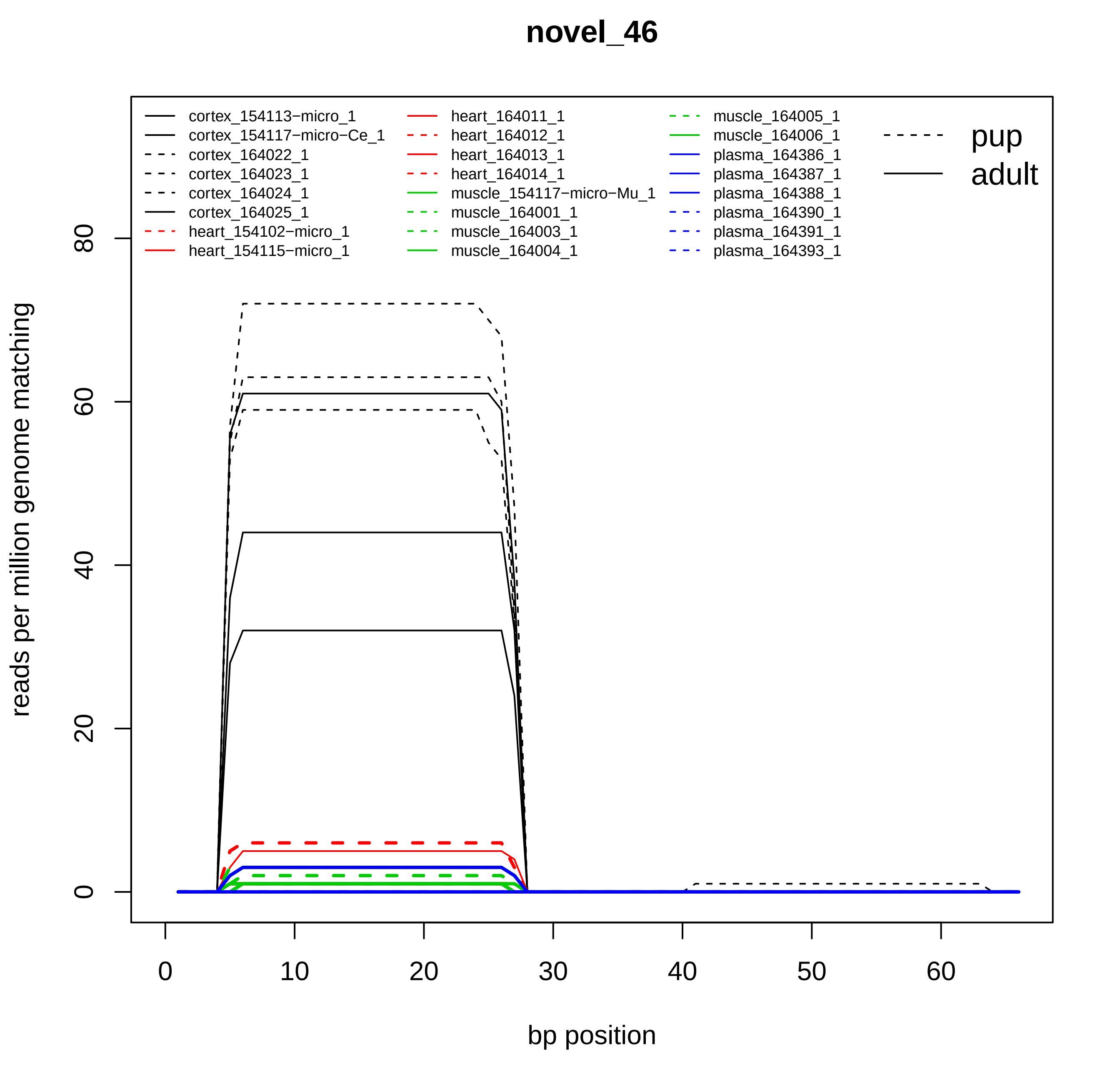

novel_46
cortex_154113−micro_1
cortex_154117−micro−Ce_1
cortex_164022_1
cortex_164023_1
cortex_164024_1
cortex_164025_1
heart_154102−micro_1
heart_154115−micro_1
heart_164011_1
heart_164012_1
heart_164013_1
heart_164014_1
muscle_154117−micro−Mu_1
muscle_164001_1
muscle_164003_1
muscle_164004_1
muscle_164005_1
muscle_164006_1
plasma_164386_1
plasma_164387_1
plasma_164388_1
plasma_164390_1
plasma_164391_1
plasma_164393_1
pup
adult
reads per million genome matching
0
20
40
60
80
0
10
20
30
40
50
60
bp position

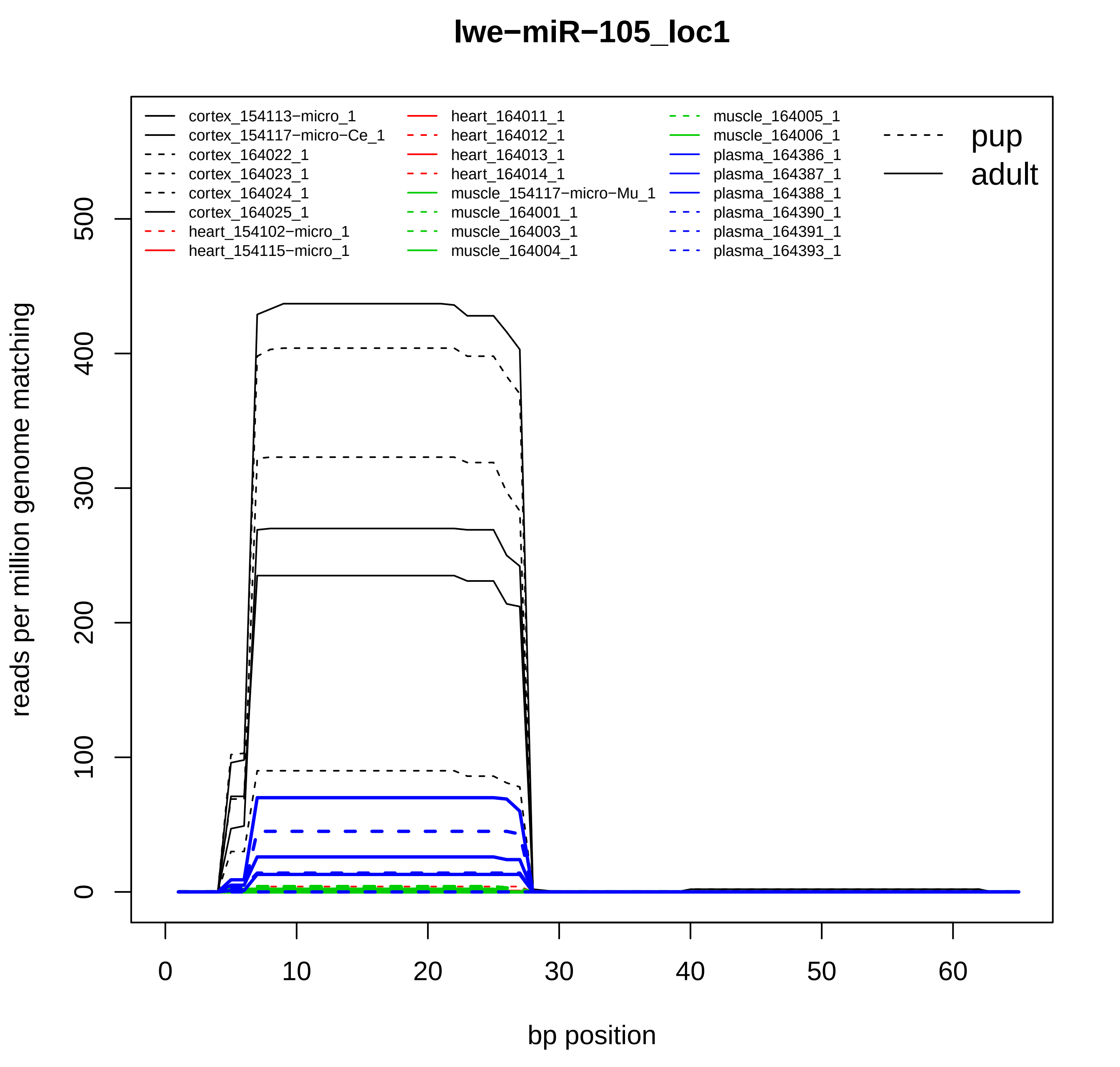

lwe−miR−105_loc1
cortex_154113−micro_1
cortex_154117−micro−Ce_1
cortex_164022_1
cortex_164023_1
cortex_164024_1
cortex_164025_1
heart_154102−micro_1
heart_154115−micro_1
heart_164011_1
heart_164012_1
heart_164013_1
heart_164014_1
muscle_154117−micro−Mu_1
muscle_164001_1
muscle_164003_1
muscle_164004_1
muscle_164005_1
muscle_164006_1
plasma_164386_1
plasma_164387_1
plasma_164388_1
plasma_164390_1
plasma_164391_1
plasma_164393_1
pup
adult
reads per million genome matching
0
100
200
300
400
500
0
10
20
30
40
50
60
bp position

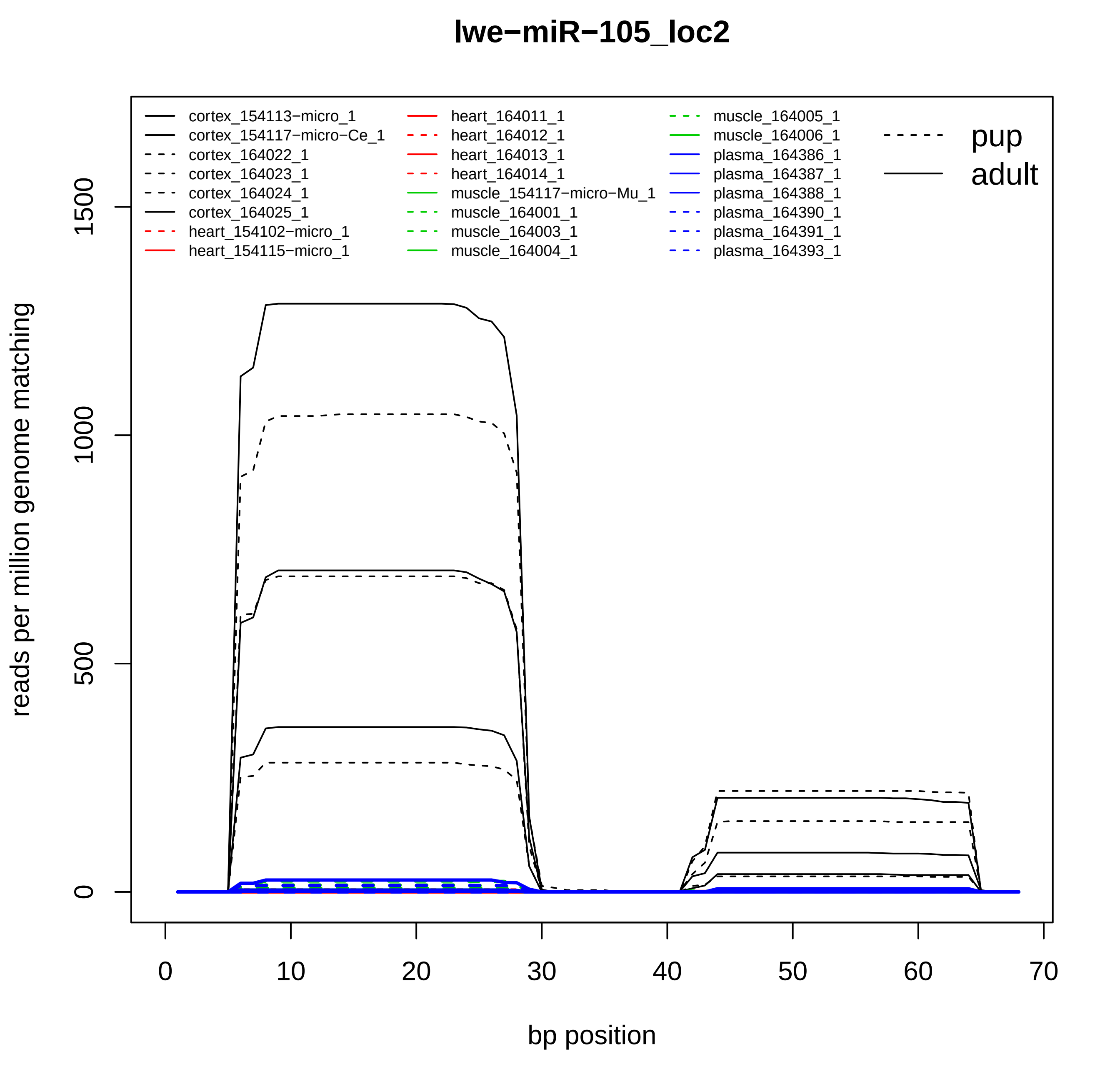
lwe−miR−105_loc2
cortex_154113−micro_1
cortex_154117−micro−Ce_1
cortex_164022_1
cortex_164023_1
cortex_164024_1
cortex_164025_1
heart_154102−micro_1
heart_154115−micro_1
heart_164011_1
heart_164012_1
heart_164013_1
heart_164014_1
muscle_154117−micro−Mu_1
muscle_164001_1
muscle_164003_1
muscle_164004_1
muscle_164005_1
muscle_164006_1
plasma_164386_1
plasma_164387_1
plasma_164388_1
plasma_164390_1
plasma_164391_1
plasma_164393_1
pup
adult
reads per million genome matching
0
500
1000
1500
bp position
0
10
20
30
40
50
60
70

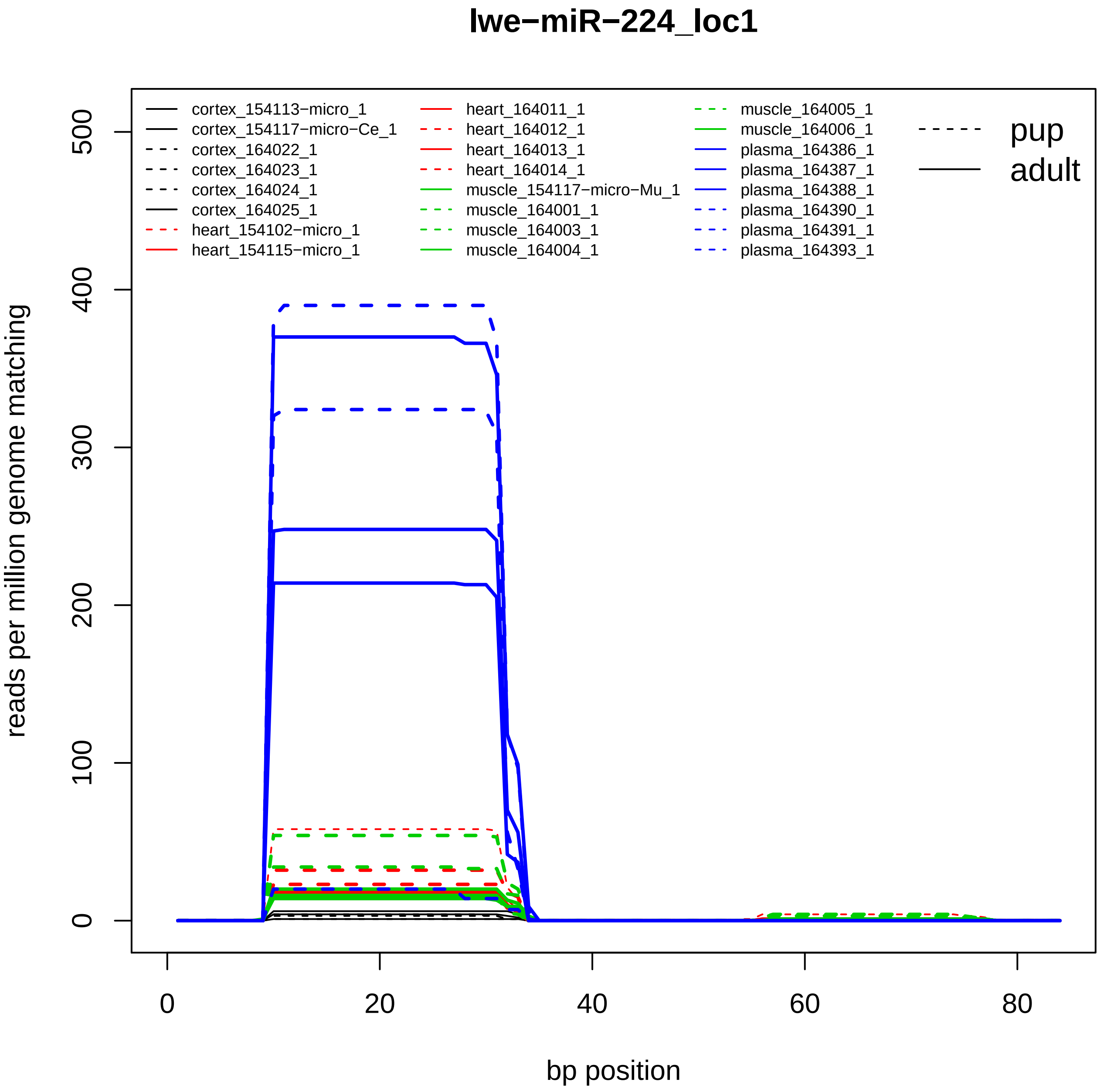

lwe−miR−224_loc1
cortex_154113−micro_1
cortex_154117−micro−Ce_1
cortex_164022_1
cortex_164023_1
cortex_164024_1
cortex_164025_1
heart_154102−micro_1
heart_154115−micro_1
heart_164011_1
heart_164012_1
heart_164013_1
heart_164014_1
muscle_154117−micro−Mu_1
muscle_164001_1
muscle_164003_1
muscle_164004_1
muscle_164005_1
muscle_164006_1
plasma_164386_1
plasma_164387_1
plasma_164388_1
plasma_164390_1
plasma_164391_1
plasma_164393_1
pup
adult
reads per million genome matching
0
100
200
300
400
500
0
20
40
60
80
bp position

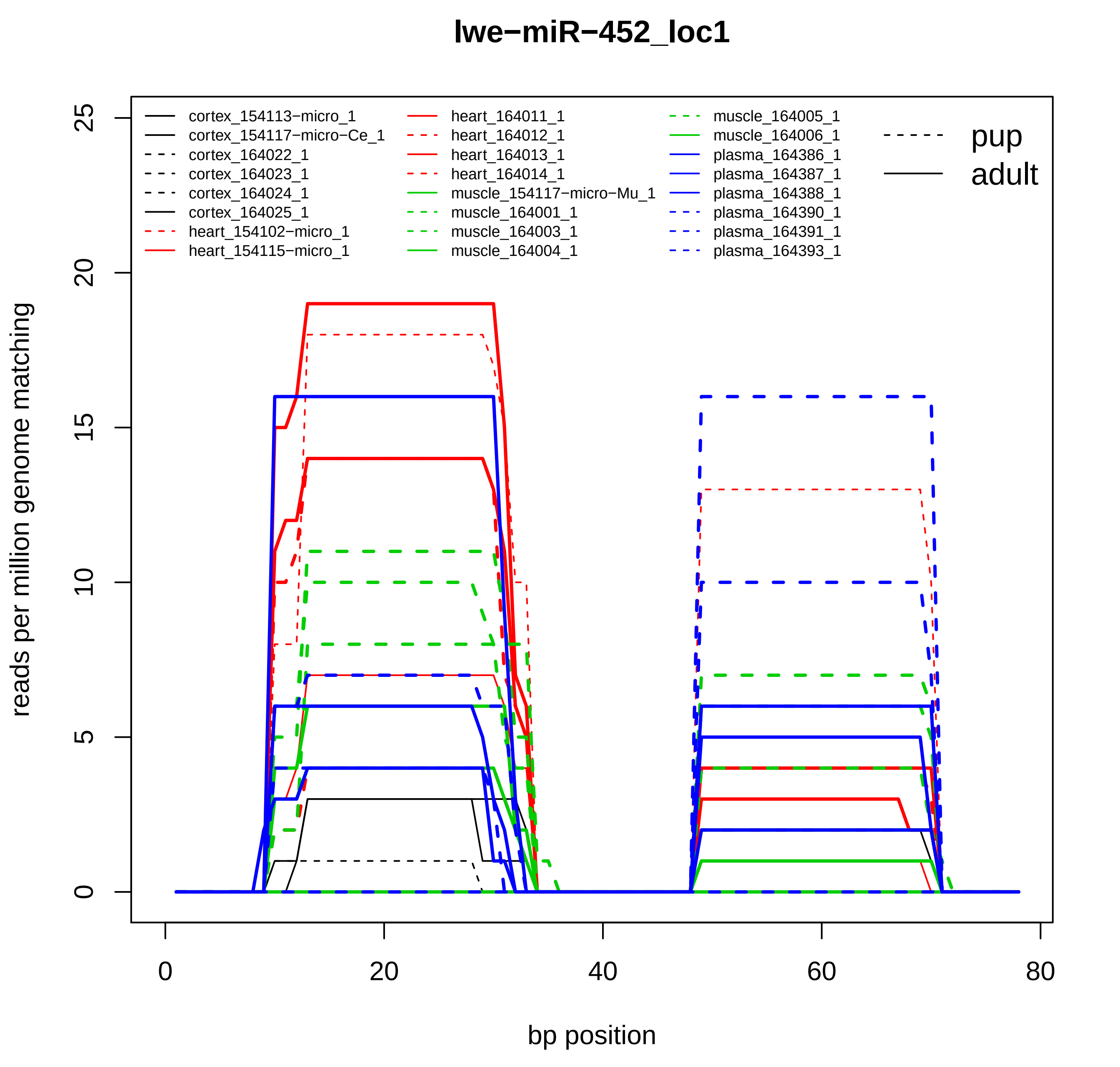

lwe−miR−452_loc1
cortex_154113−micro_1
cortex_154117−micro−Ce_1
cortex_164022_1
cortex_164023_1
cortex_164024_1
cortex_164025_1
heart_154102−micro_1
heart_154115−micro_1
heart_164011_1
heart_164012_1
heart_164013_1
heart_164014_1
muscle_154117−micro−Mu_1
muscle_164001_1
muscle_164003_1
muscle_164004_1
muscle_164005_1
muscle_164006_1
plasma_164386_1
plasma_164387_1
plasma_164388_1
plasma_164390_1
plasma_164391_1
plasma_164393_1
pup
adult
reads per million genome matching
0
5
10
15
20
25
0
20
40
60
80
bp position

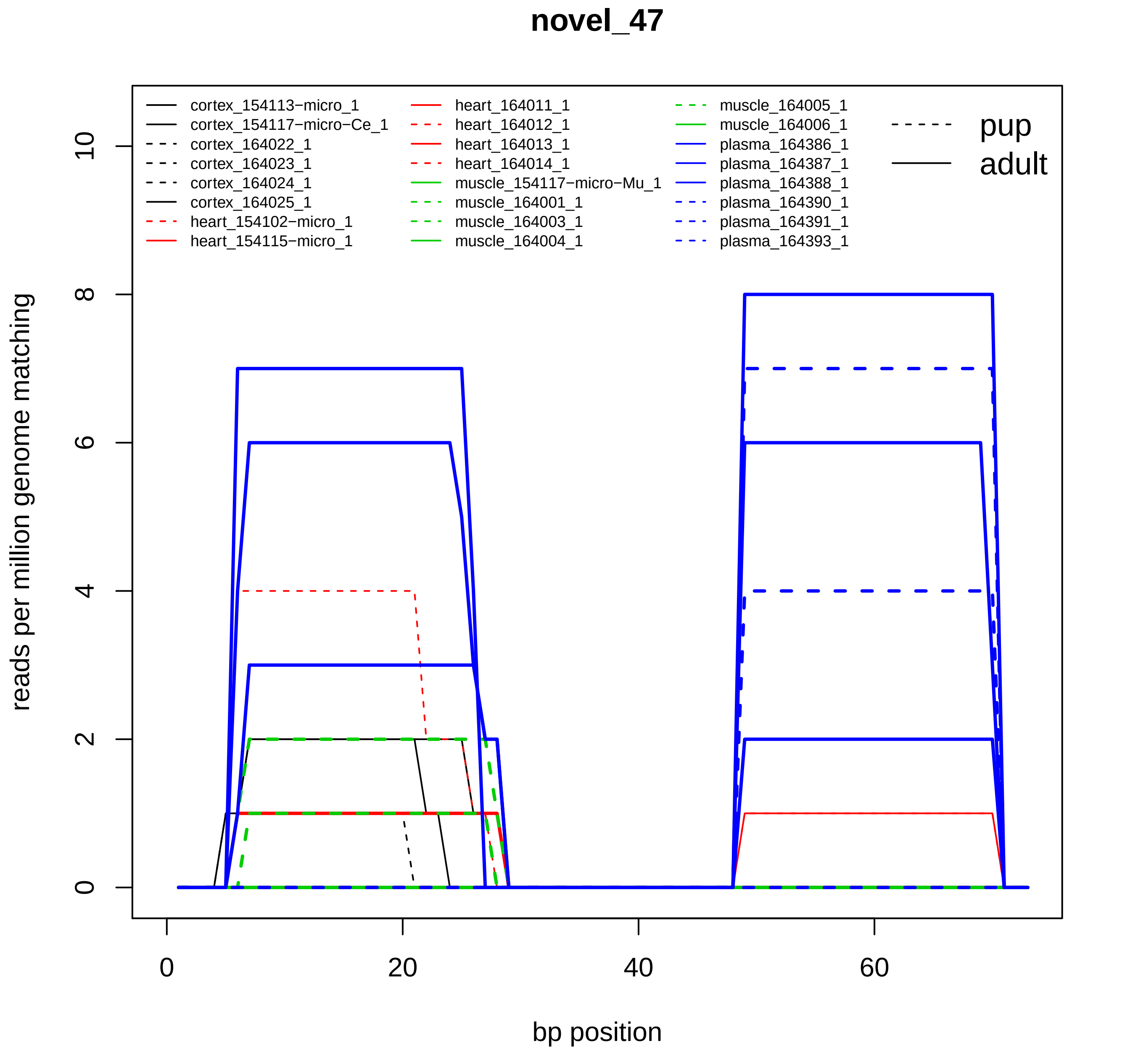
novel_47
cortex_154113−micro_1
cortex_154117−micro−Ce_1
cortex_164022_1
cortex_164023_1
cortex_164024_1
cortex_164025_1
heart_154102−micro_1
heart_154115−micro_1
heart_164011_1
heart_164012_1
heart_164013_1
heart_164014_1
muscle_154117−micro−Mu_1
muscle_164001_1
muscle_164003_1
muscle_164004_1
muscle_164005_1
muscle_164006_1
plasma_164386_1
plasma_164387_1
plasma_164388_1
plasma_164390_1
plasma_164391_1
plasma_164393_1
pup
adult
reads per million genome matching
bp position
0
2
4
6
8
10
0
20
40
60

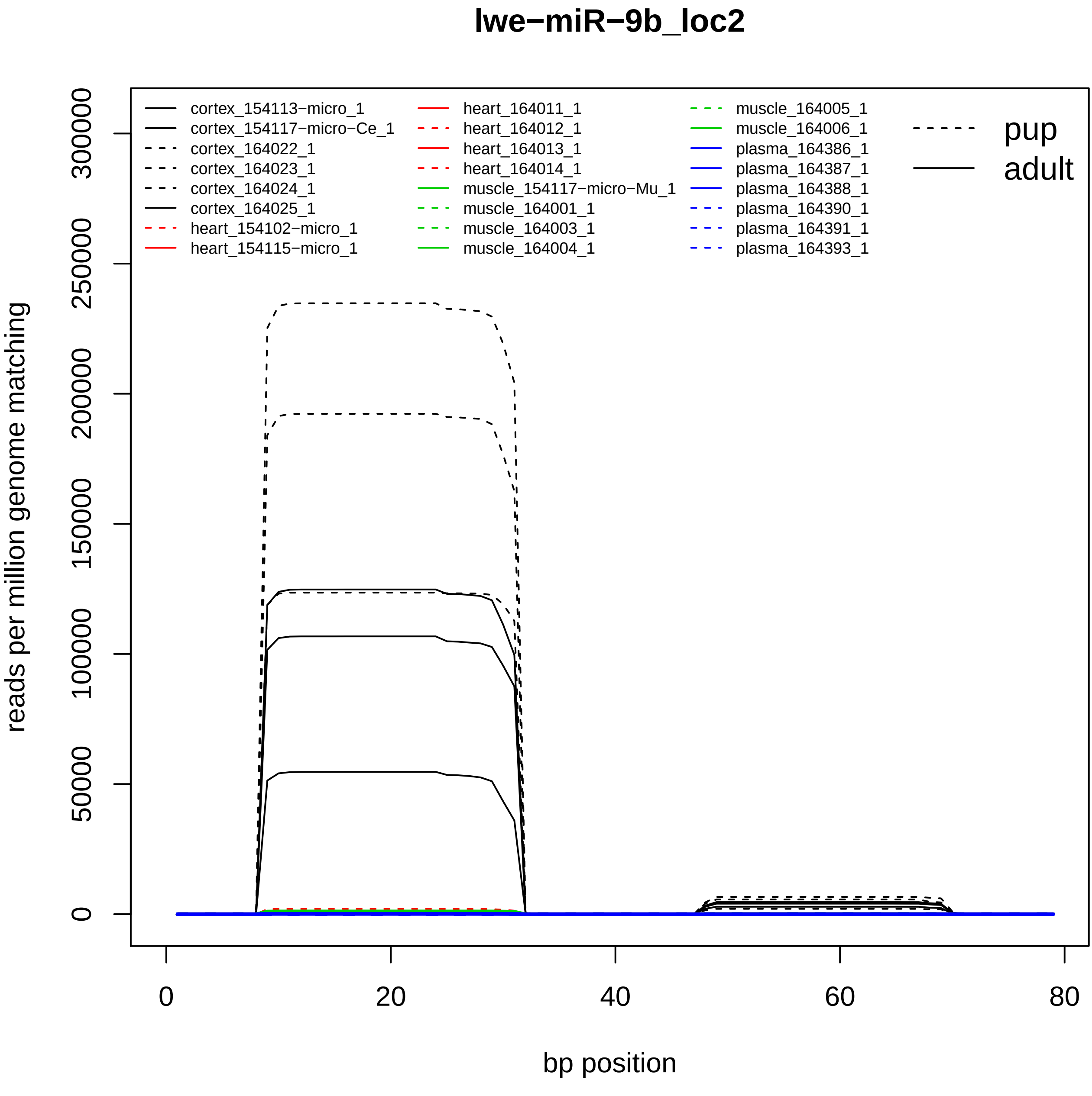
lwe−miR−9b_loc2
reads per million genome matching
bp position
0
50000
100000
150000
200000
250000
300000
0
20
40
60
80
cortex_154113−micro_1
cortex_154117−micro−Ce_1
cortex_164022_1
cortex_164023_1
cortex_164024_1
cortex_164025_1
heart_154102−micro_1
heart_154115−micro_1
heart_164011_1
heart_164012_1
heart_164013_1
heart_164014_1
muscle_154117−micro−Mu_1
muscle_164001_1
muscle_164003_1
muscle_164004_1
muscle_164005_1
muscle_164006_1
plasma_164386_1
plasma_164387_1
plasma_164388_1
plasma_164390_1
plasma_164391_1
plasma_164393_1
pup
adult

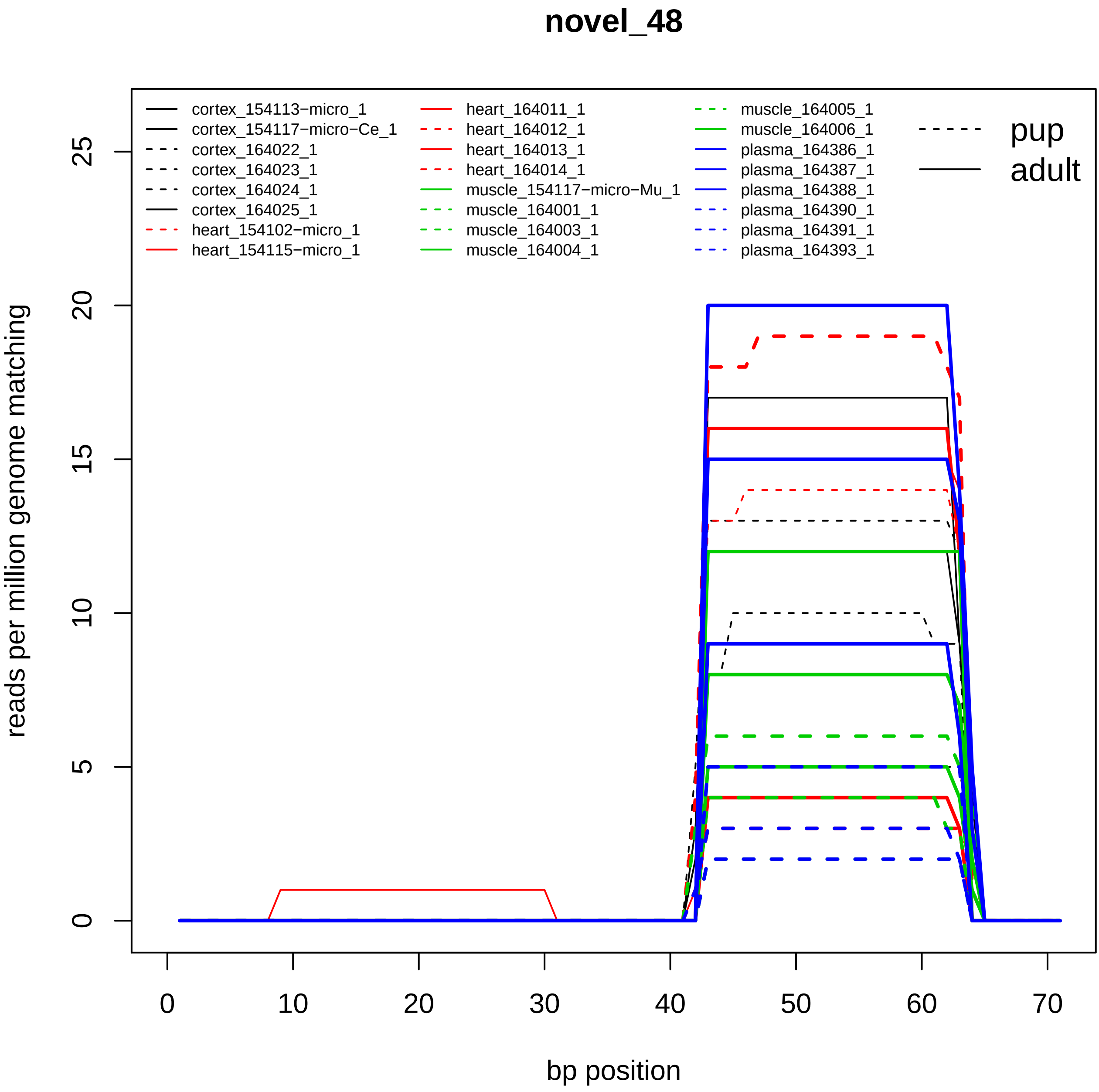
novel_48
cortex_154113−micro_1
cortex_154117−micro−Ce_1
cortex_164022_1
cortex_164023_1
cortex_164024_1
cortex_164025_1
heart_154102−micro_1
heart_154115−micro_1
heart_164011_1
heart_164012_1
heart_164013_1
heart_164014_1
muscle_154117−micro−Mu_1
muscle_164001_1
muscle_164003_1
muscle_164004_1
muscle_164005_1
muscle_164006_1
plasma_164386_1
plasma_164387_1
plasma_164388_1
plasma_164390_1
plasma_164391_1
plasma_164393_1
pup
adult
reads per million genome matching
bp position
0
5
10
15
20
25
0
10
20
30
40
50
60
70

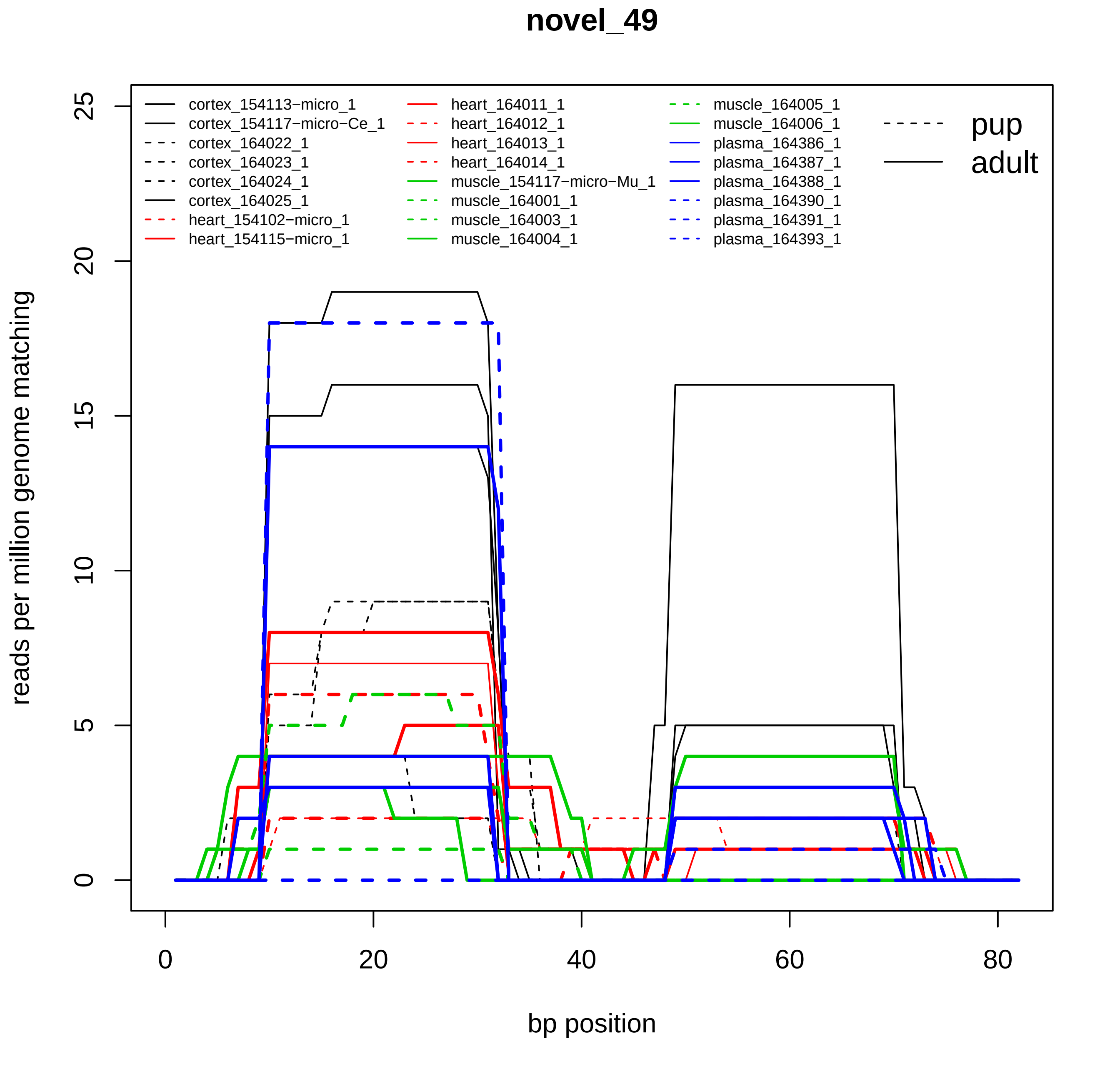

novel_49
cortex_154113−micro_1
cortex_154117−micro−Ce_1
cortex_164022_1
cortex_164023_1
cortex_164024_1
cortex_164025_1
heart_154102−micro_1
heart_154115−micro_1
heart_164011_1
heart_164012_1
heart_164013_1
heart_164014_1
muscle_154117−micro−Mu_1
muscle_164001_1
muscle_164003_1
muscle_164004_1
muscle_164005_1
muscle_164006_1
plasma_164386_1
plasma_164387_1
plasma_164388_1
plasma_164390_1
plasma_164391_1
plasma_164393_1
pup
adult
reads per million genome matching
bp position
0
5
10
15
20
25
0
20
40
60
80

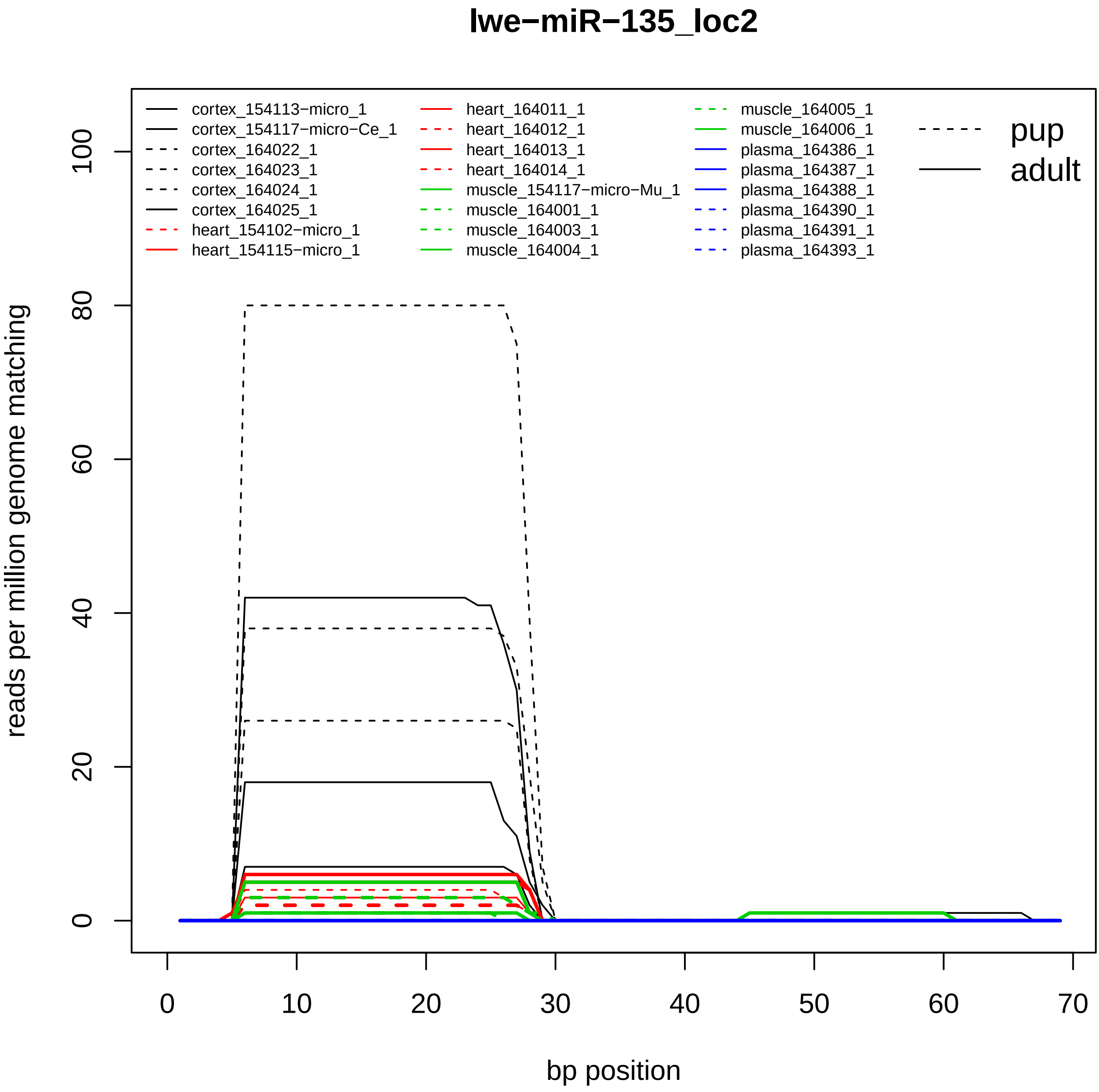

lwe−miR−135_loc2
cortex_154113−micro_1
cortex_154117−micro−Ce_1
cortex_164022_1
cortex_164023_1
cortex_164024_1
cortex_164025_1
heart_154102−micro_1
heart_154115−micro_1
heart_164011_1
heart_164012_1
heart_164013_1
heart_164014_1
muscle_154117−micro−Mu_1
muscle_164001_1
muscle_164003_1
muscle_164004_1
muscle_164005_1
muscle_164006_1
plasma_164386_1
plasma_164387_1
plasma_164388_1
plasma_164390_1
plasma_164391_1
plasma_164393_1
pup
adult
reads per million genome matching
0
20
40
60
80
100
0
10
20
30
40
50
60
70
bp position

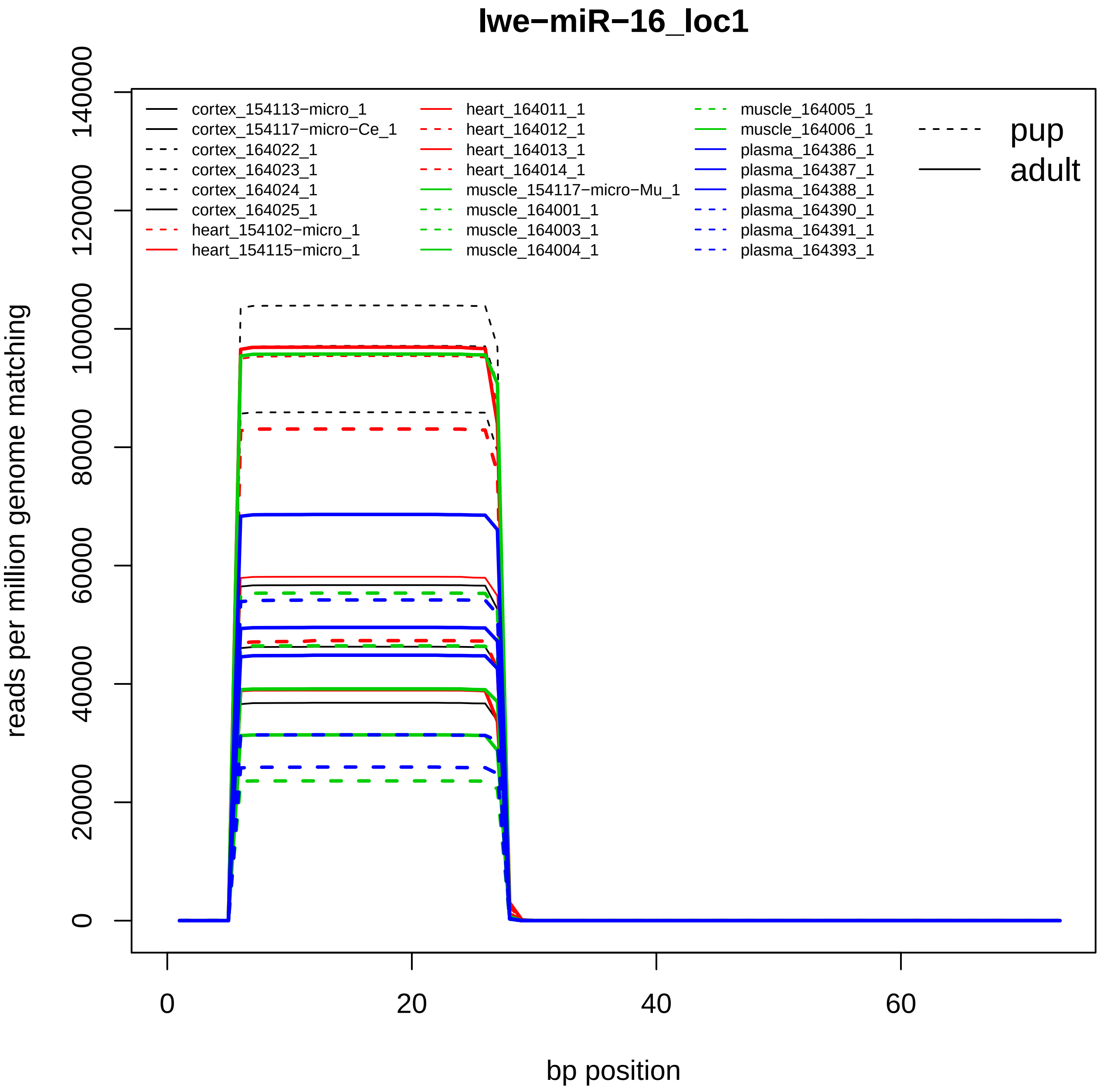

lwe−miR−16_loc1
reads per million genome matching
0
20000
40000
60000
80000
100000
120000
140000
bp position
0
20
40
60
cortex_154113−micro_1
cortex_154117−micro−Ce_1
cortex_164022_1
cortex_164023_1
cortex_164024_1
cortex_164025_1
heart_154102−micro_1
heart_154115−micro_1
heart_164011_1
heart_164012_1
heart_164013_1
heart_164014_1
muscle_154117−micro−Mu_1
muscle_164001_1
muscle_164003_1
muscle_164004_1
muscle_164005_1
muscle_164006_1
plasma_164386_1
plasma_164387_1
plasma_164388_1
plasma_164390_1
plasma_164391_1
plasma_164393_1
pup
adult

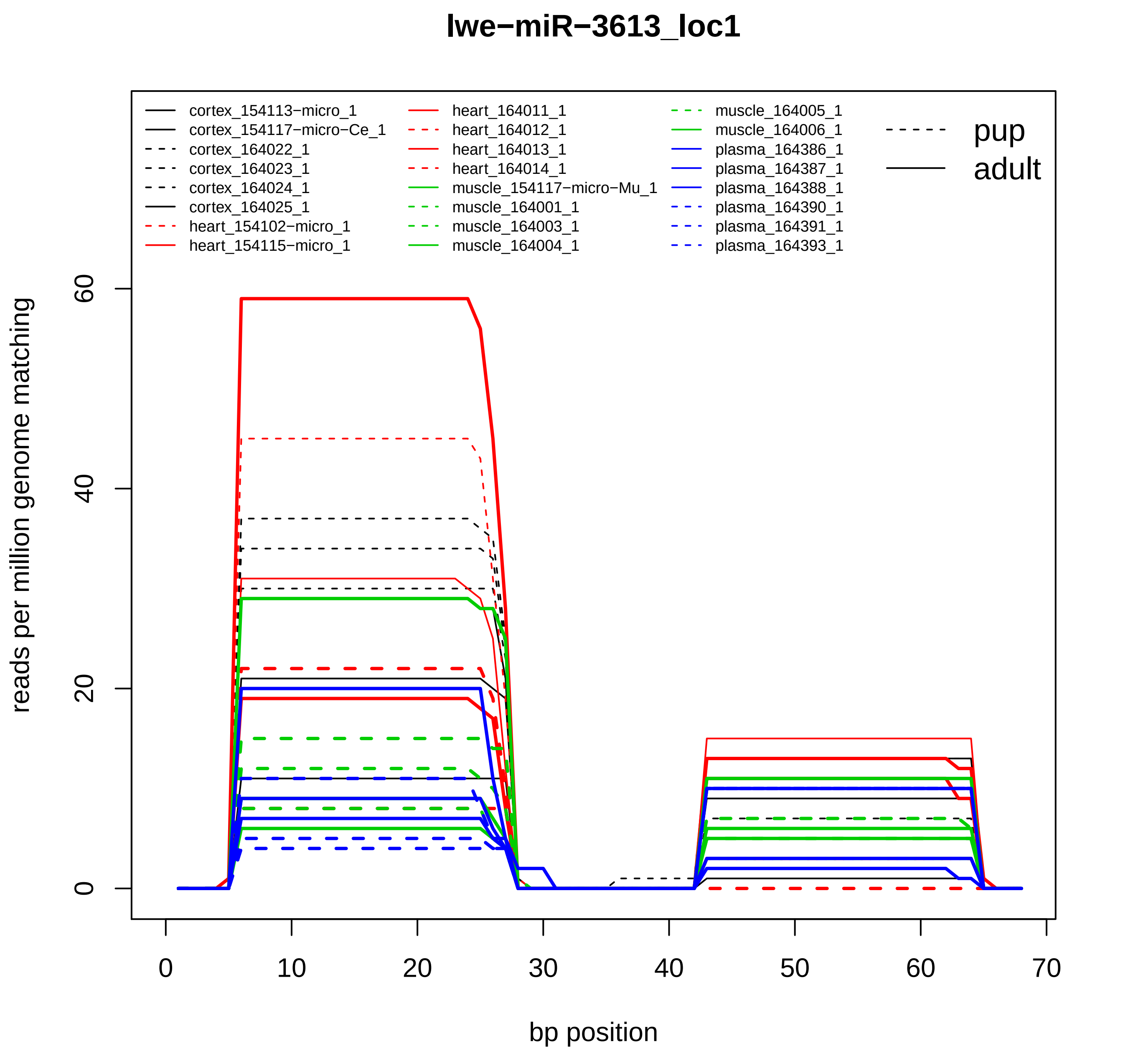

lwe−miR−3613_loc1
cortex_154113−micro_1
cortex_154117−micro−Ce_1
cortex_164022_1
cortex_164023_1
cortex_164024_1
cortex_164025_1
heart_154102−micro_1
heart_154115−micro_1
heart_164011_1
heart_164012_1
heart_164013_1
heart_164014_1
muscle_154117−micro−Mu_1
muscle_164001_1
muscle_164003_1
muscle_164004_1
muscle_164005_1
muscle_164006_1
plasma_164386_1
plasma_164387_1
plasma_164388_1
plasma_164390_1
plasma_164391_1
plasma_164393_1
pup
adult
reads per million genome matching
0
20
40
60
0
10
20
30
40
50
60
70
bp position

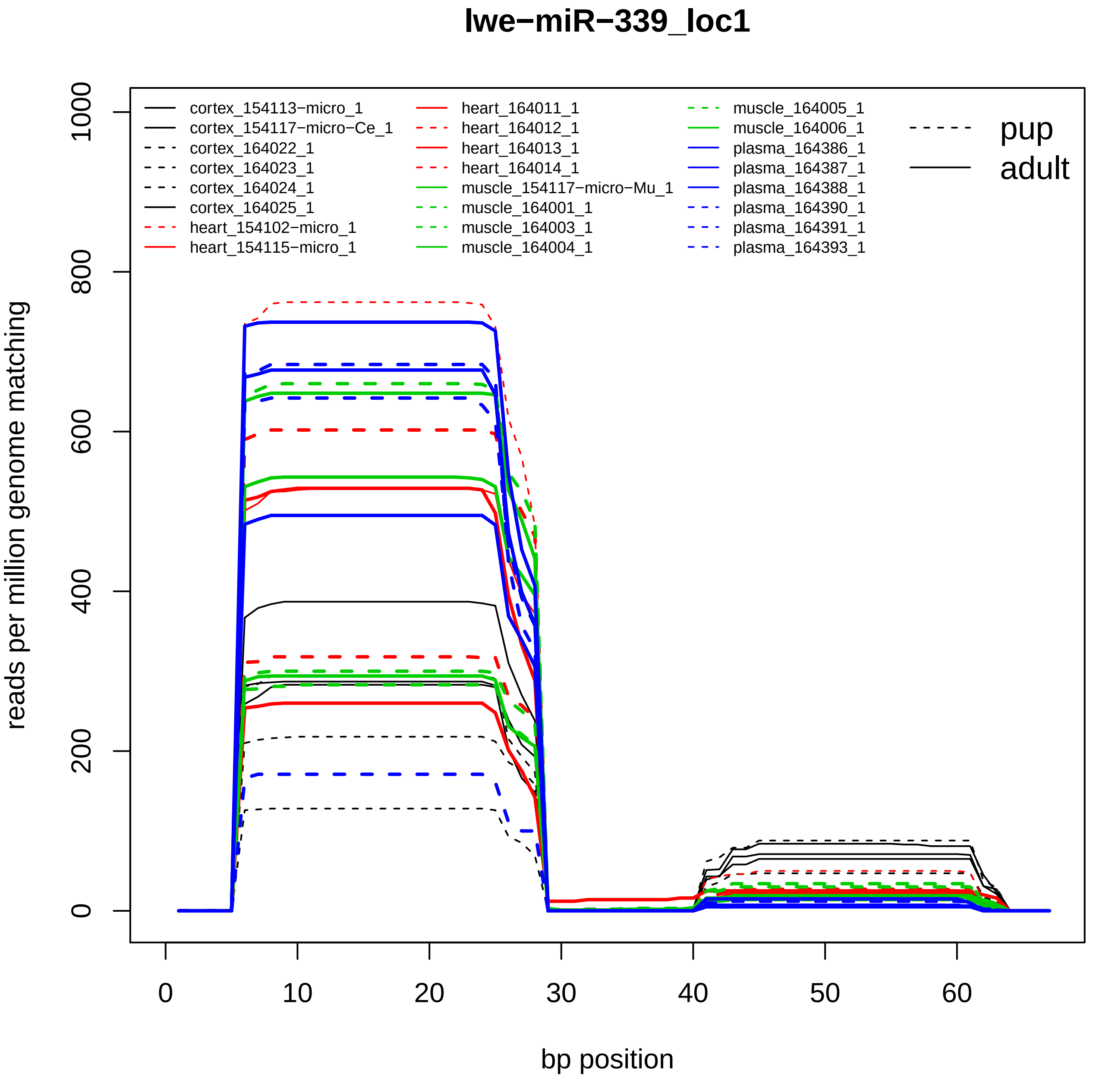
lwe−miR−339_loc1
reads per million genome matching
bp position
0
200
400
600
800
1000
0
10
20
30
40
50
60
cortex_154113−micro_1
cortex_154117−micro−Ce_1
cortex_164022_1
cortex_164023_1
cortex_164024_1
cortex_164025_1
heart_154102−micro_1
heart_154115−micro_1
heart_164011_1
heart_164012_1
heart_164013_1
heart_164014_1
muscle_154117−micro−Mu_1
muscle_164001_1
muscle_164003_1
muscle_164004_1
muscle_164005_1
muscle_164006_1
plasma_164386_1
plasma_164387_1
plasma_164388_1
plasma_164390_1
plasma_164391_1
plasma_164393_1
pup
adult

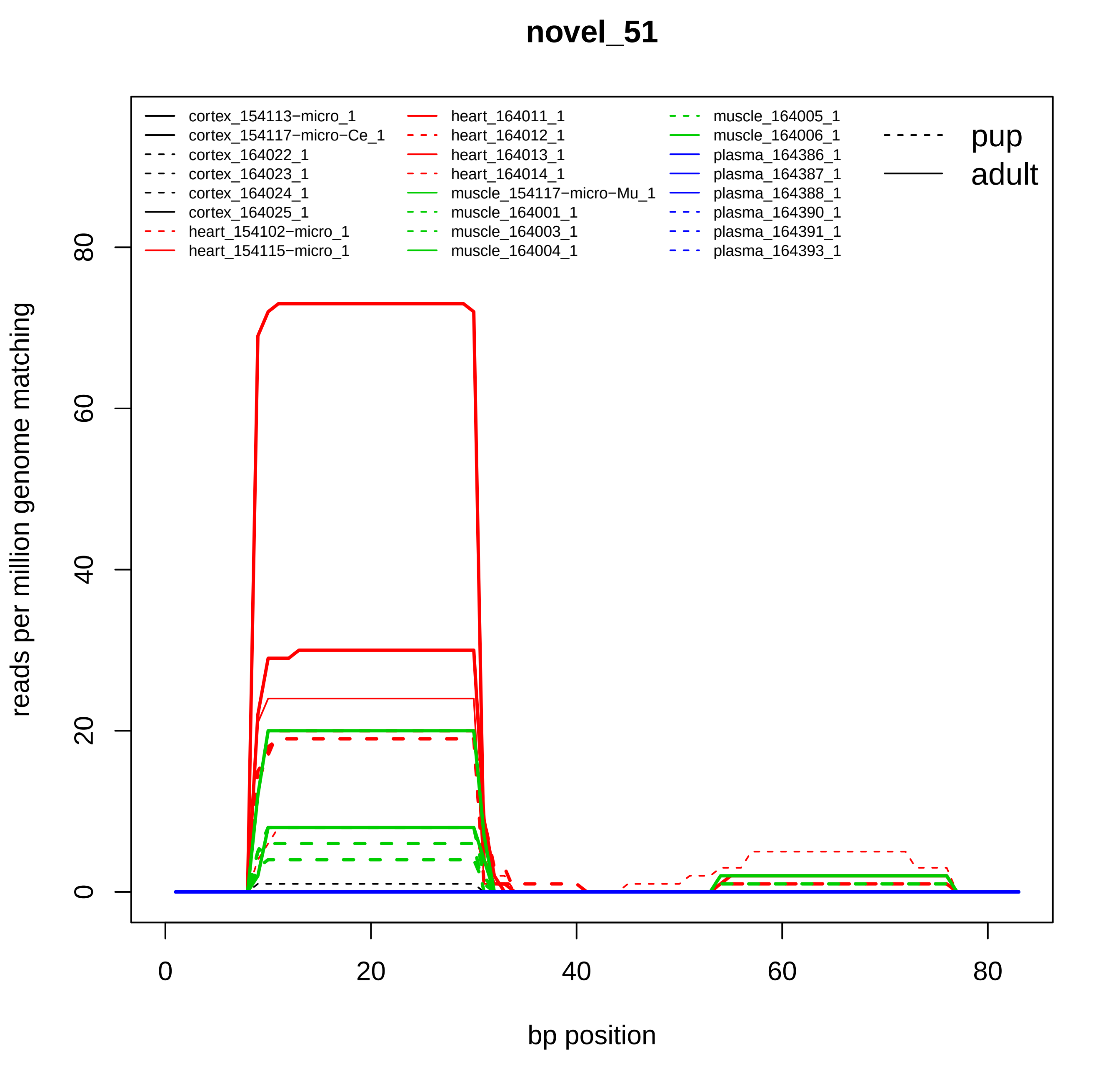
novel_51
cortex_154113−micro_1
cortex_154117−micro−Ce_1
cortex_164022_1
cortex_164023_1
cortex_164024_1
cortex_164025_1
heart_154102−micro_1
heart_154115−micro_1
heart_164011_1
heart_164012_1
heart_164013_1
heart_164014_1
muscle_154117−micro−Mu_1
muscle_164001_1
muscle_164003_1
muscle_164004_1
muscle_164005_1
muscle_164006_1
plasma_164386_1
plasma_164387_1
plasma_164388_1
plasma_164390_1
plasma_164391_1
plasma_164393_1
pup
adult
reads per million genome matching
0
20
40
60
80
bp position
0
20
40
60
80

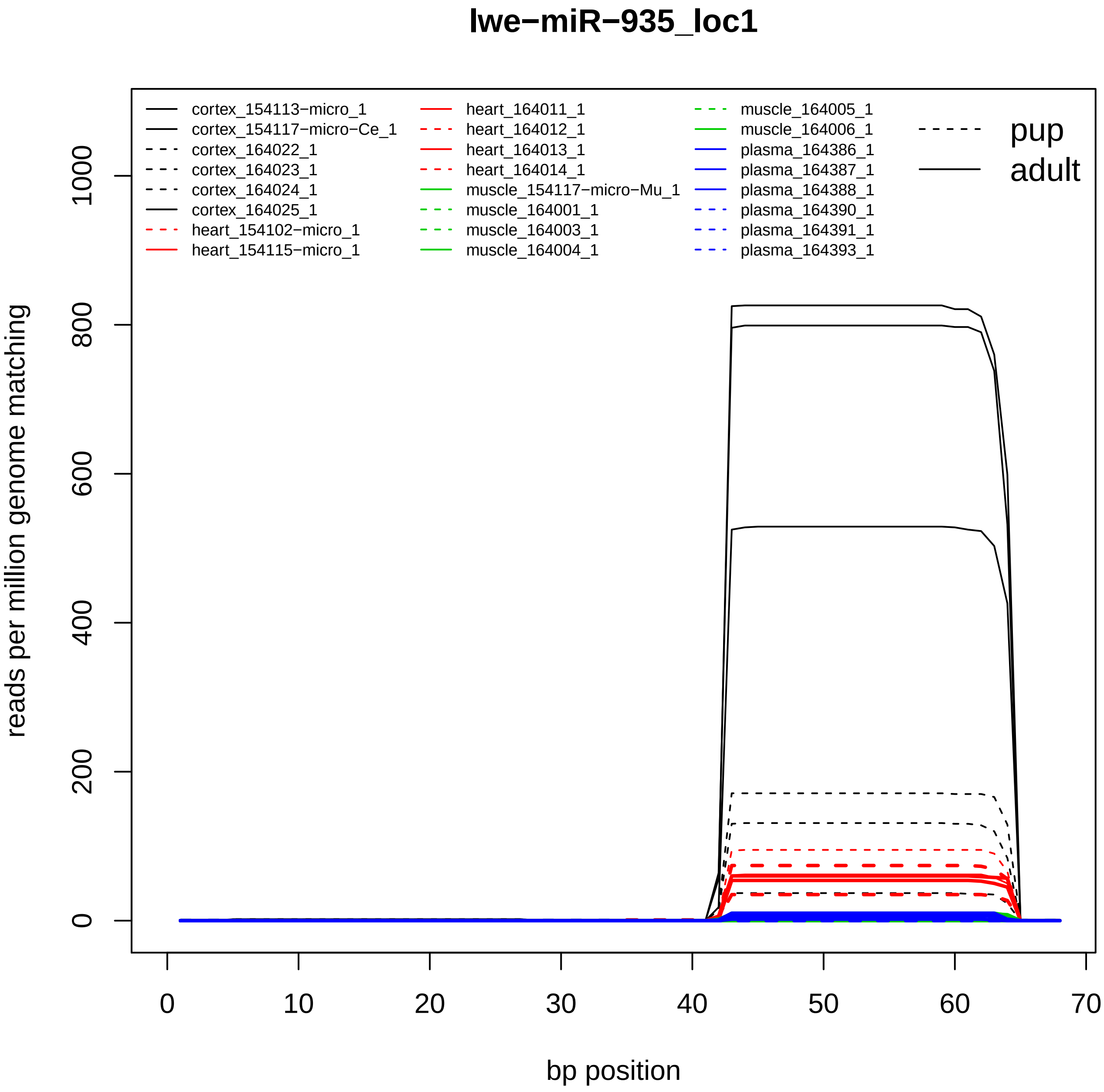

lwe−miR−935_loc1
cortex_154113−micro_1
cortex_154117−micro−Ce_1
cortex_164022_1
cortex_164023_1
cortex_164024_1
cortex_164025_1
heart_154102−micro_1
heart_154115−micro_1
heart_164011_1
heart_164012_1
heart_164013_1
heart_164014_1
muscle_154117−micro−Mu_1
muscle_164001_1
muscle_164003_1
muscle_164004_1
muscle_164005_1
muscle_164006_1
plasma_164386_1
plasma_164387_1
plasma_164388_1
plasma_164390_1
plasma_164391_1
plasma_164393_1
pup
adult
reads per million genome matching
0
200
400
600
800
1000
0
10
20
30
40
50
60
70
bp position

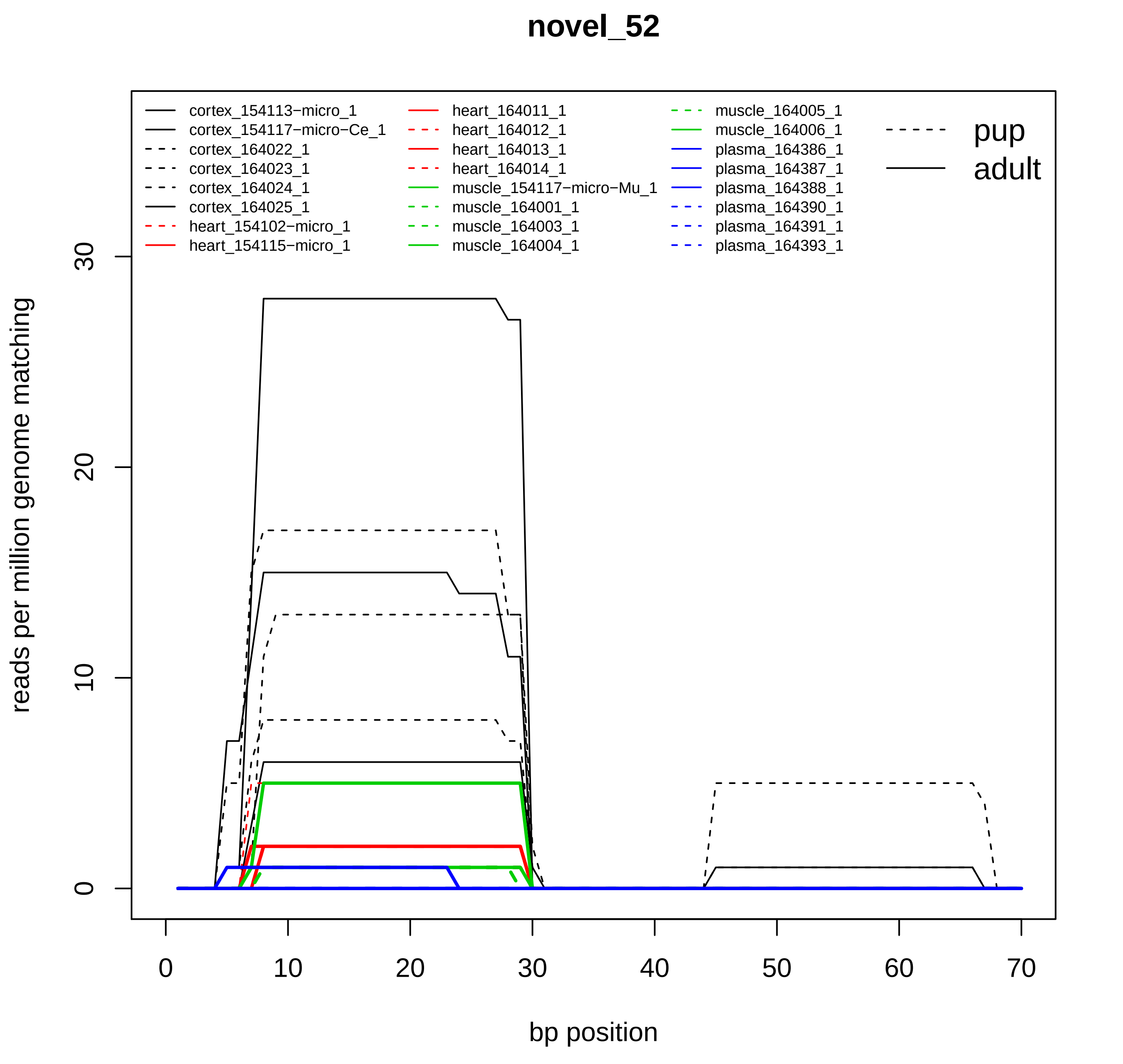

novel_52
cortex_154113−micro_1
cortex_154117−micro−Ce_1
cortex_164022_1
cortex_164023_1
cortex_164024_1
cortex_164025_1
heart_154102−micro_1
heart_154115−micro_1
heart_164011_1
heart_164012_1
heart_164013_1
heart_164014_1
muscle_154117−micro−Mu_1
muscle_164001_1
muscle_164003_1
muscle_164004_1
muscle_164005_1
muscle_164006_1
plasma_164386_1
plasma_164387_1
plasma_164388_1
plasma_164390_1
plasma_164391_1
plasma_164393_1
pup
adult
reads per million genome matching
0
10
20
30
0
10
20
30
40
50
60
70
bp position

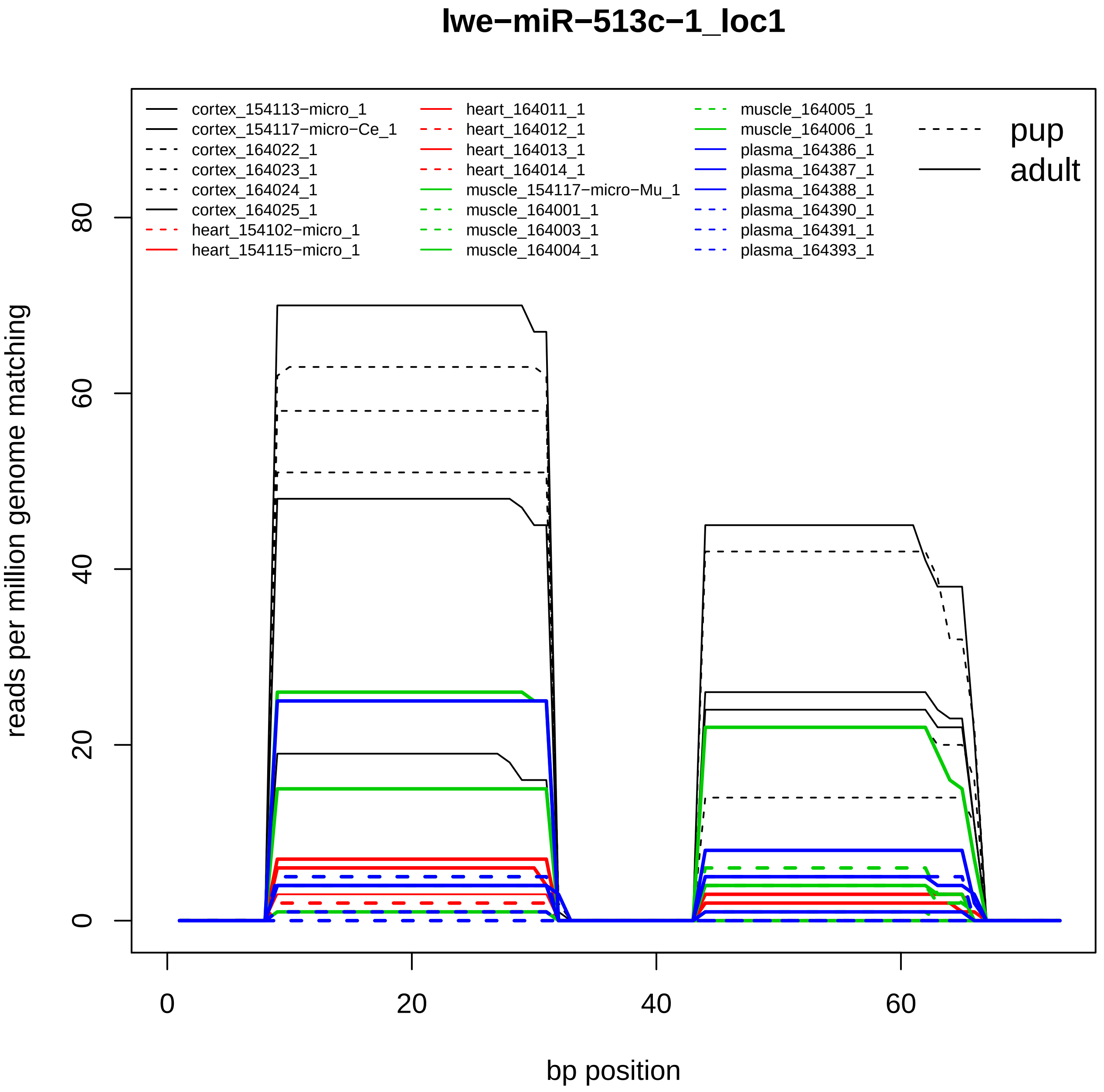
lwe−miR−513c−1_loc1
cortex_154113−micro_1
cortex_154117−micro−Ce_1
cortex_164022_1
cortex_164023_1
cortex_164024_1
cortex_164025_1
heart_154102−micro_1
heart_154115−micro_1
heart_164011_1
heart_164012_1
heart_164013_1
heart_164014_1
muscle_154117−micro−Mu_1
muscle_164001_1
muscle_164003_1
muscle_164004_1
muscle_164005_1
muscle_164006_1
plasma_164386_1
plasma_164387_1
plasma_164388_1
plasma_164390_1
plasma_164391_1
plasma_164393_1
pup
adult
reads per million genome matching
0
20
40
60
80
bp position
0
20
40
60

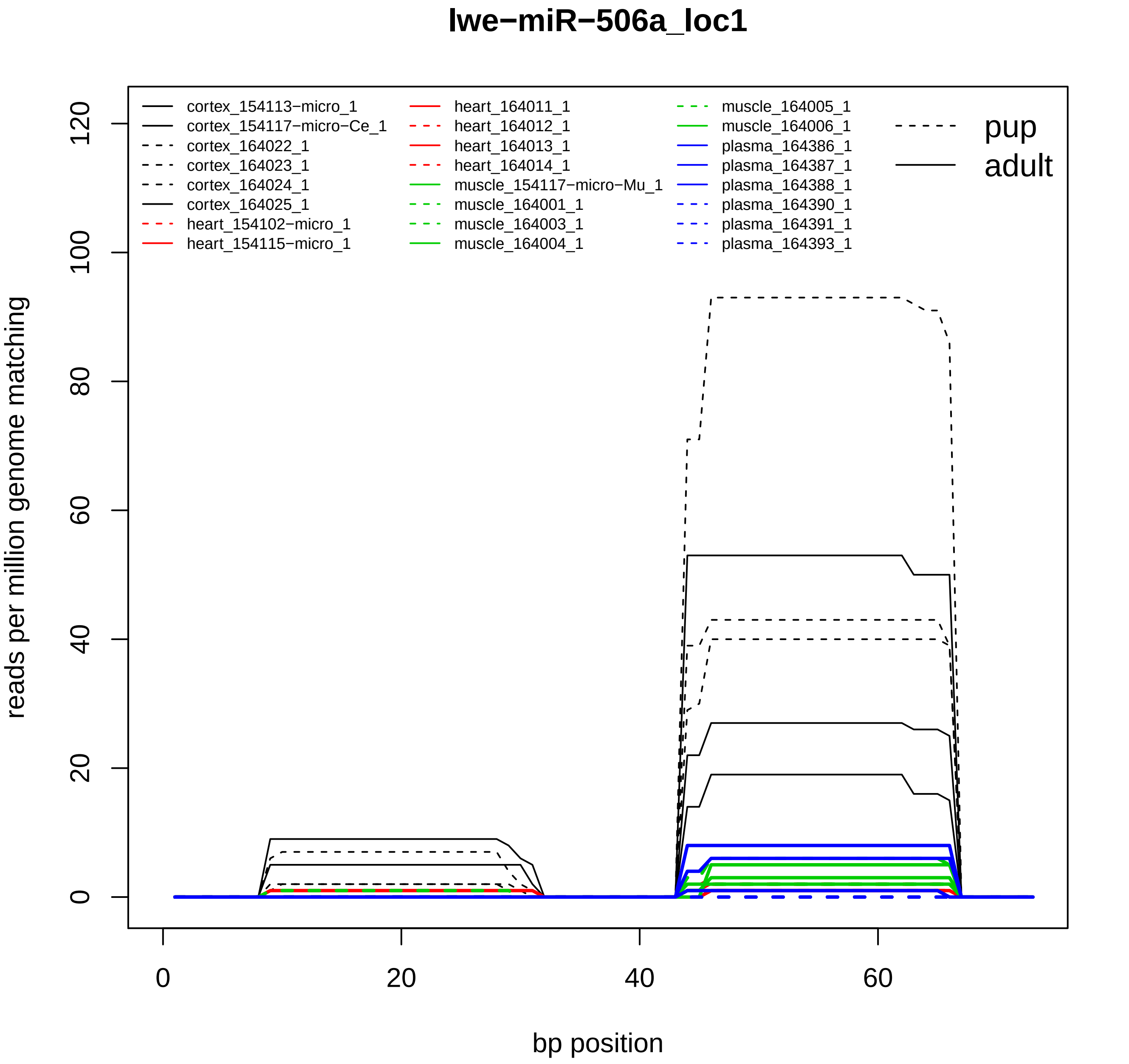
lwe−miR−506a_loc1
cortex_154113−micro_1
cortex_154117−micro−Ce_1
cortex_164022_1
cortex_164023_1
cortex_164024_1
cortex_164025_1
heart_154102−micro_1
heart_154115−micro_1
heart_164011_1
heart_164012_1
heart_164013_1
heart_164014_1
muscle_154117−micro−Mu_1
muscle_164001_1
muscle_164003_1
muscle_164004_1
muscle_164005_1
muscle_164006_1
plasma_164386_1
plasma_164387_1
plasma_164388_1
plasma_164390_1
plasma_164391_1
plasma_164393_1
pup
adult
reads per million genome matching
0
20
40
60
80
100
120
bp position
0
20
40
60

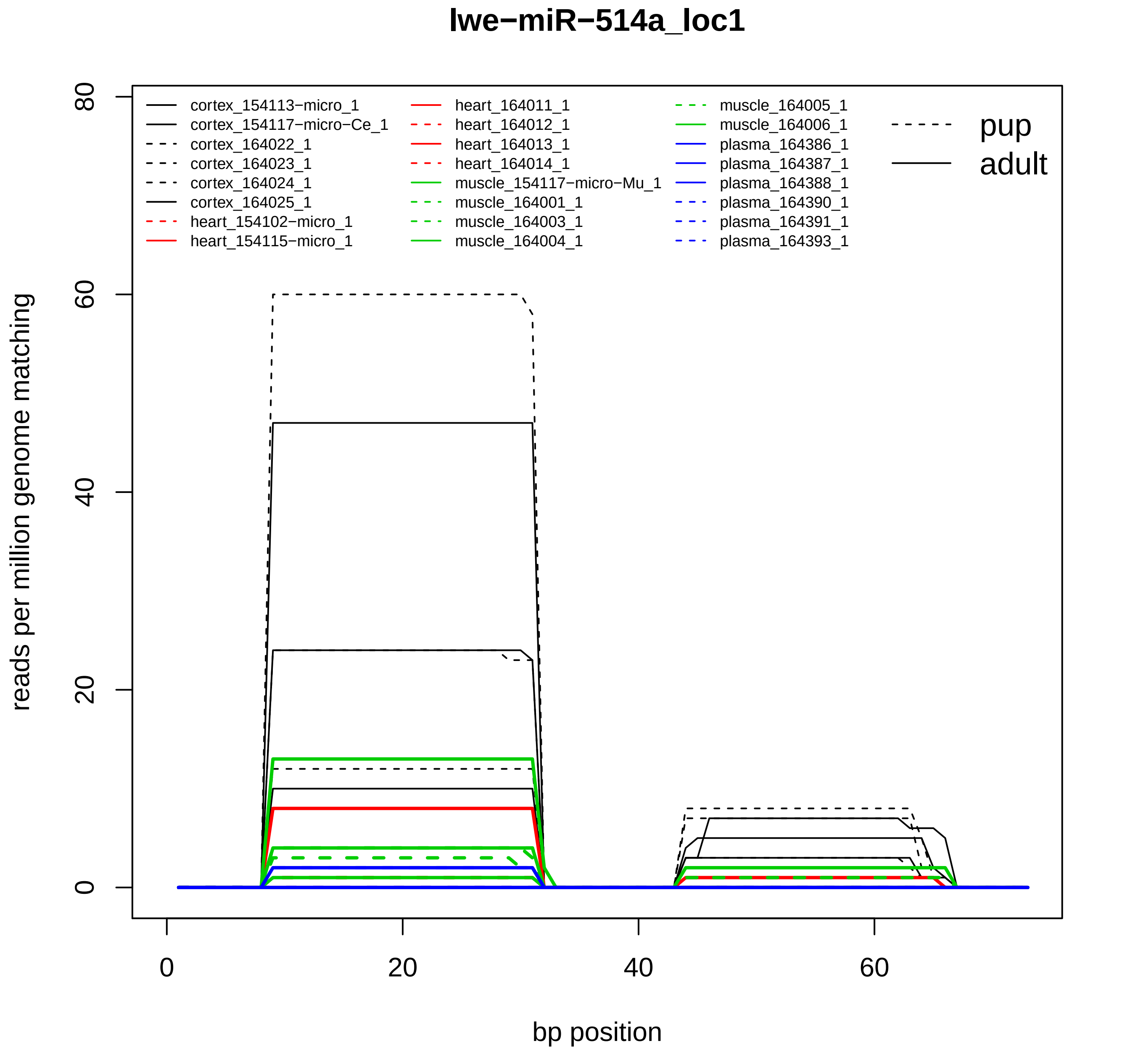
lwe−miR−514a_loc1
cortex_154113−micro_1
cortex_154117−micro−Ce_1
cortex_164022_1
cortex_164023_1
cortex_164024_1
cortex_164025_1
heart_154102−micro_1
heart_154115−micro_1
heart_164011_1
heart_164012_1
heart_164013_1
heart_164014_1
muscle_154117−micro−Mu_1
muscle_164001_1
muscle_164003_1
muscle_164004_1
muscle_164005_1
muscle_164006_1
plasma_164386_1
plasma_164387_1
plasma_164388_1
plasma_164390_1
plasma_164391_1
plasma_164393_1
pup
adult
reads per million genome matching
bp position
0
20
40
60
80

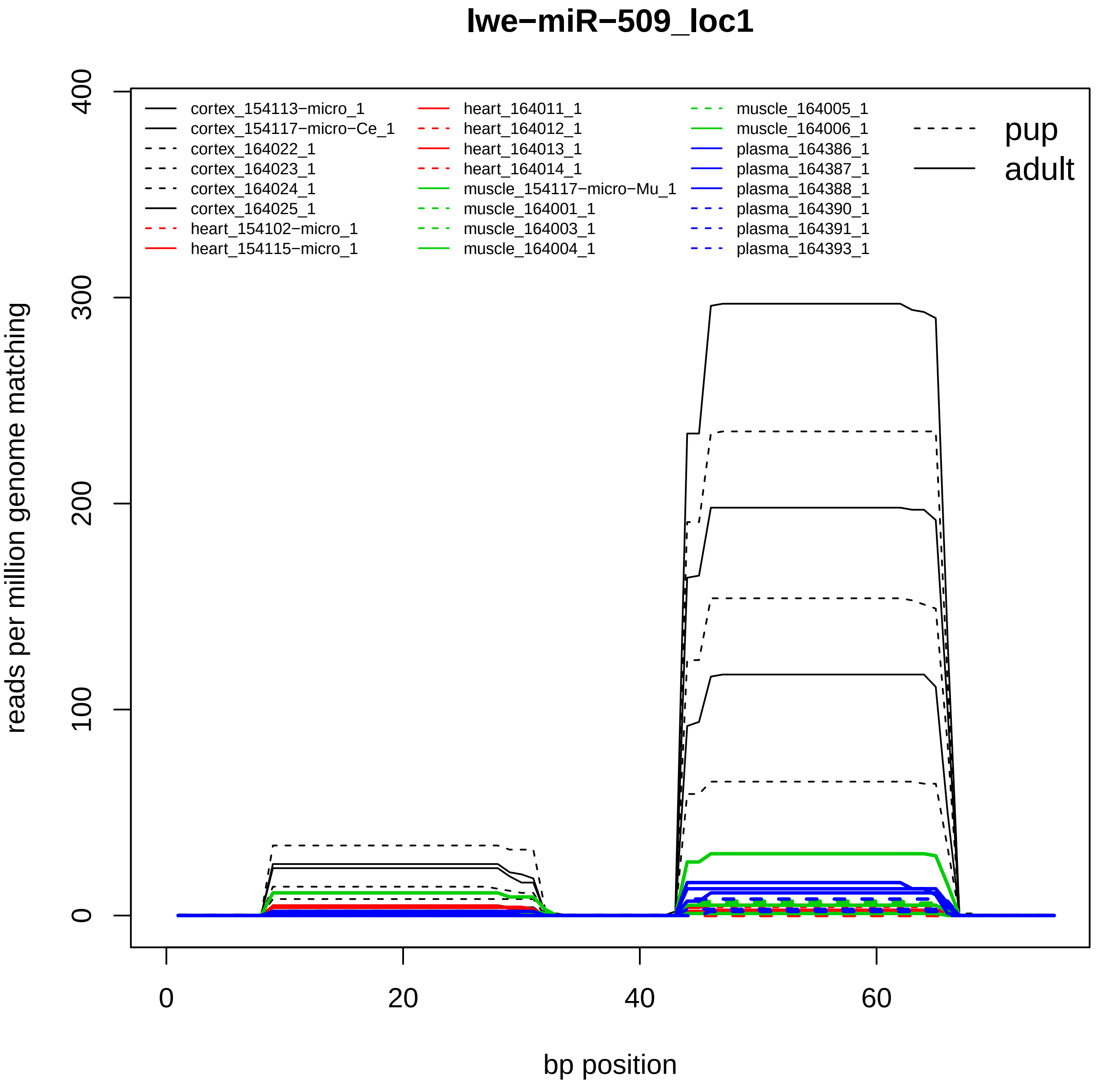
lwe−miR−509_loc1
reads per million genome matching
bp position
0
100
200
300
400
0
20
40
60
cortex_154113−micro_1
cortex_154117−micro−Ce_1
cortex_164022_1
cortex_164023_1
cortex_164024_1
cortex_164025_1
heart_154102−micro_1
heart_154115−micro_1
heart_164011_1
heart_164012_1
heart_164013_1
heart_164014_1
muscle_154117−micro−Mu_1
muscle_164001_1
muscle_164003_1
muscle_164004_1
muscle_164005_1
muscle_164006_1
plasma_164386_1
plasma_164387_1
plasma_164388_1
plasma_164390_1
plasma_164391_1
plasma_164393_1
pup
adult

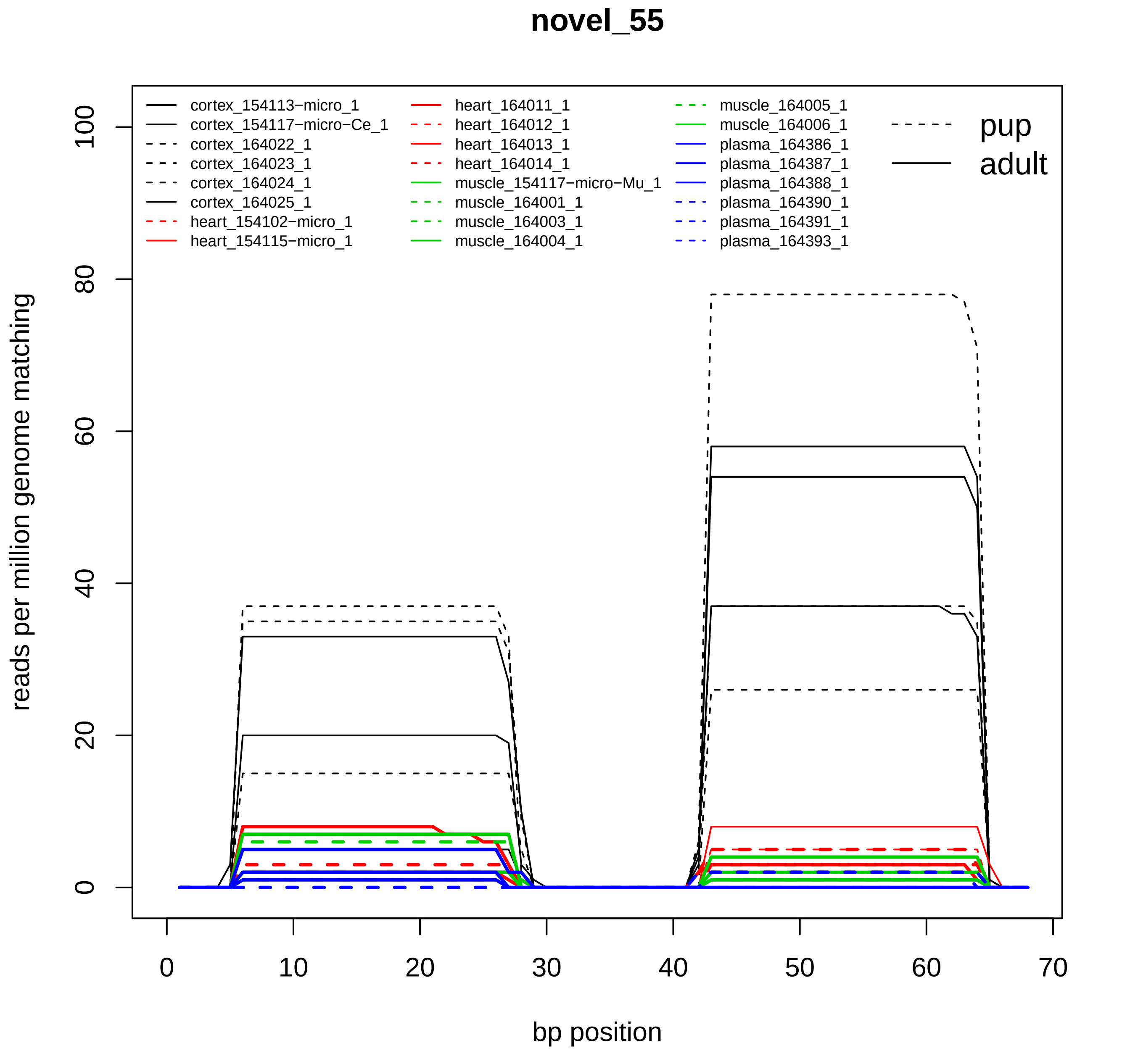

novel_55
cortex_154113−micro_1
cortex_154117−micro−Ce_1
cortex_164022_1
cortex_164023_1
cortex_164024_1
cortex_164025_1
heart_154102−micro_1
heart_154115−micro_1
heart_164011_1
heart_164012_1
heart_164013_1
heart_164014_1
muscle_154117−micro−Mu_1
muscle_164001_1
muscle_164003_1
muscle_164004_1
muscle_164005_1
muscle_164006_1
plasma_164386_1
plasma_164387_1
plasma_164388_1
plasma_164390_1
plasma_164391_1
plasma_164393_1
pup
adult
reads per million genome matching
0
20
40
60
80
100
0
10
20
30
40
50
60
70
bp position

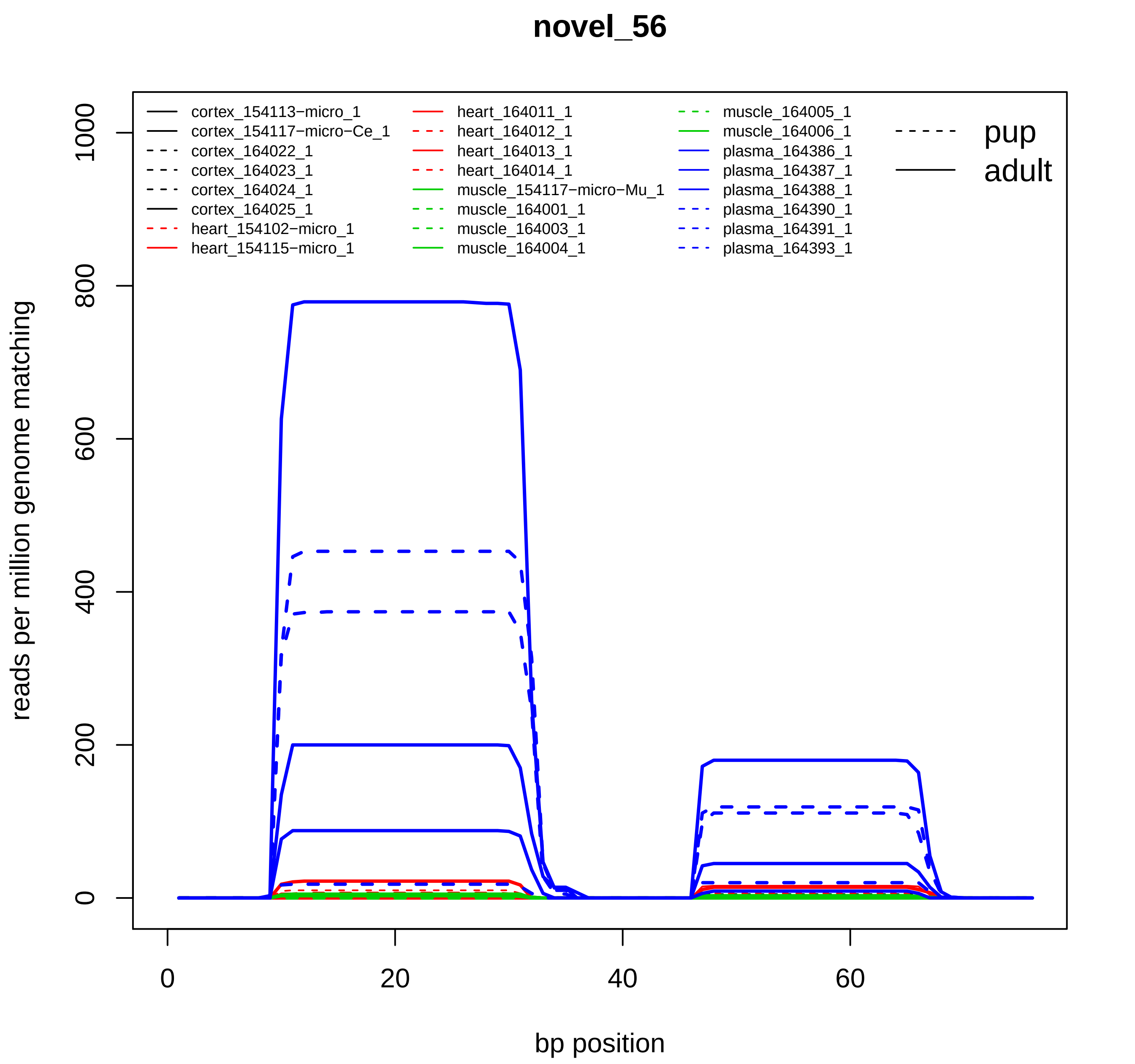
novel_56
cortex_154113−micro_1
cortex_154117−micro−Ce_1
cortex_164022_1
cortex_164023_1
cortex_164024_1
cortex_164025_1
heart_154102−micro_1
heart_154115−micro_1
heart_164011_1
heart_164012_1
heart_164013_1
heart_164014_1
muscle_154117−micro−Mu_1
muscle_164001_1
muscle_164003_1
muscle_164004_1
muscle_164005_1
muscle_164006_1
plasma_164386_1
plasma_164387_1
plasma_164388_1
plasma_164390_1
plasma_164391_1
plasma_164393_1
pup
adult
reads per million genome matching
0
200
400
600
800
1000
0
20
40
60
bp position

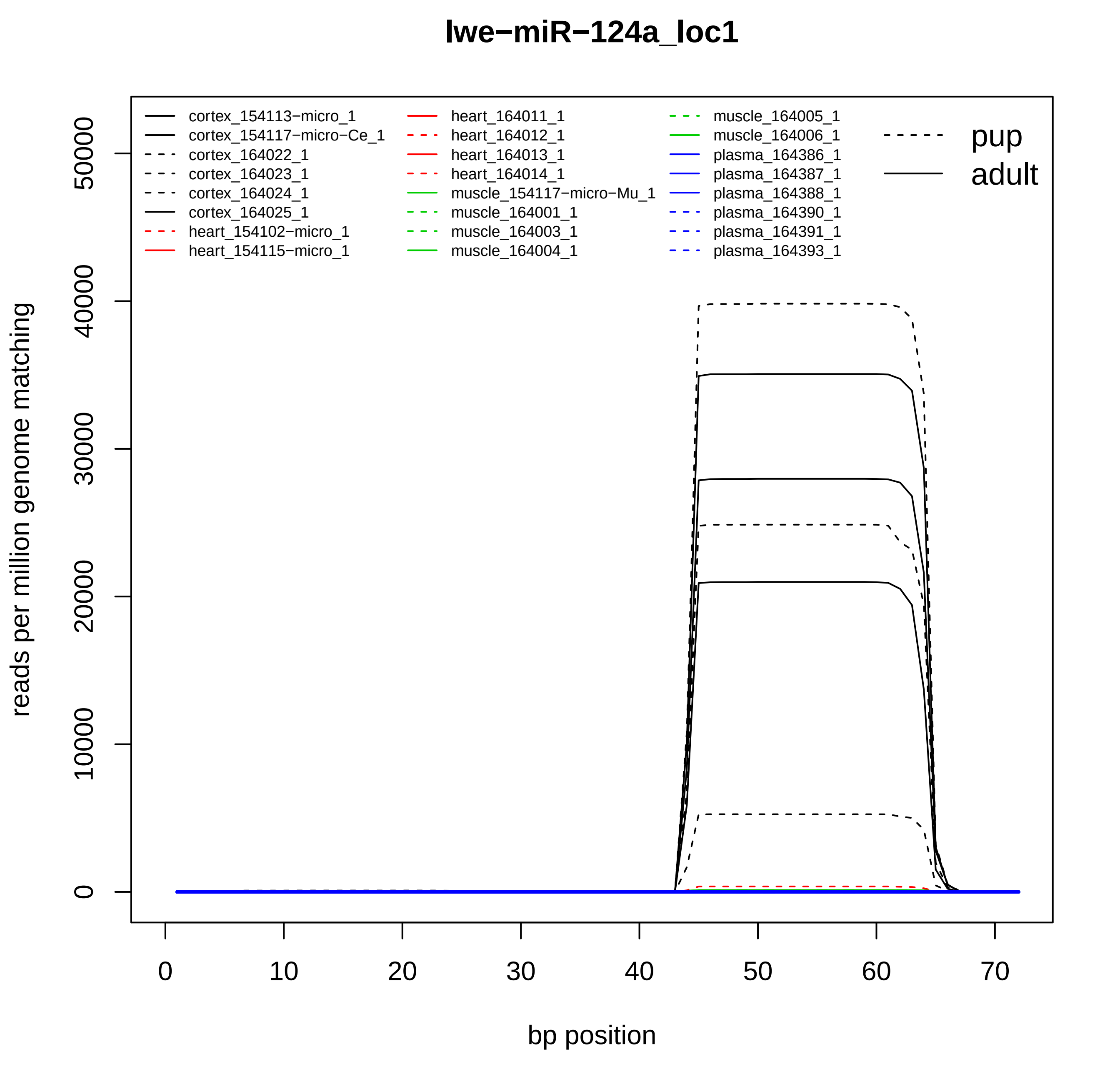

lwe−miR−124a_loc1
reads per million genome matching
bp position
0
10000
20000
30000
40000
50000
0
10
20
30
40
50
60
70
cortex_154113−micro_1
cortex_154117−micro−Ce_1
cortex_164022_1
cortex_164023_1
cortex_164024_1
cortex_164025_1
heart_154102−micro_1
heart_154115−micro_1
heart_164011_1
heart_164012_1
heart_164013_1
heart_164014_1
muscle_154117−micro−Mu_1
muscle_164001_1
muscle_164003_1
muscle_164004_1
muscle_164005_1
muscle_164006_1
plasma_164386_1
plasma_164387_1
plasma_164388_1
plasma_164390_1
plasma_164391_1
plasma_164393_1
pup
adult

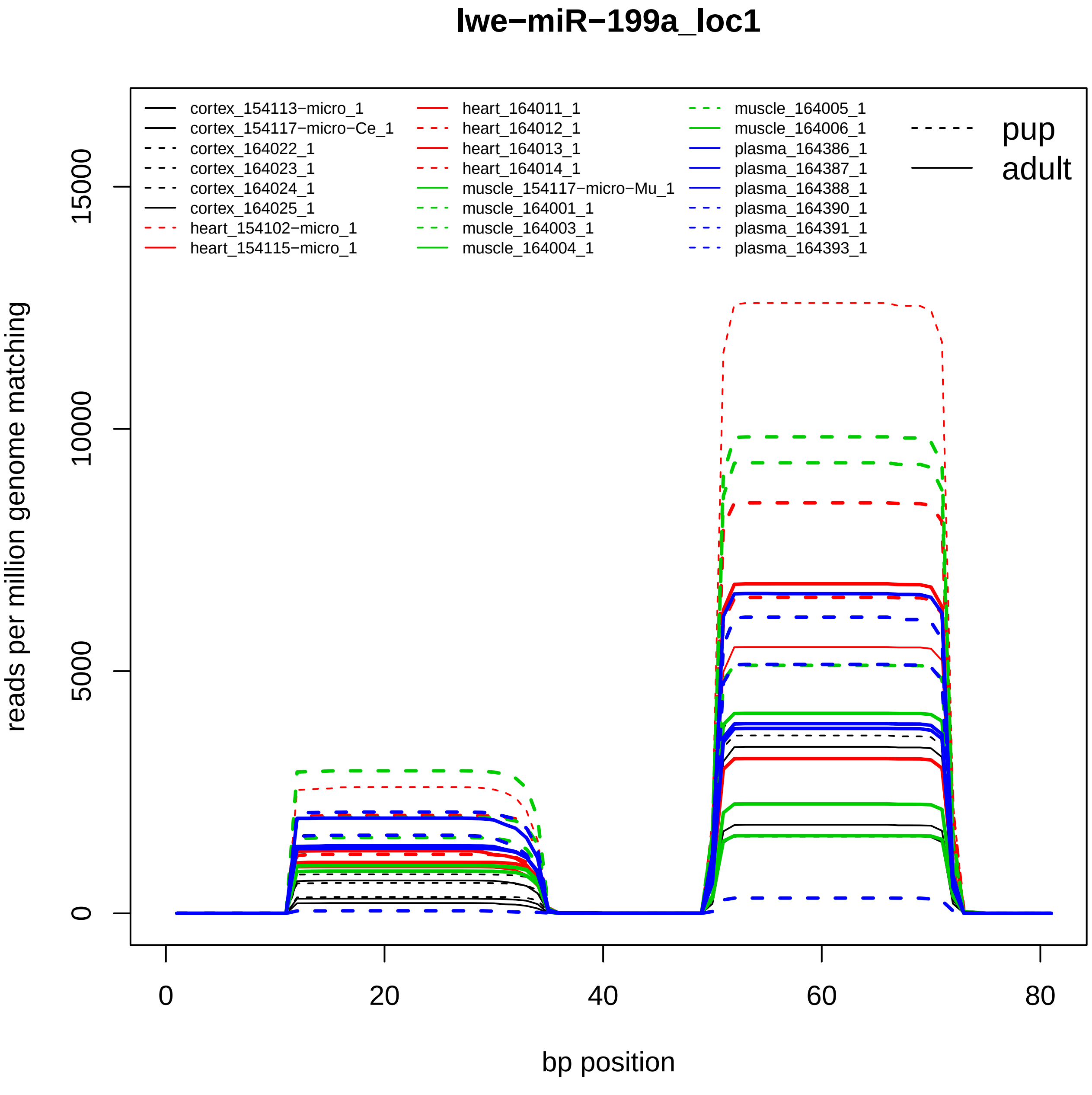
lwe−miR−199a_loc1
reads per million genome matching
bp position
0
5000
10000
15000
0
20
40
60
80
cortex_154113−micro_1
cortex_154117−micro−Ce_1
cortex_164022_1
cortex_164023_1
cortex_164024_1
cortex_164025_1
heart_154102−micro_1
heart_154115−micro_1
heart_164011_1
heart_164012_1
heart_164013_1
heart_164014_1
muscle_154117−micro−Mu_1
muscle_164001_1
muscle_164003_1
muscle_164004_1
muscle_164005_1
muscle_164006_1
plasma_164386_1
plasma_164387_1
plasma_164388_1
plasma_164390_1
plasma_164391_1
plasma_164393_1
pup
adult

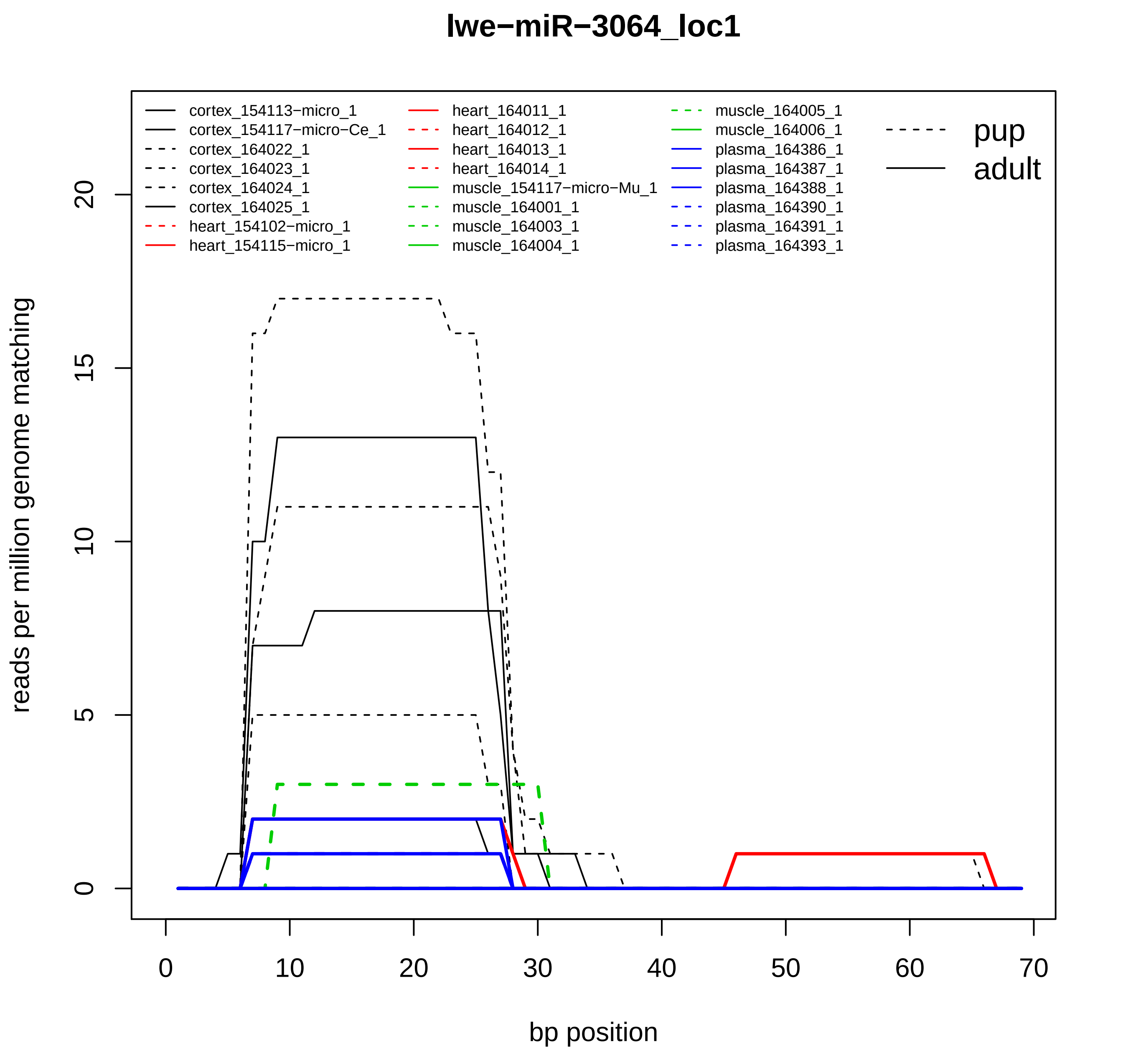
lwe−miR−3064_loc1
cortex_154113−micro_1
cortex_154117−micro−Ce_1
cortex_164022_1
cortex_164023_1
cortex_164024_1
cortex_164025_1
heart_154102−micro_1
heart_154115−micro_1
heart_164011_1
heart_164012_1
heart_164013_1
heart_164014_1
muscle_154117−micro−Mu_1
muscle_164001_1
muscle_164003_1
muscle_164004_1
muscle_164005_1
muscle_164006_1
plasma_164386_1
plasma_164387_1
plasma_164388_1
plasma_164390_1
plasma_164391_1
plasma_164393_1
pup
adult
reads per million genome matching
0
5
10
15
20
0
10
20
30
40
50
60
70
bp position

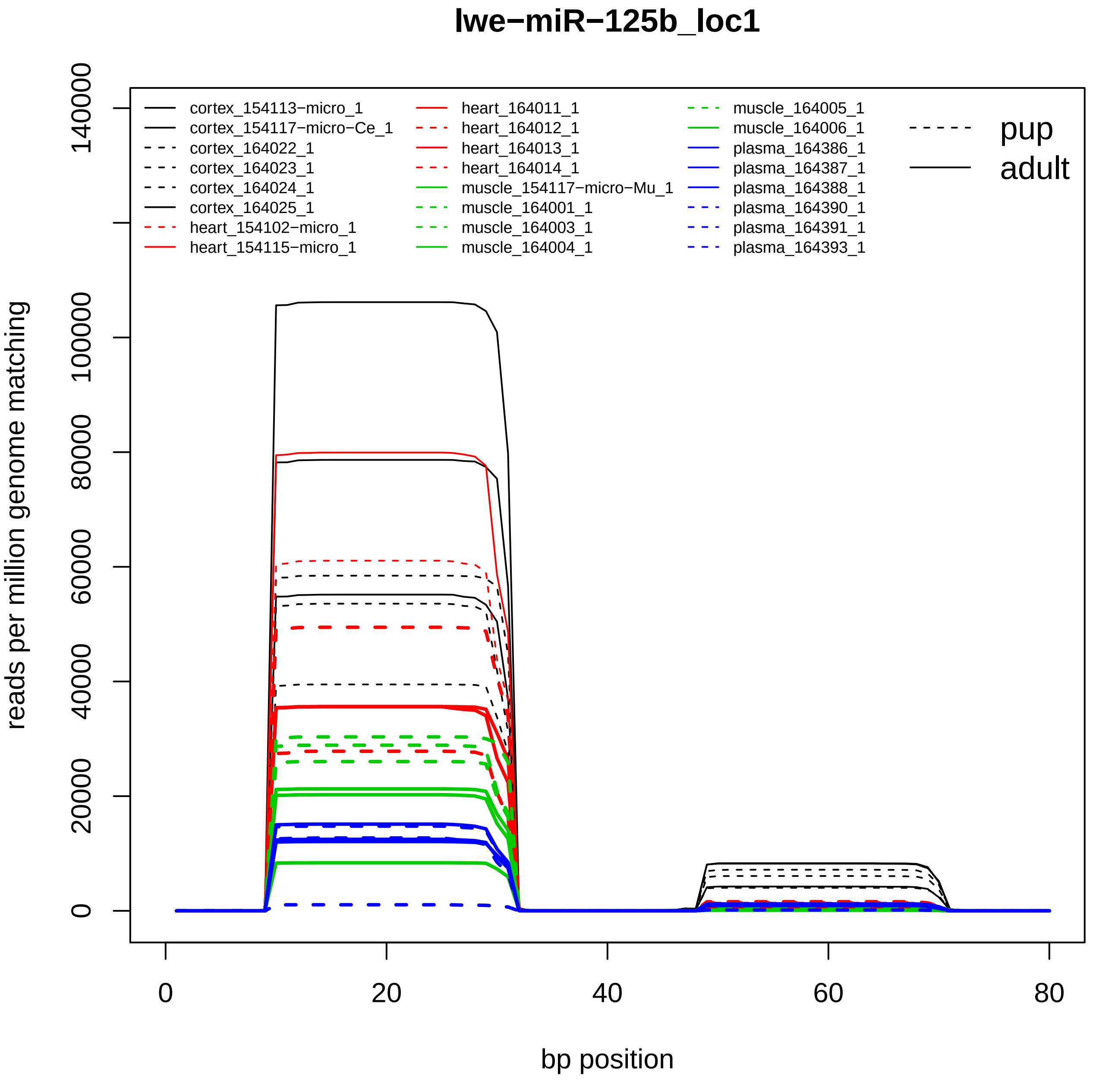

lwe−miR−125b_loc1
cortex_154113−micro_1
cortex_154117−micro−Ce_1
cortex_164022_1
cortex_164023_1
cortex_164024_1
cortex_164025_1
heart_154102−micro_1
heart_154115−micro_1
heart_164011_1
heart_164012_1
heart_164013_1
heart_164014_1
muscle_154117−micro−Mu_1
muscle_164001_1
muscle_164003_1
muscle_164004_1
muscle_164005_1
muscle_164006_1
plasma_164386_1
plasma_164387_1
plasma_164388_1
plasma_164390_1
plasma_164391_1
plasma_164393_1
pup
adult
reads per million genome matching
0
20000
40000
60000
80000
100000
140000
bp position
0
20
40
60
80

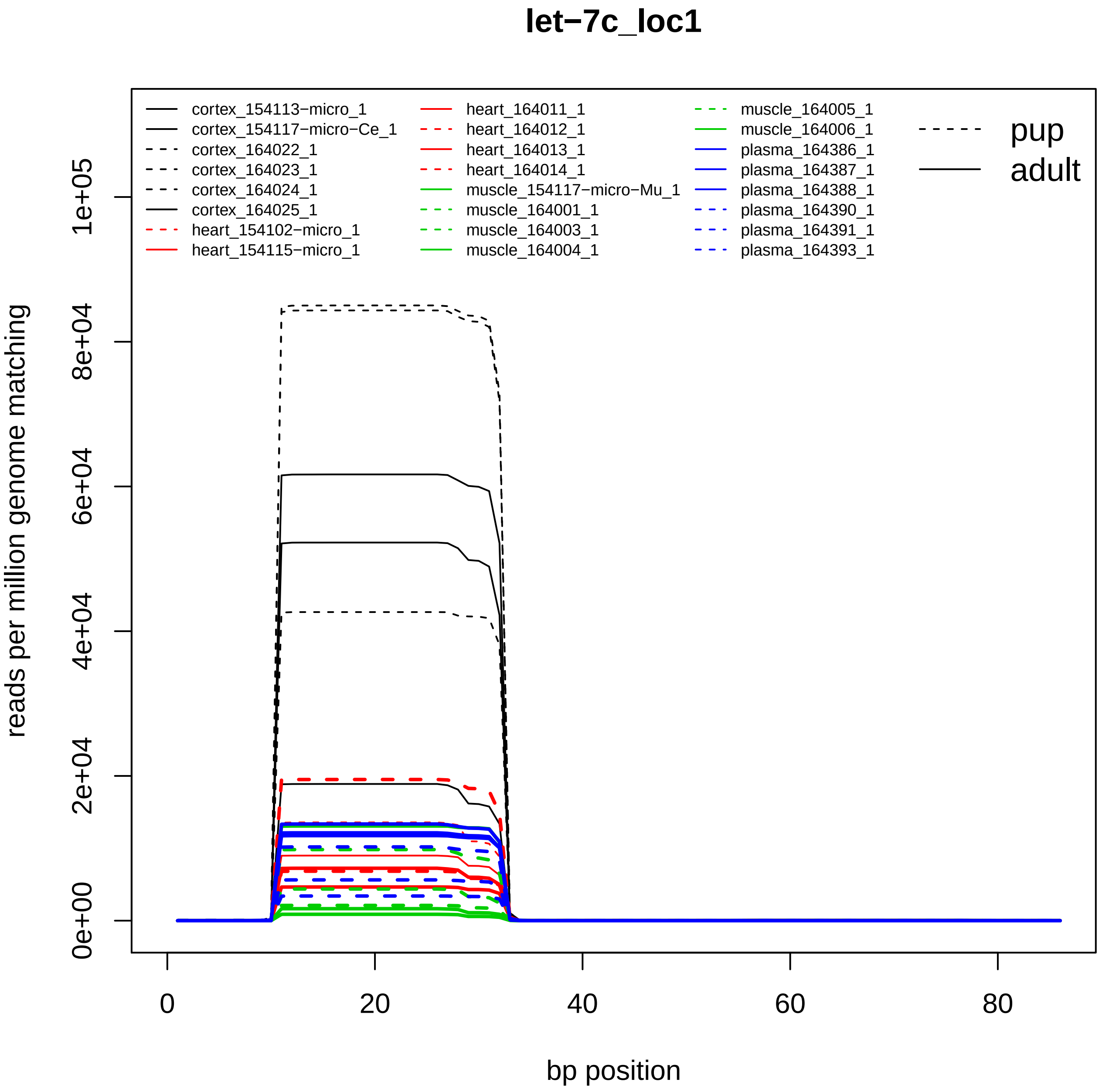

let−7c_loc1
cortex_154113−micro_1
cortex_154117−micro−Ce_1
cortex_164022_1
cortex_164023_1
cortex_164024_1
cortex_164025_1
heart_154102−micro_1
heart_154115−micro_1
heart_164011_1
heart_164012_1
heart_164013_1
heart_164014_1
muscle_154117−micro−Mu_1
muscle_164001_1
muscle_164003_1
muscle_164004_1
muscle_164005_1
muscle_164006_1
plasma_164386_1
plasma_164387_1
plasma_164388_1
plasma_164390_1
plasma_164391_1
plasma_164393_1
pup
adult
reads per million genome matching
0e+00
2e+04
4e+04
6e+04
8e+04
1e+05
0
20
40
60
80
bp position

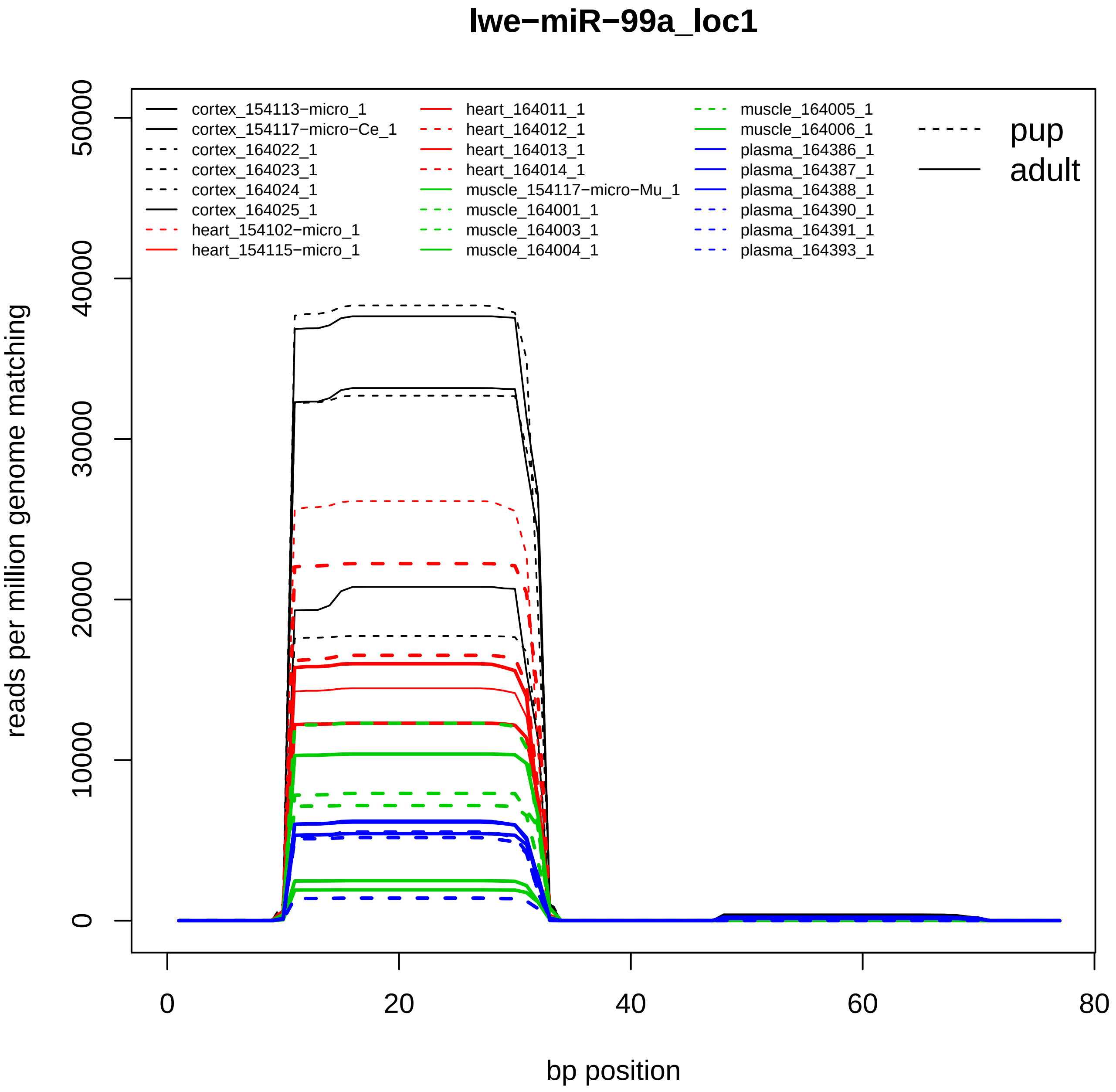
lwe−miR−99a_loc1
cortex_154113−micro_1
cortex_154117−micro−Ce_1
cortex_164022_1
cortex_164023_1
cortex_164024_1
cortex_164025_1
heart_154102−micro_1
heart_154115−micro_1
heart_164011_1
heart_164012_1
heart_164013_1
heart_164014_1
muscle_154117−micro−Mu_1
muscle_164001_1
muscle_164003_1
muscle_164004_1
muscle_164005_1
muscle_164006_1
plasma_164386_1
plasma_164387_1
plasma_164388_1
plasma_164390_1
plasma_164391_1
plasma_164393_1
pup
adult
reads per million genome matching
0
10000
20000
30000
40000
50000
0
20
40
60
80
bp position

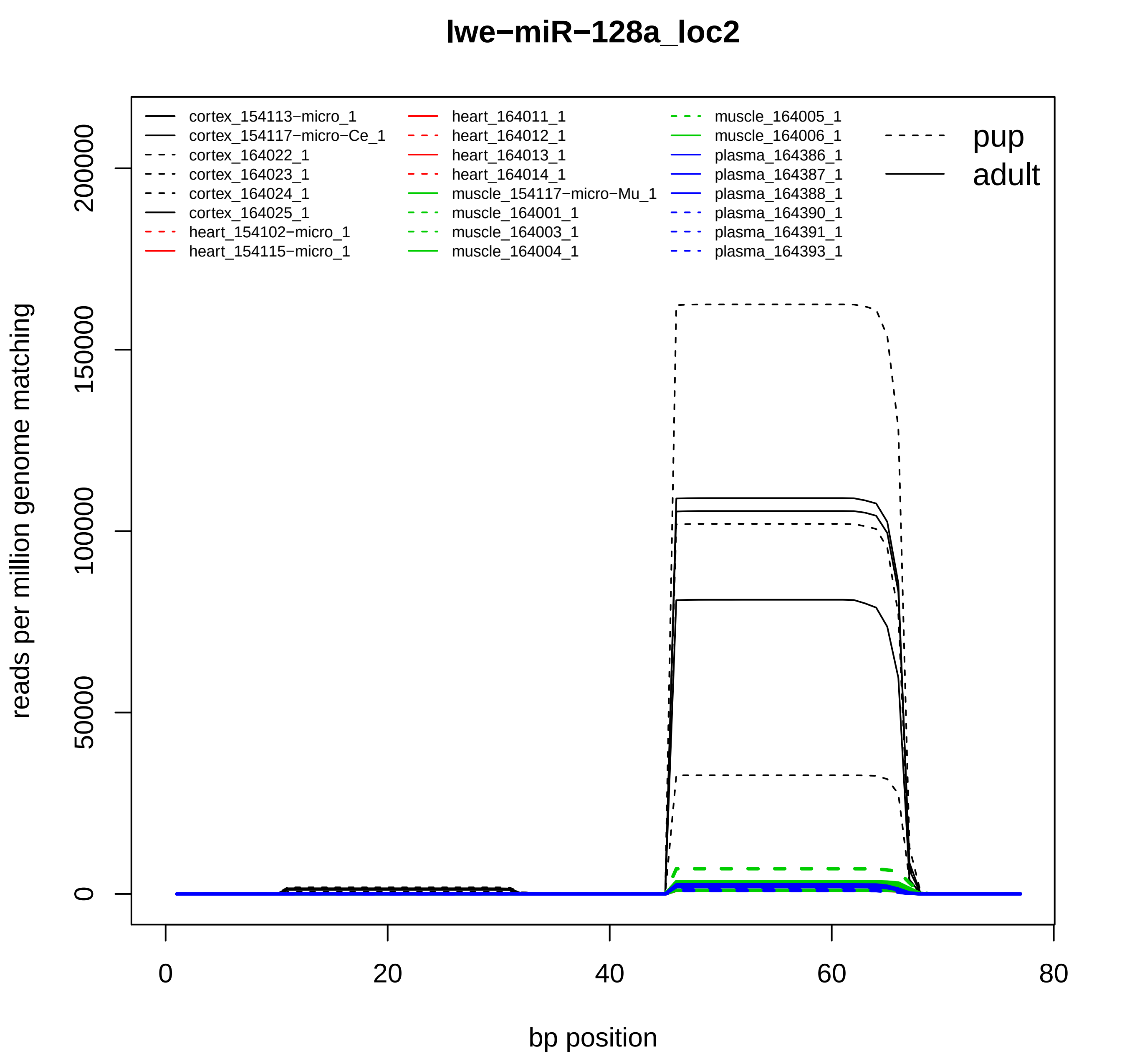

lwe−miR−128a_loc2
cortex_154113−micro_1
cortex_154117−micro−Ce_1
cortex_164022_1
cortex_164023_1
cortex_164024_1
cortex_164025_1
heart_154102−micro_1
heart_154115−micro_1
heart_164011_1
heart_164012_1
heart_164013_1
heart_164014_1
muscle_154117−micro−Mu_1
muscle_164001_1
muscle_164003_1
muscle_164004_1
muscle_164005_1
muscle_164006_1
plasma_164386_1
plasma_164387_1
plasma_164388_1
plasma_164390_1
plasma_164391_1
plasma_164393_1
pup
adult
reads per million genome matching
0
50000
100000
150000
200000
bp position
0
20
40
60
80

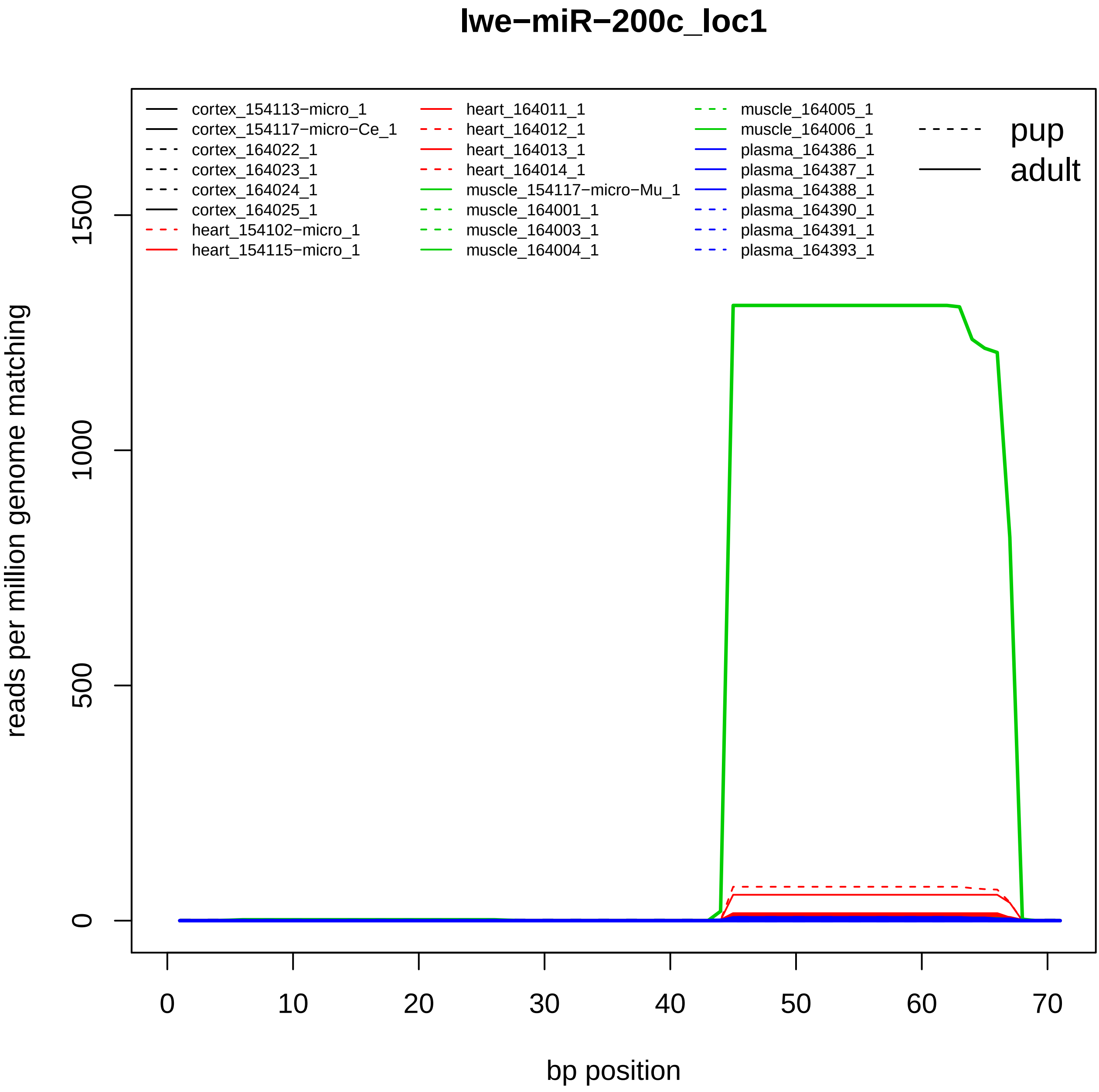

lwe−miR−200c_loc1
cortex_154113−micro_1
cortex_154117−micro−Ce_1
cortex_164022_1
cortex_164023_1
cortex_164024_1
cortex_164025_1
heart_154102−micro_1
heart_154115−micro_1
heart_164011_1
heart_164012_1
heart_164013_1
heart_164014_1
muscle_154117−micro−Mu_1
muscle_164001_1
muscle_164003_1
muscle_164004_1
muscle_164005_1
muscle_164006_1
plasma_164386_1
plasma_164387_1
plasma_164388_1
plasma_164390_1
plasma_164391_1
plasma_164393_1
pup
adult
reads per million genome matching
0
500
1000
1500
0
10
20
30
40
50
60
70
bp position

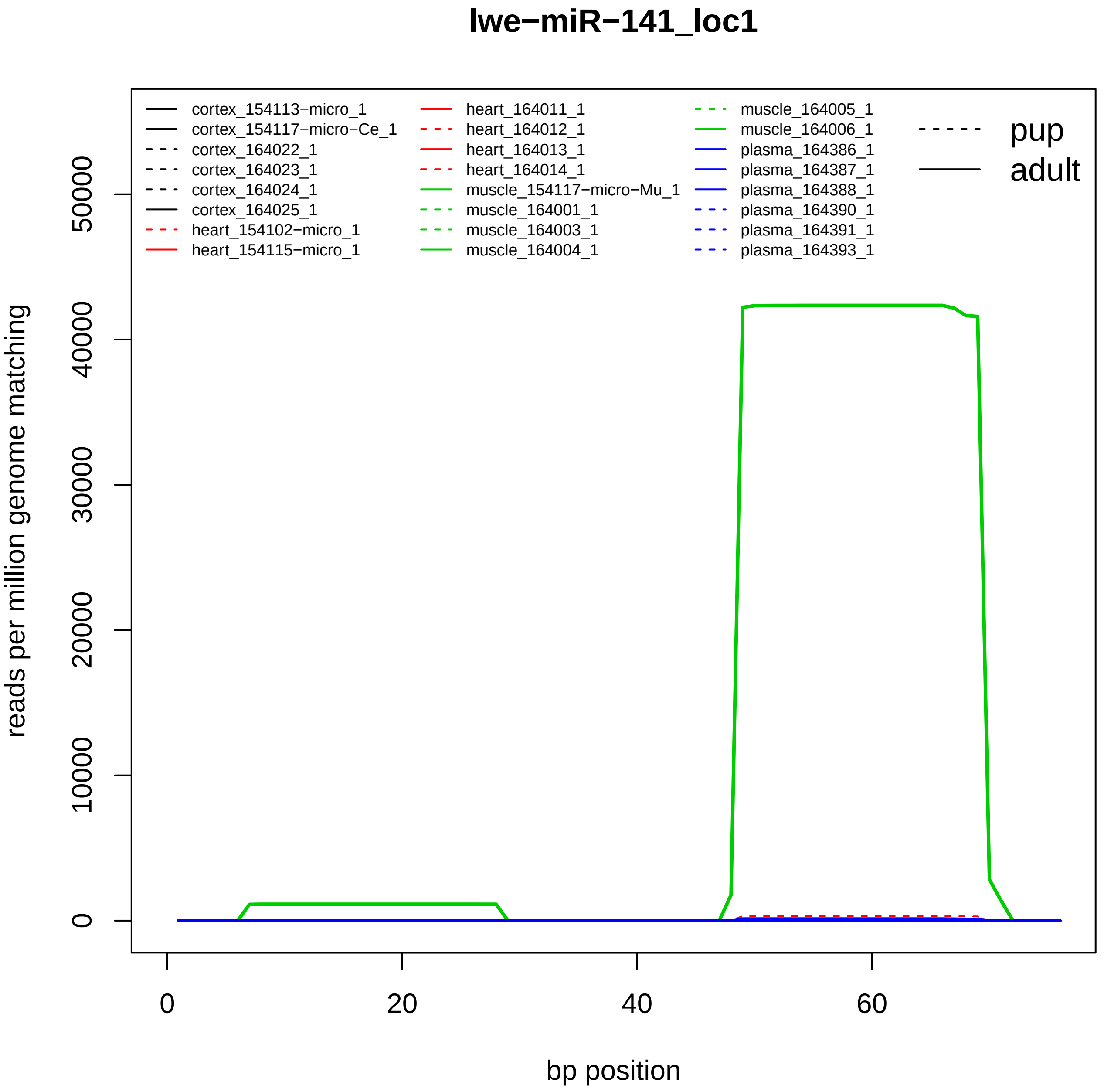
lwe−miR−141_loc1
cortex_154113−micro_1
cortex_154117−micro−Ce_1
cortex_164022_1
cortex_164023_1
cortex_164024_1
cortex_164025_1
heart_154102−micro_1
heart_154115−micro_1
heart_164011_1
heart_164012_1
heart_164013_1
heart_164014_1
muscle_154117−micro−Mu_1
muscle_164001_1
muscle_164003_1
muscle_164004_1
muscle_164005_1
muscle_164006_1
plasma_164386_1
plasma_164387_1
plasma_164388_1
plasma_164390_1
plasma_164391_1
plasma_164393_1
pup
adult
reads per million genome matching
0
10000
20000
30000
40000
50000
0
20
40
60
bp position

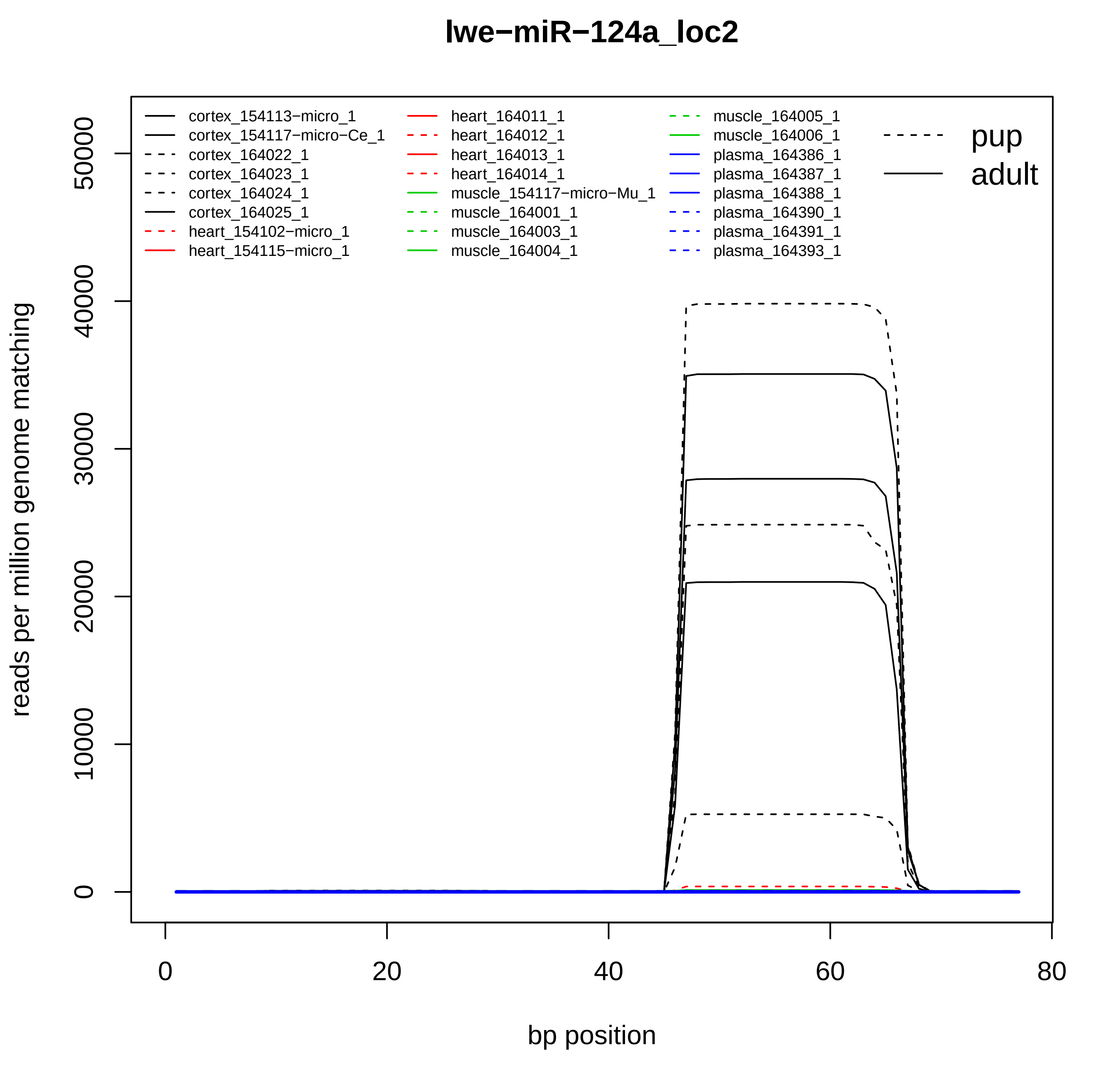
lwe−miR−124a_loc2
cortex_154113−micro_1
cortex_154117−micro−Ce_1
cortex_164022_1
cortex_164023_1
cortex_164024_1
cortex_164025_1
heart_154102−micro_1
heart_154115−micro_1
heart_164011_1
heart_164012_1
heart_164013_1
heart_164014_1
muscle_154117−micro−Mu_1
muscle_164001_1
muscle_164003_1
muscle_164004_1
muscle_164005_1
muscle_164006_1
plasma_164386_1
plasma_164387_1
plasma_164388_1
plasma_164390_1
plasma_164391_1
plasma_164393_1
pup
adult
reads per million genome matching
0
10000
20000
30000
40000
50000
bp position
0
20
40
60
80

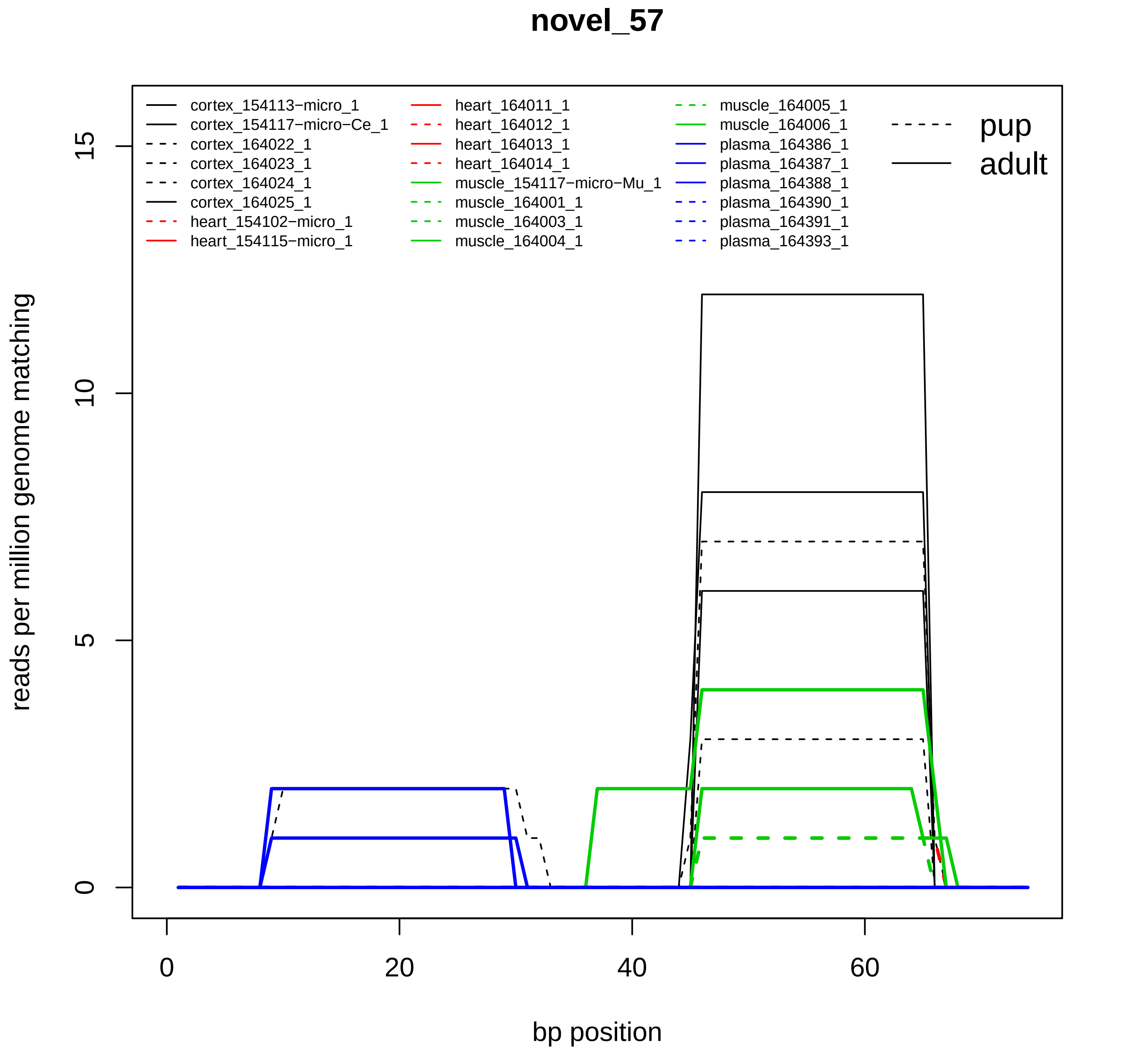

novel_57
cortex_154113−micro_1
cortex_154117−micro−Ce_1
cortex_164022_1
cortex_164023_1
cortex_164024_1
cortex_164025_1
heart_154102−micro_1
heart_154115−micro_1
heart_164011_1
heart_164012_1
heart_164013_1
heart_164014_1
muscle_154117−micro−Mu_1
muscle_164001_1
muscle_164003_1
muscle_164004_1
muscle_164005_1
muscle_164006_1
plasma_164386_1
plasma_164387_1
plasma_164388_1
plasma_164390_1
plasma_164391_1
plasma_164393_1
pup
adult
reads per million genome matching
bp position
0
5
10
15
0
20
40
60

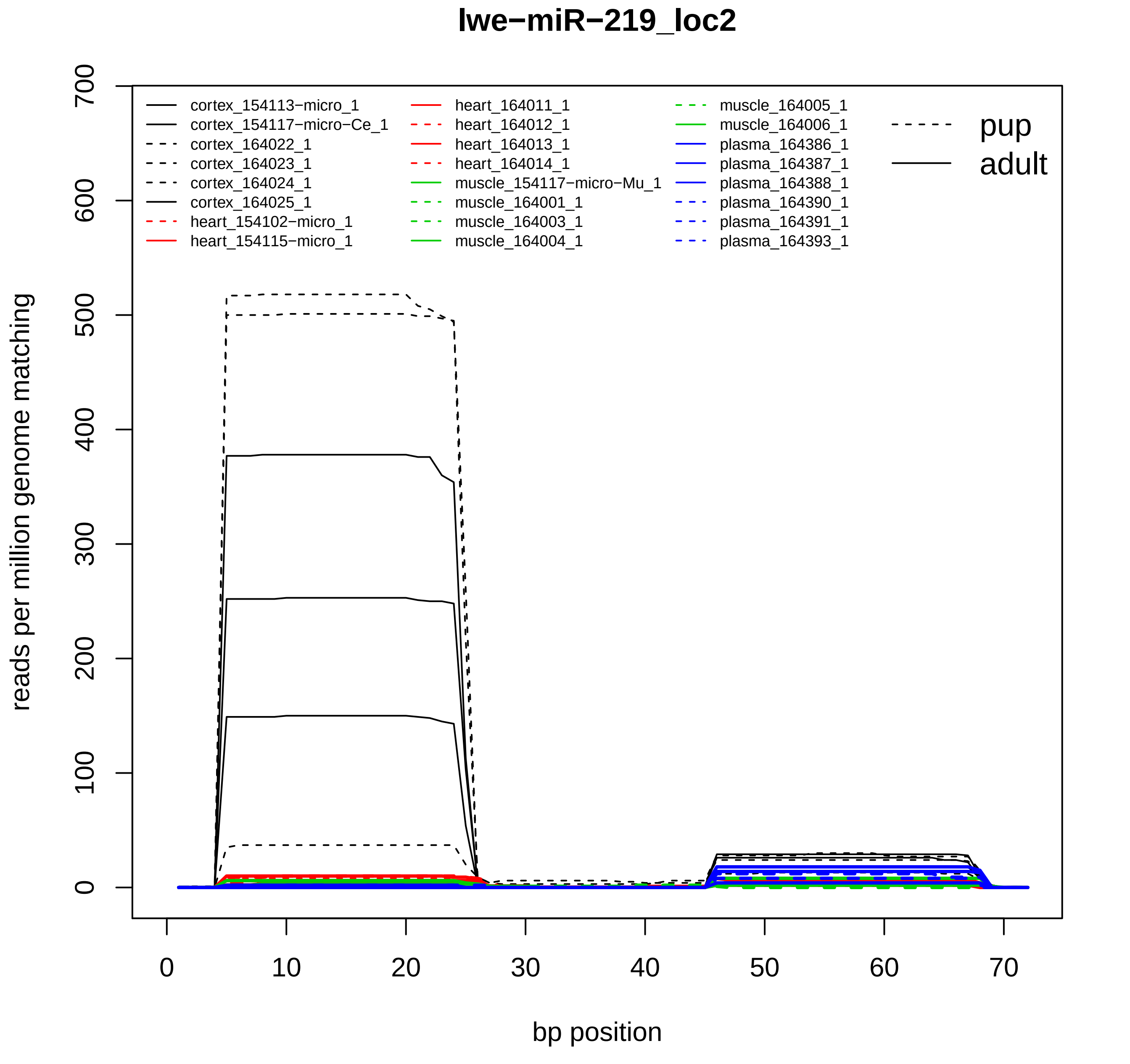

lwe−miR−219_loc2
reads per million genome matching
bp position
0
100
200
300
400
500
600
700
0
10
20
30
40
50
60
70
cortex_154113−micro_1
cortex_154117−micro−Ce_1
cortex_164022_1
cortex_164023_1
cortex_164024_1
cortex_164025_1
heart_154102−micro_1
heart_154115−micro_1
heart_164011_1
heart_164012_1
heart_164013_1
heart_164014_1
muscle_154117−micro−Mu_1
muscle_164001_1
muscle_164003_1
muscle_164004_1
muscle_164005_1
muscle_164006_1
plasma_164386_1
plasma_164387_1
plasma_164388_1
plasma_164390_1
plasma_164391_1
plasma_164393_1
pup
adult

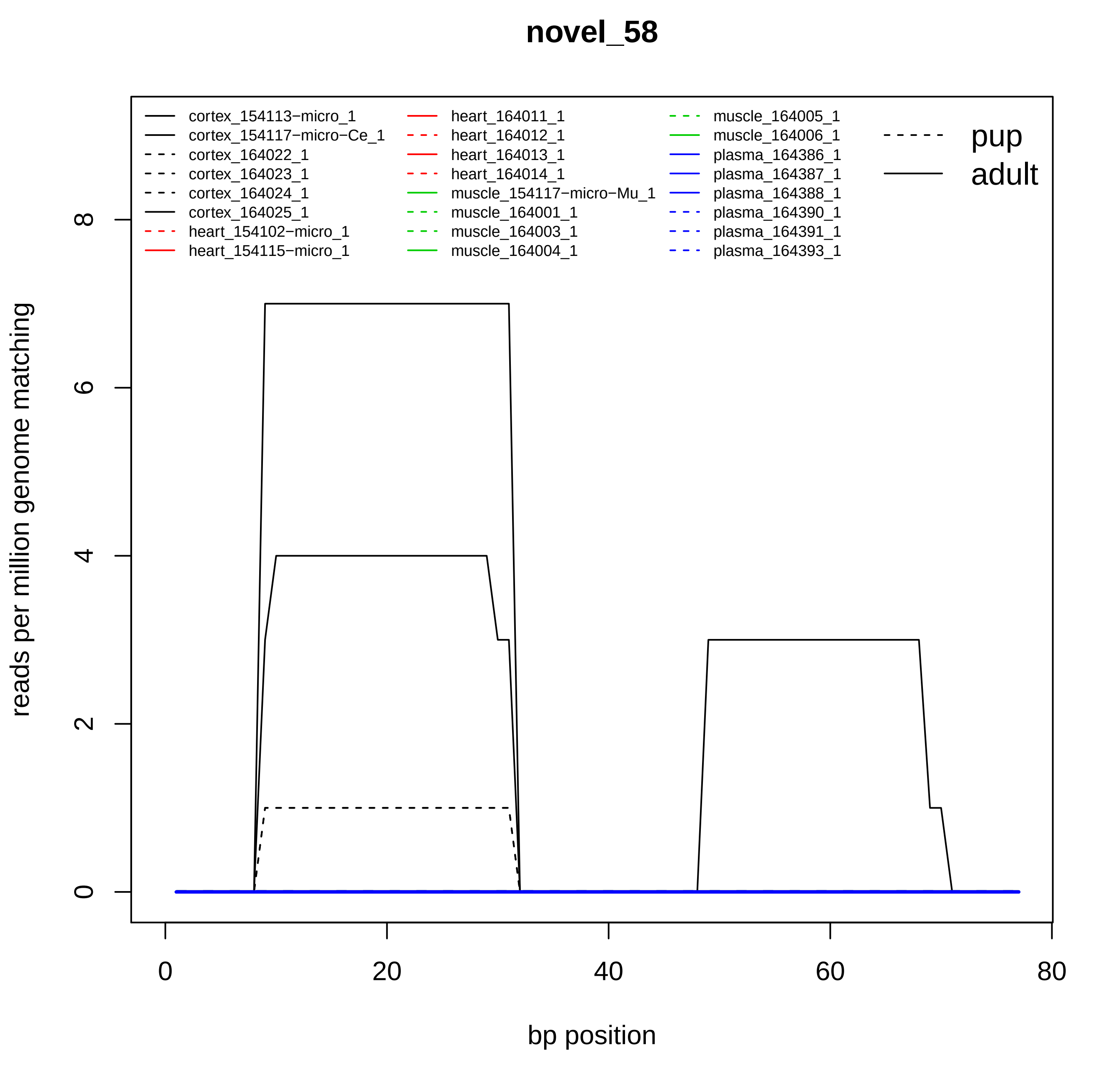

novel_58
cortex_154113−micro_1
cortex_154117−micro−Ce_1
cortex_164022_1
cortex_164023_1
cortex_164024_1
cortex_164025_1
heart_154102−micro_1
heart_154115−micro_1
heart_164011_1
heart_164012_1
heart_164013_1
heart_164014_1
muscle_154117−micro−Mu_1
muscle_164001_1
muscle_164003_1
muscle_164004_1
muscle_164005_1
muscle_164006_1
plasma_164386_1
plasma_164387_1
plasma_164388_1
plasma_164390_1
plasma_164391_1
plasma_164393_1
pup
adult
reads per million genome matching
0
2
4
6
8
0
20
40
60
80
bp position

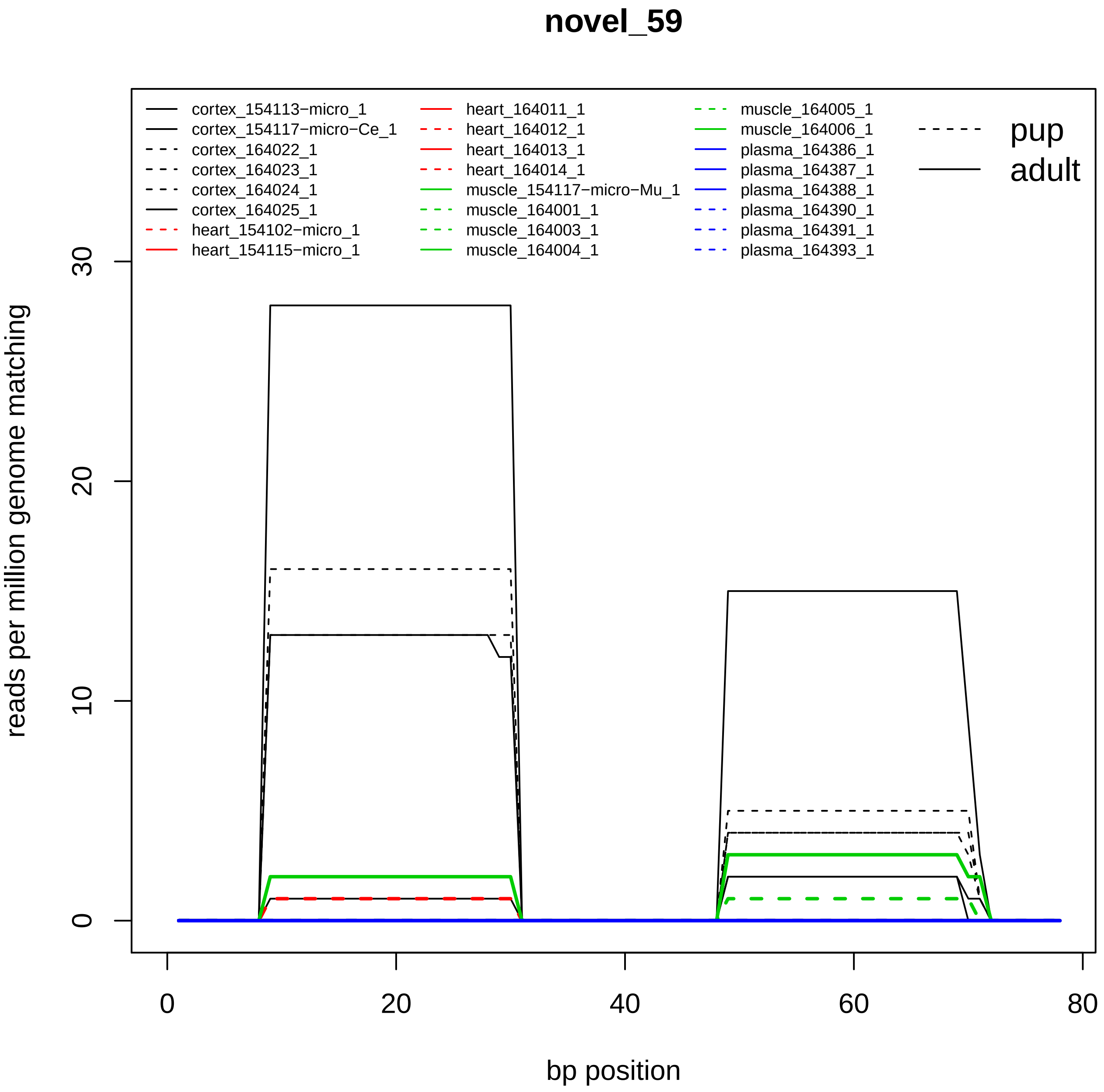

novel_59
cortex_154113−micro_1
cortex_154117−micro−Ce_1
cortex_164022_1
cortex_164023_1
cortex_164024_1
cortex_164025_1
heart_154102−micro_1
heart_154115−micro_1
heart_164011_1
heart_164012_1
heart_164013_1
heart_164014_1
muscle_154117−micro−Mu_1
muscle_164001_1
muscle_164003_1
muscle_164004_1
muscle_164005_1
muscle_164006_1
plasma_164386_1
plasma_164387_1
plasma_164388_1
plasma_164390_1
plasma_164391_1
plasma_164393_1
pup
adult
reads per million genome matching
bp position
0
10
20
30
0
20
40
60
80

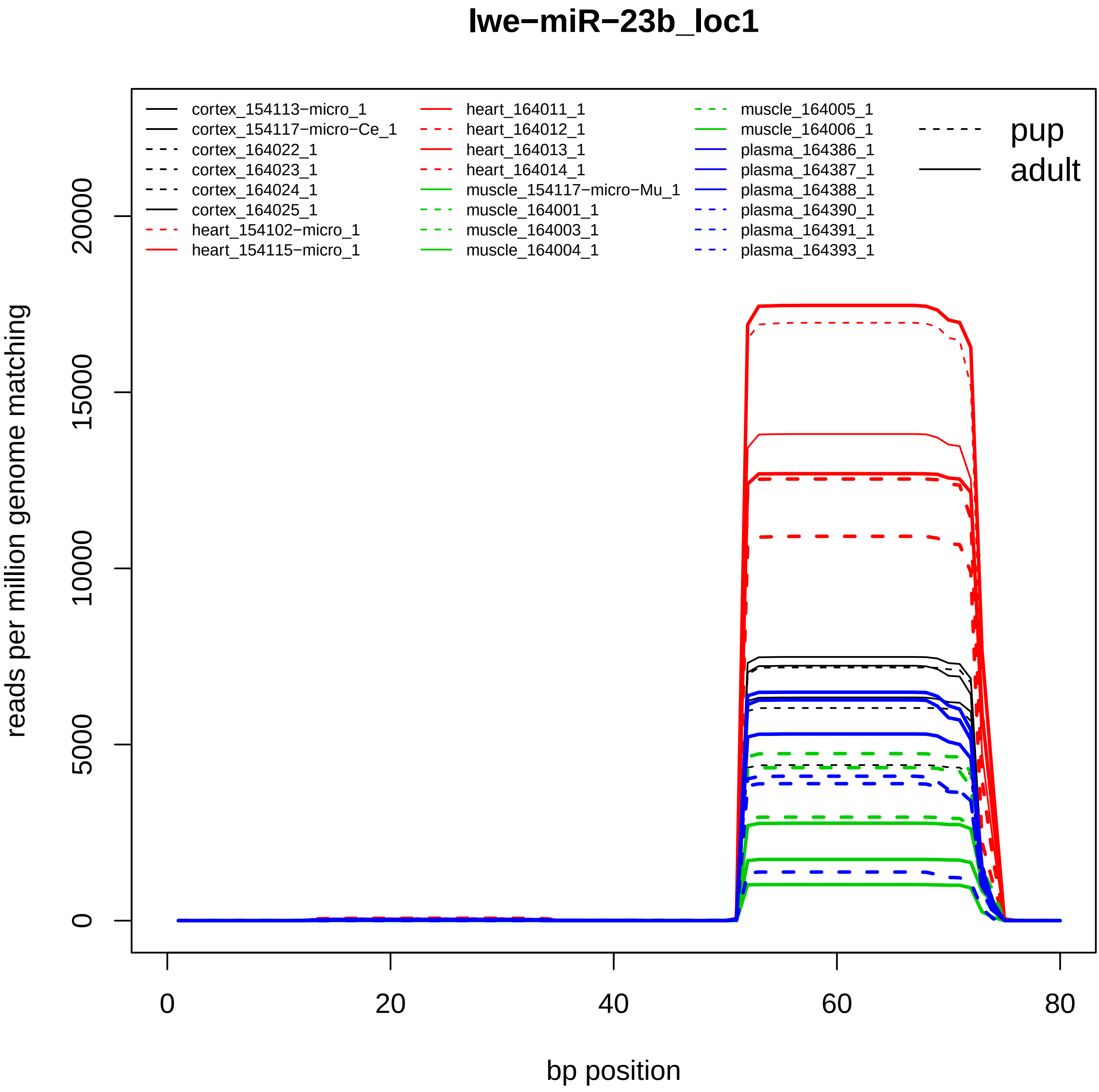
lwe−miR−23b_loc1
cortex_154113−micro_1
cortex_154117−micro−Ce_1
cortex_164022_1
cortex_164023_1
cortex_164024_1
cortex_164025_1
heart_154102−micro_1
heart_154115−micro_1
heart_164011_1
heart_164012_1
heart_164013_1
heart_164014_1
muscle_154117−micro−Mu_1
muscle_164001_1
muscle_164003_1
muscle_164004_1
muscle_164005_1
muscle_164006_1
plasma_164386_1
plasma_164387_1
plasma_164388_1
plasma_164390_1
plasma_164391_1
plasma_164393_1
pup
adult
reads per million genome matching
0
5000
10000
15000
20000
bp position
0
20
40
60
80

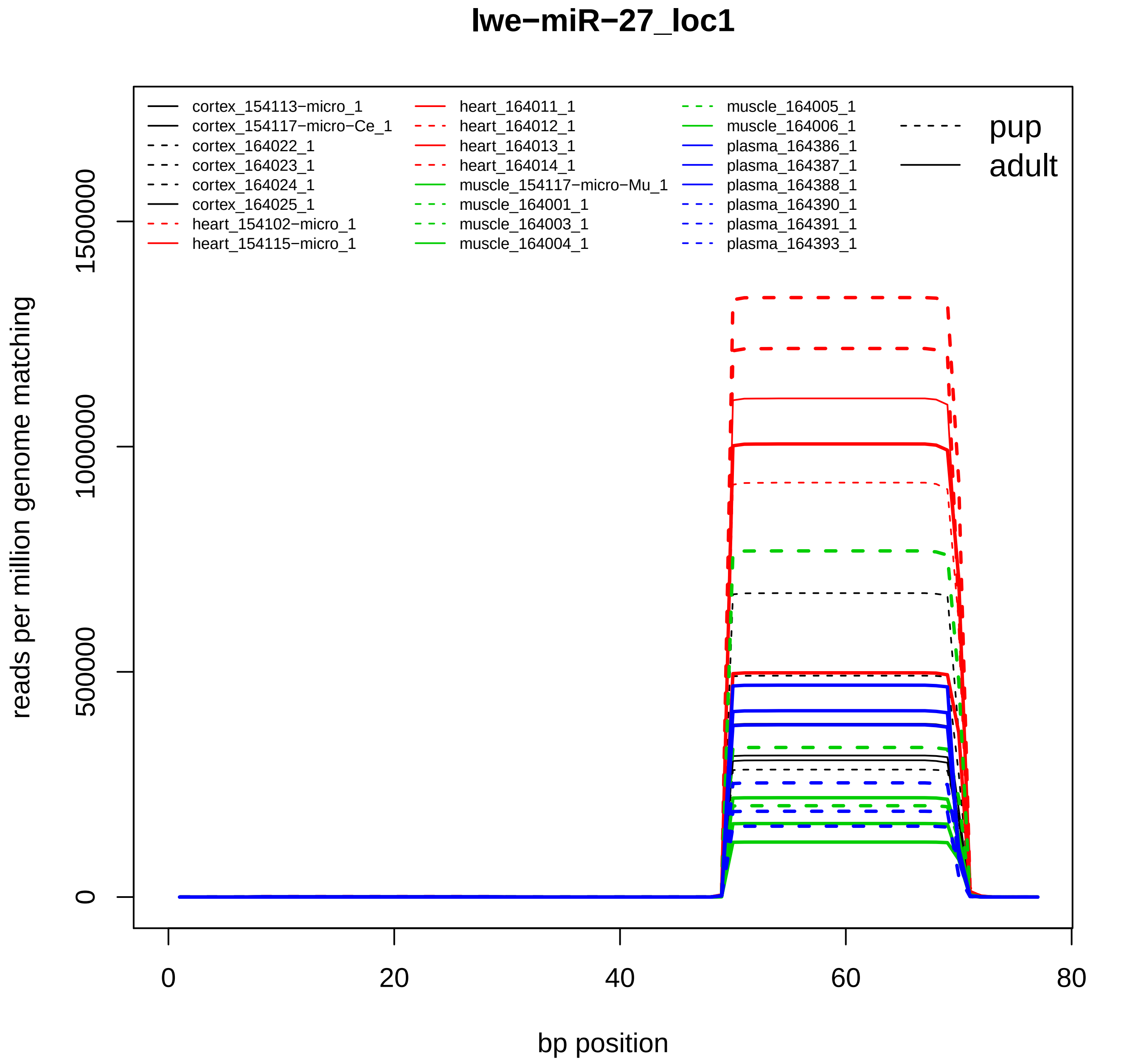
lwe−miR−27_loc1
cortex_154113−micro_1
cortex_154117−micro−Ce_1
cortex_164022_1
cortex_164023_1
cortex_164024_1
cortex_164025_1
heart_154102−micro_1
heart_154115−micro_1
heart_164011_1
heart_164012_1
heart_164013_1
heart_164014_1
muscle_154117−micro−Mu_1
muscle_164001_1
muscle_164003_1
muscle_164004_1
muscle_164005_1
muscle_164006_1
plasma_164386_1
plasma_164387_1
plasma_164388_1
plasma_164390_1
plasma_164391_1
plasma_164393_1
pup
adult
reads per million genome matching
0
500000
1000000
1500000
0
20
40
60
80
bp position

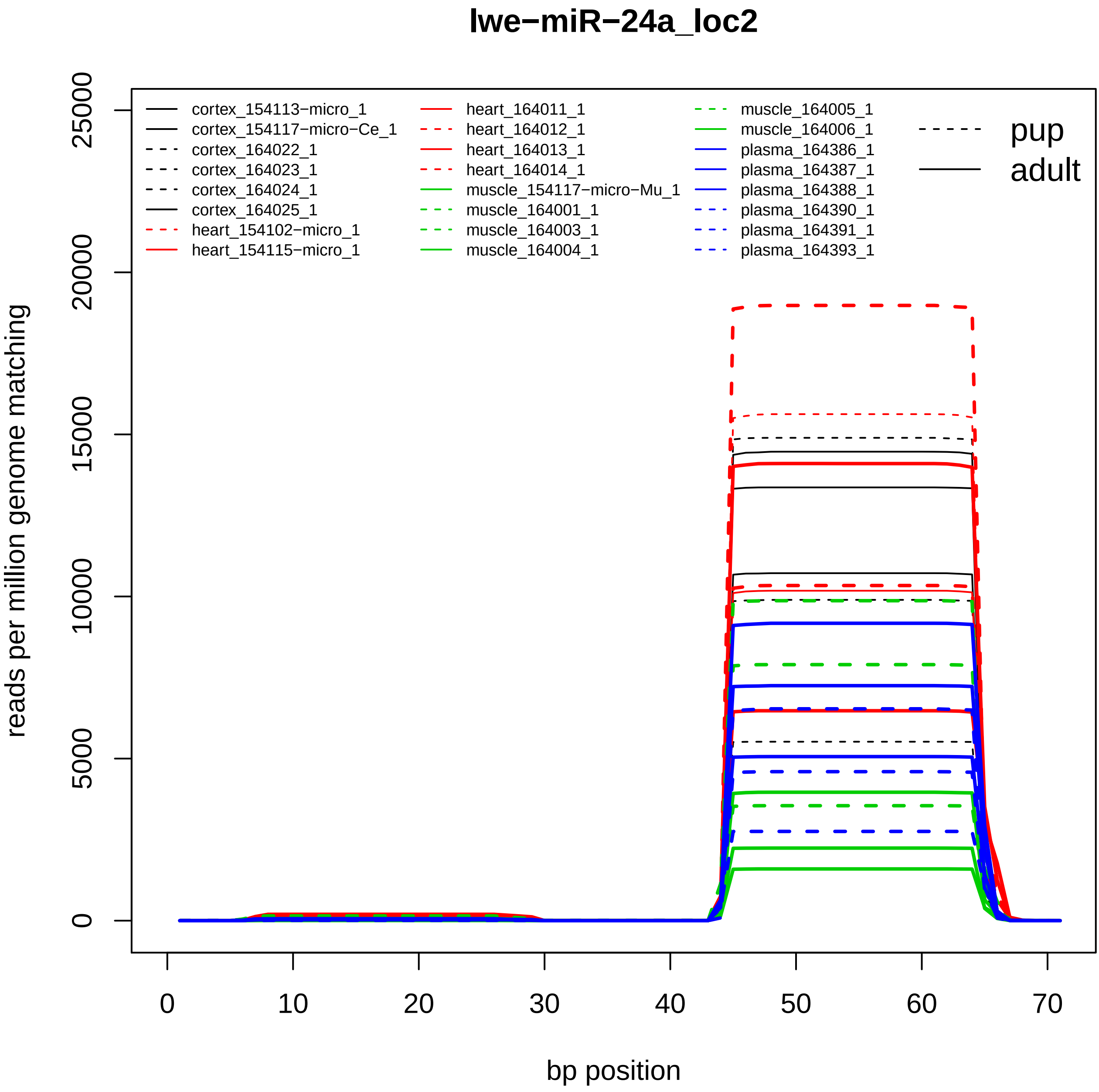

lwe−miR−24a_loc2
cortex_154113−micro_1
cortex_154117−micro−Ce_1
cortex_164022_1
cortex_164023_1
cortex_164024_1
cortex_164025_1
heart_154102−micro_1
heart_154115−micro_1
heart_164011_1
heart_164012_1
heart_164013_1
heart_164014_1
muscle_154117−micro−Mu_1
muscle_164001_1
muscle_164003_1
muscle_164004_1
muscle_164005_1
muscle_164006_1
plasma_164386_1
plasma_164387_1
plasma_164388_1
plasma_164390_1
plasma_164391_1
plasma_164393_1
pup
adult
reads per million genome matching
0
5000
10000
15000
20000
25000
bp position
0
10
20
30
40
50
60
70

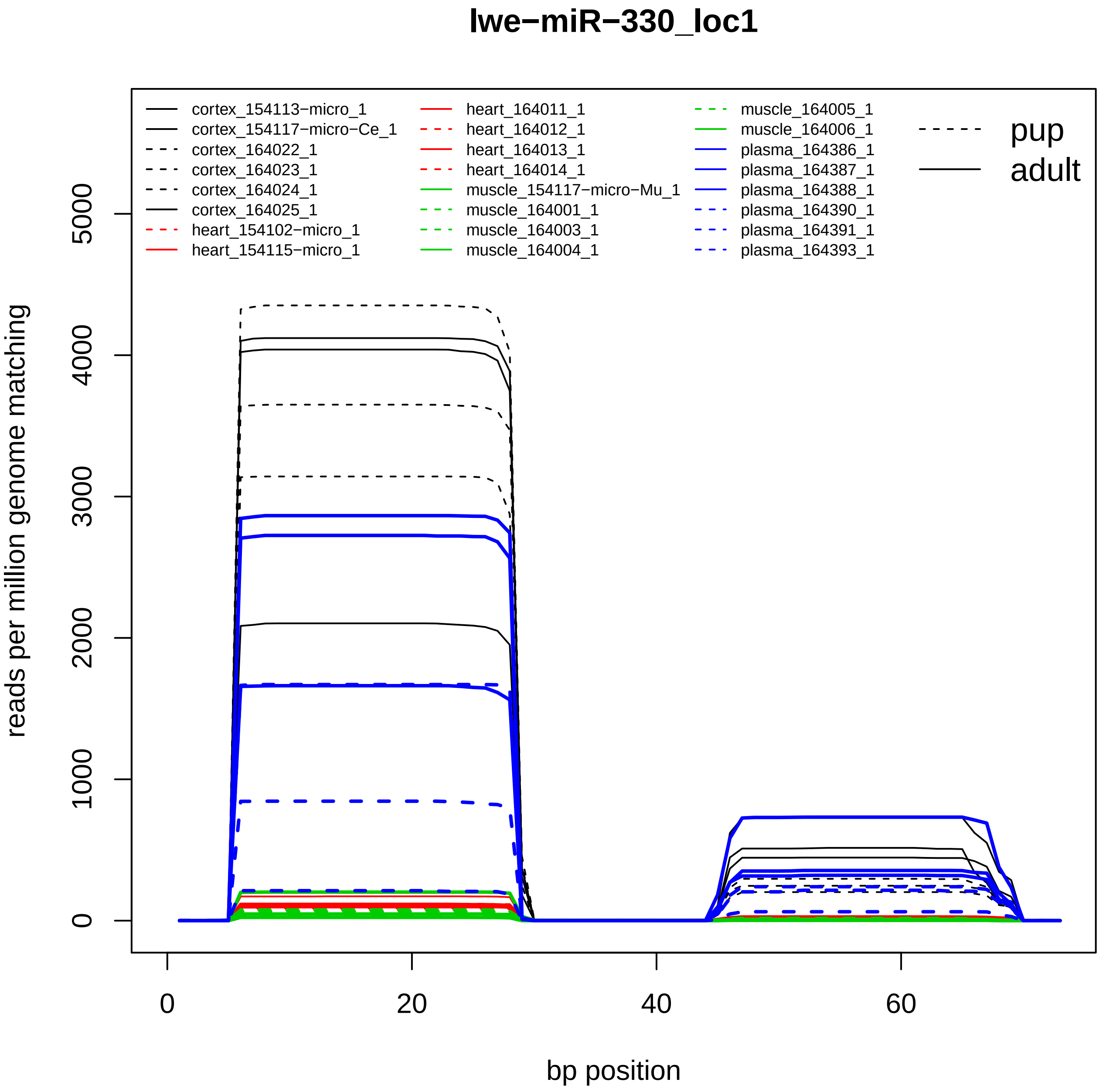

lwe−miR−330_loc1
cortex_154113−micro_1
cortex_154117−micro−Ce_1
cortex_164022_1
cortex_164023_1
cortex_164024_1
cortex_164025_1
heart_154102−micro_1
heart_154115−micro_1
heart_164011_1
heart_164012_1
heart_164013_1
heart_164014_1
muscle_154117−micro−Mu_1
muscle_164001_1
muscle_164003_1
muscle_164004_1
muscle_164005_1
muscle_164006_1
plasma_164386_1
plasma_164387_1
plasma_164388_1
plasma_164390_1
plasma_164391_1
plasma_164393_1
pup
adult
reads per million genome matching
0
1000
2000
3000
4000
5000
0
20
40
60
bp position

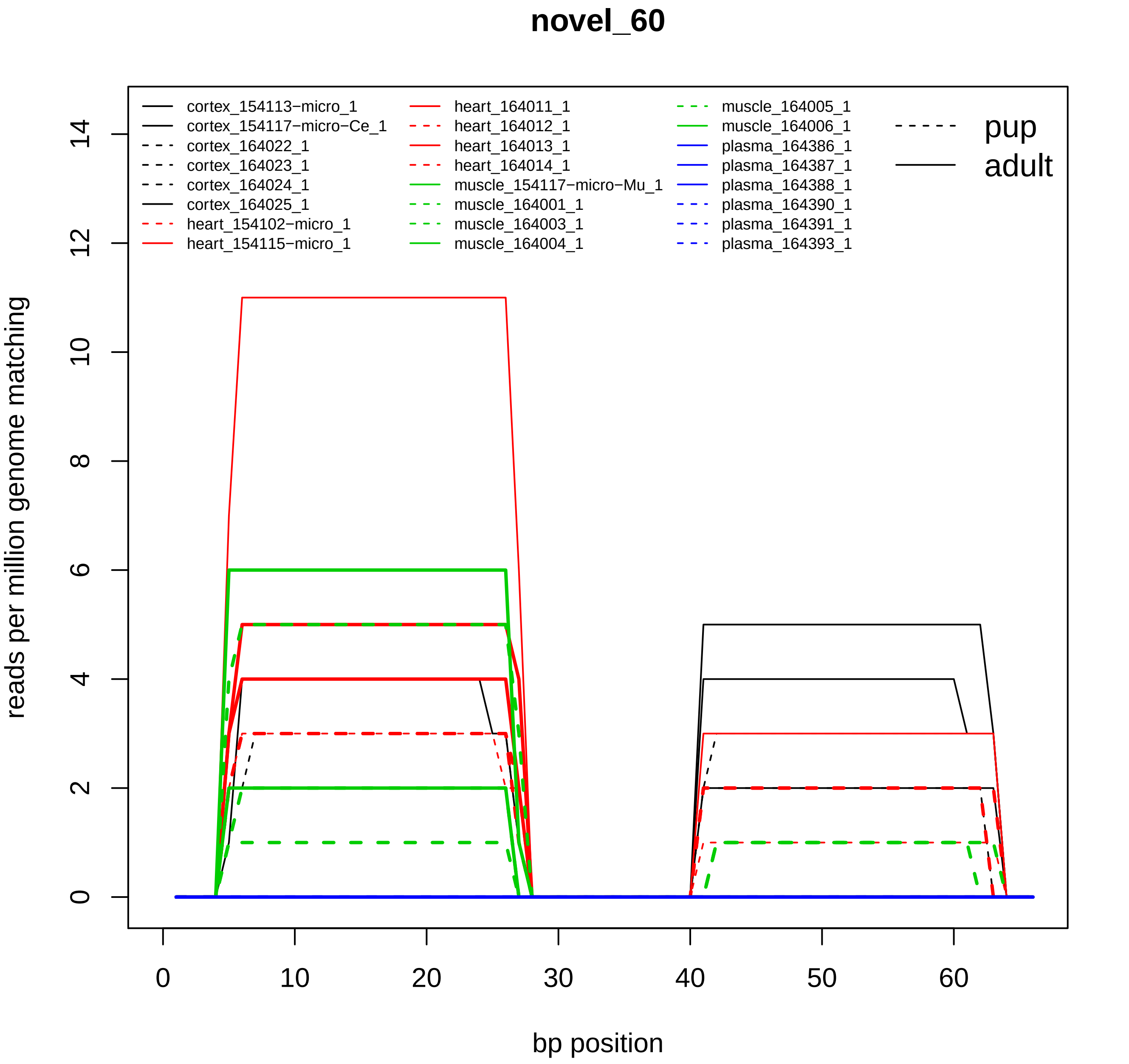

novel_60
cortex_154113−micro_1
cortex_154117−micro−Ce_1
cortex_164022_1
cortex_164023_1
cortex_164024_1
cortex_164025_1
heart_154102−micro_1
heart_154115−micro_1
heart_164011_1
heart_164012_1
heart_164013_1
heart_164014_1
muscle_154117−micro−Mu_1
muscle_164001_1
muscle_164003_1
muscle_164004_1
muscle_164005_1
muscle_164006_1
plasma_164386_1
plasma_164387_1
plasma_164388_1
plasma_164390_1
plasma_164391_1
plasma_164393_1
pup
adult
reads per million genome matching
0
2
4
6
8
10
12
14
0
10
20
30
40
50
60
bp position

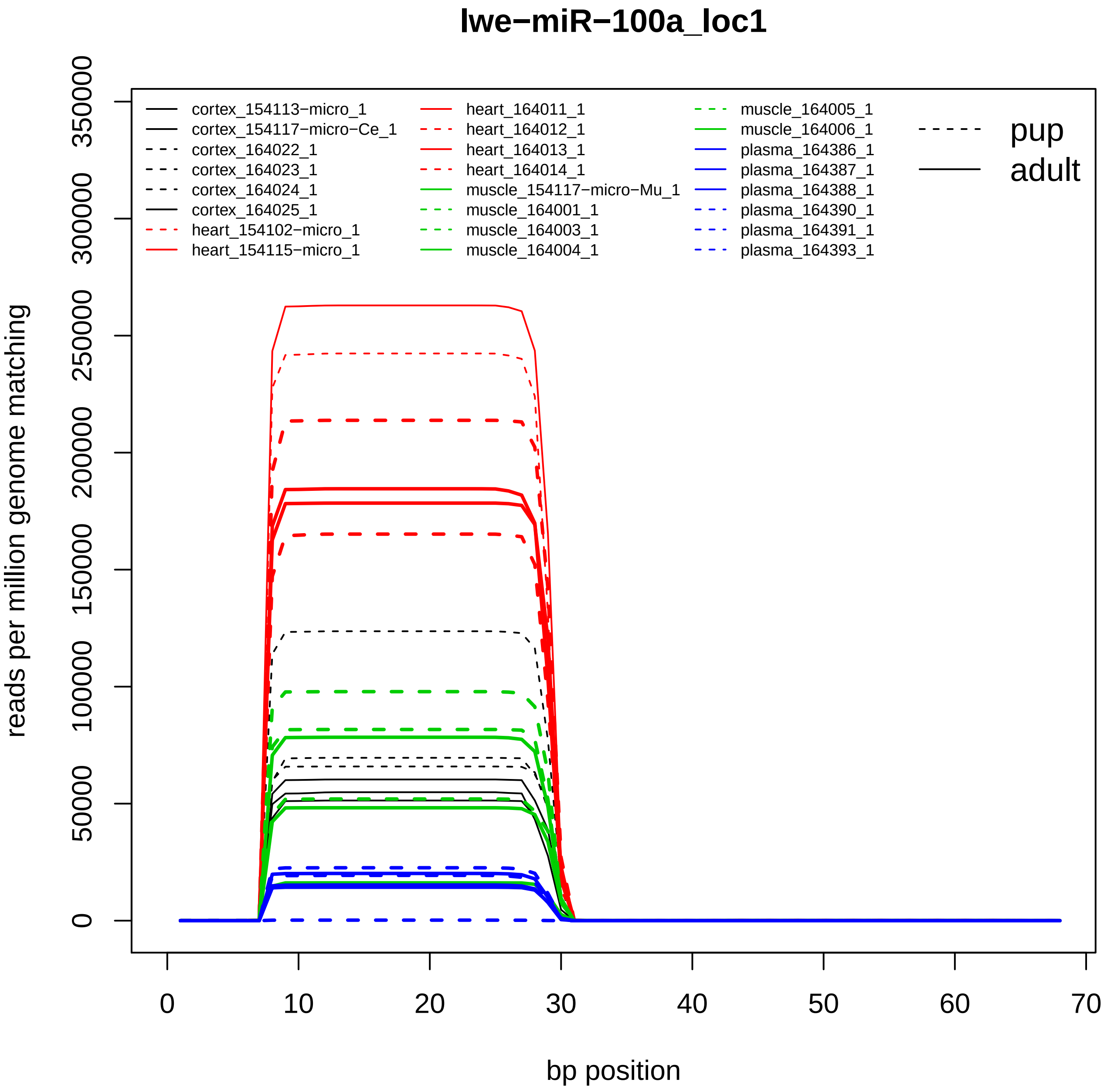

lwe−miR−100a_loc1
reads per million genome matching
bp position
0
50000
100000
150000
200000
250000
300000
350000
0
10
20
30
40
50
60
70
cortex_154113−micro_1
cortex_154117−micro−Ce_1
cortex_164022_1
cortex_164023_1
cortex_164024_1
cortex_164025_1
heart_154102−micro_1
heart_154115−micro_1
heart_164011_1
heart_164012_1
heart_164013_1
heart_164014_1
muscle_154117−micro−Mu_1
muscle_164001_1
muscle_164003_1
muscle_164004_1
muscle_164005_1
muscle_164006_1
plasma_164386_1
plasma_164387_1
plasma_164388_1
plasma_164390_1
plasma_164391_1
plasma_164393_1
pup
adult

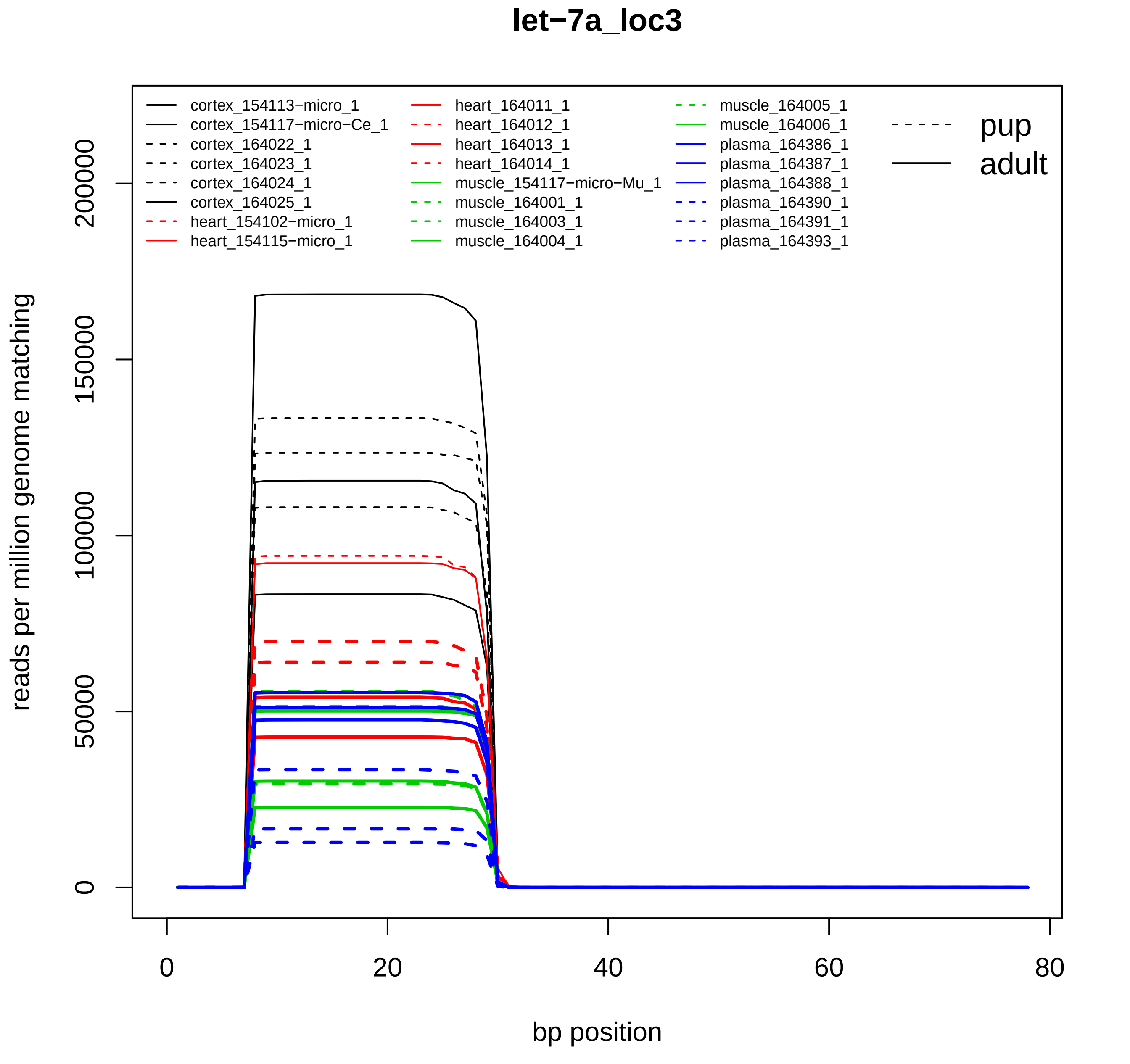
let−7a_loc3
cortex_154113−micro_1
cortex_154117−micro−Ce_1
cortex_164022_1
cortex_164023_1
cortex_164024_1
cortex_164025_1
heart_154102−micro_1
heart_154115−micro_1
heart_164011_1
heart_164012_1
heart_164013_1
heart_164014_1
muscle_154117−micro−Mu_1
muscle_164001_1
muscle_164003_1
muscle_164004_1
muscle_164005_1
muscle_164006_1
plasma_164386_1
plasma_164387_1
plasma_164388_1
plasma_164390_1
plasma_164391_1
plasma_164393_1
pup
adult
reads per million genome matching
0
50000
100000
150000
200000
bp position
0
20
40
60
80

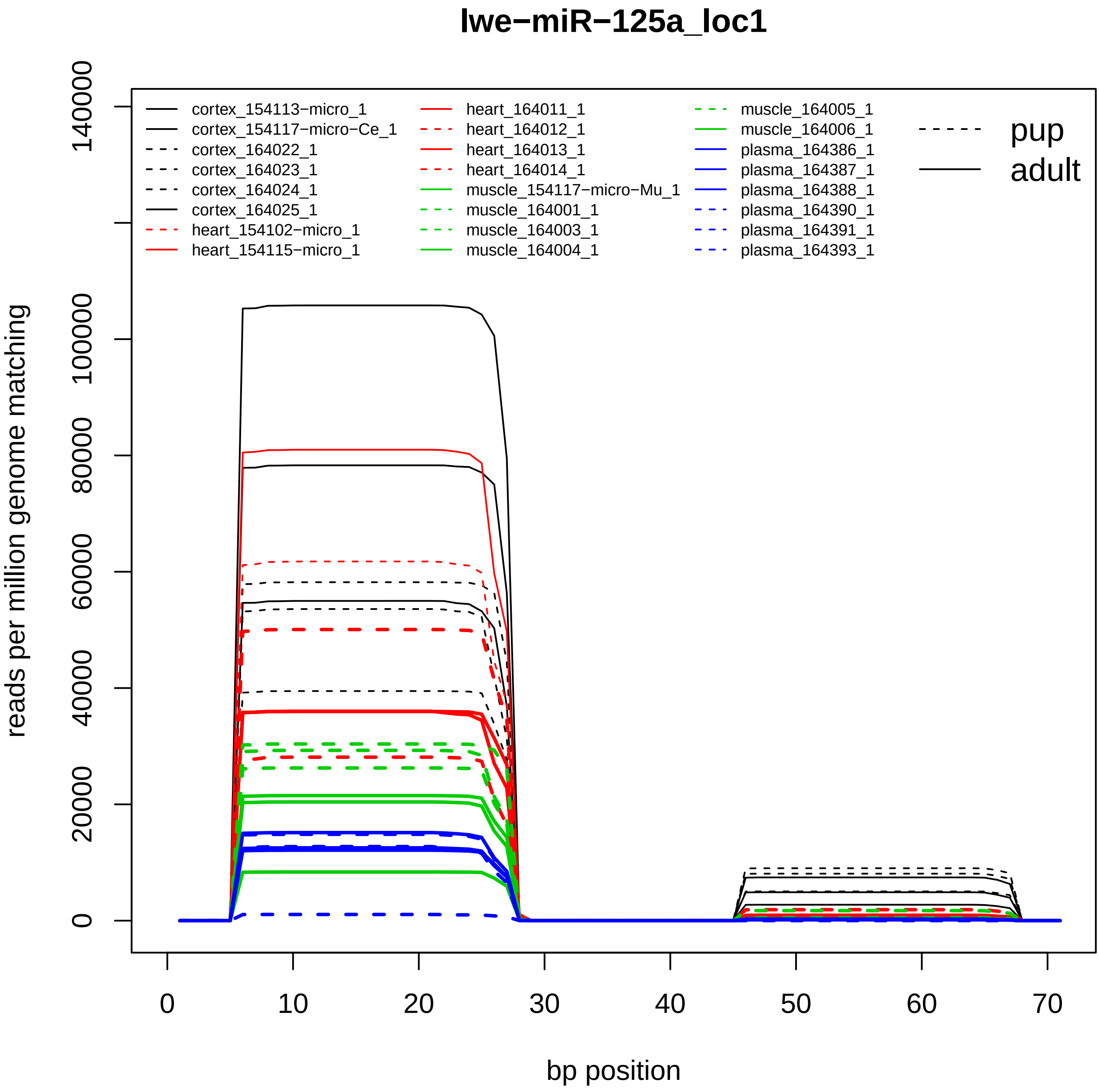
lwe−miR−125a_loc1
cortex_154113−micro_1
cortex_154117−micro−Ce_1
cortex_164022_1
cortex_164023_1
cortex_164024_1
cortex_164025_1
heart_154102−micro_1
heart_154115−micro_1
heart_164011_1
heart_164012_1
heart_164013_1
heart_164014_1
muscle_154117−micro−Mu_1
muscle_164001_1
muscle_164003_1
muscle_164004_1
muscle_164005_1
muscle_164006_1
plasma_164386_1
plasma_164387_1
plasma_164388_1
plasma_164390_1
plasma_164391_1
plasma_164393_1
pup
adult
reads per million genome matching
0
20000
40000
60000
80000
100000
140000
bp position
0
10
20
30
40
50
60
70

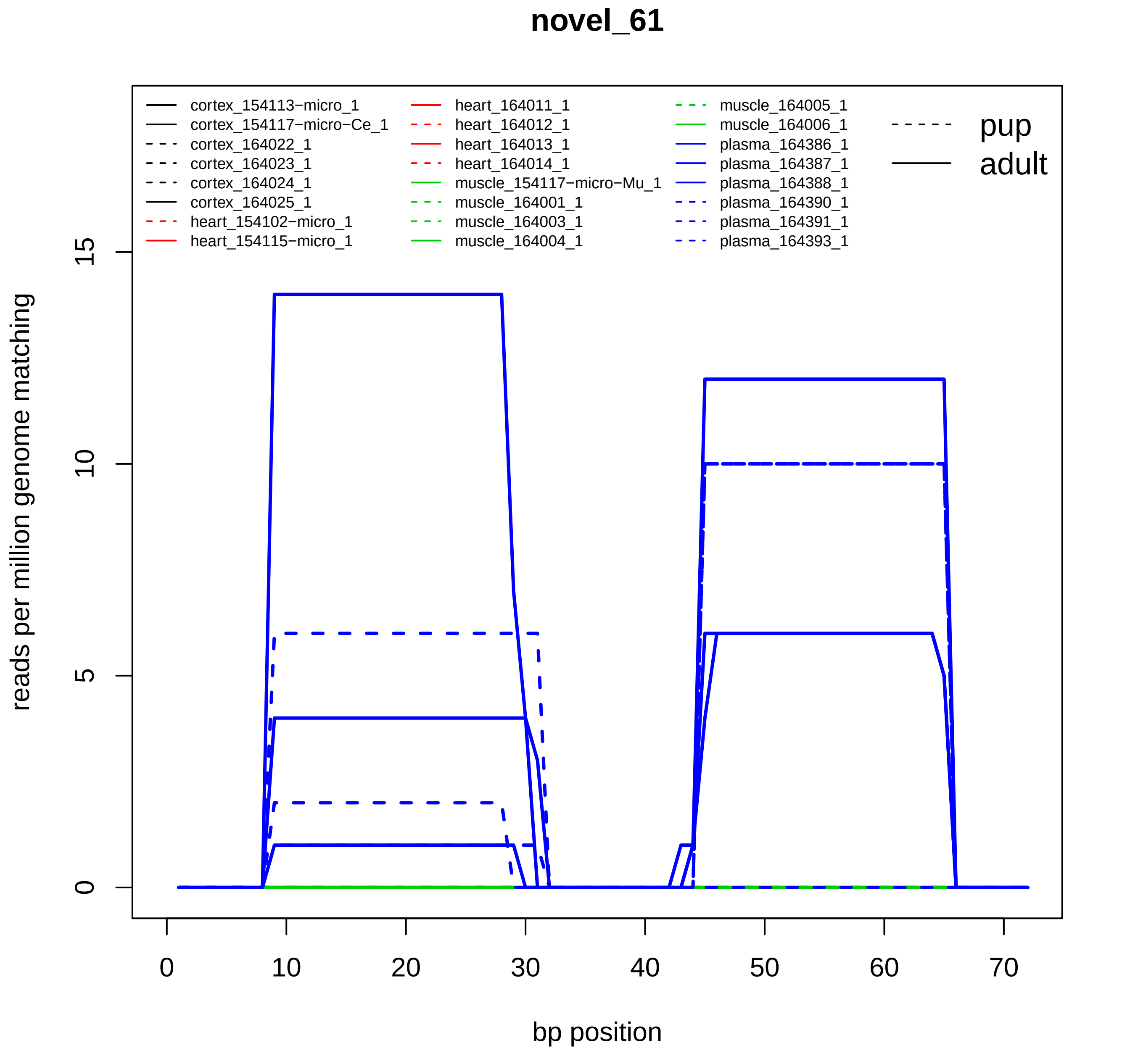
novel_61
cortex_154113−micro_1
cortex_154117−micro−Ce_1
cortex_164022_1
cortex_164023_1
cortex_164024_1
cortex_164025_1
heart_154102−micro_1
heart_154115−micro_1
heart_164011_1
heart_164012_1
heart_164013_1
heart_164014_1
muscle_154117−micro−Mu_1
muscle_164001_1
muscle_164003_1
muscle_164004_1
muscle_164005_1
muscle_164006_1
plasma_164386_1
plasma_164387_1
plasma_164388_1
plasma_164390_1
plasma_164391_1
plasma_164393_1
pup
adult
reads per million genome matching
0
5
10
15
0
10
20
30
40
50
60
70
bp position

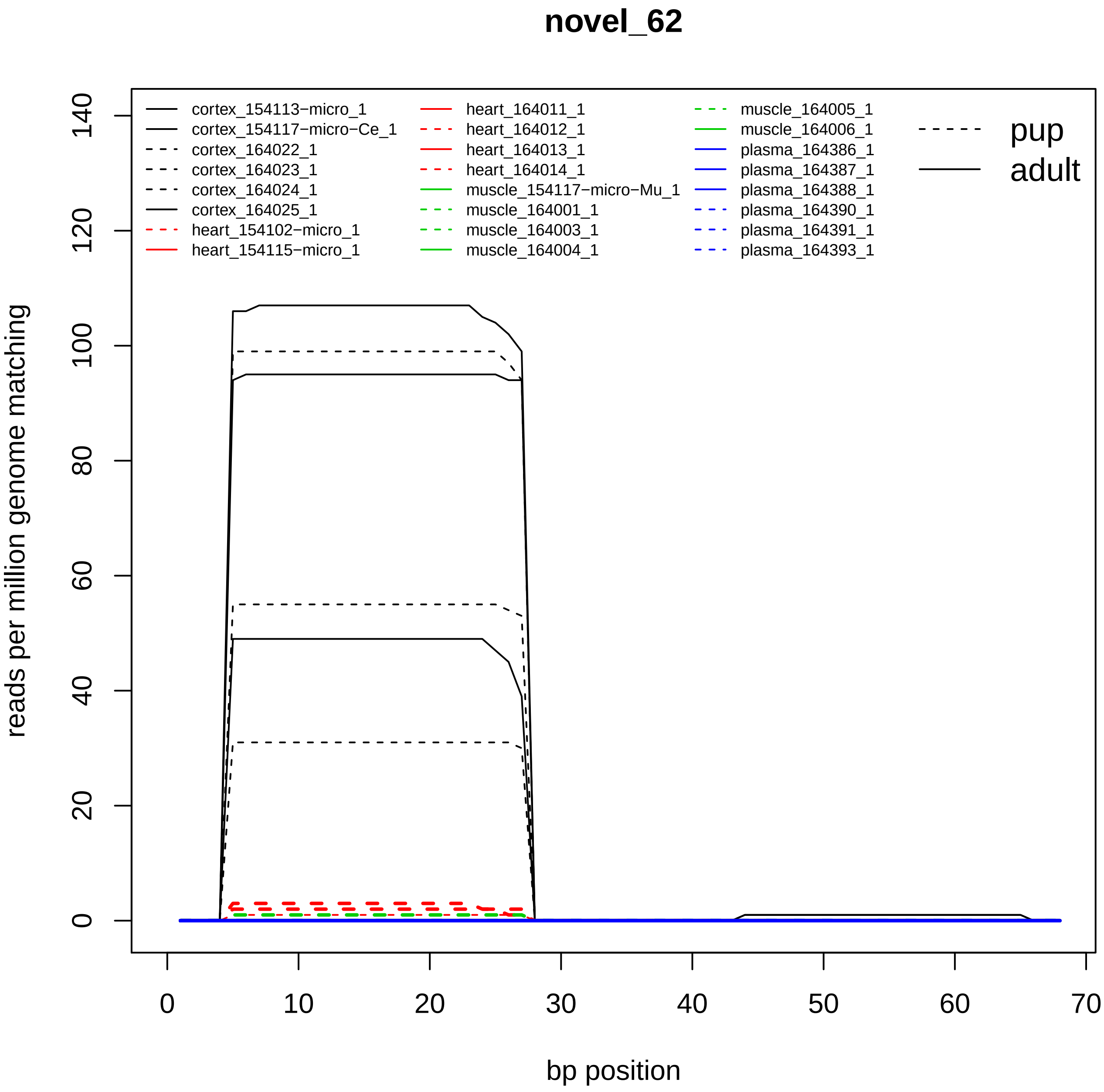

novel_62
cortex_154113−micro_1
cortex_154117−micro−Ce_1
cortex_164022_1
cortex_164023_1
cortex_164024_1
cortex_164025_1
heart_154102−micro_1
heart_154115−micro_1
heart_164011_1
heart_164012_1
heart_164013_1
heart_164014_1
muscle_154117−micro−Mu_1
muscle_164001_1
muscle_164003_1
muscle_164004_1
muscle_164005_1
muscle_164006_1
plasma_164386_1
plasma_164387_1
plasma_164388_1
plasma_164390_1
plasma_164391_1
plasma_164393_1
pup
adult
reads per million genome matching
bp position
0
20
40
60
80
100
120
140
0
10
20
30
40
50
60
70

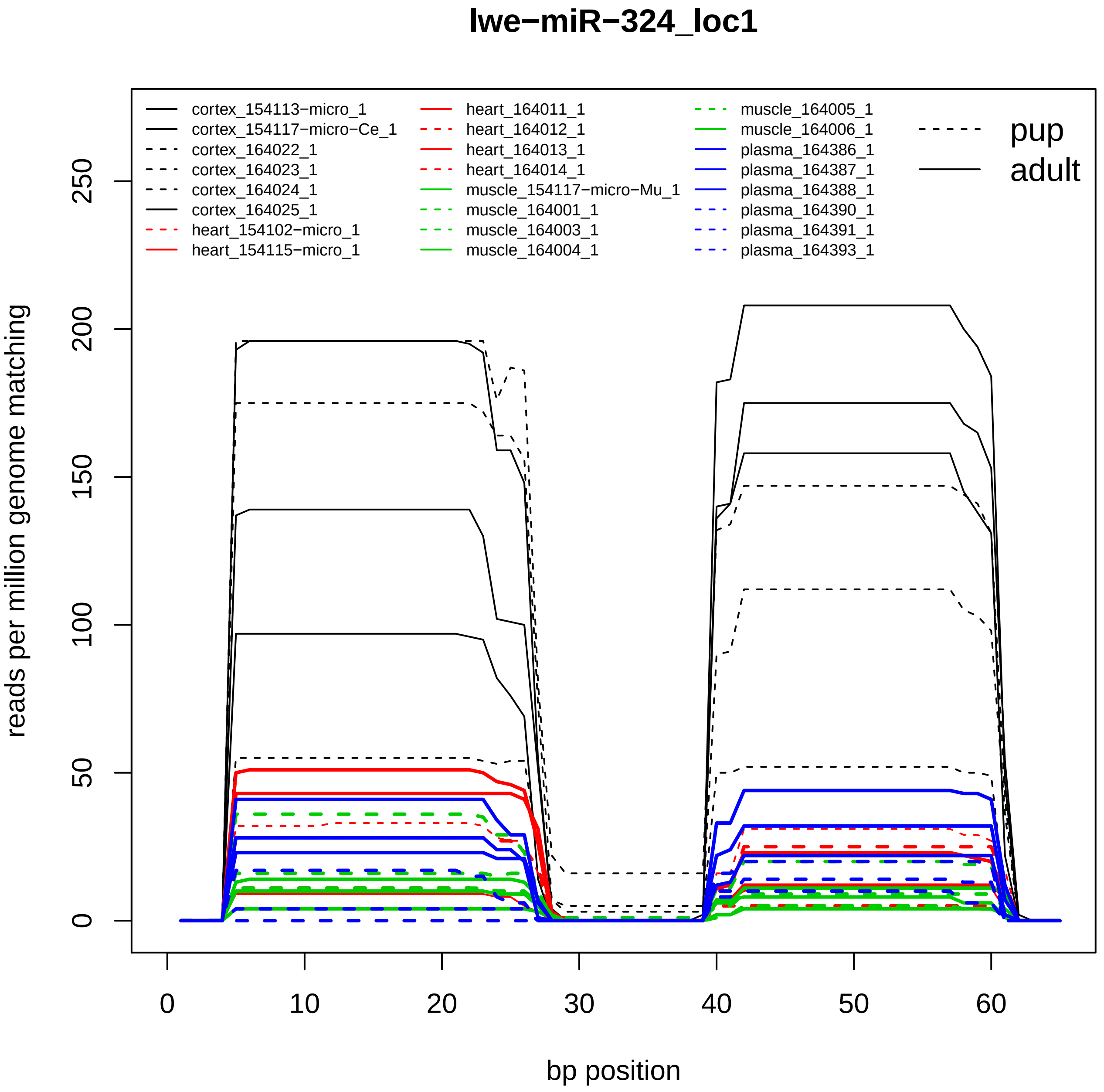
lwe−miR−324_loc1
cortex_154113−micro_1
cortex_154117−micro−Ce_1
cortex_164022_1
cortex_164023_1
cortex_164024_1
cortex_164025_1
heart_154102−micro_1
heart_154115−micro_1
heart_164011_1
heart_164012_1
heart_164013_1
heart_164014_1
muscle_154117−micro−Mu_1
muscle_164001_1
muscle_164003_1
muscle_164004_1
muscle_164005_1
muscle_164006_1
plasma_164386_1
plasma_164387_1
plasma_164388_1
plasma_164390_1
plasma_164391_1
plasma_164393_1
pup
adult
reads per million genome matching
0
50
100
150
200
250
0
10
20
30
40
50
60
bp position

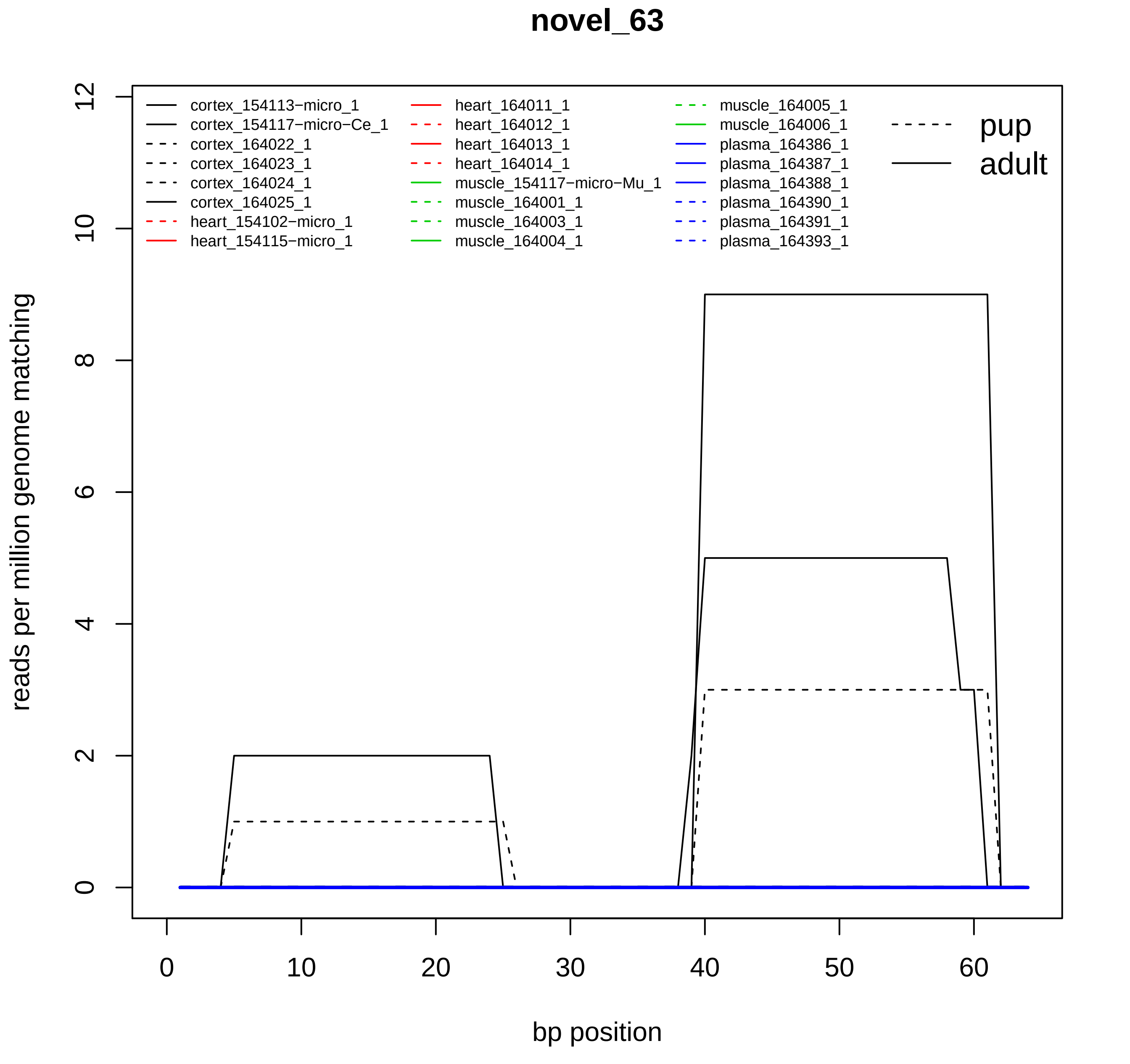
novel_63
cortex_154113−micro_1
cortex_154117−micro−Ce_1
cortex_164022_1
cortex_164023_1
cortex_164024_1
cortex_164025_1
heart_154102−micro_1
heart_154115−micro_1
heart_164011_1
heart_164012_1
heart_164013_1
heart_164014_1
muscle_154117−micro−Mu_1
muscle_164001_1
muscle_164003_1
muscle_164004_1
muscle_164005_1
muscle_164006_1
plasma_164386_1
plasma_164387_1
plasma_164388_1
plasma_164390_1
plasma_164391_1
plasma_164393_1
pup
adult
reads per million genome matching
bp position

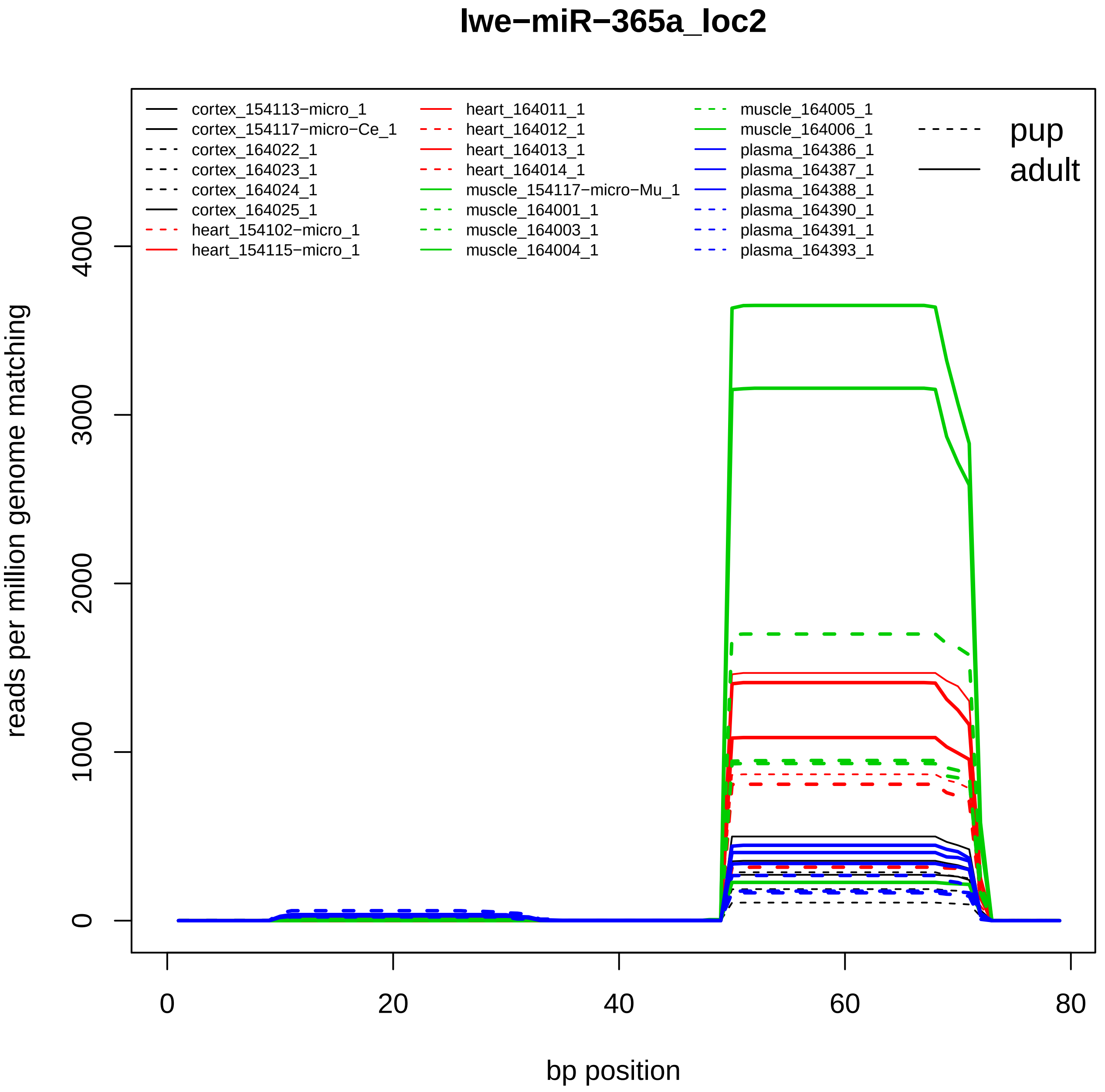
lwe−miR−365a_loc2
cortex_154113−micro_1
cortex_154117−micro−Ce_1
cortex_164022_1
cortex_164023_1
cortex_164024_1
cortex_164025_1
heart_154102−micro_1
heart_154115−micro_1
heart_164011_1
heart_164012_1
heart_164013_1
heart_164014_1
muscle_154117−micro−Mu_1
muscle_164001_1
muscle_164003_1
muscle_164004_1
muscle_164005_1
muscle_164006_1
plasma_164386_1
plasma_164387_1
plasma_164388_1
plasma_164390_1
plasma_164391_1
plasma_164393_1
pup
adult
reads per million genome matching
0
1000
2000
3000
4000
0
20
40
60
80
bp position

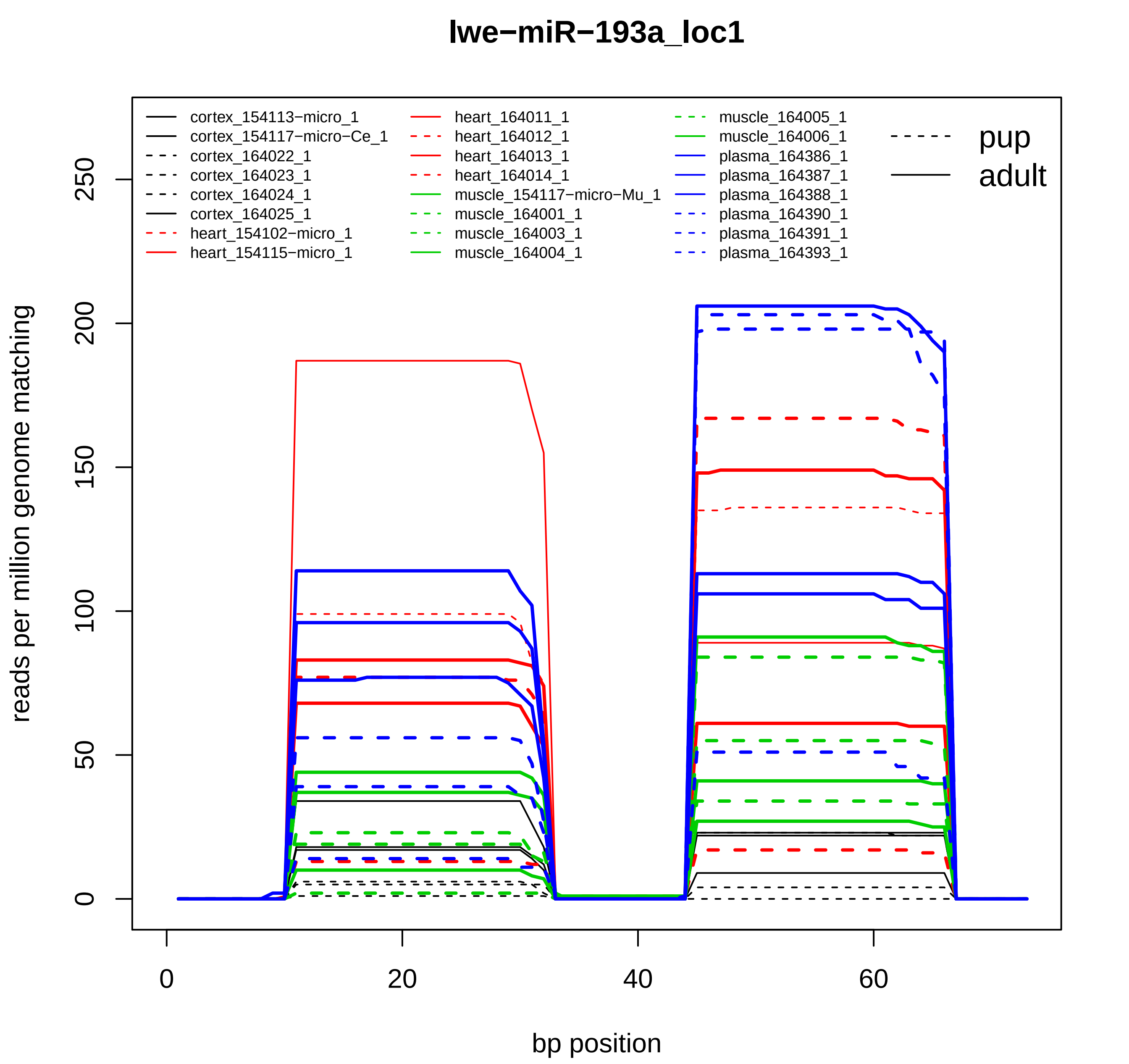
lwe−miR−193a_loc1
cortex_154113−micro_1
cortex_154117−micro−Ce_1
cortex_164022_1
cortex_164023_1
cortex_164024_1
cortex_164025_1
heart_154102−micro_1
heart_154115−micro_1
heart_164011_1
heart_164012_1
heart_164013_1
heart_164014_1
muscle_154117−micro−Mu_1
muscle_164001_1
muscle_164003_1
muscle_164004_1
muscle_164005_1
muscle_164006_1
plasma_164386_1
plasma_164387_1
plasma_164388_1
plasma_164390_1
plasma_164391_1
plasma_164393_1
pup
adult
reads per million genome matching
0
50
100
150
200
250
0
20
40
60
bp position

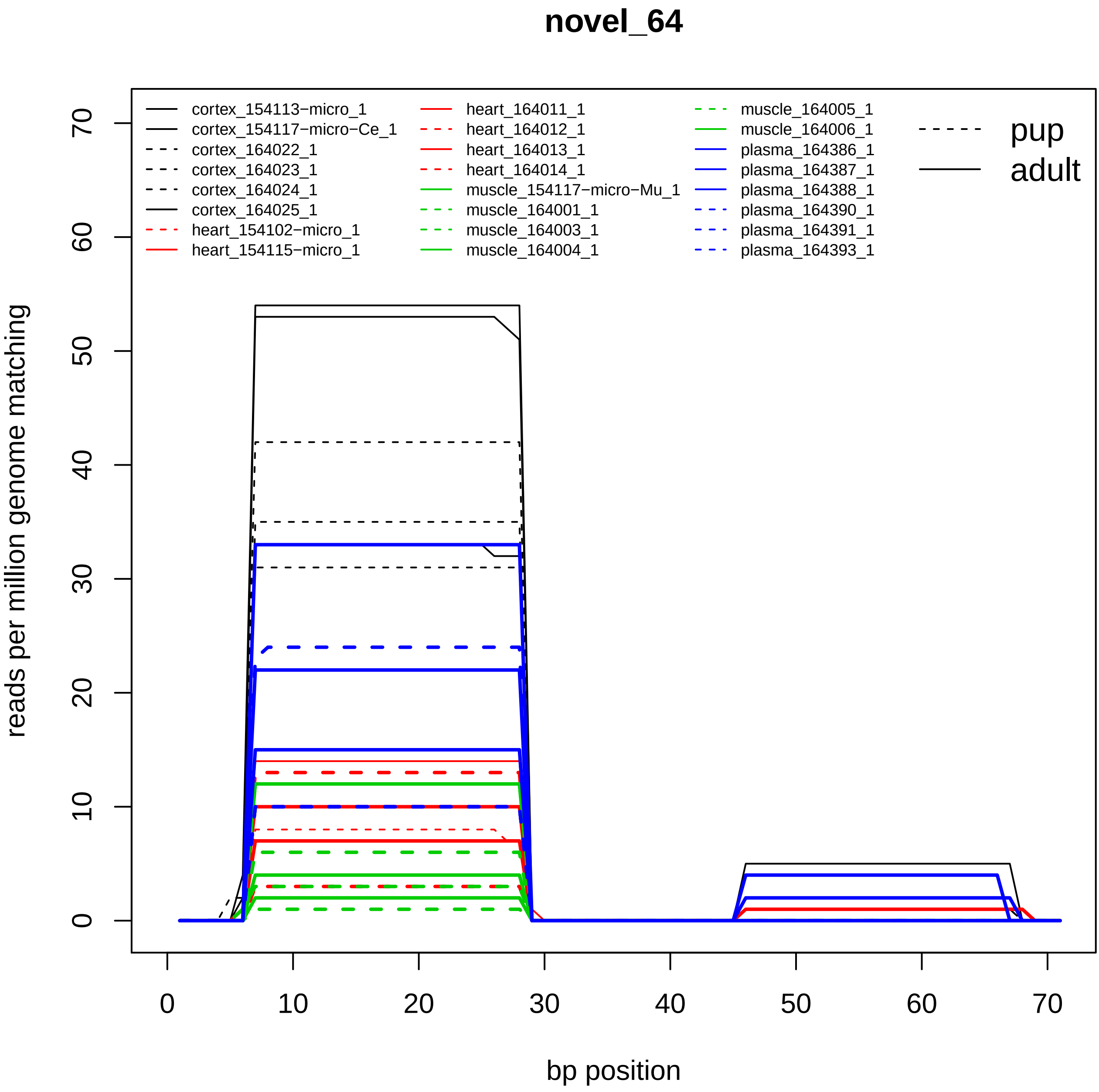
novel_64
cortex_154113−micro_1
cortex_154117−micro−Ce_1
cortex_164022_1
cortex_164023_1
cortex_164024_1
cortex_164025_1
heart_154102−micro_1
heart_154115−micro_1
heart_164011_1
heart_164012_1
heart_164013_1
heart_164014_1
muscle_154117−micro−Mu_1
muscle_164001_1
muscle_164003_1
muscle_164004_1
muscle_164005_1
muscle_164006_1
plasma_164386_1
plasma_164387_1
plasma_164388_1
plasma_164390_1
plasma_164391_1
plasma_164393_1
pup
adult
reads per million genome matching
bp position
0
10
20
30
40
50
60
70

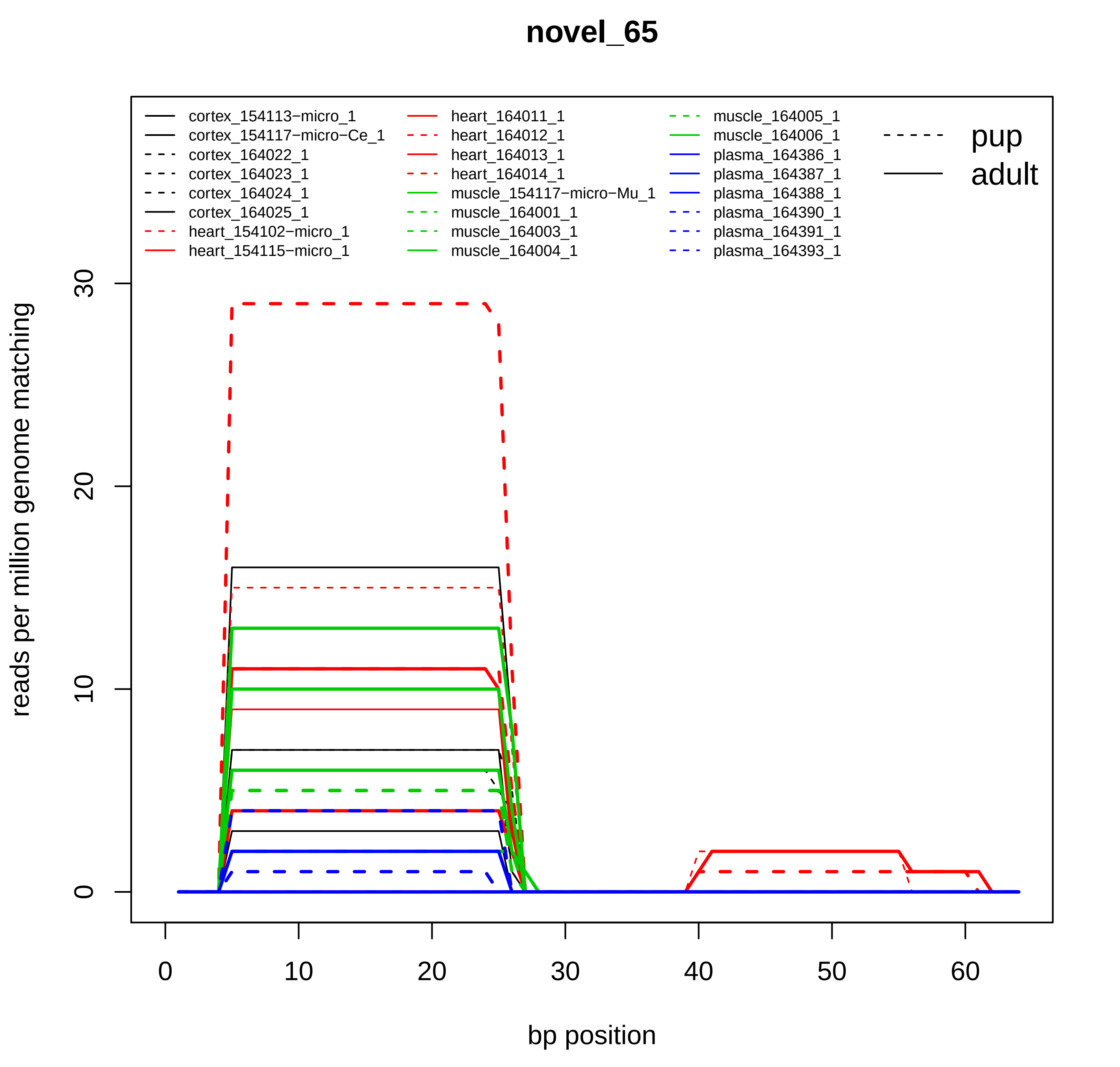
novel_65
cortex_154113−micro_1
cortex_154117−micro−Ce_1
cortex_164022_1
cortex_164023_1
cortex_164024_1
cortex_164025_1
heart_154102−micro_1
heart_154115−micro_1
heart_164011_1
heart_164012_1
heart_164013_1
heart_164014_1
muscle_154117−micro−Mu_1
muscle_164001_1
muscle_164003_1
muscle_164004_1
muscle_164005_1
muscle_164006_1
plasma_164386_1
plasma_164387_1
plasma_164388_1
plasma_164390_1
plasma_164391_1
plasma_164393_1
pup
adult
reads per million genome matching
0
10
20
30
bp position
0
10
20
30
40
50
60

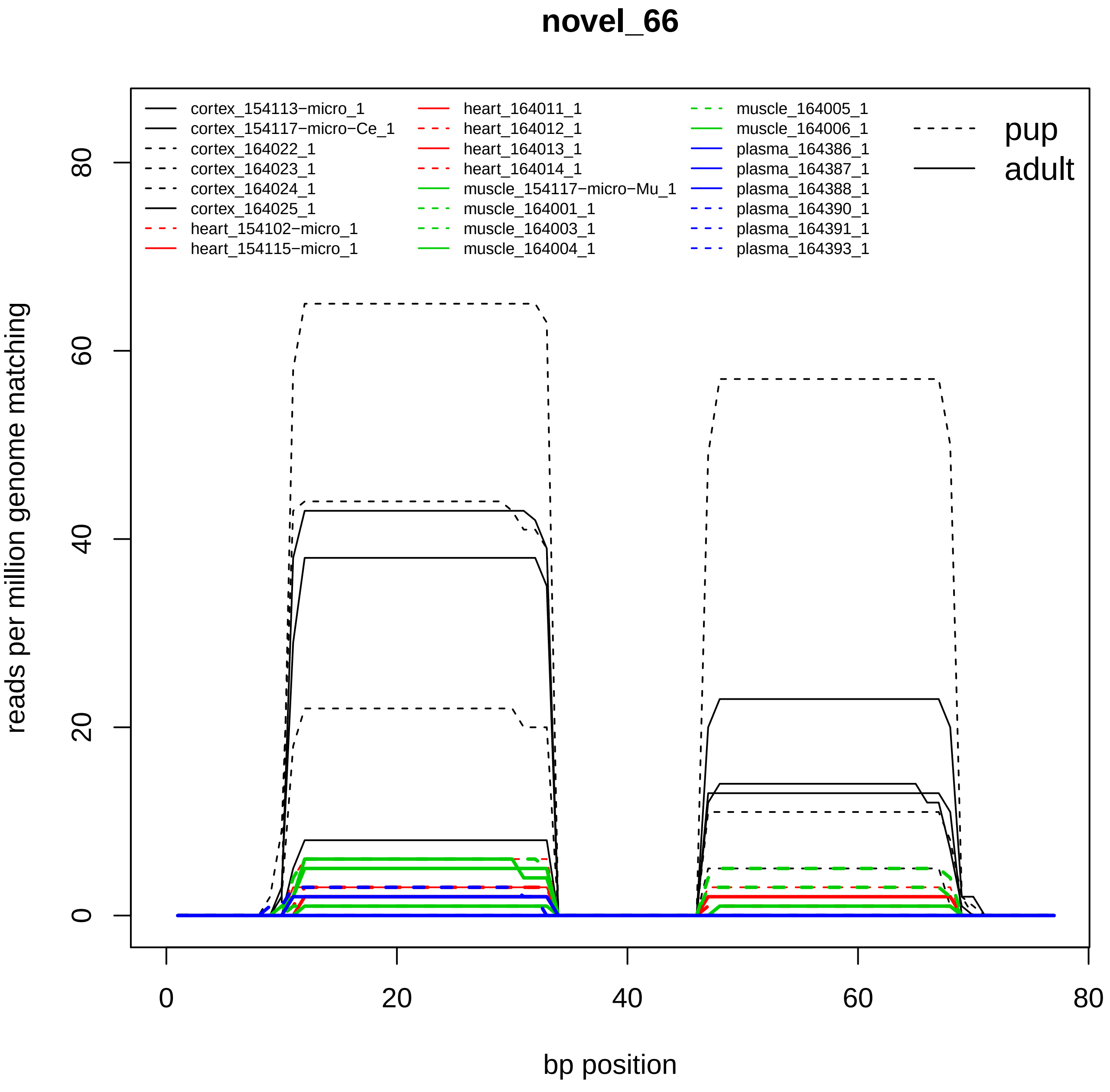
novel_66
cortex_154113−micro_1
cortex_154117−micro−Ce_1
cortex_164022_1
cortex_164023_1
cortex_164024_1
cortex_164025_1
heart_154102−micro_1
heart_154115−micro_1
heart_164011_1
heart_164012_1
heart_164013_1
heart_164014_1
muscle_154117−micro−Mu_1
muscle_164001_1
muscle_164003_1
muscle_164004_1
muscle_164005_1
muscle_164006_1
plasma_164386_1
plasma_164387_1
plasma_164388_1
plasma_164390_1
plasma_164391_1
plasma_164393_1
pup
adult
reads per million genome matching
bp position
0
20
40
60
80
0
20
40
60
80

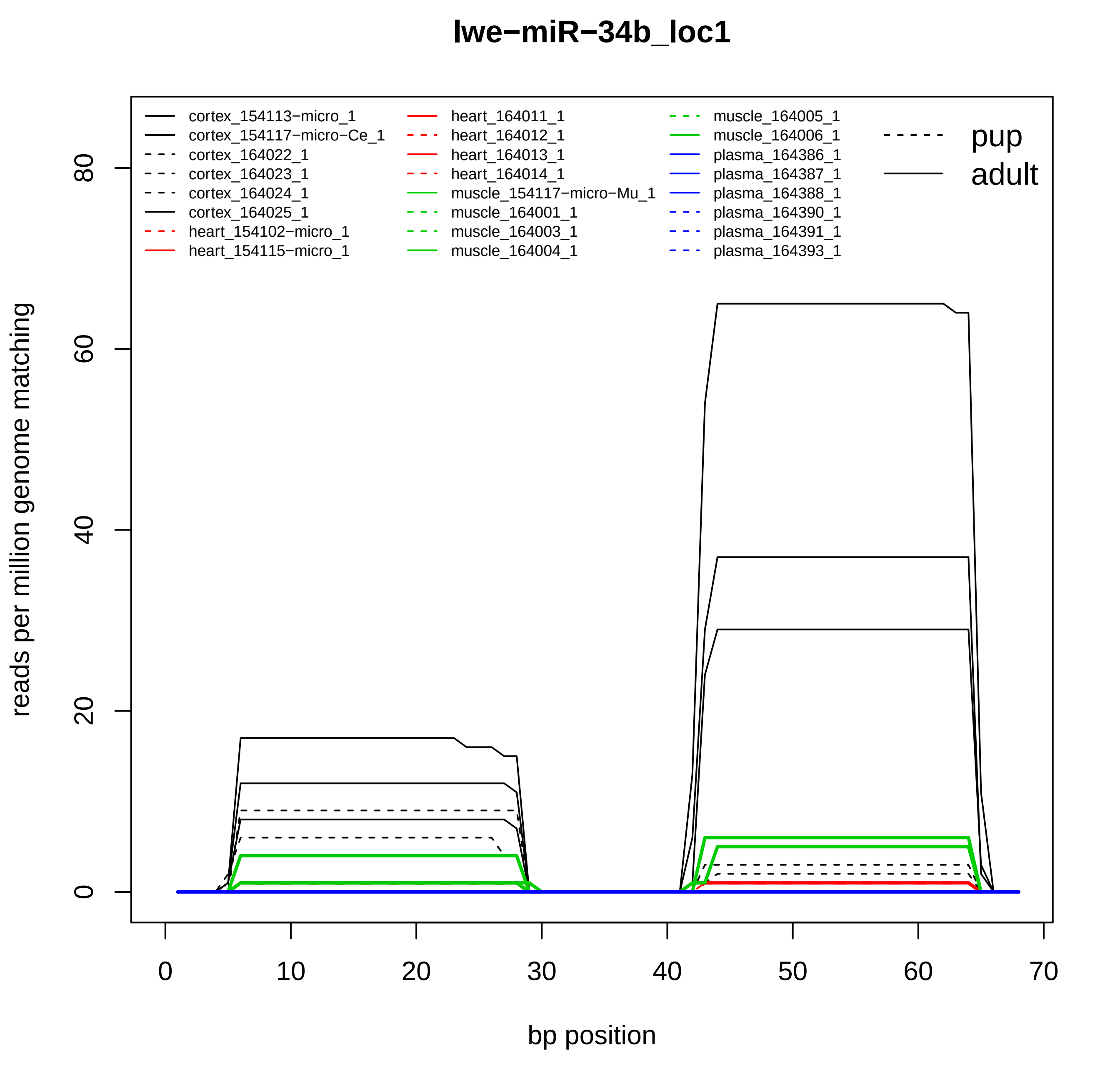

lwe−miR−34b_loc1
cortex_154113−micro_1
cortex_154117−micro−Ce_1
cortex_164022_1
cortex_164023_1
cortex_164024_1
cortex_164025_1
heart_154102−micro_1
heart_154115−micro_1
heart_164011_1
heart_164012_1
heart_164013_1
heart_164014_1
muscle_154117−micro−Mu_1
muscle_164001_1
muscle_164003_1
muscle_164004_1
muscle_164005_1
muscle_164006_1
plasma_164386_1
plasma_164387_1
plasma_164388_1
plasma_164390_1
plasma_164391_1
plasma_164393_1
pup
adult
reads per million genome matching
0
20
40
60
80
0
10
20
30
40
50
60
70
bp position

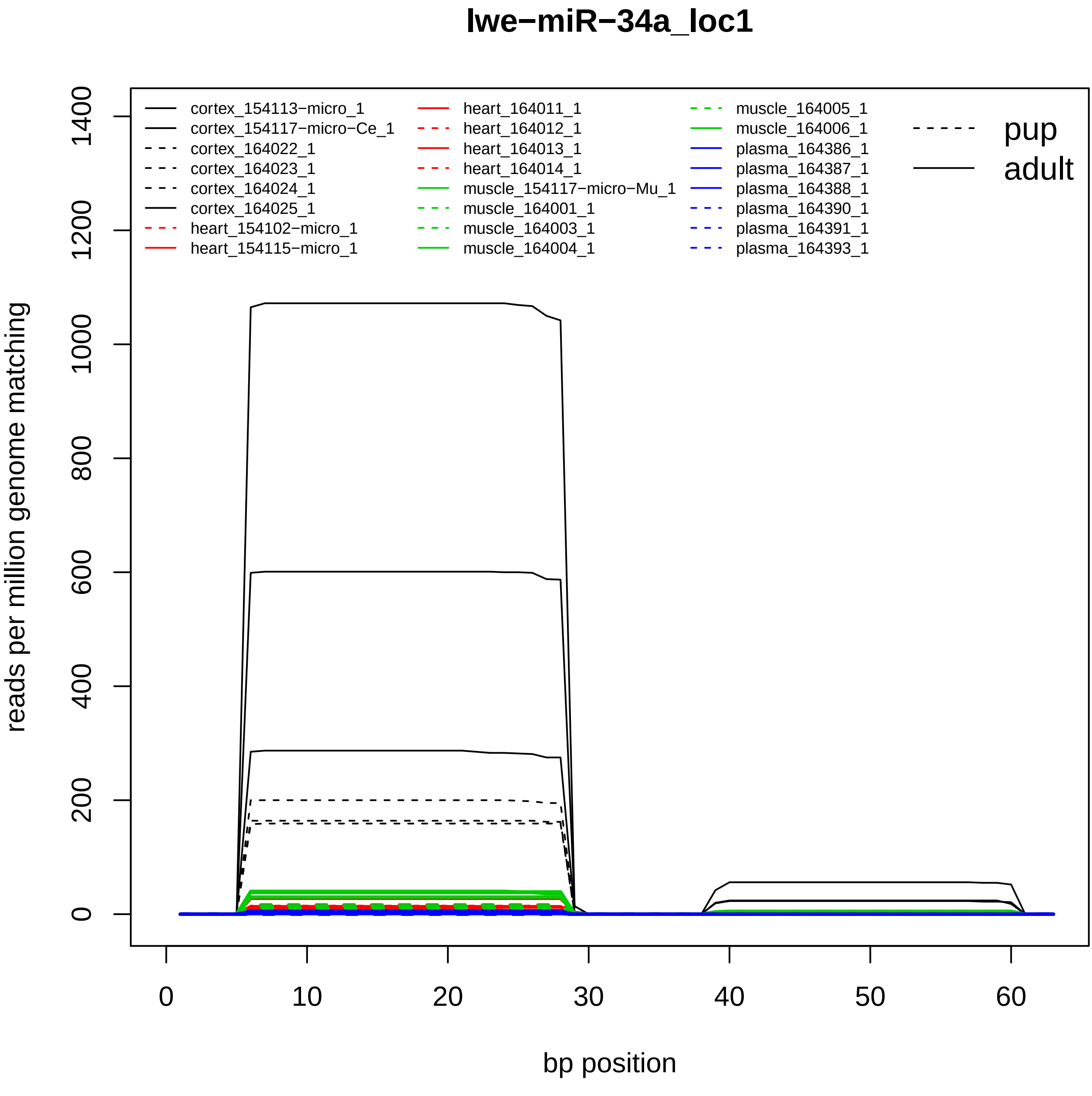

lwe−miR−34a_loc1
reads per million genome matching
bp position
0
200
400
600
800
1000
1200
1400
0
10
20
30
40
50
60
cortex_154113−micro_1
cortex_154117−micro−Ce_1
cortex_164022_1
cortex_164023_1
cortex_164024_1
cortex_164025_1
heart_154102−micro_1
heart_154115−micro_1
heart_164011_1
heart_164012_1
heart_164013_1
heart_164014_1
muscle_154117−micro−Mu_1
muscle_164001_1
muscle_164003_1
muscle_164004_1
muscle_164005_1
muscle_164006_1
plasma_164386_1
plasma_164387_1
plasma_164388_1
plasma_164390_1
plasma_164391_1
plasma_164393_1
pup
adult

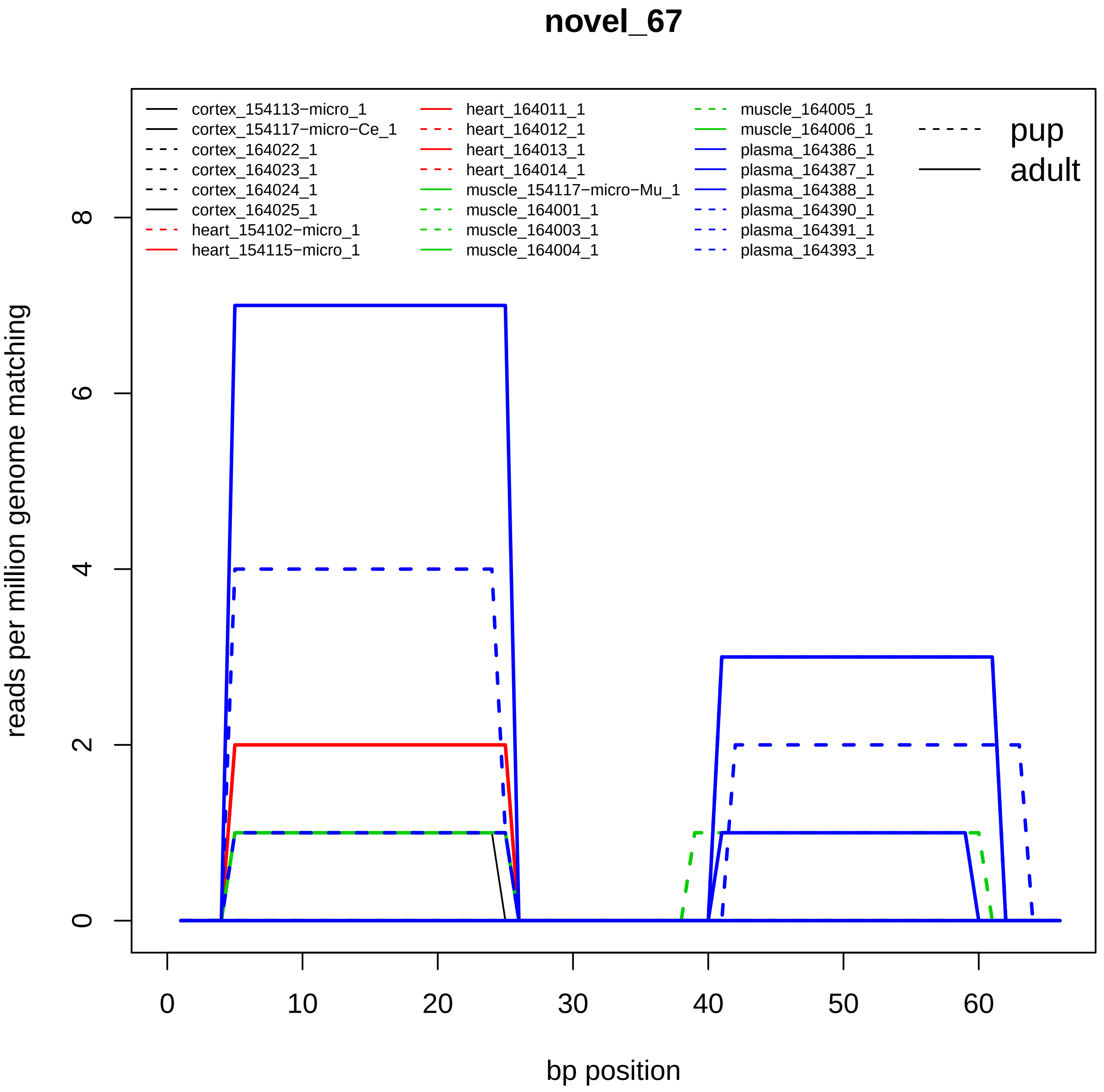
novel_67
cortex_154113−micro_1
cortex_154117−micro−Ce_1
cortex_164022_1
cortex_164023_1
cortex_164024_1
cortex_164025_1
heart_154102−micro_1
heart_154115−micro_1
heart_164011_1
heart_164012_1
heart_164013_1
heart_164014_1
muscle_154117−micro−Mu_1
muscle_164001_1
muscle_164003_1
muscle_164004_1
muscle_164005_1
muscle_164006_1
plasma_164386_1
plasma_164387_1
plasma_164388_1
plasma_164390_1
plasma_164391_1
plasma_164393_1
pup
adult
reads per million genome matching
0
2
4
6
8
0
10
20
30
40
50
60
bp position

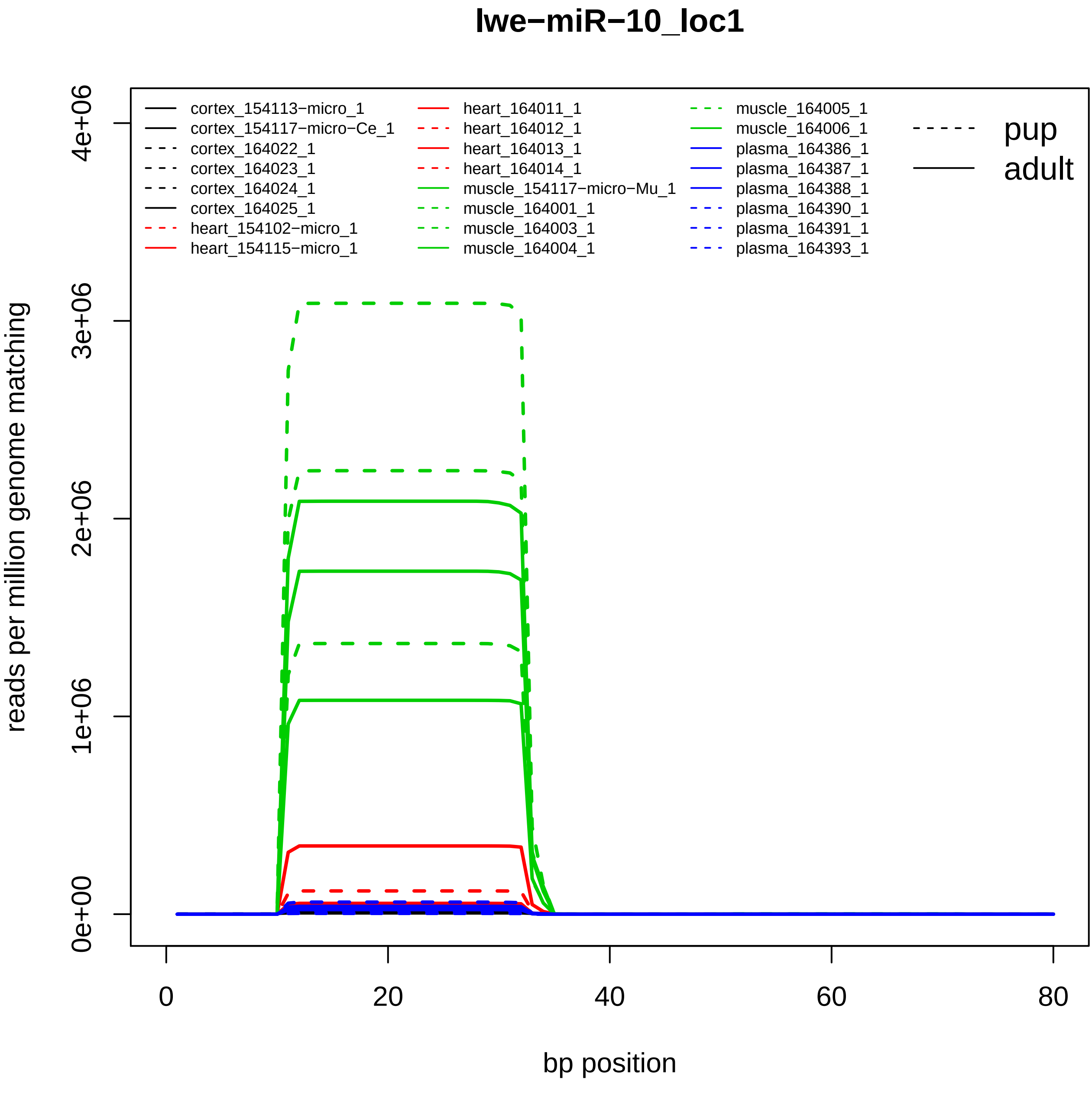
lwe−miR−10_loc1
cortex_154113−micro_1
cortex_154117−micro−Ce_1
cortex_164022_1
cortex_164023_1
cortex_164024_1
cortex_164025_1
heart_154102−micro_1
heart_154115−micro_1
heart_164011_1
heart_164012_1
heart_164013_1
heart_164014_1
muscle_154117−micro−Mu_1
muscle_164001_1
muscle_164003_1
muscle_164004_1
muscle_164005_1
muscle_164006_1
plasma_164386_1
plasma_164387_1
plasma_164388_1
plasma_164390_1
plasma_164391_1
plasma_164393_1
pup
adult
reads per million genome matching
0e+00
1e+06
2e+06
3e+06
4e+06
0
20
40
60
80
bp position

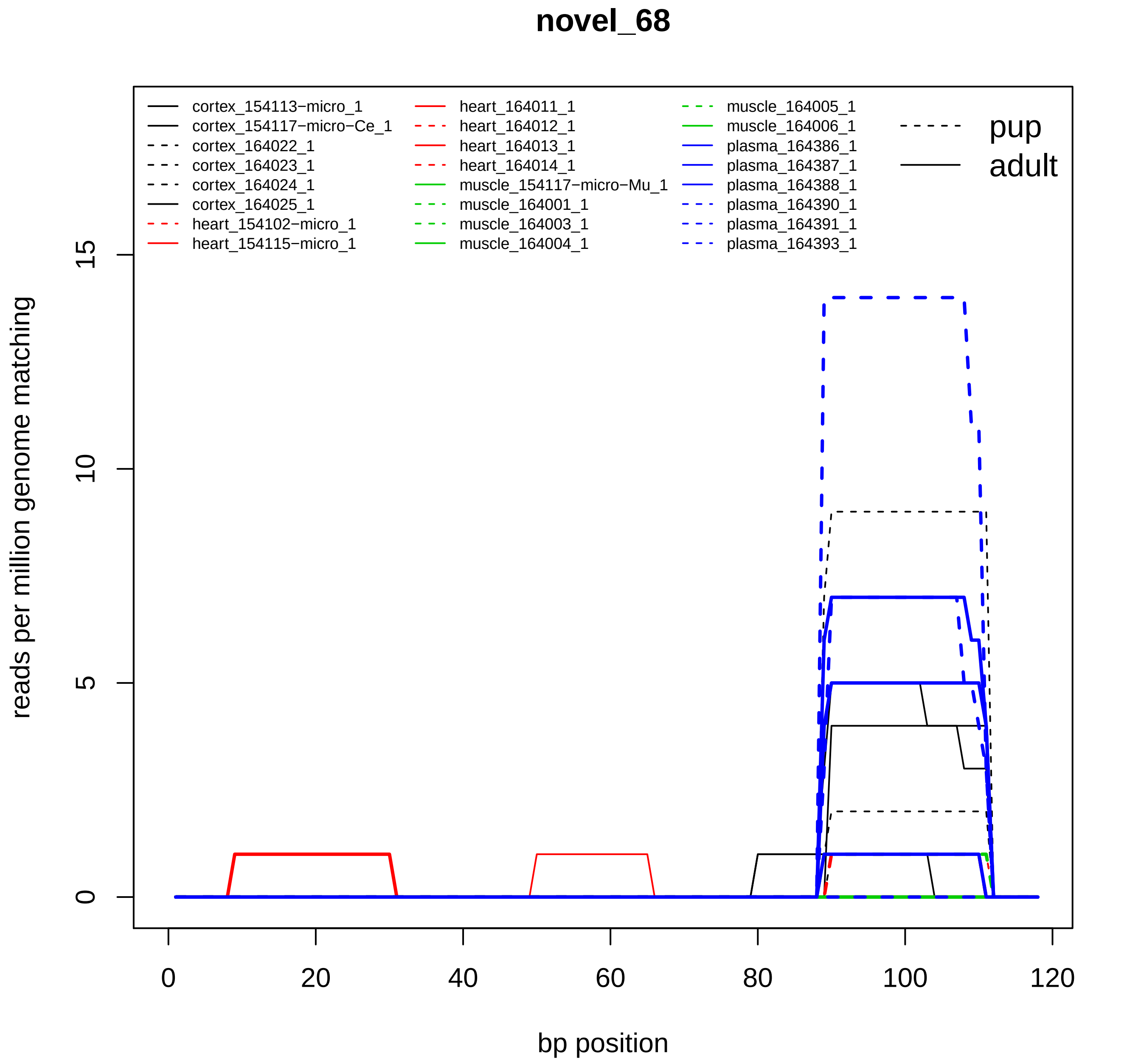
novel_68
cortex_154113−micro_1
cortex_154117−micro−Ce_1
cortex_164022_1
cortex_164023_1
cortex_164024_1
cortex_164025_1
heart_154102−micro_1
heart_154115−micro_1
heart_164011_1
heart_164012_1
heart_164013_1
heart_164014_1
muscle_154117−micro−Mu_1
muscle_164001_1
muscle_164003_1
muscle_164004_1
muscle_164005_1
muscle_164006_1
plasma_164386_1
plasma_164387_1
plasma_164388_1
plasma_164390_1
plasma_164391_1
plasma_164393_1
pup
adult
reads per million genome matching
0
5
10
15
0
20
40
60
80
100
120
bp position

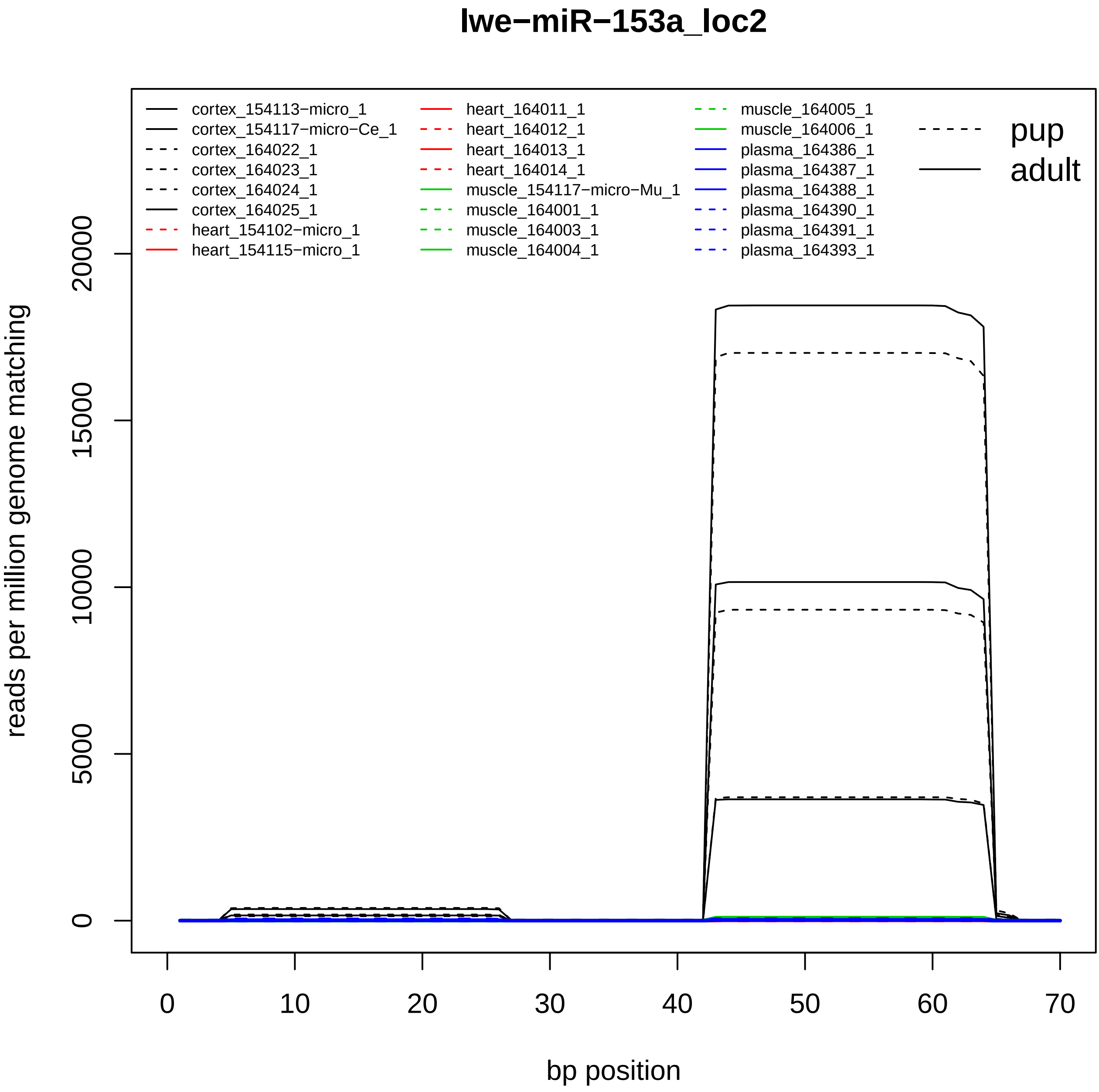
lwe−miR−153a_loc2
cortex_154113−micro_1
cortex_154117−micro−Ce_1
cortex_164022_1
cortex_164023_1
cortex_164024_1
cortex_164025_1
heart_154102−micro_1
heart_154115−micro_1
heart_164011_1
heart_164012_1
heart_164013_1
heart_164014_1
muscle_154117−micro−Mu_1
muscle_164001_1
muscle_164003_1
muscle_164004_1
muscle_164005_1
muscle_164006_1
plasma_164386_1
plasma_164387_1
plasma_164388_1
plasma_164390_1
plasma_164391_1
plasma_164393_1
pup
adult
reads per million genome matching
0
5000
10000
15000
20000
bp position
0
10
20
30
40
50
60
70

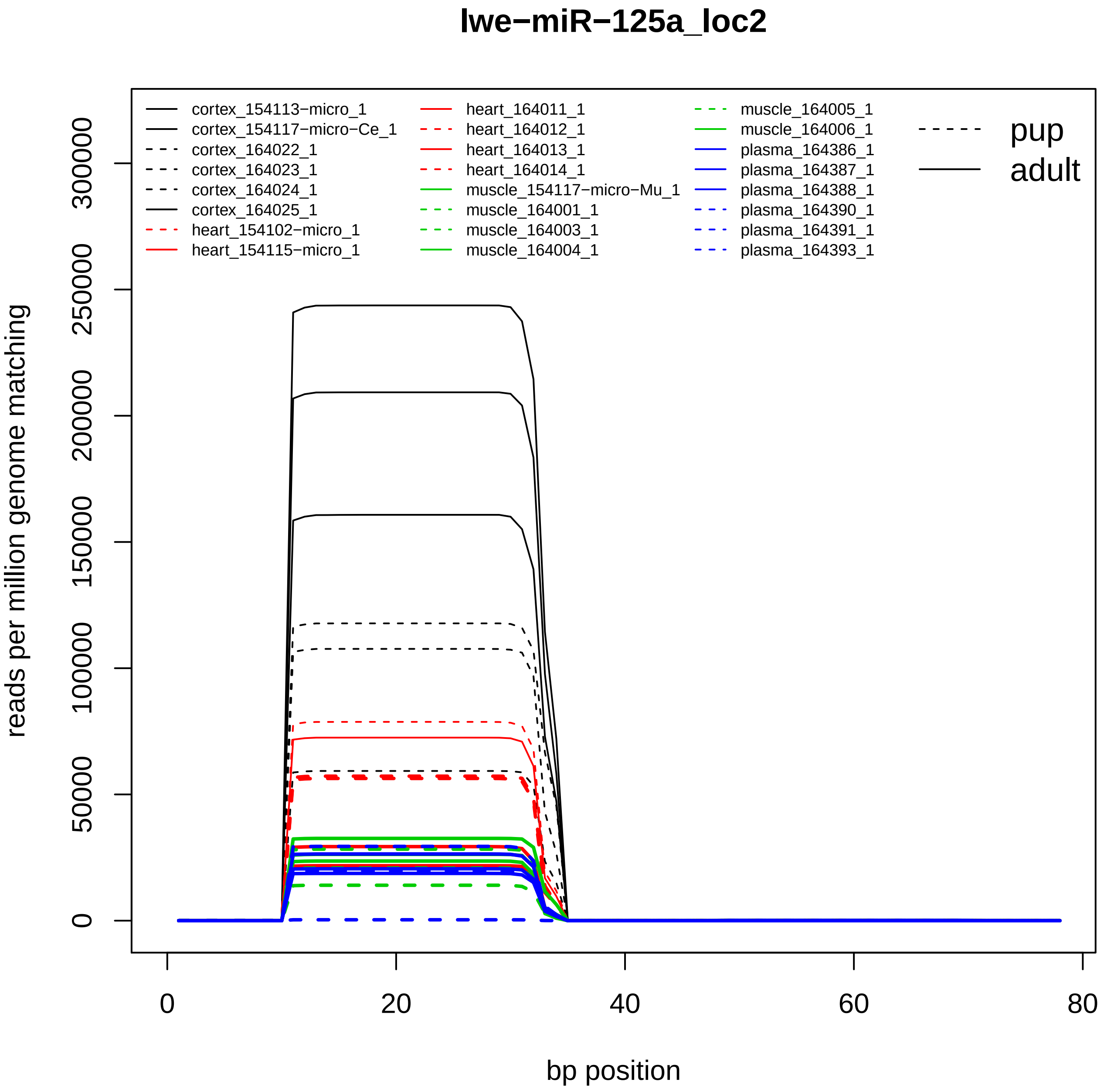
lwe−miR−125a_loc2
cortex_154113−micro_1
cortex_154117−micro−Ce_1
cortex_164022_1
cortex_164023_1
cortex_164024_1
cortex_164025_1
heart_154102−micro_1
heart_154115−micro_1
heart_164011_1
heart_164012_1
heart_164013_1
heart_164014_1
muscle_154117−micro−Mu_1
muscle_164001_1
muscle_164003_1
muscle_164004_1
muscle_164005_1
muscle_164006_1
plasma_164386_1
plasma_164387_1
plasma_164388_1
plasma_164390_1
plasma_164391_1
plasma_164393_1
pup
adult
reads per million genome matching
0
50000
100000
150000
200000
250000
300000
0
20
40
60
80
bp position

### let−7e_loc1

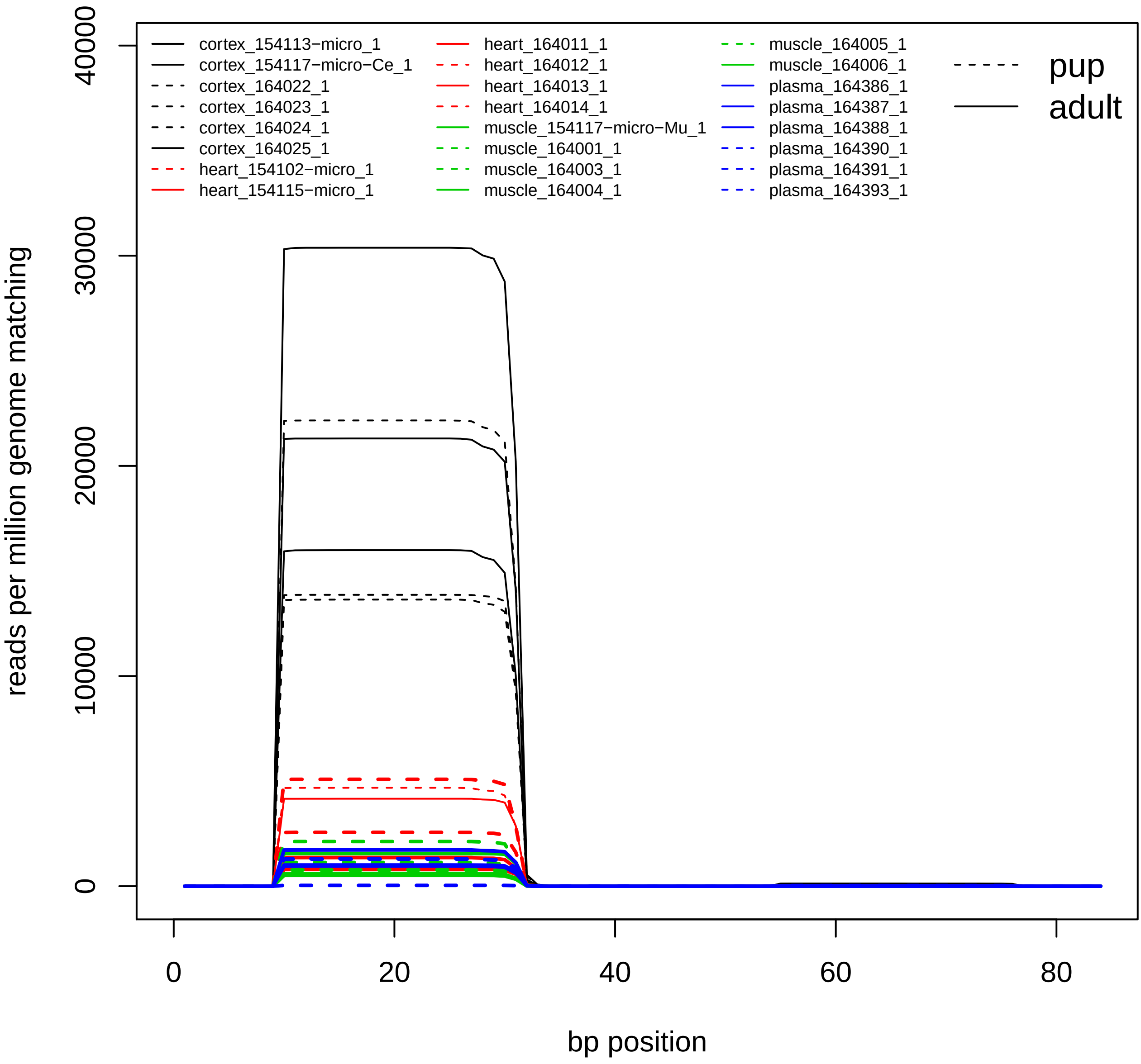

cortex_154113−micro_1
cortex_154117−micro−Ce_1
cortex_164022_1
cortex_164023_1
cortex_164024_1
cortex_164025_1
heart_154102−micro_1
heart_154115−micro_1
heart_164011_1
heart_164012_1
heart_164013_1
heart_164014_1
muscle_154117−micro−Mu_1
muscle_164001_1
muscle_164003_1
muscle_164004_1
muscle_164005_1
muscle_164006_1
plasma_164386_1
plasma_164387_1
plasma_164388_1
plasma_164390_1
plasma_164391_1
plasma_164393_1
pup
adult
reads per million genome matching
0
10000
20000
30000
40000
0
20
40
60
80
bp position

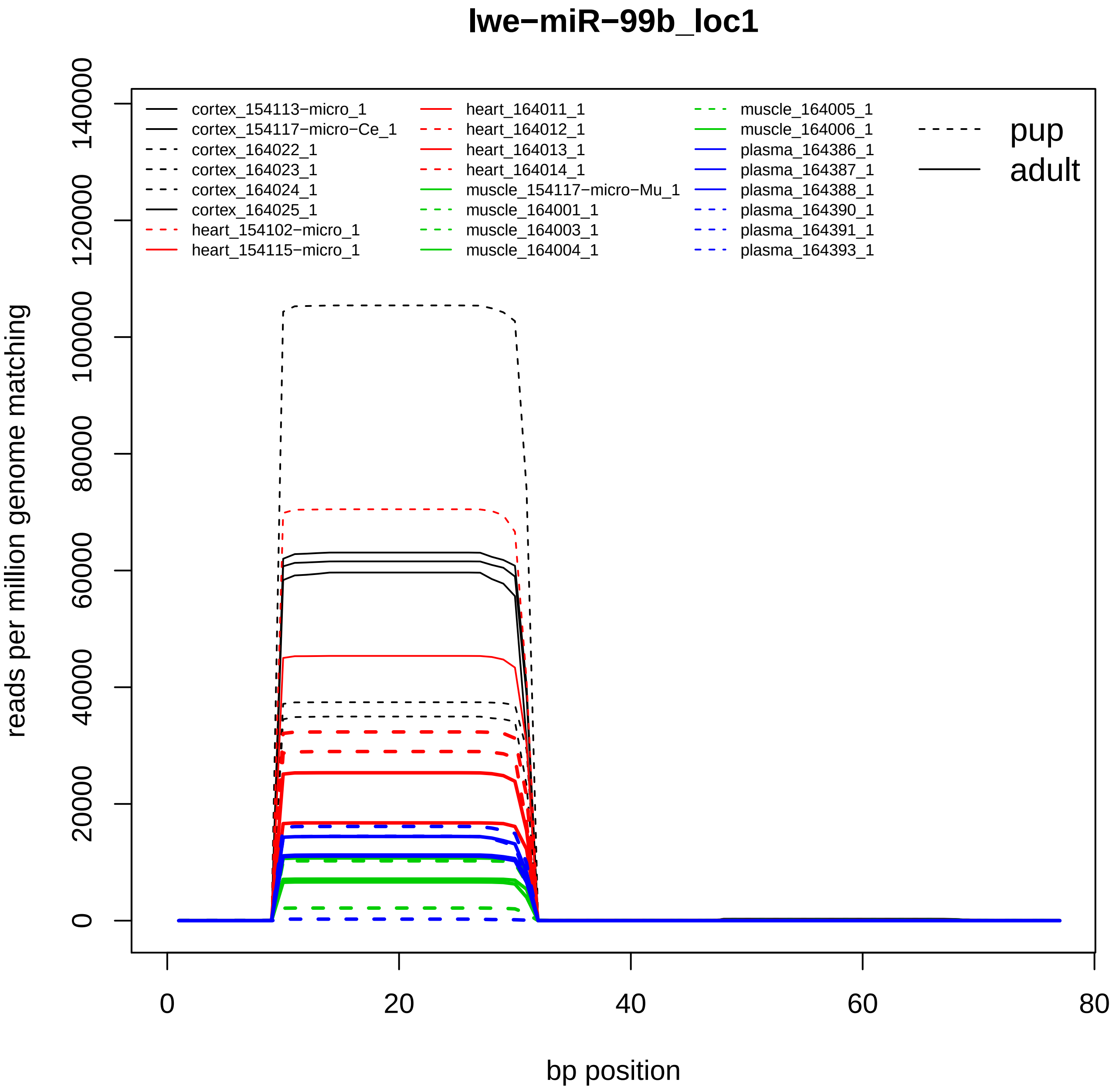

lwe−miR−99b_loc1
cortex_154113−micro_1
cortex_154117−micro−Ce_1
cortex_164022_1
cortex_164023_1
cortex_164024_1
cortex_164025_1
heart_154102−micro_1
heart_154115−micro_1
heart_164011_1
heart_164012_1
heart_164013_1
heart_164014_1
muscle_154117−micro−Mu_1
muscle_164001_1
muscle_164003_1
muscle_164004_1
muscle_164005_1
muscle_164006_1
plasma_164386_1
plasma_164387_1
plasma_164388_1
plasma_164390_1
plasma_164391_1
plasma_164393_1
pup
adult
reads per million genome matching
0
20000
40000
60000
80000
100000
120000
140000
bp position
0
20
40
60
80

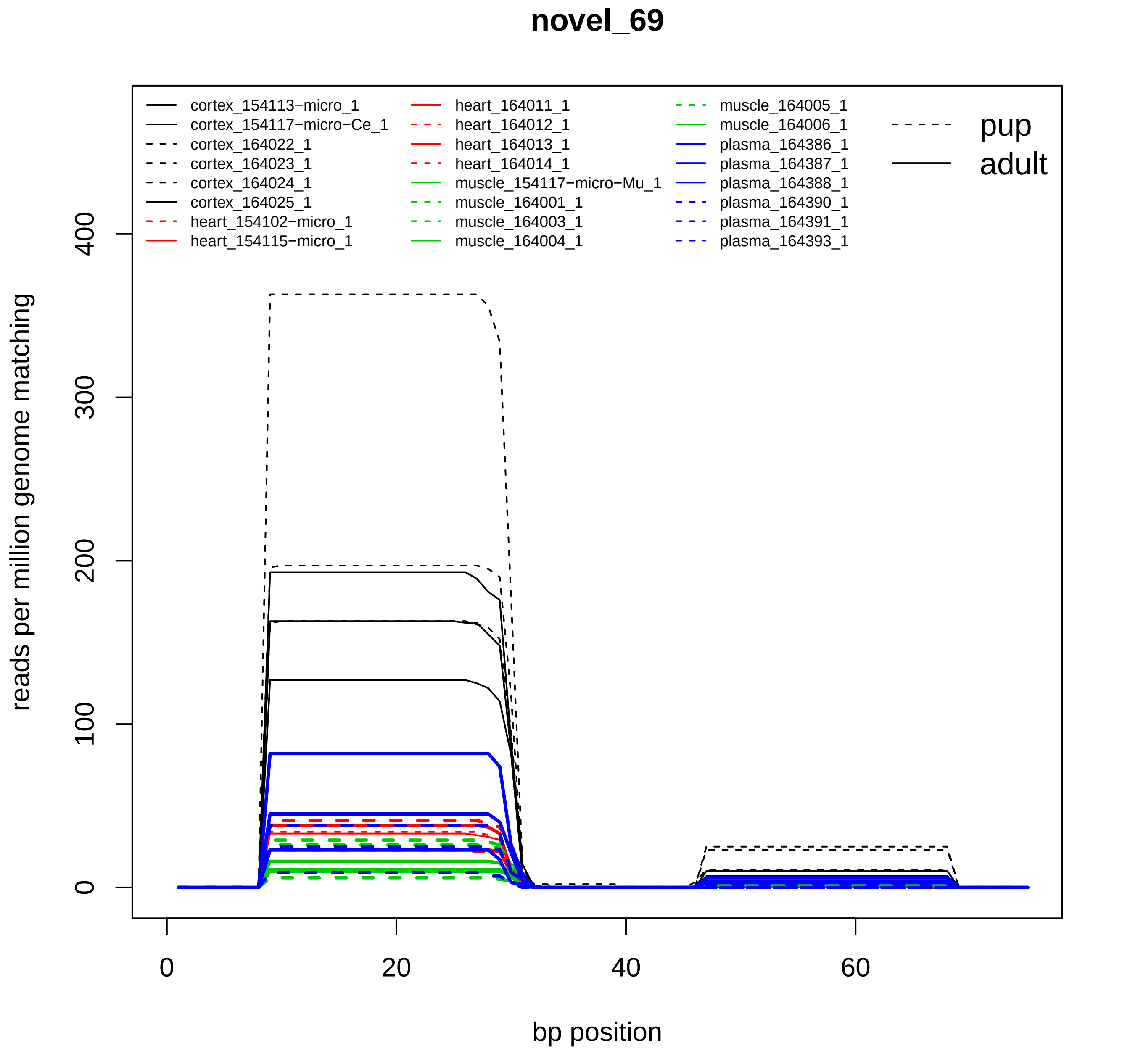
novel_69
cortex_154113−micro_1
cortex_154117−micro−Ce_1
cortex_164022_1
cortex_164023_1
cortex_164024_1
cortex_164025_1
heart_154102−micro_1
heart_154115−micro_1
heart_164011_1
heart_164012_1
heart_164013_1
heart_164014_1
muscle_154117−micro−Mu_1
muscle_164001_1
muscle_164003_1
muscle_164004_1
muscle_164005_1
muscle_164006_1
plasma_164386_1
plasma_164387_1
plasma_164388_1
plasma_164390_1
plasma_164391_1
plasma_164393_1
pup
adult
reads per million genome matching
0
100
200
300
400
bp position
0
20
40
60

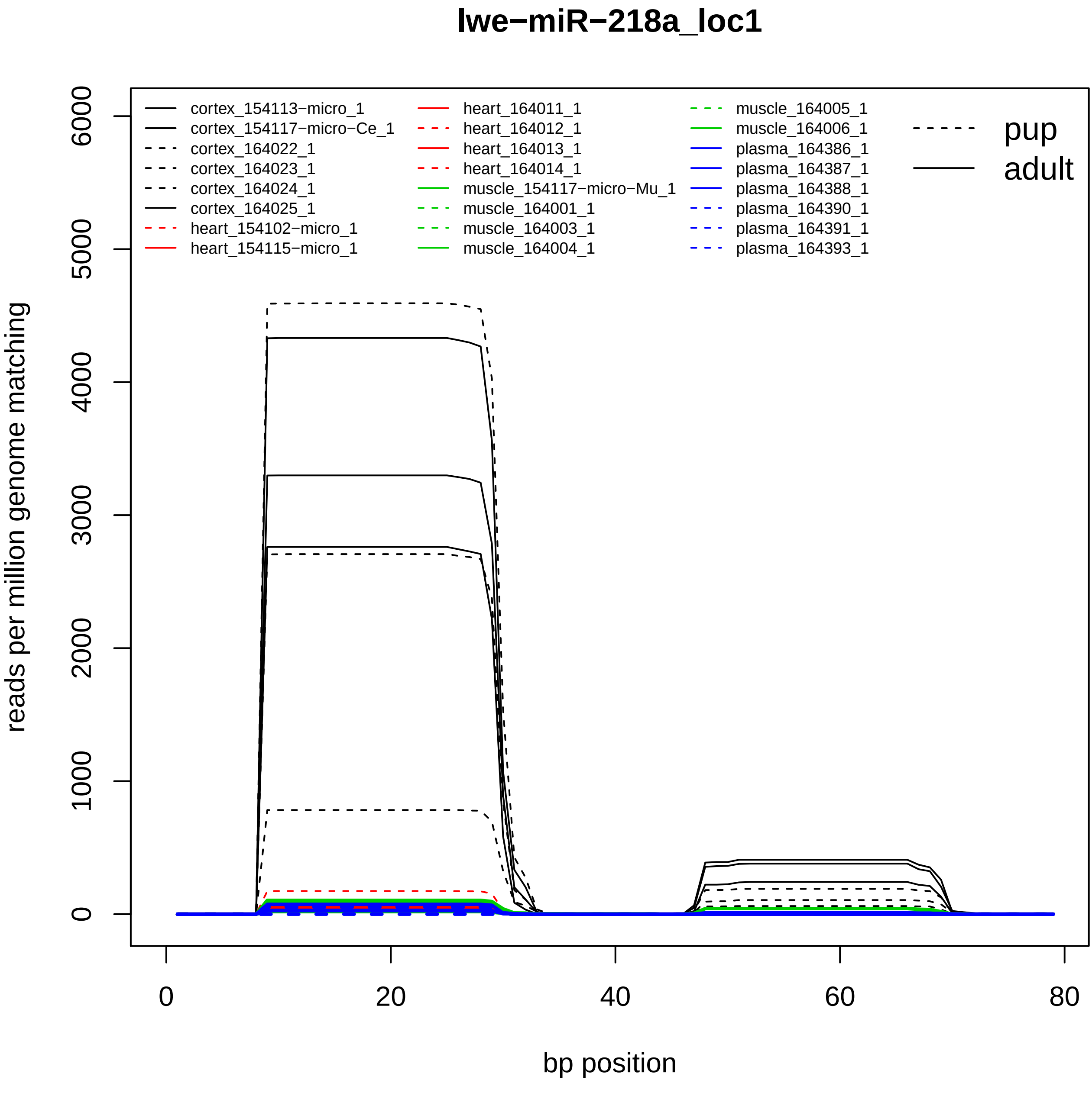
lwe−miR−218a_loc1
reads per million genome matching
bp position
0
1000
2000
3000
4000
5000
6000
0
20
40
60
80
cortex_154113−micro_1
cortex_154117−micro−Ce_1
cortex_164022_1
cortex_164023_1
cortex_164024_1
cortex_164025_1
heart_154102−micro_1
heart_154115−micro_1
heart_164011_1
heart_164012_1
heart_164013_1
heart_164014_1
muscle_154117−micro−Mu_1
muscle_164001_1
muscle_164003_1
muscle_164004_1
muscle_164005_1
muscle_164006_1
plasma_164386_1
plasma_164387_1
plasma_164388_1
plasma_164390_1
plasma_164391_1
plasma_164393_1
pup
adult

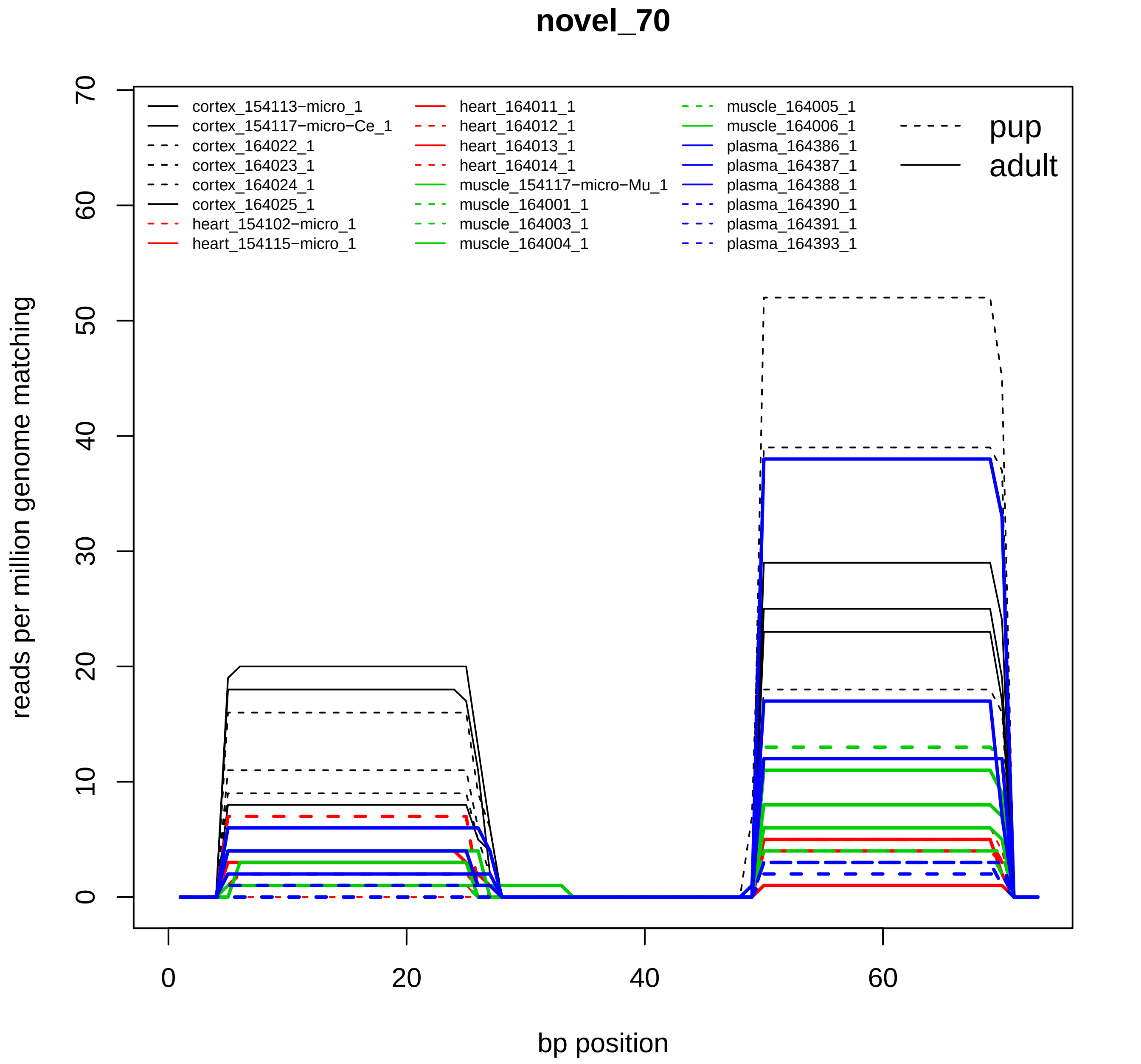
novel_70
cortex_154113−micro_1
cortex_154117−micro−Ce_1
cortex_164022_1
cortex_164023_1
cortex_164024_1
cortex_164025_1
heart_154102−micro_1
heart_154115−micro_1
heart_164011_1
heart_164012_1
heart_164013_1
heart_164014_1
muscle_154117−micro−Mu_1
muscle_164001_1
muscle_164003_1
muscle_164004_1
muscle_164005_1
muscle_164006_1
plasma_164386_1
plasma_164387_1
plasma_164388_1
plasma_164390_1
plasma_164391_1
plasma_164393_1
pup
adult
reads per million genome matching
0
10
20
30
40
50
60
70
bp position
0
20
40
60

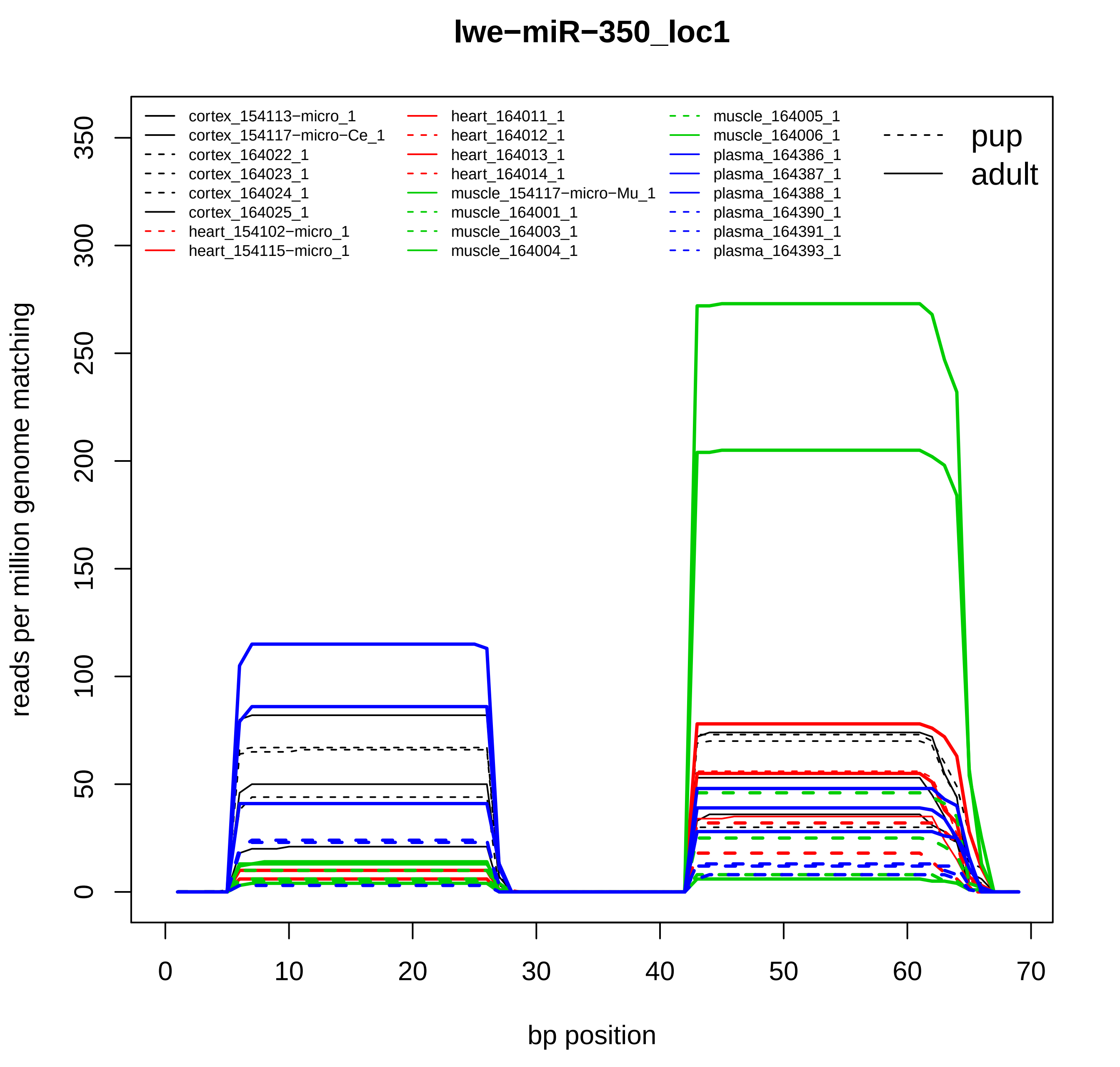
lwe−miR−350_loc1
reads per million genome matching
bp position
0
50
100
150
200
250
300
350
0
10
20
30
40
50
60
70
cortex_154113−micro_1
cortex_154117−micro−Ce_1
cortex_164022_1
cortex_164023_1
cortex_164024_1
cortex_164025_1
heart_154102−micro_1
heart_154115−micro_1
heart_164011_1
heart_164012_1
heart_164013_1
heart_164014_1
muscle_154117−micro−Mu_1
muscle_164001_1
muscle_164003_1
muscle_164004_1
muscle_164005_1
muscle_164006_1
plasma_164386_1
plasma_164387_1
plasma_164388_1
plasma_164390_1
plasma_164391_1
plasma_164393_1
pup
adult

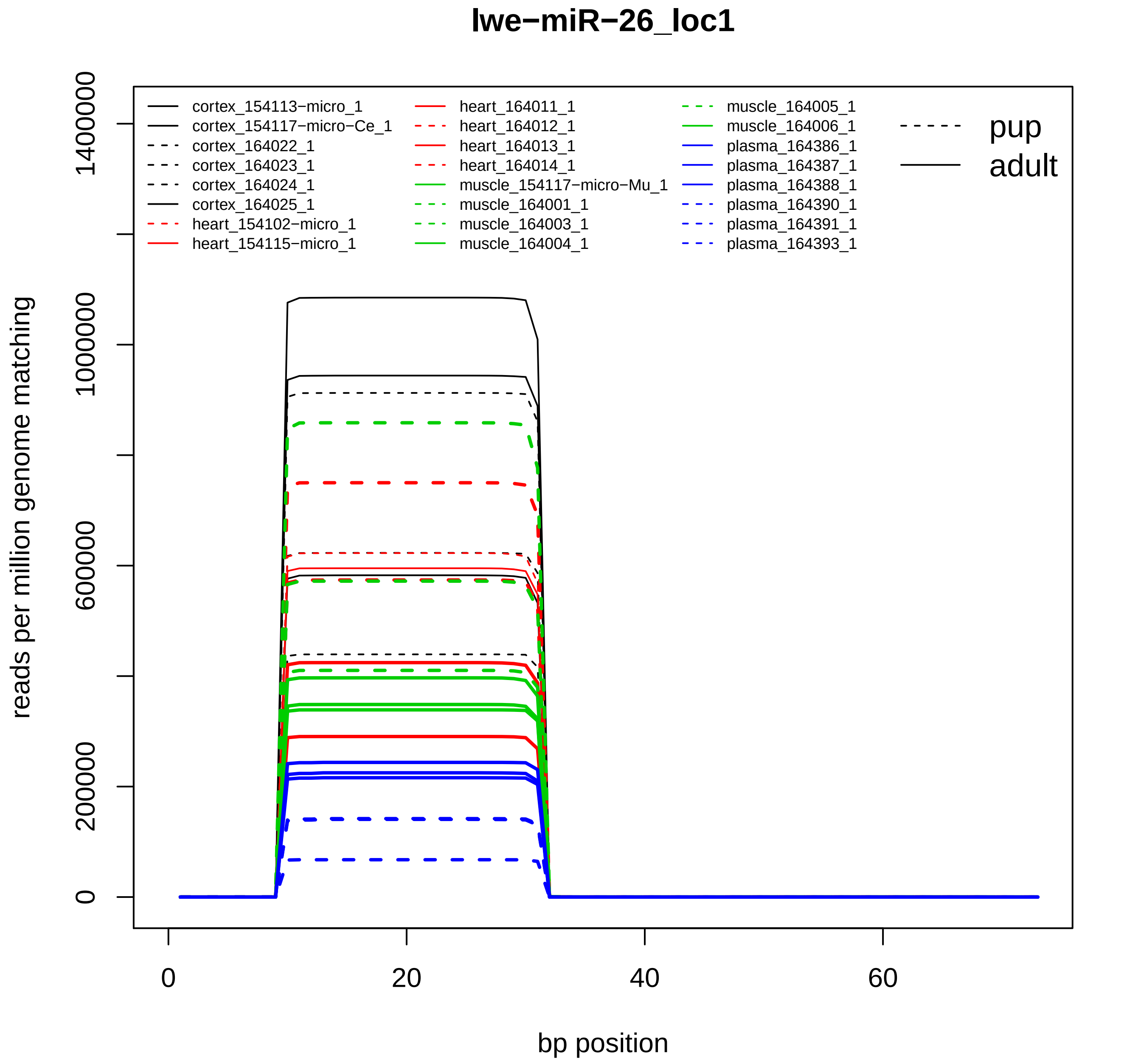

lwe−miR−26_loc1
cortex_154113−micro_1
cortex_154117−micro−Ce_1
cortex_164022_1
cortex_164023_1
cortex_164024_1
cortex_164025_1
heart_154102−micro_1
heart_154115−micro_1
heart_164011_1
heart_164012_1
heart_164013_1
heart_164014_1
muscle_154117−micro−Mu_1
muscle_164001_1
muscle_164003_1
muscle_164004_1
muscle_164005_1
muscle_164006_1
plasma_164386_1
plasma_164387_1
plasma_164388_1
plasma_164390_1
plasma_164391_1
plasma_164393_1
pup
adult
reads per million genome matching
0
200000
600000
1000000
1400000
0
20
40
60
bp position

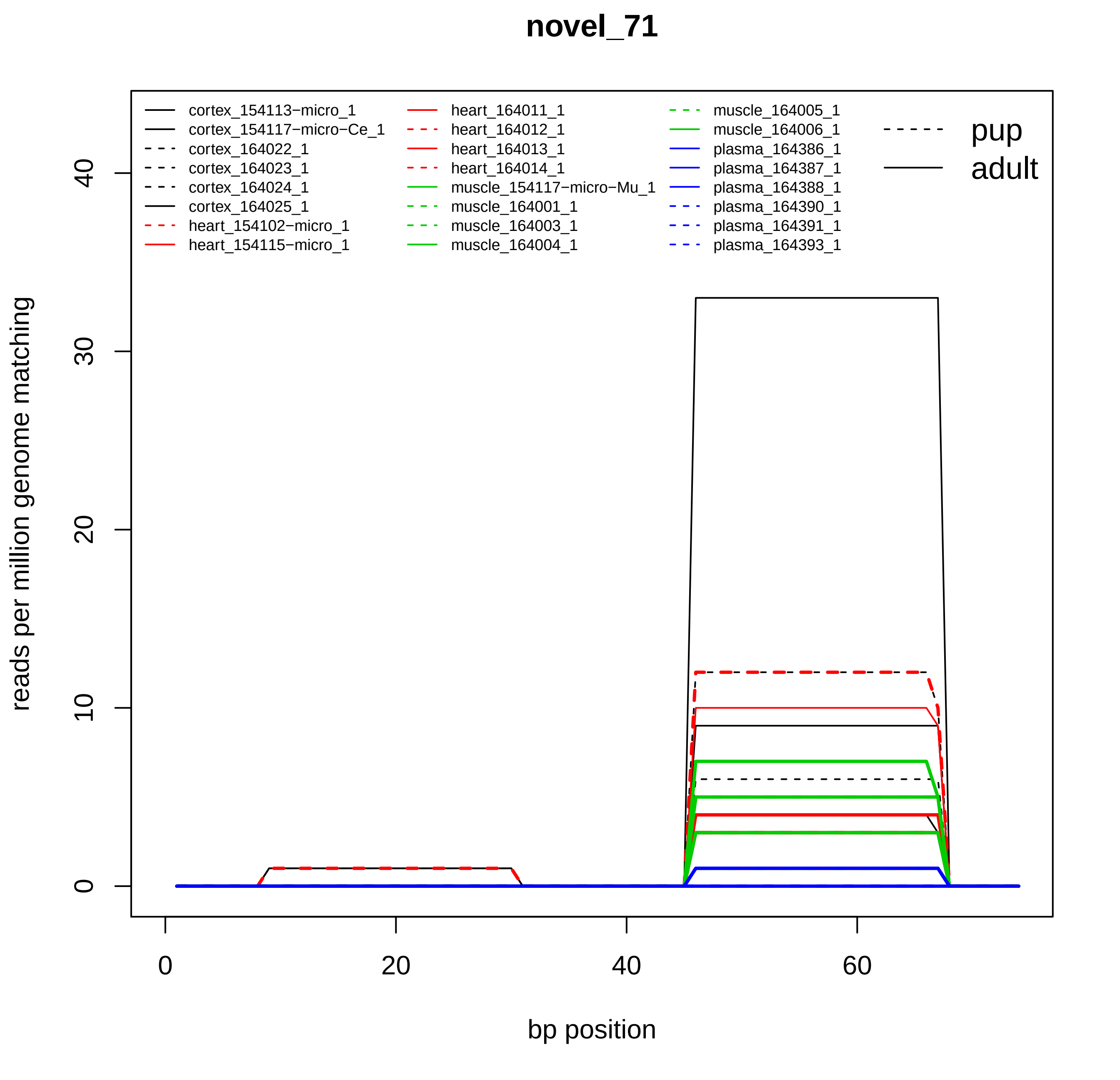
novel_71
reads per million genome matching
bp position
0
10
20
30
40
0
20
40
60
cortex_154113−micro_1
cortex_154117−micro−Ce_1
cortex_164022_1
cortex_164023_1
cortex_164024_1
cortex_164025_1
heart_154102−micro_1
heart_154115−micro_1
heart_164011_1
heart_164012_1
heart_164013_1
heart_164014_1
muscle_154117−micro−Mu_1
muscle_164001_1
muscle_164003_1
muscle_164004_1
muscle_164005_1
muscle_164006_1
plasma_164386_1
plasma_164387_1
plasma_164388_1
plasma_164390_1
plasma_164391_1
plasma_164393_1
pup
adult

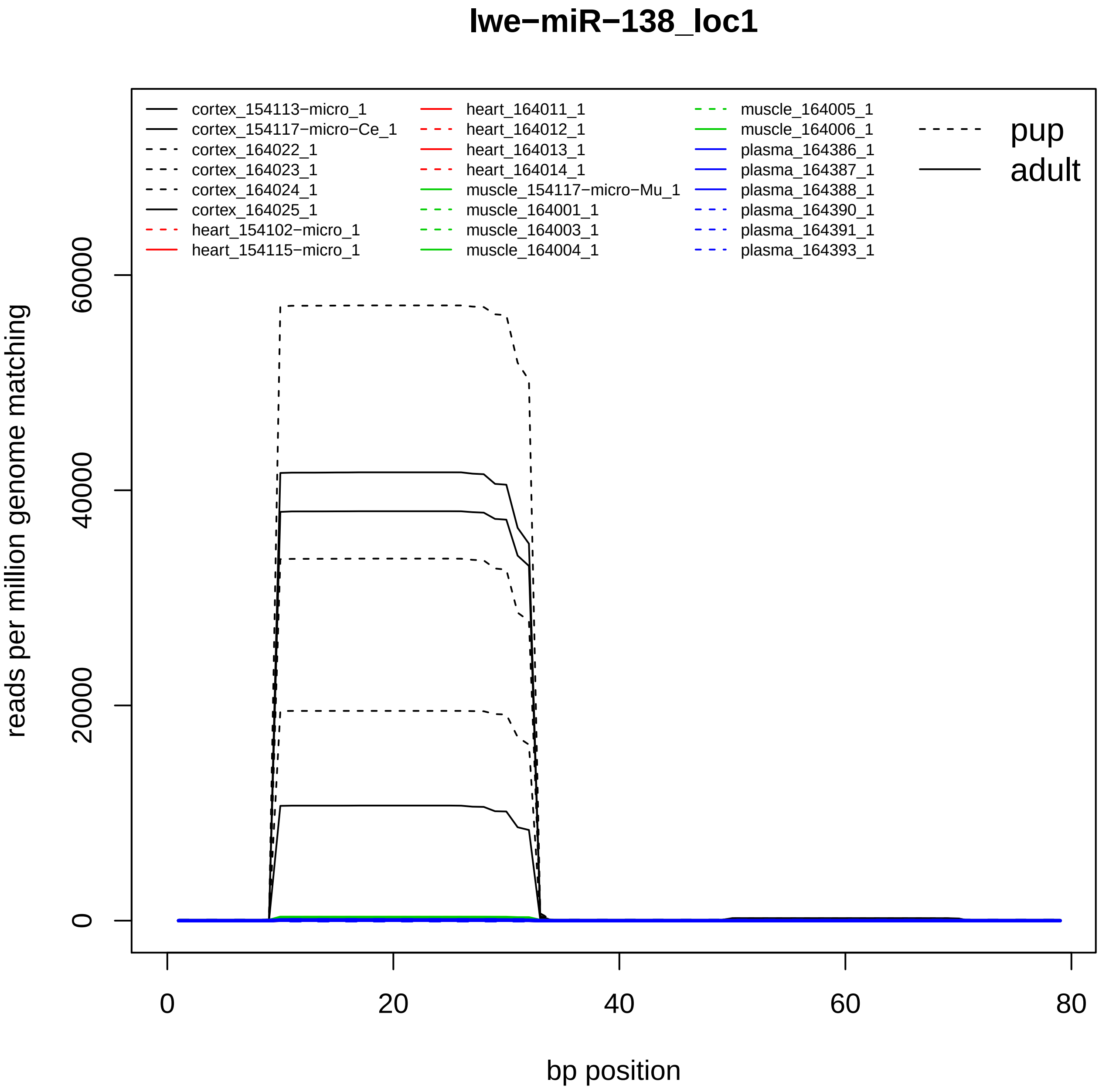
lwe−miR−138_loc1
cortex_154113−micro_1
cortex_154117−micro−Ce_1
cortex_164022_1
cortex_164023_1
cortex_164024_1
cortex_164025_1
heart_154102−micro_1
heart_154115−micro_1
heart_164011_1
heart_164012_1
heart_164013_1
heart_164014_1
muscle_154117−micro−Mu_1
muscle_164001_1
muscle_164003_1
muscle_164004_1
muscle_164005_1
muscle_164006_1
plasma_164386_1
plasma_164387_1
plasma_164388_1
plasma_164390_1
plasma_164391_1
plasma_164393_1
pup
adult
reads per million genome matching
0
20000
40000
60000
0
20
40
60
80
bp position

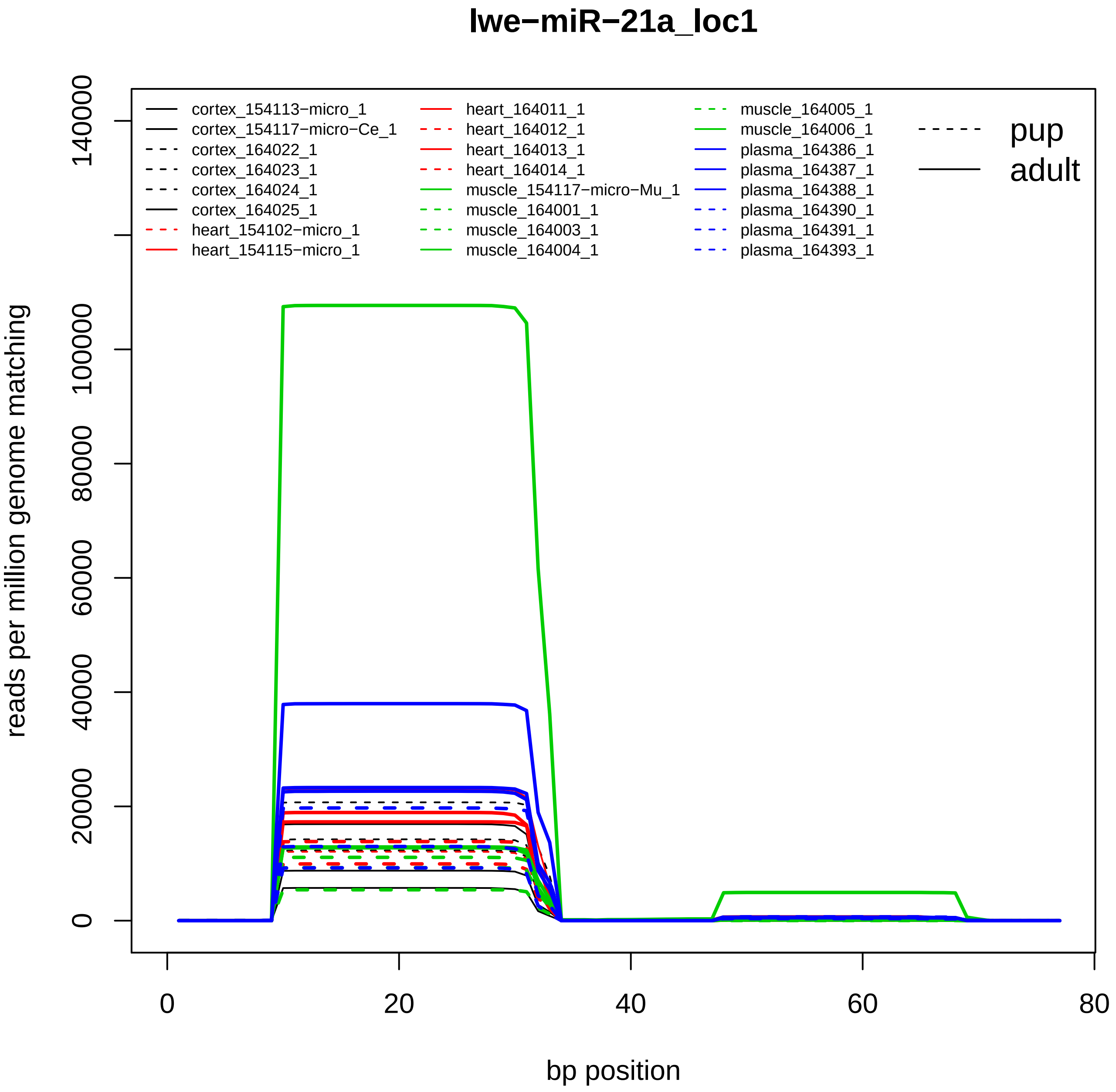
lwe−miR−21a_loc1
reads per million genome matching
bp position
0
20000
40000
60000
80000
100000
140000
0
20
40
60
80
cortex_154113−micro_1
cortex_154117−micro−Ce_1
cortex_164022_1
cortex_164023_1
cortex_164024_1
cortex_164025_1
heart_154102−micro_1
heart_154115−micro_1
heart_164011_1
heart_164012_1
heart_164013_1
heart_164014_1
muscle_154117−micro−Mu_1
muscle_164001_1
muscle_164003_1
muscle_164004_1
muscle_164005_1
muscle_164006_1
plasma_164386_1
plasma_164387_1
plasma_164388_1
plasma_164390_1
plasma_164391_1
plasma_164393_1
pup
adult

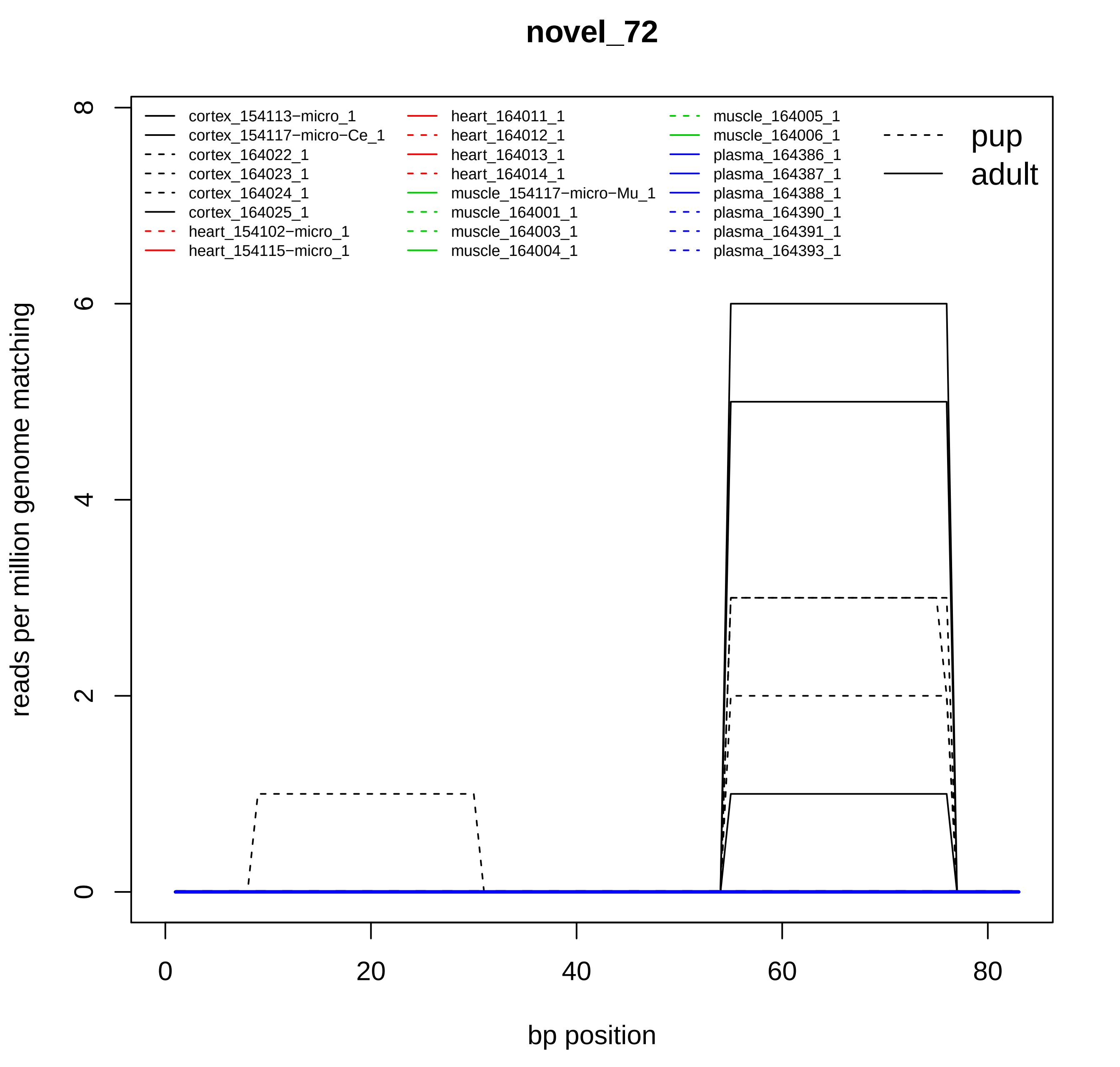
novel_72
cortex_154113−micro_1
cortex_154117−micro−Ce_1
cortex_164022_1
cortex_164023_1
cortex_164024_1
cortex_164025_1
heart_154102−micro_1
heart_154115−micro_1
heart_164011_1
heart_164012_1
heart_164013_1
heart_164014_1
muscle_154117−micro−Mu_1
muscle_164001_1
muscle_164003_1
muscle_164004_1
muscle_164005_1
muscle_164006_1
plasma_164386_1
plasma_164387_1
plasma_164388_1
plasma_164390_1
plasma_164391_1
plasma_164393_1
pup
adult
reads per million genome matching
bp position
0
2
4
6
8
0
20
40
60
80

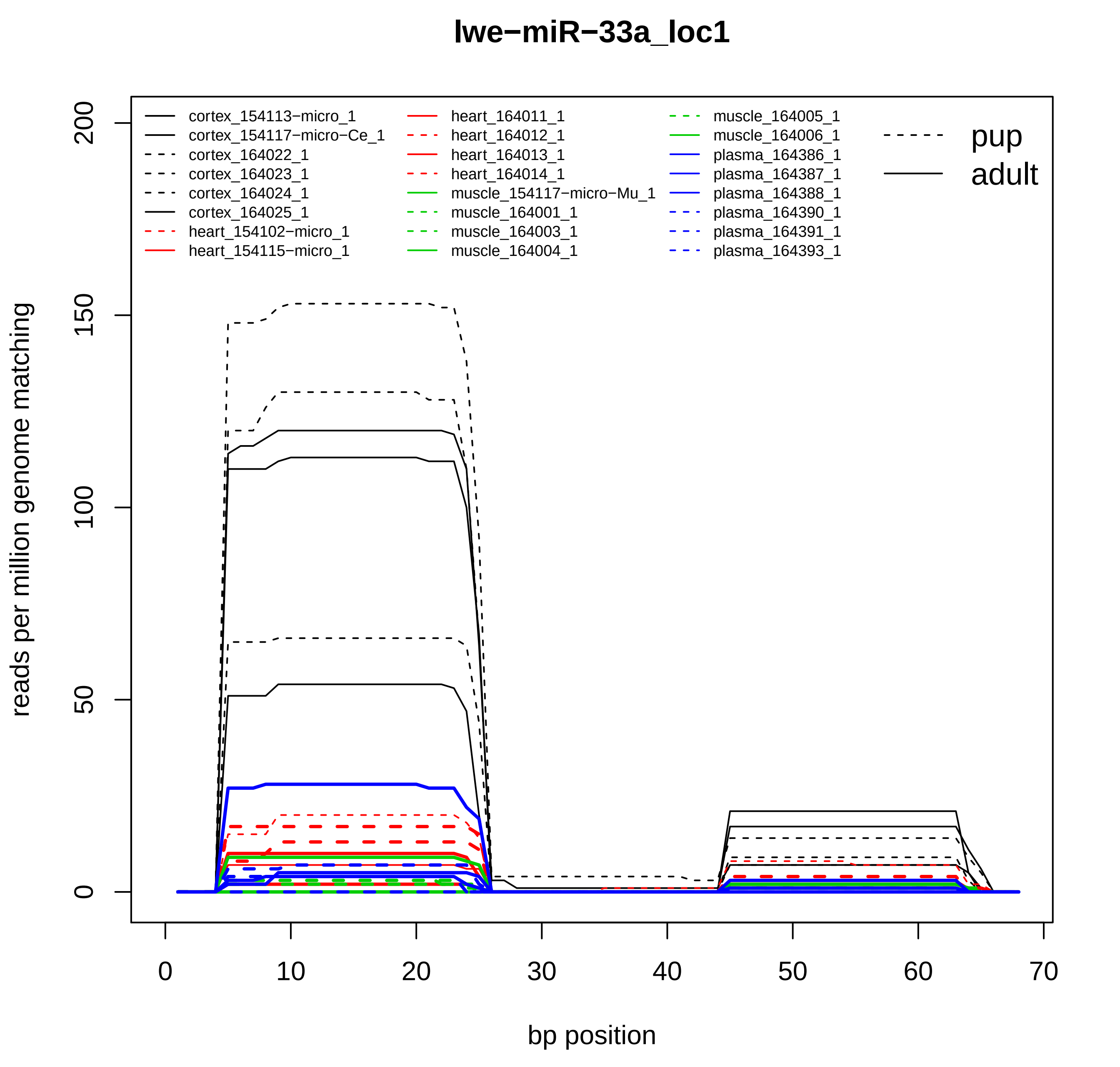
lwe−miR−33a_loc1
reads per million genome matching
bp position
0
50
100
150
200
0
10
20
30
40
50
60
70
cortex_154113−micro_1
cortex_154117−micro−Ce_1
cortex_164022_1
cortex_164023_1
cortex_164024_1
cortex_164025_1
heart_154102−micro_1
heart_154115−micro_1
heart_164011_1
heart_164012_1
heart_164013_1
heart_164014_1
muscle_154117−micro−Mu_1
muscle_164001_1
muscle_164003_1
muscle_164004_1
muscle_164005_1
muscle_164006_1
plasma_164386_1
plasma_164387_1
plasma_164388_1
plasma_164390_1
plasma_164391_1
plasma_164393_1
pup
adult

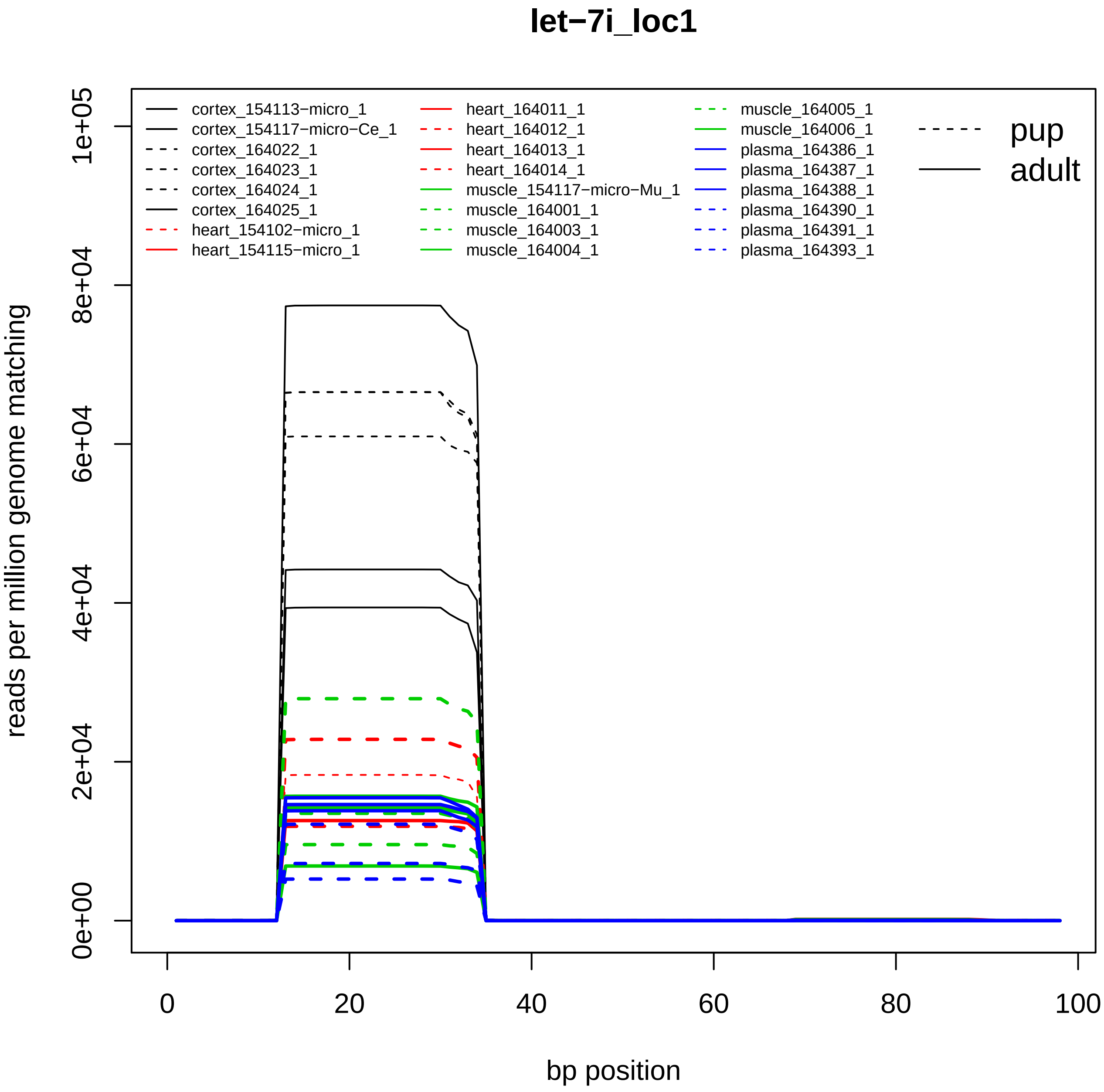

let−7i_loc1
reads per million genome matching
bp position
0e+00
2e+04
4e+04
6e+04
8e+04
1e+05
0
20
40
60
80
100
cortex_154113−micro_1
cortex_154117−micro−Ce_1
cortex_164022_1
cortex_164023_1
cortex_164024_1
cortex_164025_1
heart_154102−micro_1
heart_154115−micro_1
heart_164011_1
heart_164012_1
heart_164013_1
heart_164014_1
muscle_154117−micro−Mu_1
muscle_164001_1
muscle_164003_1
muscle_164004_1
muscle_164005_1
muscle_164006_1
plasma_164386_1
plasma_164387_1
plasma_164388_1
plasma_164390_1
plasma_164391_1
plasma_164393_1
pup
adult

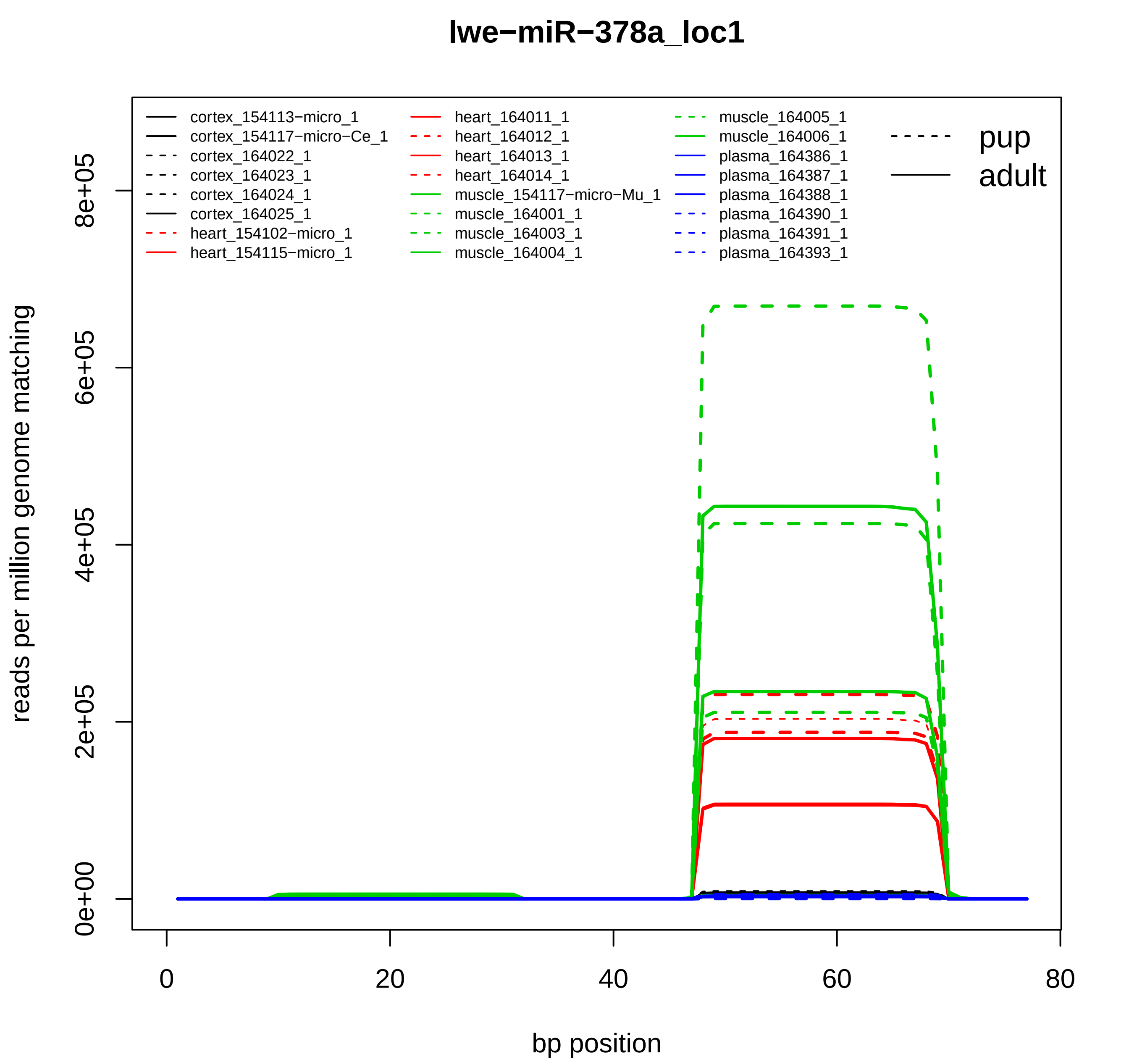
lwe−miR−378a_loc1
cortex_154113−micro_1
cortex_154117−micro−Ce_1
cortex_164022_1
cortex_164023_1
cortex_164024_1
cortex_164025_1
heart_154102−micro_1
heart_154115−micro_1
heart_164011_1
heart_164012_1
heart_164013_1
heart_164014_1
muscle_154117−micro−Mu_1
muscle_164001_1
muscle_164003_1
muscle_164004_1
muscle_164005_1
muscle_164006_1
plasma_164386_1
plasma_164387_1
plasma_164388_1
plasma_164390_1
plasma_164391_1
plasma_164393_1
pup
adult
reads per million genome matching
0e+00
2e+05
4e+05
6e+05
8e+05
0
20
40
60
80
bp position

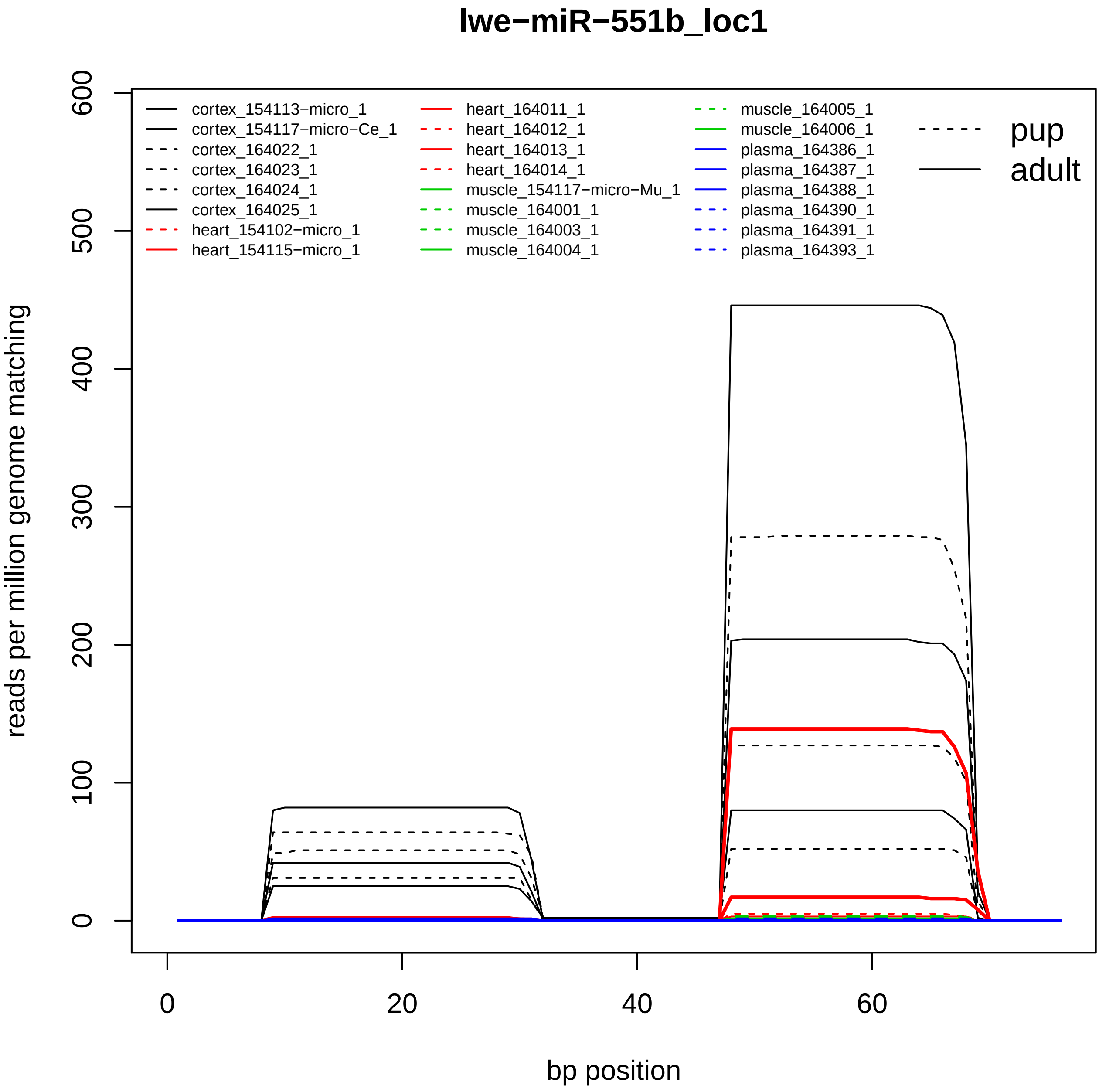
lwe−miR−551b_loc1
reads per million genome matching
bp position
0
100
200
300
400
500
600
0
20
40
60
cortex_154113−micro_1
cortex_154117−micro−Ce_1
cortex_164022_1
cortex_164023_1
cortex_164024_1
cortex_164025_1
heart_154102−micro_1
heart_154115−micro_1
heart_164011_1
heart_164012_1
heart_164013_1
heart_164014_1
muscle_154117−micro−Mu_1
muscle_164001_1
muscle_164003_1
muscle_164004_1
muscle_164005_1
muscle_164006_1
plasma_164386_1
plasma_164387_1
plasma_164388_1
plasma_164390_1
plasma_164391_1
plasma_164393_1
pup
adult

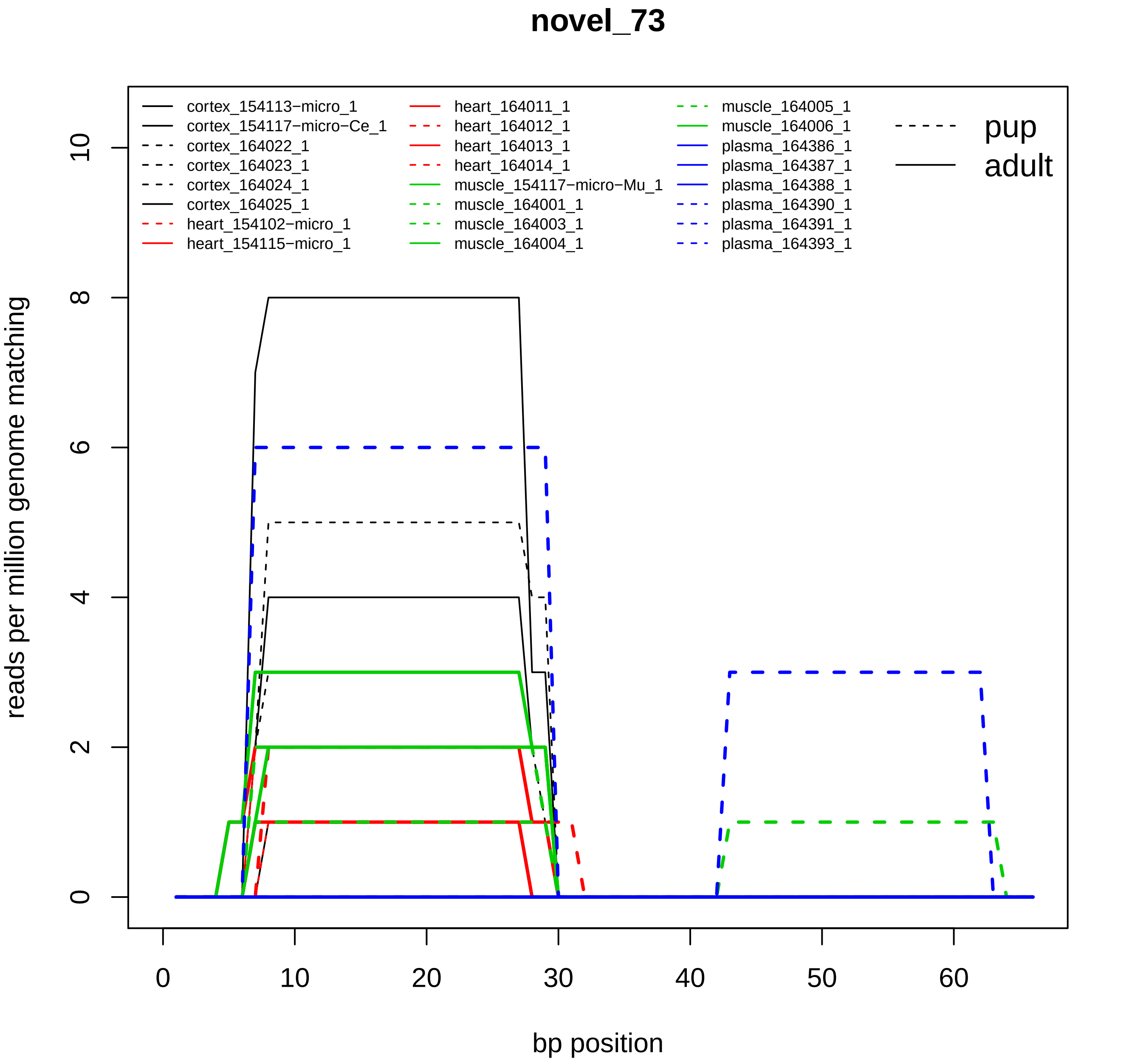
novel_73
cortex_154113−micro_1
cortex_154117−micro−Ce_1
cortex_164022_1
cortex_164023_1
cortex_164024_1
cortex_164025_1
heart_154102−micro_1
heart_154115−micro_1
heart_164011_1
heart_164012_1
heart_164013_1
heart_164014_1
muscle_154117−micro−Mu_1
muscle_164001_1
muscle_164003_1
muscle_164004_1
muscle_164005_1
muscle_164006_1
plasma_164386_1
plasma_164387_1
plasma_164388_1
plasma_164390_1
plasma_164391_1
plasma_164393_1
pup
adult
reads per million genome matching
bp position
0
2
4
6
8
10
0
10
20
30
40
50
60

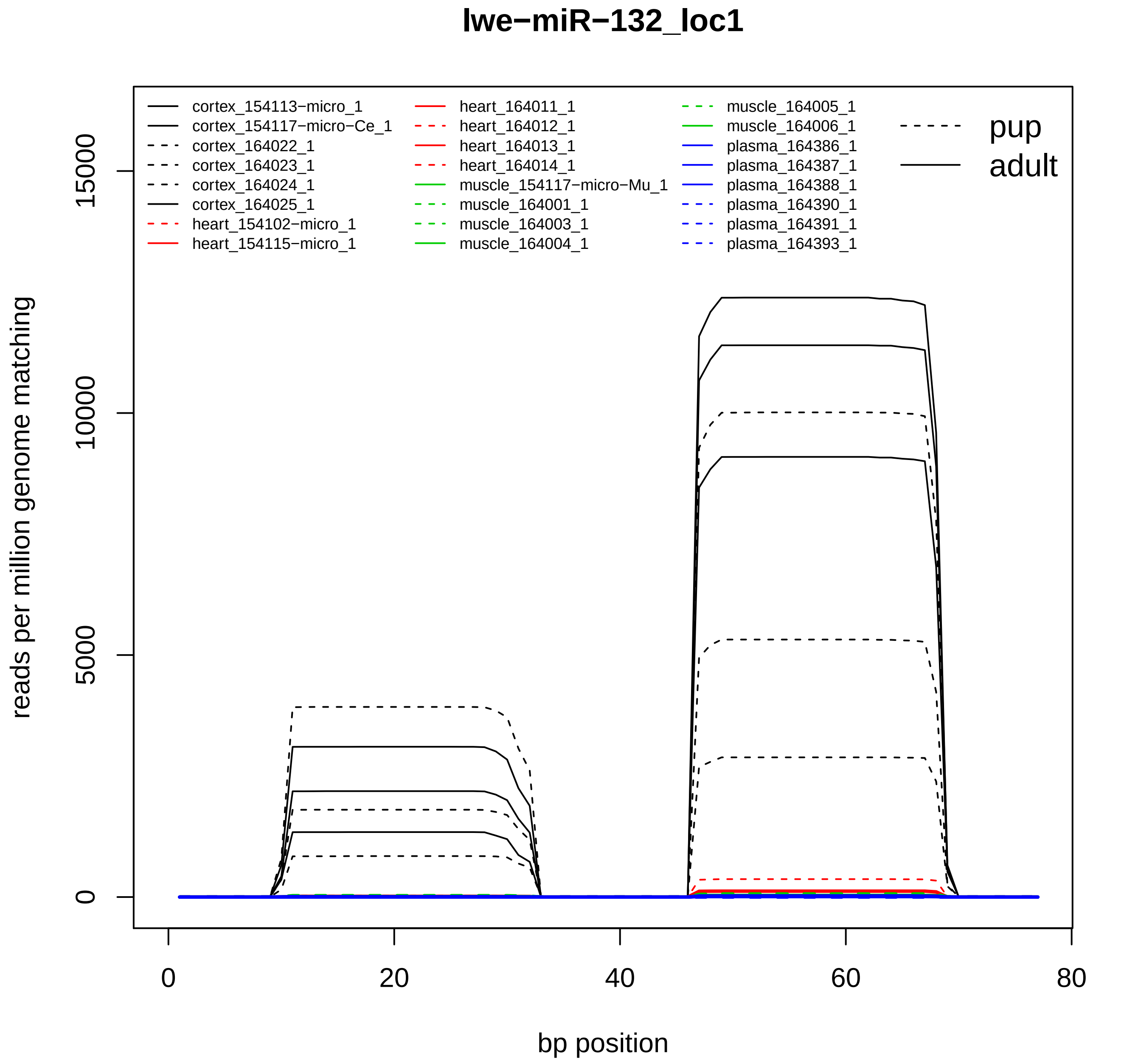

lwe−miR−132_loc1
cortex_154113−micro_1
cortex_154117−micro−Ce_1
cortex_164022_1
cortex_164023_1
cortex_164024_1
cortex_164025_1
heart_154102−micro_1
heart_154115−micro_1
heart_164011_1
heart_164012_1
heart_164013_1
heart_164014_1
muscle_154117−micro−Mu_1
muscle_164001_1
muscle_164003_1
muscle_164004_1
muscle_164005_1
muscle_164006_1
plasma_164386_1
plasma_164387_1
plasma_164388_1
plasma_164390_1
plasma_164391_1
plasma_164393_1
pup
adult
reads per million genome matching
0
5000
10000
15000
0
20
40
60
80
bp position

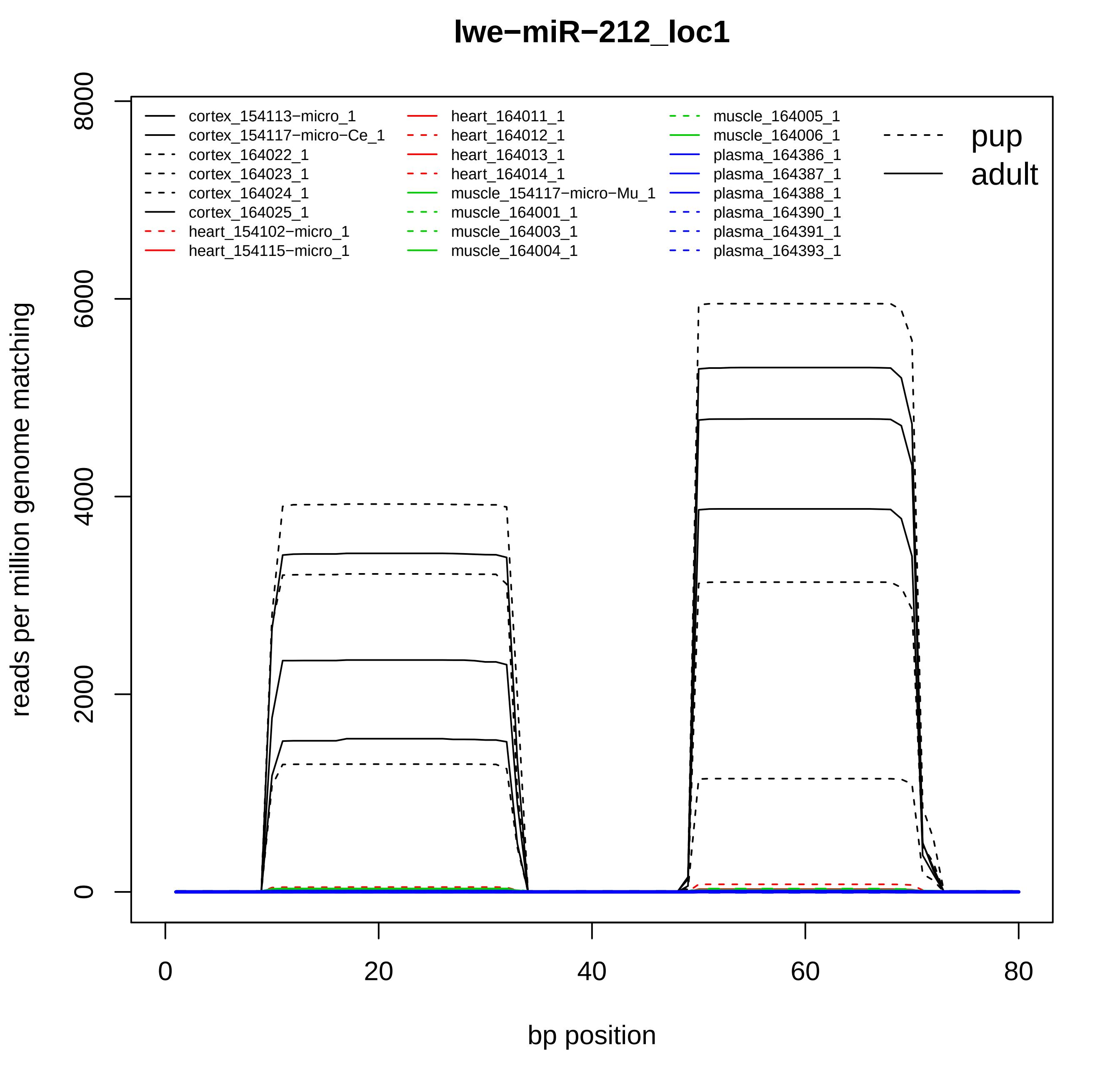

lwe−miR−212_loc1
cortex_154113−micro_1
cortex_154117−micro−Ce_1
cortex_164022_1
cortex_164023_1
cortex_164024_1
cortex_164025_1
heart_154102−micro_1
heart_154115−micro_1
heart_164011_1
heart_164012_1
heart_164013_1
heart_164014_1
muscle_154117−micro−Mu_1
muscle_164001_1
muscle_164003_1
muscle_164004_1
muscle_164005_1
muscle_164006_1
plasma_164386_1
plasma_164387_1
plasma_164388_1
plasma_164390_1
plasma_164391_1
plasma_164393_1
pup
adult
reads per million genome matching
0
2000
4000
6000
8000
bp position
0
20
40
60
80

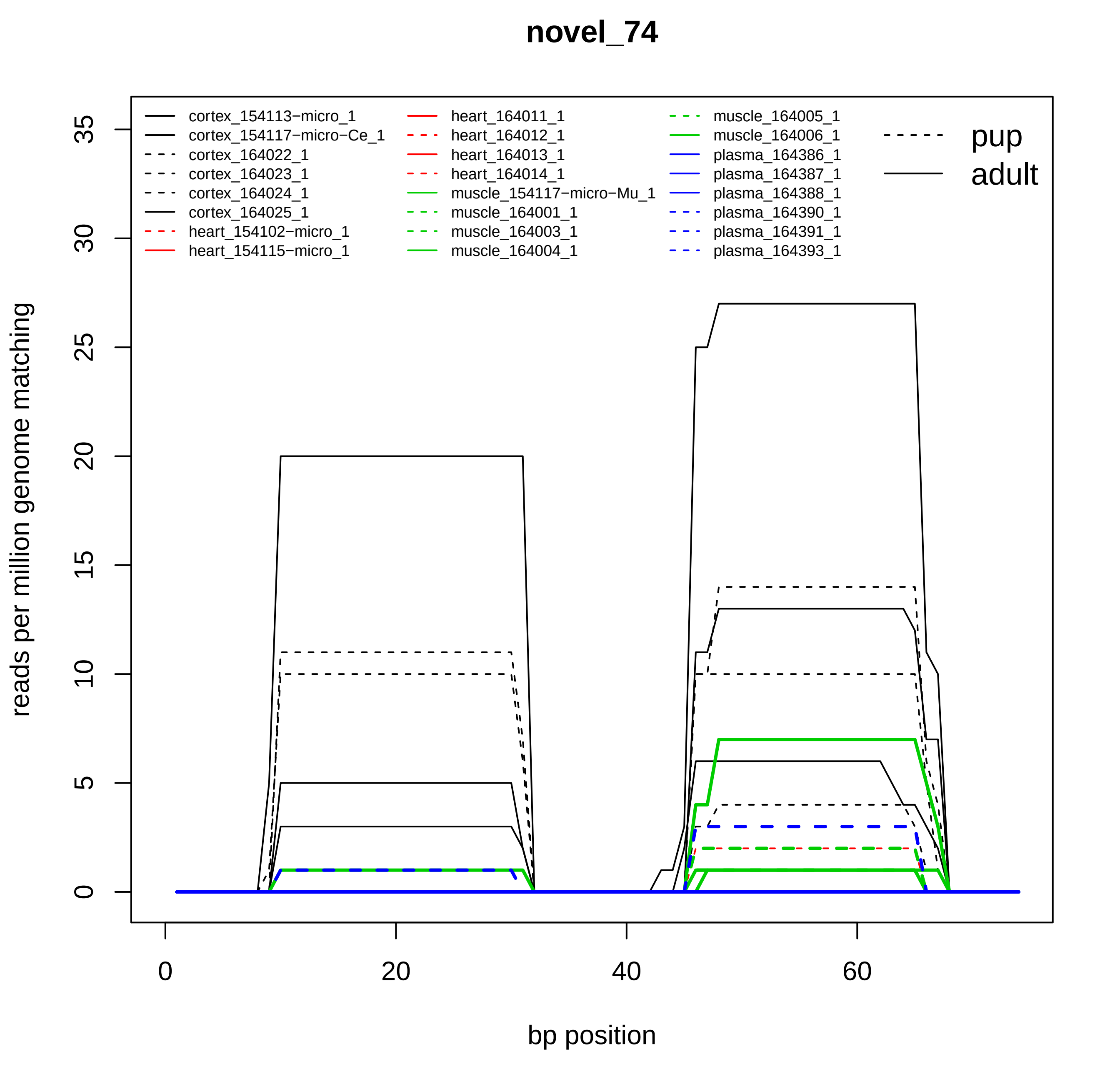
novel_74
cortex_154113−micro_1
cortex_154117−micro−Ce_1
cortex_164022_1
cortex_164023_1
cortex_164024_1
cortex_164025_1
heart_154102−micro_1
heart_154115−micro_1
heart_164011_1
heart_164012_1
heart_164013_1
heart_164014_1
muscle_154117−micro−Mu_1
muscle_164001_1
muscle_164003_1
muscle_164004_1
muscle_164005_1
muscle_164006_1
plasma_164386_1
plasma_164387_1
plasma_164388_1
plasma_164390_1
plasma_164391_1
plasma_164393_1
pup
adult
reads per million genome matching
bp position
0
5
10
15
20
25
30
35
0
20
40
60

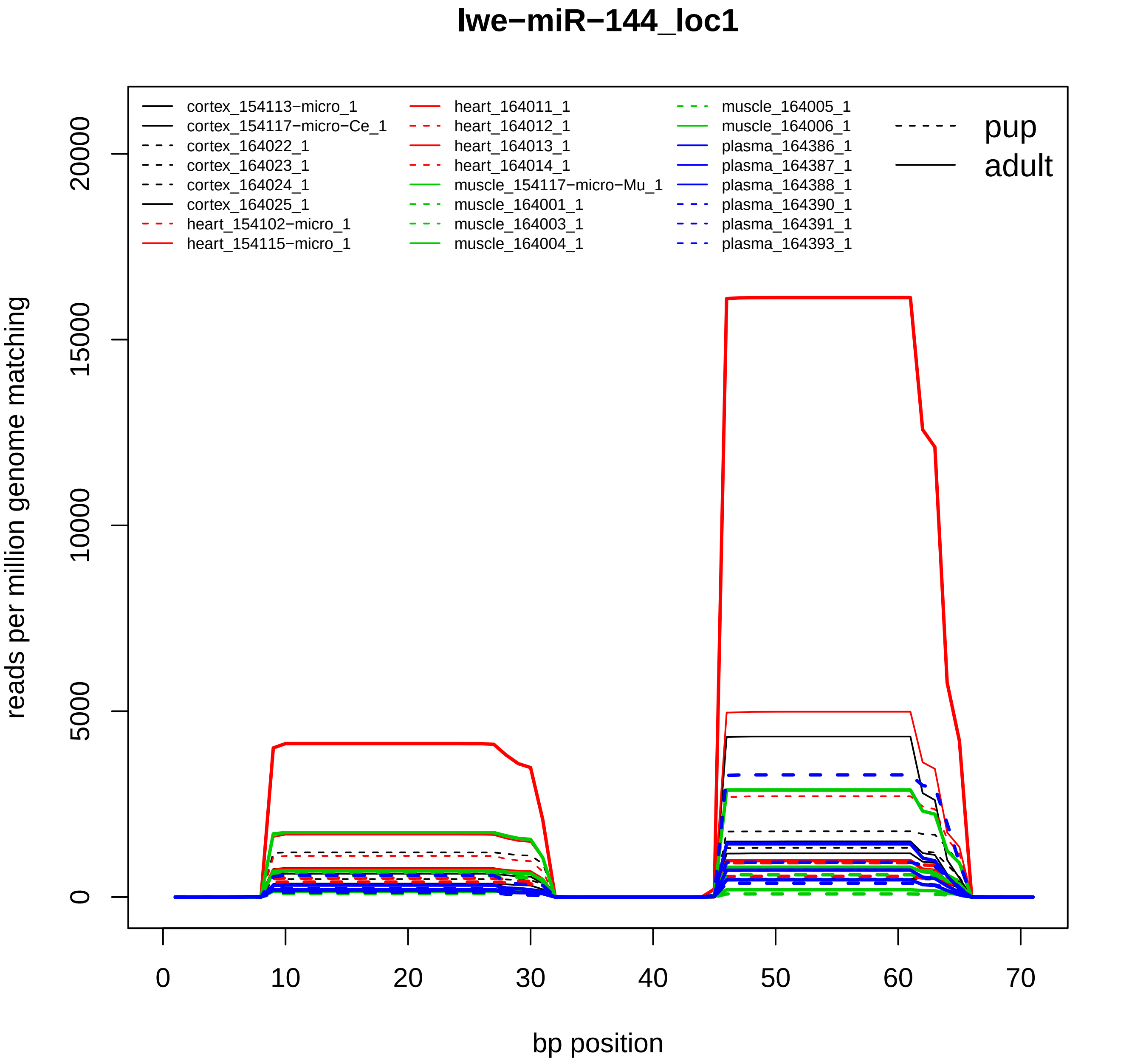
lwe−miR−144_loc1
cortex_154113−micro_1
cortex_154117−micro−Ce_1
cortex_164022_1
cortex_164023_1
cortex_164024_1
cortex_164025_1
heart_154102−micro_1
heart_154115−micro_1
heart_164011_1
heart_164012_1
heart_164013_1
heart_164014_1
muscle_154117−micro−Mu_1
muscle_164001_1
muscle_164003_1
muscle_164004_1
muscle_164005_1
muscle_164006_1
plasma_164386_1
plasma_164387_1
plasma_164388_1
plasma_164390_1
plasma_164391_1
plasma_164393_1
pup
adult
reads per million genome matching
0
5000
10000
15000
20000
0
10
20
30
40
50
60
70
bp position

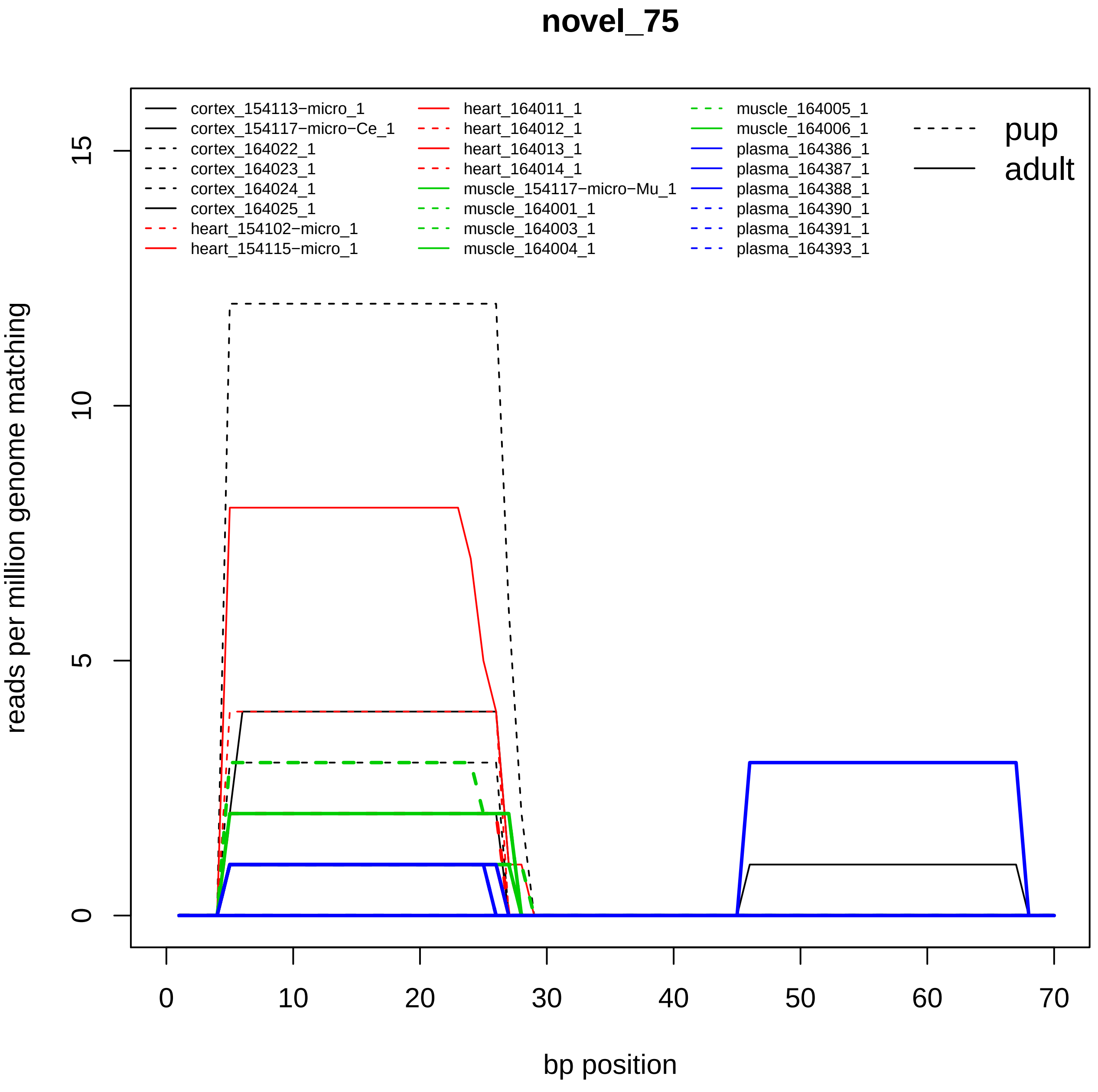

novel_75
cortex_154113−micro_1
cortex_154117−micro−Ce_1
cortex_164022_1
cortex_164023_1
cortex_164024_1
cortex_164025_1
heart_154102−micro_1
heart_154115−micro_1
heart_164011_1
heart_164012_1
heart_164013_1
heart_164014_1
muscle_154117−micro−Mu_1
muscle_164001_1
muscle_164003_1
muscle_164004_1
muscle_164005_1
muscle_164006_1
plasma_164386_1
plasma_164387_1
plasma_164388_1
plasma_164390_1
plasma_164391_1
plasma_164393_1
pup
adult
reads per million genome matching
bp position
0
5
10
15
0
10
20
30
40
50
60
70

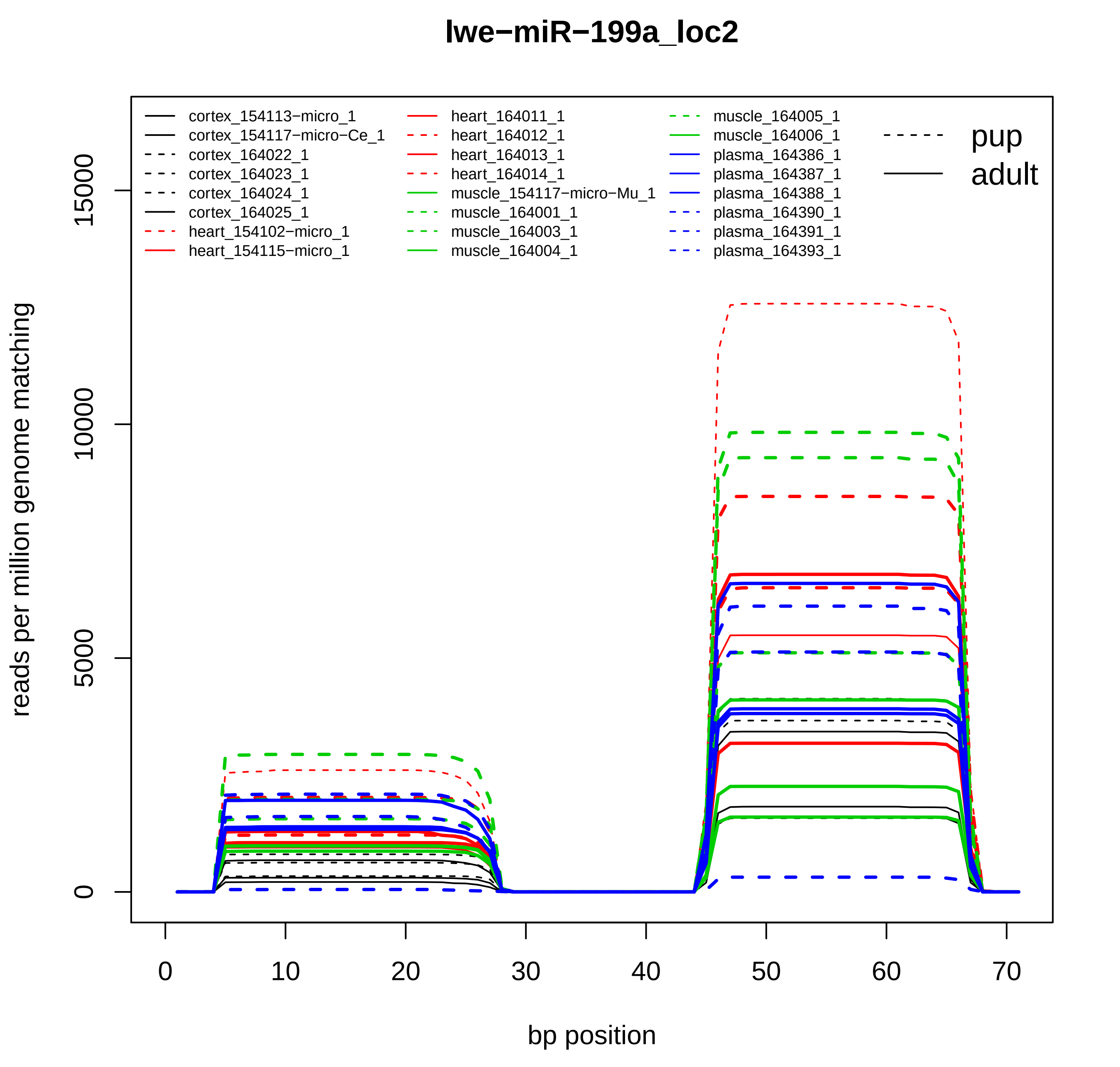

lwe−miR−199a_loc2
cortex_154113−micro_1
cortex_154117−micro−Ce_1
cortex_164022_1
cortex_164023_1
cortex_164024_1
cortex_164025_1
heart_154102−micro_1
heart_154115−micro_1
heart_164011_1
heart_164012_1
heart_164013_1
heart_164014_1
muscle_154117−micro−Mu_1
muscle_164001_1
muscle_164003_1
muscle_164004_1
muscle_164005_1
muscle_164006_1
plasma_164386_1
plasma_164387_1
plasma_164388_1
plasma_164390_1
plasma_164391_1
plasma_164393_1
pup
adult
reads per million genome matching
0
5000
10000
15000
bp position
0
10
20
30
40
50
60
70

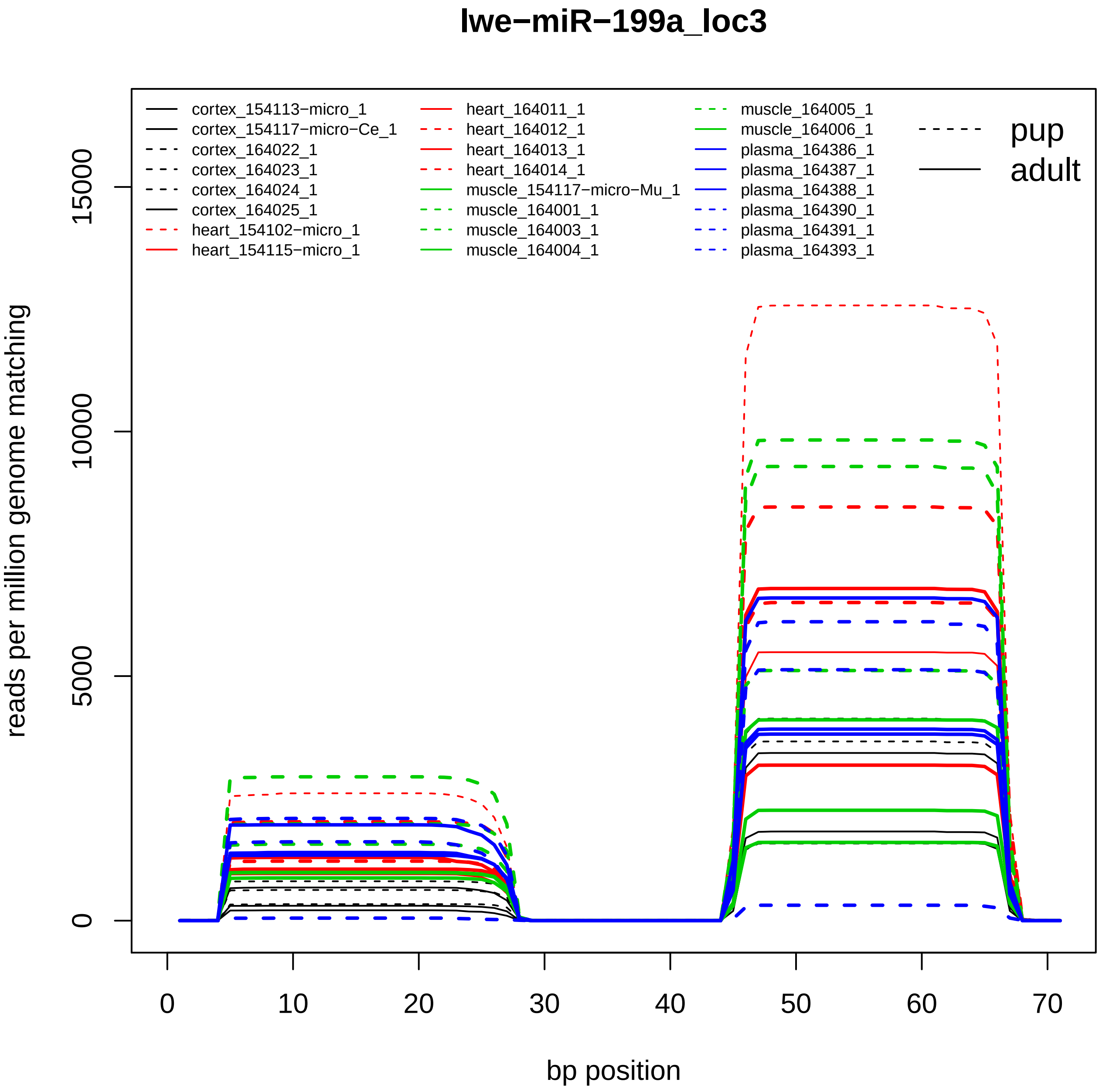
lwe−miR−199a_loc3
reads per million genome matching
bp position
0
5000
10000
15000
0
10
20
30
40
50
60
70
cortex_154113−micro_1
cortex_154117−micro−Ce_1
cortex_164022_1
cortex_164023_1
cortex_164024_1
cortex_164025_1
heart_154102−micro_1
heart_154115−micro_1
heart_164011_1
heart_164012_1
heart_164013_1
heart_164014_1
muscle_154117−micro−Mu_1
muscle_164001_1
muscle_164003_1
muscle_164004_1
muscle_164005_1
muscle_164006_1
plasma_164386_1
plasma_164387_1
plasma_164388_1
plasma_164390_1
plasma_164391_1
plasma_164393_1
pup
adult

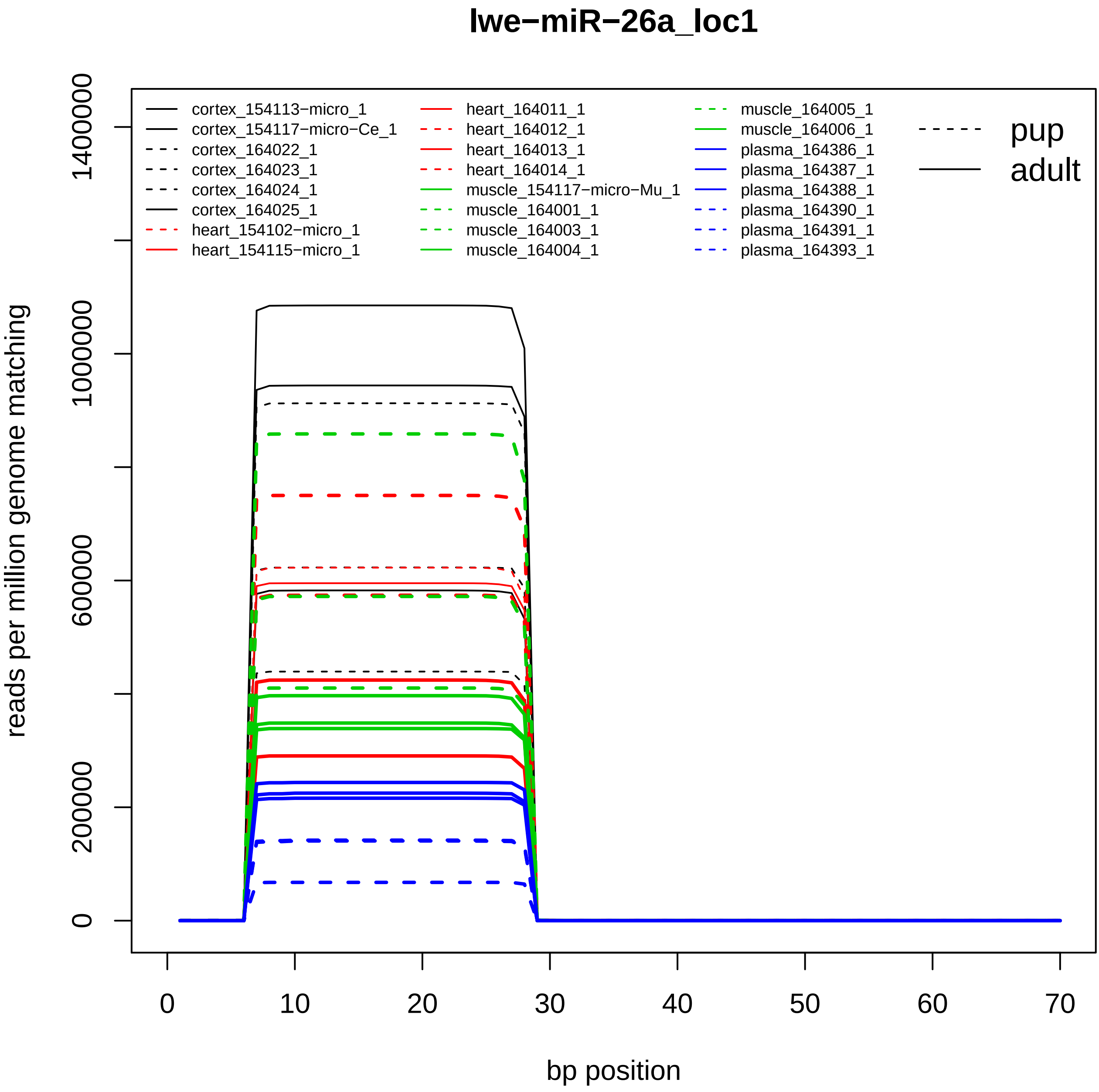

lwe−miR−26a_loc1
cortex_154113−micro_1
cortex_154117−micro−Ce_1
cortex_164022_1
cortex_164023_1
cortex_164024_1
cortex_164025_1
heart_154102−micro_1
heart_154115−micro_1
heart_164011_1
heart_164012_1
heart_164013_1
heart_164014_1
muscle_154117−micro−Mu_1
muscle_164001_1
muscle_164003_1
muscle_164004_1
muscle_164005_1
muscle_164006_1
plasma_164386_1
plasma_164387_1
plasma_164388_1
plasma_164390_1
plasma_164391_1
plasma_164393_1
pup
adult
reads per million genome matching
0
200000
600000
1000000
1400000
bp position
0
10
20
30
40
50
60
70

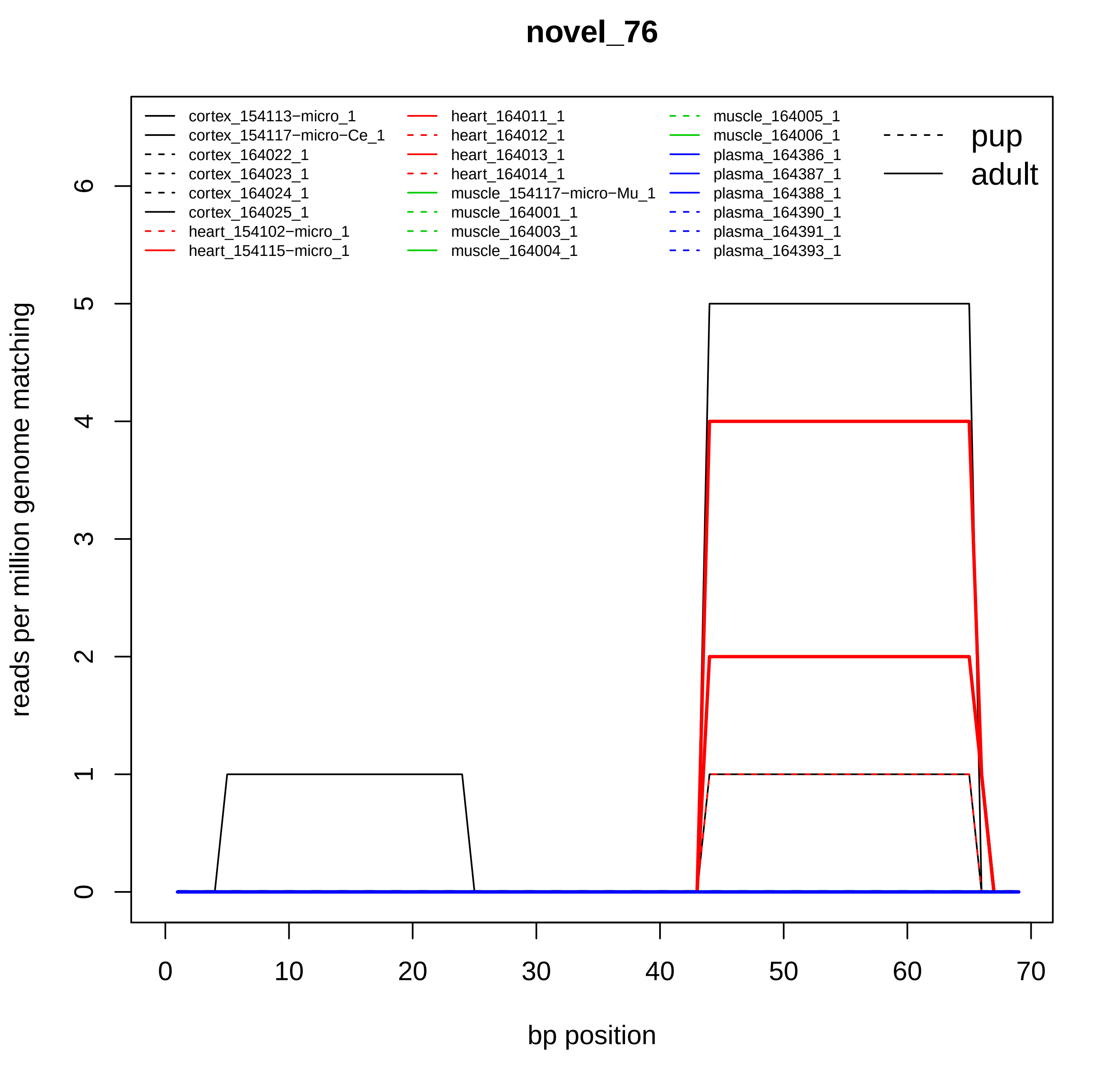

novel_76
cortex_154113−micro_1
cortex_154117−micro−Ce_1
cortex_164022_1
cortex_164023_1
cortex_164024_1
cortex_164025_1
heart_154102−micro_1
heart_154115−micro_1
heart_164011_1
heart_164012_1
heart_164013_1
heart_164014_1
muscle_154117−micro−Mu_1
muscle_164001_1
muscle_164003_1
muscle_164004_1
muscle_164005_1
muscle_164006_1
plasma_164386_1
plasma_164387_1
plasma_164388_1
plasma_164390_1
plasma_164391_1
plasma_164393_1
pup
adult
reads per million genome matching
bp position
0
1
2
3
4
5
6
0
10
20
30
40
50
60
70

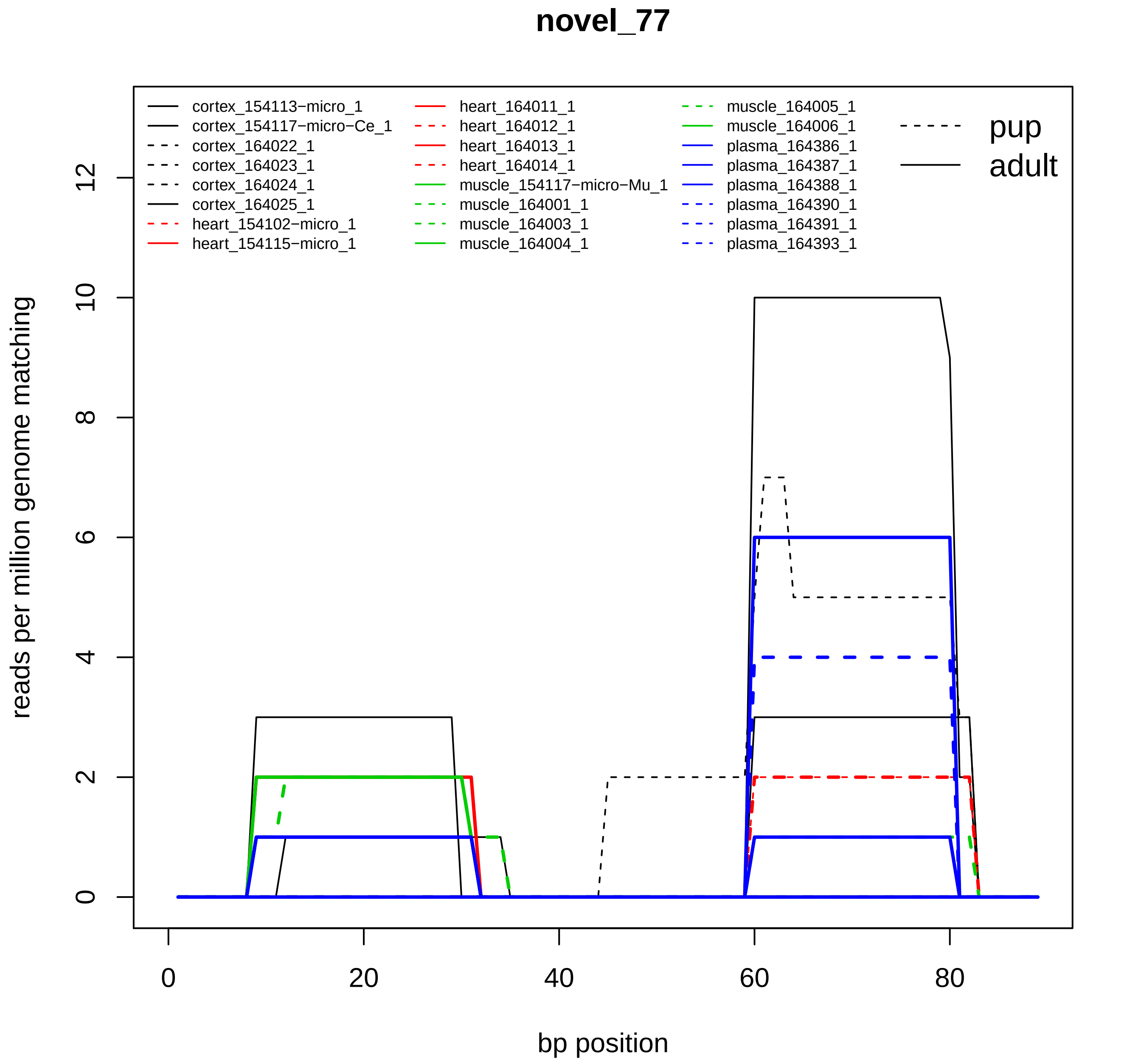

novel_77
cortex_154113−micro_1
cortex_154117−micro−Ce_1
cortex_164022_1
cortex_164023_1
cortex_164024_1
cortex_164025_1
heart_154102−micro_1
heart_154115−micro_1
heart_164011_1
heart_164012_1
heart_164013_1
heart_164014_1
muscle_154117−micro−Mu_1
muscle_164001_1
muscle_164003_1
muscle_164004_1
muscle_164005_1
muscle_164006_1
plasma_164386_1
plasma_164387_1
plasma_164388_1
plasma_164390_1
plasma_164391_1
plasma_164393_1
pup
adult
reads per million genome matching
0
2
4
6
8
10
12
0
20
40
60
80
bp position

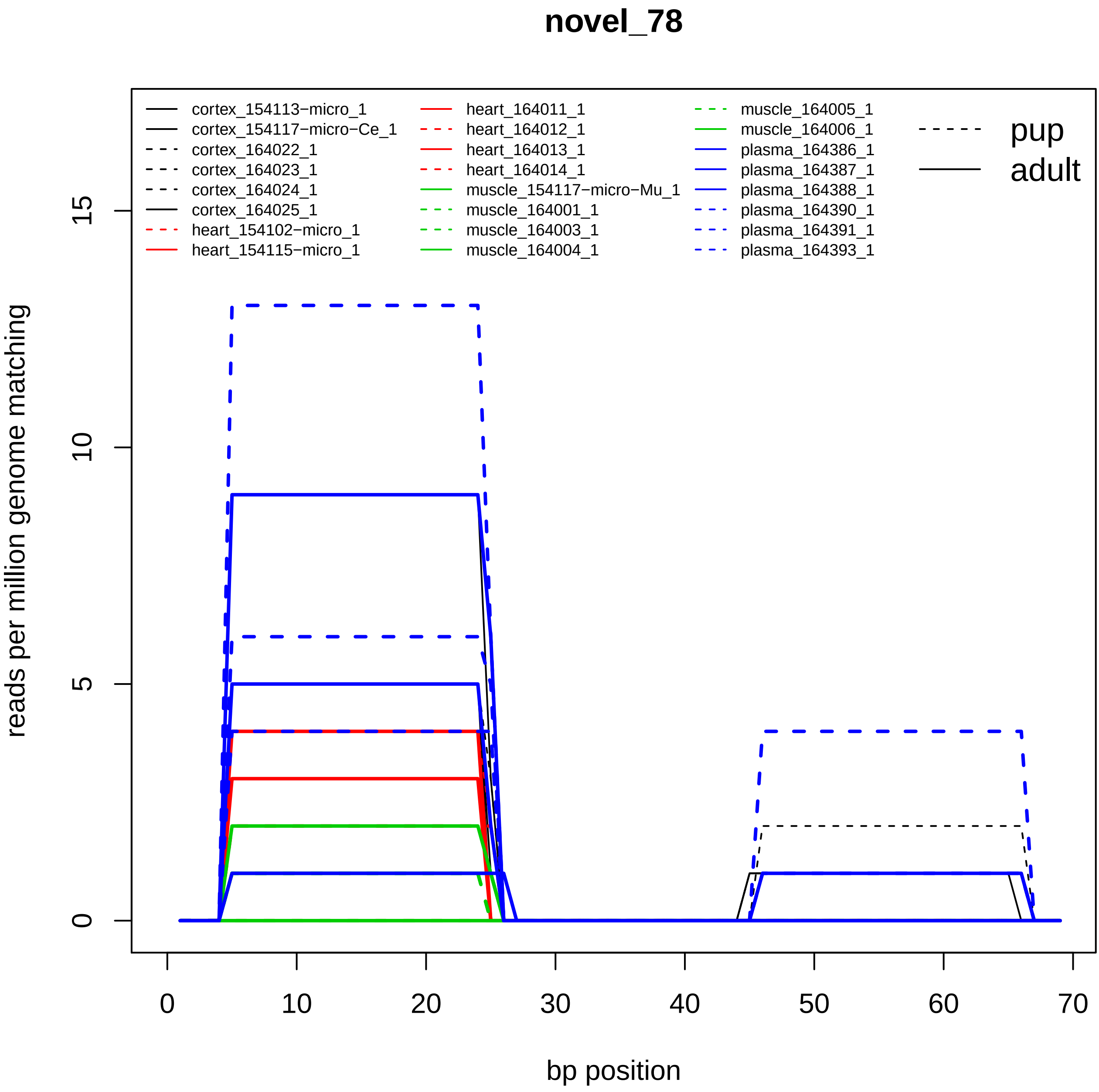

novel_78
cortex_154113−micro_1
cortex_154117−micro−Ce_1
cortex_164022_1
cortex_164023_1
cortex_164024_1
cortex_164025_1
heart_154102−micro_1
heart_154115−micro_1
heart_164011_1
heart_164012_1
heart_164013_1
heart_164014_1
muscle_154117−micro−Mu_1
muscle_164001_1
muscle_164003_1
muscle_164004_1
muscle_164005_1
muscle_164006_1
plasma_164386_1
plasma_164387_1
plasma_164388_1
plasma_164390_1
plasma_164391_1
plasma_164393_1
pup
adult
reads per million genome matching
0
5
10
15
0
10
20
30
40
50
60
70
bp position

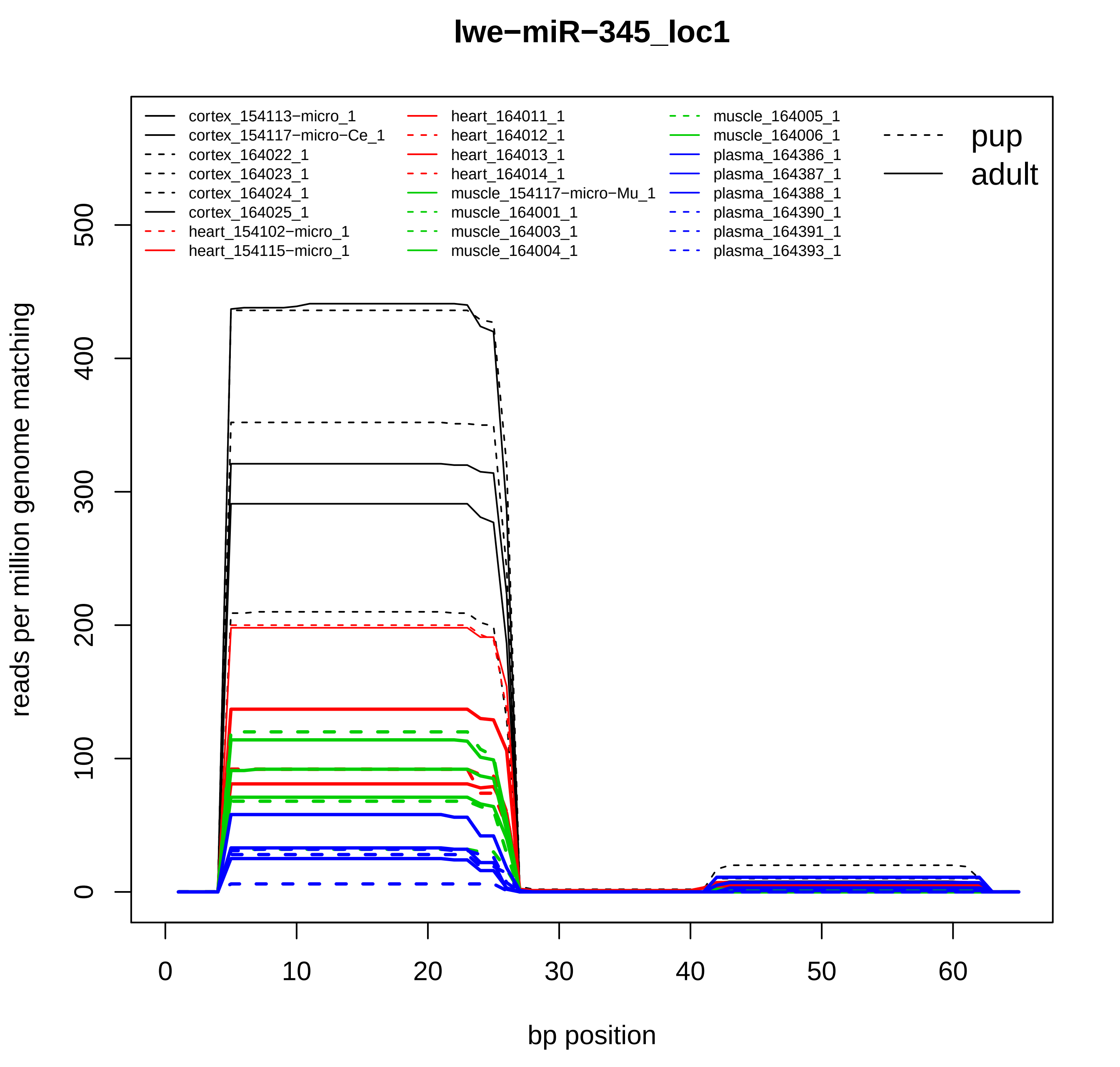
lwe−miR−345_loc1
cortex_154113−micro_1
cortex_154117−micro−Ce_1
cortex_164022_1
cortex_164023_1
cortex_164024_1
cortex_164025_1
heart_154102−micro_1
heart_154115−micro_1
heart_164011_1
heart_164012_1
heart_164013_1
heart_164014_1
muscle_154117−micro−Mu_1
muscle_164001_1
muscle_164003_1
muscle_164004_1
muscle_164005_1
muscle_164006_1
plasma_164386_1
plasma_164387_1
plasma_164388_1
plasma_164390_1
plasma_164391_1
plasma_164393_1
pup
adult
reads per million genome matching
0
100
200
300
400
500
0
10
20
30
40
50
60
bp position

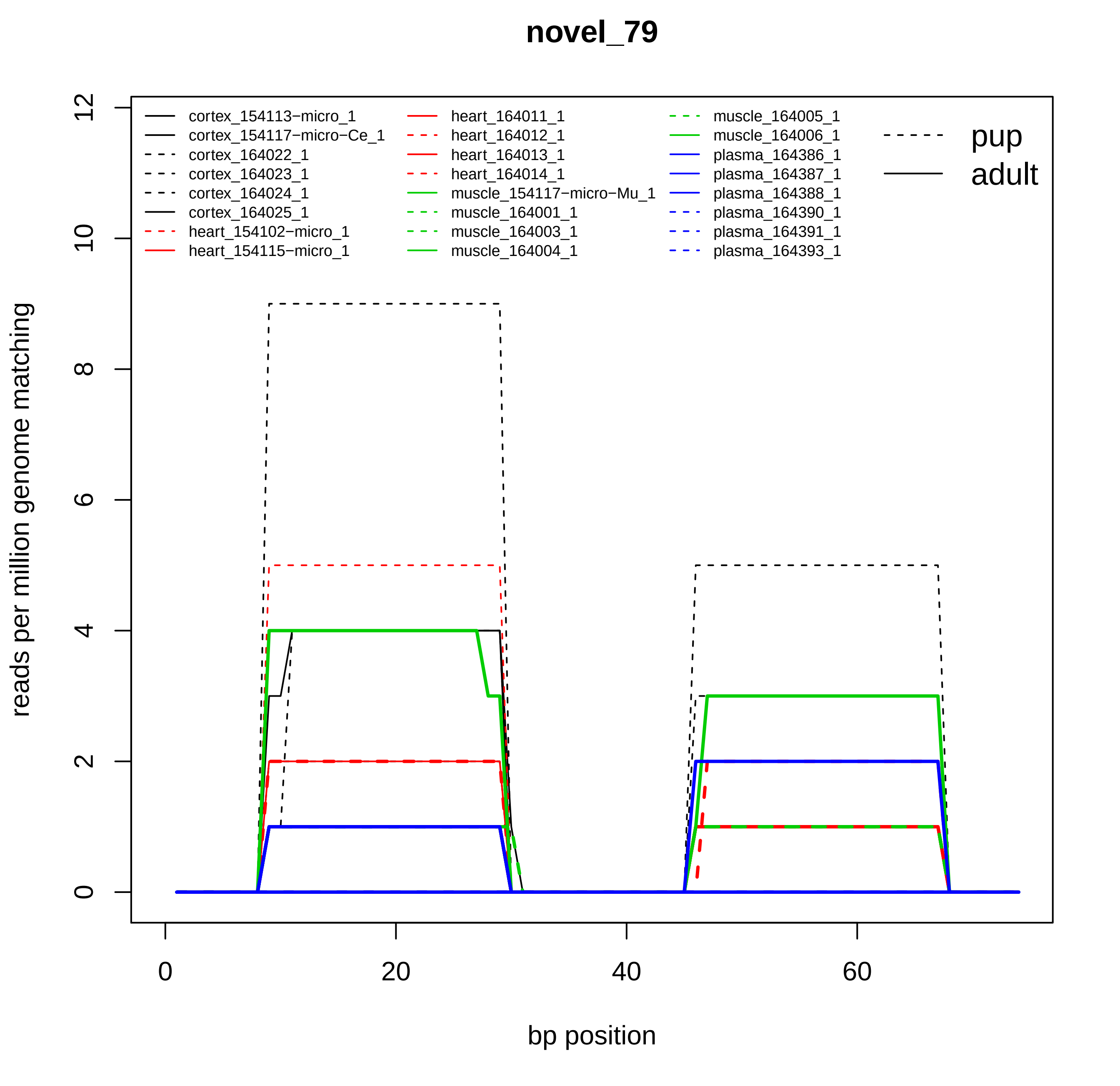

novel_79
cortex_154113−micro_1
cortex_154117−micro−Ce_1
cortex_164022_1
cortex_164023_1
cortex_164024_1
cortex_164025_1
heart_154102−micro_1
heart_154115−micro_1
heart_164011_1
heart_164012_1
heart_164013_1
heart_164014_1
muscle_154117−micro−Mu_1
muscle_164001_1
muscle_164003_1
muscle_164004_1
muscle_164005_1
muscle_164006_1
plasma_164386_1
plasma_164387_1
plasma_164388_1
plasma_164390_1
plasma_164391_1
plasma_164393_1
pup
adult
reads per million genome matching
bp position
0
2
4
6
8
10
12
0
20
40
60

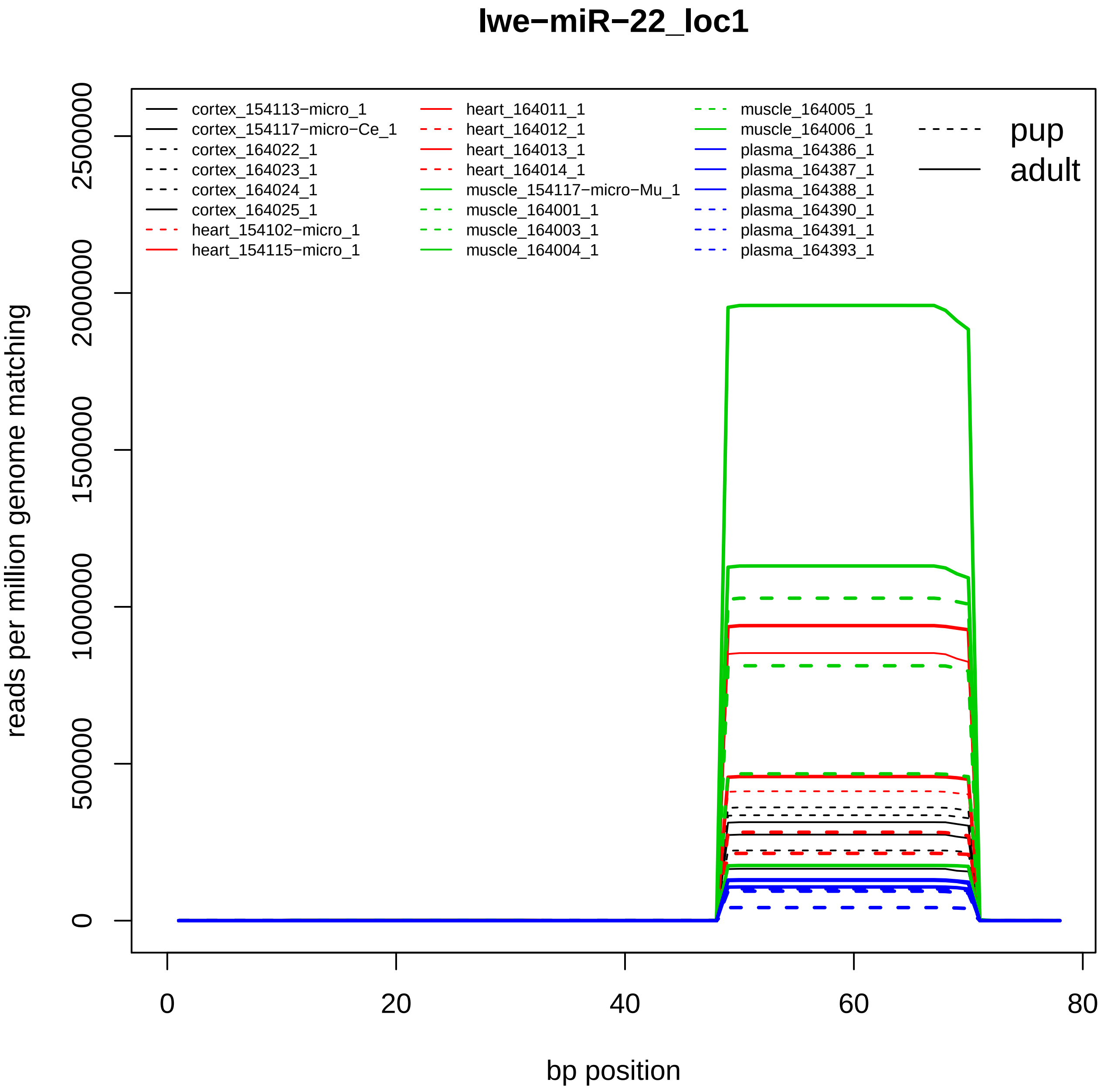
lwe−miR−22_loc1
reads per million genome matching
0
500000
1000000
1500000
2000000
2500000
0
20
40
60
80
bp position
cortex_154113−micro_1
cortex_154117−micro−Ce_1
cortex_164022_1
cortex_164023_1
cortex_164024_1
cortex_164025_1
heart_154102−micro_1
heart_154115−micro_1
heart_164011_1
heart_164012_1
heart_164013_1
heart_164014_1
muscle_154117−micro−Mu_1
muscle_164001_1
muscle_164003_1
muscle_164004_1
muscle_164005_1
muscle_164006_1
plasma_164386_1
plasma_164387_1
plasma_164388_1
plasma_164390_1
plasma_164391_1
plasma_164393_1
pup
adult

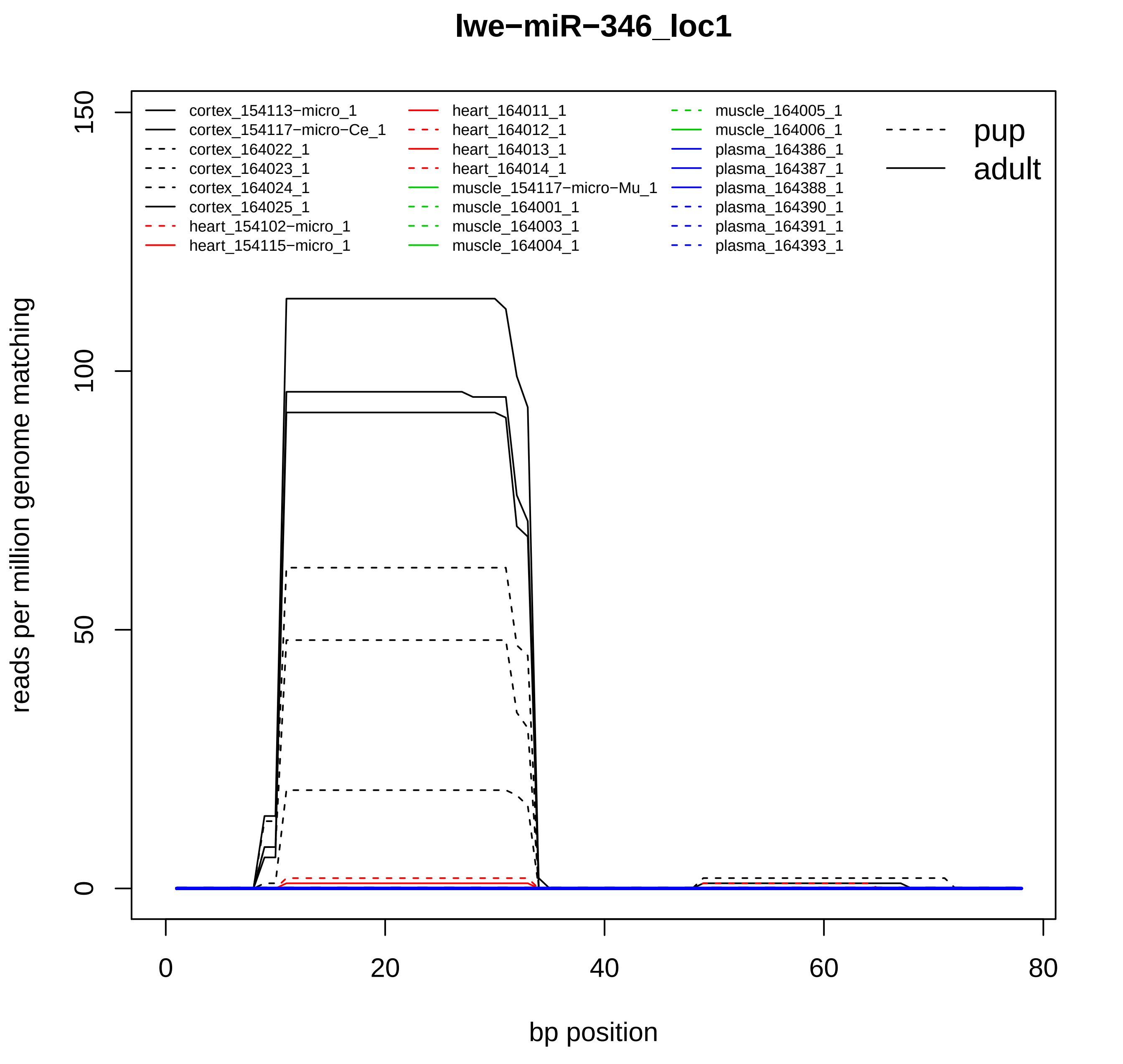
lwe−miR−346_loc1
cortex_154113−micro_1
cortex_154117−micro−Ce_1
cortex_164022_1
cortex_164023_1
cortex_164024_1
cortex_164025_1
heart_154102−micro_1
heart_154115−micro_1
heart_164011_1
heart_164012_1
heart_164013_1
heart_164014_1
muscle_154117−micro−Mu_1
muscle_164001_1
muscle_164003_1
muscle_164004_1
muscle_164005_1
muscle_164006_1
plasma_164386_1
plasma_164387_1
plasma_164388_1
plasma_164390_1
plasma_164391_1
plasma_164393_1
pup
adult
reads per million genome matching
bp position
0
50
100
150
0
20
40
60
80

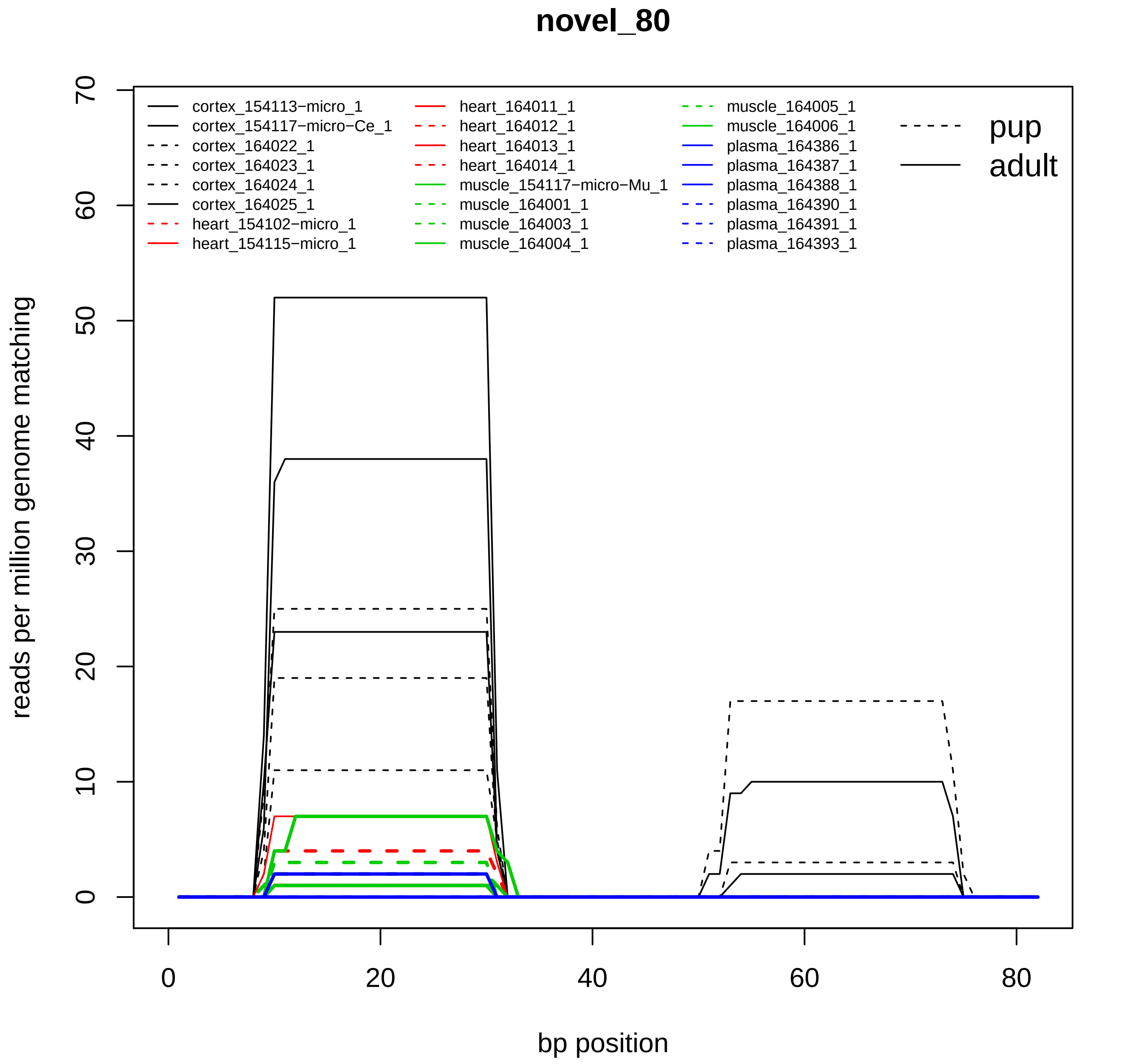

novel_80
reads per million genome matching
bp position
0
10
20
30
40
50
60
70
0
20
40
60
80
cortex_154113−micro_1
cortex_154117−micro−Ce_1
cortex_164022_1
cortex_164023_1
cortex_164024_1
cortex_164025_1
heart_154102−micro_1
heart_154115−micro_1
heart_164011_1
heart_164012_1
heart_164013_1
heart_164014_1
muscle_154117−micro−Mu_1
muscle_164001_1
muscle_164003_1
muscle_164004_1
muscle_164005_1
muscle_164006_1
plasma_164386_1
plasma_164387_1
plasma_164388_1
plasma_164390_1
plasma_164391_1
plasma_164393_1
pup
adult

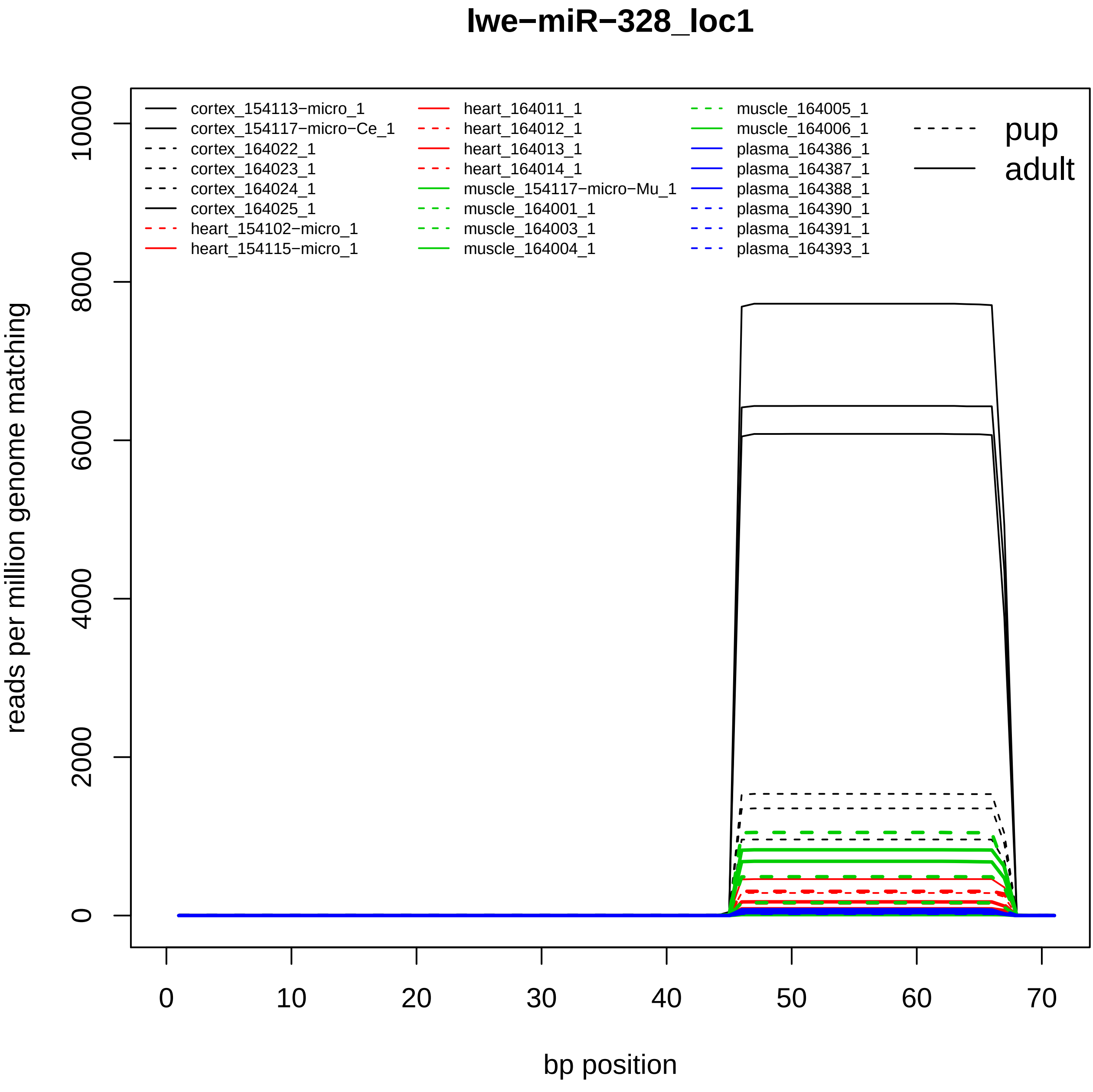
lwe−miR−328_loc1
cortex_154113−micro_1
cortex_154117−micro−Ce_1
cortex_164022_1
cortex_164023_1
cortex_164024_1
cortex_164025_1
heart_154102−micro_1
heart_154115−micro_1
heart_164011_1
heart_164012_1
heart_164013_1
heart_164014_1
muscle_154117−micro−Mu_1
muscle_164001_1
muscle_164003_1
muscle_164004_1
muscle_164005_1
muscle_164006_1
plasma_164386_1
plasma_164387_1
plasma_164388_1
plasma_164390_1
plasma_164391_1
plasma_164393_1
pup
adult
reads per million genome matching
0
2000
4000
6000
8000
10000
0
10
20
30
40
50
60
70
bp position

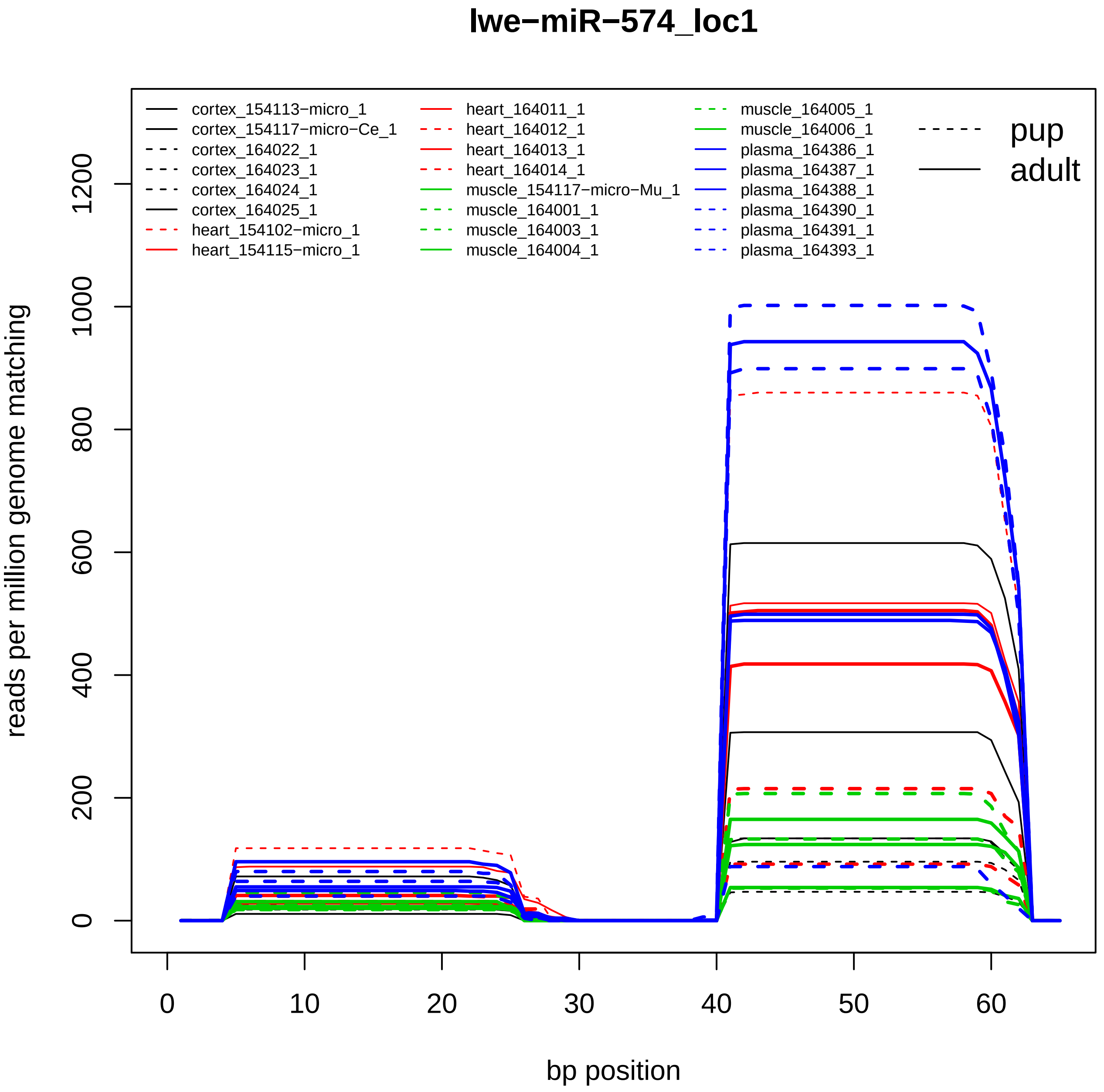

lwe−miR−574_loc1
reads per million genome matching
bp position
0
200
400
600
800
1000
1200
0
10
20
30
40
50
60
cortex_154113−micro_1
cortex_154117−micro−Ce_1
cortex_164022_1
cortex_164023_1
cortex_164024_1
cortex_164025_1
heart_154102−micro_1
heart_154115−micro_1
heart_164011_1
heart_164012_1
heart_164013_1
heart_164014_1
muscle_154117−micro−Mu_1
muscle_164001_1
muscle_164003_1
muscle_164004_1
muscle_164005_1
muscle_164006_1
plasma_164386_1
plasma_164387_1
plasma_164388_1
plasma_164390_1
plasma_164391_1
plasma_164393_1
pup
adult

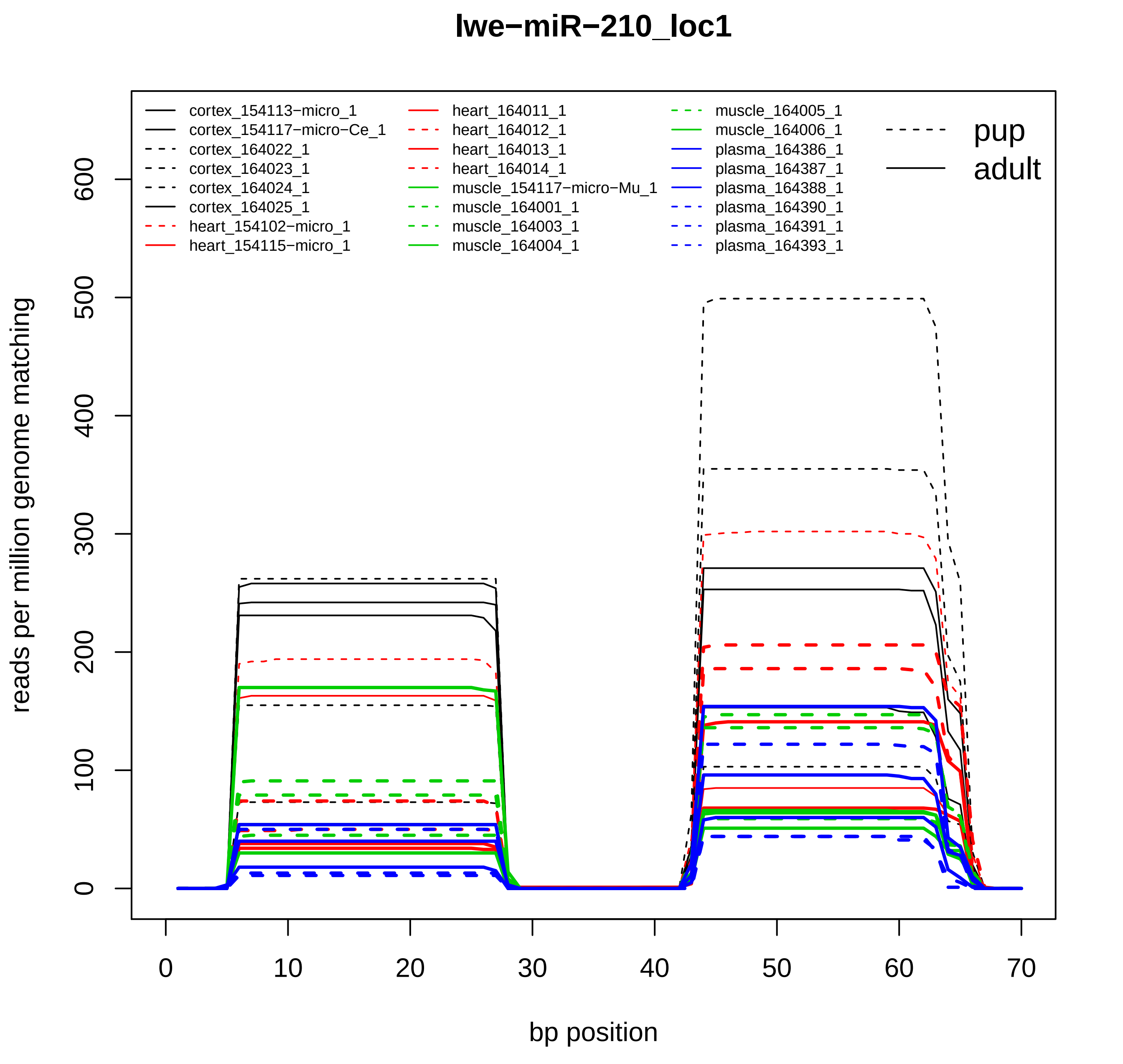
lwe−miR−210_loc1
cortex_154113−micro_1
cortex_154117−micro−Ce_1
cortex_164022_1
cortex_164023_1
cortex_164024_1
cortex_164025_1
heart_154102−micro_1
heart_154115−micro_1
heart_164011_1
heart_164012_1
heart_164013_1
heart_164014_1
muscle_154117−micro−Mu_1
muscle_164001_1
muscle_164003_1
muscle_164004_1
muscle_164005_1
muscle_164006_1
plasma_164386_1
plasma_164387_1
plasma_164388_1
plasma_164390_1
plasma_164391_1
plasma_164393_1
pup
adult
reads per million genome matching
0
100
200
300
400
500
600
0
10
20
30
40
50
60
70
bp position

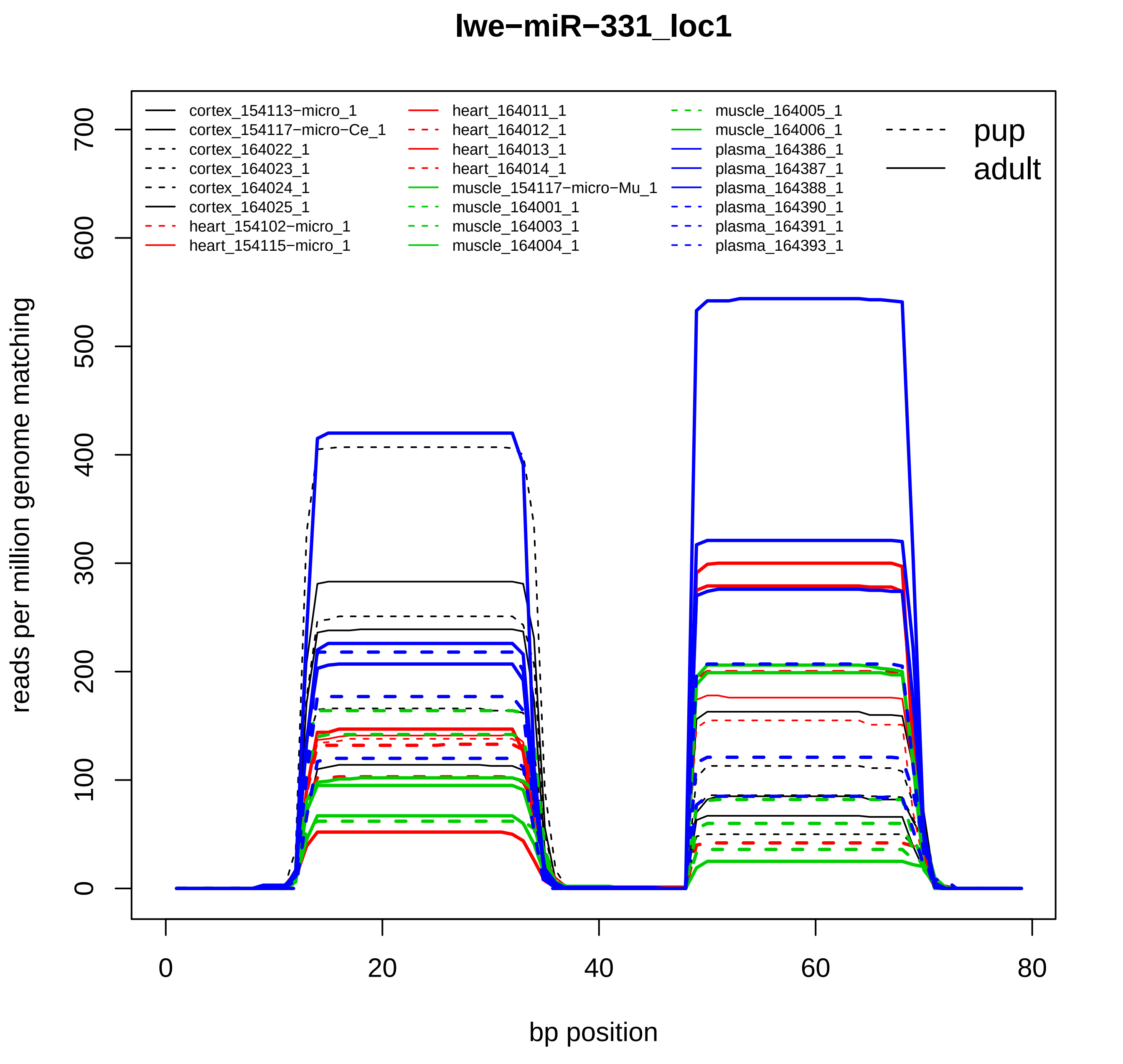

lwe−miR−331_loc1
cortex_154113−micro_1
cortex_154117−micro−Ce_1
cortex_164022_1
cortex_164023_1
cortex_164024_1
cortex_164025_1
heart_154102−micro_1
heart_154115−micro_1
heart_164011_1
heart_164012_1
heart_164013_1
heart_164014_1
muscle_154117−micro−Mu_1
muscle_164001_1
muscle_164003_1
muscle_164004_1
muscle_164005_1
muscle_164006_1
plasma_164386_1
plasma_164387_1
plasma_164388_1
plasma_164390_1
plasma_164391_1
plasma_164393_1
pup
adult
reads per million genome matching
0
100
200
300
400
500
600
700
0
20
40
60
80
bp position

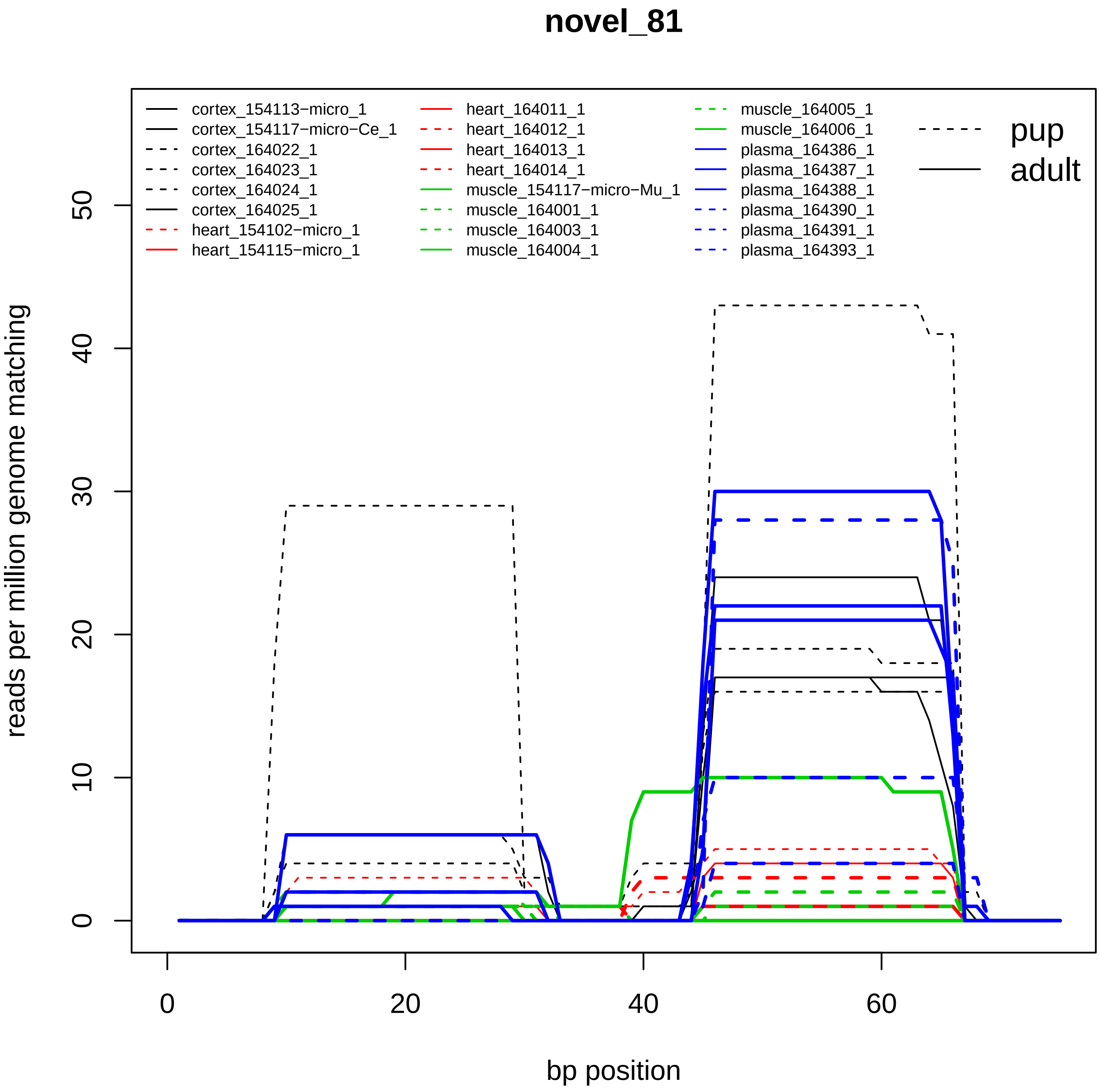
novel_81
cortex_154113−micro_1
cortex_154117−micro−Ce_1
cortex_164022_1
cortex_164023_1
cortex_164024_1
cortex_164025_1
heart_154102−micro_1
heart_154115−micro_1
heart_164011_1
heart_164012_1
heart_164013_1
heart_164014_1
muscle_154117−micro−Mu_1
muscle_164001_1
muscle_164003_1
muscle_164004_1
muscle_164005_1
muscle_164006_1
plasma_164386_1
plasma_164387_1
plasma_164388_1
plasma_164390_1
plasma_164391_1
plasma_164393_1
pup
adult
reads per million genome matching
0
10
20
30
40
50
0
20
40
60
bp position

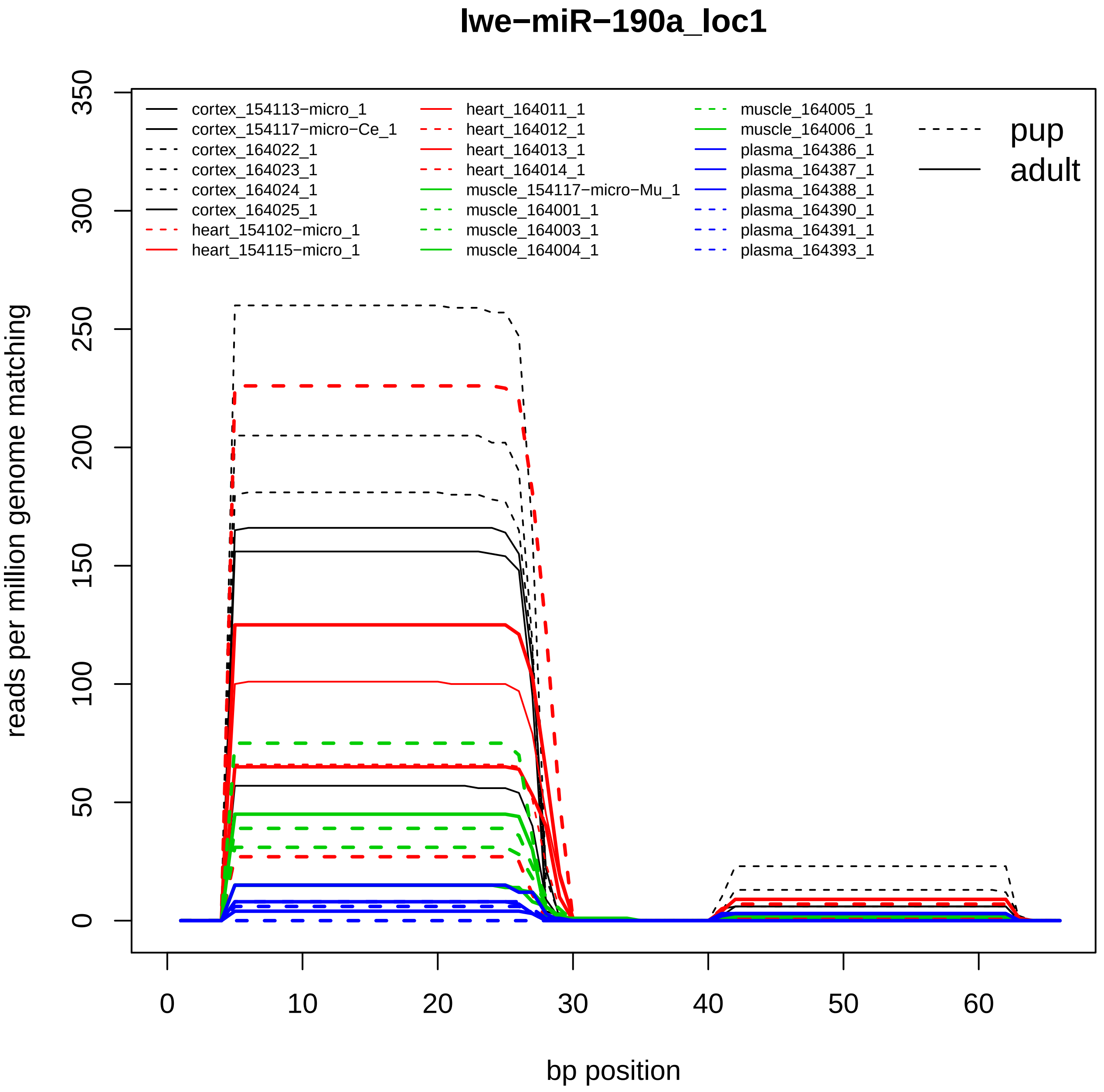
lwe−miR−190a_loc1
reads per million genome matching
bp position
0
50
100
150
200
250
300
350
0
10
20
30
40
50
60
cortex_154113−micro_1
cortex_154117−micro−Ce_1
cortex_164022_1
cortex_164023_1
cortex_164024_1
cortex_164025_1
heart_154102−micro_1
heart_154115−micro_1
heart_164011_1
heart_164012_1
heart_164013_1
heart_164014_1
muscle_154117−micro−Mu_1
muscle_164001_1
muscle_164003_1
muscle_164004_1
muscle_164005_1
muscle_164006_1
plasma_164386_1
plasma_164387_1
plasma_164388_1
plasma_164390_1
plasma_164391_1
plasma_164393_1
pup
adult

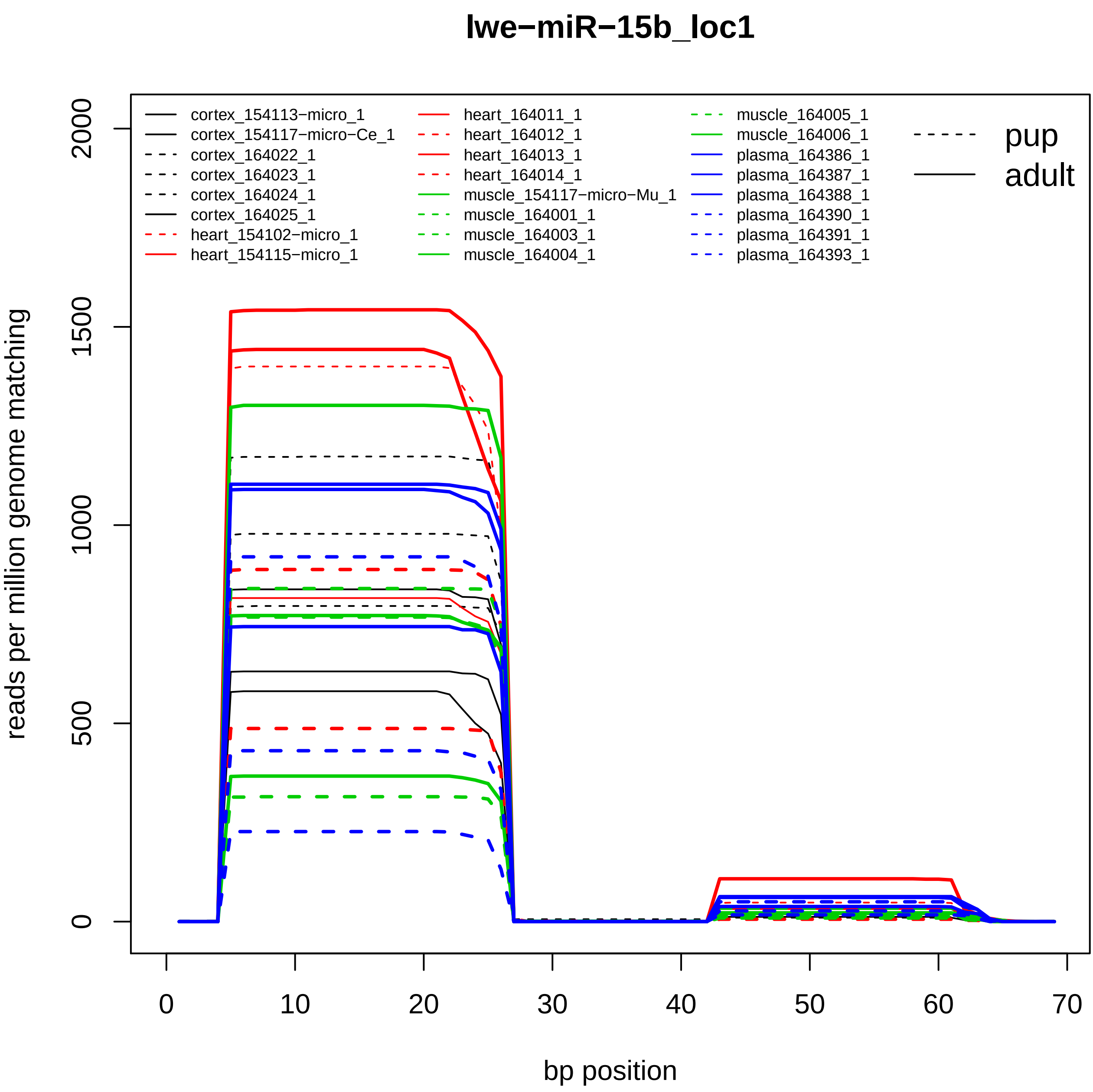
lwe−miR−15b_loc1
cortex_154113−micro_1
cortex_154117−micro−Ce_1
cortex_164022_1
cortex_164023_1
cortex_164024_1
cortex_164025_1
heart_154102−micro_1
heart_154115−micro_1
heart_164011_1
heart_164012_1
heart_164013_1
heart_164014_1
muscle_154117−micro−Mu_1
muscle_164001_1
muscle_164003_1
muscle_164004_1
muscle_164005_1
muscle_164006_1
plasma_164386_1
plasma_164387_1
plasma_164388_1
plasma_164390_1
plasma_164391_1
plasma_164393_1
pup
adult
reads per million genome matching
bp position
0
500
1000
1500
2000
0
10
20
30
40
50
60
70

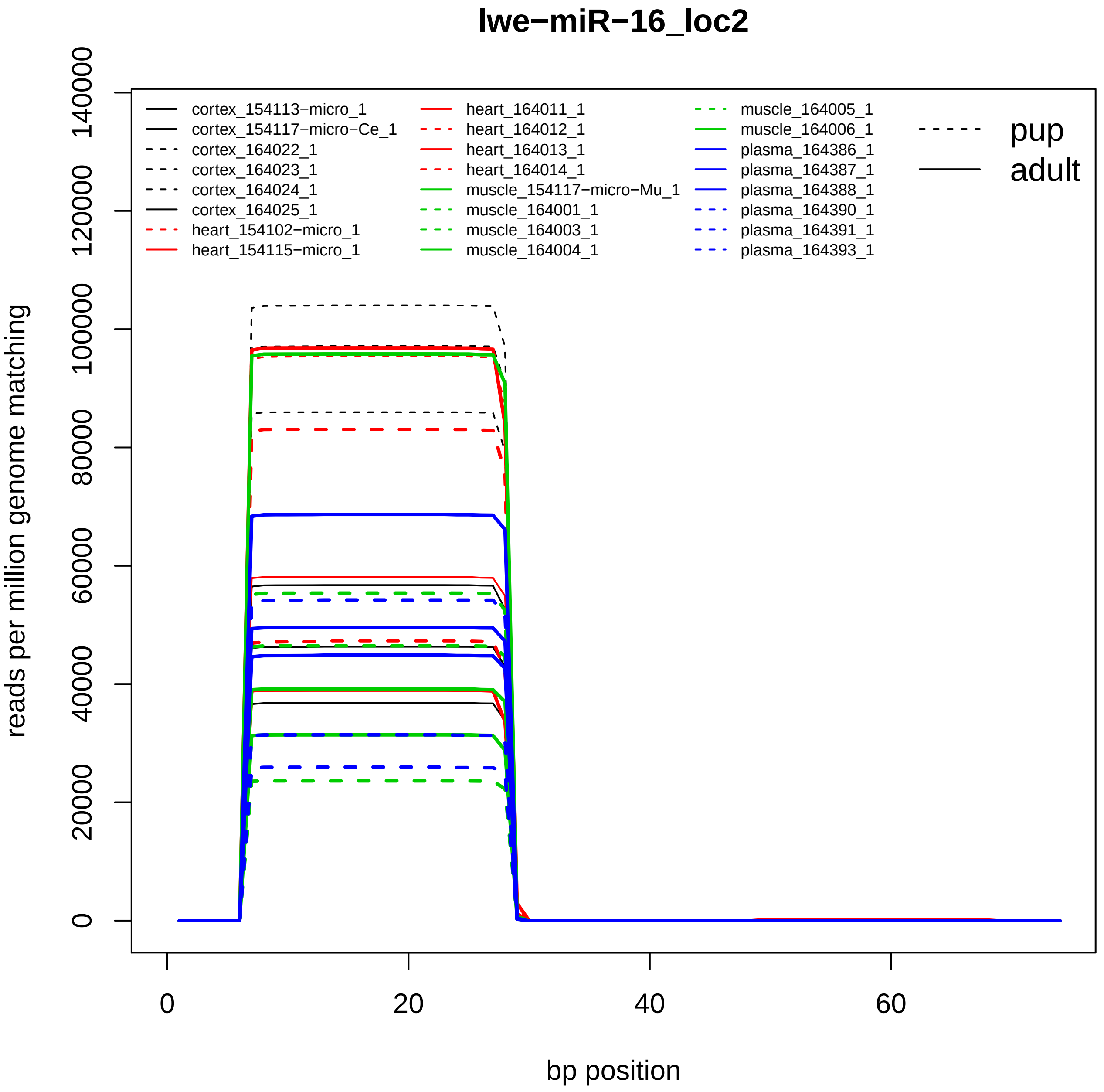
lwe−miR−16_loc2
reads per million genome matching
bp position
0
20000
40000
60000
80000
100000
120000
140000
0
20
40
60
cortex_154113−micro_1
cortex_154117−micro−Ce_1
cortex_164022_1
cortex_164023_1
cortex_164024_1
cortex_164025_1
heart_154102−micro_1
heart_154115−micro_1
heart_164011_1
heart_164012_1
heart_164013_1
heart_164014_1
muscle_154117−micro−Mu_1
muscle_164001_1
muscle_164003_1
muscle_164004_1
muscle_164005_1
muscle_164006_1
plasma_164386_1
plasma_164387_1
plasma_164388_1
plasma_164390_1
plasma_164391_1
plasma_164393_1
pup
adult

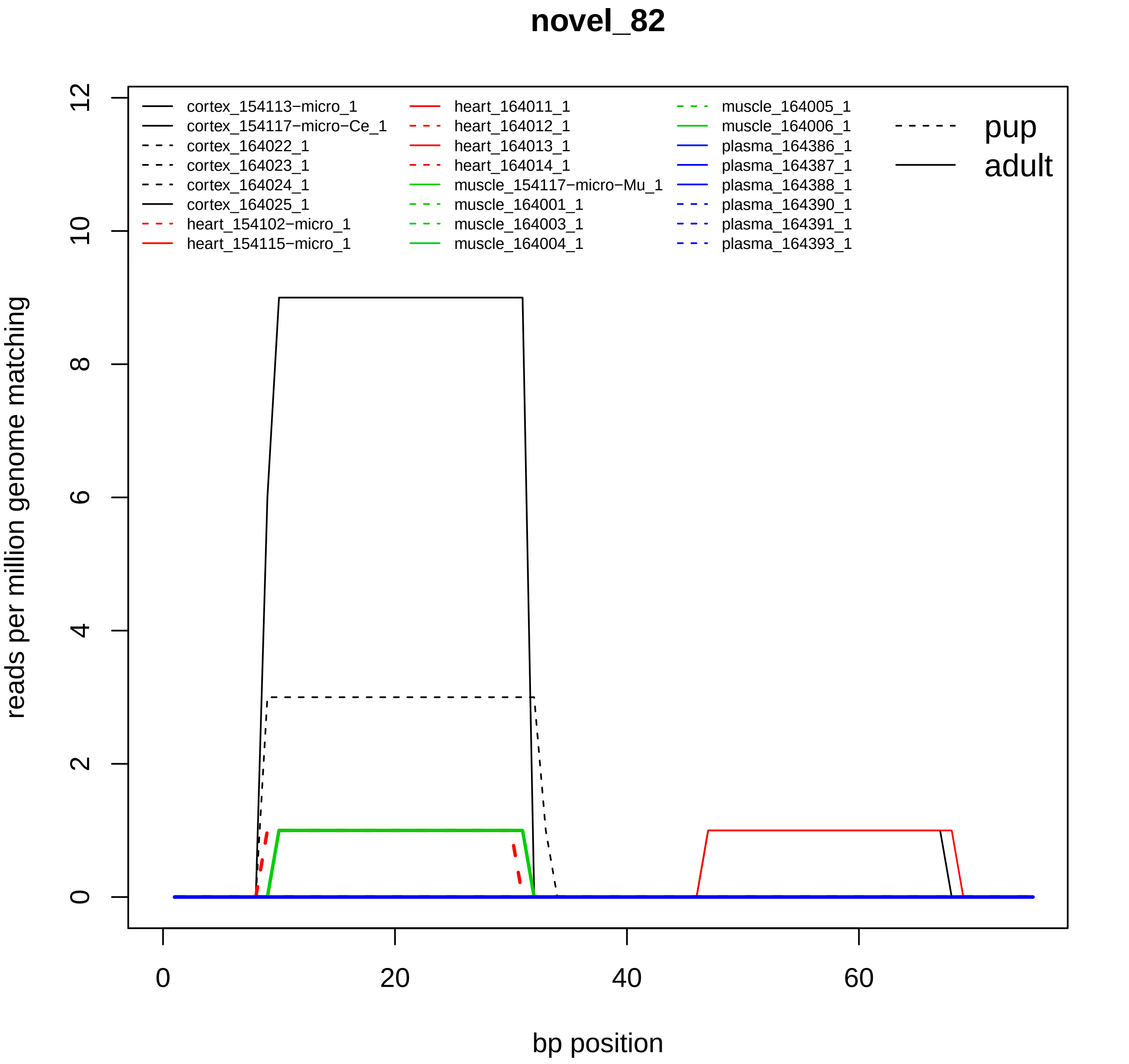
novel_82
cortex_154113−micro_1
cortex_154117−micro−Ce_1
cortex_164022_1
cortex_164023_1
cortex_164024_1
cortex_164025_1
heart_154102−micro_1
heart_154115−micro_1
heart_164011_1
heart_164012_1
heart_164013_1
heart_164014_1
muscle_154117−micro−Mu_1
muscle_164001_1
muscle_164003_1
muscle_164004_1
muscle_164005_1
muscle_164006_1
plasma_164386_1
plasma_164387_1
plasma_164388_1
plasma_164390_1
plasma_164391_1
plasma_164393_1
pup
adult
reads per million genome matching
bp position
0
2
4
6
8
10
12
0
20
40
60

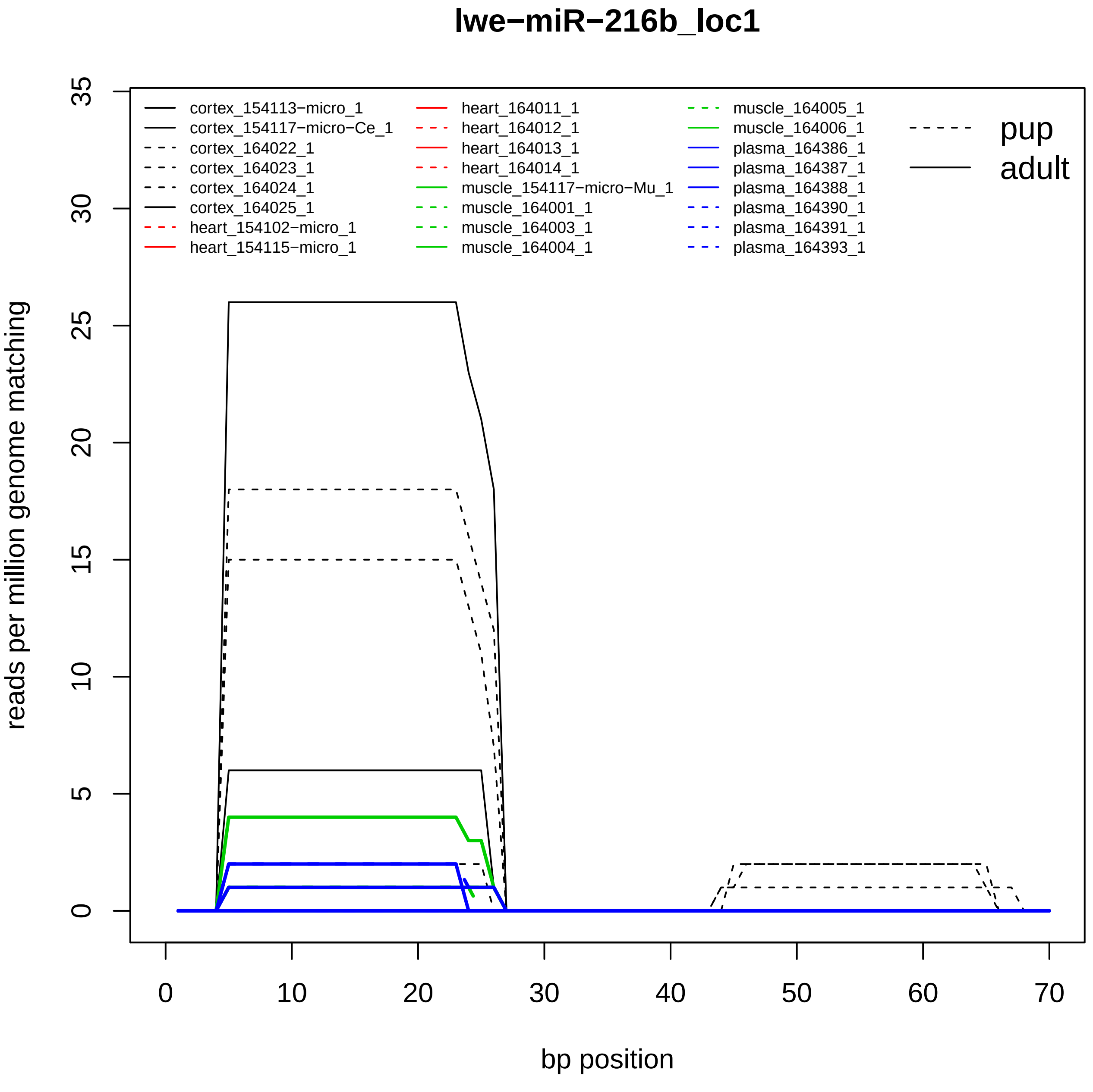
lwe−miR−216b_loc1
cortex_154113−micro_1
cortex_154117−micro−Ce_1
cortex_164022_1
cortex_164023_1
cortex_164024_1
cortex_164025_1
heart_154102−micro_1
heart_154115−micro_1
heart_164011_1
heart_164012_1
heart_164013_1
heart_164014_1
muscle_154117−micro−Mu_1
muscle_164001_1
muscle_164003_1
muscle_164004_1
muscle_164005_1
muscle_164006_1
plasma_164386_1
plasma_164387_1
plasma_164388_1
plasma_164390_1
plasma_164391_1
plasma_164393_1
pup
adult
reads per million genome matching
bp position
0
5
10
15
20
25
30
35
0
10
20
30
40
50
60
70

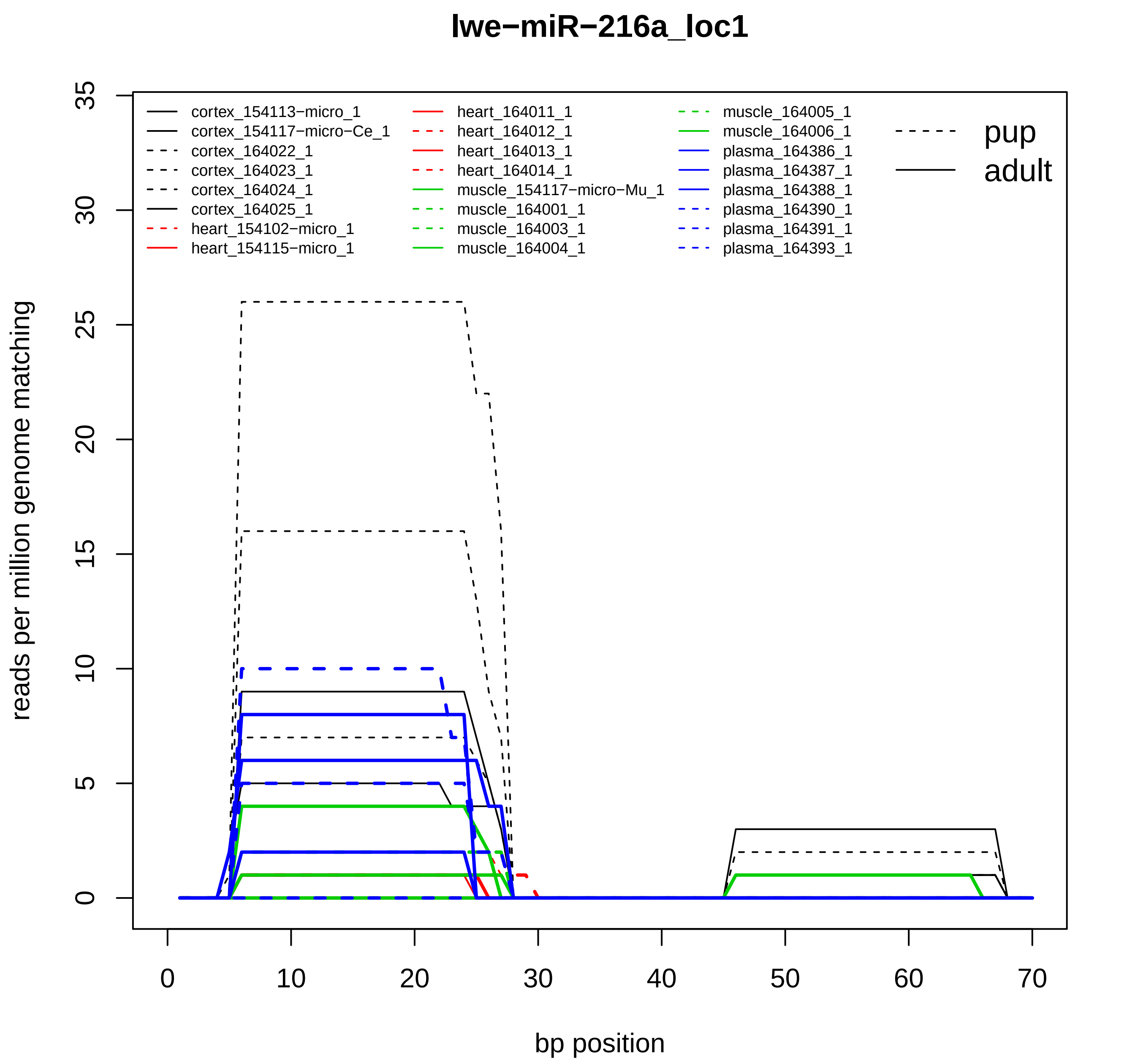

lwe−miR−216a_loc1
reads per million genome matching
bp position
0
5
10
15
20
25
30
35
0
10
20
30
40
50
60
70
cortex_154113−micro_1
cortex_154117−micro−Ce_1
cortex_164022_1
cortex_164023_1
cortex_164024_1
cortex_164025_1
heart_154102−micro_1
heart_154115−micro_1
heart_164011_1
heart_164012_1
heart_164013_1
heart_164014_1
muscle_154117−micro−Mu_1
muscle_164001_1
muscle_164003_1
muscle_164004_1
muscle_164005_1
muscle_164006_1
plasma_164386_1
plasma_164387_1
plasma_164388_1
plasma_164390_1
plasma_164391_1
plasma_164393_1
pup
adult

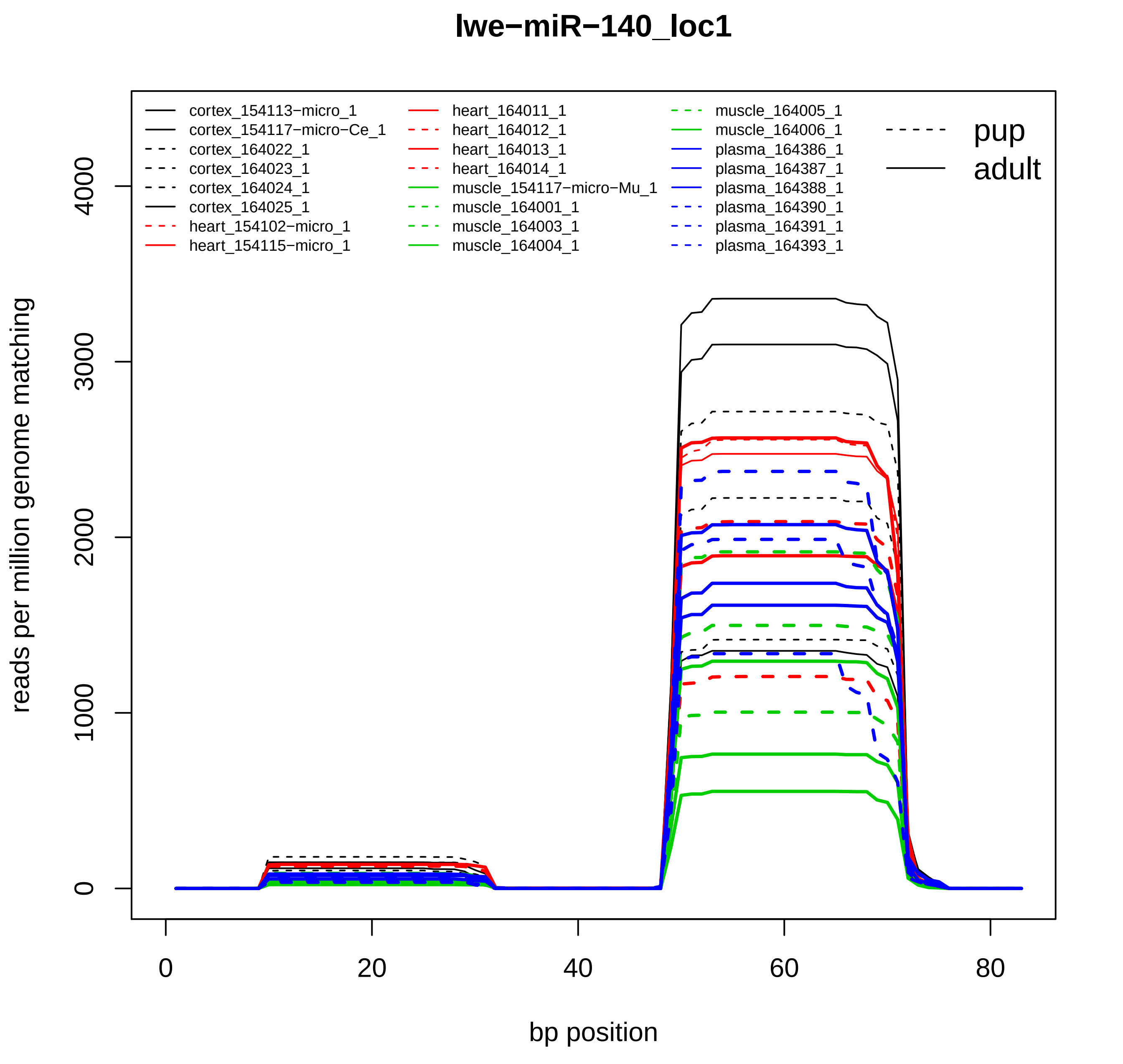

lwe−miR−140_loc1
cortex_154113−micro_1
cortex_154117−micro−Ce_1
cortex_164022_1
cortex_164023_1
cortex_164024_1
cortex_164025_1
heart_154102−micro_1
heart_154115−micro_1
heart_164011_1
heart_164012_1
heart_164013_1
heart_164014_1
muscle_154117−micro−Mu_1
muscle_164001_1
muscle_164003_1
muscle_164004_1
muscle_164005_1
muscle_164006_1
plasma_164386_1
plasma_164387_1
plasma_164388_1
plasma_164390_1
plasma_164391_1
plasma_164393_1
pup
adult
reads per million genome matching
0
1000
2000
3000
4000
0
20
40
60
80
bp position

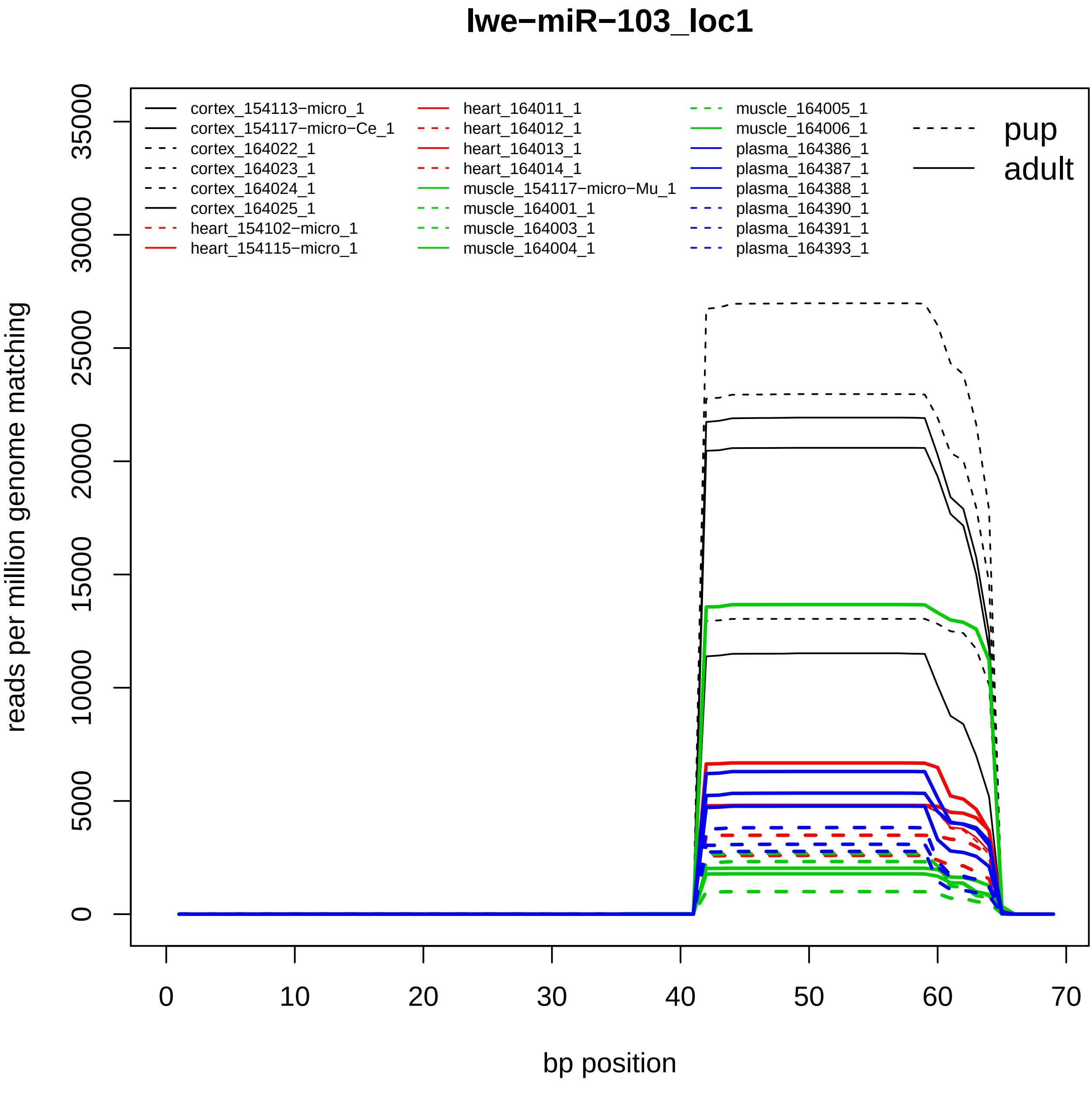

lwe−miR−103_loc1
cortex_154113−micro_1
cortex_154117−micro−Ce_1
cortex_164022_1
cortex_164023_1
cortex_164024_1
cortex_164025_1
heart_154102−micro_1
heart_154115−micro_1
heart_164011_1
heart_164012_1
heart_164013_1
heart_164014_1
muscle_154117−micro−Mu_1
muscle_164001_1
muscle_164003_1
muscle_164004_1
muscle_164005_1
muscle_164006_1
plasma_164386_1
plasma_164387_1
plasma_164388_1
plasma_164390_1
plasma_164391_1
plasma_164393_1
pup
adult
reads per million genome matching
0
5000
10000
15000
20000
25000
30000
35000
bp position
0
10
20
30
40
50
60
70

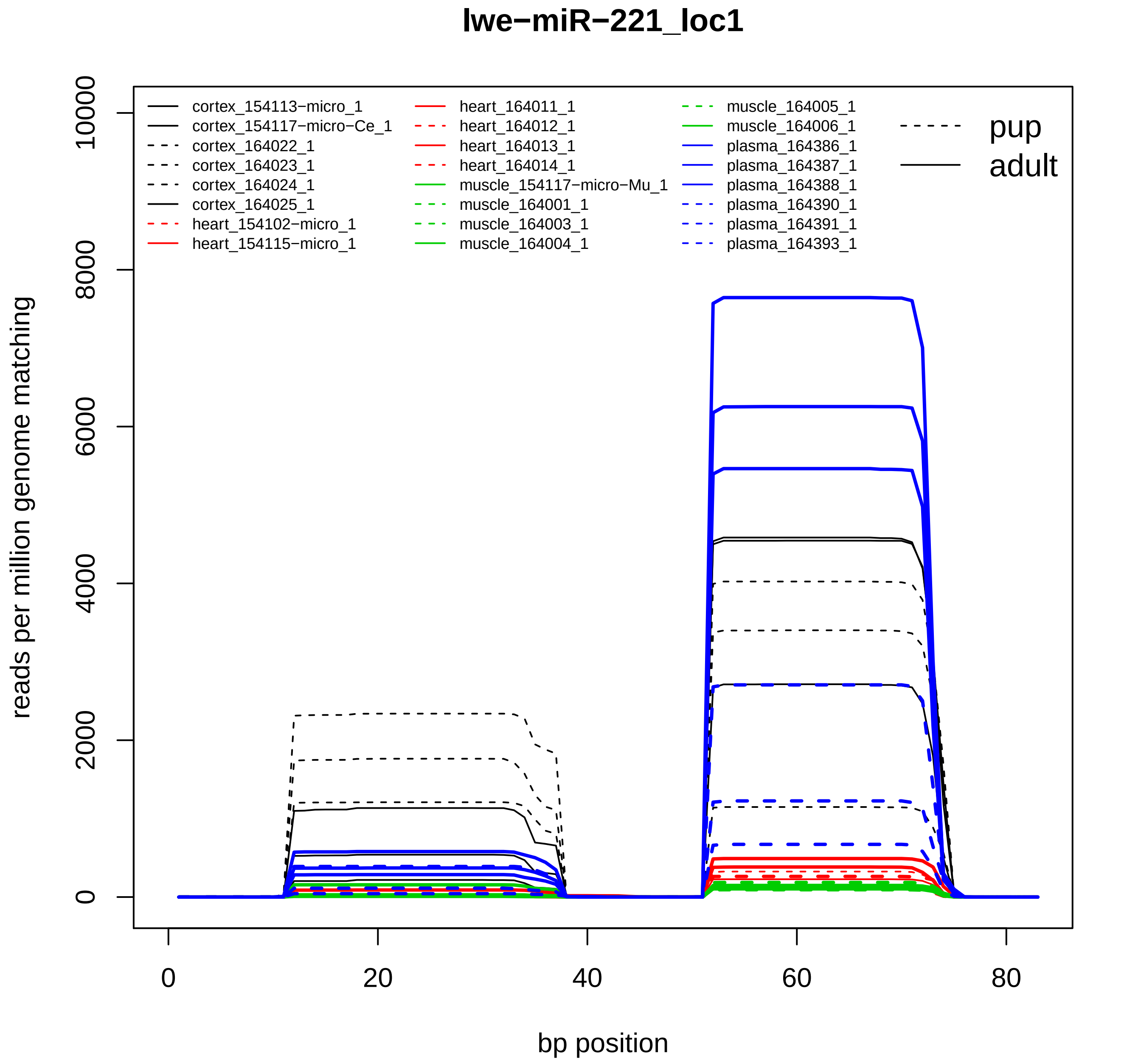
lwe−miR−221_loc1
cortex_154113−micro_1
cortex_154117−micro−Ce_1
cortex_164022_1
cortex_164023_1
cortex_164024_1
cortex_164025_1
heart_154102−micro_1
heart_154115−micro_1
heart_164011_1
heart_164012_1
heart_164013_1
heart_164014_1
muscle_154117−micro−Mu_1
muscle_164001_1
muscle_164003_1
muscle_164004_1
muscle_164005_1
muscle_164006_1
plasma_164386_1
plasma_164387_1
plasma_164388_1
plasma_164390_1
plasma_164391_1
plasma_164393_1
pup
adult
reads per million genome matching
0
2000
4000
6000
8000
10000
0
20
40
60
80
bp position

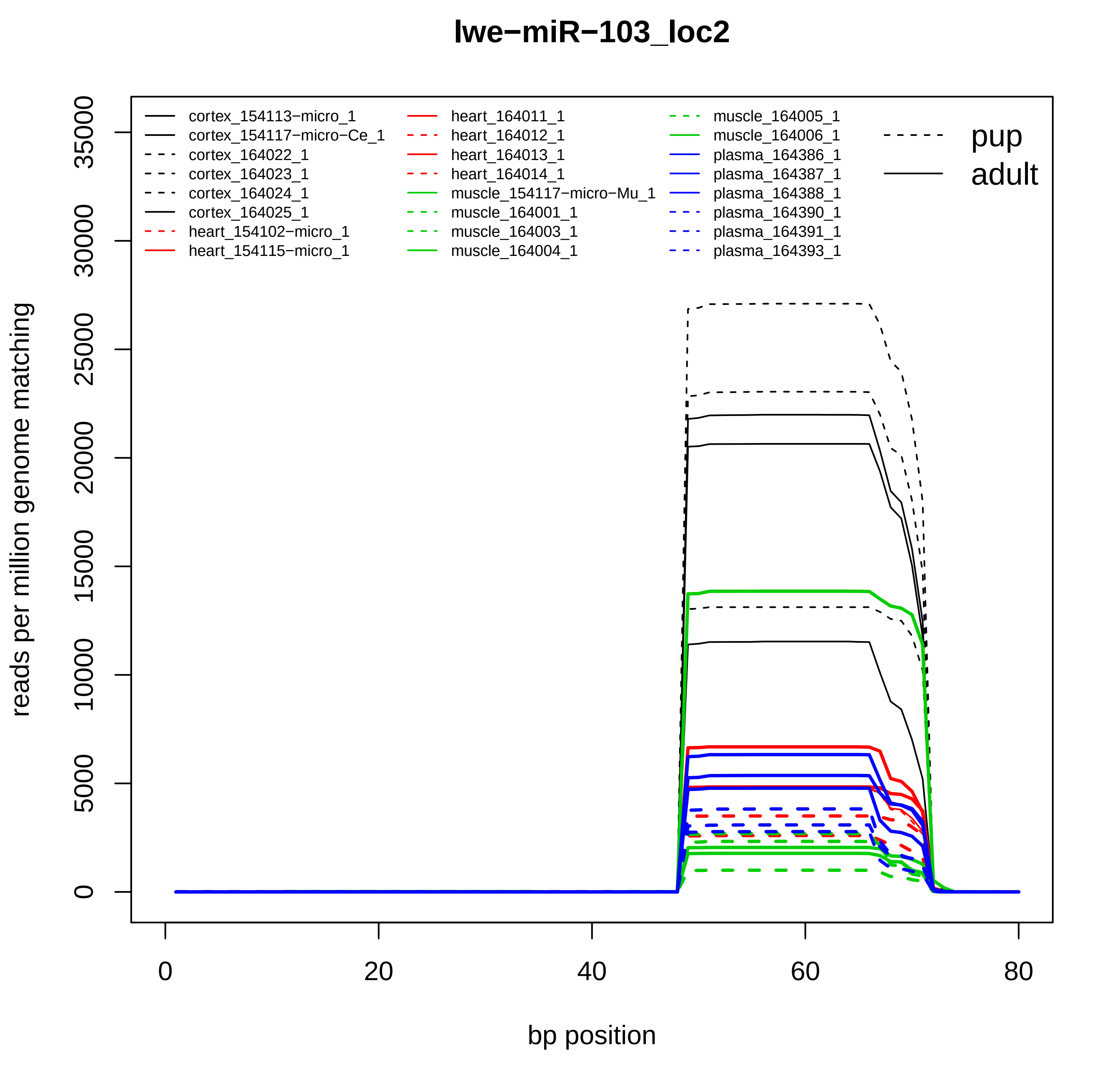
lwe−miR−103_loc2
reads per million genome matching
bp position
0
5000
10000
15000
20000
25000
30000
35000
0
20
40
60
80
cortex_154113−micro_1
cortex_154117−micro−Ce_1
cortex_164022_1
cortex_164023_1
cortex_164024_1
cortex_164025_1
heart_154102−micro_1
heart_154115−micro_1
heart_164011_1
heart_164012_1
heart_164013_1
heart_164014_1
muscle_154117−micro−Mu_1
muscle_164001_1
muscle_164003_1
muscle_164004_1
muscle_164005_1
muscle_164006_1
plasma_164386_1
plasma_164387_1
plasma_164388_1
plasma_164390_1
plasma_164391_1
plasma_164393_1
pup
adult

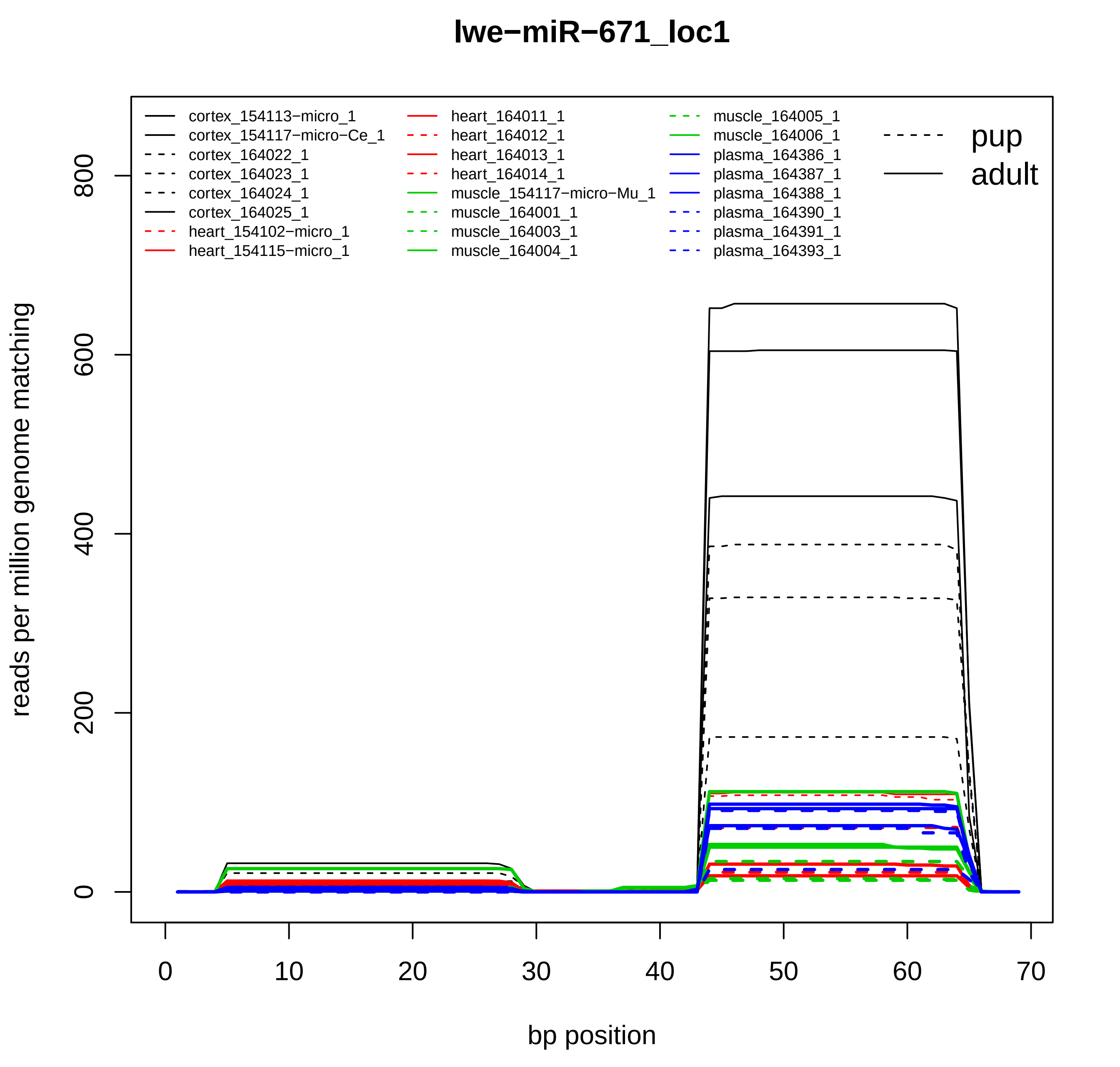

lwe−miR−671_loc1
cortex_154113−micro_1
cortex_154117−micro−Ce_1
cortex_164022_1
cortex_164023_1
cortex_164024_1
cortex_164025_1
heart_154102−micro_1
heart_154115−micro_1
heart_164011_1
heart_164012_1
heart_164013_1
heart_164014_1
muscle_154117−micro−Mu_1
muscle_164001_1
muscle_164003_1
muscle_164004_1
muscle_164005_1
muscle_164006_1
plasma_164386_1
plasma_164387_1
plasma_164388_1
plasma_164390_1
plasma_164391_1
plasma_164393_1
pup
adult
reads per million genome matching
0
200
400
600
800
bp position
0
10
20
30
40
50
60
70

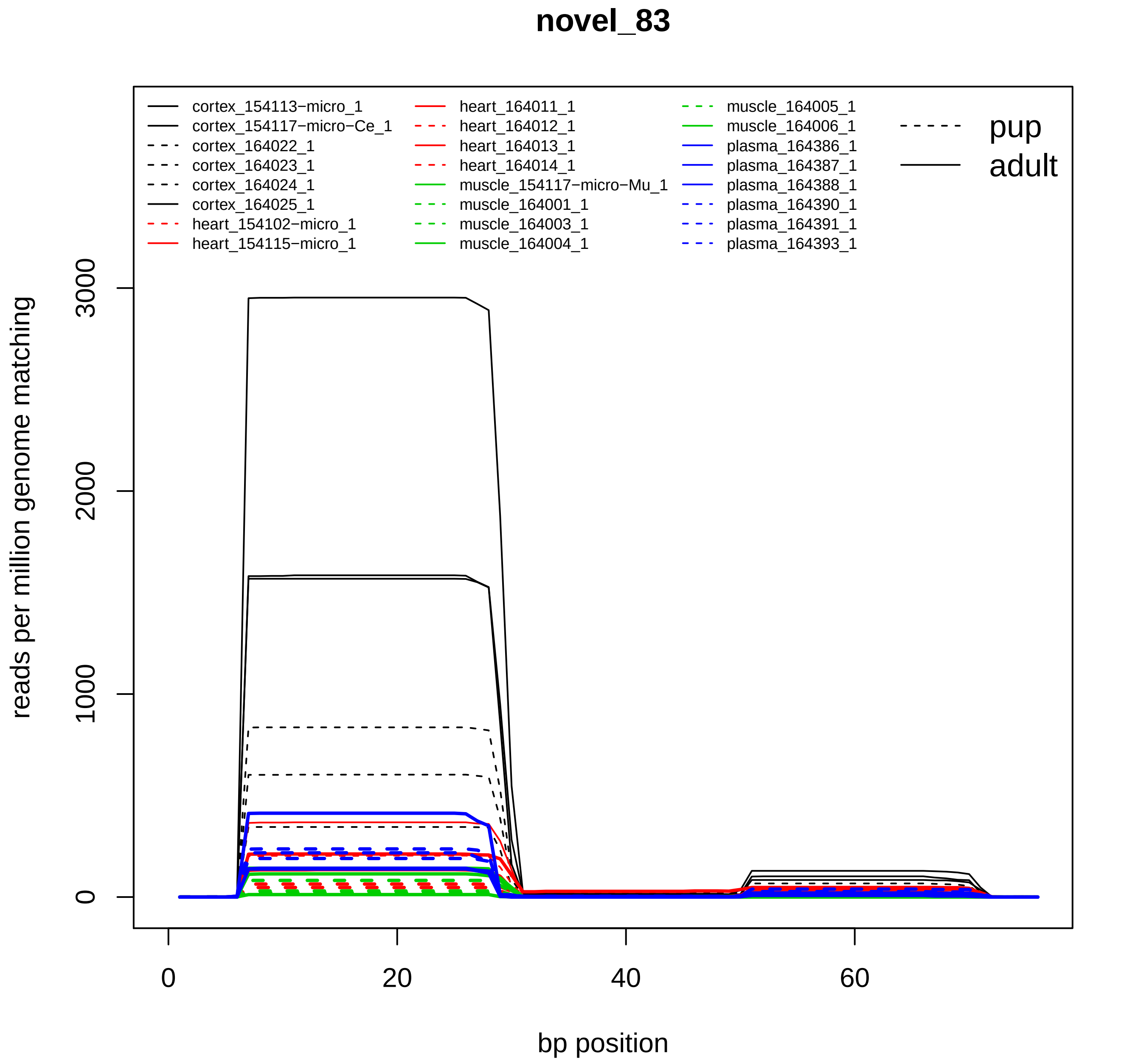
novel_83
cortex_154113−micro_1
cortex_154117−micro−Ce_1
cortex_164022_1
cortex_164023_1
cortex_164024_1
cortex_164025_1
heart_154102−micro_1
heart_154115−micro_1
heart_164011_1
heart_164012_1
heart_164013_1
heart_164014_1
muscle_154117−micro−Mu_1
muscle_164001_1
muscle_164003_1
muscle_164004_1
muscle_164005_1
muscle_164006_1
plasma_164386_1
plasma_164387_1
plasma_164388_1
plasma_164390_1
plasma_164391_1
plasma_164393_1
pup
adult
reads per million genome matching
0
1000
2000
3000
0
20
40
60
bp position

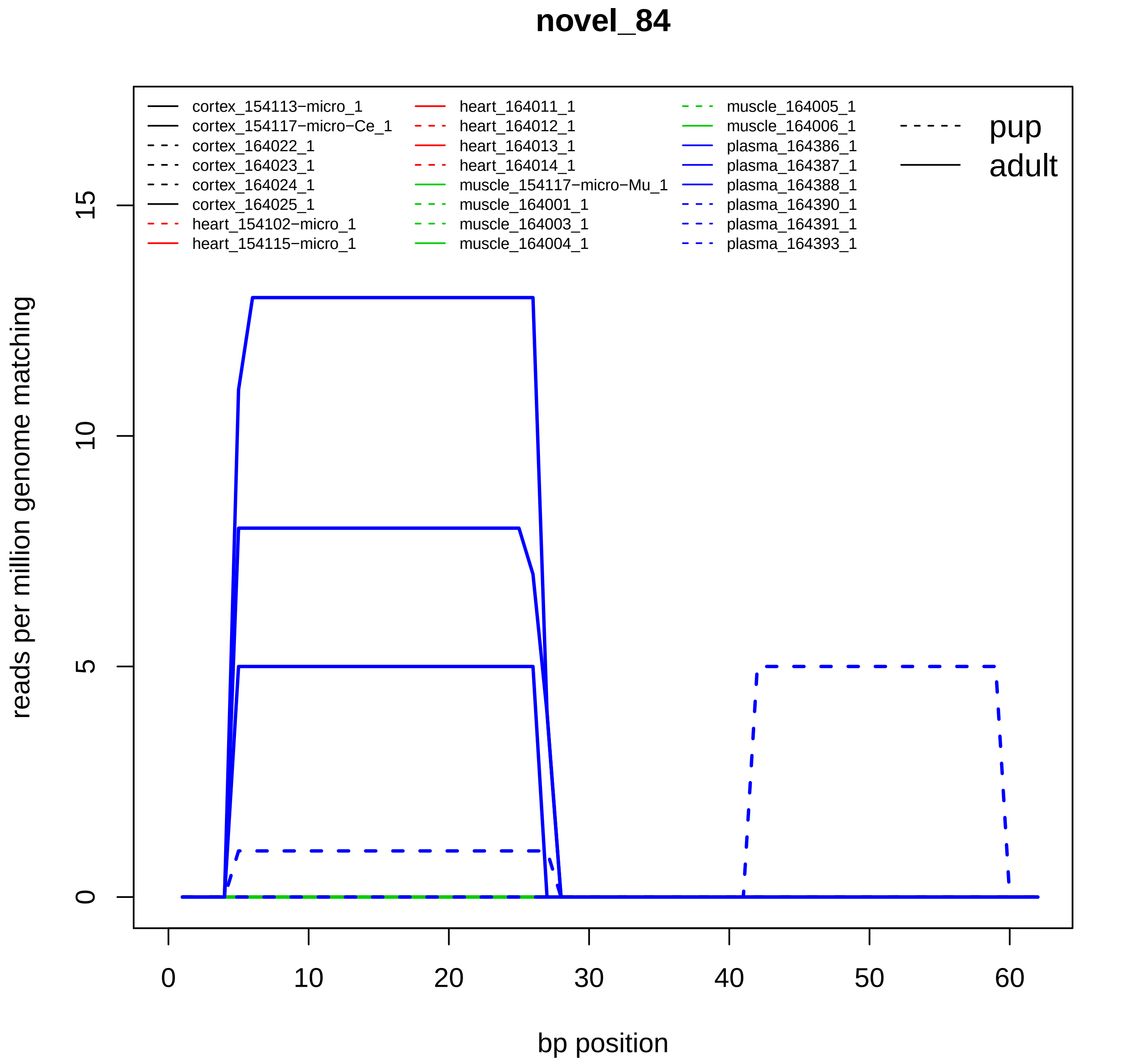
novel_84
cortex_154113−micro_1
cortex_154117−micro−Ce_1
cortex_164022_1
cortex_164023_1
cortex_164024_1
cortex_164025_1
heart_154102−micro_1
heart_154115−micro_1
heart_164011_1
heart_164012_1
heart_164013_1
heart_164014_1
muscle_154117−micro−Mu_1
muscle_164001_1
muscle_164003_1
muscle_164004_1
muscle_164005_1
muscle_164006_1
plasma_164386_1
plasma_164387_1
plasma_164388_1
plasma_164390_1
plasma_164391_1
plasma_164393_1
pup
adult
reads per million genome matching
0
5
10
15
bp position
0
10
20
30
40
50
60

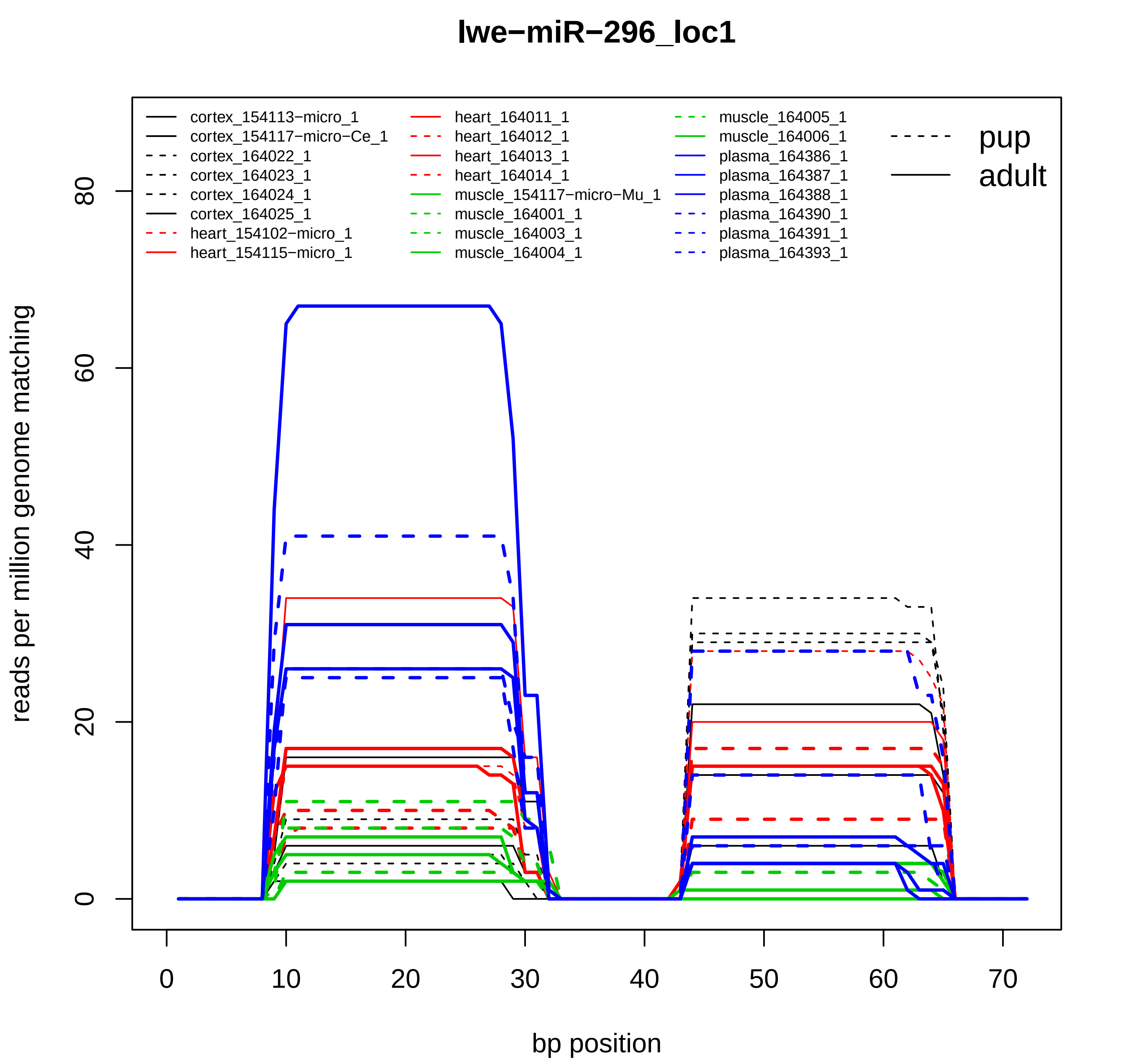
lwe−miR−296_loc1
cortex_154113−micro_1
cortex_154117−micro−Ce_1
cortex_164022_1
cortex_164023_1
cortex_164024_1
cortex_164025_1
heart_154102−micro_1
heart_154115−micro_1
heart_164011_1
heart_164012_1
heart_164013_1
heart_164014_1
muscle_154117−micro−Mu_1
muscle_164001_1
muscle_164003_1
muscle_164004_1
muscle_164005_1
muscle_164006_1
plasma_164386_1
plasma_164387_1
plasma_164388_1
plasma_164390_1
plasma_164391_1
plasma_164393_1
pup
adult
reads per million genome matching
0
20
40
60
80
0
10
20
30
40
50
60
70
bp position

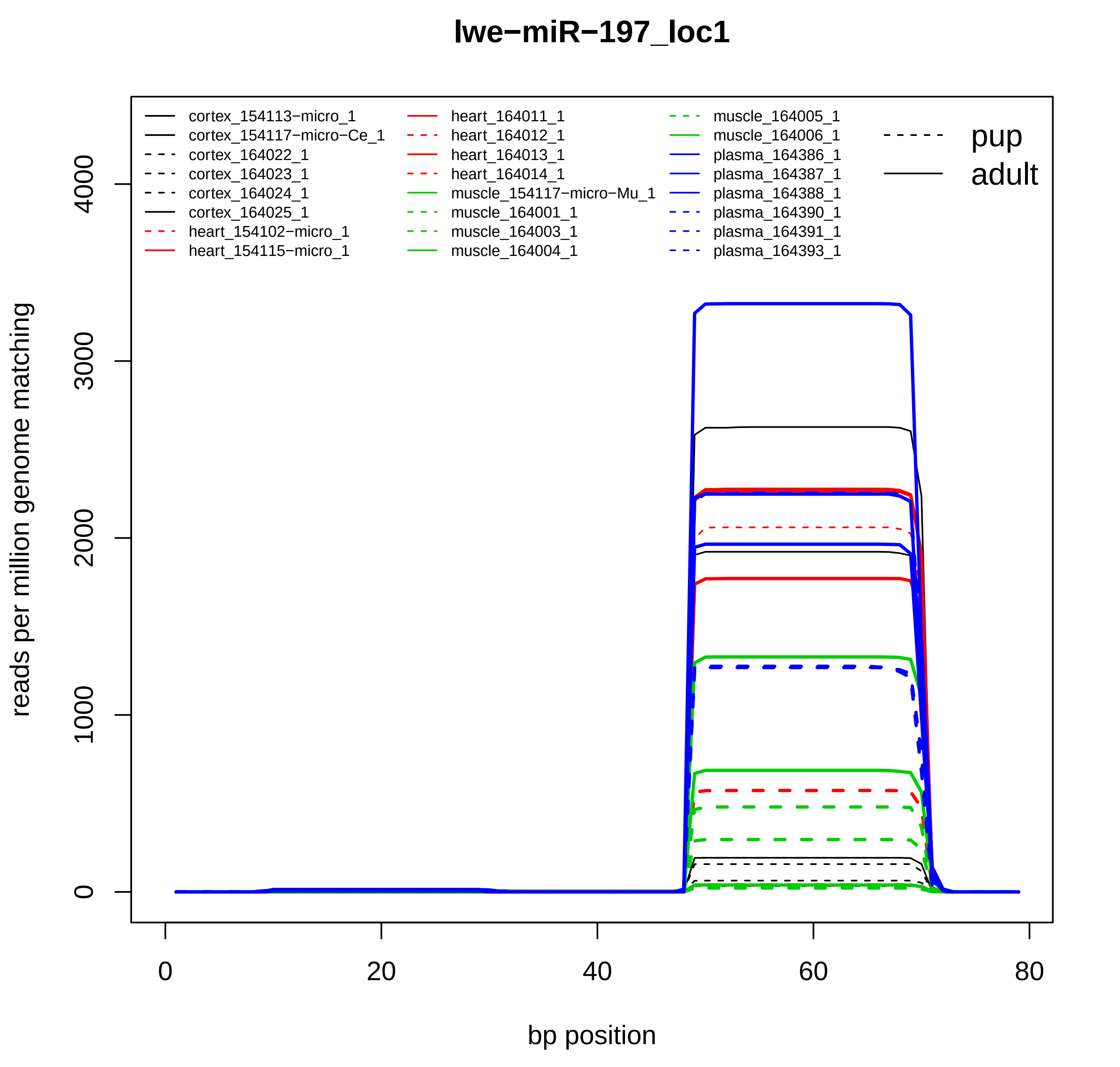
lwe−miR−197_loc1
cortex_154113−micro_1
cortex_154117−micro−Ce_1
cortex_164022_1
cortex_164023_1
cortex_164024_1
cortex_164025_1
heart_154102−micro_1
heart_154115−micro_1
heart_164011_1
heart_164012_1
heart_164013_1
heart_164014_1
muscle_154117−micro−Mu_1
muscle_164001_1
muscle_164003_1
muscle_164004_1
muscle_164005_1
muscle_164006_1
plasma_164386_1
plasma_164387_1
plasma_164388_1
plasma_164390_1
plasma_164391_1
plasma_164393_1
pup
adult
reads per million genome matching
0
1000
2000
3000
4000
0
20
40
60
80
bp position

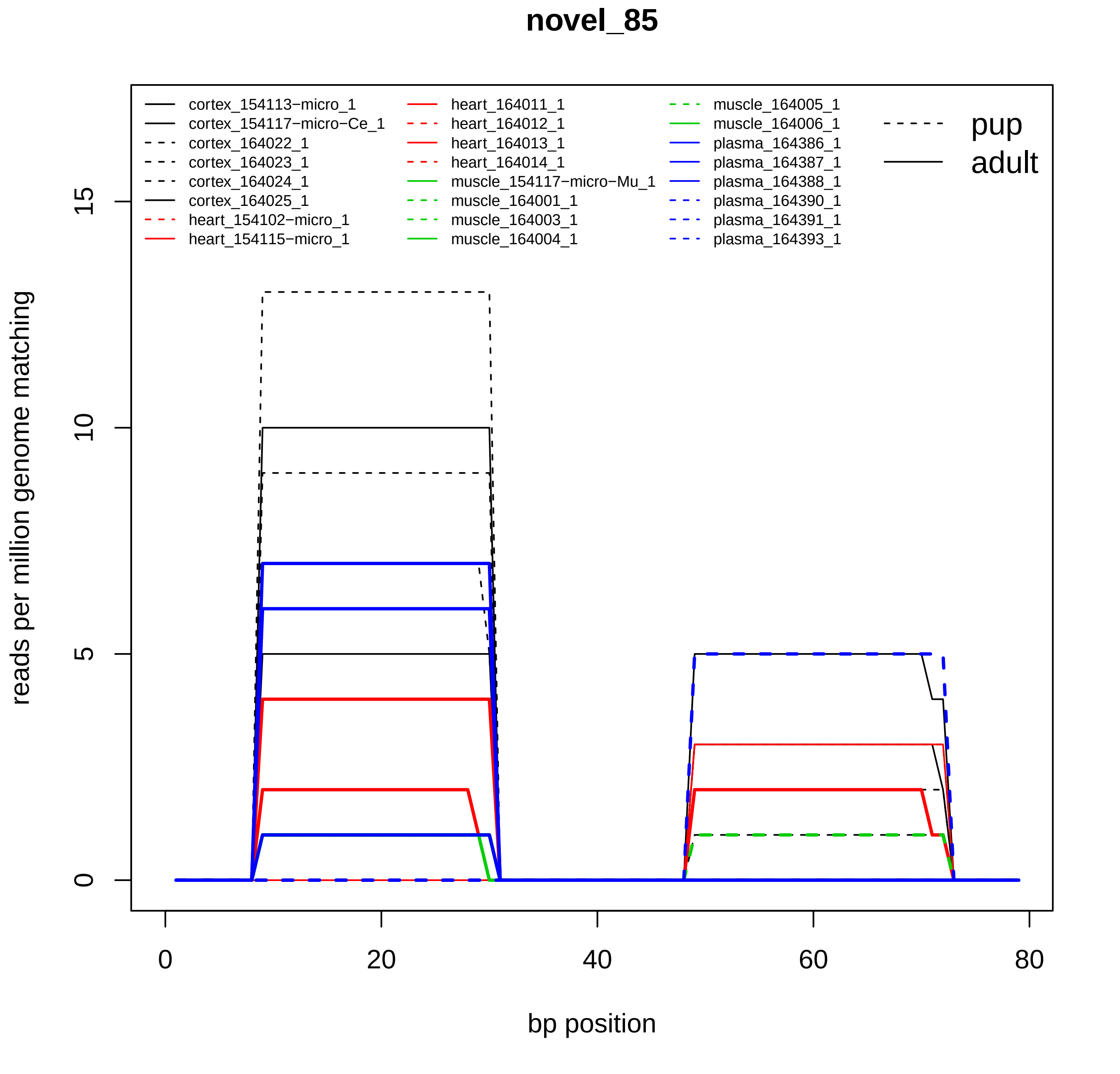

novel_85
cortex_154113−micro_1
cortex_154117−micro−Ce_1
cortex_164022_1
cortex_164023_1
cortex_164024_1
cortex_164025_1
heart_154102−micro_1
heart_154115−micro_1
heart_164011_1
heart_164012_1
heart_164013_1
heart_164014_1
muscle_154117−micro−Mu_1
muscle_164001_1
muscle_164003_1
muscle_164004_1
muscle_164005_1
muscle_164006_1
plasma_164386_1
plasma_164387_1
plasma_164388_1
plasma_164390_1
plasma_164391_1
plasma_164393_1
pup
adult
reads per million genome matching
0
5
10
15
0
20
40
60
80
bp position

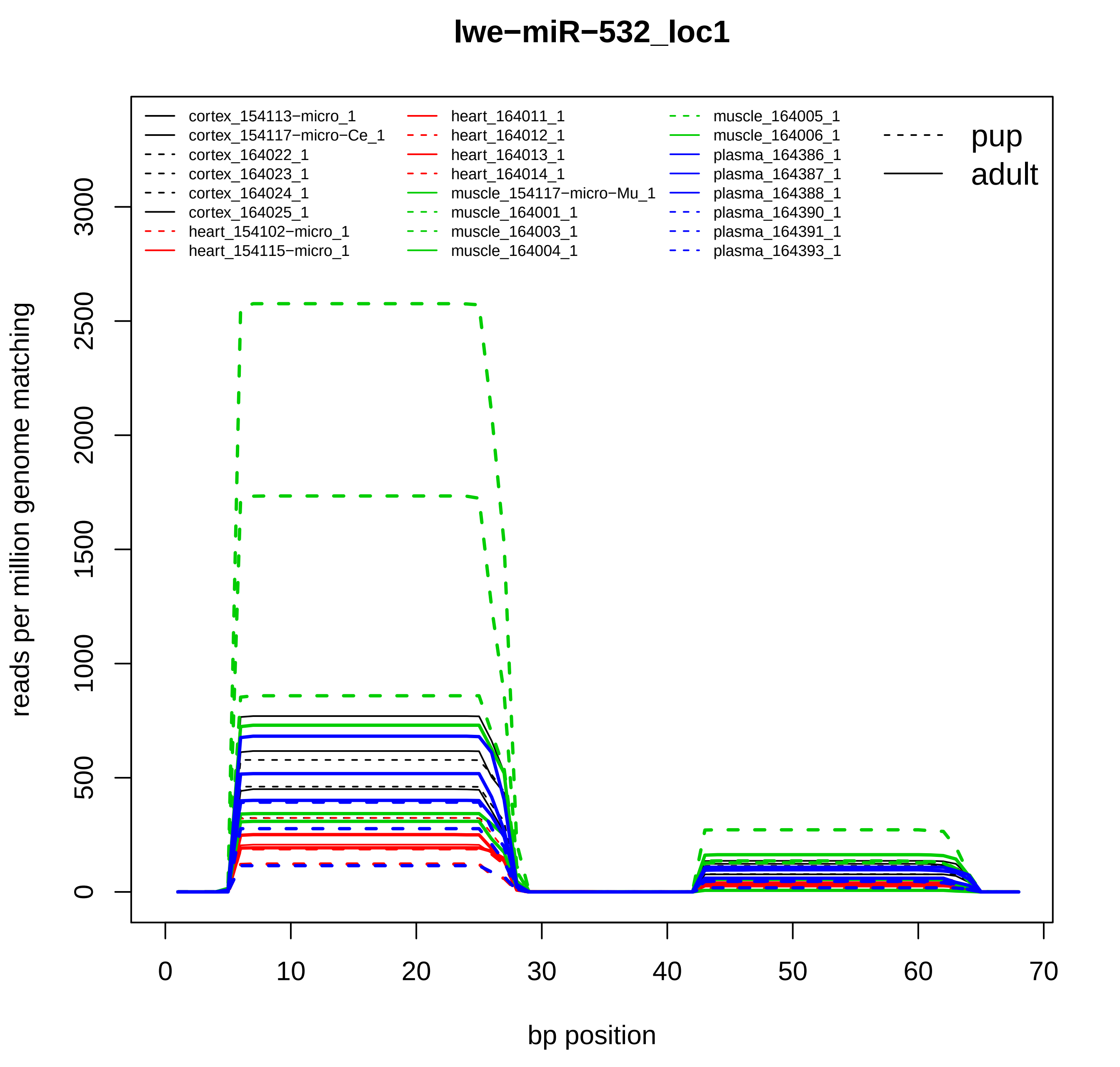

lwe−miR−532_loc1
reads per million genome matching
bp position
0
500
1000
1500
2000
2500
3000
0
10
20
30
40
50
60
70
cortex_154113−micro_1
cortex_154117−micro−Ce_1
cortex_164022_1
cortex_164023_1
cortex_164024_1
cortex_164025_1
heart_154102−micro_1
heart_154115−micro_1
heart_164011_1
heart_164012_1
heart_164013_1
heart_164014_1
muscle_154117−micro−Mu_1
muscle_164001_1
muscle_164003_1
muscle_164004_1
muscle_164005_1
muscle_164006_1
plasma_164386_1
plasma_164387_1
plasma_164388_1
plasma_164390_1
plasma_164391_1
plasma_164393_1
pup
adult

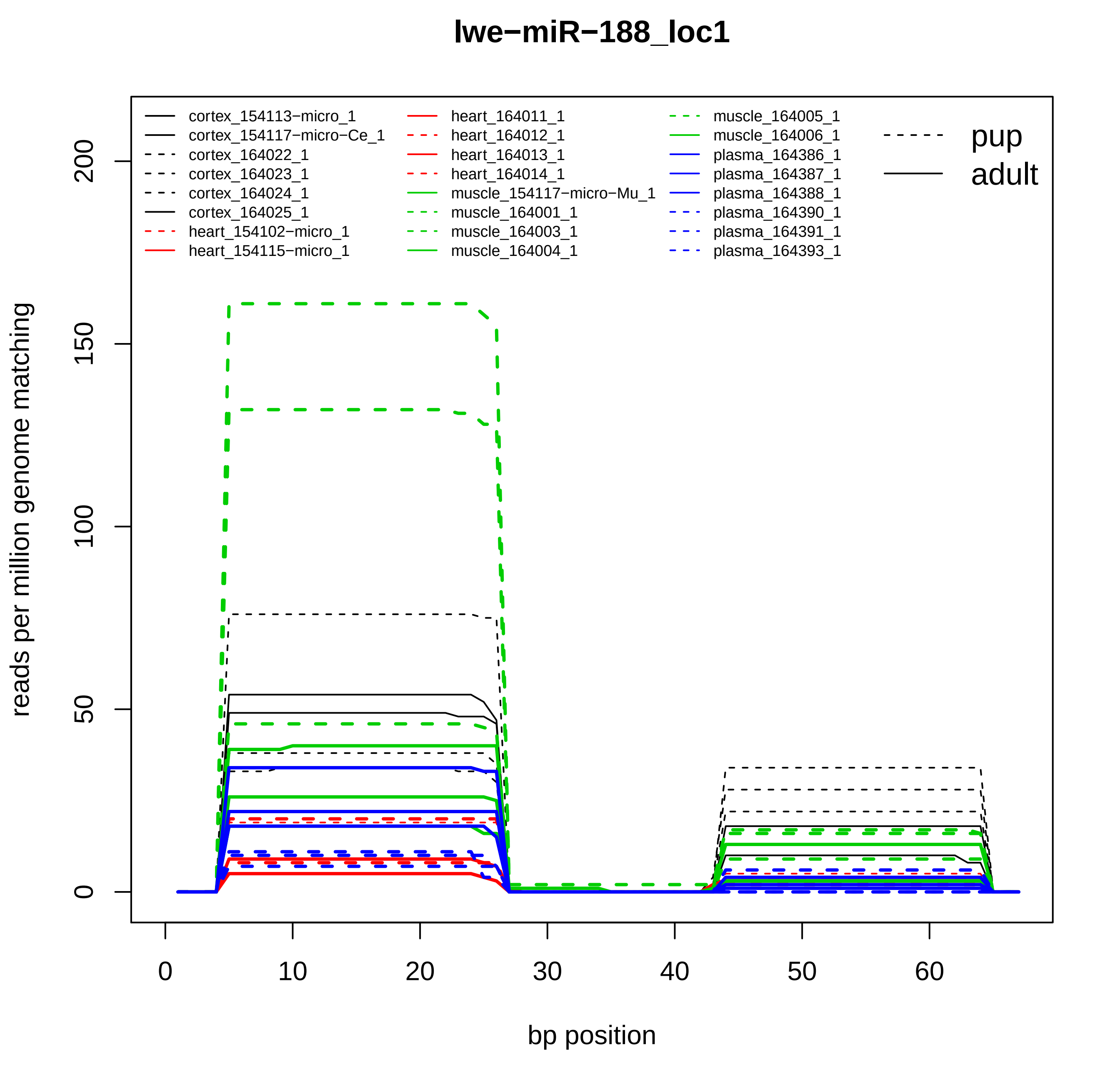

lwe−miR−188_loc1
cortex_154113−micro_1
cortex_154117−micro−Ce_1
cortex_164022_1
cortex_164023_1
cortex_164024_1
cortex_164025_1
heart_154102−micro_1
heart_154115−micro_1
heart_164011_1
heart_164012_1
heart_164013_1
heart_164014_1
muscle_154117−micro−Mu_1
muscle_164001_1
muscle_164003_1
muscle_164004_1
muscle_164005_1
muscle_164006_1
plasma_164386_1
plasma_164387_1
plasma_164388_1
plasma_164390_1
plasma_164391_1
plasma_164393_1
pup
adult
reads per million genome matching
bp position
0
50
100
150
200
0
10
20
30
40
50
60

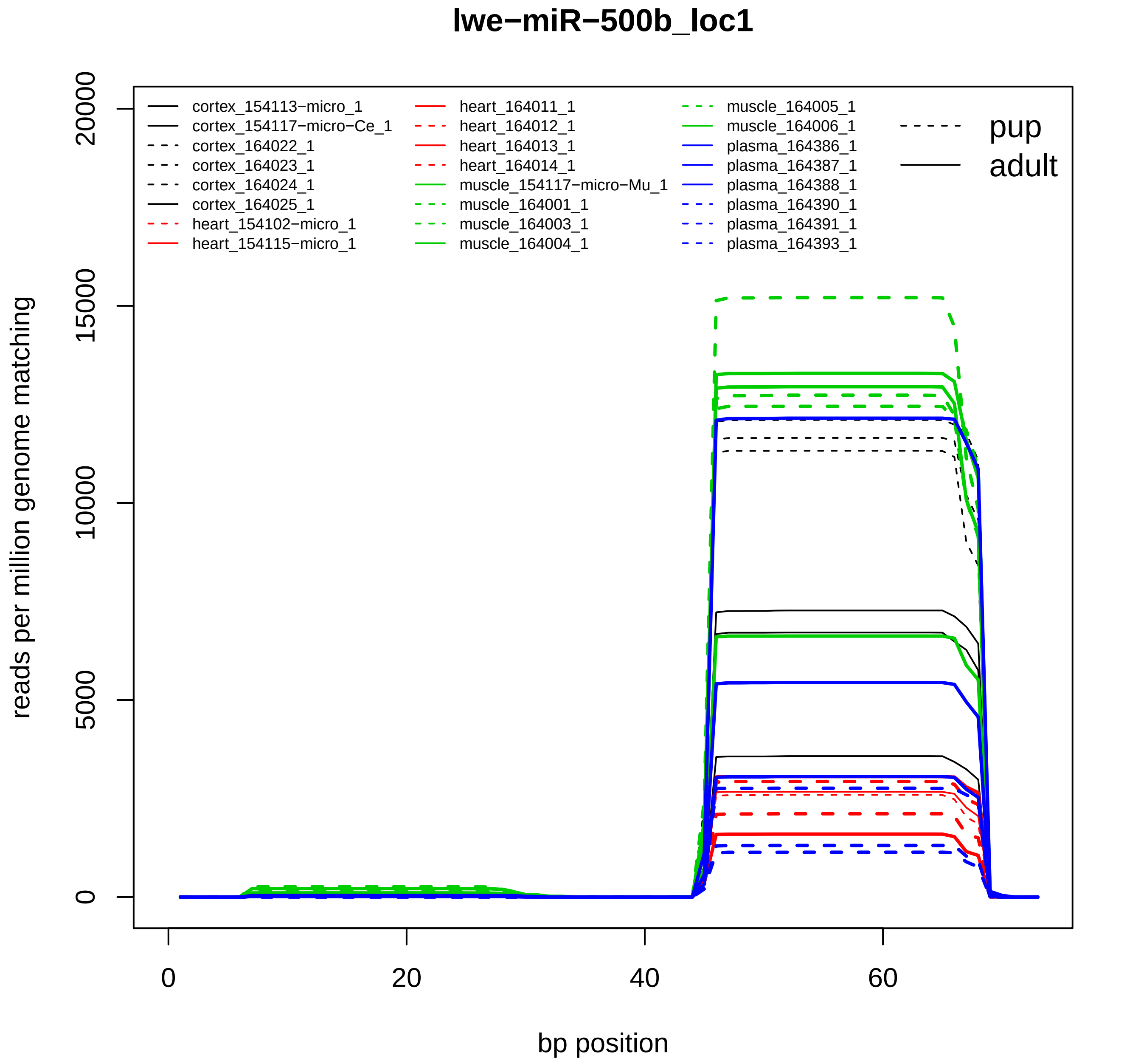
lwe−miR−500b_loc1
cortex_154113−micro_1
cortex_154117−micro−Ce_1
cortex_164022_1
cortex_164023_1
cortex_164024_1
cortex_164025_1
heart_154102−micro_1
heart_154115−micro_1
heart_164011_1
heart_164012_1
heart_164013_1
heart_164014_1
muscle_154117−micro−Mu_1
muscle_164001_1
muscle_164003_1
muscle_164004_1
muscle_164005_1
muscle_164006_1
plasma_164386_1
plasma_164387_1
plasma_164388_1
plasma_164390_1
plasma_164391_1
plasma_164393_1
pup
adult
reads per million genome matching
20000
15000
10000
5000
0
0
20
40
60
bp position

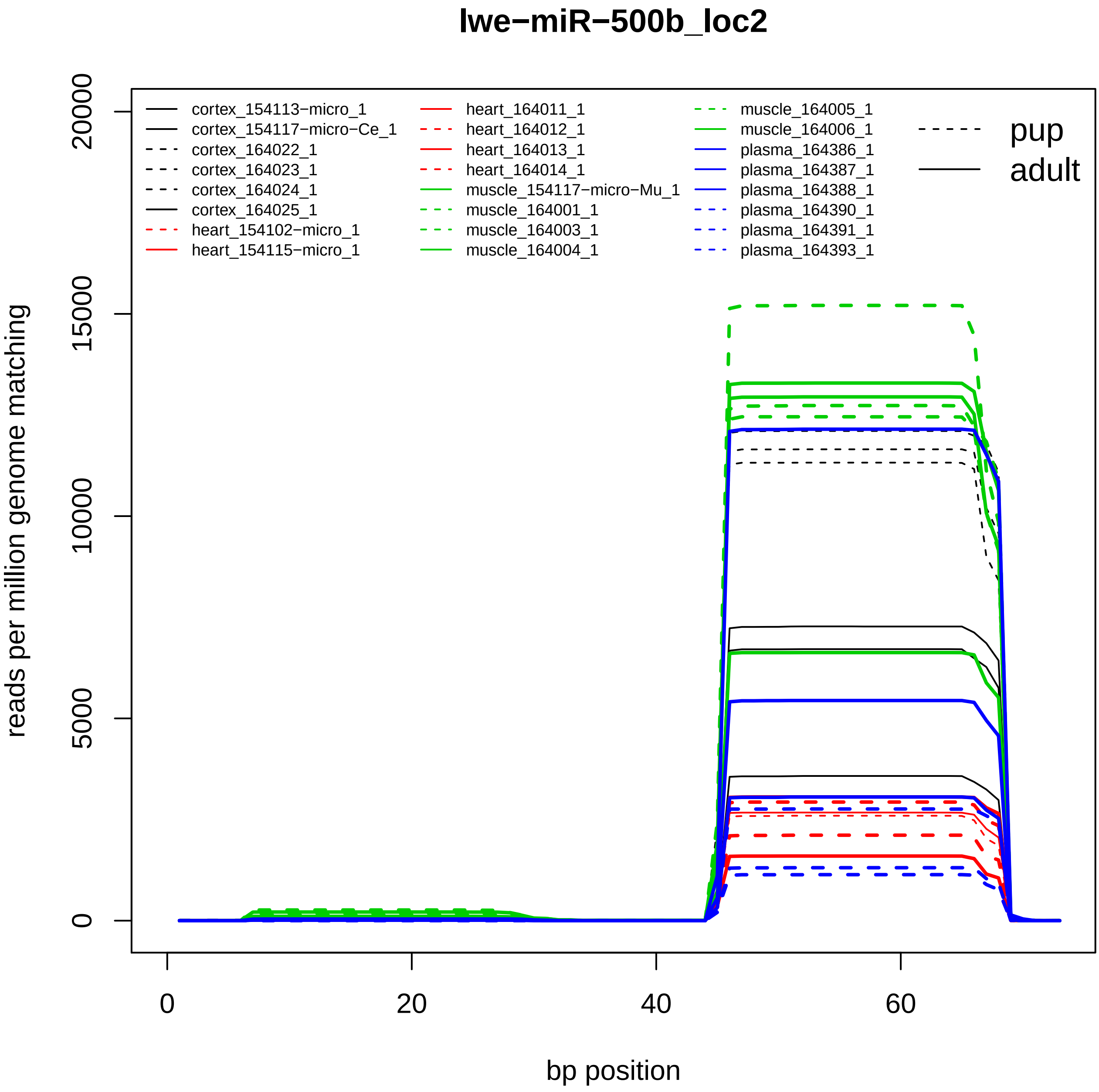
lwe−miR−500b_loc2
reads per million genome matching
bp position
0
5000
10000
15000
20000
0
20
40
60
cortex_154113−micro_1
cortex_154117−micro−Ce_1
cortex_164022_1
cortex_164023_1
cortex_164024_1
cortex_164025_1
heart_154102−micro_1
heart_154115−micro_1
heart_164011_1
heart_164012_1
heart_164013_1
heart_164014_1
muscle_154117−micro−Mu_1
muscle_164001_1
muscle_164003_1
muscle_164004_1
muscle_164005_1
muscle_164006_1
plasma_164386_1
plasma_164387_1
plasma_164388_1
plasma_164390_1
plasma_164391_1
plasma_164393_1
pup
adult

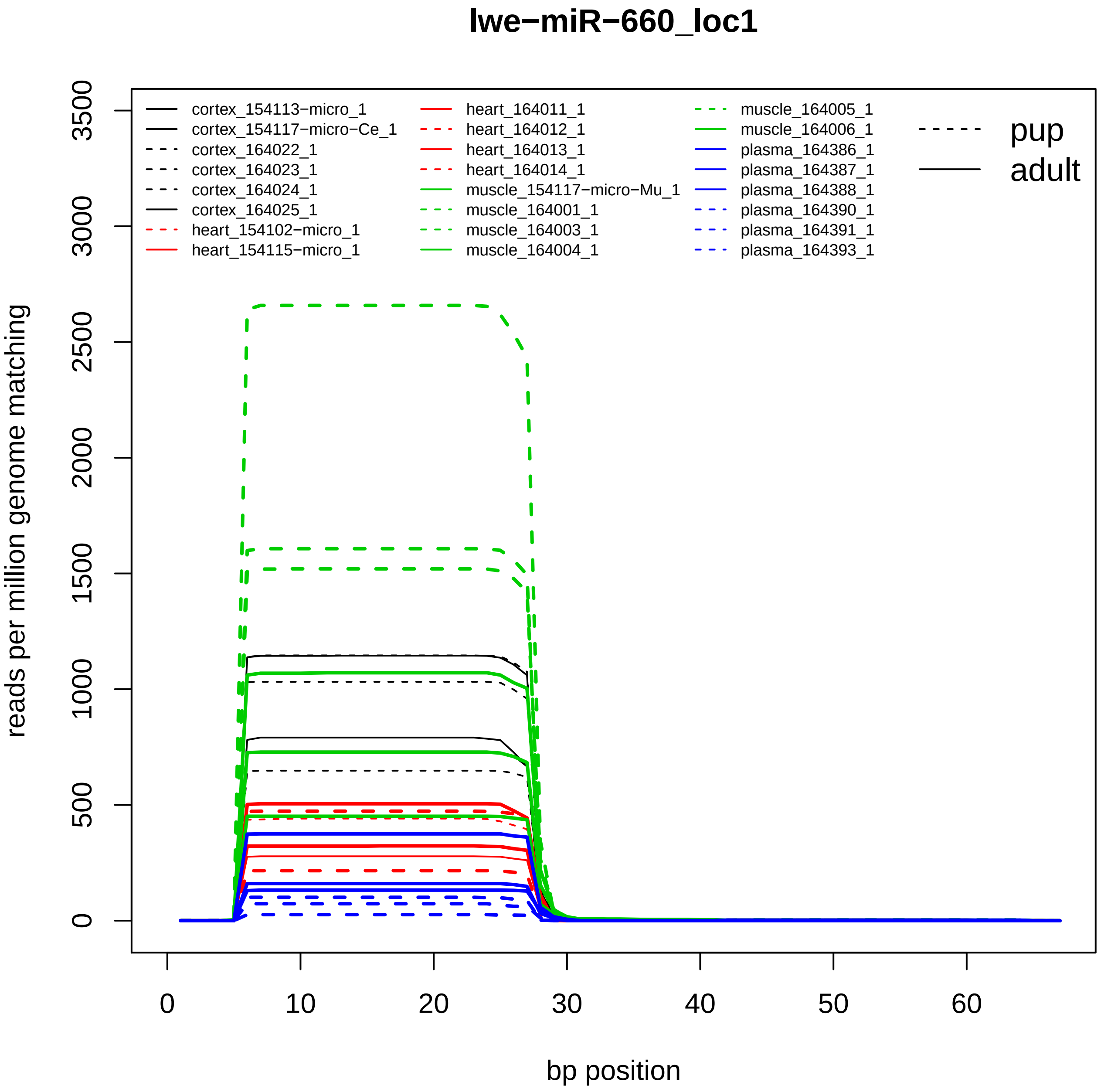

lwe−miR−660_loc1
reads per million genome matching
bp position
0
500
1000
1500
2000
2500
3000
3500
0
10
20
30
40
50
60
cortex_154113−micro_1
cortex_154117−micro−Ce_1
cortex_164022_1
cortex_164023_1
cortex_164024_1
cortex_164025_1
heart_154102−micro_1
heart_154115−micro_1
heart_164011_1
heart_164012_1
heart_164013_1
heart_164014_1
muscle_154117−micro−Mu_1
muscle_164001_1
muscle_164003_1
muscle_164004_1
muscle_164005_1
muscle_164006_1
plasma_164386_1
plasma_164387_1
plasma_164388_1
plasma_164390_1
plasma_164391_1
plasma_164393_1
pup
adult

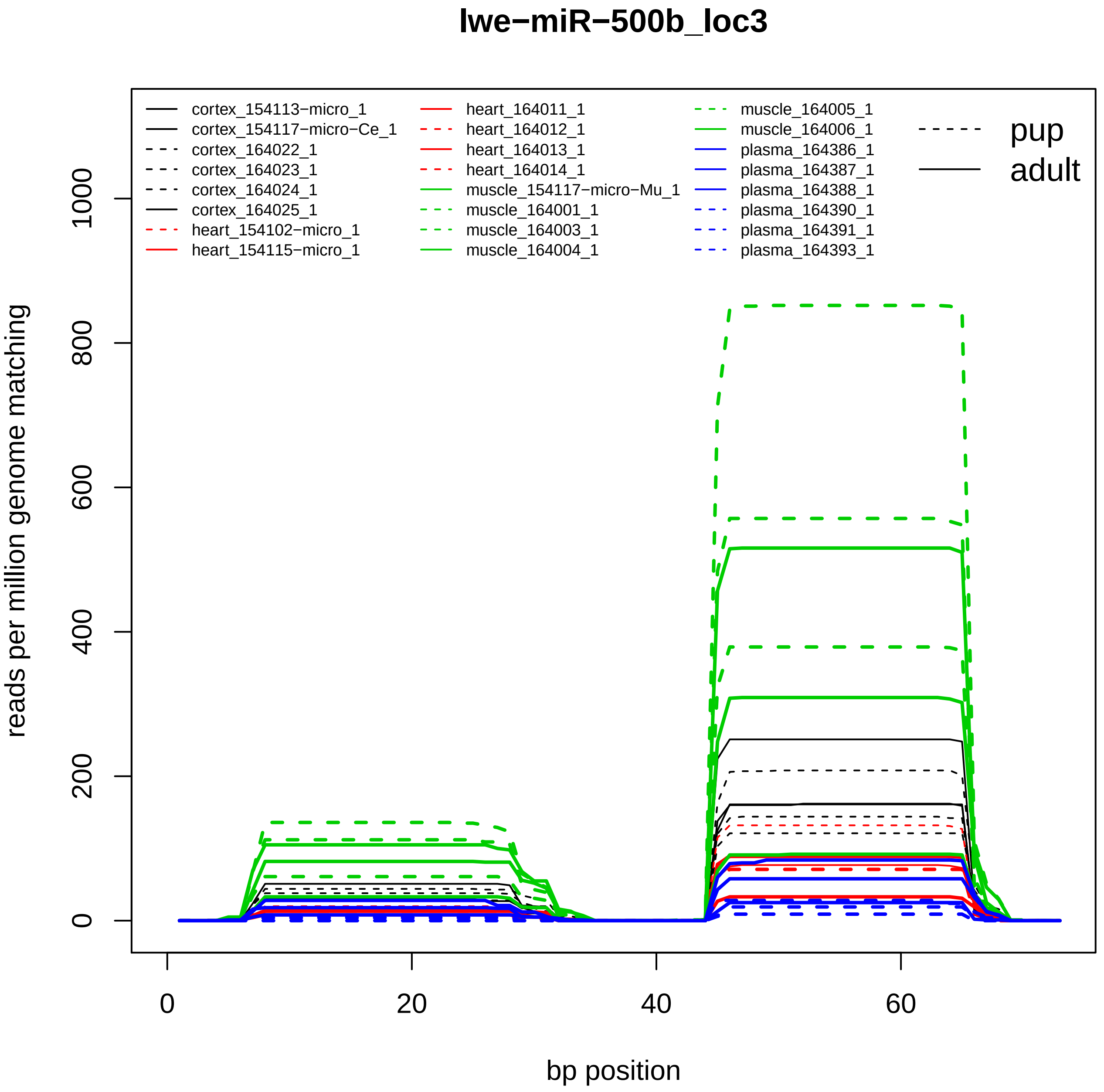

lwe−miR−500b_loc3
cortex_154113−micro_1
cortex_154117−micro−Ce_1
cortex_164022_1
cortex_164023_1
cortex_164024_1
cortex_164025_1
heart_154102−micro_1
heart_154115−micro_1
heart_164011_1
heart_164012_1
heart_164013_1
heart_164014_1
muscle_154117−micro−Mu_1
muscle_164001_1
muscle_164003_1
muscle_164004_1
muscle_164005_1
muscle_164006_1
plasma_164386_1
plasma_164387_1
plasma_164388_1
plasma_164390_1
plasma_164391_1
plasma_164393_1
pup
adult
reads per million genome matching
0
200
400
600
800
1000
0
20
40
60
bp position

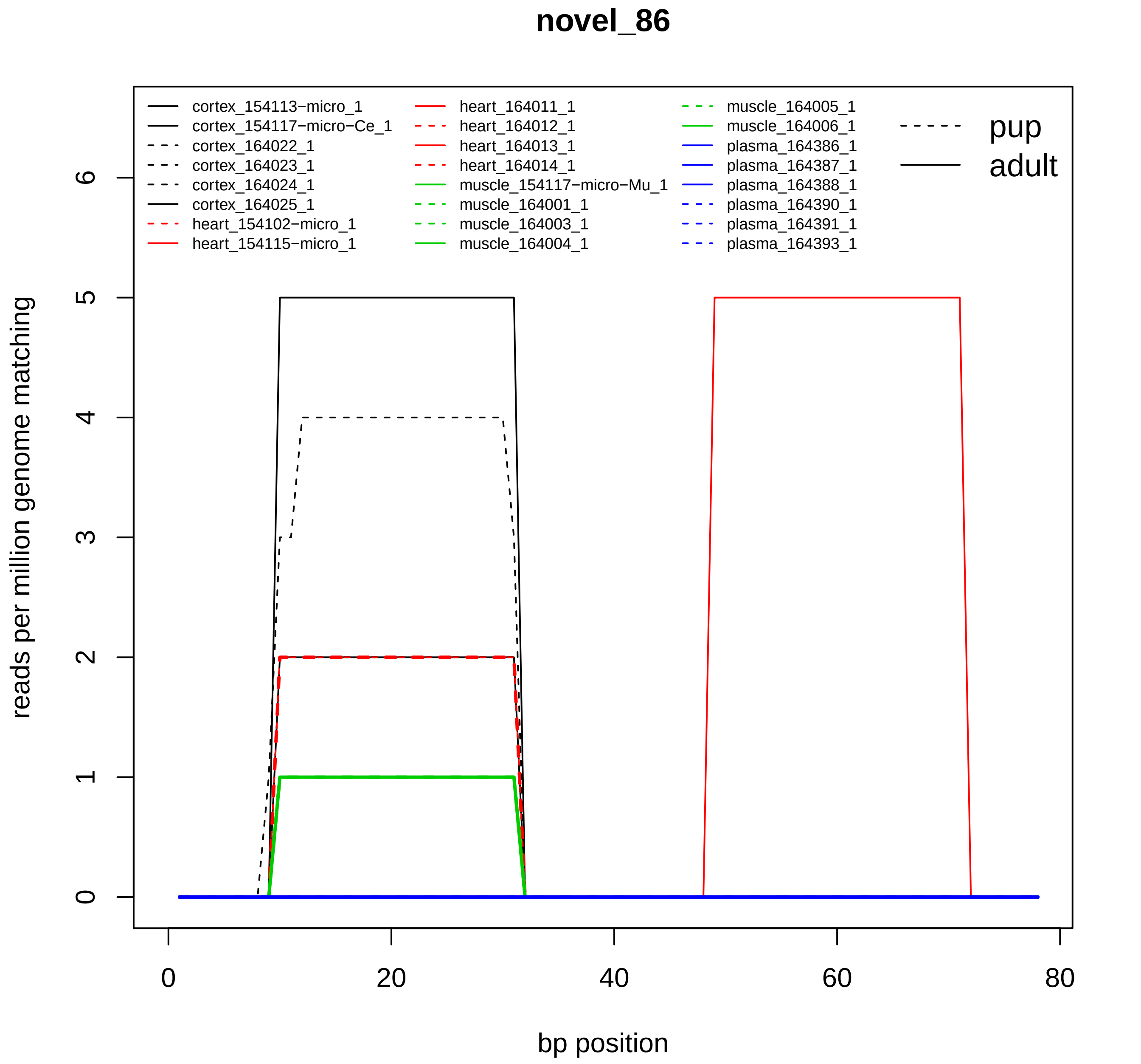
novel_86
reads per million genome matching
bp position
0
1
2
3
4
5
6
0
20
40
60
80
cortex_154113−micro_1
cortex_154117−micro−Ce_1
cortex_164022_1
cortex_164023_1
cortex_164024_1
cortex_164025_1
heart_154102−micro_1
heart_154115−micro_1
heart_164011_1
heart_164012_1
heart_164013_1
heart_164014_1
muscle_154117−micro−Mu_1
muscle_164001_1
muscle_164003_1
muscle_164004_1
muscle_164005_1
muscle_164006_1
plasma_164386_1
plasma_164387_1
plasma_164388_1
plasma_164390_1
plasma_164391_1
plasma_164393_1
pup
adult

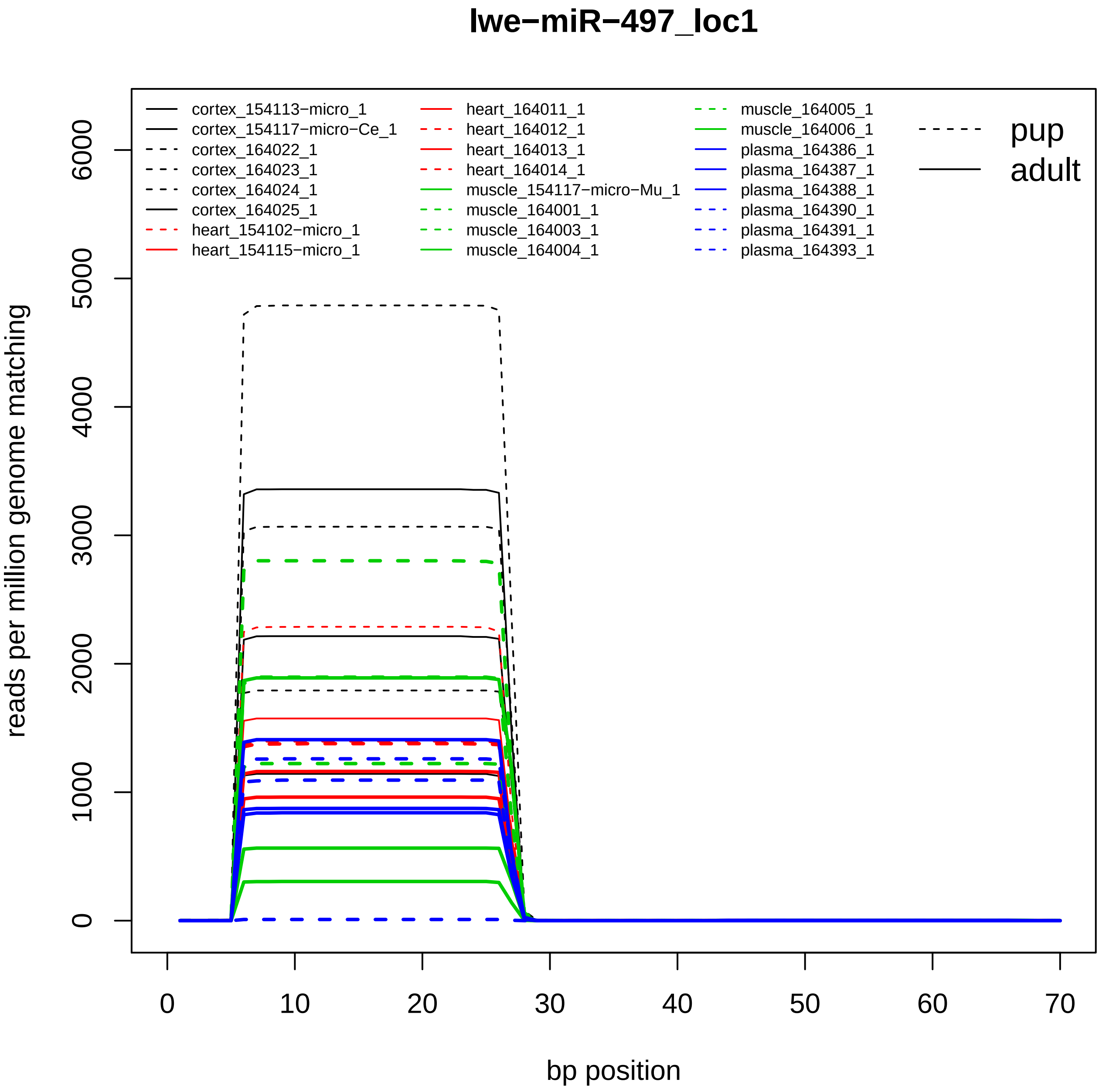
lwe−miR−497_loc1
cortex_154113−micro_1
cortex_154117−micro−Ce_1
cortex_164022_1
cortex_164023_1
cortex_164024_1
cortex_164025_1
heart_154102−micro_1
heart_154115−micro_1
heart_164011_1
heart_164012_1
heart_164013_1
heart_164014_1
muscle_154117−micro−Mu_1
muscle_164001_1
muscle_164003_1
muscle_164004_1
muscle_164005_1
muscle_164006_1
plasma_164386_1
plasma_164387_1
plasma_164388_1
plasma_164390_1
plasma_164391_1
plasma_164393_1
pup
adult
reads per million genome matching
0
1000
2000
3000
4000
5000
6000
bp position
0
10
20
30
40
50
60
70

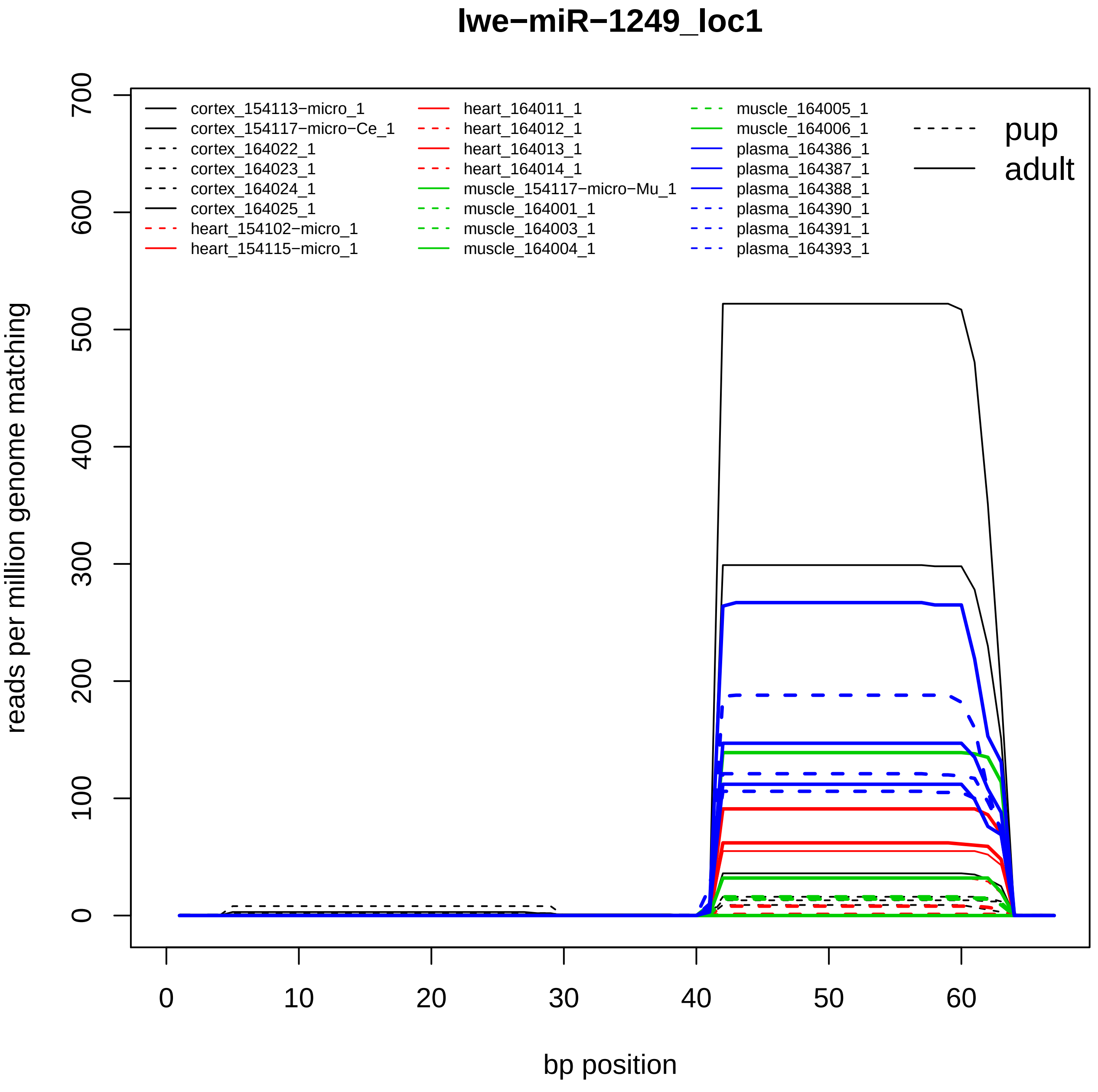
lwe−miR−1249_loc1
reads per million genome matching
bp position
0
100
200
300
400
500
600
700
0
10
20
30
40
50
60
cortex_154113−micro_1
cortex_154117−micro−Ce_1
cortex_164022_1
cortex_164023_1
cortex_164024_1
cortex_164025_1
heart_154102−micro_1
heart_154115−micro_1
heart_164011_1
heart_164012_1
heart_164013_1
heart_164014_1
muscle_154117−micro−Mu_1
muscle_164001_1
muscle_164003_1
muscle_164004_1
muscle_164005_1
muscle_164006_1
plasma_164386_1
plasma_164387_1
plasma_164388_1
plasma_164390_1
plasma_164391_1
plasma_164393_1
pup
adult

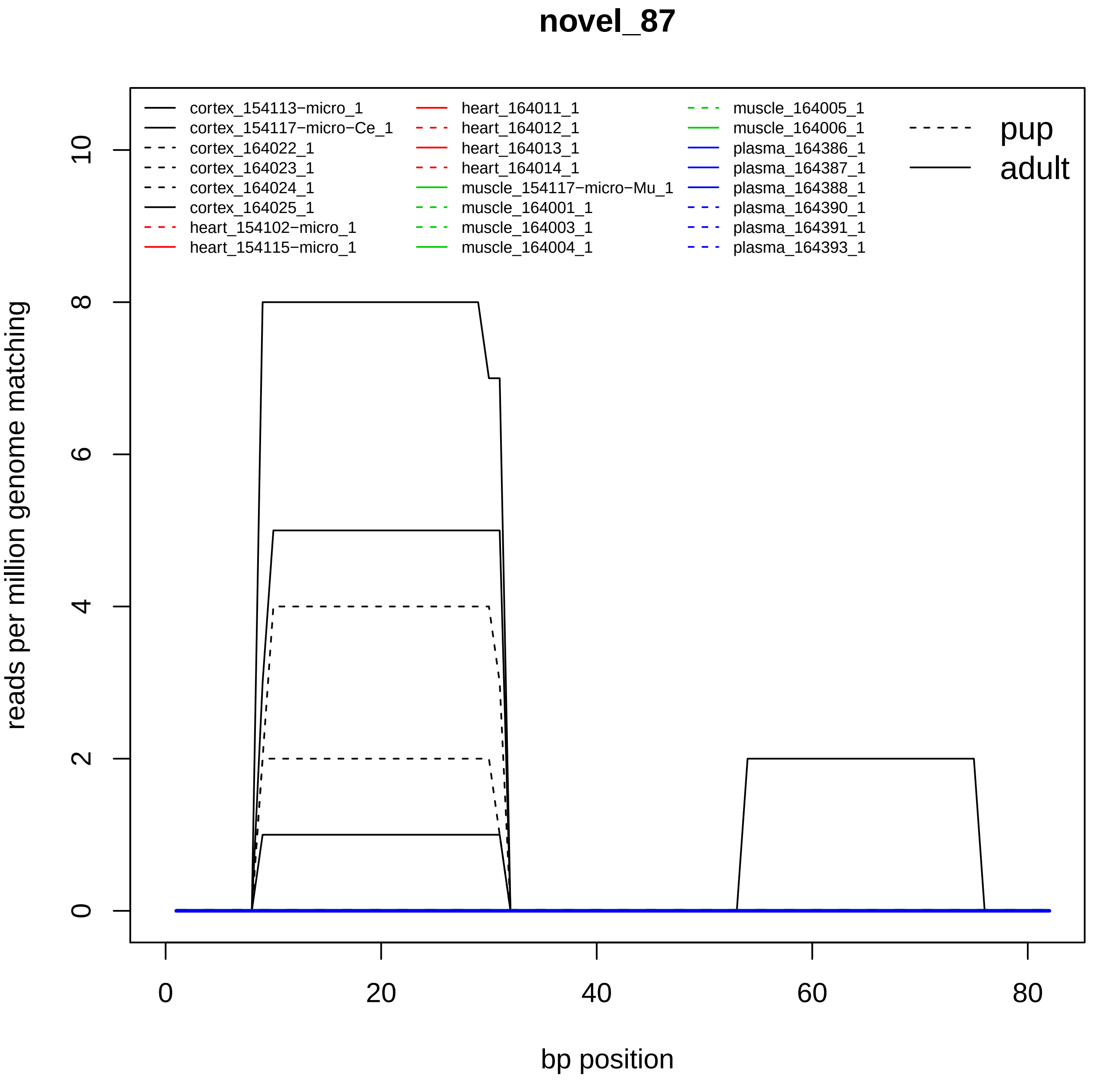

novel_87
cortex_154113−micro_1
cortex_154117−micro−Ce_1
cortex_164022_1
cortex_164023_1
cortex_164024_1
cortex_164025_1
heart_154102−micro_1
heart_154115−micro_1
heart_164011_1
heart_164012_1
heart_164013_1
heart_164014_1
muscle_154117−micro−Mu_1
muscle_164001_1
muscle_164003_1
muscle_164004_1
muscle_164005_1
muscle_164006_1
plasma_164386_1
plasma_164387_1
plasma_164388_1
plasma_164390_1
plasma_164391_1
plasma_164393_1
pup
adult
reads per million genome matching
0
2
4
6
8
10
bp position
0
20
40
60
80

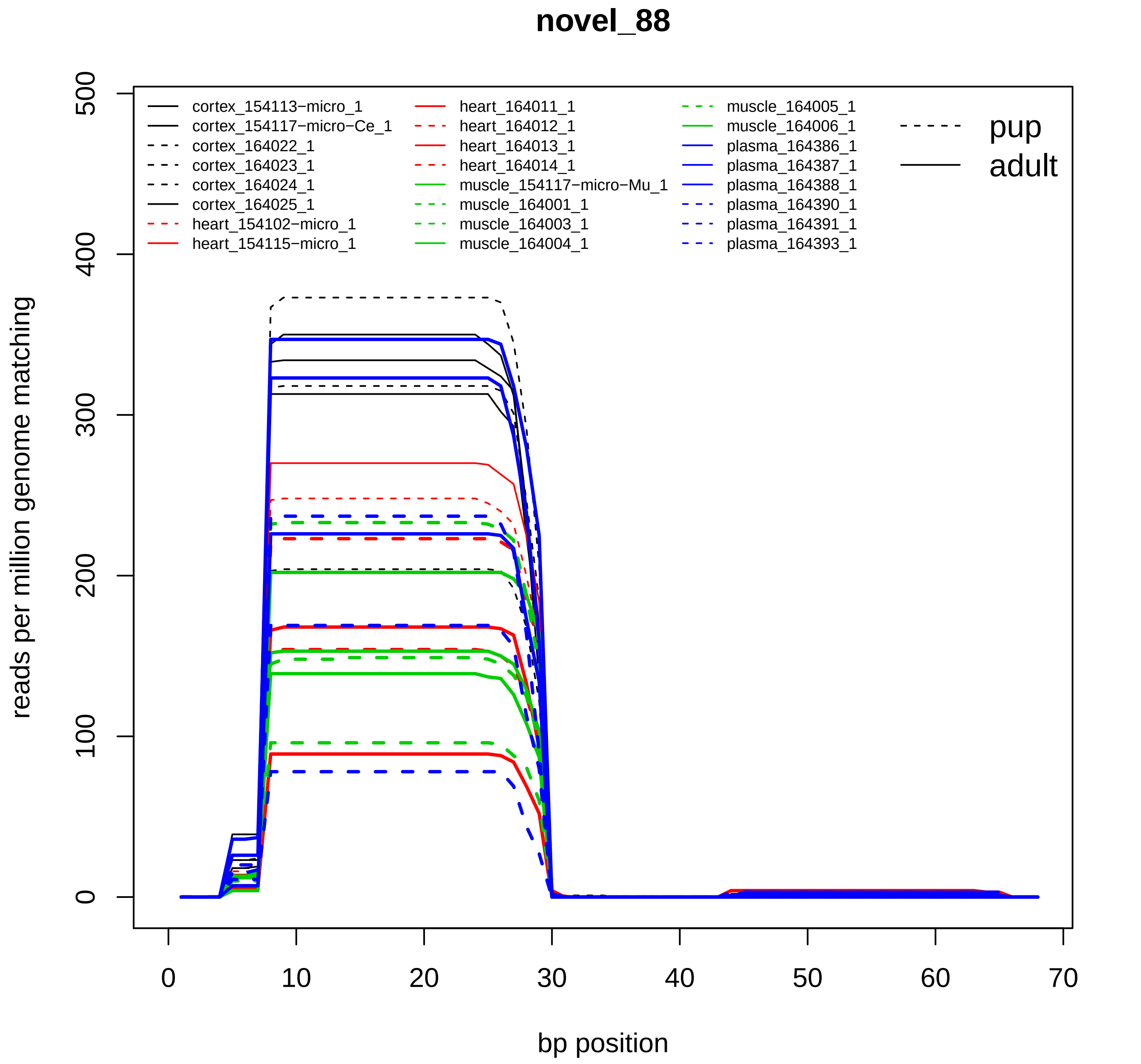

novel_88
cortex_154113−micro_1
cortex_154117−micro−Ce_1
cortex_164022_1
cortex_164023_1
cortex_164024_1
cortex_164025_1
heart_154102−micro_1
heart_154115−micro_1
heart_164011_1
heart_164012_1
heart_164013_1
heart_164014_1
muscle_154117−micro−Mu_1
muscle_164001_1
muscle_164003_1
muscle_164004_1
muscle_164005_1
muscle_164006_1
plasma_164386_1
plasma_164387_1
plasma_164388_1
plasma_164390_1
plasma_164391_1
plasma_164393_1
pup
adult
reads per million genome matching
bp position
0
100
200
300
400
500
0
10
20
30
40
50
60
70

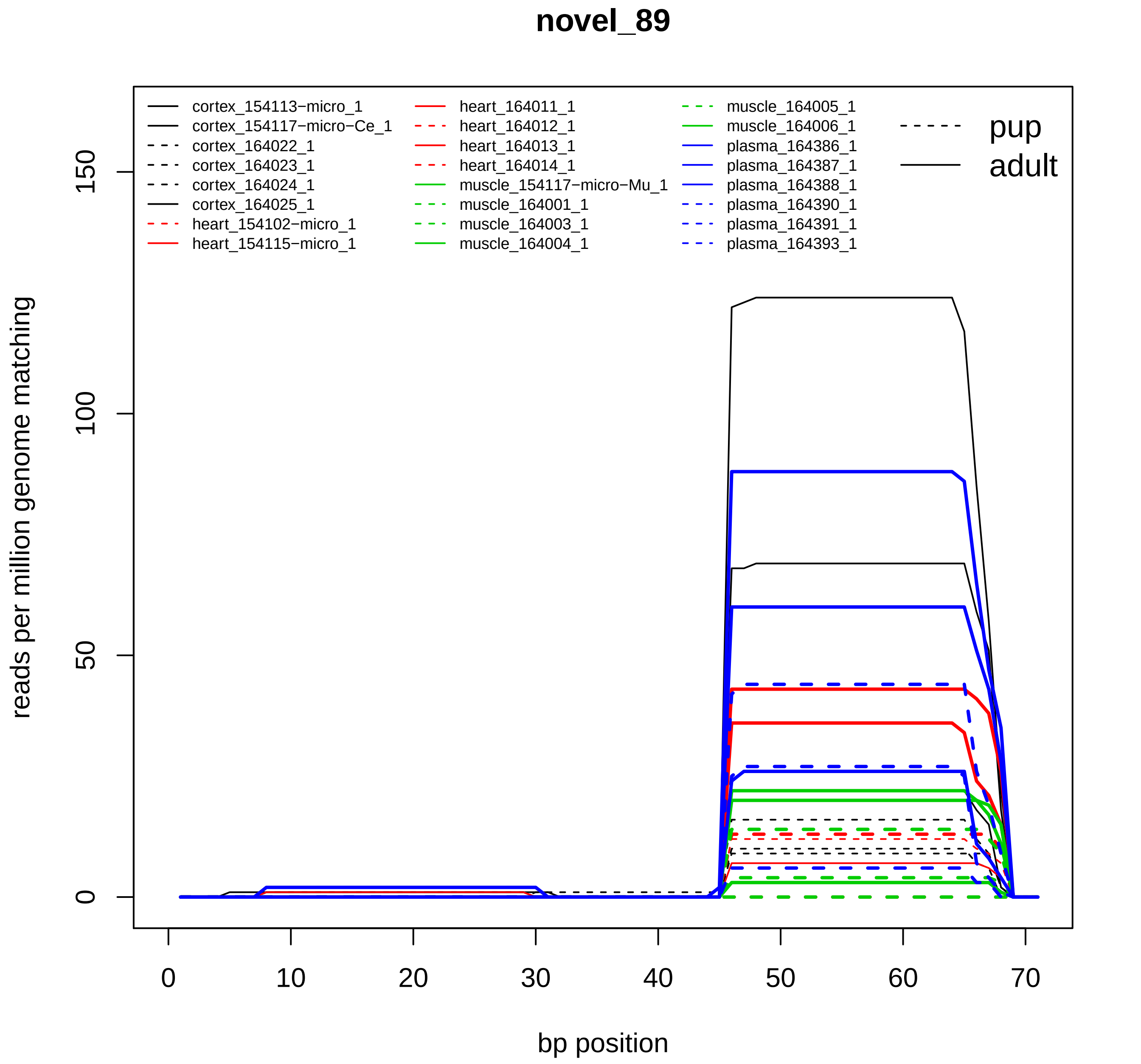

novel_89
cortex_154113−micro_1
cortex_154117−micro−Ce_1
cortex_164022_1
cortex_164023_1
cortex_164024_1
cortex_164025_1
heart_154102−micro_1
heart_154115−micro_1
heart_164011_1
heart_164012_1
heart_164013_1
heart_164014_1
muscle_154117−micro−Mu_1
muscle_164001_1
muscle_164003_1
muscle_164004_1
muscle_164005_1
muscle_164006_1
plasma_164386_1
plasma_164387_1
plasma_164388_1
plasma_164390_1
plasma_164391_1
plasma_164393_1
pup
adult
reads per million genome matching
bp position
0
50
100
150
0
10
20
30
40
50
60
70

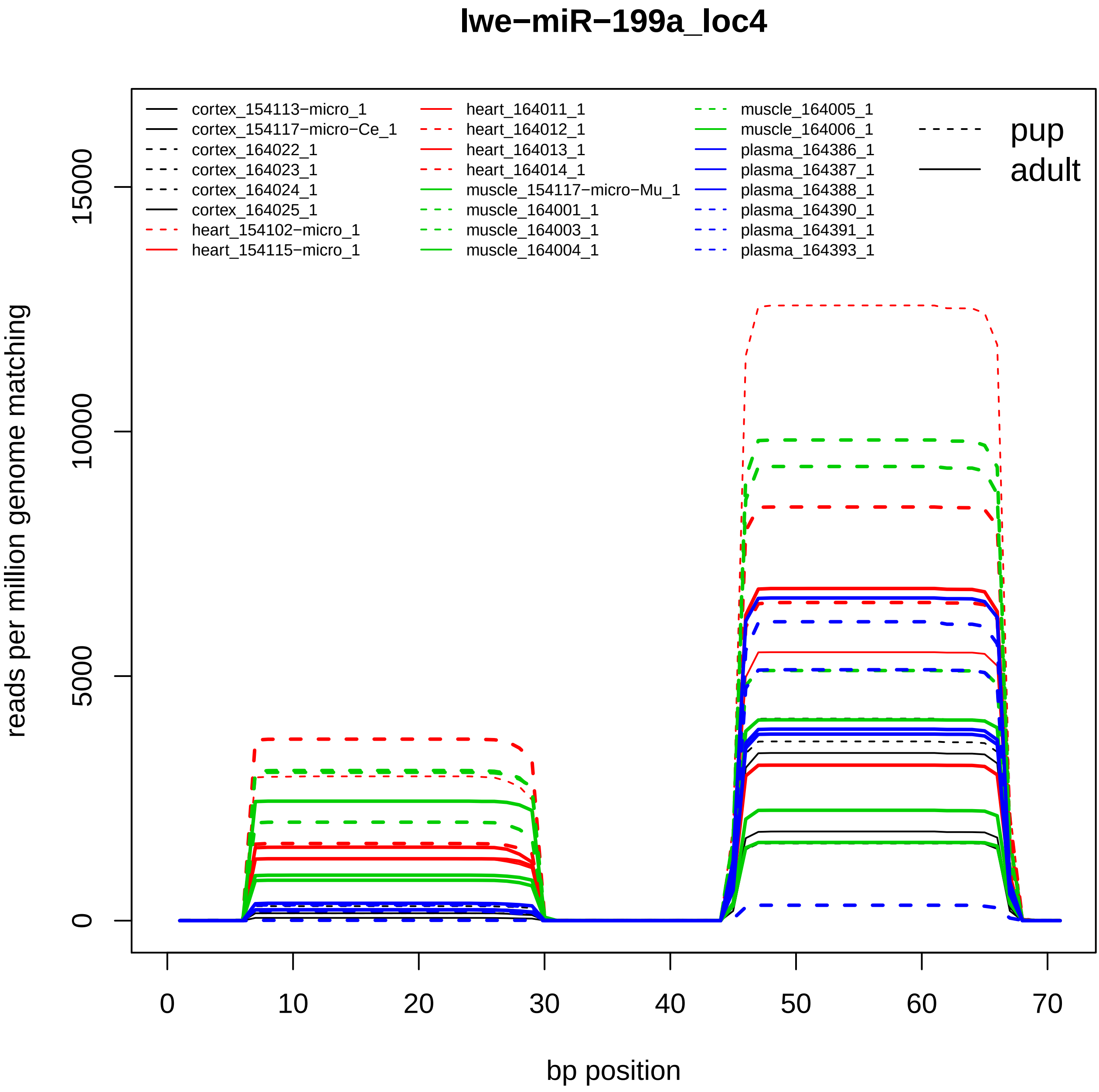
lwe−miR−199a_loc4
cortex_154113−micro_1
cortex_154117−micro−Ce_1
cortex_164022_1
cortex_164023_1
cortex_164024_1
cortex_164025_1
heart_154102−micro_1
heart_154115−micro_1
heart_164011_1
heart_164012_1
heart_164013_1
heart_164014_1
muscle_154117−micro−Mu_1
muscle_164001_1
muscle_164003_1
muscle_164004_1
muscle_164005_1
muscle_164006_1
plasma_164386_1
plasma_164387_1
plasma_164388_1
plasma_164390_1
plasma_164391_1
plasma_164393_1
pup
adult
reads per million genome matching
0
5000
10000
15000
bp position
0
10
20
30
40
50
60
70

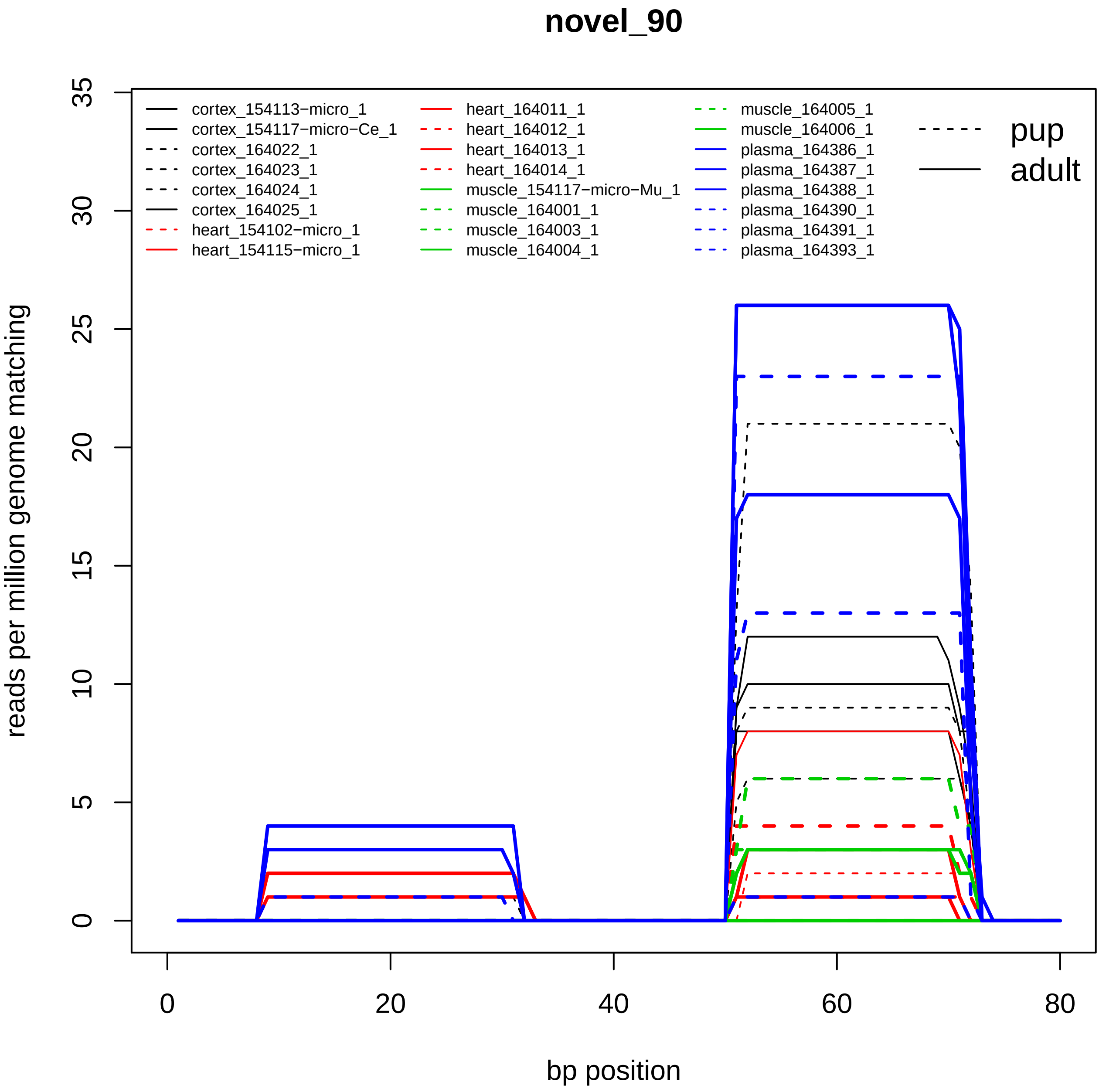
novel_90
cortex_154113−micro_1
cortex_154117−micro−Ce_1
cortex_164022_1
cortex_164023_1
cortex_164024_1
cortex_164025_1
heart_154102−micro_1
heart_154115−micro_1
heart_164011_1
heart_164012_1
heart_164013_1
heart_164014_1
muscle_154117−micro−Mu_1
muscle_164001_1
muscle_164003_1
muscle_164004_1
muscle_164005_1
muscle_164006_1
plasma_164386_1
plasma_164387_1
plasma_164388_1
plasma_164390_1
plasma_164391_1
plasma_164393_1
pup
adult
reads per million genome matching
bp position
0
5
10
15
20
25
30
35
0
20
40
60
80

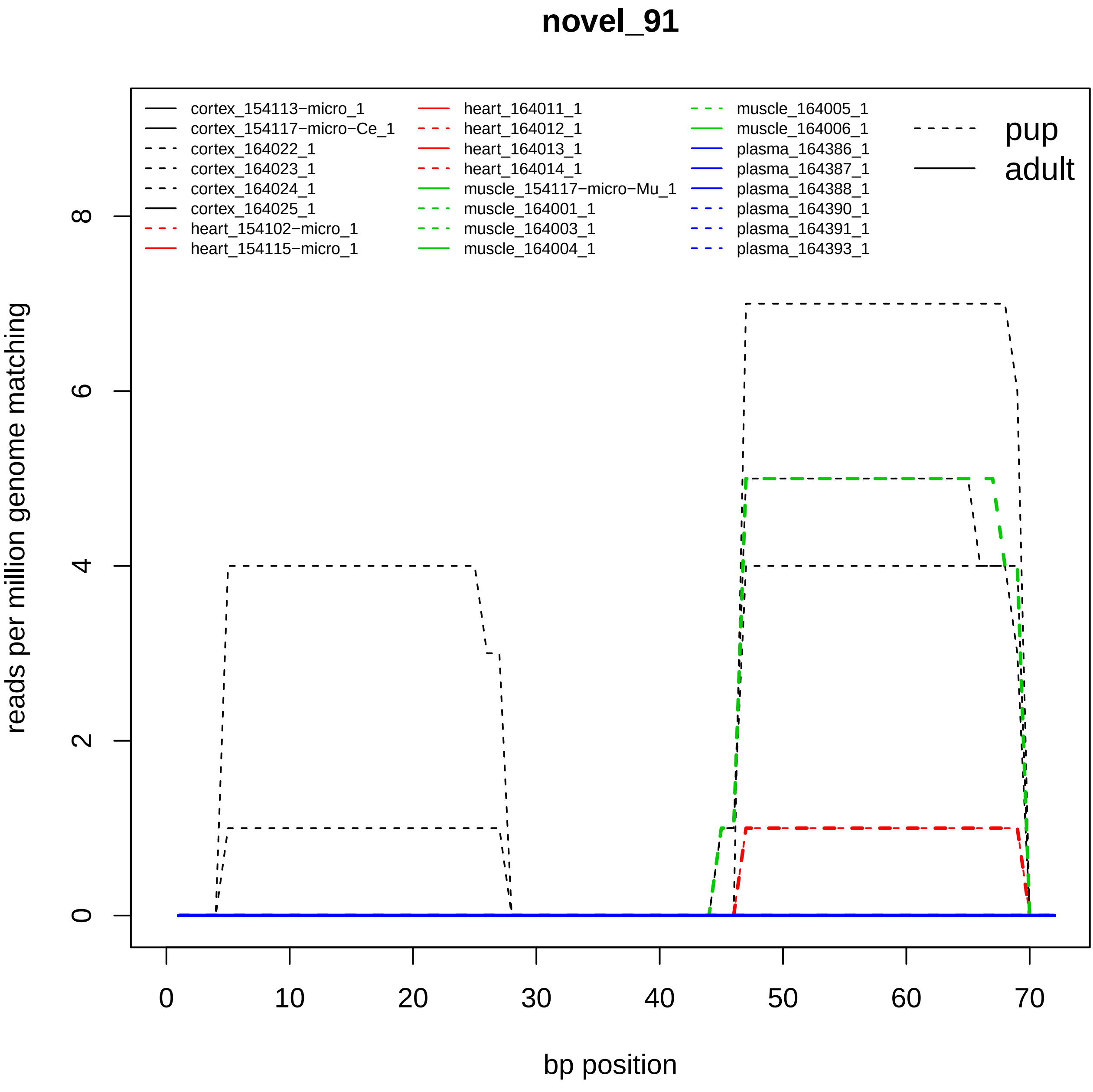
novel_91
cortex_154113−micro_1
cortex_154117−micro−Ce_1
cortex_164022_1
cortex_164023_1
cortex_164024_1
cortex_164025_1
heart_154102−micro_1
heart_154115−micro_1
heart_164011_1
heart_164012_1
heart_164013_1
heart_164014_1
muscle_154117−micro−Mu_1
muscle_164001_1
muscle_164003_1
muscle_164004_1
muscle_164005_1
muscle_164006_1
plasma_164386_1
plasma_164387_1
plasma_164388_1
plasma_164390_1
plasma_164391_1
plasma_164393_1
pup
adult
reads per million genome matching
0
2
4
6
8
0
10
20
30
40
50
60
70
bp position

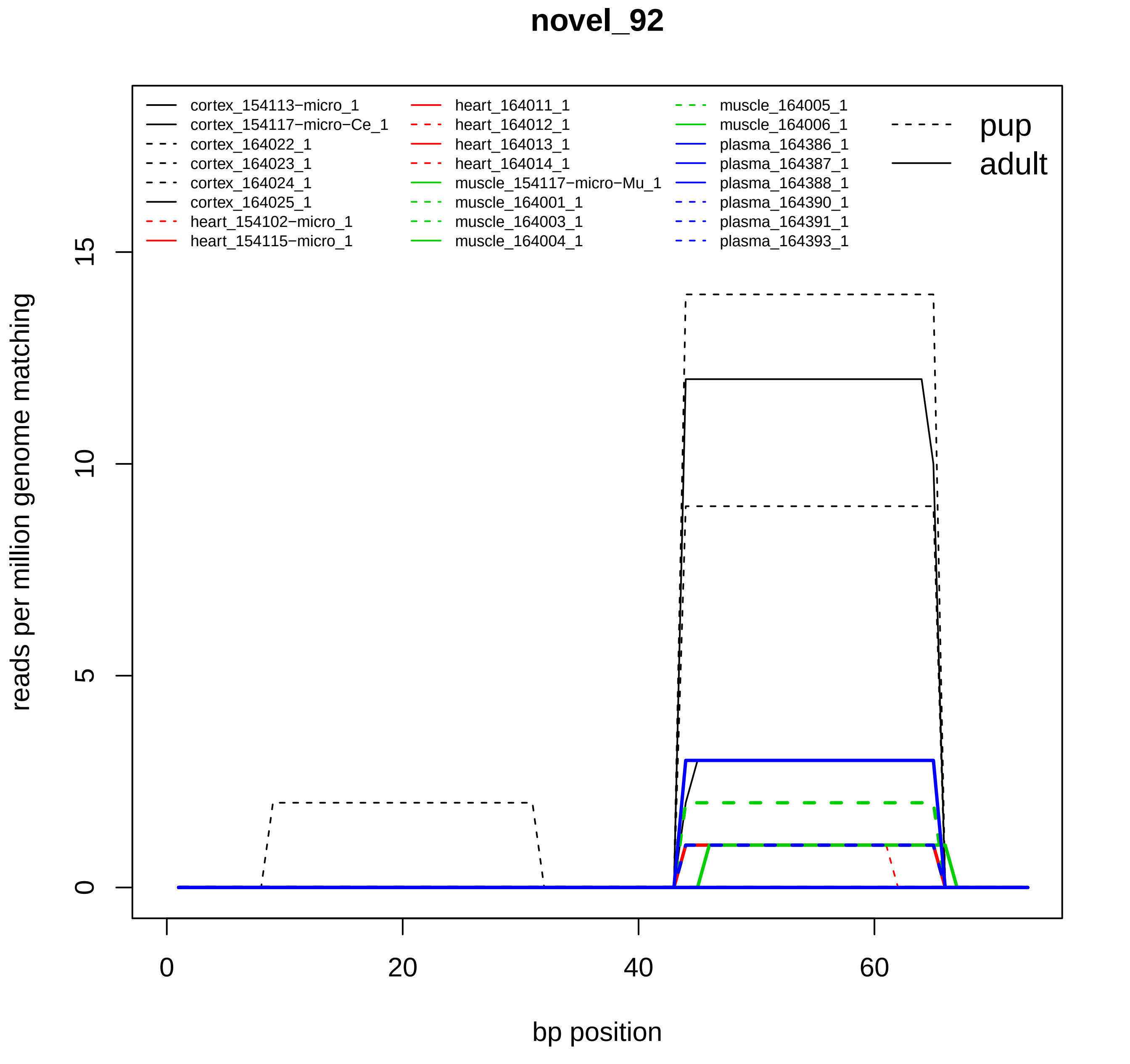
novel_92
cortex_154113−micro_1
cortex_154117−micro−Ce_1
cortex_164022_1
cortex_164023_1
cortex_164024_1
cortex_164025_1
heart_154102−micro_1
heart_154115−micro_1
heart_164011_1
heart_164012_1
heart_164013_1
heart_164014_1
muscle_154117−micro−Mu_1
muscle_164001_1
muscle_164003_1
muscle_164004_1
muscle_164005_1
muscle_164006_1
plasma_164386_1
plasma_164387_1
plasma_164388_1
plasma_164390_1
plasma_164391_1
plasma_164393_1
pup
adult
reads per million genome matching
0
5
10
15
0
20
40
60
bp position

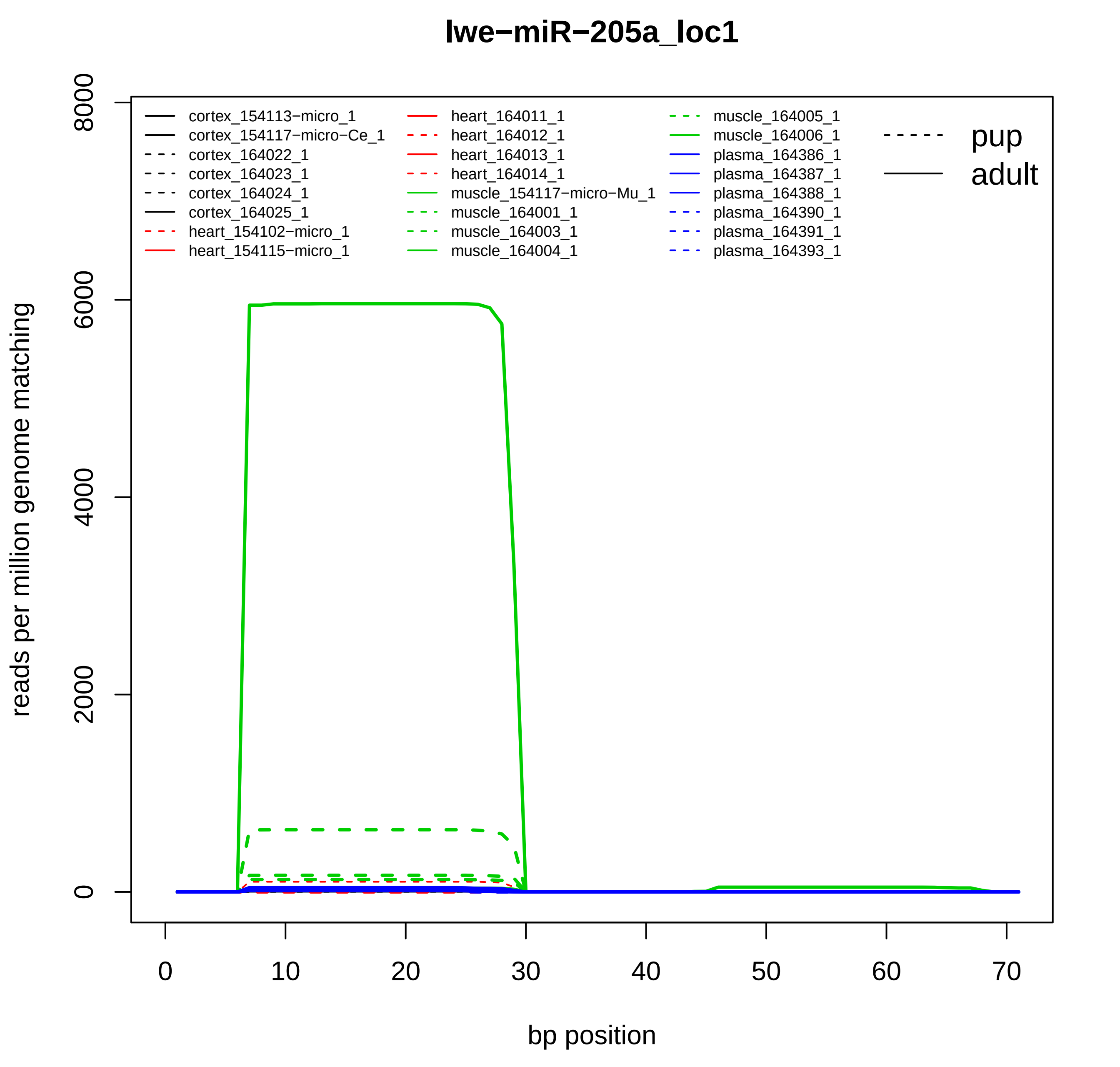

lwe−miR−205a_loc1
cortex_154113−micro_1
cortex_154117−micro−Ce_1
cortex_164022_1
cortex_164023_1
cortex_164024_1
cortex_164025_1
heart_154102−micro_1
heart_154115−micro_1
heart_164011_1
heart_164012_1
heart_164013_1
heart_164014_1
muscle_154117−micro−Mu_1
muscle_164001_1
muscle_164003_1
muscle_164004_1
muscle_164005_1
muscle_164006_1
plasma_164386_1
plasma_164387_1
plasma_164388_1
plasma_164390_1
plasma_164391_1
plasma_164393_1
pup
adult
reads per million genome matching
0
2000
4000
6000
8000
bp position
0
10
20
30
40
50
60
70

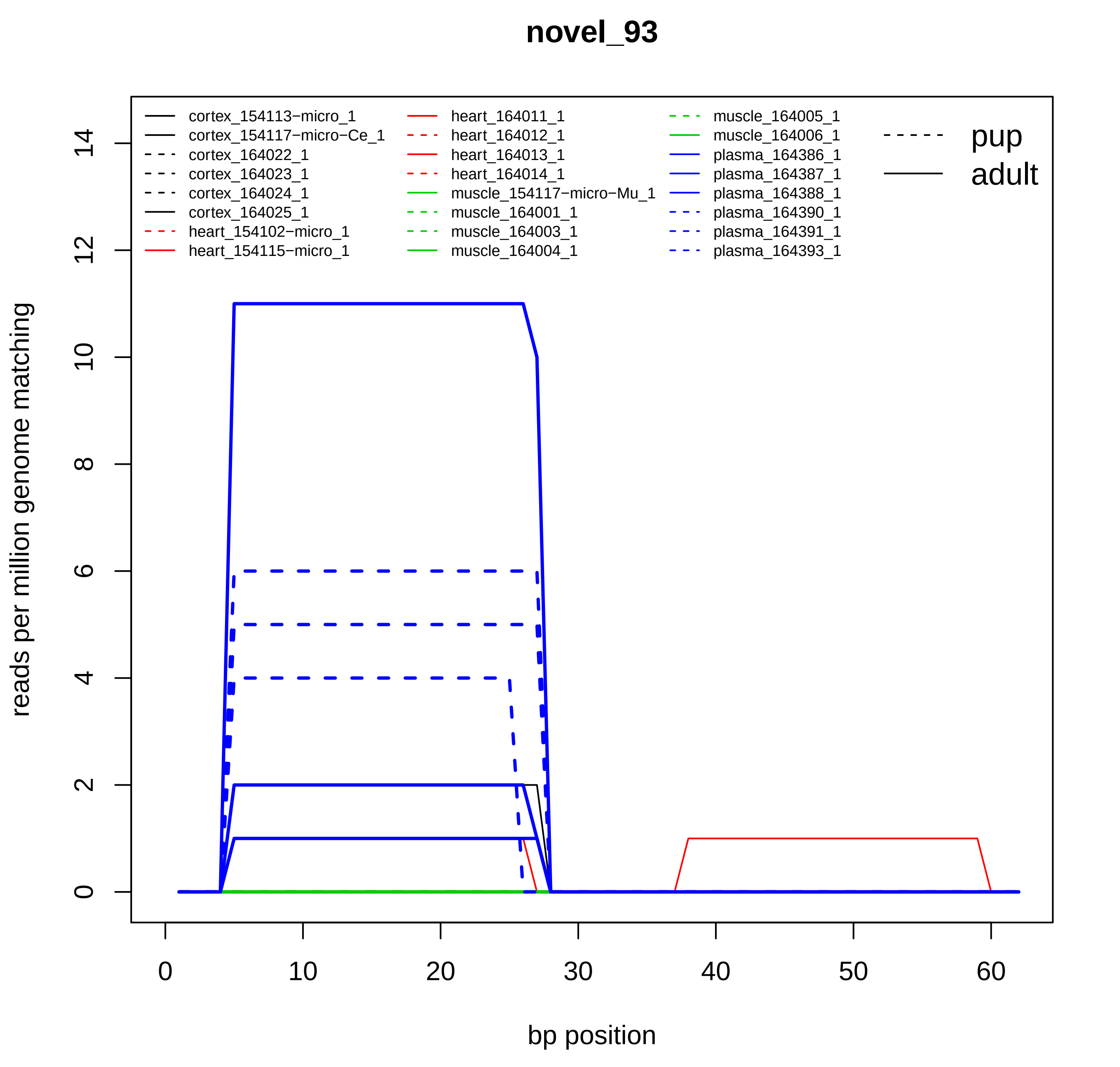
novel_93
cortex_154113−micro_1
cortex_154117−micro−Ce_1
cortex_164022_1
cortex_164023_1
cortex_164024_1
cortex_164025_1
heart_154102−micro_1
heart_154115−micro_1
heart_164011_1
heart_164012_1
heart_164013_1
heart_164014_1
muscle_154117−micro−Mu_1
muscle_164001_1
muscle_164003_1
muscle_164004_1
muscle_164005_1
muscle_164006_1
plasma_164386_1
plasma_164387_1
plasma_164388_1
plasma_164390_1
plasma_164391_1
plasma_164393_1
pup
adult
reads per million genome matching
bp position
0
2
4
6
8
10
12
14
0
10
20
30
40
50
60

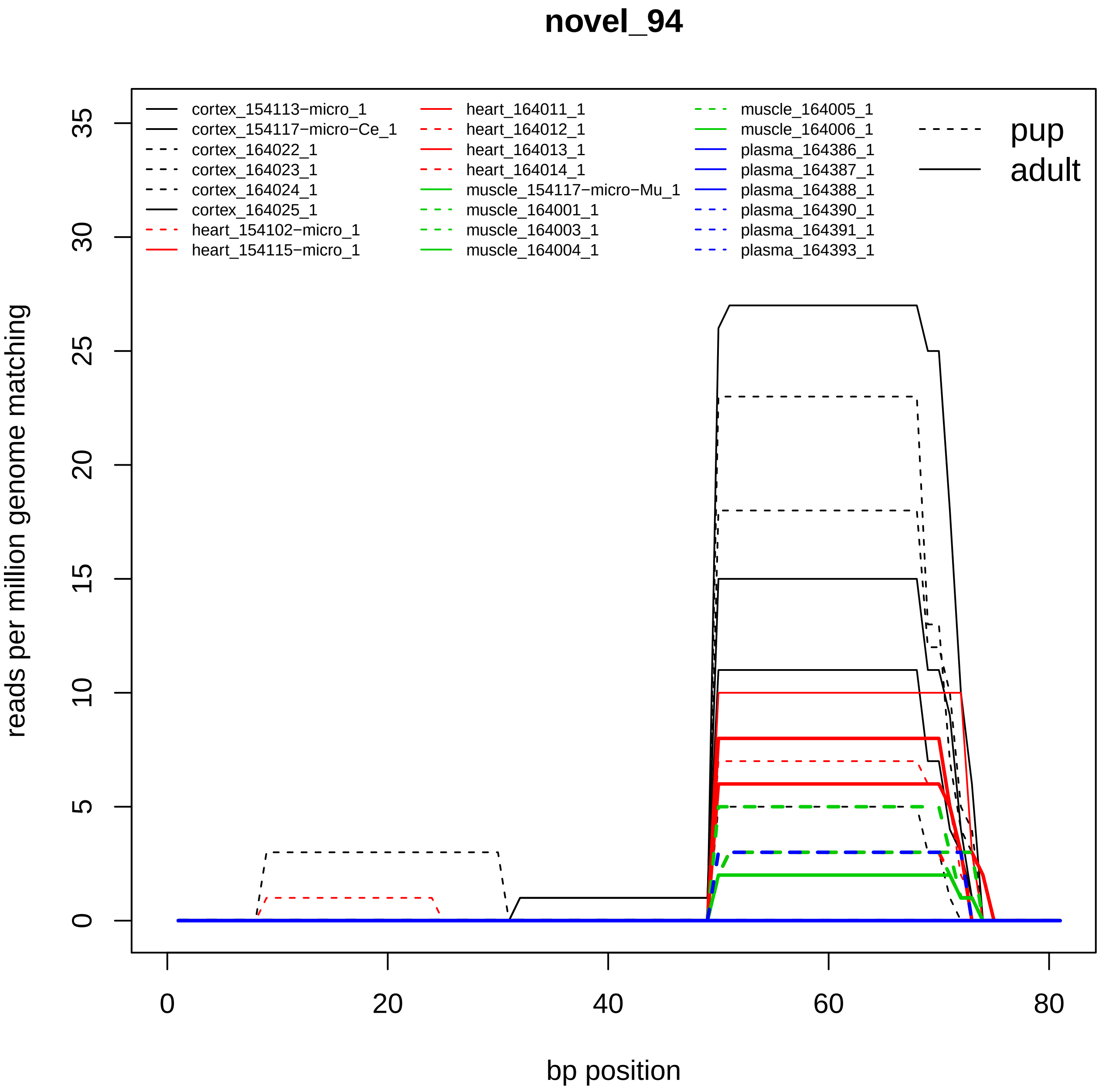

novel_94
cortex_154113−micro_1
cortex_154117−micro−Ce_1
cortex_164022_1
cortex_164023_1
cortex_164024_1
cortex_164025_1
heart_154102−micro_1
heart_154115−micro_1
heart_164011_1
heart_164012_1
heart_164013_1
heart_164014_1
muscle_154117−micro−Mu_1
muscle_164001_1
muscle_164003_1
muscle_164004_1
muscle_164005_1
muscle_164006_1
plasma_164386_1
plasma_164387_1
plasma_164388_1
plasma_164390_1
plasma_164391_1
plasma_164393_1
pup
adult
reads per million genome matching
0
5
10
15
20
25
30
35
0
20
40
60
80
bp position

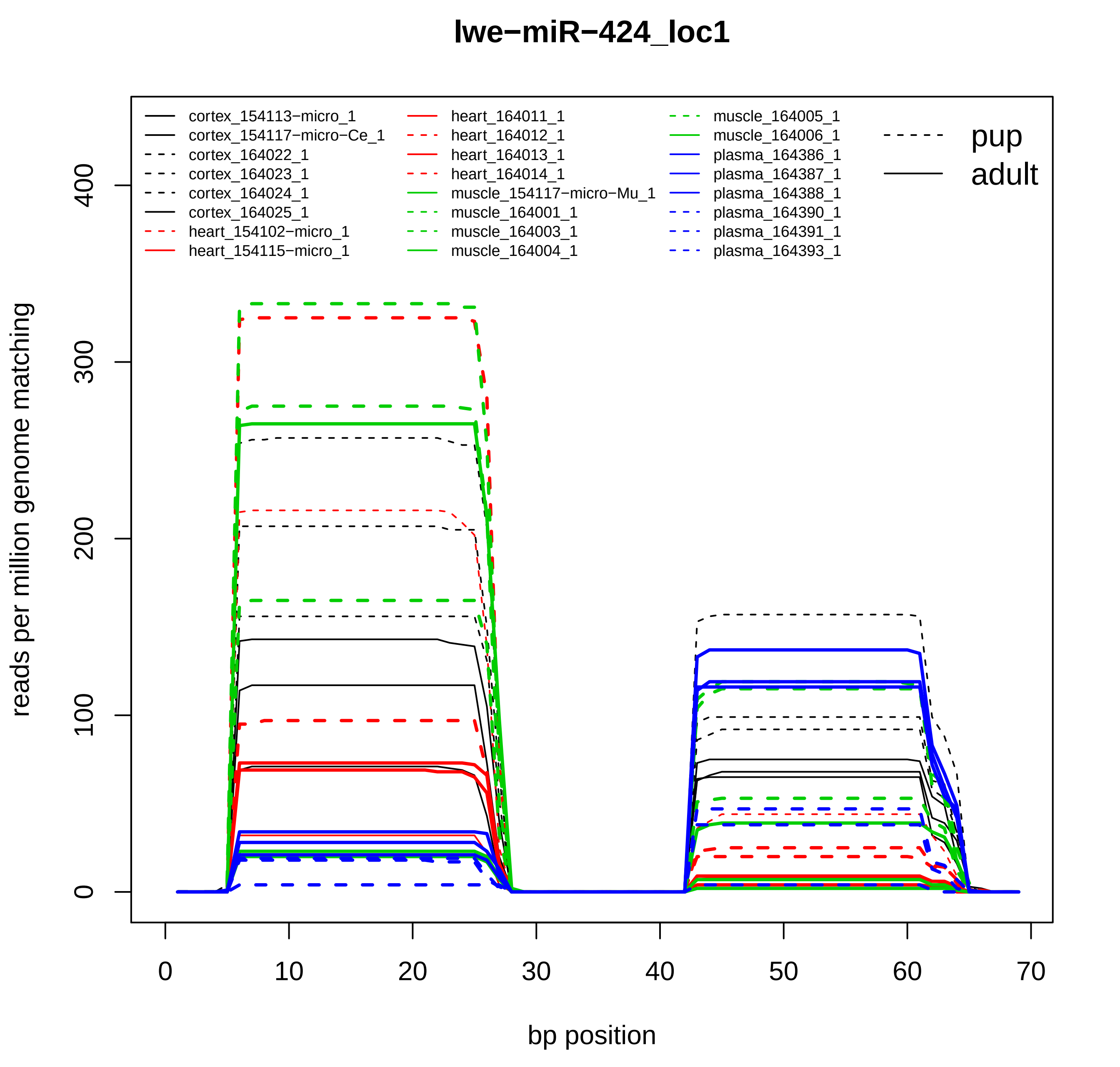
lwe−miR−424_loc1
reads per million genome matching
bp position
0
100
200
300
400
0
10
20
30
40
50
60
70
cortex_154113−micro_1
cortex_154117−micro−Ce_1
cortex_164022_1
cortex_164023_1
cortex_164024_1
cortex_164025_1
heart_154102−micro_1
heart_154115−micro_1
heart_164011_1
heart_164012_1
heart_164013_1
heart_164014_1
muscle_154117−micro−Mu_1
muscle_164001_1
muscle_164003_1
muscle_164004_1
muscle_164005_1
muscle_164006_1
plasma_164386_1
plasma_164387_1
plasma_164388_1
plasma_164390_1
plasma_164391_1
plasma_164393_1
pup
adult

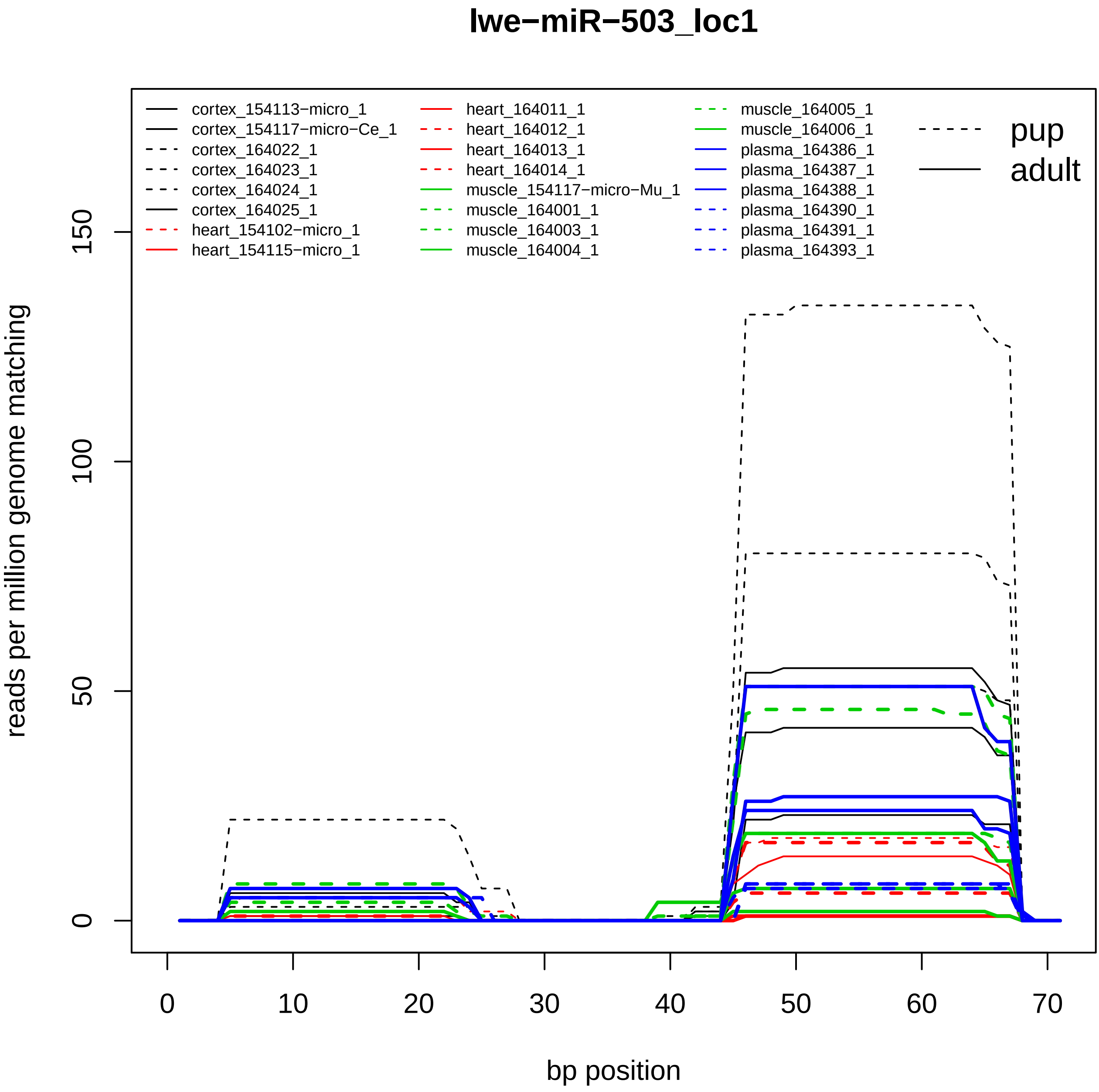
lwe−miR−503_loc1
cortex_154113−micro_1
cortex_154117−micro−Ce_1
cortex_164022_1
cortex_164023_1
cortex_164024_1
cortex_164025_1
heart_154102−micro_1
heart_154115−micro_1
heart_164011_1
heart_164012_1
heart_164013_1
heart_164014_1
muscle_154117−micro−Mu_1
muscle_164001_1
muscle_164003_1
muscle_164004_1
muscle_164005_1
muscle_164006_1
plasma_164386_1
plasma_164387_1
plasma_164388_1
plasma_164390_1
plasma_164391_1
plasma_164393_1
pup
adult
reads per million genome matching
0
50
100
150
0
10
20
30
40
50
60
70
bp position

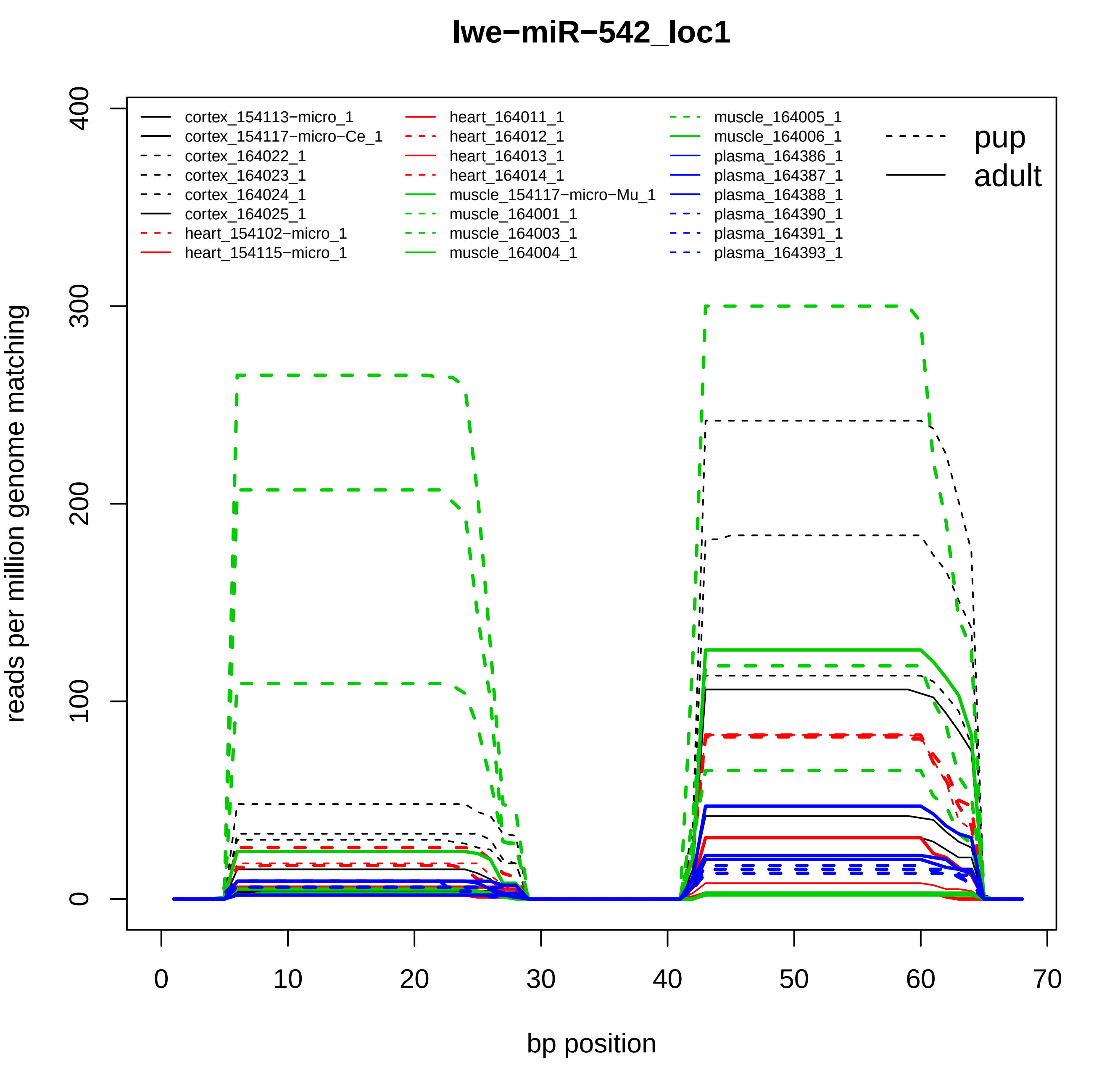
lwe−miR−542_loc1
cortex_154113−micro_1
cortex_154117−micro−Ce_1
cortex_164022_1
cortex_164023_1
cortex_164024_1
cortex_164025_1
heart_154102−micro_1
heart_154115−micro_1
heart_164011_1
heart_164012_1
heart_164013_1
heart_164014_1
muscle_154117−micro−Mu_1
muscle_164001_1
muscle_164003_1
muscle_164004_1
muscle_164005_1
muscle_164006_1
plasma_164386_1
plasma_164387_1
plasma_164388_1
plasma_164390_1
plasma_164391_1
plasma_164393_1
pup
adult
reads per million genome matching
0
100
200
300
400
bp position
0
10
20
30
40
50
60
70

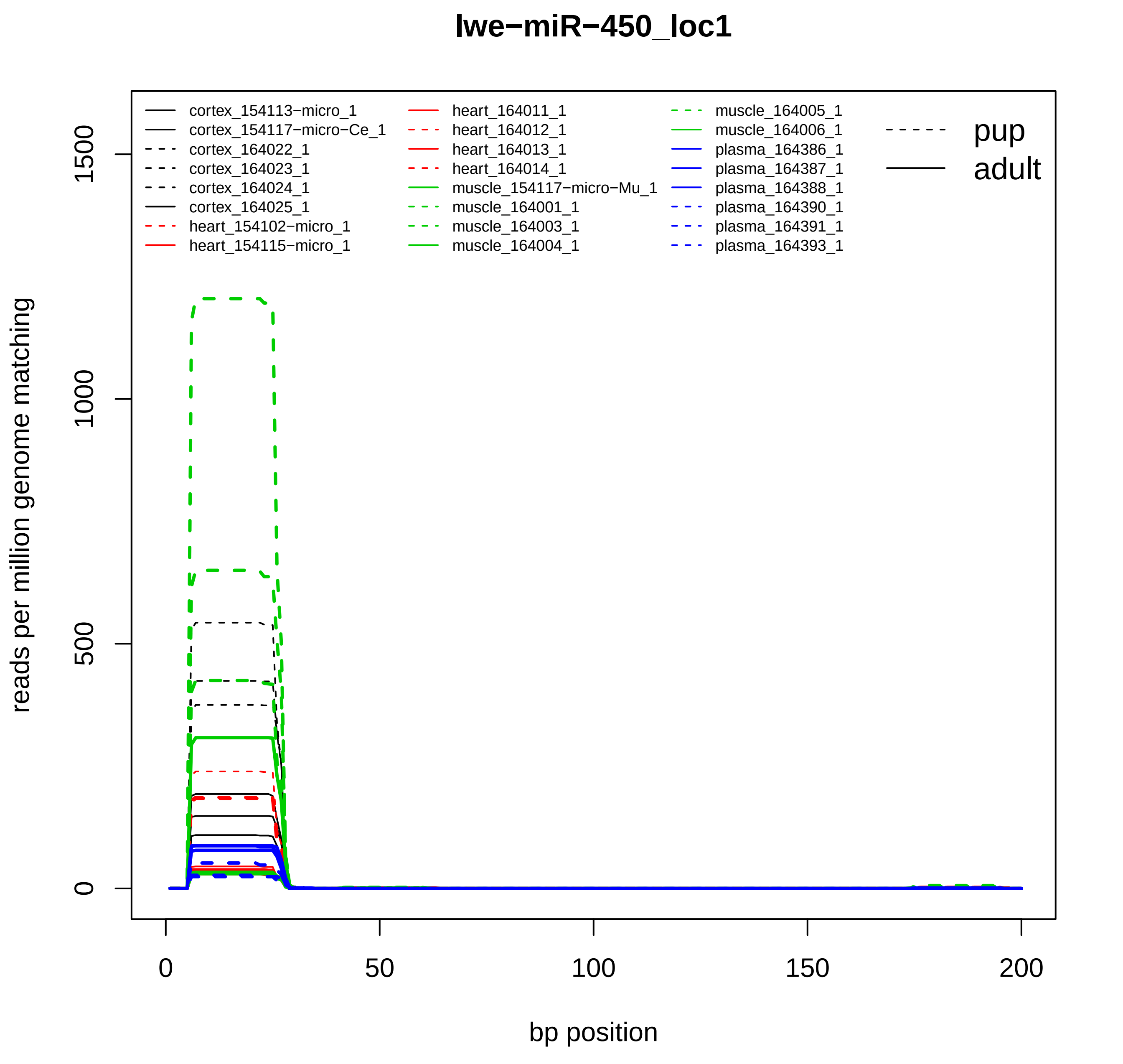

lwe−miR−450_loc1
cortex_154113−micro_1
cortex_154117−micro−Ce_1
cortex_164022_1
cortex_164023_1
cortex_164024_1
cortex_164025_1
heart_154102−micro_1
heart_154115−micro_1
heart_164011_1
heart_164012_1
heart_164013_1
heart_164014_1
muscle_154117−micro−Mu_1
muscle_164001_1
muscle_164003_1
muscle_164004_1
muscle_164005_1
muscle_164006_1
plasma_164386_1
plasma_164387_1
plasma_164388_1
plasma_164390_1
plasma_164391_1
plasma_164393_1
pup
adult
reads per million genome matching
0
500
1000
1500
bp position
0
50
100
150
200

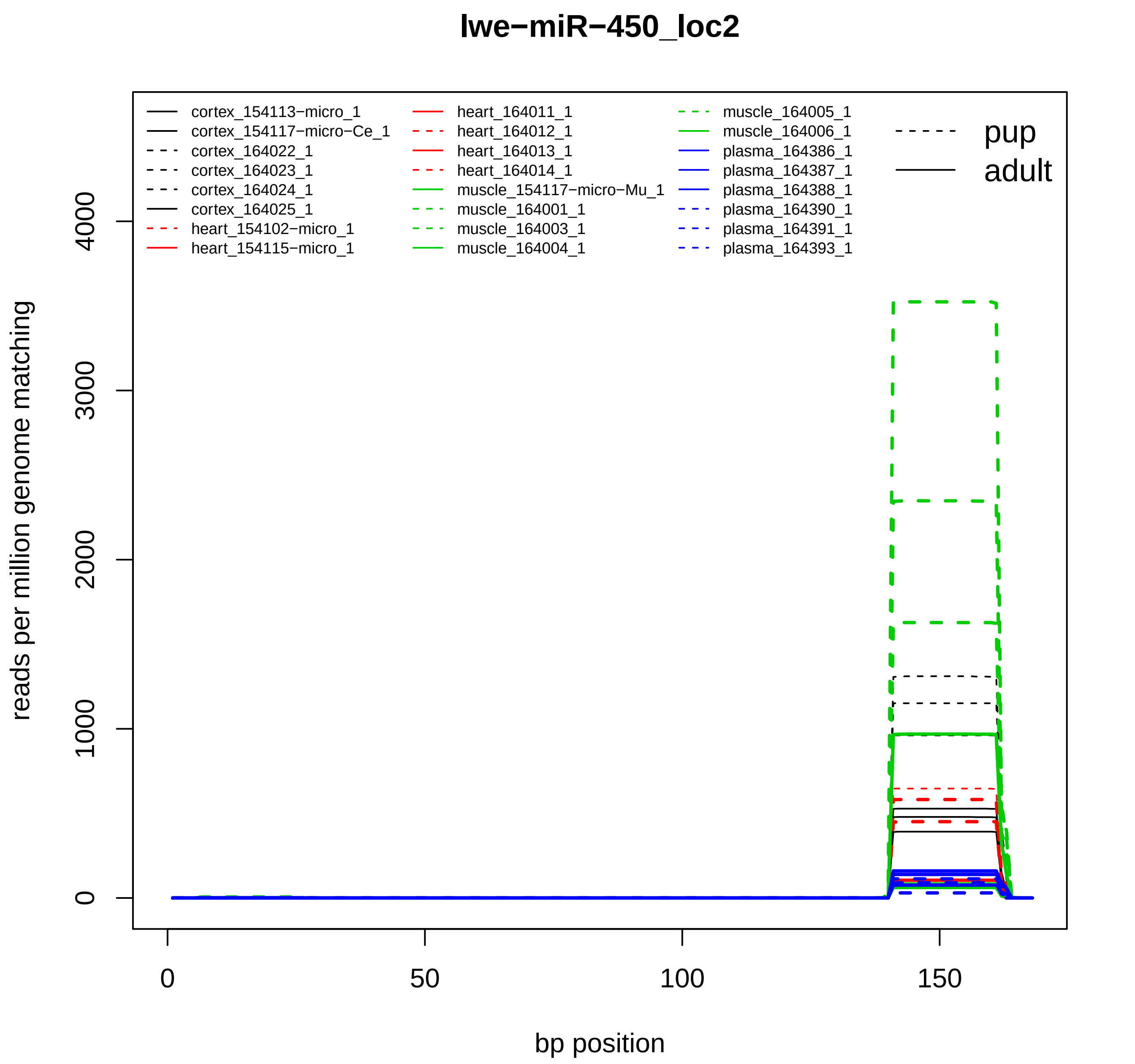

lwe−miR−450_loc2
cortex_154113−micro_1
cortex_154117−micro−Ce_1
cortex_164022_1
cortex_164023_1
cortex_164024_1
cortex_164025_1
heart_154102−micro_1
heart_154115−micro_1
heart_164011_1
heart_164012_1
heart_164013_1
heart_164014_1
muscle_154117−micro−Mu_1
muscle_164001_1
muscle_164003_1
muscle_164004_1
muscle_164005_1
muscle_164006_1
plasma_164386_1
plasma_164387_1
plasma_164388_1
plasma_164390_1
plasma_164391_1
plasma_164393_1
pup
adult
reads per million genome matching
0
1000
2000
3000
4000
0
50
100
150
bp position

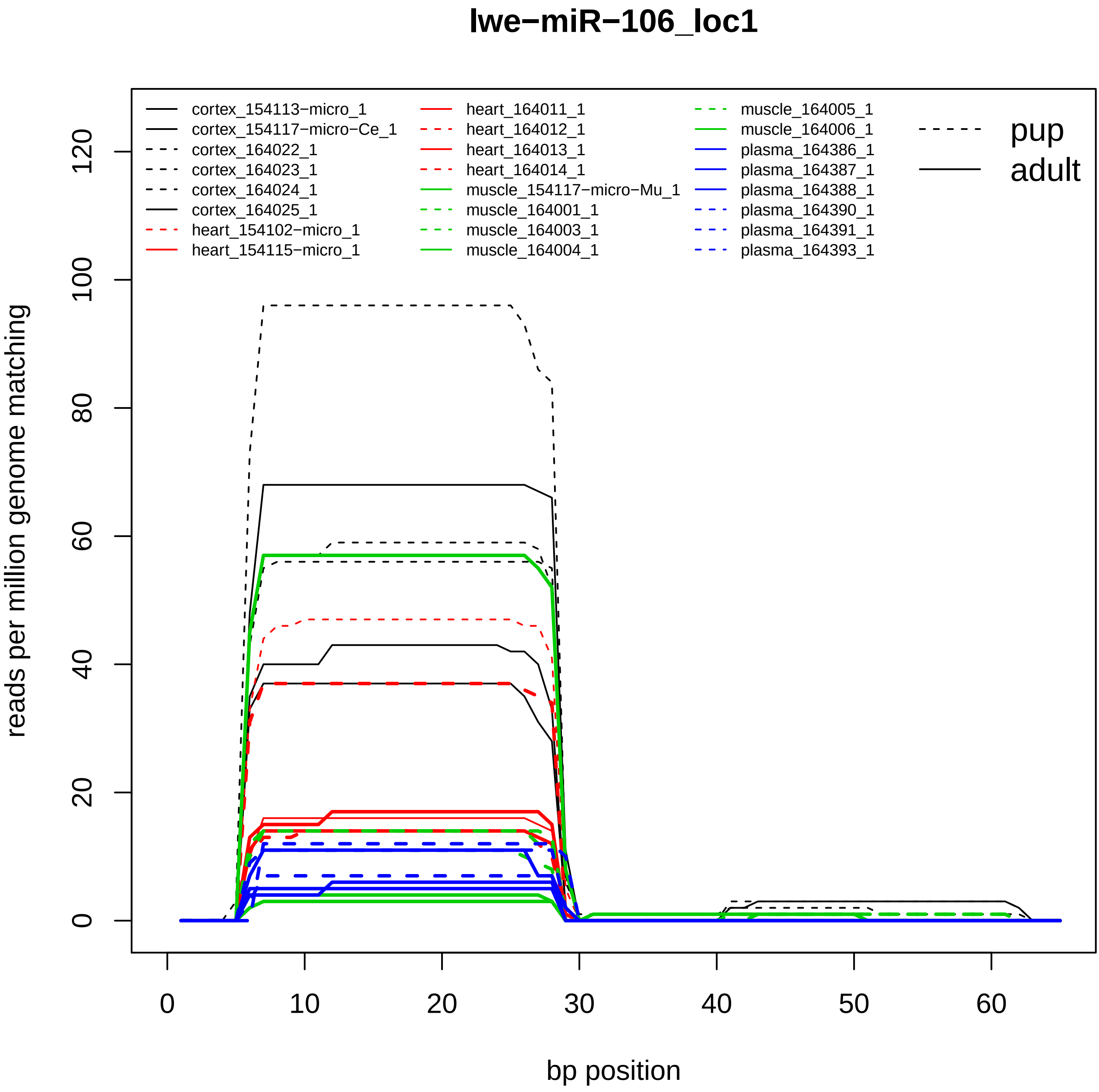
lwe−miR−106_loc1
cortex_154113−micro_1
cortex_154117−micro−Ce_1
cortex_164022_1
cortex_164023_1
cortex_164024_1
cortex_164025_1
heart_154102−micro_1
heart_154115−micro_1
heart_164011_1
heart_164012_1
heart_164013_1
heart_164014_1
muscle_154117−micro−Mu_1
muscle_164001_1
muscle_164003_1
muscle_164004_1
muscle_164005_1
muscle_164006_1
plasma_164386_1
plasma_164387_1
plasma_164388_1
plasma_164390_1
plasma_164391_1
plasma_164393_1
pup
adult
reads per million genome matching
0
20
40
60
80
100
120
0
10
20
30
40
50
60
bp position

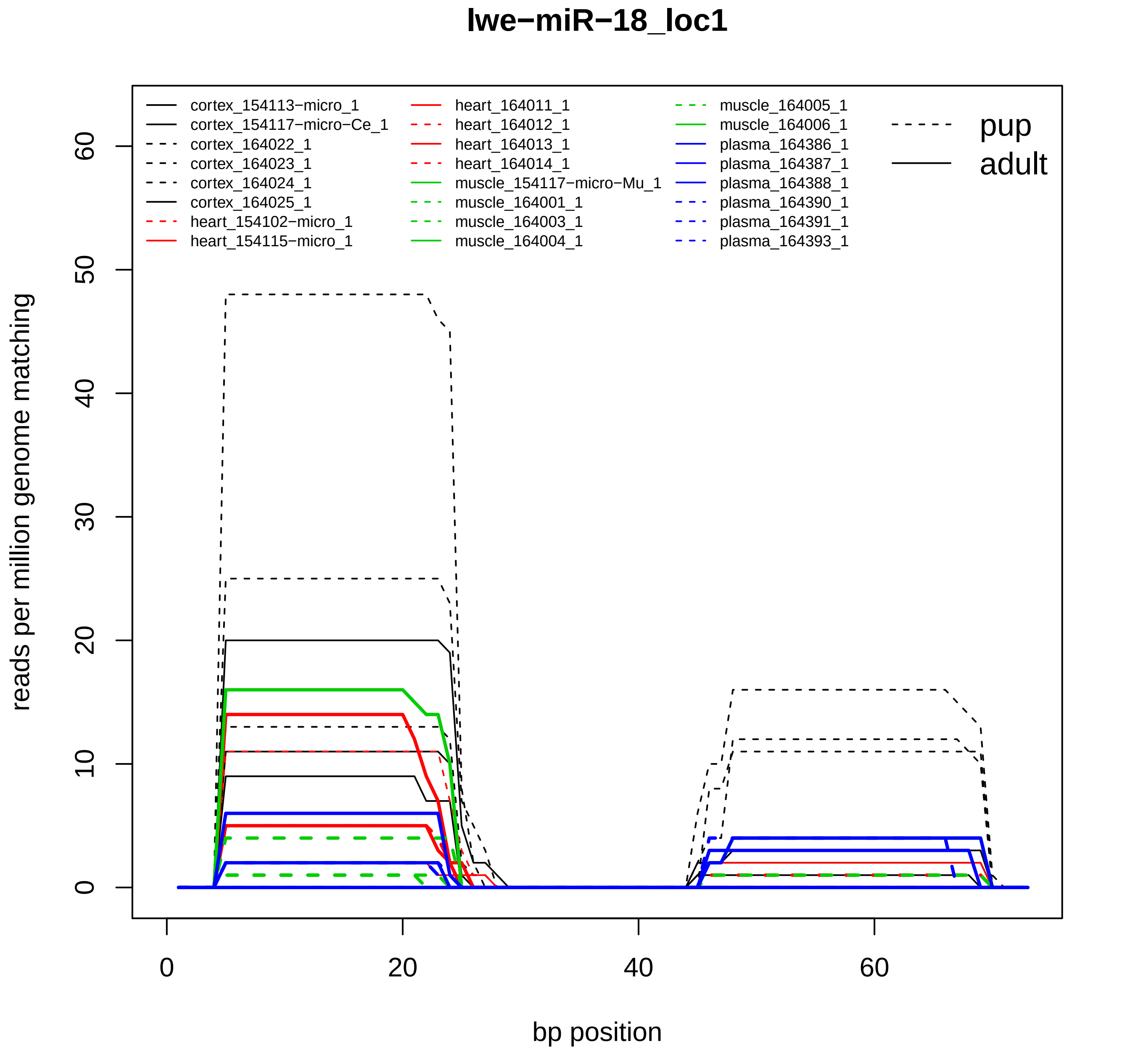
lwe−miR−18_loc1
cortex_154113−micro_1
cortex_154117−micro−Ce_1
cortex_164022_1
cortex_164023_1
cortex_164024_1
cortex_164025_1
heart_154102−micro_1
heart_154115−micro_1
heart_164011_1
heart_164012_1
heart_164013_1
heart_164014_1
muscle_154117−micro−Mu_1
muscle_164001_1
muscle_164003_1
muscle_164004_1
muscle_164005_1
muscle_164006_1
plasma_164386_1
plasma_164387_1
plasma_164388_1
plasma_164390_1
plasma_164391_1
plasma_164393_1
pup
adult
reads per million genome matching
0
10
20
30
40
50
60
0
20
40
60
bp position

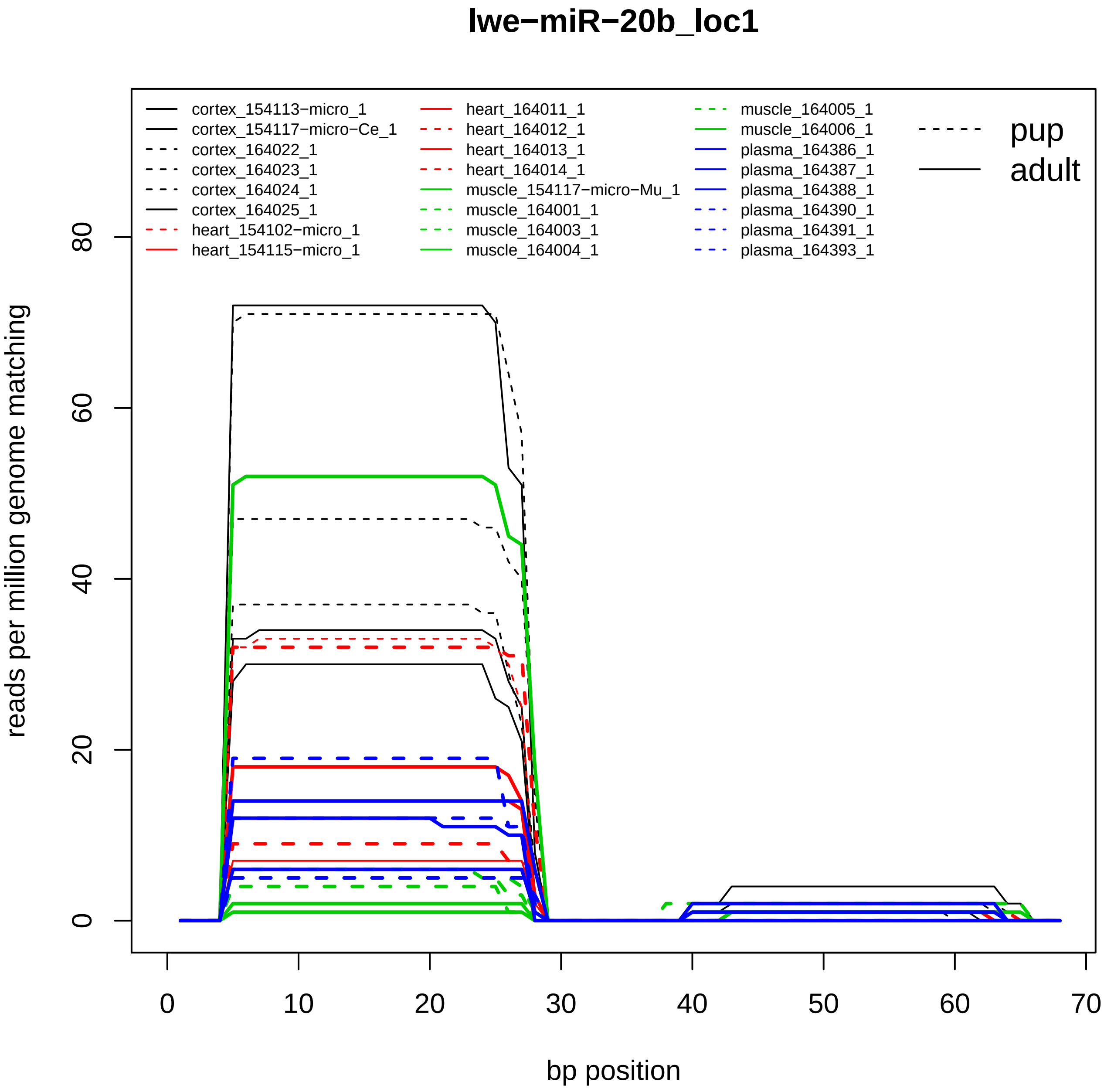
lwe−miR−20b_loc1
cortex_154113−micro_1
cortex_154117−micro−Ce_1
cortex_164022_1
cortex_164023_1
cortex_164024_1
cortex_164025_1
heart_154102−micro_1
heart_154115−micro_1
heart_164011_1
heart_164012_1
heart_164013_1
heart_164014_1
muscle_154117−micro−Mu_1
muscle_164001_1
muscle_164003_1
muscle_164004_1
muscle_164005_1
muscle_164006_1
plasma_164386_1
plasma_164387_1
plasma_164388_1
plasma_164390_1
plasma_164391_1
plasma_164393_1
pup
adult
reads per million genome matching
bp position
0
20
40
60
80
0
10
20
30
40
50
60
70

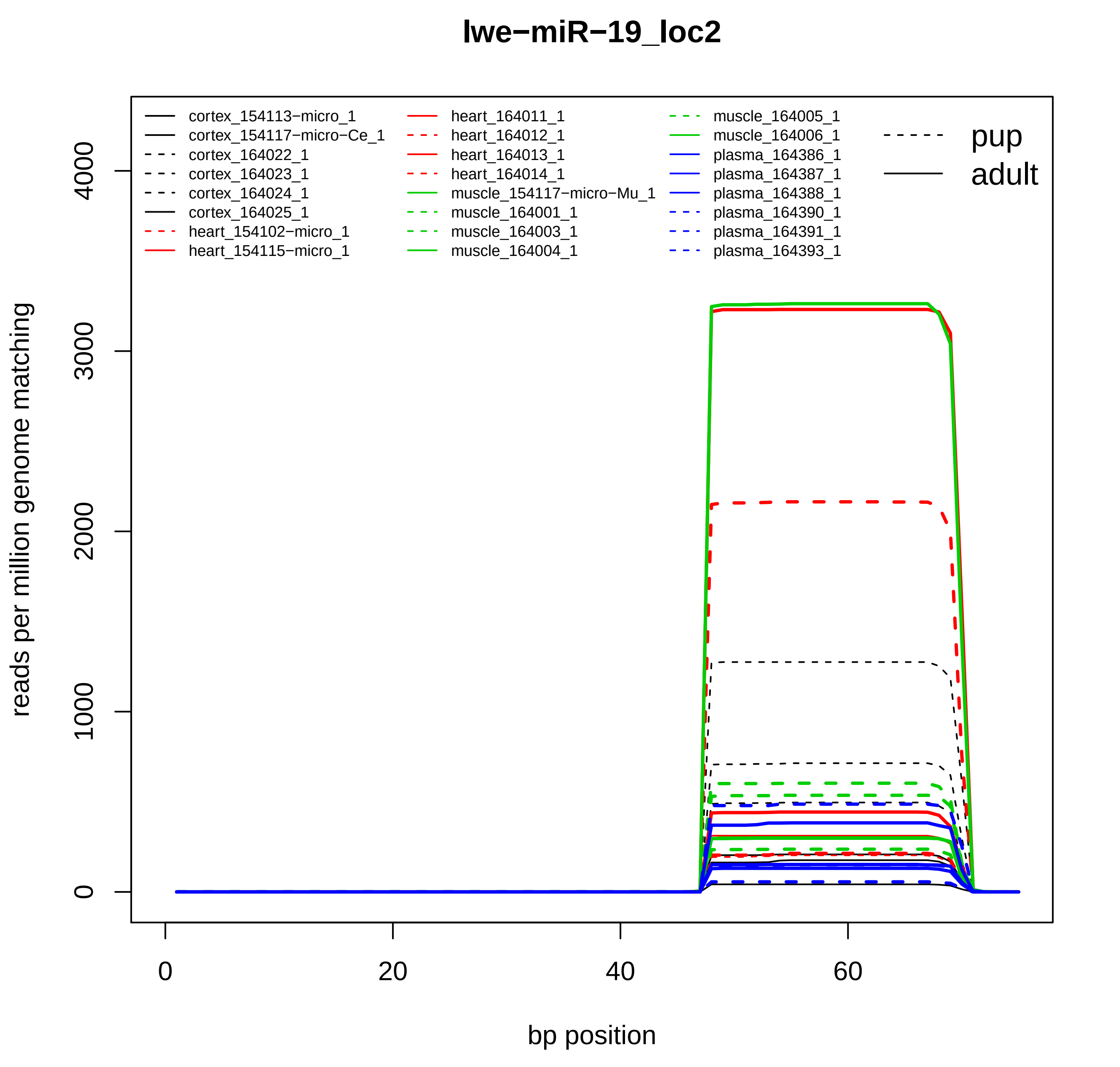
lwe−miR−19_loc2
cortex_154113−micro_1
cortex_154117−micro−Ce_1
cortex_164022_1
cortex_164023_1
cortex_164024_1
cortex_164025_1
heart_154102−micro_1
heart_154115−micro_1
heart_164011_1
heart_164012_1
heart_164013_1
heart_164014_1
muscle_154117−micro−Mu_1
muscle_164001_1
muscle_164003_1
muscle_164004_1
muscle_164005_1
muscle_164006_1
plasma_164386_1
plasma_164387_1
plasma_164388_1
plasma_164390_1
plasma_164391_1
plasma_164393_1
pup
adult
reads per million genome matching
0
1000
2000
3000
4000
0
20
40
60
bp position

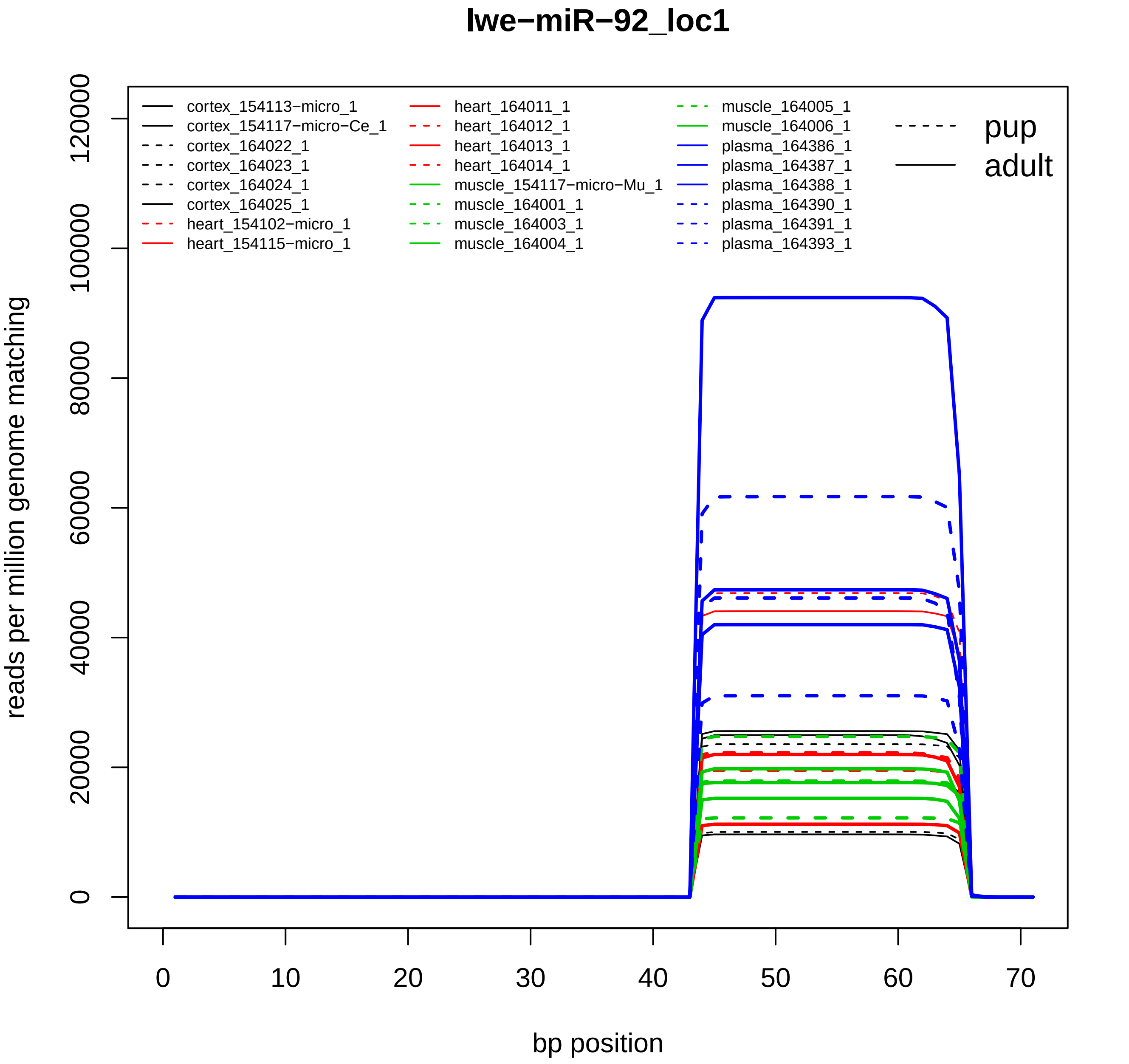

lwe−miR−92_loc1
reads per million genome matching
bp position
0
20000
40000
60000
80000
100000
120000
0
10
20
30
40
50
60
70
cortex_154113−micro_1
cortex_154117−micro−Ce_1
cortex_164022_1
cortex_164023_1
cortex_164024_1
cortex_164025_1
heart_154102−micro_1
heart_154115−micro_1
heart_164011_1
heart_164012_1
heart_164013_1
heart_164014_1
muscle_154117−micro−Mu_1
muscle_164001_1
muscle_164003_1
muscle_164004_1
muscle_164005_1
muscle_164006_1
plasma_164386_1
plasma_164387_1
plasma_164388_1
plasma_164390_1
plasma_164391_1
plasma_164393_1
pup
adult

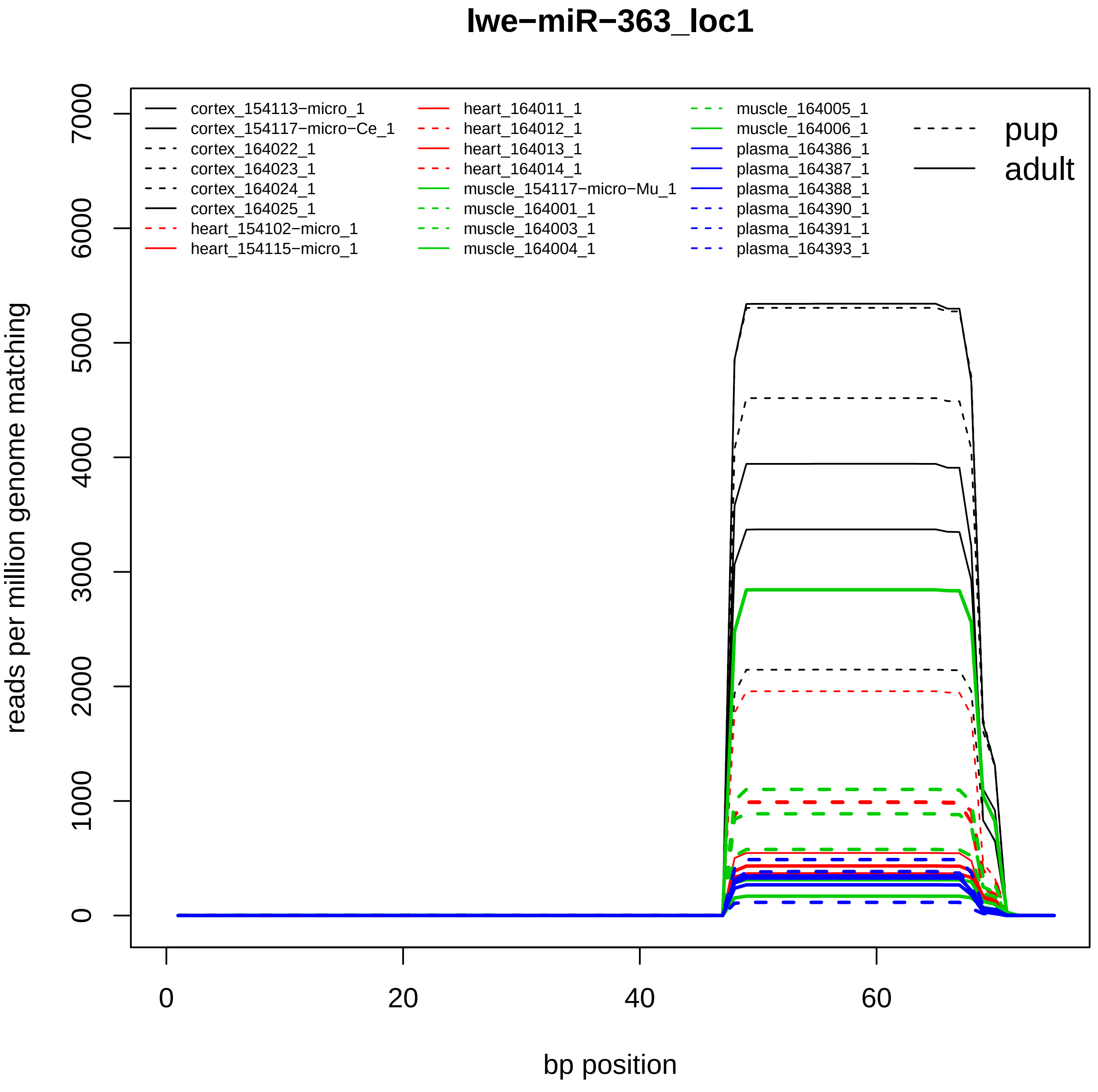
lwe−miR−363_loc1
cortex_154113−micro_1
cortex_154117−micro−Ce_1
cortex_164022_1
cortex_164023_1
cortex_164024_1
cortex_164025_1
heart_154102−micro_1
heart_154115−micro_1
heart_164011_1
heart_164012_1
heart_164013_1
heart_164014_1
muscle_154117−micro−Mu_1
muscle_164001_1
muscle_164003_1
muscle_164004_1
muscle_164005_1
muscle_164006_1
plasma_164386_1
plasma_164387_1
plasma_164388_1
plasma_164390_1
plasma_164391_1
plasma_164393_1
pup
adult
reads per million genome matching
bp position
0
1000
2000
3000
4000
5000
6000
7000
0
20
40
60

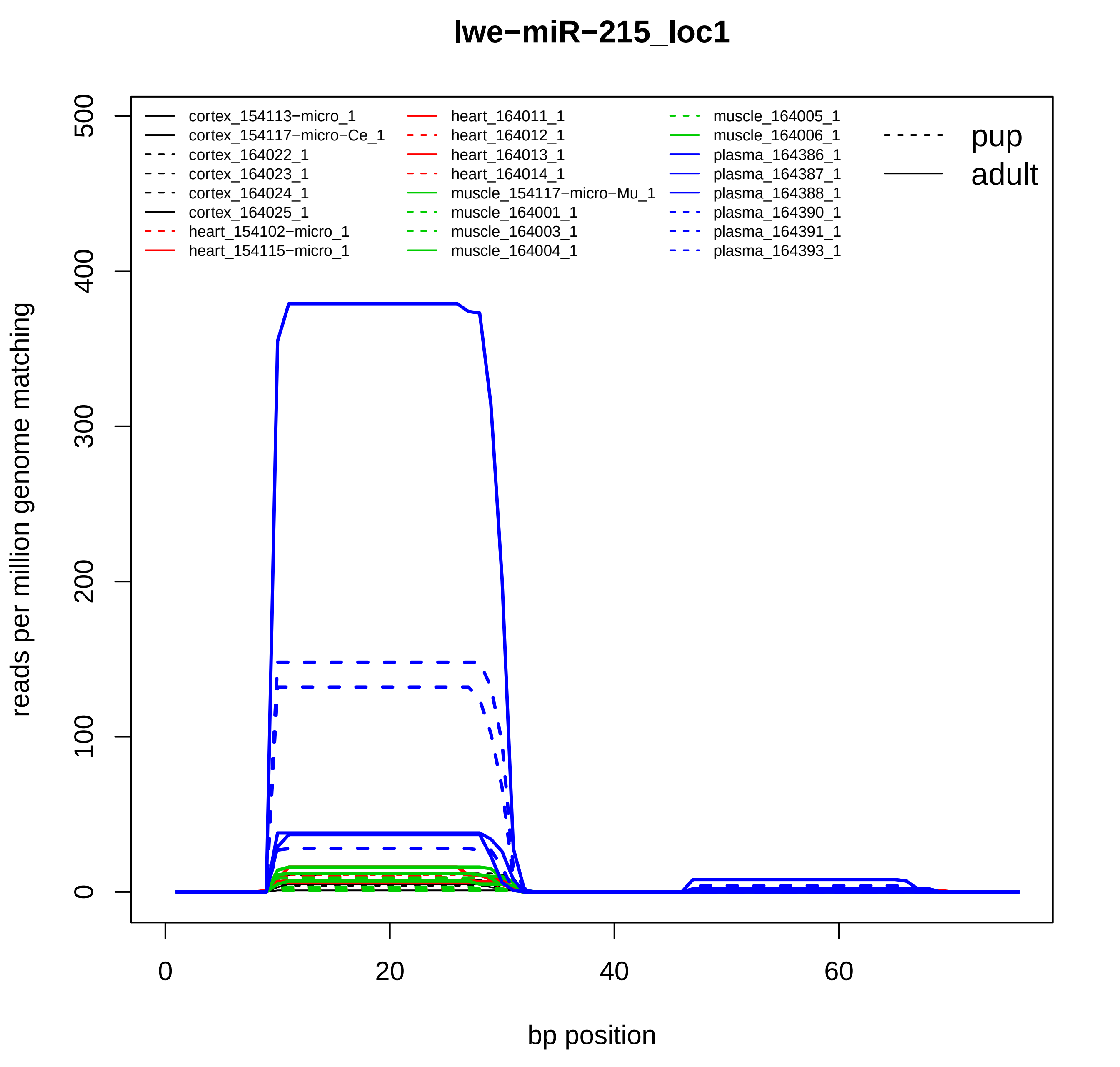

lwe−miR−215_loc1
reads per million genome matching
bp position
0
100
200
300
400
500
0
20
40
60
cortex_154113−micro_1
cortex_154117−micro−Ce_1
cortex_164022_1
cortex_164023_1
cortex_164024_1
cortex_164025_1
heart_154102−micro_1
heart_154115−micro_1
heart_164011_1
heart_164012_1
heart_164013_1
heart_164014_1
muscle_154117−micro−Mu_1
muscle_164001_1
muscle_164003_1
muscle_164004_1
muscle_164005_1
muscle_164006_1
plasma_164386_1
plasma_164387_1
plasma_164388_1
plasma_164390_1
plasma_164391_1
plasma_164393_1
pup
adult

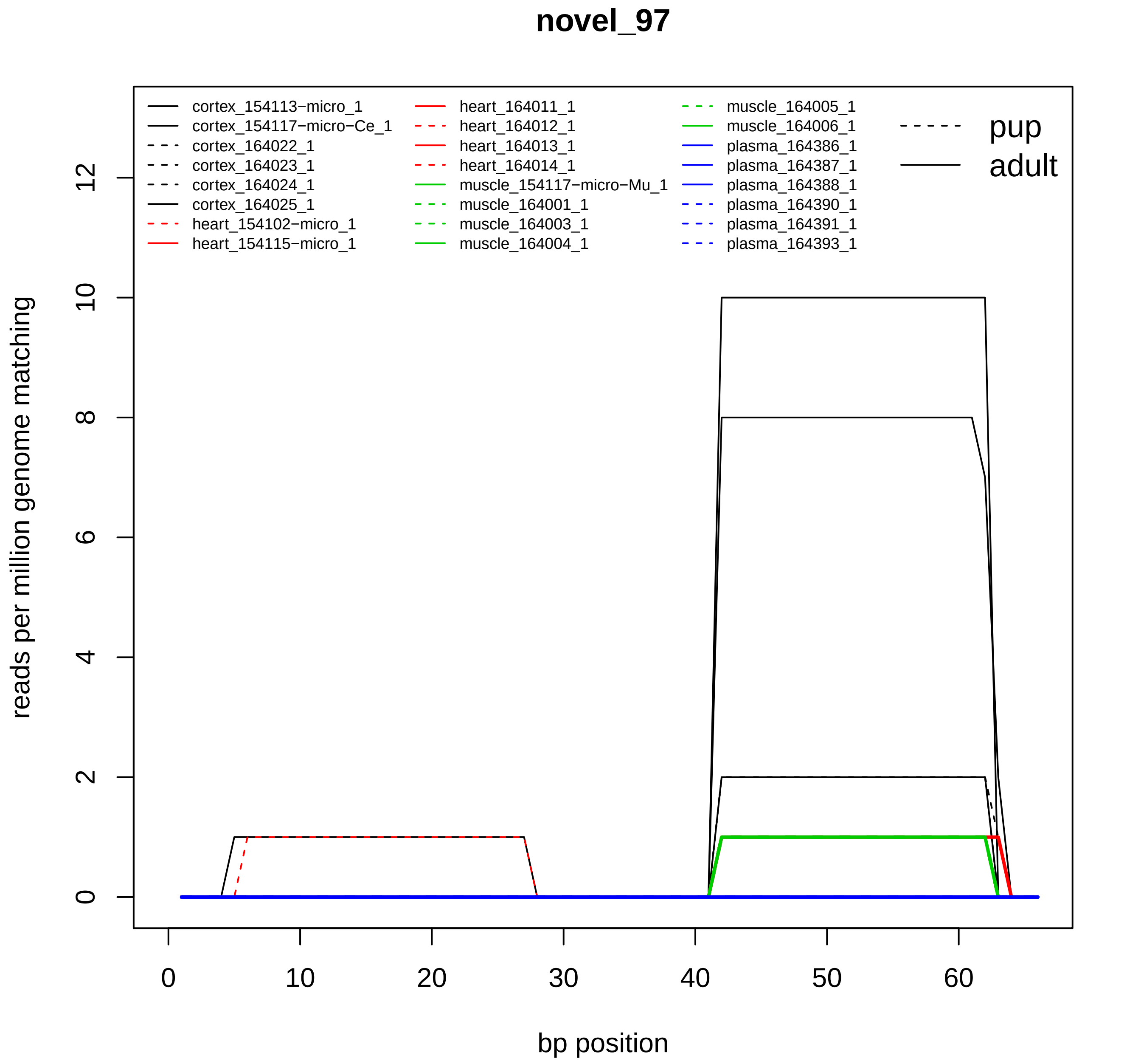

novel_97
cortex_154113−micro_1
cortex_154117−micro−Ce_1
cortex_164022_1
cortex_164023_1
cortex_164024_1
cortex_164025_1
heart_154102−micro_1
heart_154115−micro_1
heart_164011_1
heart_164012_1
heart_164013_1
heart_164014_1
muscle_154117−micro−Mu_1
muscle_164001_1
muscle_164003_1
muscle_164004_1
muscle_164005_1
muscle_164006_1
plasma_164386_1
plasma_164387_1
plasma_164388_1
plasma_164390_1
plasma_164391_1
plasma_164393_1
pup
adult
reads per million genome matching
bp position
0
2
4
6
8
10
12
0
10
20
30
40
50
60

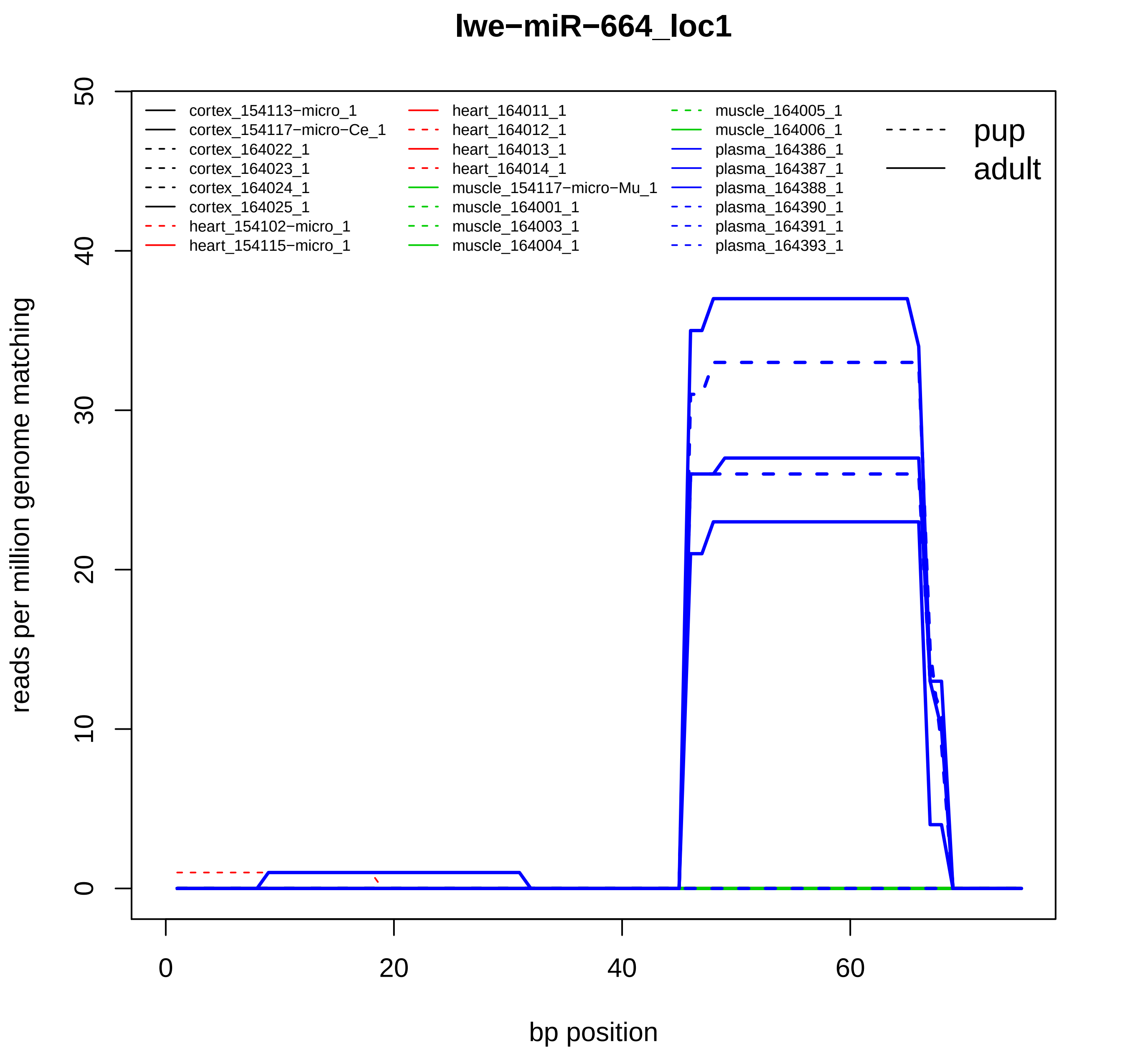
lwe−miR−664_loc1
cortex_154113−micro_1
cortex_154117−micro−Ce_1
cortex_164022_1
cortex_164023_1
cortex_164024_1
cortex_164025_1
heart_154102−micro_1
heart_154115−micro_1
heart_164011_1
heart_164012_1
heart_164013_1
heart_164014_1
muscle_154117−micro−Mu_1
muscle_164001_1
muscle_164003_1
muscle_164004_1
muscle_164005_1
muscle_164006_1
plasma_164386_1
plasma_164387_1
plasma_164388_1
plasma_164390_1
plasma_164391_1
plasma_164393_1
pup
adult
reads per million genome matching
bp position
0
10
20
30
40
50
0
20
40
60

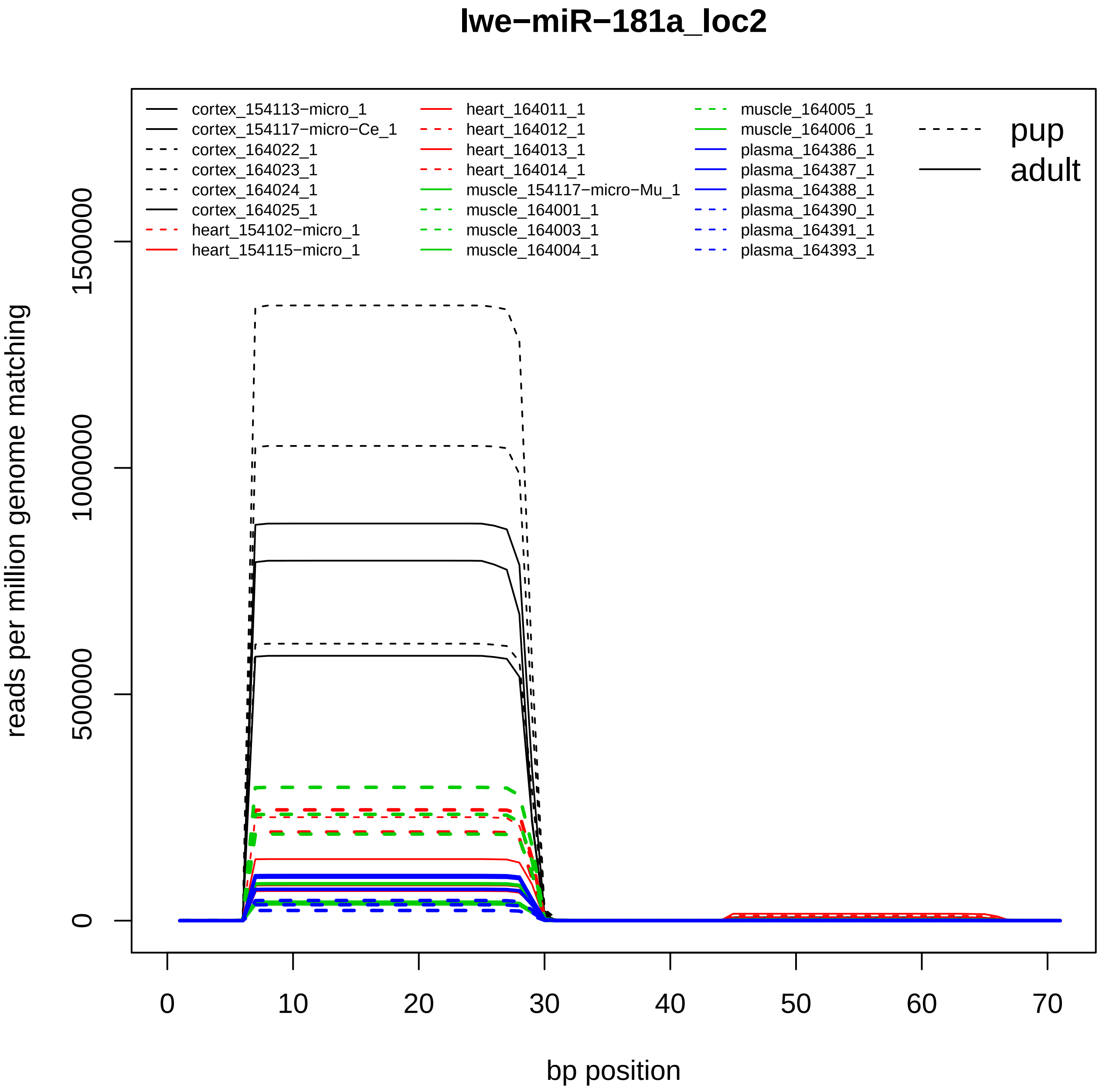

lwe−miR−181a_loc2
cortex_154113−micro_1
cortex_154117−micro−Ce_1
cortex_164022_1
cortex_164023_1
cortex_164024_1
cortex_164025_1
heart_154102−micro_1
heart_154115−micro_1
heart_164011_1
heart_164012_1
heart_164013_1
heart_164014_1
muscle_154117−micro−Mu_1
muscle_164001_1
muscle_164003_1
muscle_164004_1
muscle_164005_1
muscle_164006_1
plasma_164386_1
plasma_164387_1
plasma_164388_1
plasma_164390_1
plasma_164391_1
plasma_164393_1
pup
adult
reads per million genome matching
0
500000
1000000
1500000
bp position
0
10
20
30
40
50
60
70

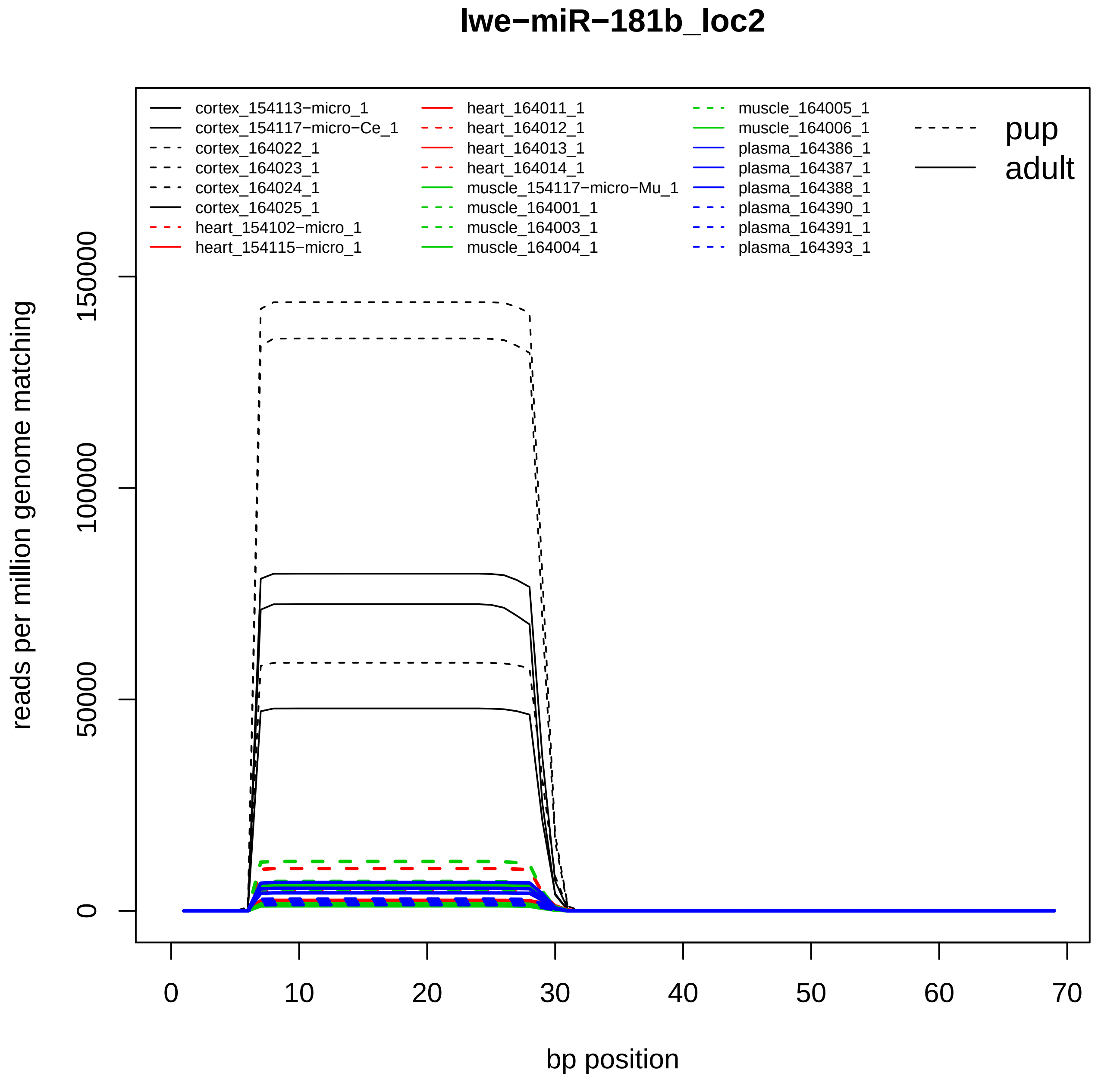
lwe−miR−181b_loc2
cortex_154113−micro_1
cortex_154117−micro−Ce_1
cortex_164022_1
cortex_164023_1
cortex_164024_1
cortex_164025_1
heart_154102−micro_1
heart_154115−micro_1
heart_164011_1
heart_164012_1
heart_164013_1
heart_164014_1
muscle_154117−micro−Mu_1
muscle_164001_1
muscle_164003_1
muscle_164004_1
muscle_164005_1
muscle_164006_1
plasma_164386_1
plasma_164387_1
plasma_164388_1
plasma_164390_1
plasma_164391_1
plasma_164393_1
pup
adult
reads per million genome matching
0
50000
100000
150000
0
10
20
30
40
50
60
70
bp position

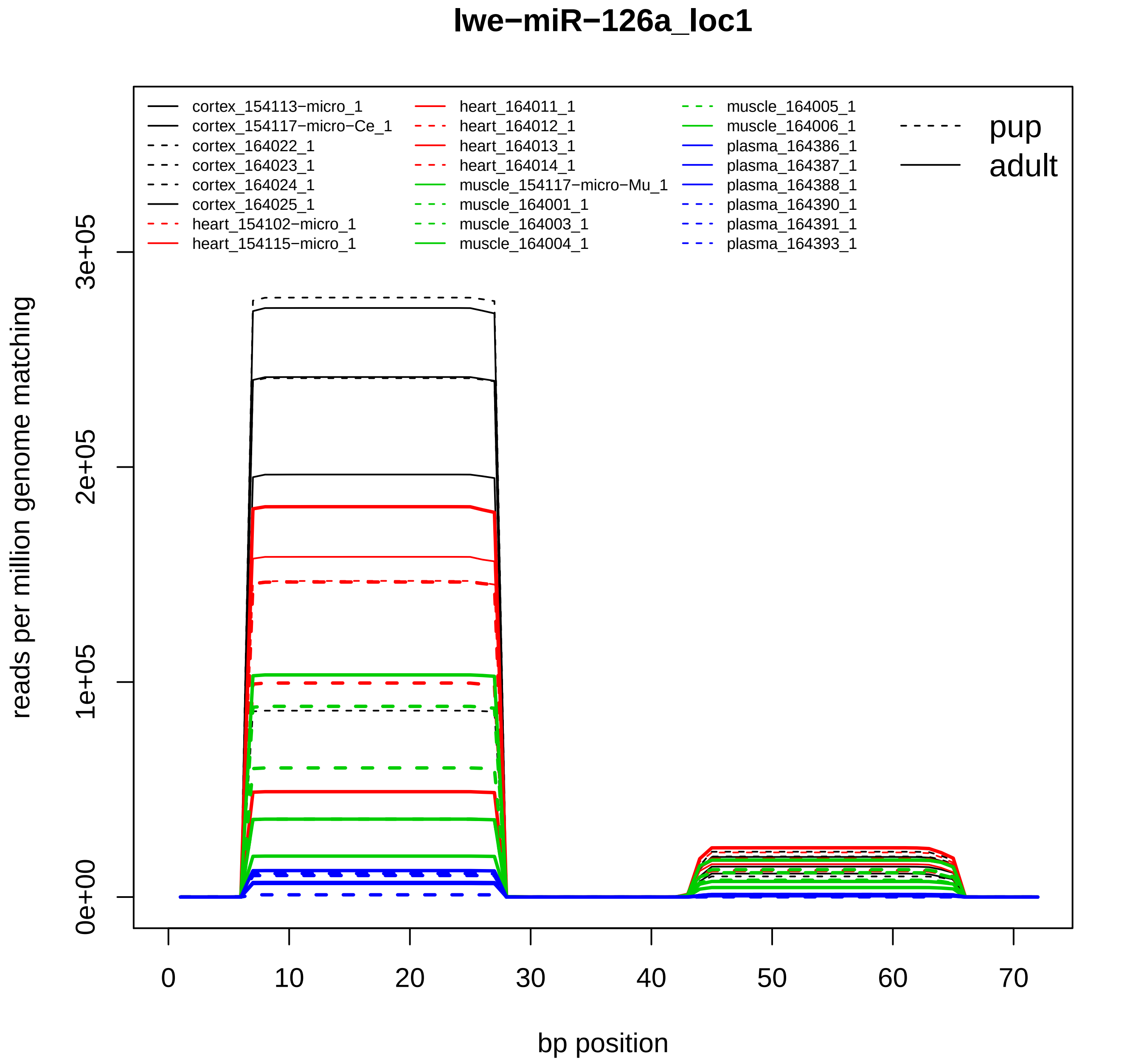
lwe−miR−126a_loc1
cortex_154113−micro_1
cortex_154117−micro−Ce_1
cortex_164022_1
cortex_164023_1
cortex_164024_1
cortex_164025_1
heart_154102−micro_1
heart_154115−micro_1
heart_164011_1
heart_164012_1
heart_164013_1
heart_164014_1
muscle_154117−micro−Mu_1
muscle_164001_1
muscle_164003_1
muscle_164004_1
muscle_164005_1
muscle_164006_1
plasma_164386_1
plasma_164387_1
plasma_164388_1
plasma_164390_1
plasma_164391_1
plasma_164393_1
pup
adult
reads per million genome matching
0e+00
1e+05
2e+05
3e+05
0
10
20
30
40
50
60
70
bp position

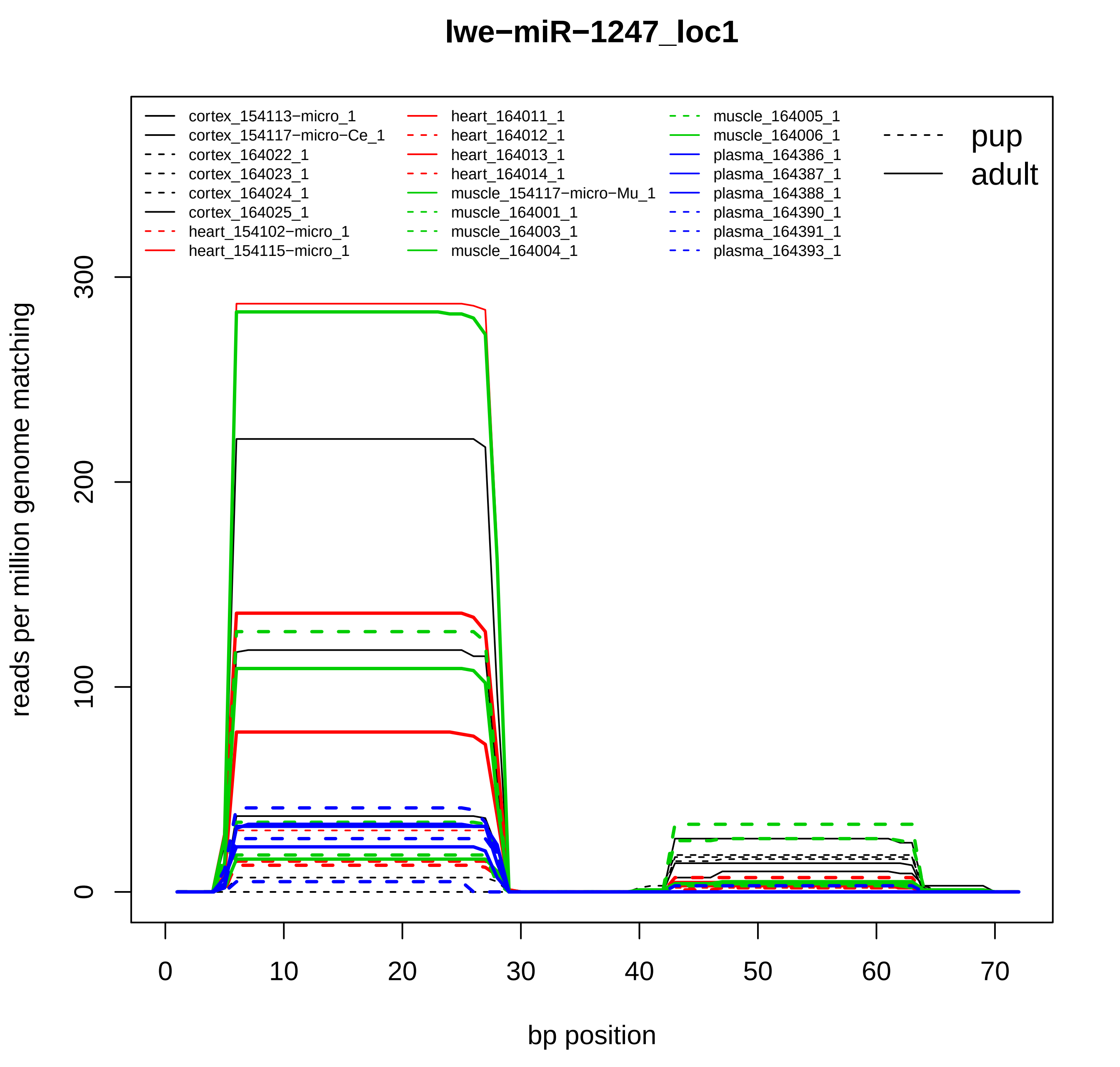
lwe−miR−1247_loc1
cortex_154113−micro_1
cortex_154117−micro−Ce_1
cortex_164022_1
cortex_164023_1
cortex_164024_1
cortex_164025_1
heart_154102−micro_1
heart_154115−micro_1
heart_164011_1
heart_164012_1
heart_164013_1
heart_164014_1
muscle_154117−micro−Mu_1
muscle_164001_1
muscle_164003_1
muscle_164004_1
muscle_164005_1
muscle_164006_1
plasma_164386_1
plasma_164387_1
plasma_164388_1
plasma_164390_1
plasma_164391_1
plasma_164393_1
pup
adult
reads per million genome matching
0
100
200
300
bp position
0
10
20
30
40
50
60
70

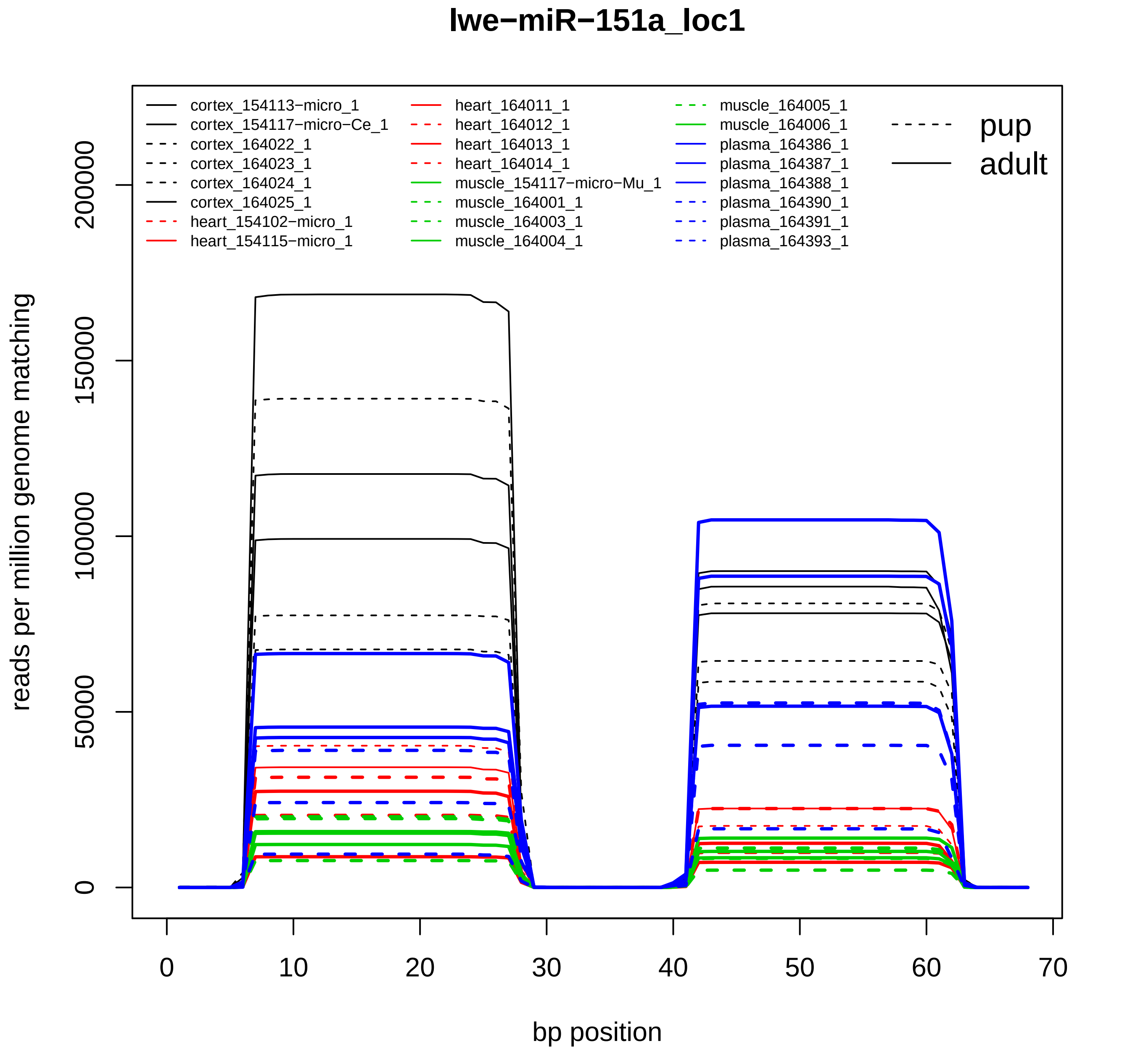
lwe−miR−151a_loc1
cortex_154113−micro_1
cortex_154117−micro−Ce_1
cortex_164022_1
cortex_164023_1
cortex_164024_1
cortex_164025_1
heart_154102−micro_1
heart_154115−micro_1
heart_164011_1
heart_164012_1
heart_164013_1
heart_164014_1
muscle_154117−micro−Mu_1
muscle_164001_1
muscle_164003_1
muscle_164004_1
muscle_164005_1
muscle_164006_1
plasma_164386_1
plasma_164387_1
plasma_164388_1
plasma_164390_1
plasma_164391_1
plasma_164393_1
pup
adult
reads per million genome matching
0
50000
100000
150000
200000
bp position
0
10
20
30
40
50
60
70

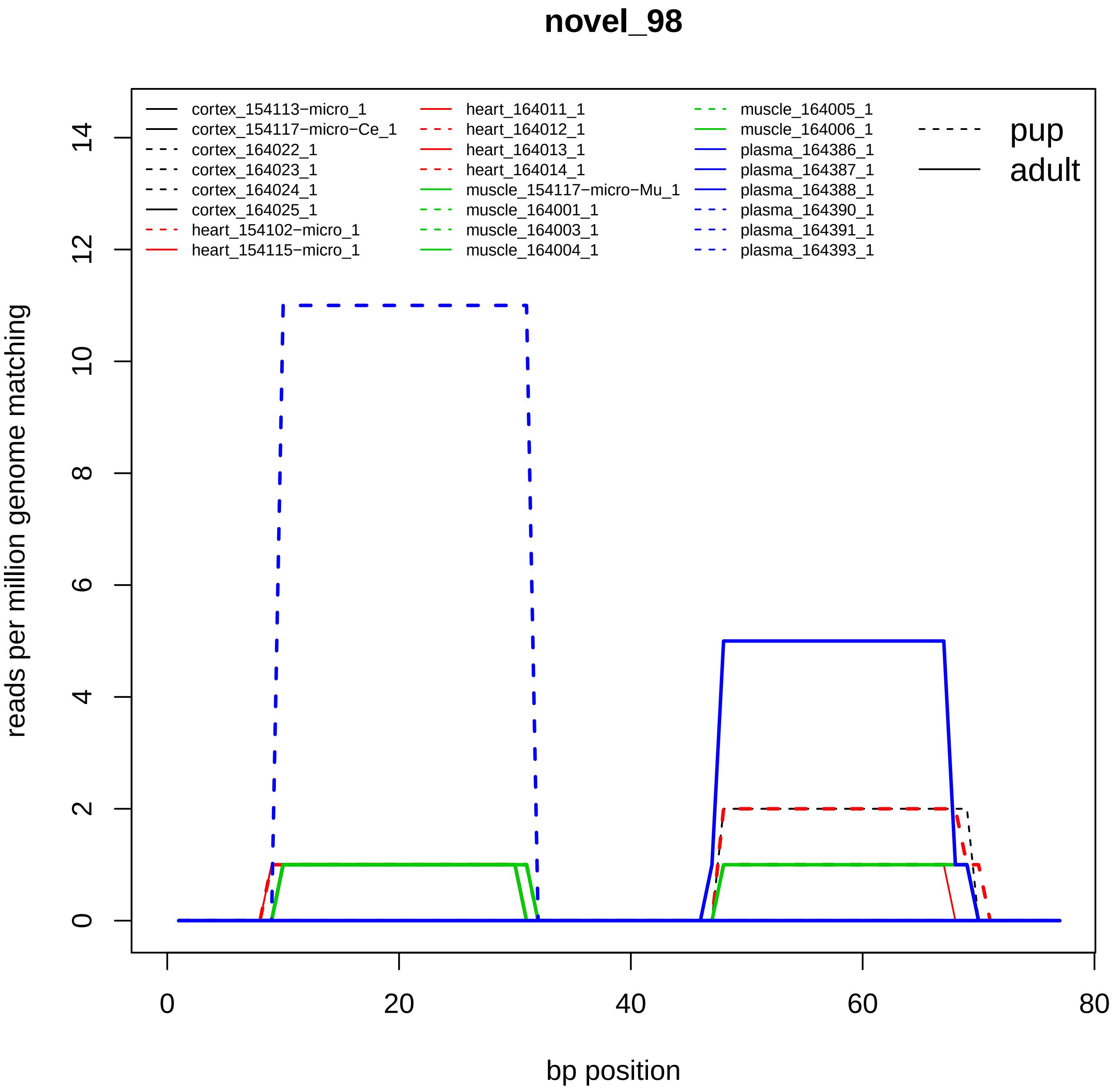
novel_98
cortex_154113−micro_1
cortex_154117−micro−Ce_1
cortex_164022_1
cortex_164023_1
cortex_164024_1
cortex_164025_1
heart_154102−micro_1
heart_154115−micro_1
heart_164011_1
heart_164012_1
heart_164013_1
heart_164014_1
muscle_154117−micro−Mu_1
muscle_164001_1
muscle_164003_1
muscle_164004_1
muscle_164005_1
muscle_164006_1
plasma_164386_1
plasma_164387_1
plasma_164388_1
plasma_164390_1
plasma_164391_1
plasma_164393_1
pup
adult
reads per million genome matching
0
2
4
6
8
10
12
14
0
20
40
60
80
bp position

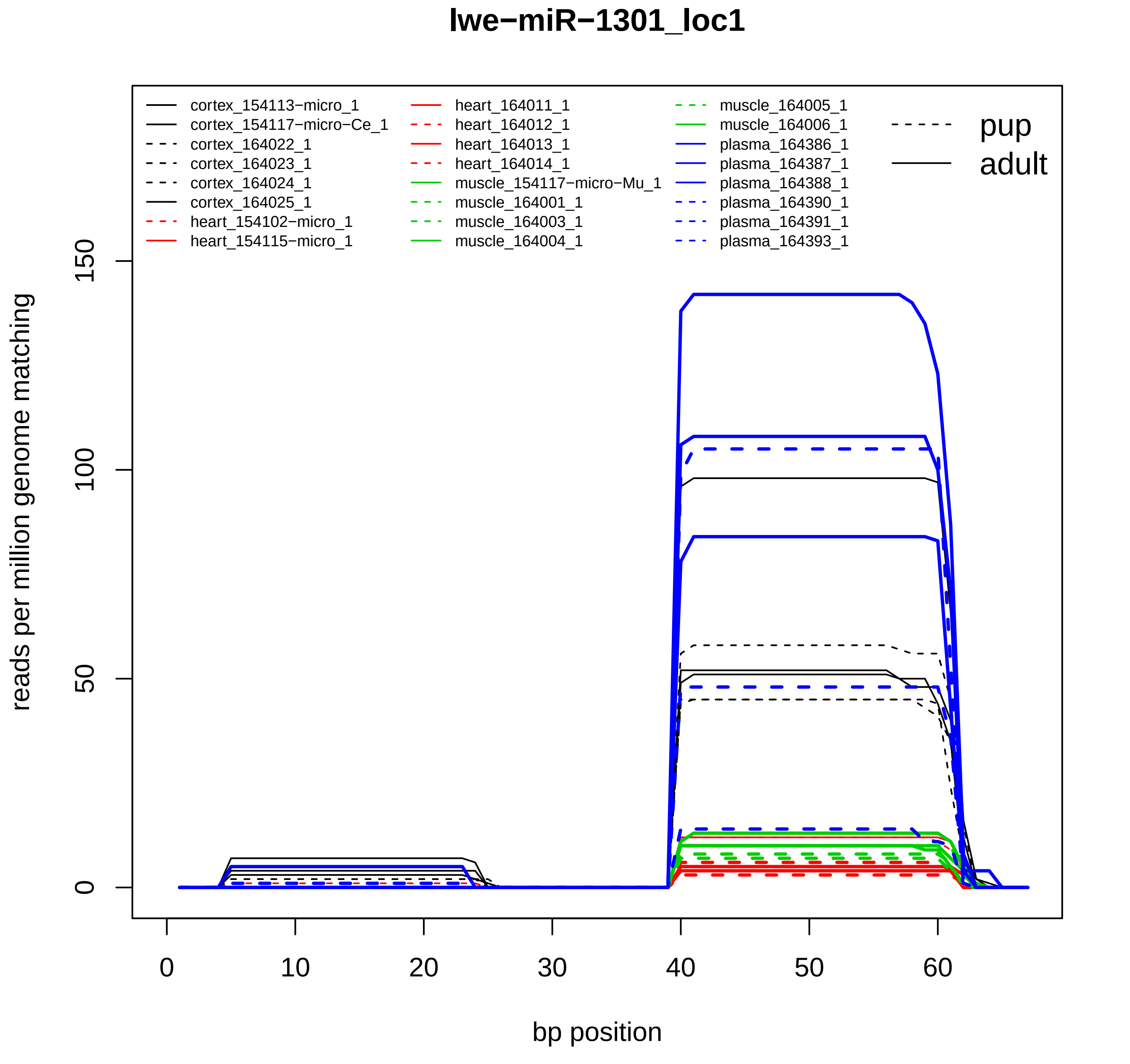
lwe−miR−1301_loc1
cortex_154113−micro_1
cortex_154117−micro−Ce_1
cortex_164022_1
cortex_164023_1
cortex_164024_1
cortex_164025_1
heart_154102−micro_1
heart_154115−micro_1
heart_164011_1
heart_164012_1
heart_164013_1
heart_164014_1
muscle_154117−micro−Mu_1
muscle_164001_1
muscle_164003_1
muscle_164004_1
muscle_164005_1
muscle_164006_1
plasma_164386_1
plasma_164387_1
plasma_164388_1
plasma_164390_1
plasma_164391_1
plasma_164393_1
pup
adult
reads per million genome matching
0
50
100
150
0
10
20
30
40
50
60
bp position

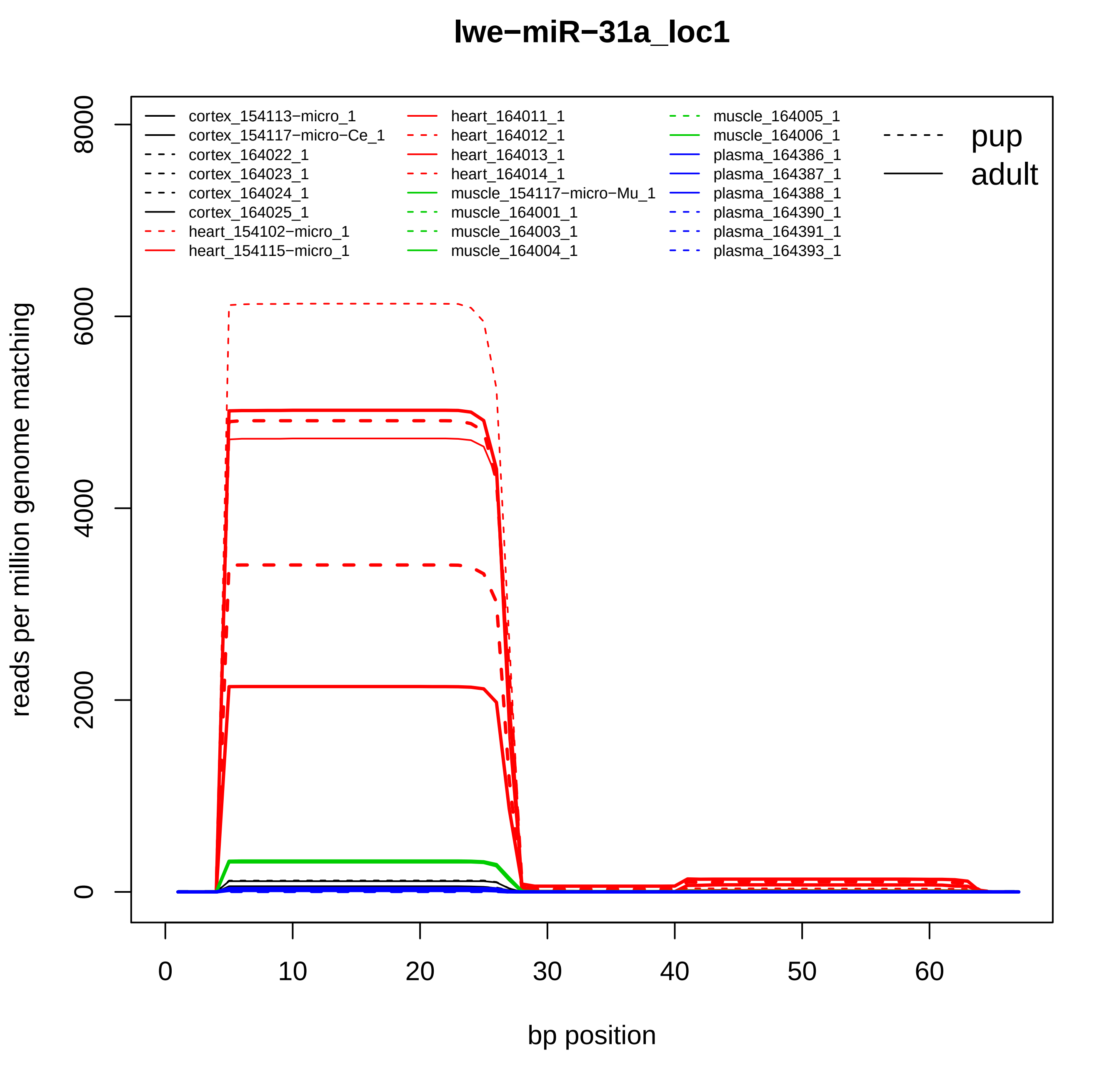

lwe−miR−31a_loc1
reads per million genome matching
bp position
0
2000
4000
6000
8000
0
10
20
30
40
50
60
cortex_154113−micro_1
cortex_154117−micro−Ce_1
cortex_164022_1
cortex_164023_1
cortex_164024_1
cortex_164025_1
heart_154102−micro_1
heart_154115−micro_1
heart_164011_1
heart_164012_1
heart_164013_1
heart_164014_1
muscle_154117−micro−Mu_1
muscle_164001_1
muscle_164003_1
muscle_164004_1
muscle_164005_1
muscle_164006_1
plasma_164386_1
plasma_164387_1
plasma_164388_1
plasma_164390_1
plasma_164391_1
plasma_164393_1
pup
adult

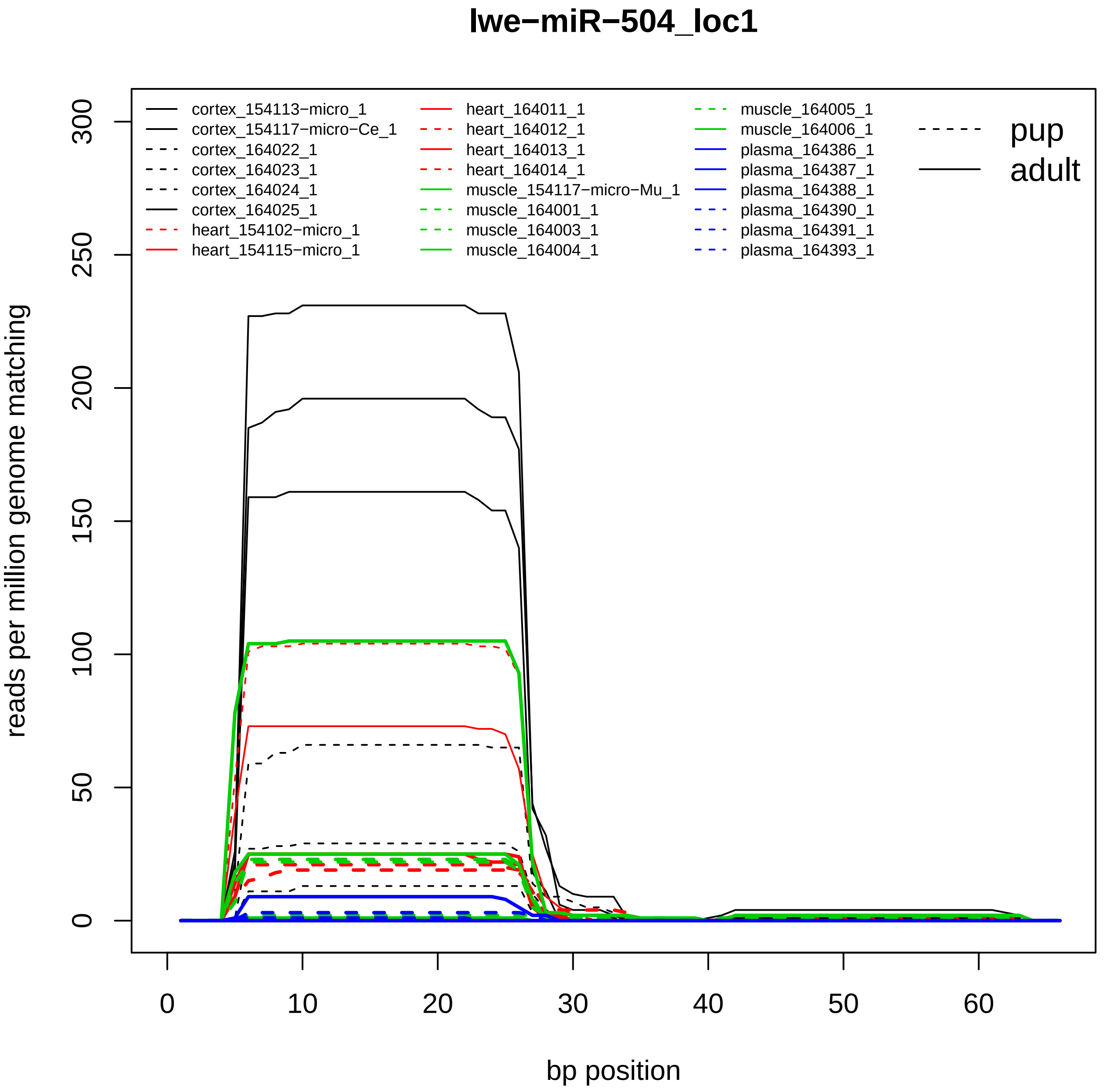

lwe−miR−504_loc1
cortex_154113−micro_1
cortex_154117−micro−Ce_1
cortex_164022_1
cortex_164023_1
cortex_164024_1
cortex_164025_1
heart_154102−micro_1
heart_154115−micro_1
heart_164011_1
heart_164012_1
heart_164013_1
heart_164014_1
muscle_154117−micro−Mu_1
muscle_164001_1
muscle_164003_1
muscle_164004_1
muscle_164005_1
muscle_164006_1
plasma_164386_1
plasma_164387_1
plasma_164388_1
plasma_164390_1
plasma_164391_1
plasma_164393_1
pup
adult
reads per million genome matching
bp position
0
50
100
150
200
250
300
0
10
20
30
40
50
60

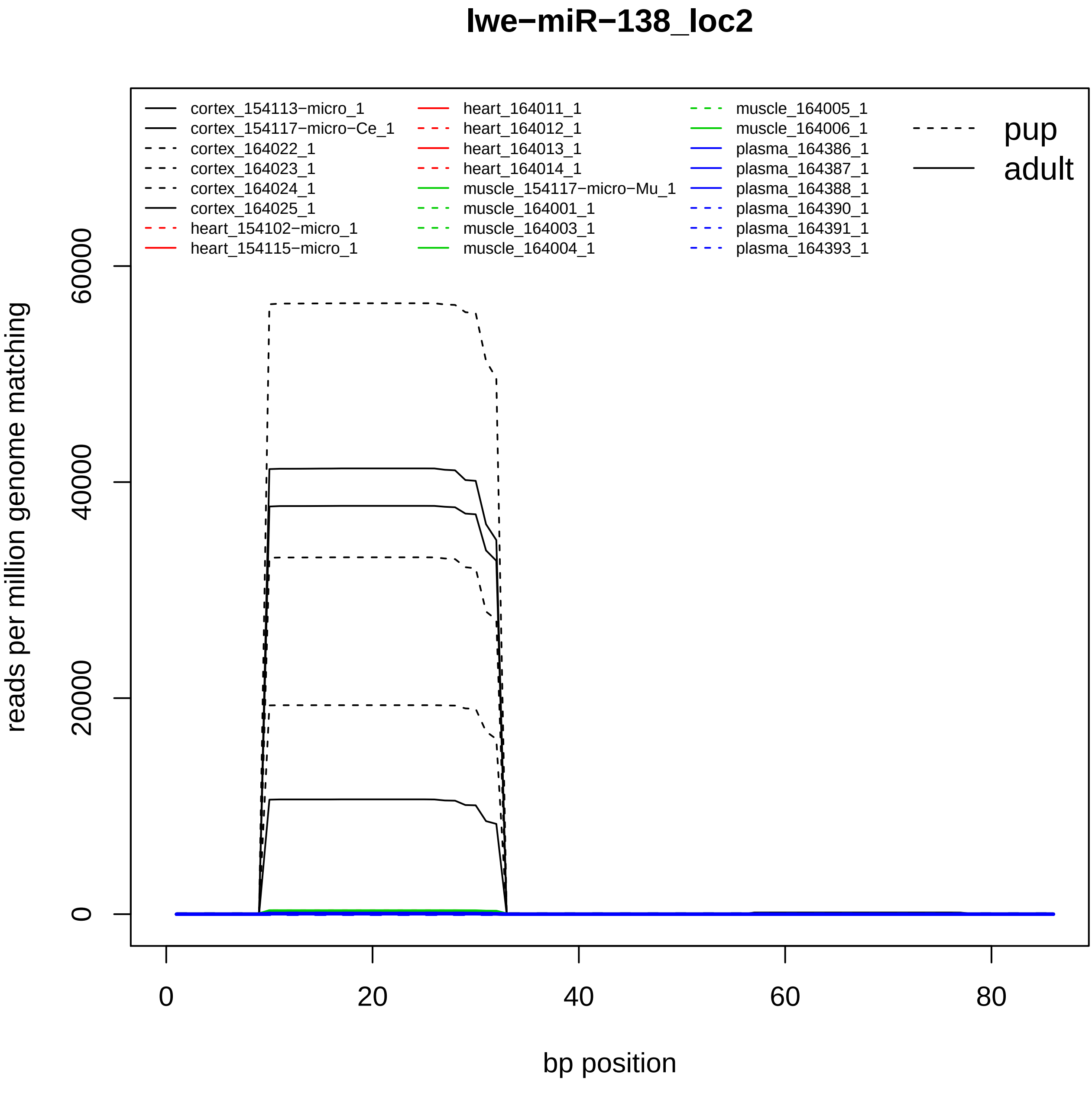
lwe−miR−138_loc2
cortex_154113−micro_1
cortex_154117−micro−Ce_1
cortex_164022_1
cortex_164023_1
cortex_164024_1
cortex_164025_1
heart_154102−micro_1
heart_154115−micro_1
heart_164011_1
heart_164012_1
heart_164013_1
heart_164014_1
muscle_154117−micro−Mu_1
muscle_164001_1
muscle_164003_1
muscle_164004_1
muscle_164005_1
muscle_164006_1
plasma_164386_1
plasma_164387_1
plasma_164388_1
plasma_164390_1
plasma_164391_1
plasma_164393_1
pup
adult
reads per million genome matching
0
20000
40000
60000
0
20
40
60
80
bp position

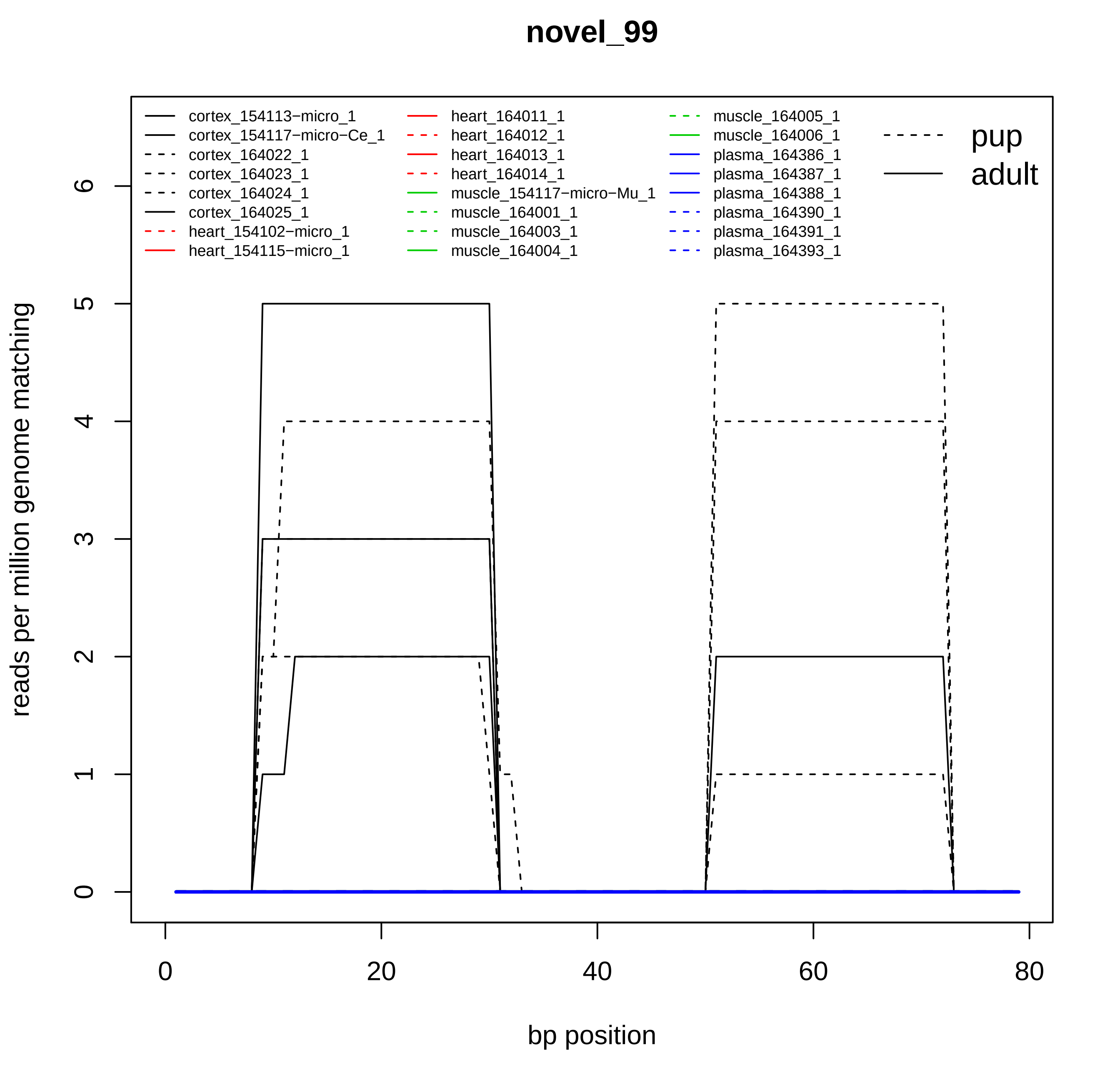
novel_99
reads per million genome matching
bp position
0
1
2
3
4
5
6
0
20
40
60
80
cortex_154113−micro_1
cortex_154117−micro−Ce_1
cortex_164022_1
cortex_164023_1
cortex_164024_1
cortex_164025_1
heart_154102−micro_1
heart_154115−micro_1
heart_164011_1
heart_164012_1
heart_164013_1
heart_164014_1
muscle_154117−micro−Mu_1
muscle_164001_1
muscle_164003_1
muscle_164004_1
muscle_164005_1
muscle_164006_1
plasma_164386_1
plasma_164387_1
plasma_164388_1
plasma_164390_1
plasma_164391_1
plasma_164393_1
pup
adult

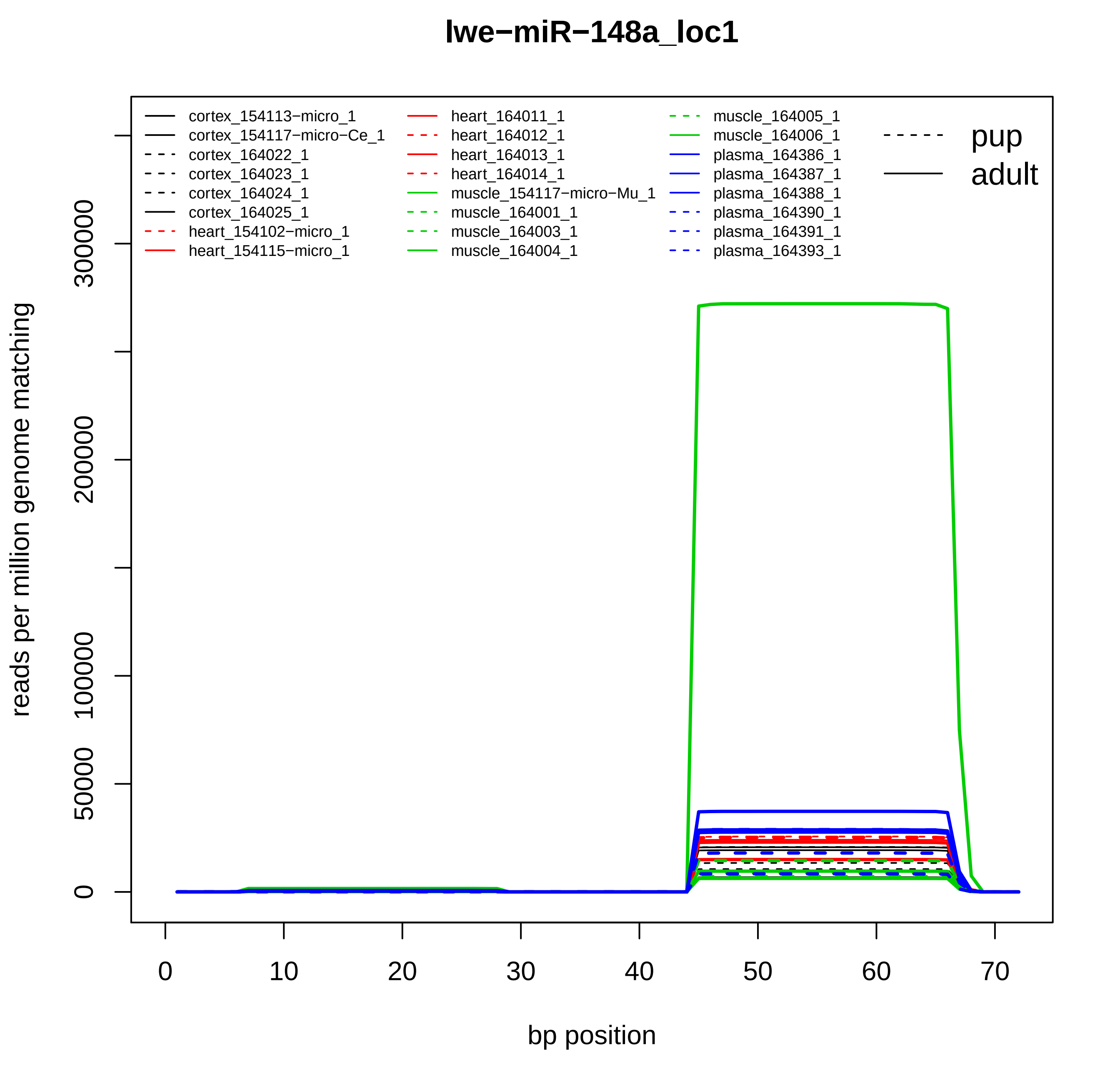

lwe−miR−148a_loc1
cortex_154113−micro_1
cortex_154117−micro−Ce_1
cortex_164022_1
cortex_164023_1
cortex_164024_1
cortex_164025_1
heart_154102−micro_1
heart_154115−micro_1
heart_164011_1
heart_164012_1
heart_164013_1
heart_164014_1
muscle_154117−micro−Mu_1
muscle_164001_1
muscle_164003_1
muscle_164004_1
muscle_164005_1
muscle_164006_1
plasma_164386_1
plasma_164387_1
plasma_164388_1
plasma_164390_1
plasma_164391_1
plasma_164393_1
pup
adult
reads per million genome matching
0
50000
100000
200000
300000
bp position
0
10
20
30
40
50
60
70

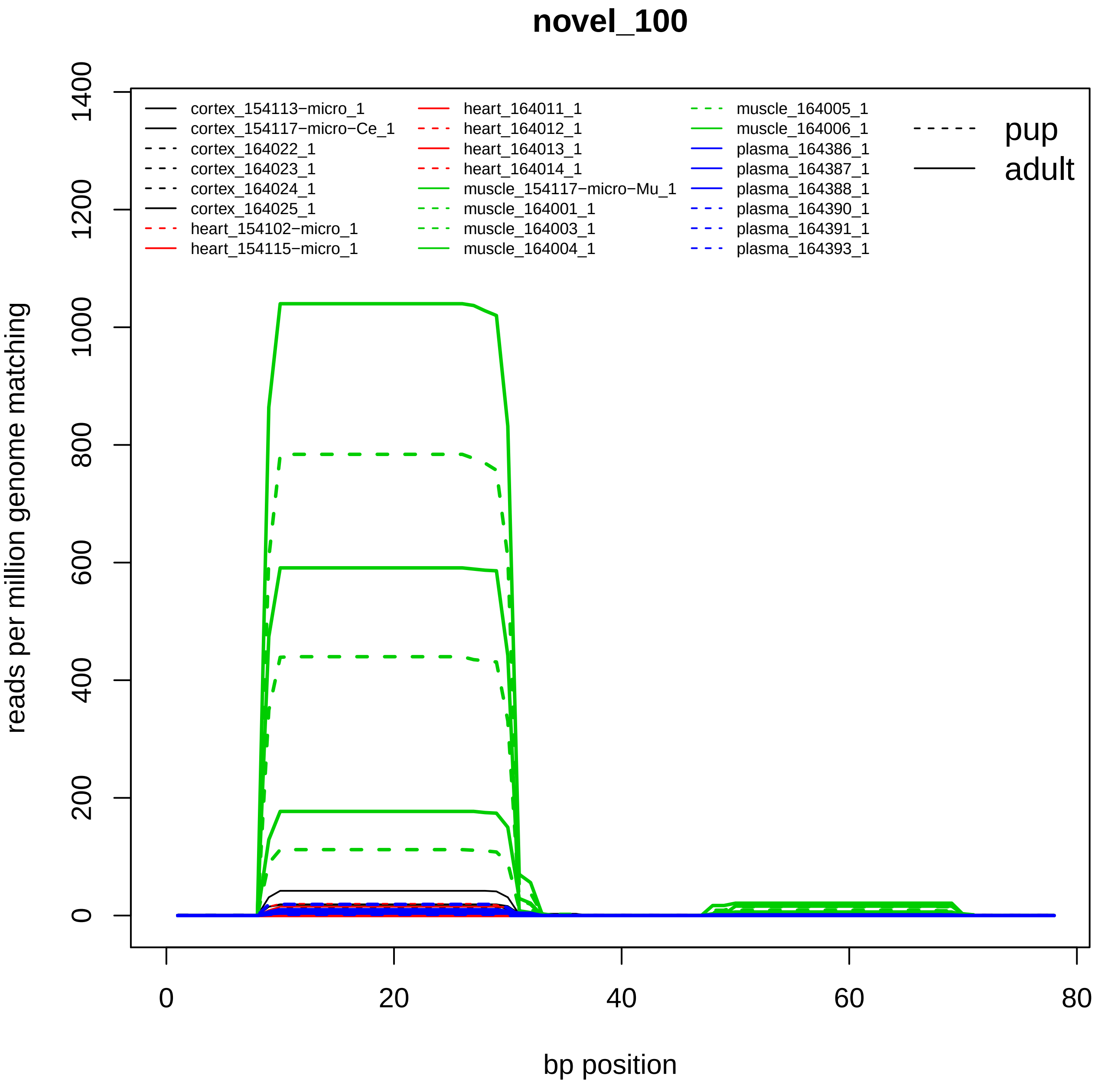

novel_100
cortex_154113−micro_1
cortex_154117−micro−Ce_1
cortex_164022_1
cortex_164023_1
cortex_164024_1
cortex_164025_1
heart_154102−micro_1
heart_154115−micro_1
heart_164011_1
heart_164012_1
heart_164013_1
heart_164014_1
muscle_154117−micro−Mu_1
muscle_164001_1
muscle_164003_1
muscle_164004_1
muscle_164005_1
muscle_164006_1
plasma_164386_1
plasma_164387_1
plasma_164388_1
plasma_164390_1
plasma_164391_1
plasma_164393_1
pup
adult
reads per million genome matching
0
200
400
600
800
1000
1200
1400
0
20
40
60
80
bp position

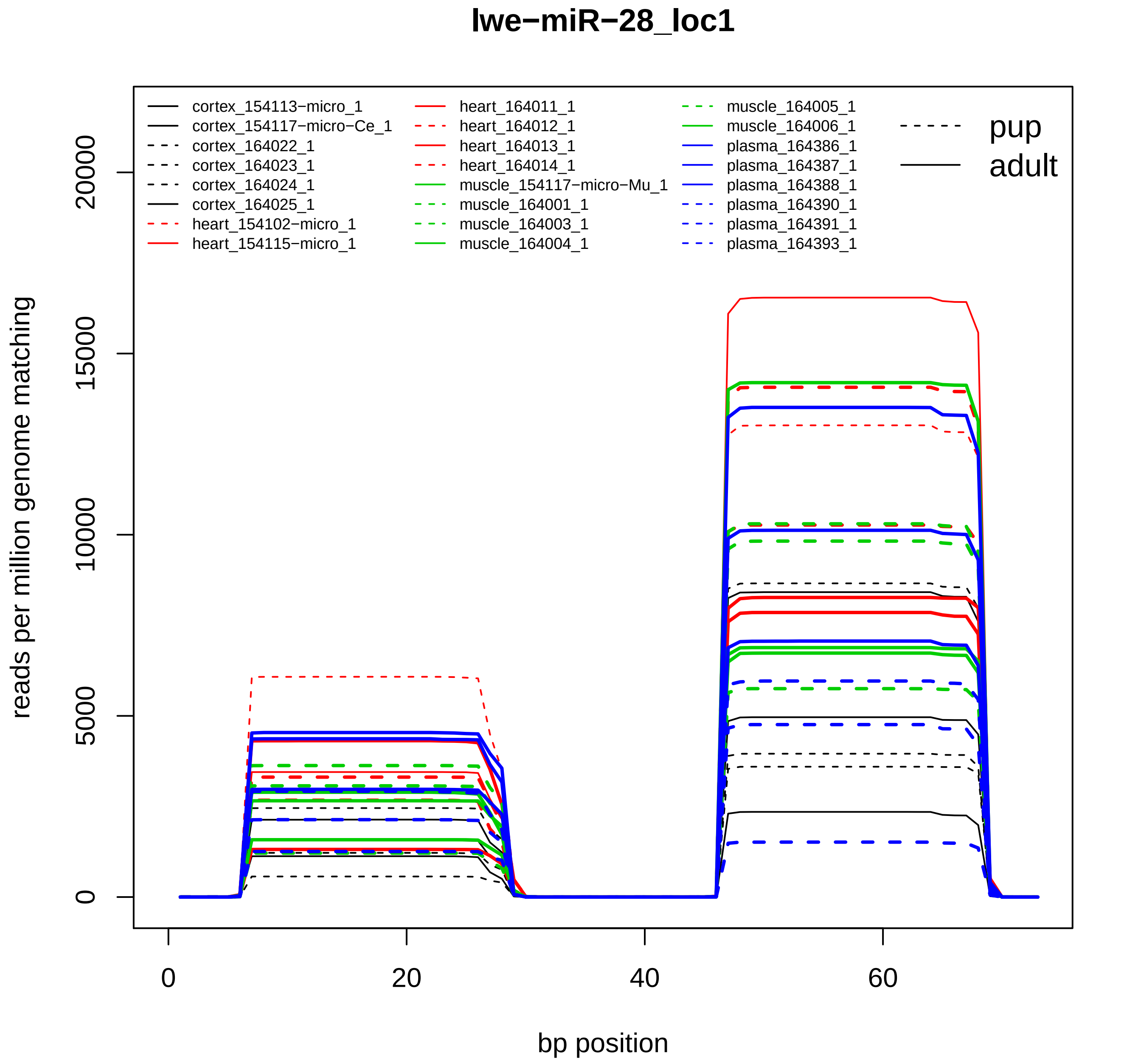
lwe−miR−28_loc1
cortex_154113−micro_1
cortex_154117−micro−Ce_1
cortex_164022_1
cortex_164023_1
cortex_164024_1
cortex_164025_1
heart_154102−micro_1
heart_154115−micro_1
heart_164011_1
heart_164012_1
heart_164013_1
heart_164014_1
muscle_154117−micro−Mu_1
muscle_164001_1
muscle_164003_1
muscle_164004_1
muscle_164005_1
muscle_164006_1
plasma_164386_1
plasma_164387_1
plasma_164388_1
plasma_164390_1
plasma_164391_1
plasma_164393_1
pup
adult
reads per million genome matching
0
5000
10000
15000
20000
bp position
0
20
40
60

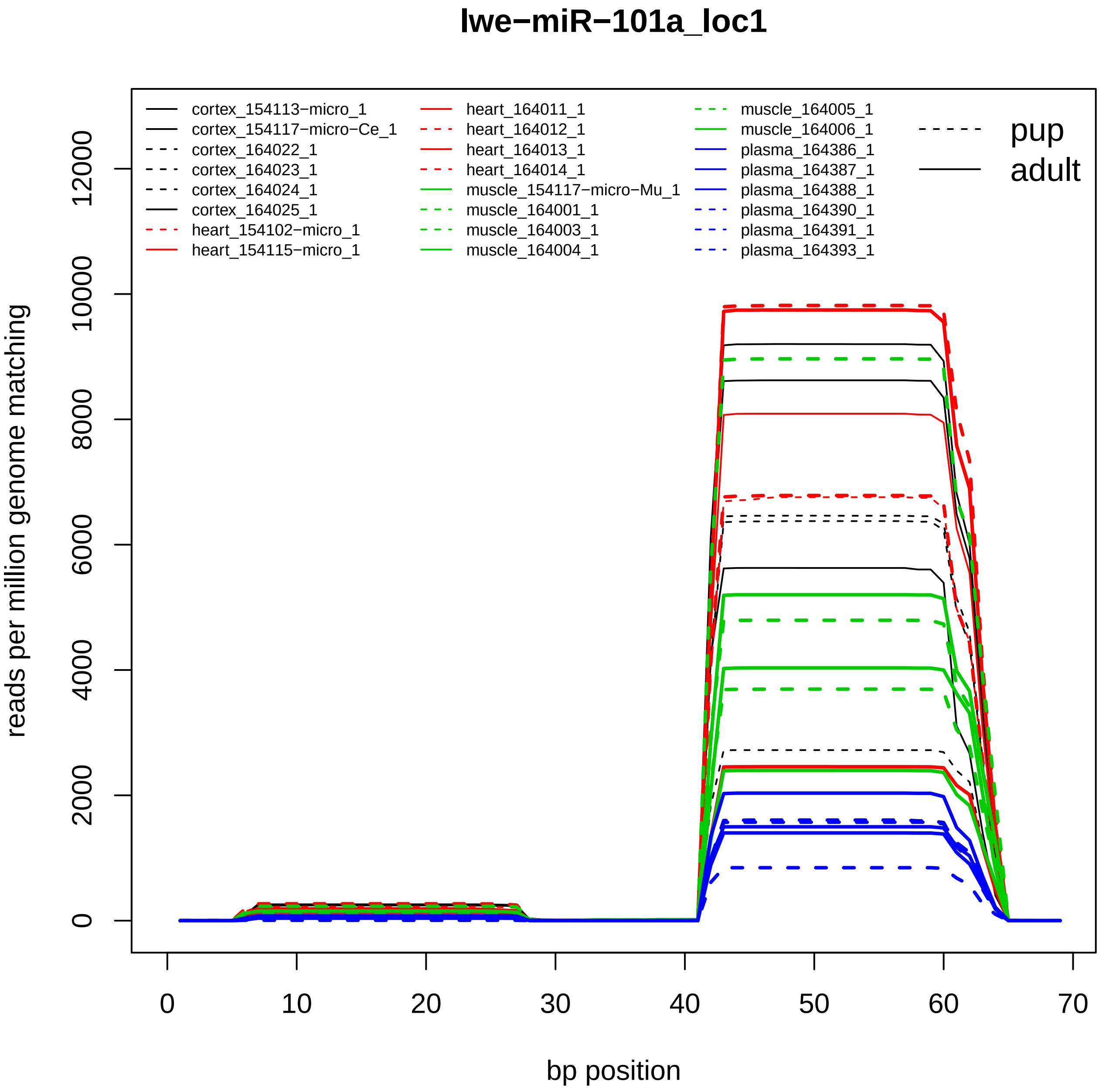

lwe−miR−101a_loc1
cortex_154113−micro_1
cortex_154117−micro−Ce_1
cortex_164022_1
cortex_164023_1
cortex_164024_1
cortex_164025_1
heart_154102−micro_1
heart_154115−micro_1
heart_164011_1
heart_164012_1
heart_164013_1
heart_164014_1
muscle_154117−micro−Mu_1
muscle_164001_1
muscle_164003_1
muscle_164004_1
muscle_164005_1
muscle_164006_1
plasma_164386_1
plasma_164387_1
plasma_164388_1
plasma_164390_1
plasma_164391_1
plasma_164393_1
pup
adult
reads per million genome matching
0
2000
4000
6000
8000
10000
12000
bp position
0
10
20
30
40
50
60
70

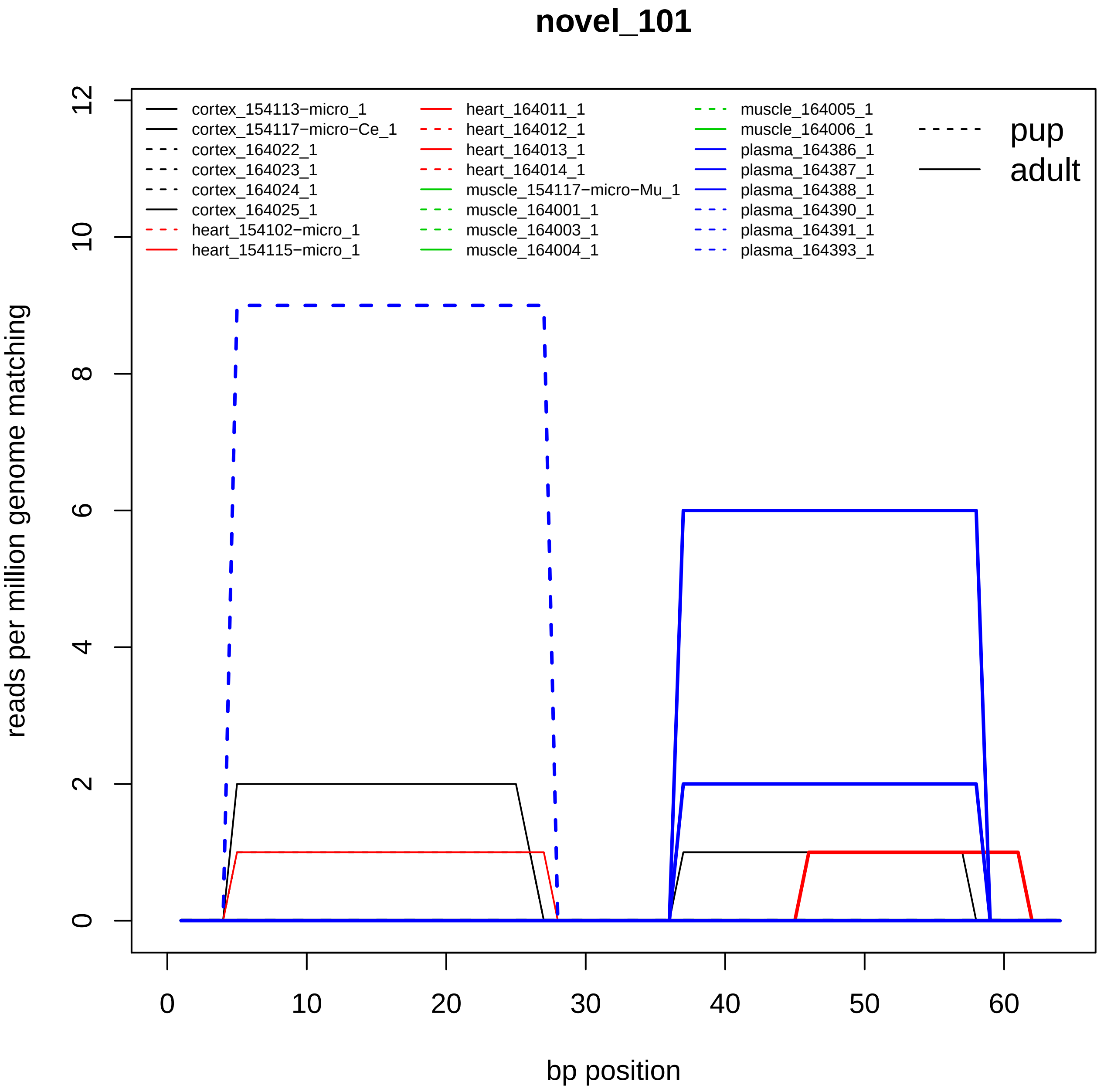
novel_101
cortex_154113−micro_1
cortex_154117−micro−Ce_1
cortex_164022_1
cortex_164023_1
cortex_164024_1
cortex_164025_1
heart_154102−micro_1
heart_154115−micro_1
heart_164011_1
heart_164012_1
heart_164013_1
heart_164014_1
muscle_154117−micro−Mu_1
muscle_164001_1
muscle_164003_1
muscle_164004_1
muscle_164005_1
muscle_164006_1
plasma_164386_1
plasma_164387_1
plasma_164388_1
plasma_164390_1
plasma_164391_1
plasma_164393_1
pup
adult
reads per million genome matching
bp position
0
2
4
6
8
10
12
0
10
20
30
40
50
60

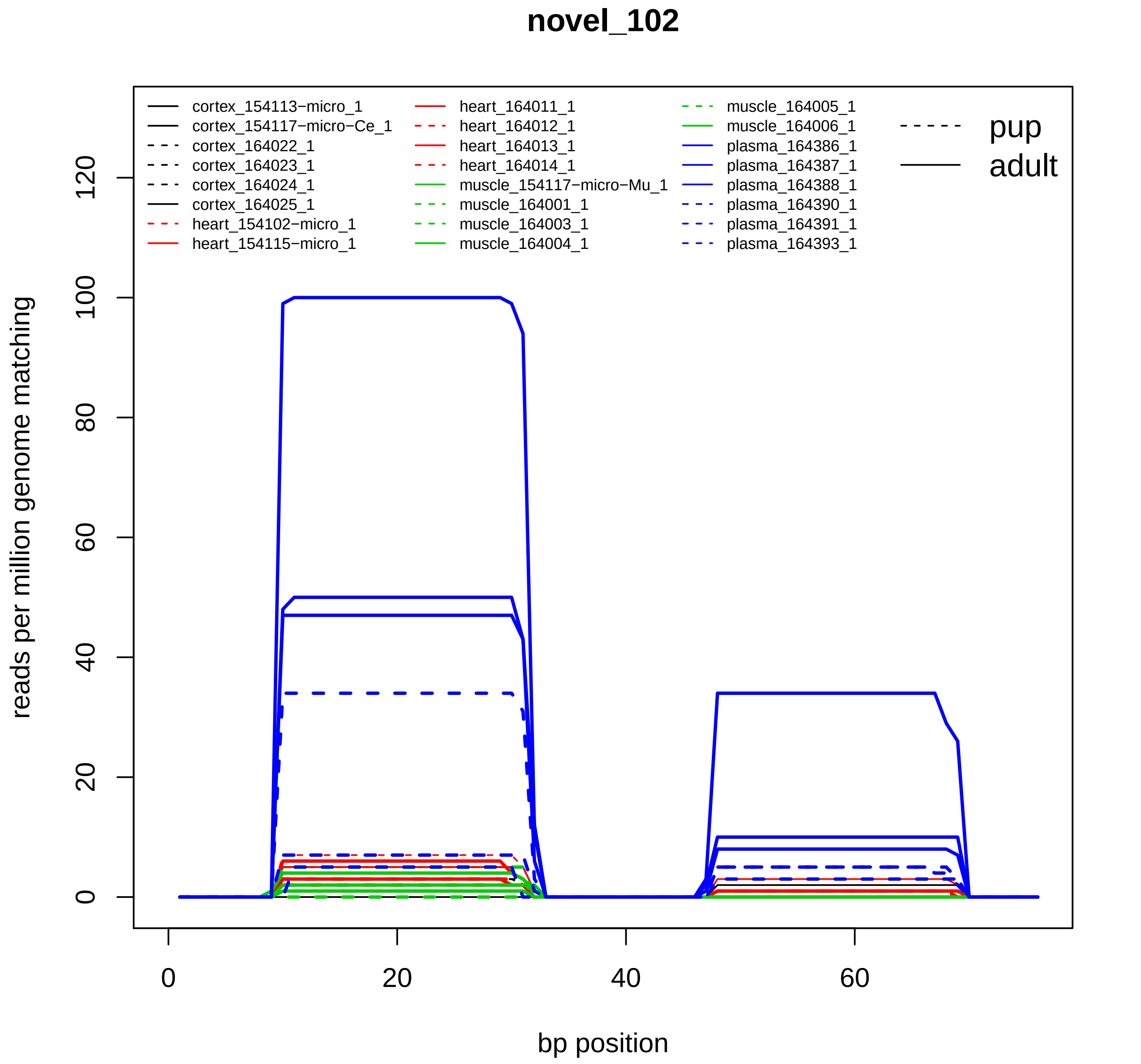

novel_102
cortex_154113−micro_1
cortex_154117−micro−Ce_1
cortex_164022_1
cortex_164023_1
cortex_164024_1
cortex_164025_1
heart_154102−micro_1
heart_154115−micro_1
heart_164011_1
heart_164012_1
heart_164013_1
heart_164014_1
muscle_154117−micro−Mu_1
muscle_164001_1
muscle_164003_1
muscle_164004_1
muscle_164005_1
muscle_164006_1
plasma_164386_1
plasma_164387_1
plasma_164388_1
plasma_164390_1
plasma_164391_1
plasma_164393_1
pup
adult
reads per million genome matching
0
20
40
60
80
100
120
bp position
0
20
40
60

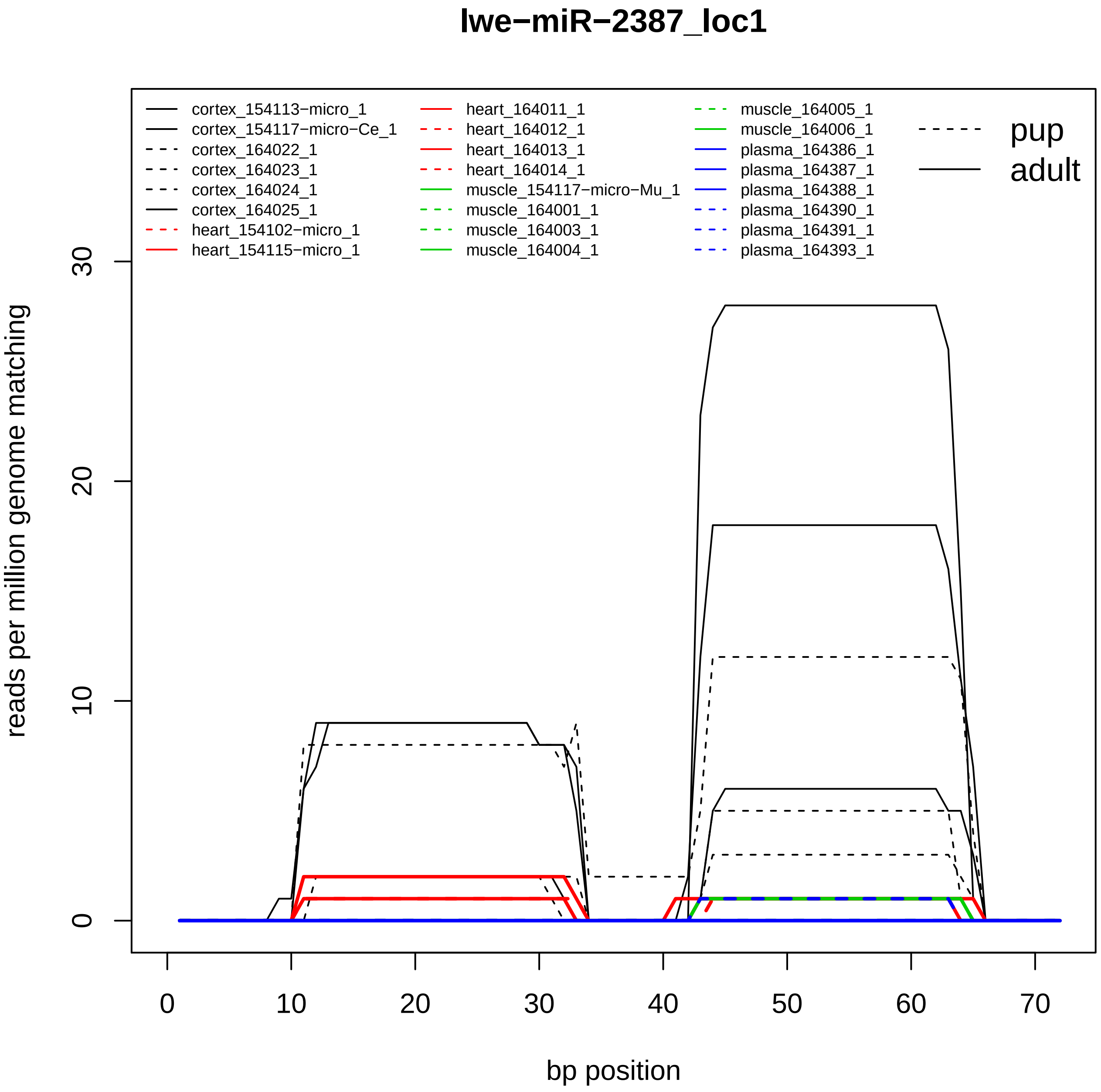
lwe−miR−2387_loc1
cortex_154113−micro_1
cortex_154117−micro−Ce_1
cortex_164022_1
cortex_164023_1
cortex_164024_1
cortex_164025_1
heart_154102−micro_1
heart_154115−micro_1
heart_164011_1
heart_164012_1
heart_164013_1
heart_164014_1
muscle_154117−micro−Mu_1
muscle_164001_1
muscle_164003_1
muscle_164004_1
muscle_164005_1
muscle_164006_1
plasma_164386_1
plasma_164387_1
plasma_164388_1
plasma_164390_1
plasma_164391_1
plasma_164393_1
pup
adult
reads per million genome matching
0
10
20
30
0
10
20
30
40
50
60
70
bp position

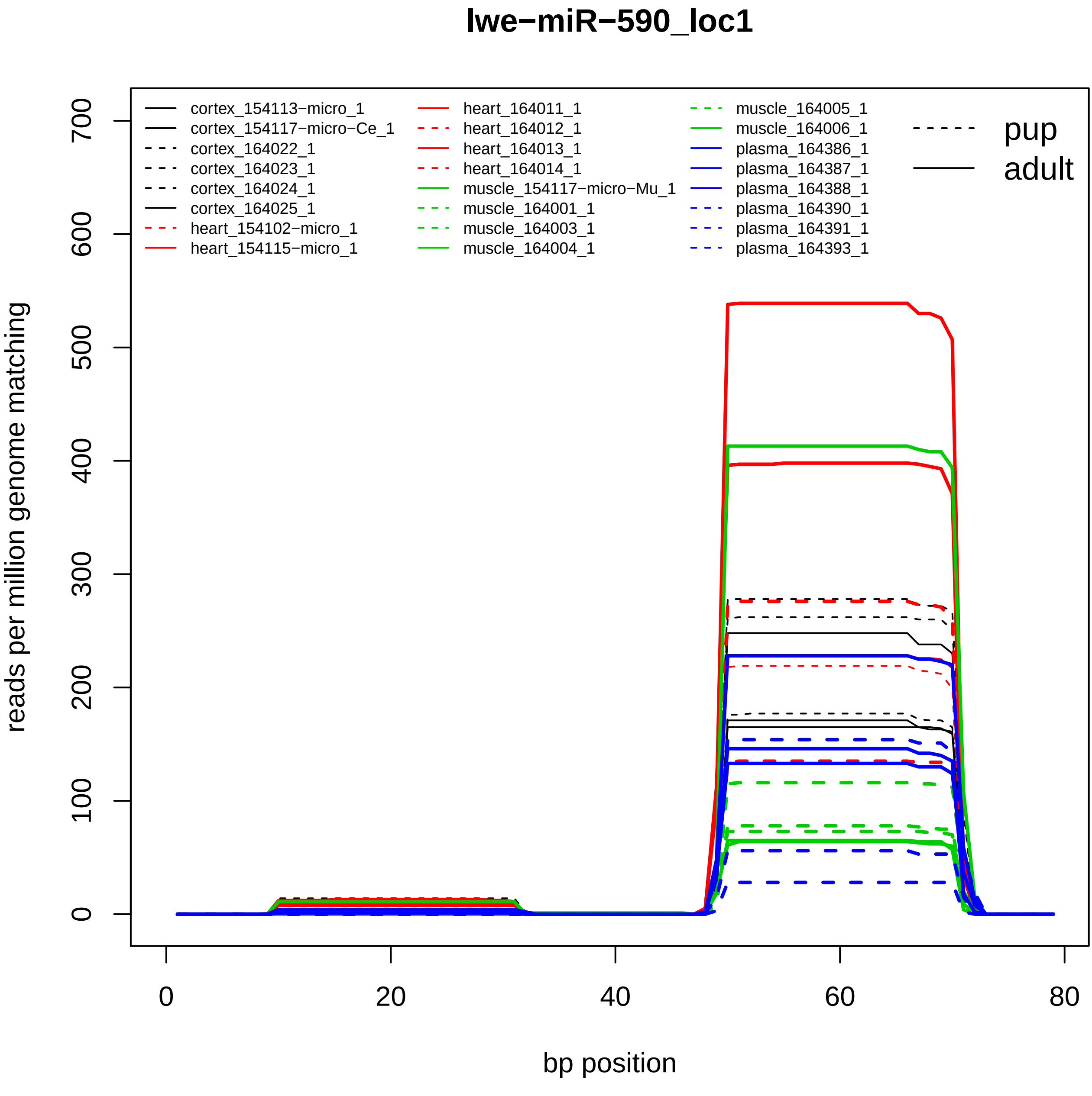

lwe−miR−590_loc1
reads per million genome matching
bp position
0
100
200
300
400
500
600
700
0
20
40
60
80
cortex_154113−micro_1
cortex_154117−micro−Ce_1
cortex_164022_1
cortex_164023_1
cortex_164024_1
cortex_164025_1
heart_154102−micro_1
heart_154115−micro_1
heart_164011_1
heart_164012_1
heart_164013_1
heart_164014_1
muscle_154117−micro−Mu_1
muscle_164001_1
muscle_164003_1
muscle_164004_1
muscle_164005_1
muscle_164006_1
plasma_164386_1
plasma_164387_1
plasma_164388_1
plasma_164390_1
plasma_164391_1
plasma_164393_1
pup
adult

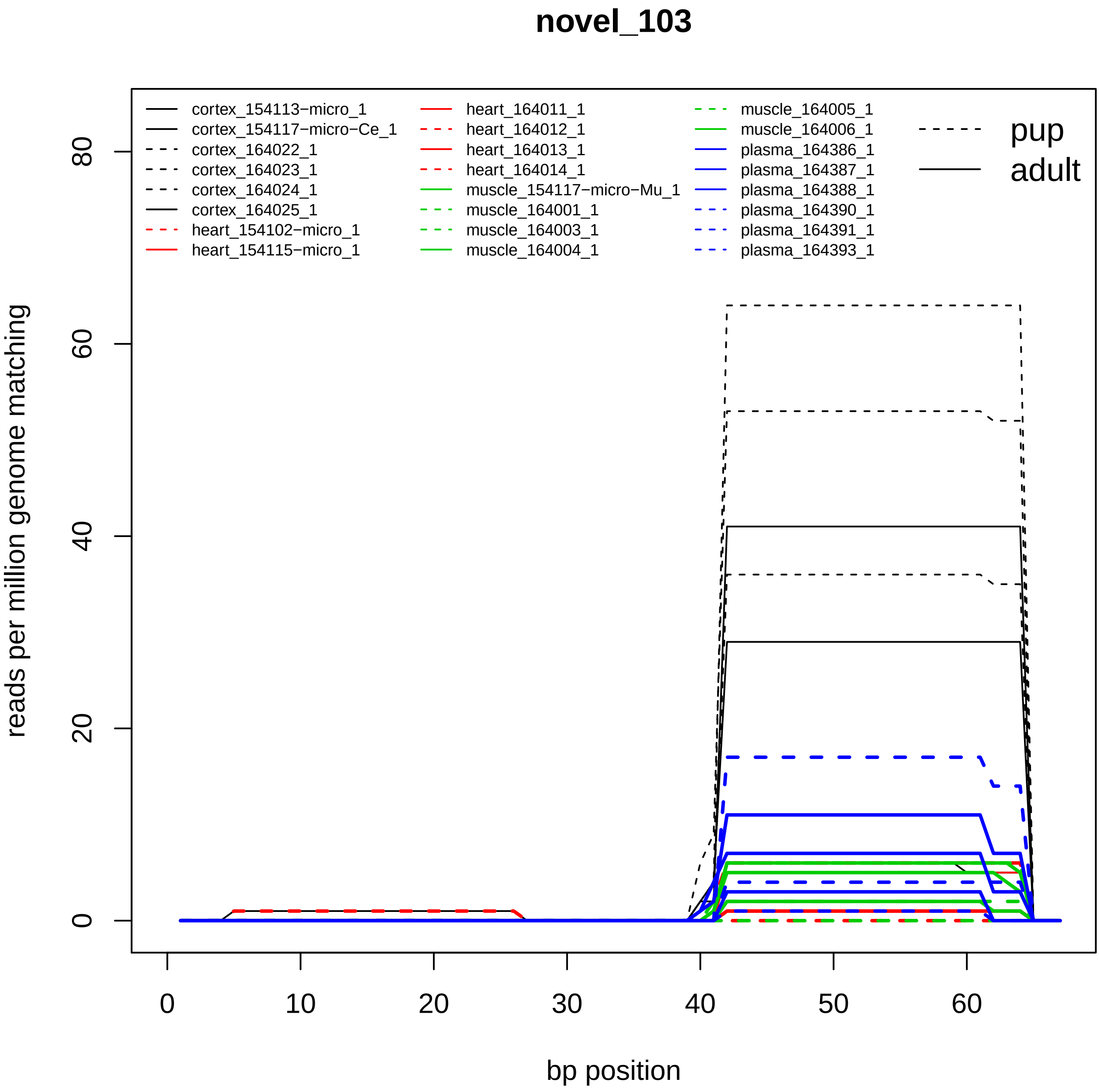

novel_103
cortex_154113−micro_1
cortex_154117−micro−Ce_1
cortex_164022_1
cortex_164023_1
cortex_164024_1
cortex_164025_1
heart_154102−micro_1
heart_154115−micro_1
heart_164011_1
heart_164012_1
heart_164013_1
heart_164014_1
muscle_154117−micro−Mu_1
muscle_164001_1
muscle_164003_1
muscle_164004_1
muscle_164005_1
muscle_164006_1
plasma_164386_1
plasma_164387_1
plasma_164388_1
plasma_164390_1
plasma_164391_1
plasma_164393_1
pup
adult
reads per million genome matching
bp position
0
20
40
60
80
0
10
20
30
40
50
60

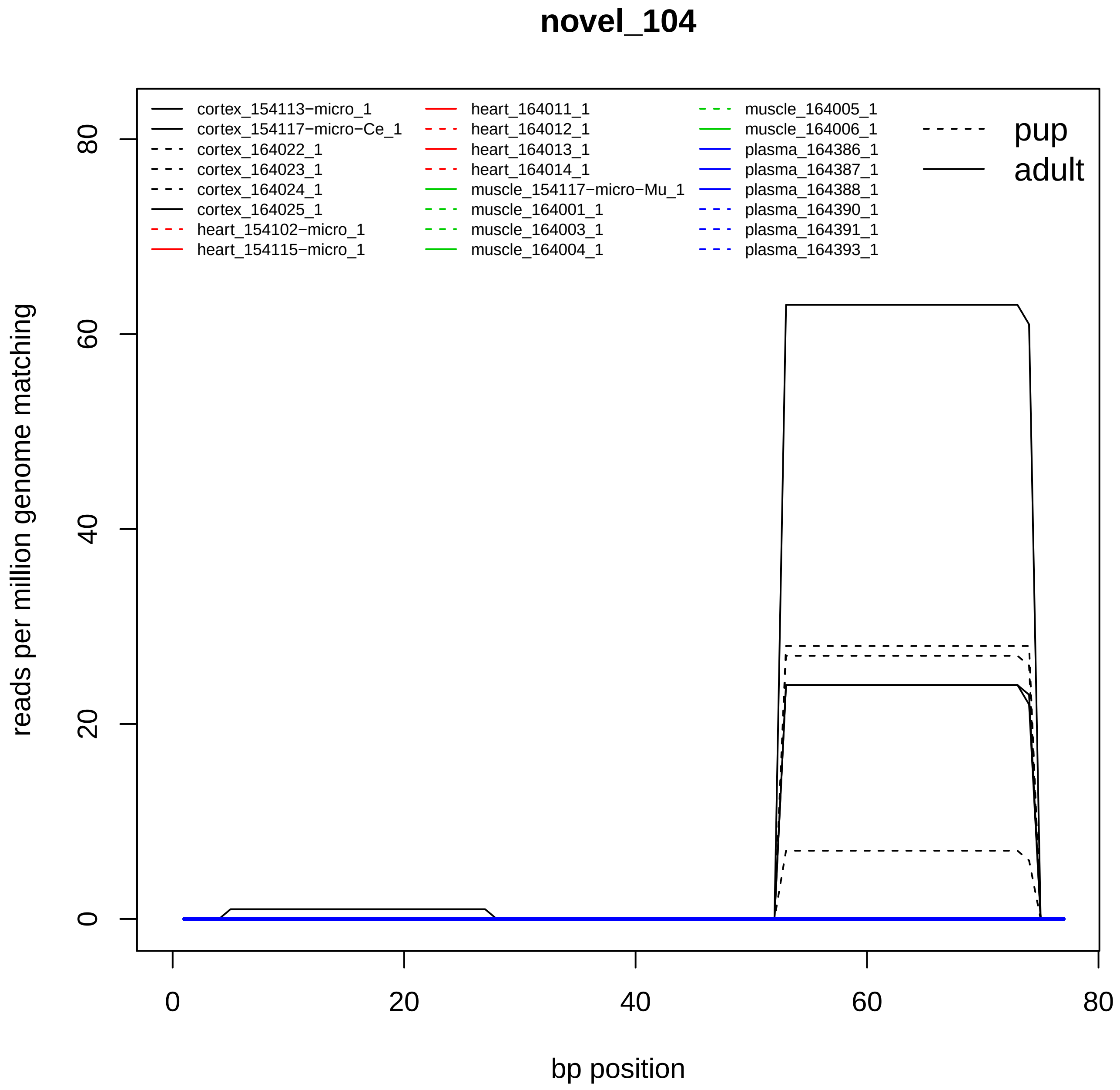

novel_104
reads per million genome matching
bp position
0
20
40
60
80
0
20
40
60
80
cortex_154113−micro_1
cortex_154117−micro−Ce_1
cortex_164022_1
cortex_164023_1
cortex_164024_1
cortex_164025_1
heart_154102−micro_1
heart_154115−micro_1
heart_164011_1
heart_164012_1
heart_164013_1
heart_164014_1
muscle_154117−micro−Mu_1
muscle_164001_1
muscle_164003_1
muscle_164004_1
muscle_164005_1
muscle_164006_1
plasma_164386_1
plasma_164387_1
plasma_164388_1
plasma_164390_1
plasma_164391_1
plasma_164393_1
pup
adult

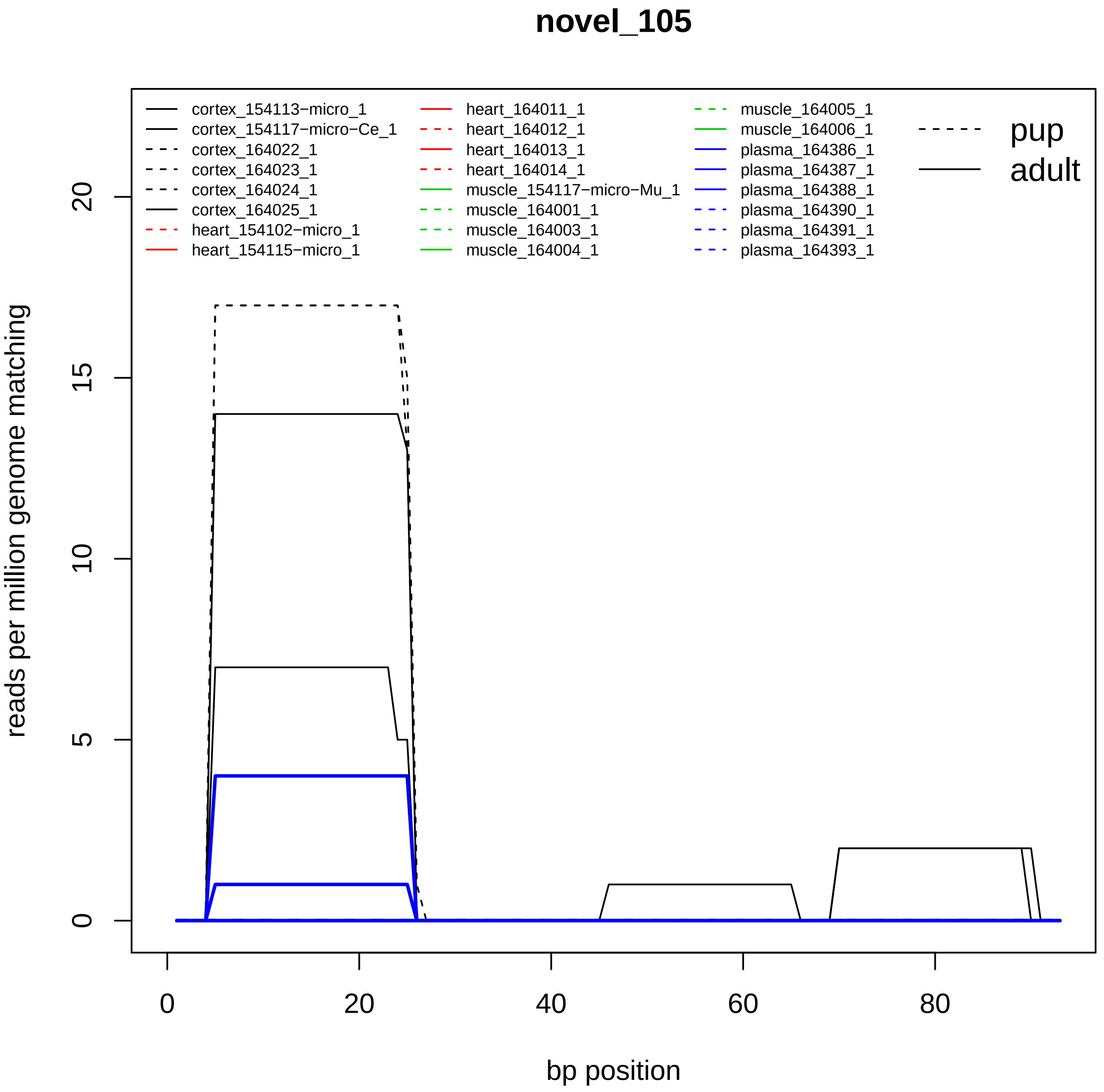

novel_105
cortex_154113−micro_1
cortex_154117−micro−Ce_1
cortex_164022_1
cortex_164023_1
cortex_164024_1
cortex_164025_1
heart_154102−micro_1
heart_154115−micro_1
heart_164011_1
heart_164012_1
heart_164013_1
heart_164014_1
muscle_154117−micro−Mu_1
muscle_164001_1
muscle_164003_1
muscle_164004_1
muscle_164005_1
muscle_164006_1
plasma_164386_1
plasma_164387_1
plasma_164388_1
plasma_164390_1
plasma_164391_1
plasma_164393_1
pup
adult
reads per million genome matching
0
5
10
15
20
0
20
40
60
80
bp position

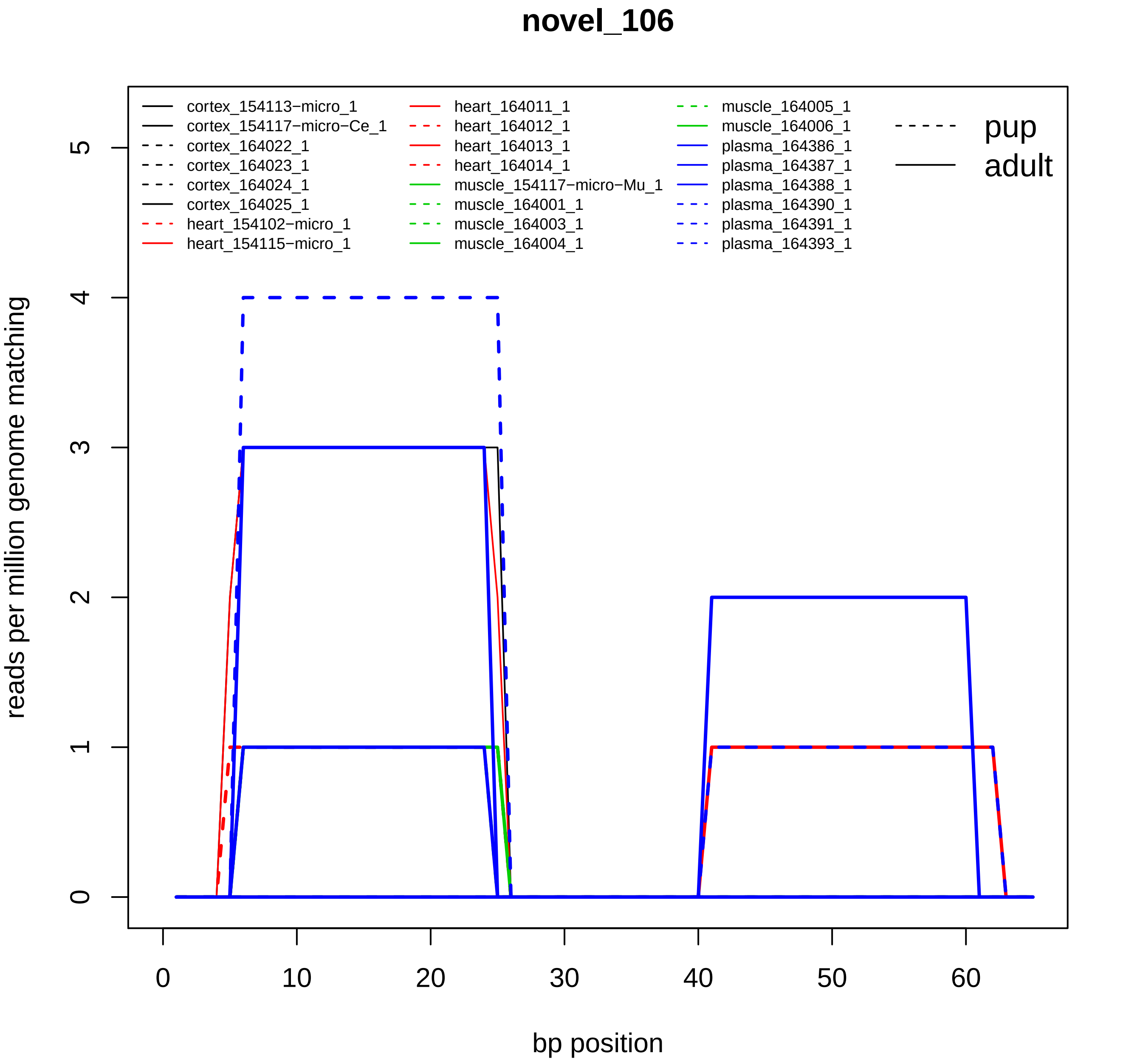
novel_106
cortex_154113−micro_1
cortex_154117−micro−Ce_1
cortex_164022_1
cortex_164023_1
cortex_164024_1
cortex_164025_1
heart_154102−micro_1
heart_154115−micro_1
heart_164011_1
heart_164012_1
heart_164013_1
heart_164014_1
muscle_154117−micro−Mu_1
muscle_164001_1
muscle_164003_1
muscle_164004_1
muscle_164005_1
muscle_164006_1
plasma_164386_1
plasma_164387_1
plasma_164388_1
plasma_164390_1
plasma_164391_1
plasma_164393_1
pup
adult
reads per million genome matching
bp position
0
1
2
3
4
5
0
10
20
30
40
50
60

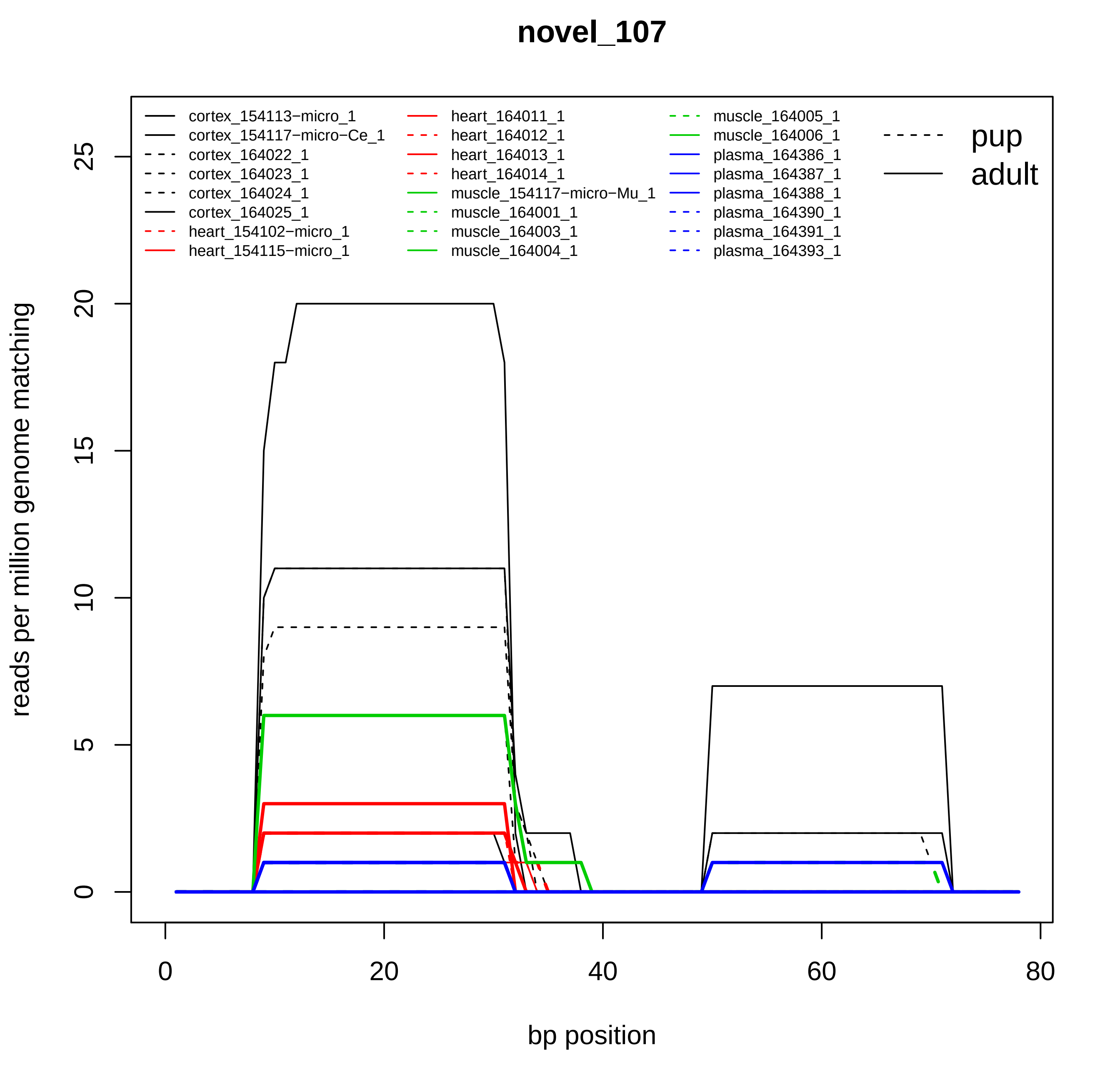

novel_107
cortex_154113−micro_1
cortex_154117−micro−Ce_1
cortex_164022_1
cortex_164023_1
cortex_164024_1
cortex_164025_1
heart_154102−micro_1
heart_154115−micro_1
heart_164011_1
heart_164012_1
heart_164013_1
heart_164014_1
muscle_154117−micro−Mu_1
muscle_164001_1
muscle_164003_1
muscle_164004_1
muscle_164005_1
muscle_164006_1
plasma_164386_1
plasma_164387_1
plasma_164388_1
plasma_164390_1
plasma_164391_1
plasma_164393_1
pup
adult
reads per million genome matching
bp position
0
5
10
15
20
25
0
20
40
60
80

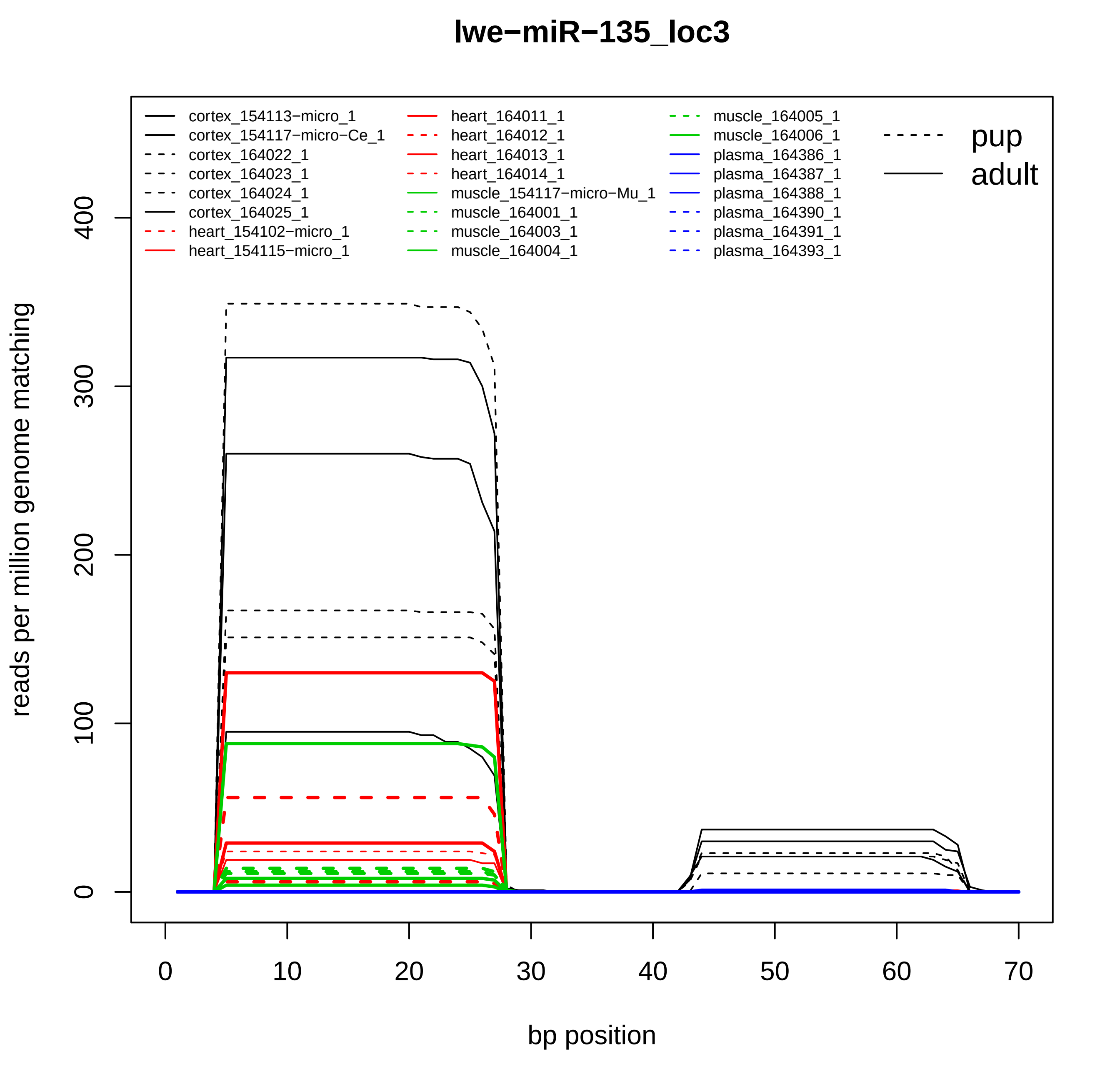

lwe−miR−135_loc3
cortex_154113−micro_1
cortex_154117−micro−Ce_1
cortex_164022_1
cortex_164023_1
cortex_164024_1
cortex_164025_1
heart_154102−micro_1
heart_154115−micro_1
heart_164011_1
heart_164012_1
heart_164013_1
heart_164014_1
muscle_154117−micro−Mu_1
muscle_164001_1
muscle_164003_1
muscle_164004_1
muscle_164005_1
muscle_164006_1
plasma_164386_1
plasma_164387_1
plasma_164388_1
plasma_164390_1
plasma_164391_1
plasma_164393_1
pup
adult
reads per million genome matching
0
100
200
300
400
0
10
20
30
40
50
60
70
bp position

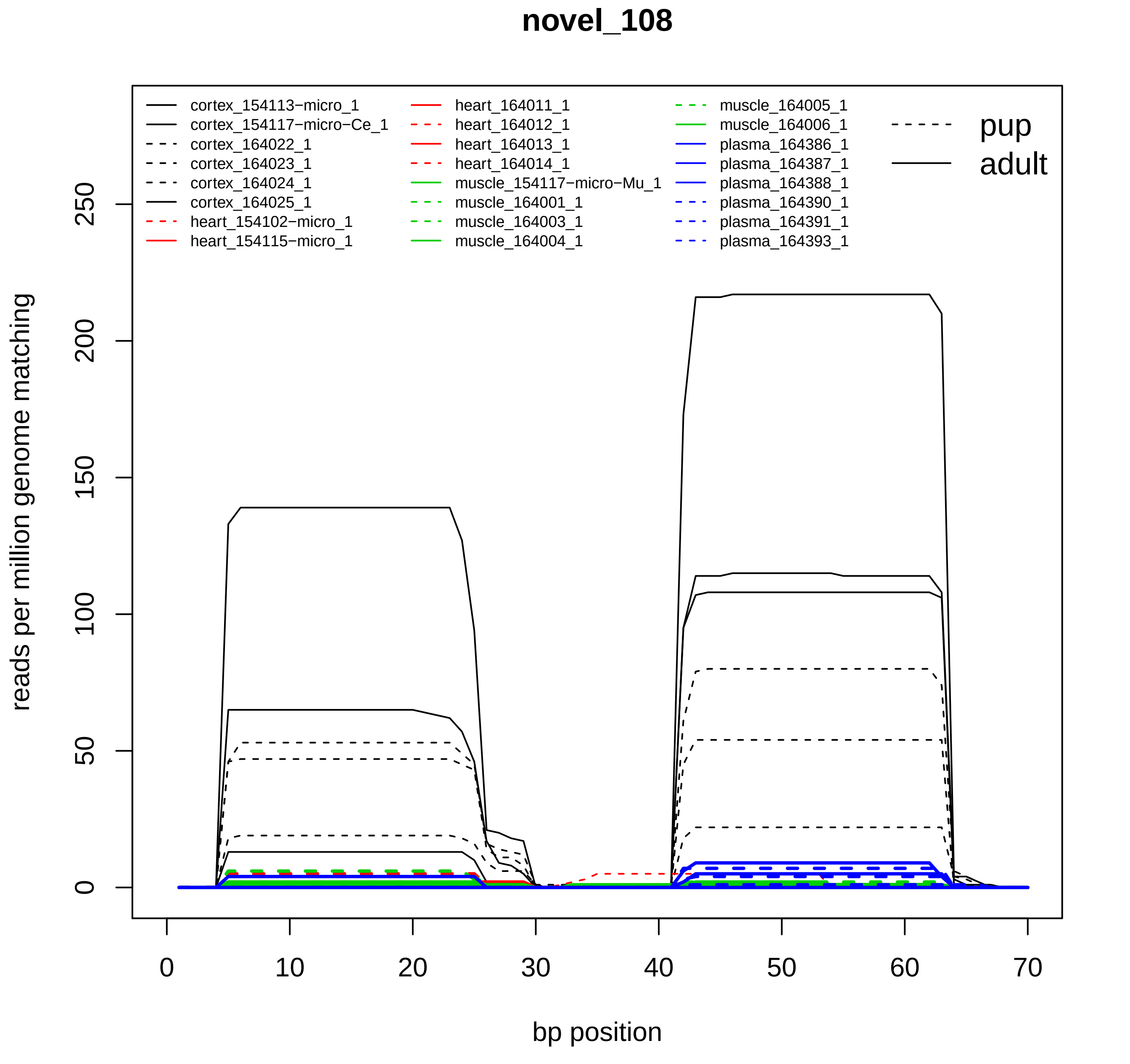

novel_108
reads per million genome matching
bp position
0
50
100
150
200
250
0
10
20
30
40
50
60
70
cortex_154113−micro_1
cortex_154117−micro−Ce_1
cortex_164022_1
cortex_164023_1
cortex_164024_1
cortex_164025_1
heart_154102−micro_1
heart_154115−micro_1
heart_164011_1
heart_164012_1
heart_164013_1
heart_164014_1
muscle_154117−micro−Mu_1
muscle_164001_1
muscle_164003_1
muscle_164004_1
muscle_164005_1
muscle_164006_1
plasma_164386_1
plasma_164387_1
plasma_164388_1
plasma_164390_1
plasma_164391_1
plasma_164393_1
pup
adult

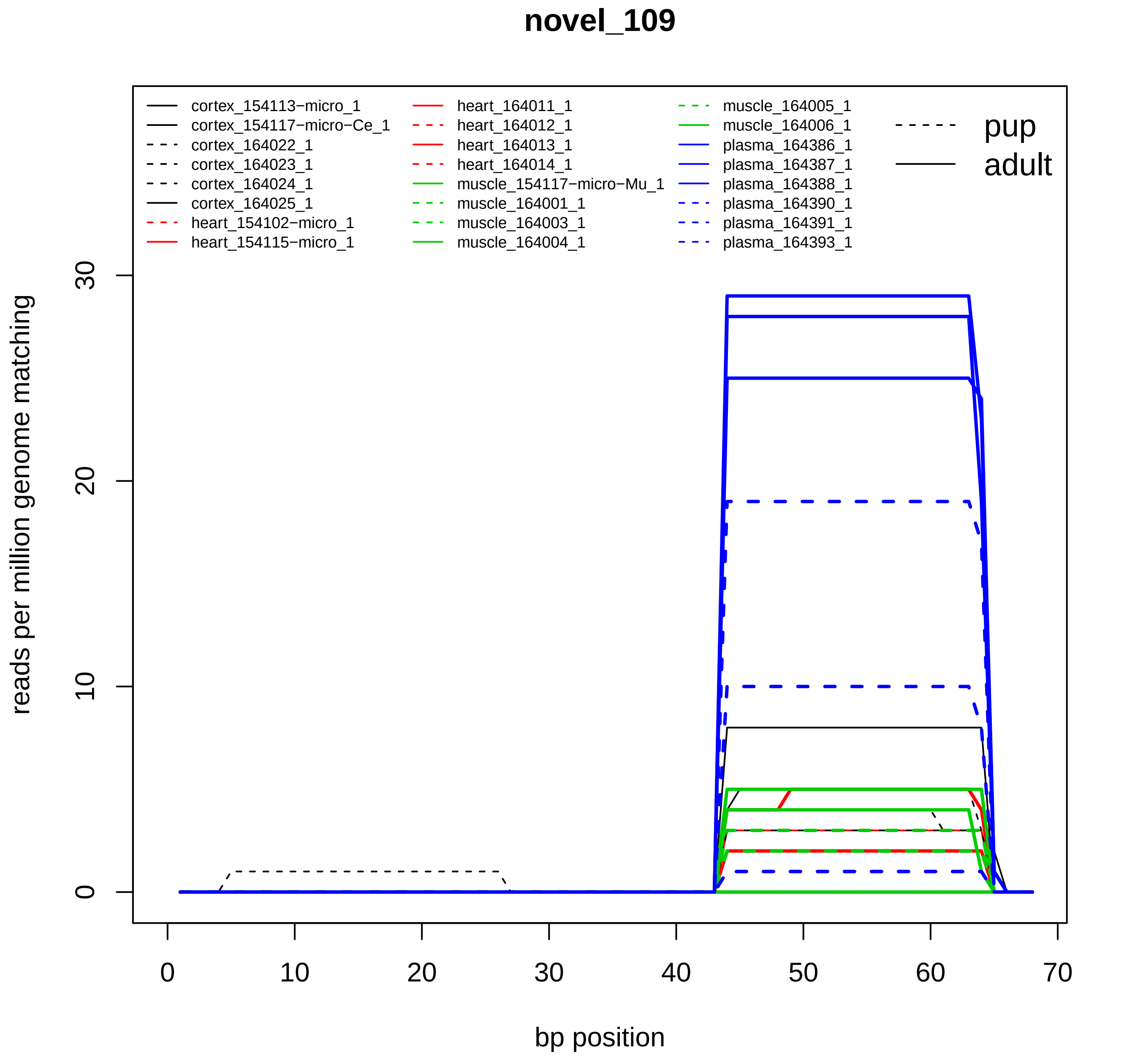

novel_109
cortex_154113−micro_1
cortex_154117−micro−Ce_1
cortex_164022_1
cortex_164023_1
cortex_164024_1
cortex_164025_1
heart_154102−micro_1
heart_154115−micro_1
heart_164011_1
heart_164012_1
heart_164013_1
heart_164014_1
muscle_154117−micro−Mu_1
muscle_164001_1
muscle_164003_1
muscle_164004_1
muscle_164005_1
muscle_164006_1
plasma_164386_1
plasma_164387_1
plasma_164388_1
plasma_164390_1
plasma_164391_1
plasma_164393_1
pup
adult
reads per million genome matching
0
10
20
30
bp position
0
10
20
30
40
50
60
70

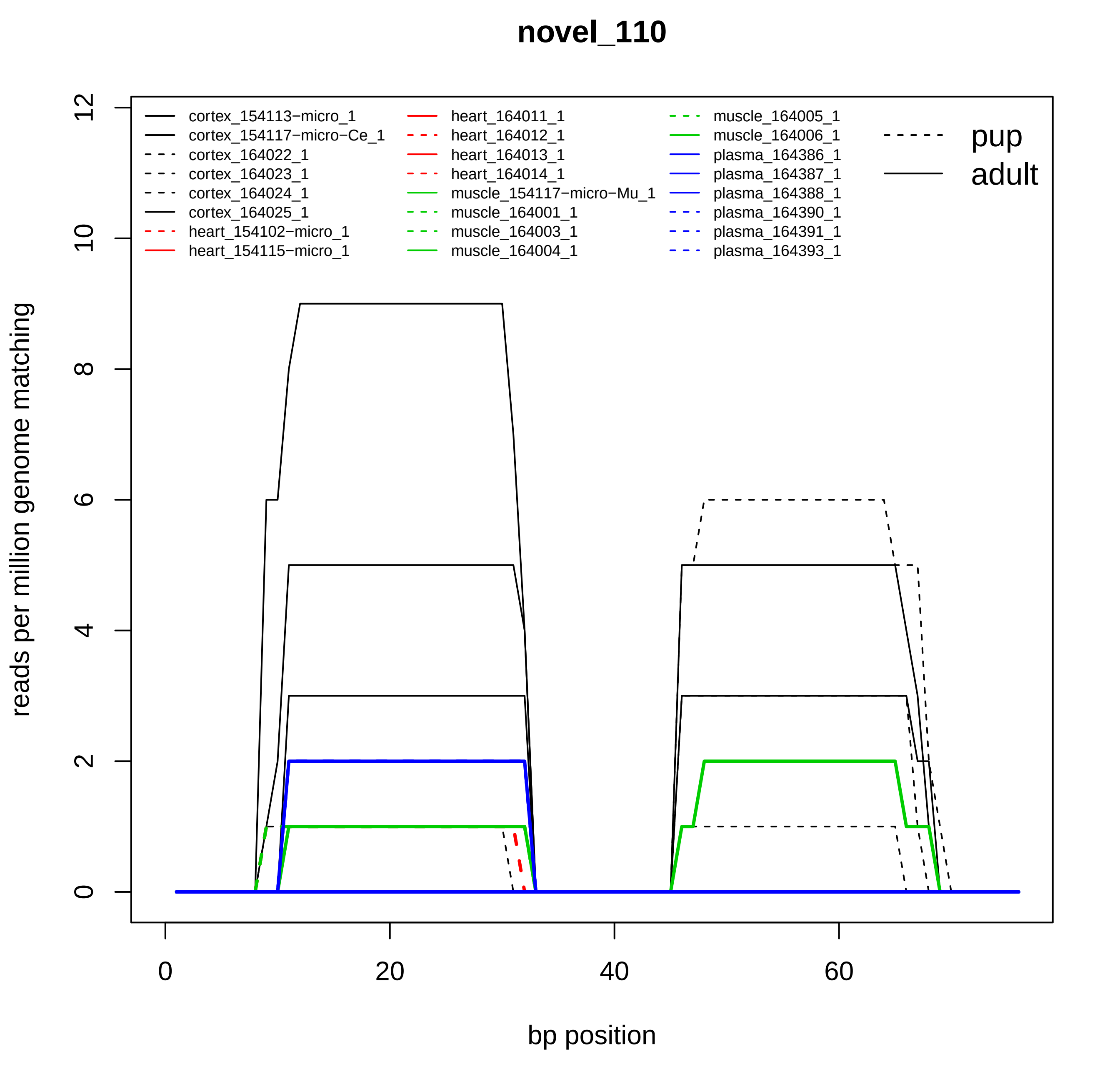
novel_110
cortex_154113−micro_1
cortex_154117−micro−Ce_1
cortex_164022_1
cortex_164023_1
cortex_164024_1
cortex_164025_1
heart_154102−micro_1
heart_154115−micro_1
heart_164011_1
heart_164012_1
heart_164013_1
heart_164014_1
muscle_154117−micro−Mu_1
muscle_164001_1
muscle_164003_1
muscle_164004_1
muscle_164005_1
muscle_164006_1
plasma_164386_1
plasma_164387_1
plasma_164388_1
plasma_164390_1
plasma_164391_1
plasma_164393_1
pup
adult
reads per million genome matching
bp position
0
2
4
6
8
10
12
0
20
40
60

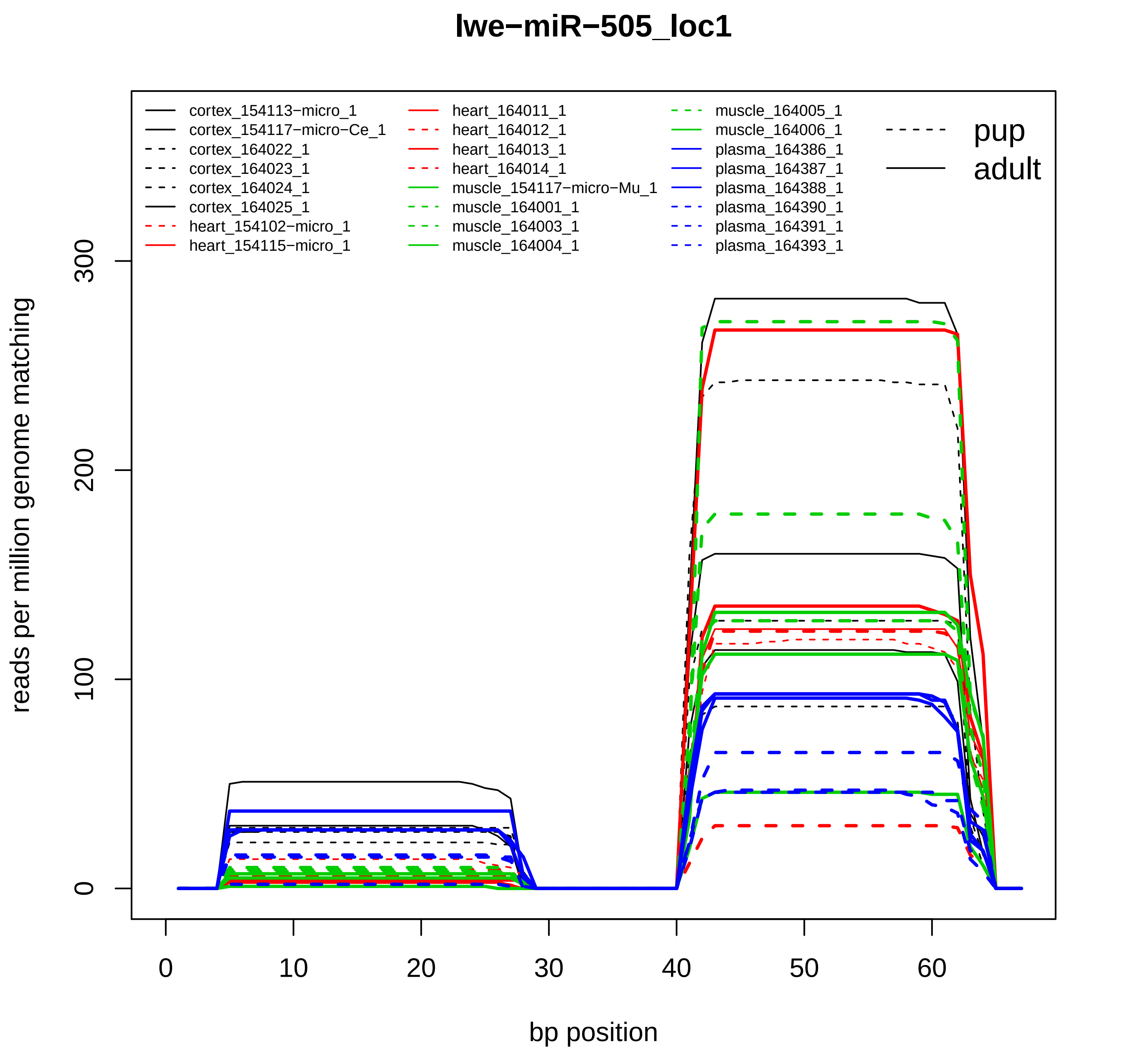
lwe−miR−505_loc1
cortex_154113−micro_1
cortex_154117−micro−Ce_1
cortex_164022_1
cortex_164023_1
cortex_164024_1
cortex_164025_1
heart_154102−micro_1
heart_154115−micro_1
heart_164011_1
heart_164012_1
heart_164013_1
heart_164014_1
muscle_154117−micro−Mu_1
muscle_164001_1
muscle_164003_1
muscle_164004_1
muscle_164005_1
muscle_164006_1
plasma_164386_1
plasma_164387_1
plasma_164388_1
plasma_164390_1
plasma_164391_1
plasma_164393_1
pup
adult
reads per million genome matching
bp position
0
100
200
300
0
10
20
30
40
50
60

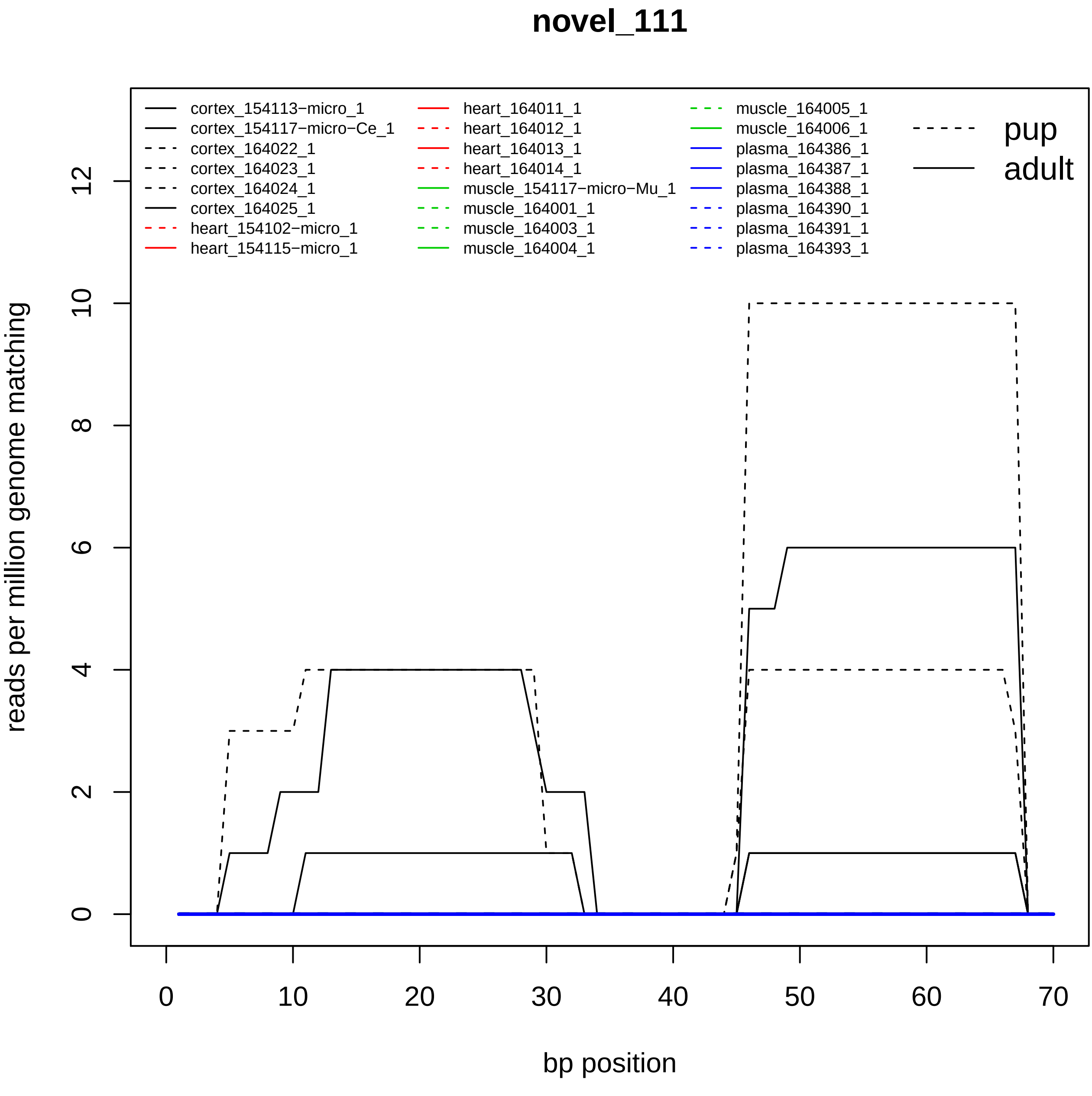
novel_111
cortex_154113−micro_1
cortex_154117−micro−Ce_1
cortex_164022_1
cortex_164023_1
cortex_164024_1
cortex_164025_1
heart_154102−micro_1
heart_154115−micro_1
heart_164011_1
heart_164012_1
heart_164013_1
heart_164014_1
muscle_154117−micro−Mu_1
muscle_164001_1
muscle_164003_1
muscle_164004_1
muscle_164005_1
muscle_164006_1
plasma_164386_1
plasma_164387_1
plasma_164388_1
plasma_164390_1
plasma_164391_1
plasma_164393_1
pup
adult
reads per million genome matching
bp position
0
2
4
6
8
10
12
0
10
20
30
40
50
60
70

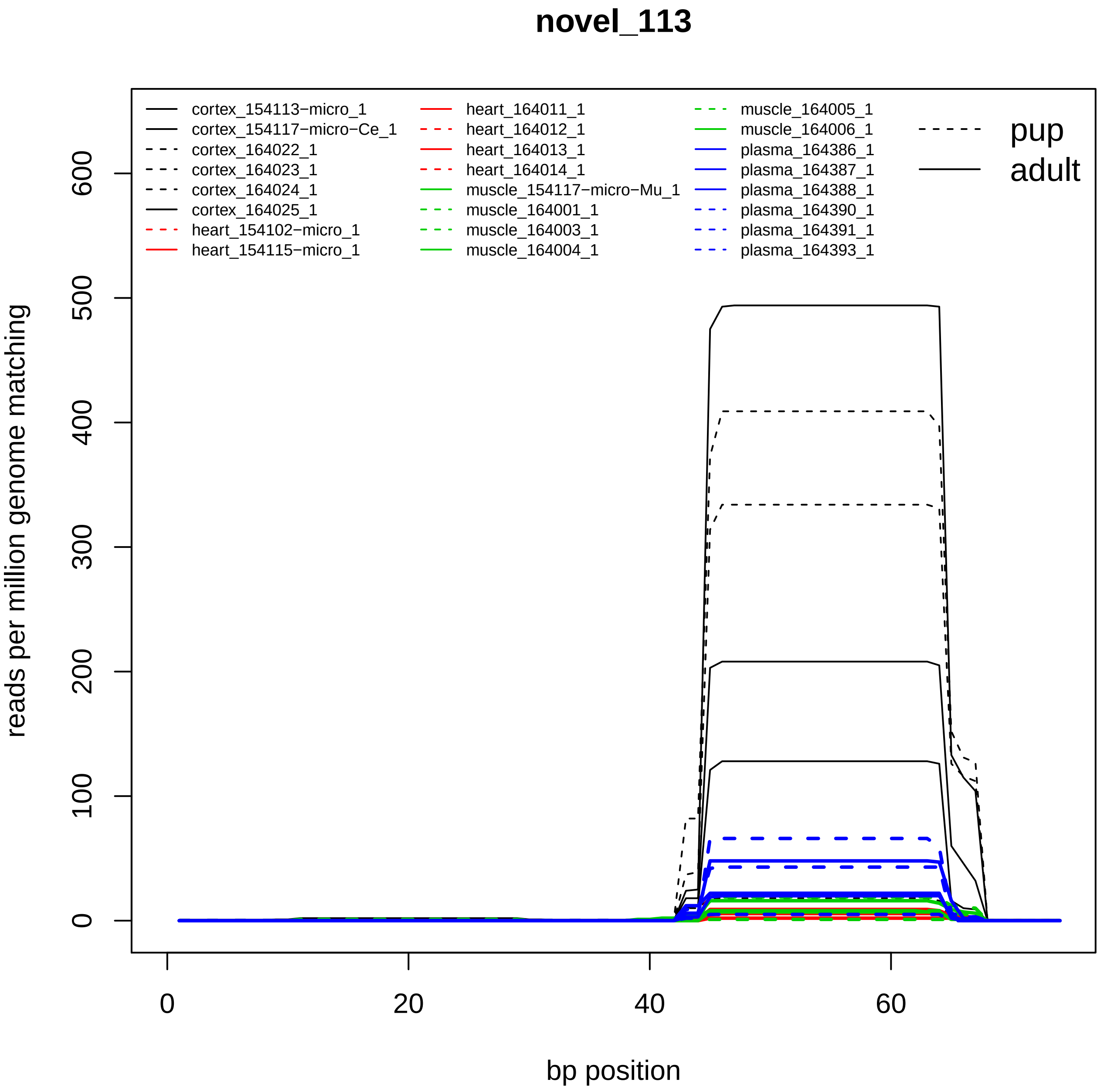
novel_113
cortex_154113−micro_1
cortex_154117−micro−Ce_1
cortex_164022_1
cortex_164023_1
cortex_164024_1
cortex_164025_1
heart_154102−micro_1
heart_154115−micro_1
heart_164011_1
heart_164012_1
heart_164013_1
heart_164014_1
muscle_154117−micro−Mu_1
muscle_164001_1
muscle_164003_1
muscle_164004_1
muscle_164005_1
muscle_164006_1
plasma_164386_1
plasma_164387_1
plasma_164388_1
plasma_164390_1
plasma_164391_1
plasma_164393_1
pup
adult
reads per million genome matching
0
100
200
300
400
500
600
0
20
40
60
bp position

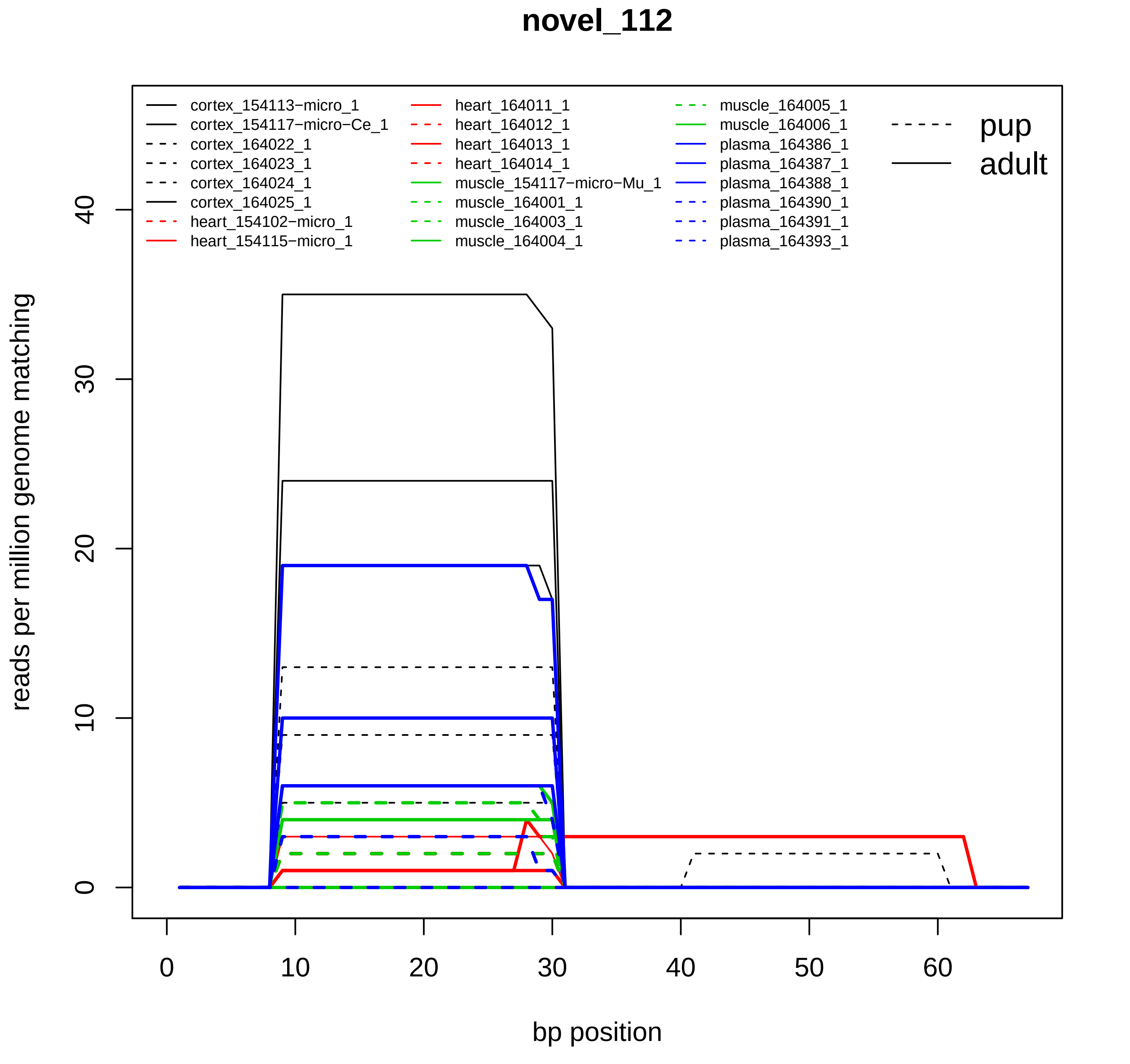

novel_112
cortex_154113−micro_1
cortex_154117−micro−Ce_1
cortex_164022_1
cortex_164023_1
cortex_164024_1
cortex_164025_1
heart_154102−micro_1
heart_154115−micro_1
heart_164011_1
heart_164012_1
heart_164013_1
heart_164014_1
muscle_154117−micro−Mu_1
muscle_164001_1
muscle_164003_1
muscle_164004_1
muscle_164005_1
muscle_164006_1
plasma_164386_1
plasma_164387_1
plasma_164388_1
plasma_164390_1
plasma_164391_1
plasma_164393_1
pup
adult
reads per million genome matching
0
10
20
30
40
0
10
20
30
40
50
60
bp position

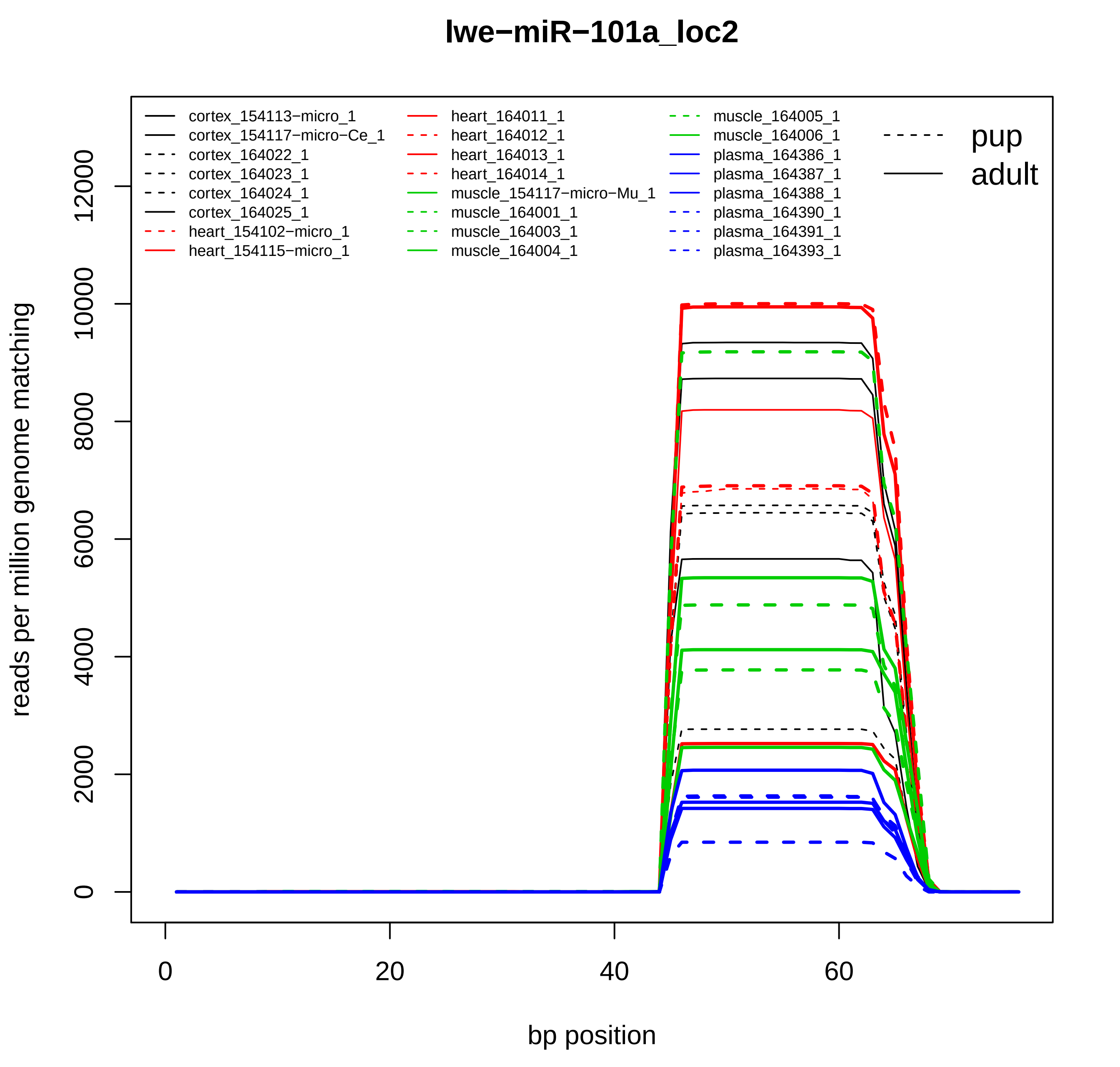

lwe−miR−101a_loc2
cortex_154113−micro_1
cortex_154117−micro−Ce_1
cortex_164022_1
cortex_164023_1
cortex_164024_1
cortex_164025_1
heart_154102−micro_1
heart_154115−micro_1
heart_164011_1
heart_164012_1
heart_164013_1
heart_164014_1
muscle_154117−micro−Mu_1
muscle_164001_1
muscle_164003_1
muscle_164004_1
muscle_164005_1
muscle_164006_1
plasma_164386_1
plasma_164387_1
plasma_164388_1
plasma_164390_1
plasma_164391_1
plasma_164393_1
pup
adult
reads per million genome matching
0
2000
4000
6000
8000
10000
12000
0
20
40
60
bp position

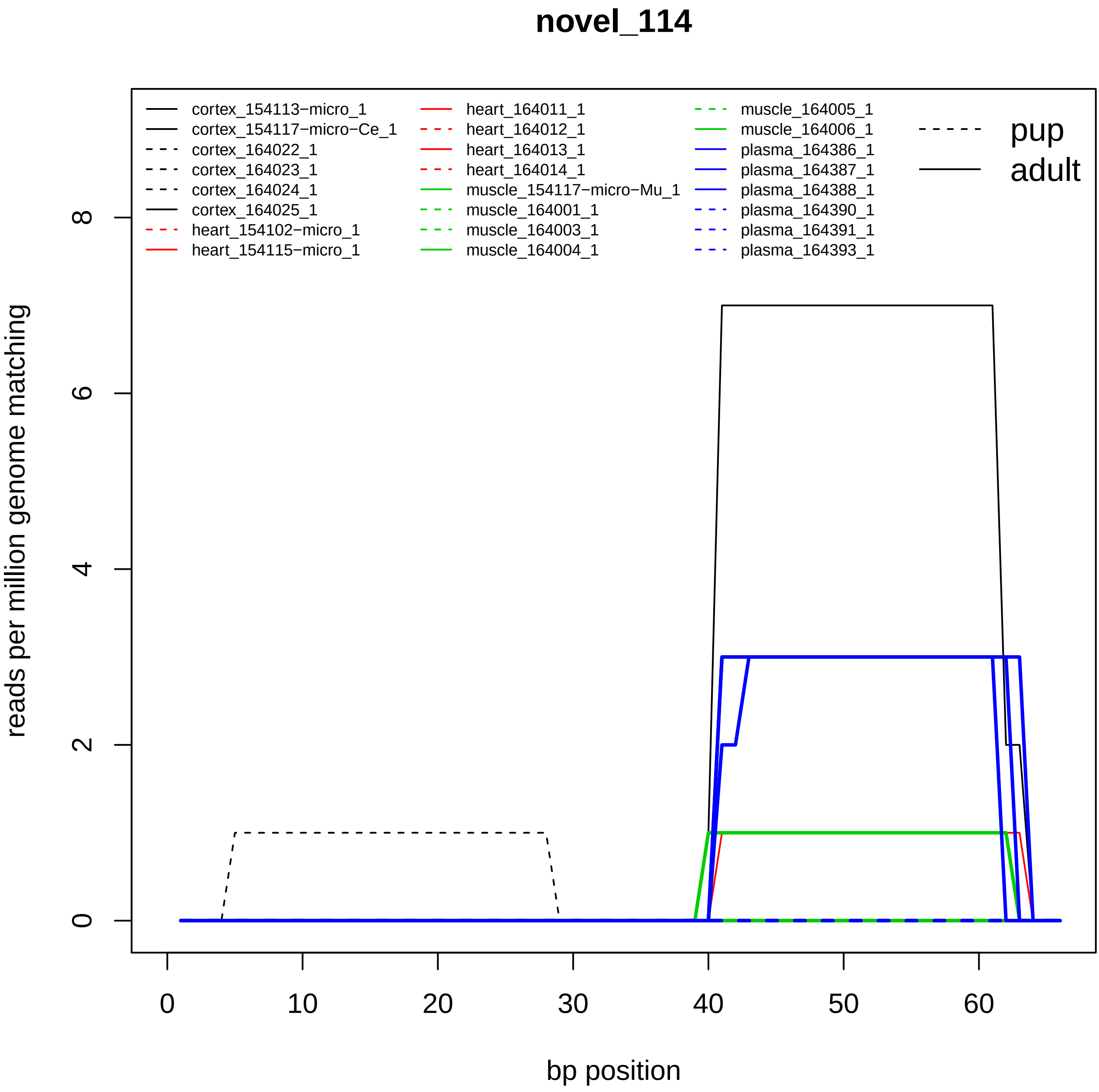
novel_114
cortex_154113−micro_1
cortex_154117−micro−Ce_1
cortex_164022_1
cortex_164023_1
cortex_164024_1
cortex_164025_1
heart_154102−micro_1
heart_154115−micro_1
heart_164011_1
heart_164012_1
heart_164013_1
heart_164014_1
muscle_154117−micro−Mu_1
muscle_164001_1
muscle_164003_1
muscle_164004_1
muscle_164005_1
muscle_164006_1
plasma_164386_1
plasma_164387_1
plasma_164388_1
plasma_164390_1
plasma_164391_1
plasma_164393_1
pup
adult
reads per million genome matching
bp position
0
2
4
6
8
0
10
20
30
40
50
60

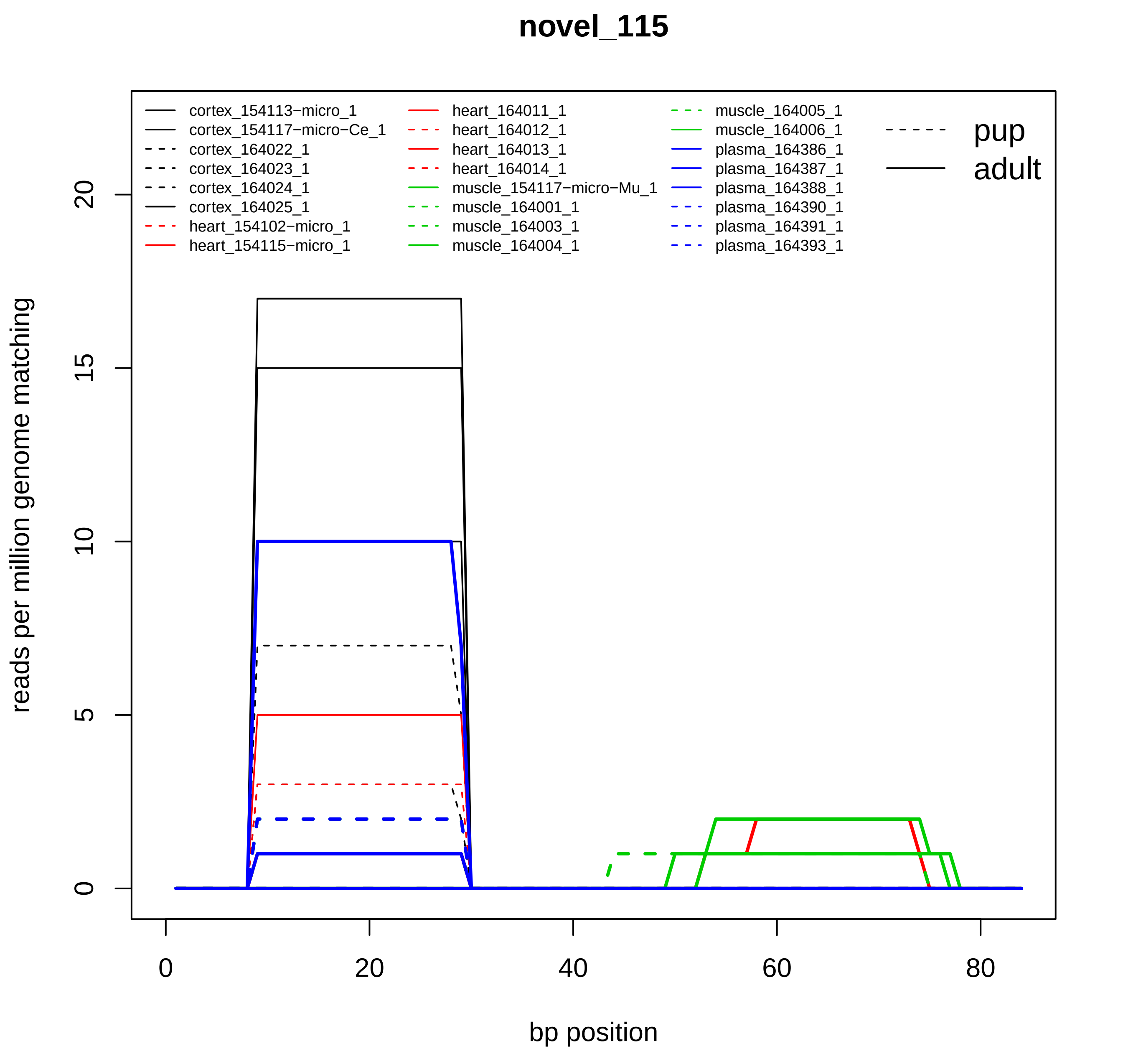

novel_115
cortex_154113−micro_1
cortex_154117−micro−Ce_1
cortex_164022_1
cortex_164023_1
cortex_164024_1
cortex_164025_1
heart_154102−micro_1
heart_154115−micro_1
heart_164011_1
heart_164012_1
heart_164013_1
heart_164014_1
muscle_154117−micro−Mu_1
muscle_164001_1
muscle_164003_1
muscle_164004_1
muscle_164005_1
muscle_164006_1
plasma_164386_1
plasma_164387_1
plasma_164388_1
plasma_164390_1
plasma_164391_1
plasma_164393_1
pup
adult
reads per million genome matching
0
5
10
15
20
0
20
40
60
80
bp position

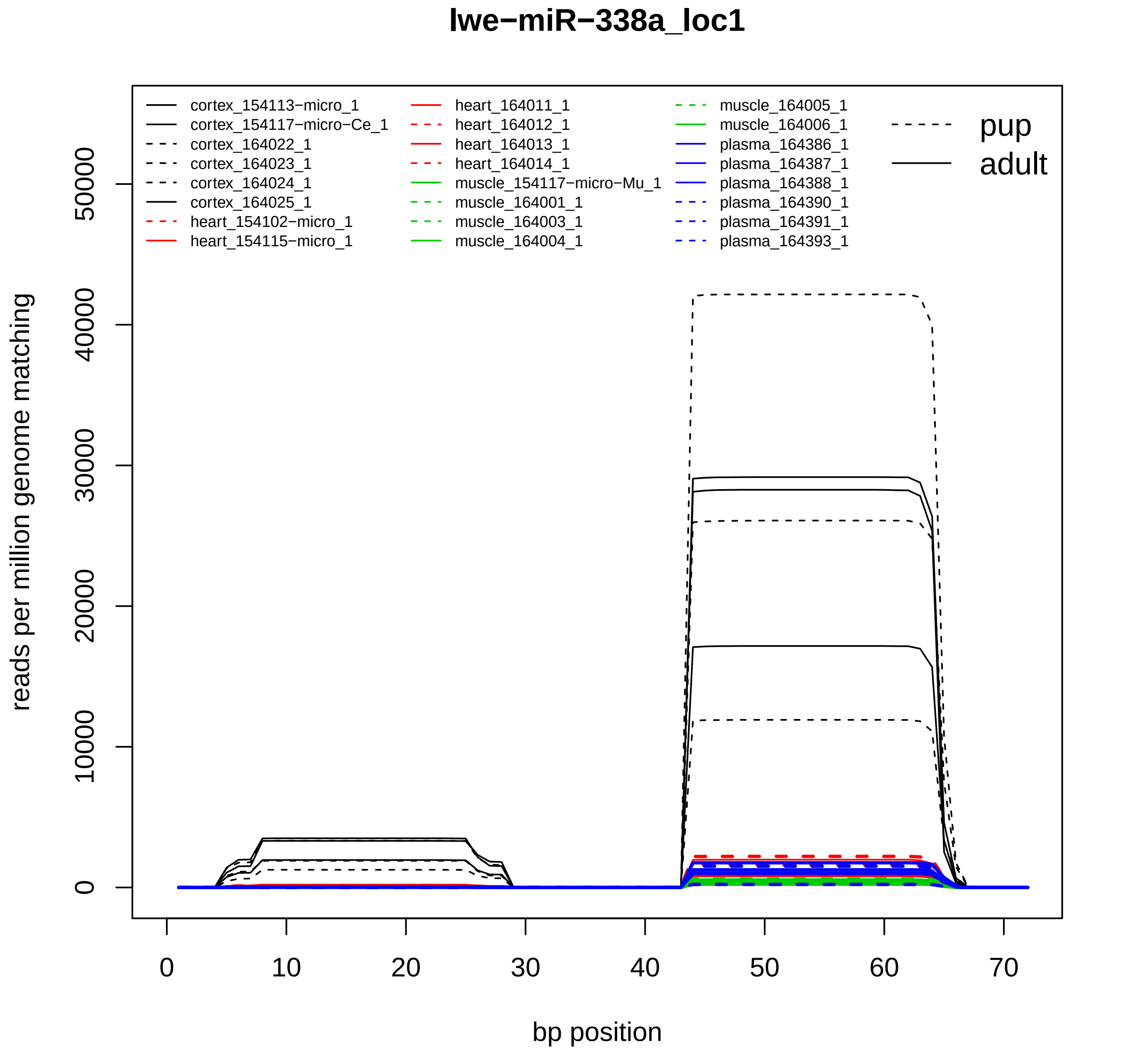
lwe−miR−338a_loc1
cortex_154113−micro_1
cortex_154117−micro−Ce_1
cortex_164022_1
cortex_164023_1
cortex_164024_1
cortex_164025_1
heart_154102−micro_1
heart_154115−micro_1
heart_164011_1
heart_164012_1
heart_164013_1
heart_164014_1
muscle_154117−micro−Mu_1
muscle_164001_1
muscle_164003_1
muscle_164004_1
muscle_164005_1
muscle_164006_1
plasma_164386_1
plasma_164387_1
plasma_164388_1
plasma_164390_1
plasma_164391_1
plasma_164393_1
pup
adult
reads per million genome matching
0
10000
20000
30000
40000
50000
bp position
0
10
20
30
40
50
60
70

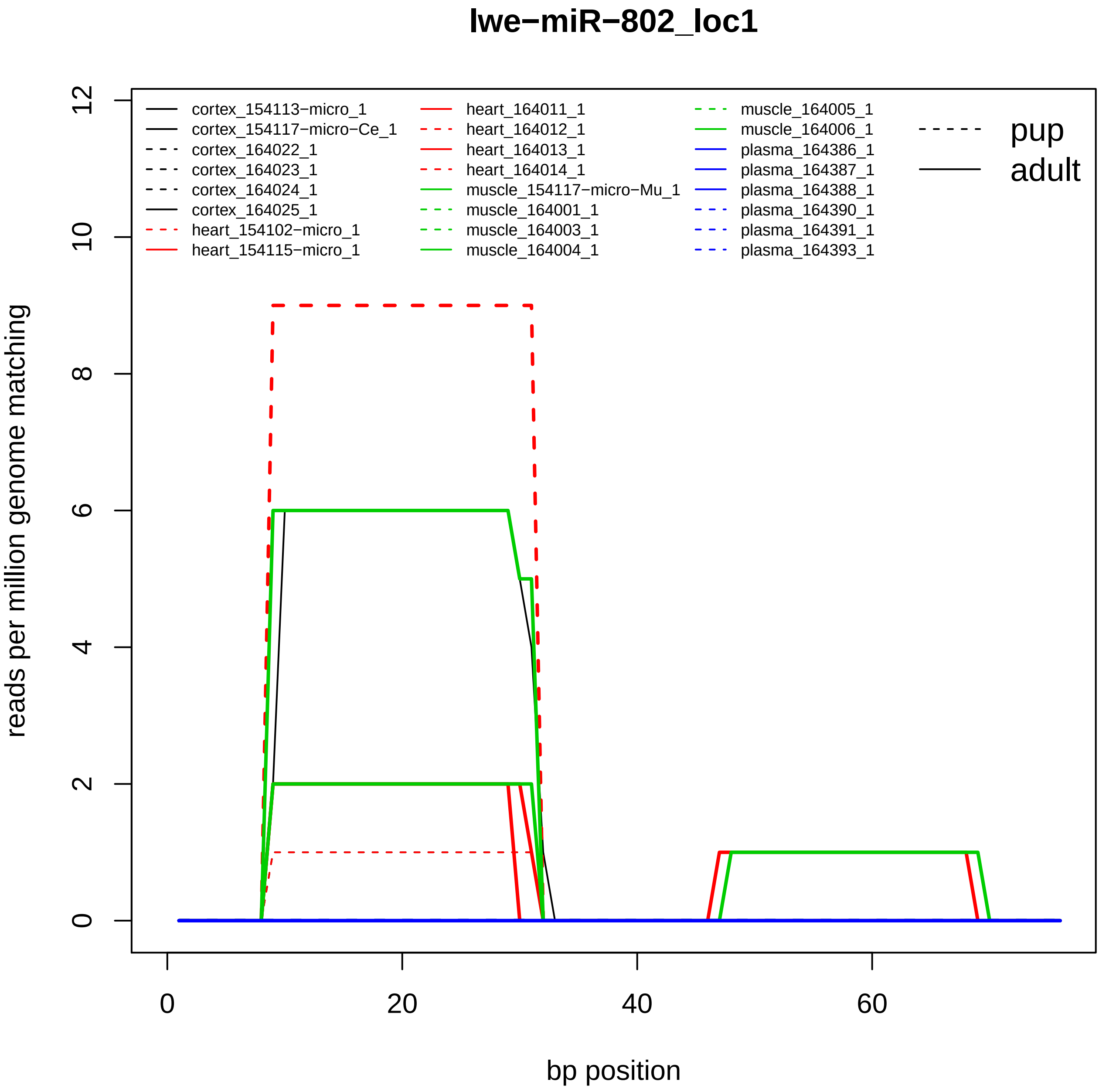

lwe−miR−802_loc1
reads per million genome matching
bp position
0
2
4
6
8
10
12
0
20
40
60
cortex_154113−micro_1
cortex_154117−micro−Ce_1
cortex_164022_1
cortex_164023_1
cortex_164024_1
cortex_164025_1
heart_154102−micro_1
heart_154115−micro_1
heart_164011_1
heart_164012_1
heart_164013_1
heart_164014_1
muscle_154117−micro−Mu_1
muscle_164001_1
muscle_164003_1
muscle_164004_1
muscle_164005_1
muscle_164006_1
plasma_164386_1
plasma_164387_1
plasma_164388_1
plasma_164390_1
plasma_164391_1
plasma_164393_1
pup
adult

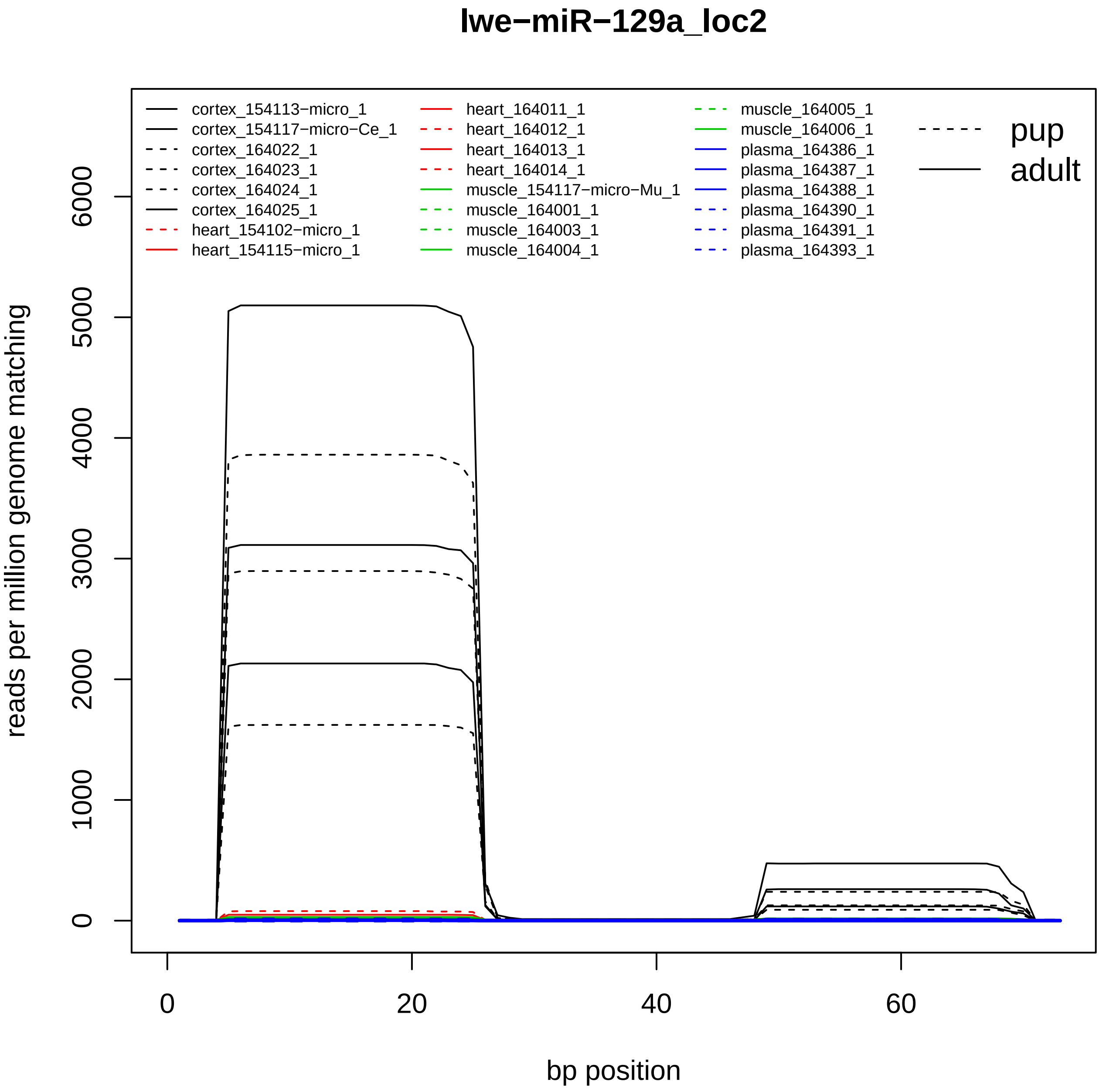

lwe−miR−129a_loc2
cortex_154113−micro_1
cortex_154117−micro−Ce_1
cortex_164022_1
cortex_164023_1
cortex_164024_1
cortex_164025_1
heart_154102−micro_1
heart_154115−micro_1
heart_164011_1
heart_164012_1
heart_164013_1
heart_164014_1
muscle_154117−micro−Mu_1
muscle_164001_1
muscle_164003_1
muscle_164004_1
muscle_164005_1
muscle_164006_1
plasma_164386_1
plasma_164387_1
plasma_164388_1
plasma_164390_1
plasma_164391_1
plasma_164393_1
pup
adult
reads per million genome matching
0
1000
2000
3000
4000
5000
6000
0
20
40
60
bp position

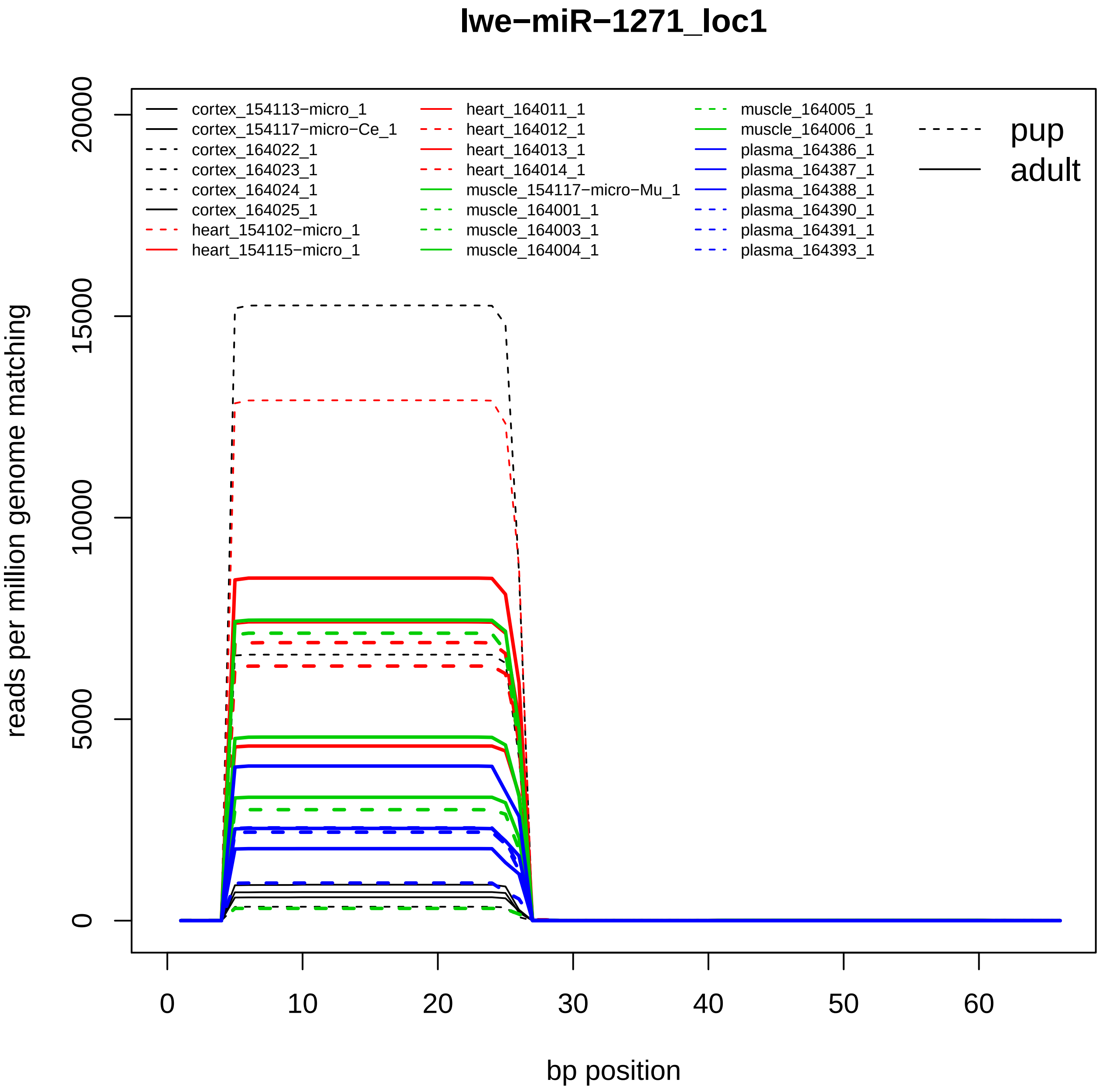
lwe−miR−1271_loc1
cortex_154113−micro_1
cortex_154117−micro−Ce_1
cortex_164022_1
cortex_164023_1
cortex_164024_1
cortex_164025_1
heart_154102−micro_1
heart_154115−micro_1
heart_164011_1
heart_164012_1
heart_164013_1
heart_164014_1
muscle_154117−micro−Mu_1
muscle_164001_1
muscle_164003_1
muscle_164004_1
muscle_164005_1
muscle_164006_1
plasma_164386_1
plasma_164387_1
plasma_164388_1
plasma_164390_1
plasma_164391_1
plasma_164393_1
pup
adult
reads per million genome matching
0
5000
10000
15000
20000
bp position
0
10
20
30
40
50
60

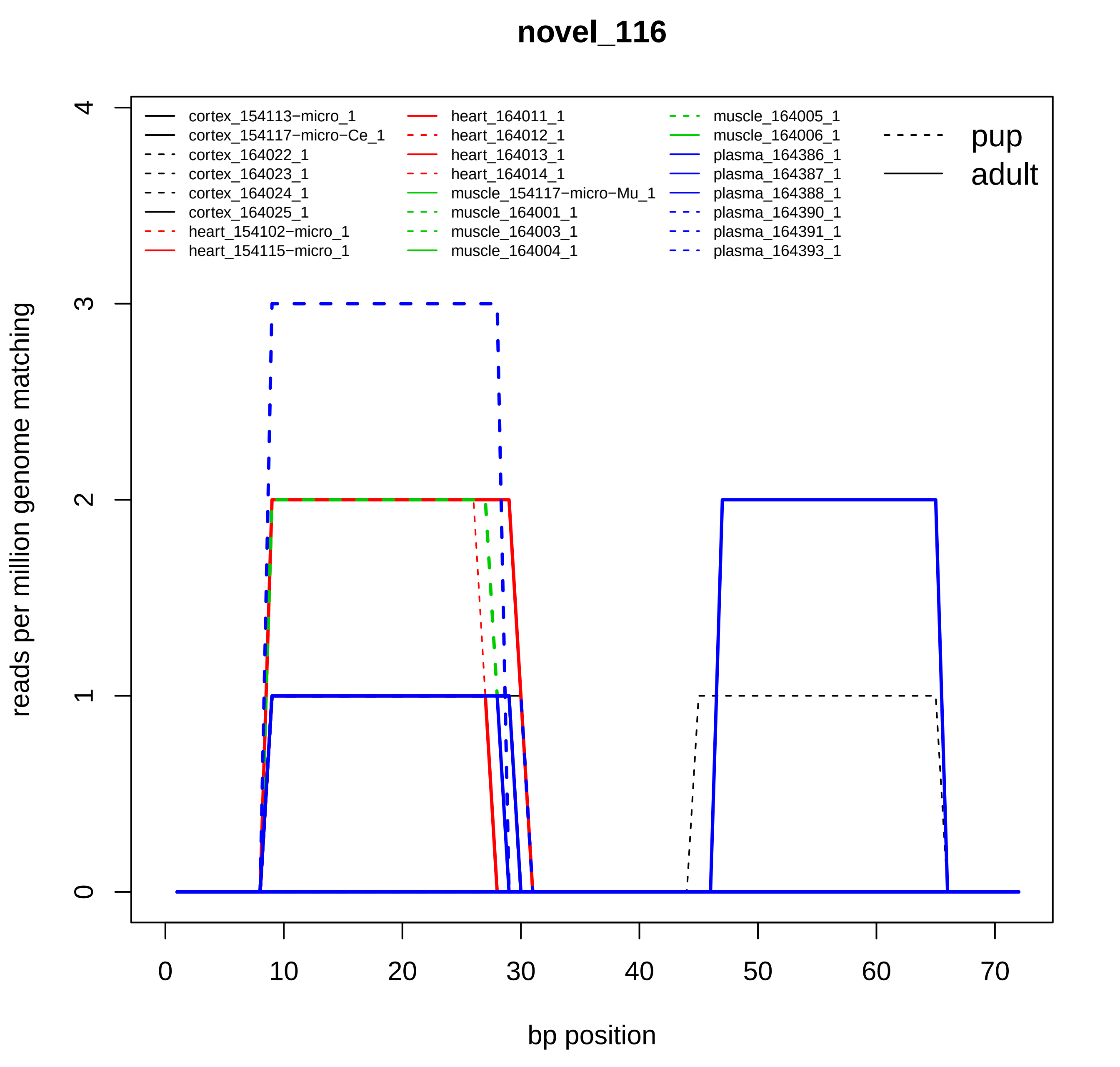

novel_116
cortex_154113−micro_1
cortex_154117−micro−Ce_1
cortex_164022_1
cortex_164023_1
cortex_164024_1
cortex_164025_1
heart_154102−micro_1
heart_154115−micro_1
heart_164011_1
heart_164012_1
heart_164013_1
heart_164014_1
muscle_154117−micro−Mu_1
muscle_164001_1
muscle_164003_1
muscle_164004_1
muscle_164005_1
muscle_164006_1
plasma_164386_1
plasma_164387_1
plasma_164388_1
plasma_164390_1
plasma_164391_1
plasma_164393_1
pup
adult
reads per million genome matching
bp position
0
1
2
3
4
0
10
20
30
40
50
60
70

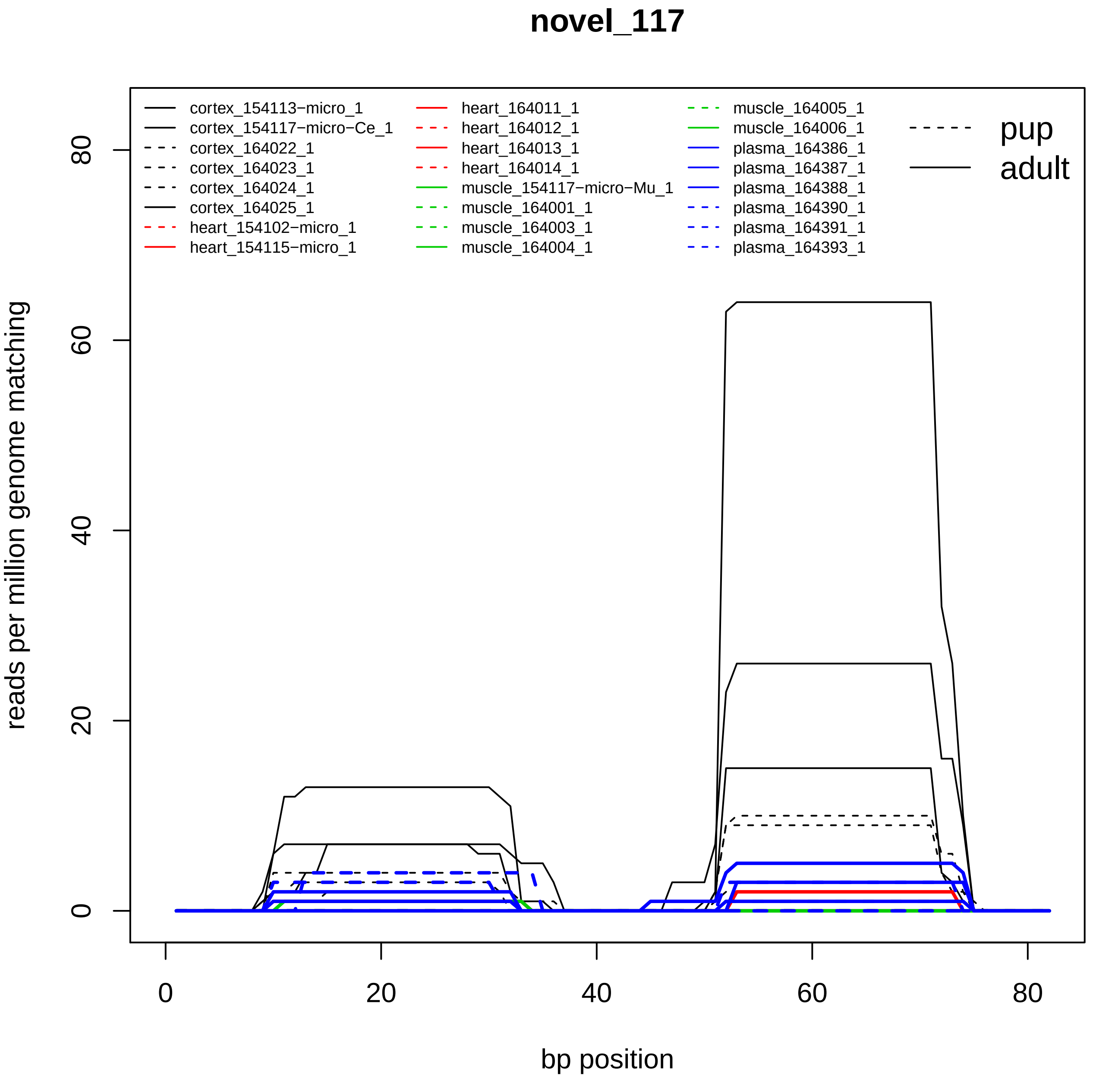

novel_117
cortex_154113−micro_1
cortex_154117−micro−Ce_1
cortex_164022_1
cortex_164023_1
cortex_164024_1
cortex_164025_1
heart_154102−micro_1
heart_154115−micro_1
heart_164011_1
heart_164012_1
heart_164013_1
heart_164014_1
muscle_154117−micro−Mu_1
muscle_164001_1
muscle_164003_1
muscle_164004_1
muscle_164005_1
muscle_164006_1
plasma_164386_1
plasma_164387_1
plasma_164388_1
plasma_164390_1
plasma_164391_1
plasma_164393_1
pup
adult
reads per million genome matching
0
20
40
60
80
bp position

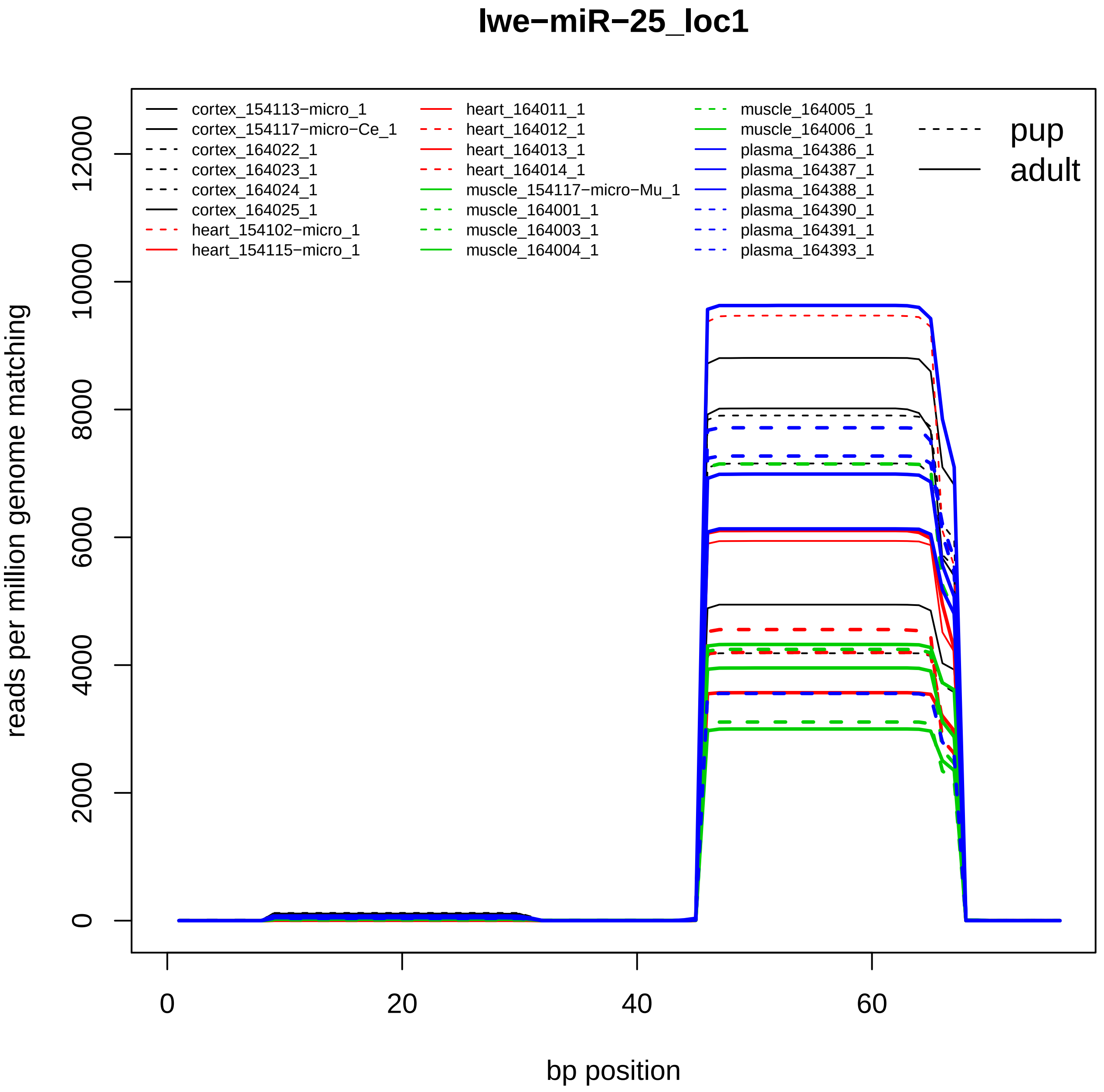
lwe−miR−25_loc1
reads per million genome matching
bp position
0
2000
4000
6000
8000
10000
12000
0
20
40
60
cortex_154113−micro_1
cortex_154117−micro−Ce_1
cortex_164022_1
cortex_164023_1
cortex_164024_1
cortex_164025_1
heart_154102−micro_1
heart_154115−micro_1
heart_164011_1
heart_164012_1
heart_164013_1
heart_164014_1
muscle_154117−micro−Mu_1
muscle_164001_1
muscle_164003_1
muscle_164004_1
muscle_164005_1
muscle_164006_1
plasma_164386_1
plasma_164387_1
plasma_164388_1
plasma_164390_1
plasma_164391_1
plasma_164393_1
pup
adult

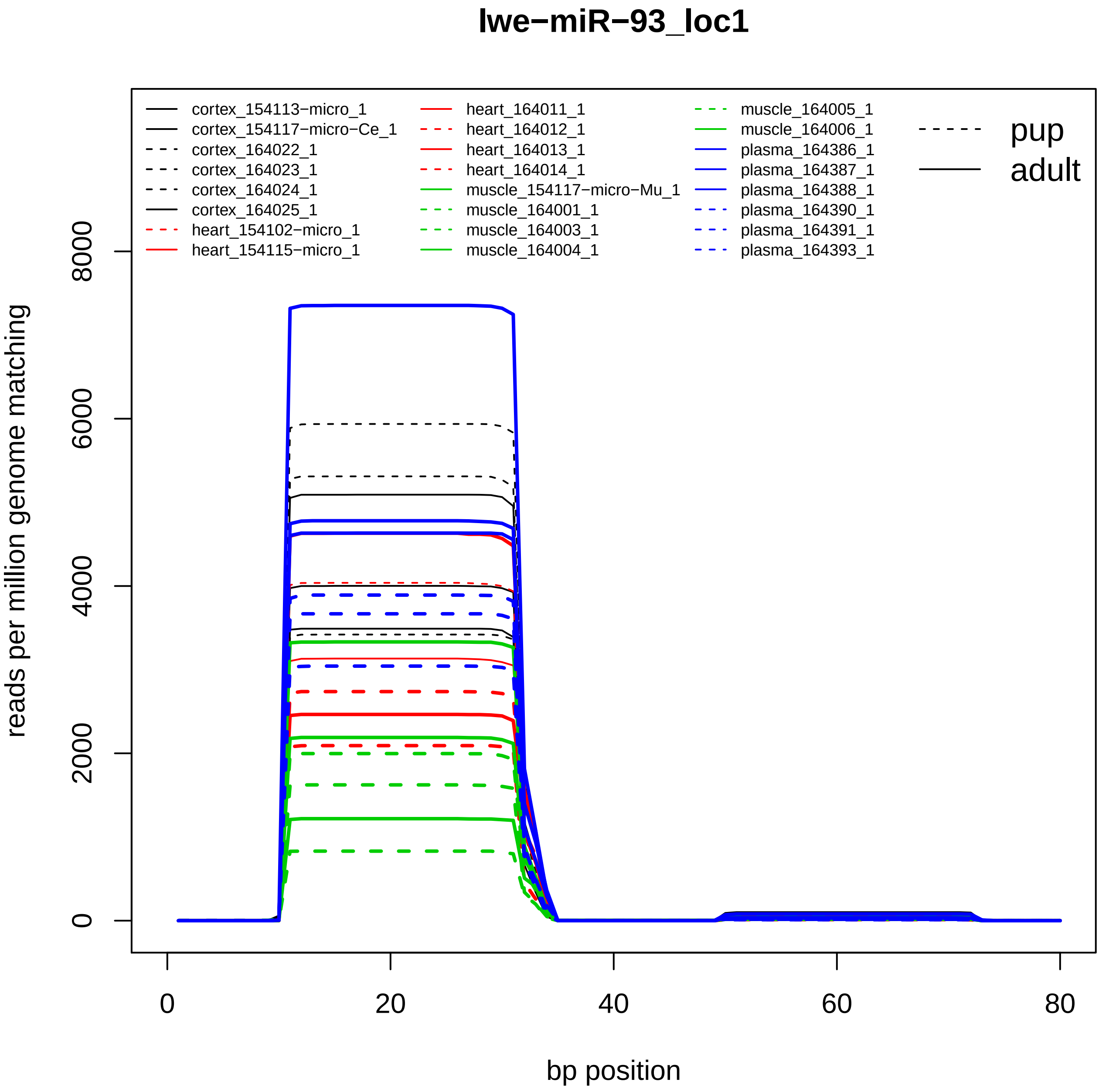
lwe−miR−93_loc1
cortex_154113−micro_1
cortex_154117−micro−Ce_1
cortex_164022_1
cortex_164023_1
cortex_164024_1
cortex_164025_1
heart_154102−micro_1
heart_154115−micro_1
heart_164011_1
heart_164012_1
heart_164013_1
heart_164014_1
muscle_154117−micro−Mu_1
muscle_164001_1
muscle_164003_1
muscle_164004_1
muscle_164005_1
muscle_164006_1
plasma_164386_1
plasma_164387_1
plasma_164388_1
plasma_164390_1
plasma_164391_1
plasma_164393_1
pup
adult
reads per million genome matching
0
2000
4000
6000
8000
0
20
40
60
80
bp position

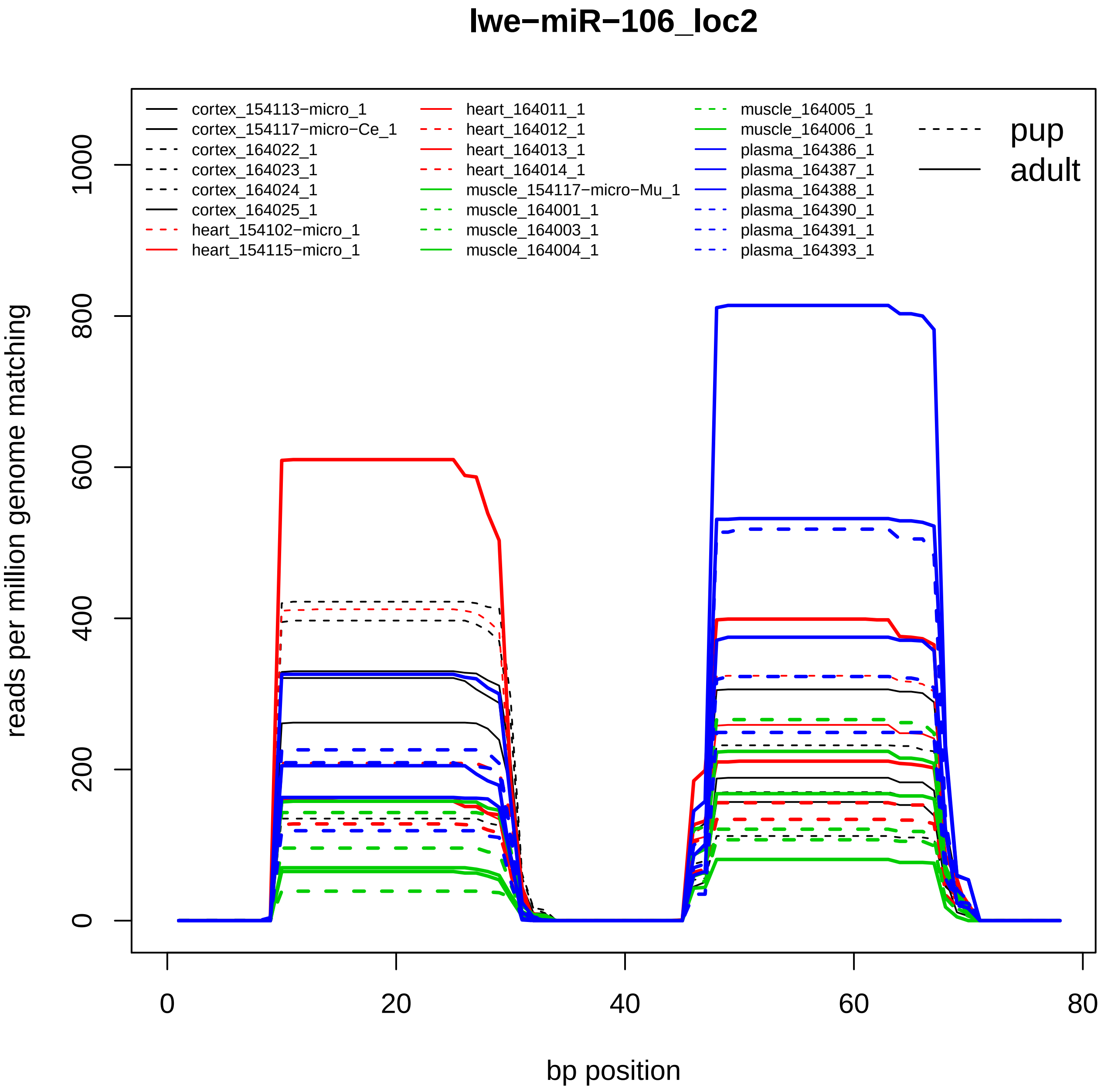
lwe−miR−106_loc2
cortex_154113−micro_1
cortex_154117−micro−Ce_1
cortex_164022_1
cortex_164023_1
cortex_164024_1
cortex_164025_1
heart_154102−micro_1
heart_154115−micro_1
heart_164011_1
heart_164012_1
heart_164013_1
heart_164014_1
muscle_154117−micro−Mu_1
muscle_164001_1
muscle_164003_1
muscle_164004_1
muscle_164005_1
muscle_164006_1
plasma_164386_1
plasma_164387_1
plasma_164388_1
plasma_164390_1
plasma_164391_1
plasma_164393_1
pup
adult
reads per million genome matching
0
200
400
600
800
1000
bp position
0
20
40
60
80

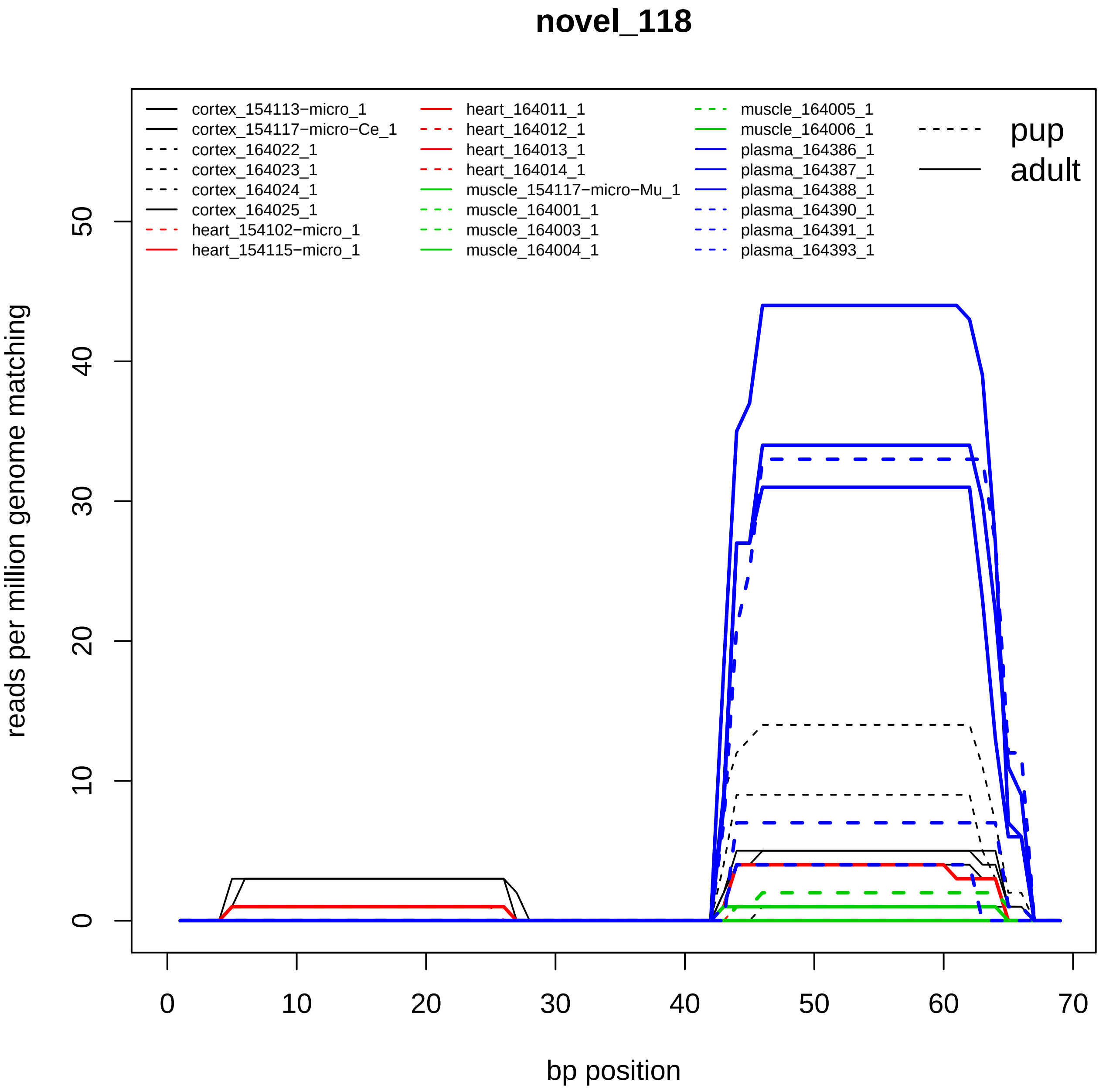

novel_118
cortex_154113−micro_1
cortex_154117−micro−Ce_1
cortex_164022_1
cortex_164023_1
cortex_164024_1
cortex_164025_1
heart_154102−micro_1
heart_154115−micro_1
heart_164011_1
heart_164012_1
heart_164013_1
heart_164014_1
muscle_154117−micro−Mu_1
muscle_164001_1
muscle_164003_1
muscle_164004_1
muscle_164005_1
muscle_164006_1
plasma_164386_1
plasma_164387_1
plasma_164388_1
plasma_164390_1
plasma_164391_1
plasma_164393_1
pup
adult
reads per million genome matching
bp position
0
10
20
30
40
50
60
70

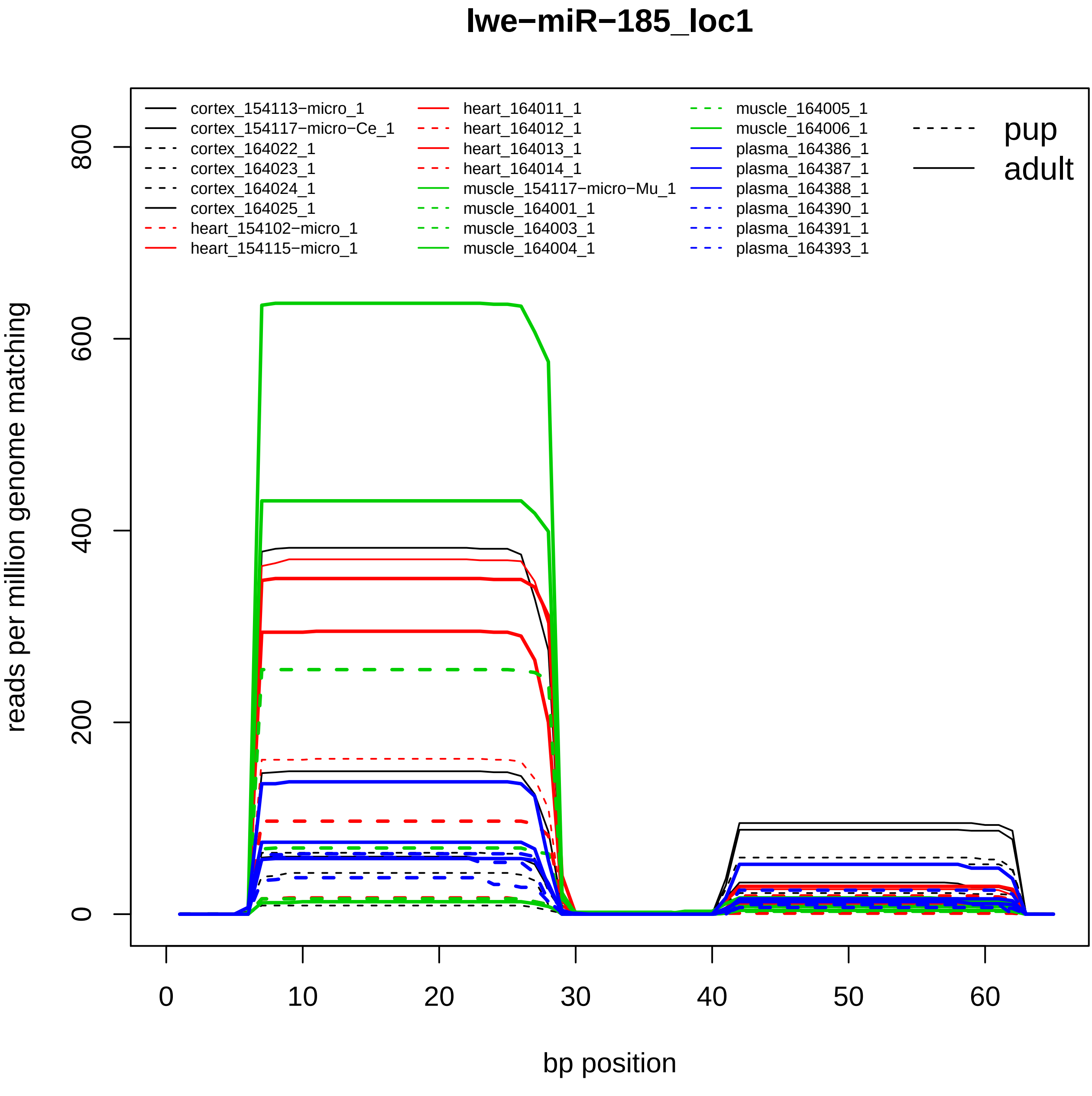

lwe−miR−185_loc1
reads per million genome matching
bp position
0
200
400
600
800
0
10
20
30
40
50
60
cortex_154113−micro_1
cortex_154117−micro−Ce_1
cortex_164022_1
cortex_164023_1
cortex_164024_1
cortex_164025_1
heart_154102−micro_1
heart_154115−micro_1
heart_164011_1
heart_164012_1
heart_164013_1
heart_164014_1
muscle_154117−micro−Mu_1
muscle_164001_1
muscle_164003_1
muscle_164004_1
muscle_164005_1
muscle_164006_1
plasma_164386_1
plasma_164387_1
plasma_164388_1
plasma_164390_1
plasma_164391_1
plasma_164393_1
pup
adult

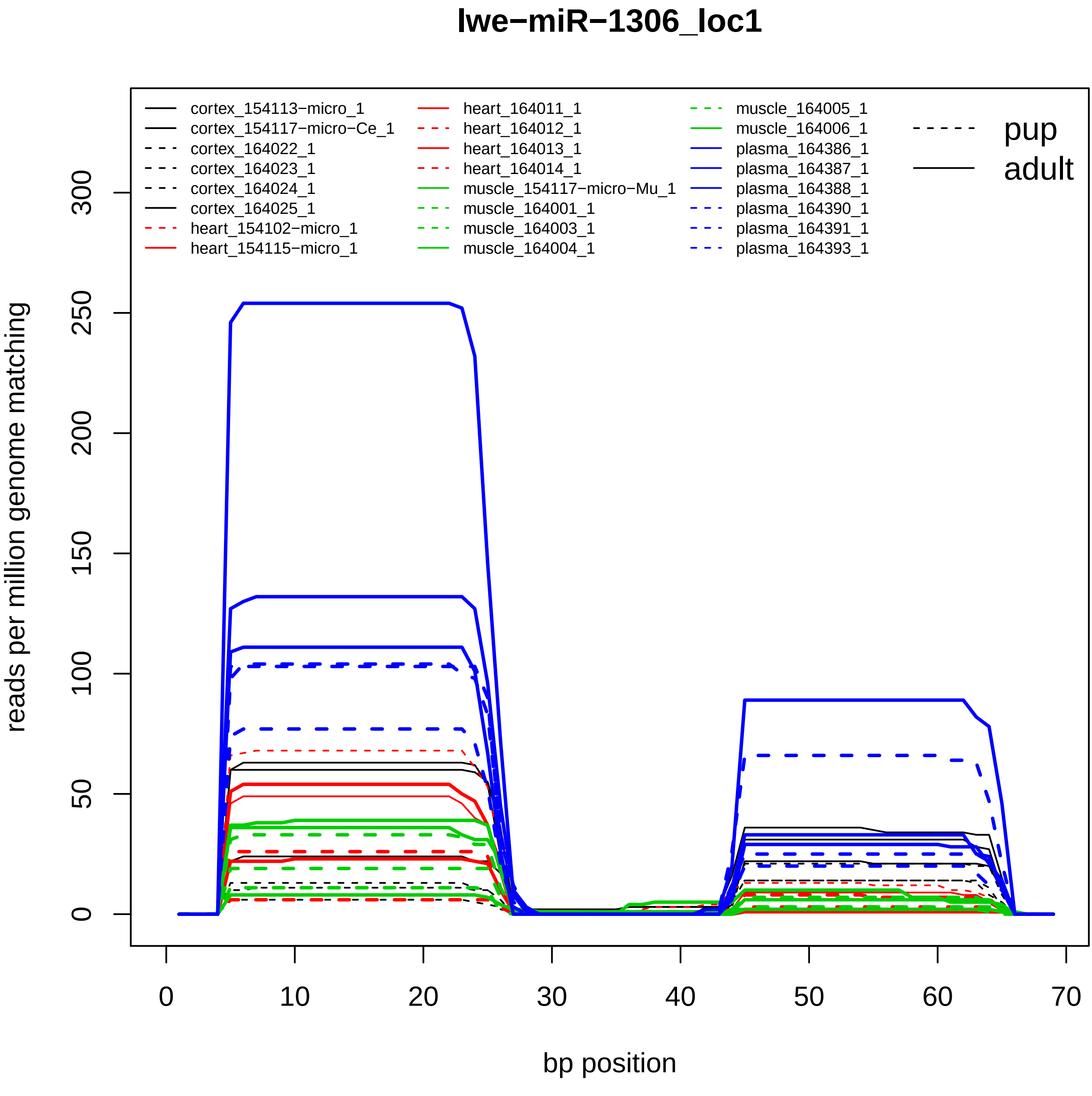
lwe−miR−1306_loc1
cortex_154113−micro_1
cortex_154117−micro−Ce_1
cortex_164022_1
cortex_164023_1
cortex_164024_1
cortex_164025_1
heart_154102−micro_1
heart_154115−micro_1
heart_164011_1
heart_164012_1
heart_164013_1
heart_164014_1
muscle_154117−micro−Mu_1
muscle_164001_1
muscle_164003_1
muscle_164004_1
muscle_164005_1
muscle_164006_1
plasma_164386_1
plasma_164387_1
plasma_164388_1
plasma_164390_1
plasma_164391_1
plasma_164393_1
pup
adult
reads per million genome matching
0
50
100
150
200
250
300
0
10
20
30
40
50
60
70
bp position

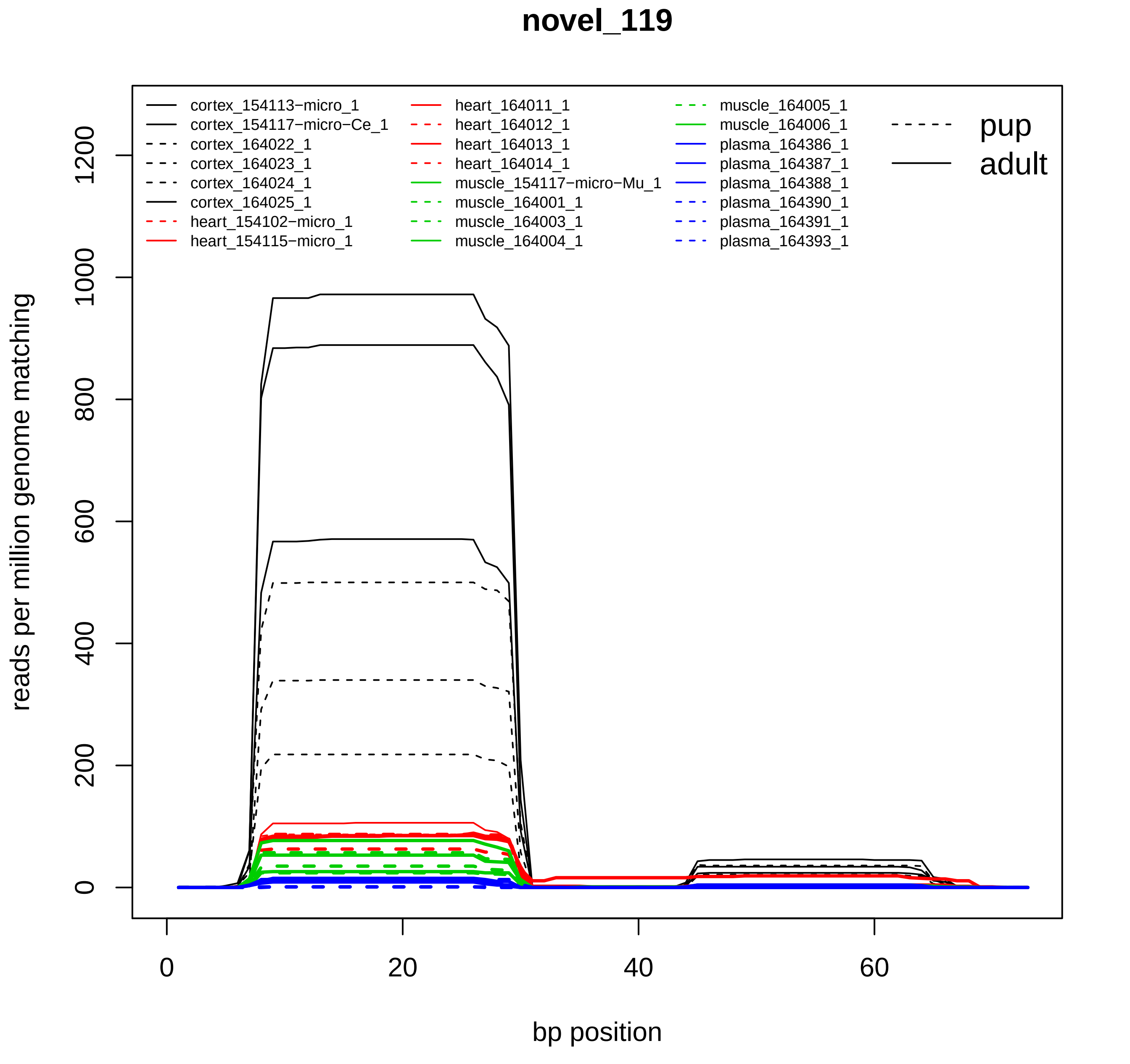

novel_119
cortex_154113−micro_1
cortex_154117−micro−Ce_1
cortex_164022_1
cortex_164023_1
cortex_164024_1
cortex_164025_1
heart_154102−micro_1
heart_154115−micro_1
heart_164011_1
heart_164012_1
heart_164013_1
heart_164014_1
muscle_154117−micro−Mu_1
muscle_164001_1
muscle_164003_1
muscle_164004_1
muscle_164005_1
muscle_164006_1
plasma_164386_1
plasma_164387_1
plasma_164388_1
plasma_164390_1
plasma_164391_1
plasma_164393_1
pup
adult
reads per million genome matching
0
200
400
600
800
1000
1200
bp position
0
20
40
60

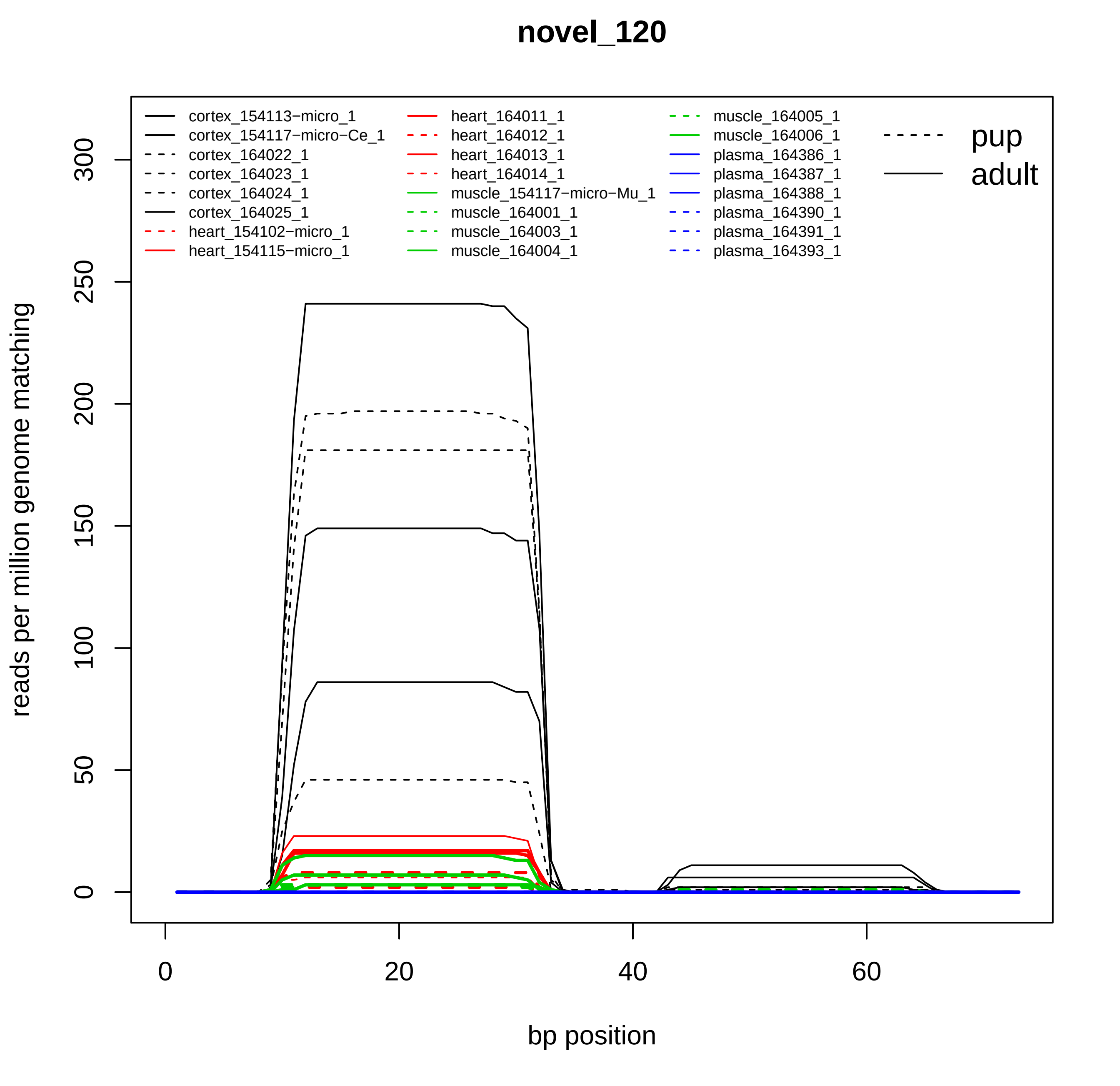

novel_120
cortex_154113−micro_1
cortex_154117−micro−Ce_1
cortex_164022_1
cortex_164023_1
cortex_164024_1
cortex_164025_1
heart_154102−micro_1
heart_154115−micro_1
heart_164011_1
heart_164012_1
heart_164013_1
heart_164014_1
muscle_154117−micro−Mu_1
muscle_164001_1
muscle_164003_1
muscle_164004_1
muscle_164005_1
muscle_164006_1
plasma_164386_1
plasma_164387_1
plasma_164388_1
plasma_164390_1
plasma_164391_1
plasma_164393_1
pup
adult
reads per million genome matching
0
50
100
150
200
250
300
0
20
40
60
bp position

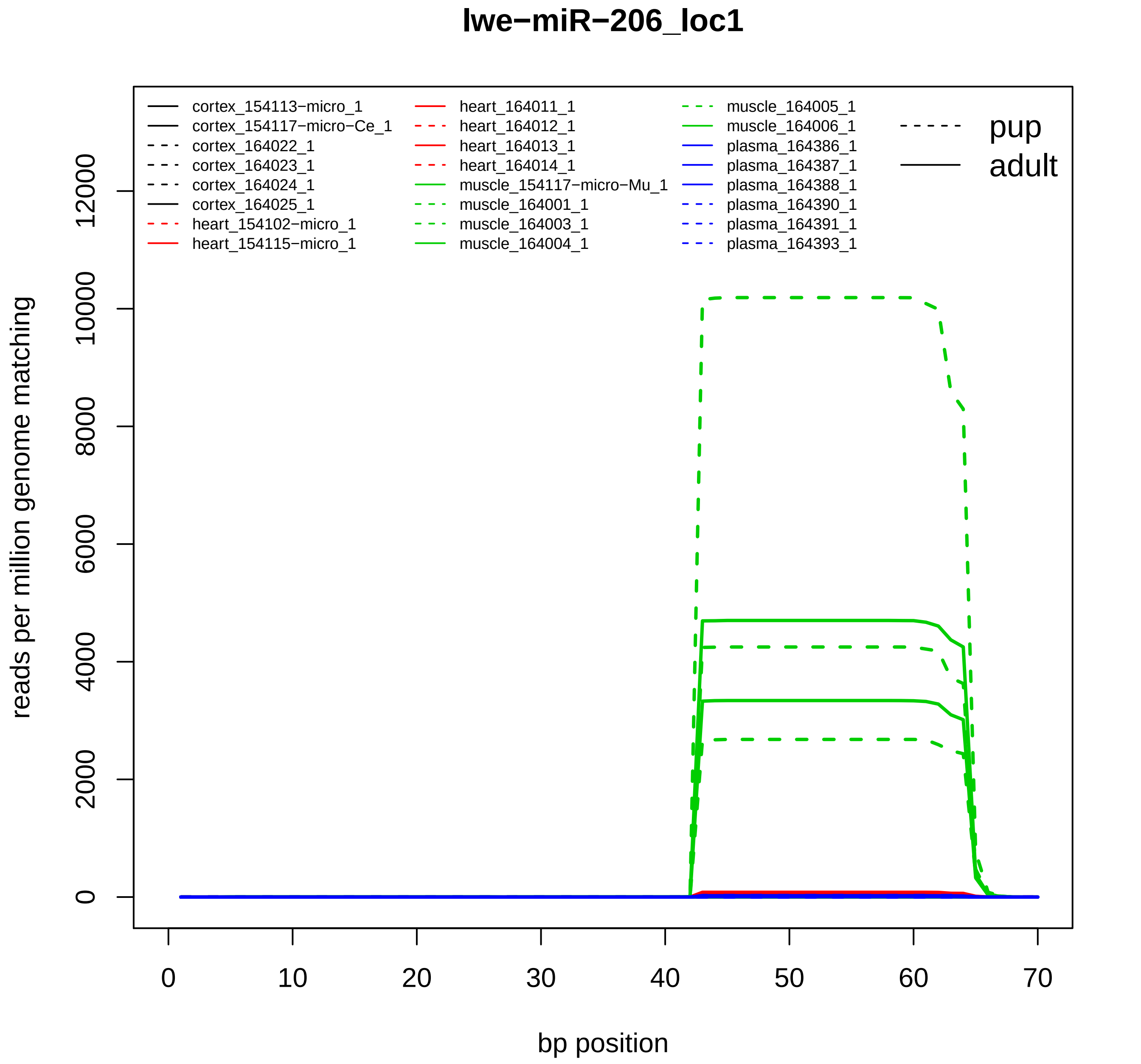
lwe−miR−206_loc1
cortex_154113−micro_1
cortex_154117−micro−Ce_1
cortex_164022_1
cortex_164023_1
cortex_164024_1
cortex_164025_1
heart_154102−micro_1
heart_154115−micro_1
heart_164011_1
heart_164012_1
heart_164013_1
heart_164014_1
muscle_154117−micro−Mu_1
muscle_164001_1
muscle_164003_1
muscle_164004_1
muscle_164005_1
muscle_164006_1
plasma_164386_1
plasma_164387_1
plasma_164388_1
plasma_164390_1
plasma_164391_1
plasma_164393_1
pup
adult
reads per million genome matching
0
2000
4000
6000
8000
10000
12000
0
10
20
30
40
50
60
70
bp position

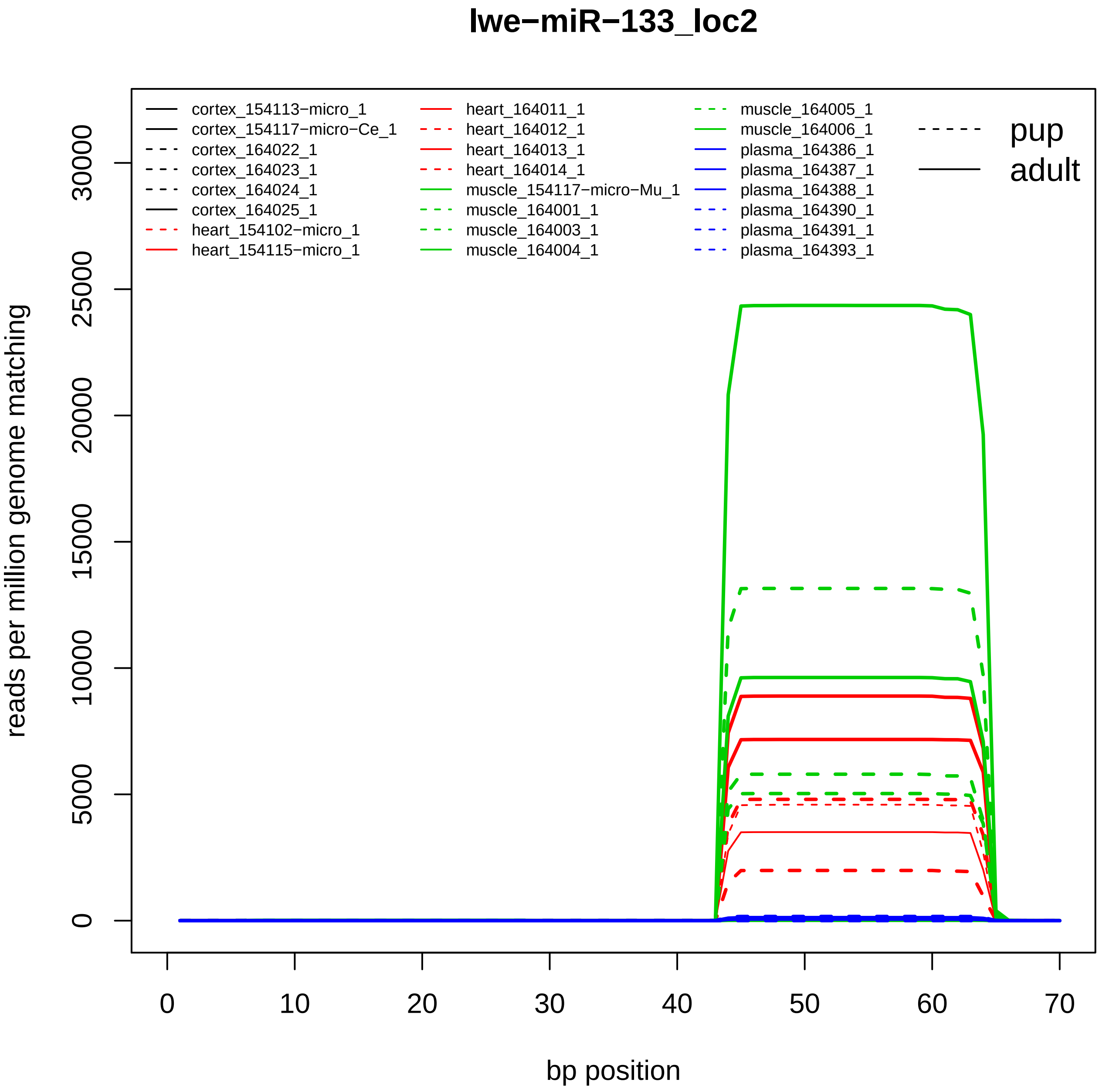
lwe−miR−133_loc2
cortex_154113−micro_1
cortex_154117−micro−Ce_1
cortex_164022_1
cortex_164023_1
cortex_164024_1
cortex_164025_1
heart_154102−micro_1
heart_154115−micro_1
heart_164011_1
heart_164012_1
heart_164013_1
heart_164014_1
muscle_154117−micro−Mu_1
muscle_164001_1
muscle_164003_1
muscle_164004_1
muscle_164005_1
muscle_164006_1
plasma_164386_1
plasma_164387_1
plasma_164388_1
plasma_164390_1
plasma_164391_1
plasma_164393_1
pup
adult
reads per million genome matching
0
5000
10000
15000
20000
25000
30000
0
10
20
30
40
50
60
70
bp position

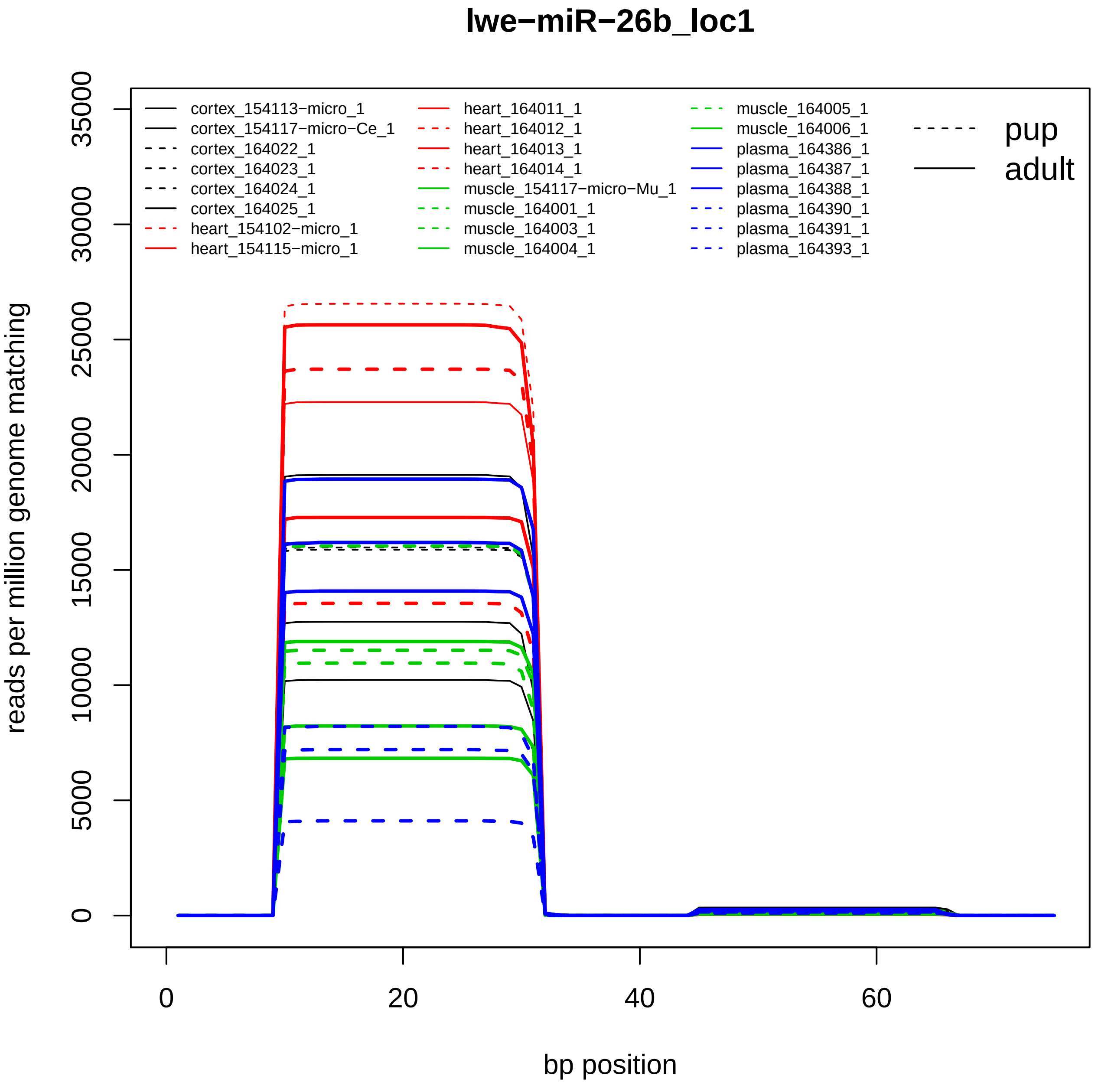

lwe−miR−26b_loc1
cortex_154113−micro_1
cortex_154117−micro−Ce_1
cortex_164022_1
cortex_164023_1
cortex_164024_1
cortex_164025_1
heart_154102−micro_1
heart_154115−micro_1
heart_164011_1
heart_164012_1
heart_164013_1
heart_164014_1
muscle_154117−micro−Mu_1
muscle_164001_1
muscle_164003_1
muscle_164004_1
muscle_164005_1
muscle_164006_1
plasma_164386_1
plasma_164387_1
plasma_164388_1
plasma_164390_1
plasma_164391_1
plasma_164393_1
pup
adult
reads per million genome matching
0
5000
10000
15000
20000
25000
30000
35000
bp position
0
20
40
60

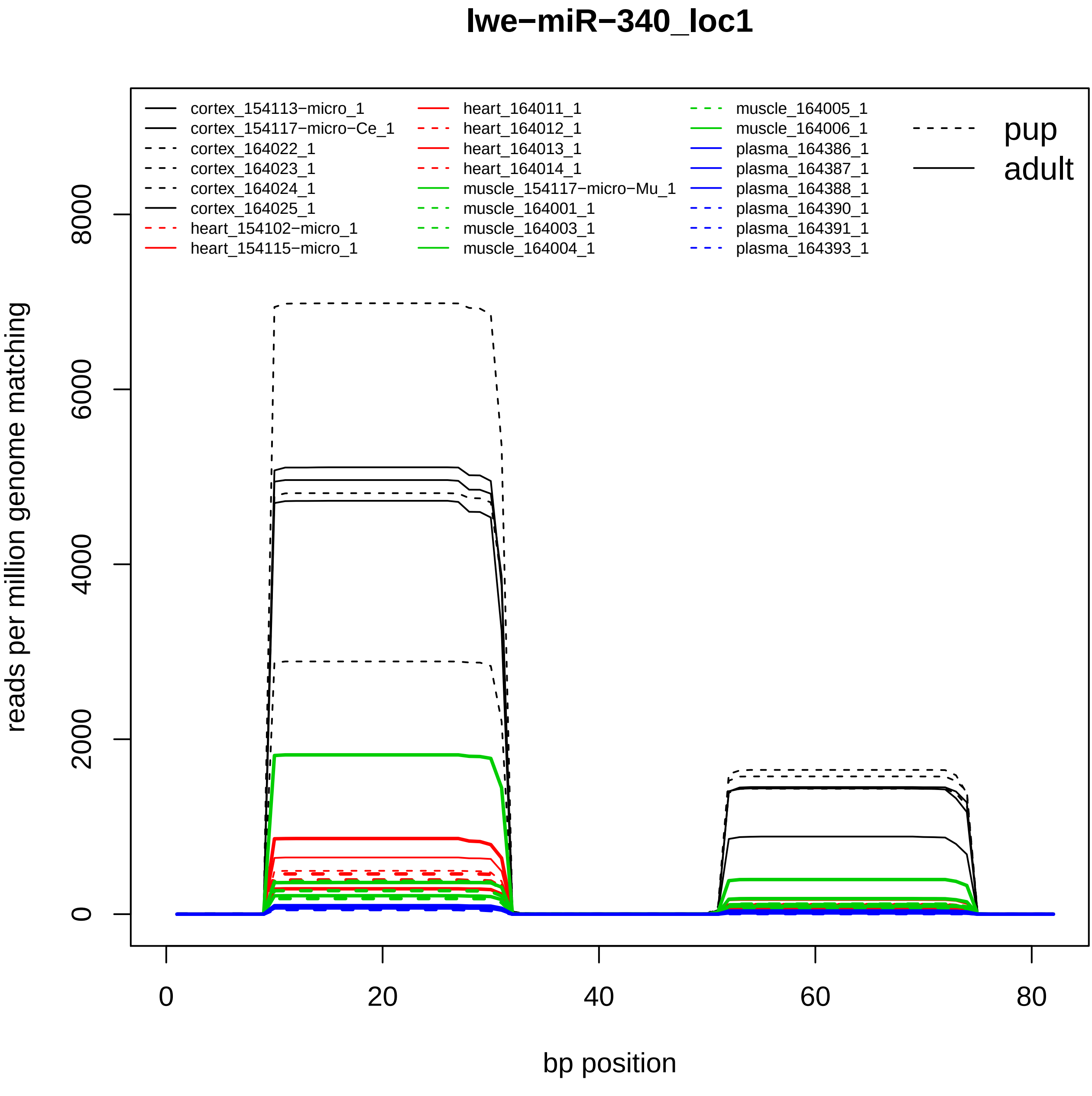

lwe−miR−340_loc1
cortex_154113−micro_1
cortex_154117−micro−Ce_1
cortex_164022_1
cortex_164023_1
cortex_164024_1
cortex_164025_1
heart_154102−micro_1
heart_154115−micro_1
heart_164011_1
heart_164012_1
heart_164013_1
heart_164014_1
muscle_154117−micro−Mu_1
muscle_164001_1
muscle_164003_1
muscle_164004_1
muscle_164005_1
muscle_164006_1
plasma_164386_1
plasma_164387_1
plasma_164388_1
plasma_164390_1
plasma_164391_1
plasma_164393_1
pup
adult
reads per million genome matching
0
2000
4000
6000
8000
0
20
40
60
80
bp position

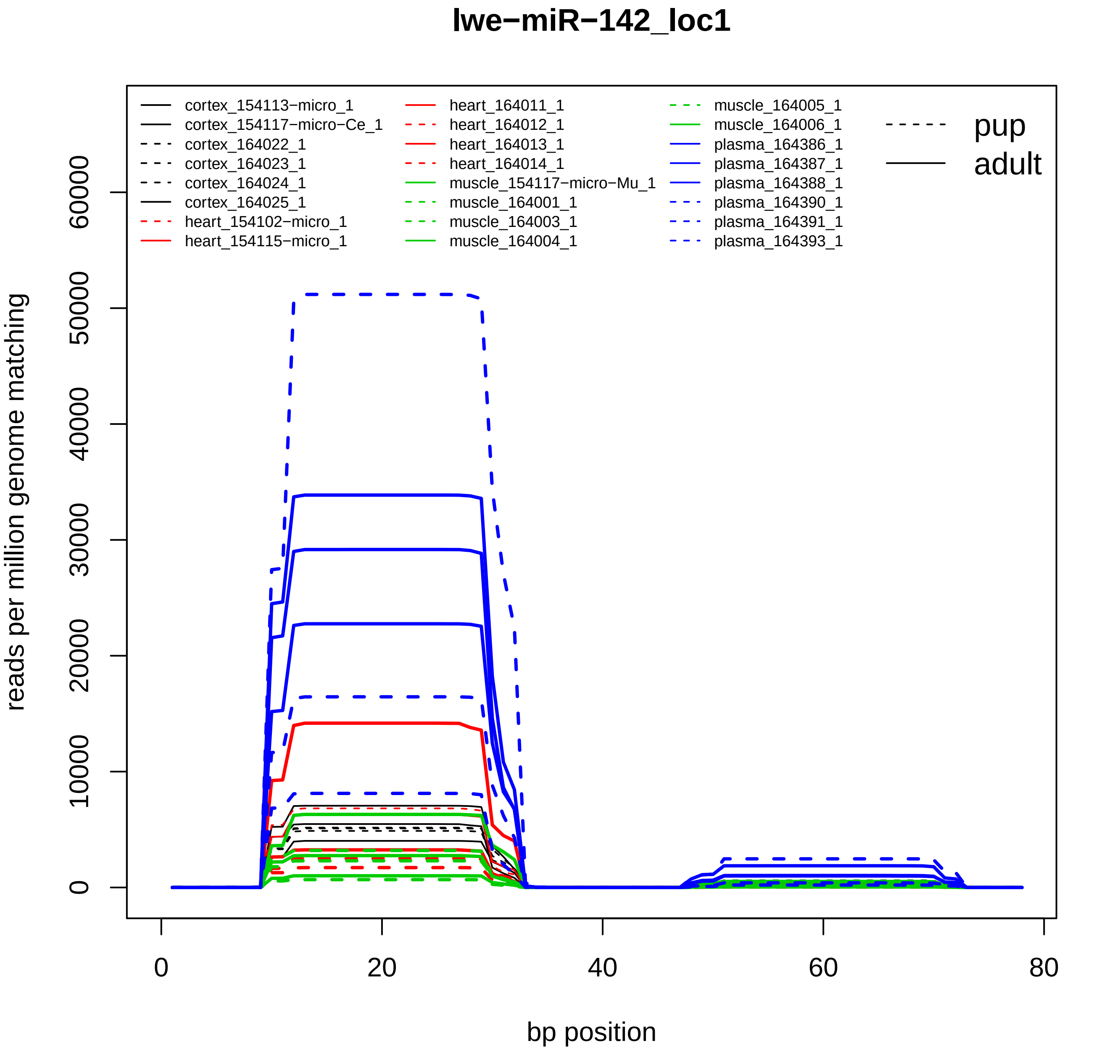

lwe−miR−142_loc1
cortex_154113−micro_1
cortex_154117−micro−Ce_1
cortex_164022_1
cortex_164023_1
cortex_164024_1
cortex_164025_1
heart_154102−micro_1
heart_154115−micro_1
heart_164011_1
heart_164012_1
heart_164013_1
heart_164014_1
muscle_154117−micro−Mu_1
muscle_164001_1
muscle_164003_1
muscle_164004_1
muscle_164005_1
muscle_164006_1
plasma_164386_1
plasma_164387_1
plasma_164388_1
plasma_164390_1
plasma_164391_1
plasma_164393_1
pup
adult
reads per million genome matching
0
10000
20000
30000
40000
50000
60000
0
20
40
60
80
bp position

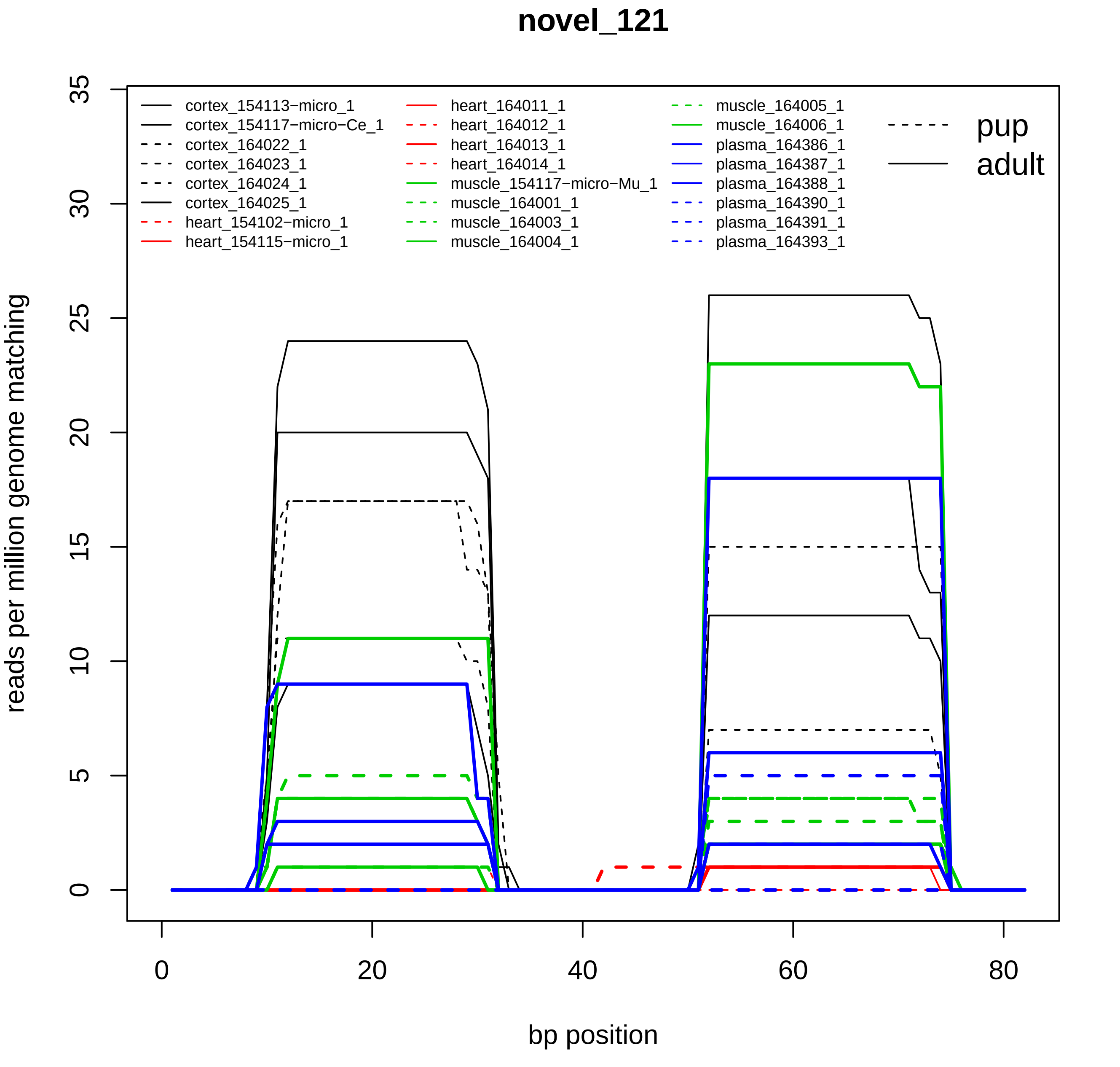
novel_121
cortex_154113−micro_1
cortex_154117−micro−Ce_1
cortex_164022_1
cortex_164023_1
cortex_164024_1
cortex_164025_1
heart_154102−micro_1
heart_154115−micro_1
heart_164011_1
heart_164012_1
heart_164013_1
heart_164014_1
muscle_154117−micro−Mu_1
muscle_164001_1
muscle_164003_1
muscle_164004_1
muscle_164005_1
muscle_164006_1
plasma_164386_1
plasma_164387_1
plasma_164388_1
plasma_164390_1
plasma_164391_1
plasma_164393_1
pup
adult
reads per million genome matching
bp position
0
5
10
15
20
25
30
35
0
20
40
60
80

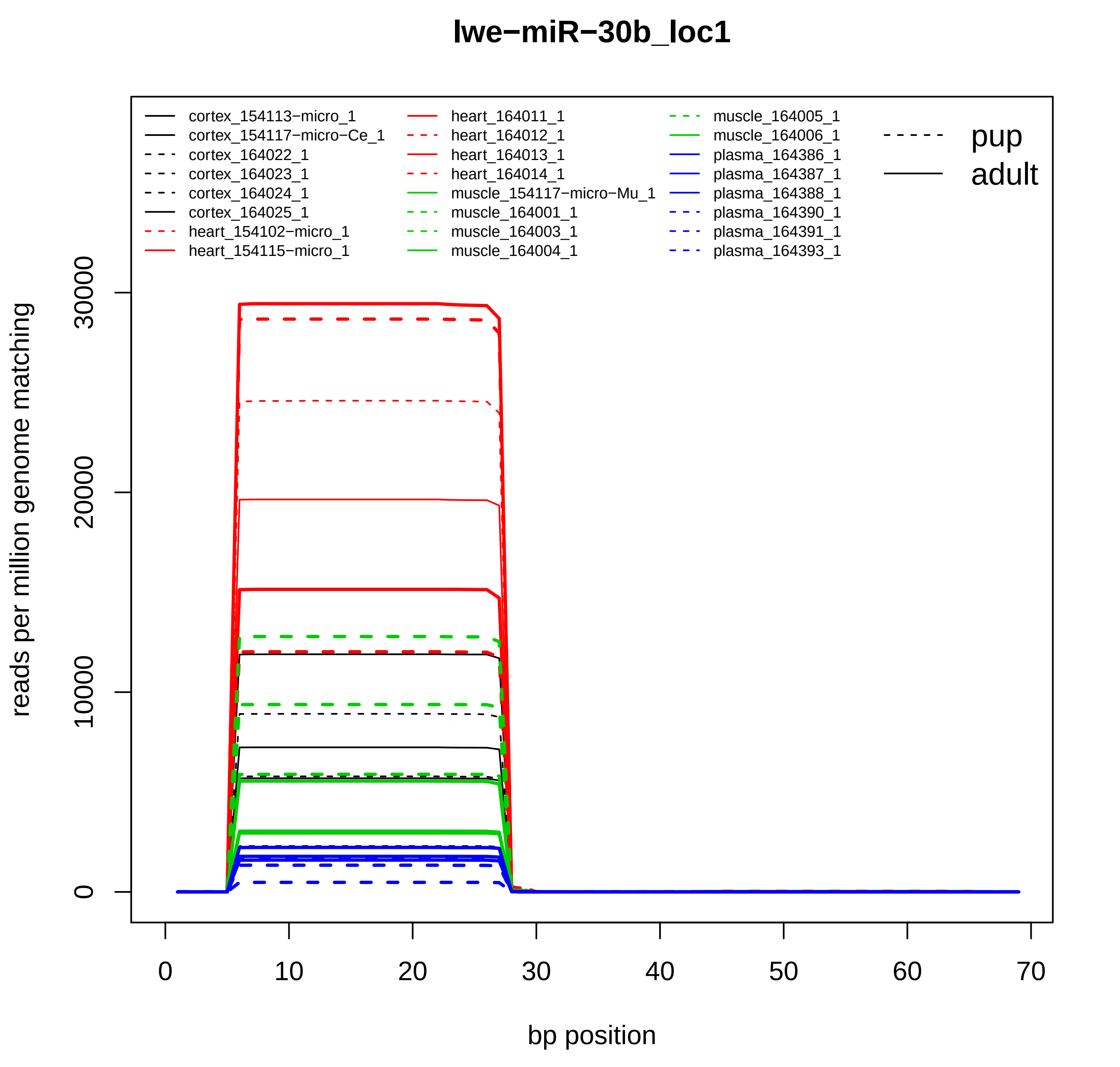
lwe−miR−30b_loc1
cortex_154113−micro_1
cortex_154117−micro−Ce_1
cortex_164022_1
cortex_164023_1
cortex_164024_1
cortex_164025_1
heart_154102−micro_1
heart_154115−micro_1
heart_164011_1
heart_164012_1
heart_164013_1
heart_164014_1
muscle_154117−micro−Mu_1
muscle_164001_1
muscle_164003_1
muscle_164004_1
muscle_164005_1
muscle_164006_1
plasma_164386_1
plasma_164387_1
plasma_164388_1
plasma_164390_1
plasma_164391_1
plasma_164393_1
pup
adult
reads per million genome matching
0
10000
20000
30000
0
10
20
30
40
50
60
70
bp position

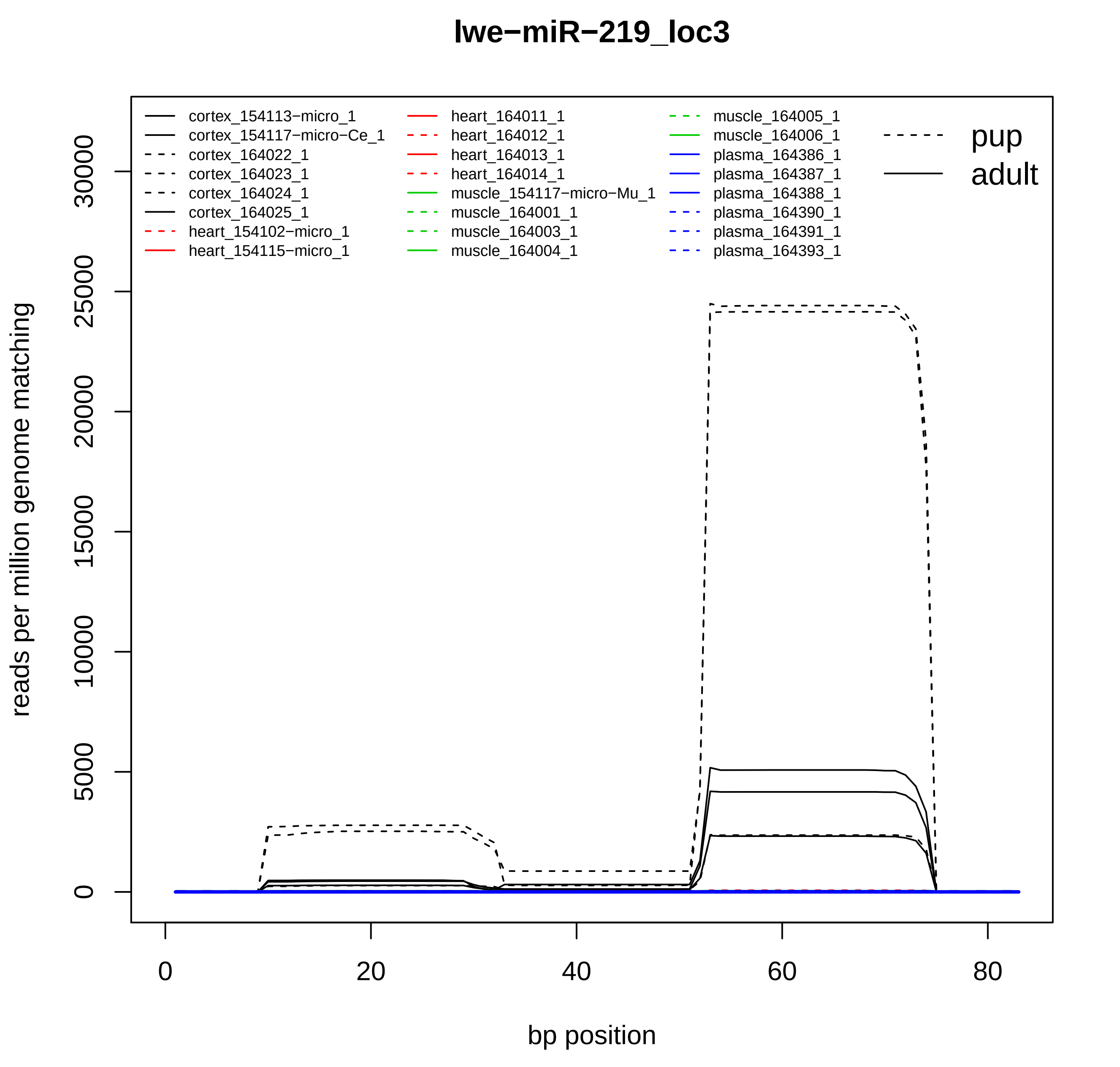
lwe−miR−219_loc3
cortex_154113−micro_1
cortex_154117−micro−Ce_1
cortex_164022_1
cortex_164023_1
cortex_164024_1
cortex_164025_1
heart_154102−micro_1
heart_154115−micro_1
heart_164011_1
heart_164012_1
heart_164013_1
heart_164014_1
muscle_154117−micro−Mu_1
muscle_164001_1
muscle_164003_1
muscle_164004_1
muscle_164005_1
muscle_164006_1
plasma_164386_1
plasma_164387_1
plasma_164388_1
plasma_164390_1
plasma_164391_1
plasma_164393_1
pup
adult
reads per million genome matching
0
5000
10000
15000
20000
25000
30000
0
20
40
60
80
bp position

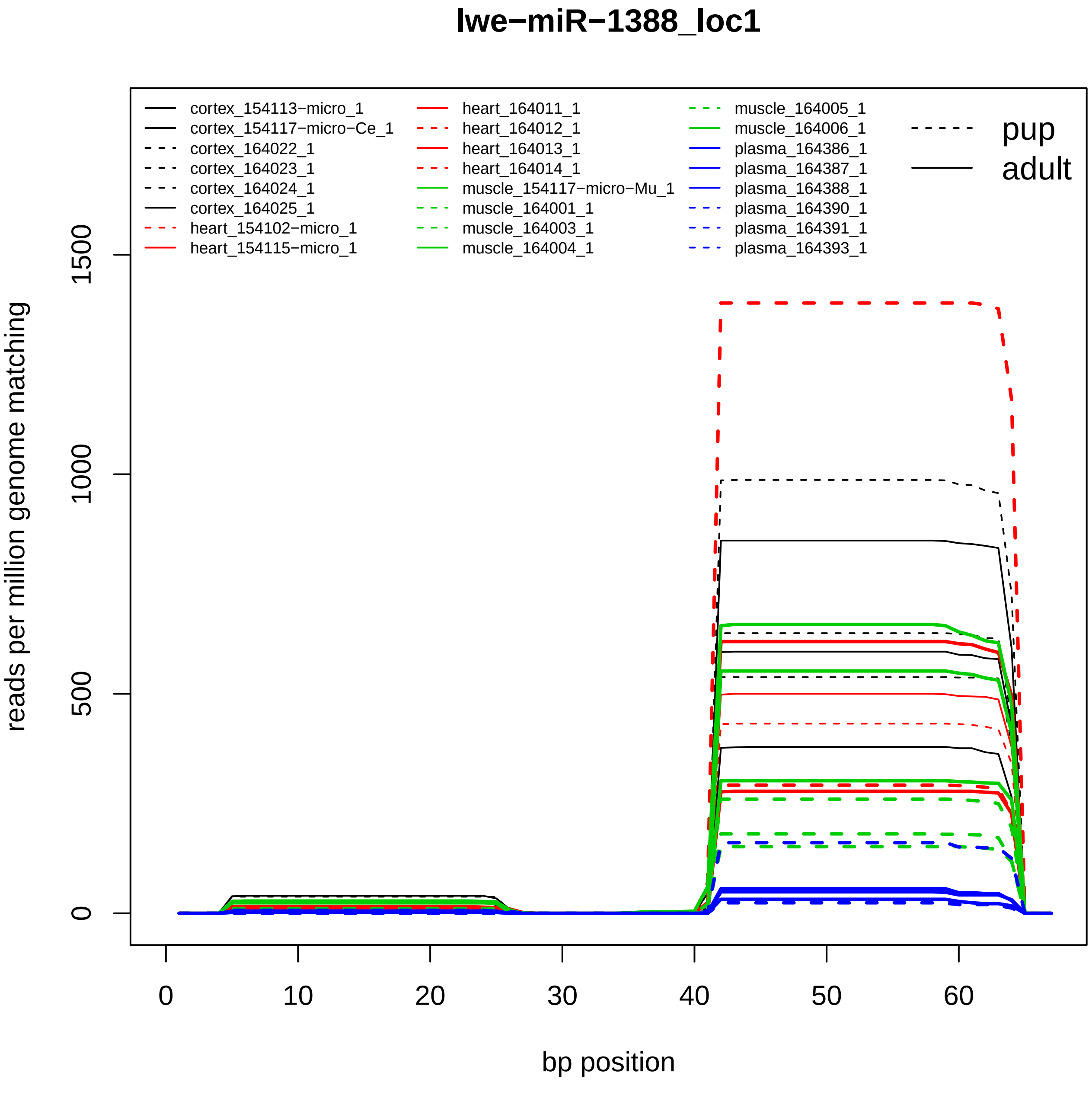
lwe−miR−1388_loc1
cortex_154113−micro_1
cortex_154117−micro−Ce_1
cortex_164022_1
cortex_164023_1
cortex_164024_1
cortex_164025_1
heart_154102−micro_1
heart_154115−micro_1
heart_164011_1
heart_164012_1
heart_164013_1
heart_164014_1
muscle_154117−micro−Mu_1
muscle_164001_1
muscle_164003_1
muscle_164004_1
muscle_164005_1
muscle_164006_1
plasma_164386_1
plasma_164387_1
plasma_164388_1
plasma_164390_1
plasma_164391_1
plasma_164393_1
pup
adult
reads per million genome matching
0
500
1000
1500
0
10
20
30
40
50
60
bp position

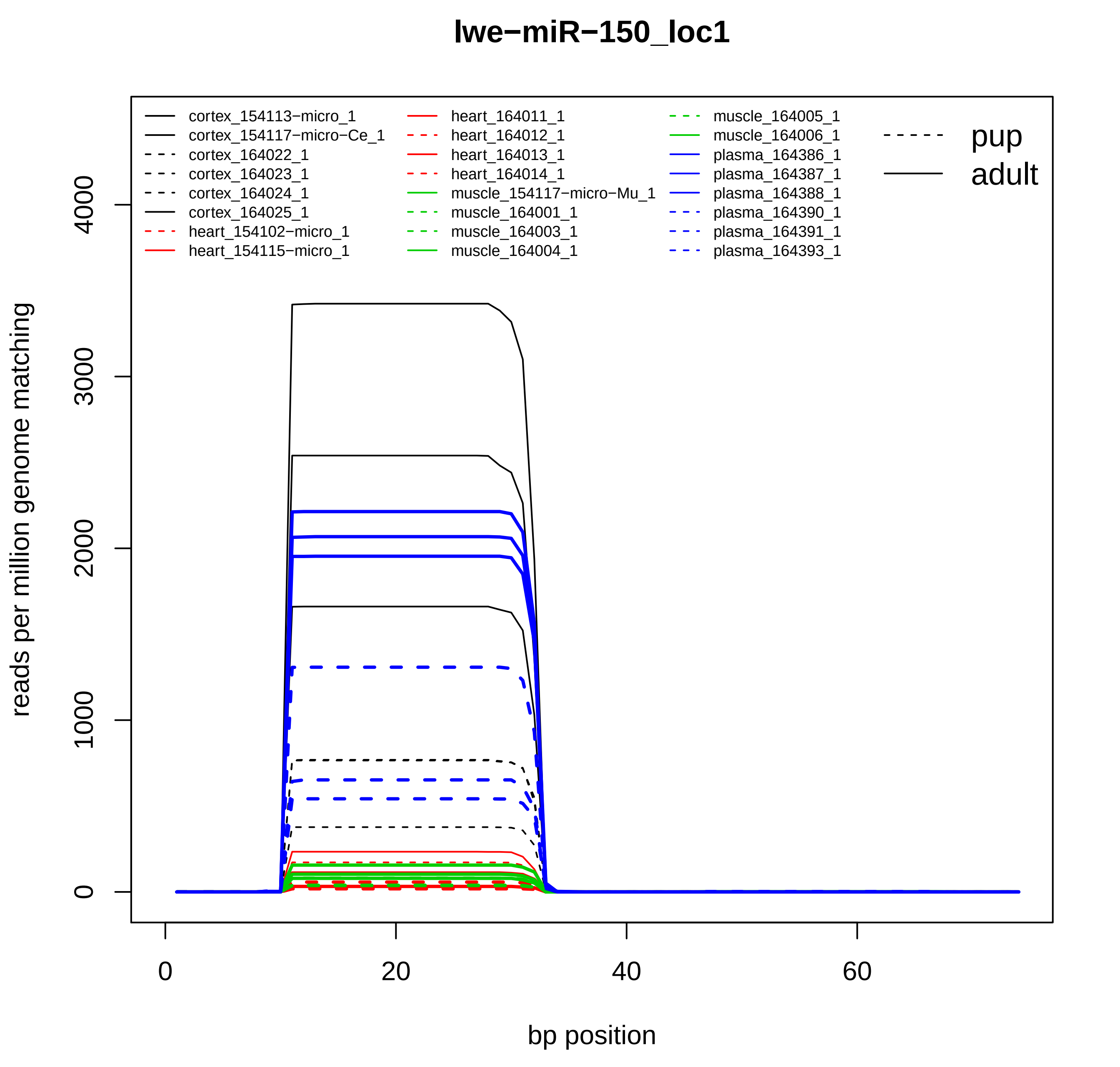
lwe−miR−150_loc1
cortex_154113−micro_1
cortex_154117−micro−Ce_1
cortex_164022_1
cortex_164023_1
cortex_164024_1
cortex_164025_1
heart_154102−micro_1
heart_154115−micro_1
heart_164011_1
heart_164012_1
heart_164013_1
heart_164014_1
muscle_154117−micro−Mu_1
muscle_164001_1
muscle_164003_1
muscle_164004_1
muscle_164005_1
muscle_164006_1
plasma_164386_1
plasma_164387_1
plasma_164388_1
plasma_164390_1
plasma_164391_1
plasma_164393_1
pup
adult
reads per million genome matching
0
1000
2000
3000
4000
0
20
40
60
bp position

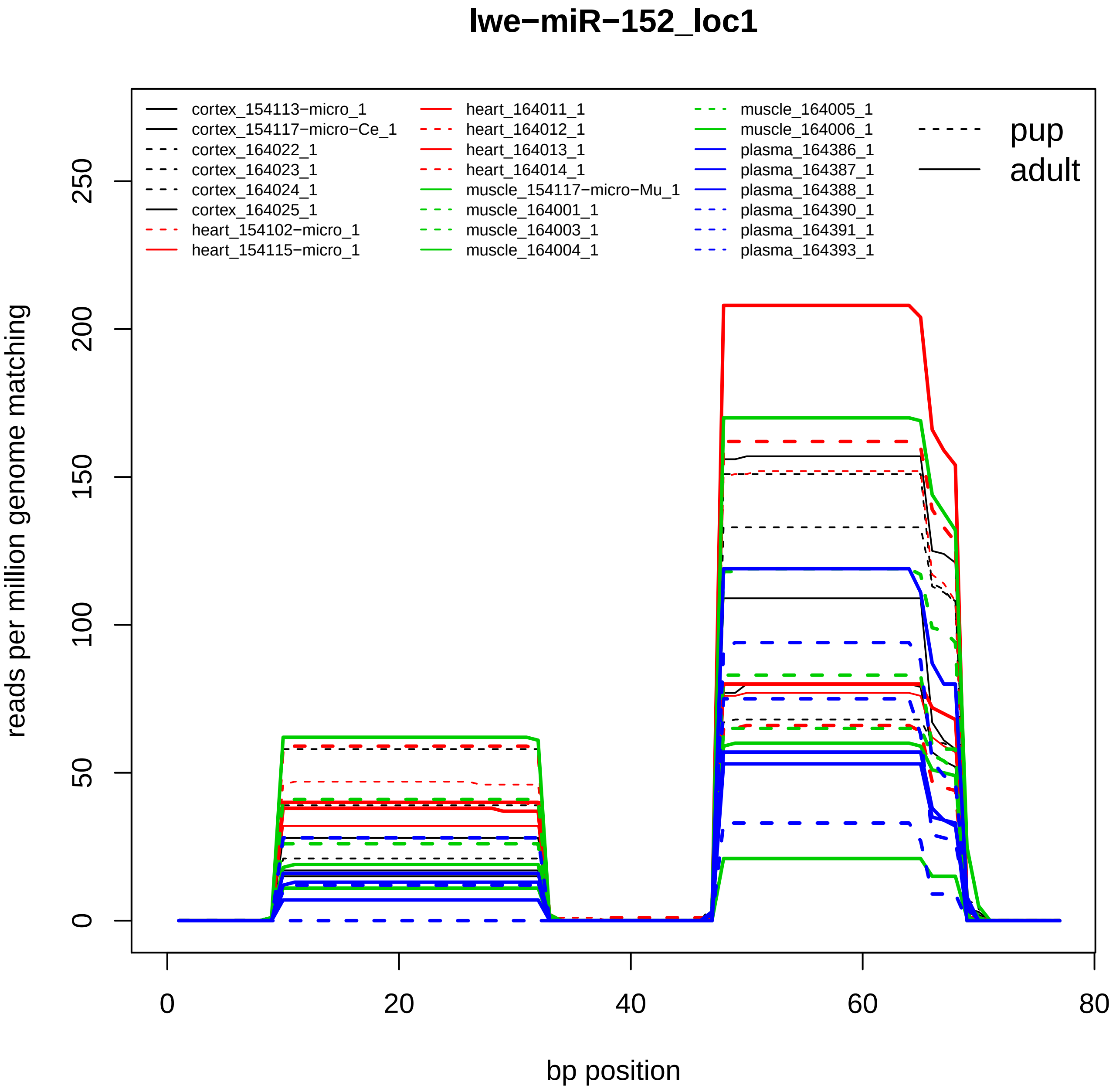
lwe−miR−152_loc1
cortex_154113−micro_1
cortex_154117−micro−Ce_1
cortex_164022_1
cortex_164023_1
cortex_164024_1
cortex_164025_1
heart_154102−micro_1
heart_154115−micro_1
heart_164011_1
heart_164012_1
heart_164013_1
heart_164014_1
muscle_154117−micro−Mu_1
muscle_164001_1
muscle_164003_1
muscle_164004_1
muscle_164005_1
muscle_164006_1
plasma_164386_1
plasma_164387_1
plasma_164388_1
plasma_164390_1
plasma_164391_1
plasma_164393_1
pup
adult
reads per million genome matching
0
50
100
150
200
250
0
20
40
60
80
bp position

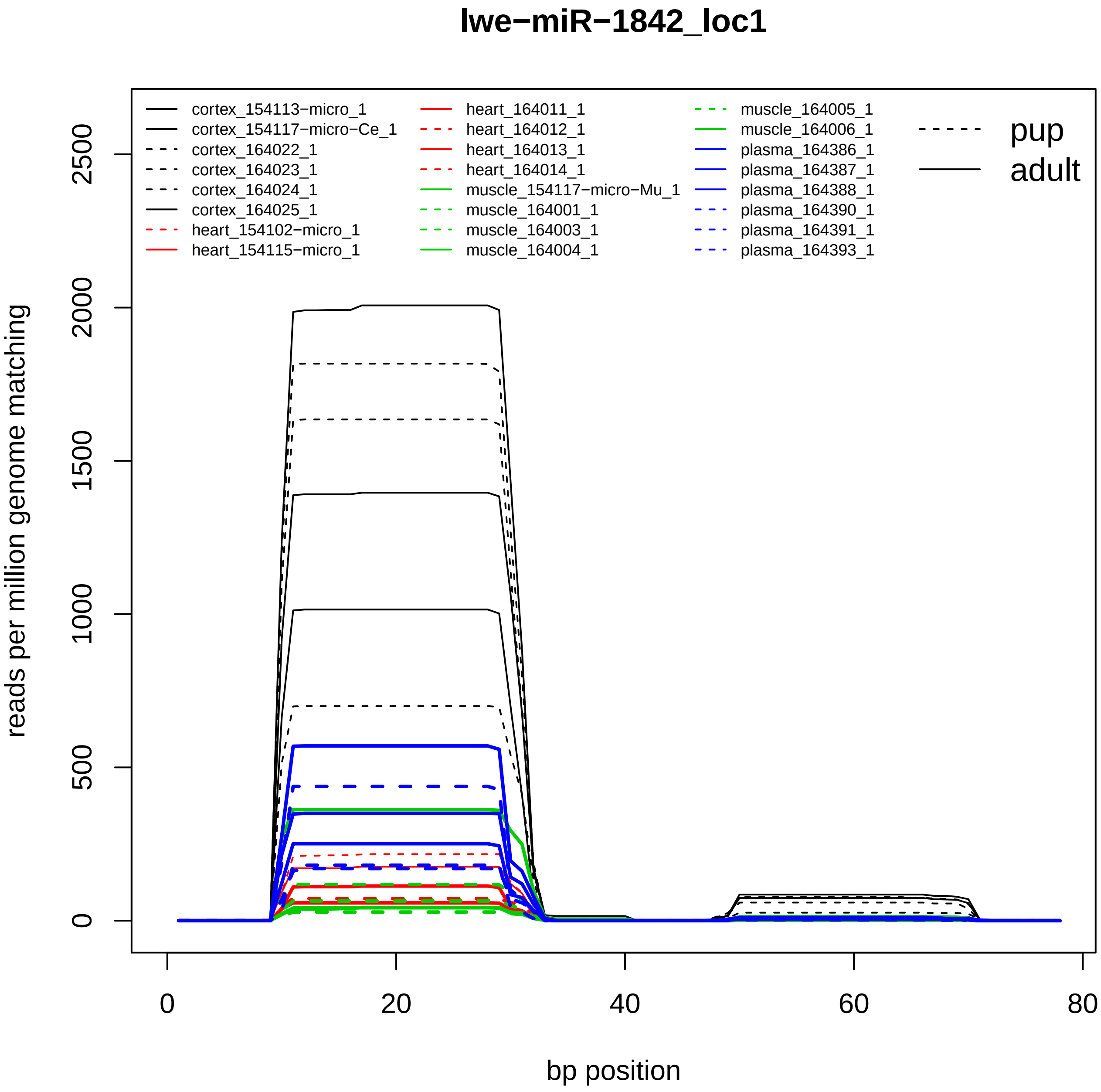
lwe−miR−1842_loc1
cortex_154113−micro_1
cortex_154117−micro−Ce_1
cortex_164022_1
cortex_164023_1
cortex_164024_1
cortex_164025_1
heart_154102−micro_1
heart_154115−micro_1
heart_164011_1
heart_164012_1
heart_164013_1
heart_164014_1
muscle_154117−micro−Mu_1
muscle_164001_1
muscle_164003_1
muscle_164004_1
muscle_164005_1
muscle_164006_1
plasma_164386_1
plasma_164387_1
plasma_164388_1
plasma_164390_1
plasma_164391_1
plasma_164393_1
pup
adult
reads per million genome matching
0
500
1000
1500
2000
2500
bp position
0
20
40
60
80

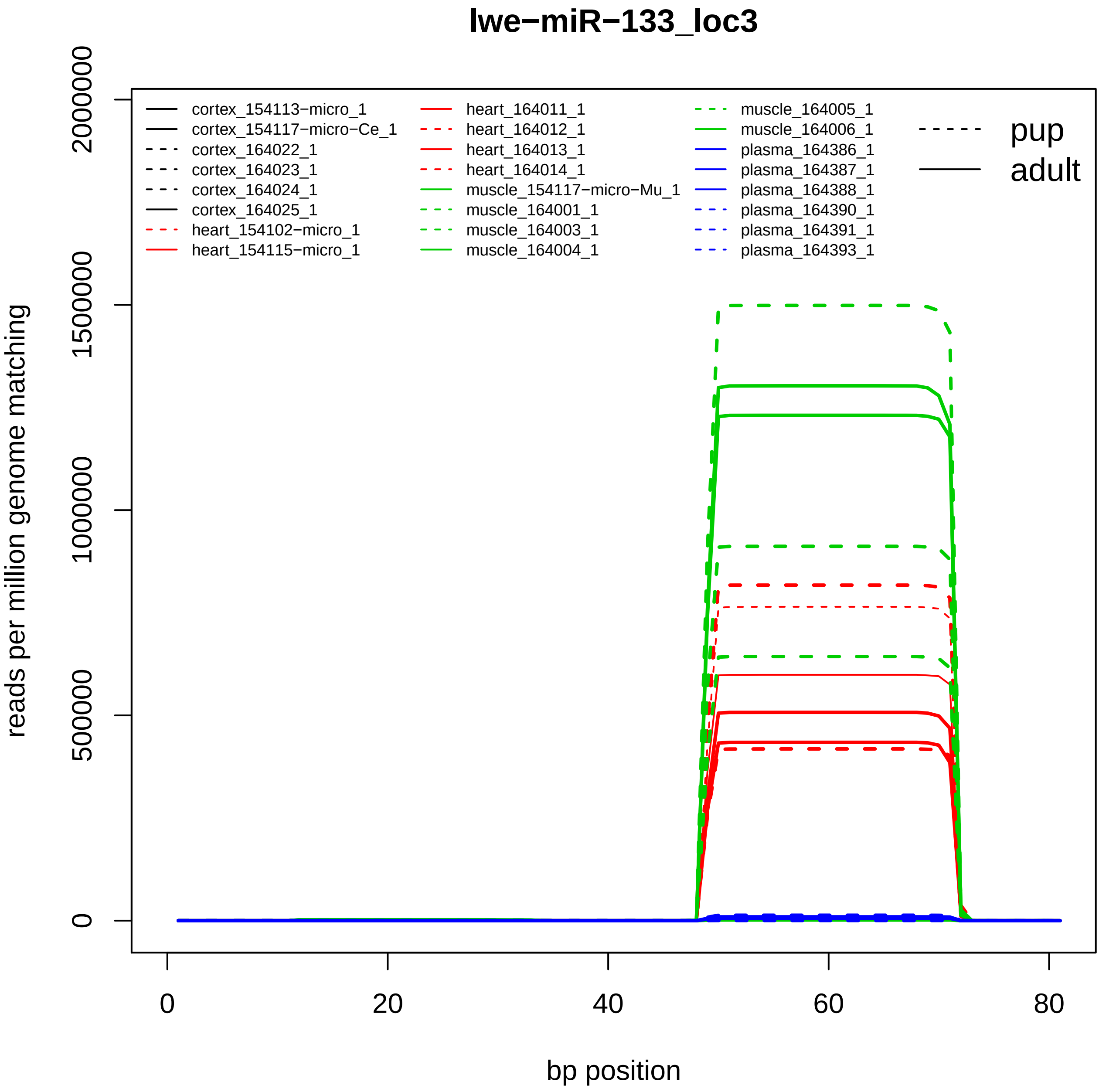
lwe−miR−133_loc3
reads per million genome matching
bp position
0
500000
1000000
1500000
2000000
0
20
40
60
80
cortex_154113−micro_1
cortex_154117−micro−Ce_1
cortex_164022_1
cortex_164023_1
cortex_164024_1
cortex_164025_1
heart_154102−micro_1
heart_154115−micro_1
heart_164011_1
heart_164012_1
heart_164013_1
heart_164014_1
muscle_154117−micro−Mu_1
muscle_164001_1
muscle_164003_1
muscle_164004_1
muscle_164005_1
muscle_164006_1
plasma_164386_1
plasma_164387_1
plasma_164388_1
plasma_164390_1
plasma_164391_1
plasma_164393_1
pup
adult

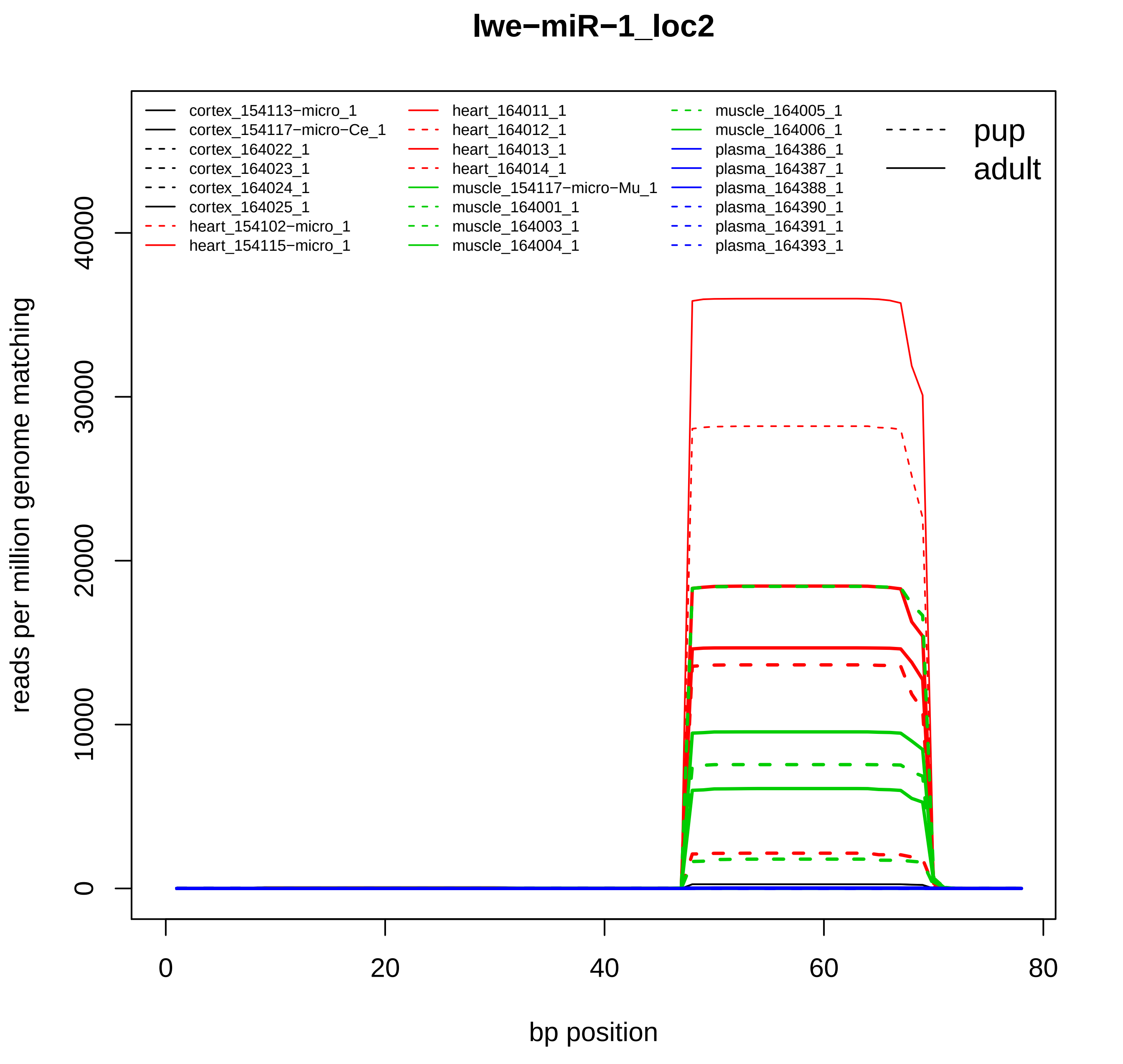

lwe−miR−1_loc2
cortex_154113−micro_1
cortex_154117−micro−Ce_1
cortex_164022_1
cortex_164023_1
cortex_164024_1
cortex_164025_1
heart_154102−micro_1
heart_154115−micro_1
heart_164011_1
heart_164012_1
heart_164013_1
heart_164014_1
muscle_154117−micro−Mu_1
muscle_164001_1
muscle_164003_1
muscle_164004_1
muscle_164005_1
muscle_164006_1
plasma_164386_1
plasma_164387_1
plasma_164388_1
plasma_164390_1
plasma_164391_1
plasma_164393_1
pup
adult
reads per million genome matching
0
10000
20000
30000
40000
0
20
40
60
80
bp position

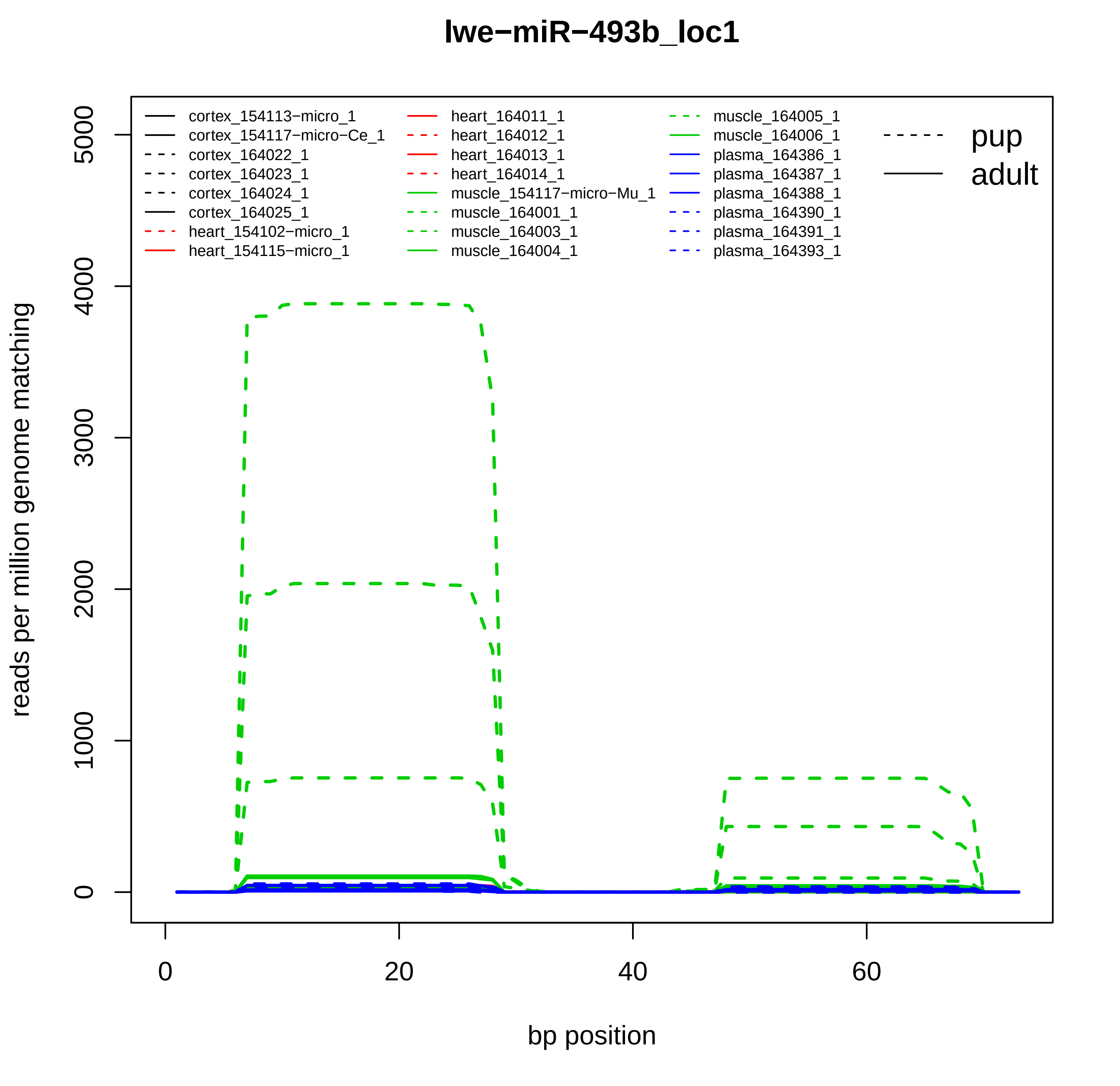
lwe−miR−493b_loc1
cortex_154113−micro_1
cortex_154117−micro−Ce_1
cortex_164022_1
cortex_164023_1
cortex_164024_1
cortex_164025_1
heart_154102−micro_1
heart_154115−micro_1
heart_164011_1
heart_164012_1
heart_164013_1
heart_164014_1
muscle_154117−micro−Mu_1
muscle_164001_1
muscle_164003_1
muscle_164004_1
muscle_164005_1
muscle_164006_1
plasma_164386_1
plasma_164387_1
plasma_164388_1
plasma_164390_1
plasma_164391_1
plasma_164393_1
pup
adult
reads per million genome matching
0
1000
2000
3000
4000
5000
0
20
40
60
bp position

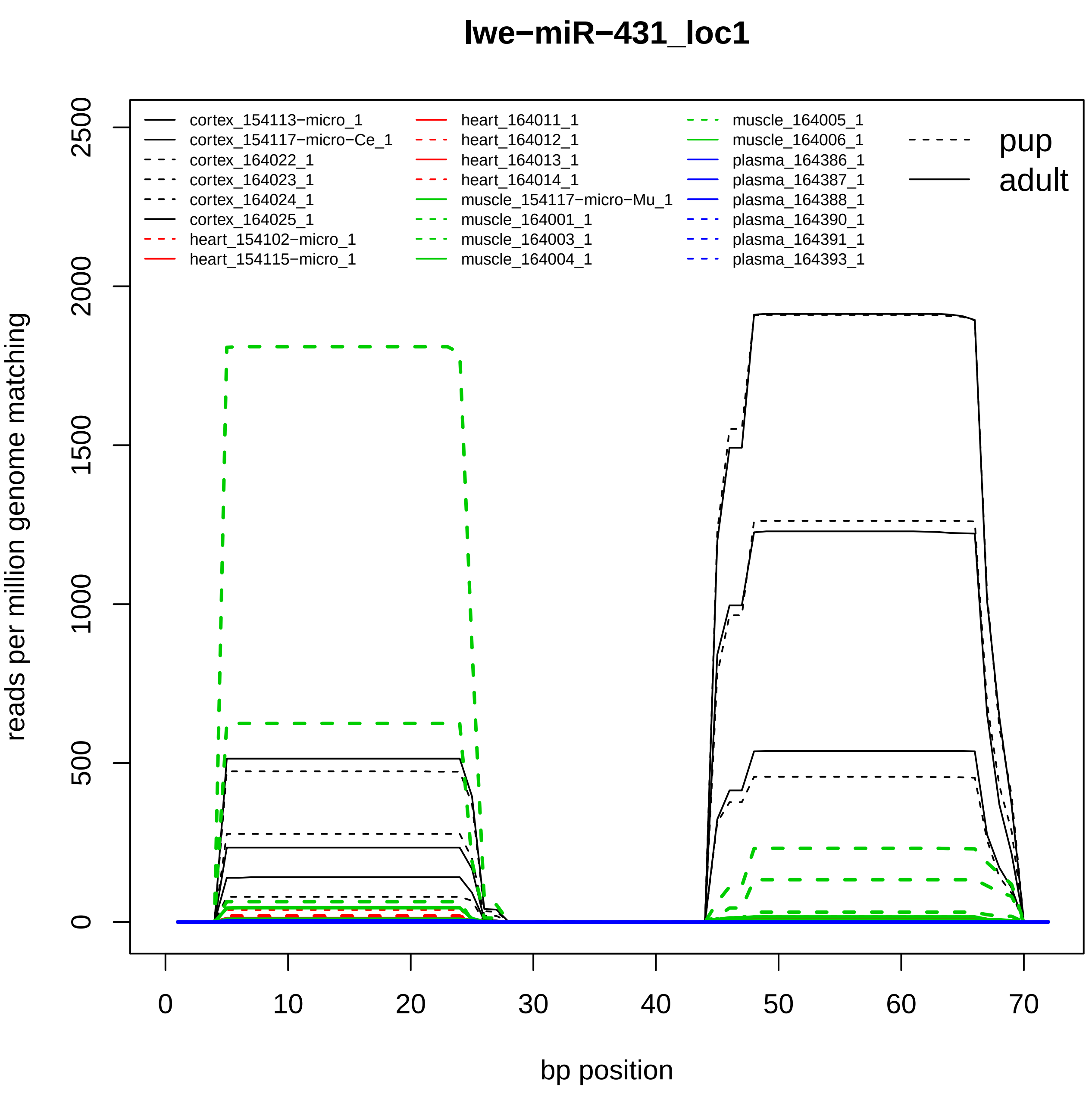
lwe−miR−431_loc1
cortex_154113−micro_1
cortex_154117−micro−Ce_1
cortex_164022_1
cortex_164023_1
cortex_164024_1
cortex_164025_1
heart_154102−micro_1
heart_154115−micro_1
heart_164011_1
heart_164012_1
heart_164013_1
heart_164014_1
muscle_154117−micro−Mu_1
muscle_164001_1
muscle_164003_1
muscle_164004_1
muscle_164005_1
muscle_164006_1
plasma_164386_1
plasma_164387_1
plasma_164388_1
plasma_164390_1
plasma_164391_1
plasma_164393_1
pup
adult
reads per million genome matching
0
500
1000
1500
2000
2500
0
10
20
30
40
50
60
70
bp position

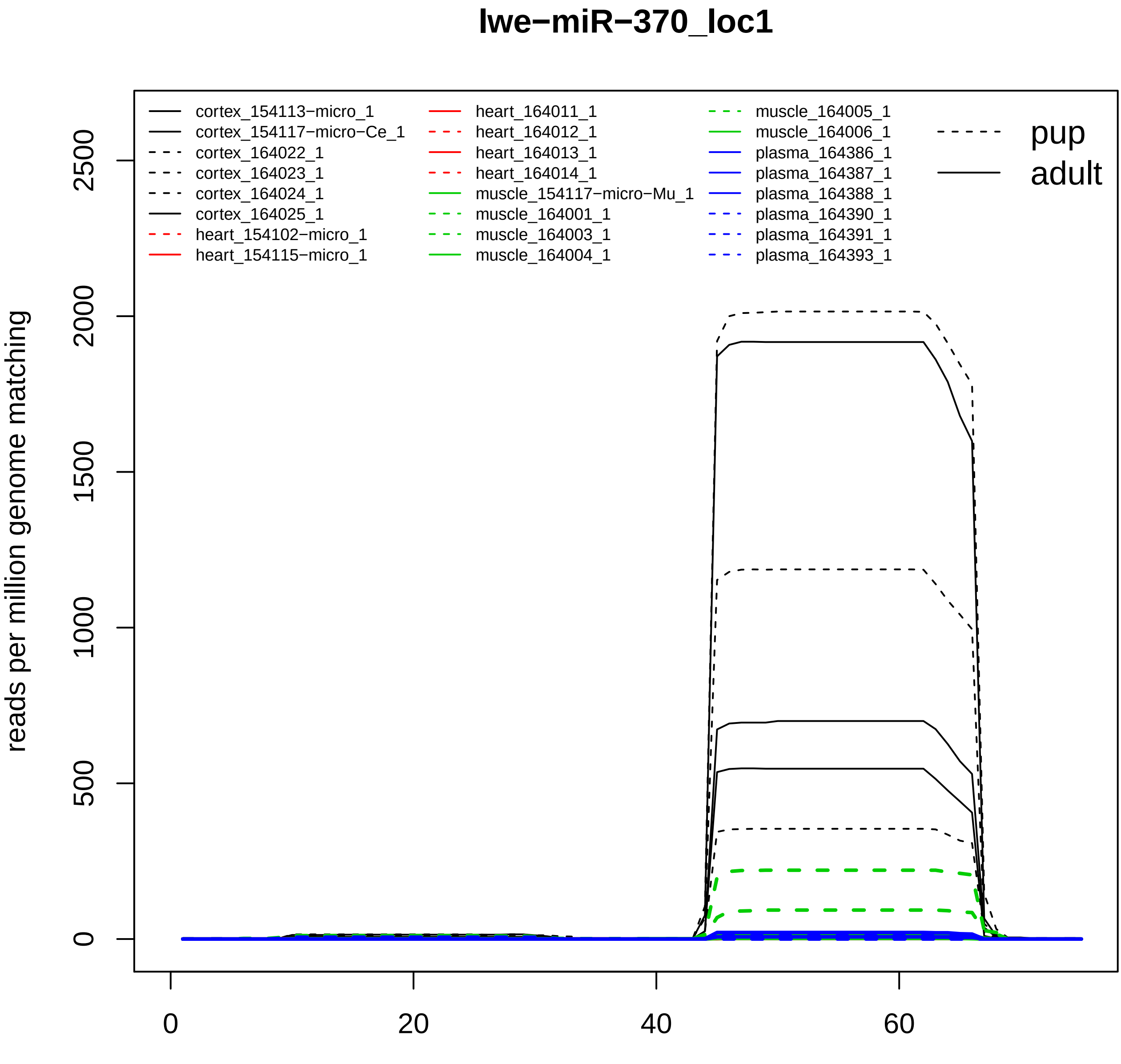
lwe−miR−370_loc1
cortex_154113−micro_1
cortex_154117−micro−Ce_1
cortex_164022_1
cortex_164023_1
cortex_164024_1
cortex_164025_1
heart_154102−micro_1
heart_154115−micro_1
heart_164011_1
heart_164012_1
heart_164013_1
heart_164014_1
muscle_154117−micro−Mu_1
muscle_164001_1
muscle_164003_1
muscle_164004_1
muscle_164005_1
muscle_164006_1
plasma_164386_1
plasma_164387_1
plasma_164388_1
plasma_164390_1
plasma_164391_1
plasma_164393_1
pup
adult
reads per million genome matching
0
500
1000
1500
2000
2500
bp position
0
20
40
60

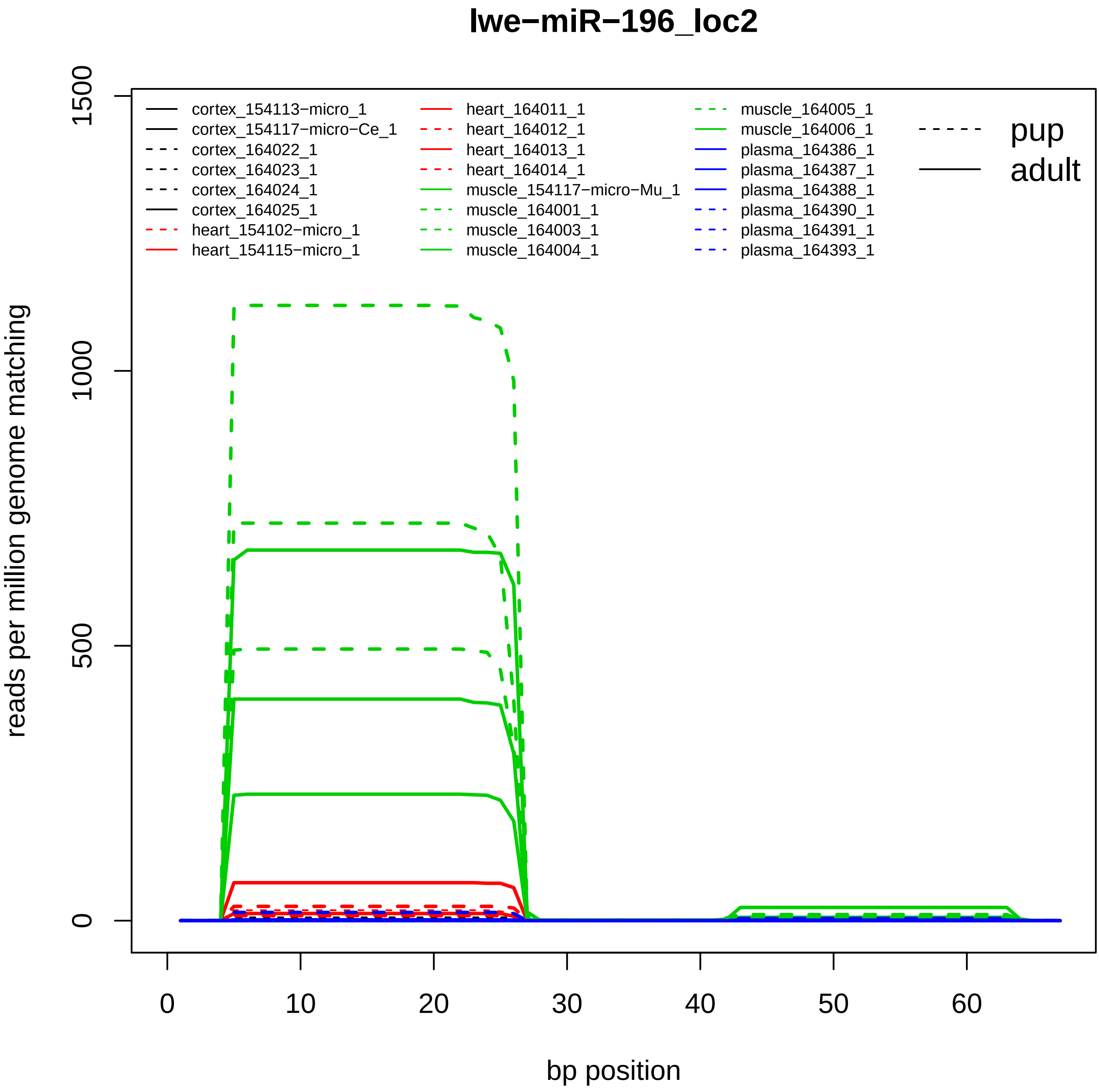

lwe−miR−196_loc2
reads per million genome matching
bp position
0
500
1000
1500
0
10
20
30
40
50
60
cortex_154113−micro_1
cortex_154117−micro−Ce_1
cortex_164022_1
cortex_164023_1
cortex_164024_1
cortex_164025_1
heart_154102−micro_1
heart_154115−micro_1
heart_164011_1
heart_164012_1
heart_164013_1
heart_164014_1
muscle_154117−micro−Mu_1
muscle_164001_1
muscle_164003_1
muscle_164004_1
muscle_164005_1
muscle_164006_1
plasma_164386_1
plasma_164387_1
plasma_164388_1
plasma_164390_1
plasma_164391_1
plasma_164393_1
pup
adult

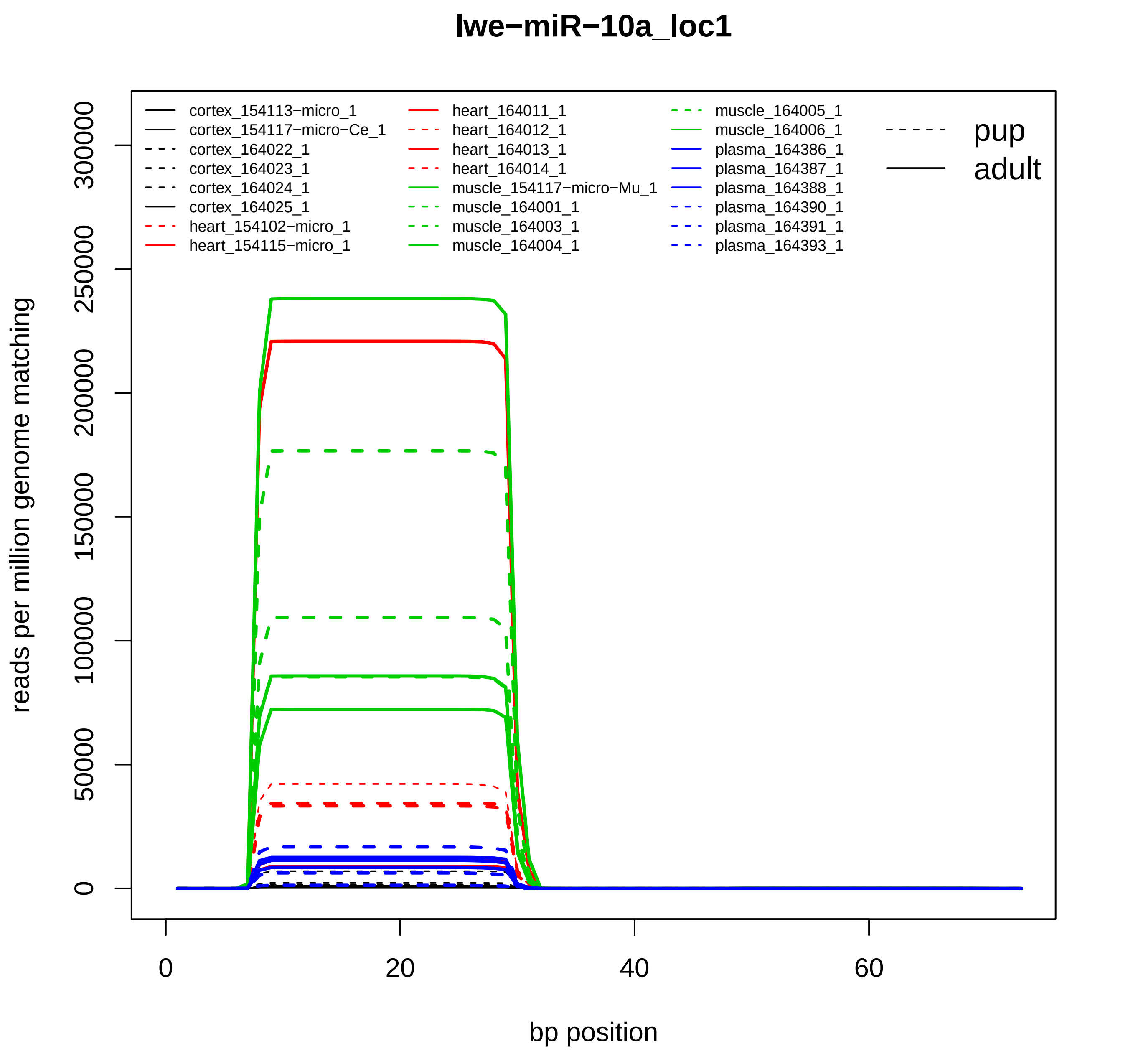
lwe−miR−10a_loc1
reads per million genome matching
bp position
0
50000
100000
150000
200000
250000
300000
0
20
40
60
cortex_154113−micro_1
cortex_154117−micro−Ce_1
cortex_164022_1
cortex_164023_1
cortex_164024_1
cortex_164025_1
heart_154102−micro_1
heart_154115−micro_1
heart_164011_1
heart_164012_1
heart_164013_1
heart_164014_1
muscle_154117−micro−Mu_1
muscle_164001_1
muscle_164003_1
muscle_164004_1
muscle_164005_1
muscle_164006_1
plasma_164386_1
plasma_164387_1
plasma_164388_1
plasma_164390_1
plasma_164391_1
plasma_164393_1
pup
adult

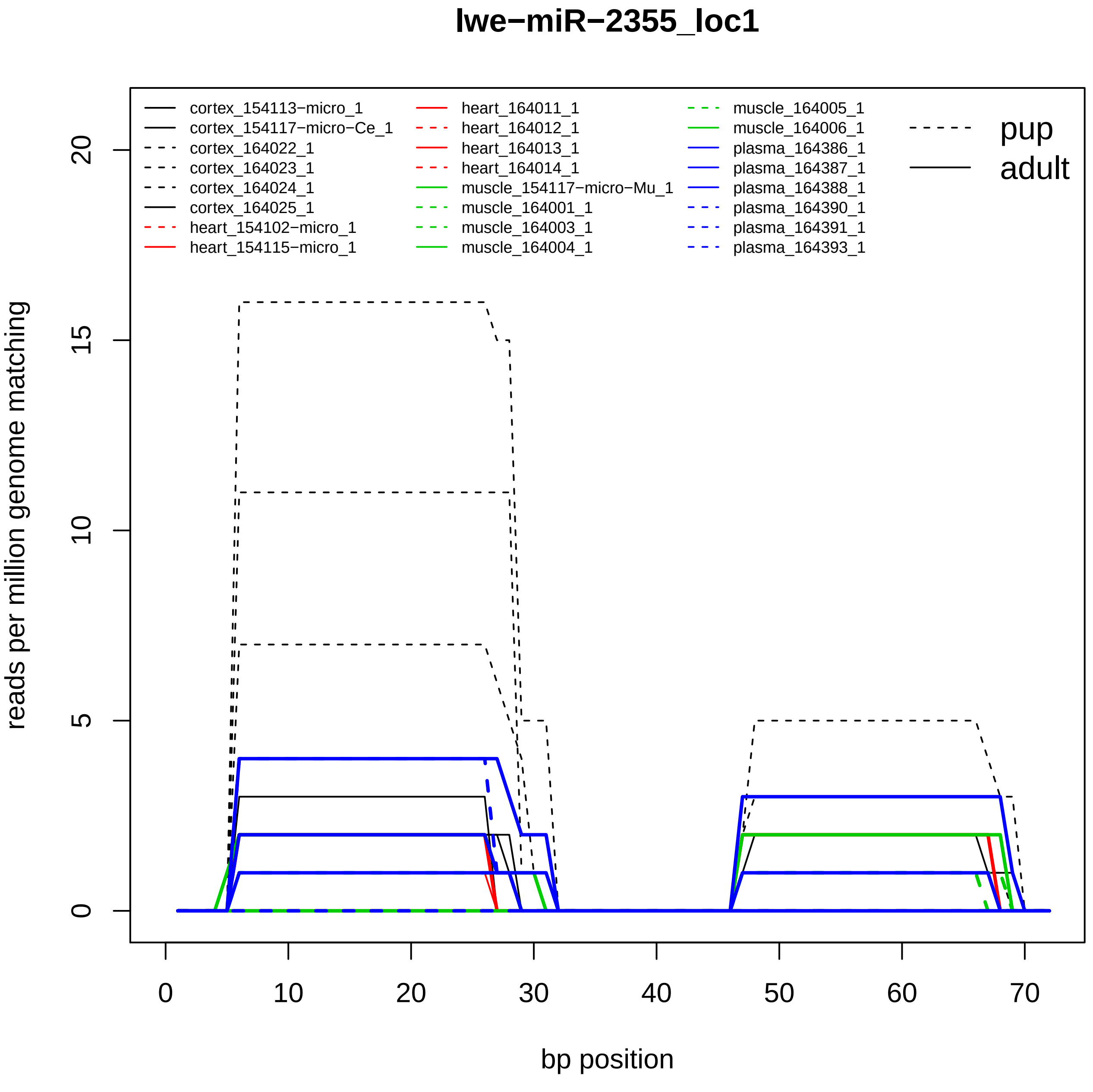
lwe−miR−2355_loc1
cortex_154113−micro_1
cortex_154117−micro−Ce_1
cortex_164022_1
cortex_164023_1
cortex_164024_1
cortex_164025_1
heart_154102−micro_1
heart_154115−micro_1
heart_164011_1
heart_164012_1
heart_164013_1
heart_164014_1
muscle_154117−micro−Mu_1
muscle_164001_1
muscle_164003_1
muscle_164004_1
muscle_164005_1
muscle_164006_1
plasma_164386_1
plasma_164387_1
plasma_164388_1
plasma_164390_1
plasma_164391_1
plasma_164393_1
pup
adult
reads per million genome matching
bp position
0
5
10
15
20
0
10
20
30
40
50
60
70

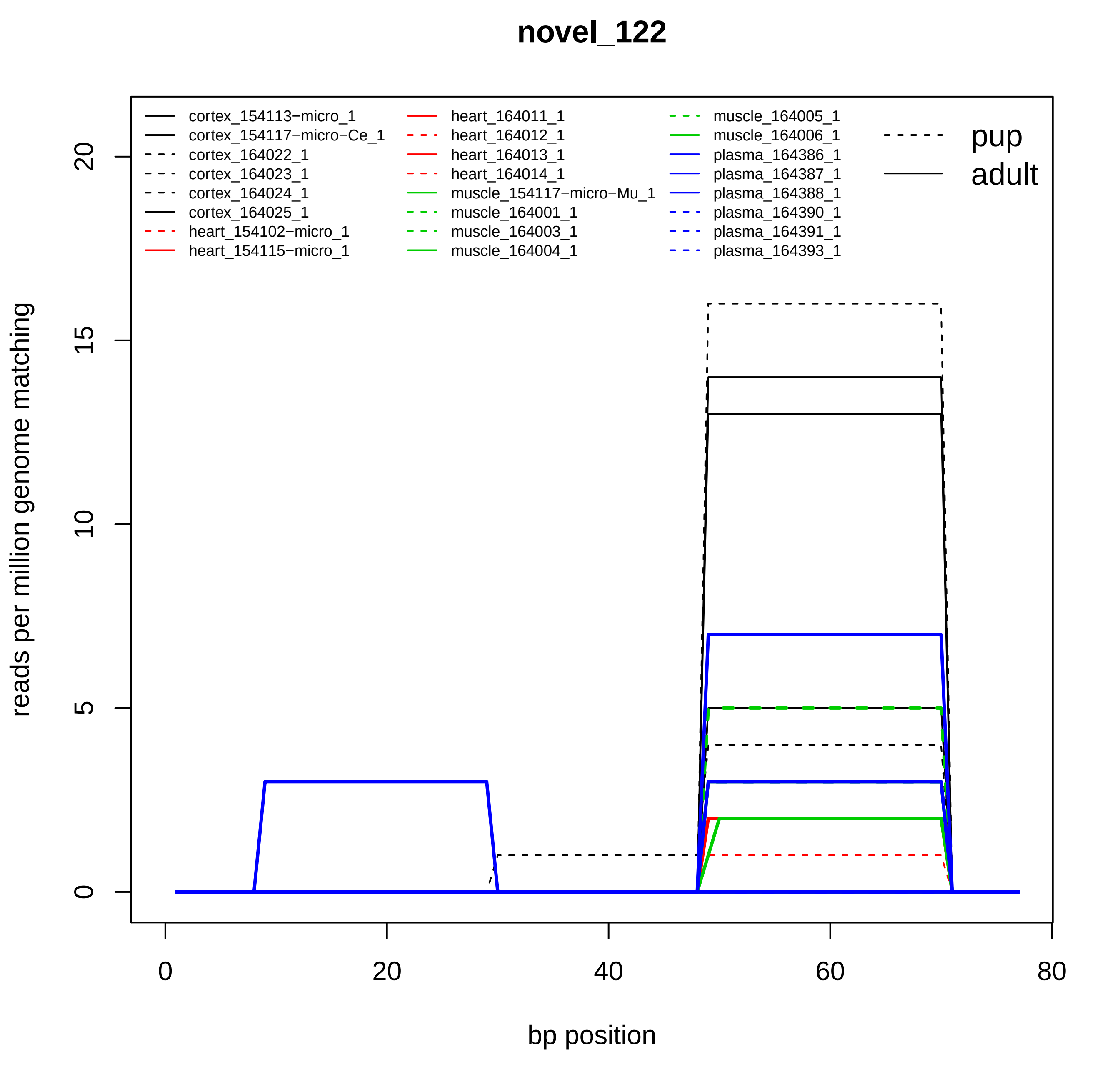
novel_122
cortex_154113−micro_1
cortex_154117−micro−Ce_1
cortex_164022_1
cortex_164023_1
cortex_164024_1
cortex_164025_1
heart_154102−micro_1
heart_154115−micro_1
heart_164011_1
heart_164012_1
heart_164013_1
heart_164014_1
muscle_154117−micro−Mu_1
muscle_164001_1
muscle_164003_1
muscle_164004_1
muscle_164005_1
muscle_164006_1
plasma_164386_1
plasma_164387_1
plasma_164388_1
plasma_164390_1
plasma_164391_1
plasma_164393_1
pup
adult
reads per million genome matching
0
5
10
15
20
bp position
0
20
40
60
80

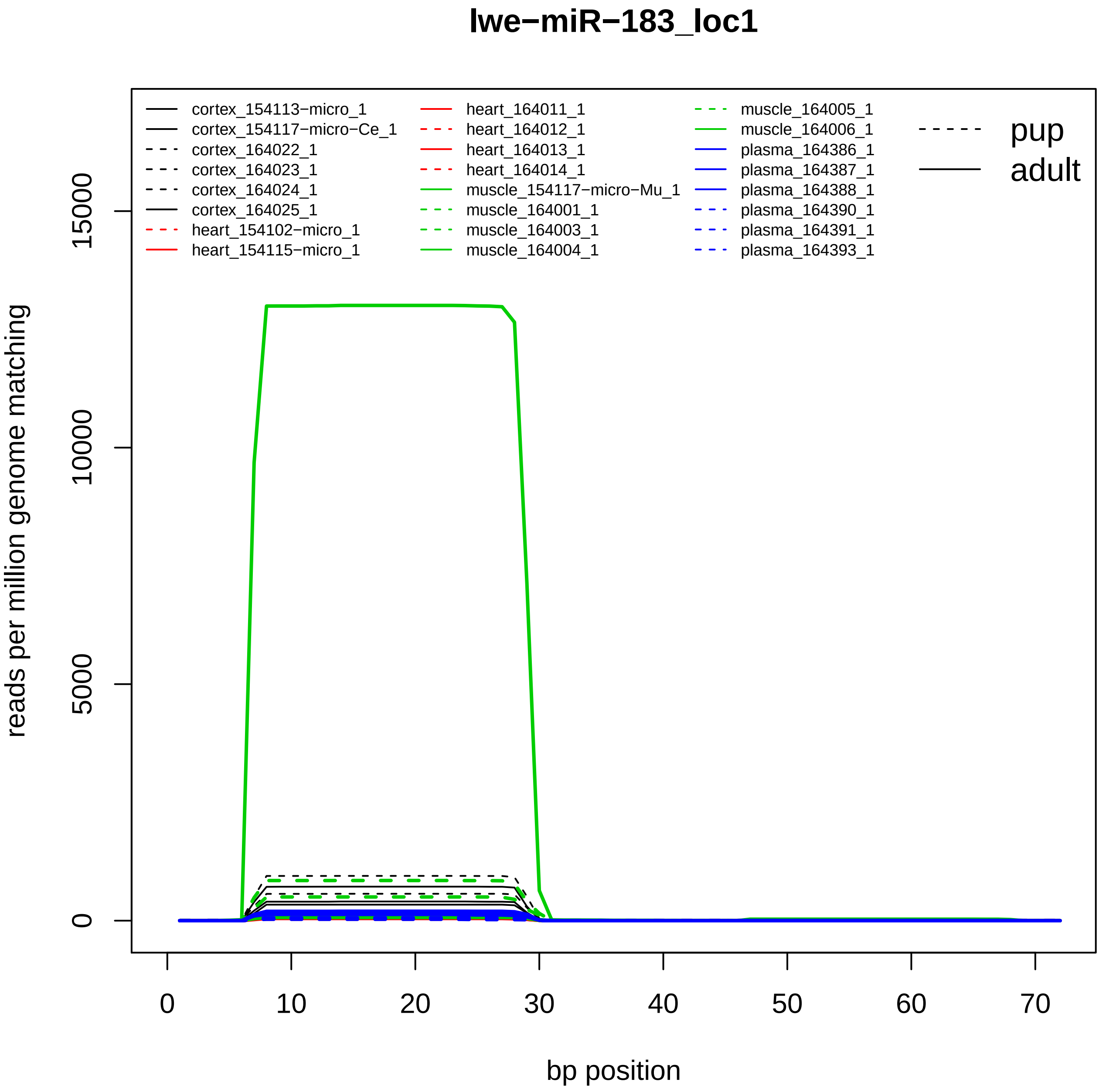
lwe−miR−183_loc1
cortex_154113−micro_1
cortex_154117−micro−Ce_1
cortex_164022_1
cortex_164023_1
cortex_164024_1
cortex_164025_1
heart_154102−micro_1
heart_154115−micro_1
heart_164011_1
heart_164012_1
heart_164013_1
heart_164014_1
muscle_154117−micro−Mu_1
muscle_164001_1
muscle_164003_1
muscle_164004_1
muscle_164005_1
muscle_164006_1
plasma_164386_1
plasma_164387_1
plasma_164388_1
plasma_164390_1
plasma_164391_1
plasma_164393_1
pup
adult
reads per million genome matching
0
5000
10000
15000
0
10
20
30
40
50
60
70
bp position

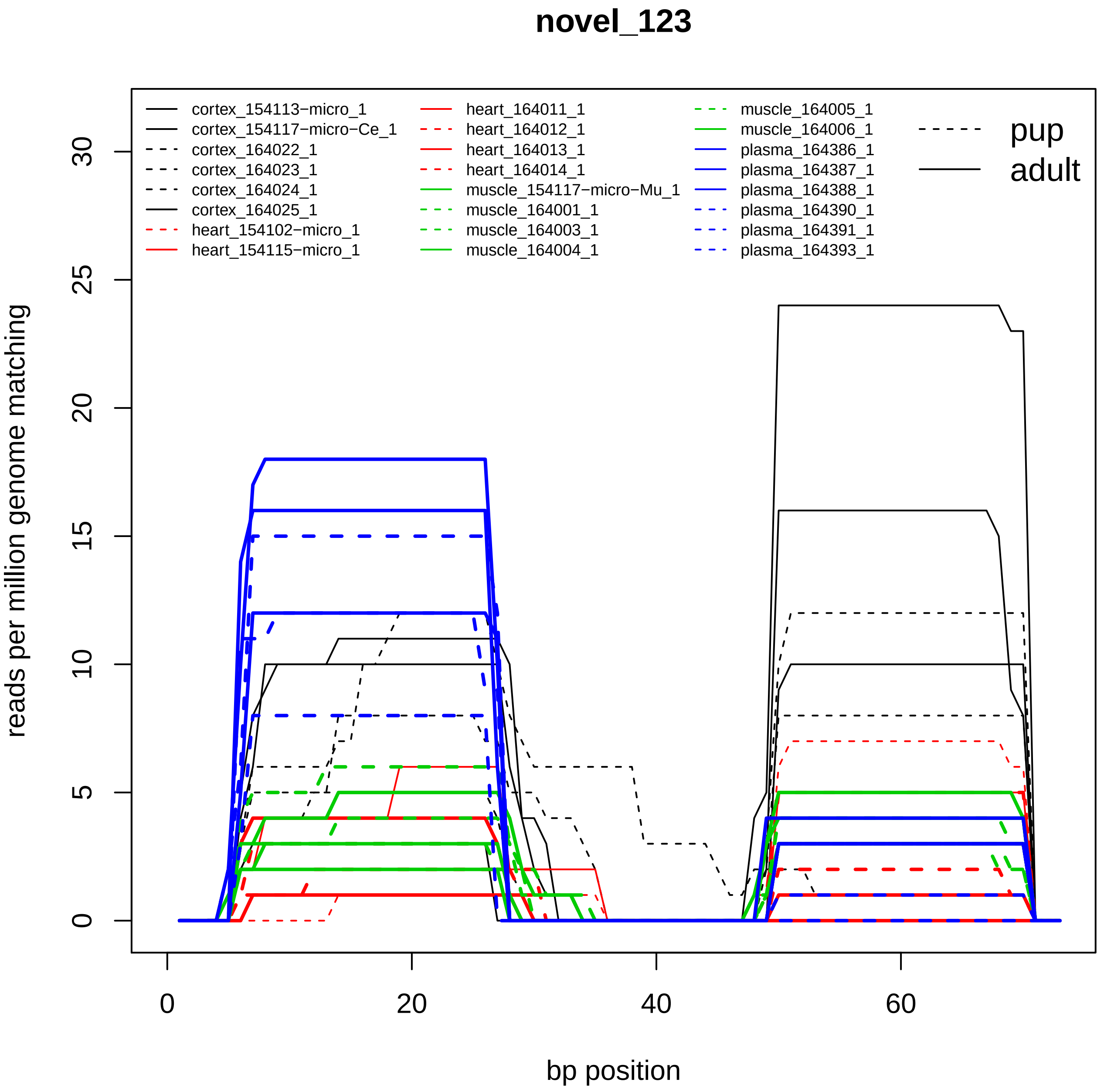

novel_123
cortex_154113−micro_1
cortex_154117−micro−Ce_1
cortex_164022_1
cortex_164023_1
cortex_164024_1
cortex_164025_1
heart_154102−micro_1
heart_154115−micro_1
heart_164011_1
heart_164012_1
heart_164013_1
heart_164014_1
muscle_154117−micro−Mu_1
muscle_164001_1
muscle_164003_1
muscle_164004_1
muscle_164005_1
muscle_164006_1
plasma_164386_1
plasma_164387_1
plasma_164388_1
plasma_164390_1
plasma_164391_1
plasma_164393_1
pup
adult
reads per million genome matching
0
5
10
15
20
25
30
0
20
40
60
bp position

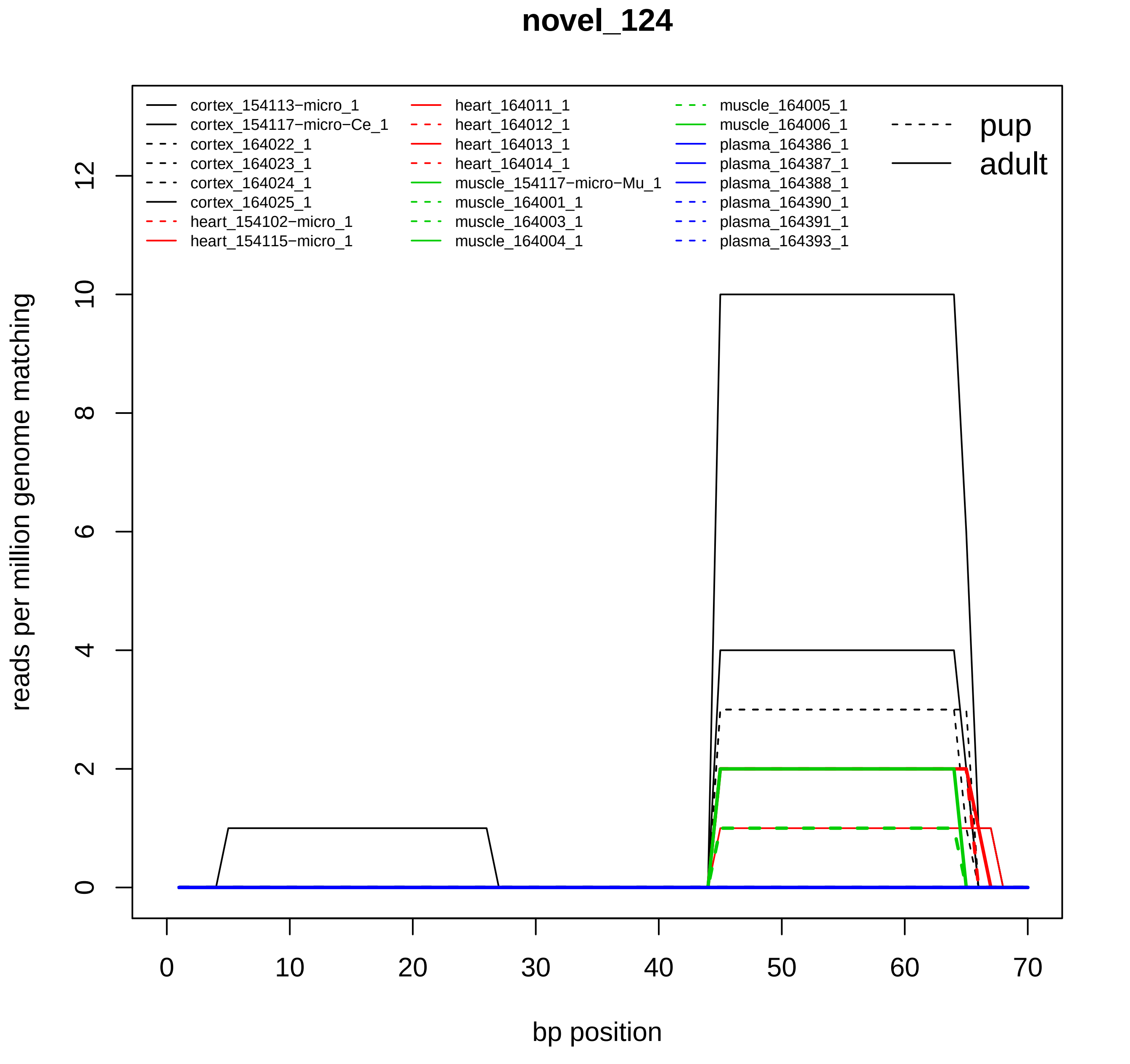
novel_124
cortex_154113−micro_1
cortex_154117−micro−Ce_1
cortex_164022_1
cortex_164023_1
cortex_164024_1
cortex_164025_1
heart_154102−micro_1
heart_154115−micro_1
heart_164011_1
heart_164012_1
heart_164013_1
heart_164014_1
muscle_154117−micro−Mu_1
muscle_164001_1
muscle_164003_1
muscle_164004_1
muscle_164005_1
muscle_164006_1
plasma_164386_1
plasma_164387_1
plasma_164388_1
plasma_164390_1
plasma_164391_1
plasma_164393_1
pup
adult
reads per million genome matching
bp position
0
2
4
6
8
10
12
0
10
20
30
40
50
60
70

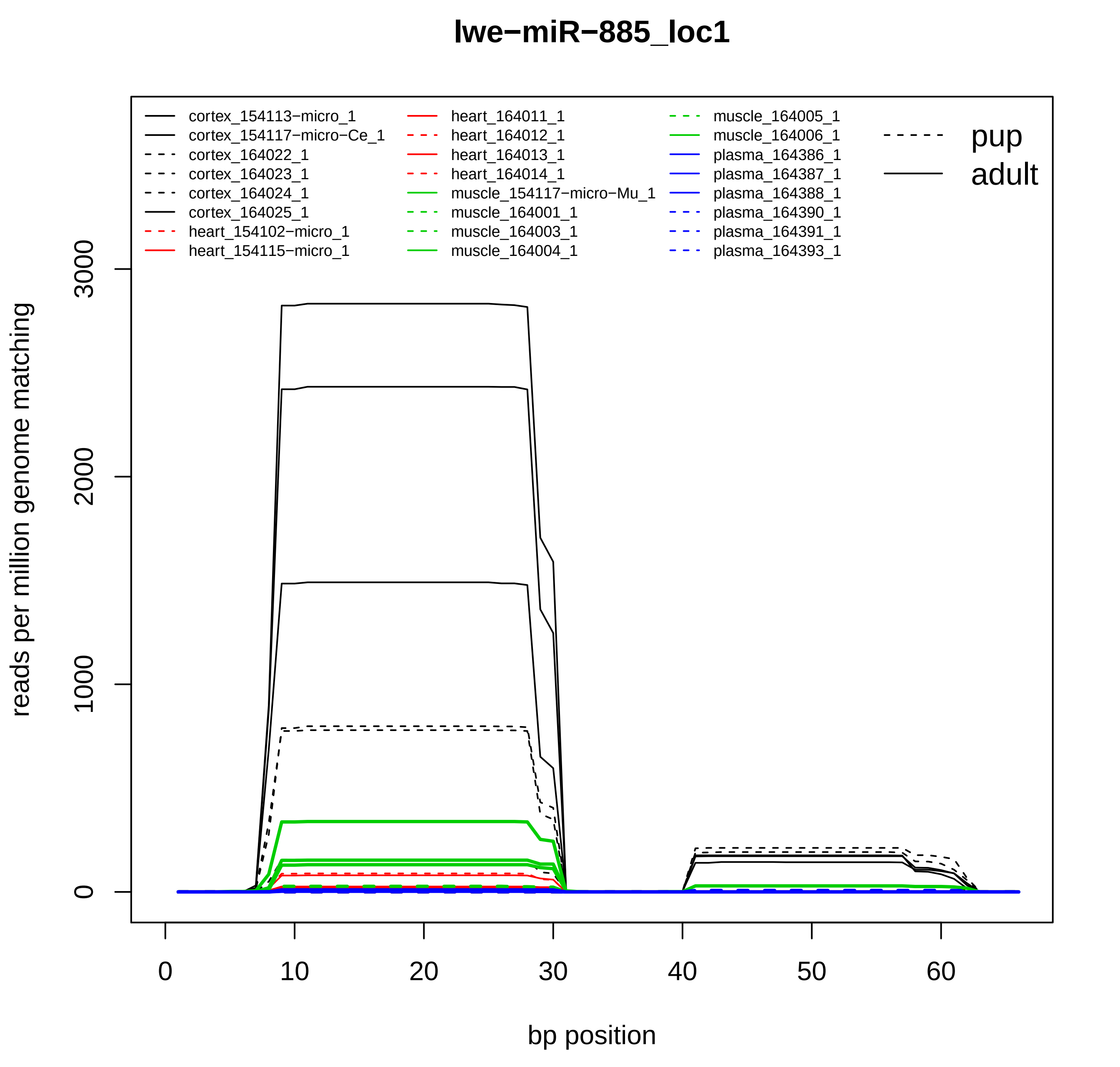
lwe−miR−885_loc1
cortex_154113−micro_1
cortex_154117−micro−Ce_1
cortex_164022_1
cortex_164023_1
cortex_164024_1
cortex_164025_1
heart_154102−micro_1
heart_154115−micro_1
heart_164011_1
heart_164012_1
heart_164013_1
heart_164014_1
muscle_154117−micro−Mu_1
muscle_164001_1
muscle_164003_1
muscle_164004_1
muscle_164005_1
muscle_164006_1
plasma_164386_1
plasma_164387_1
plasma_164388_1
plasma_164390_1
plasma_164391_1
plasma_164393_1
pup
adult
reads per million genome matching
0
1000
2000
3000
bp position
0
10
20
30
40
50
60

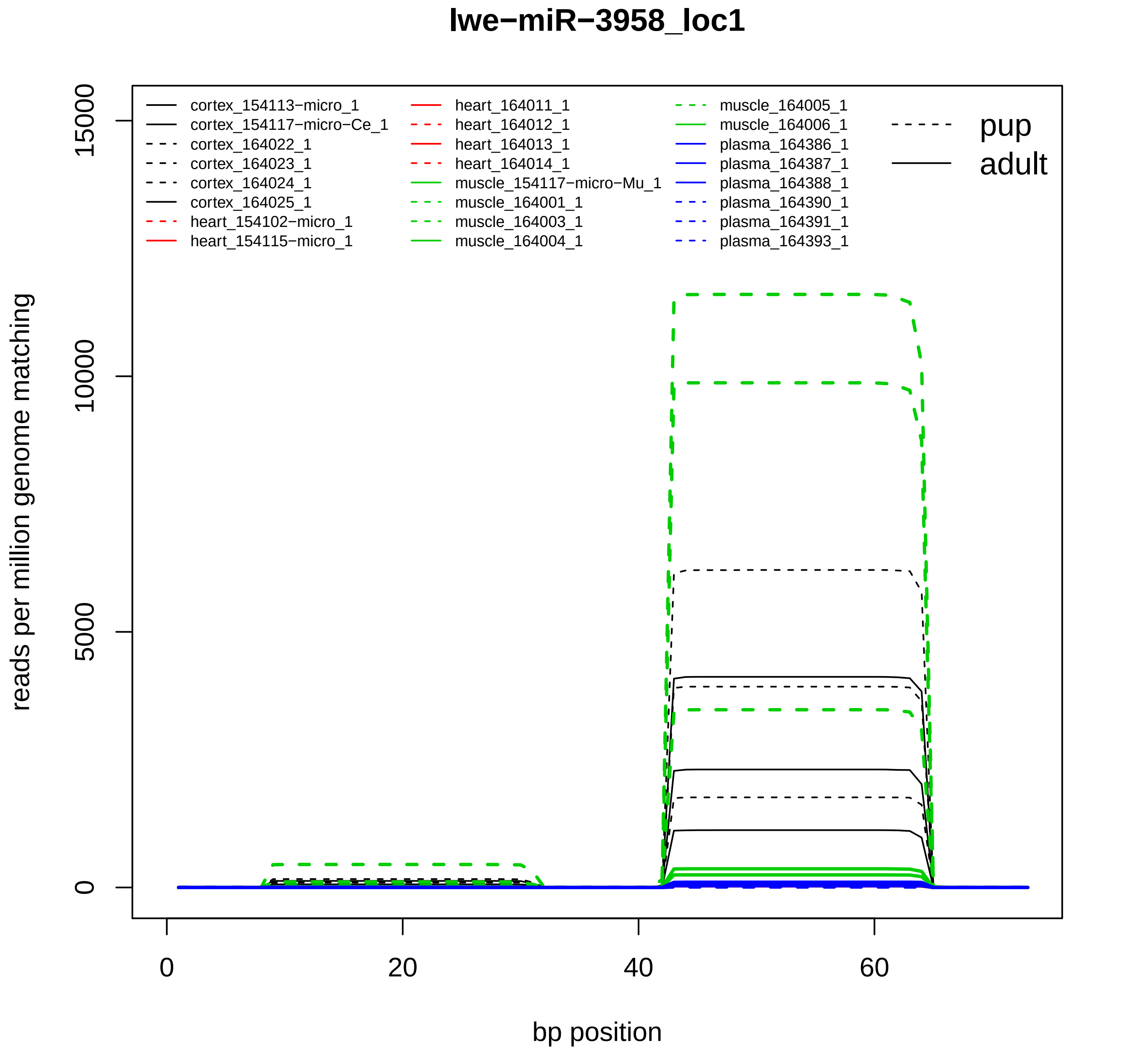
lwe−miR−3958_loc1
cortex_154113−micro_1
cortex_154117−micro−Ce_1
cortex_164022_1
cortex_164023_1
cortex_164024_1
cortex_164025_1
heart_154102−micro_1
heart_154115−micro_1
heart_164011_1
heart_164012_1
heart_164013_1
heart_164014_1
muscle_154117−micro−Mu_1
muscle_164001_1
muscle_164003_1
muscle_164004_1
muscle_164005_1
muscle_164006_1
plasma_164386_1
plasma_164387_1
plasma_164388_1
plasma_164390_1
plasma_164391_1
plasma_164393_1
pup
adult
reads per million genome matching
0
5000
10000
15000
0
20
40
60
bp position

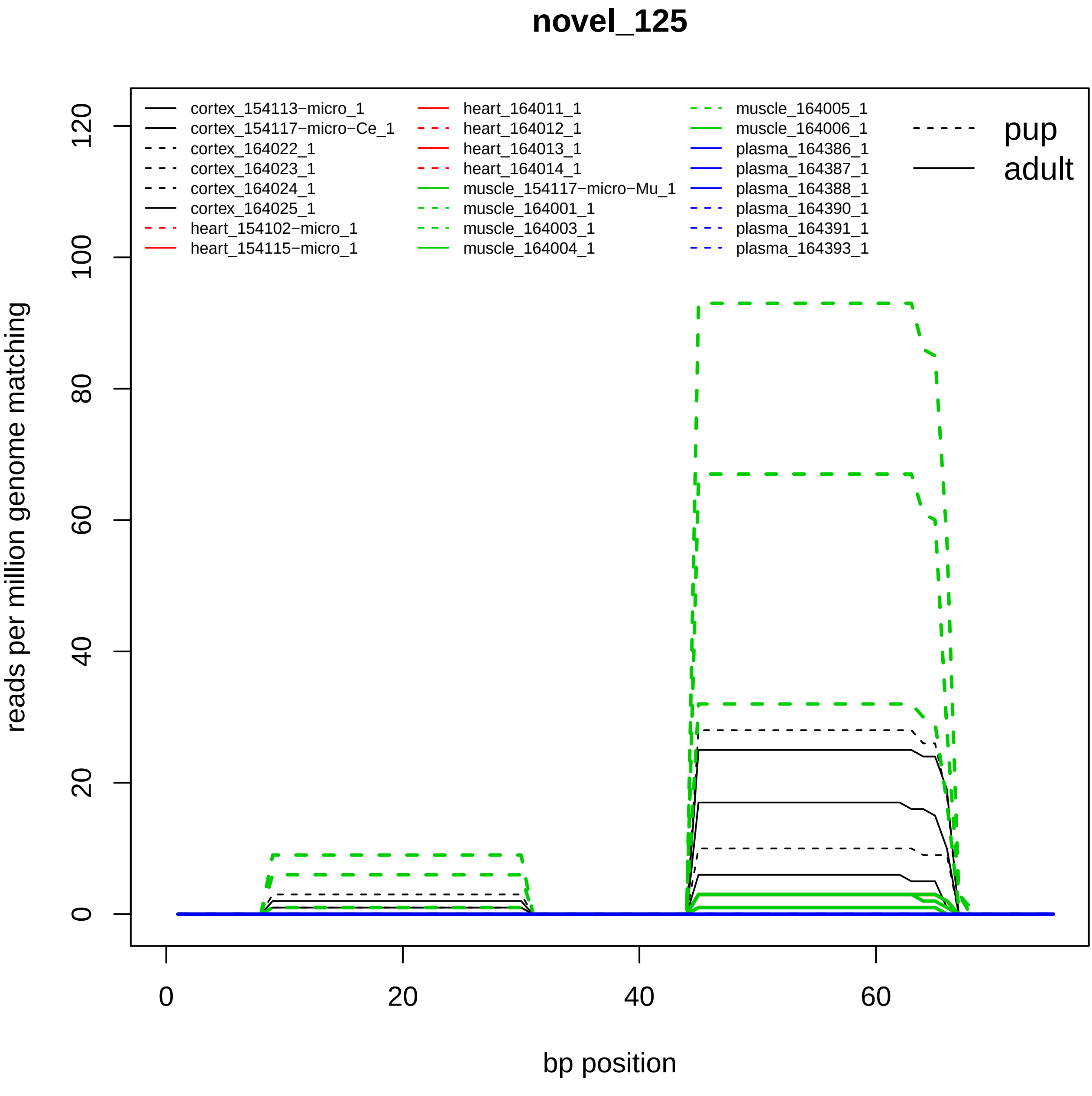

novel_125
cortex_154113−micro_1
cortex_154117−micro−Ce_1
cortex_164022_1
cortex_164023_1
cortex_164024_1
cortex_164025_1
heart_154102−micro_1
heart_154115−micro_1
heart_164011_1
heart_164012_1
heart_164013_1
heart_164014_1
muscle_154117−micro−Mu_1
muscle_164001_1
muscle_164003_1
muscle_164004_1
muscle_164005_1
muscle_164006_1
plasma_164386_1
plasma_164387_1
plasma_164388_1
plasma_164390_1
plasma_164391_1
plasma_164393_1
pup
adult
reads per million genome matching
0
20
40
60
80
100
120
bp position
0
20
40
60

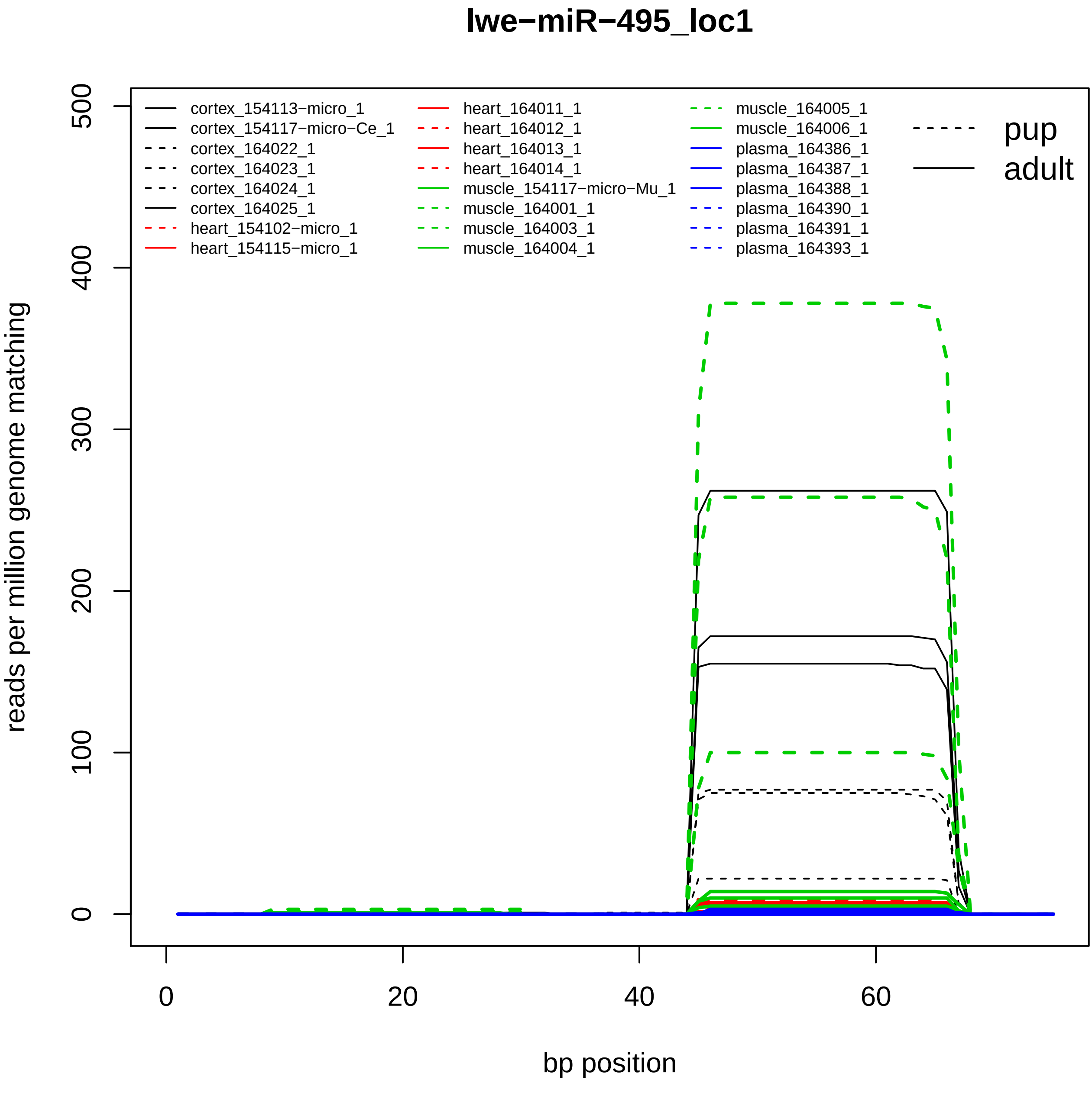
lwe−miR−495_loc1
reads per million genome matching
bp position
0
100
200
300
400
500
0
20
40
60
cortex_154113−micro_1
cortex_154117−micro−Ce_1
cortex_164022_1
cortex_164023_1
cortex_164024_1
cortex_164025_1
heart_154102−micro_1
heart_154115−micro_1
heart_164011_1
heart_164012_1
heart_164013_1
heart_164014_1
muscle_154117−micro−Mu_1
muscle_164001_1
muscle_164003_1
muscle_164004_1
muscle_164005_1
muscle_164006_1
plasma_164386_1
plasma_164387_1
plasma_164388_1
plasma_164390_1
plasma_164391_1
plasma_164393_1
pup
adult

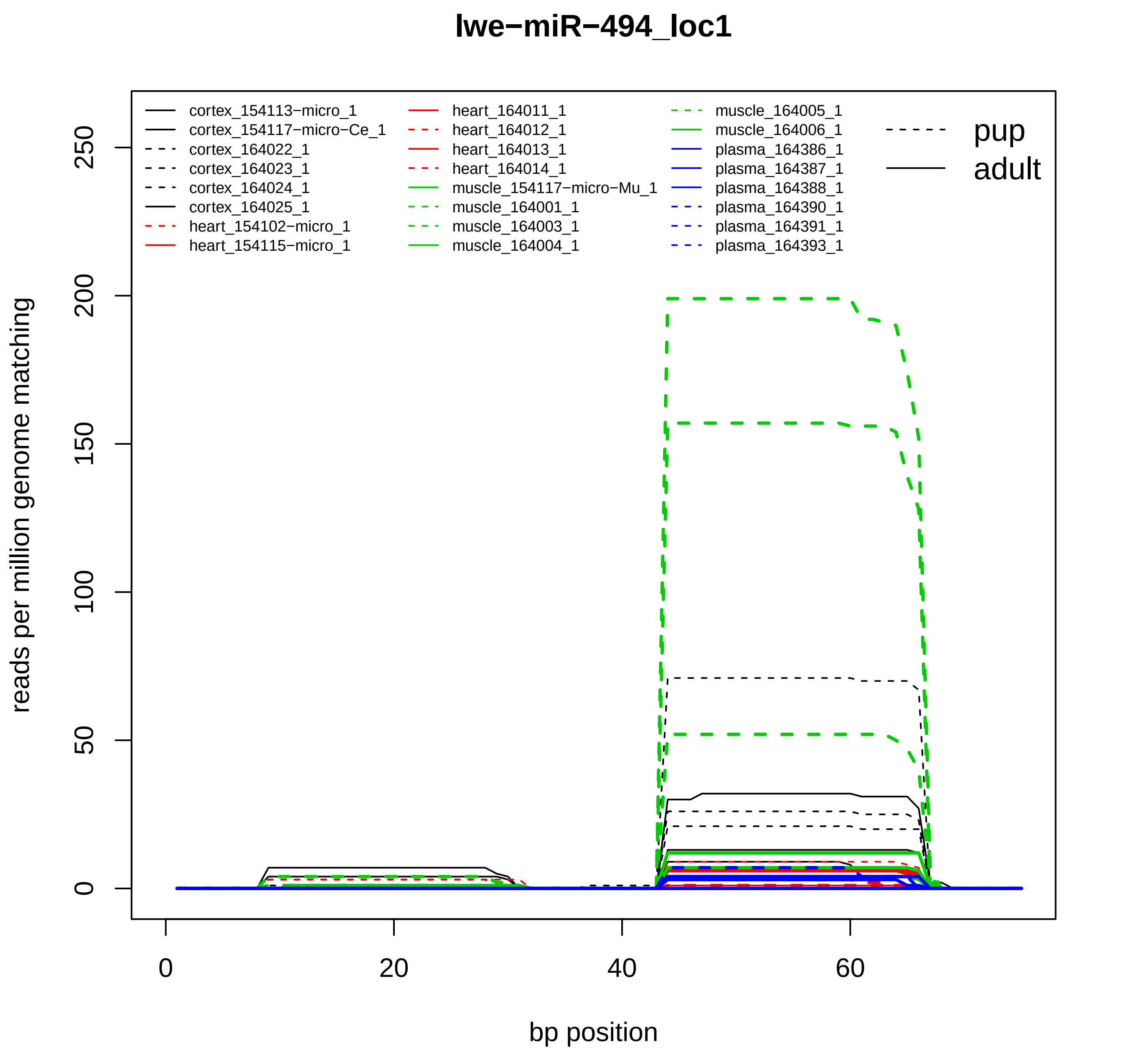
lwe−miR−494_loc1
cortex_154113−micro_1
cortex_154117−micro−Ce_1
cortex_164022_1
cortex_164023_1
cortex_164024_1
cortex_164025_1
heart_154102−micro_1
heart_154115−micro_1
heart_164011_1
heart_164012_1
heart_164013_1
heart_164014_1
muscle_154117−micro−Mu_1
muscle_164001_1
muscle_164003_1
muscle_164004_1
muscle_164005_1
muscle_164006_1
plasma_164386_1
plasma_164387_1
plasma_164388_1
plasma_164390_1
plasma_164391_1
plasma_164393_1
pup
adult
reads per million genome matching
0
50
100
150
200
250
0
20
40
60
bp position

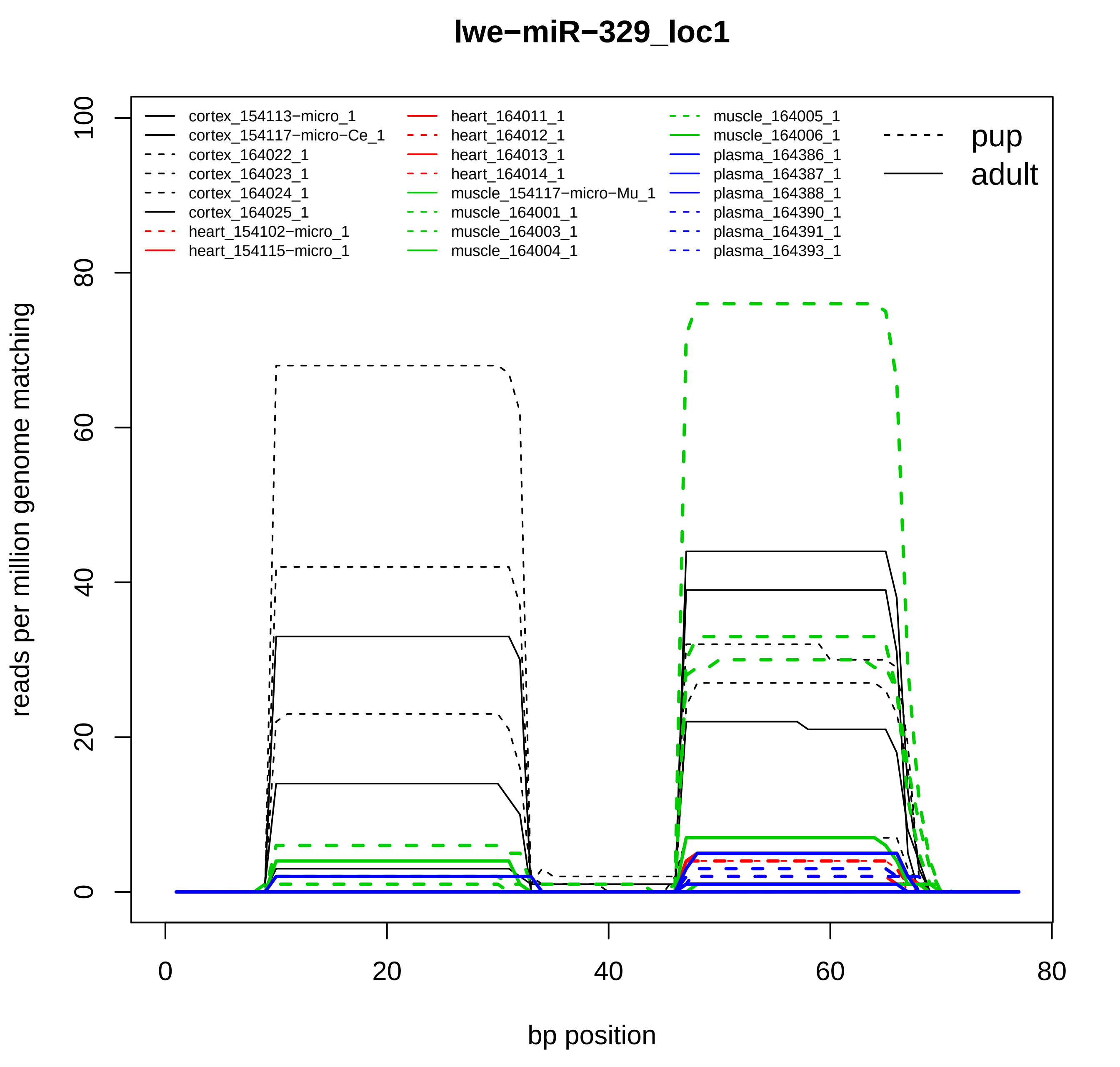
lwe−miR−329_loc1
cortex_154113−micro_1
cortex_154117−micro−Ce_1
cortex_164022_1
cortex_164023_1
cortex_164024_1
cortex_164025_1
heart_154102−micro_1
heart_154115−micro_1
heart_164011_1
heart_164012_1
heart_164013_1
heart_164014_1
muscle_154117−micro−Mu_1
muscle_164001_1
muscle_164003_1
muscle_164004_1
muscle_164005_1
muscle_164006_1
plasma_164386_1
plasma_164387_1
plasma_164388_1
plasma_164390_1
plasma_164391_1
plasma_164393_1
pup
adult
reads per million genome matching
0
20
40
60
80
100
bp position
0
20
40
60
80

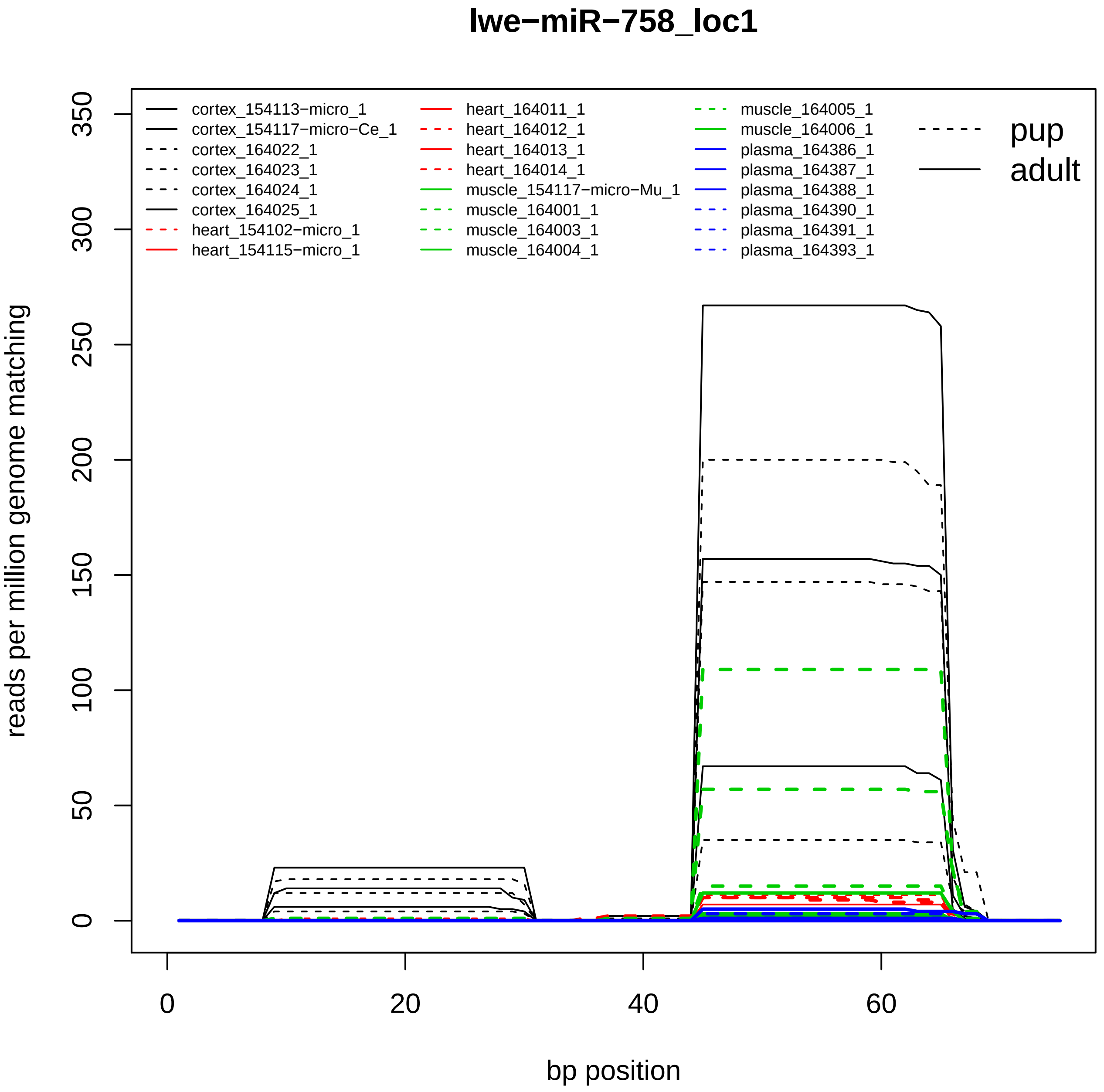

lwe−miR−758_loc1
cortex_154113−micro_1
cortex_154117−micro−Ce_1
cortex_164022_1
cortex_164023_1
cortex_164024_1
cortex_164025_1
heart_154102−micro_1
heart_154115−micro_1
heart_164011_1
heart_164012_1
heart_164013_1
heart_164014_1
muscle_154117−micro−Mu_1
muscle_164001_1
muscle_164003_1
muscle_164004_1
muscle_164005_1
muscle_164006_1
plasma_164386_1
plasma_164387_1
plasma_164388_1
plasma_164390_1
plasma_164391_1
plasma_164393_1
pup
adult
reads per million genome matching
0
50
100
150
200
250
300
350
bp position
0
20
40
60

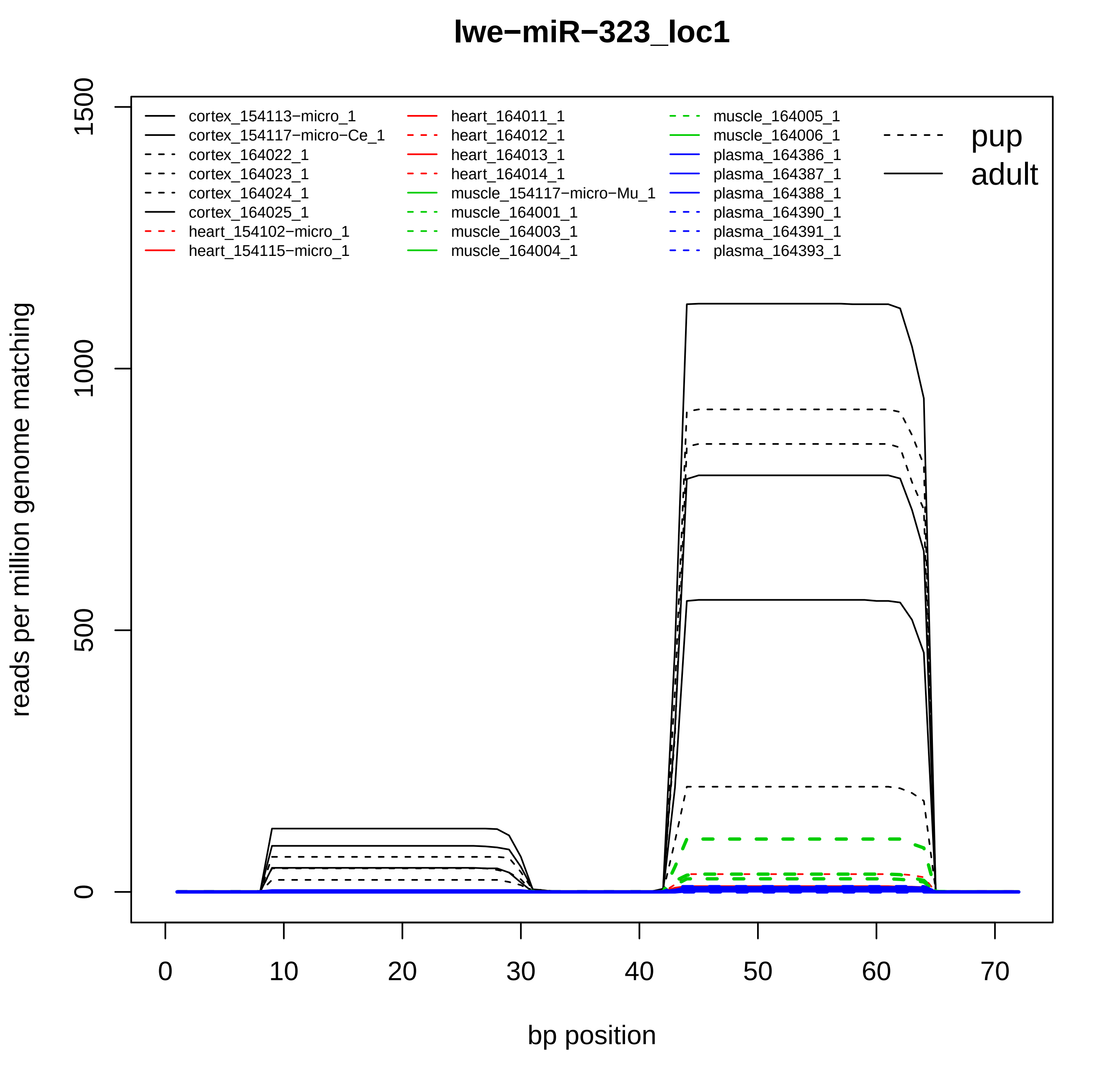

lwe−miR−323_loc1
reads per million genome matching
bp position
0
500
1000
1500
0
10
20
30
40
50
60
70
cortex_154113−micro_1
cortex_154117−micro−Ce_1
cortex_164022_1
cortex_164023_1
cortex_164024_1
cortex_164025_1
heart_154102−micro_1
heart_154115−micro_1
heart_164011_1
heart_164012_1
heart_164013_1
heart_164014_1
muscle_154117−micro−Mu_1
muscle_164001_1
muscle_164003_1
muscle_164004_1
muscle_164005_1
muscle_164006_1
plasma_164386_1
plasma_164387_1
plasma_164388_1
plasma_164390_1
plasma_164391_1
plasma_164393_1
pup
adult

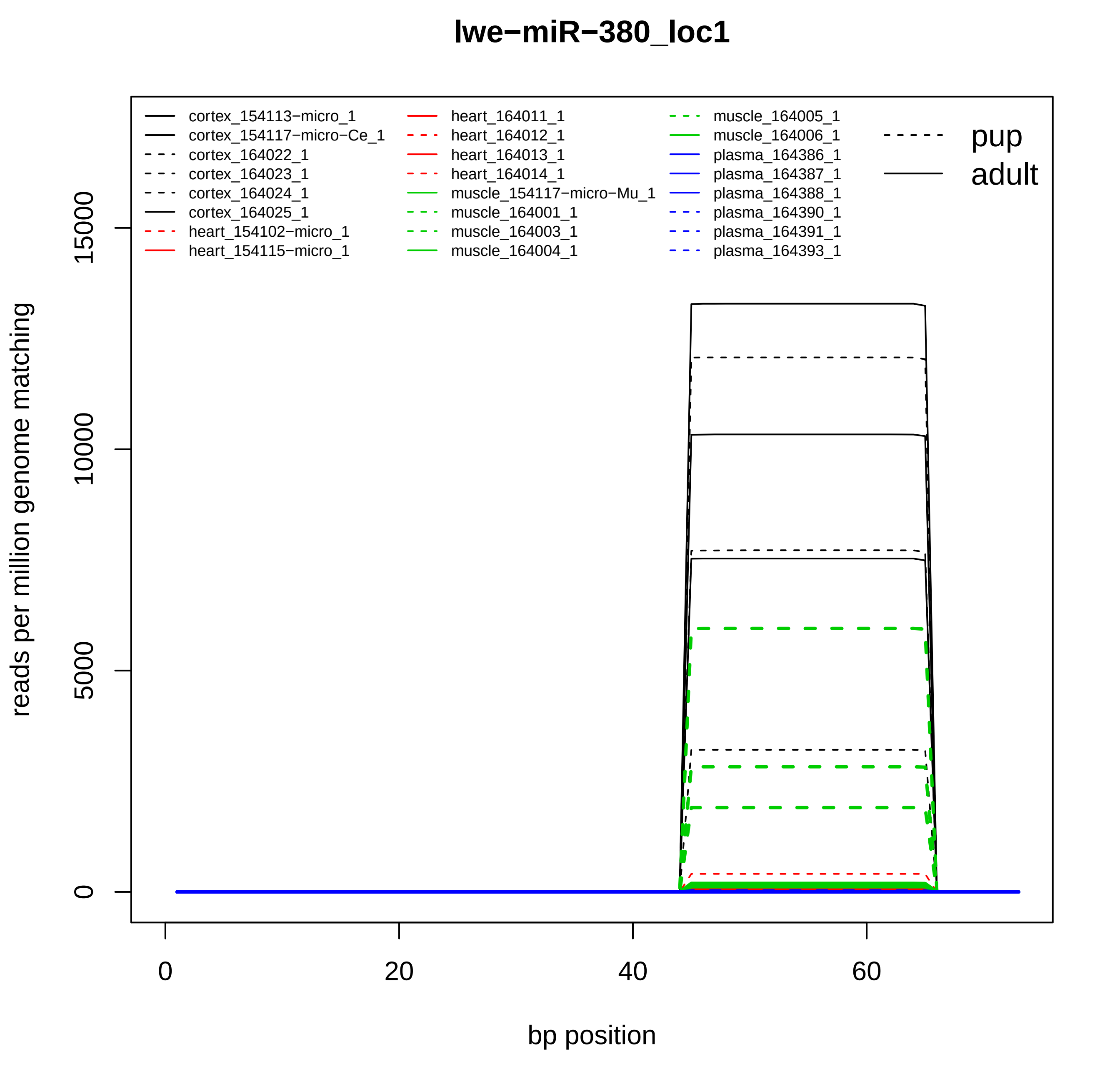
lwe−miR−380_loc1
cortex_154113−micro_1
cortex_154117−micro−Ce_1
cortex_164022_1
cortex_164023_1
cortex_164024_1
cortex_164025_1
heart_154102−micro_1
heart_154115−micro_1
heart_164011_1
heart_164012_1
heart_164013_1
heart_164014_1
muscle_154117−micro−Mu_1
muscle_164001_1
muscle_164003_1
muscle_164004_1
muscle_164005_1
muscle_164006_1
plasma_164386_1
plasma_164387_1
plasma_164388_1
plasma_164390_1
plasma_164391_1
plasma_164393_1
pup
adult
reads per million genome matching
0
5000
10000
15000
bp position
0
20
40
60

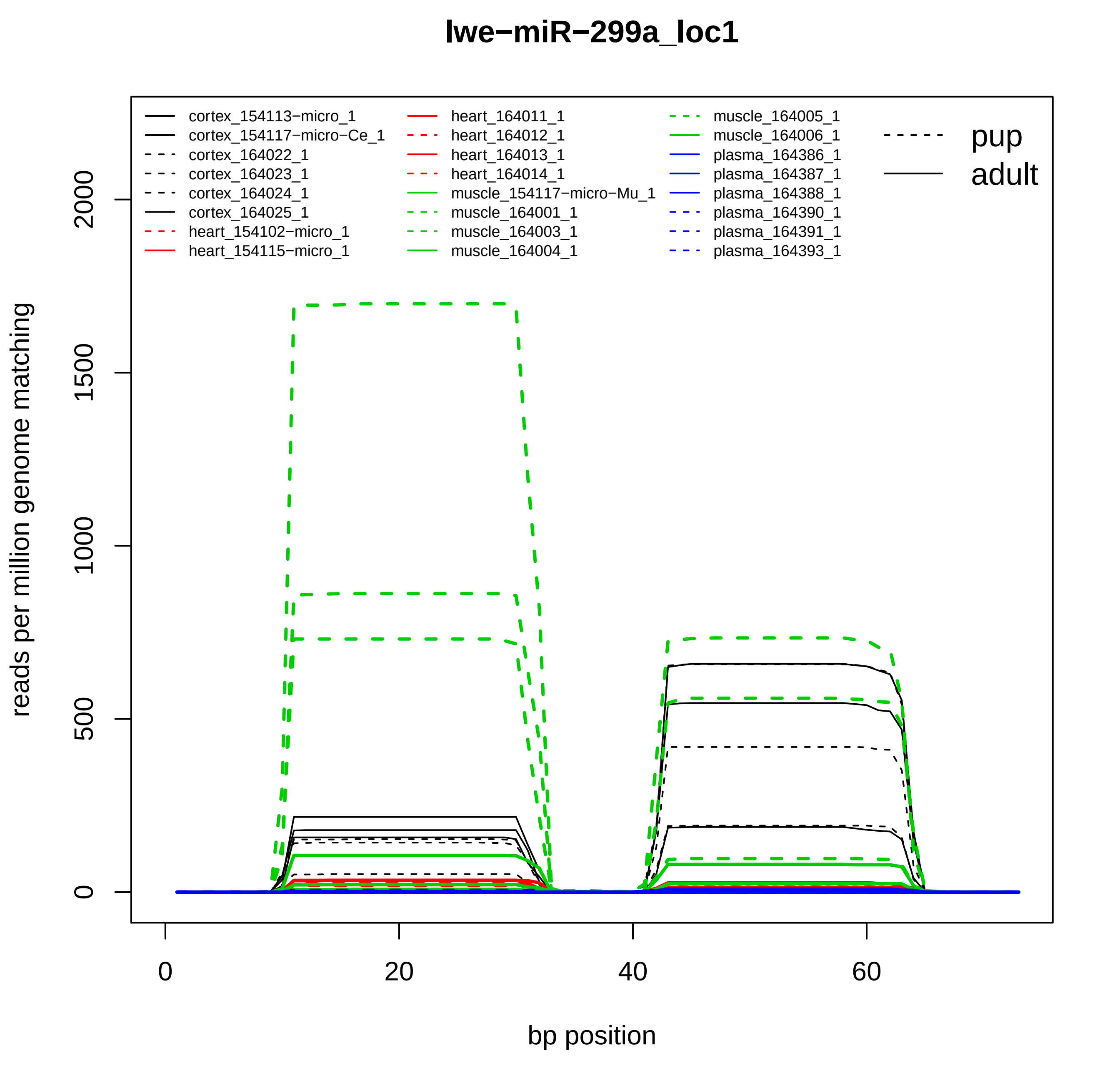

lwe−miR−299a_loc1
cortex_154113−micro_1
cortex_154117−micro−Ce_1
cortex_164022_1
cortex_164023_1
cortex_164024_1
cortex_164025_1
heart_154102−micro_1
heart_154115−micro_1
heart_164011_1
heart_164012_1
heart_164013_1
heart_164014_1
muscle_154117−micro−Mu_1
muscle_164001_1
muscle_164003_1
muscle_164004_1
muscle_164005_1
muscle_164006_1
plasma_164386_1
plasma_164387_1
plasma_164388_1
plasma_164390_1
plasma_164391_1
plasma_164393_1
pup
adult
reads per million genome matching
0
500
1000
1500
2000
0
20
40
60
bp position

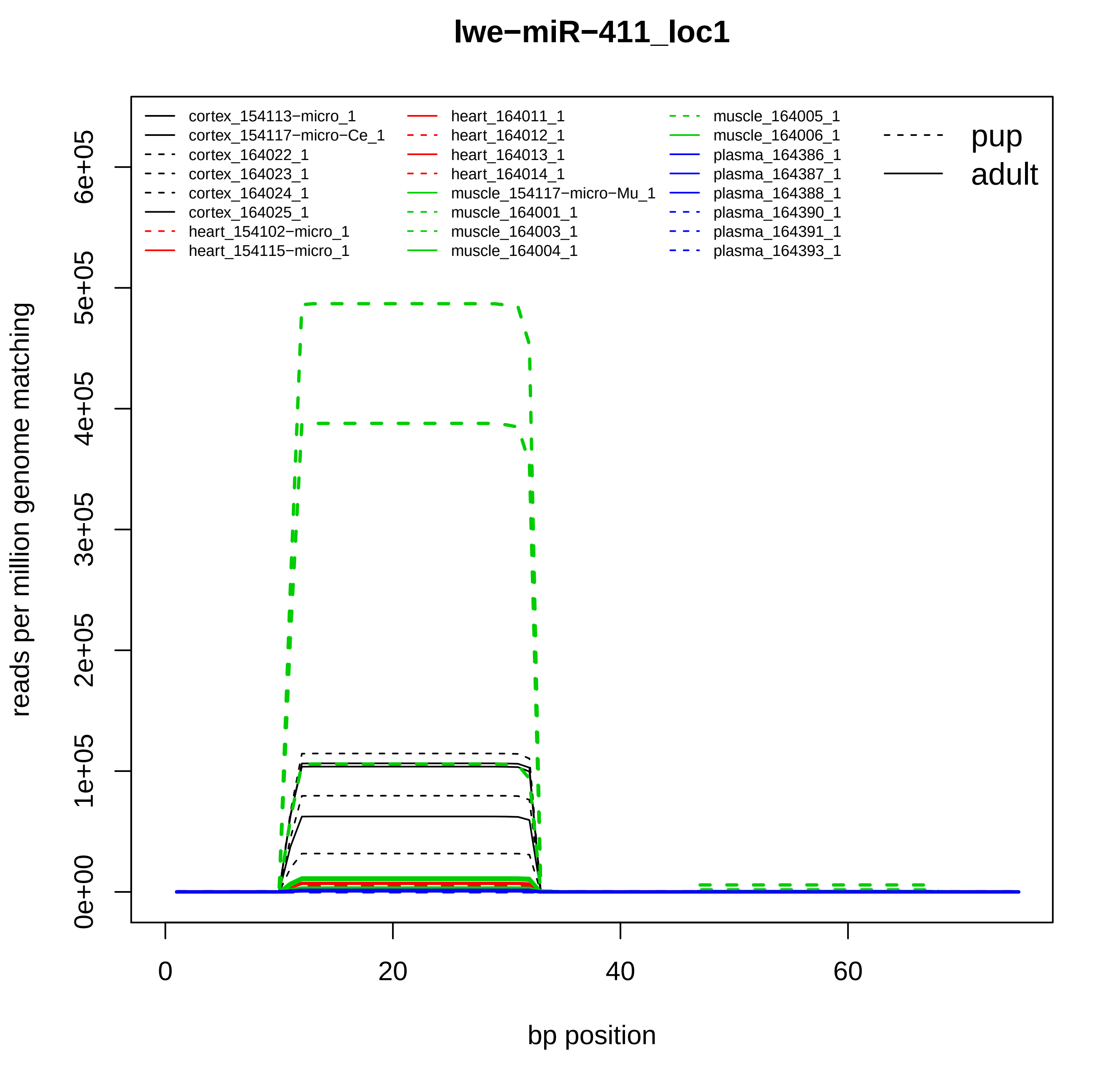

lwe−miR−411_loc1
cortex_154113−micro_1
cortex_154117−micro−Ce_1
cortex_164022_1
cortex_164023_1
cortex_164024_1
cortex_164025_1
heart_154102−micro_1
heart_154115−micro_1
heart_164011_1
heart_164012_1
heart_164013_1
heart_164014_1
muscle_154117−micro−Mu_1
muscle_164001_1
muscle_164003_1
muscle_164004_1
muscle_164005_1
muscle_164006_1
plasma_164386_1
plasma_164387_1
plasma_164388_1
plasma_164390_1
plasma_164391_1
plasma_164393_1
pup
adult
reads per million genome matching
0e+00
1e+05
2e+05
3e+05
4e+05
5e+05
6e+05
0
20
40
60
bp position

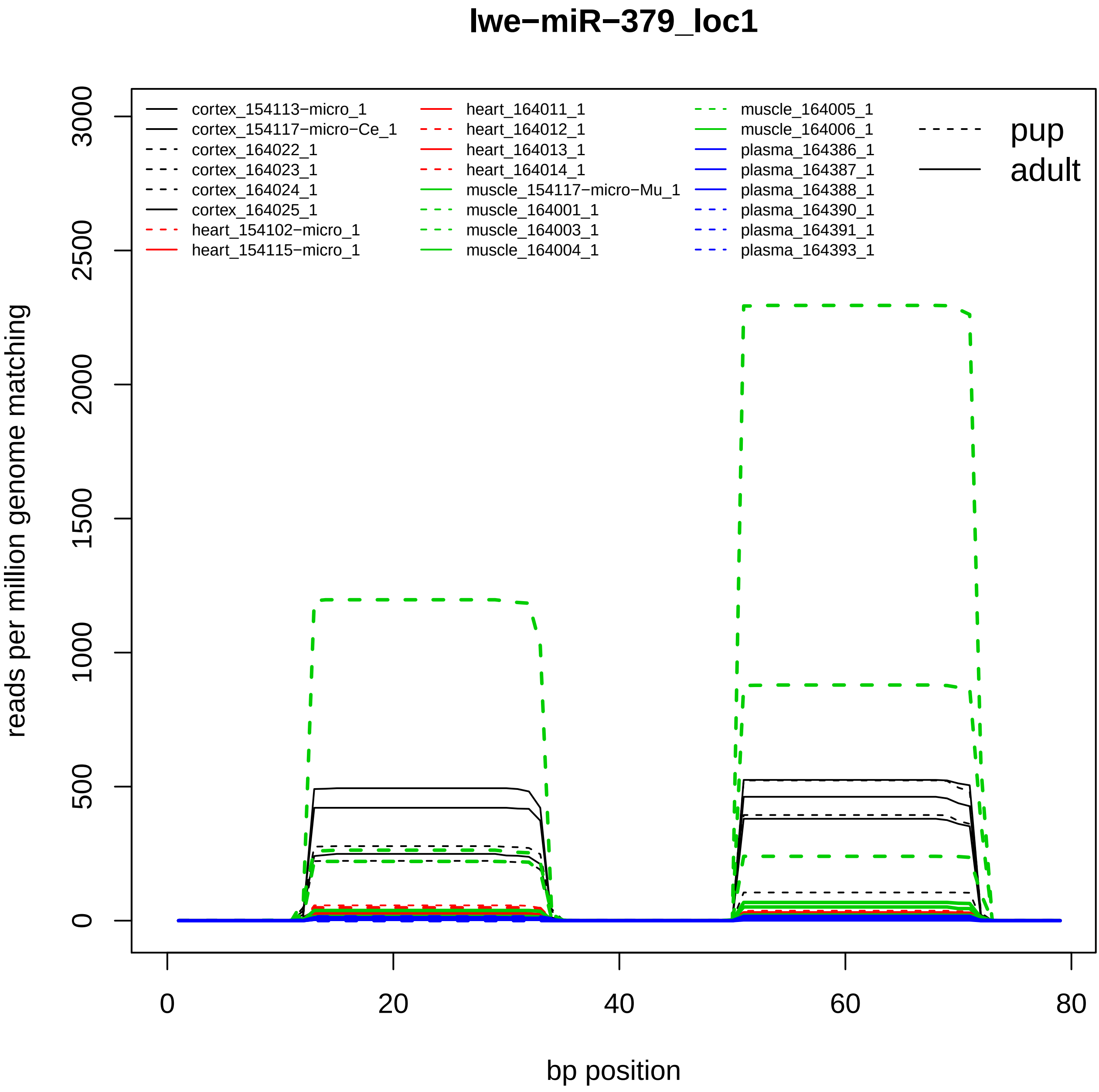
lwe−miR−379_loc1
reads per million genome matching
bp position
0
20
40
60
80
0
500
1000
1500
2000
2500
3000
cortex_154113−micro_1
cortex_154117−micro−Ce_1
cortex_164022_1
cortex_164023_1
cortex_164024_1
cortex_164025_1
heart_154102−micro_1
heart_154115−micro_1
heart_164011_1
heart_164012_1
heart_164013_1
heart_164014_1
muscle_154117−micro−Mu_1
muscle_164001_1
muscle_164003_1
muscle_164004_1
muscle_164005_1
muscle_164006_1
plasma_164386_1
plasma_164387_1
plasma_164388_1
plasma_164390_1
plasma_164391_1
plasma_164393_1
pup
adult

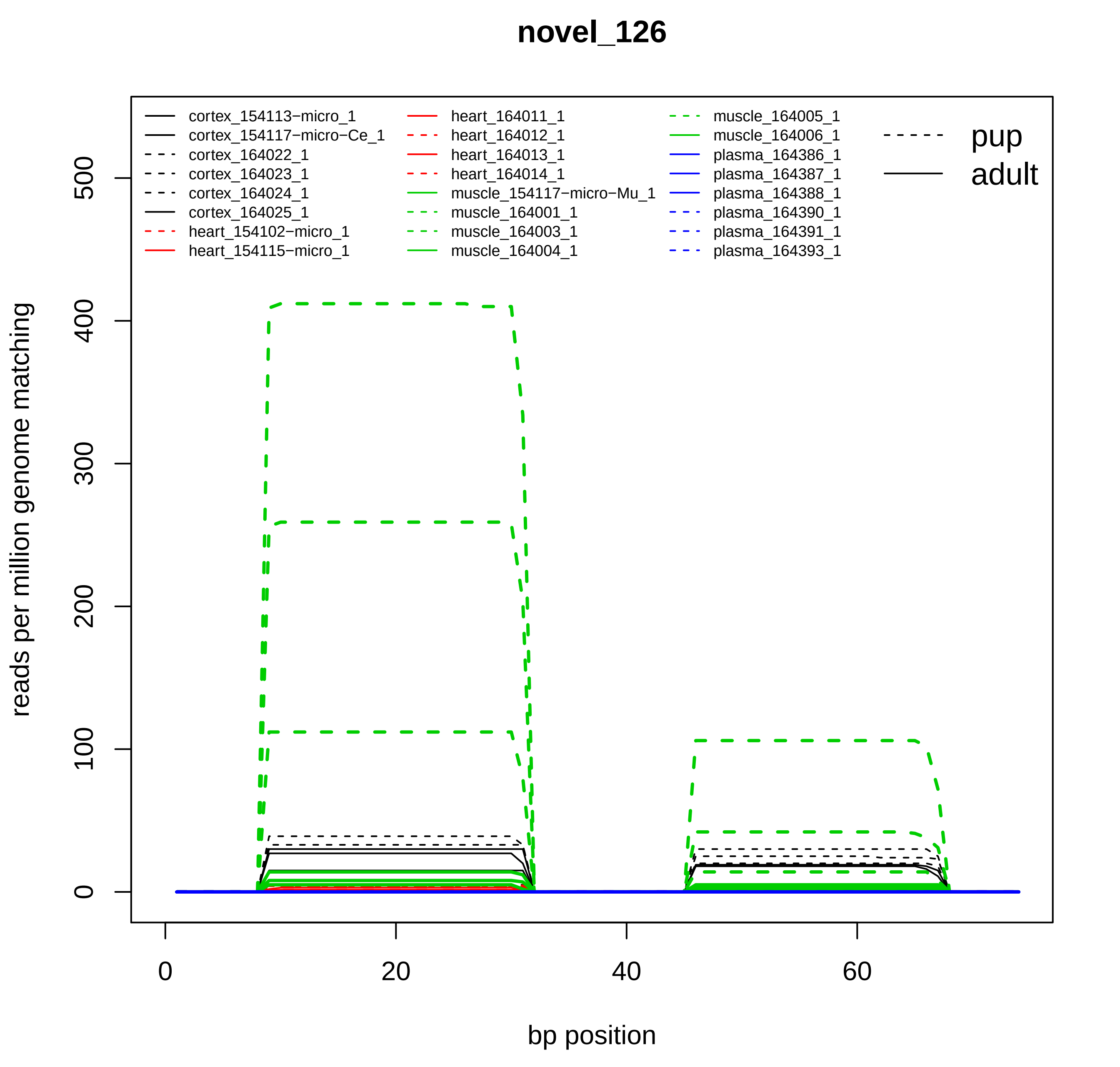

novel_126
cortex_154113−micro_1
cortex_154117−micro−Ce_1
cortex_164022_1
cortex_164023_1
cortex_164024_1
cortex_164025_1
heart_154102−micro_1
heart_154115−micro_1
heart_164011_1
heart_164012_1
heart_164013_1
heart_164014_1
muscle_154117−micro−Mu_1
muscle_164001_1
muscle_164003_1
muscle_164004_1
muscle_164005_1
muscle_164006_1
plasma_164386_1
plasma_164387_1
plasma_164388_1
plasma_164390_1
plasma_164391_1
plasma_164393_1
pup
adult
reads per million genome matching
0
100
200
300
400
500
0
20
40
60
bp position

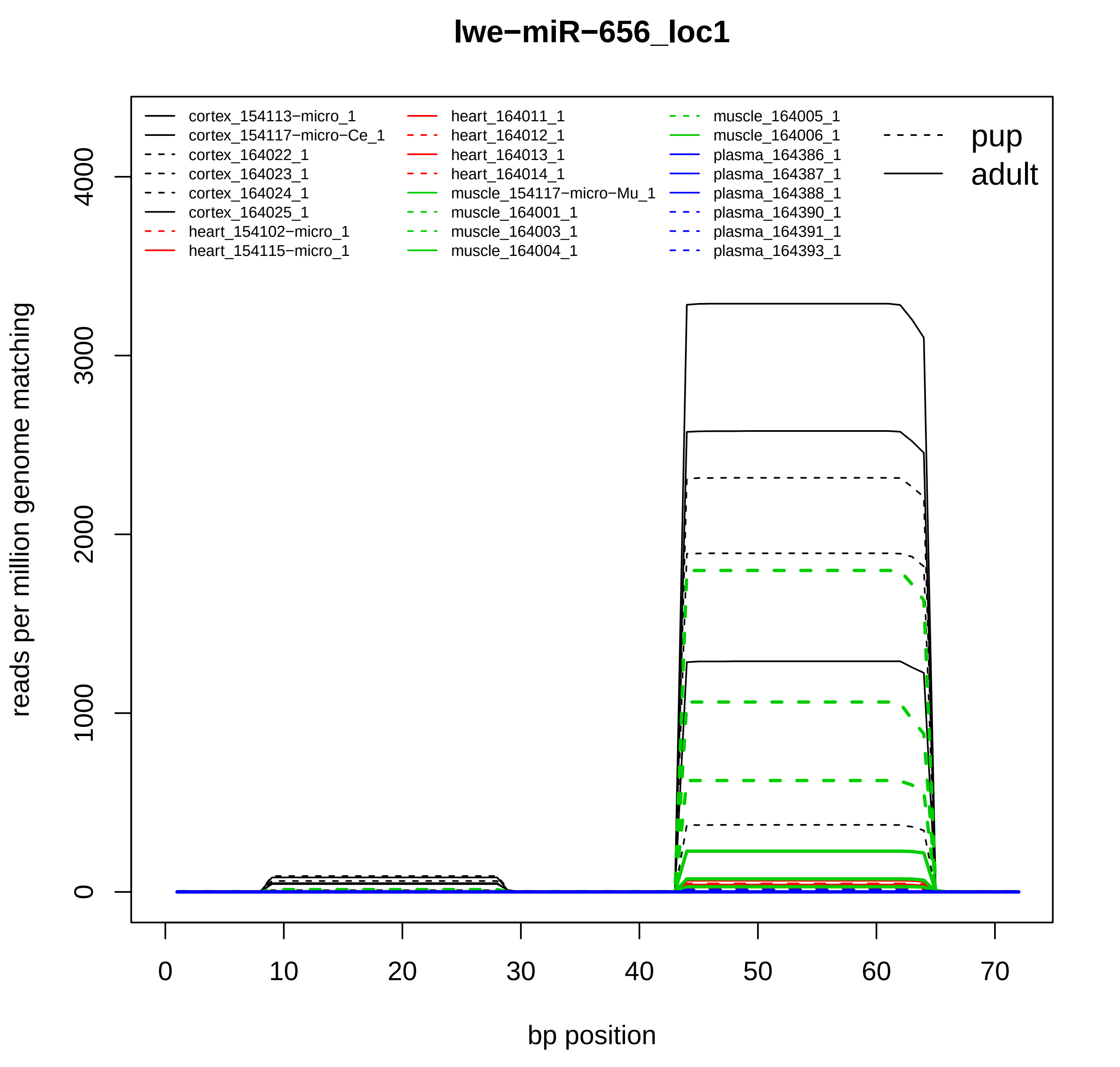

lwe−miR−656_loc1
cortex_154113−micro_1
cortex_154117−micro−Ce_1
cortex_164022_1
cortex_164023_1
cortex_164024_1
cortex_164025_1
heart_154102−micro_1
heart_154115−micro_1
heart_164011_1
heart_164012_1
heart_164013_1
heart_164014_1
muscle_154117−micro−Mu_1
muscle_164001_1
muscle_164003_1
muscle_164004_1
muscle_164005_1
muscle_164006_1
plasma_164386_1
plasma_164387_1
plasma_164388_1
plasma_164390_1
plasma_164391_1
plasma_164393_1
pup
adult
reads per million genome matching
0
1000
2000
3000
4000
bp position
0
10
20
30
40
50
60
70

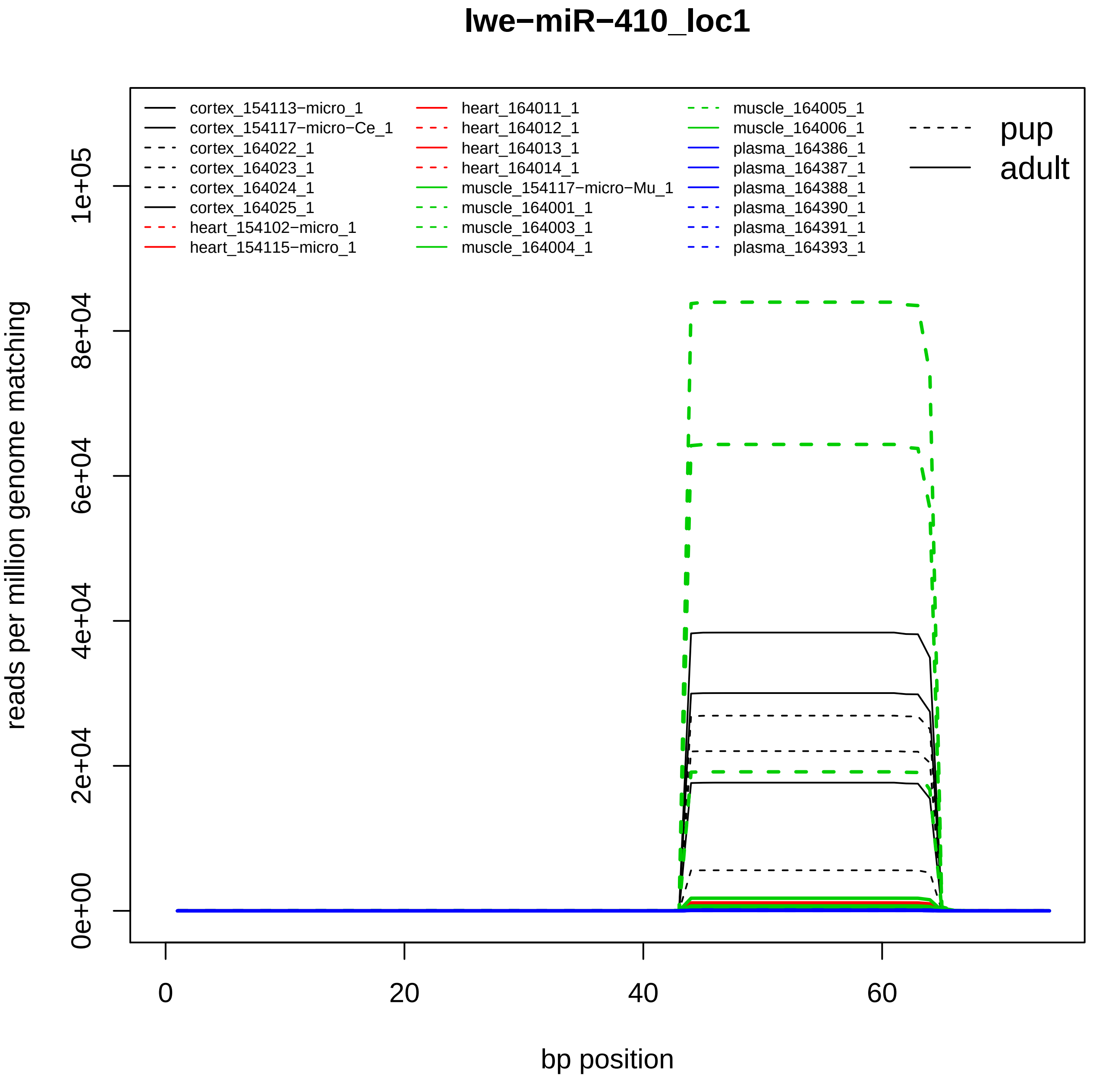
lwe−miR−410_loc1
cortex_154113−micro_1
cortex_154117−micro−Ce_1
cortex_164022_1
cortex_164023_1
cortex_164024_1
cortex_164025_1
heart_154102−micro_1
heart_154115−micro_1
heart_164011_1
heart_164012_1
heart_164013_1
heart_164014_1
muscle_154117−micro−Mu_1
muscle_164001_1
muscle_164003_1
muscle_164004_1
muscle_164005_1
muscle_164006_1
plasma_164386_1
plasma_164387_1
plasma_164388_1
plasma_164390_1
plasma_164391_1
plasma_164393_1
pup
adult
reads per million genome matching
0e+00
2e+04
4e+04
6e+04
8e+04
1e+05
0
20
40
60
bp position

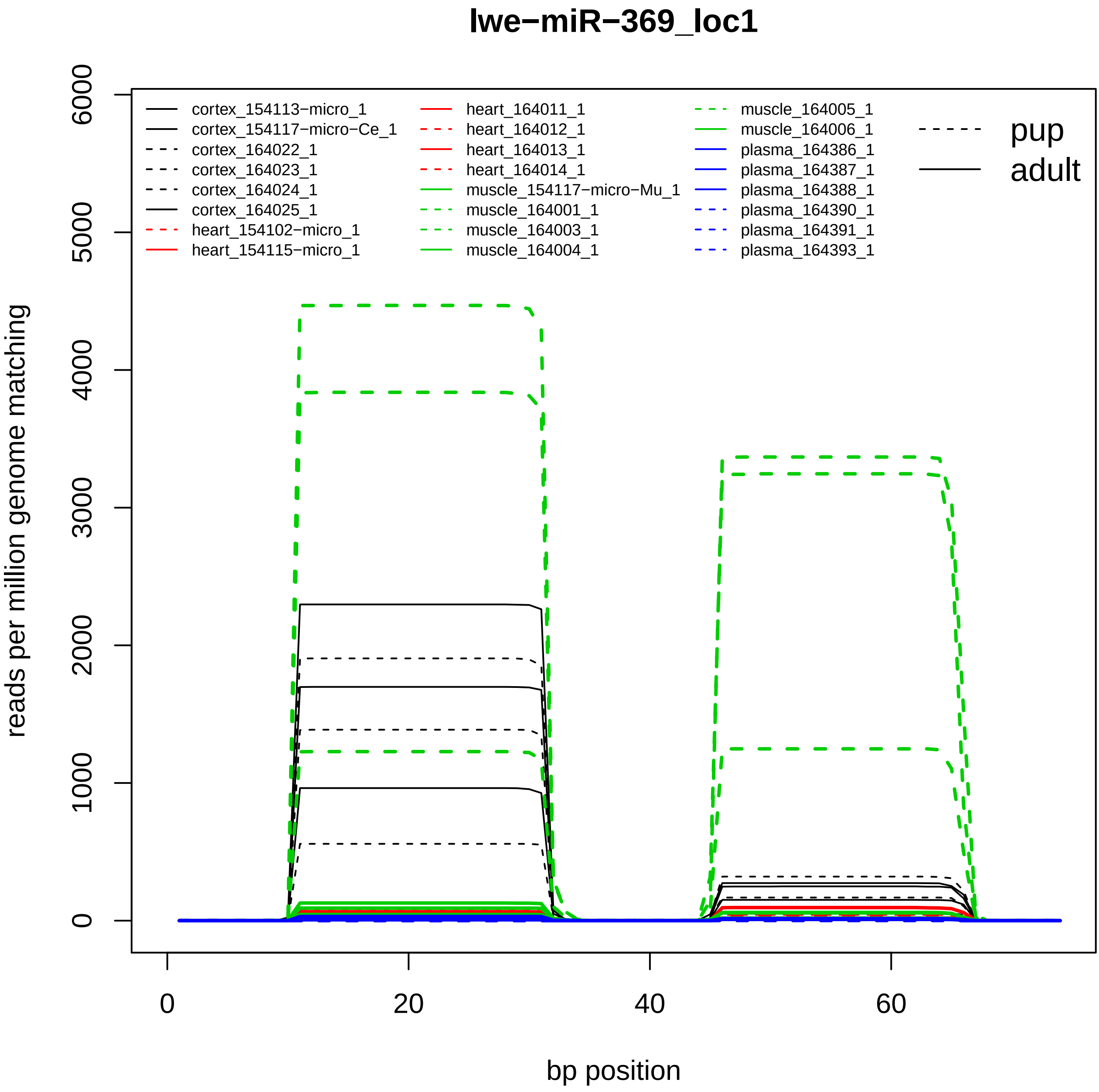
lwe−miR−369_loc1
cortex_154113−micro_1
cortex_154117−micro−Ce_1
cortex_164022_1
cortex_164023_1
cortex_164024_1
cortex_164025_1
heart_154102−micro_1
heart_154115−micro_1
heart_164011_1
heart_164012_1
heart_164013_1
heart_164014_1
muscle_154117−micro−Mu_1
muscle_164001_1
muscle_164003_1
muscle_164004_1
muscle_164005_1
muscle_164006_1
plasma_164386_1
plasma_164387_1
plasma_164388_1
plasma_164390_1
plasma_164391_1
plasma_164393_1
pup
adult
reads per million genome matching
bp position
0
1000
2000
3000
4000
5000
6000
0
20
40
60

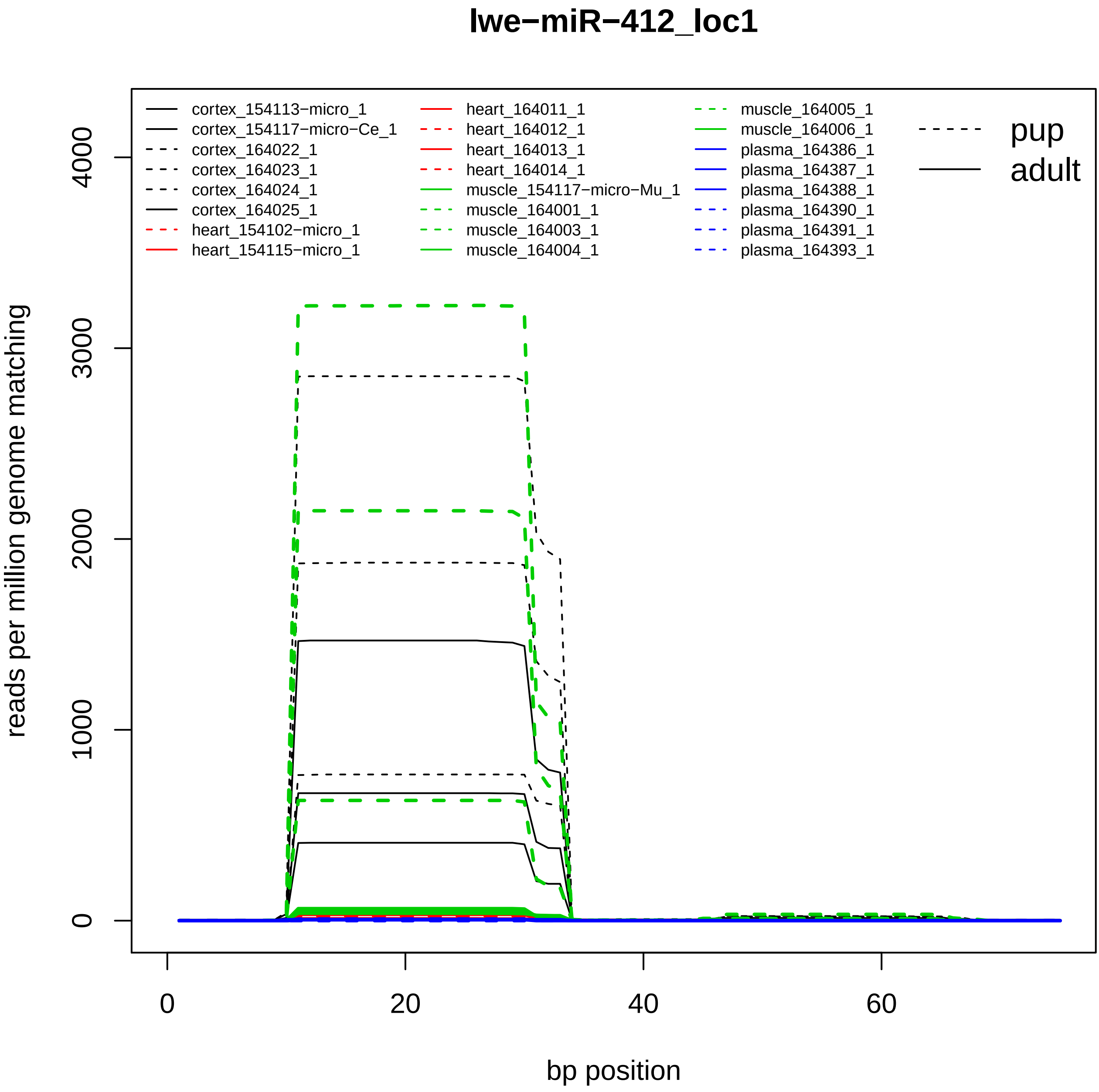

lwe−miR−412_loc1
cortex_154113−micro_1
cortex_154117−micro−Ce_1
cortex_164022_1
cortex_164023_1
cortex_164024_1
cortex_164025_1
heart_154102−micro_1
heart_154115−micro_1
heart_164011_1
heart_164012_1
heart_164013_1
heart_164014_1
muscle_154117−micro−Mu_1
muscle_164001_1
muscle_164003_1
muscle_164004_1
muscle_164005_1
muscle_164006_1
plasma_164386_1
plasma_164387_1
plasma_164388_1
plasma_164390_1
plasma_164391_1
plasma_164393_1
pup
adult
reads per million genome matching
0
1000
2000
3000
4000
0
20
40
60
bp position

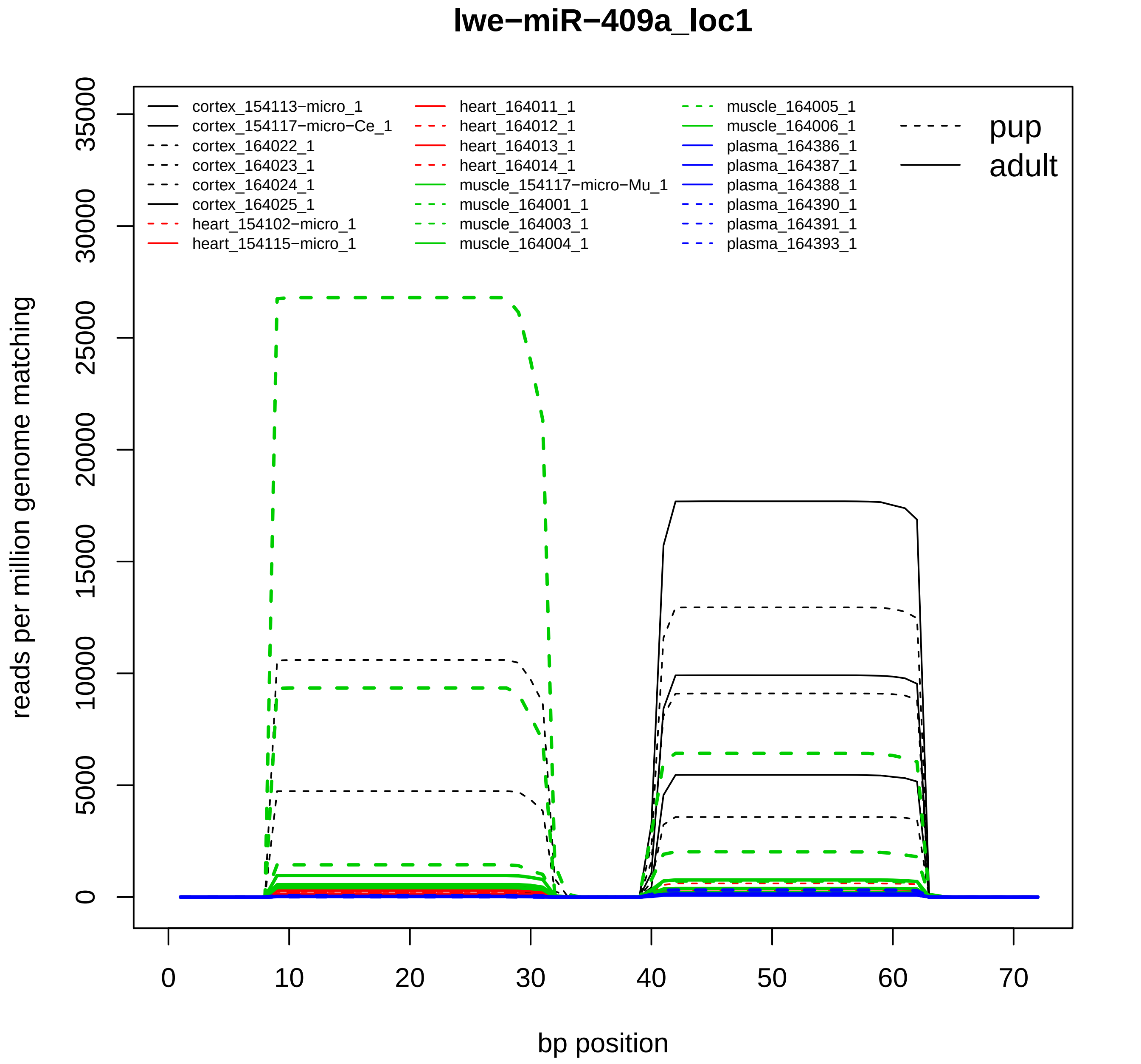
lwe−miR−409a_loc1
cortex_154113−micro_1
cortex_154117−micro−Ce_1
cortex_164022_1
cortex_164023_1
cortex_164024_1
cortex_164025_1
heart_154102−micro_1
heart_154115−micro_1
heart_164011_1
heart_164012_1
heart_164013_1
heart_164014_1
muscle_154117−micro−Mu_1
muscle_164001_1
muscle_164003_1
muscle_164004_1
muscle_164005_1
muscle_164006_1
plasma_164386_1
plasma_164387_1
plasma_164388_1
plasma_164390_1
plasma_164391_1
plasma_164393_1
pup
adult
reads per million genome matching
0
5000
10000
15000
20000
25000
30000
35000
bp position
0
10
20
30
40
50
60
70

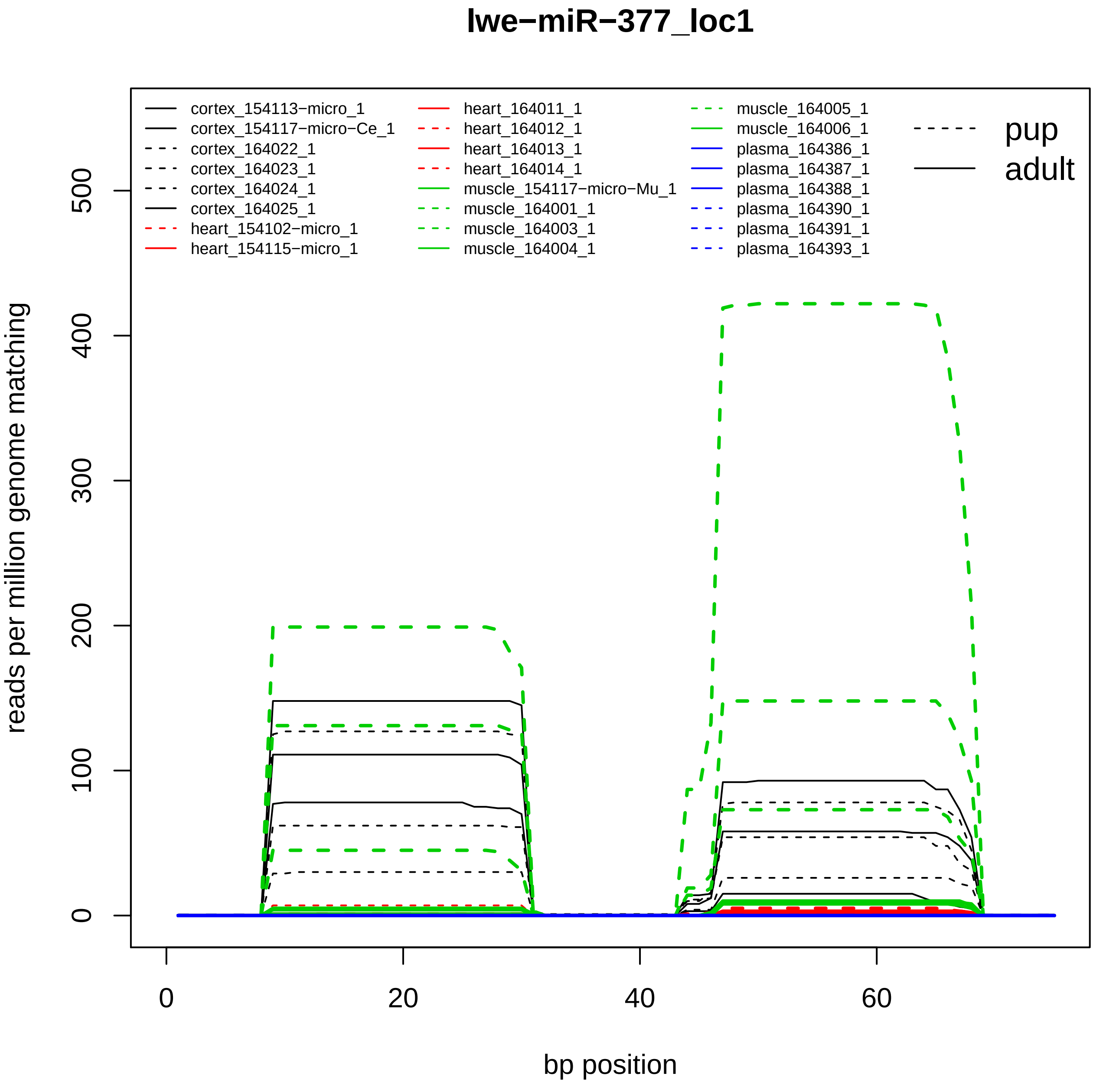

lwe−miR−377_loc1
cortex_154113−micro_1
cortex_154117−micro−Ce_1
cortex_164022_1
cortex_164023_1
cortex_164024_1
cortex_164025_1
heart_154102−micro_1
heart_154115−micro_1
heart_164011_1
heart_164012_1
heart_164013_1
heart_164014_1
muscle_154117−micro−Mu_1
muscle_164001_1
muscle_164003_1
muscle_164004_1
muscle_164005_1
muscle_164006_1
plasma_164386_1
plasma_164387_1
plasma_164388_1
plasma_164390_1
plasma_164391_1
plasma_164393_1
pup
adult
reads per million genome matching
0
100
200
300
400
500
0
20
40
60
bp position

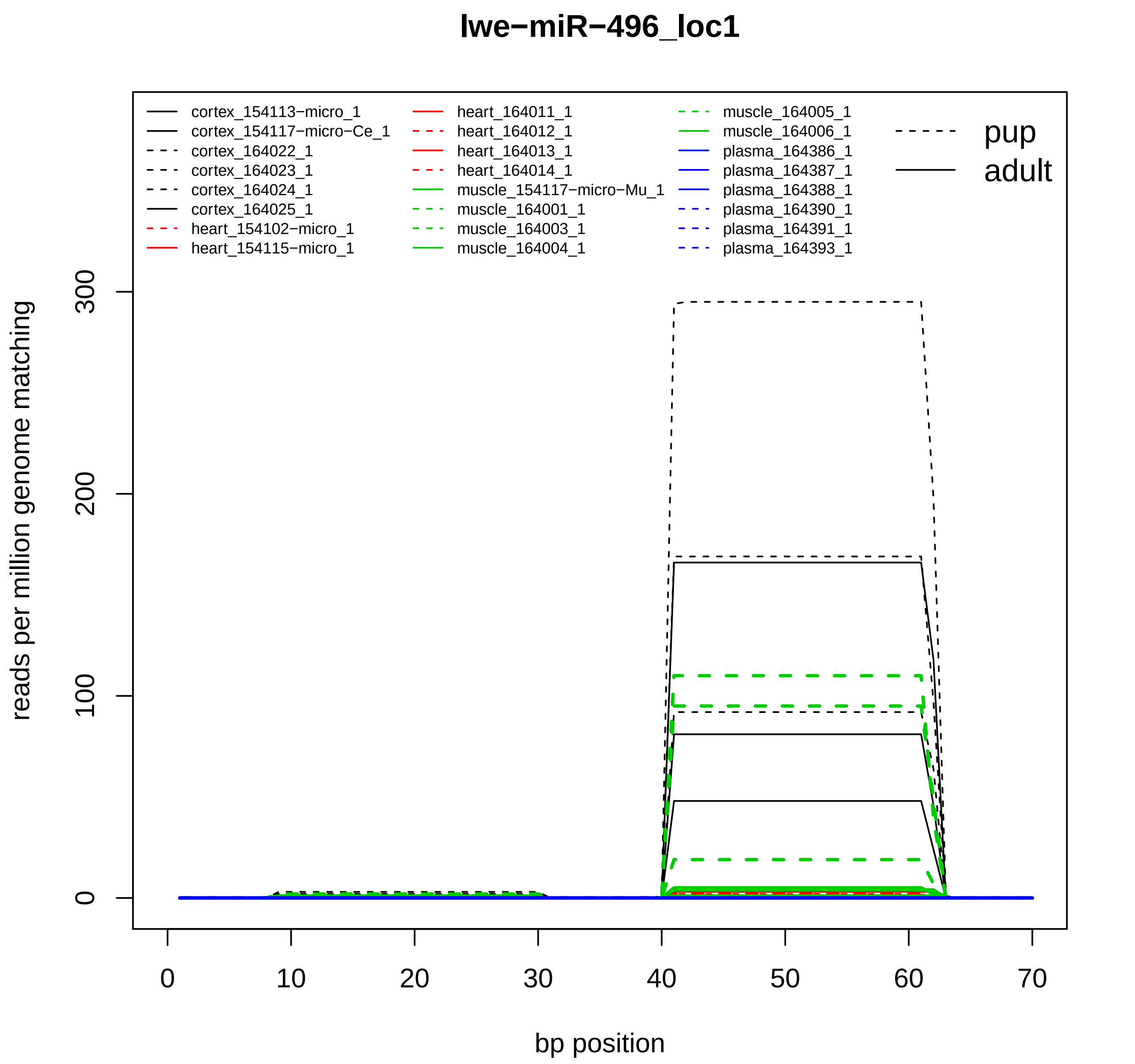
lwe−miR−496_loc1
cortex_154113−micro_1
cortex_154117−micro−Ce_1
cortex_164022_1
cortex_164023_1
cortex_164024_1
cortex_164025_1
heart_154102−micro_1
heart_154115−micro_1
heart_164011_1
heart_164012_1
heart_164013_1
heart_164014_1
muscle_154117−micro−Mu_1
muscle_164001_1
muscle_164003_1
muscle_164004_1
muscle_164005_1
muscle_164006_1
plasma_164386_1
plasma_164387_1
plasma_164388_1
plasma_164390_1
plasma_164391_1
plasma_164393_1
pup
adult
reads per million genome matching
0
100
200
300
0
10
20
30
40
50
60
70
bp position

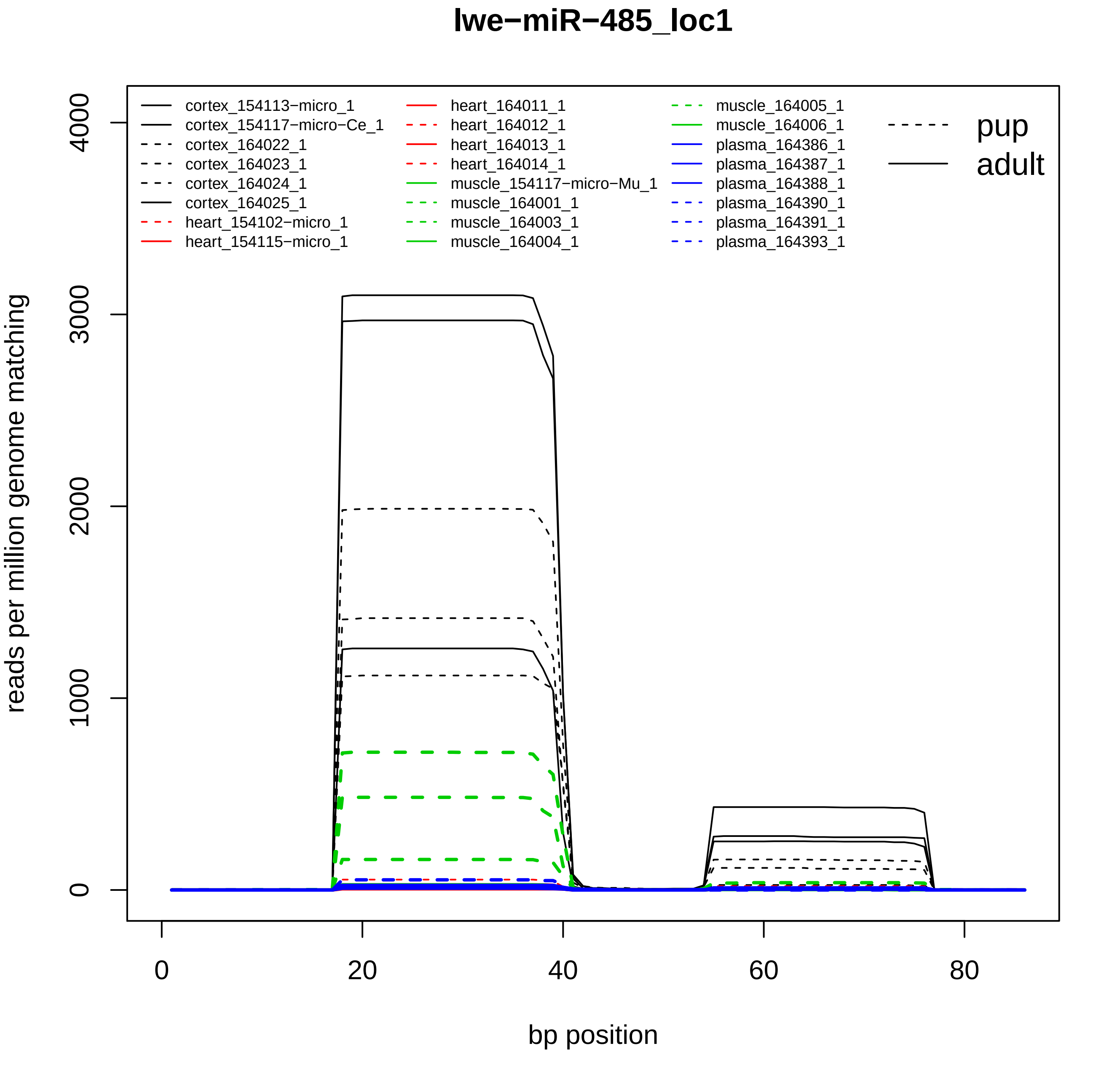
lwe−miR−485_loc1
reads per million genome matching
bp position
0
1000
2000
3000
4000
0
20
40
60
80
cortex_154113−micro_1
cortex_154117−micro−Ce_1
cortex_164022_1
cortex_164023_1
cortex_164024_1
cortex_164025_1
heart_154102−micro_1
heart_154115−micro_1
heart_164011_1
heart_164012_1
heart_164013_1
heart_164014_1
muscle_154117−micro−Mu_1
muscle_164001_1
muscle_164003_1
muscle_164004_1
muscle_164005_1
muscle_164006_1
plasma_164386_1
plasma_164387_1
plasma_164388_1
plasma_164390_1
plasma_164391_1
plasma_164393_1
pup
adult

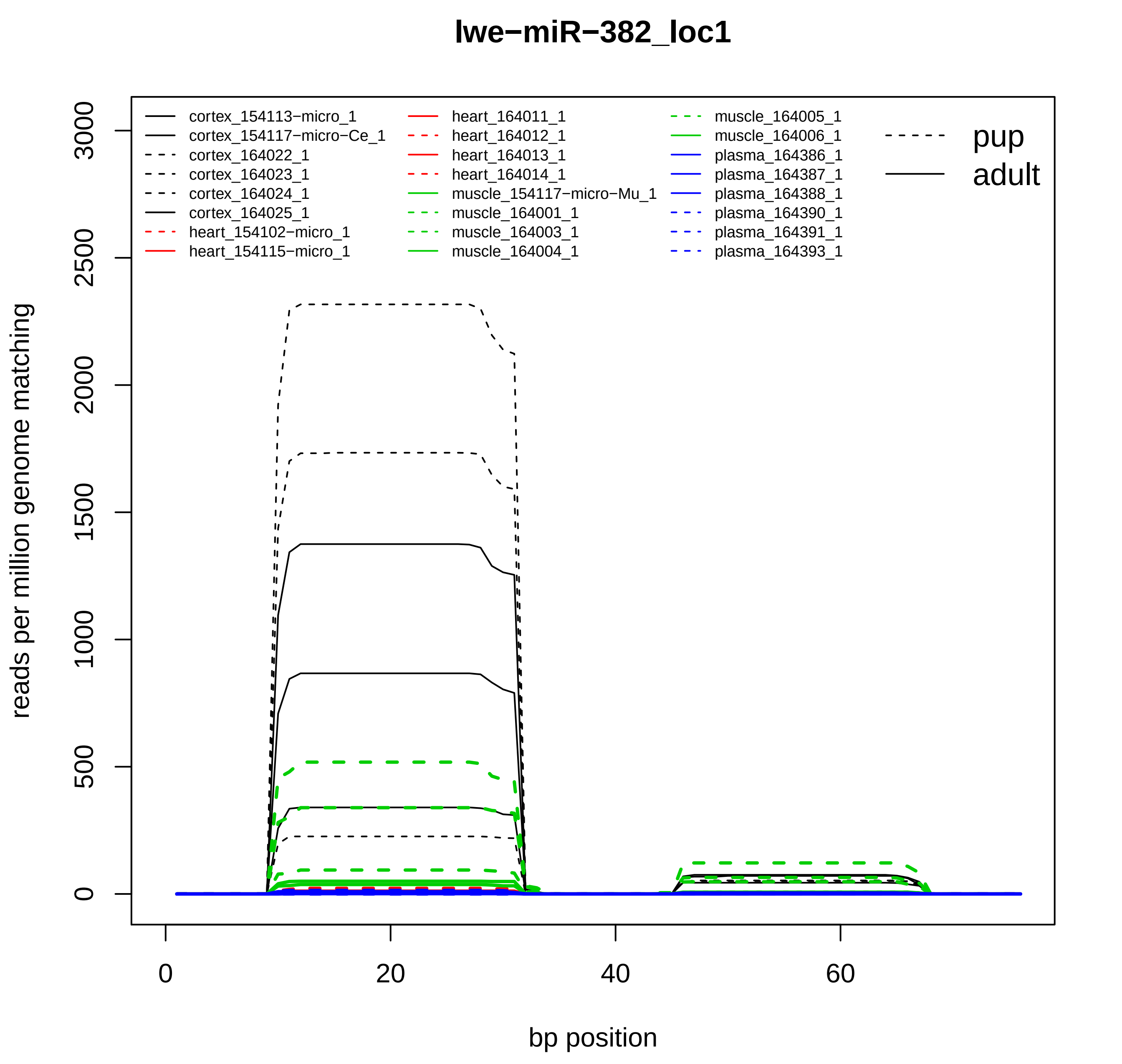
lwe−miR−382_loc1
reads per million genome matching
bp position
0
500
1000
1500
2000
2500
3000
0
20
40
60
cortex_154113−micro_1
cortex_154117−micro−Ce_1
cortex_164022_1
cortex_164023_1
cortex_164024_1
cortex_164025_1
heart_154102−micro_1
heart_154115−micro_1
heart_164011_1
heart_164012_1
heart_164013_1
heart_164014_1
muscle_154117−micro−Mu_1
muscle_164001_1
muscle_164003_1
muscle_164004_1
muscle_164005_1
muscle_164006_1
plasma_164386_1
plasma_164387_1
plasma_164388_1
plasma_164390_1
plasma_164391_1
plasma_164393_1
pup
adult

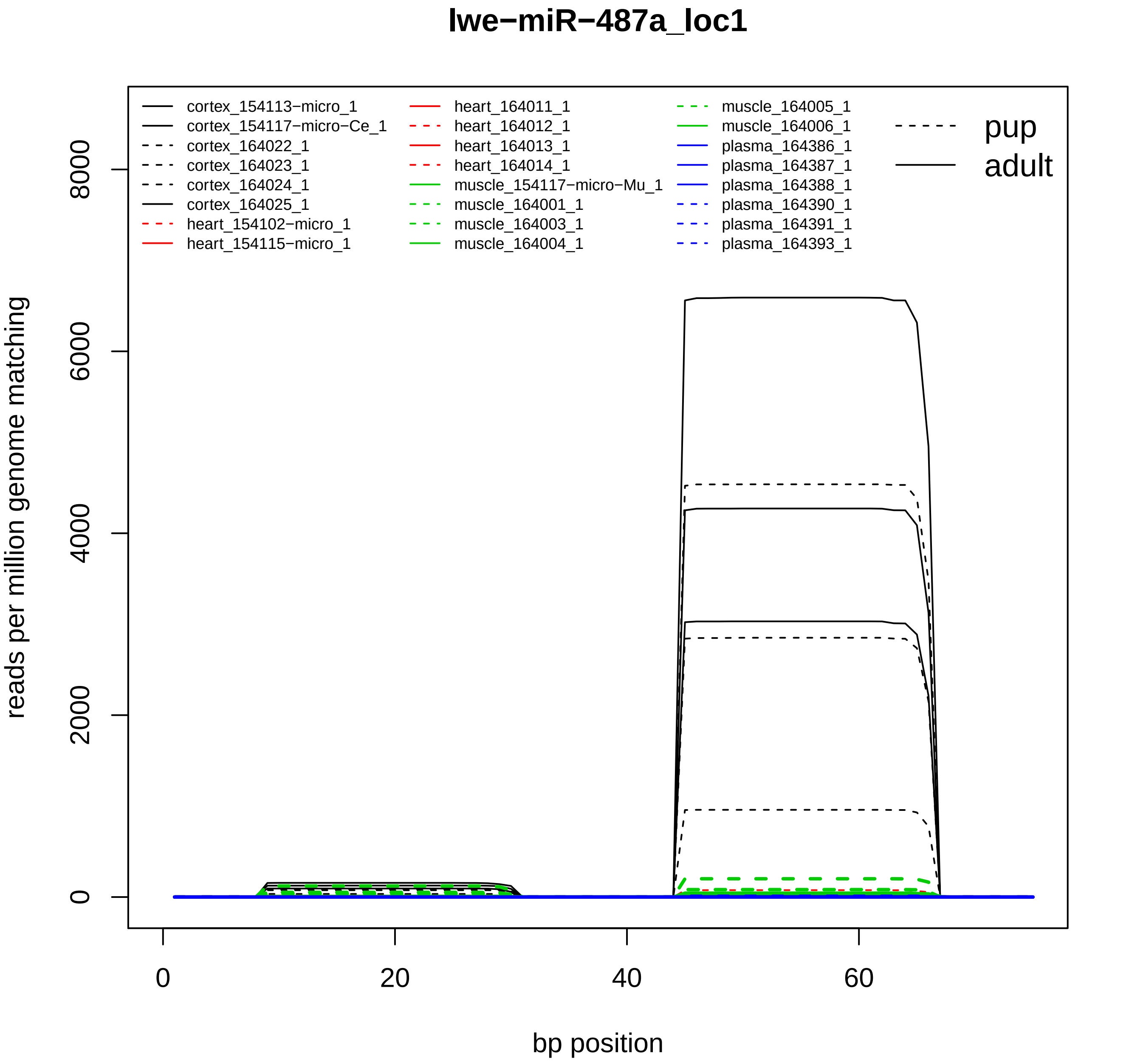
lwe−miR−487a_loc1
cortex_154113−micro_1
cortex_154117−micro−Ce_1
cortex_164022_1
cortex_164023_1
cortex_164024_1
cortex_164025_1
heart_154102−micro_1
heart_154115−micro_1
heart_164011_1
heart_164012_1
heart_164013_1
heart_164014_1
muscle_154117−micro−Mu_1
muscle_164001_1
muscle_164003_1
muscle_164004_1
muscle_164005_1
muscle_164006_1
plasma_164386_1
plasma_164387_1
plasma_164388_1
plasma_164390_1
plasma_164391_1
plasma_164393_1
pup
adult
reads per million genome matching
0
2000
4000
6000
8000
0
20
40
60
bp position

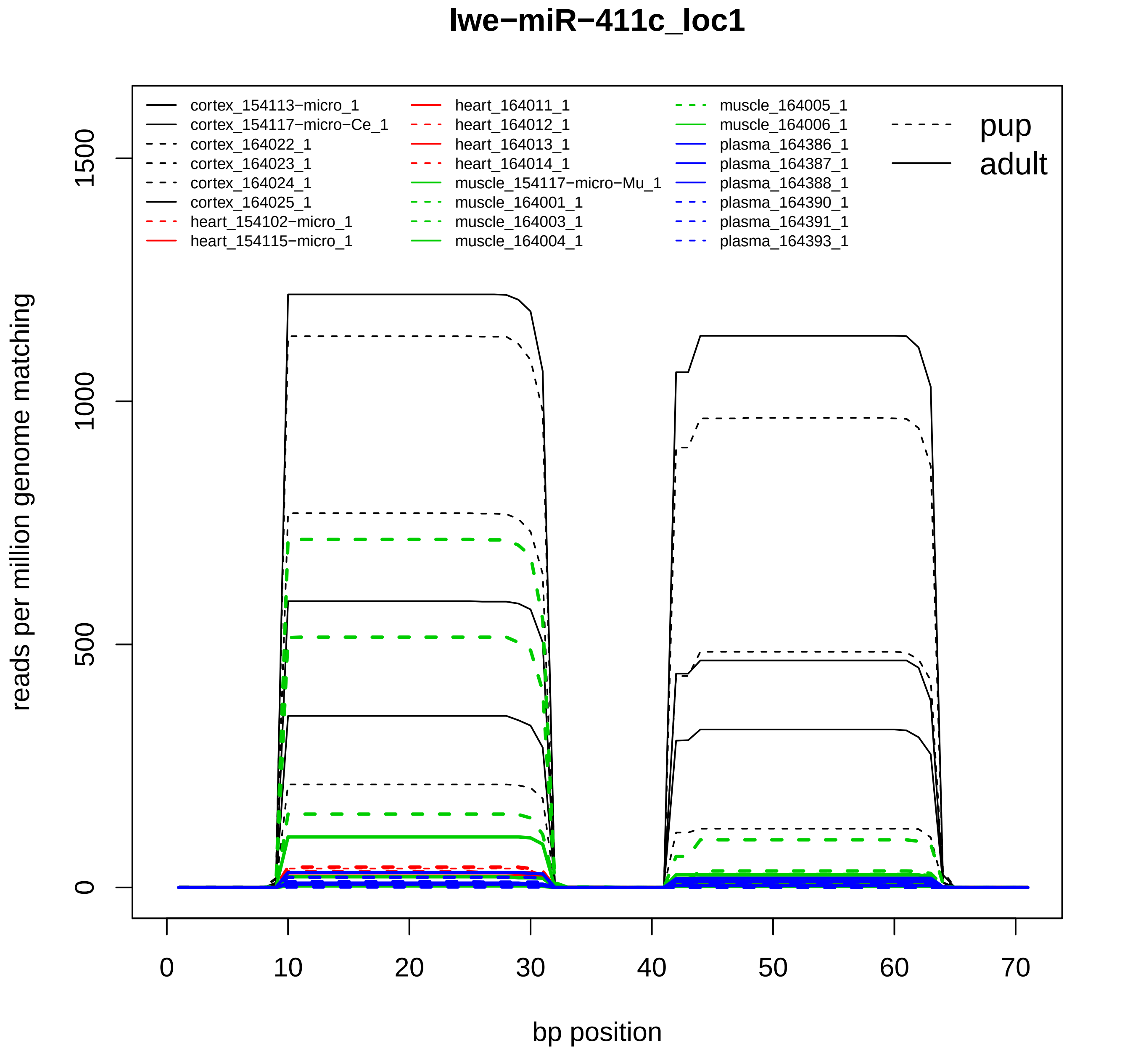

lwe−miR−411c_loc1
cortex_154113−micro_1
cortex_154117−micro−Ce_1
cortex_164022_1
cortex_164023_1
cortex_164024_1
cortex_164025_1
heart_154102−micro_1
heart_154115−micro_1
heart_164011_1
heart_164012_1
heart_164013_1
heart_164014_1
muscle_154117−micro−Mu_1
muscle_164001_1
muscle_164003_1
muscle_164004_1
muscle_164005_1
muscle_164006_1
plasma_164386_1
plasma_164387_1
plasma_164388_1
plasma_164390_1
plasma_164391_1
plasma_164393_1
pup
adult
reads per million genome matching
0
500
1000
1500
bp position
0
10
20
30
40
50
60
70

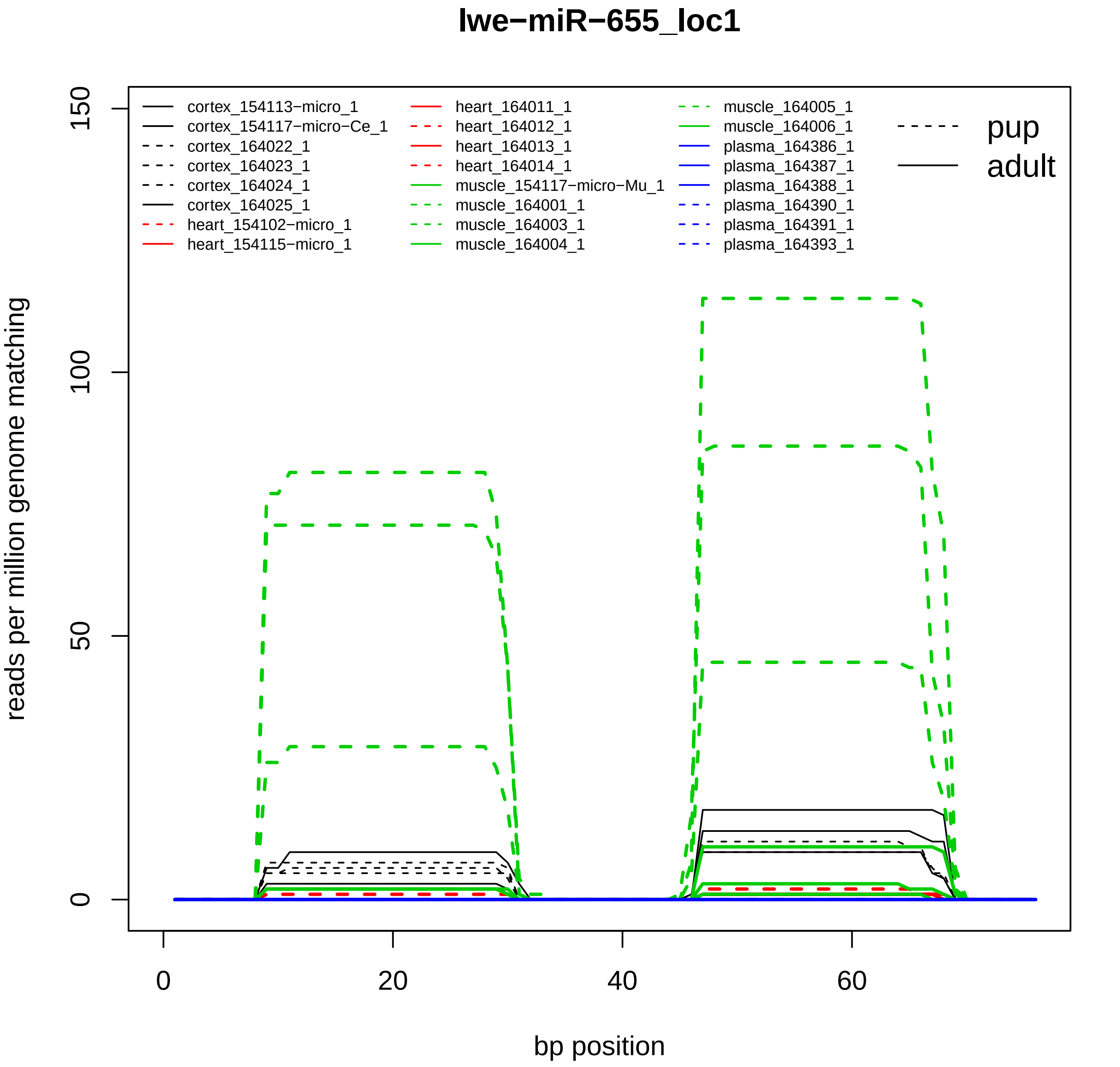

lwe−miR−655_loc1
reads per million genome matching
bp position
0
50
100
150
0
20
40
60
cortex_154113−micro_1
cortex_154117−micro−Ce_1
cortex_164022_1
cortex_164023_1
cortex_164024_1
cortex_164025_1
heart_154102−micro_1
heart_154115−micro_1
heart_164011_1
heart_164012_1
heart_164013_1
heart_164014_1
muscle_154117−micro−Mu_1
muscle_164001_1
muscle_164003_1
muscle_164004_1
muscle_164005_1
muscle_164006_1
plasma_164386_1
plasma_164387_1
plasma_164388_1
plasma_164390_1
plasma_164391_1
plasma_164393_1
pup
adult

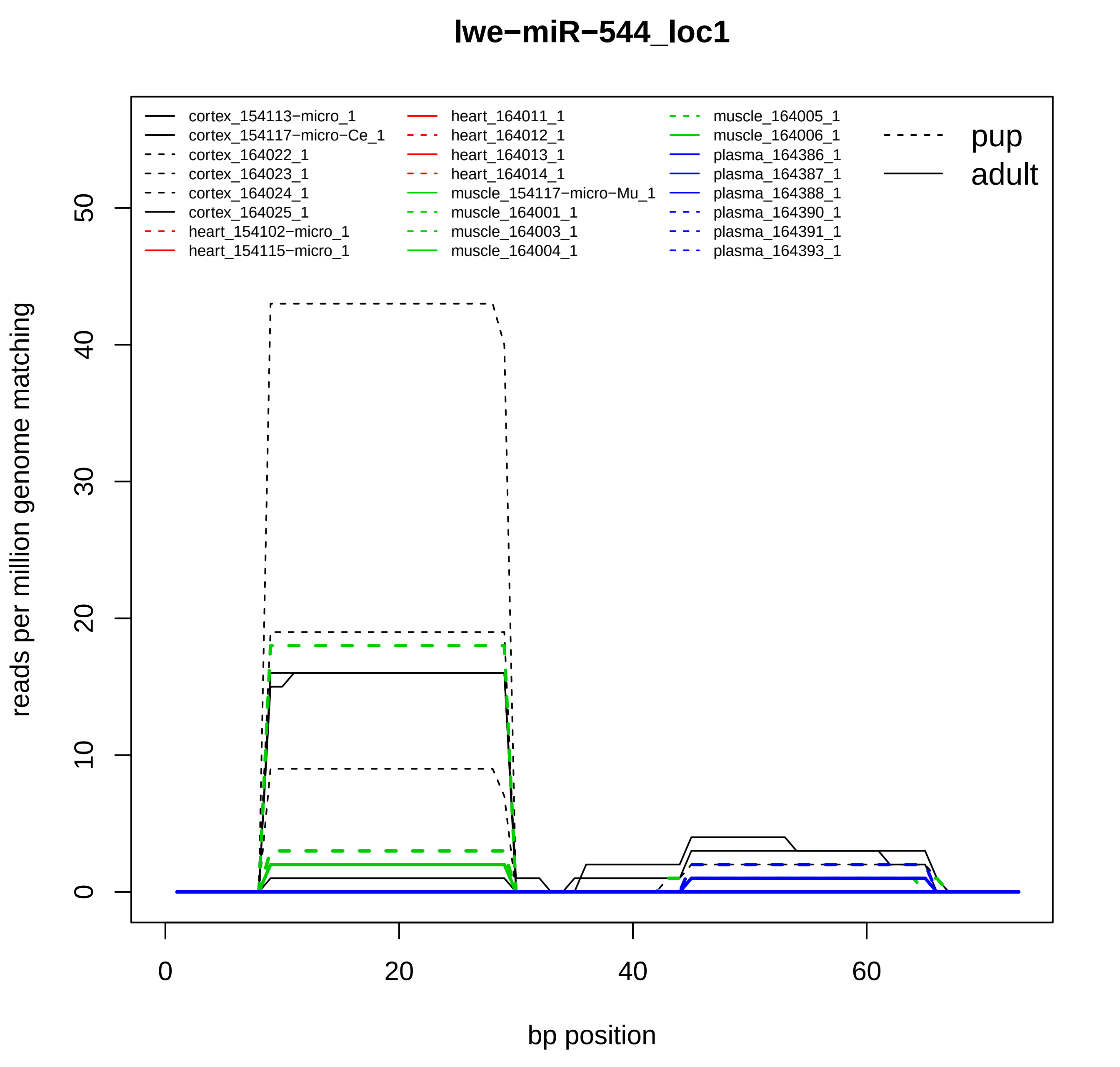

lwe−miR−544_loc1
cortex_154113−micro_1
cortex_154117−micro−Ce_1
cortex_164022_1
cortex_164023_1
cortex_164024_1
cortex_164025_1
heart_154102−micro_1
heart_154115−micro_1
heart_164011_1
heart_164012_1
heart_164013_1
heart_164014_1
muscle_154117−micro−Mu_1
muscle_164001_1
muscle_164003_1
muscle_164004_1
muscle_164005_1
muscle_164006_1
plasma_164386_1
plasma_164387_1
plasma_164388_1
plasma_164390_1
plasma_164391_1
plasma_164393_1
pup
adult
reads per million genome matching
0
10
20
30
40
50
0
20
40
60
bp position

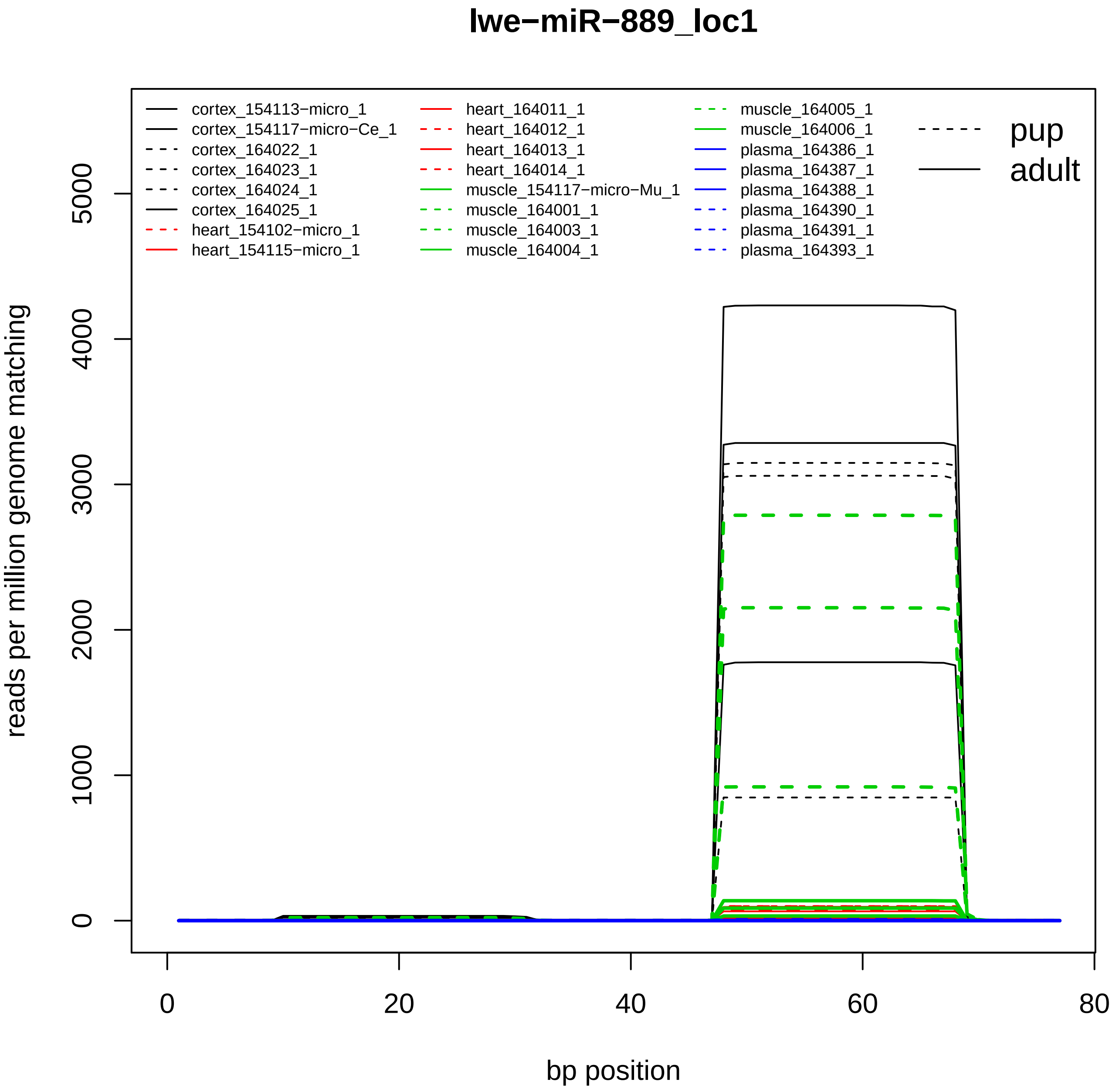

lwe−miR−889_loc1
cortex_154113−micro_1
cortex_154117−micro−Ce_1
cortex_164022_1
cortex_164023_1
cortex_164024_1
cortex_164025_1
heart_154102−micro_1
heart_154115−micro_1
heart_164011_1
heart_164012_1
heart_164013_1
heart_164014_1
muscle_154117−micro−Mu_1
muscle_164001_1
muscle_164003_1
muscle_164004_1
muscle_164005_1
muscle_164006_1
plasma_164386_1
plasma_164387_1
plasma_164388_1
plasma_164390_1
plasma_164391_1
plasma_164393_1
pup
adult
reads per million genome matching
0
1000
2000
3000
4000
5000
0
20
40
60
80
bp position

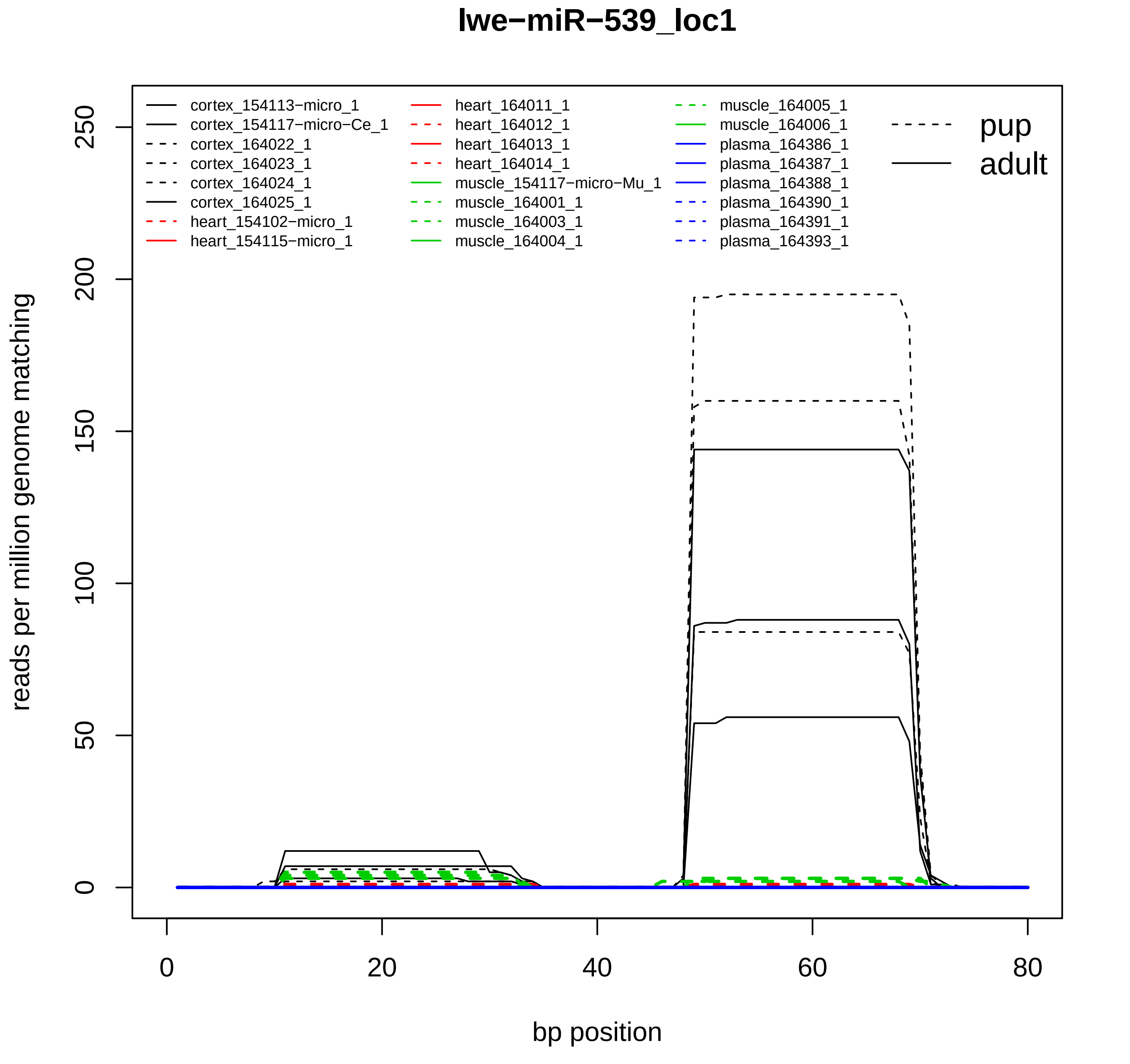

lwe−miR−539_loc1
cortex_154113−micro_1
cortex_154117−micro−Ce_1
cortex_164022_1
cortex_164023_1
cortex_164024_1
cortex_164025_1
heart_154102−micro_1
heart_154115−micro_1
heart_164011_1
heart_164012_1
heart_164013_1
heart_164014_1
muscle_154117−micro−Mu_1
muscle_164001_1
muscle_164003_1
muscle_164004_1
muscle_164005_1
muscle_164006_1
plasma_164386_1
plasma_164387_1
plasma_164388_1
plasma_164390_1
plasma_164391_1
plasma_164393_1
pup
adult
0
50
100
150
200
250
reads per million genome matching
0
20
40
60
80
bp position

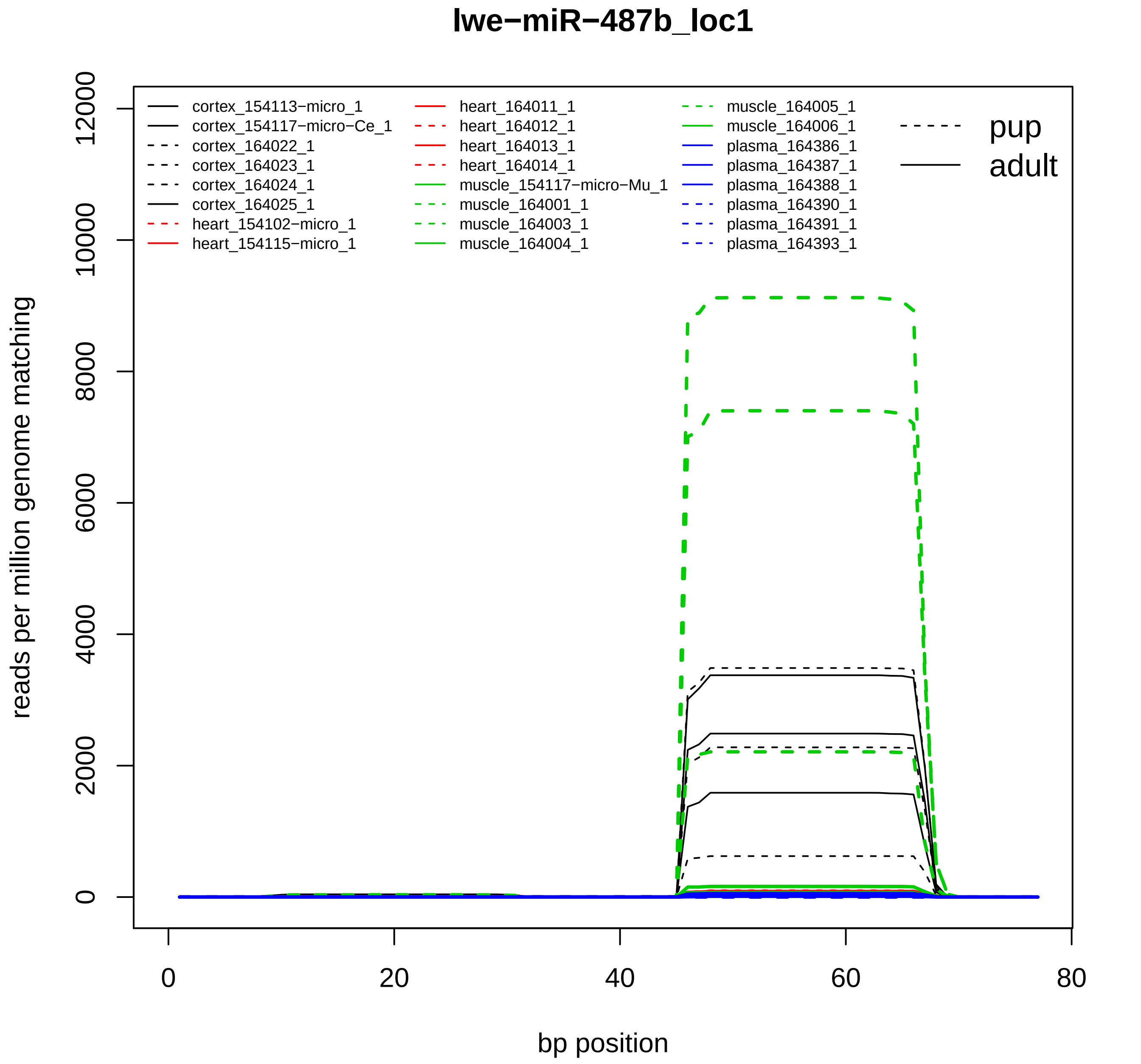
lwe−miR−487b_loc1
cortex_154113−micro_1
cortex_154117−micro−Ce_1
cortex_164022_1
cortex_164023_1
cortex_164024_1
cortex_164025_1
heart_154102−micro_1
heart_154115−micro_1
heart_164011_1
heart_164012_1
heart_164013_1
heart_164014_1
muscle_154117−micro−Mu_1
muscle_164001_1
muscle_164003_1
muscle_164004_1
muscle_164005_1
muscle_164006_1
plasma_164386_1
plasma_164387_1
plasma_164388_1
plasma_164390_1
plasma_164391_1
plasma_164393_1
pup
adult
reads per million genome matching
0
2000
4000
6000
8000
10000
12000
0
20
40
60
80
bp position

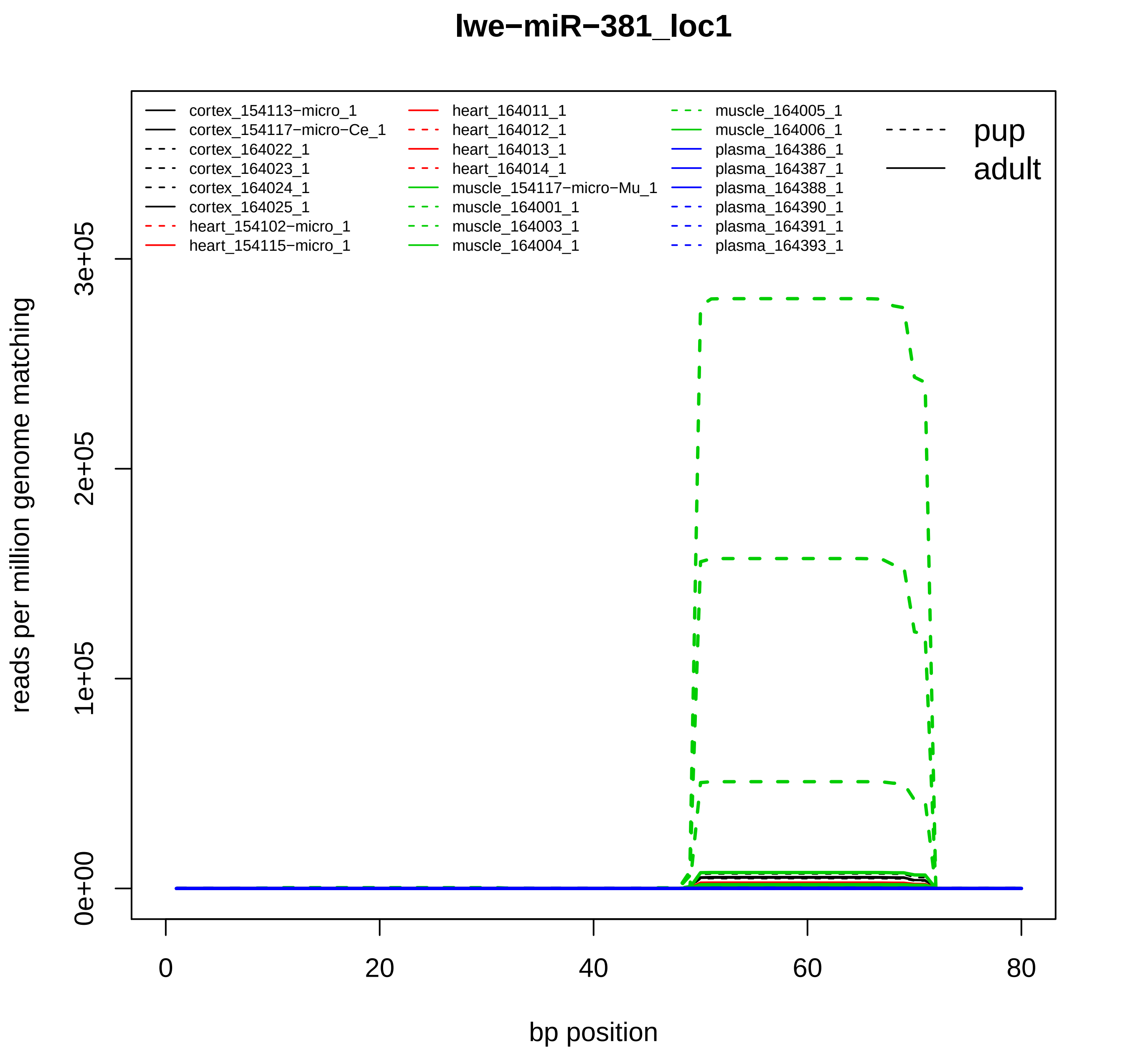
lwe−miR−381_loc1
cortex_154113−micro_1
cortex_154117−micro−Ce_1
cortex_164022_1
cortex_164023_1
cortex_164024_1
cortex_164025_1
heart_154102−micro_1
heart_154115−micro_1
heart_164011_1
heart_164012_1
heart_164013_1
heart_164014_1
muscle_154117−micro−Mu_1
muscle_164001_1
muscle_164003_1
muscle_164004_1
muscle_164005_1
muscle_164006_1
plasma_164386_1
plasma_164387_1
plasma_164388_1
plasma_164390_1
plasma_164391_1
plasma_164393_1
pup
adult
reads per million genome matching
0e+00
1e+05
2e+05
3e+05
0
20
40
60
80
bp position

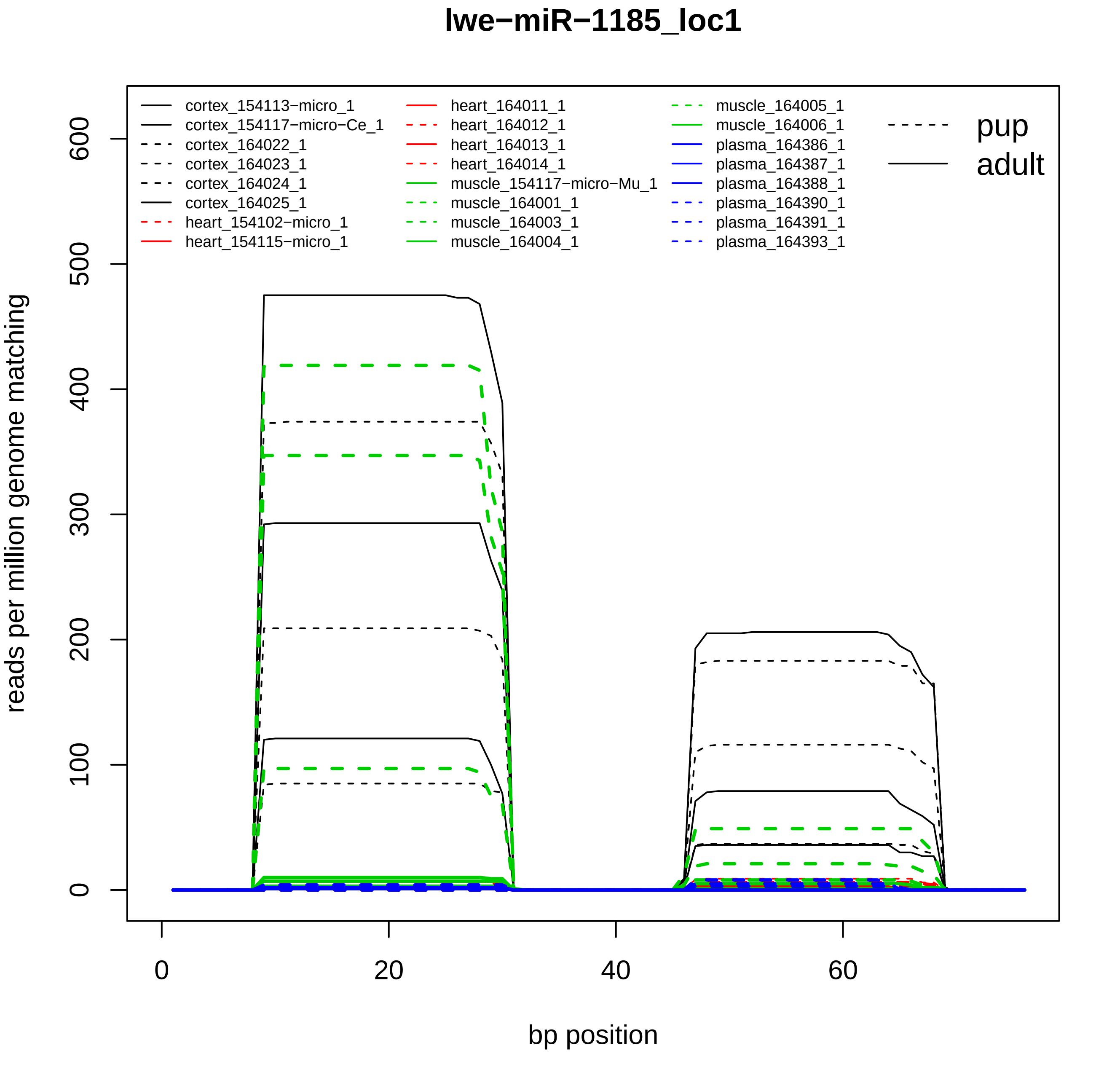
lwe−miR−1185_loc1
cortex_154113−micro_1
cortex_154117−micro−Ce_1
cortex_164022_1
cortex_164023_1
cortex_164024_1
cortex_164025_1
heart_154102−micro_1
heart_154115−micro_1
heart_164011_1
heart_164012_1
heart_164013_1
heart_164014_1
muscle_154117−micro−Mu_1
muscle_164001_1
muscle_164003_1
muscle_164004_1
muscle_164005_1
muscle_164006_1
plasma_164386_1
plasma_164387_1
plasma_164388_1
plasma_164390_1
plasma_164391_1
plasma_164393_1
pup
adult
reads per million genome matching
0
100
200
300
400
500
600
0
20
40
60
bp position

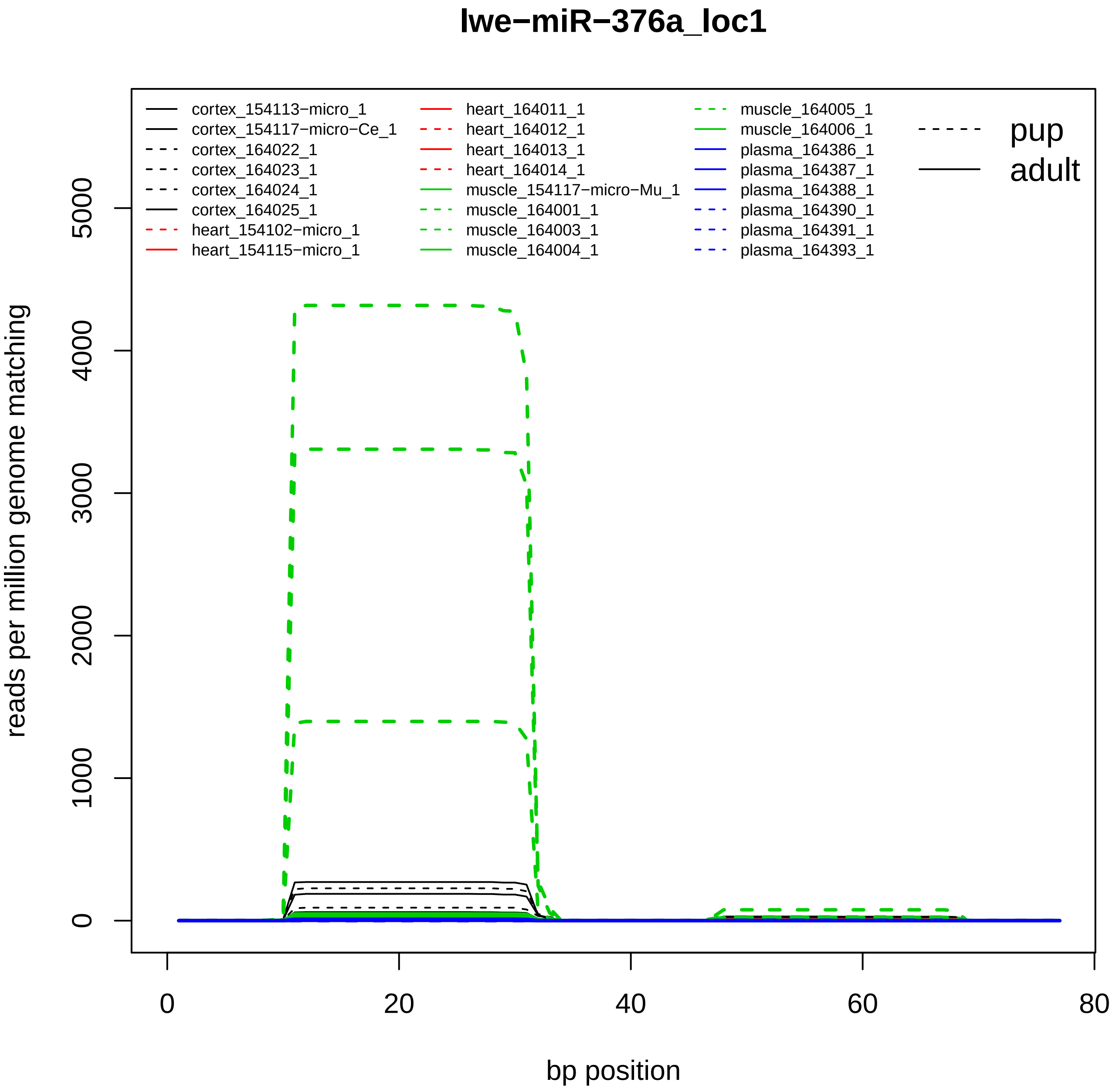
lwe−miR−376a_loc1
cortex_154113−micro_1
cortex_154117−micro−Ce_1
cortex_164022_1
cortex_164023_1
cortex_164024_1
cortex_164025_1
heart_154102−micro_1
heart_154115−micro_1
heart_164011_1
heart_164012_1
heart_164013_1
heart_164014_1
muscle_154117−micro−Mu_1
muscle_164001_1
muscle_164003_1
muscle_164004_1
muscle_164005_1
muscle_164006_1
plasma_164386_1
plasma_164387_1
plasma_164388_1
plasma_164390_1
plasma_164391_1
plasma_164393_1
pup
adult
reads per million genome matching
0
1000
2000
3000
4000
5000
bp position
0
20
40
60
80

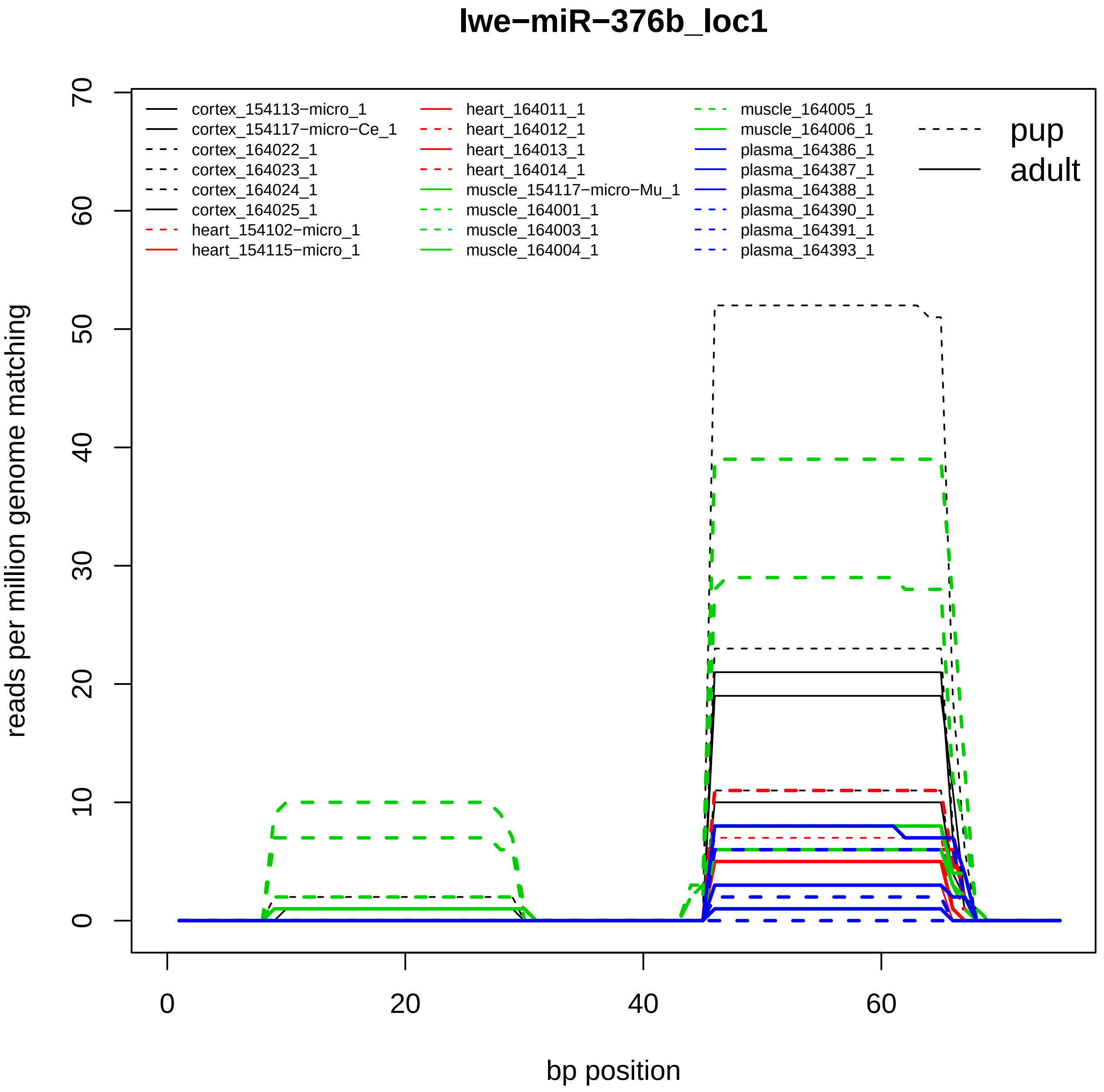

lwe−miR−376b_loc1
cortex_154113−micro_1
cortex_154117−micro−Ce_1
cortex_164022_1
cortex_164023_1
cortex_164024_1
cortex_164025_1
heart_154102−micro_1
heart_154115−micro_1
heart_164011_1
heart_164012_1
heart_164013_1
heart_164014_1
muscle_154117−micro−Mu_1
muscle_164001_1
muscle_164003_1
muscle_164004_1
muscle_164005_1
muscle_164006_1
plasma_164386_1
plasma_164387_1
plasma_164388_1
plasma_164390_1
plasma_164391_1
plasma_164393_1
pup
adult
reads per million genome matching
0
10
20
30
40
50
60
70
0
20
40
60
bp position

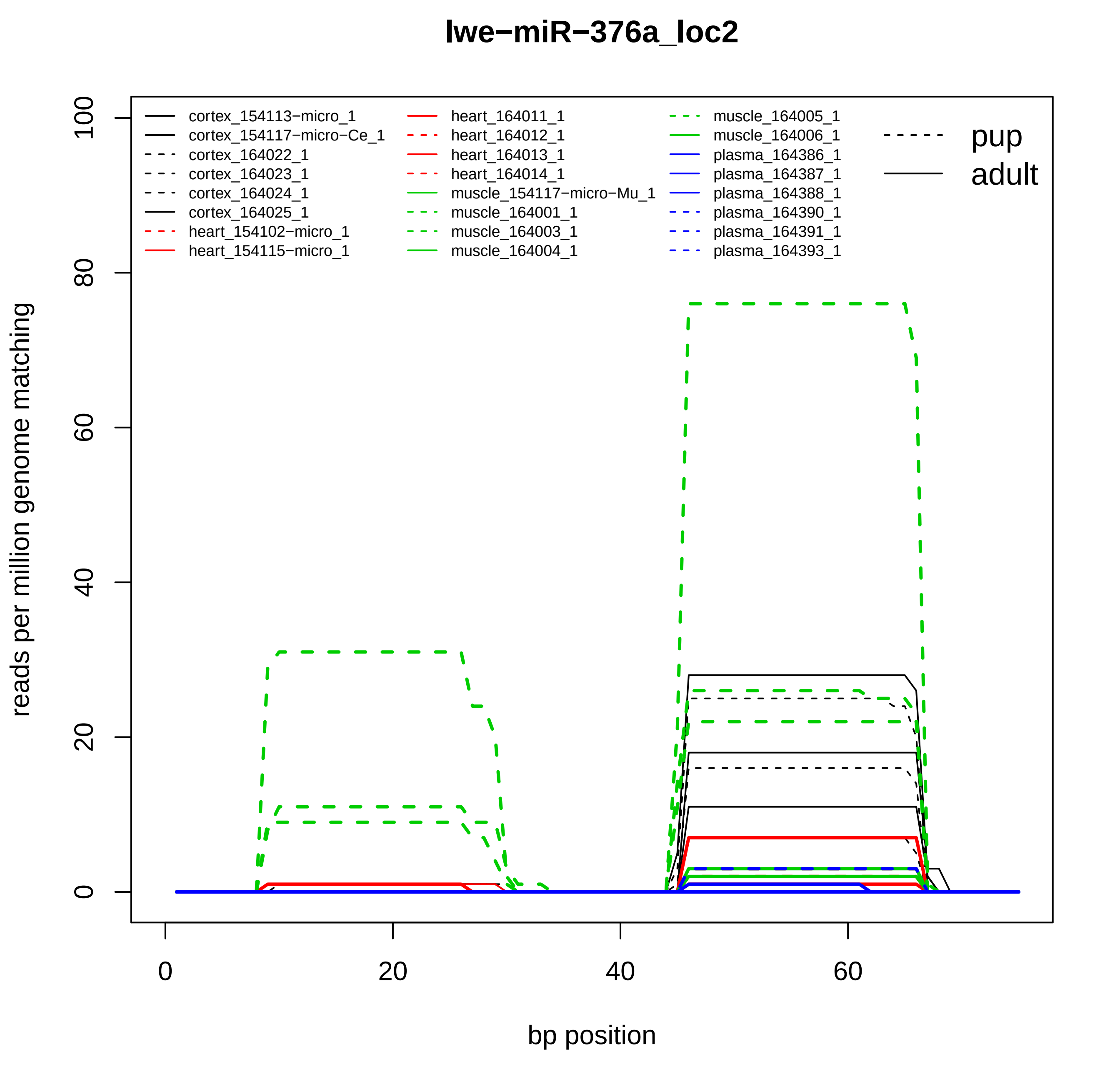

lwe−miR−376a_loc2
reads per million genome matching
bp position
0
20
40
60
80
100
0
20
40
60
cortex_154113−micro_1
cortex_154117−micro−Ce_1
cortex_164022_1
cortex_164023_1
cortex_164024_1
cortex_164025_1
heart_154102−micro_1
heart_154115−micro_1
heart_164011_1
heart_164012_1
heart_164013_1
heart_164014_1
muscle_154117−micro−Mu_1
muscle_164001_1
muscle_164003_1
muscle_164004_1
muscle_164005_1
muscle_164006_1
plasma_164386_1
plasma_164387_1
plasma_164388_1
plasma_164390_1
plasma_164391_1
plasma_164393_1
pup
adult

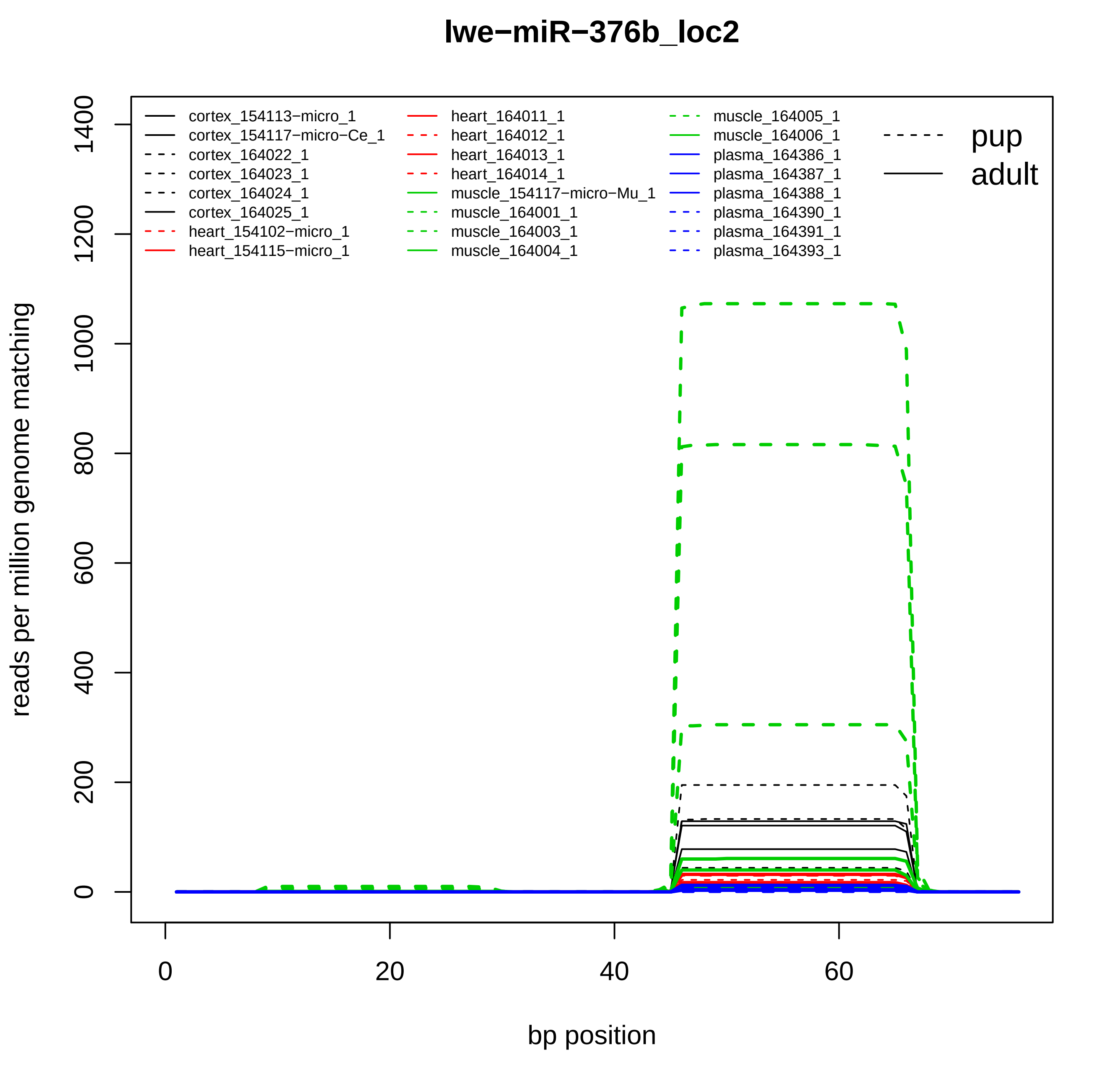
lwe−miR−376b_loc2
reads per million genome matching
bp position
0
200
400
600
800
1000
1200
1400
0
20
40
60
cortex_154113−micro_1
cortex_154117−micro−Ce_1
cortex_164022_1
cortex_164023_1
cortex_164024_1
cortex_164025_1
heart_154102−micro_1
heart_154115−micro_1
heart_164011_1
heart_164012_1
heart_164013_1
heart_164014_1
muscle_154117−micro−Mu_1
muscle_164001_1
muscle_164003_1
muscle_164004_1
muscle_164005_1
muscle_164006_1
plasma_164386_1
plasma_164387_1
plasma_164388_1
plasma_164390_1
plasma_164391_1
plasma_164393_1
pup
adult

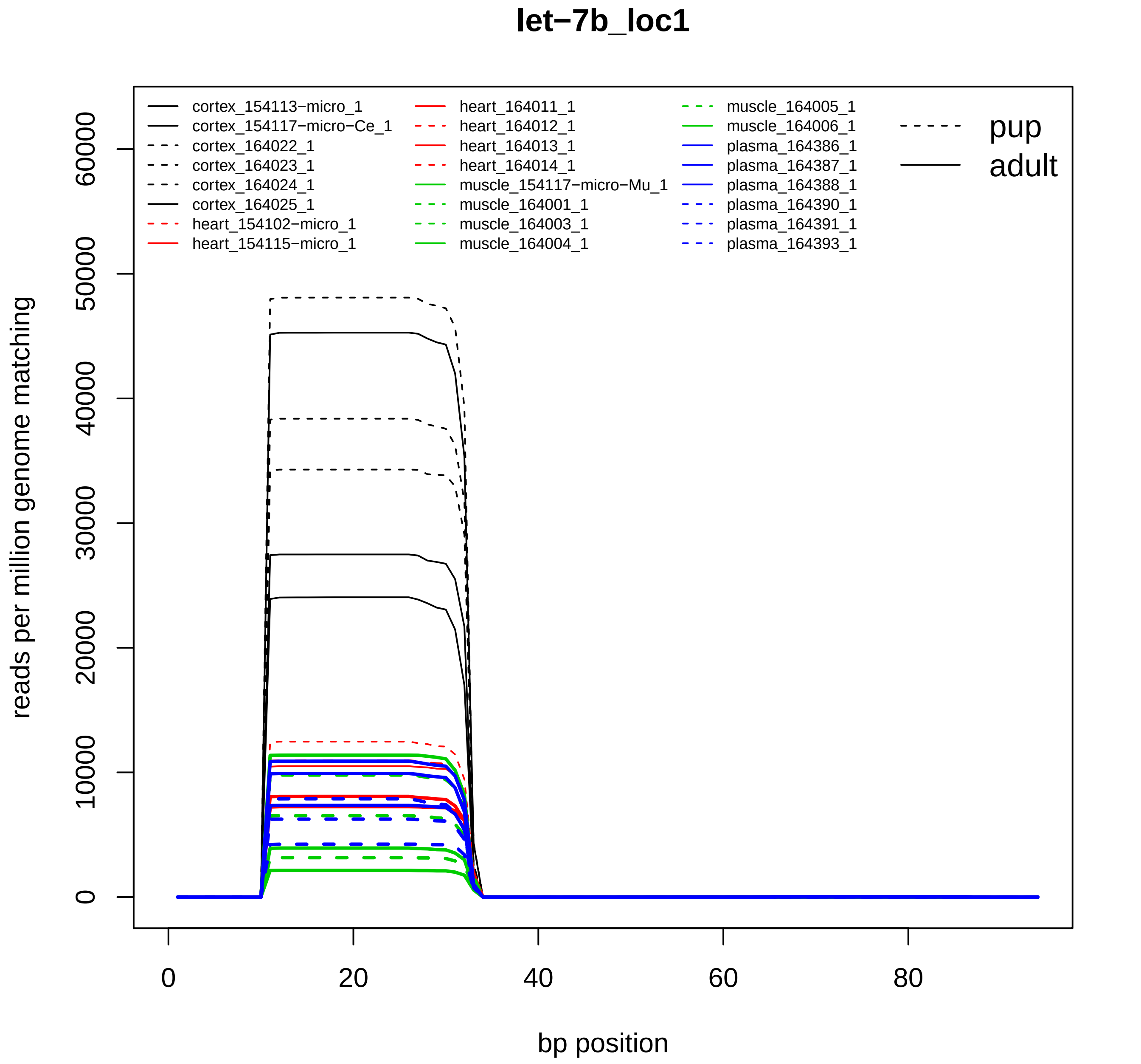

let−7b_loc1
cortex_154113−micro_1
cortex_154117−micro−Ce_1
cortex_164022_1
cortex_164023_1
cortex_164024_1
cortex_164025_1
heart_154102−micro_1
heart_154115−micro_1
heart_164011_1
heart_164012_1
heart_164013_1
heart_164014_1
muscle_154117−micro−Mu_1
muscle_164001_1
muscle_164003_1
muscle_164004_1
muscle_164005_1
muscle_164006_1
plasma_164386_1
plasma_164387_1
plasma_164388_1
plasma_164390_1
plasma_164391_1
plasma_164393_1
pup
adult
reads per million genome matching
0
10000
20000
30000
40000
50000
60000
0
20
40
60
80
bp position

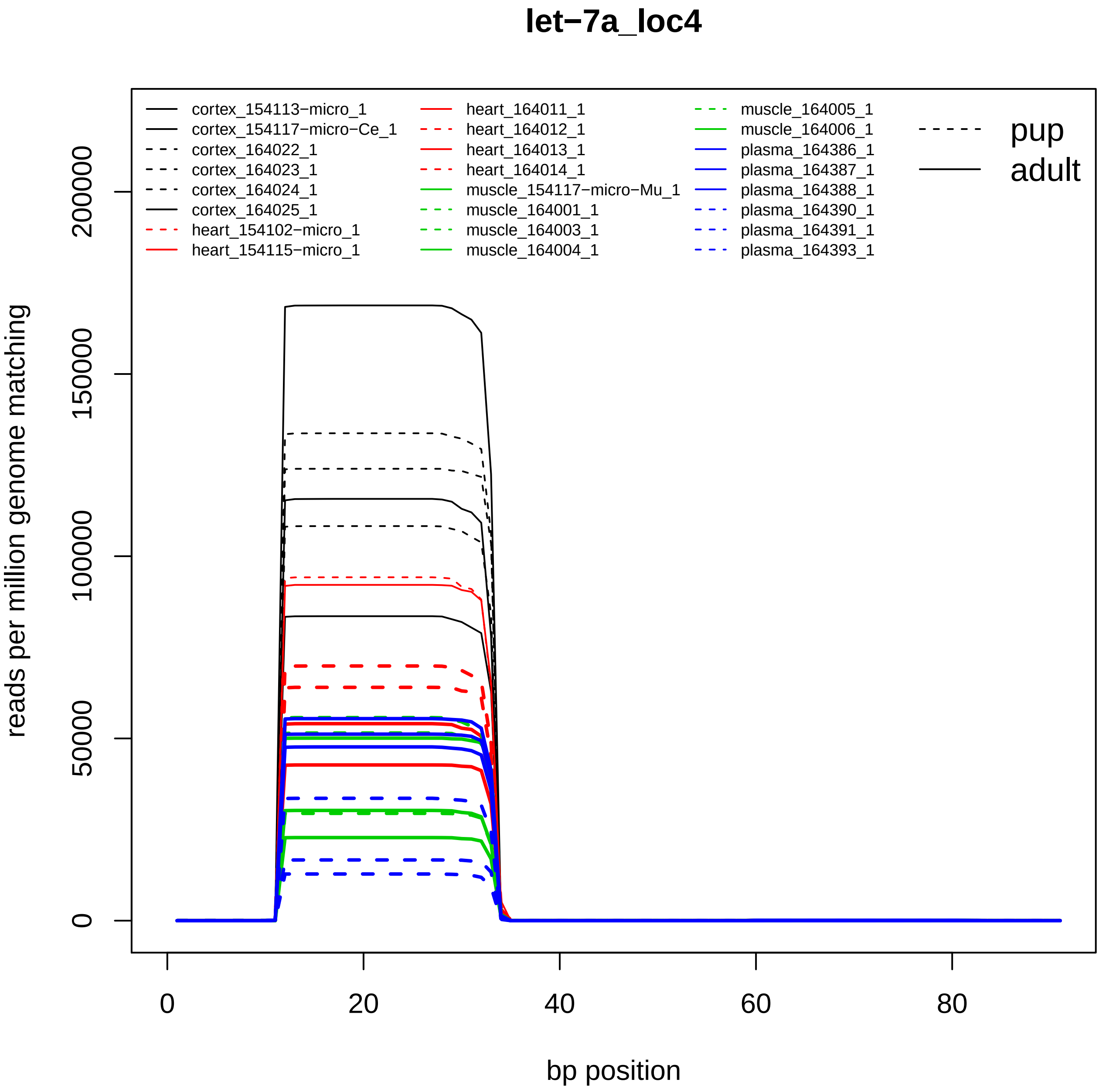
let−7a_loc4
cortex_154113−micro_1
cortex_154117−micro−Ce_1
cortex_164022_1
cortex_164023_1
cortex_164024_1
cortex_164025_1
heart_154102−micro_1
heart_154115−micro_1
heart_164011_1
heart_164012_1
heart_164013_1
heart_164014_1
muscle_154117−micro−Mu_1
muscle_164001_1
muscle_164003_1
muscle_164004_1
muscle_164005_1
muscle_164006_1
plasma_164386_1
plasma_164387_1
plasma_164388_1
plasma_164390_1
plasma_164391_1
plasma_164393_1
pup
adult
reads per million genome matching
0
50000
100000
150000
200000
0
20
40
60
80
bp position

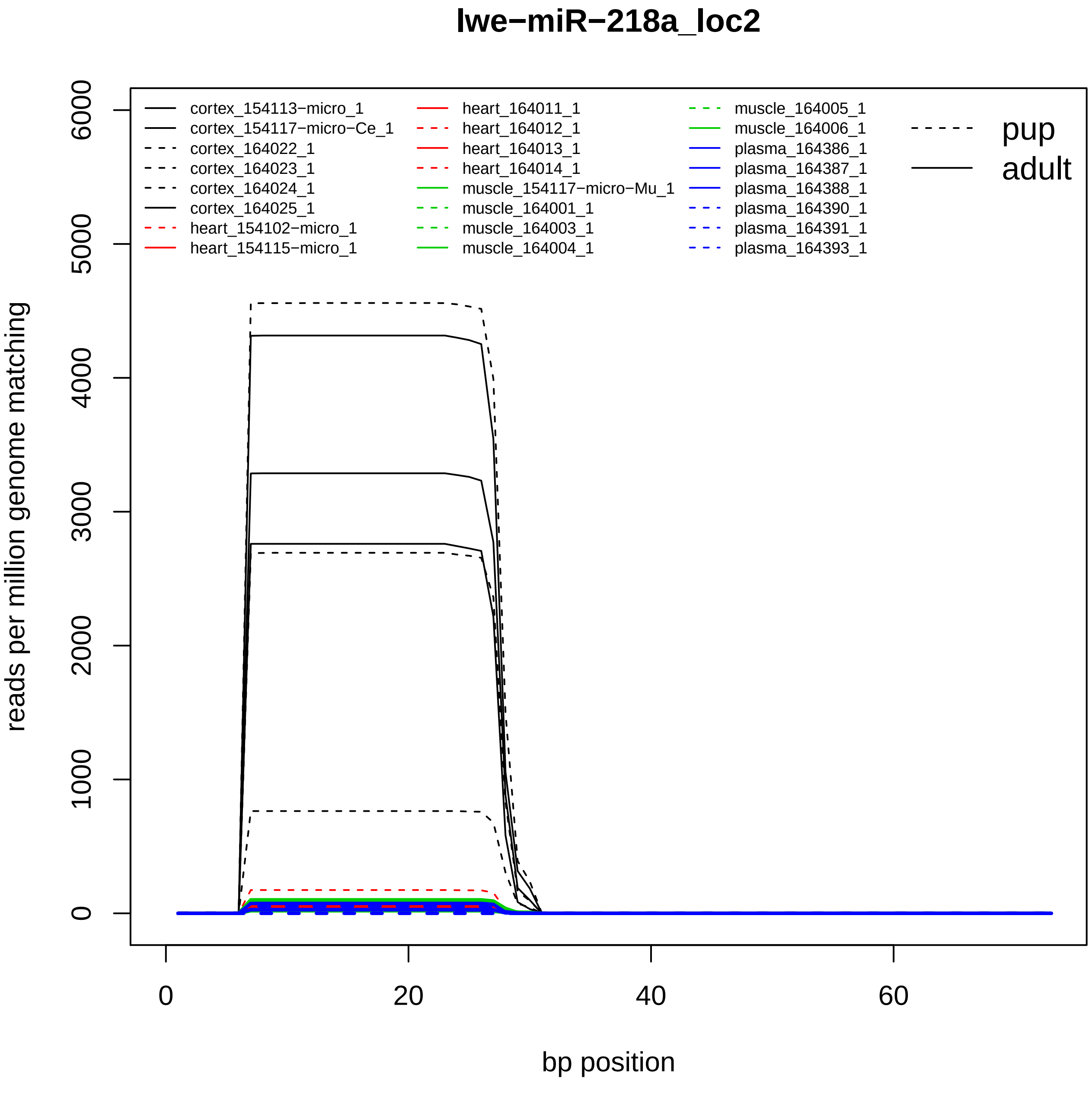
lwe−miR−218a_loc2
reads per million genome matching
bp position
0
1000
2000
3000
4000
5000
6000
0
20
40
60
cortex_154113−micro_1
cortex_154117−micro−Ce_1
cortex_164022_1
cortex_164023_1
cortex_164024_1
cortex_164025_1
heart_154102−micro_1
heart_154115−micro_1
heart_164011_1
heart_164012_1
heart_164013_1
heart_164014_1
muscle_154117−micro−Mu_1
muscle_164001_1
muscle_164003_1
muscle_164004_1
muscle_164005_1
muscle_164006_1
plasma_164386_1
plasma_164387_1
plasma_164388_1
plasma_164390_1
plasma_164391_1
plasma_164393_1
pup
adult

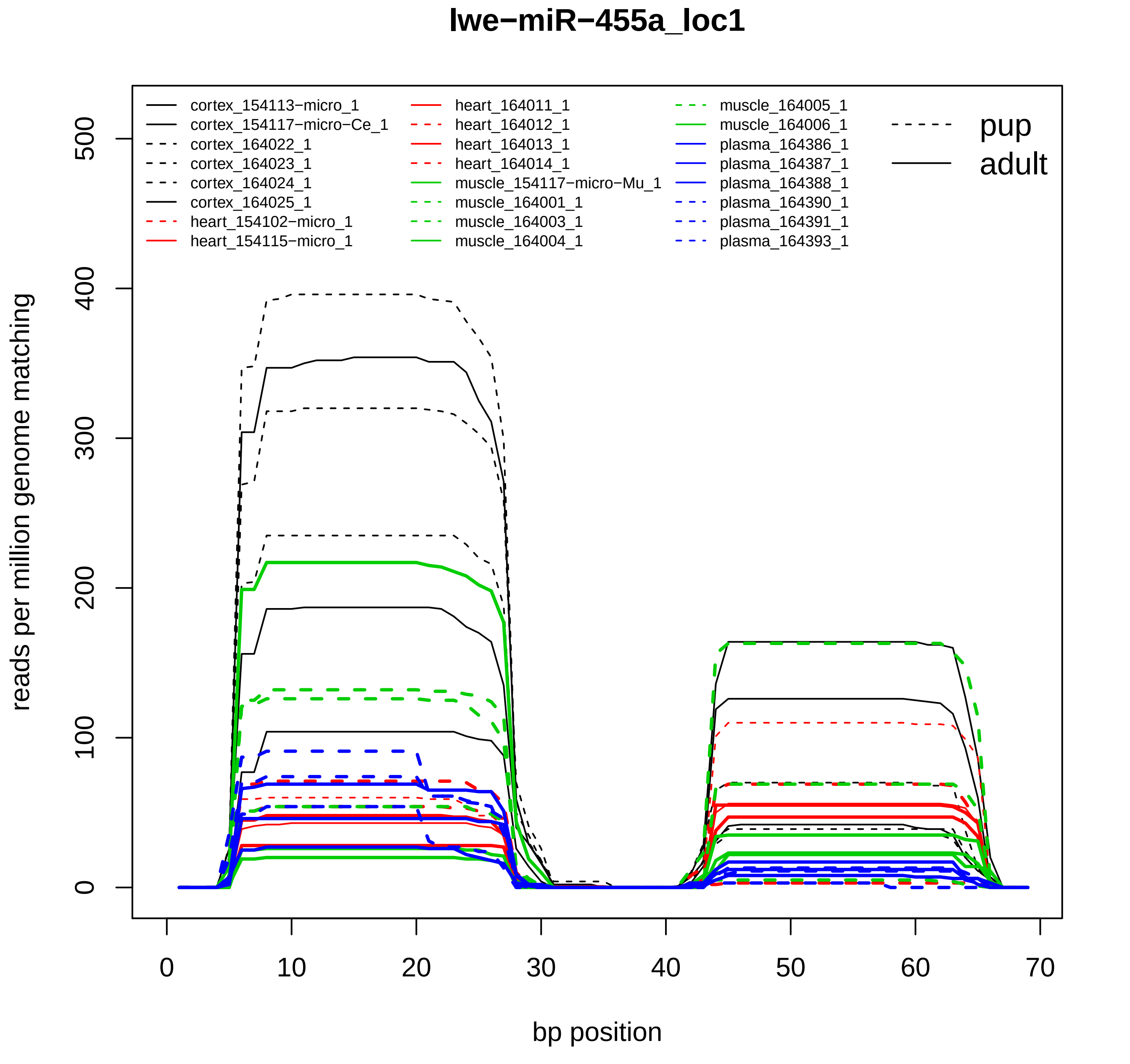
lwe−miR−455a_loc1
reads per million genome matching
bp position
0
100
200
300
400
500
0
10
20
30
40
50
60
70
cortex_154113−micro_1
cortex_154117−micro−Ce_1
cortex_164022_1
cortex_164023_1
cortex_164024_1
cortex_164025_1
heart_154102−micro_1
heart_154115−micro_1
heart_164011_1
heart_164012_1
heart_164013_1
heart_164014_1
muscle_154117−micro−Mu_1
muscle_164001_1
muscle_164003_1
muscle_164004_1
muscle_164005_1
muscle_164006_1
plasma_164386_1
plasma_164387_1
plasma_164388_1
plasma_164390_1
plasma_164391_1
plasma_164393_1
pup
adult

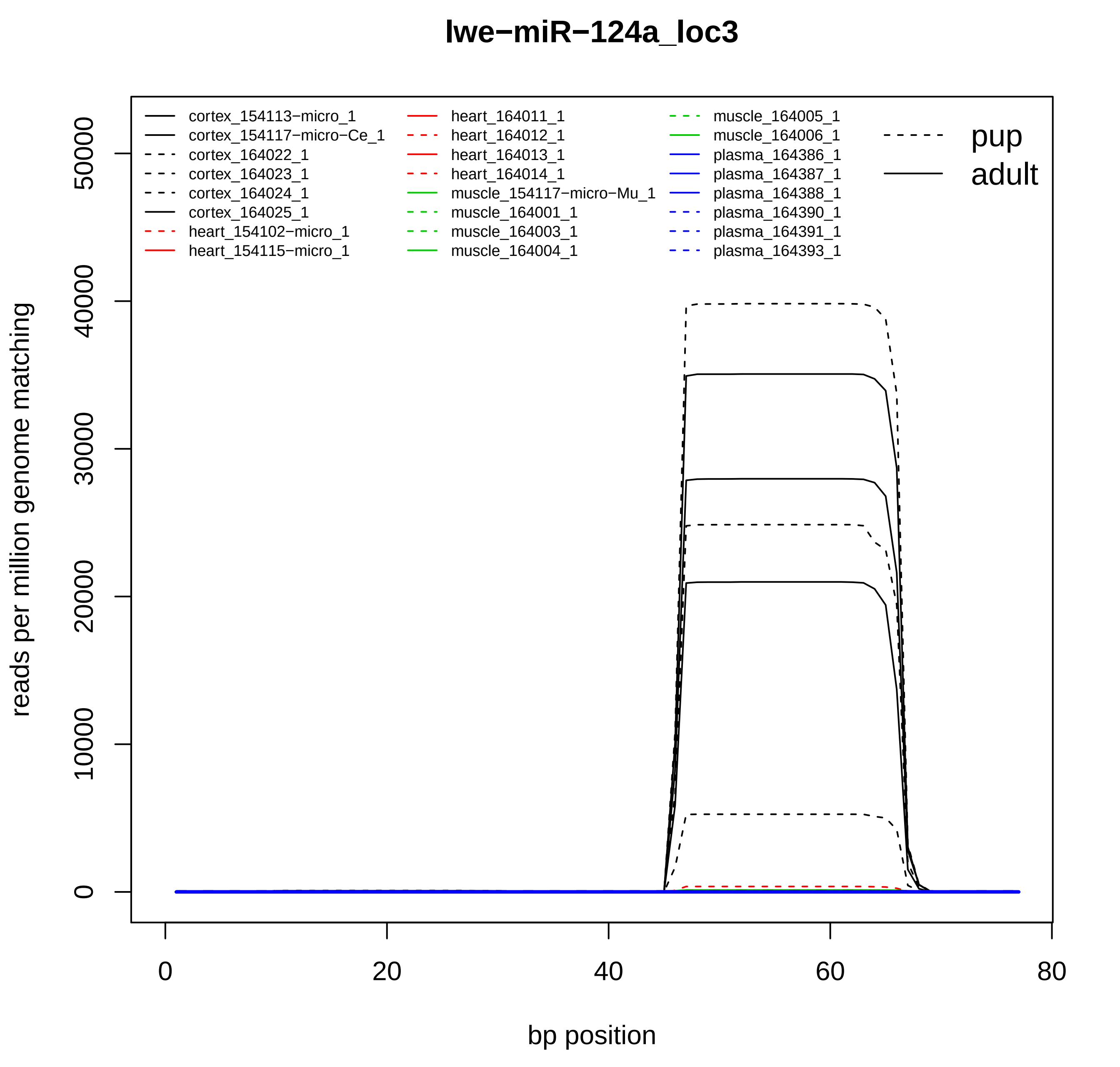
lwe−miR−124a_loc3
cortex_154113−micro_1
cortex_154117−micro−Ce_1
cortex_164022_1
cortex_164023_1
cortex_164024_1
cortex_164025_1
heart_154102−micro_1
heart_154115−micro_1
heart_164011_1
heart_164012_1
heart_164013_1
heart_164014_1
muscle_154117−micro−Mu_1
muscle_164001_1
muscle_164003_1
muscle_164004_1
muscle_164005_1
muscle_164006_1
plasma_164386_1
plasma_164387_1
plasma_164388_1
plasma_164390_1
plasma_164391_1
plasma_164393_1
pup
adult
reads per million genome matching
0
10000
20000
30000
40000
50000
bp position
0
20
40
60
80

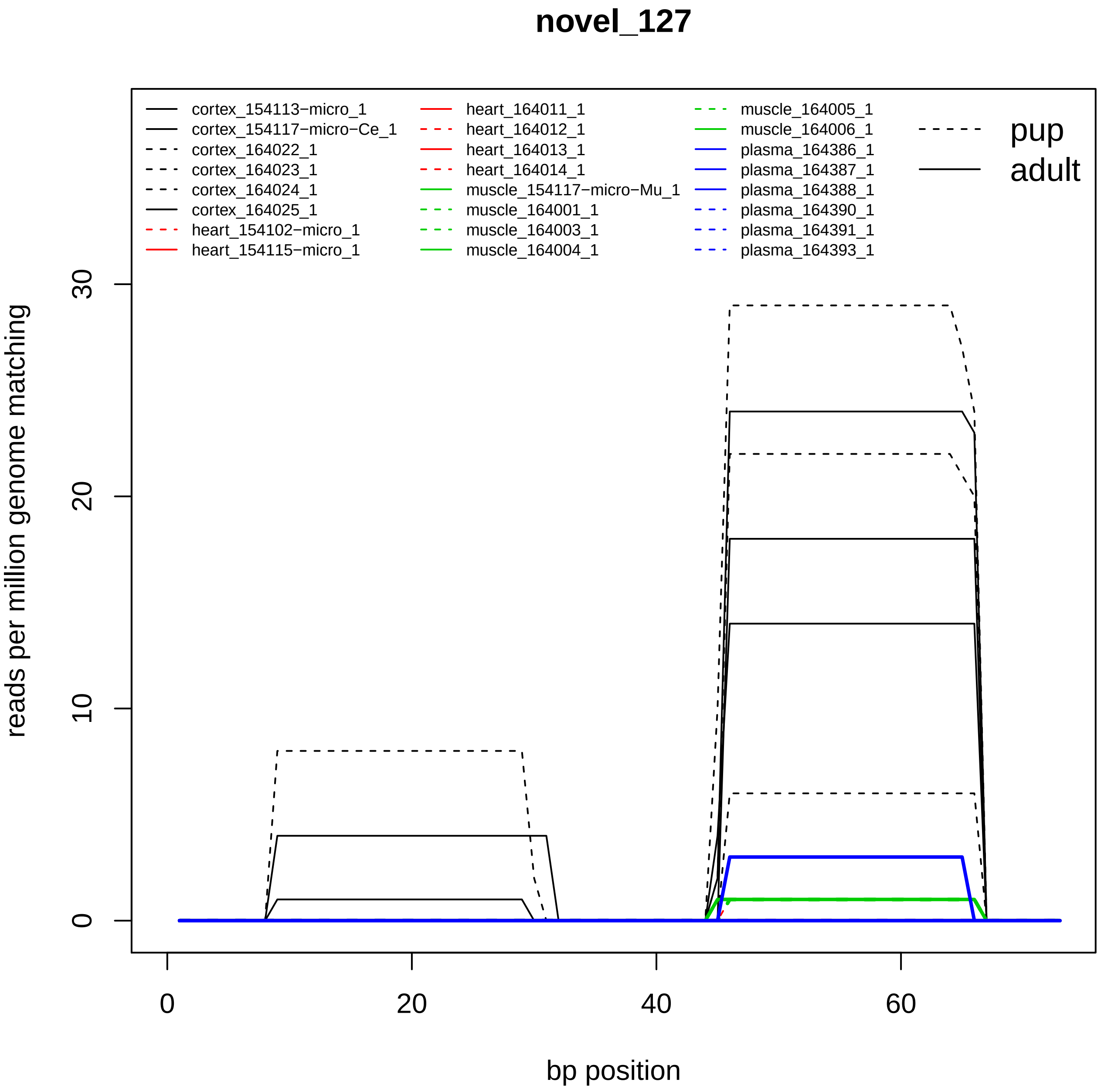
novel_127
cortex_154113−micro_1
cortex_154117−micro−Ce_1
cortex_164022_1
cortex_164023_1
cortex_164024_1
cortex_164025_1
heart_154102−micro_1
heart_154115−micro_1
heart_164011_1
heart_164012_1
heart_164013_1
heart_164014_1
muscle_154117−micro−Mu_1
muscle_164001_1
muscle_164003_1
muscle_164004_1
muscle_164005_1
muscle_164006_1
plasma_164386_1
plasma_164387_1
plasma_164388_1
plasma_164390_1
plasma_164391_1
plasma_164393_1
pup
adult
reads per million genome matching
0
10
20
30
0
20
40
60
bp position

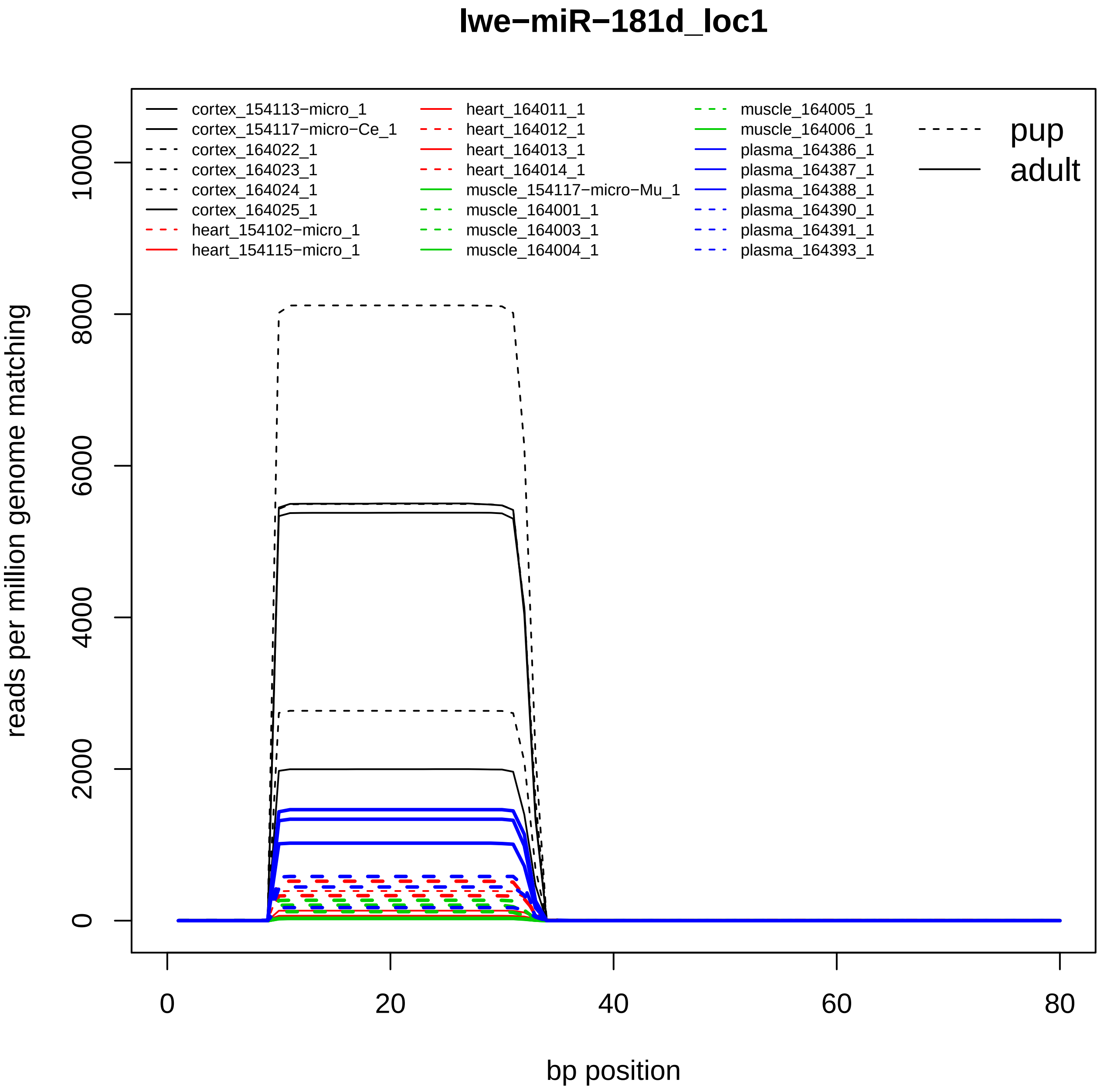
lwe−miR−181d_loc1
cortex_154113−micro_1
cortex_154117−micro−Ce_1
cortex_164022_1
cortex_164023_1
cortex_164024_1
cortex_164025_1
heart_154102−micro_1
heart_154115−micro_1
heart_164011_1
heart_164012_1
heart_164013_1
heart_164014_1
muscle_154117−micro−Mu_1
muscle_164001_1
muscle_164003_1
muscle_164004_1
muscle_164005_1
muscle_164006_1
plasma_164386_1
plasma_164387_1
plasma_164388_1
plasma_164390_1
plasma_164391_1
plasma_164393_1
pup
adult
reads per million genome matching
0
2000
4000
6000
8000
10000
0
20
40
60
80
bp position

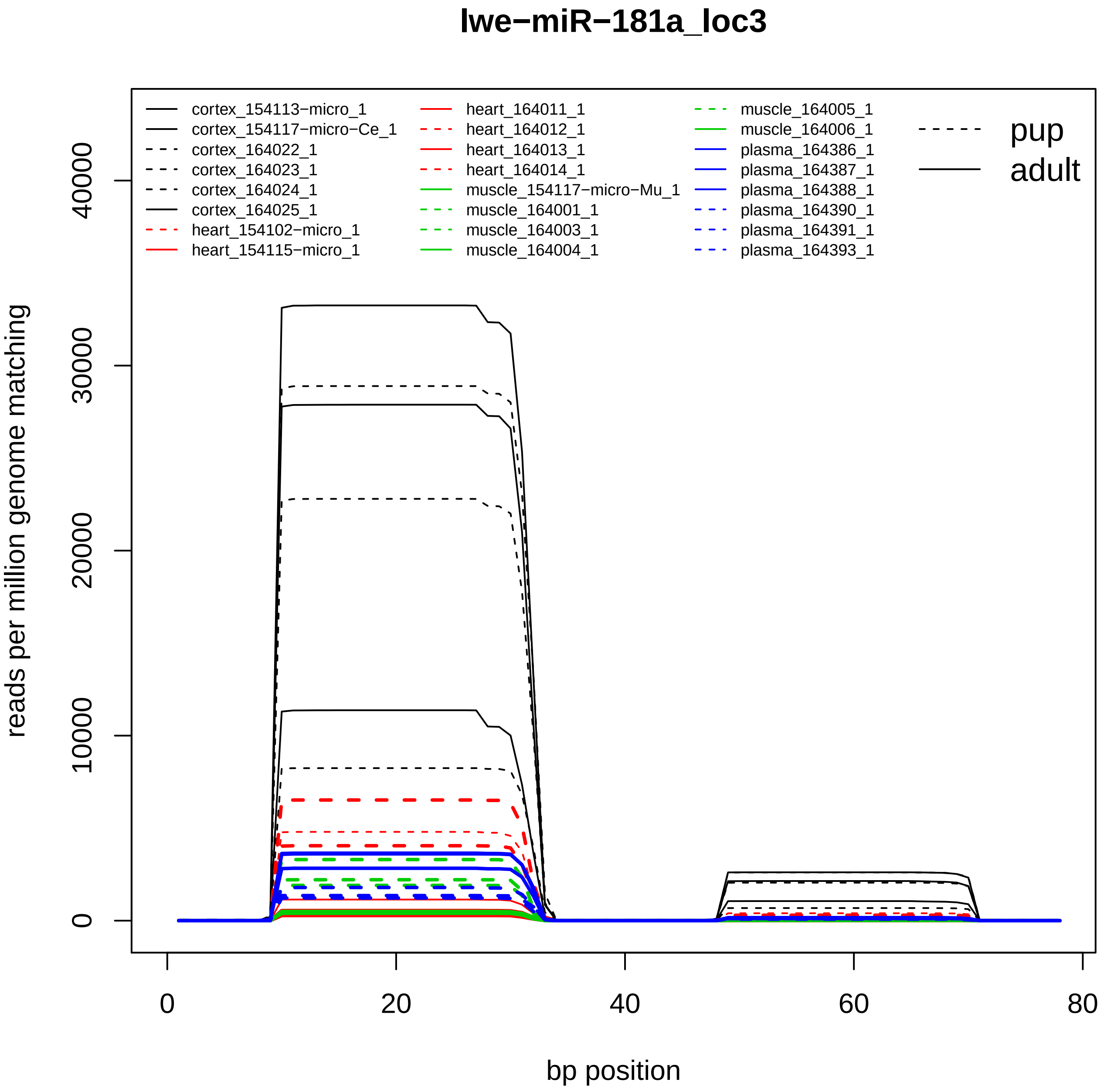
lwe−miR−181a_loc3
cortex_154113−micro_1
cortex_154117−micro−Ce_1
cortex_164022_1
cortex_164023_1
cortex_164024_1
cortex_164025_1
heart_154102−micro_1
heart_154115−micro_1
heart_164011_1
heart_164012_1
heart_164013_1
heart_164014_1
muscle_154117−micro−Mu_1
muscle_164001_1
muscle_164003_1
muscle_164004_1
muscle_164005_1
muscle_164006_1
plasma_164386_1
plasma_164387_1
plasma_164388_1
plasma_164390_1
plasma_164391_1
plasma_164393_1
pup
adult
reads per million genome matching
0
10000
20000
30000
40000
0
20
40
60
80
bp position

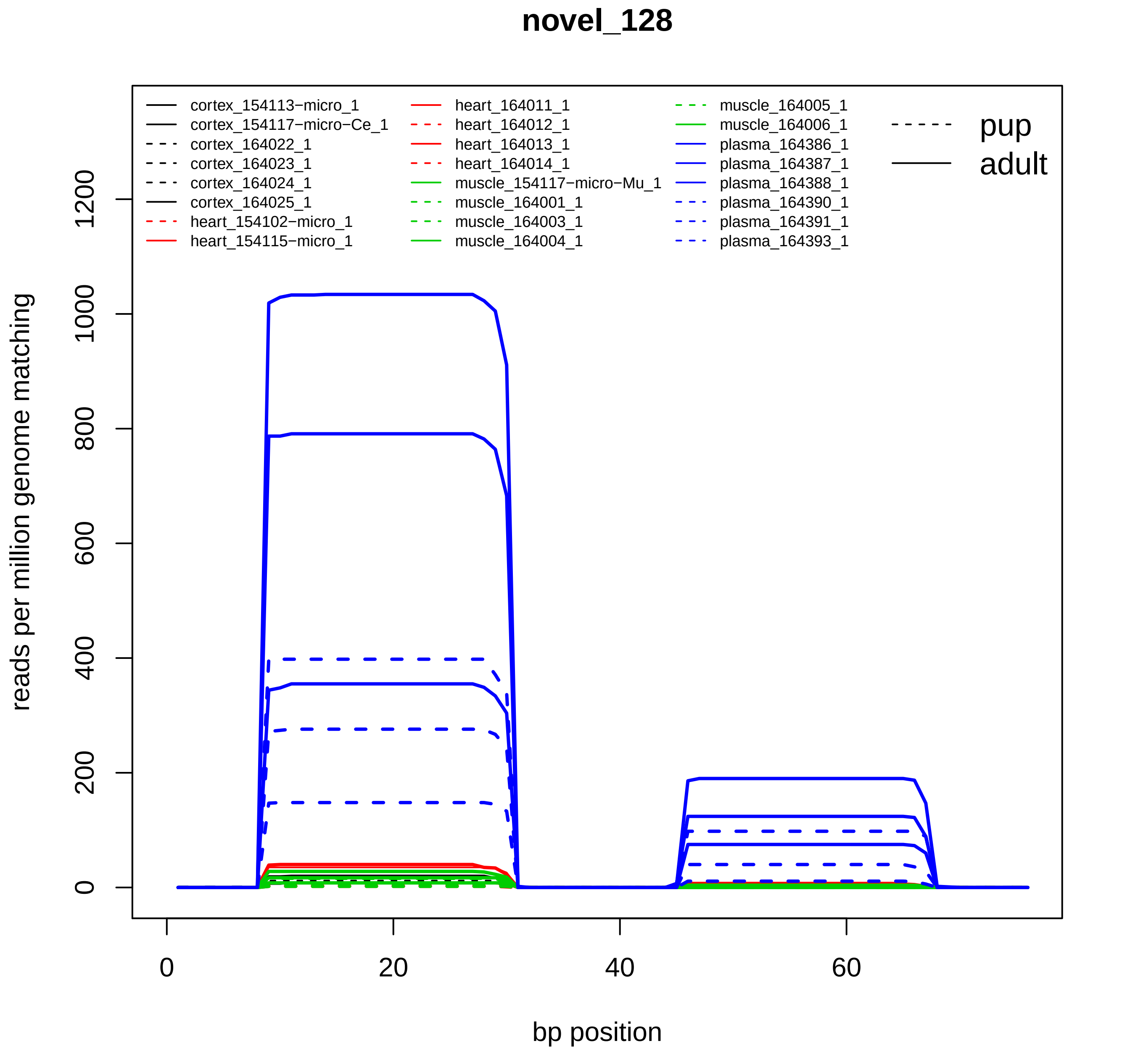
novel_128
cortex_154113−micro_1
cortex_154117−micro−Ce_1
cortex_164022_1
cortex_164023_1
cortex_164024_1
cortex_164025_1
heart_154102−micro_1
heart_154115−micro_1
heart_164011_1
heart_164012_1
heart_164013_1
heart_164014_1
muscle_154117−micro−Mu_1
muscle_164001_1
muscle_164003_1
muscle_164004_1
muscle_164005_1
muscle_164006_1
plasma_164386_1
plasma_164387_1
plasma_164388_1
plasma_164390_1
plasma_164391_1
plasma_164393_1
pup
adult
reads per million genome matching
0
200
400
600
800
1000
1200
0
20
40
60
bp position

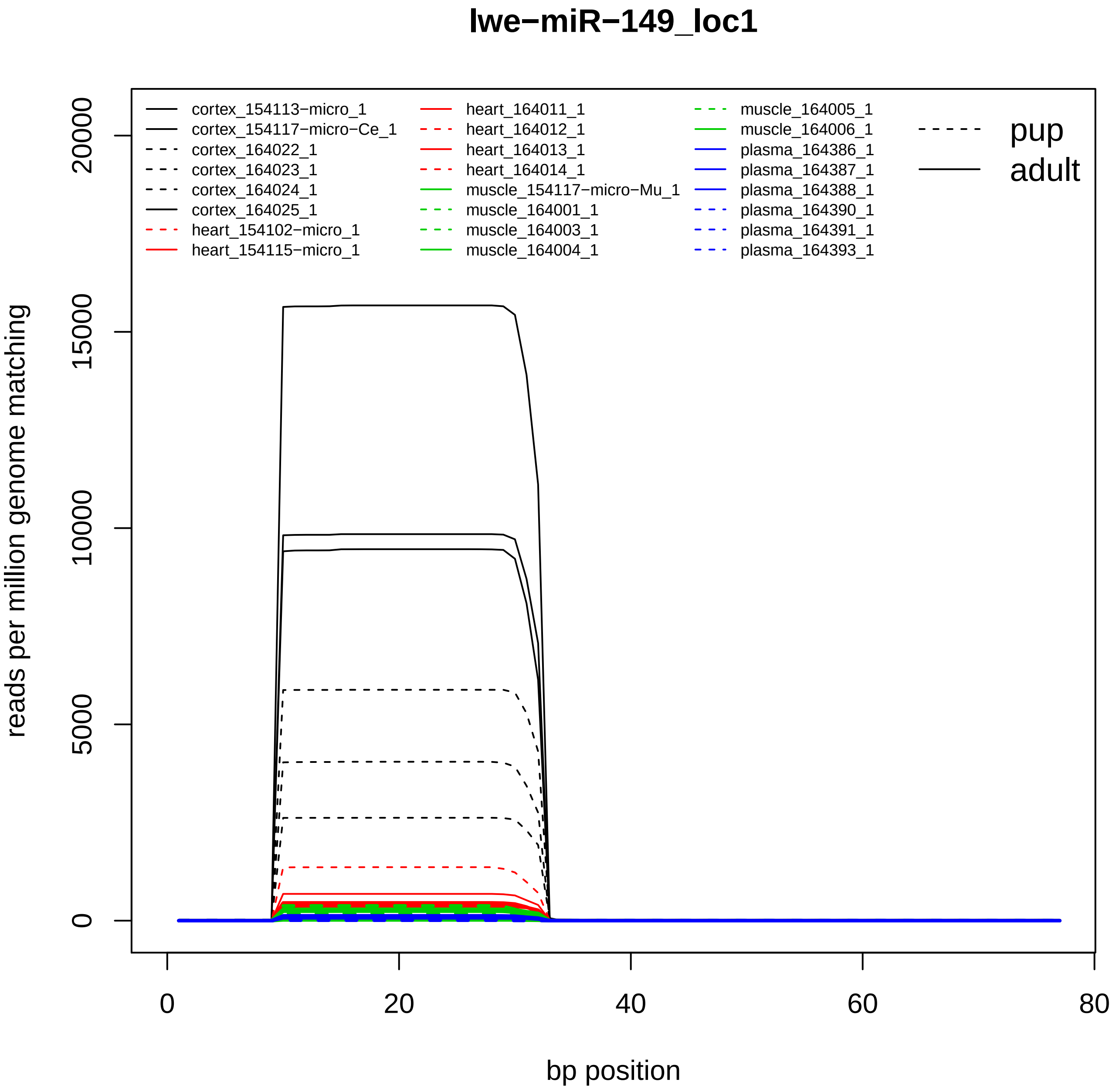
lwe−miR−149_loc1
reads per million genome matching
bp position
0
5000
10000
15000
20000
0
20
40
60
80
cortex_154113−micro_1
cortex_154117−micro−Ce_1
cortex_164022_1
cortex_164023_1
cortex_164024_1
cortex_164025_1
heart_154102−micro_1
heart_154115−micro_1
heart_164011_1
heart_164012_1
heart_164013_1
heart_164014_1
muscle_154117−micro−Mu_1
muscle_164001_1
muscle_164003_1
muscle_164004_1
muscle_164005_1
muscle_164006_1
plasma_164386_1
plasma_164387_1
plasma_164388_1
plasma_164390_1
plasma_164391_1
plasma_164393_1
pup
adult

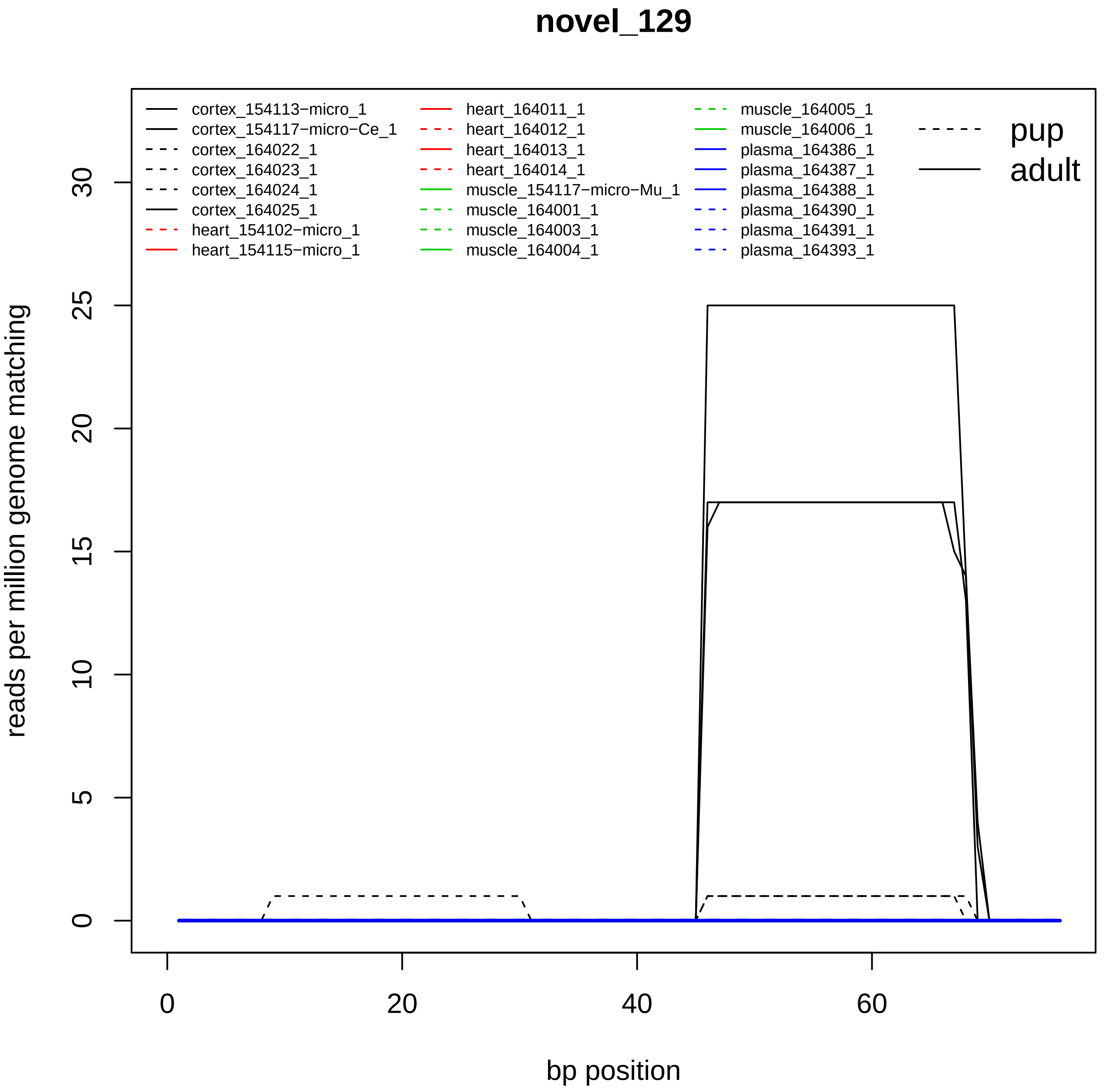
novel_129
cortex_154113−micro_1
cortex_154117−micro−Ce_1
cortex_164022_1
cortex_164023_1
cortex_164024_1
cortex_164025_1
heart_154102−micro_1
heart_154115−micro_1
heart_164011_1
heart_164012_1
heart_164013_1
heart_164014_1
muscle_154117−micro−Mu_1
muscle_164001_1
muscle_164003_1
muscle_164004_1
muscle_164005_1
muscle_164006_1
plasma_164386_1
plasma_164387_1
plasma_164388_1
plasma_164390_1
plasma_164391_1
plasma_164393_1
pup
adult
reads per million genome matching
bp position
0
5
10
15
20
25
30
0
20
40
60

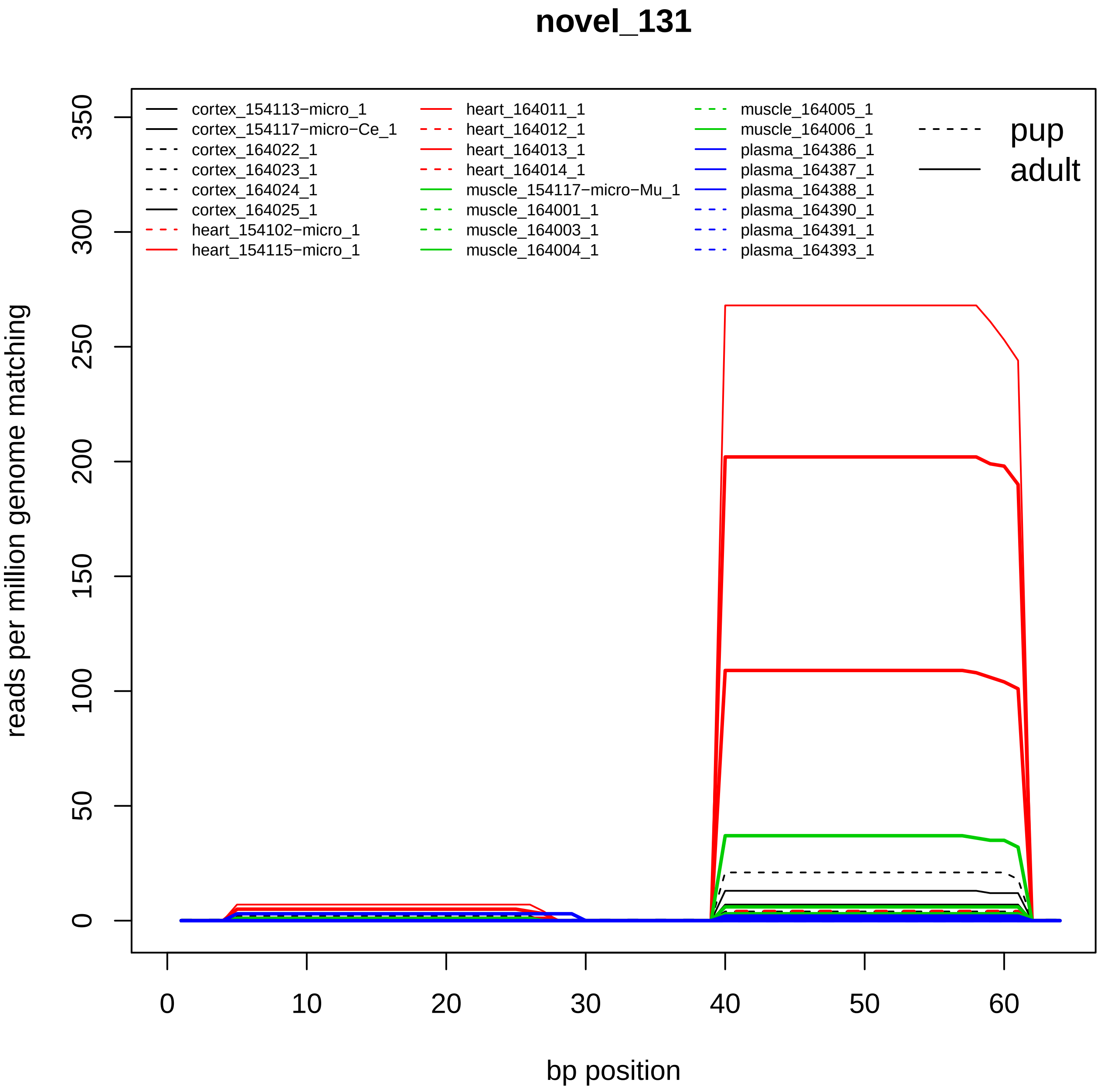
novel_131
cortex_154113−micro_1
cortex_154117−micro−Ce_1
cortex_164022_1
cortex_164023_1
cortex_164024_1
cortex_164025_1
heart_154102−micro_1
heart_154115−micro_1
heart_164011_1
heart_164012_1
heart_164013_1
heart_164014_1
muscle_154117−micro−Mu_1
muscle_164001_1
muscle_164003_1
muscle_164004_1
muscle_164005_1
muscle_164006_1
plasma_164386_1
plasma_164387_1
plasma_164388_1
plasma_164390_1
plasma_164391_1
plasma_164393_1
pup
adult
reads per million genome matching
bp position
0
50
100
150
200
250
300
350
0
10
20
30
40
50
60

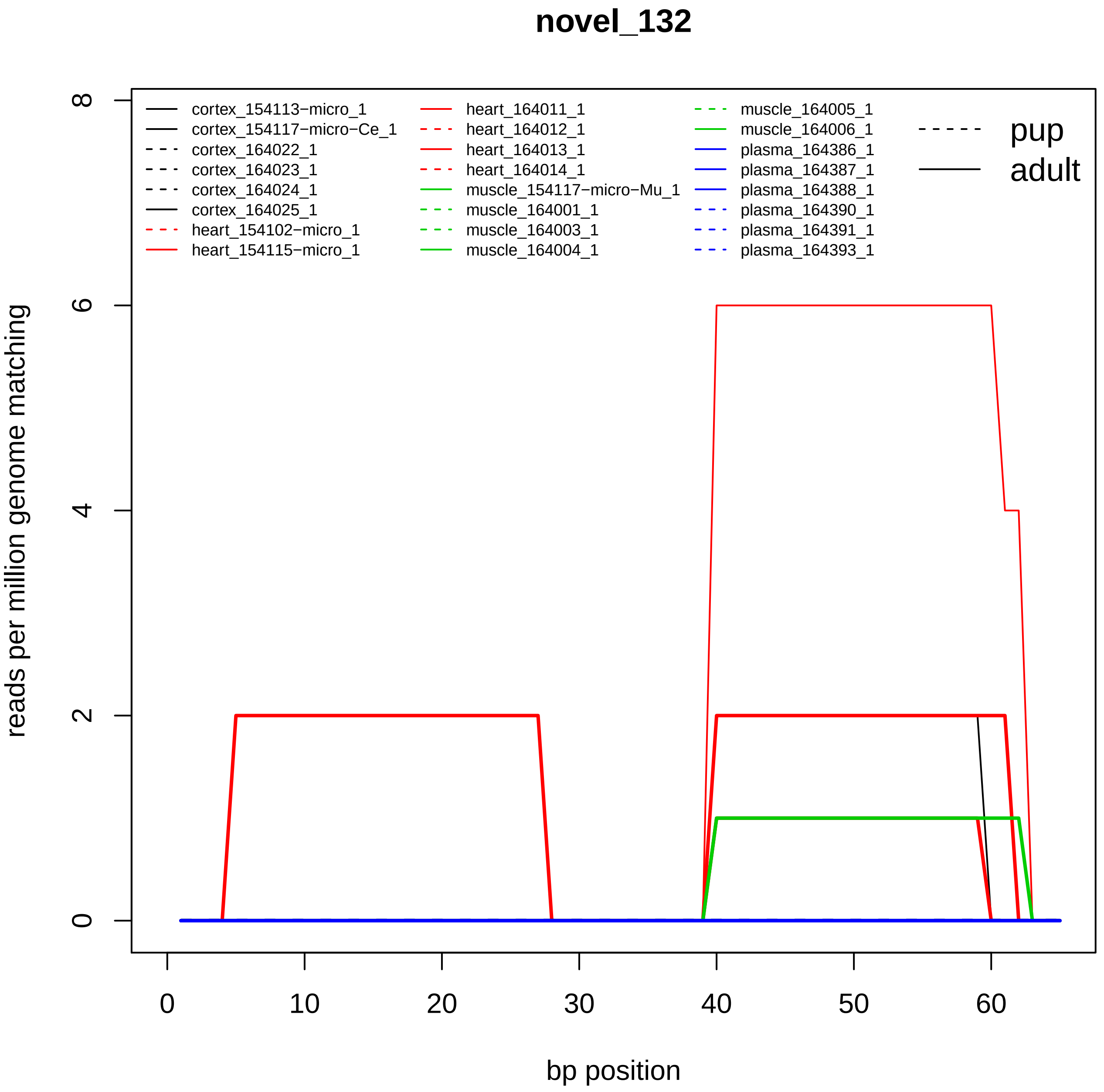
novel_132
cortex_154113−micro_1
cortex_154117−micro−Ce_1
cortex_164022_1
cortex_164023_1
cortex_164024_1
cortex_164025_1
heart_154102−micro_1
heart_154115−micro_1
heart_164011_1
heart_164012_1
heart_164013_1
heart_164014_1
muscle_154117−micro−Mu_1
muscle_164001_1
muscle_164003_1
muscle_164004_1
muscle_164005_1
muscle_164006_1
plasma_164386_1
plasma_164387_1
plasma_164388_1
plasma_164390_1
plasma_164391_1
plasma_164393_1
pup
adult
reads per million genome matching
0
2
4
6
8
0
10
20
30
40
50
60
bp position

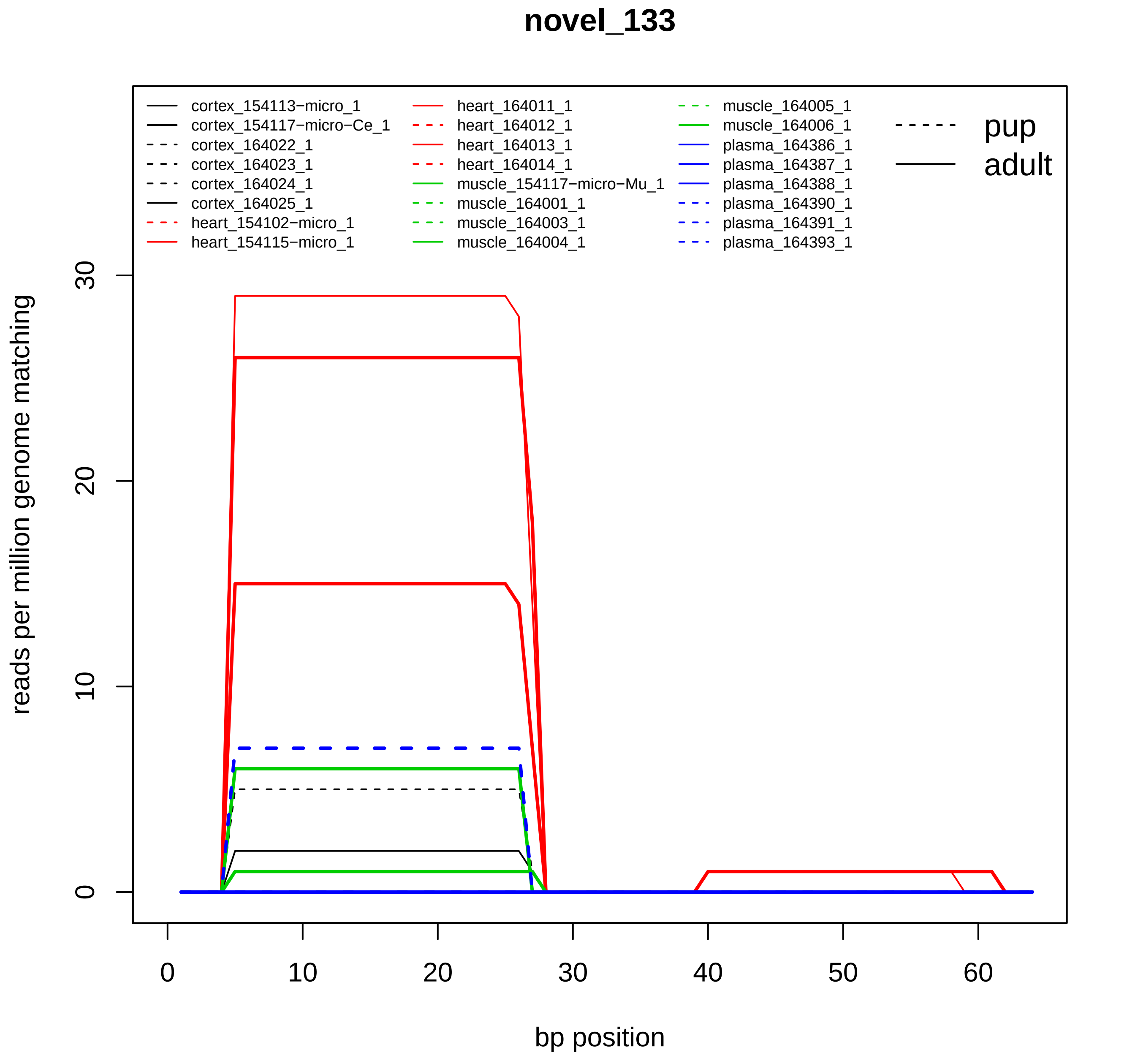

novel_133
cortex_154113−micro_1
cortex_154117−micro−Ce_1
cortex_164022_1
cortex_164023_1
cortex_164024_1
cortex_164025_1
heart_154102−micro_1
heart_154115−micro_1
heart_164011_1
heart_164012_1
heart_164013_1
heart_164014_1
muscle_154117−micro−Mu_1
muscle_164001_1
muscle_164003_1
muscle_164004_1
muscle_164005_1
muscle_164006_1
plasma_164386_1
plasma_164387_1
plasma_164388_1
plasma_164390_1
plasma_164391_1
plasma_164393_1
pup
adult
reads per million genome matching
0
10
20
30
0
10
20
30
40
50
60
bp position

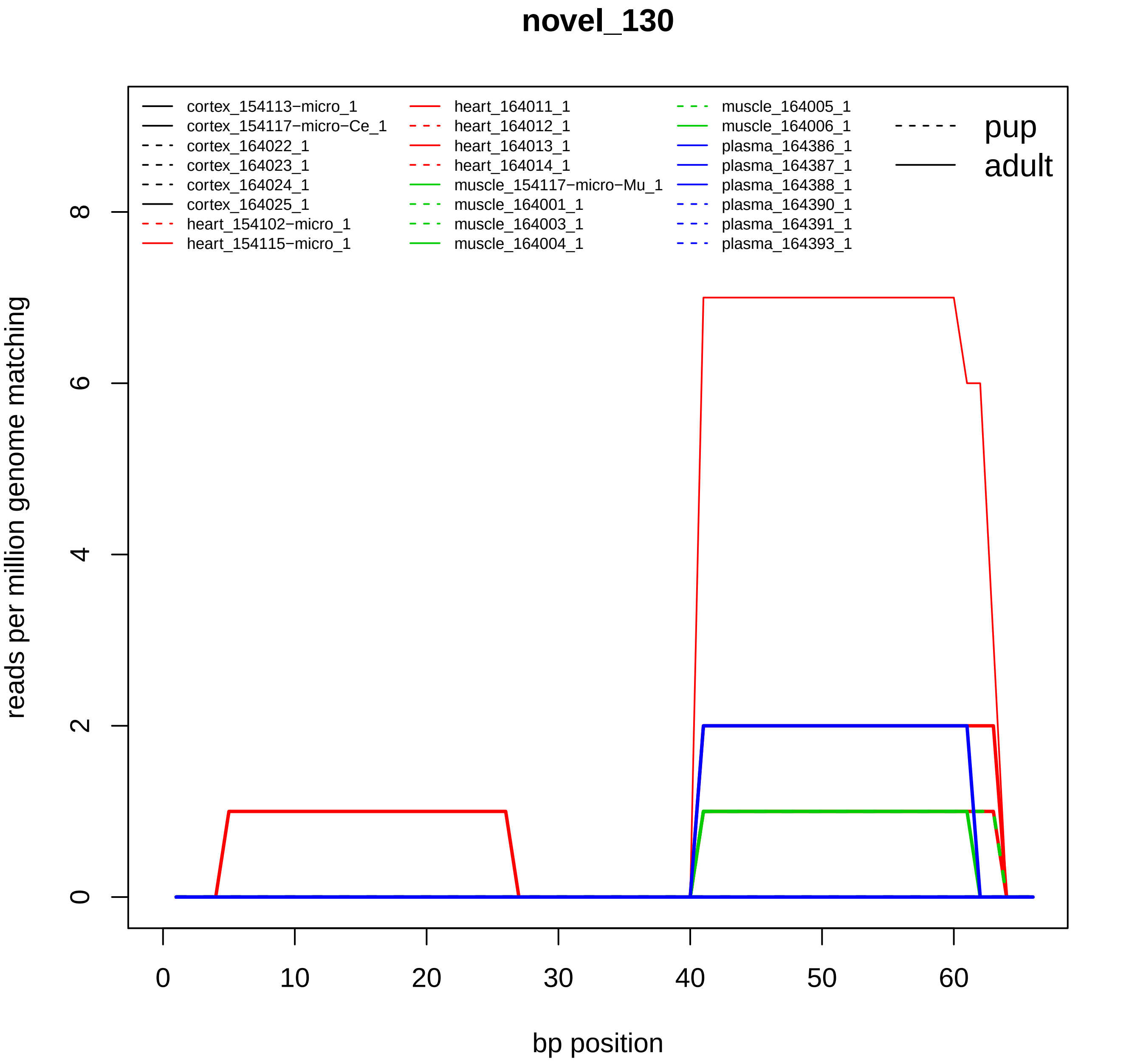

novel_130
cortex_154113−micro_1
cortex_154117−micro−Ce_1
cortex_164022_1
cortex_164023_1
cortex_164024_1
cortex_164025_1
heart_154102−micro_1
heart_154115−micro_1
heart_164011_1
heart_164012_1
heart_164013_1
heart_164014_1
muscle_154117−micro−Mu_1
muscle_164001_1
muscle_164003_1
muscle_164004_1
muscle_164005_1
muscle_164006_1
plasma_164386_1
plasma_164387_1
plasma_164388_1
plasma_164390_1
plasma_164391_1
plasma_164393_1
pup
adult
reads per million genome matching
0
2
4
6
8
0
10
20
30
40
50
60
bp position

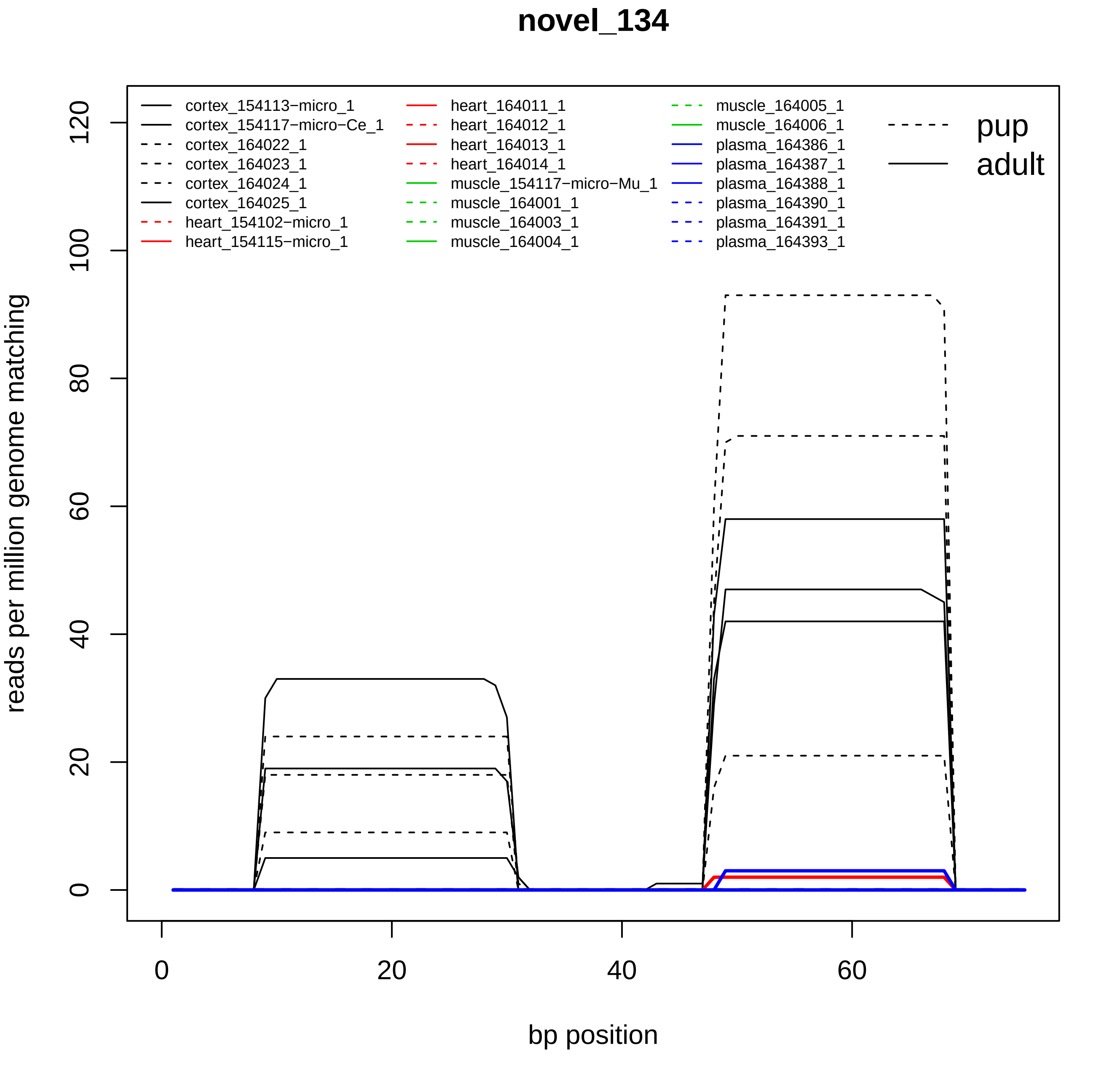

novel_134
cortex_154113−micro_1
cortex_154117−micro−Ce_1
cortex_164022_1
cortex_164023_1
cortex_164024_1
cortex_164025_1
heart_154102−micro_1
heart_154115−micro_1
heart_164011_1
heart_164012_1
heart_164013_1
heart_164014_1
muscle_154117−micro−Mu_1
muscle_164001_1
muscle_164003_1
muscle_164004_1
muscle_164005_1
muscle_164006_1
plasma_164386_1
plasma_164387_1
plasma_164388_1
plasma_164390_1
plasma_164391_1
plasma_164393_1
pup
adult
reads per million genome matching
bp position
0
20
40
60
80
100
120

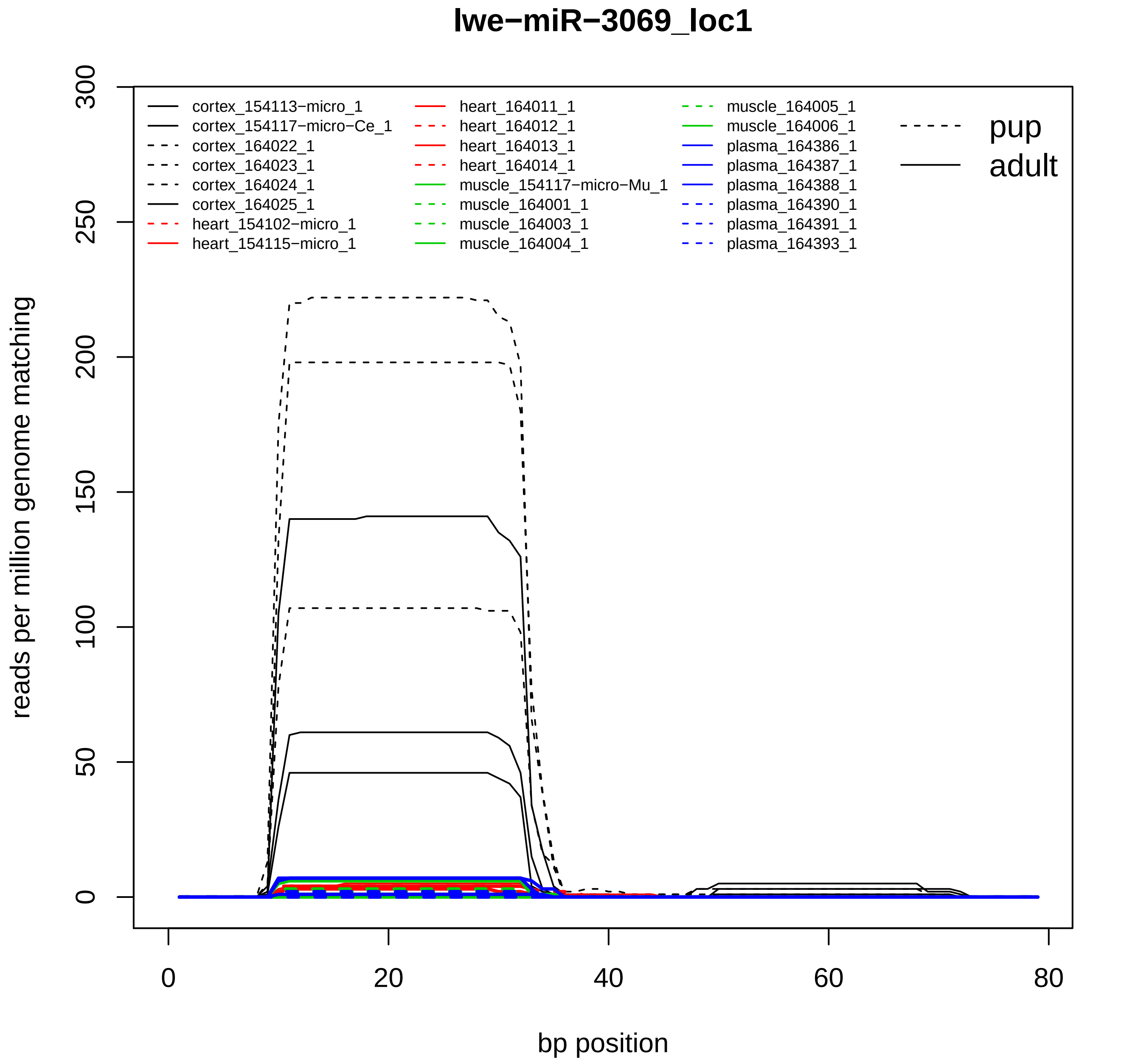

lwe−miR−3069_loc1
reads per million genome matching
bp position
0
50
100
150
200
250
300
0
20
40
60
80
cortex_154113−micro_1
cortex_154117−micro−Ce_1
cortex_164022_1
cortex_164023_1
cortex_164024_1
cortex_164025_1
heart_154102−micro_1
heart_154115−micro_1
heart_164011_1
heart_164012_1
heart_164013_1
heart_164014_1
muscle_154117−micro−Mu_1
muscle_164001_1
muscle_164003_1
muscle_164004_1
muscle_164005_1
muscle_164006_1
plasma_164386_1
plasma_164387_1
plasma_164388_1
plasma_164390_1
plasma_164391_1
plasma_164393_1
pup
adult

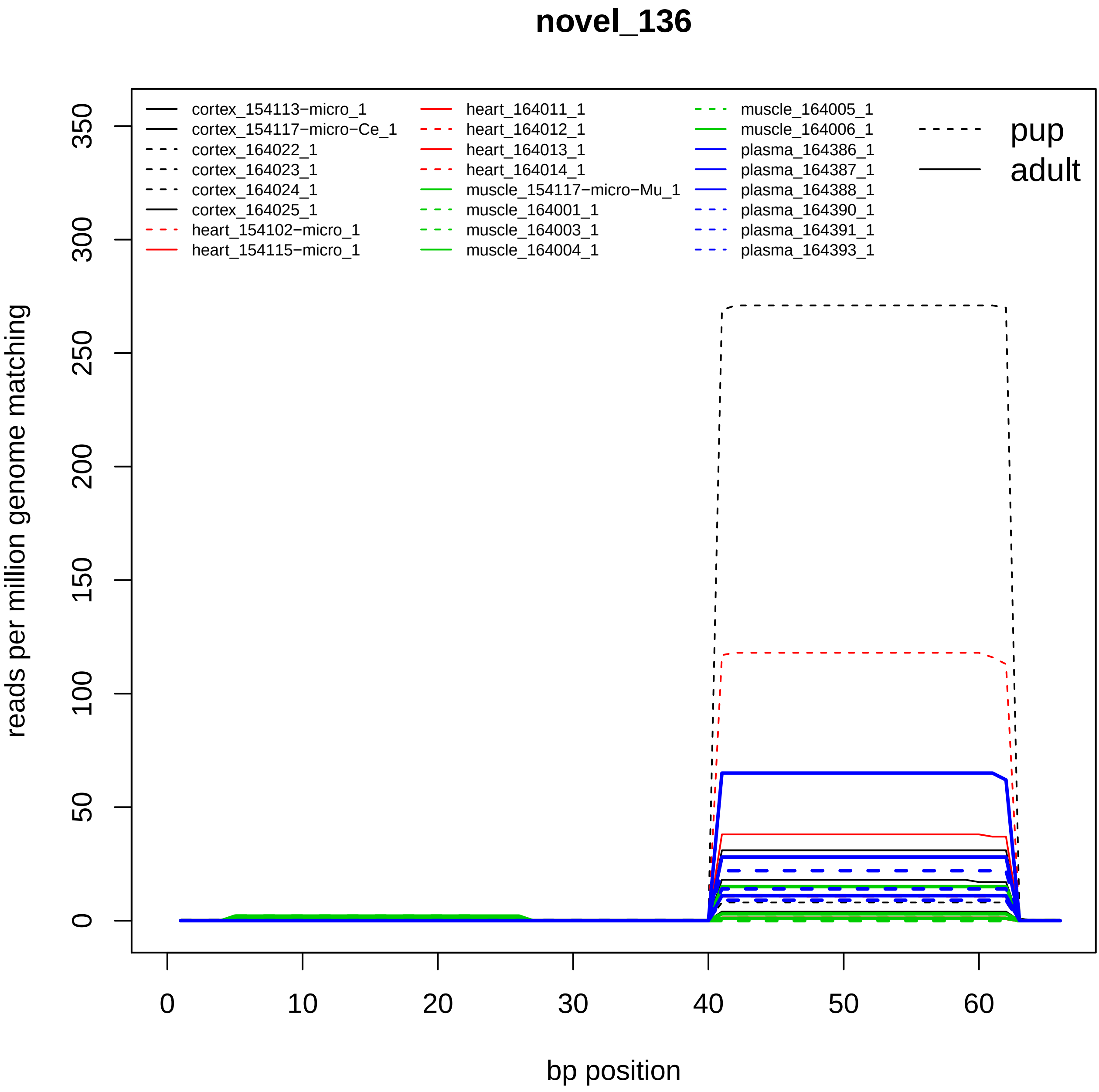

novel_136
cortex_154113−micro_1
cortex_154117−micro−Ce_1
cortex_164022_1
cortex_164023_1
cortex_164024_1
cortex_164025_1
heart_154102−micro_1
heart_154115−micro_1
heart_164011_1
heart_164012_1
heart_164013_1
heart_164014_1
muscle_154117−micro−Mu_1
muscle_164001_1
muscle_164003_1
muscle_164004_1
muscle_164005_1
muscle_164006_1
plasma_164386_1
plasma_164387_1
plasma_164388_1
plasma_164390_1
plasma_164391_1
plasma_164393_1
pup
adult
reads per million genome matching
0
50
100
150
200
250
300
350
0
10
20
30
40
50
60
bp position

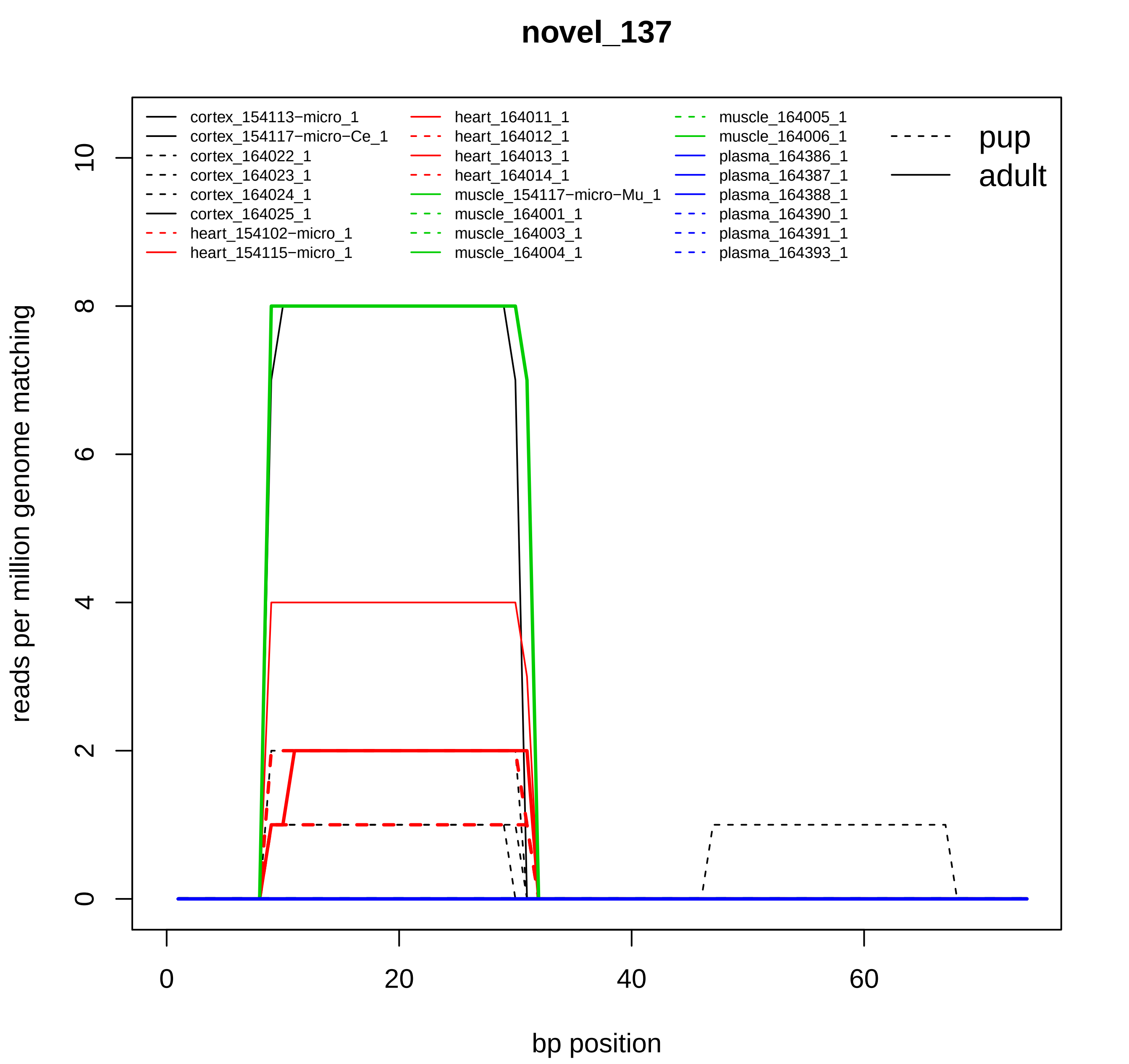

novel_137
cortex_154113−micro_1
cortex_154117−micro−Ce_1
cortex_164022_1
cortex_164023_1
cortex_164024_1
cortex_164025_1
heart_154102−micro_1
heart_154115−micro_1
heart_164011_1
heart_164012_1
heart_164013_1
heart_164014_1
muscle_154117−micro−Mu_1
muscle_164001_1
muscle_164003_1
muscle_164004_1
muscle_164005_1
muscle_164006_1
plasma_164386_1
plasma_164387_1
plasma_164388_1
plasma_164390_1
plasma_164391_1
plasma_164393_1
pup
adult
reads per million genome matching
bp position
0
2
4
6
8
10
0
20
40
60

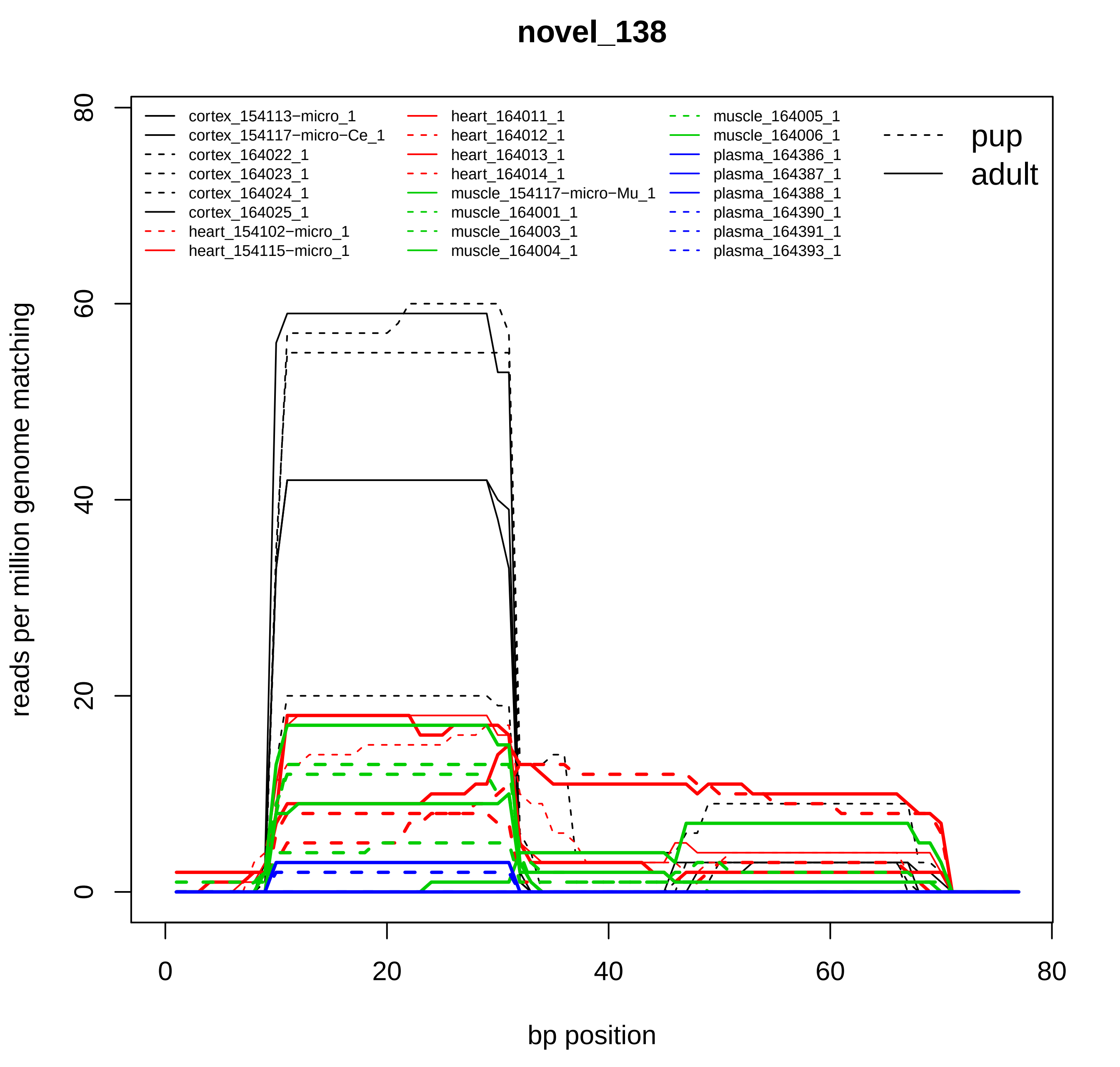
novel_138
cortex_154113−micro_1
cortex_154117−micro−Ce_1
cortex_164022_1
cortex_164023_1
cortex_164024_1
cortex_164025_1
heart_154102−micro_1
heart_154115−micro_1
heart_164011_1
heart_164012_1
heart_164013_1
heart_164014_1
muscle_154117−micro−Mu_1
muscle_164001_1
muscle_164003_1
muscle_164004_1
muscle_164005_1
muscle_164006_1
plasma_164386_1
plasma_164387_1
plasma_164388_1
plasma_164390_1
plasma_164391_1
plasma_164393_1
pup
adult
reads per million genome matching
bp position
0
20
40
60
80

### lwe−miR−200b_loc1

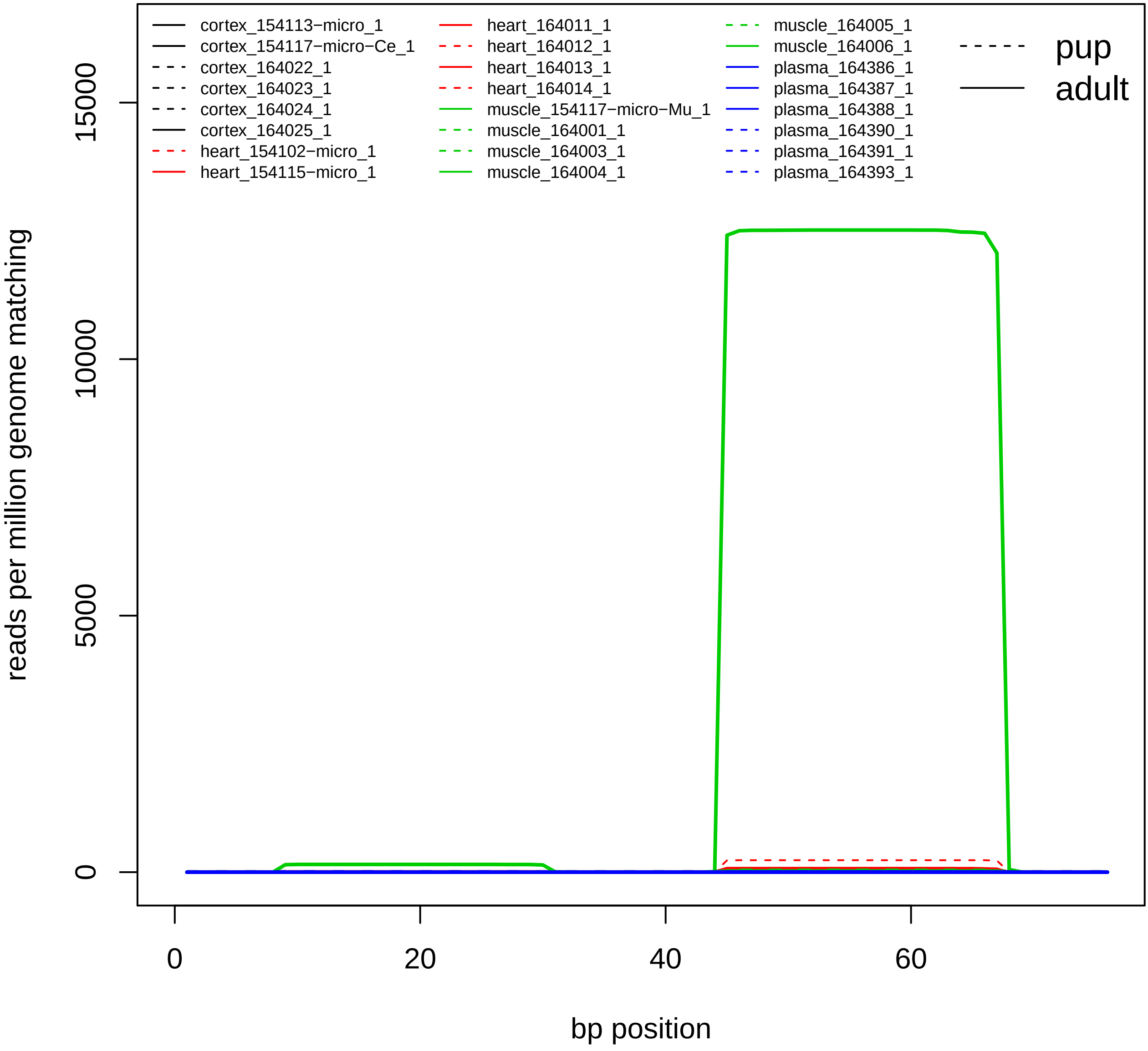

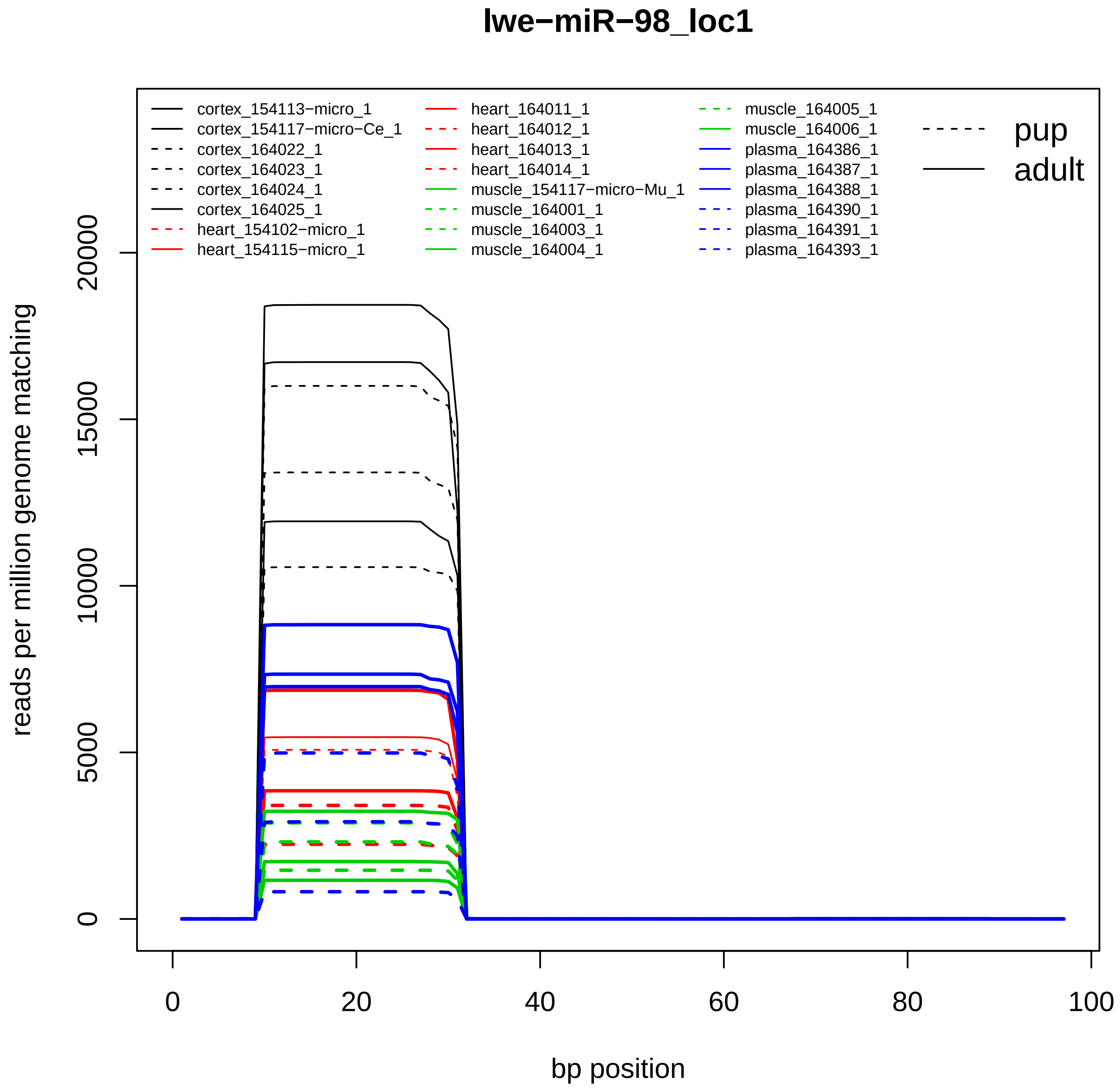
lwe−miR−98_loc1
cortex_154113−micro_1
cortex_154117−micro−Ce_1
cortex_164022_1
cortex_164023_1
cortex_164024_1
cortex_164025_1
heart_154102−micro_1
heart_154115−micro_1
heart_164011_1
heart_164012_1
heart_164013_1
heart_164014_1
muscle_154117−micro−Mu_1
muscle_164001_1
muscle_164003_1
muscle_164004_1
muscle_164005_1
muscle_164006_1
plasma_164386_1
plasma_164387_1
plasma_164388_1
plasma_164390_1
plasma_164391_1
plasma_164393_1
pup
adult
reads per million genome matching
0
5000
10000
15000
20000
0
20
40
60
80
100
bp position

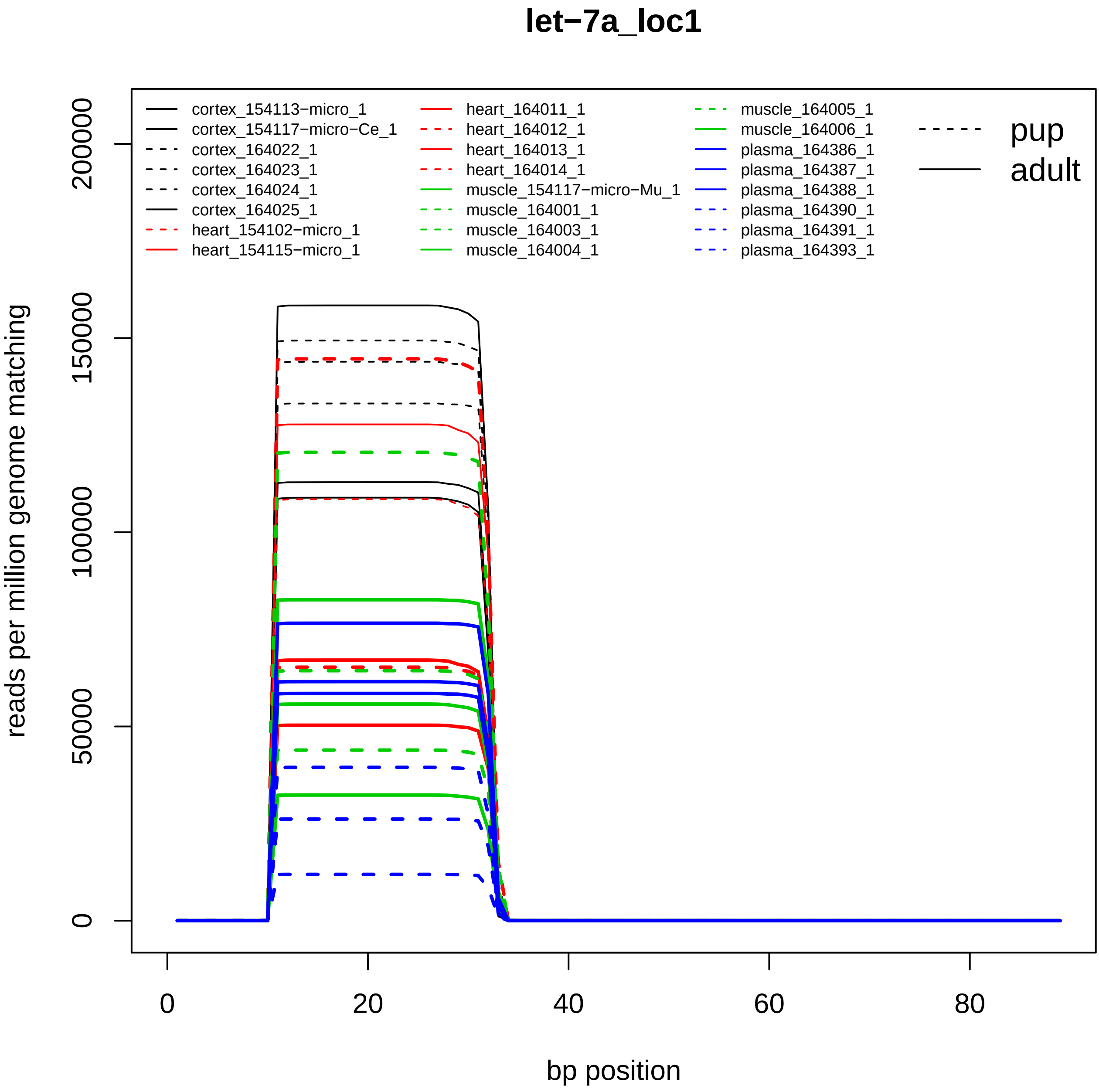
let−7a_loc1
reads per million genome matching
bp position
0
50000
100000
150000
200000
0
20
40
60
80
cortex_154113−micro_1
cortex_154117−micro−Ce_1
cortex_164022_1
cortex_164023_1
cortex_164024_1
cortex_164025_1
heart_154102−micro_1
heart_154115−micro_1
heart_164011_1
heart_164012_1
heart_164013_1
heart_164014_1
muscle_154117−micro−Mu_1
muscle_164001_1
muscle_164003_1
muscle_164004_1
muscle_164005_1
muscle_164006_1
plasma_164386_1
plasma_164387_1
plasma_164388_1
plasma_164390_1
plasma_164391_1
plasma_164393_1
pup
adult

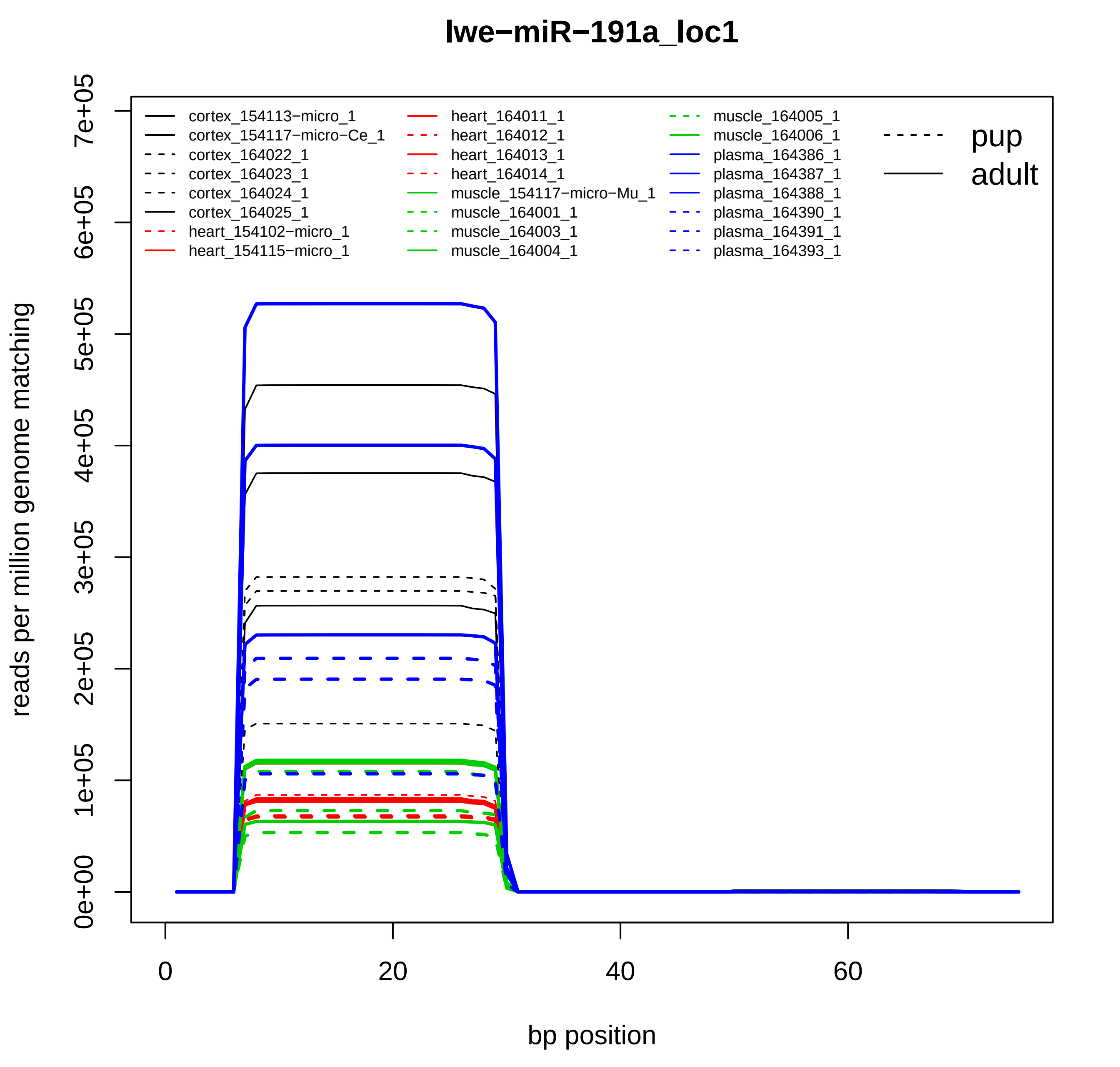
lwe−miR−191a_loc1
cortex_154113−micro_1
cortex_154117−micro−Ce_1
cortex_164022_1
cortex_164023_1
cortex_164024_1
cortex_164025_1
heart_154102−micro_1
heart_154115−micro_1
heart_164011_1
heart_164012_1
heart_164013_1
heart_164014_1
muscle_154117−micro−Mu_1
muscle_164001_1
muscle_164003_1
muscle_164004_1
muscle_164005_1
muscle_164006_1
plasma_164386_1
plasma_164387_1
plasma_164388_1
plasma_164390_1
plasma_164391_1
plasma_164393_1
pup
adult
reads per million genome matching
0e+00
1e+05
2e+05
3e+05
4e+05
5e+05
6e+05
7e+05
0
20
40
60
bp position

### lwe−miR−425_loc1

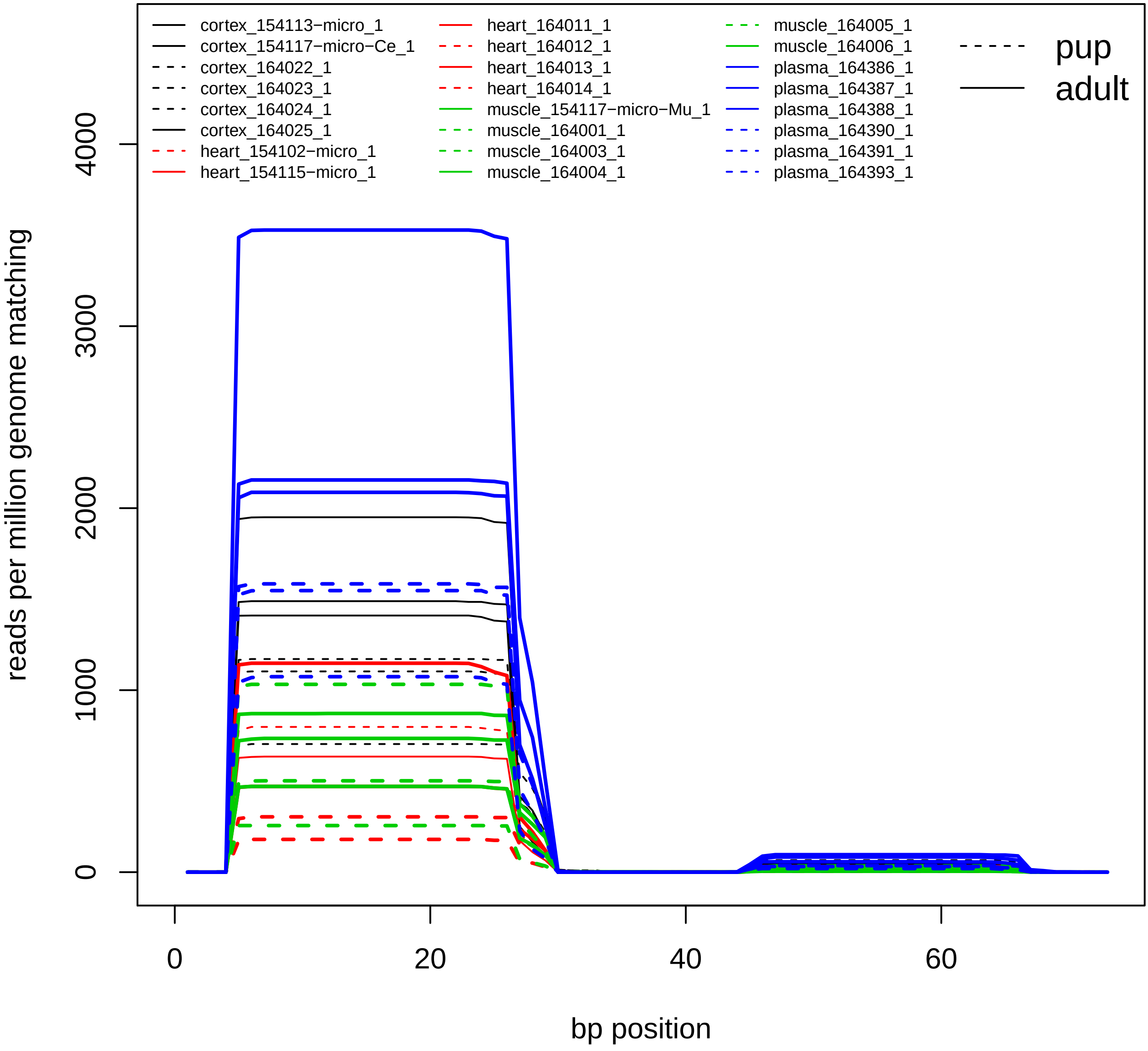

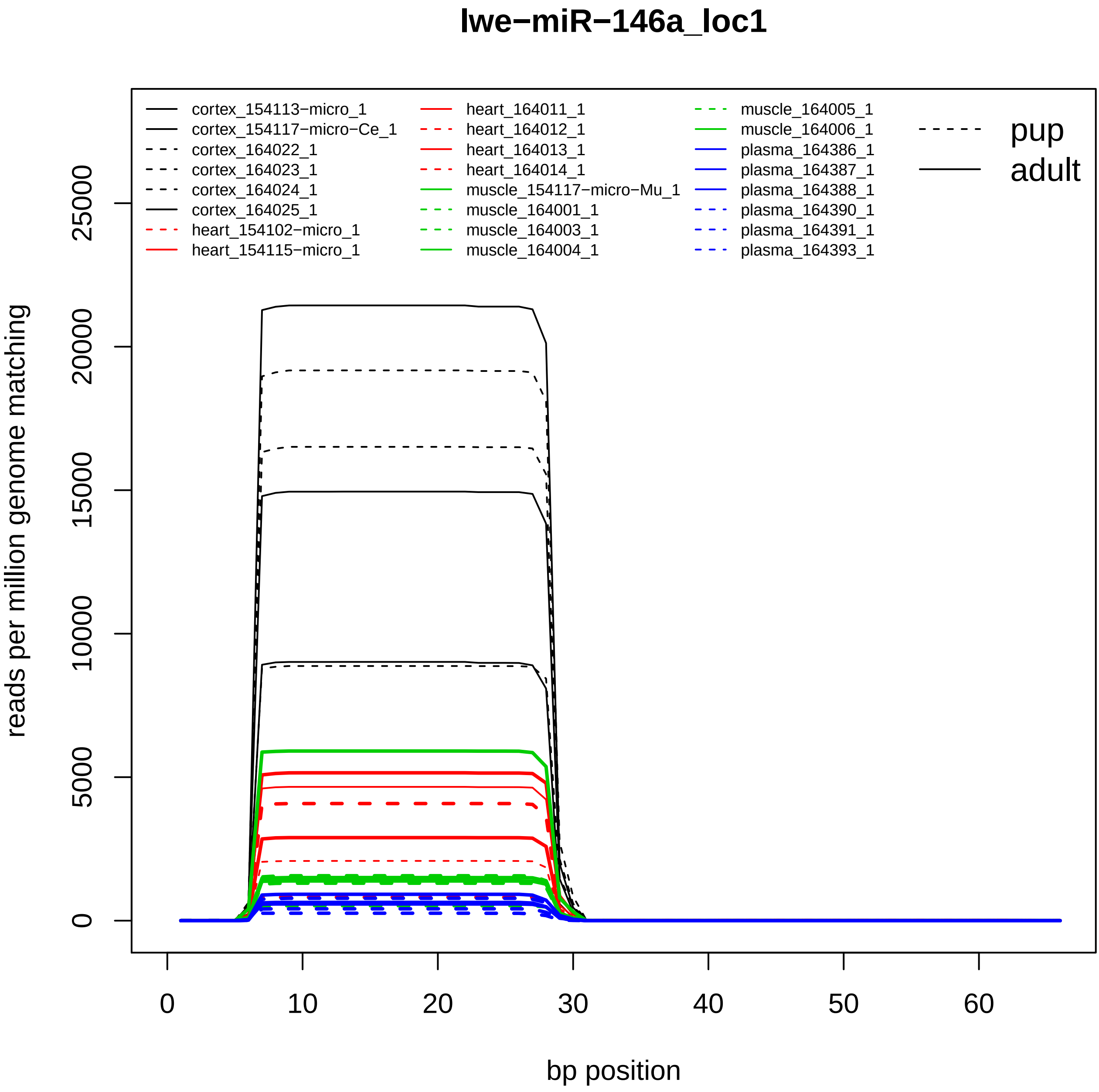
lwe−miR−146a_loc1
cortex_154113−micro_1
cortex_154117−micro−Ce_1
cortex_164022_1
cortex_164023_1
cortex_164024_1
cortex_164025_1
heart_154102−micro_1
heart_154115−micro_1
heart_164011_1
heart_164012_1
heart_164013_1
heart_164014_1
muscle_154117−micro−Mu_1
muscle_164001_1
muscle_164003_1
muscle_164004_1
muscle_164005_1
muscle_164006_1
plasma_164386_1
plasma_164387_1
plasma_164388_1
plasma_164390_1
plasma_164391_1
plasma_164393_1
pup
adult
reads per million genome matching
0
5000
10000
15000
20000
25000
0
10
20
30
40
50
60
bp position

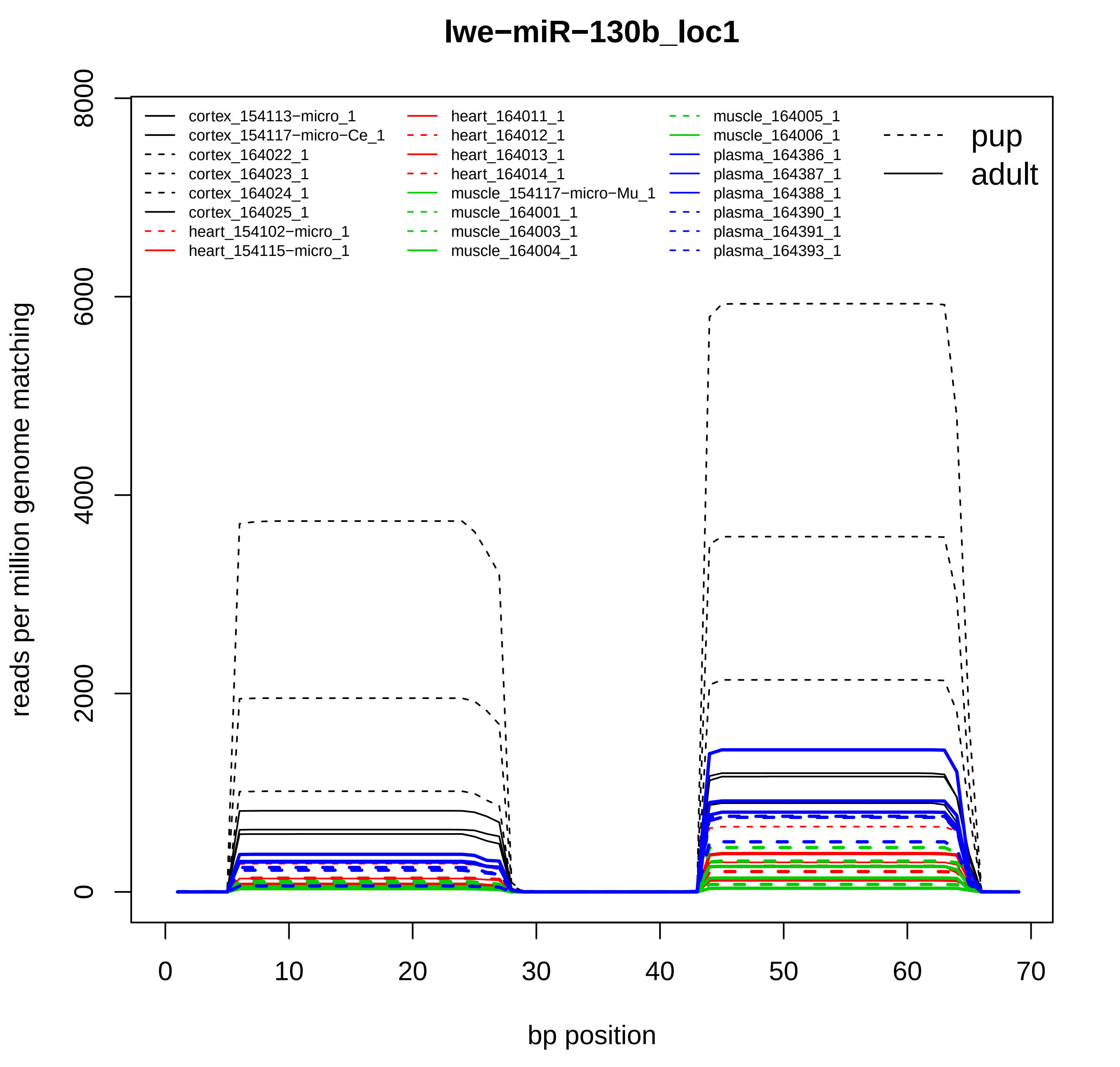
lwe−miR−130b_loc1
reads per million genome matching
bp position
0
2000
4000
6000
8000
0
10
20
30
40
50
60
70
cortex_154113−micro_1
cortex_154117−micro−Ce_1
cortex_164022_1
cortex_164023_1
cortex_164024_1
cortex_164025_1
heart_154102−micro_1
heart_154115−micro_1
heart_164011_1
heart_164012_1
heart_164013_1
heart_164014_1
muscle_154117−micro−Mu_1
muscle_164001_1
muscle_164003_1
muscle_164004_1
muscle_164005_1
muscle_164006_1
plasma_164386_1
plasma_164387_1
plasma_164388_1
plasma_164390_1
plasma_164391_1
plasma_164393_1
pup
adult

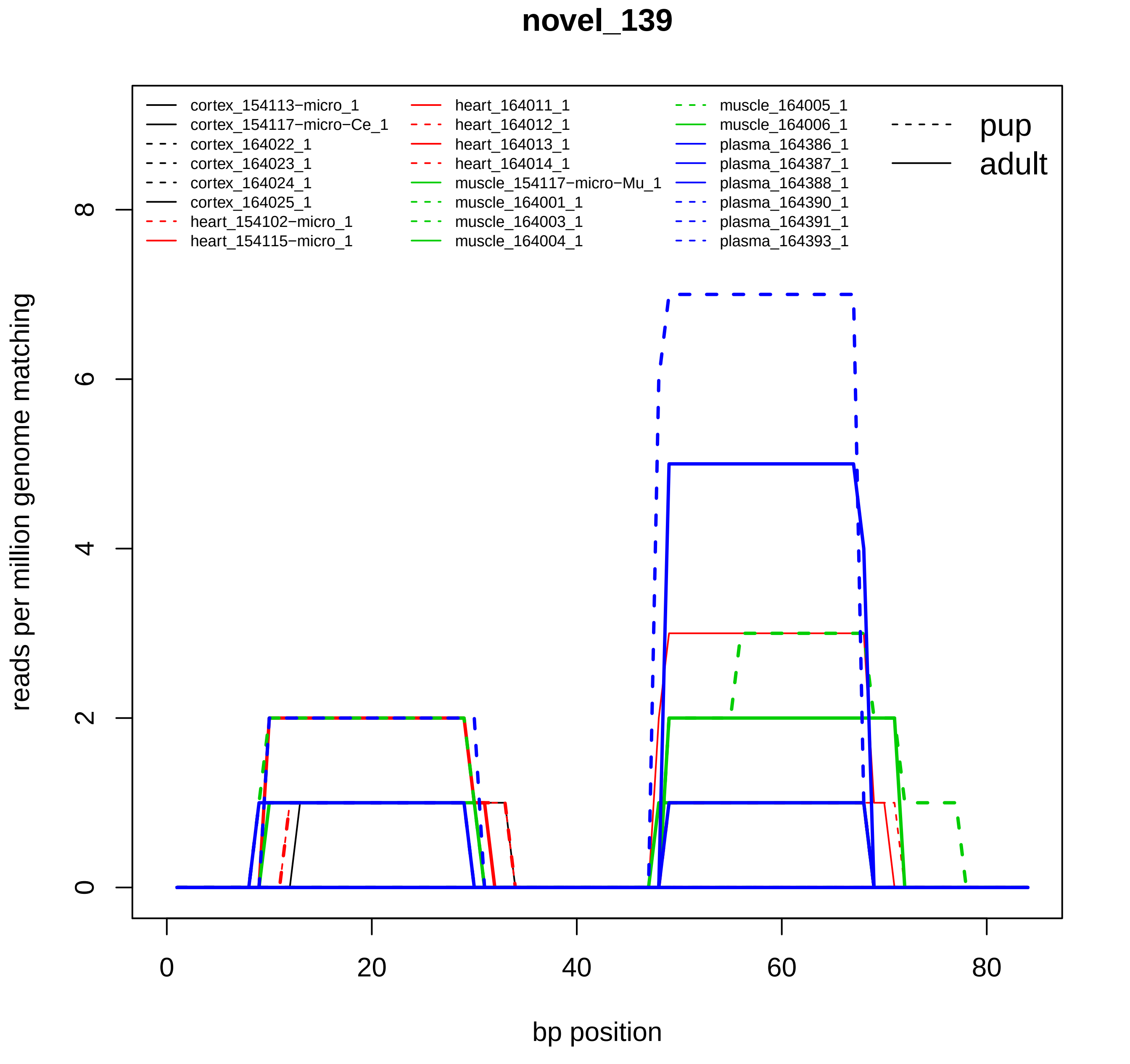
novel_139
cortex_154113−micro_1
cortex_154117−micro−Ce_1
cortex_164022_1
cortex_164023_1
cortex_164024_1
cortex_164025_1
heart_154102−micro_1
heart_154115−micro_1
heart_164011_1
heart_164012_1
heart_164013_1
heart_164014_1
muscle_154117−micro−Mu_1
muscle_164001_1
muscle_164003_1
muscle_164004_1
muscle_164005_1
muscle_164006_1
plasma_164386_1
plasma_164387_1
plasma_164388_1
plasma_164390_1
plasma_164391_1
plasma_164393_1
pup
adult
reads per million genome matching
0
2
4
6
8
0
20
40
60
80
bp position

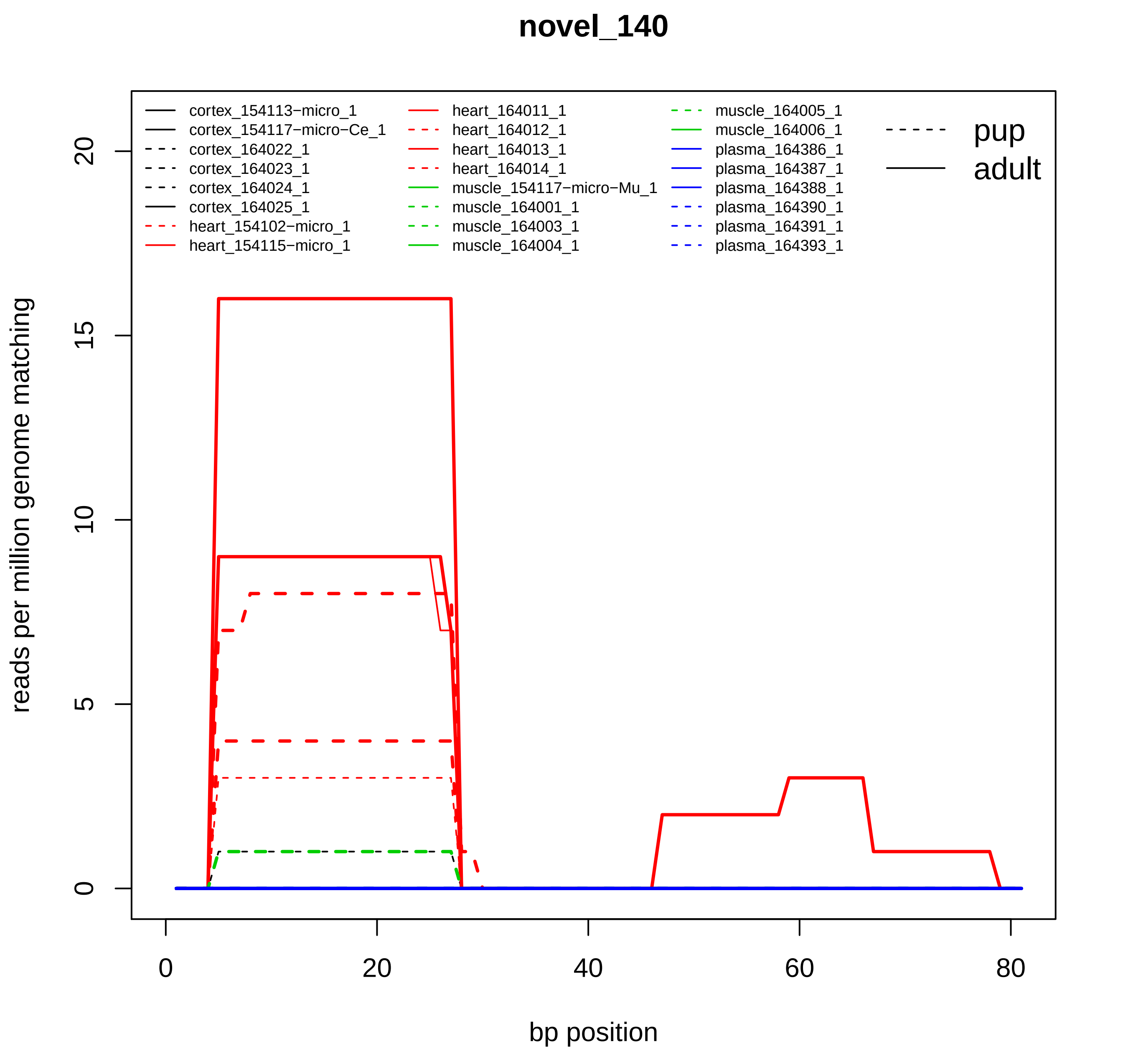
novel_140
cortex_154113−micro_1
cortex_154117−micro−Ce_1
cortex_164022_1
cortex_164023_1
cortex_164024_1
cortex_164025_1
heart_154102−micro_1
heart_154115−micro_1
heart_164011_1
heart_164012_1
heart_164013_1
heart_164014_1
muscle_154117−micro−Mu_1
muscle_164001_1
muscle_164003_1
muscle_164004_1
muscle_164005_1
muscle_164006_1
plasma_164386_1
plasma_164387_1
plasma_164388_1
plasma_164390_1
plasma_164391_1
plasma_164393_1
pup
adult
reads per million genome matching
0
5
10
15
20
bp position
0
20
40
60
80

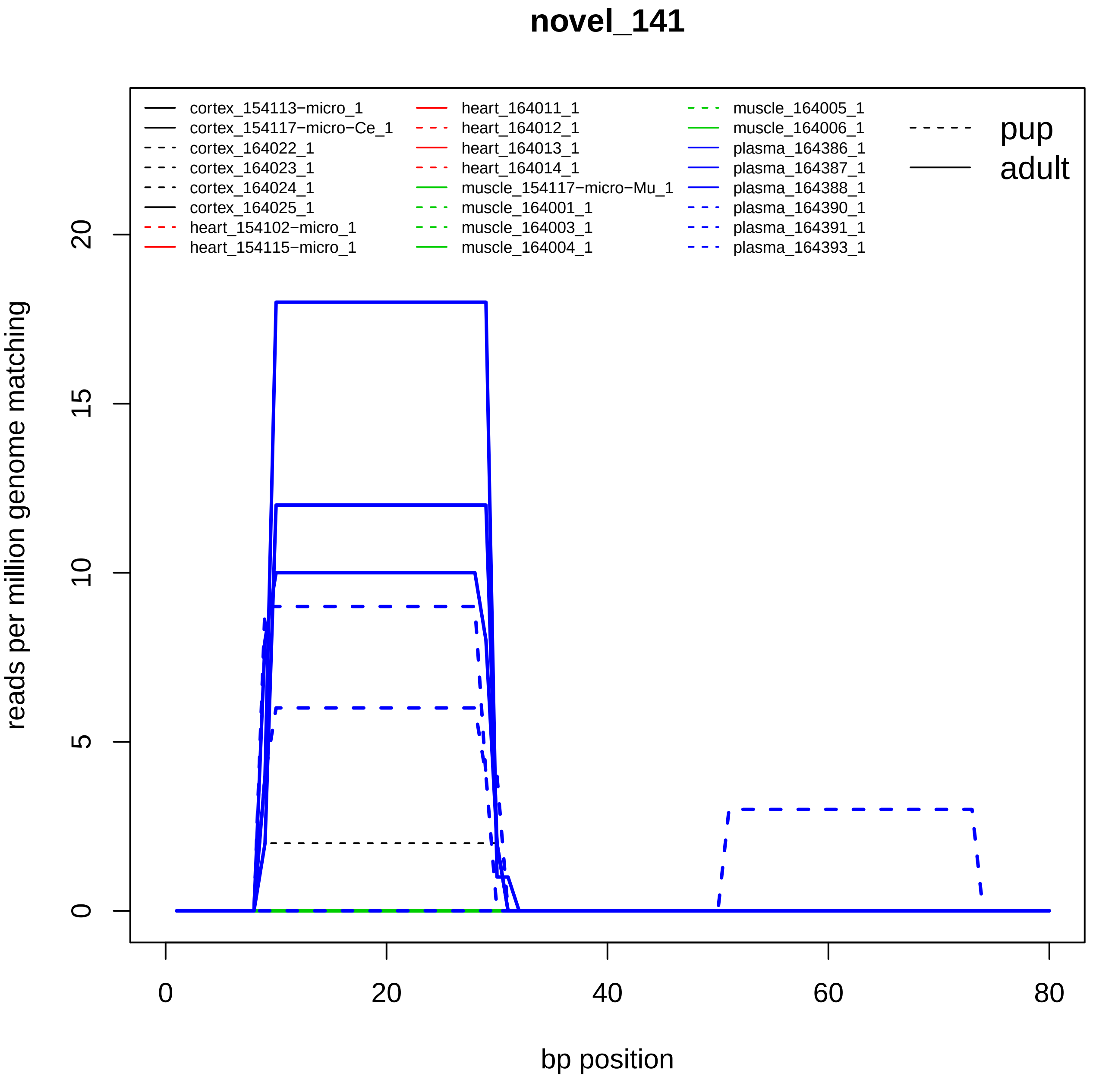
novel_141
cortex_154113−micro_1
cortex_154117−micro−Ce_1
cortex_164022_1
cortex_164023_1
cortex_164024_1
cortex_164025_1
heart_154102−micro_1
heart_154115−micro_1
heart_164011_1
heart_164012_1
heart_164013_1
heart_164014_1
muscle_154117−micro−Mu_1
muscle_164001_1
muscle_164003_1
muscle_164004_1
muscle_164005_1
muscle_164006_1
plasma_164386_1
plasma_164387_1
plasma_164388_1
plasma_164390_1
plasma_164391_1
plasma_164393_1
pup
adult
reads per million genome matching
0
5
10
15
20
0
20
40
60
80
bp position

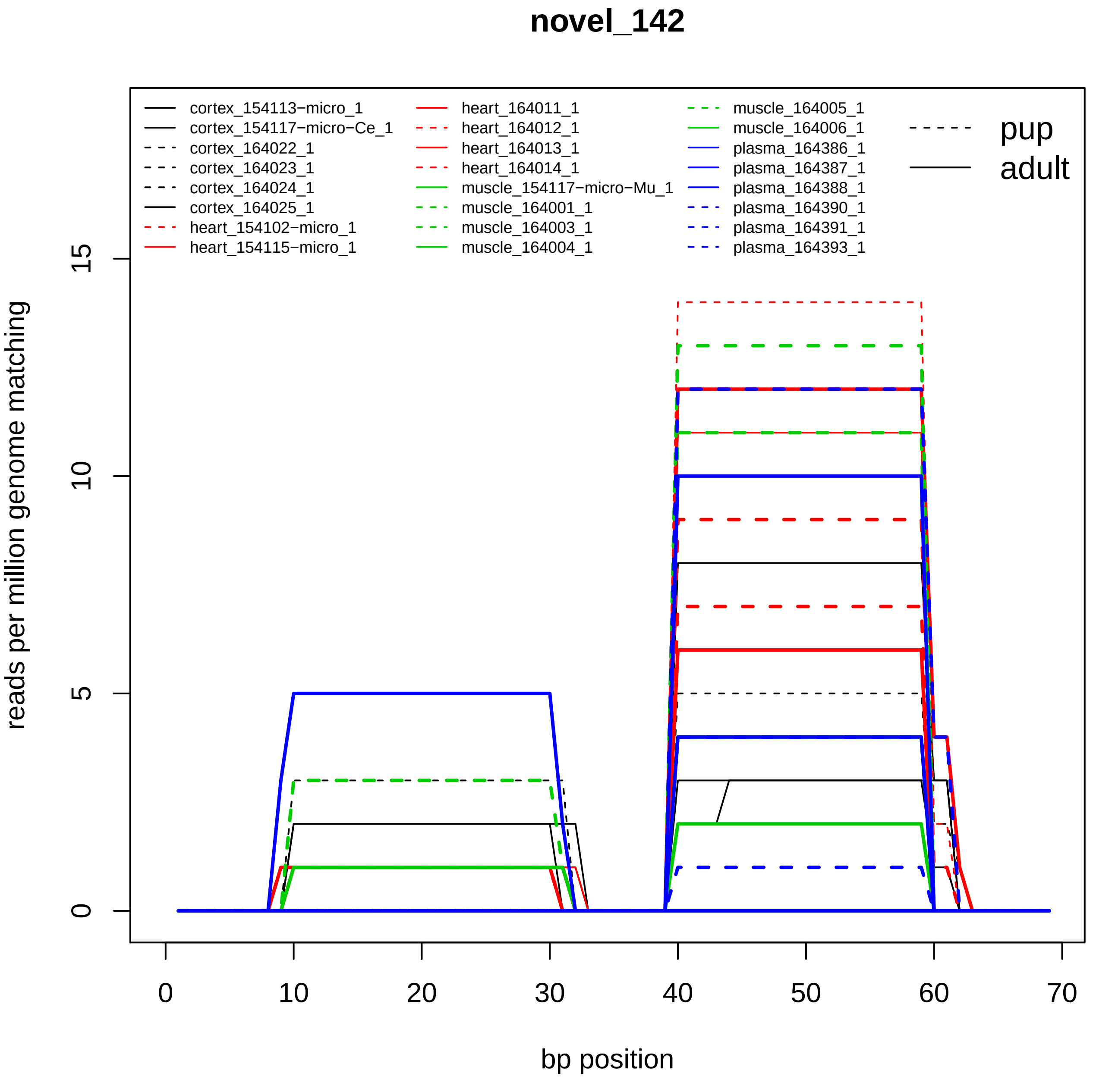
novel_142
cortex_154113−micro_1
cortex_154117−micro−Ce_1
cortex_164022_1
cortex_164023_1
cortex_164024_1
cortex_164025_1
heart_154102−micro_1
heart_154115−micro_1
heart_164011_1
heart_164012_1
heart_164013_1
heart_164014_1
muscle_154117−micro−Mu_1
muscle_164001_1
muscle_164003_1
muscle_164004_1
muscle_164005_1
muscle_164006_1
plasma_164386_1
plasma_164387_1
plasma_164388_1
plasma_164390_1
plasma_164391_1
plasma_164393_1
pup
adult
reads per million genome matching
0
5
10
15
bp position
0
10
20
30
40
50
60
70

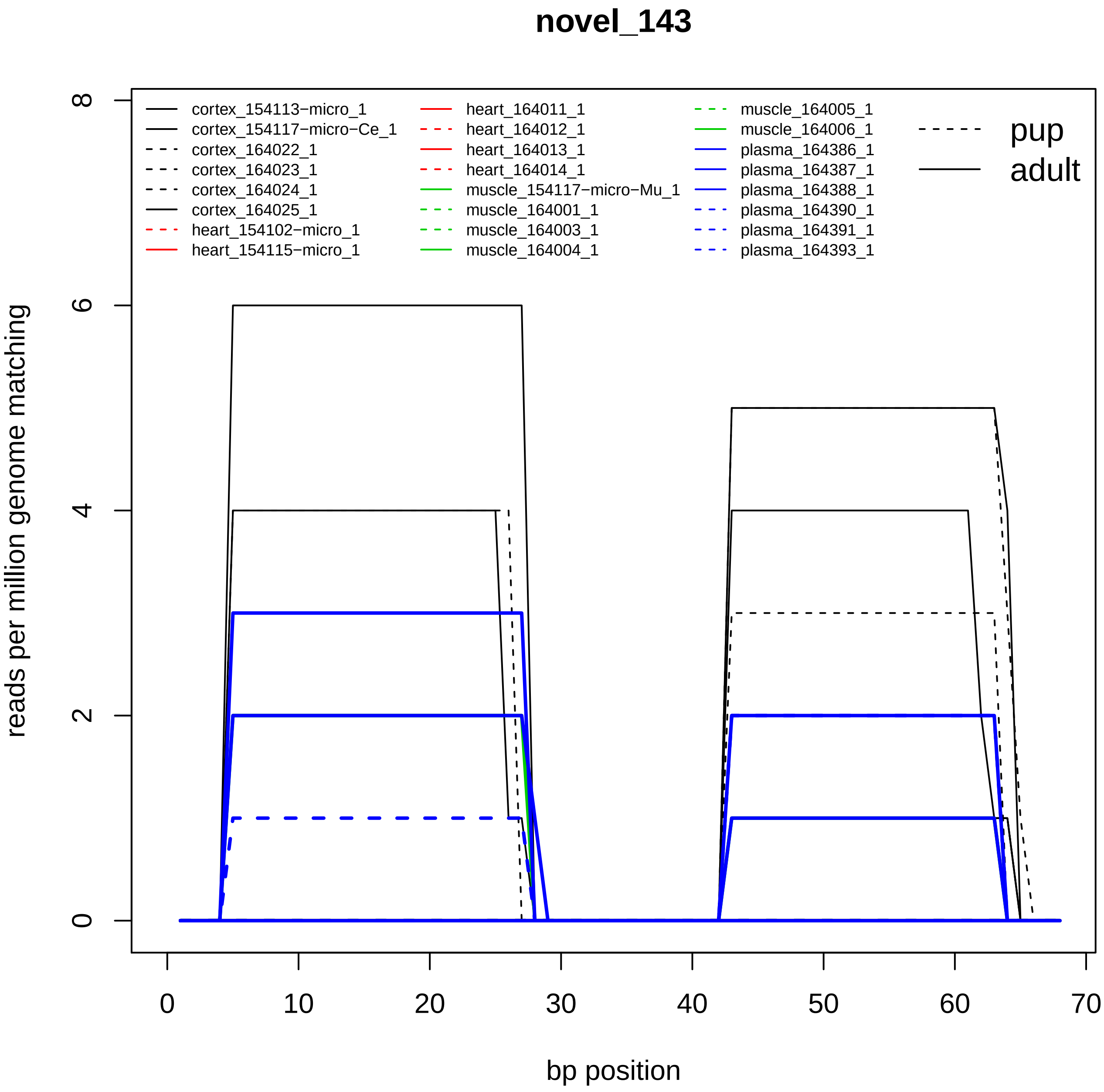

novel_143
cortex_154113−micro_1
cortex_154117−micro−Ce_1
cortex_164022_1
cortex_164023_1
cortex_164024_1
cortex_164025_1
heart_154102−micro_1
heart_154115−micro_1
heart_164011_1
heart_164012_1
heart_164013_1
heart_164014_1
muscle_154117−micro−Mu_1
muscle_164001_1
muscle_164003_1
muscle_164004_1
muscle_164005_1
muscle_164006_1
plasma_164386_1
plasma_164387_1
plasma_164388_1
plasma_164390_1
plasma_164391_1
plasma_164393_1
pup
adult
reads per million genome matching
bp position
0
2
4
6
8
0
10
20
30
40
50
60
70

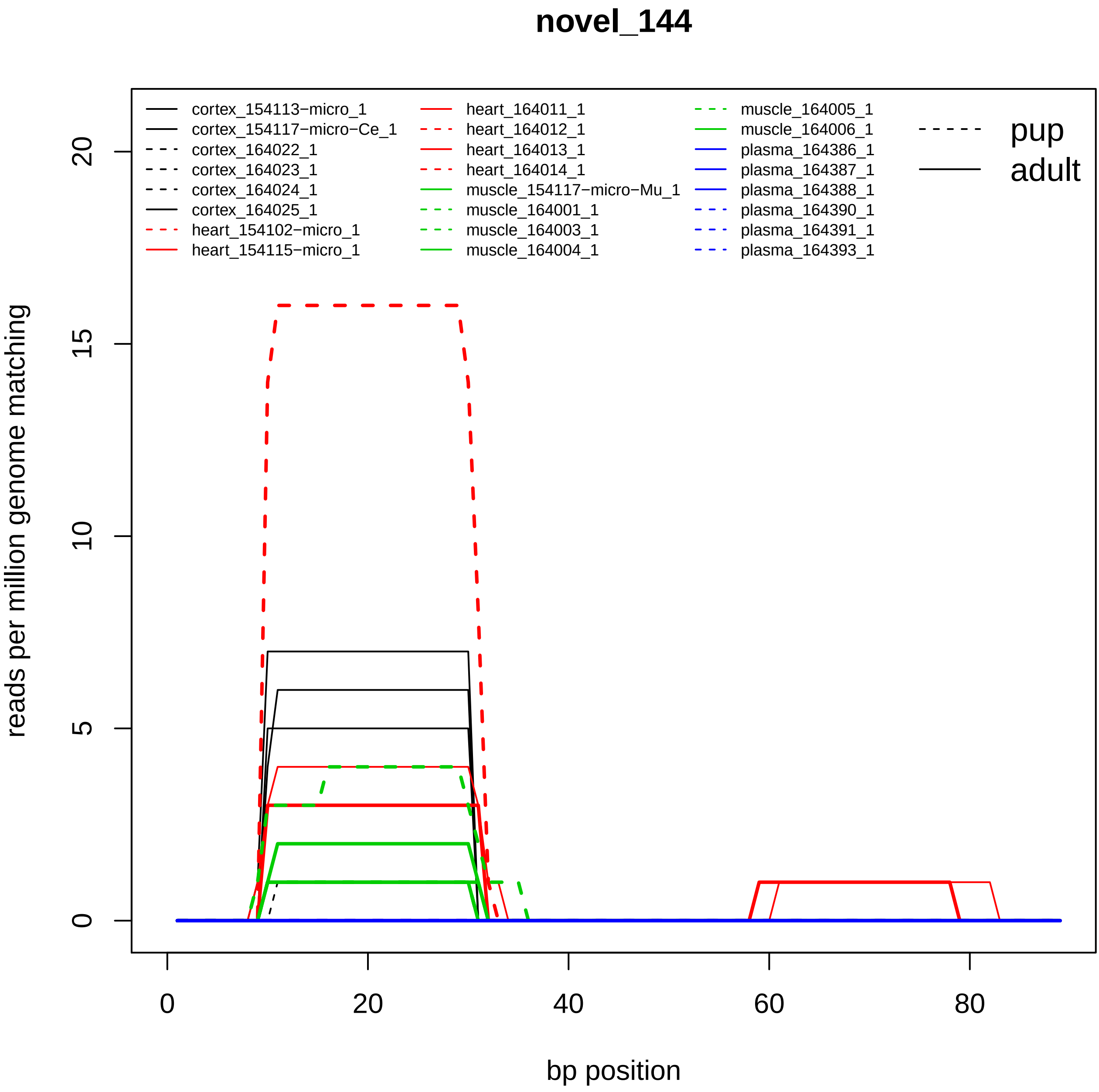
novel_144
cortex_154113−micro_1
cortex_154117−micro−Ce_1
cortex_164022_1
cortex_164023_1
cortex_164024_1
cortex_164025_1
heart_154102−micro_1
heart_154115−micro_1
heart_164011_1
heart_164012_1
heart_164013_1
heart_164014_1
muscle_154117−micro−Mu_1
muscle_164001_1
muscle_164003_1
muscle_164004_1
muscle_164005_1
muscle_164006_1
plasma_164386_1
plasma_164387_1
plasma_164388_1
plasma_164390_1
plasma_164391_1
plasma_164393_1
pup
adult
reads per million genome matching
0
5
10
15
20
0
20
40
60
80
bp position

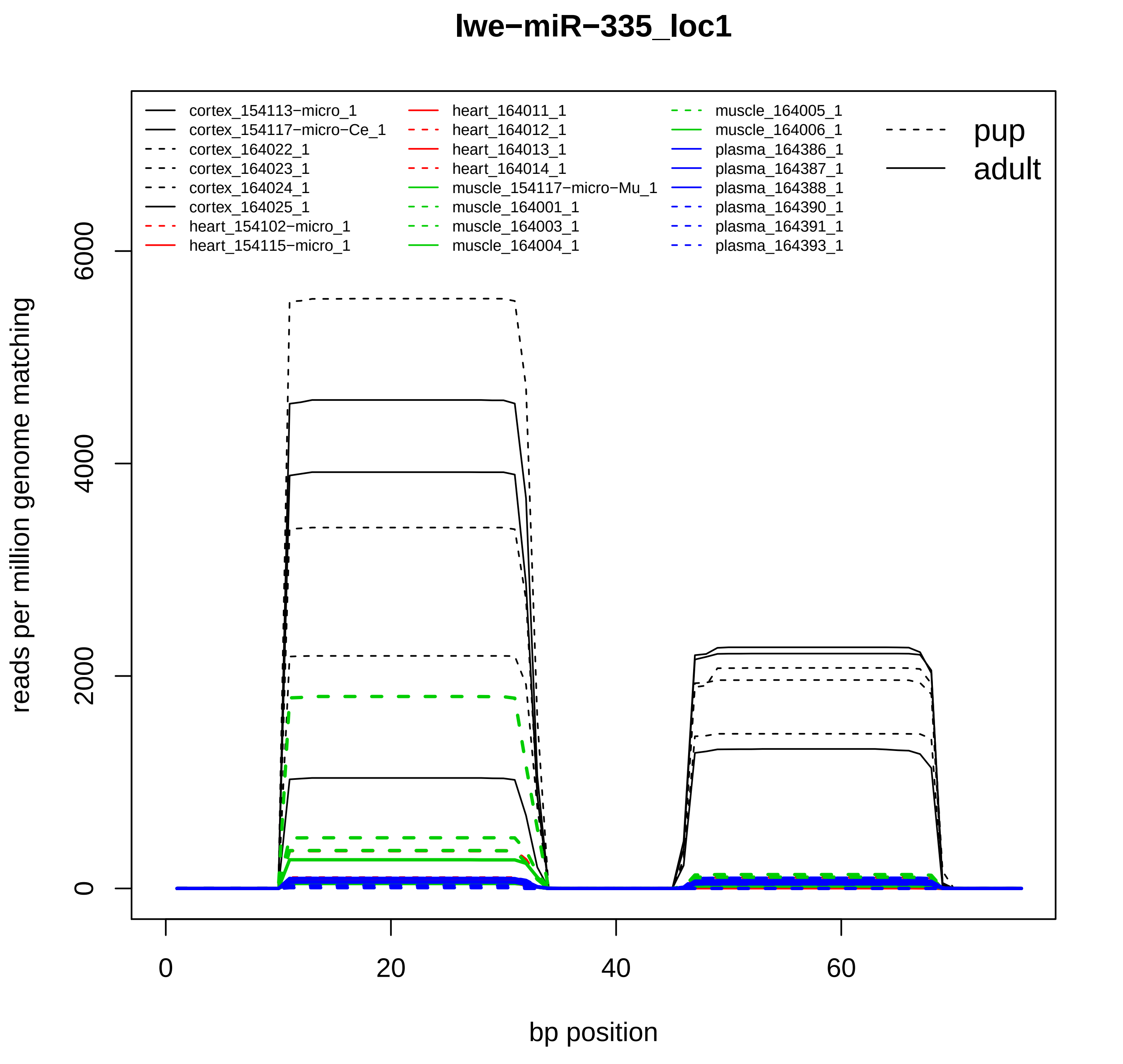
lwe−miR−335_loc1
cortex_154113−micro_1
cortex_154117−micro−Ce_1
cortex_164022_1
cortex_164023_1
cortex_164024_1
cortex_164025_1
heart_154102−micro_1
heart_154115−micro_1
heart_164011_1
heart_164012_1
heart_164013_1
heart_164014_1
muscle_154117−micro−Mu_1
muscle_164001_1
muscle_164003_1
muscle_164004_1
muscle_164005_1
muscle_164006_1
plasma_164386_1
plasma_164387_1
plasma_164388_1
plasma_164390_1
plasma_164391_1
plasma_164393_1
pup
adult
reads per million genome matching
0
2000
4000
6000
0
20
40
60
bp position

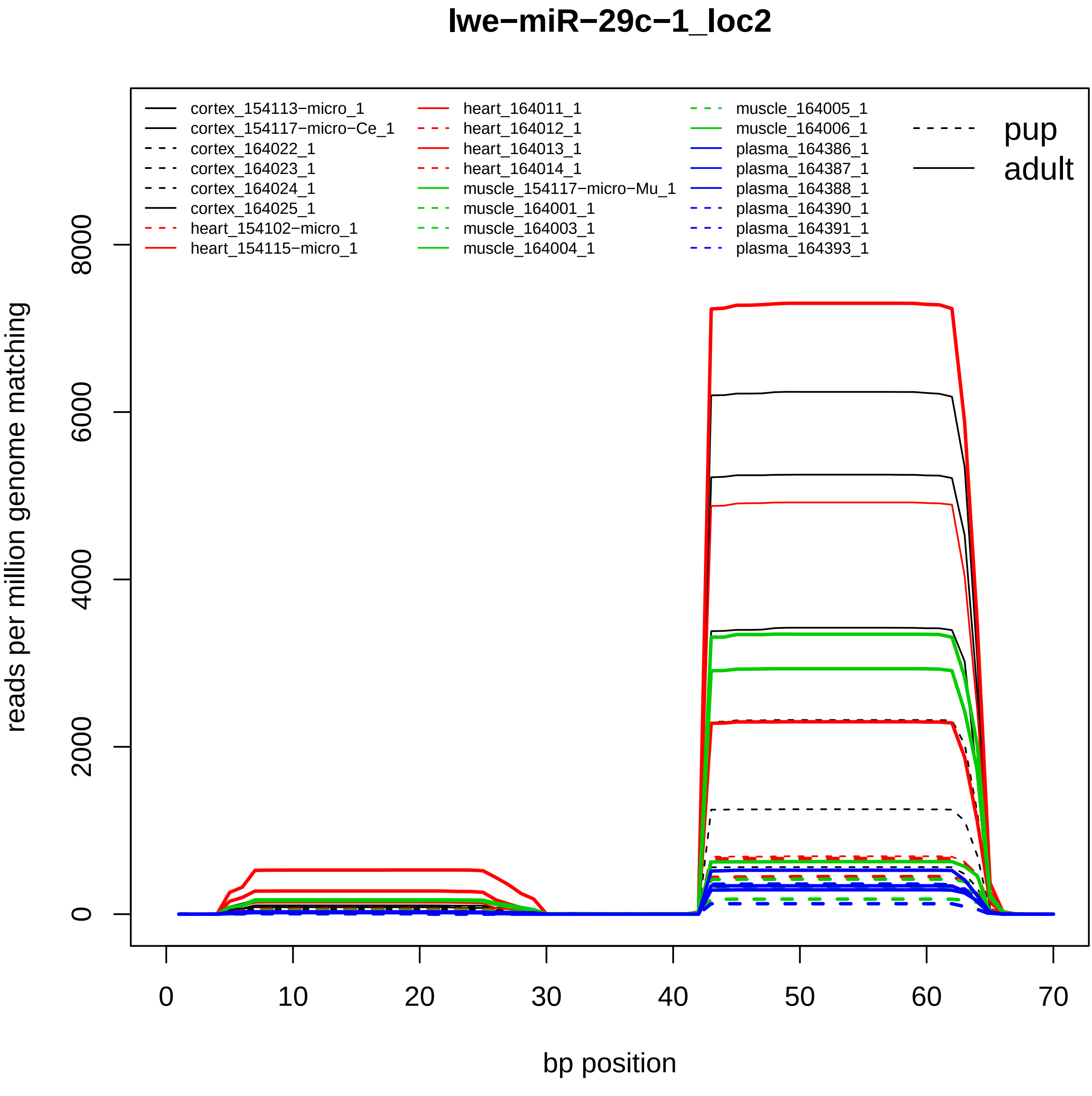

lwe−miR−29c−1_loc2
cortex_154113−micro_1
cortex_154117−micro−Ce_1
cortex_164022_1
cortex_164023_1
cortex_164024_1
cortex_164025_1
heart_154102−micro_1
heart_154115−micro_1
heart_164011_1
heart_164012_1
heart_164013_1
heart_164014_1
muscle_154117−micro−Mu_1
muscle_164001_1
muscle_164003_1
muscle_164004_1
muscle_164005_1
muscle_164006_1
plasma_164386_1
plasma_164387_1
plasma_164388_1
plasma_164390_1
plasma_164391_1
plasma_164393_1
pup
adult
reads per million genome matching
0
2000
4000
6000
8000
bp position
0
10
20
30
40
50
60
70

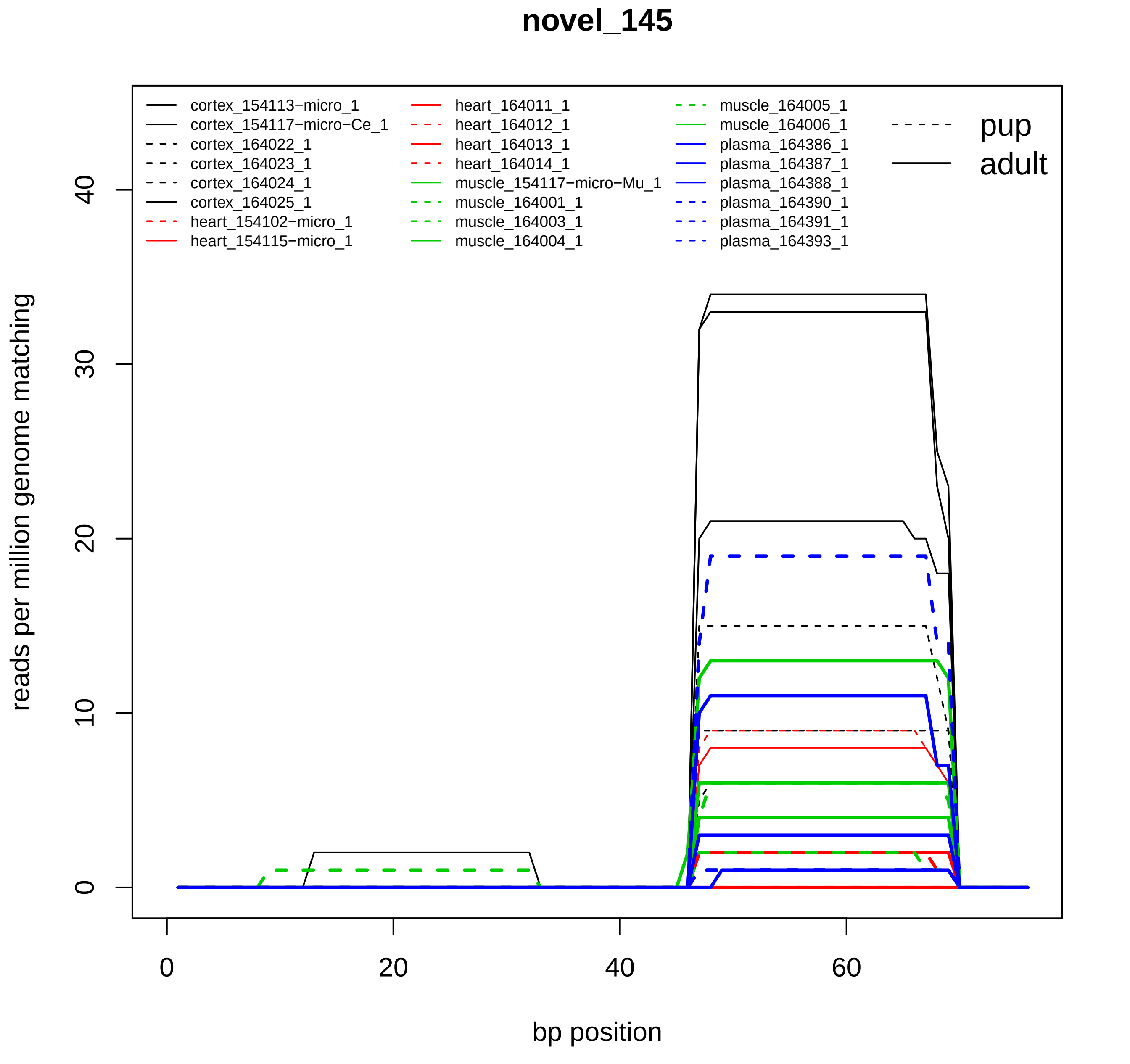
novel_145
cortex_154113−micro_1
cortex_154117−micro−Ce_1
cortex_164022_1
cortex_164023_1
cortex_164024_1
cortex_164025_1
heart_154102−micro_1
heart_154115−micro_1
heart_164011_1
heart_164012_1
heart_164013_1
heart_164014_1
muscle_154117−micro−Mu_1
muscle_164001_1
muscle_164003_1
muscle_164004_1
muscle_164005_1
muscle_164006_1
plasma_164386_1
plasma_164387_1
plasma_164388_1
plasma_164390_1
plasma_164391_1
plasma_164393_1
pup
adult
reads per million genome matching
0
10
20
30
40
0
20
40
60
bp position

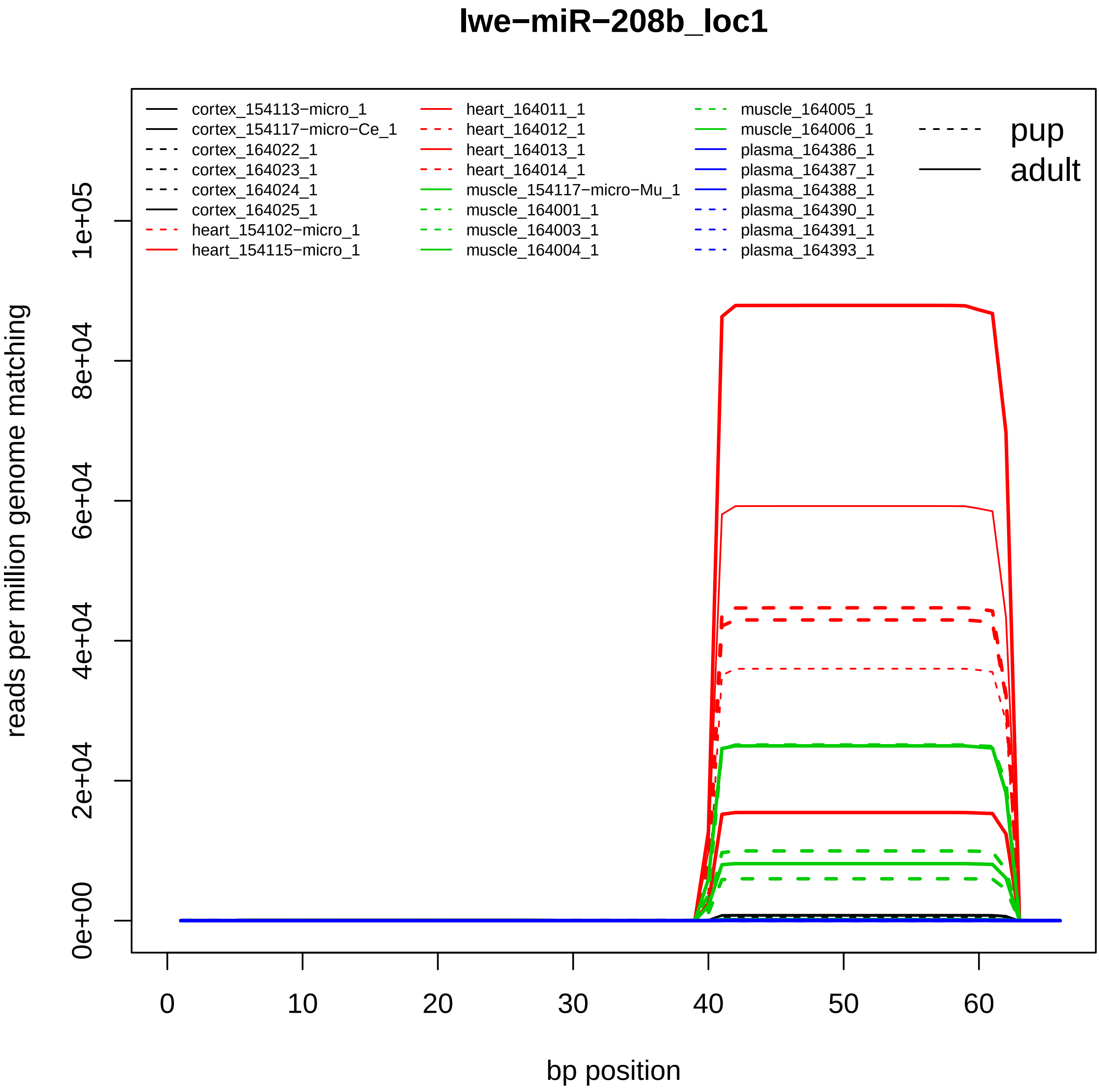
lwe−miR−208b_loc1
cortex_154113−micro_1
cortex_154117−micro−Ce_1
cortex_164022_1
cortex_164023_1
cortex_164024_1
cortex_164025_1
heart_154102−micro_1
heart_154115−micro_1
heart_164011_1
heart_164012_1
heart_164013_1
heart_164014_1
muscle_154117−micro−Mu_1
muscle_164001_1
muscle_164003_1
muscle_164004_1
muscle_164005_1
muscle_164006_1
plasma_164386_1
plasma_164387_1
plasma_164388_1
plasma_164390_1
plasma_164391_1
plasma_164393_1
pup
adult
reads per million genome matching
0e+00
2e+04
4e+04
6e+04
8e+04
1e+05
0
10
20
30
40
50
60
bp position

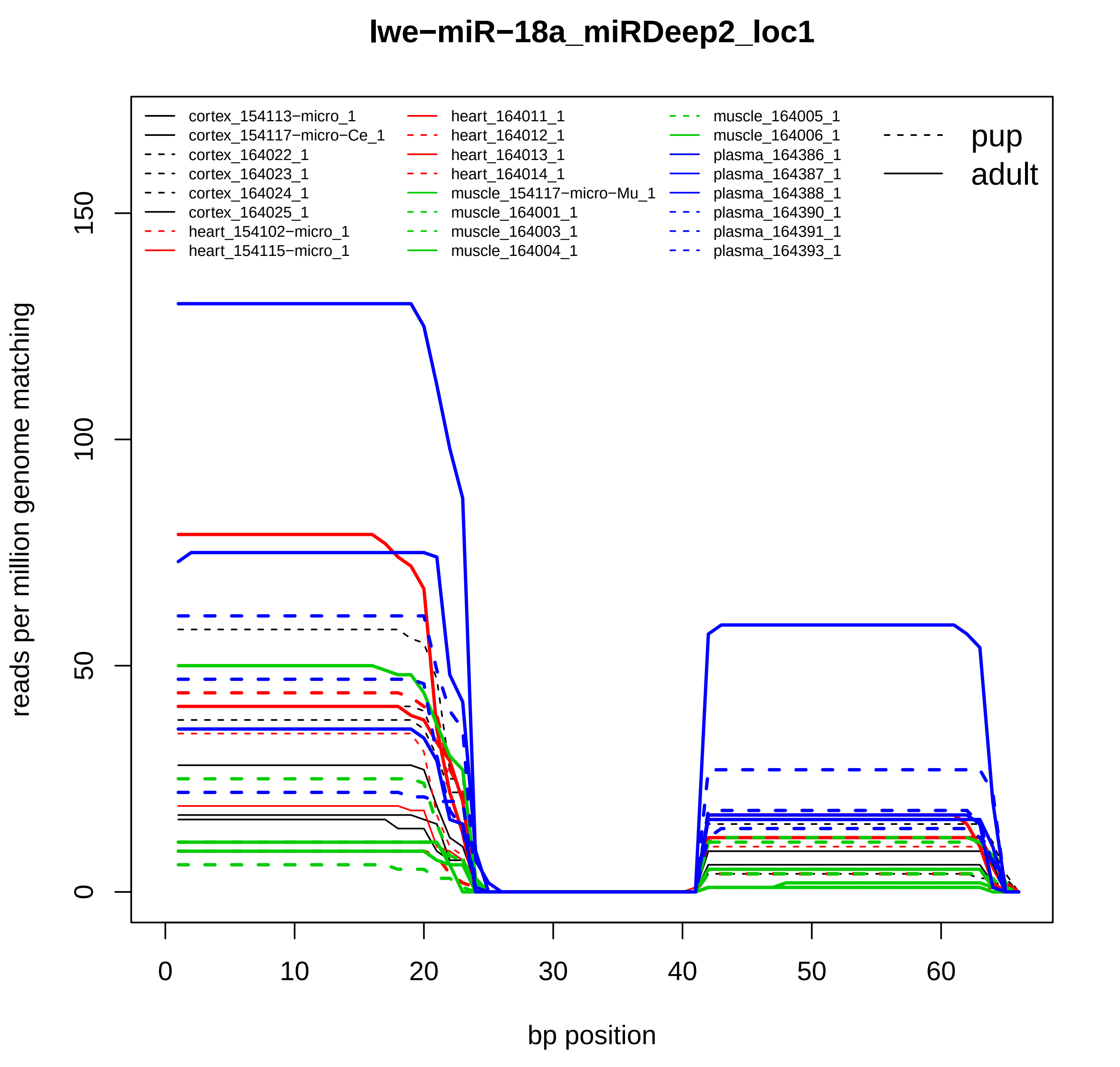
lwe−miR−18a_miRDeep2_loc1
cortex_154113−micro_1
cortex_154117−micro−Ce_1
cortex_164022_1
cortex_164023_1
cortex_164024_1
cortex_164025_1
heart_154102−micro_1
heart_154115−micro_1
heart_164011_1
heart_164012_1
heart_164013_1
heart_164014_1
muscle_154117−micro−Mu_1
muscle_164001_1
muscle_164003_1
muscle_164004_1
muscle_164005_1
muscle_164006_1
plasma_164386_1
plasma_164387_1
plasma_164388_1
plasma_164390_1
plasma_164391_1
plasma_164393_1
pup
adult
reads per million genome matching
0
50
100
150
0
10
20
30
40
50
60
bp position

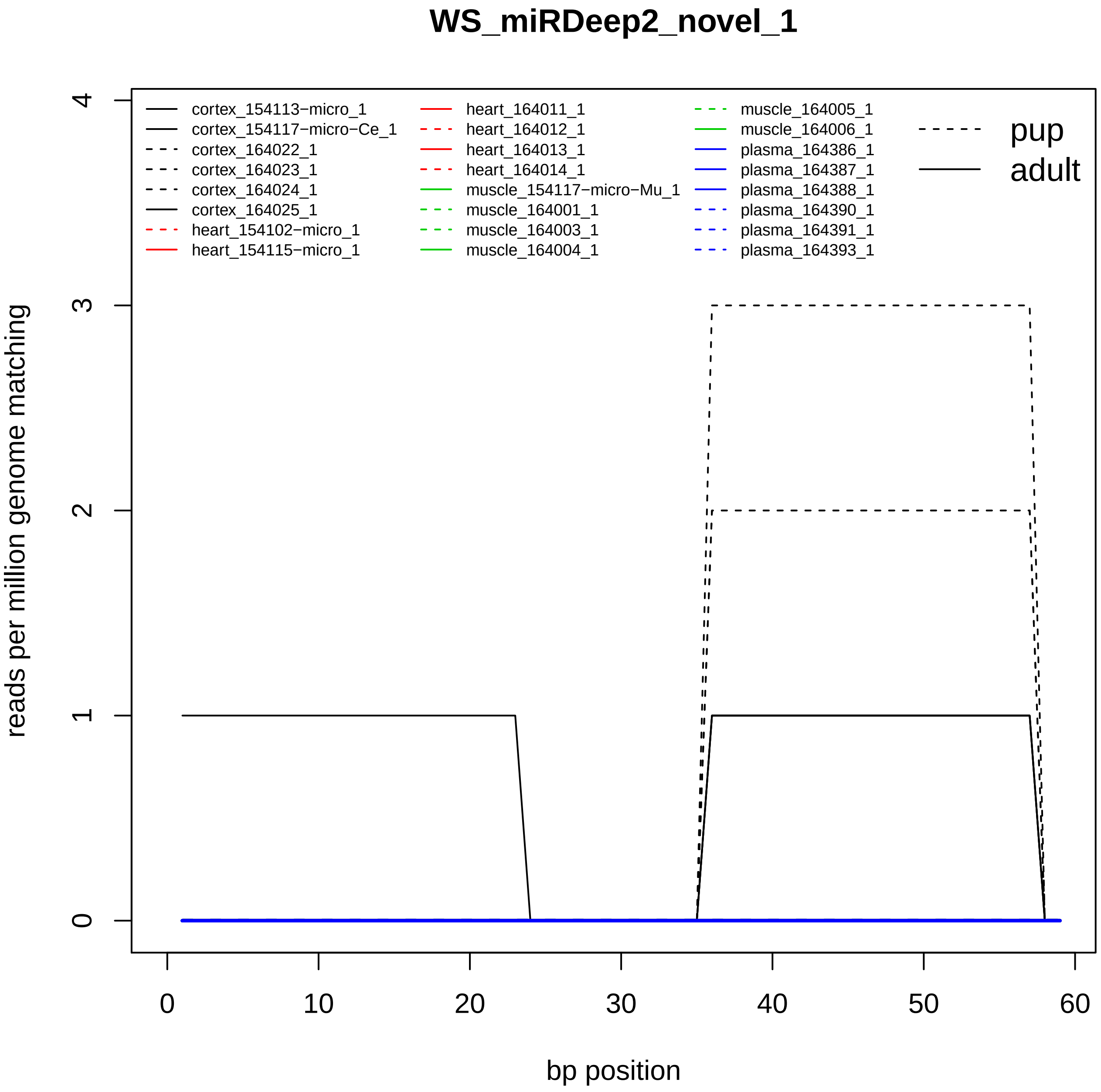

WS_miRDeep2_novel_1
cortex_154113−micro_1
cortex_154117−micro−Ce_1
cortex_164022_1
cortex_164023_1
cortex_164024_1
cortex_164025_1
heart_154102−micro_1
heart_154115−micro_1
heart_164011_1
heart_164012_1
heart_164013_1
heart_164014_1
muscle_154117−micro−Mu_1
muscle_164001_1
muscle_164003_1
muscle_164004_1
muscle_164005_1
muscle_164006_1
plasma_164386_1
plasma_164387_1
plasma_164388_1
plasma_164390_1
plasma_164391_1
plasma_164393_1
pup
adult
reads per million genome matching
0
1
2
3
4
0
10
20
30
40
50
60
bp position

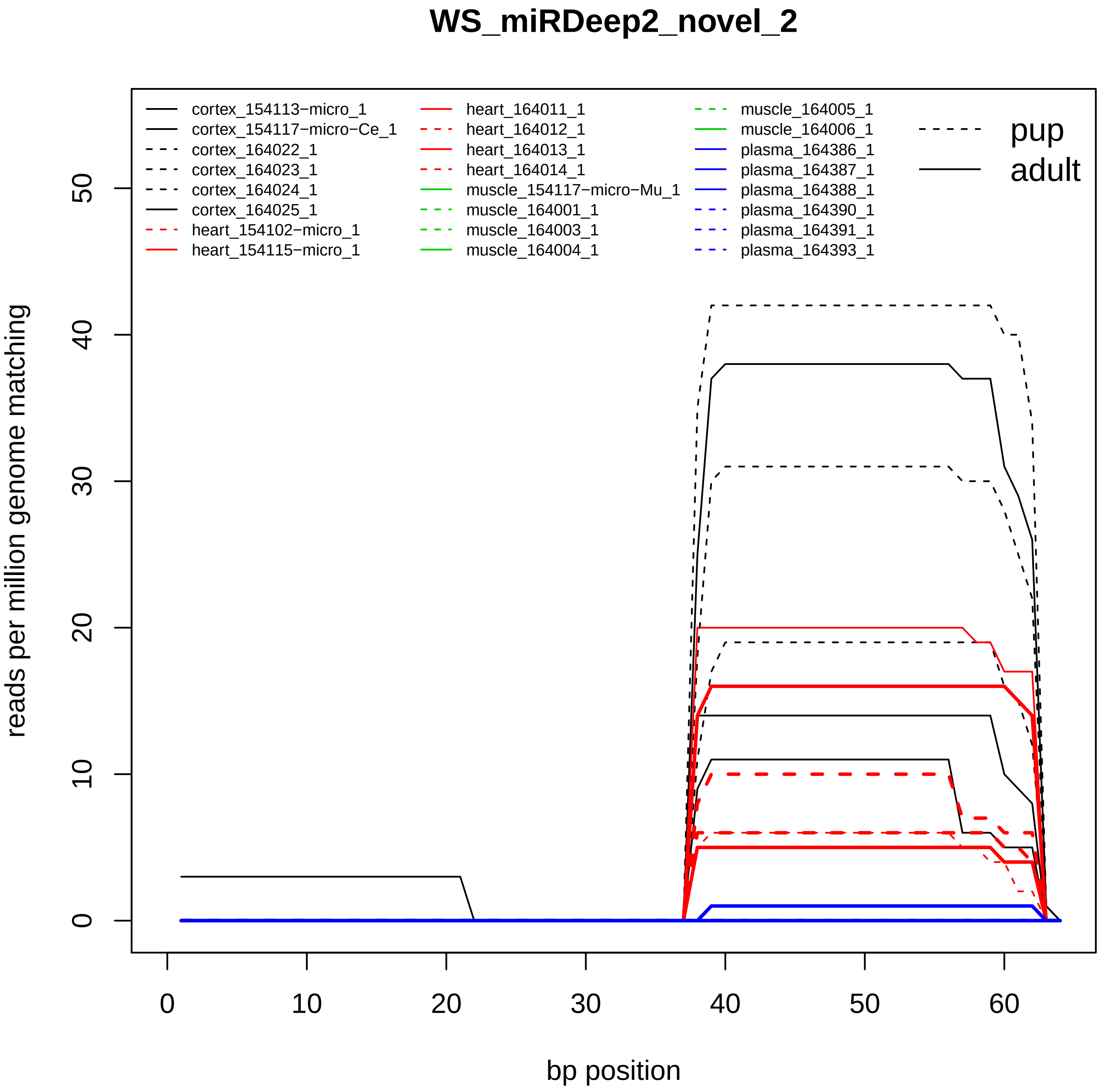
WS_miRDeep2_novel_2
cortex_154113−micro_1
cortex_154117−micro−Ce_1
cortex_164022_1
cortex_164023_1
cortex_164024_1
cortex_164025_1
heart_154102−micro_1
heart_154115−micro_1
heart_164011_1
heart_164012_1
heart_164013_1
heart_164014_1
muscle_154117−micro−Mu_1
muscle_164001_1
muscle_164003_1
muscle_164004_1
muscle_164005_1
muscle_164006_1
plasma_164386_1
plasma_164387_1
plasma_164388_1
plasma_164390_1
plasma_164391_1
plasma_164393_1
pup
adult
reads per million genome matching
bp position
0
10
20
30
40
50
60

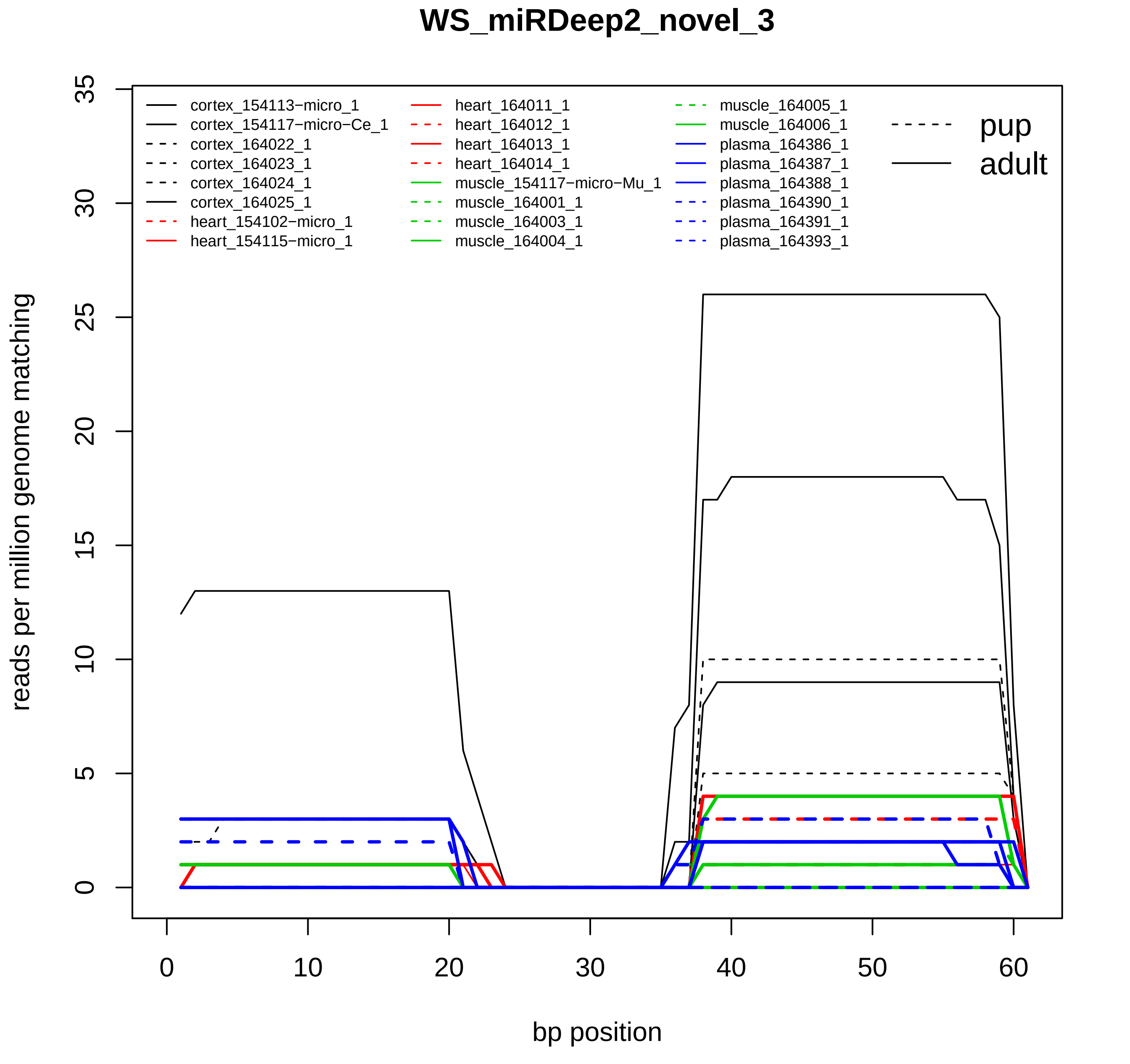

WS_miRDeep2_novel_3
cortex_154113−micro_1
cortex_154117−micro−Ce_1
cortex_164022_1
cortex_164023_1
cortex_164024_1
cortex_164025_1
heart_154102−micro_1
heart_154115−micro_1
heart_164011_1
heart_164012_1
heart_164013_1
heart_164014_1
muscle_154117−micro−Mu_1
muscle_164001_1
muscle_164003_1
muscle_164004_1
muscle_164005_1
muscle_164006_1
plasma_164386_1
plasma_164387_1
plasma_164388_1
plasma_164390_1
plasma_164391_1
plasma_164393_1
pup
adult
reads per million genome matching
bp position
0
5
10
15
20
25
30
35
0
10
20
30
40
50
60

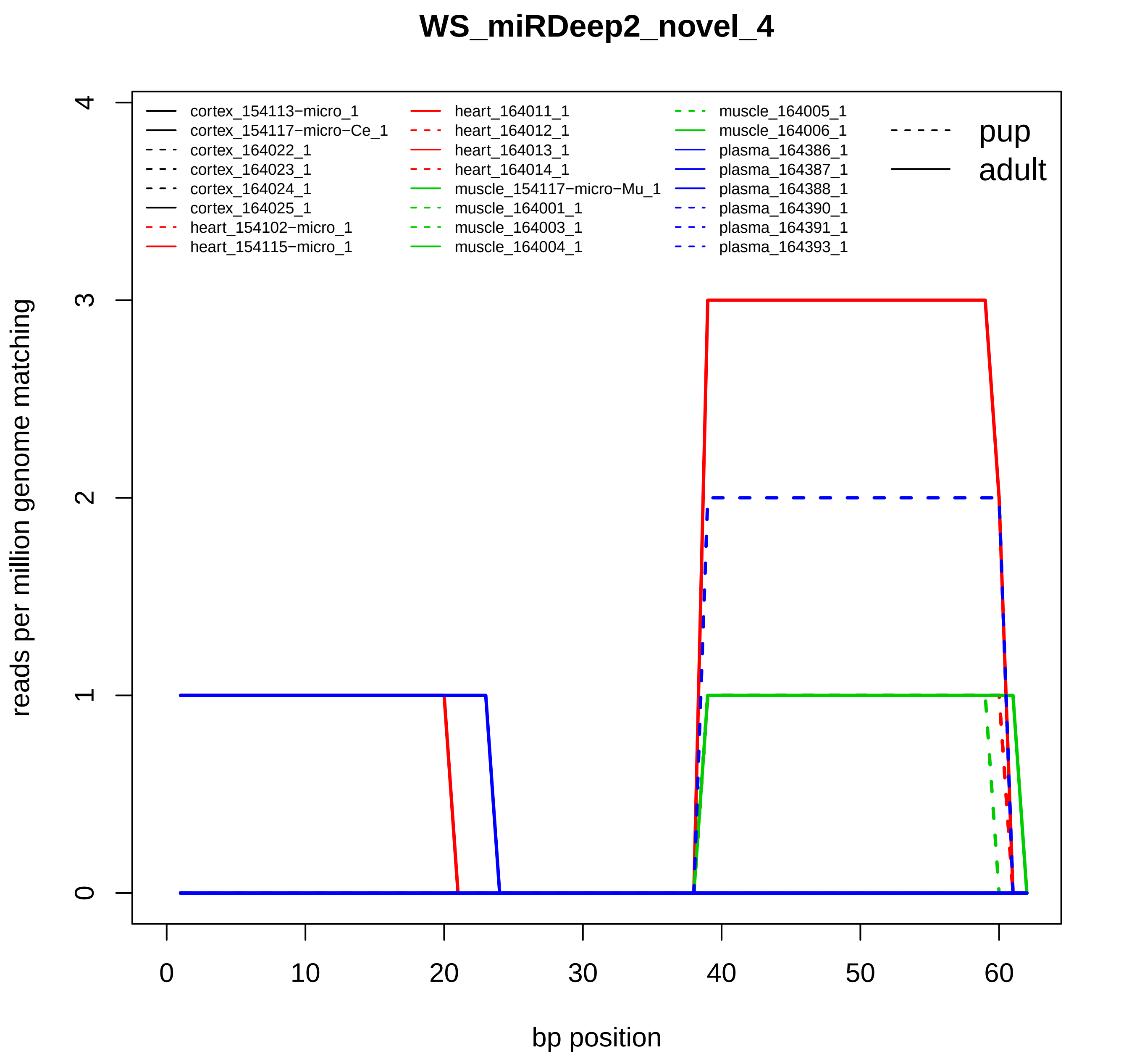

WS_miRDeep2_novel_4
cortex_154113−micro_1
cortex_154117−micro−Ce_1
cortex_164022_1
cortex_164023_1
cortex_164024_1
cortex_164025_1
heart_154102−micro_1
heart_154115−micro_1
heart_164011_1
heart_164012_1
heart_164013_1
heart_164014_1
muscle_154117−micro−Mu_1
muscle_164001_1
muscle_164003_1
muscle_164004_1
muscle_164005_1
muscle_164006_1
plasma_164386_1
plasma_164387_1
plasma_164388_1
plasma_164390_1
plasma_164391_1
plasma_164393_1
pup
adult
reads per million genome matching
bp position
0
1
2
3
4
0
10
20
30
40
50
60

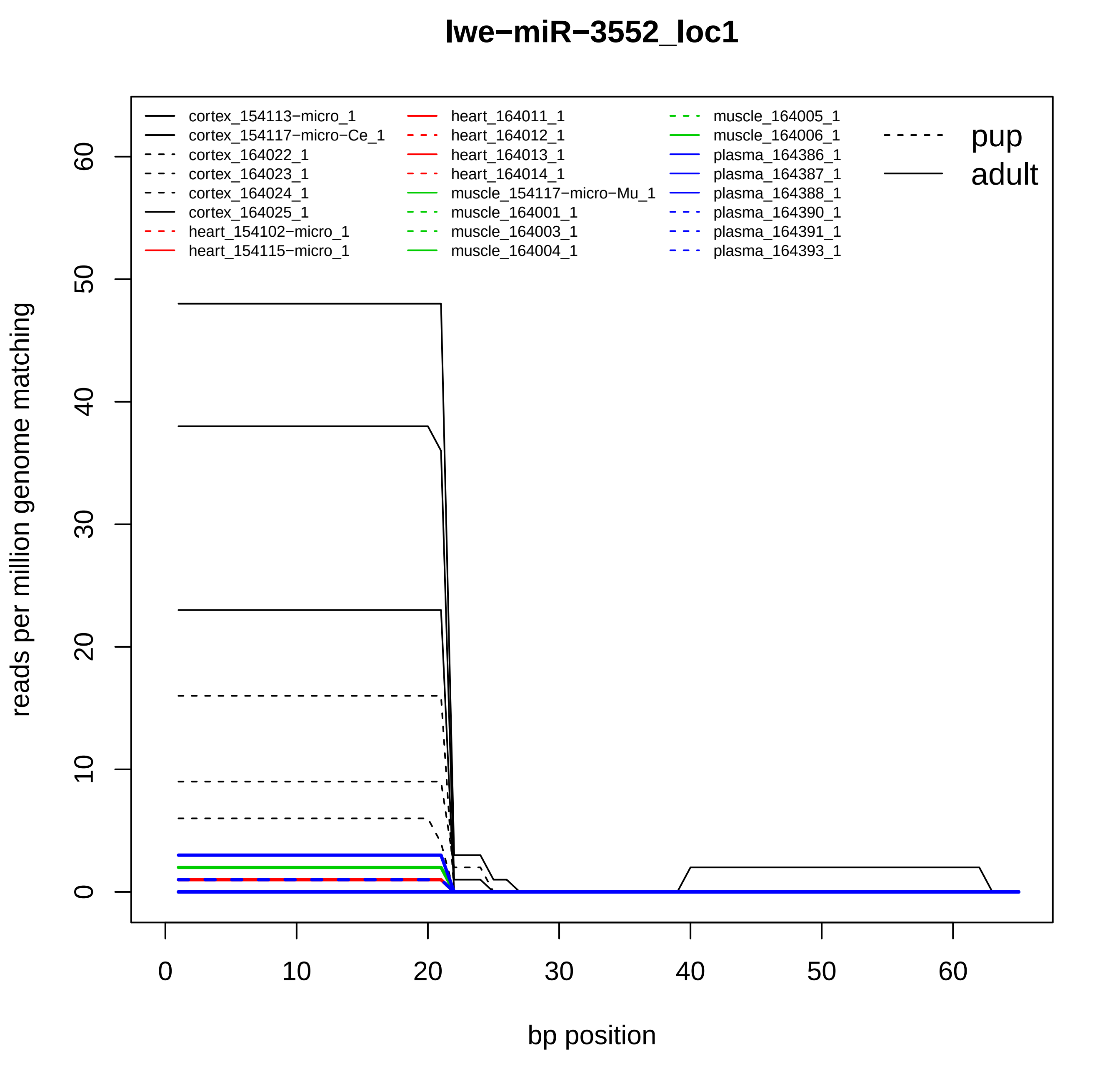

lwe−miR−3552_loc1
cortex_154113−micro_1
cortex_154117−micro−Ce_1
cortex_164022_1
cortex_164023_1
cortex_164024_1
cortex_164025_1
heart_154102−micro_1
heart_154115−micro_1
heart_164011_1
heart_164012_1
heart_164013_1
heart_164014_1
muscle_154117−micro−Mu_1
muscle_164001_1
muscle_164003_1
muscle_164004_1
muscle_164005_1
muscle_164006_1
plasma_164386_1
plasma_164387_1
plasma_164388_1
plasma_164390_1
plasma_164391_1
plasma_164393_1
pup
adult
reads per million genome matching
bp position
0
10
20
30
40
50
60

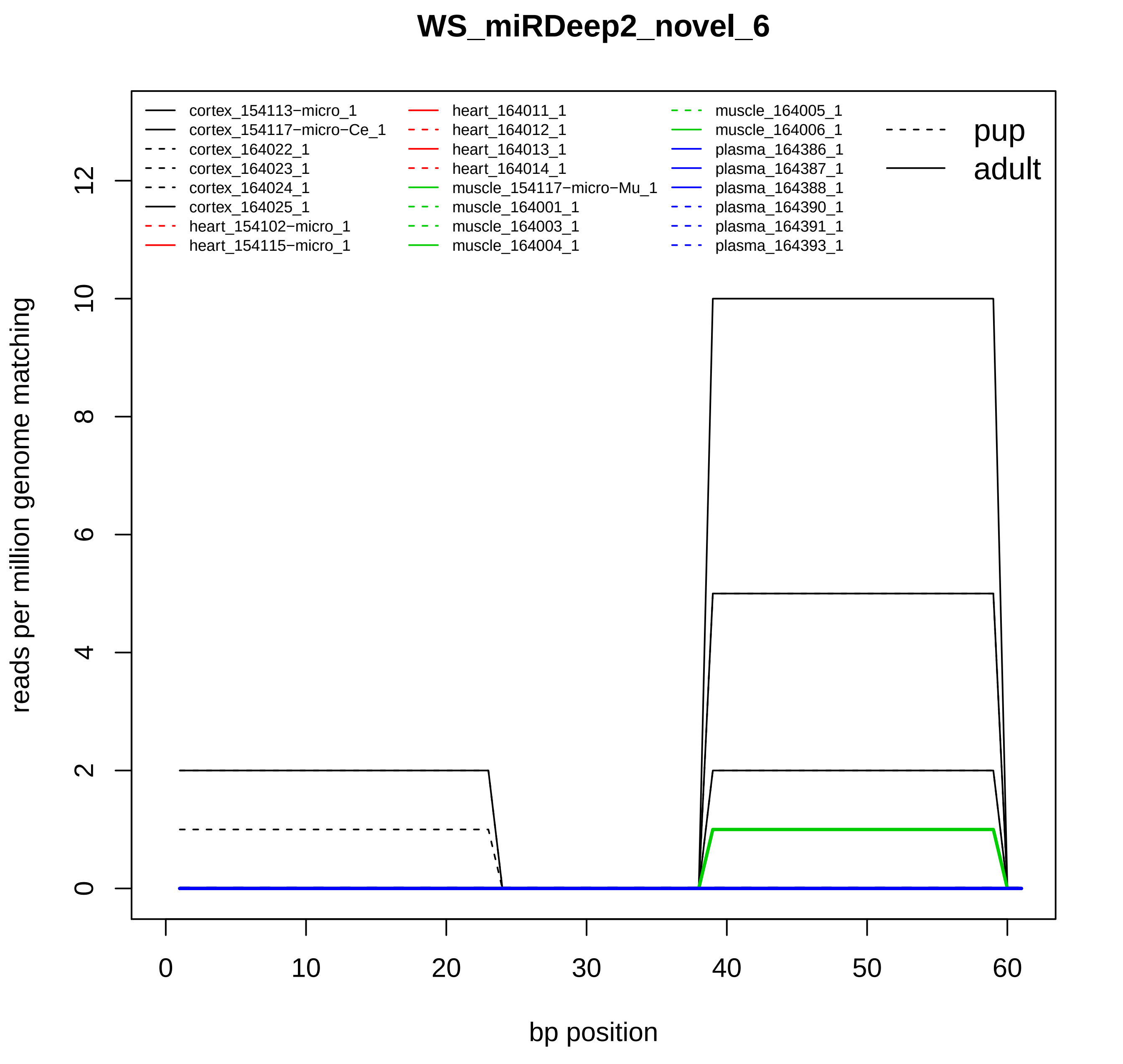
WS_miRDeep2_novel_6
cortex_154113−micro_1
cortex_154117−micro−Ce_1
cortex_164022_1
cortex_164023_1
cortex_164024_1
cortex_164025_1
heart_154102−micro_1
heart_154115−micro_1
heart_164011_1
heart_164012_1
heart_164013_1
heart_164014_1
muscle_154117−micro−Mu_1
muscle_164001_1
muscle_164003_1
muscle_164004_1
muscle_164005_1
muscle_164006_1
plasma_164386_1
plasma_164387_1
plasma_164388_1
plasma_164390_1
plasma_164391_1
plasma_164393_1
pup
adult
reads per million genome matching
0
2
4
6
8
10
12
0
10
20
30
40
50
60
bp position

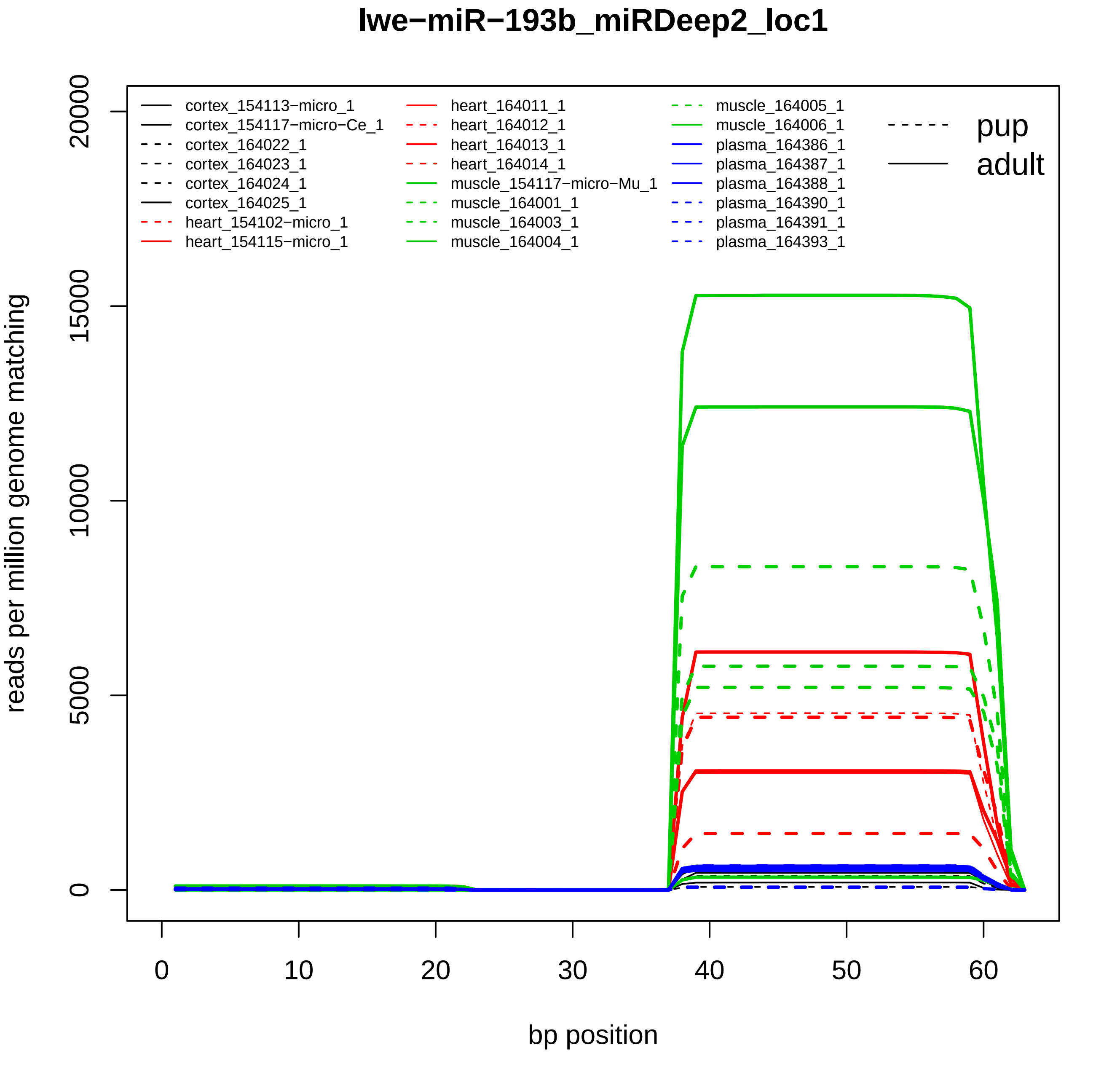
lwe−miR−193b_miRDeep2_loc1
reads per million genome matching
bp position
0
5000
10000
15000
20000
0
10
20
30
40
50
60
cortex_154113−micro_1
cortex_154117−micro−Ce_1
cortex_164022_1
cortex_164023_1
cortex_164024_1
cortex_164025_1
heart_154102−micro_1
heart_154115−micro_1
heart_164011_1
heart_164012_1
heart_164013_1
heart_164014_1
muscle_154117−micro−Mu_1
muscle_164001_1
muscle_164003_1
muscle_164004_1
muscle_164005_1
muscle_164006_1
plasma_164386_1
plasma_164387_1
plasma_164388_1
plasma_164390_1
plasma_164391_1
plasma_164393_1
pup
adult

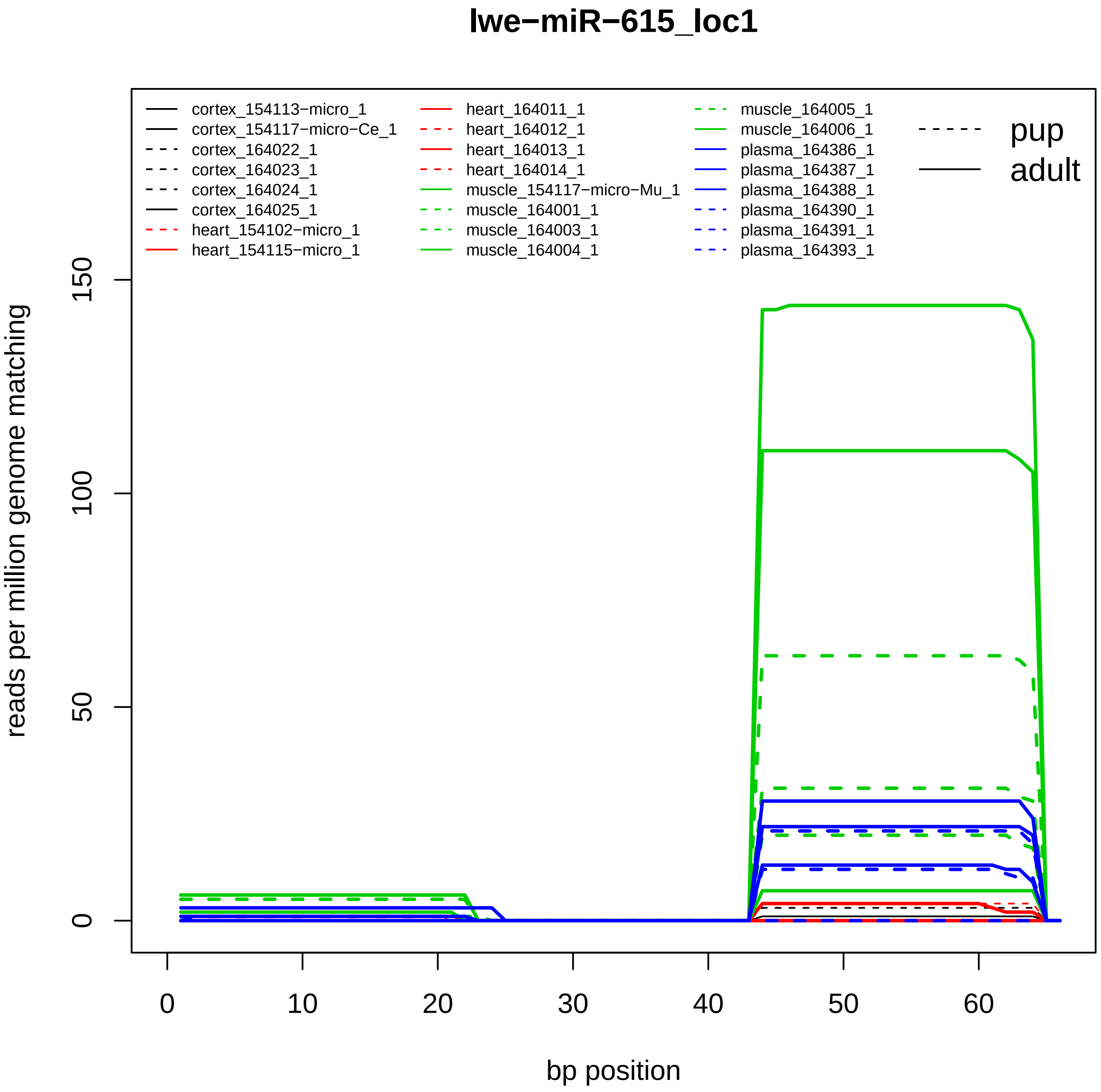
lwe−miR−615_loc1
cortex_154113−micro_1
cortex_154117−micro−Ce_1
cortex_164022_1
cortex_164023_1
cortex_164024_1
cortex_164025_1
heart_154102−micro_1
heart_154115−micro_1
heart_164011_1
heart_164012_1
heart_164013_1
heart_164014_1
muscle_154117−micro−Mu_1
muscle_164001_1
muscle_164003_1
muscle_164004_1
muscle_164005_1
muscle_164006_1
plasma_164386_1
plasma_164387_1
plasma_164388_1
plasma_164390_1
plasma_164391_1
plasma_164393_1
pup
adult
reads per million genome matching
0
50
100
150
0
10
20
30
40
50
60
bp position

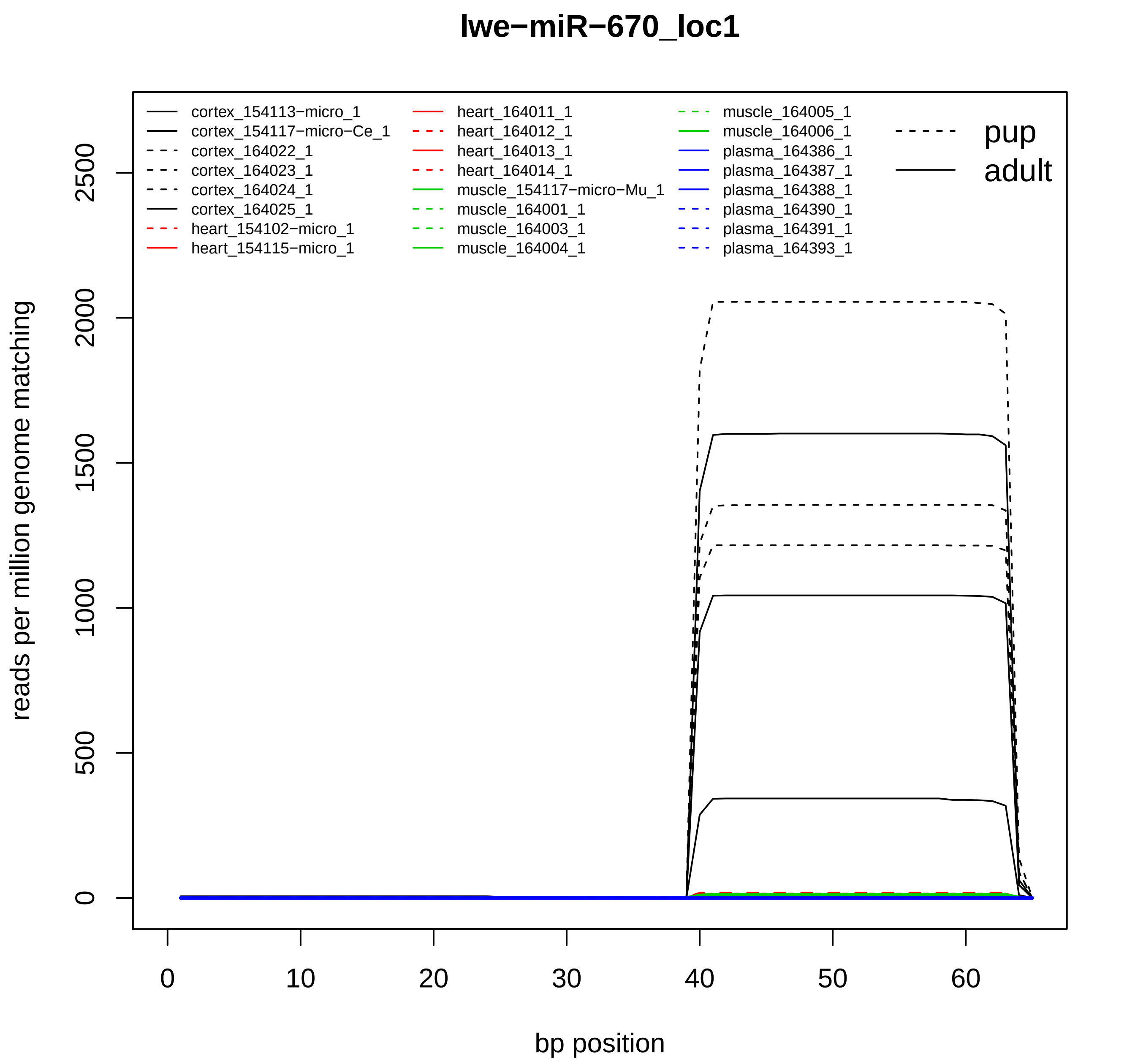

lwe−miR−670_loc1
cortex_154113−micro_1
cortex_154117−micro−Ce_1
cortex_164022_1
cortex_164023_1
cortex_164024_1
cortex_164025_1
heart_154102−micro_1
heart_154115−micro_1
heart_164011_1
heart_164012_1
heart_164013_1
heart_164014_1
muscle_154117−micro−Mu_1
muscle_164001_1
muscle_164003_1
muscle_164004_1
muscle_164005_1
muscle_164006_1
plasma_164386_1
plasma_164387_1
plasma_164388_1
plasma_164390_1
plasma_164391_1
plasma_164393_1
pup
adult
reads per million genome matching
0
500
1000
1500
2000
2500
0
10
20
30
40
50
60
bp position

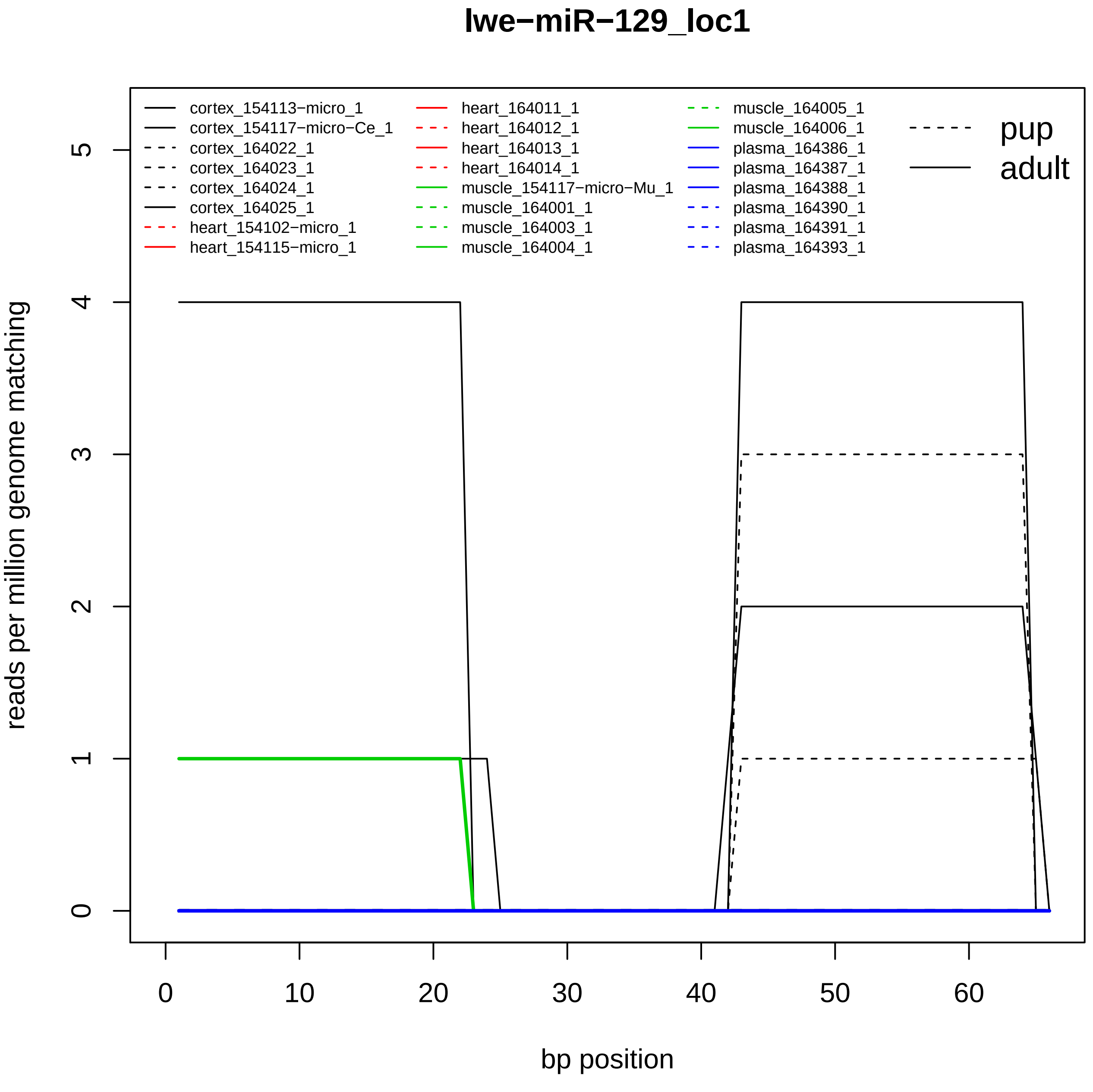
lwe−miR−129_loc1
cortex_154113−micro_1
cortex_154117−micro−Ce_1
cortex_164022_1
cortex_164023_1
cortex_164024_1
cortex_164025_1
heart_154102−micro_1
heart_154115−micro_1
heart_164011_1
heart_164012_1
heart_164013_1
heart_164014_1
muscle_154117−micro−Mu_1
muscle_164001_1
muscle_164003_1
muscle_164004_1
muscle_164005_1
muscle_164006_1
plasma_164386_1
plasma_164387_1
plasma_164388_1
plasma_164390_1
plasma_164391_1
plasma_164393_1
pup
adult
reads per million genome matching
0
1
2
3
4
5
0
10
20
30
40
50
60
bp position

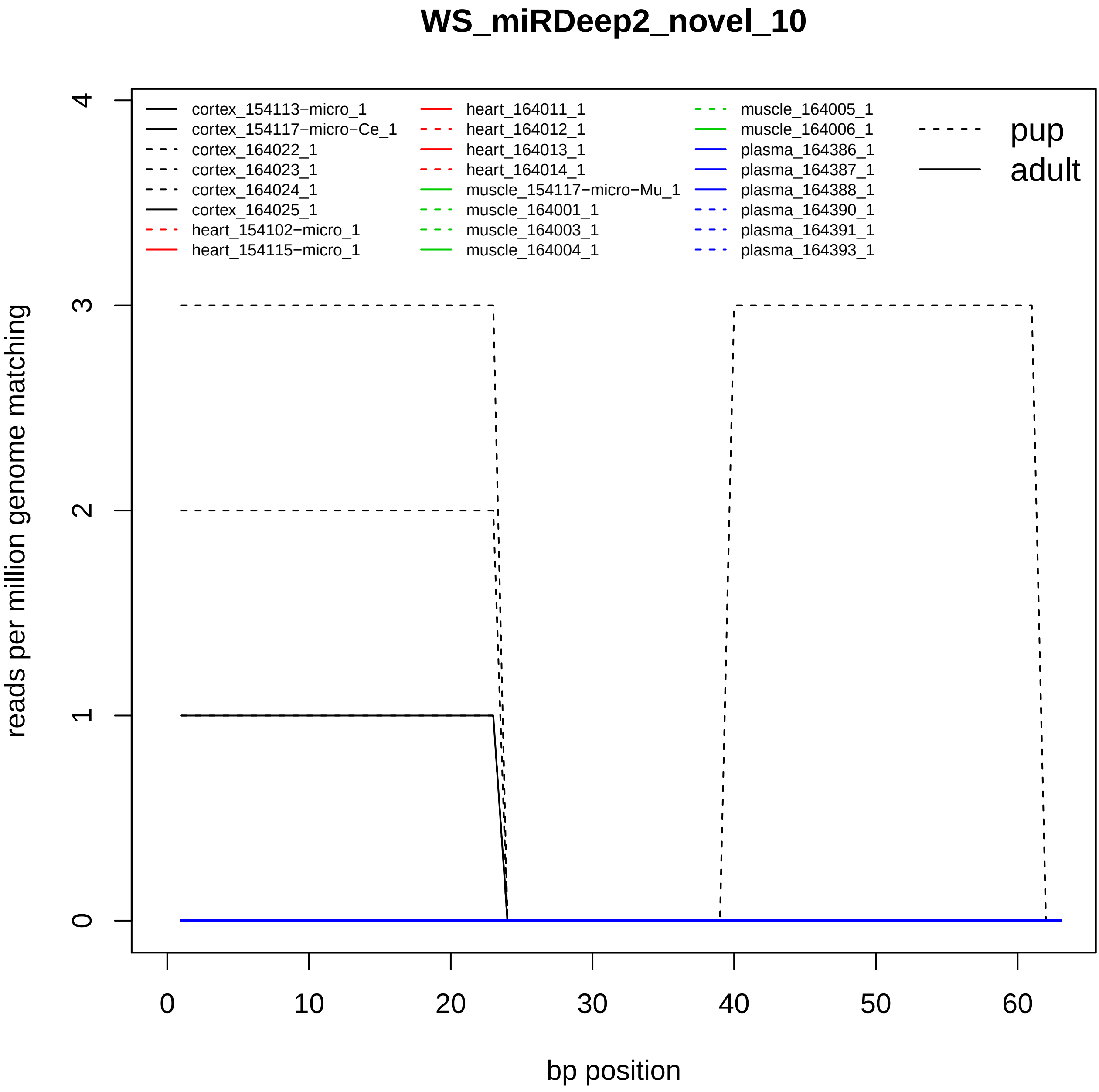

WS_miRDeep2_novel_10
cortex_154113−micro_1
cortex_154117−micro−Ce_1
cortex_164022_1
cortex_164023_1
cortex_164024_1
cortex_164025_1
heart_154102−micro_1
heart_154115−micro_1
heart_164011_1
heart_164012_1
heart_164013_1
heart_164014_1
muscle_154117−micro−Mu_1
muscle_164001_1
muscle_164003_1
muscle_164004_1
muscle_164005_1
muscle_164006_1
plasma_164386_1
plasma_164387_1
plasma_164388_1
plasma_164390_1
plasma_164391_1
plasma_164393_1
pup
adult
reads per million genome matching
bp position
0
1
2
3
4
0
10
20
30
40
50
60

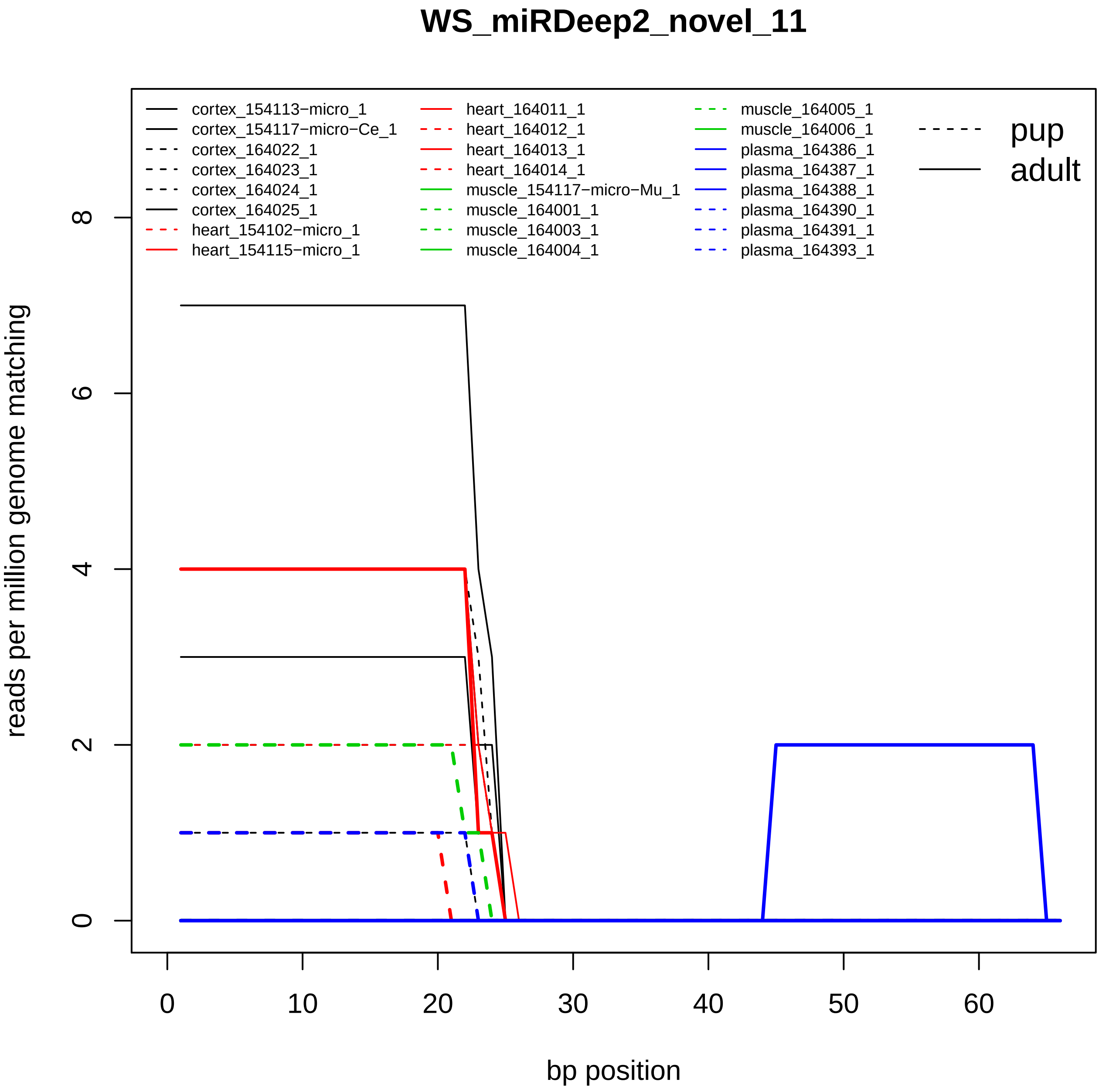

WS_miRDeep2_novel_11
cortex_154113−micro_1
cortex_154117−micro−Ce_1
cortex_164022_1
cortex_164023_1
cortex_164024_1
cortex_164025_1
heart_154102−micro_1
heart_154115−micro_1
heart_164011_1
heart_164012_1
heart_164013_1
heart_164014_1
muscle_154117−micro−Mu_1
muscle_164001_1
muscle_164003_1
muscle_164004_1
muscle_164005_1
muscle_164006_1
plasma_164386_1
plasma_164387_1
plasma_164388_1
plasma_164390_1
plasma_164391_1
plasma_164393_1
pup
adult
reads per million genome matching
0
2
4
6
8
0
10
20
30
40
50
60
bp position

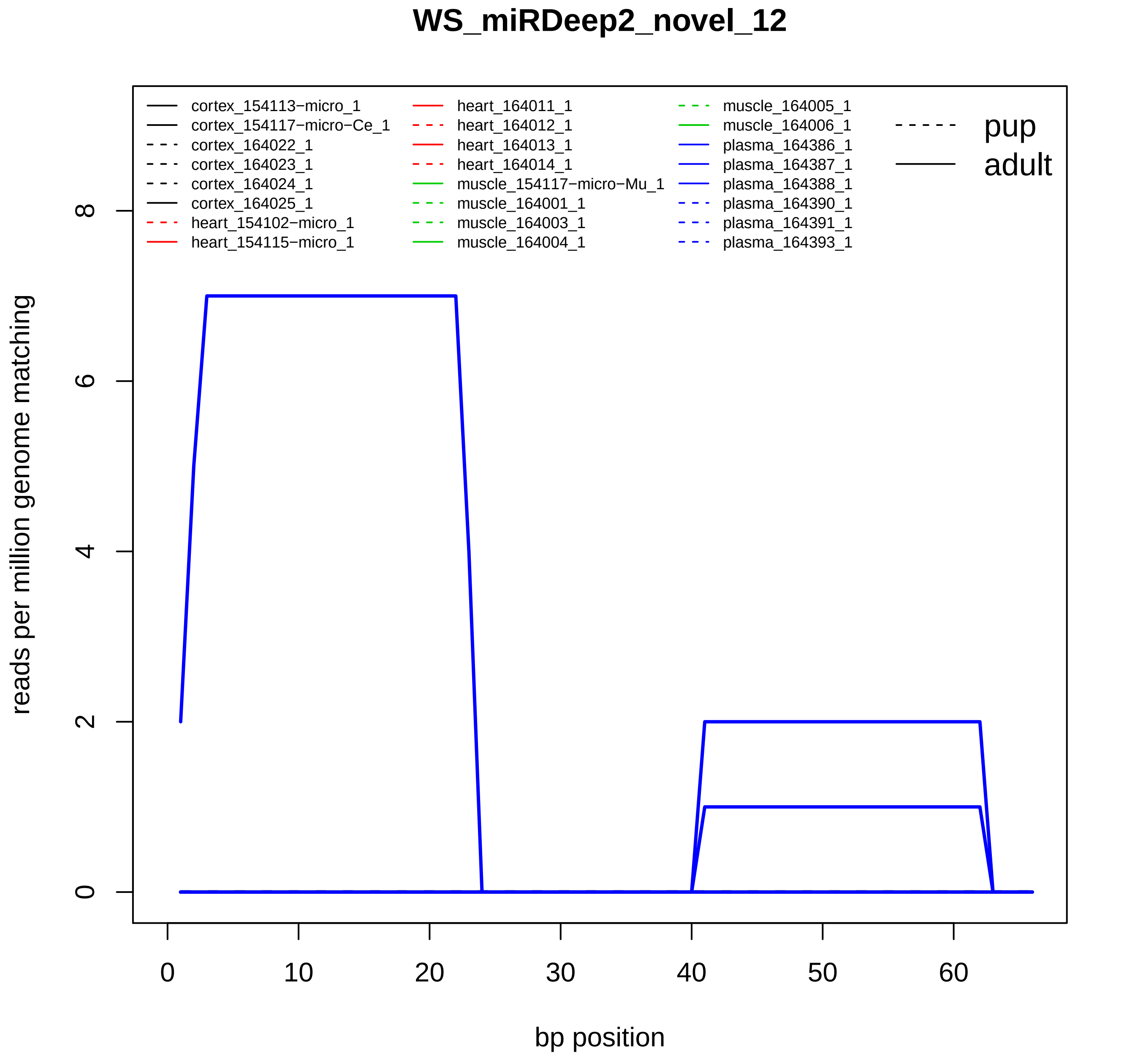

WS_miRDeep2_novel_12
cortex_154113−micro_1
cortex_154117−micro−Ce_1
cortex_164022_1
cortex_164023_1
cortex_164024_1
cortex_164025_1
heart_154102−micro_1
heart_154115−micro_1
heart_164011_1
heart_164012_1
heart_164013_1
heart_164014_1
muscle_154117−micro−Mu_1
muscle_164001_1
muscle_164003_1
muscle_164004_1
muscle_164005_1
muscle_164006_1
plasma_164386_1
plasma_164387_1
plasma_164388_1
plasma_164390_1
plasma_164391_1
plasma_164393_1
pup
adult
reads per million genome matching
0
2
4
6
8
0
10
20
30
40
50
60
bp position

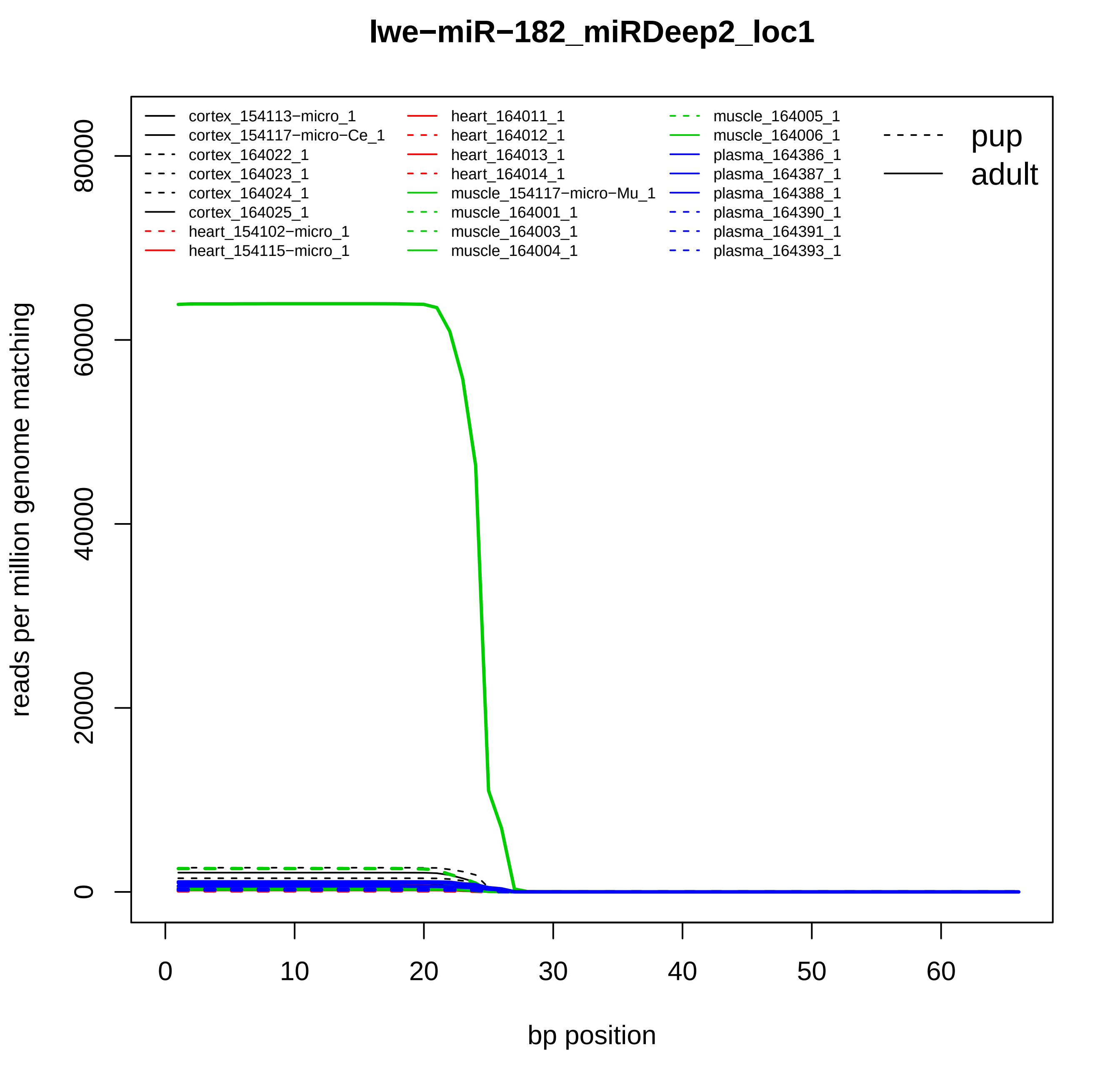
lwe−miR−182_miRDeep2_loc1
cortex_154113−micro_1
cortex_154117−micro−Ce_1
cortex_164022_1
cortex_164023_1
cortex_164024_1
cortex_164025_1
heart_154102−micro_1
heart_154115−micro_1
heart_164011_1
heart_164012_1
heart_164013_1
heart_164014_1
muscle_154117−micro−Mu_1
muscle_164001_1
muscle_164003_1
muscle_164004_1
muscle_164005_1
muscle_164006_1
plasma_164386_1
plasma_164387_1
plasma_164388_1
plasma_164390_1
plasma_164391_1
plasma_164393_1
pup
adult
reads per million genome matching
0
20000
40000
60000
80000
bp position
0
10
20
30
40
50
60

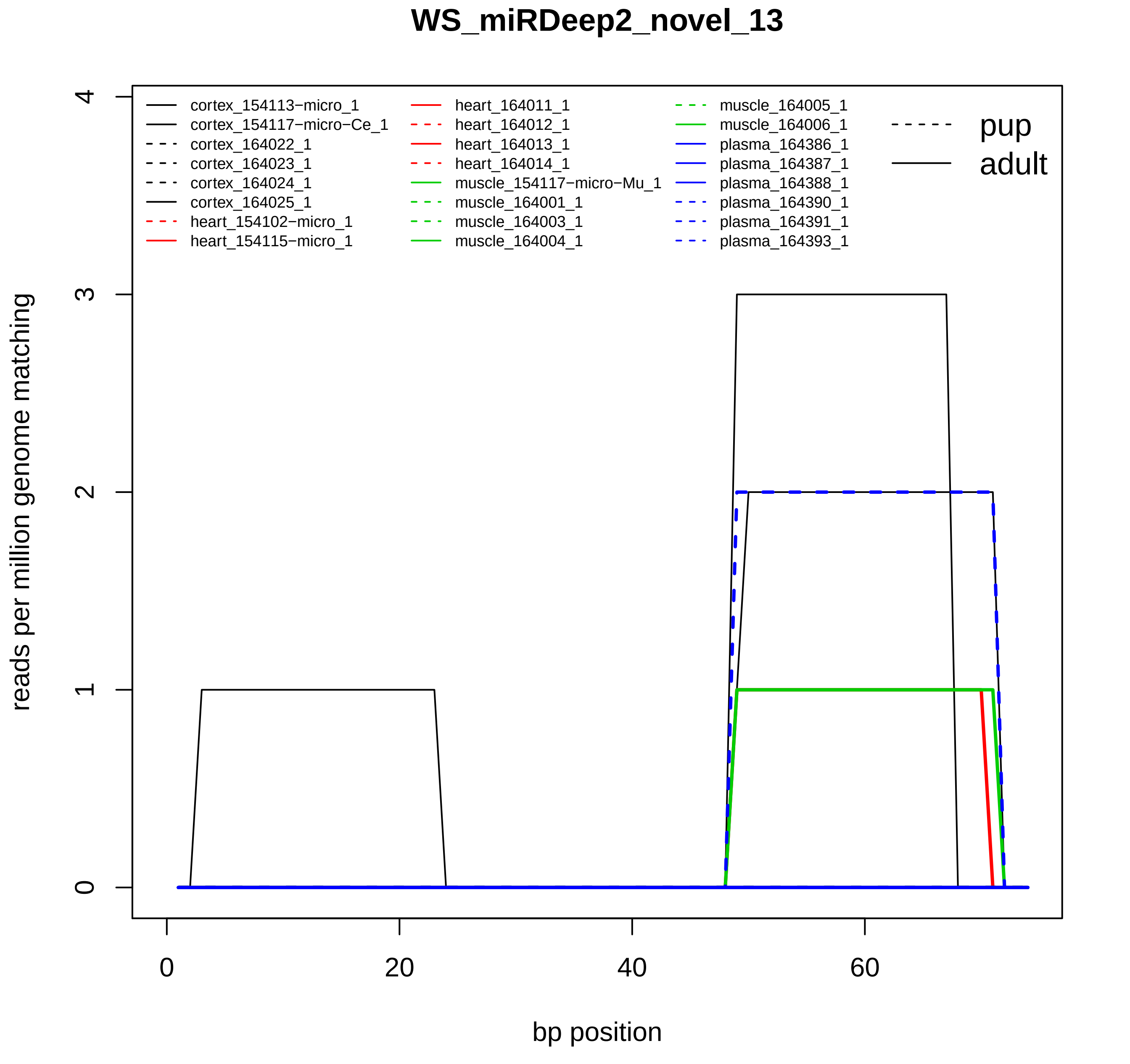
WS_miRDeep2_novel_13
reads per million genome matching
bp position
0
1
2
3
4
0
20
40
60
cortex_154113−micro_1
cortex_154117−micro−Ce_1
cortex_164022_1
cortex_164023_1
cortex_164024_1
cortex_164025_1
heart_154102−micro_1
heart_154115−micro_1
heart_164011_1
heart_164012_1
heart_164013_1
heart_164014_1
muscle_154117−micro−Mu_1
muscle_164001_1
muscle_164003_1
muscle_164004_1
muscle_164005_1
muscle_164006_1
plasma_164386_1
plasma_164387_1
plasma_164388_1
plasma_164390_1
plasma_164391_1
plasma_164393_1
pup
adult

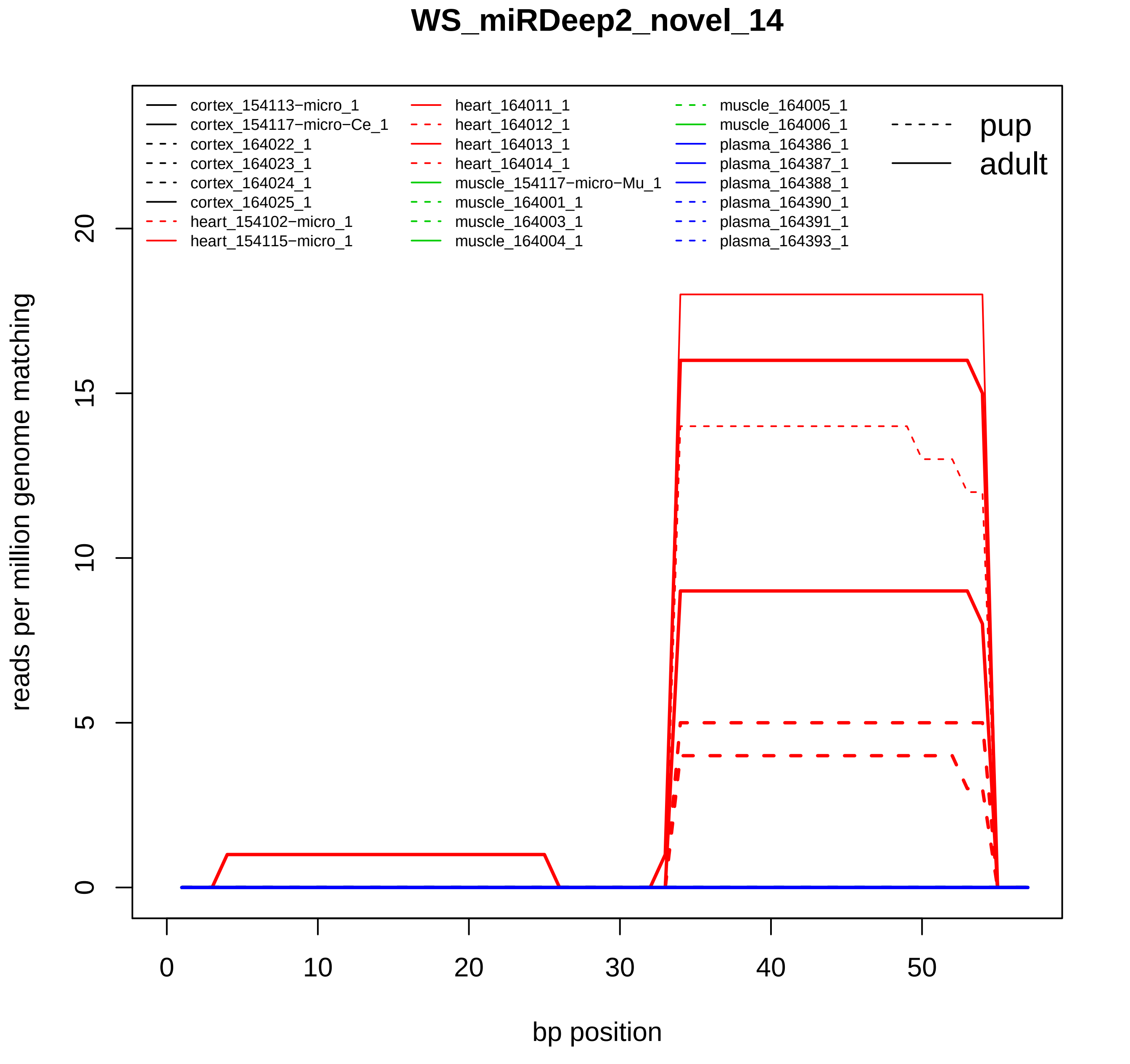
WS_miRDeep2_novel_14
cortex_154113−micro_1
cortex_154117−micro−Ce_1
cortex_164022_1
cortex_164023_1
cortex_164024_1
cortex_164025_1
heart_154102−micro_1
heart_154115−micro_1
heart_164011_1
heart_164012_1
heart_164013_1
heart_164014_1
muscle_154117−micro−Mu_1
muscle_164001_1
muscle_164003_1
muscle_164004_1
muscle_164005_1
muscle_164006_1
plasma_164386_1
plasma_164387_1
plasma_164388_1
plasma_164390_1
plasma_164391_1
plasma_164393_1
pup
adult
reads per million genome matching
0
5
10
15
20
0
10
20
30
40
50
bp position

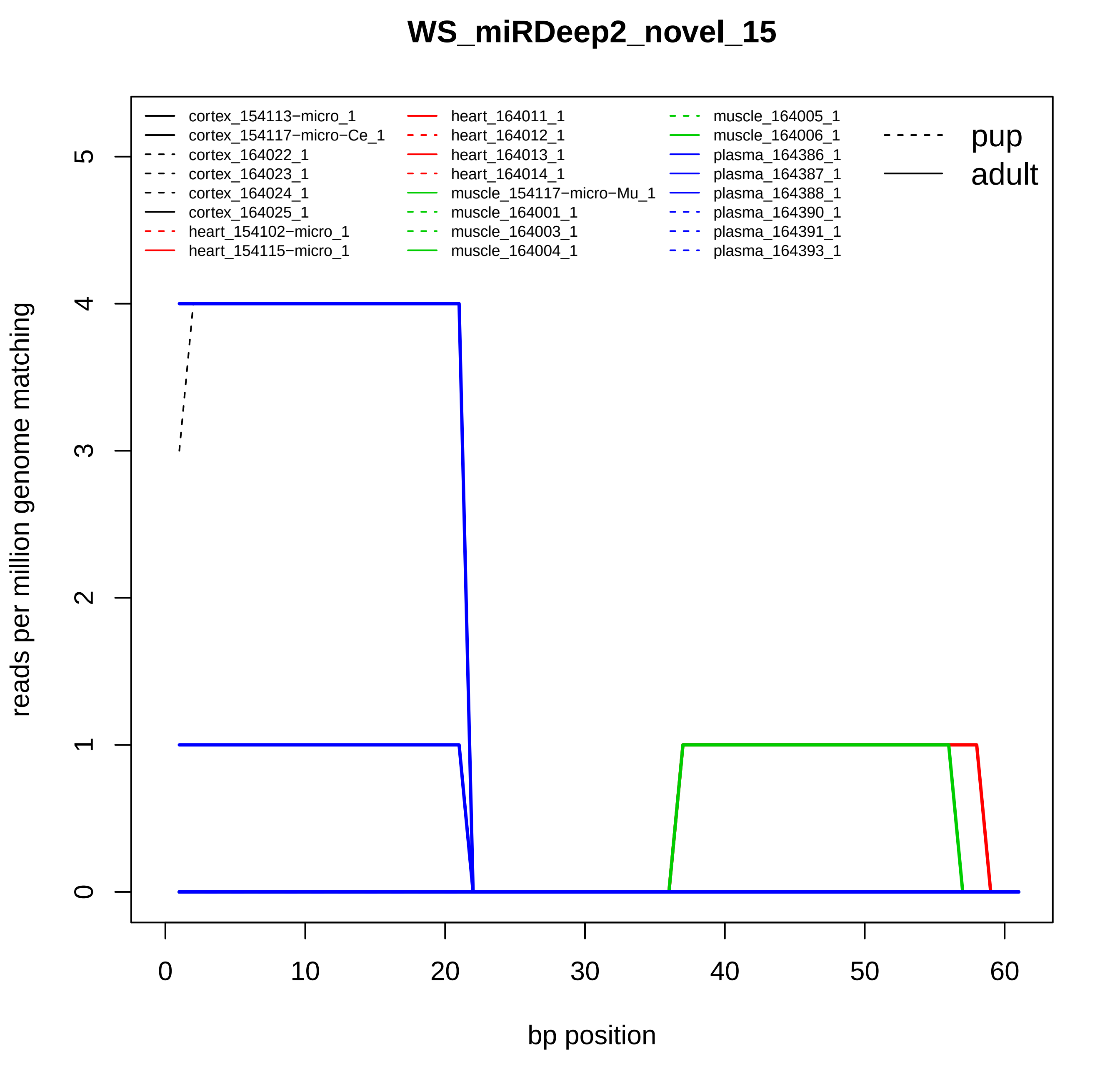

WS_miRDeep2_novel_15
cortex_154113−micro_1
cortex_154117−micro−Ce_1
cortex_164022_1
cortex_164023_1
cortex_164024_1
cortex_164025_1
heart_154102−micro_1
heart_154115−micro_1
heart_164011_1
heart_164012_1
heart_164013_1
heart_164014_1
muscle_154117−micro−Mu_1
muscle_164001_1
muscle_164003_1
muscle_164004_1
muscle_164005_1
muscle_164006_1
plasma_164386_1
plasma_164387_1
plasma_164388_1
plasma_164390_1
plasma_164391_1
plasma_164393_1
pup
adult
reads per million genome matching
0
1
2
3
4
5
0
10
20
30
40
50
60
bp position

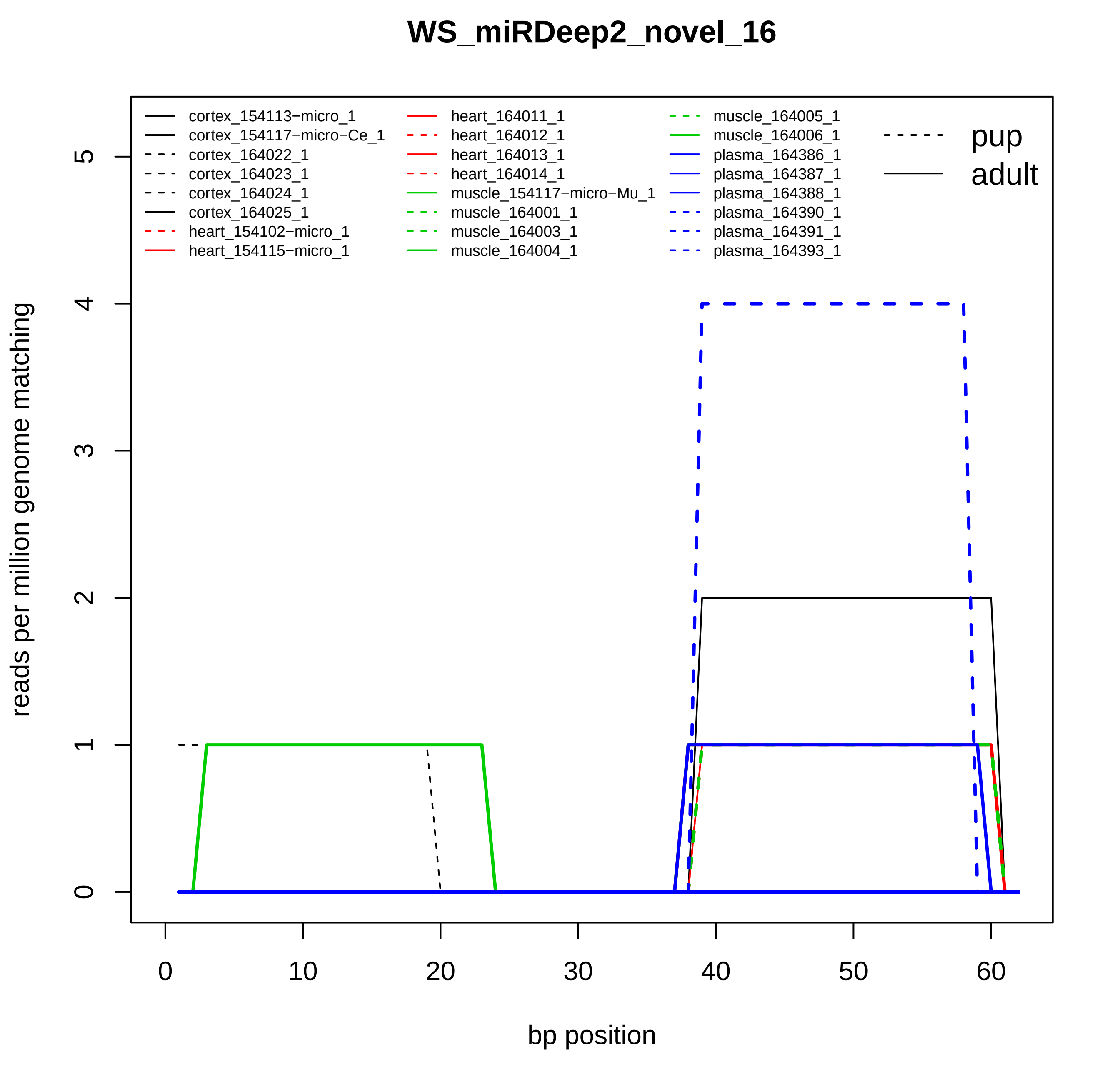
WS_miRDeep2_novel_16
cortex_154113−micro_1
cortex_154117−micro−Ce_1
cortex_164022_1
cortex_164023_1
cortex_164024_1
cortex_164025_1
heart_154102−micro_1
heart_154115−micro_1
heart_164011_1
heart_164012_1
heart_164013_1
heart_164014_1
muscle_154117−micro−Mu_1
muscle_164001_1
muscle_164003_1
muscle_164004_1
muscle_164005_1
muscle_164006_1
plasma_164386_1
plasma_164387_1
plasma_164388_1
plasma_164390_1
plasma_164391_1
plasma_164393_1
pup
adult
reads per million genome matching
0
1
2
3
4
5
0
10
20
30
40
50
60
bp position

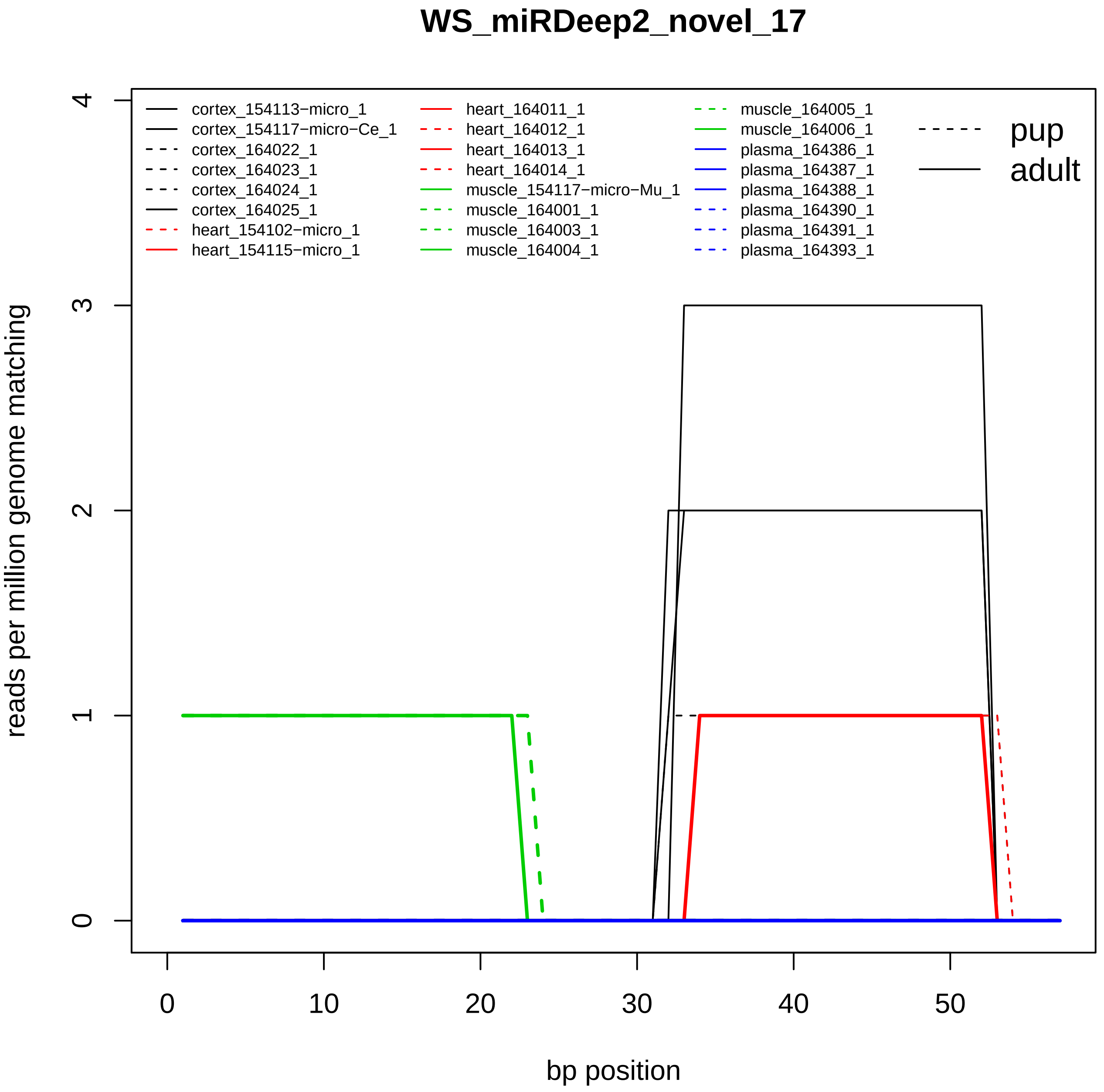

WS_miRDeep2_novel_17
cortex_154113−micro_1
cortex_154117−micro−Ce_1
cortex_164022_1
cortex_164023_1
cortex_164024_1
cortex_164025_1
heart_154102−micro_1
heart_154115−micro_1
heart_164011_1
heart_164012_1
heart_164013_1
heart_164014_1
muscle_154117−micro−Mu_1
muscle_164001_1
muscle_164003_1
muscle_164004_1
muscle_164005_1
muscle_164006_1
plasma_164386_1
plasma_164387_1
plasma_164388_1
plasma_164390_1
plasma_164391_1
plasma_164393_1
pup
adult
reads per million genome matching
0
1
2
3
4
bp position
0
10
20
30
40
50

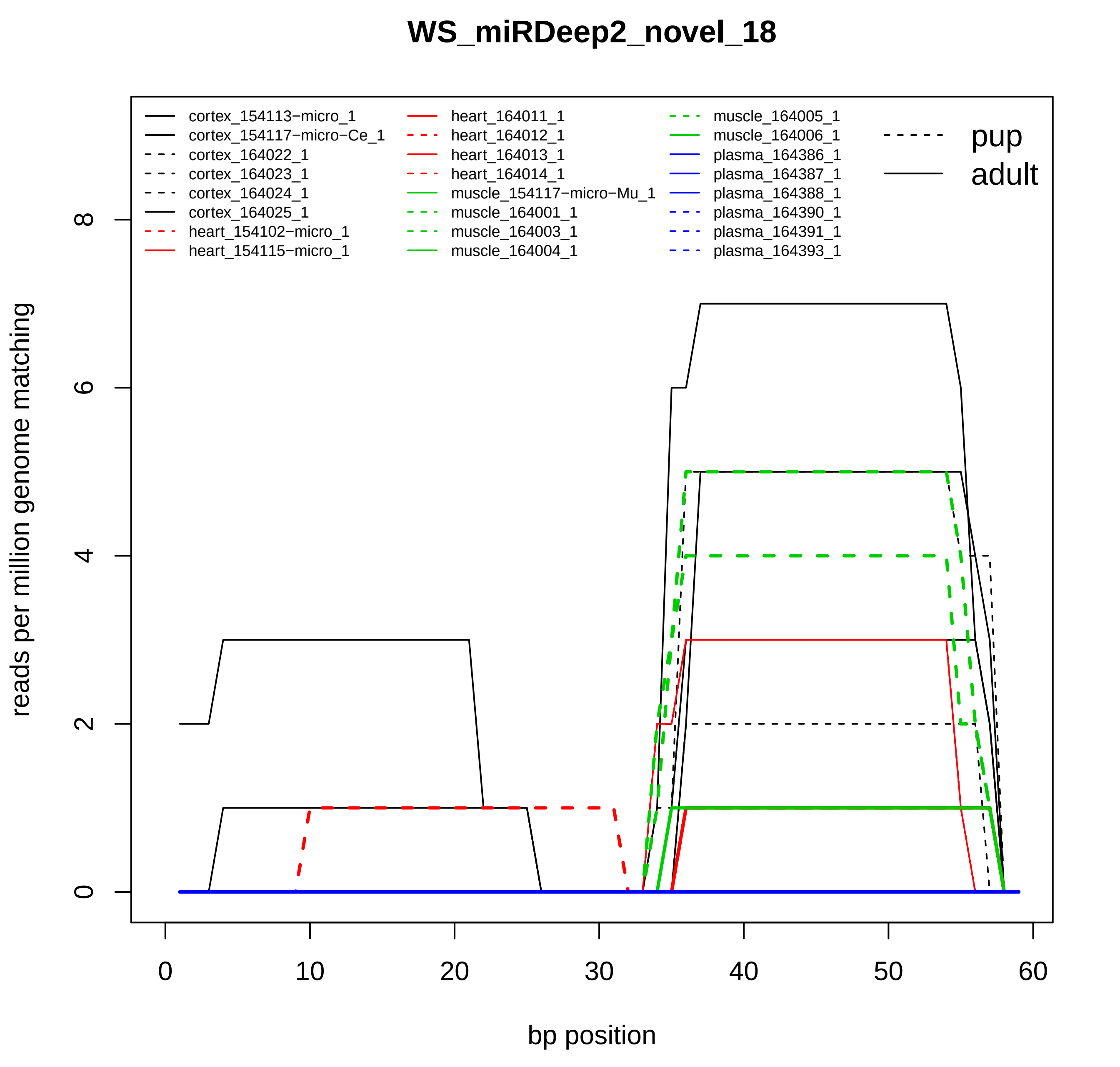
WS_miRDeep2_novel_18
cortex_154113−micro_1
cortex_154117−micro−Ce_1
cortex_164022_1
cortex_164023_1
cortex_164024_1
cortex_164025_1
heart_154102−micro_1
heart_154115−micro_1
heart_164011_1
heart_164012_1
heart_164013_1
heart_164014_1
muscle_154117−micro−Mu_1
muscle_164001_1
muscle_164003_1
muscle_164004_1
muscle_164005_1
muscle_164006_1
plasma_164386_1
plasma_164387_1
plasma_164388_1
plasma_164390_1
plasma_164391_1
plasma_164393_1
pup
adult
reads per million genome matching
0
2
4
6
8
bp position
0
10
20
30
40
50
60

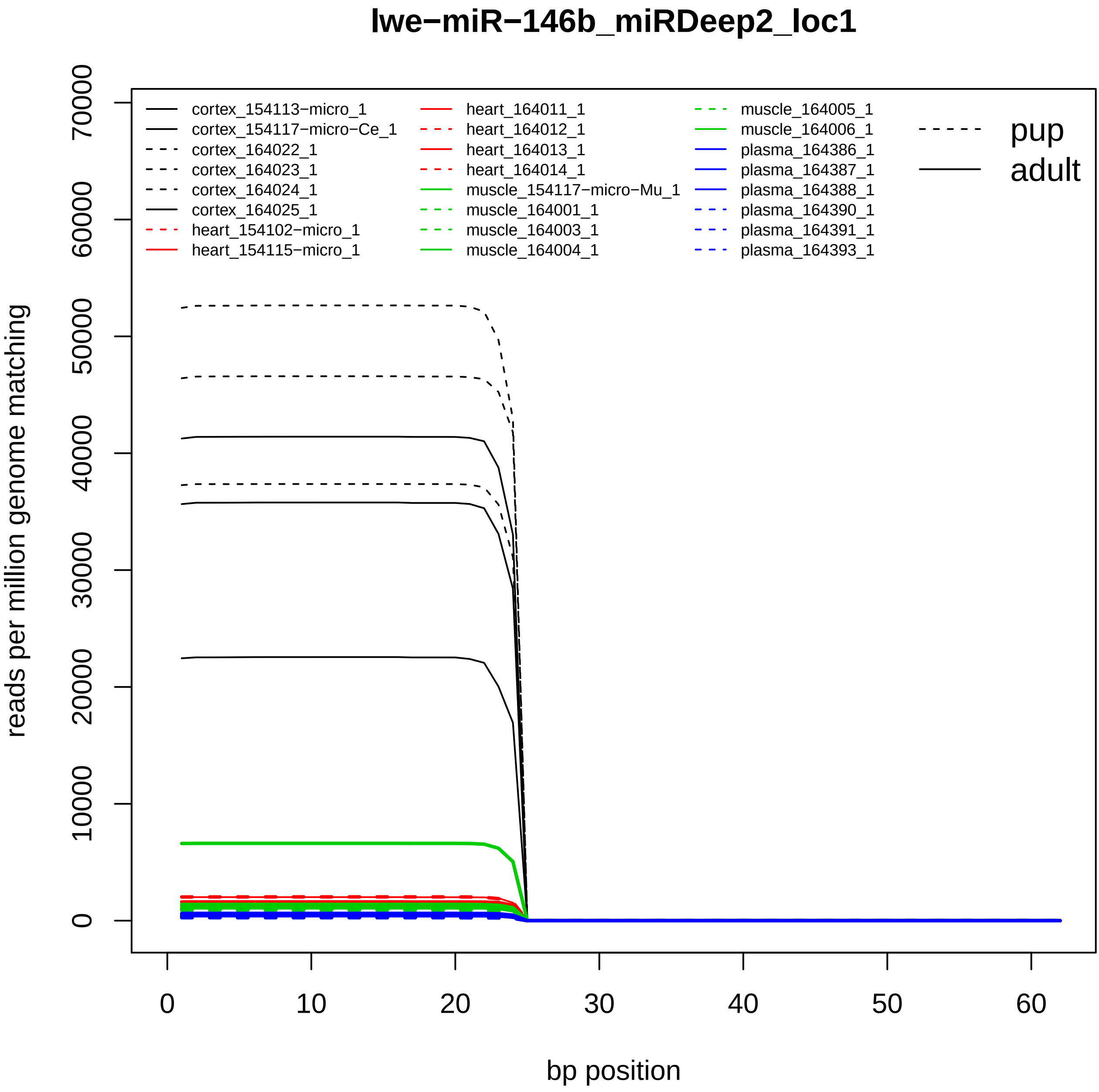

lwe−miR−146b_miRDeep2_loc1
reads per million genome matching
bp position
0
10000
20000
30000
40000
50000
60000
70000
0
10
20
30
40
50
60
cortex_154113−micro_1
cortex_154117−micro−Ce_1
cortex_164022_1
cortex_164023_1
cortex_164024_1
cortex_164025_1
heart_154102−micro_1
heart_154115−micro_1
heart_164011_1
heart_164012_1
heart_164013_1
heart_164014_1
muscle_154117−micro−Mu_1
muscle_164001_1
muscle_164003_1
muscle_164004_1
muscle_164005_1
muscle_164006_1
plasma_164386_1
plasma_164387_1
plasma_164388_1
plasma_164390_1
plasma_164391_1
plasma_164393_1
pup
adult

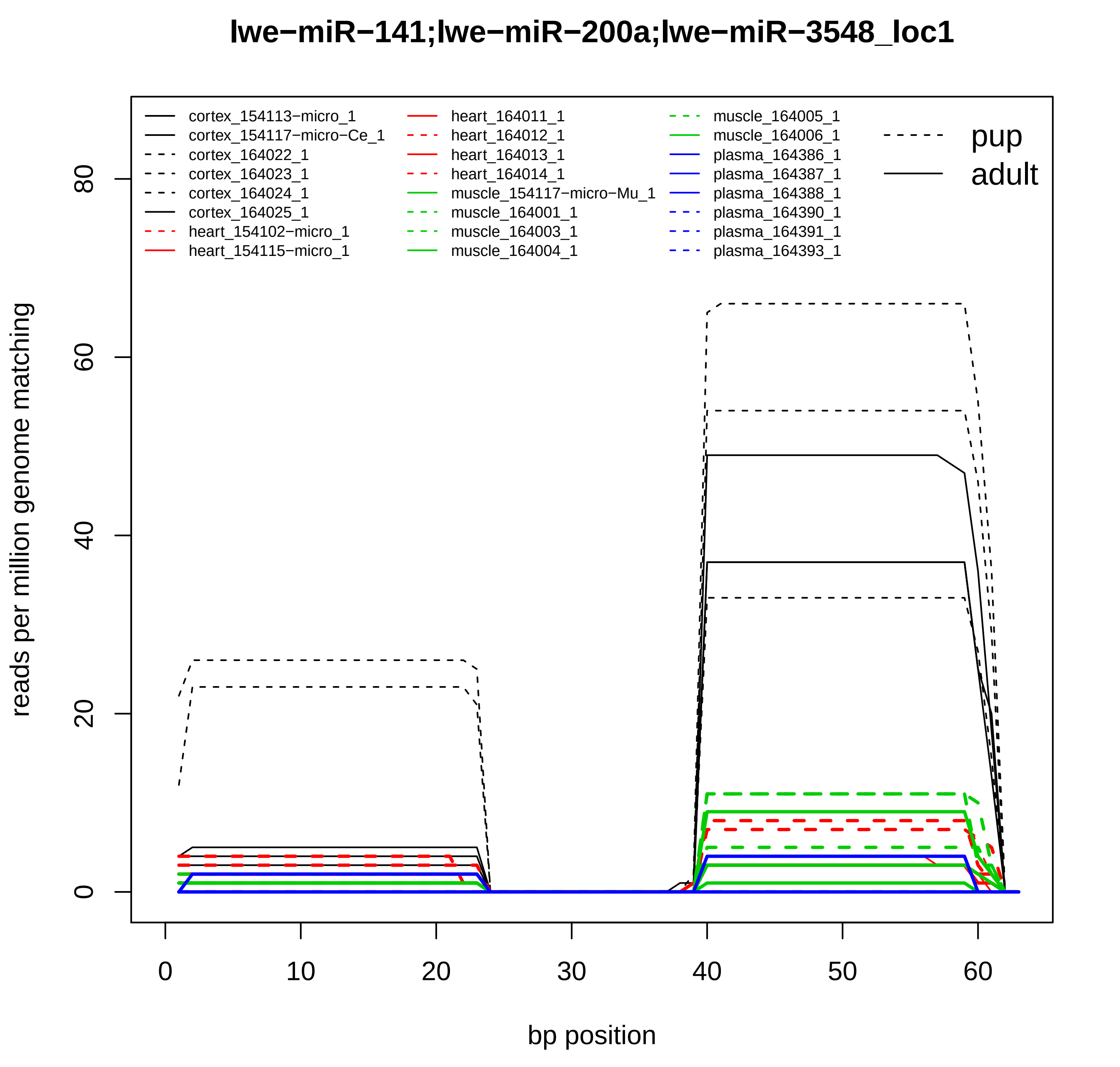
lwe−miR−141;lwe−miR−200a;lwe−miR−3548_loc1
cortex_154113−micro_1
cortex_154117−micro−Ce_1
cortex_164022_1
cortex_164023_1
cortex_164024_1
cortex_164025_1
heart_154102−micro_1
heart_154115−micro_1
heart_164011_1
heart_164012_1
heart_164013_1
heart_164014_1
muscle_154117−micro−Mu_1
muscle_164001_1
muscle_164003_1
muscle_164004_1
muscle_164005_1
muscle_164006_1
plasma_164386_1
plasma_164387_1
plasma_164388_1
plasma_164390_1
plasma_164391_1
plasma_164393_1
pup
adult
reads per million genome matching
0
20
40
60
80
0
10
20
30
40
50
60
bp position

### lwe−miR−302a_miRDeep2_loc1

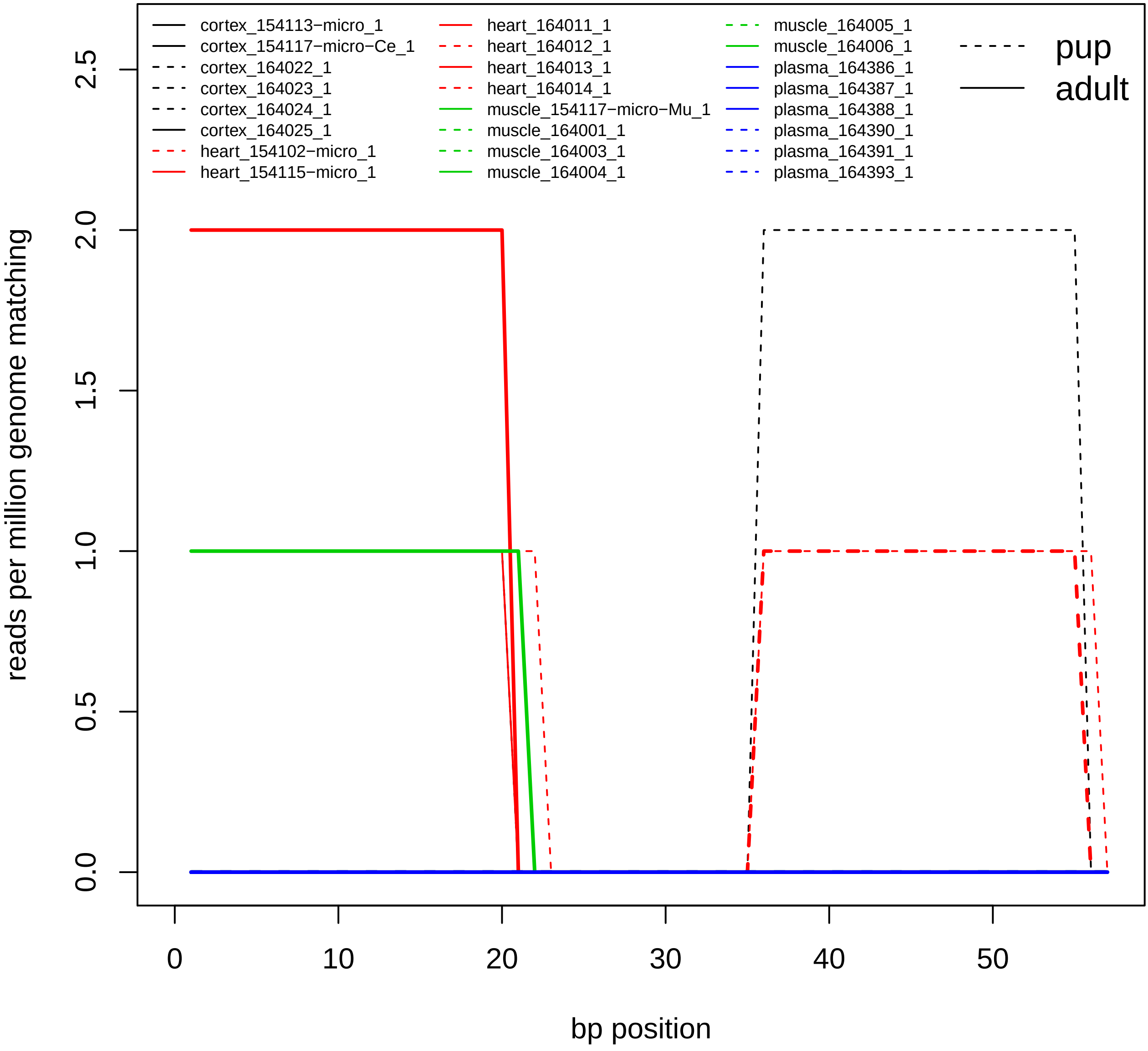

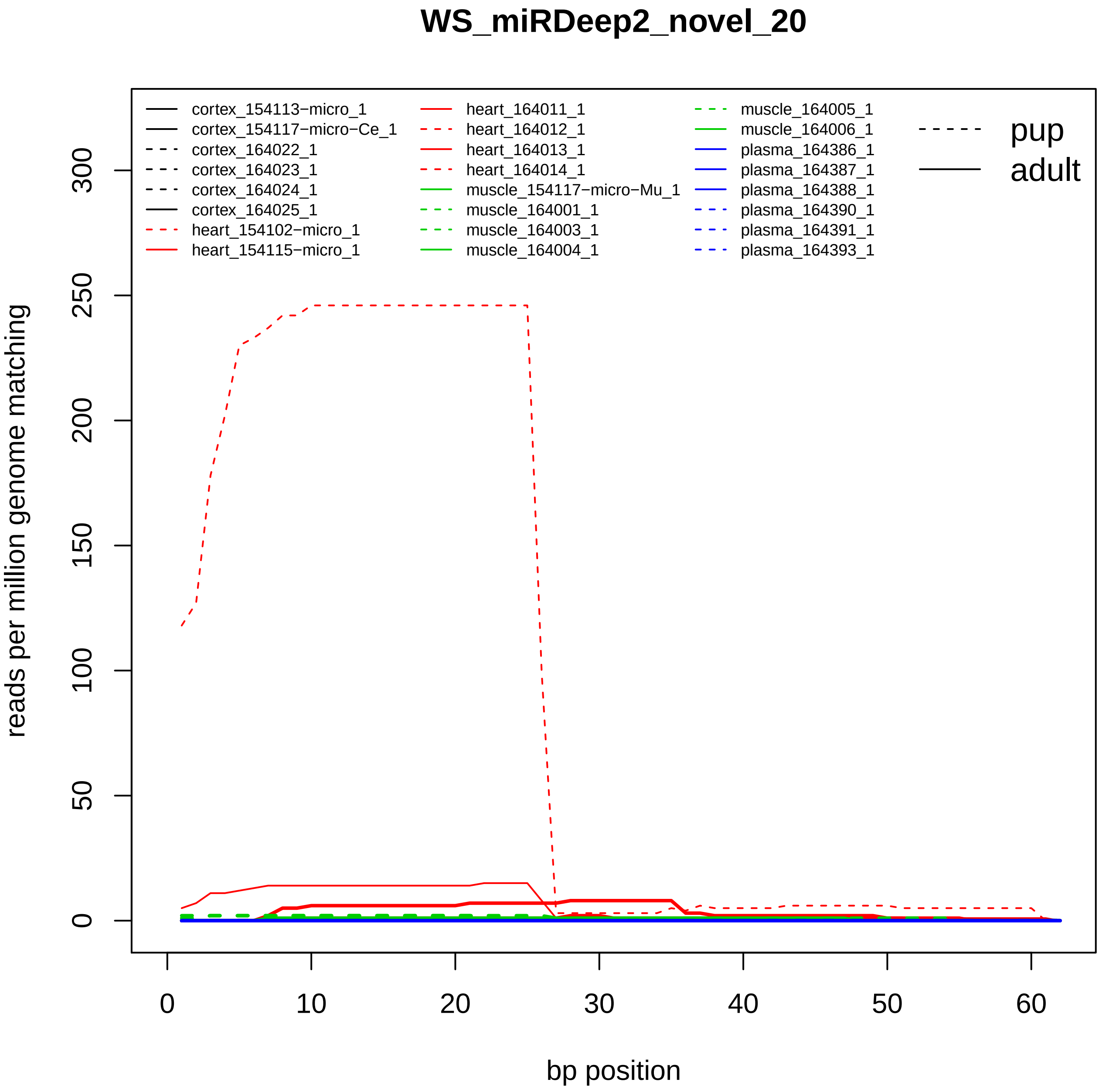

WS_miRDeep2_novel_20
cortex_154113−micro_1
cortex_154117−micro−Ce_1
cortex_164022_1
cortex_164023_1
cortex_164024_1
cortex_164025_1
heart_154102−micro_1
heart_154115−micro_1
heart_164011_1
heart_164012_1
heart_164013_1
heart_164014_1
muscle_154117−micro−Mu_1
muscle_164001_1
muscle_164003_1
muscle_164004_1
muscle_164005_1
muscle_164006_1
plasma_164386_1
plasma_164387_1
plasma_164388_1
plasma_164390_1
plasma_164391_1
plasma_164393_1
pup
adult
reads per million genome matching
0
50
100
150
200
250
300
bp position
0
10
20
30
40
50
60

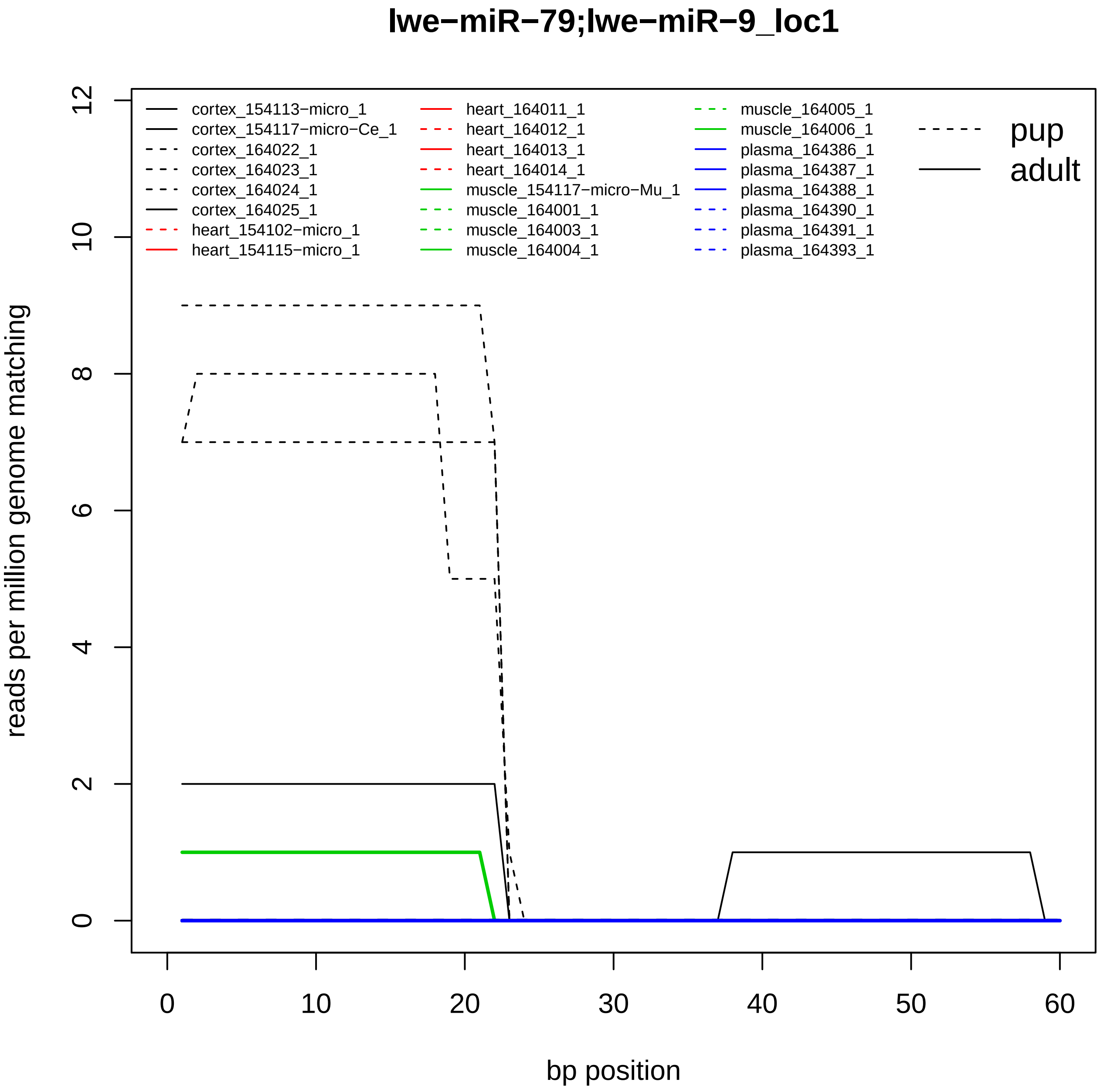

lwe−miR−79;lwe−miR−9_loc1
cortex_154113−micro_1
cortex_154117−micro−Ce_1
cortex_164022_1
cortex_164023_1
cortex_164024_1
cortex_164025_1
heart_154102−micro_1
heart_154115−micro_1
heart_164011_1
heart_164012_1
heart_164013_1
heart_164014_1
muscle_154117−micro−Mu_1
muscle_164001_1
muscle_164003_1
muscle_164004_1
muscle_164005_1
muscle_164006_1
plasma_164386_1
plasma_164387_1
plasma_164388_1
plasma_164390_1
plasma_164391_1
plasma_164393_1
pup
adult
reads per million genome matching
bp position
0
2
4
6
8
10
12
0
10
20
30
40
50
60

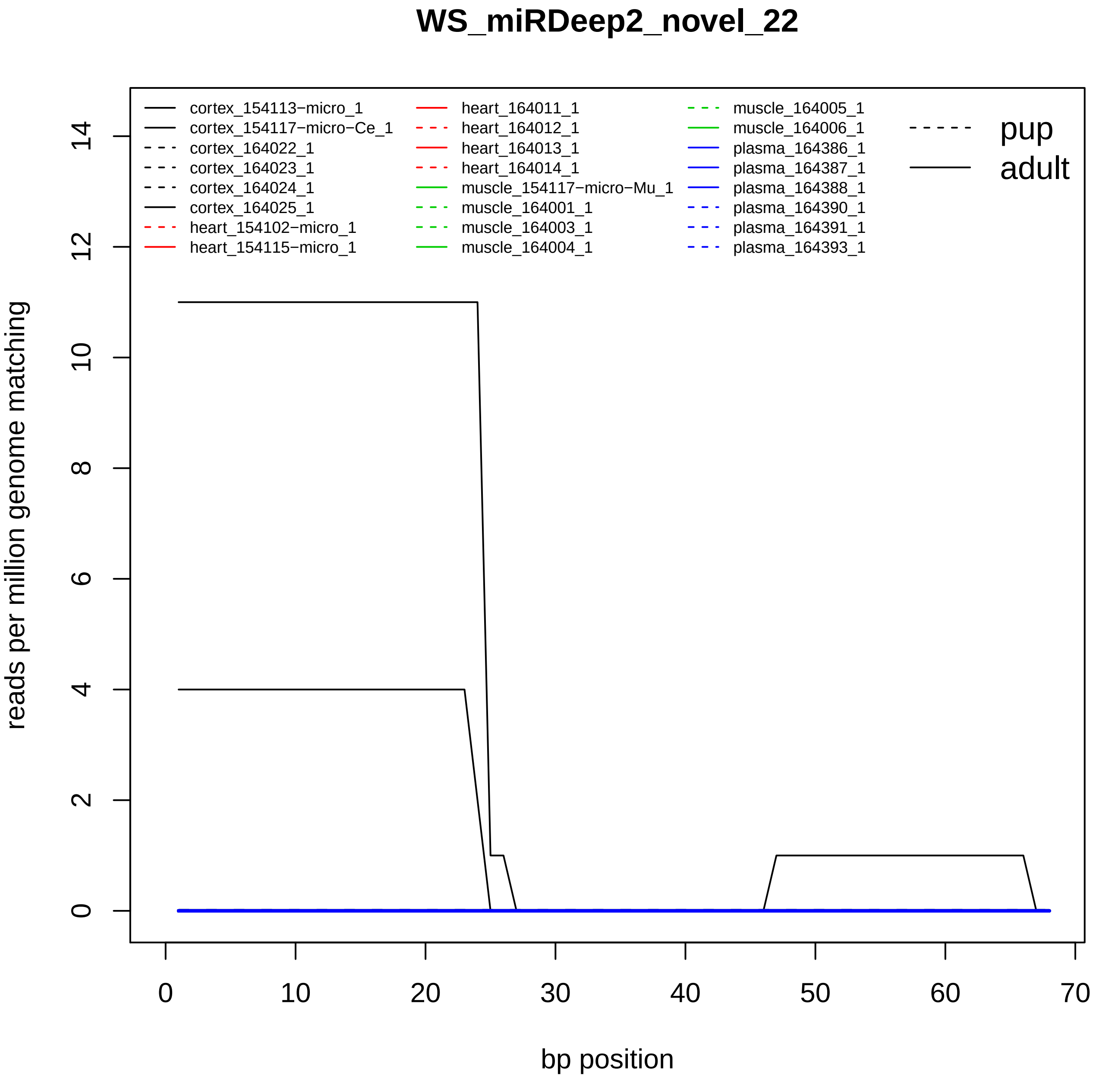

WS_miRDeep2_novel_22
cortex_154113−micro_1
cortex_154117−micro−Ce_1
cortex_164022_1
cortex_164023_1
cortex_164024_1
cortex_164025_1
heart_154102−micro_1
heart_154115−micro_1
heart_164011_1
heart_164012_1
heart_164013_1
heart_164014_1
muscle_154117−micro−Mu_1
muscle_164001_1
muscle_164003_1
muscle_164004_1
muscle_164005_1
muscle_164006_1
plasma_164386_1
plasma_164387_1
plasma_164388_1
plasma_164390_1
plasma_164391_1
plasma_164393_1
pup
adult
reads per million genome matching
0
2
4
6
8
10
12
14
bp position
0
10
20
30
40
50
60
70

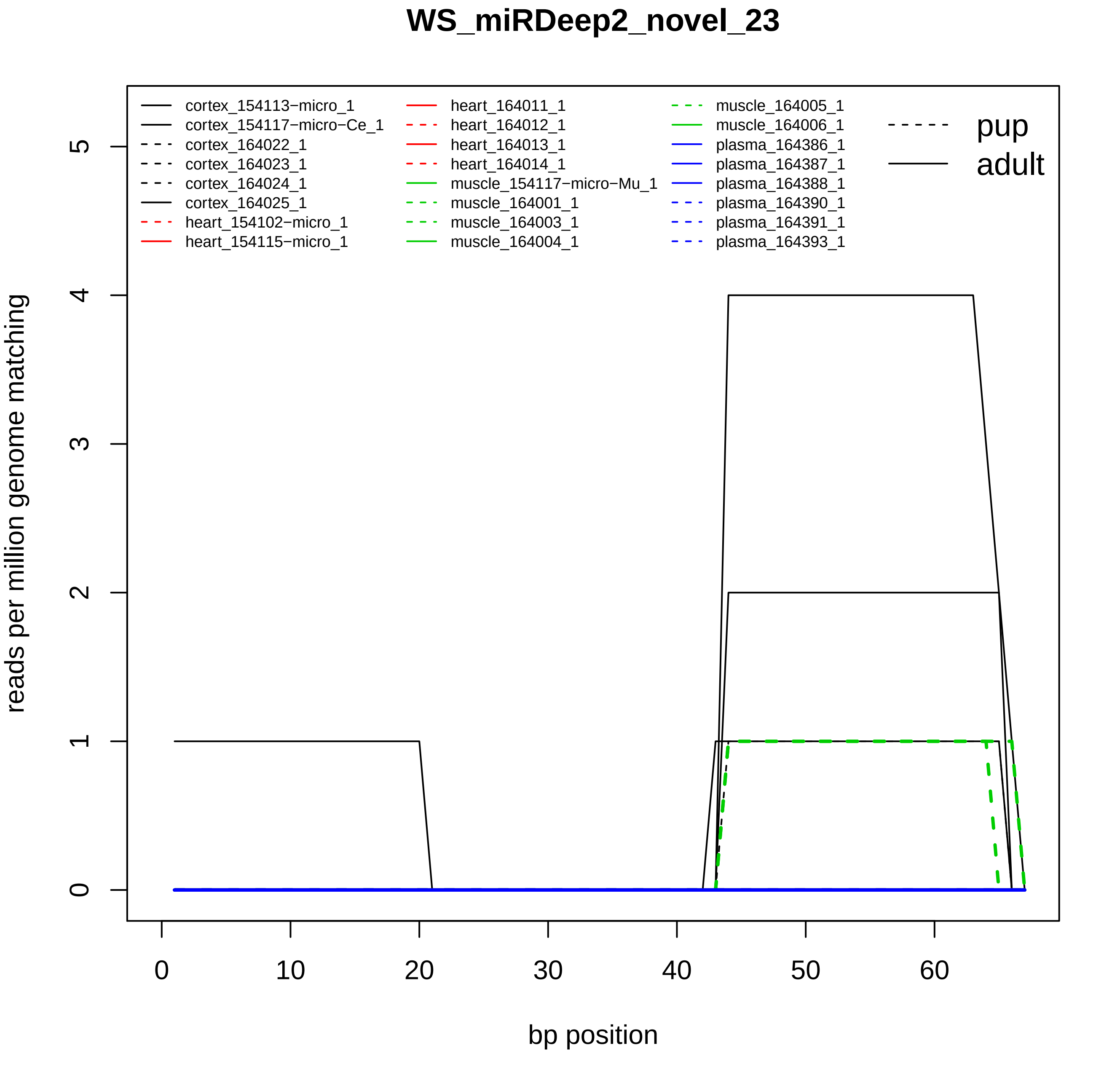
WS_miRDeep2_novel_23
cortex_154113−micro_1
cortex_154117−micro−Ce_1
cortex_164022_1
cortex_164023_1
cortex_164024_1
cortex_164025_1
heart_154102−micro_1
heart_154115−micro_1
heart_164011_1
heart_164012_1
heart_164013_1
heart_164014_1
muscle_154117−micro−Mu_1
muscle_164001_1
muscle_164003_1
muscle_164004_1
muscle_164005_1
muscle_164006_1
plasma_164386_1
plasma_164387_1
plasma_164388_1
plasma_164390_1
plasma_164391_1
plasma_164393_1
pup
adult
reads per million genome matching
0
1
2
3
4
5
0
10
20
30
40
50
60
bp position

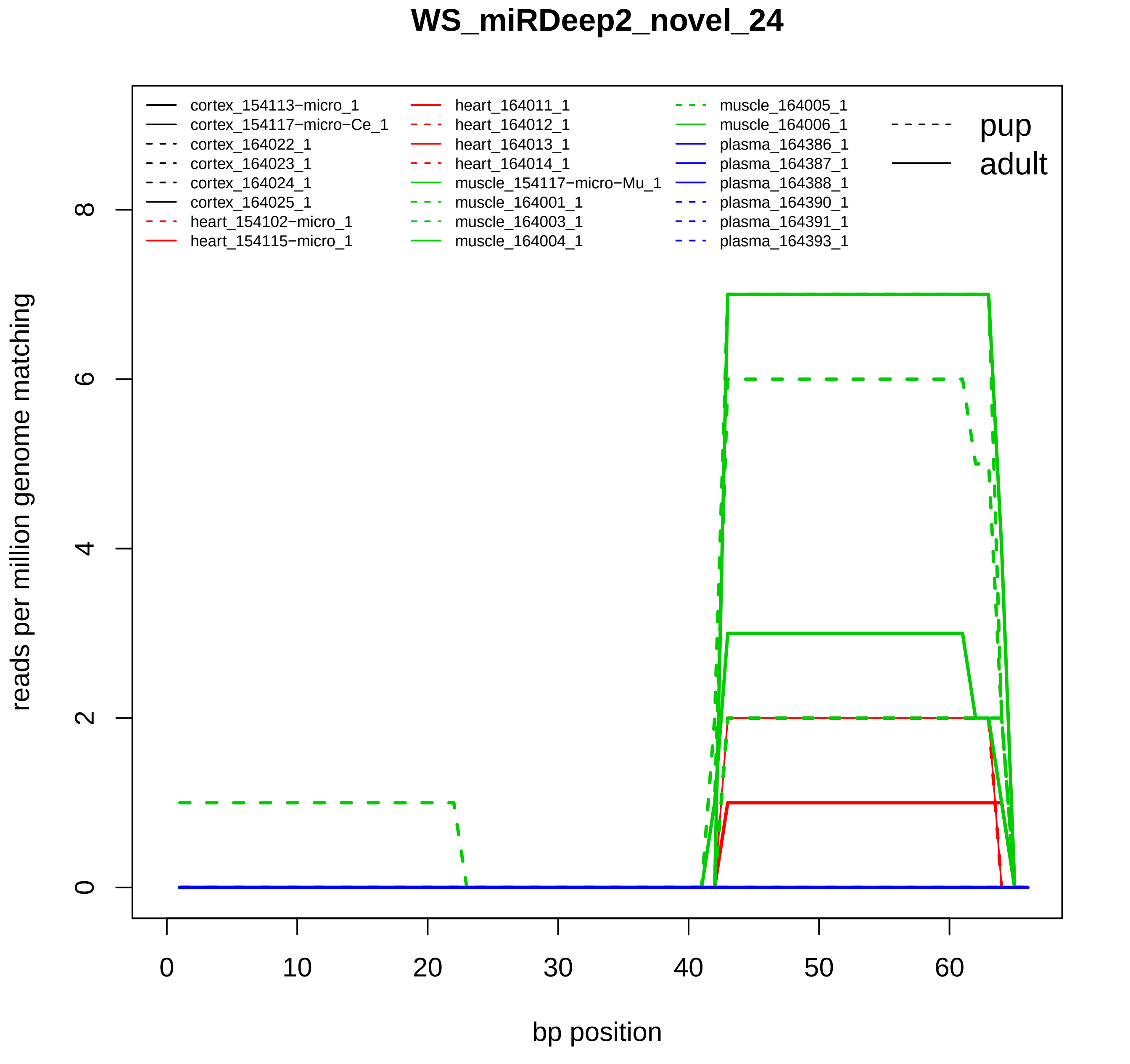
WS_miRDeep2_novel_24
cortex_154113−micro_1
cortex_154117−micro−Ce_1
cortex_164022_1
cortex_164023_1
cortex_164024_1
cortex_164025_1
heart_154102−micro_1
heart_154115−micro_1
heart_164011_1
heart_164012_1
heart_164013_1
heart_164014_1
muscle_154117−micro−Mu_1
muscle_164001_1
muscle_164003_1
muscle_164004_1
muscle_164005_1
muscle_164006_1
plasma_164386_1
plasma_164387_1
plasma_164388_1
plasma_164390_1
plasma_164391_1
plasma_164393_1
pup
adult
reads per million genome matching
0
2
4
6
8
0
10
20
30
40
50
60
bp position

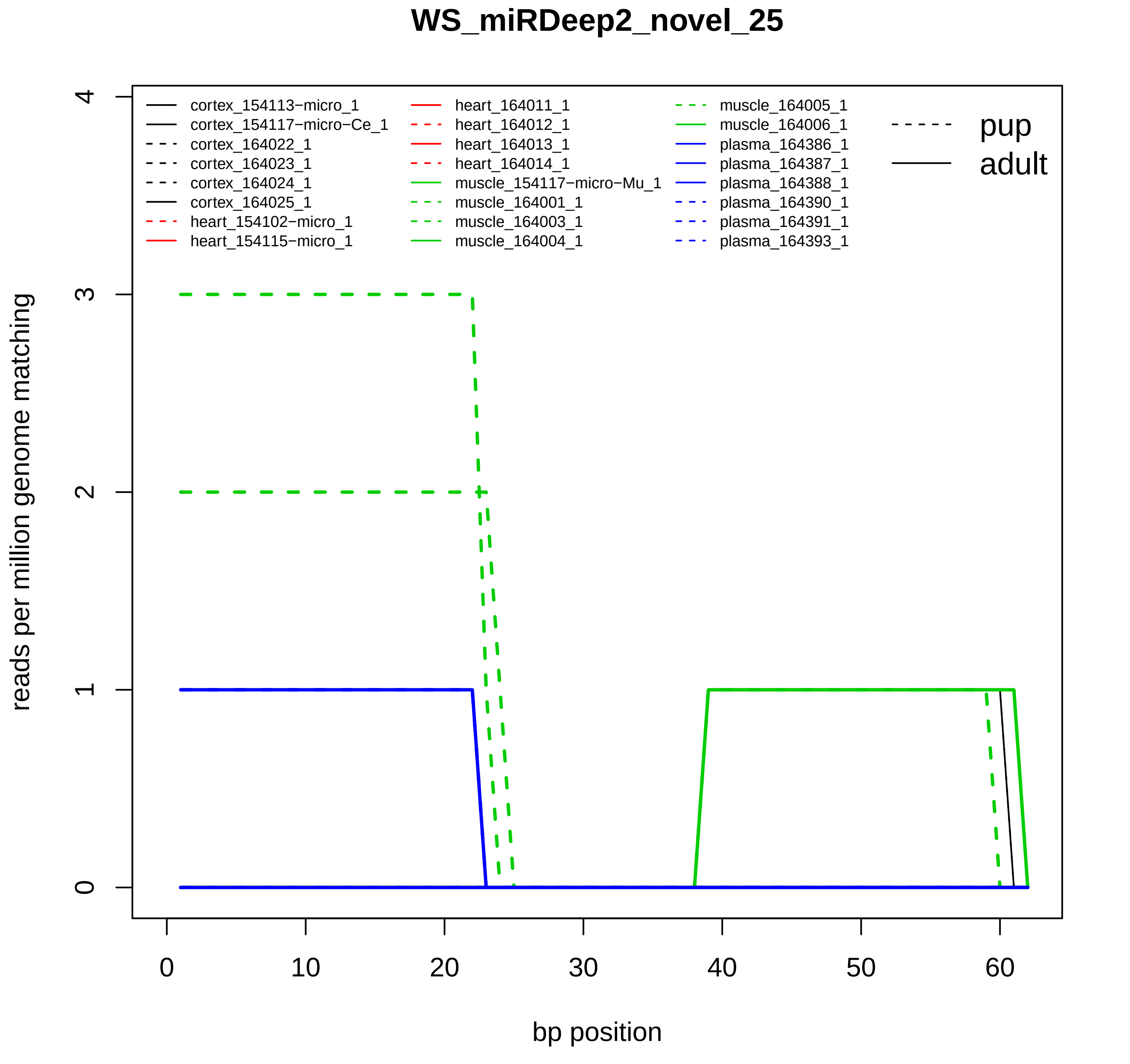

WS_miRDeep2_novel_25
cortex_154113−micro_1
cortex_154117−micro−Ce_1
cortex_164022_1
cortex_164023_1
cortex_164024_1
cortex_164025_1
heart_154102−micro_1
heart_154115−micro_1
heart_164011_1
heart_164012_1
heart_164013_1
heart_164014_1
muscle_154117−micro−Mu_1
muscle_164001_1
muscle_164003_1
muscle_164004_1
muscle_164005_1
muscle_164006_1
plasma_164386_1
plasma_164387_1
plasma_164388_1
plasma_164390_1
plasma_164391_1
plasma_164393_1
pup
adult
reads per million genome matching
0
1
2
3
4
bp position
0
10
20
30
40
50
60

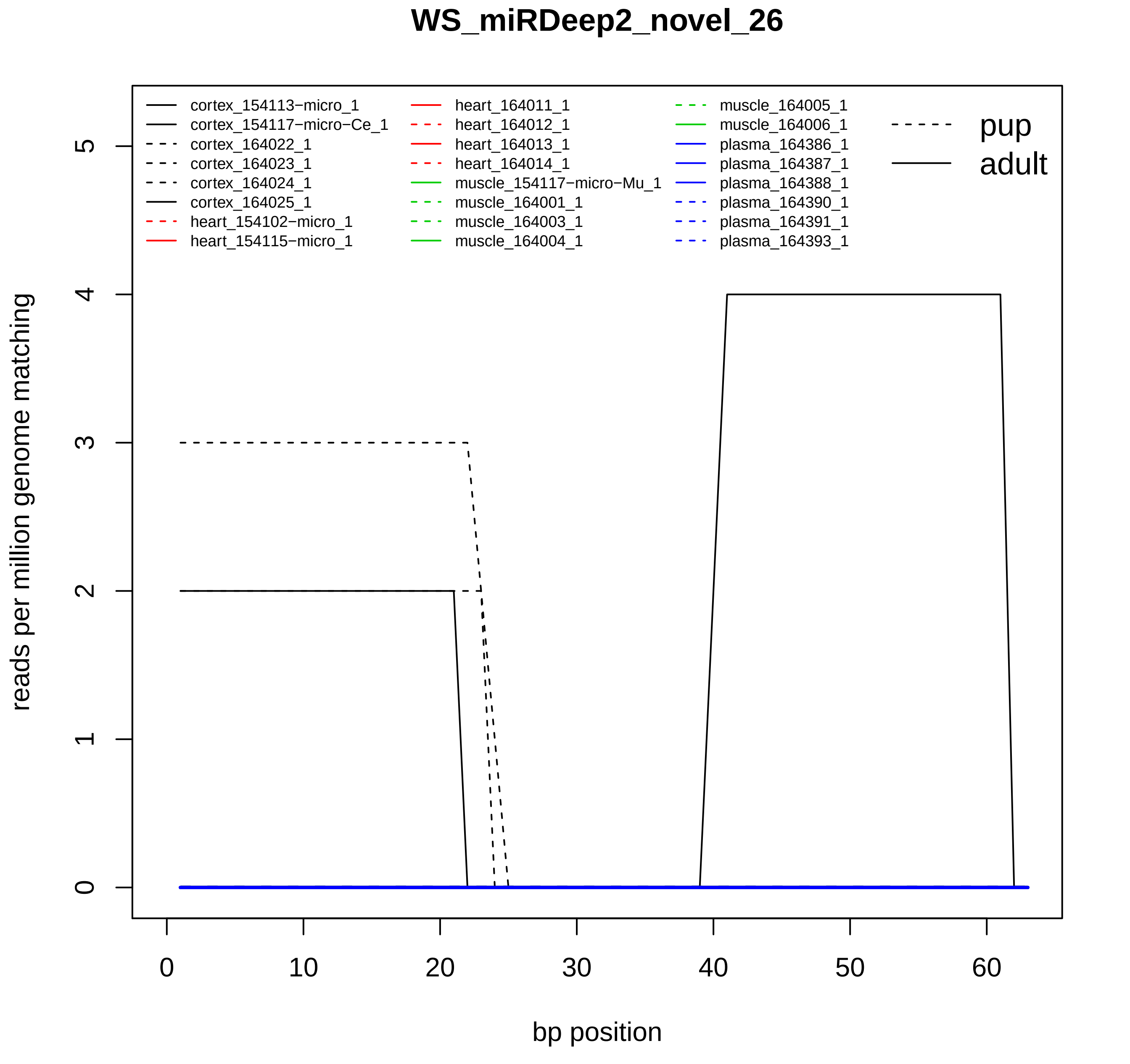

WS_miRDeep2_novel_26
cortex_154113−micro_1
cortex_154117−micro−Ce_1
cortex_164022_1
cortex_164023_1
cortex_164024_1
cortex_164025_1
heart_154102−micro_1
heart_154115−micro_1
heart_164011_1
heart_164012_1
heart_164013_1
heart_164014_1
muscle_154117−micro−Mu_1
muscle_164001_1
muscle_164003_1
muscle_164004_1
muscle_164005_1
muscle_164006_1
plasma_164386_1
plasma_164387_1
plasma_164388_1
plasma_164390_1
plasma_164391_1
plasma_164393_1
pup
adult
reads per million genome matching
0
1
2
3
4
5
0
10
20
30
40
50
60
bp position

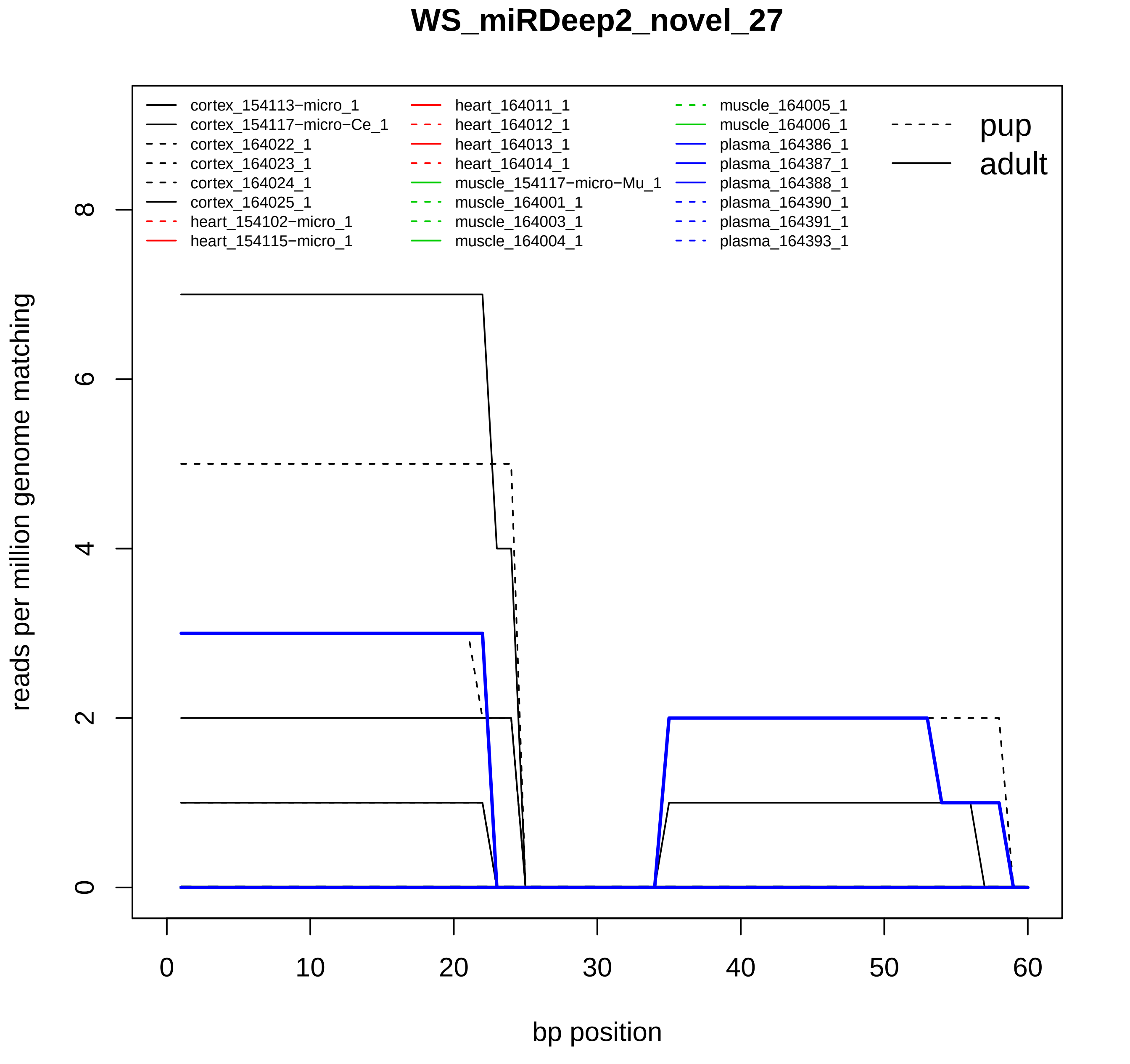
WS_miRDeep2_novel_27
cortex_154113−micro_1
cortex_154117−micro−Ce_1
cortex_164022_1
cortex_164023_1
cortex_164024_1
cortex_164025_1
heart_154102−micro_1
heart_154115−micro_1
heart_164011_1
heart_164012_1
heart_164013_1
heart_164014_1
muscle_154117−micro−Mu_1
muscle_164001_1
muscle_164003_1
muscle_164004_1
muscle_164005_1
muscle_164006_1
plasma_164386_1
plasma_164387_1
plasma_164388_1
plasma_164390_1
plasma_164391_1
plasma_164393_1
pup
adult
reads per million genome matching
0
2
4
6
8
0
10
20
30
40
50
60
bp position

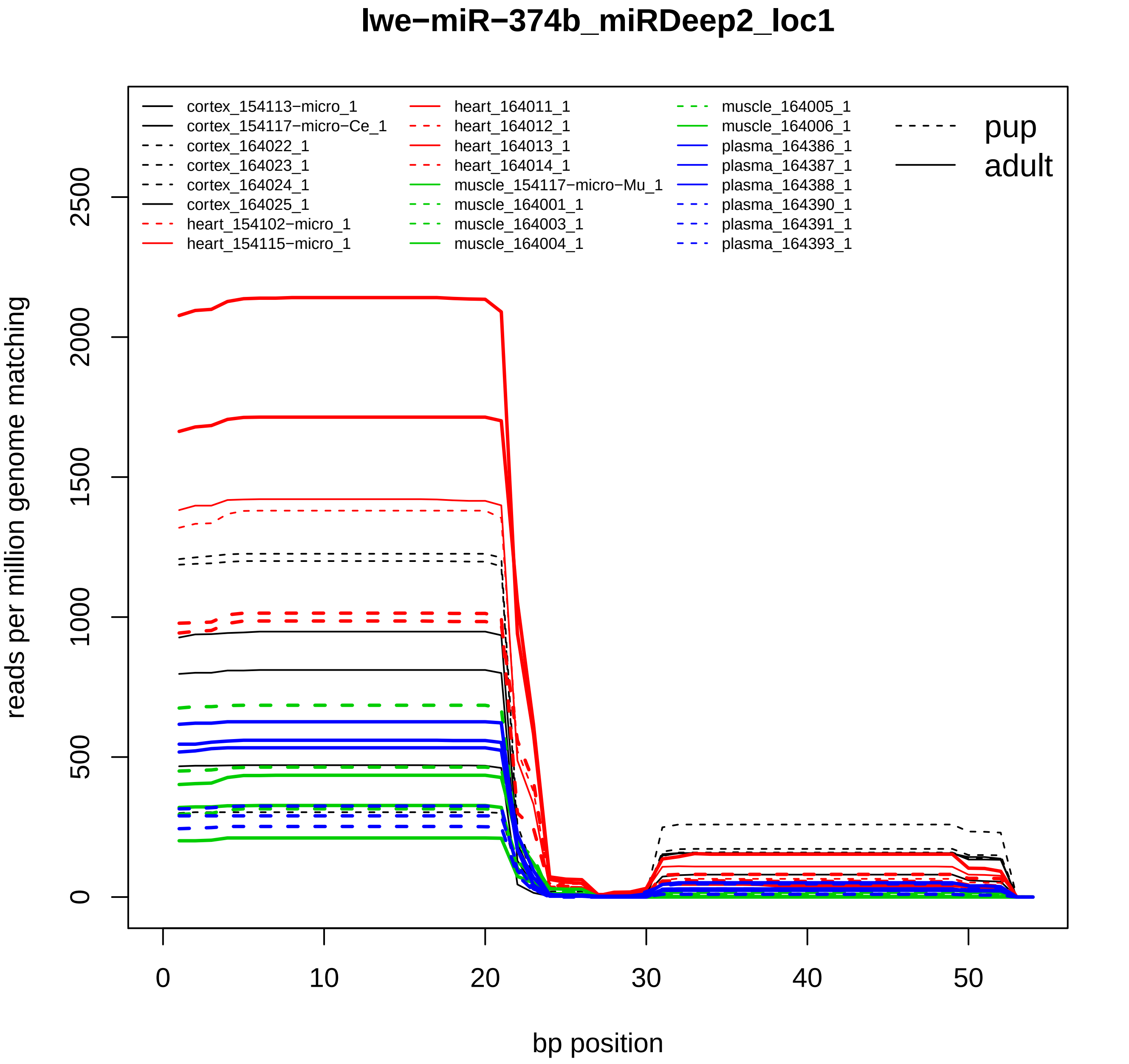
lwe−miR−374b_miRDeep2_loc1
reads per million genome matching
bp position
0
500
1000
1500
2000
2500
0
10
20
30
40
50
cortex_154113−micro_1
cortex_154117−micro−Ce_1
cortex_164022_1
cortex_164023_1
cortex_164024_1
cortex_164025_1
heart_154102−micro_1
heart_154115−micro_1
heart_164011_1
heart_164012_1
heart_164013_1
heart_164014_1
muscle_154117−micro−Mu_1
muscle_164001_1
muscle_164003_1
muscle_164004_1
muscle_164005_1
muscle_164006_1
plasma_164386_1
plasma_164387_1
plasma_164388_1
plasma_164390_1
plasma_164391_1
plasma_164393_1
pup
adult

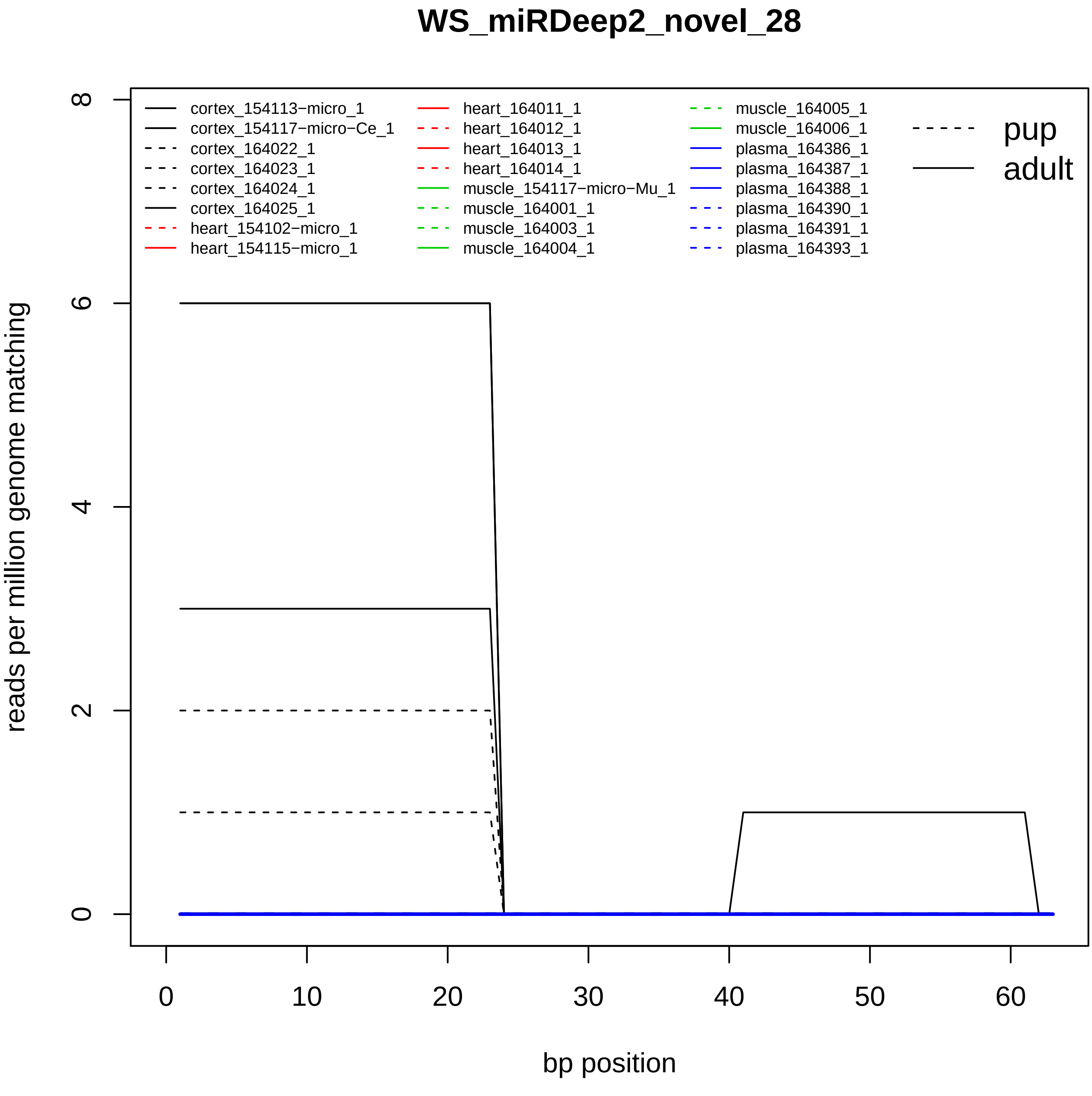
WS_miRDeep2_novel_28
cortex_154113−micro_1
cortex_154117−micro−Ce_1
cortex_164022_1
cortex_164023_1
cortex_164024_1
cortex_164025_1
heart_154102−micro_1
heart_154115−micro_1
heart_164011_1
heart_164012_1
heart_164013_1
heart_164014_1
muscle_154117−micro−Mu_1
muscle_164001_1
muscle_164003_1
muscle_164004_1
muscle_164005_1
muscle_164006_1
plasma_164386_1
plasma_164387_1
plasma_164388_1
plasma_164390_1
plasma_164391_1
plasma_164393_1
pup
adult
reads per million genome matching
bp position
0
2
4
6
8
0
10
20
30
40
50
60

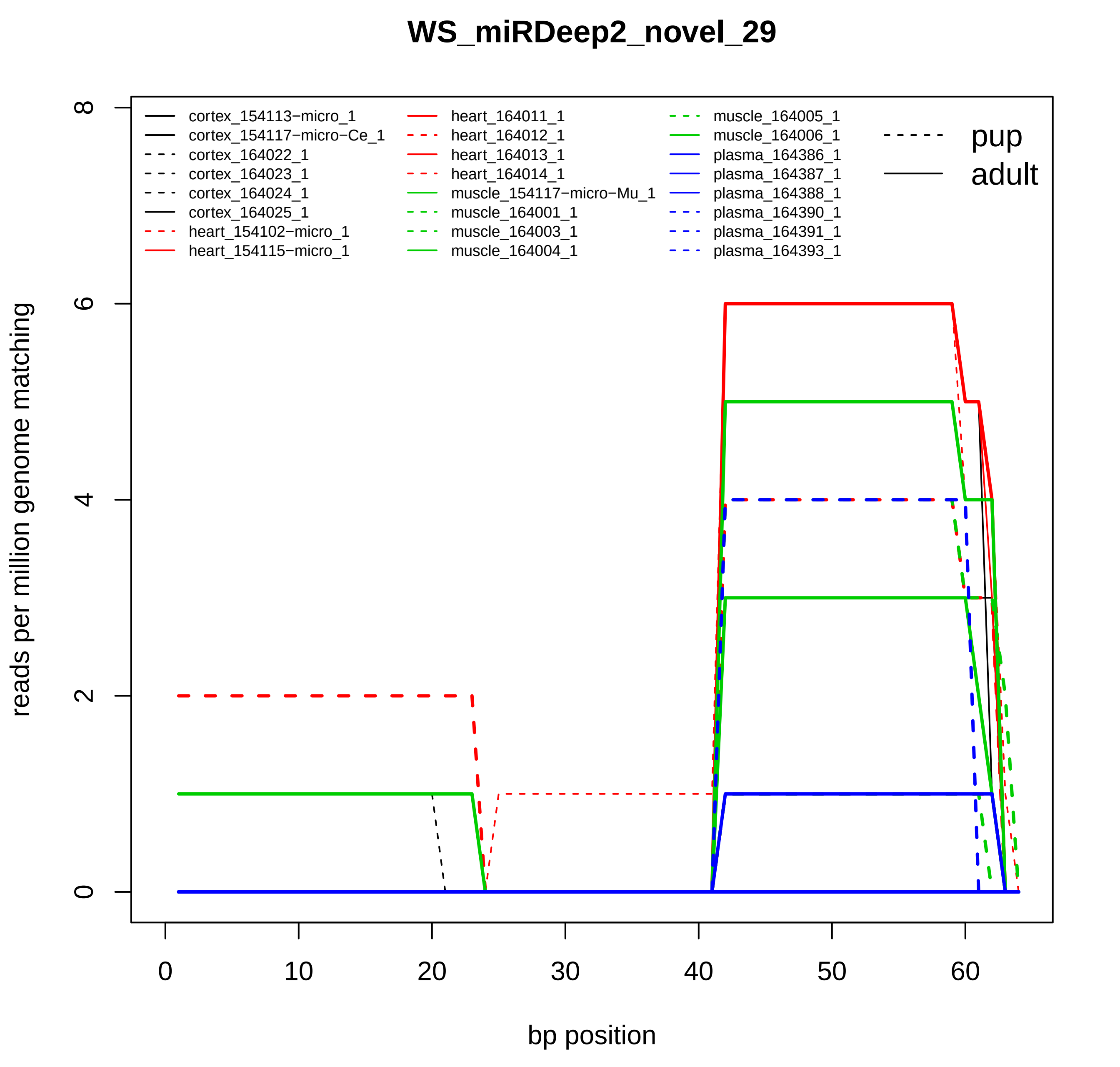

WS_miRDeep2_novel_29
cortex_154113−micro_1
cortex_154117−micro−Ce_1
cortex_164022_1
cortex_164023_1
cortex_164024_1
cortex_164025_1
heart_154102−micro_1
heart_154115−micro_1
heart_164011_1
heart_164012_1
heart_164013_1
heart_164014_1
muscle_154117−micro−Mu_1
muscle_164001_1
muscle_164003_1
muscle_164004_1
muscle_164005_1
muscle_164006_1
plasma_164386_1
plasma_164387_1
plasma_164388_1
plasma_164390_1
plasma_164391_1
plasma_164393_1
pup
adult
reads per million genome matching
bp position
0
2
4
6
8
0
10
20
30
40
50
60

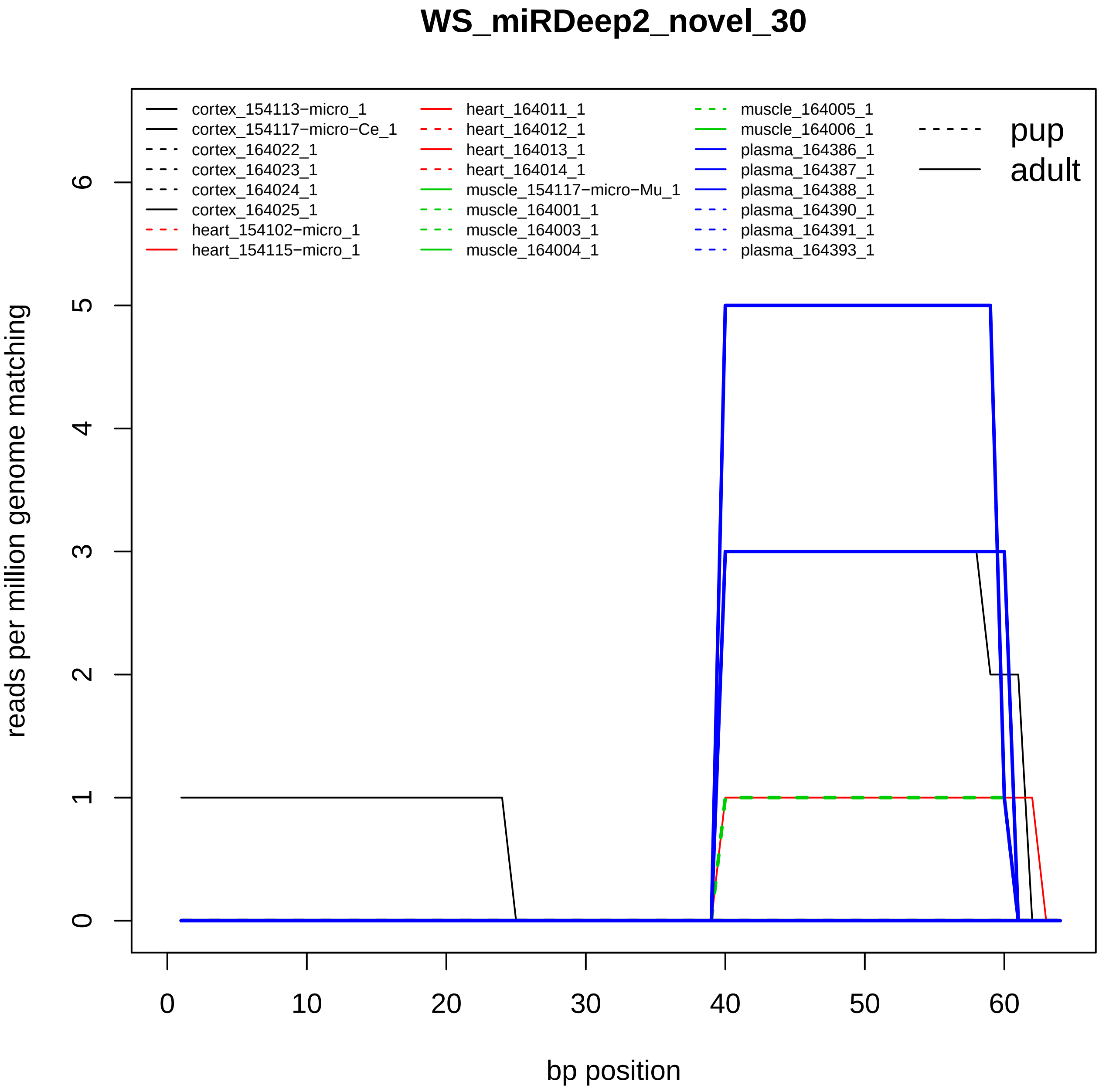
WS_miRDeep2_novel_30
cortex_154113−micro_1
cortex_154117−micro−Ce_1
cortex_164022_1
cortex_164023_1
cortex_164024_1
cortex_164025_1
heart_154102−micro_1
heart_154115−micro_1
heart_164011_1
heart_164012_1
heart_164013_1
heart_164014_1
muscle_154117−micro−Mu_1
muscle_164001_1
muscle_164003_1
muscle_164004_1
muscle_164005_1
muscle_164006_1
plasma_164386_1
plasma_164387_1
plasma_164388_1
plasma_164390_1
plasma_164391_1
plasma_164393_1
pup
adult
reads per million genome matching
0
1
2
3
4
5
6
0
10
20
30
40
50
60
bp position

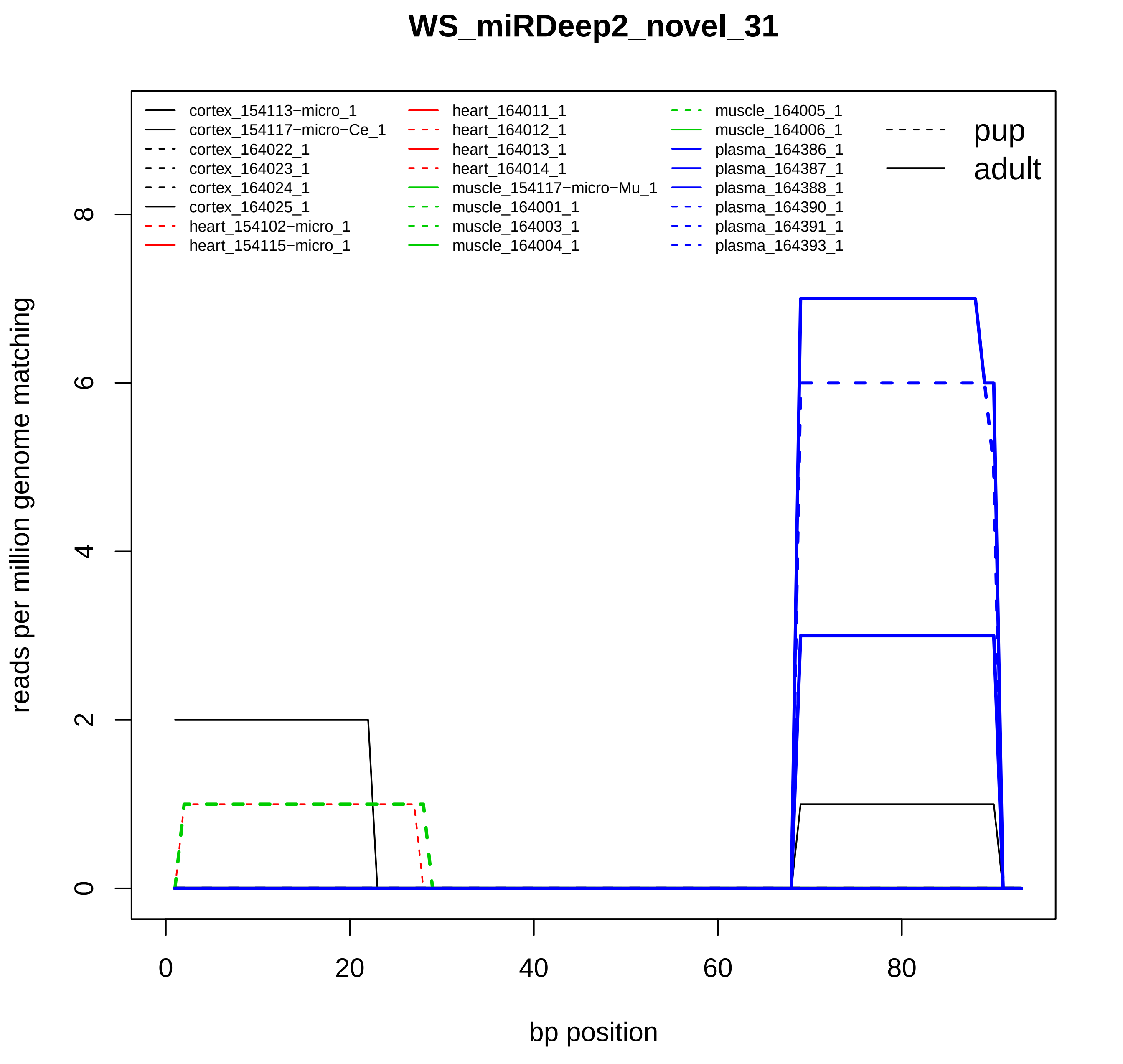

WS_miRDeep2_novel_31
cortex_154113−micro_1
cortex_154117−micro−Ce_1
cortex_164022_1
cortex_164023_1
cortex_164024_1
cortex_164025_1
heart_154102−micro_1
heart_154115−micro_1
heart_164011_1
heart_164012_1
heart_164013_1
heart_164014_1
muscle_154117−micro−Mu_1
muscle_164001_1
muscle_164003_1
muscle_164004_1
muscle_164005_1
muscle_164006_1
plasma_164386_1
plasma_164387_1
plasma_164388_1
plasma_164390_1
plasma_164391_1
plasma_164393_1
pup
adult
reads per million genome matching
0
2
4
6
8
0
20
40
60
80
bp position

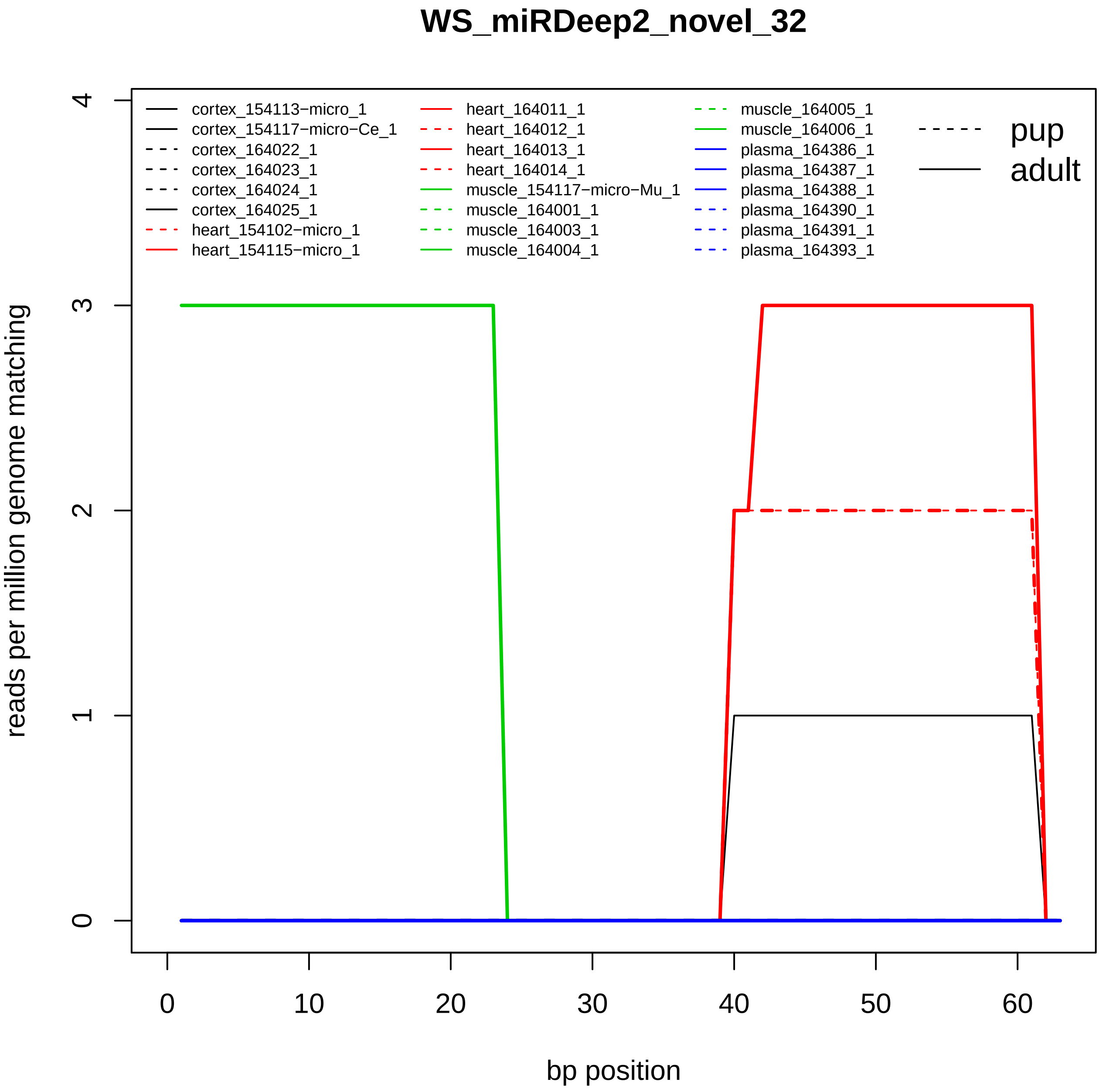

WS_miRDeep2_novel_32
cortex_154113−micro_1
cortex_154117−micro−Ce_1
cortex_164022_1
cortex_164023_1
cortex_164024_1
cortex_164025_1
heart_154102−micro_1
heart_154115−micro_1
heart_164011_1
heart_164012_1
heart_164013_1
heart_164014_1
muscle_154117−micro−Mu_1
muscle_164001_1
muscle_164003_1
muscle_164004_1
muscle_164005_1
muscle_164006_1
plasma_164386_1
plasma_164387_1
plasma_164388_1
plasma_164390_1
plasma_164391_1
plasma_164393_1
pup
adult
reads per million genome matching
0
1
2
3
4
0
10
20
30
40
50
60
bp position

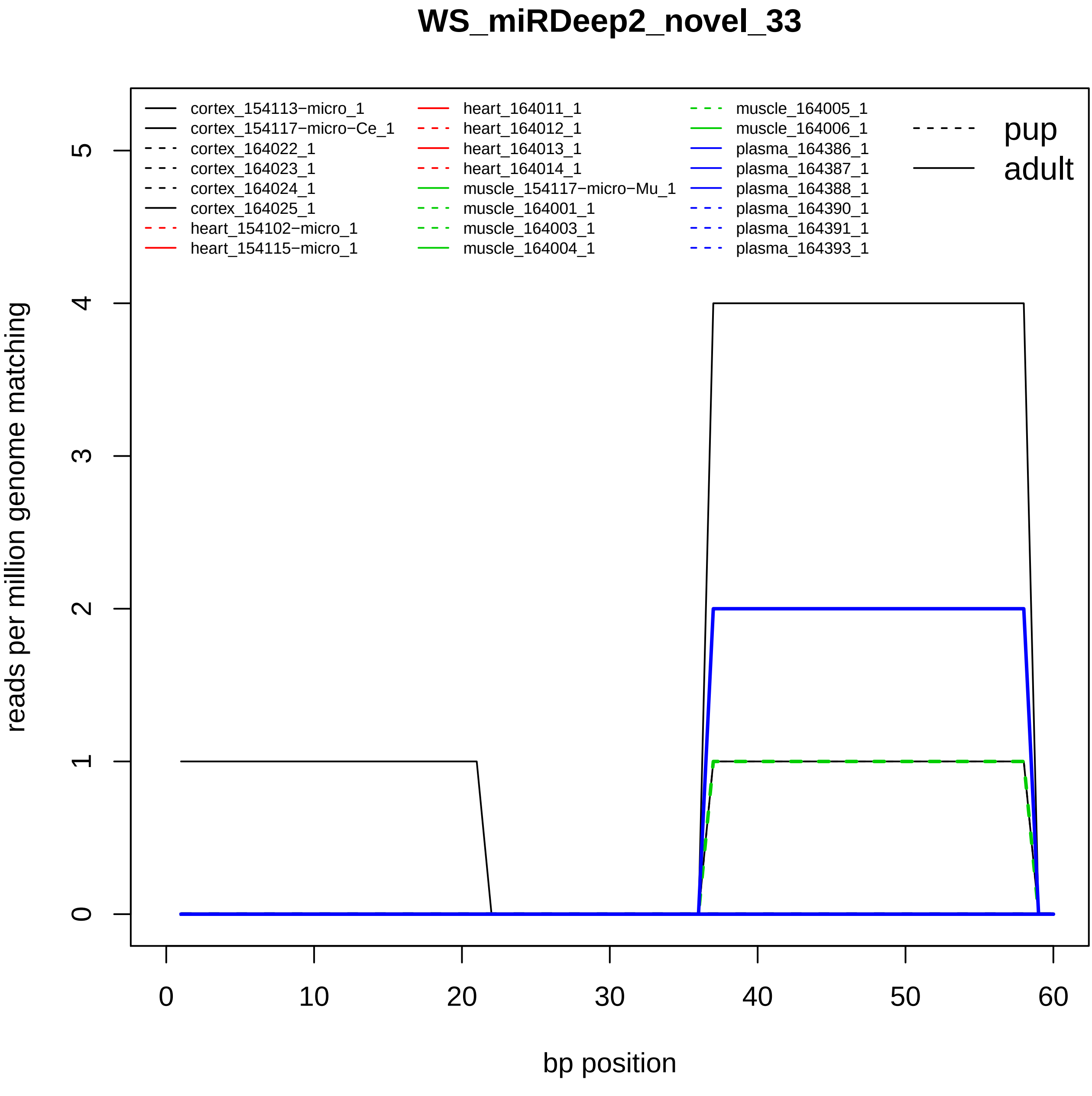

WS_miRDeep2_novel_33
cortex_154113−micro_1
cortex_154117−micro−Ce_1
cortex_164022_1
cortex_164023_1
cortex_164024_1
cortex_164025_1
heart_154102−micro_1
heart_154115−micro_1
heart_164011_1
heart_164012_1
heart_164013_1
heart_164014_1
muscle_154117−micro−Mu_1
muscle_164001_1
muscle_164003_1
muscle_164004_1
muscle_164005_1
muscle_164006_1
plasma_164386_1
plasma_164387_1
plasma_164388_1
plasma_164390_1
plasma_164391_1
plasma_164393_1
pup
adult
reads per million genome matching
bp position
0
1
2
3
4
5
0
10
20
30
40
50
60

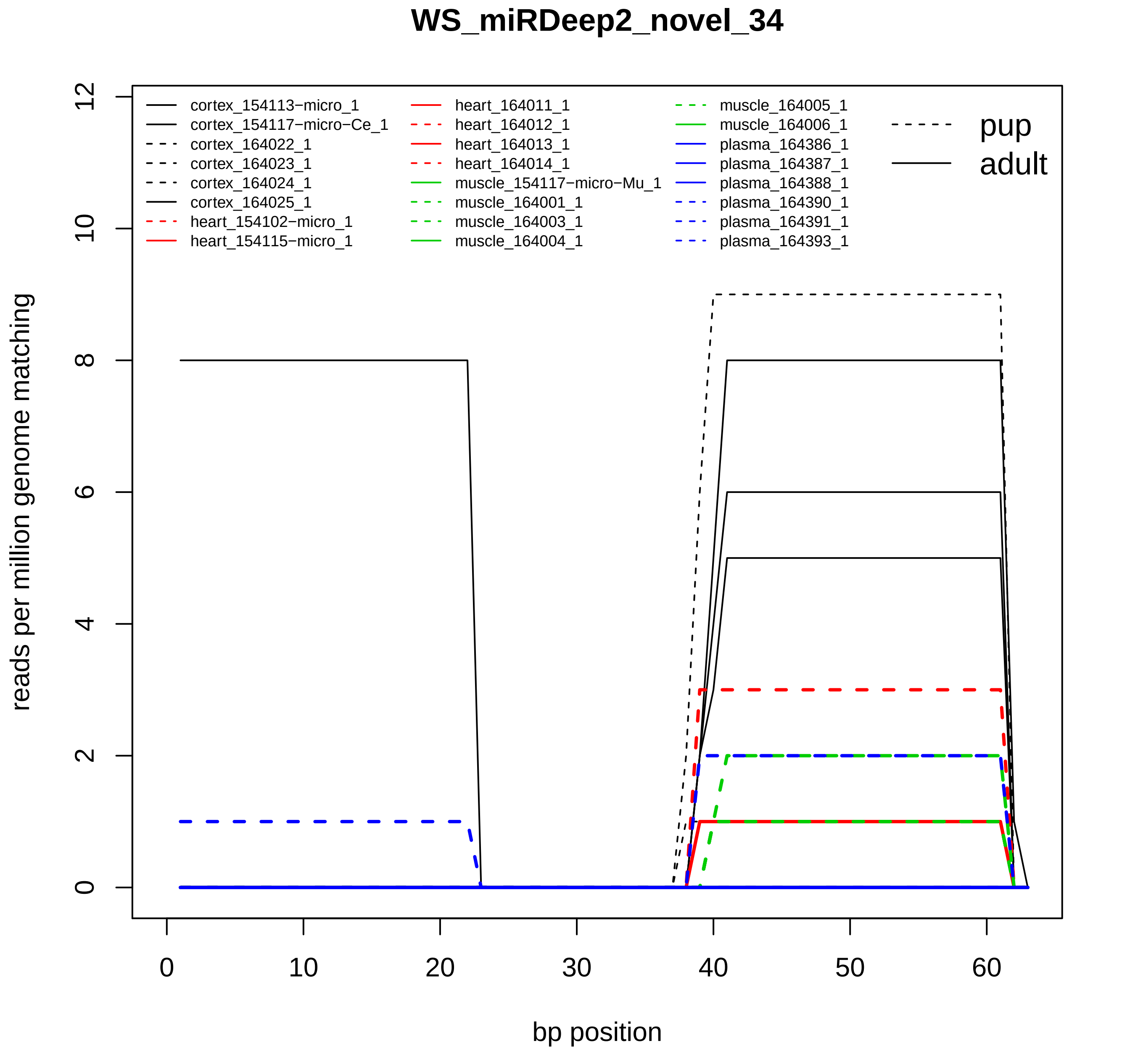

WS_miRDeep2_novel_34
cortex_154113−micro_1
cortex_154117−micro−Ce_1
cortex_164022_1
cortex_164023_1
cortex_164024_1
cortex_164025_1
heart_154102−micro_1
heart_154115−micro_1
heart_164011_1
heart_164012_1
heart_164013_1
heart_164014_1
muscle_154117−micro−Mu_1
muscle_164001_1
muscle_164003_1
muscle_164004_1
muscle_164005_1
muscle_164006_1
plasma_164386_1
plasma_164387_1
plasma_164388_1
plasma_164390_1
plasma_164391_1
plasma_164393_1
pup
adult
reads per million genome matching
0
2
4
6
8
10
12
bp position
0
10
20
30
40
50
60

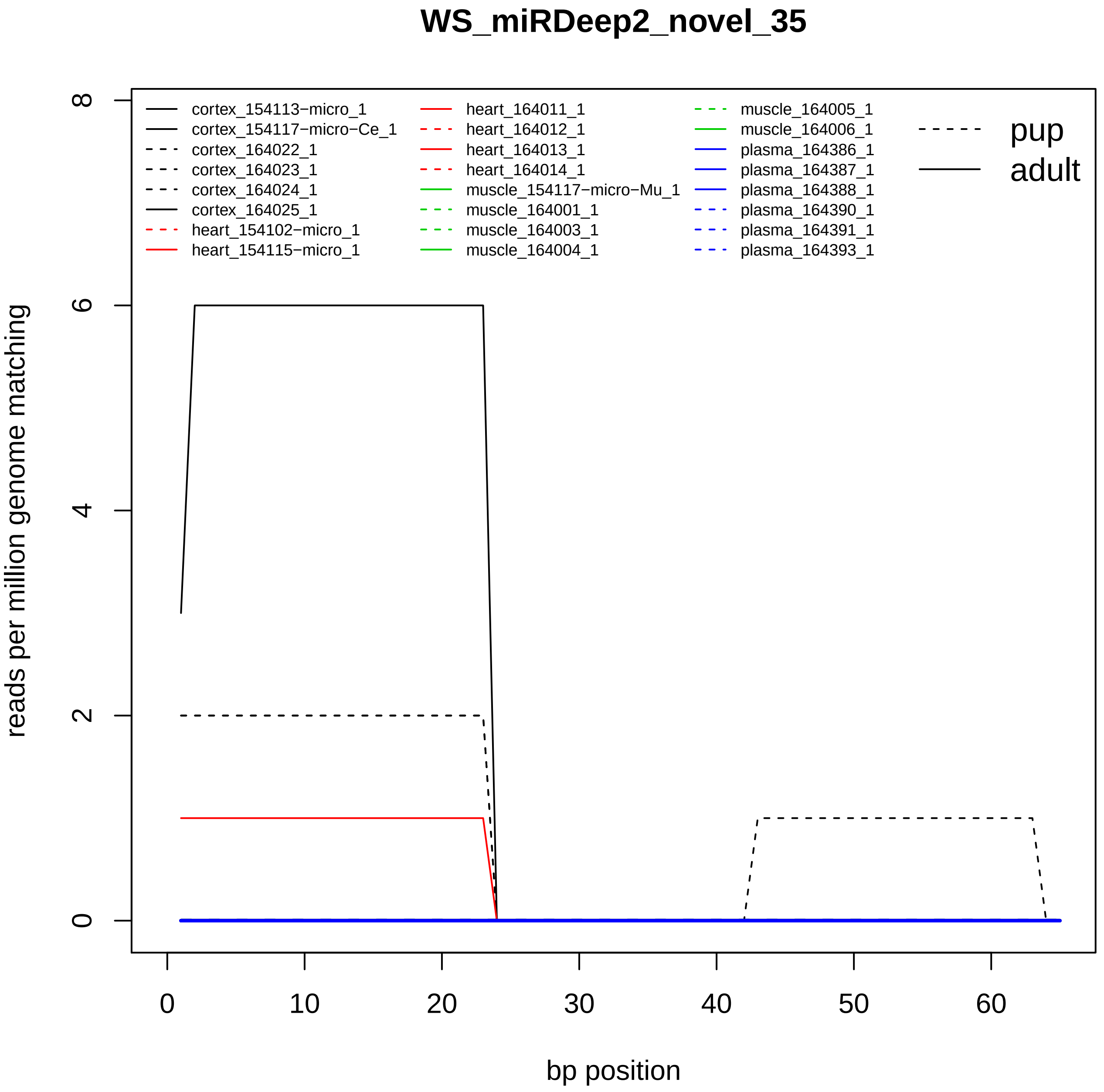

WS_miRDeep2_novel_35
reads per million genome matching
bp position
0
2
4
6
8
0
10
20
30
40
50
60
cortex_154113−micro_1
cortex_154117−micro−Ce_1
cortex_164022_1
cortex_164023_1
cortex_164024_1
cortex_164025_1
heart_154102−micro_1
heart_154115−micro_1
heart_164011_1
heart_164012_1
heart_164013_1
heart_164014_1
muscle_154117−micro−Mu_1
muscle_164001_1
muscle_164003_1
muscle_164004_1
muscle_164005_1
muscle_164006_1
plasma_164386_1
plasma_164387_1
plasma_164388_1
plasma_164390_1
plasma_164391_1
plasma_164393_1
pup
adult

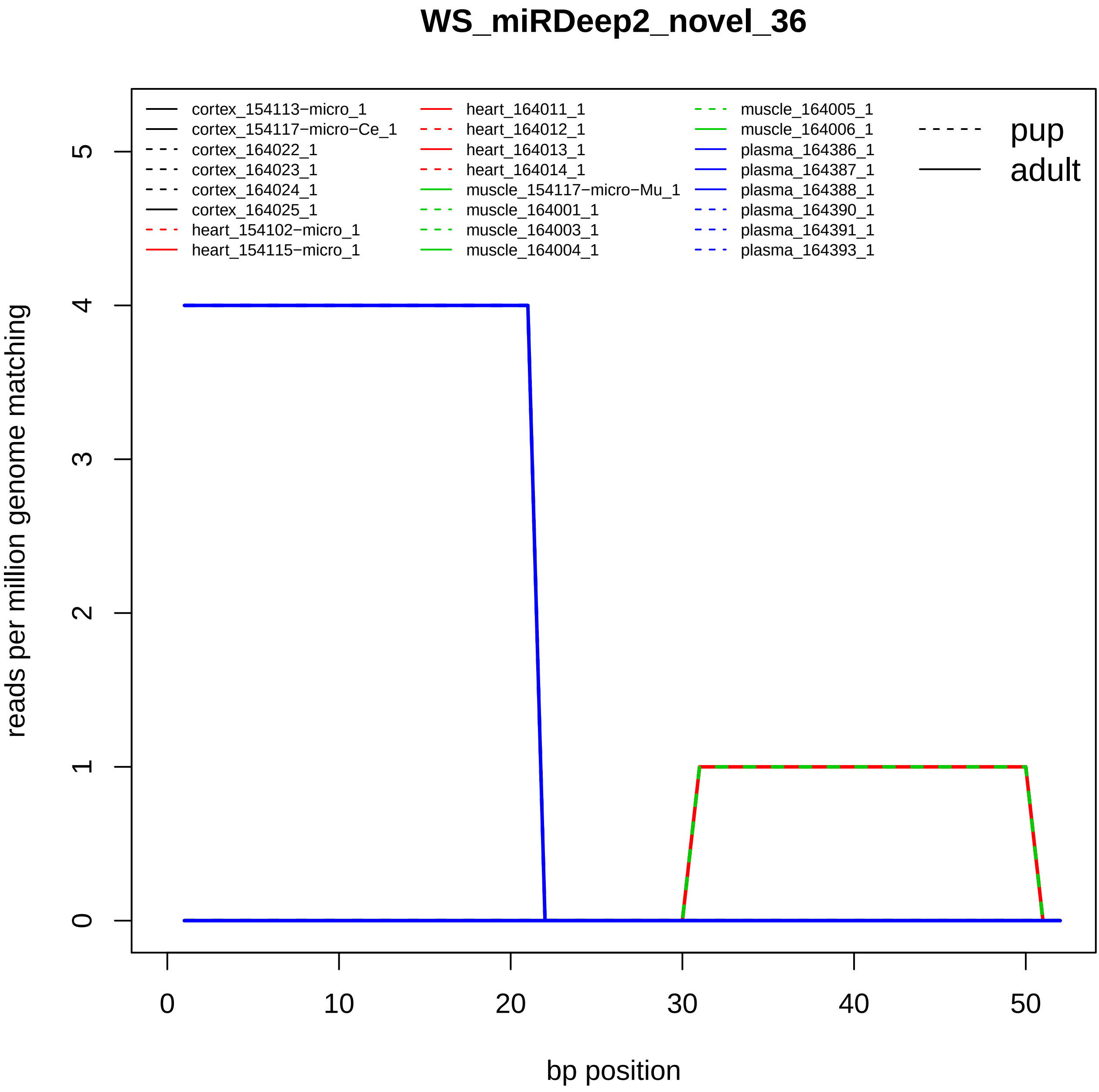
WS_miRDeep2_novel_36
cortex_154113−micro_1
cortex_154117−micro−Ce_1
cortex_164022_1
cortex_164023_1
cortex_164024_1
cortex_164025_1
heart_154102−micro_1
heart_154115−micro_1
heart_164011_1
heart_164012_1
heart_164013_1
heart_164014_1
muscle_154117−micro−Mu_1
muscle_164001_1
muscle_164003_1
muscle_164004_1
muscle_164005_1
muscle_164006_1
plasma_164386_1
plasma_164387_1
plasma_164388_1
plasma_164390_1
plasma_164391_1
plasma_164393_1
pup
adult
reads per million genome matching
0
1
2
3
4
5
0
10
20
30
40
50
bp position

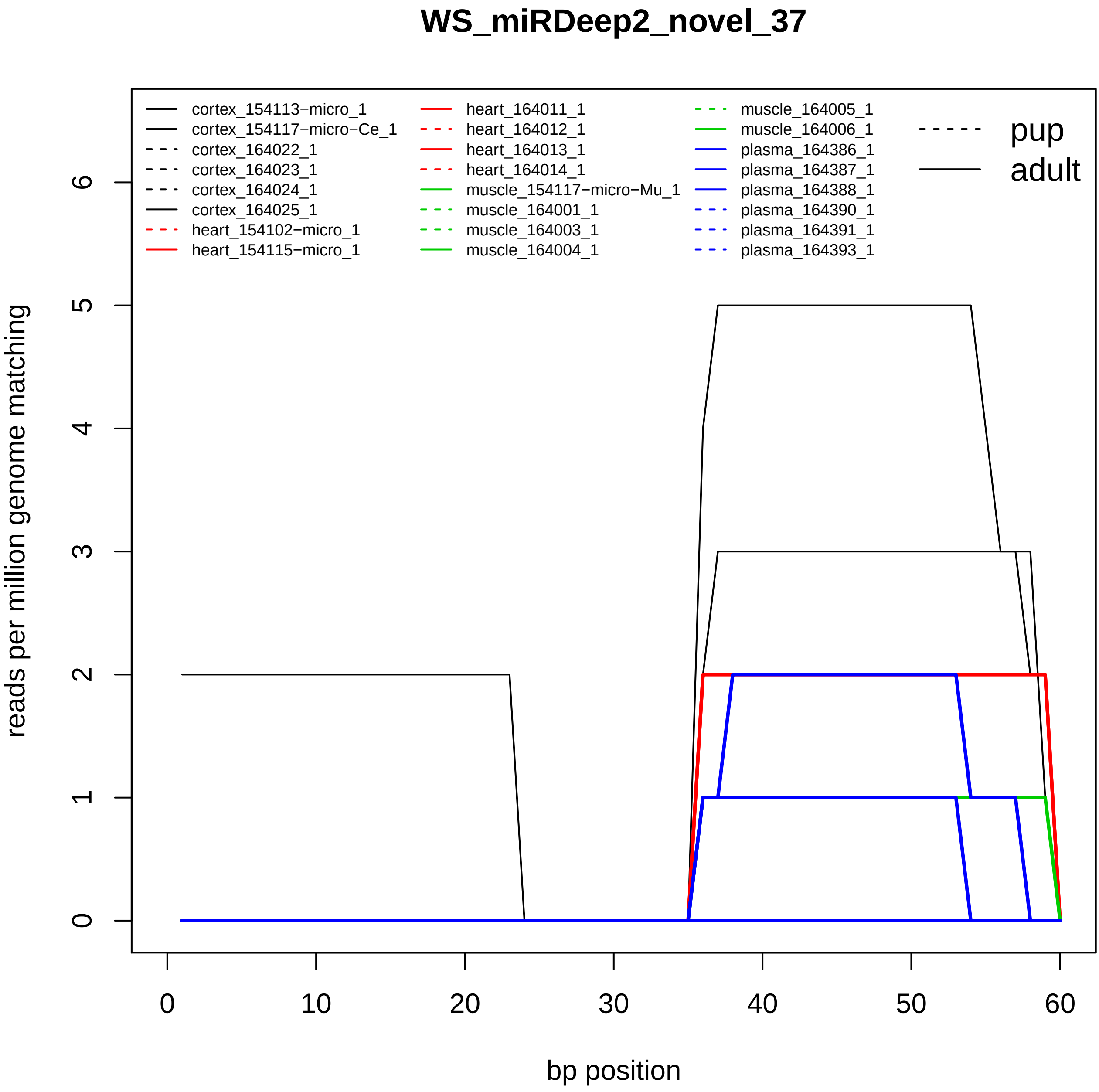

WS_miRDeep2_novel_37
cortex_154113−micro_1
cortex_154117−micro−Ce_1
cortex_164022_1
cortex_164023_1
cortex_164024_1
cortex_164025_1
heart_154102−micro_1
heart_154115−micro_1
heart_164011_1
heart_164012_1
heart_164013_1
heart_164014_1
muscle_154117−micro−Mu_1
muscle_164001_1
muscle_164003_1
muscle_164004_1
muscle_164005_1
muscle_164006_1
plasma_164386_1
plasma_164387_1
plasma_164388_1
plasma_164390_1
plasma_164391_1
plasma_164393_1
pup
adult
reads per million genome matching
bp position
0
1
2
3
4
5
6
0
10
20
30
40
50
60

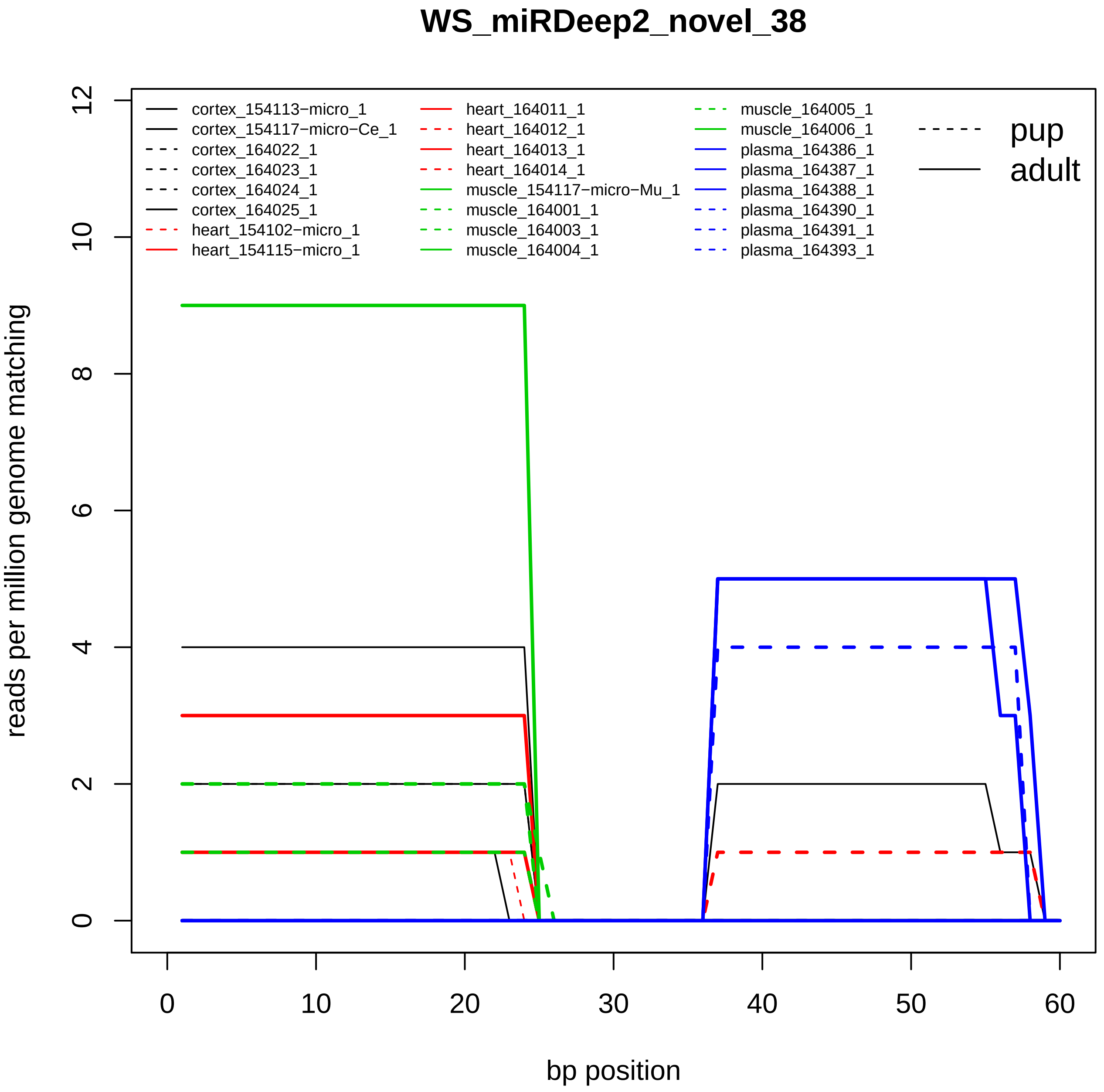

WS_miRDeep2_novel_38
cortex_154113−micro_1
cortex_154117−micro−Ce_1
cortex_164022_1
cortex_164023_1
cortex_164024_1
cortex_164025_1
heart_154102−micro_1
heart_154115−micro_1
heart_164011_1
heart_164012_1
heart_164013_1
heart_164014_1
muscle_154117−micro−Mu_1
muscle_164001_1
muscle_164003_1
muscle_164004_1
muscle_164005_1
muscle_164006_1
plasma_164386_1
plasma_164387_1
plasma_164388_1
plasma_164390_1
plasma_164391_1
plasma_164393_1
pup
adult
reads per million genome matching
bp position
0
2
4
6
8
10
12
0
10
20
30
40
50
60

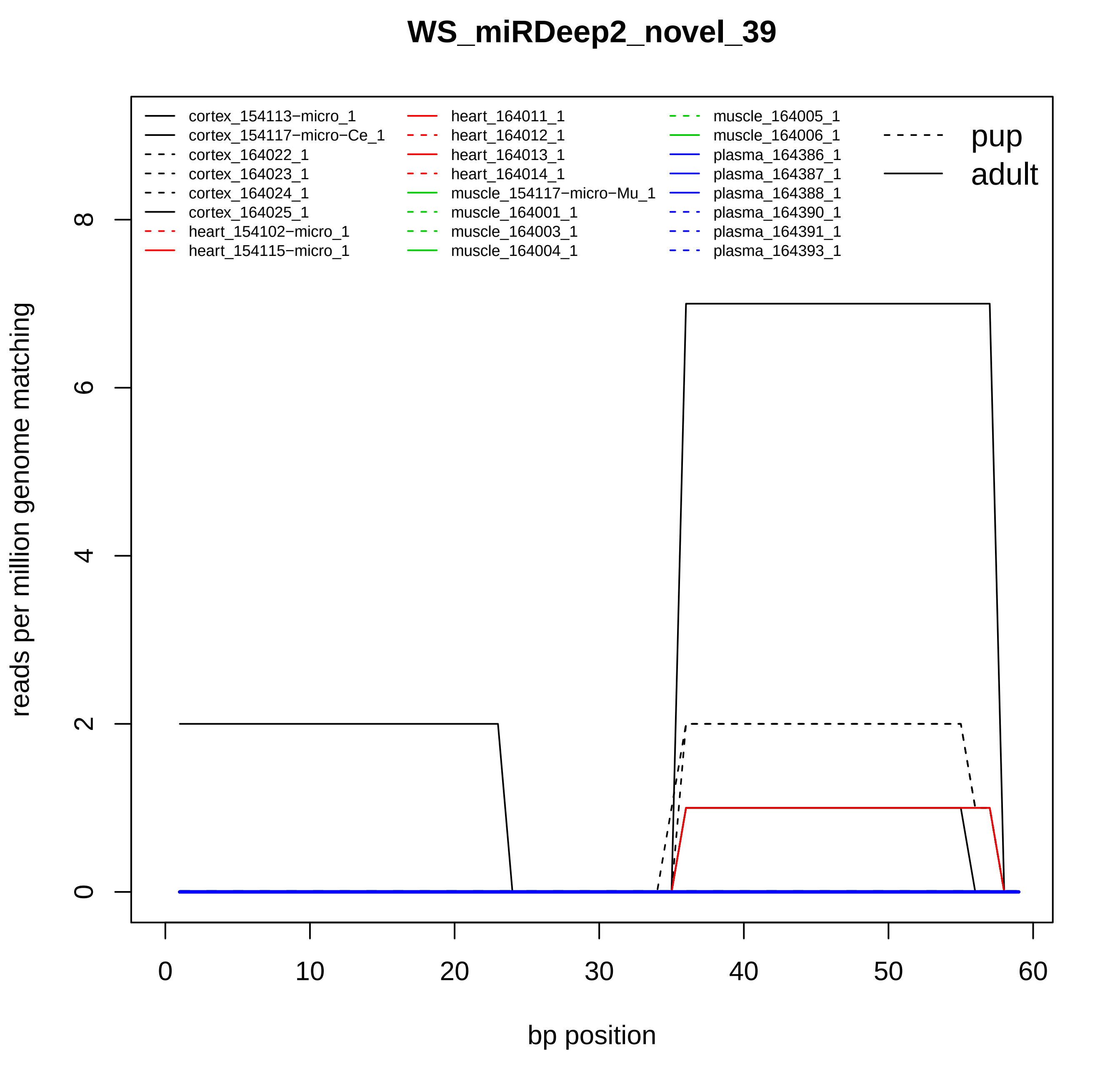
WS_miRDeep2_novel_39
cortex_154113−micro_1
cortex_154117−micro−Ce_1
cortex_164022_1
cortex_164023_1
cortex_164024_1
cortex_164025_1
heart_154102−micro_1
heart_154115−micro_1
heart_164011_1
heart_164012_1
heart_164013_1
heart_164014_1
muscle_154117−micro−Mu_1
muscle_164001_1
muscle_164003_1
muscle_164004_1
muscle_164005_1
muscle_164006_1
plasma_164386_1
plasma_164387_1
plasma_164388_1
plasma_164390_1
plasma_164391_1
plasma_164393_1
pup
adult
reads per million genome matching
0
2
4
6
8
0
10
20
30
40
50
60
bp position

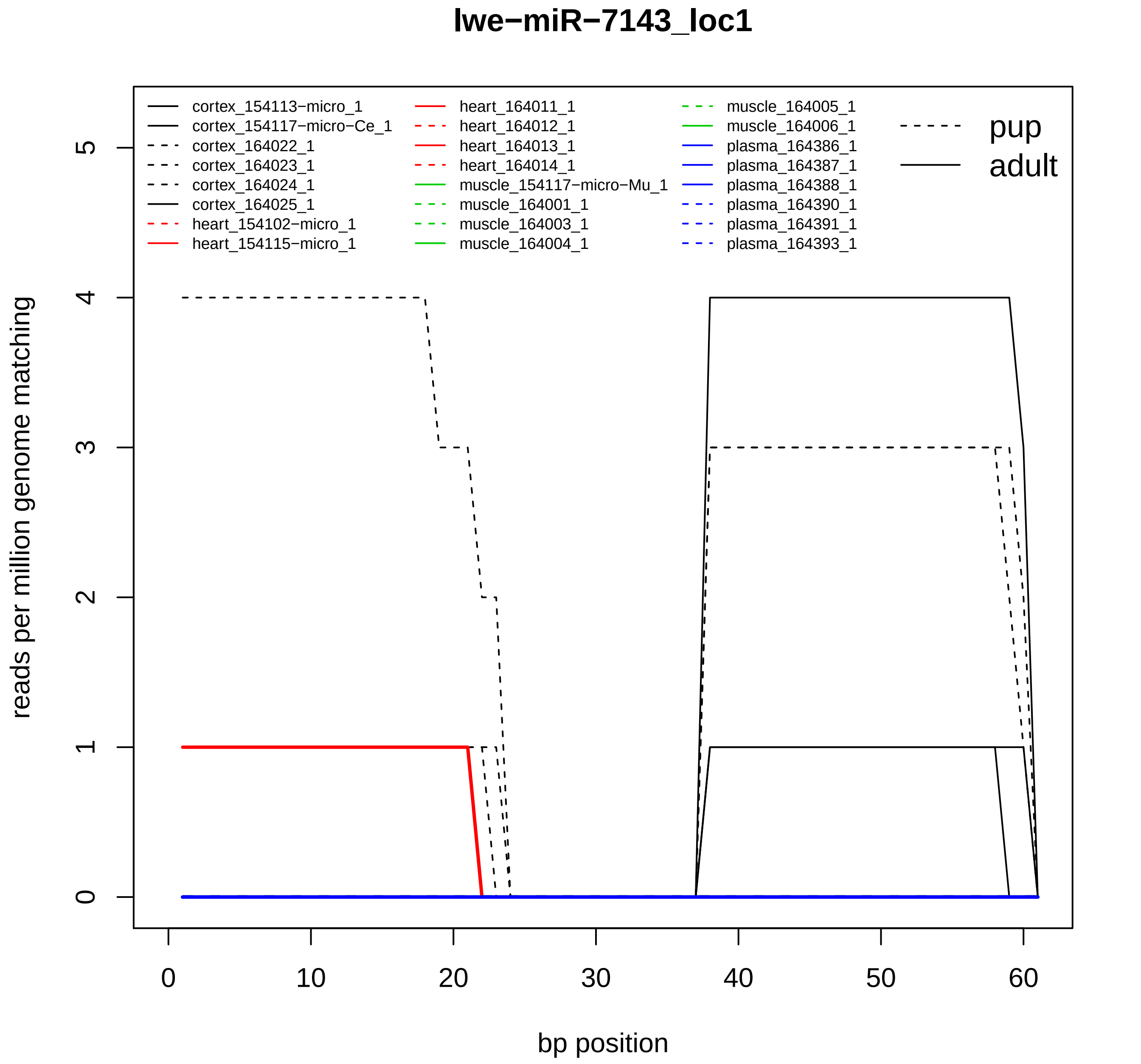
lwe−miR−7143_loc1
cortex_154113−micro_1
cortex_154117−micro−Ce_1
cortex_164022_1
cortex_164023_1
cortex_164024_1
cortex_164025_1
heart_154102−micro_1
heart_154115−micro_1
heart_164011_1
heart_164012_1
heart_164013_1
heart_164014_1
muscle_154117−micro−Mu_1
muscle_164001_1
muscle_164003_1
muscle_164004_1
muscle_164005_1
muscle_164006_1
plasma_164386_1
plasma_164387_1
plasma_164388_1
plasma_164390_1
plasma_164391_1
plasma_164393_1
pup
adult
reads per million genome matching
bp position
0
1
2
3
4
5
0
10
20
30
40
50
60

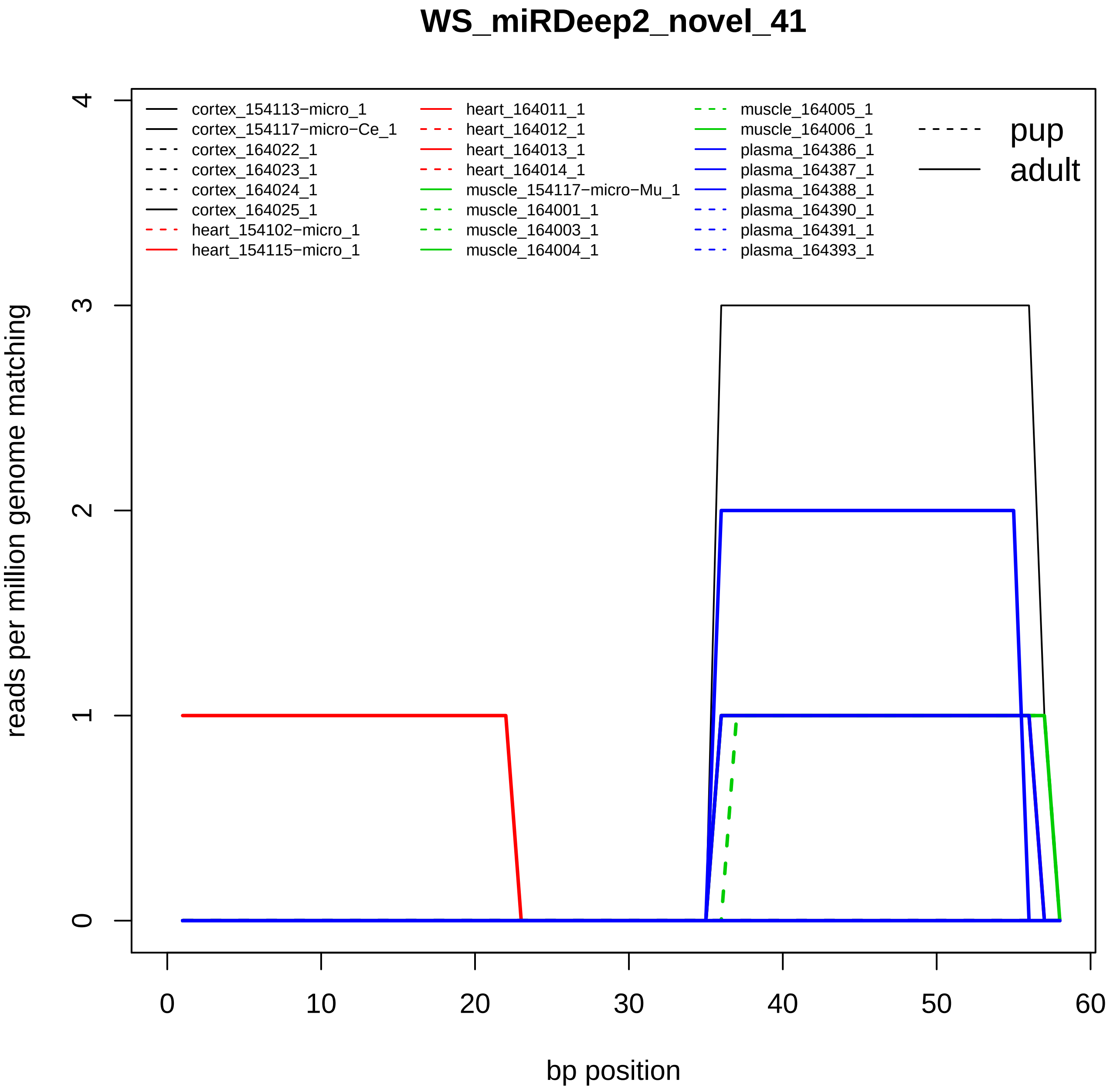
WS_miRDeep2_novel_41
cortex_154113−micro_1
cortex_154117−micro−Ce_1
cortex_164022_1
cortex_164023_1
cortex_164024_1
cortex_164025_1
heart_154102−micro_1
heart_154115−micro_1
heart_164011_1
heart_164012_1
heart_164013_1
heart_164014_1
muscle_154117−micro−Mu_1
muscle_164001_1
muscle_164003_1
muscle_164004_1
muscle_164005_1
muscle_164006_1
plasma_164386_1
plasma_164387_1
plasma_164388_1
plasma_164390_1
plasma_164391_1
plasma_164393_1
pup
adult
reads per million genome matching
0
1
2
3
4
0
10
20
30
40
50
60
bp position

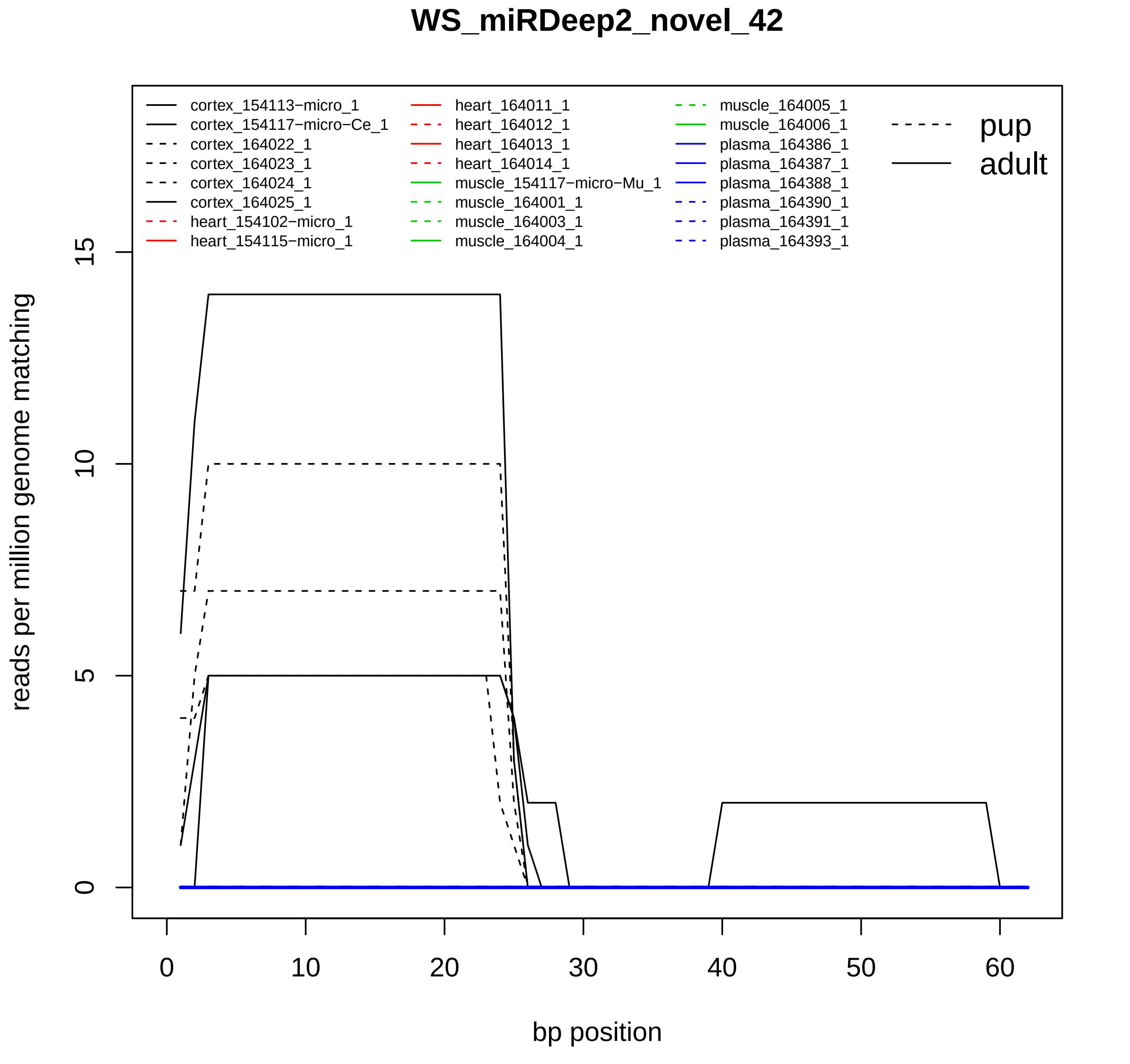
WS_miRDeep2_novel_42
cortex_154113−micro_1
cortex_154117−micro−Ce_1
cortex_164022_1
cortex_164023_1
cortex_164024_1
cortex_164025_1
heart_154102−micro_1
heart_154115−micro_1
heart_164011_1
heart_164012_1
heart_164013_1
heart_164014_1
muscle_154117−micro−Mu_1
muscle_164001_1
muscle_164003_1
muscle_164004_1
muscle_164005_1
muscle_164006_1
plasma_164386_1
plasma_164387_1
plasma_164388_1
plasma_164390_1
plasma_164391_1
plasma_164393_1
pup
adult
reads per million genome matching
0
5
10
15
0
10
20
30
40
50
60
bp position

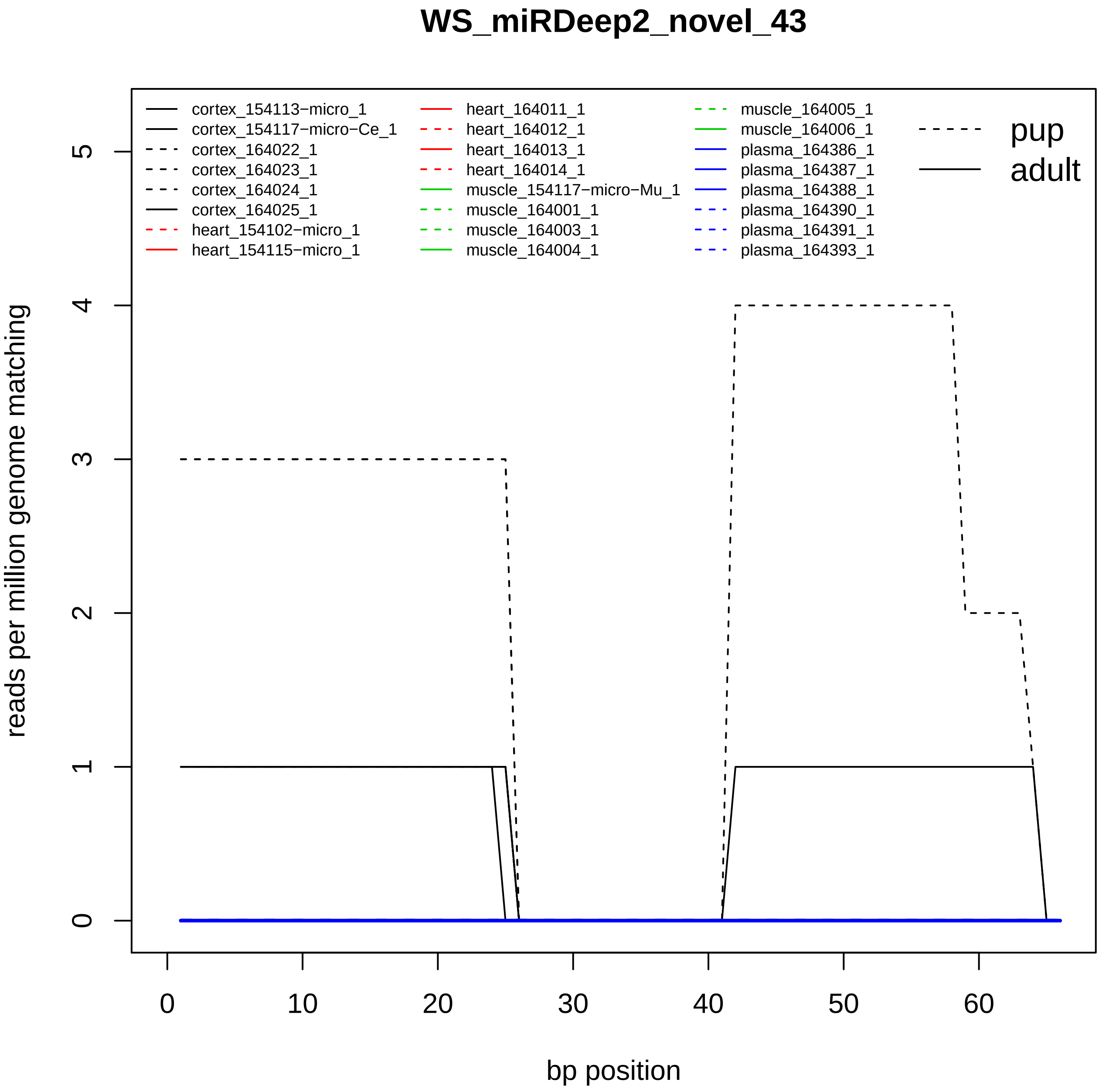

WS_miRDeep2_novel_43
cortex_154113−micro_1
cortex_154117−micro−Ce_1
cortex_164022_1
cortex_164023_1
cortex_164024_1
cortex_164025_1
heart_154102−micro_1
heart_154115−micro_1
heart_164011_1
heart_164012_1
heart_164013_1
heart_164014_1
muscle_154117−micro−Mu_1
muscle_164001_1
muscle_164003_1
muscle_164004_1
muscle_164005_1
muscle_164006_1
plasma_164386_1
plasma_164387_1
plasma_164388_1
plasma_164390_1
plasma_164391_1
plasma_164393_1
pup
adult
reads per million genome matching
bp position
0
1
2
3
4
5
0
10
20
30
40
50
60

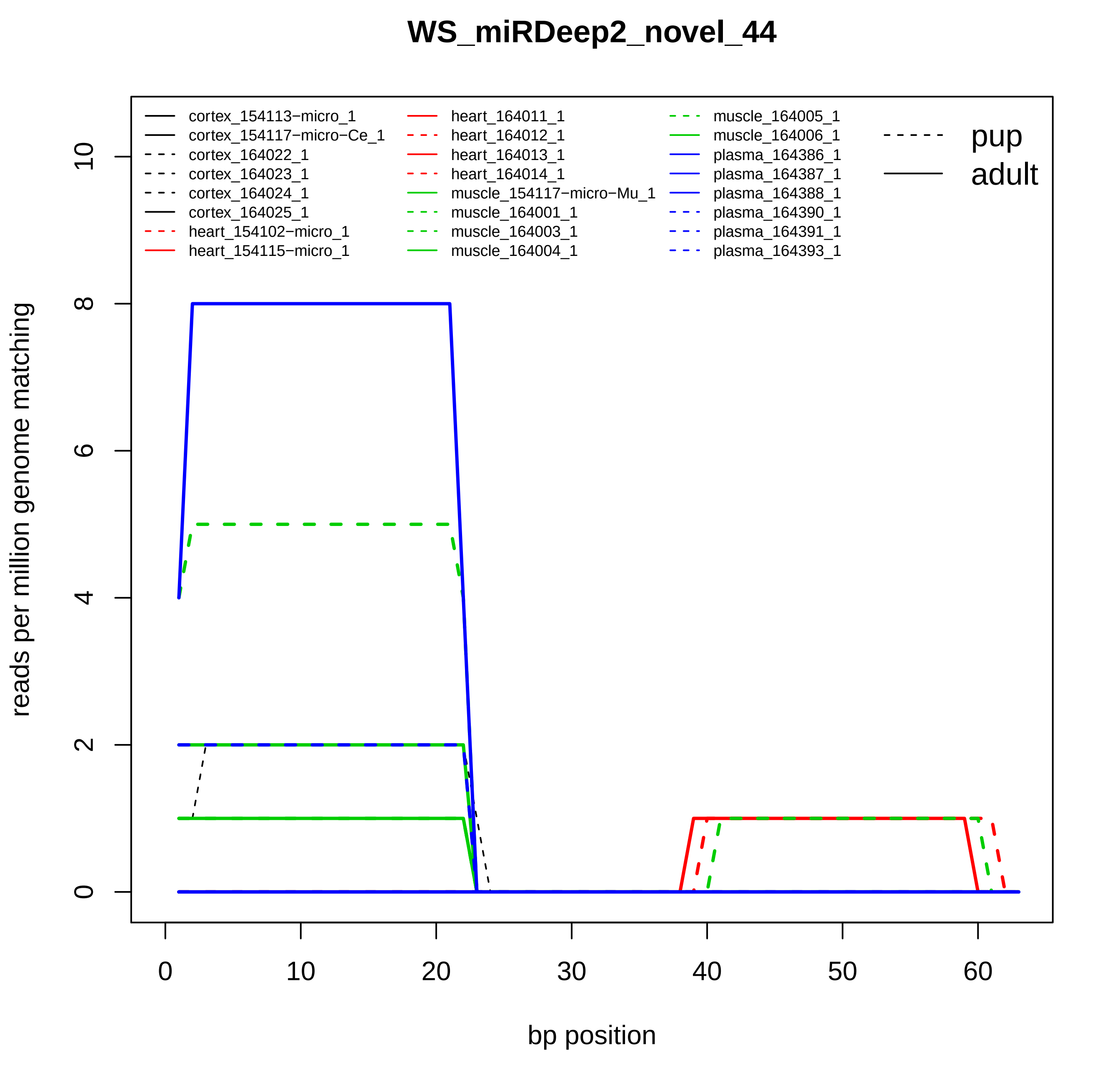
WS_miRDeep2_novel_44
cortex_154113−micro_1
cortex_154117−micro−Ce_1
cortex_164022_1
cortex_164023_1
cortex_164024_1
cortex_164025_1
heart_154102−micro_1
heart_154115−micro_1
heart_164011_1
heart_164012_1
heart_164013_1
heart_164014_1
muscle_154117−micro−Mu_1
muscle_164001_1
muscle_164003_1
muscle_164004_1
muscle_164005_1
muscle_164006_1
plasma_164386_1
plasma_164387_1
plasma_164388_1
plasma_164390_1
plasma_164391_1
plasma_164393_1
pup
adult
reads per million genome matching
0
2
4
6
8
10
0
10
20
30
40
50
60
bp position

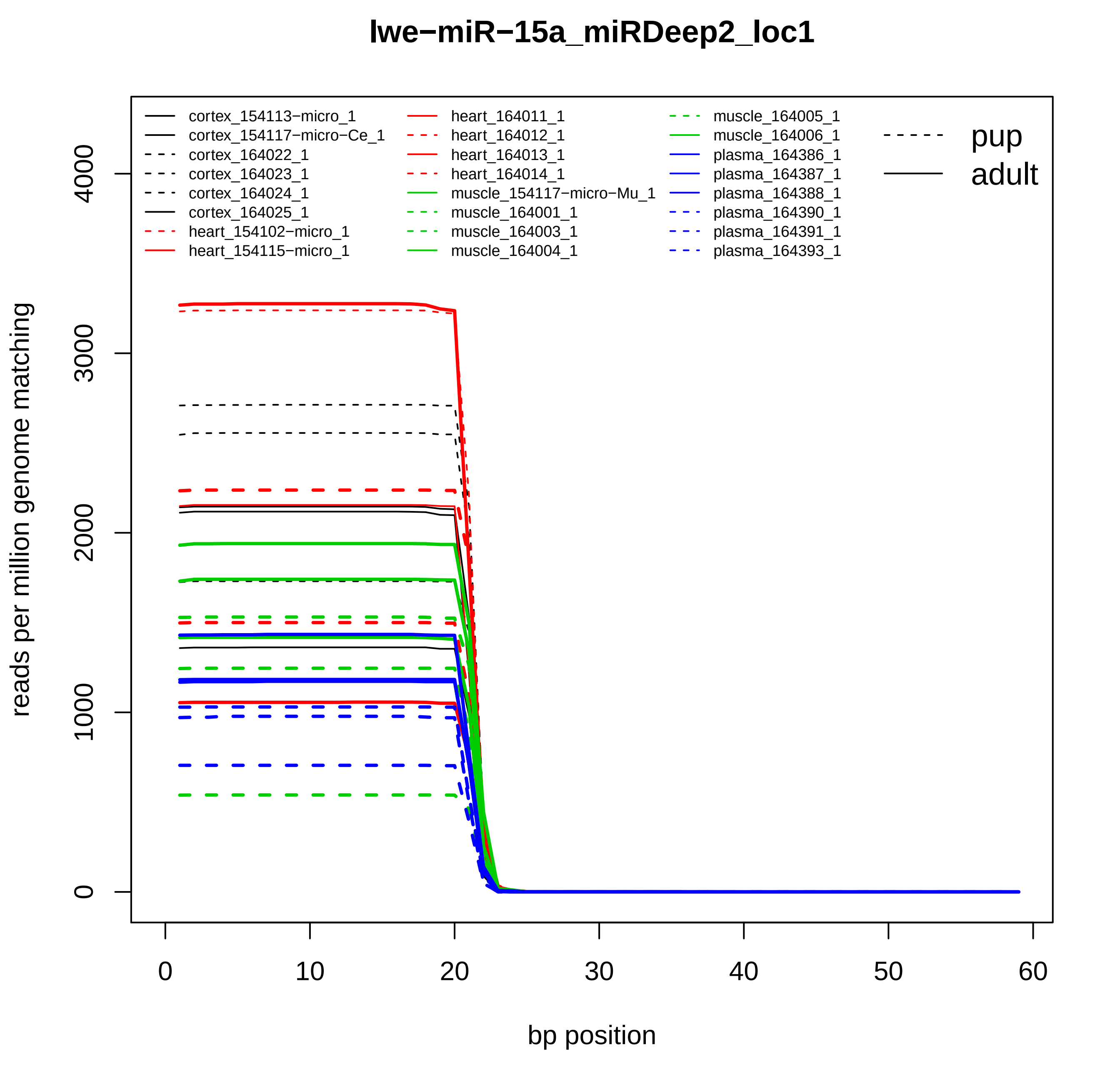
lwe−miR−15a_miRDeep2_loc1
cortex_154113−micro_1
cortex_154117−micro−Ce_1
cortex_164022_1
cortex_164023_1
cortex_164024_1
cortex_164025_1
heart_154102−micro_1
heart_154115−micro_1
heart_164011_1
heart_164012_1
heart_164013_1
heart_164014_1
muscle_154117−micro−Mu_1
muscle_164001_1
muscle_164003_1
muscle_164004_1
muscle_164005_1
muscle_164006_1
plasma_164386_1
plasma_164387_1
plasma_164388_1
plasma_164390_1
plasma_164391_1
plasma_164393_1
pup
adult
reads per million genome matching
0
1000
2000
3000
4000
bp position
0
10
20
30
40
50
60

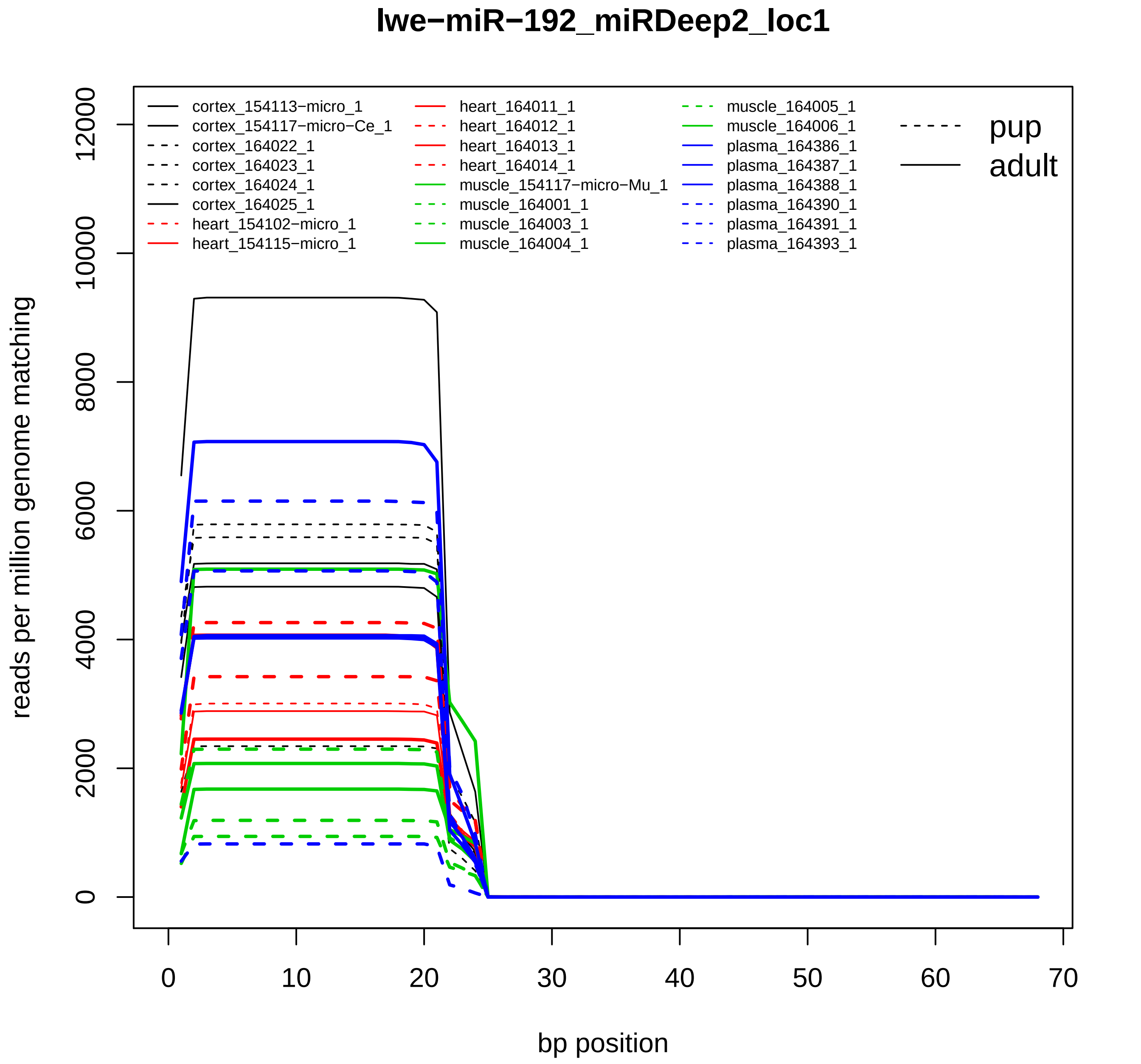

lwe−miR−192_miRDeep2_loc1
reads per million genome matching
bp position
0
2000
4000
6000
8000
10000
12000
0
10
20
30
40
50
60
70
cortex_154113−micro_1
cortex_154117−micro−Ce_1
cortex_164022_1
cortex_164023_1
cortex_164024_1
cortex_164025_1
heart_154102−micro_1
heart_154115−micro_1
heart_164011_1
heart_164012_1
heart_164013_1
heart_164014_1
muscle_154117−micro−Mu_1
muscle_164001_1
muscle_164003_1
muscle_164004_1
muscle_164005_1
muscle_164006_1
plasma_164386_1
plasma_164387_1
plasma_164388_1
plasma_164390_1
plasma_164391_1
plasma_164393_1
pup
adult

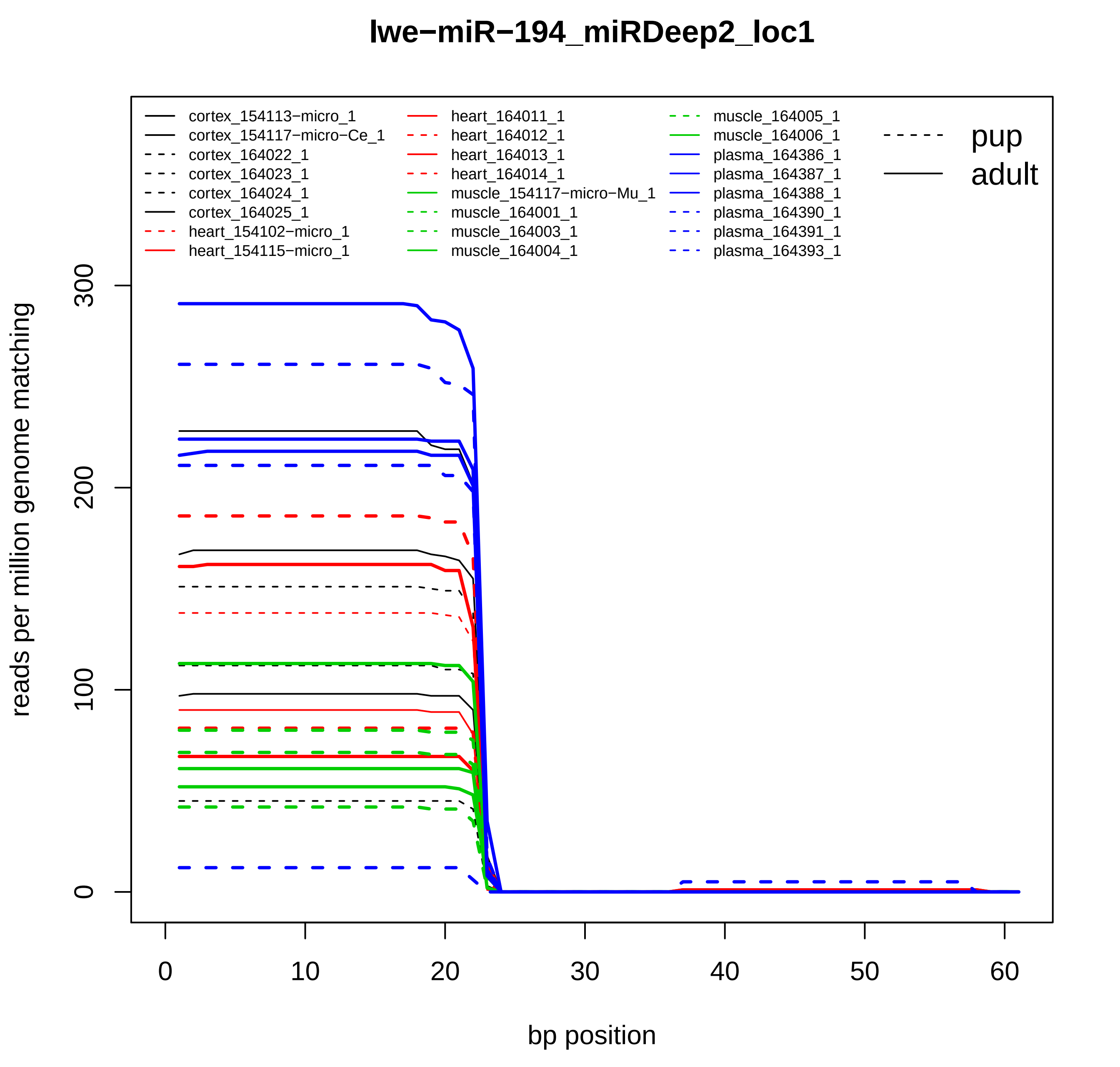

lwe−miR−194_miRDeep2_loc1
cortex_154113−micro_1
cortex_154117−micro−Ce_1
cortex_164022_1
cortex_164023_1
cortex_164024_1
cortex_164025_1
heart_154102−micro_1
heart_154115−micro_1
heart_164011_1
heart_164012_1
heart_164013_1
heart_164014_1
muscle_154117−micro−Mu_1
muscle_164001_1
muscle_164003_1
muscle_164004_1
muscle_164005_1
muscle_164006_1
plasma_164386_1
plasma_164387_1
plasma_164388_1
plasma_164390_1
plasma_164391_1
plasma_164393_1
pup
adult
reads per million genome matching
0
100
200
300
bp position
0
10
20
30
40
50
60

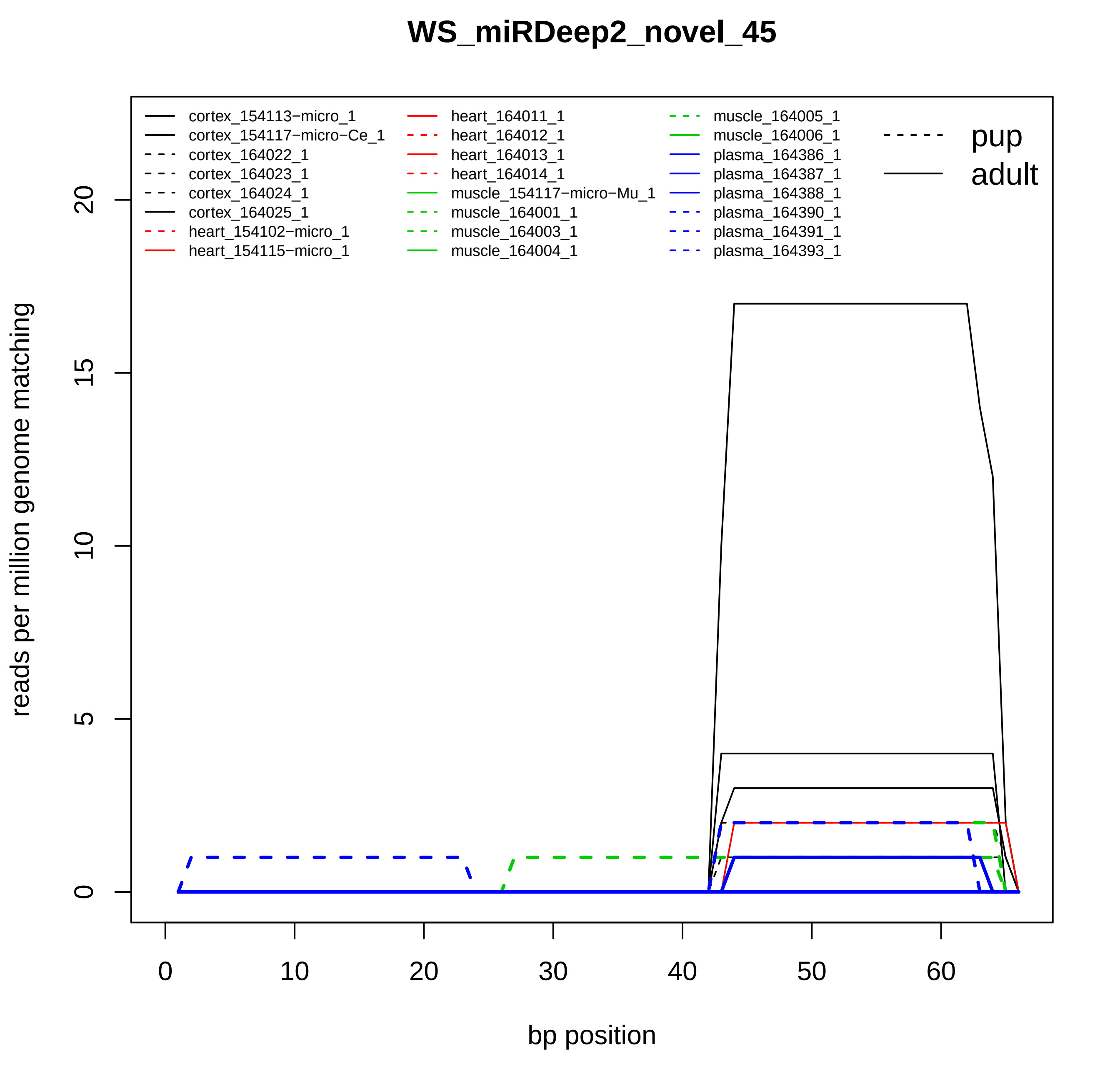
WS_miRDeep2_novel_45
cortex_154113−micro_1
cortex_154117−micro−Ce_1
cortex_164022_1
cortex_164023_1
cortex_164024_1
cortex_164025_1
heart_154102−micro_1
heart_154115−micro_1
heart_164011_1
heart_164012_1
heart_164013_1
heart_164014_1
muscle_154117−micro−Mu_1
muscle_164001_1
muscle_164003_1
muscle_164004_1
muscle_164005_1
muscle_164006_1
plasma_164386_1
plasma_164387_1
plasma_164388_1
plasma_164390_1
plasma_164391_1
plasma_164393_1
pup
adult
reads per million genome matching
0
5
10
15
20
bp position
0
10
20
30
40
50
60

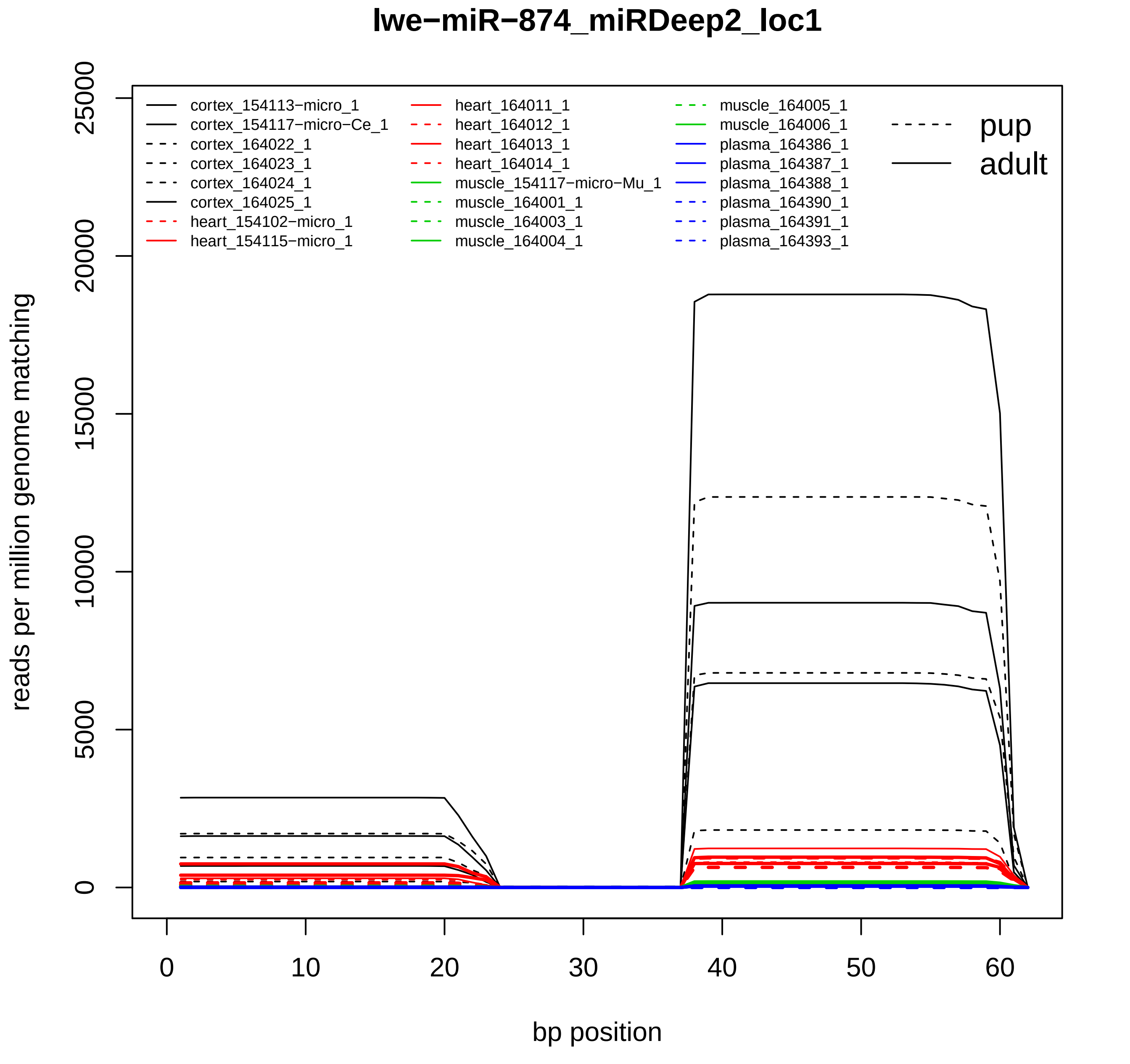
lwe−miR−874_miRDeep2_loc1
reads per million genome matching
bp position
0
5000
10000
15000
20000
25000
0
10
20
30
40
50
60
cortex_154113−micro_1
cortex_154117−micro−Ce_1
cortex_164022_1
cortex_164023_1
cortex_164024_1
cortex_164025_1
heart_154102−micro_1
heart_154115−micro_1
heart_164011_1
heart_164012_1
heart_164013_1
heart_164014_1
muscle_154117−micro−Mu_1
muscle_164001_1
muscle_164003_1
muscle_164004_1
muscle_164005_1
muscle_164006_1
plasma_164386_1
plasma_164387_1
plasma_164388_1
plasma_164390_1
plasma_164391_1
plasma_164393_1
pup
adult

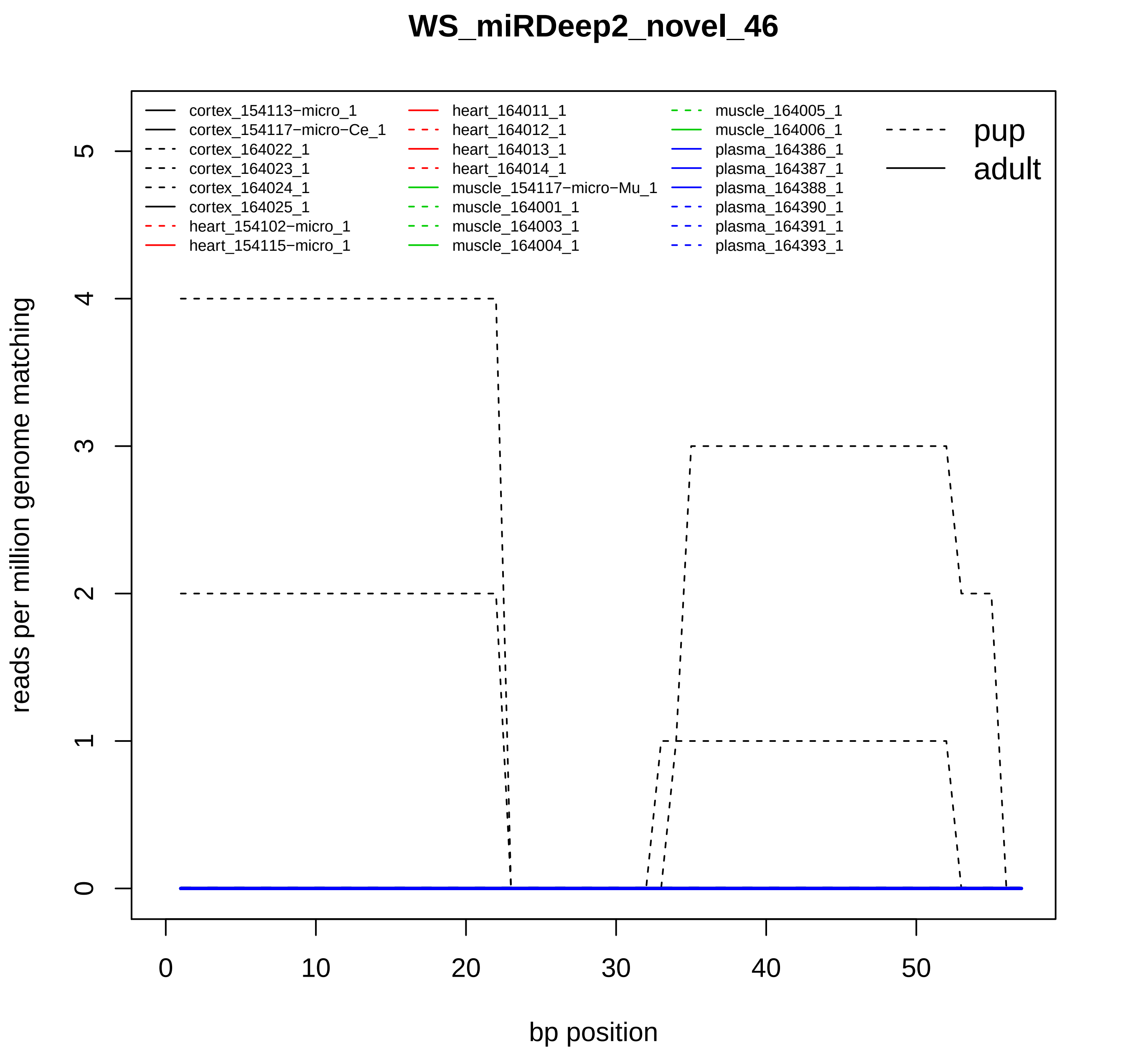

WS_miRDeep2_novel_46
cortex_154113−micro_1
cortex_154117−micro−Ce_1
cortex_164022_1
cortex_164023_1
cortex_164024_1
cortex_164025_1
heart_154102−micro_1
heart_154115−micro_1
heart_164011_1
heart_164012_1
heart_164013_1
heart_164014_1
muscle_154117−micro−Mu_1
muscle_164001_1
muscle_164003_1
muscle_164004_1
muscle_164005_1
muscle_164006_1
plasma_164386_1
plasma_164387_1
plasma_164388_1
plasma_164390_1
plasma_164391_1
plasma_164393_1
pup
adult
reads per million genome matching
bp position
0
1
2
3
4
5
0
10
20
30
40
50

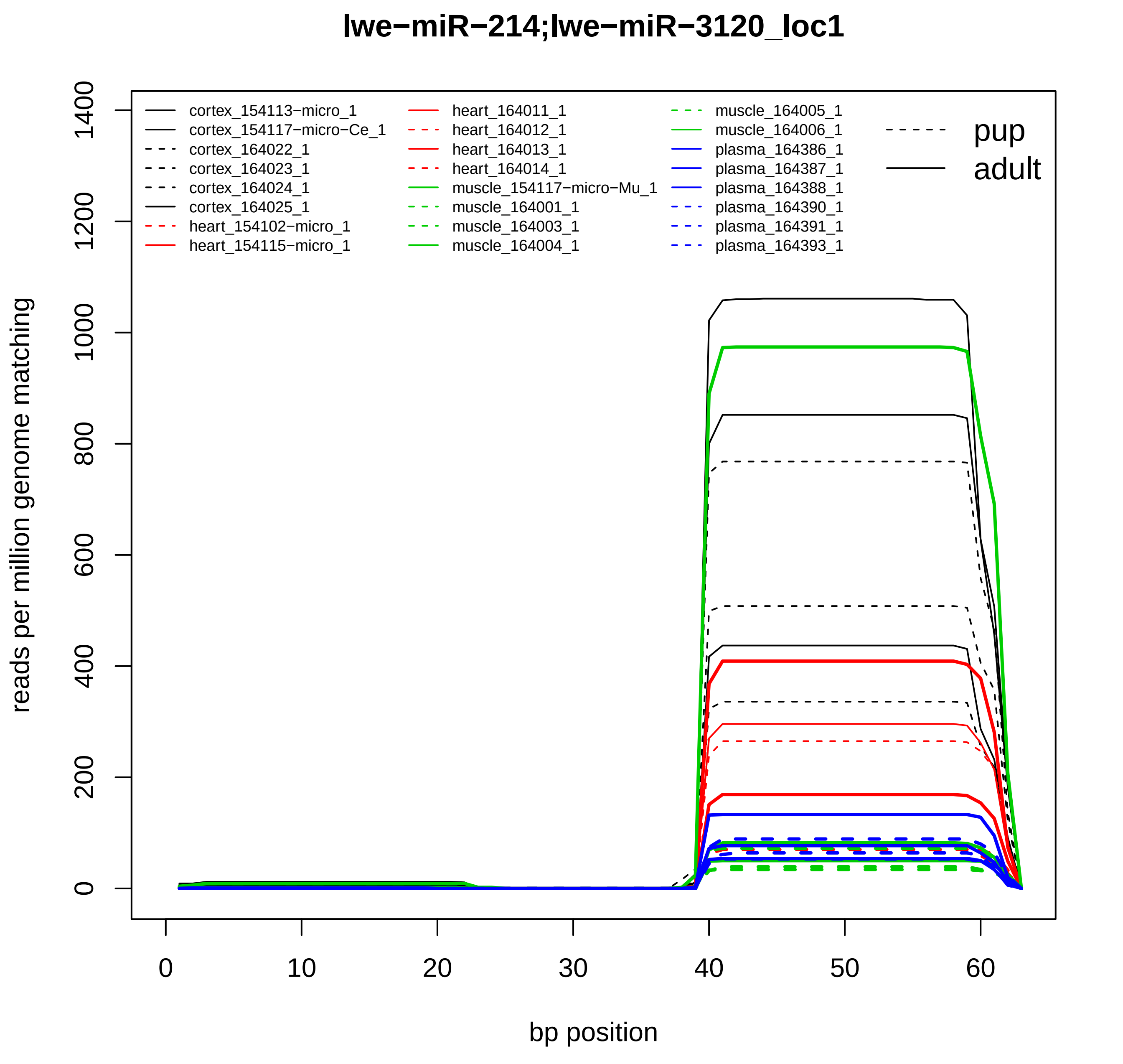

lwe−miR−214;lwe−miR−3120_loc1
cortex_154113−micro_1
cortex_154117−micro−Ce_1
cortex_164022_1
cortex_164023_1
cortex_164024_1
cortex_164025_1
heart_154102−micro_1
heart_154115−micro_1
heart_164011_1
heart_164012_1
heart_164013_1
heart_164014_1
muscle_154117−micro−Mu_1
muscle_164001_1
muscle_164003_1
muscle_164004_1
muscle_164005_1
muscle_164006_1
plasma_164386_1
plasma_164387_1
plasma_164388_1
plasma_164390_1
plasma_164391_1
plasma_164393_1
pup
adult
reads per million genome matching
0
200
400
600
800
1000
1200
1400
0
10
20
30
40
50
60
bp position

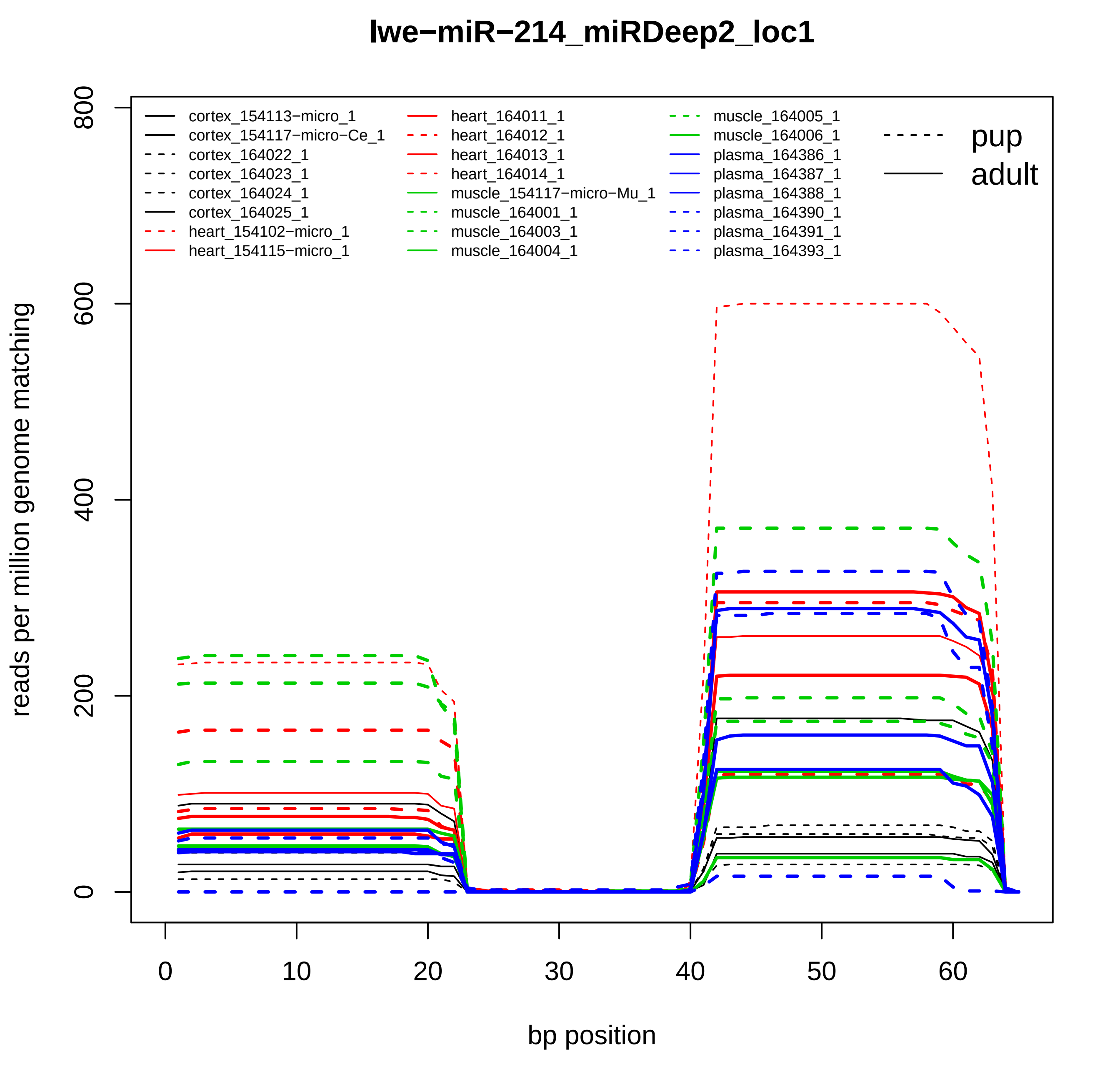

lwe−miR−214_miRDeep2_loc1
cortex_154113−micro_1
cortex_154117−micro−Ce_1
cortex_164022_1
cortex_164023_1
cortex_164024_1
cortex_164025_1
heart_154102−micro_1
heart_154115−micro_1
heart_164011_1
heart_164012_1
heart_164013_1
heart_164014_1
muscle_154117−micro−Mu_1
muscle_164001_1
muscle_164003_1
muscle_164004_1
muscle_164005_1
muscle_164006_1
plasma_164386_1
plasma_164387_1
plasma_164388_1
plasma_164390_1
plasma_164391_1
plasma_164393_1
pup
adult
reads per million genome matching
0
200
400
600
800
bp position
0
10
20
30
40
50
60

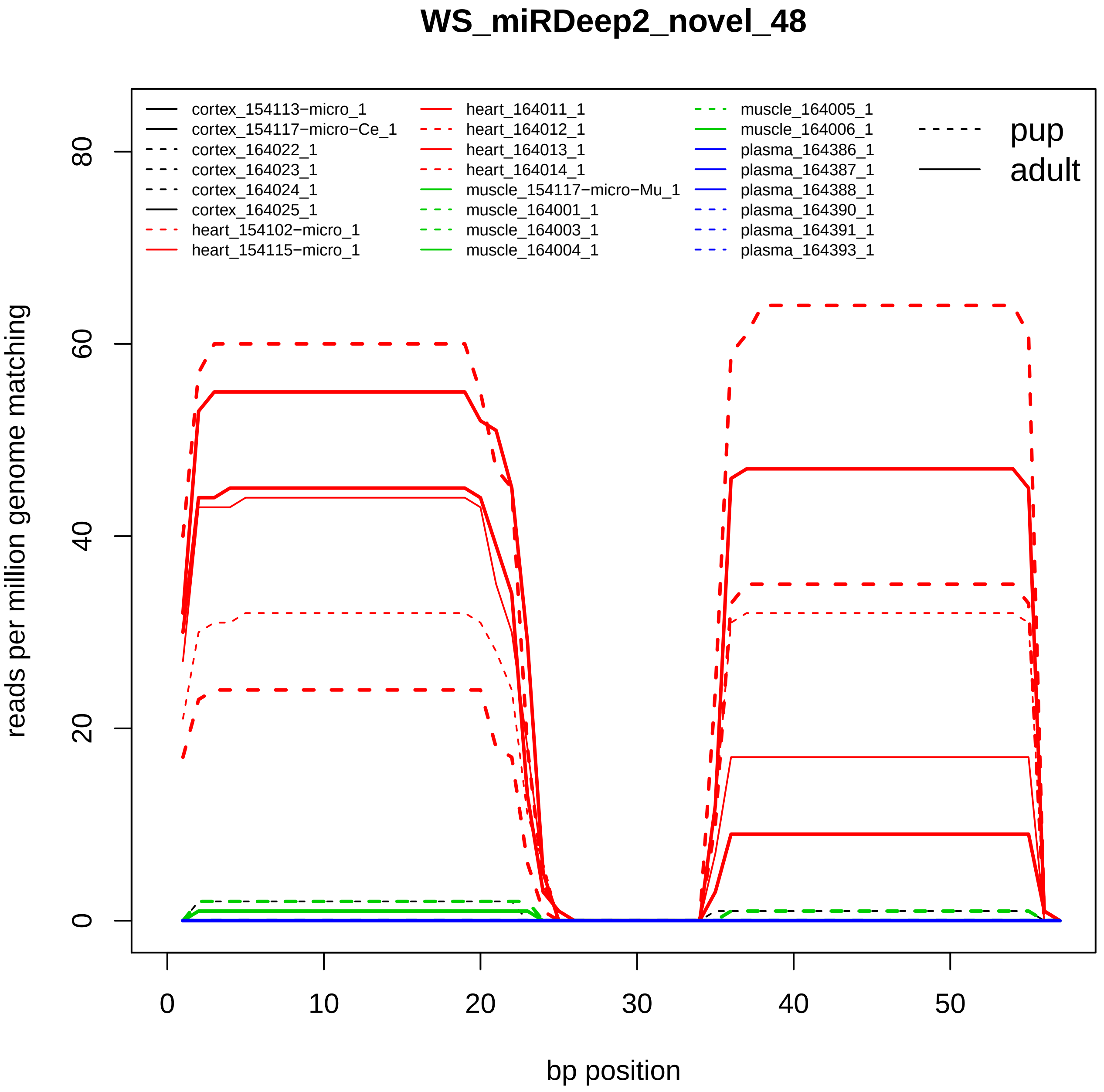

WS_miRDeep2_novel_48
cortex_154113−micro_1
cortex_154117−micro−Ce_1
cortex_164022_1
cortex_164023_1
cortex_164024_1
cortex_164025_1
heart_154102−micro_1
heart_154115−micro_1
heart_164011_1
heart_164012_1
heart_164013_1
heart_164014_1
muscle_154117−micro−Mu_1
muscle_164001_1
muscle_164003_1
muscle_164004_1
muscle_164005_1
muscle_164006_1
plasma_164386_1
plasma_164387_1
plasma_164388_1
plasma_164390_1
plasma_164391_1
plasma_164393_1
pup
adult
reads per million genome matching
bp position
0
20
40
60
80
0
10
20
30
40
50

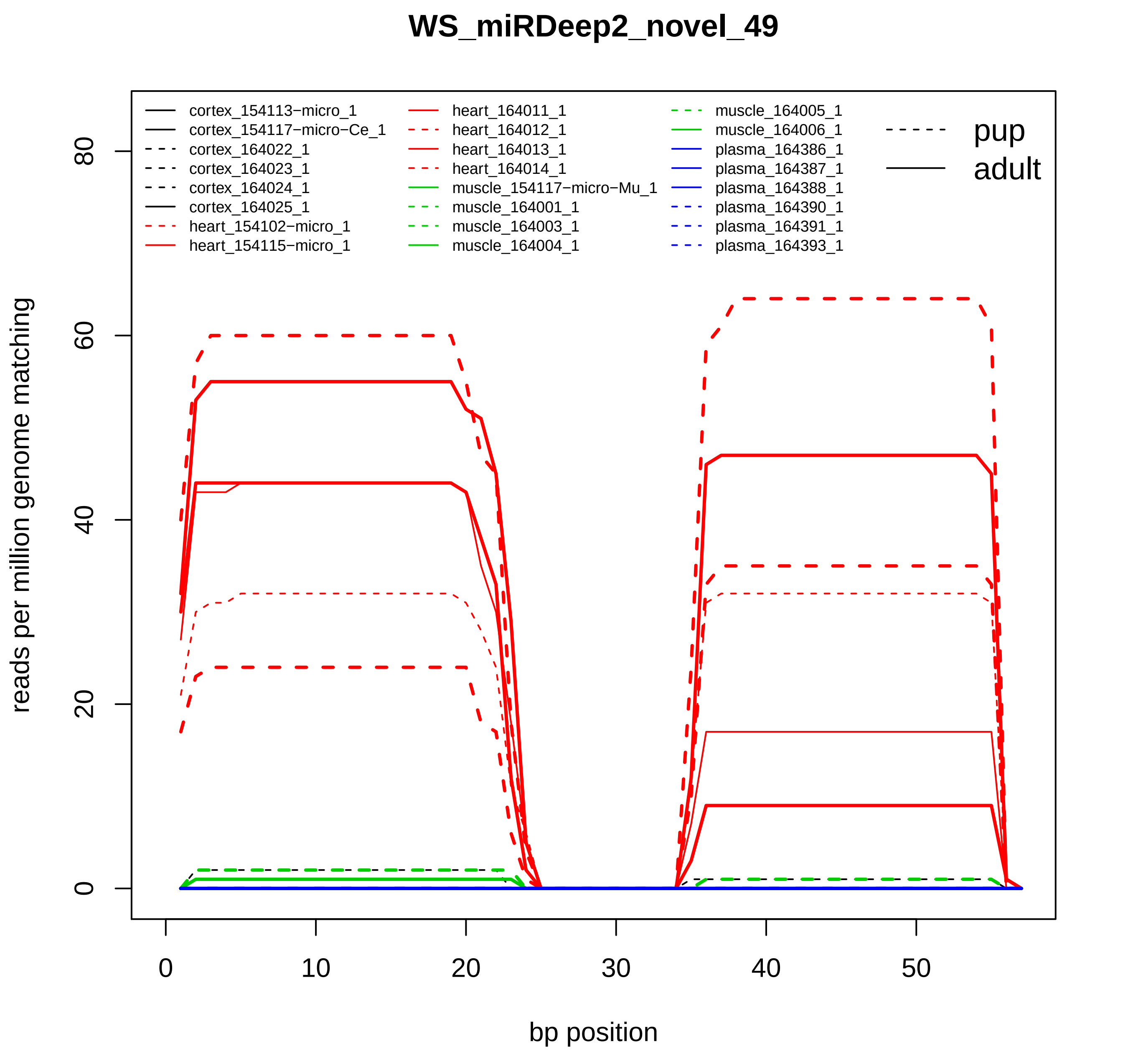

WS_miRDeep2_novel_49
cortex_154113−micro_1
cortex_154117−micro−Ce_1
cortex_164022_1
cortex_164023_1
cortex_164024_1
cortex_164025_1
heart_154102−micro_1
heart_154115−micro_1
heart_164011_1
heart_164012_1
heart_164013_1
heart_164014_1
muscle_154117−micro−Mu_1
muscle_164001_1
muscle_164003_1
muscle_164004_1
muscle_164005_1
muscle_164006_1
plasma_164386_1
plasma_164387_1
plasma_164388_1
plasma_164390_1
plasma_164391_1
plasma_164393_1
pup
adult
reads per million genome matching
0
20
40
60
80
0
10
20
30
40
50
bp position

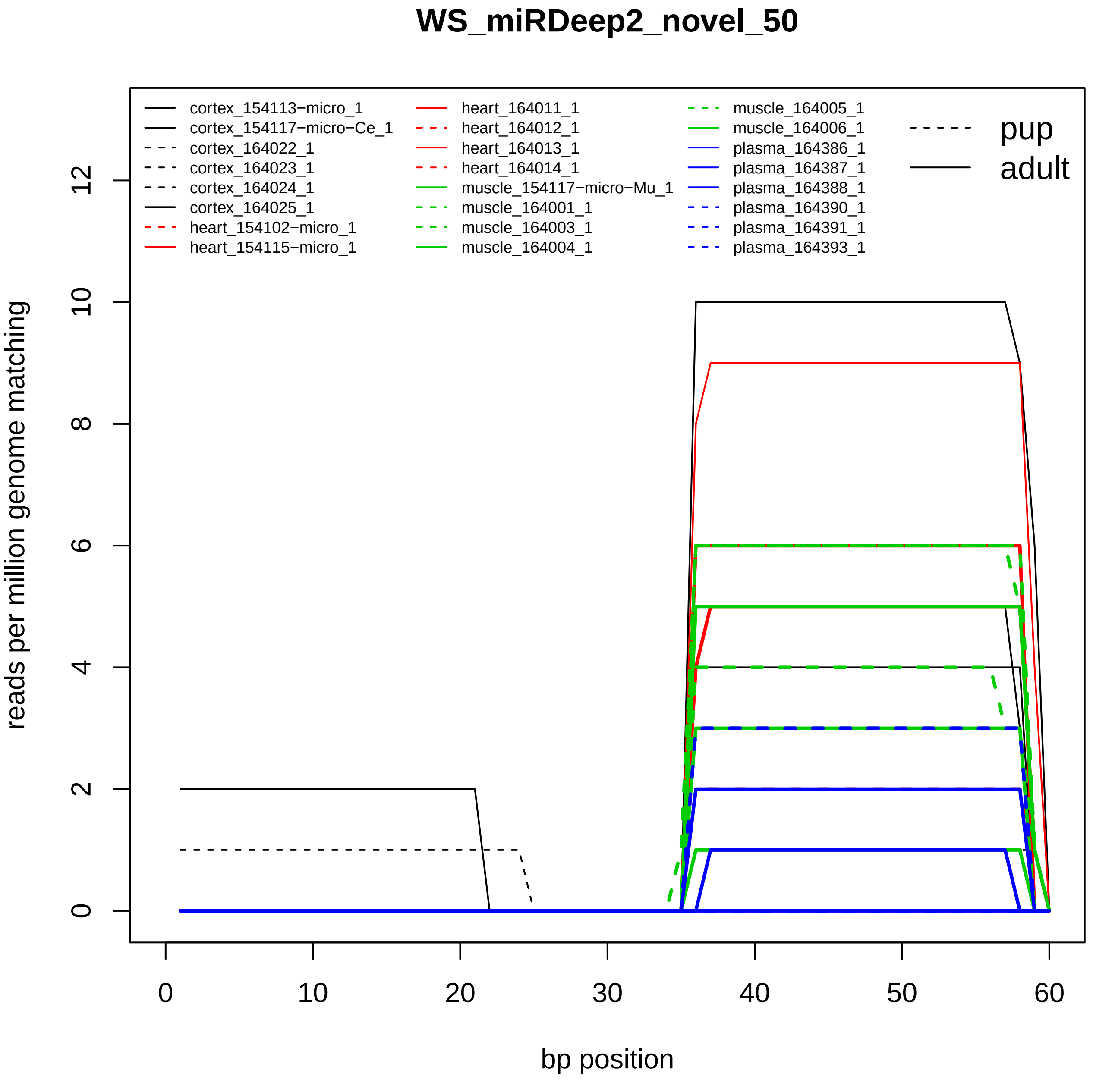

WS_miRDeep2_novel_50
cortex_154113−micro_1
cortex_154117−micro−Ce_1
cortex_164022_1
cortex_164023_1
cortex_164024_1
cortex_164025_1
heart_154102−micro_1
heart_154115−micro_1
heart_164011_1
heart_164012_1
heart_164013_1
heart_164014_1
muscle_154117−micro−Mu_1
muscle_164001_1
muscle_164003_1
muscle_164004_1
muscle_164005_1
muscle_164006_1
plasma_164386_1
plasma_164387_1
plasma_164388_1
plasma_164390_1
plasma_164391_1
plasma_164393_1
pup
adult
reads per million genome matching
bp position
0
2
4
6
8
10
12
0
10
20
30
40
50
60

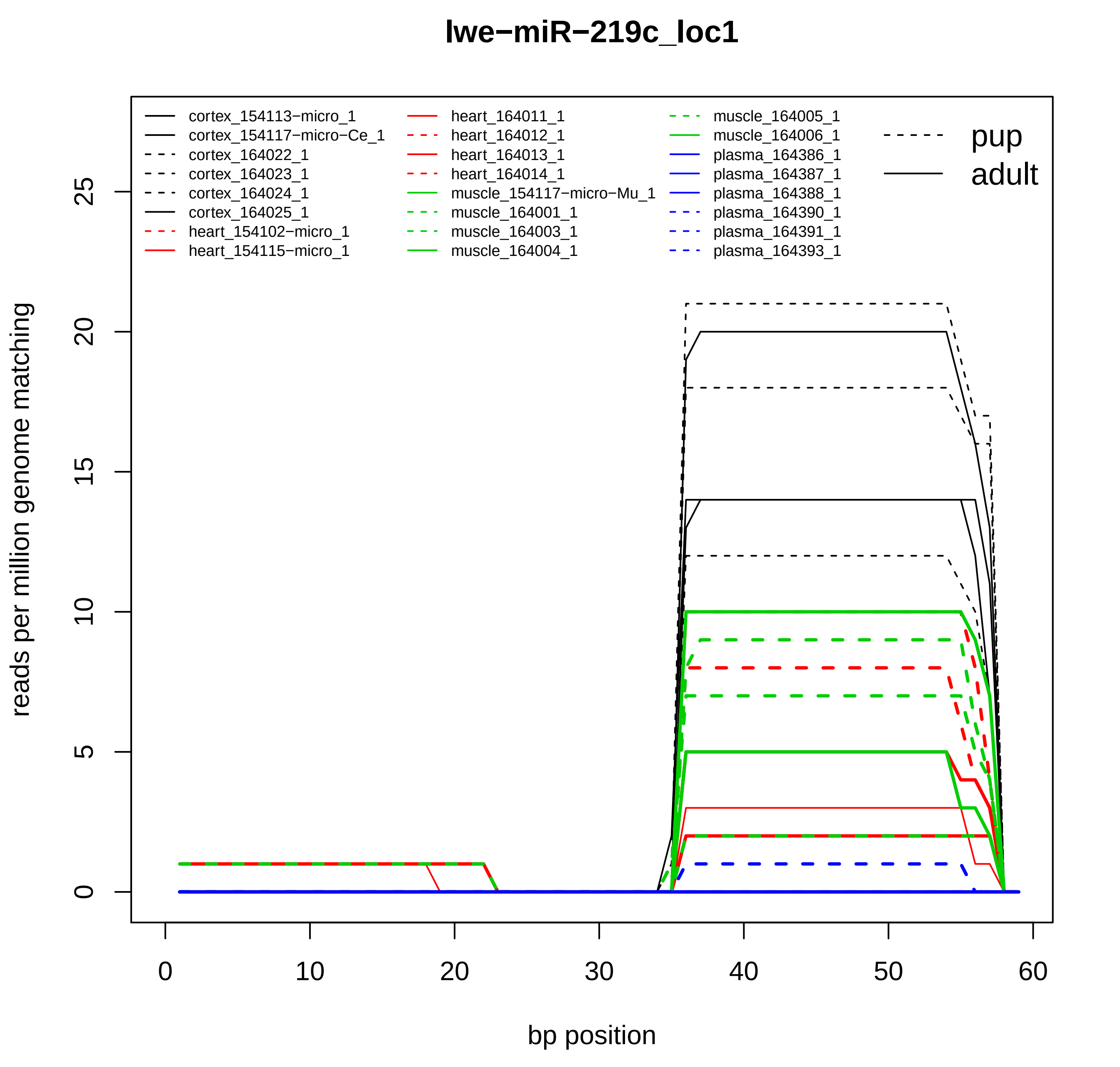

lwe−miR−219c_loc1
cortex_154113−micro_1
cortex_154117−micro−Ce_1
cortex_164022_1
cortex_164023_1
cortex_164024_1
cortex_164025_1
heart_154102−micro_1
heart_154115−micro_1
heart_164011_1
heart_164012_1
heart_164013_1
heart_164014_1
muscle_154117−micro−Mu_1
muscle_164001_1
muscle_164003_1
muscle_164004_1
muscle_164005_1
muscle_164006_1
plasma_164386_1
plasma_164387_1
plasma_164388_1
plasma_164390_1
plasma_164391_1
plasma_164393_1
pup
adult
reads per million genome matching
bp position
0
5
10
15
20
25
0
10
20
30
40
50
60

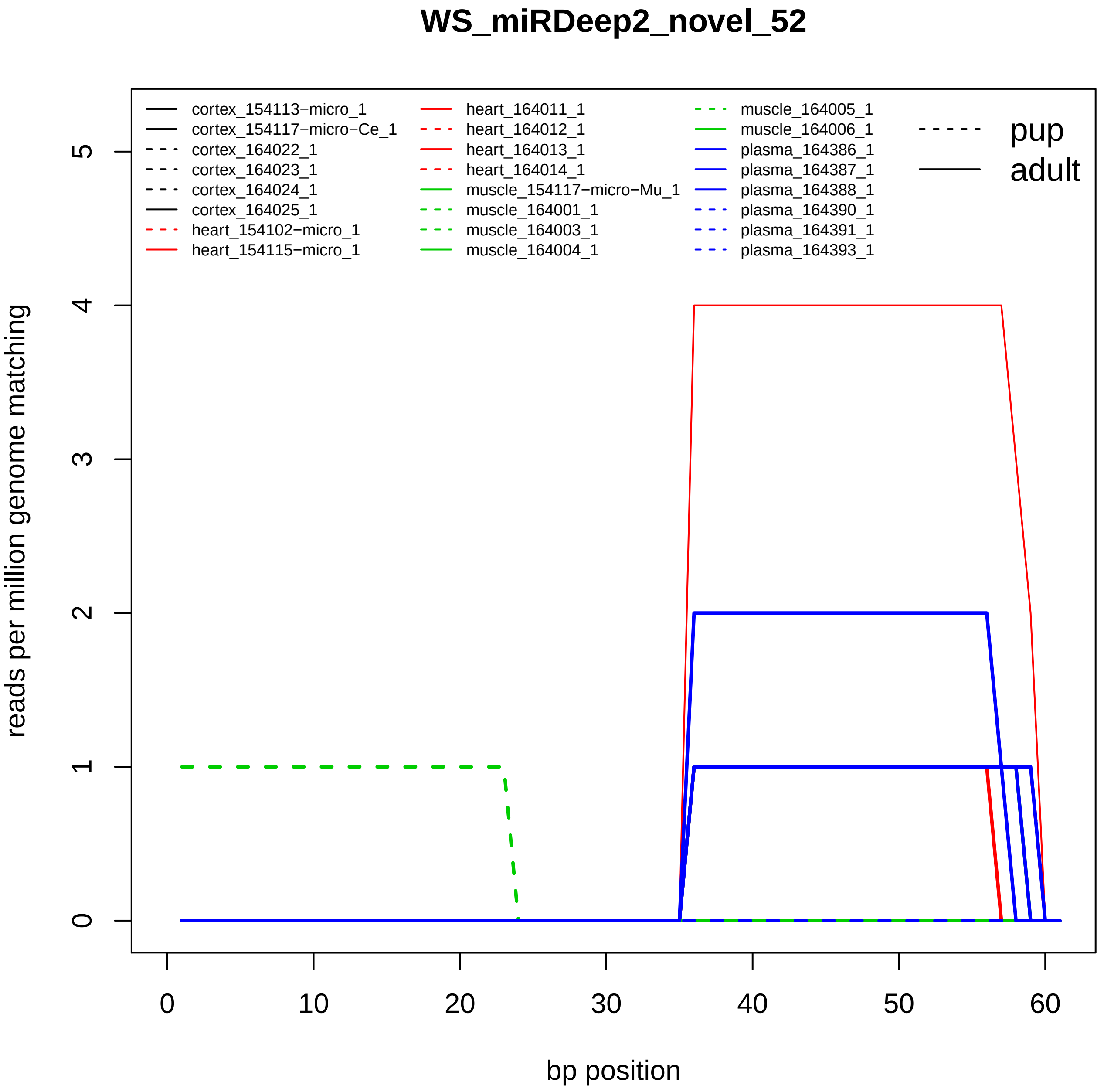
WS_miRDeep2_novel_52
cortex_154113−micro_1
cortex_154117−micro−Ce_1
cortex_164022_1
cortex_164023_1
cortex_164024_1
cortex_164025_1
heart_154102−micro_1
heart_154115−micro_1
heart_164011_1
heart_164012_1
heart_164013_1
heart_164014_1
muscle_154117−micro−Mu_1
muscle_164001_1
muscle_164003_1
muscle_164004_1
muscle_164005_1
muscle_164006_1
plasma_164386_1
plasma_164387_1
plasma_164388_1
plasma_164390_1
plasma_164391_1
plasma_164393_1
pup
adult
reads per million genome matching
bp position
0
1
2
3
4
5
0
10
20
30
40
50
60

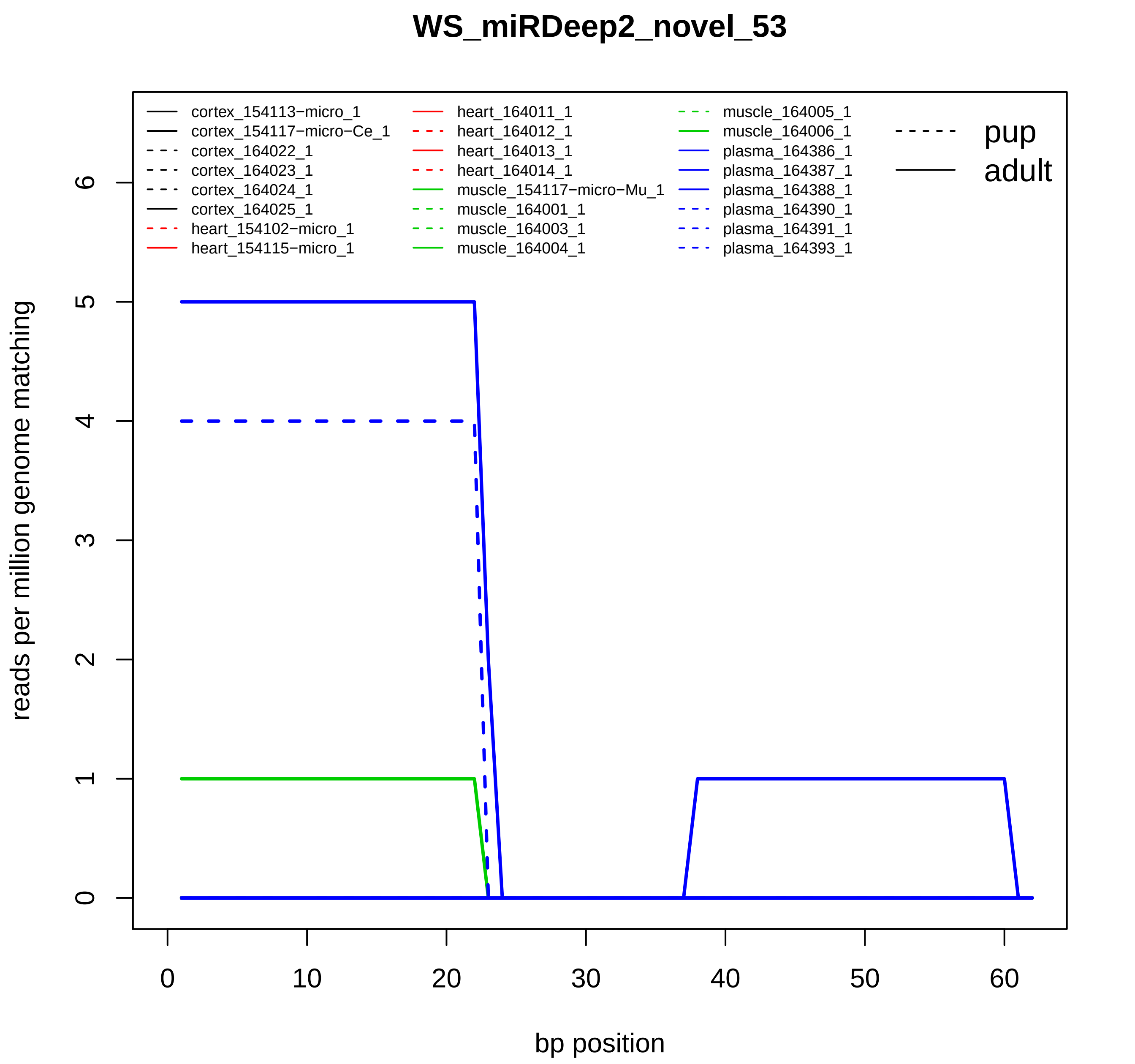

WS_miRDeep2_novel_53
cortex_154113−micro_1
cortex_154117−micro−Ce_1
cortex_164022_1
cortex_164023_1
cortex_164024_1
cortex_164025_1
heart_154102−micro_1
heart_154115−micro_1
heart_164011_1
heart_164012_1
heart_164013_1
heart_164014_1
muscle_154117−micro−Mu_1
muscle_164001_1
muscle_164003_1
muscle_164004_1
muscle_164005_1
muscle_164006_1
plasma_164386_1
plasma_164387_1
plasma_164388_1
plasma_164390_1
plasma_164391_1
plasma_164393_1
pup
adult
reads per million genome matching
0
1
2
3
4
5
6
0
10
20
30
40
50
60
bp position

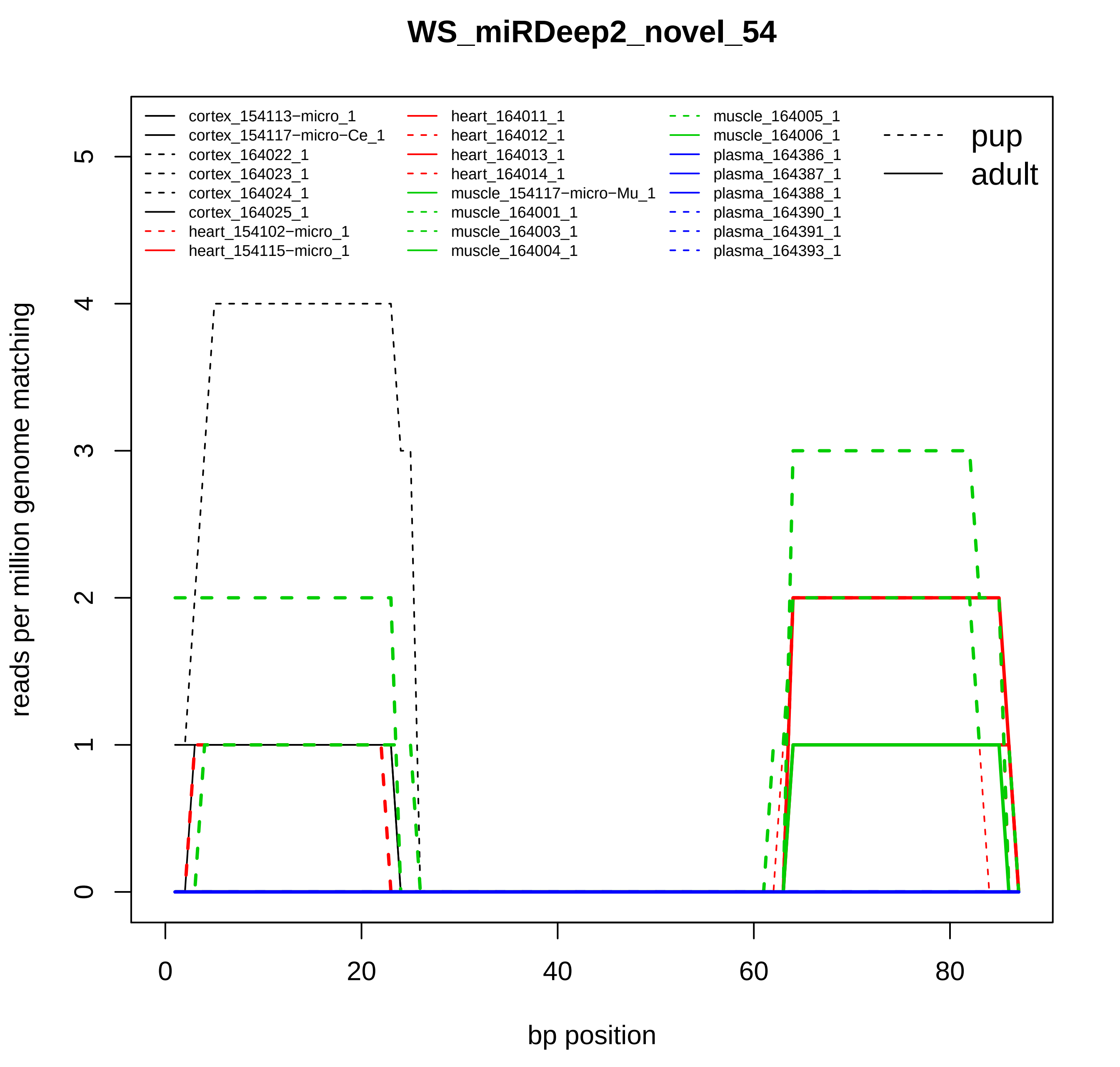

WS_miRDeep2_novel_54
cortex_154113−micro_1
cortex_154117−micro−Ce_1
cortex_164022_1
cortex_164023_1
cortex_164024_1
cortex_164025_1
heart_154102−micro_1
heart_154115−micro_1
heart_164011_1
heart_164012_1
heart_164013_1
heart_164014_1
muscle_154117−micro−Mu_1
muscle_164001_1
muscle_164003_1
muscle_164004_1
muscle_164005_1
muscle_164006_1
plasma_164386_1
plasma_164387_1
plasma_164388_1
plasma_164390_1
plasma_164391_1
plasma_164393_1
pup
adult
reads per million genome matching
0
1
2
3
4
5
0
20
40
60
80
bp position

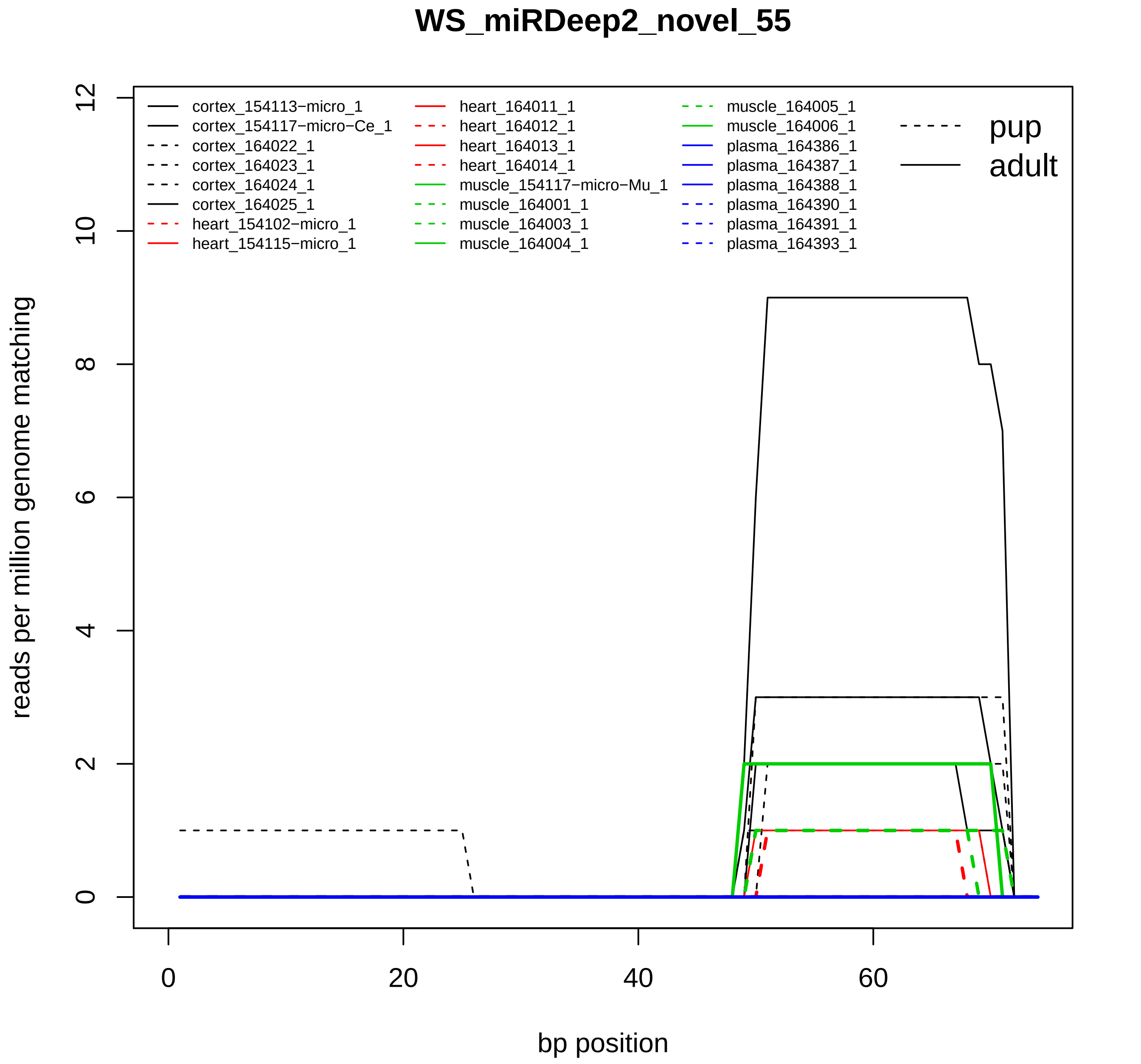

WS_miRDeep2_novel_55
reads per million genome matching
bp position
0
2
4
6
8
10
12
0
20
40
60
cortex_154113−micro_1
cortex_154117−micro−Ce_1
cortex_164022_1
cortex_164023_1
cortex_164024_1
cortex_164025_1
heart_154102−micro_1
heart_154115−micro_1
heart_164011_1
heart_164012_1
heart_164013_1
heart_164014_1
muscle_154117−micro−Mu_1
muscle_164001_1
muscle_164003_1
muscle_164004_1
muscle_164005_1
muscle_164006_1
plasma_164386_1
plasma_164387_1
plasma_164388_1
plasma_164390_1
plasma_164391_1
plasma_164393_1
pup
adult

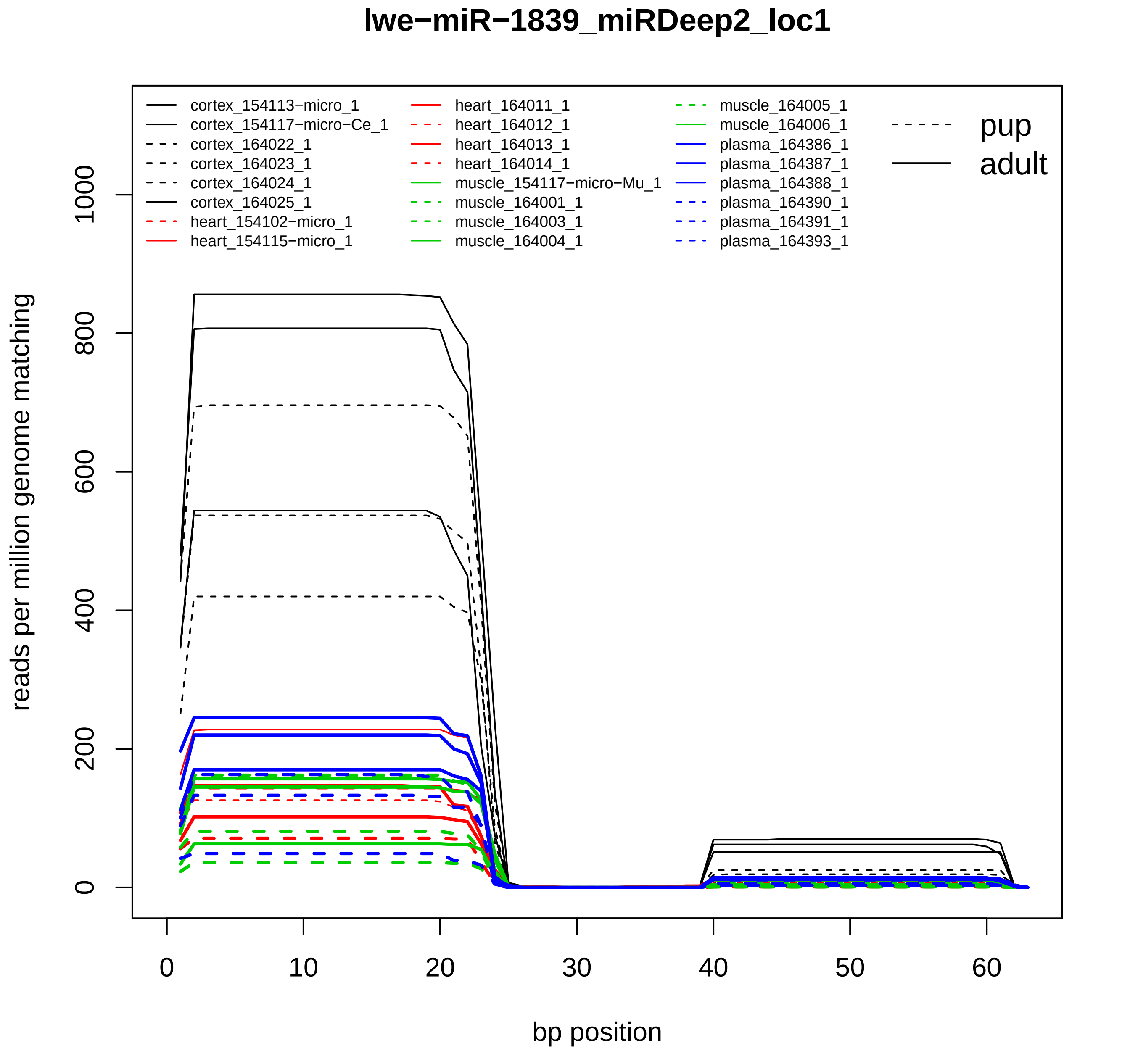
lwe−miR−1839_miRDeep2_loc1
cortex_154113−micro_1
cortex_154117−micro−Ce_1
cortex_164022_1
cortex_164023_1
cortex_164024_1
cortex_164025_1
heart_154102−micro_1
heart_154115−micro_1
heart_164011_1
heart_164012_1
heart_164013_1
heart_164014_1
muscle_154117−micro−Mu_1
muscle_164001_1
muscle_164003_1
muscle_164004_1
muscle_164005_1
muscle_164006_1
plasma_164386_1
plasma_164387_1
plasma_164388_1
plasma_164390_1
plasma_164391_1
plasma_164393_1
pup
adult
reads per million genome matching
0
200
400
600
800
1000
bp position
0
10
20
30
40
50
60

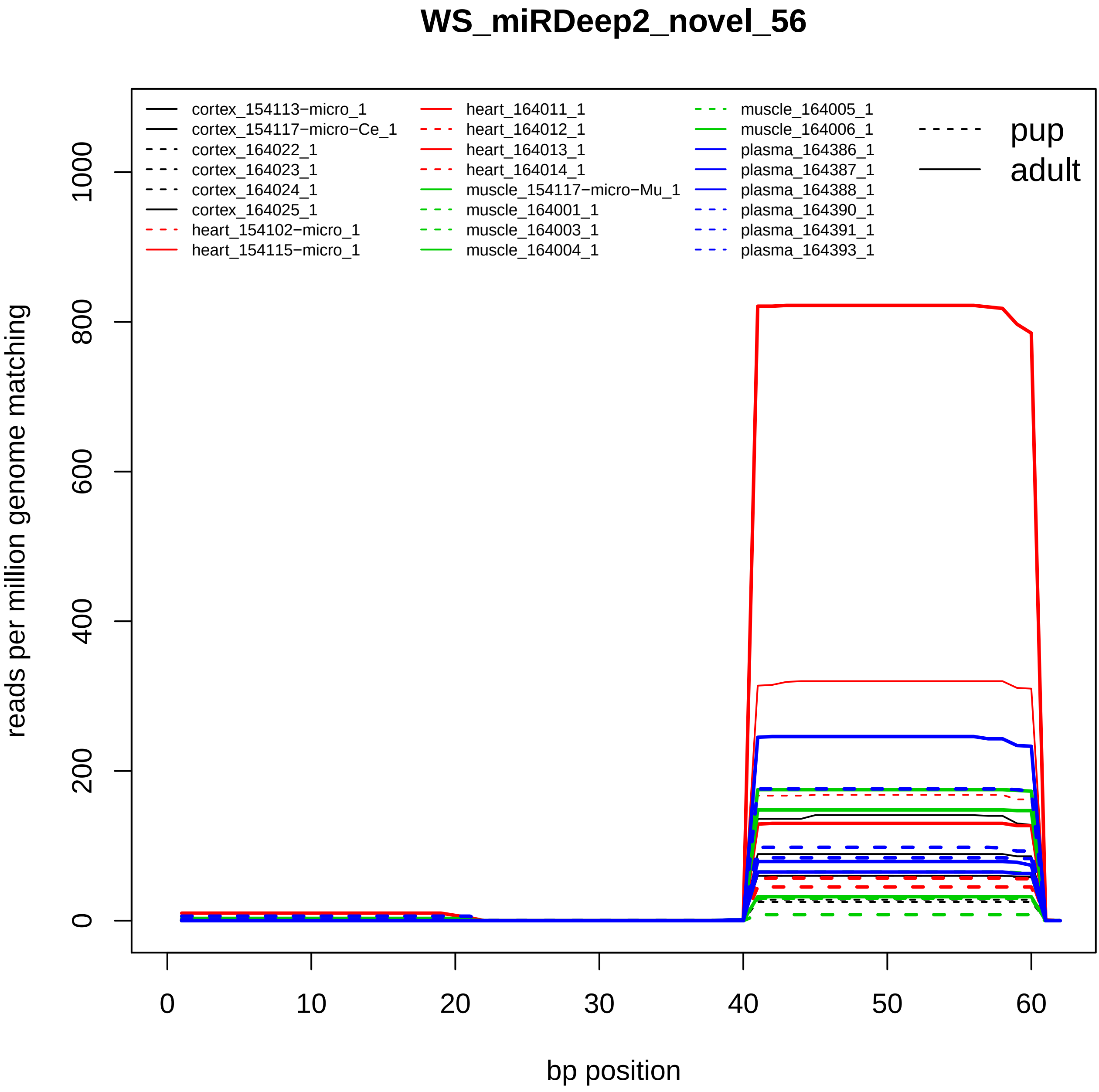

WS_miRDeep2_novel_56
cortex_154113−micro_1
cortex_154117−micro−Ce_1
cortex_164022_1
cortex_164023_1
cortex_164024_1
cortex_164025_1
heart_154102−micro_1
heart_154115−micro_1
heart_164011_1
heart_164012_1
heart_164013_1
heart_164014_1
muscle_154117−micro−Mu_1
muscle_164001_1
muscle_164003_1
muscle_164004_1
muscle_164005_1
muscle_164006_1
plasma_164386_1
plasma_164387_1
plasma_164388_1
plasma_164390_1
plasma_164391_1
plasma_164393_1
pup
adult
reads per million genome matching
0
200
400
600
800
1000
bp position
0
10
20
30
40
50
60

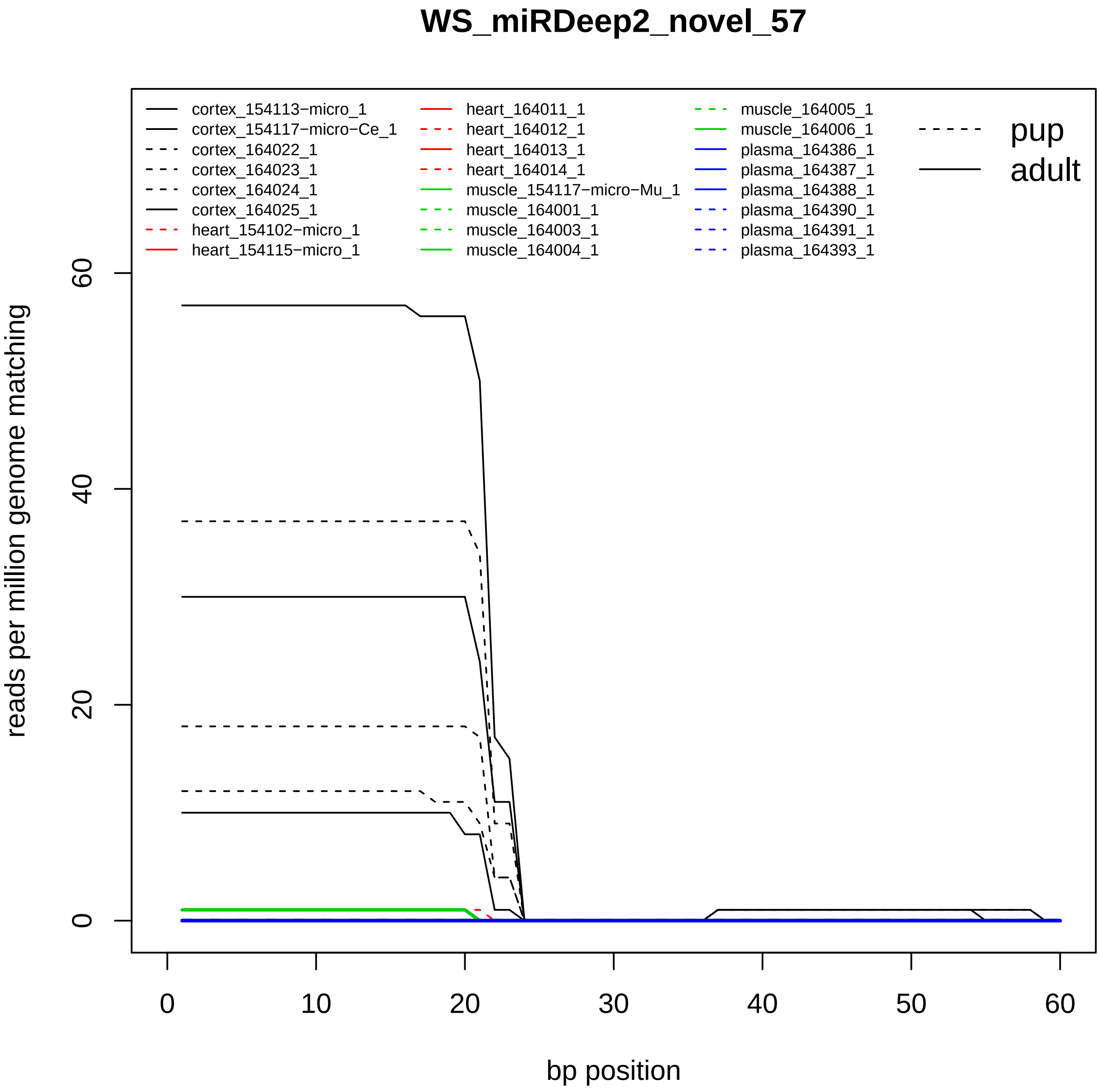

WS_miRDeep2_novel_57
cortex_154113−micro_1
cortex_154117−micro−Ce_1
cortex_164022_1
cortex_164023_1
cortex_164024_1
cortex_164025_1
heart_154102−micro_1
heart_154115−micro_1
heart_164011_1
heart_164012_1
heart_164013_1
heart_164014_1
muscle_154117−micro−Mu_1
muscle_164001_1
muscle_164003_1
muscle_164004_1
muscle_164005_1
muscle_164006_1
plasma_164386_1
plasma_164387_1
plasma_164388_1
plasma_164390_1
plasma_164391_1
plasma_164393_1
pup
adult
reads per million genome matching
0
20
40
60
0
10
20
30
40
50
60
bp position

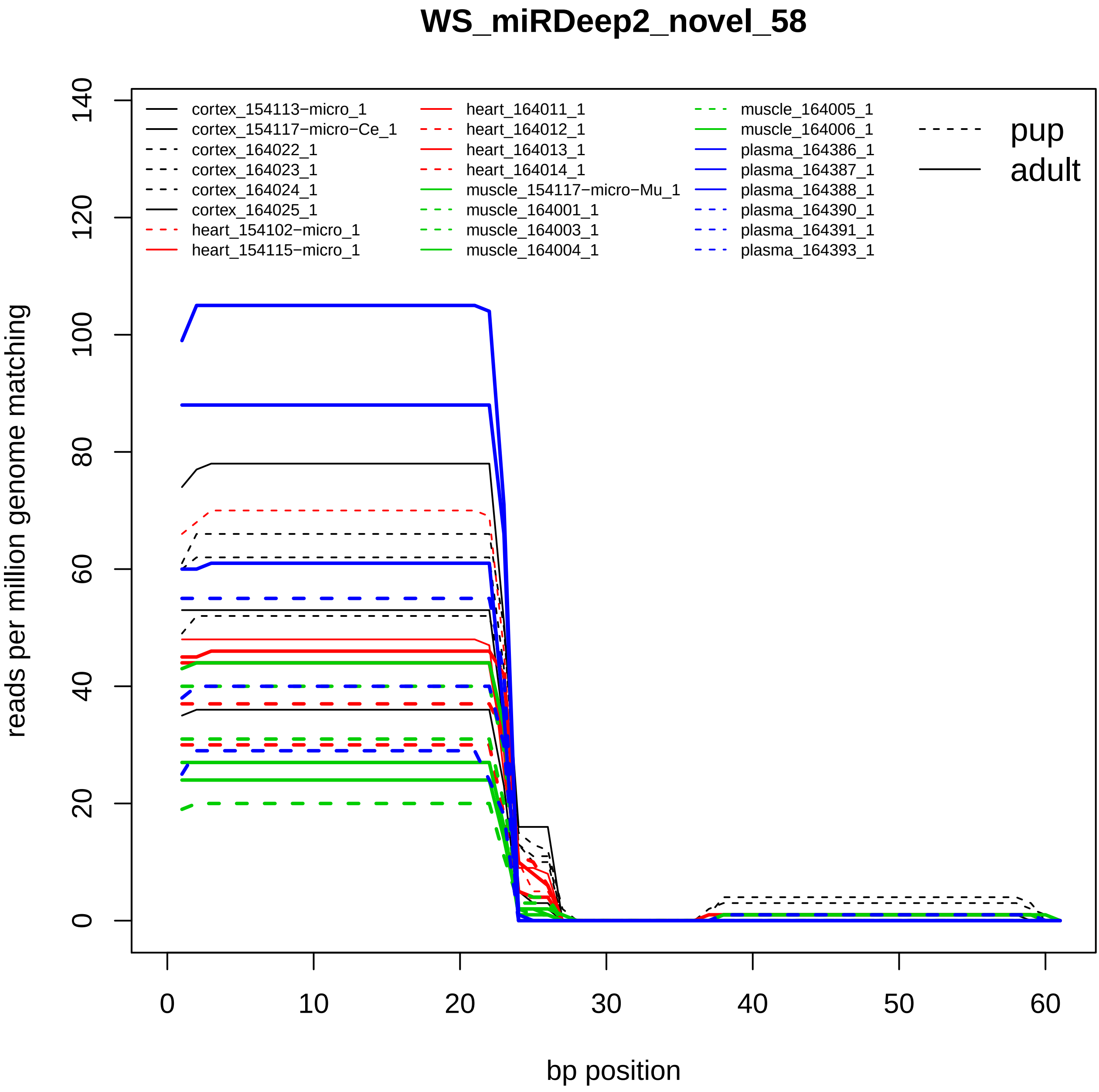

WS_miRDeep2_novel_58
cortex_154113−micro_1
cortex_154117−micro−Ce_1
cortex_164022_1
cortex_164023_1
cortex_164024_1
cortex_164025_1
heart_154102−micro_1
heart_154115−micro_1
heart_164011_1
heart_164012_1
heart_164013_1
heart_164014_1
muscle_154117−micro−Mu_1
muscle_164001_1
muscle_164003_1
muscle_164004_1
muscle_164005_1
muscle_164006_1
plasma_164386_1
plasma_164387_1
plasma_164388_1
plasma_164390_1
plasma_164391_1
plasma_164393_1
pup
adult
reads per million genome matching
bp position
0
20
40
60
80
100
120
140
0
10
20
30
40
50
60

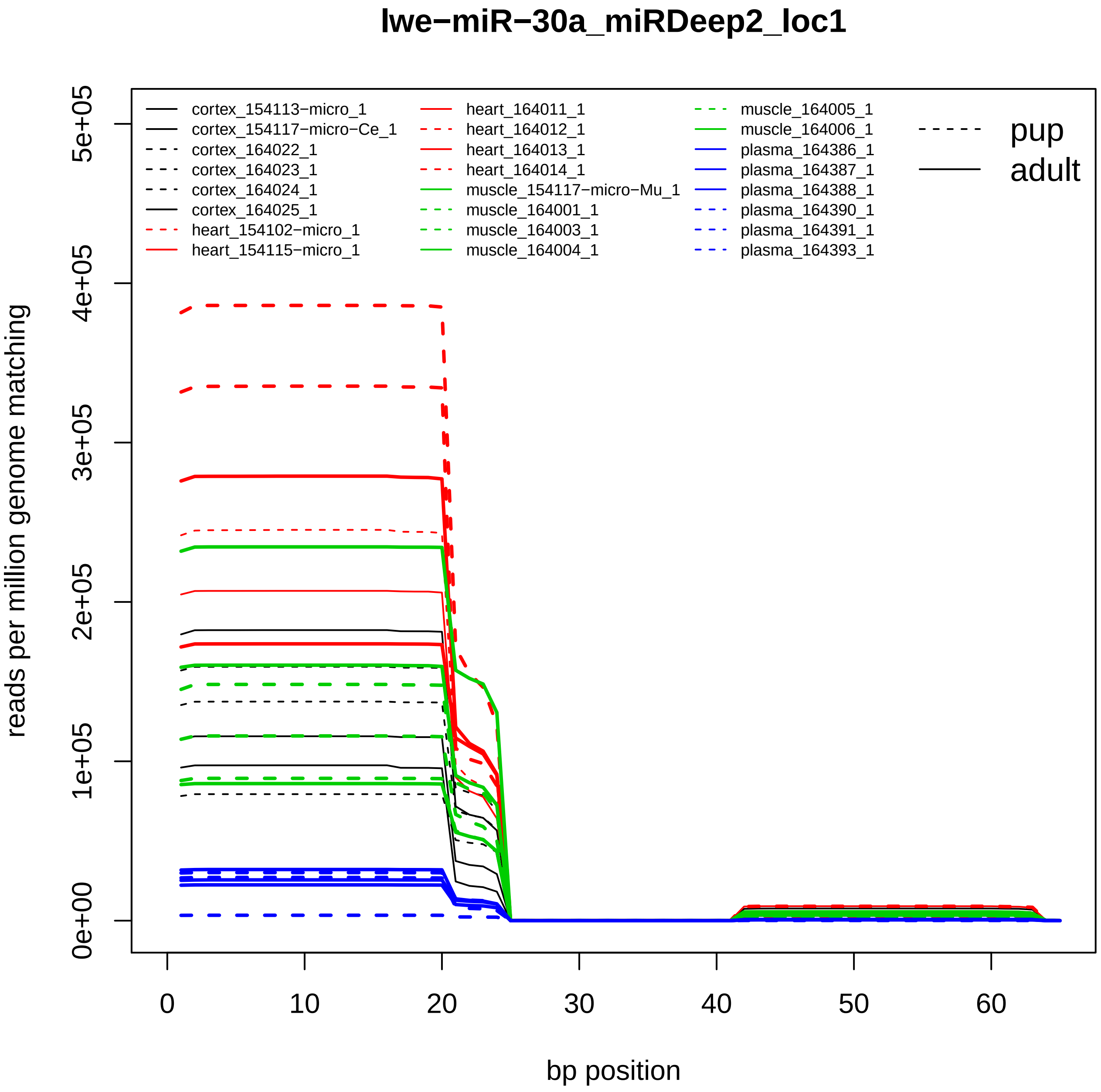

lwe−miR−30a_miRDeep2_loc1
cortex_154113−micro_1
cortex_154117−micro−Ce_1
cortex_164022_1
cortex_164023_1
cortex_164024_1
cortex_164025_1
heart_154102−micro_1
heart_154115−micro_1
heart_164011_1
heart_164012_1
heart_164013_1
heart_164014_1
muscle_154117−micro−Mu_1
muscle_164001_1
muscle_164003_1
muscle_164004_1
muscle_164005_1
muscle_164006_1
plasma_164386_1
plasma_164387_1
plasma_164388_1
plasma_164390_1
plasma_164391_1
plasma_164393_1
pup
adult
reads per million genome matching
0e+00
1e+05
2e+05
3e+05
4e+05
5e+05
0
10
20
30
40
50
60
bp position

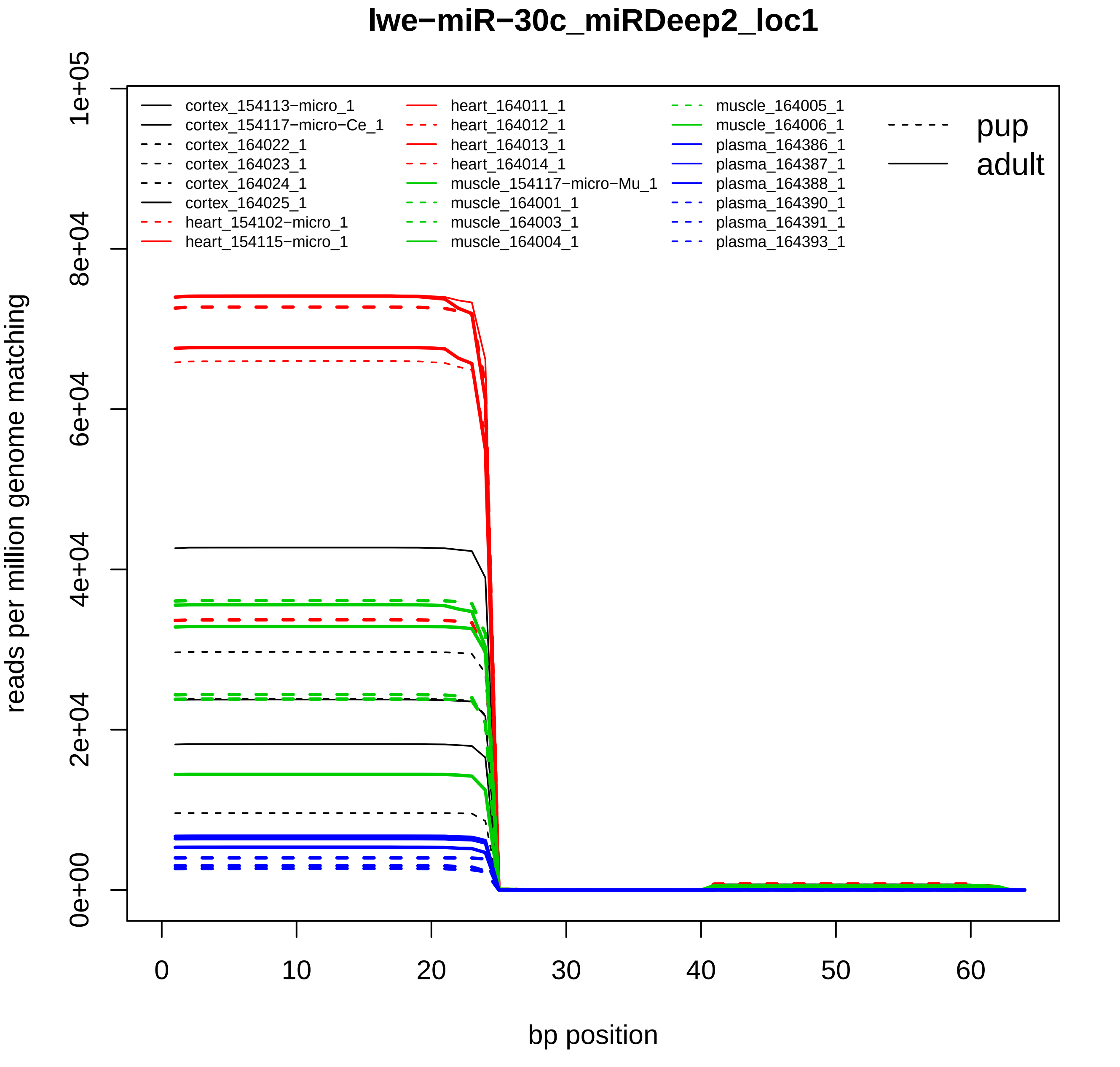

lwe−miR−30c_miRDeep2_loc1
reads per million genome matching
bp position
0e+00
2e+04
4e+04
6e+04
8e+04
1e+05
0
10
20
30
40
50
60
cortex_154113−micro_1
cortex_154117−micro−Ce_1
cortex_164022_1
cortex_164023_1
cortex_164024_1
cortex_164025_1
heart_154102−micro_1
heart_154115−micro_1
heart_164011_1
heart_164012_1
heart_164013_1
heart_164014_1
muscle_154117−micro−Mu_1
muscle_164001_1
muscle_164003_1
muscle_164004_1
muscle_164005_1
muscle_164006_1
plasma_164386_1
plasma_164387_1
plasma_164388_1
plasma_164390_1
plasma_164391_1
plasma_164393_1
pup
adult

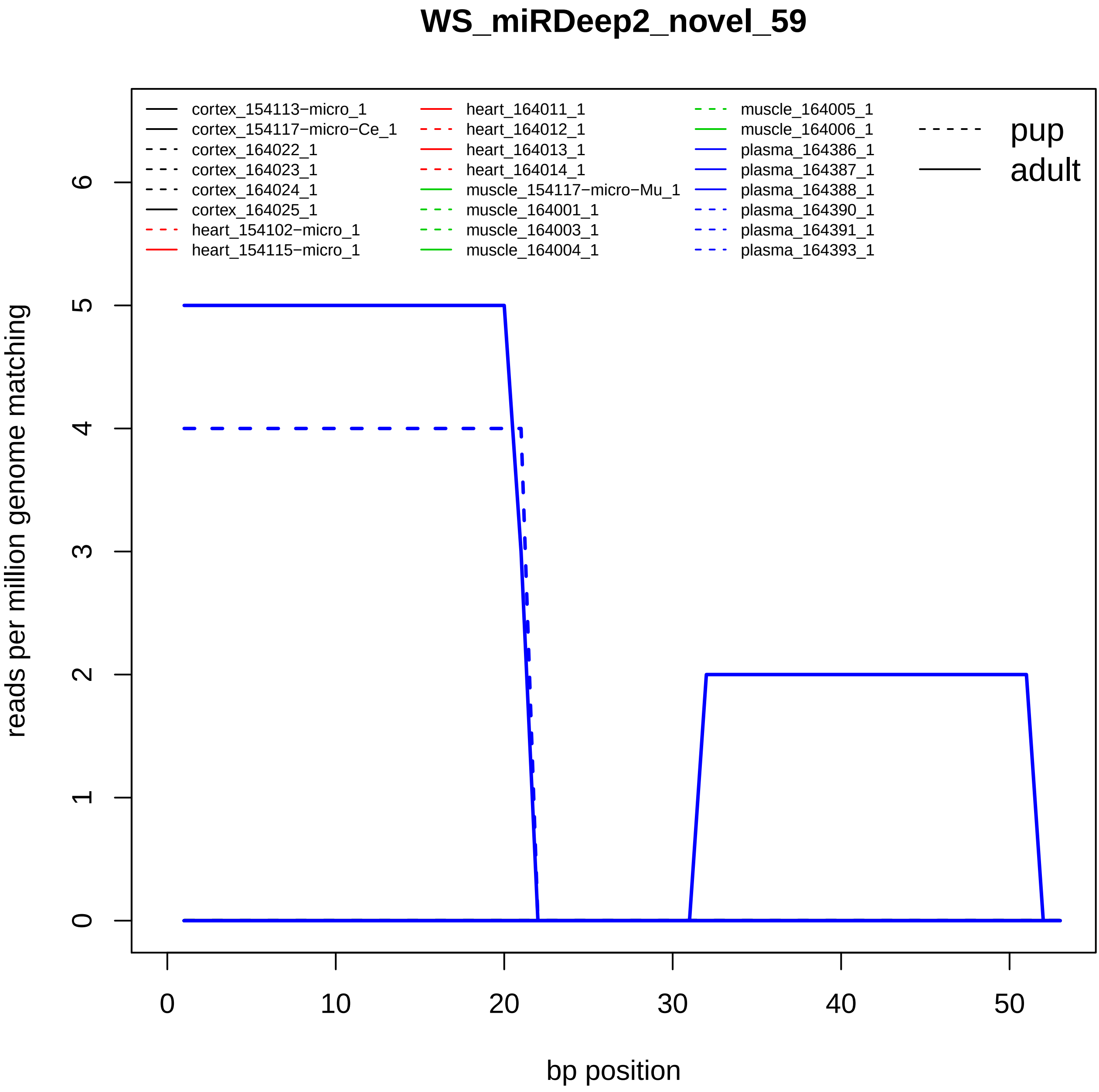
WS_miRDeep2_novel_59
cortex_154113−micro_1
cortex_154117−micro−Ce_1
cortex_164022_1
cortex_164023_1
cortex_164024_1
cortex_164025_1
heart_154102−micro_1
heart_154115−micro_1
heart_164011_1
heart_164012_1
heart_164013_1
heart_164014_1
muscle_154117−micro−Mu_1
muscle_164001_1
muscle_164003_1
muscle_164004_1
muscle_164005_1
muscle_164006_1
plasma_164386_1
plasma_164387_1
plasma_164388_1
plasma_164390_1
plasma_164391_1
plasma_164393_1
pup
adult
reads per million genome matching
0
1
2
3
4
5
6
0
10
20
30
40
50
bp position

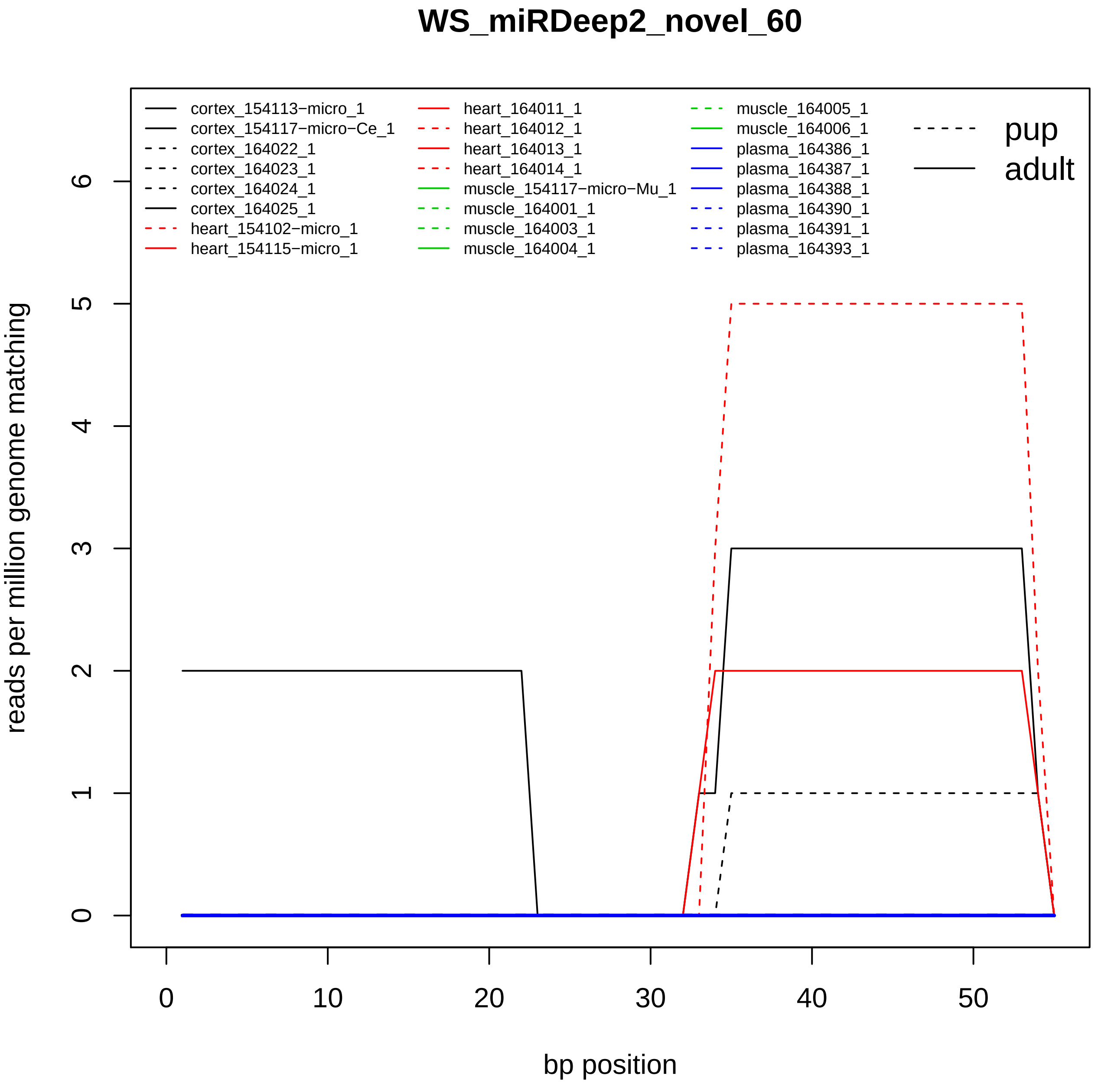

WS_miRDeep2_novel_60
cortex_154113−micro_1
cortex_154117−micro−Ce_1
cortex_164022_1
cortex_164023_1
cortex_164024_1
cortex_164025_1
heart_154102−micro_1
heart_154115−micro_1
heart_164011_1
heart_164012_1
heart_164013_1
heart_164014_1
muscle_154117−micro−Mu_1
muscle_164001_1
muscle_164003_1
muscle_164004_1
muscle_164005_1
muscle_164006_1
plasma_164386_1
plasma_164387_1
plasma_164388_1
plasma_164390_1
plasma_164391_1
plasma_164393_1
pup
adult
reads per million genome matching
0
1
2
3
4
5
6
0
10
20
30
40
50
bp position

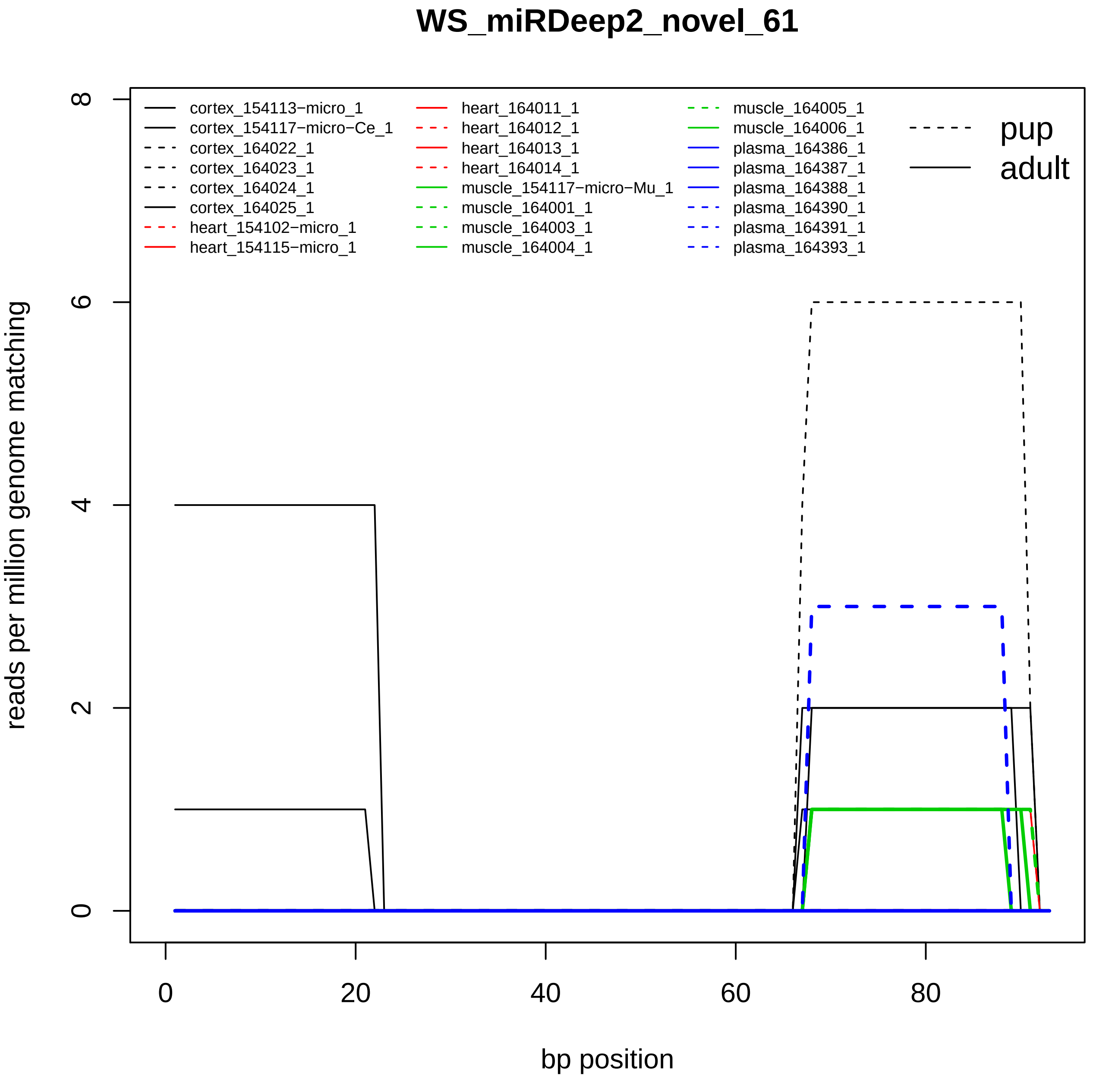
WS_miRDeep2_novel_61
cortex_154113−micro_1
cortex_154117−micro−Ce_1
cortex_164022_1
cortex_164023_1
cortex_164024_1
cortex_164025_1
heart_154102−micro_1
heart_154115−micro_1
heart_164011_1
heart_164012_1
heart_164013_1
heart_164014_1
muscle_154117−micro−Mu_1
muscle_164001_1
muscle_164003_1
muscle_164004_1
muscle_164005_1
muscle_164006_1
plasma_164386_1
plasma_164387_1
plasma_164388_1
plasma_164390_1
plasma_164391_1
plasma_164393_1
pup
adult
reads per million genome matching
bp position
0
2
4
6
8
0
20
40
60
80

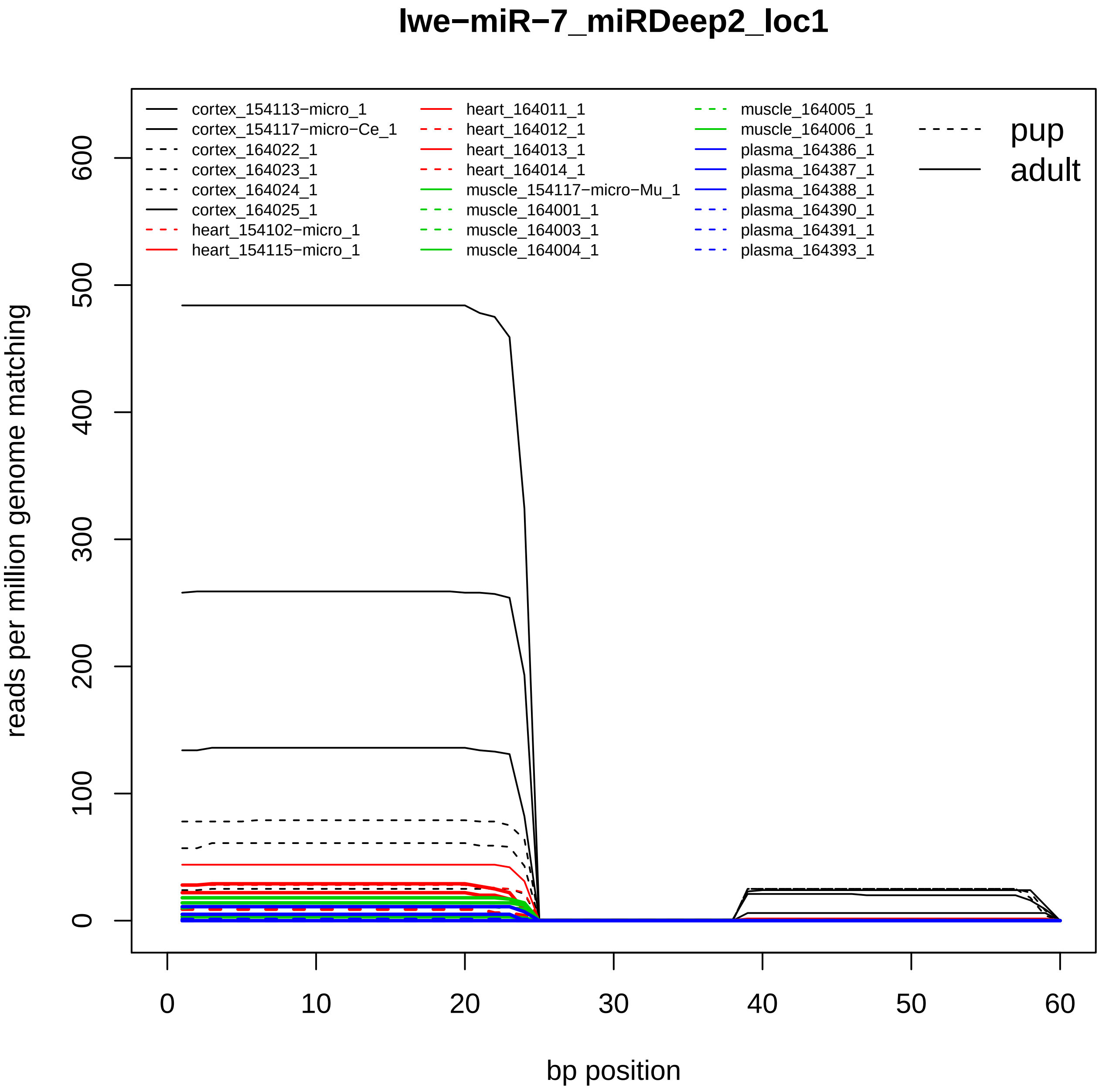

lwe−miR−7_miRDeep2_loc1
cortex_154113−micro_1
cortex_154117−micro−Ce_1
cortex_164022_1
cortex_164023_1
cortex_164024_1
cortex_164025_1
heart_154102−micro_1
heart_154115−micro_1
heart_164011_1
heart_164012_1
heart_164013_1
heart_164014_1
muscle_154117−micro−Mu_1
muscle_164001_1
muscle_164003_1
muscle_164004_1
muscle_164005_1
muscle_164006_1
plasma_164386_1
plasma_164387_1
plasma_164388_1
plasma_164390_1
plasma_164391_1
plasma_164393_1
pup
adult
reads per million genome matching
0
100
200
300
400
500
600
0
10
20
30
40
50
60
bp position

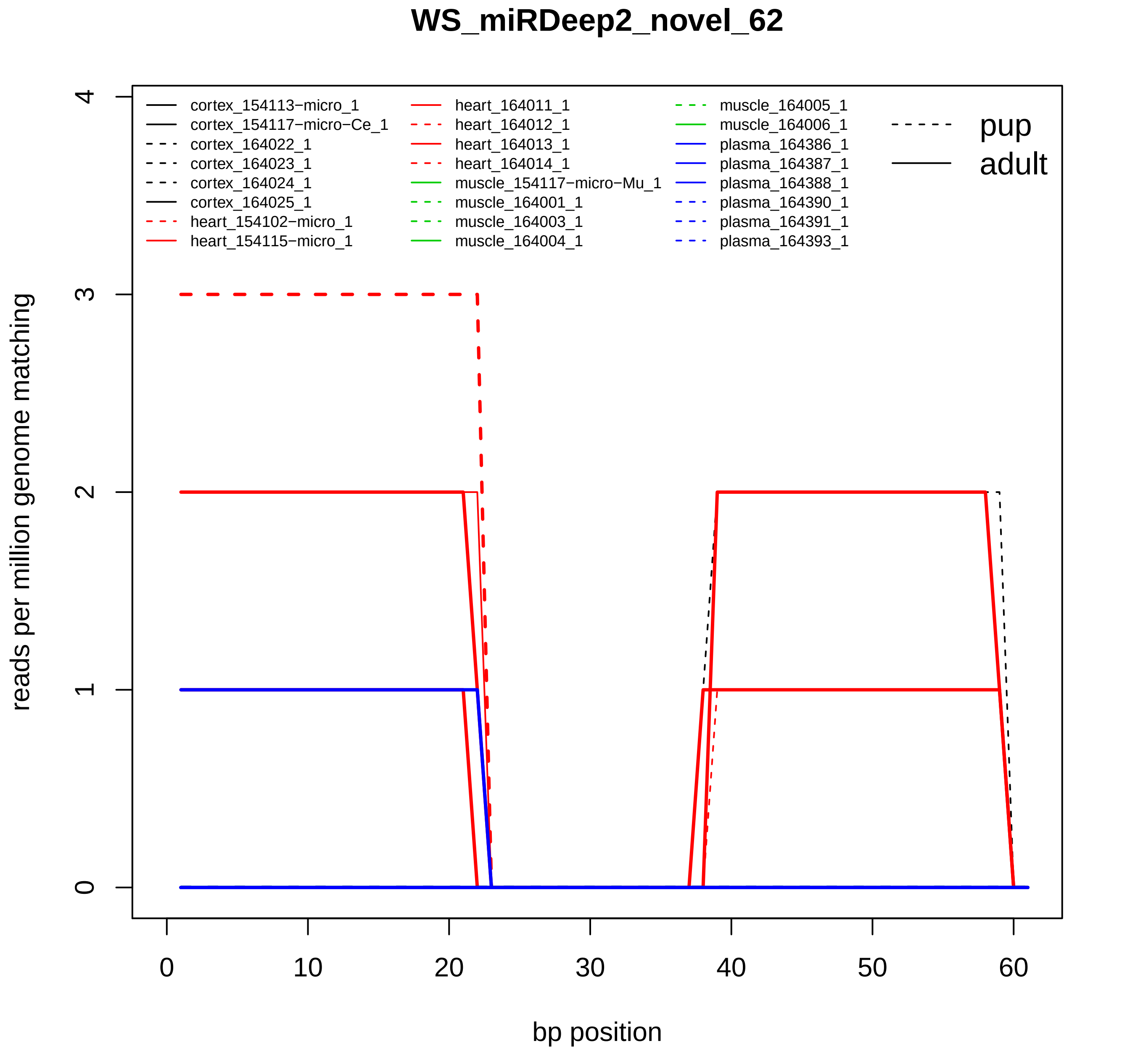

WS_miRDeep2_novel_62
cortex_154113−micro_1
cortex_154117−micro−Ce_1
cortex_164022_1
cortex_164023_1
cortex_164024_1
cortex_164025_1
heart_154102−micro_1
heart_154115−micro_1
heart_164011_1
heart_164012_1
heart_164013_1
heart_164014_1
muscle_154117−micro−Mu_1
muscle_164001_1
muscle_164003_1
muscle_164004_1
muscle_164005_1
muscle_164006_1
plasma_164386_1
plasma_164387_1
plasma_164388_1
plasma_164390_1
plasma_164391_1
plasma_164393_1
pup
adult
reads per million genome matching
bp position
0
1
2
3
4
0
10
20
30
40
50
60

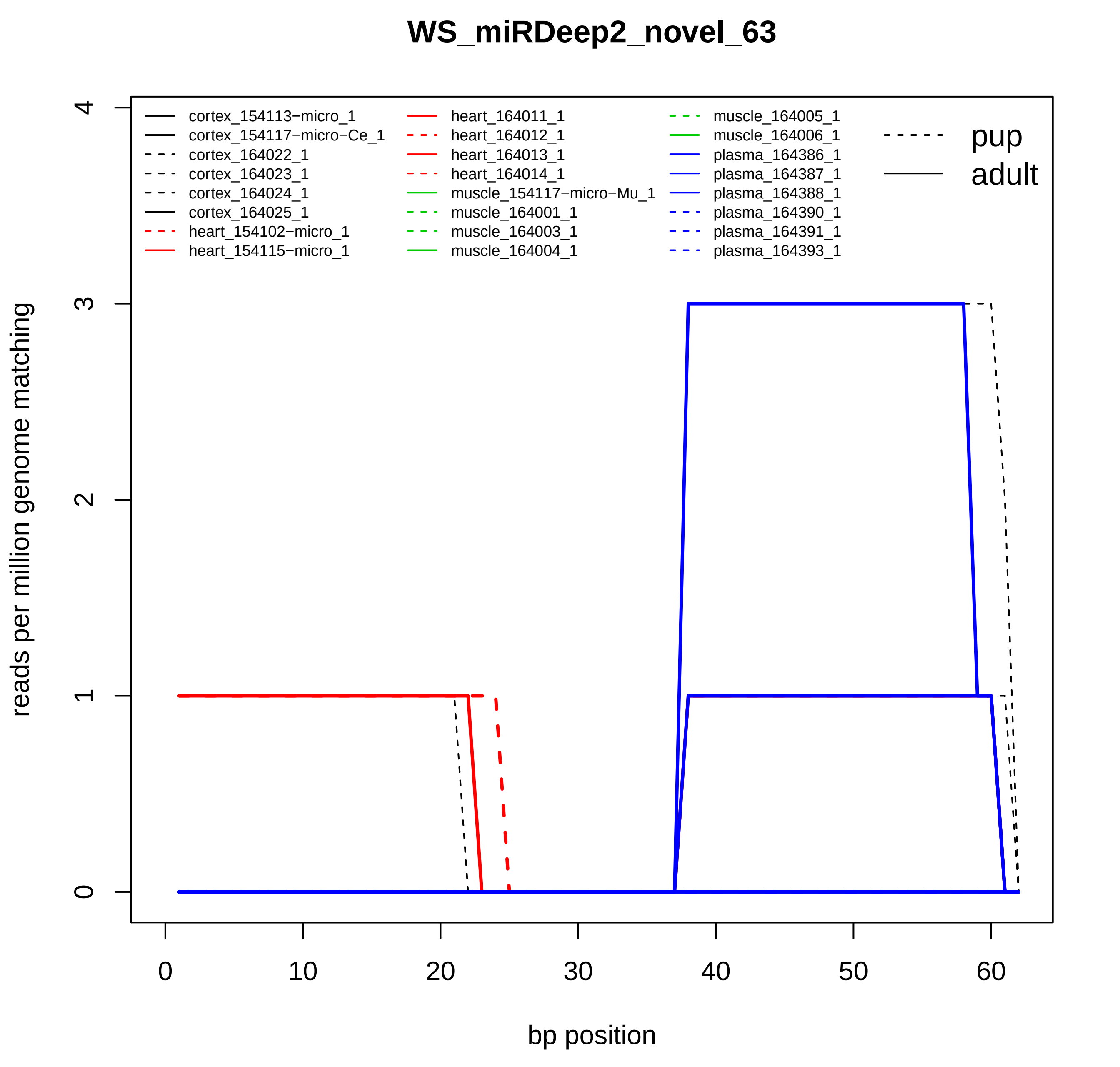

WS_miRDeep2_novel_63
cortex_154113−micro_1
cortex_154117−micro−Ce_1
cortex_164022_1
cortex_164023_1
cortex_164024_1
cortex_164025_1
heart_154102−micro_1
heart_154115−micro_1
heart_164011_1
heart_164012_1
heart_164013_1
heart_164014_1
muscle_154117−micro−Mu_1
muscle_164001_1
muscle_164003_1
muscle_164004_1
muscle_164005_1
muscle_164006_1
plasma_164386_1
plasma_164387_1
plasma_164388_1
plasma_164390_1
plasma_164391_1
plasma_164393_1
pup
adult
reads per million genome matching
bp position
0
1
2
3
4
0
10
20
30
40
50
60

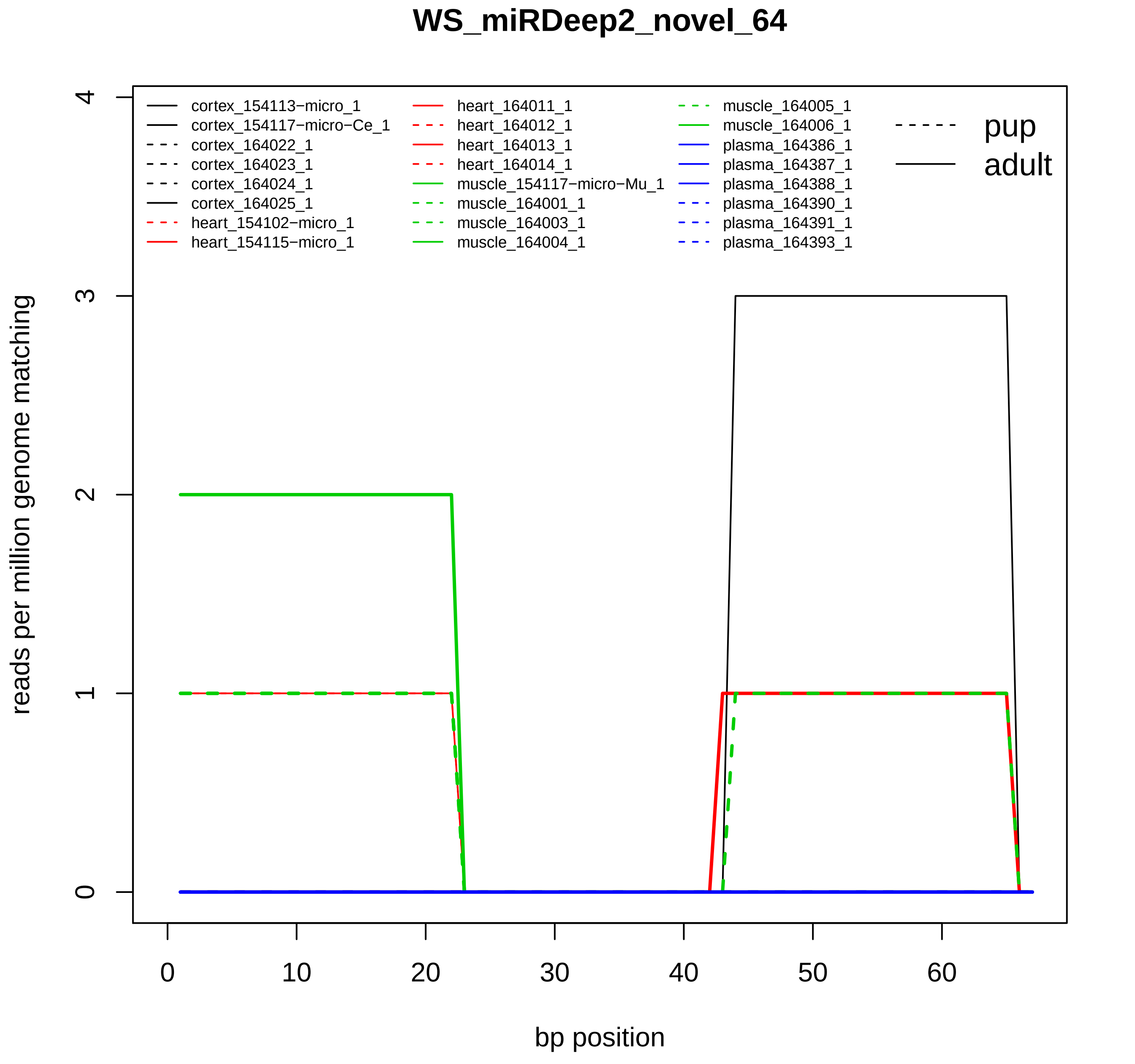

WS_miRDeep2_novel_64
cortex_154113−micro_1
cortex_154117−micro−Ce_1
cortex_164022_1
cortex_164023_1
cortex_164024_1
cortex_164025_1
heart_154102−micro_1
heart_154115−micro_1
heart_164011_1
heart_164012_1
heart_164013_1
heart_164014_1
muscle_154117−micro−Mu_1
muscle_164001_1
muscle_164003_1
muscle_164004_1
muscle_164005_1
muscle_164006_1
plasma_164386_1
plasma_164387_1
plasma_164388_1
plasma_164390_1
plasma_164391_1
plasma_164393_1
pup
adult
reads per million genome matching
bp position
0
1
2
3
4
0
10
20
30
40
50
60

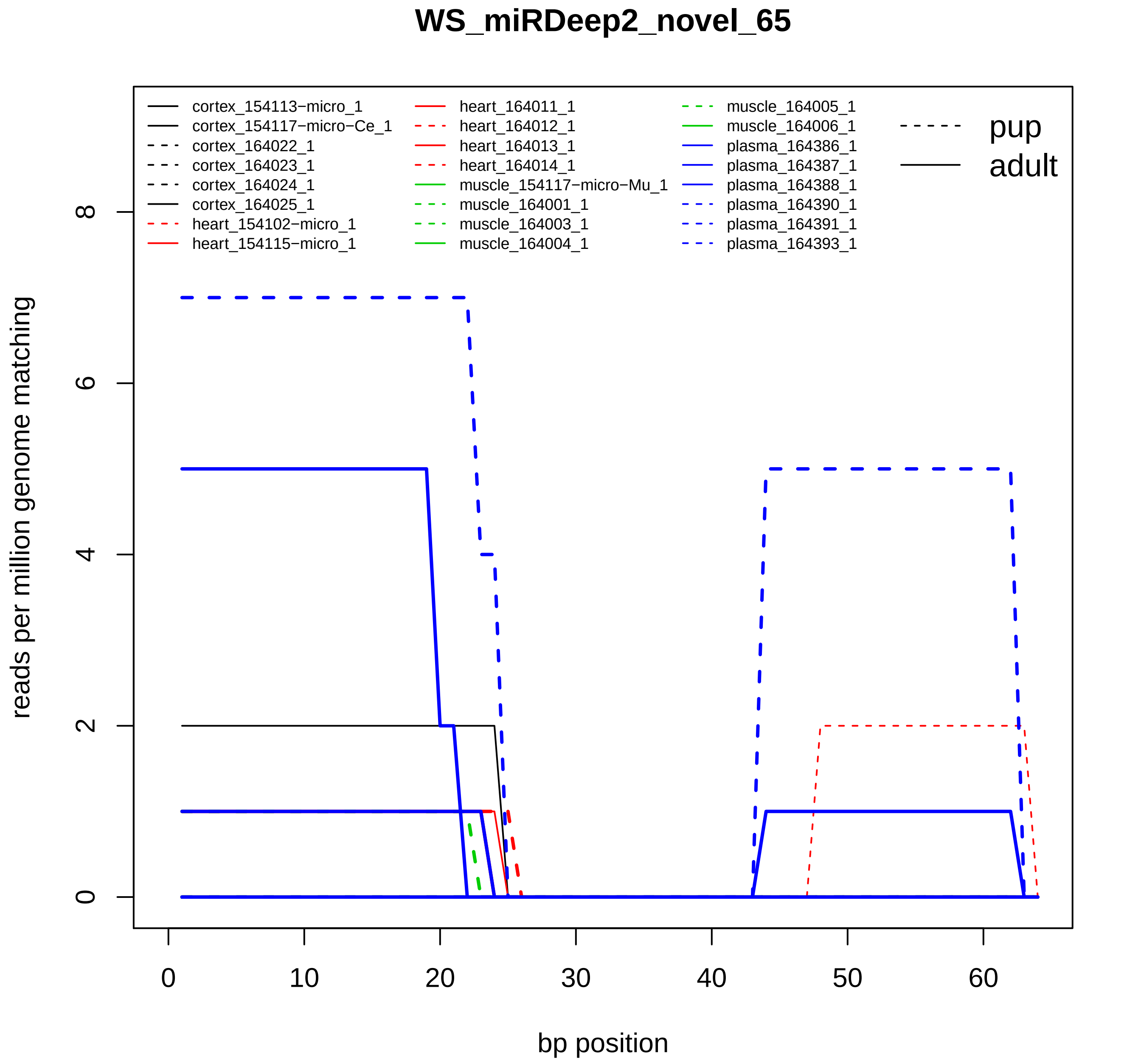

WS_miRDeep2_novel_65
cortex_154113−micro_1
cortex_154117−micro−Ce_1
cortex_164022_1
cortex_164023_1
cortex_164024_1
cortex_164025_1
heart_154102−micro_1
heart_154115−micro_1
heart_164011_1
heart_164012_1
heart_164013_1
heart_164014_1
muscle_154117−micro−Mu_1
muscle_164001_1
muscle_164003_1
muscle_164004_1
muscle_164005_1
muscle_164006_1
plasma_164386_1
plasma_164387_1
plasma_164388_1
plasma_164390_1
plasma_164391_1
plasma_164393_1
pup
adult
reads per million genome matching
0
2
4
6
8
0
10
20
30
40
50
60
bp position

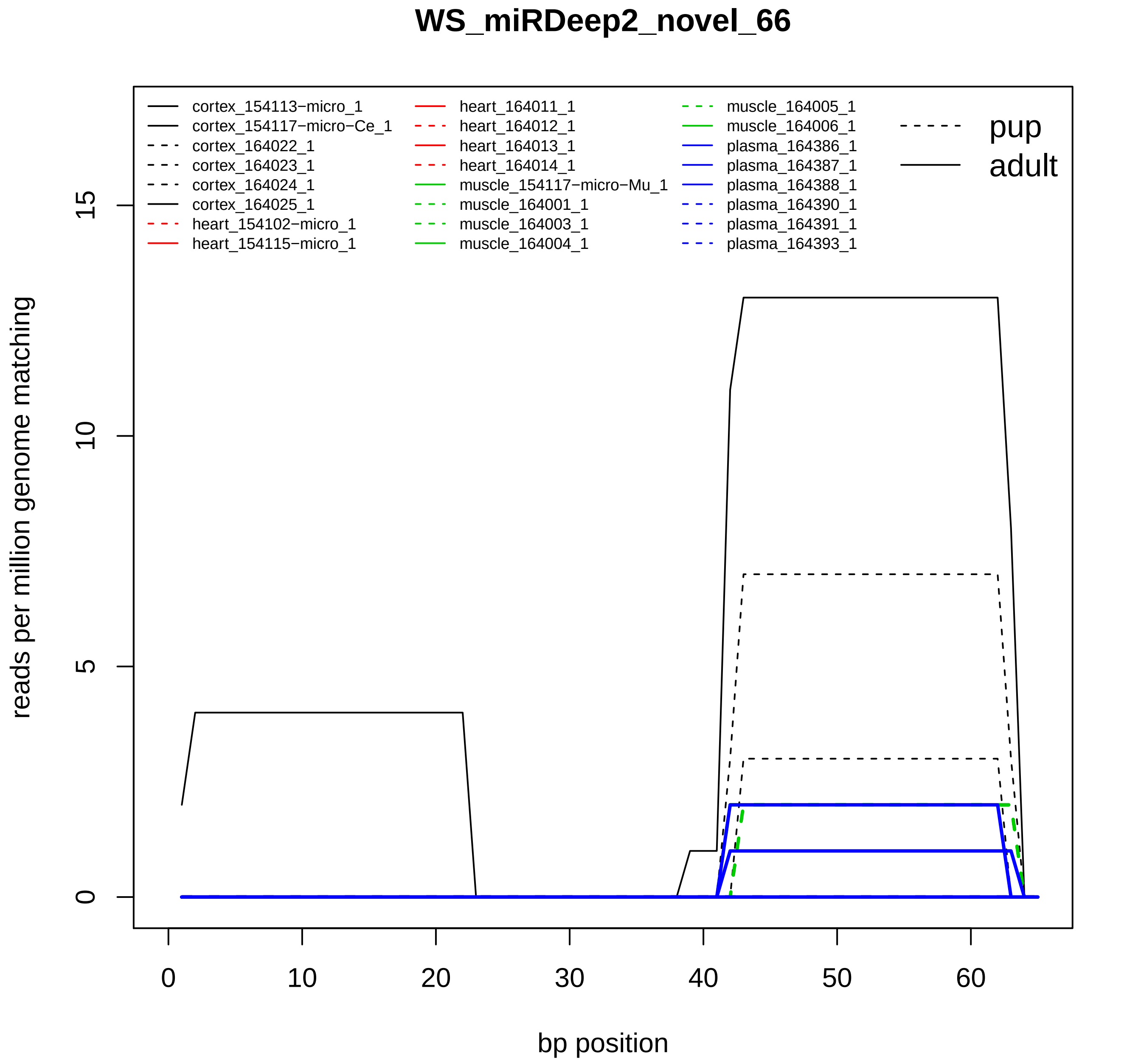
WS_miRDeep2_novel_66
cortex_154113−micro_1
cortex_154117−micro−Ce_1
cortex_164022_1
cortex_164023_1
cortex_164024_1
cortex_164025_1
heart_154102−micro_1
heart_154115−micro_1
heart_164011_1
heart_164012_1
heart_164013_1
heart_164014_1
muscle_154117−micro−Mu_1
muscle_164001_1
muscle_164003_1
muscle_164004_1
muscle_164005_1
muscle_164006_1
plasma_164386_1
plasma_164387_1
plasma_164388_1
plasma_164390_1
plasma_164391_1
plasma_164393_1
pup
adult
reads per million genome matching
0
5
10
15
0
10
20
30
40
50
60
bp position

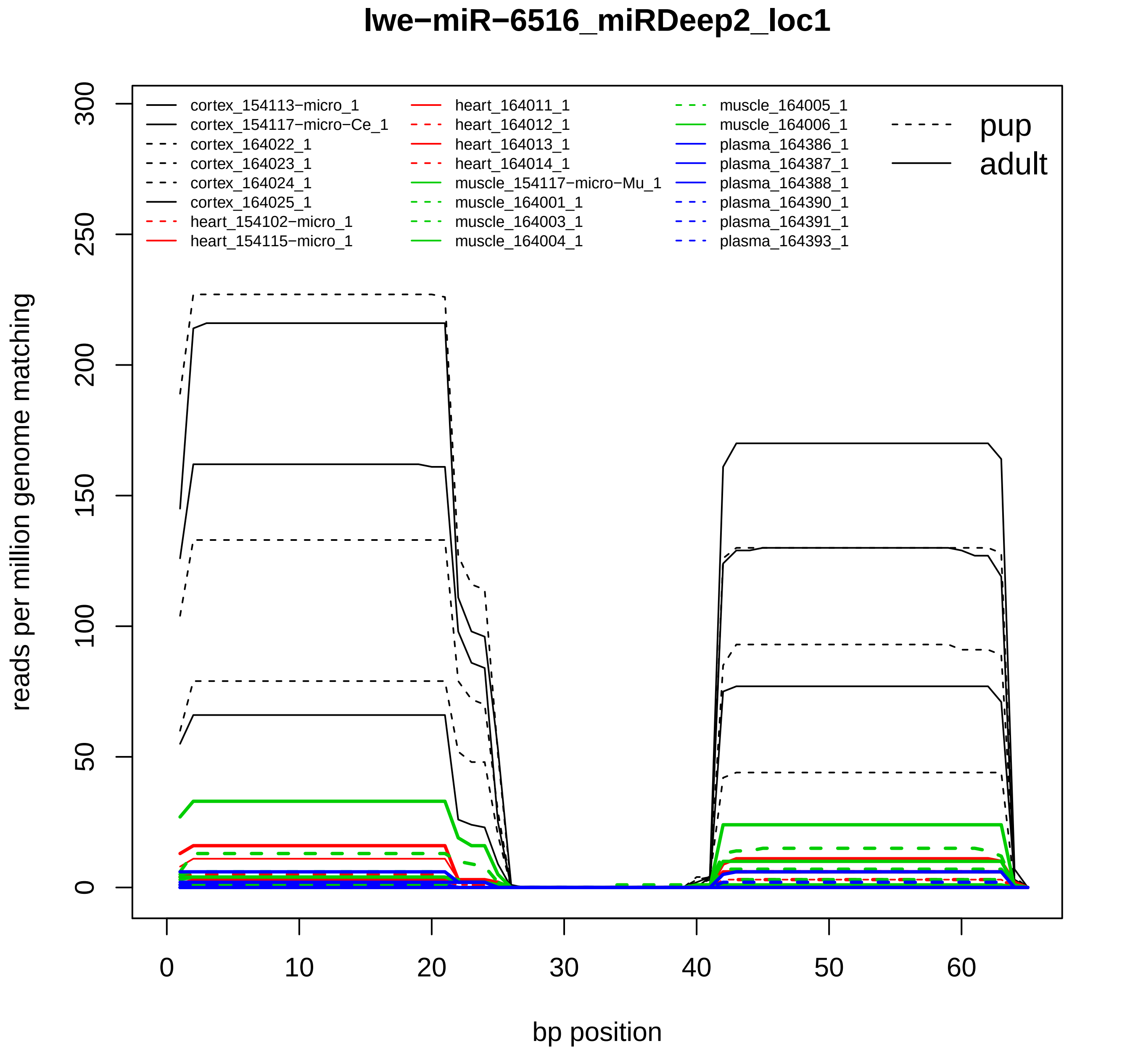
lwe−miR−6516_miRDeep2_loc1
reads per million genome matching
bp position
0
50
100
150
200
250
300
0
10
20
30
40
50
60
cortex_154113−micro_1
cortex_154117−micro−Ce_1
cortex_164022_1
cortex_164023_1
cortex_164024_1
cortex_164025_1
heart_154102−micro_1
heart_154115−micro_1
heart_164011_1
heart_164012_1
heart_164013_1
heart_164014_1
muscle_154117−micro−Mu_1
muscle_164001_1
muscle_164003_1
muscle_164004_1
muscle_164005_1
muscle_164006_1
plasma_164386_1
plasma_164387_1
plasma_164388_1
plasma_164390_1
plasma_164391_1
plasma_164393_1
pup
adult

### WS_miRDeep2_novel_67

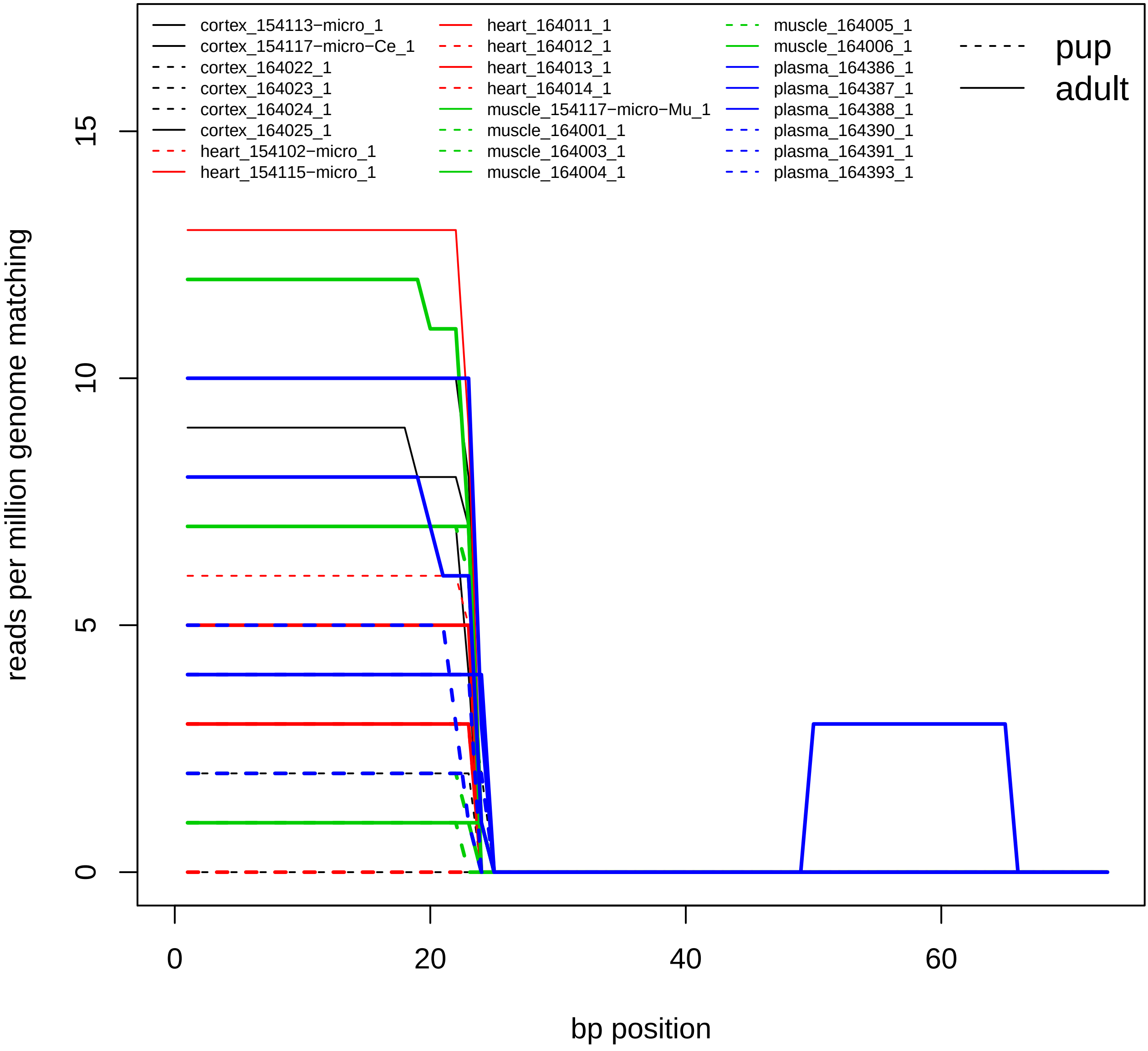

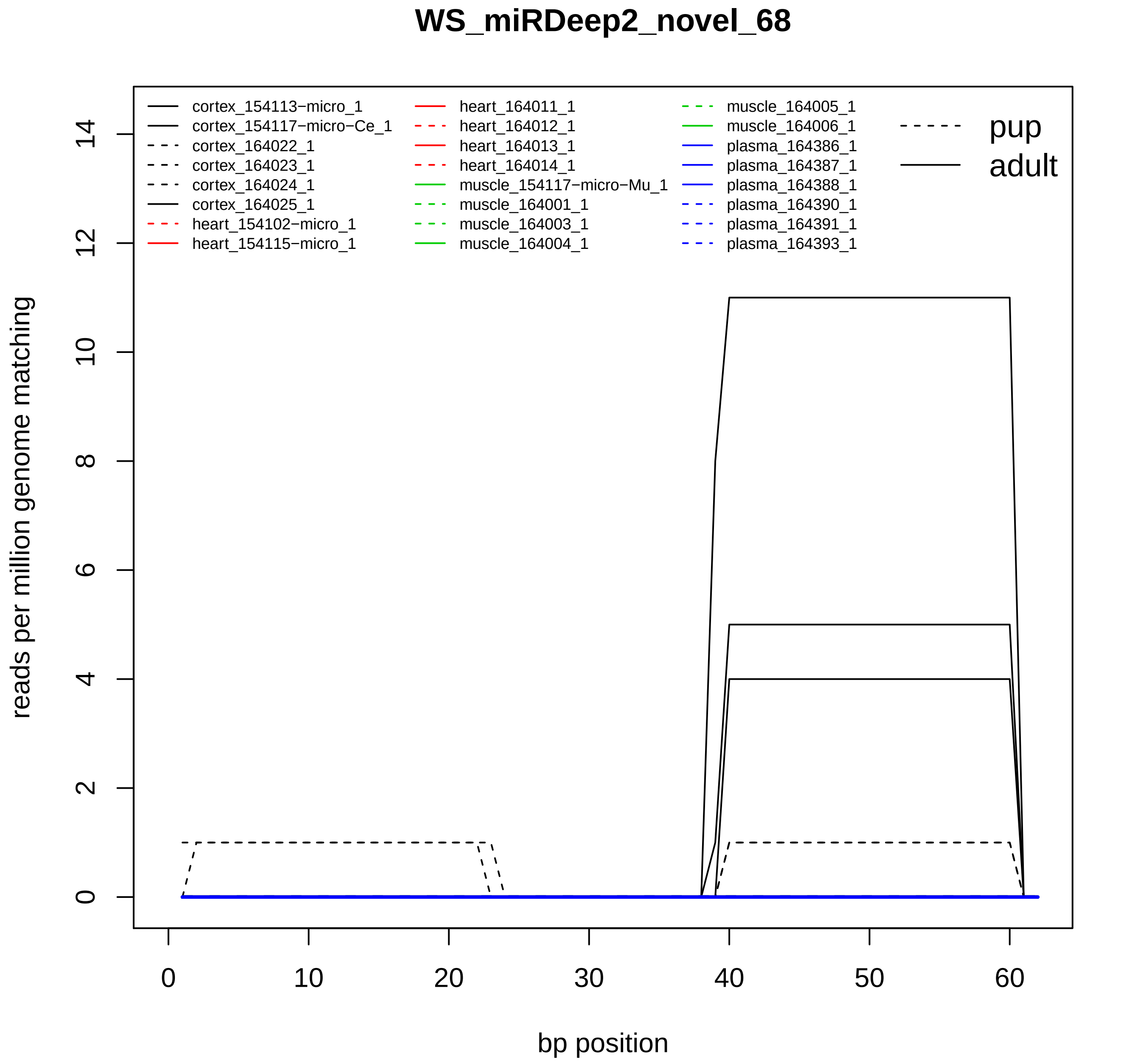
WS_miRDeep2_novel_68
cortex_154113−micro_1
cortex_154117−micro−Ce_1
cortex_164022_1
cortex_164023_1
cortex_164024_1
cortex_164025_1
heart_154102−micro_1
heart_154115−micro_1
heart_164011_1
heart_164012_1
heart_164013_1
heart_164014_1
muscle_154117−micro−Mu_1
muscle_164001_1
muscle_164003_1
muscle_164004_1
muscle_164005_1
muscle_164006_1
plasma_164386_1
plasma_164387_1
plasma_164388_1
plasma_164390_1
plasma_164391_1
plasma_164393_1
pup
adult
reads per million genome matching
bp position
0
2
4
6
8
10
12
14
0
10
20
30
40
50
60

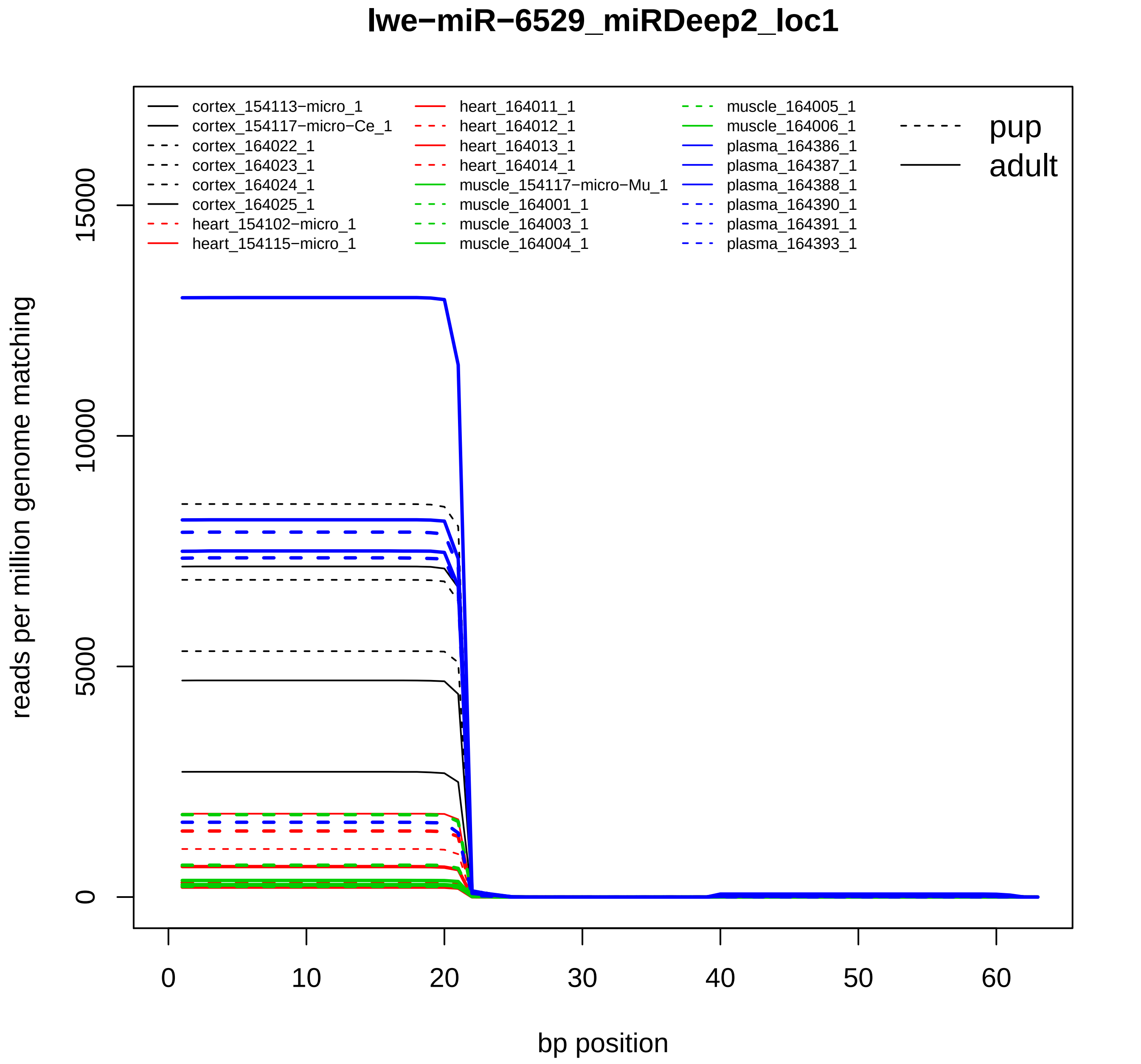

lwe−miR−6529_miRDeep2_loc1
cortex_154113−micro_1
cortex_154117−micro−Ce_1
cortex_164022_1
cortex_164023_1
cortex_164024_1
cortex_164025_1
heart_154102−micro_1
heart_154115−micro_1
heart_164011_1
heart_164012_1
heart_164013_1
heart_164014_1
muscle_154117−micro−Mu_1
muscle_164001_1
muscle_164003_1
muscle_164004_1
muscle_164005_1
muscle_164006_1
plasma_164386_1
plasma_164387_1
plasma_164388_1
plasma_164390_1
plasma_164391_1
plasma_164393_1
pup
adult
reads per million genome matching
0
5000
10000
15000
0
10
20
30
40
50
60
bp position

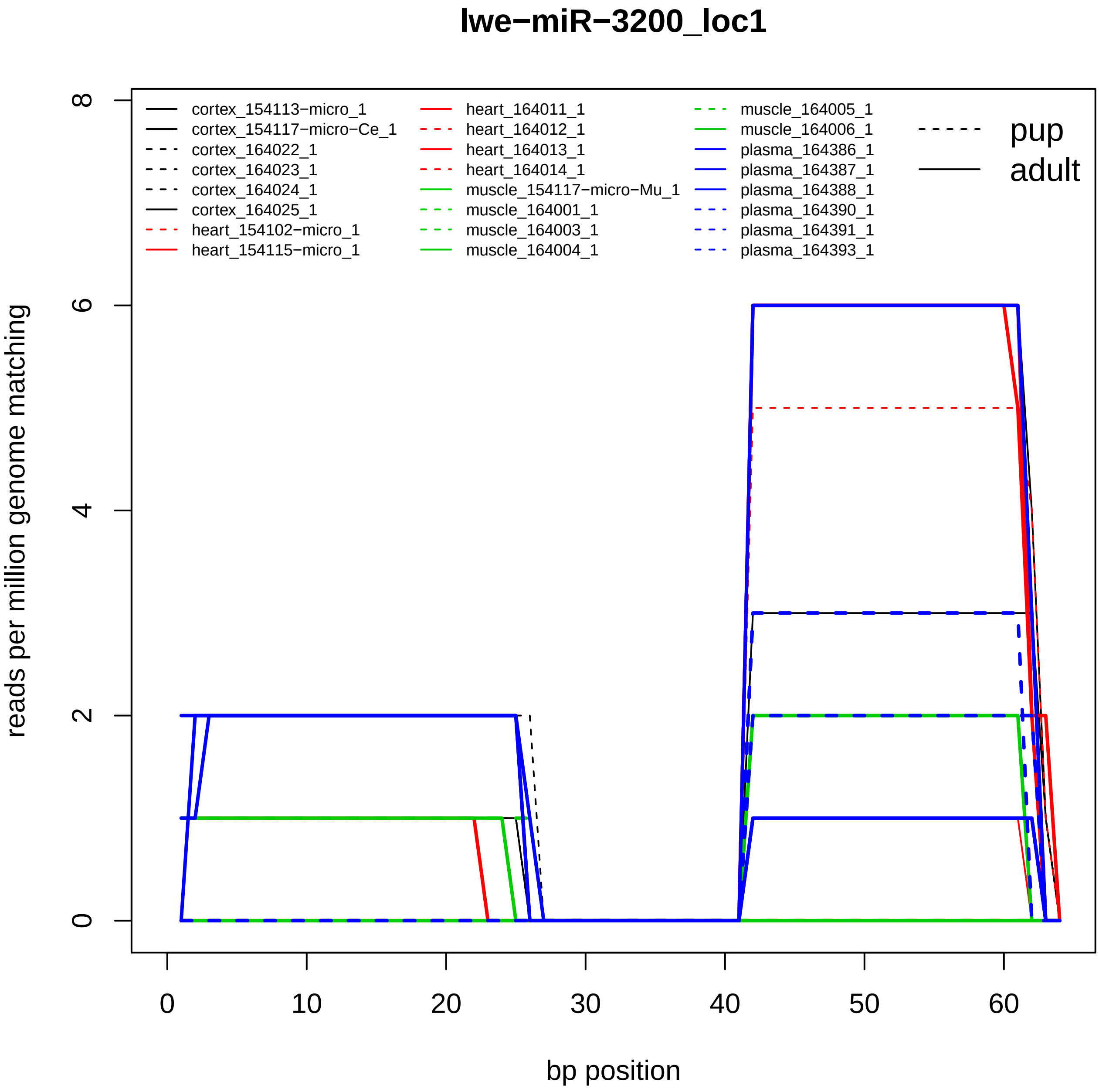
lwe−miR−3200_loc1
reads per million genome matching
bp position
0
2
4
6
8
0
10
20
30
40
50
60
cortex_154113−micro_1
cortex_154117−micro−Ce_1
cortex_164022_1
cortex_164023_1
cortex_164024_1
cortex_164025_1
heart_154102−micro_1
heart_154115−micro_1
heart_164011_1
heart_164012_1
heart_164013_1
heart_164014_1
muscle_154117−micro−Mu_1
muscle_164001_1
muscle_164003_1
muscle_164004_1
muscle_164005_1
muscle_164006_1
plasma_164386_1
plasma_164387_1
plasma_164388_1
plasma_164390_1
plasma_164391_1
plasma_164393_1
pup
adult

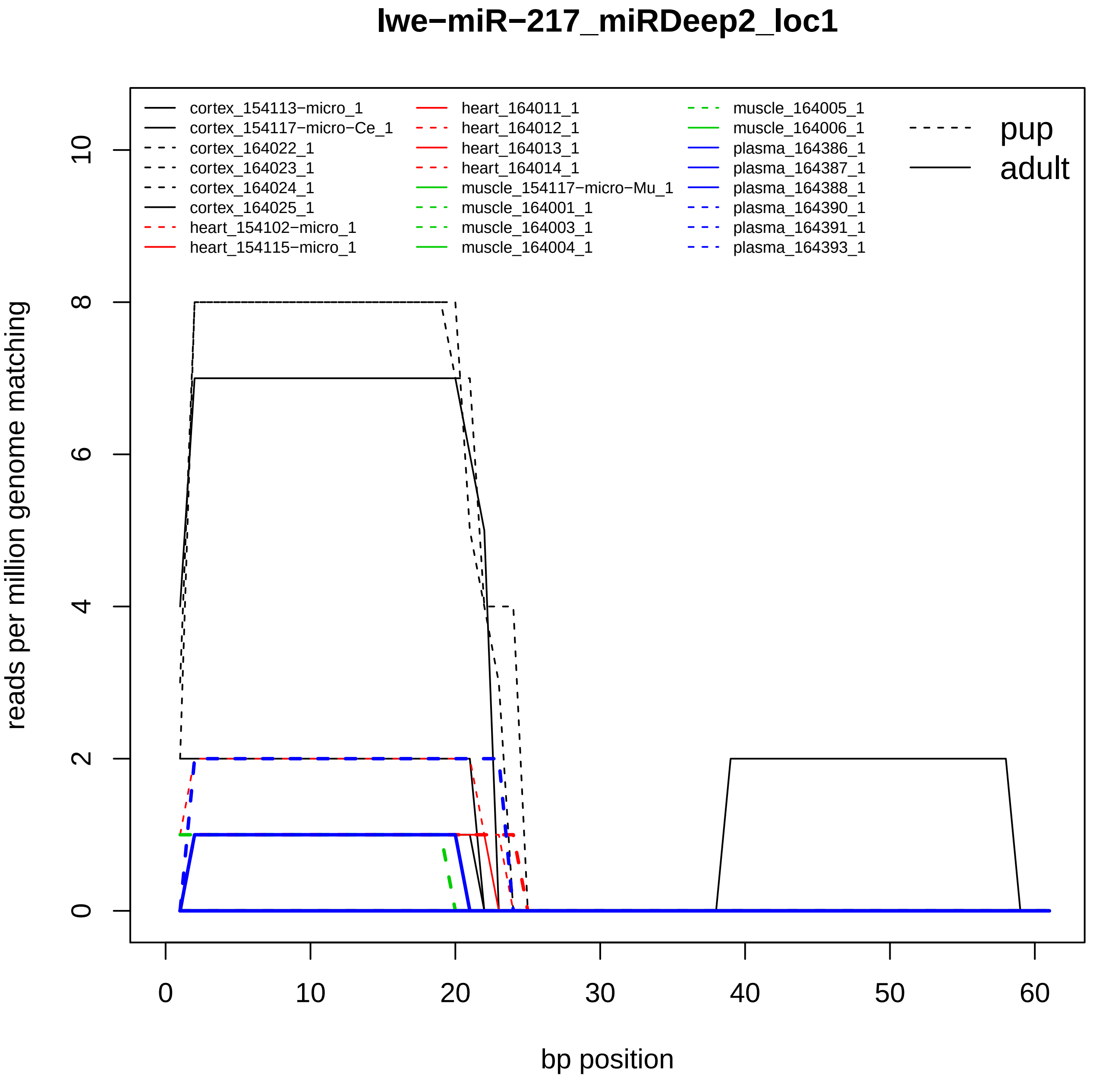

lwe−miR−217_miRDeep2_loc1
cortex_154113−micro_1
cortex_154117−micro−Ce_1
cortex_164022_1
cortex_164023_1
cortex_164024_1
cortex_164025_1
heart_154102−micro_1
heart_154115−micro_1
heart_164011_1
heart_164012_1
heart_164013_1
heart_164014_1
muscle_154117−micro−Mu_1
muscle_164001_1
muscle_164003_1
muscle_164004_1
muscle_164005_1
muscle_164006_1
plasma_164386_1
plasma_164387_1
plasma_164388_1
plasma_164390_1
plasma_164391_1
plasma_164393_1
pup
adult
reads per million genome matching
bp position
0
2
4
6
8
10
0
10
20
30
40
50
60

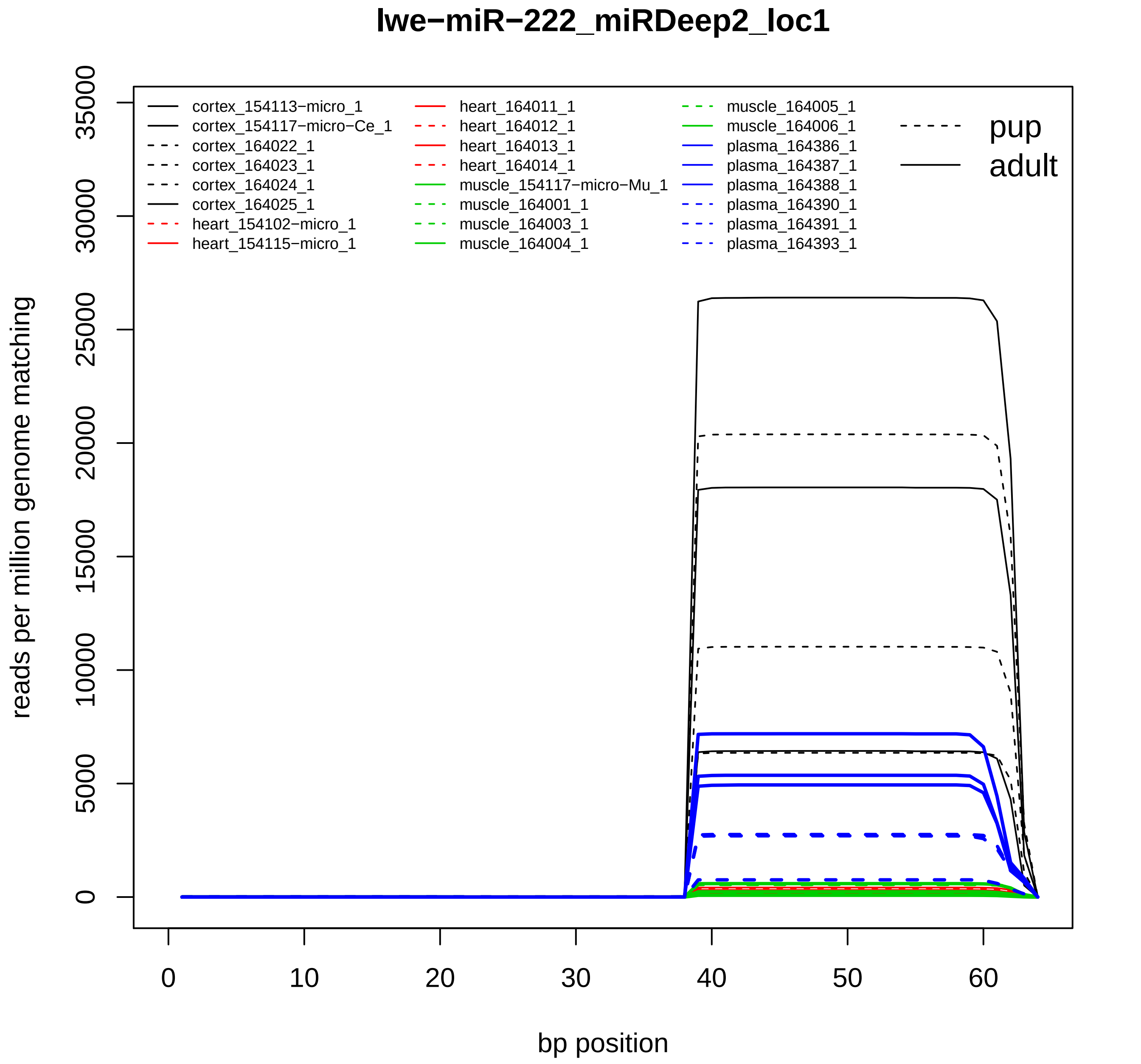
lwe−miR−222_miRDeep2_loc1
cortex_154113−micro_1
cortex_154117−micro−Ce_1
cortex_164022_1
cortex_164023_1
cortex_164024_1
cortex_164025_1
heart_154102−micro_1
heart_154115−micro_1
heart_164011_1
heart_164012_1
heart_164013_1
heart_164014_1
muscle_154117−micro−Mu_1
muscle_164001_1
muscle_164003_1
muscle_164004_1
muscle_164005_1
muscle_164006_1
plasma_164386_1
plasma_164387_1
plasma_164388_1
plasma_164390_1
plasma_164391_1
plasma_164393_1
pup
adult
reads per million genome matching
0
5000
10000
15000
20000
25000
30000
35000
0
10
20
30
40
50
60
bp position

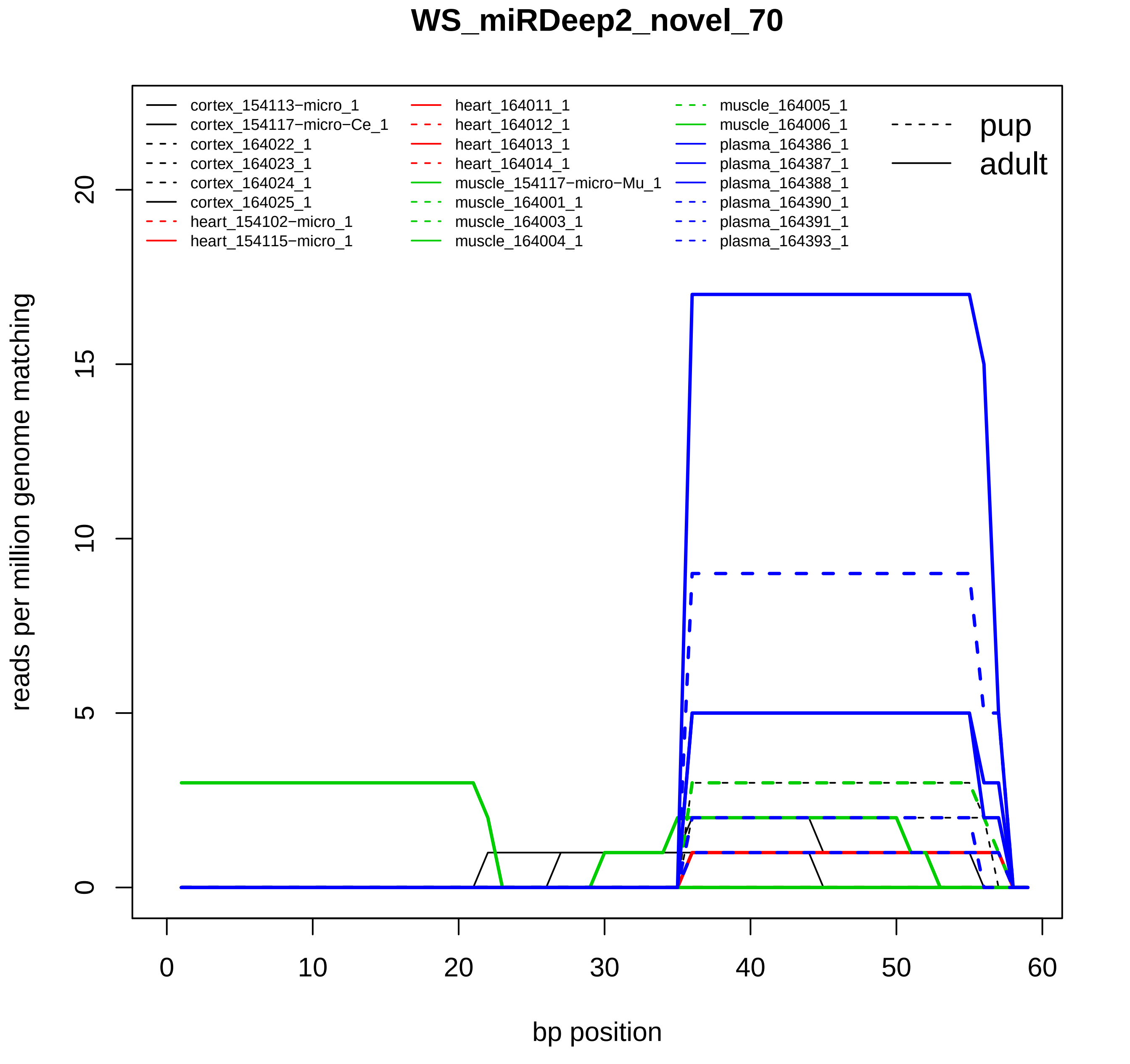
WS_miRDeep2_novel_70
cortex_154113−micro_1
cortex_154117−micro−Ce_1
cortex_164022_1
cortex_164023_1
cortex_164024_1
cortex_164025_1
heart_154102−micro_1
heart_154115−micro_1
heart_164011_1
heart_164012_1
heart_164013_1
heart_164014_1
muscle_154117−micro−Mu_1
muscle_164001_1
muscle_164003_1
muscle_164004_1
muscle_164005_1
muscle_164006_1
plasma_164386_1
plasma_164387_1
plasma_164388_1
plasma_164390_1
plasma_164391_1
plasma_164393_1
pup
adult
reads per million genome matching
bp position
0
5
10
15
20
0
10
20
30
40
50
60

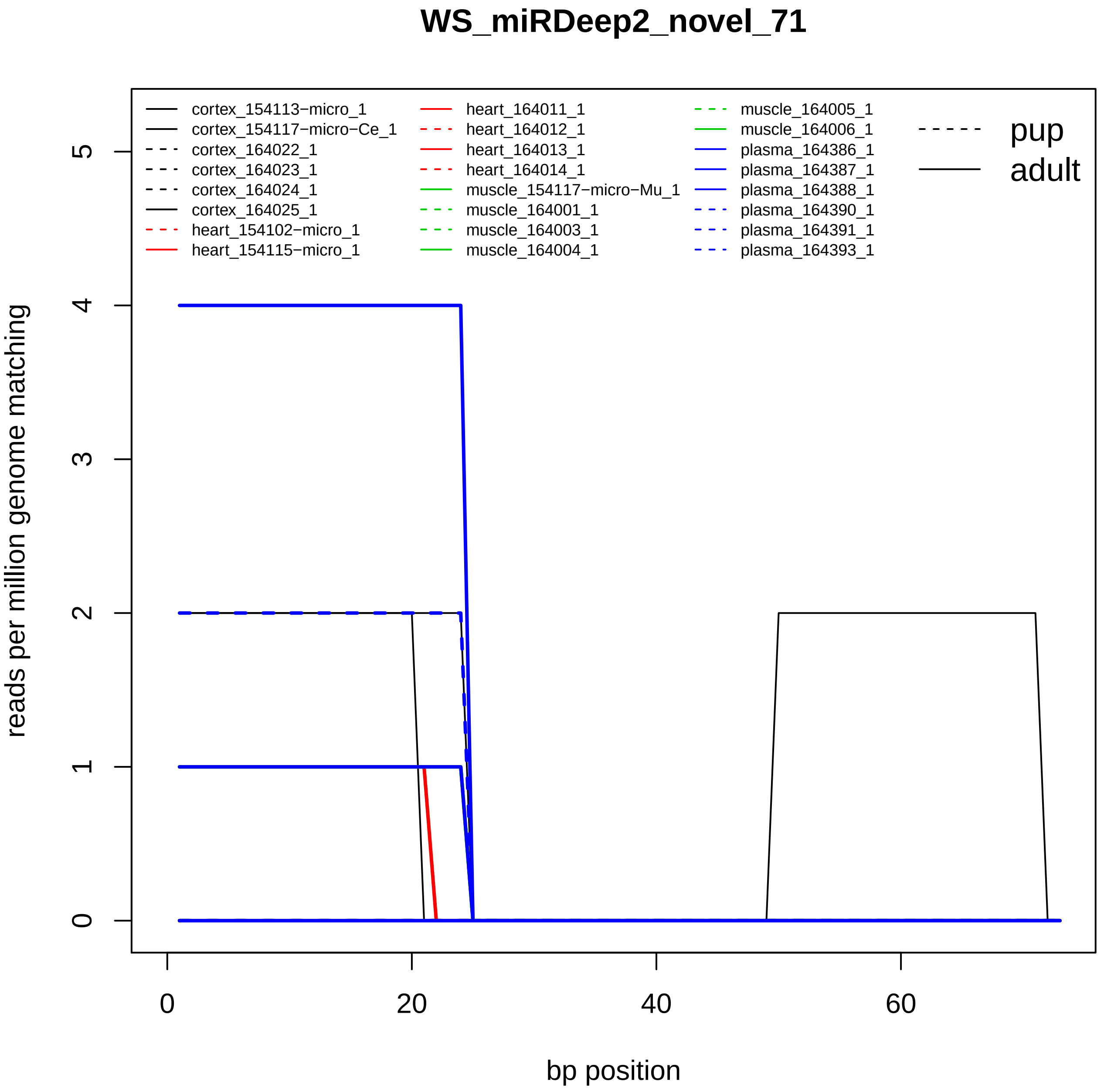
WS_miRDeep2_novel_71
cortex_154113−micro_1
cortex_154117−micro−Ce_1
cortex_164022_1
cortex_164023_1
cortex_164024_1
cortex_164025_1
heart_154102−micro_1
heart_154115−micro_1
heart_164011_1
heart_164012_1
heart_164013_1
heart_164014_1
muscle_154117−micro−Mu_1
muscle_164001_1
muscle_164003_1
muscle_164004_1
muscle_164005_1
muscle_164006_1
plasma_164386_1
plasma_164387_1
plasma_164388_1
plasma_164390_1
plasma_164391_1
plasma_164393_1
pup
adult
reads per million genome matching
0
1
2
3
4
5
0
20
40
60
bp position

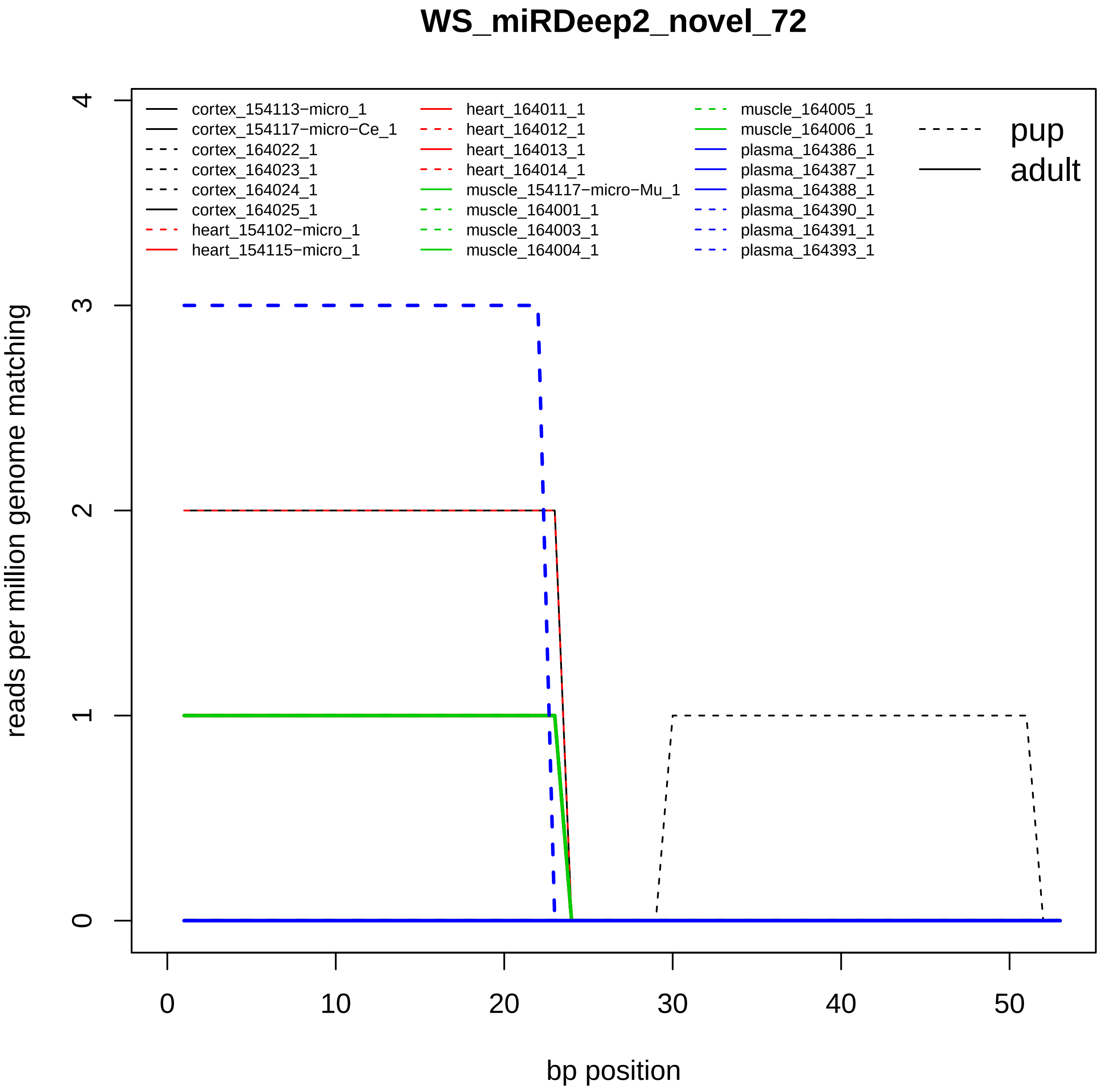

WS_miRDeep2_novel_72
cortex_154113−micro_1
cortex_154117−micro−Ce_1
cortex_164022_1
cortex_164023_1
cortex_164024_1
cortex_164025_1
heart_154102−micro_1
heart_154115−micro_1
heart_164011_1
heart_164012_1
heart_164013_1
heart_164014_1
muscle_154117−micro−Mu_1
muscle_164001_1
muscle_164003_1
muscle_164004_1
muscle_164005_1
muscle_164006_1
plasma_164386_1
plasma_164387_1
plasma_164388_1
plasma_164390_1
plasma_164391_1
plasma_164393_1
pup
adult
reads per million genome matching
bp position
0
1
2
3
4
0
10
20
30
40
50

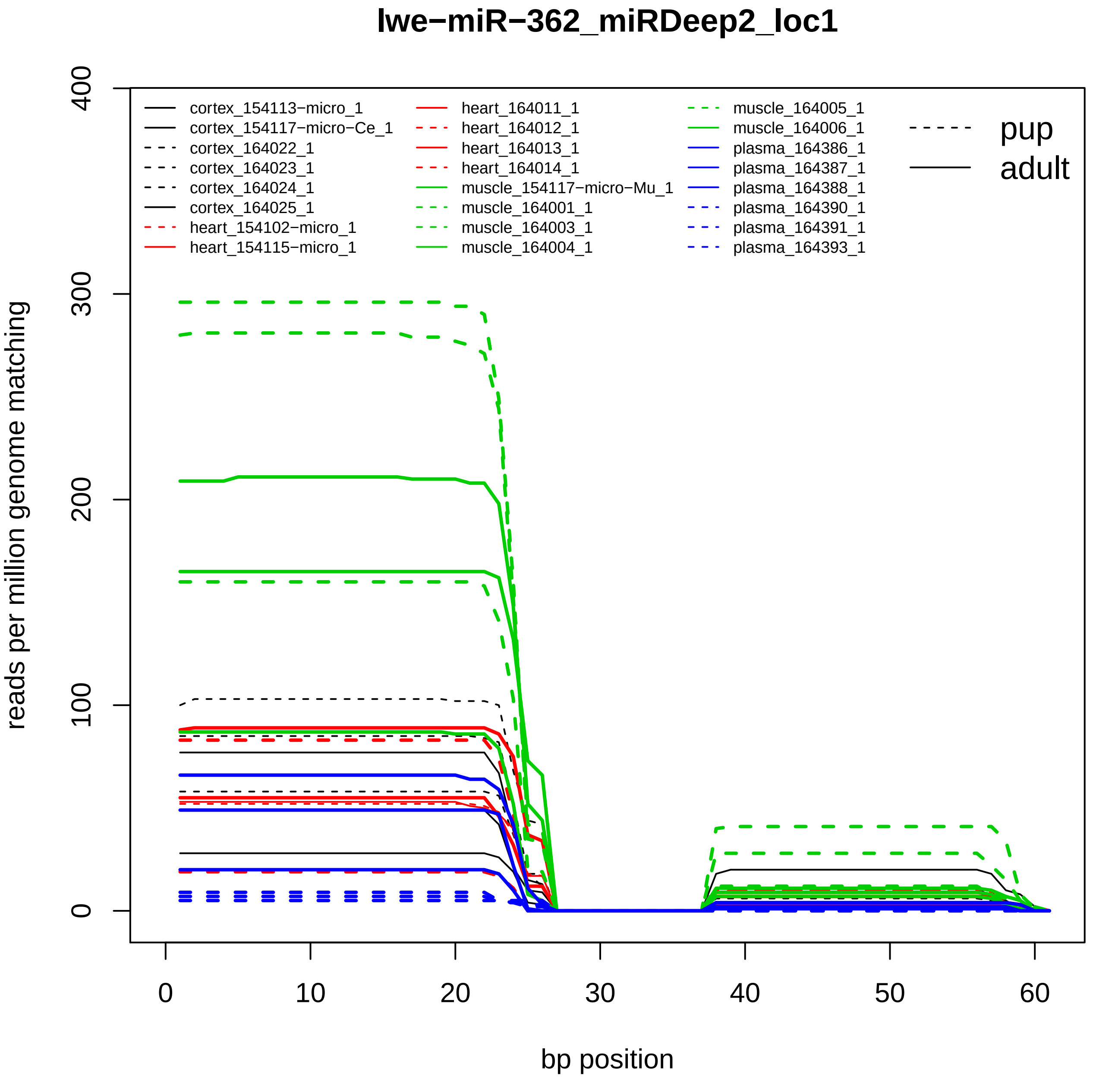
lwe−miR−362_miRDeep2_loc1
reads per million genome matching
bp position
0
100
200
300
400
0
10
20
30
40
50
60
cortex_154113−micro_1
cortex_154117−micro−Ce_1
cortex_164022_1
cortex_164023_1
cortex_164024_1
cortex_164025_1
heart_154102−micro_1
heart_154115−micro_1
heart_164011_1
heart_164012_1
heart_164013_1
heart_164014_1
muscle_154117−micro−Mu_1
muscle_164001_1
muscle_164003_1
muscle_164004_1
muscle_164005_1
muscle_164006_1
plasma_164386_1
plasma_164387_1
plasma_164388_1
plasma_164390_1
plasma_164391_1
plasma_164393_1
pup
adult

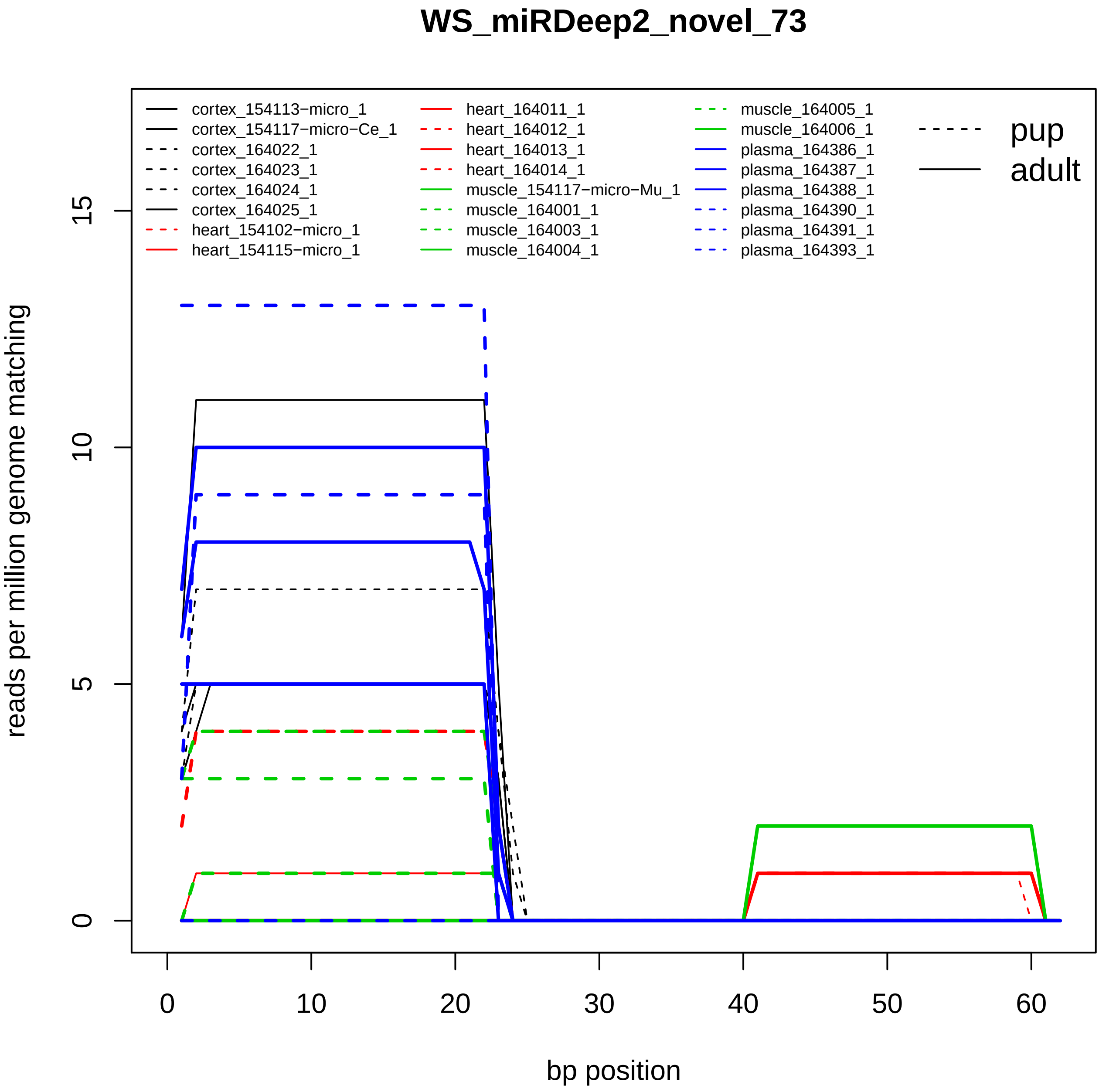

WS_miRDeep2_novel_73
cortex_154113−micro_1
cortex_154117−micro−Ce_1
cortex_164022_1
cortex_164023_1
cortex_164024_1
cortex_164025_1
heart_154102−micro_1
heart_154115−micro_1
heart_164011_1
heart_164012_1
heart_164013_1
heart_164014_1
muscle_154117−micro−Mu_1
muscle_164001_1
muscle_164003_1
muscle_164004_1
muscle_164005_1
muscle_164006_1
plasma_164386_1
plasma_164387_1
plasma_164388_1
plasma_164390_1
plasma_164391_1
plasma_164393_1
pup
adult
reads per million genome matching
0
5
10
15
0
10
20
30
40
50
60
bp position

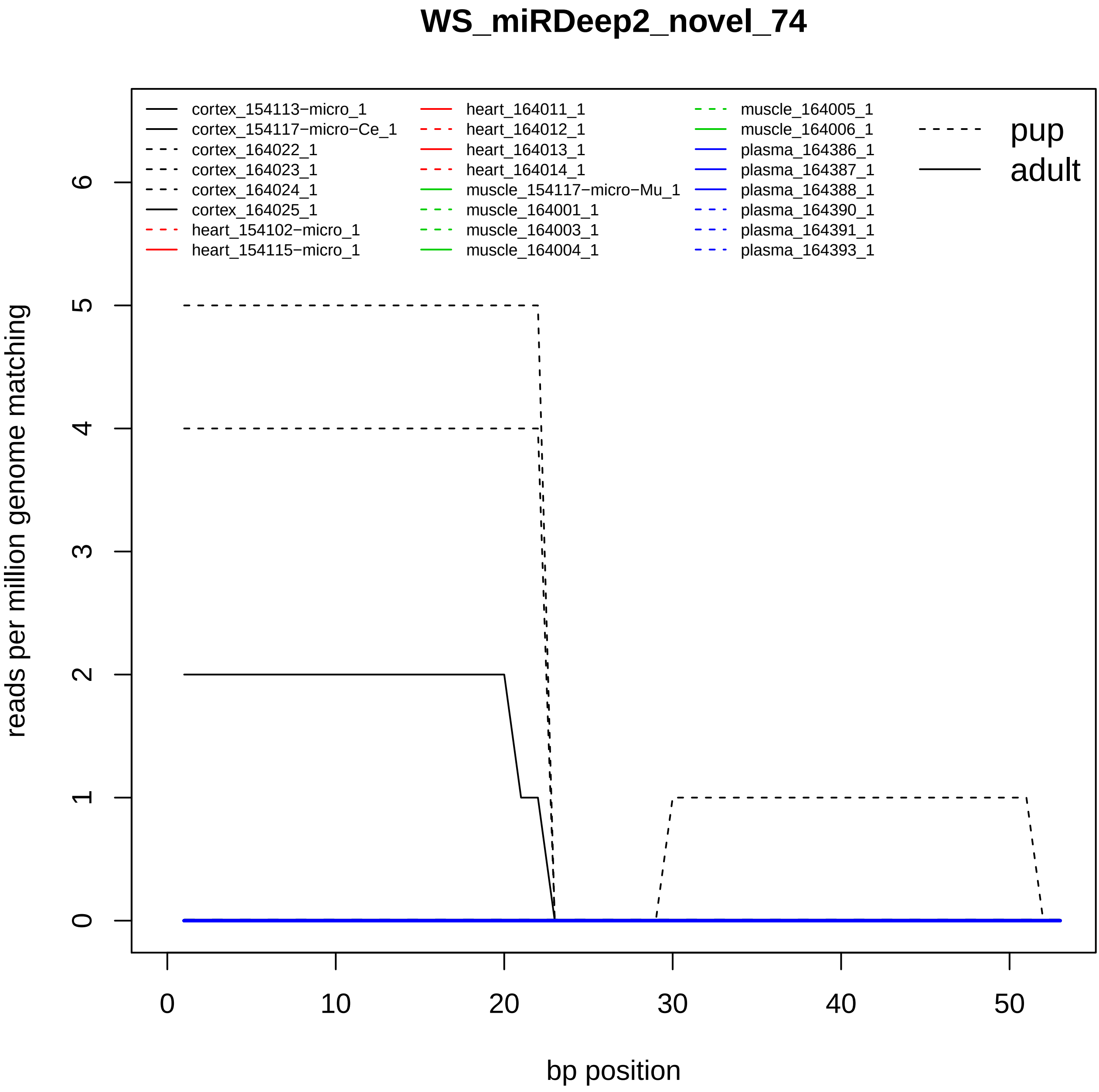

WS_miRDeep2_novel_74
cortex_154113−micro_1
cortex_154117−micro−Ce_1
cortex_164022_1
cortex_164023_1
cortex_164024_1
cortex_164025_1
heart_154102−micro_1
heart_154115−micro_1
heart_164011_1
heart_164012_1
heart_164013_1
heart_164014_1
muscle_154117−micro−Mu_1
muscle_164001_1
muscle_164003_1
muscle_164004_1
muscle_164005_1
muscle_164006_1
plasma_164386_1
plasma_164387_1
plasma_164388_1
plasma_164390_1
plasma_164391_1
plasma_164393_1
pup
adult
reads per million genome matching
0
1
2
3
4
5
6
0
10
20
30
40
50
bp position

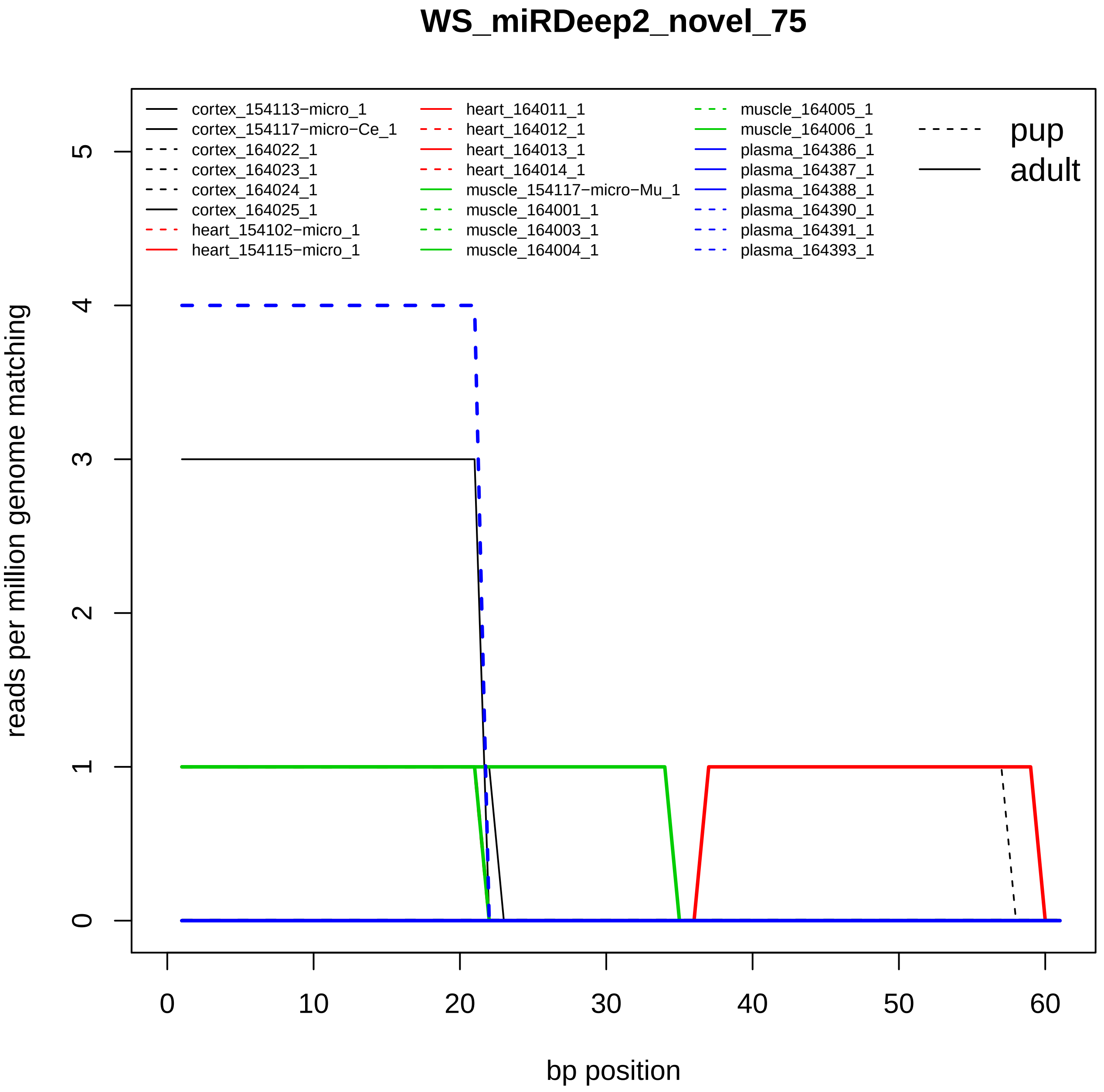

WS_miRDeep2_novel_75
cortex_154113−micro_1
cortex_154117−micro−Ce_1
cortex_164022_1
cortex_164023_1
cortex_164024_1
cortex_164025_1
heart_154102−micro_1
heart_154115−micro_1
heart_164011_1
heart_164012_1
heart_164013_1
heart_164014_1
muscle_154117−micro−Mu_1
muscle_164001_1
muscle_164003_1
muscle_164004_1
muscle_164005_1
muscle_164006_1
plasma_164386_1
plasma_164387_1
plasma_164388_1
plasma_164390_1
plasma_164391_1
plasma_164393_1
pup
adult
reads per million genome matching
bp position
0
1
2
3
4
5
0
10
20
30
40
50
60

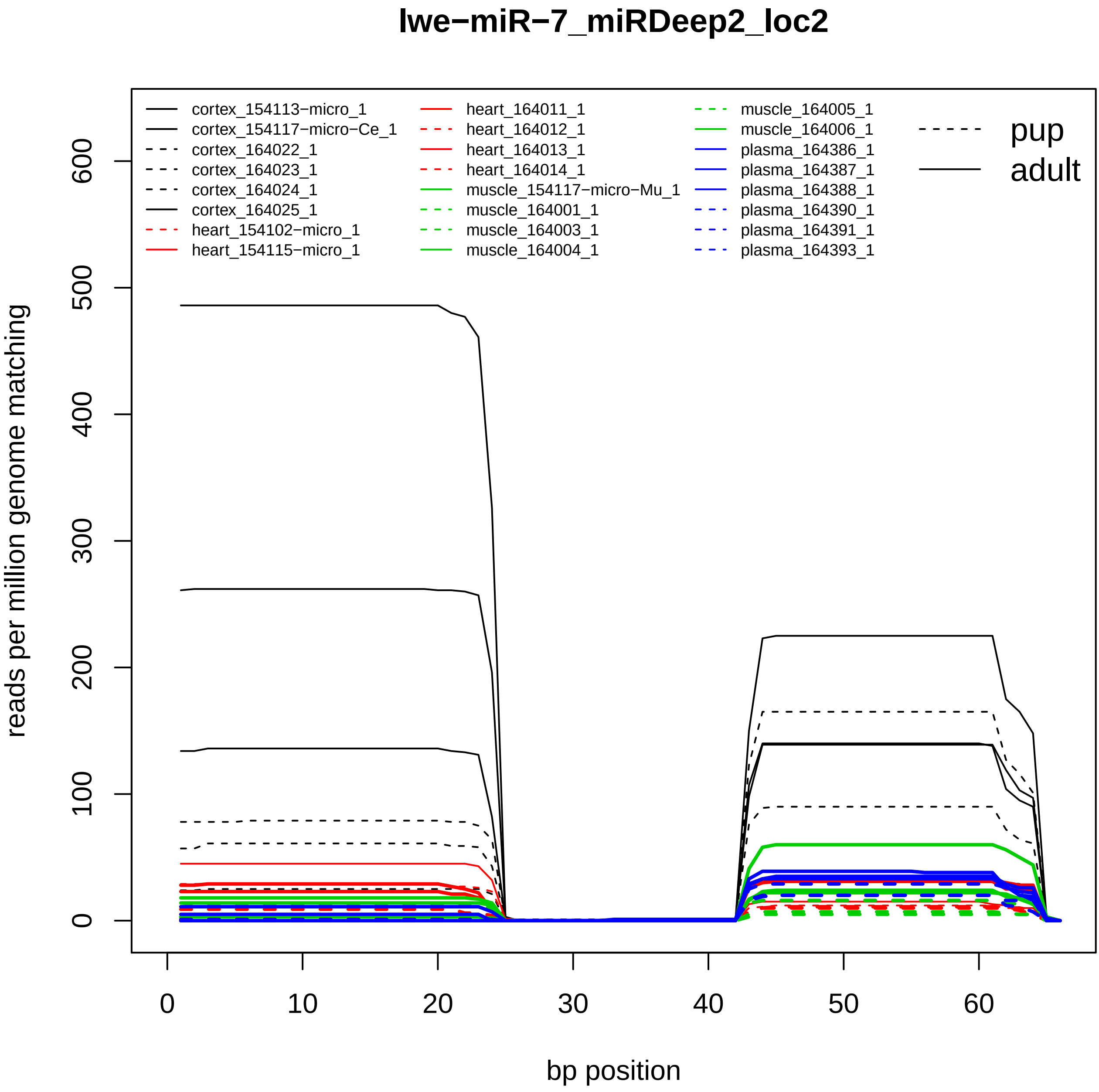

lwe−miR−7_miRDeep2_loc2
reads per million genome matching
bp position
0
100
200
300
400
500
600
0
10
20
30
40
50
60
cortex_154113−micro_1
cortex_154117−micro−Ce_1
cortex_164022_1
cortex_164023_1
cortex_164024_1
cortex_164025_1
heart_154102−micro_1
heart_154115−micro_1
heart_164011_1
heart_164012_1
heart_164013_1
heart_164014_1
muscle_154117−micro−Mu_1
muscle_164001_1
muscle_164003_1
muscle_164004_1
muscle_164005_1
muscle_164006_1
plasma_164386_1
plasma_164387_1
plasma_164388_1
plasma_164390_1
plasma_164391_1
plasma_164393_1
pup
adult

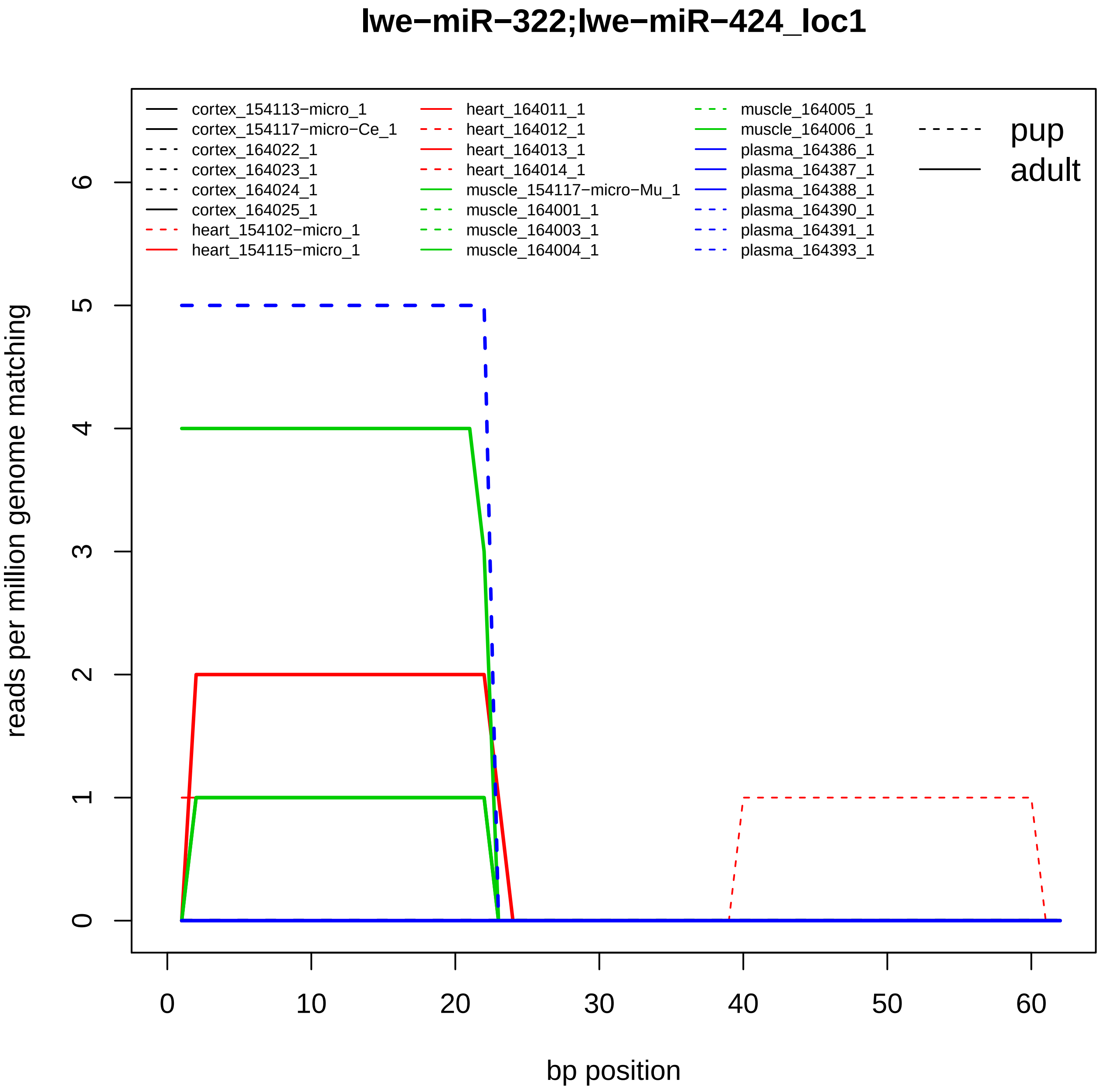
lwe−miR−322;lwe−miR−424_loc1
cortex_154113−micro_1
cortex_154117−micro−Ce_1
cortex_164022_1
cortex_164023_1
cortex_164024_1
cortex_164025_1
heart_154102−micro_1
heart_154115−micro_1
heart_164011_1
heart_164012_1
heart_164013_1
heart_164014_1
muscle_154117−micro−Mu_1
muscle_164001_1
muscle_164003_1
muscle_164004_1
muscle_164005_1
muscle_164006_1
plasma_164386_1
plasma_164387_1
plasma_164388_1
plasma_164390_1
plasma_164391_1
plasma_164393_1
pup
adult
reads per million genome matching
0
1
2
3
4
5
6
0
10
20
30
40
50
60
bp position

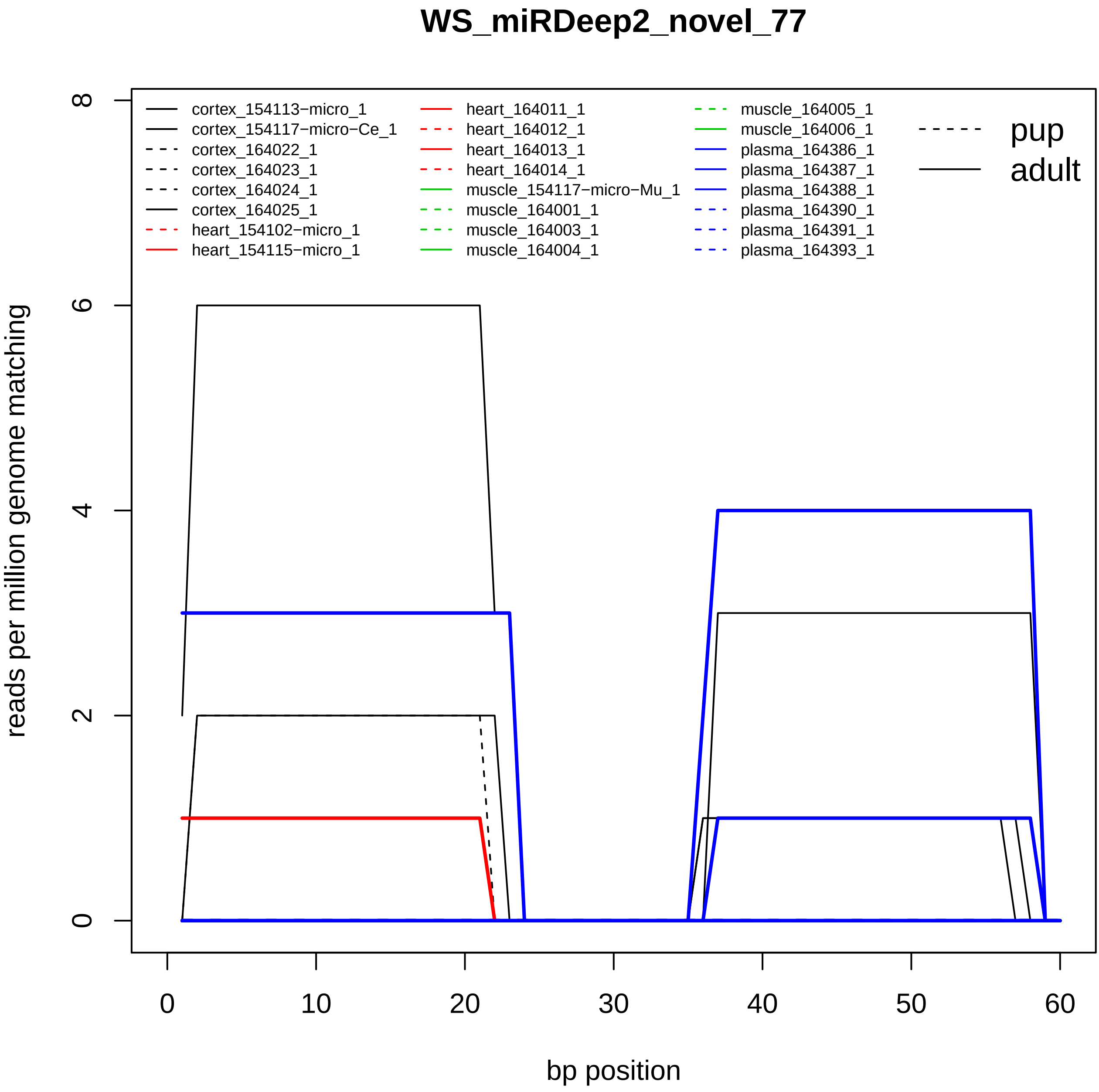

WS_miRDeep2_novel_77
cortex_154113−micro_1
cortex_154117−micro−Ce_1
cortex_164022_1
cortex_164023_1
cortex_164024_1
cortex_164025_1
heart_154102−micro_1
heart_154115−micro_1
heart_164011_1
heart_164012_1
heart_164013_1
heart_164014_1
muscle_154117−micro−Mu_1
muscle_164001_1
muscle_164003_1
muscle_164004_1
muscle_164005_1
muscle_164006_1
plasma_164386_1
plasma_164387_1
plasma_164388_1
plasma_164390_1
plasma_164391_1
plasma_164393_1
pup
adult
reads per million genome matching
bp position
0
2
4
6
8
0
10
20
30
40
50
60

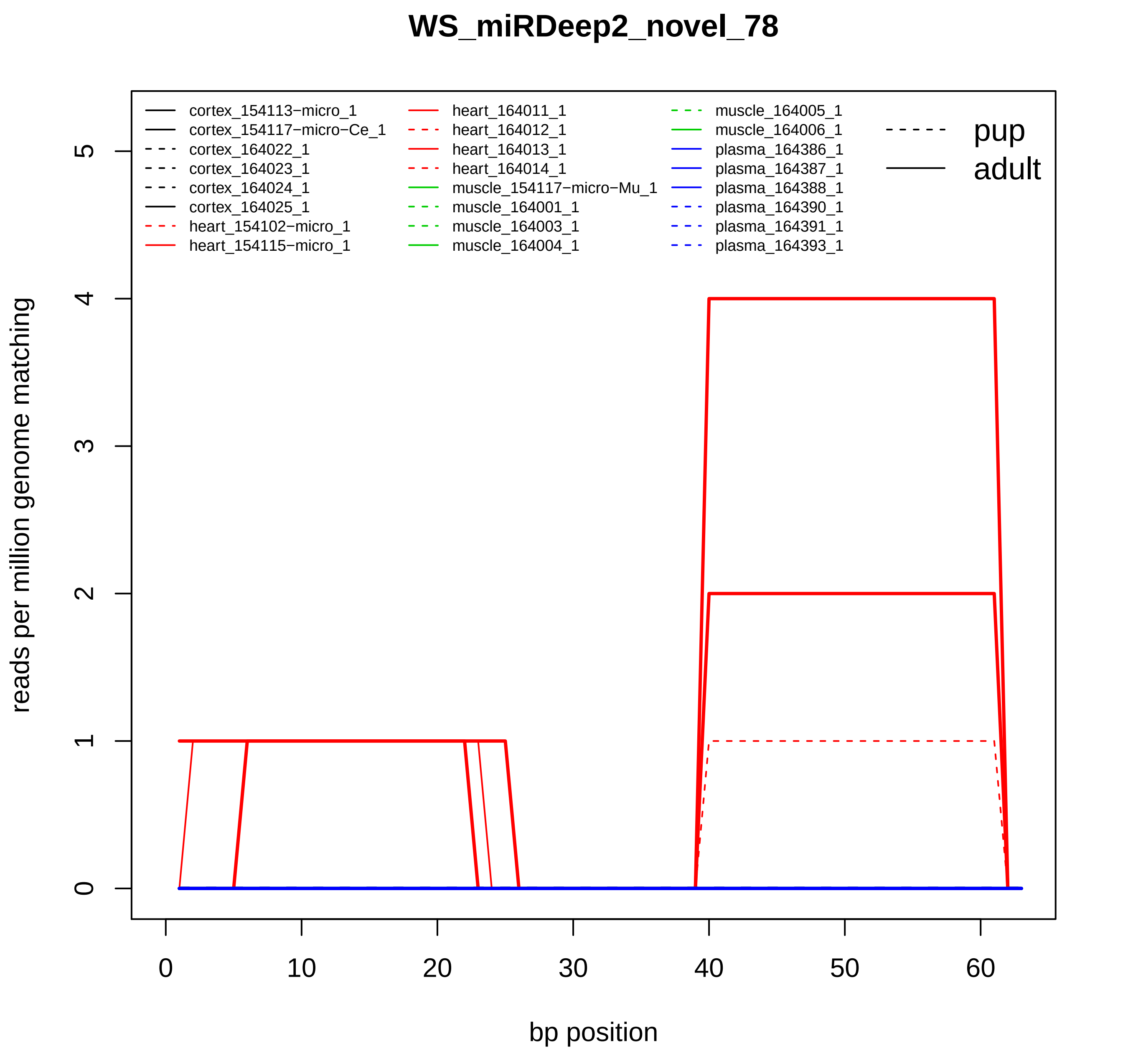

WS_miRDeep2_novel_78
cortex_154113−micro_1
cortex_154117−micro−Ce_1
cortex_164022_1
cortex_164023_1
cortex_164024_1
cortex_164025_1
heart_154102−micro_1
heart_154115−micro_1
heart_164011_1
heart_164012_1
heart_164013_1
heart_164014_1
muscle_154117−micro−Mu_1
muscle_164001_1
muscle_164003_1
muscle_164004_1
muscle_164005_1
muscle_164006_1
plasma_164386_1
plasma_164387_1
plasma_164388_1
plasma_164390_1
plasma_164391_1
plasma_164393_1
pup
adult
reads per million genome matching
0
1
2
3
4
5
0
10
20
30
40
50
60
bp position

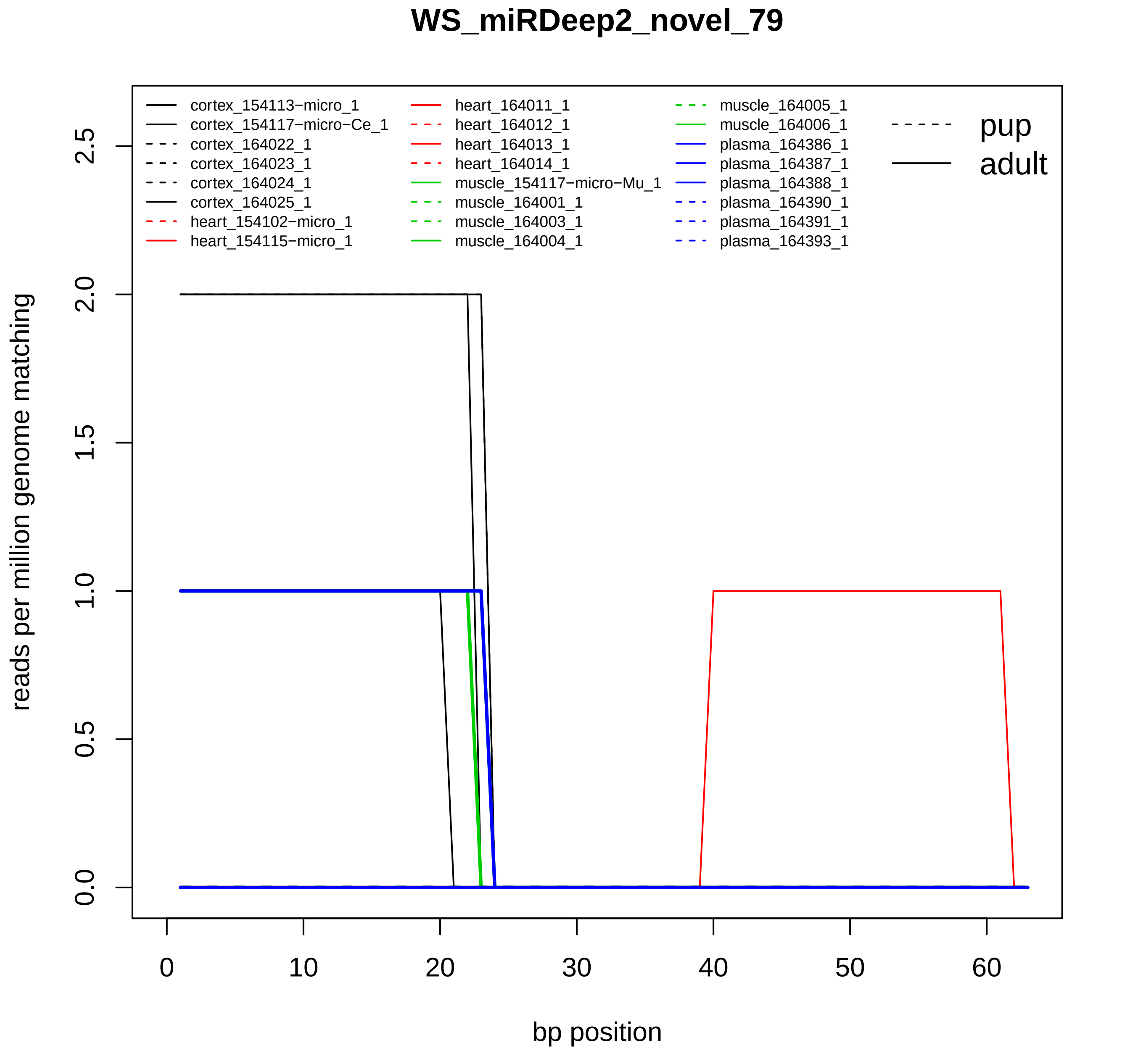
WS_miRDeep2_novel_79
cortex_154113−micro_1
cortex_154117−micro−Ce_1
cortex_164022_1
cortex_164023_1
cortex_164024_1
cortex_164025_1
heart_154102−micro_1
heart_154115−micro_1
heart_164011_1
heart_164012_1
heart_164013_1
heart_164014_1
muscle_154117−micro−Mu_1
muscle_164001_1
muscle_164003_1
muscle_164004_1
muscle_164005_1
muscle_164006_1
plasma_164386_1
plasma_164387_1
plasma_164388_1
plasma_164390_1
plasma_164391_1
plasma_164393_1
pup
adult
reads per million genome matching
0.0
0.5
1.0
1.5
2.0
2.5
0
10
20
30
40
50
60
bp position

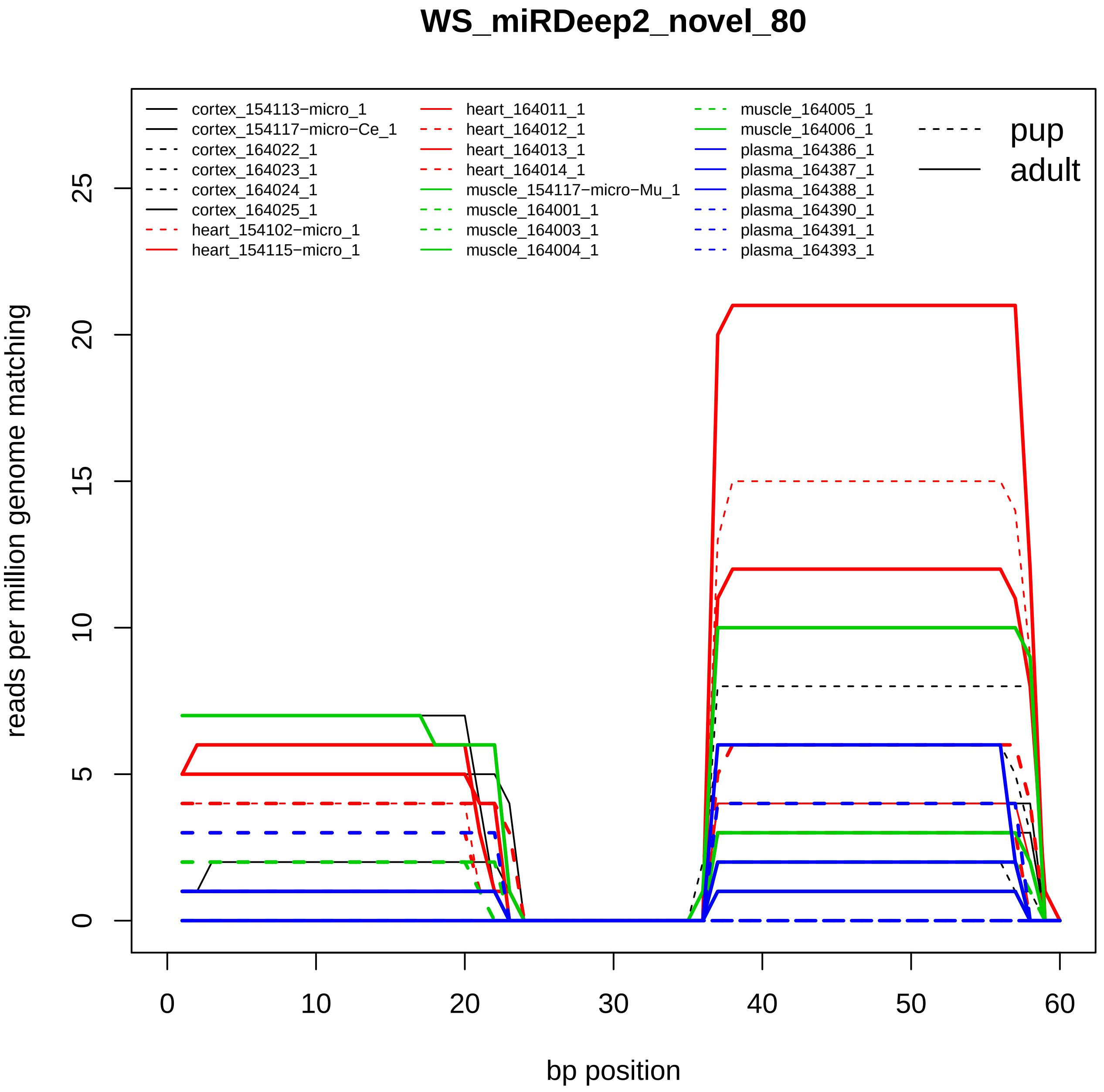

WS_miRDeep2_novel_80
cortex_154113−micro_1
cortex_154117−micro−Ce_1
cortex_164022_1
cortex_164023_1
cortex_164024_1
cortex_164025_1
heart_154102−micro_1
heart_154115−micro_1
heart_164011_1
heart_164012_1
heart_164013_1
heart_164014_1
muscle_154117−micro−Mu_1
muscle_164001_1
muscle_164003_1
muscle_164004_1
muscle_164005_1
muscle_164006_1
plasma_164386_1
plasma_164387_1
plasma_164388_1
plasma_164390_1
plasma_164391_1
plasma_164393_1
pup
adult
reads per million genome matching
0
5
10
15
20
25
bp position
0
10
20
30
40
50
60

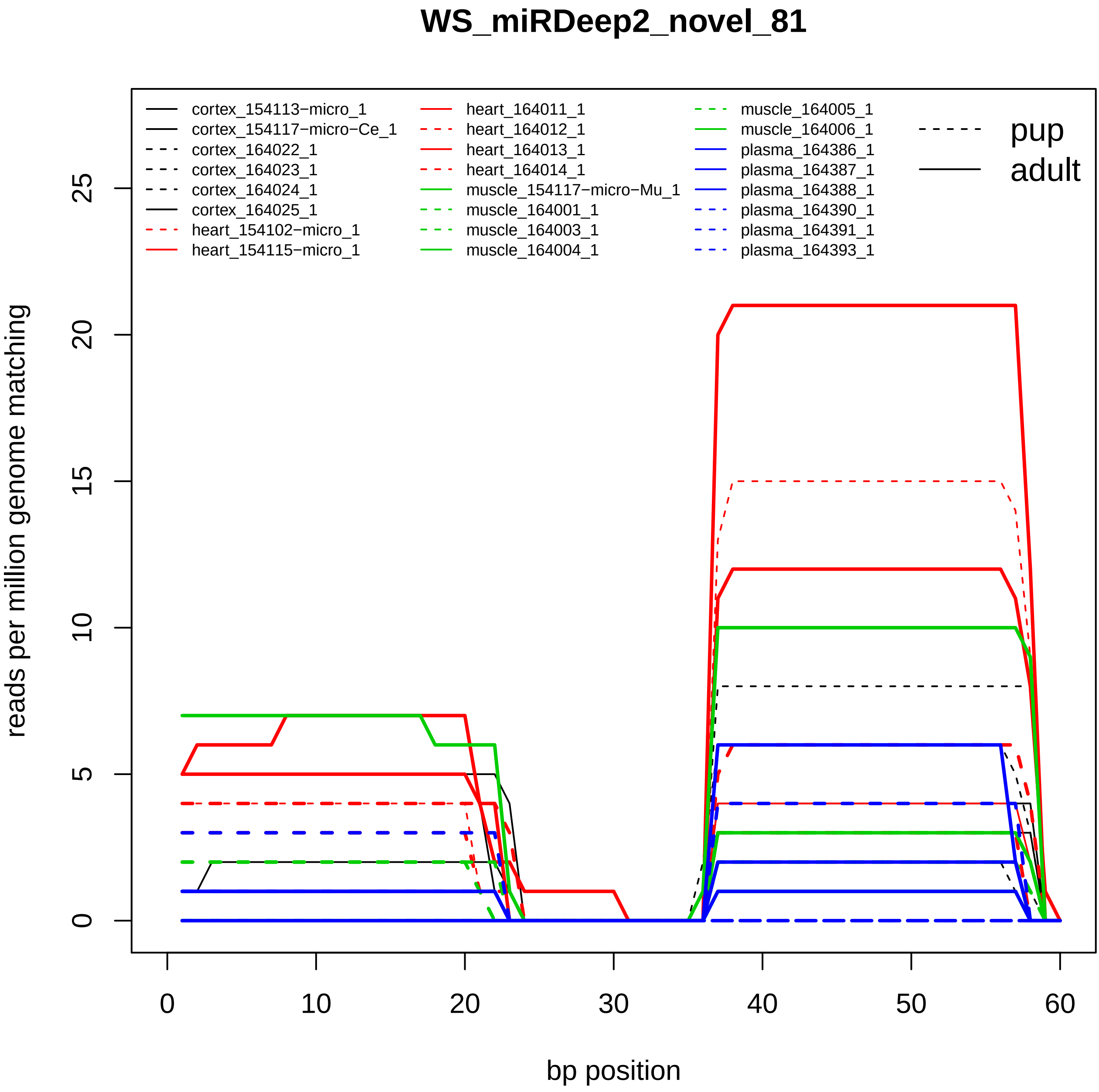

WS_miRDeep2_novel_81
cortex_154113−micro_1
cortex_154117−micro−Ce_1
cortex_164022_1
cortex_164023_1
cortex_164024_1
cortex_164025_1
heart_154102−micro_1
heart_154115−micro_1
heart_164011_1
heart_164012_1
heart_164013_1
heart_164014_1
muscle_154117−micro−Mu_1
muscle_164001_1
muscle_164003_1
muscle_164004_1
muscle_164005_1
muscle_164006_1
plasma_164386_1
plasma_164387_1
plasma_164388_1
plasma_164390_1
plasma_164391_1
plasma_164393_1
pup
adult
reads per million genome matching
bp position
0
5
10
15
20
25
0
10
20
30
40
50
60

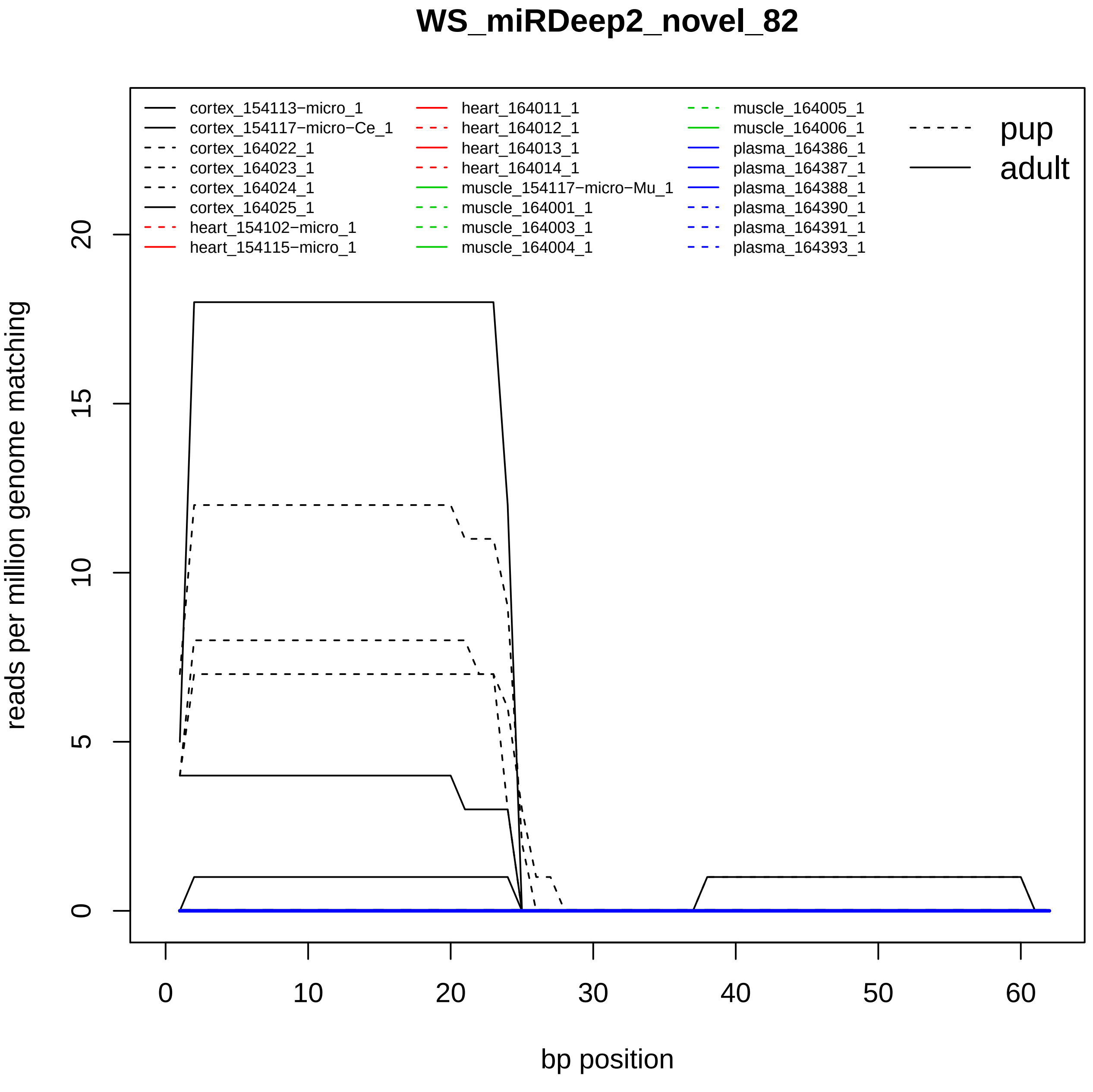

WS_miRDeep2_novel_82
cortex_154113−micro_1
cortex_154117−micro−Ce_1
cortex_164022_1
cortex_164023_1
cortex_164024_1
cortex_164025_1
heart_154102−micro_1
heart_154115−micro_1
heart_164011_1
heart_164012_1
heart_164013_1
heart_164014_1
muscle_154117−micro−Mu_1
muscle_164001_1
muscle_164003_1
muscle_164004_1
muscle_164005_1
muscle_164006_1
plasma_164386_1
plasma_164387_1
plasma_164388_1
plasma_164390_1
plasma_164391_1
plasma_164393_1
pup
adult
reads per million genome matching
0
5
10
15
20
bp position
0
10
20
30
40
50
60

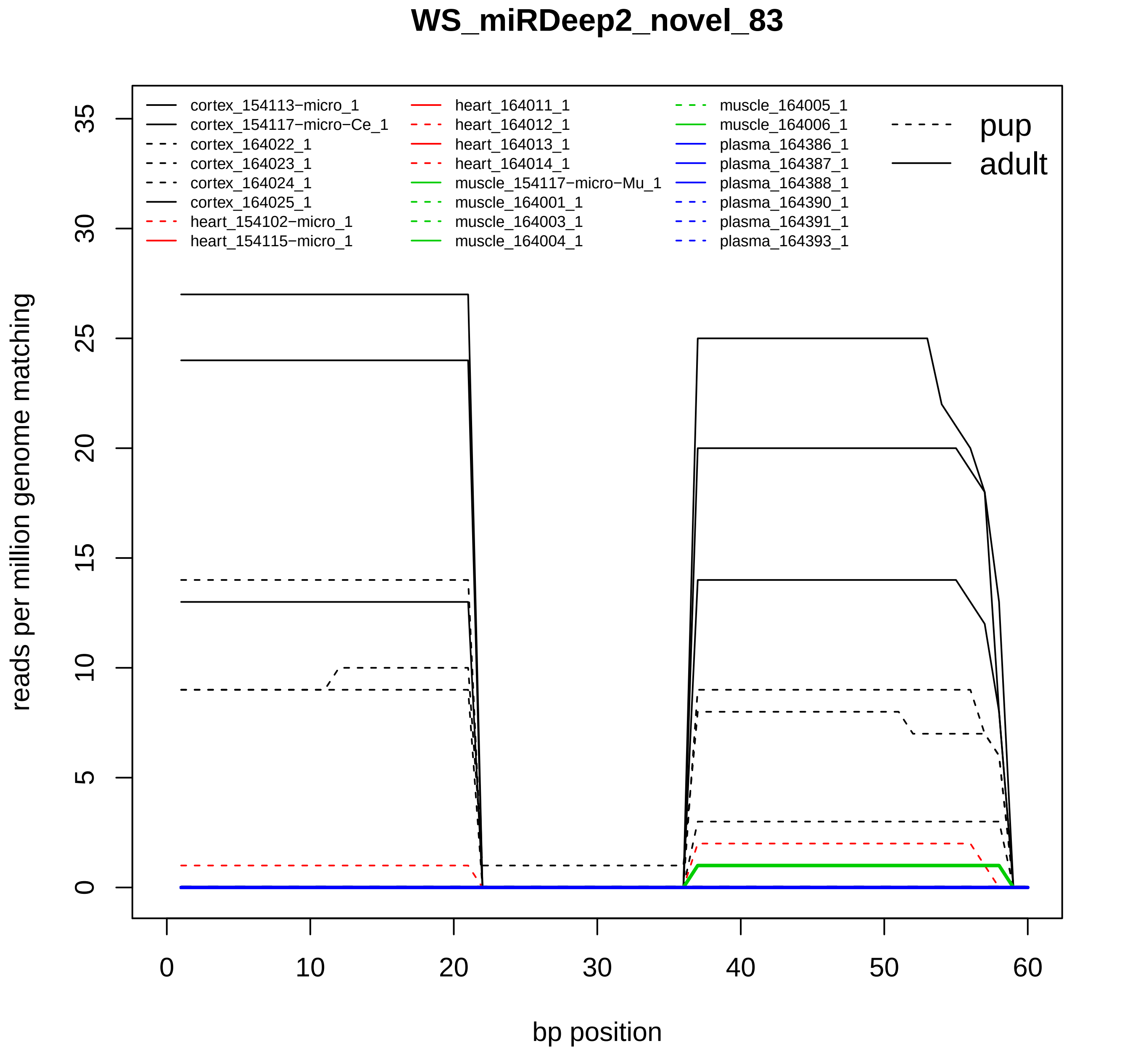

WS_miRDeep2_novel_83
cortex_154113−micro_1
cortex_154117−micro−Ce_1
cortex_164022_1
cortex_164023_1
cortex_164024_1
cortex_164025_1
heart_154102−micro_1
heart_154115−micro_1
heart_164011_1
heart_164012_1
heart_164013_1
heart_164014_1
muscle_154117−micro−Mu_1
muscle_164001_1
muscle_164003_1
muscle_164004_1
muscle_164005_1
muscle_164006_1
plasma_164386_1
plasma_164387_1
plasma_164388_1
plasma_164390_1
plasma_164391_1
plasma_164393_1
pup
adult
reads per million genome matching
0
5
10
15
20
25
30
35
0
10
20
30
40
50
60
bp position

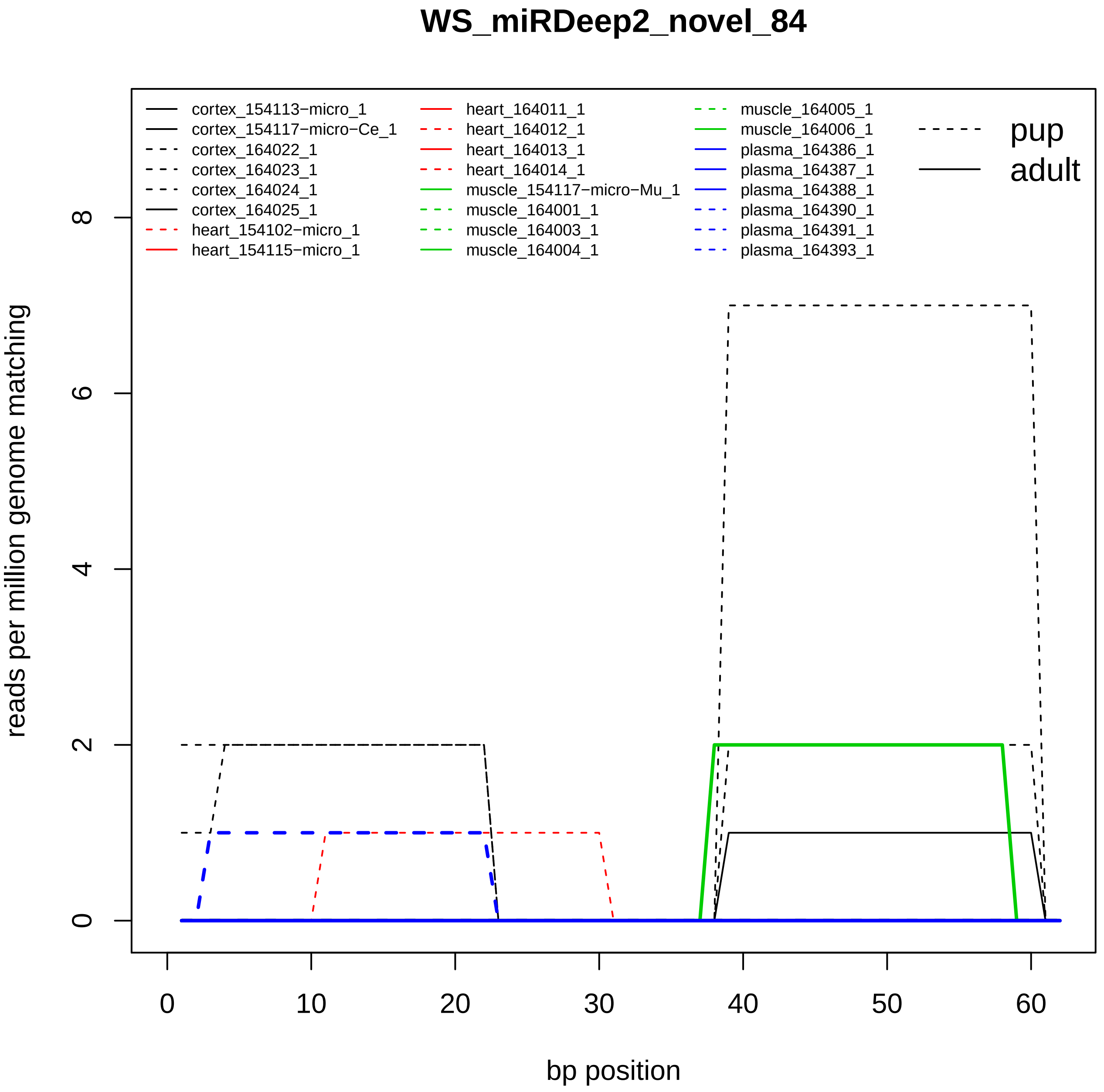
WS_miRDeep2_novel_84
cortex_154113−micro_1
cortex_154117−micro−Ce_1
cortex_164022_1
cortex_164023_1
cortex_164024_1
cortex_164025_1
heart_154102−micro_1
heart_154115−micro_1
heart_164011_1
heart_164012_1
heart_164013_1
heart_164014_1
muscle_154117−micro−Mu_1
muscle_164001_1
muscle_164003_1
muscle_164004_1
muscle_164005_1
muscle_164006_1
plasma_164386_1
plasma_164387_1
plasma_164388_1
plasma_164390_1
plasma_164391_1
plasma_164393_1
pup
adult
reads per million genome matching
0
2
4
6
8
bp position
0
10
20
30
40
50
60

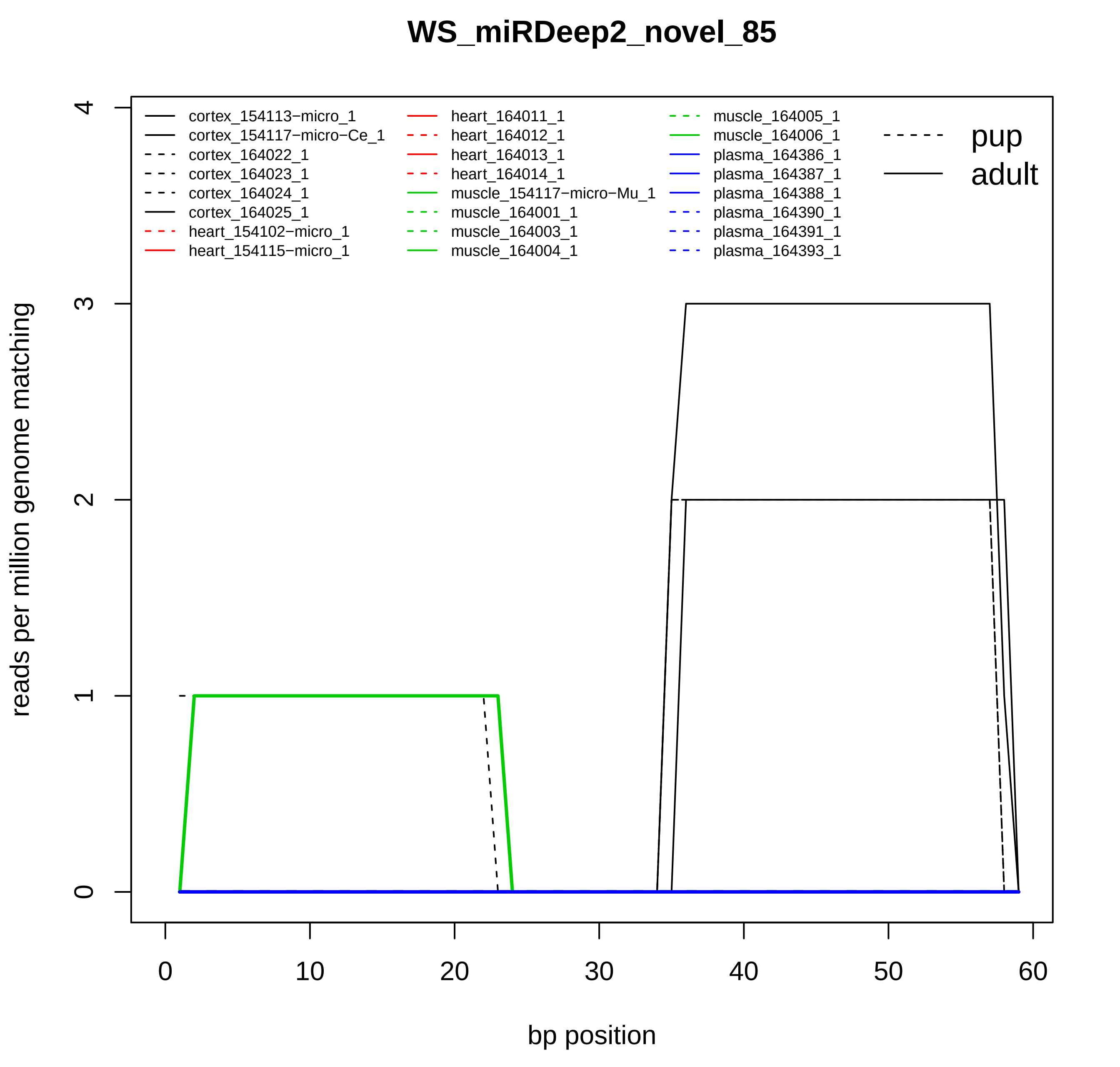

WS_miRDeep2_novel_85
cortex_154113−micro_1
cortex_154117−micro−Ce_1
cortex_164022_1
cortex_164023_1
cortex_164024_1
cortex_164025_1
heart_154102−micro_1
heart_154115−micro_1
heart_164011_1
heart_164012_1
heart_164013_1
heart_164014_1
muscle_154117−micro−Mu_1
muscle_164001_1
muscle_164003_1
muscle_164004_1
muscle_164005_1
muscle_164006_1
plasma_164386_1
plasma_164387_1
plasma_164388_1
plasma_164390_1
plasma_164391_1
plasma_164393_1
pup
adult
reads per million genome matching
0
1
2
3
4
bp position
0
10
20
30
40
50
60

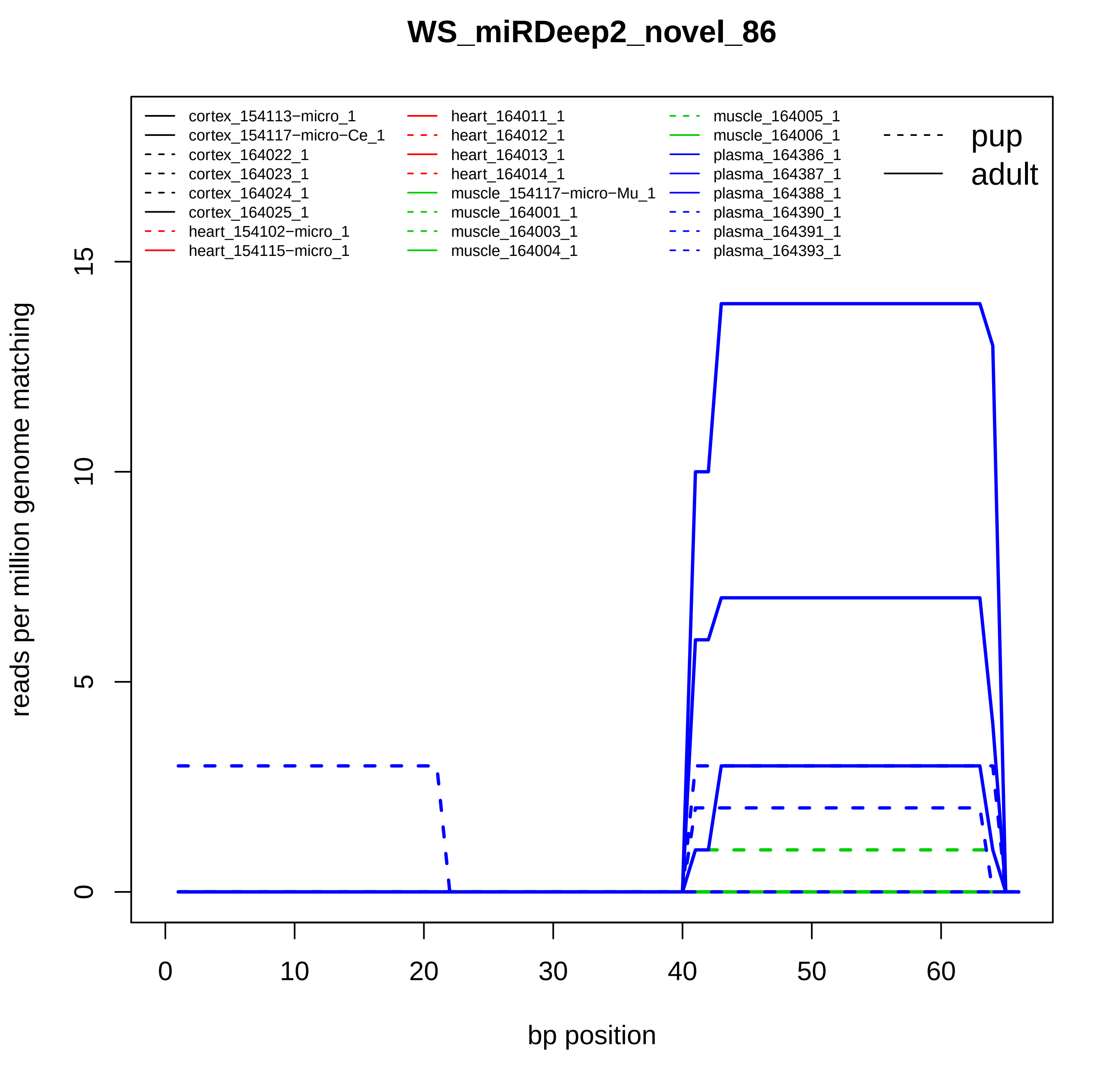
WS_miRDeep2_novel_86
cortex_154113−micro_1
cortex_154117−micro−Ce_1
cortex_164022_1
cortex_164023_1
cortex_164024_1
cortex_164025_1
heart_154102−micro_1
heart_154115−micro_1
heart_164011_1
heart_164012_1
heart_164013_1
heart_164014_1
muscle_154117−micro−Mu_1
muscle_164001_1
muscle_164003_1
muscle_164004_1
muscle_164005_1
muscle_164006_1
plasma_164386_1
plasma_164387_1
plasma_164388_1
plasma_164390_1
plasma_164391_1
plasma_164393_1
pup
adult
reads per million genome matching
0
5
10
15
bp position
0
10
20
30
40
50
60

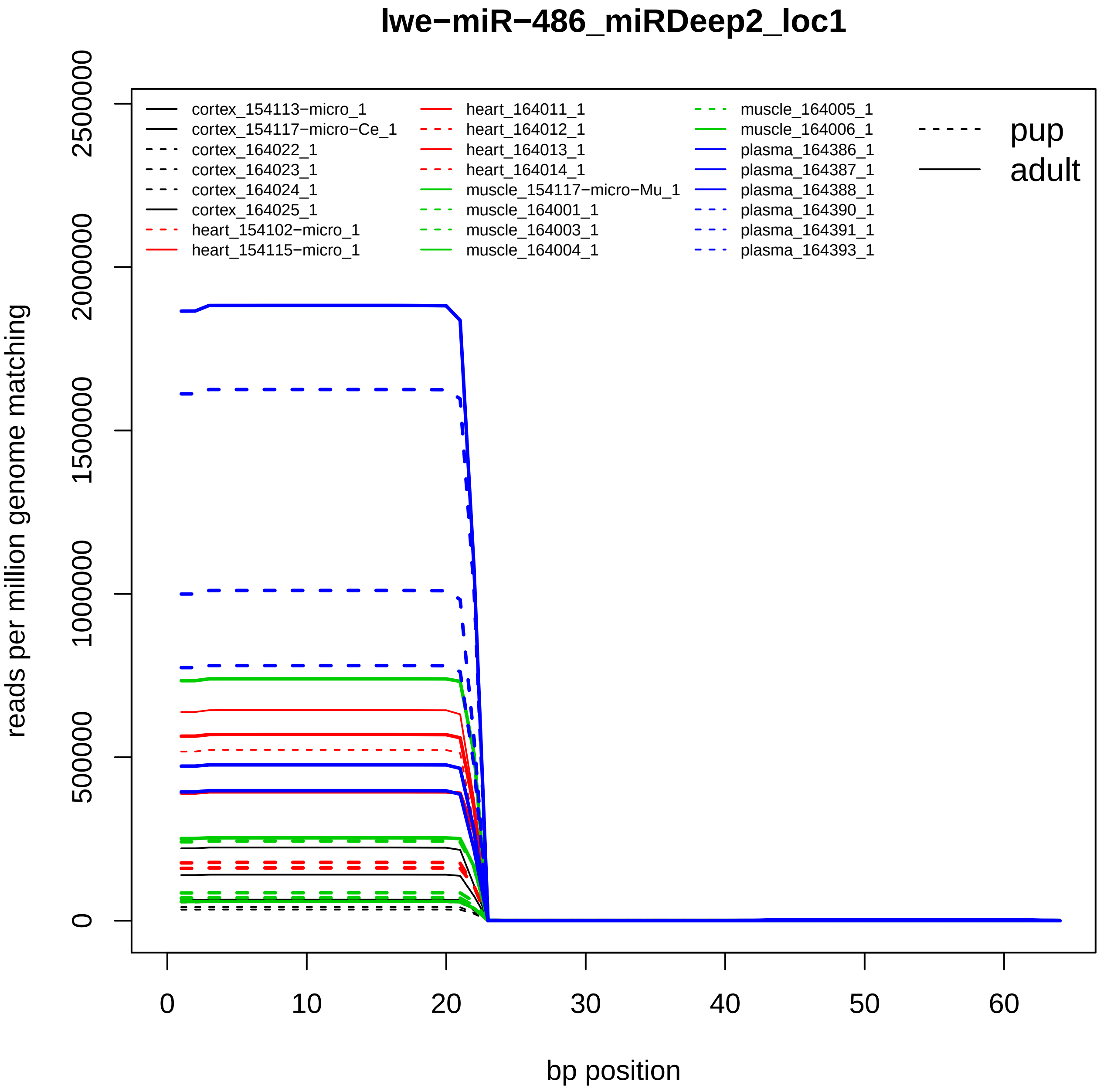
lwe−miR−486_miRDeep2_loc1
reads per million genome matching
bp position
0
500000
1000000
1500000
2000000
2500000
0
10
20
30
40
50
60
cortex_154113−micro_1
cortex_154117−micro−Ce_1
cortex_164022_1
cortex_164023_1
cortex_164024_1
cortex_164025_1
heart_154102−micro_1
heart_154115−micro_1
heart_164011_1
heart_164012_1
heart_164013_1
heart_164014_1
muscle_154117−micro−Mu_1
muscle_164001_1
muscle_164003_1
muscle_164004_1
muscle_164005_1
muscle_164006_1
plasma_164386_1
plasma_164387_1
plasma_164388_1
plasma_164390_1
plasma_164391_1
plasma_164393_1
pup
adult

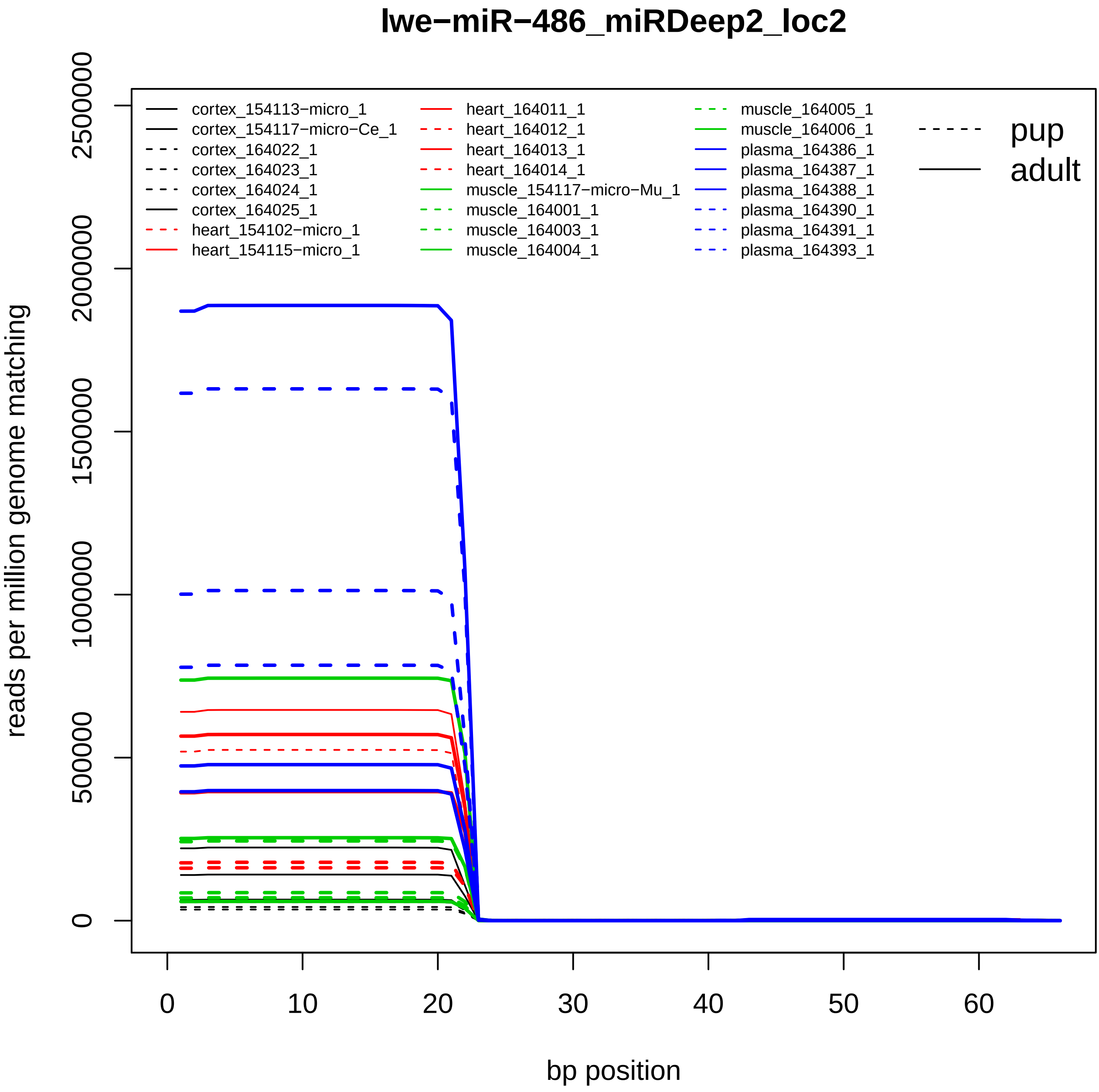

lwe−miR−486_miRDeep2_loc2
reads per million genome matching
bp position
0
500000
1000000
1500000
2000000
2500000
0
10
20
30
40
50
60
cortex_154113−micro_1
cortex_154117−micro−Ce_1
cortex_164022_1
cortex_164023_1
cortex_164024_1
cortex_164025_1
heart_154102−micro_1
heart_154115−micro_1
heart_164011_1
heart_164012_1
heart_164013_1
heart_164014_1
muscle_154117−micro−Mu_1
muscle_164001_1
muscle_164003_1
muscle_164004_1
muscle_164005_1
muscle_164006_1
plasma_164386_1
plasma_164387_1
plasma_164388_1
plasma_164390_1
plasma_164391_1
plasma_164393_1
pup
adult

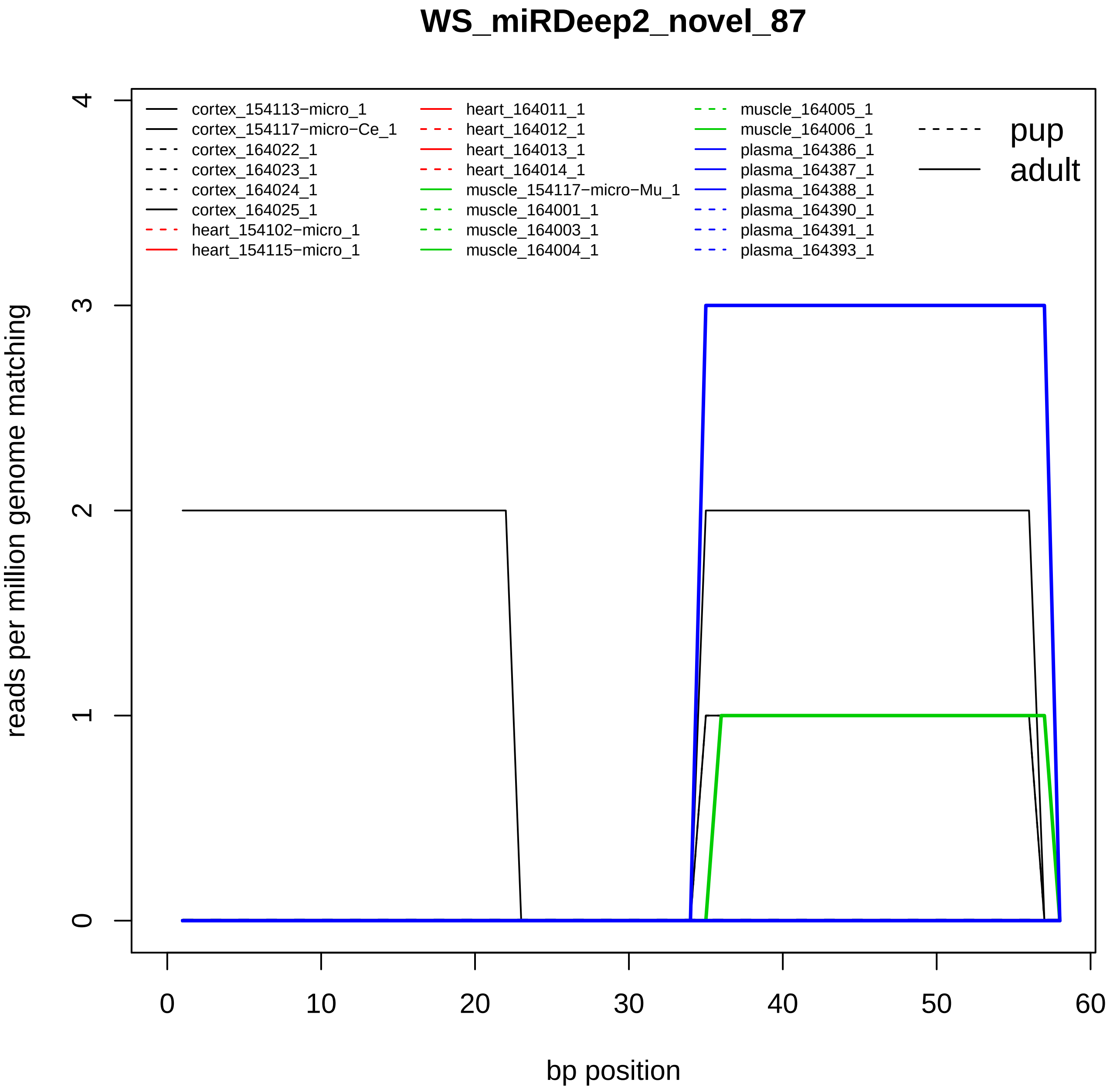

WS_miRDeep2_novel_87
cortex_154113−micro_1
cortex_154117−micro−Ce_1
cortex_164022_1
cortex_164023_1
cortex_164024_1
cortex_164025_1
heart_154102−micro_1
heart_154115−micro_1
heart_164011_1
heart_164012_1
heart_164013_1
heart_164014_1
muscle_154117−micro−Mu_1
muscle_164001_1
muscle_164003_1
muscle_164004_1
muscle_164005_1
muscle_164006_1
plasma_164386_1
plasma_164387_1
plasma_164388_1
plasma_164390_1
plasma_164391_1
plasma_164393_1
pup
adult
reads per million genome matching
0
1
2
3
4
0
10
20
30
40
50
60
bp position

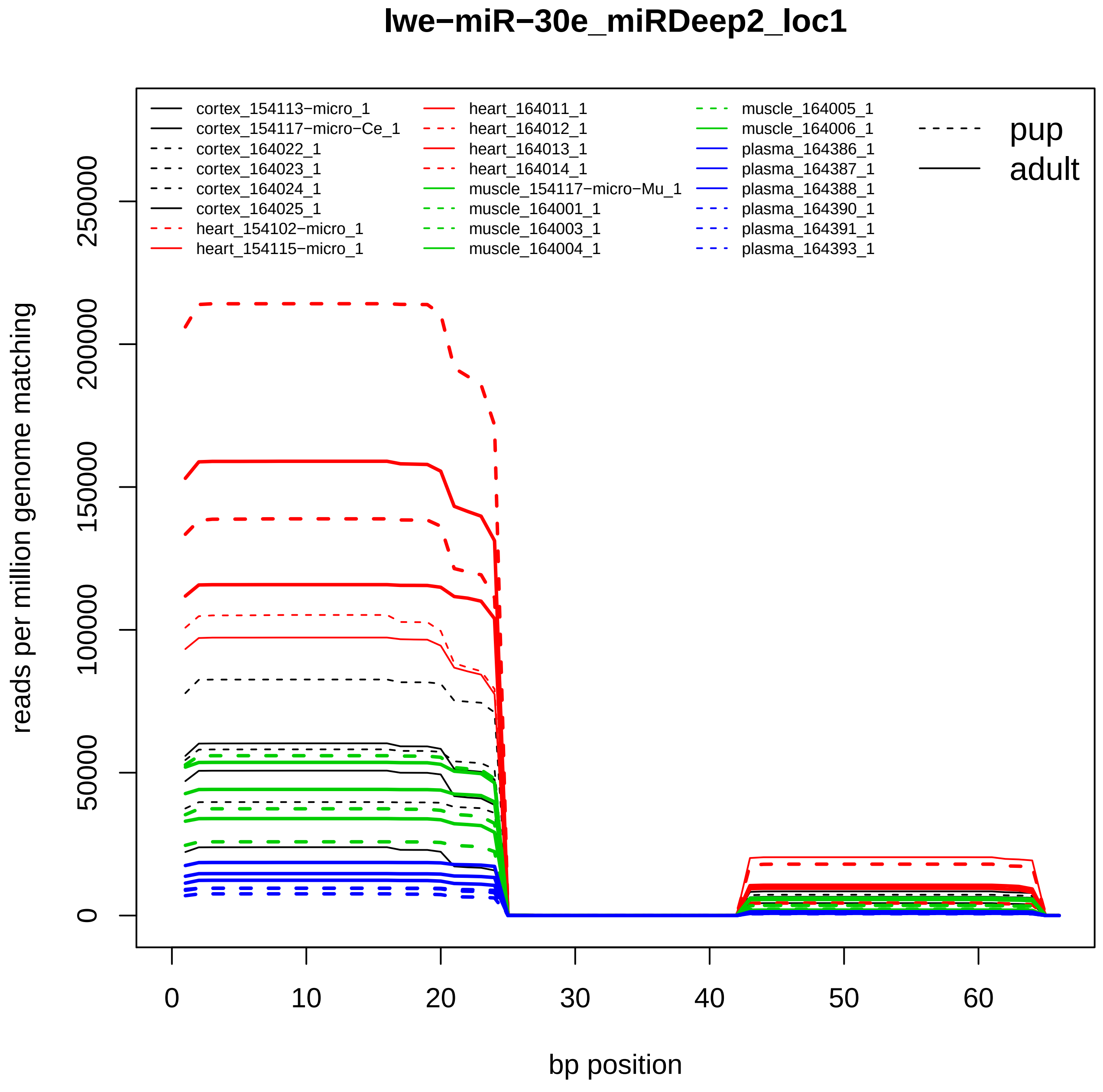
lwe−miR−30e_miRDeep2_loc1
cortex_154113−micro_1
cortex_154117−micro−Ce_1
cortex_164022_1
cortex_164023_1
cortex_164024_1
cortex_164025_1
heart_154102−micro_1
heart_154115−micro_1
heart_164011_1
heart_164012_1
heart_164013_1
heart_164014_1
muscle_154117−micro−Mu_1
muscle_164001_1
muscle_164003_1
muscle_164004_1
muscle_164005_1
muscle_164006_1
plasma_164386_1
plasma_164387_1
plasma_164388_1
plasma_164390_1
plasma_164391_1
plasma_164393_1
pup
adult
reads per million genome matching
0
50000
100000
150000
200000
250000
0
10
20
30
40
50
60
bp position

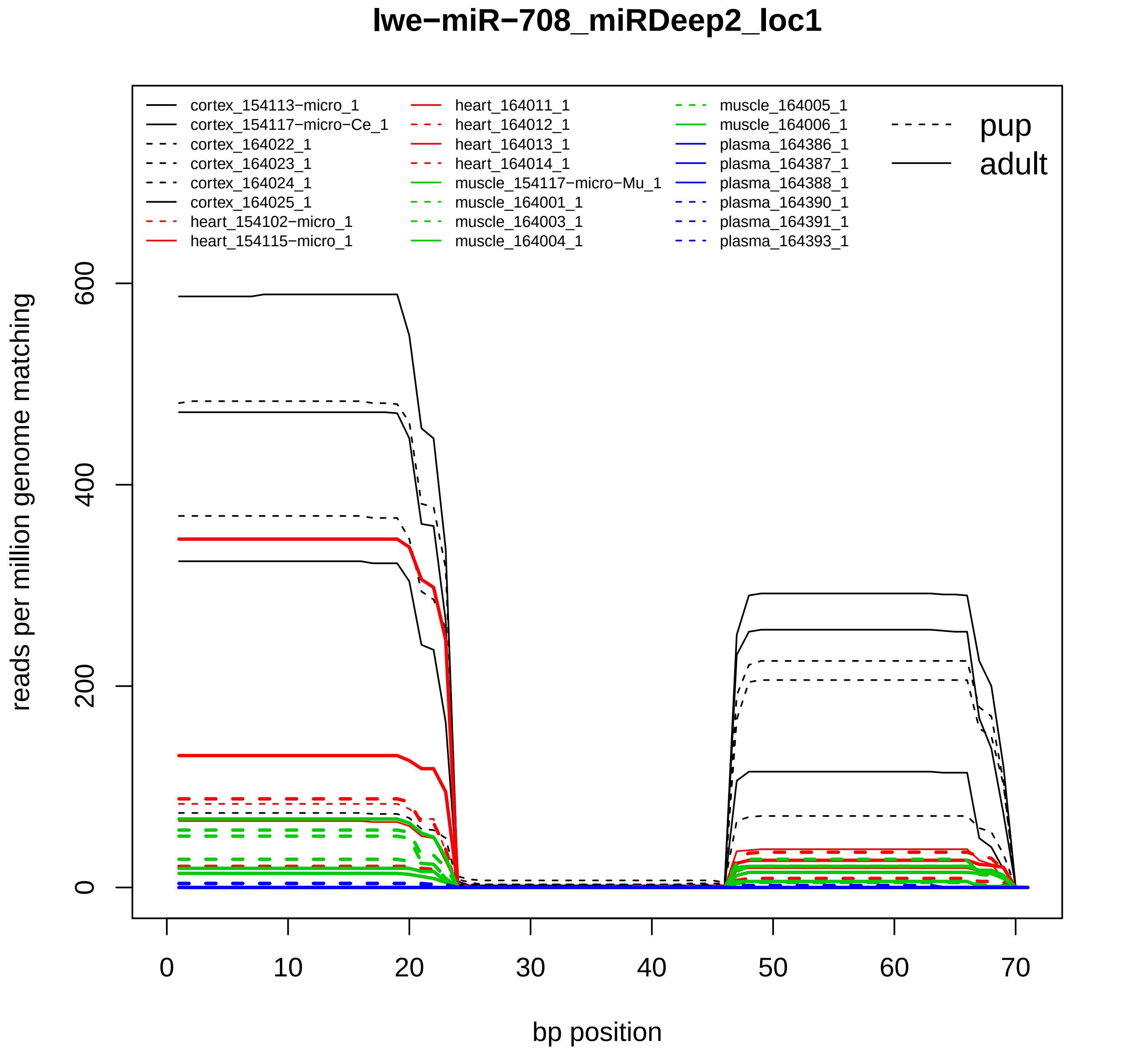

lwe−miR−708_miRDeep2_loc1
cortex_154113−micro_1
cortex_154117−micro−Ce_1
cortex_164022_1
cortex_164023_1
cortex_164024_1
cortex_164025_1
heart_154102−micro_1
heart_154115−micro_1
heart_164011_1
heart_164012_1
heart_164013_1
heart_164014_1
muscle_154117−micro−Mu_1
muscle_164001_1
muscle_164003_1
muscle_164004_1
muscle_164005_1
muscle_164006_1
plasma_164386_1
plasma_164387_1
plasma_164388_1
plasma_164390_1
plasma_164391_1
plasma_164393_1
pup
adult
reads per million genome matching
0
200
400
600
0
10
20
30
40
50
60
70
bp position

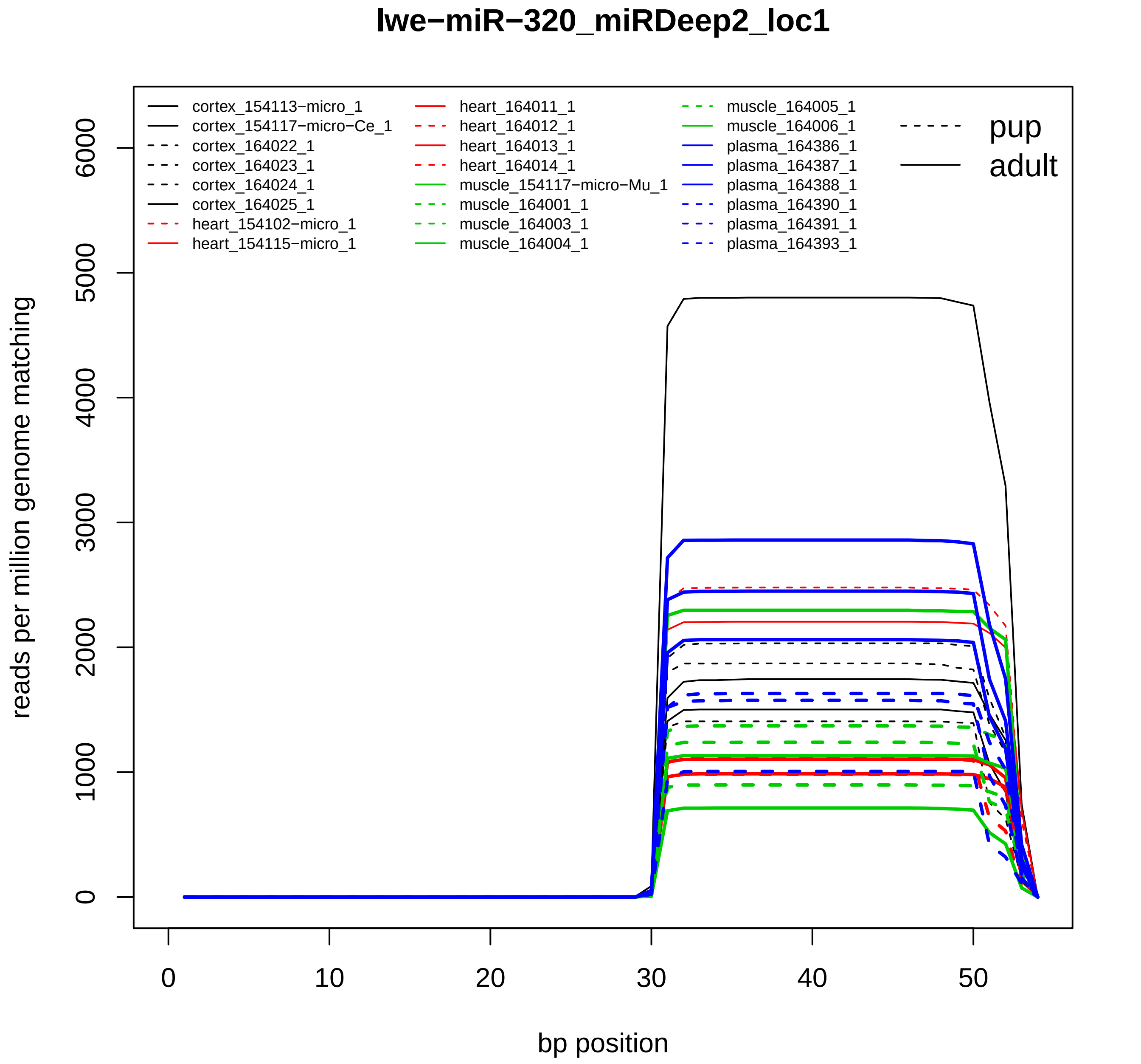
lwe−miR−320_miRDeep2_loc1
cortex_154113−micro_1
cortex_154117−micro−Ce_1
cortex_164022_1
cortex_164023_1
cortex_164024_1
cortex_164025_1
heart_154102−micro_1
heart_154115−micro_1
heart_164011_1
heart_164012_1
heart_164013_1
heart_164014_1
muscle_154117−micro−Mu_1
muscle_164001_1
muscle_164003_1
muscle_164004_1
muscle_164005_1
muscle_164006_1
plasma_164386_1
plasma_164387_1
plasma_164388_1
plasma_164390_1
plasma_164391_1
plasma_164393_1
pup
adult
reads per million genome matching
0
1000
2000
3000
4000
5000
6000
0
10
20
30
40
50
bp position

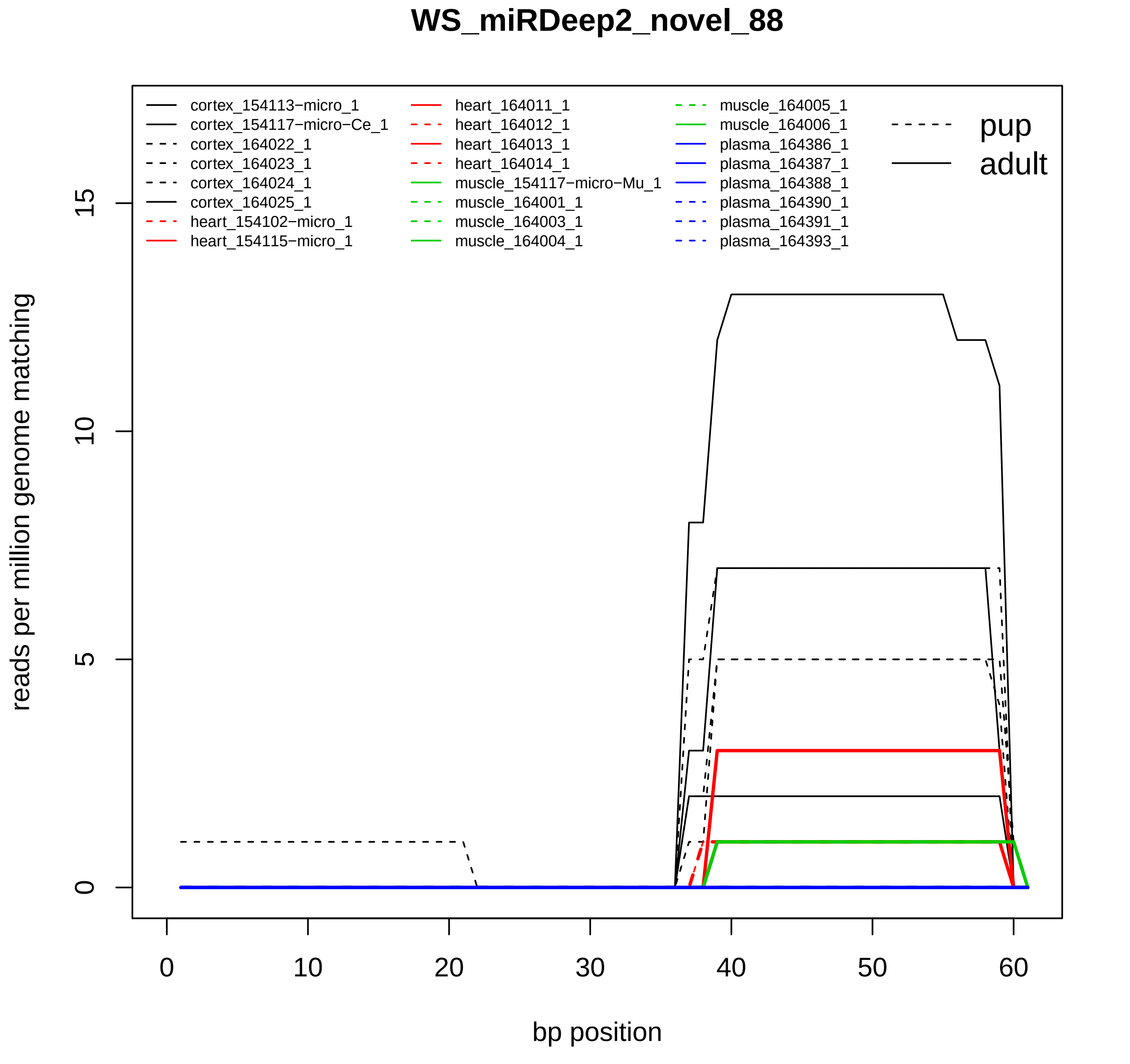

WS_miRDeep2_novel_88
cortex_154113−micro_1
cortex_154117−micro−Ce_1
cortex_164022_1
cortex_164023_1
cortex_164024_1
cortex_164025_1
heart_154102−micro_1
heart_154115−micro_1
heart_164011_1
heart_164012_1
heart_164013_1
heart_164014_1
muscle_154117−micro−Mu_1
muscle_164001_1
muscle_164003_1
muscle_164004_1
muscle_164005_1
muscle_164006_1
plasma_164386_1
plasma_164387_1
plasma_164388_1
plasma_164390_1
plasma_164391_1
plasma_164393_1
pup
adult
reads per million genome matching
0
5
10
15
0
10
20
30
40
50
60
bp position

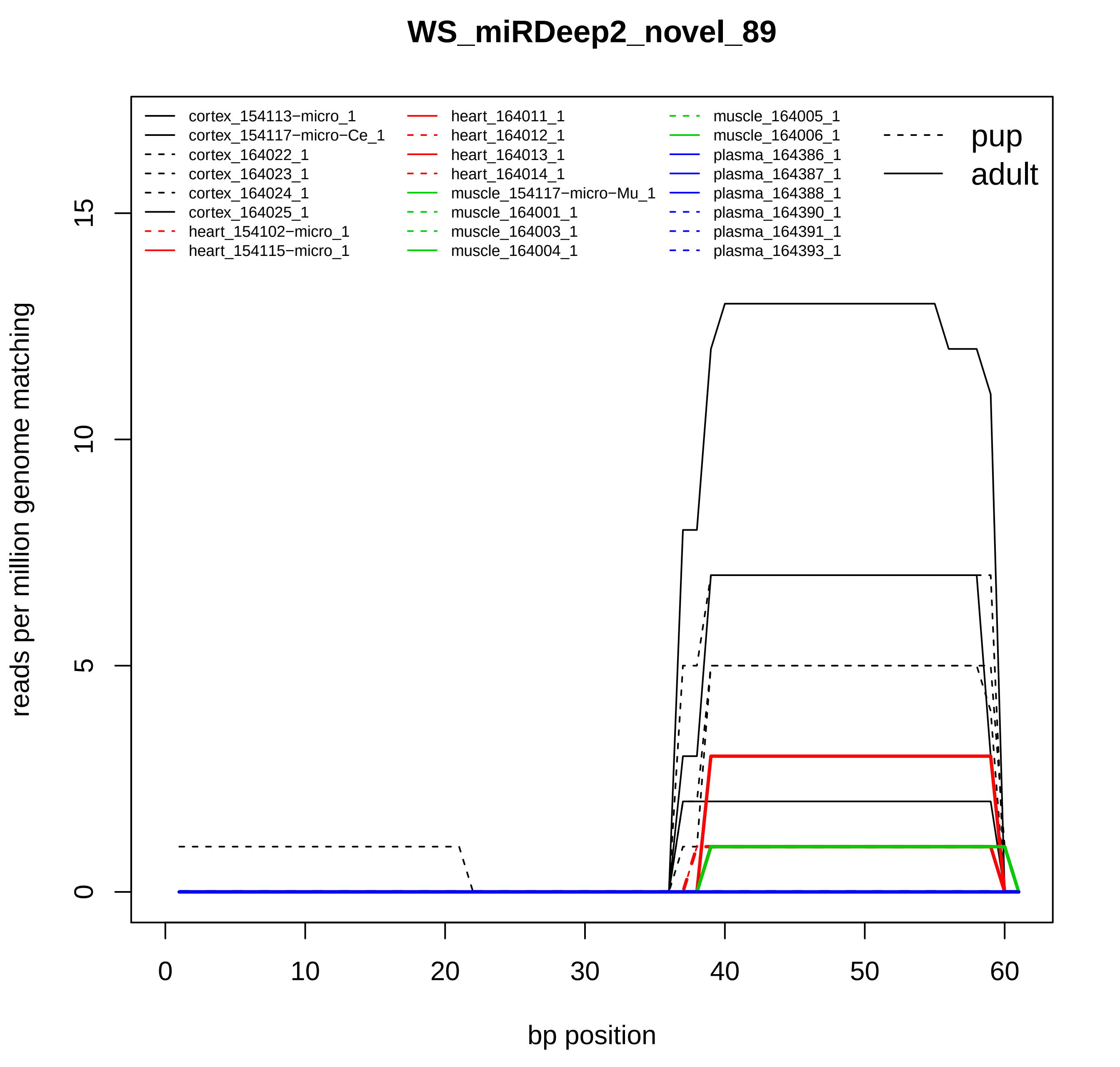

WS_miRDeep2_novel_89
cortex_154113−micro_1
cortex_154117−micro−Ce_1
cortex_164022_1
cortex_164023_1
cortex_164024_1
cortex_164025_1
heart_154102−micro_1
heart_154115−micro_1
heart_164011_1
heart_164012_1
heart_164013_1
heart_164014_1
muscle_154117−micro−Mu_1
muscle_164001_1
muscle_164003_1
muscle_164004_1
muscle_164005_1
muscle_164006_1
plasma_164386_1
plasma_164387_1
plasma_164388_1
plasma_164390_1
plasma_164391_1
plasma_164393_1
pup
adult
reads per million genome matching
0
5
10
15
0
10
20
30
40
50
60
bp position

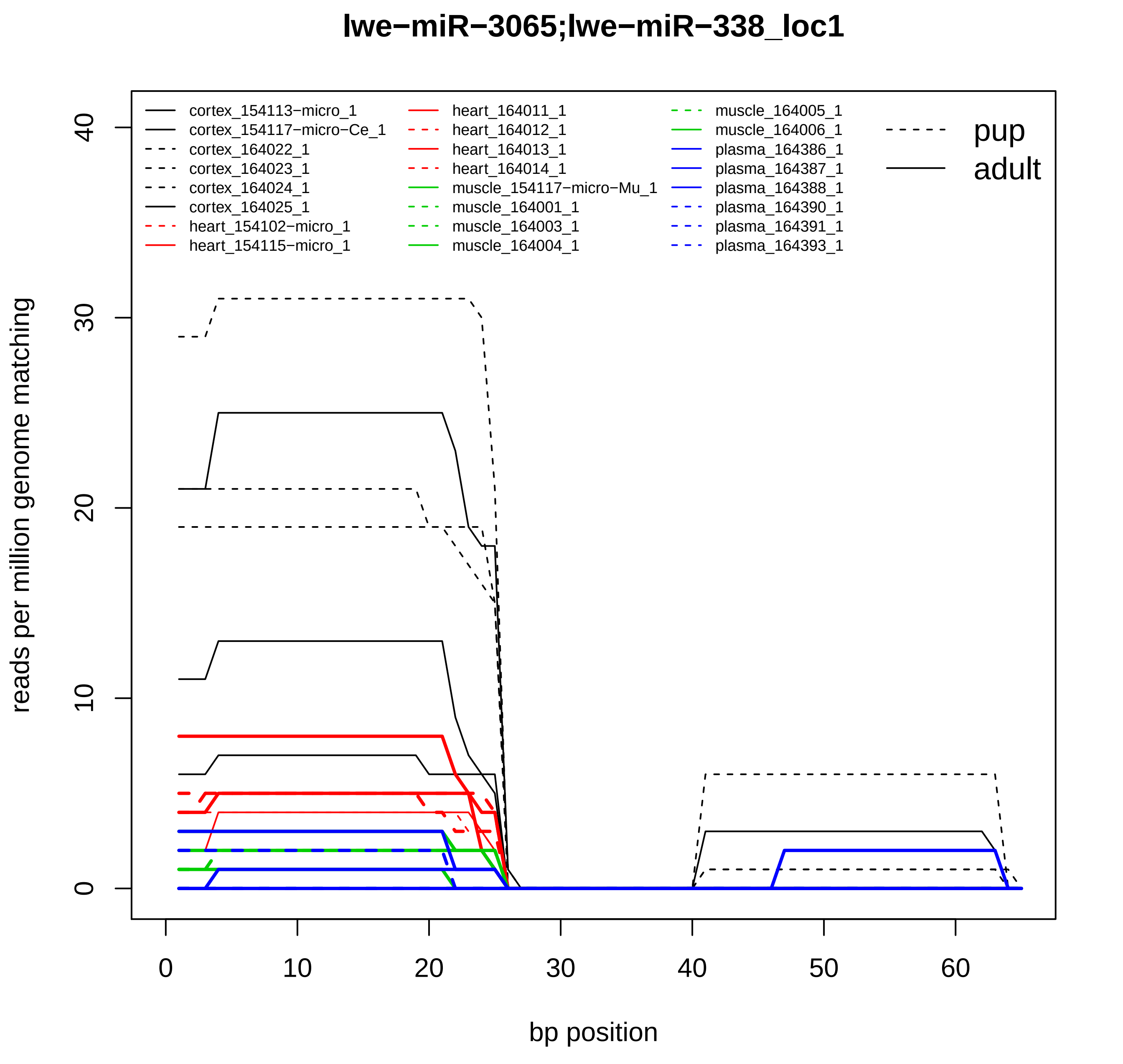
lwe−miR−3065;lwe−miR−338_loc1
cortex_154113−micro_1
cortex_154117−micro−Ce_1
cortex_164022_1
cortex_164023_1
cortex_164024_1
cortex_164025_1
heart_154102−micro_1
heart_154115−micro_1
heart_164011_1
heart_164012_1
heart_164013_1
heart_164014_1
muscle_154117−micro−Mu_1
muscle_164001_1
muscle_164003_1
muscle_164004_1
muscle_164005_1
muscle_164006_1
plasma_164386_1
plasma_164387_1
plasma_164388_1
plasma_164390_1
plasma_164391_1
plasma_164393_1
pup
adult
reads per million genome matching
0
10
20
30
40
0
10
20
30
40
50
60
bp position

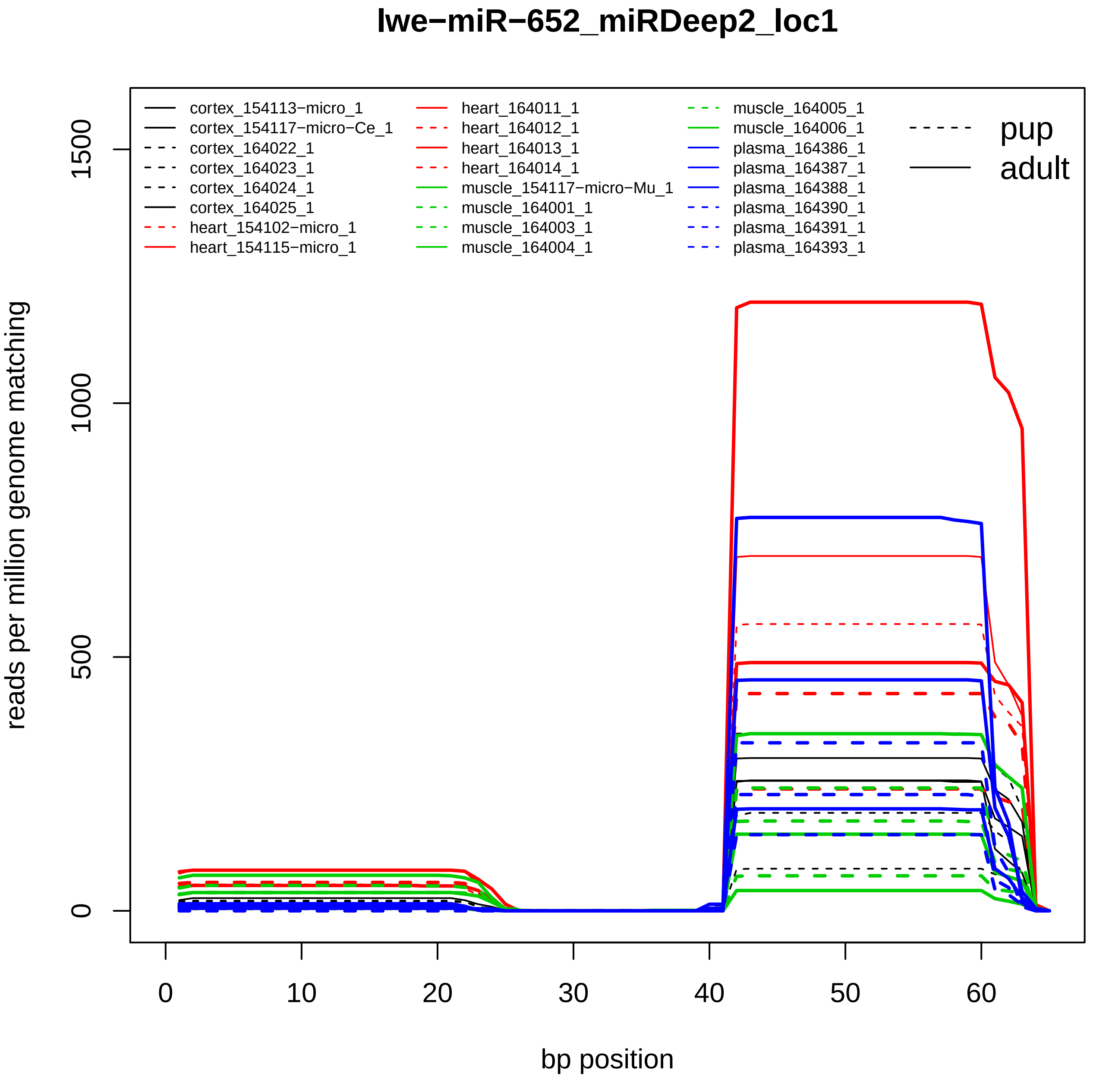

lwe−miR−652_miRDeep2_loc1
cortex_154113−micro_1
cortex_154117−micro−Ce_1
cortex_164022_1
cortex_164023_1
cortex_164024_1
cortex_164025_1
heart_154102−micro_1
heart_154115−micro_1
heart_164011_1
heart_164012_1
heart_164013_1
heart_164014_1
muscle_154117−micro−Mu_1
muscle_164001_1
muscle_164003_1
muscle_164004_1
muscle_164005_1
muscle_164006_1
plasma_164386_1
plasma_164387_1
plasma_164388_1
plasma_164390_1
plasma_164391_1
plasma_164393_1
pup
adult
reads per million genome matching
0
500
1000
1500
0
10
20
30
40
50
60
bp position

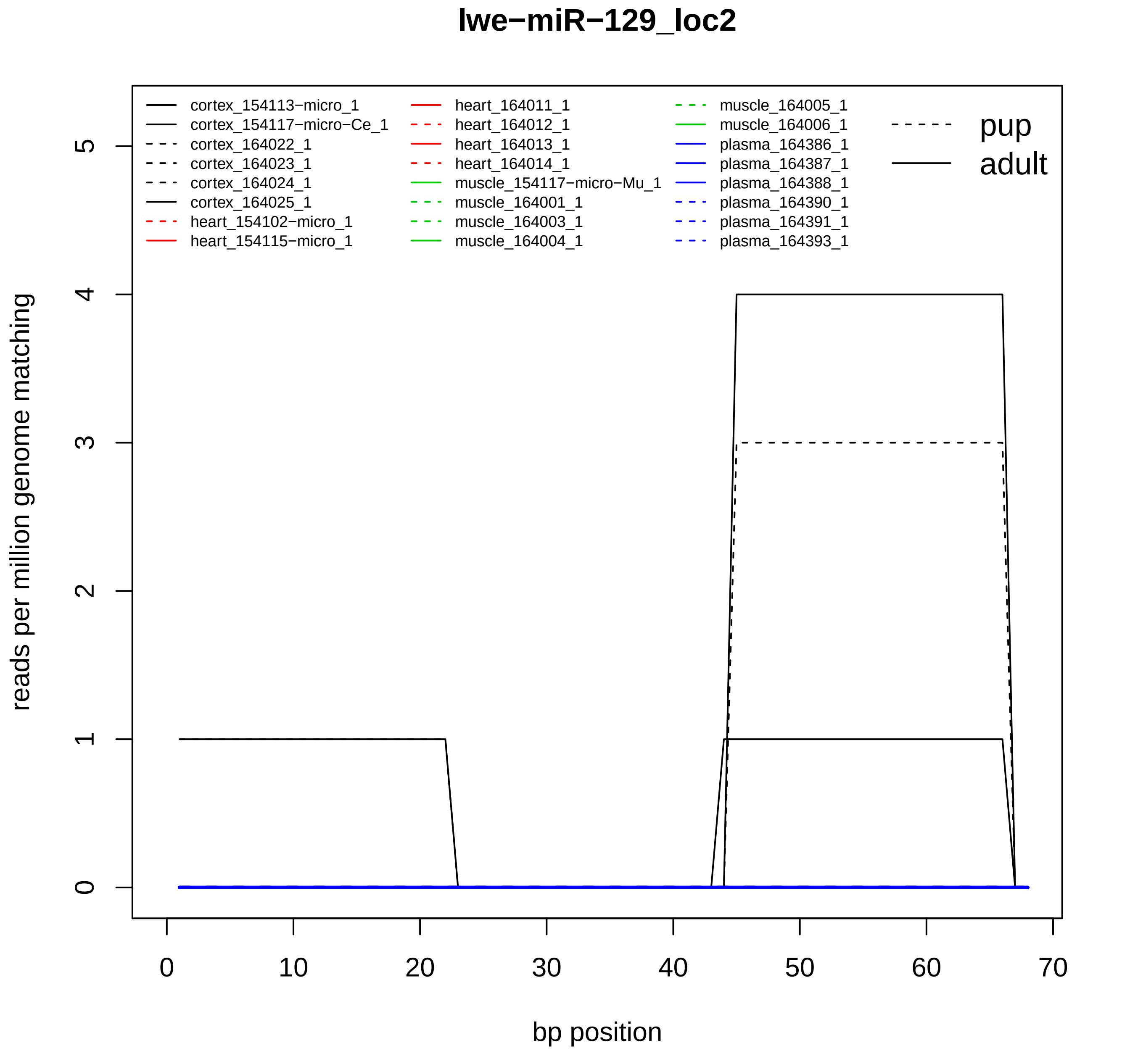

lwe−miR−129_loc2
cortex_154113−micro_1
cortex_154117−micro−Ce_1
cortex_164022_1
cortex_164023_1
cortex_164024_1
cortex_164025_1
heart_154102−micro_1
heart_154115−micro_1
heart_164011_1
heart_164012_1
heart_164013_1
heart_164014_1
muscle_154117−micro−Mu_1
muscle_164001_1
muscle_164003_1
muscle_164004_1
muscle_164005_1
muscle_164006_1
plasma_164386_1
plasma_164387_1
plasma_164388_1
plasma_164390_1
plasma_164391_1
plasma_164393_1
pup
adult
reads per million genome matching
0
1
2
3
4
5
0
10
20
30
40
50
60
70
bp position

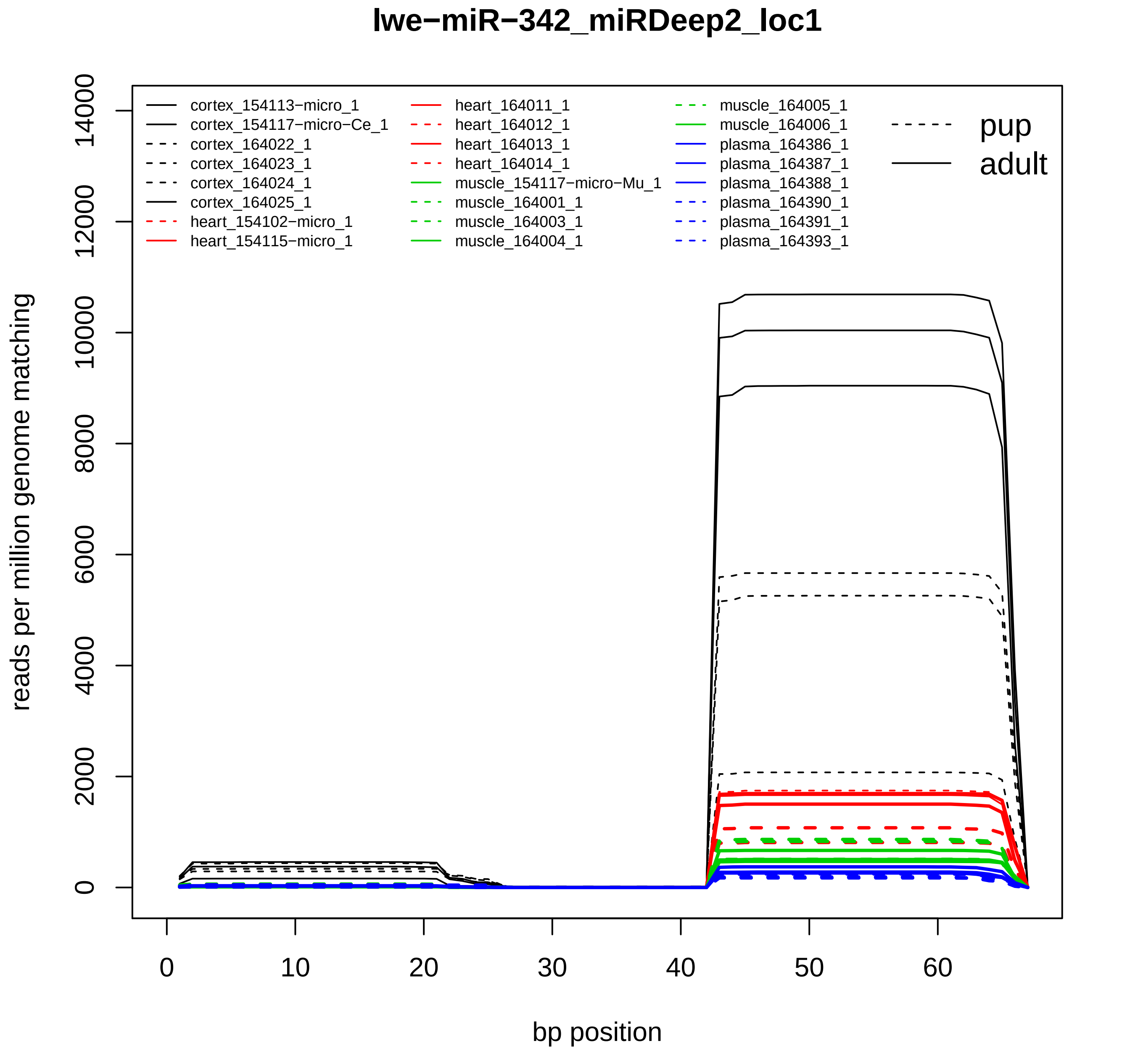

lwe−miR−342_miRDeep2_loc1
reads per million genome matching
bp position
0
2000
4000
6000
8000
10000
12000
14000
0
10
20
30
40
50
60
cortex_154113−micro_1
cortex_154117−micro−Ce_1
cortex_164022_1
cortex_164023_1
cortex_164024_1
cortex_164025_1
heart_154102−micro_1
heart_154115−micro_1
heart_164011_1
heart_164012_1
heart_164013_1
heart_164014_1
muscle_154117−micro−Mu_1
muscle_164001_1
muscle_164003_1
muscle_164004_1
muscle_164005_1
muscle_164006_1
plasma_164386_1
plasma_164387_1
plasma_164388_1
plasma_164390_1
plasma_164391_1
plasma_164393_1
pup
adult

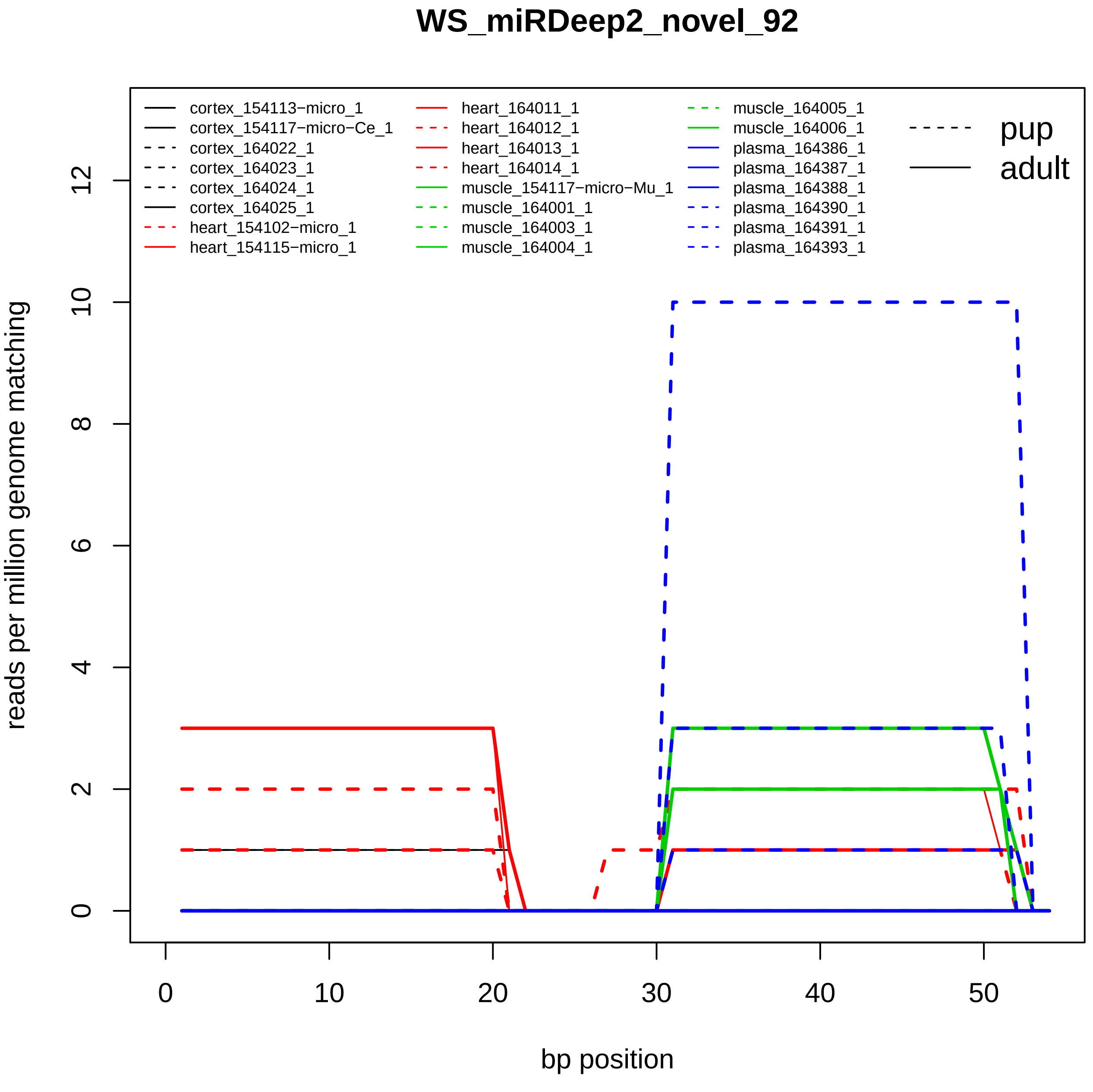

WS_miRDeep2_novel_92
cortex_154113−micro_1
cortex_154117−micro−Ce_1
cortex_164022_1
cortex_164023_1
cortex_164024_1
cortex_164025_1
heart_154102−micro_1
heart_154115−micro_1
heart_164011_1
heart_164012_1
heart_164013_1
heart_164014_1
muscle_154117−micro−Mu_1
muscle_164001_1
muscle_164003_1
muscle_164004_1
muscle_164005_1
muscle_164006_1
plasma_164386_1
plasma_164387_1
plasma_164388_1
plasma_164390_1
plasma_164391_1
plasma_164393_1
pup
adult
reads per million genome matching
0
2
4
6
8
10
12
0
10
20
30
40
50
bp position

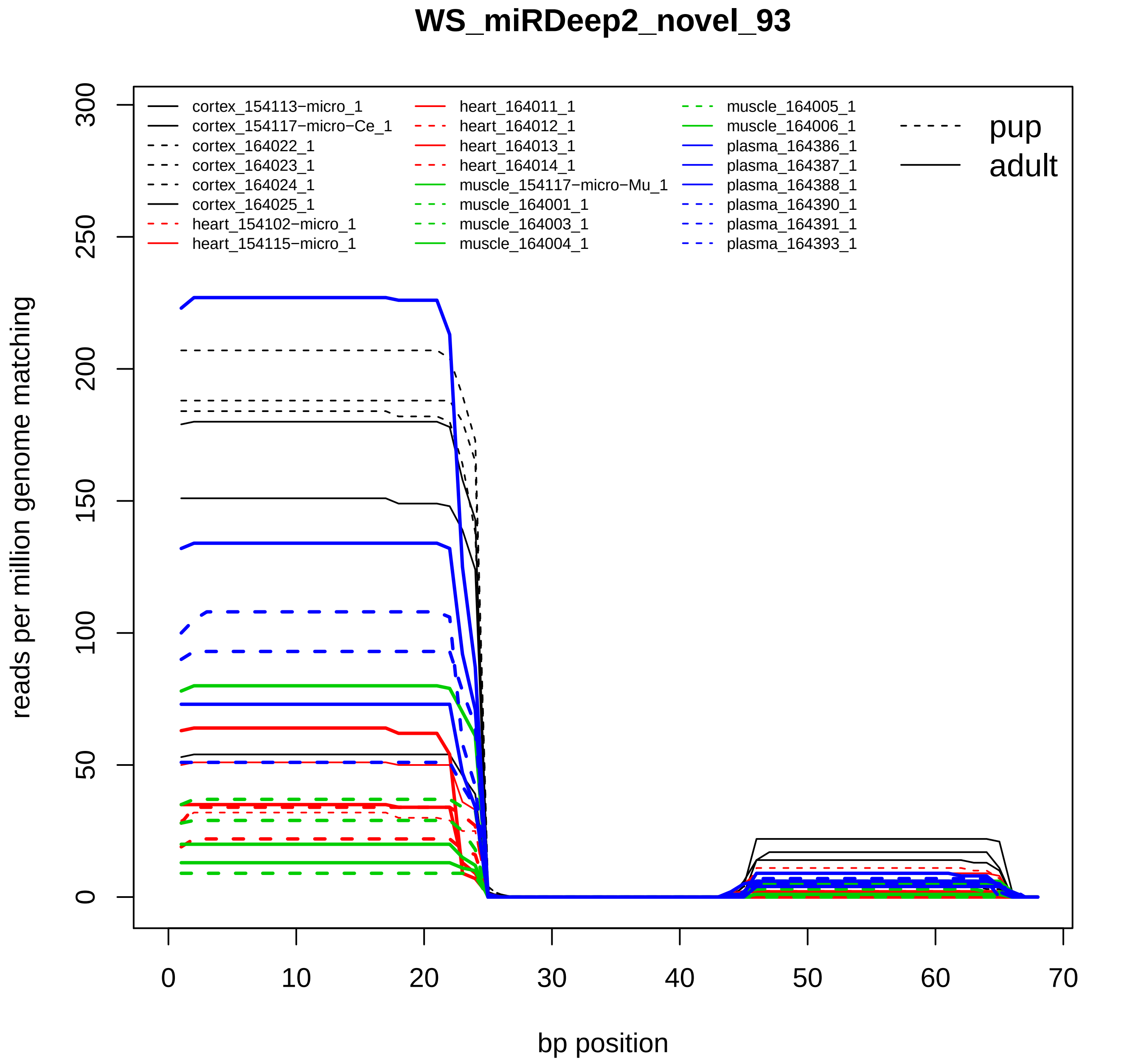

WS_miRDeep2_novel_93
cortex_154113−micro_1
cortex_154117−micro−Ce_1
cortex_164022_1
cortex_164023_1
cortex_164024_1
cortex_164025_1
heart_154102−micro_1
heart_154115−micro_1
heart_164011_1
heart_164012_1
heart_164013_1
heart_164014_1
muscle_154117−micro−Mu_1
muscle_164001_1
muscle_164003_1
muscle_164004_1
muscle_164005_1
muscle_164006_1
plasma_164386_1
plasma_164387_1
plasma_164388_1
plasma_164390_1
plasma_164391_1
plasma_164393_1
pup
adult
reads per million genome matching
0
50
100
150
200
250
300
bp position
0
10
20
30
40
50
60
70

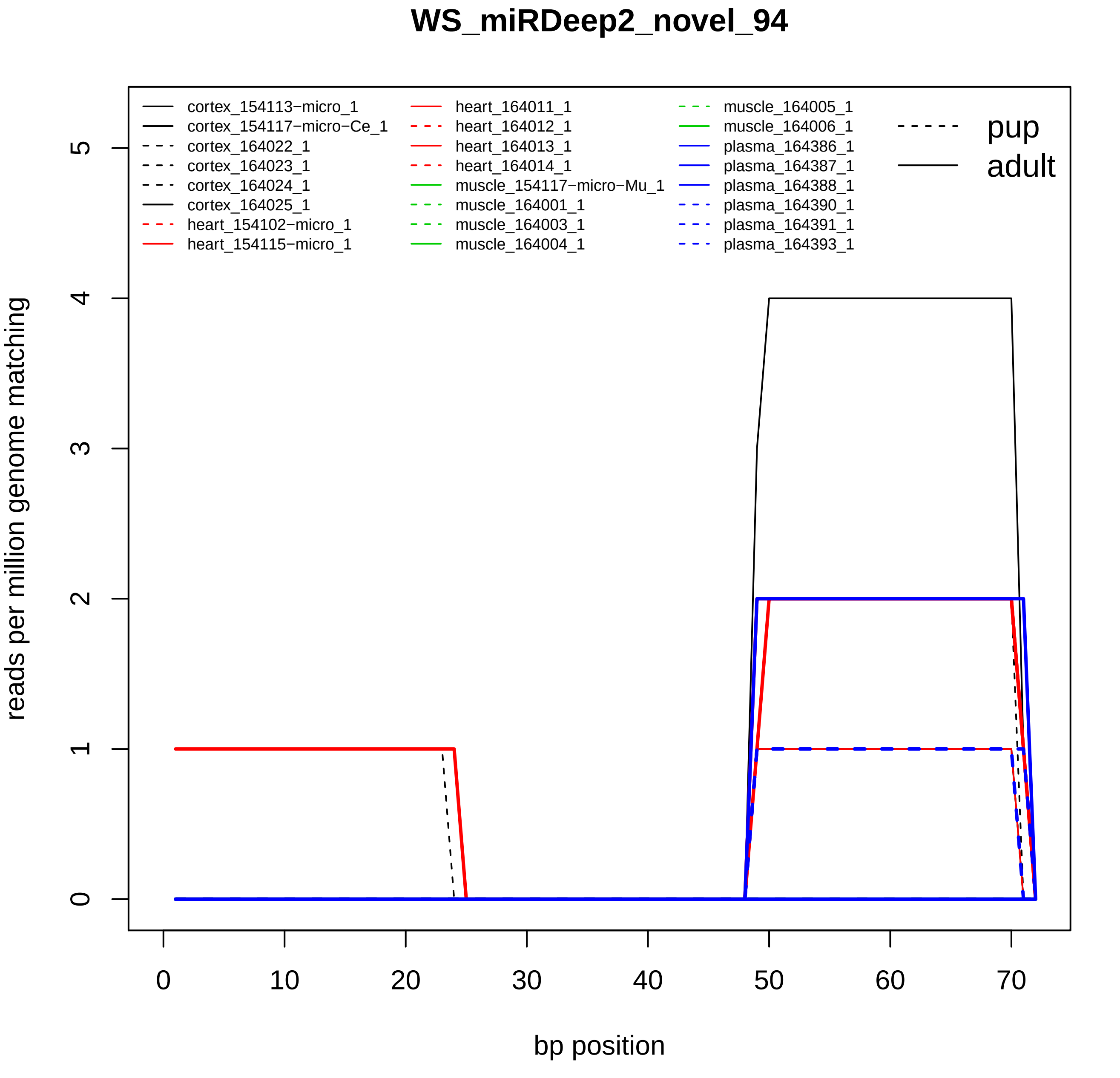

WS_miRDeep2_novel_94
cortex_154113−micro_1
cortex_154117−micro−Ce_1
cortex_164022_1
cortex_164023_1
cortex_164024_1
cortex_164025_1
heart_154102−micro_1
heart_154115−micro_1
heart_164011_1
heart_164012_1
heart_164013_1
heart_164014_1
muscle_154117−micro−Mu_1
muscle_164001_1
muscle_164003_1
muscle_164004_1
muscle_164005_1
muscle_164006_1
plasma_164386_1
plasma_164387_1
plasma_164388_1
plasma_164390_1
plasma_164391_1
plasma_164393_1
pup
adult
reads per million genome matching
0
1
2
3
4
5
bp position
0
10
20
30
40
50
60
70

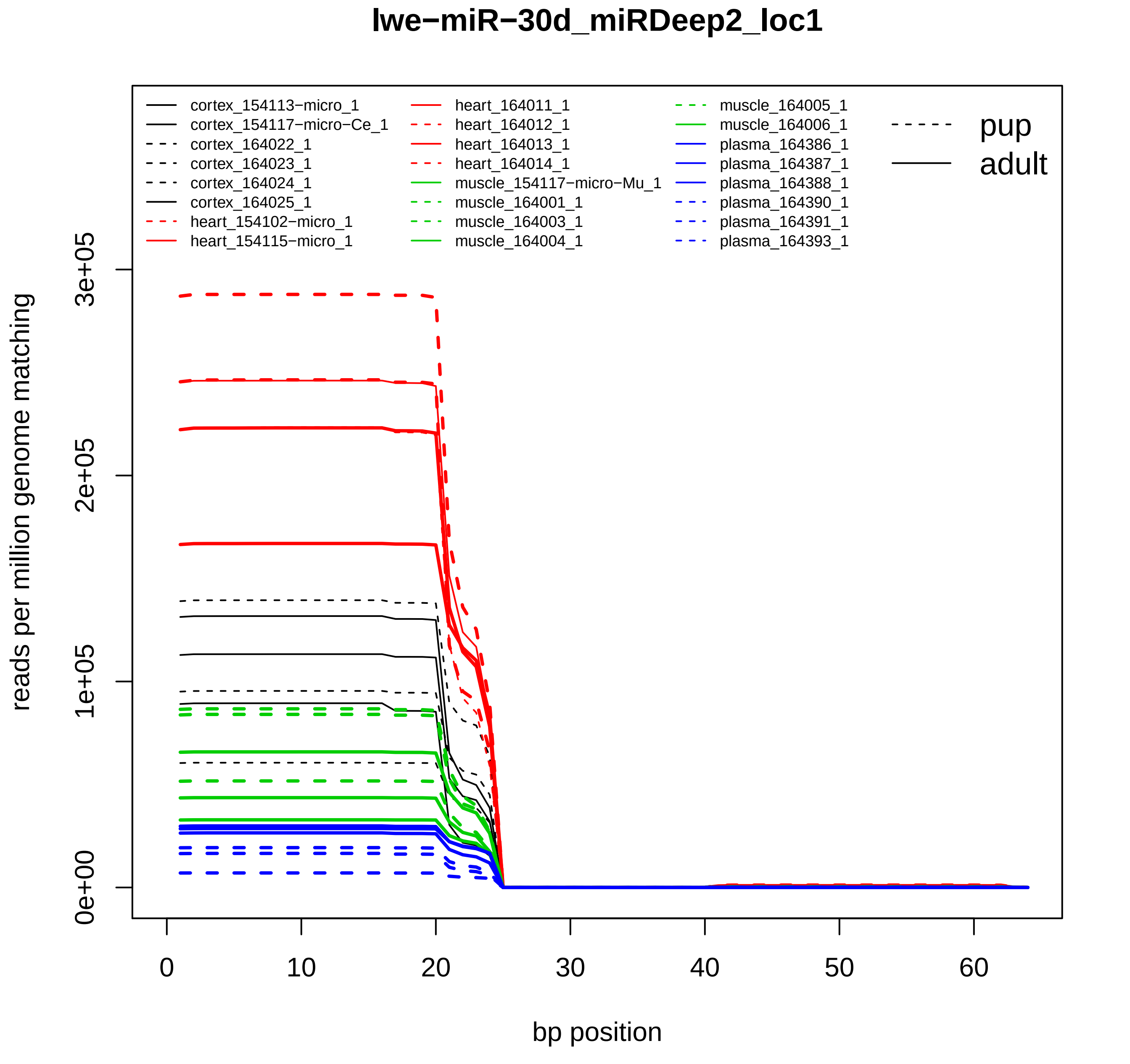

lwe−miR−30d_miRDeep2_loc1
cortex_154113−micro_1
cortex_154117−micro−Ce_1
cortex_164022_1
cortex_164023_1
cortex_164024_1
cortex_164025_1
heart_154102−micro_1
heart_154115−micro_1
heart_164011_1
heart_164012_1
heart_164013_1
heart_164014_1
muscle_154117−micro−Mu_1
muscle_164001_1
muscle_164003_1
muscle_164004_1
muscle_164005_1
muscle_164006_1
plasma_164386_1
plasma_164387_1
plasma_164388_1
plasma_164390_1
plasma_164391_1
plasma_164393_1
pup
adult
reads per million genome matching
0e+00
1e+05
2e+05
3e+05
0
10
20
30
40
50
60
bp position

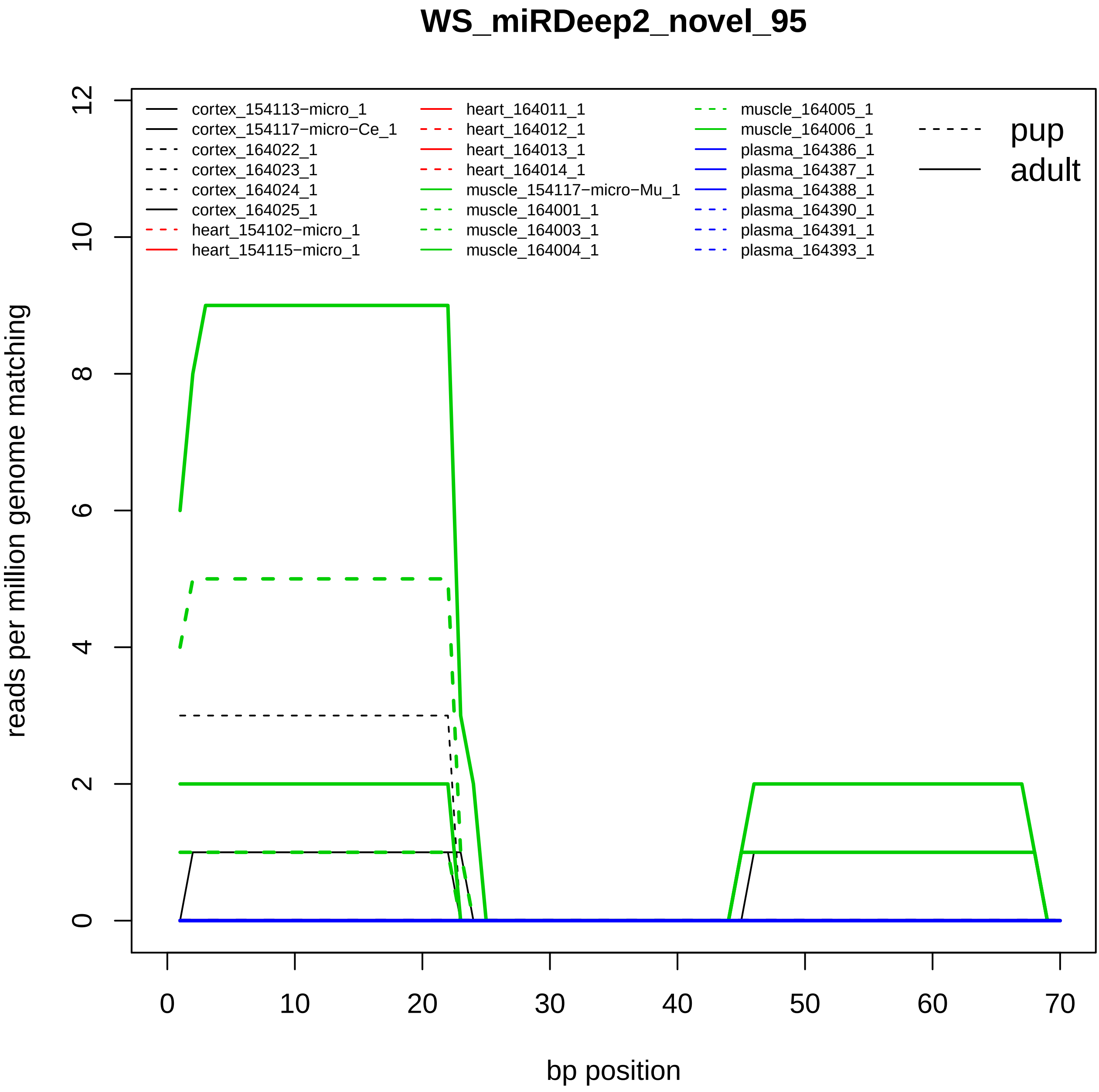

WS_miRDeep2_novel_95
cortex_154113−micro_1
cortex_154117−micro−Ce_1
cortex_164022_1
cortex_164023_1
cortex_164024_1
cortex_164025_1
heart_154102−micro_1
heart_154115−micro_1
heart_164011_1
heart_164012_1
heart_164013_1
heart_164014_1
muscle_154117−micro−Mu_1
muscle_164001_1
muscle_164003_1
muscle_164004_1
muscle_164005_1
muscle_164006_1
plasma_164386_1
plasma_164387_1
plasma_164388_1
plasma_164390_1
plasma_164391_1
plasma_164393_1
pup
adult
reads per million genome matching
0
2
4
6
8
10
12
0
10
20
30
40
50
60
70
bp position

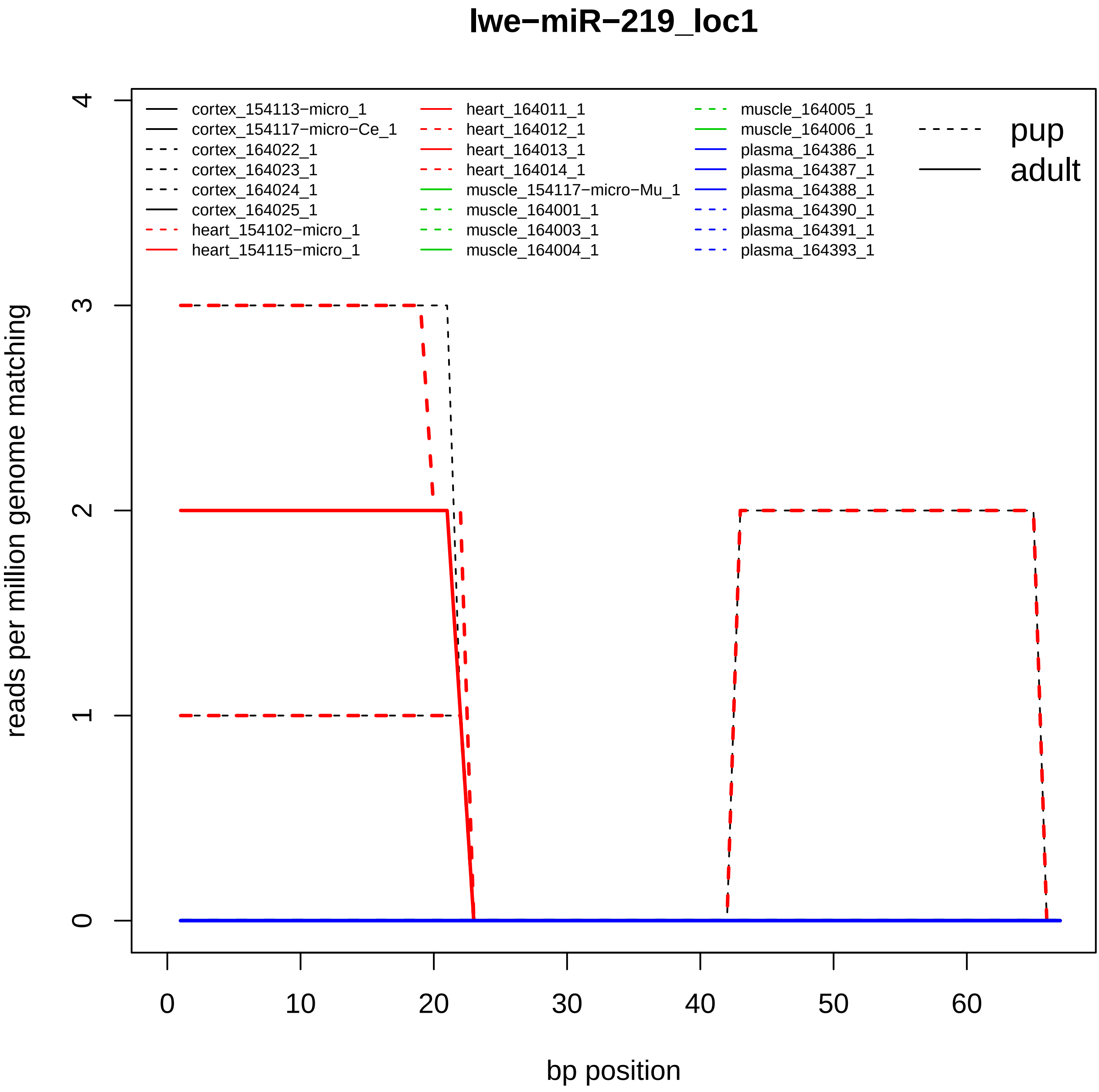

lwe−miR−219_loc1
reads per million genome matching
bp position
0
1
2
3
4
0
10
20
30
40
50
60
cortex_154113−micro_1
cortex_154117−micro−Ce_1
cortex_164022_1
cortex_164023_1
cortex_164024_1
cortex_164025_1
heart_154102−micro_1
heart_154115−micro_1
heart_164011_1
heart_164012_1
heart_164013_1
heart_164014_1
muscle_154117−micro−Mu_1
muscle_164001_1
muscle_164003_1
muscle_164004_1
muscle_164005_1
muscle_164006_1
plasma_164386_1
plasma_164387_1
plasma_164388_1
plasma_164390_1
plasma_164391_1
plasma_164393_1
pup
adult

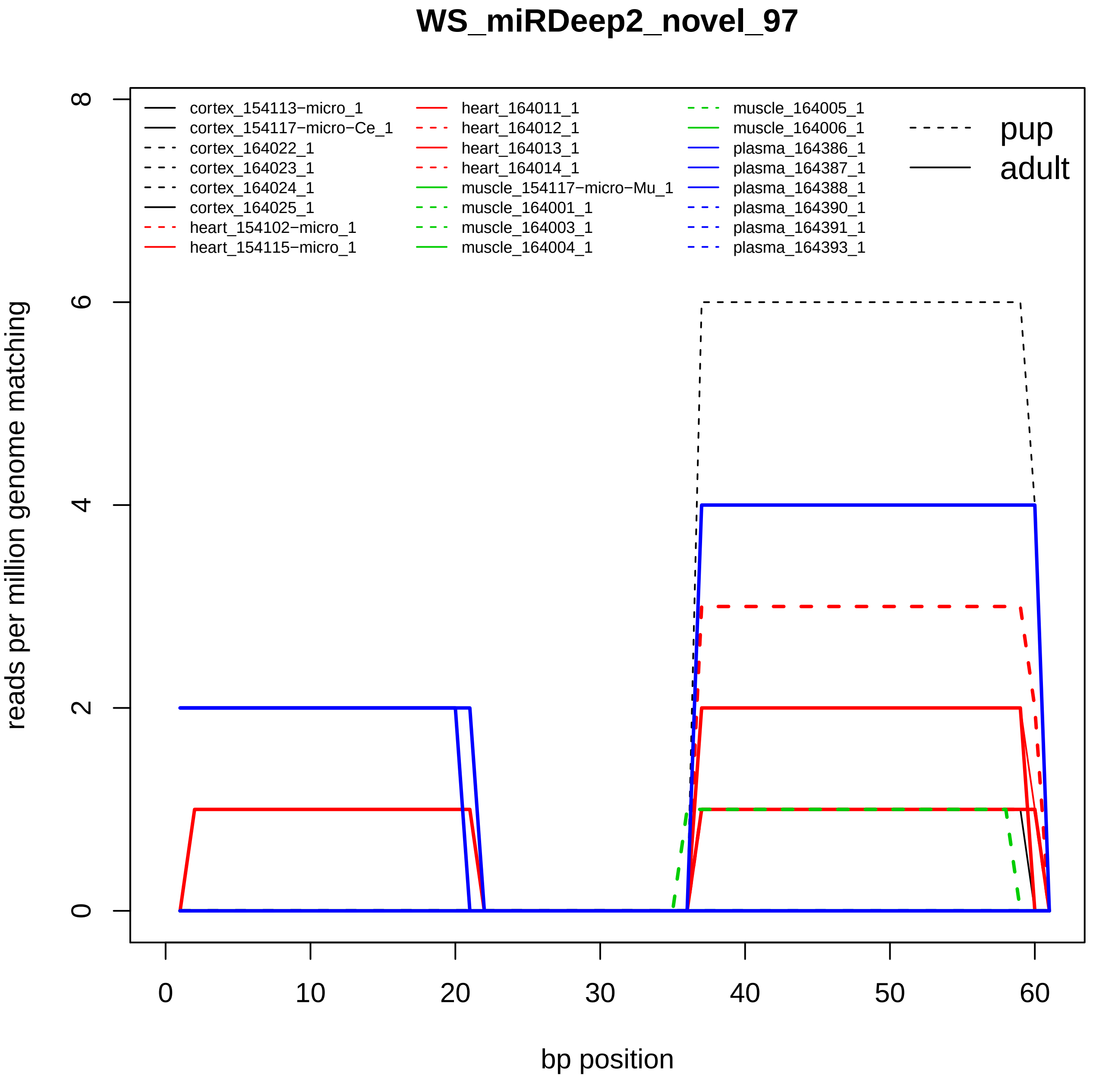

WS_miRDeep2_novel_97
cortex_154113−micro_1
cortex_154117−micro−Ce_1
cortex_164022_1
cortex_164023_1
cortex_164024_1
cortex_164025_1
heart_154102−micro_1
heart_154115−micro_1
heart_164011_1
heart_164012_1
heart_164013_1
heart_164014_1
muscle_154117−micro−Mu_1
muscle_164001_1
muscle_164003_1
muscle_164004_1
muscle_164005_1
muscle_164006_1
plasma_164386_1
plasma_164387_1
plasma_164388_1
plasma_164390_1
plasma_164391_1
plasma_164393_1
pup
adult
reads per million genome matching
0
2
4
6
8
0
10
20
30
40
50
60
bp position

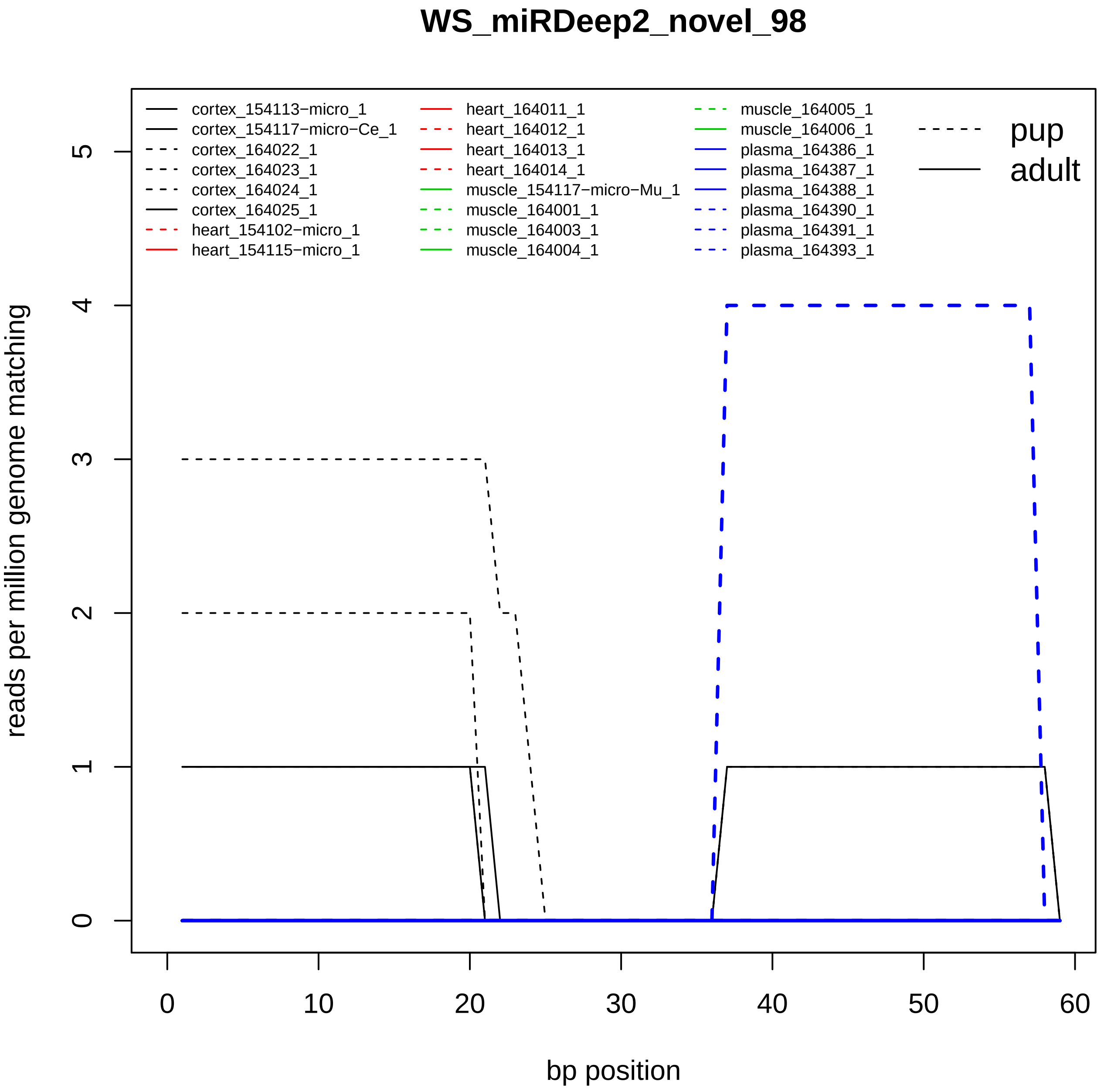
WS_miRDeep2_novel_98
cortex_154113−micro_1
cortex_154117−micro−Ce_1
cortex_164022_1
cortex_164023_1
cortex_164024_1
cortex_164025_1
heart_154102−micro_1
heart_154115−micro_1
heart_164011_1
heart_164012_1
heart_164013_1
heart_164014_1
muscle_154117−micro−Mu_1
muscle_164001_1
muscle_164003_1
muscle_164004_1
muscle_164005_1
muscle_164006_1
plasma_164386_1
plasma_164387_1
plasma_164388_1
plasma_164390_1
plasma_164391_1
plasma_164393_1
pup
adult
reads per million genome matching
bp position
0
1
2
3
4
5
0
10
20
30
40
50
60

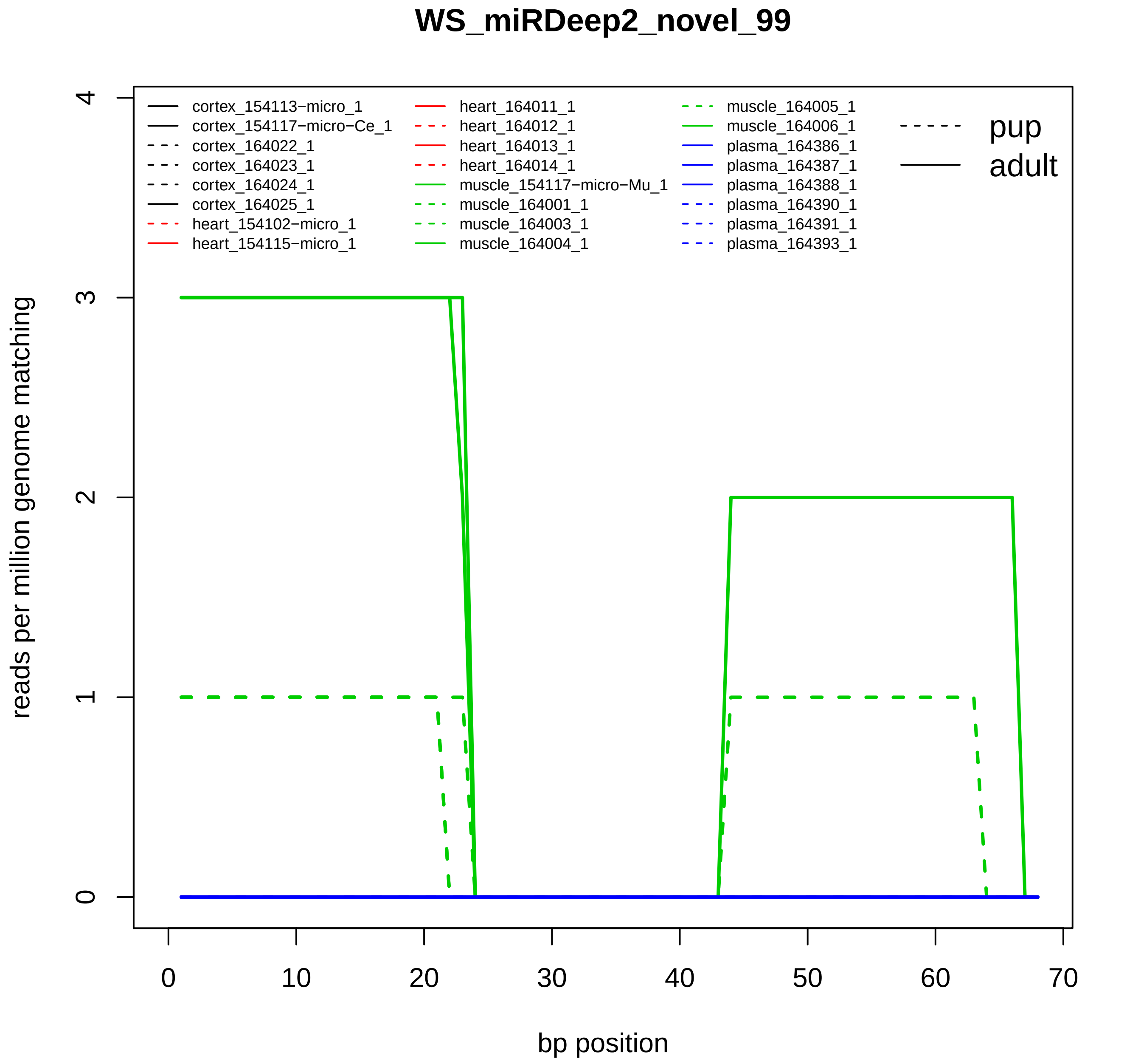
WS_miRDeep2_novel_99
cortex_154113−micro_1
cortex_154117−micro−Ce_1
cortex_164022_1
cortex_164023_1
cortex_164024_1
cortex_164025_1
heart_154102−micro_1
heart_154115−micro_1
heart_164011_1
heart_164012_1
heart_164013_1
heart_164014_1
muscle_154117−micro−Mu_1
muscle_164001_1
muscle_164003_1
muscle_164004_1
muscle_164005_1
muscle_164006_1
plasma_164386_1
plasma_164387_1
plasma_164388_1
plasma_164390_1
plasma_164391_1
plasma_164393_1
pup
adult
reads per million genome matching
bp position
0
1
2
3
4
0
10
20
30
40
50
60
70

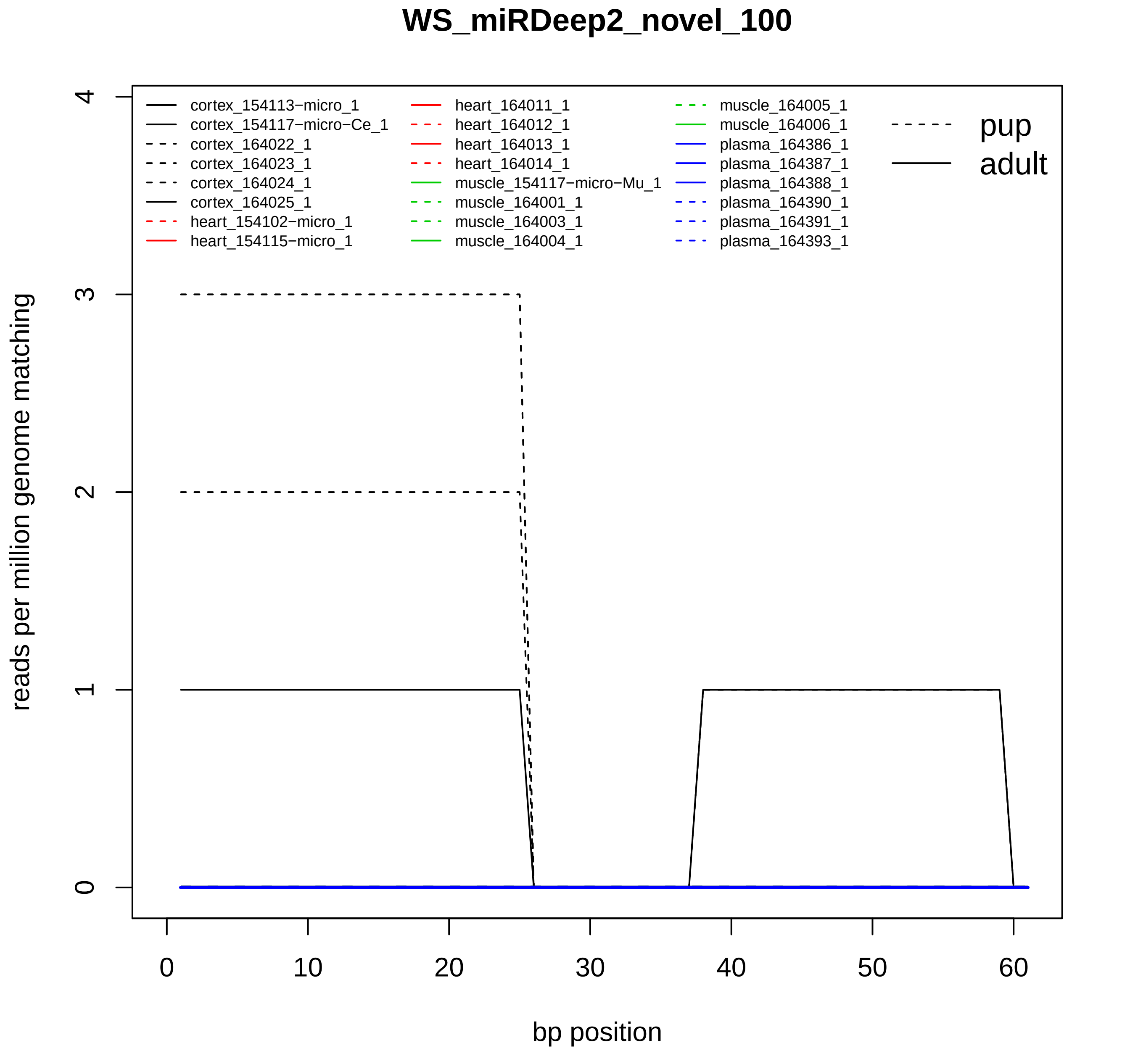

WS_miRDeep2_novel_100
cortex_154113−micro_1
cortex_154117−micro−Ce_1
cortex_164022_1
cortex_164023_1
cortex_164024_1
cortex_164025_1
heart_154102−micro_1
heart_154115−micro_1
heart_164011_1
heart_164012_1
heart_164013_1
heart_164014_1
muscle_154117−micro−Mu_1
muscle_164001_1
muscle_164003_1
muscle_164004_1
muscle_164005_1
muscle_164006_1
plasma_164386_1
plasma_164387_1
plasma_164388_1
plasma_164390_1
plasma_164391_1
plasma_164393_1
pup
adult
reads per million genome matching
0
1
2
3
4
0
10
20
30
40
50
60
bp position

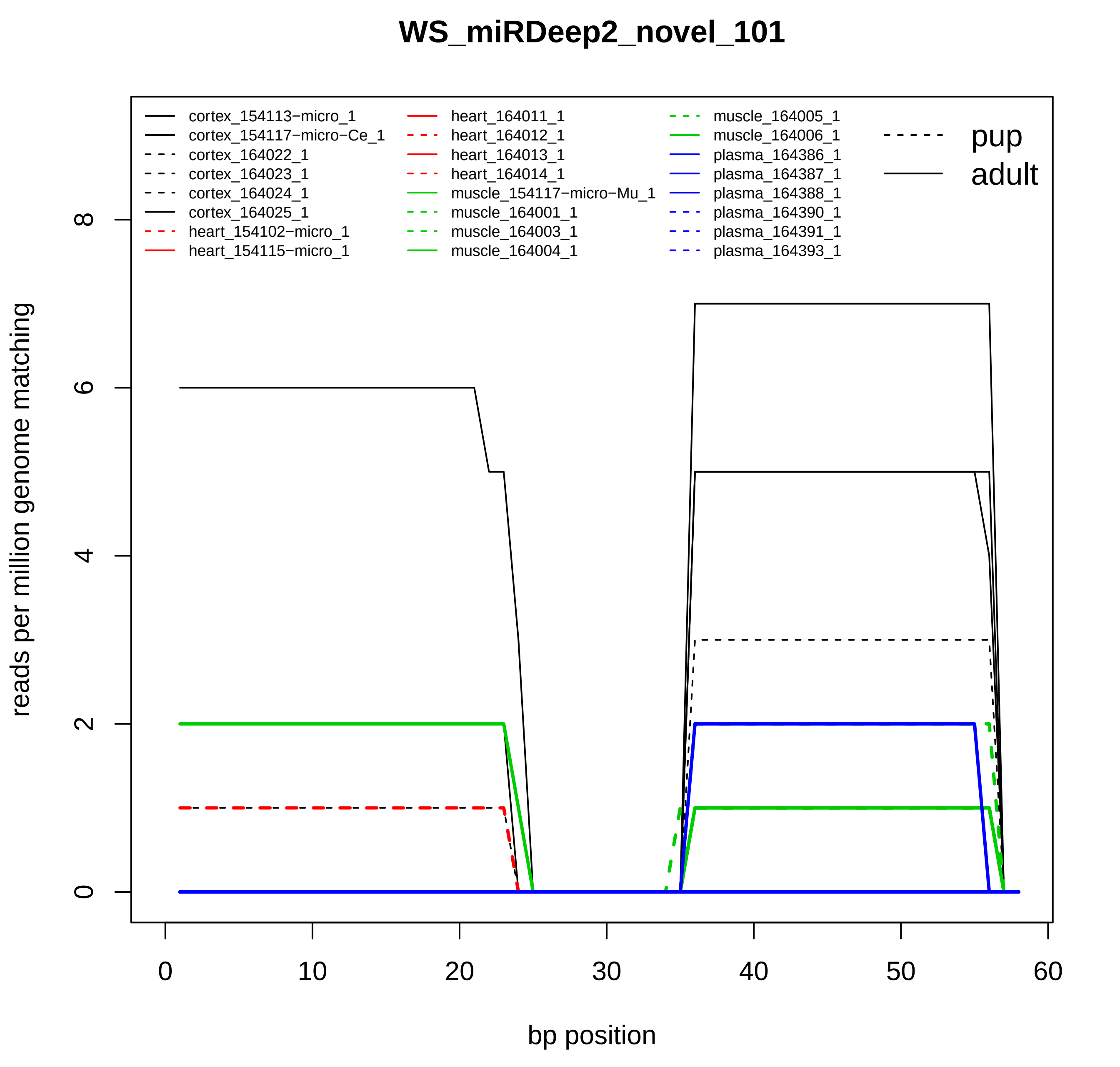
WS_miRDeep2_novel_101
cortex_154113−micro_1
cortex_154117−micro−Ce_1
cortex_164022_1
cortex_164023_1
cortex_164024_1
cortex_164025_1
heart_154102−micro_1
heart_154115−micro_1
heart_164011_1
heart_164012_1
heart_164013_1
heart_164014_1
muscle_154117−micro−Mu_1
muscle_164001_1
muscle_164003_1
muscle_164004_1
muscle_164005_1
muscle_164006_1
plasma_164386_1
plasma_164387_1
plasma_164388_1
plasma_164390_1
plasma_164391_1
plasma_164393_1
pup
adult
reads per million genome matching
0
2
4
6
8
bp position
0
10
20
30
40
50
60

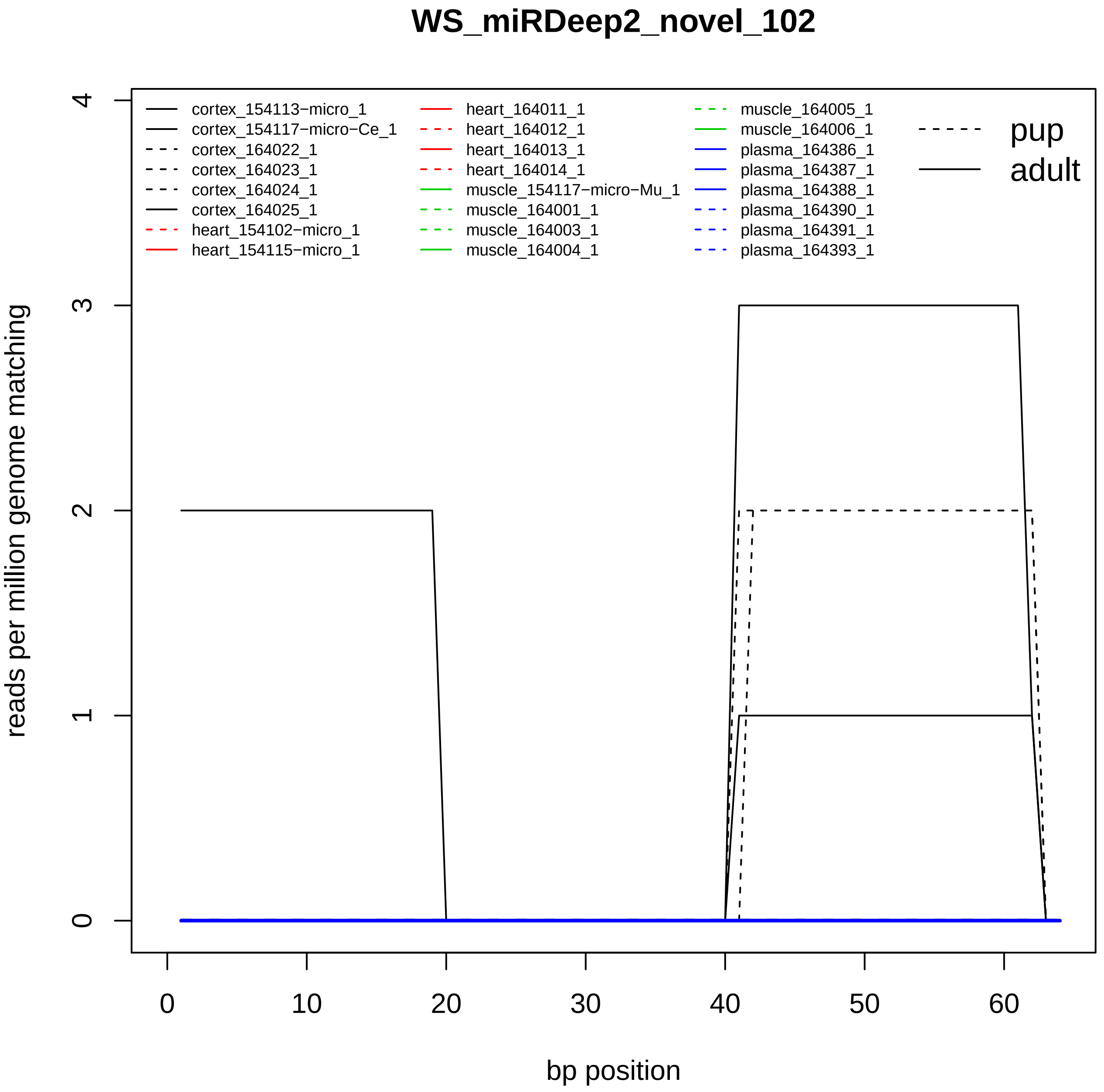

WS_miRDeep2_novel_102
cortex_154113−micro_1
cortex_154117−micro−Ce_1
cortex_164022_1
cortex_164023_1
cortex_164024_1
cortex_164025_1
heart_154102−micro_1
heart_154115−micro_1
heart_164011_1
heart_164012_1
heart_164013_1
heart_164014_1
muscle_154117−micro−Mu_1
muscle_164001_1
muscle_164003_1
muscle_164004_1
muscle_164005_1
muscle_164006_1
plasma_164386_1
plasma_164387_1
plasma_164388_1
plasma_164390_1
plasma_164391_1
plasma_164393_1
pup
adult
reads per million genome matching
bp position
0
1
2
3
4
0
10
20
30
40
50
60

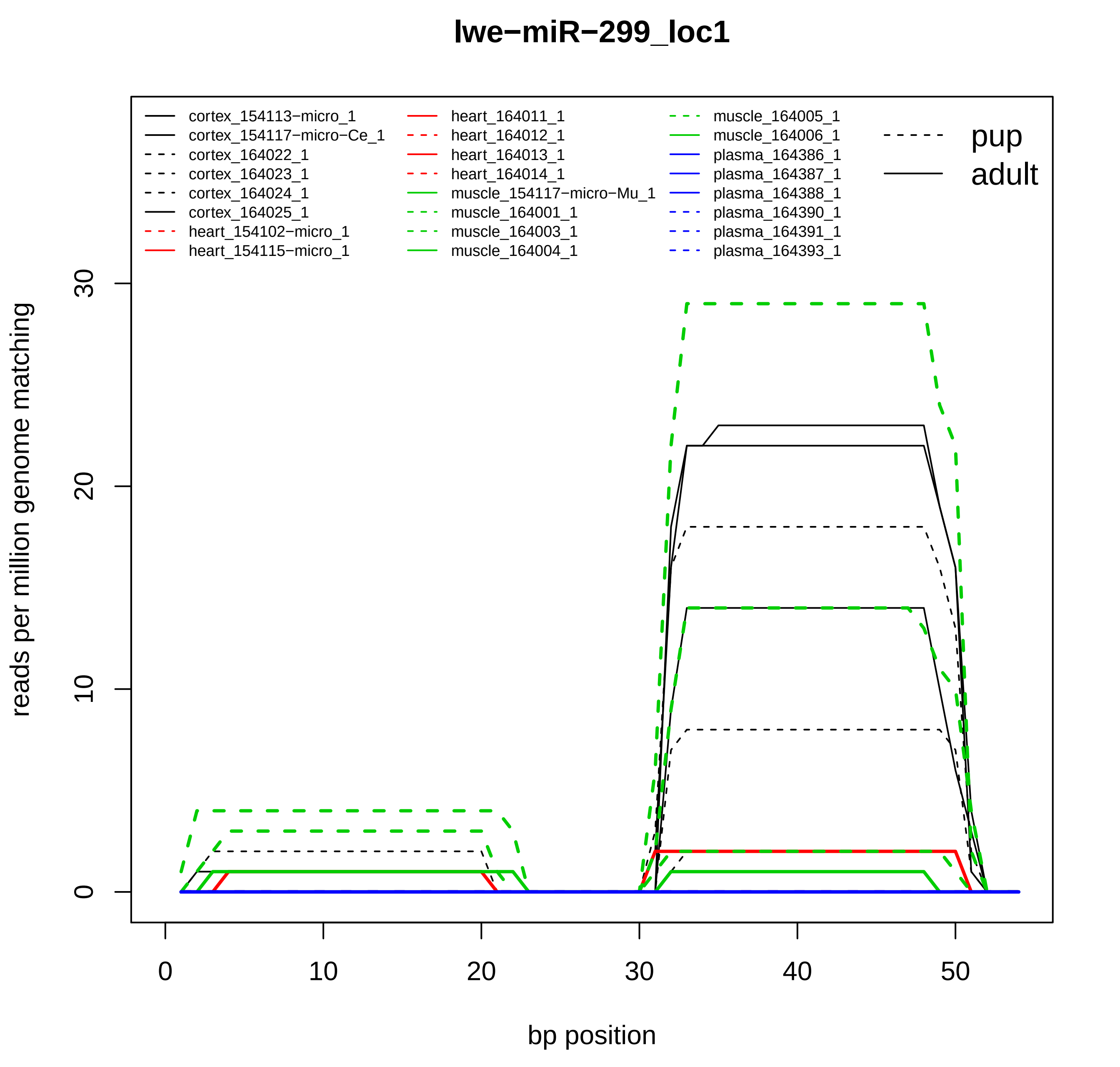
lwe−miR−299_loc1
cortex_154113−micro_1
cortex_154117−micro−Ce_1
cortex_164022_1
cortex_164023_1
cortex_164024_1
cortex_164025_1
heart_154102−micro_1
heart_154115−micro_1
heart_164011_1
heart_164012_1
heart_164013_1
heart_164014_1
muscle_154117−micro−Mu_1
muscle_164001_1
muscle_164003_1
muscle_164004_1
muscle_164005_1
muscle_164006_1
plasma_164386_1
plasma_164387_1
plasma_164388_1
plasma_164390_1
plasma_164391_1
plasma_164393_1
pup
adult
reads per million genome matching
0
10
20
30
0
10
20
30
40
50
bp position

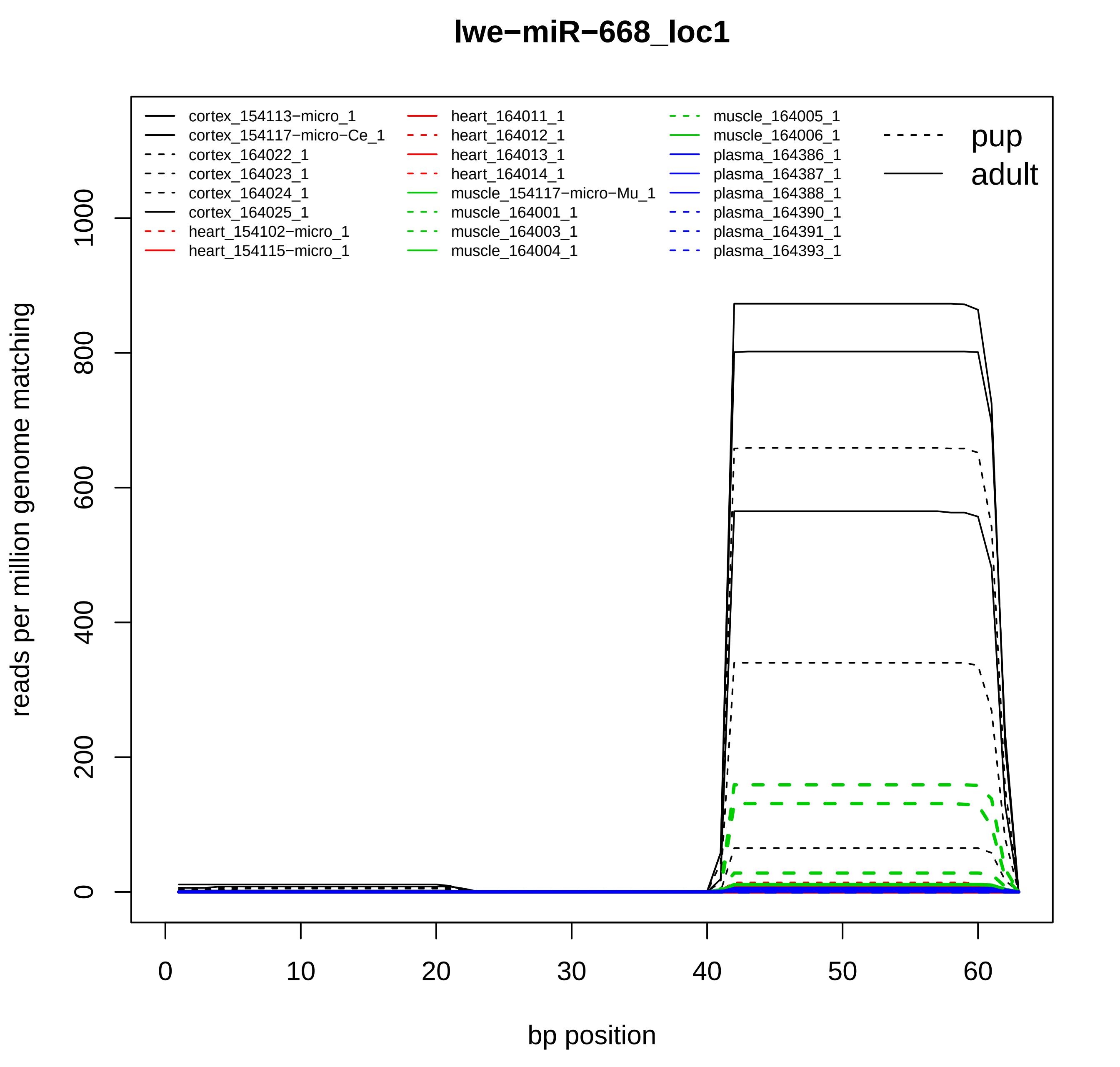
lwe−miR−668_loc1
cortex_154113−micro_1
cortex_154117−micro−Ce_1
cortex_164022_1
cortex_164023_1
cortex_164024_1
cortex_164025_1
heart_154102−micro_1
heart_154115−micro_1
heart_164011_1
heart_164012_1
heart_164013_1
heart_164014_1
muscle_154117−micro−Mu_1
muscle_164001_1
muscle_164003_1
muscle_164004_1
muscle_164005_1
muscle_164006_1
plasma_164386_1
plasma_164387_1
plasma_164388_1
plasma_164390_1
plasma_164391_1
plasma_164393_1
pup
adult
reads per million genome matching
bp position
0
200
400
600
800
1000
0
10
20
30
40
50
60

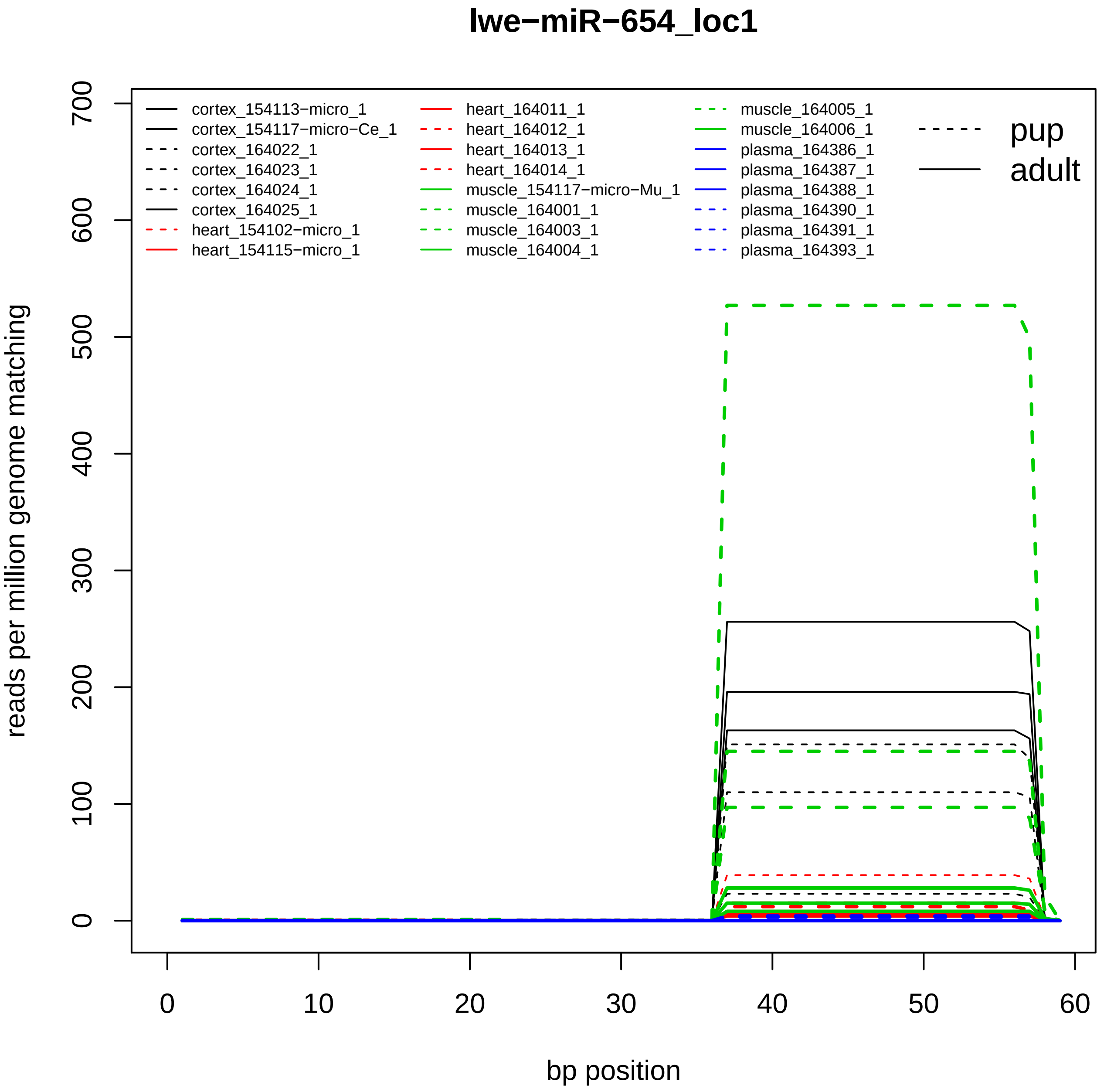
lwe−miR−654_loc1
reads per million genome matching
bp position
0
100
200
300
400
500
600
700
0
10
20
30
40
50
60
cortex_154113−micro_1
cortex_154117−micro−Ce_1
cortex_164022_1
cortex_164023_1
cortex_164024_1
cortex_164025_1
heart_154102−micro_1
heart_154115−micro_1
heart_164011_1
heart_164012_1
heart_164013_1
heart_164014_1
muscle_154117−micro−Mu_1
muscle_164001_1
muscle_164003_1
muscle_164004_1
muscle_164005_1
muscle_164006_1
plasma_164386_1
plasma_164387_1
plasma_164388_1
plasma_164390_1
plasma_164391_1
plasma_164393_1
pup
adult

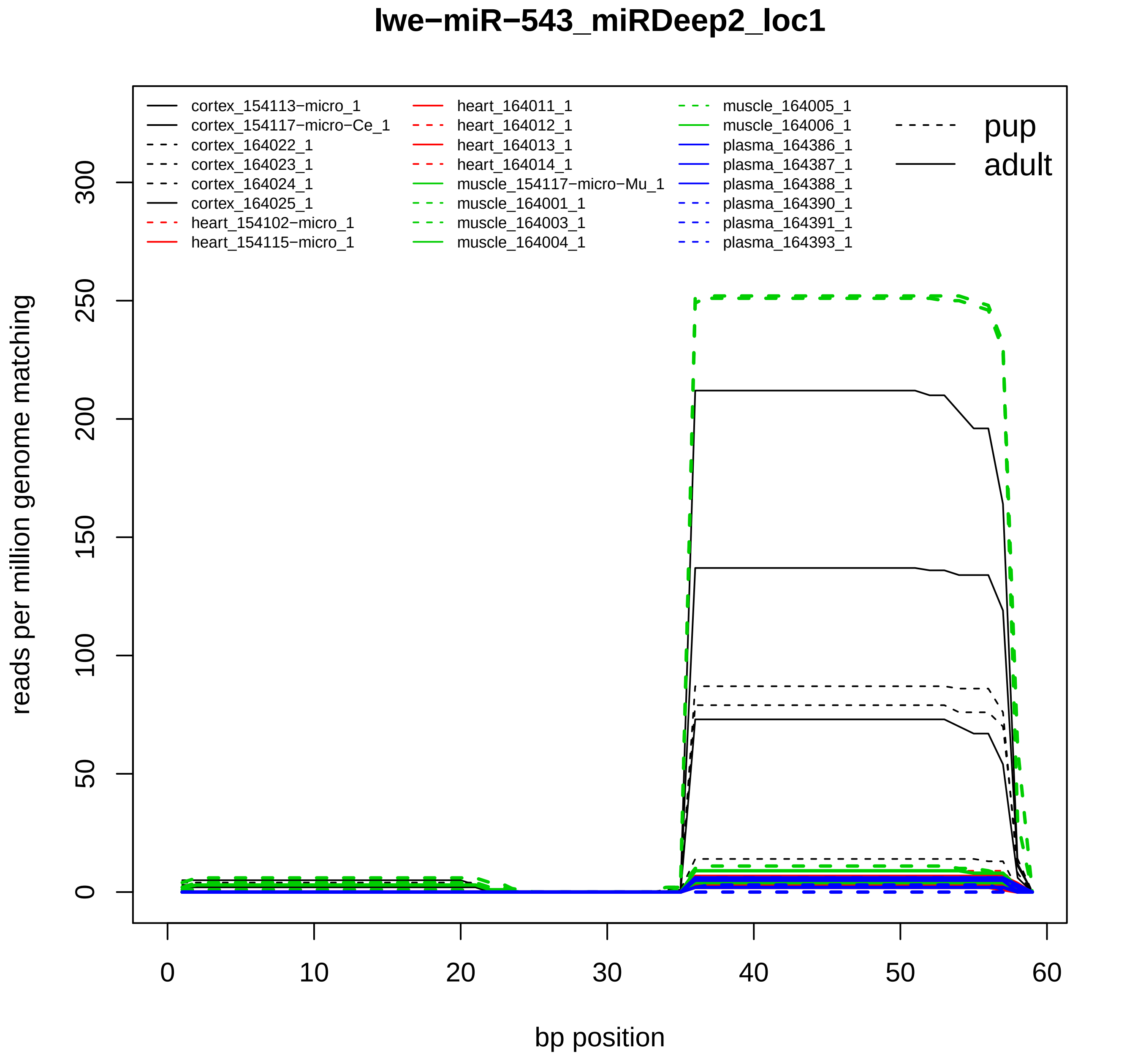

lwe−miR−543_miRDeep2_loc1
cortex_154113−micro_1
cortex_154117−micro−Ce_1
cortex_164022_1
cortex_164023_1
cortex_164024_1
cortex_164025_1
heart_154102−micro_1
heart_154115−micro_1
heart_164011_1
heart_164012_1
heart_164013_1
heart_164014_1
muscle_154117−micro−Mu_1
muscle_164001_1
muscle_164003_1
muscle_164004_1
muscle_164005_1
muscle_164006_1
plasma_164386_1
plasma_164387_1
plasma_164388_1
plasma_164390_1
plasma_164391_1
plasma_164393_1
pup
adult
reads per million genome matching
0
50
100
150
200
250
300
bp position
0
10
20
30
40
50
60

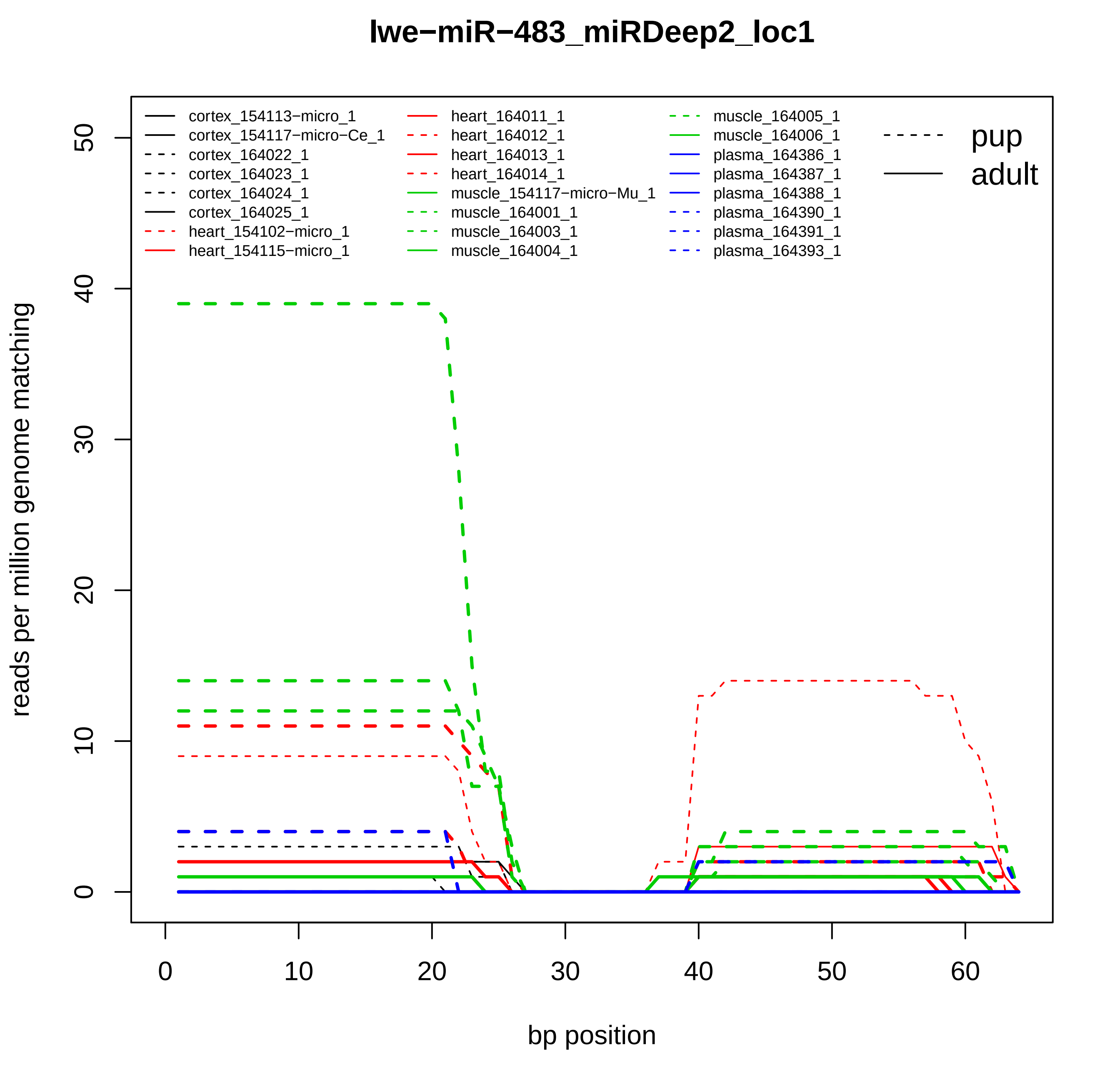

lwe−miR−483_miRDeep2_loc1
cortex_154113−micro_1
cortex_154117−micro−Ce_1
cortex_164022_1
cortex_164023_1
cortex_164024_1
cortex_164025_1
heart_154102−micro_1
heart_154115−micro_1
heart_164011_1
heart_164012_1
heart_164013_1
heart_164014_1
muscle_154117−micro−Mu_1
muscle_164001_1
muscle_164003_1
muscle_164004_1
muscle_164005_1
muscle_164006_1
plasma_164386_1
plasma_164387_1
plasma_164388_1
plasma_164390_1
plasma_164391_1
plasma_164393_1
pup
adult
reads per million genome matching
bp position
0
10
20
30
40
50
60

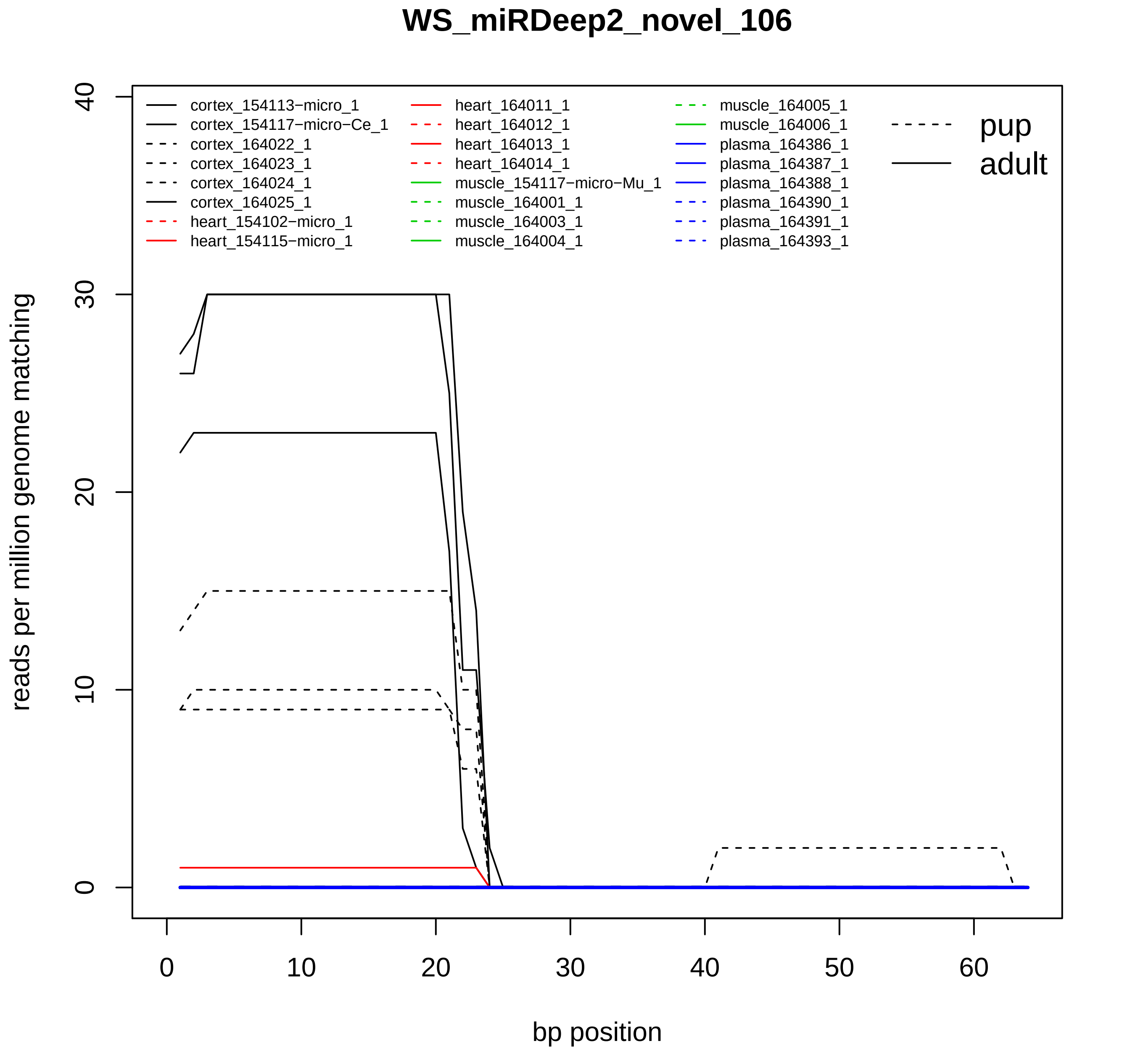
WS_miRDeep2_novel_106
cortex_154113−micro_1
cortex_154117−micro−Ce_1
cortex_164022_1
cortex_164023_1
cortex_164024_1
cortex_164025_1
heart_154102−micro_1
heart_154115−micro_1
heart_164011_1
heart_164012_1
heart_164013_1
heart_164014_1
muscle_154117−micro−Mu_1
muscle_164001_1
muscle_164003_1
muscle_164004_1
muscle_164005_1
muscle_164006_1
plasma_164386_1
plasma_164387_1
plasma_164388_1
plasma_164390_1
plasma_164391_1
plasma_164393_1
pup
adult
reads per million genome matching
0
10
20
30
40
bp position
0
10
20
30
40
50
60

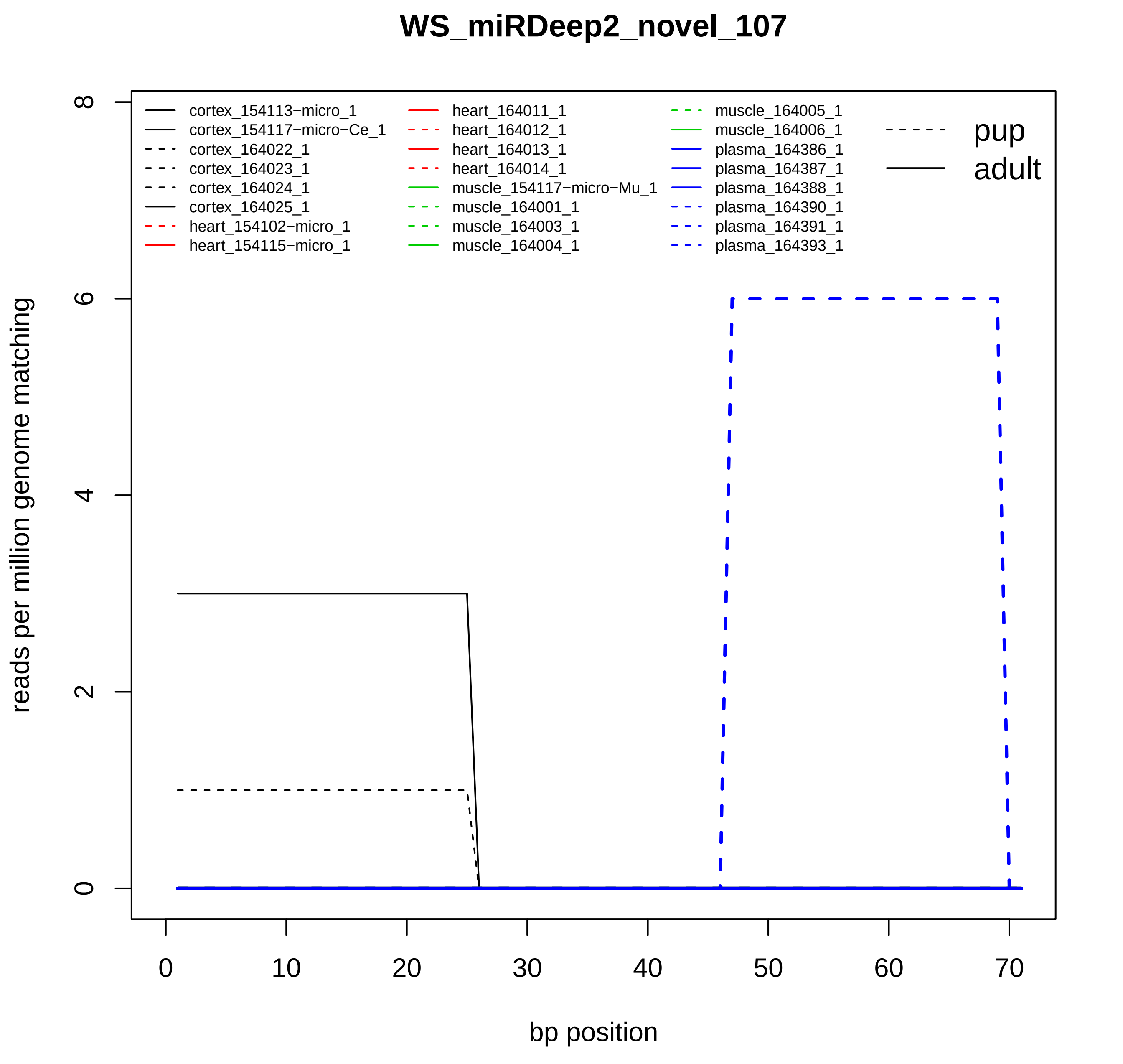
WS_miRDeep2_novel_107
reads per million genome matching
bp position
0
2
4
6
8
0
10
20
30
40
50
60
70
cortex_154113−micro_1
cortex_154117−micro−Ce_1
cortex_164022_1
cortex_164023_1
cortex_164024_1
cortex_164025_1
heart_154102−micro_1
heart_154115−micro_1
heart_164011_1
heart_164012_1
heart_164013_1
heart_164014_1
muscle_154117−micro−Mu_1
muscle_164001_1
muscle_164003_1
muscle_164004_1
muscle_164005_1
muscle_164006_1
plasma_164386_1
plasma_164387_1
plasma_164388_1
plasma_164390_1
plasma_164391_1
plasma_164393_1
pup
adult

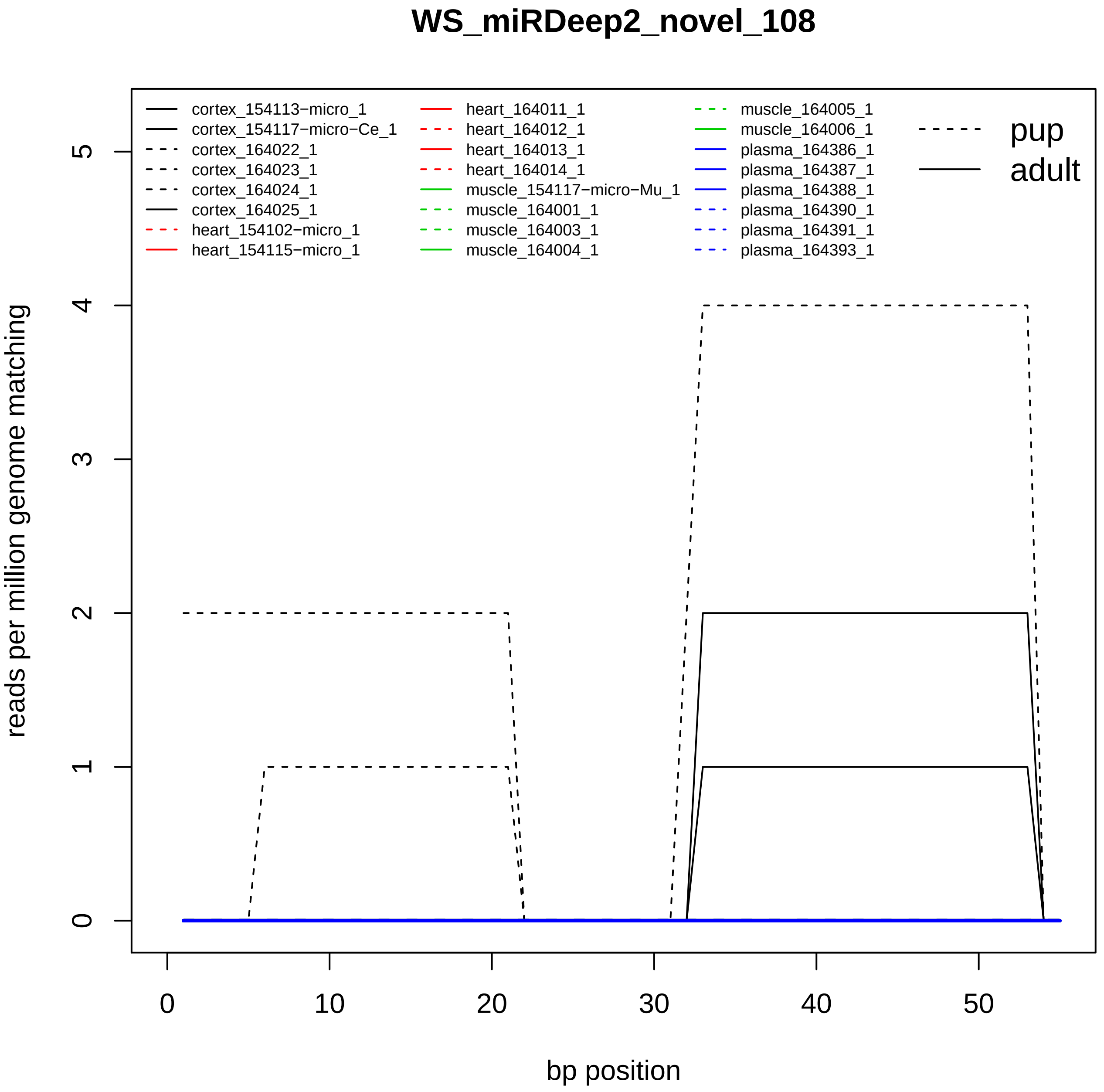

WS_miRDeep2_novel_108
cortex_154113−micro_1
cortex_154117−micro−Ce_1
cortex_164022_1
cortex_164023_1
cortex_164024_1
cortex_164025_1
heart_154102−micro_1
heart_154115−micro_1
heart_164011_1
heart_164012_1
heart_164013_1
heart_164014_1
muscle_154117−micro−Mu_1
muscle_164001_1
muscle_164003_1
muscle_164004_1
muscle_164005_1
muscle_164006_1
plasma_164386_1
plasma_164387_1
plasma_164388_1
plasma_164390_1
plasma_164391_1
plasma_164393_1
pup
adult
reads per million genome matching
bp position
0
1
2
3
4
5
0
10
20
30
40
50

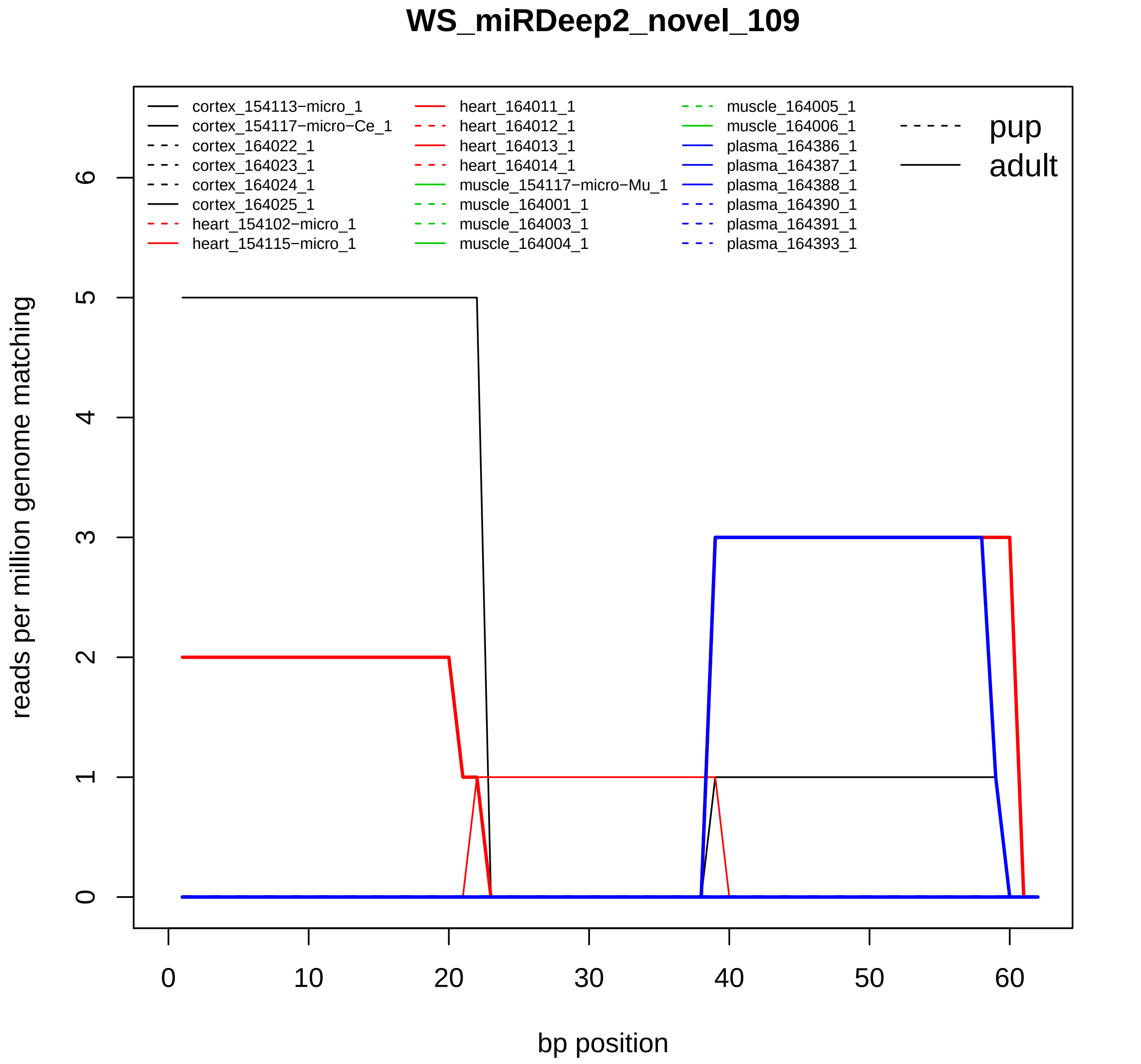
WS_miRDeep2_novel_109
cortex_154113−micro_1
cortex_154117−micro−Ce_1
cortex_164022_1
cortex_164023_1
cortex_164024_1
cortex_164025_1
heart_154102−micro_1
heart_154115−micro_1
heart_164011_1
heart_164012_1
heart_164013_1
heart_164014_1
muscle_154117−micro−Mu_1
muscle_164001_1
muscle_164003_1
muscle_164004_1
muscle_164005_1
muscle_164006_1
plasma_164386_1
plasma_164387_1
plasma_164388_1
plasma_164390_1
plasma_164391_1
plasma_164393_1
pup
adult
reads per million genome matching
0
1
2
3
4
5
6
0
10
20
30
40
50
60
bp position

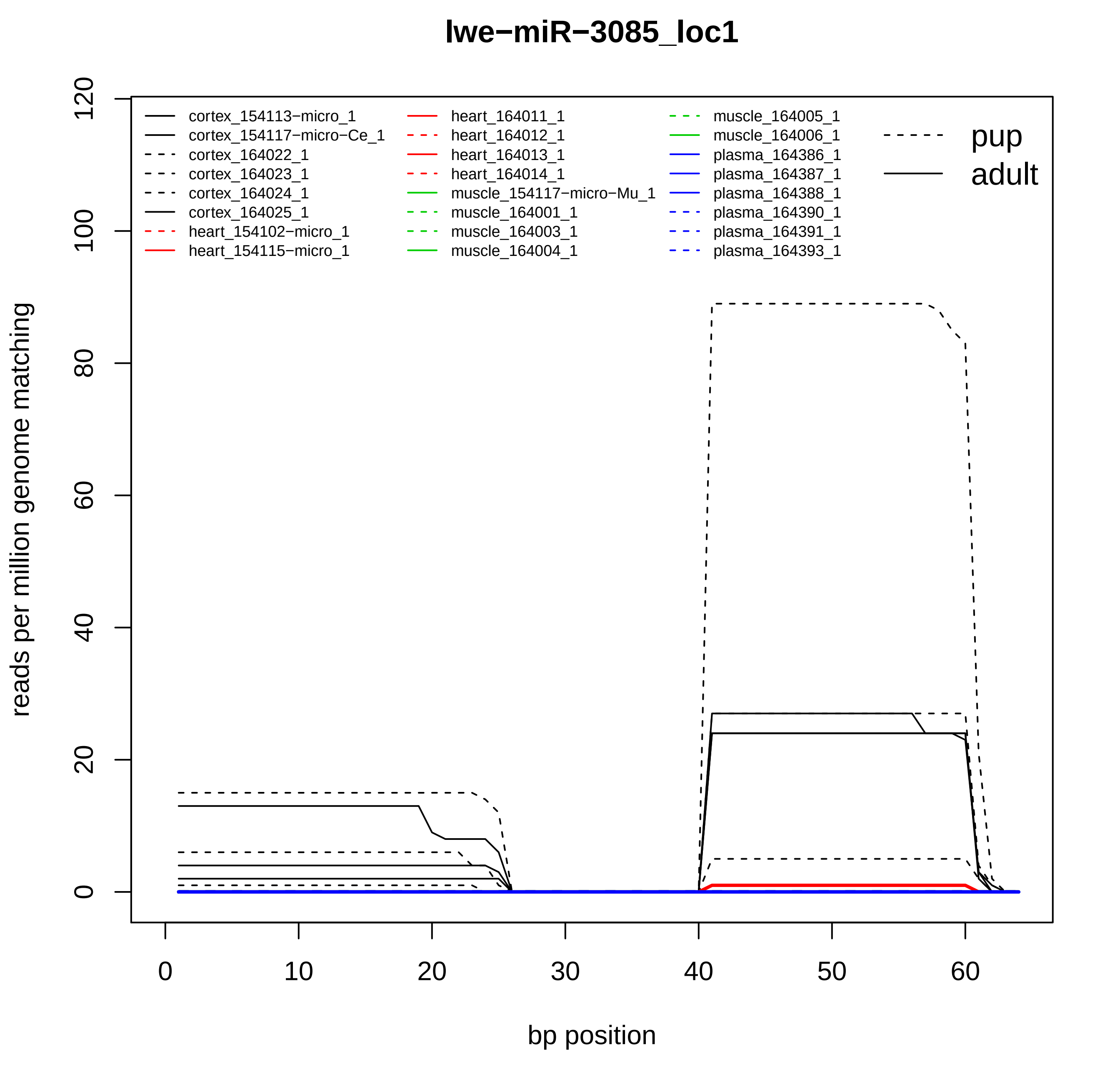
lwe−miR−3085_loc1
cortex_154113−micro_1
cortex_154117−micro−Ce_1
cortex_164022_1
cortex_164023_1
cortex_164024_1
cortex_164025_1
heart_154102−micro_1
heart_154115−micro_1
heart_164011_1
heart_164012_1
heart_164013_1
heart_164014_1
muscle_154117−micro−Mu_1
muscle_164001_1
muscle_164003_1
muscle_164004_1
muscle_164005_1
muscle_164006_1
plasma_164386_1
plasma_164387_1
plasma_164388_1
plasma_164390_1
plasma_164391_1
plasma_164393_1
pup
adult
reads per million genome matching
bp position
0
20
40
60
80
100
120
0
10
20
30
40
50
60

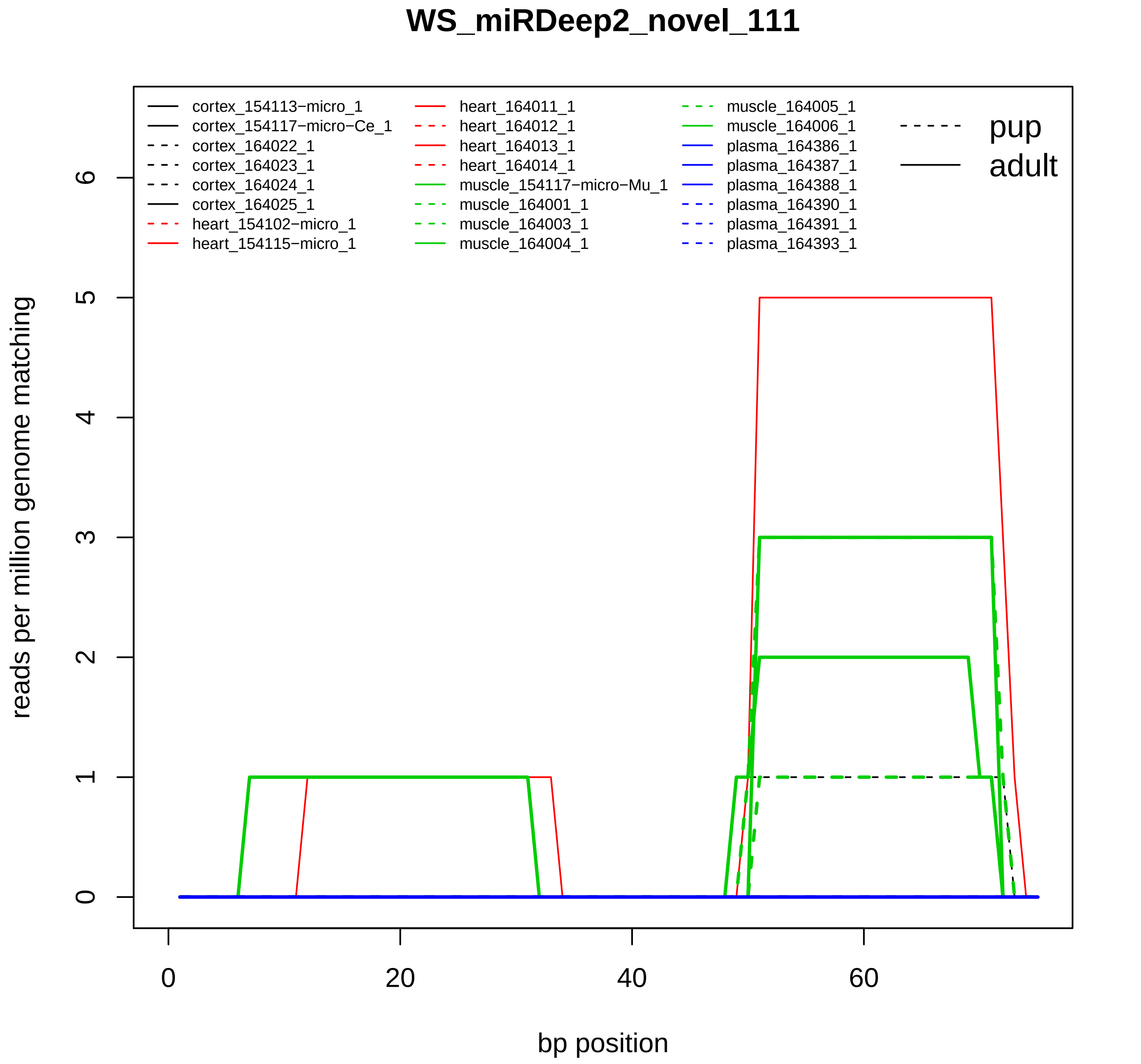

WS_miRDeep2_novel_111
cortex_154113−micro_1
cortex_154117−micro−Ce_1
cortex_164022_1
cortex_164023_1
cortex_164024_1
cortex_164025_1
heart_154102−micro_1
heart_154115−micro_1
heart_164011_1
heart_164012_1
heart_164013_1
heart_164014_1
muscle_154117−micro−Mu_1
muscle_164001_1
muscle_164003_1
muscle_164004_1
muscle_164005_1
muscle_164006_1
plasma_164386_1
plasma_164387_1
plasma_164388_1
plasma_164390_1
plasma_164391_1
plasma_164393_1
pup
adult
reads per million genome matching
0
1
2
3
4
5
6
0
20
40
60
bp position

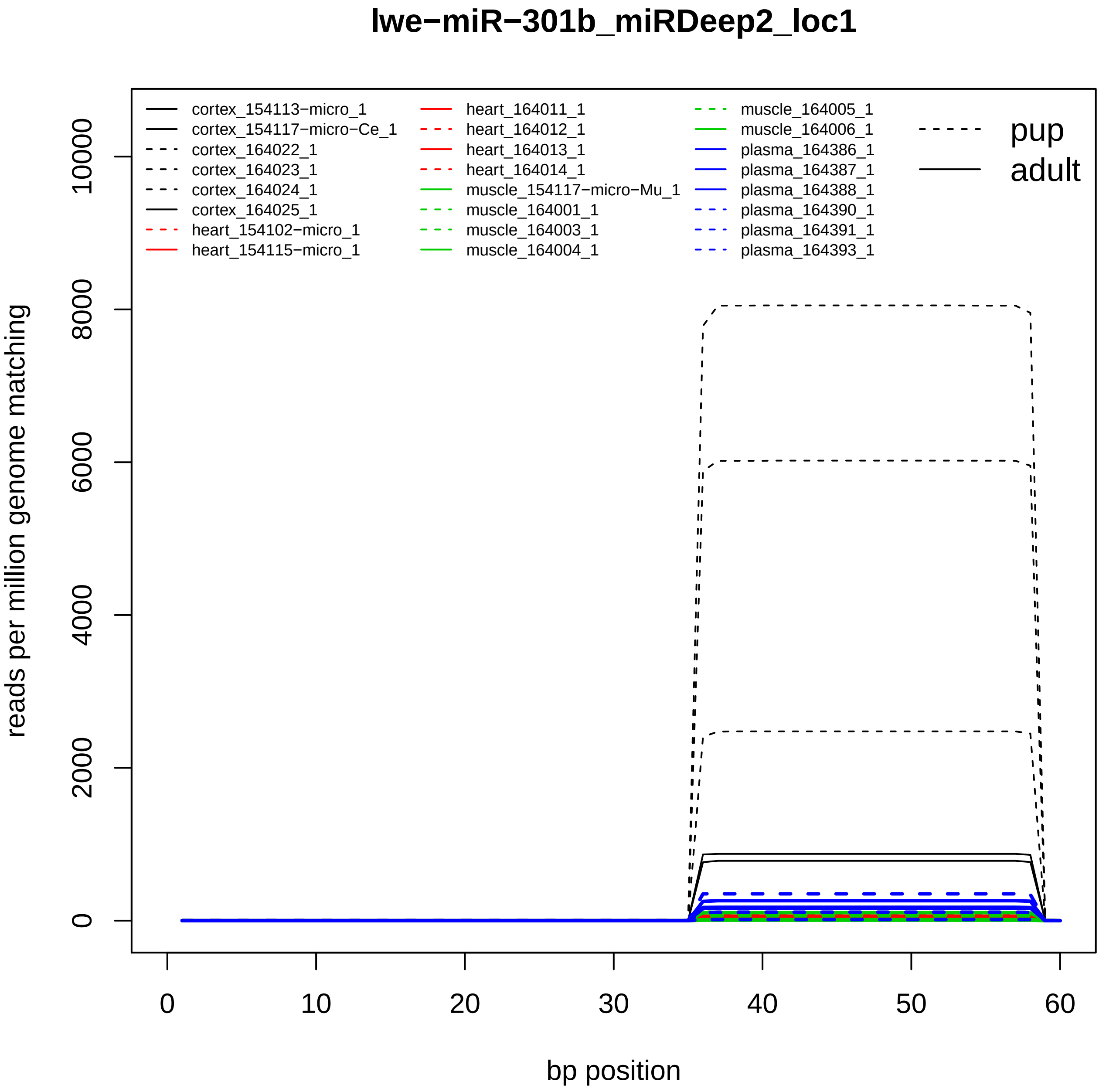
lwe−miR−301b_miRDeep2_loc1
reads per million genome matching
bp position
0
2000
4000
6000
8000
10000
0
10
20
30
40
50
60
cortex_154113−micro_1
cortex_154117−micro−Ce_1
cortex_164022_1
cortex_164023_1
cortex_164024_1
cortex_164025_1
heart_154102−micro_1
heart_154115−micro_1
heart_164011_1
heart_164012_1
heart_164013_1
heart_164014_1
muscle_154117−micro−Mu_1
muscle_164001_1
muscle_164003_1
muscle_164004_1
muscle_164005_1
muscle_164006_1
plasma_164386_1
plasma_164387_1
plasma_164388_1
plasma_164390_1
plasma_164391_1
plasma_164393_1
pup
adult

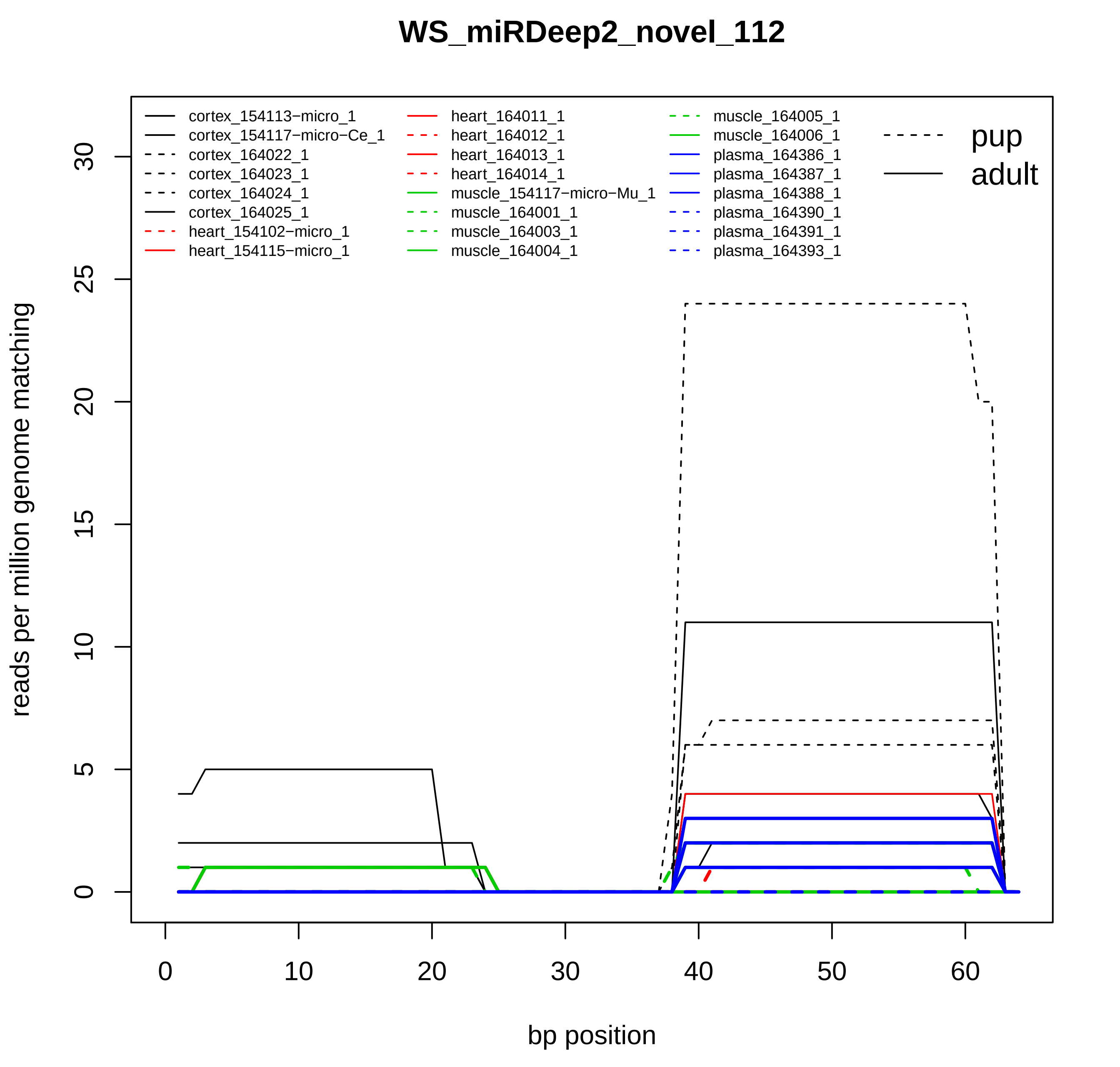

WS_miRDeep2_novel_112
cortex_154113−micro_1
cortex_154117−micro−Ce_1
cortex_164022_1
cortex_164023_1
cortex_164024_1
cortex_164025_1
heart_154102−micro_1
heart_154115−micro_1
heart_164011_1
heart_164012_1
heart_164013_1
heart_164014_1
muscle_154117−micro−Mu_1
muscle_164001_1
muscle_164003_1
muscle_164004_1
muscle_164005_1
muscle_164006_1
plasma_164386_1
plasma_164387_1
plasma_164388_1
plasma_164390_1
plasma_164391_1
plasma_164393_1
pup
adult
reads per million genome matching
0
5
10
15
20
25
30
0
10
20
30
40
50
60
bp position

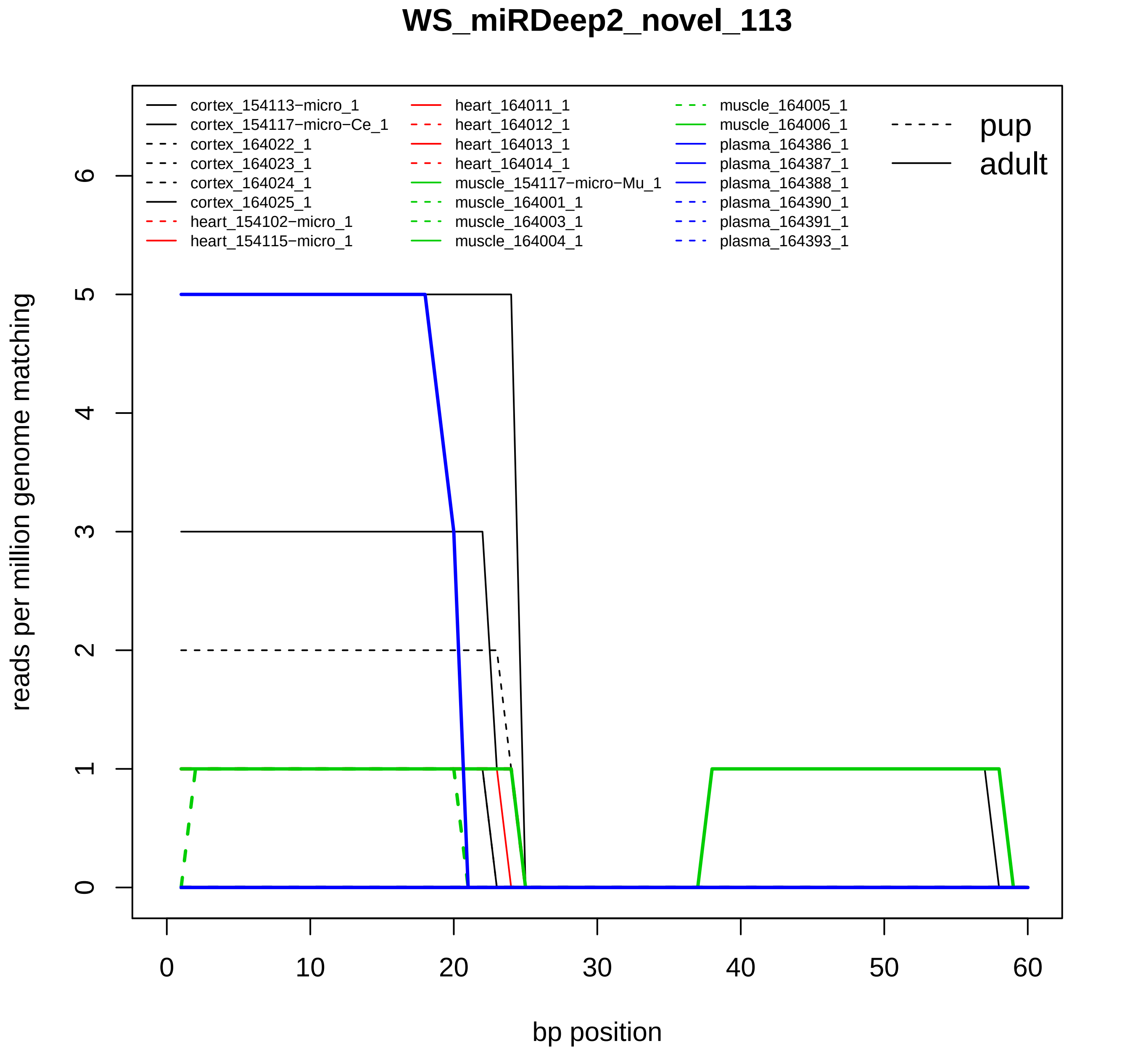

WS_miRDeep2_novel_113
cortex_154113−micro_1
cortex_154117−micro−Ce_1
cortex_164022_1
cortex_164023_1
cortex_164024_1
cortex_164025_1
heart_154102−micro_1
heart_154115−micro_1
heart_164011_1
heart_164012_1
heart_164013_1
heart_164014_1
muscle_154117−micro−Mu_1
muscle_164001_1
muscle_164003_1
muscle_164004_1
muscle_164005_1
muscle_164006_1
plasma_164386_1
plasma_164387_1
plasma_164388_1
plasma_164390_1
plasma_164391_1
plasma_164393_1
pup
adult
reads per million genome matching
0
1
2
3
4
5
6
0
10
20
30
40
50
60
bp position

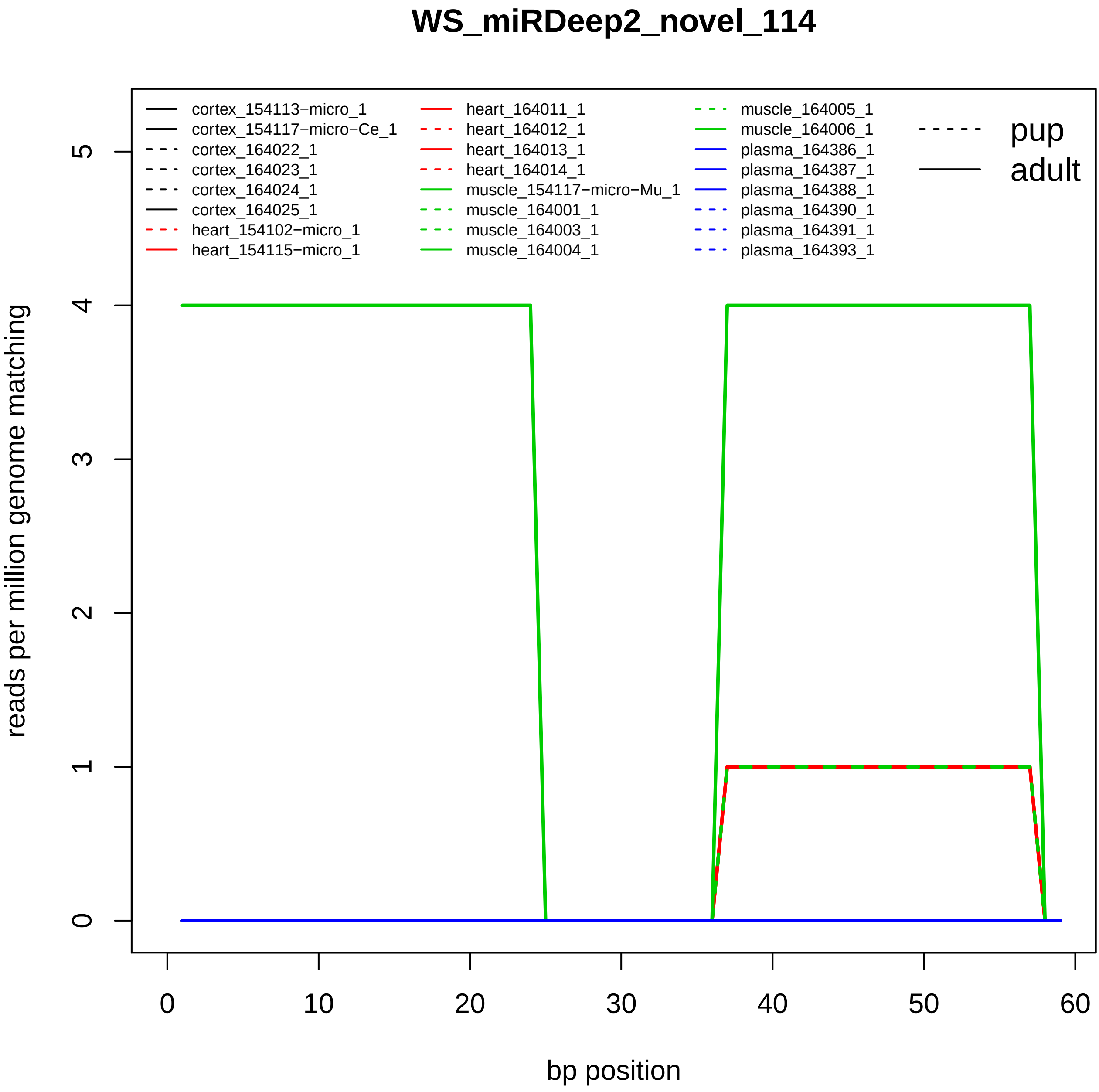
WS_miRDeep2_novel_114
cortex_154113−micro_1
cortex_154117−micro−Ce_1
cortex_164022_1
cortex_164023_1
cortex_164024_1
cortex_164025_1
heart_154102−micro_1
heart_154115−micro_1
heart_164011_1
heart_164012_1
heart_164013_1
heart_164014_1
muscle_154117−micro−Mu_1
muscle_164001_1
muscle_164003_1
muscle_164004_1
muscle_164005_1
muscle_164006_1
plasma_164386_1
plasma_164387_1
plasma_164388_1
plasma_164390_1
plasma_164391_1
plasma_164393_1
pup
adult
reads per million genome matching
0
1
2
3
4
5
0
10
20
30
40
50
60
bp position

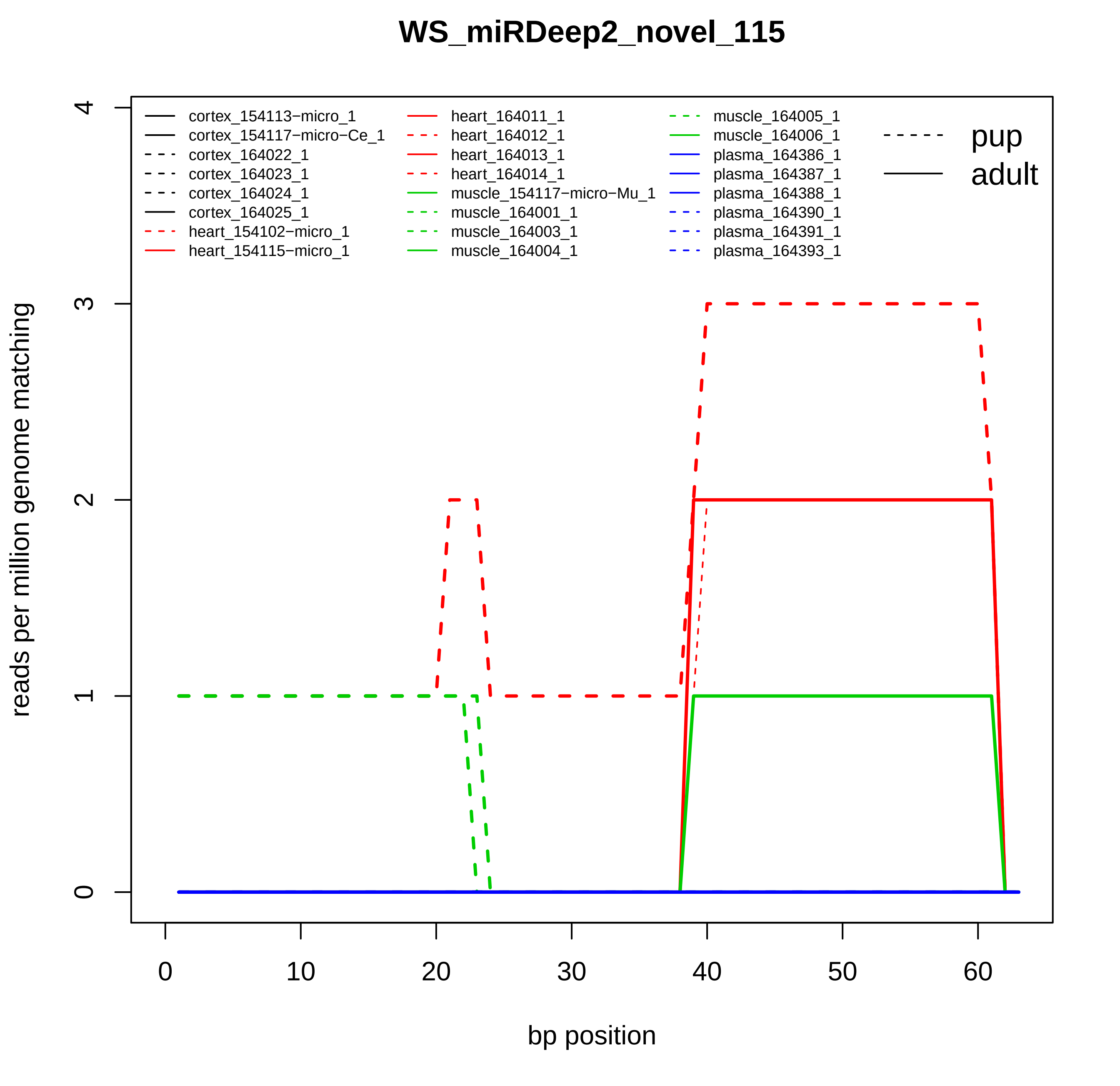

WS_miRDeep2_novel_115
reads per million genome matching
bp position
0
1
2
3
4
0
10
20
30
40
50
60
cortex_154113−micro_1
cortex_154117−micro−Ce_1
cortex_164022_1
cortex_164023_1
cortex_164024_1
cortex_164025_1
heart_154102−micro_1
heart_154115−micro_1
heart_164011_1
heart_164012_1
heart_164013_1
heart_164014_1
muscle_154117−micro−Mu_1
muscle_164001_1
muscle_164003_1
muscle_164004_1
muscle_164005_1
muscle_164006_1
plasma_164386_1
plasma_164387_1
plasma_164388_1
plasma_164390_1
plasma_164391_1
plasma_164393_1
pup
adult

Supplementary Figure S6. Comparison of miRNA-5p and miRNA-3p abundance (reads per million genome matching) for all 559 annotated miRNA loci. Colors separate tissues, while different shapes separate pup and adult individuals.

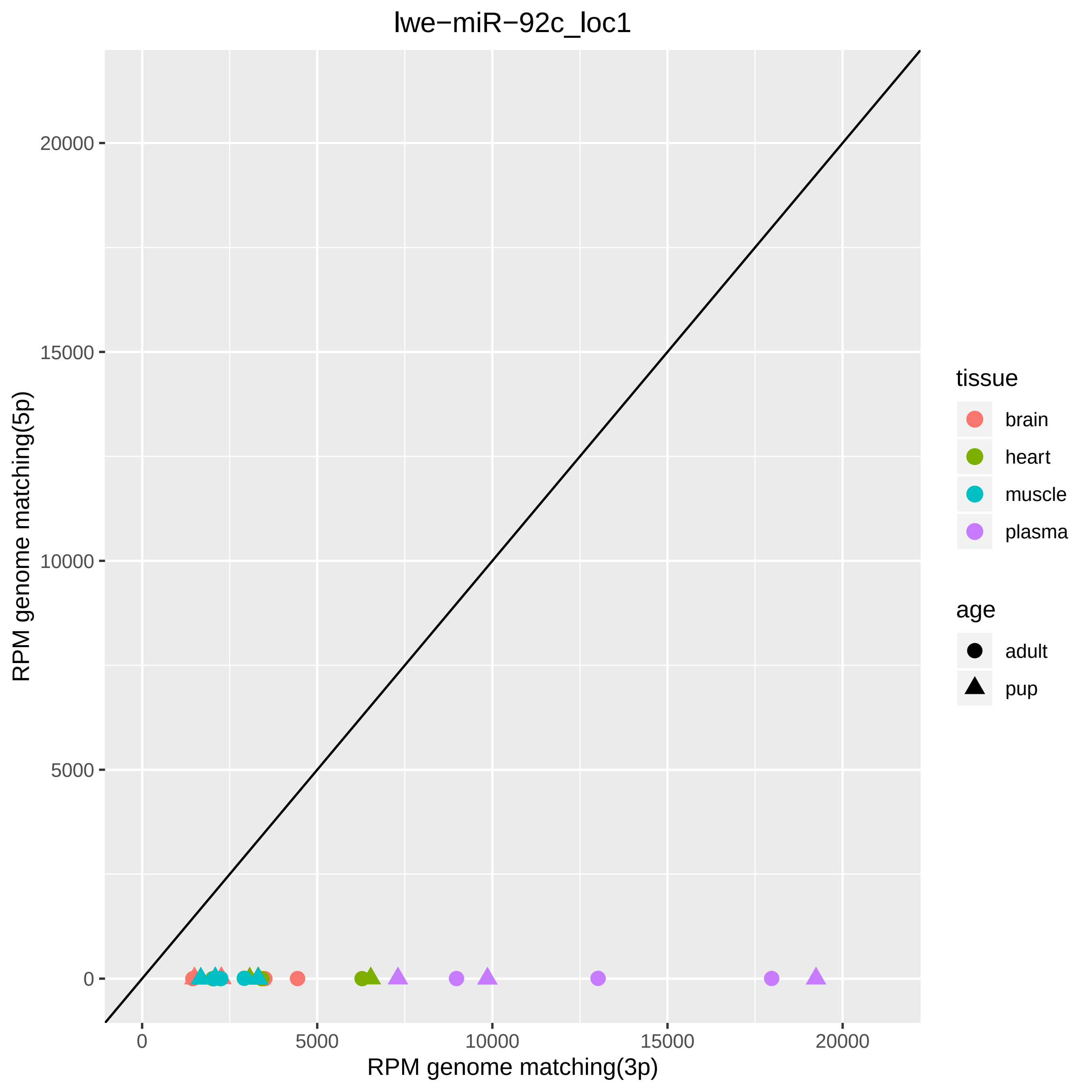

lwe−miR−92c_loc1
RPM genome matching(5p)
RPM genome matching(3p)
0
5000
10000
15000
20000
tissue
brain
heart
muscle
plasma
age
adult
pup

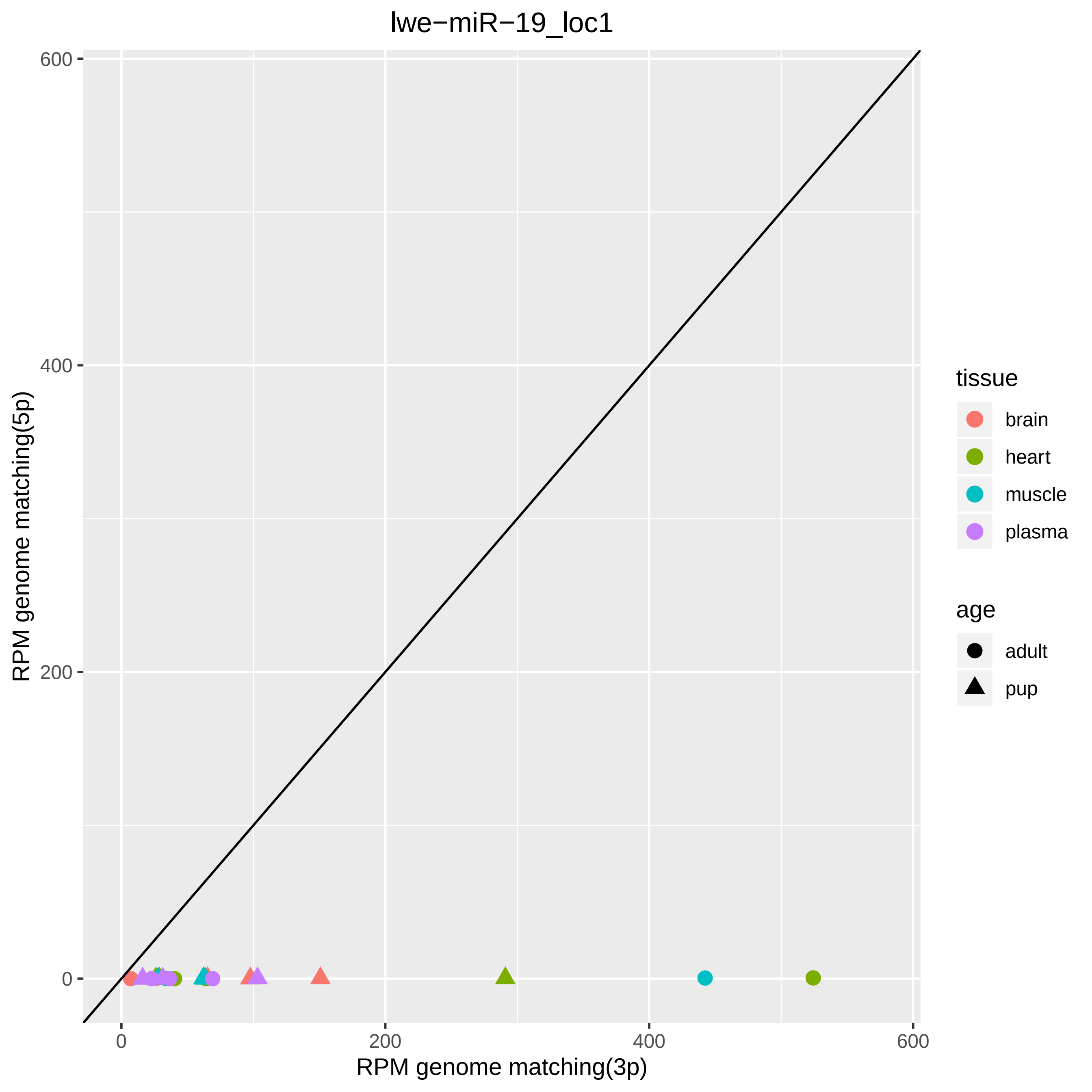

lwe−miR−19_loc1
RPM genome matching(5p)
RPM genome matching(3p)
0
200
400
600
tissue
brain
heart
muscle
plasma
age
adult
pup

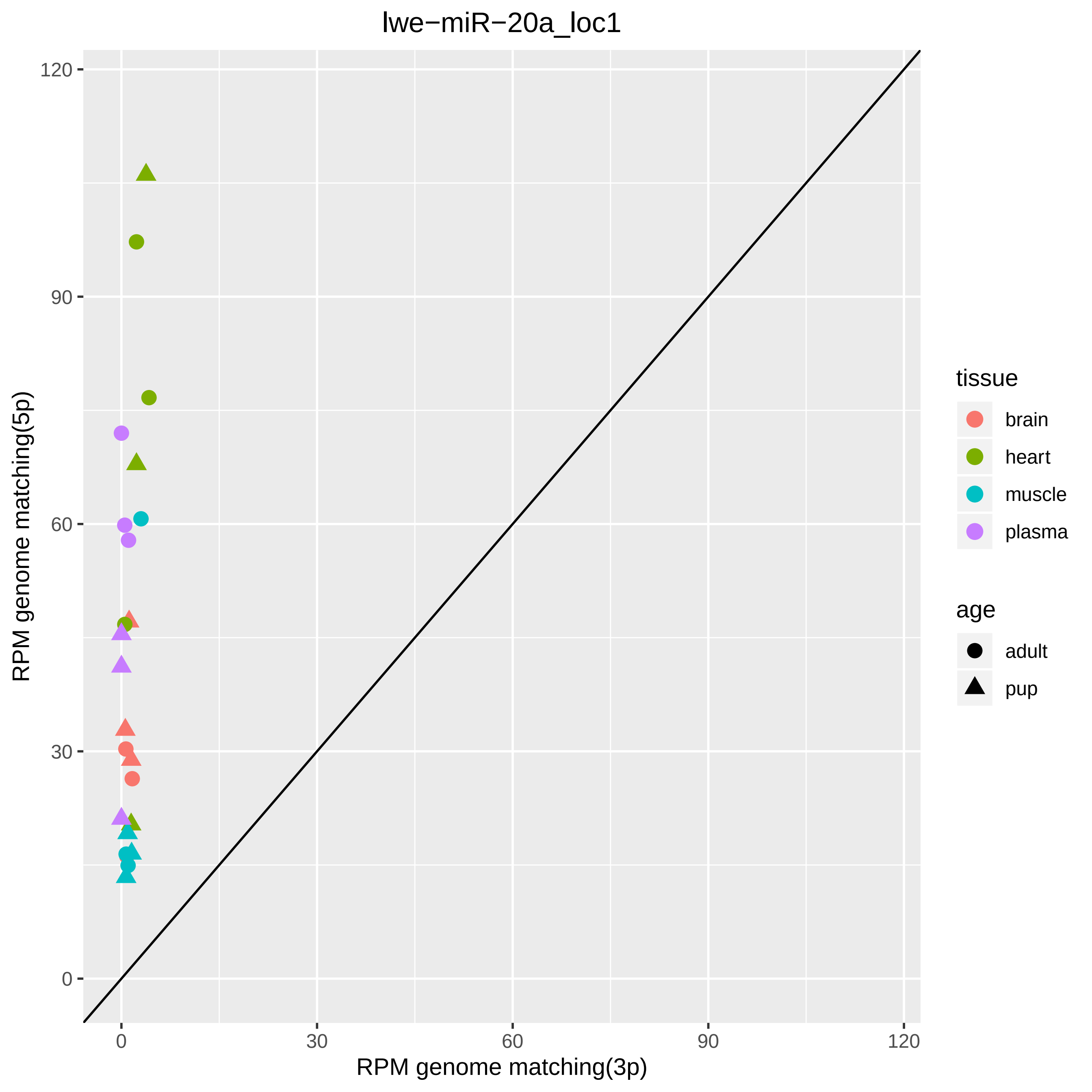

lwe−miR−20a_loc1
RPM genome matching(5p)
RPM genome matching(3p)
120
90
60
30
0
0
30
60
90
120
tissue
brain
heart
muscle
plasma
age
adult
pup

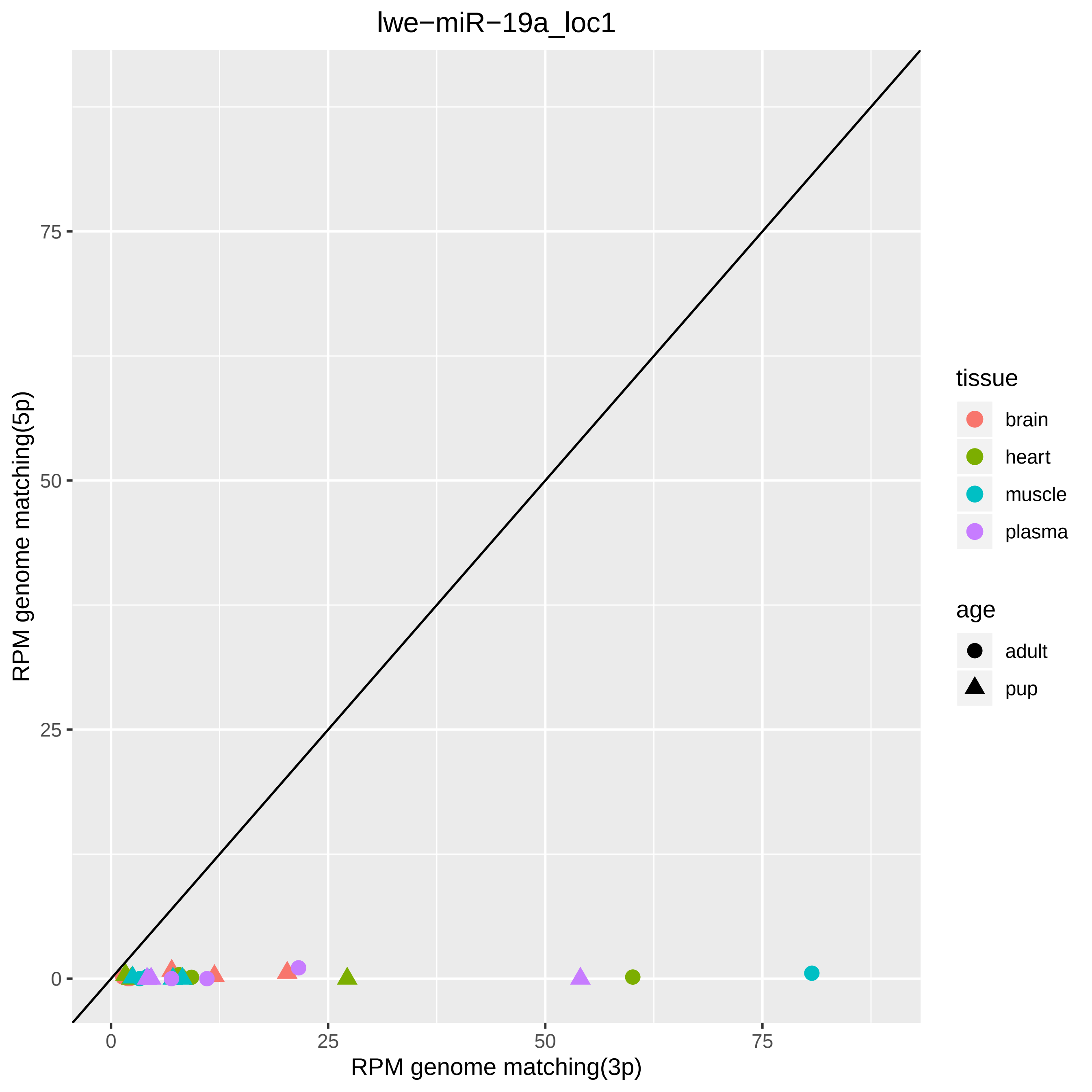

lwe−miR−19a_loc1
RPM genome matching(5p)
RPM genome matching(3p)
0
25
50
75
tissue
brain
heart
muscle
plasma
age
adult
pup

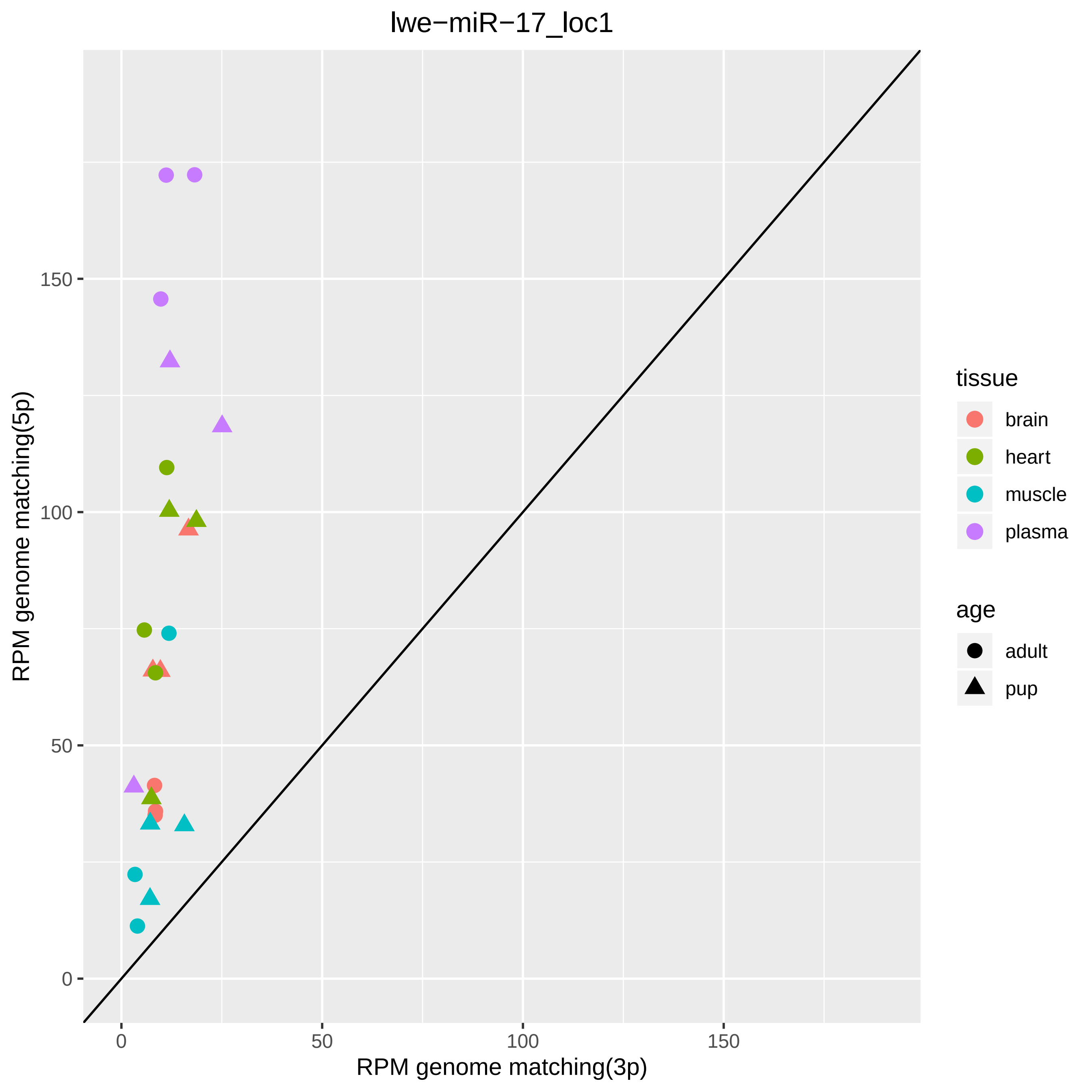
lwe−miR−17_loc1
RPM genome matching(5p)
RPM genome matching(3p)
0
50
100
150
tissue
brain
heart
muscle
plasma
age
adult
pup

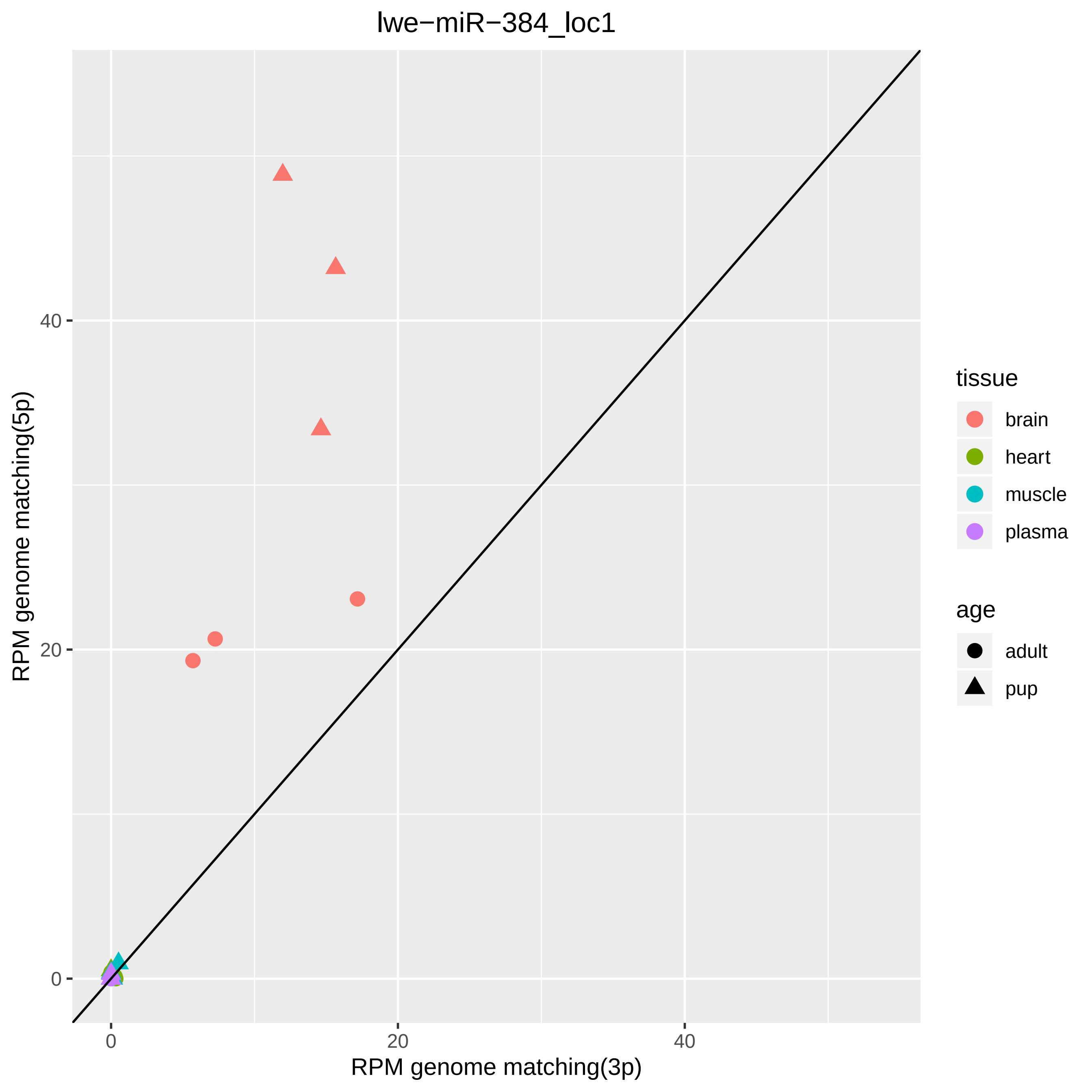

lwe−miR−384_loc1
RPM genome matching(5p)
RPM genome matching(3p)
0
20
40
0
20
40
tissue
brain
heart
muscle
plasma
age
adult
pup

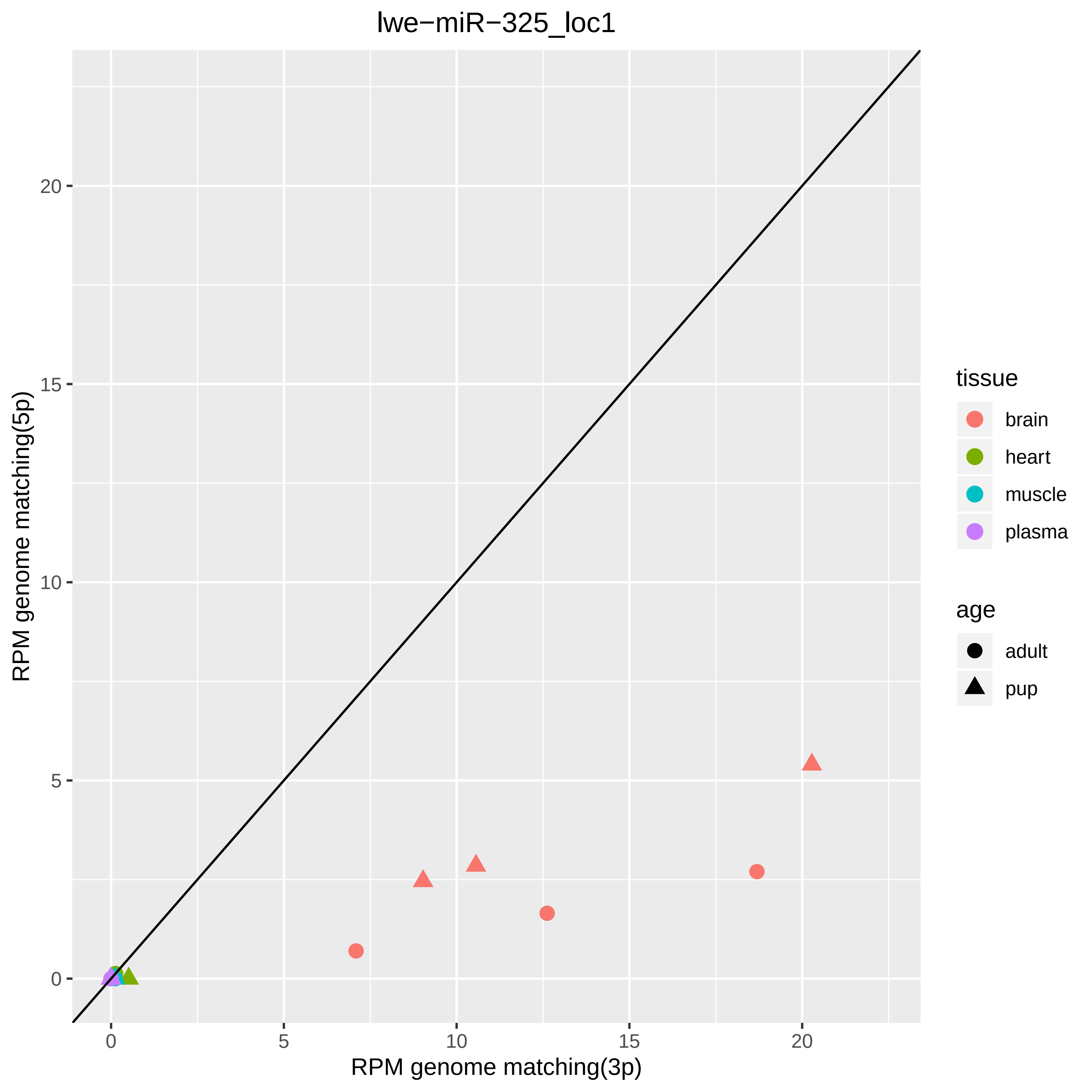

lwe−miR−325_loc1
RPM genome matching(5p)
RPM genome matching(3p)
0
5
10
15
20
tissue
brain
heart
muscle
plasma
age
adult
pup

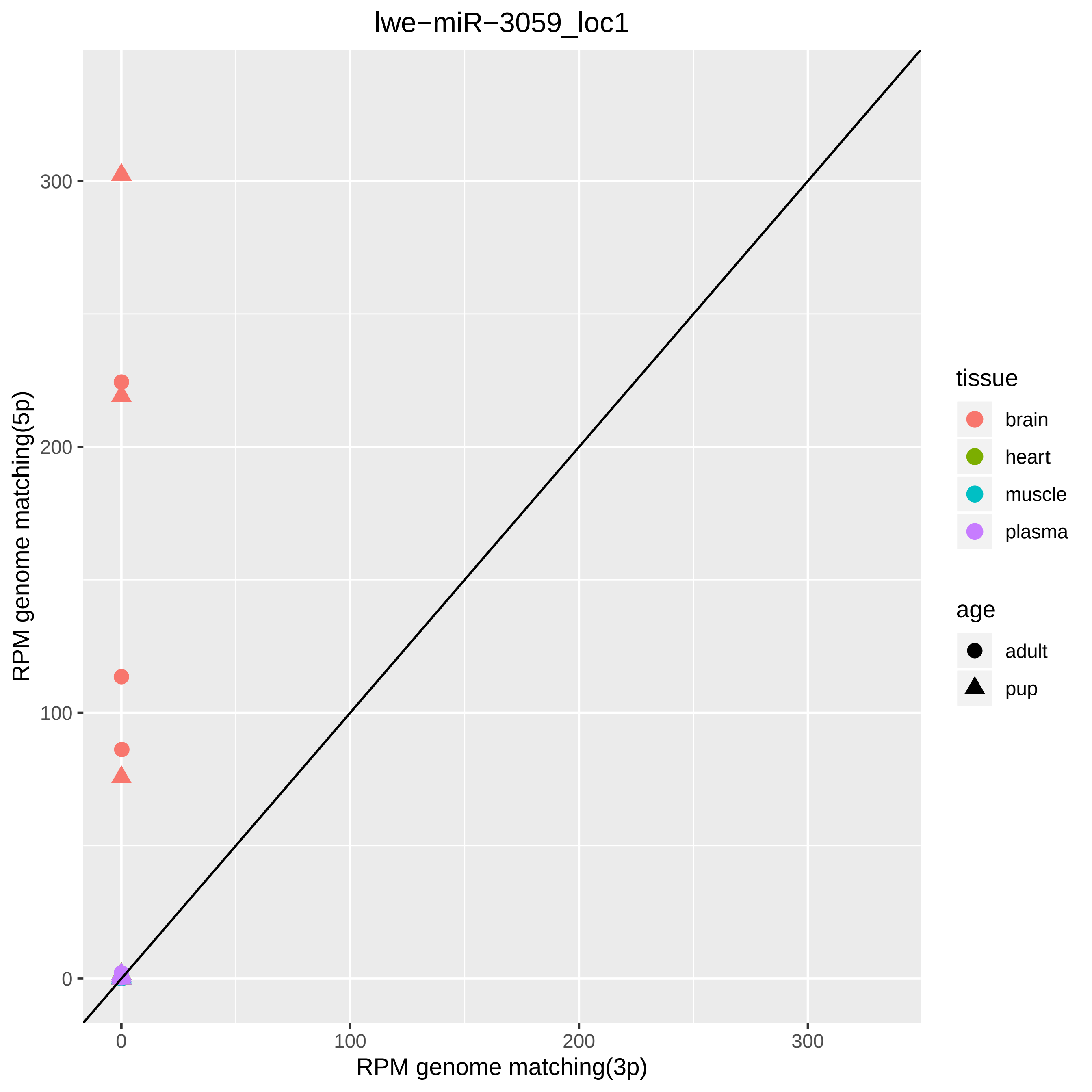

lwe−miR−3059_loc1
RPM genome matching(5p)
RPM genome matching(3p)
0
100
200
300
tissue
brain
heart
muscle
plasma
age
adult
pup

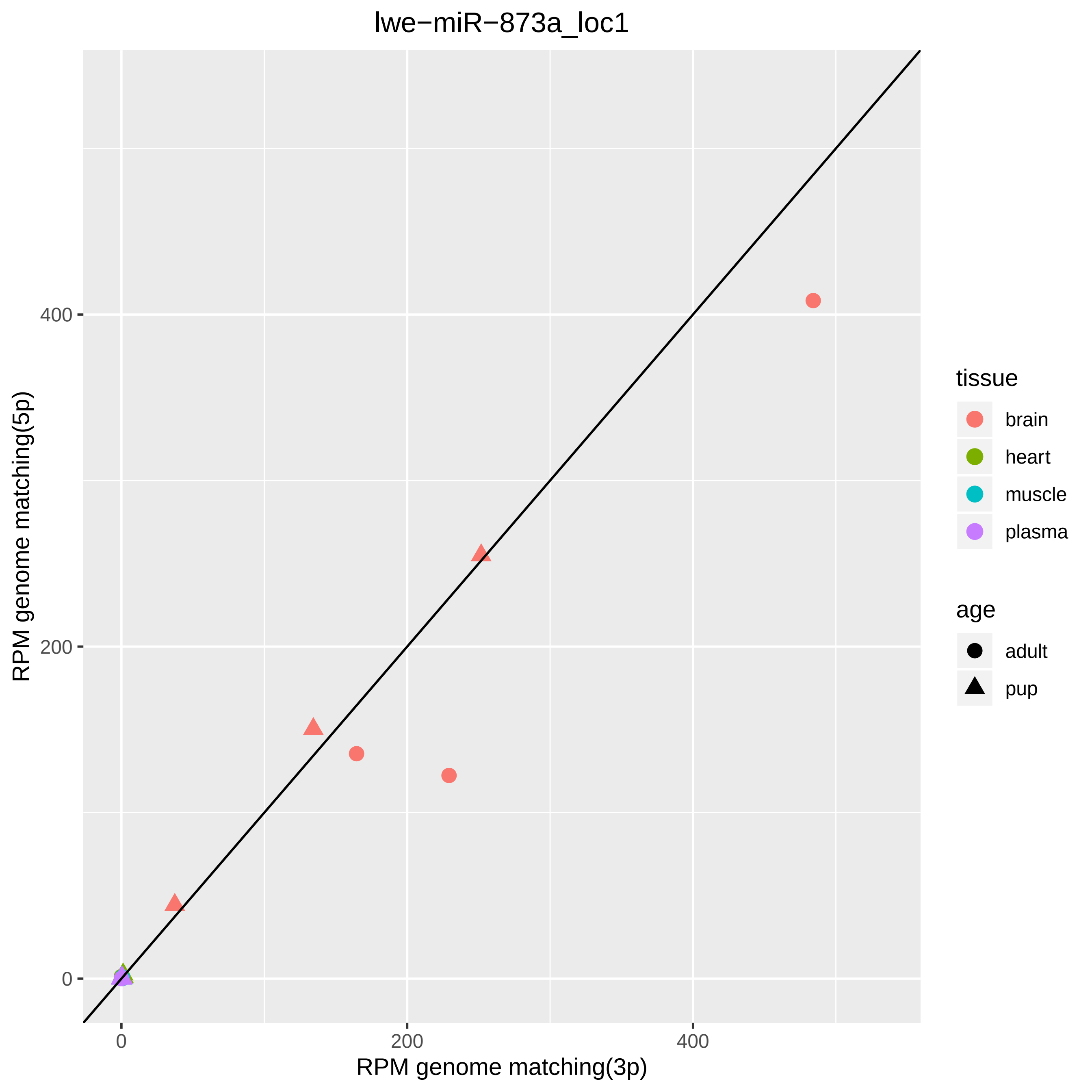
lwe−miR−873a_loc1
RPM genome matching(5p)
RPM genome matching(3p)
0
200
400
0
200
400
tissue
brain
heart
muscle
plasma
age
adult
pup

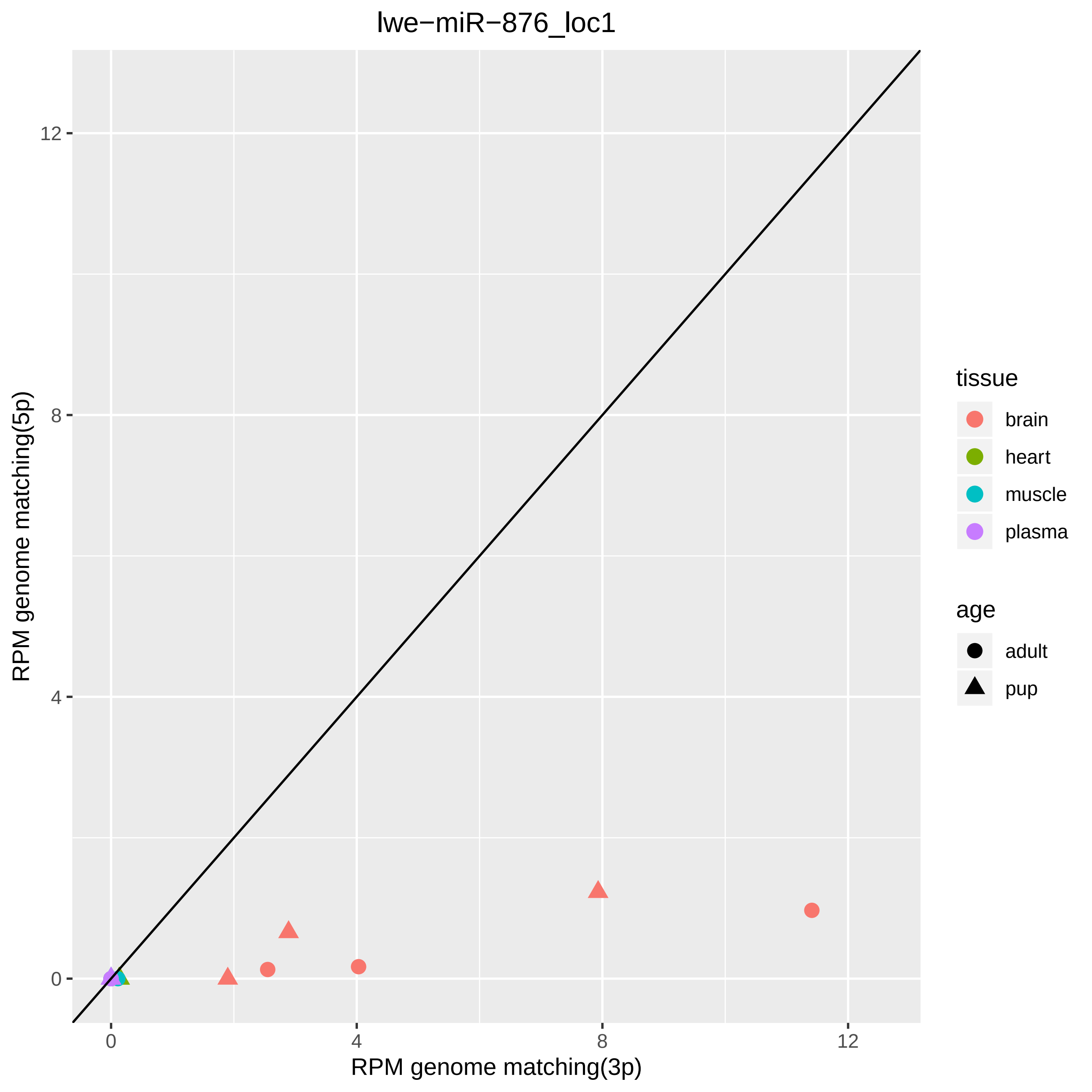
lwe−miR−876_loc1
RPM genome matching(5p)
RPM genome matching(3p)
0
4
8
12
0
4
8
12
tissue
brain
heart
muscle
plasma
age
adult
pup

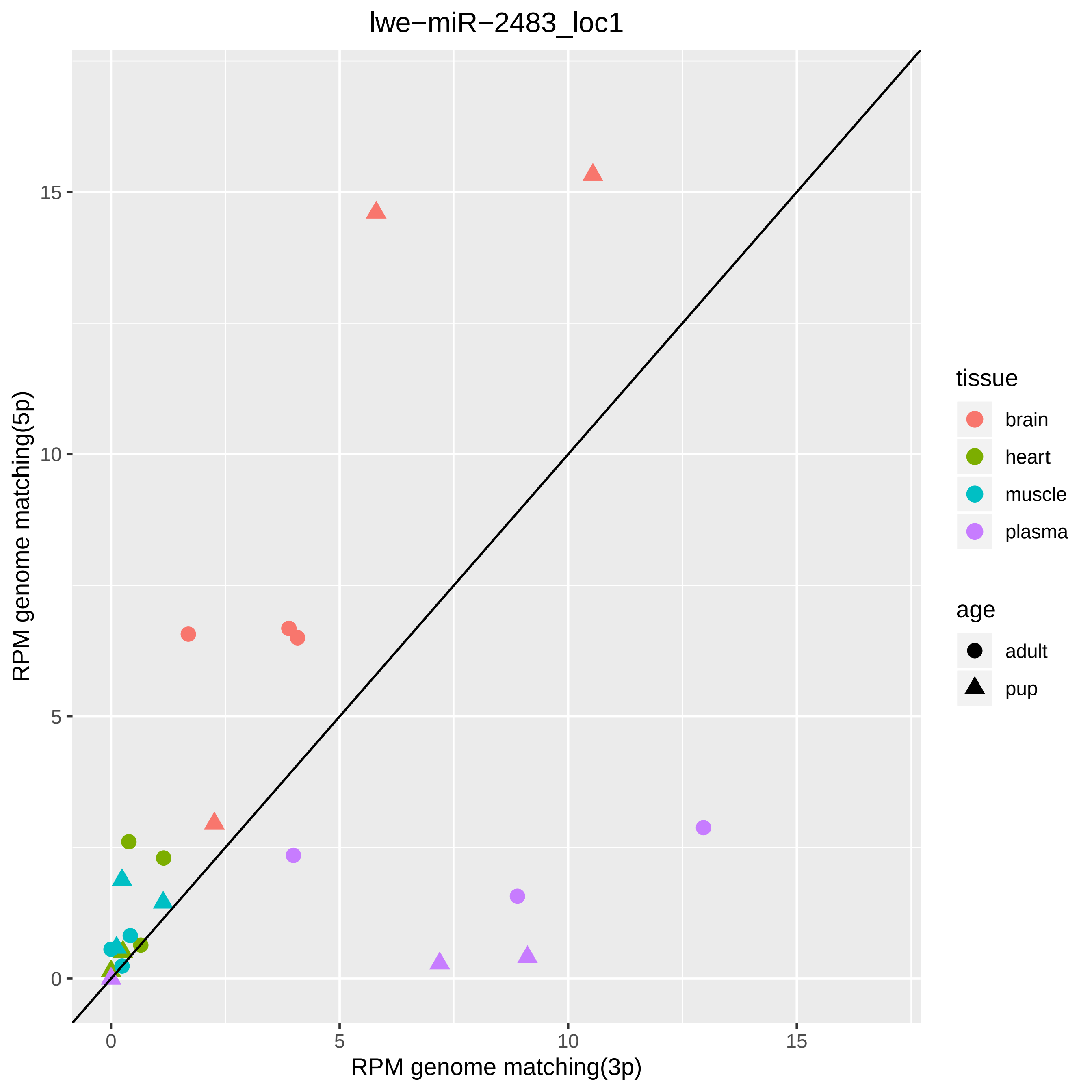
lwe−miR−2483_loc1
RPM genome matching(5p)
RPM genome matching(3p)
0
5
10
15
tissue
brain
heart
muscle
plasma
age
adult
pup

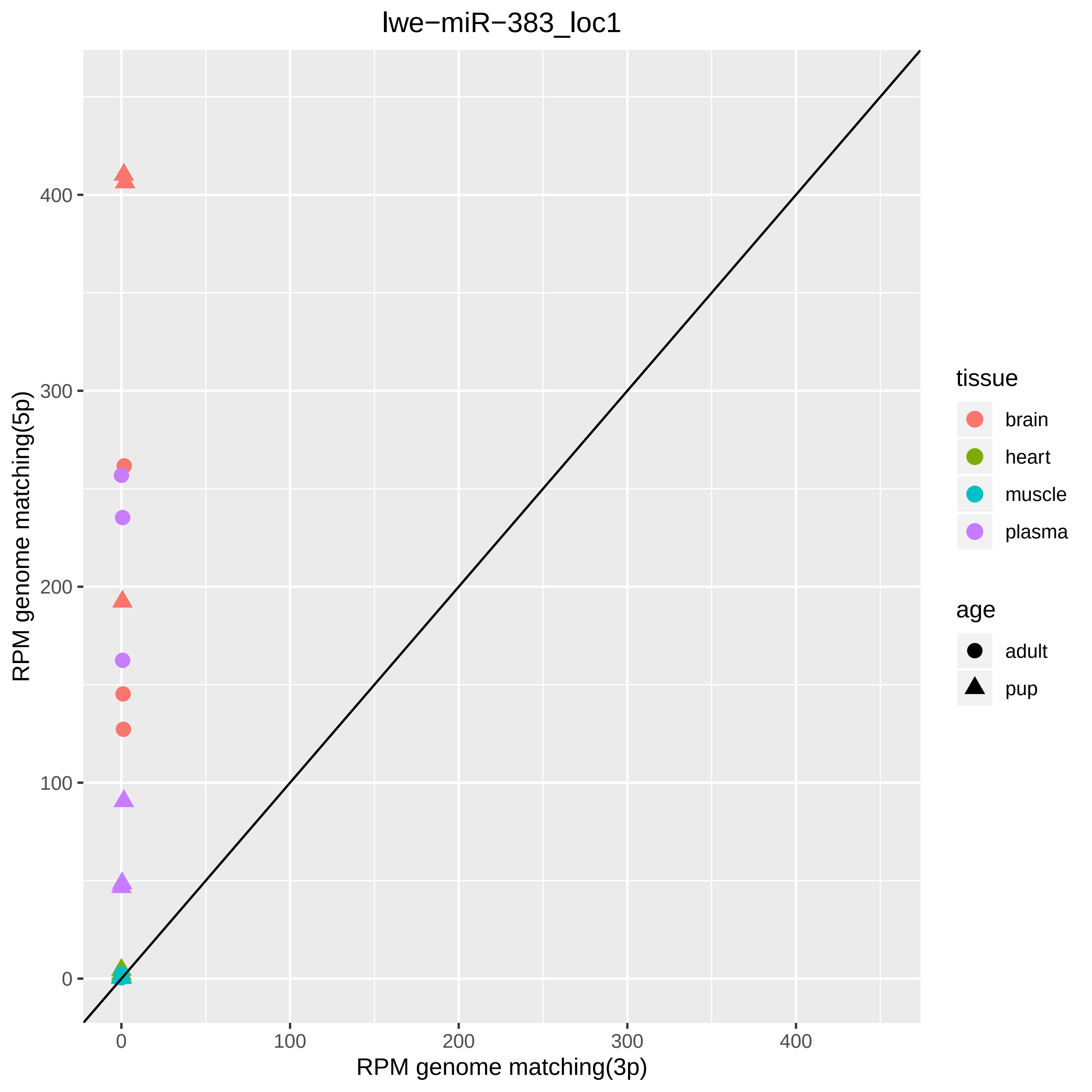
lwe−miR−383_loc1
RPM genome matching(5p)
RPM genome matching(3p)
0
100
200
300
400
0
100
200
300
400
tissue
brain
heart
muscle
plasma
age
adult
pup

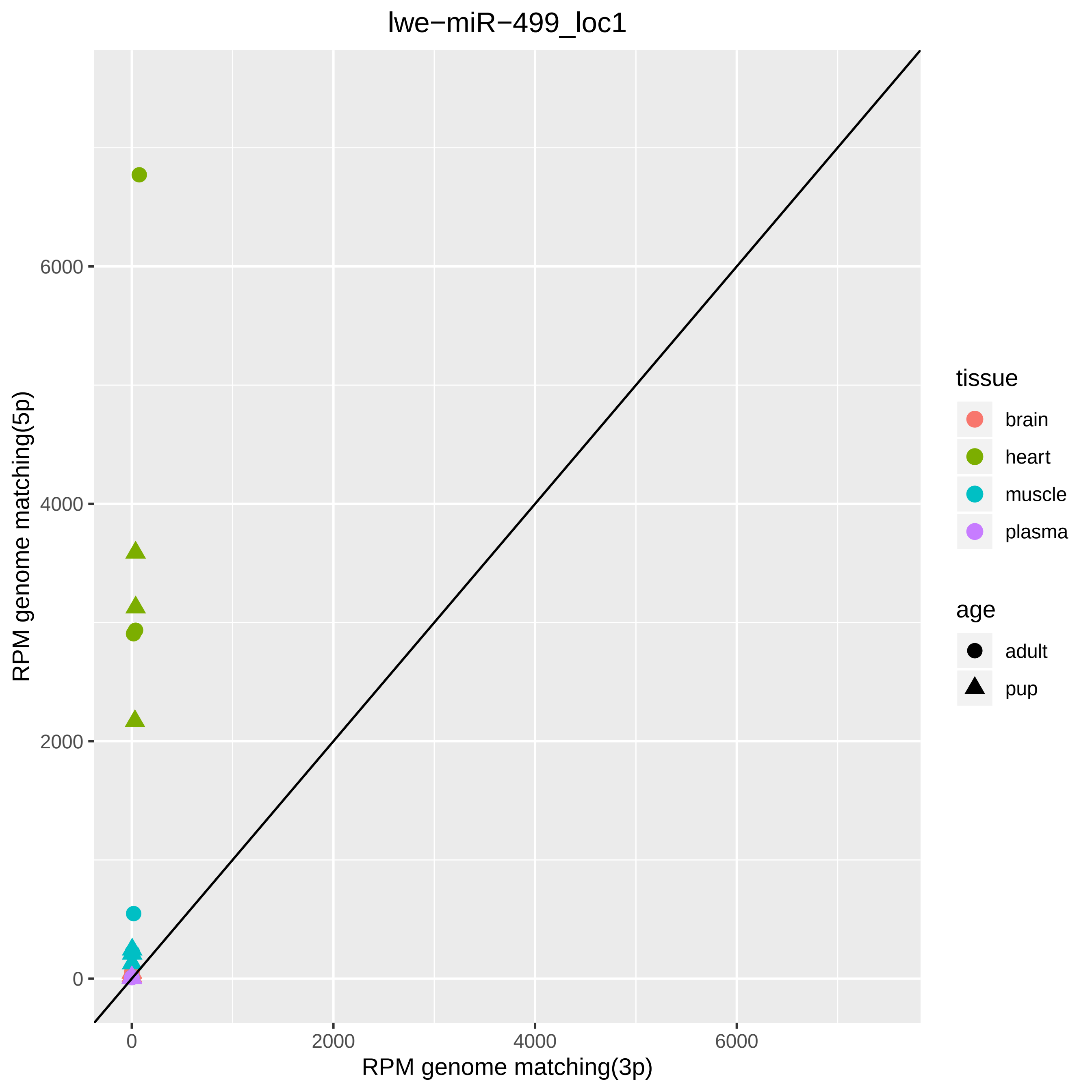

lwe−miR−499_loc1
RPM genome matching(5p)
6000
4000
2000
0
0
2000
4000
6000
RPM genome matching(3p)
tissue
brain
heart
muscle
plasma
age
adult
pup

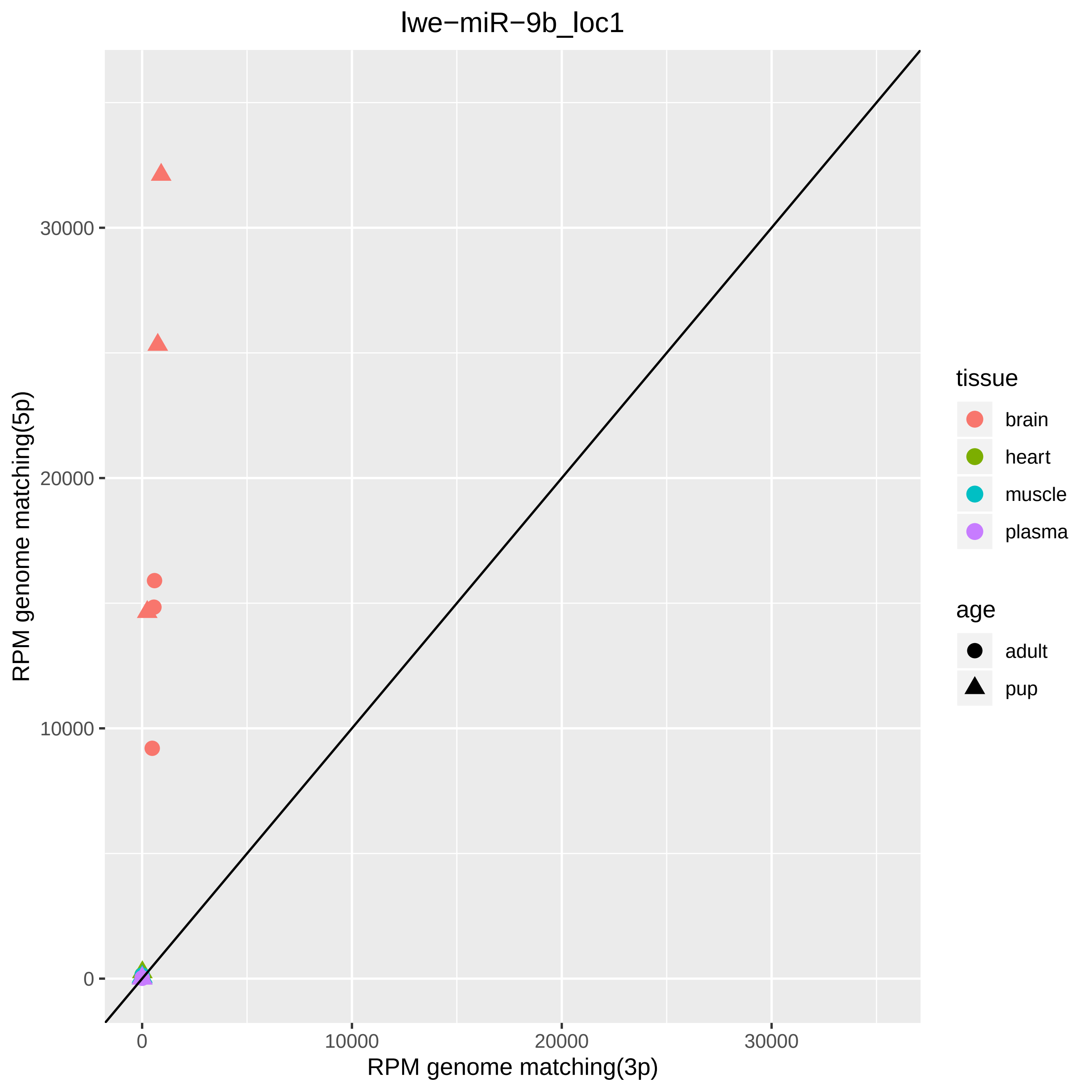
lwe−miR−9b_loc1
RPM genome matching(5p)
RPM genome matching(3p)
0
10000
20000
30000
0
10000
20000
30000
tissue
brain
heart
muscle
plasma
age
adult
pup

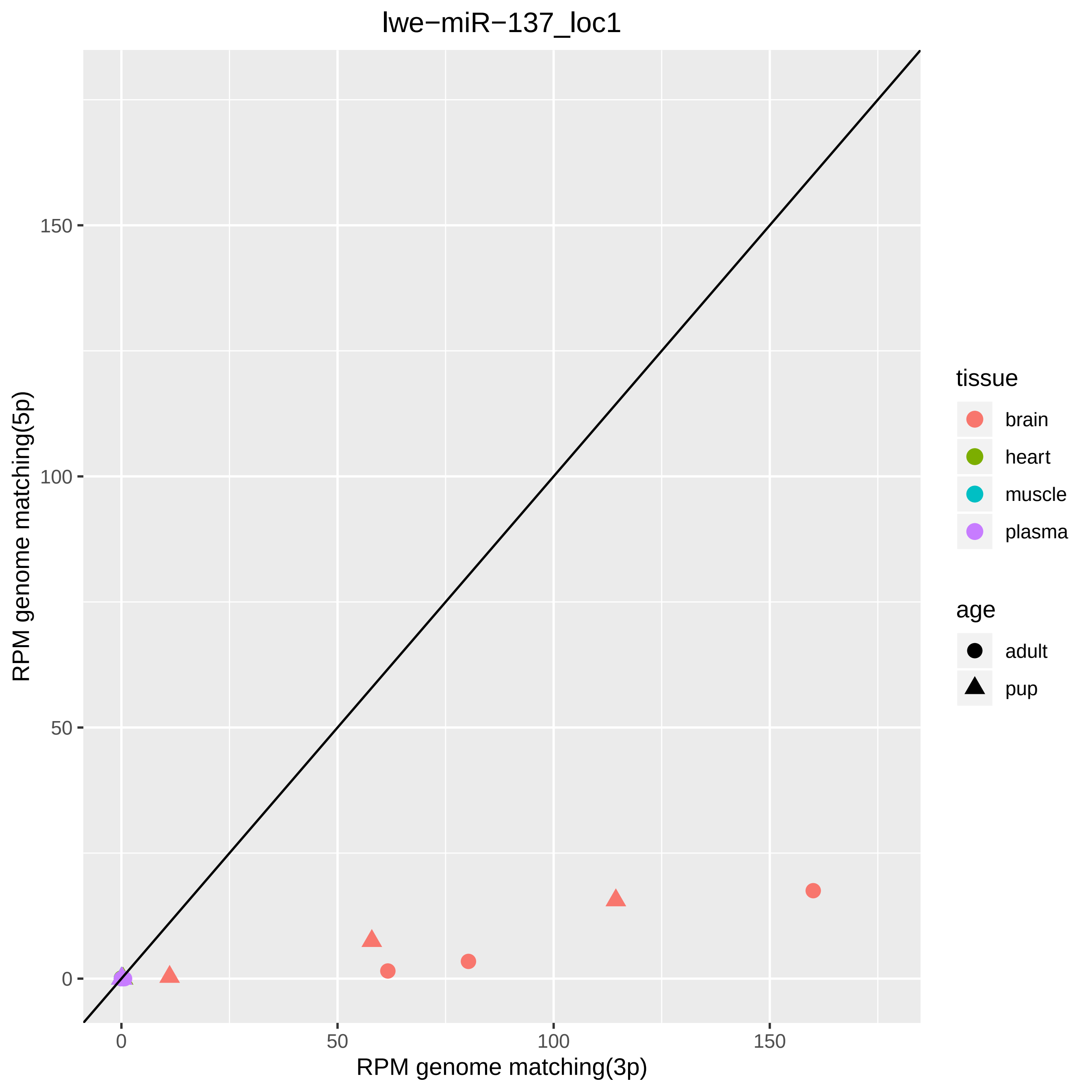
lwe−miR−137_loc1
RPM genome matching(5p)
RPM genome matching(3p)
0
50
100
150
tissue
brain
heart
muscle
plasma
age
adult
pup

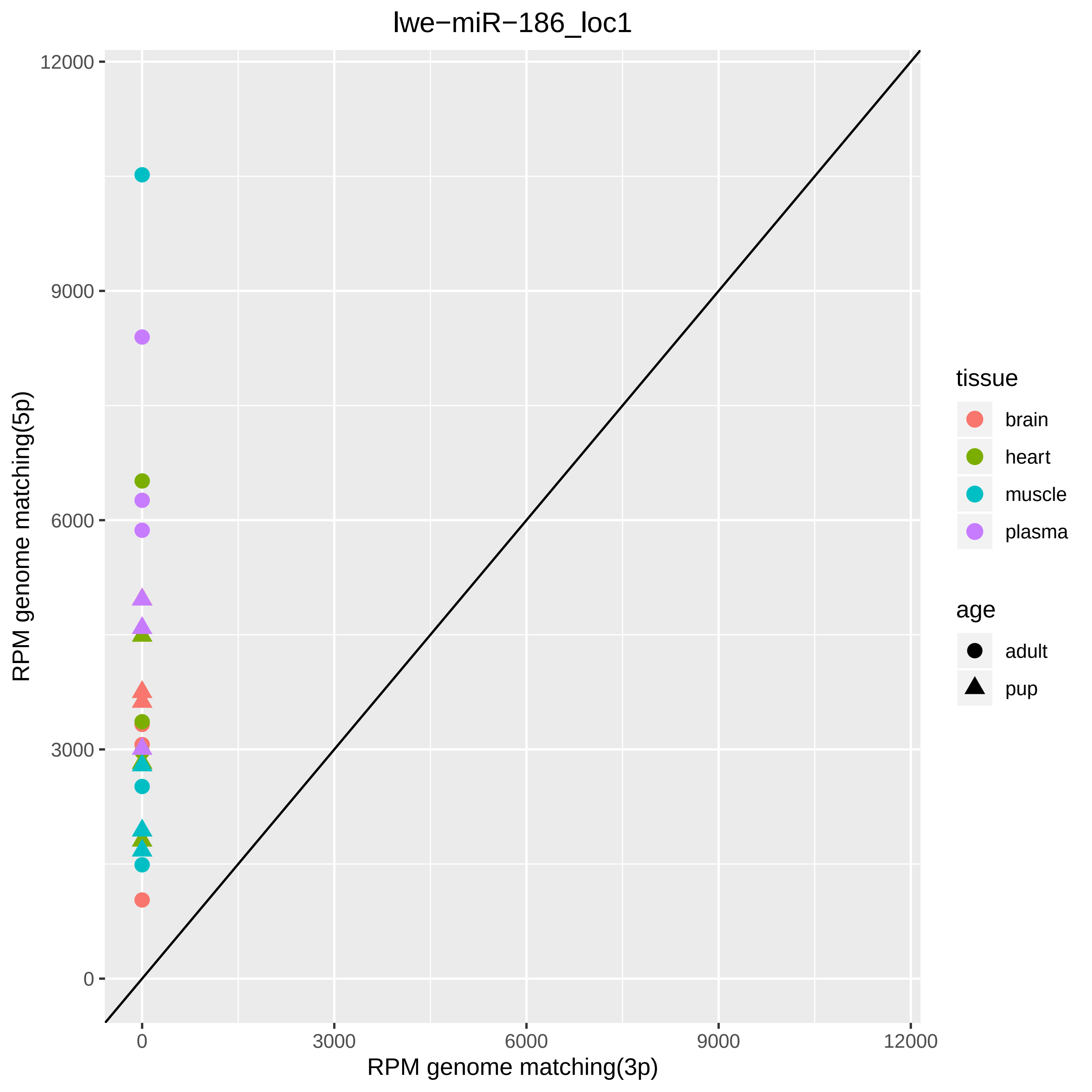
lwe−miR−186_loc1
RPM genome matching(5p)
RPM genome matching(3p)
0
3000
6000
9000
12000
tissue
brain
heart
muscle
plasma
age
adult
pup

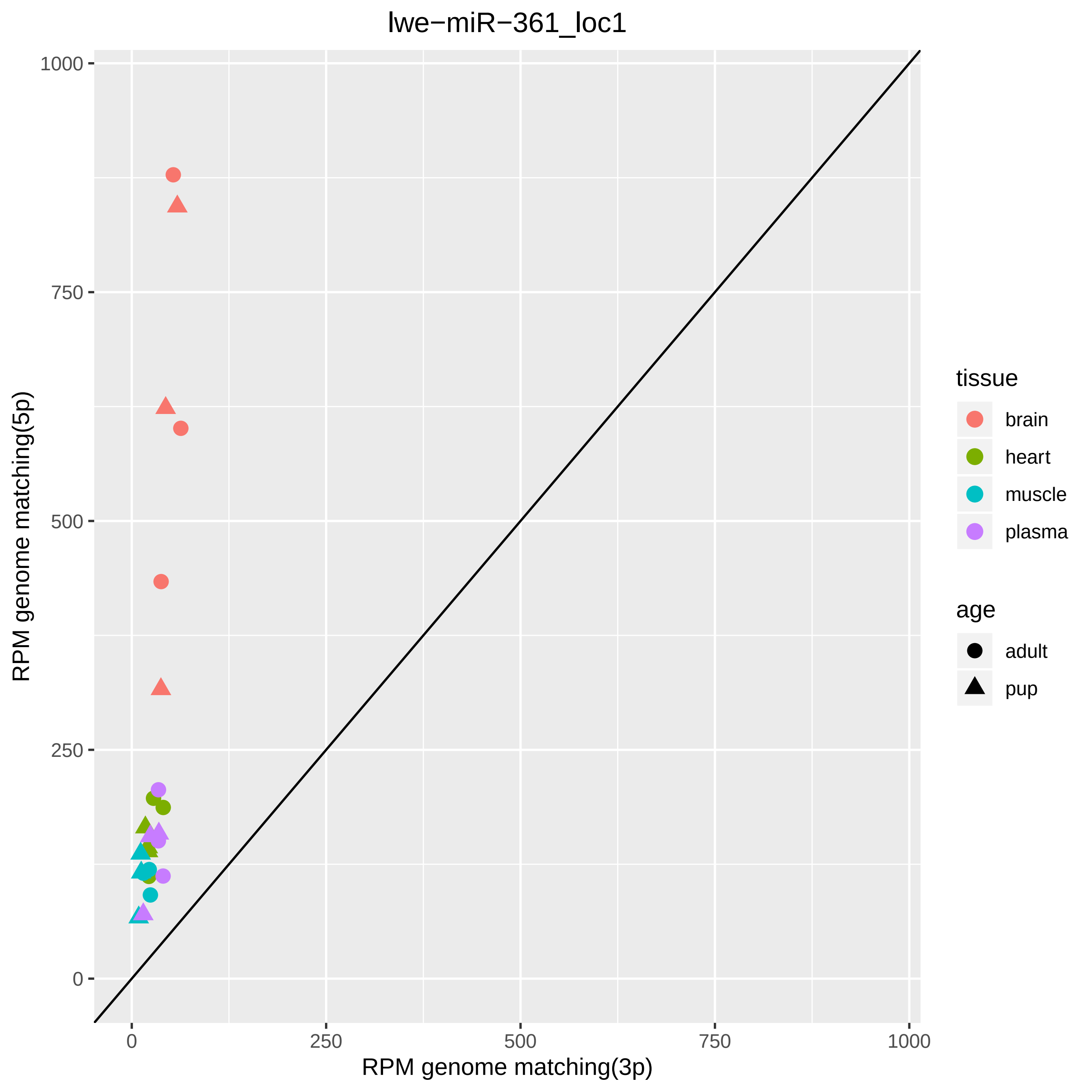

lwe−miR−361_loc1
1000
750
500
250
0
0
250
500
750
1000
RPM genome matching(5p)
RPM genome matching(3p)
tissue
brain
heart
muscle
plasma
age
adult
pup

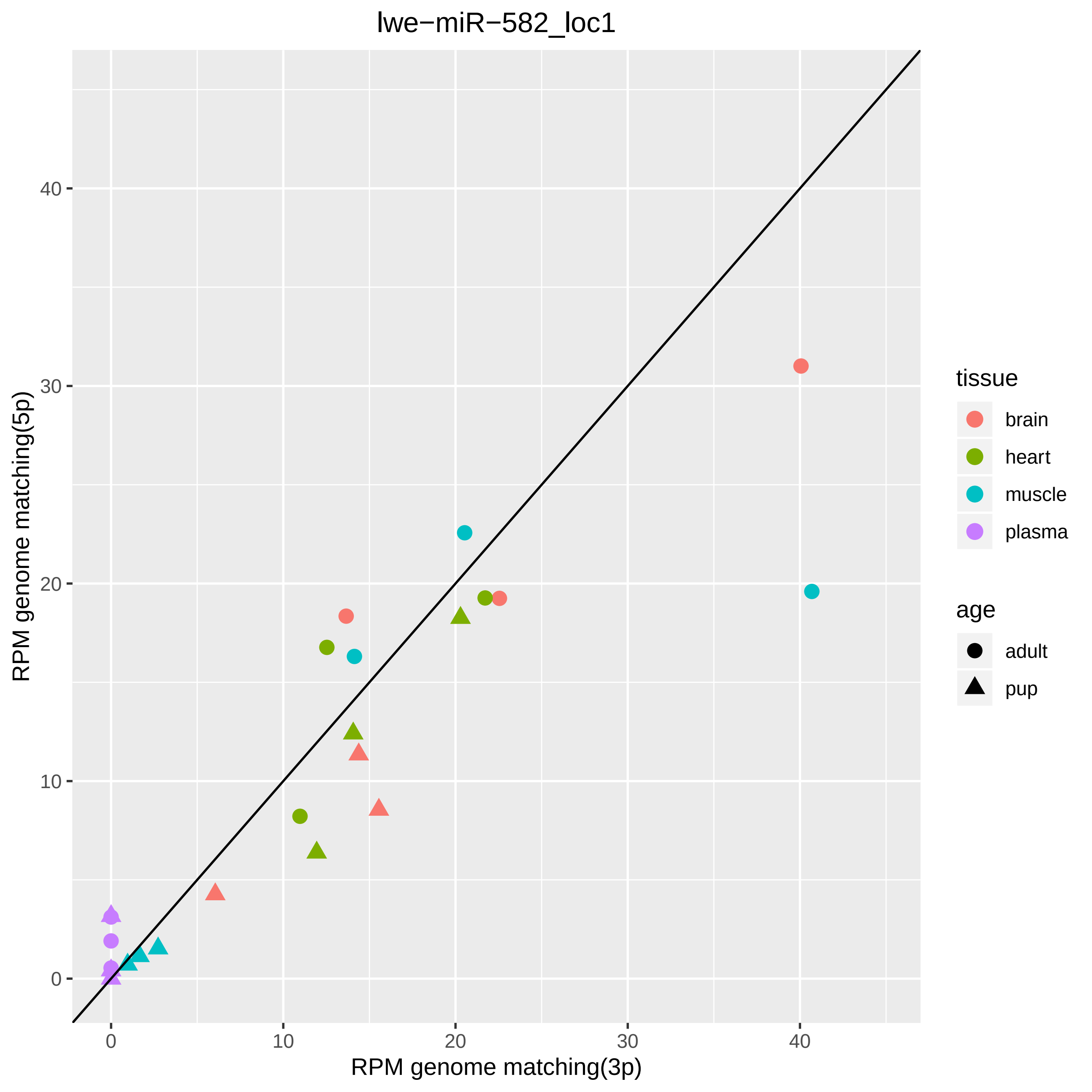
lwe−miR−582_loc1
RPM genome matching(5p)
RPM genome matching(3p)
0
10
20
30
40
0
10
20
30
40
tissue
brain
heart
muscle
plasma
age
adult
pup

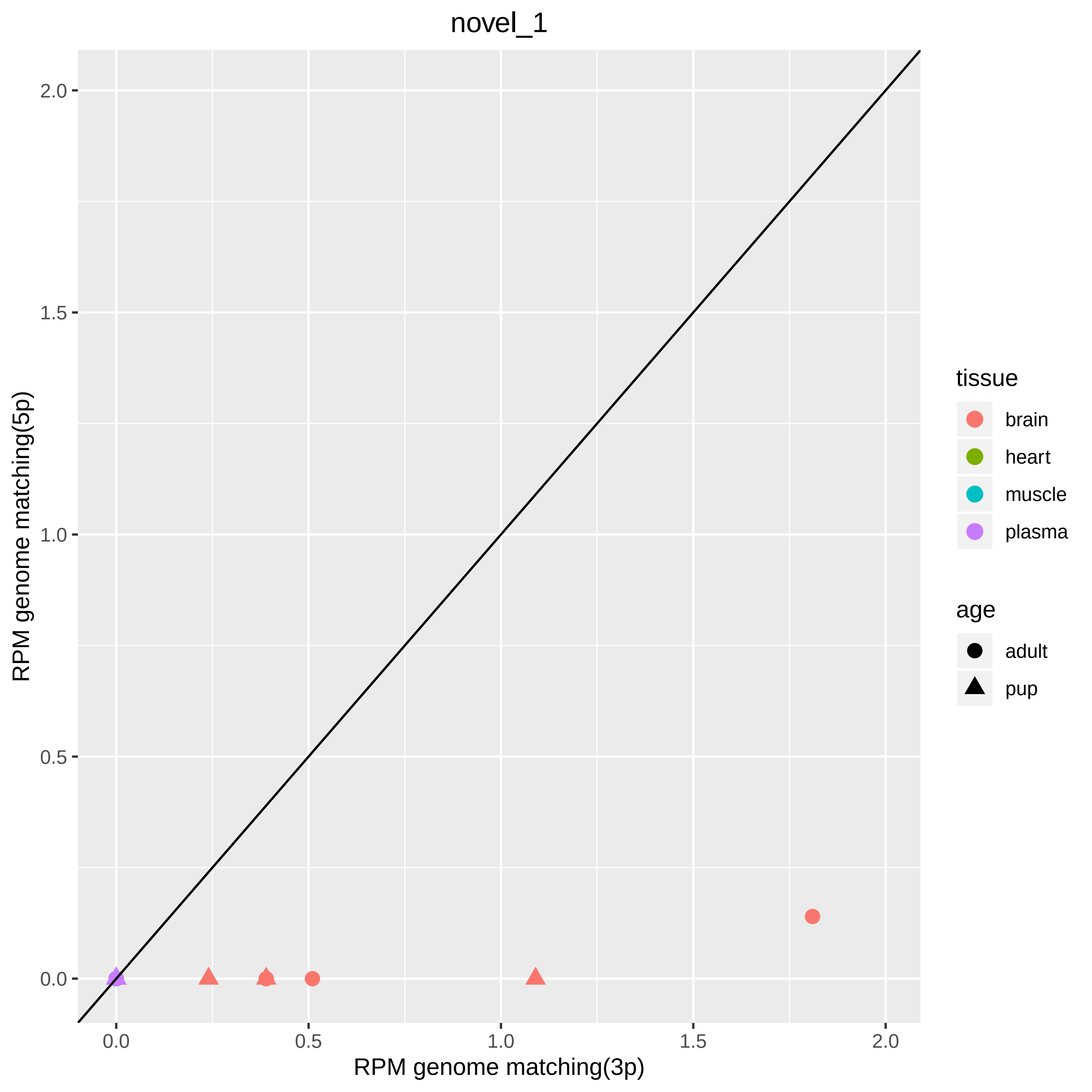
novel_1
RPM genome matching(5p)
RPM genome matching(3p)
0.0
0.5
1.0
1.5
2.0
tissue
brain
heart
muscle
plasma
age
adult
pup

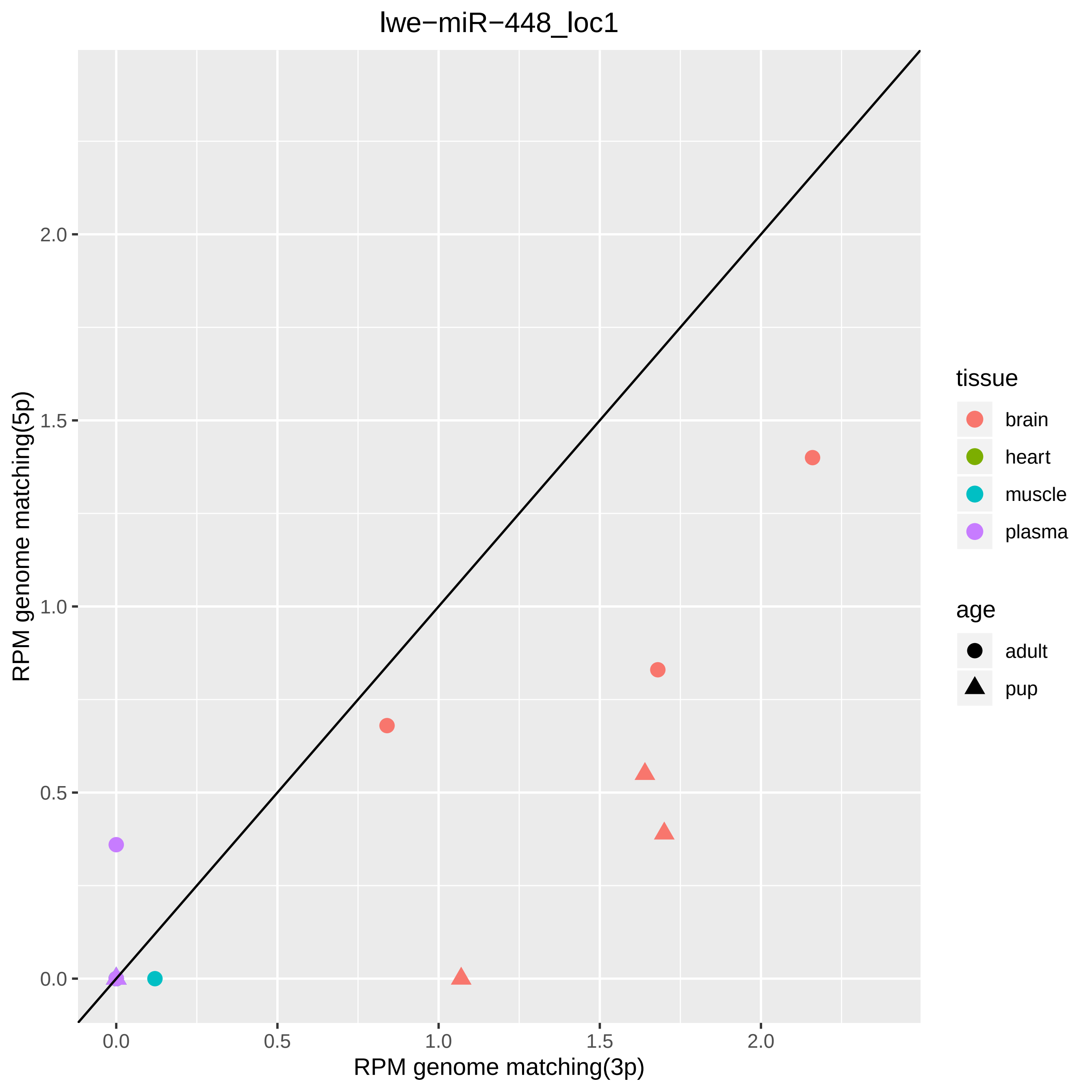
lwe−miR−448_loc1
RPM genome matching(5p)
RPM genome matching(3p)
0.0
0.5
1.0
1.5
2.0
tissue
brain
heart
muscle
plasma
age
adult
pup

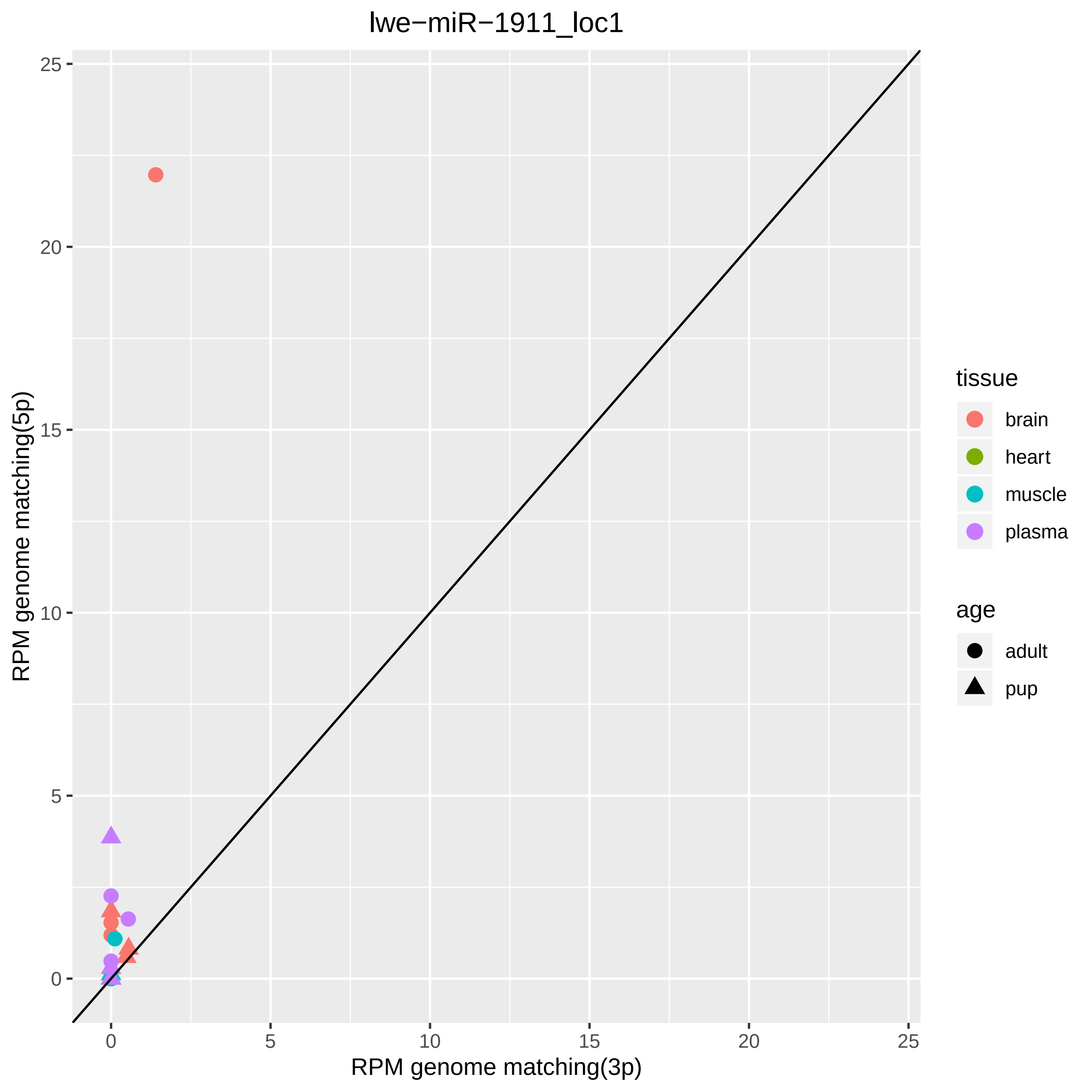
lwe−miR−1911_loc1
RPM genome matching(5p)
RPM genome matching(3p)
0
5
10
15
20
25
tissue
brain
heart
muscle
plasma
age
adult
pup

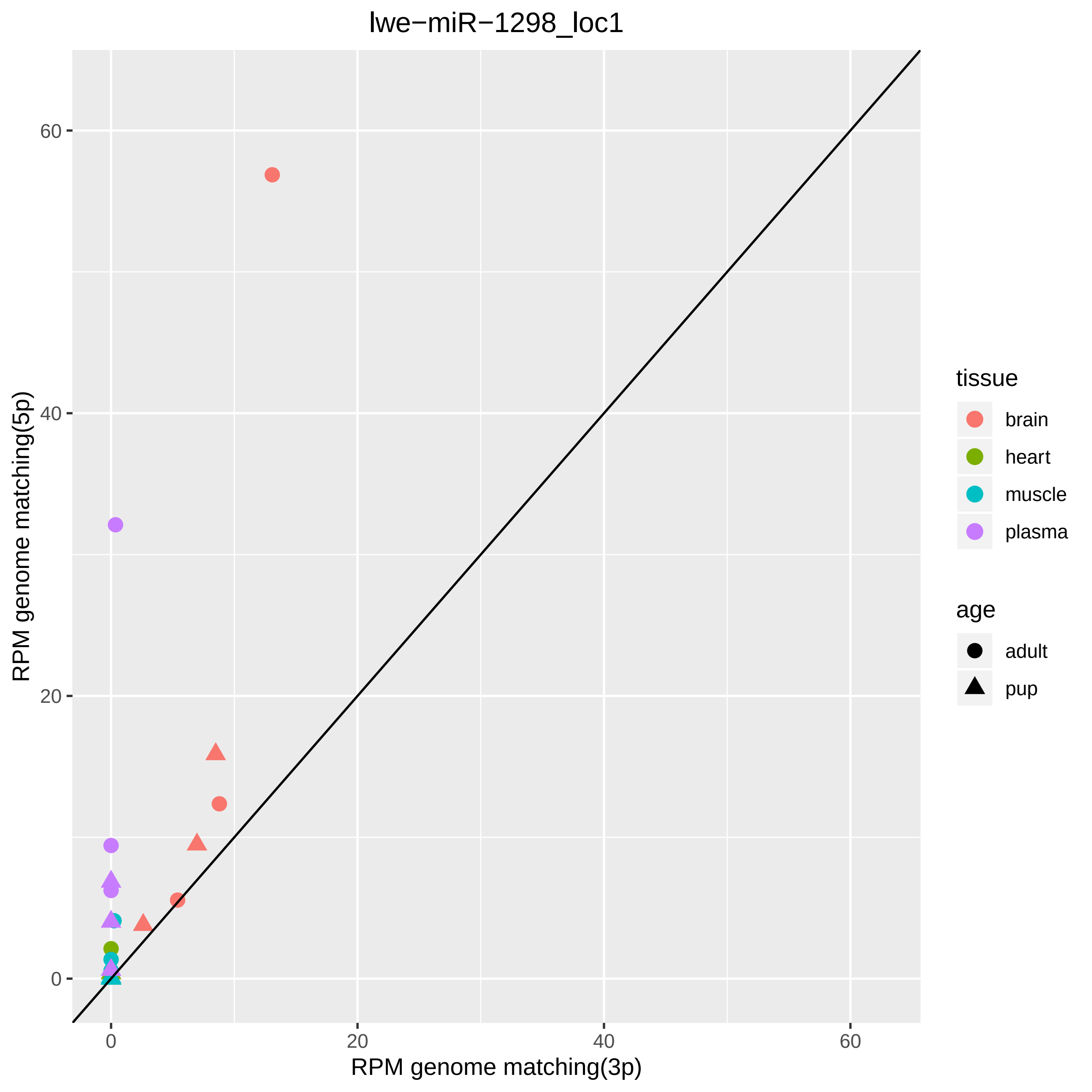
lwe−miR−1298_loc1
RPM genome matching(5p)
RPM genome matching(3p)
0
20
40
60
0
20
40
60
tissue
brain
heart
muscle
plasma
age
adult
pup

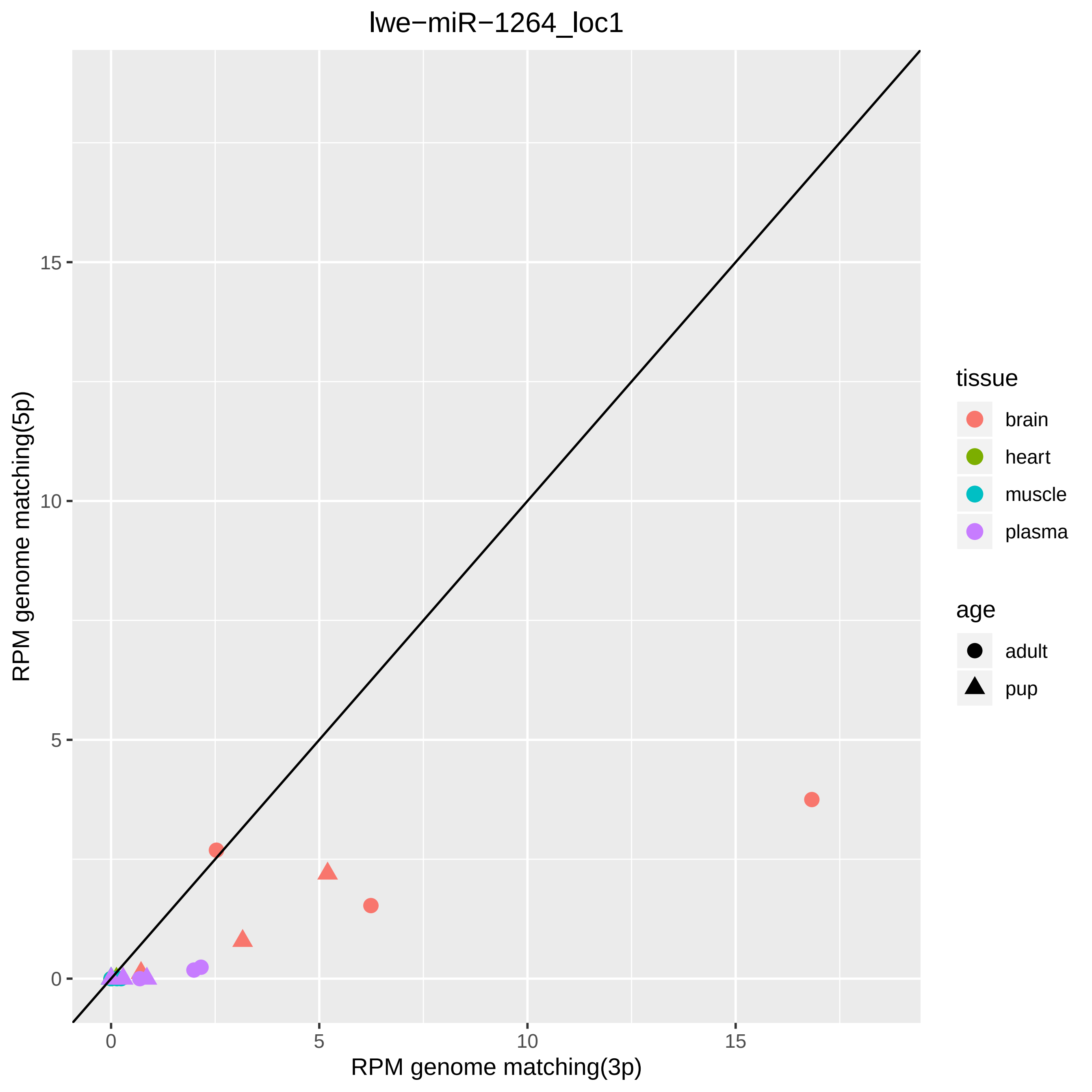

lwe−miR−1264_loc1
RPM genome matching(5p)
RPM genome matching(3p)
0
5
10
15
0
5
10
15
tissue
brain
heart
muscle
plasma
age
adult
pup

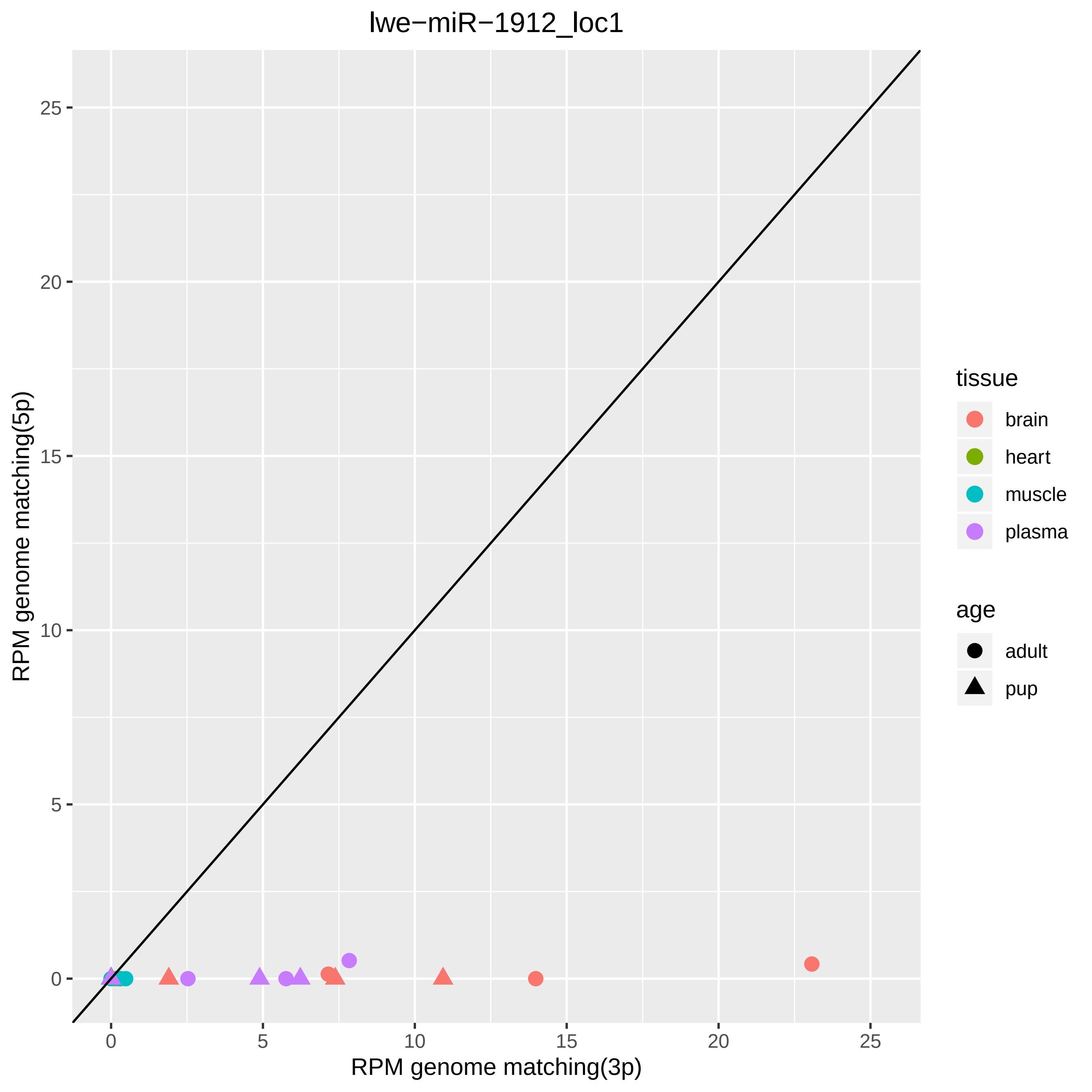

lwe−miR−1912_loc1
RPM genome matching(5p)
RPM genome matching(3p)
0
5
10
15
20
25
tissue
brain
heart
muscle
plasma
age
adult
pup

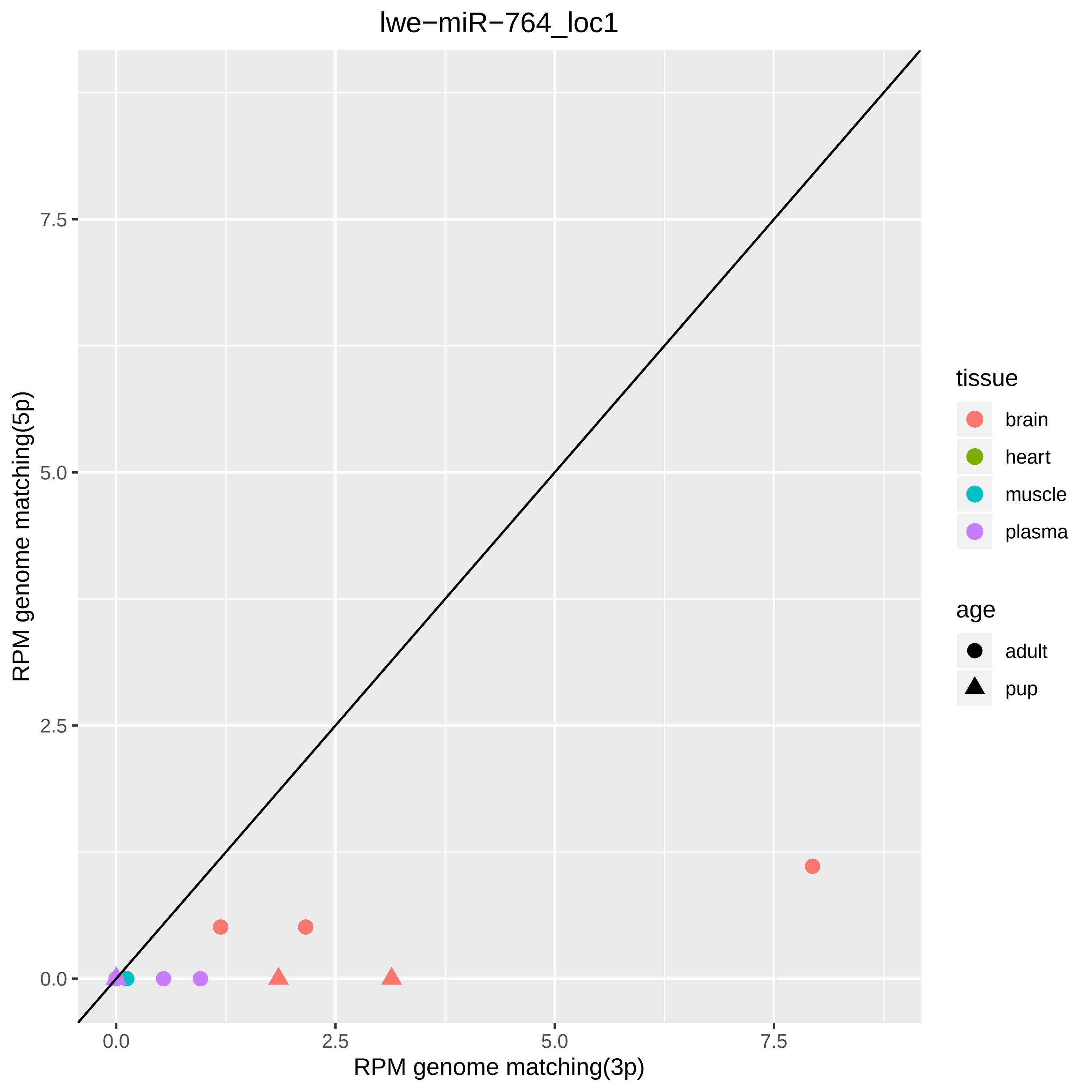

lwe−miR−764_loc1
RPM genome matching(5p)
RPM genome matching(3p)
0.0
2.5
5.0
7.5
tissue
brain
heart
muscle
plasma
age
adult
pup

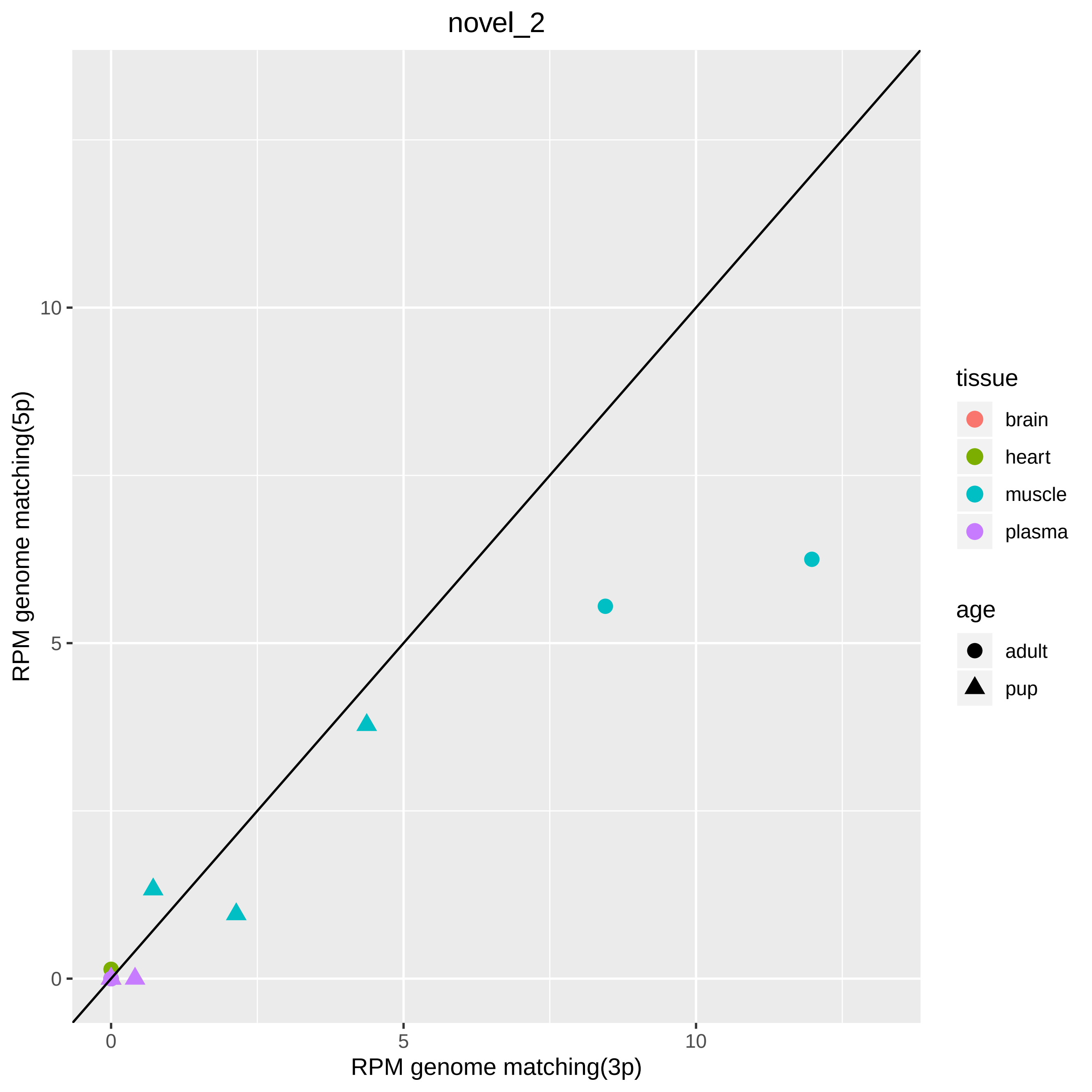
novel_2
RPM genome matching(5p)
RPM genome matching(3p)
0
5
10
0
5
10
tissue
brain
heart
muscle
plasma
age
adult
pup

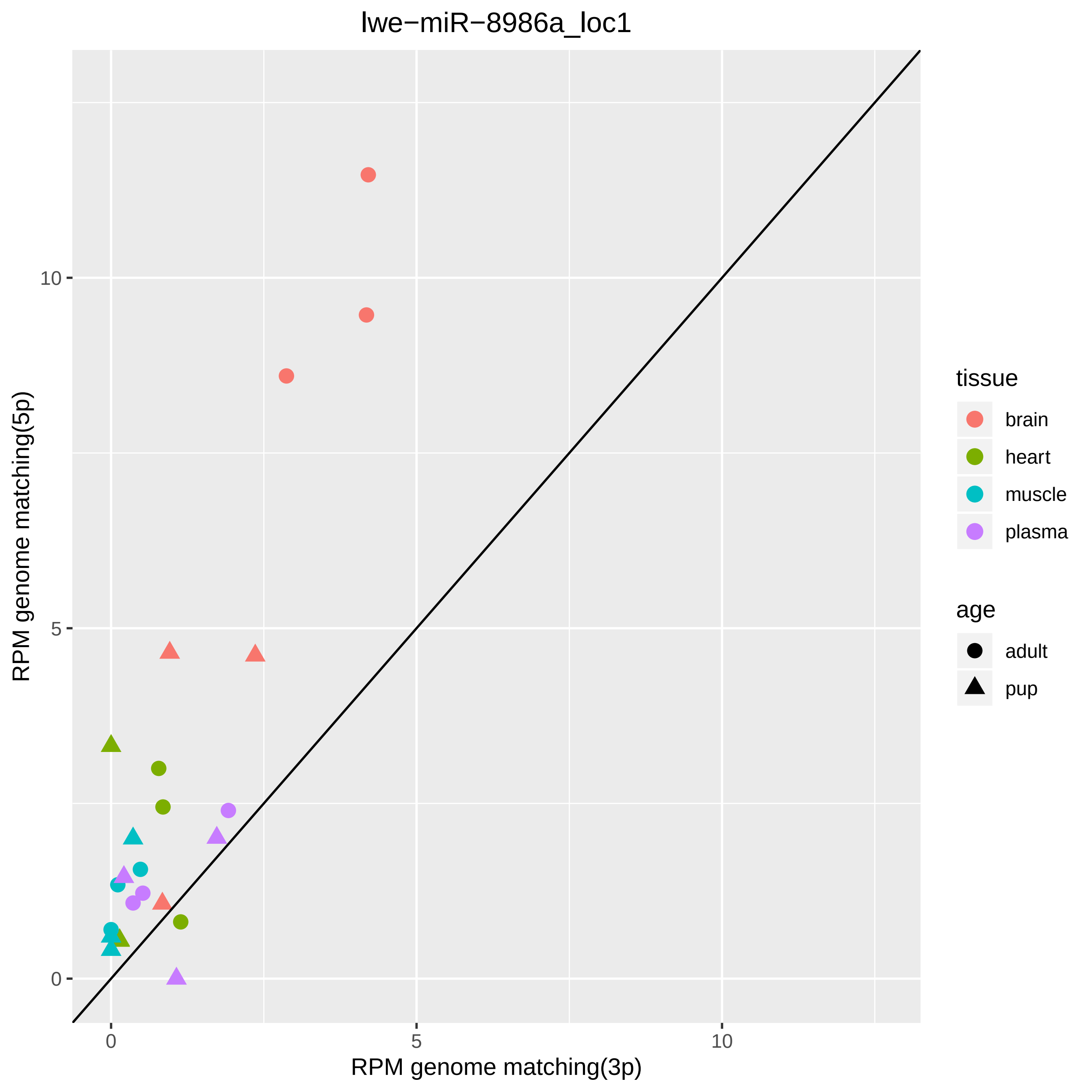

lwe−miR−8986a_loc1
RPM genome matching(5p)
RPM genome matching(3p)
0
5
10
0
5
10
tissue
brain
heart
muscle
plasma
age
adult
pup

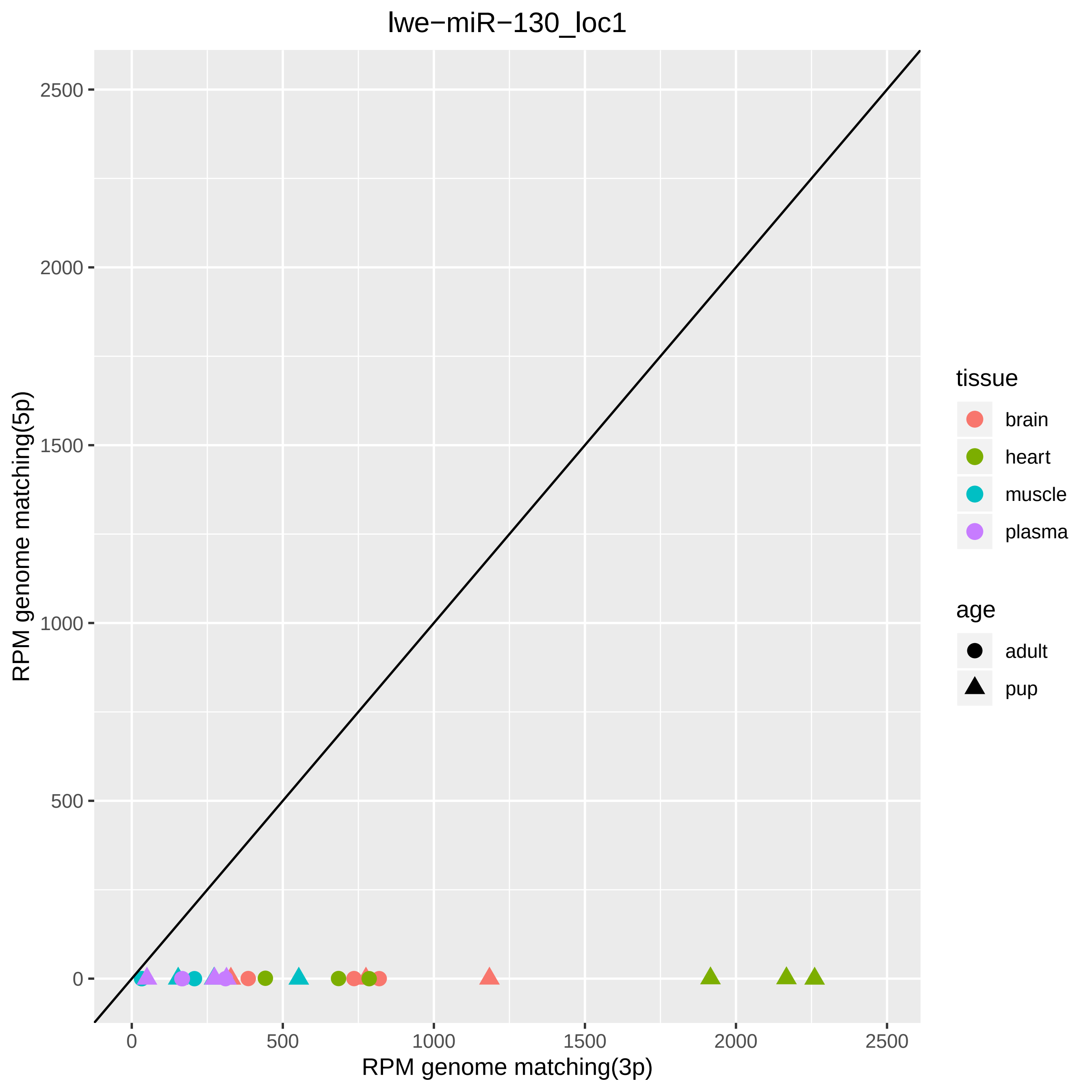

lwe−miR−130_loc1
RPM genome matching(5p)
RPM genome matching(3p)
0
500
1000
1500
2000
2500
tissue
brain
heart
muscle
plasma
age
adult
pup

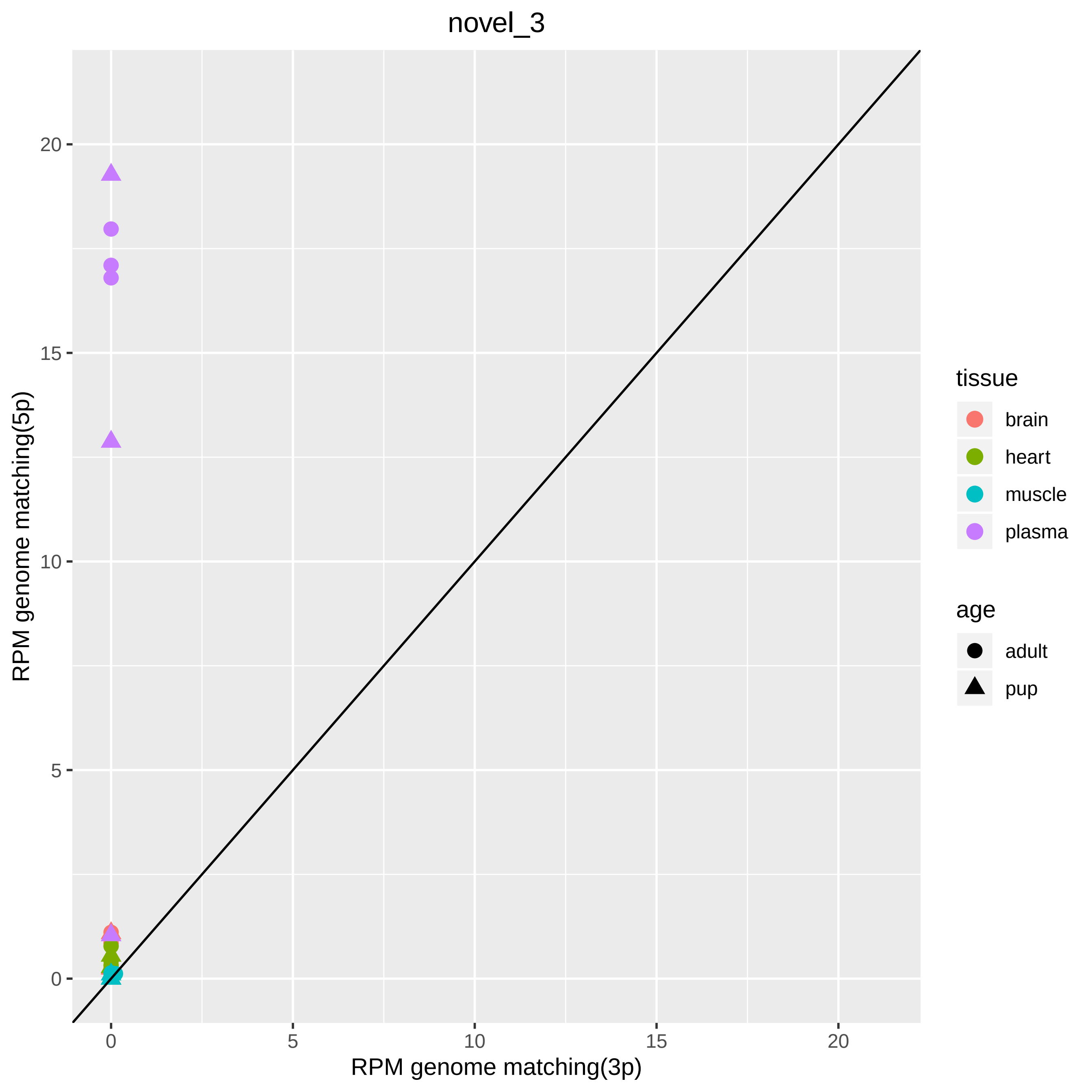

novel_3
RPM genome matching(5p)
RPM genome matching(3p)
0
5
10
15
20
tissue
brain
heart
muscle
plasma
age
adult
pup

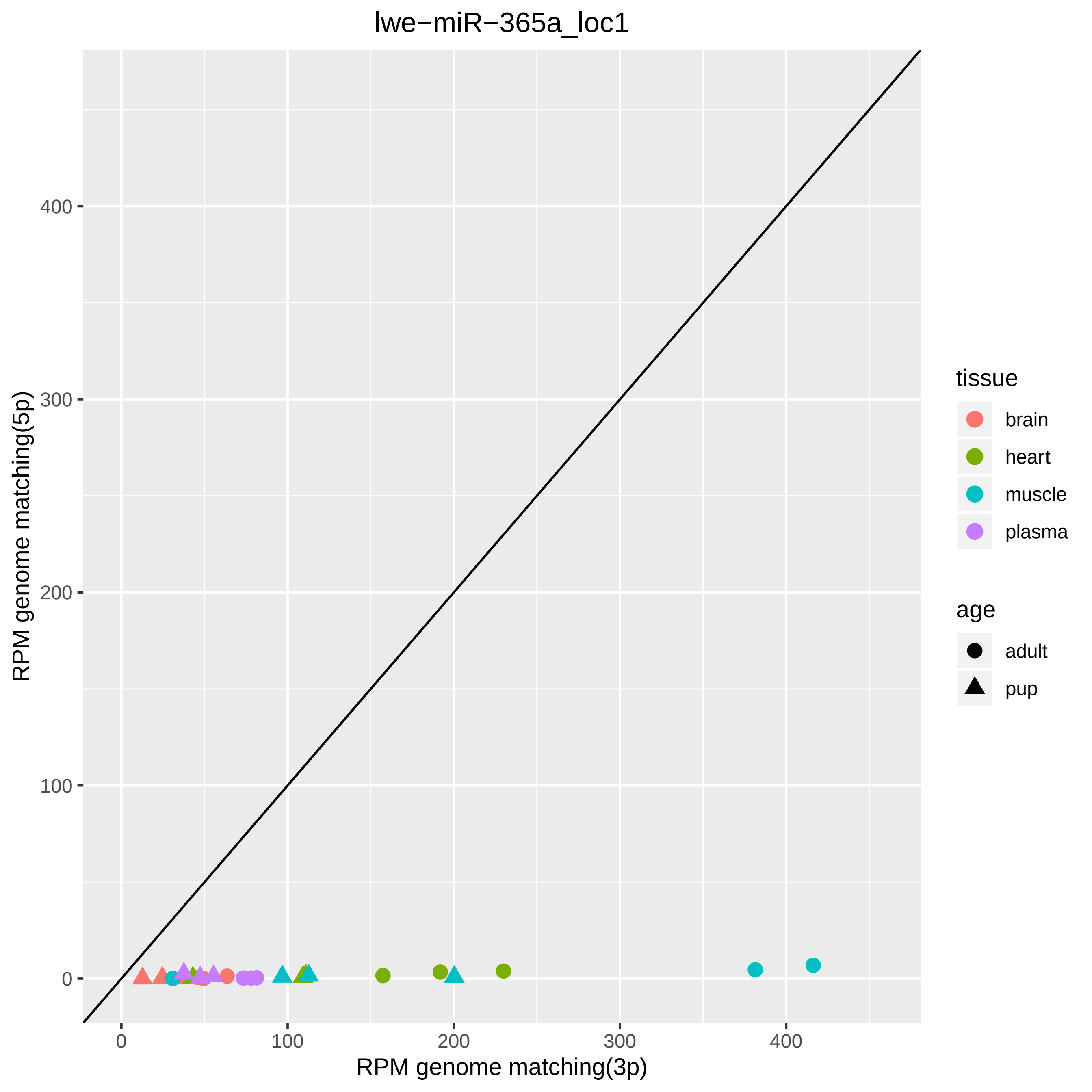

lwe−miR−365a_loc1
RPM genome matching(5p)
RPM genome matching(3p)
0
100
200
300
400
tissue
brain
heart
muscle
plasma
age
adult
pup

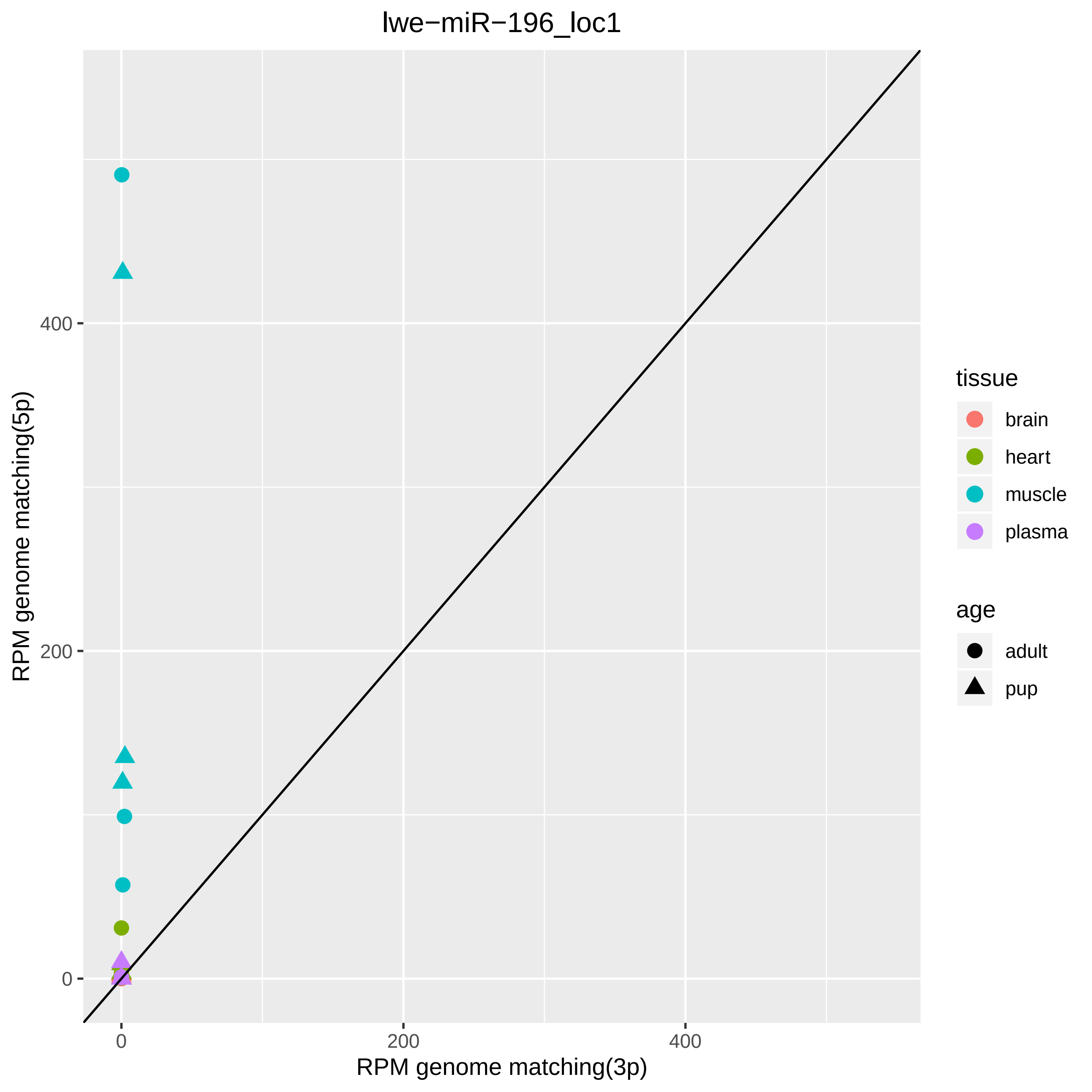
lwe−miR−196_loc1
RPM genome matching(5p)
RPM genome matching(3p)
0
200
400
0
200
400
tissue
brain
heart
muscle
plasma
age
adult
pup

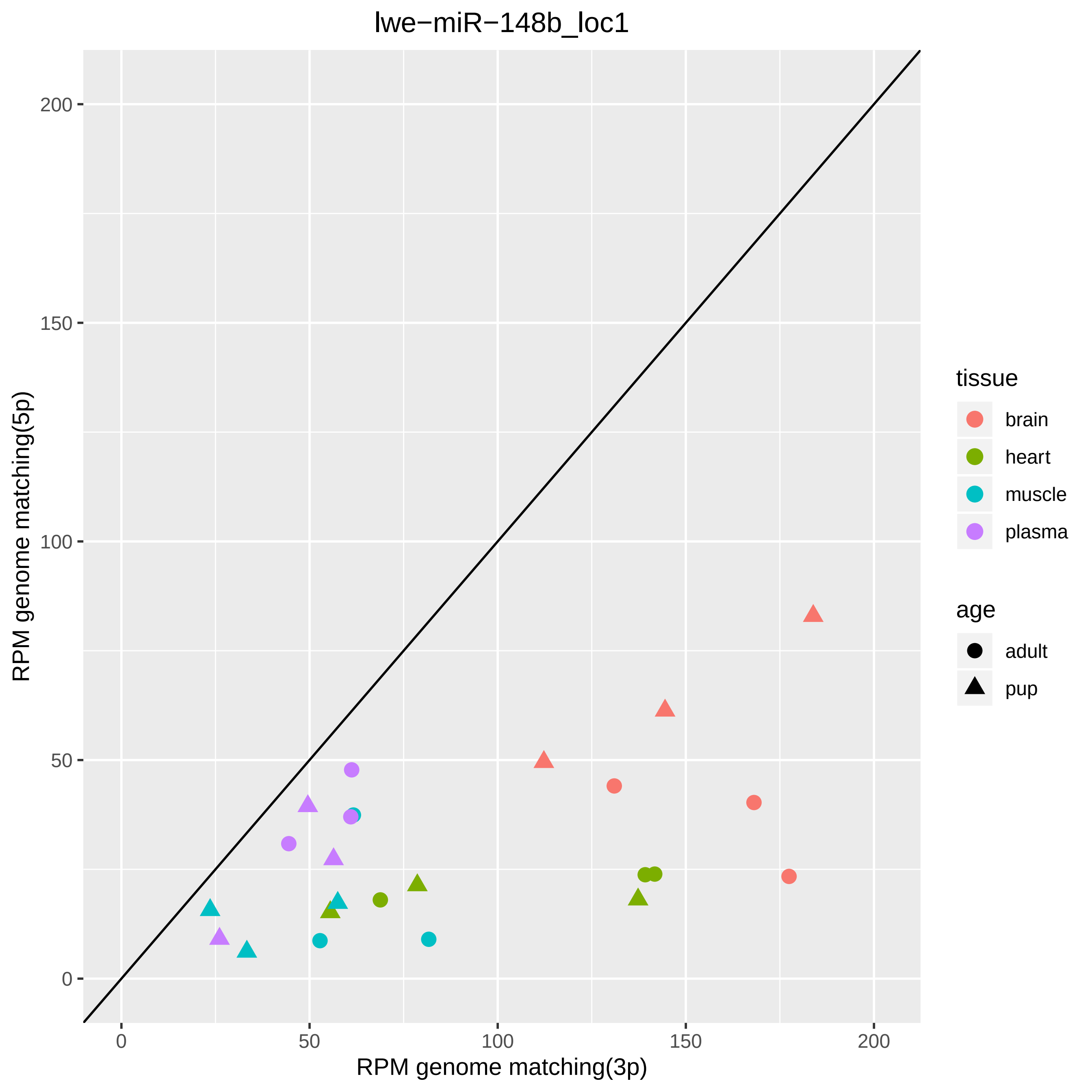
lwe−miR−148b_loc1
RPM genome matching(5p)
RPM genome matching(3p)
0
50
100
150
200
tissue
brain
heart
muscle
plasma
age
adult
pup

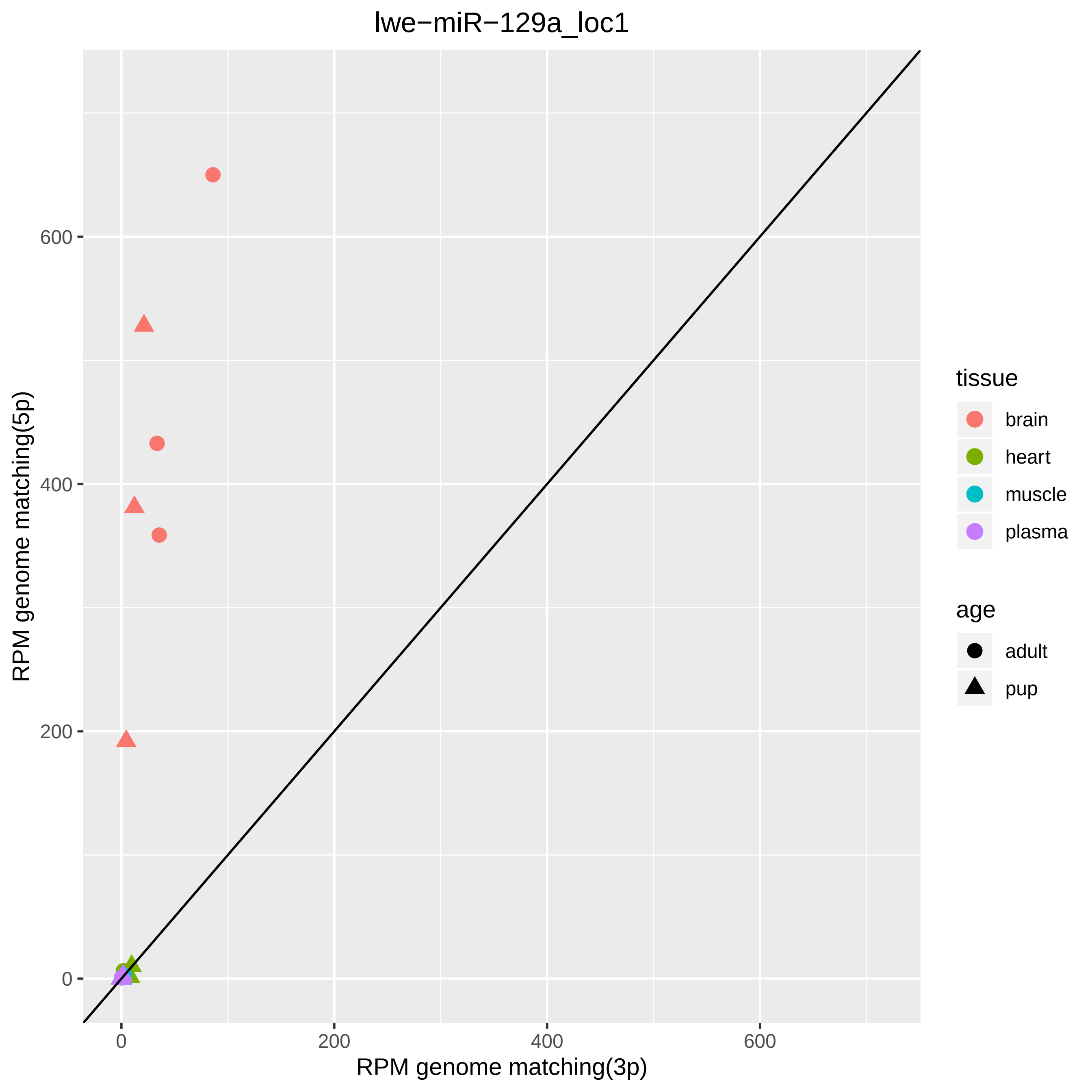
lwe−miR−129a_loc1
RPM genome matching(5p)
RPM genome matching(3p)
0
200
400
600
tissue
brain
heart
muscle
plasma
age
adult
pup

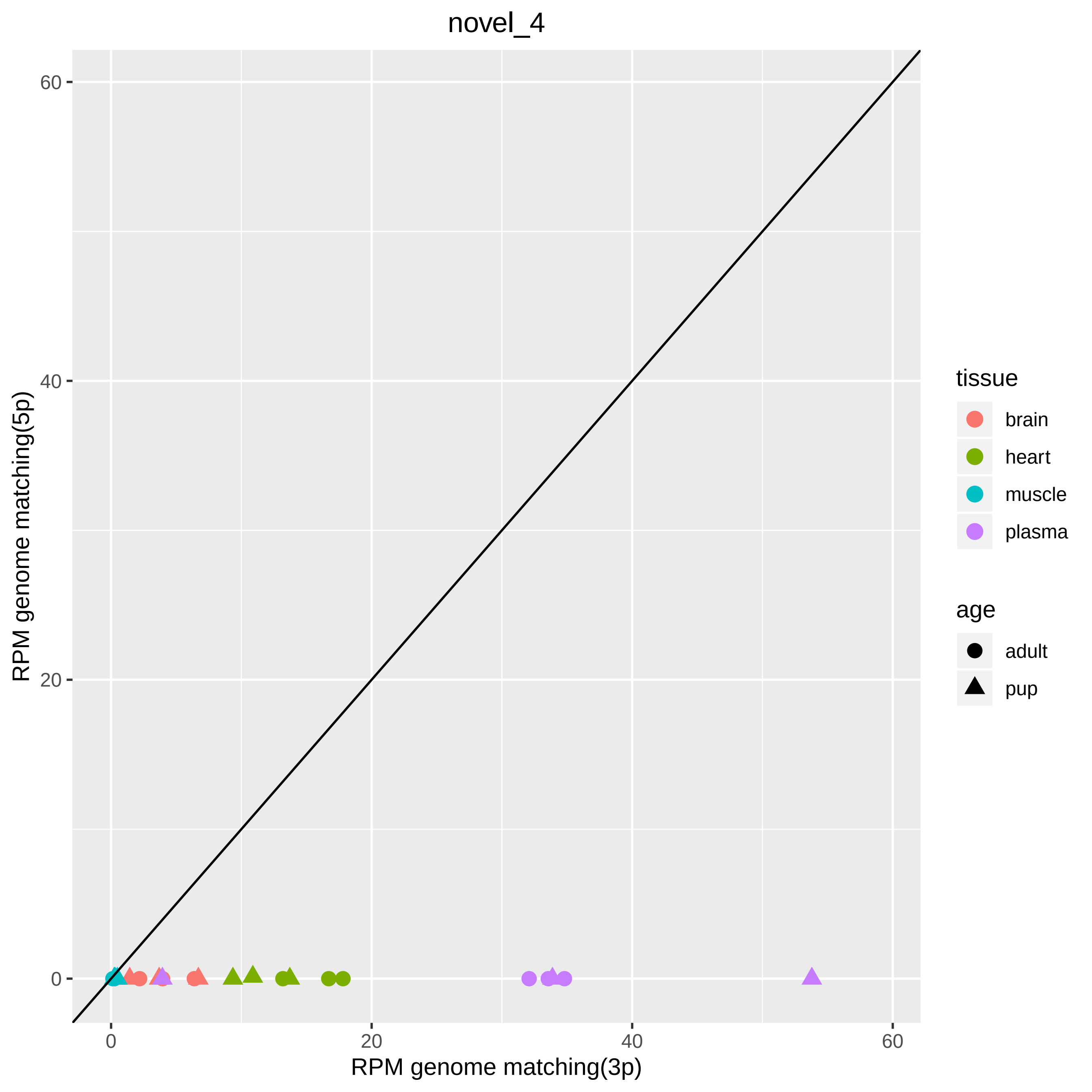

novel_4
RPM genome matching(5p)
RPM genome matching(3p)
0
20
40
60
0
20
40
60
tissue
brain
heart
muscle
plasma
age
adult
pup

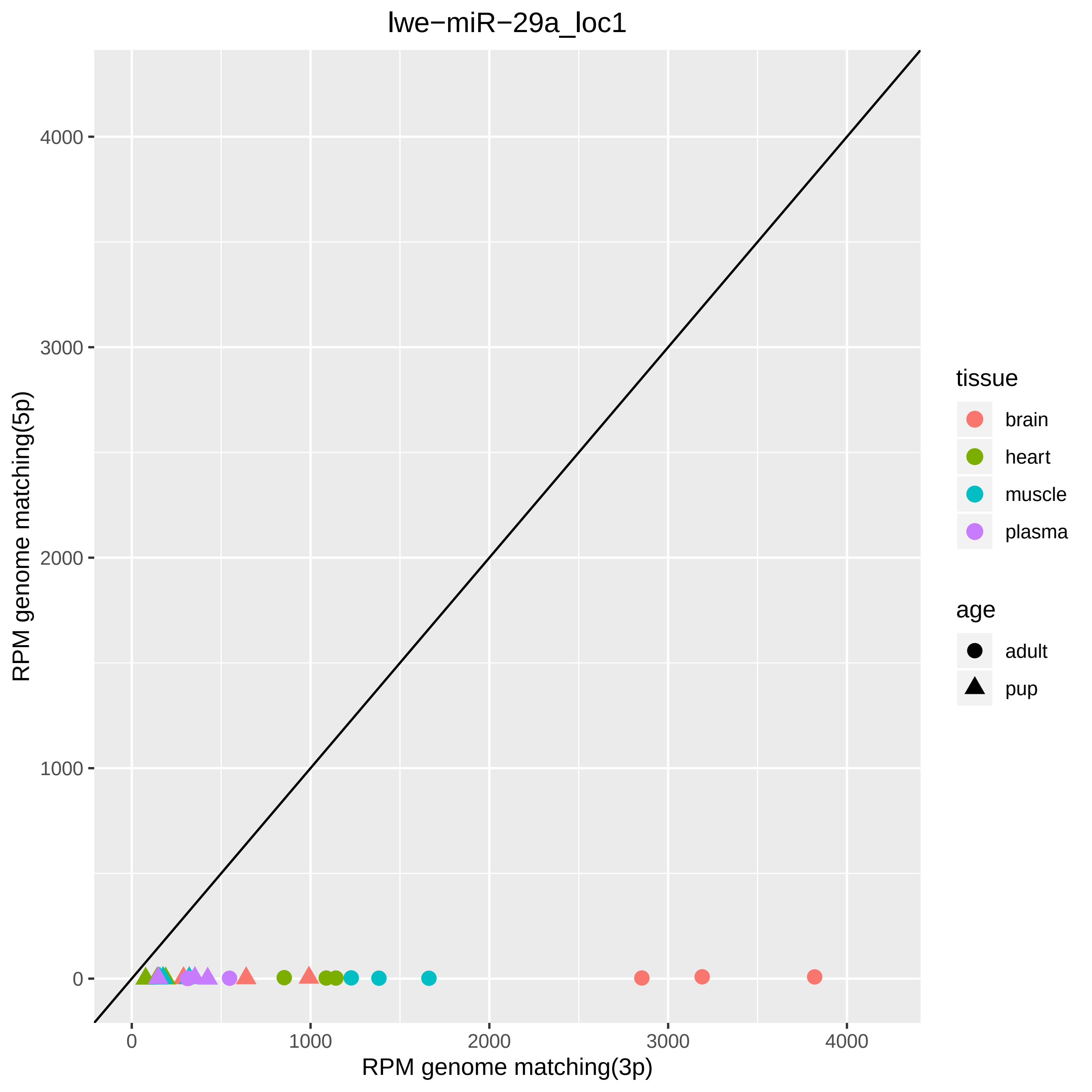

lwe−miR−29a_loc1
RPM genome matching(5p)
RPM genome matching(3p)
0
1000
2000
3000
4000
tissue
brain
heart
muscle
plasma
age
adult
pup

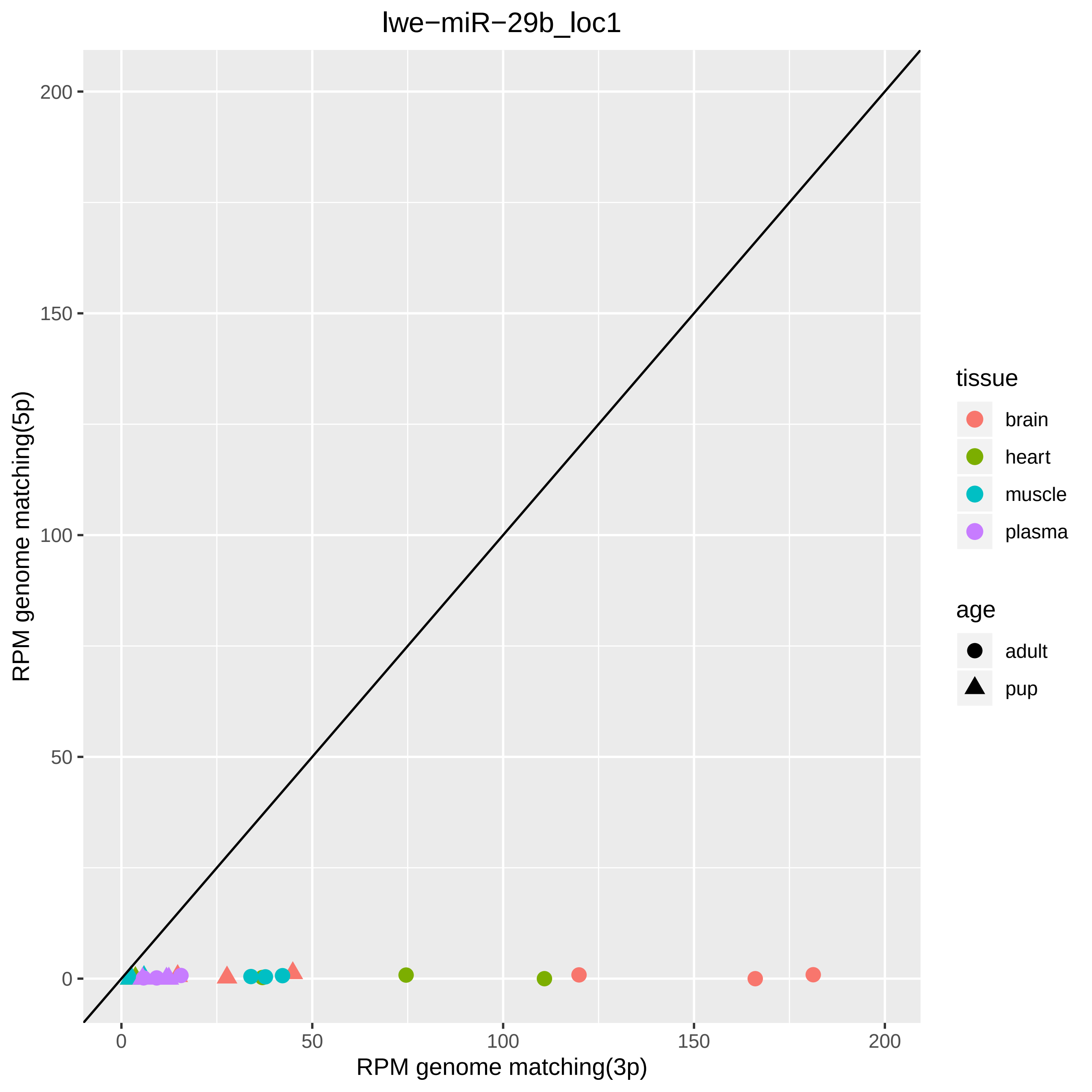

lwe−miR−29b_loc1
RPM genome matching(5p)
RPM genome matching(3p)
0
50
100
150
200
tissue
brain
heart
muscle
plasma
age
adult
pup

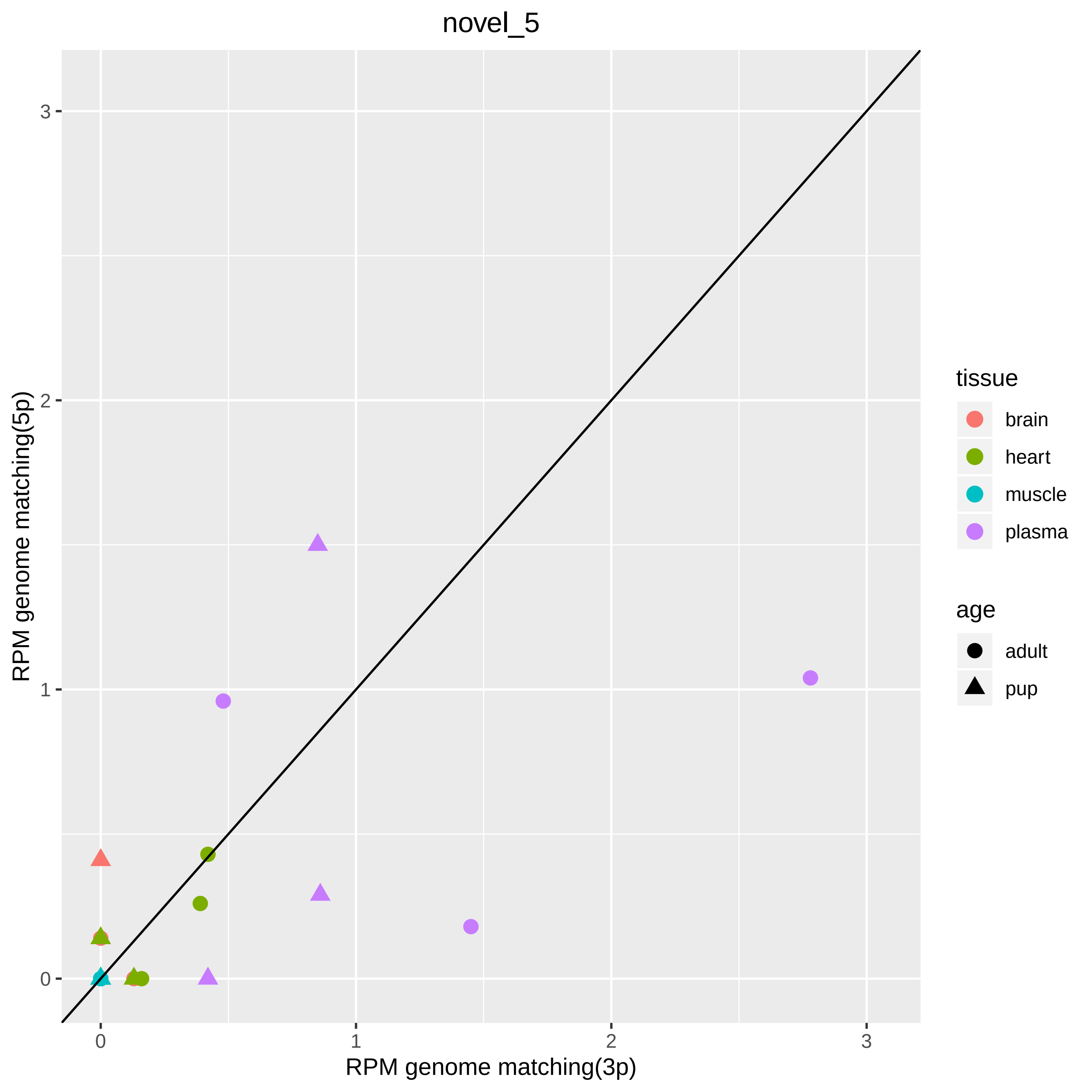

novel_5
RPM genome matching(5p)
RPM genome matching(3p)
0
1
2
3
0
1
2
3
tissue
brain
heart
muscle
plasma
age
adult
pup

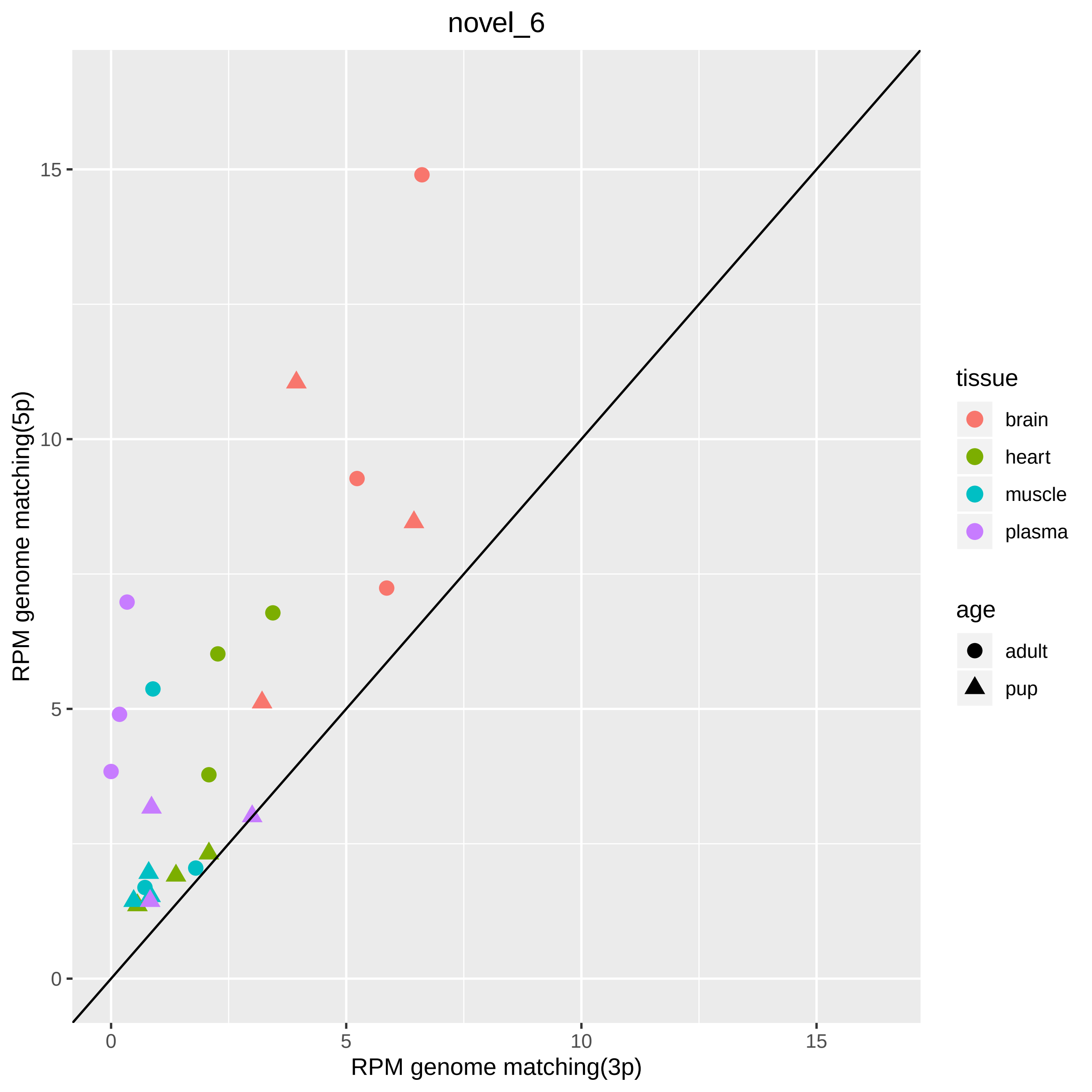
novel_6
RPM genome matching(5p)
RPM genome matching(3p)
0
5
10
15
0
5
10
15
tissue
brain
heart
muscle
plasma
age
adult
pup

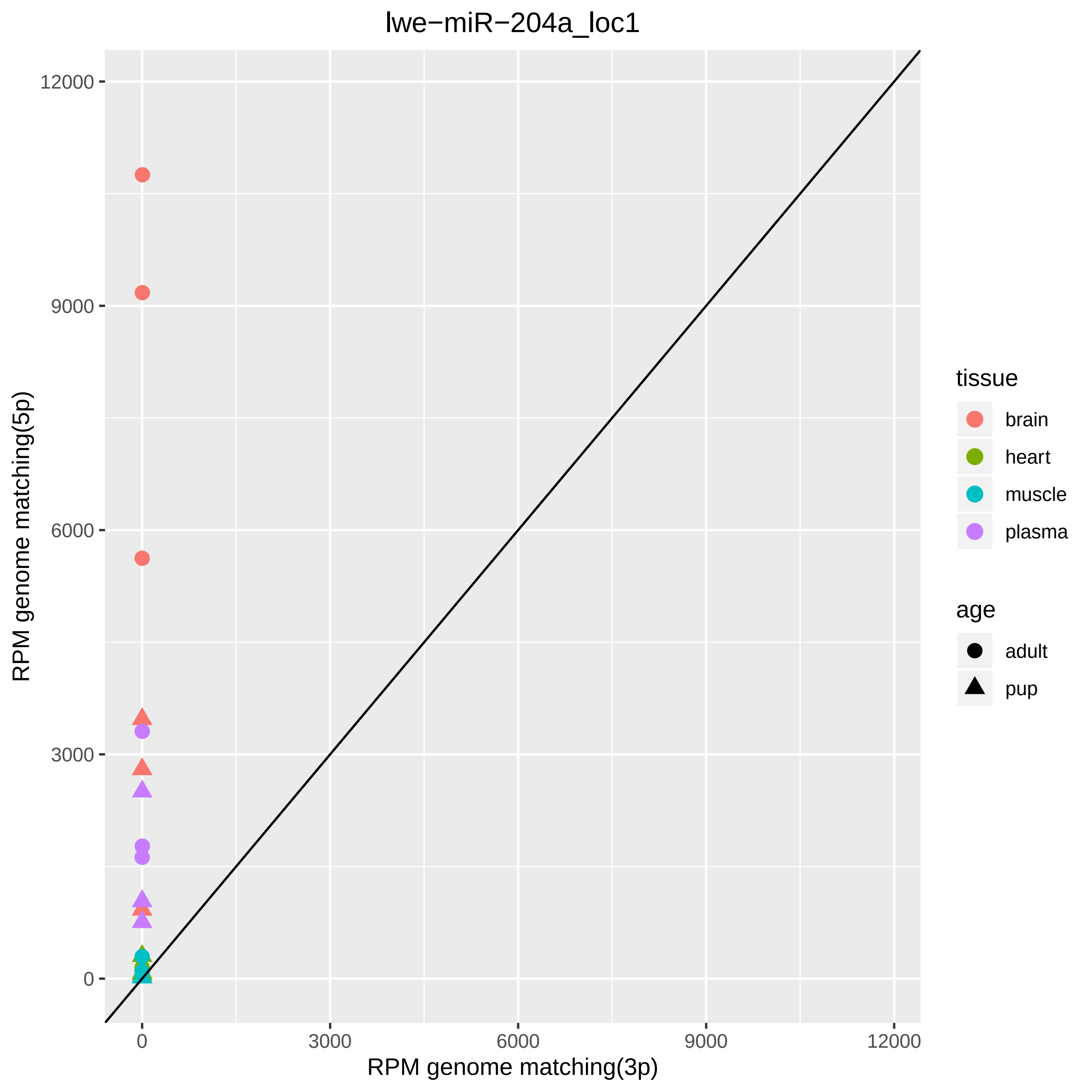
lwe−miR−204a_loc1
RPM genome matching(5p)
RPM genome matching(3p)
0
3000
6000
9000
12000
tissue
brain
heart
muscle
plasma
age
adult
pup

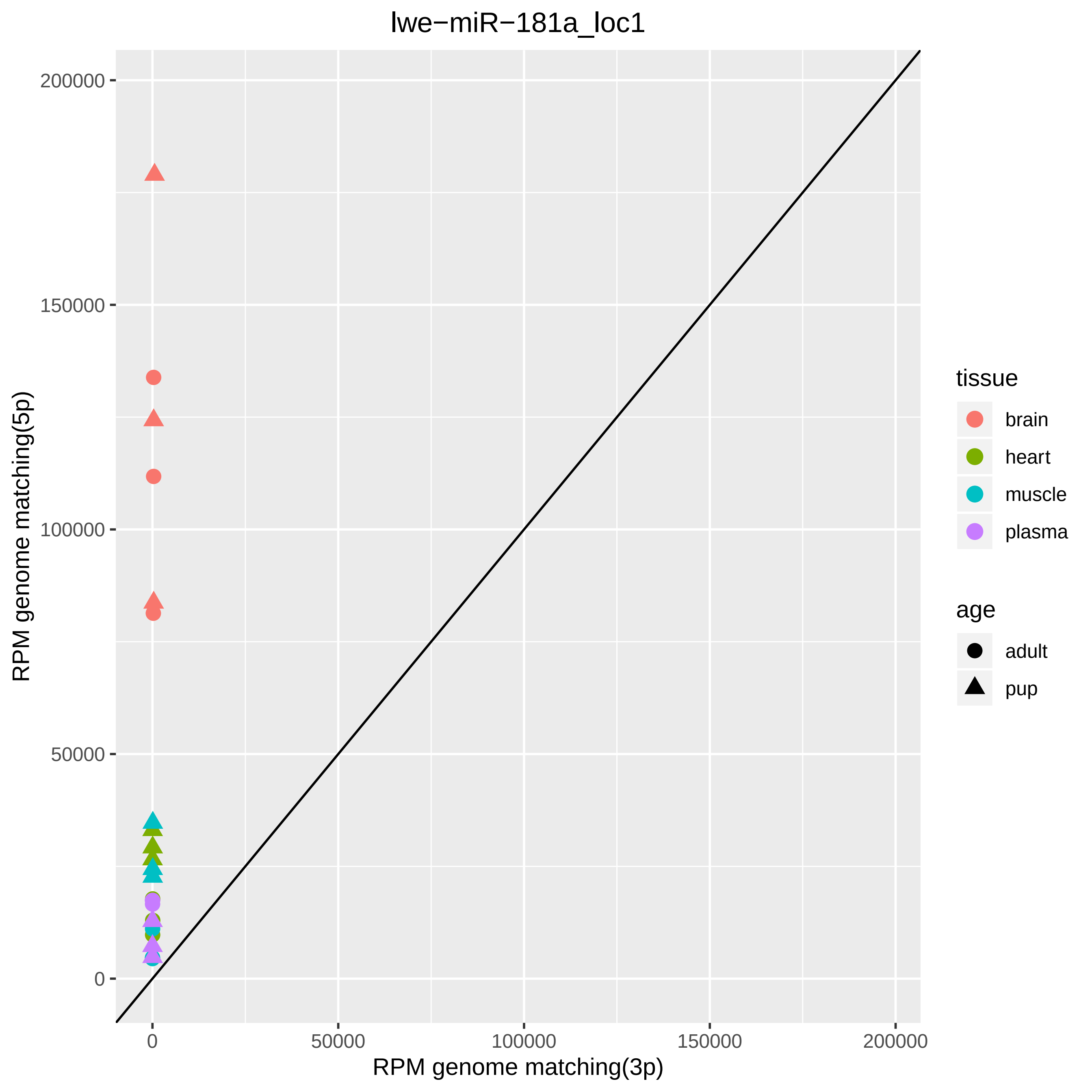
lwe−miR−181a_loc1
RPM genome matching(5p)
RPM genome matching(3p)
0
50000
100000
150000
200000
tissue
brain
heart
muscle
plasma
age
adult
pup

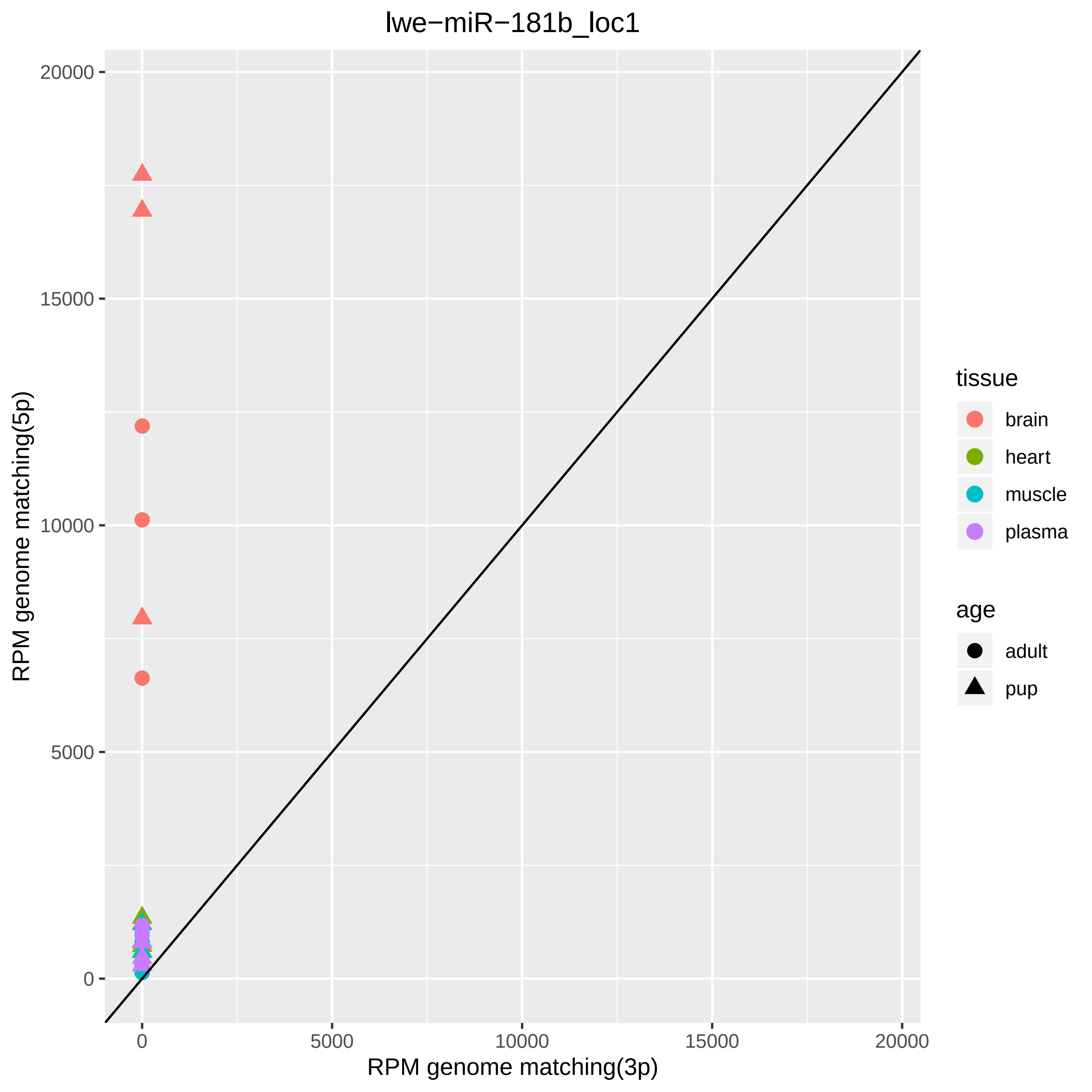
lwe−miR−181b_loc1
RPM genome matching(5p)
RPM genome matching(3p)
20000
15000
10000
5000
0
0
5000
10000
15000
20000
tissue
brain
heart
muscle
plasma
age
adult
pup

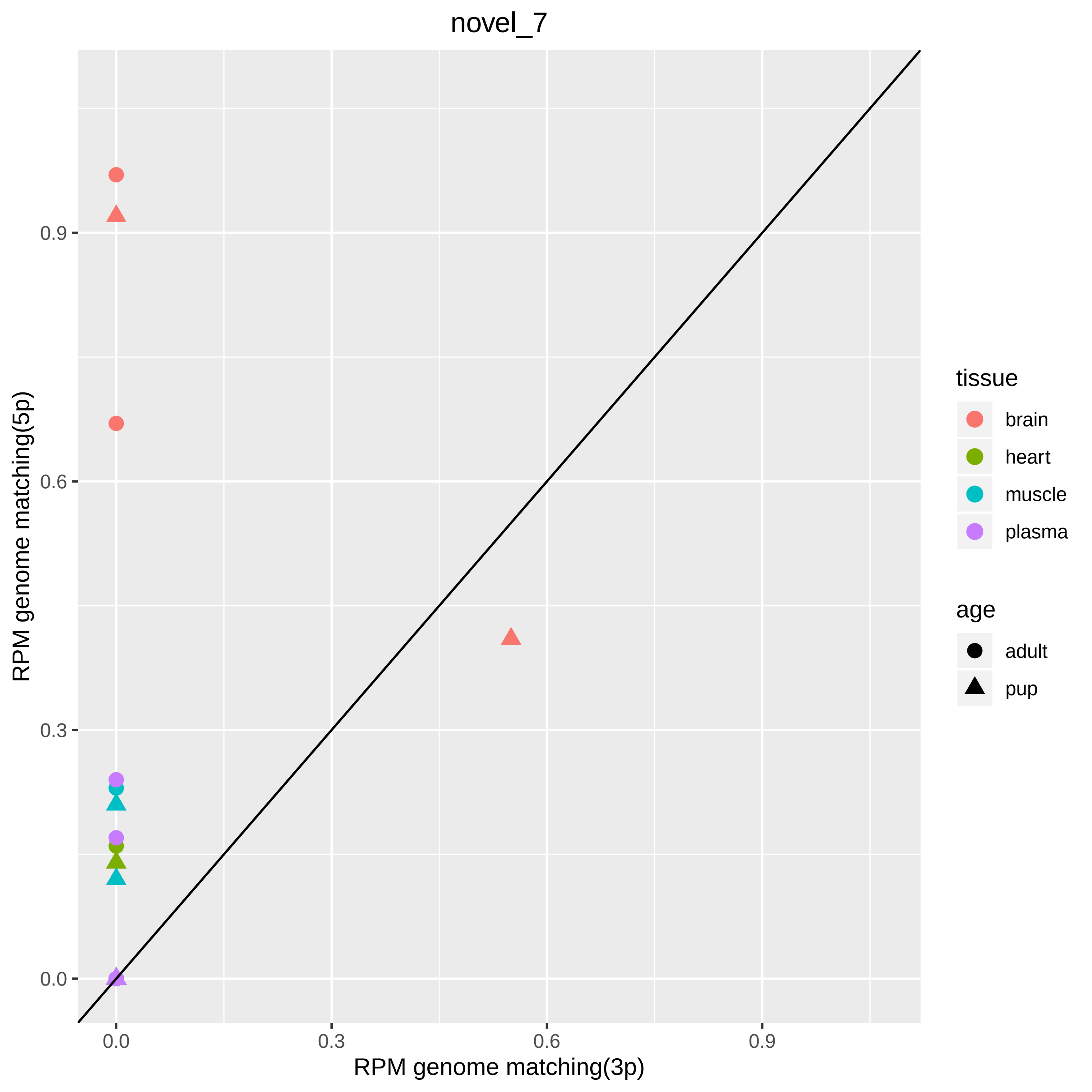
novel_7
RPM genome matching(5p)
RPM genome matching(3p)
0.0
0.3
0.6
0.9
tissue
brain
heart
muscle
plasma
age
adult
pup

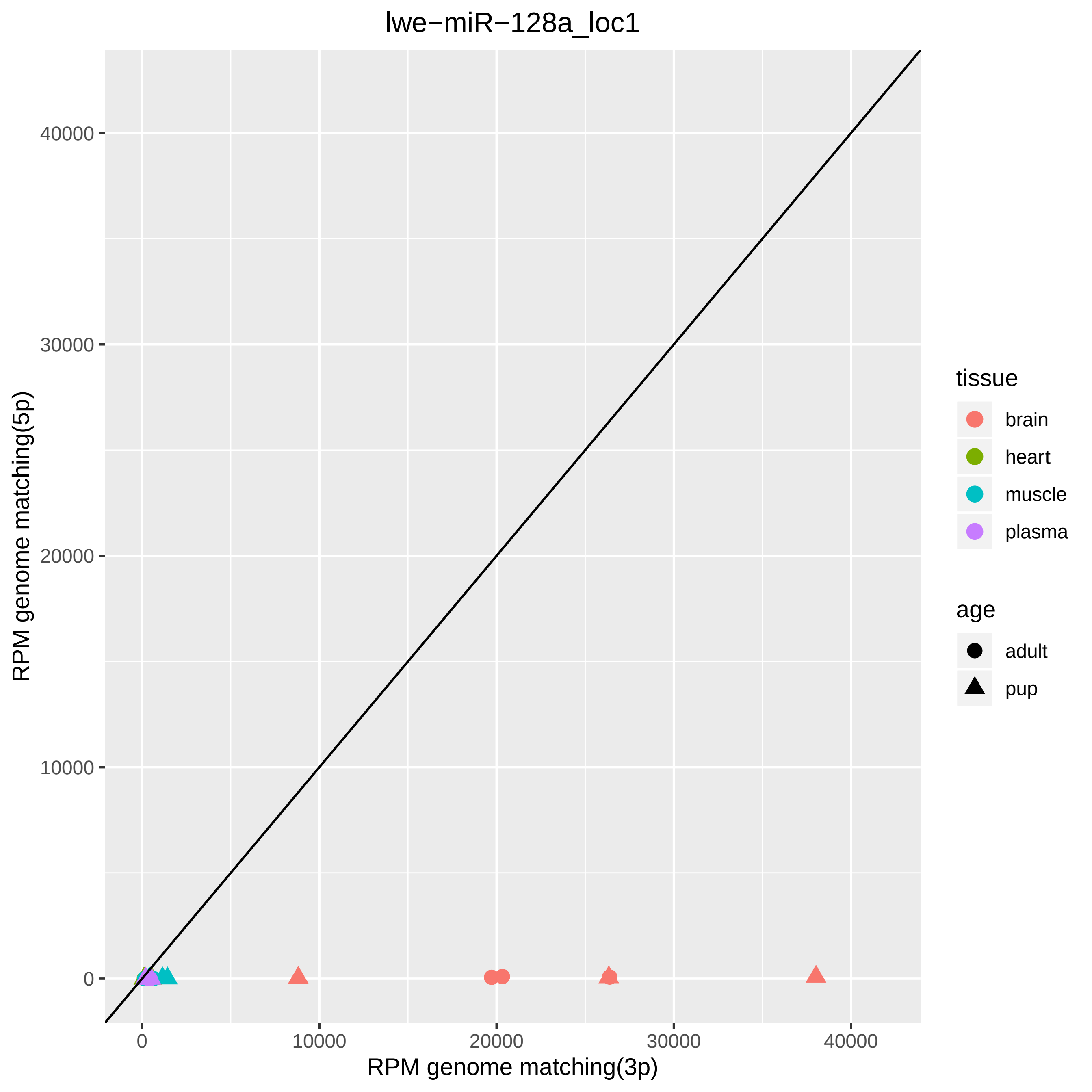
lwe−miR−128a_loc1
RPM genome matching(5p)
RPM genome matching(3p)
0
10000
20000
30000
40000
tissue
brain
heart
muscle
plasma
age
adult
pup

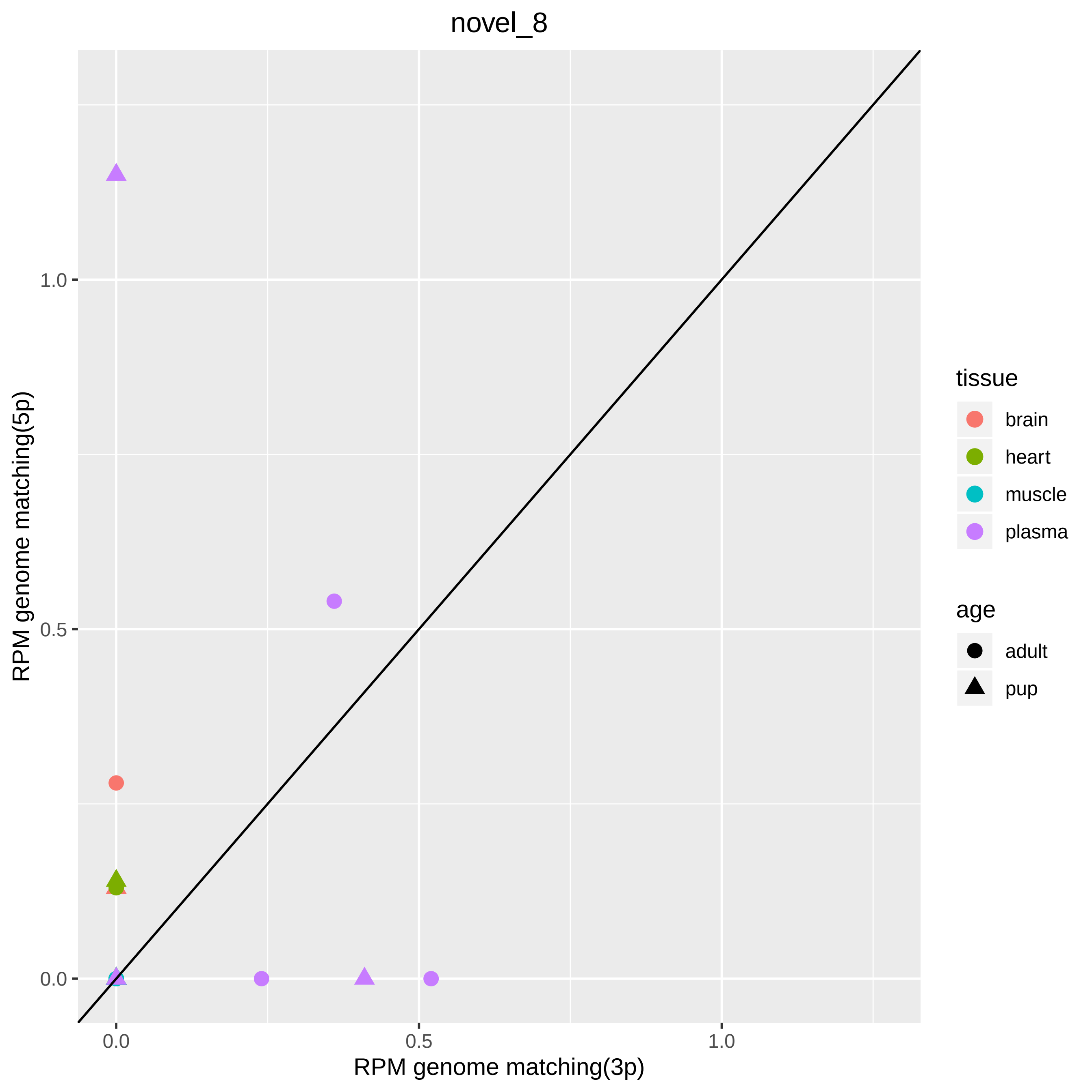

novel_8
RPM genome matching(5p)
RPM genome matching(3p)
0.0
0.5
1.0
0.0
0.5
1.0
tissue
brain
heart
muscle
plasma
age
adult
pup

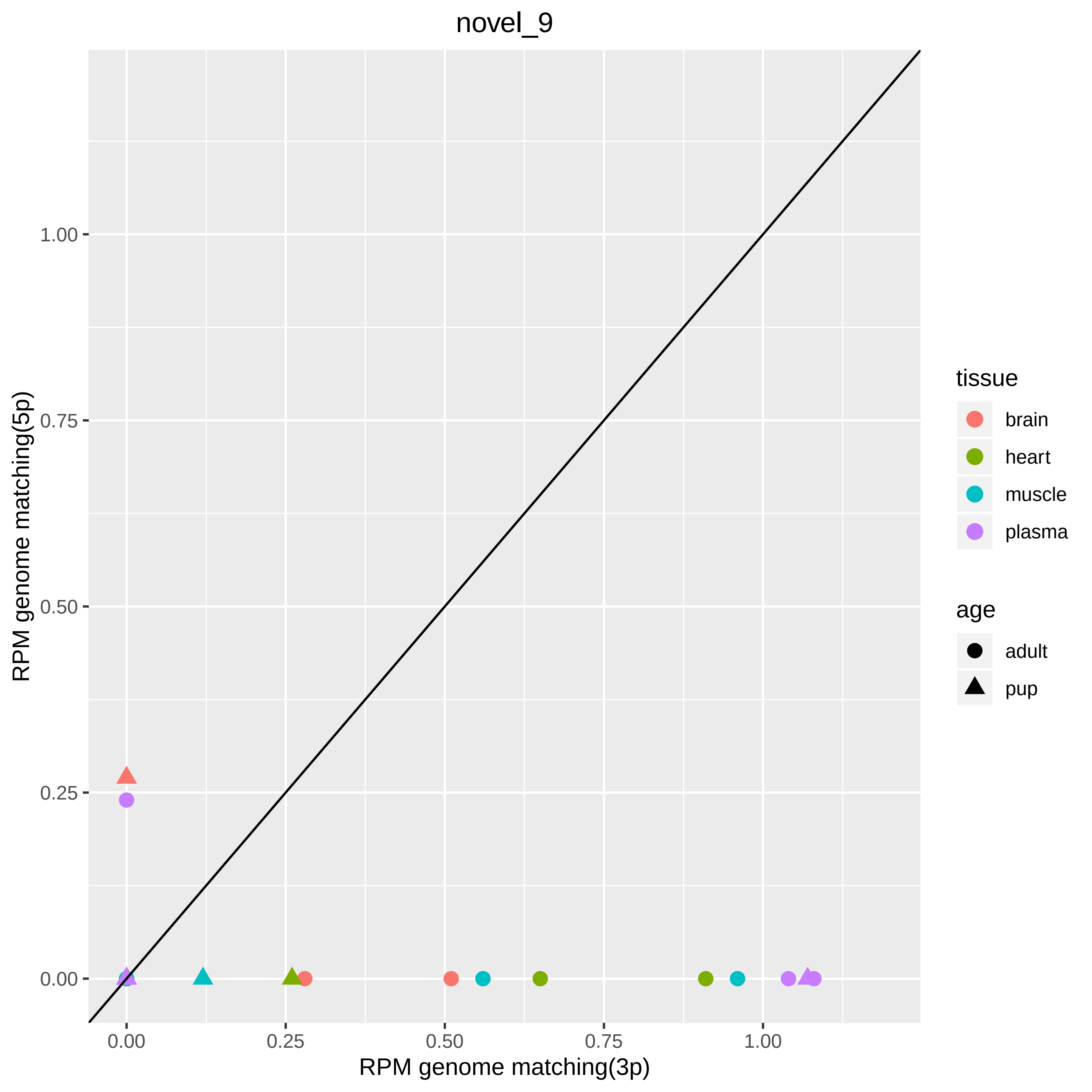
novel_9
RPM genome matching(5p)
RPM genome matching(3p)
1.00
0.75
0.50
0.25
0.00
0.00
0.25
0.50
0.75
1.00
tissue
brain
heart
muscle
plasma
age
adult
pup

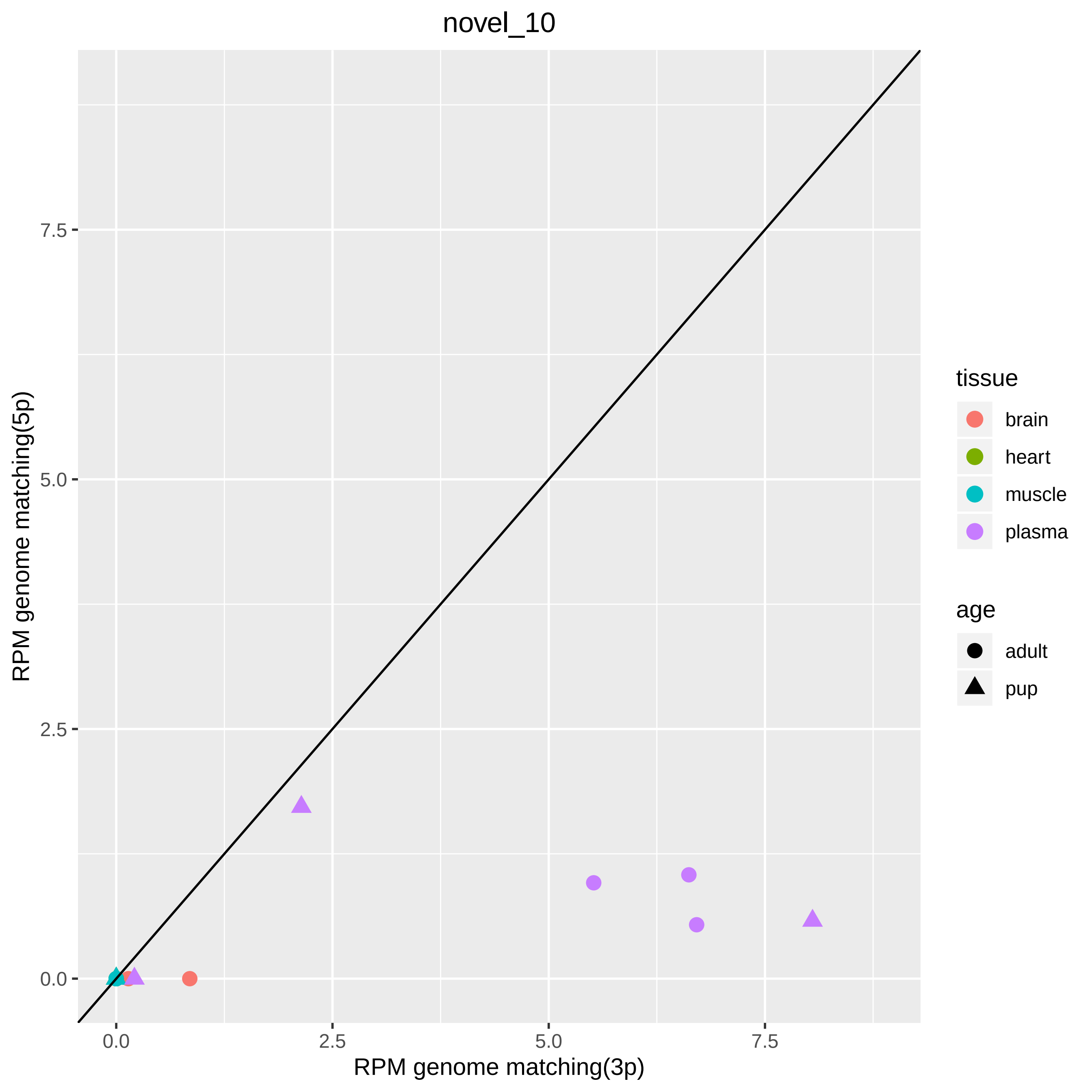
novel_10
RPM genome matching(5p)
RPM genome matching(3p)
0.0
2.5
5.0
7.5
tissue
brain
heart
muscle
plasma
age
adult
pup

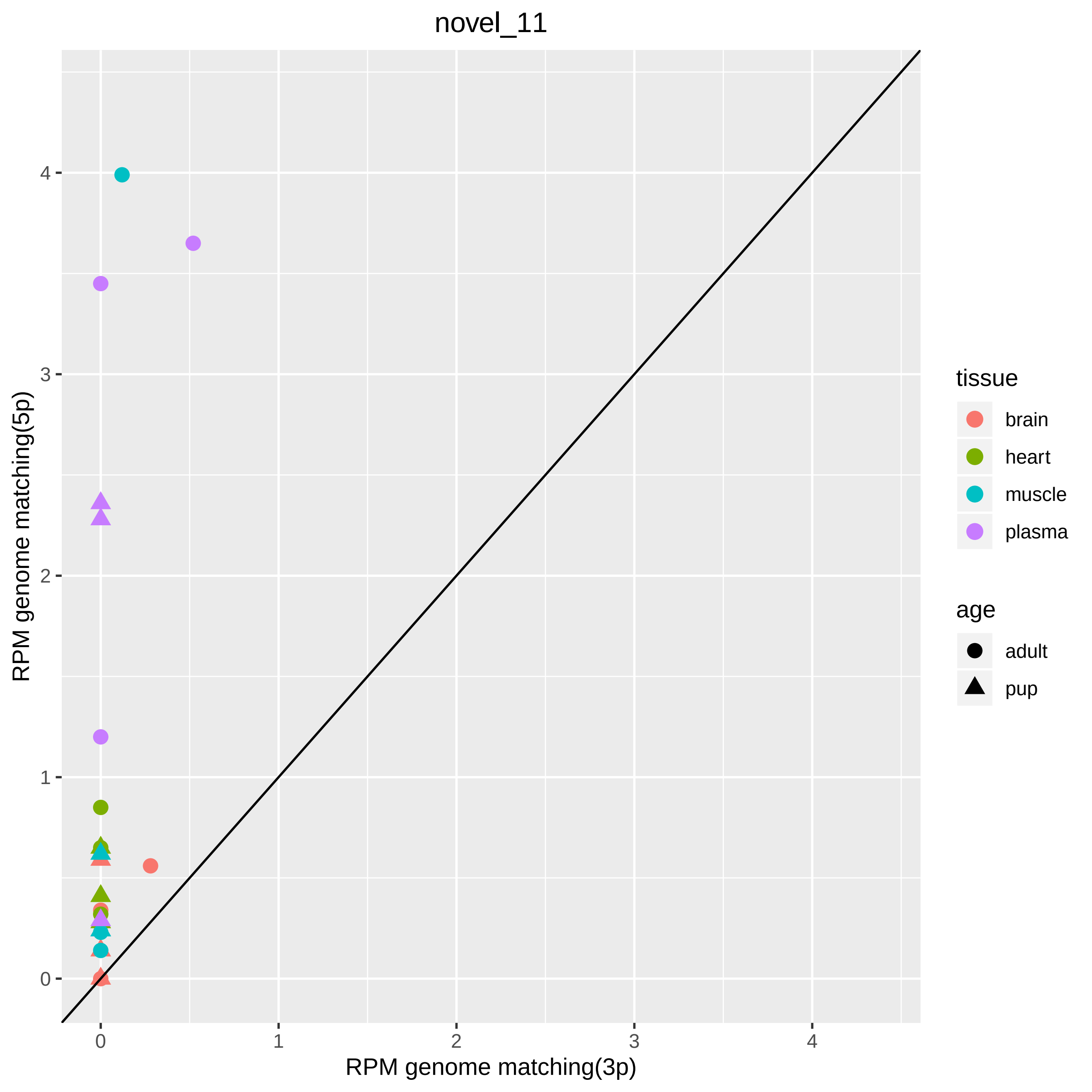

novel_11
RPM genome matching(5p)
RPM genome matching(3p)
0
1
2
3
4
0
1
2
3
4
tissue
brain
heart
muscle
plasma
age
adult
pup

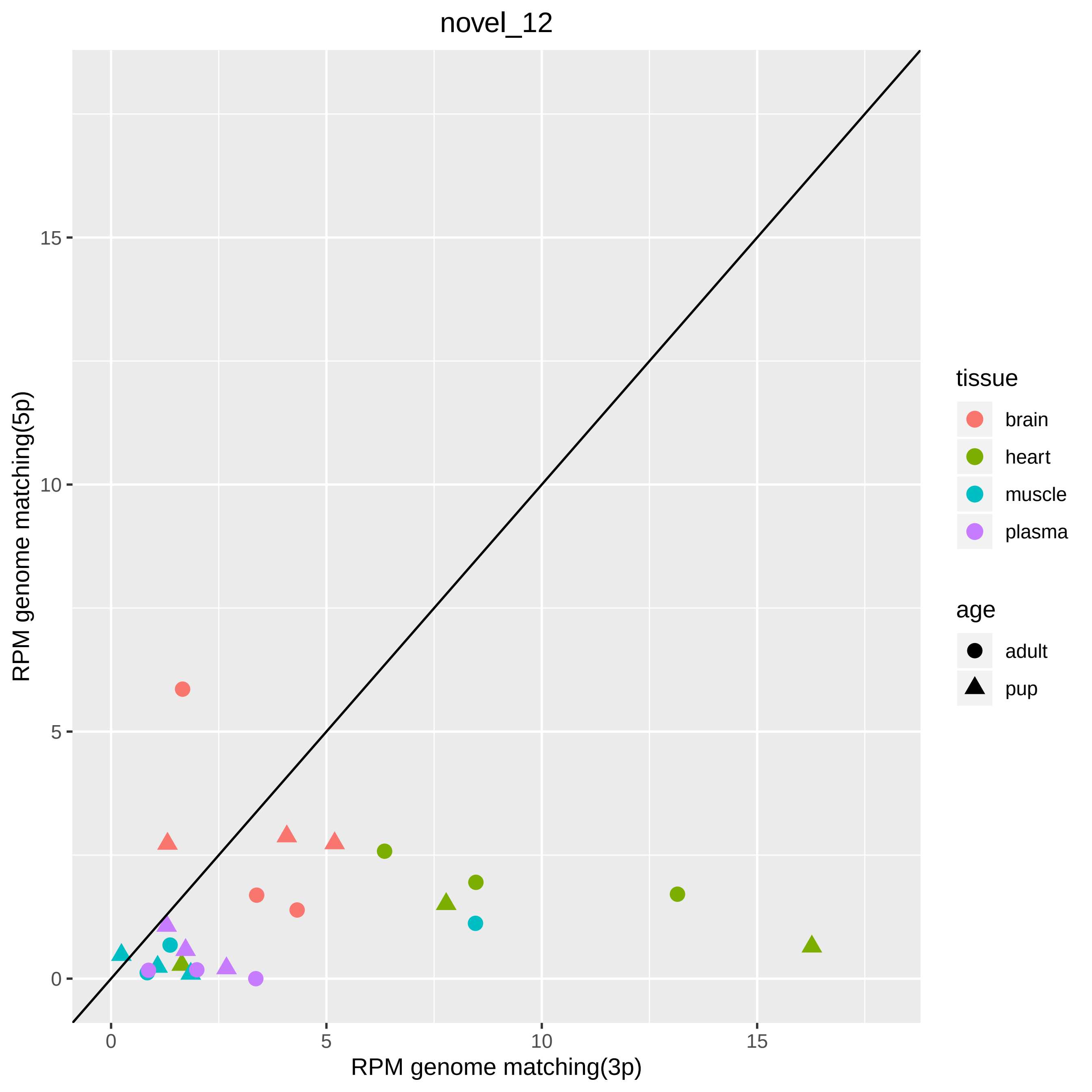

novel_12
RPM genome matching(5p)
RPM genome matching(3p)
0
5
10
15
tissue
brain
heart
muscle
plasma
age
adult
pup

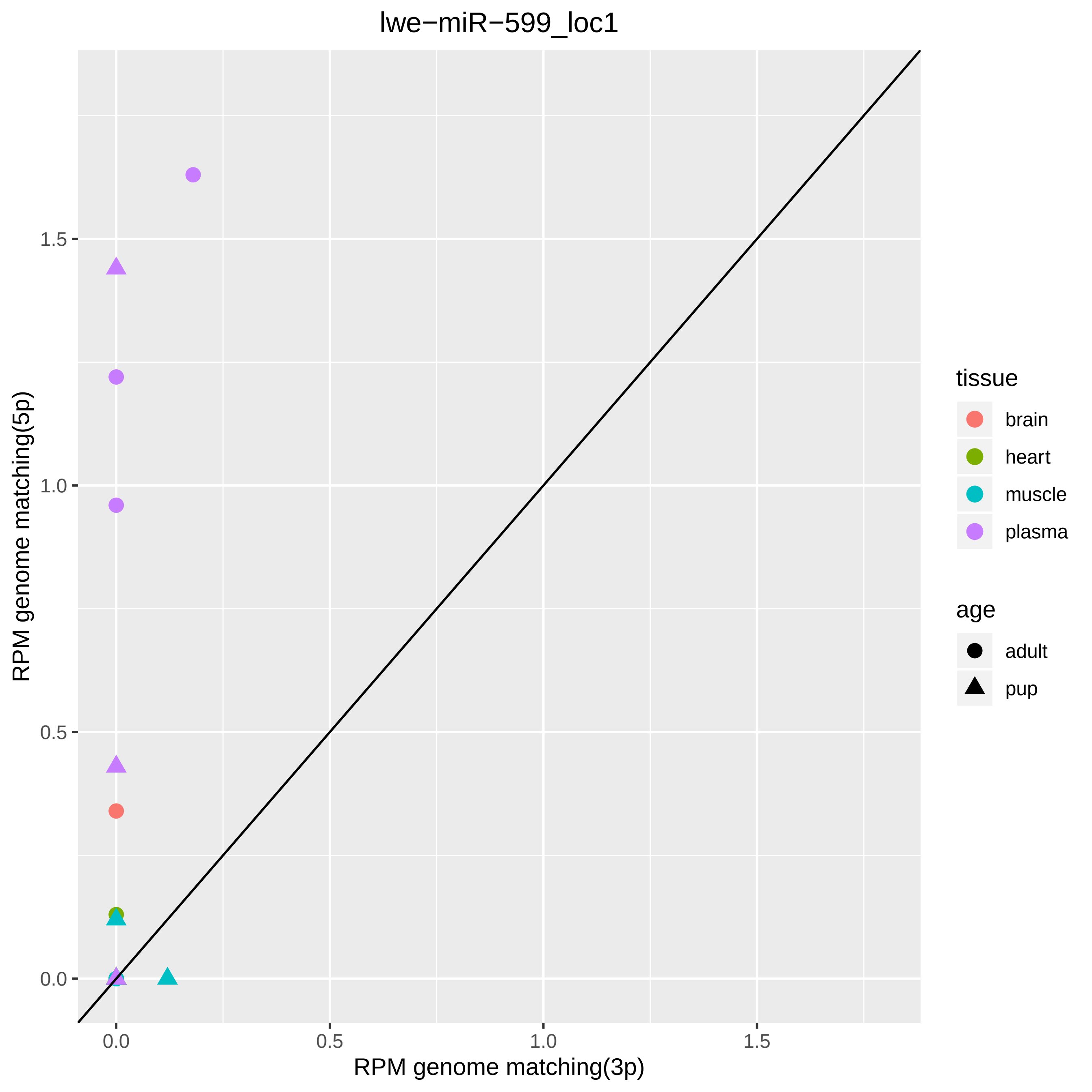
lwe−miR−599_loc1
RPM genome matching(5p)
RPM genome matching(3p)
0.0
0.5
1.0
1.5
0.0
0.5
1.0
1.5
tissue
brain
heart
muscle
plasma
age
adult
pup

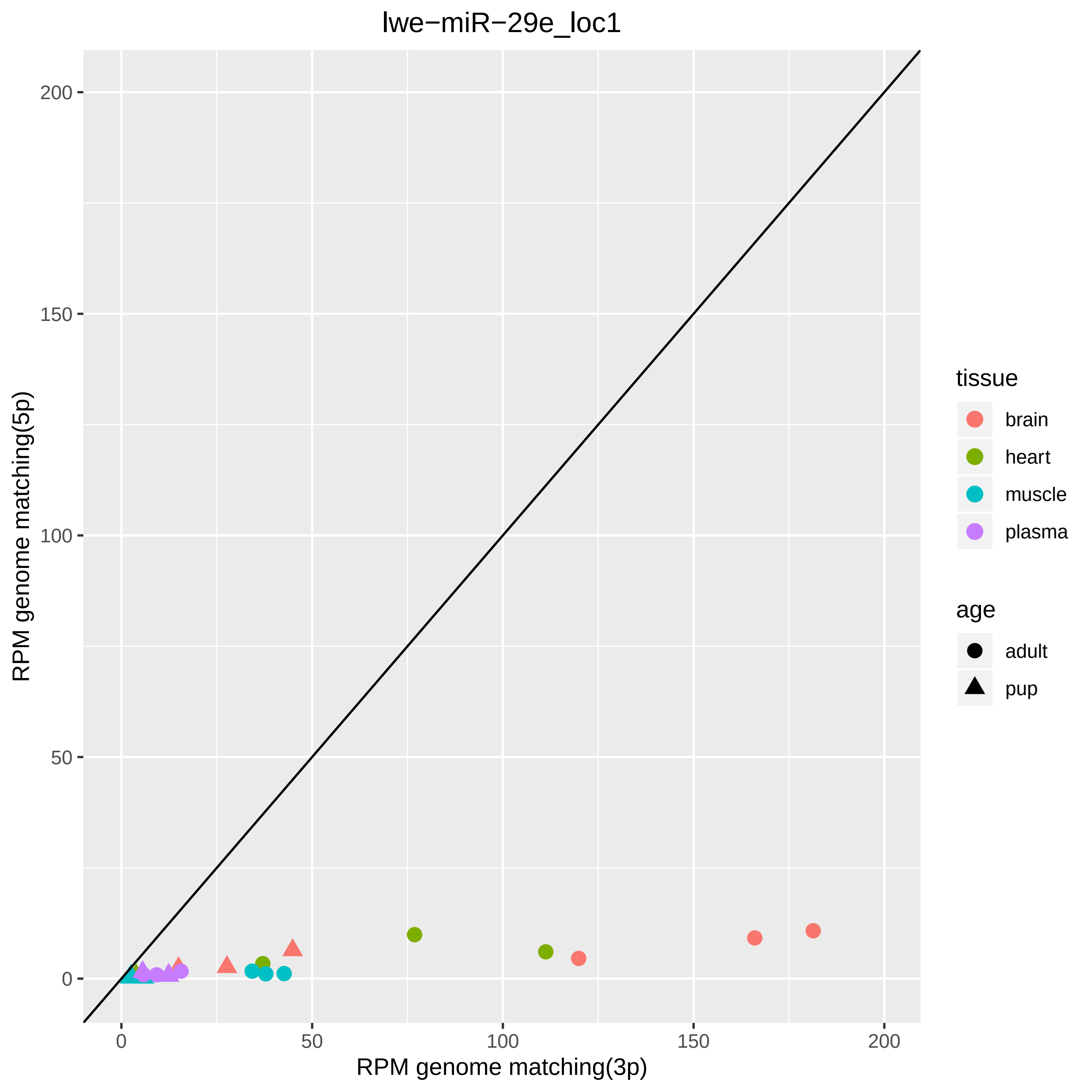
lwe−miR−29e_loc1
RPM genome matching(5p)
RPM genome matching(3p)
0
50
100
150
200
0
50
100
150
200
tissue
brain
heart
muscle
plasma
age
adult
pup

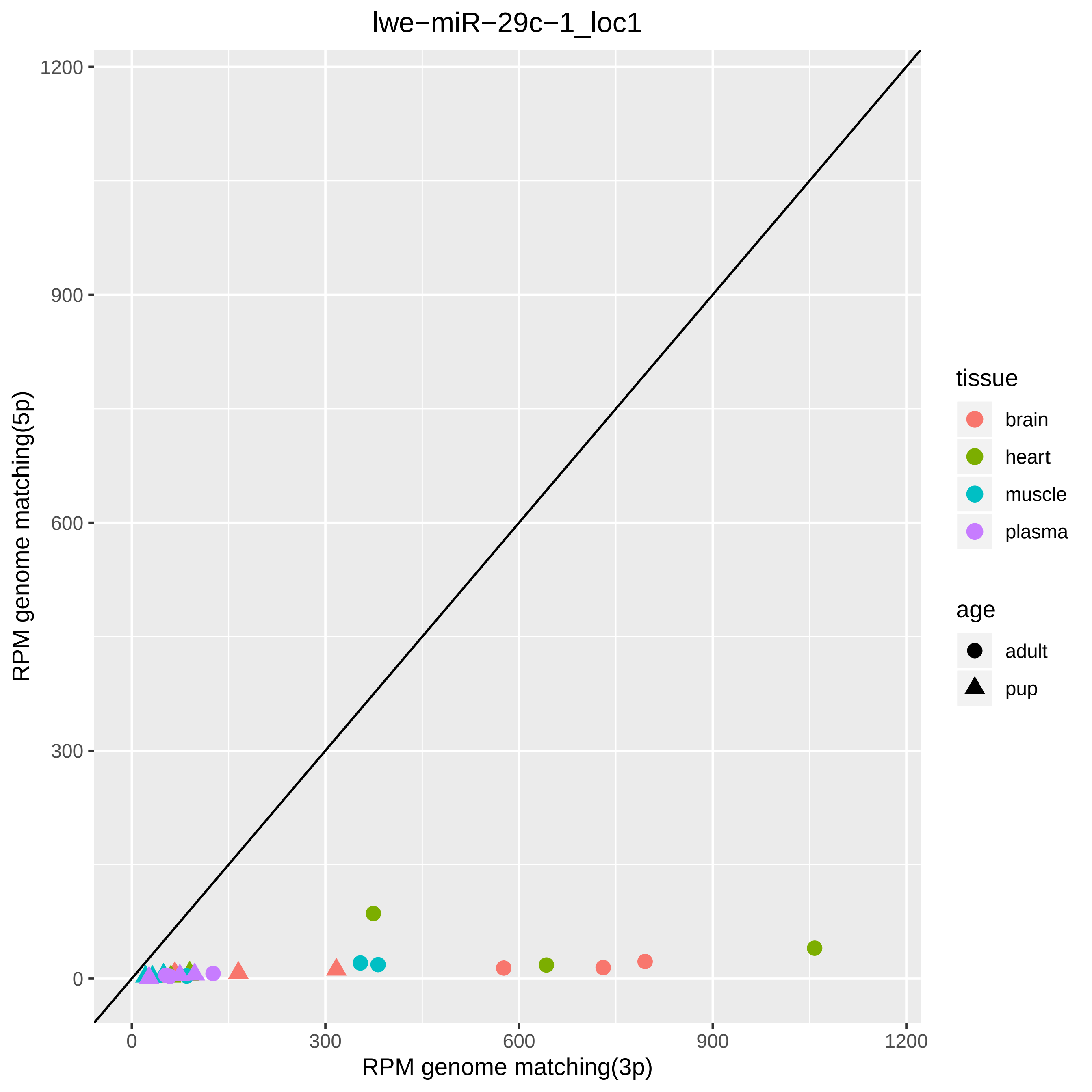
lwe−miR−29c−1_loc1
RPM genome matching(5p)
RPM genome matching(3p)
0
300
600
900
1200
0
300
600
900
1200
tissue
brain
heart
muscle
plasma
age
adult
pup

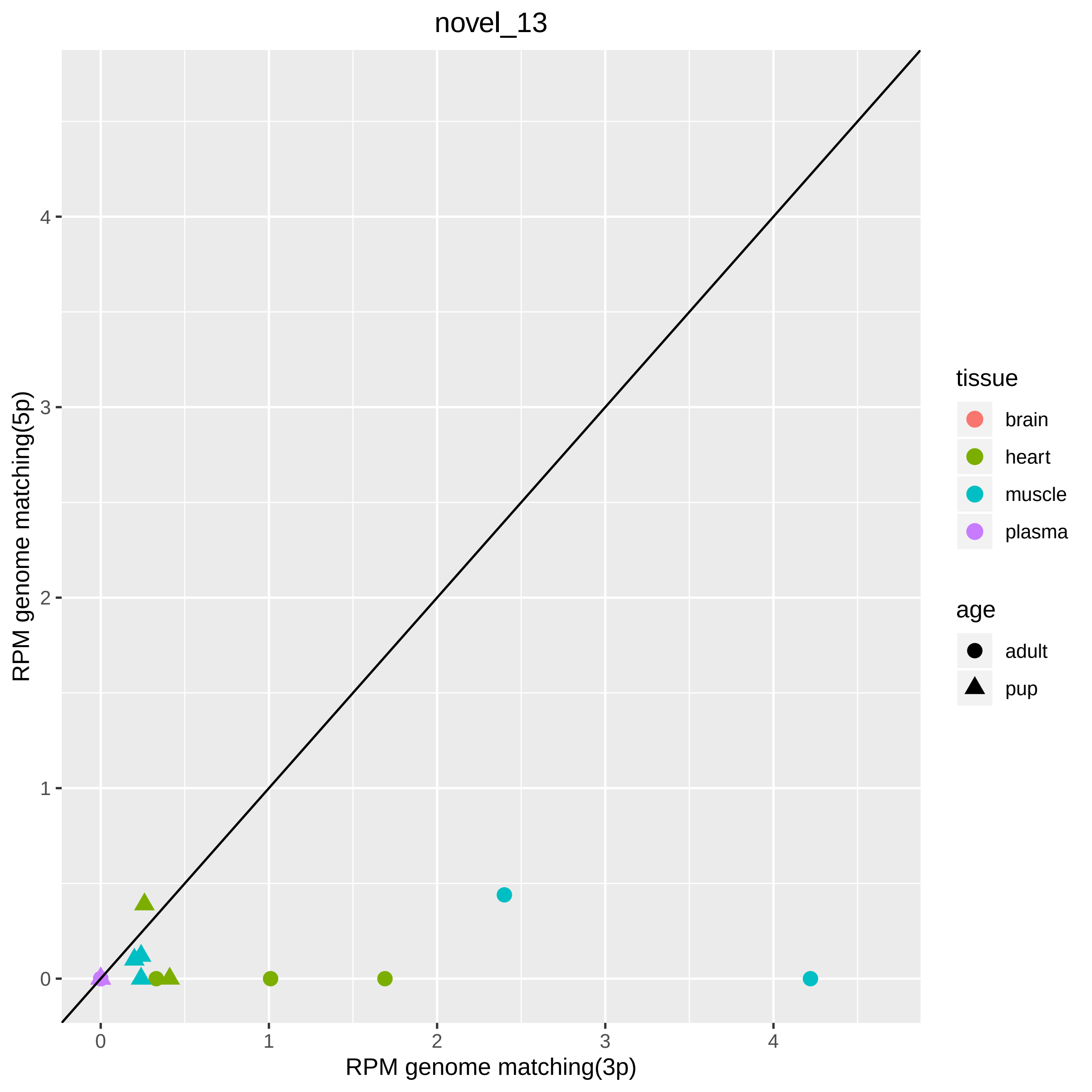
novel_13
RPM genome matching(5p)
RPM genome matching(3p)
0
1
2
3
4
0
1
2
3
4
tissue
brain
heart
muscle
plasma
age
adult
pup

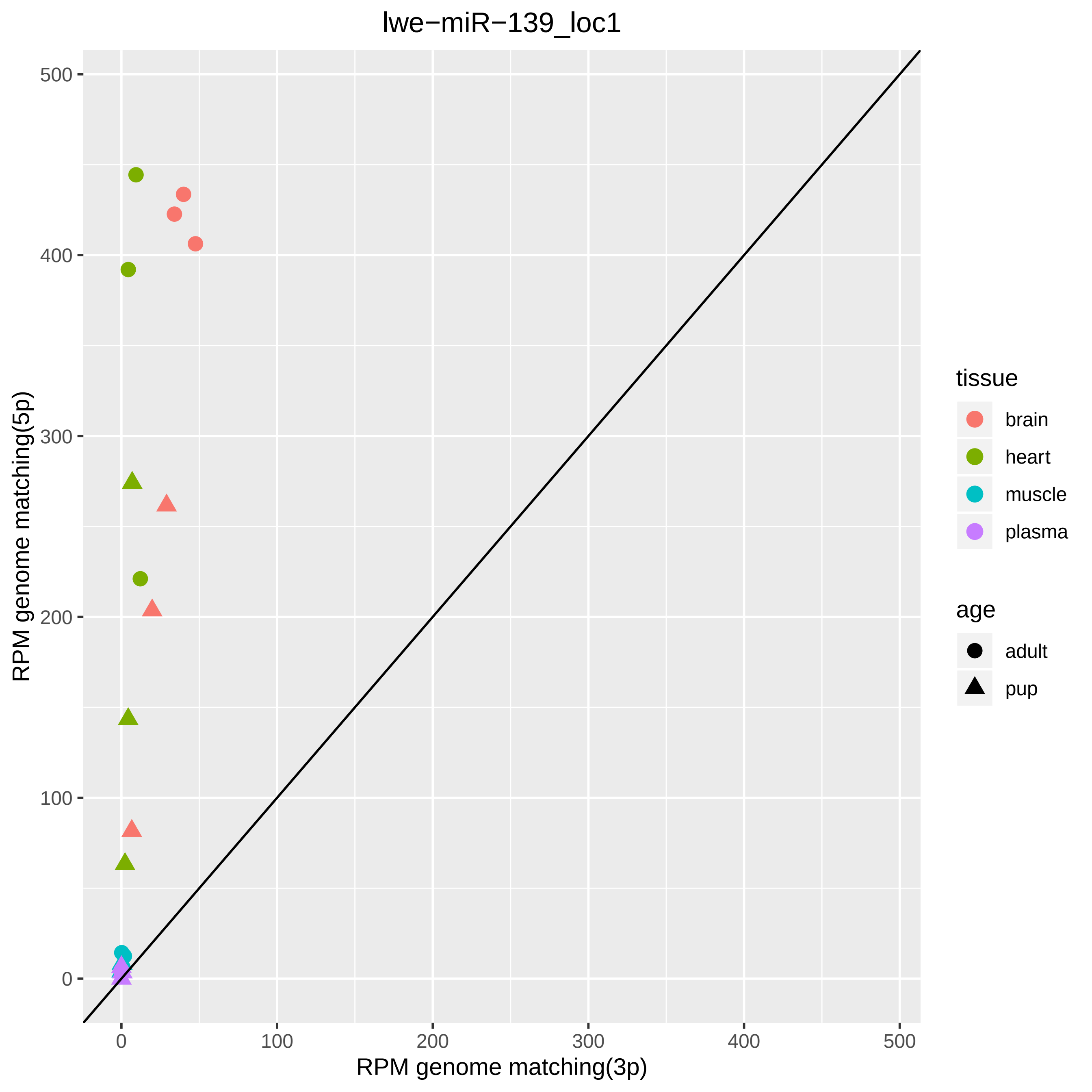
lwe−miR−139_loc1
RPM genome matching(5p)
RPM genome matching(3p)
0
100
200
300
400
500
tissue
brain
heart
muscle
plasma
age
adult
pup

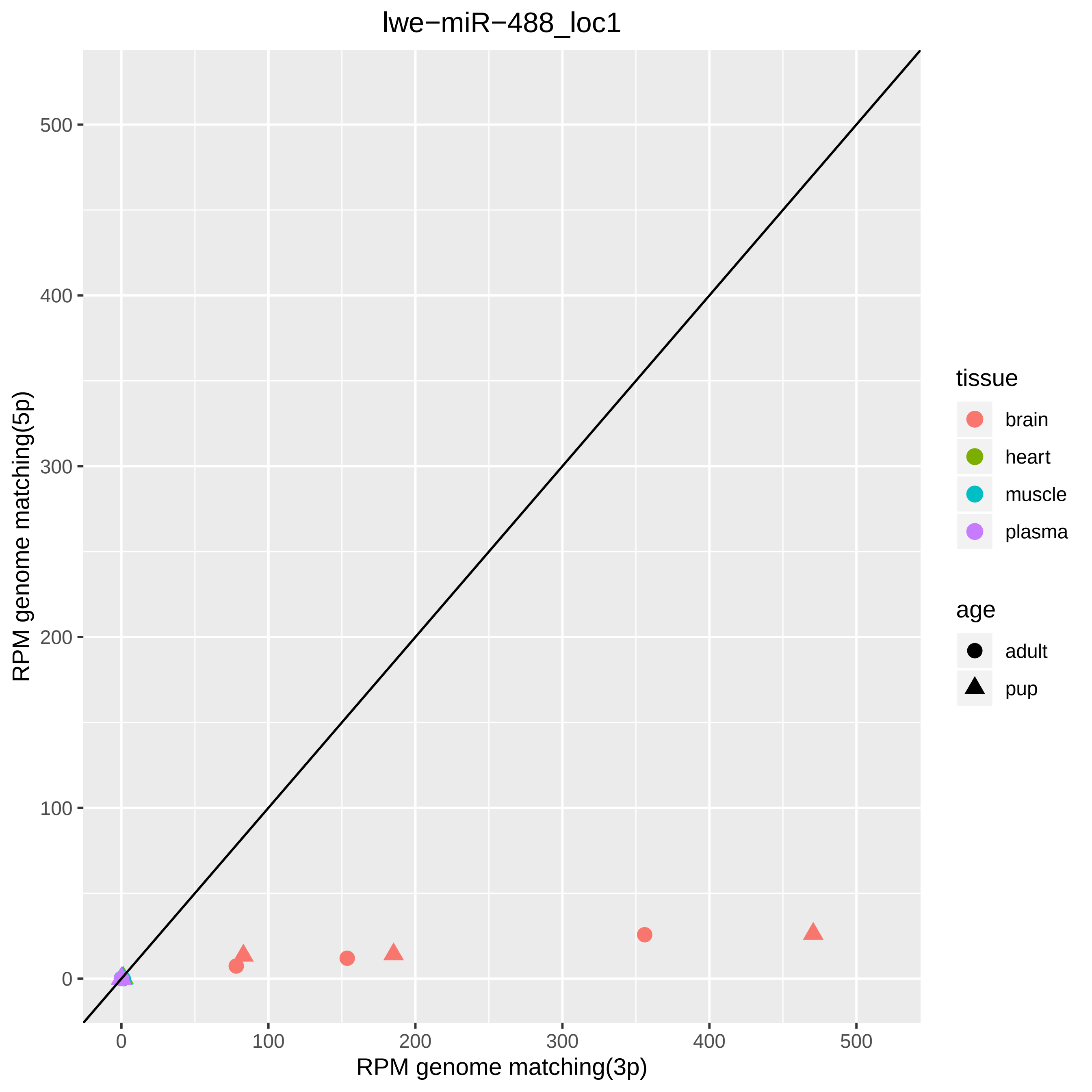
lwe−miR−488_loc1
500
400
300
200
100
0
0
100
200
300
400
500
RPM genome matching(5p)
RPM genome matching(3p)
tissue
brain
heart
muscle
plasma
age
adult
pup

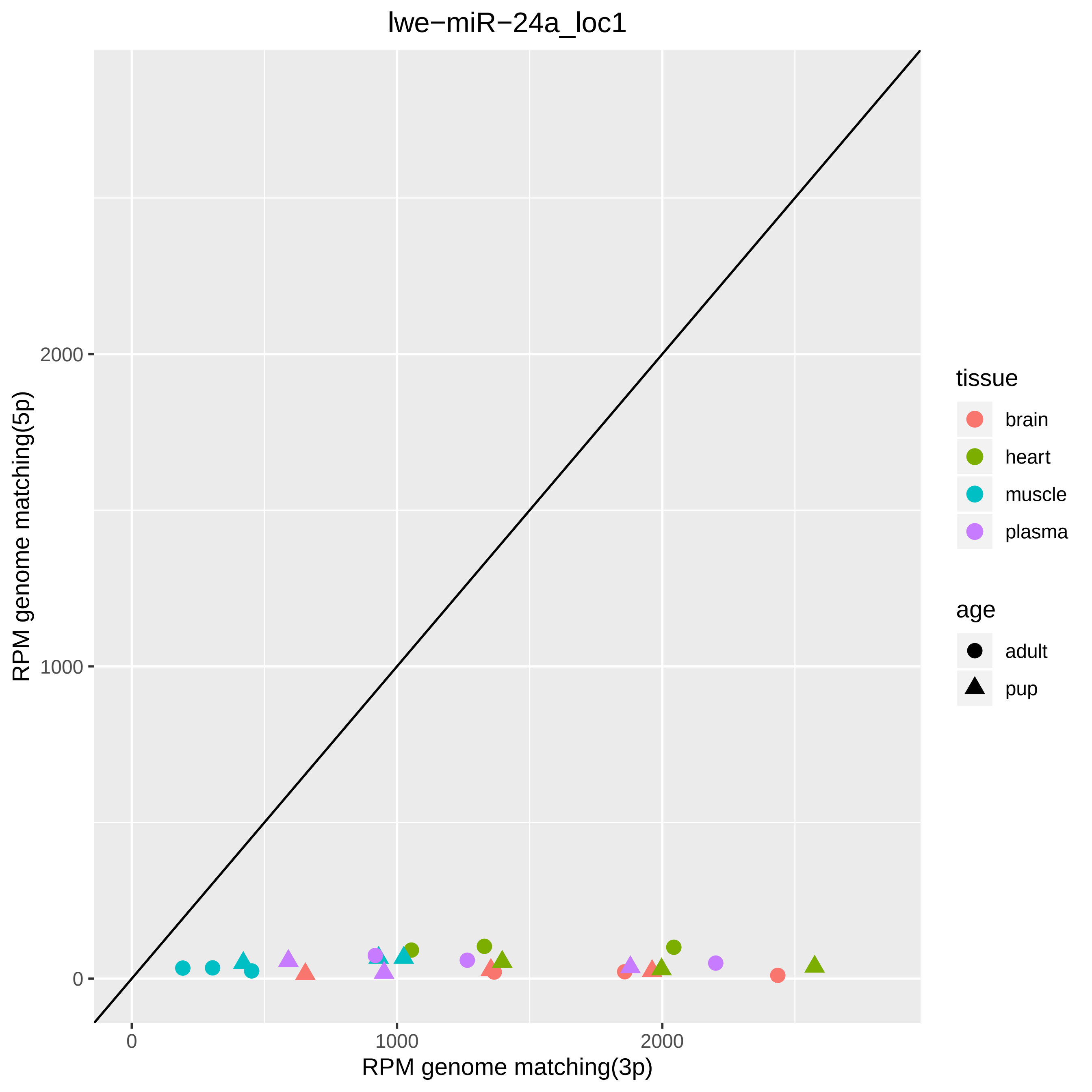
lwe−miR−24a_loc1
RPM genome matching(5p)
RPM genome matching(3p)
0
1000
2000
tissue
brain
heart
muscle
plasma
age
adult
pup

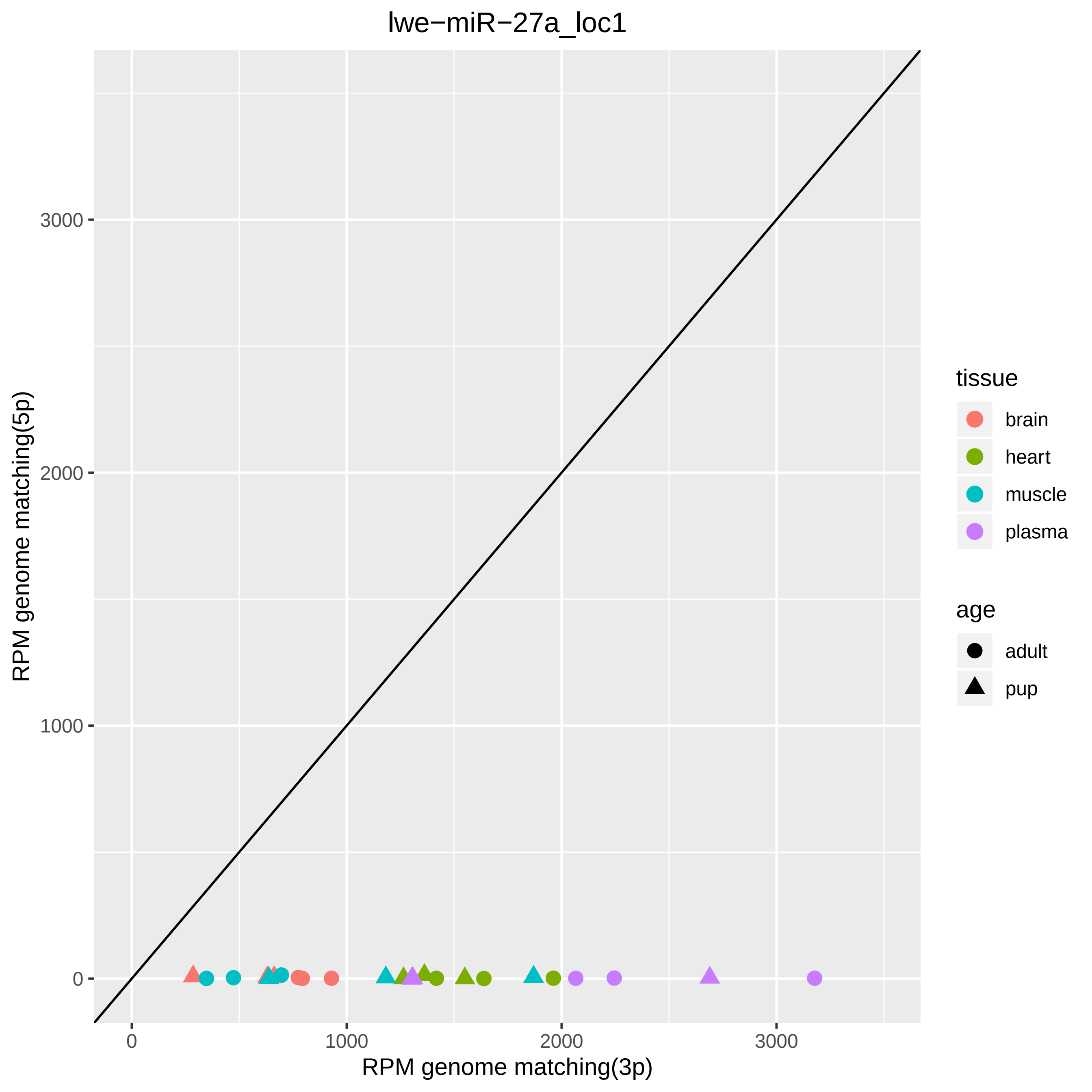
lwe−miR−27a_loc1
RPM genome matching(5p)
RPM genome matching(3p)
0
1000
2000
3000
0
1000
2000
3000
tissue
brain
heart
muscle
plasma
age
adult
pup

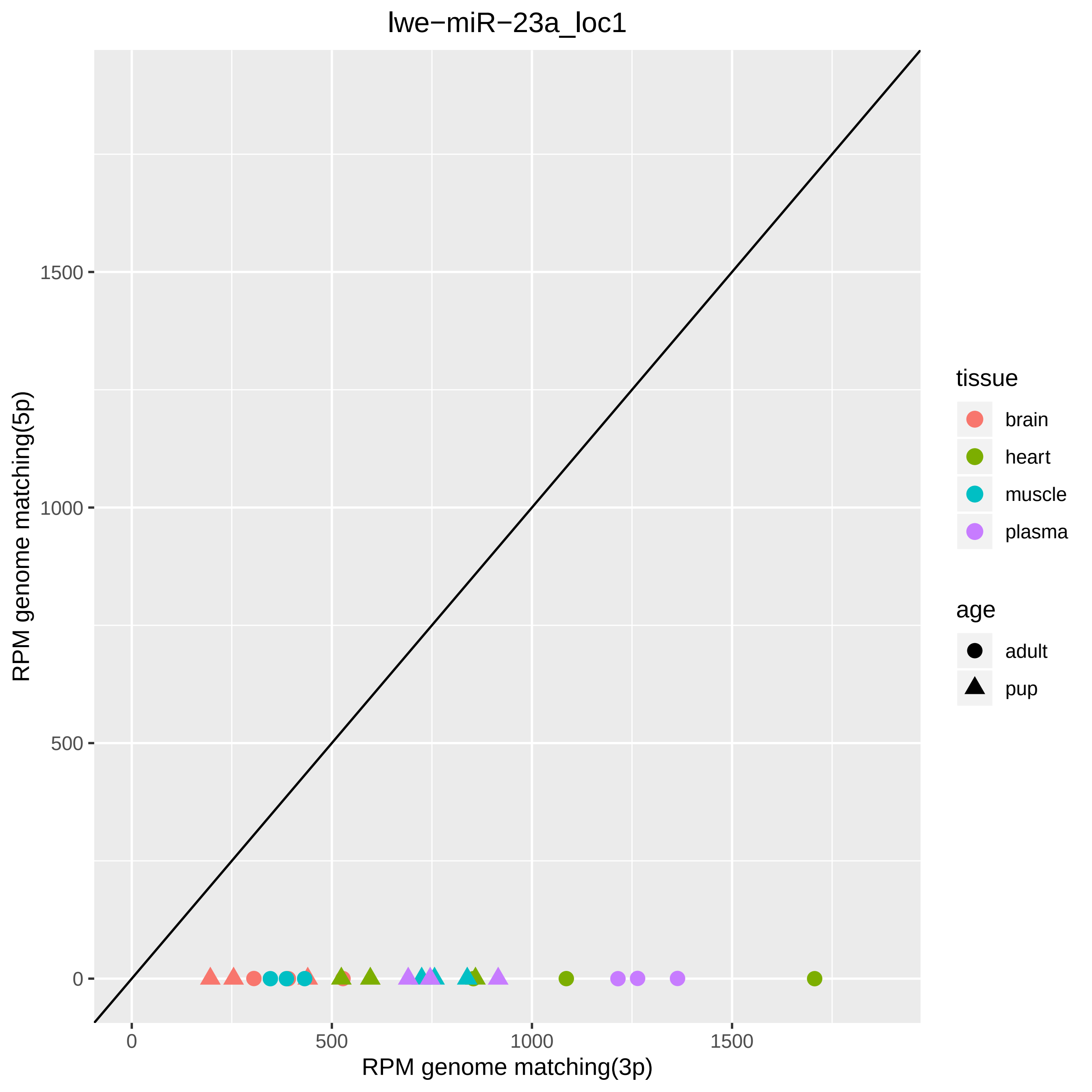

lwe−miR−23a_loc1
RPM genome matching(5p)
RPM genome matching(3p)
0
500
1000
1500
0
500
1000
1500
tissue
brain
heart
muscle
plasma
age
adult
pup

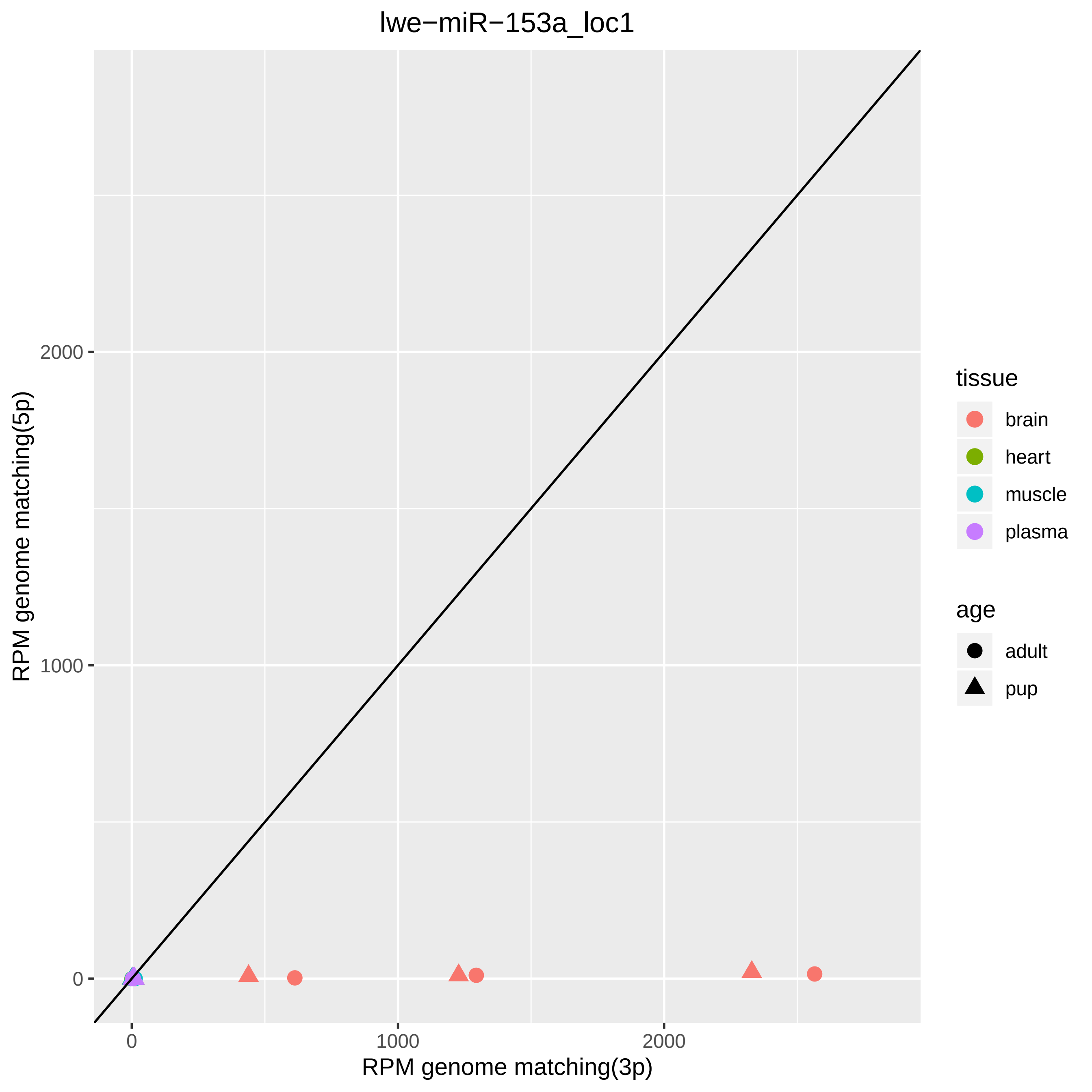

lwe−miR−153a_loc1
RPM genome matching(5p)
2000
1000
0
0
1000
2000
RPM genome matching(3p)
tissue
brain
heart
muscle
plasma
age
adult
pup

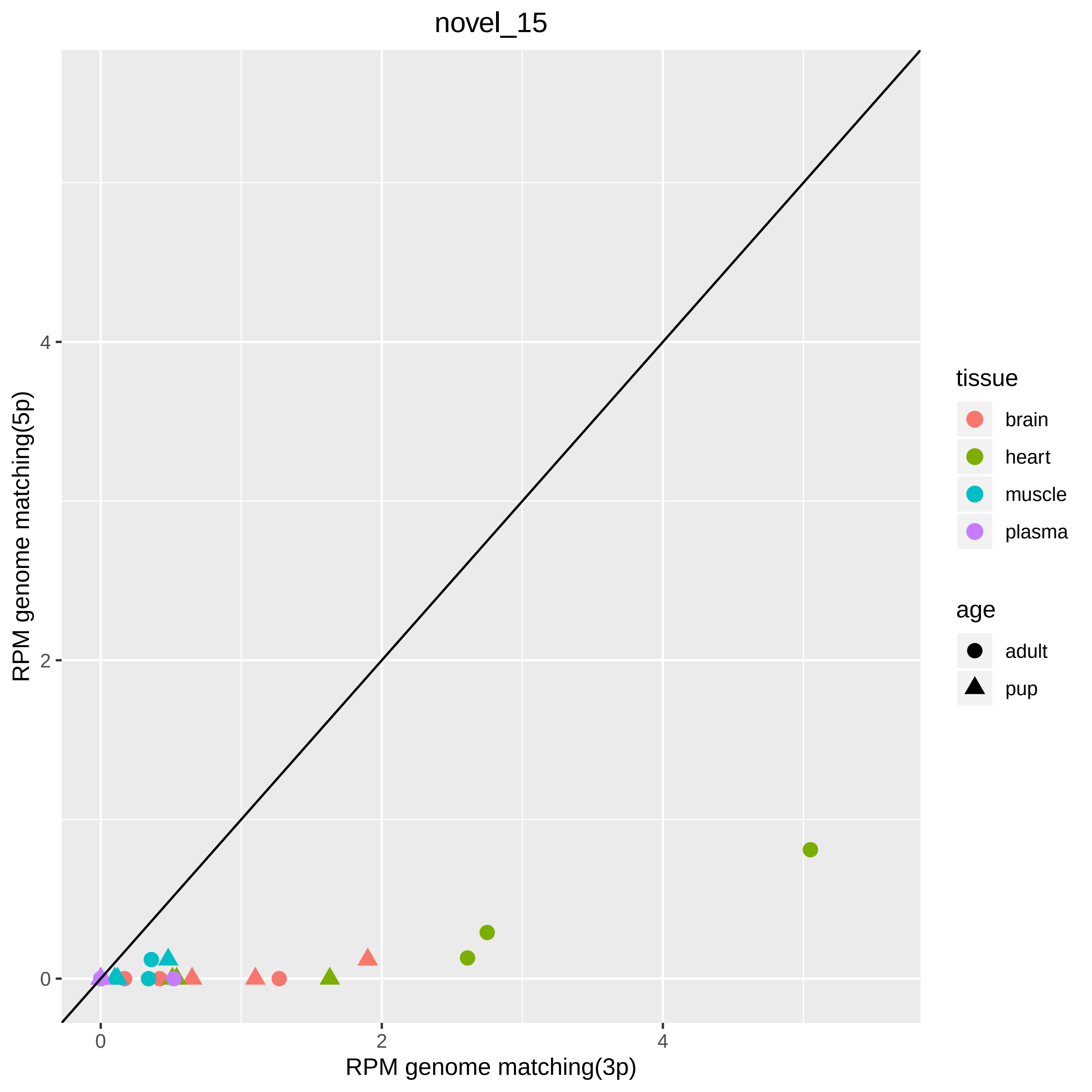

novel_15
RPM genome matching(5p)
RPM genome matching(3p)
0
2
4
0
2
4
tissue
brain
heart
muscle
plasma
age
adult
pup

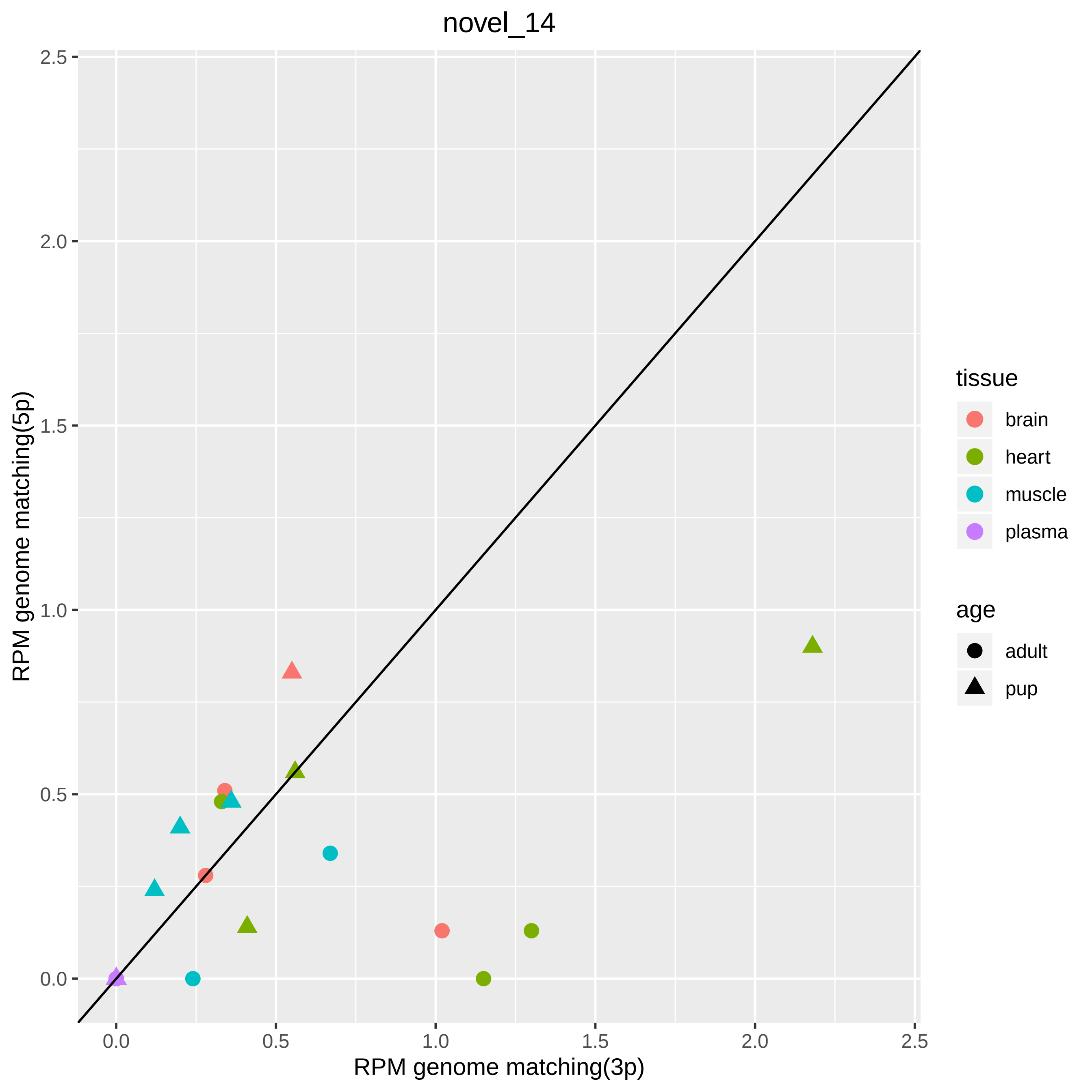
novel_14
RPM genome matching(5p)
RPM genome matching(3p)
0.0
0.5
1.0
1.5
2.0
2.5
tissue
brain
heart
muscle
plasma
age
adult
pup

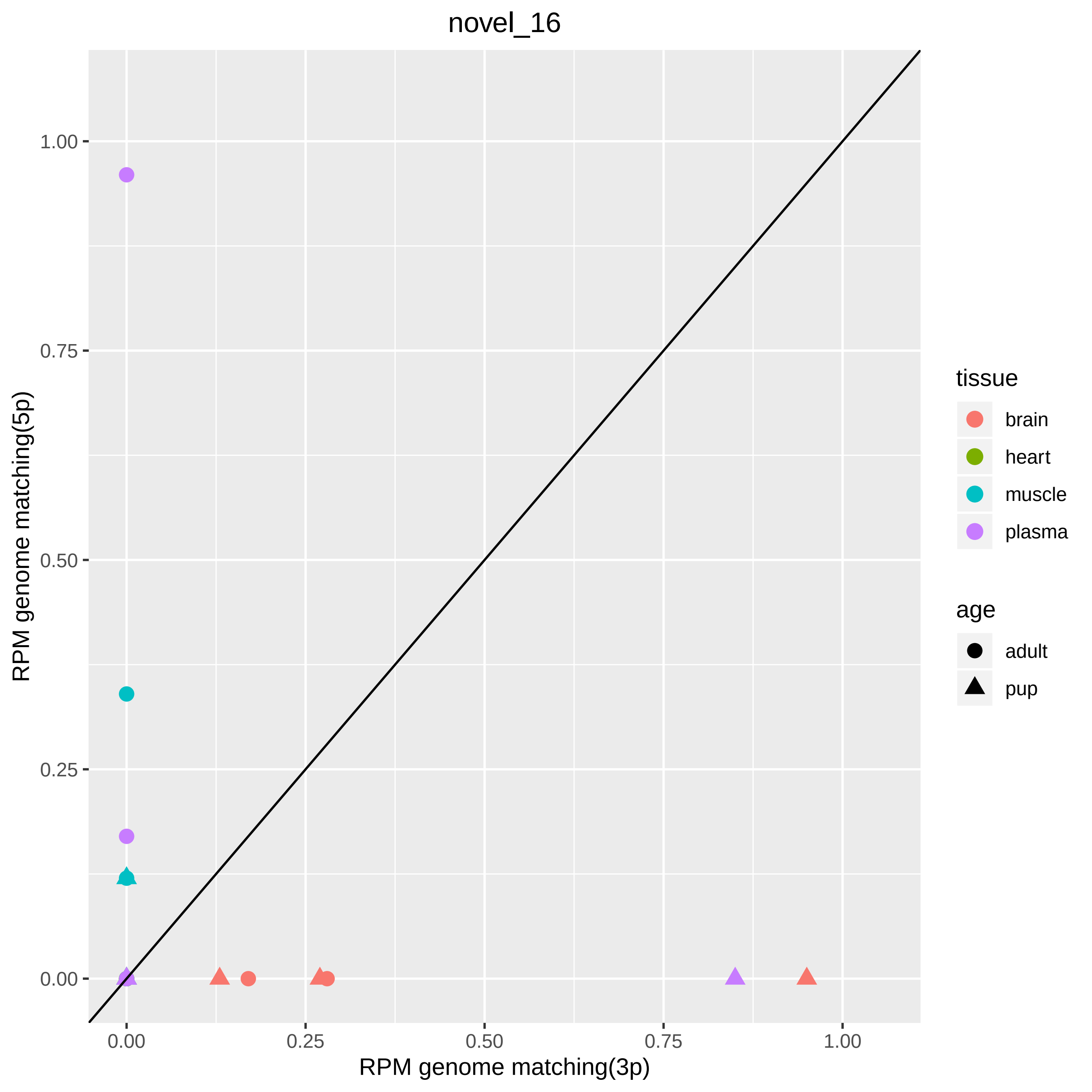

novel_16
RPM genome matching(5p)
RPM genome matching(3p)
0.00
0.25
0.50
0.75
1.00
tissue
brain
heart
muscle
plasma
age
adult
pup

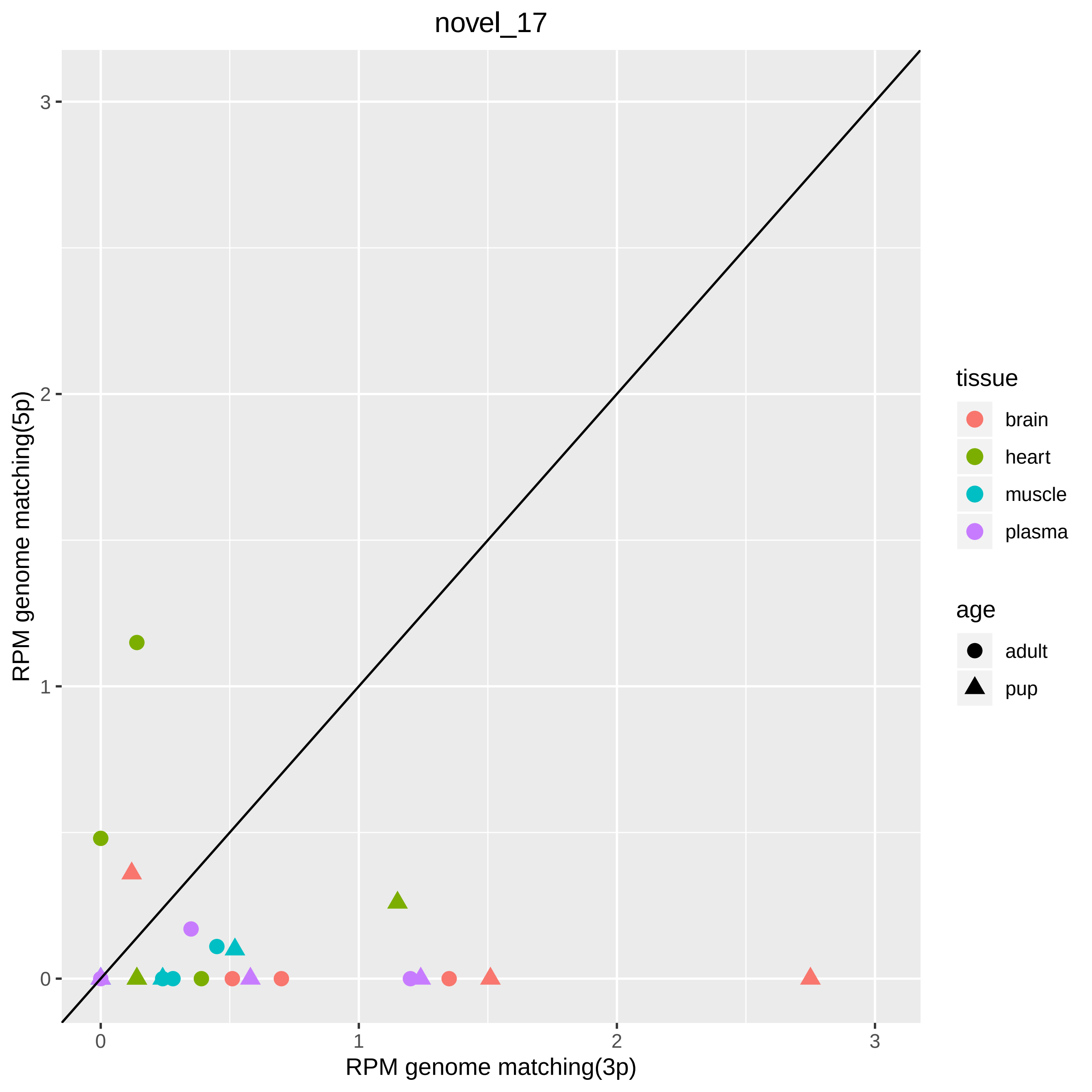

novel_17
RPM genome matching(5p)
RPM genome matching(3p)
0
1
2
3
tissue
brain
heart
muscle
plasma
age
adult
pup

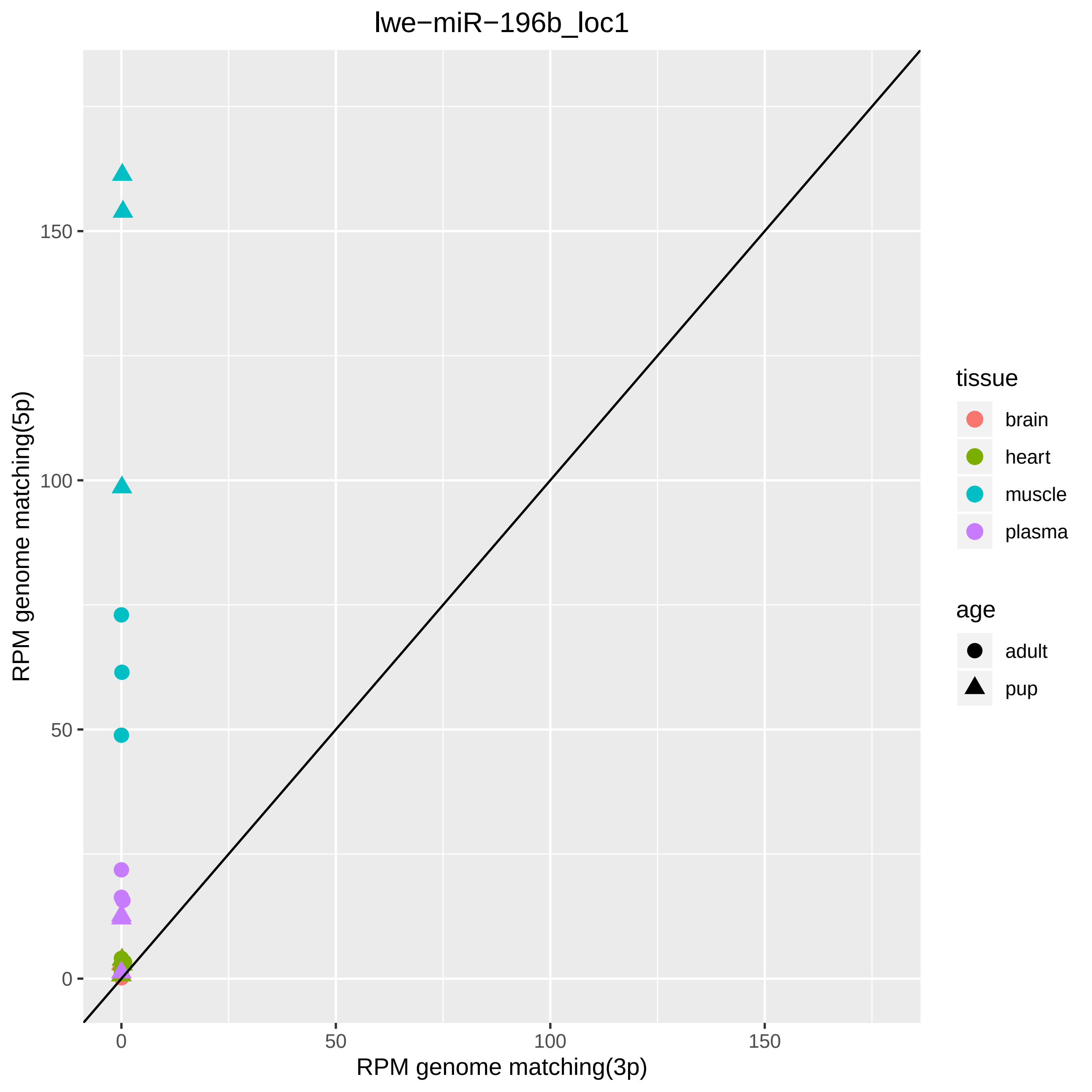

lwe−miR−196b_loc1
RPM genome matching(5p)
RPM genome matching(3p)
0
50
100
150
0
50
100
150
tissue
brain
heart
muscle
plasma
age
adult
pup

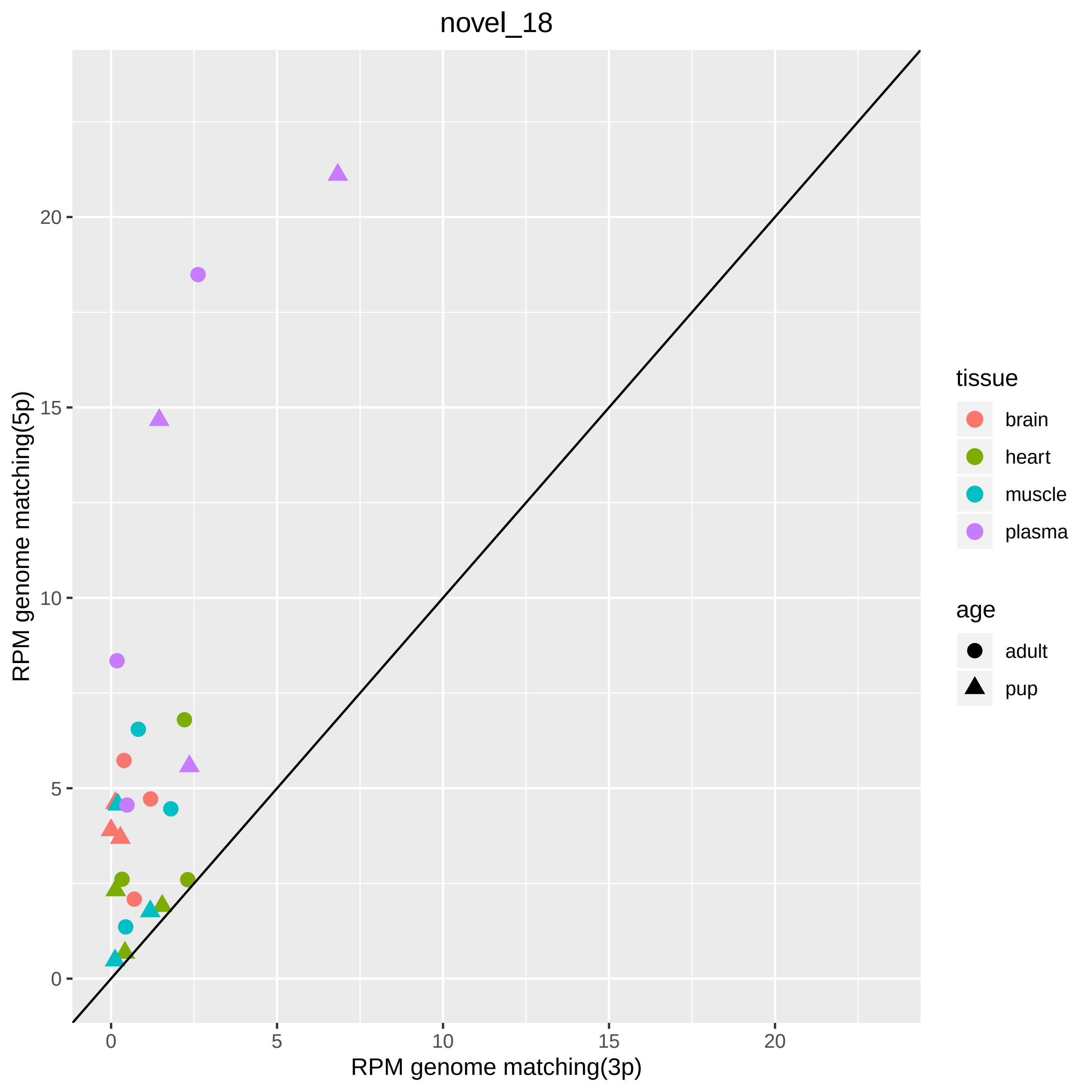

novel_18
RPM genome matching(5p)
RPM genome matching(3p)
0
5
10
15
20
tissue
brain
heart
muscle
plasma
age
adult
pup

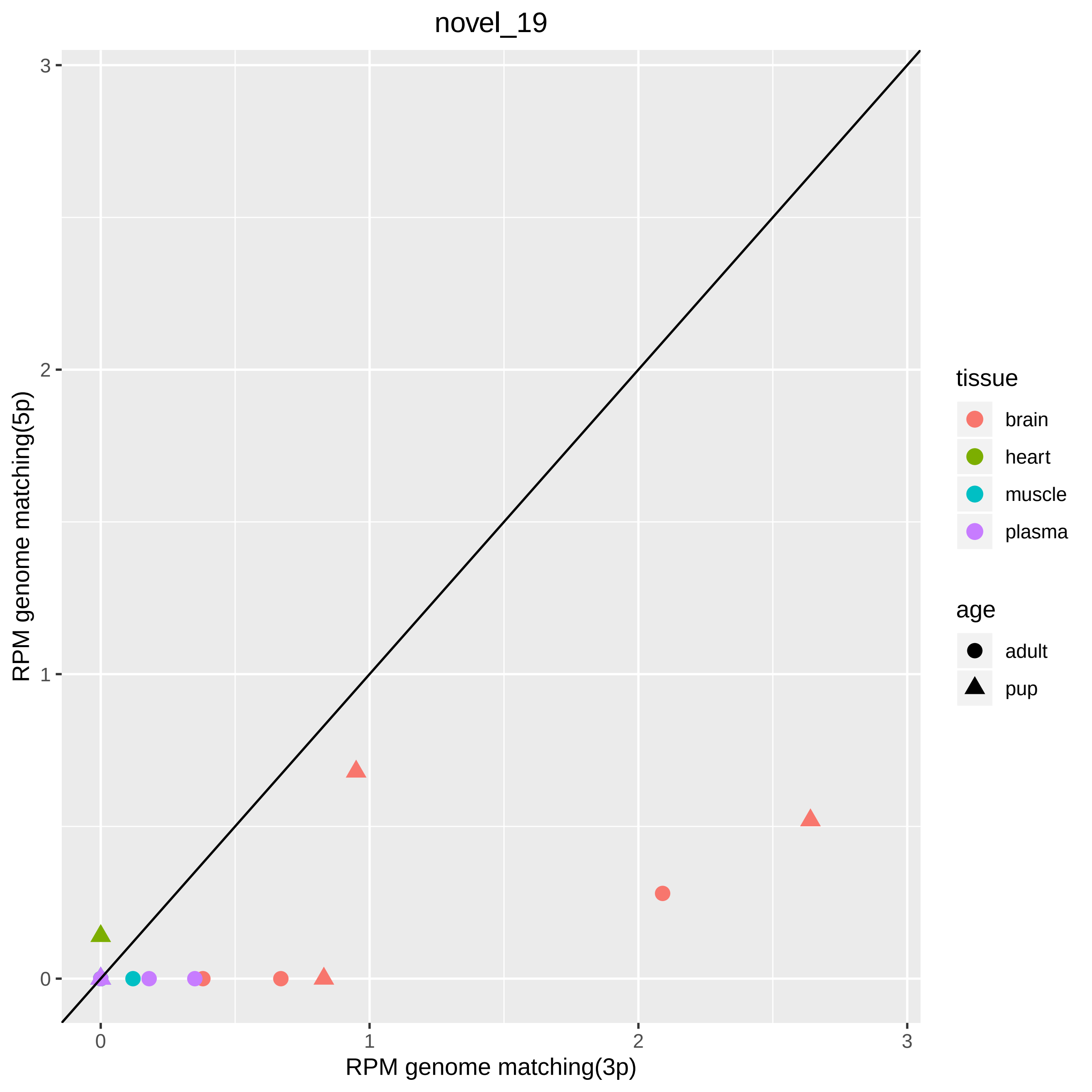

novel_19
RPM genome matching(5p)
RPM genome matching(3p)
0
1
2
3
0
1
2
3
tissue
brain
heart
muscle
plasma
age
adult
pup

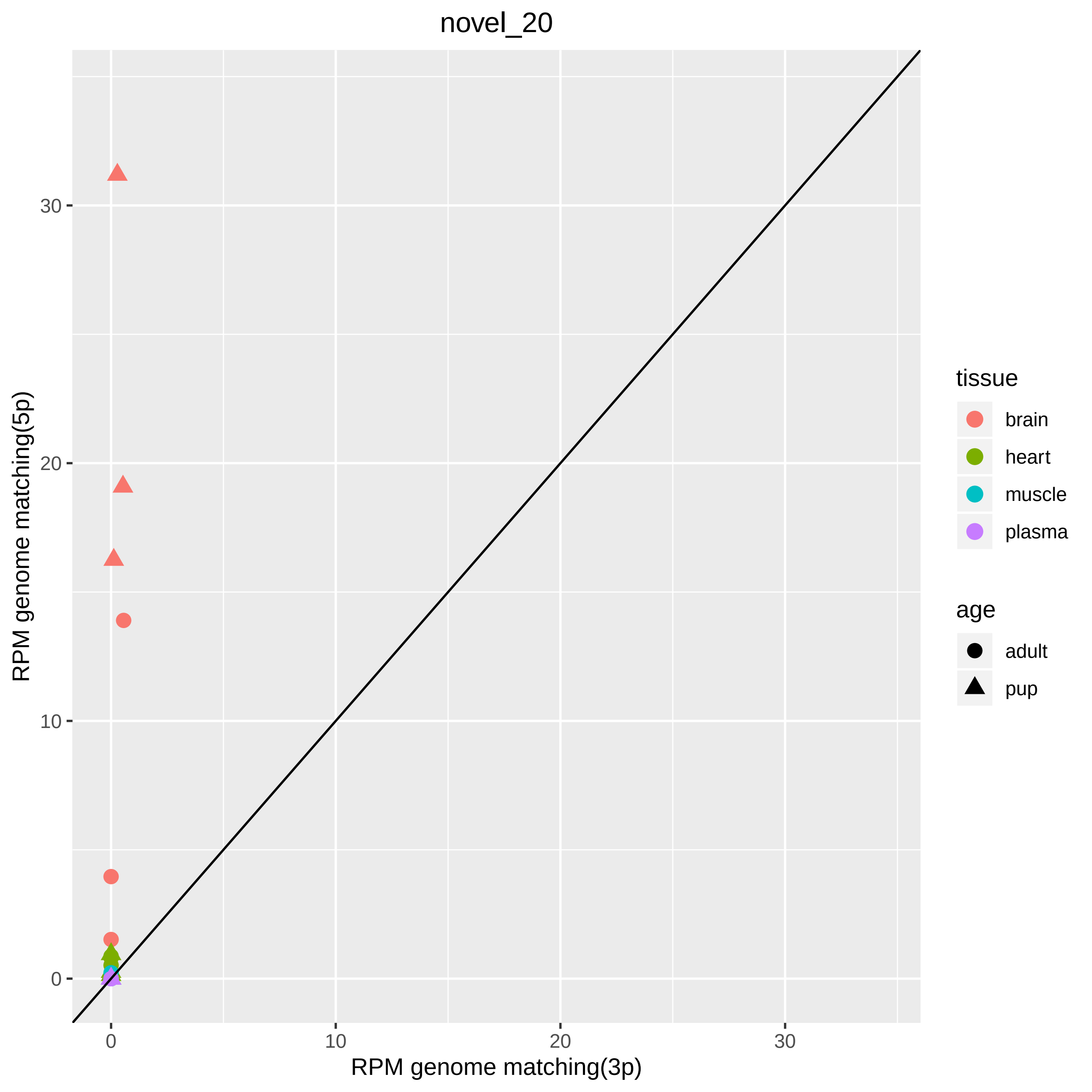
novel_20
RPM genome matching(5p)
RPM genome matching(3p)
0
10
20
30
tissue
brain
heart
muscle
plasma
age
adult
pup

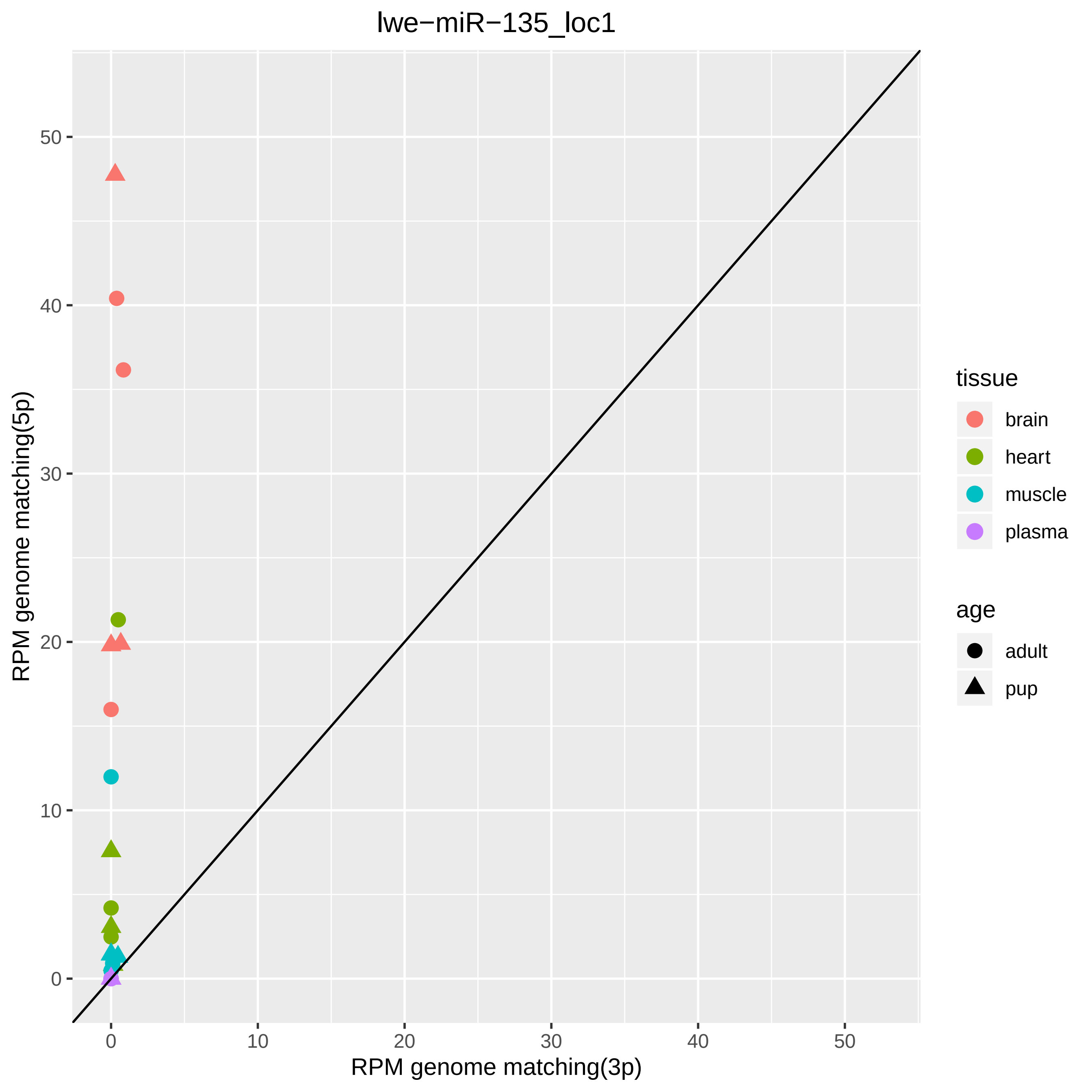
lwe−miR−135_loc1
RPM genome matching(5p)
RPM genome matching(3p)
0
10
20
30
40
50
tissue
brain
heart
muscle
plasma
age
adult
pup

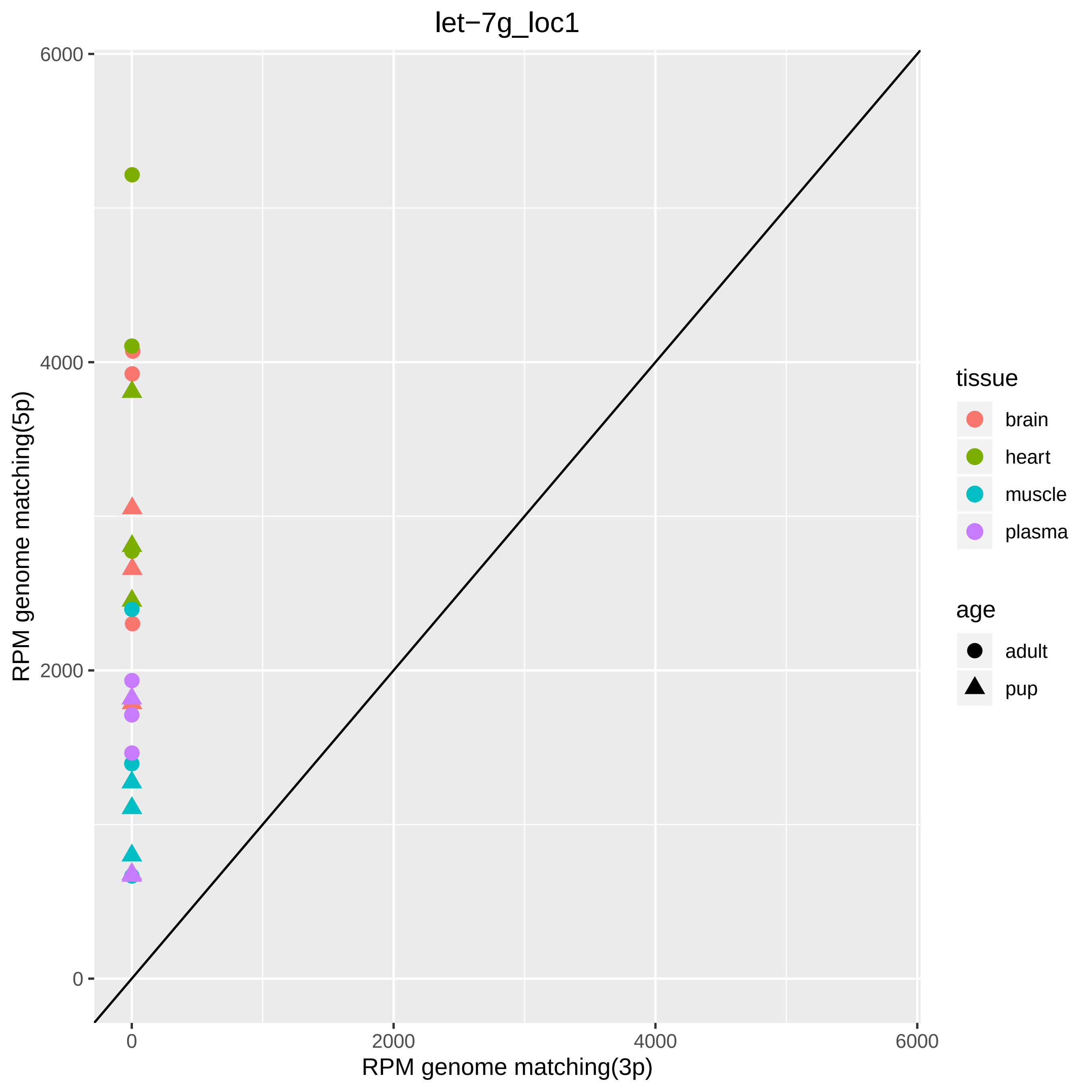

let−7g_loc1
RPM genome matching(5p)
RPM genome matching(3p)
0
2000
4000
6000
0
2000
4000
6000
tissue
brain
heart
muscle
plasma
age
adult
pup

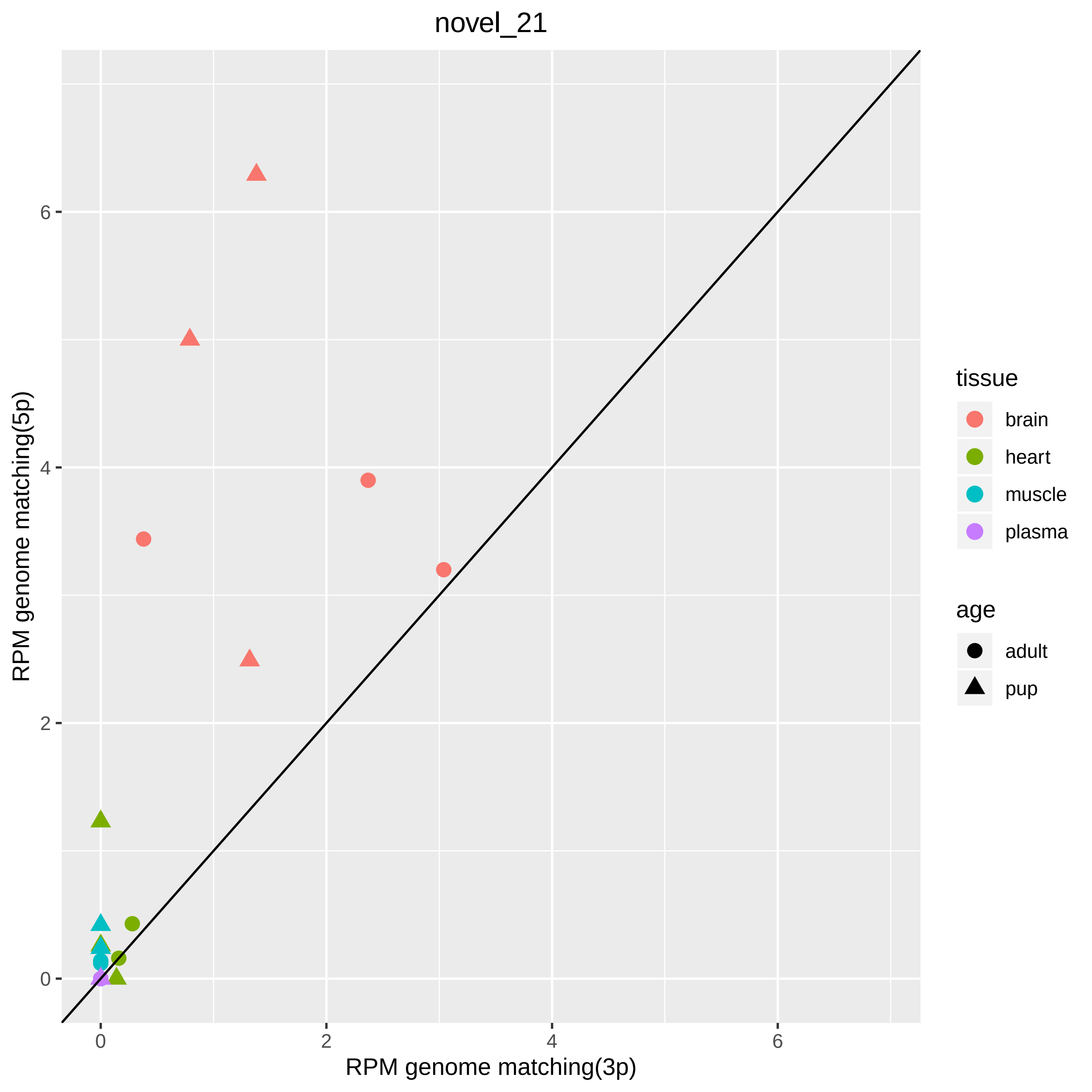

novel_21
RPM genome matching(5p)
RPM genome matching(3p)
0
2
4
6
tissue
brain
heart
muscle
plasma
age
adult
pup

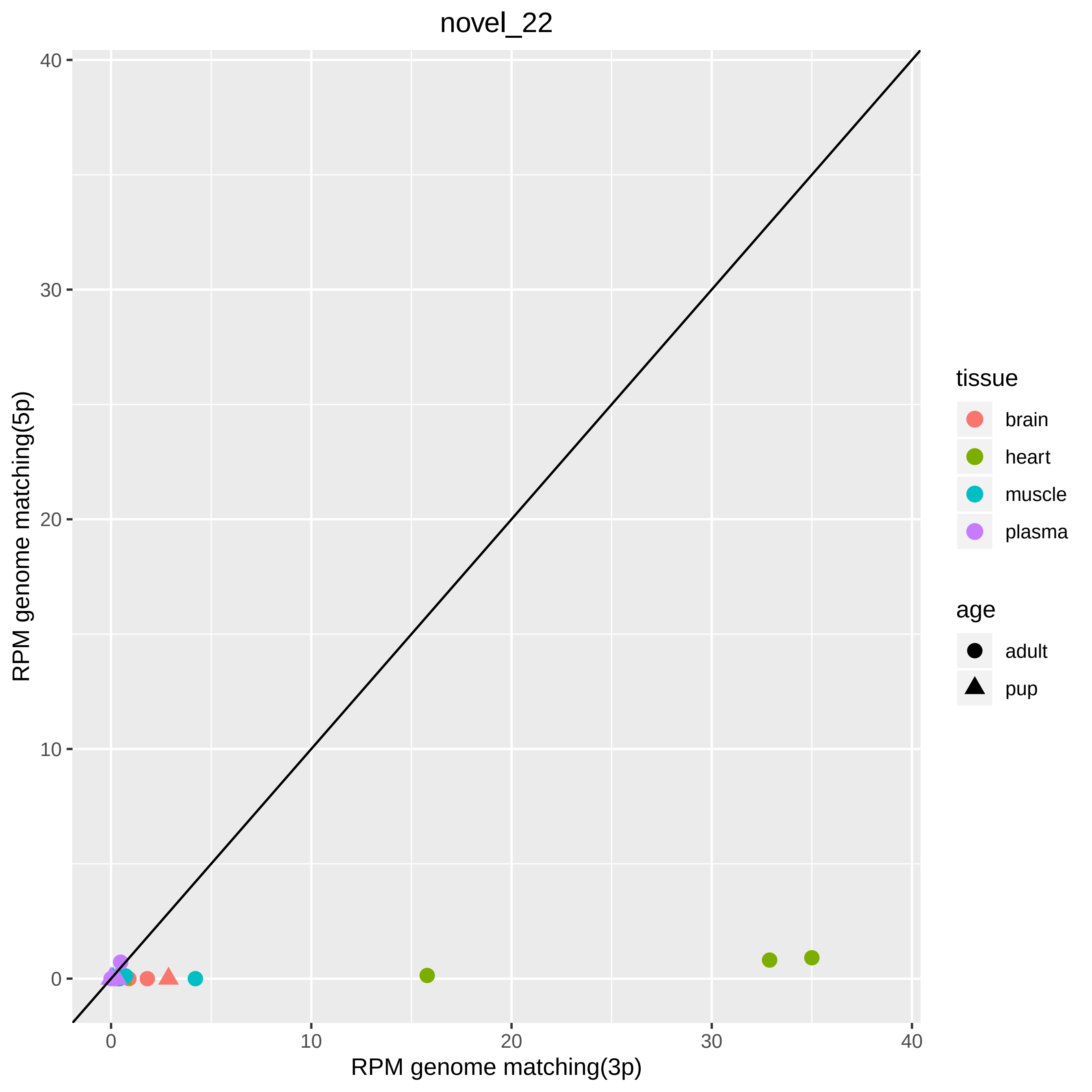

novel_22
RPM genome matching(5p)
RPM genome matching(3p)
0
10
20
30
40
0
10
20
30
40
tissue
brain
heart
muscle
plasma
age
adult
pup

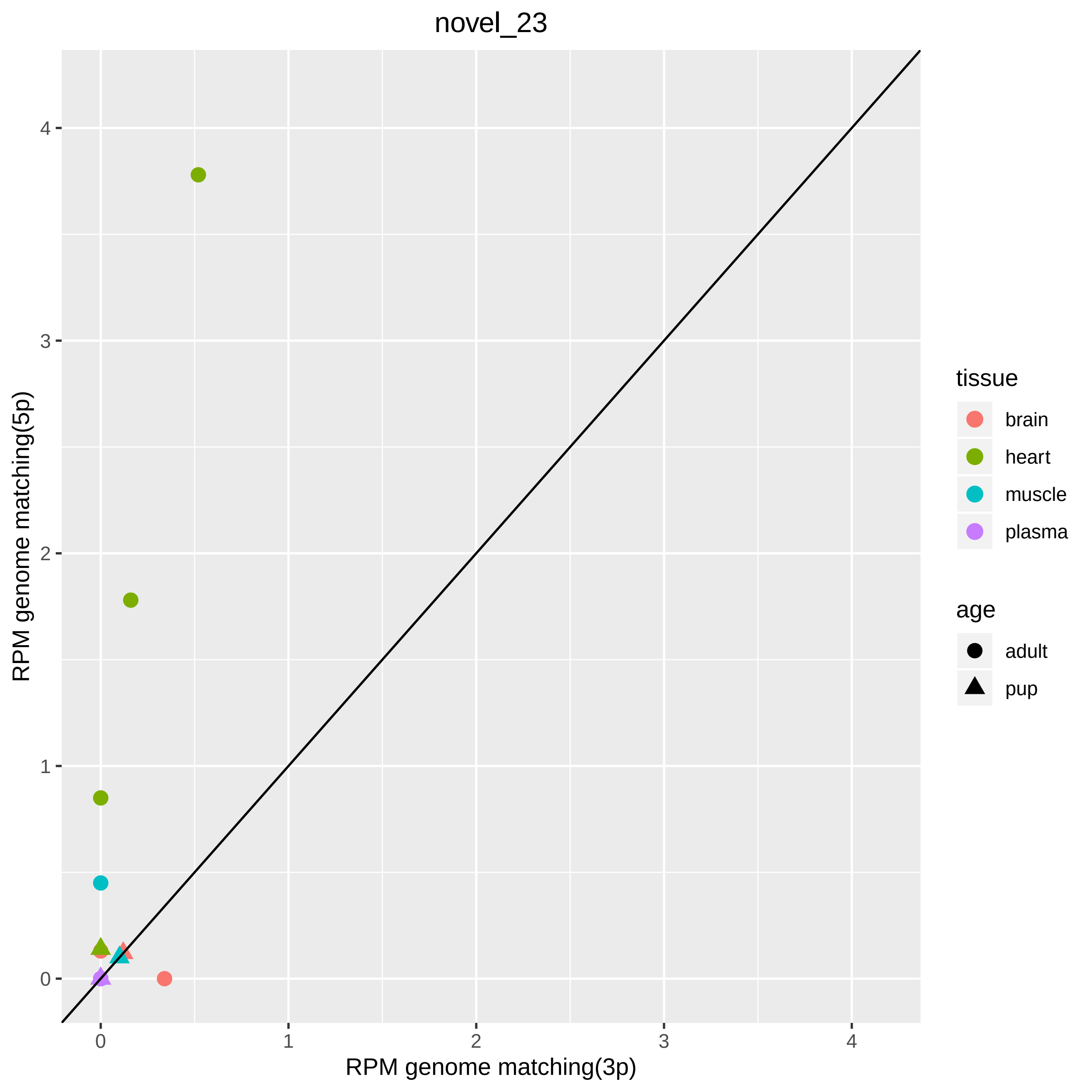
novel_23
RPM genome matching(5p)
RPM genome matching(3p)
0
1
2
3
4
0
1
2
3
4
tissue
brain
heart
muscle
plasma
age
adult
pup

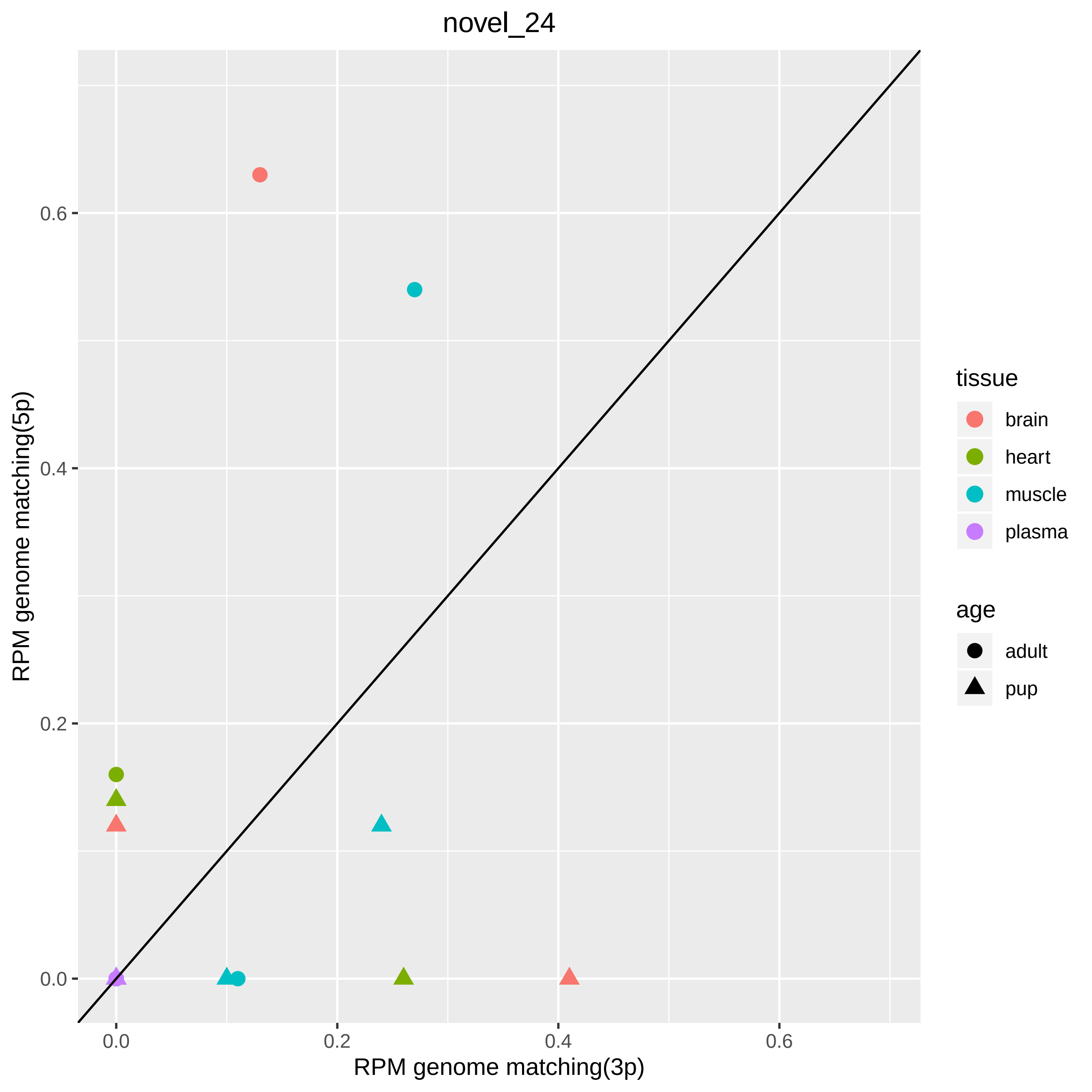

novel_24
RPM genome matching(5p)
RPM genome matching(3p)
0.0
0.2
0.4
0.6
0.0
0.2
0.4
0.6
tissue
brain
heart
muscle
plasma
age
adult
pup

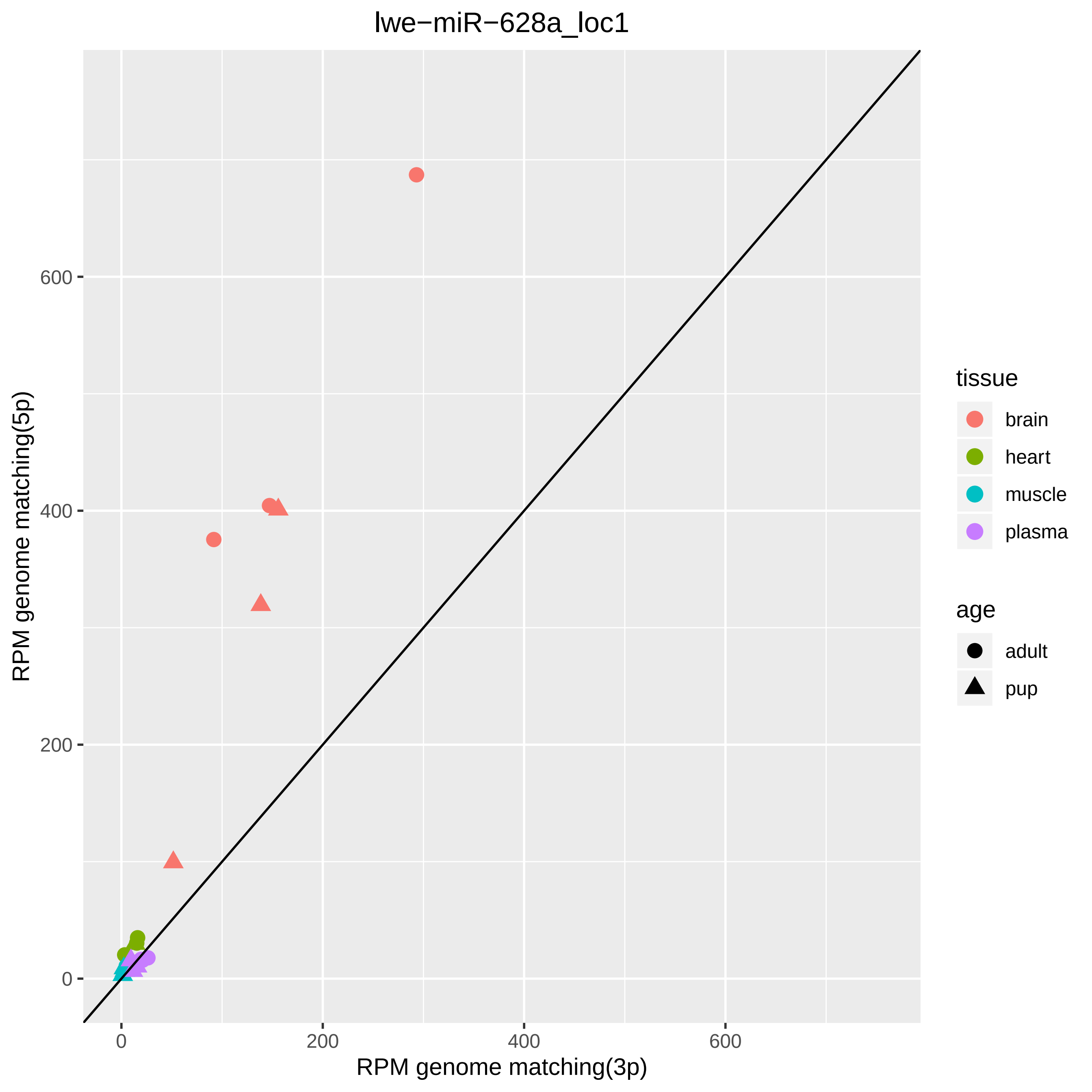

lwe−miR−628a_loc1
RPM genome matching(5p)
RPM genome matching(3p)
0
200
400
600
0
200
400
600
tissue
brain
heart
muscle
plasma
age
adult
pup

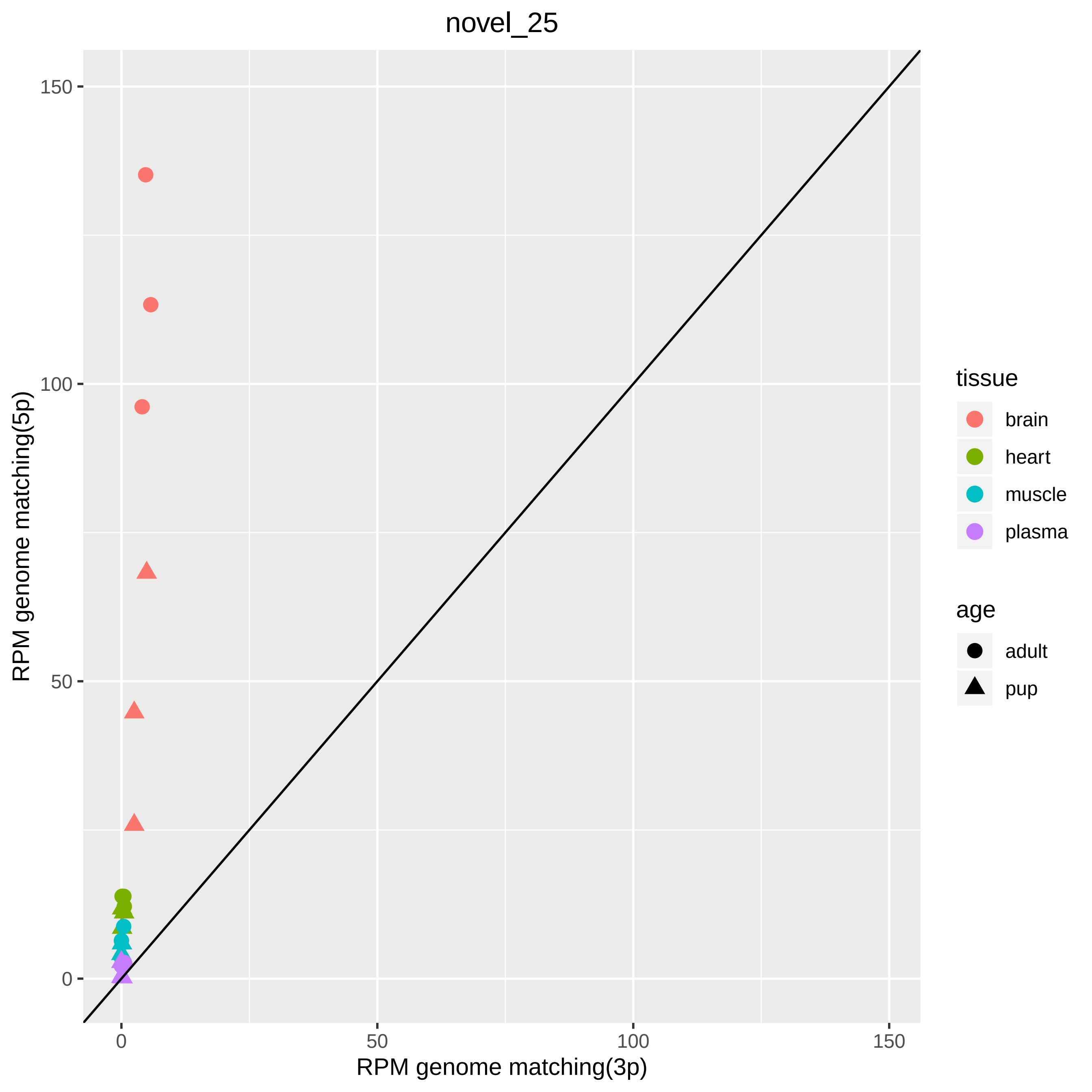
novel_25
RPM genome matching(5p)
RPM genome matching(3p)
0
50
100
150
0
50
100
150
tissue
brain
heart
muscle
plasma
age
adult
pup

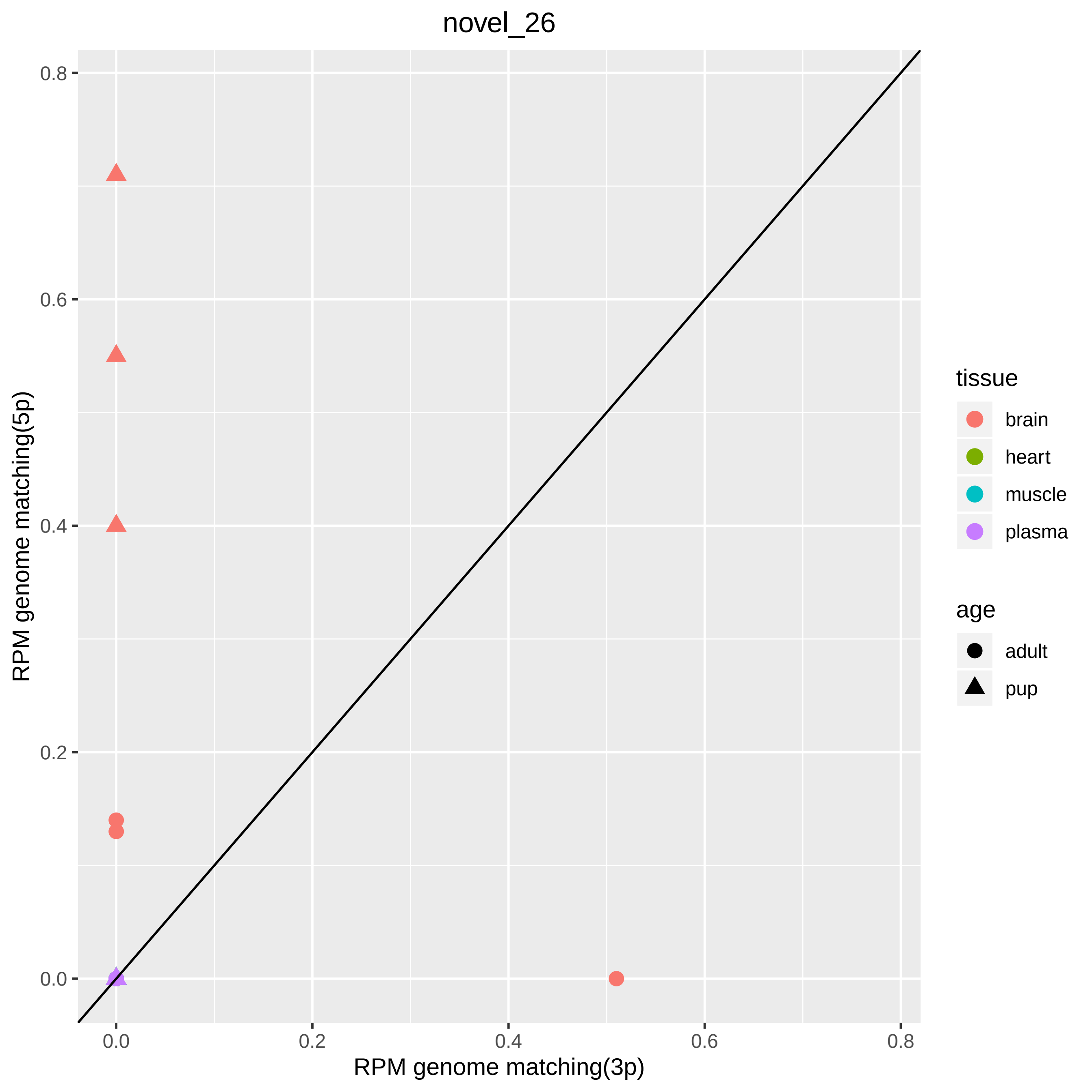

novel_26
0.8
0.6
0.4
0.2
0.0
0.0
0.2
0.4
0.6
0.8
RPM genome matching(5p)
RPM genome matching(3p)
tissue
brain
heart
muscle
plasma
age
adult
pup

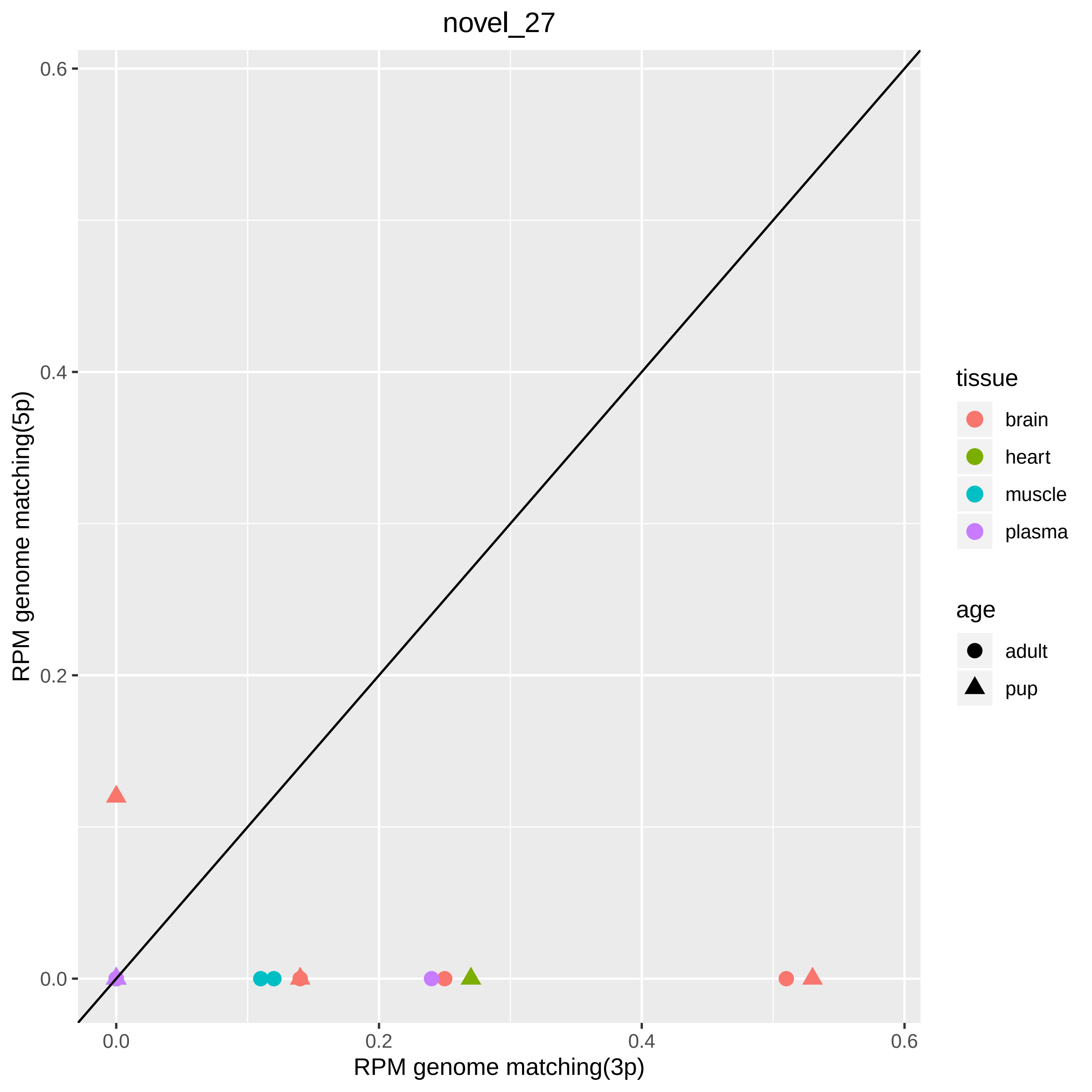
novel_27
RPM genome matching(5p)
RPM genome matching(3p)
0.0
0.2
0.4
0.6
tissue
brain
heart
muscle
plasma
age
adult
pup

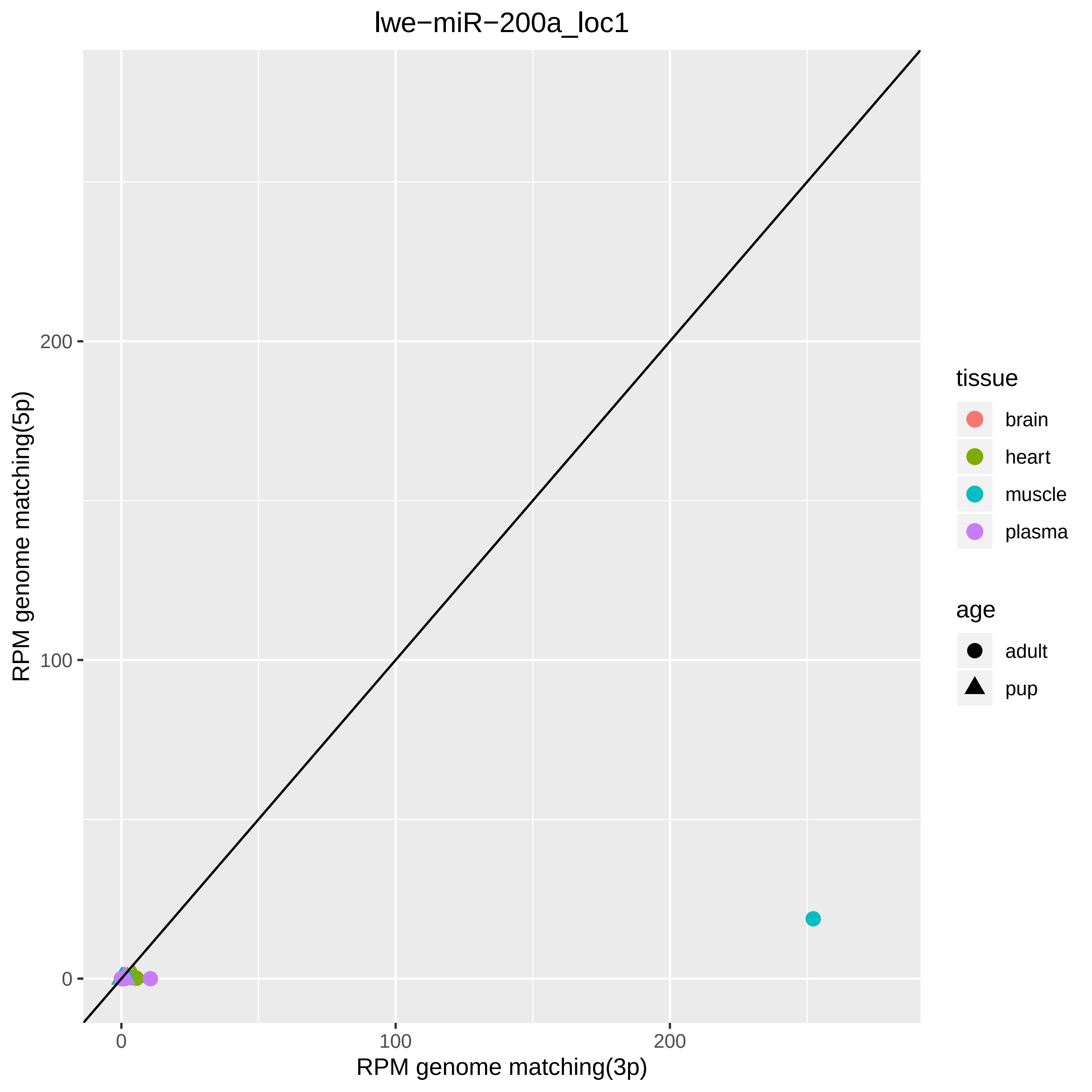
lwe−miR−200a_loc1
RPM genome matching(5p)
RPM genome matching(3p)
0
100
200
0
100
200
tissue
brain
heart
muscle
plasma
age
adult
pup

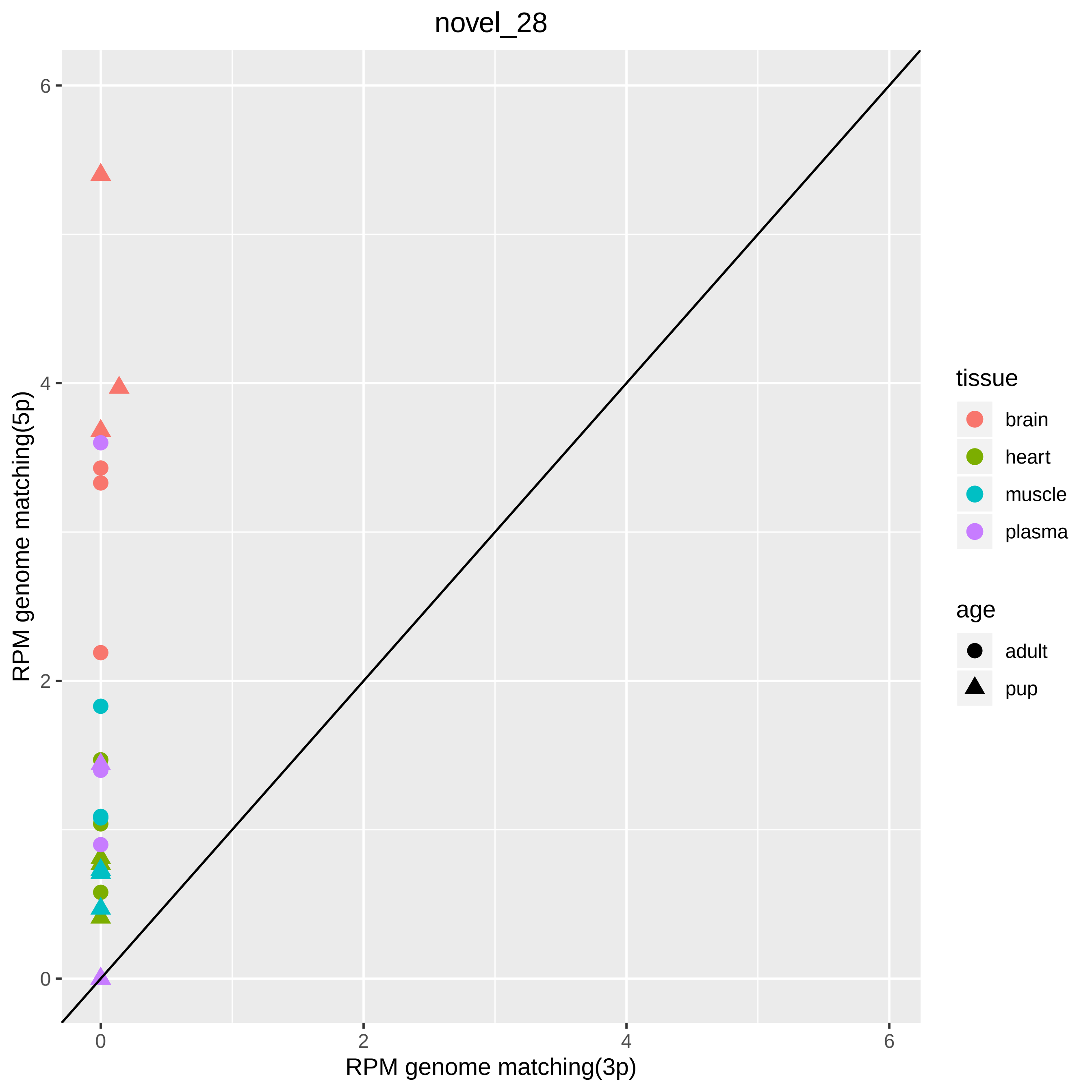
novel_28
RPM genome matching(5p)
RPM genome matching(3p)
0
2
4
6
0
2
4
6
tissue
brain
heart
muscle
plasma
age
adult
pup

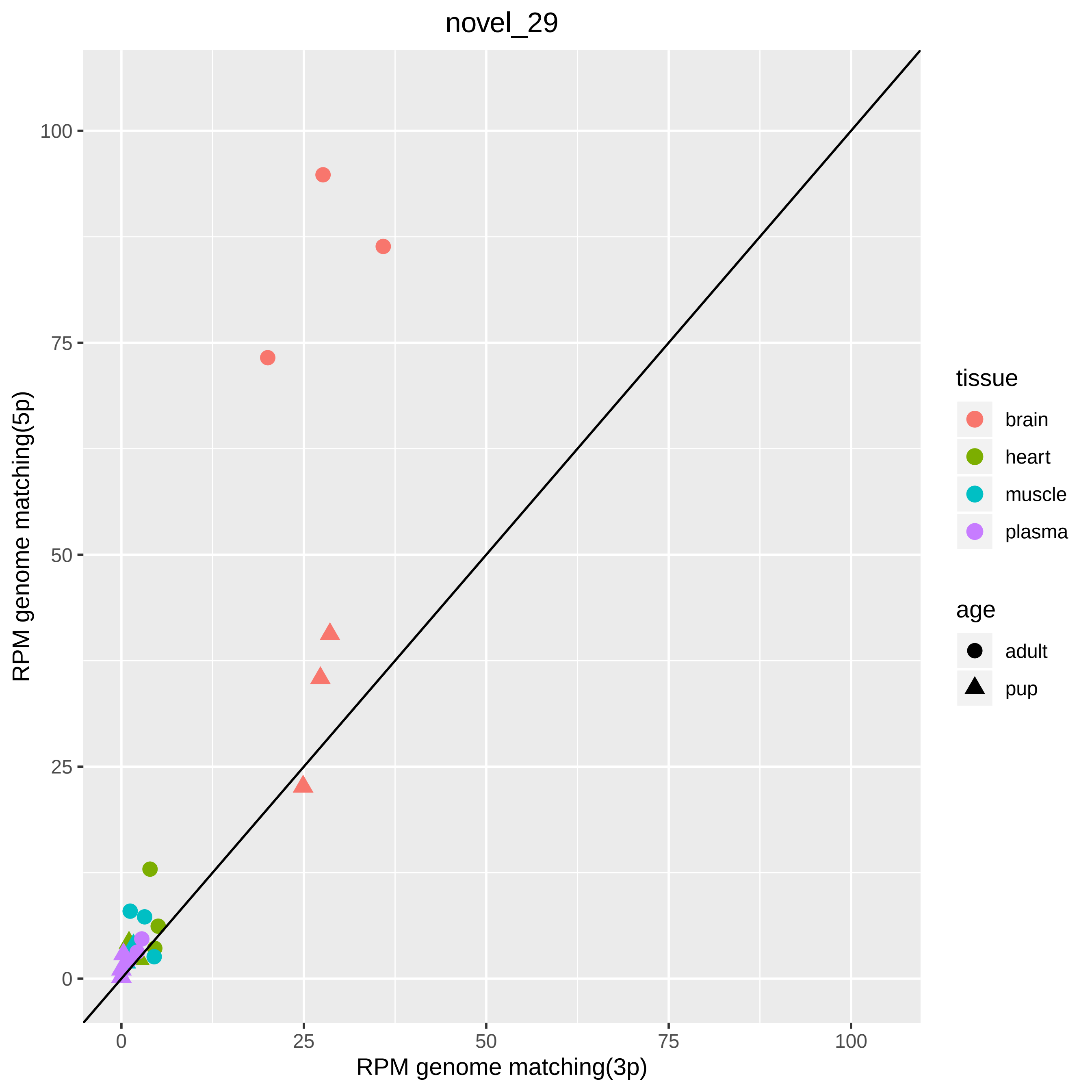
novel_29
RPM genome matching(5p)
RPM genome matching(3p)
0
25
50
75
100
tissue
brain
heart
muscle
plasma
age
adult
pup

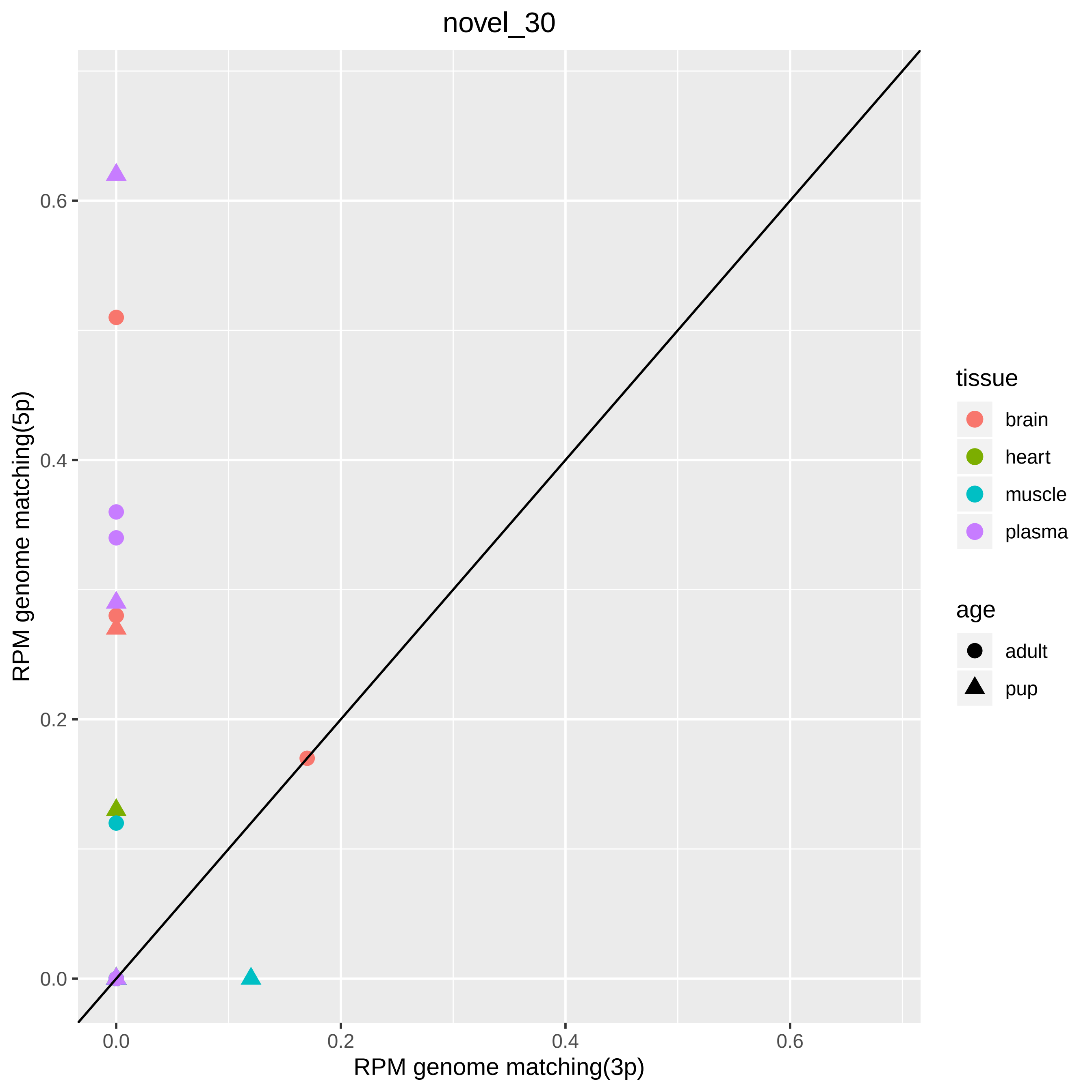
novel_30
RPM genome matching(5p)
RPM genome matching(3p)
0.0
0.2
0.4
0.6
tissue
brain
heart
muscle
plasma
age
adult
pup

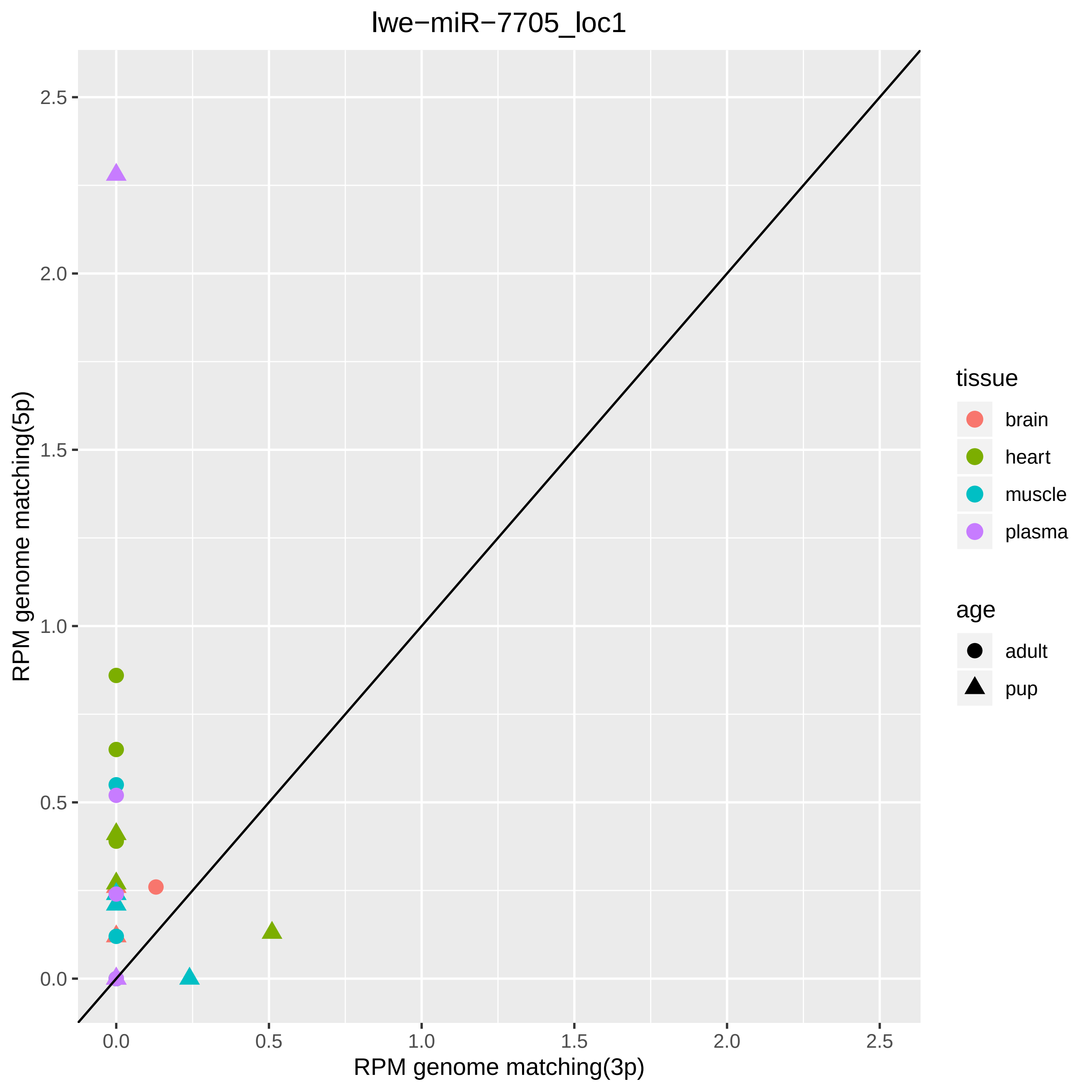
lwe−miR−7705_loc1
RPM genome matching(5p)
RPM genome matching(3p)
0.0
0.5
1.0
1.5
2.0
2.5
tissue
brain
heart
muscle
plasma
age
adult
pup

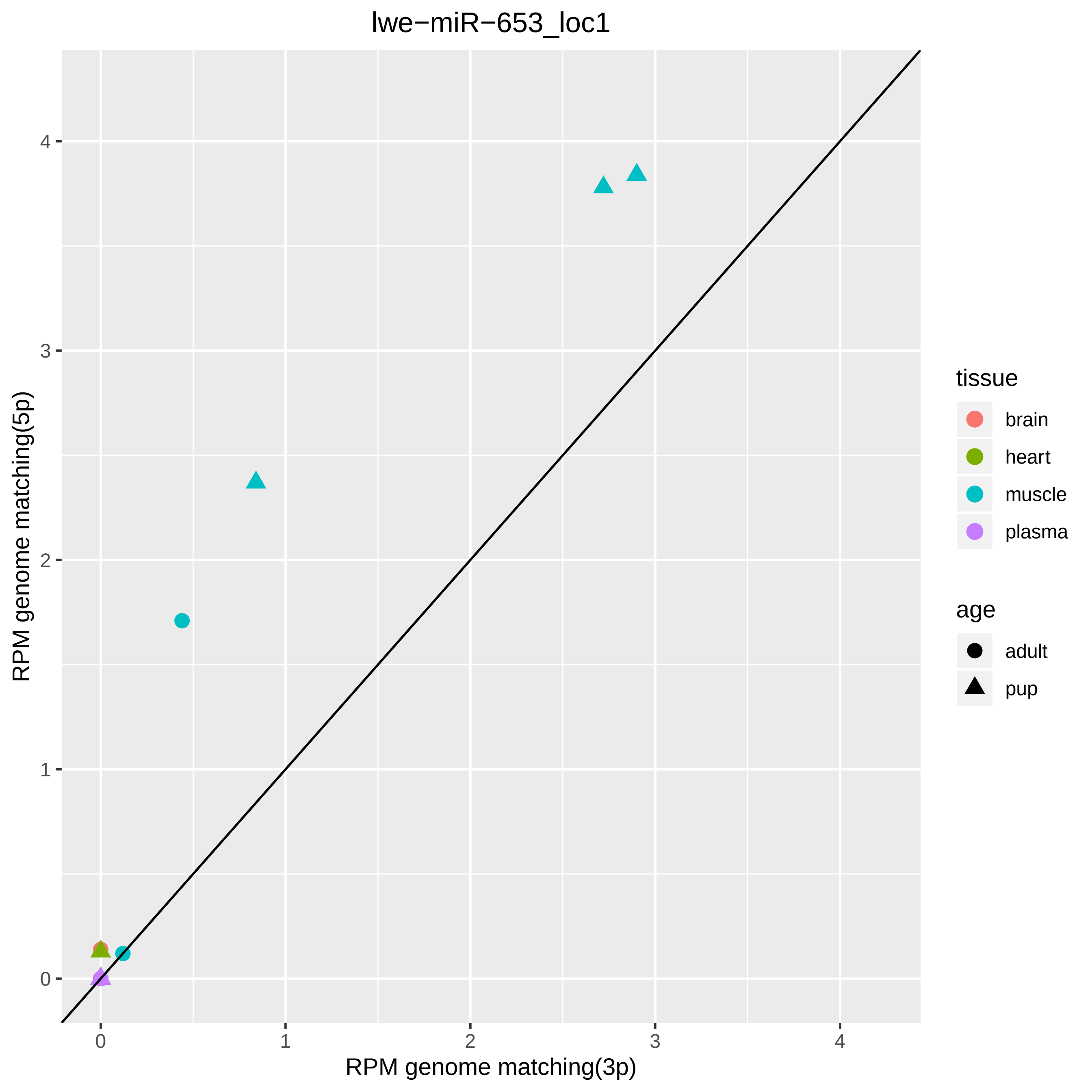
lwe−miR−653_loc1
RPM genome matching(5p)
RPM genome matching(3p)
0
1
2
3
4
tissue
brain
heart
muscle
plasma
age
adult
pup

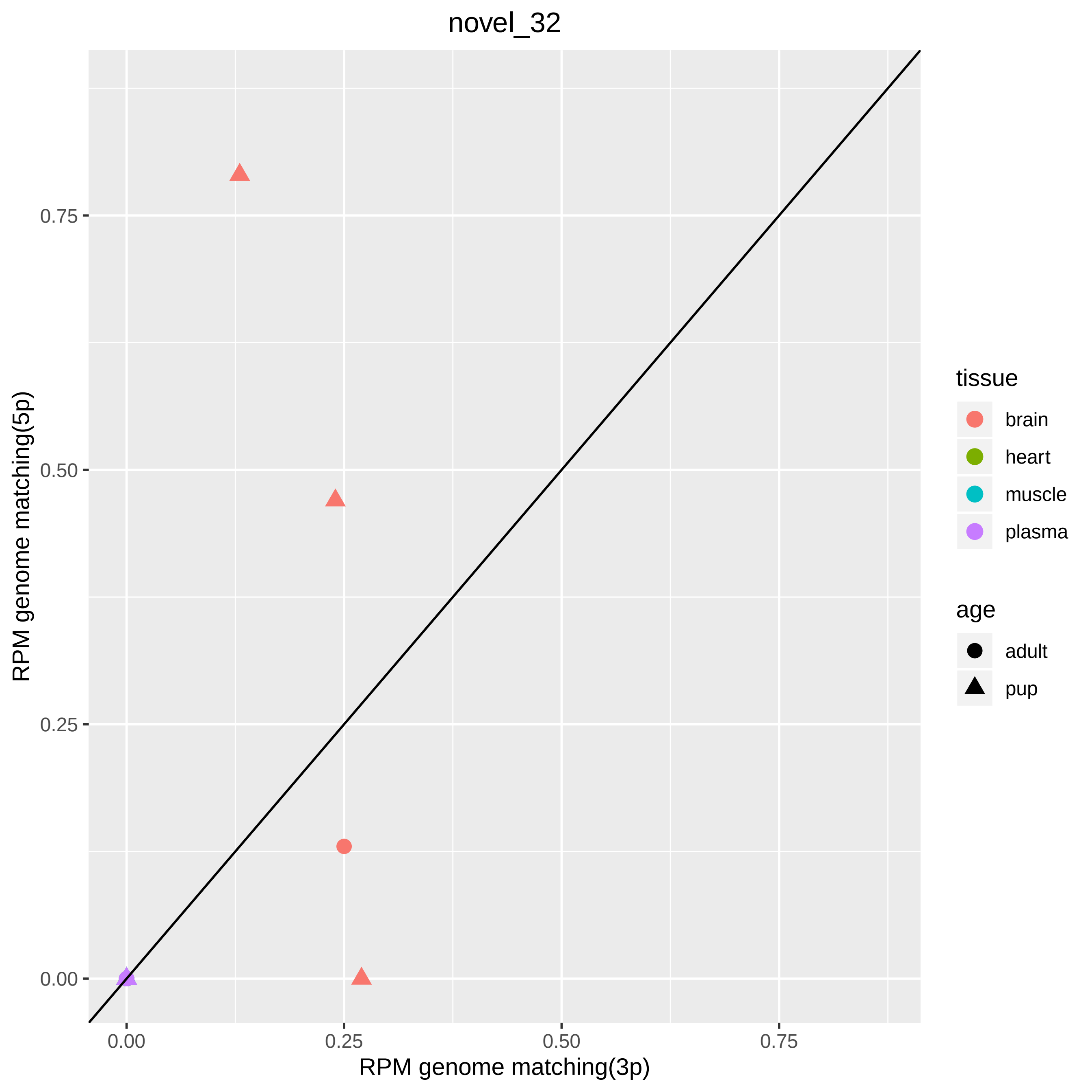

novel_32
RPM genome matching(5p)
RPM genome matching(3p)
0.00
0.25
0.50
0.75
tissue
brain
heart
muscle
plasma
age
adult
pup

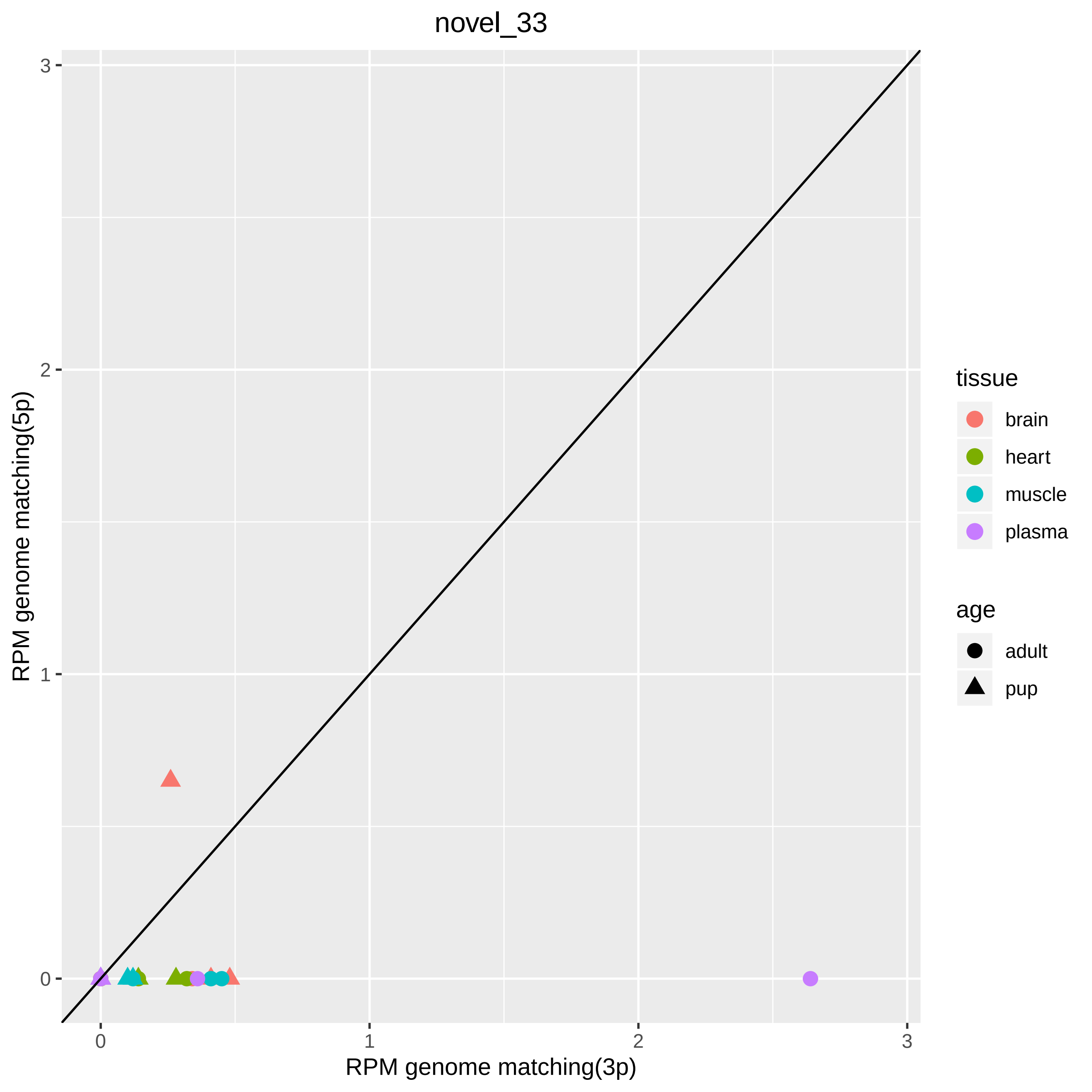
novel_33
RPM genome matching(5p)
RPM genome matching(3p)
0
1
2
3
tissue
brain
heart
muscle
plasma
age
adult
pup

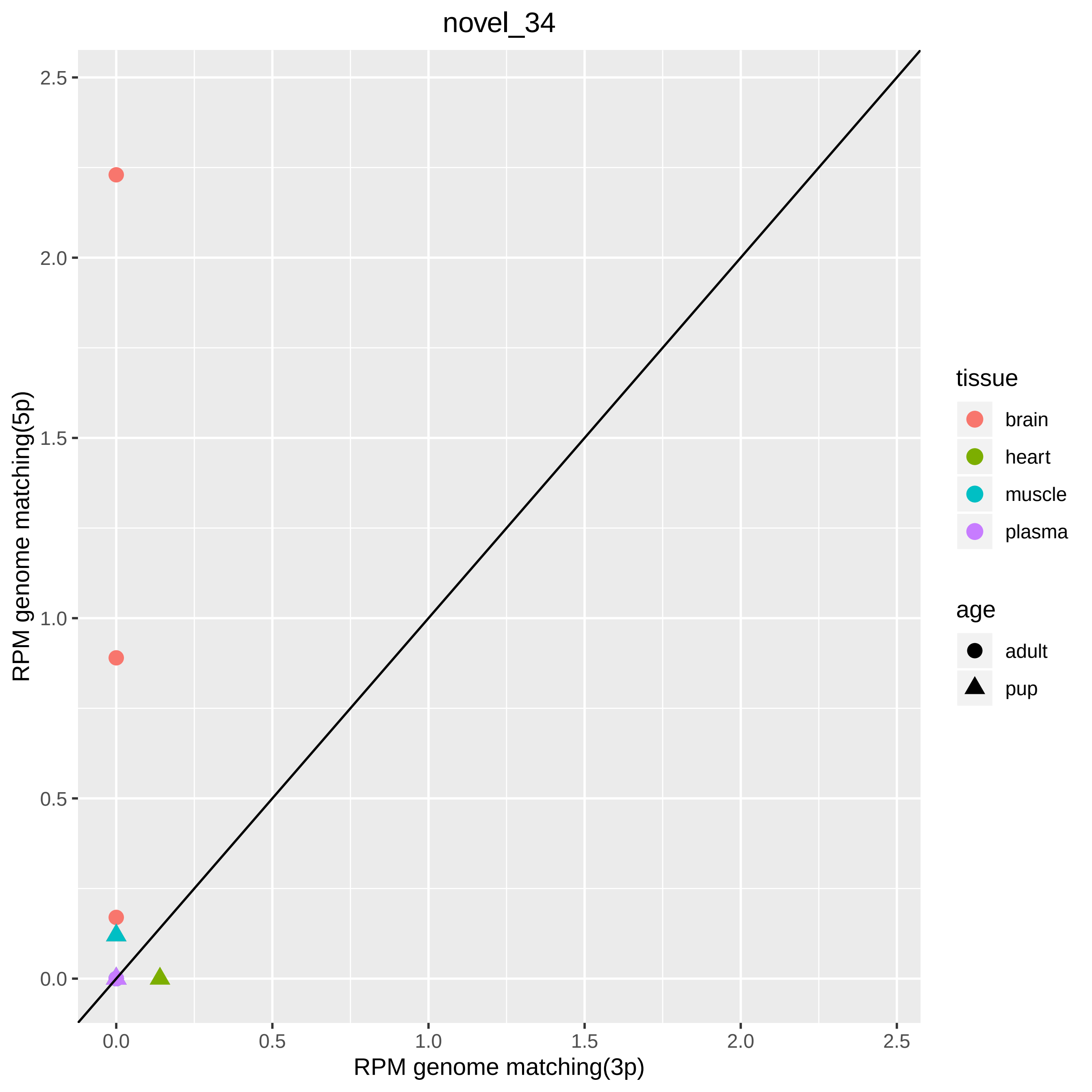

novel_34
RPM genome matching(5p)
RPM genome matching(3p)
0.0
0.5
1.0
1.5
2.0
2.5
tissue
brain
heart
muscle
plasma
age
adult
pup

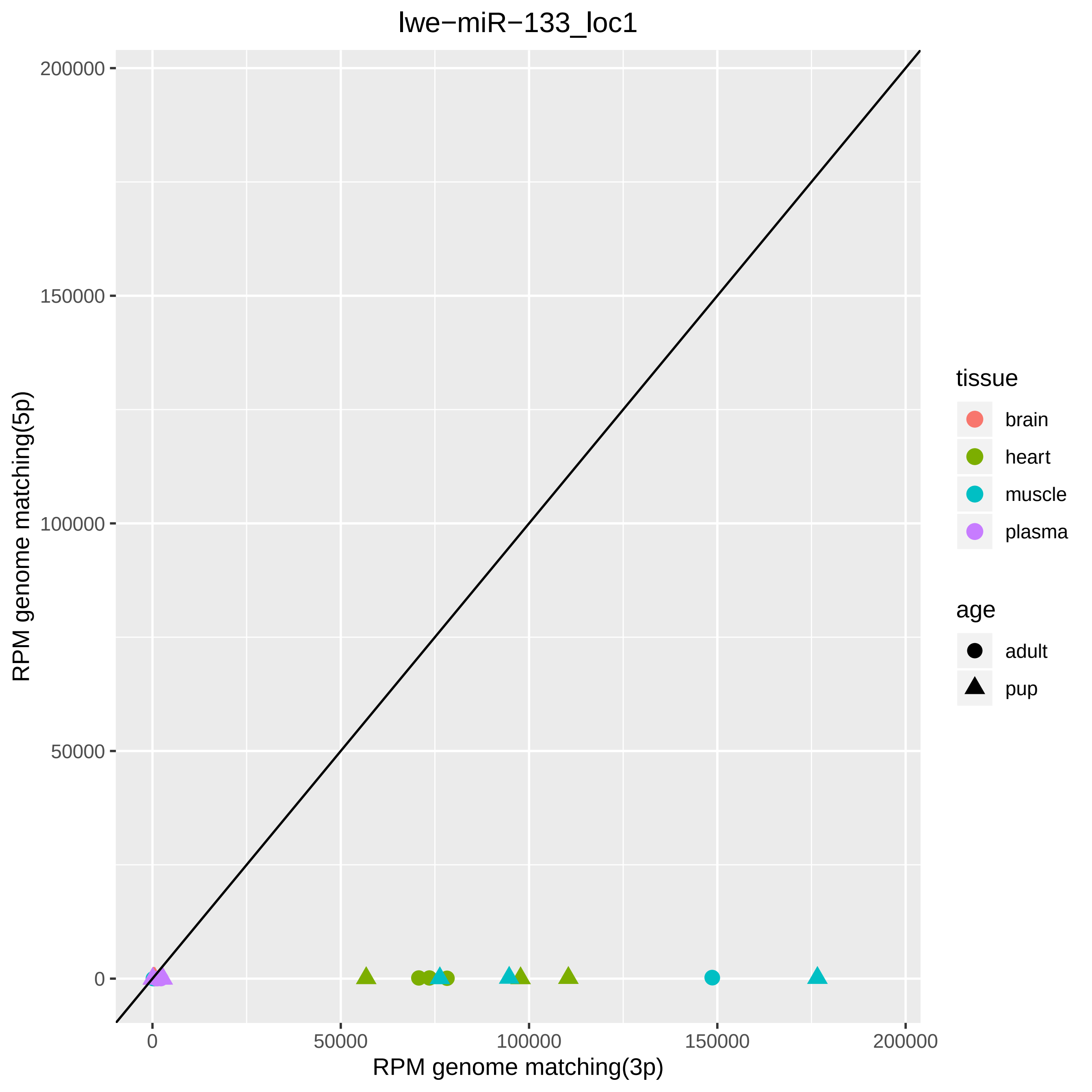

lwe−miR−133_loc1
RPM genome matching(5p)
RPM genome matching(3p)
200000
150000
100000
50000
0
0
50000
100000
150000
200000
tissue
brain
heart
muscle
plasma
age
adult
pup

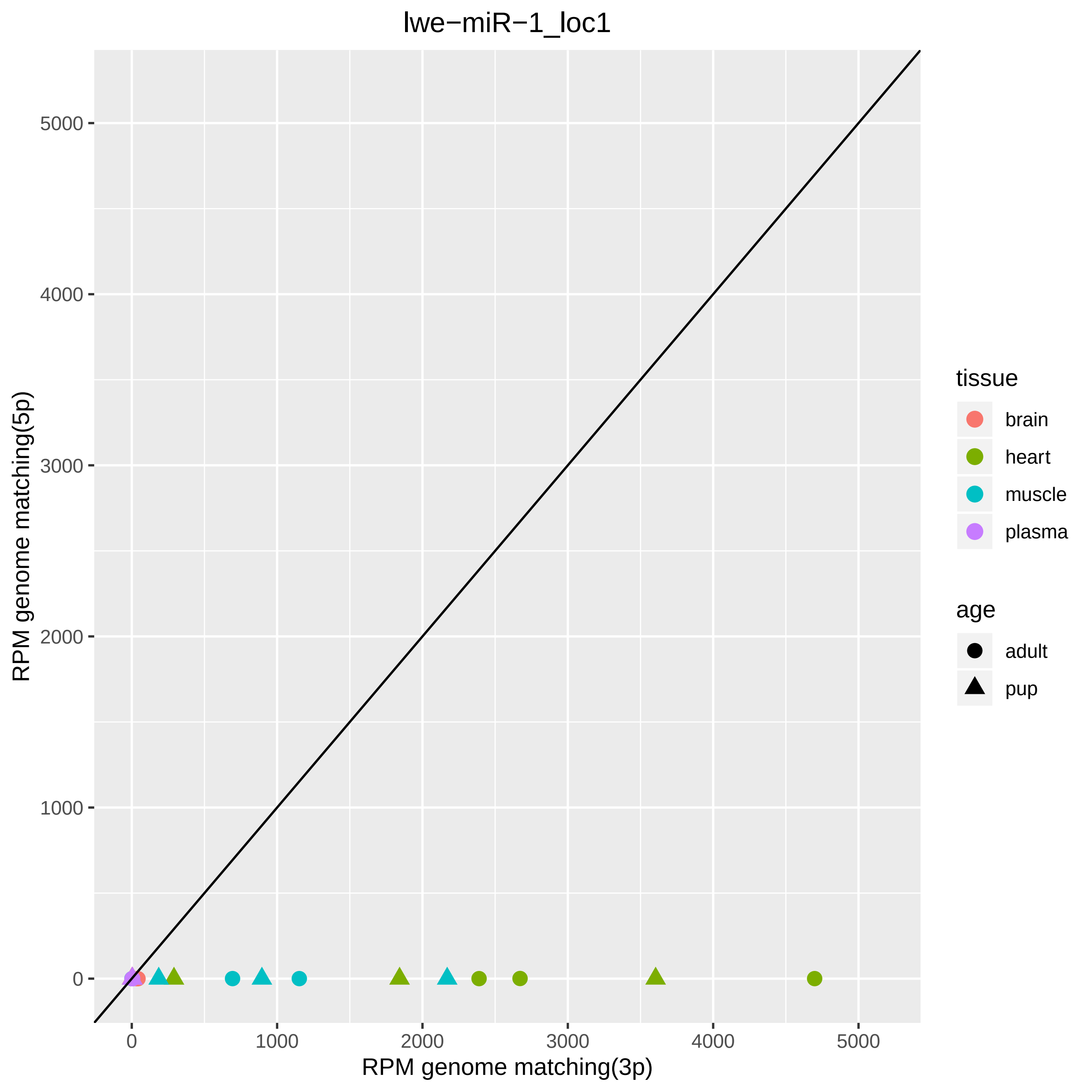
lwe−miR−1_loc1
RPM genome matching(5p)
RPM genome matching(3p)
0
1000
2000
3000
4000
5000
tissue
brain
heart
muscle
plasma
age
adult
pup

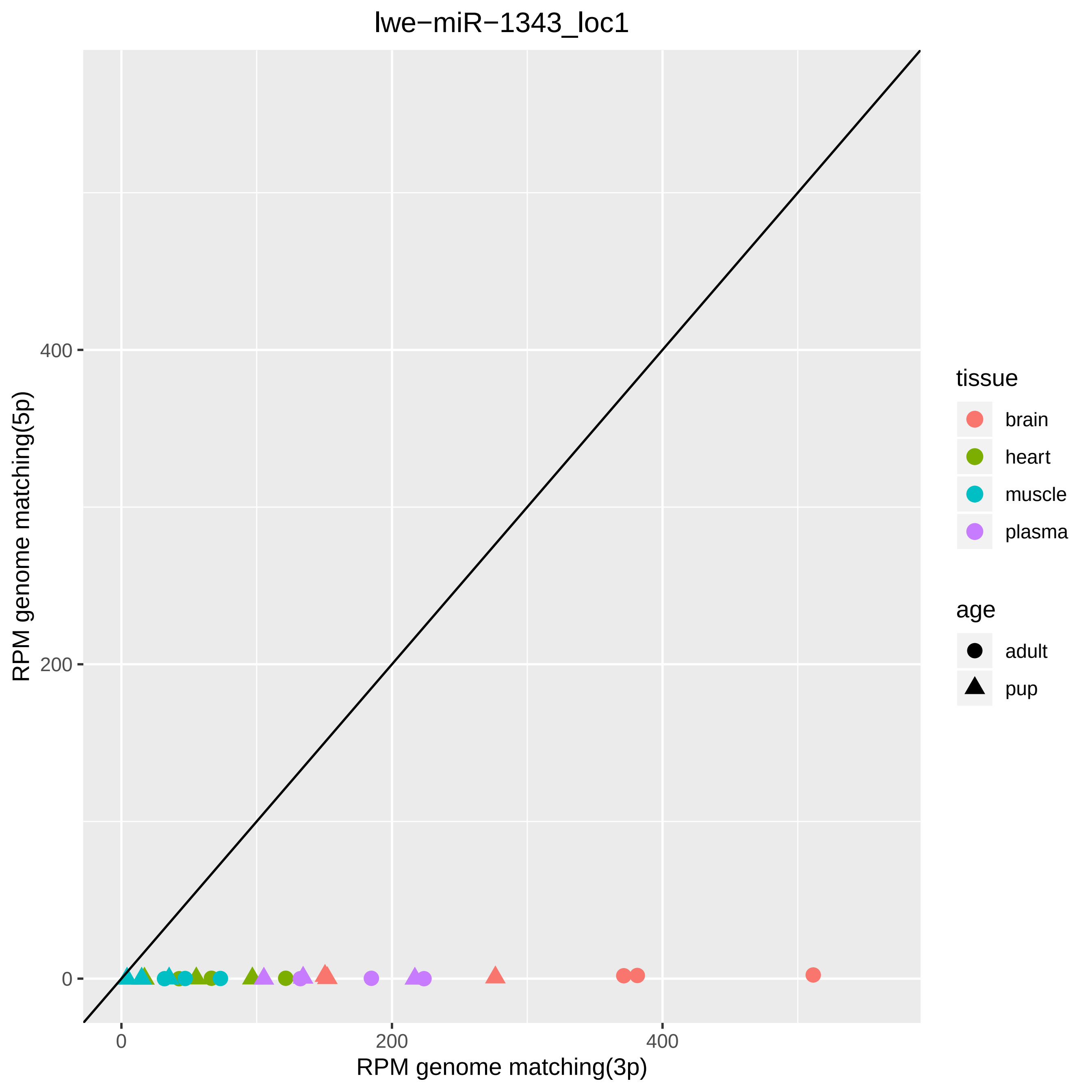

lwe−miR−1343_loc1
RPM genome matching(5p)
RPM genome matching(3p)
0
200
400
0
200
400
tissue
brain
heart
muscle
plasma
age
adult
pup

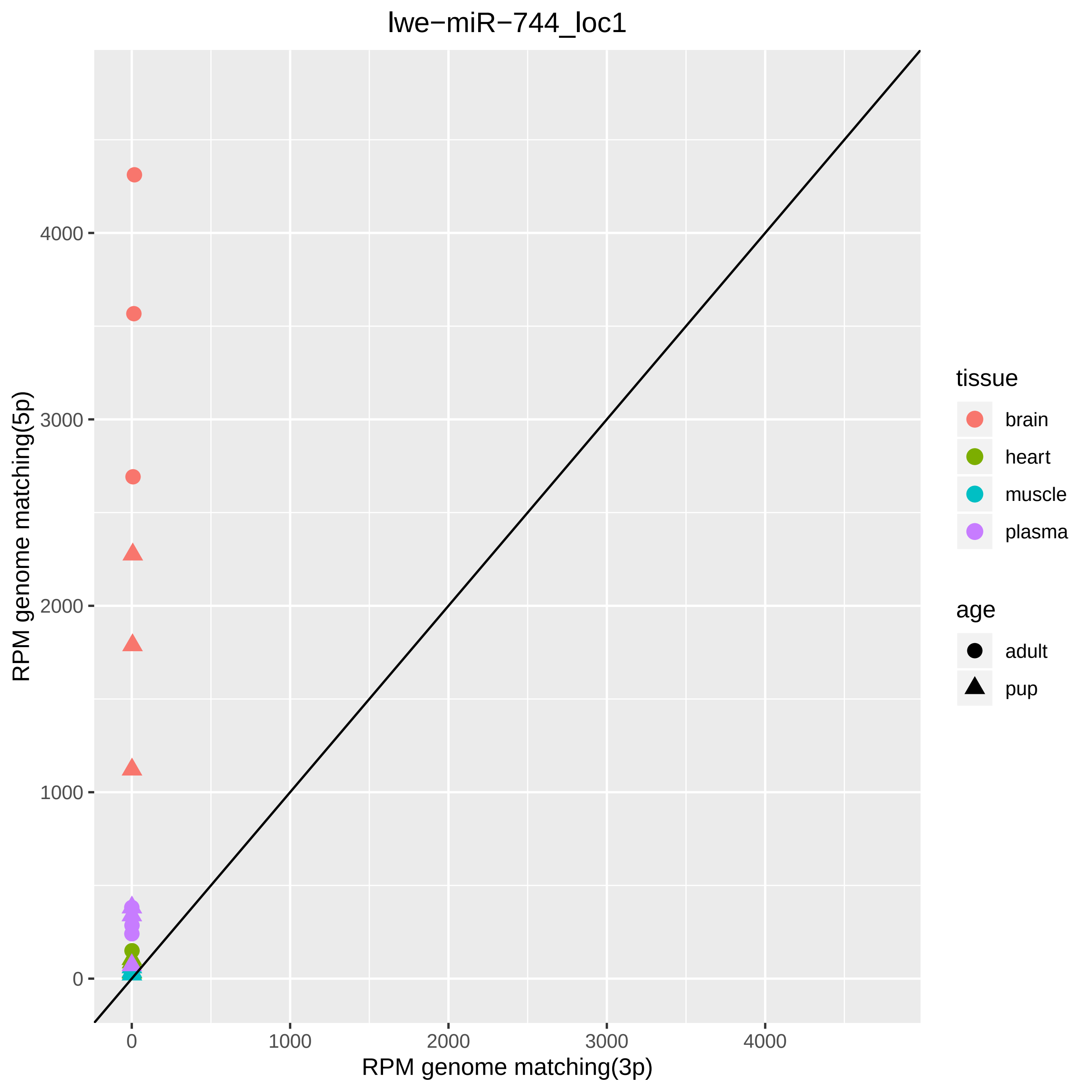

lwe−miR−744_loc1
RPM genome matching(5p)
RPM genome matching(3p)
0
1000
2000
3000
4000
tissue
brain
heart
muscle
plasma
age
adult
pup

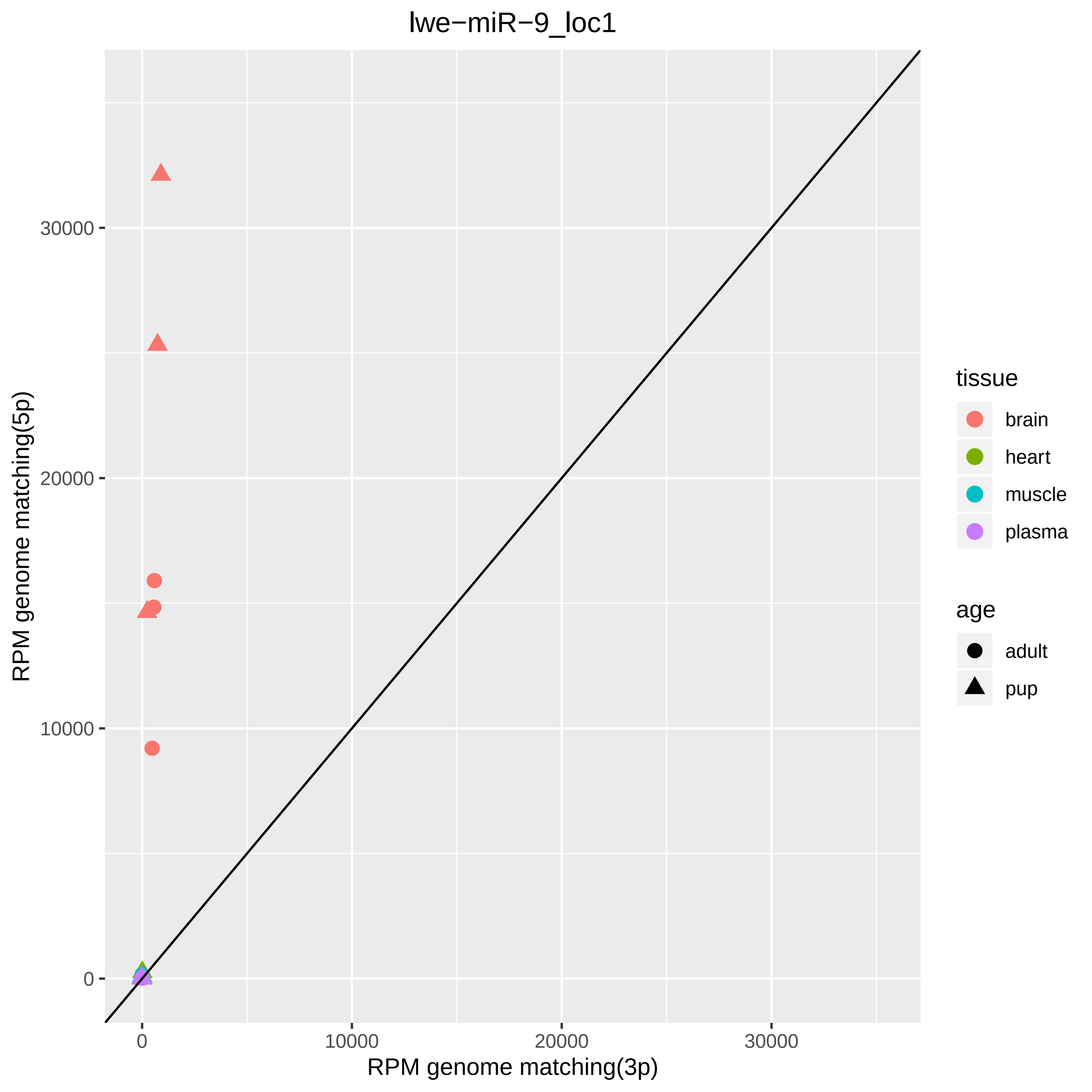
lwe−miR−9_loc1
RPM genome matching(5p)
RPM genome matching(3p)
30000
20000
10000
0
0
10000
20000
30000
tissue
brain
heart
muscle
plasma
age
adult
pup

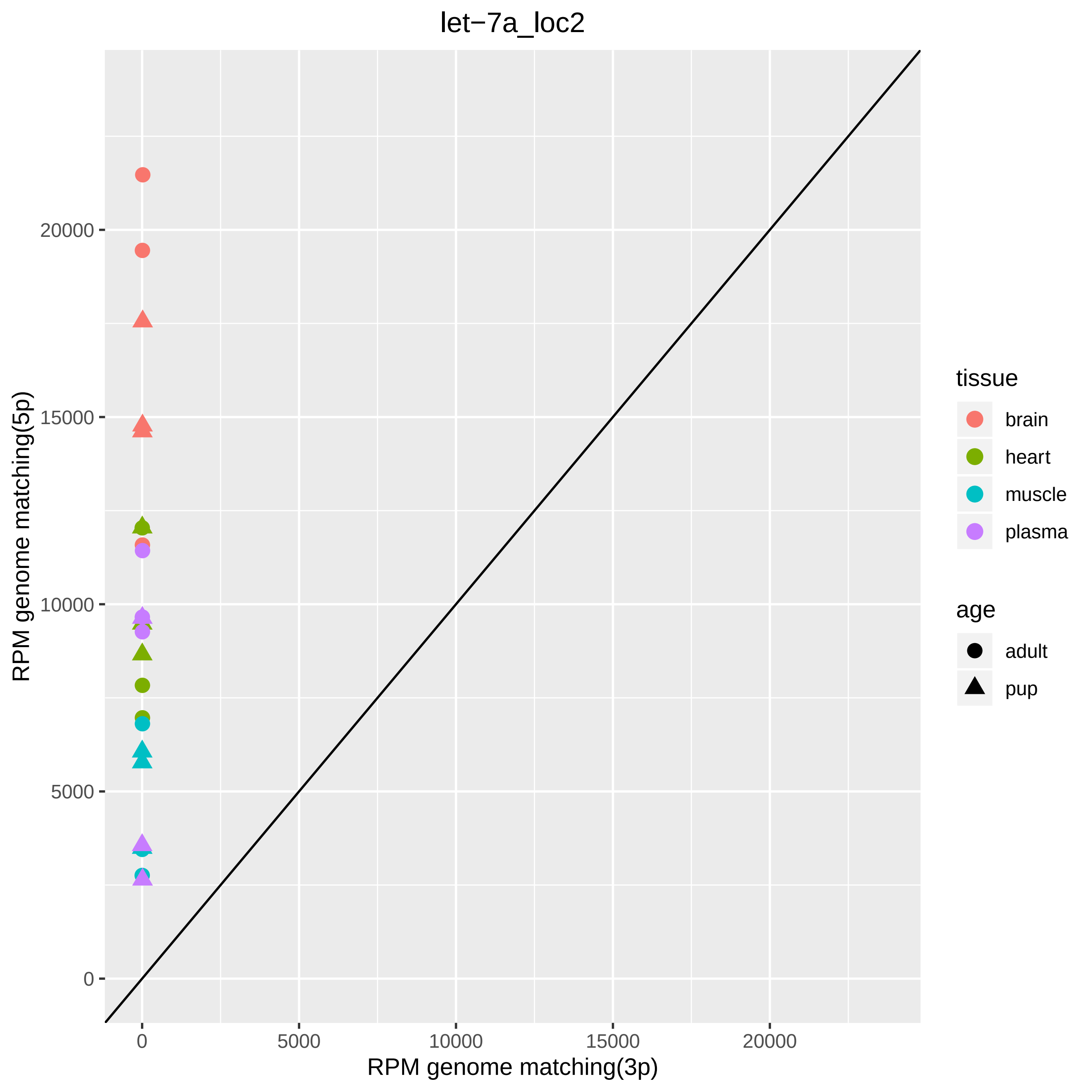
let−7a_loc2
RPM genome matching(5p)
RPM genome matching(3p)
0
5000
10000
15000
20000
tissue
brain
heart
muscle
plasma
age
adult
pup

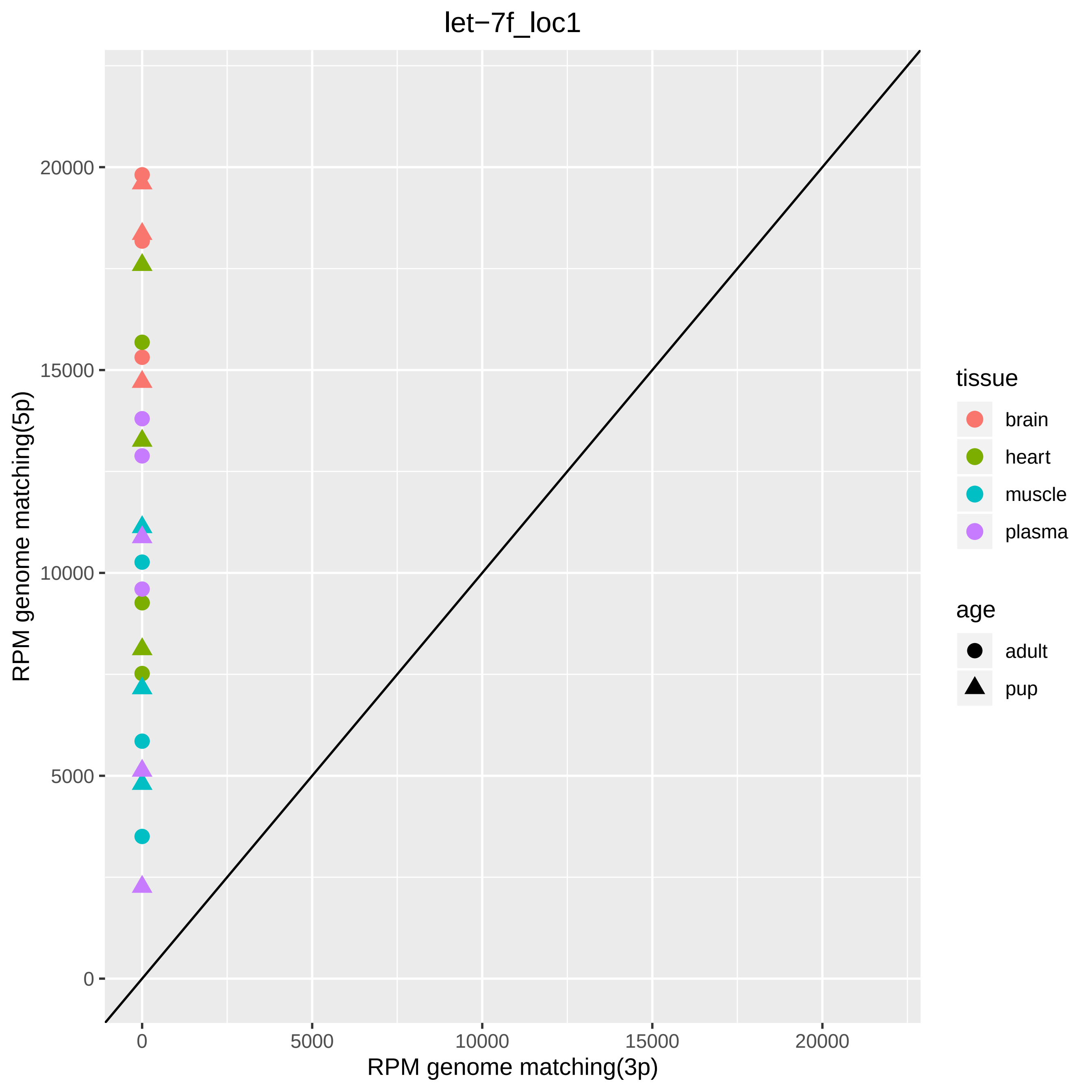

let−7f_loc1
RPM genome matching(5p)
RPM genome matching(3p)
0
5000
10000
15000
20000
tissue
brain
heart
muscle
plasma
age
adult
pup

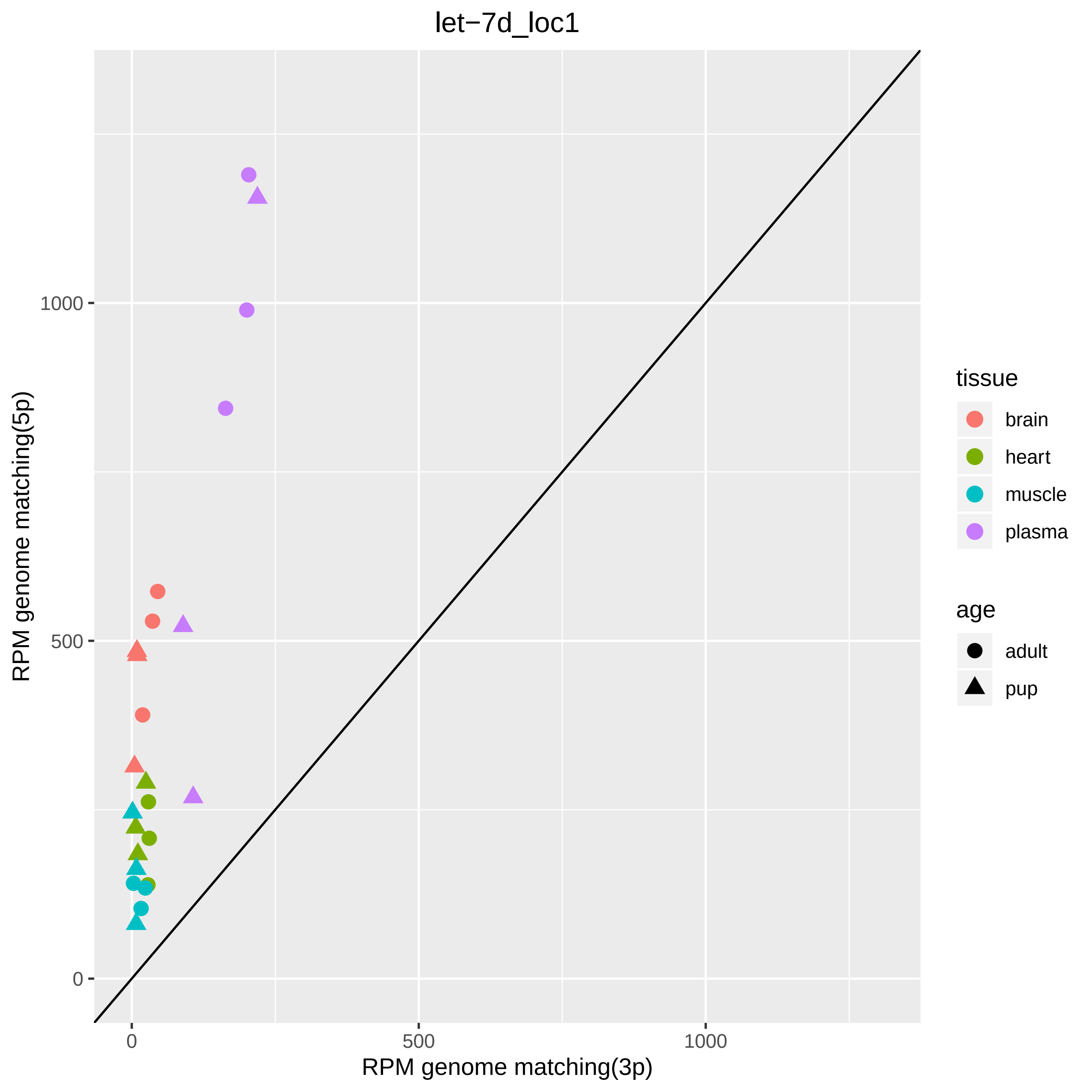

let−7d_loc1
RPM genome matching(5p)
1000
500
0
0
500
1000
RPM genome matching(3p)
tissue
brain
heart
muscle
plasma
age
adult
pup

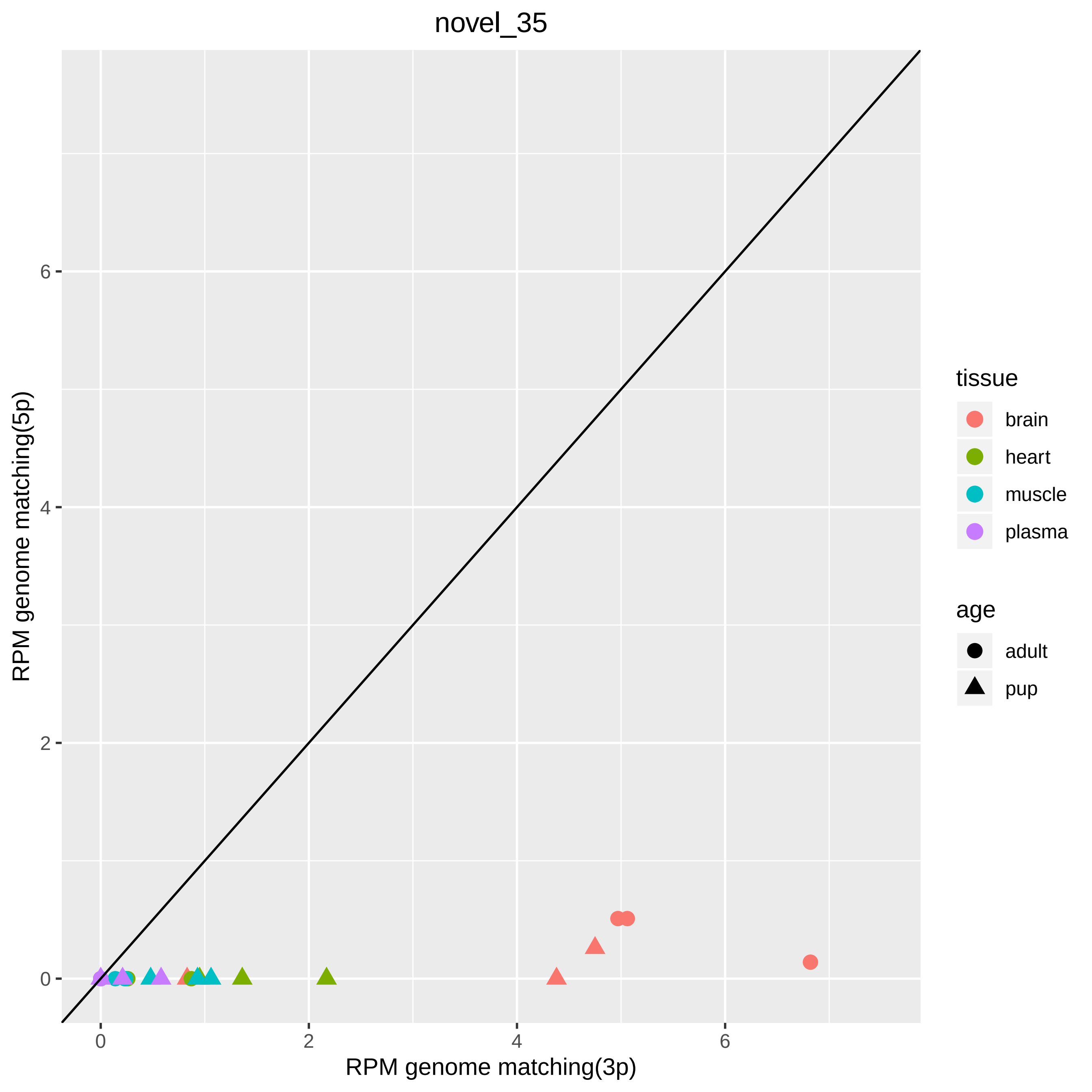
novel_35
RPM genome matching(5p)
RPM genome matching(3p)
0
2
4
6
0
2
4
6
tissue
brain
heart
muscle
plasma
age
adult
pup

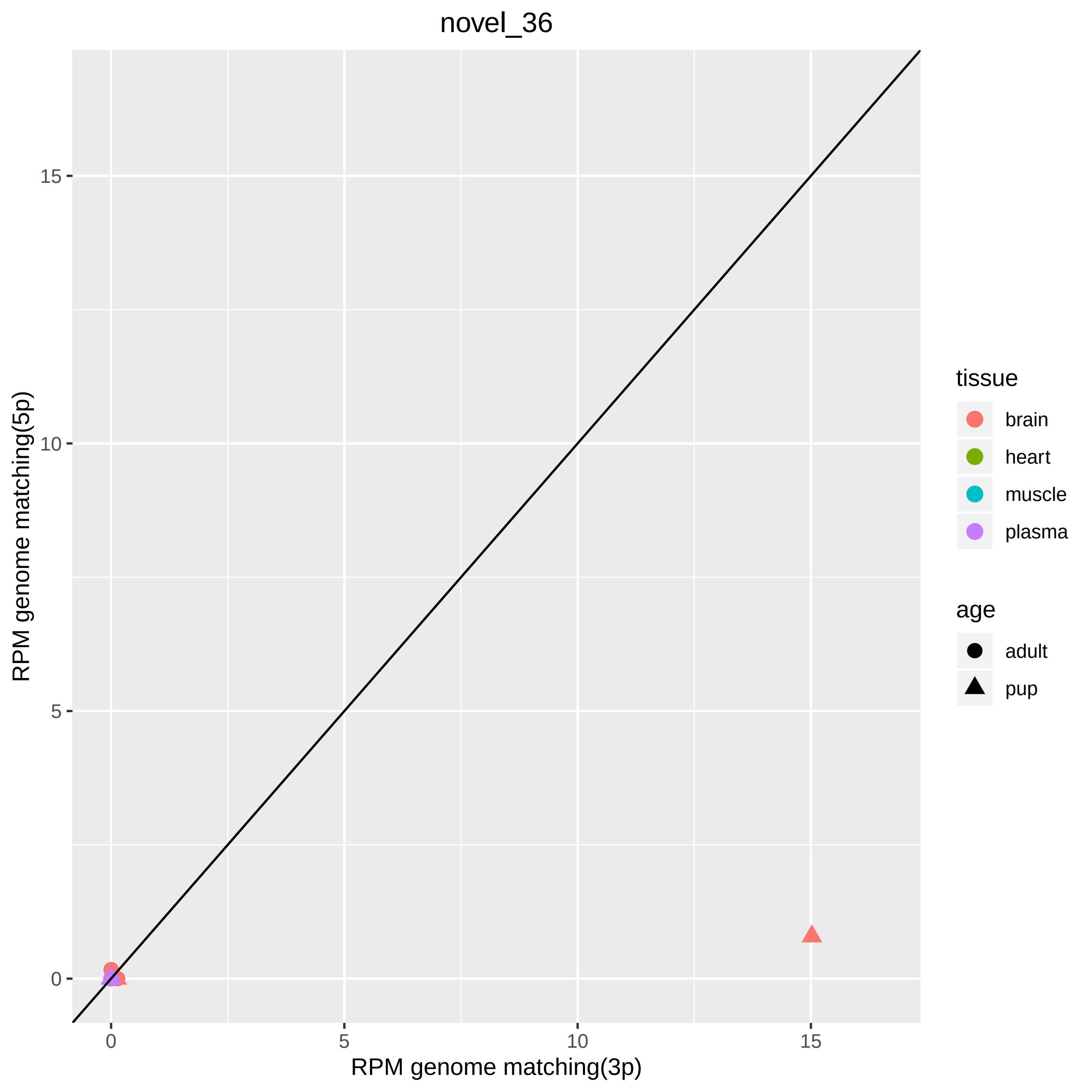
novel_36
RPM genome matching(5p)
RPM genome matching(3p)
0
5
10
15
0
5
10
15
tissue
brain
heart
muscle
plasma
age
adult
pup

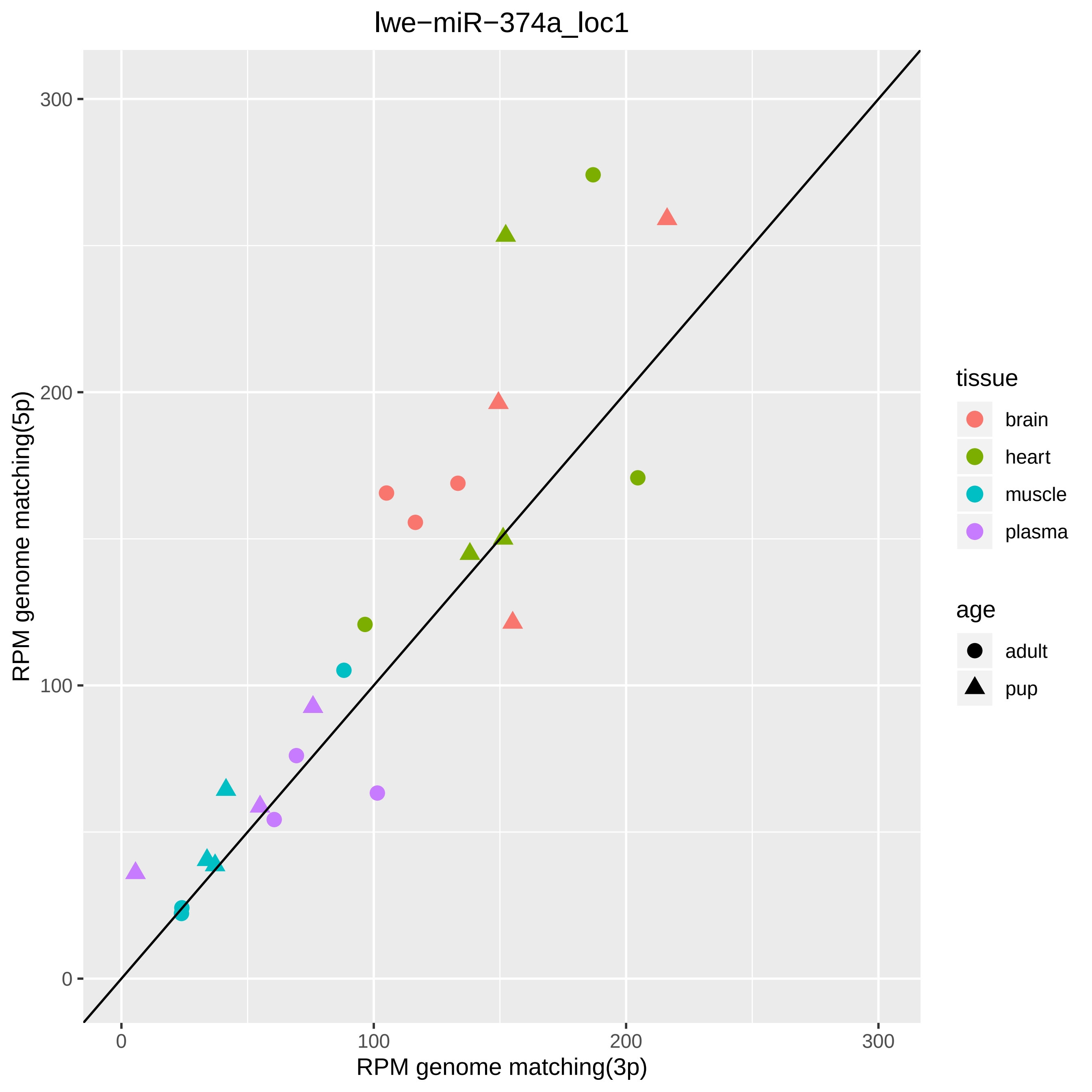

lwe−miR−374a_loc1
RPM genome matching(5p)
RPM genome matching(3p)
0
100
200
300
0
100
200
300
tissue
brain
heart
muscle
plasma
age
adult
pup

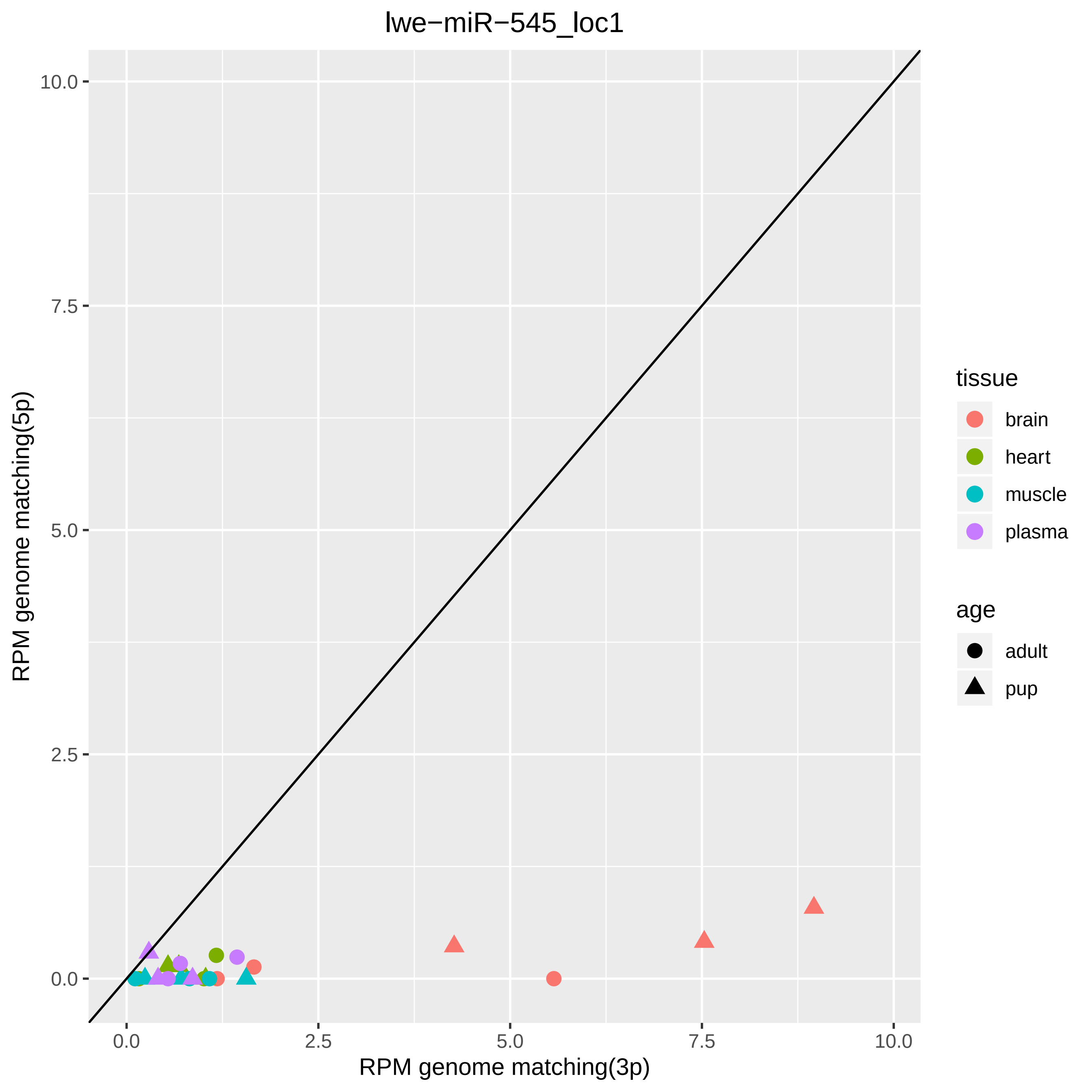

lwe−miR−545_loc1
RPM genome matching(5p)
RPM genome matching(3p)
10.0
7.5
5.0
2.5
0.0
0.0
2.5
5.0
7.5
10.0
tissue
brain
heart
muscle
plasma
age
adult
pup

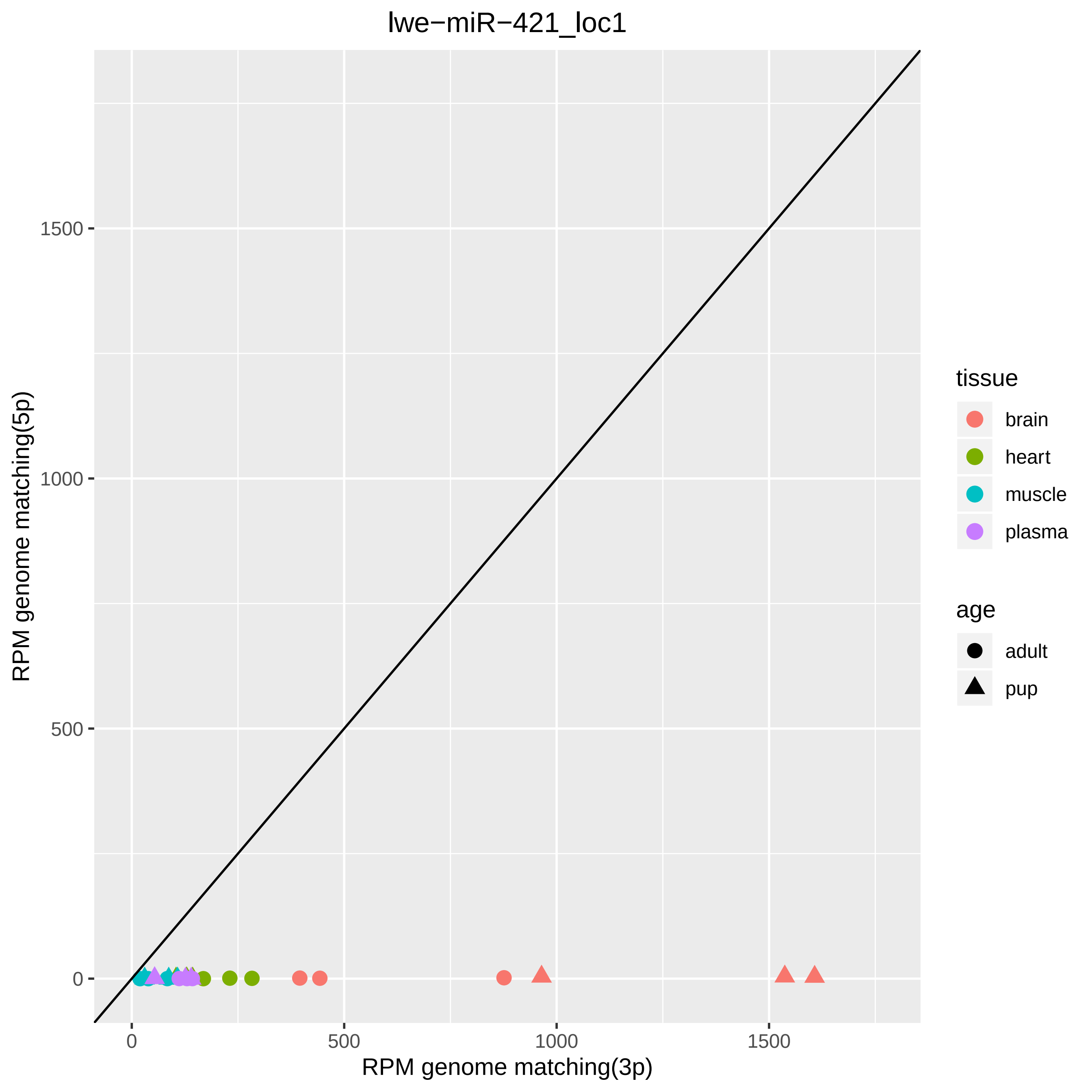

lwe−miR−421_loc1
RPM genome matching(5p)
RPM genome matching(3p)
0
500
1000
1500
tissue
brain
heart
muscle
plasma
age
adult
pup

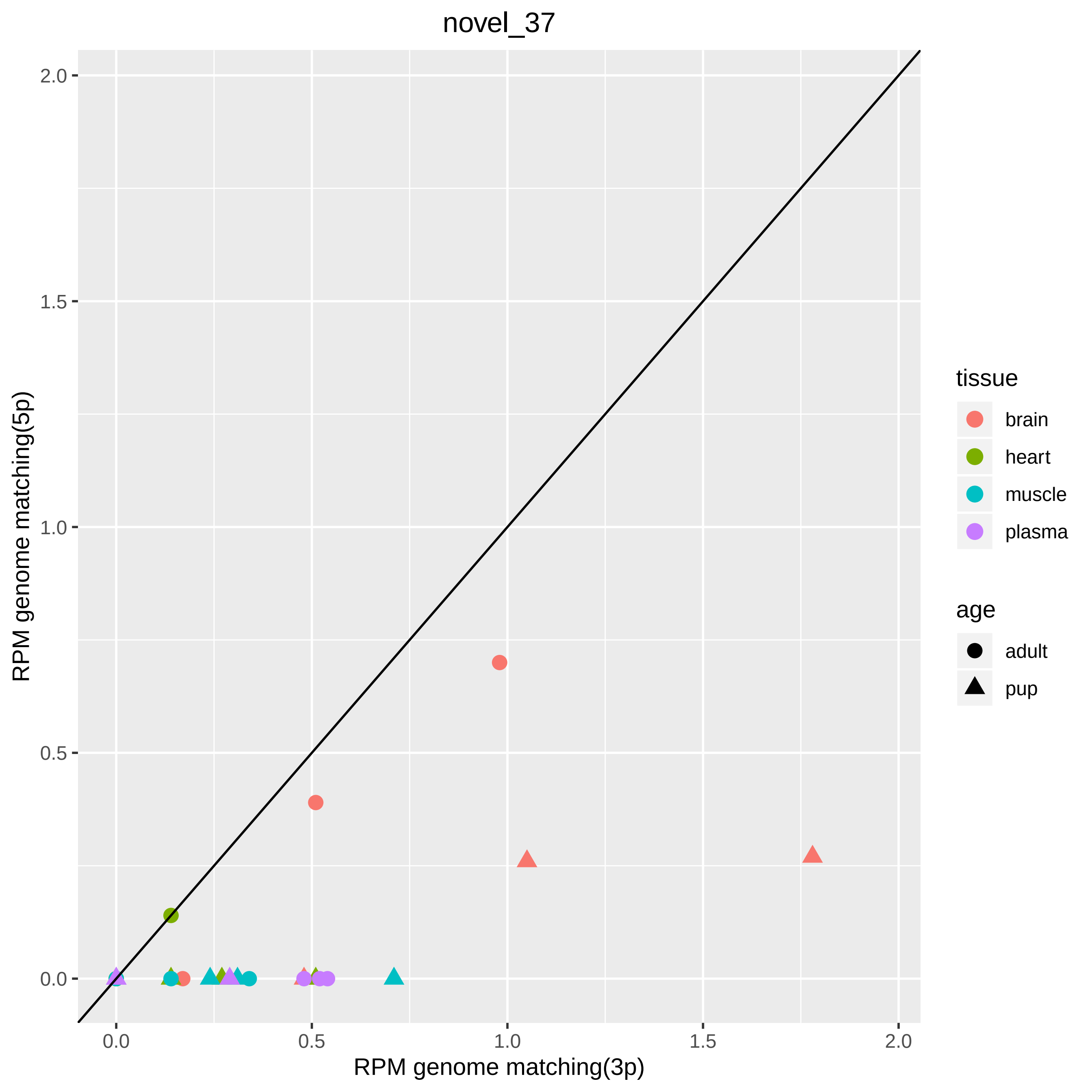

novel_37
RPM genome matching(5p)
RPM genome matching(3p)
0.0
0.5
1.0
1.5
2.0
tissue
brain
heart
muscle
plasma
age
adult
pup

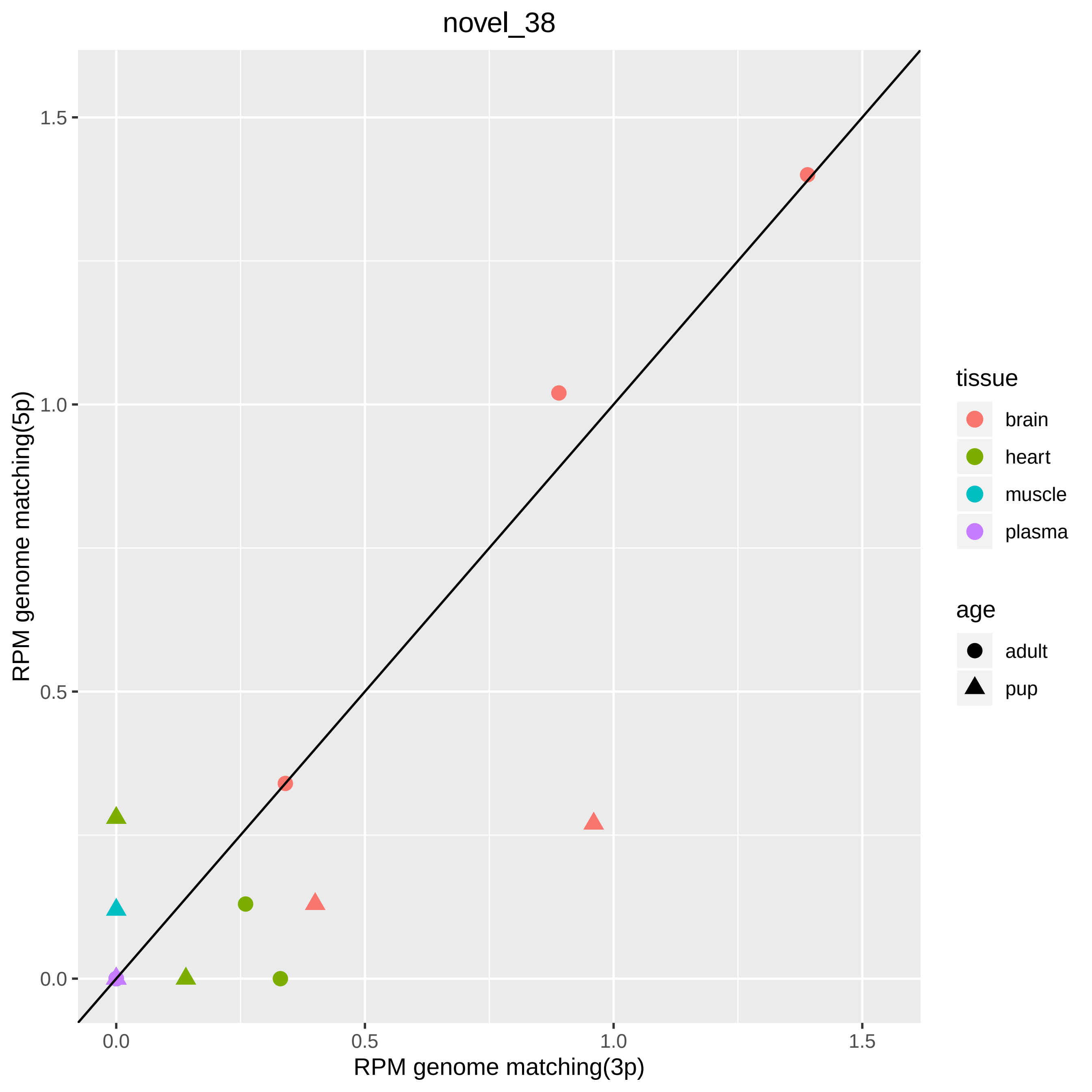

novel_38
1.5
1.0
0.5
0.0
0.0
0.5
1.0
1.5
RPM genome matching(5p)
RPM genome matching(3p)
tissue
brain
heart
muscle
plasma
age
adult
pup

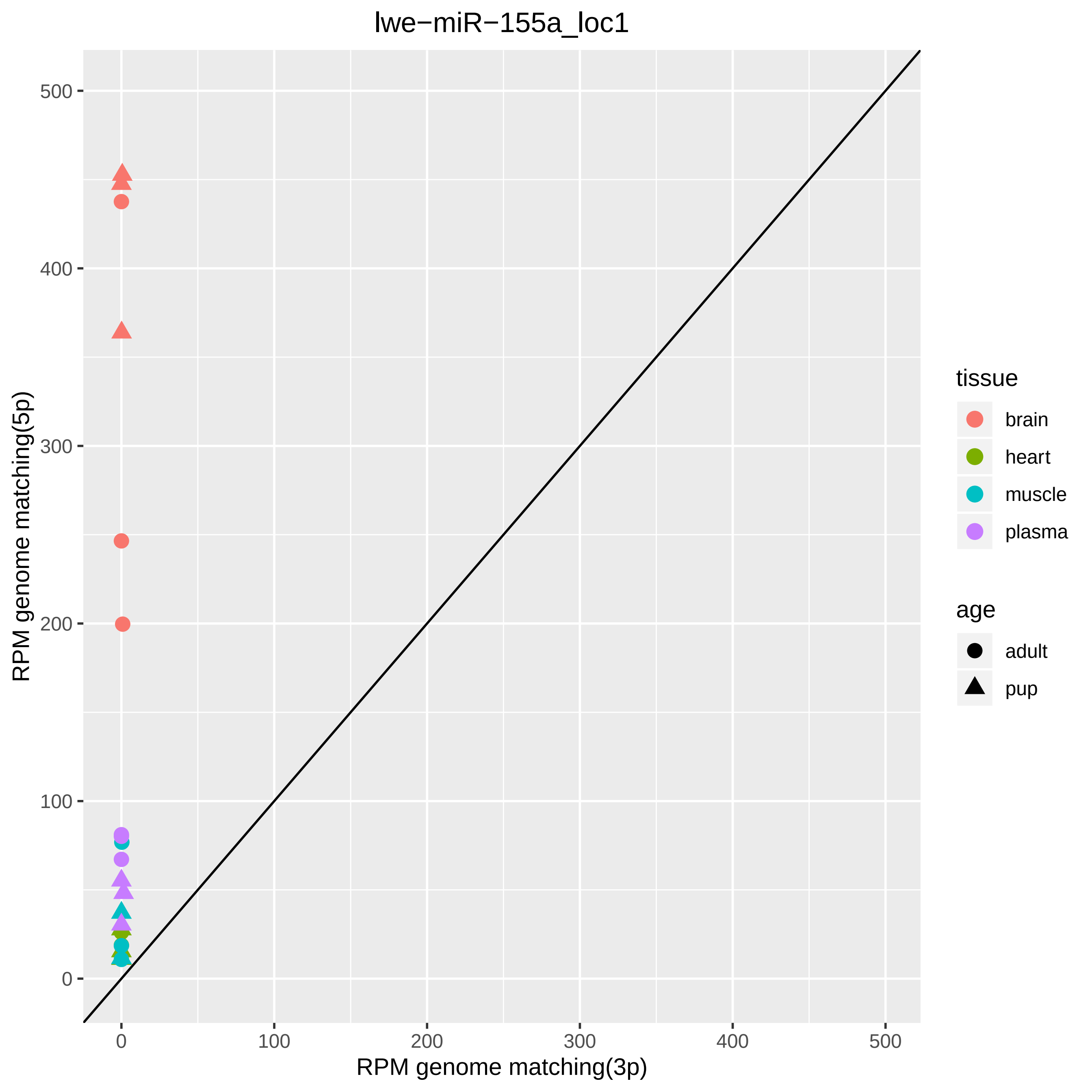
lwe−miR−155a_loc1
RPM genome matching(5p)
RPM genome matching(3p)
0
100
200
300
400
500
tissue
brain
heart
muscle
plasma
age
adult
pup

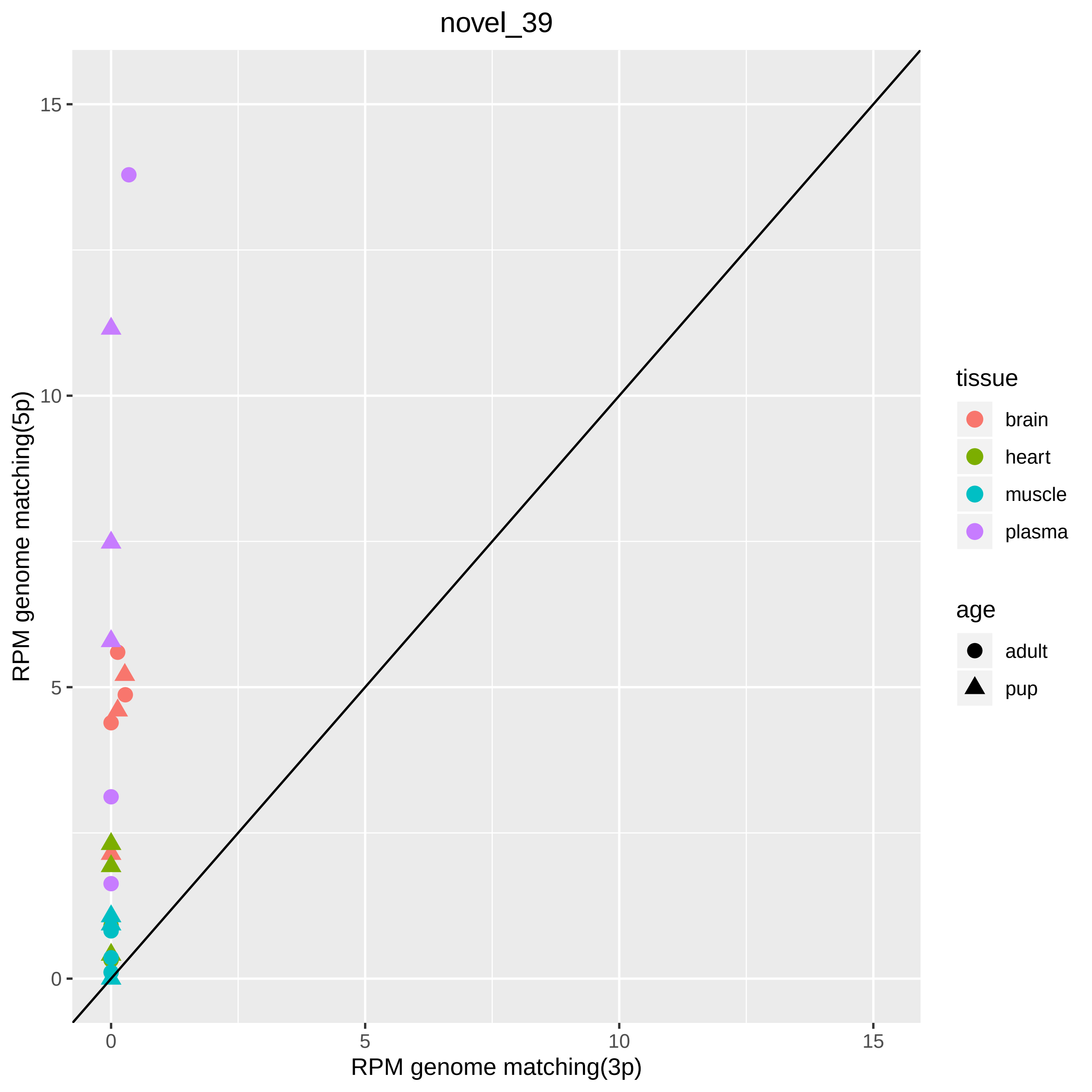

novel_39
RPM genome matching(5p)
RPM genome matching(3p)
0
5
10
15
0
5
10
15
tissue
brain
heart
muscle
plasma
age
adult
pup

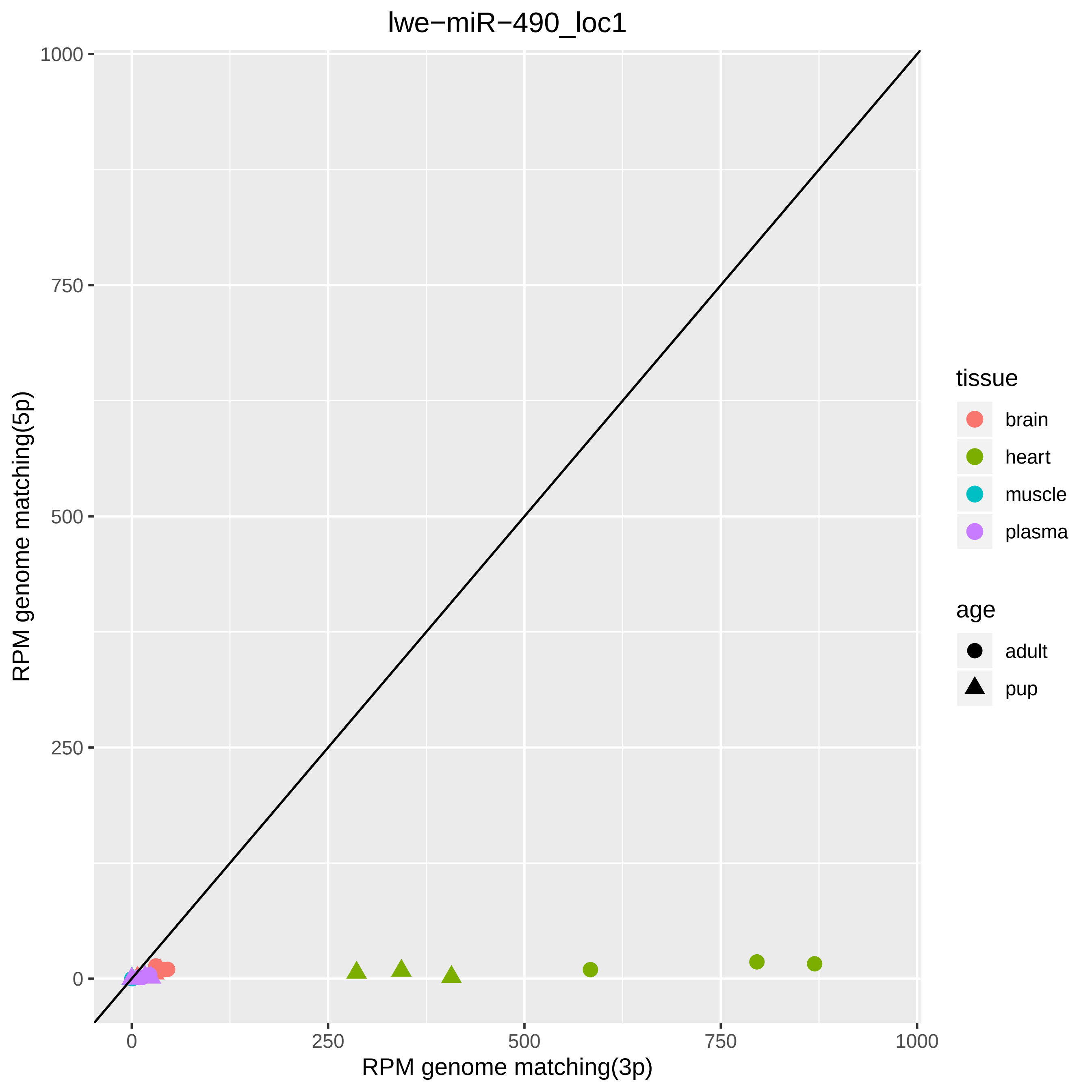

lwe−miR−490_loc1
RPM genome matching(5p)
RPM genome matching(3p)
0
250
500
750
1000
tissue
brain
heart
muscle
plasma
age
adult
pup

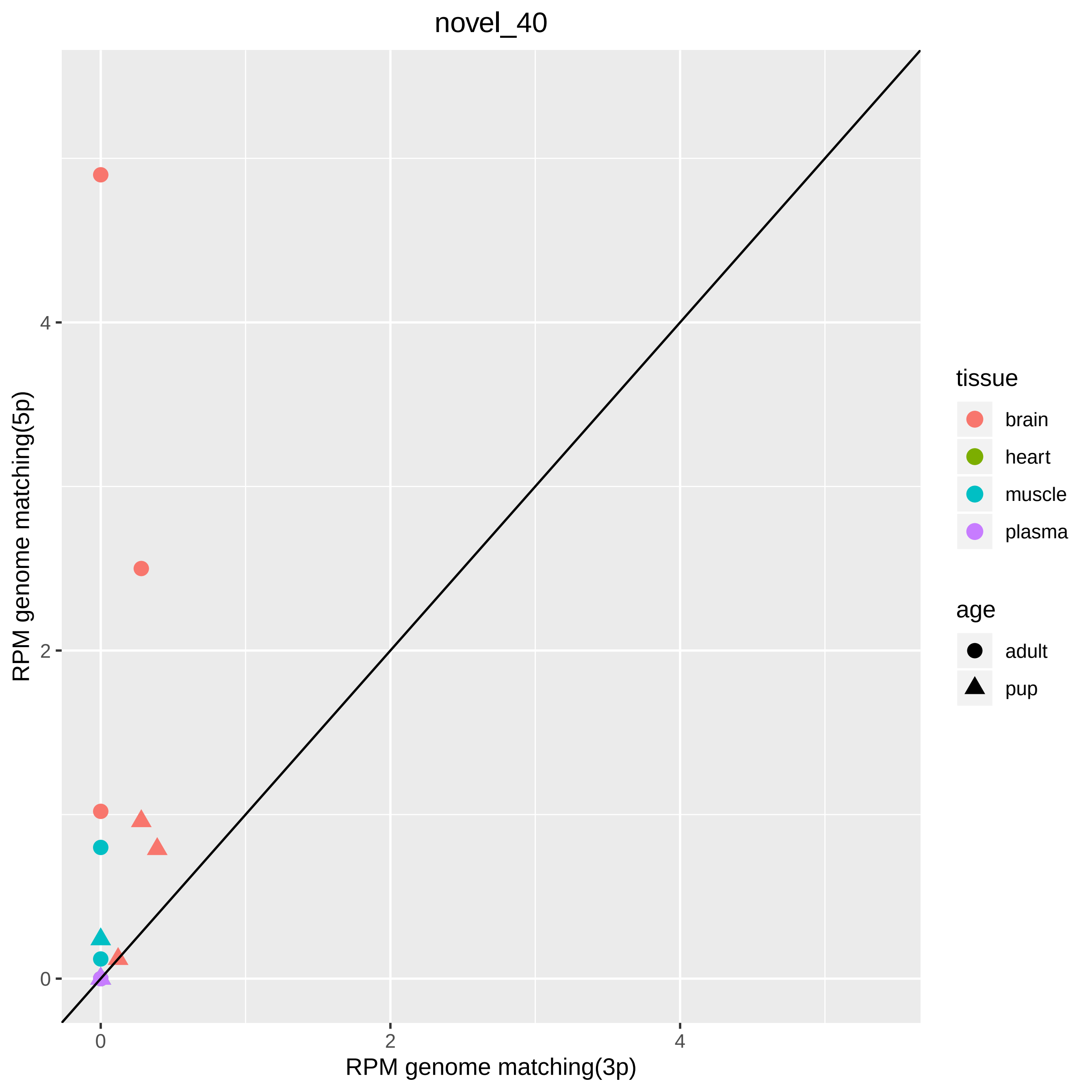

novel_40
RPM genome matching(5p)
RPM genome matching(3p)
0
2
4
tissue
brain
heart
muscle
plasma
age
adult
pup

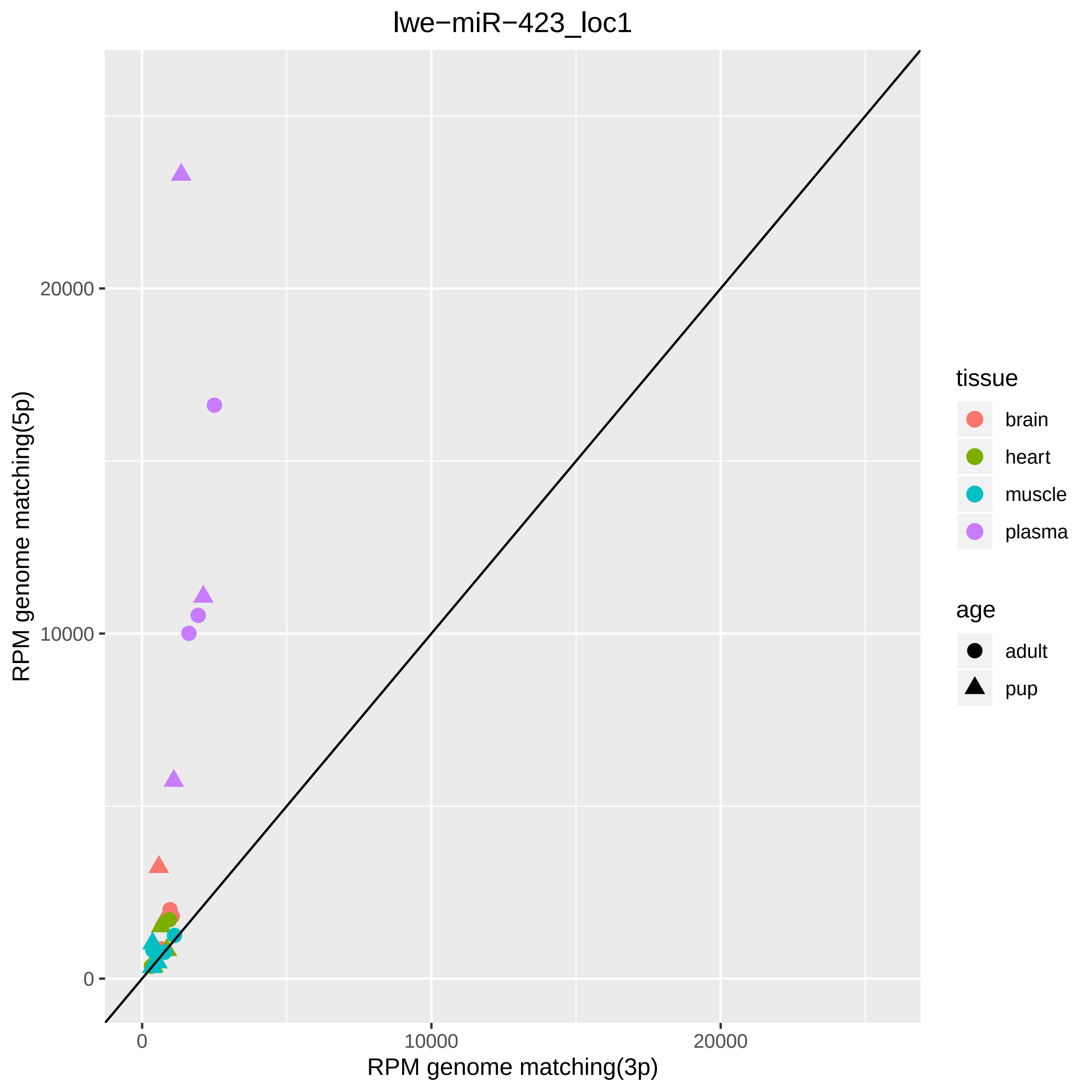
lwe−miR−423_loc1
RPM genome matching(5p)
RPM genome matching(3p)
0
10000
20000
0
10000
20000
tissue
brain
heart
muscle
plasma
age
adult
pup

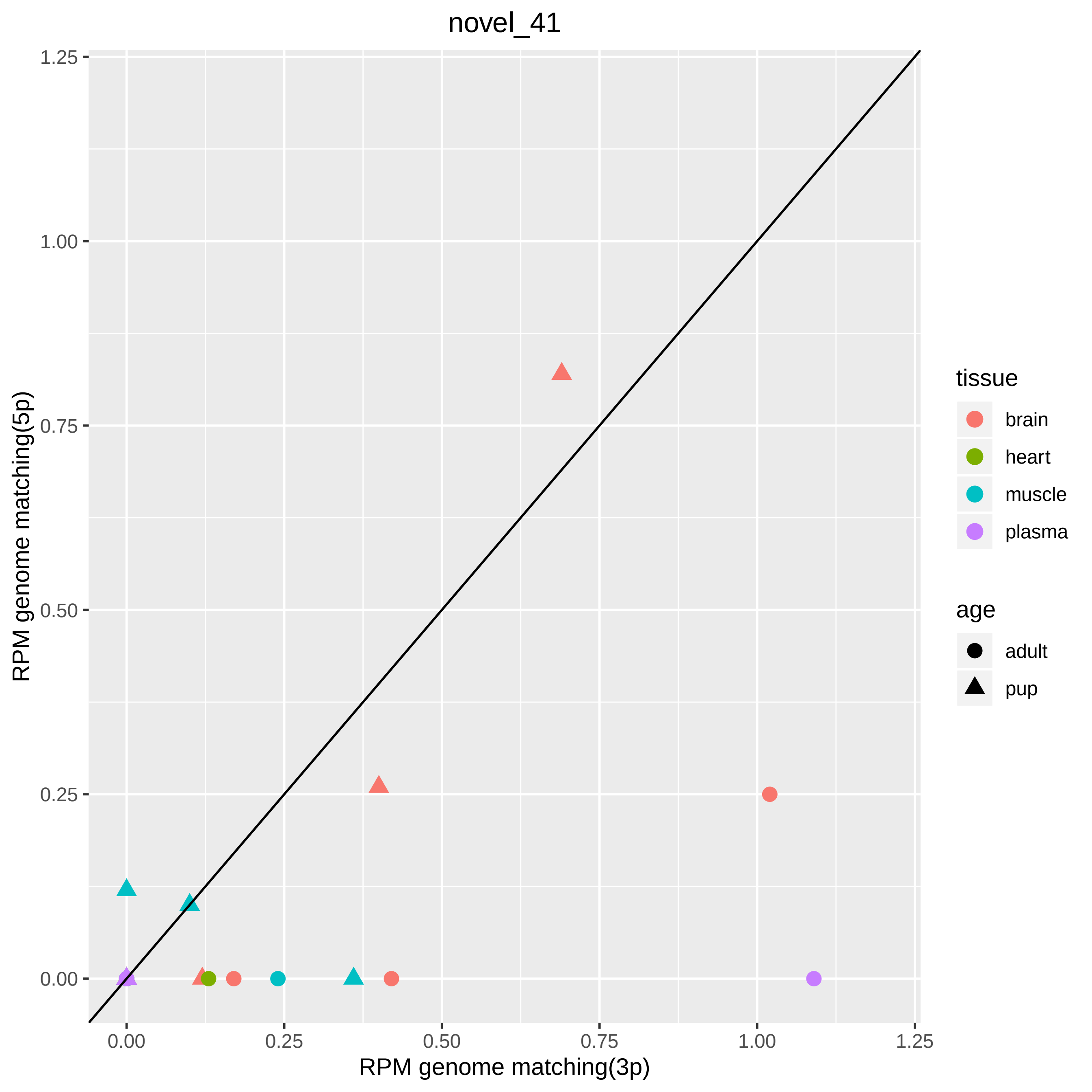

novel_41
RPM genome matching(5p)
RPM genome matching(3p)
1.25
1.00
0.75
0.50
0.25
0.00
0.00
0.25
0.50
0.75
1.00
1.25
tissue
brain
heart
muscle
plasma
age
adult
pup

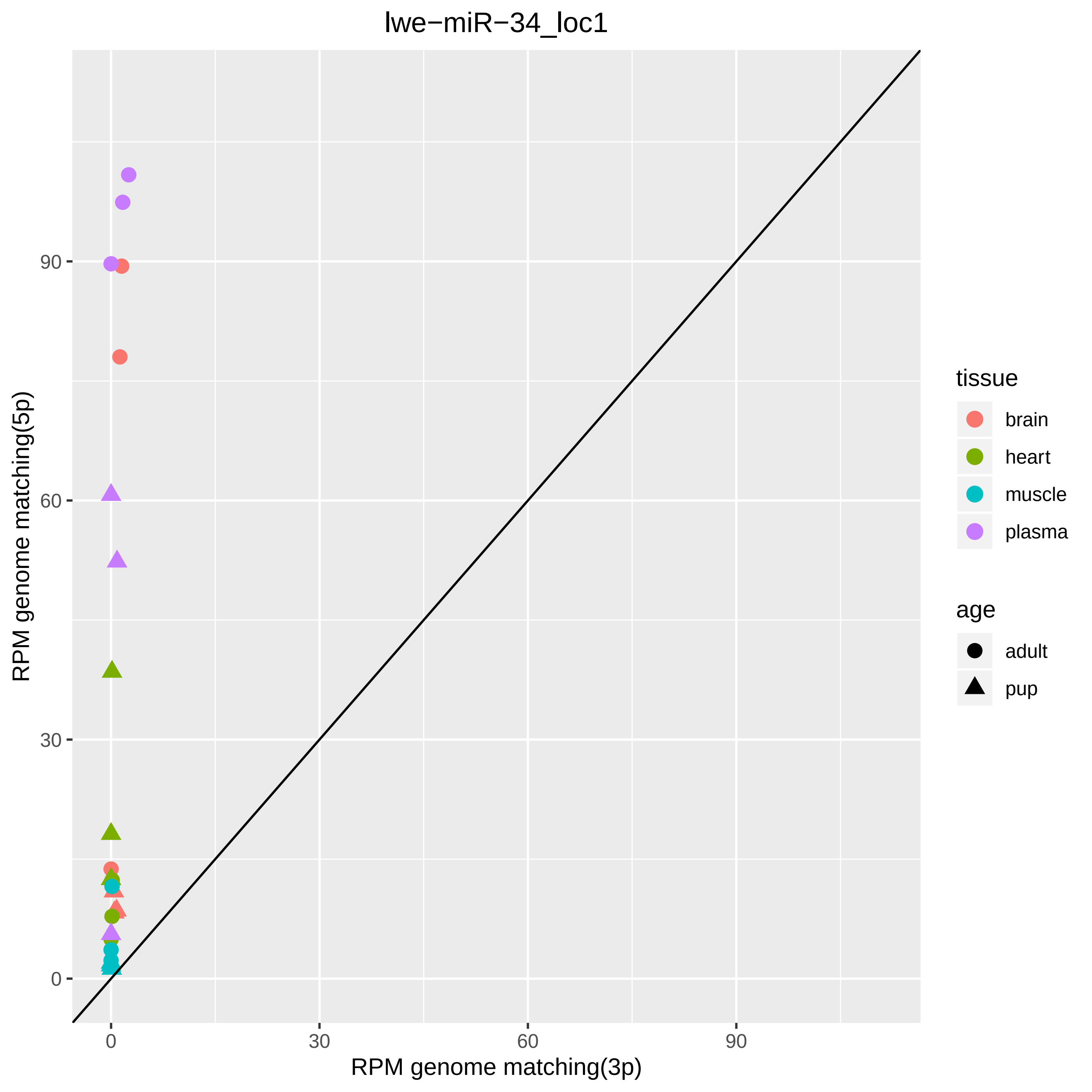

lwe−miR−34_loc1
RPM genome matching(5p)
RPM genome matching(3p)
0
30
60
90
0
30
60
90
tissue
brain
heart
muscle
plasma
age
adult
pup

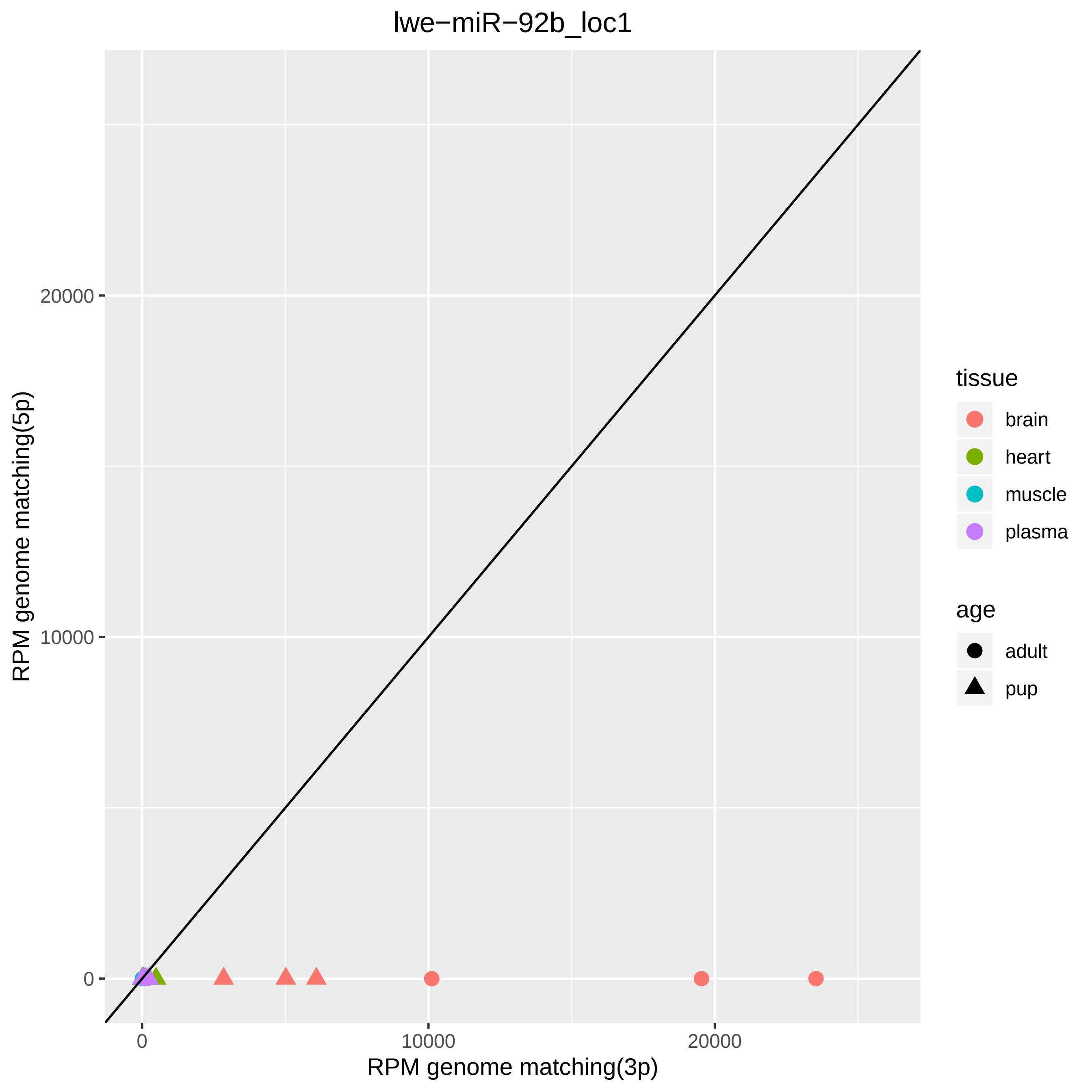

lwe−miR−92b_loc1
RPM genome matching(5p)
RPM genome matching(3p)
20000
10000
0
0
10000
20000
tissue
brain
heart
muscle
plasma
age
adult
pup

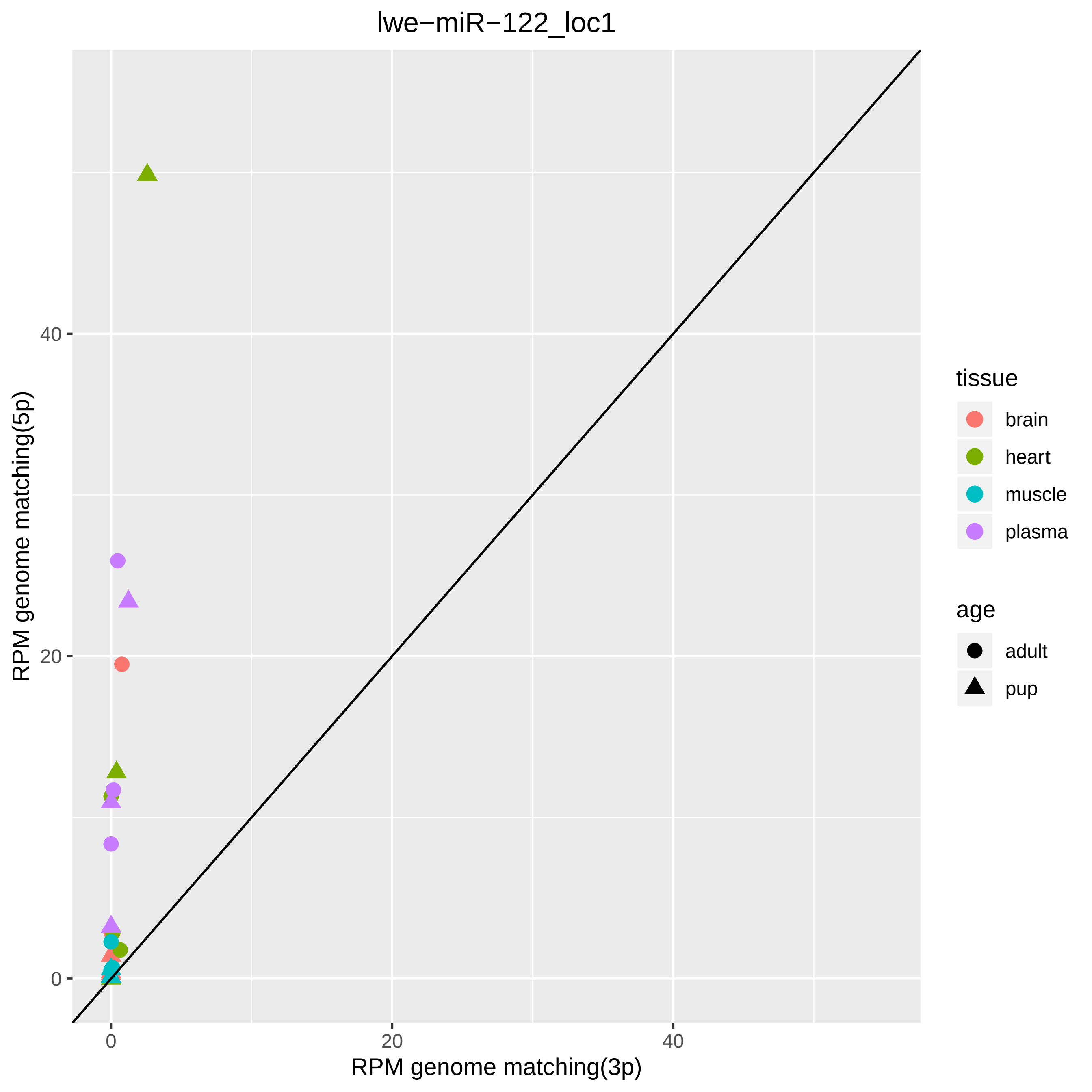
lwe−miR−122_loc1
RPM genome matching(5p)
RPM genome matching(3p)
0
20
40
0
20
40
tissue
brain
heart
muscle
plasma
age
adult
pup

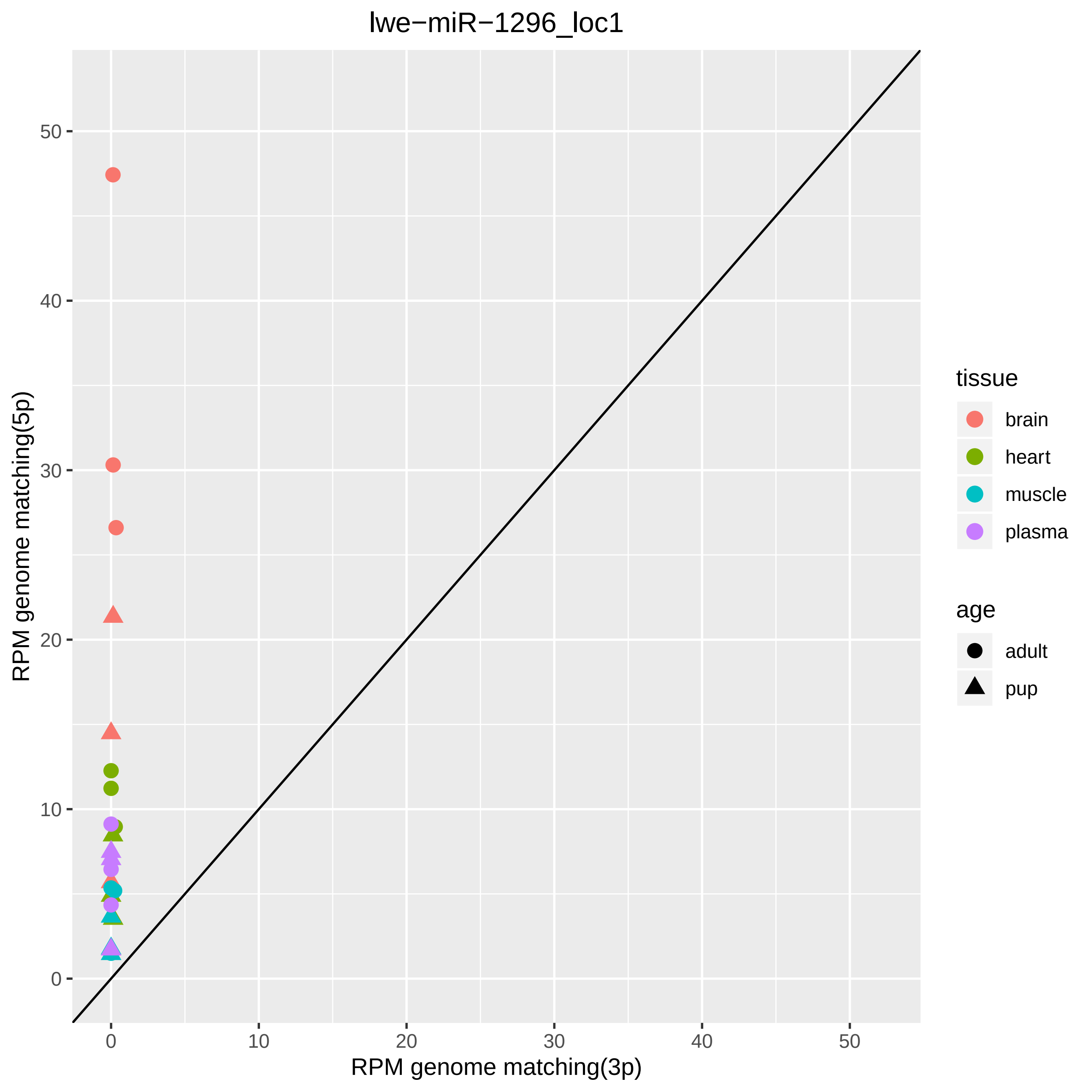
lwe−miR−1296_loc1
RPM genome matching(5p)
RPM genome matching(3p)
0
10
20
30
40
50
tissue
brain
heart
muscle
plasma
age
adult
pup

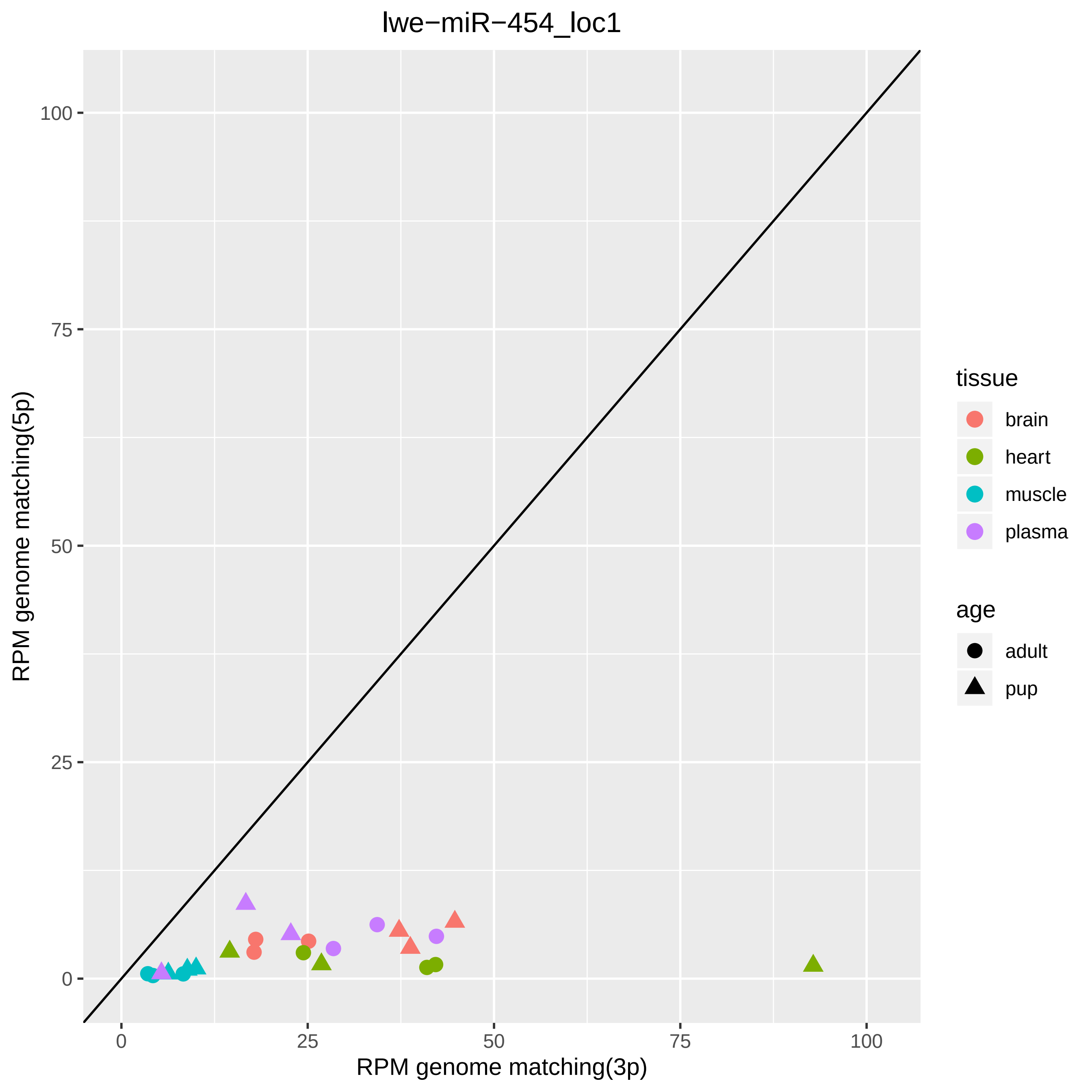

lwe−miR−454_loc1
RPM genome matching(5p)
RPM genome matching(3p)
0
25
50
75
100
0
25
50
75
100
tissue
brain
heart
muscle
plasma
age
adult
pup

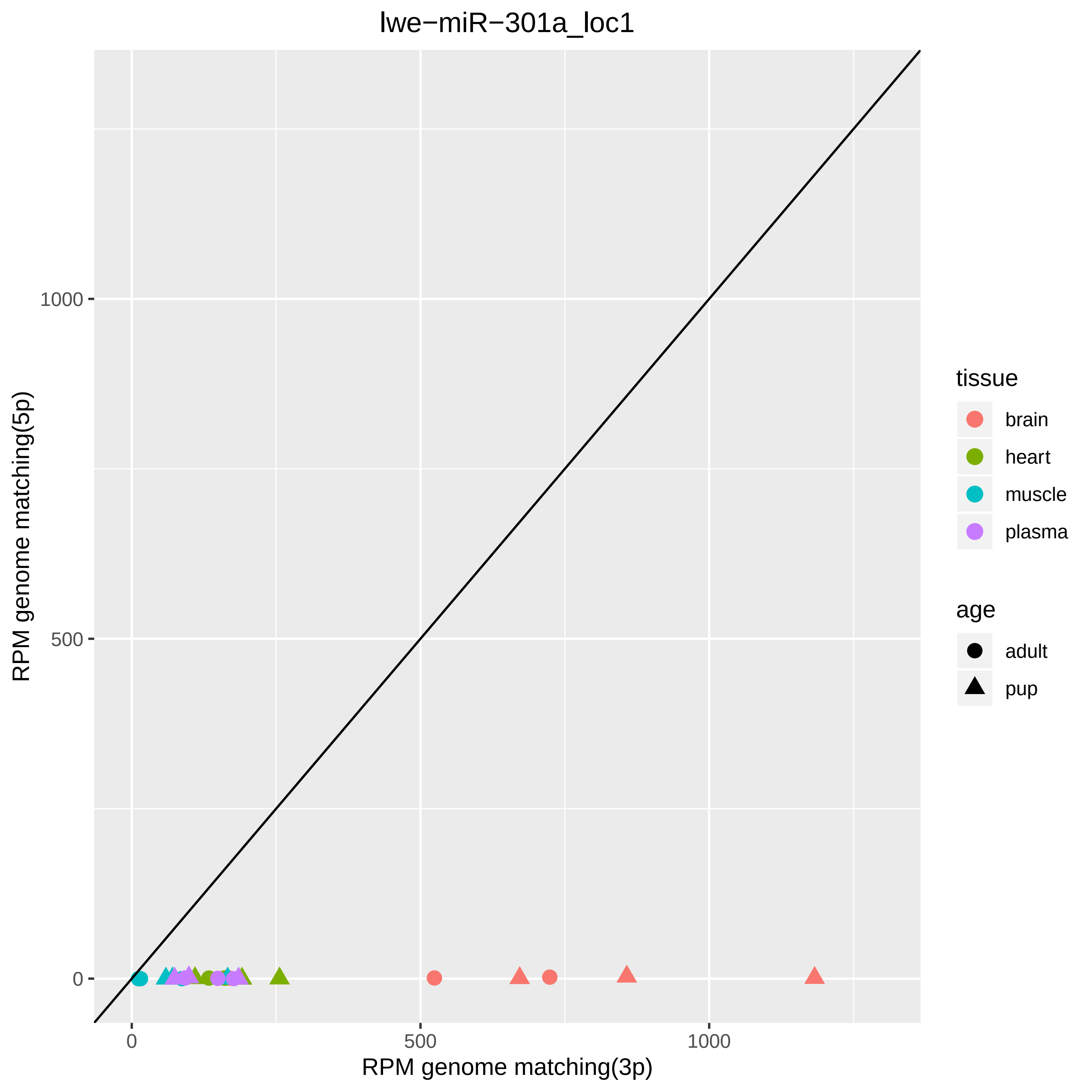
lwe−miR−301a_loc1
RPM genome matching(5p)
RPM genome matching(3p)
0
500
1000
0
500
1000
tissue
brain
heart
muscle
plasma
age
adult
pup

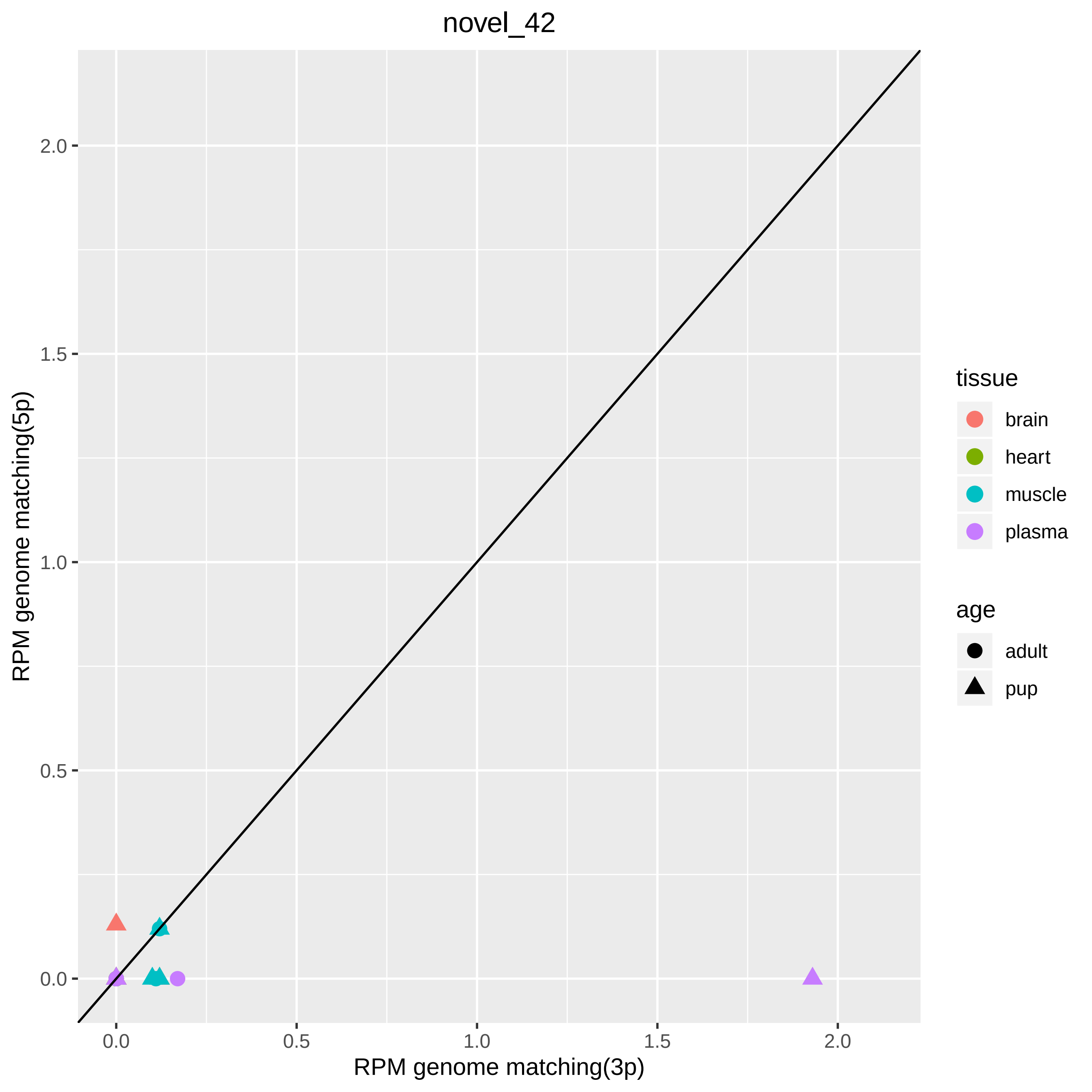

novel_42
RPM genome matching(5p)
RPM genome matching(3p)
0.0
0.5
1.0
1.5
2.0
tissue
brain
heart
muscle
plasma
age
adult
pup

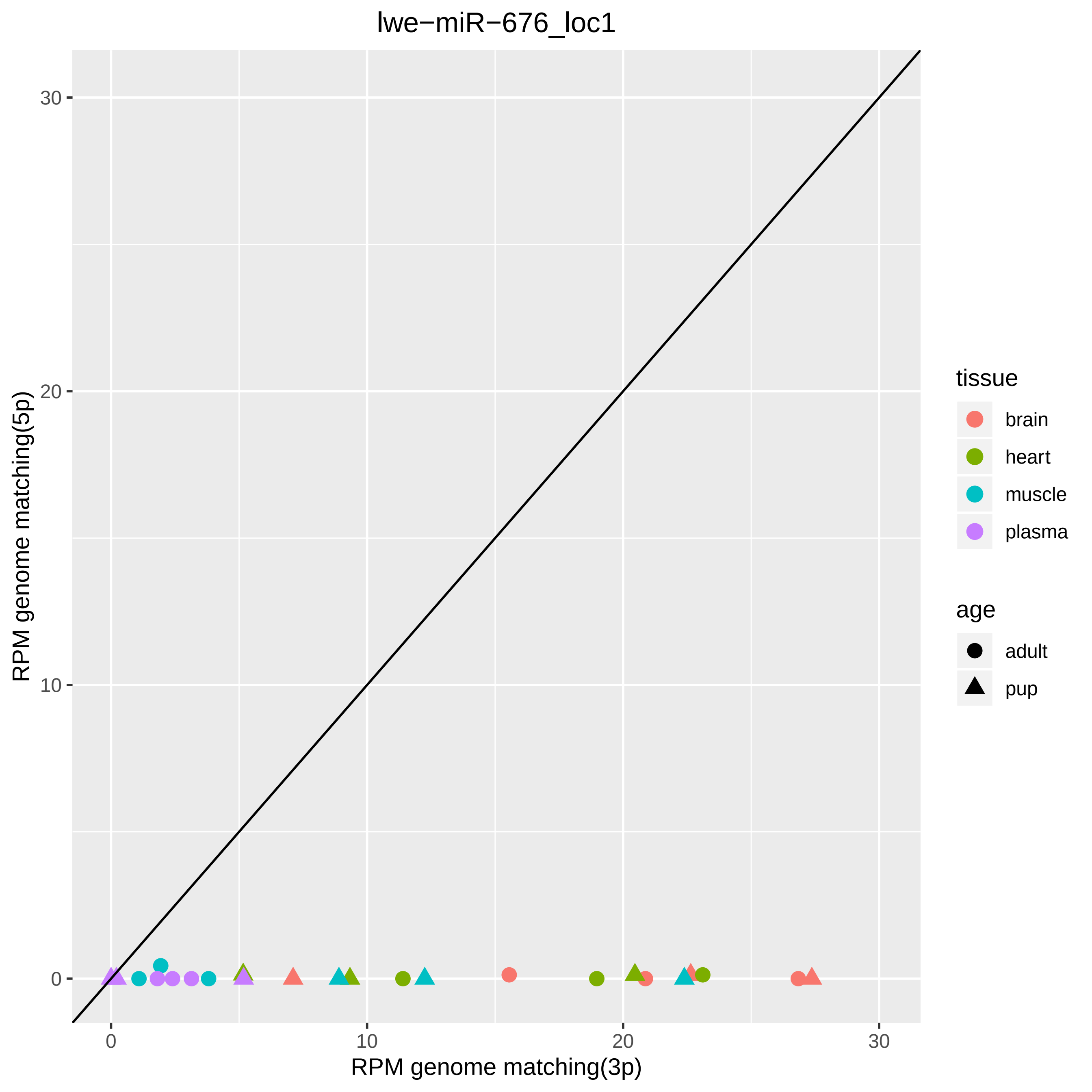
lwe−miR−676_loc1
RPM genome matching(5p)
RPM genome matching(3p)
0
10
20
30
0
10
20
30
tissue
brain
heart
muscle
plasma
age
adult
pup

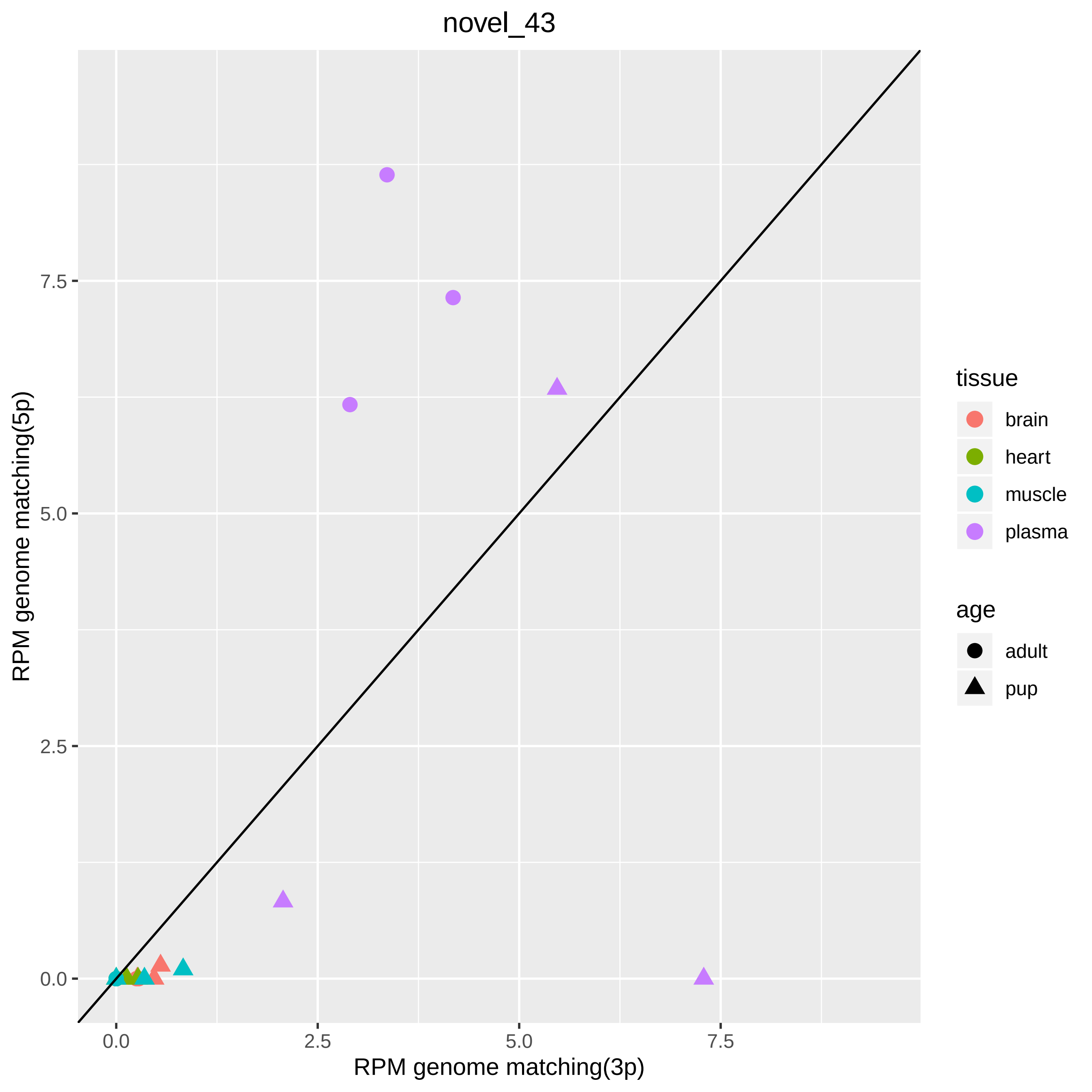

novel_43
RPM genome matching(5p)
RPM genome matching(3p)
0.0
2.5
5.0
7.5
0.0
2.5
5.0
7.5
tissue
brain
heart
muscle
plasma
age
adult
pup

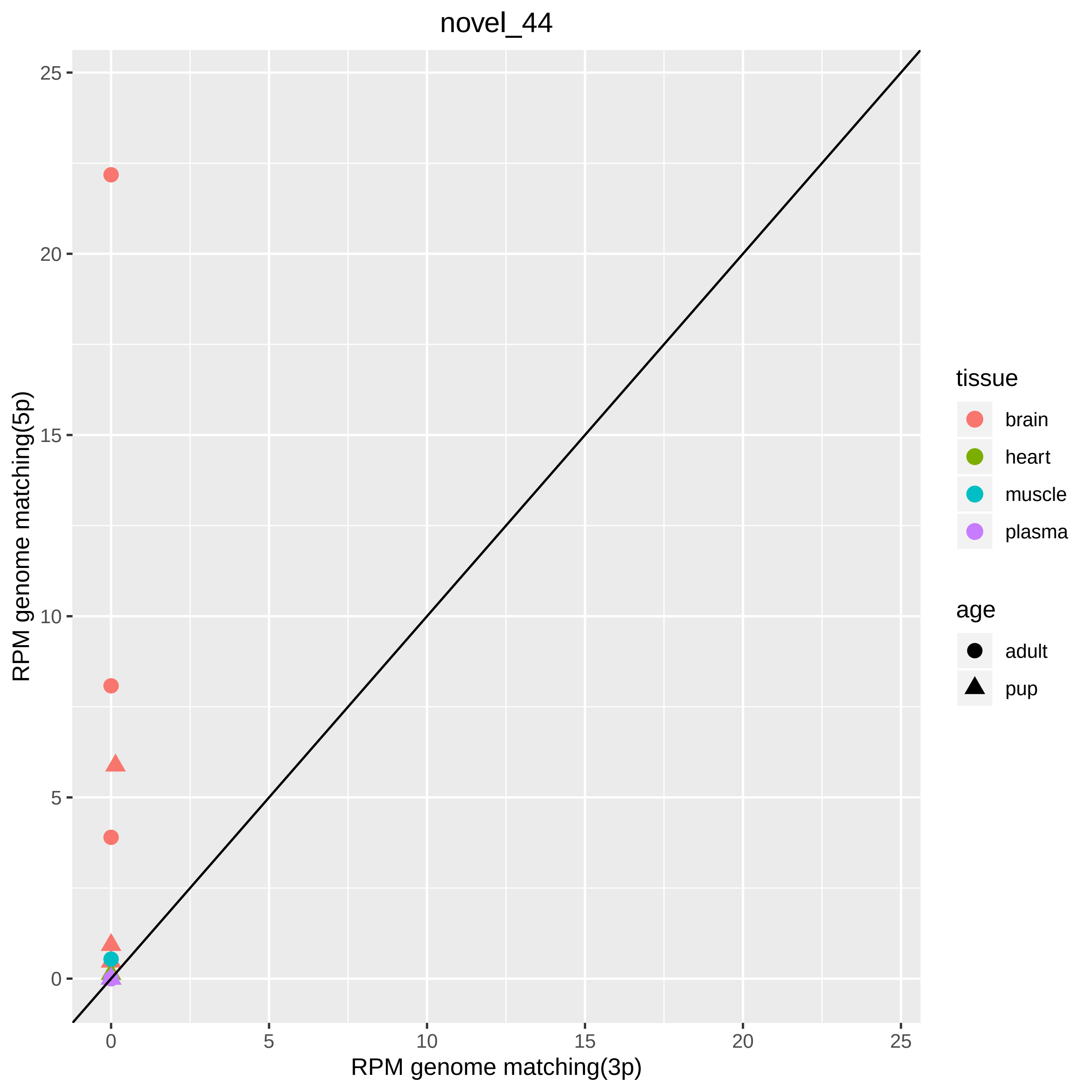
novel_44
RPM genome matching(5p)
RPM genome matching(3p)
0
5
10
15
20
25
tissue
brain
heart
muscle
plasma
age
adult
pup

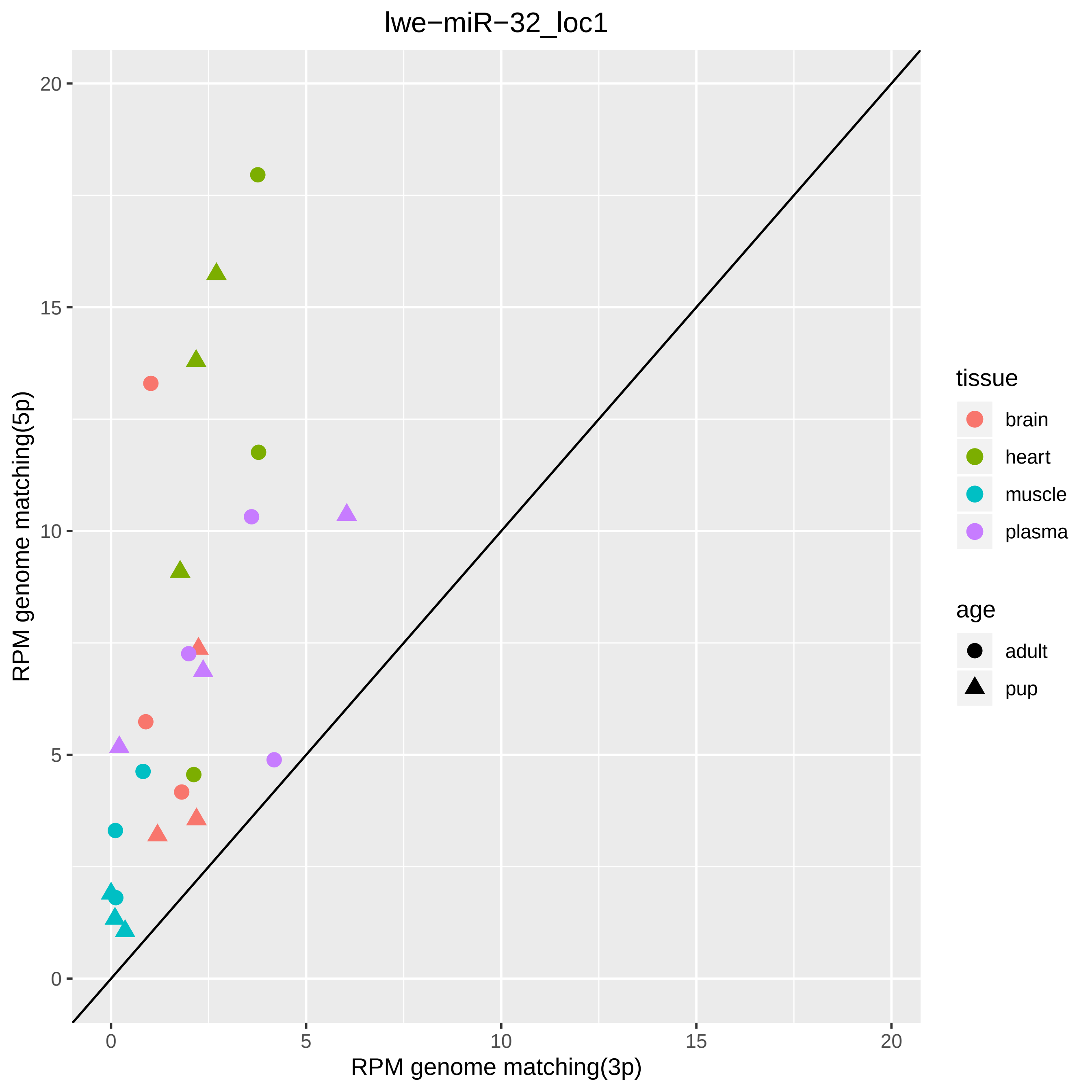
lwe−miR−32_loc1
RPM genome matching(5p)
RPM genome matching(3p)
0
5
10
15
20
tissue
brain
heart
muscle
plasma
age
adult
pup

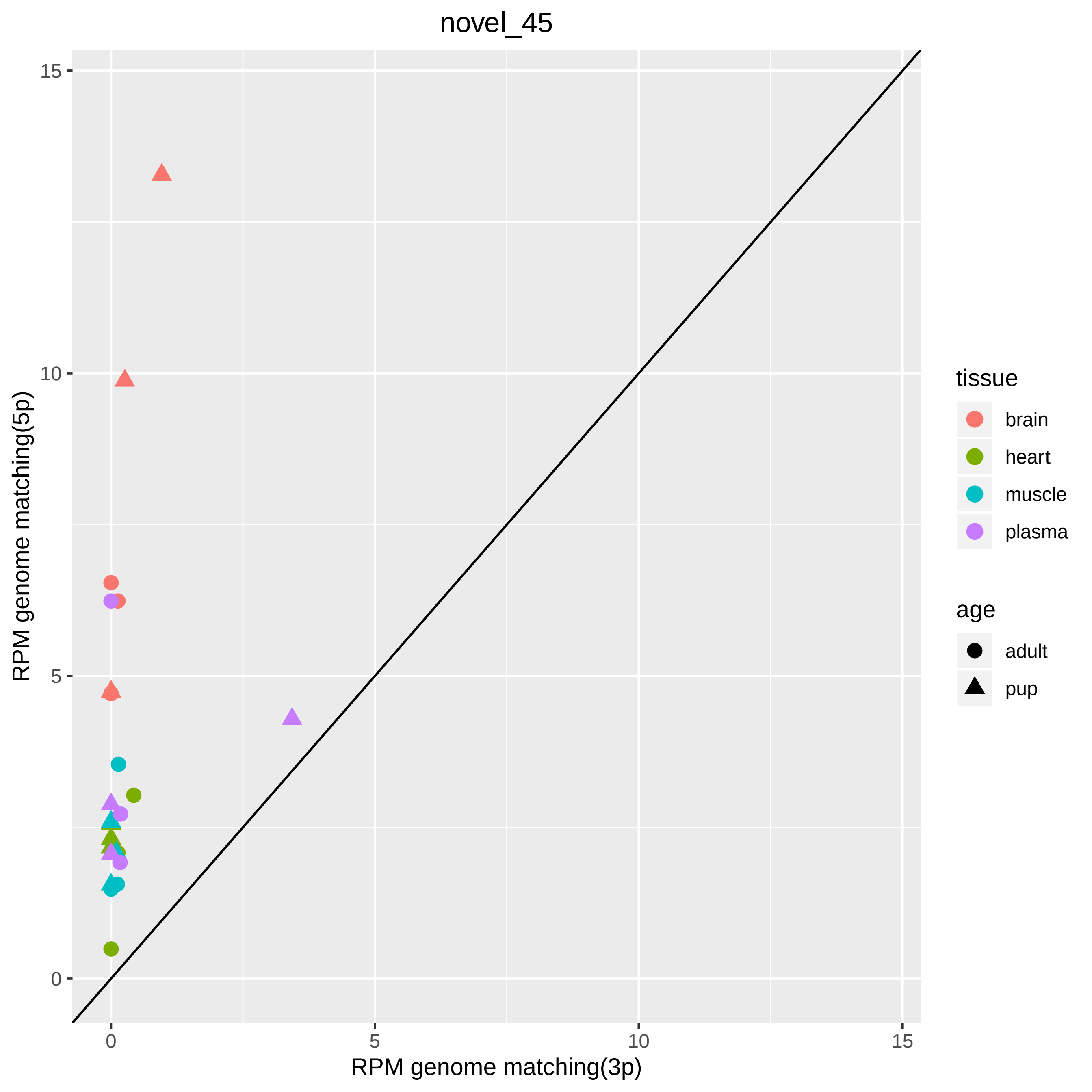
novel_45
RPM genome matching(5p)
RPM genome matching(3p)
0
5
10
15
tissue
brain
heart
muscle
plasma
age
adult
pup

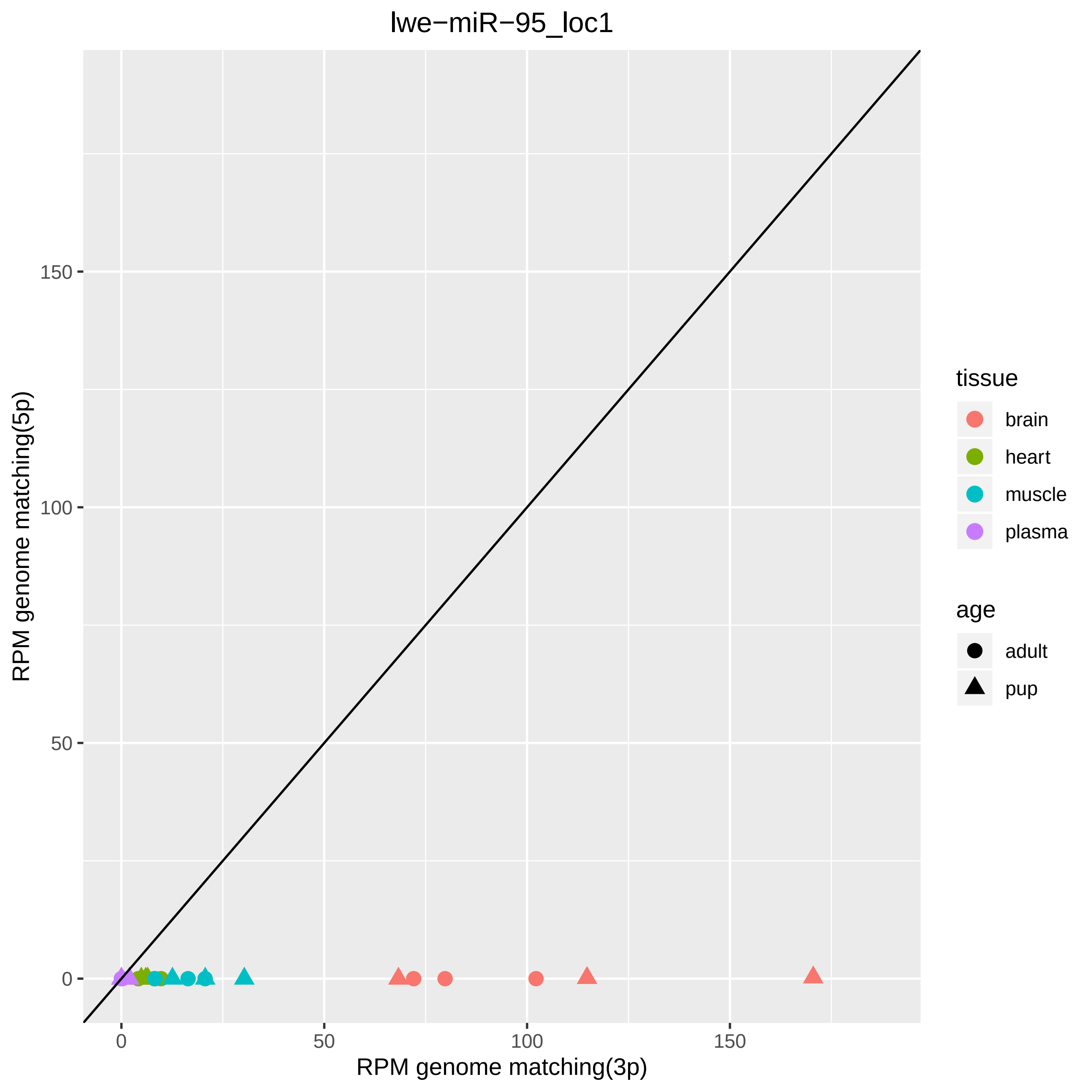

lwe−miR−95_loc1
RPM genome matching(5p)
RPM genome matching(3p)
0
50
100
150
0
50
100
150
tissue
brain
heart
muscle
plasma
age
adult
pup

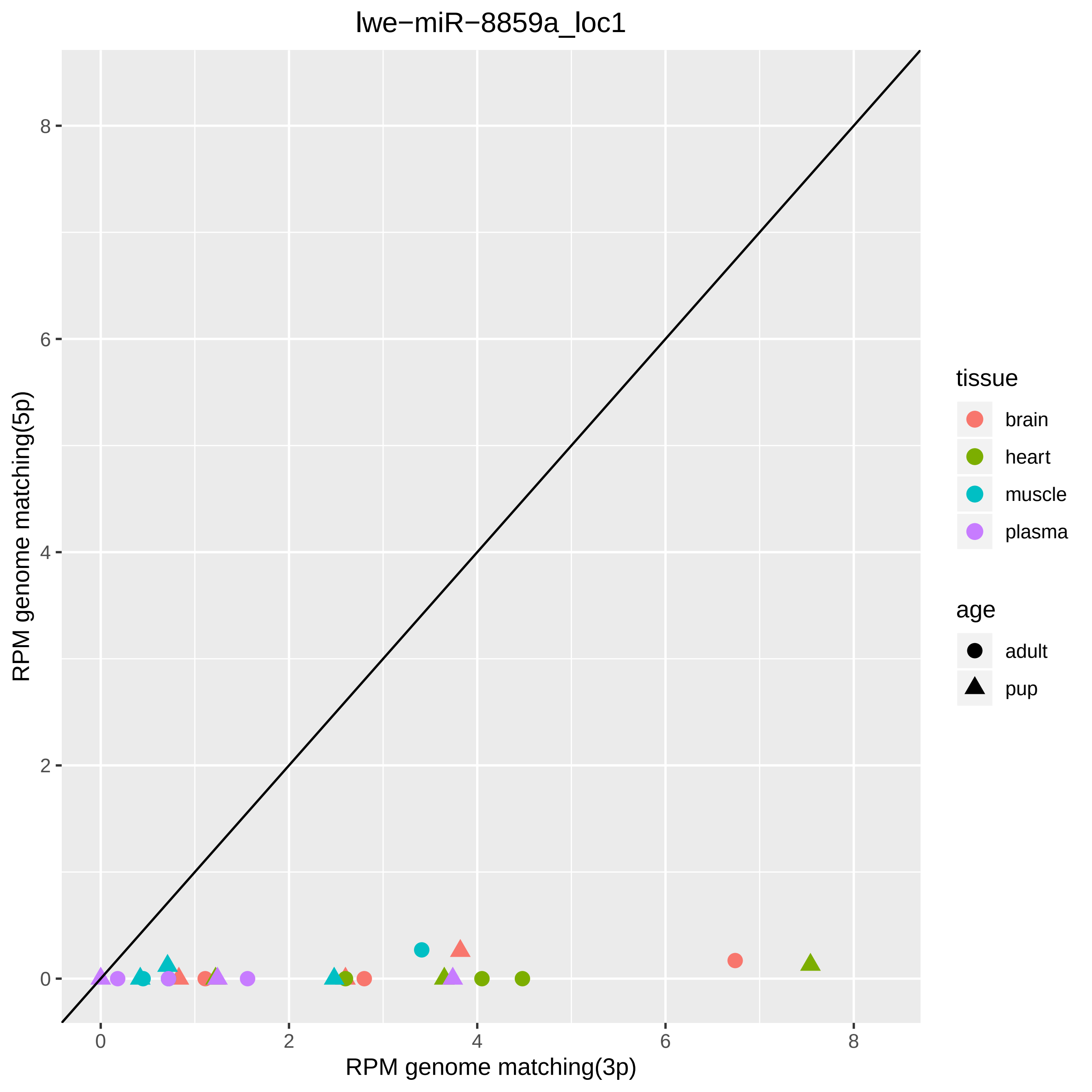
lwe−miR−8859a_loc1
RPM genome matching(5p)
RPM genome matching(3p)
0
2
4
6
8
tissue
brain
heart
muscle
plasma
age
adult
pup

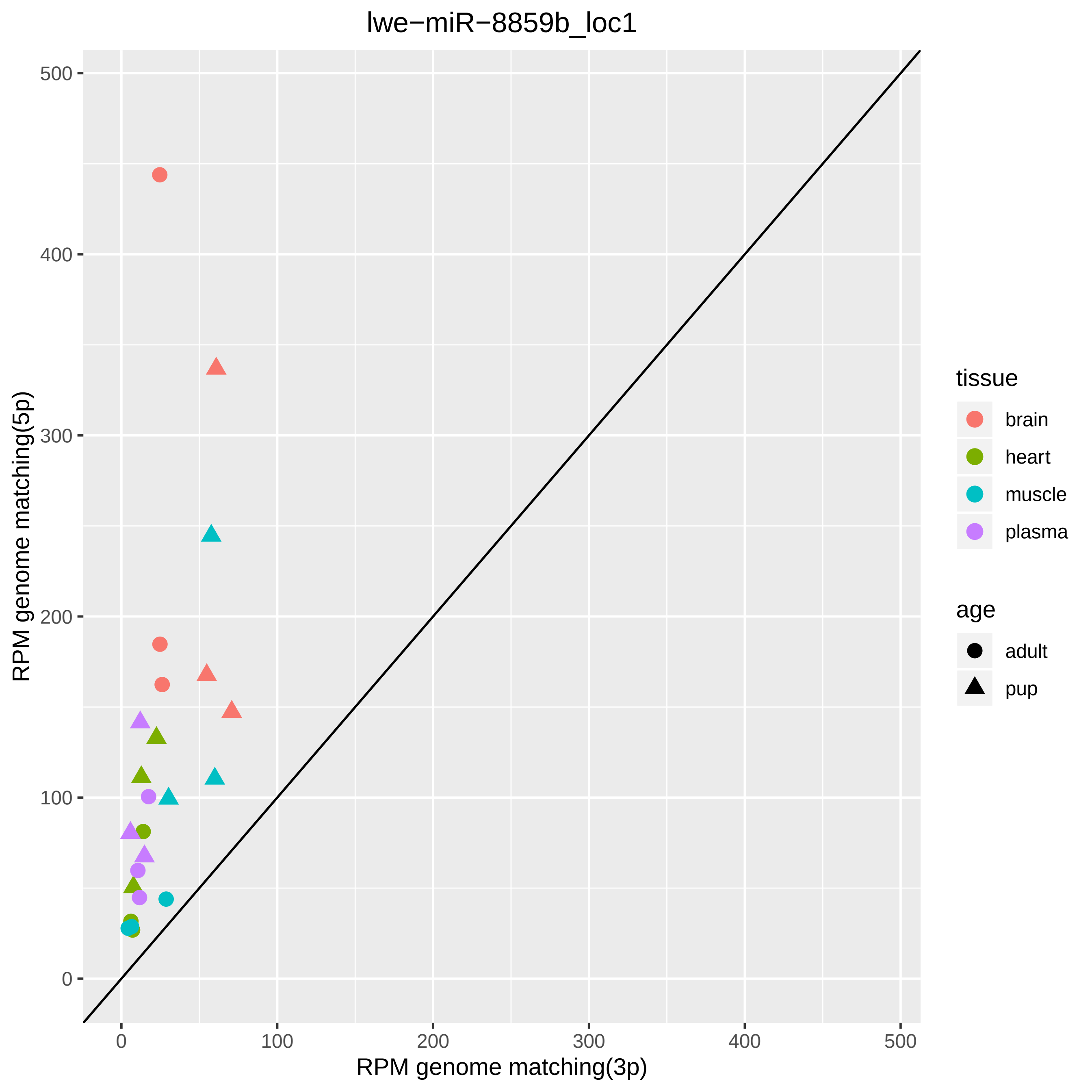
lwe−miR−8859b_loc1
RPM genome matching(5p)
RPM genome matching(3p)
0
100
200
300
400
500
tissue
brain
heart
muscle
plasma
age
adult
pup

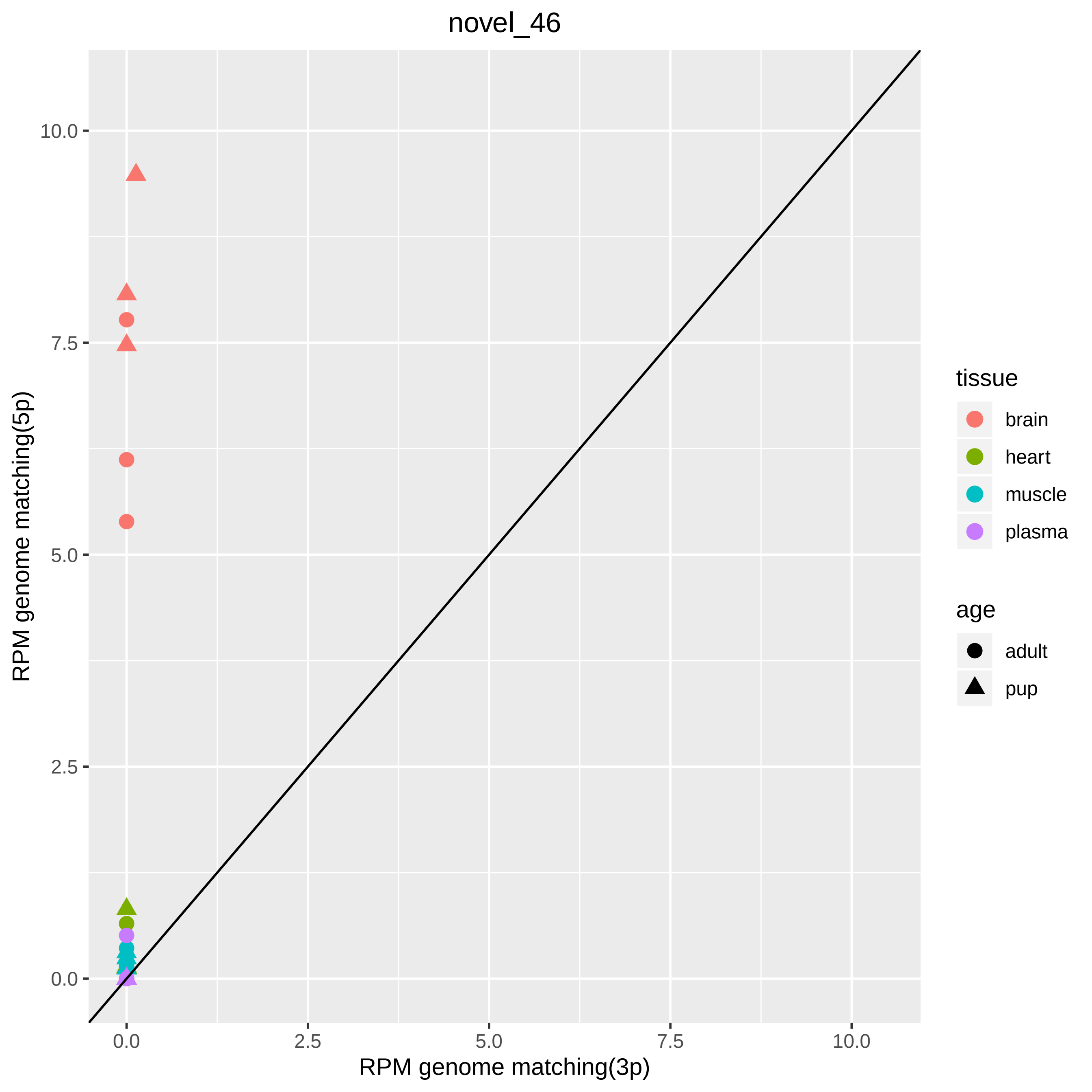

novel_46
RPM genome matching(5p)
RPM genome matching(3p)
10.0
7.5
5.0
2.5
0.0
0.0
2.5
5.0
7.5
10.0
tissue
brain
heart
muscle
plasma
age
adult
pup

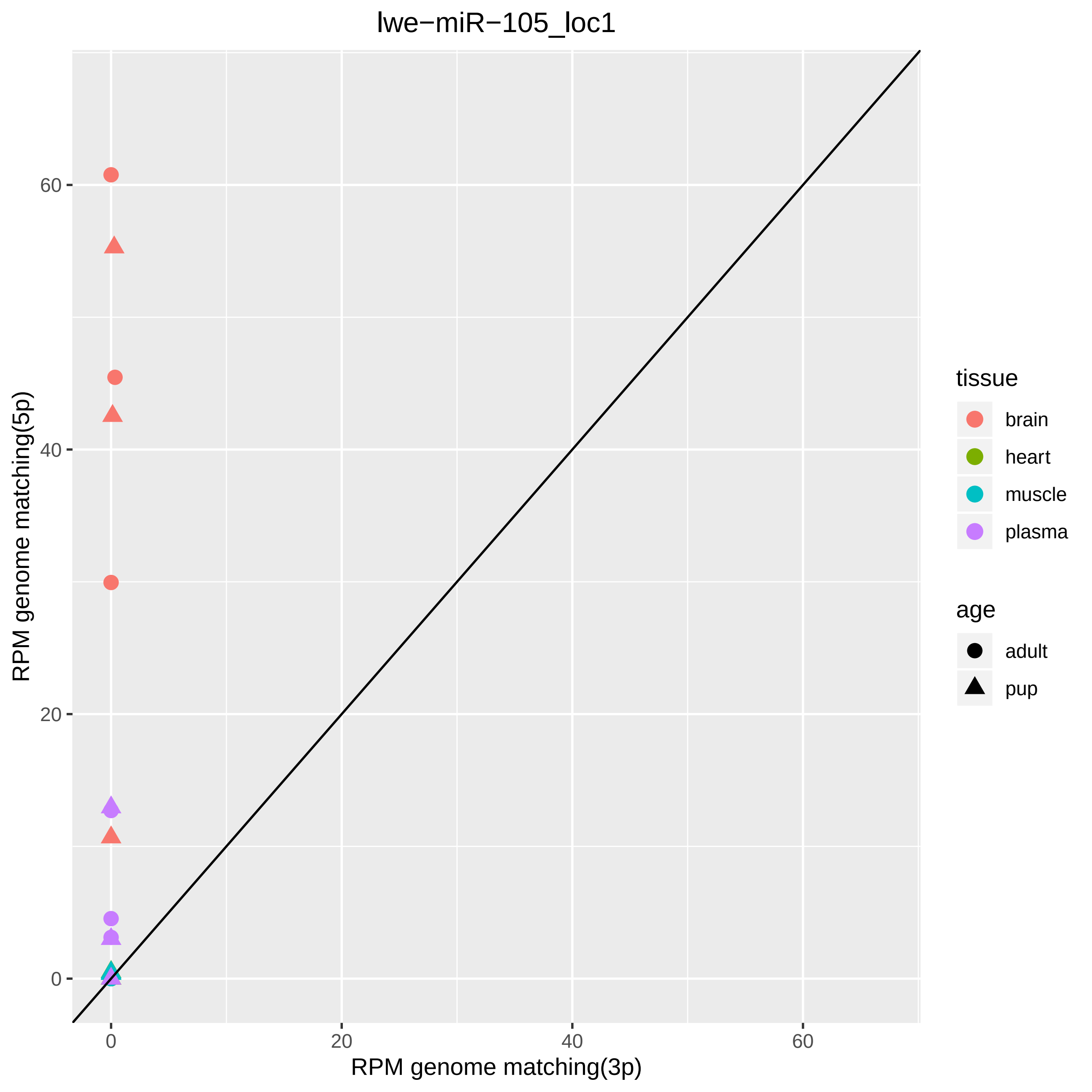

lwe−miR−105_loc1
RPM genome matching(5p)
RPM genome matching(3p)
0
20
40
60
0
20
40
60
tissue
brain
heart
muscle
plasma
age
adult
pup

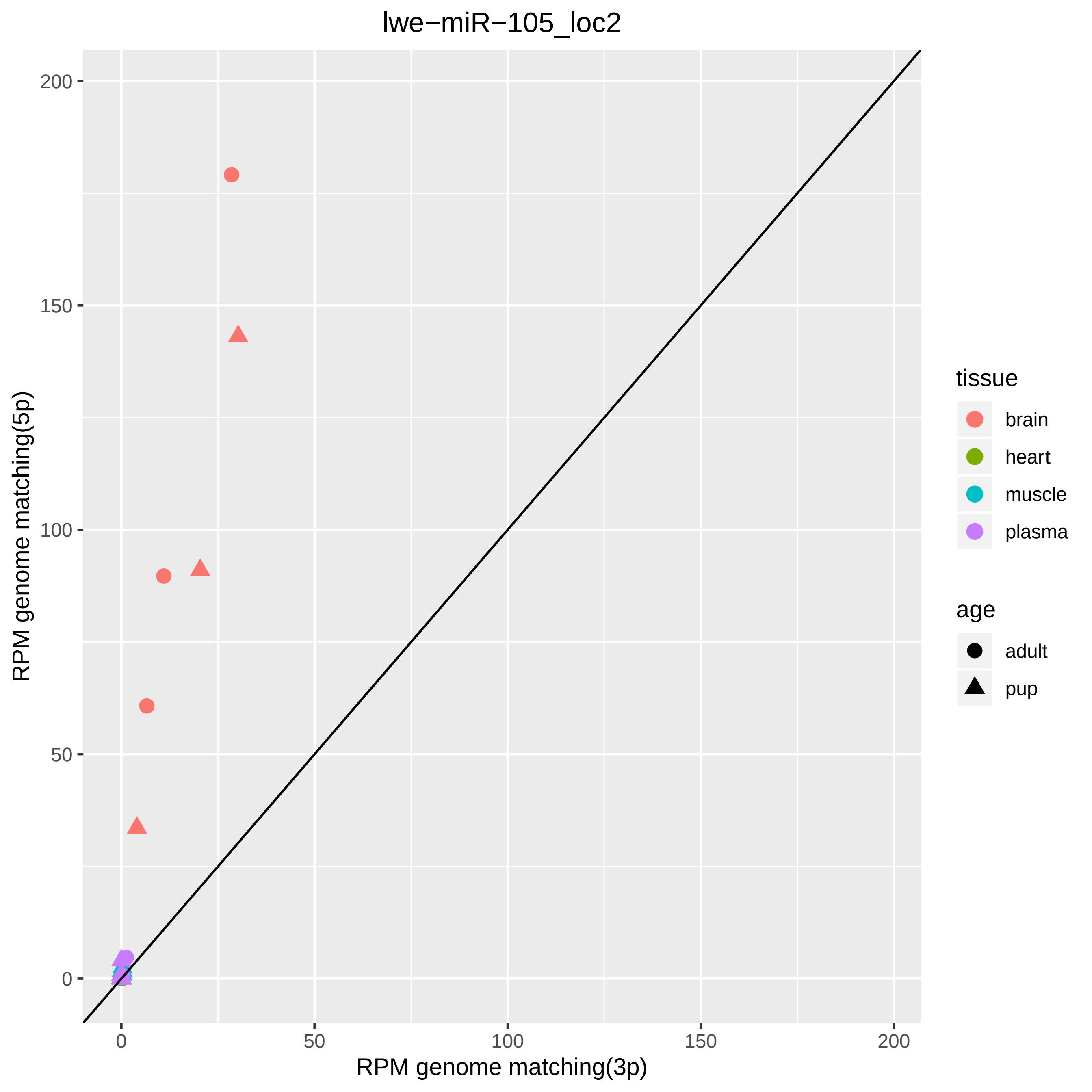

lwe−miR−105_loc2
RPM genome matching(5p)
RPM genome matching(3p)
200
150
100
50
0
0
50
100
150
200
tissue
brain
heart
muscle
plasma
age
adult
pup

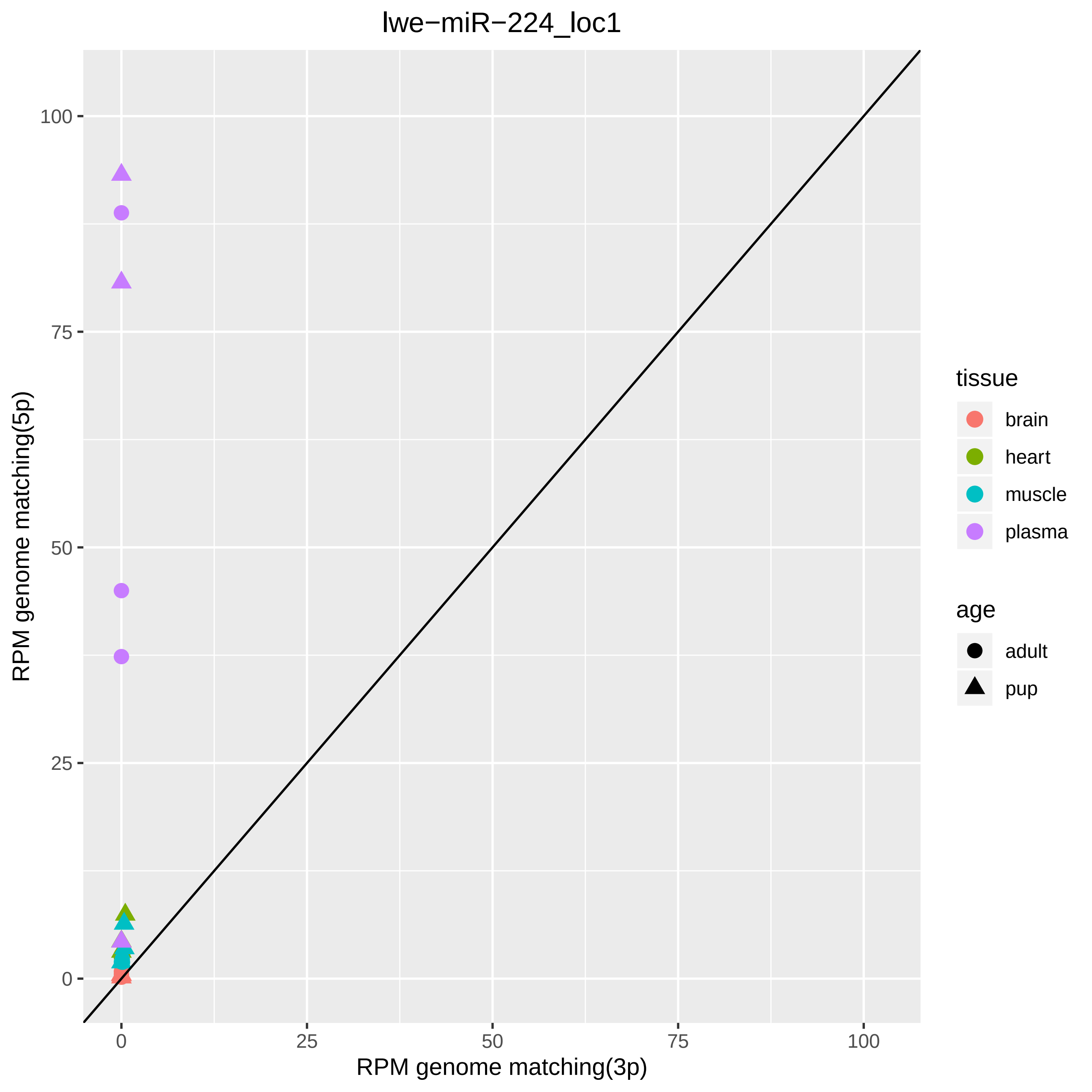

lwe−miR−224_loc1
RPM genome matching(5p)
RPM genome matching(3p)
0
25
50
75
100
tissue
brain
heart
muscle
plasma
age
adult
pup

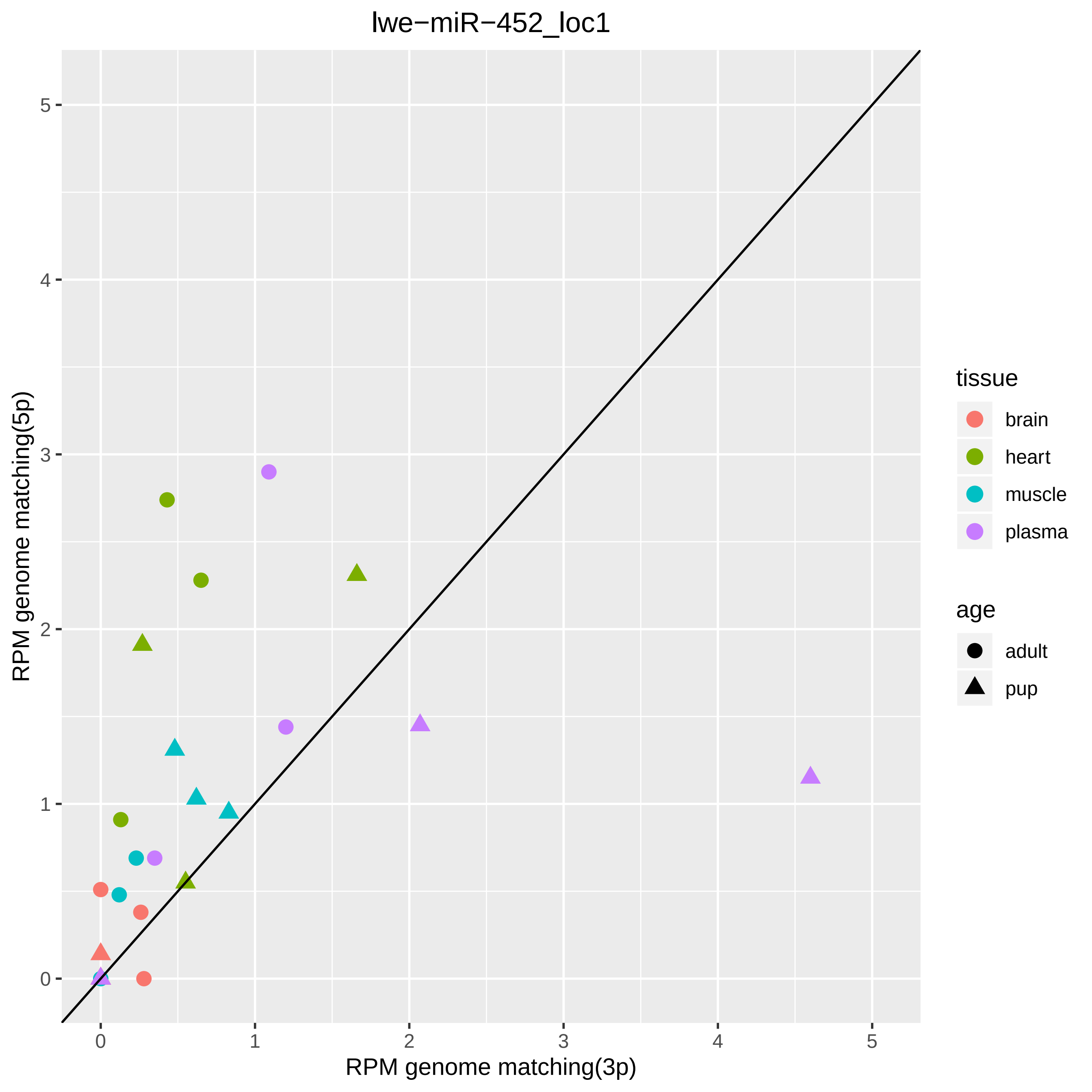

lwe−miR−452_loc1
RPM genome matching(5p)
RPM genome matching(3p)
0
1
2
3
4
5
tissue
brain
heart
muscle
plasma
age
adult
pup

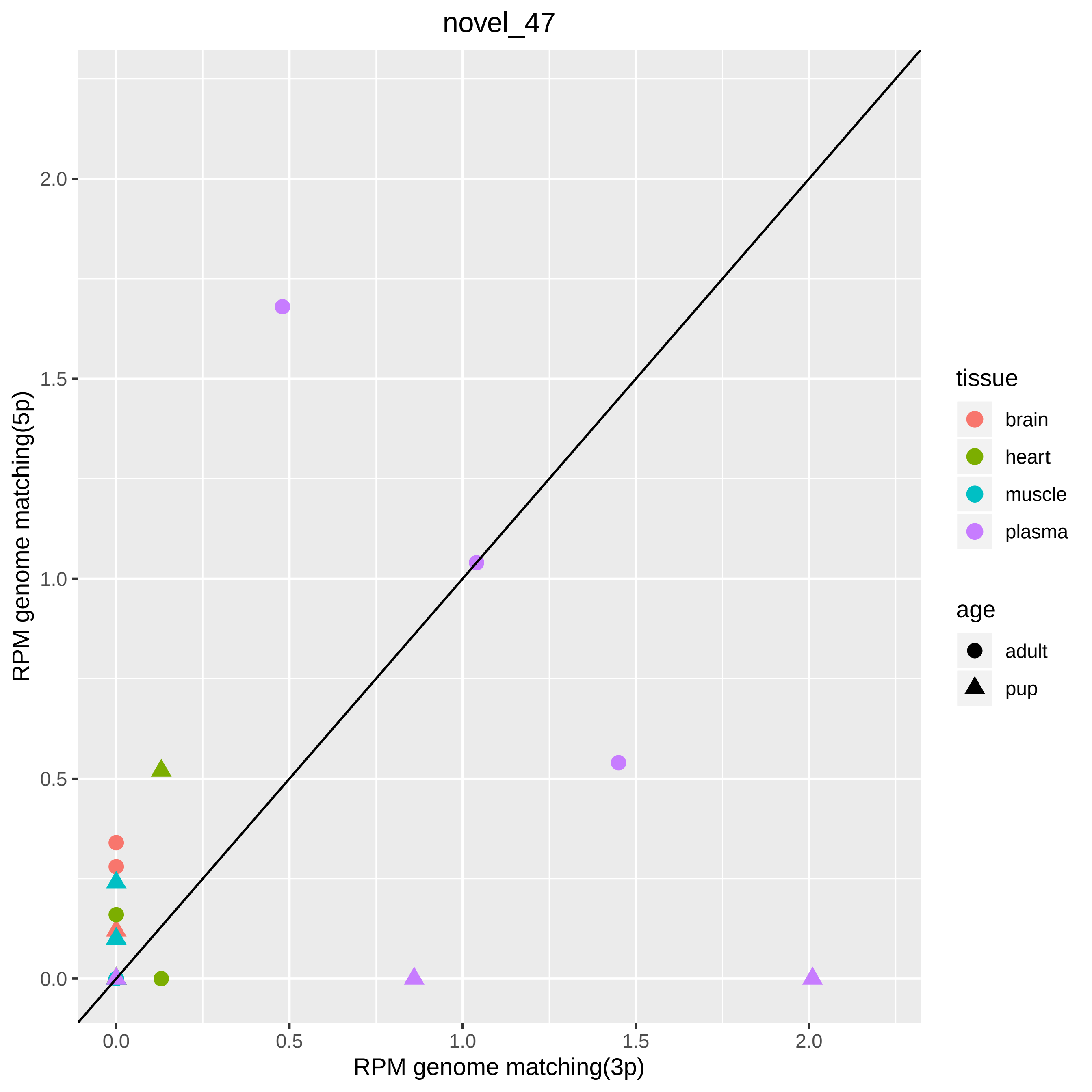
novel_47
RPM genome matching(5p)
RPM genome matching(3p)
0.0
0.5
1.0
1.5
2.0
tissue
brain
heart
muscle
plasma
age
adult
pup

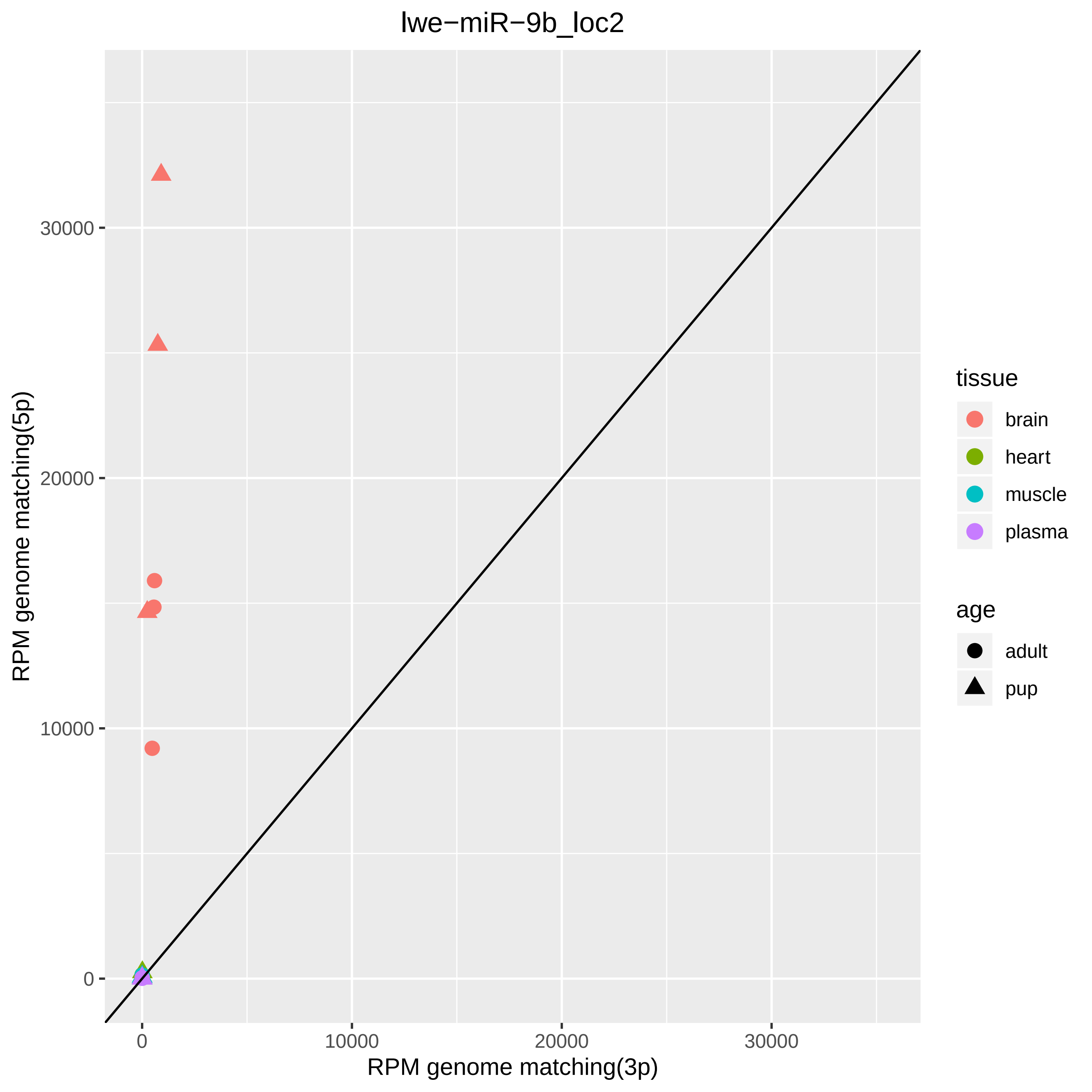

lwe−miR−9b_loc2
RPM genome matching(5p)
RPM genome matching(3p)
0
10000
20000
30000
0
10000
20000
30000
tissue
brain
heart
muscle
plasma
age
adult
pup

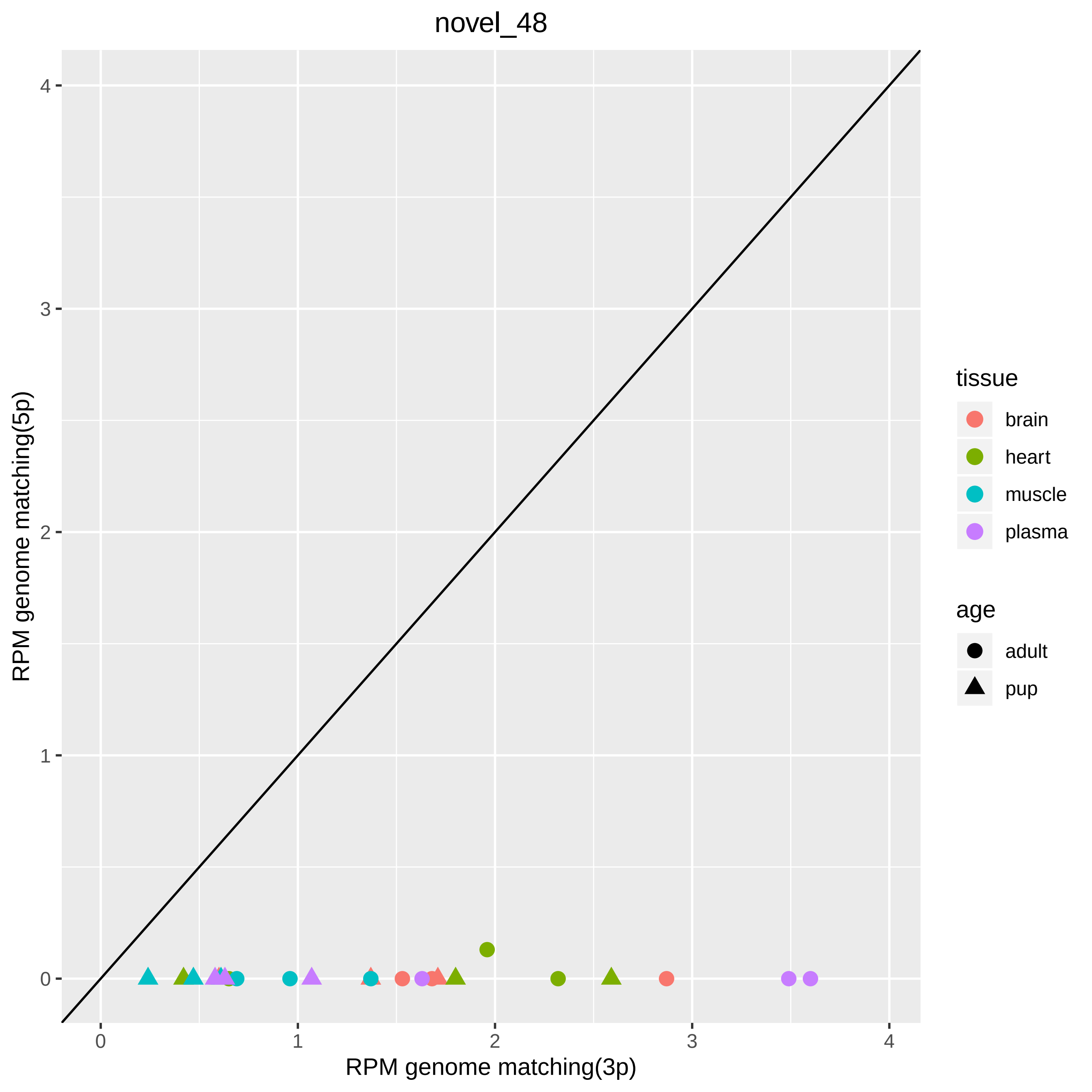
novel_48
RPM genome matching(5p)
RPM genome matching(3p)
0
1
2
3
4
0
1
2
3
4
tissue
brain
heart
muscle
plasma
age
adult
pup

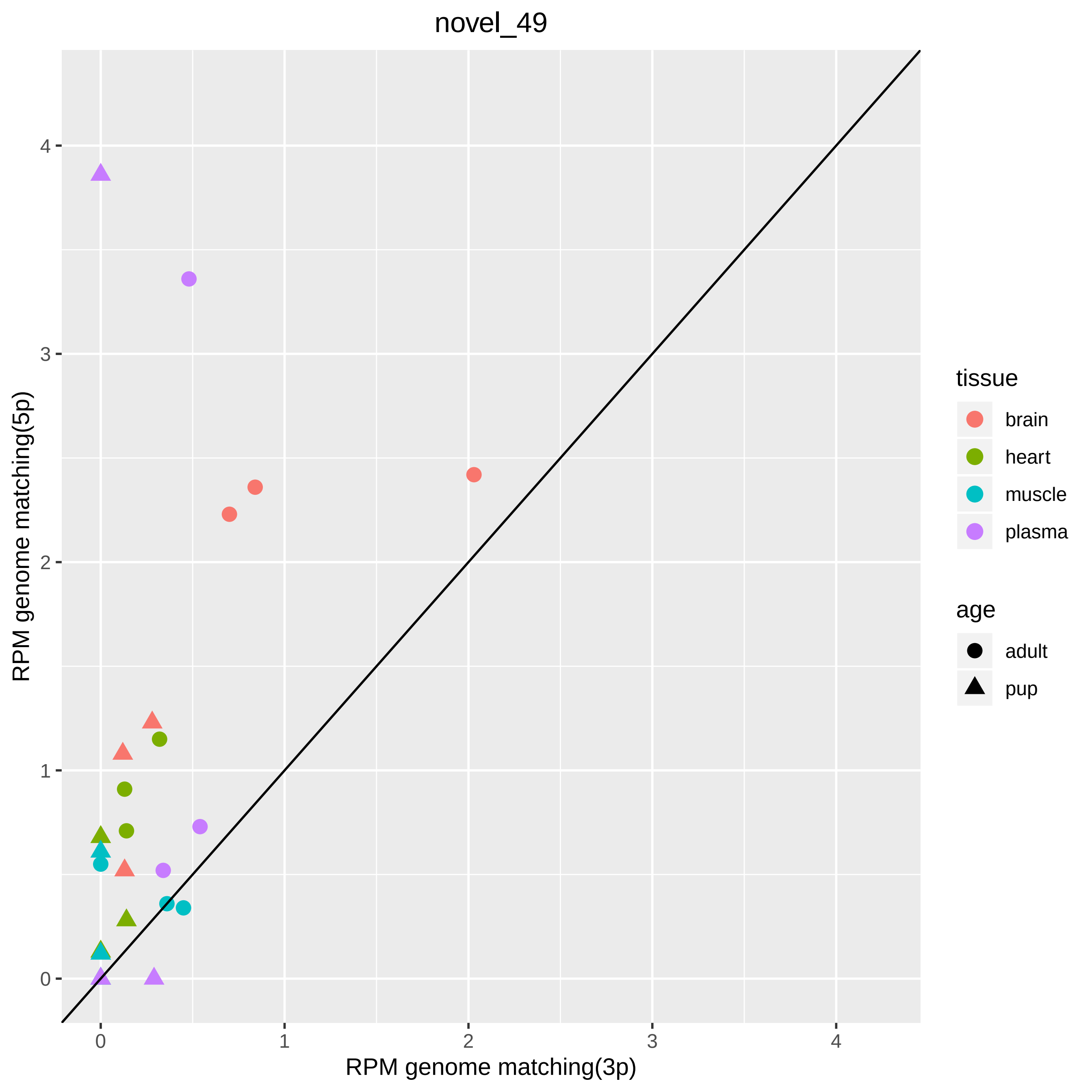

novel_49
RPM genome matching(5p)
RPM genome matching(3p)
0
1
2
3
4
0
1
2
3
4
tissue
brain
heart
muscle
plasma
age
adult
pup

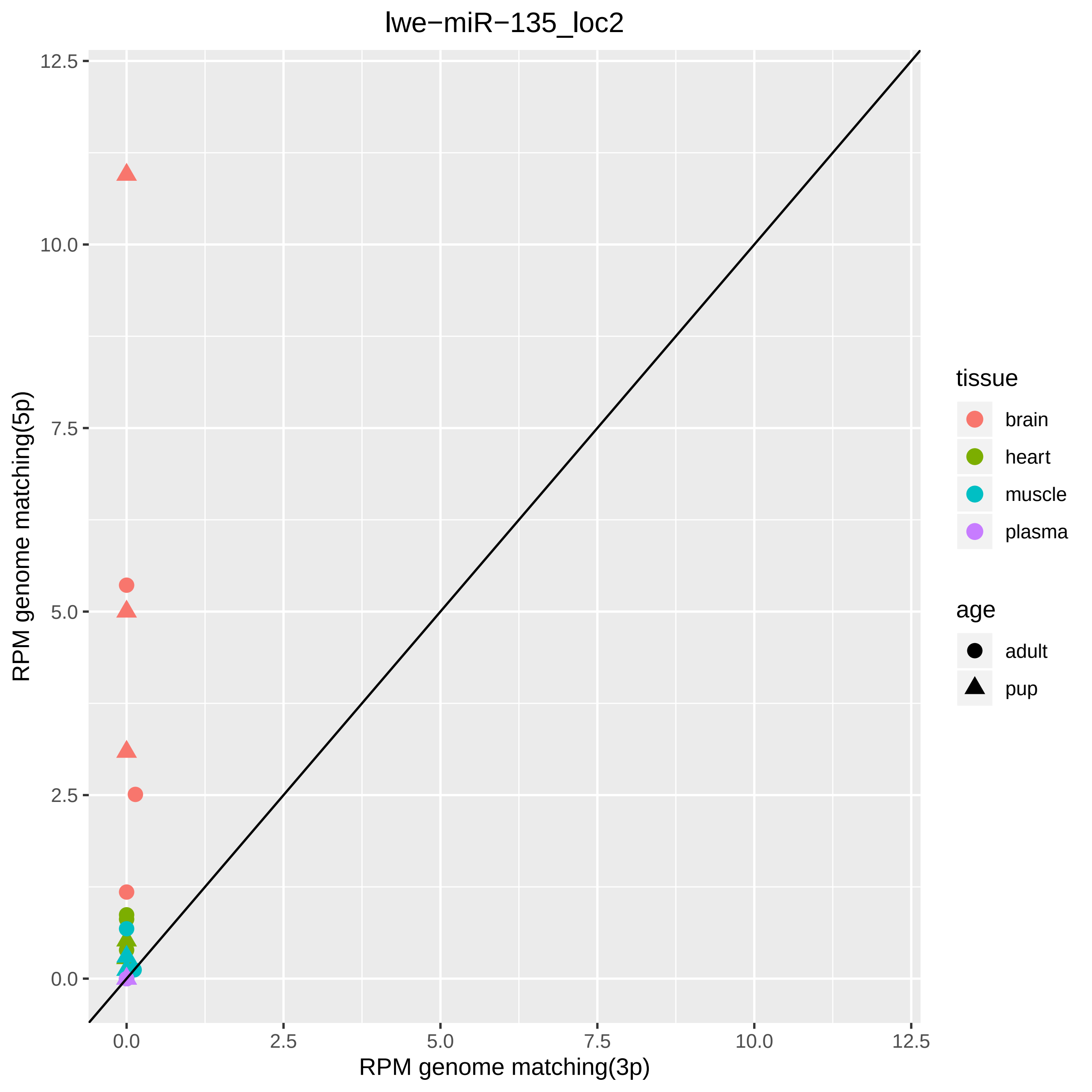
lwe−miR−135_loc2
RPM genome matching(5p)
RPM genome matching(3p)
0.0
2.5
5.0
7.5
10.0
12.5
tissue
brain
heart
muscle
plasma
age
adult
pup

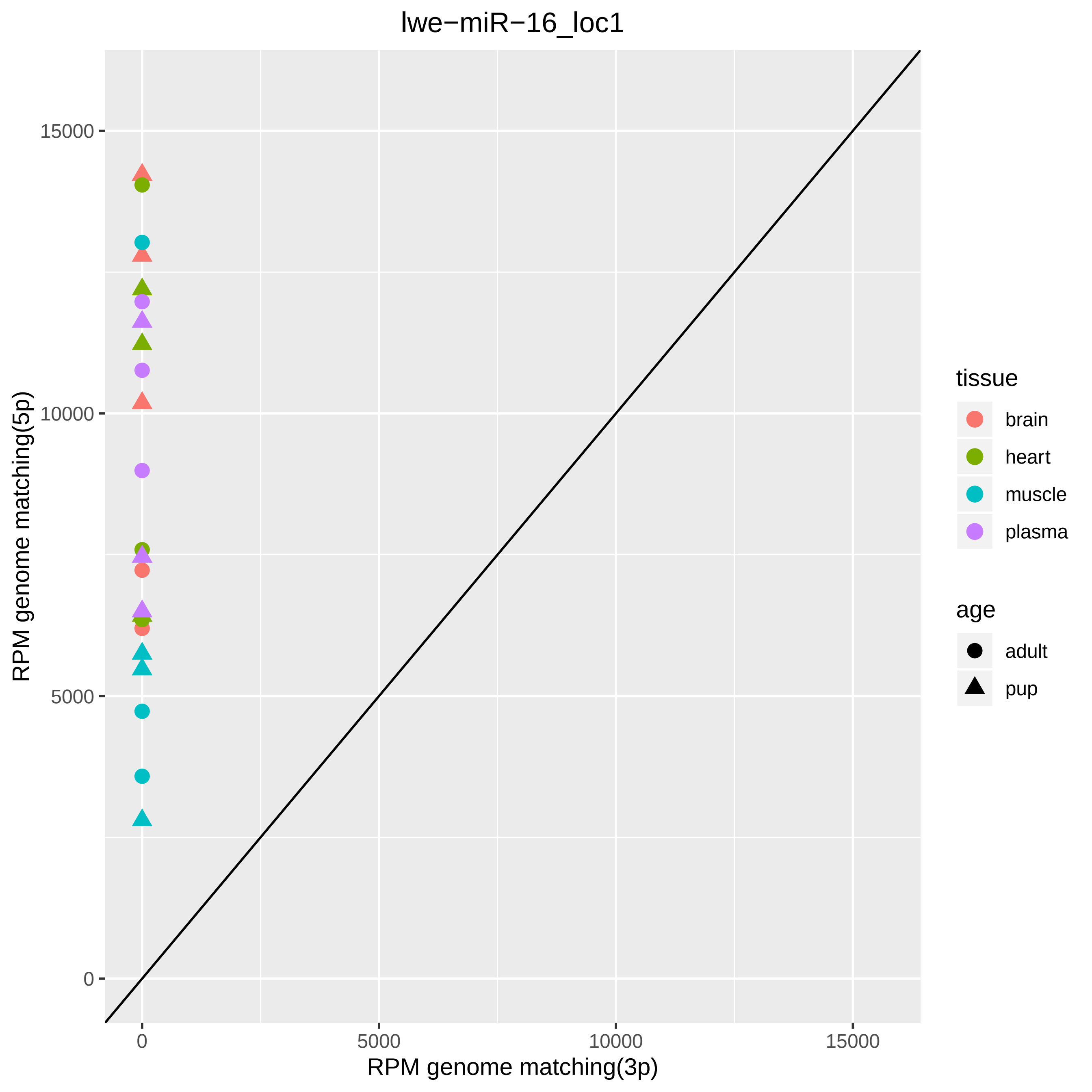
lwe−miR−16_loc1
RPM genome matching(5p)
RPM genome matching(3p)
0
5000
10000
15000
0
5000
10000
15000
tissue
brain
heart
muscle
plasma
age
adult
pup

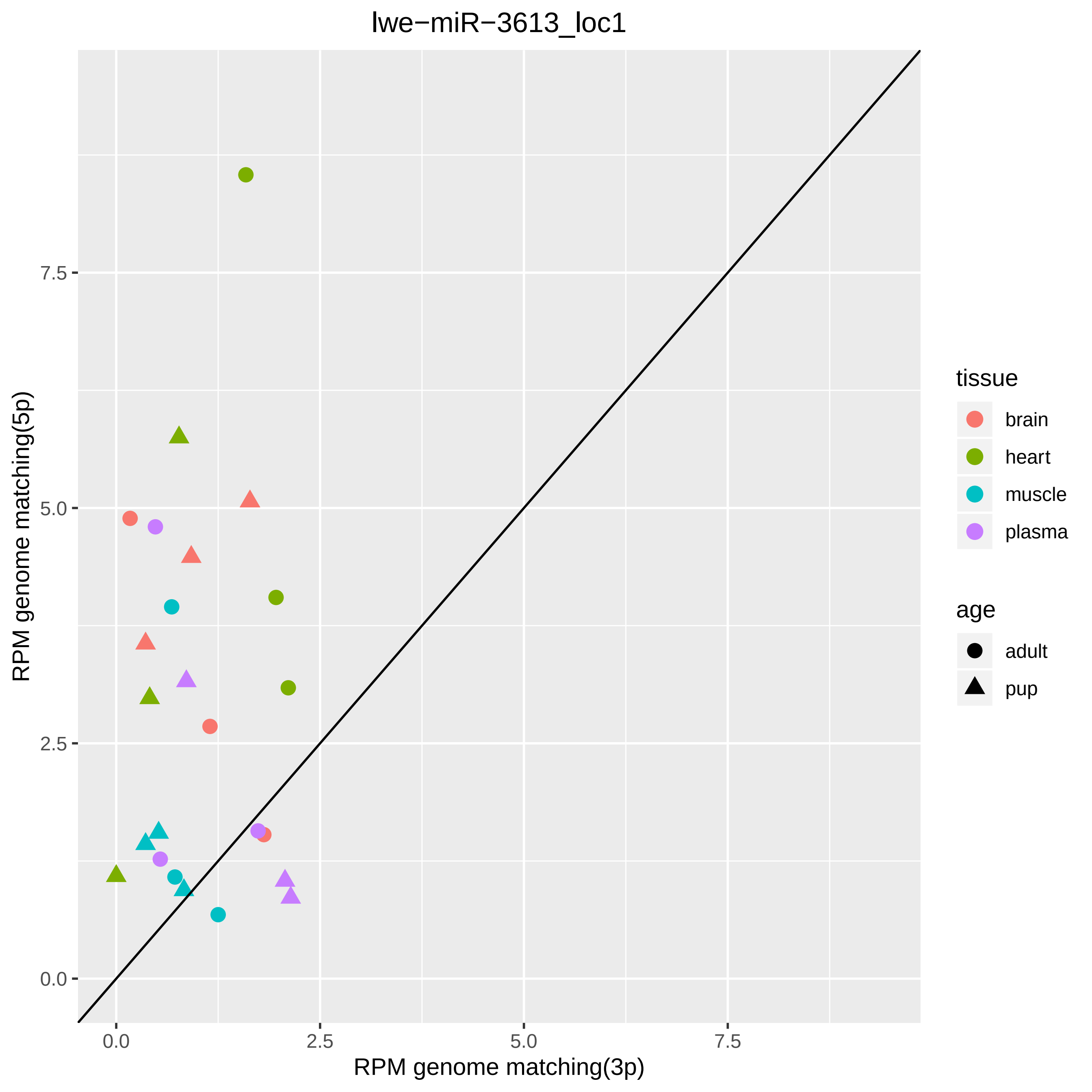
lwe−miR−3613_loc1
RPM genome matching(5p)
RPM genome matching(3p)
0.0
2.5
5.0
7.5
0.0
2.5
5.0
7.5
tissue
brain
heart
muscle
plasma
age
adult
pup

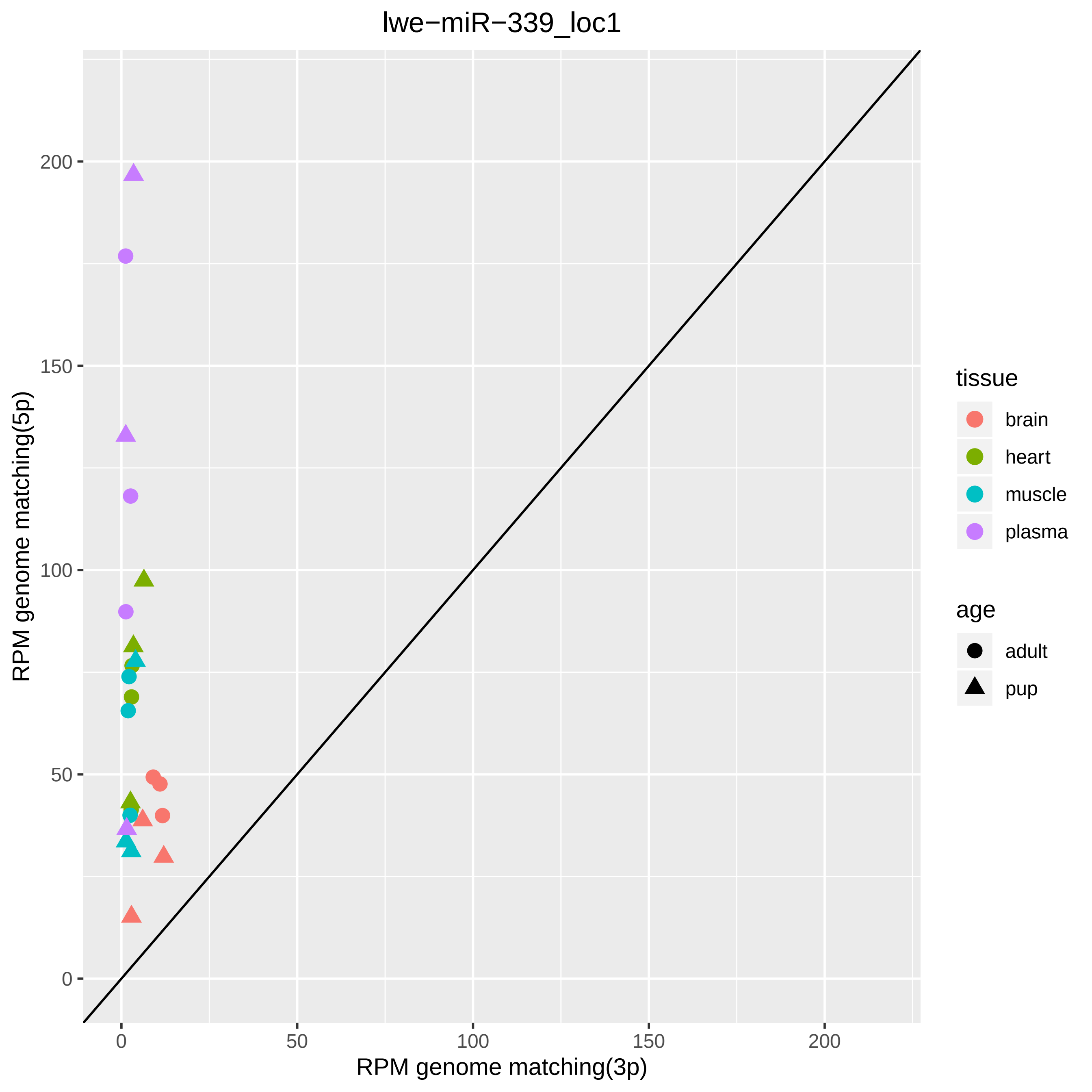
lwe−miR−339_loc1
RPM genome matching(5p)
RPM genome matching(3p)
0
50
100
150
200
tissue
brain
heart
muscle
plasma
age
adult
pup

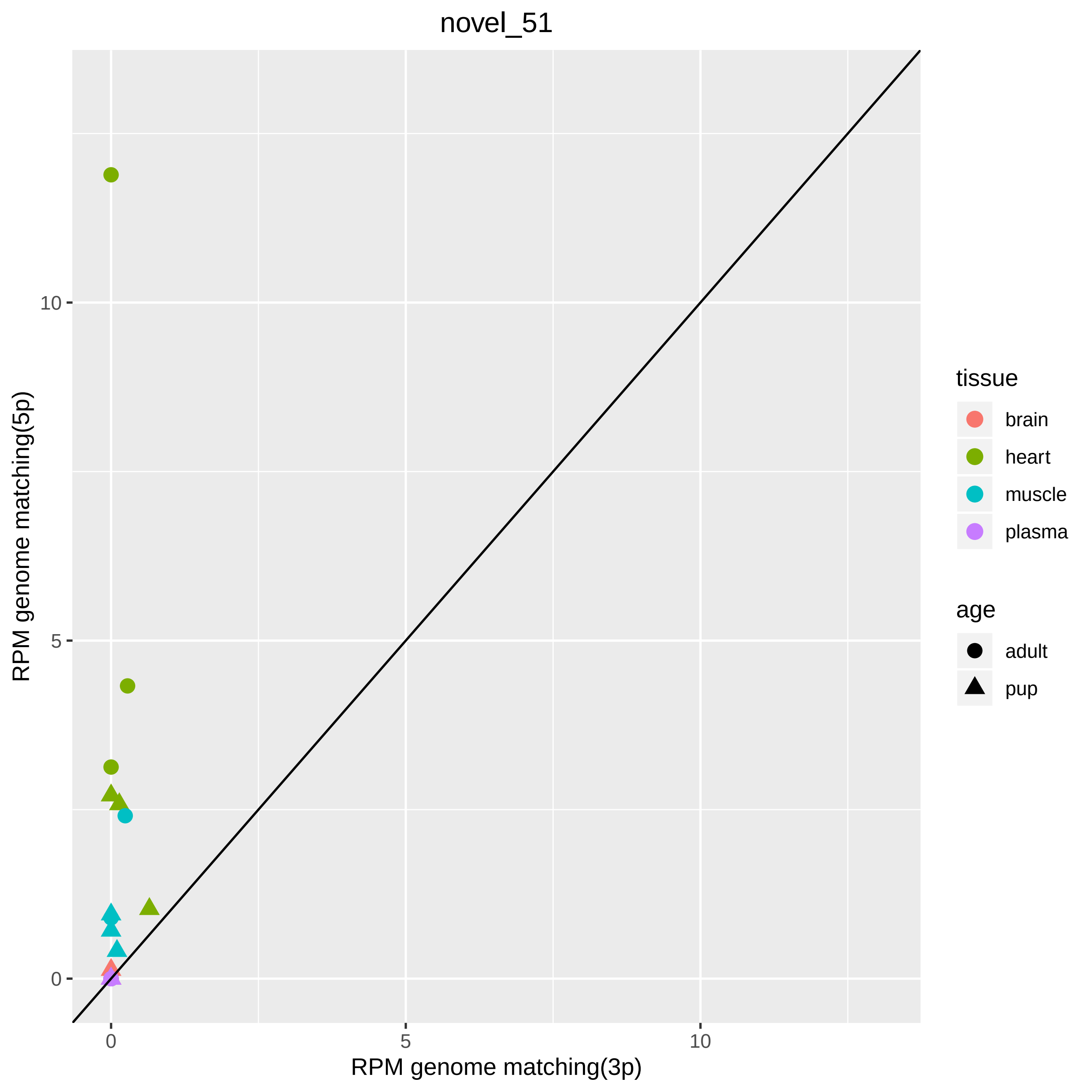
novel_51
RPM genome matching(5p)
RPM genome matching(3p)
0
5
10
0
5
10
tissue
brain
heart
muscle
plasma
age
adult
pup

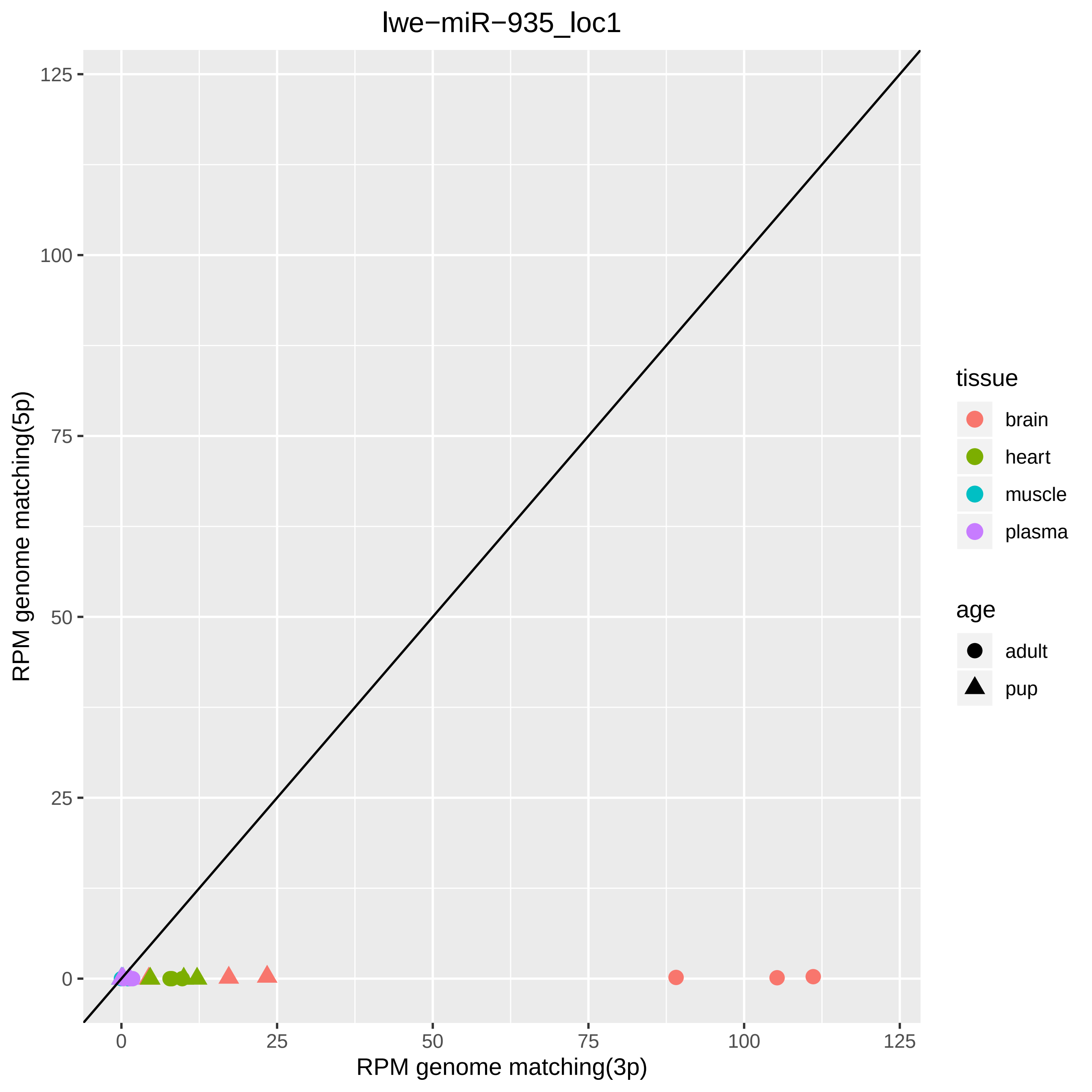

lwe−miR−935_loc1
RPM genome matching(5p)
RPM genome matching(3p)
0
25
50
75
100
125
tissue
brain
heart
muscle
plasma
age
adult
pup

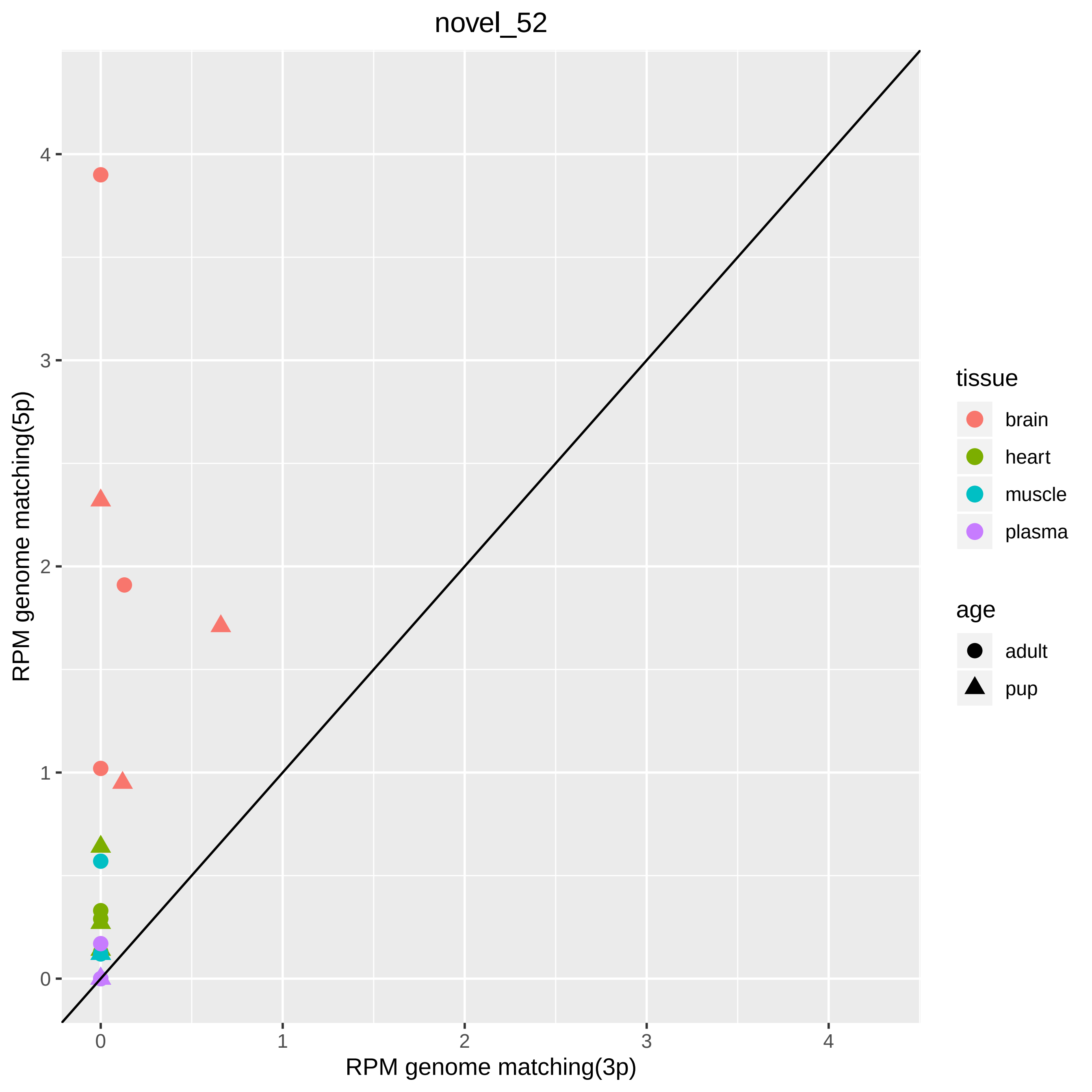

novel_52
RPM genome matching(5p)
RPM genome matching(3p)
0
1
2
3
4
0
1
2
3
4
tissue
brain
heart
muscle
plasma
age
adult
pup

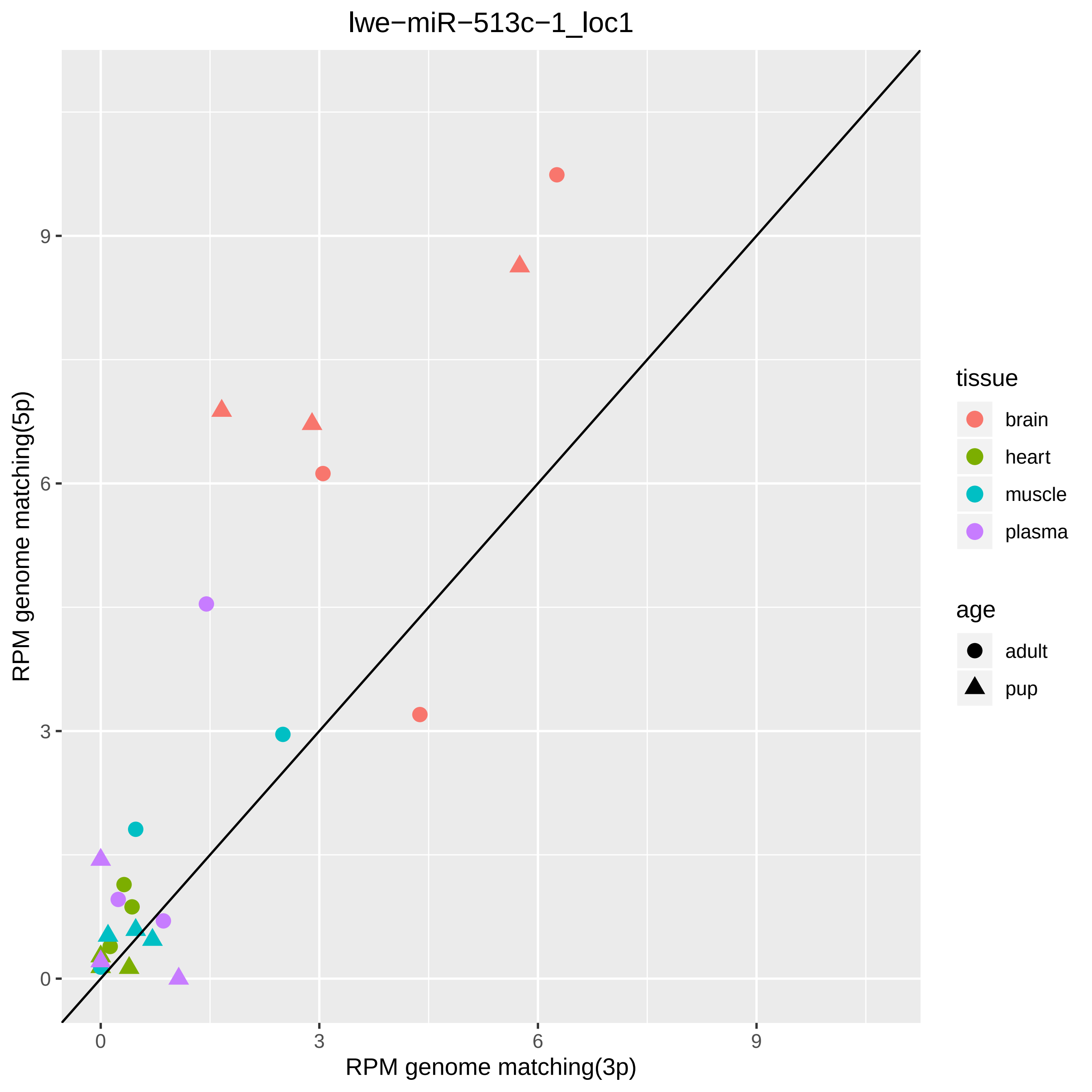
lwe−miR−513c−1_loc1
RPM genome matching(5p)
RPM genome matching(3p)
0
3
6
9
0
3
6
9
tissue
brain
heart
muscle
plasma
age
adult
pup

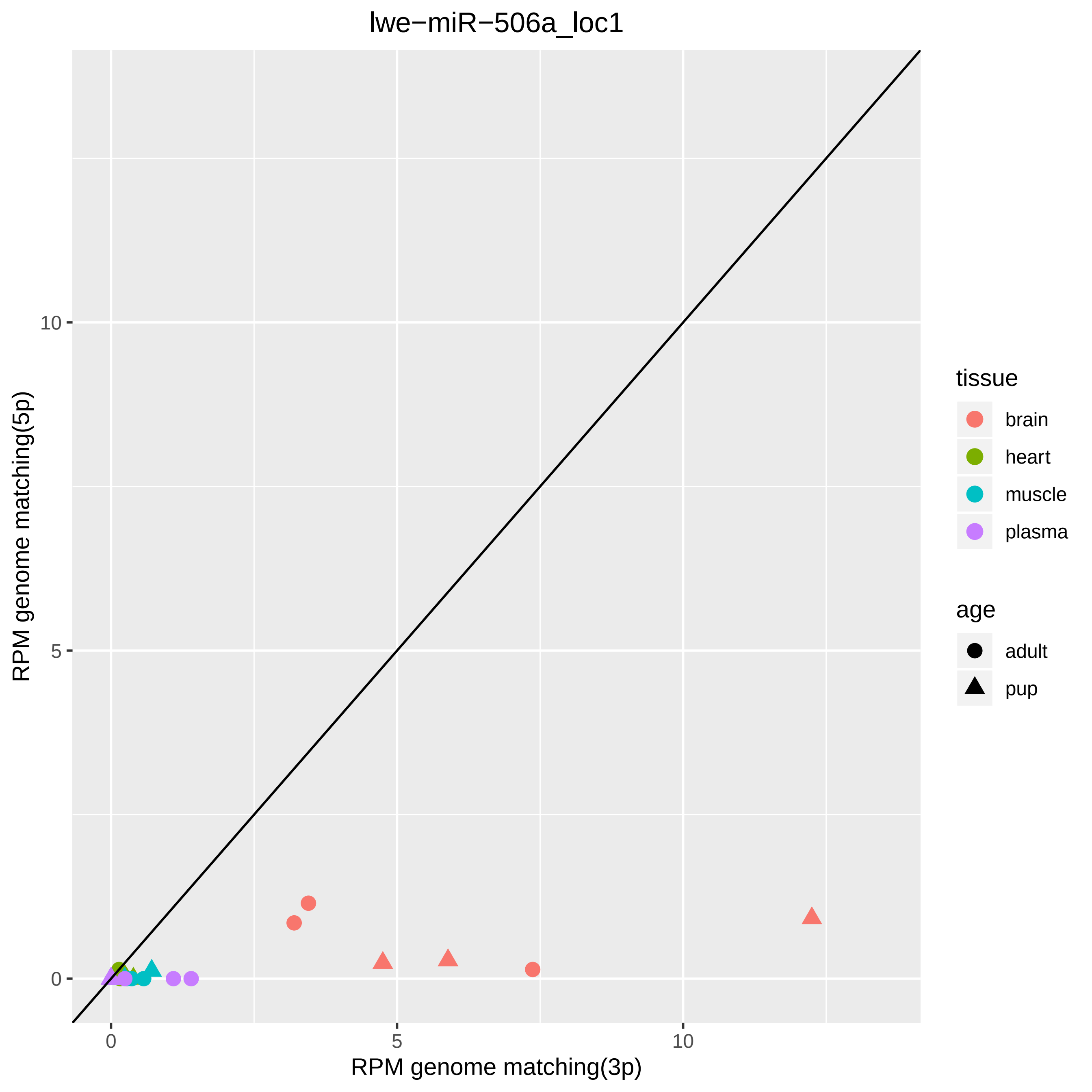
lwe−miR−506a_loc1
RPM genome matching(5p)
RPM genome matching(3p)
0
5
10
0
5
10
tissue
brain
heart
muscle
plasma
age
adult
pup

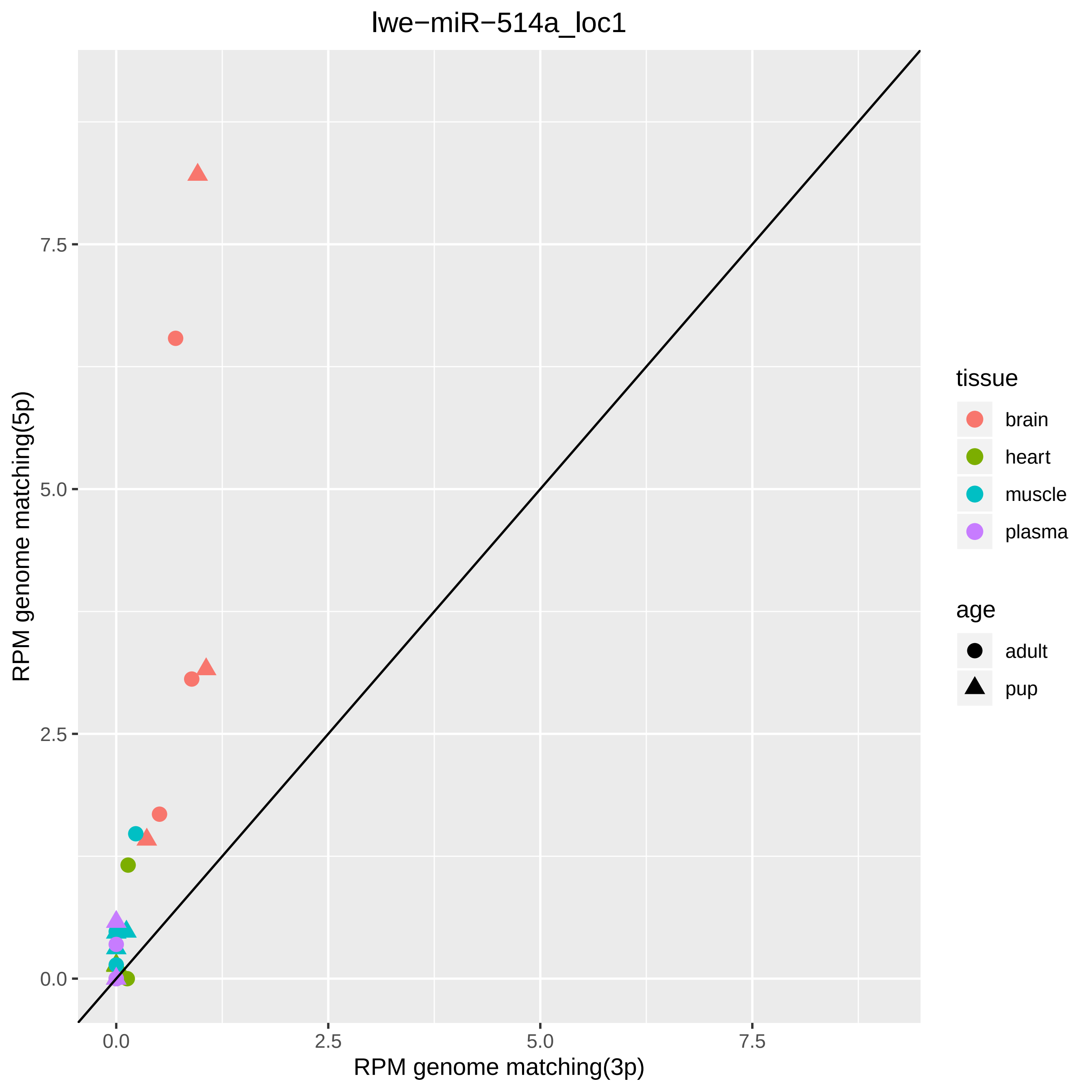

lwe−miR−514a_loc1
RPM genome matching(5p)
RPM genome matching(3p)
0.0
2.5
5.0
7.5
tissue
brain
heart
muscle
plasma
age
adult
pup

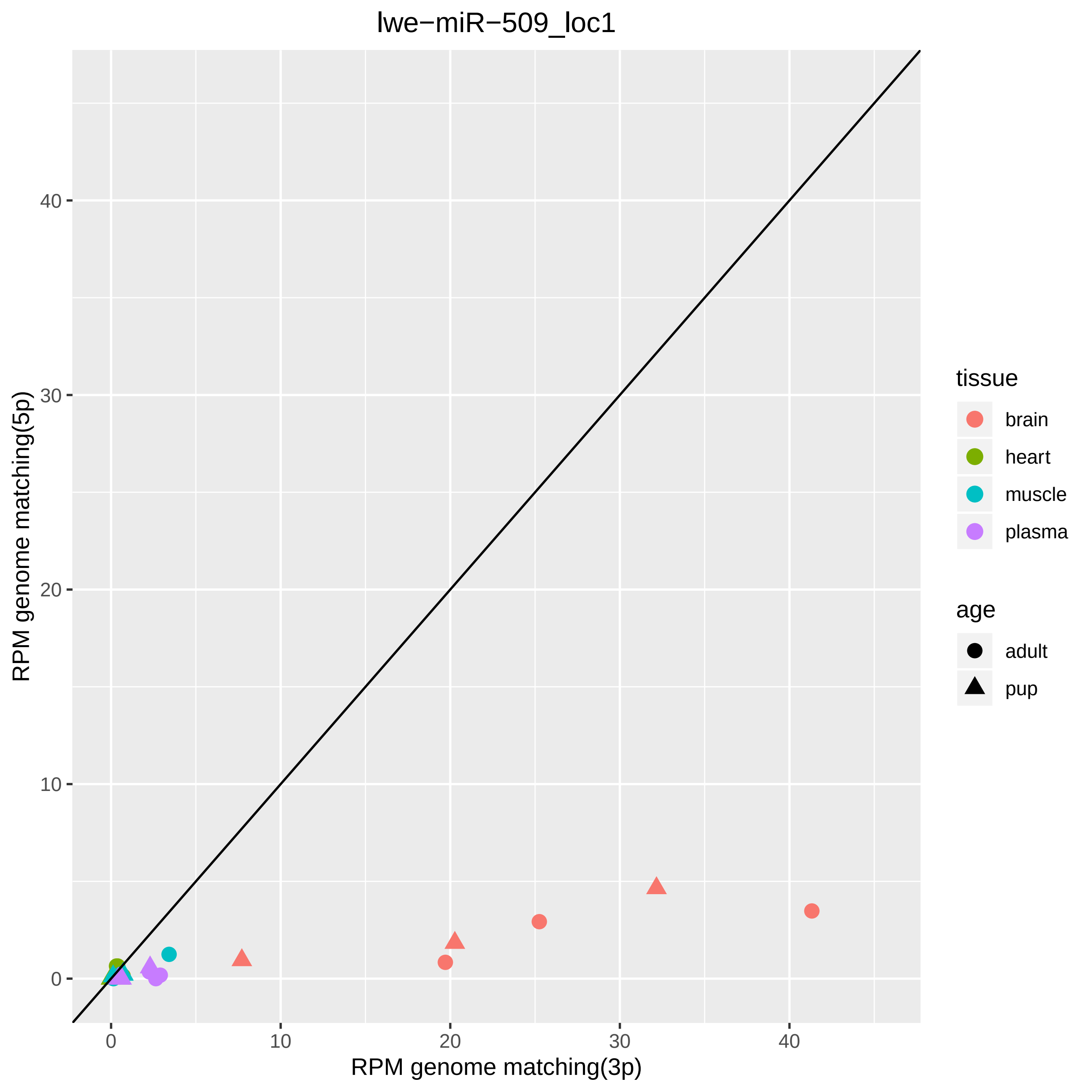
lwe−miR−509_loc1
RPM genome matching(5p)
RPM genome matching(3p)
0
10
20
30
40
0
10
20
30
40
tissue
brain
heart
muscle
plasma
age
adult
pup

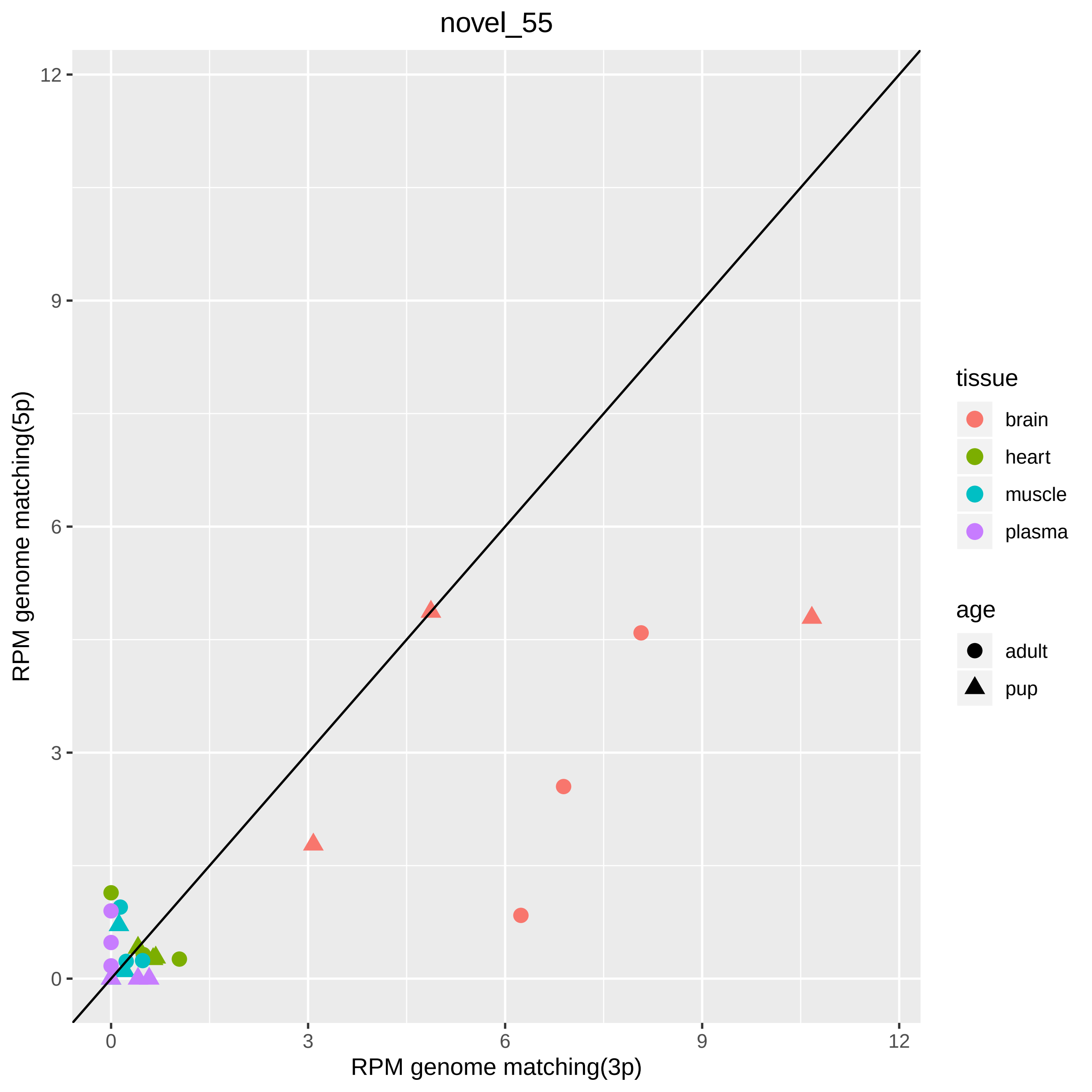
novel_55
RPM genome matching(5p)
RPM genome matching(3p)
0
3
6
9
12
tissue
brain
heart
muscle
plasma
age
adult
pup

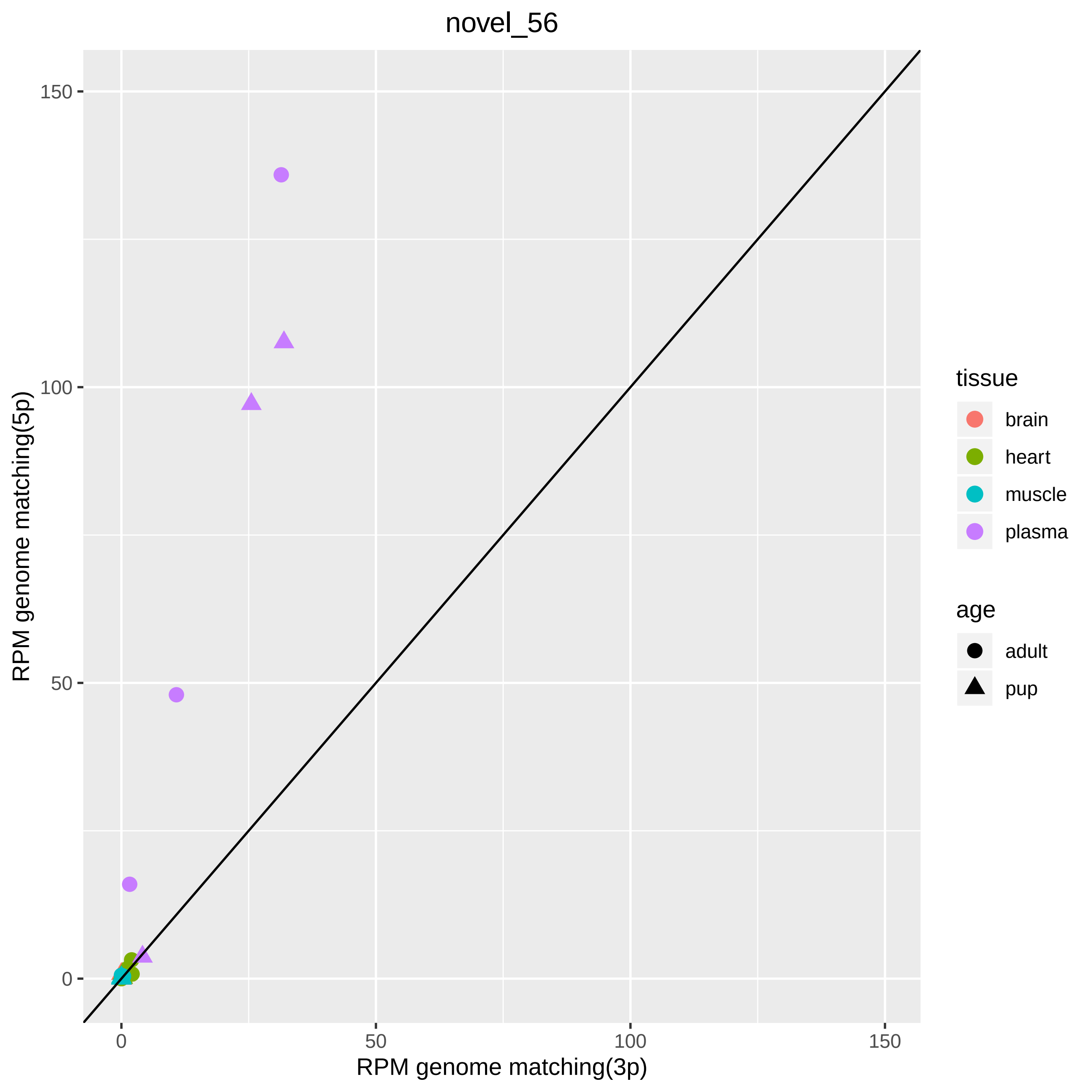

novel_56
RPM genome matching(5p)
RPM genome matching(3p)
150
100
50
0
0
50
100
150
tissue
brain
heart
muscle
plasma
age
adult
pup

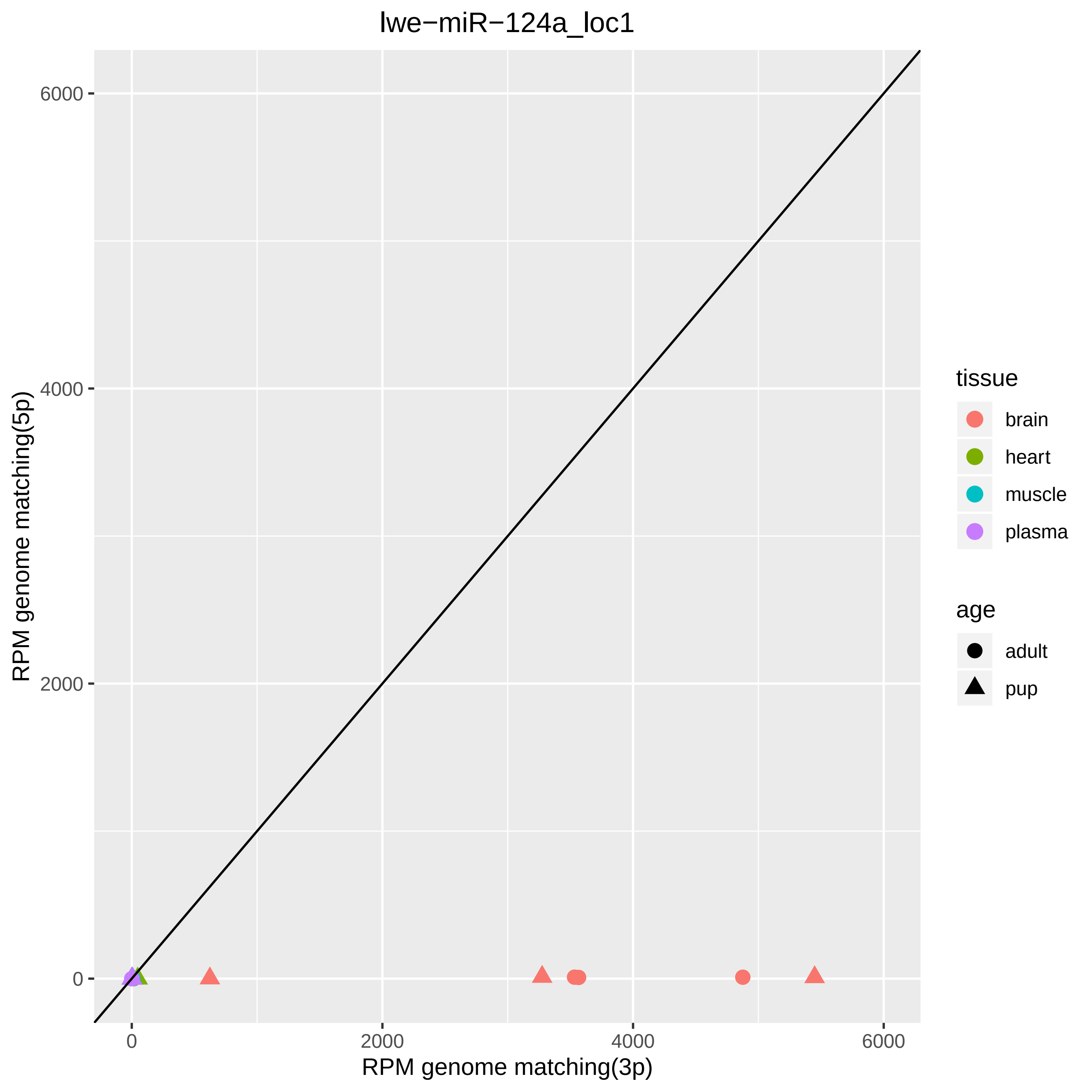

lwe−miR−124a_loc1
RPM genome matching(5p)
RPM genome matching(3p)
0
2000
4000
6000
0
2000
4000
6000
tissue
brain
heart
muscle
plasma
age
adult
pup

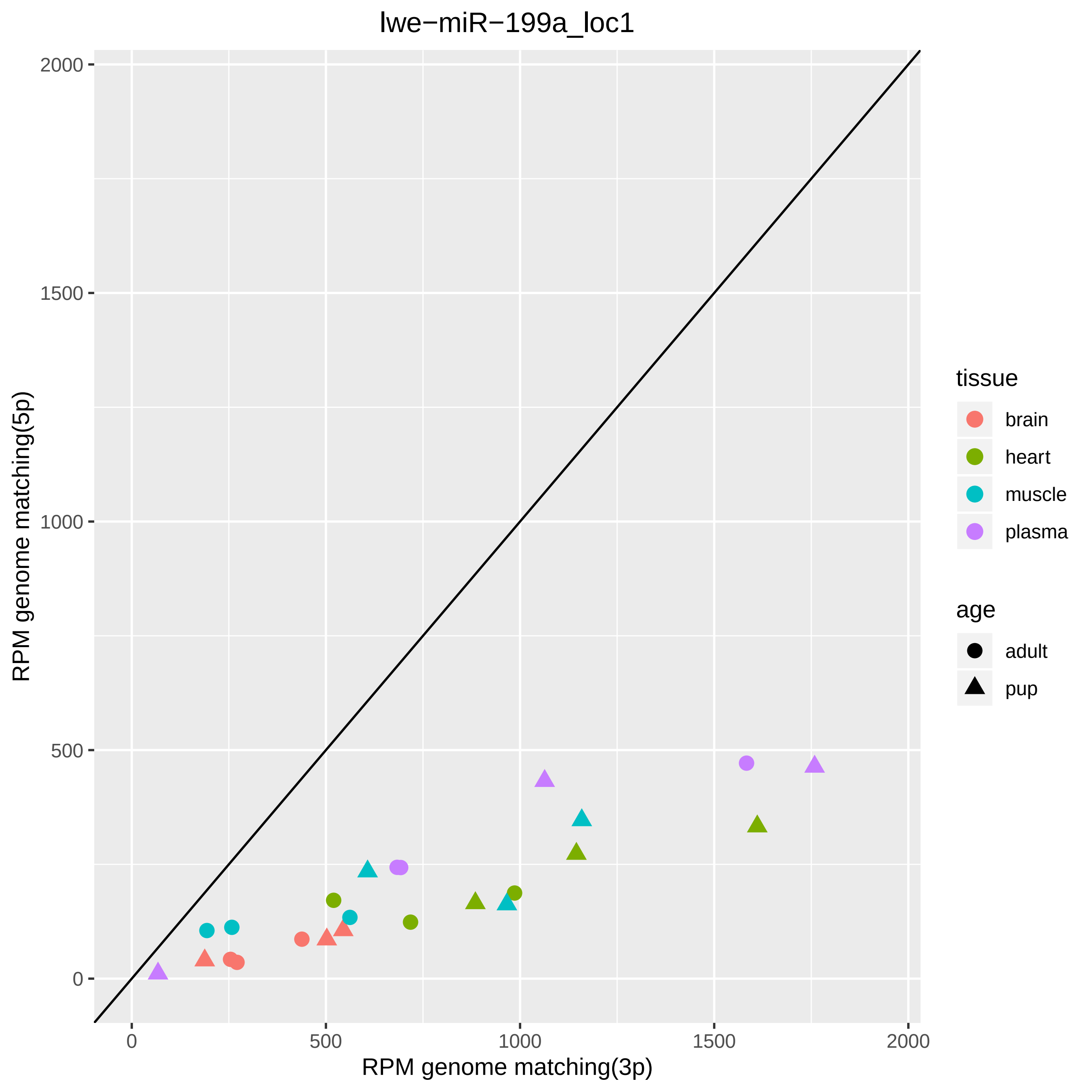
lwe−miR−199a_loc1
RPM genome matching(5p)
RPM genome matching(3p)
0
500
1000
1500
2000
tissue
brain
heart
muscle
plasma
age
adult
pup

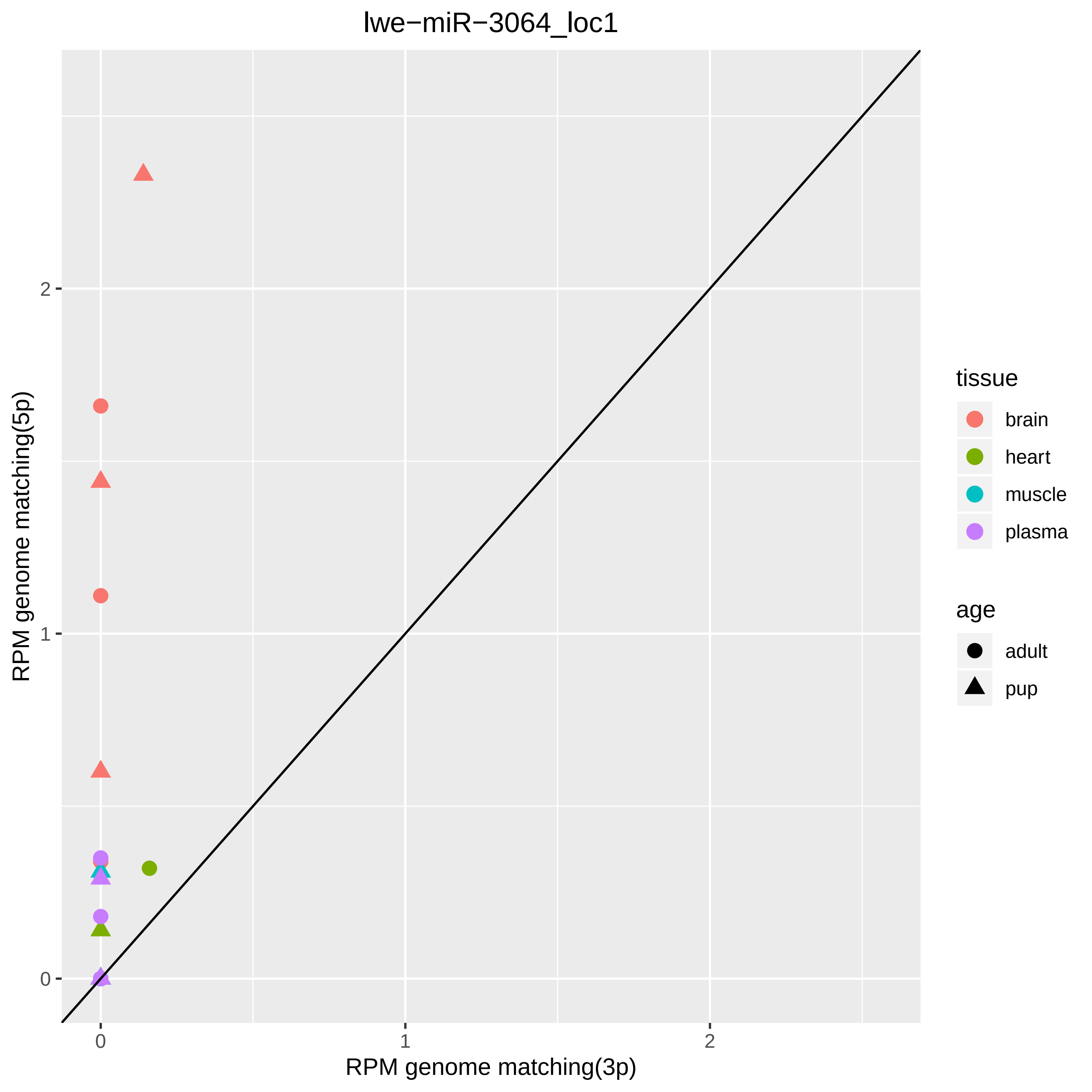
lwe−miR−3064_loc1
RPM genome matching(5p)
RPM genome matching(3p)
0
1
2
0
1
2
tissue
brain
heart
muscle
plasma
age
adult
pup

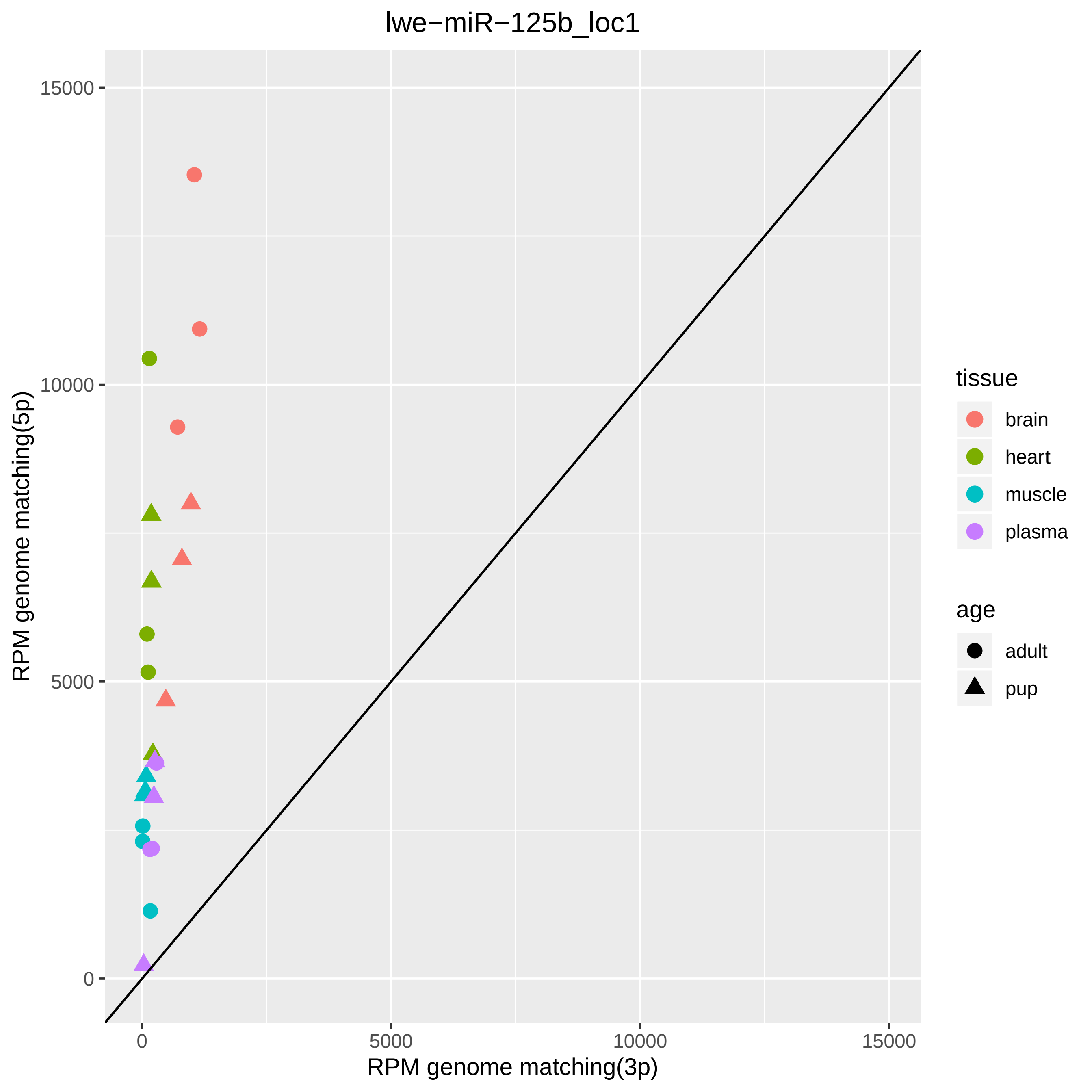

lwe−miR−125b_loc1
15000
10000
5000
0
0
5000
10000
15000
RPM genome matching(5p)
RPM genome matching(3p)
tissue
brain
heart
muscle
plasma
age
adult
pup

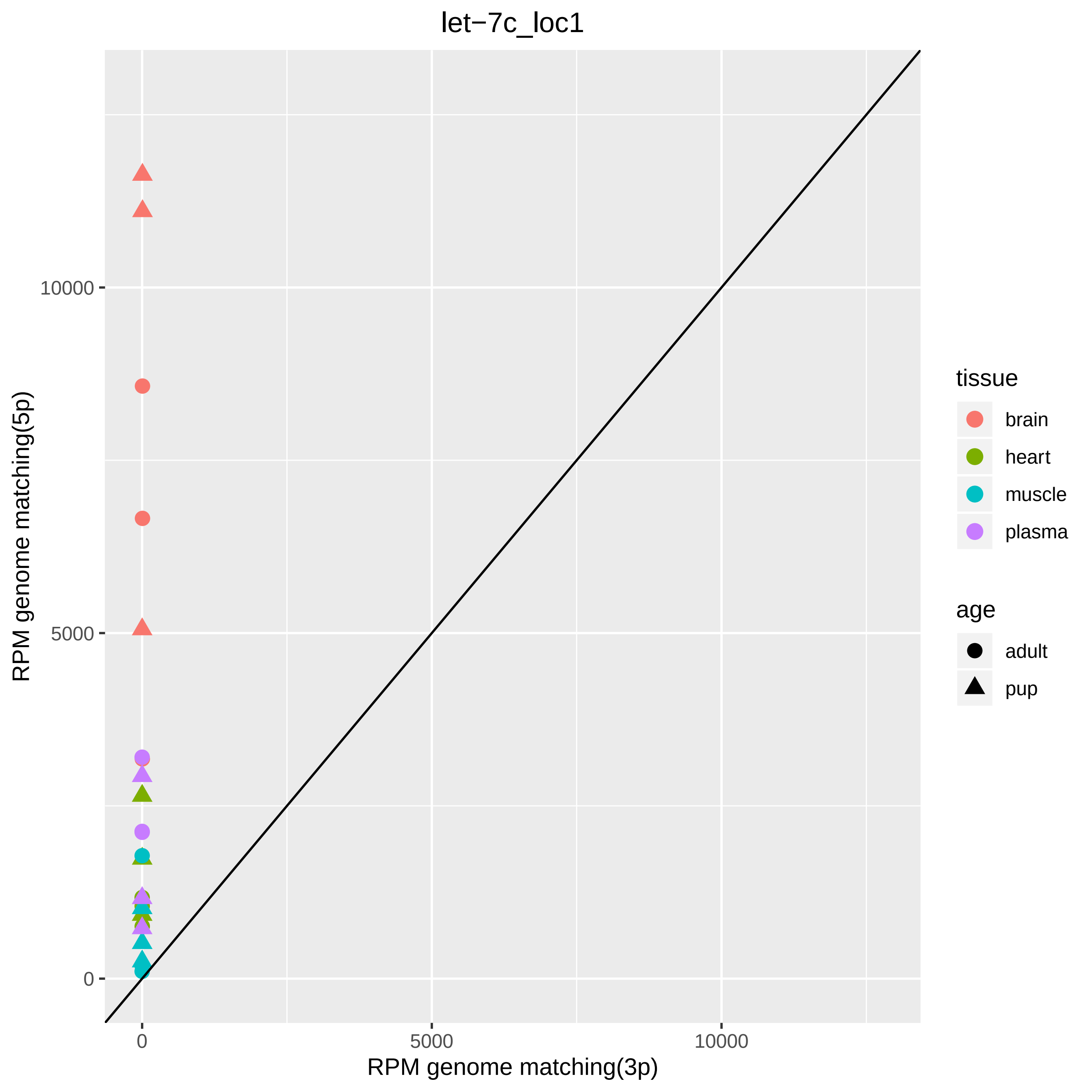
let−7c_loc1
RPM genome matching(5p)
RPM genome matching(3p)
10000
5000
0
0
5000
10000
tissue
brain
heart
muscle
plasma
age
adult
pup

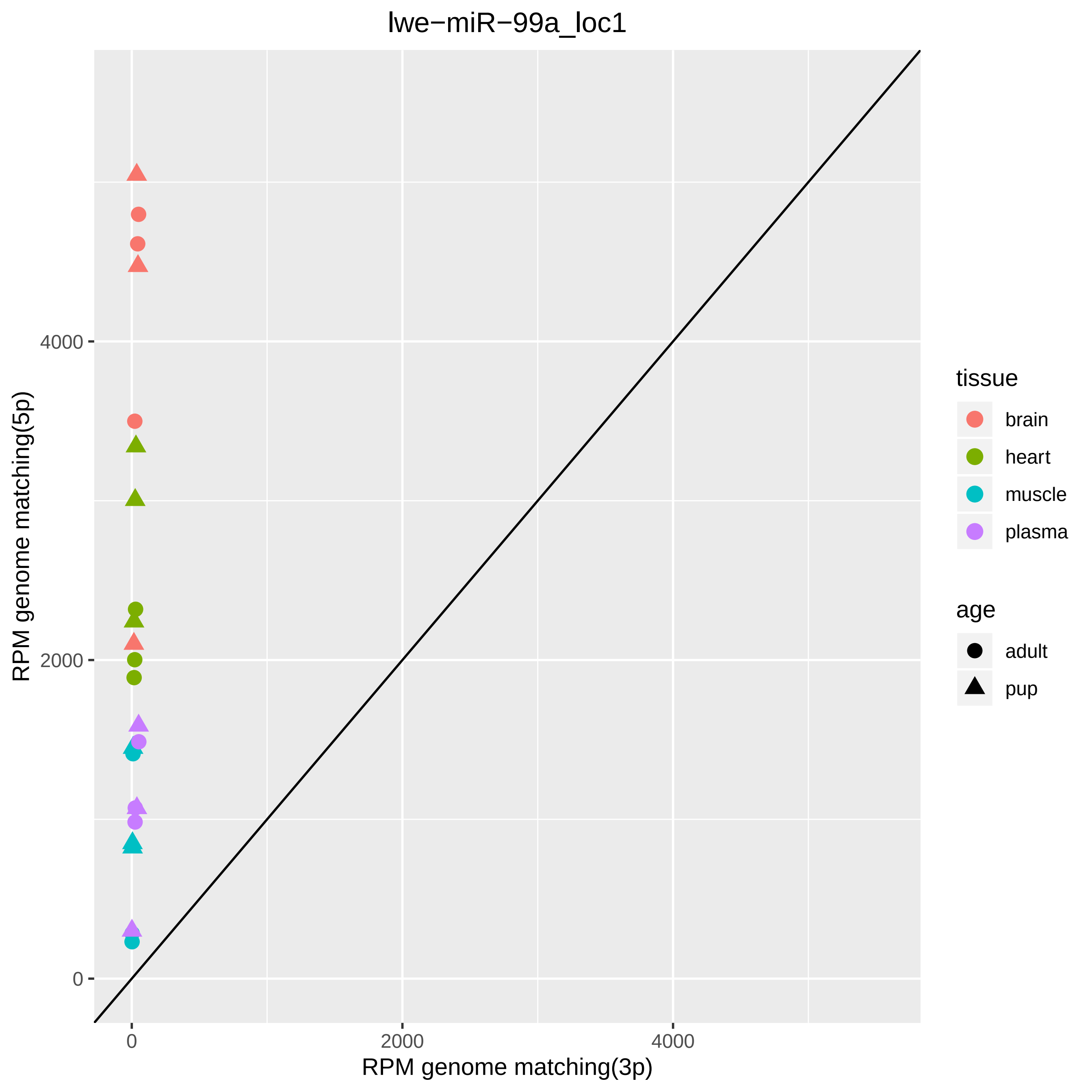

lwe−miR−99a_loc1
RPM genome matching(5p)
4000
2000
0
RPM genome matching(3p)
0
2000
4000
tissue
brain
heart
muscle
plasma
age
adult
pup

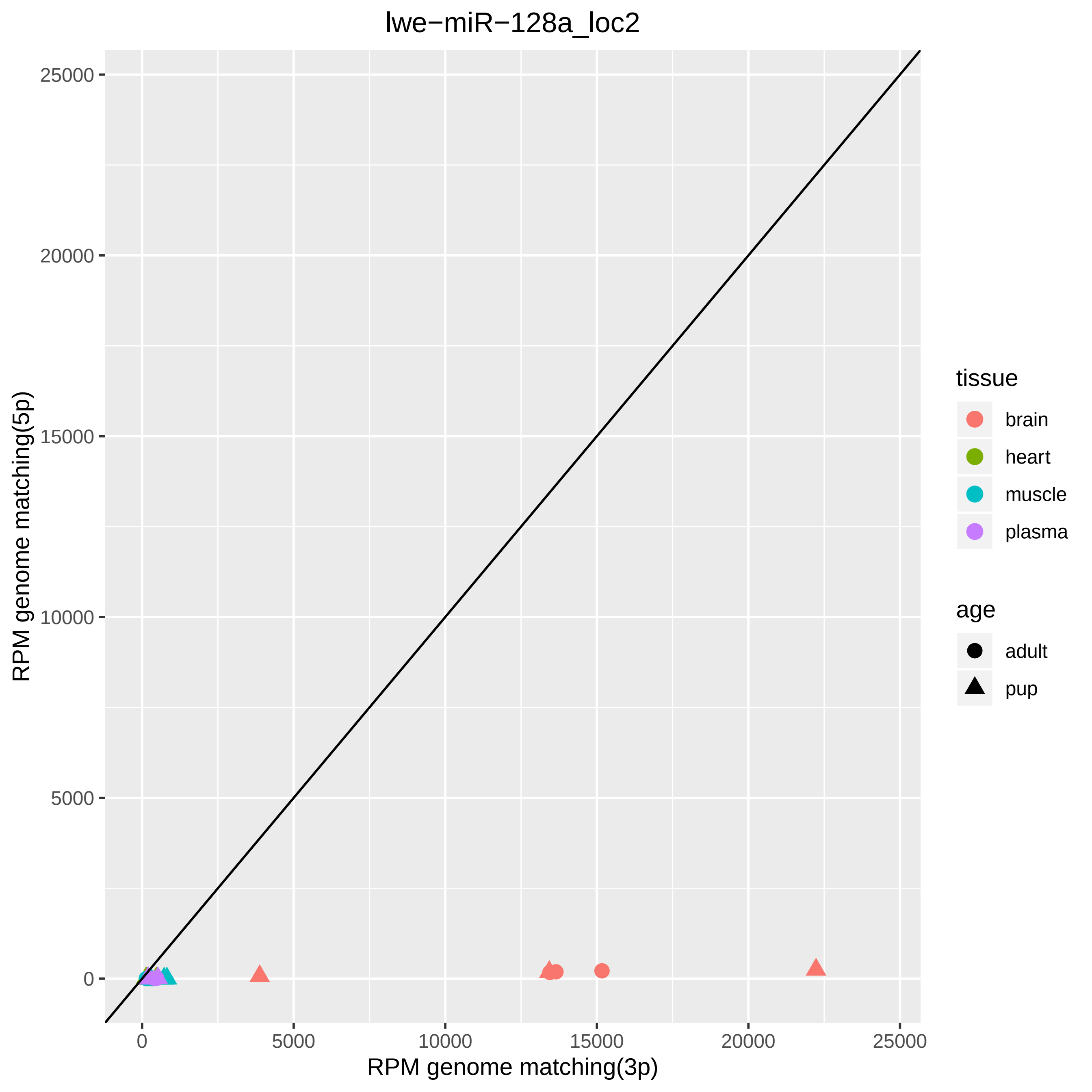
lwe−miR−128a_loc2
RPM genome matching(5p)
RPM genome matching(3p)
0
5000
10000
15000
20000
25000
tissue
brain
heart
muscle
plasma
age
adult
pup

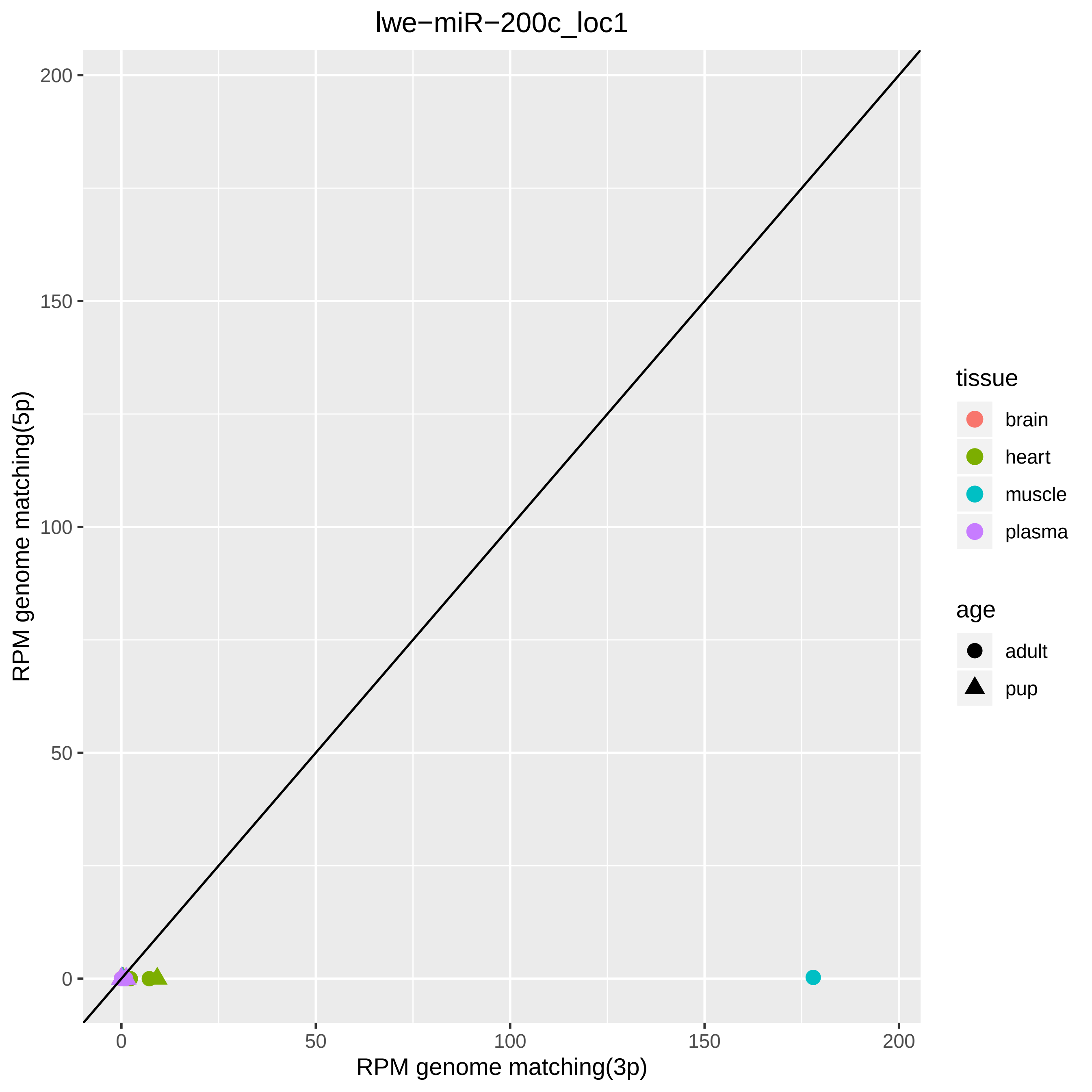

lwe−miR−200c_loc1
RPM genome matching(5p)
RPM genome matching(3p)
0
50
100
150
200
tissue
brain
heart
muscle
plasma
age
adult
pup

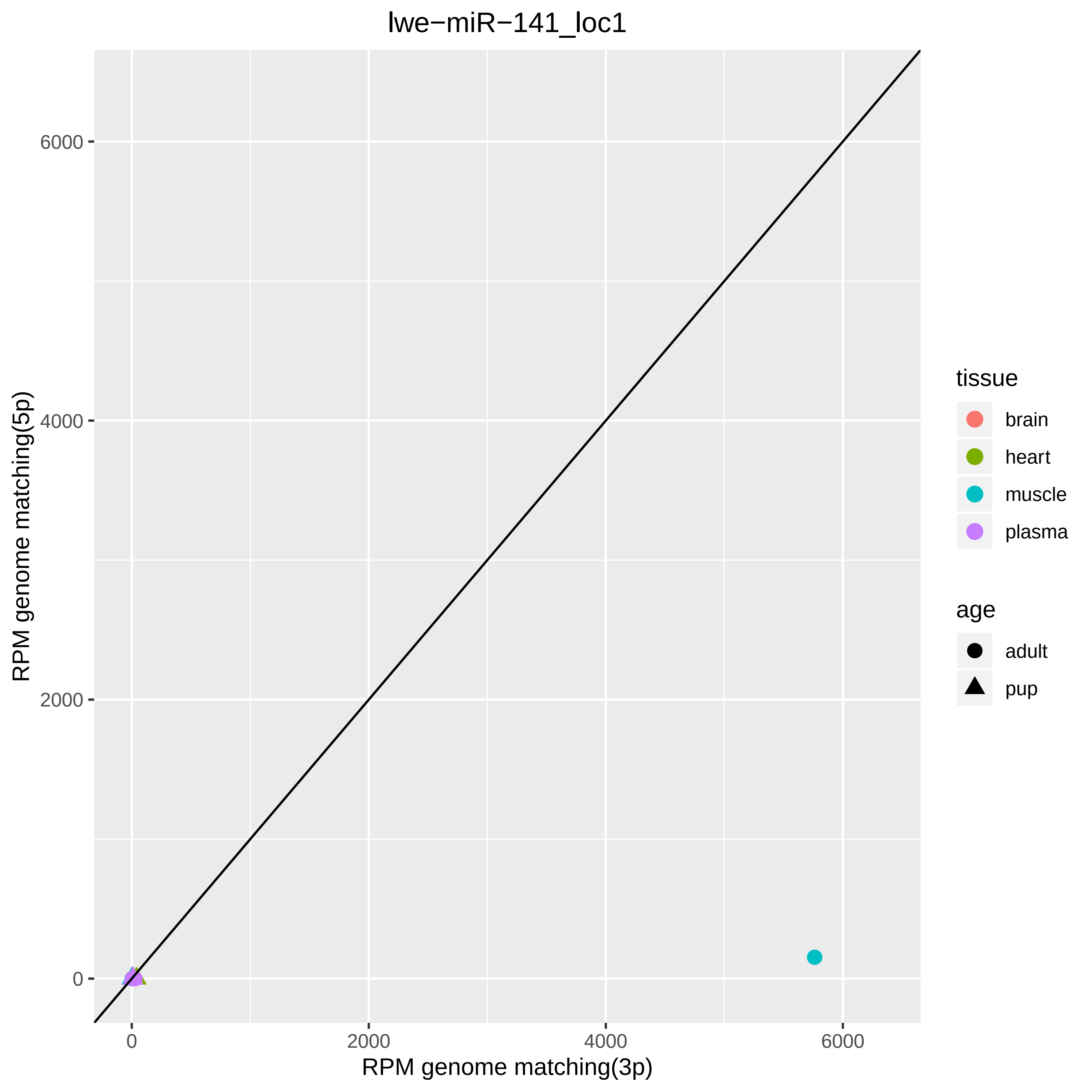
lwe−miR−141_loc1
RPM genome matching(5p)
RPM genome matching(3p)
0
2000
4000
6000
0
2000
4000
6000
tissue
brain
heart
muscle
plasma
age
adult
pup

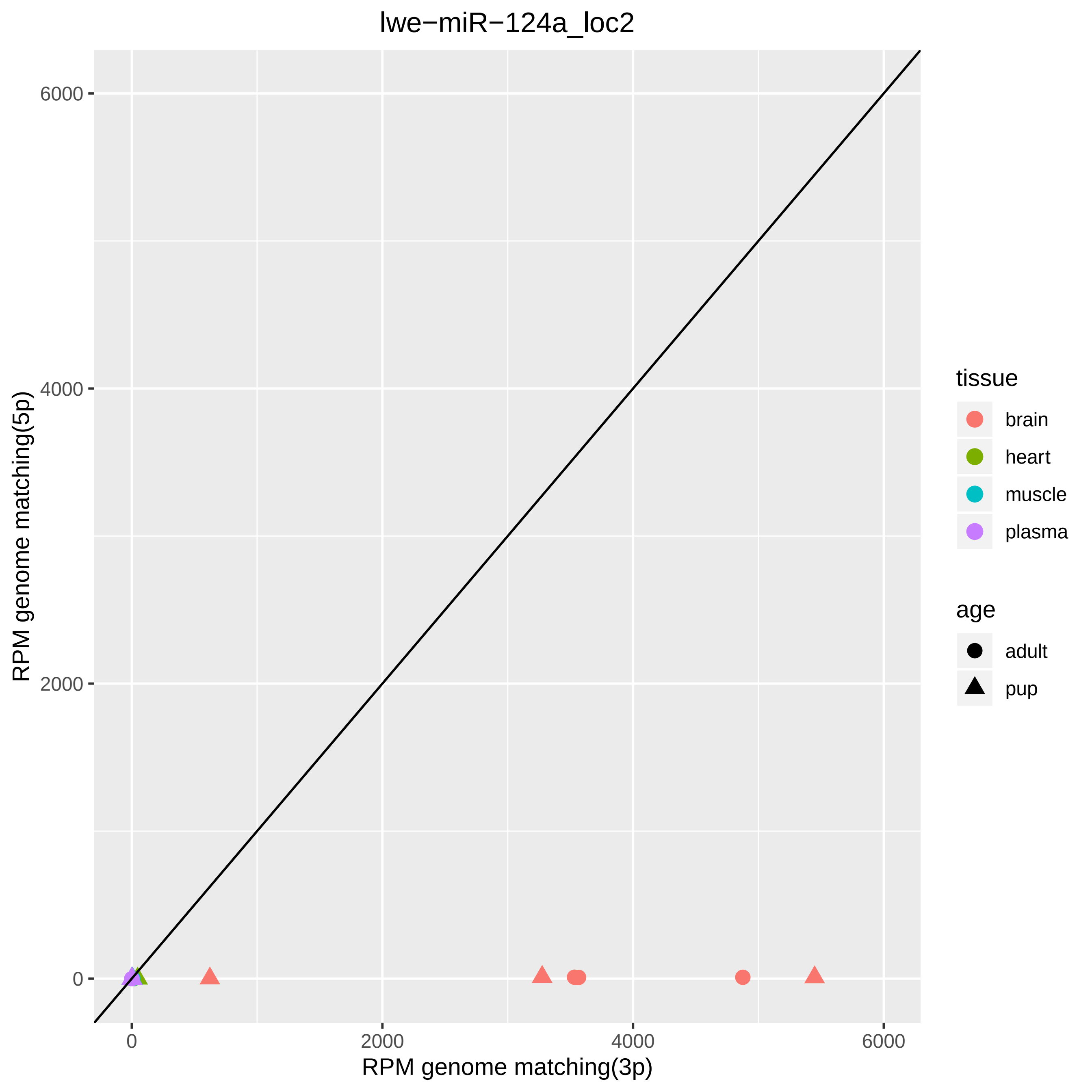

lwe−miR−124a_loc2
RPM genome matching(5p)
RPM genome matching(3p)
0
2000
4000
6000
0
2000
4000
6000
tissue
brain
heart
muscle
plasma
age
adult
pup

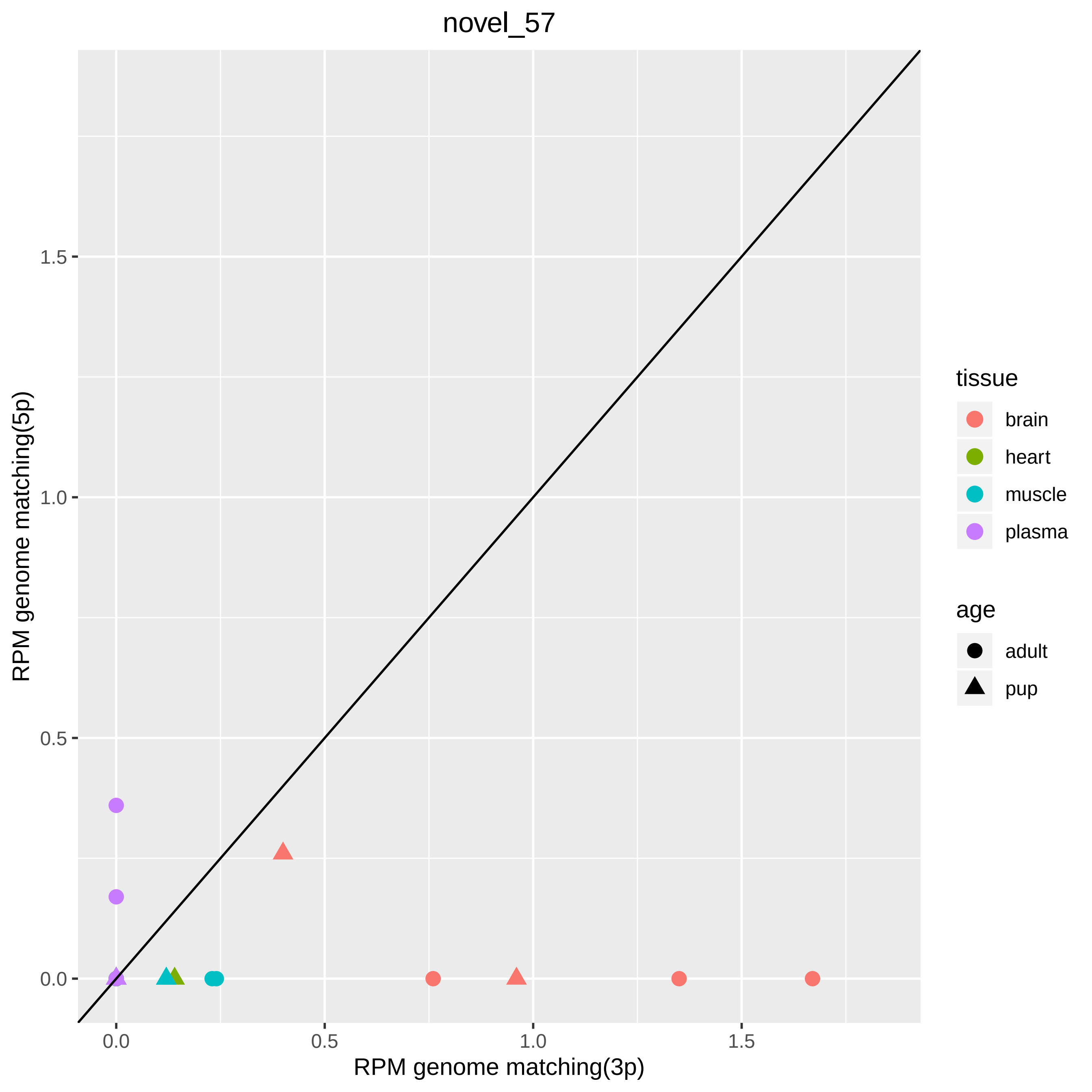
novel_57
RPM genome matching(5p)
RPM genome matching(3p)
0.0
0.5
1.0
1.5
tissue
brain
heart
muscle
plasma
age
adult
pup

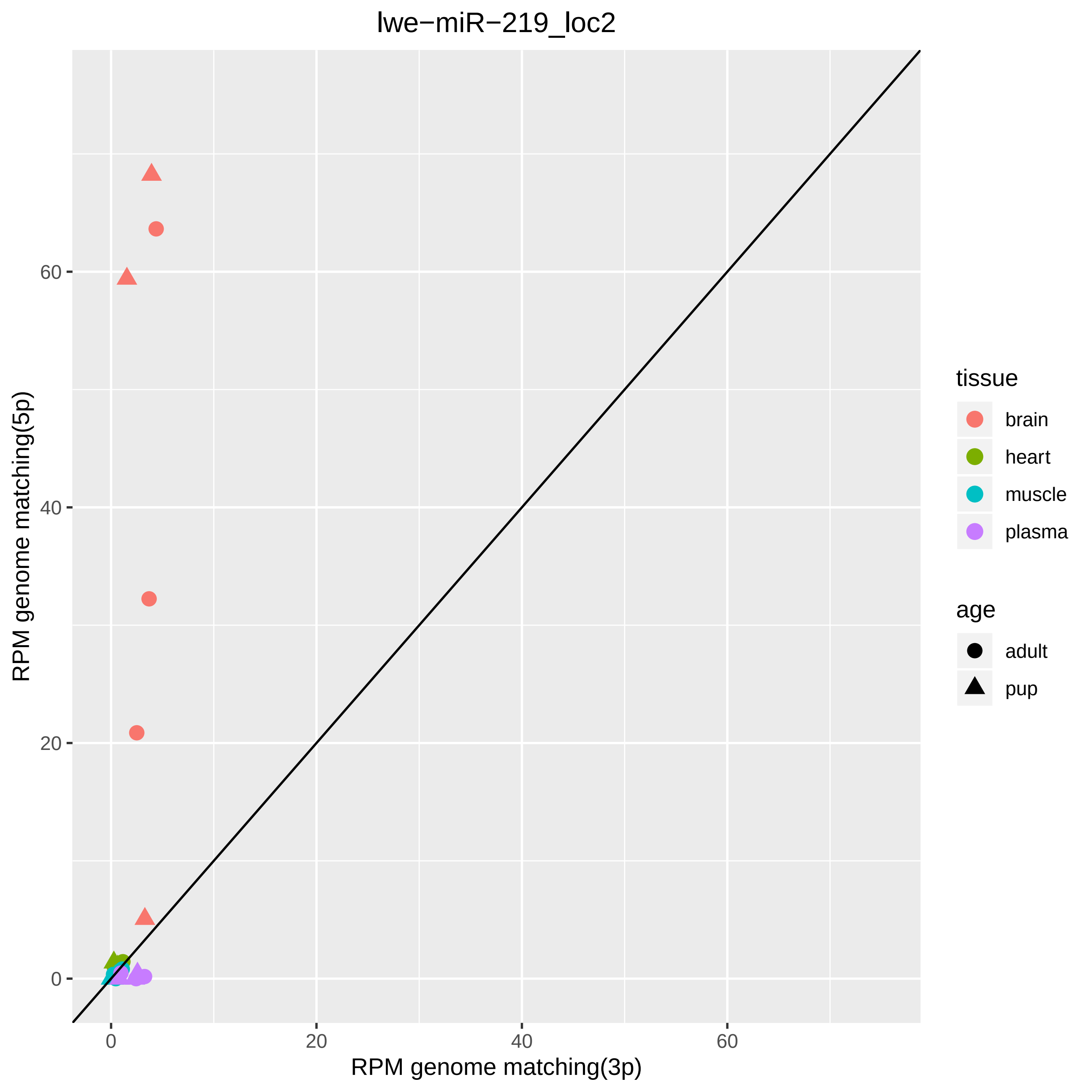
lwe−miR−219_loc2
RPM genome matching(5p)
RPM genome matching(3p)
0
20
40
60
tissue
brain
heart
muscle
plasma
age
adult
pup

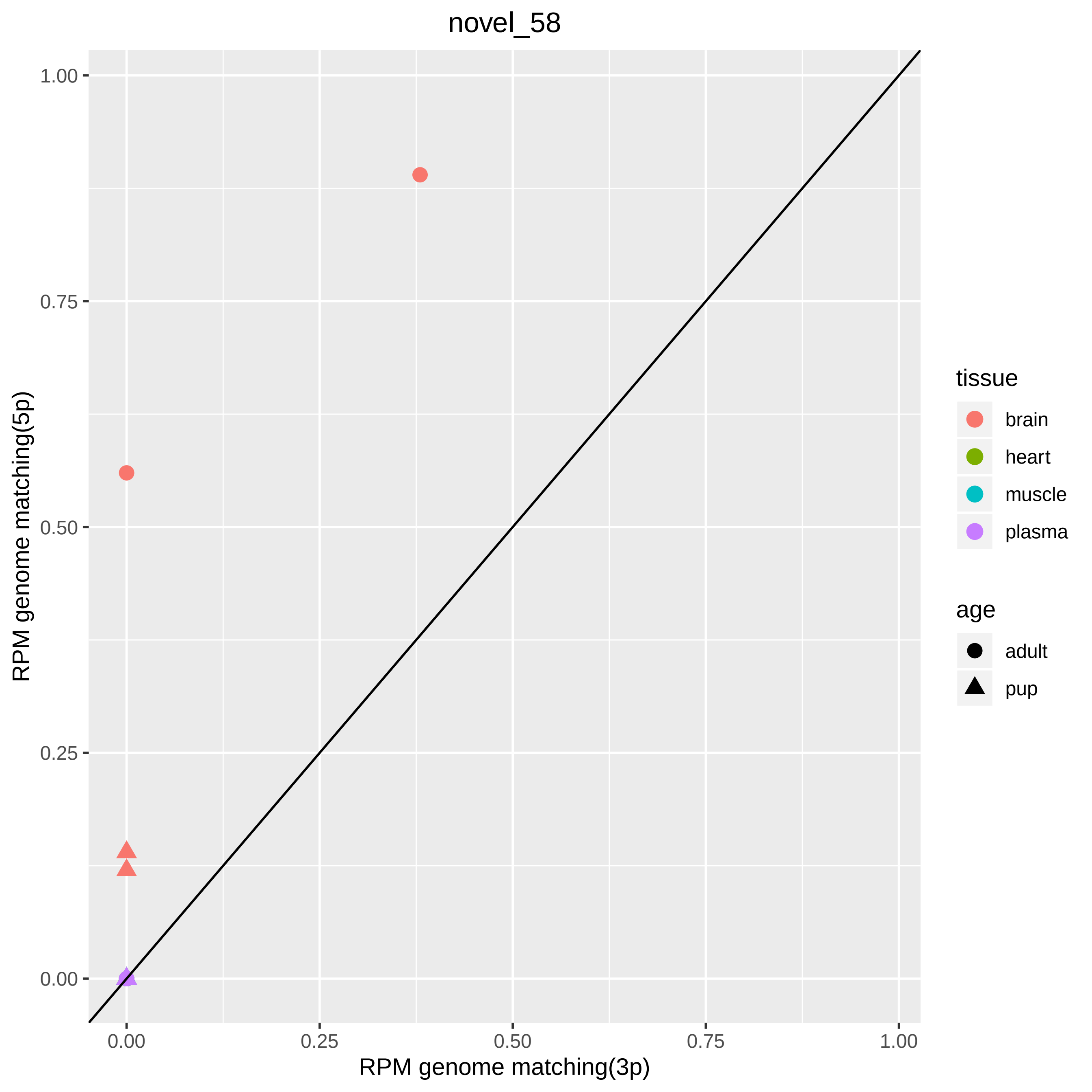
novel_58
RPM genome matching(5p)
RPM genome matching(3p)
1.00
0.75
0.50
0.25
0.00
0.00
0.25
0.50
0.75
1.00
tissue
brain
heart
muscle
plasma
age
adult
pup

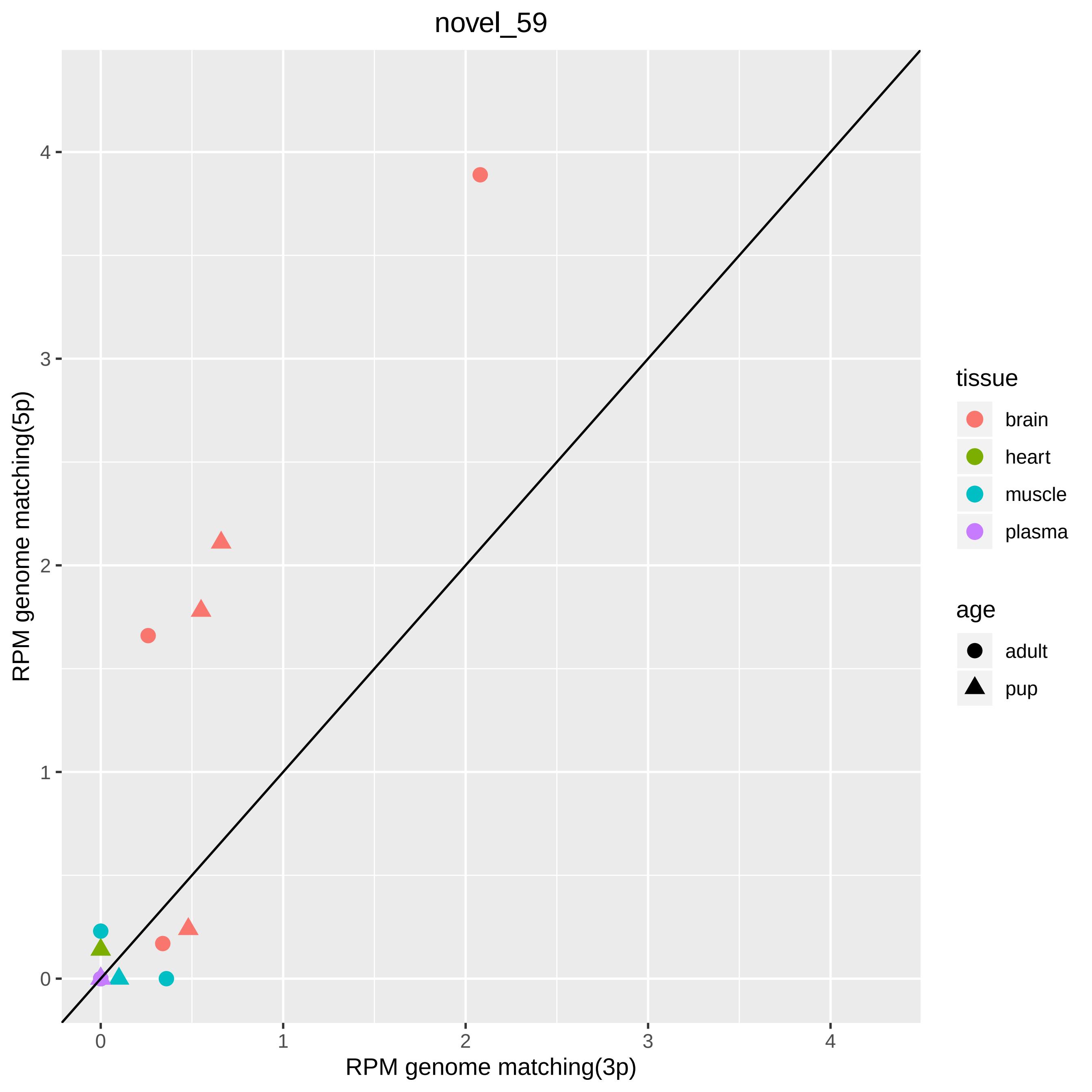
novel_59
RPM genome matching(5p)
RPM genome matching(3p)
0
1
2
3
4
0
1
2
3
4
tissue
brain
heart
muscle
plasma
age
adult
pup

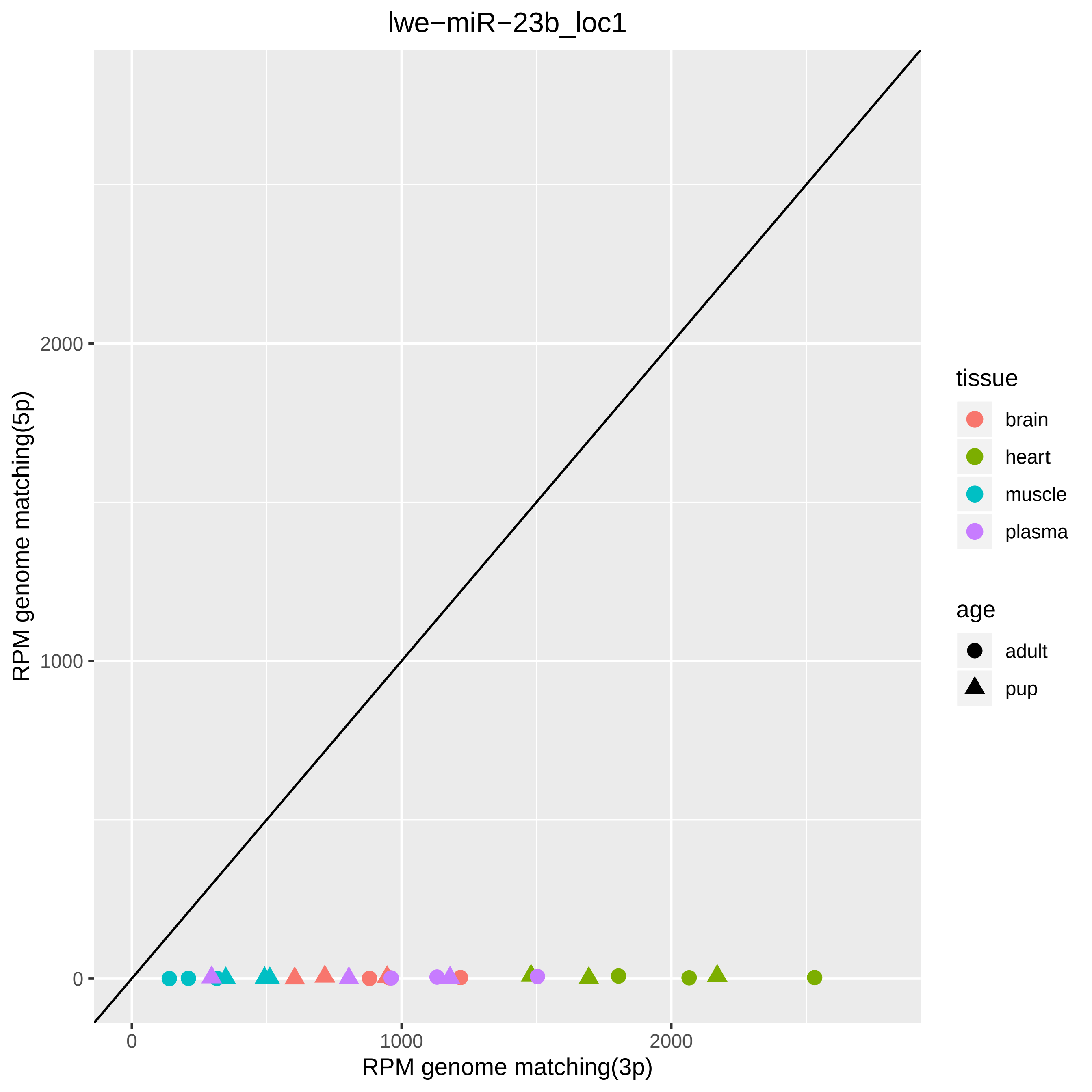
lwe−miR−23b_loc1
RPM genome matching(5p)
RPM genome matching(3p)
0
1000
2000
0
1000
2000
tissue
brain
heart
muscle
plasma
age
adult
pup

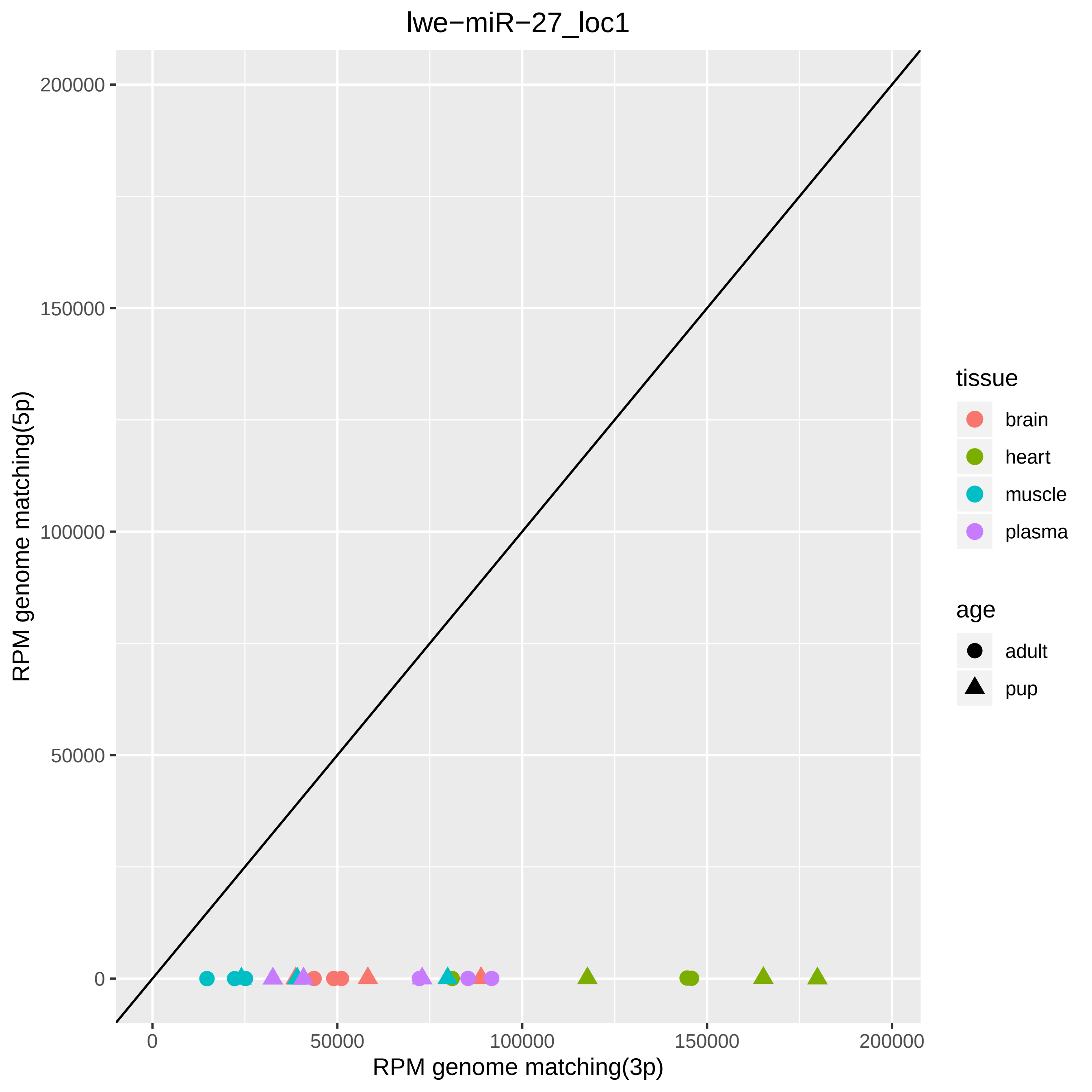

lwe−miR−27_loc1
200000
150000
100000
50000
0
RPM genome matching(5p)
0
50000
100000
150000
200000
RPM genome matching(3p)
tissue
brain
heart
muscle
plasma
age
adult
pup

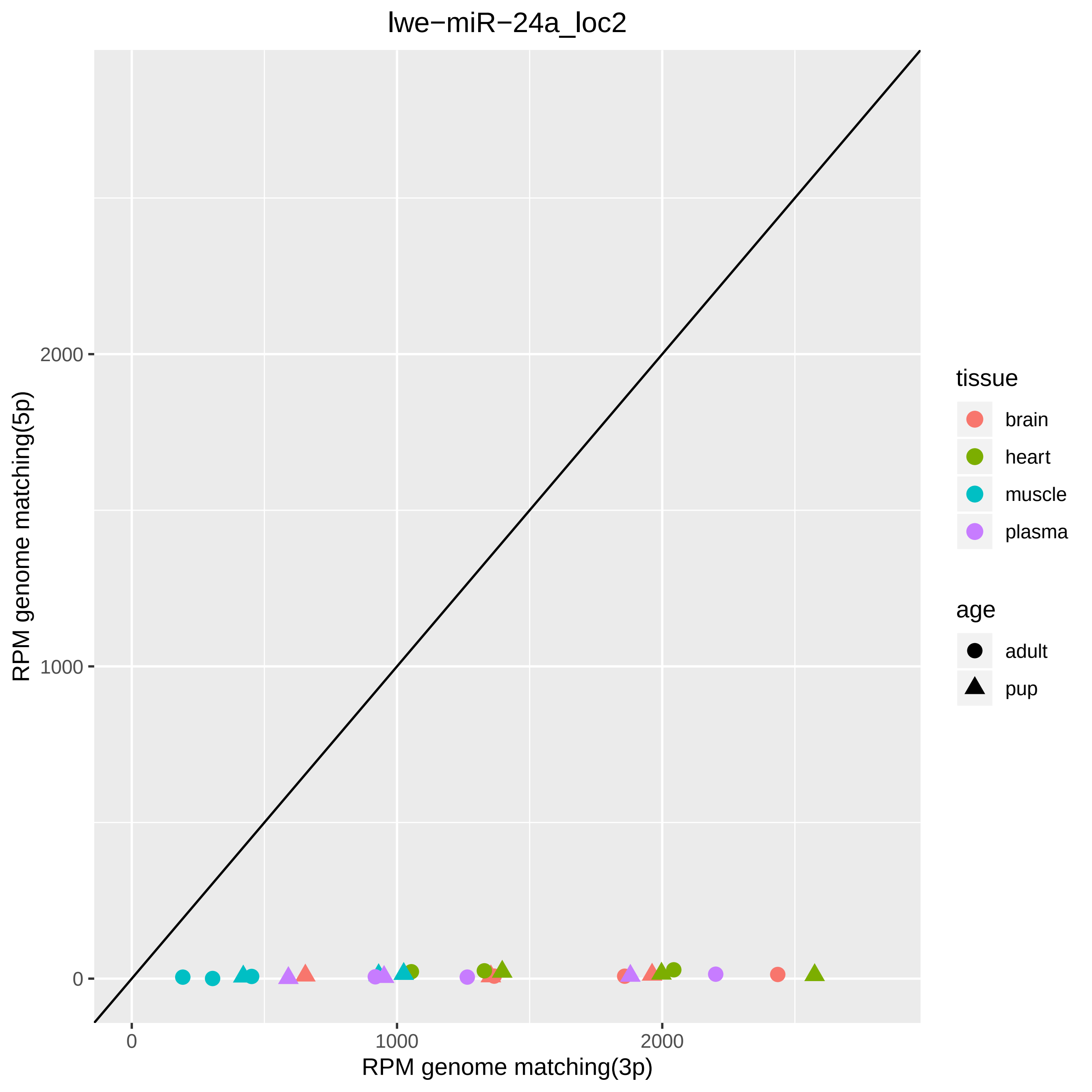
lwe−miR−24a_loc2
RPM genome matching(5p)
RPM genome matching(3p)
0
1000
2000
0
1000
2000
tissue
brain
heart
muscle
plasma
age
adult
pup

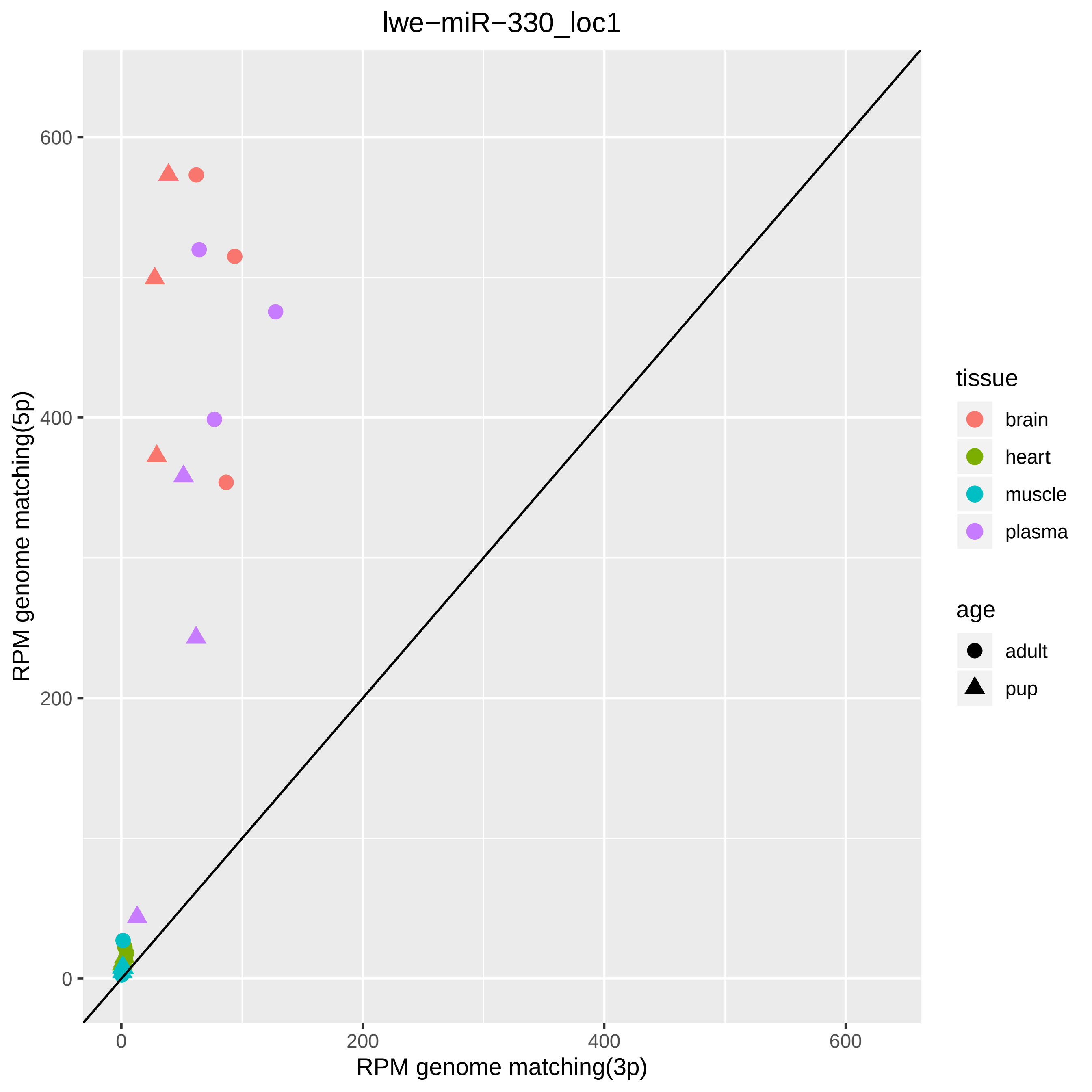
lwe−miR−330_loc1
RPM genome matching(5p)
RPM genome matching(3p)
0
200
400
600
0
200
400
600
tissue
brain
heart
muscle
plasma
age
adult
pup

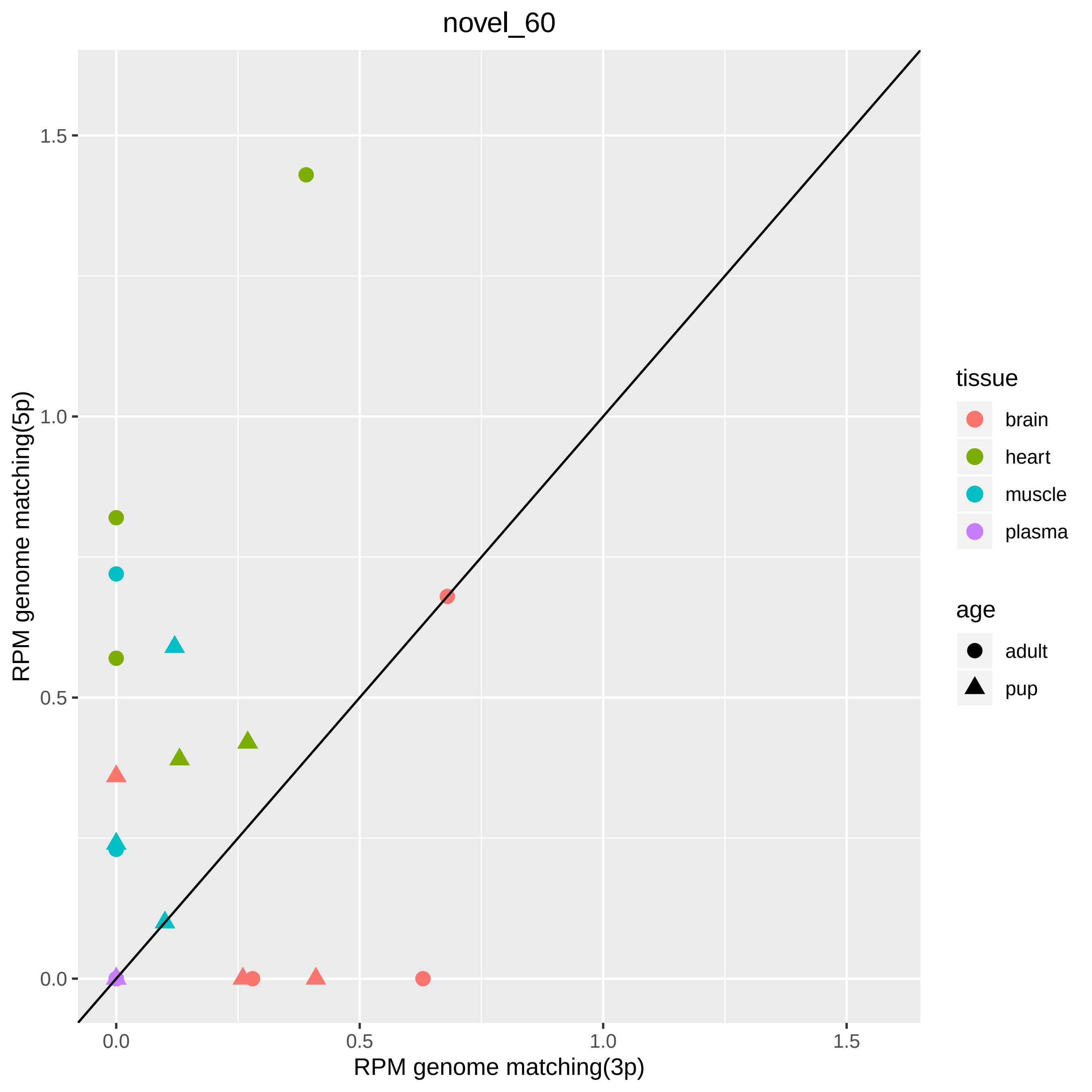
novel_60
RPM genome matching(5p)
RPM genome matching(3p)
0.0
0.5
1.0
1.5
tissue
brain
heart
muscle
plasma
age
adult
pup

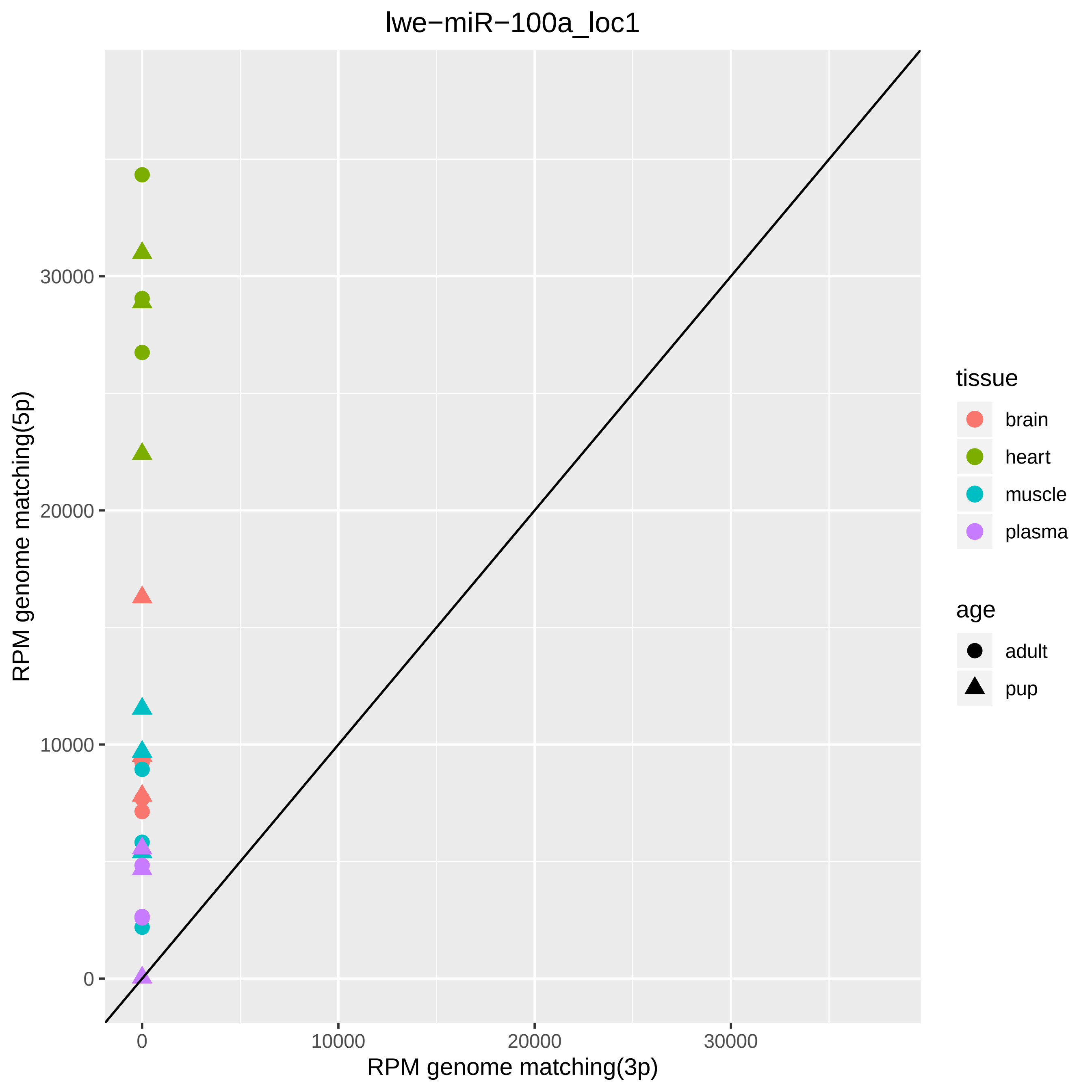
lwe−miR−100a_loc1
RPM genome matching(5p)
RPM genome matching(3p)
0
10000
20000
30000
0
10000
20000
30000
tissue
brain
heart
muscle
plasma
age
adult
pup

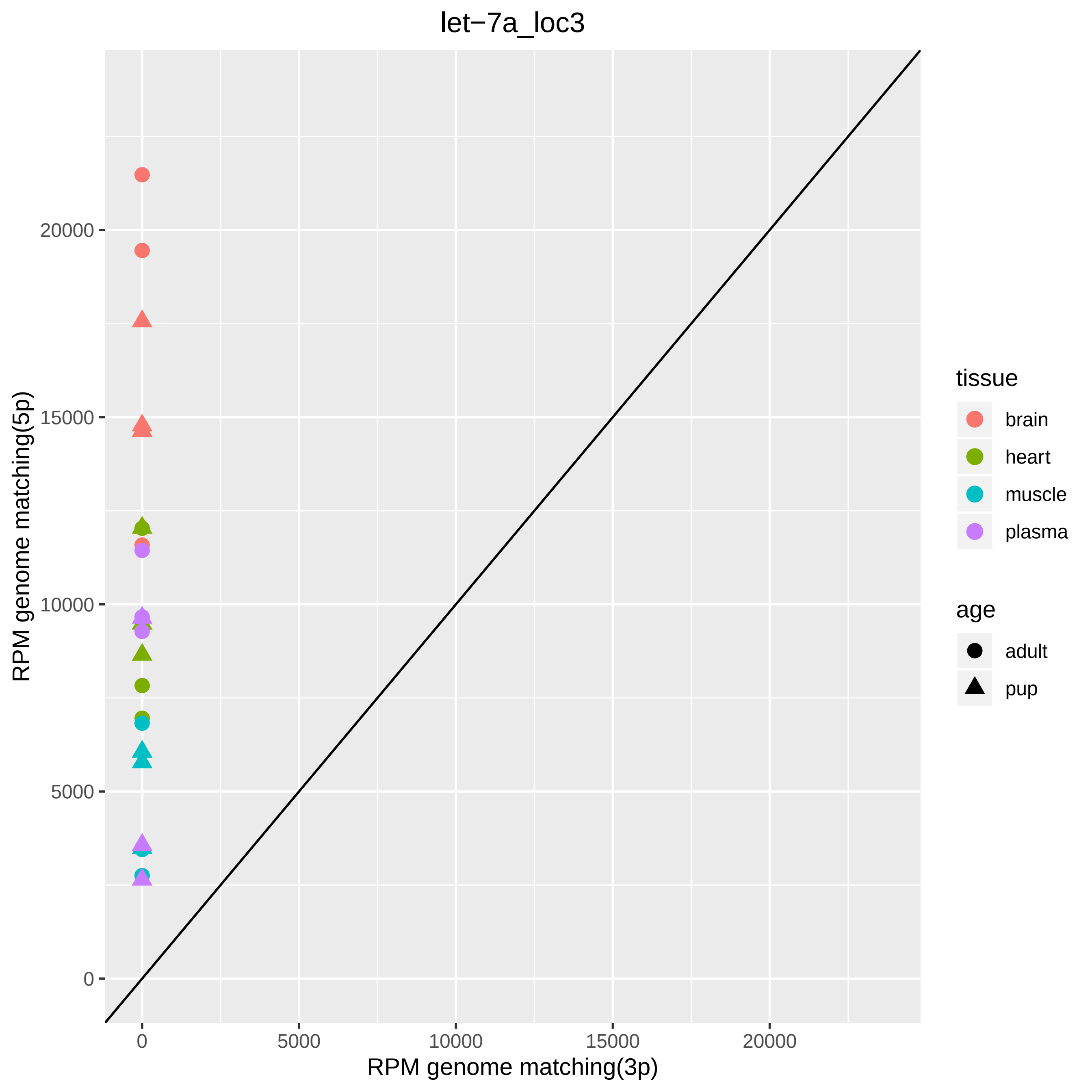

let−7a_loc3
RPM genome matching(5p)
RPM genome matching(3p)
0
5000
10000
15000
20000
tissue
brain
heart
muscle
plasma
age
adult
pup

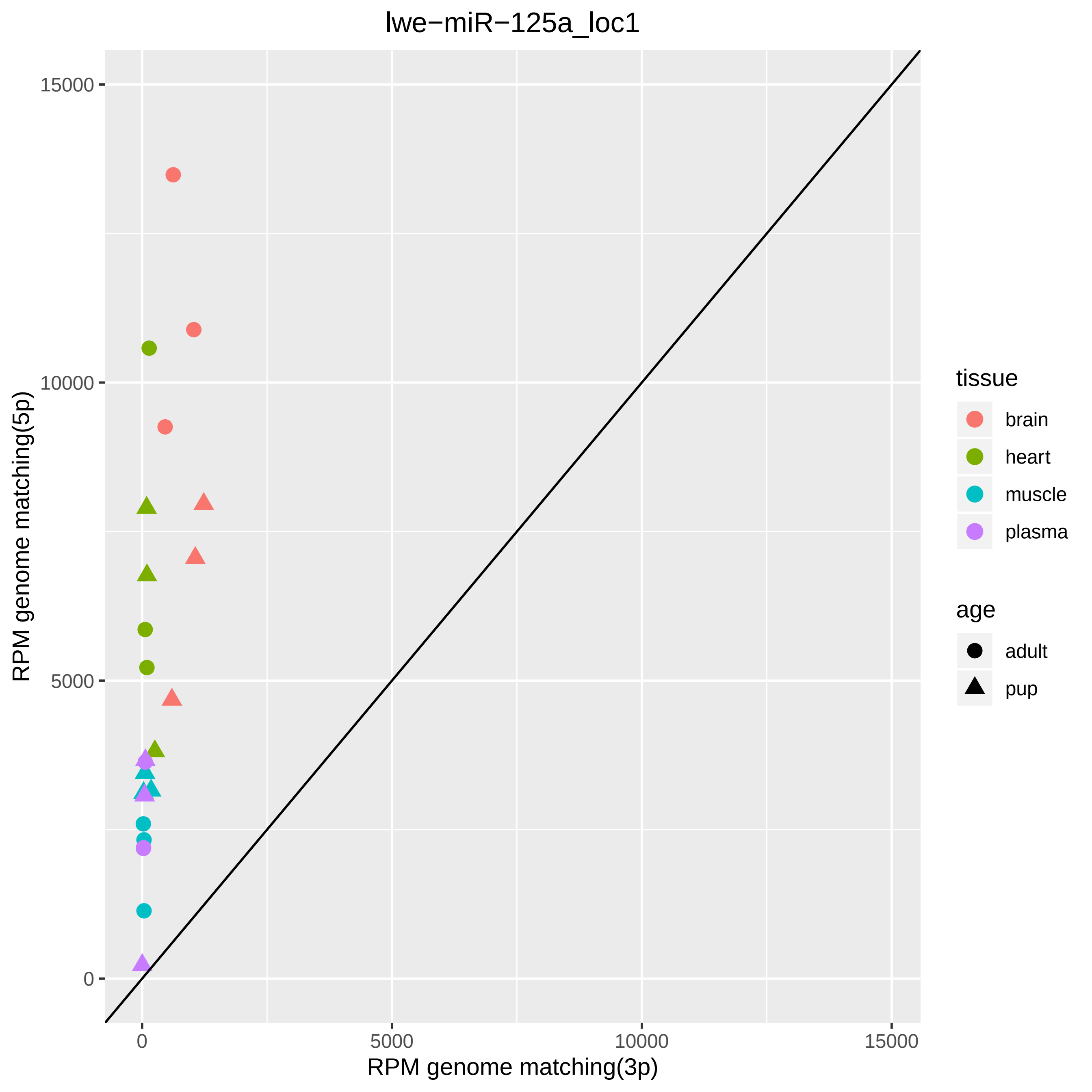
lwe−miR−125a_loc1
15000
10000
5000
0
0
5000
10000
15000
RPM genome matching(5p)
RPM genome matching(3p)
tissue
brain
heart
muscle
plasma
age
adult
pup

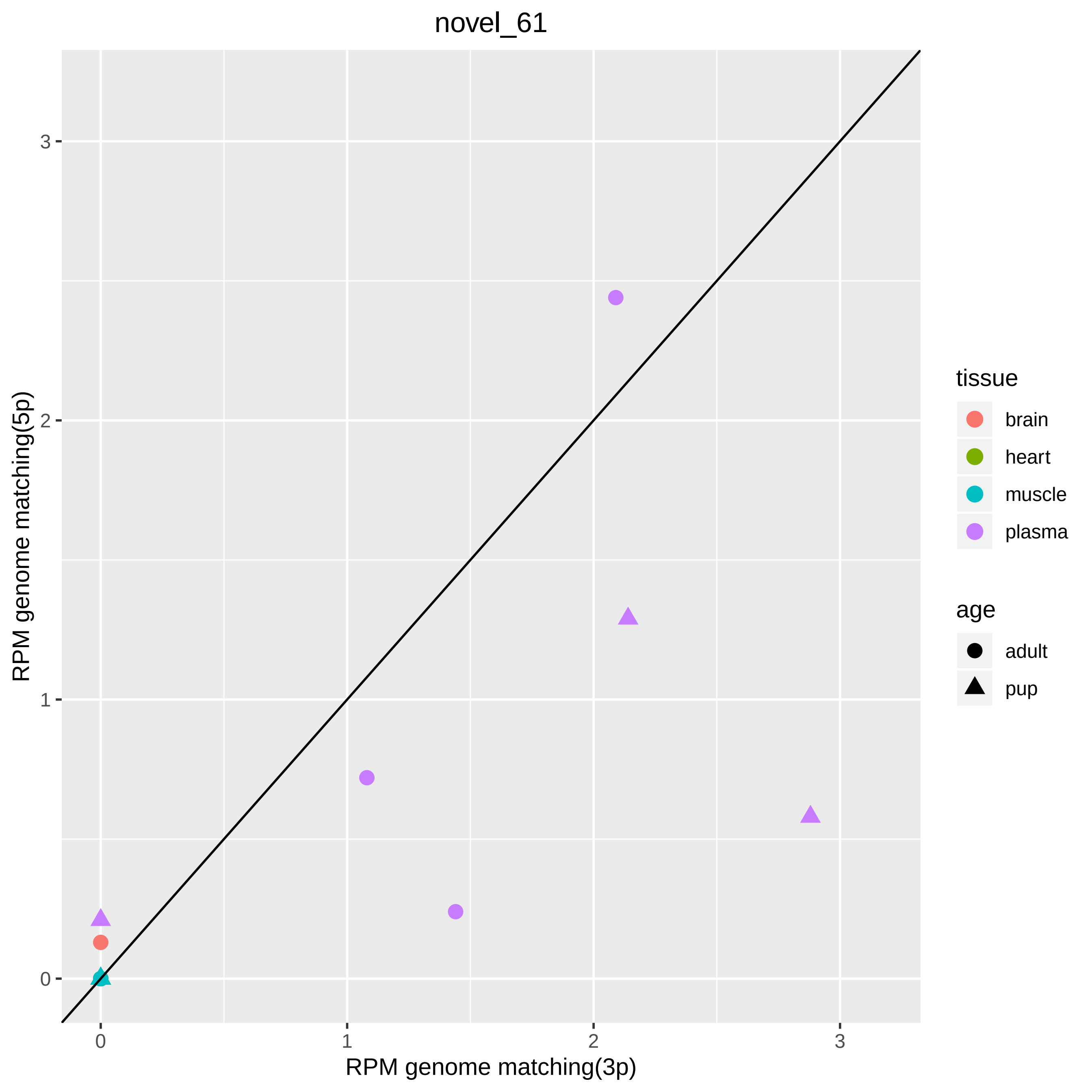
novel_61
RPM genome matching(5p)
RPM genome matching(3p)
0
1
2
3
0
1
2
3
tissue
brain
heart
muscle
plasma
age
adult
pup

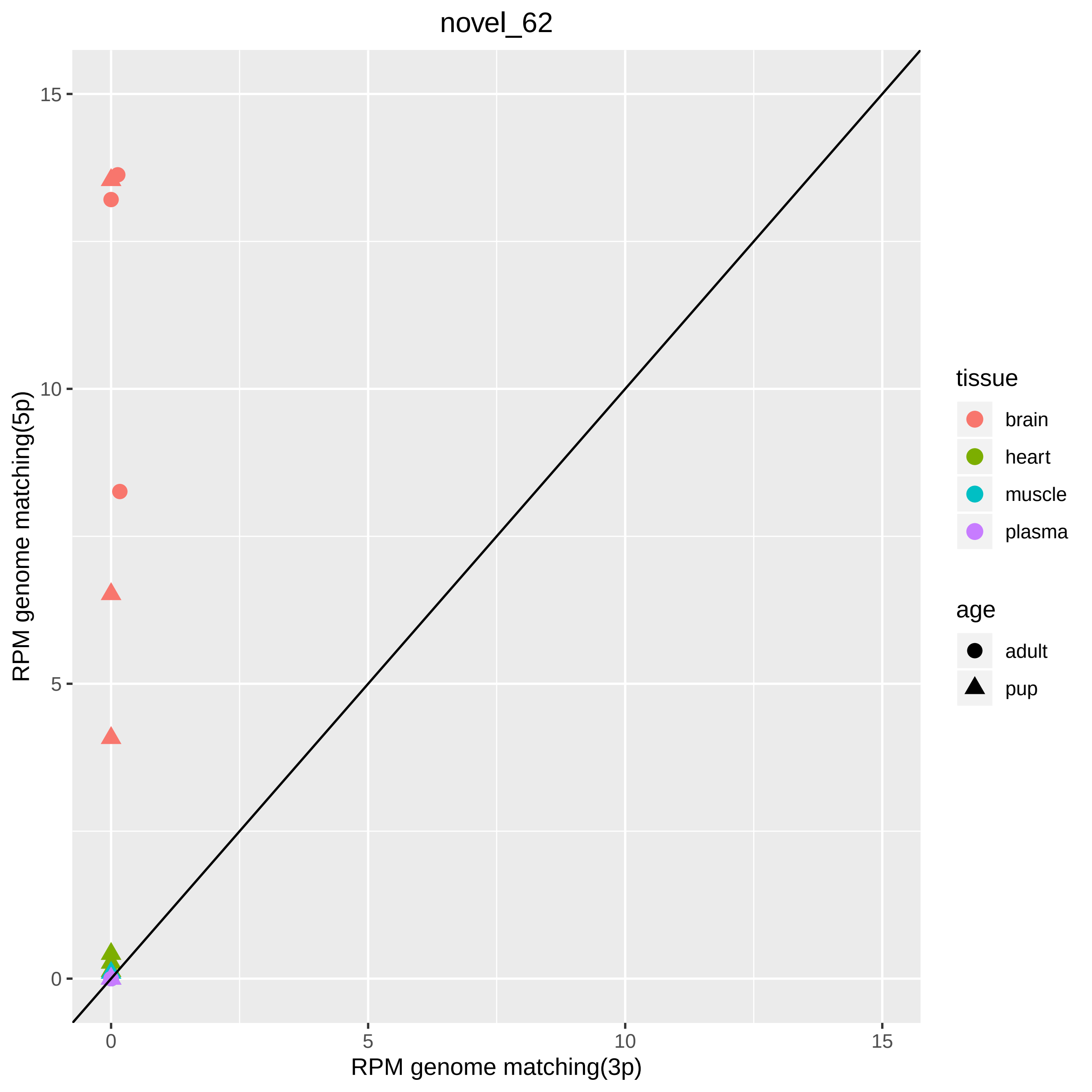

novel_62
RPM genome matching(5p)
RPM genome matching(3p)
0
5
10
15
tissue
brain
heart
muscle
plasma
age
adult
pup

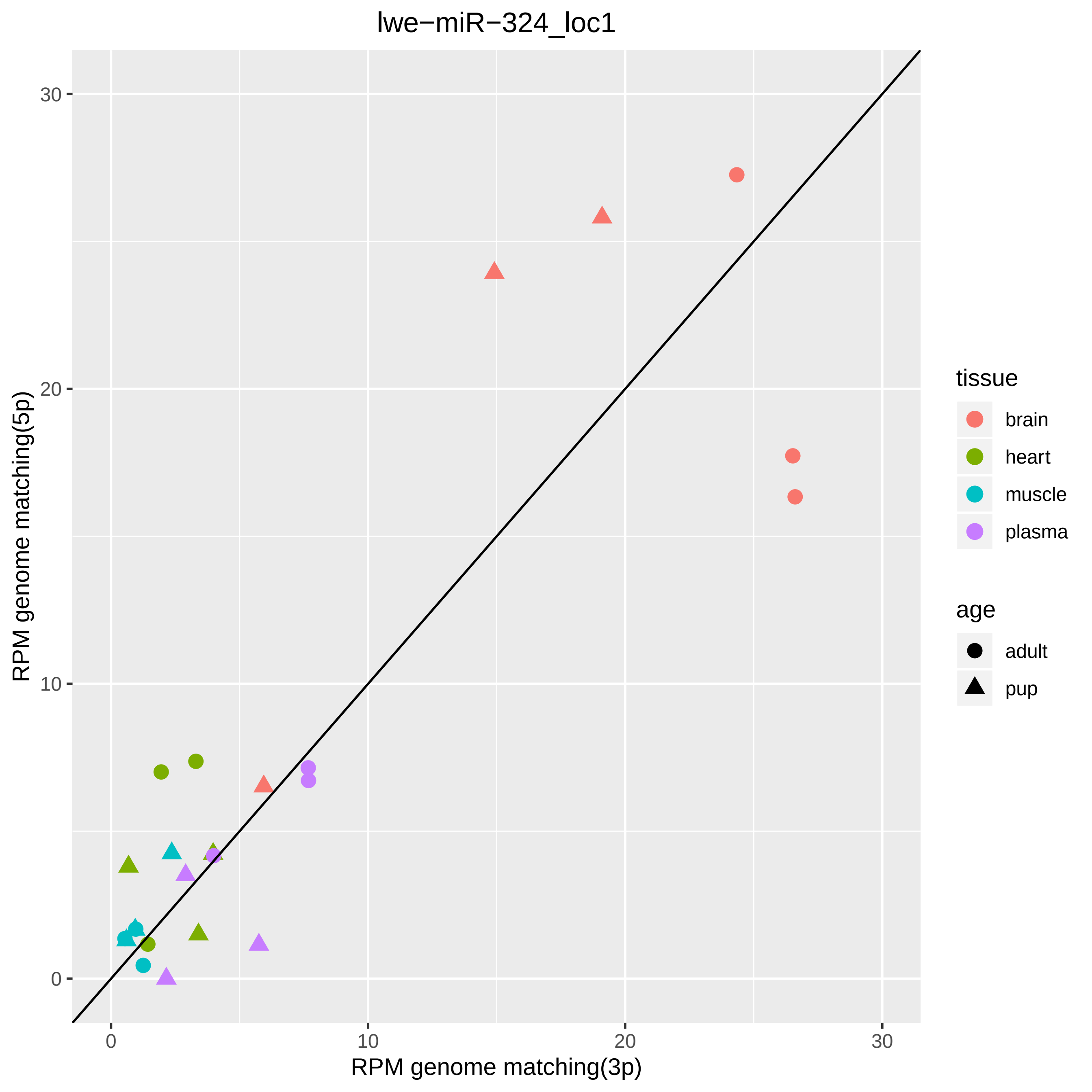
lwe−miR−324_loc1
RPM genome matching(5p)
RPM genome matching(3p)
0
10
20
30
0
10
20
30
tissue
brain
heart
muscle
plasma
age
adult
pup

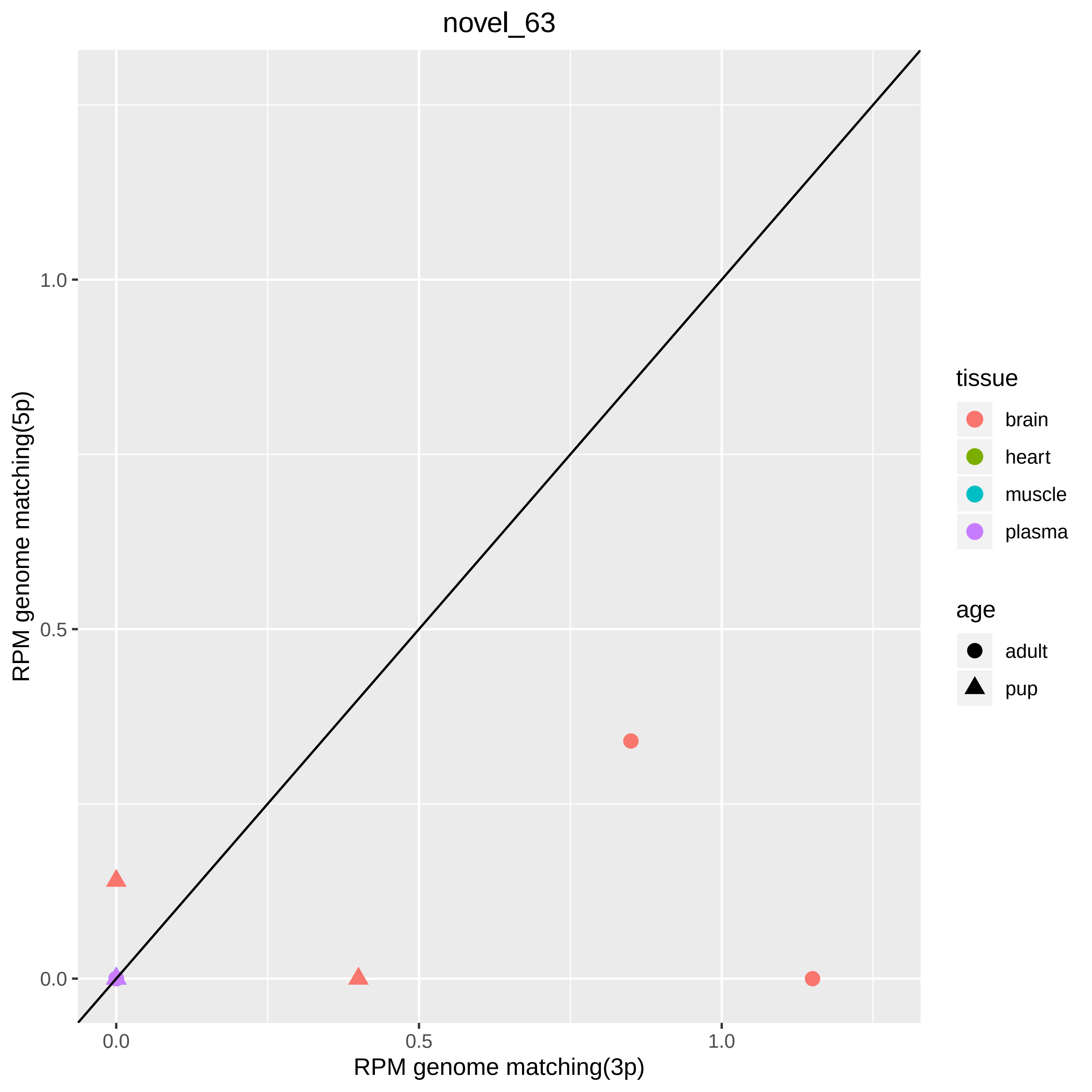
novel_63
RPM genome matching(5p)
RPM genome matching(3p)
0.0
0.5
1.0
0.0
0.5
1.0
tissue
brain
heart
muscle
plasma
age
adult
pup

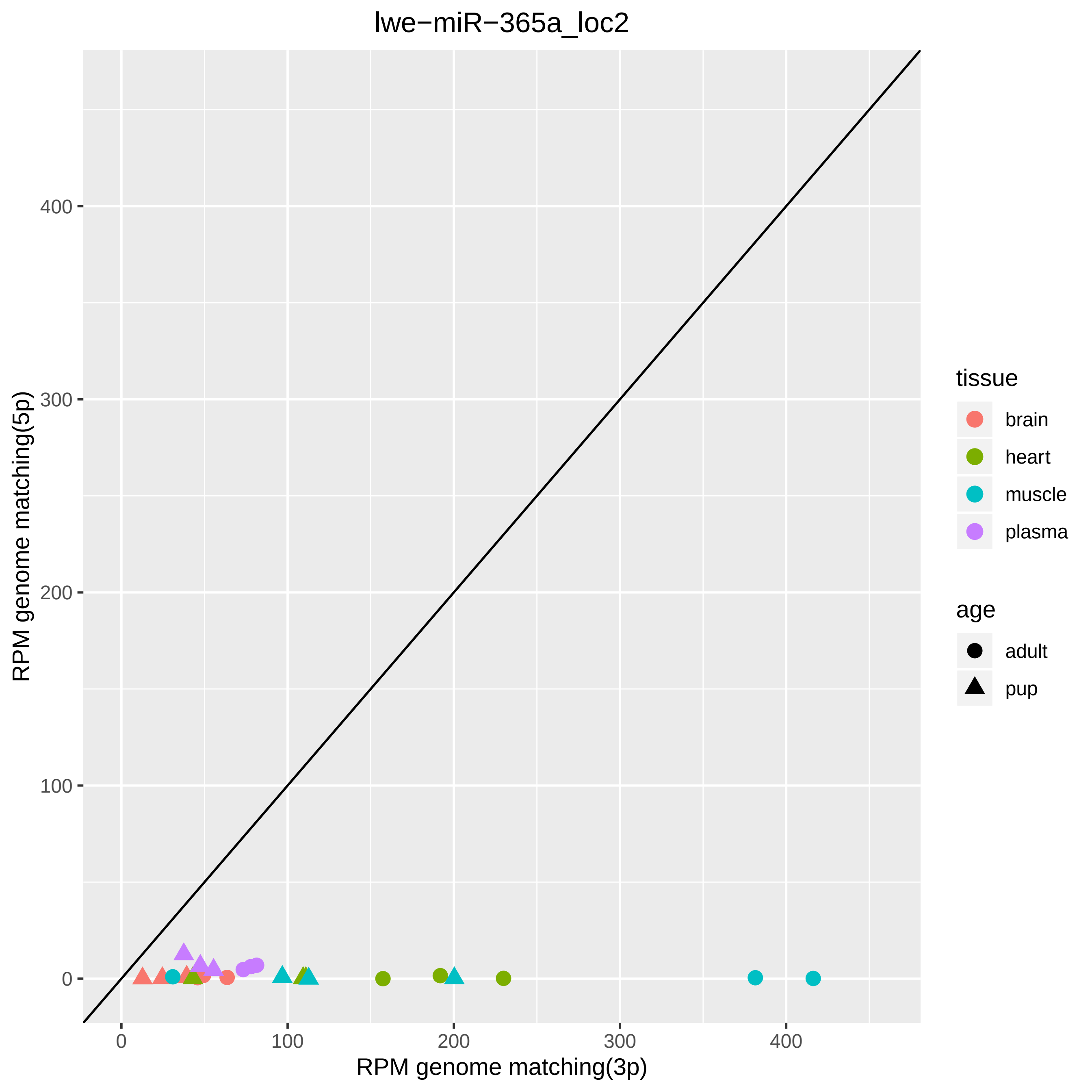

lwe−miR−365a_loc2
RPM genome matching(5p)
RPM genome matching(3p)
0
100
200
300
400
0
100
200
300
400
tissue
brain
heart
muscle
plasma
age
adult
pup

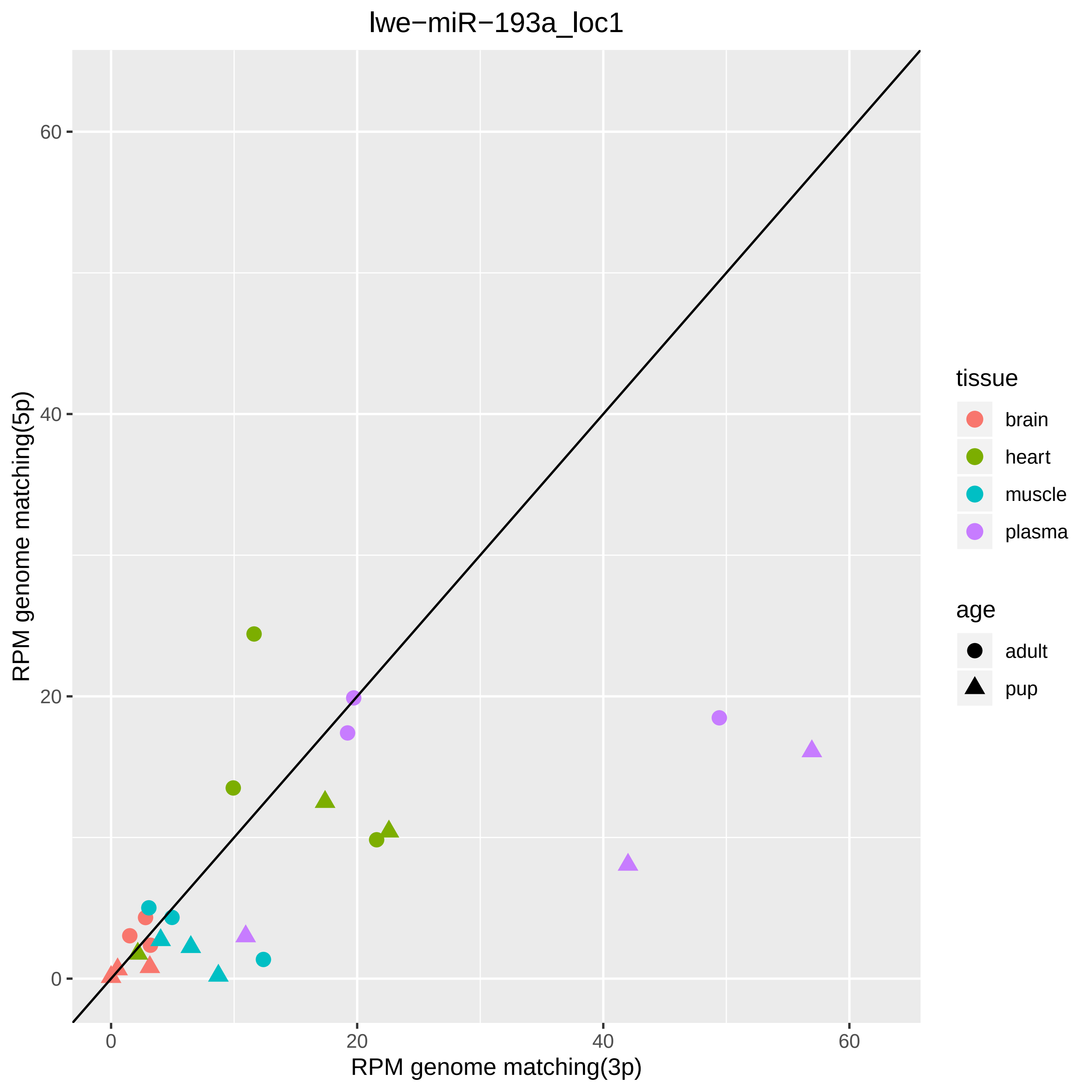
lwe−miR−193a_loc1
RPM genome matching(5p)
RPM genome matching(3p)
0
20
40
60
0
20
40
60
tissue
brain
heart
muscle
plasma
age
adult
pup

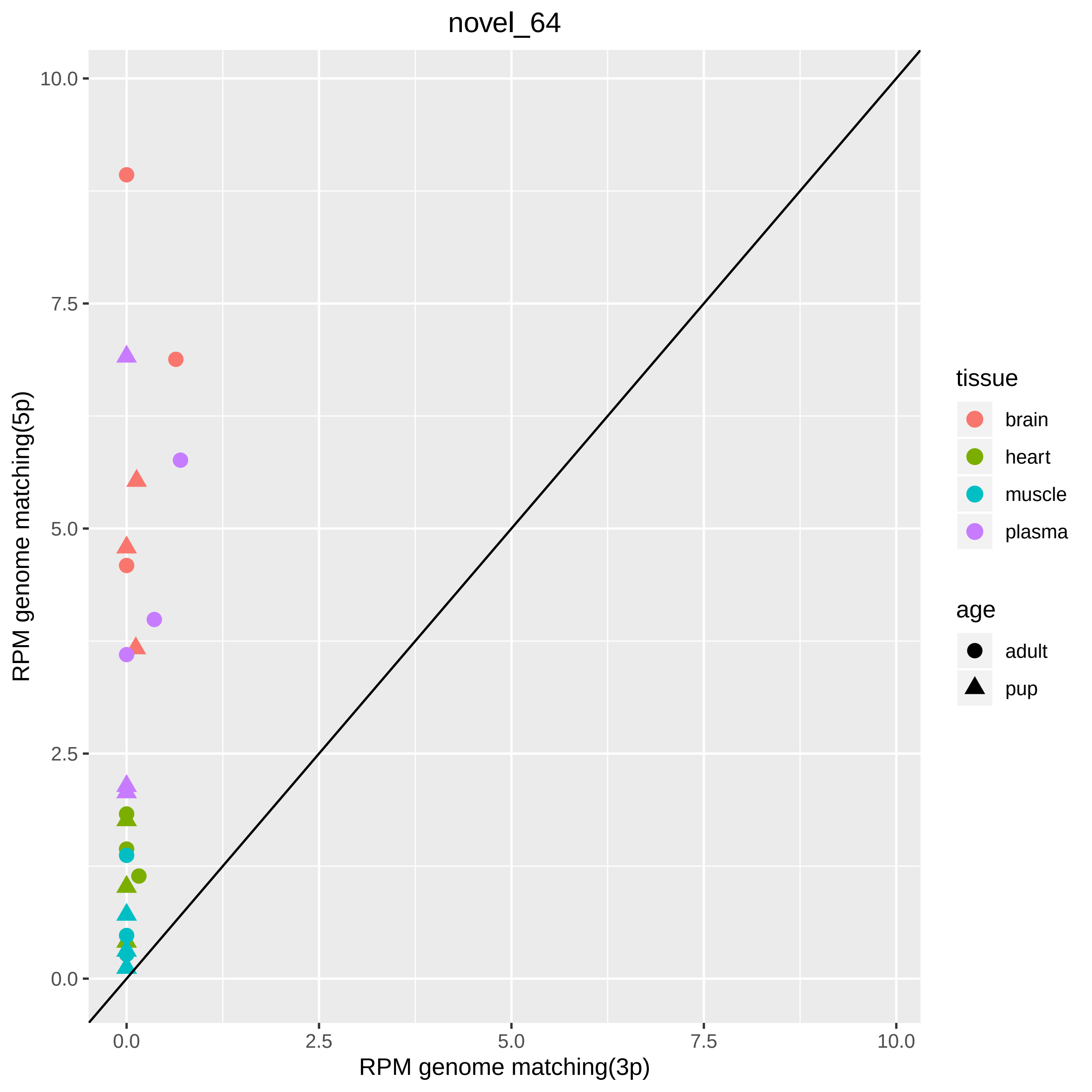
novel_64
RPM genome matching(5p)
RPM genome matching(3p)
10.0
7.5
5.0
2.5
0.0
0.0
2.5
5.0
7.5
10.0
tissue
brain
heart
muscle
plasma
age
adult
pup

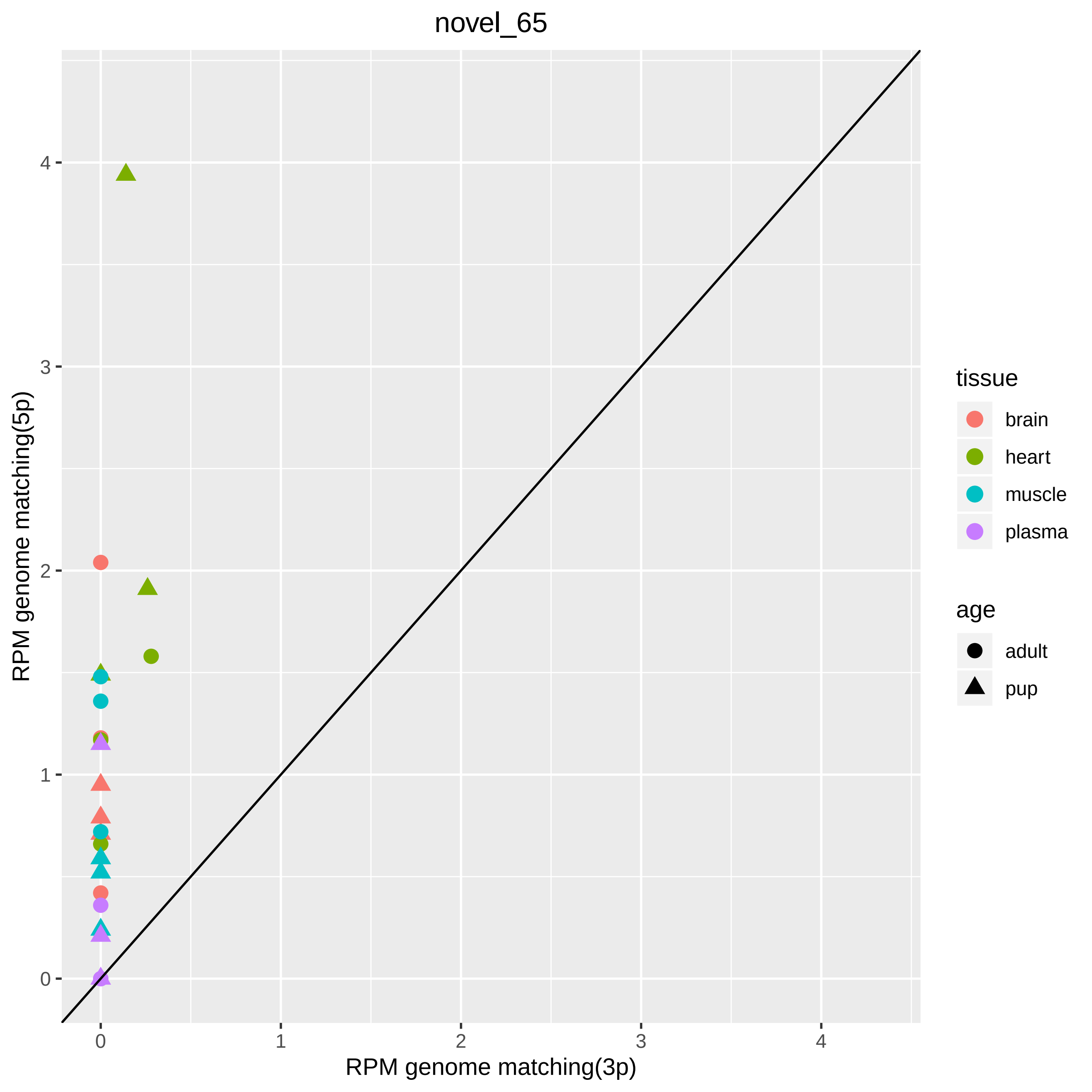

novel_65
RPM genome matching(5p)
RPM genome matching(3p)
0
1
2
3
4
0
1
2
3
4
tissue
brain
heart
muscle
plasma
age
adult
pup

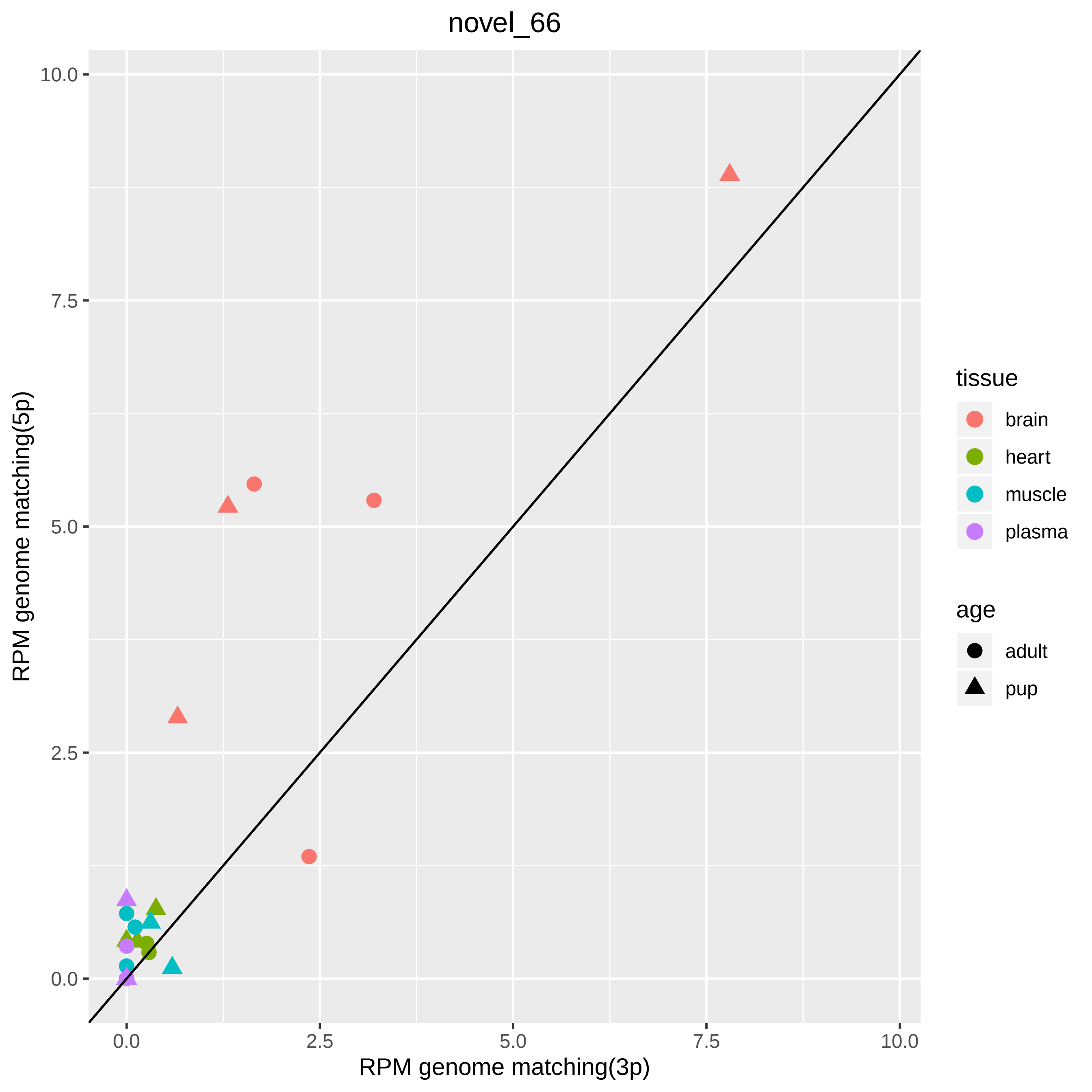

novel_66
RPM genome matching(5p)
RPM genome matching(3p)
10.0
7.5
5.0
2.5
0.0
0.0
2.5
5.0
7.5
10.0
tissue
brain
heart
muscle
plasma
age
adult
pup

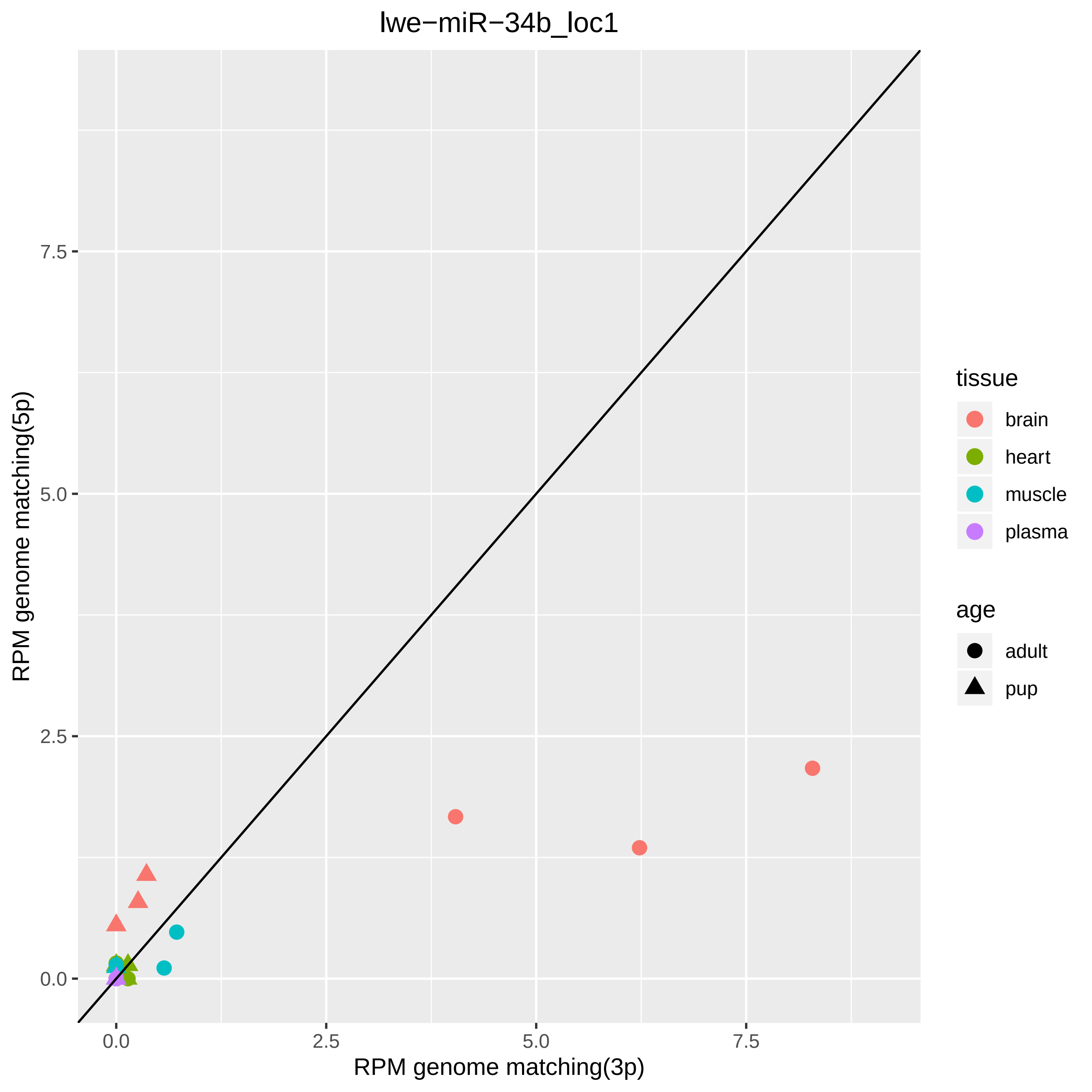

lwe−miR−34b_loc1
RPM genome matching(5p)
RPM genome matching(3p)
0.0
2.5
5.0
7.5
tissue
brain
heart
muscle
plasma
age
adult
pup

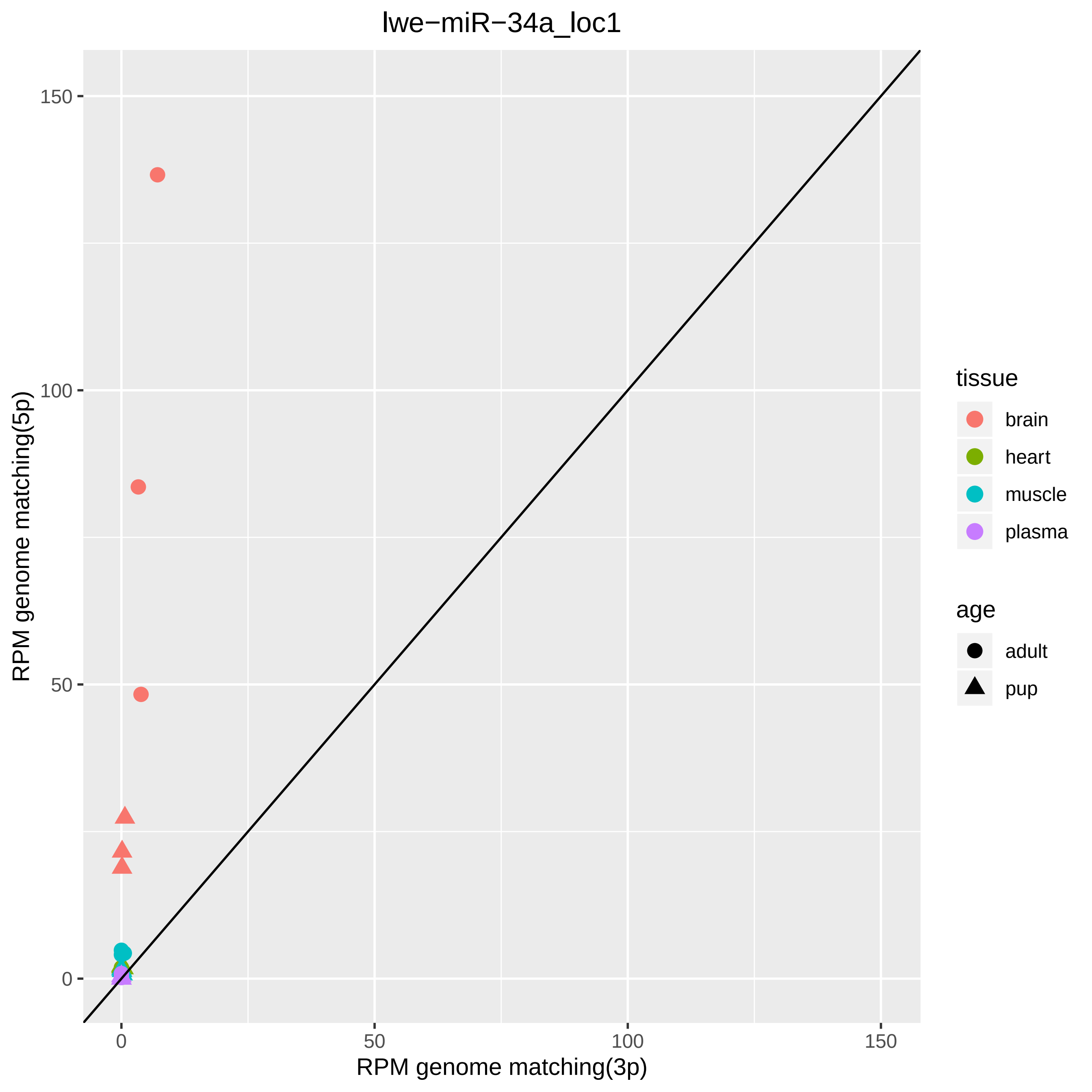

lwe−miR−34a_loc1
RPM genome matching(5p)
RPM genome matching(3p)
150
100
50
0
0
50
100
150
tissue
brain
heart
muscle
plasma
age
adult
pup

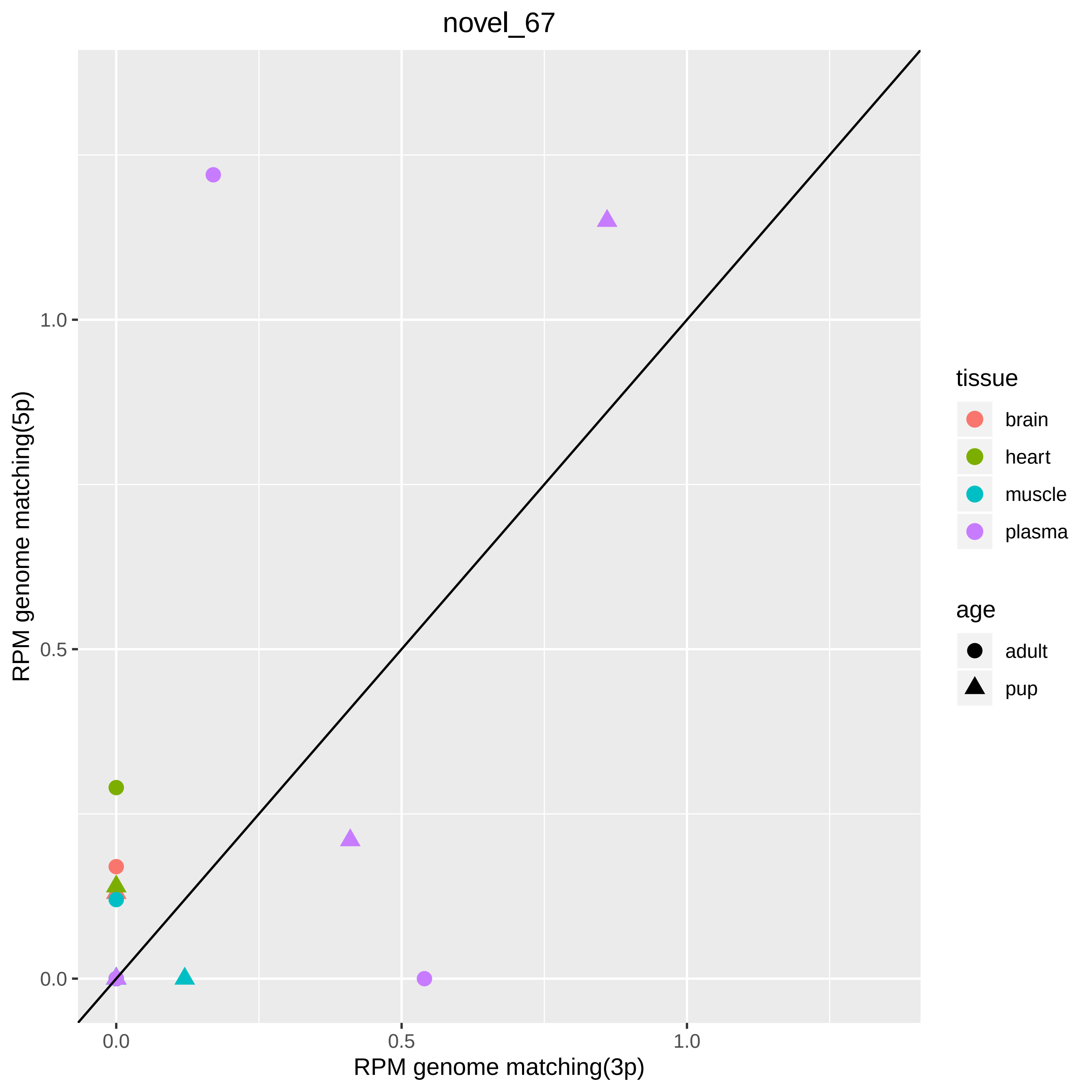

novel_67
RPM genome matching(5p)
RPM genome matching(3p)
0.0
0.5
1.0
tissue
brain
heart
muscle
plasma
age
adult
pup

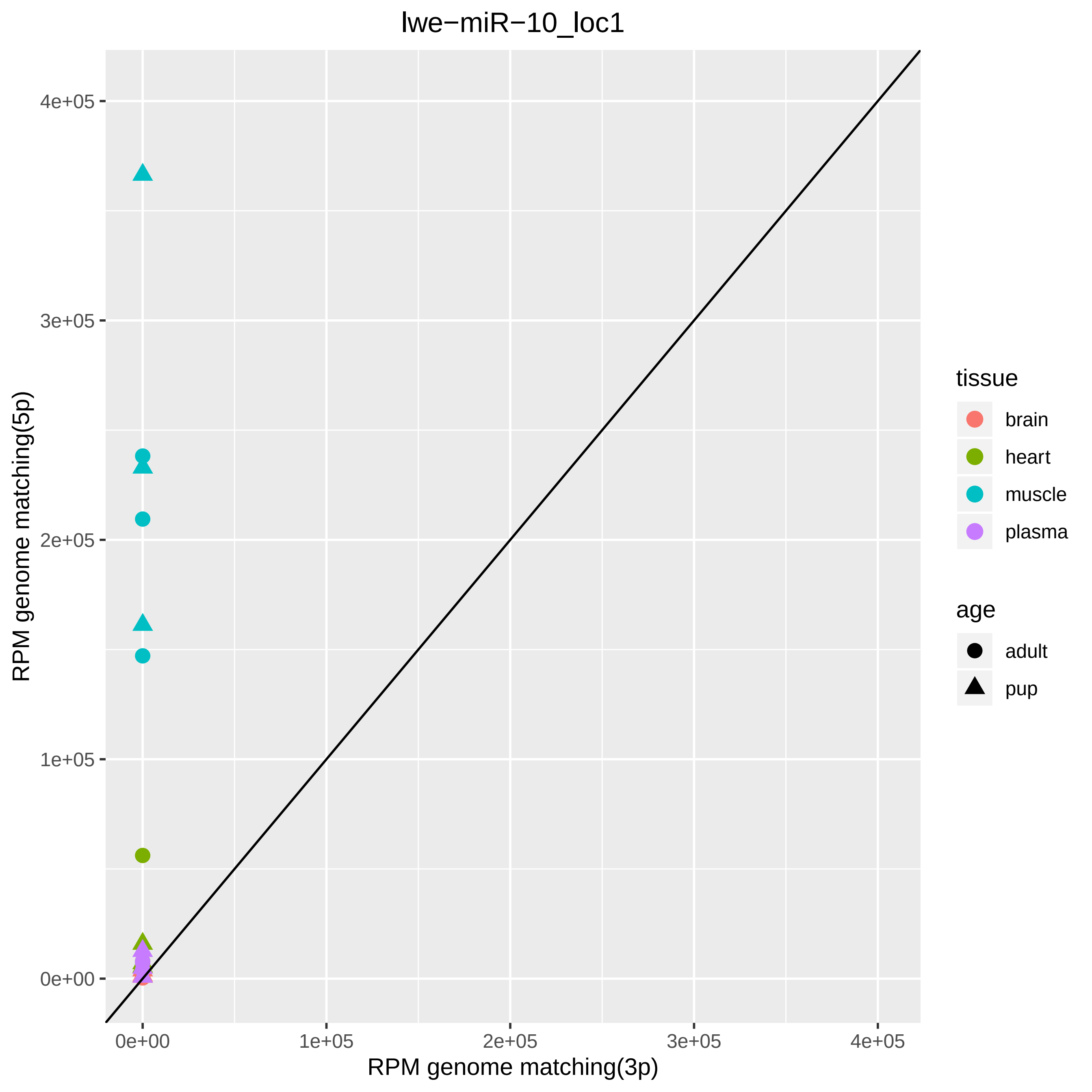

lwe−miR−10_loc1
4e+05
3e+05
2e+05
1e+05
0e+00
RPM genome matching(5p)
0e+00
1e+05
2e+05
3e+05
4e+05
RPM genome matching(3p)
tissue
brain
heart
muscle
plasma
age
adult
pup

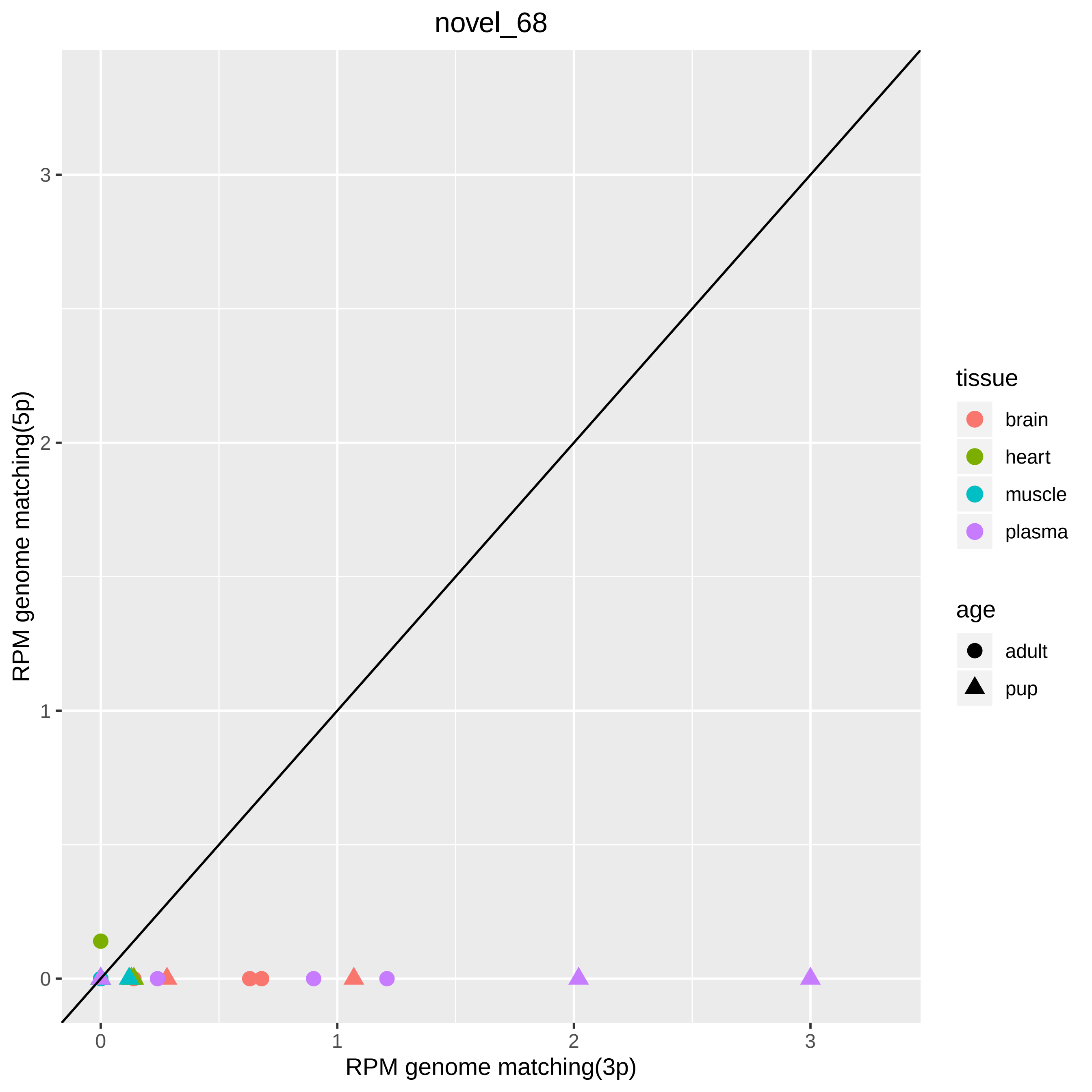

novel_68
RPM genome matching(5p)
RPM genome matching(3p)
0
1
2
3
0
1
2
3
tissue
brain
heart
muscle
plasma
age
adult
pup

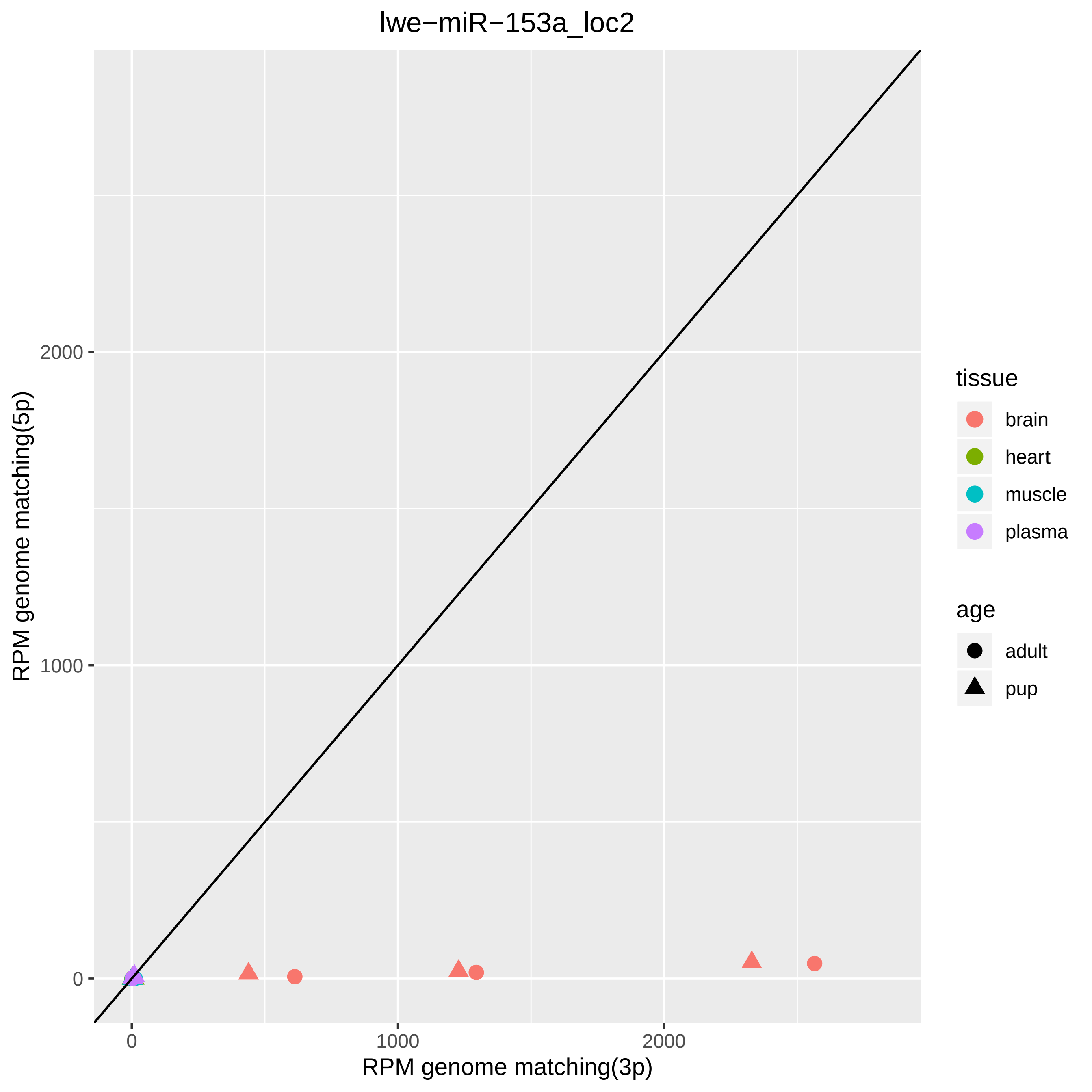

lwe−miR−153a_loc2
RPM genome matching(5p)
RPM genome matching(3p)
0
1000
2000
0
1000
2000
tissue
brain
heart
muscle
plasma
age
adult
pup

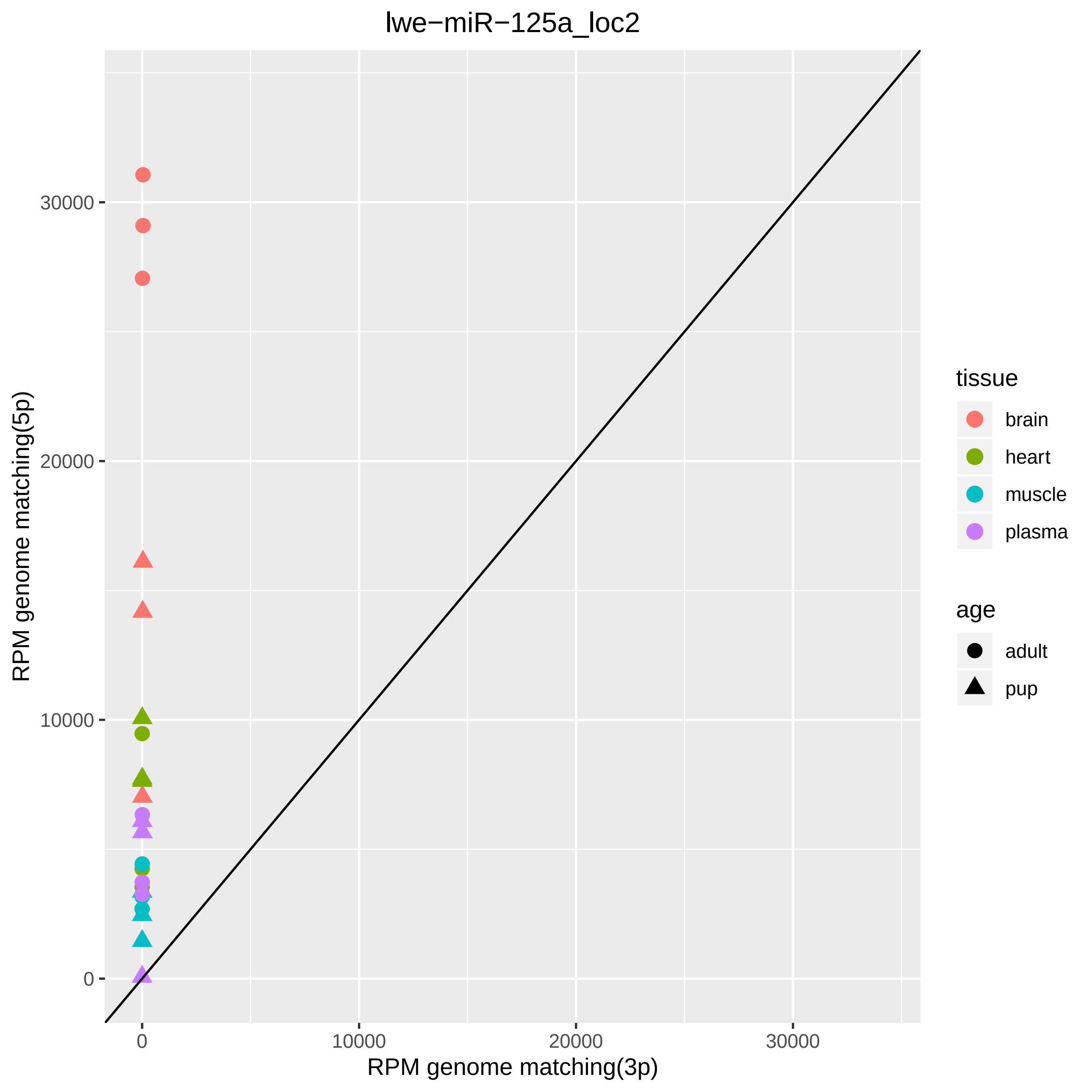

lwe−miR−125a_loc2
RPM genome matching(5p)
RPM genome matching(3p)
0
10000
20000
30000
0
10000
20000
30000
tissue
brain
heart
muscle
plasma
age
adult
pup

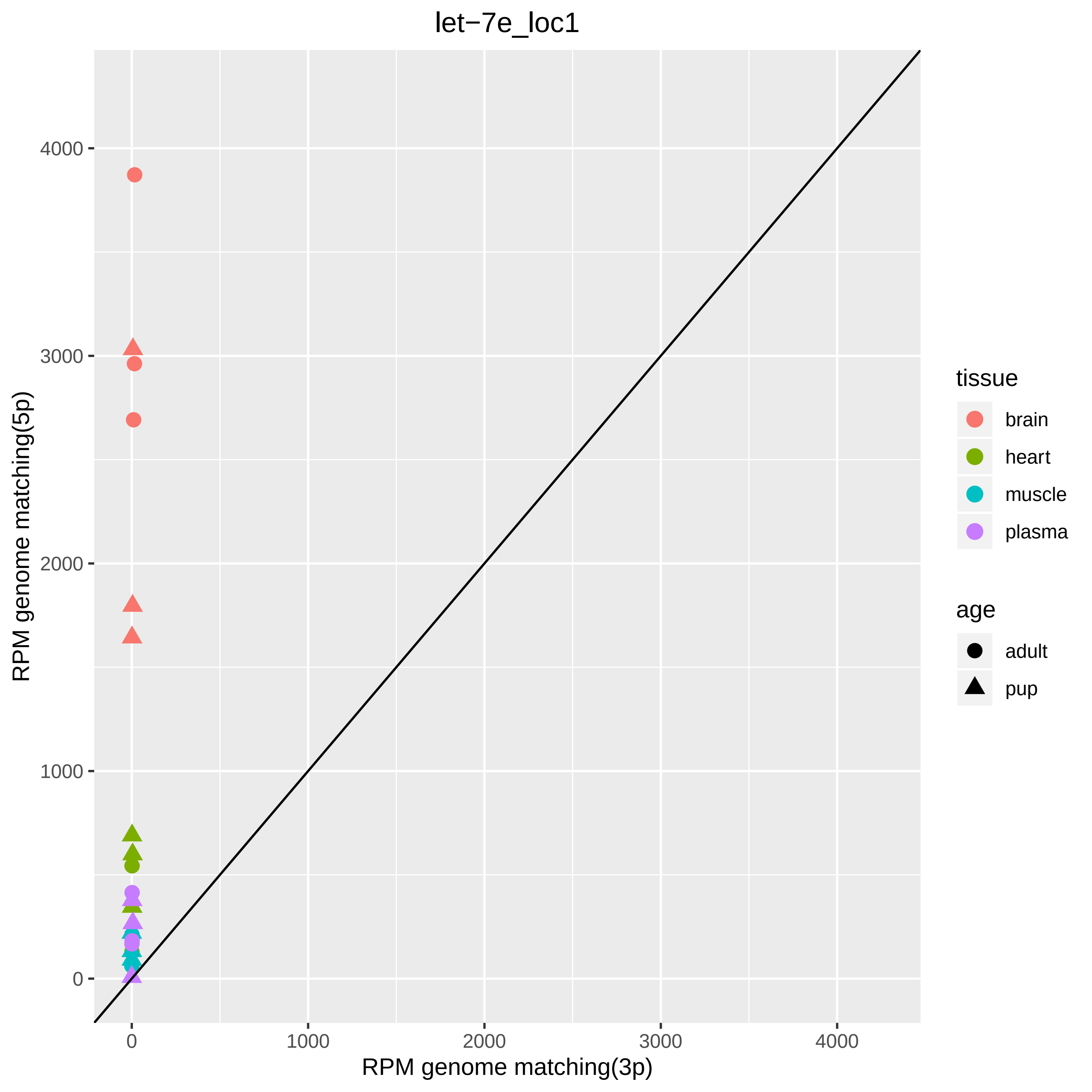
let−7e_loc1
RPM genome matching(5p)
RPM genome matching(3p)
4000
3000
2000
1000
0
0
1000
2000
3000
4000
tissue
brain
heart
muscle
plasma
age
adult
pup

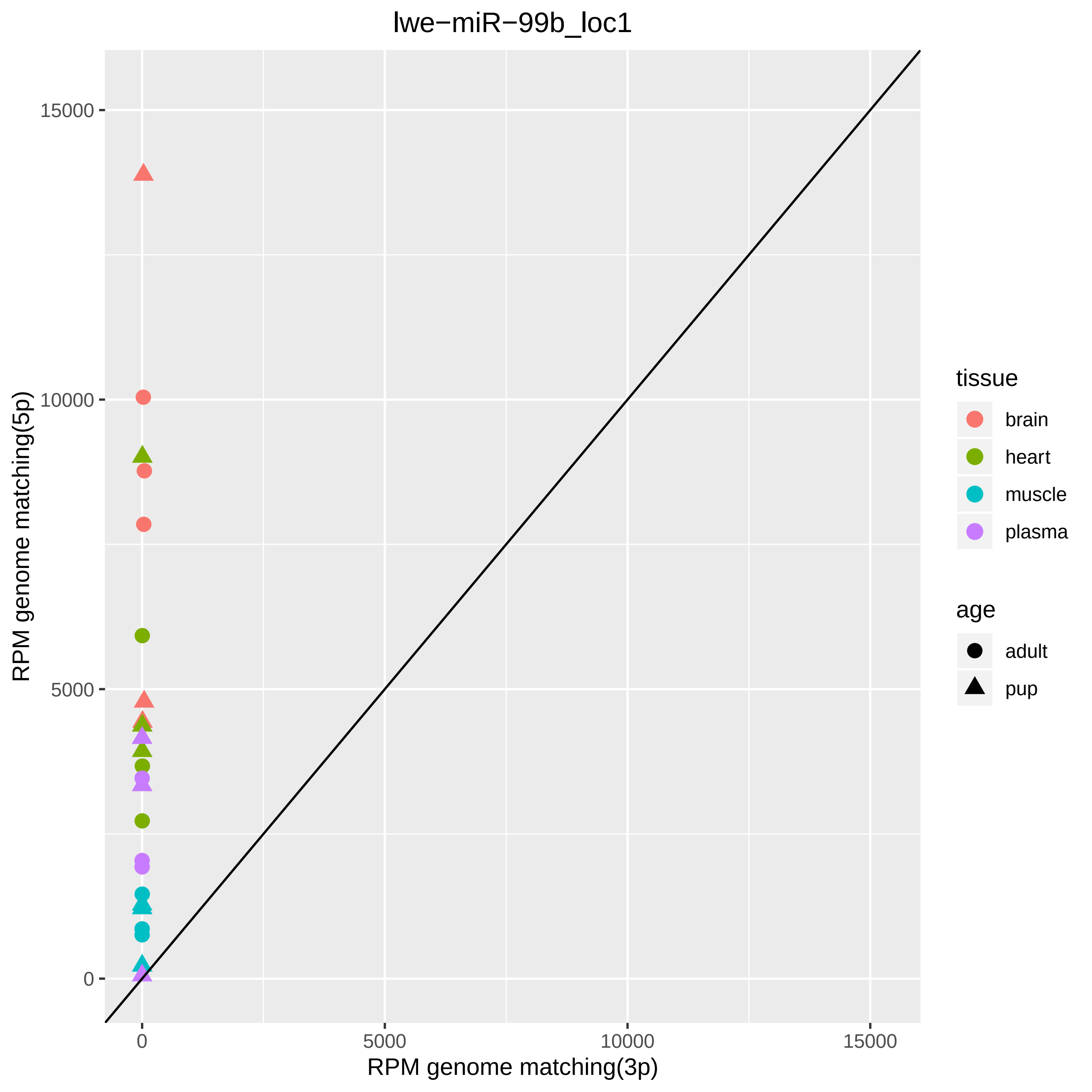
lwe−miR−99b_loc1
RPM genome matching(5p)
RPM genome matching(3p)
0
5000
10000
15000
0
5000
10000
15000
tissue
brain
heart
muscle
plasma
age
adult
pup

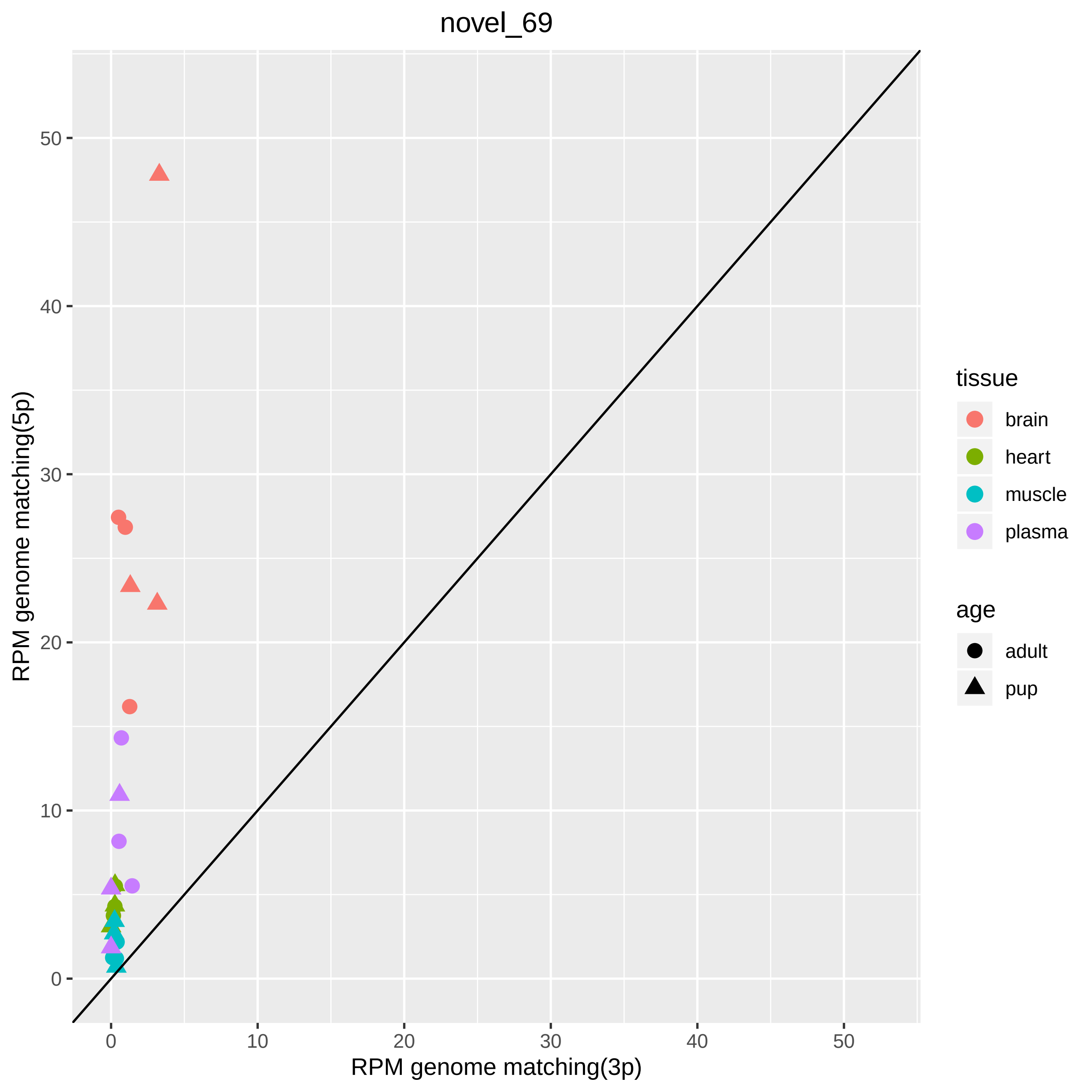

novel_69
RPM genome matching(5p)
RPM genome matching(3p)
0
10
20
30
40
50
0
10
20
30
40
50
tissue
brain
heart
muscle
plasma
age
adult
pup

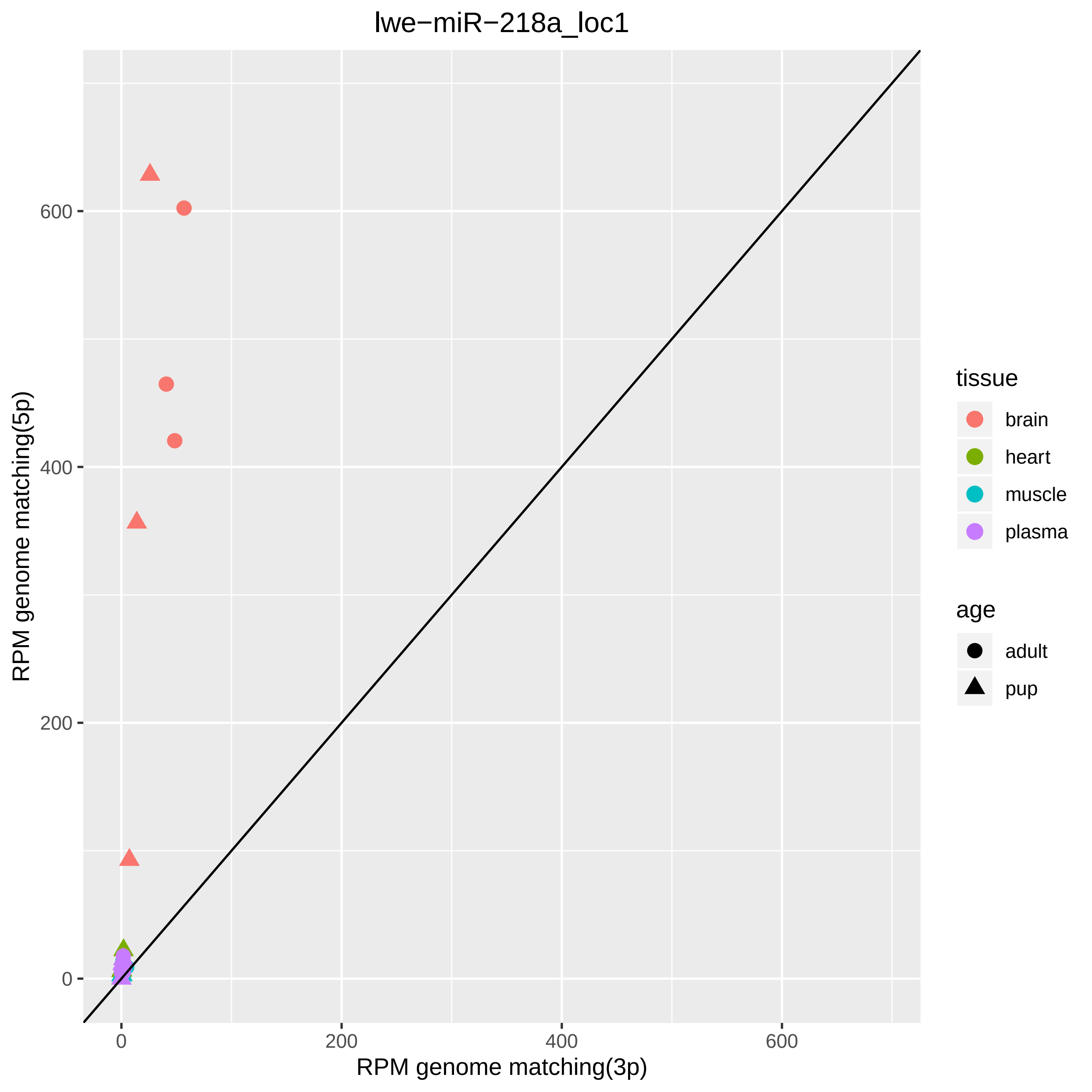
lwe−miR−218a_loc1
RPM genome matching(5p)
RPM genome matching(3p)
0
200
400
600
0
200
400
600
tissue
brain
heart
muscle
plasma
age
adult
pup

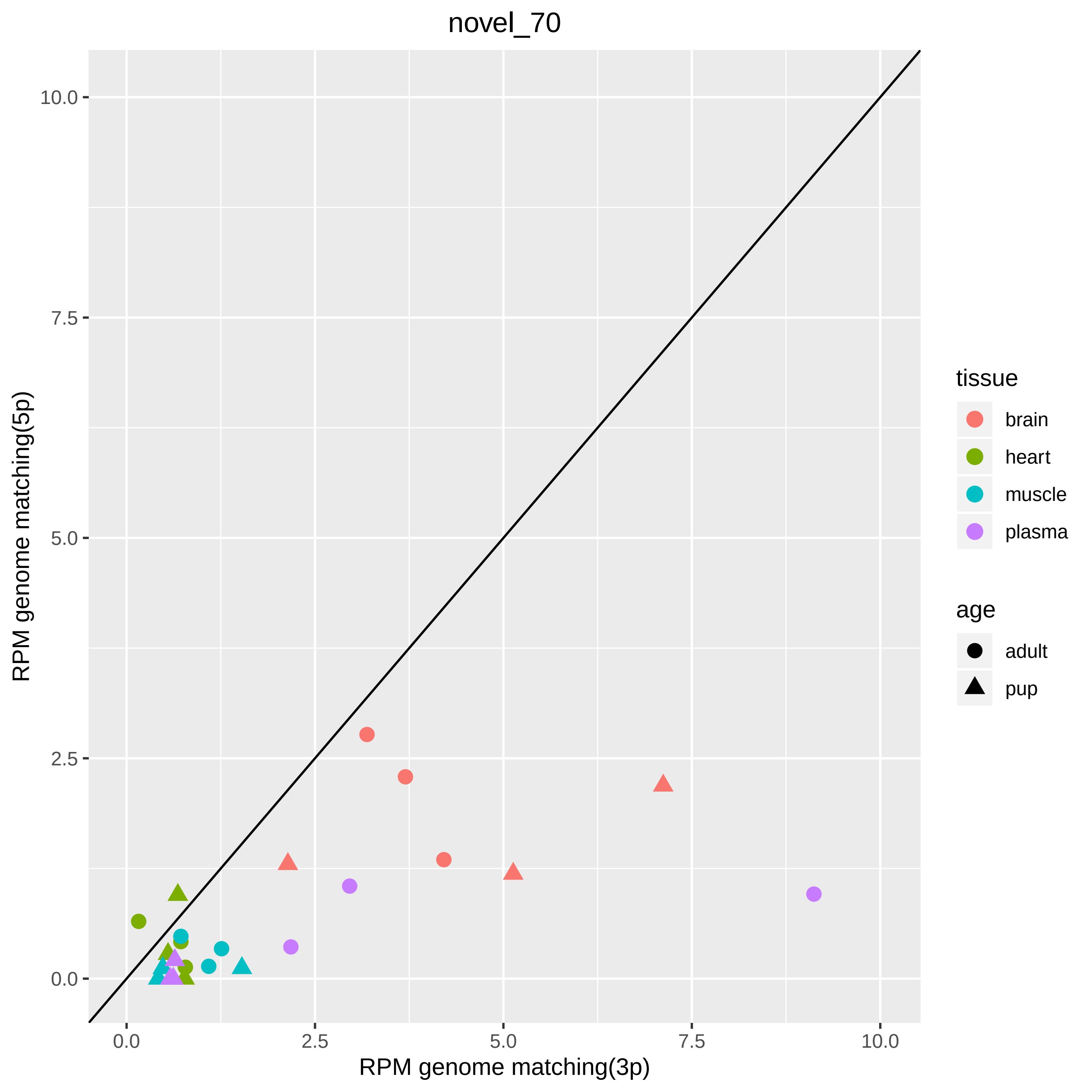
novel_70
RPM genome matching(5p)
RPM genome matching(3p)
0.0
2.5
5.0
7.5
10.0
tissue
brain
heart
muscle
plasma
age
adult
pup

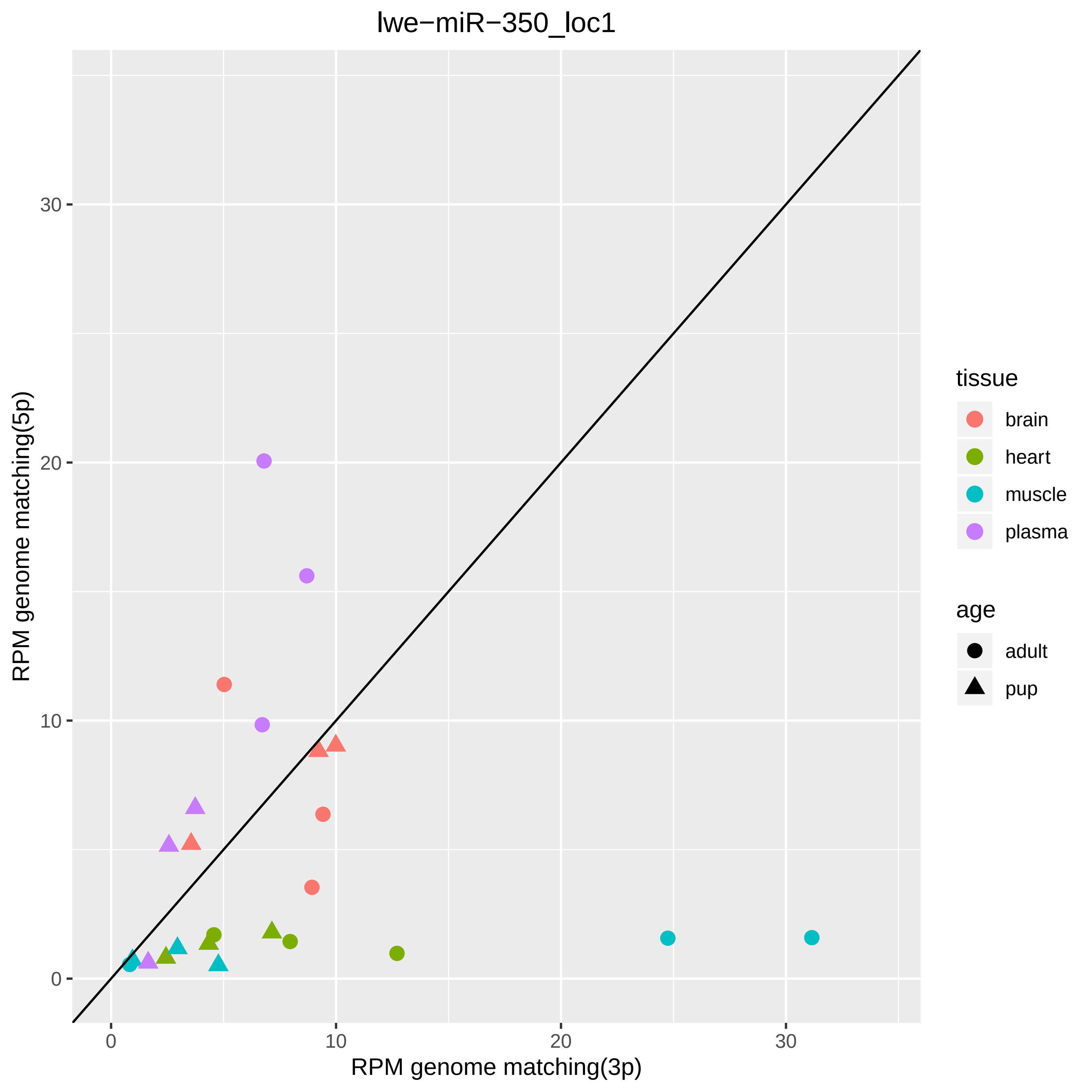

lwe−miR−350_loc1
RPM genome matching(5p)
RPM genome matching(3p)
0
10
20
30
tissue
brain
heart
muscle
plasma
age
adult
pup

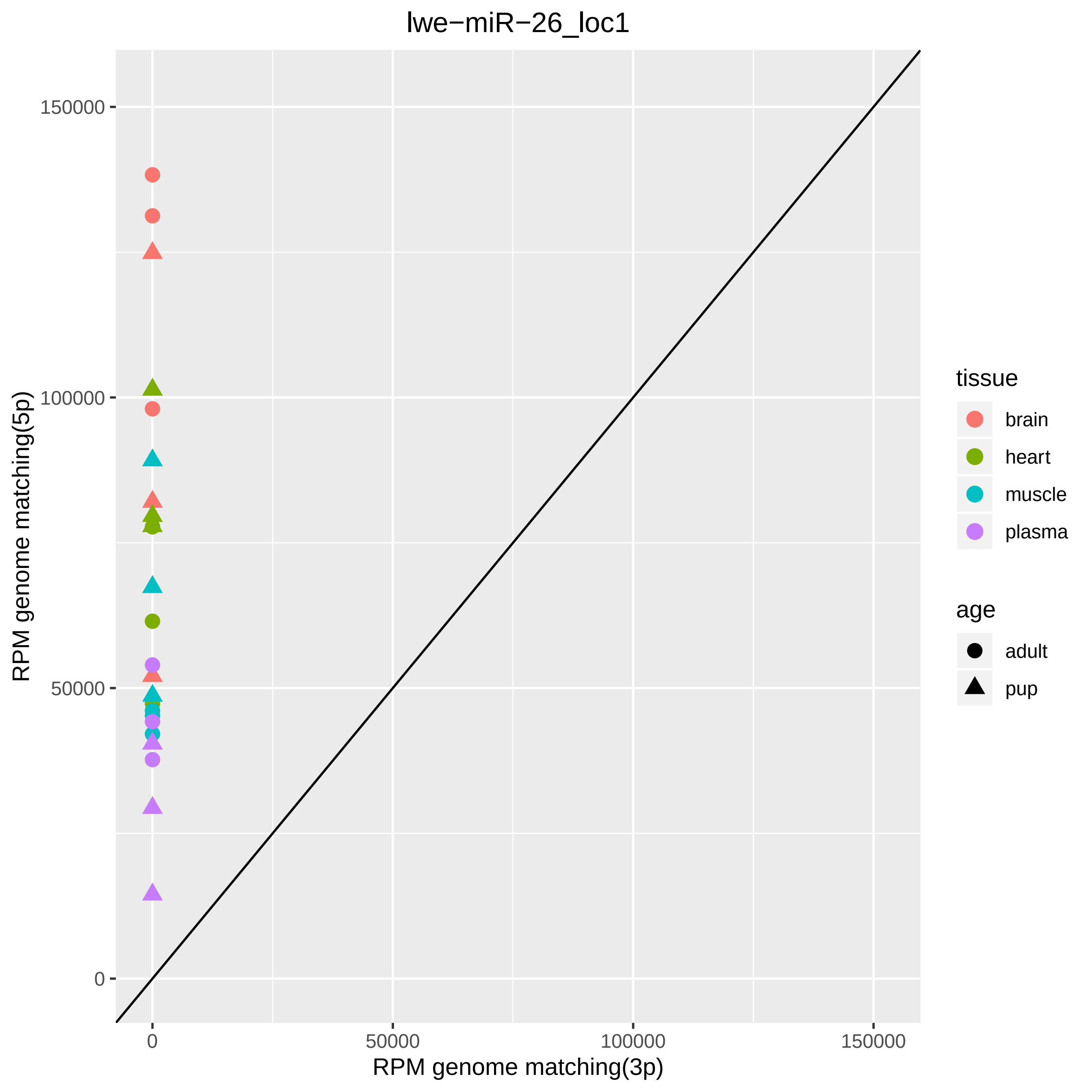
lwe−miR−26_loc1
RPM genome matching(5p)
RPM genome matching(3p)
0
50000
100000
150000
0
50000
100000
150000
tissue
brain
heart
muscle
plasma
age
adult
pup

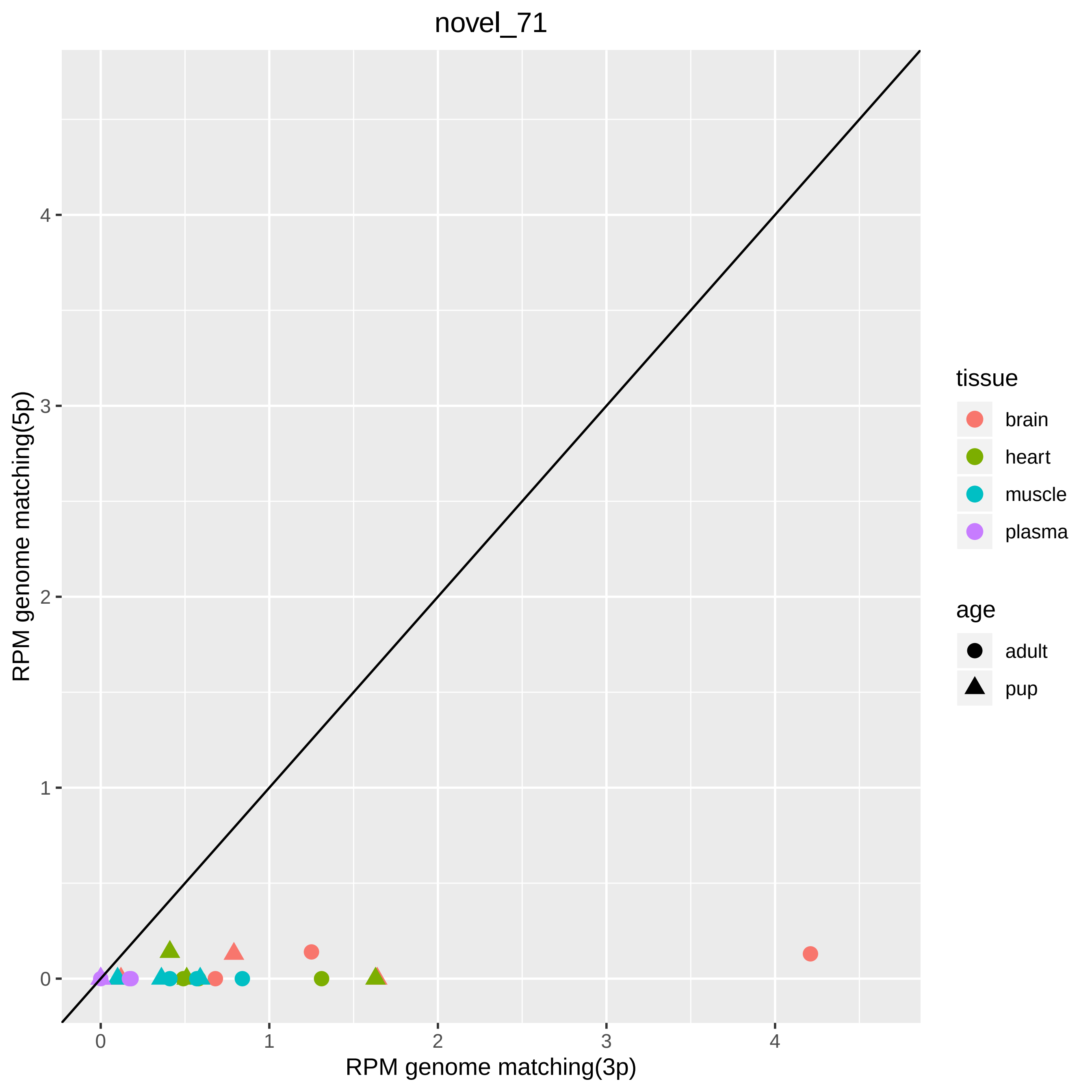

novel_71
RPM genome matching(5p)
RPM genome matching(3p)
0
1
2
3
4
tissue
brain
heart
muscle
plasma
age
adult
pup

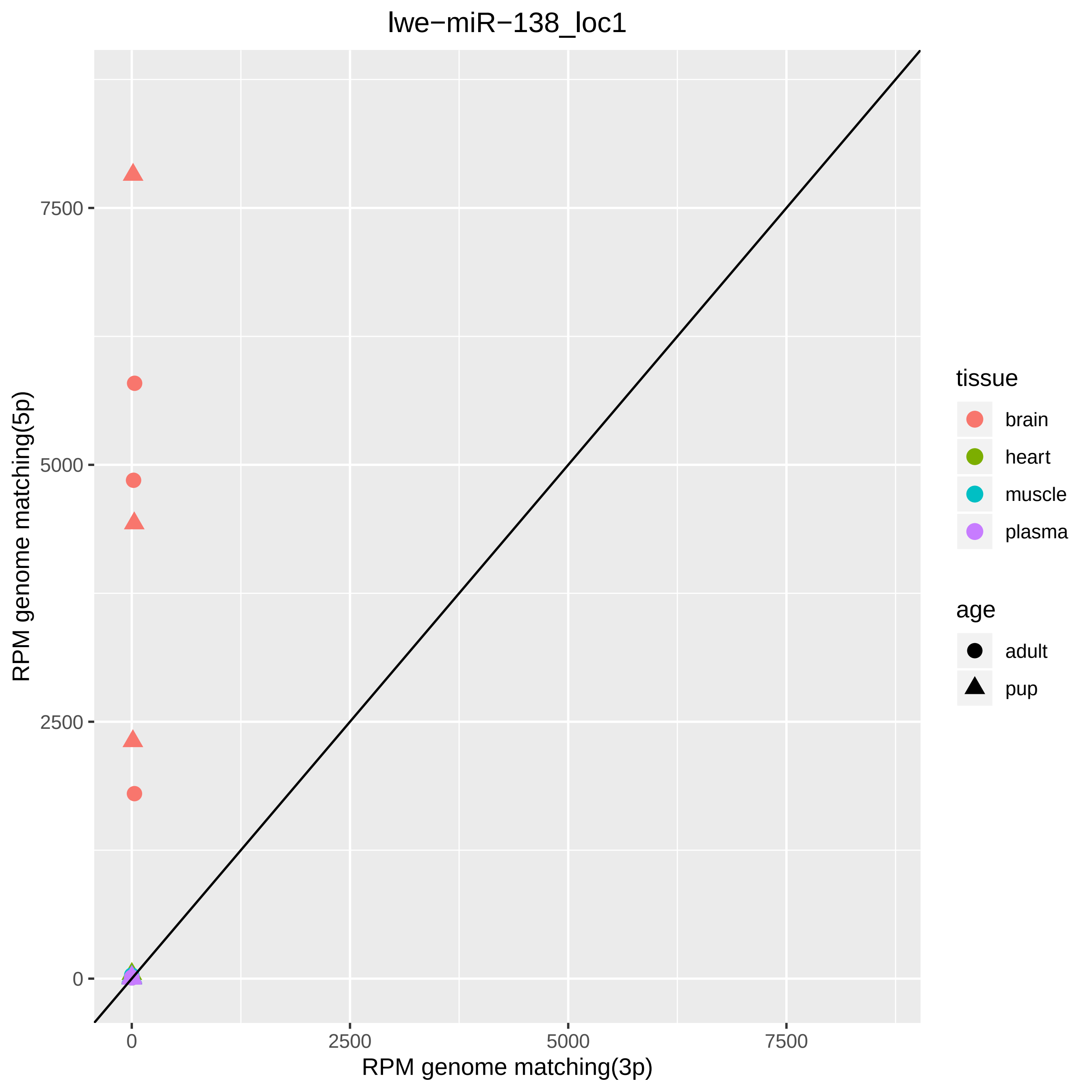
lwe−miR−138_loc1
RPM genome matching(5p)
RPM genome matching(3p)
0
2500
5000
7500
0
2500
5000
7500
tissue
brain
heart
muscle
plasma
age
adult
pup

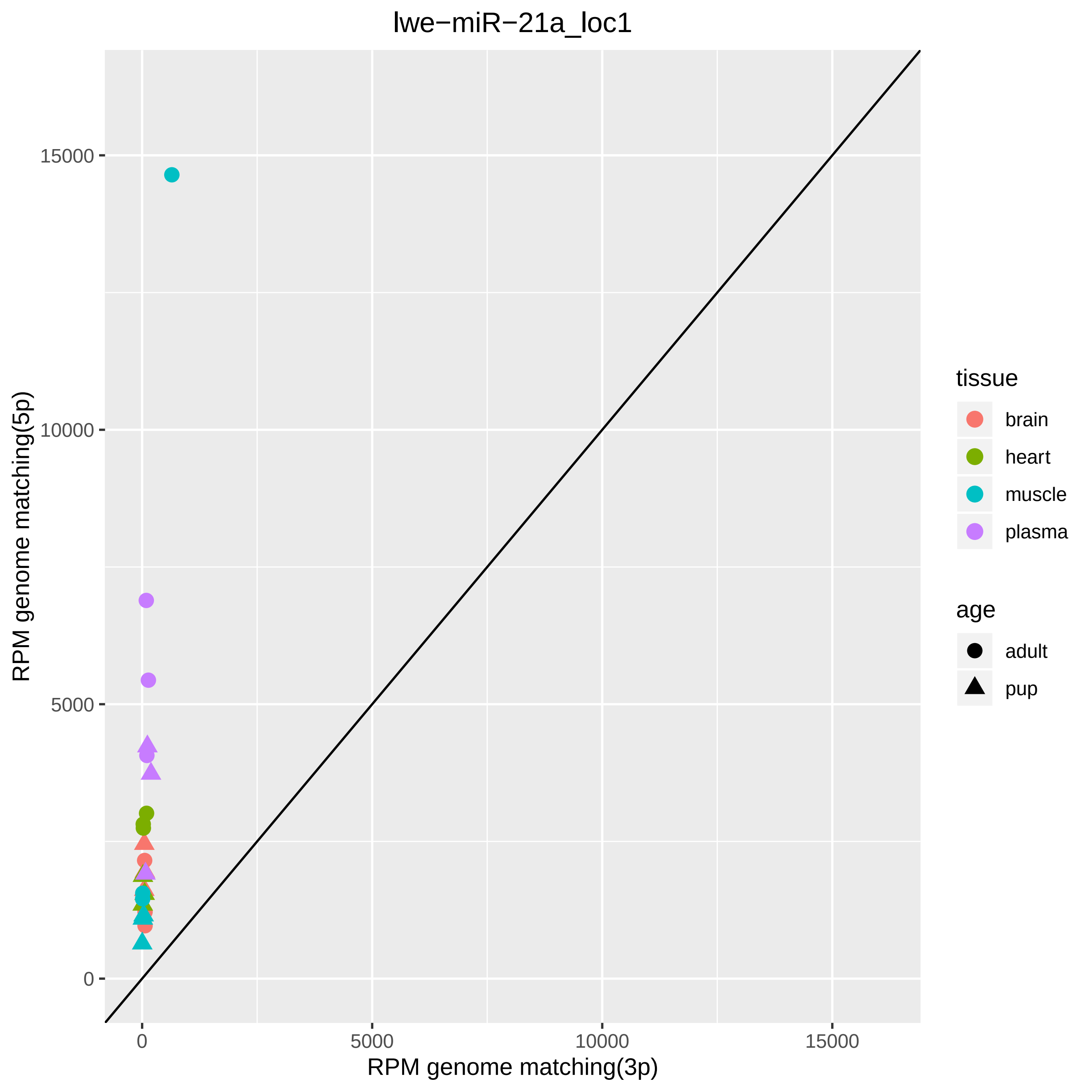
lwe−miR−21a_loc1
RPM genome matching(5p)
RPM genome matching(3p)
0
5000
10000
15000
0
5000
10000
15000
tissue
brain
heart
muscle
plasma
age
adult
pup

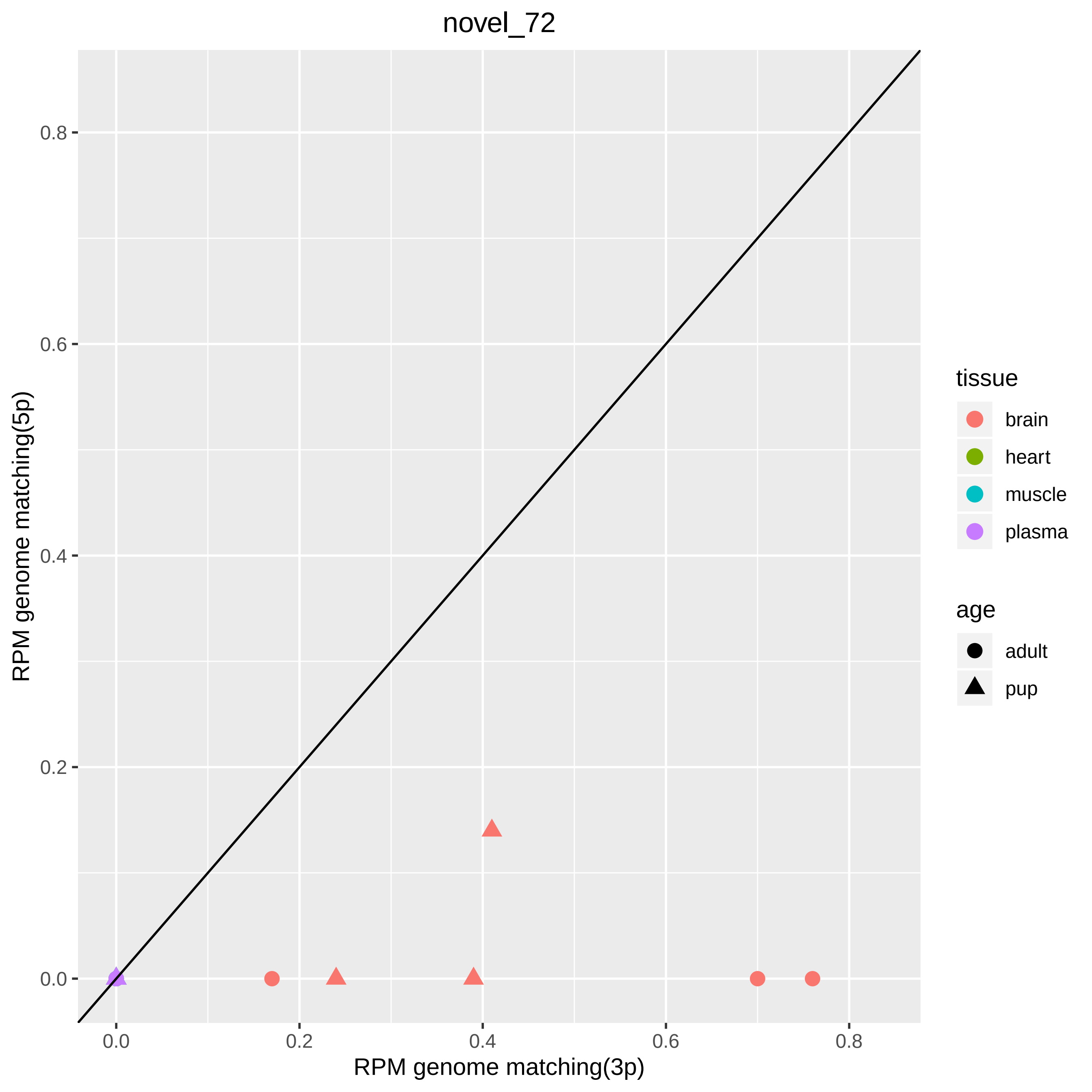
novel_72
RPM genome matching(5p)
RPM genome matching(3p)
0.0
0.2
0.4
0.6
0.8
tissue
brain
heart
muscle
plasma
age
adult
pup

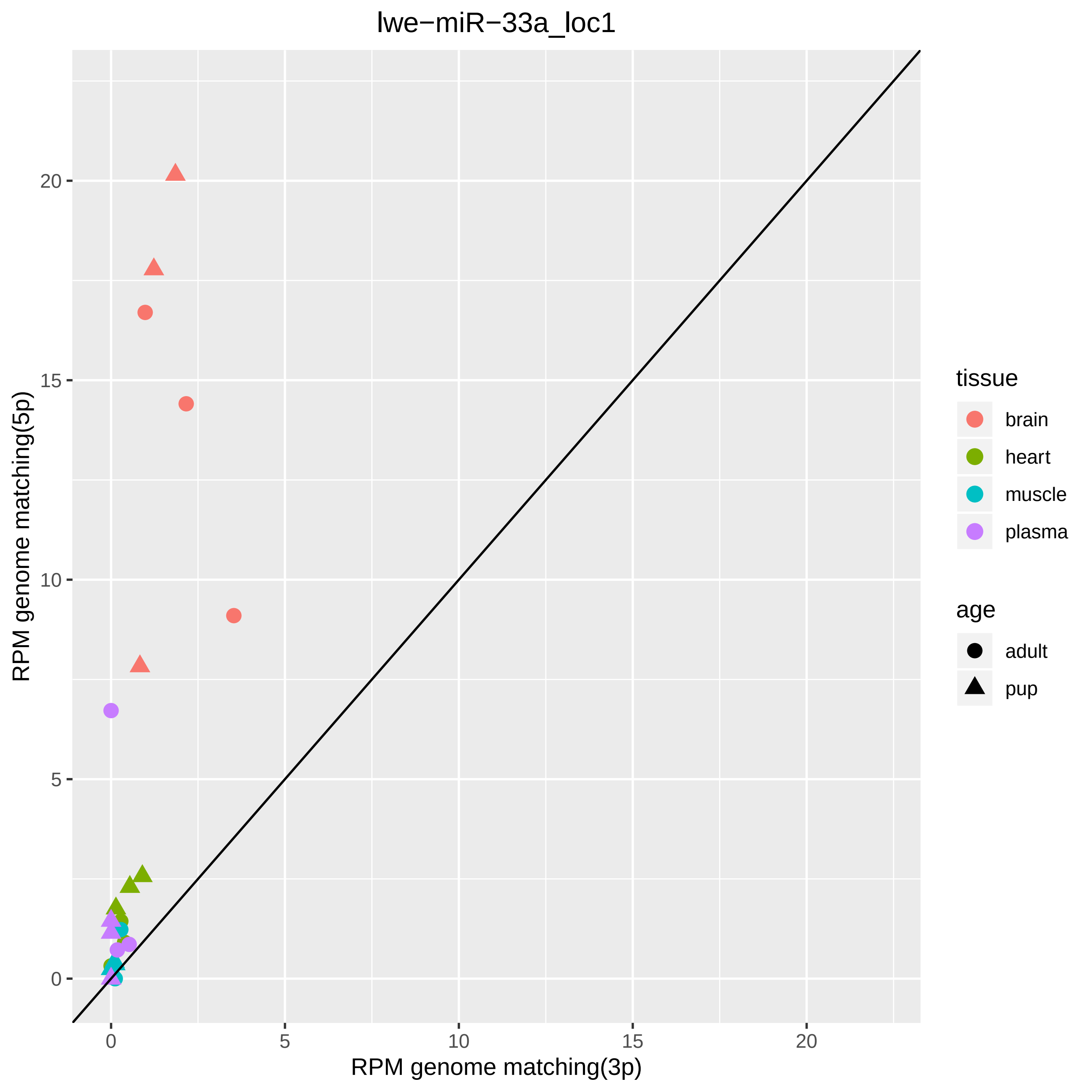

lwe−miR−33a_loc1
RPM genome matching(5p)
RPM genome matching(3p)
0
5
10
15
20
tissue
brain
heart
muscle
plasma
age
adult
pup

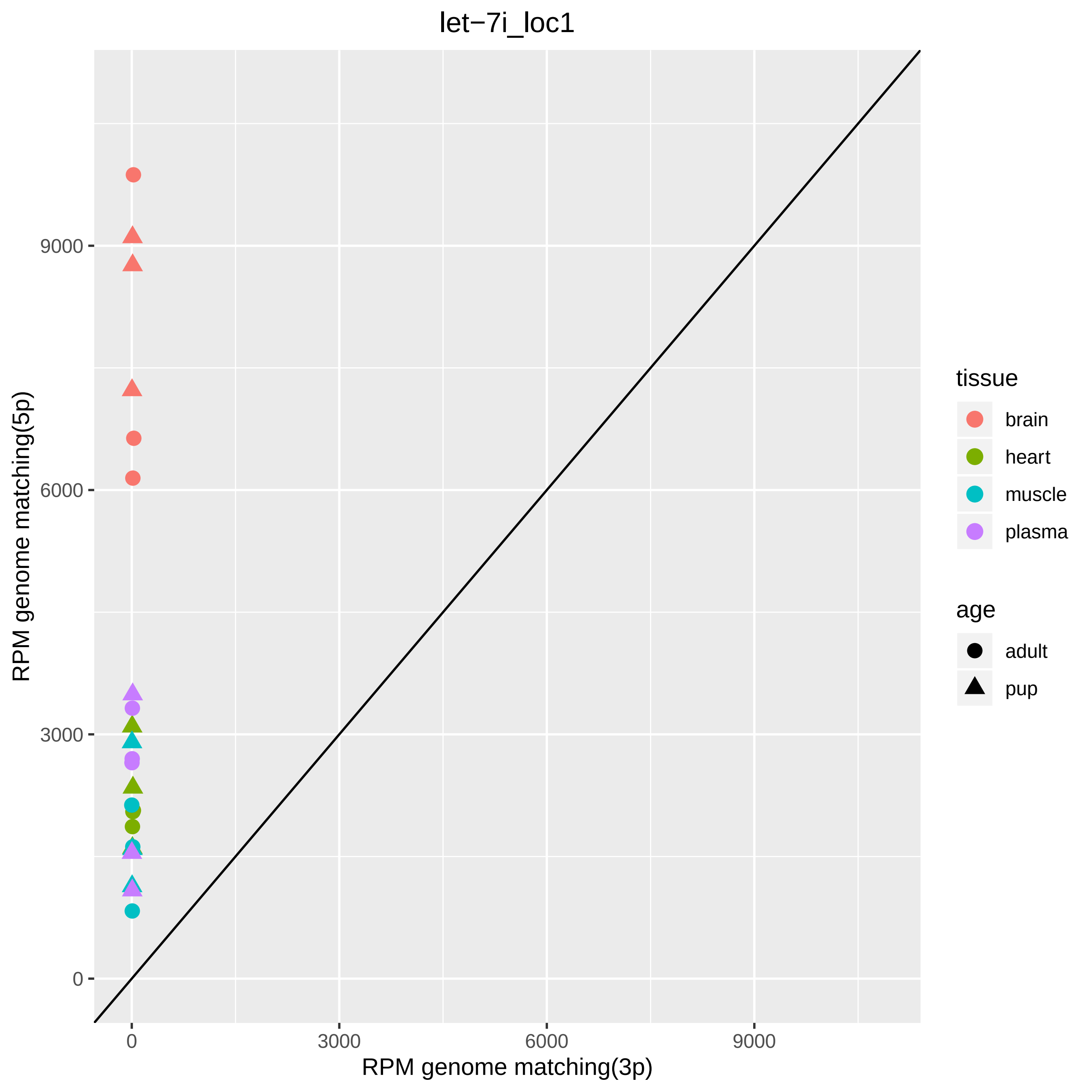
let−7i_loc1
RPM genome matching(5p)
RPM genome matching(3p)
9000
6000
3000
0
0
3000
6000
9000
tissue
brain
heart
muscle
plasma
age
adult
pup

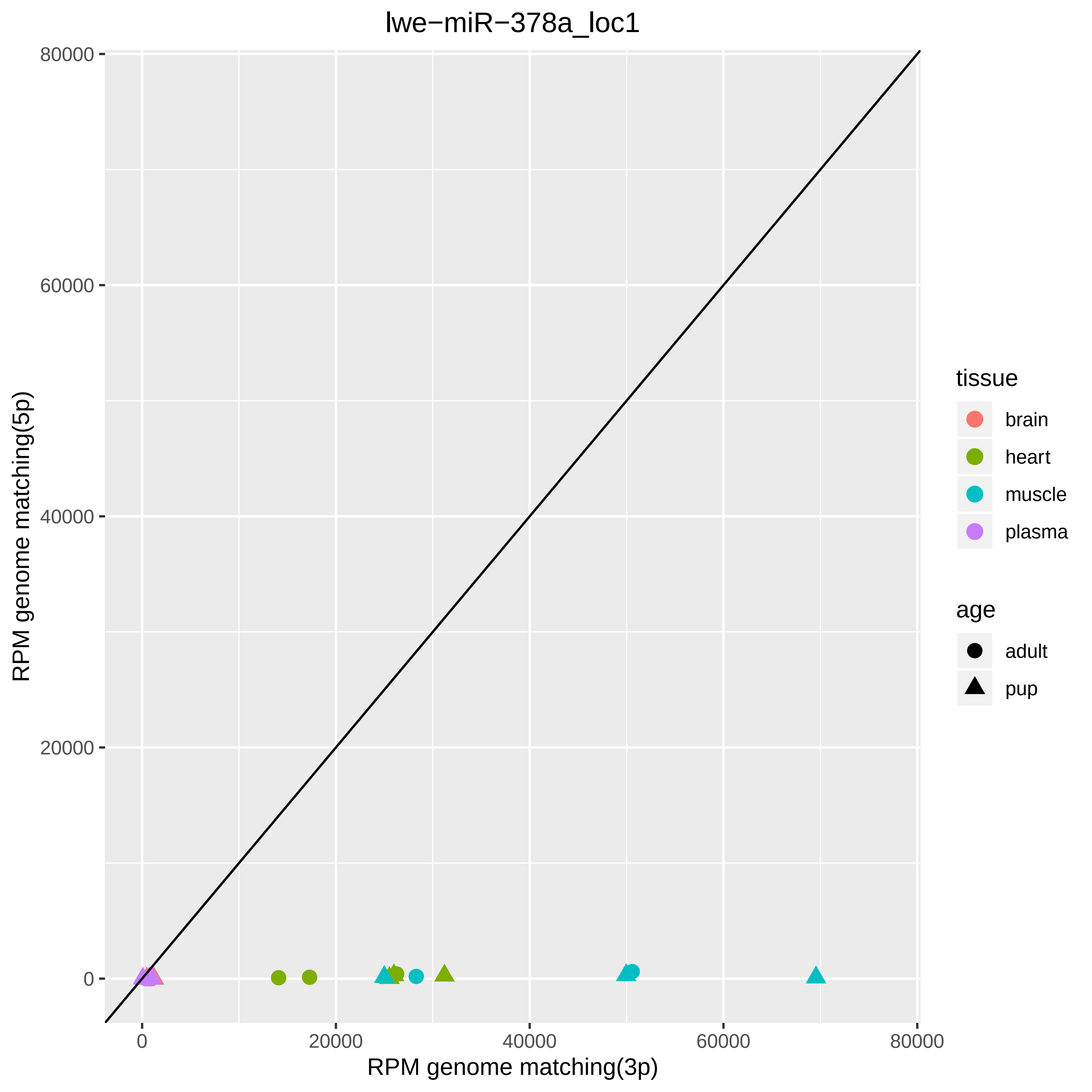

lwe−miR−378a_loc1
RPM genome matching(5p)
RPM genome matching(3p)
0
20000
40000
60000
80000
tissue
brain
heart
muscle
plasma
age
adult
pup

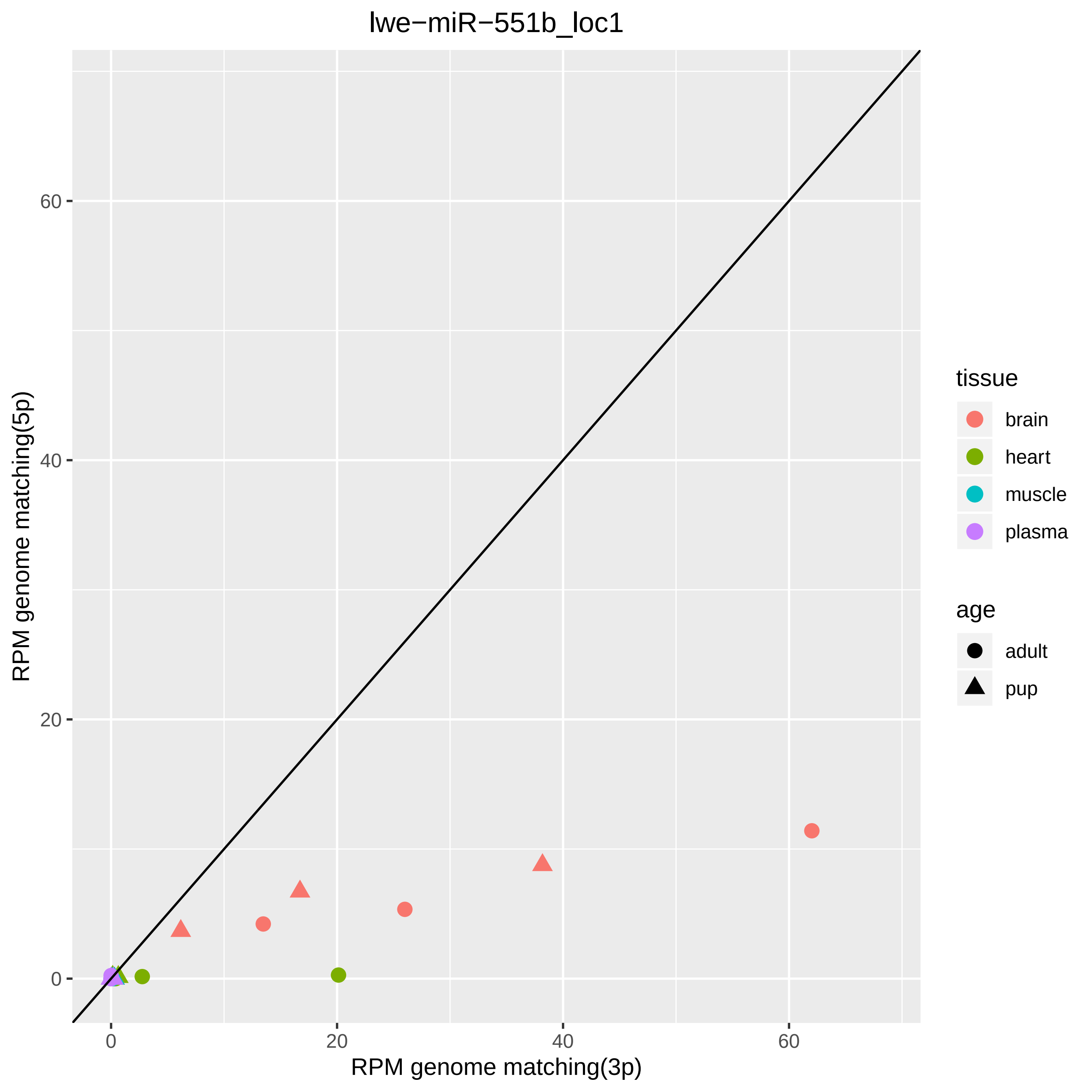

lwe−miR−551b_loc1
RPM genome matching(5p)
RPM genome matching(3p)
0
20
40
60
0
20
40
60
tissue
brain
heart
muscle
plasma
age
adult
pup

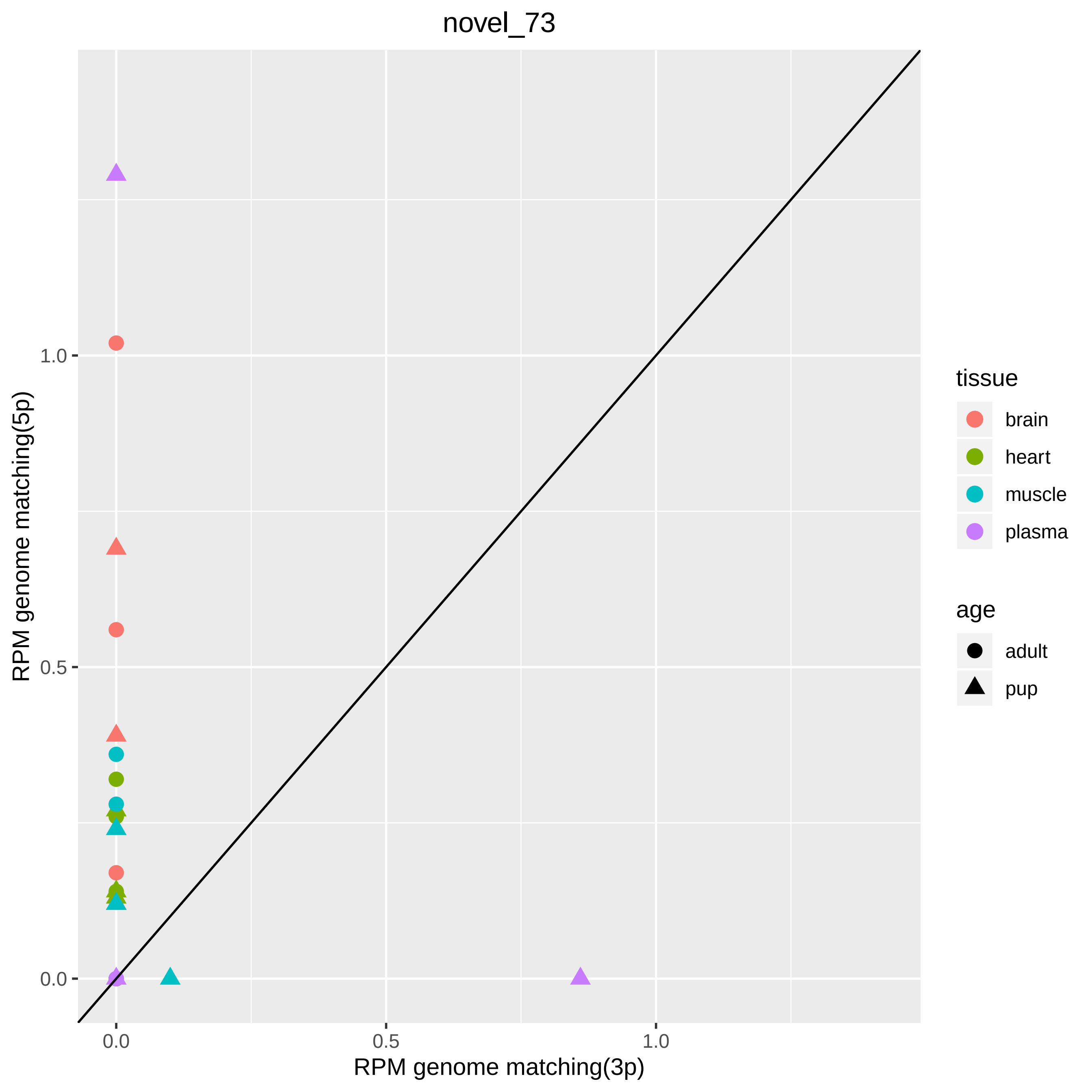

novel_73
RPM genome matching(5p)
RPM genome matching(3p)
0.0
0.5
1.0
0.0
0.5
1.0
tissue
brain
heart
muscle
plasma
age
adult
pup

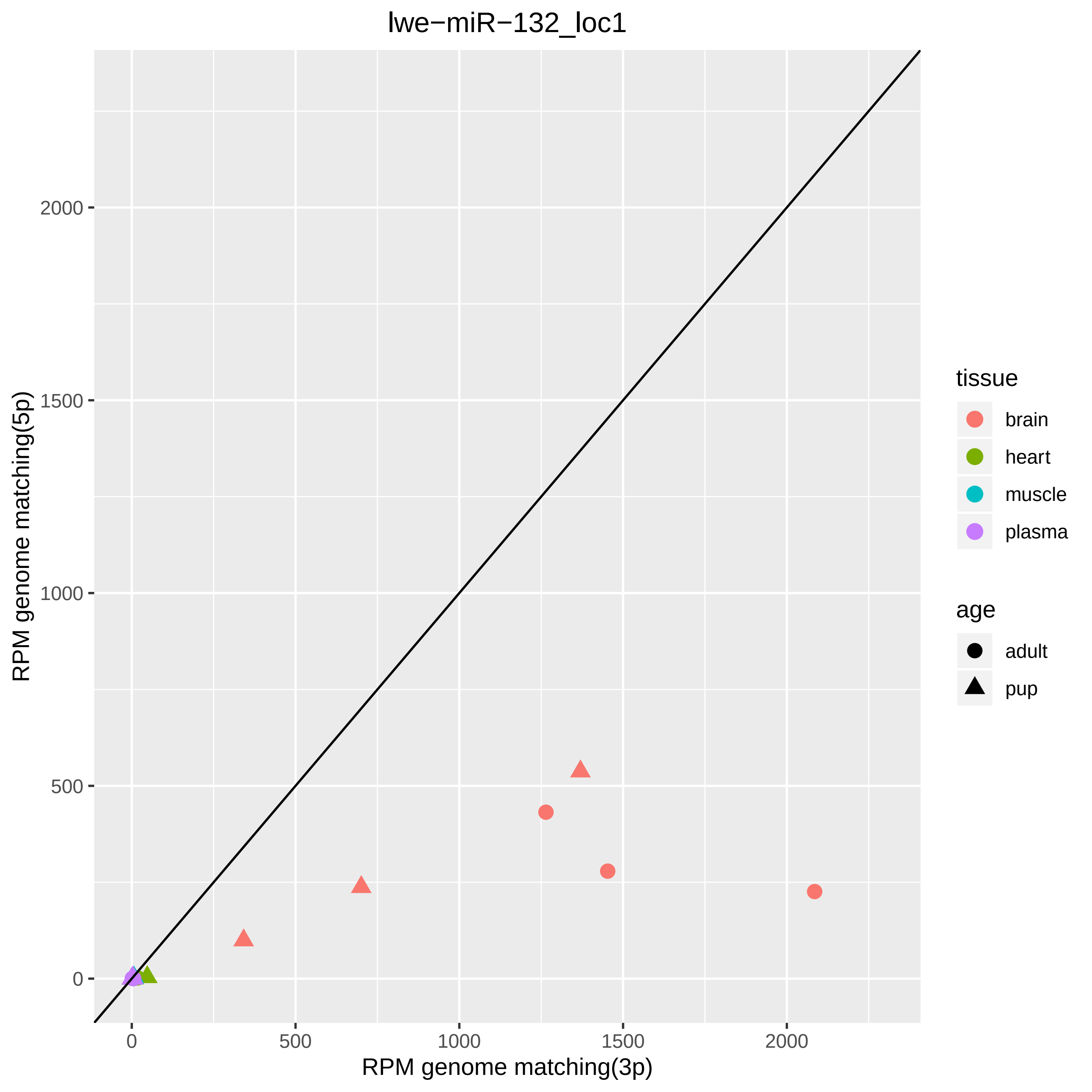

lwe−miR−132_loc1
RPM genome matching(5p)
RPM genome matching(3p)
0
500
1000
1500
2000
tissue
brain
heart
muscle
plasma
age
adult
pup

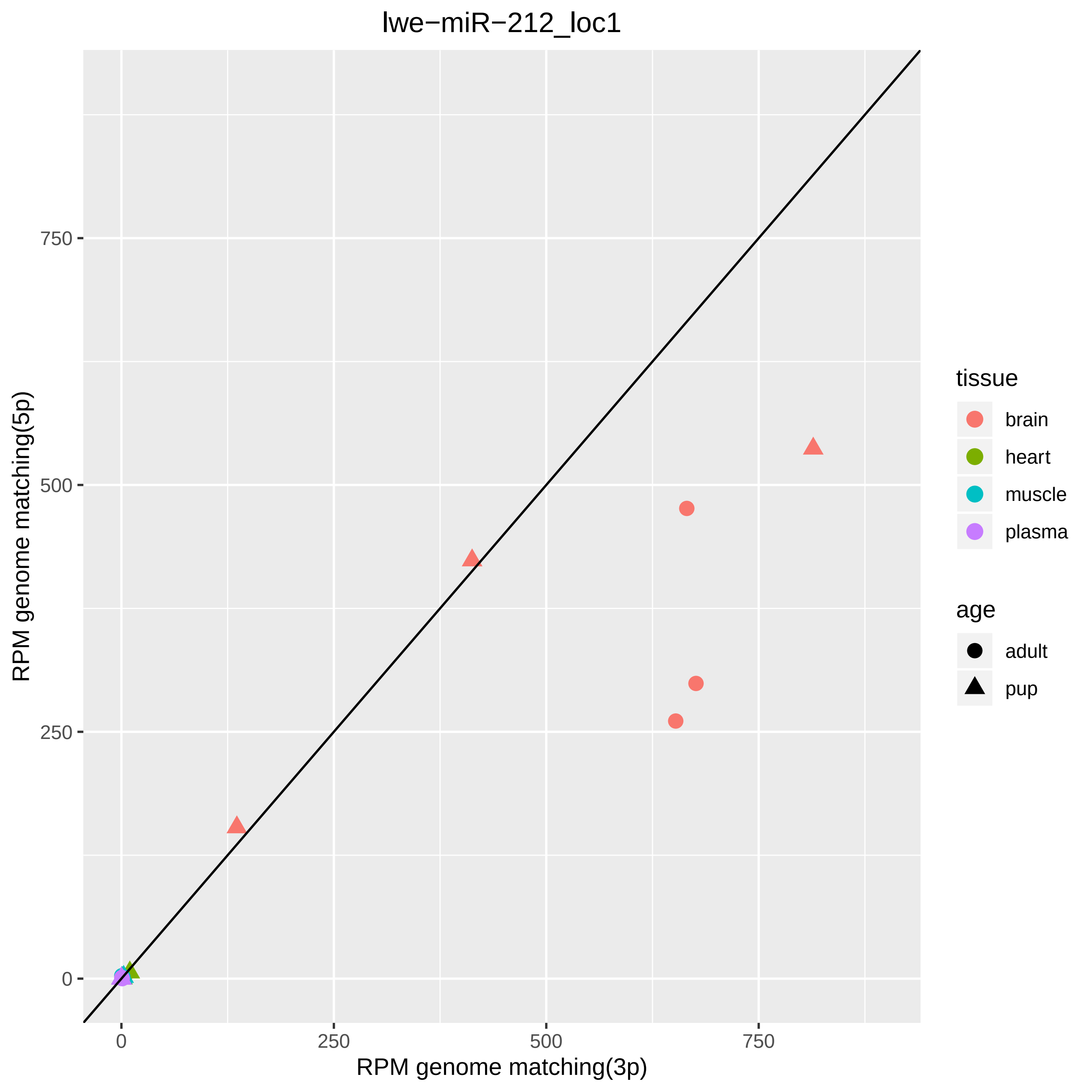

lwe−miR−212_loc1
RPM genome matching(5p)
RPM genome matching(3p)
0
250
500
750
0
250
500
750
tissue
brain
heart
muscle
plasma
age
adult
pup

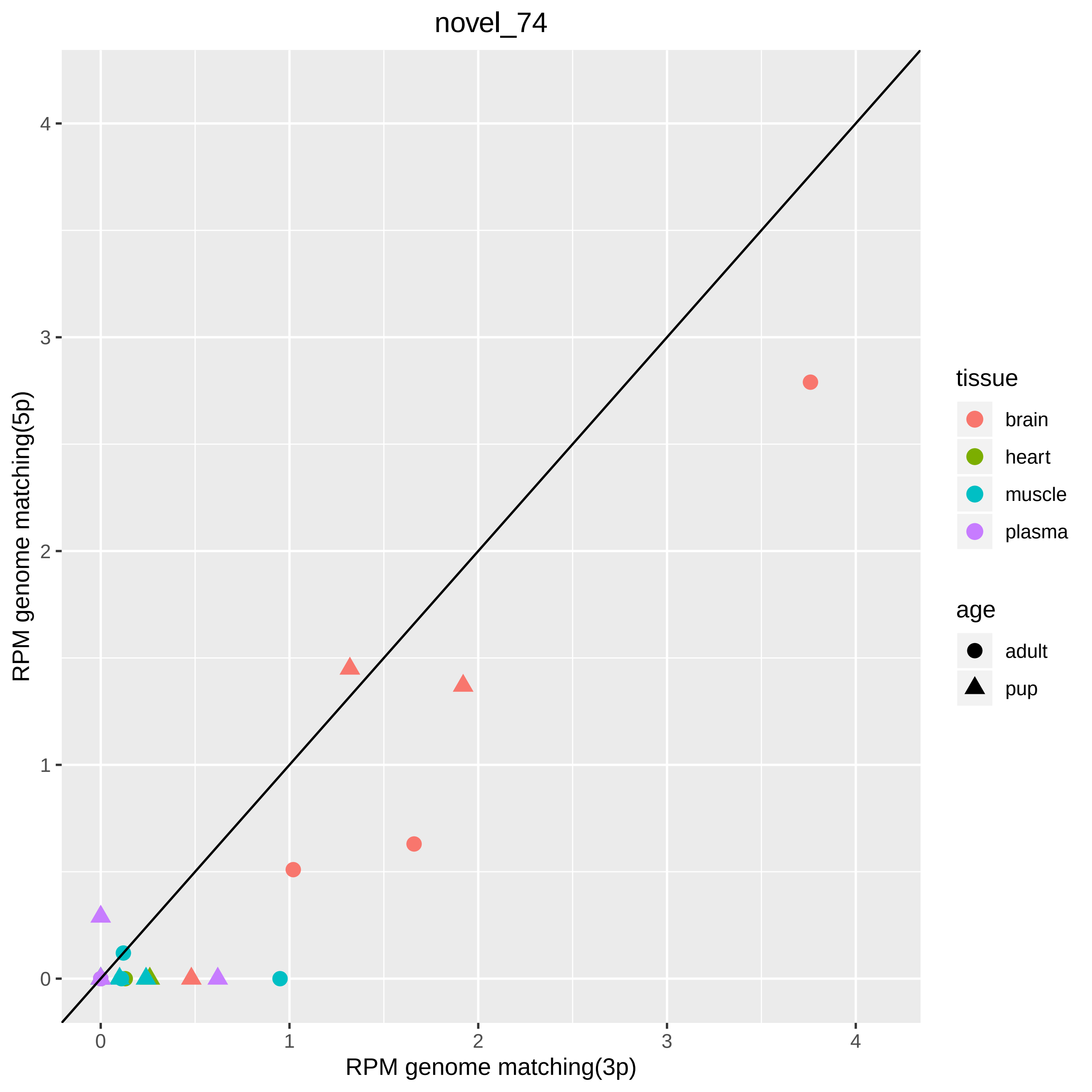

novel_74
RPM genome matching(5p)
RPM genome matching(3p)
0
1
2
3
4
0
1
2
3
4
tissue
brain
heart
muscle
plasma
age
adult
pup

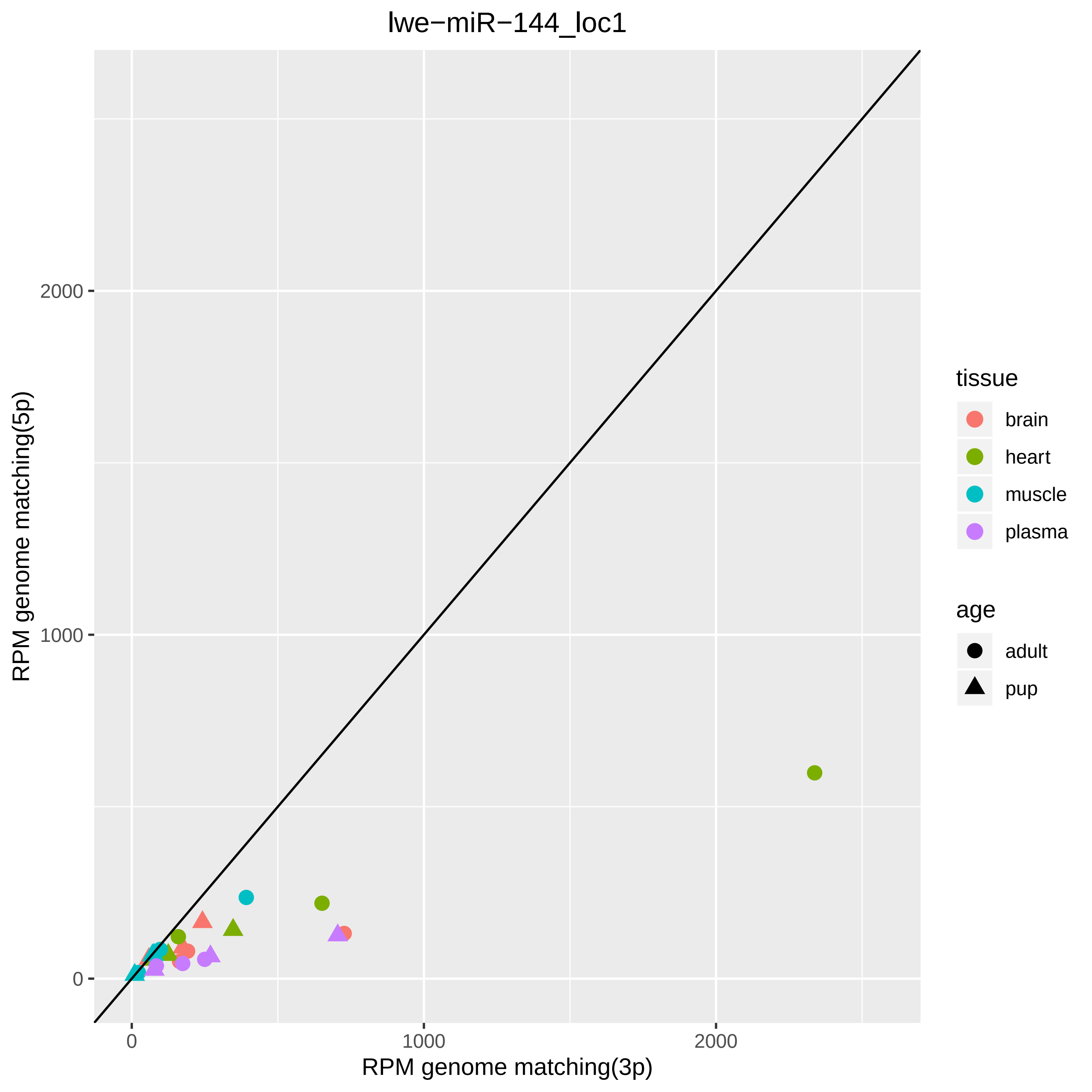
lwe−miR−144_loc1
RPM genome matching(5p)
RPM genome matching(3p)
2000
1000
0
0
1000
2000
tissue
brain
heart
muscle
plasma
age
adult
pup

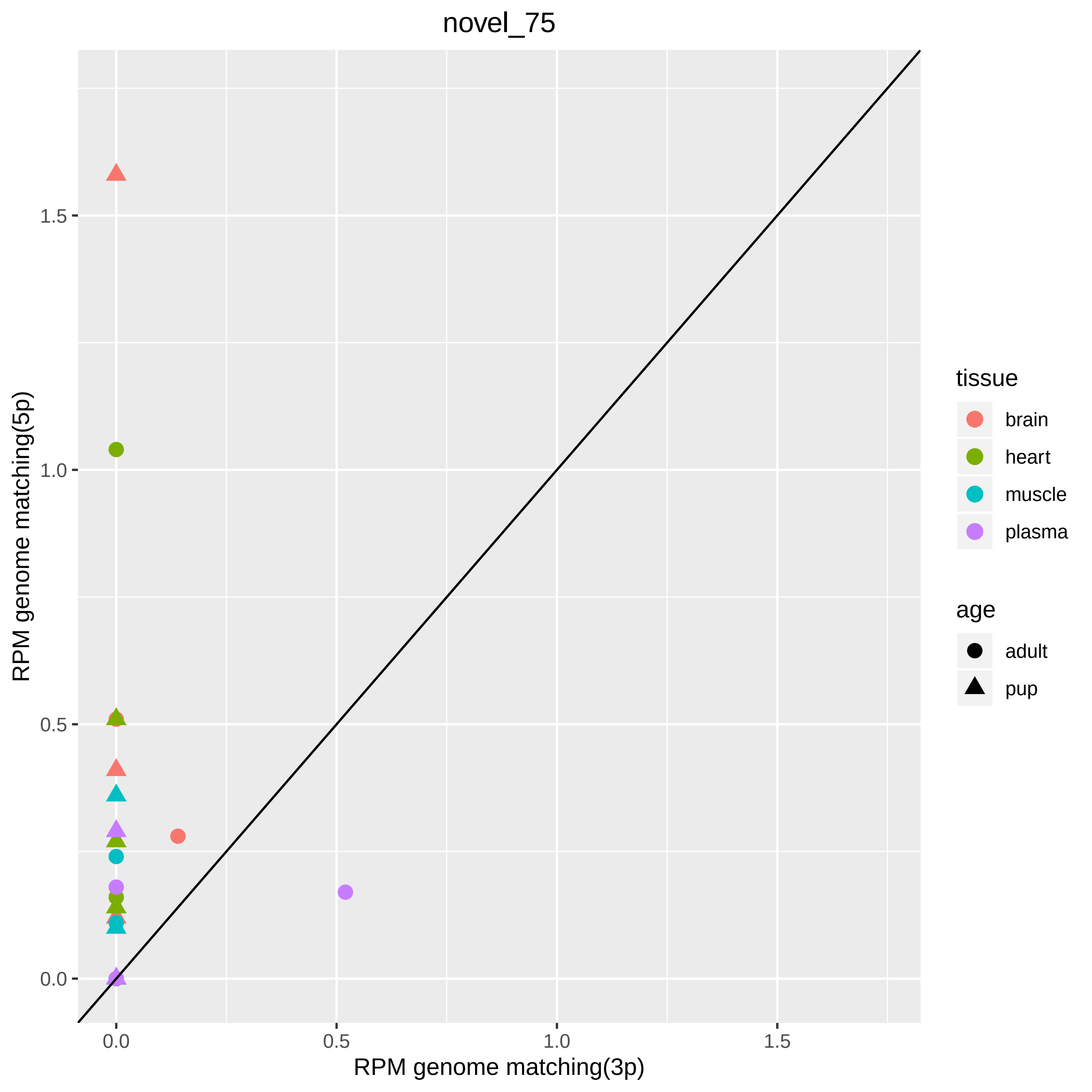

novel_75
RPM genome matching(5p)
RPM genome matching(3p)
0.0
0.5
1.0
1.5
tissue
brain
heart
muscle
plasma
age
adult
pup

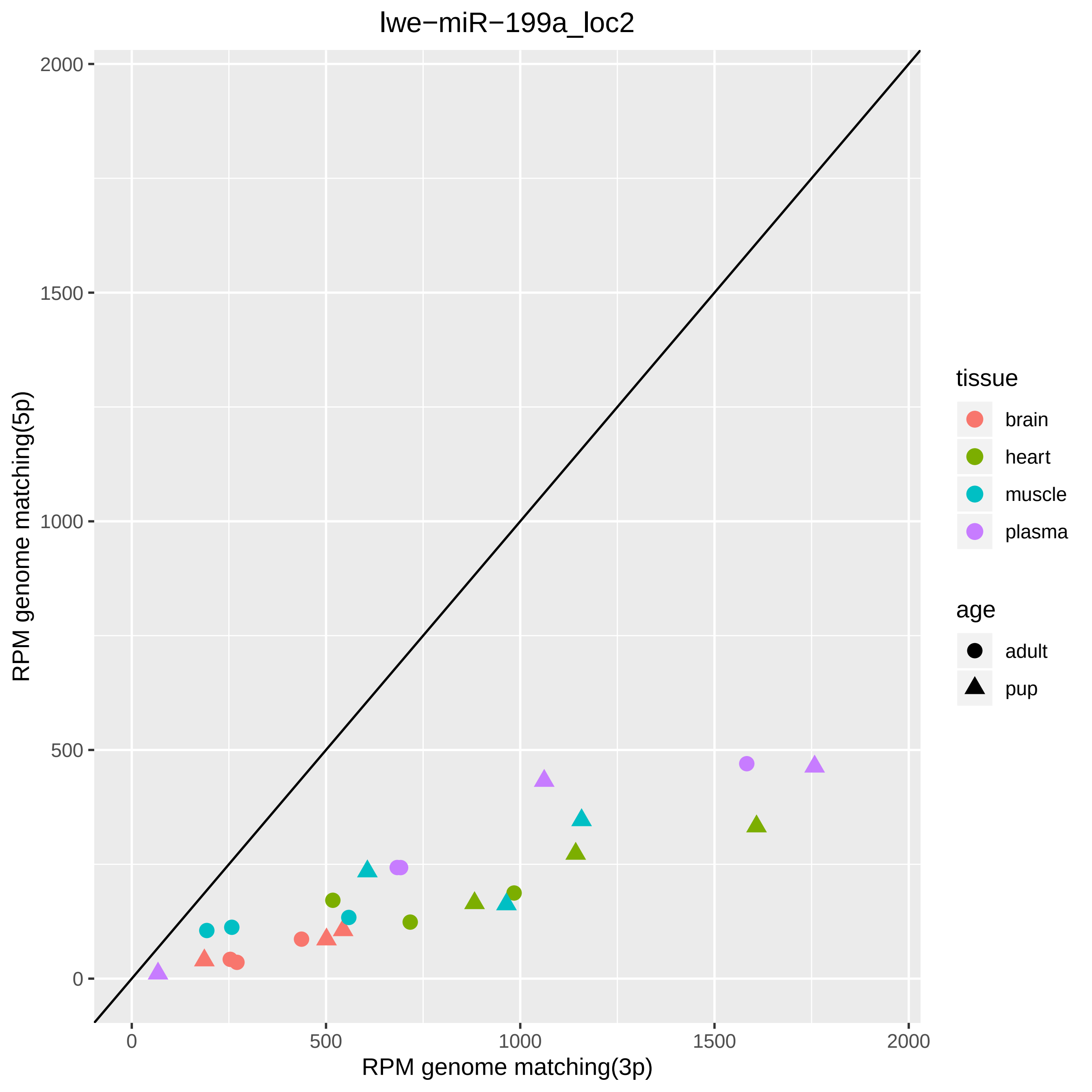
lwe−miR−199a_loc2
RPM genome matching(5p)
2000
1500
1000
500
0
0
500
1000
1500
2000
RPM genome matching(3p)
tissue
brain
heart
muscle
plasma
age
adult
pup

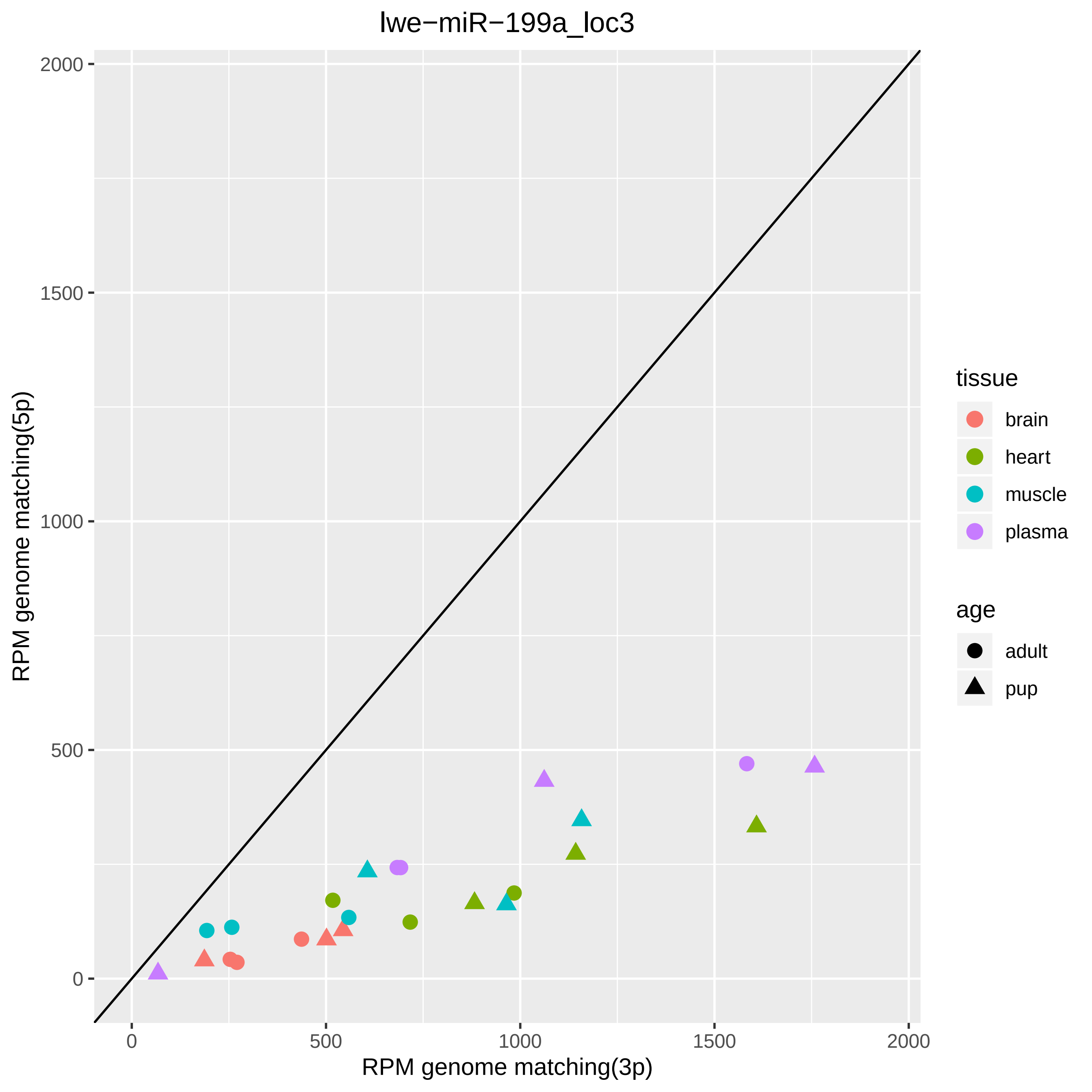

lwe−miR−199a_loc3
RPM genome matching(5p)
RPM genome matching(3p)
0
500
1000
1500
2000
tissue
brain
heart
muscle
plasma
age
adult
pup

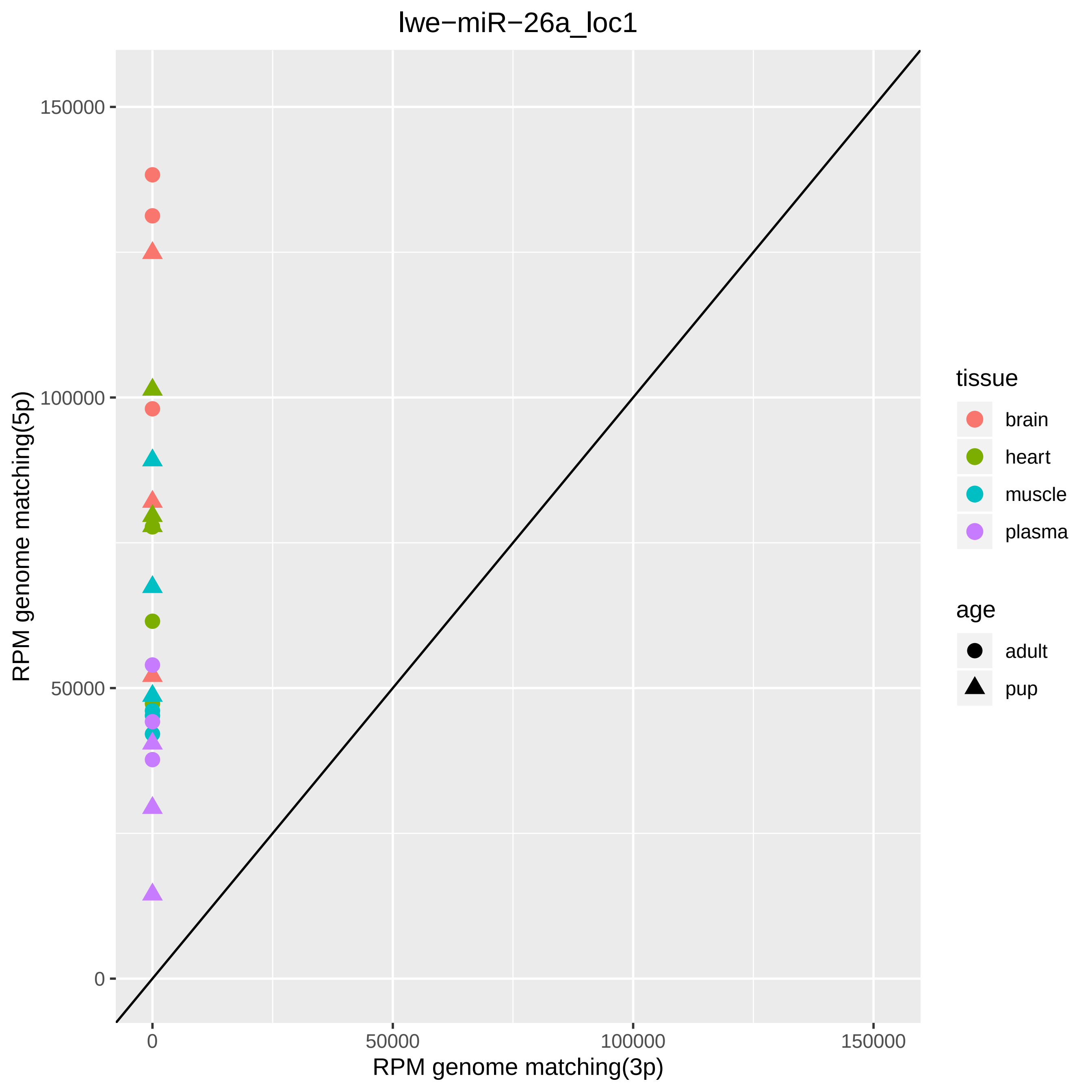

lwe−miR−26a_loc1
RPM genome matching(5p)
RPM genome matching(3p)
150000
100000
50000
0
0
50000
100000
150000
tissue
brain
heart
muscle
plasma
age
adult
pup

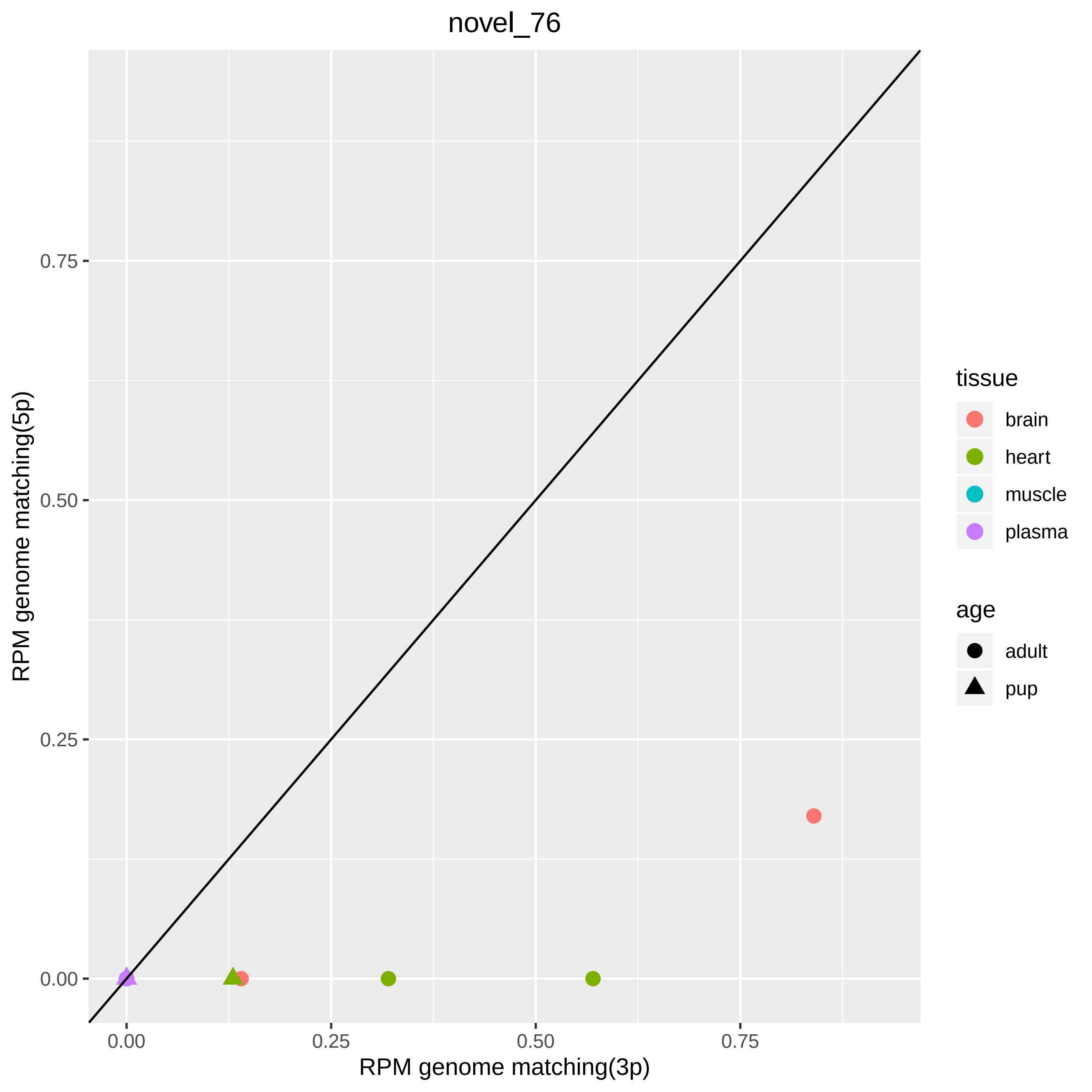

novel_76
RPM genome matching(5p)
RPM genome matching(3p)
0.00
0.25
0.50
0.75
tissue
brain
heart
muscle
plasma
age
adult
pup

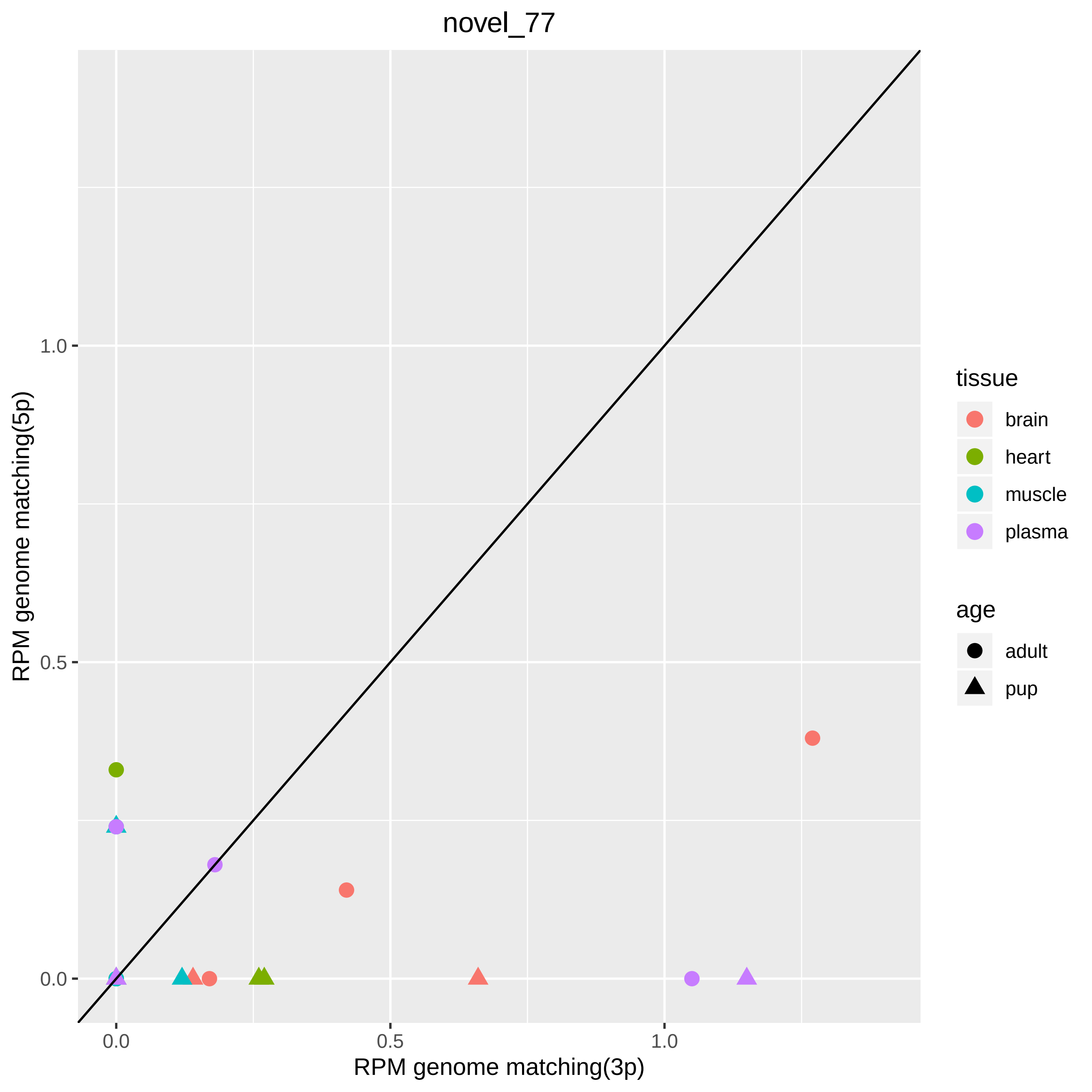

novel_77
RPM genome matching(5p)
RPM genome matching(3p)
0.0
0.5
1.0
tissue
brain
heart
muscle
plasma
age
adult
pup

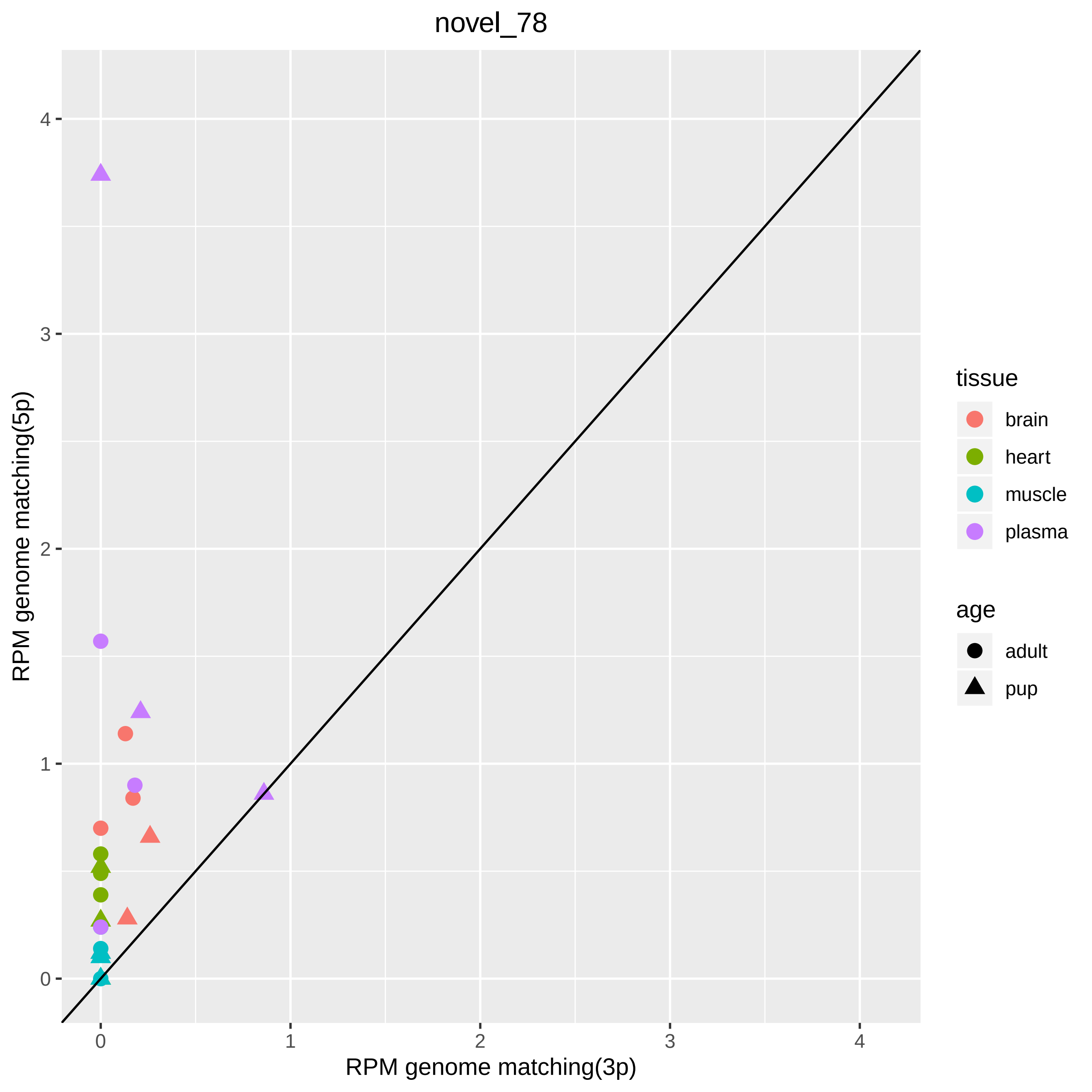

novel_78
RPM genome matching(5p)
RPM genome matching(3p)
0
1
2
3
4
tissue
brain
heart
muscle
plasma
age
adult
pup

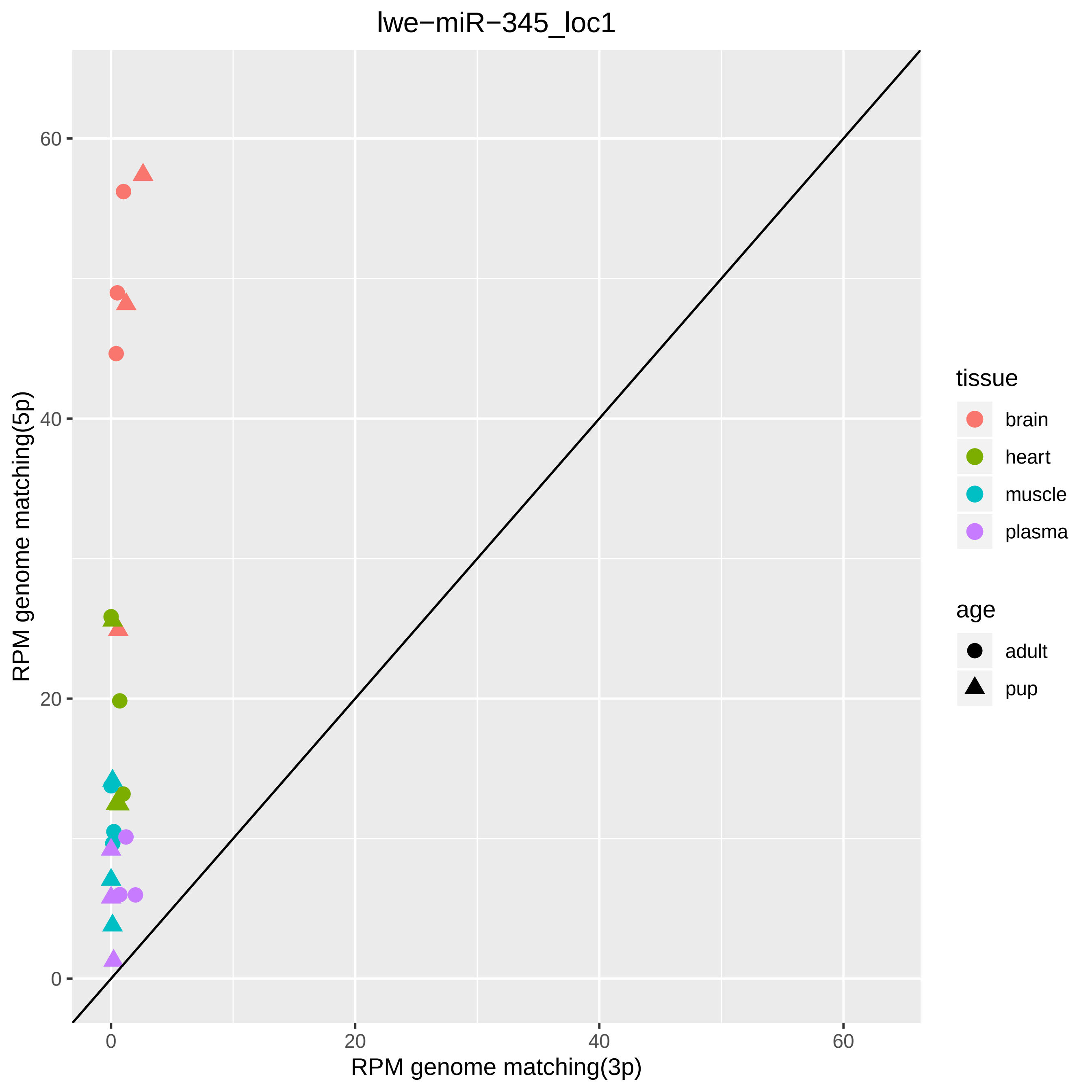
lwe−miR−345_loc1
RPM genome matching(5p)
RPM genome matching(3p)
0
20
40
60
0
20
40
60
tissue
brain
heart
muscle
plasma
age
adult
pup

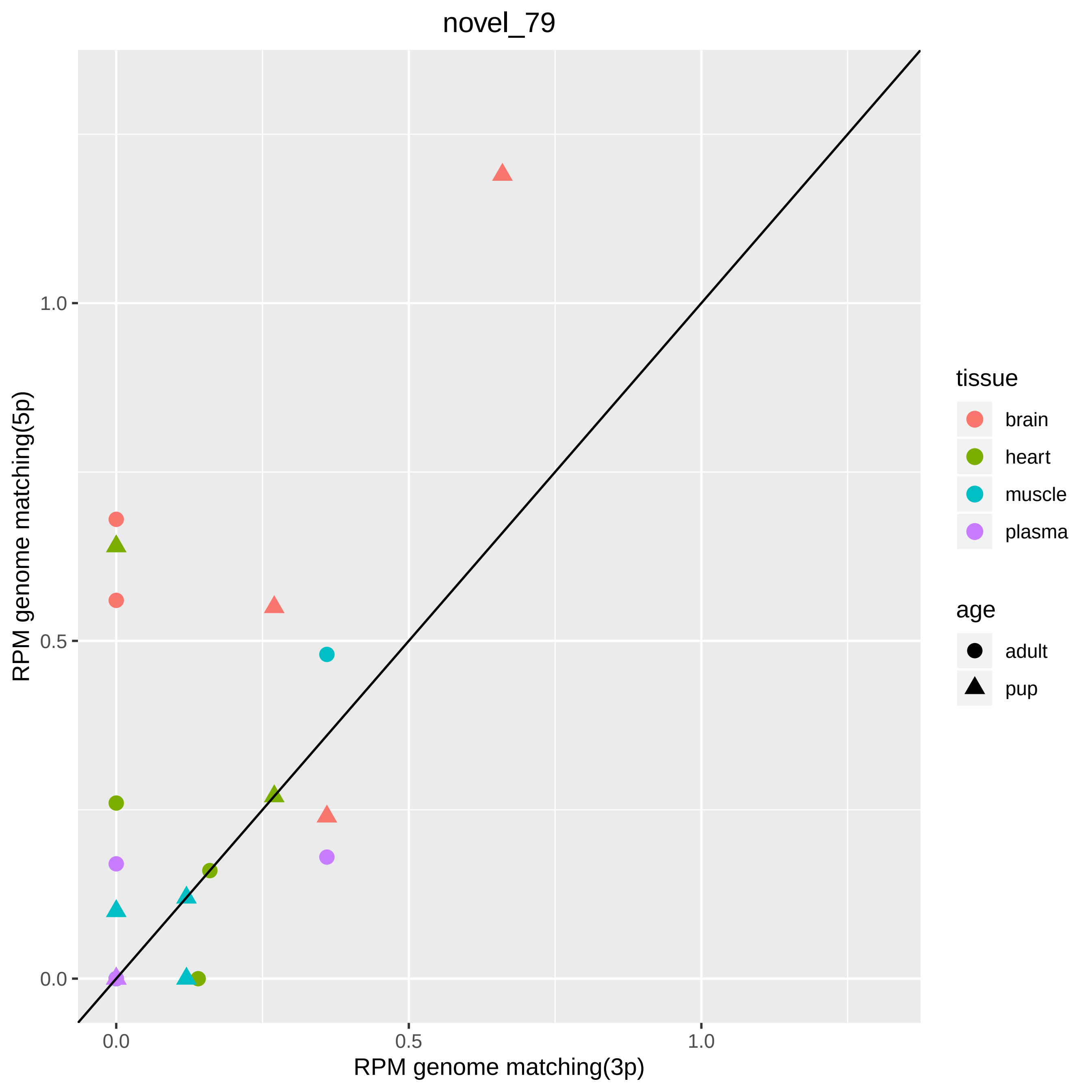

novel_79
RPM genome matching(5p)
RPM genome matching(3p)
0.0
0.5
1.0
tissue
brain
heart
muscle
plasma
age
adult
pup

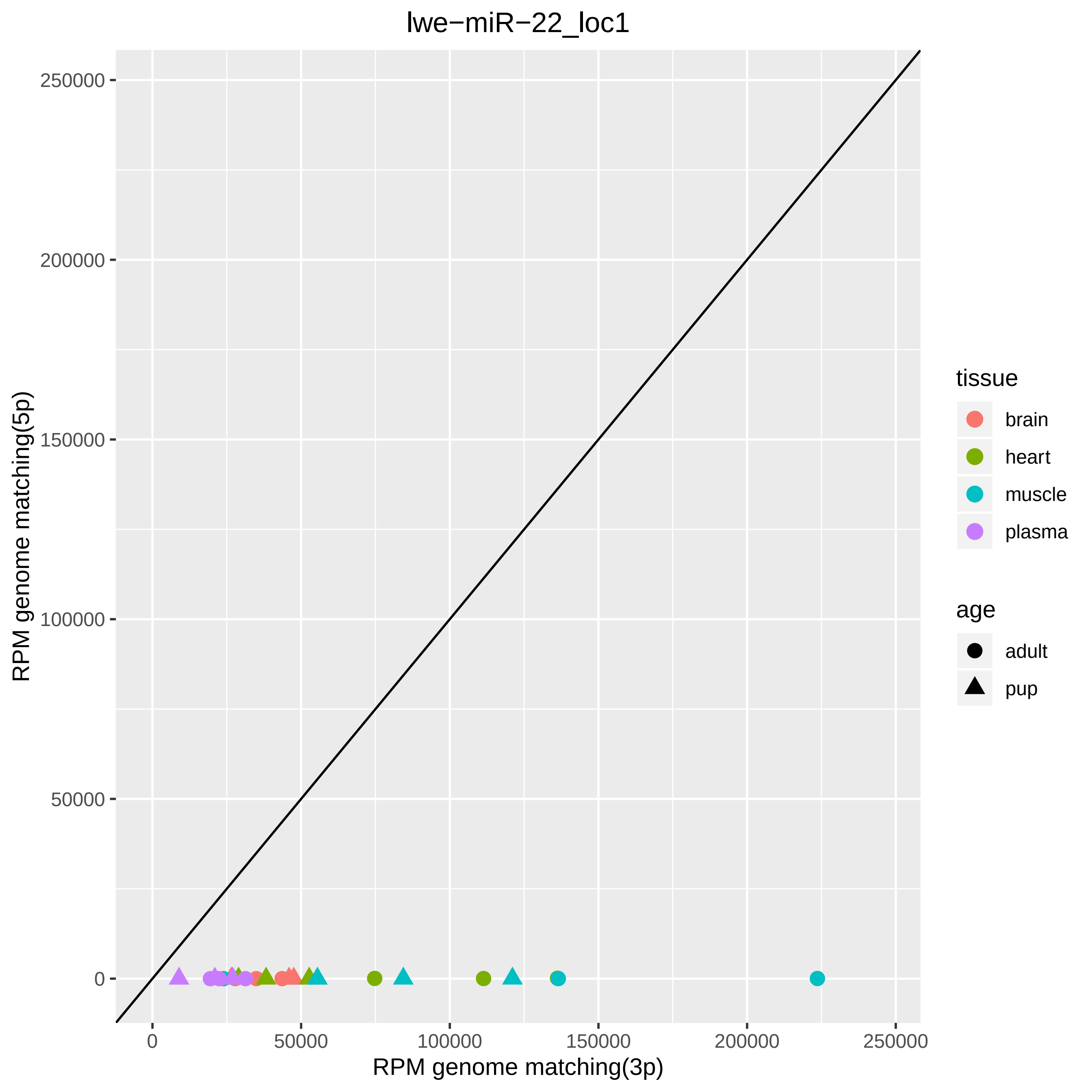
lwe−miR−22_loc1
RPM genome matching(5p)
RPM genome matching(3p)
0
50000
100000
150000
200000
250000
tissue
brain
heart
muscle
plasma
age
adult
pup

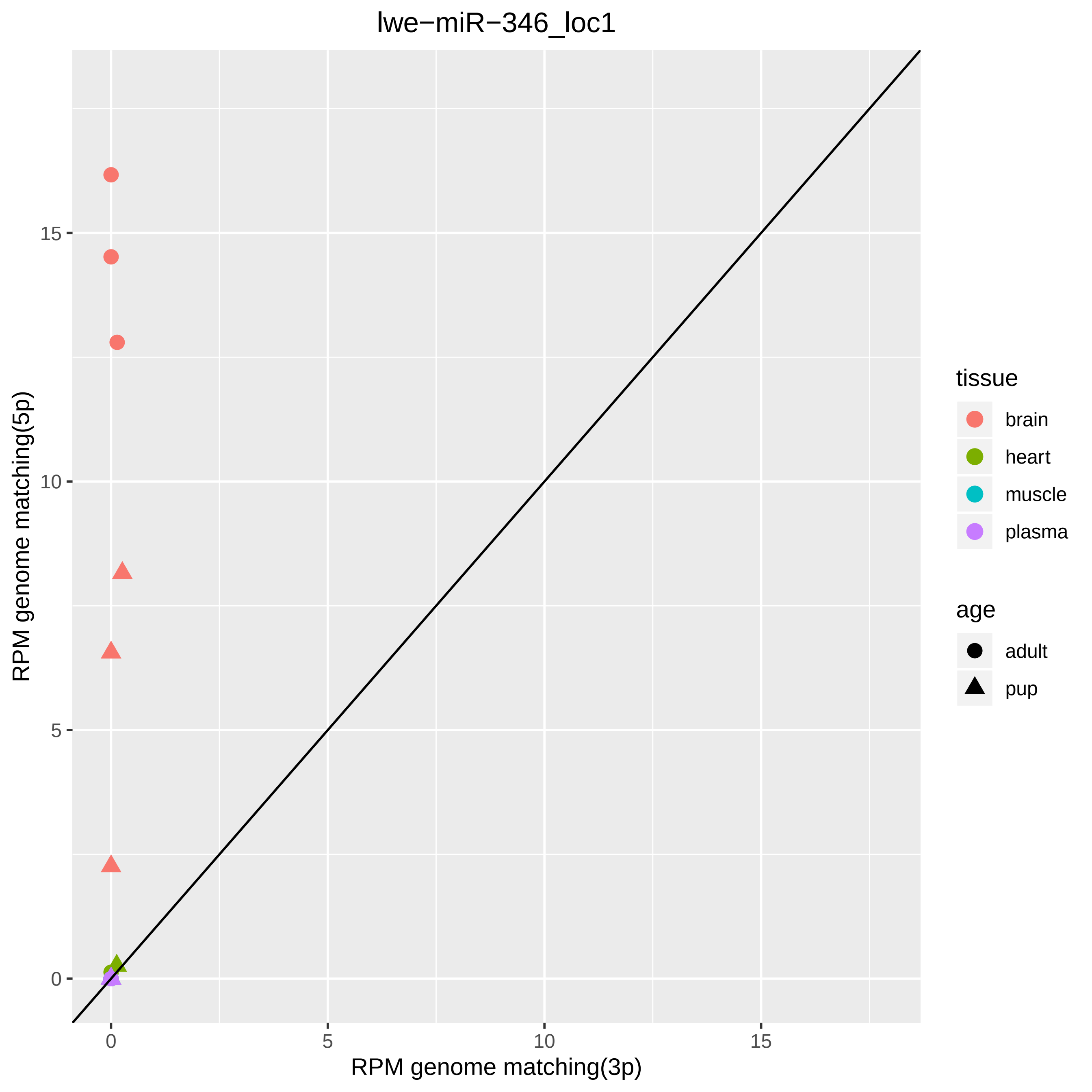

lwe−miR−346_loc1
RPM genome matching(5p)
RPM genome matching(3p)
0
5
10
15
tissue
brain
heart
muscle
plasma
age
adult
pup

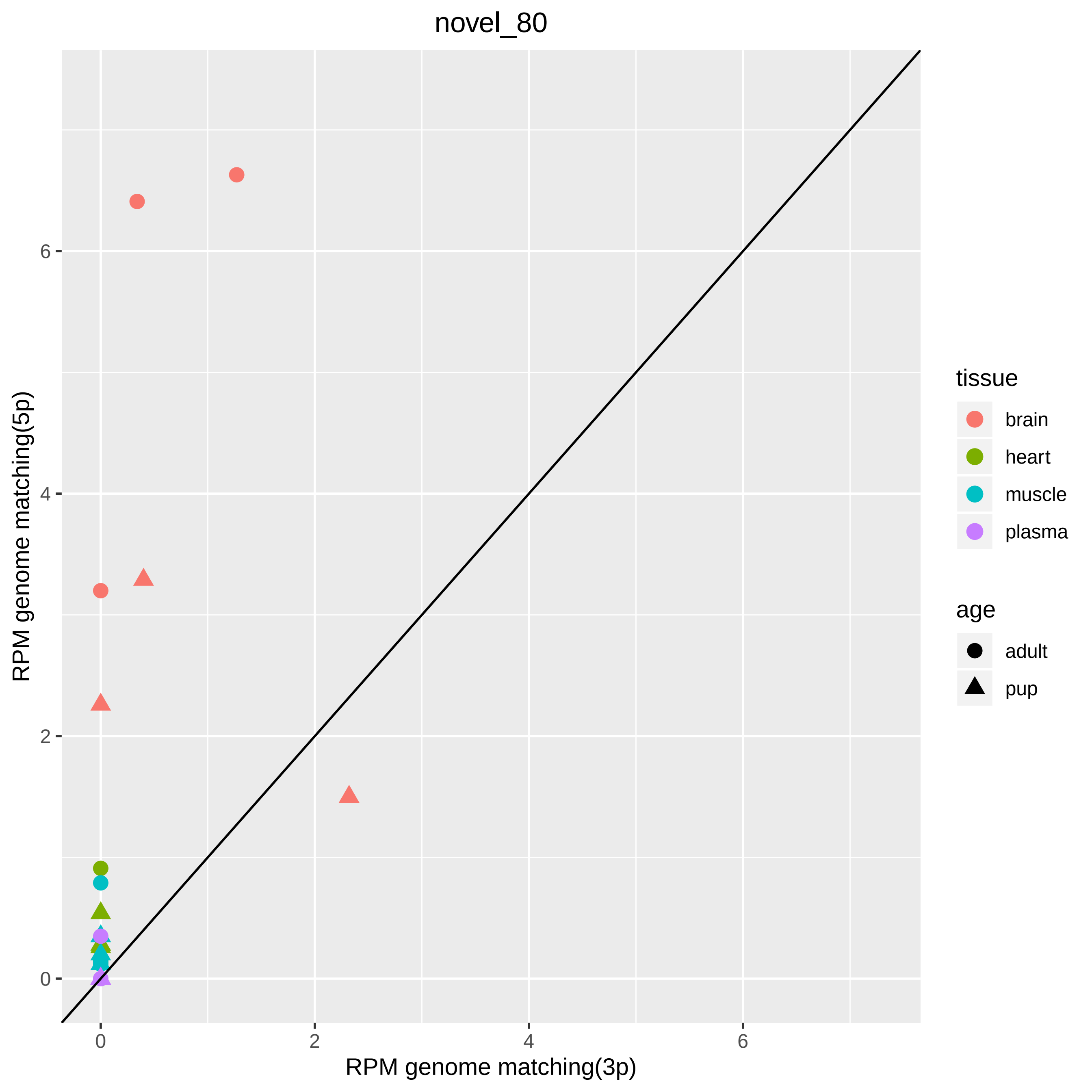

novel_80
RPM genome matching(5p)
RPM genome matching(3p)
0
2
4
6
tissue
brain
heart
muscle
plasma
age
adult
pup

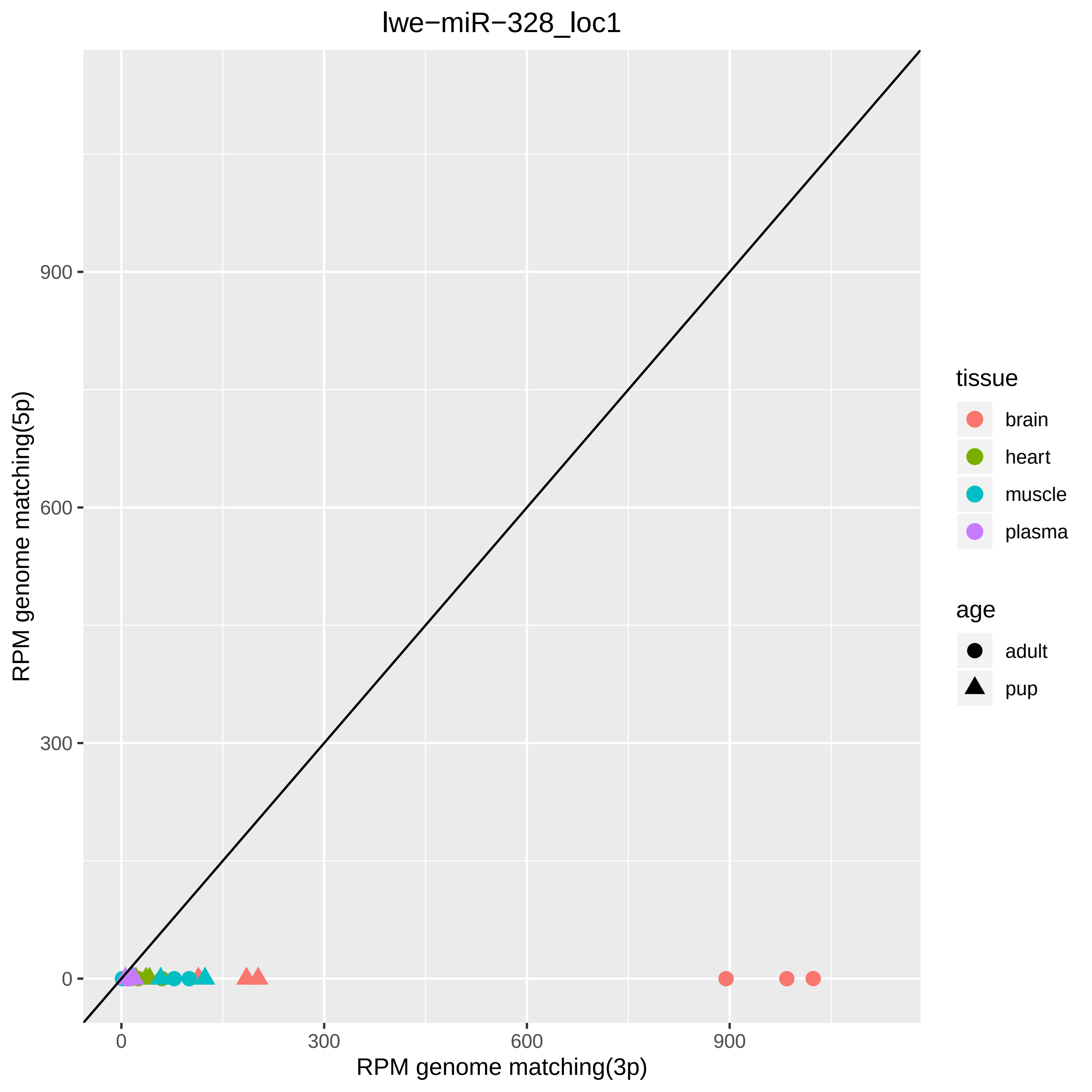

lwe−miR−328_loc1
RPM genome matching(5p)
RPM genome matching(3p)
0
300
600
900
0
300
600
900
tissue
brain
heart
muscle
plasma
age
adult
pup

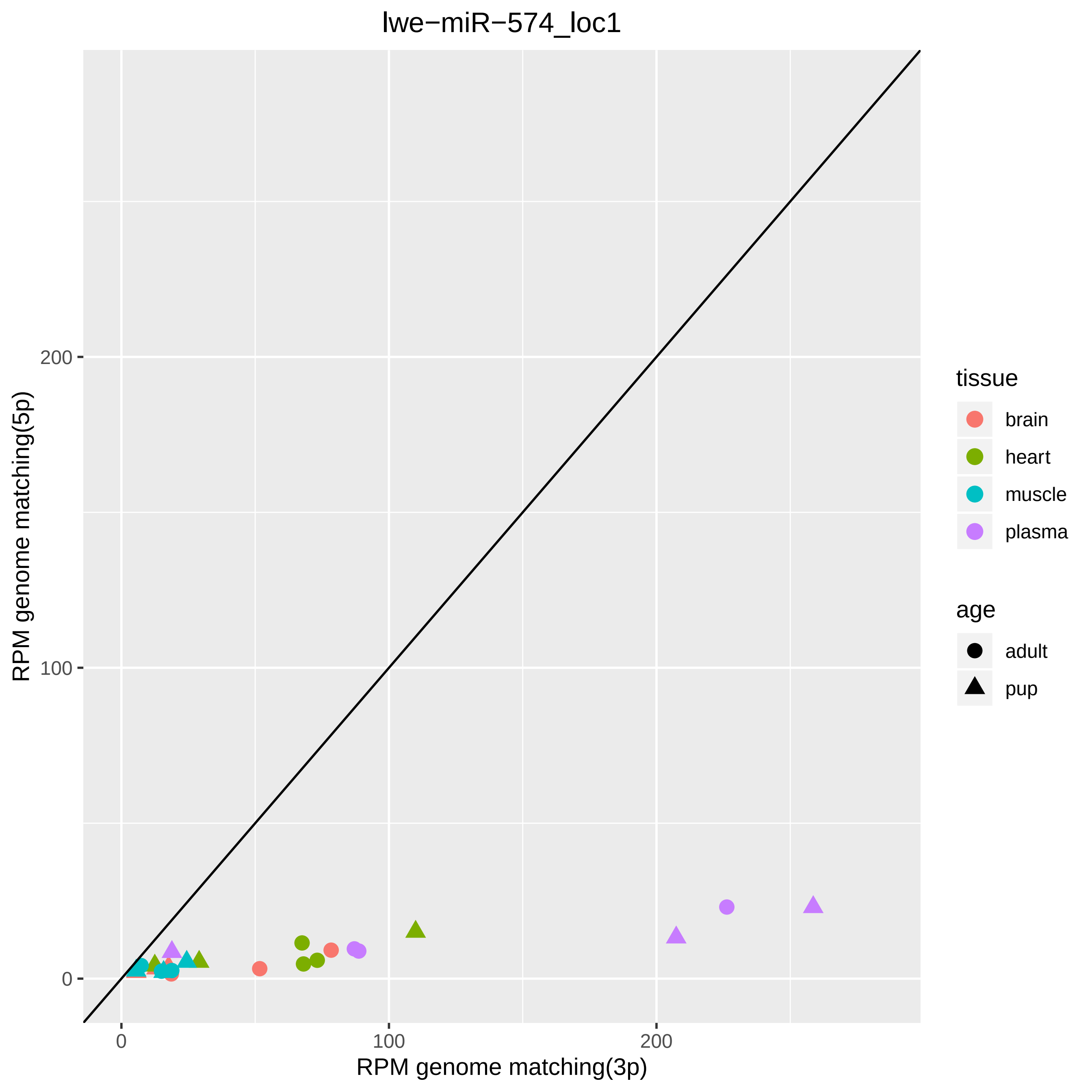
lwe−miR−574_loc1
RPM genome matching(5p)
RPM genome matching(3p)
0
100
200
0
100
200
tissue
brain
heart
muscle
plasma
age
adult
pup

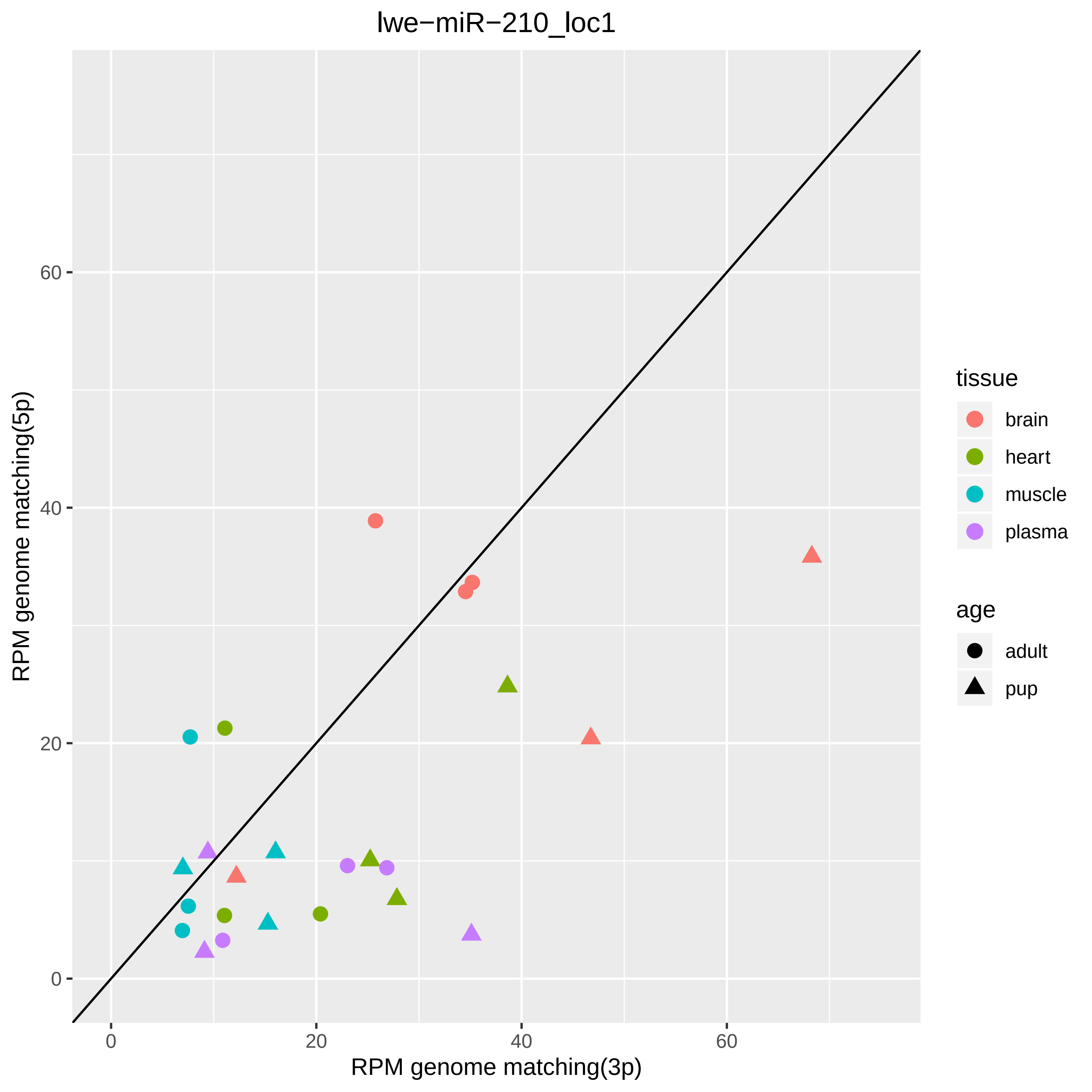
lwe−miR−210_loc1
RPM genome matching(5p)
RPM genome matching(3p)
0
20
40
60
0
20
40
60
tissue
brain
heart
muscle
plasma
age
adult
pup

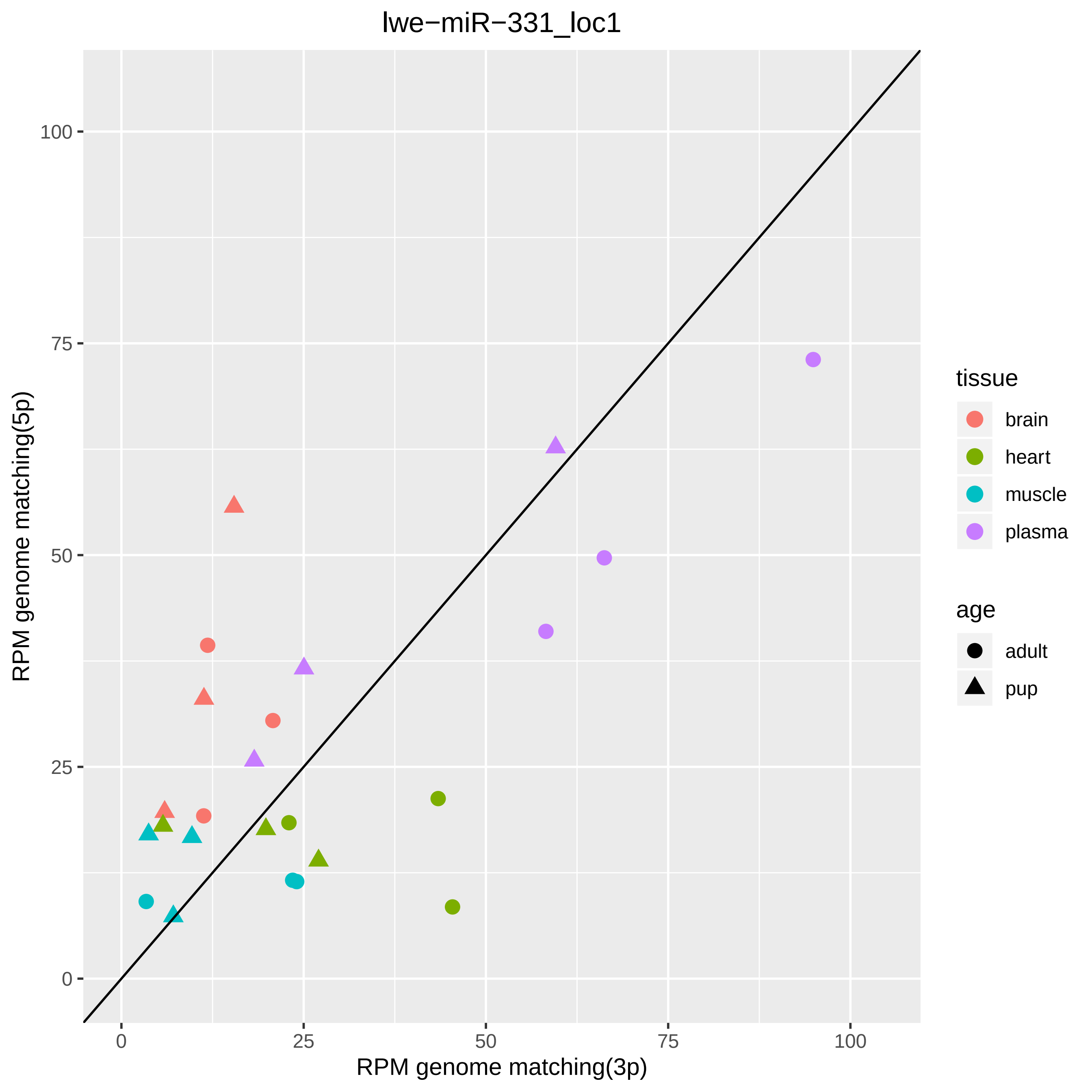
lwe−miR−331_loc1
RPM genome matching(5p)
RPM genome matching(3p)
0
25
50
75
100
tissue
brain
heart
muscle
plasma
age
adult
pup

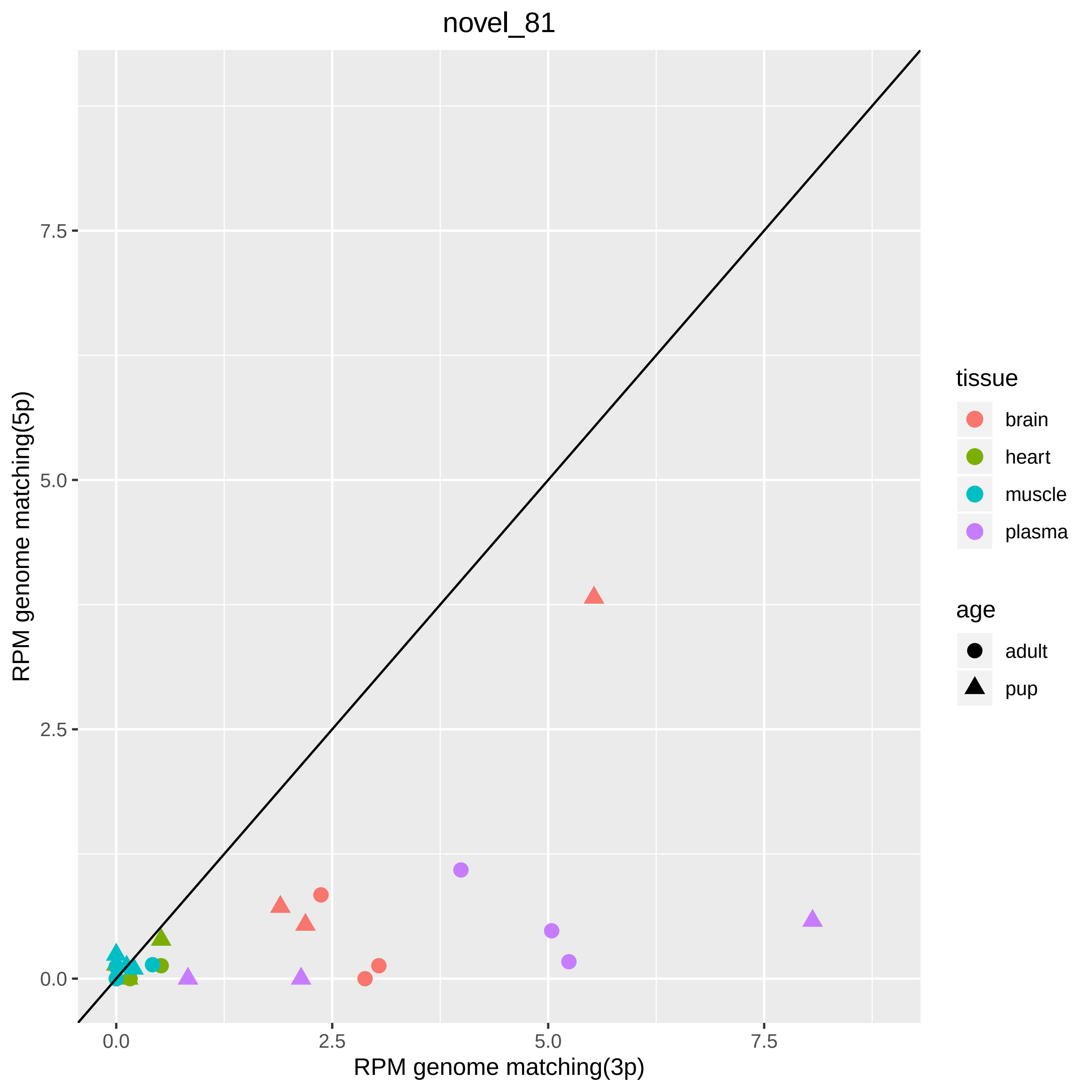
novel_81
RPM genome matching(5p)
RPM genome matching(3p)
0.0
2.5
5.0
7.5
tissue
brain
heart
muscle
plasma
age
adult
pup

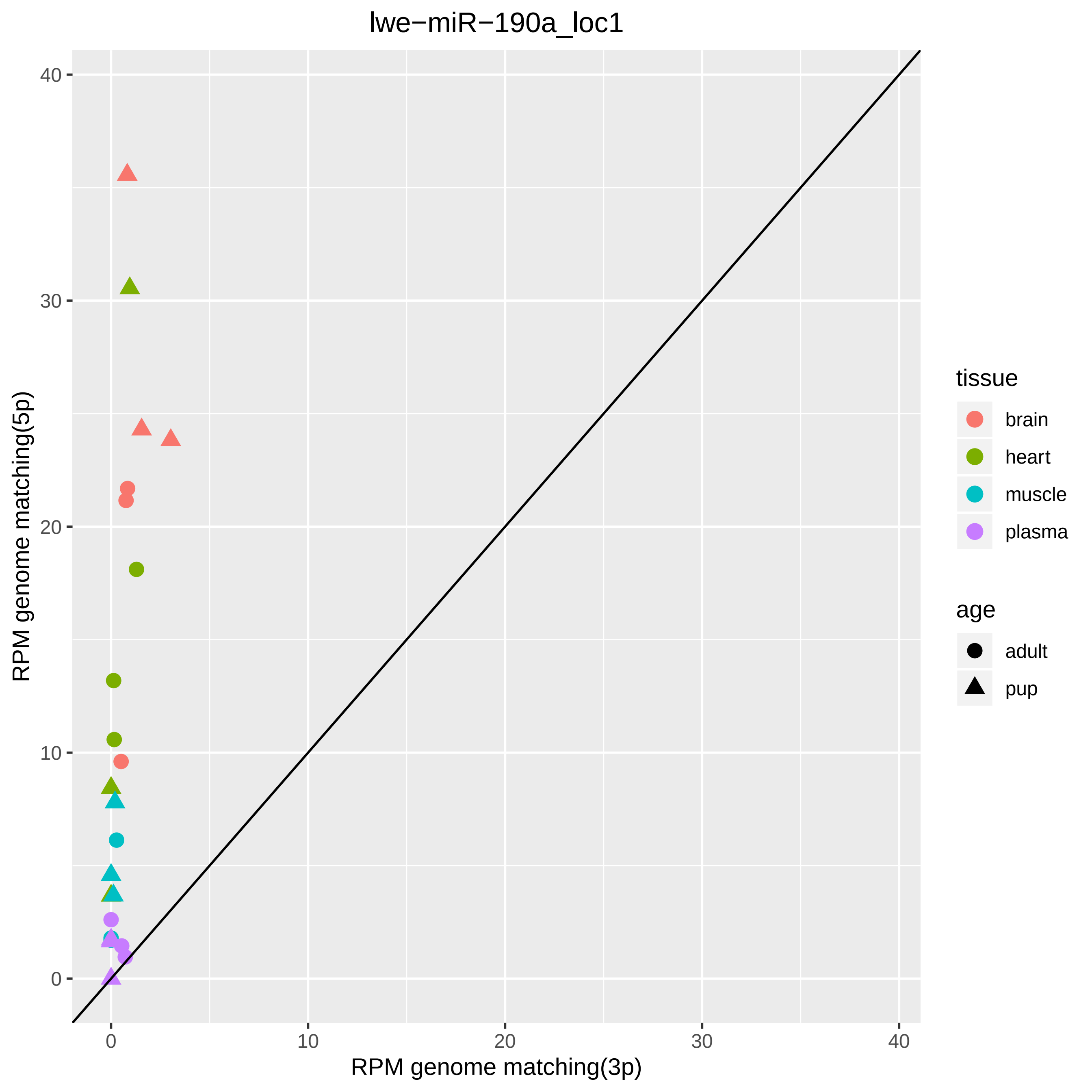

lwe−miR−190a_loc1
RPM genome matching(5p)
RPM genome matching(3p)
0
10
20
30
40
0
10
20
30
40
tissue
brain
heart
muscle
plasma
age
adult
pup

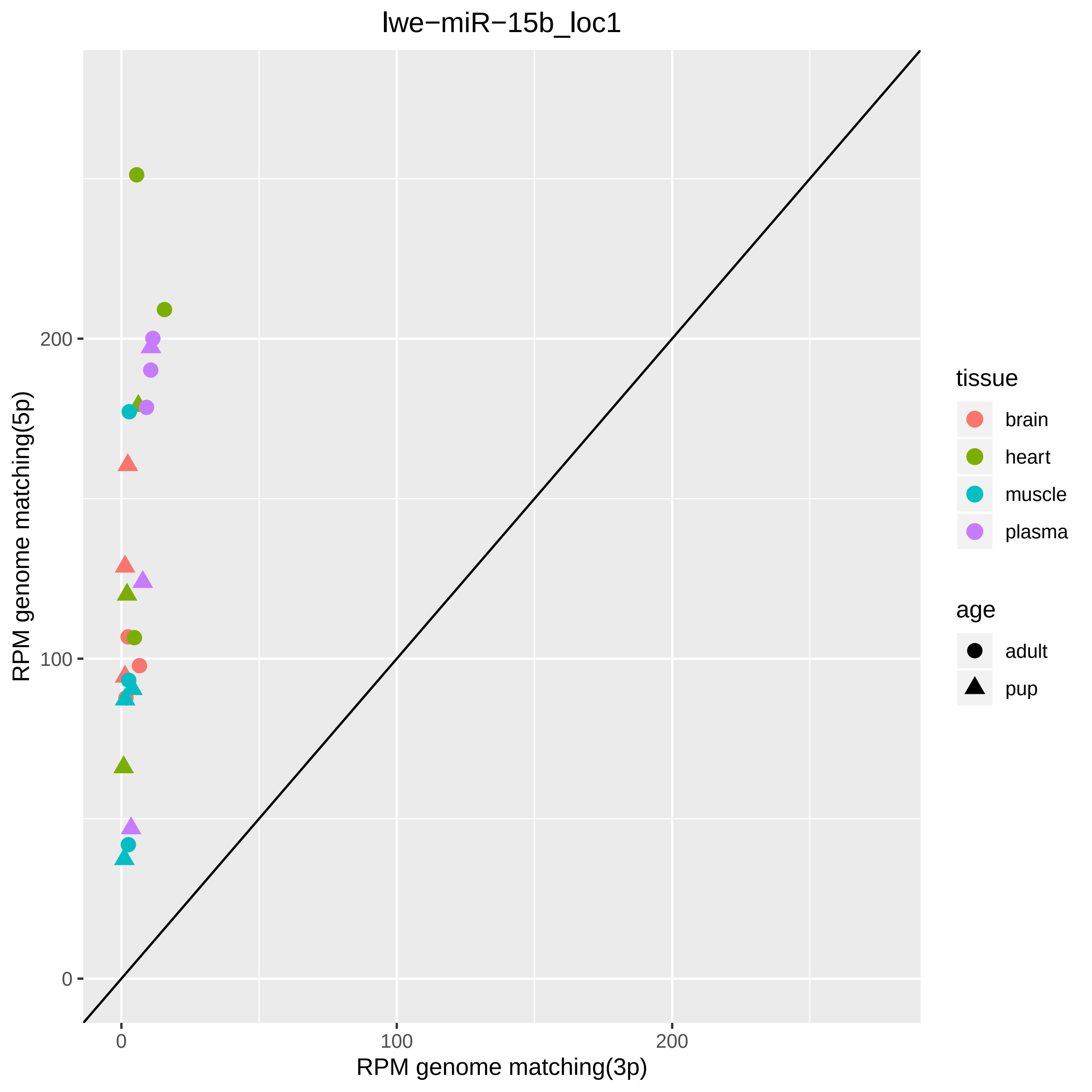

lwe−miR−15b_loc1
RPM genome matching(5p)
RPM genome matching(3p)
0
100
200
0
100
200
tissue
brain
heart
muscle
plasma
age
adult
pup

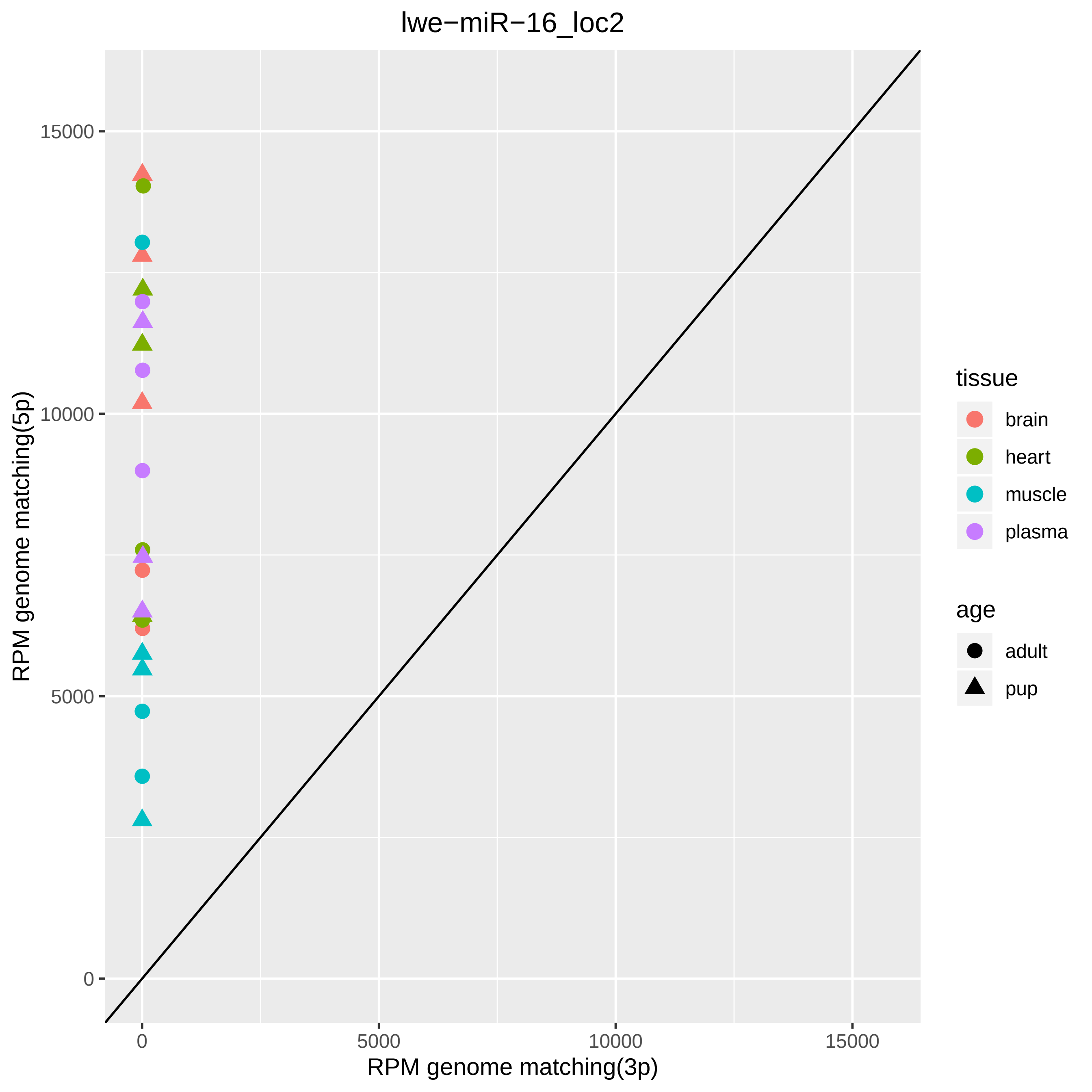

lwe−miR−16_loc2
15000
10000
5000
0
0
5000
10000
15000
RPM genome matching(5p)
RPM genome matching(3p)
tissue
brain
heart
muscle
plasma
age
adult
pup

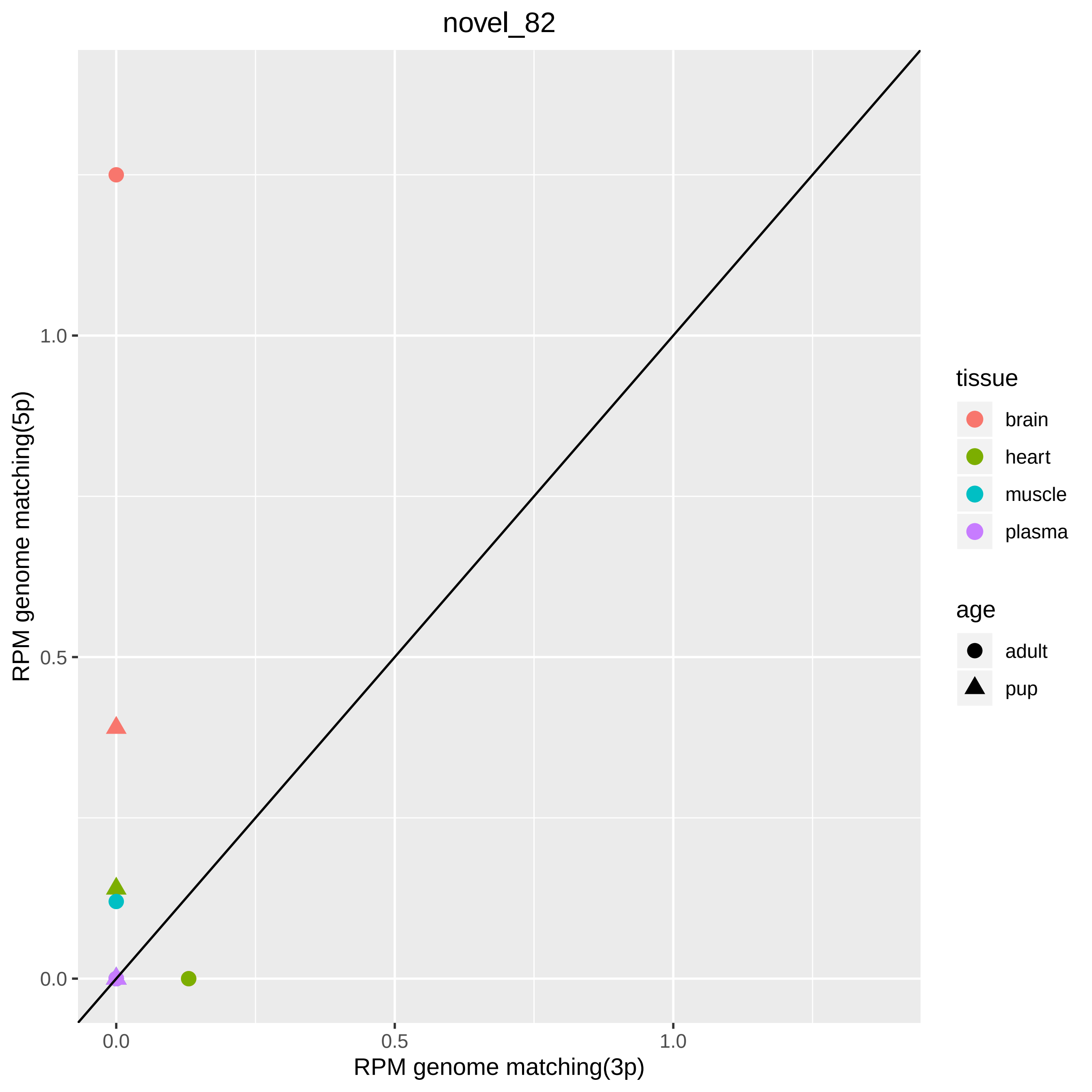
novel_82
RPM genome matching(5p)
RPM genome matching(3p)
0.0
0.5
1.0
tissue
brain
heart
muscle
plasma
age
adult
pup

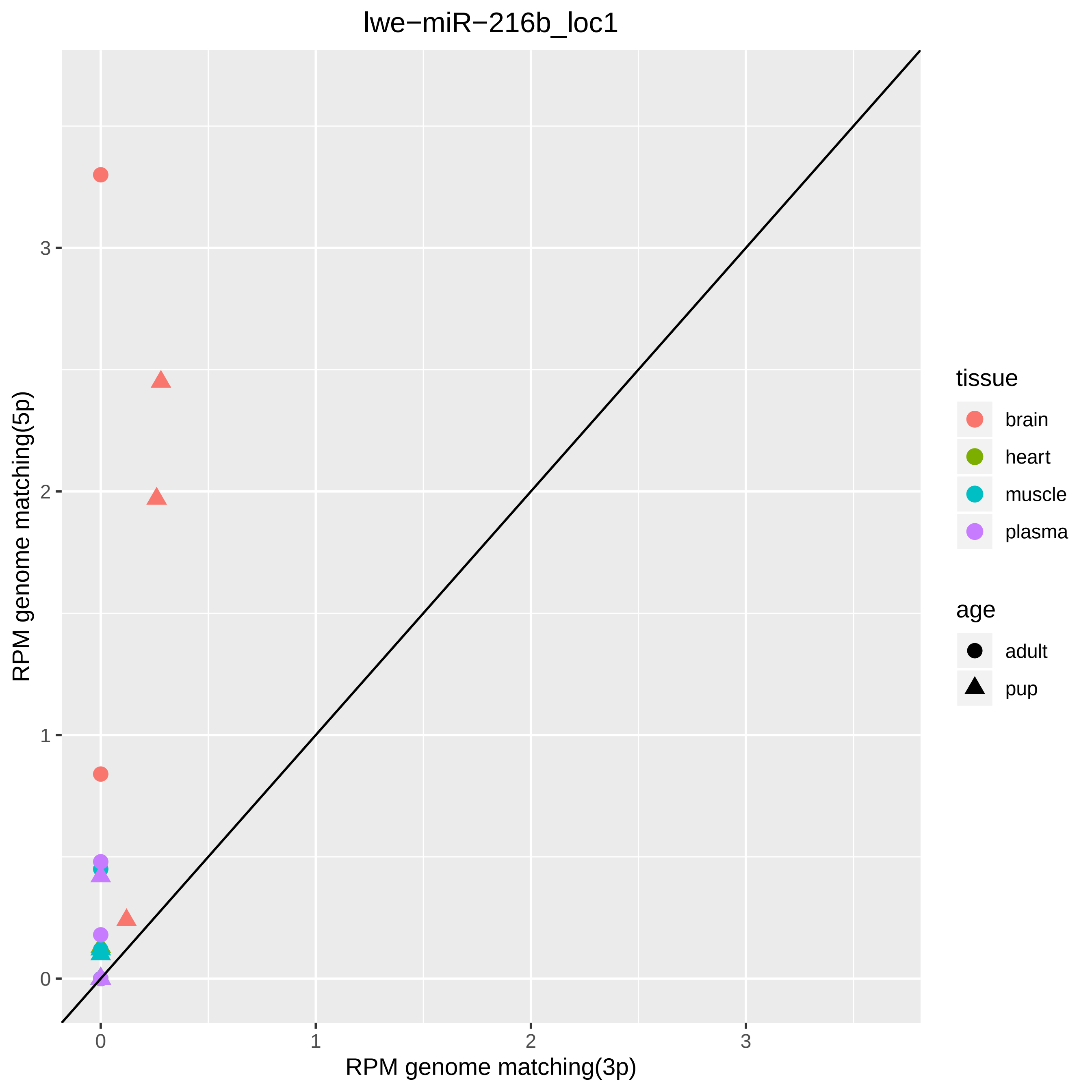

lwe−miR−216b_loc1
RPM genome matching(5p)
RPM genome matching(3p)
0
1
2
3
tissue
brain
heart
muscle
plasma
age
adult
pup

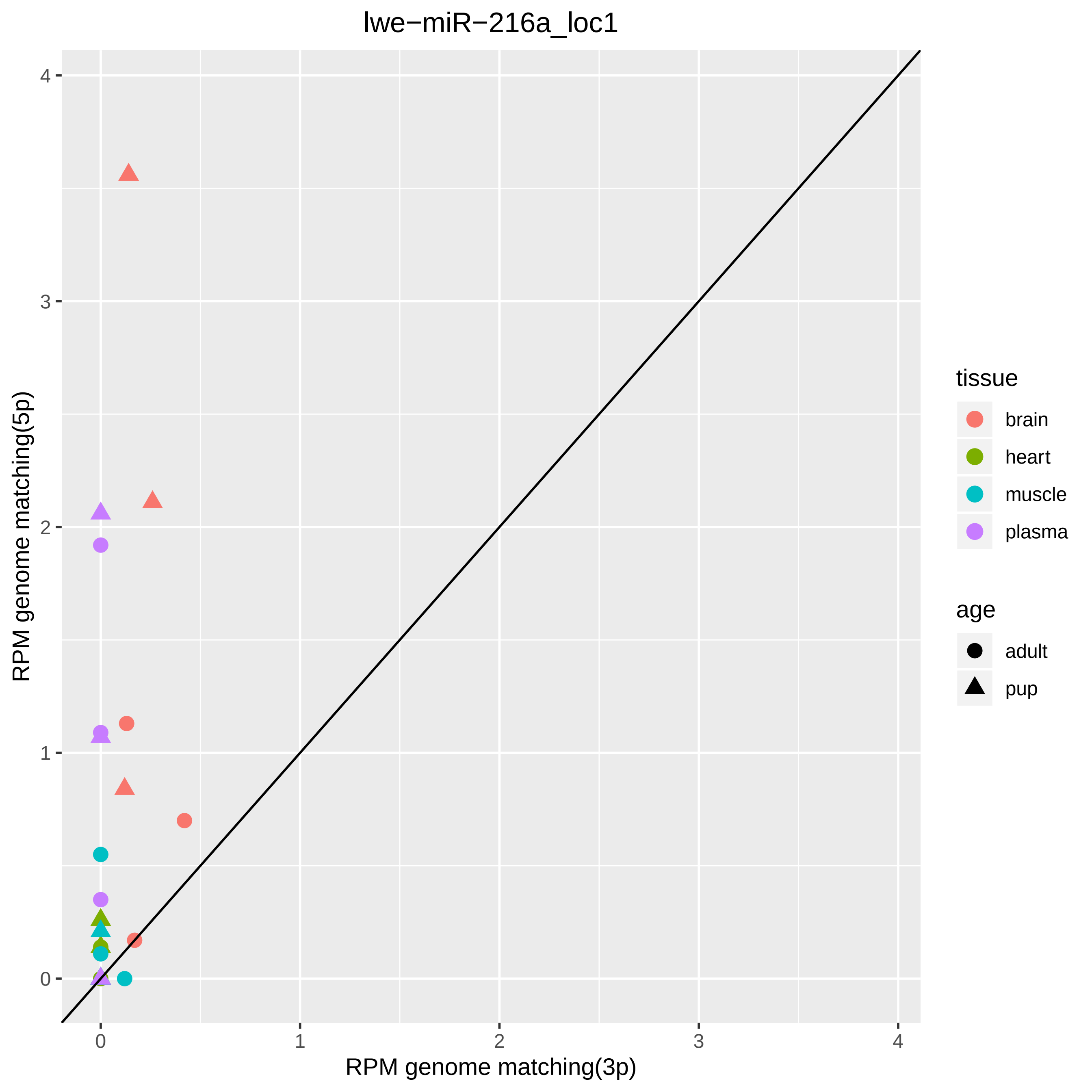

lwe−miR−216a_loc1
RPM genome matching(5p)
RPM genome matching(3p)
0
1
2
3
4
0
1
2
3
4
tissue
brain
heart
muscle
plasma
age
adult
pup

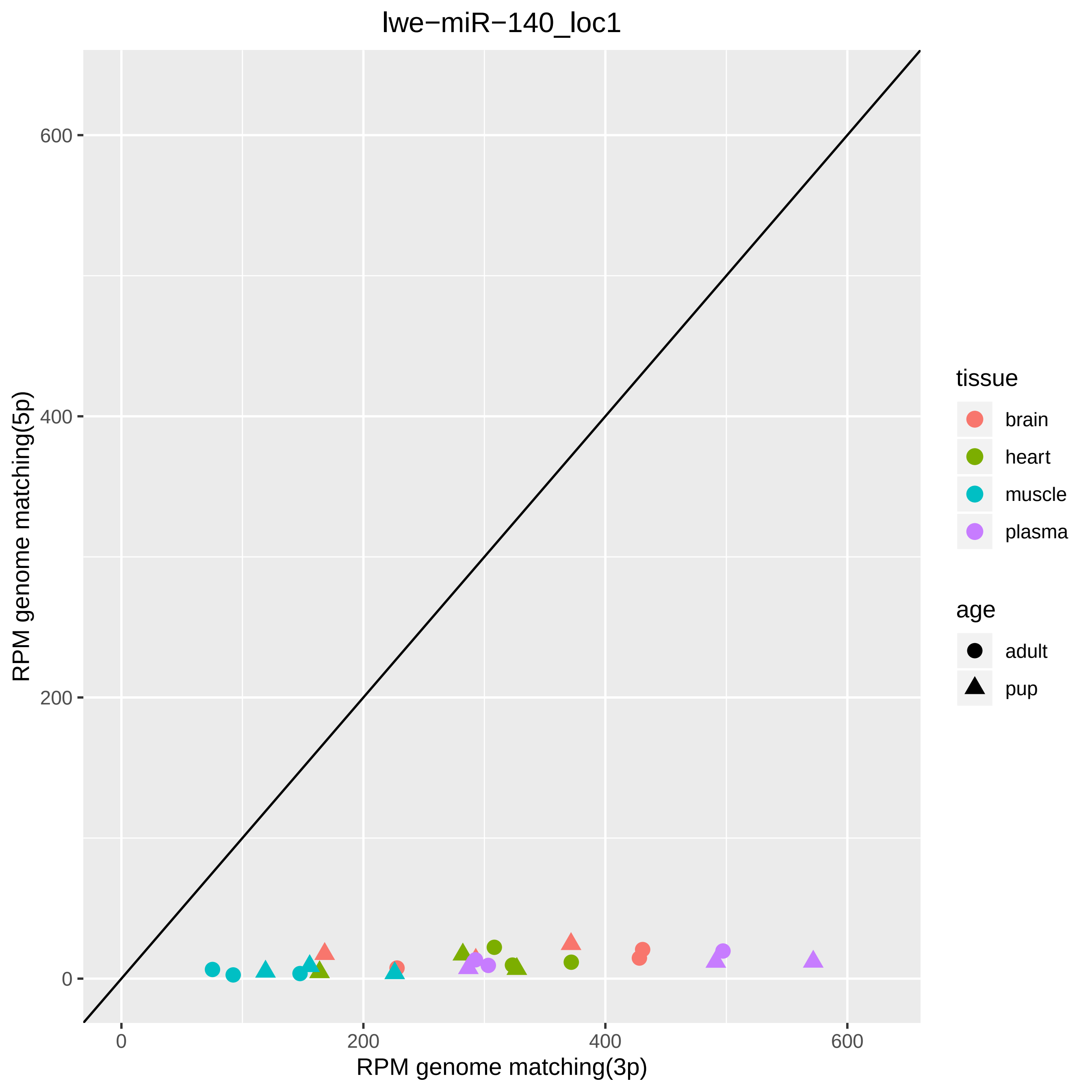
lwe−miR−140_loc1
RPM genome matching(5p)
RPM genome matching(3p)
0
200
400
600
0
200
400
600
tissue
brain
heart
muscle
plasma
age
adult
pup

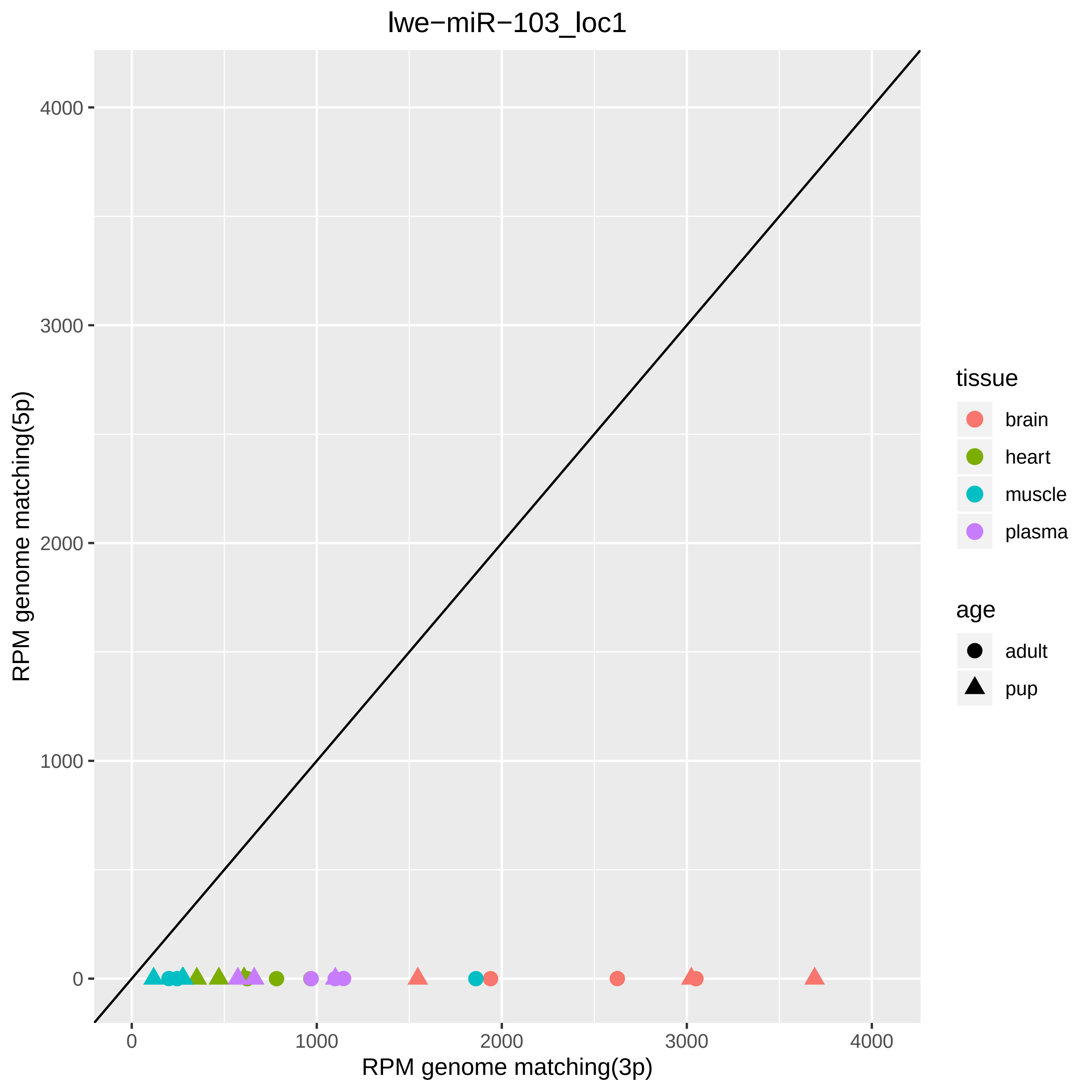

lwe−miR−103_loc1
RPM genome matching(5p)
RPM genome matching(3p)
0
1000
2000
3000
4000
tissue
brain
heart
muscle
plasma
age
adult
pup

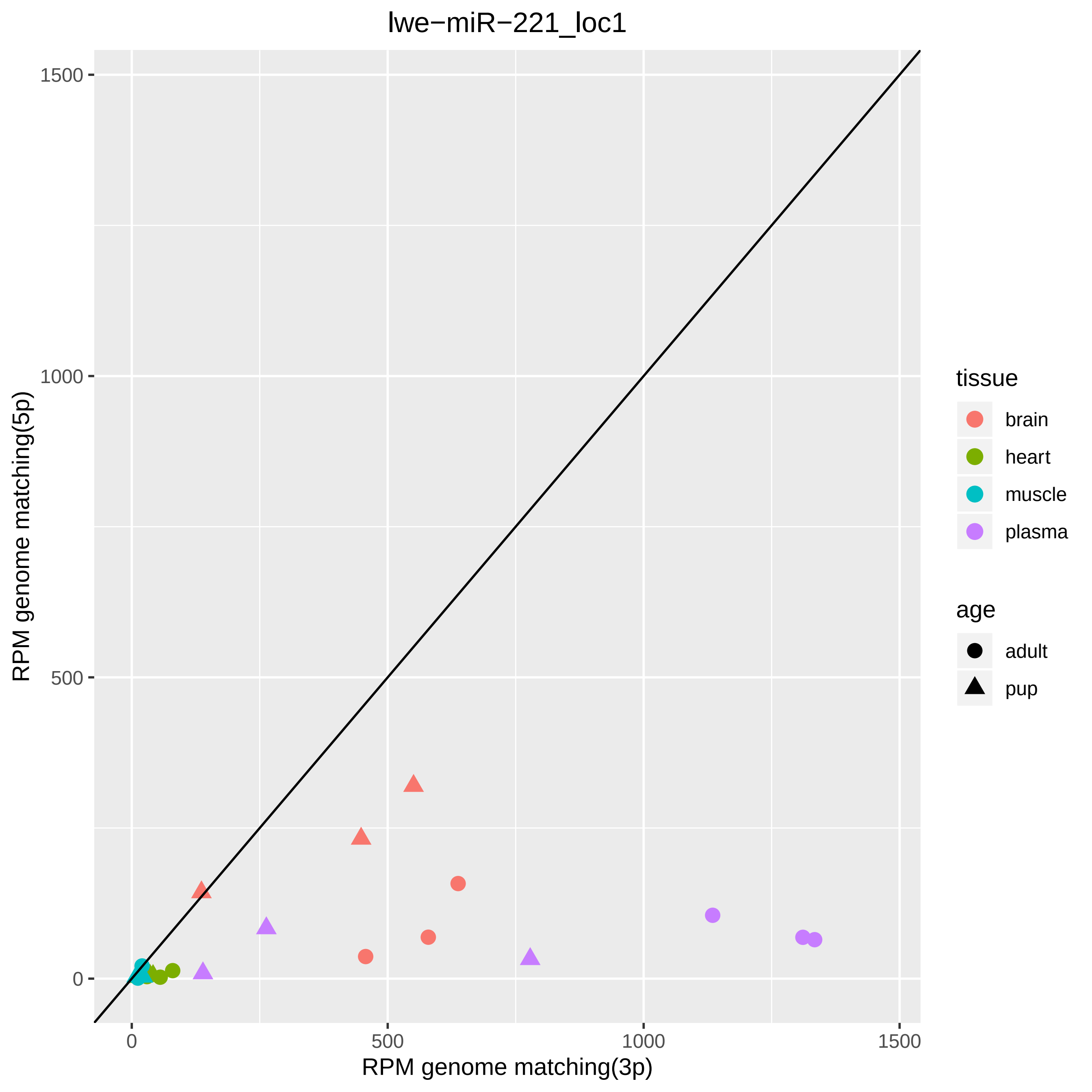
lwe−miR−221_loc1
RPM genome matching(5p)
RPM genome matching(3p)
0
500
1000
1500
tissue
brain
heart
muscle
plasma
age
adult
pup

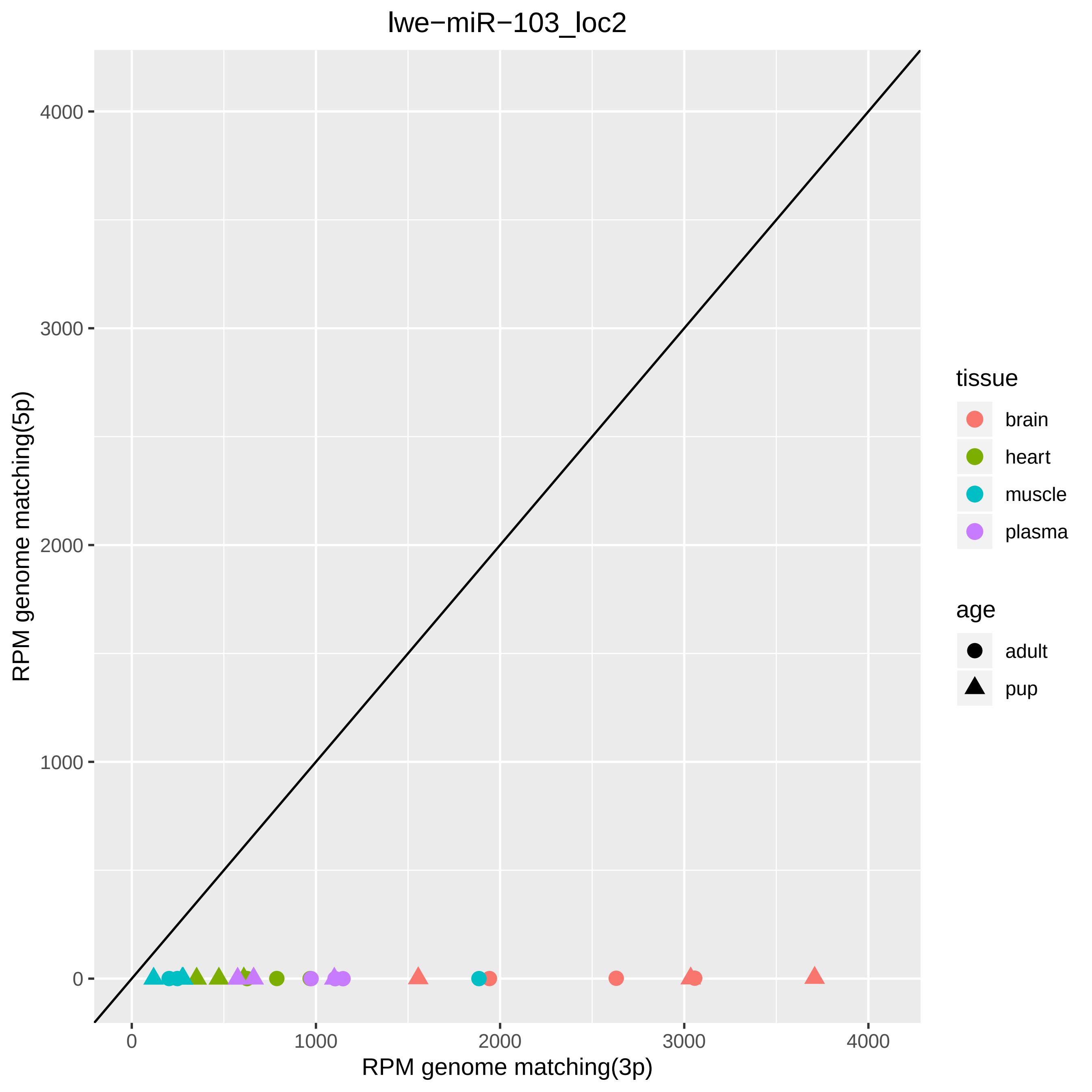

lwe−miR−103_loc2
RPM genome matching(5p)
RPM genome matching(3p)
0
1000
2000
3000
4000
tissue
brain
heart
muscle
plasma
age
adult
pup

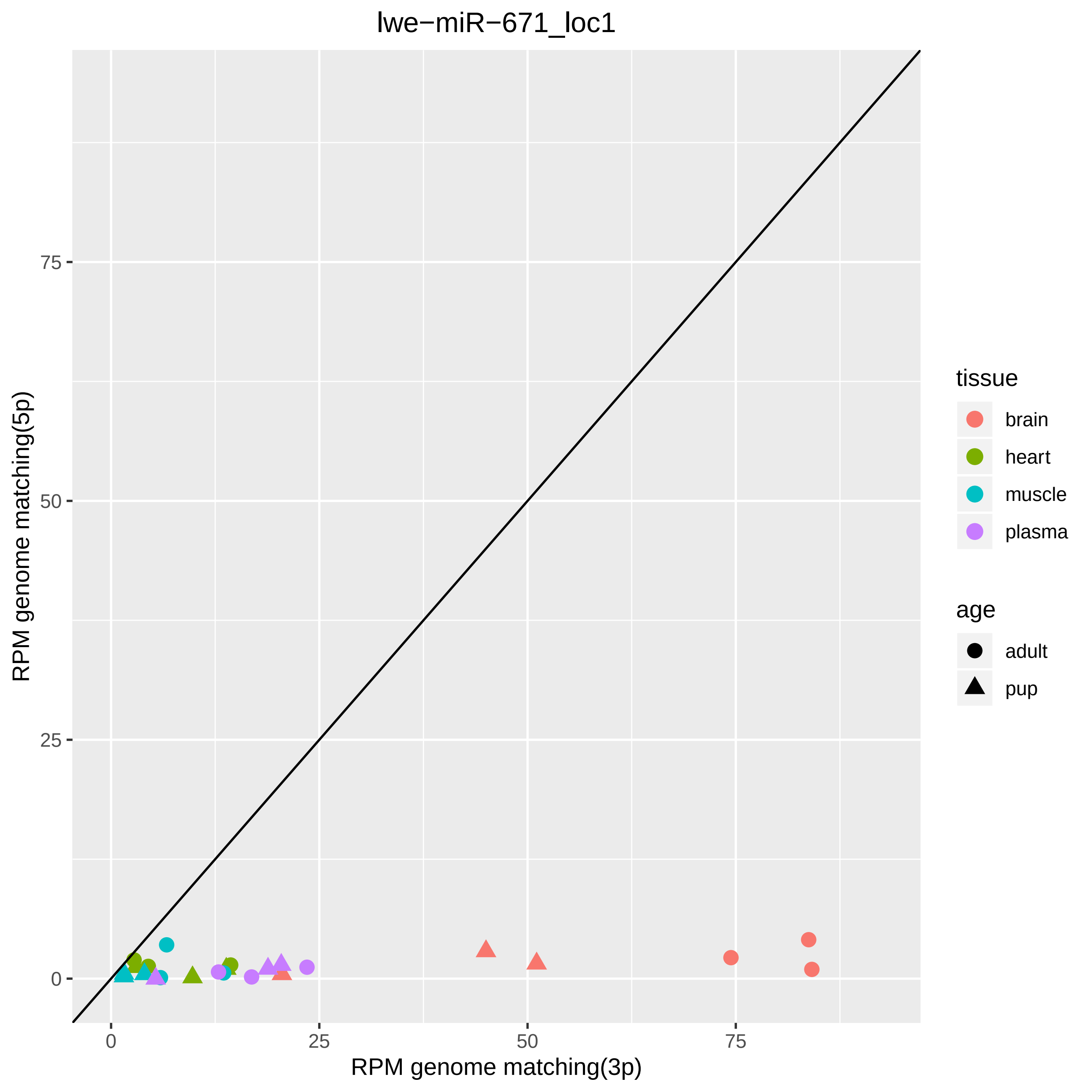
lwe−miR−671_loc1
RPM genome matching(5p)
RPM genome matching(3p)
0
25
50
75
tissue
brain
heart
muscle
plasma
age
adult
pup

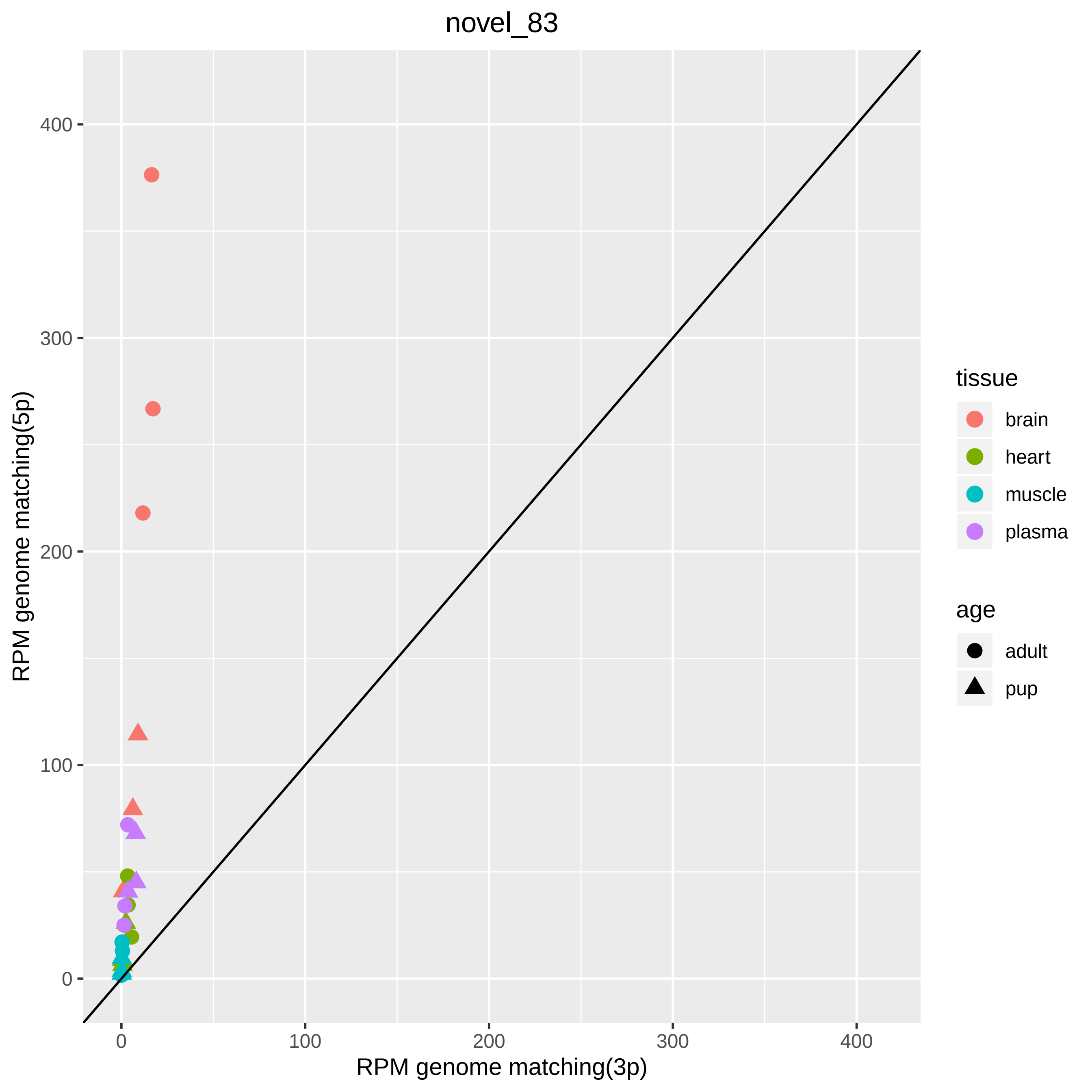
novel_83
RPM genome matching(5p)
RPM genome matching(3p)
0
100
200
300
400
tissue
brain
heart
muscle
plasma
age
adult
pup

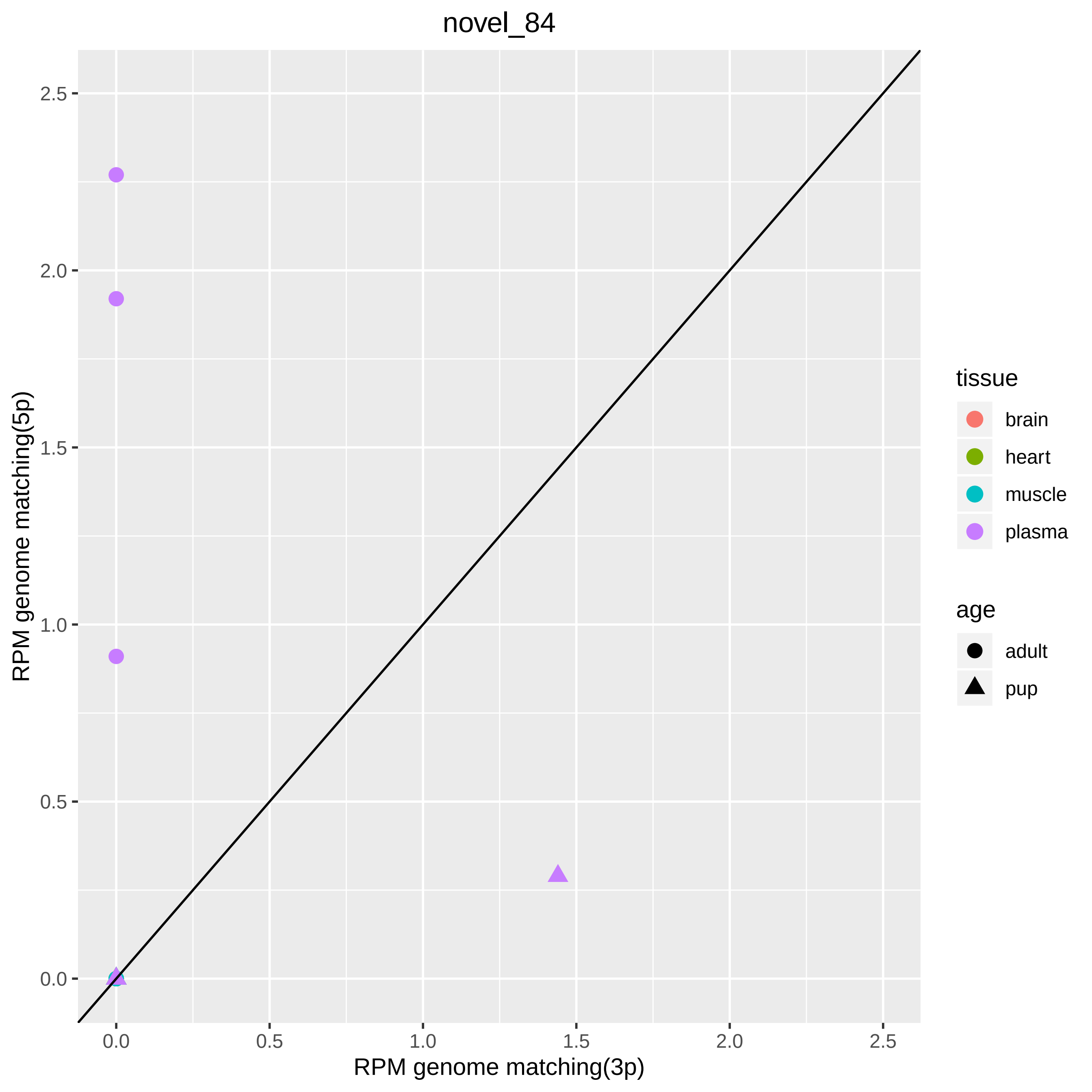

novel_84
RPM genome matching(5p)
RPM genome matching(3p)
0.0
0.5
1.0
1.5
2.0
2.5
tissue
brain
heart
muscle
plasma
age
adult
pup

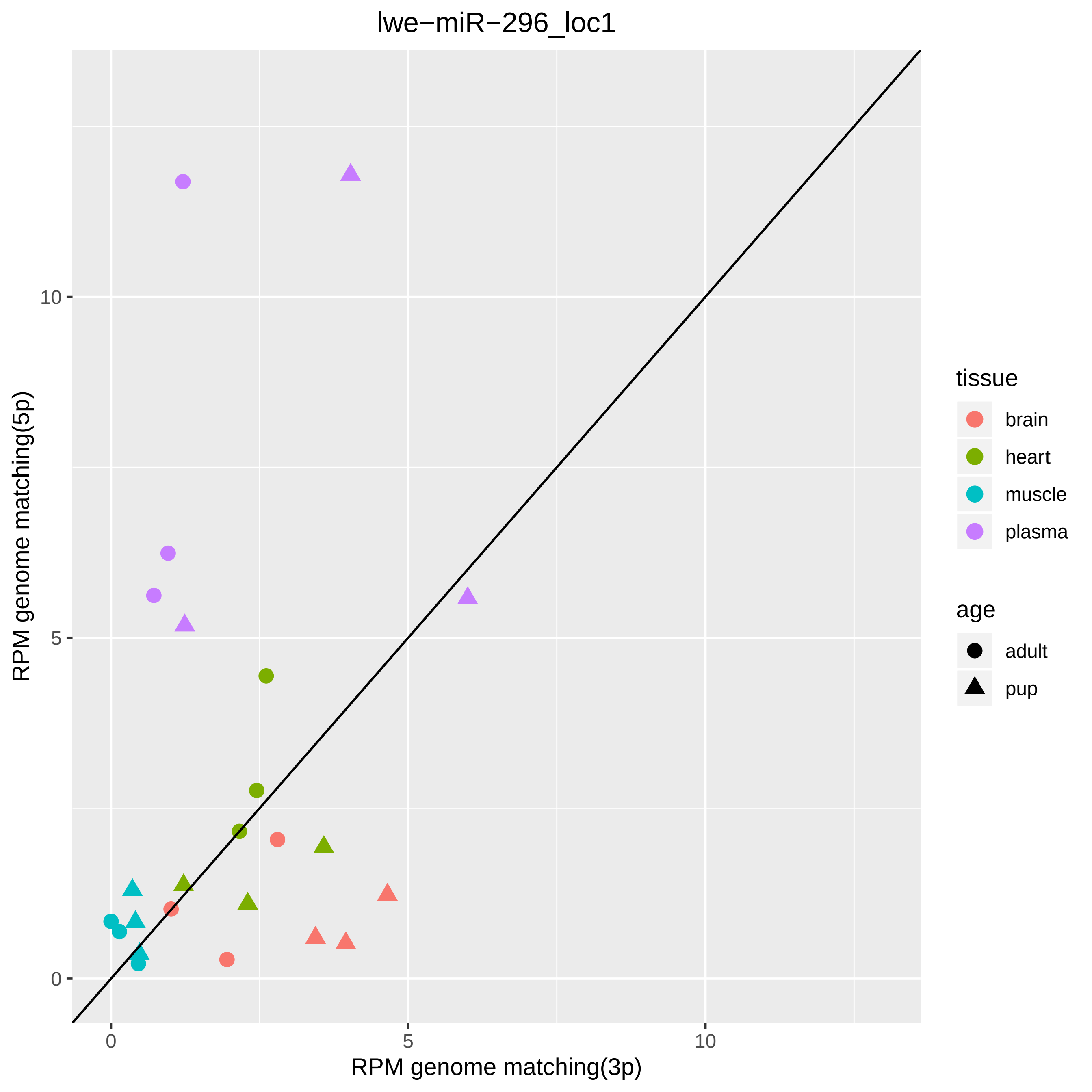

lwe−miR−296_loc1
RPM genome matching(5p)
RPM genome matching(3p)
0
5
10
tissue
brain
heart
muscle
plasma
age
adult
pup

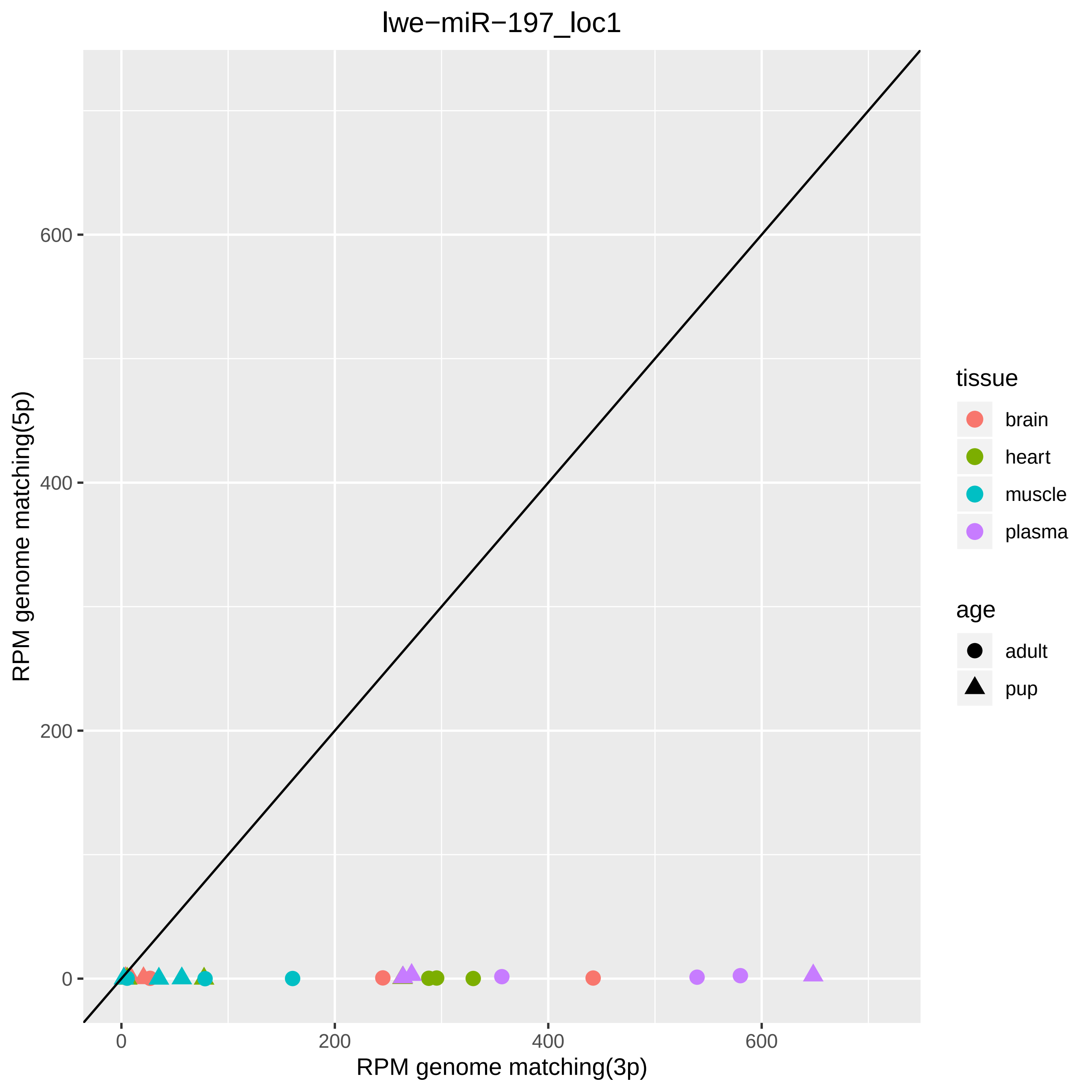

lwe−miR−197_loc1
RPM genome matching(5p)
RPM genome matching(3p)
600
400
200
0
0
200
400
600
tissue
brain
heart
muscle
plasma
age
adult
pup

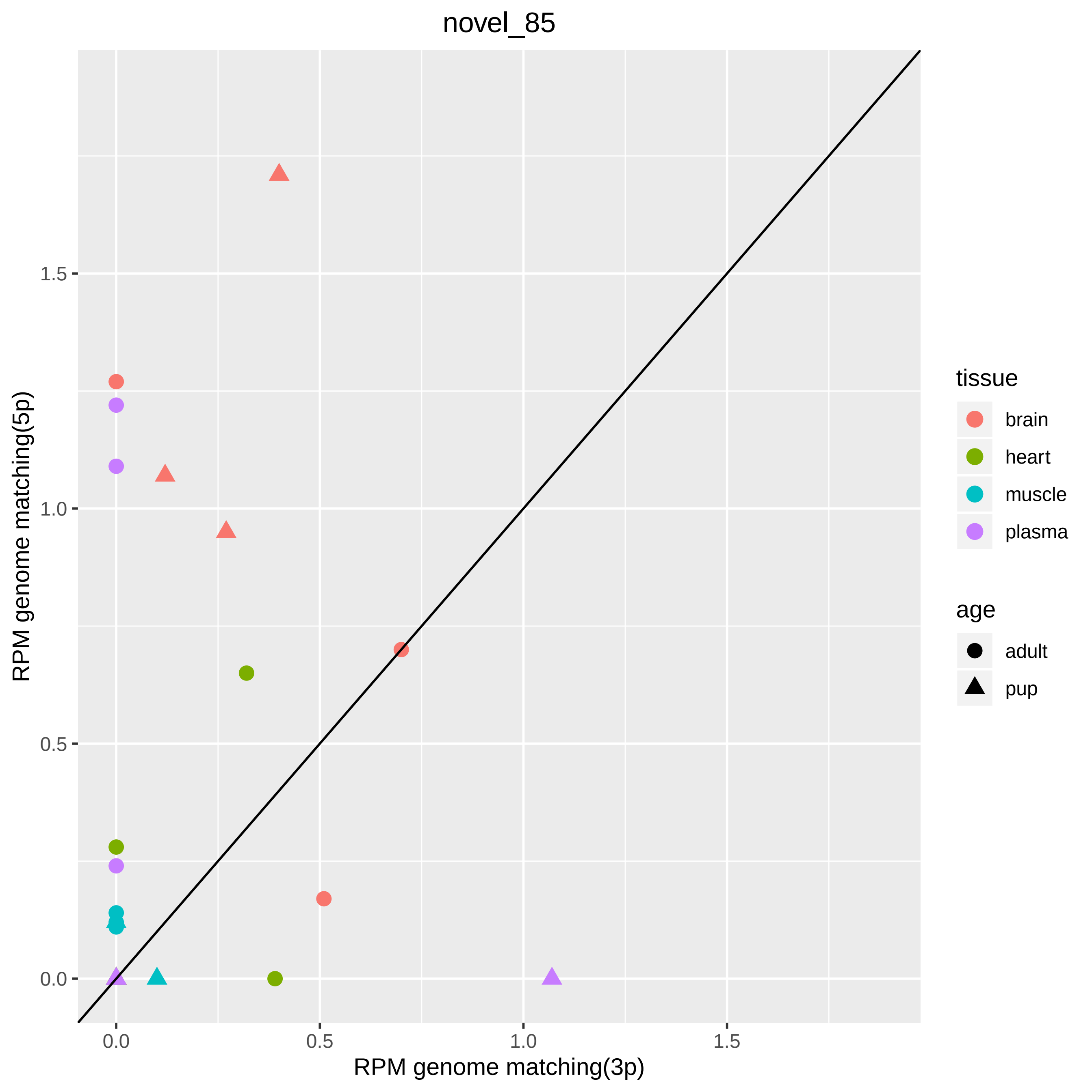

novel_85
RPM genome matching(5p)
RPM genome matching(3p)
0.0
0.5
1.0
1.5
0.0
0.5
1.0
1.5
tissue
brain
heart
muscle
plasma
age
adult
pup

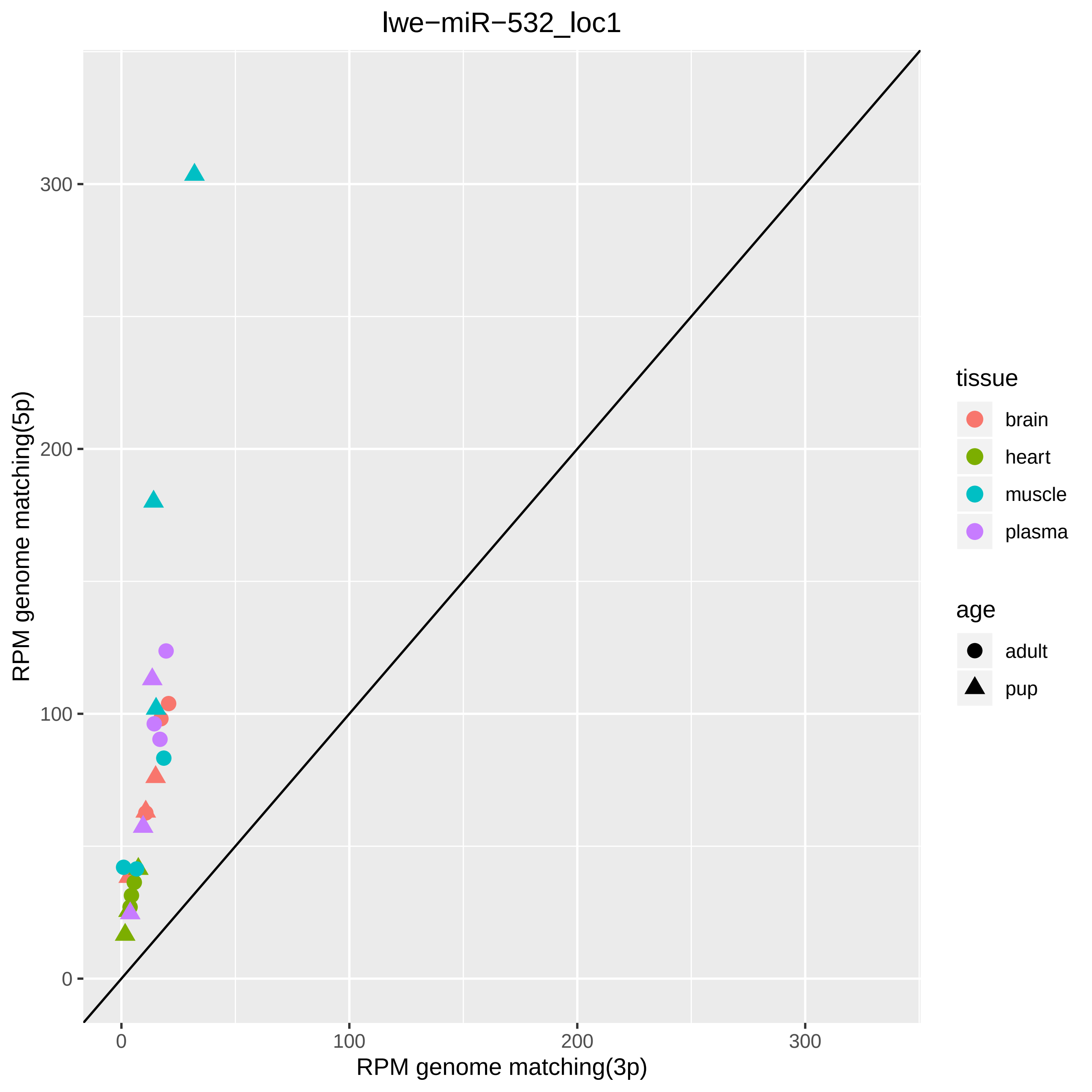

lwe−miR−532_loc1
RPM genome matching(5p)
RPM genome matching(3p)
0
100
200
300
0
100
200
300
tissue
brain
heart
muscle
plasma
age
adult
pup

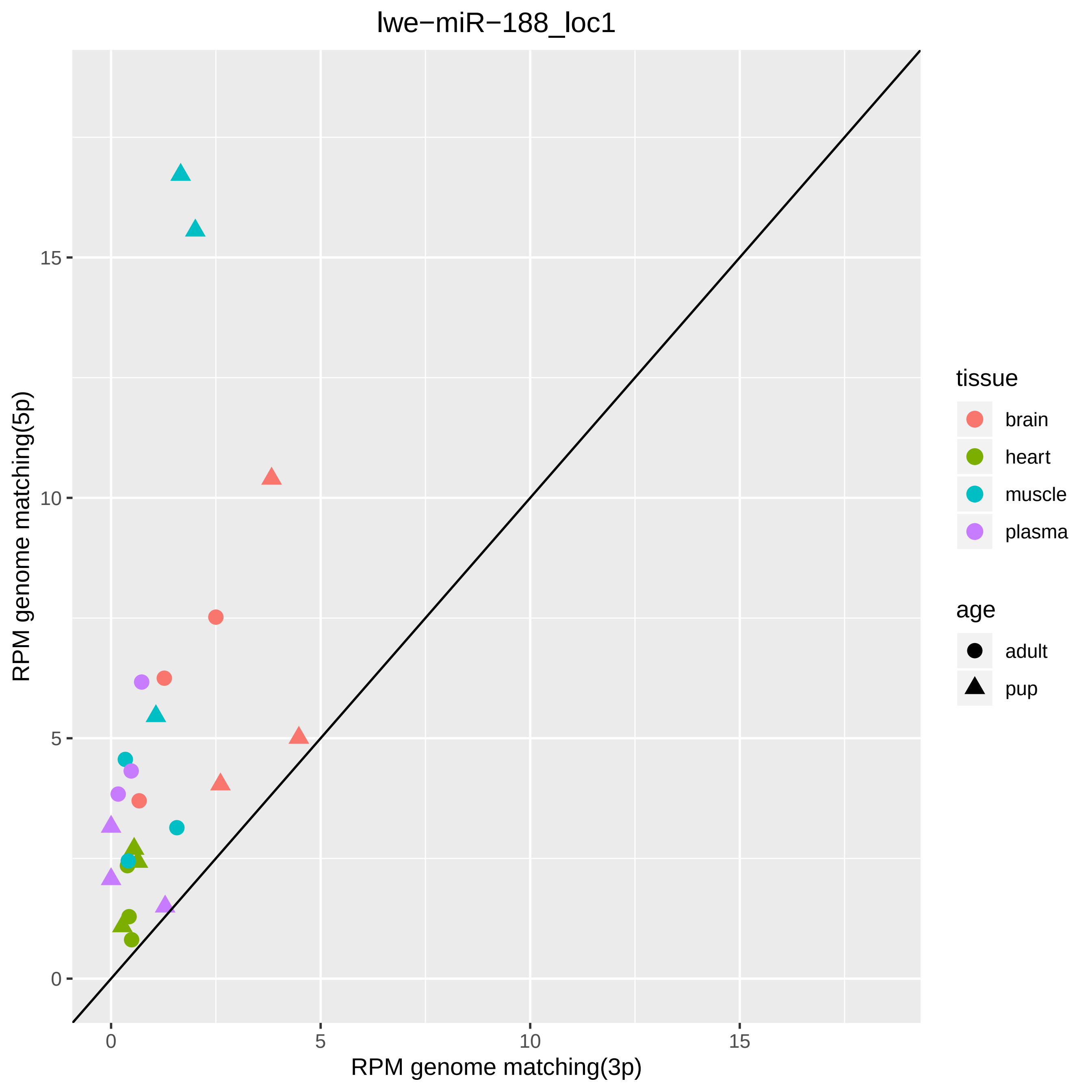

lwe−miR−188_loc1
RPM genome matching(5p)
RPM genome matching(3p)
0
5
10
15
0
5
10
15
tissue
brain
heart
muscle
plasma
age
adult
pup

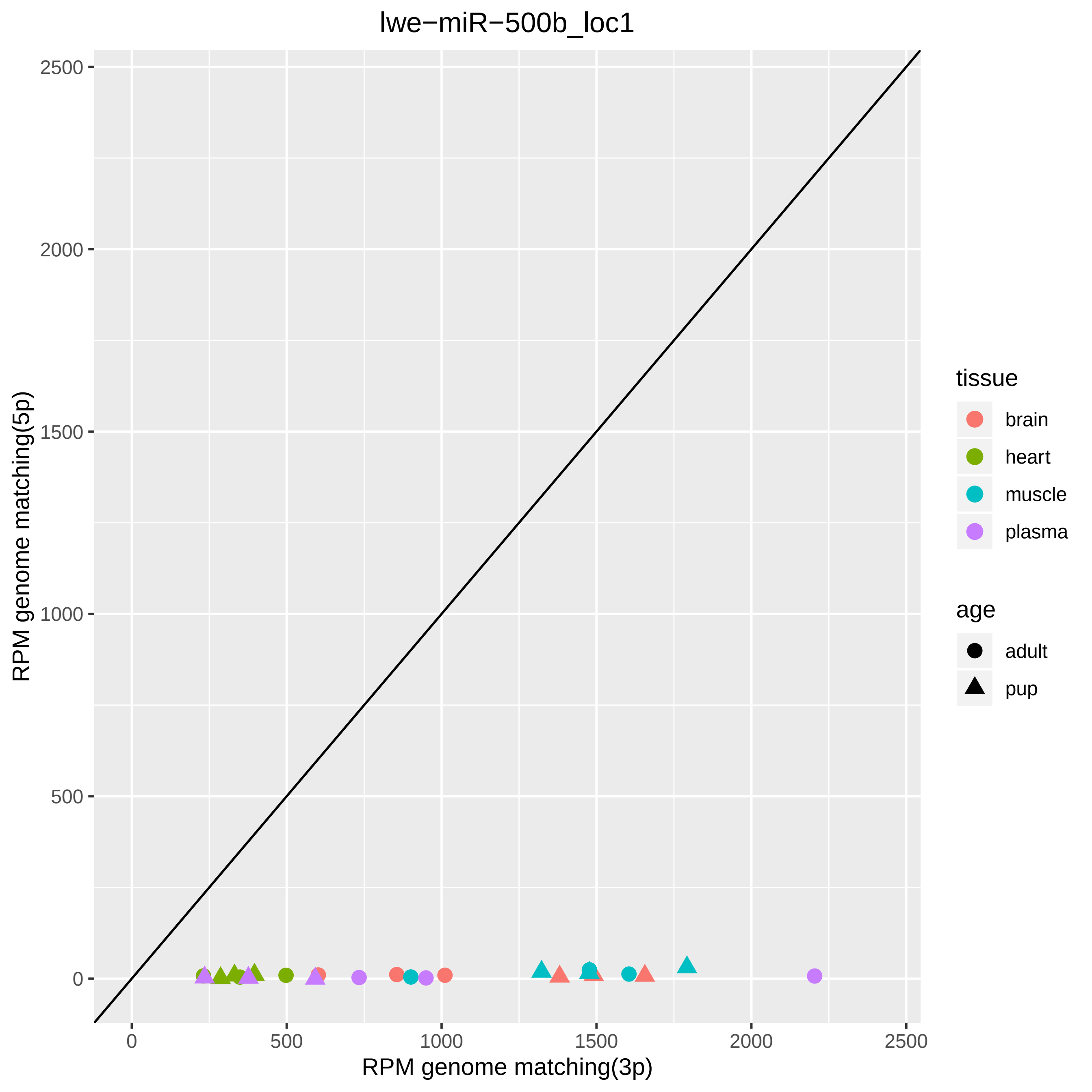

lwe−miR−500b_loc1
RPM genome matching(5p)
RPM genome matching(3p)
0
500
1000
1500
2000
2500
tissue
brain
heart
muscle
plasma
age
adult
pup

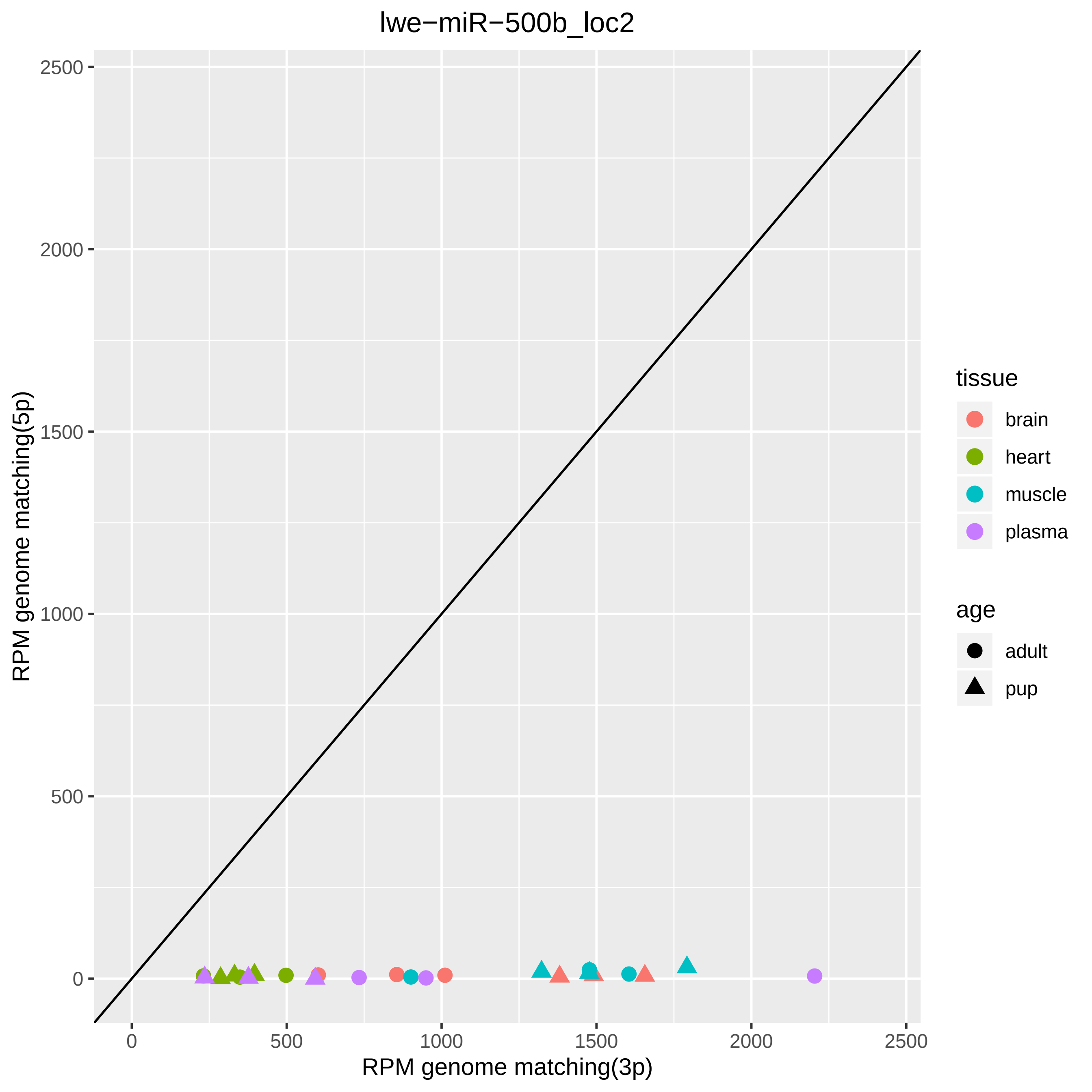

lwe−miR−500b_loc2
RPM genome matching(5p)
RPM genome matching(3p)
0
500
1000
1500
2000
2500
tissue
brain
heart
muscle
plasma
age
adult
pup

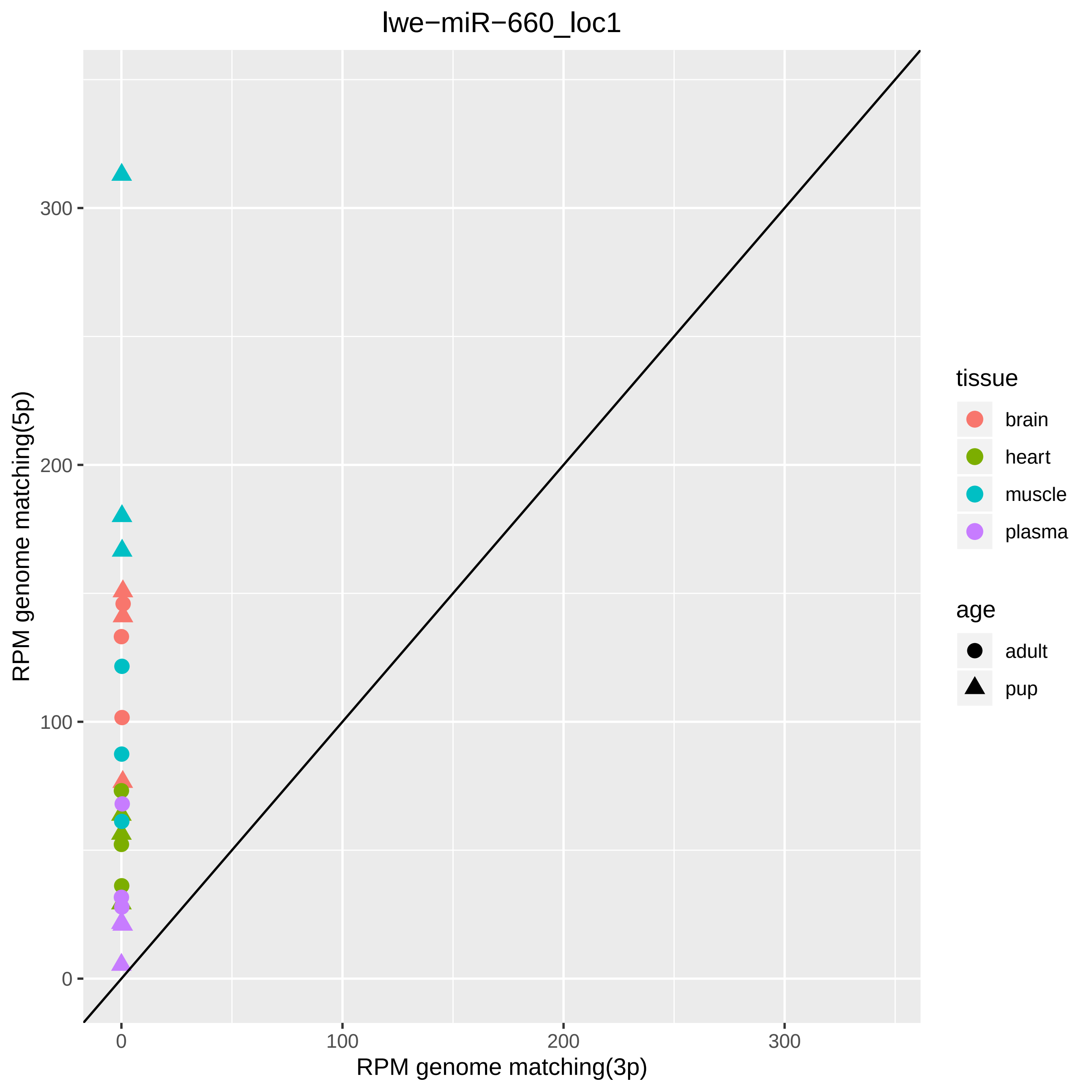
lwe−miR−660_loc1
RPM genome matching(5p)
RPM genome matching(3p)
300
200
100
0
0
100
200
300
tissue
brain
heart
muscle
plasma
age
adult
pup

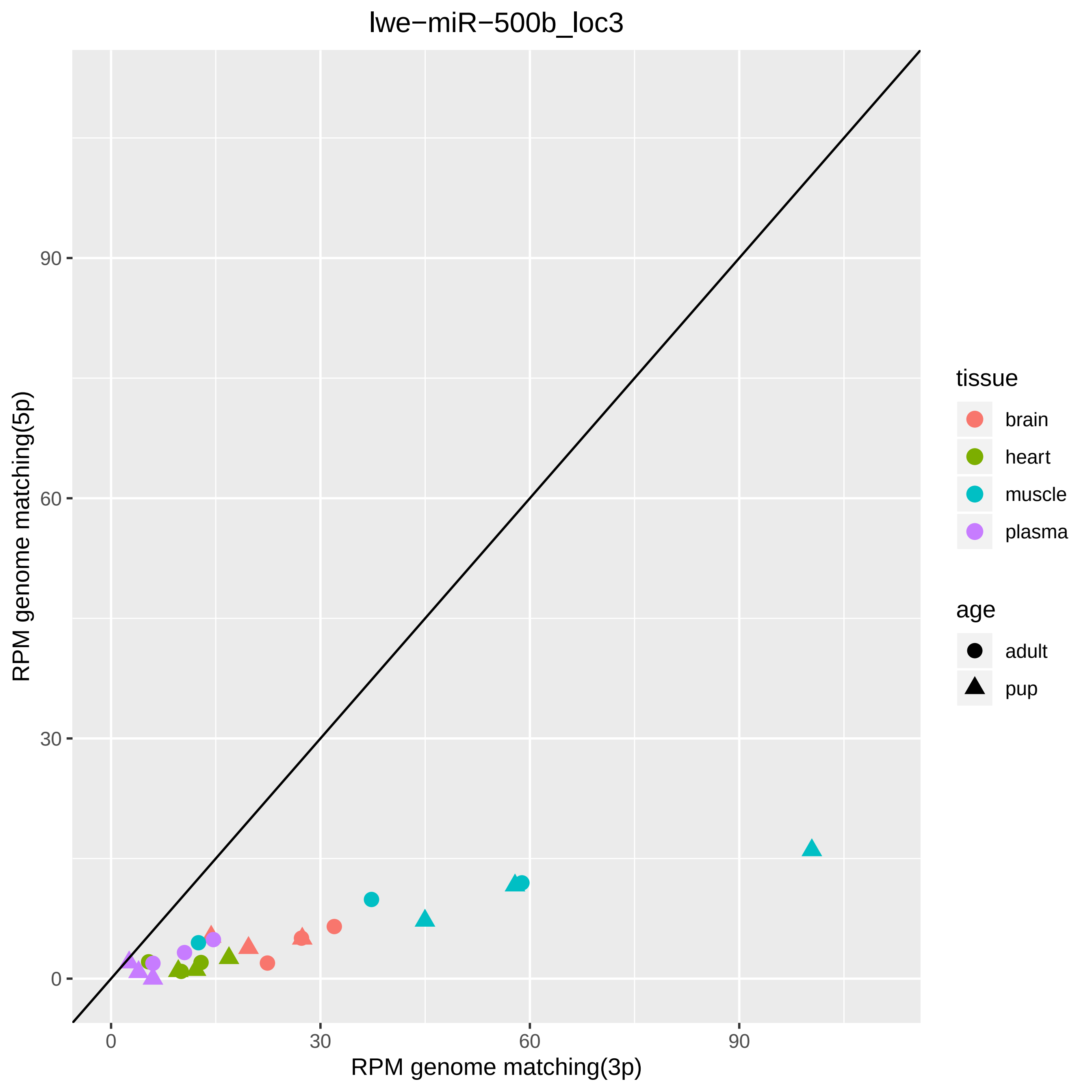

lwe−miR−500b_loc3
RPM genome matching(5p)
RPM genome matching(3p)
0
30
60
90
tissue
brain
heart
muscle
plasma
age
adult
pup

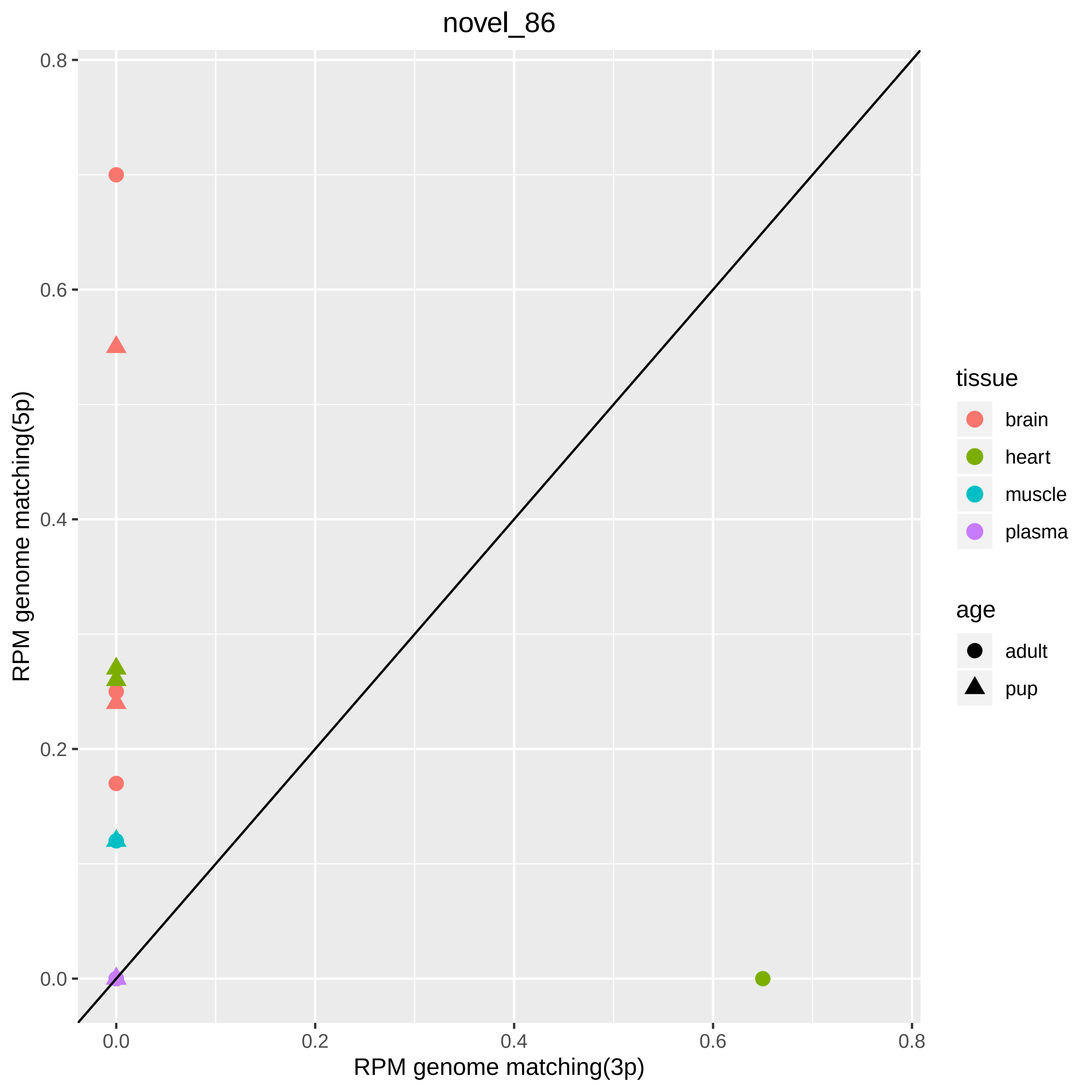
novel_86
RPM genome matching(5p)
RPM genome matching(3p)
0.0
0.2
0.4
0.6
0.8
tissue
brain
heart
muscle
plasma
age
adult
pup

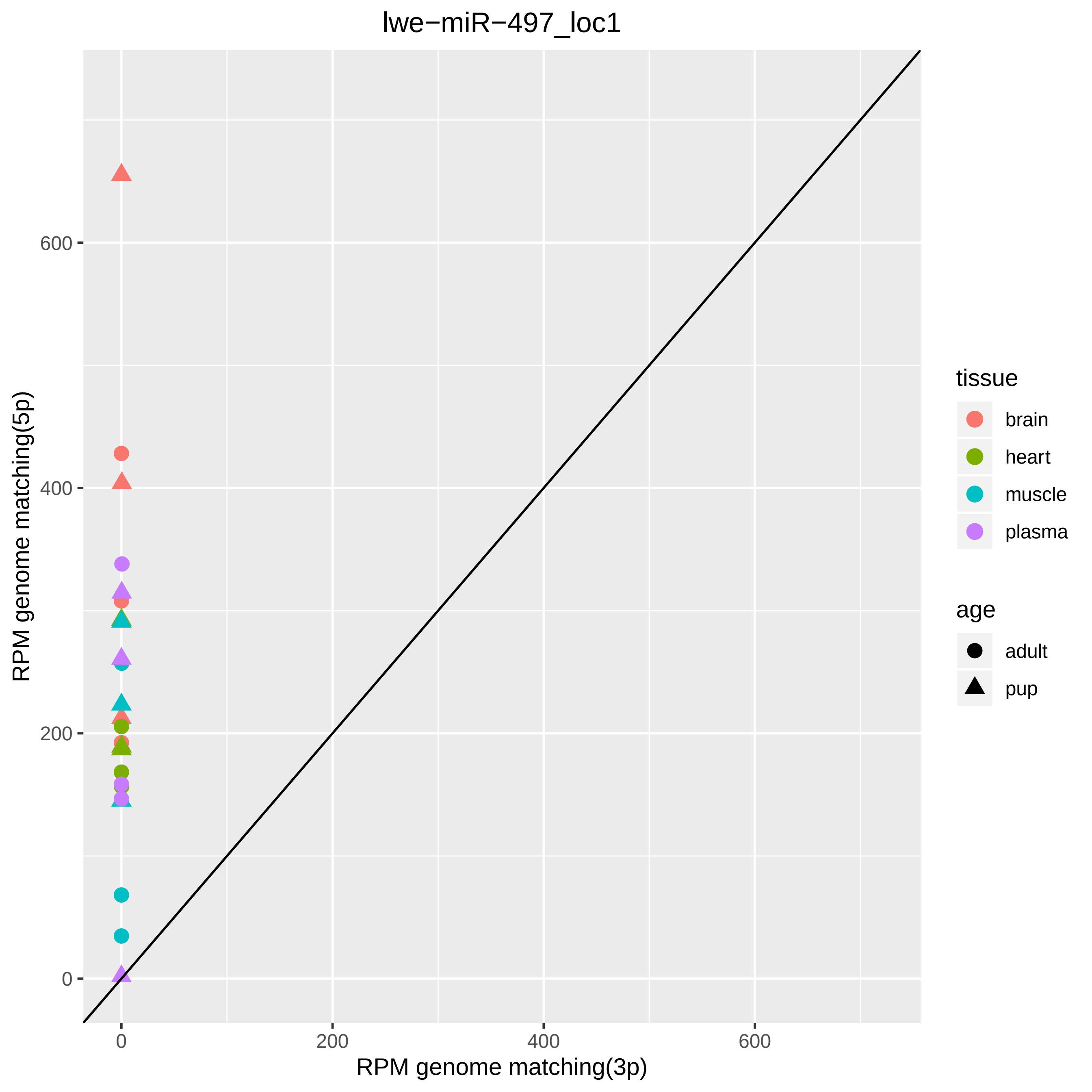

lwe−miR−497_loc1
RPM genome matching(5p)
RPM genome matching(3p)
0
200
400
600
tissue
brain
heart
muscle
plasma
age
adult
pup

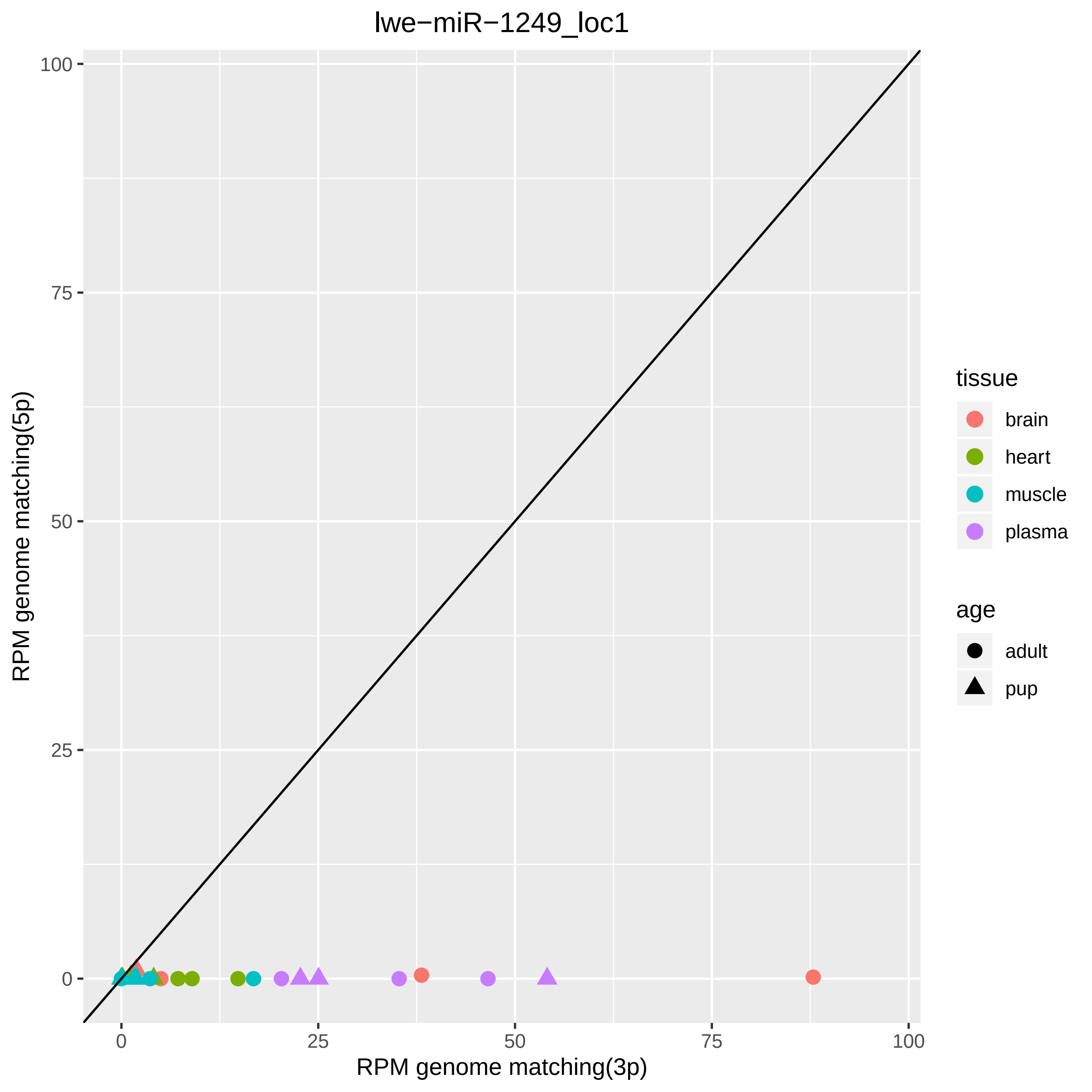

lwe−miR−1249_loc1
RPM genome matching(5p)
RPM genome matching(3p)
0
25
50
75
100
0
25
50
75
100
tissue
brain
heart
muscle
plasma
age
adult
pup

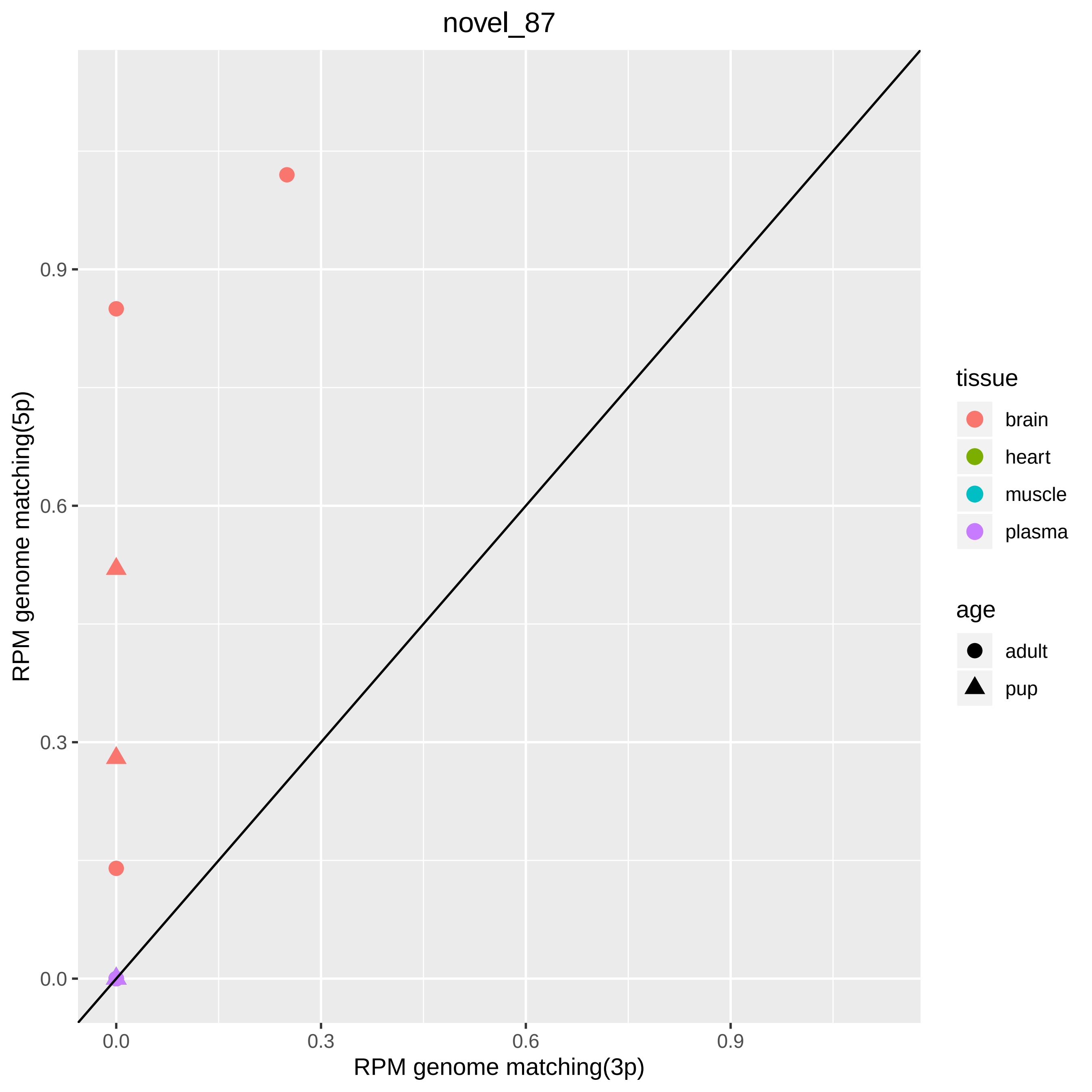
novel_87
RPM genome matching(5p)
RPM genome matching(3p)
0.0
0.3
0.6
0.9
tissue
brain
heart
muscle
plasma
age
adult
pup

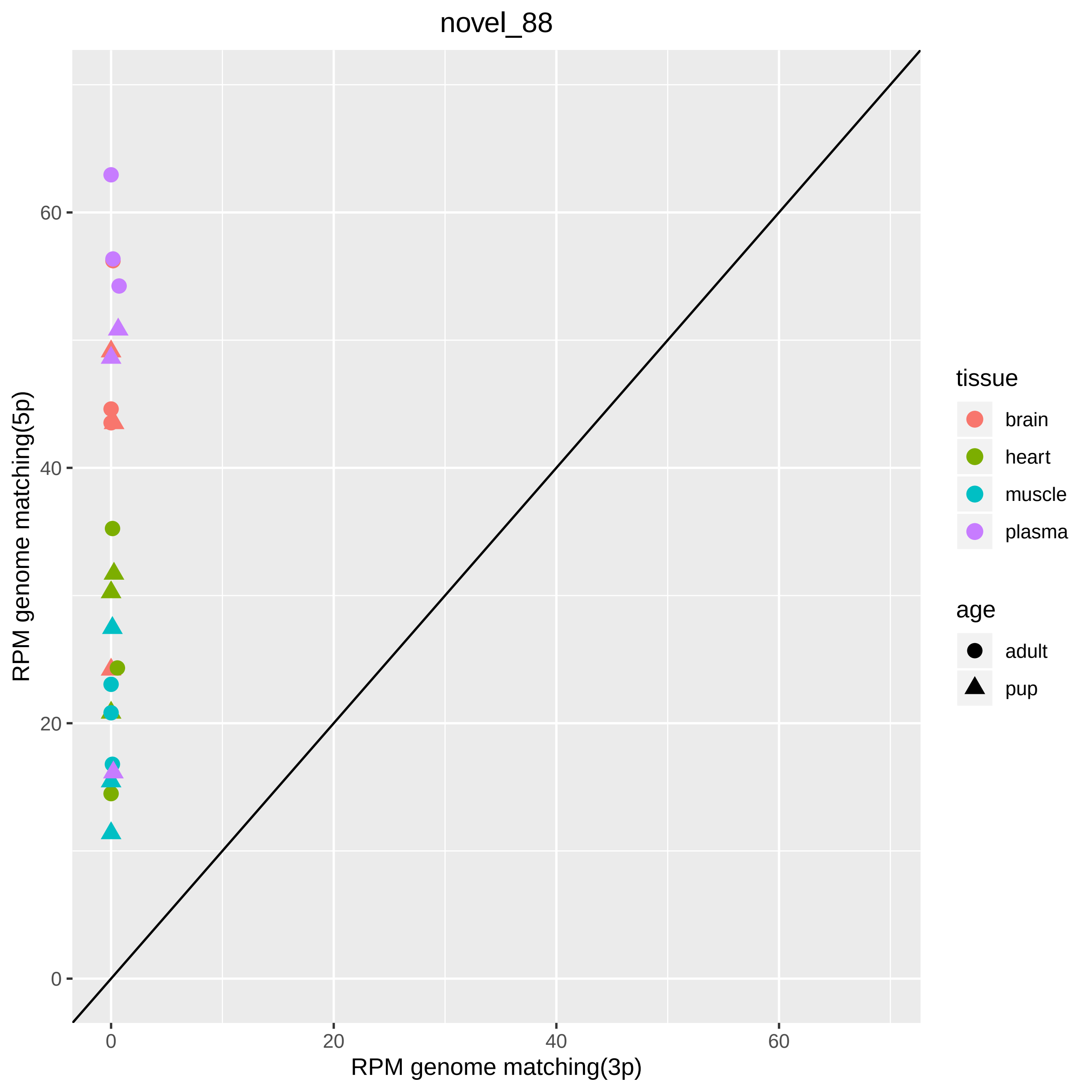
novel_88
RPM genome matching(5p)
RPM genome matching(3p)
0
20
40
60
tissue
brain
heart
muscle
plasma
age
adult
pup

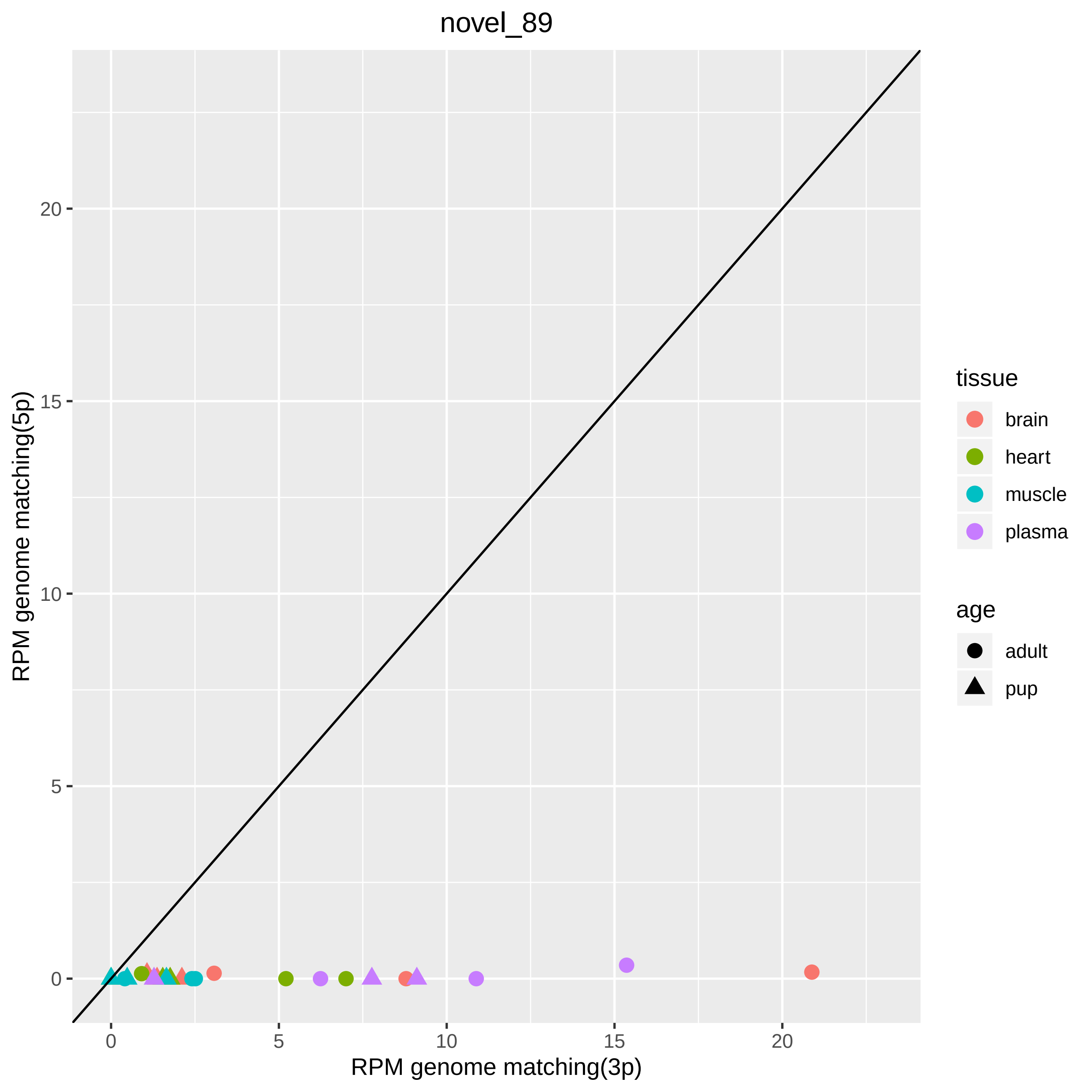
novel_89
RPM genome matching(5p)
RPM genome matching(3p)
0
5
10
15
20
tissue
brain
heart
muscle
plasma
age
adult
pup

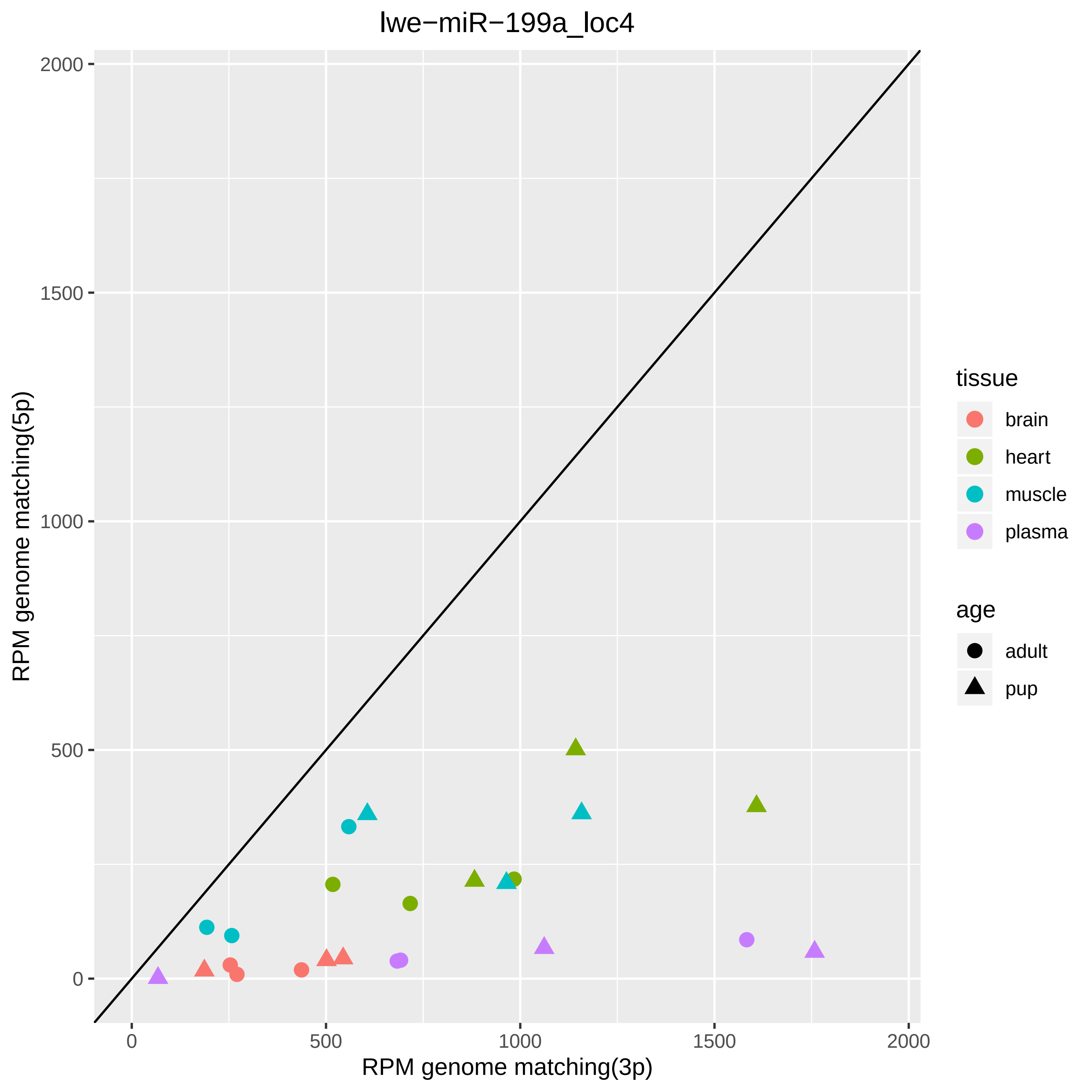
lwe−miR−199a_loc4
RPM genome matching(5p)
RPM genome matching(3p)
0
500
1000
1500
2000
0
500
1000
1500
2000
tissue
brain
heart
muscle
plasma
age
adult
pup

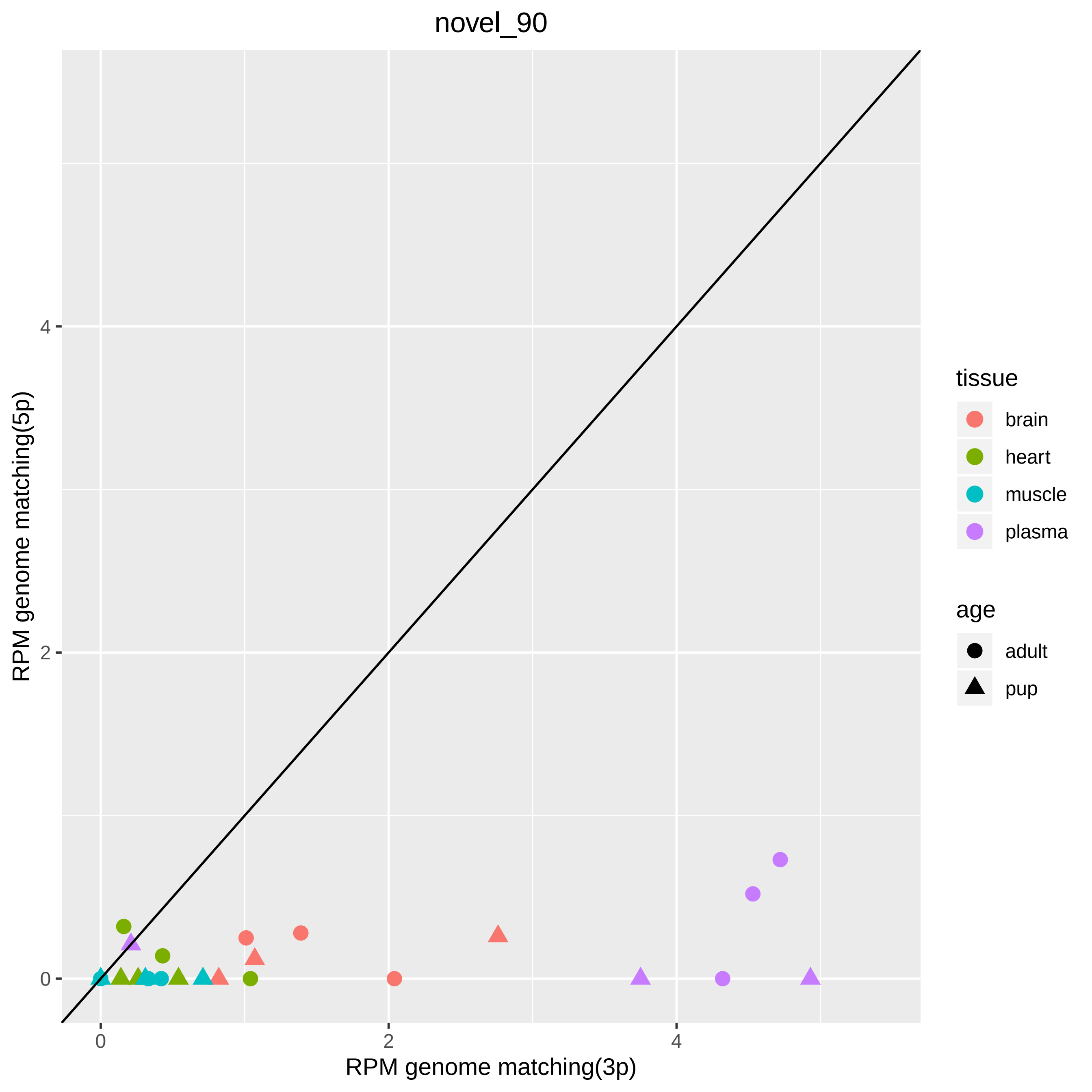

novel_90
RPM genome matching(5p)
RPM genome matching(3p)
0
2
4
tissue
brain
heart
muscle
plasma
age
adult
pup

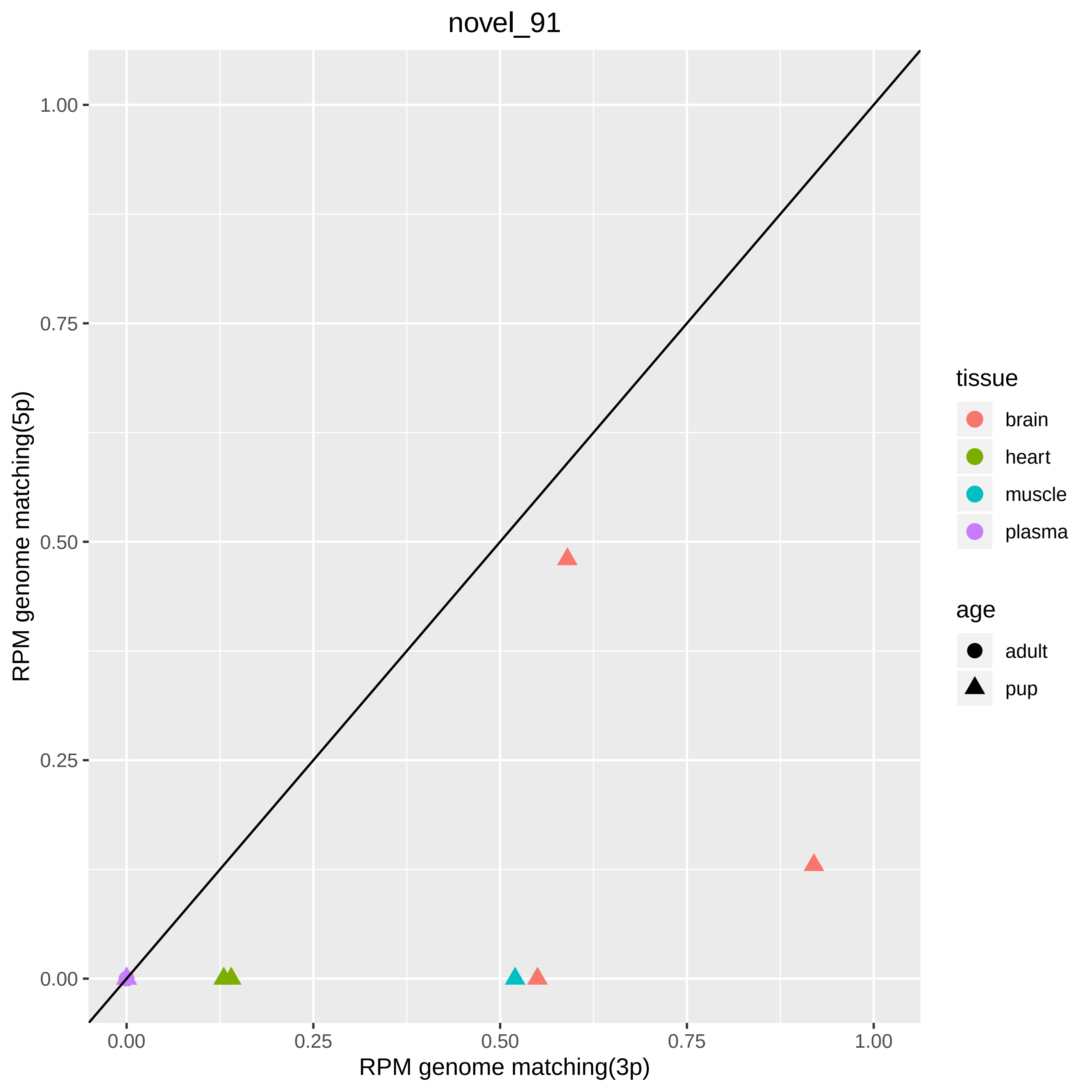

novel_91
RPM genome matching(5p)
RPM genome matching(3p)
0.00
0.25
0.50
0.75
1.00
tissue
brain
heart
muscle
plasma
age
adult
pup

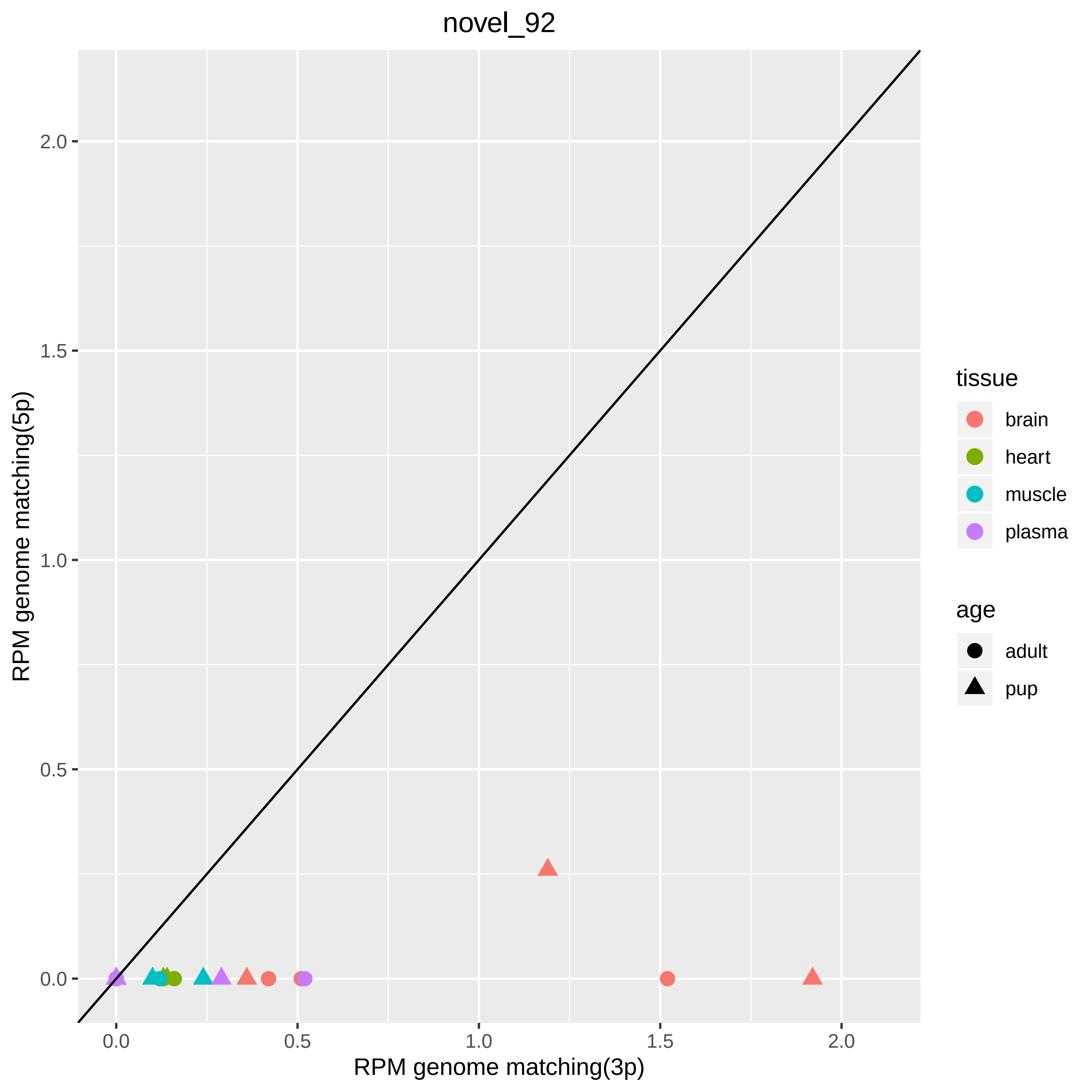
novel_92
RPM genome matching(5p)
RPM genome matching(3p)
0.0
0.5
1.0
1.5
2.0
tissue
brain
heart
muscle
plasma
age
adult
pup

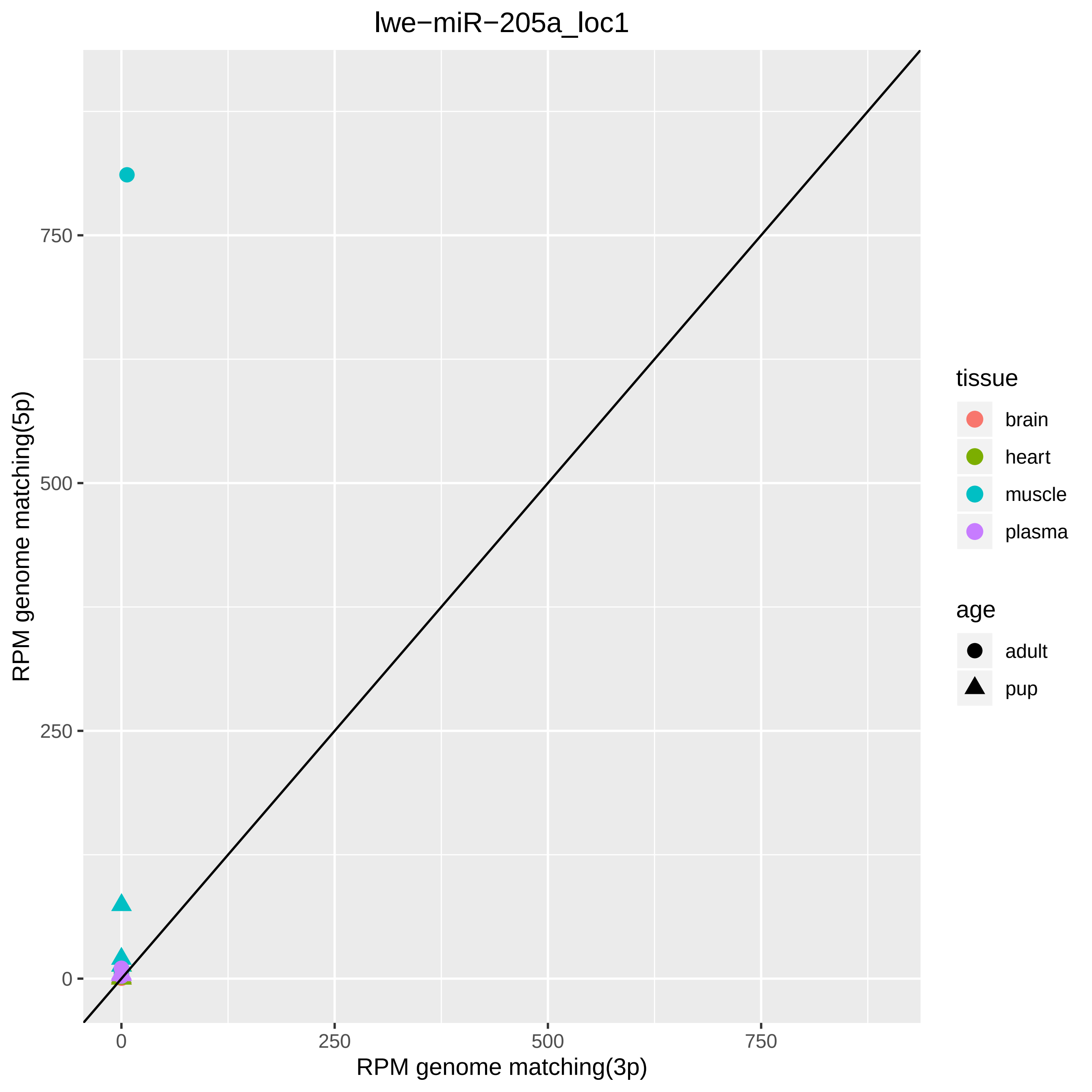
lwe−miR−205a_loc1
RPM genome matching(5p)
RPM genome matching(3p)
0
250
500
750
tissue
brain
heart
muscle
plasma
age
adult
pup

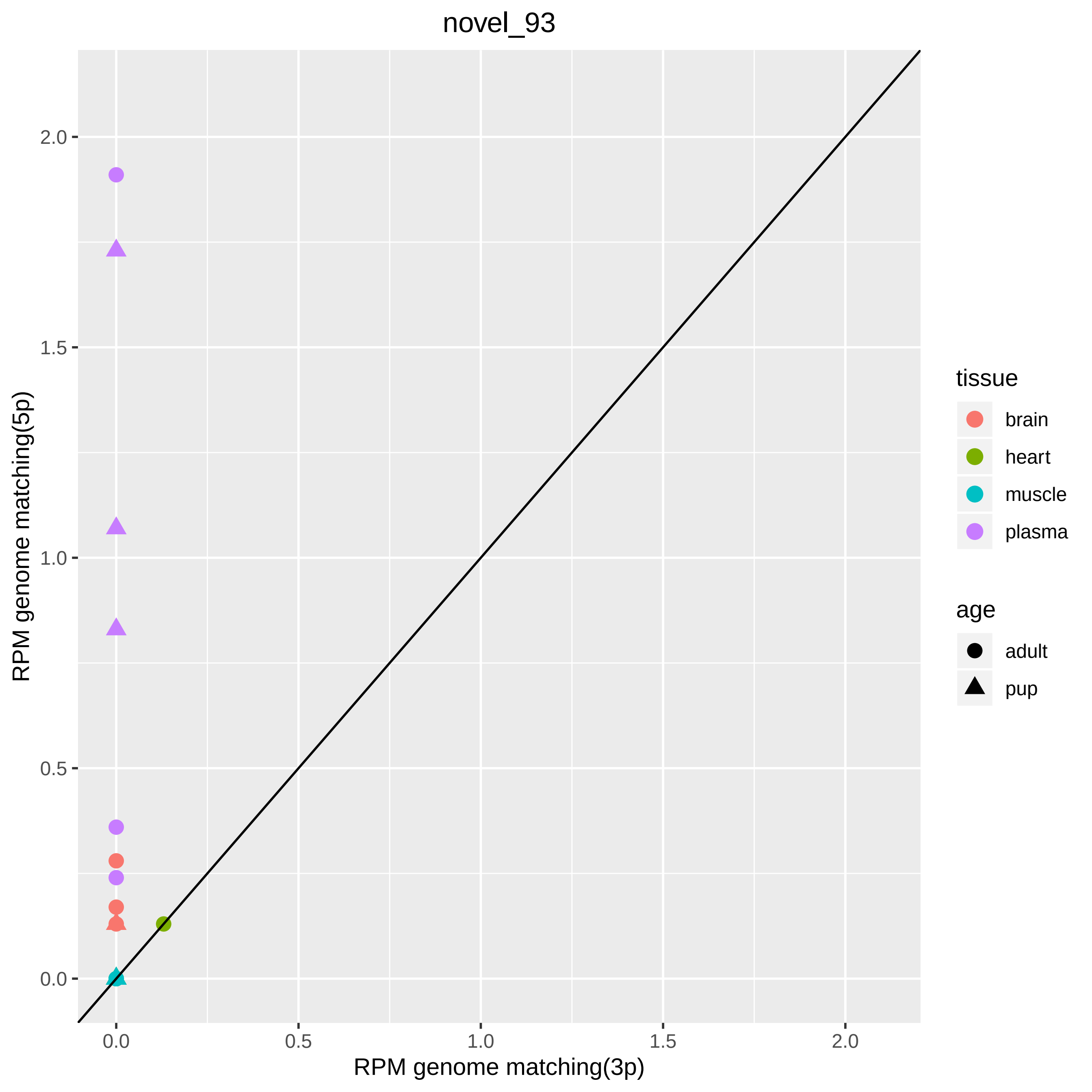

novel_93
RPM genome matching(5p)
RPM genome matching(3p)
0.0
0.5
1.0
1.5
2.0
tissue
brain
heart
muscle
plasma
age
adult
pup

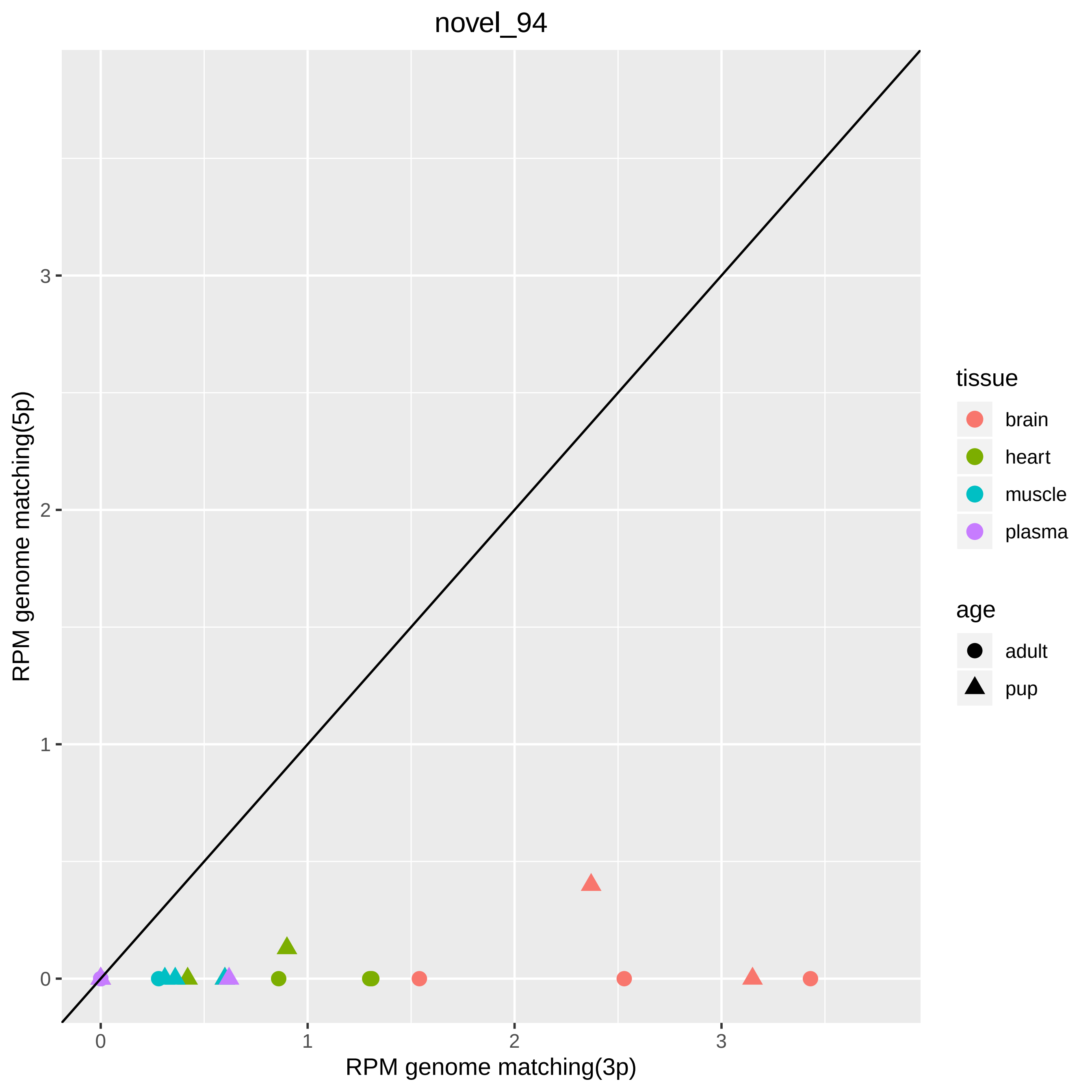

novel_94
RPM genome matching(5p)
RPM genome matching(3p)
0
1
2
3
0
1
2
3
tissue
brain
heart
muscle
plasma
age
adult
pup

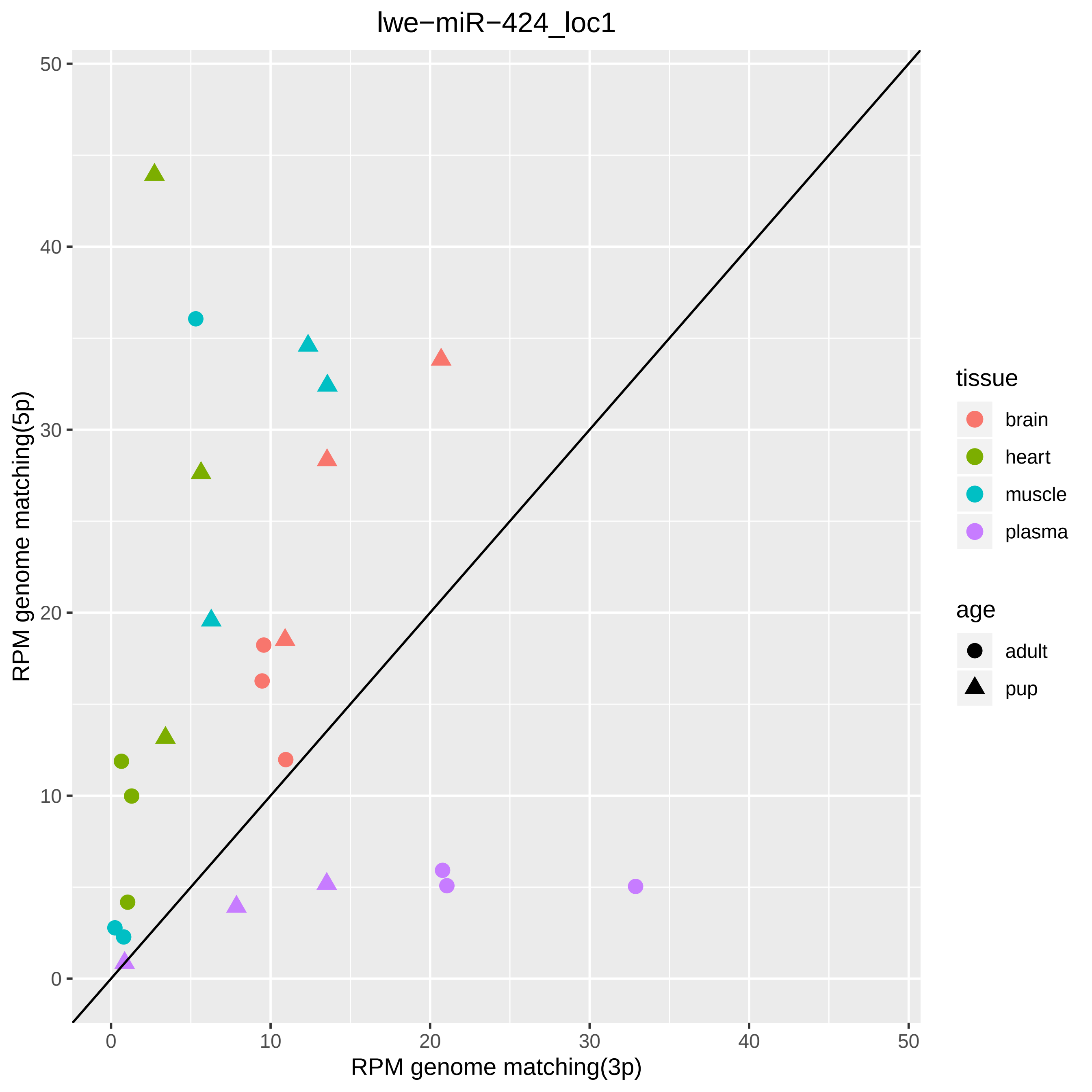
lwe−miR−424_loc1
RPM genome matching(5p)
RPM genome matching(3p)
0
10
20
30
40
50
tissue
brain
heart
muscle
plasma
age
adult
pup

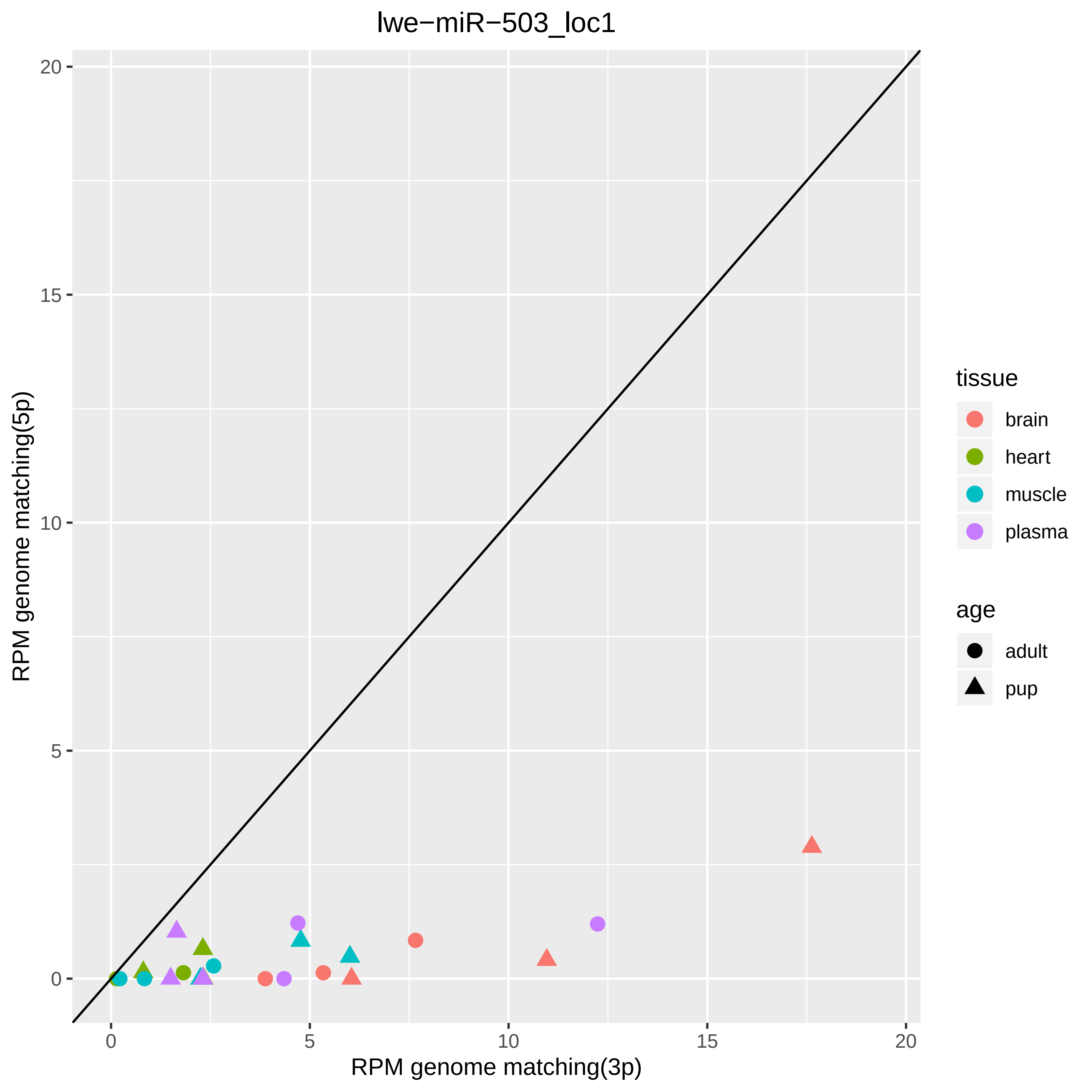

lwe−miR−503_loc1
RPM genome matching(5p)
RPM genome matching(3p)
0
5
10
15
20
0
5
10
15
20
tissue
brain
heart
muscle
plasma
age
adult
pup

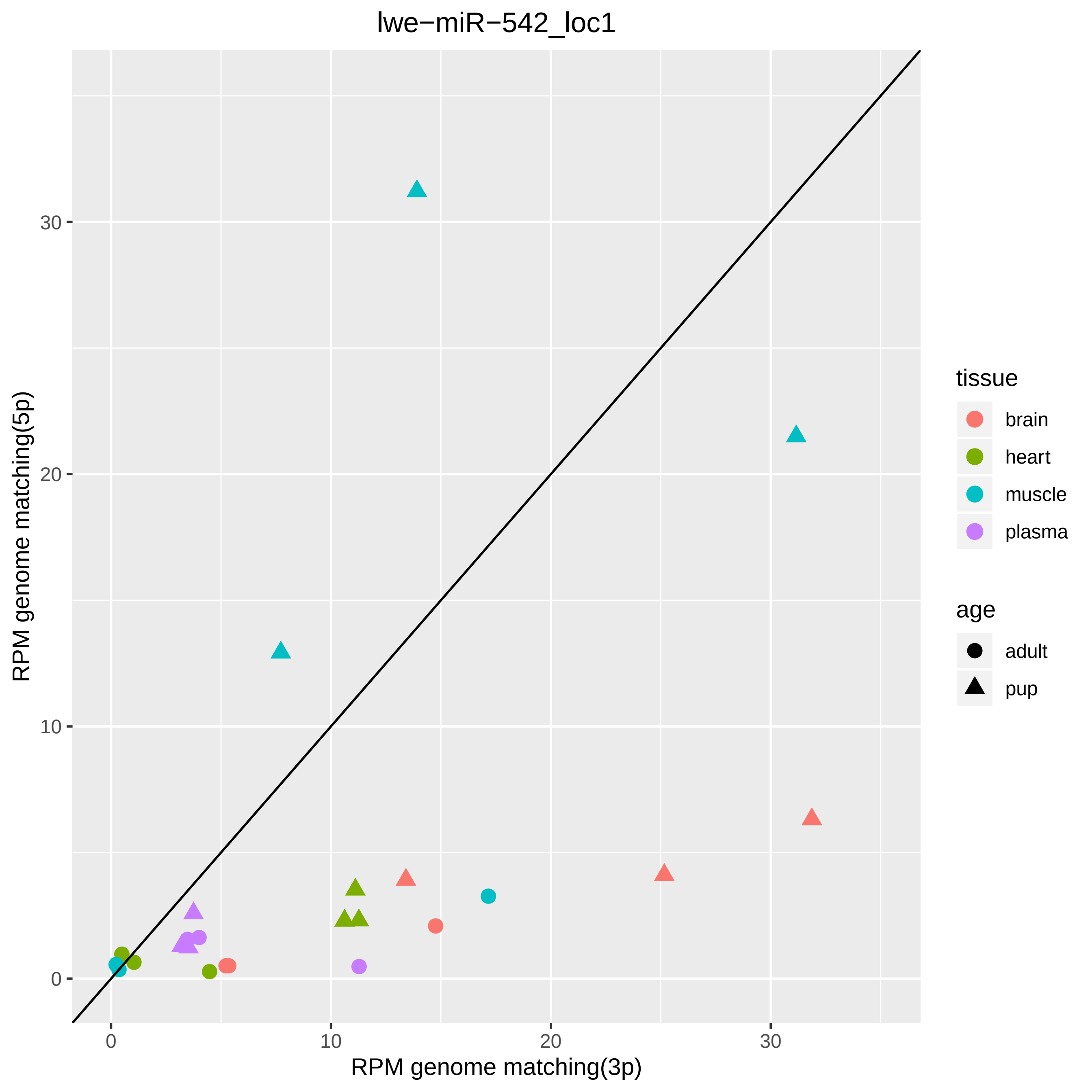

lwe−miR−542_loc1
RPM genome matching(5p)
RPM genome matching(3p)
0
10
20
30
tissue
brain
heart
muscle
plasma
age
adult
pup

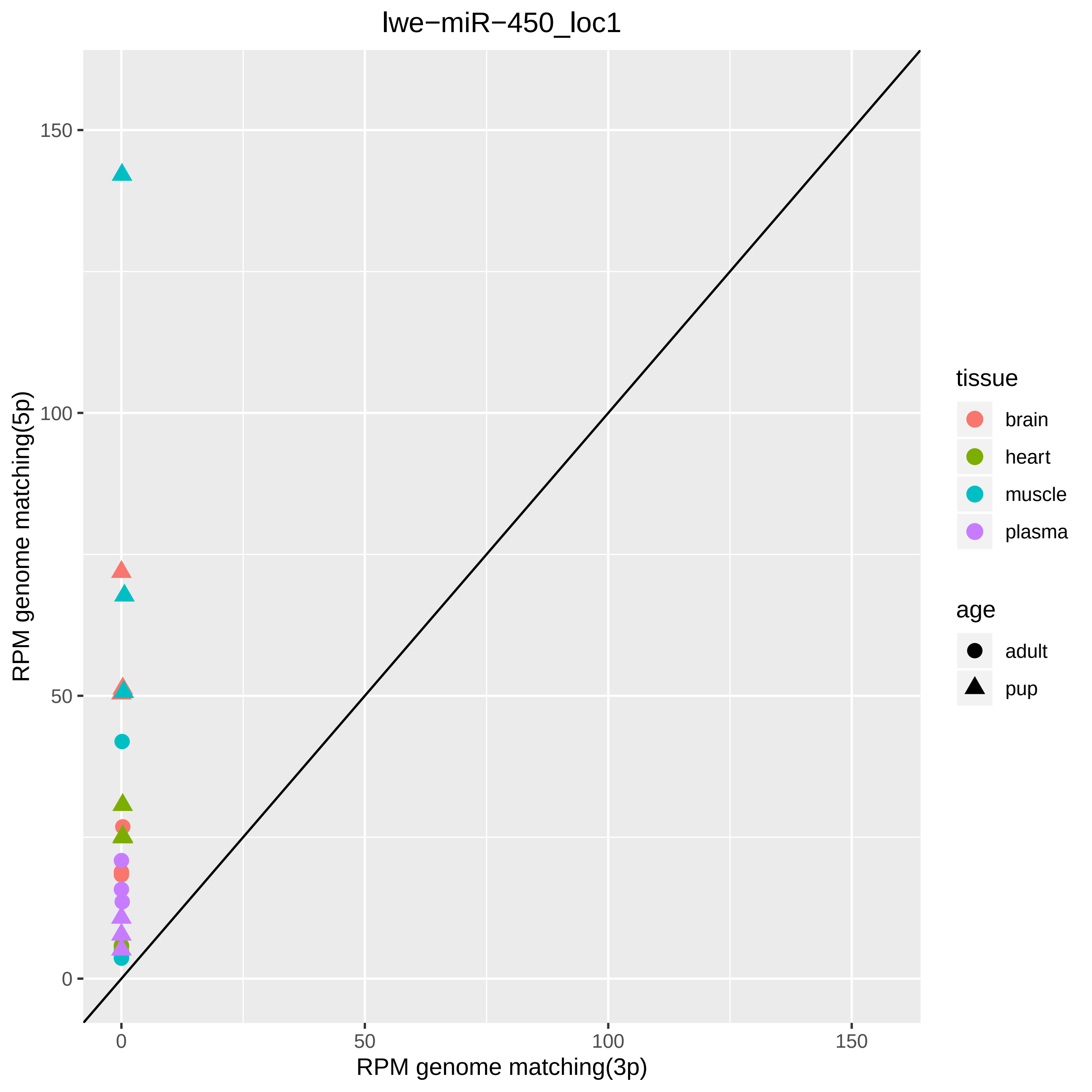
lwe−miR−450_loc1
RPM genome matching(5p)
RPM genome matching(3p)
0
50
100
150
tissue
brain
heart
muscle
plasma
age
adult
pup

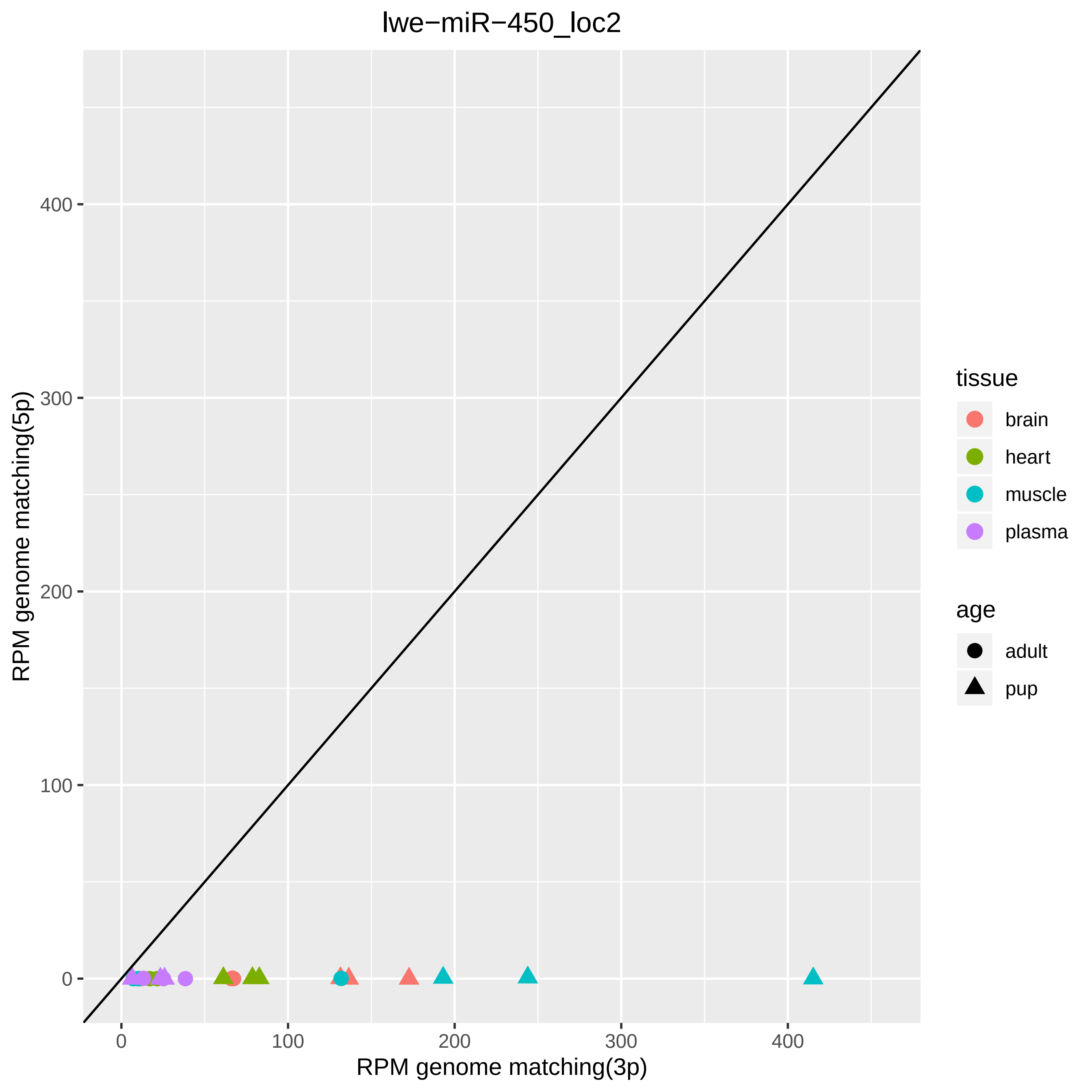

lwe−miR−450_loc2
RPM genome matching(5p)
RPM genome matching(3p)
0
100
200
300
400
tissue
brain
heart
muscle
plasma
age
adult
pup

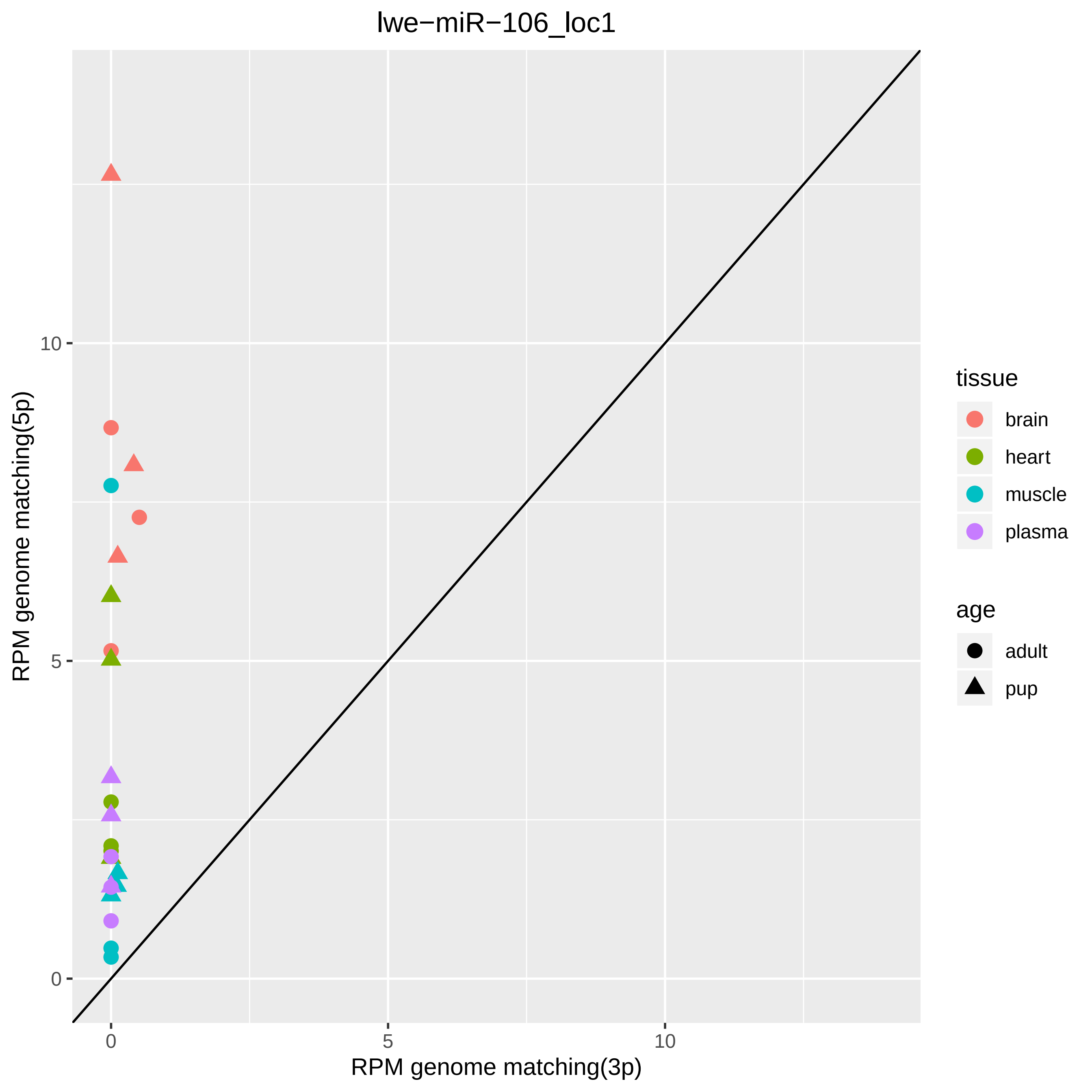

lwe−miR−106_loc1
RPM genome matching(5p)
RPM genome matching(3p)
0
5
10
0
5
10
tissue
brain
heart
muscle
plasma
age
adult
pup

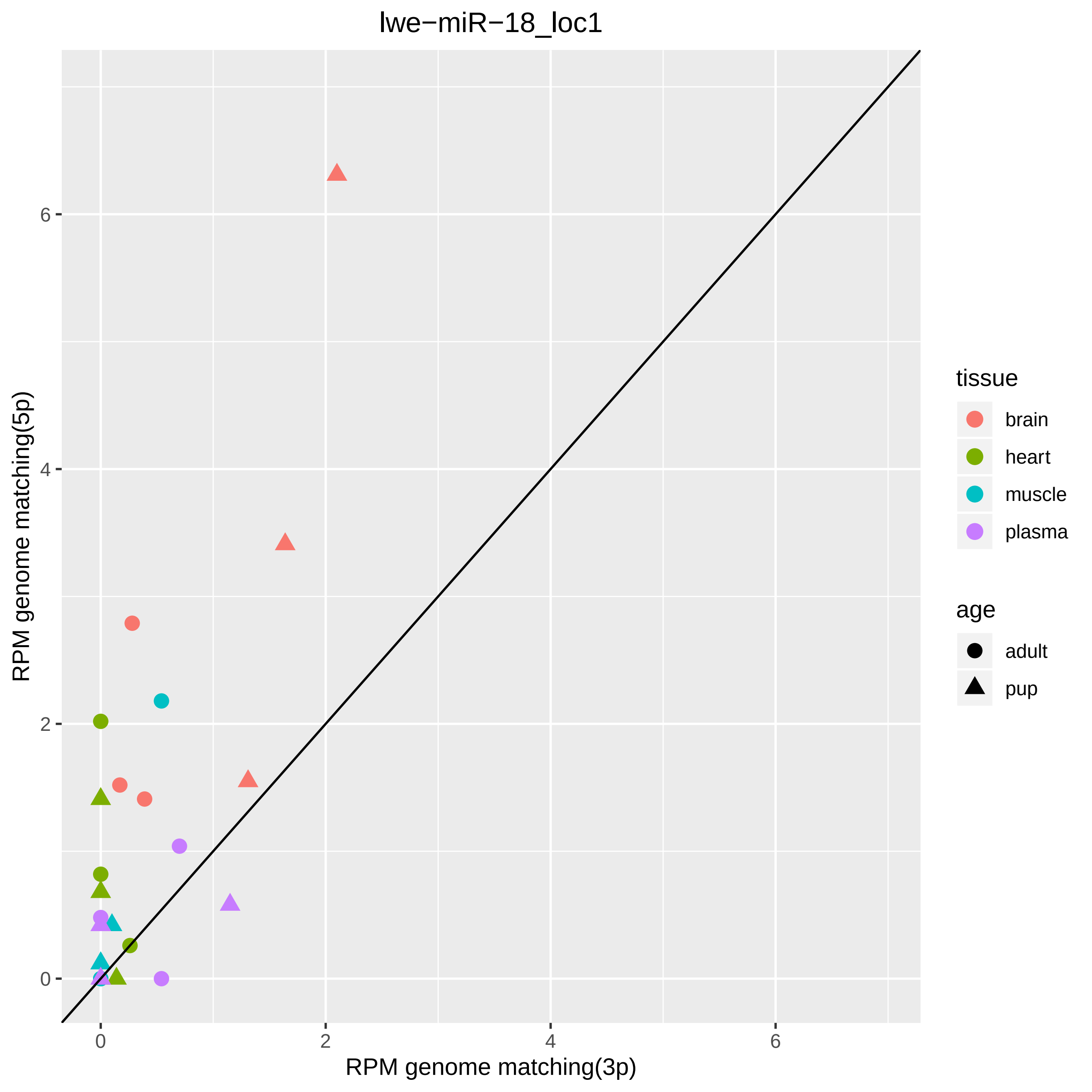

lwe−miR−18_loc1
RPM genome matching(5p)
RPM genome matching(3p)
0
2
4
6
tissue
brain
heart
muscle
plasma
age
adult
pup

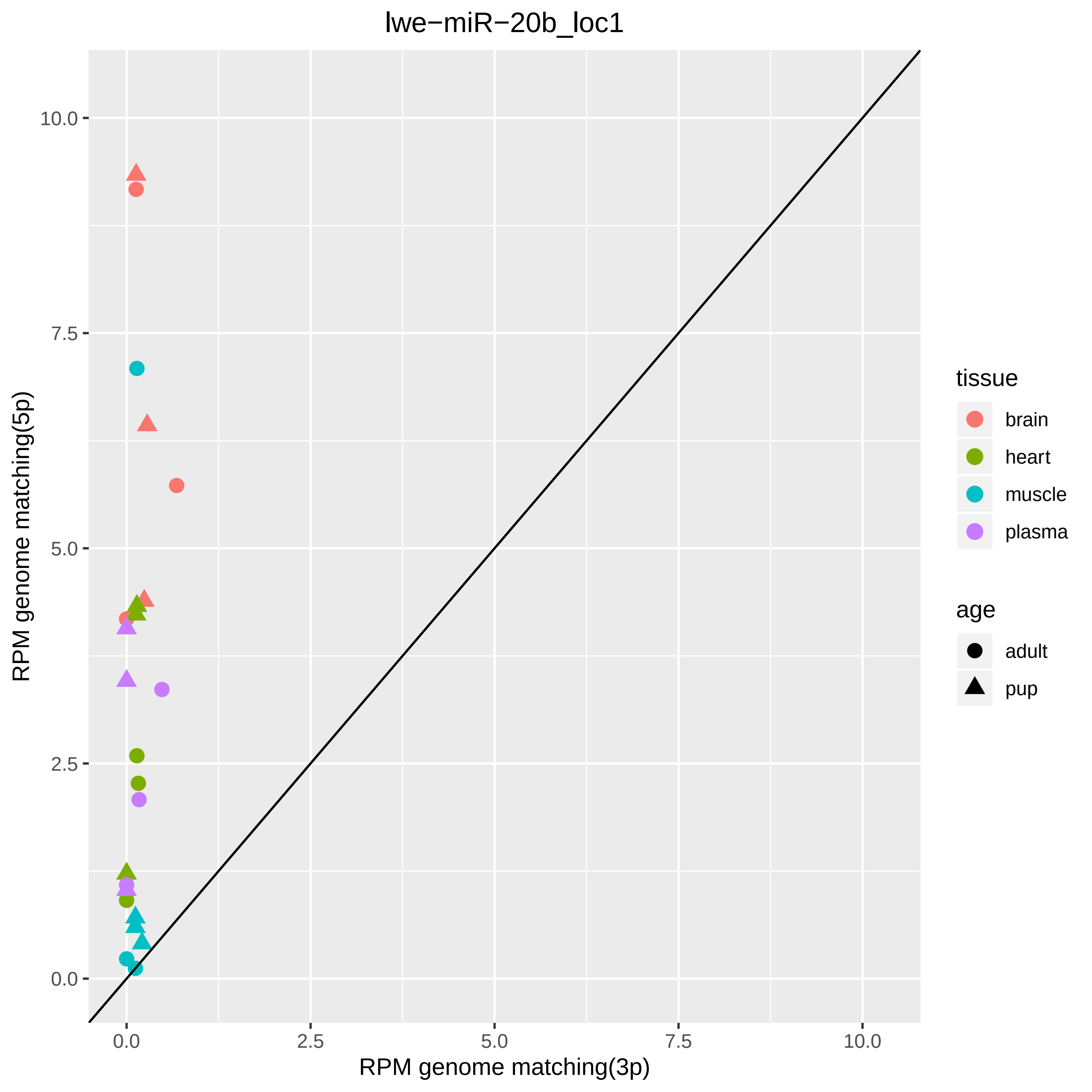

lwe−miR−20b_loc1
RPM genome matching(5p)
RPM genome matching(3p)
10.0
7.5
5.0
2.5
0.0
0.0
2.5
5.0
7.5
10.0
tissue
brain
heart
muscle
plasma
age
adult
pup

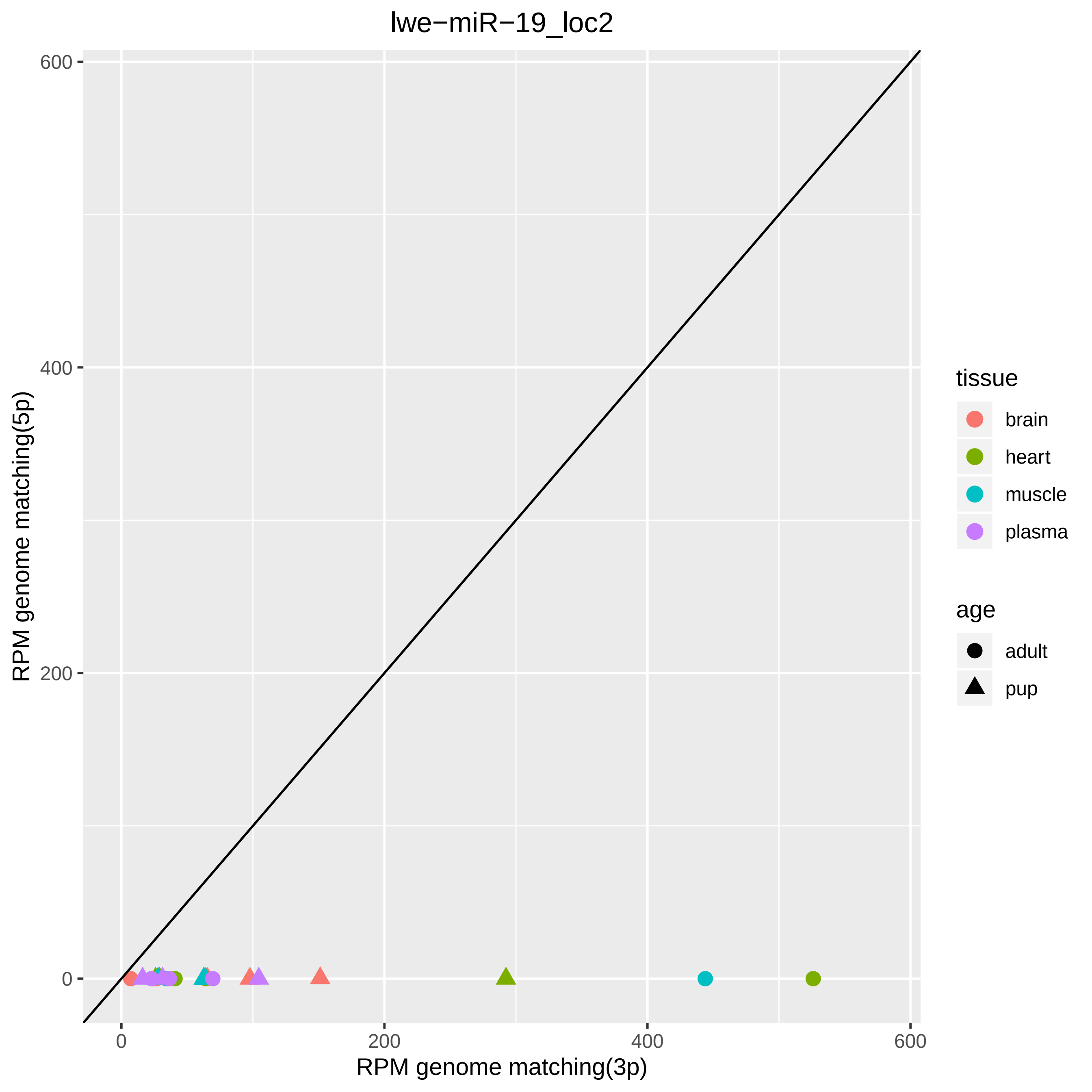

lwe−miR−19_loc2
RPM genome matching(5p)
RPM genome matching(3p)
0
200
400
600
0
200
400
600
tissue
brain
heart
muscle
plasma
age
adult
pup

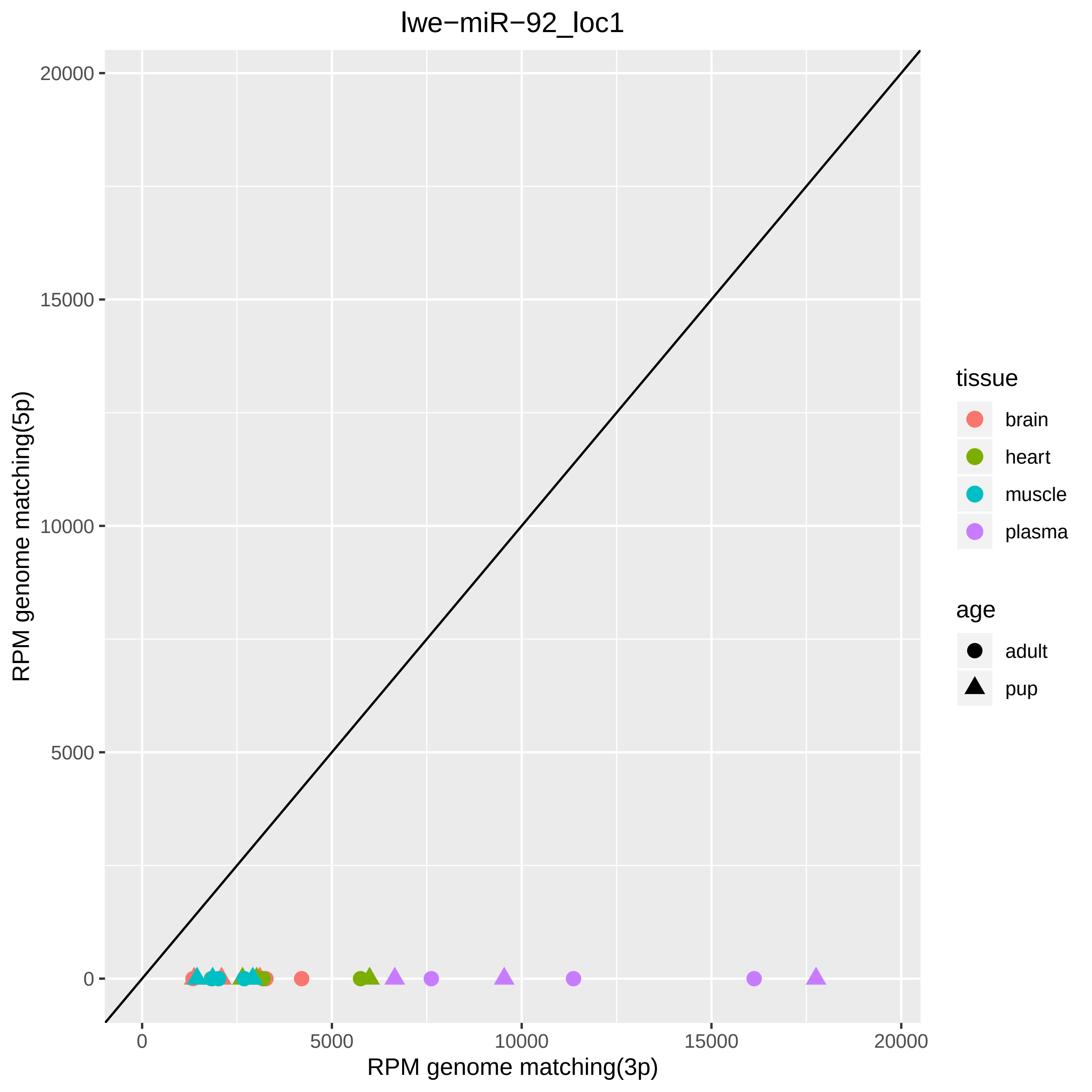
lwe−miR−92_loc1
RPM genome matching(5p)
RPM genome matching(3p)
20000
15000
10000
5000
0
0
5000
10000
15000
20000
tissue
brain
heart
muscle
plasma
age
adult
pup

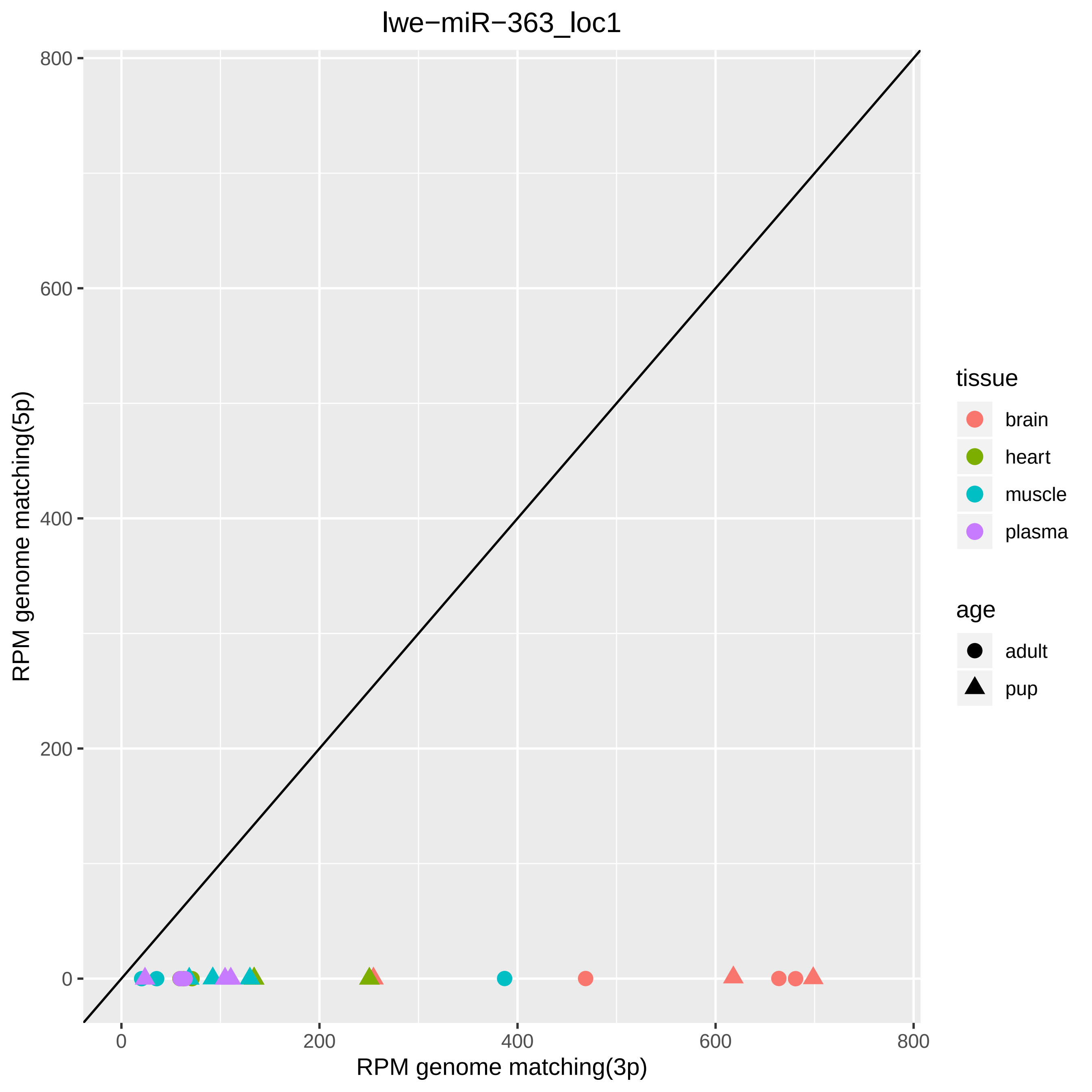
lwe−miR−363_loc1
RPM genome matching(5p)
RPM genome matching(3p)
0
200
400
600
800
0
200
400
600
800
tissue
brain
heart
muscle
plasma
age
adult
pup

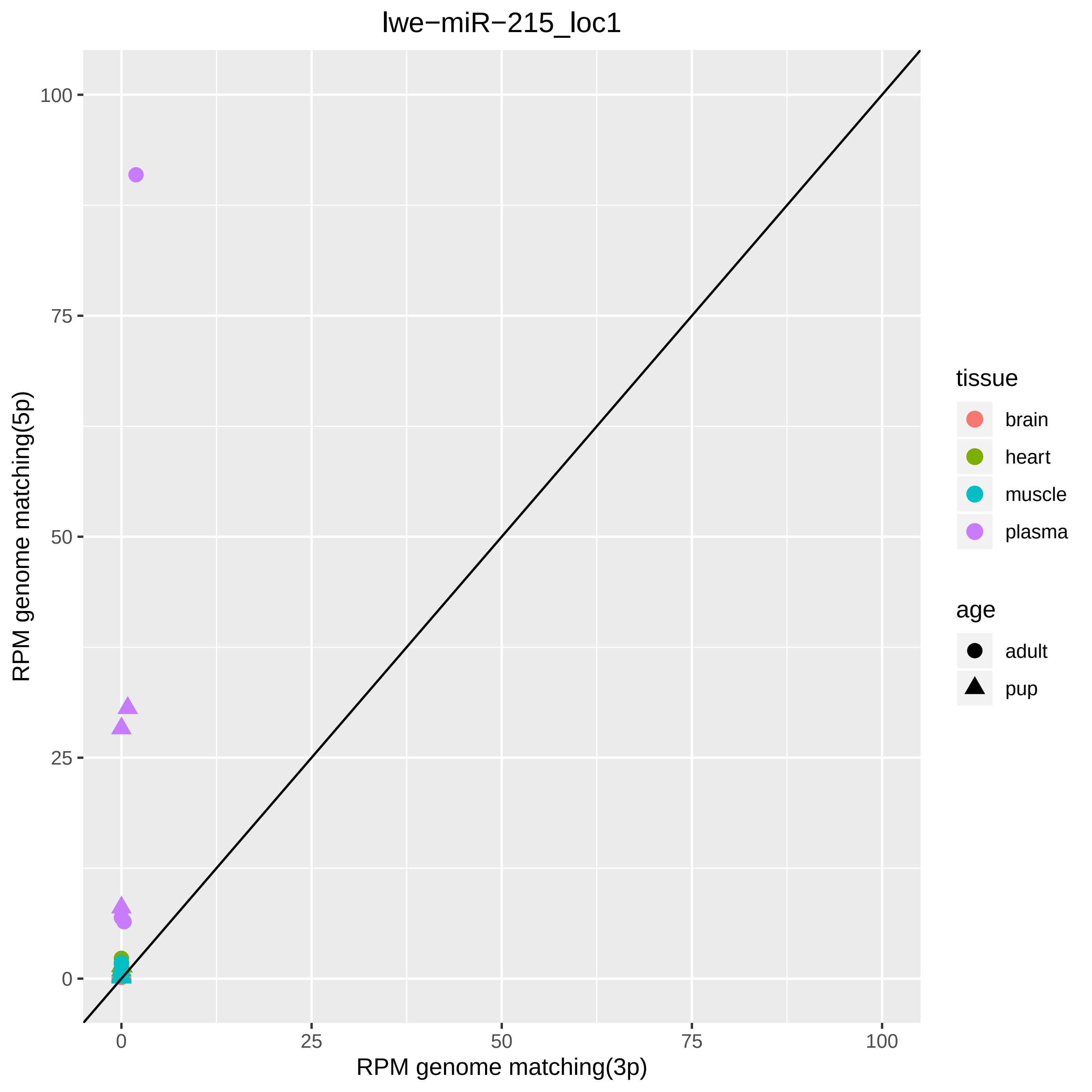
lwe−miR−215_loc1
RPM genome matching(5p)
RPM genome matching(3p)
0
25
50
75
100
0
25
50
75
100
tissue
brain
heart
muscle
plasma
age
adult
pup

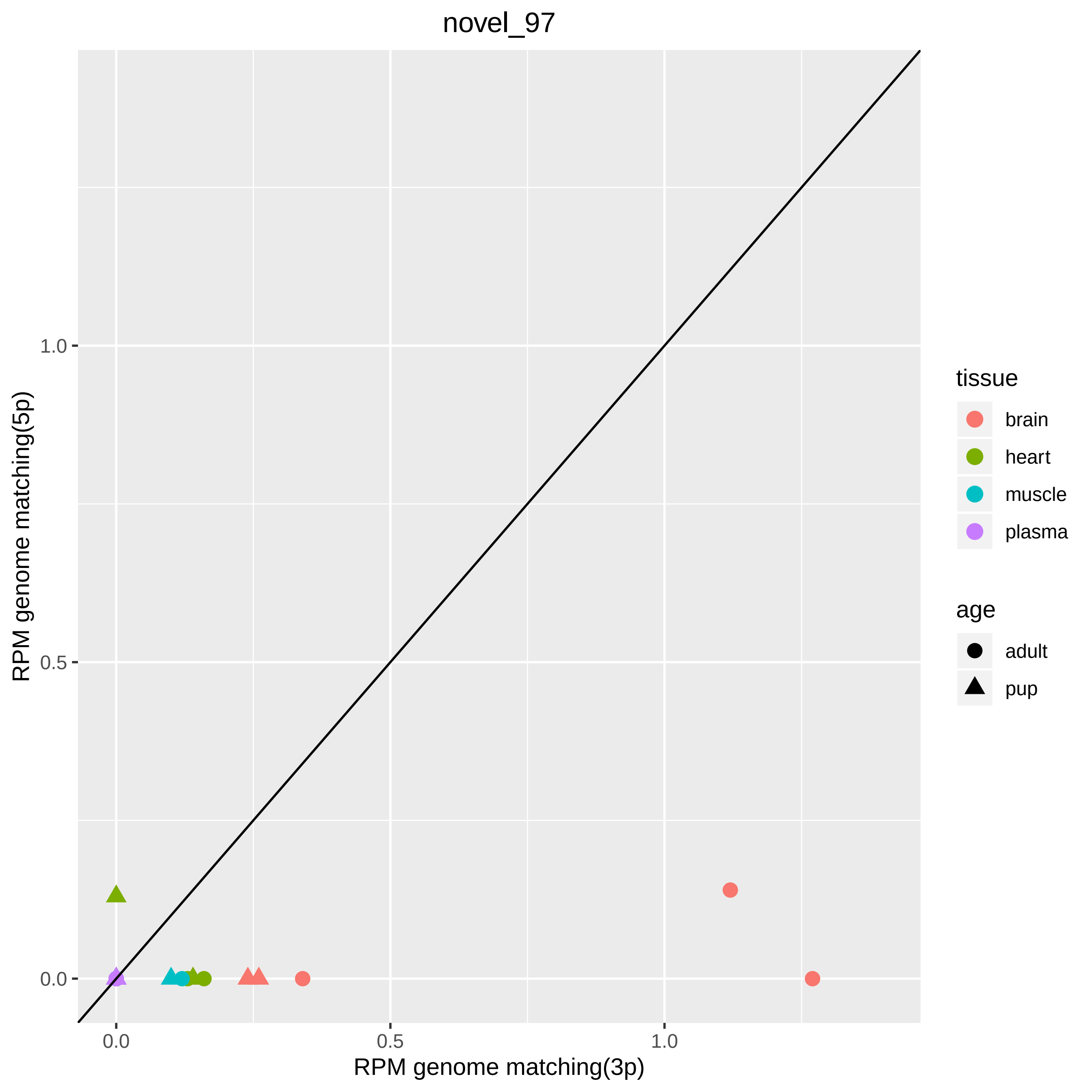
novel_97
RPM genome matching(5p)
RPM genome matching(3p)
1.0
0.5
0.0
0.0
0.5
1.0
tissue
brain
heart
muscle
plasma
age
adult
pup

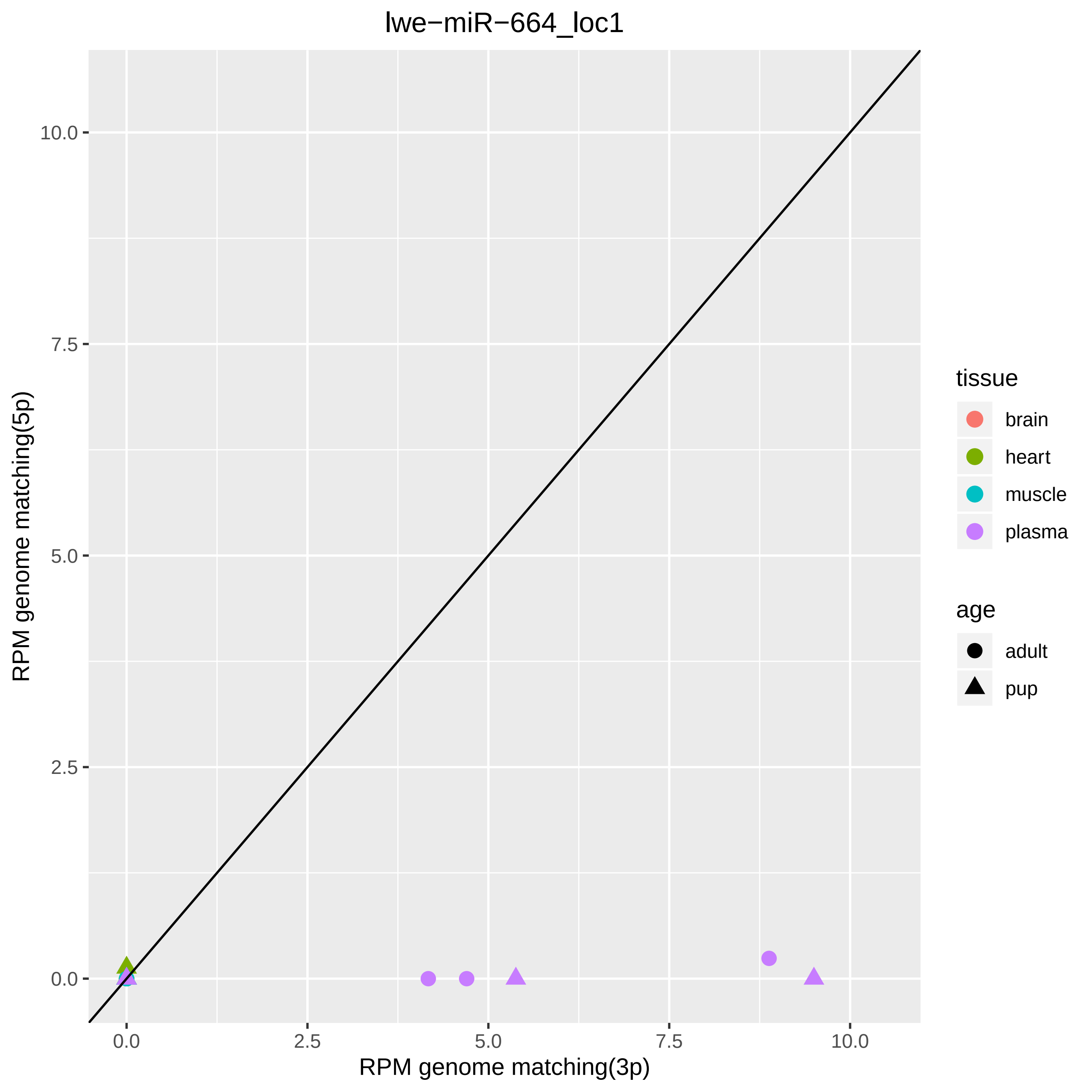
lwe−miR−664_loc1
RPM genome matching(5p)
RPM genome matching(3p)
10.0
7.5
5.0
2.5
0.0
0.0
2.5
5.0
7.5
10.0
tissue
brain
heart
muscle
plasma
age
adult
pup

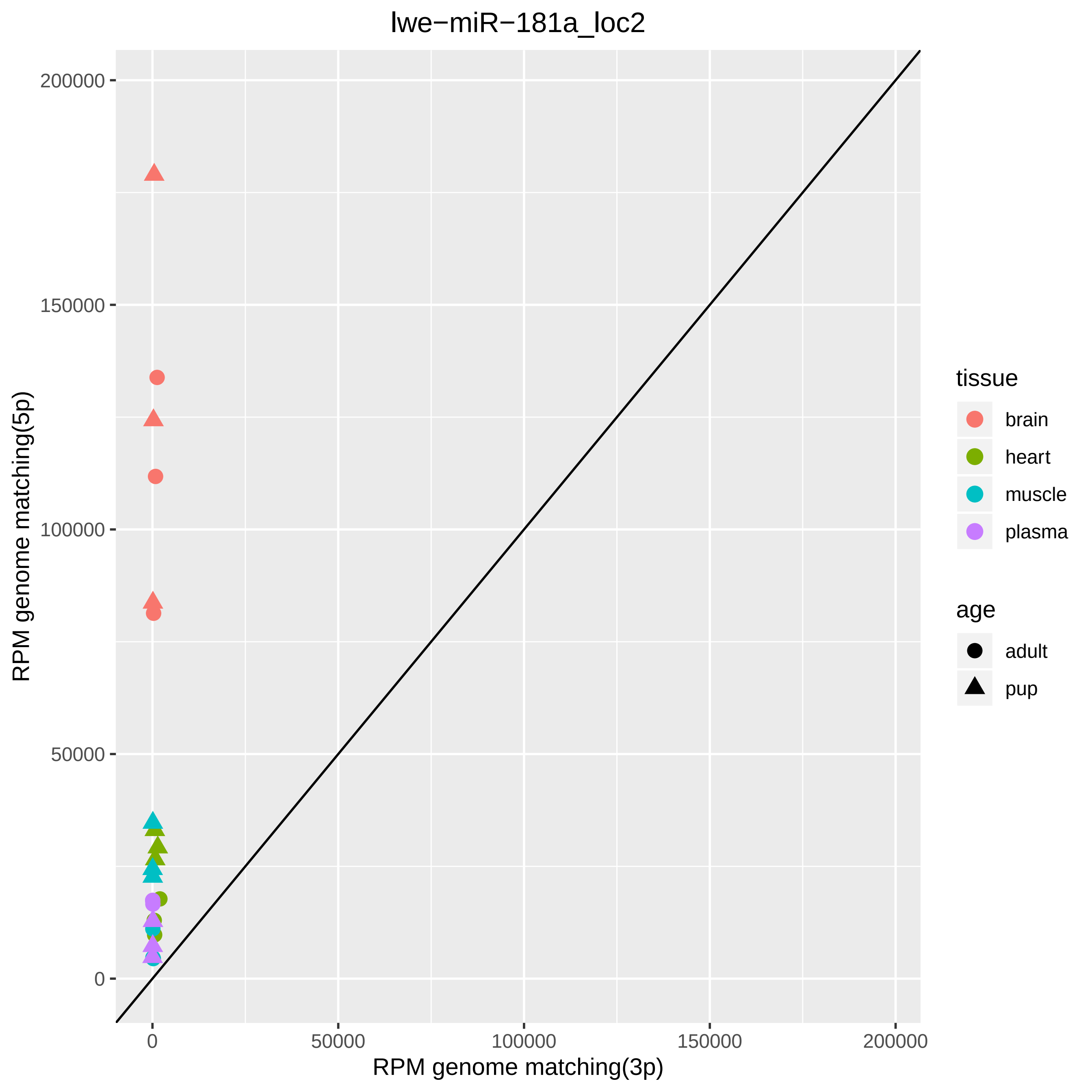

lwe−miR−181a_loc2
RPM genome matching(5p)
RPM genome matching(3p)
200000
150000
100000
50000
0
0
50000
100000
150000
200000
tissue
brain
heart
muscle
plasma
age
adult
pup

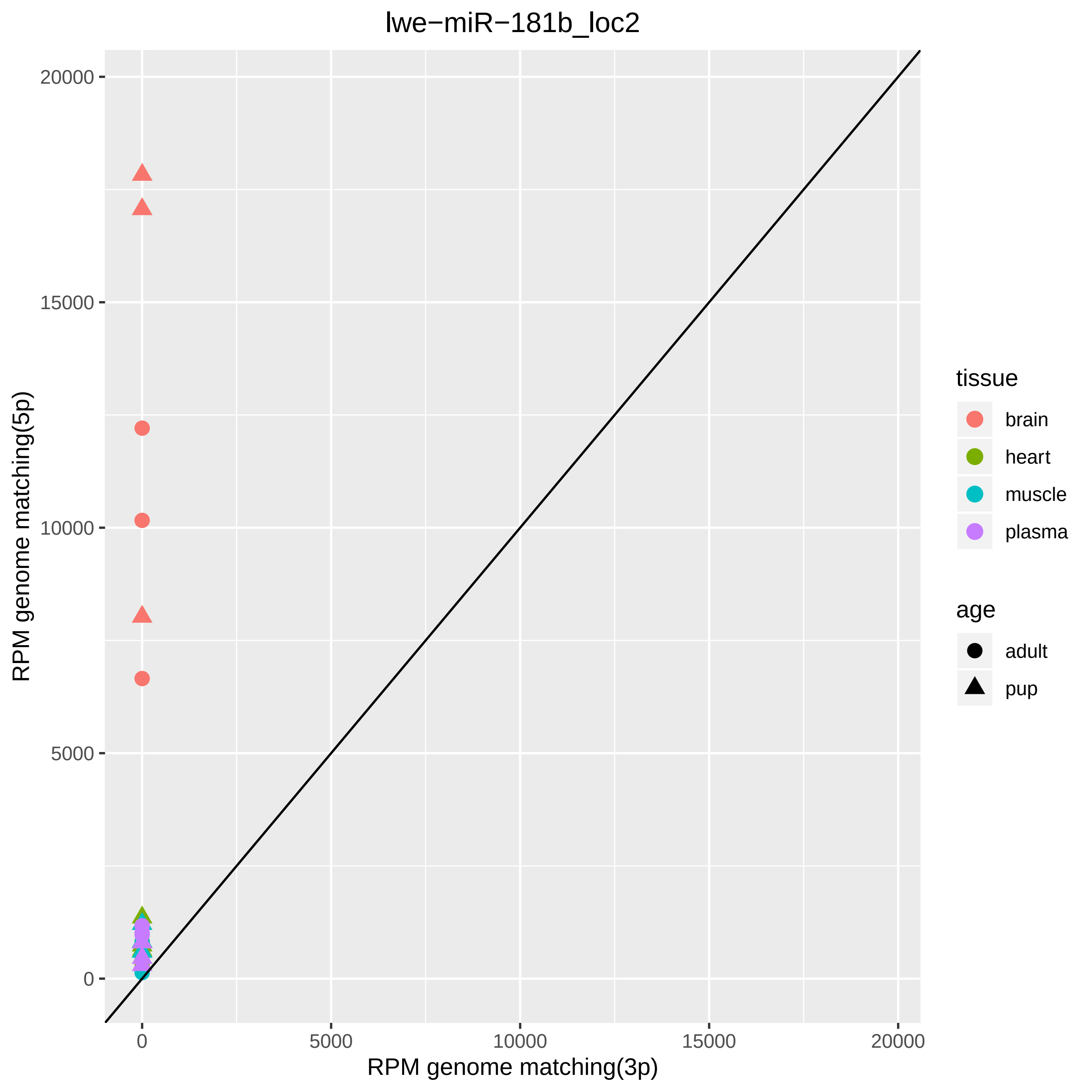
lwe−miR−181b_loc2
RPM genome matching(5p)
RPM genome matching(3p)
0
5000
10000
15000
20000
tissue
brain
heart
muscle
plasma
age
adult
pup

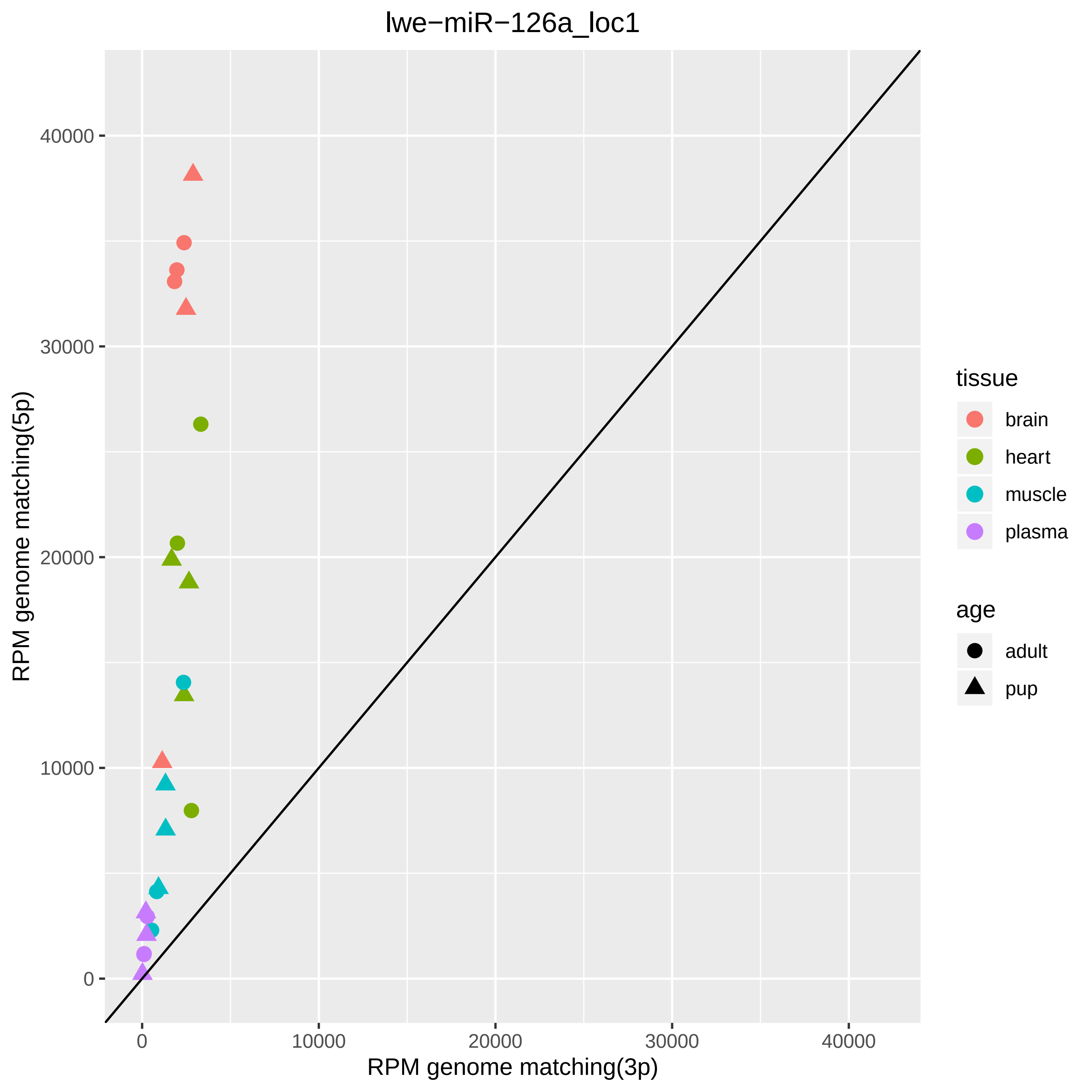
lwe−miR−126a_loc1
RPM genome matching(5p)
RPM genome matching(3p)
0
10000
20000
30000
40000
0
10000
20000
30000
40000
tissue
brain
heart
muscle
plasma
age
adult
pup

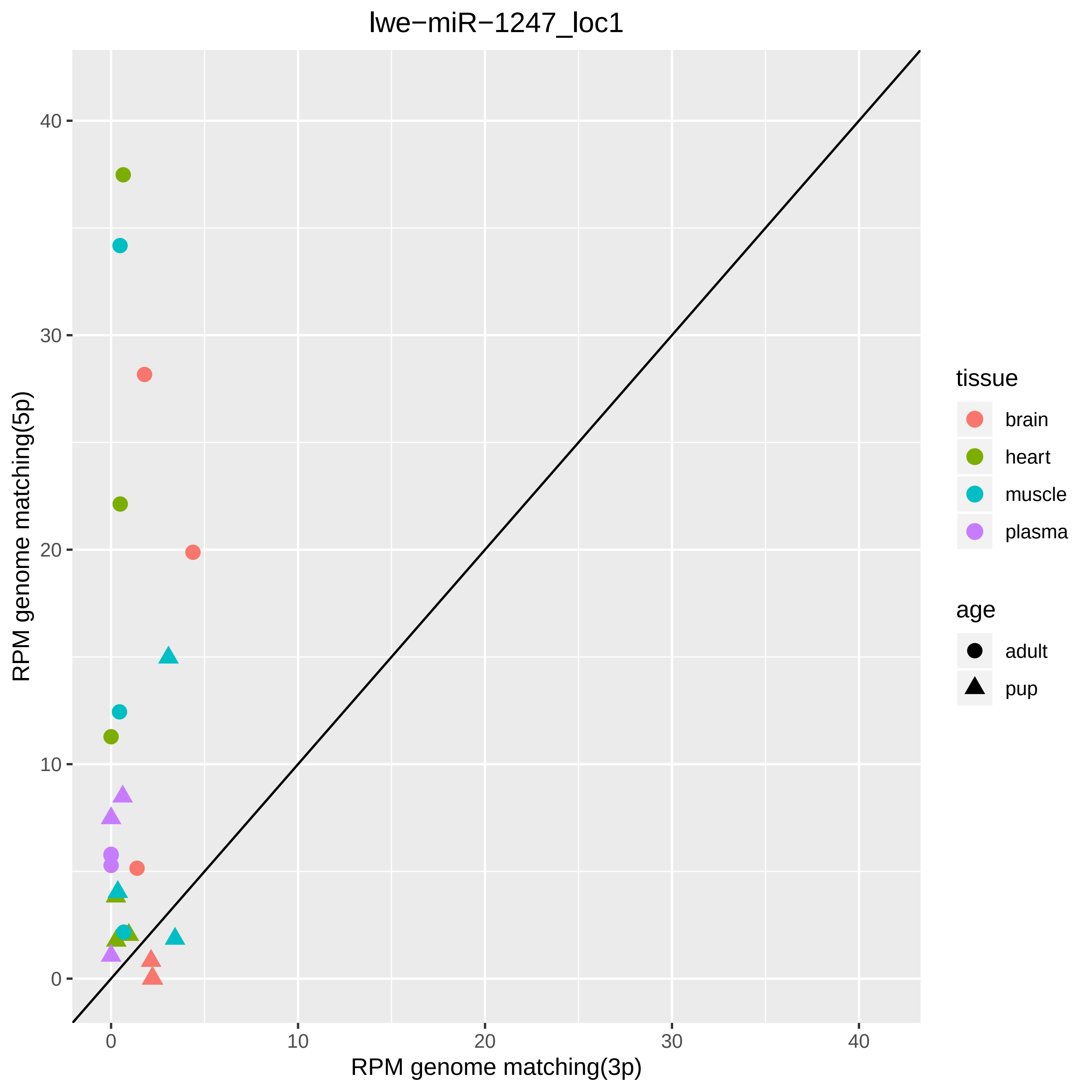

lwe−miR−1247_loc1
RPM genome matching(5p)
RPM genome matching(3p)
0
10
20
30
40
0
10
20
30
40
tissue
brain
heart
muscle
plasma
age
adult
pup

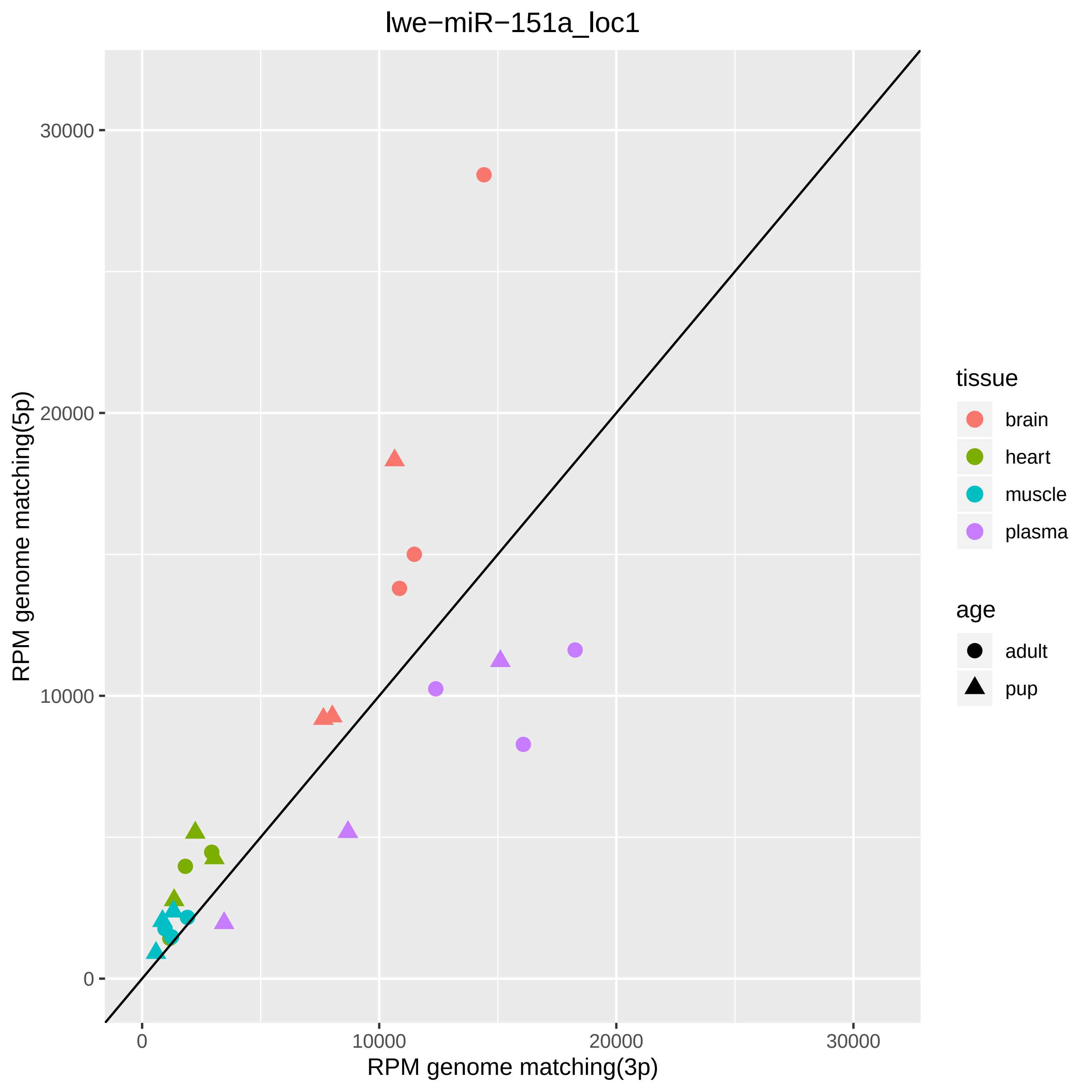
lwe−miR−151a_loc1
RPM genome matching(5p)
RPM genome matching(3p)
0
10000
20000
30000
tissue
brain
heart
muscle
plasma
age
adult
pup

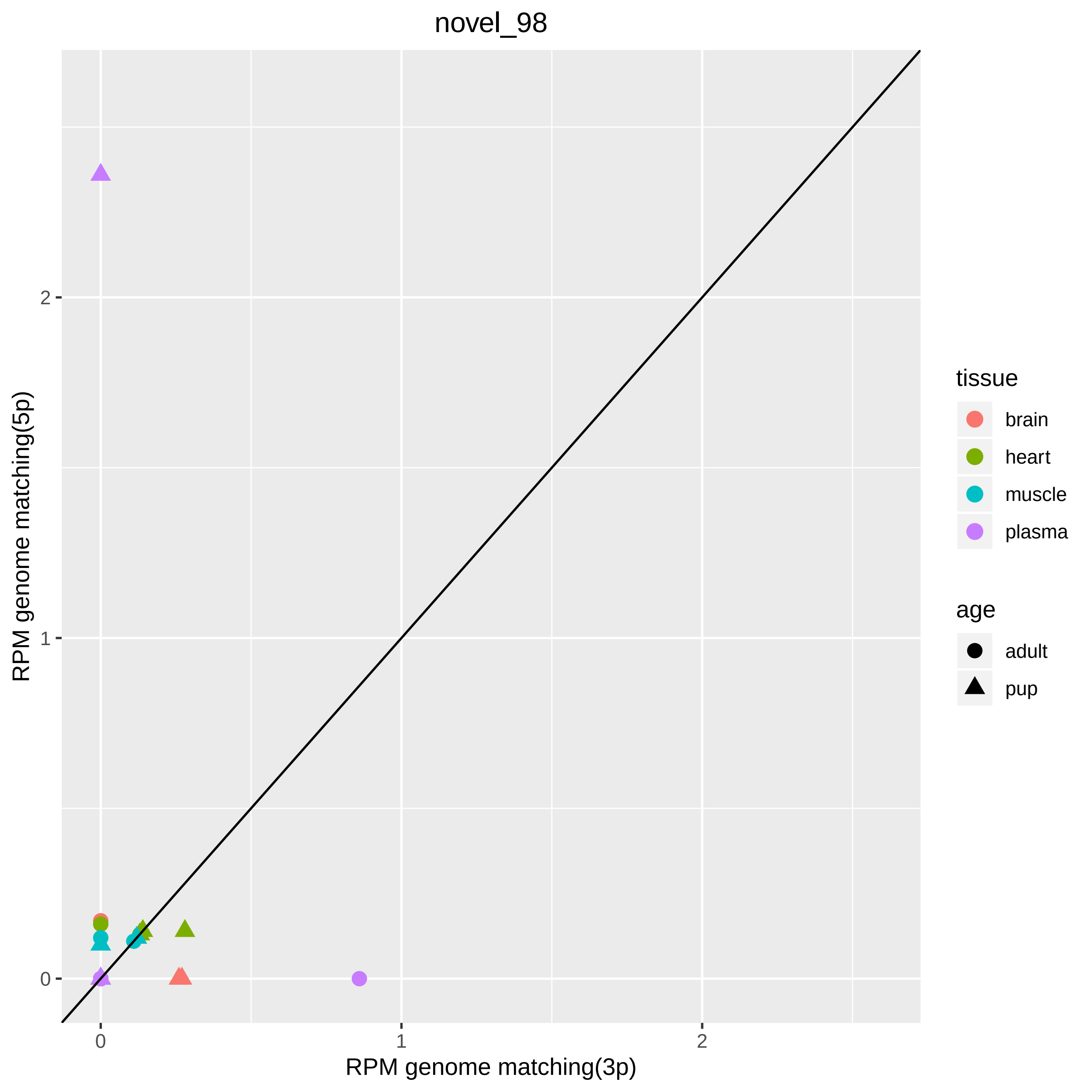

novel_98
RPM genome matching(5p)
RPM genome matching(3p)
0
1
2
0
1
2
tissue
brain
heart
muscle
plasma
age
adult
pup

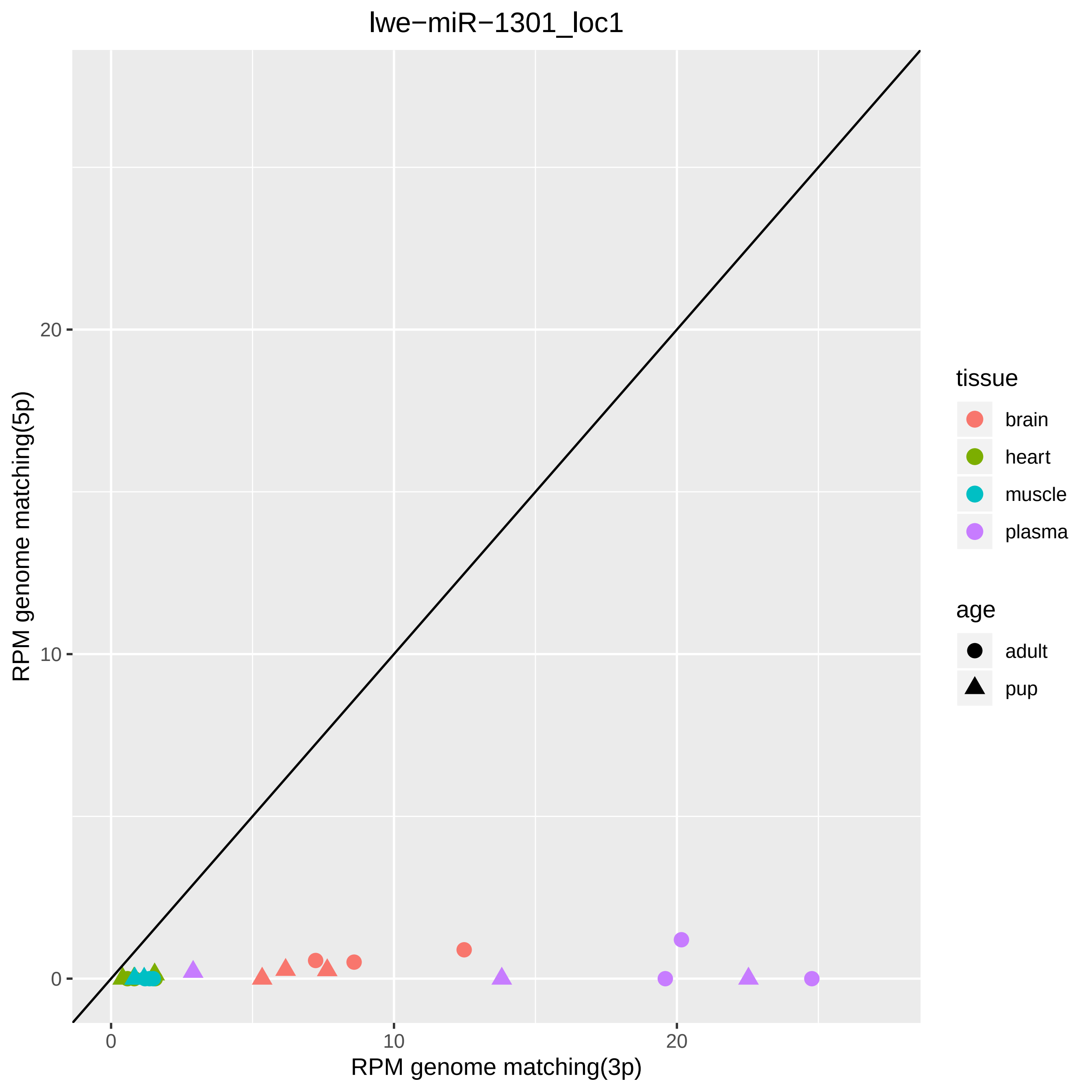

lwe−miR−1301_loc1
RPM genome matching(5p)
RPM genome matching(3p)
0
10
20
0
10
20
tissue
brain
heart
muscle
plasma
age
adult
pup

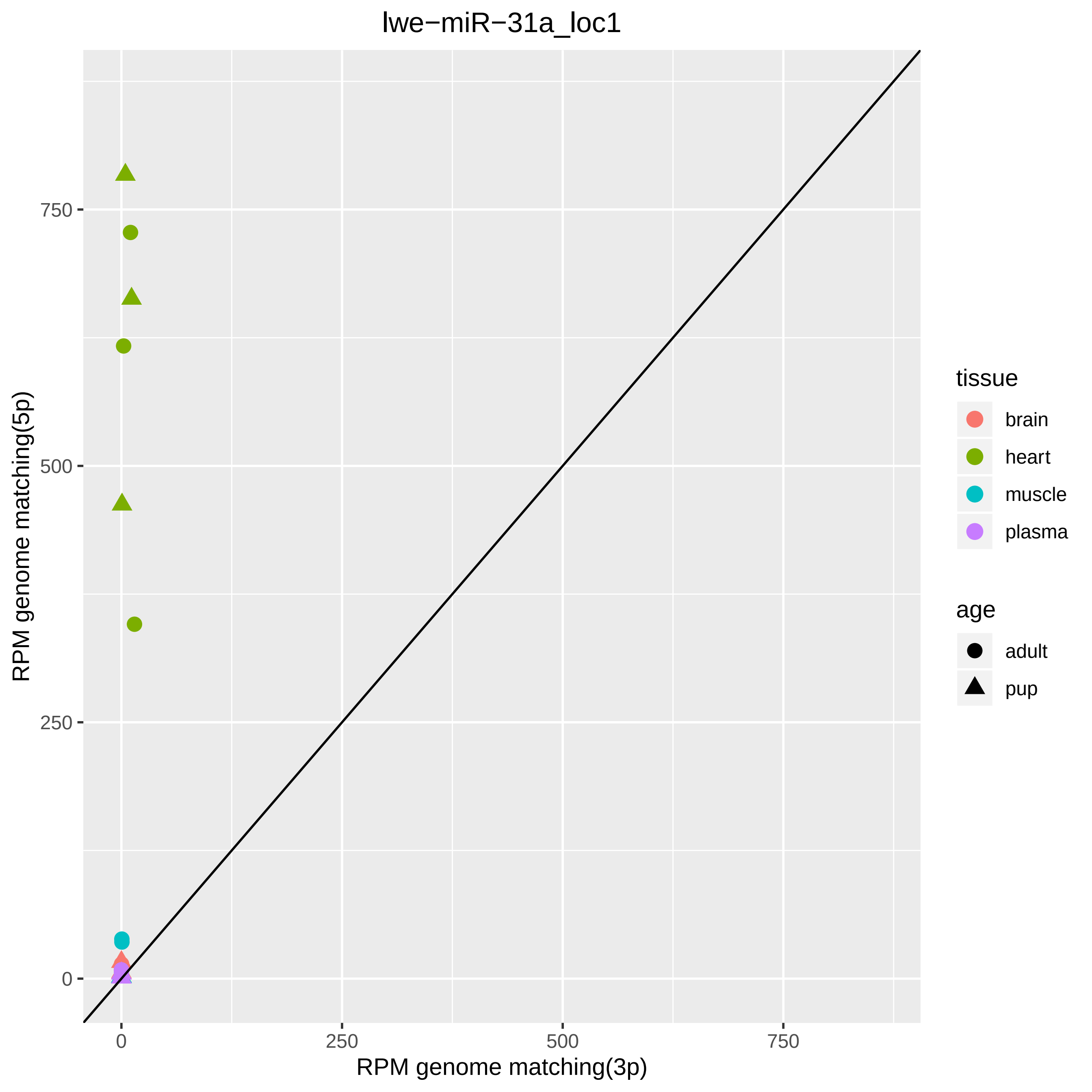

lwe−miR−31a_loc1
RPM genome matching(5p)
RPM genome matching(3p)
0
250
500
750
0
250
500
750
tissue
brain
heart
muscle
plasma
age
adult
pup

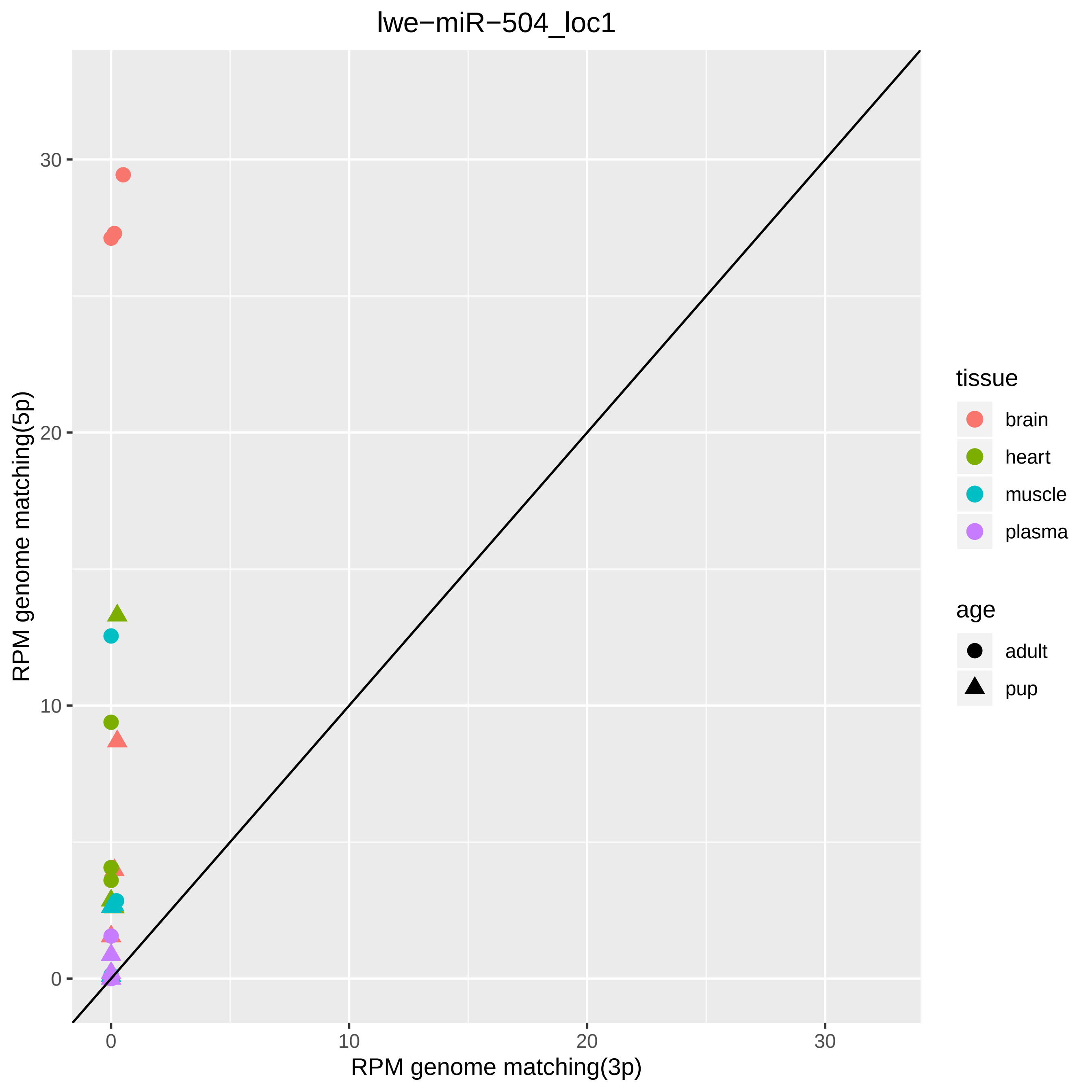

lwe−miR−504_loc1
RPM genome matching(5p)
RPM genome matching(3p)
0
10
20
30
0
10
20
30
tissue
brain
heart
muscle
plasma
age
adult
pup

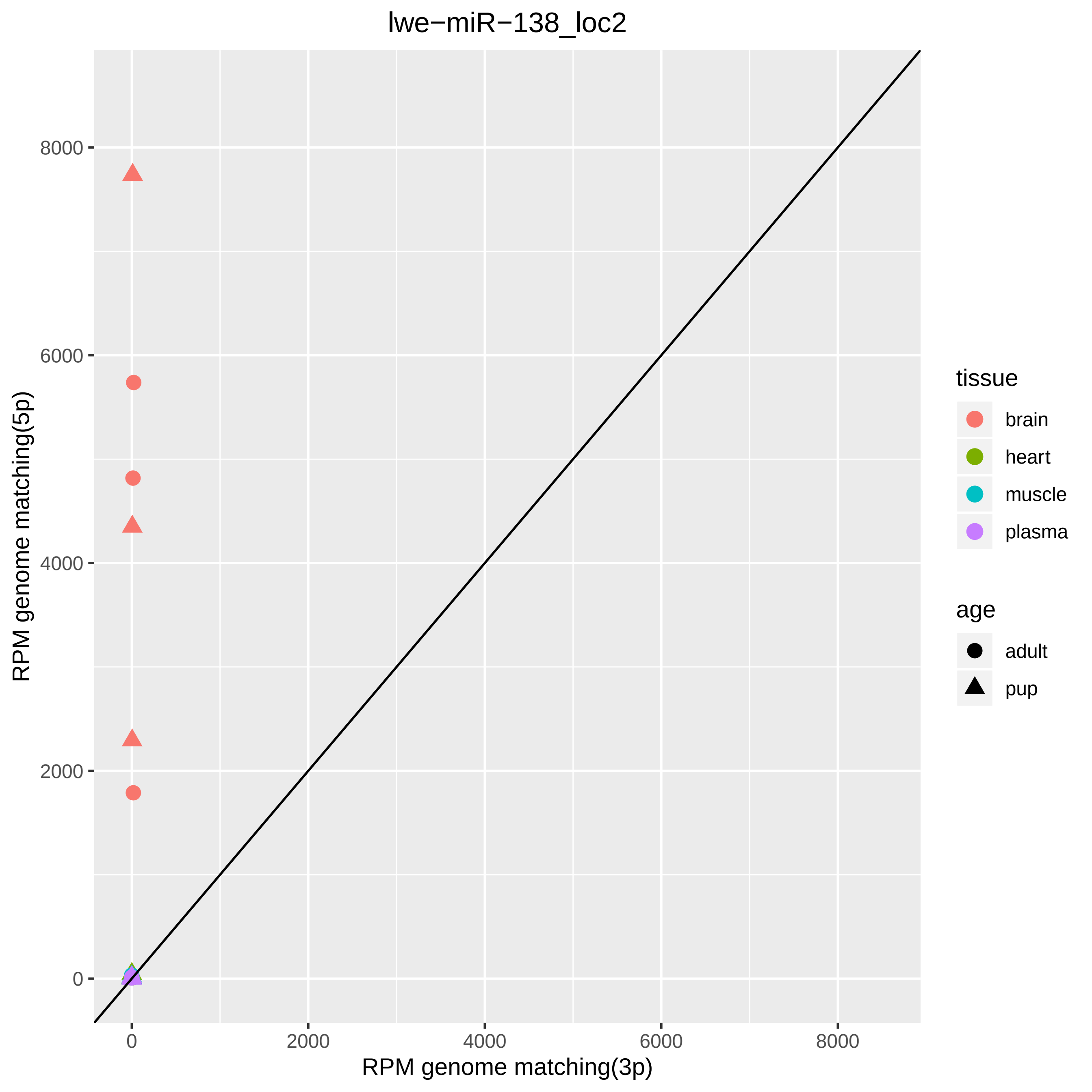
lwe−miR−138_loc2
RPM genome matching(5p)
RPM genome matching(3p)
0
2000
4000
6000
8000
0
2000
4000
6000
8000
tissue
brain
heart
muscle
plasma
age
adult
pup

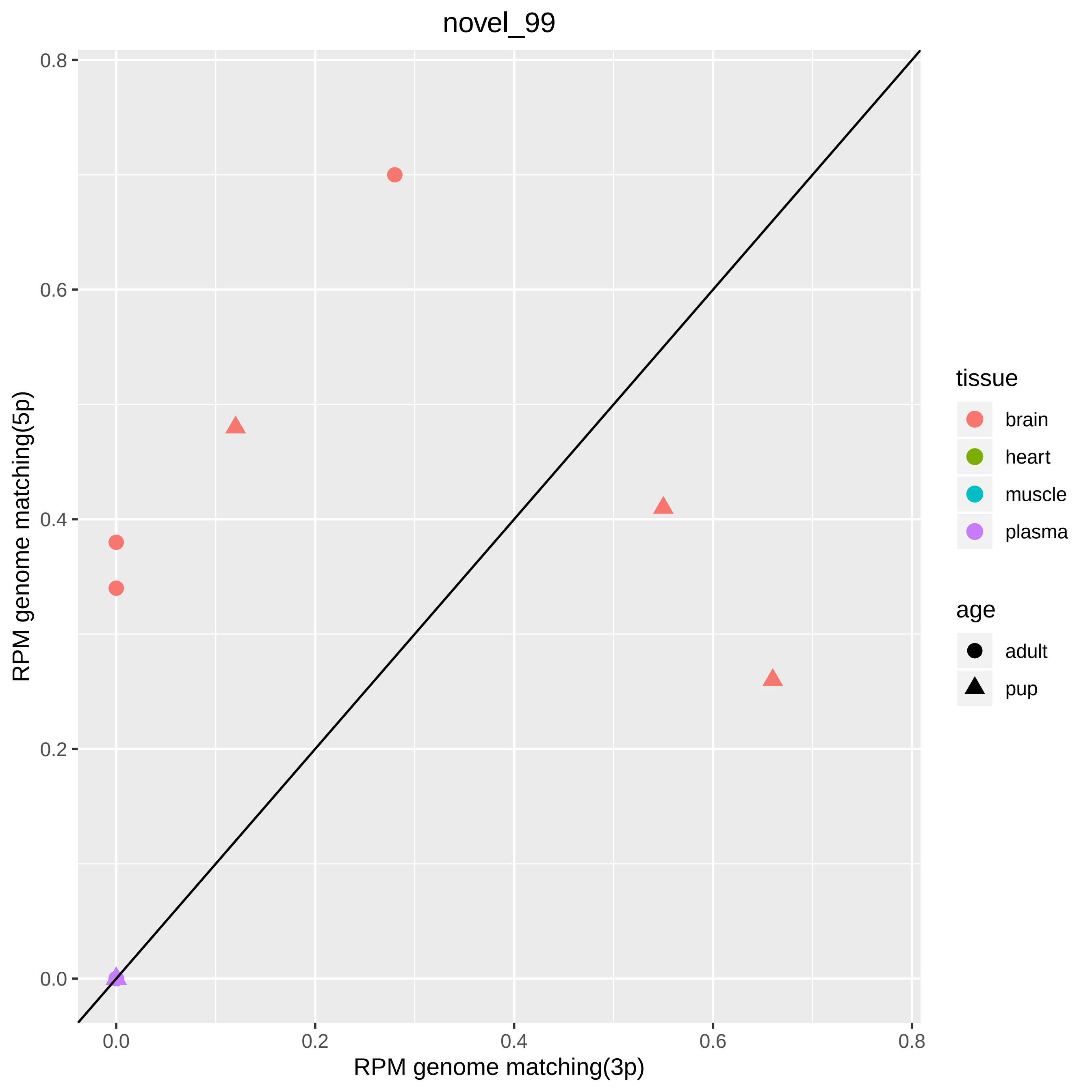

novel_99
RPM genome matching(5p)
RPM genome matching(3p)
0.0
0.2
0.4
0.6
0.8
0.0
0.2
0.4
0.6
0.8
tissue
brain
heart
muscle
plasma
age
adult
pup

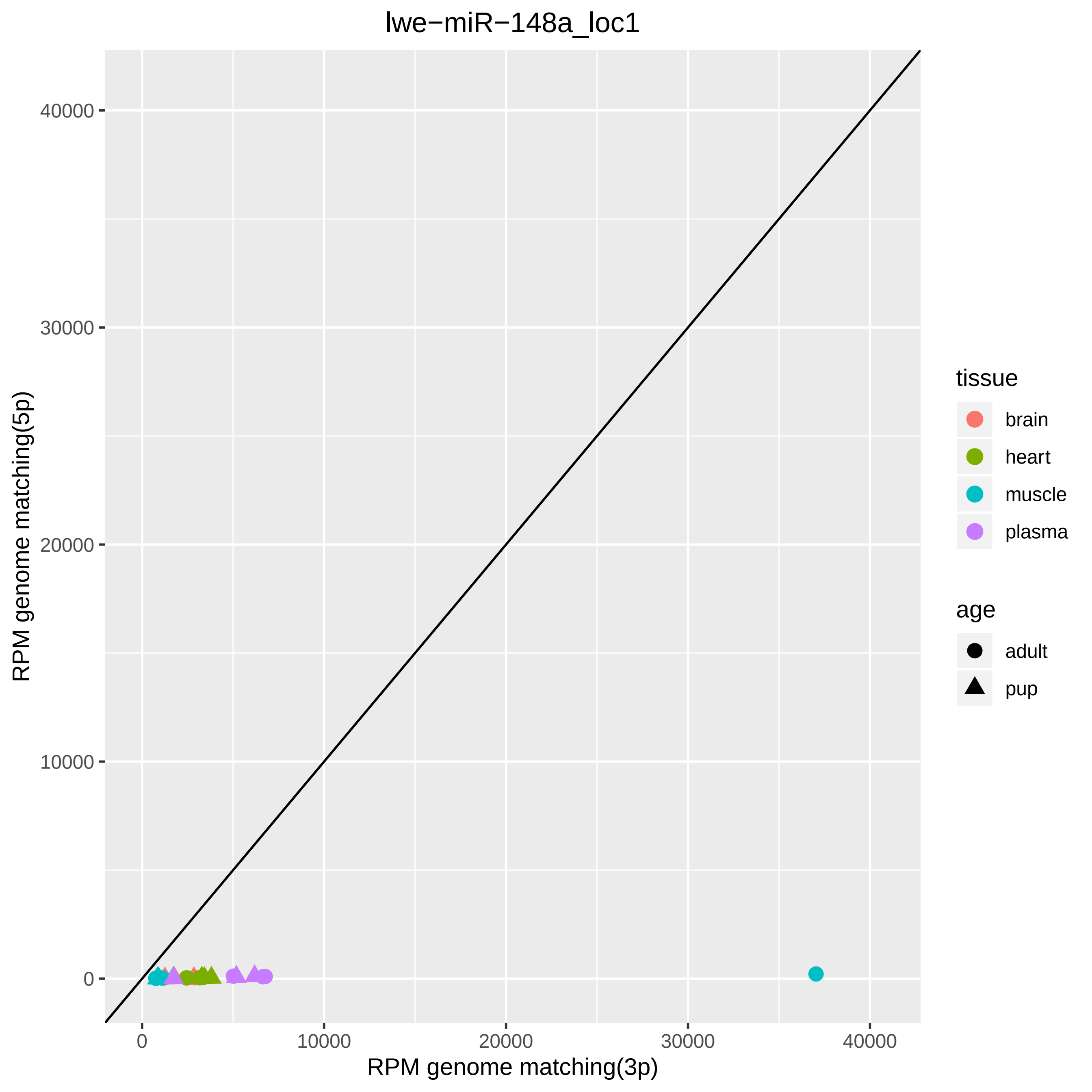

lwe−miR−148a_loc1
RPM genome matching(5p)
RPM genome matching(3p)
0
10000
20000
30000
40000
0
10000
20000
30000
40000
tissue
brain
heart
muscle
plasma
age
adult
pup

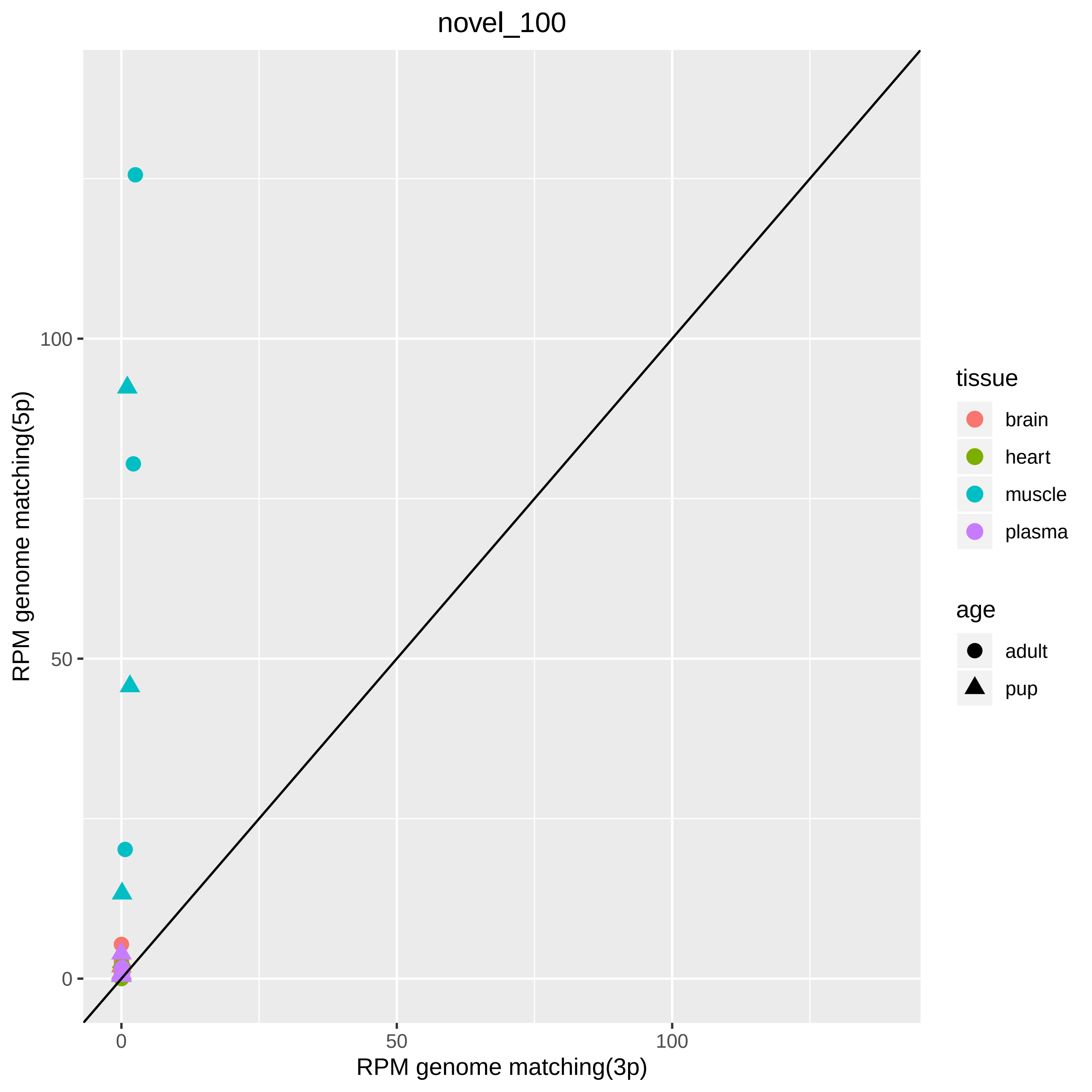

novel_100
RPM genome matching(5p)
RPM genome matching(3p)
100
50
0
0
50
100
tissue
brain
heart
muscle
plasma
age
adult
pup

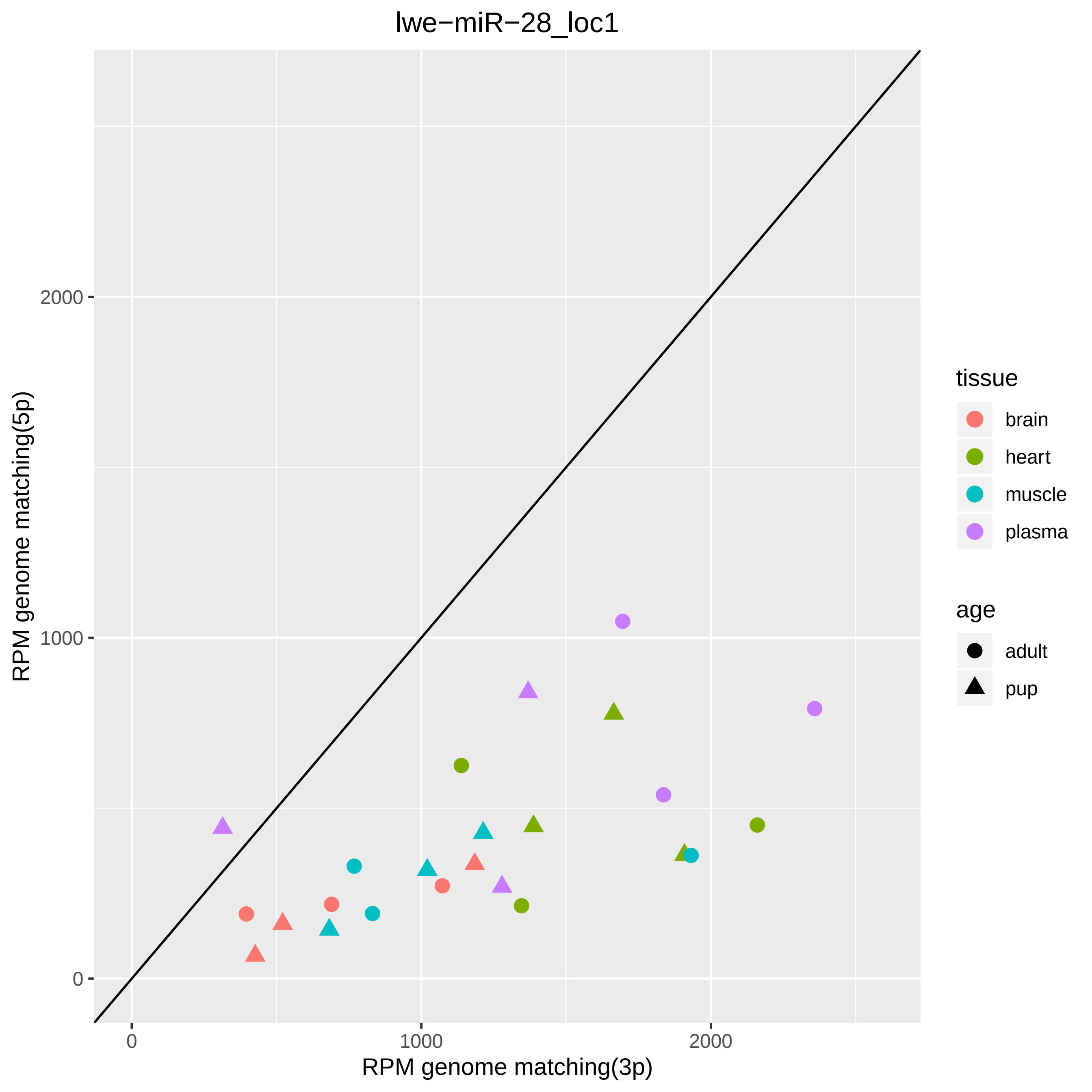

lwe−miR−28_loc1
RPM genome matching(5p)
RPM genome matching(3p)
0
1000
2000
0
1000
2000
tissue
brain
heart
muscle
plasma
age
adult
pup

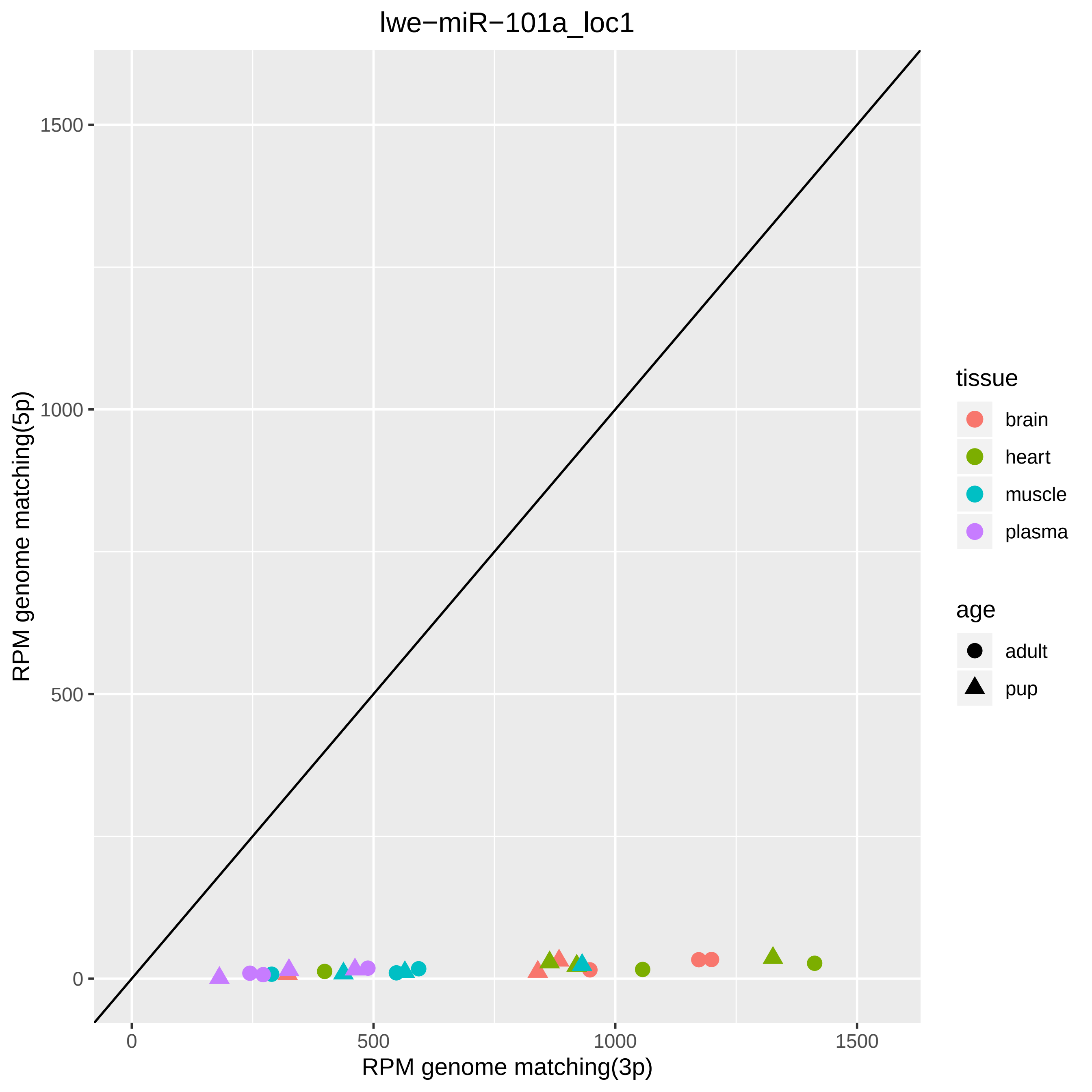

lwe−miR−101a_loc1
RPM genome matching(5p)
RPM genome matching(3p)
0
500
1000
1500
tissue
brain
heart
muscle
plasma
age
adult
pup

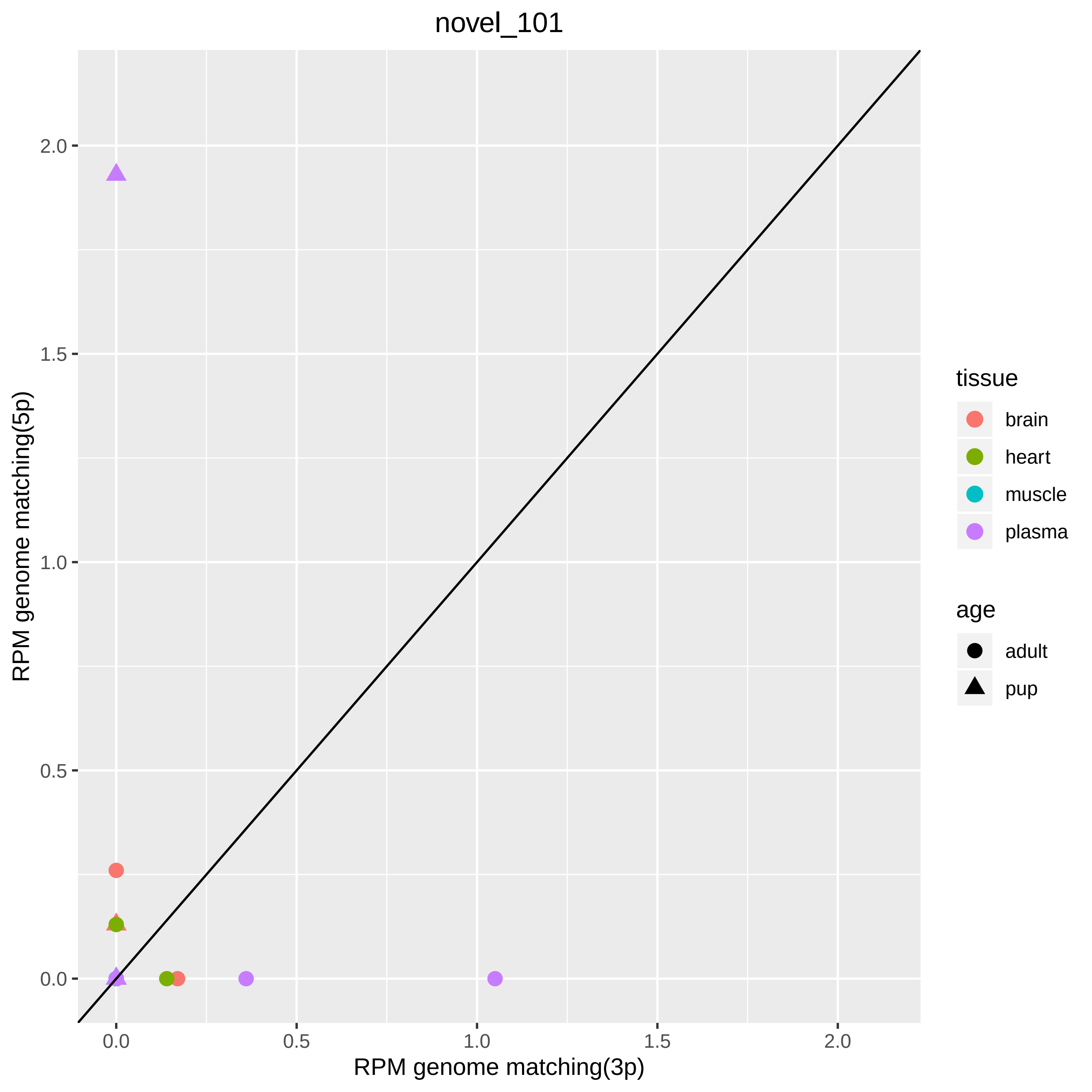
novel_101
RPM genome matching(5p)
RPM genome matching(3p)
0.0
0.5
1.0
1.5
2.0
tissue
brain
heart
muscle
plasma
age
adult
pup

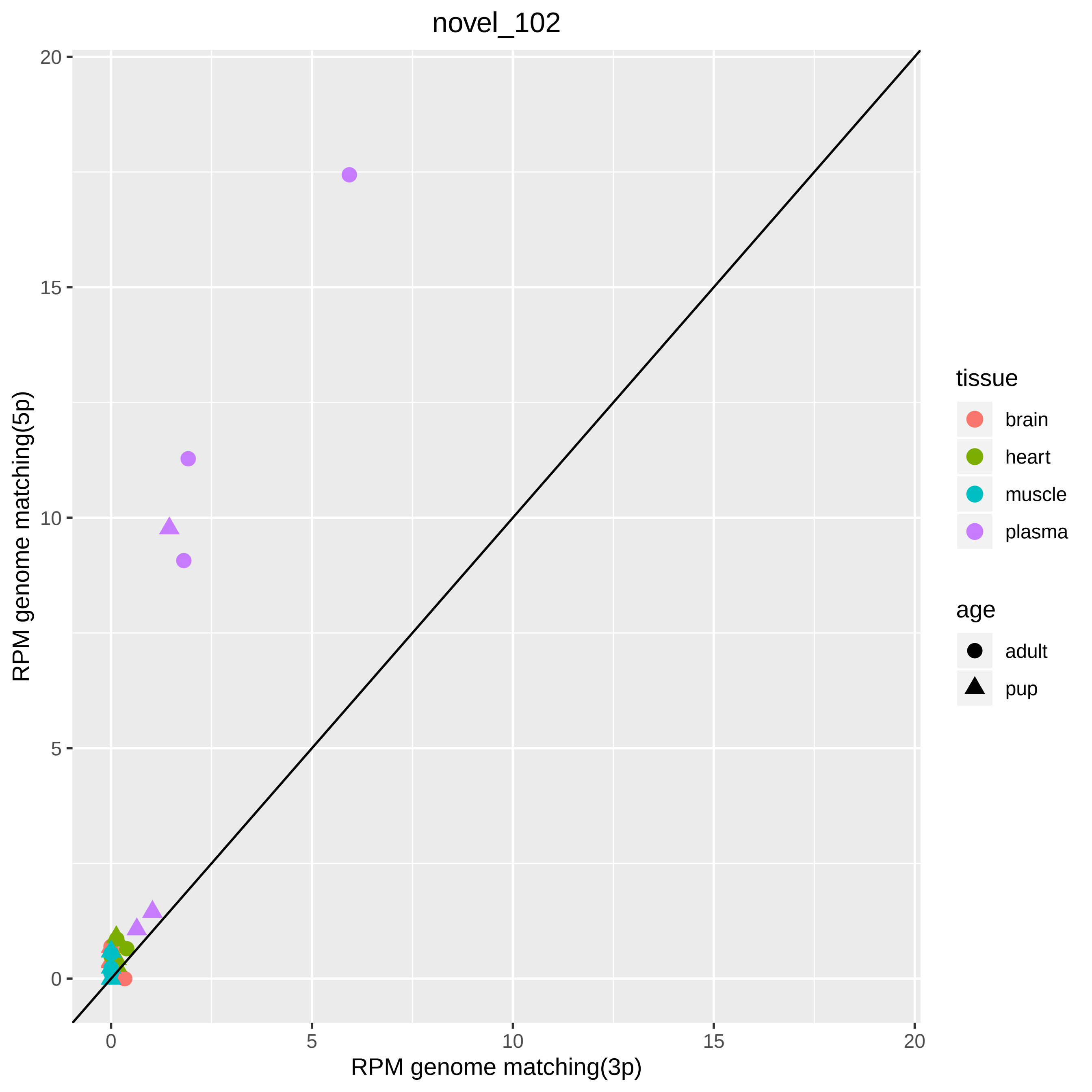

novel_102
RPM genome matching(5p)
RPM genome matching(3p)
0
5
10
15
20
0
5
10
15
20
tissue
brain
heart
muscle
plasma
age
adult
pup

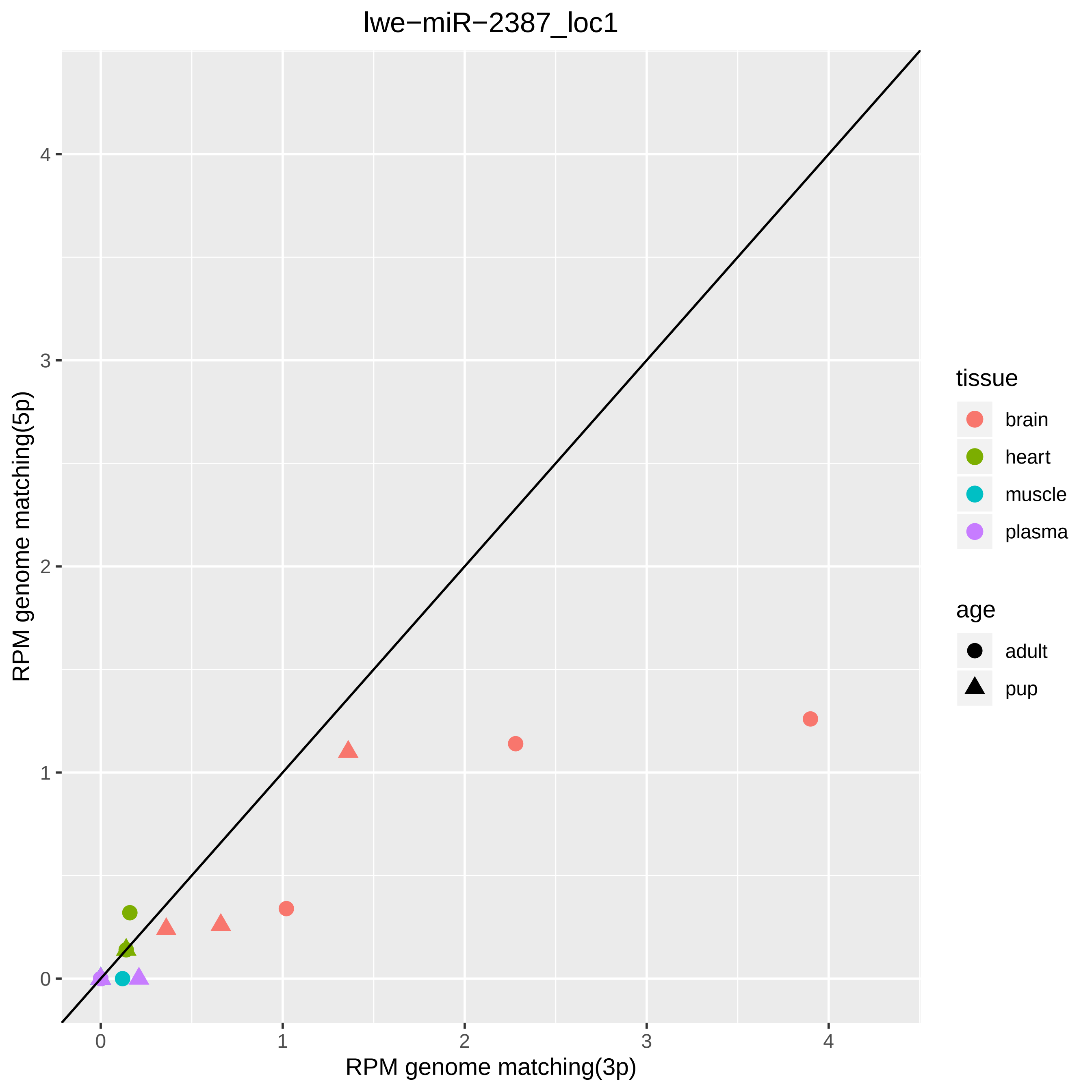
lwe−miR−2387_loc1
RPM genome matching(5p)
RPM genome matching(3p)
0
1
2
3
4
tissue
brain
heart
muscle
plasma
age
adult
pup

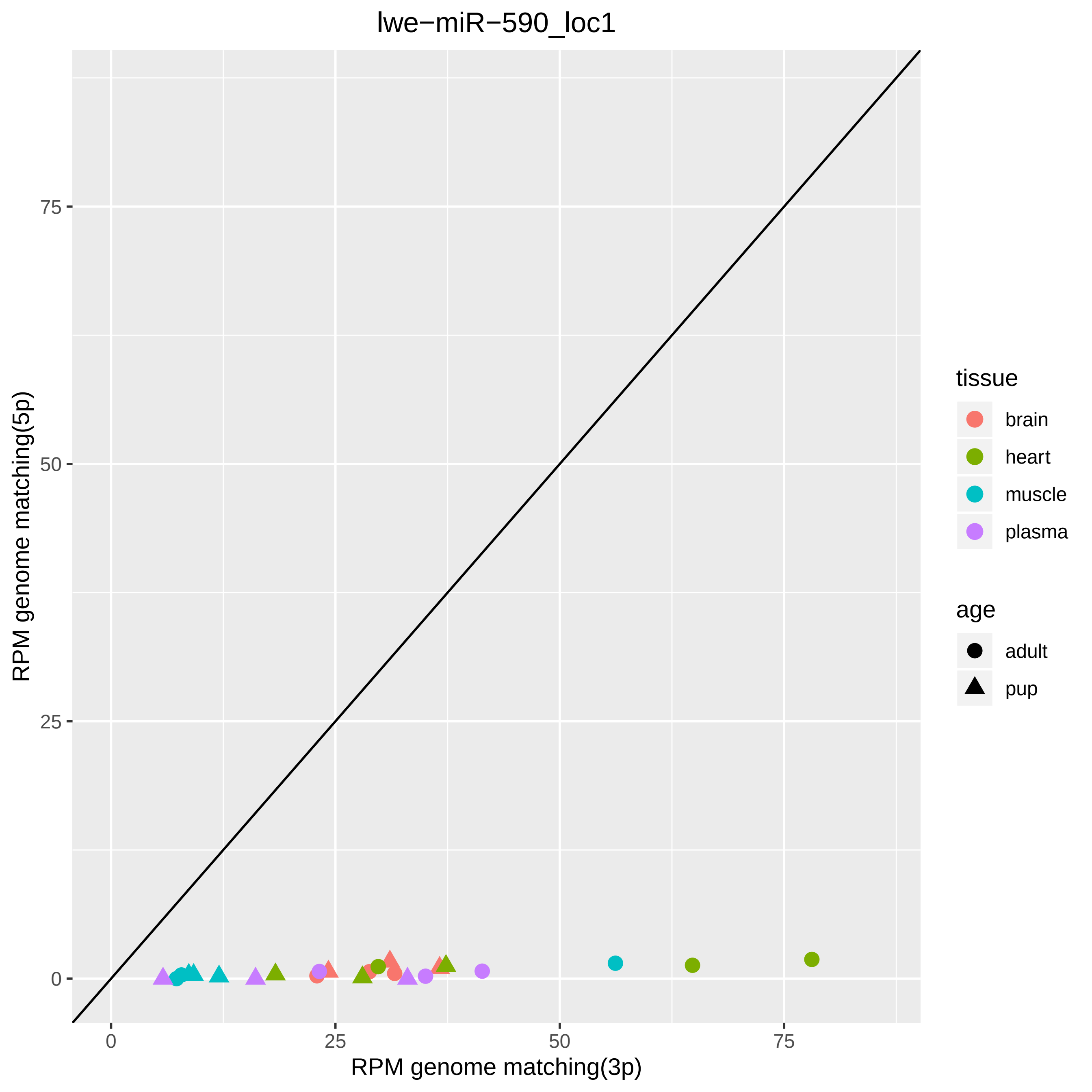

lwe−miR−590_loc1
RPM genome matching(5p)
RPM genome matching(3p)
0
25
50
75
tissue
brain
heart
muscle
plasma
age
adult
pup

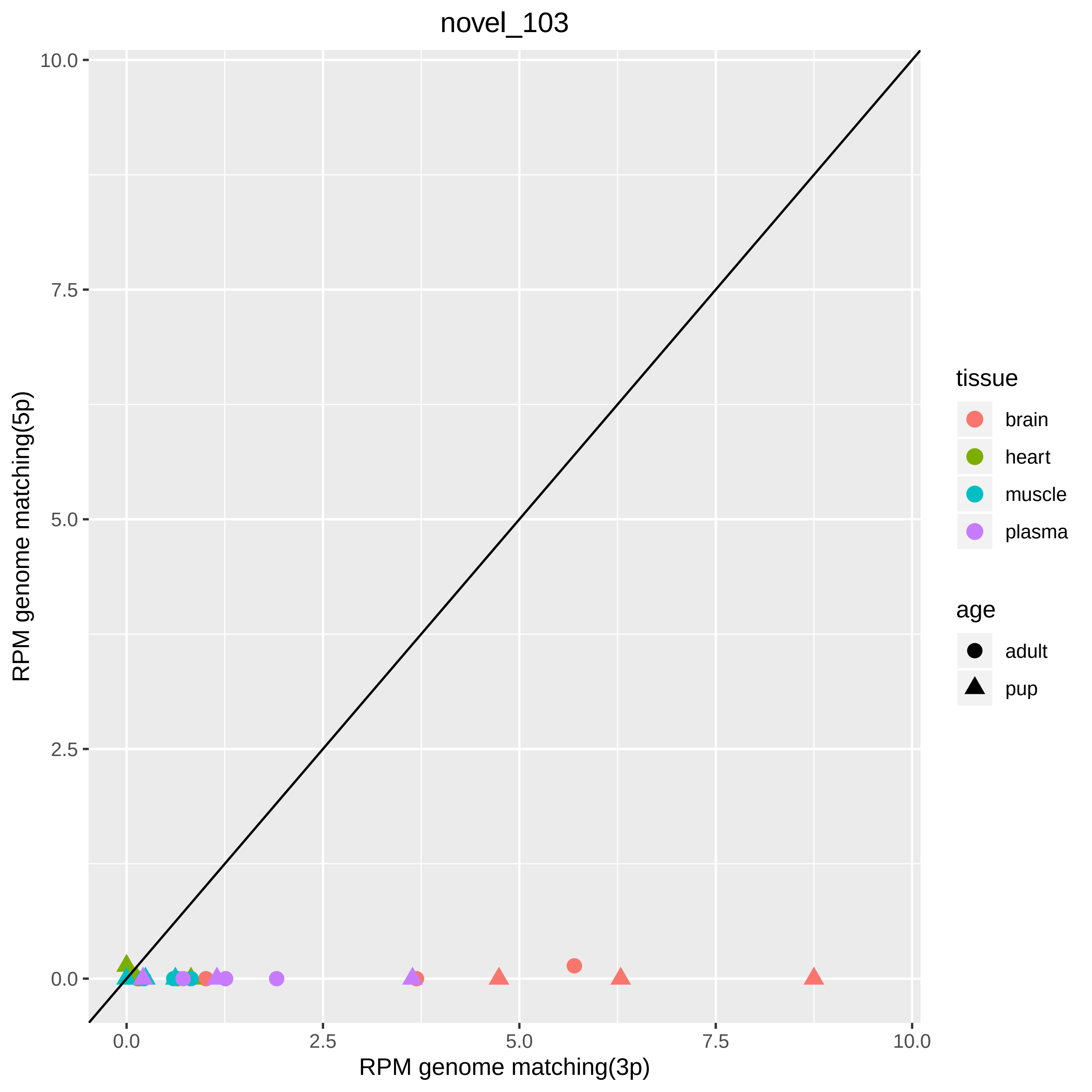

novel_103
RPM genome matching(5p)
RPM genome matching(3p)
0.0
2.5
5.0
7.5
10.0
tissue
brain
heart
muscle
plasma
age
adult
pup

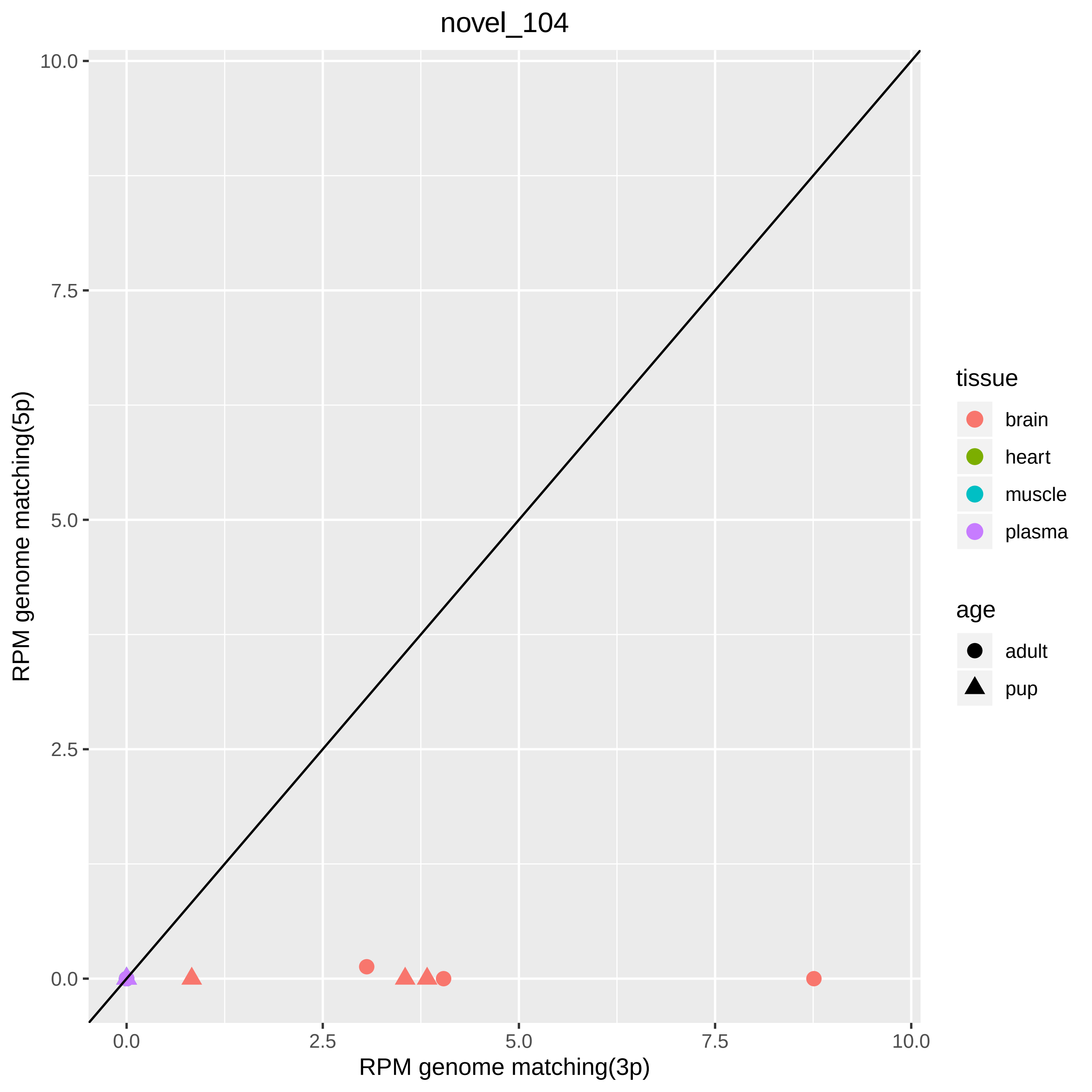
novel_104
RPM genome matching(5p)
RPM genome matching(3p)
0.0
2.5
5.0
7.5
10.0
0.0
2.5
5.0
7.5
10.0
tissue
brain
heart
muscle
plasma
age
adult
pup

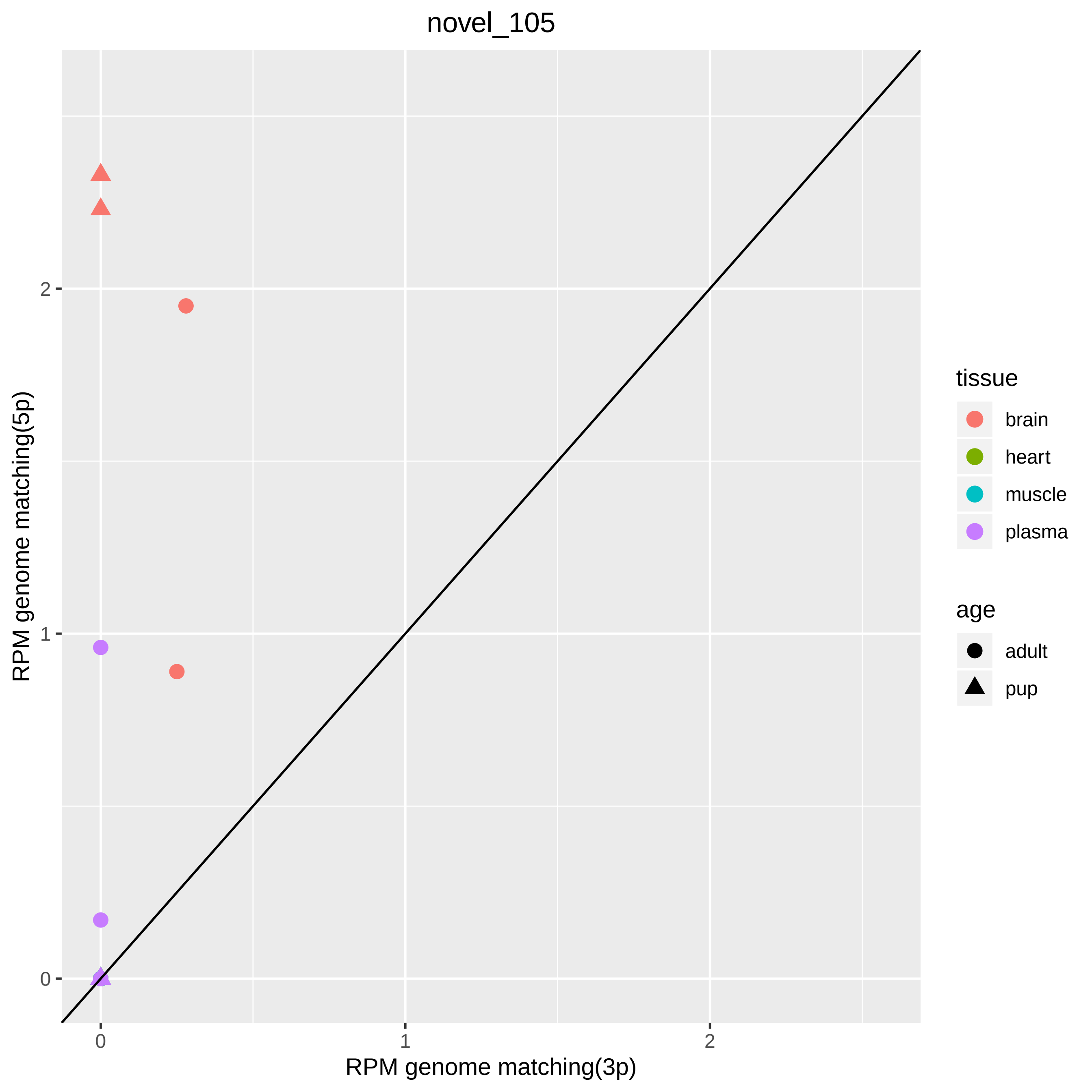
novel_105
RPM genome matching(5p)
RPM genome matching(3p)
0
1
2
tissue
brain
heart
muscle
plasma
age
adult
pup

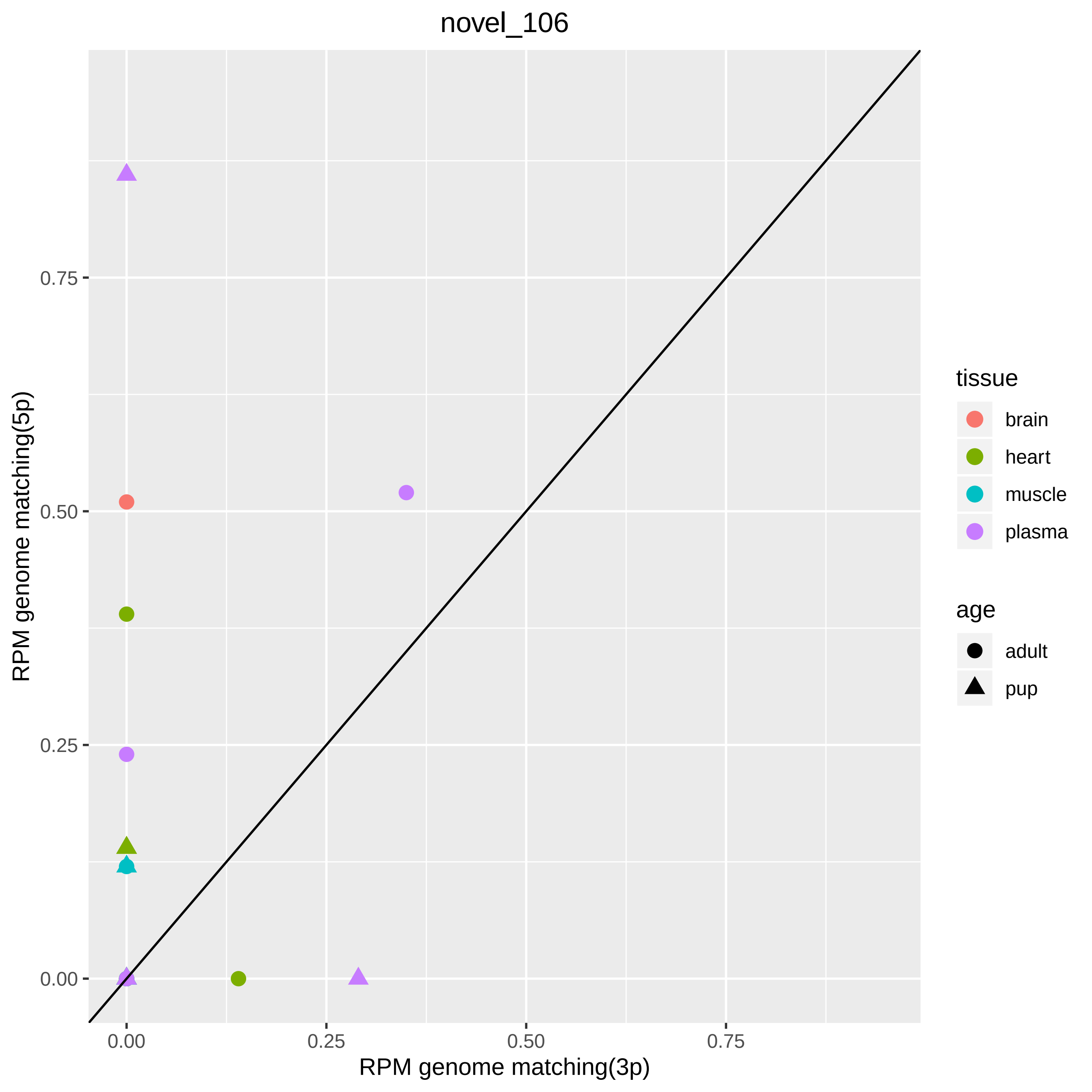

novel_106
RPM genome matching(5p)
RPM genome matching(3p)
0.00
0.25
0.50
0.75
tissue
brain
heart
muscle
plasma
age
adult
pup

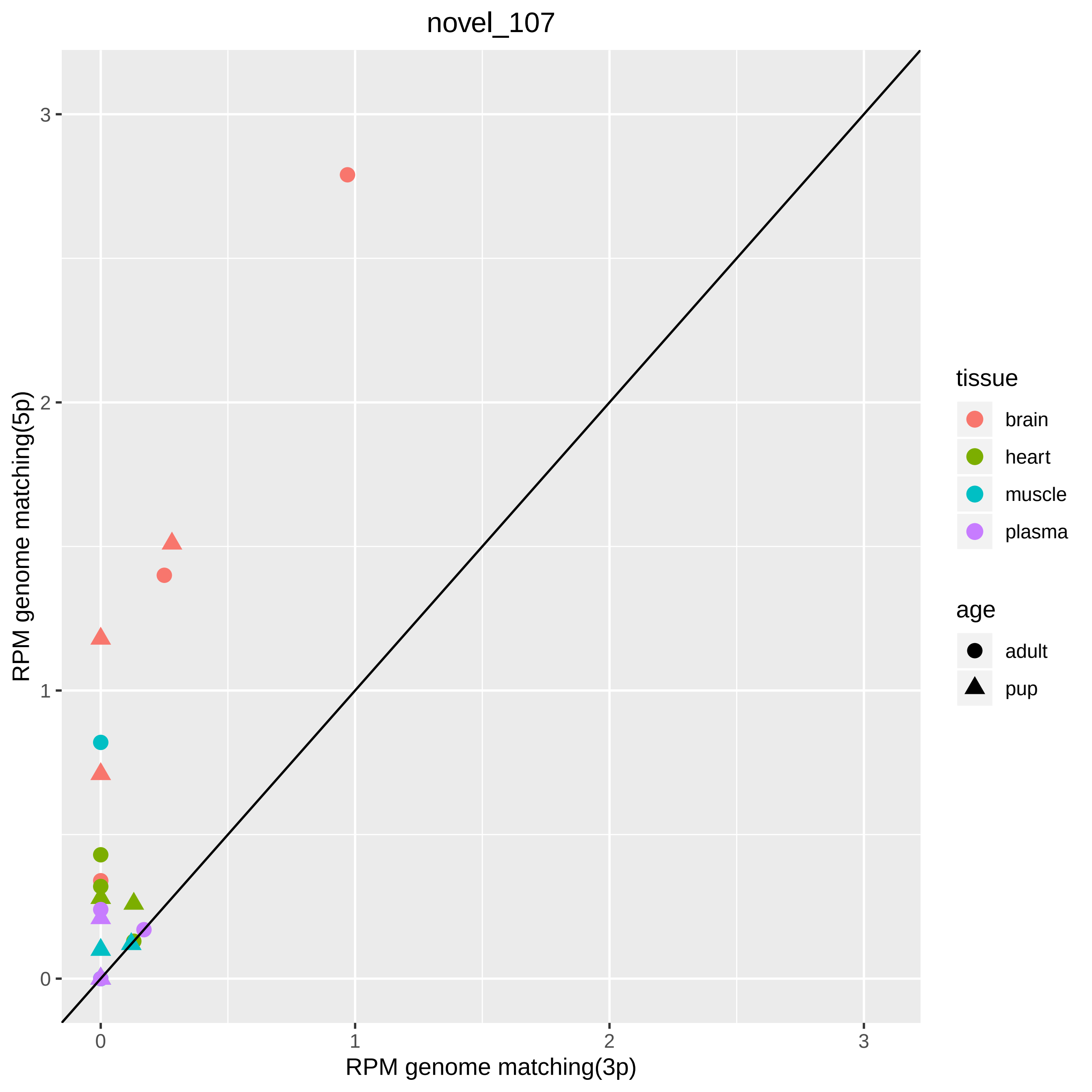

novel_107
RPM genome matching(5p)
RPM genome matching(3p)
0
1
2
3
0
1
2
3
tissue
brain
heart
muscle
plasma
age
adult
pup

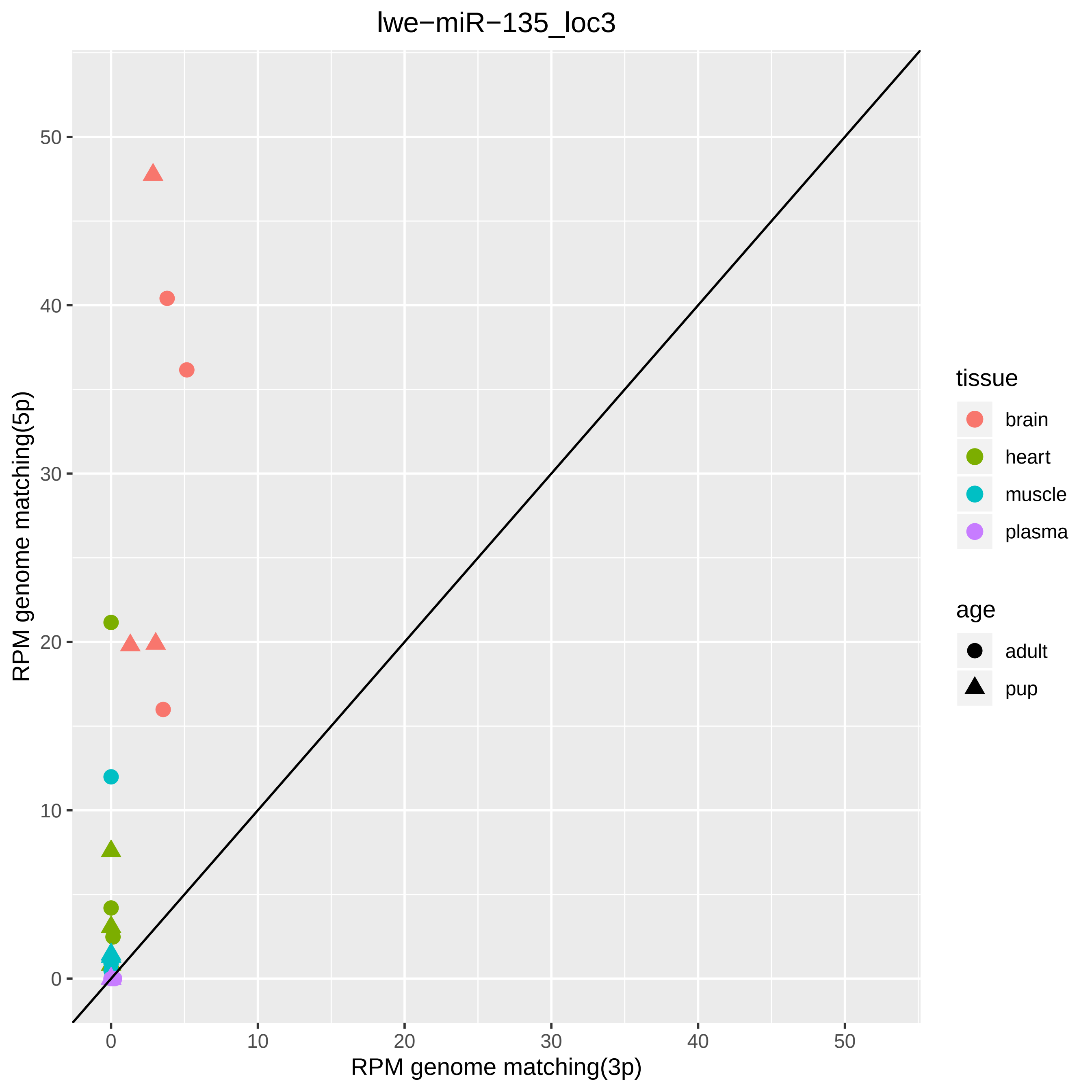

lwe−miR−135_loc3
RPM genome matching(5p)
RPM genome matching(3p)
0
10
20
30
40
50
tissue
brain
heart
muscle
plasma
age
adult
pup

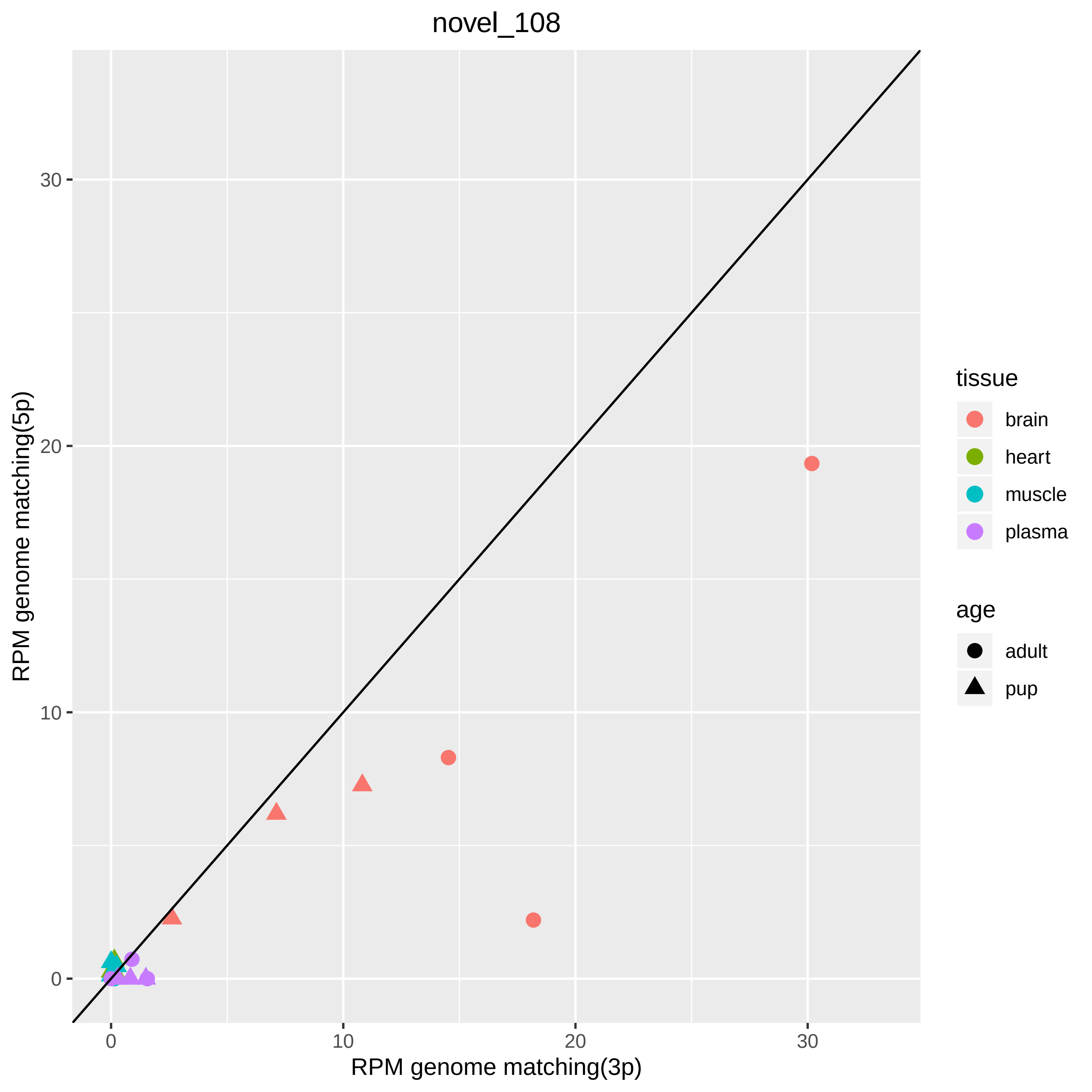

novel_108
RPM genome matching(5p)
RPM genome matching(3p)
0
10
20
30
0
10
20
30
tissue
brain
heart
muscle
plasma
age
adult
pup

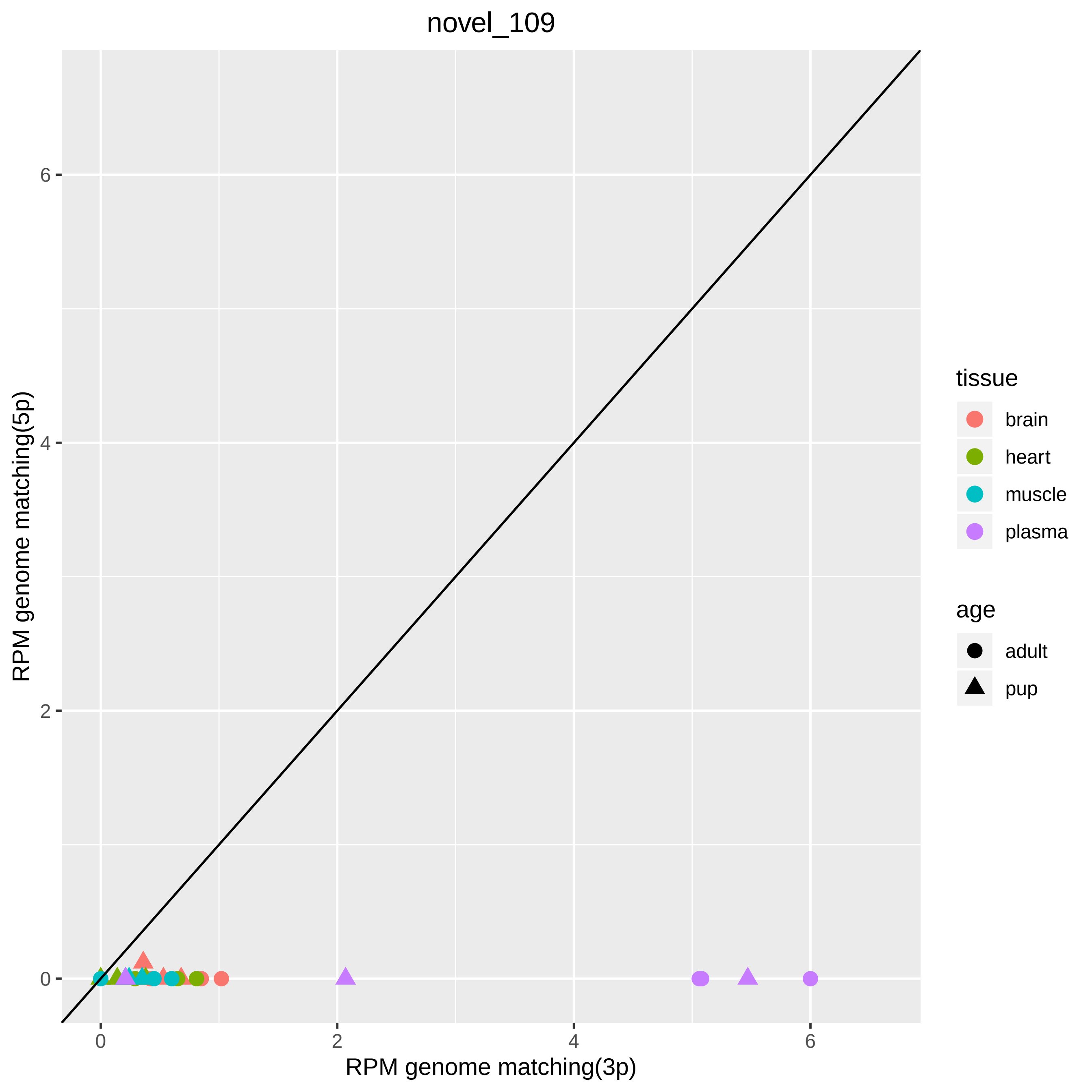

novel_109
RPM genome matching(5p)
RPM genome matching(3p)
0
2
4
6
0
2
4
6
tissue
brain
heart
muscle
plasma
age
adult
pup

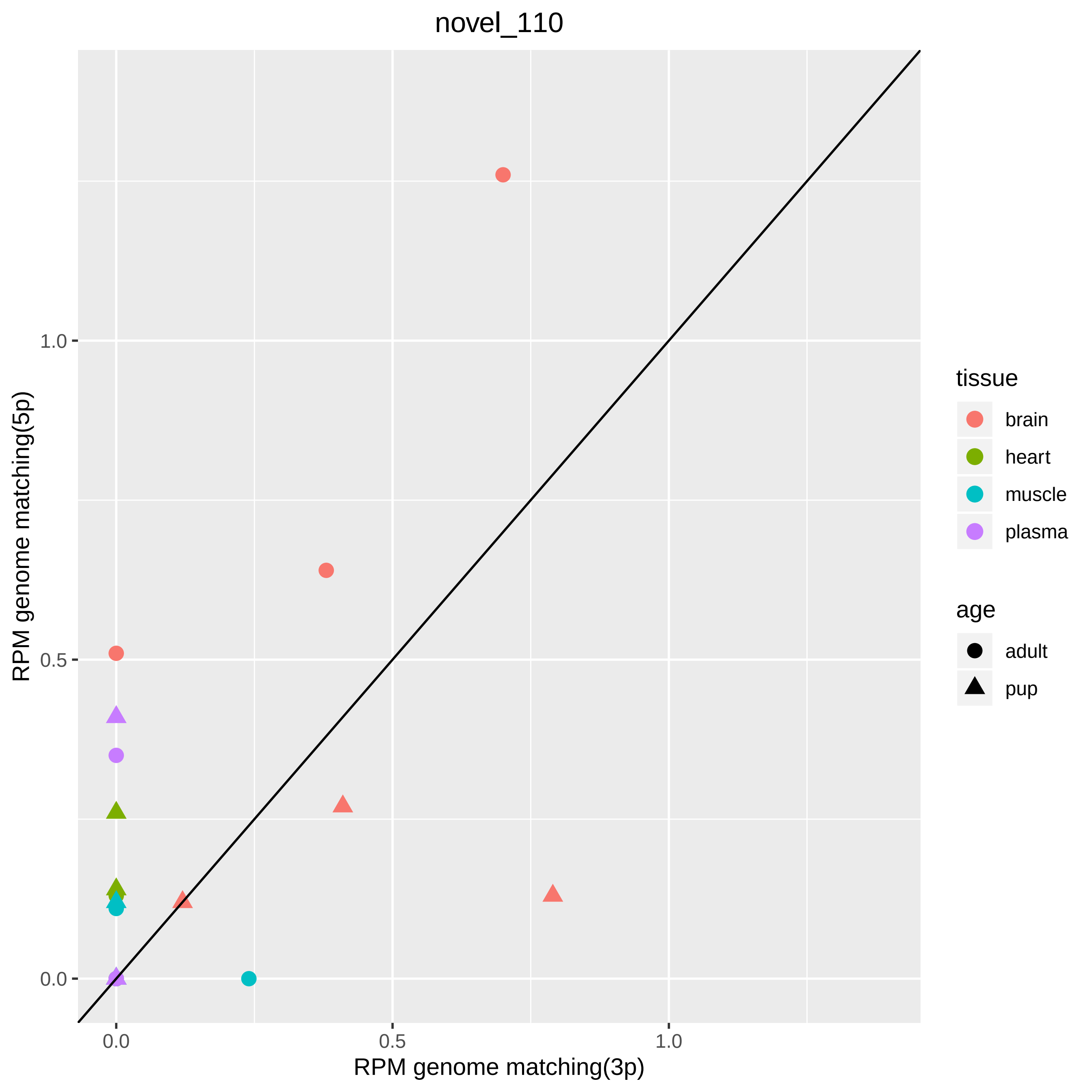

novel_110
RPM genome matching(5p)
RPM genome matching(3p)
0.0
0.5
1.0
tissue
brain
heart
muscle
plasma
age
adult
pup

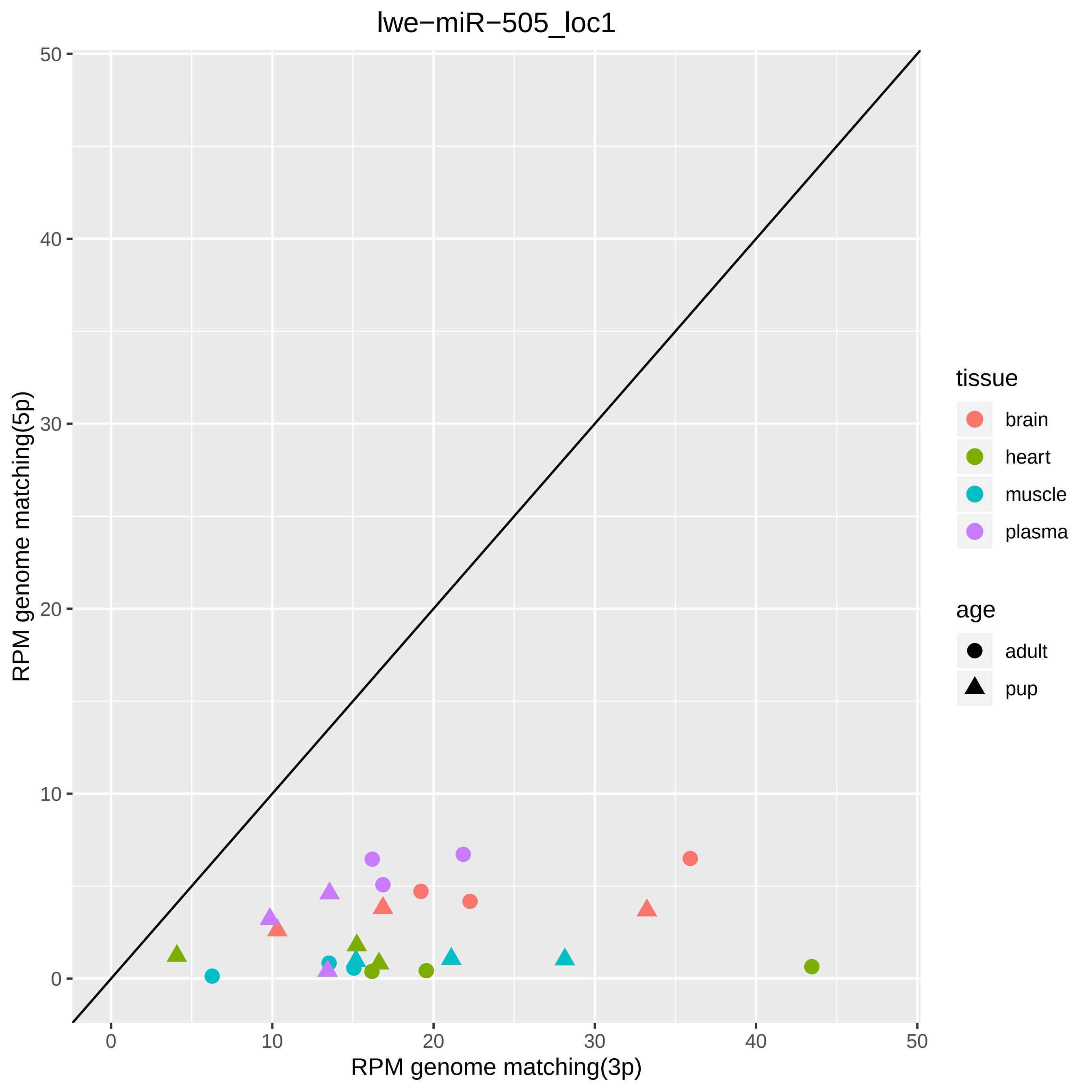
lwe−miR−505_loc1
RPM genome matching(5p)
RPM genome matching(3p)
0
10
20
30
40
50
0
10
20
30
40
50
tissue
brain
heart
muscle
plasma
age
adult
pup

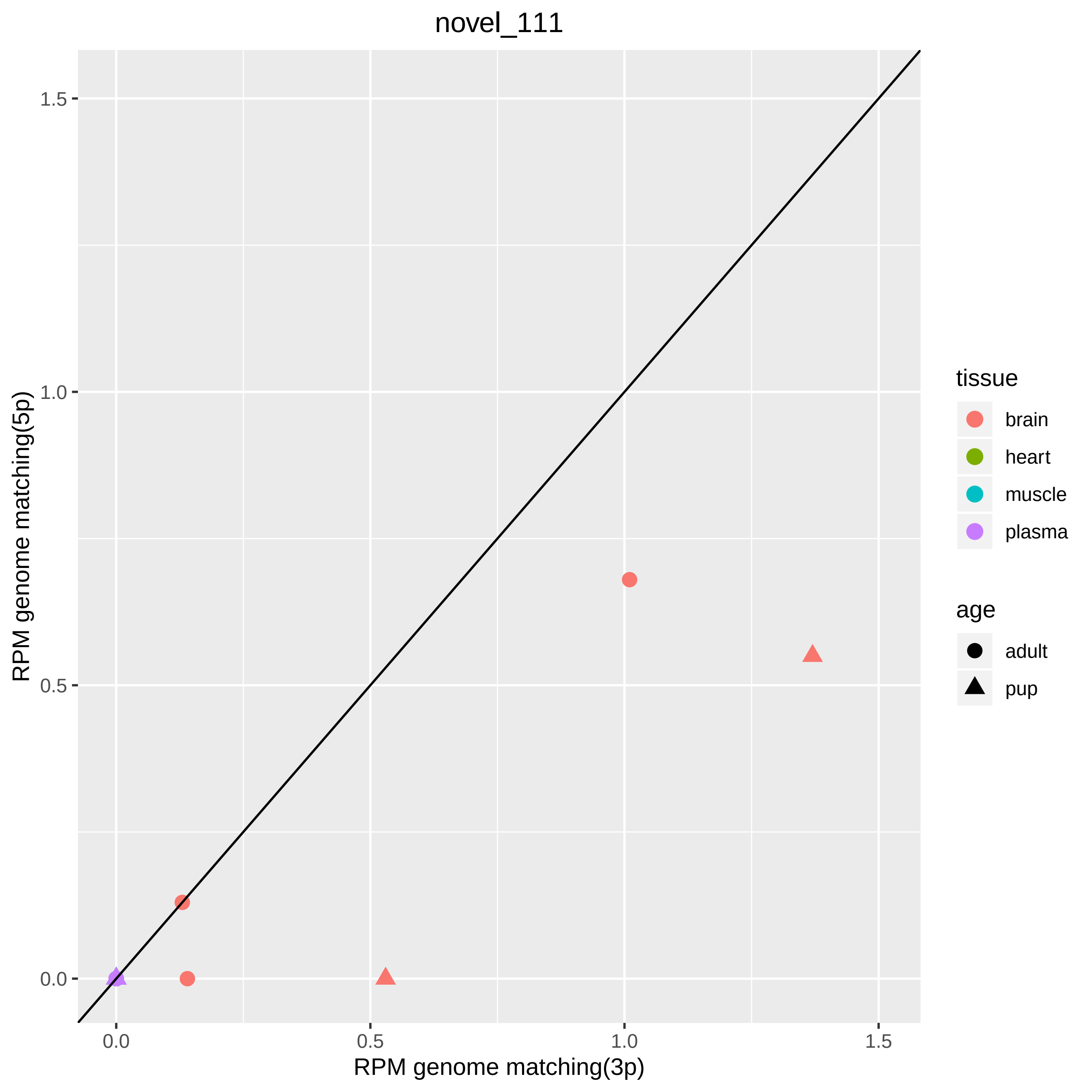

novel_111
RPM genome matching(5p)
RPM genome matching(3p)
0.0
0.5
1.0
1.5
0.0
0.5
1.0
1.5
tissue
brain
heart
muscle
plasma
age
adult
pup

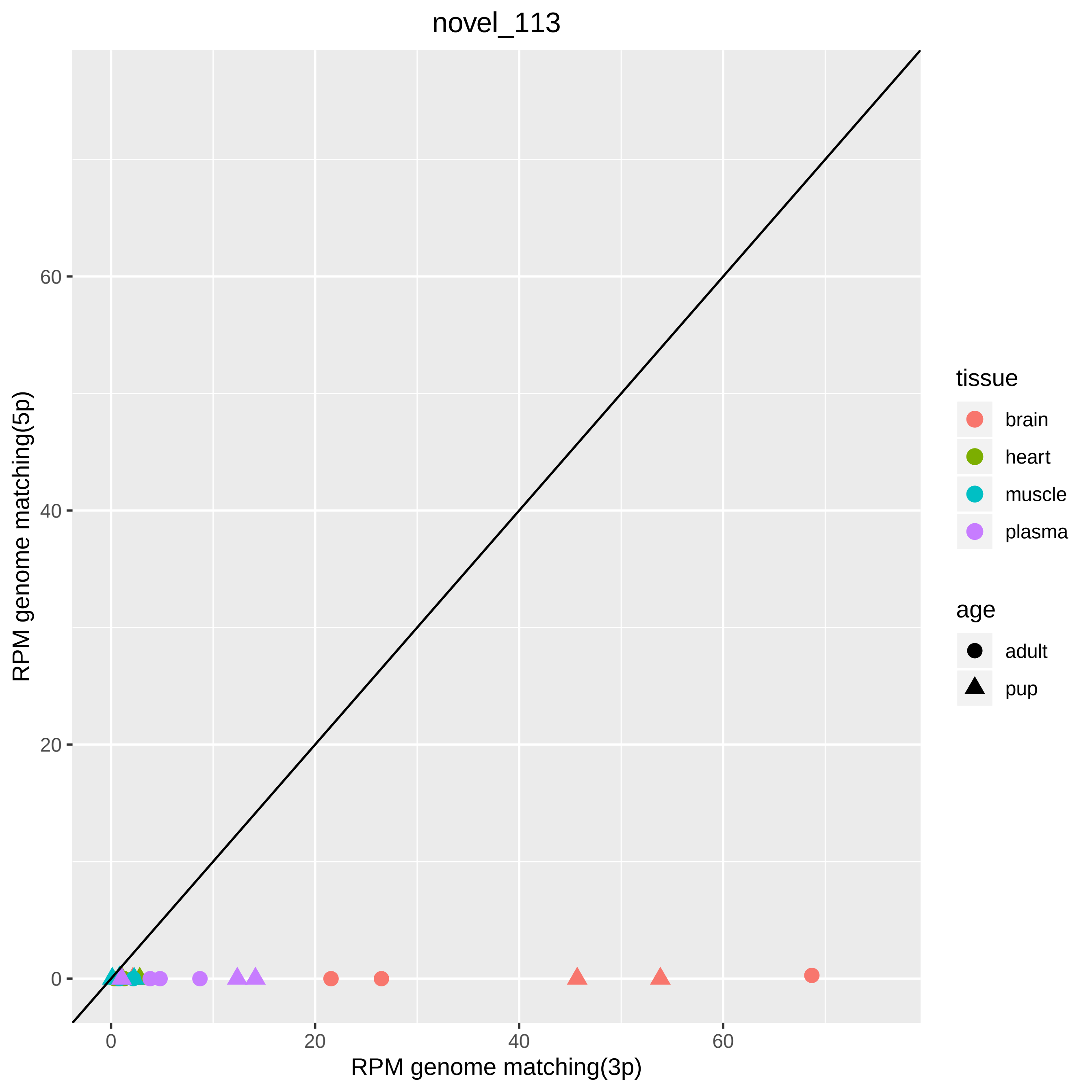
novel_113
RPM genome matching(5p)
RPM genome matching(3p)
0
20
40
60
0
20
40
60
tissue
brain
heart
muscle
plasma
age
adult
pup

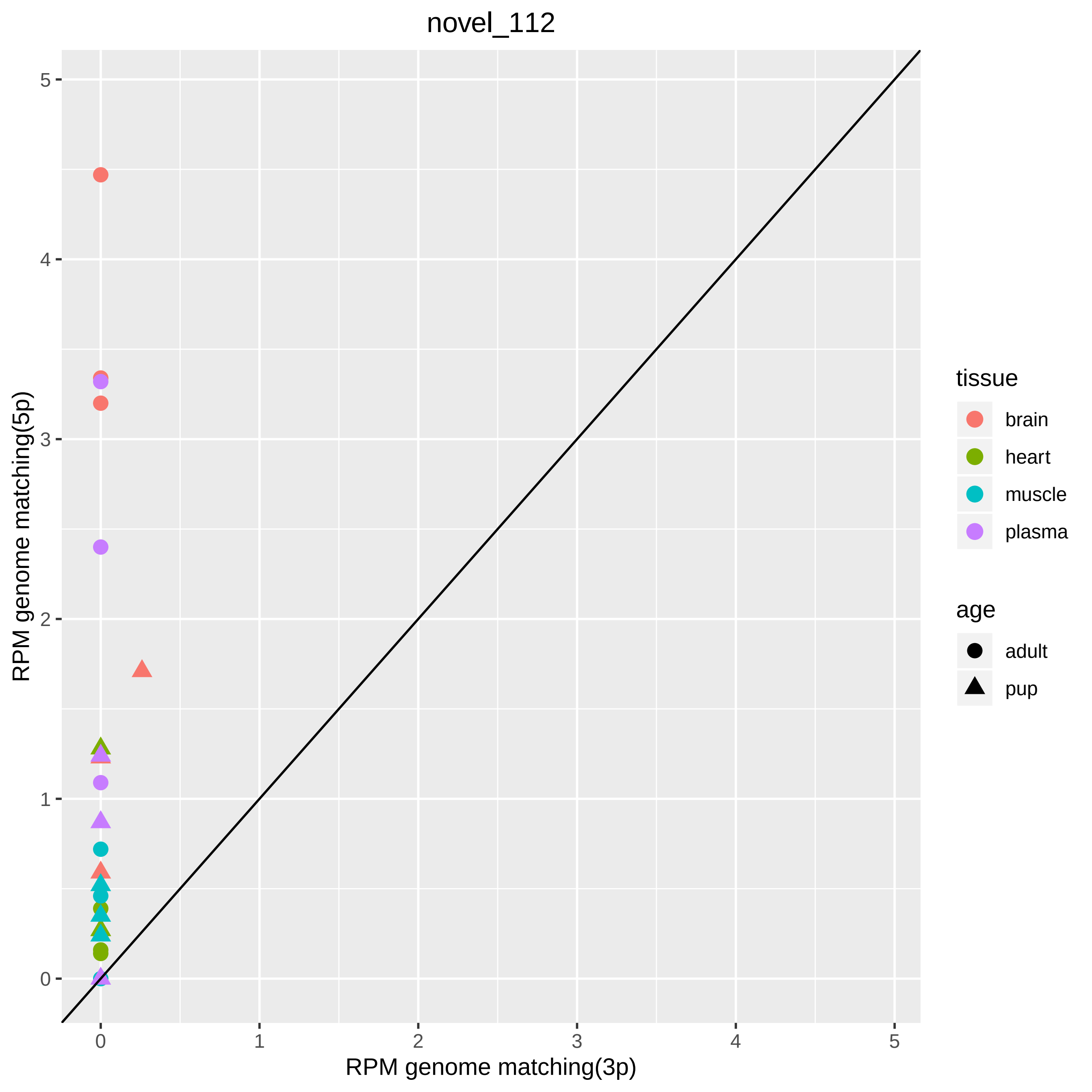

novel_112
RPM genome matching(5p)
RPM genome matching(3p)
0
1
2
3
4
5
0
1
2
3
4
5
tissue
brain
heart
muscle
plasma
age
adult
pup

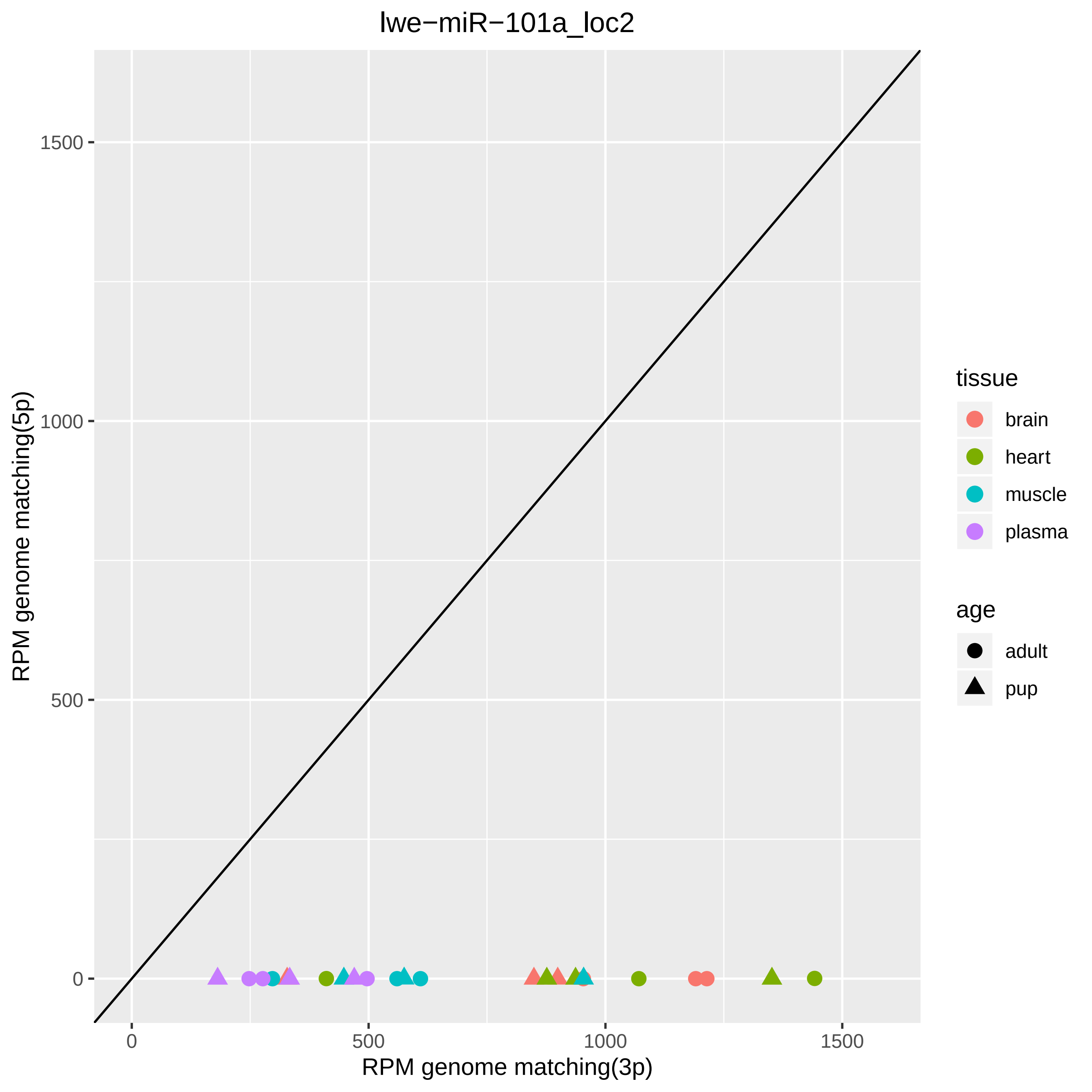
lwe−miR−101a_loc2
RPM genome matching(5p)
RPM genome matching(3p)
1500
1000
500
0
0
500
1000
1500
tissue
brain
heart
muscle
plasma
age
adult
pup

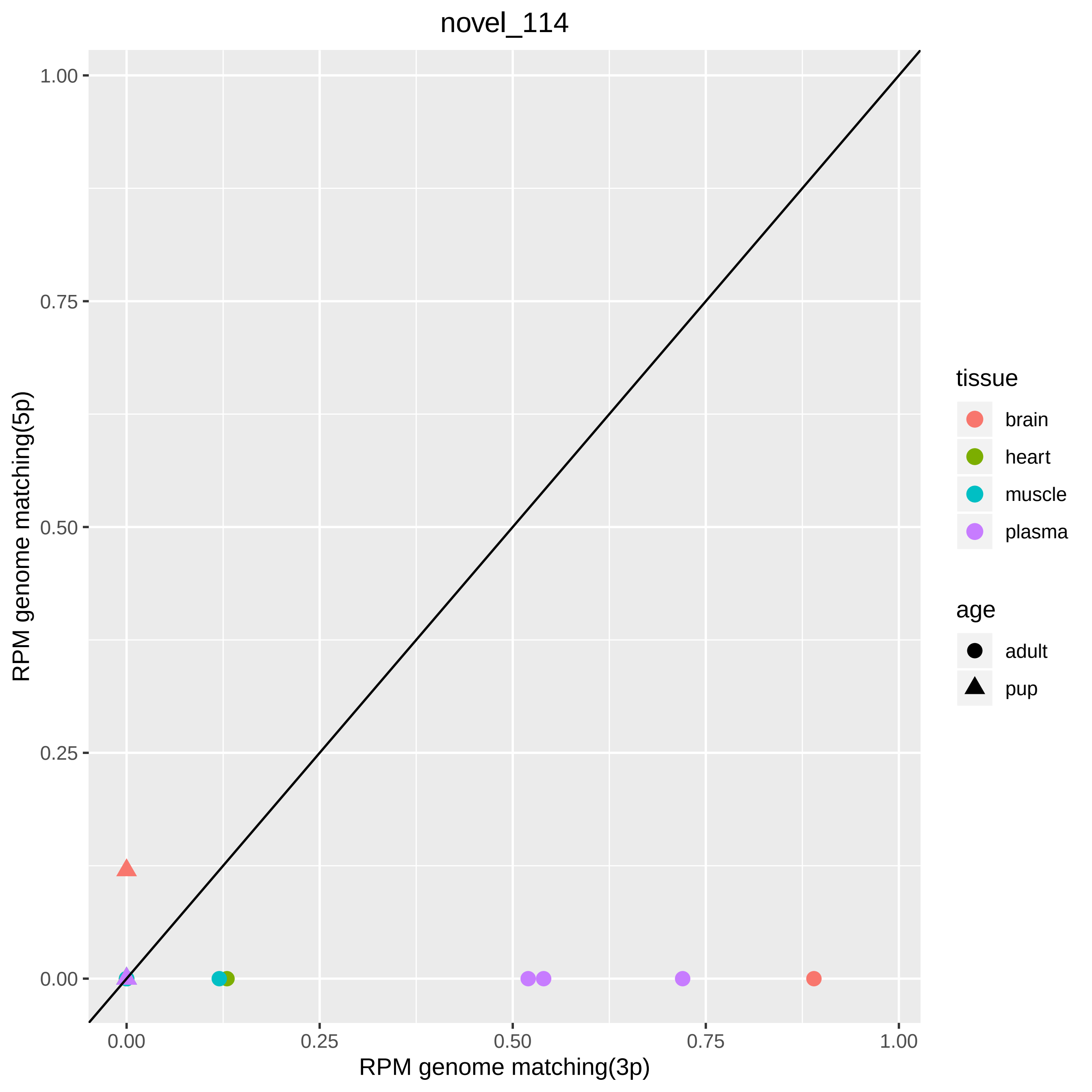
novel_114
RPM genome matching(5p)
RPM genome matching(3p)
1.00
0.75
0.50
0.25
0.00
0.00
0.25
0.50
0.75
1.00
tissue
brain
heart
muscle
plasma
age
adult
pup

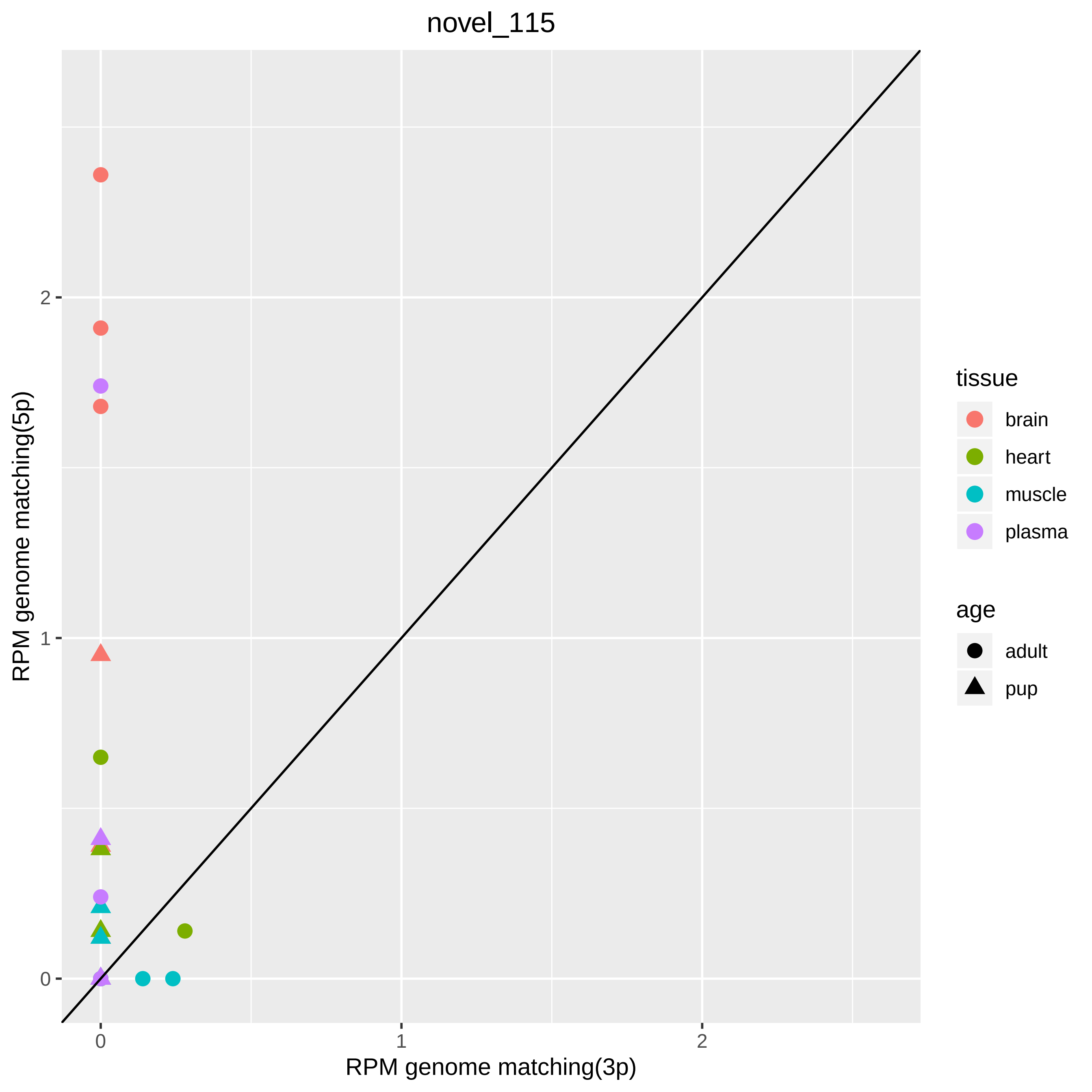
novel_115
RPM genome matching(5p)
RPM genome matching(3p)
0
1
2
0
1
2
tissue
brain
heart
muscle
plasma
age
adult
pup

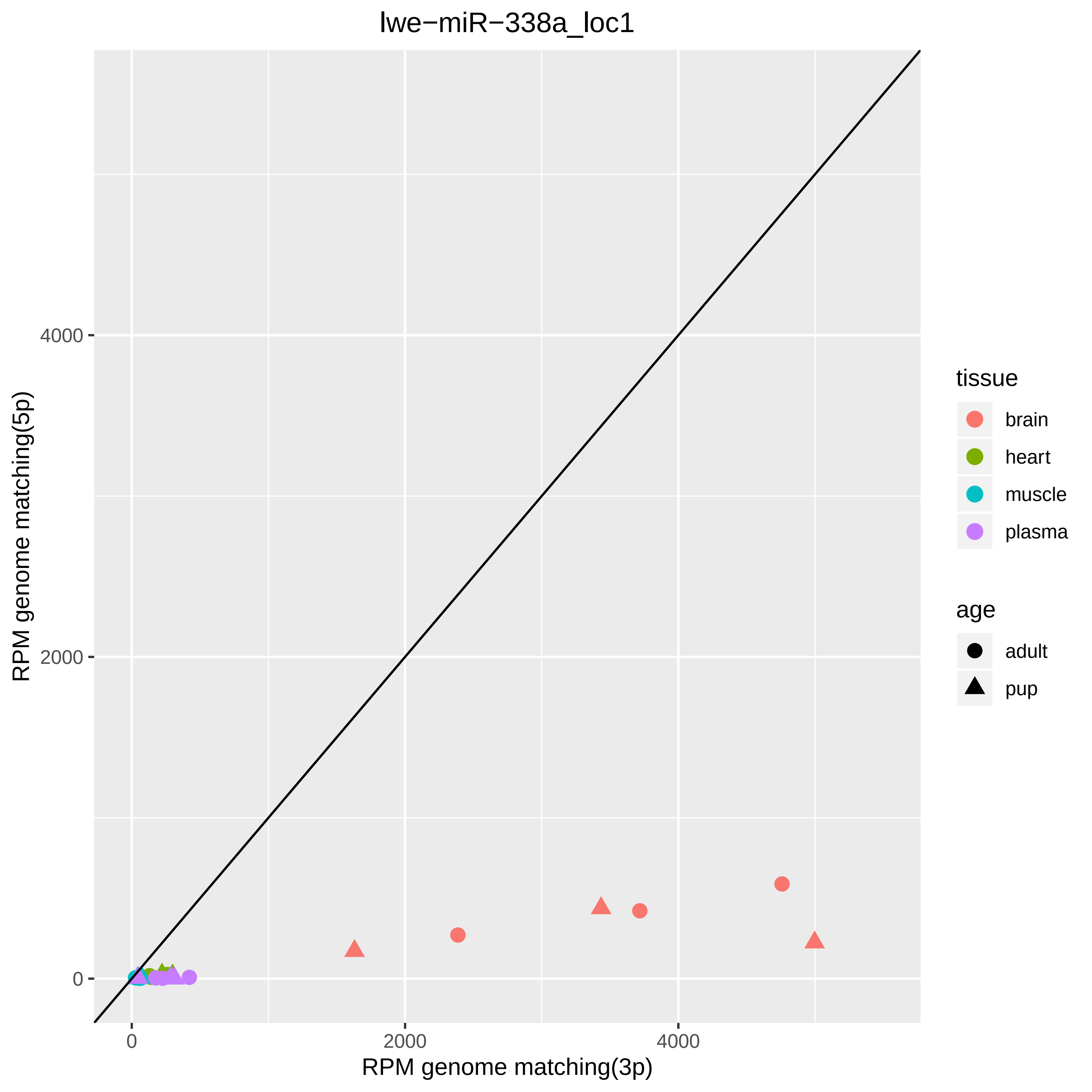

lwe−miR−338a_loc1
RPM genome matching(5p)
RPM genome matching(3p)
0
2000
4000
0
2000
4000
tissue
brain
heart
muscle
plasma
age
adult
pup

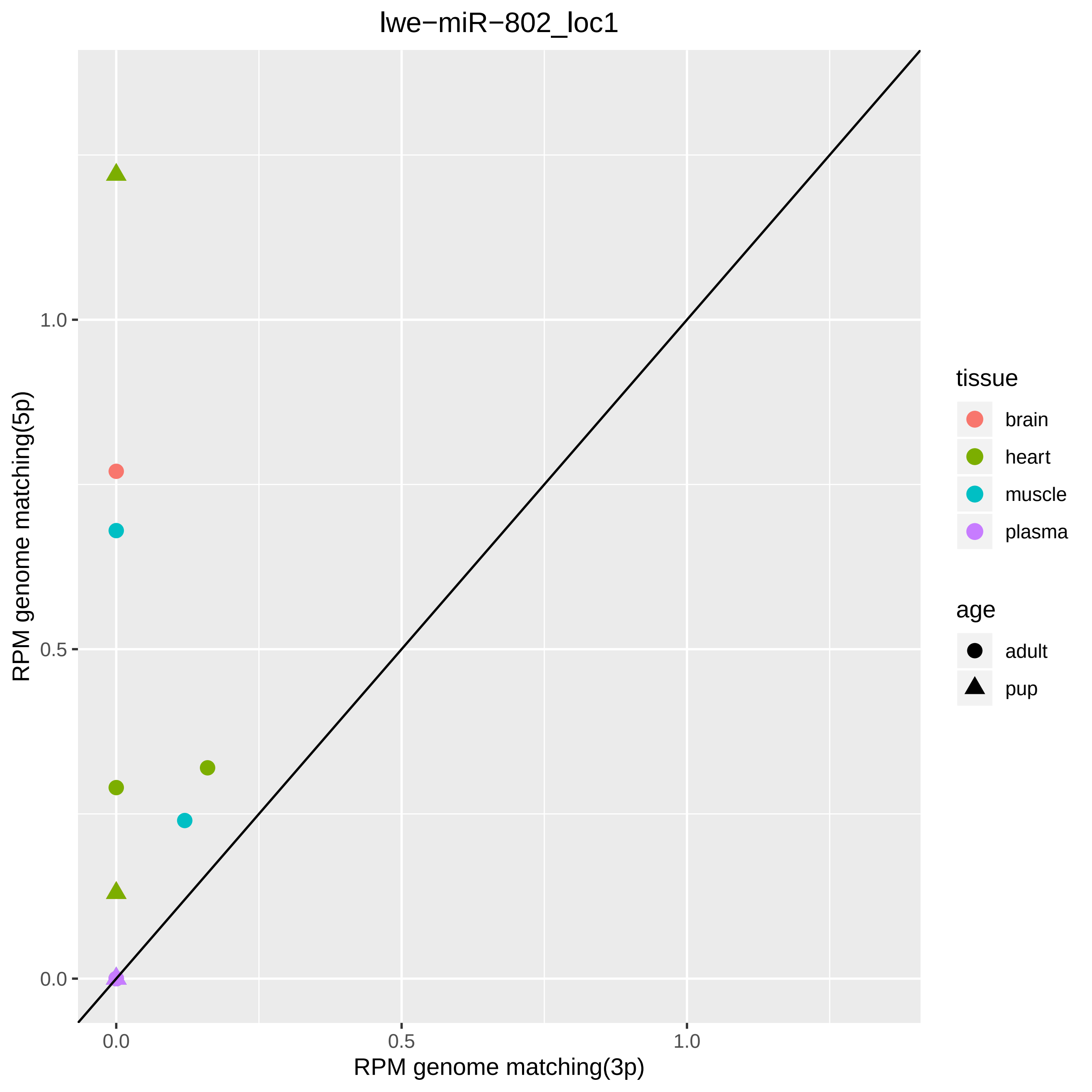
lwe−miR−802_loc1
RPM genome matching(5p)
RPM genome matching(3p)
0.0
0.5
1.0
0.0
0.5
1.0
tissue
brain
heart
muscle
plasma
age
adult
pup

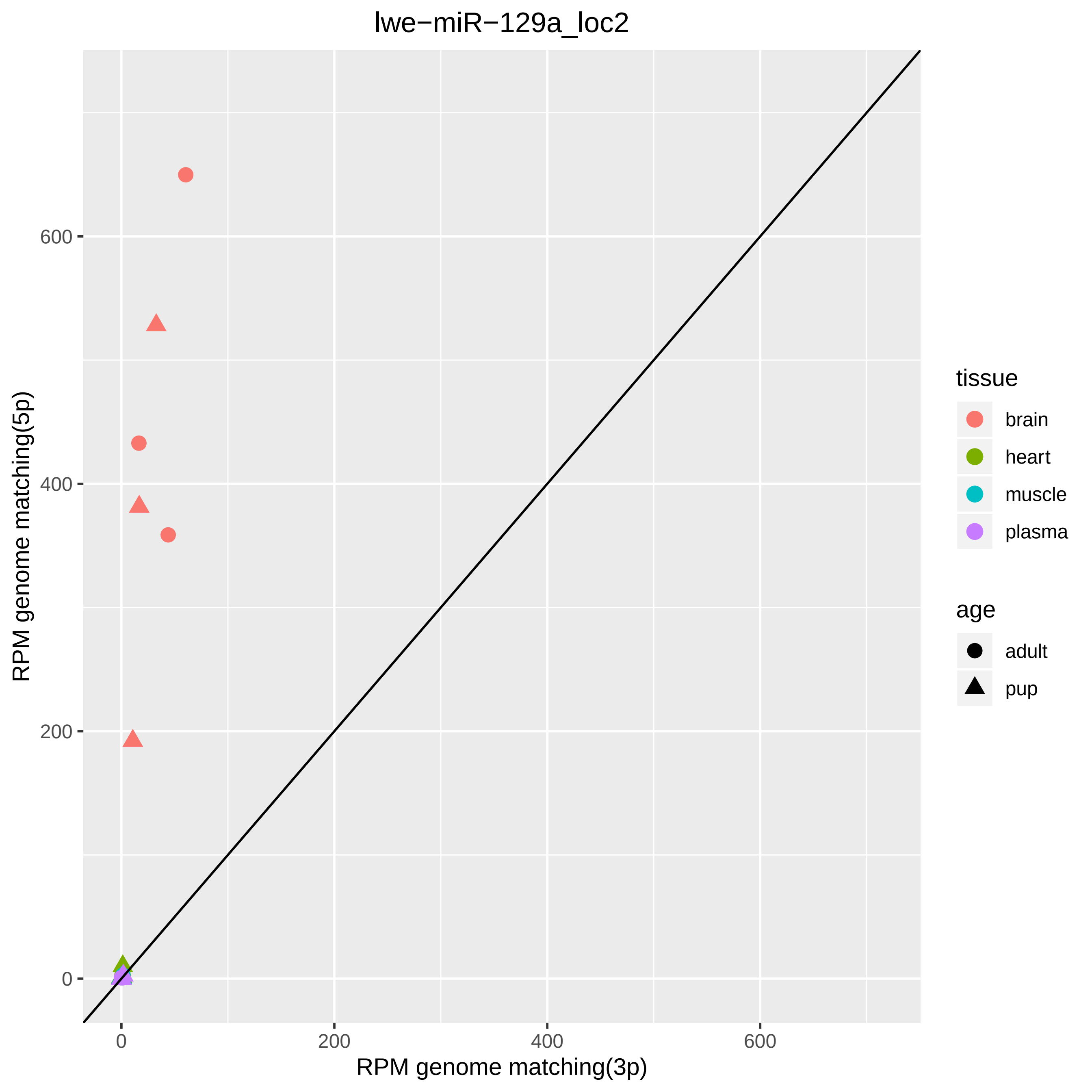

lwe−miR−129a_loc2
RPM genome matching(5p)
RPM genome matching(3p)
0
200
400
600
tissue
brain
heart
muscle
plasma
age
adult
pup

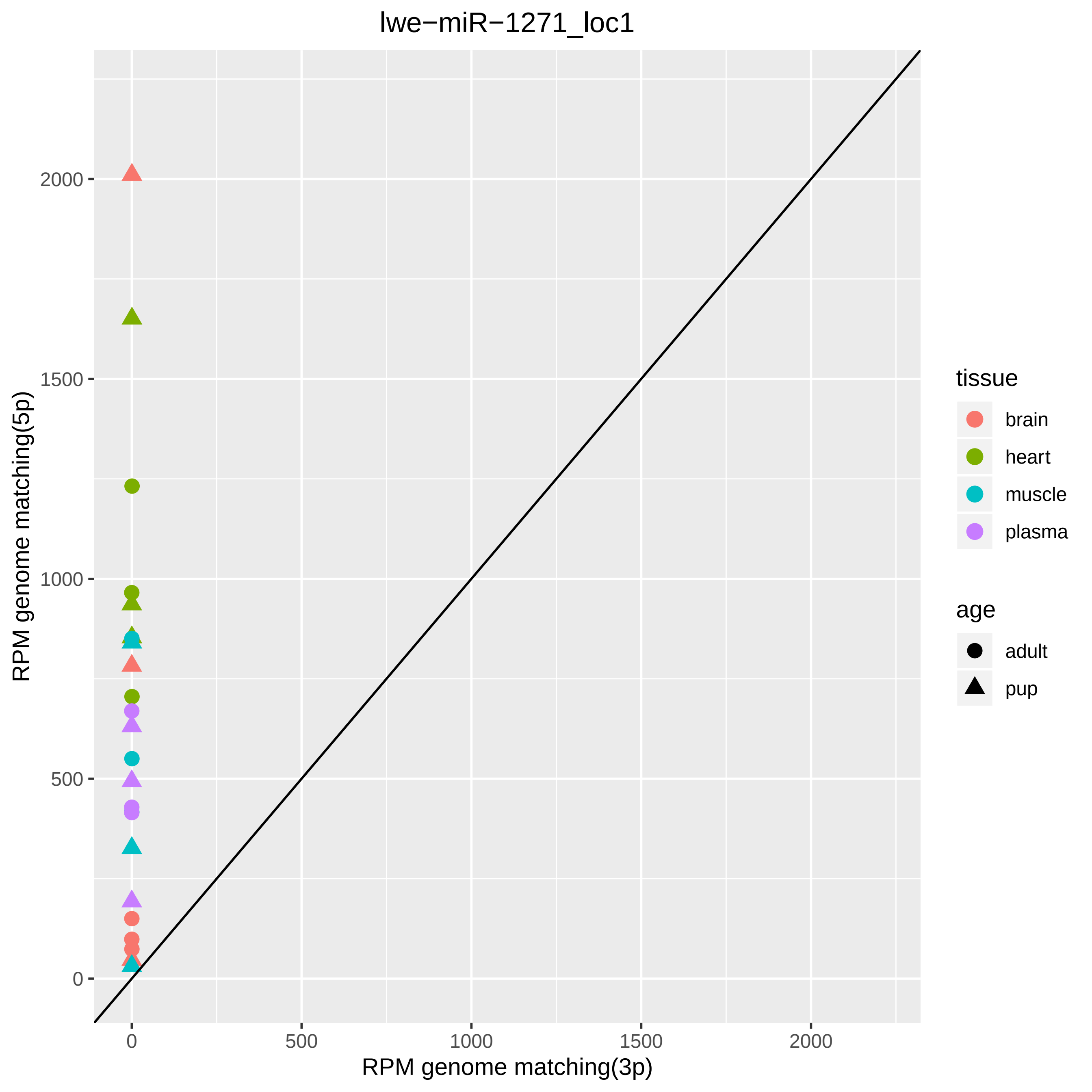
lwe−miR−1271_loc1
RPM genome matching(5p)
RPM genome matching(3p)
0
500
1000
1500
2000
tissue
brain
heart
muscle
plasma
age
adult
pup

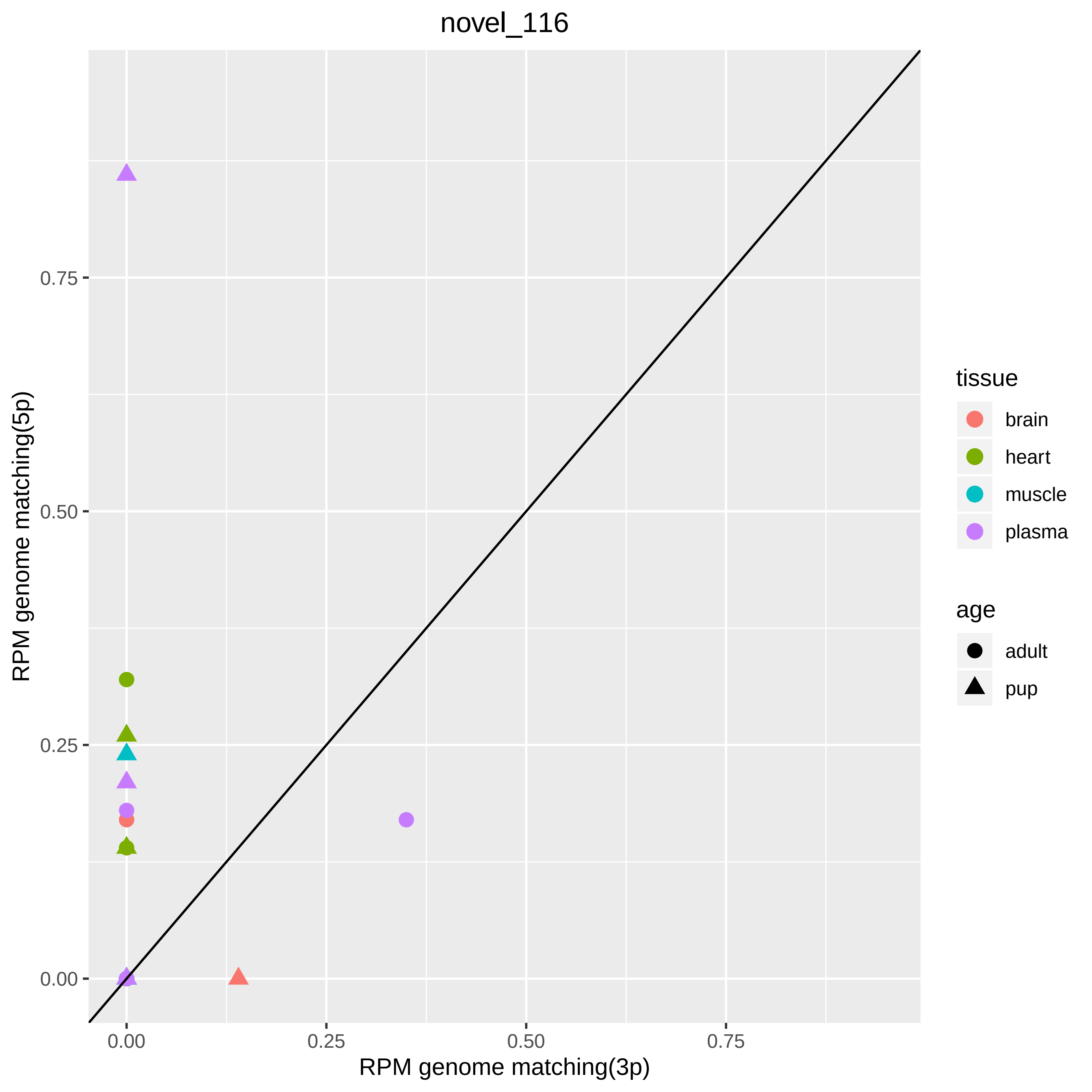

novel_116
RPM genome matching(5p)
RPM genome matching(3p)
0.00
0.25
0.50
0.75
tissue
brain
heart
muscle
plasma
age
adult
pup

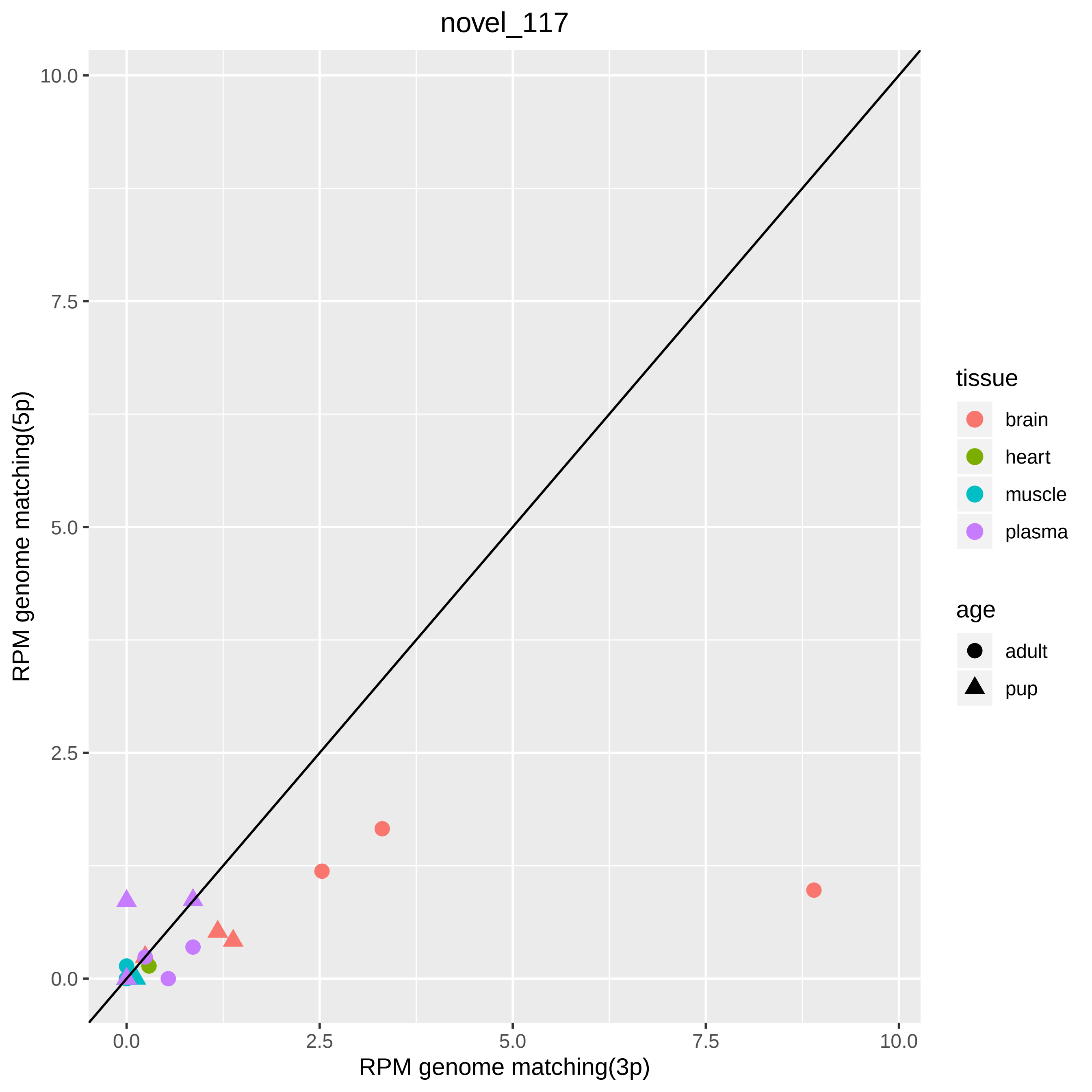

novel_117
RPM genome matching(5p)
RPM genome matching(3p)
0.0
2.5
5.0
7.5
10.0
tissue
brain
heart
muscle
plasma
age
adult
pup

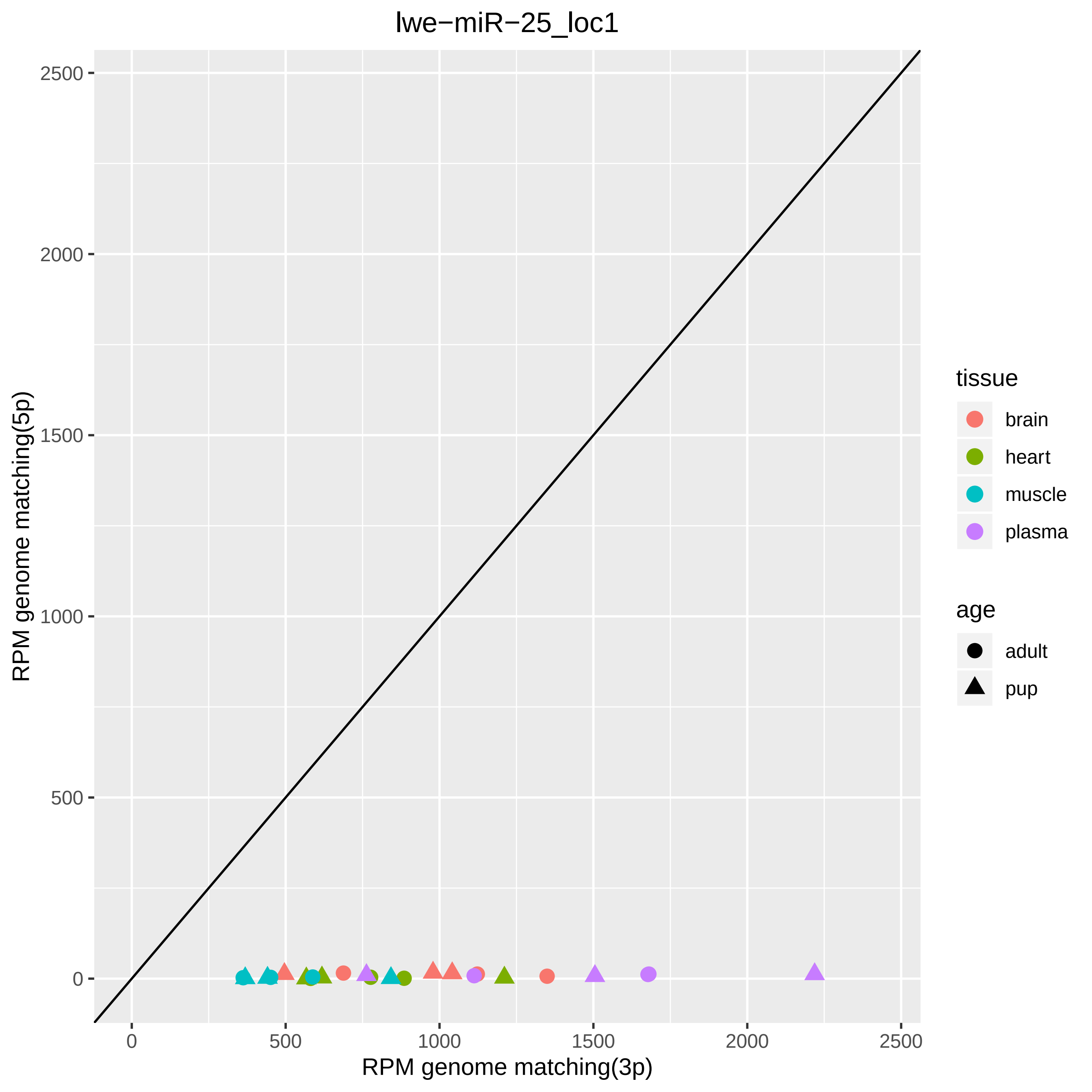

lwe−miR−25_loc1
RPM genome matching(5p)
RPM genome matching(3p)
0
500
1000
1500
2000
2500
tissue
brain
heart
muscle
plasma
age
adult
pup

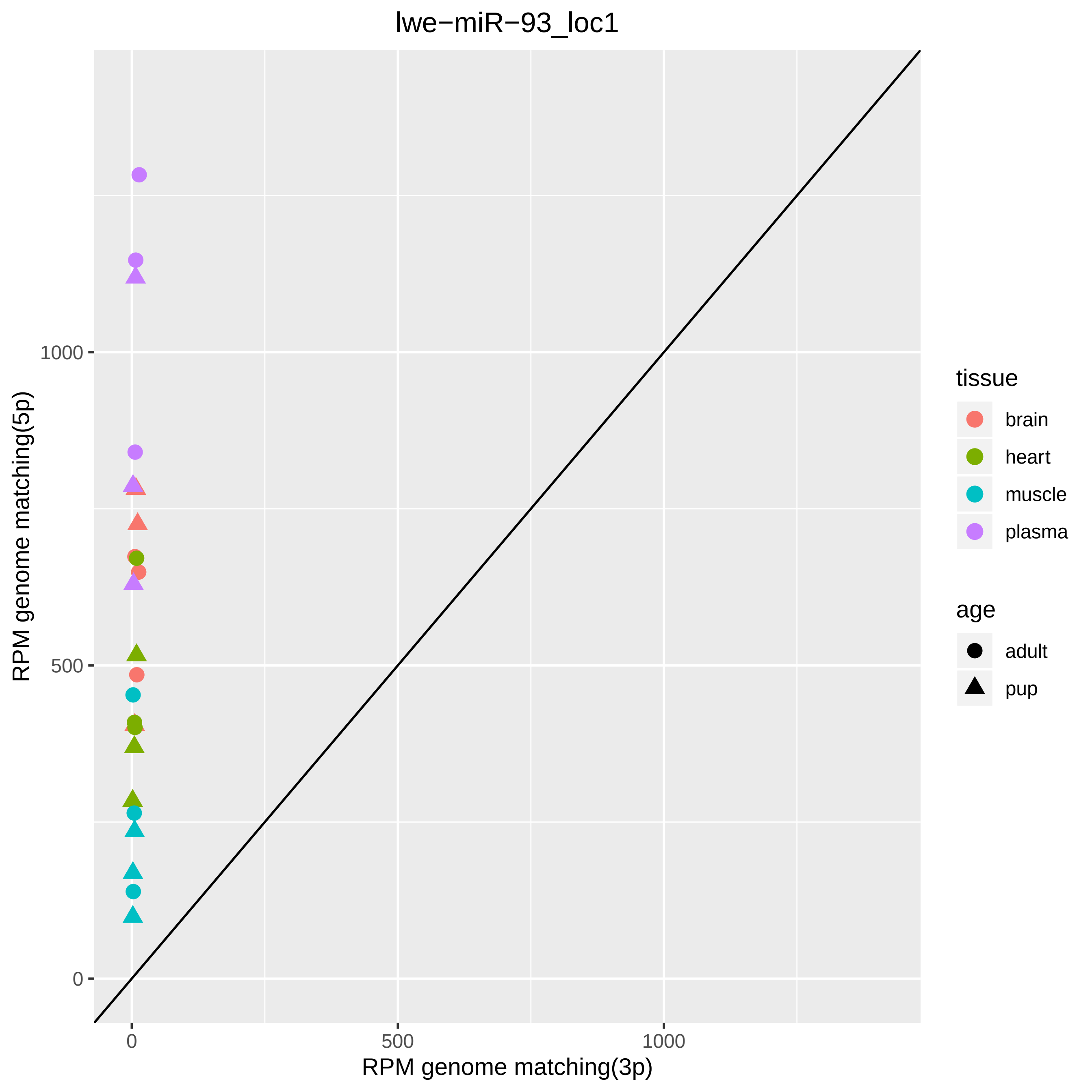
lwe−miR−93_loc1
RPM genome matching(5p)
1000
500
0
RPM genome matching(3p)
0
500
1000
tissue
brain
heart
muscle
plasma
age
adult
pup

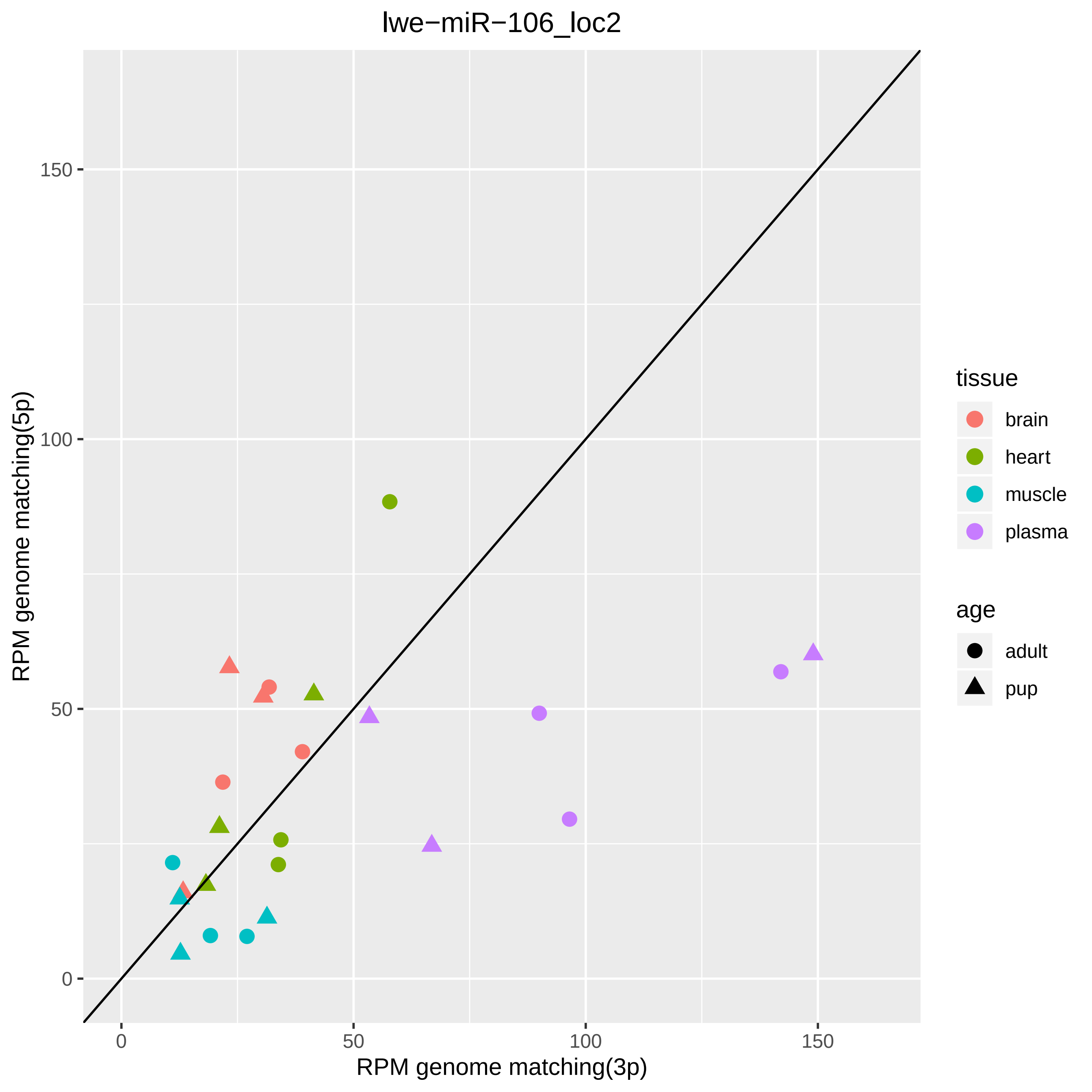

lwe−miR−106_loc2
RPM genome matching(5p)
RPM genome matching(3p)
0
50
100
150
tissue
brain
heart
muscle
plasma
age
adult
pup

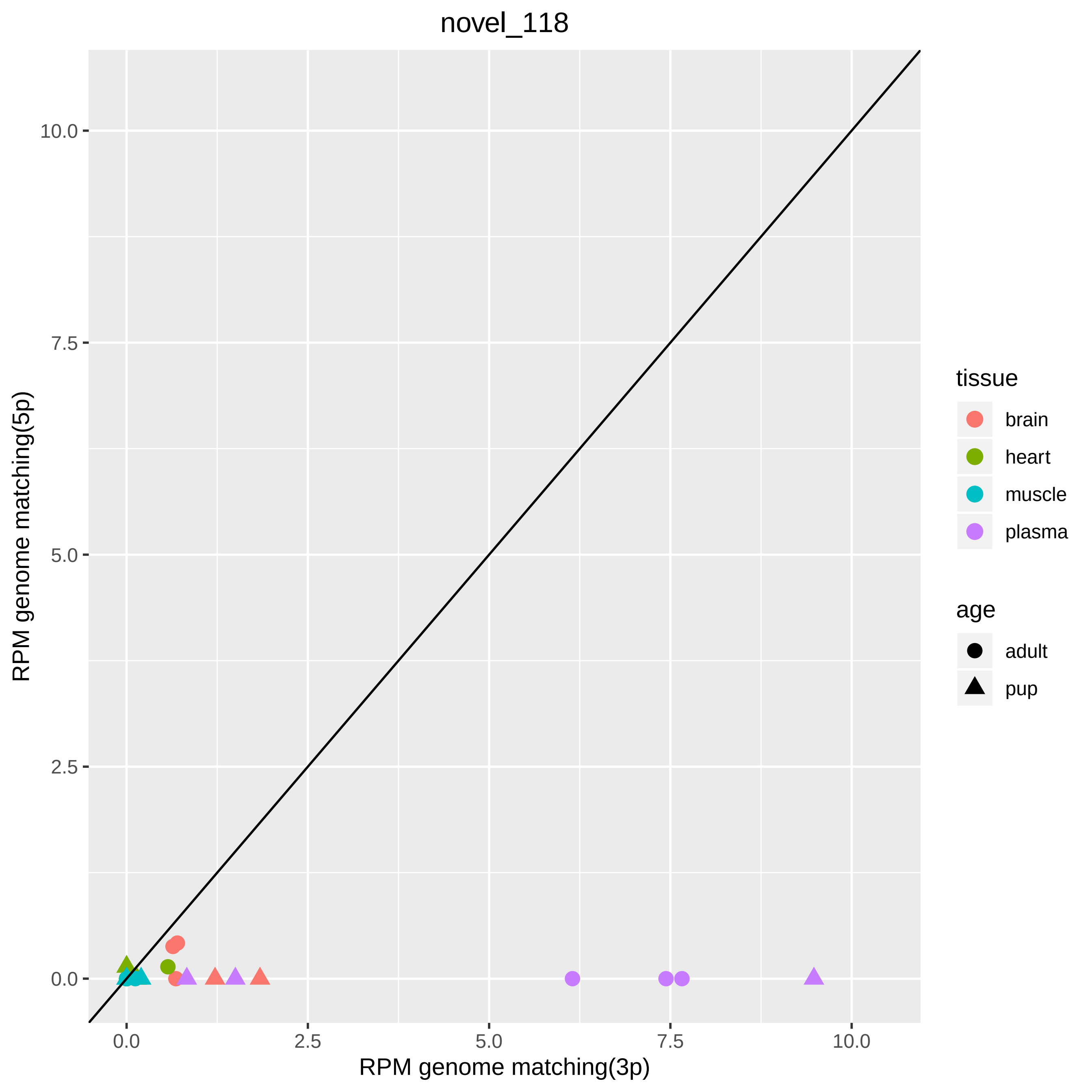
novel_118
RPM genome matching(5p)
RPM genome matching(3p)
0.0
2.5
5.0
7.5
10.0
tissue
brain
heart
muscle
plasma
age
adult
pup

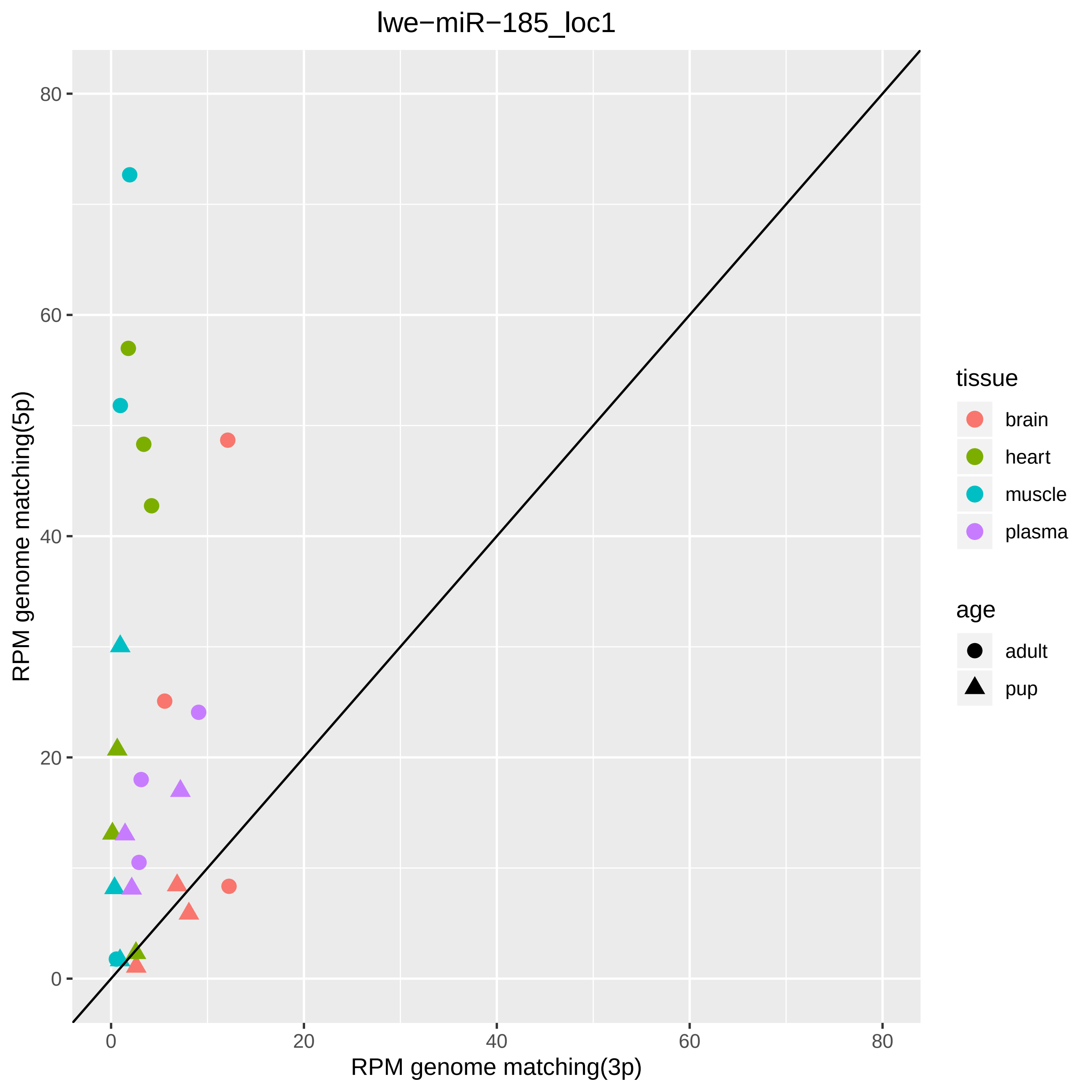
lwe−miR−185_loc1
RPM genome matching(5p)
RPM genome matching(3p)
0
20
40
60
80
0
20
40
60
80
tissue
brain
heart
muscle
plasma
age
adult
pup

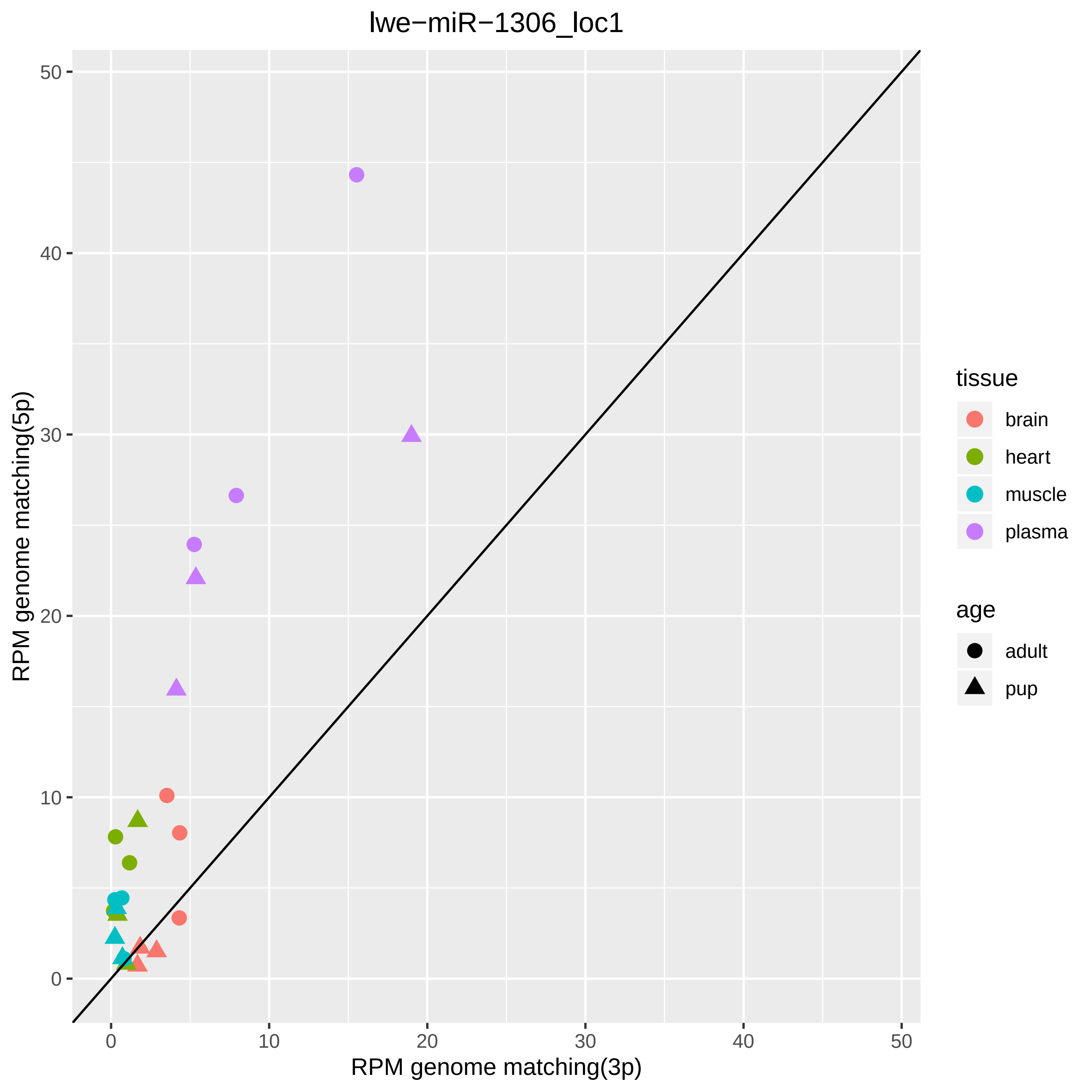
lwe−miR−1306_loc1
RPM genome matching(5p)
RPM genome matching(3p)
0
10
20
30
40
50
0
10
20
30
40
50
tissue
brain
heart
muscle
plasma
age
adult
pup

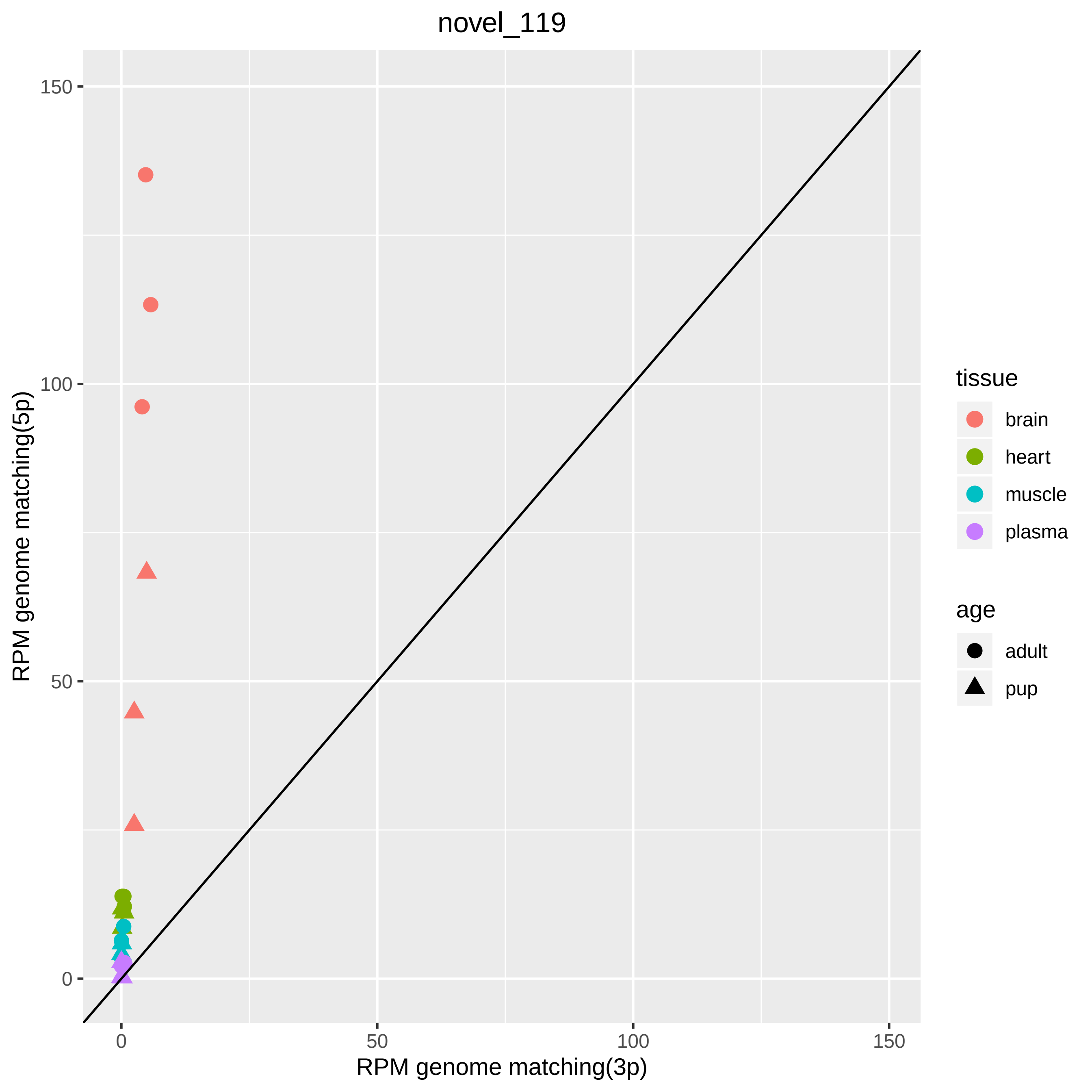

novel_119
150
100
50
0
0
50
100
150
RPM genome matching(5p)
RPM genome matching(3p)
tissue
brain
heart
muscle
plasma
age
adult
pup

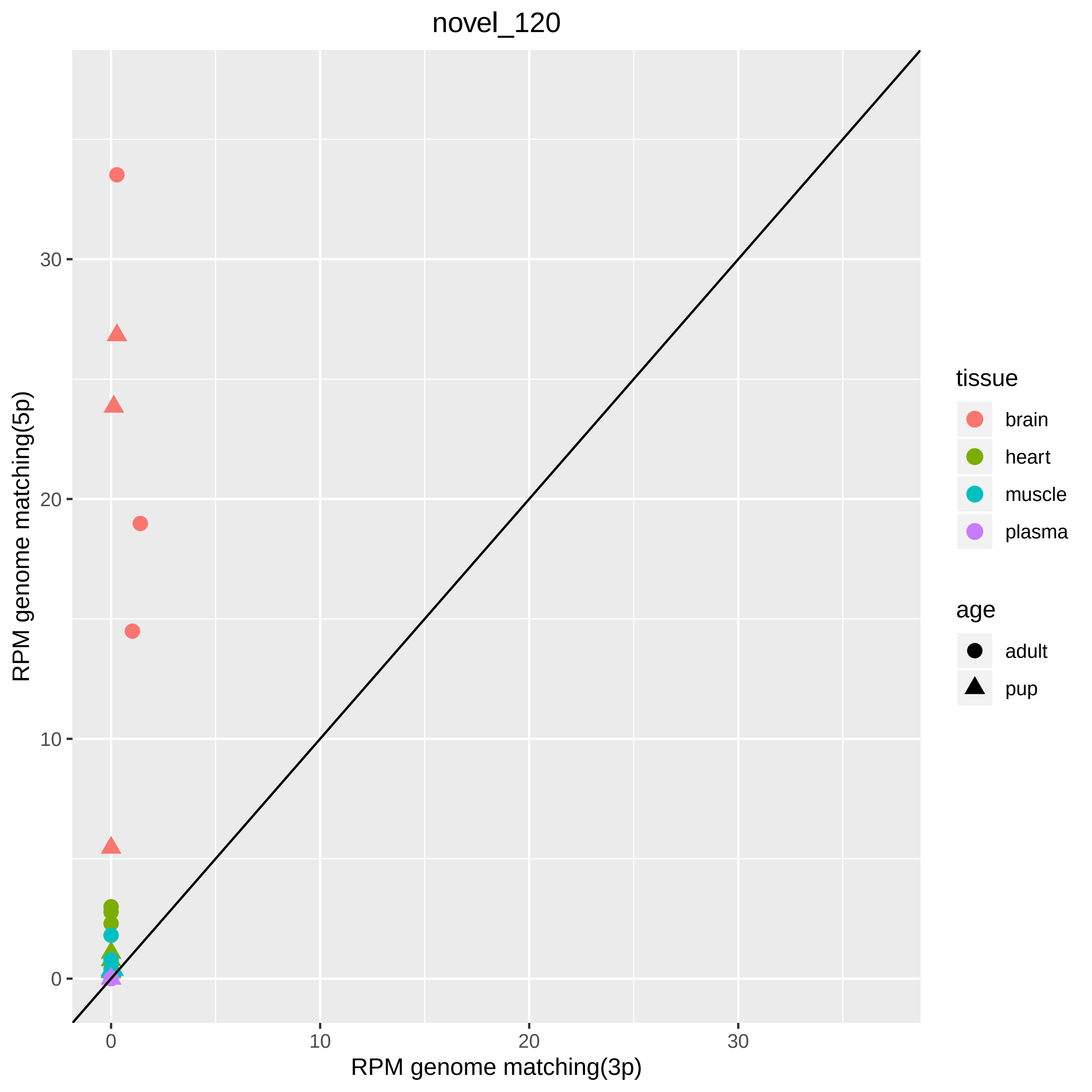

novel_120
RPM genome matching(5p)
RPM genome matching(3p)
0
10
20
30
0
10
20
30
tissue
brain
heart
muscle
plasma
age
adult
pup

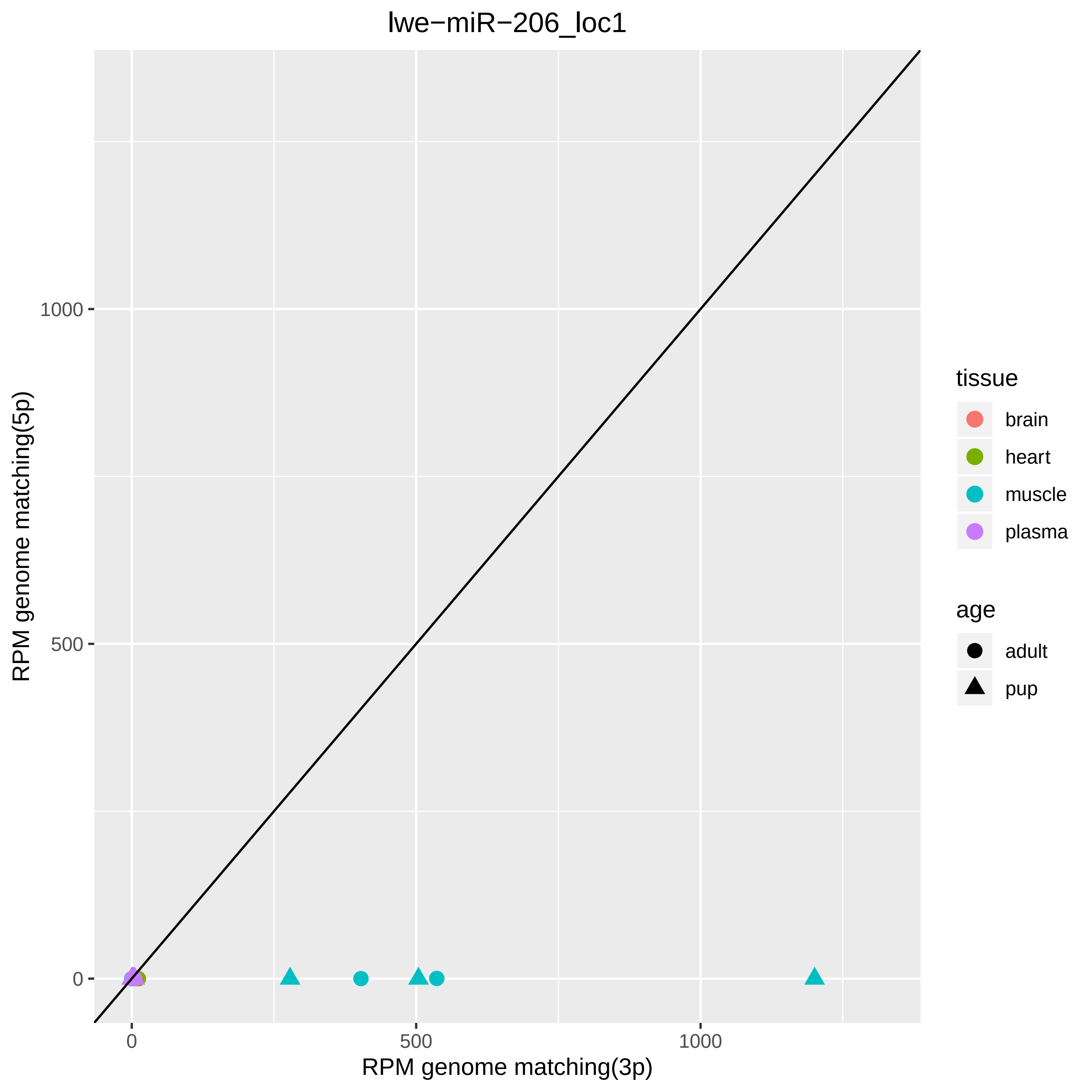

lwe−miR−206_loc1
RPM genome matching(5p)
1000
500
0
0
500
1000
RPM genome matching(3p)
tissue
brain
heart
muscle
plasma
age
adult
pup

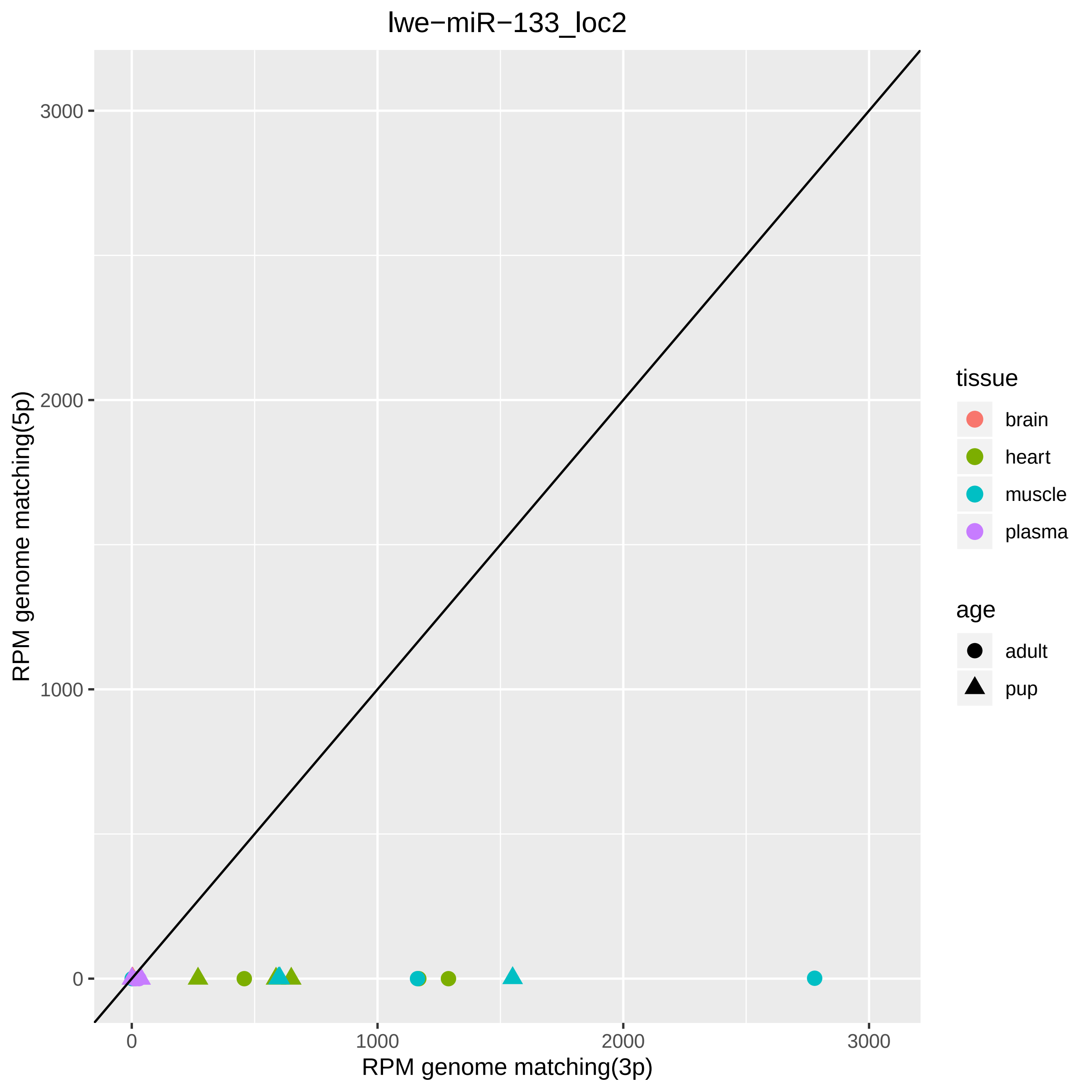

lwe−miR−133_loc2
RPM genome matching(5p)
3000
2000
1000
0
0
1000
2000
3000
RPM genome matching(3p)
tissue
brain
heart
muscle
plasma
age
adult
pup

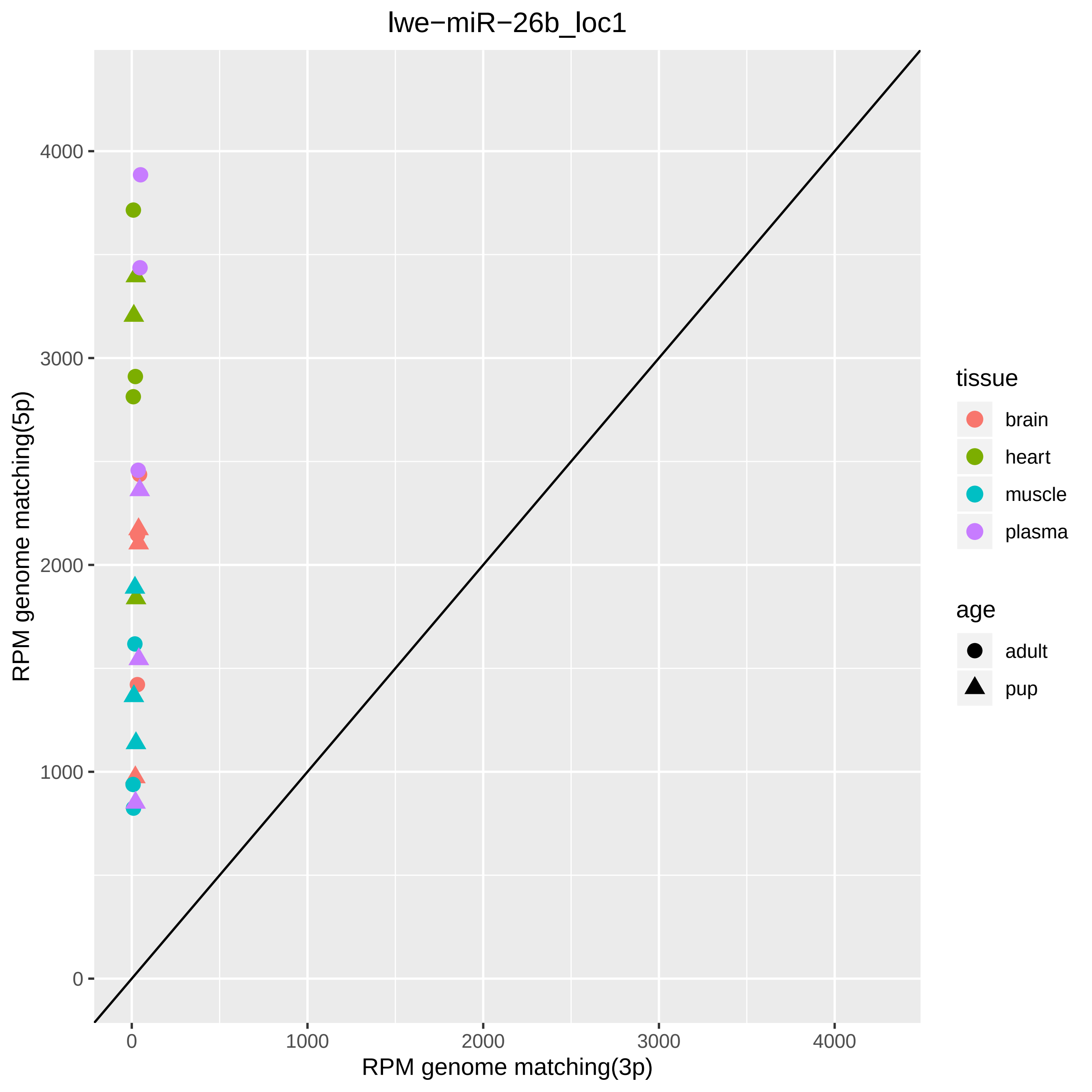

lwe−miR−26b_loc1
RPM genome matching(5p)
RPM genome matching(3p)
0
1000
2000
3000
4000
0
1000
2000
3000
4000
tissue
brain
heart
muscle
plasma
age
adult
pup

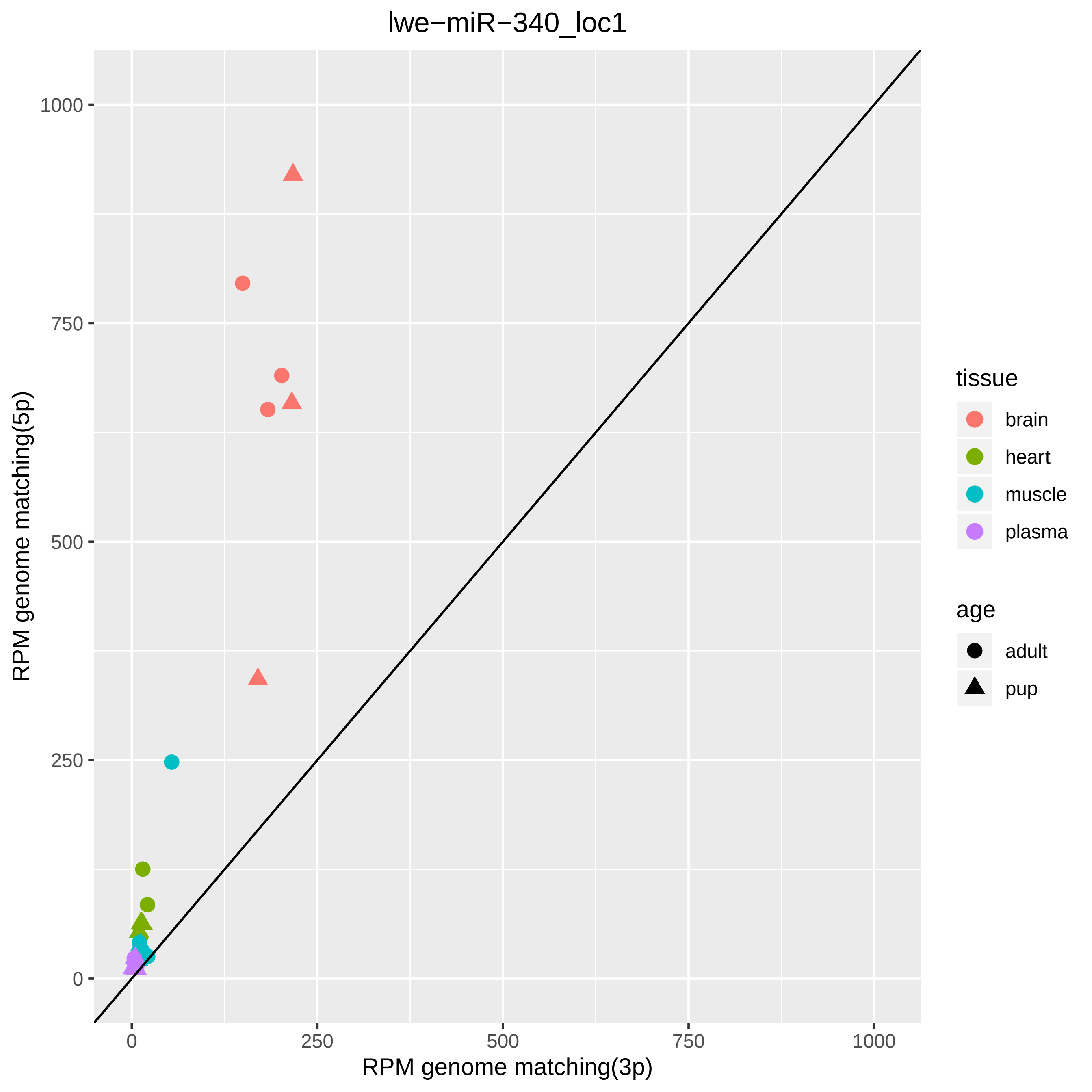

lwe−miR−340_loc1
RPM genome matching(5p)
RPM genome matching(3p)
1000
750
500
250
0
0
250
500
750
1000
tissue
brain
heart
muscle
plasma
age
adult
pup

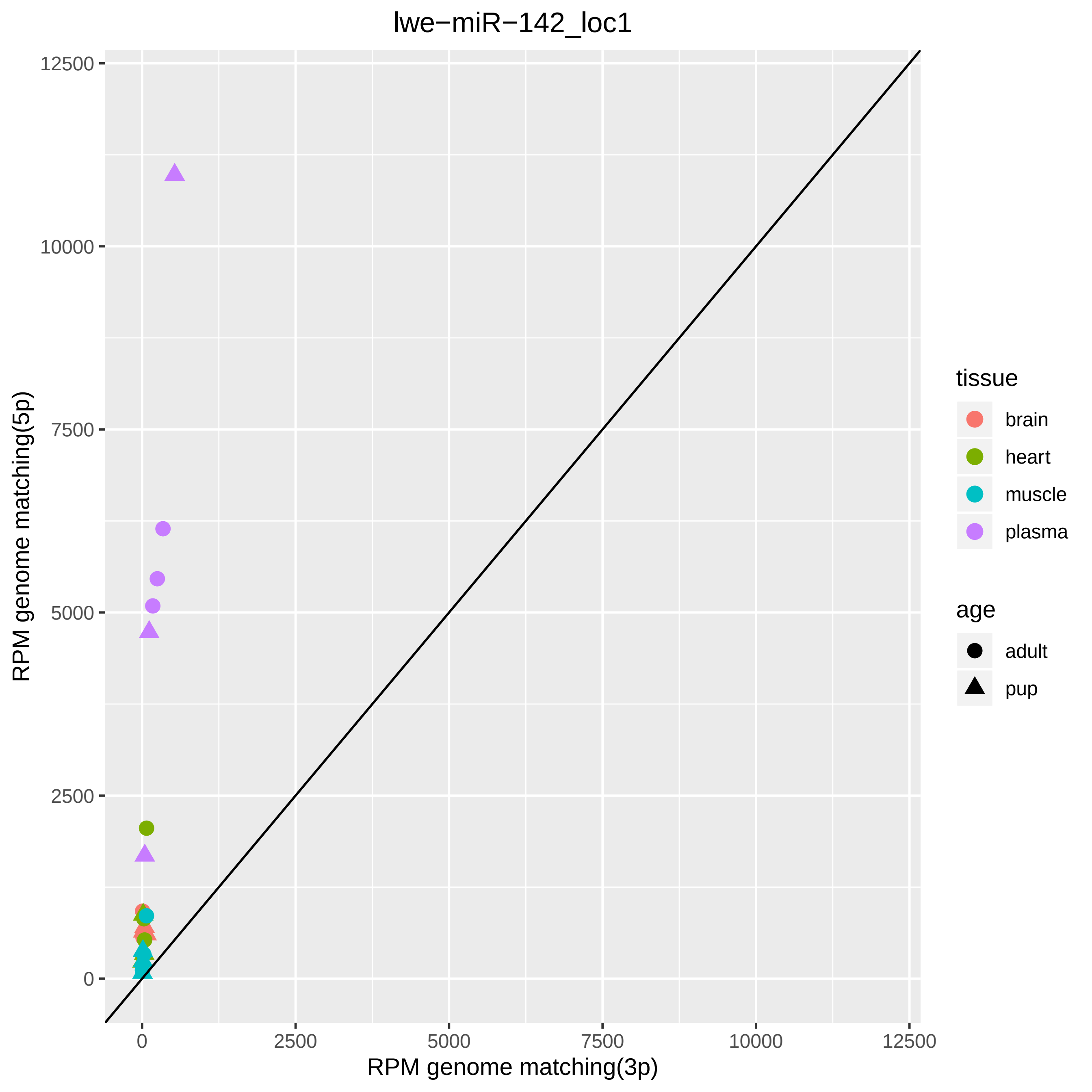

lwe−miR−142_loc1
RPM genome matching(5p)
RPM genome matching(3p)
12500
10000
7500
5000
2500
0
0
2500
5000
7500
10000
12500
tissue
brain
heart
muscle
plasma
age
adult
pup

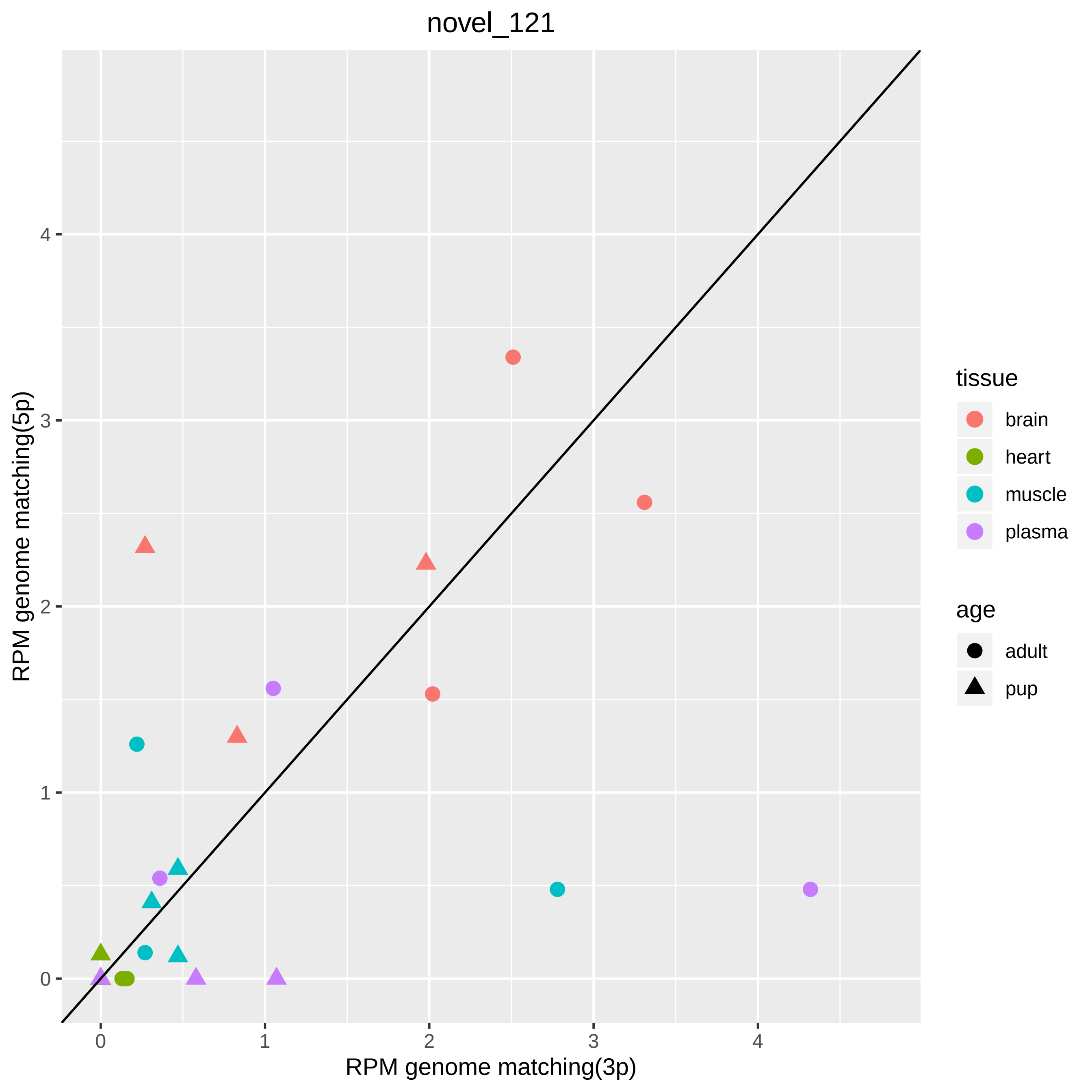
novel_121
RPM genome matching(5p)
RPM genome matching(3p)
0
1
2
3
4
tissue
brain
heart
muscle
plasma
age
adult
pup

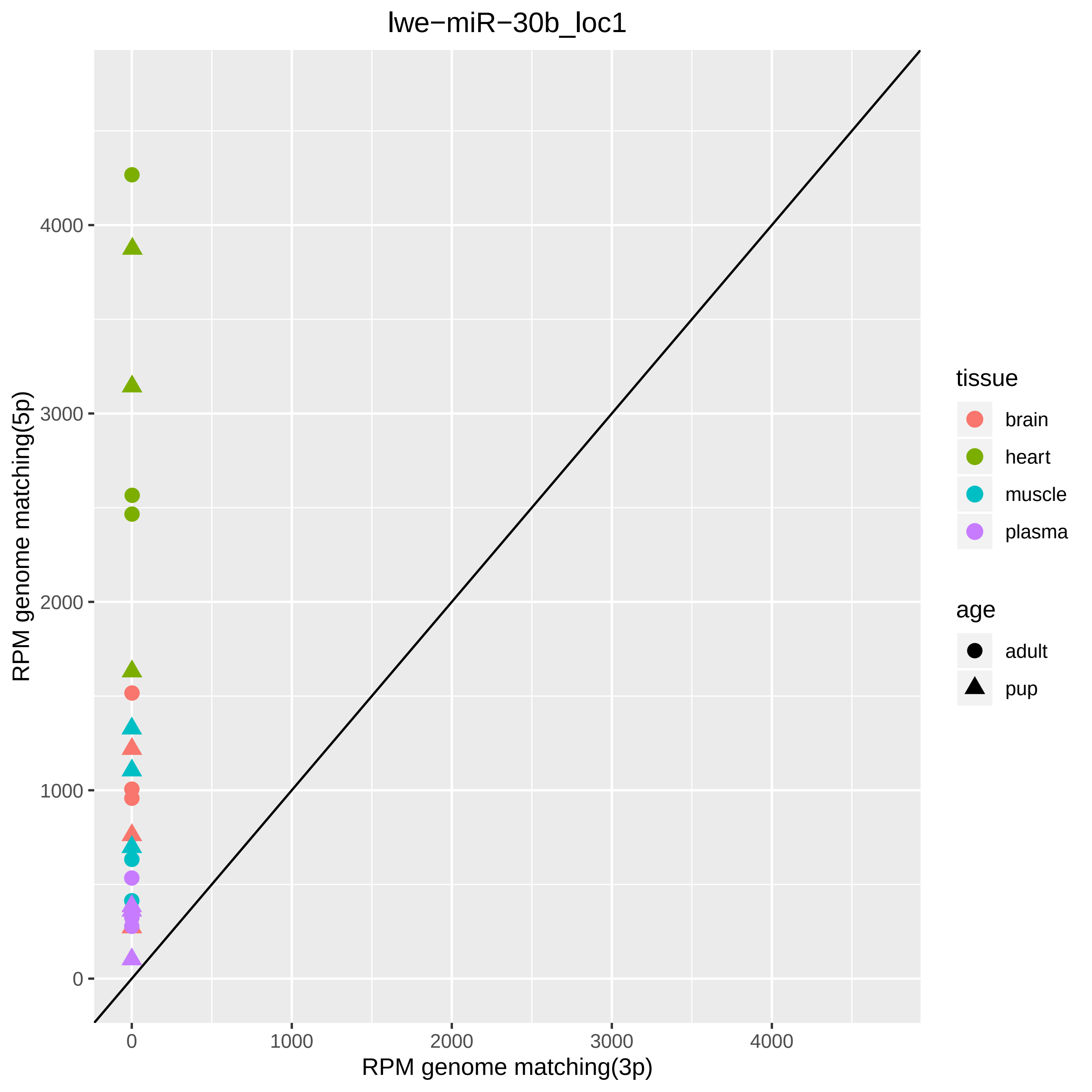

lwe−miR−30b_loc1
RPM genome matching(5p)
RPM genome matching(3p)
0
1000
2000
3000
4000
tissue
brain
heart
muscle
plasma
age
adult
pup

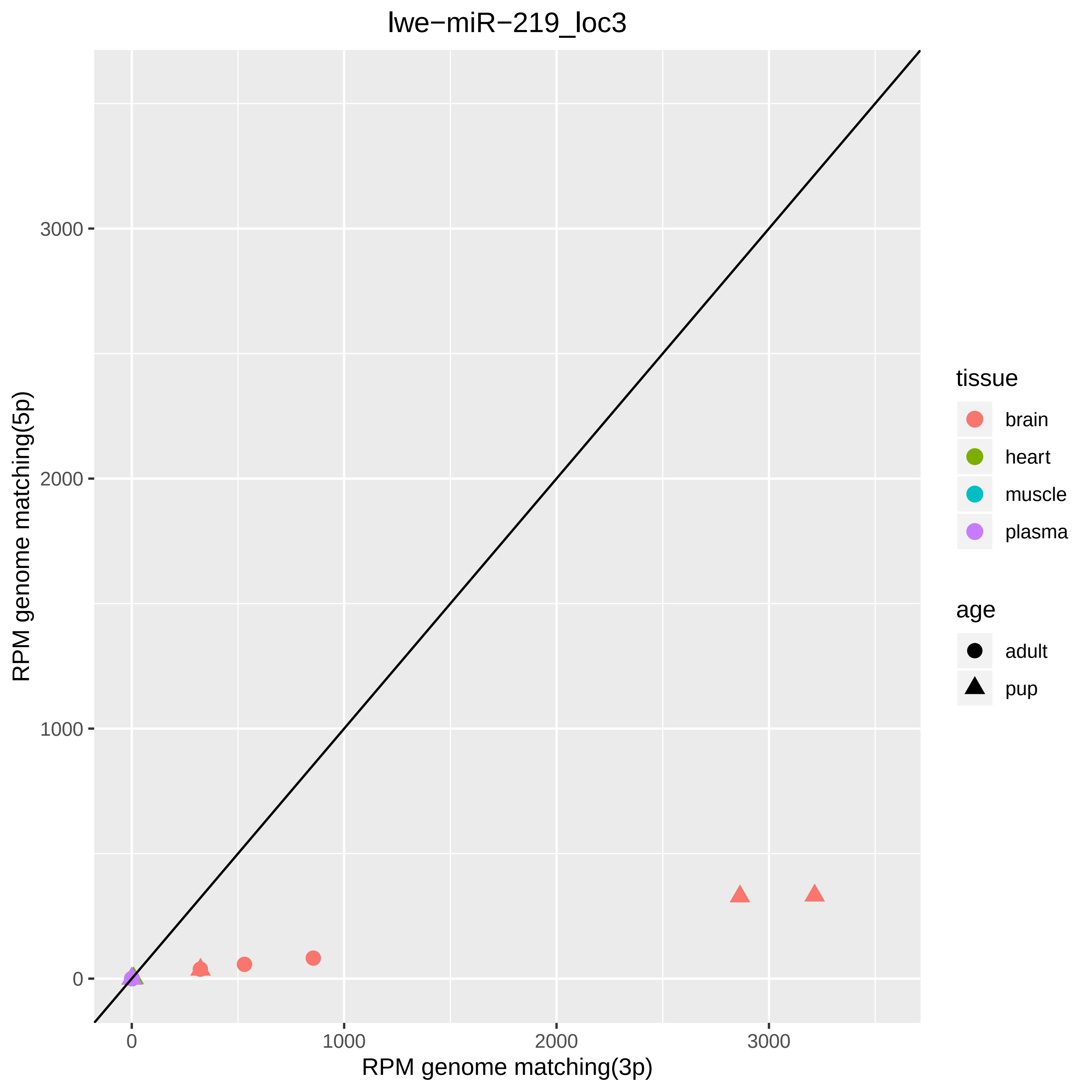

lwe−miR−219_loc3
RPM genome matching(5p)
3000
2000
1000
0
0
1000
2000
3000
RPM genome matching(3p)
tissue
brain
heart
muscle
plasma
age
adult
pup

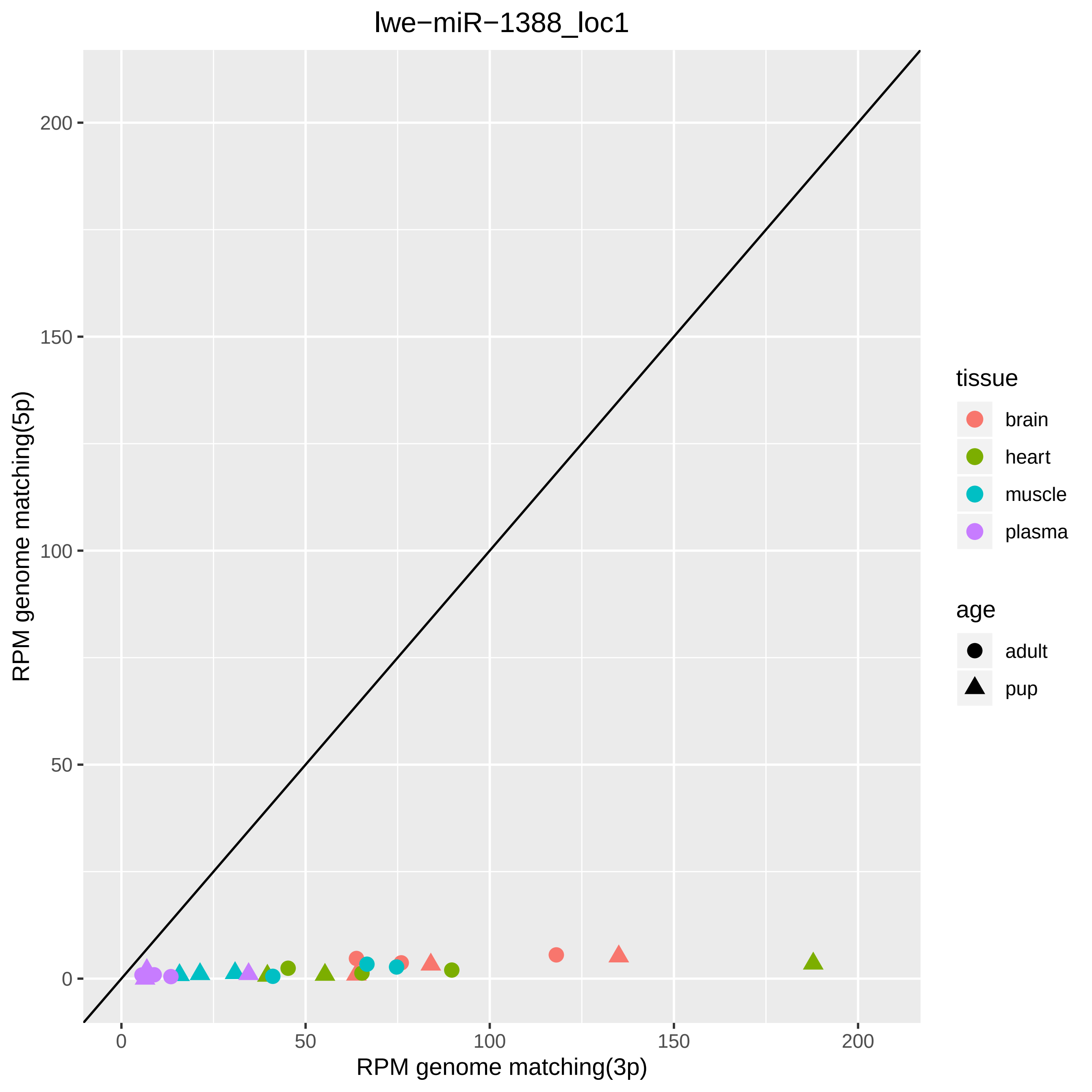

lwe−miR−1388_loc1
RPM genome matching(5p)
RPM genome matching(3p)
0
50
100
150
200
0
50
100
150
200
tissue
brain
heart
muscle
plasma
age
adult
pup

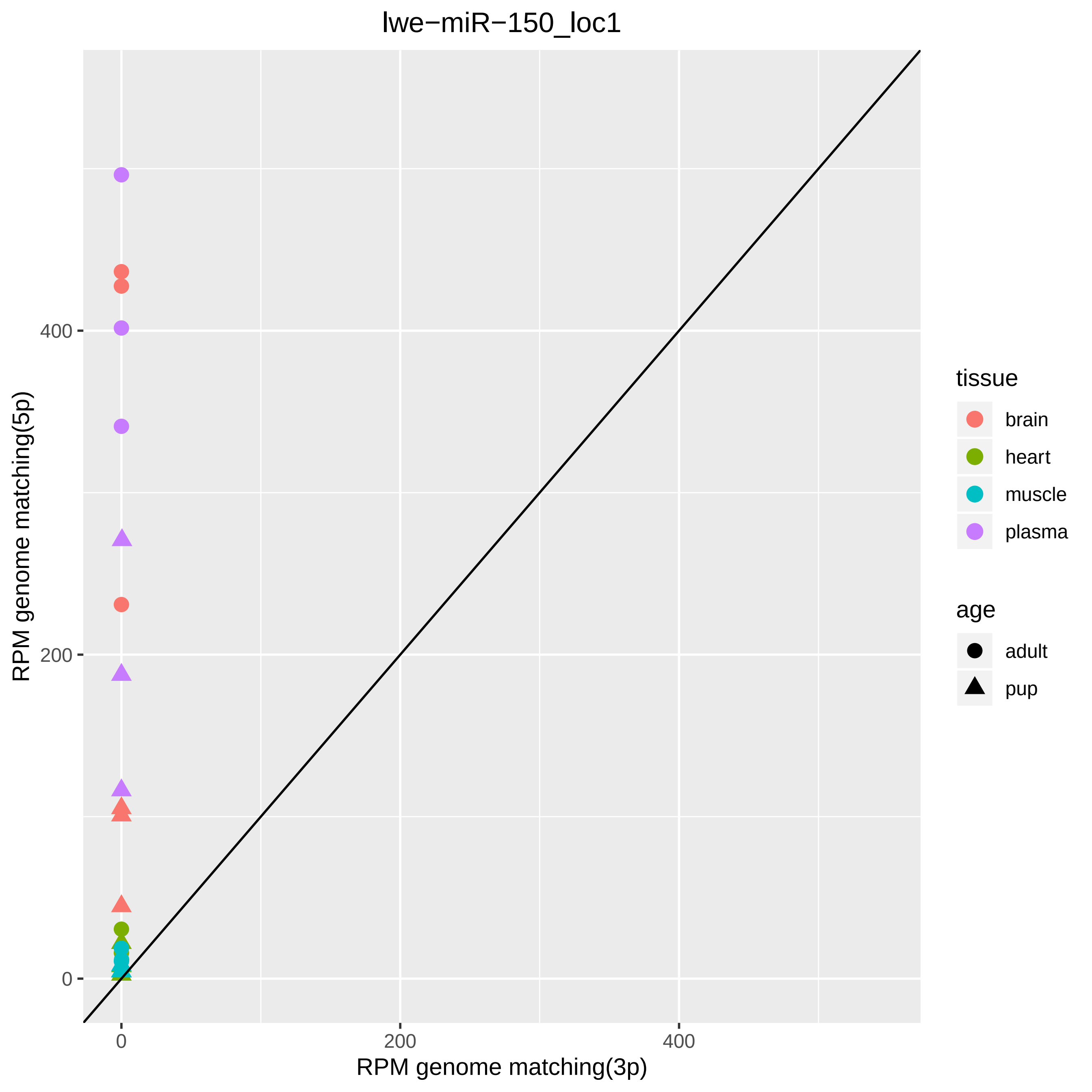

lwe−miR−150_loc1
RPM genome matching(5p)
RPM genome matching(3p)
400
200
0
0
200
400
tissue
brain
heart
muscle
plasma
age
adult
pup

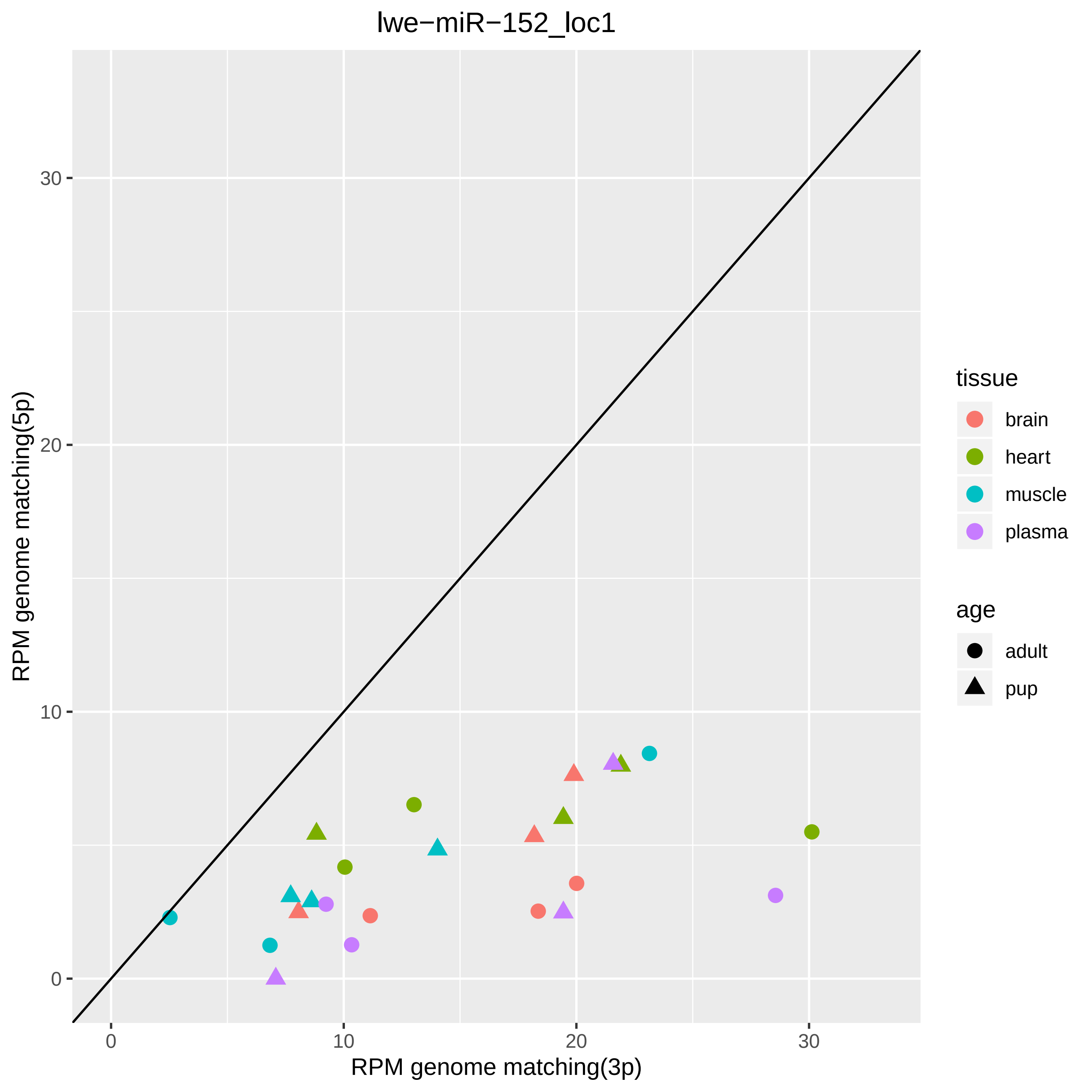

lwe−miR−152_loc1
RPM genome matching(5p)
RPM genome matching(3p)
0
10
20
30
0
10
20
30
tissue
brain
heart
muscle
plasma
age
adult
pup

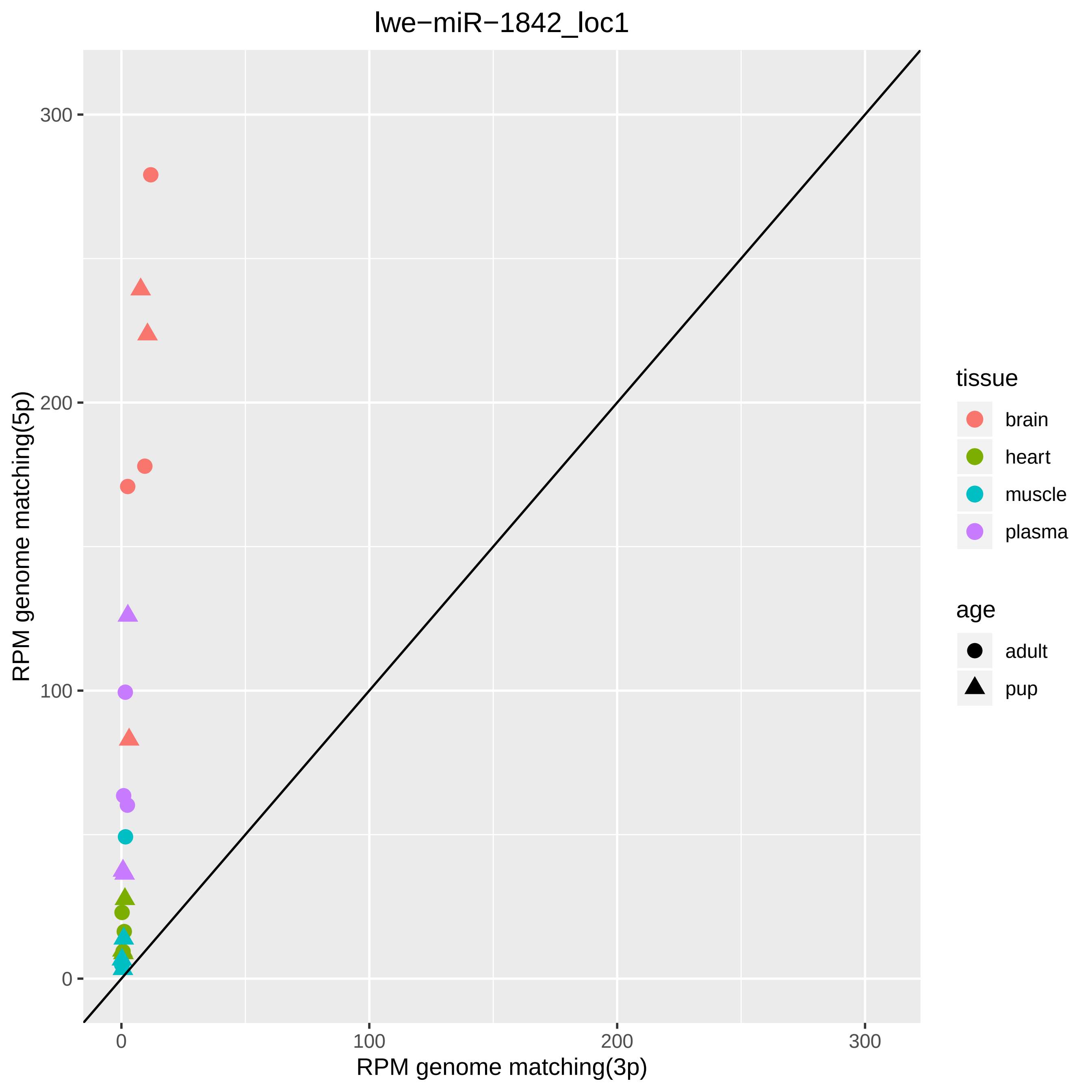
lwe−miR−1842_loc1
RPM genome matching(5p)
RPM genome matching(3p)
0
100
200
300
0
100
200
300
tissue
brain
heart
muscle
plasma
age
adult
pup

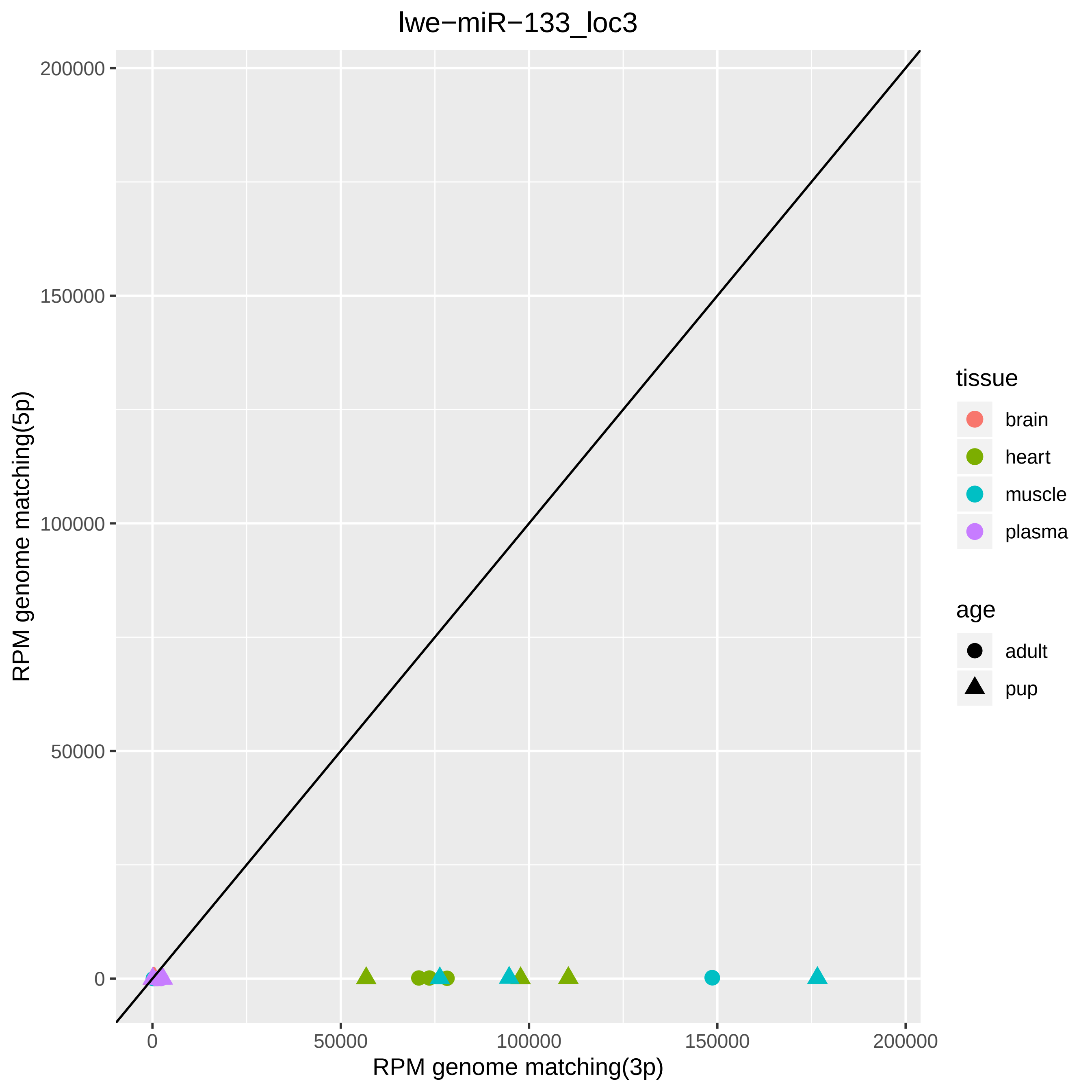
lwe−miR−133_loc3
RPM genome matching(5p)
RPM genome matching(3p)
200000
150000
100000
50000
0
tissue
brain
heart
muscle
plasma
age
adult
pup

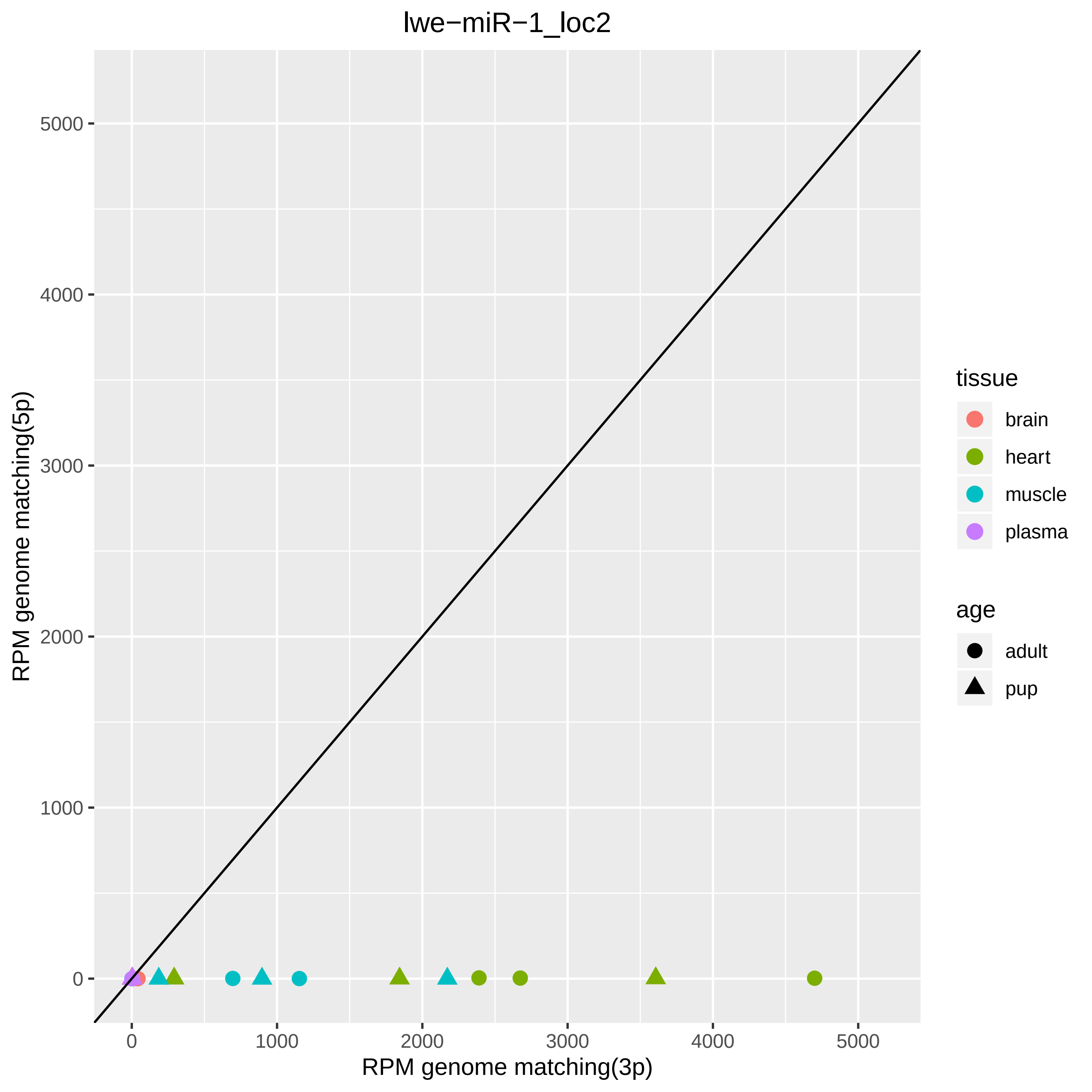
lwe−miR−1_loc2
RPM genome matching(5p)
RPM genome matching(3p)
0
1000
2000
3000
4000
5000
0
1000
2000
3000
4000
5000
tissue
brain
heart
muscle
plasma
age
adult
pup

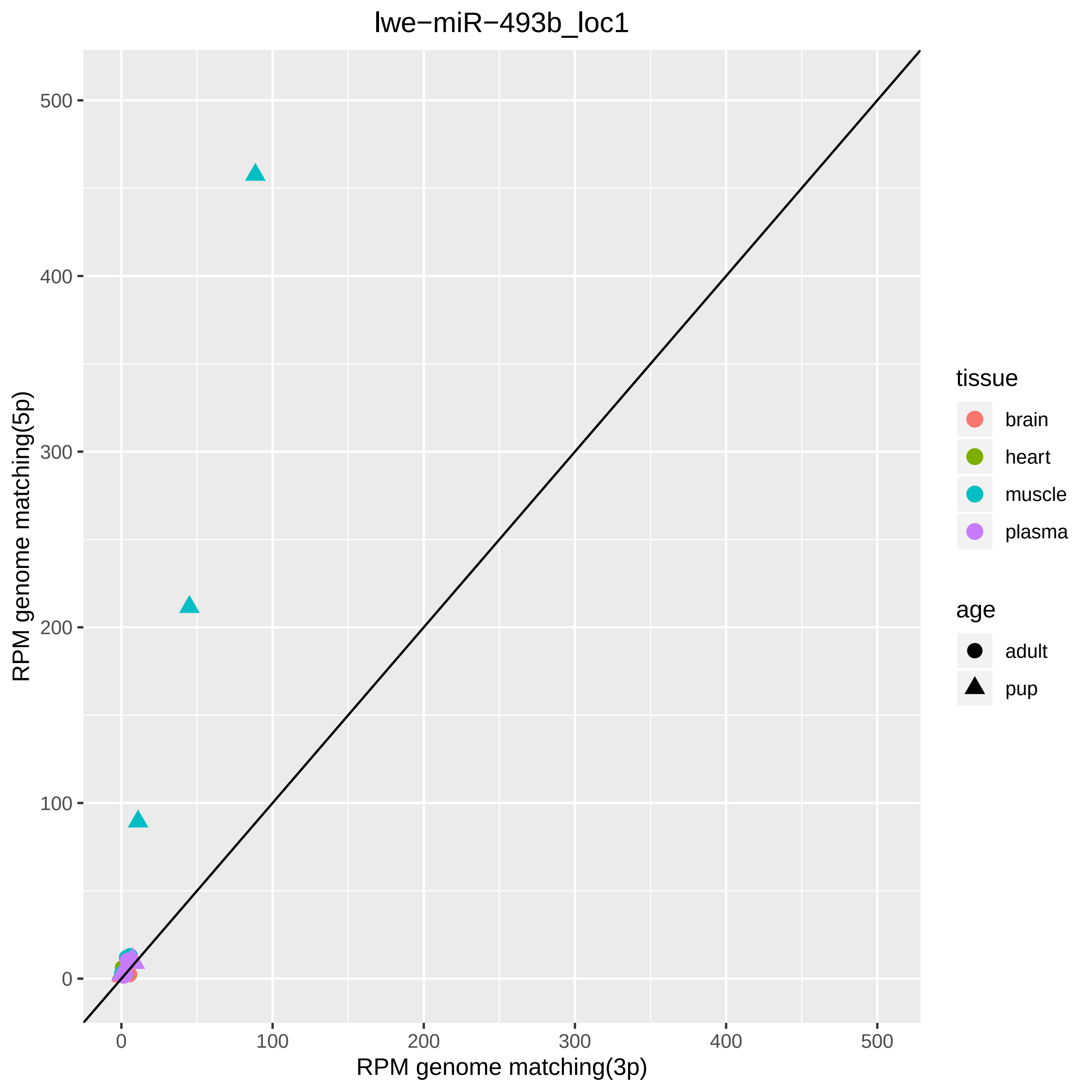
lwe−miR−493b_loc1
RPM genome matching(5p)
RPM genome matching(3p)
0
100
200
300
400
500
0
100
200
300
400
500
tissue
brain
heart
muscle
plasma
age
adult
pup

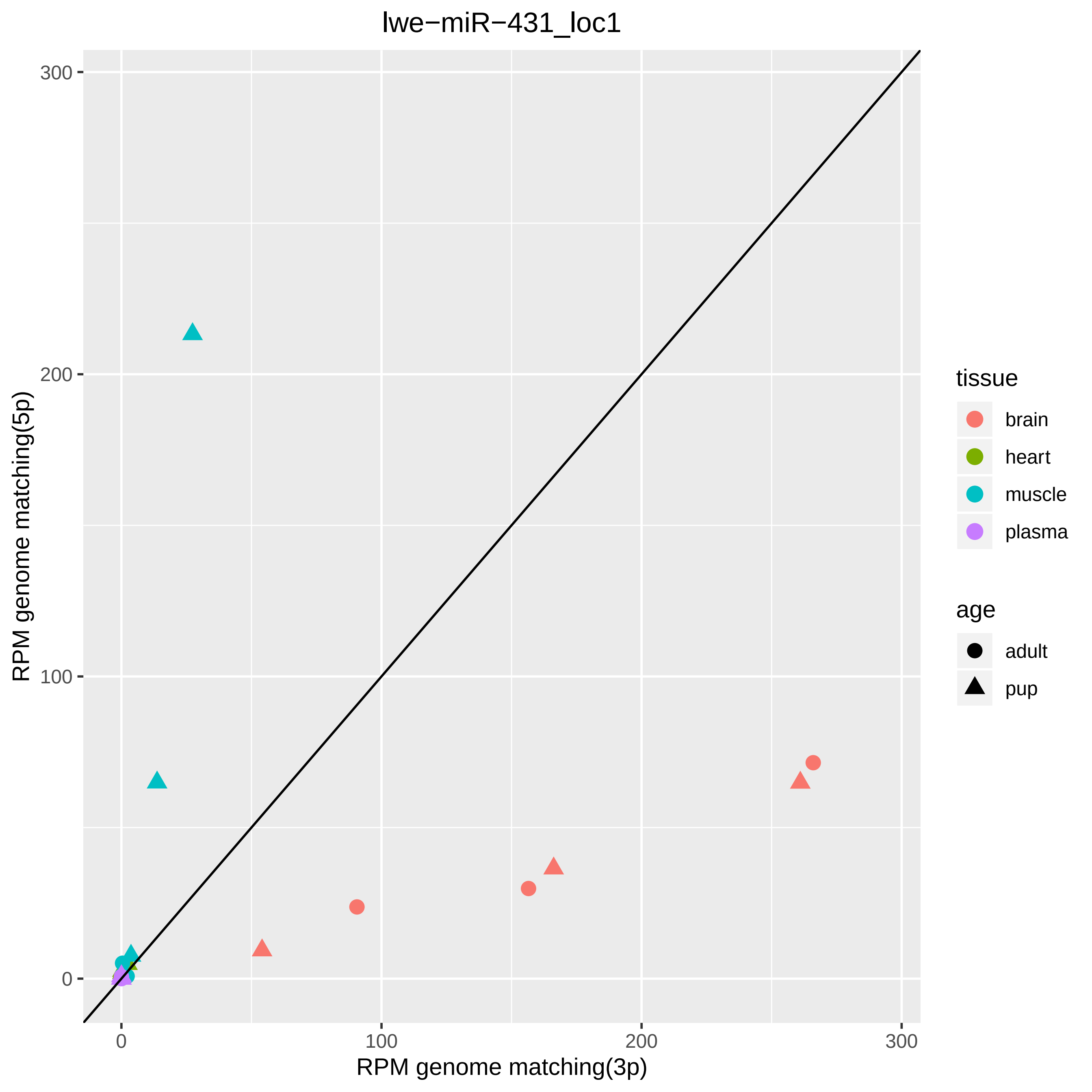

lwe−miR−431_loc1
300
200
100
0
0
100
200
300
RPM genome matching(5p)
RPM genome matching(3p)
tissue
brain
heart
muscle
plasma
age
adult
pup

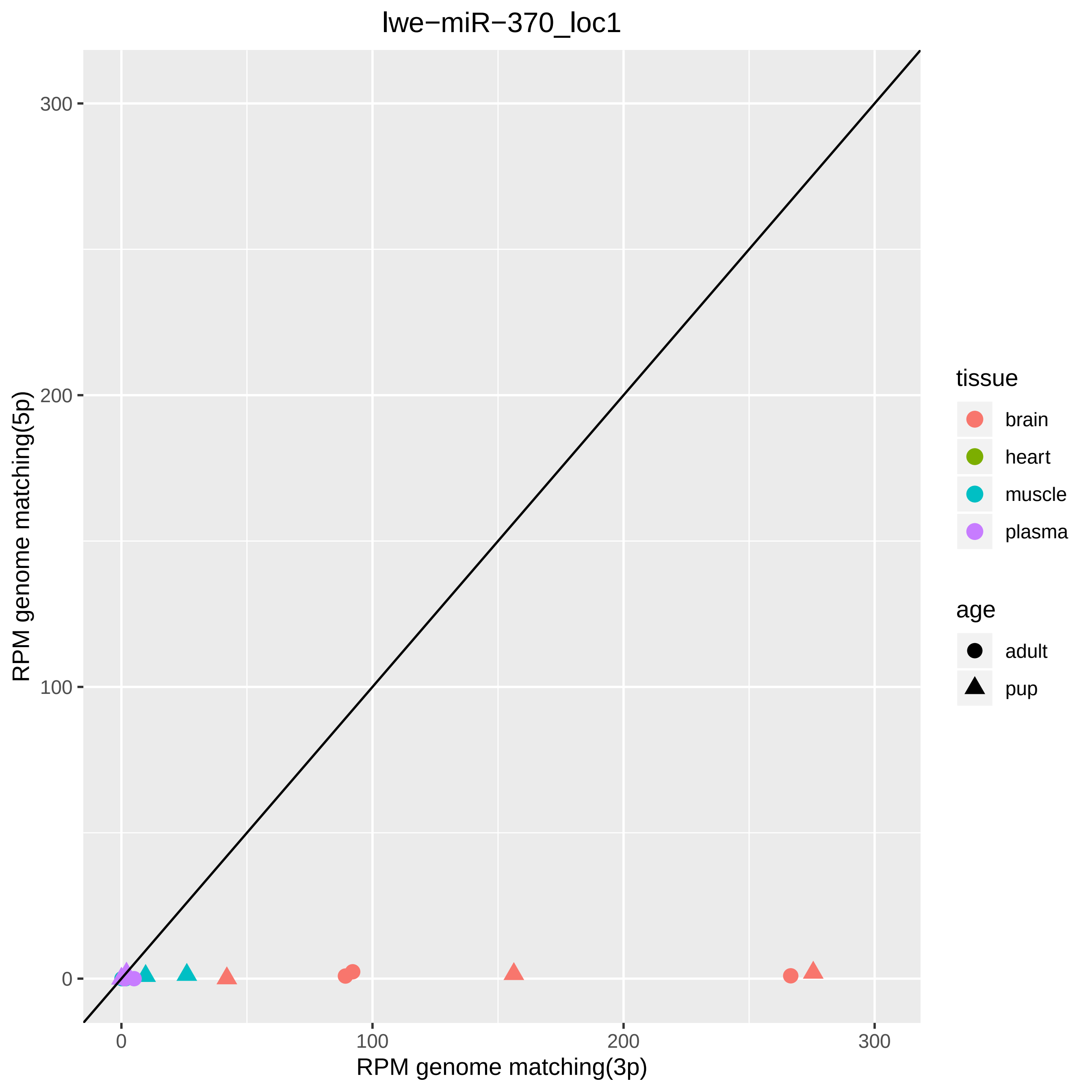
lwe−miR−370_loc1
RPM genome matching(5p)
300
200
100
0
0
100
200
300
RPM genome matching(3p)
tissue
brain
heart
muscle
plasma
age
adult
pup

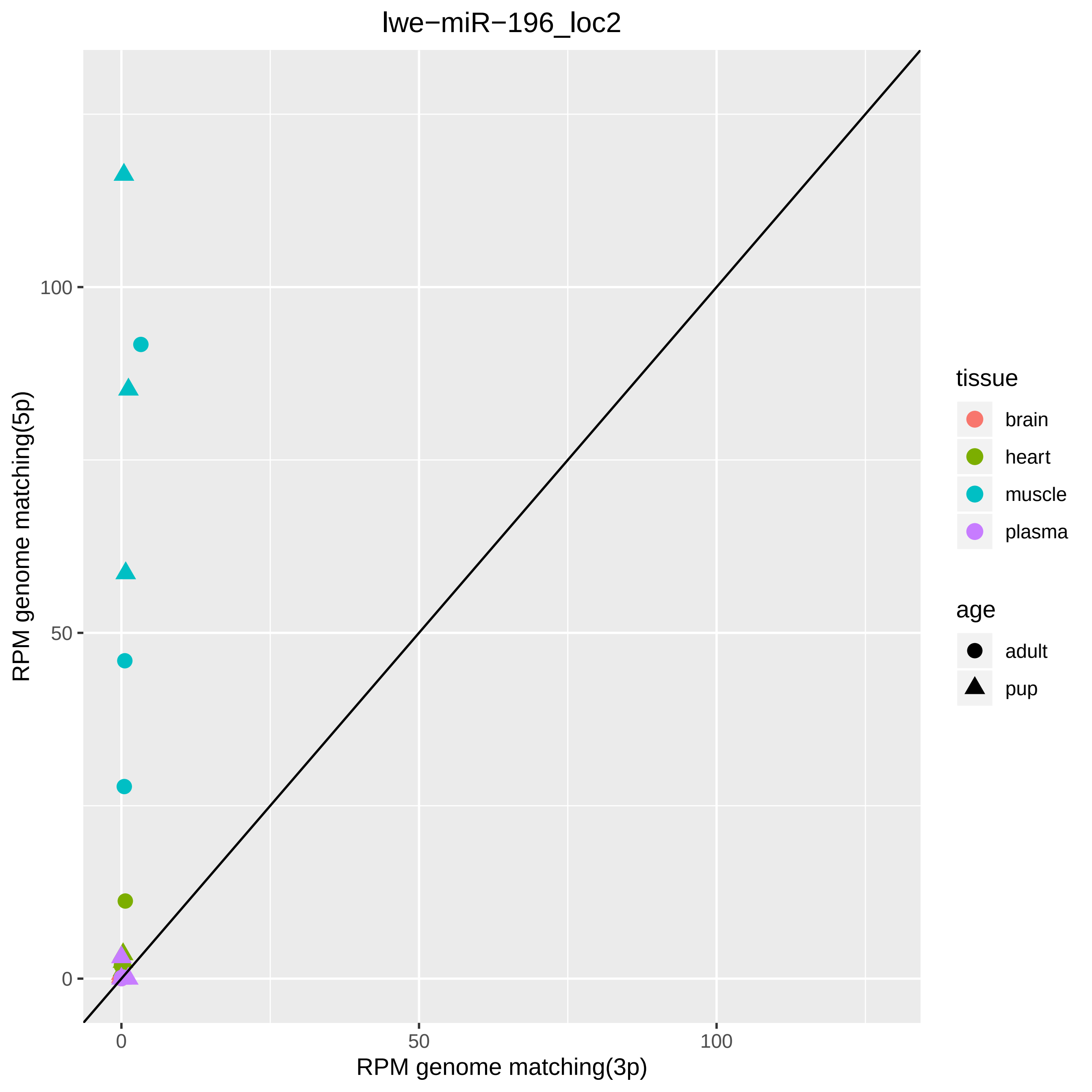
lwe−miR−196_loc2
RPM genome matching(5p)
RPM genome matching(3p)
0
50
100
0
50
100
tissue
brain
heart
muscle
plasma
age
adult
pup

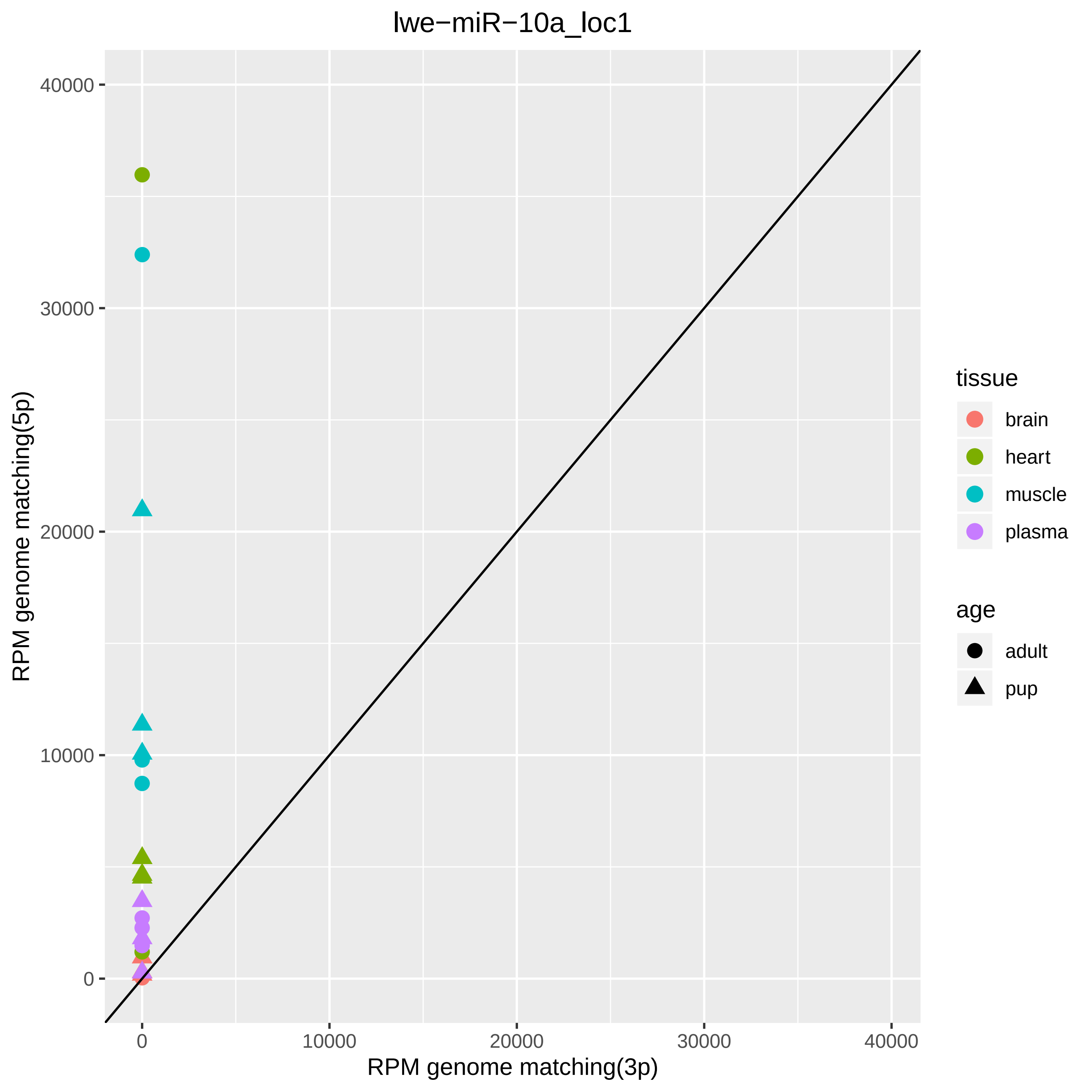
lwe−miR−10a_loc1
RPM genome matching(5p)
RPM genome matching(3p)
0
10000
20000
30000
40000
tissue
brain
heart
muscle
plasma
age
adult
pup

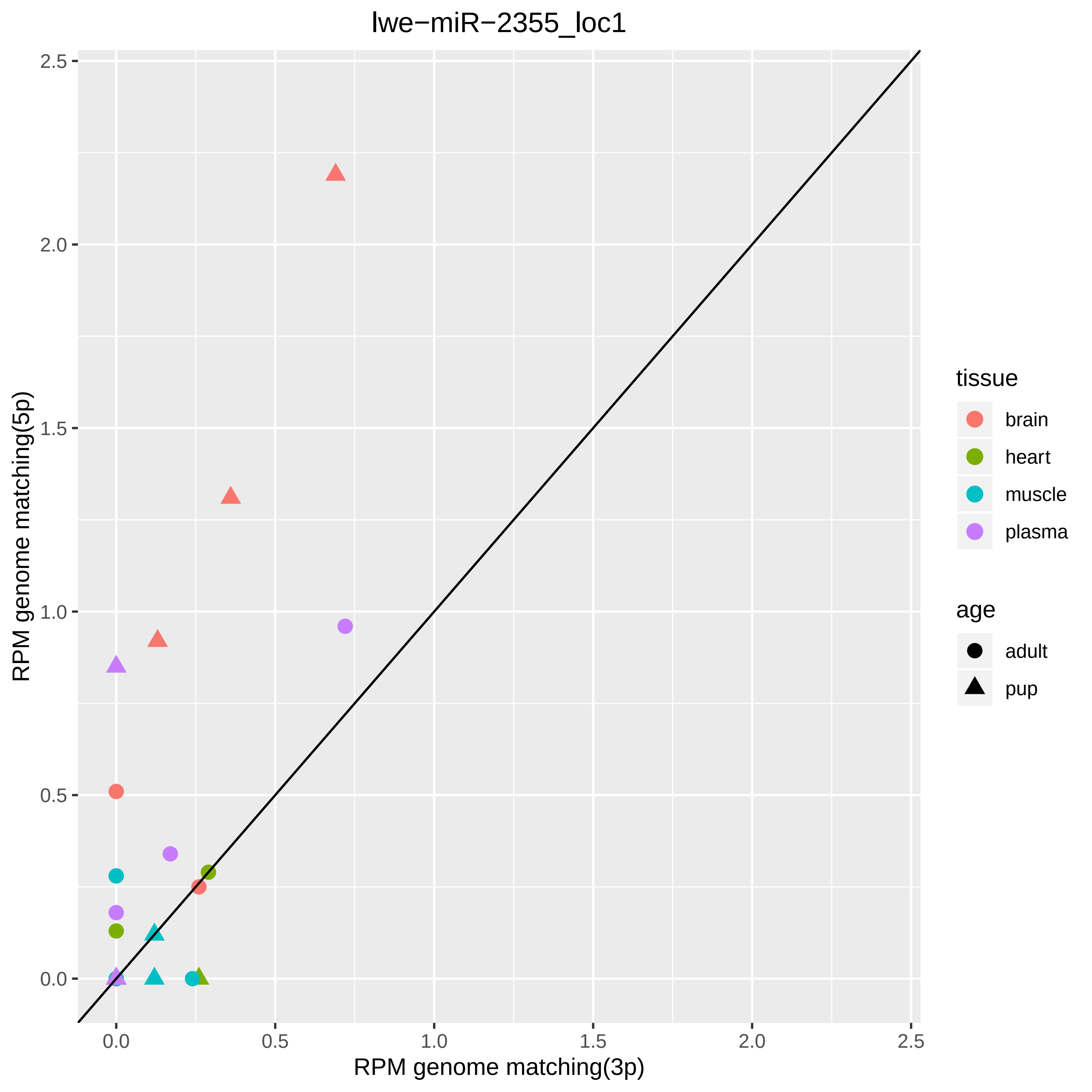
lwe−miR−2355_loc1
RPM genome matching(5p)
RPM genome matching(3p)
0.0
0.5
1.0
1.5
2.0
2.5
tissue
brain
heart
muscle
plasma
age
adult
pup

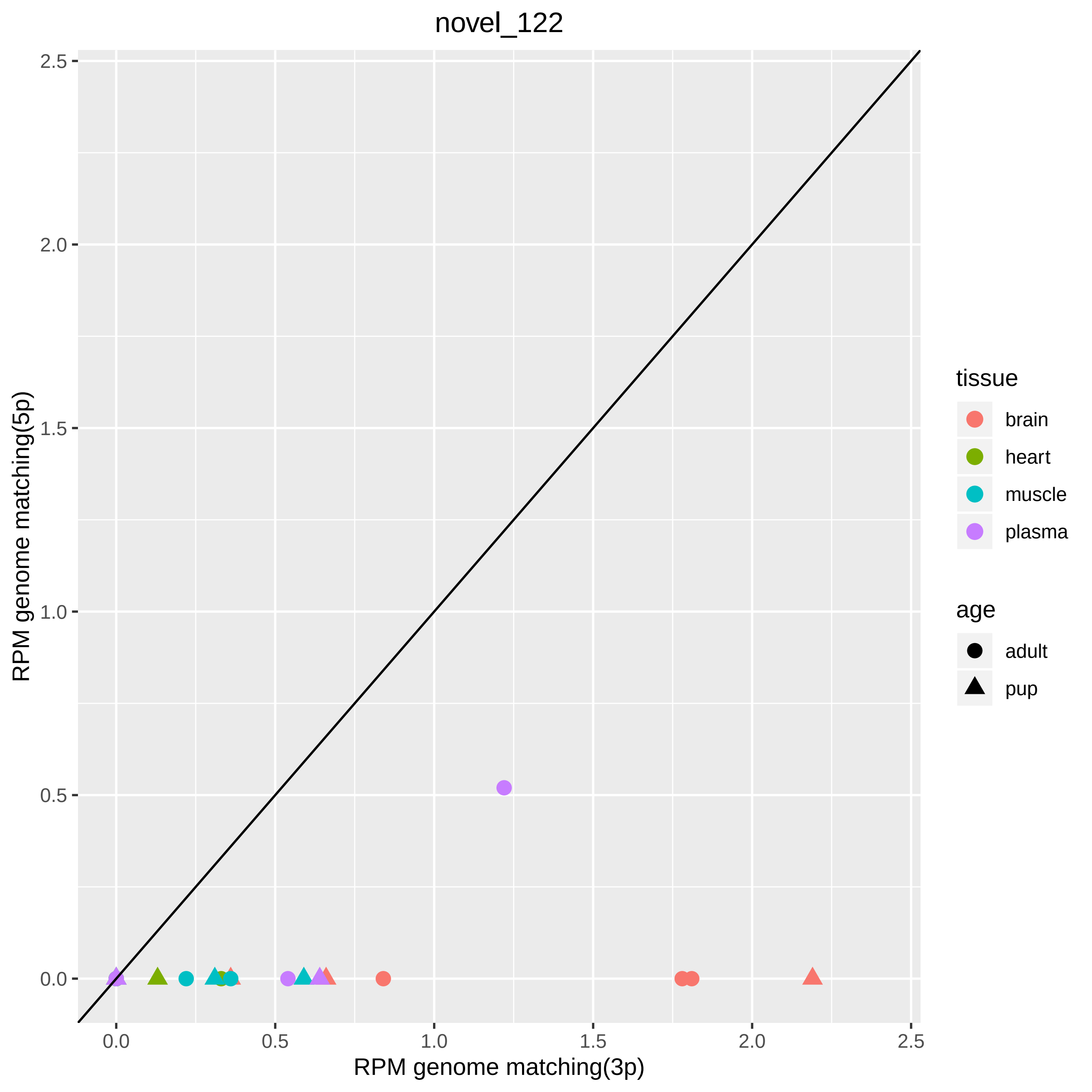
novel_122
RPM genome matching(5p)
RPM genome matching(3p)
0.0
0.5
1.0
1.5
2.0
2.5
tissue
brain
heart
muscle
plasma
age
adult
pup

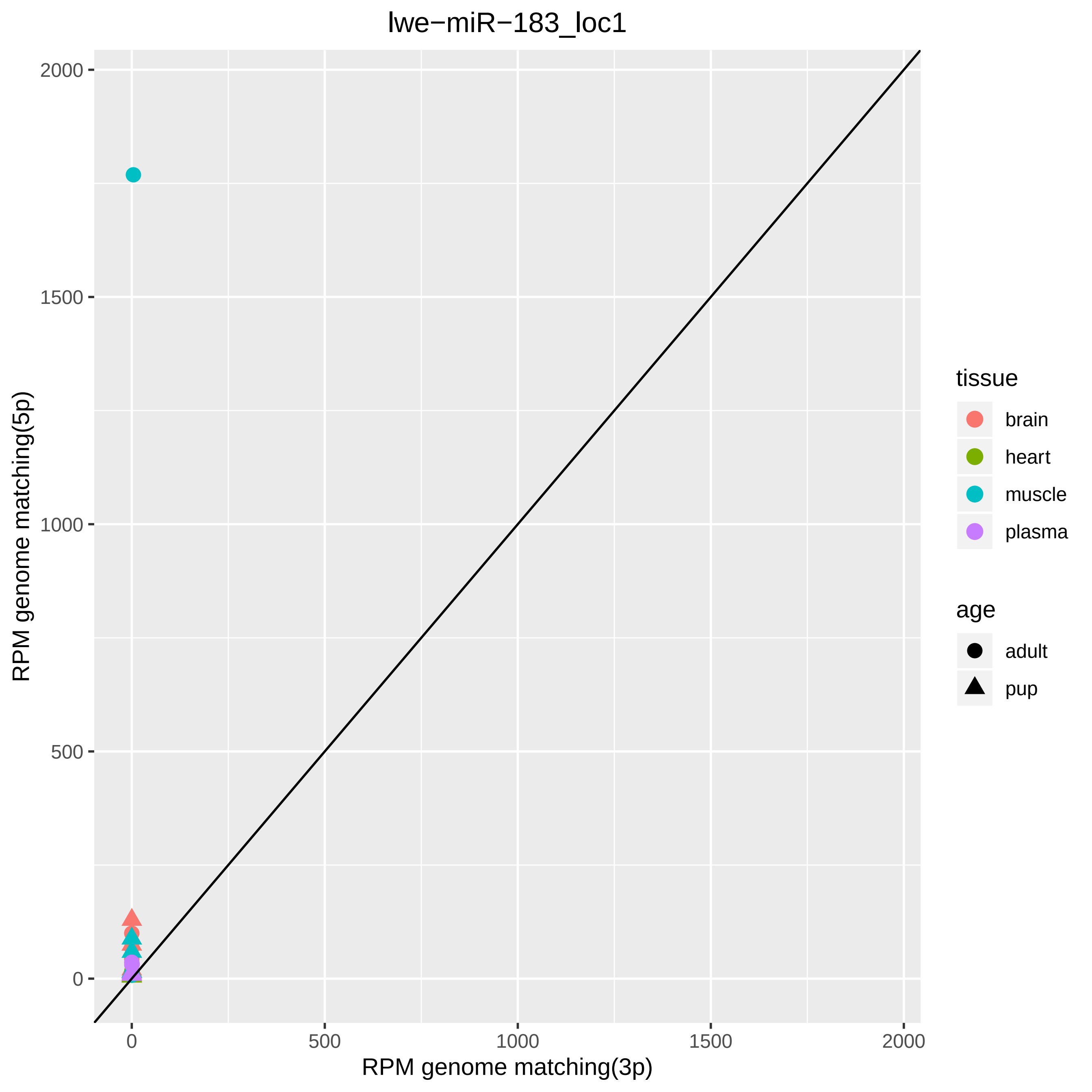

lwe−miR−183_loc1
RPM genome matching(5p)
2000
1500
1000
500
0
0
500
1000
1500
2000
RPM genome matching(3p)
tissue
brain
heart
muscle
plasma
age
adult
pup

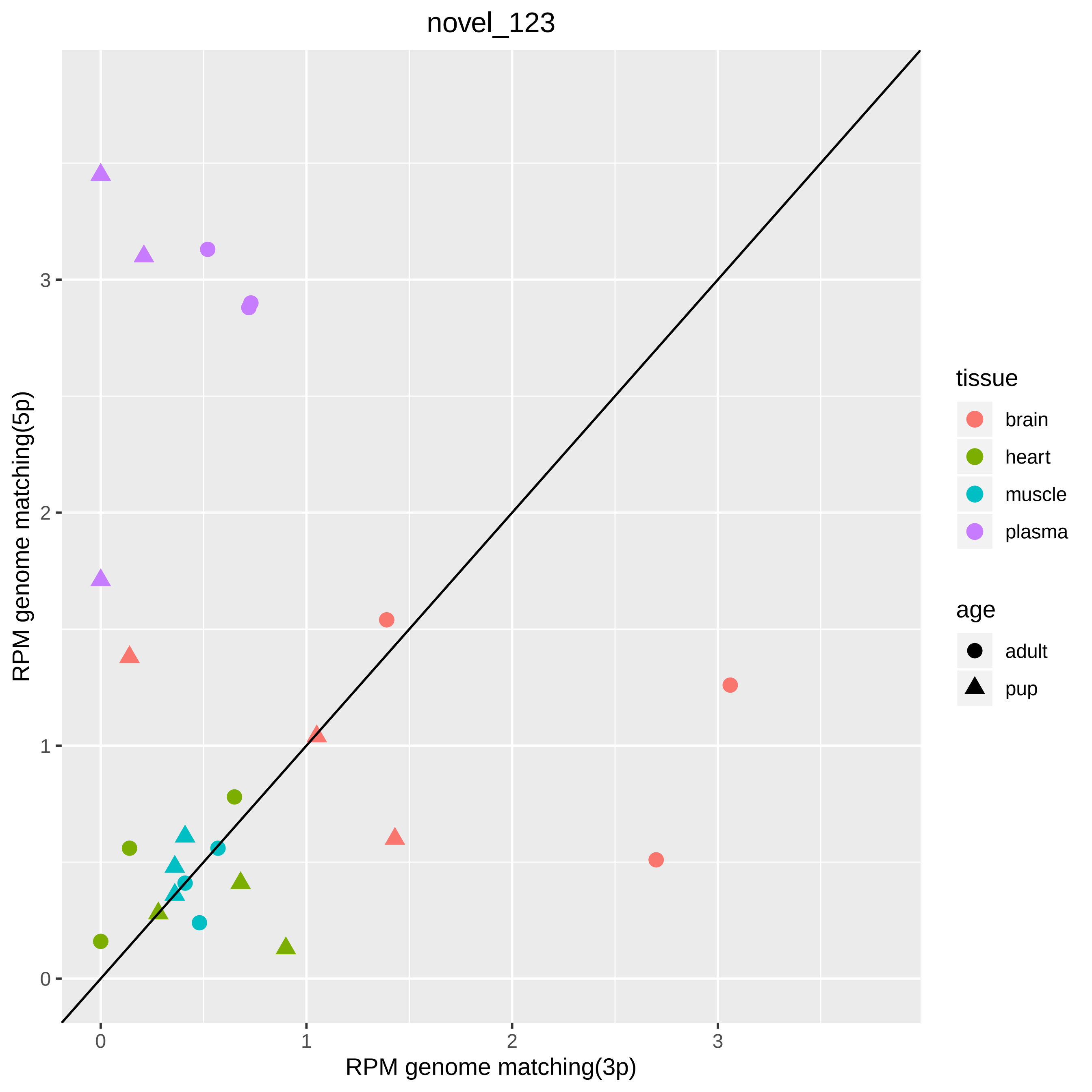

novel_123
RPM genome matching(5p)
RPM genome matching(3p)
0
1
2
3
0
1
2
3
tissue
brain
heart
muscle
plasma
age
adult
pup

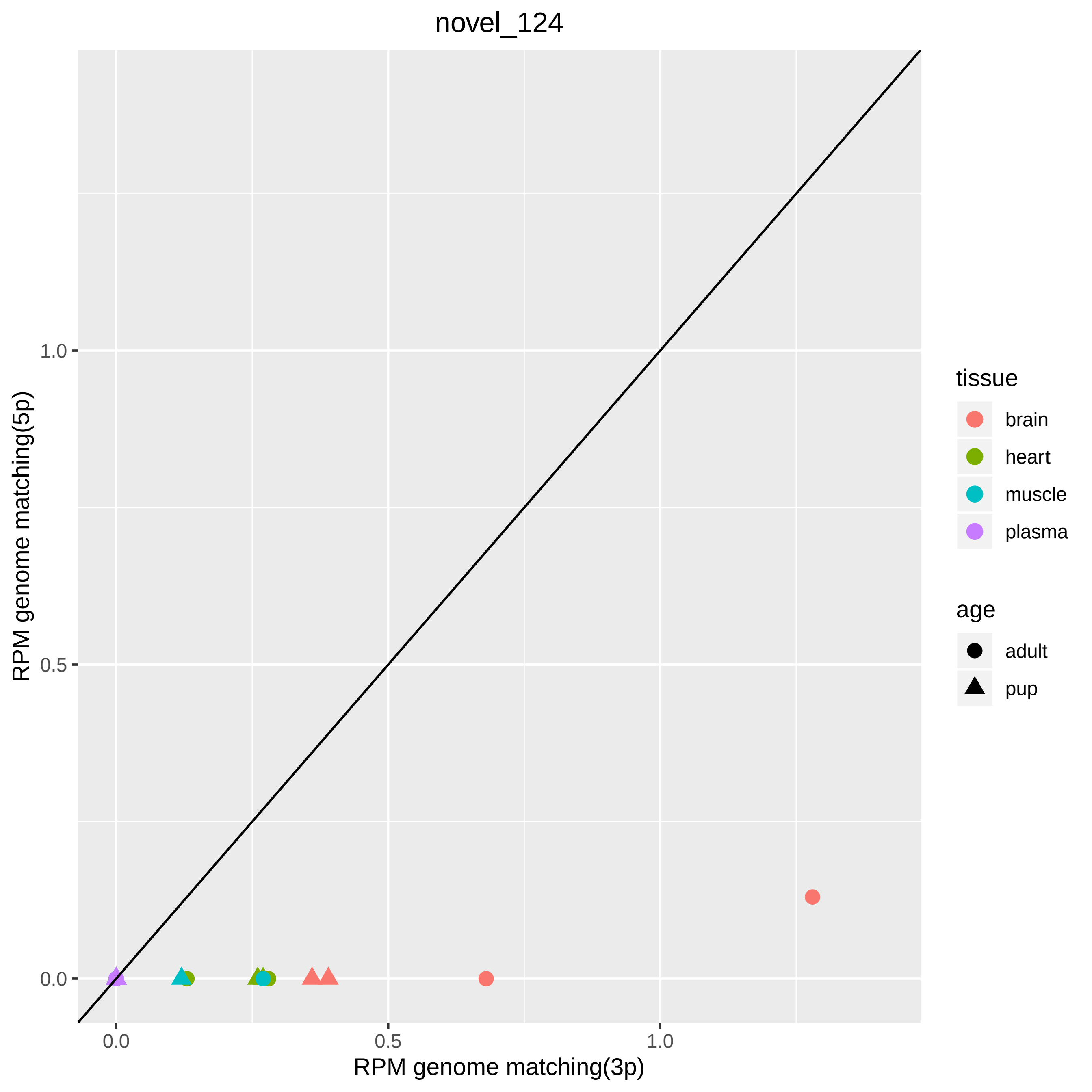

novel_124
RPM genome matching(5p)
RPM genome matching(3p)
0.0
0.5
1.0
tissue
brain
heart
muscle
plasma
age
adult
pup

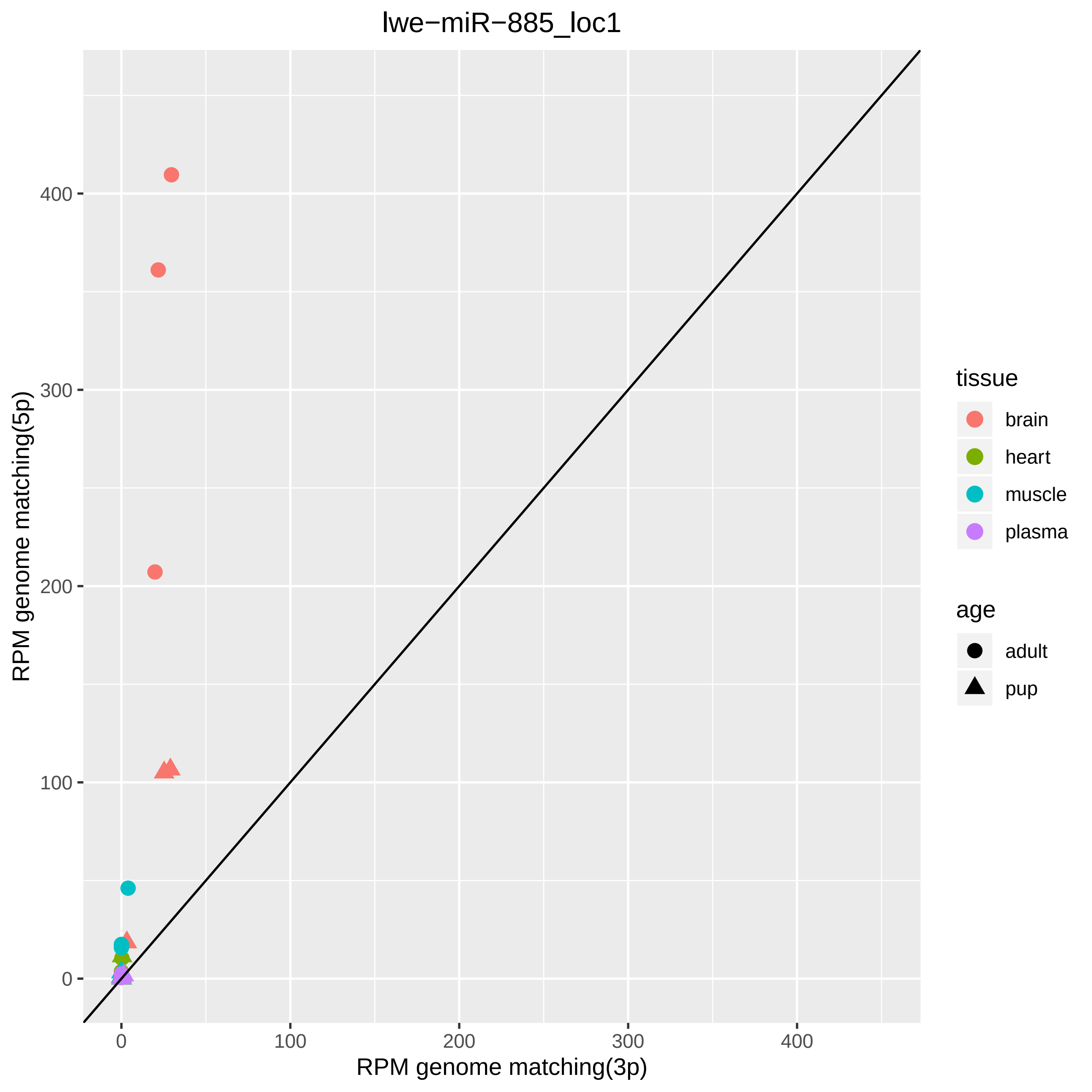
lwe−miR−885_loc1
RPM genome matching(5p)
RPM genome matching(3p)
0
100
200
300
400
0
100
200
300
400
tissue
brain
heart
muscle
plasma
age
adult
pup

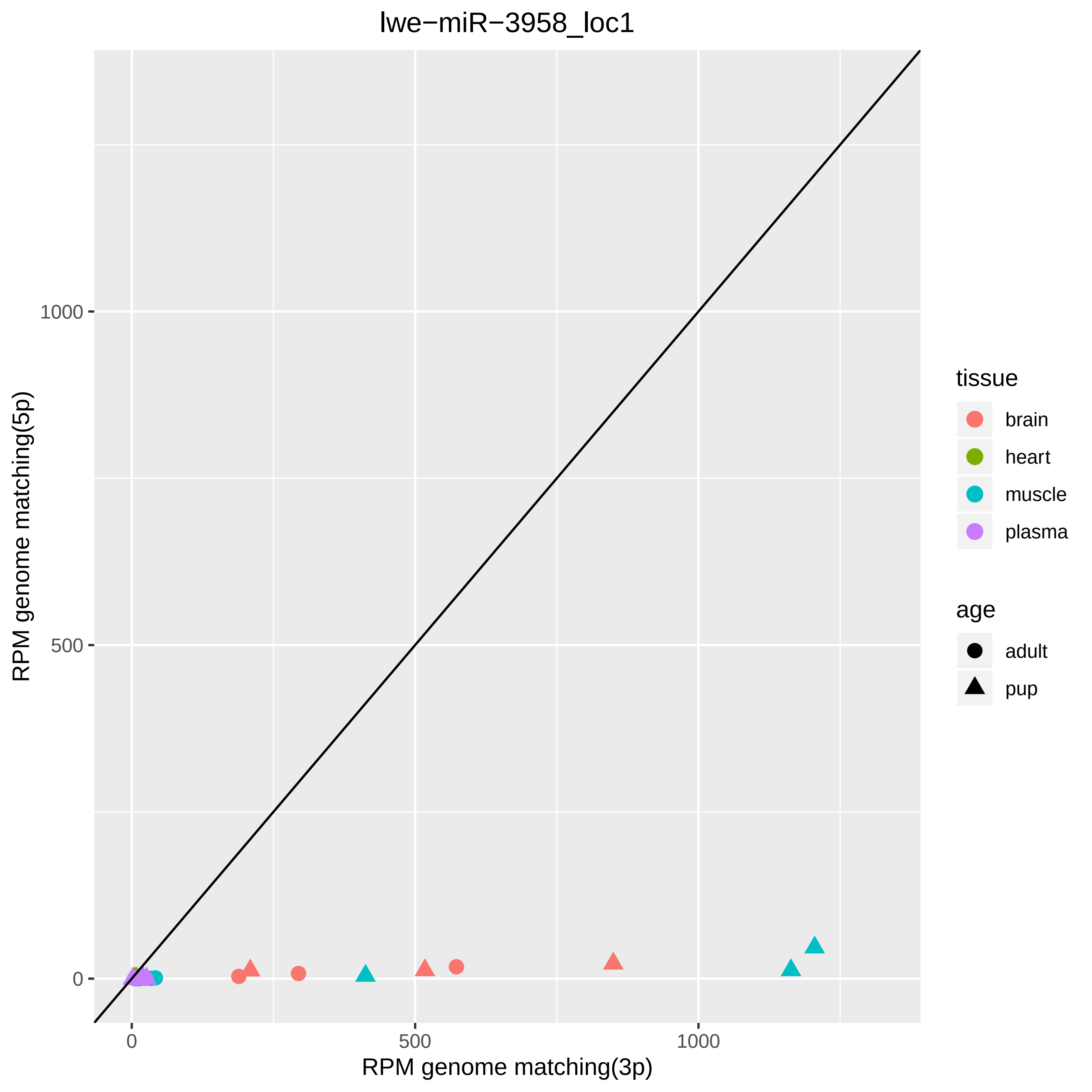
lwe−miR−3958_loc1
RPM genome matching(5p)
RPM genome matching(3p)
1000
500
0
0
500
1000
tissue
brain
heart
muscle
plasma
age
adult
pup

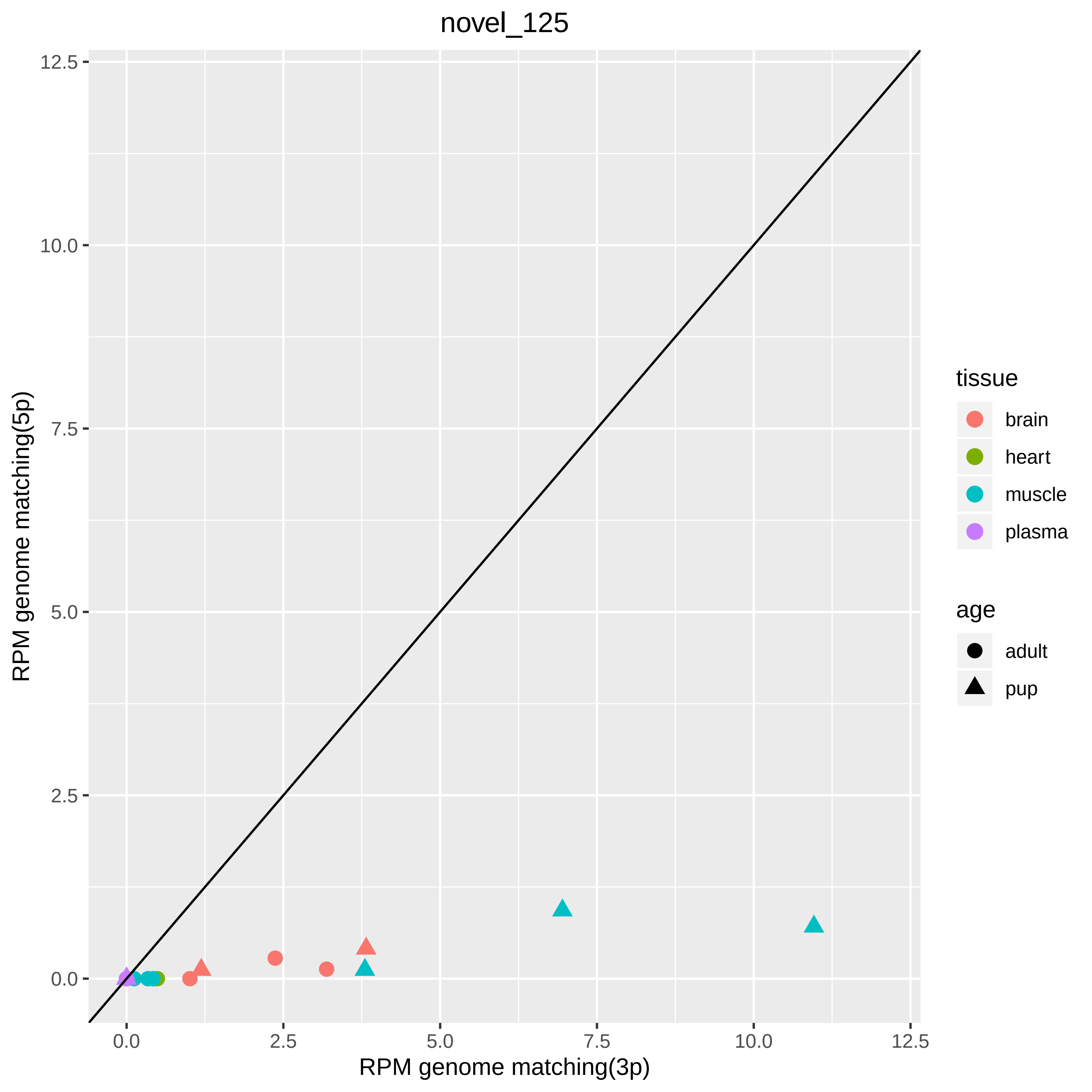
novel_125
RPM genome matching(5p)
RPM genome matching(3p)
0.0
2.5
5.0
7.5
10.0
12.5
tissue
brain
heart
muscle
plasma
age
adult
pup

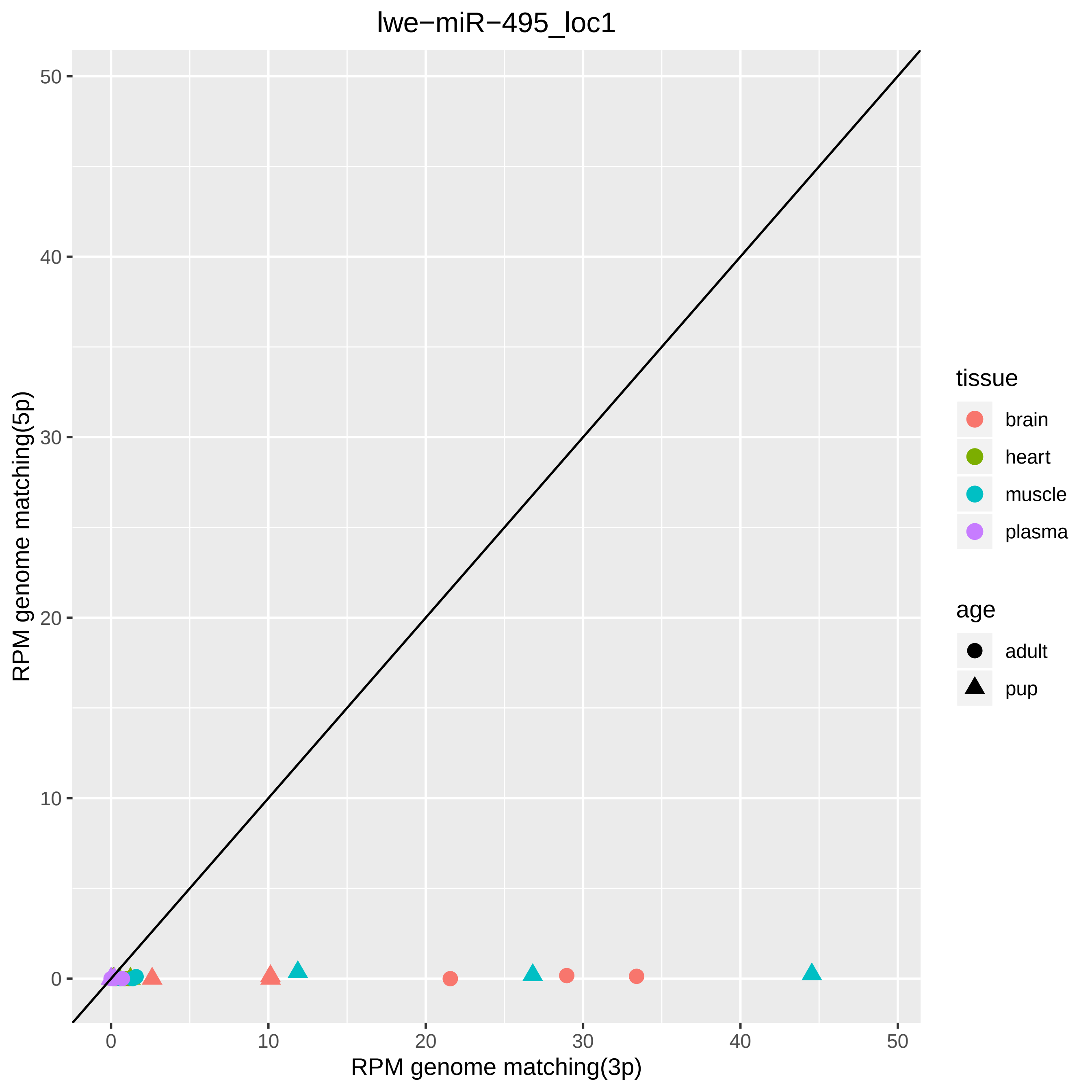

lwe−miR−495_loc1
RPM genome matching(5p)
RPM genome matching(3p)
0
10
20
30
40
50
tissue
brain
heart
muscle
plasma
age
adult
pup

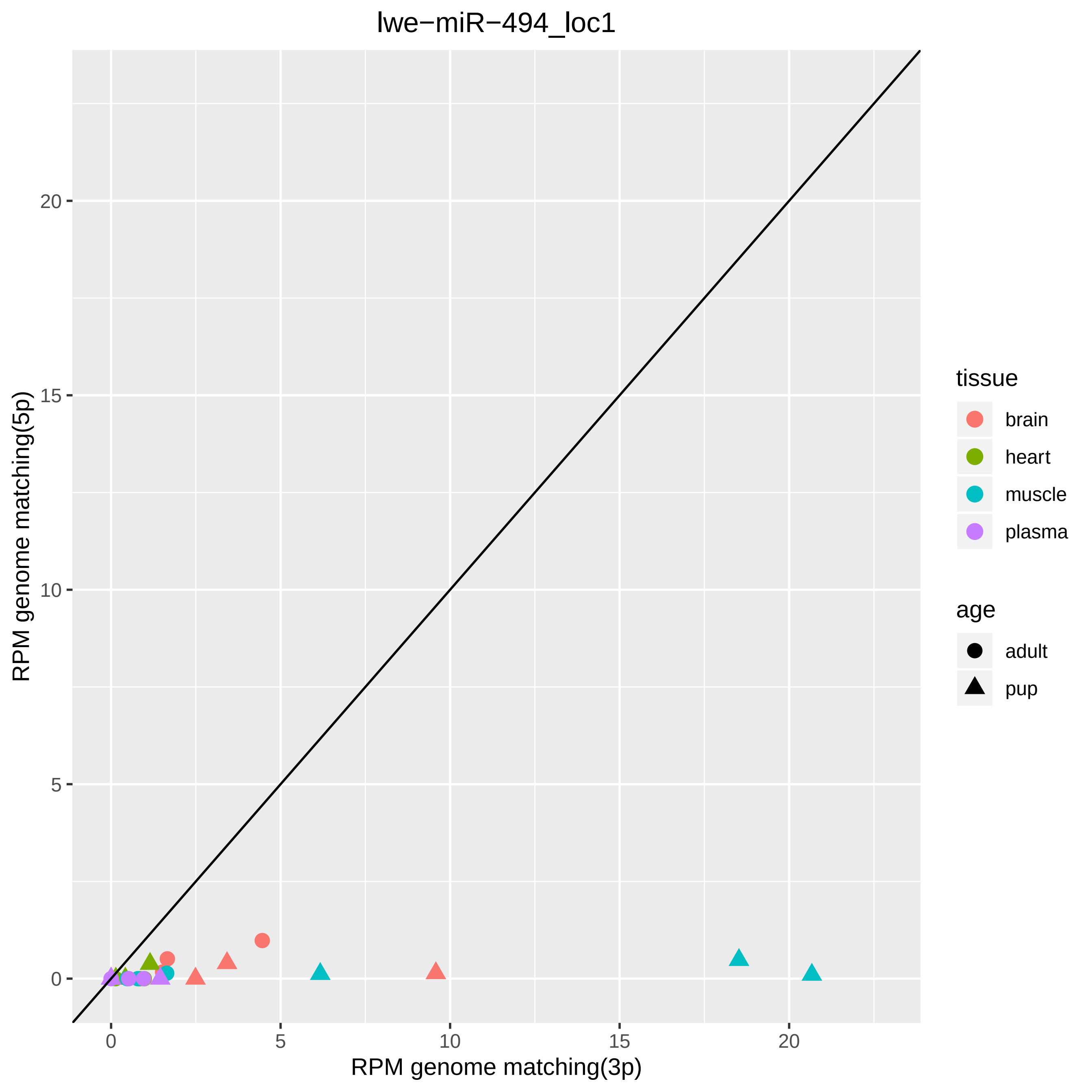
lwe−miR−494_loc1
RPM genome matching(5p)
RPM genome matching(3p)
0
5
10
15
20
0
5
10
15
20
tissue
brain
heart
muscle
plasma
age
adult
pup

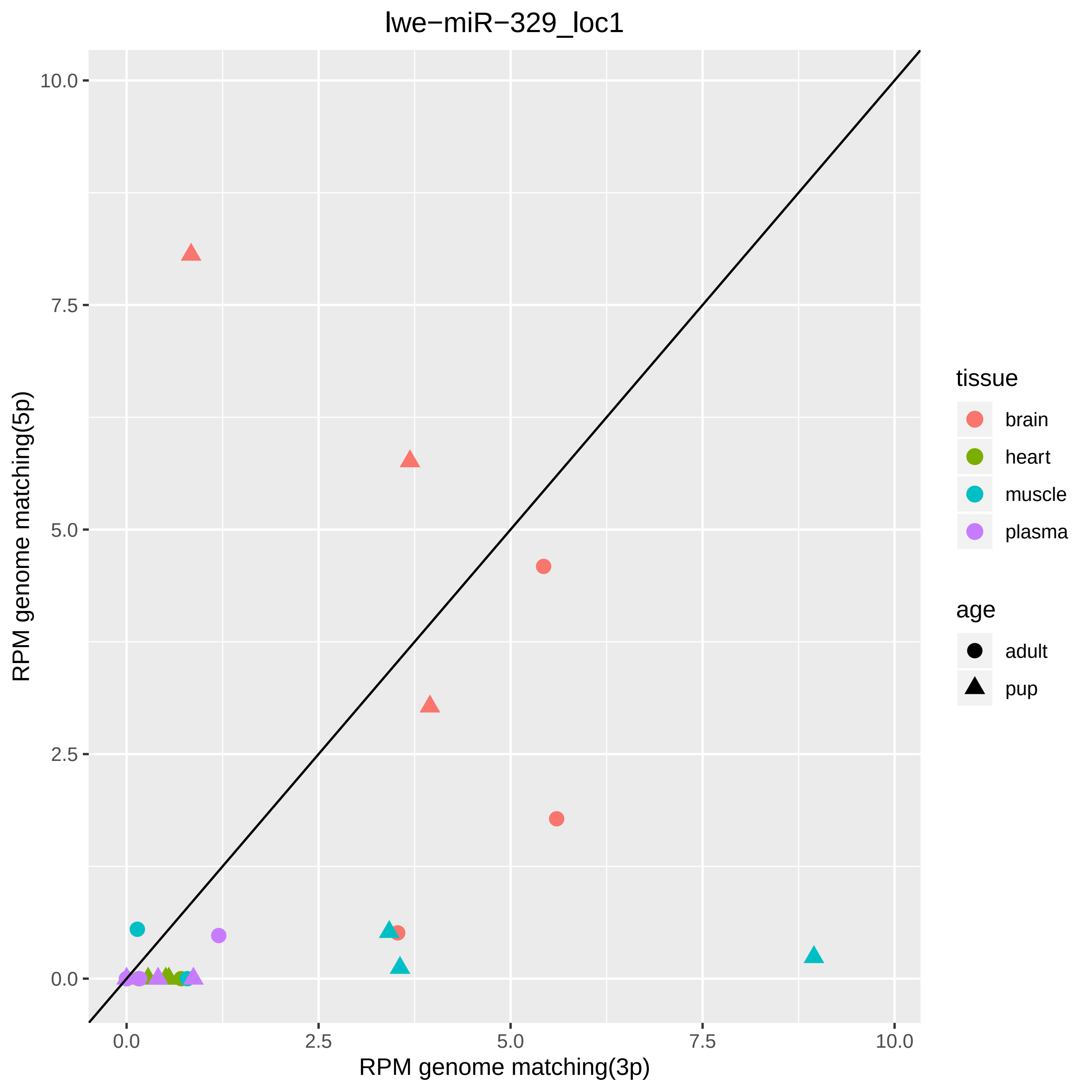
lwe−miR−329_loc1
RPM genome matching(5p)
RPM genome matching(3p)
10.0
7.5
5.0
2.5
0.0
0.0
2.5
5.0
7.5
10.0
tissue
brain
heart
muscle
plasma
age
adult
pup

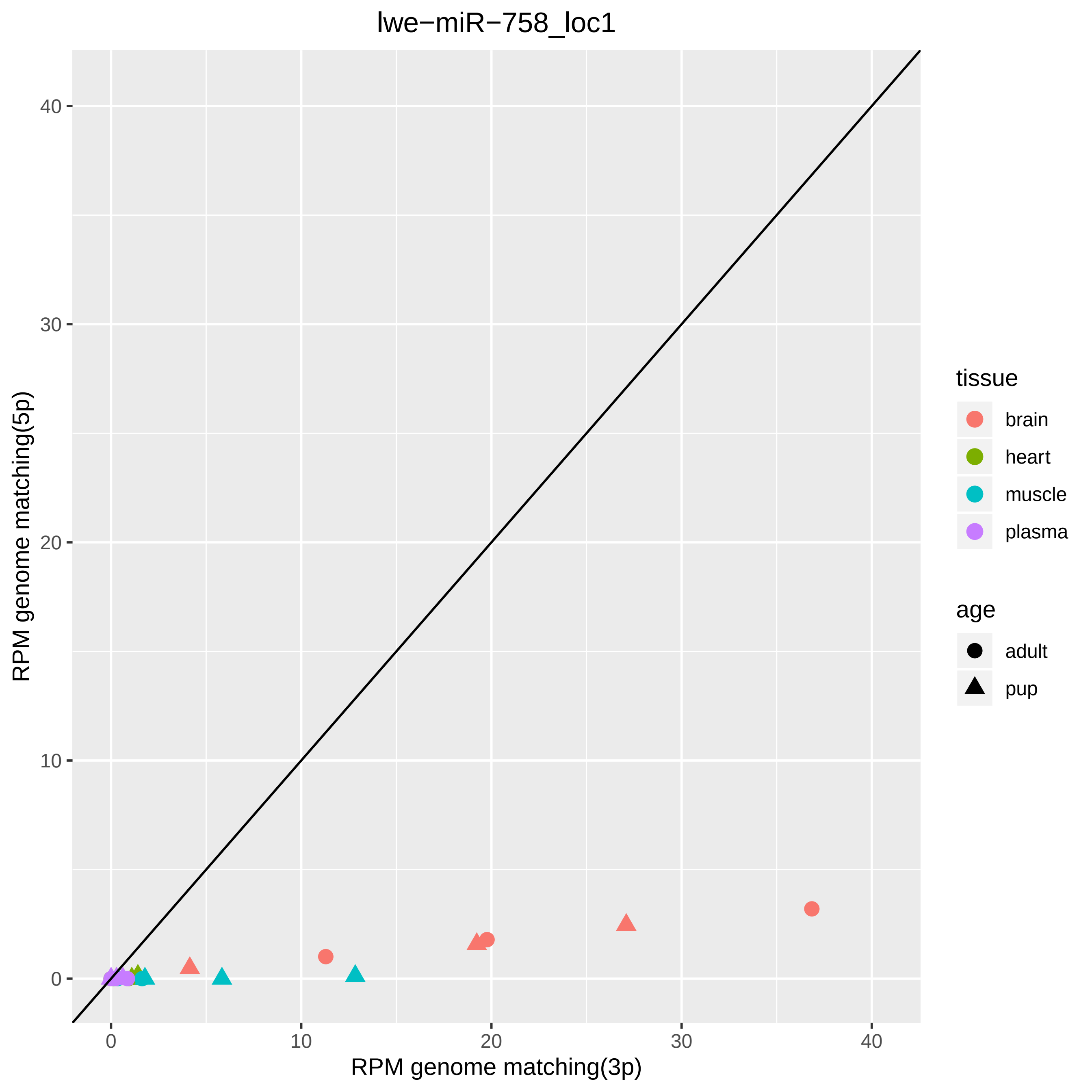
lwe−miR−758_loc1
RPM genome matching(5p)
RPM genome matching(3p)
0
10
20
30
40
tissue
brain
heart
muscle
plasma
age
adult
pup

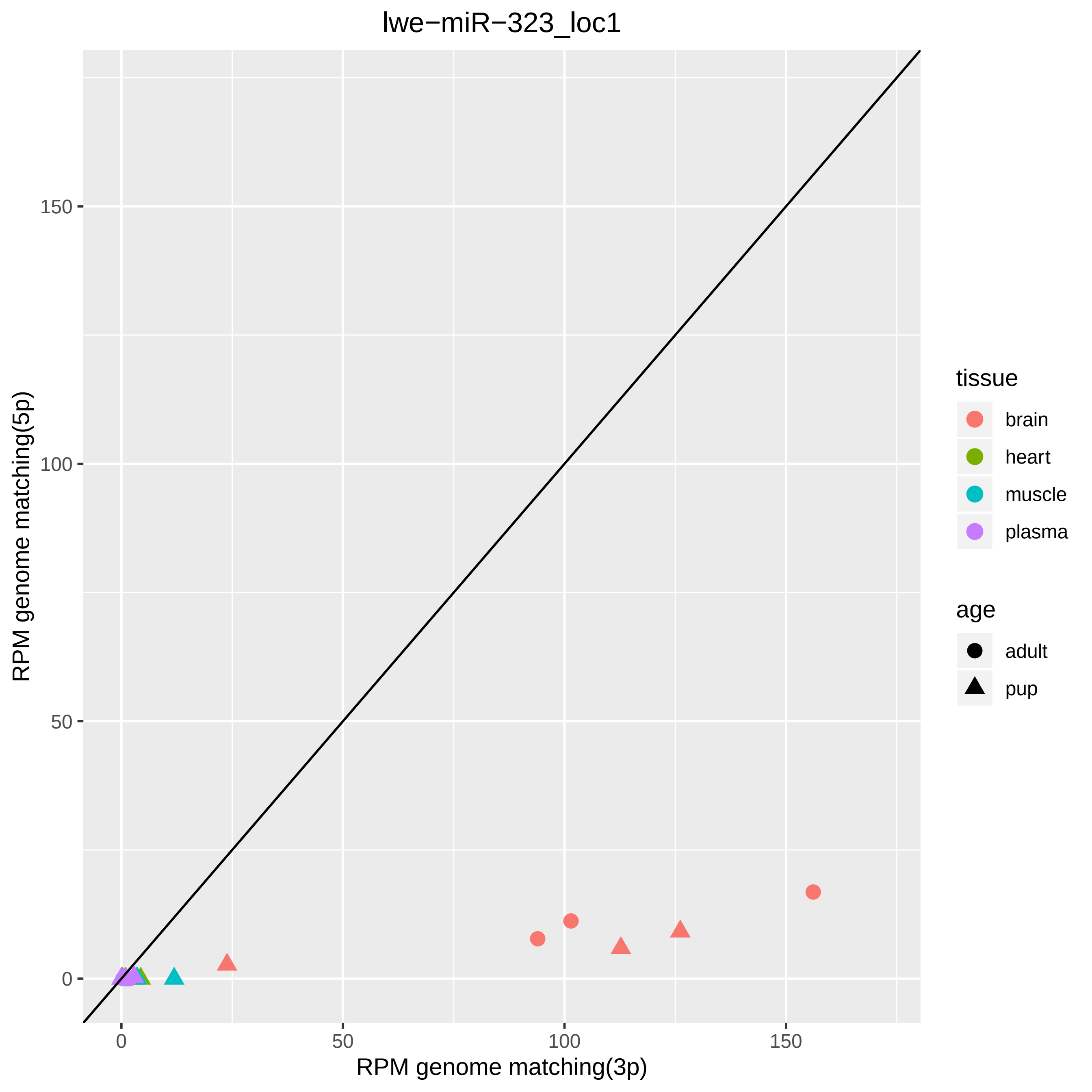

lwe−miR−323_loc1
RPM genome matching(5p)
RPM genome matching(3p)
0
50
100
150
tissue
brain
heart
muscle
plasma
age
adult
pup

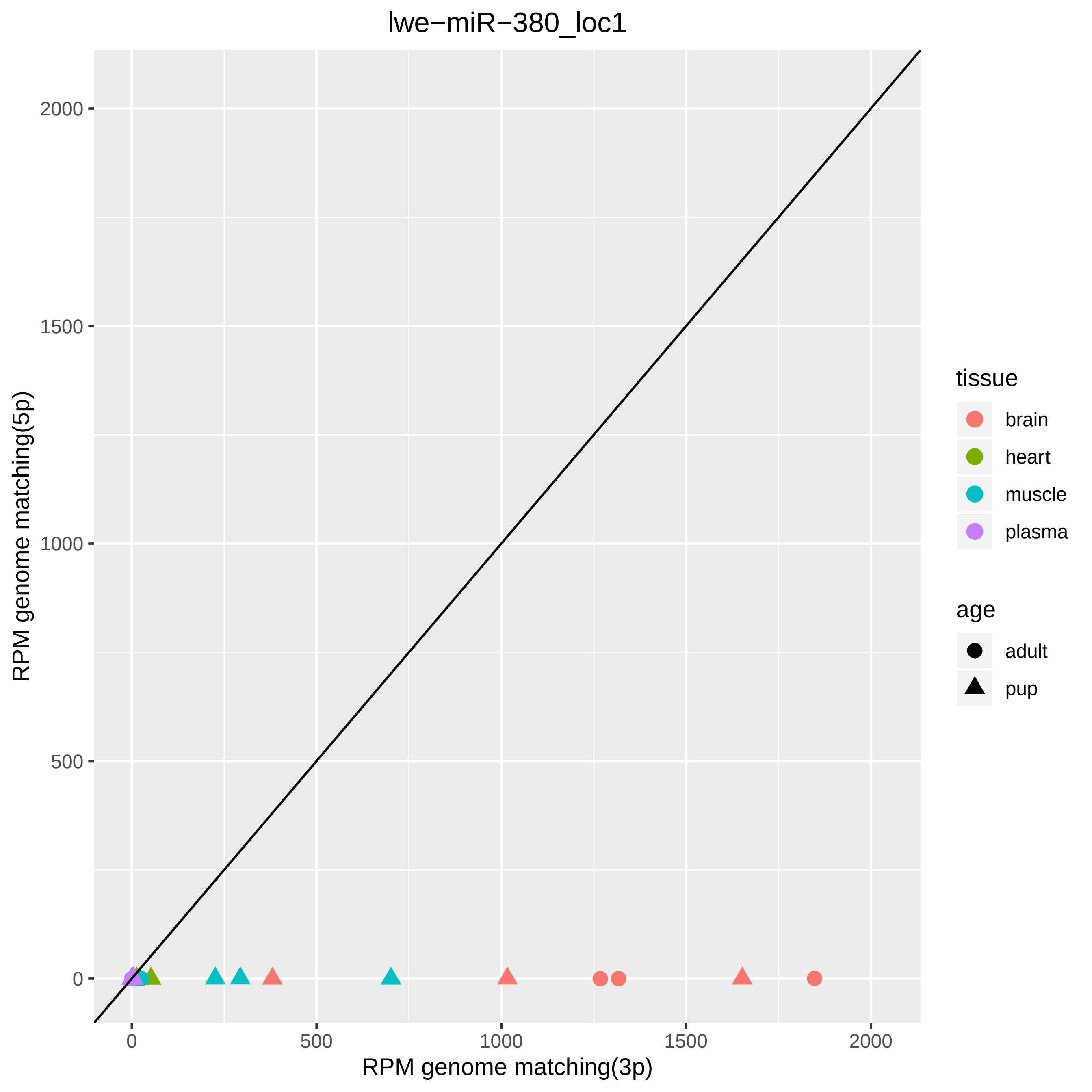

lwe−miR−380_loc1
RPM genome matching(5p)
RPM genome matching(3p)
0
500
1000
1500
2000
tissue
brain
heart
muscle
plasma
age
adult
pup

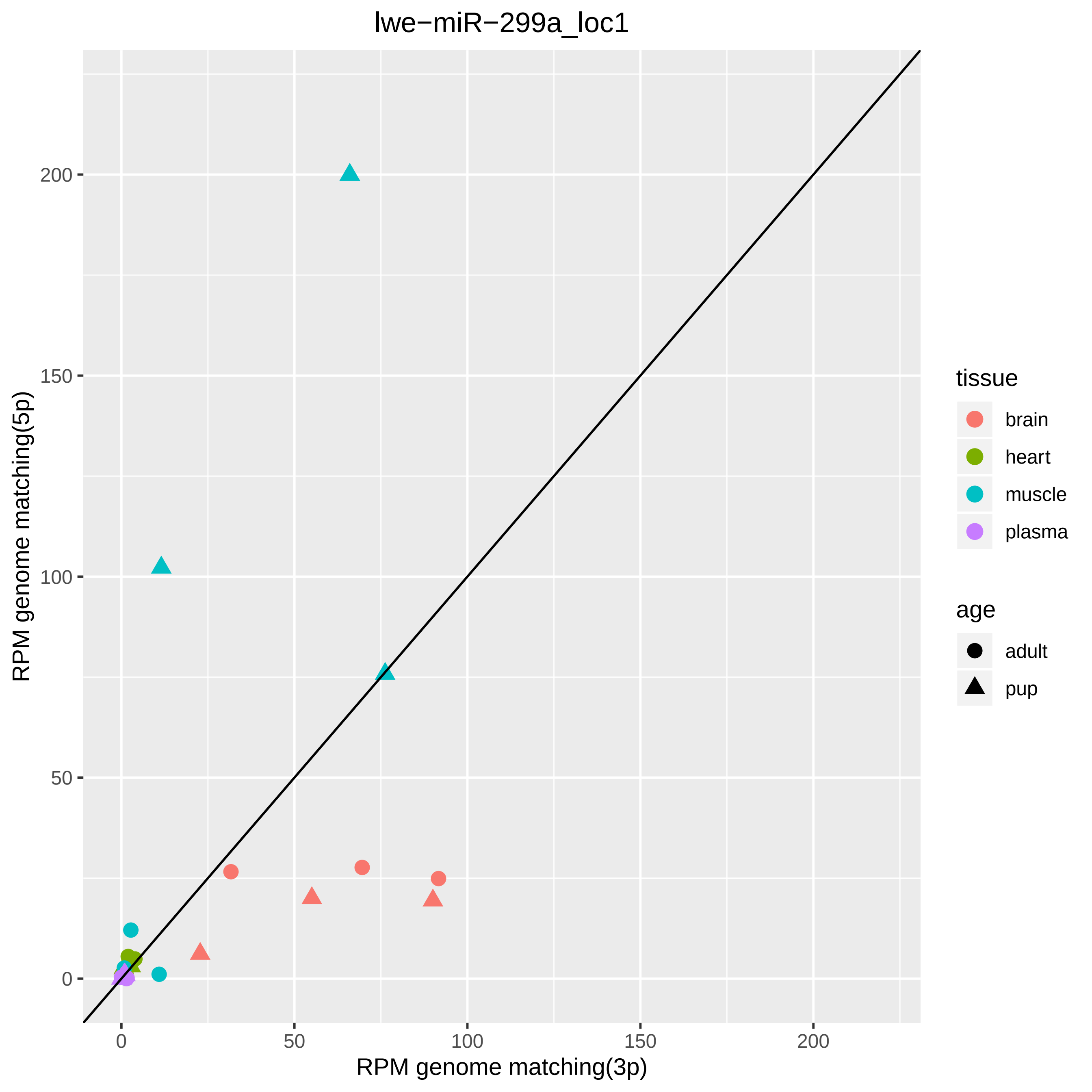
lwe−miR−299a_loc1
RPM genome matching(5p)
RPM genome matching(3p)
0
50
100
150
200
tissue
brain
heart
muscle
plasma
age
adult
pup

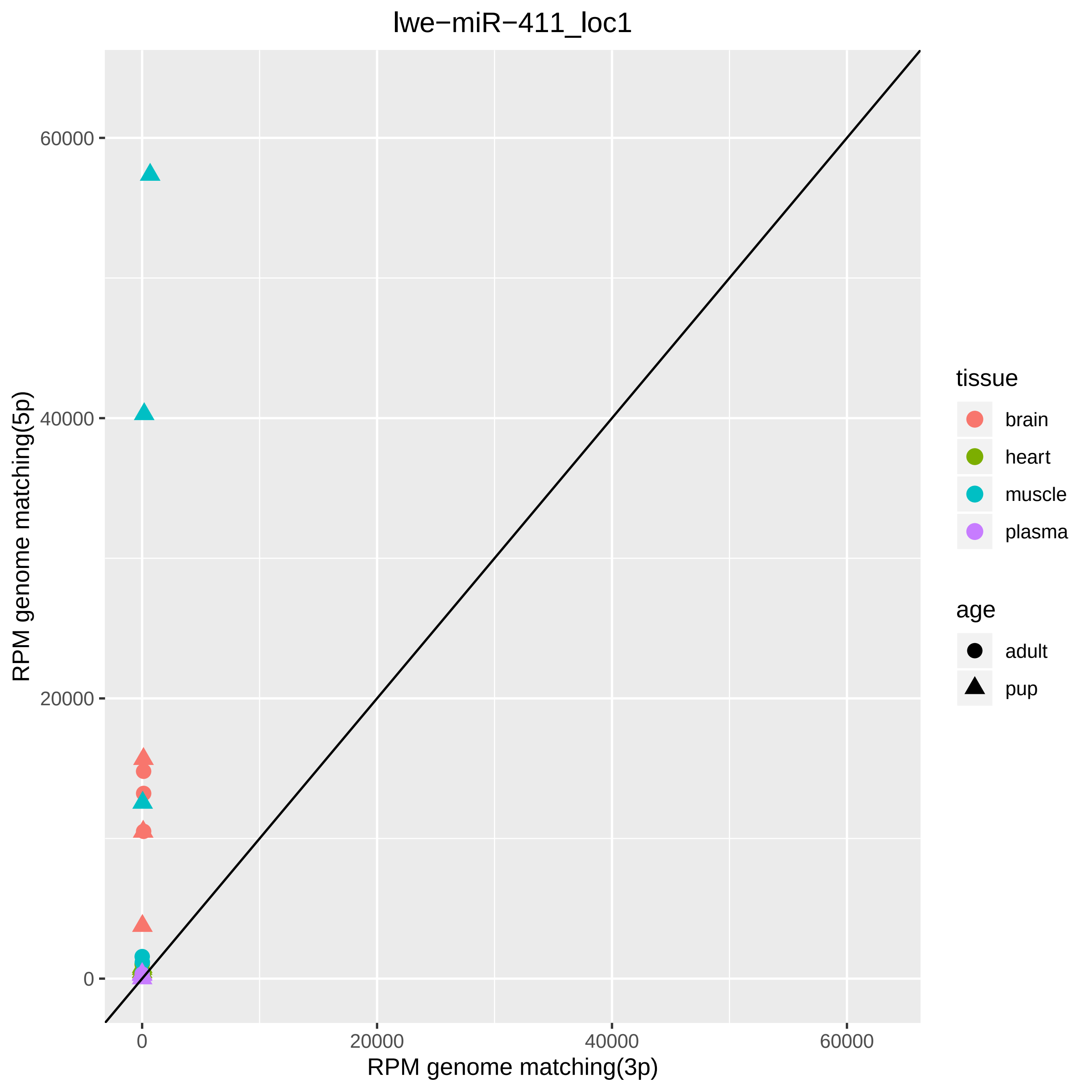

lwe−miR−411_loc1
60000
40000
20000
0
0
20000
40000
60000
RPM genome matching(5p)
RPM genome matching(3p)
tissue
brain
heart
muscle
plasma
age
adult
pup

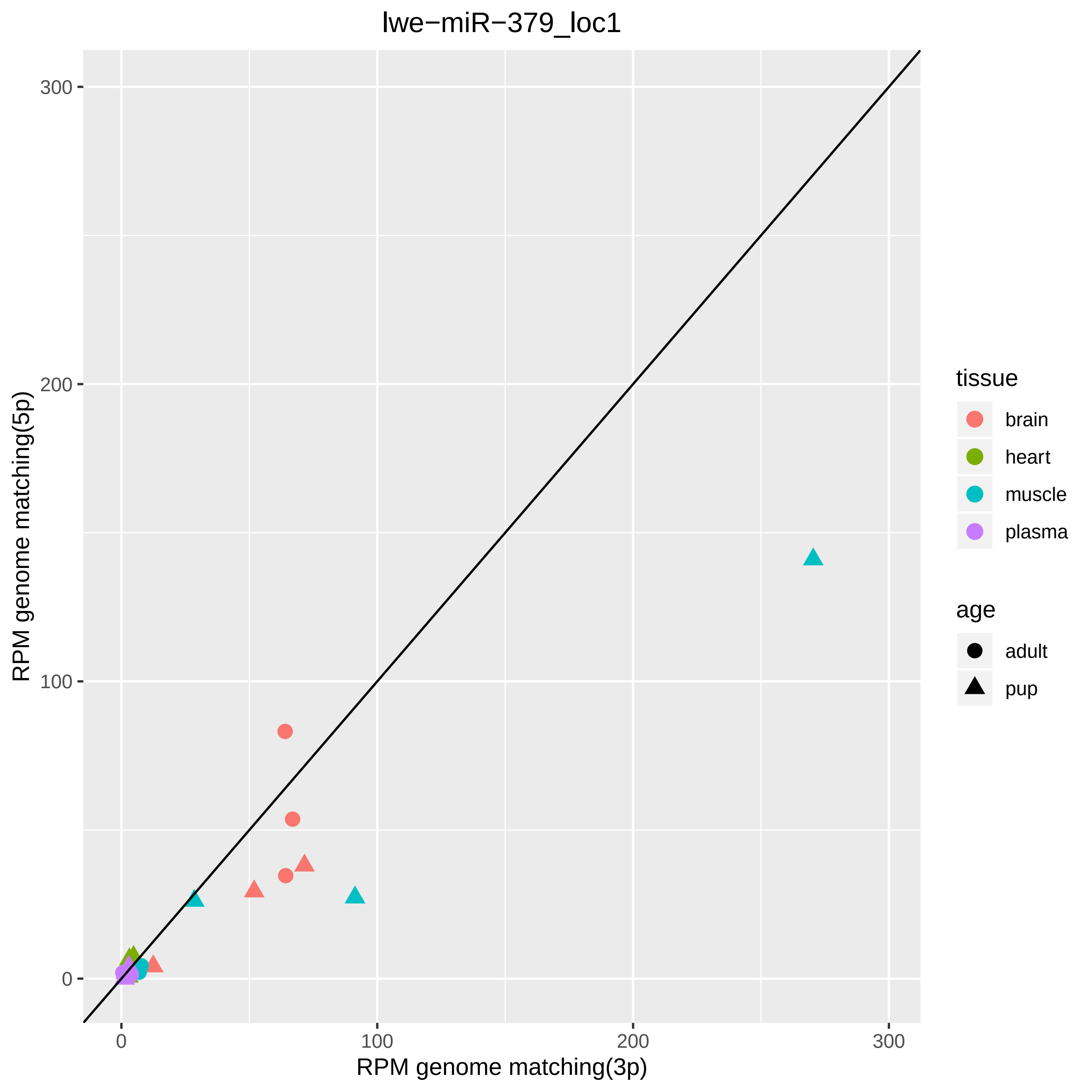
lwe−miR−379_loc1
RPM genome matching(5p)
RPM genome matching(3p)
0
100
200
300
0
100
200
300
tissue
brain
heart
muscle
plasma
age
adult
pup

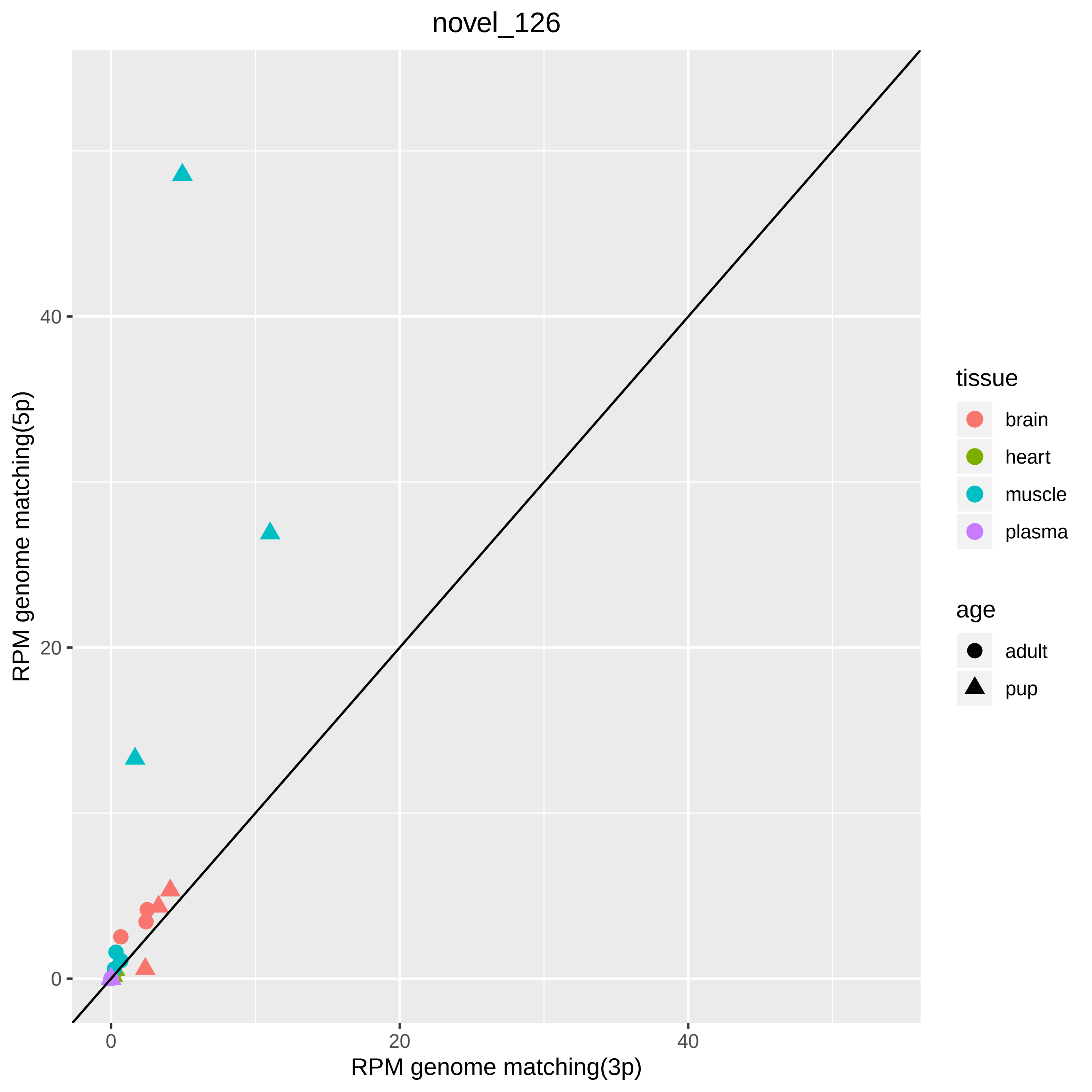

novel_126
RPM genome matching(5p)
RPM genome matching(3p)
0
20
40
0
20
40
tissue
brain
heart
muscle
plasma
age
adult
pup

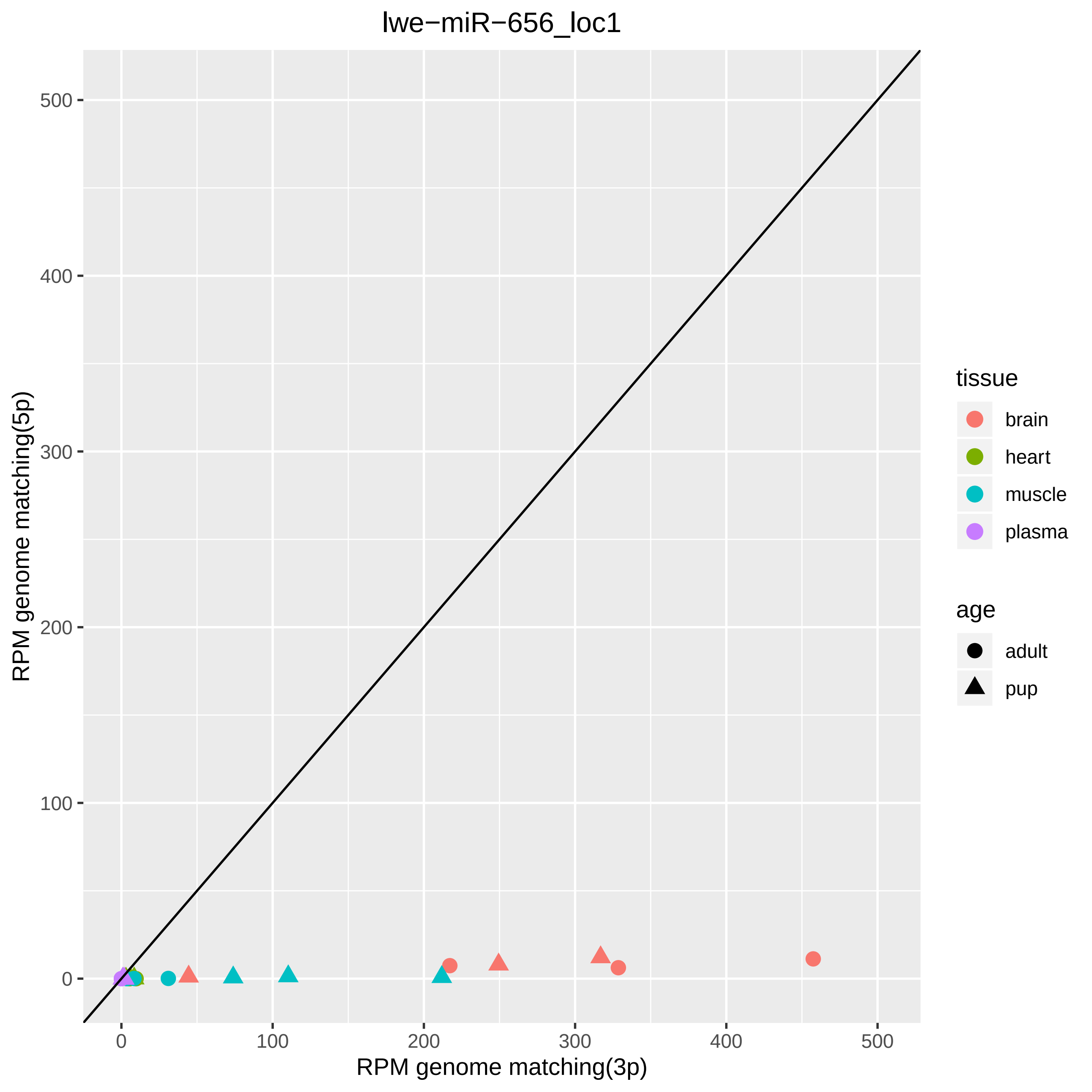
lwe−miR−656_loc1
RPM genome matching(5p)
RPM genome matching(3p)
0
100
200
300
400
500
tissue
brain
heart
muscle
plasma
age
adult
pup

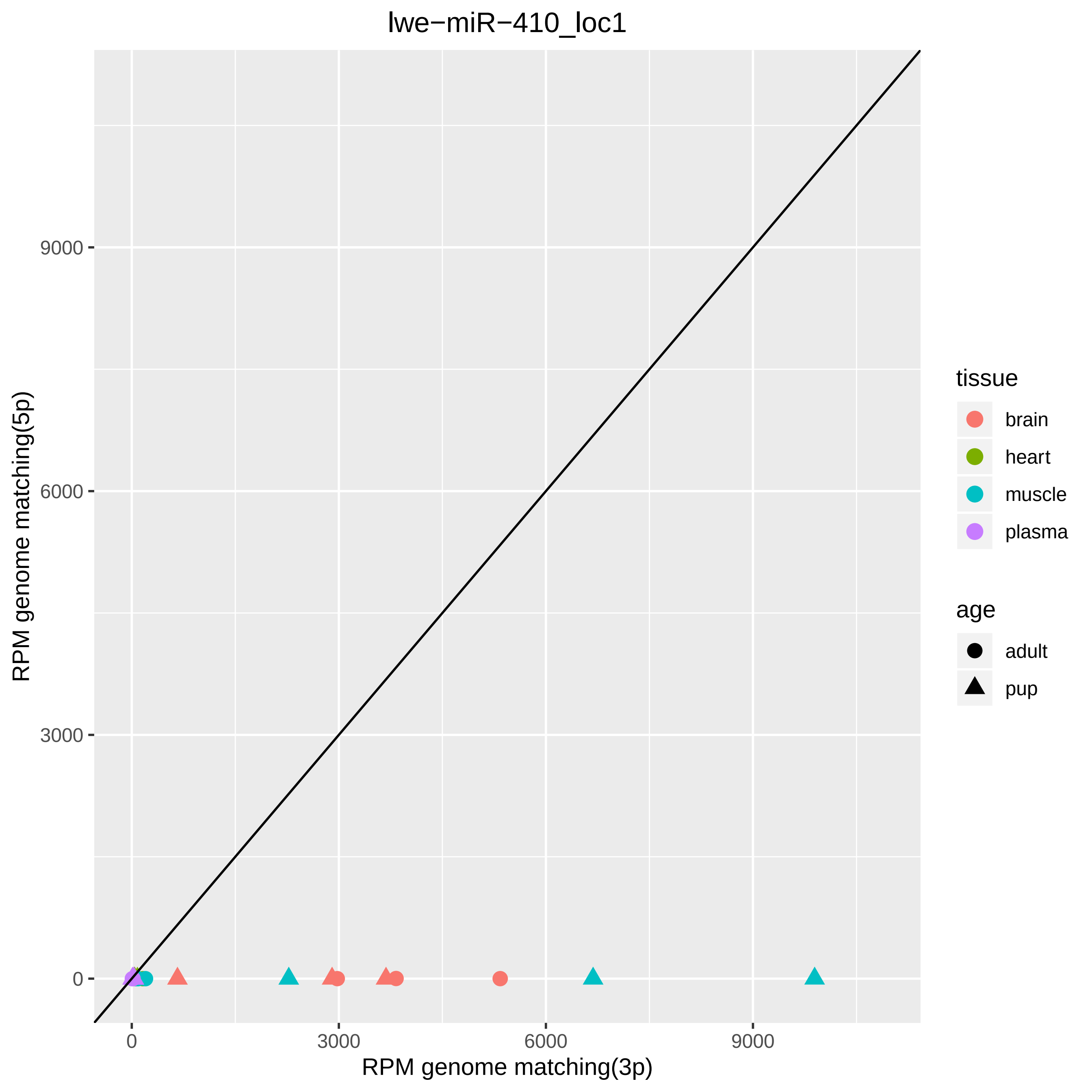
lwe−miR−410_loc1
RPM genome matching(5p)
RPM genome matching(3p)
0
3000
6000
9000
tissue
brain
heart
muscle
plasma
age
adult
pup

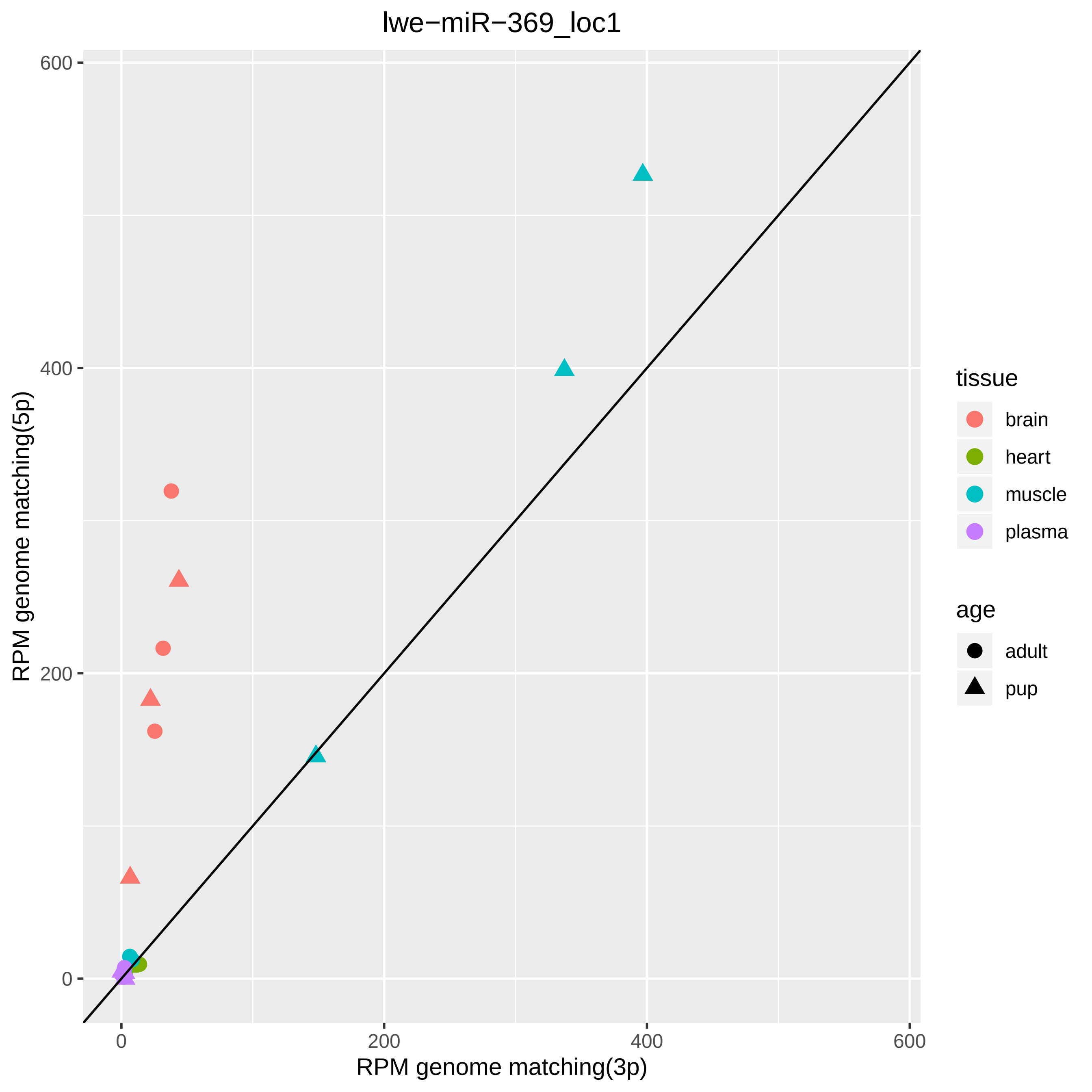

lwe−miR−369_loc1
RPM genome matching(5p)
RPM genome matching(3p)
0
200
400
600
0
200
400
600
tissue
brain
heart
muscle
plasma
age
adult
pup

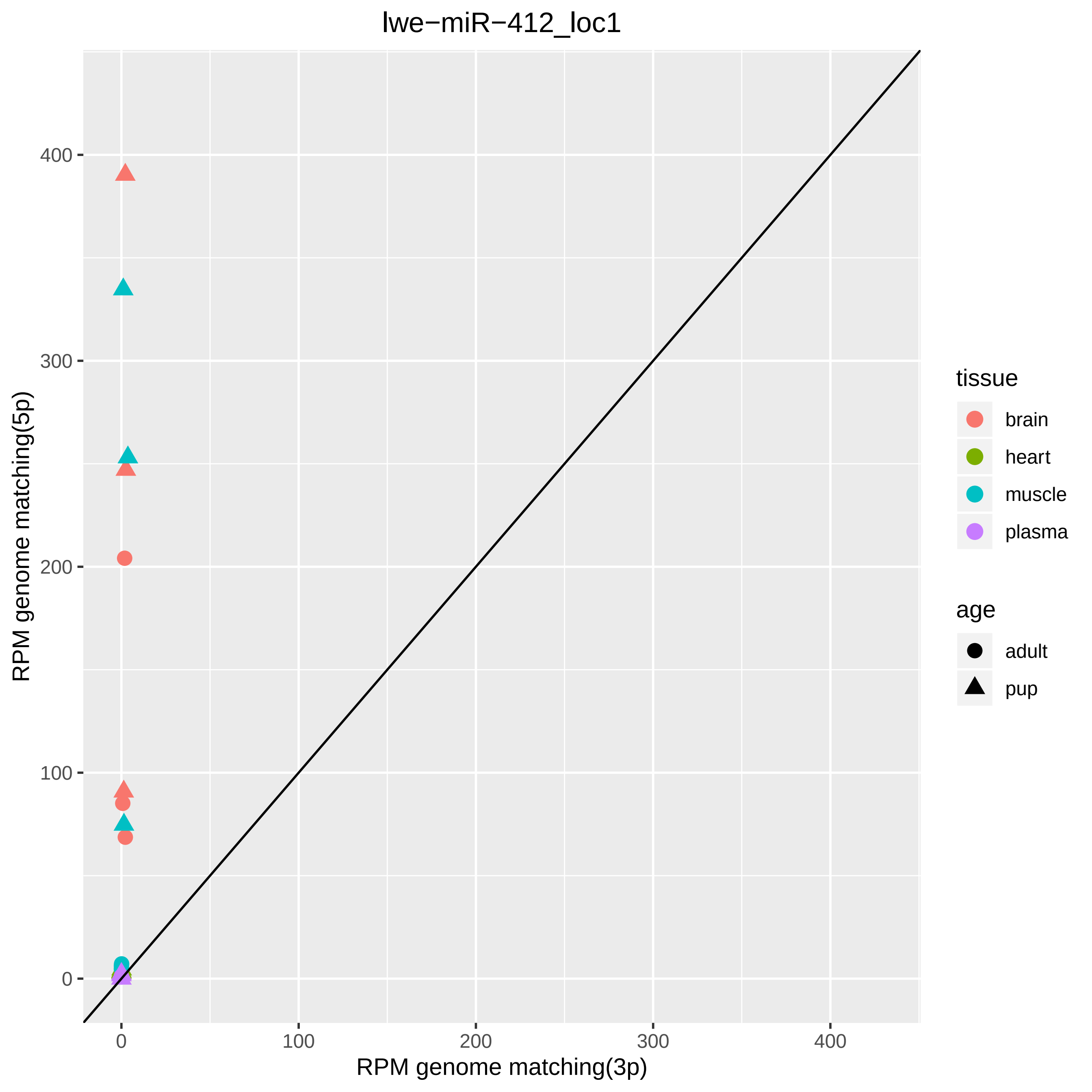

lwe−miR−412_loc1
RPM genome matching(5p)
RPM genome matching(3p)
0
100
200
300
400
tissue
brain
heart
muscle
plasma
age
adult
pup

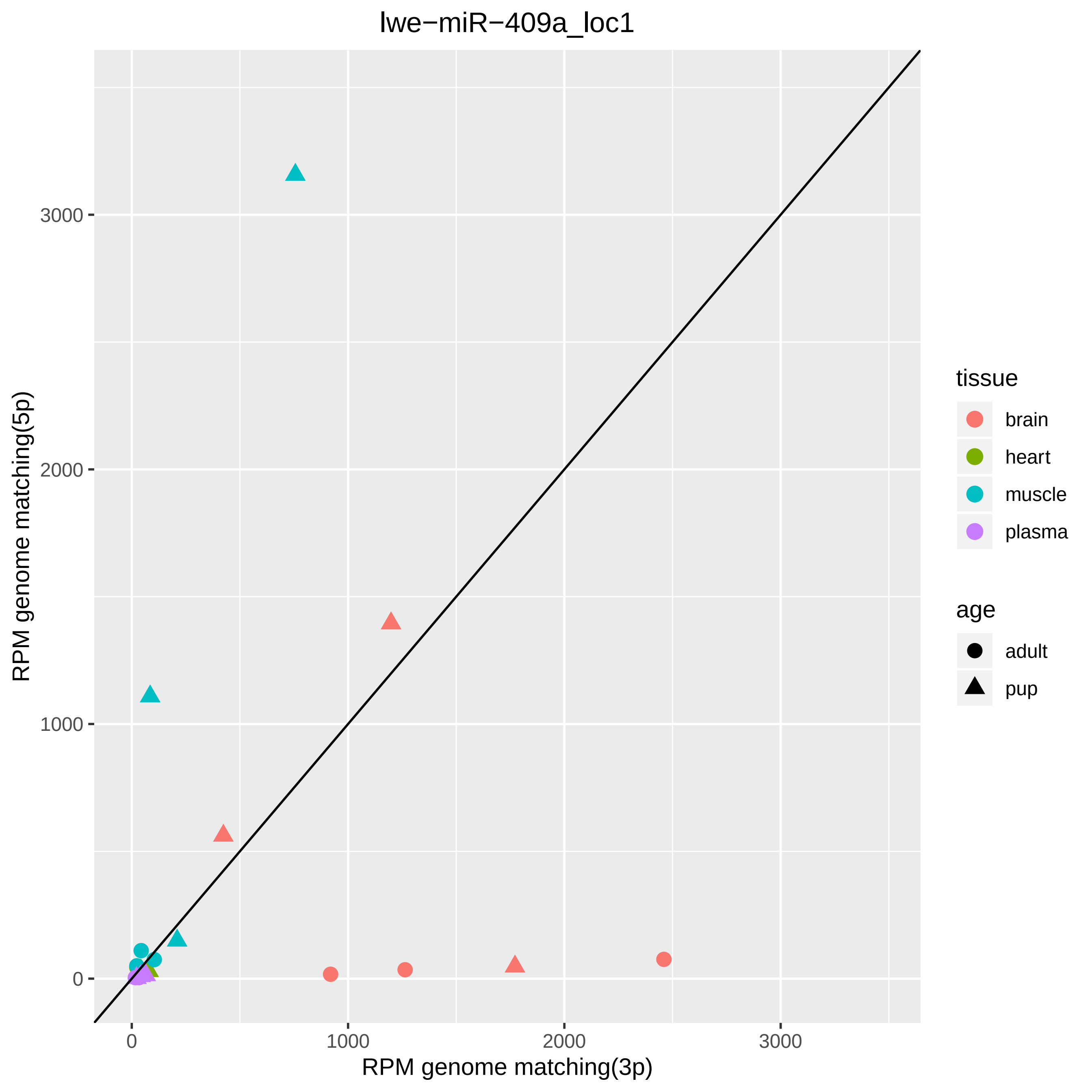

lwe−miR−409a_loc1
RPM genome matching(5p)
RPM genome matching(3p)
0
1000
2000
3000
0
1000
2000
3000
tissue
brain
heart
muscle
plasma
age
adult
pup

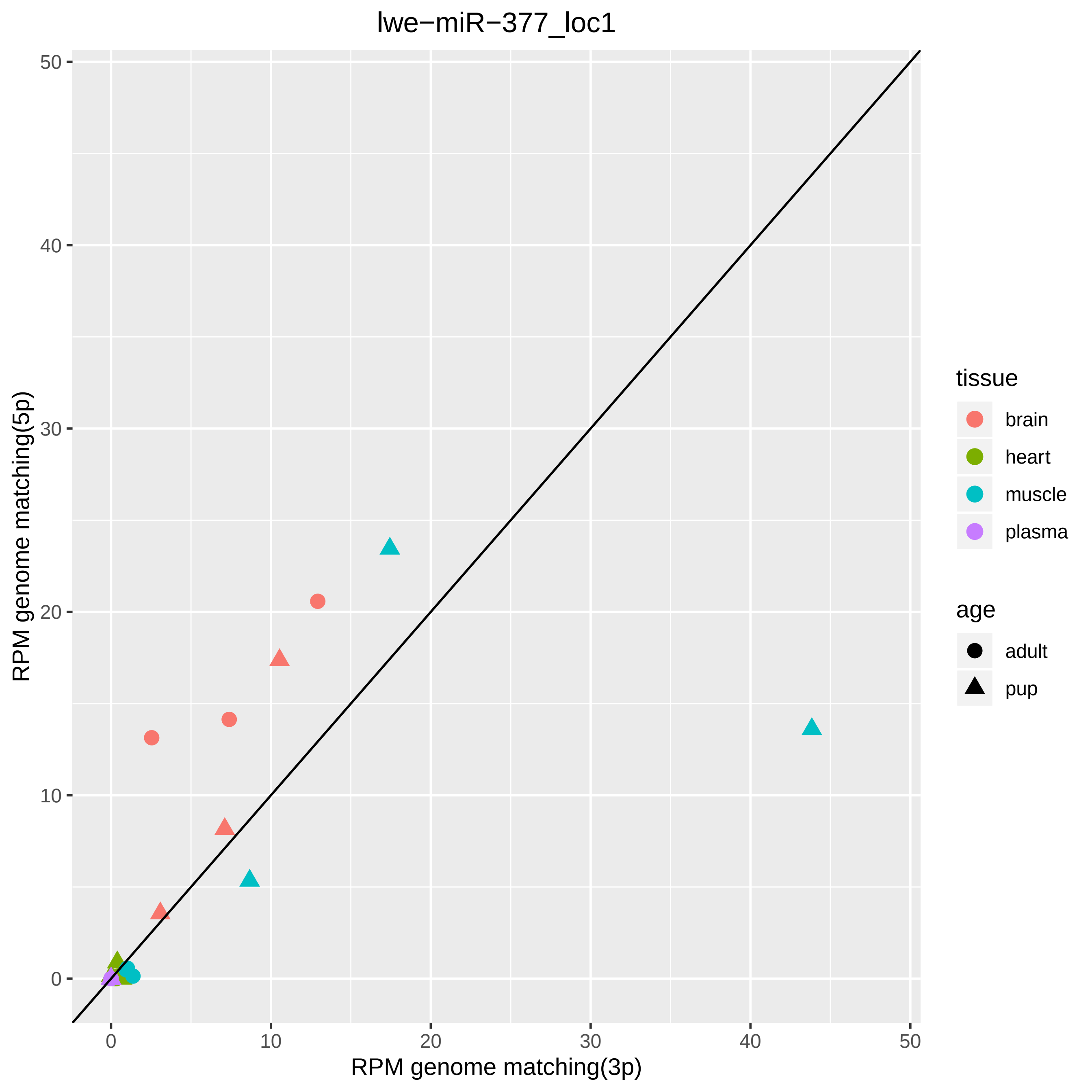

lwe−miR−377_loc1
RPM genome matching(5p)
RPM genome matching(3p)
0
10
20
30
40
50
0
10
20
30
40
50
tissue
brain
heart
muscle
plasma
age
adult
pup

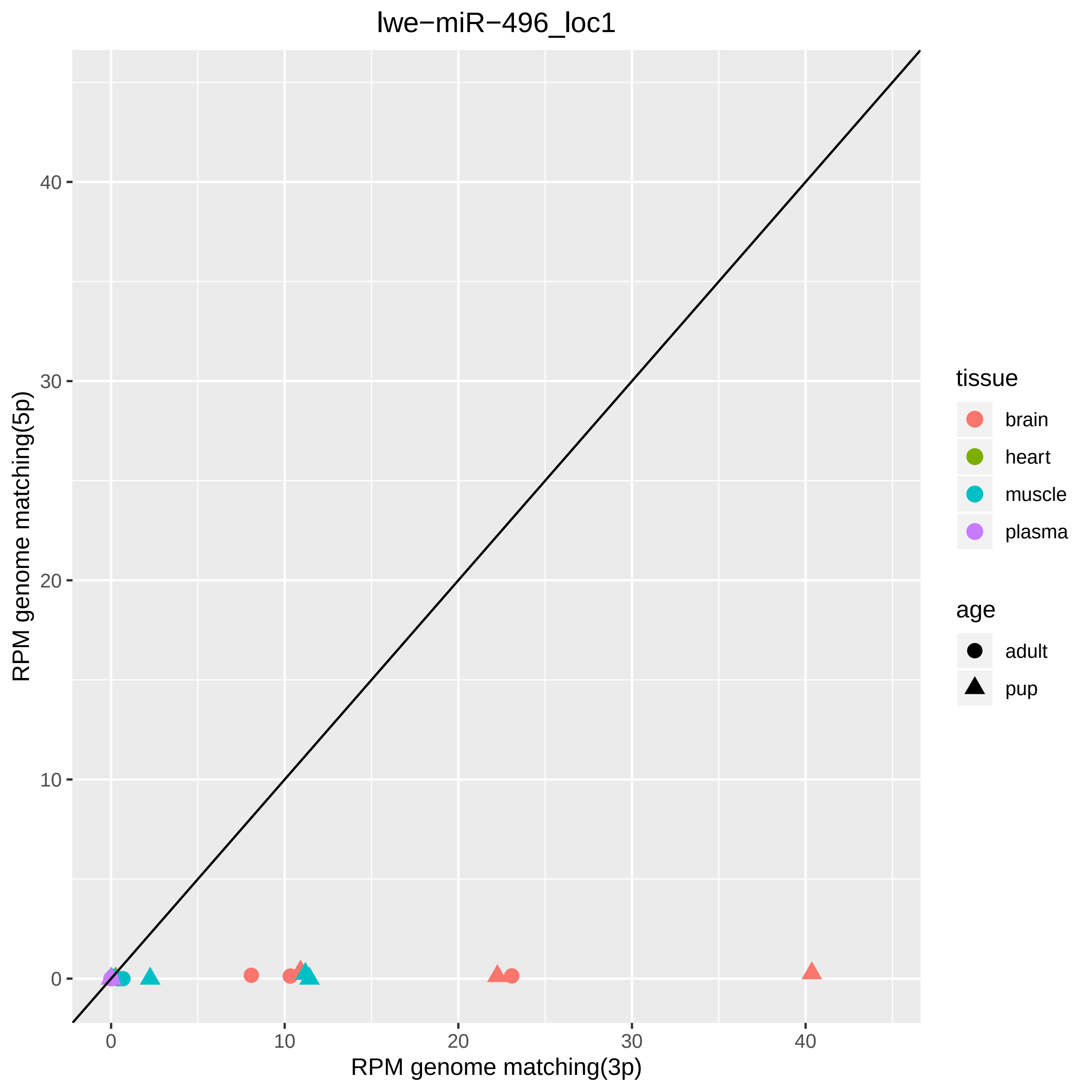
lwe−miR−496_loc1
RPM genome matching(5p)
RPM genome matching(3p)
0
10
20
30
40
0
10
20
30
40
tissue
brain
heart
muscle
plasma
age
adult
pup

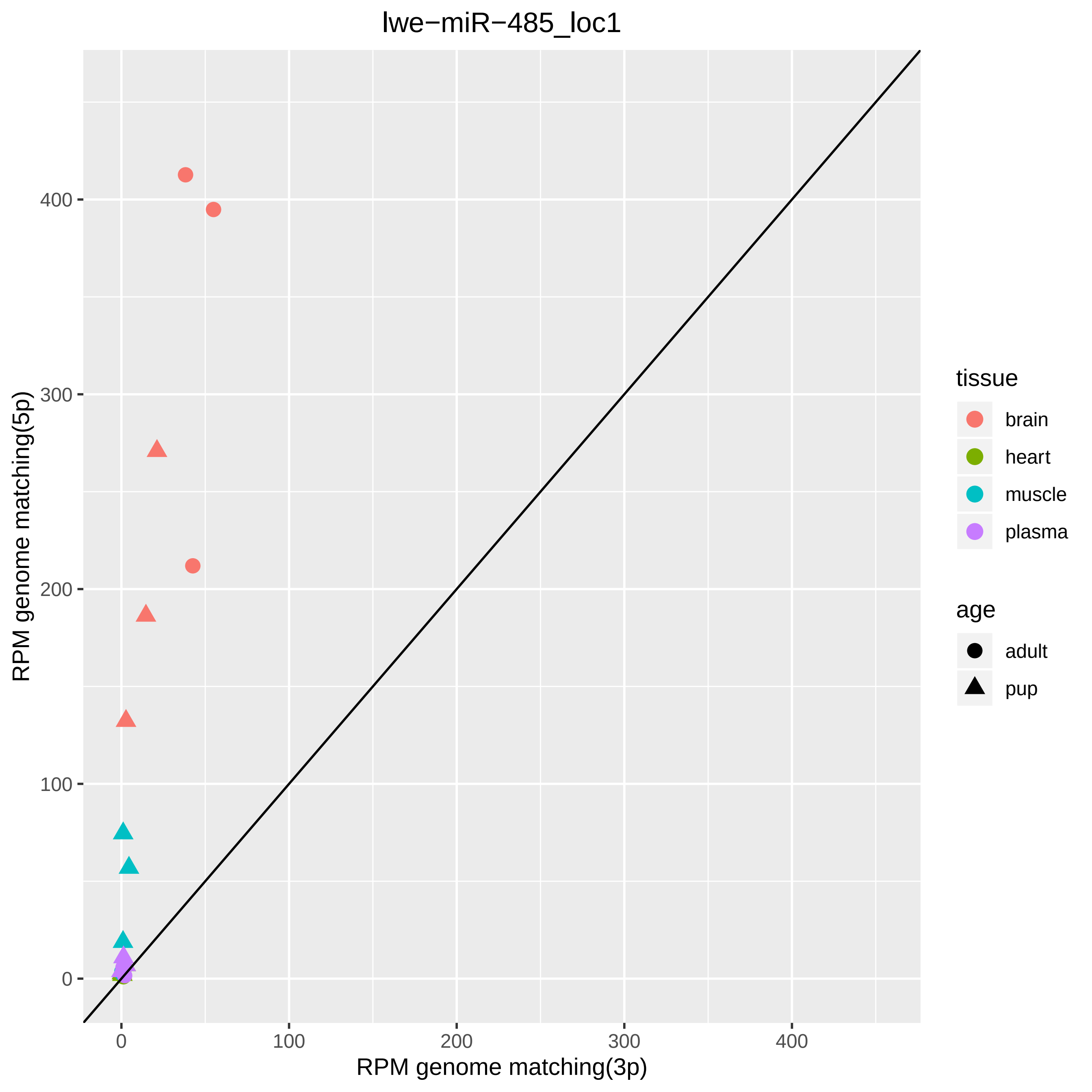
lwe−miR−485_loc1
RPM genome matching(5p)
RPM genome matching(3p)
0
100
200
300
400
tissue
brain
heart
muscle
plasma
age
adult
pup

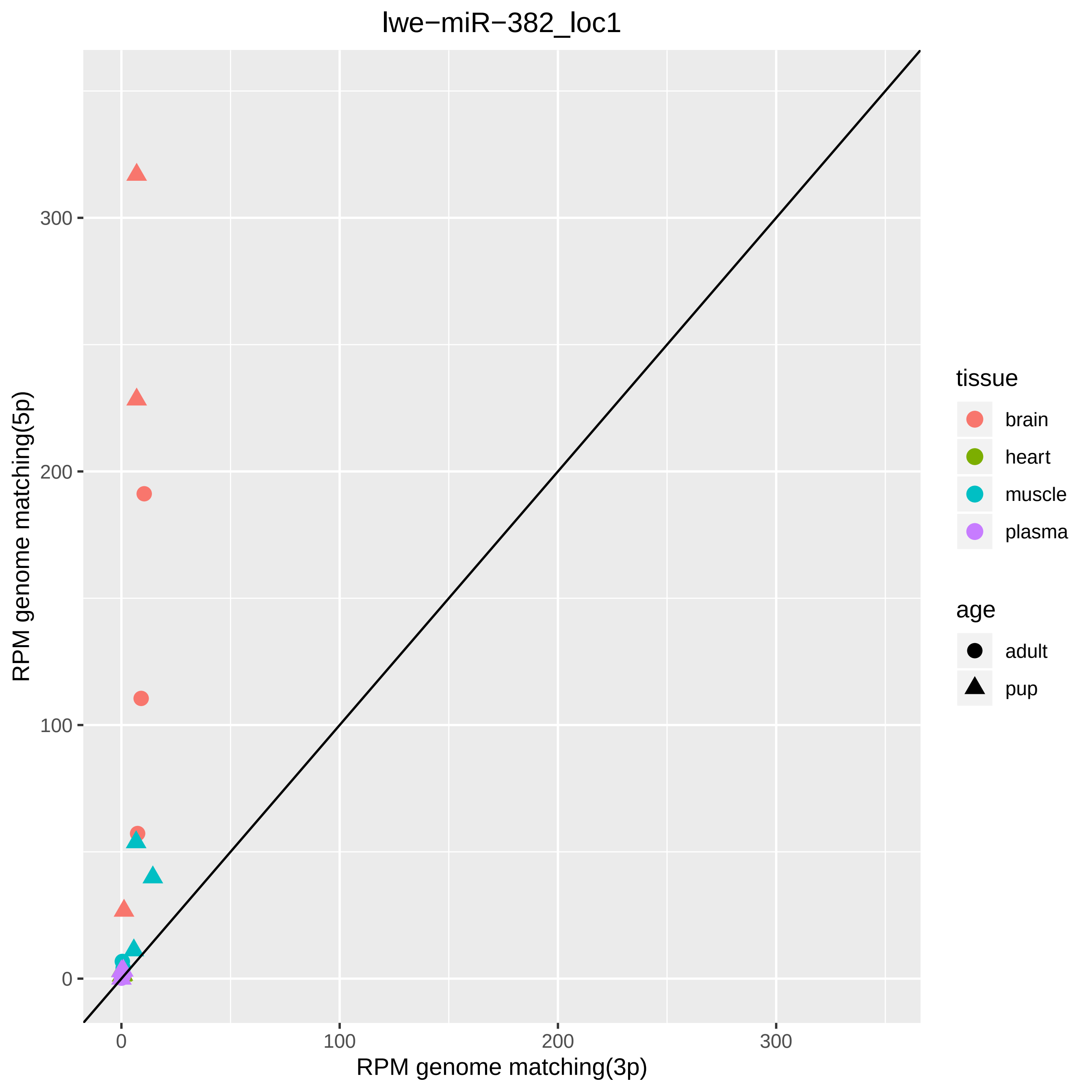
lwe−miR−382_loc1
300
200
100
0
0
100
200
300
RPM genome matching(5p)
RPM genome matching(3p)
tissue
brain
heart
muscle
plasma
age
adult
pup

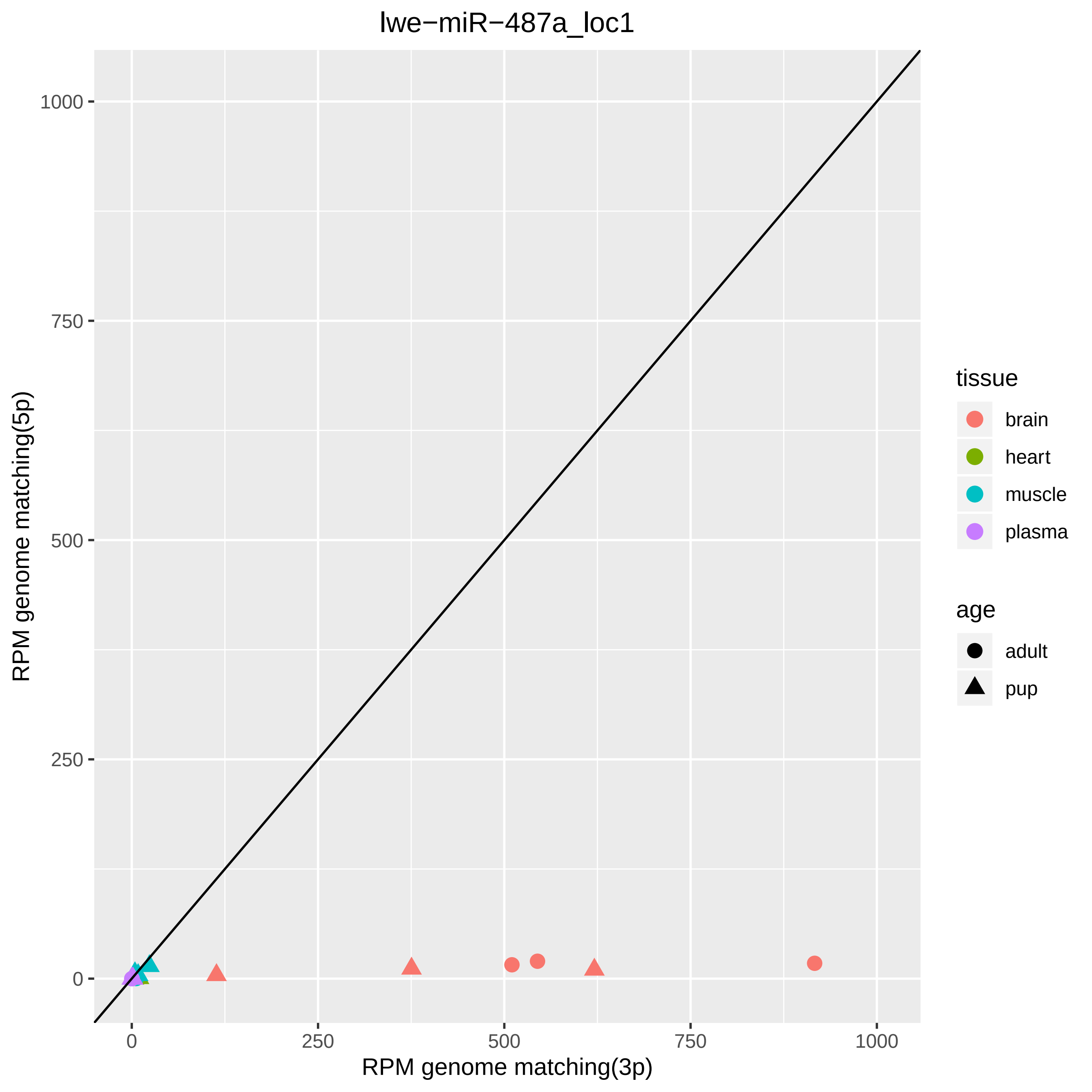

lwe−miR−487a_loc1
RPM genome matching(5p)
RPM genome matching(3p)
0
250
500
750
1000
tissue
brain
heart
muscle
plasma
age
adult
pup

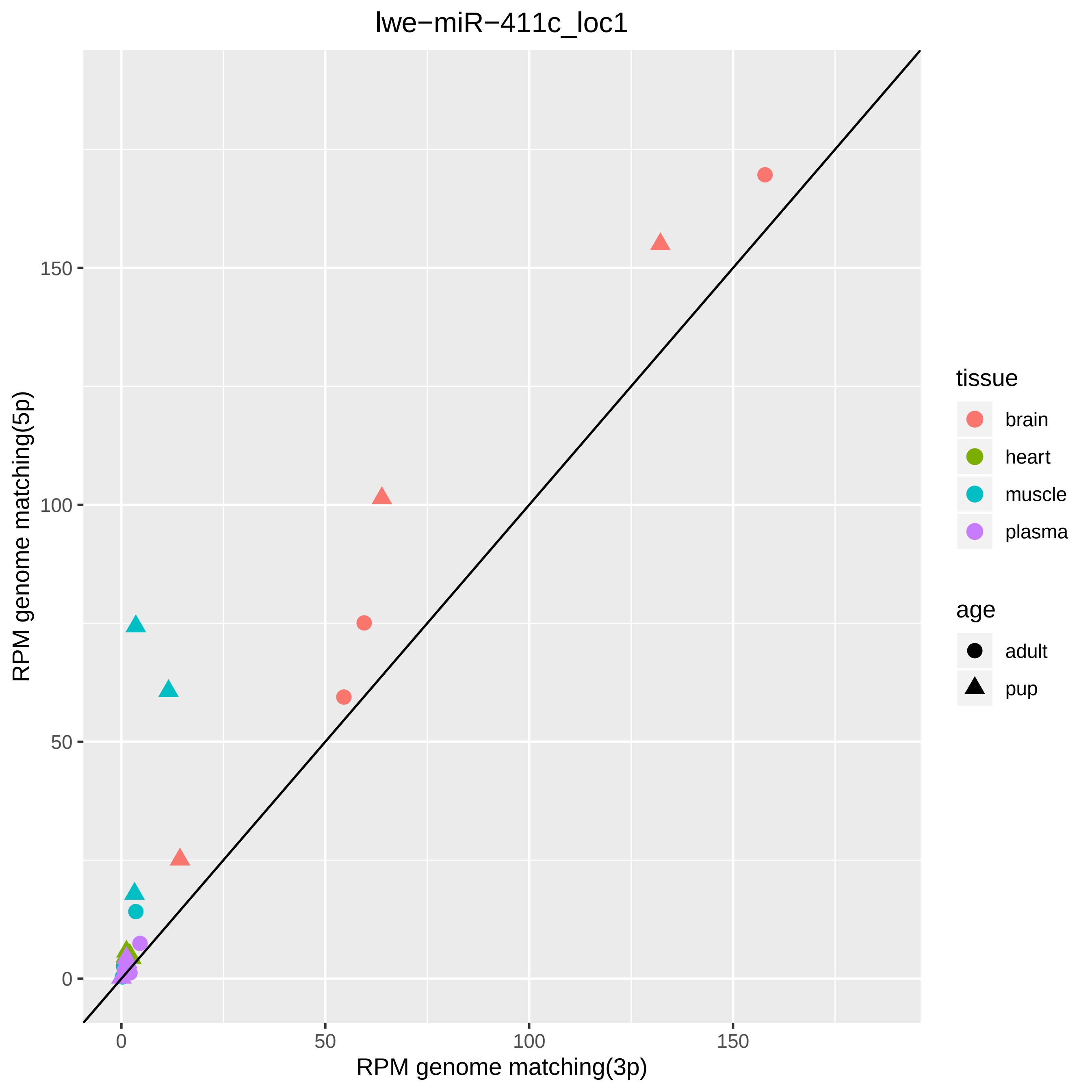
lwe−miR−411c_loc1
RPM genome matching(5p)
RPM genome matching(3p)
0
50
100
150
tissue
brain
heart
muscle
plasma
age
adult
pup

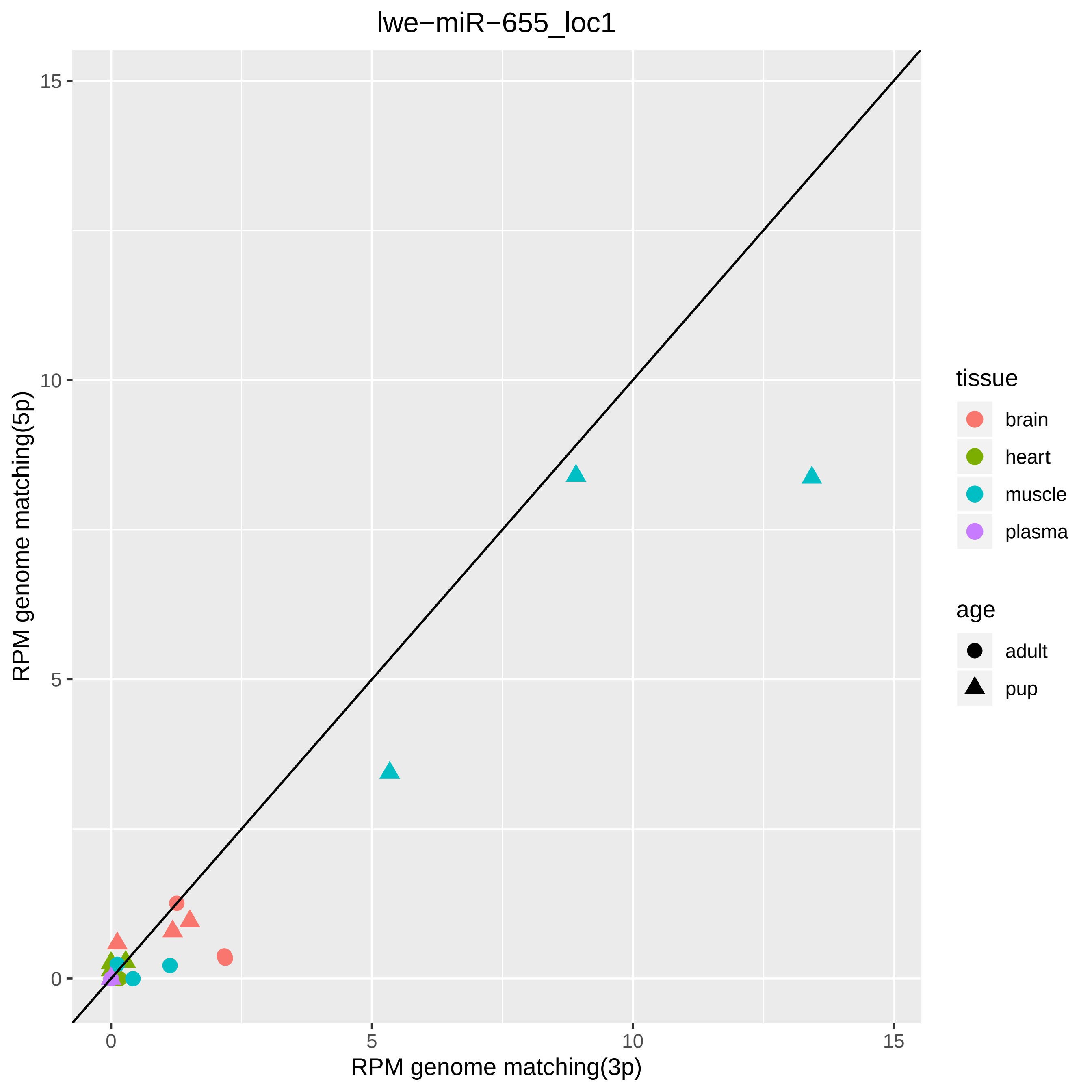

lwe−miR−655_loc1
RPM genome matching(5p)
RPM genome matching(3p)
0
5
10
15
0
5
10
15
tissue
brain
heart
muscle
plasma
age
adult
pup

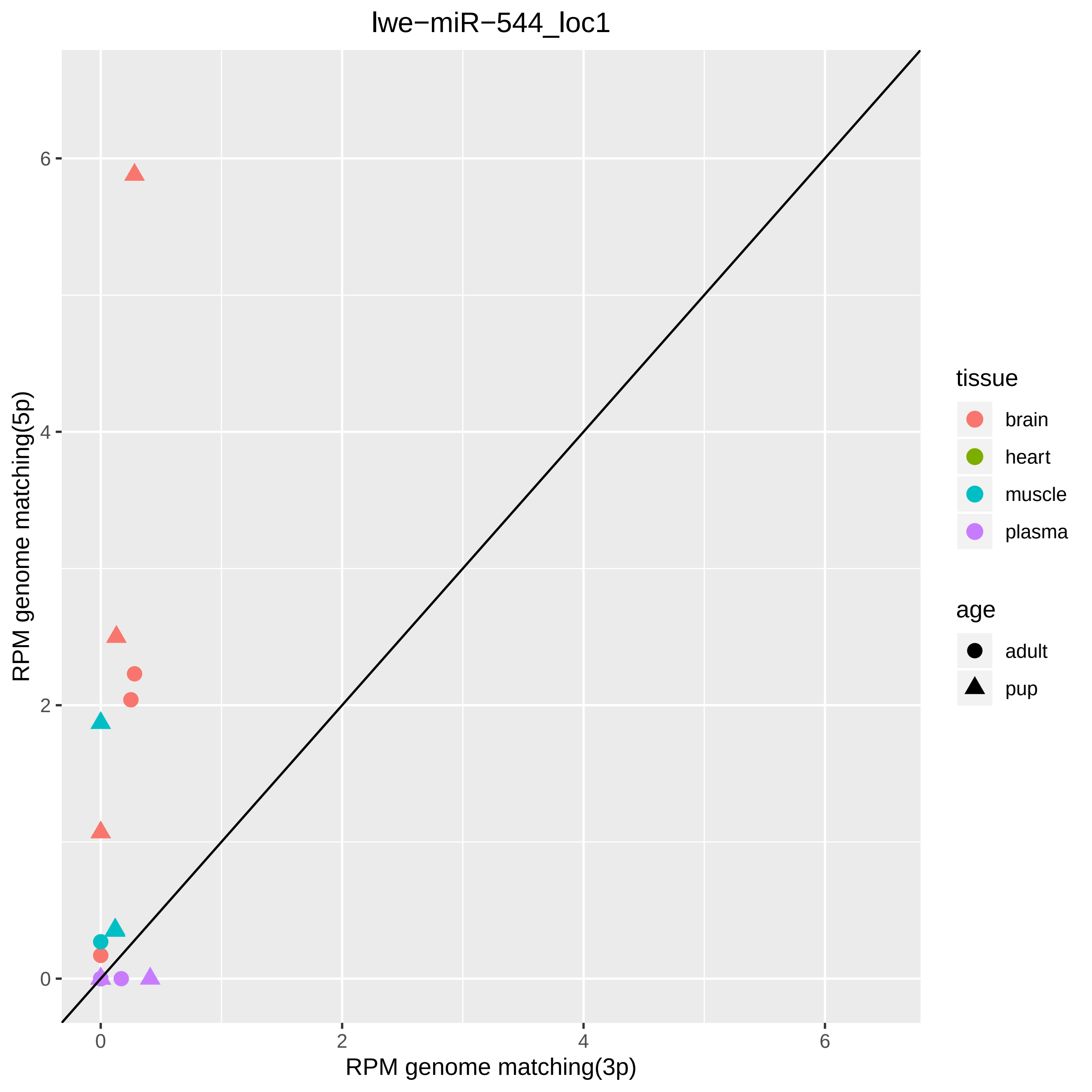

lwe−miR−544_loc1
RPM genome matching(5p)
RPM genome matching(3p)
0
2
4
6
tissue
brain
heart
muscle
plasma
age
adult
pup

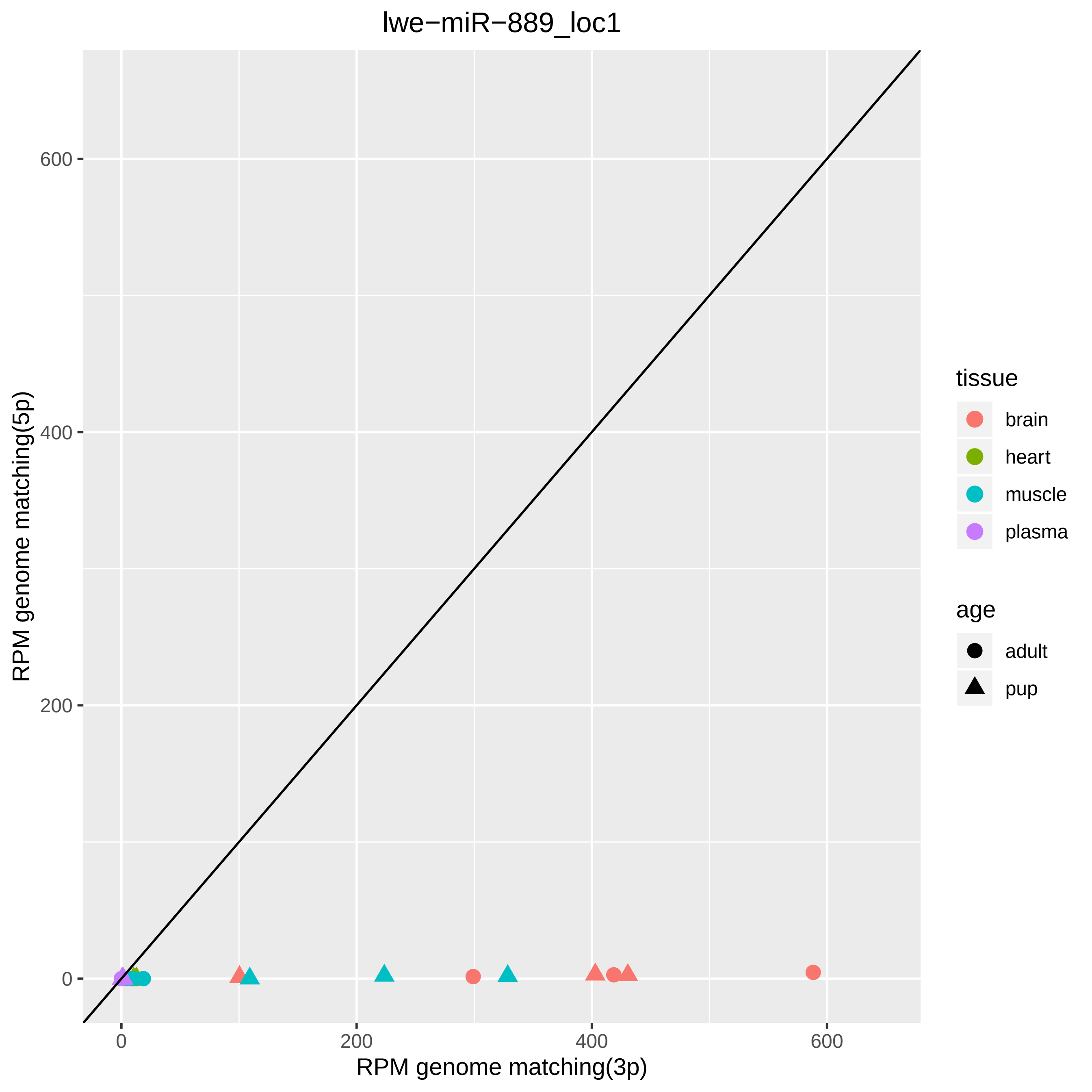
lwe−miR−889_loc1
RPM genome matching(5p)
RPM genome matching(3p)
0
200
400
600
tissue
brain
heart
muscle
plasma
age
adult
pup

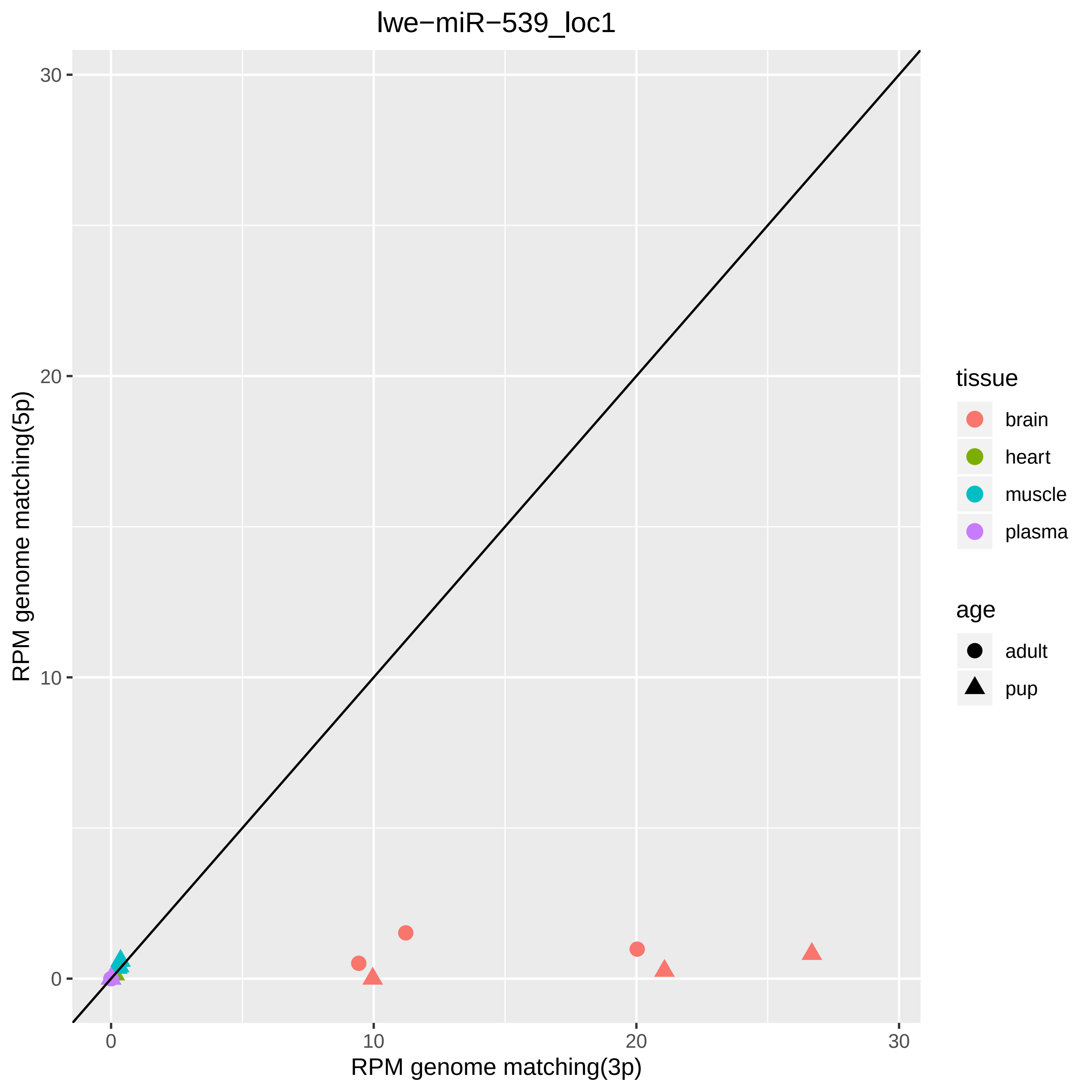
lwe−miR−539_loc1
RPM genome matching(5p)
RPM genome matching(3p)
0
10
20
30
0
10
20
30
tissue
brain
heart
muscle
plasma
age
adult
pup

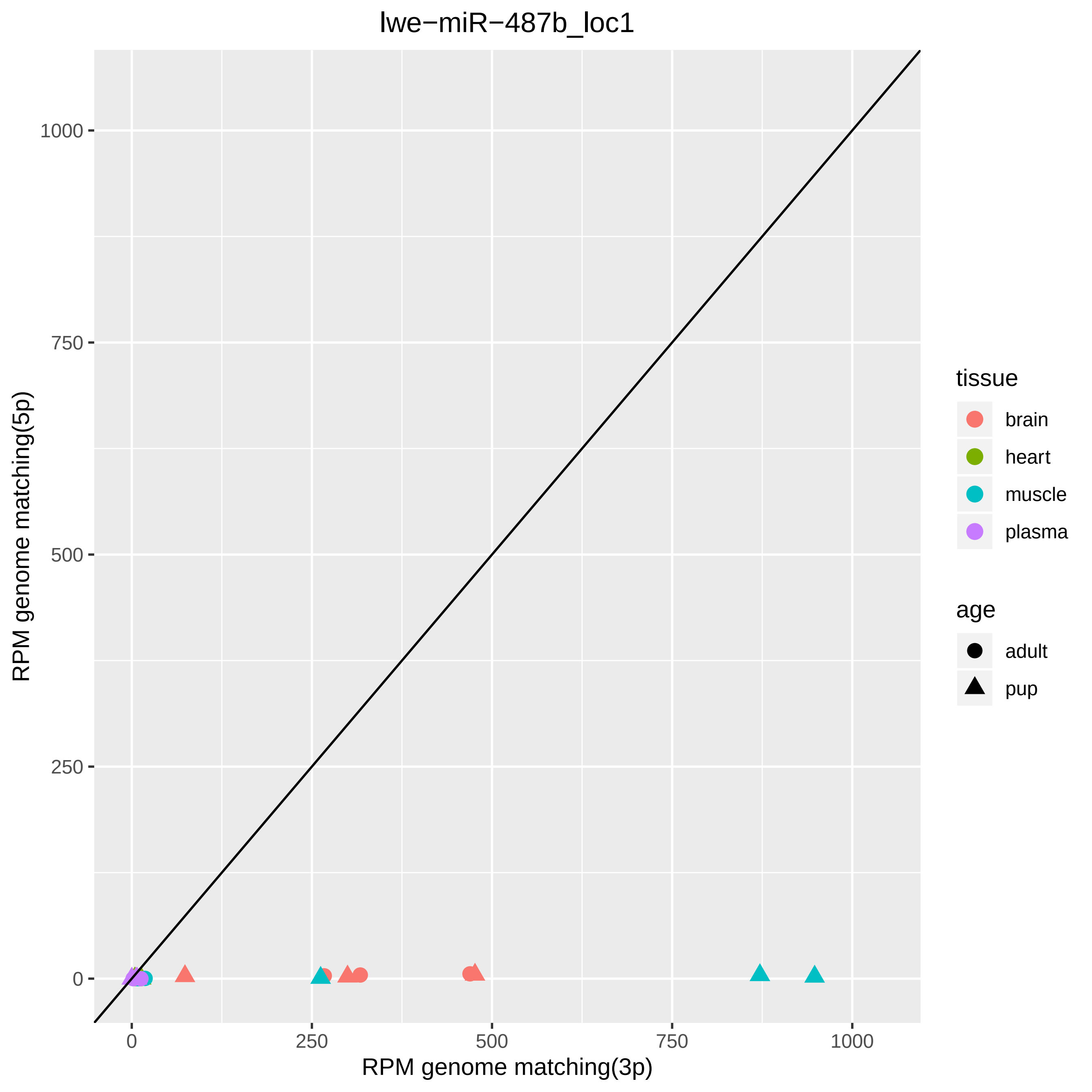
lwe−miR−487b_loc1
RPM genome matching(5p)
RPM genome matching(3p)
1000
750
500
250
0
0
250
500
750
1000
tissue
brain
heart
muscle
plasma
age
adult
pup

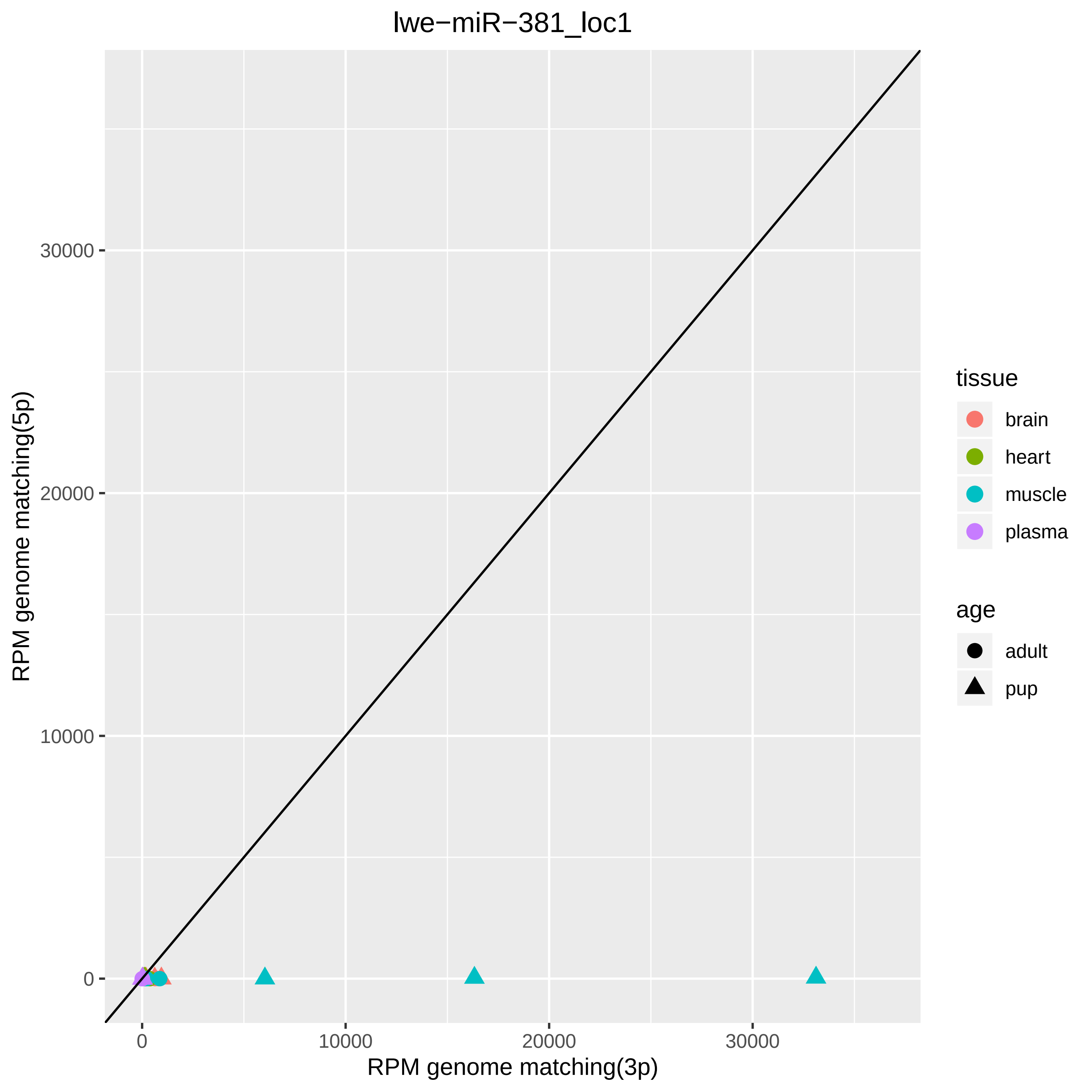

lwe−miR−381_loc1
RPM genome matching(5p)
RPM genome matching(3p)
0
10000
20000
30000
tissue
brain
heart
muscle
plasma
age
adult
pup

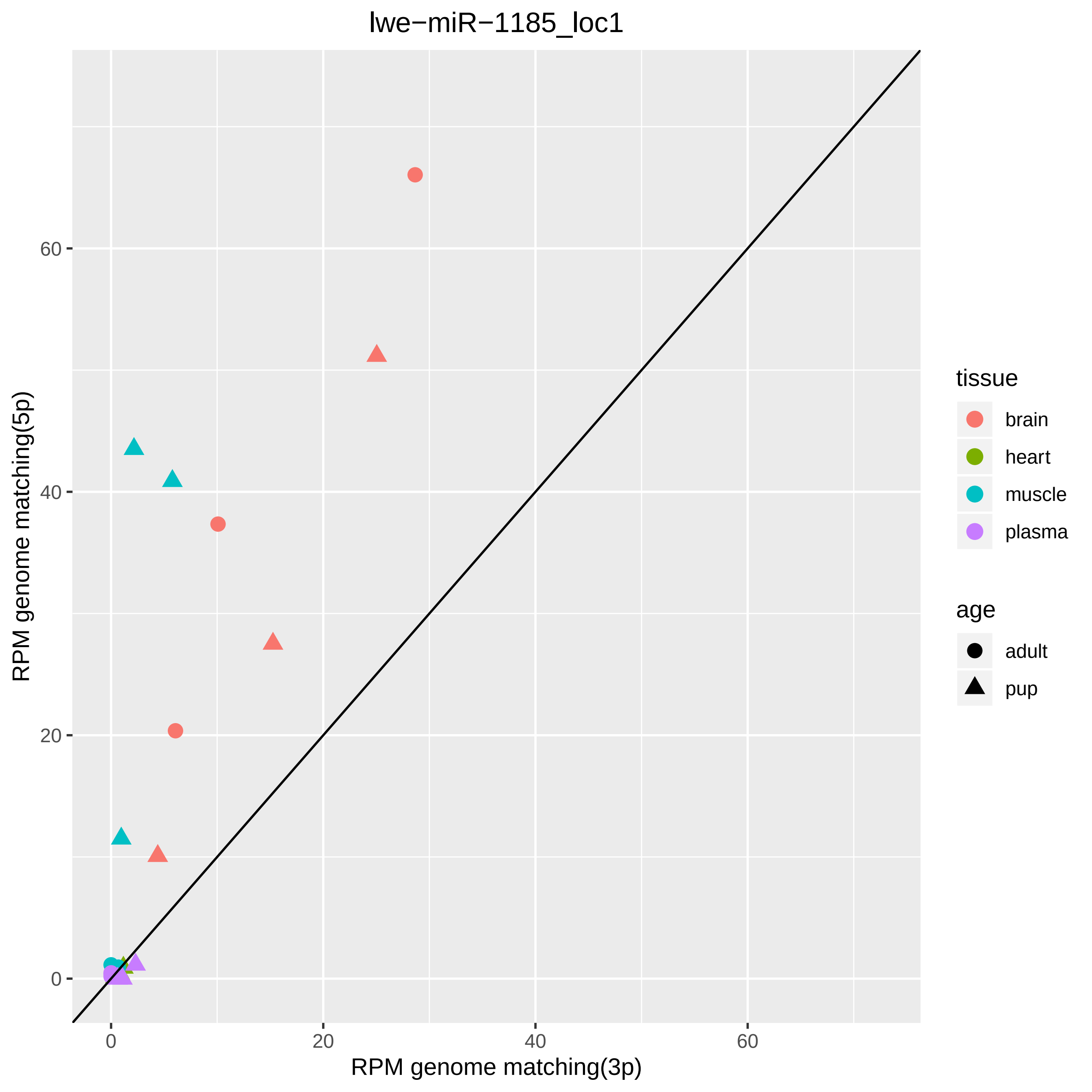
lwe−miR−1185_loc1
RPM genome matching(5p)
RPM genome matching(3p)
0
20
40
60
tissue
brain
heart
muscle
plasma
age
adult
pup

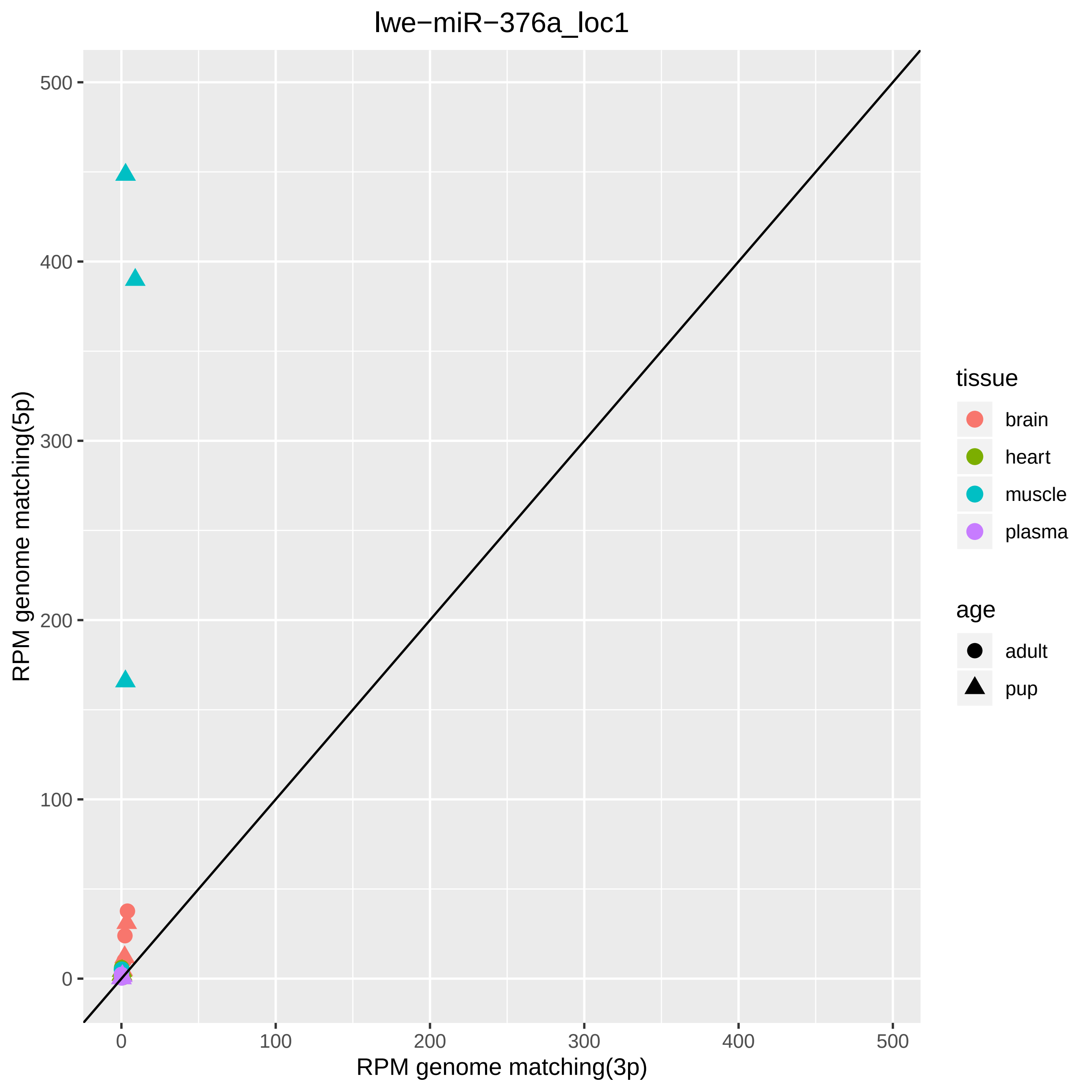
lwe−miR−376a_loc1
RPM genome matching(5p)
RPM genome matching(3p)
0
100
200
300
400
500
0
100
200
300
400
500
tissue
brain
heart
muscle
plasma
age
adult
pup

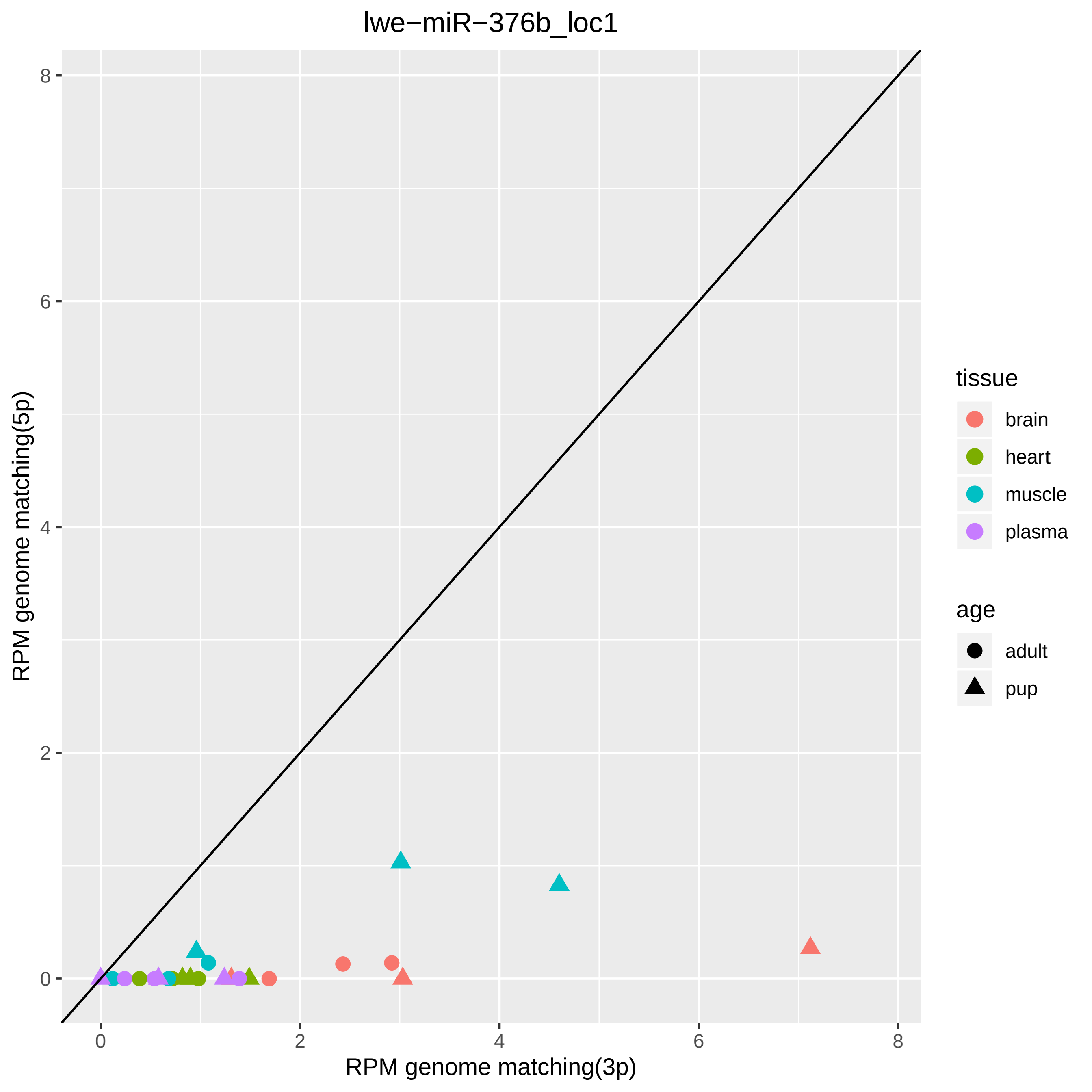
lwe−miR−376b_loc1
RPM genome matching(5p)
RPM genome matching(3p)
0
2
4
6
8
tissue
brain
heart
muscle
plasma
age
adult
pup

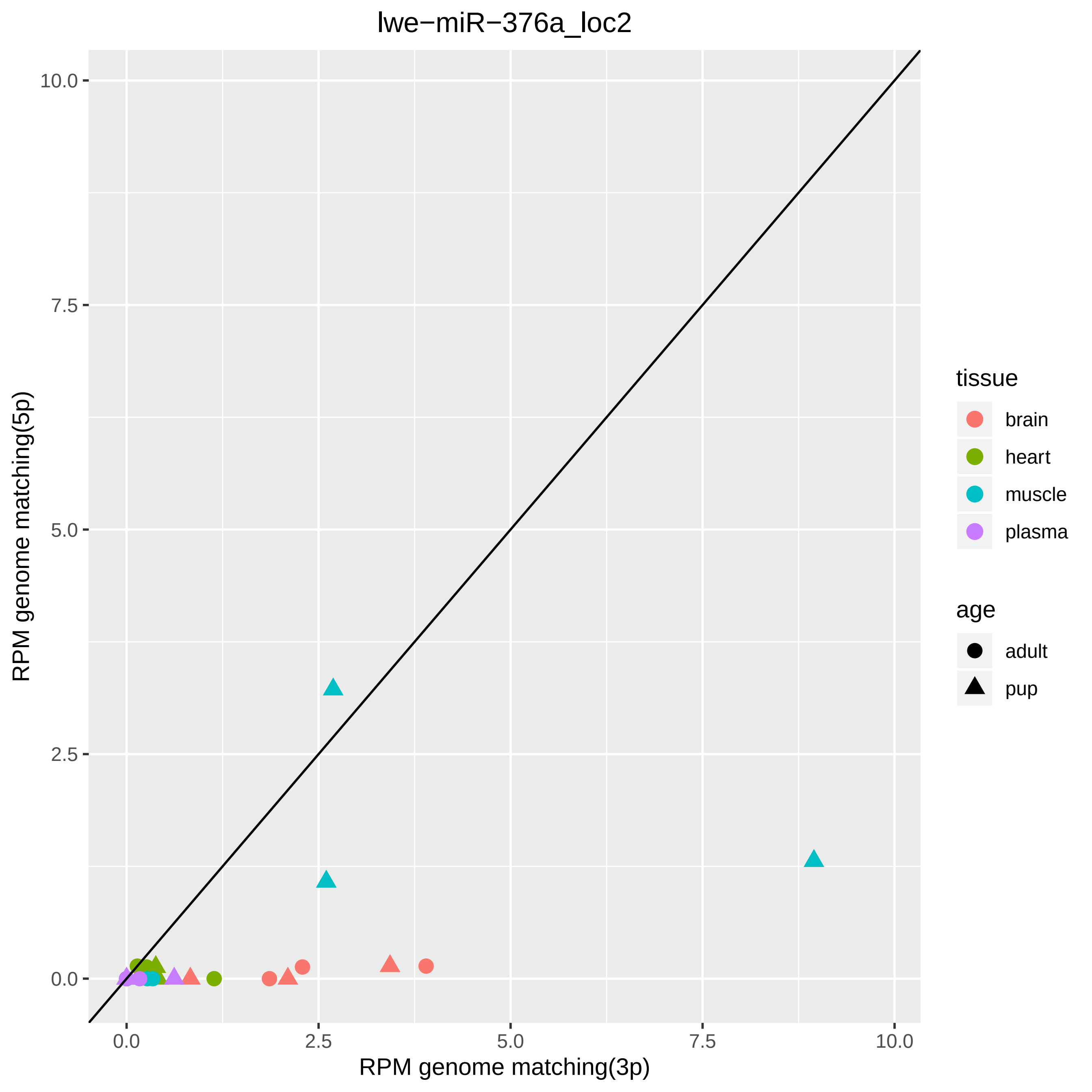
lwe−miR−376a_loc2
RPM genome matching(5p)
RPM genome matching(3p)
0.0
2.5
5.0
7.5
10.0
tissue
brain
heart
muscle
plasma
age
adult
pup

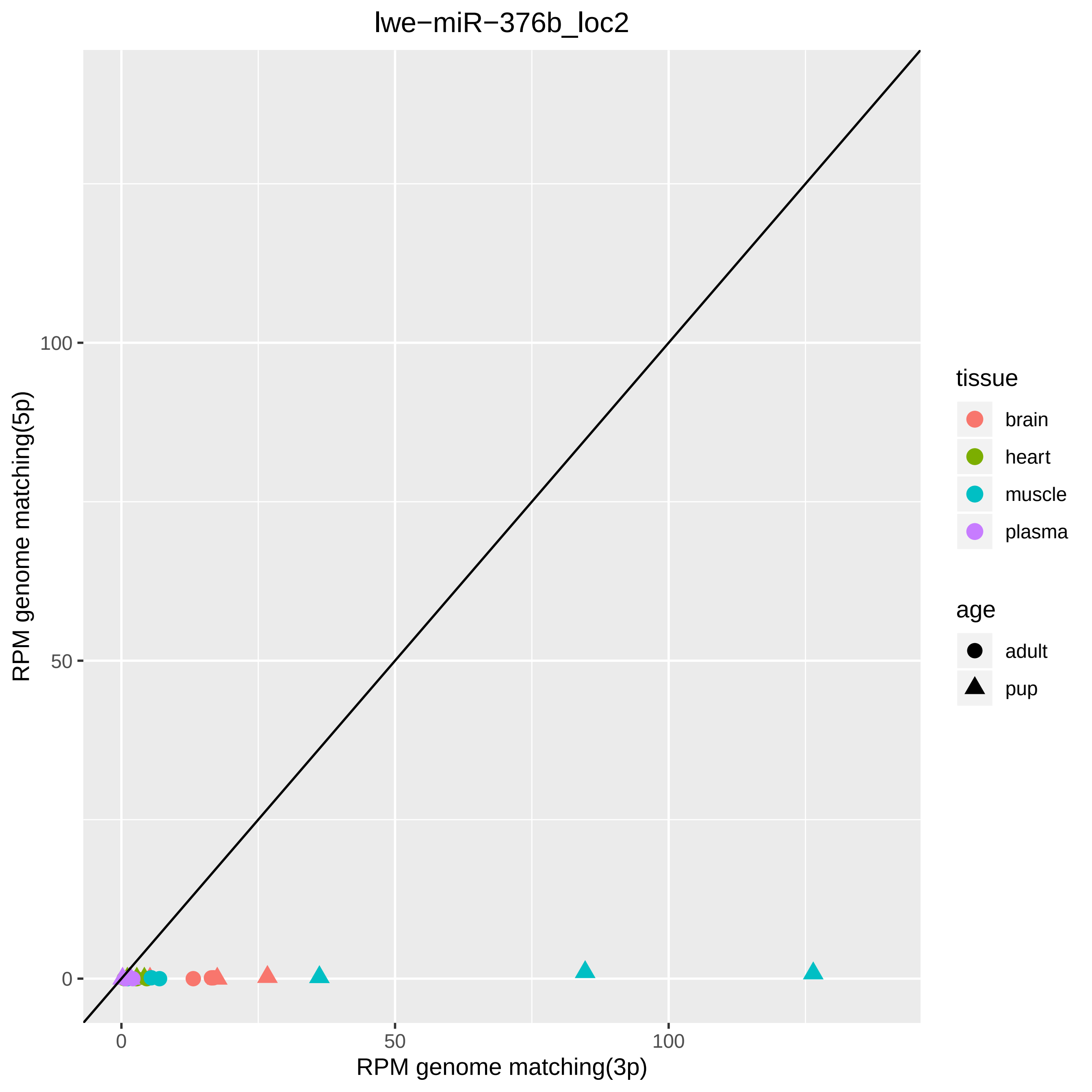

lwe−miR−376b_loc2
RPM genome matching(5p)
RPM genome matching(3p)
0
50
100
0
50
100
tissue
brain
heart
muscle
plasma
age
adult
pup

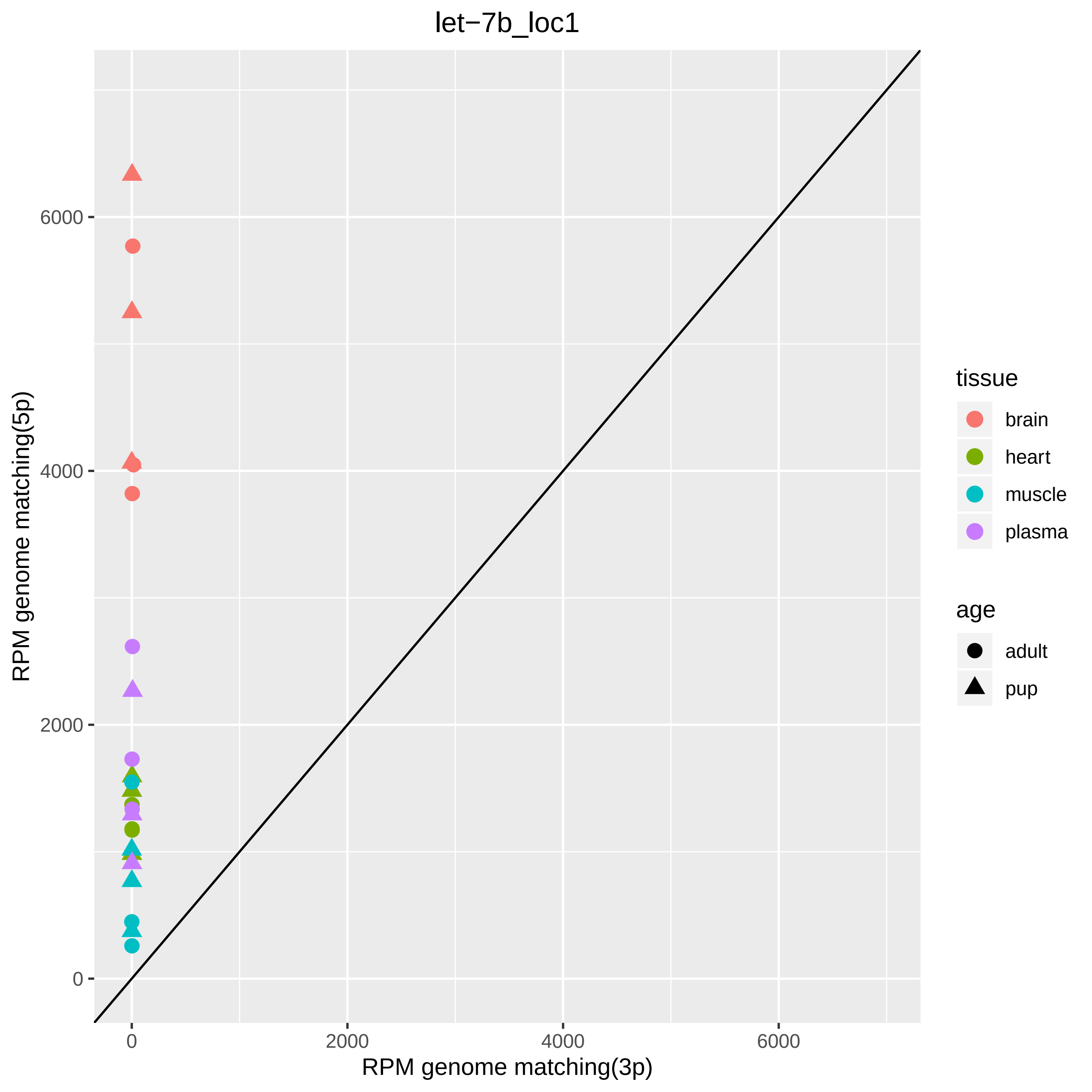

let−7b_loc1
RPM genome matching(5p)
RPM genome matching(3p)
0
2000
4000
6000
tissue
brain
heart
muscle
plasma
age
adult
pup

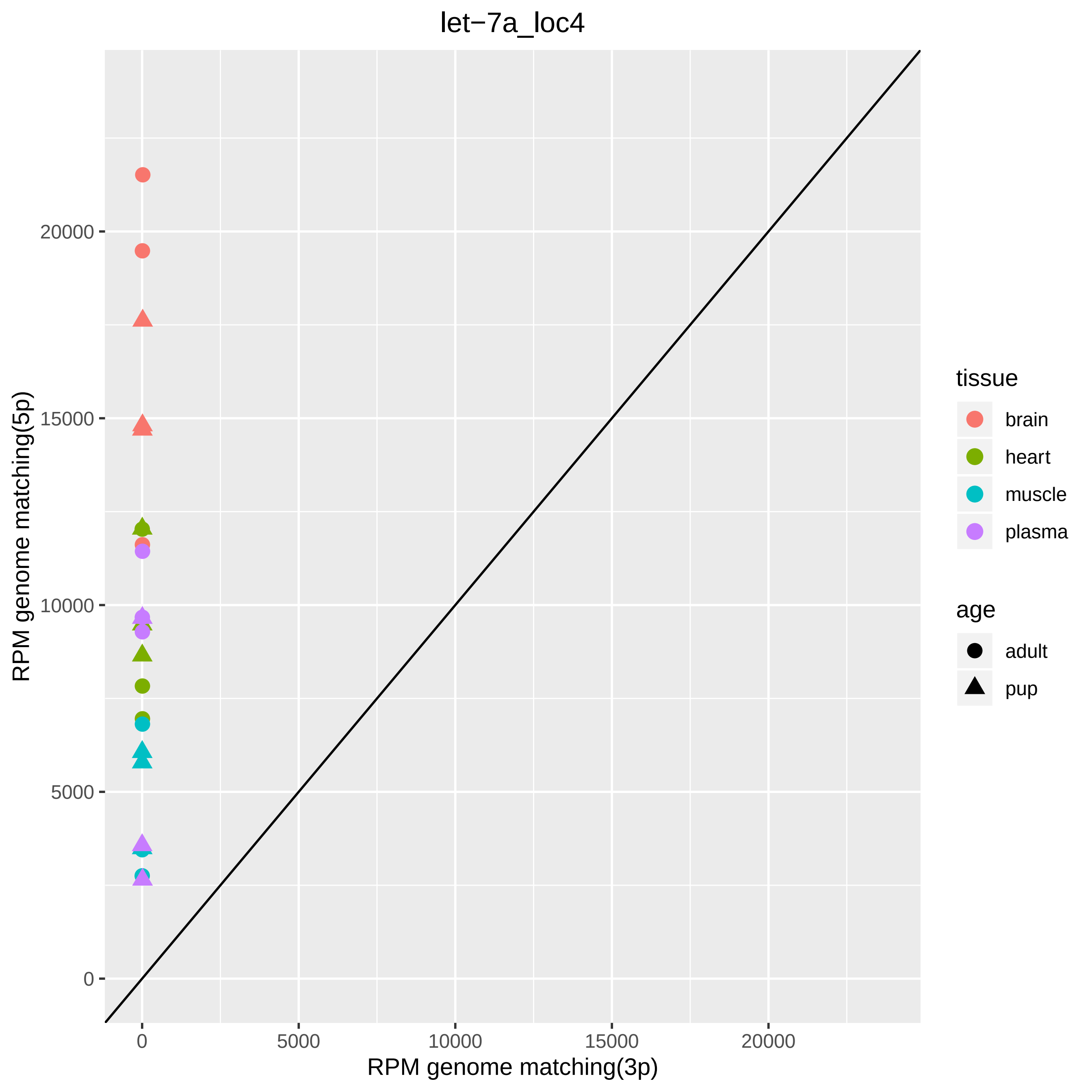

let−7a_loc4
RPM genome matching(5p)
RPM genome matching(3p)
0
5000
10000
15000
20000
tissue
brain
heart
muscle
plasma
age
adult
pup

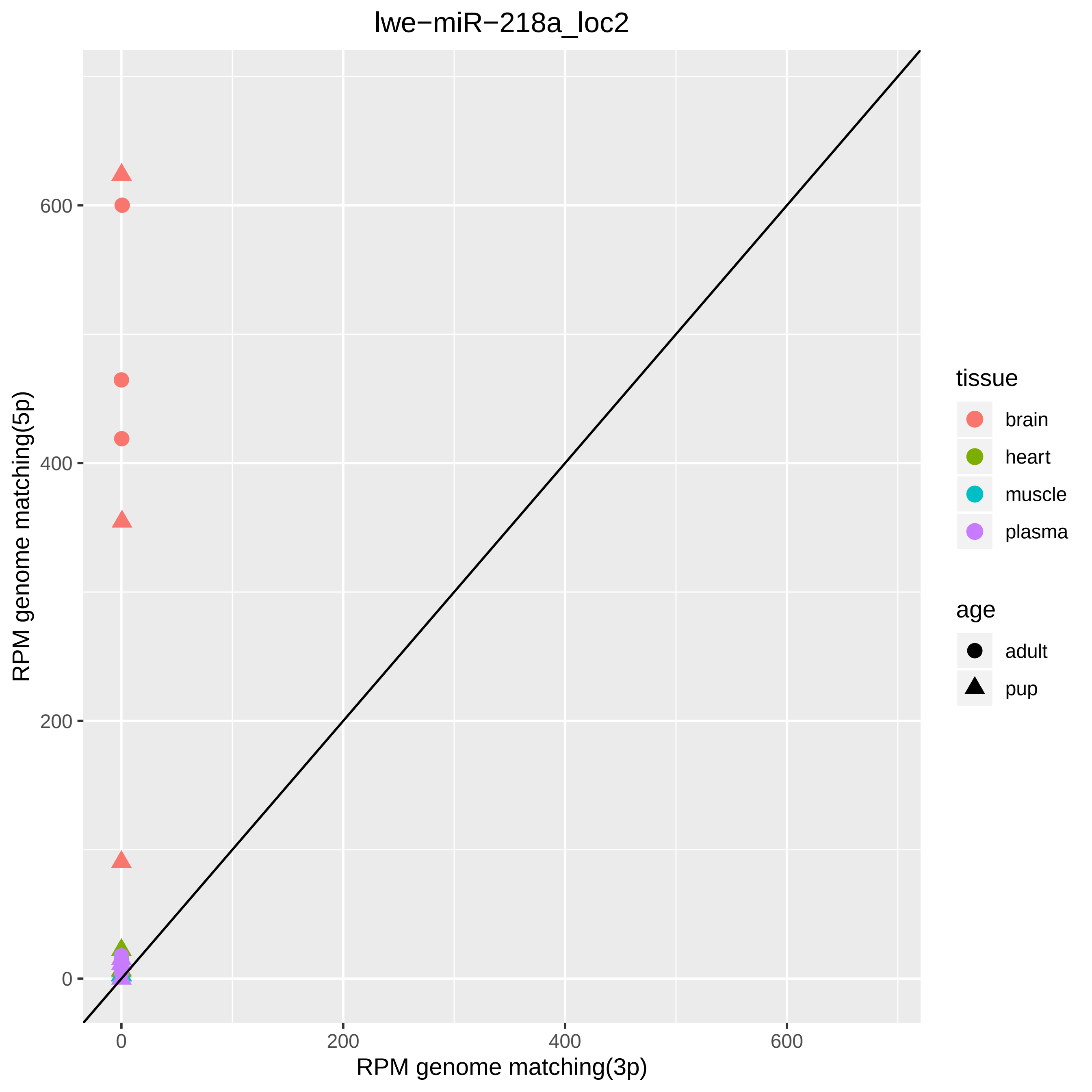

lwe−miR−218a_loc2
RPM genome matching(5p)
RPM genome matching(3p)
0
200
400
600
0
200
400
600
tissue
brain
heart
muscle
plasma
age
adult
pup

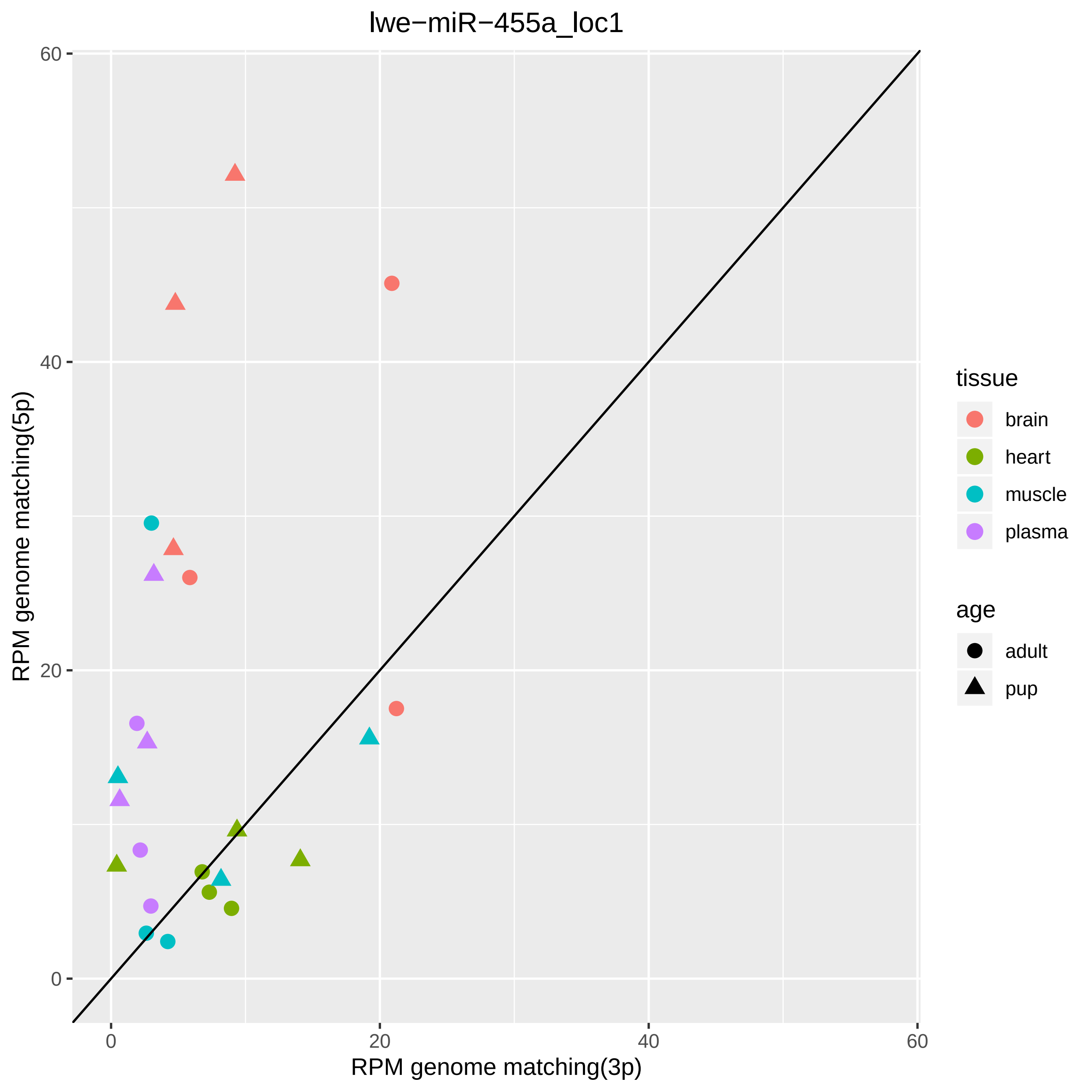
lwe−miR−455a_loc1
RPM genome matching(5p)
RPM genome matching(3p)
0
20
40
60
0
20
40
60
tissue
brain
heart
muscle
plasma
age
adult
pup

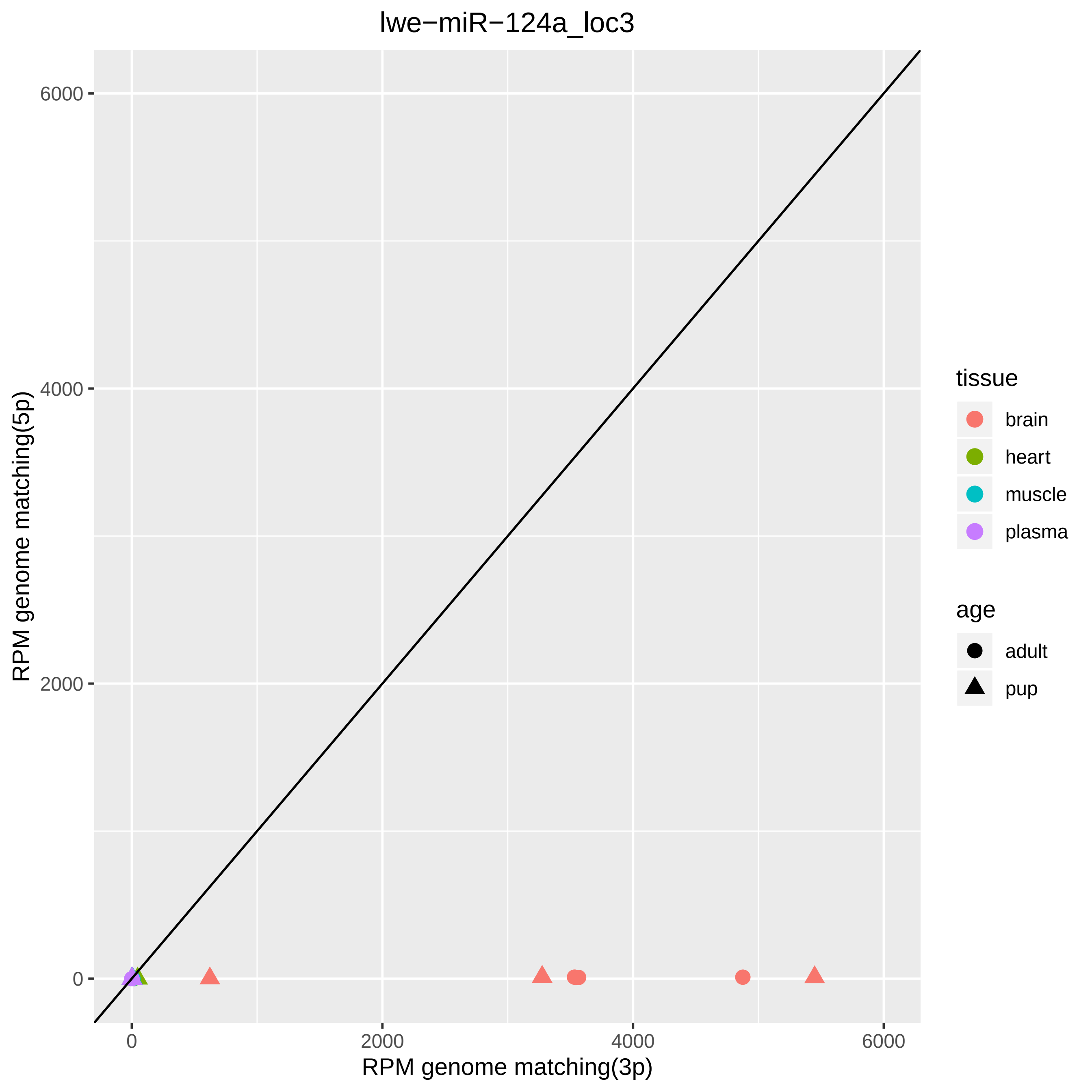
lwe−miR−124a_loc3
RPM genome matching(5p)
6000
4000
2000
0
0
2000
4000
6000
RPM genome matching(3p)
tissue
brain
heart
muscle
plasma
age
adult
pup

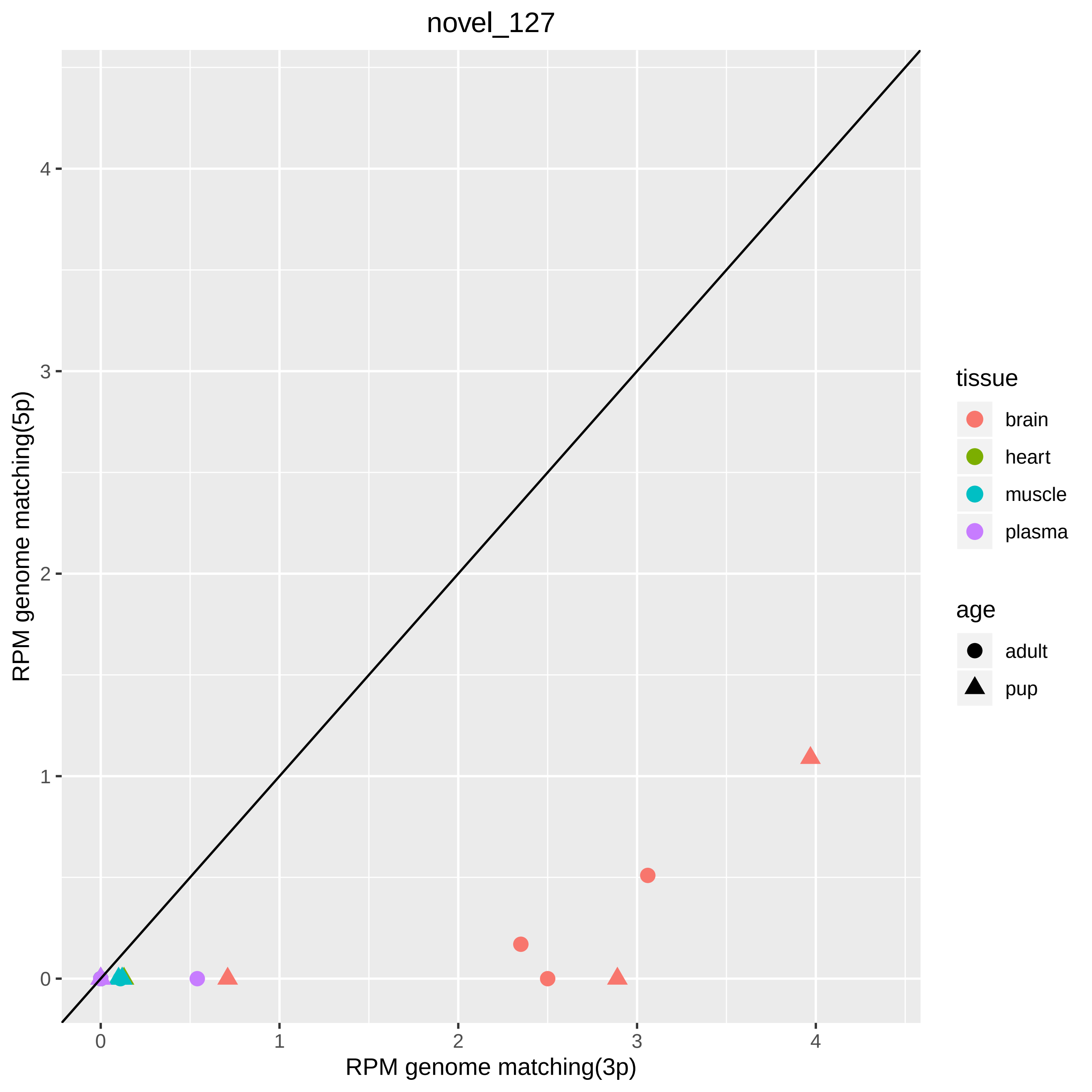

novel_127
RPM genome matching(5p)
RPM genome matching(3p)
0
1
2
3
4
0
1
2
3
4
tissue
brain
heart
muscle
plasma
age
adult
pup

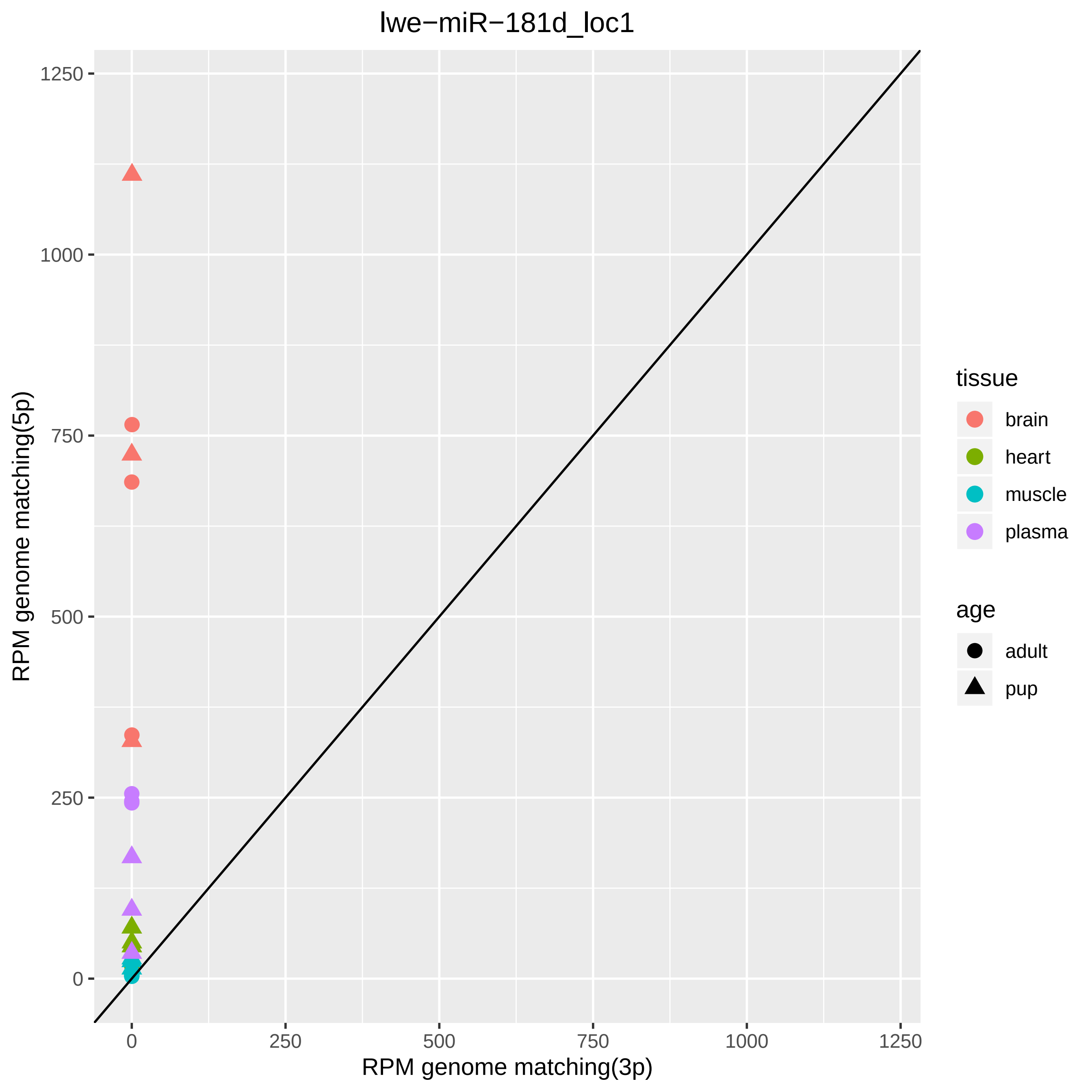

lwe−miR−181d_loc1
RPM genome matching(5p)
RPM genome matching(3p)
0
250
500
750
1000
1250
tissue
brain
heart
muscle
plasma
age
adult
pup

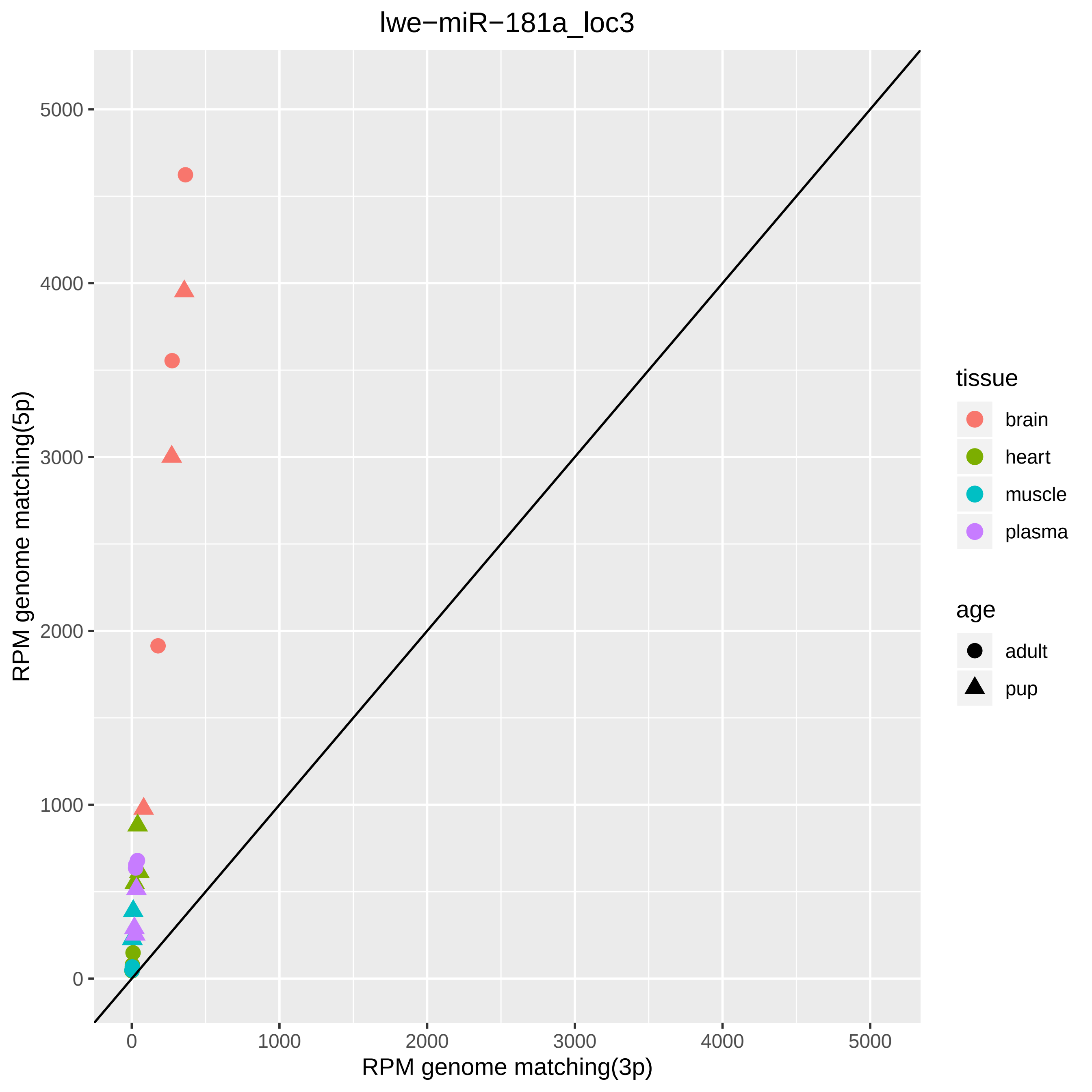
lwe−miR−181a_loc3
RPM genome matching(5p)
RPM genome matching(3p)
0
1000
2000
3000
4000
5000
tissue
brain
heart
muscle
plasma
age
adult
pup

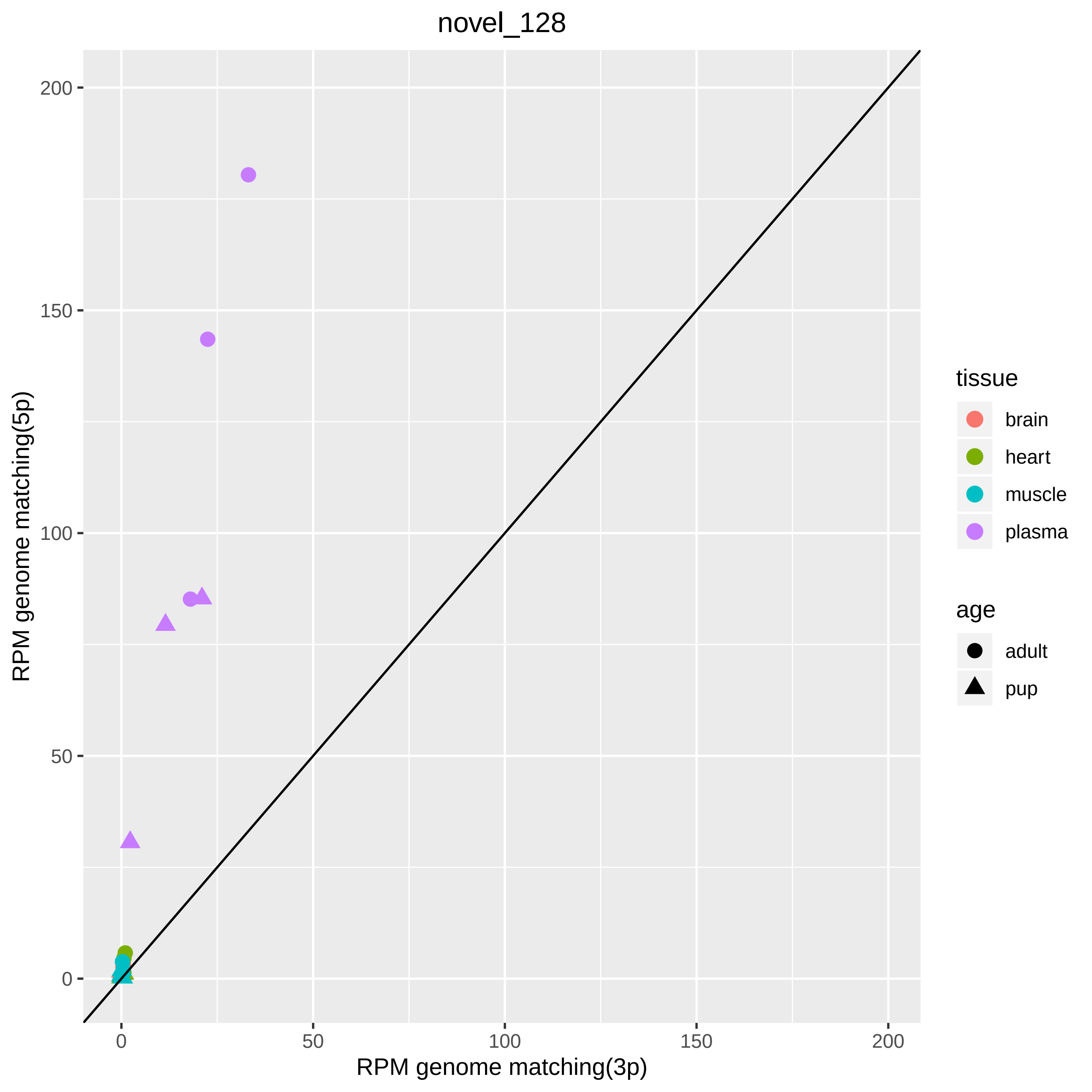

novel_128
RPM genome matching(5p)
RPM genome matching(3p)
200
150
100
50
0
0
50
100
150
200
tissue
brain
heart
muscle
plasma
age
adult
pup

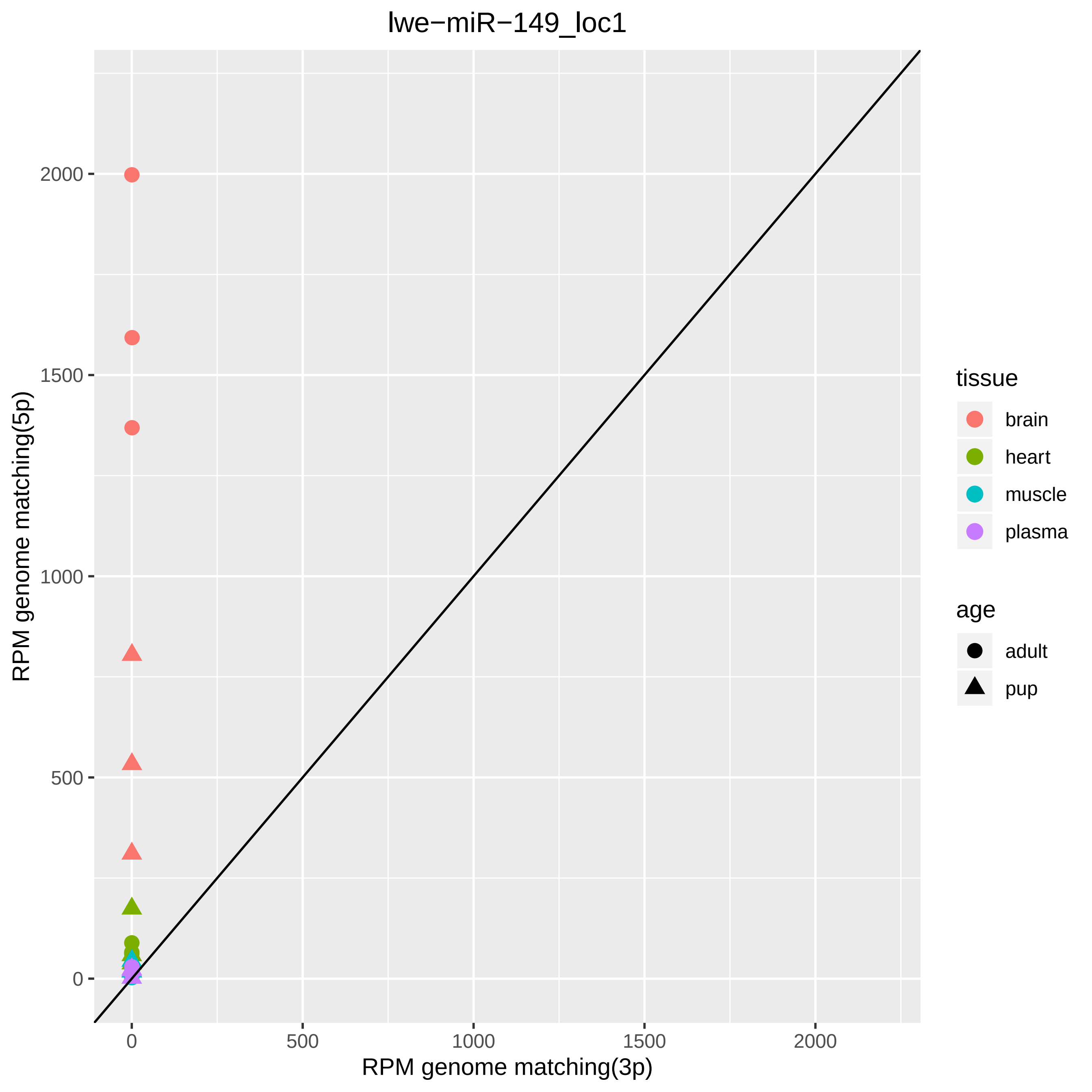
lwe−miR−149_loc1
RPM genome matching(5p)
RPM genome matching(3p)
0
500
1000
1500
2000
tissue
brain
heart
muscle
plasma
age
adult
pup

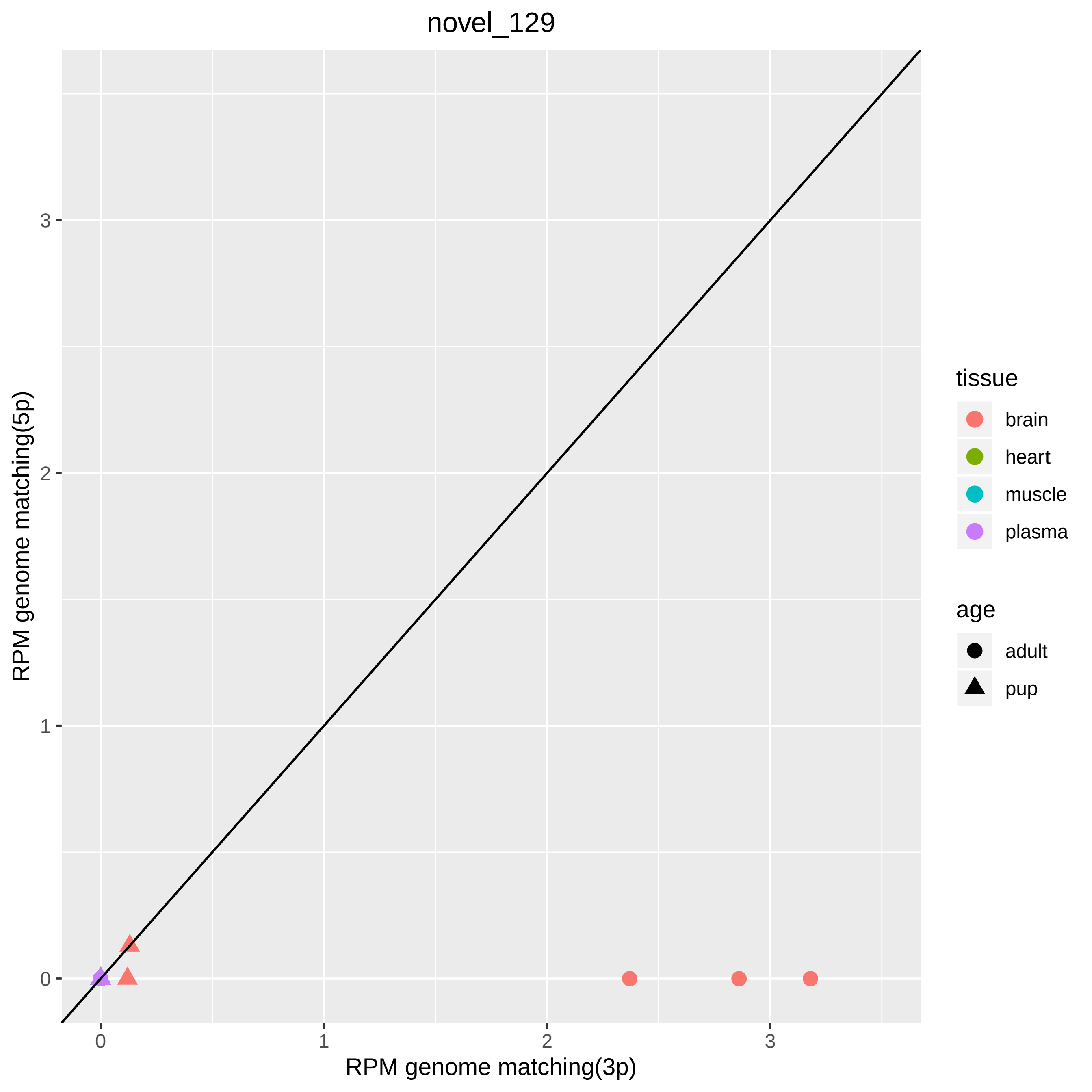

novel_129
RPM genome matching(5p)
RPM genome matching(3p)
0
1
2
3
0
1
2
3
tissue
brain
heart
muscle
plasma
age
adult
pup

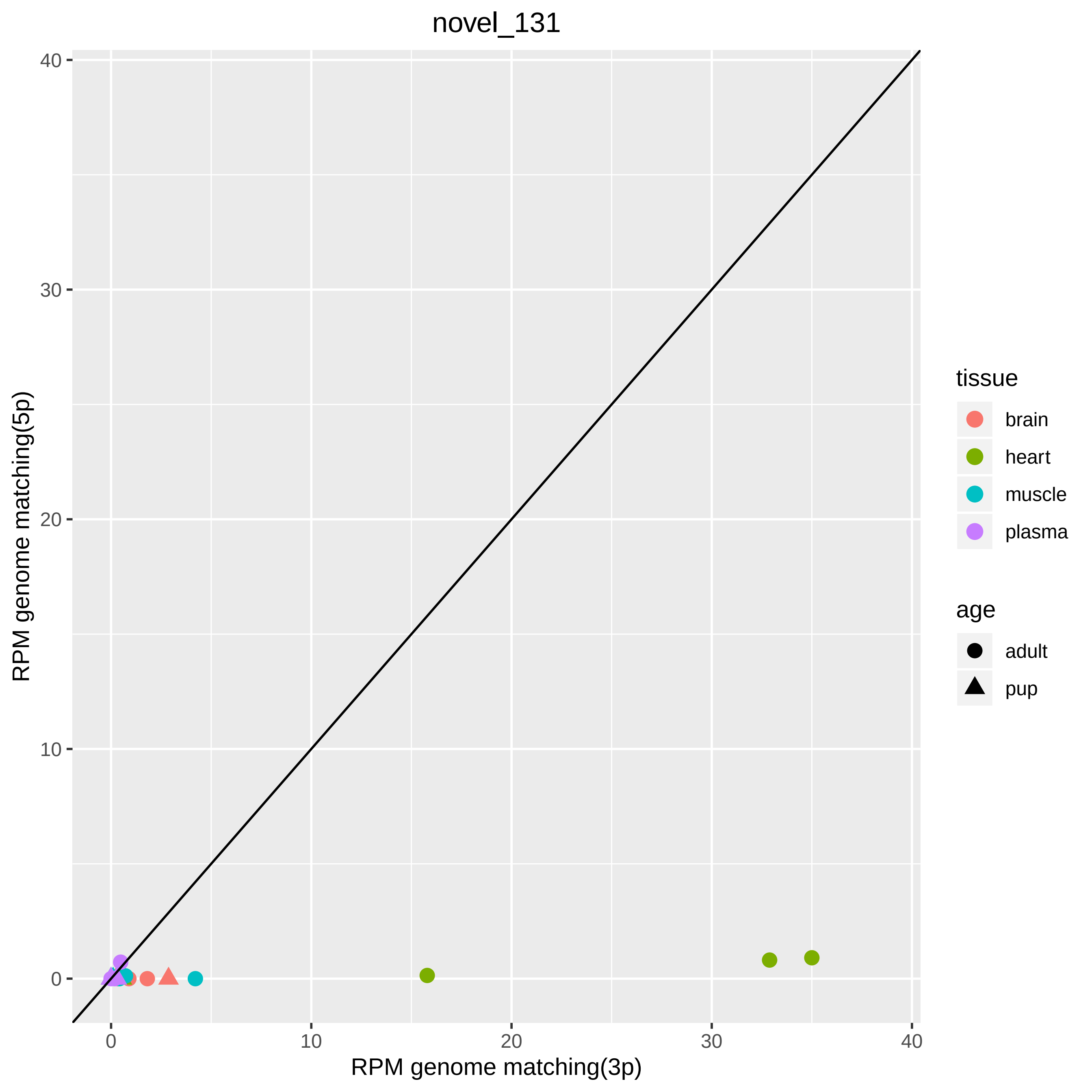
novel_131
RPM genome matching(5p)
RPM genome matching(3p)
0
10
20
30
40
0
10
20
30
40
tissue
brain
heart
muscle
plasma
age
adult
pup

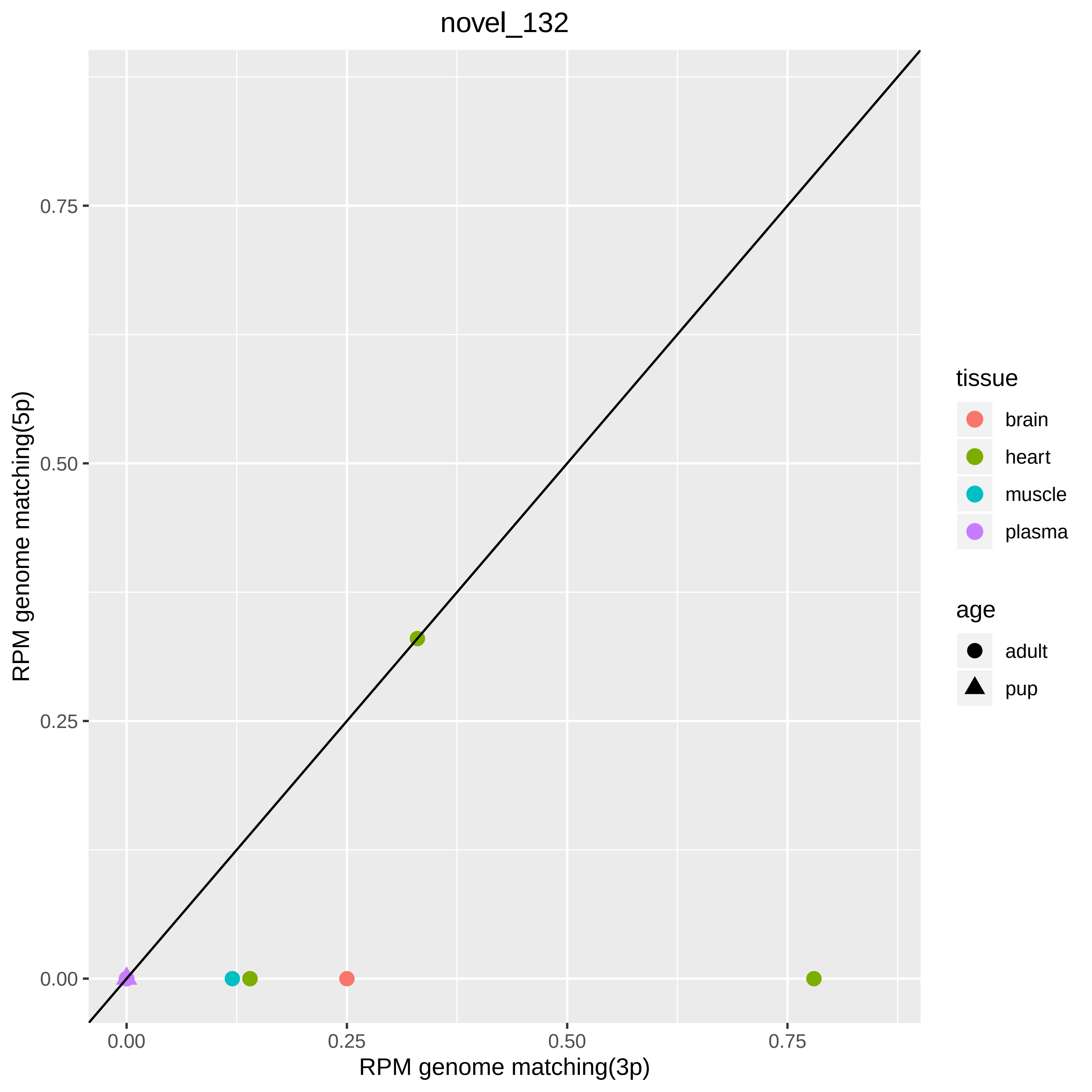
novel_132
RPM genome matching(5p)
RPM genome matching(3p)
0.00
0.25
0.50
0.75
tissue
brain
heart
muscle
plasma
age
adult
pup

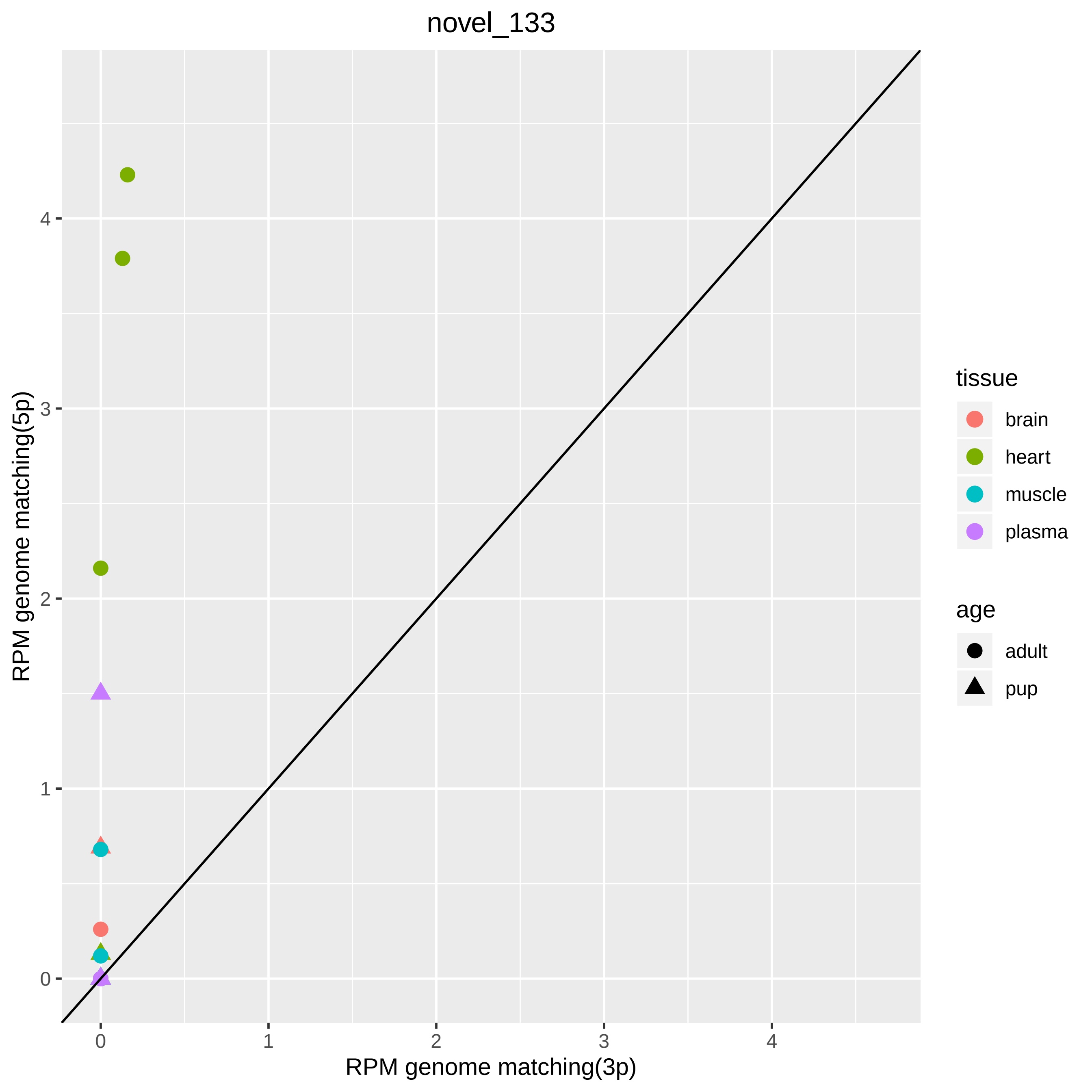

novel_133
RPM genome matching(5p)
RPM genome matching(3p)
0
1
2
3
4
0
1
2
3
4
tissue
brain
heart
muscle
plasma
age
adult
pup

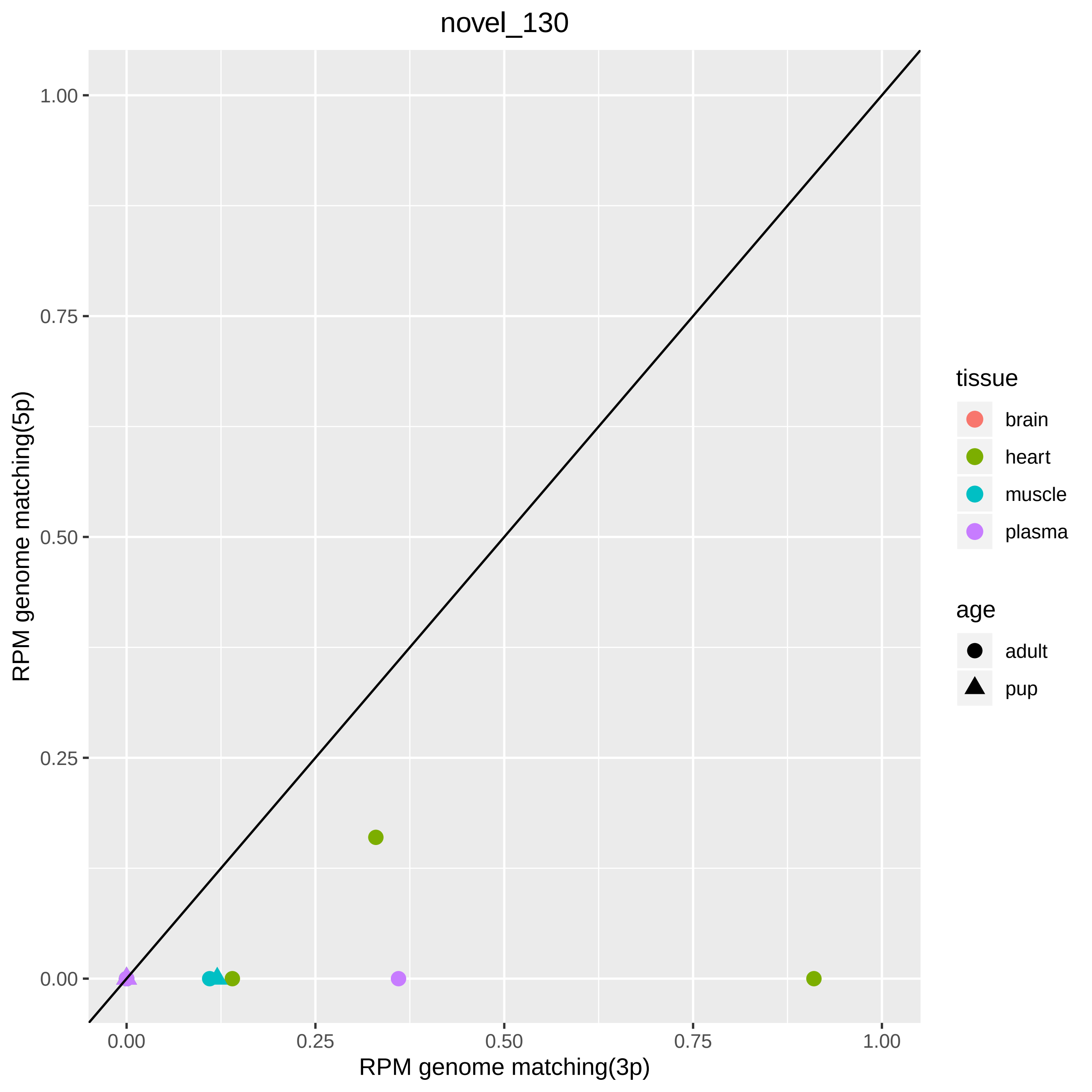
novel_130
RPM genome matching(5p)
RPM genome matching(3p)
1.00
0.75
0.50
0.25
0.00
0.00
0.25
0.50
0.75
1.00
tissue
brain
heart
muscle
plasma
age
adult
pup

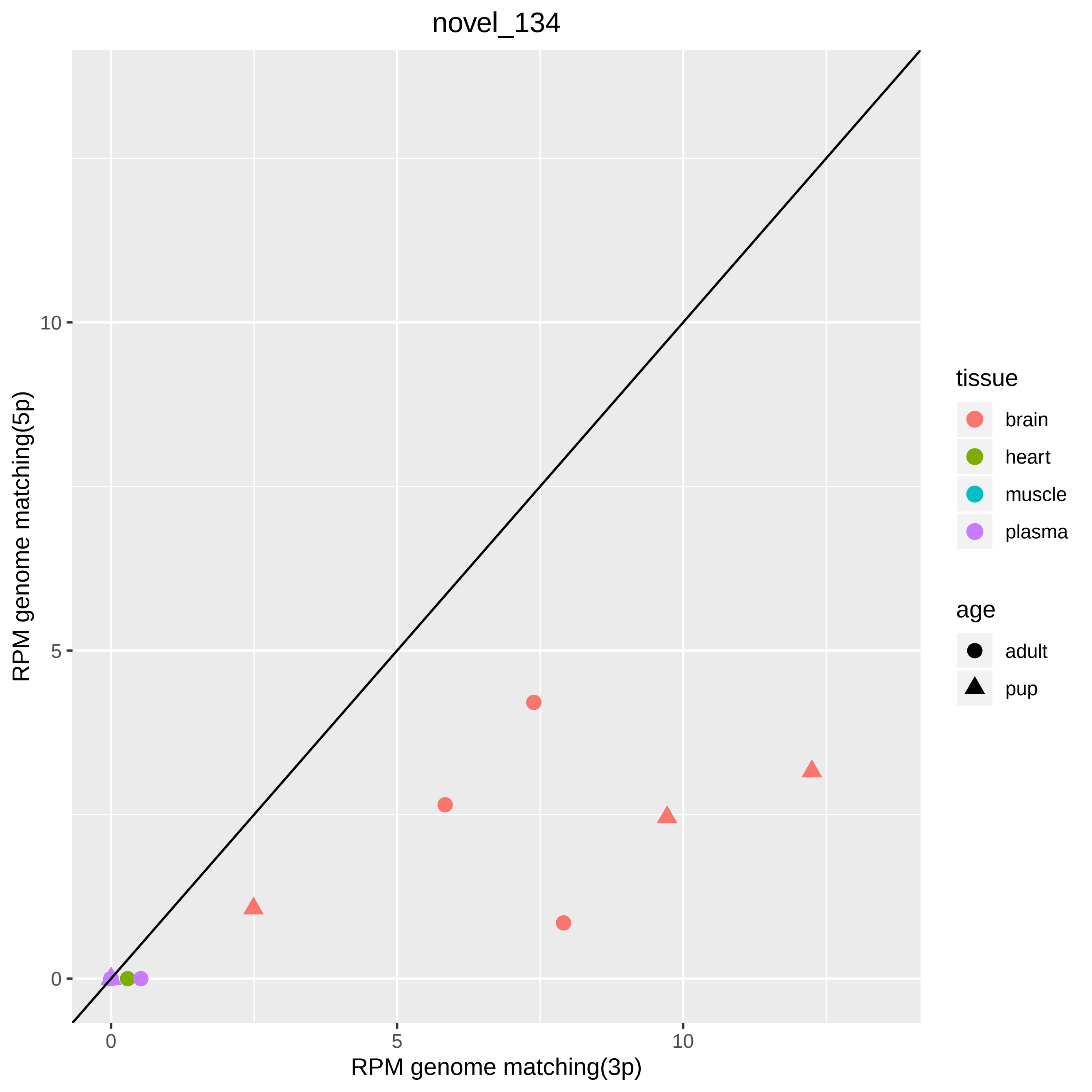
novel_134
RPM genome matching(5p)
RPM genome matching(3p)
0
5
10
0
5
10
tissue
brain
heart
muscle
plasma
age
adult
pup

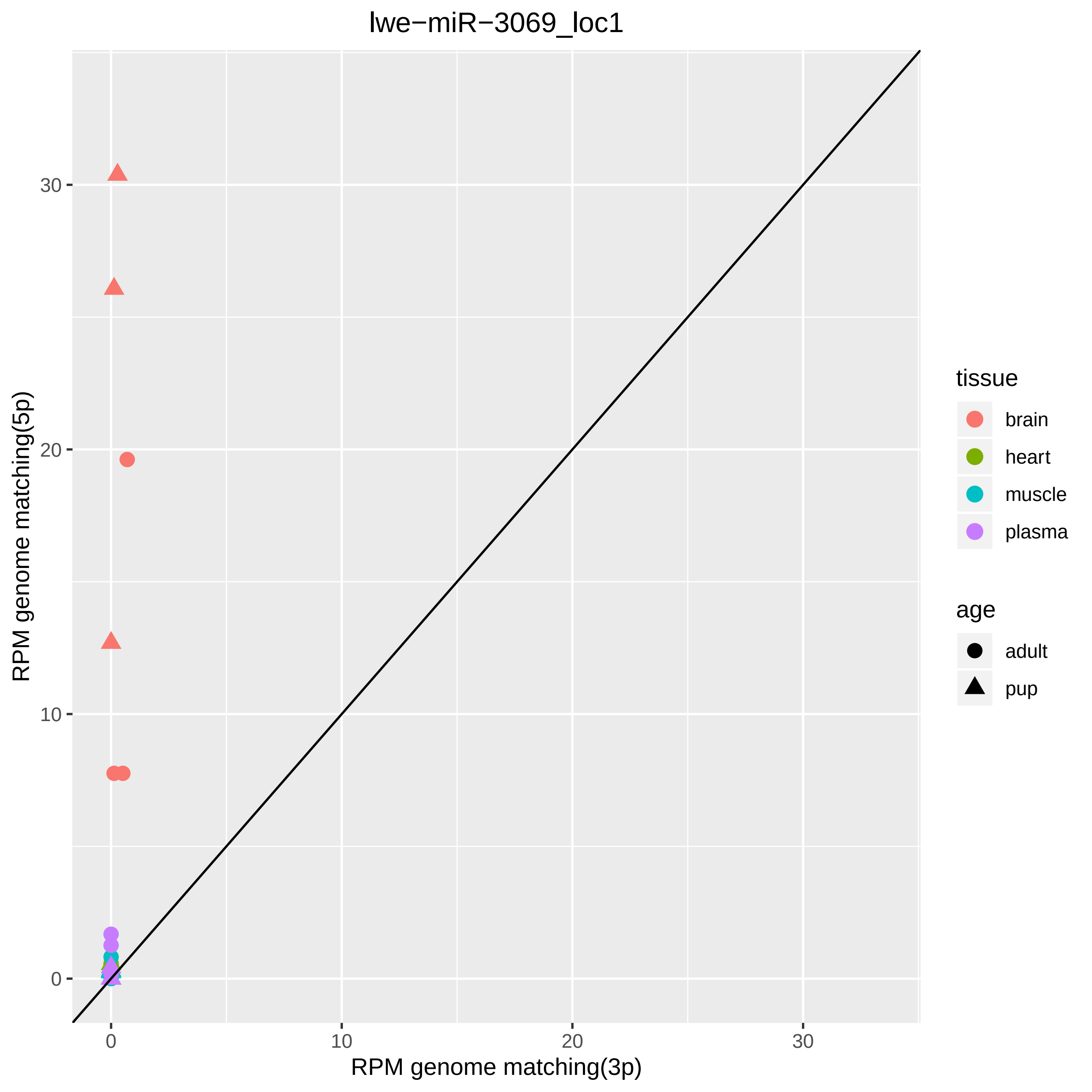
lwe−miR−3069_loc1
RPM genome matching(5p)
RPM genome matching(3p)
0
10
20
30
0
10
20
30
tissue
brain
heart
muscle
plasma
age
adult
pup

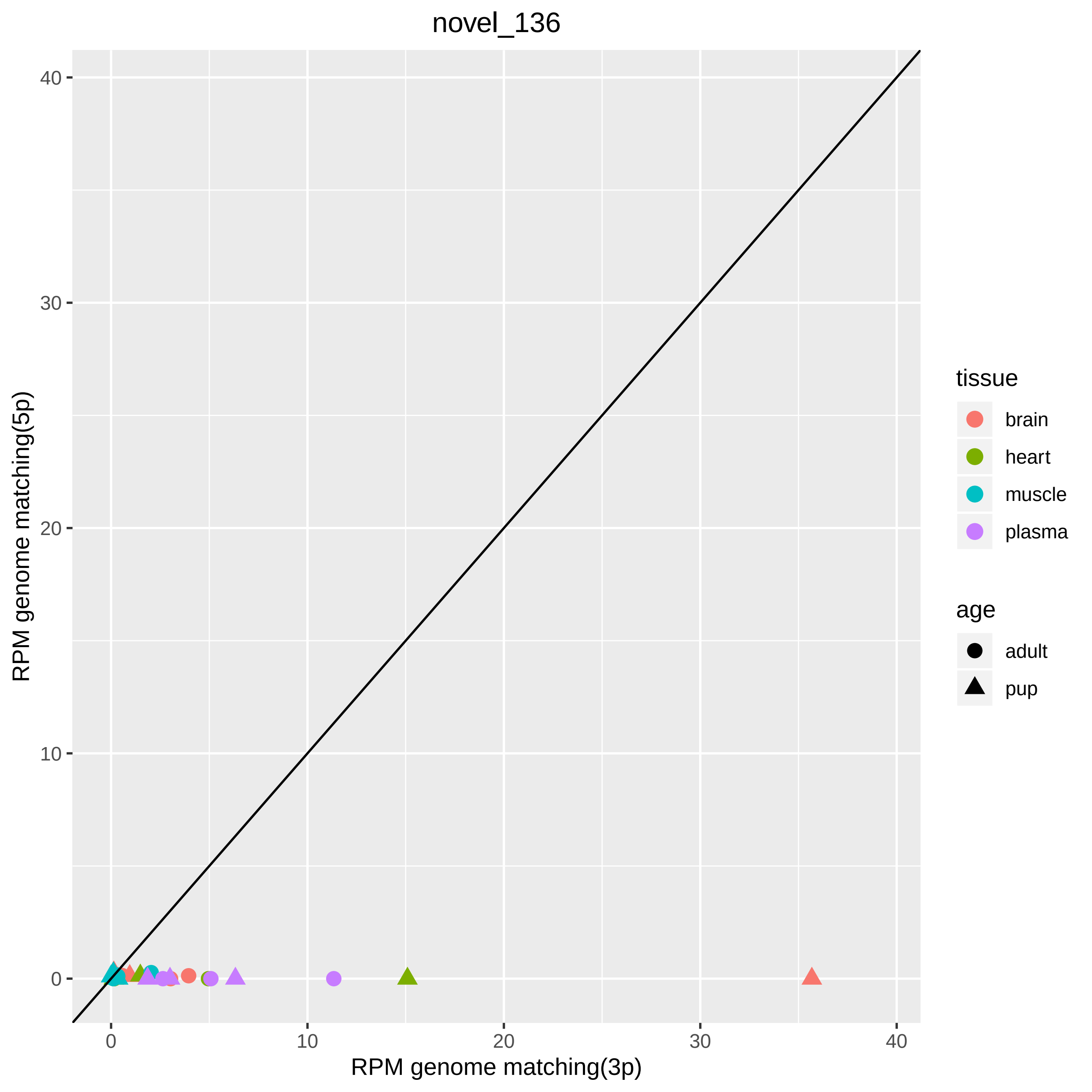

novel_136
RPM genome matching(5p)
RPM genome matching(3p)
0
10
20
30
40
tissue
brain
heart
muscle
plasma
age
adult
pup

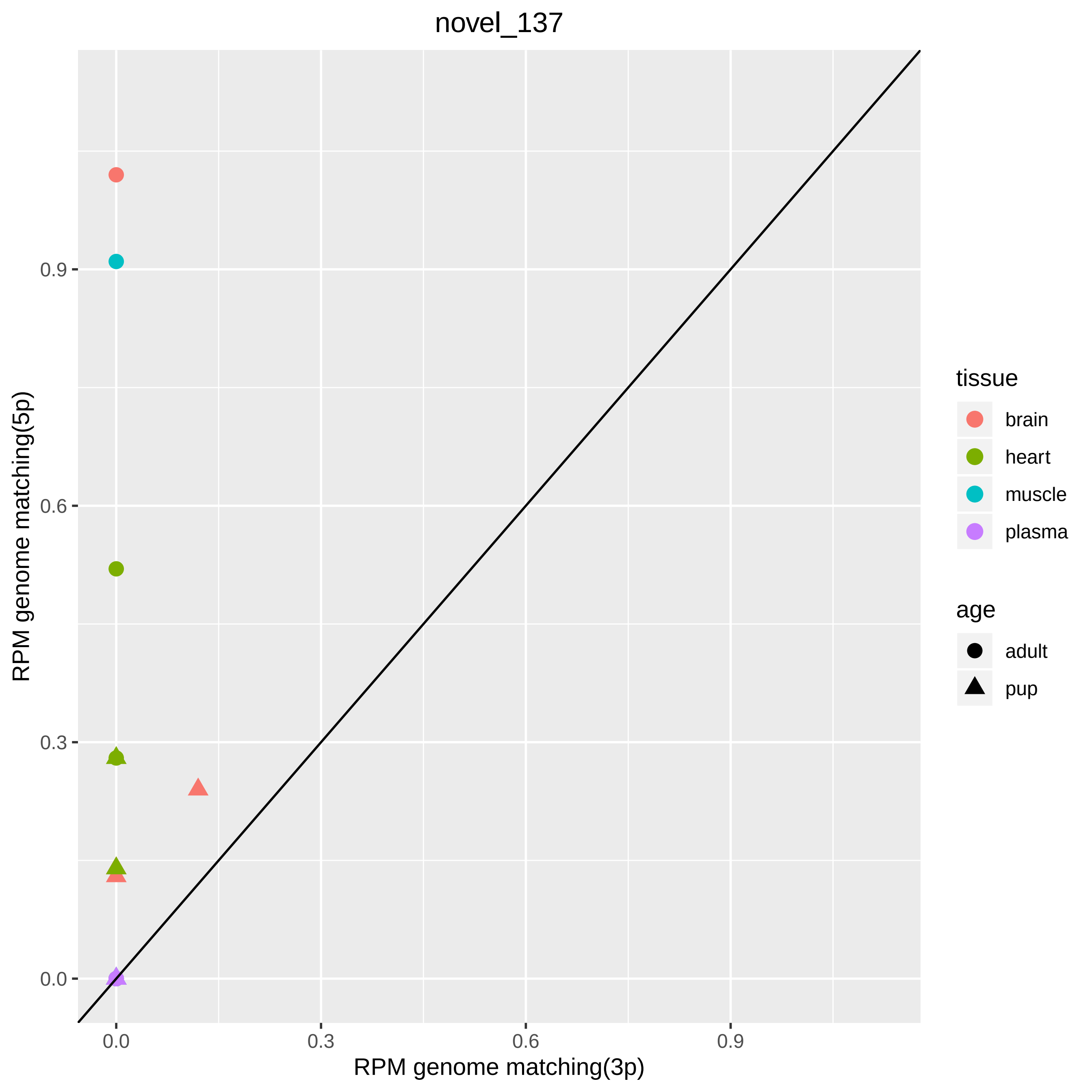
novel_137
RPM genome matching(5p)
0.9
0.6
0.3
0.0
0.0
0.3
0.6
0.9
RPM genome matching(3p)
tissue
brain
heart
muscle
plasma
age
adult
pup

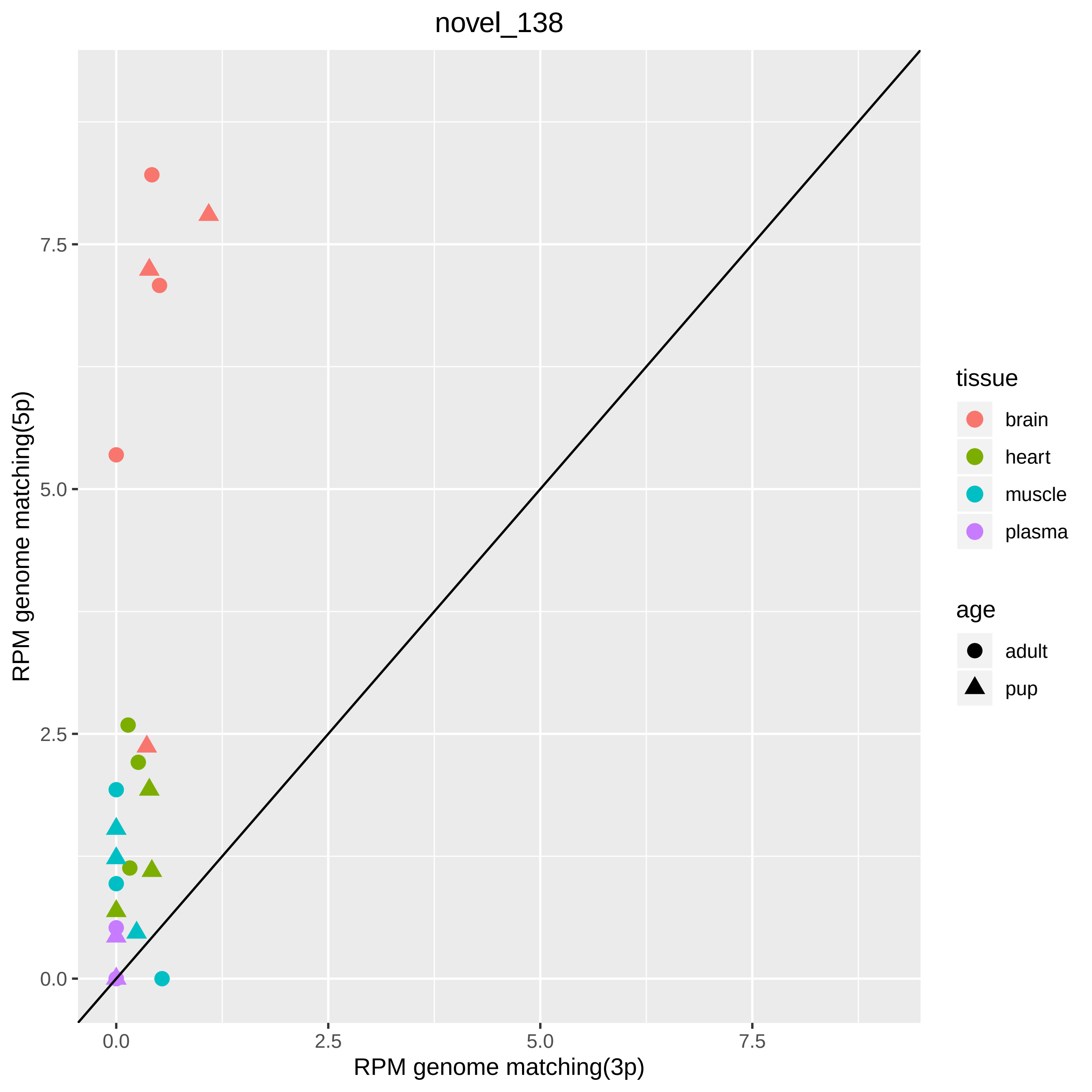
novel_138
RPM genome matching(5p)
RPM genome matching(3p)
0.0
2.5
5.0
7.5
0.0
2.5
5.0
7.5
tissue
brain
heart
muscle
plasma
age
adult
pup

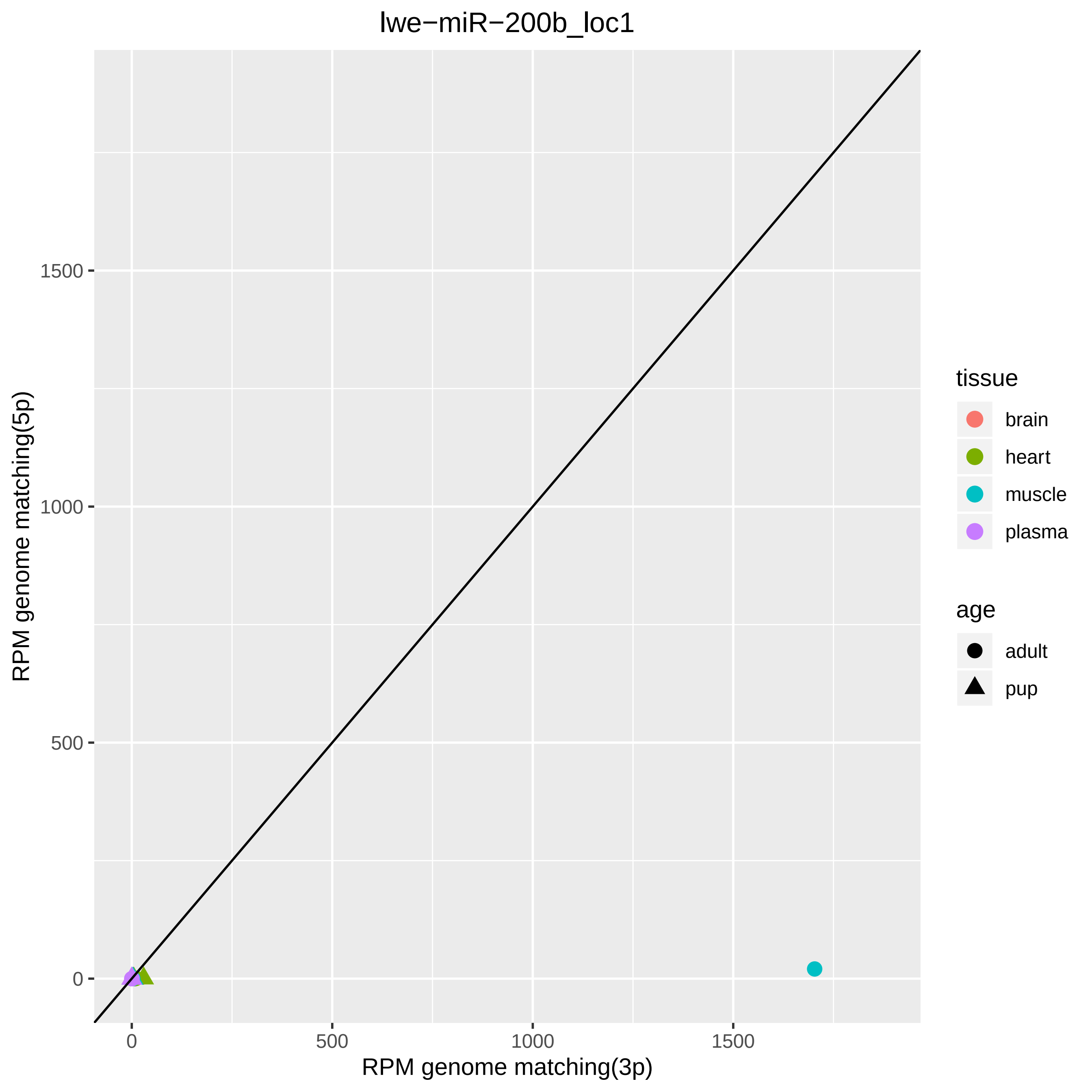

lwe−miR−200b_loc1
RPM genome matching(5p)
RPM genome matching(3p)
0
500
1000
1500
tissue
brain
heart
muscle
plasma
age
adult
pup

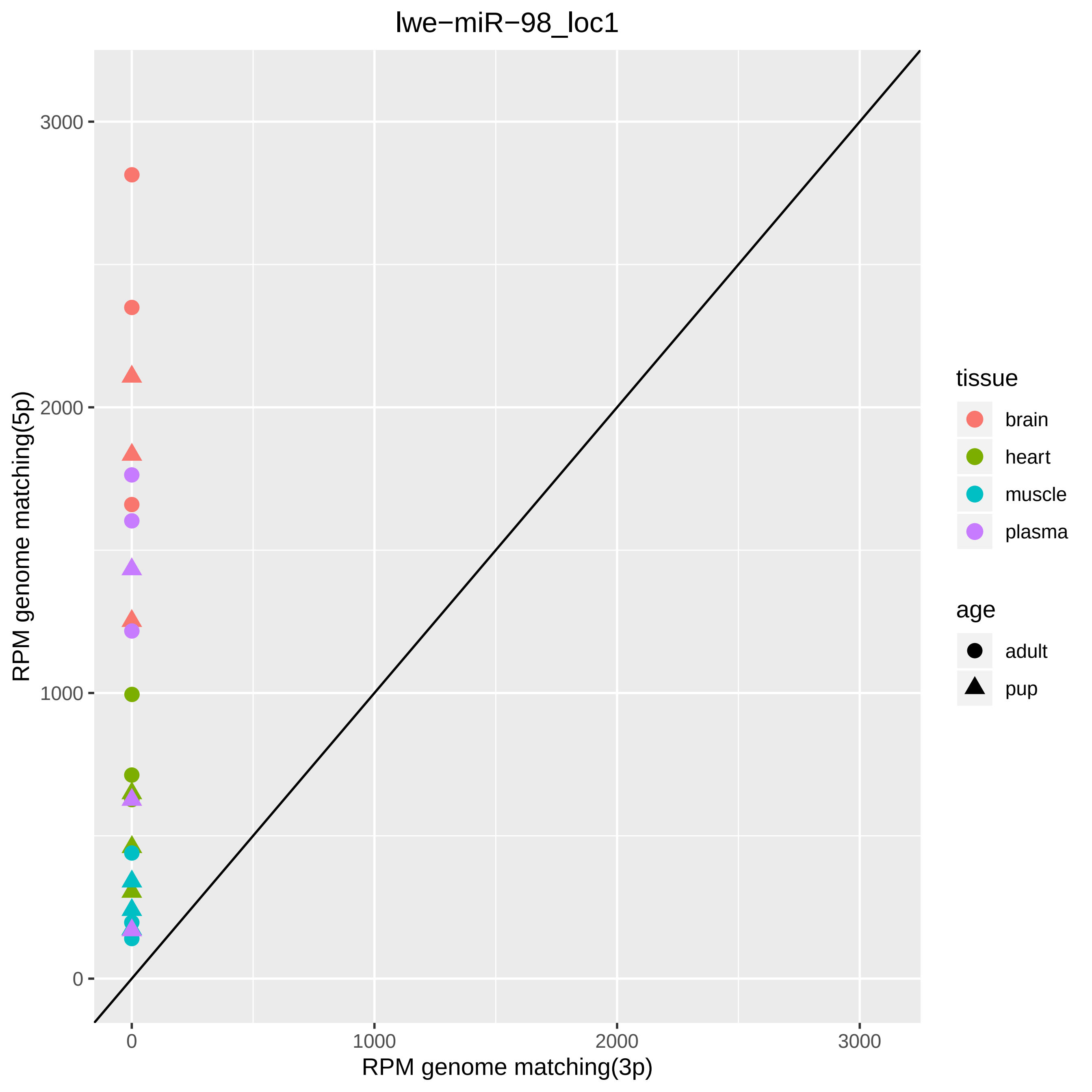

lwe−miR−98_loc1
RPM genome matching(5p)
3000
2000
1000
0
RPM genome matching(3p)
0
1000
2000
3000
tissue
brain
heart
muscle
plasma
age
adult
pup

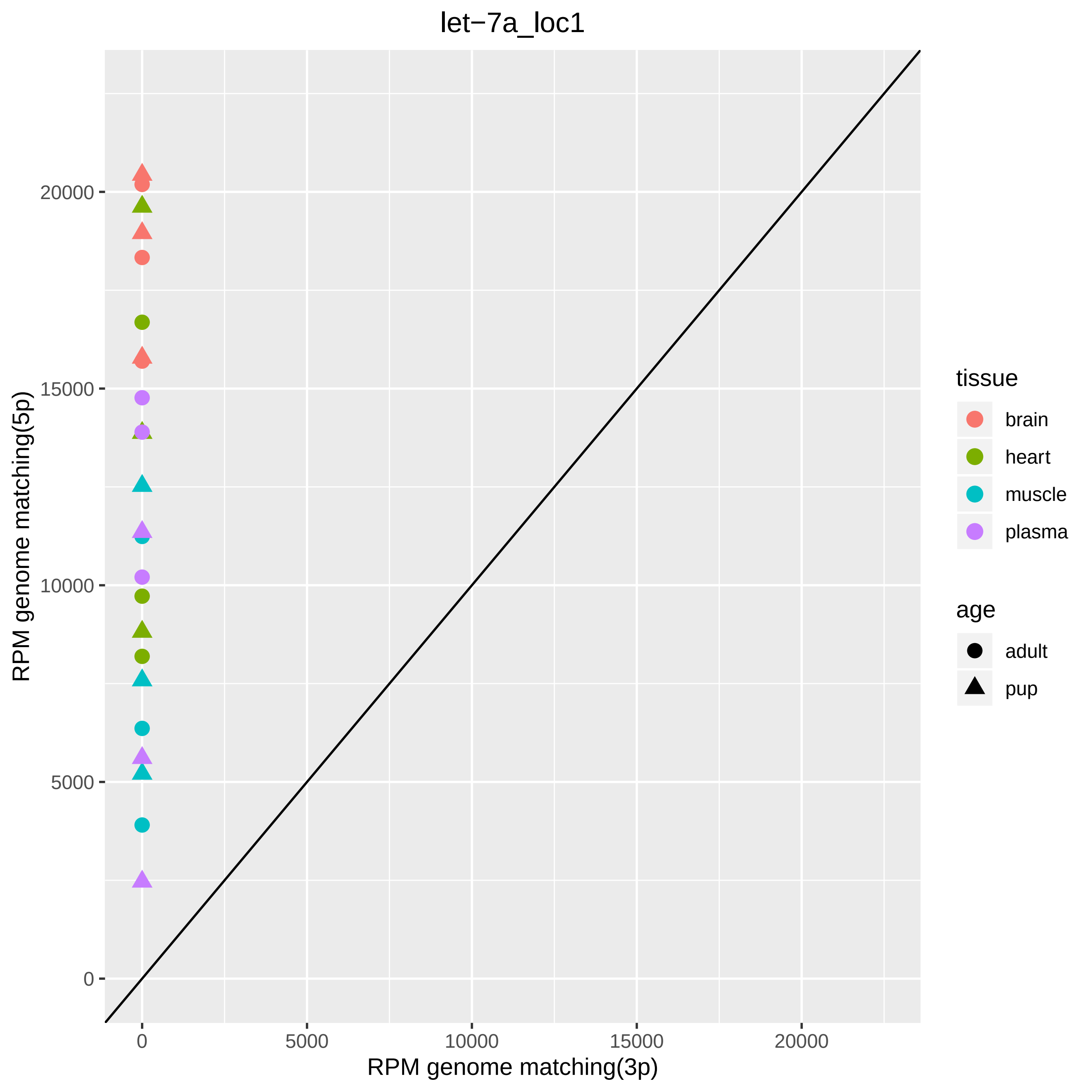
let−7a_loc1
RPM genome matching(5p)
RPM genome matching(3p)
0
5000
10000
15000
20000
0
5000
10000
15000
20000
tissue
brain
heart
muscle
plasma
age
adult
pup

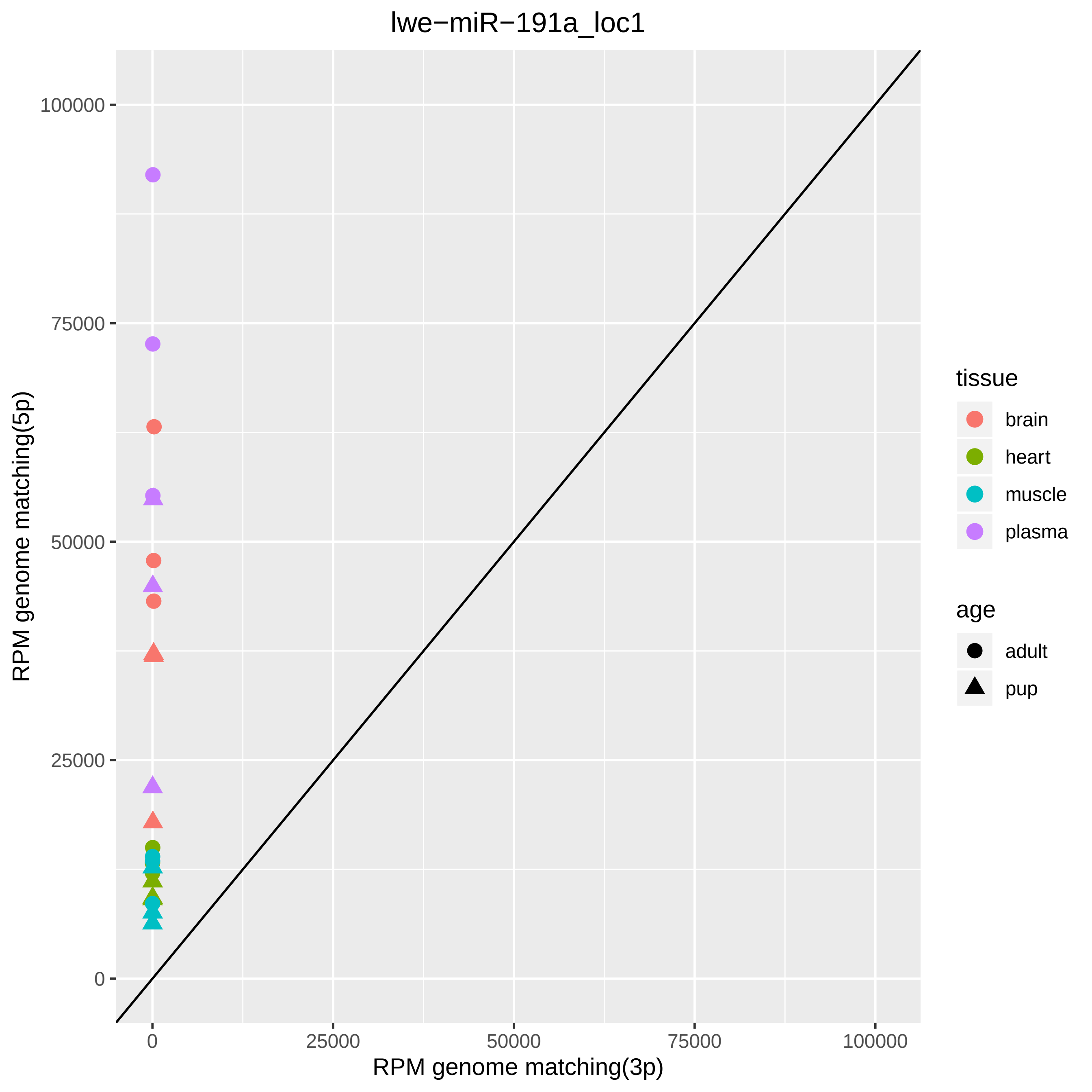
lwe−miR−191a_loc1
RPM genome matching(5p)
RPM genome matching(3p)
0
25000
50000
75000
100000
tissue
brain
heart
muscle
plasma
age
adult
pup

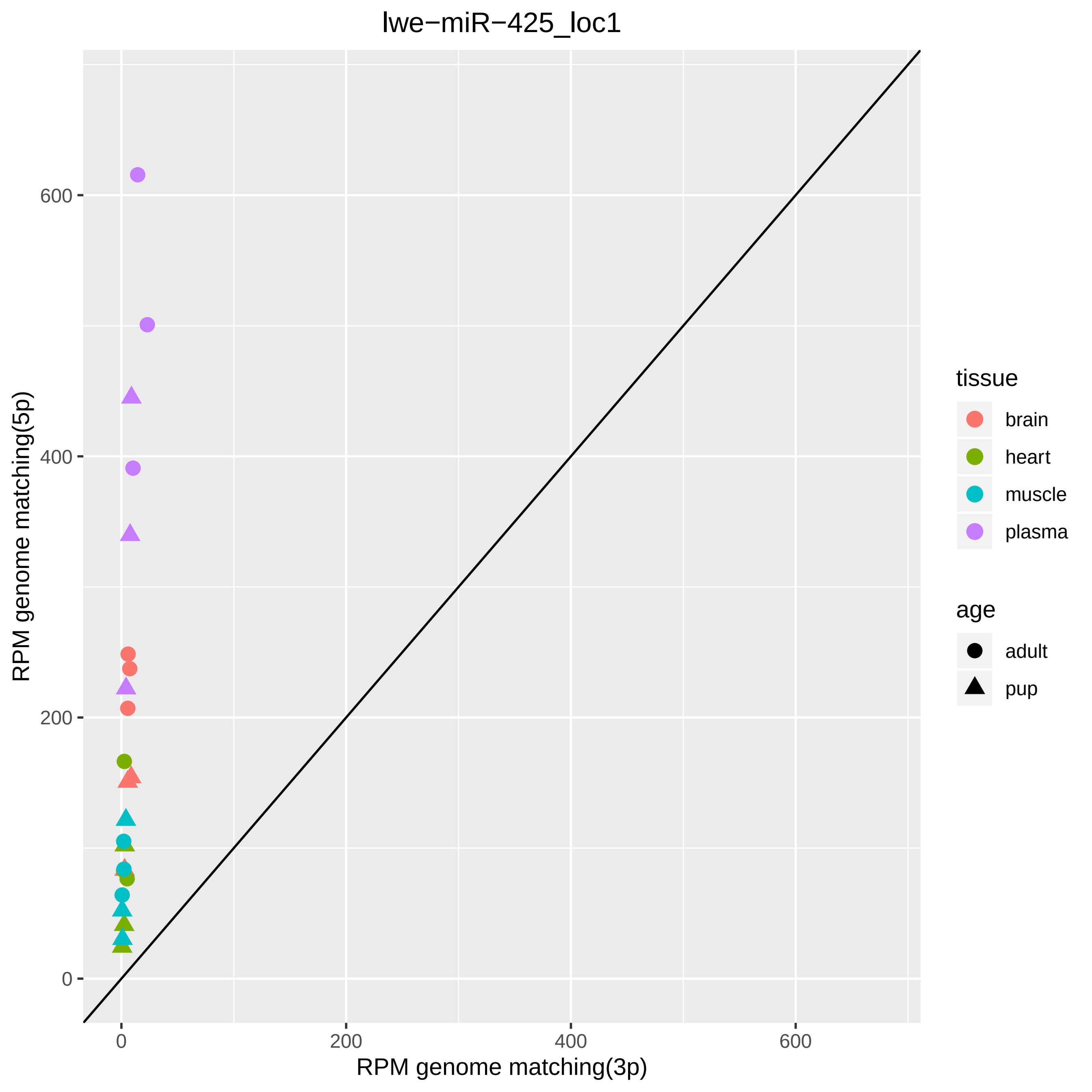

lwe−miR−425_loc1
RPM genome matching(5p)
RPM genome matching(3p)
0
200
400
600
tissue
brain
heart
muscle
plasma
age
adult
pup

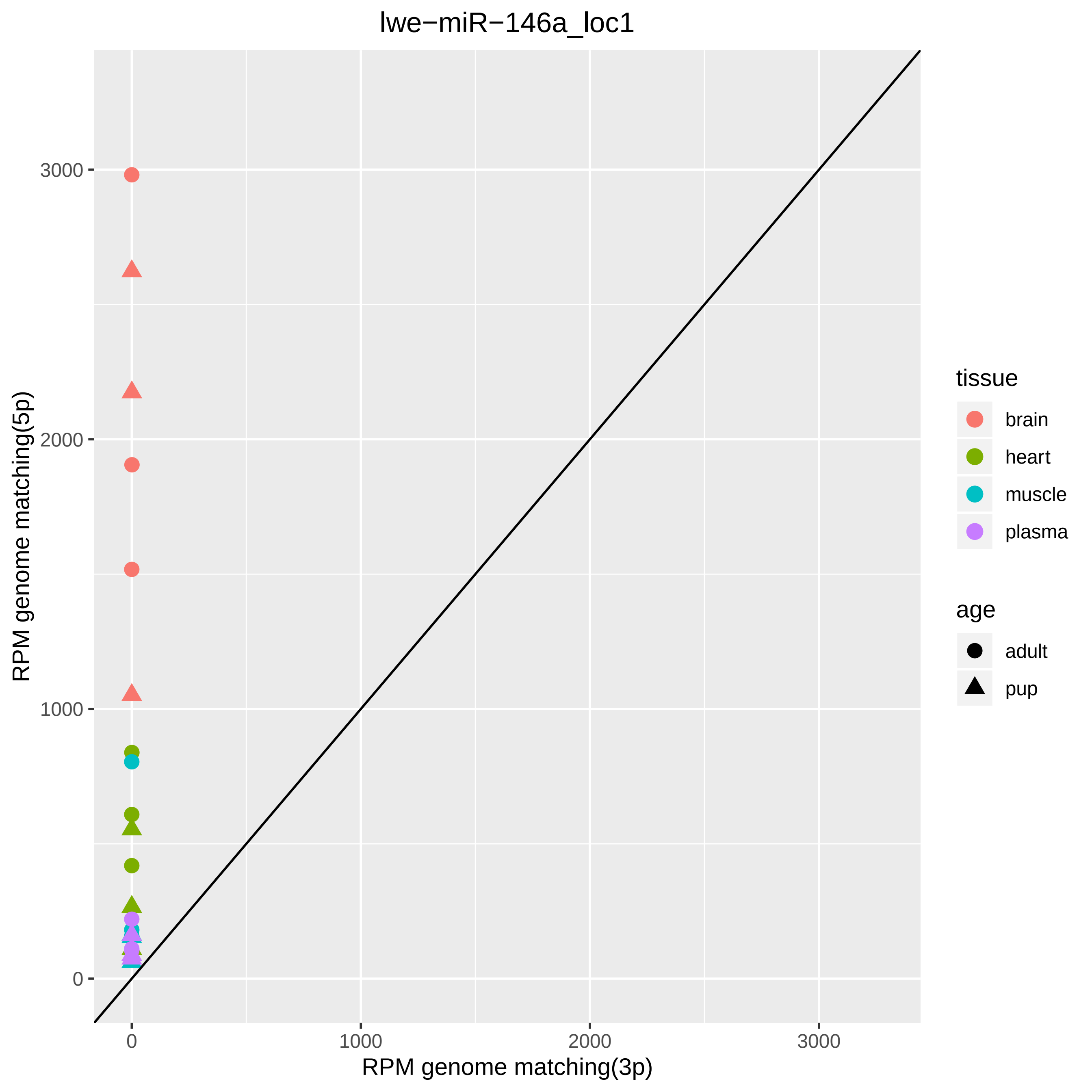
lwe−miR−146a_loc1
RPM genome matching(5p)
3000
2000
1000
0
0
1000
2000
3000
RPM genome matching(3p)
tissue
brain
heart
muscle
plasma
age
adult
pup

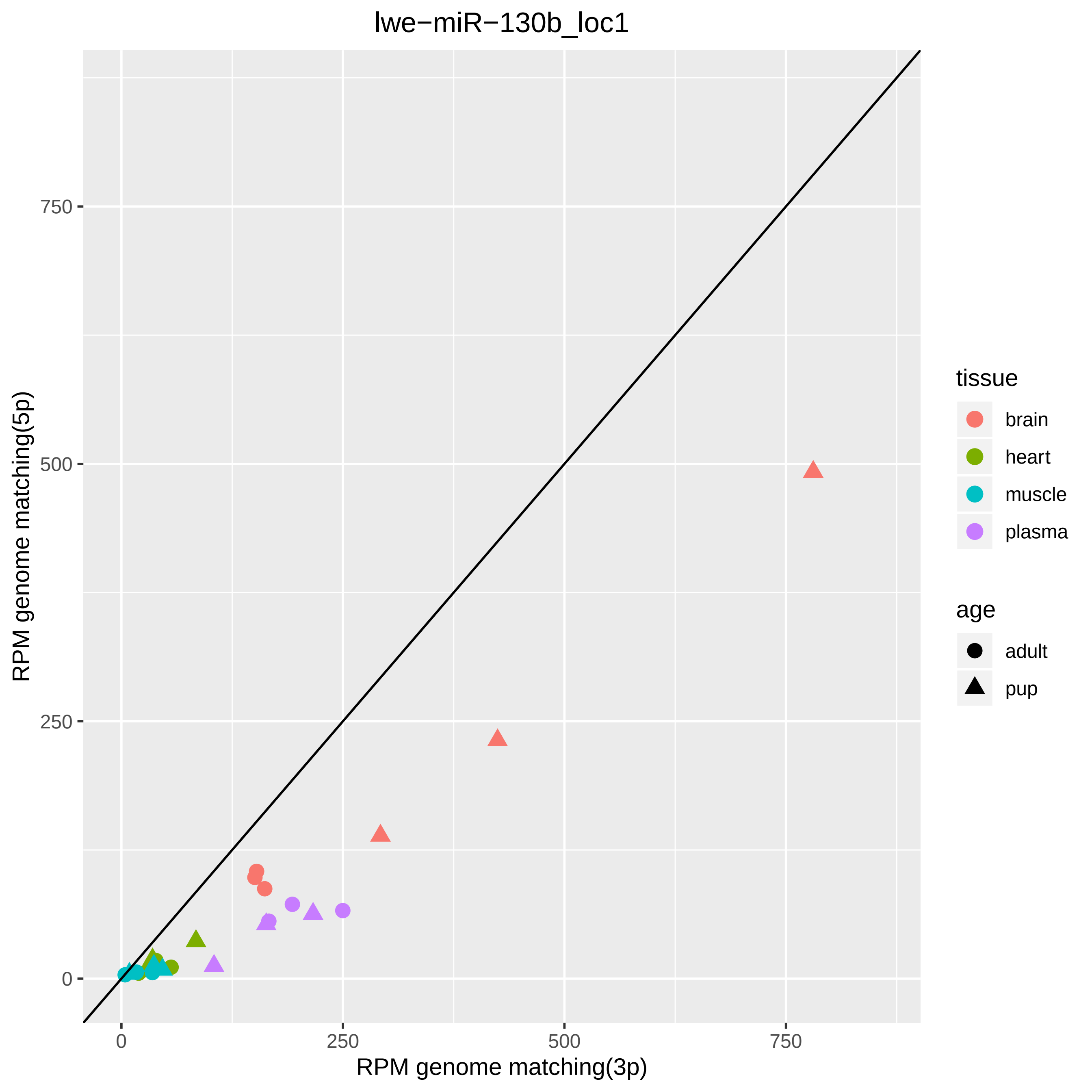
lwe−miR−130b_loc1
RPM genome matching(5p)
RPM genome matching(3p)
0
250
500
750
0
250
500
750
tissue
brain
heart
muscle
plasma
age
adult
pup

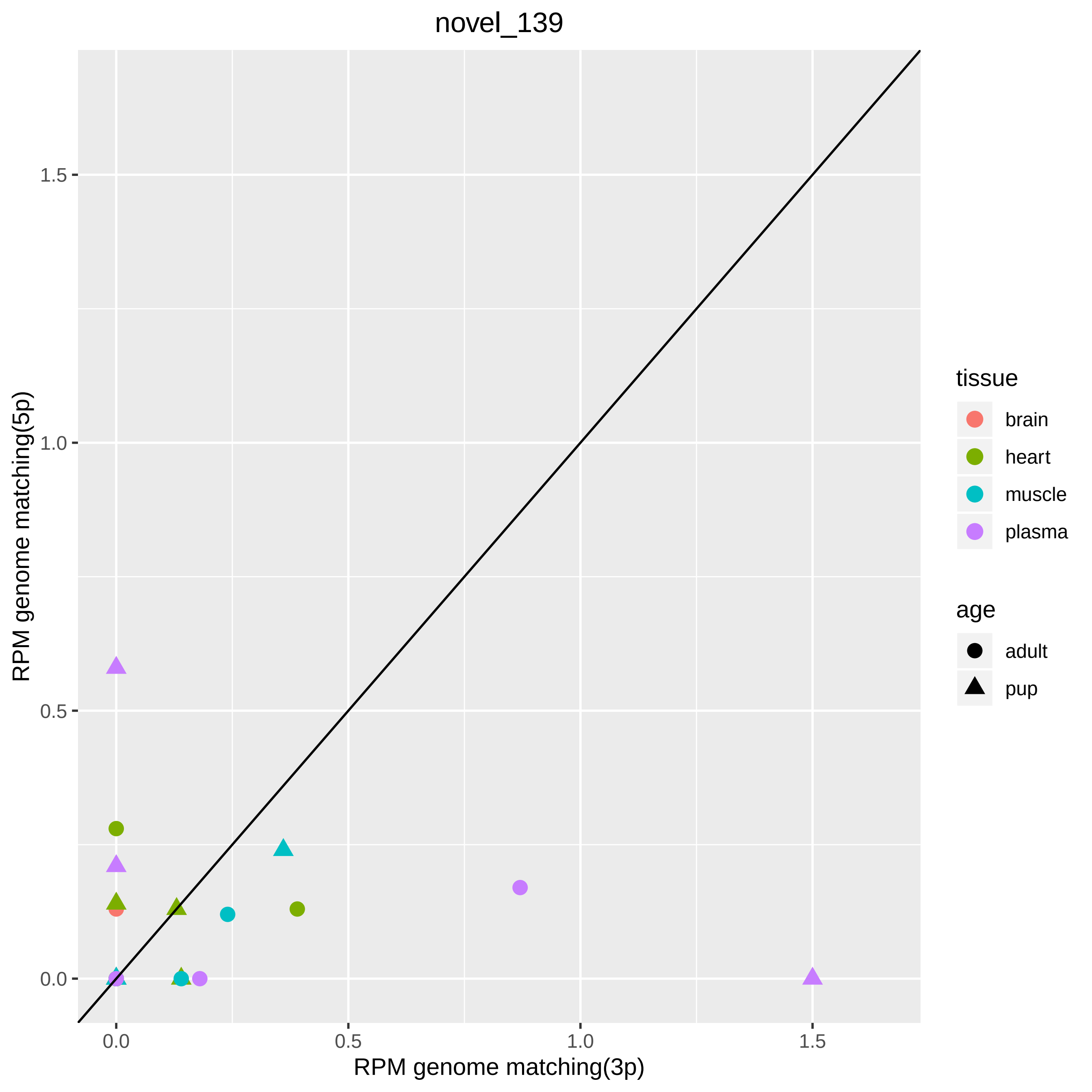

novel_139
RPM genome matching(5p)
RPM genome matching(3p)
0.0
0.5
1.0
1.5
tissue
brain
heart
muscle
plasma
age
adult
pup

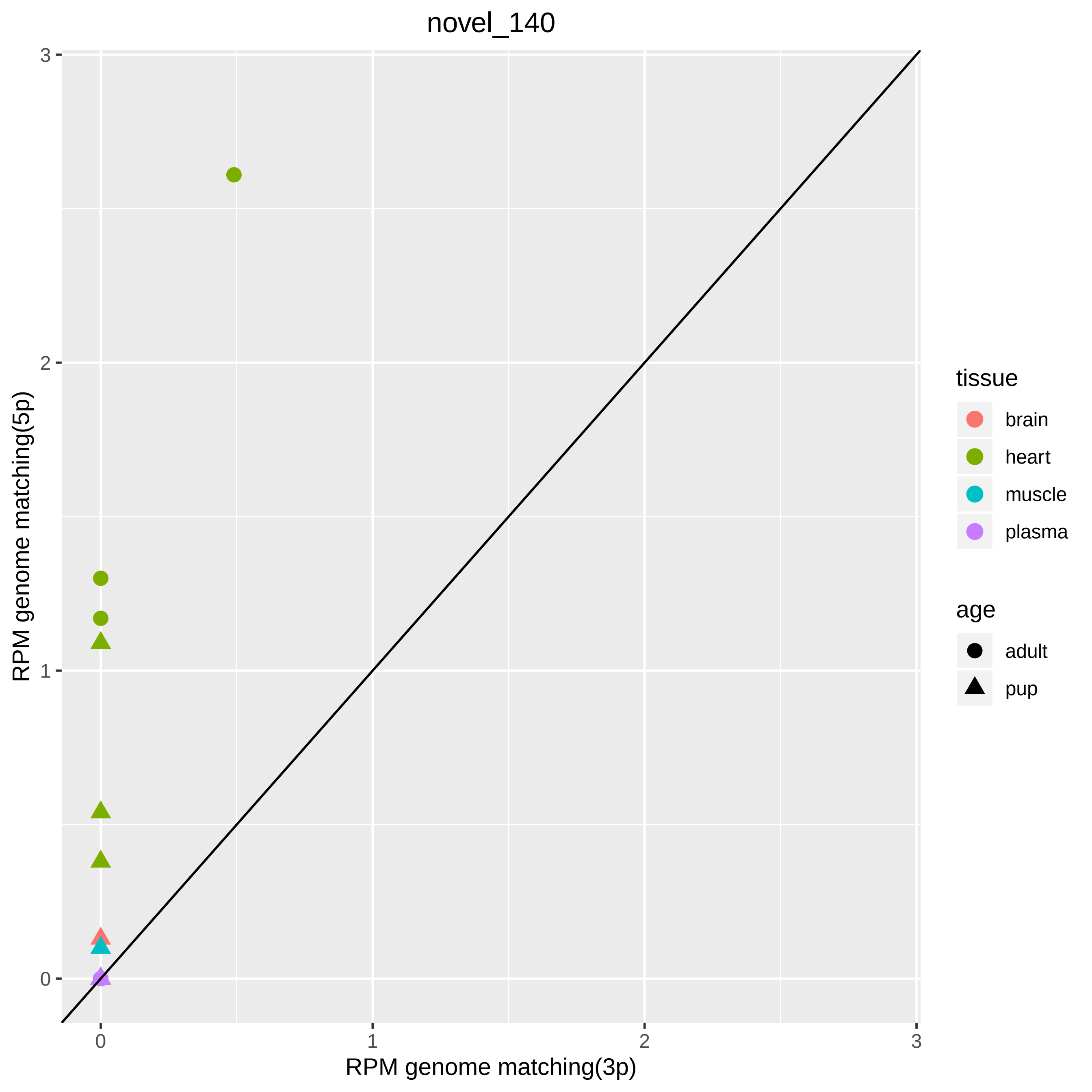
novel_140
RPM genome matching(5p)
RPM genome matching(3p)
0
1
2
3
0
1
2
3
tissue
brain
heart
muscle
plasma
age
adult
pup

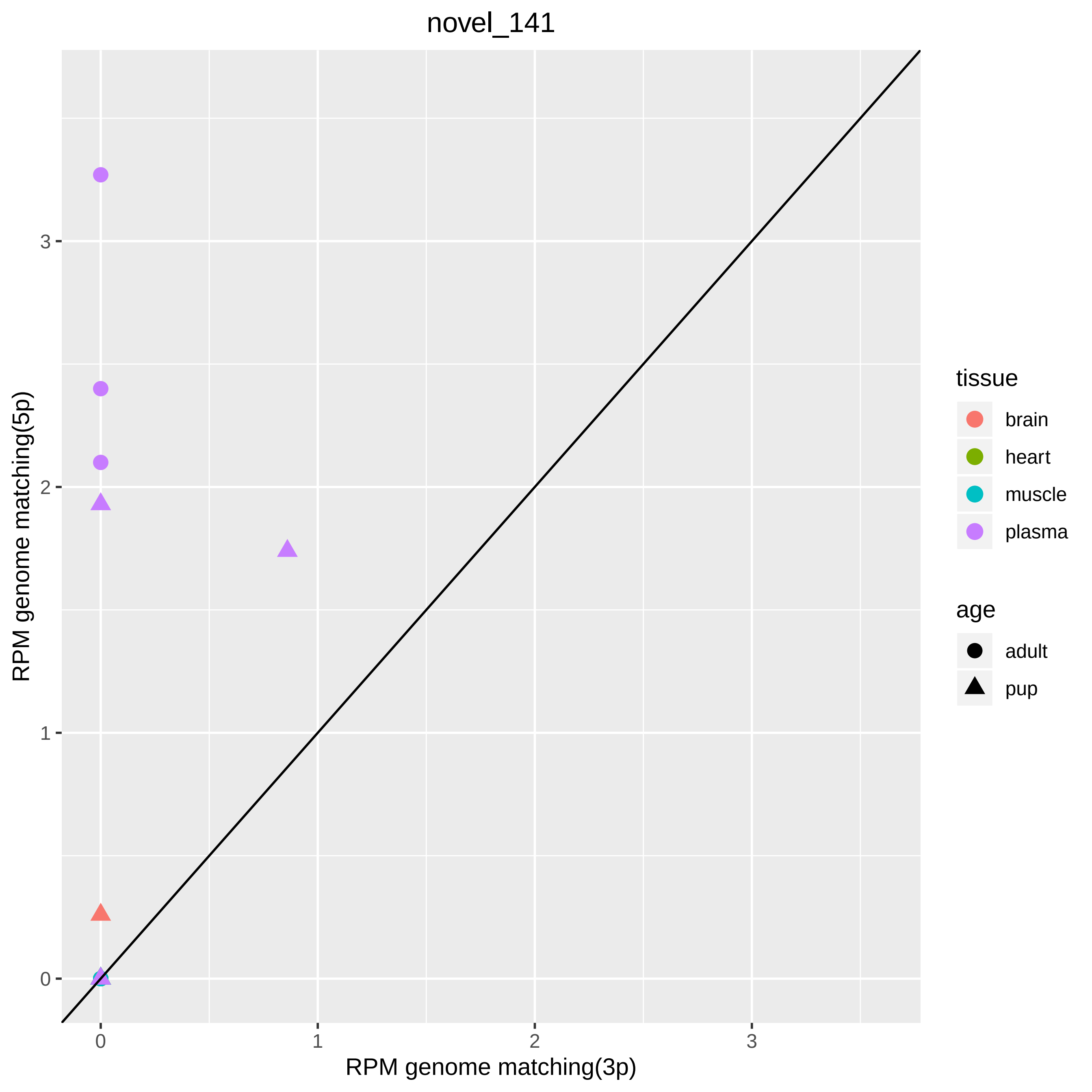
novel_141
RPM genome matching(5p)
RPM genome matching(3p)
0
1
2
3
0
1
2
3
tissue
brain
heart
muscle
plasma
age
adult
pup

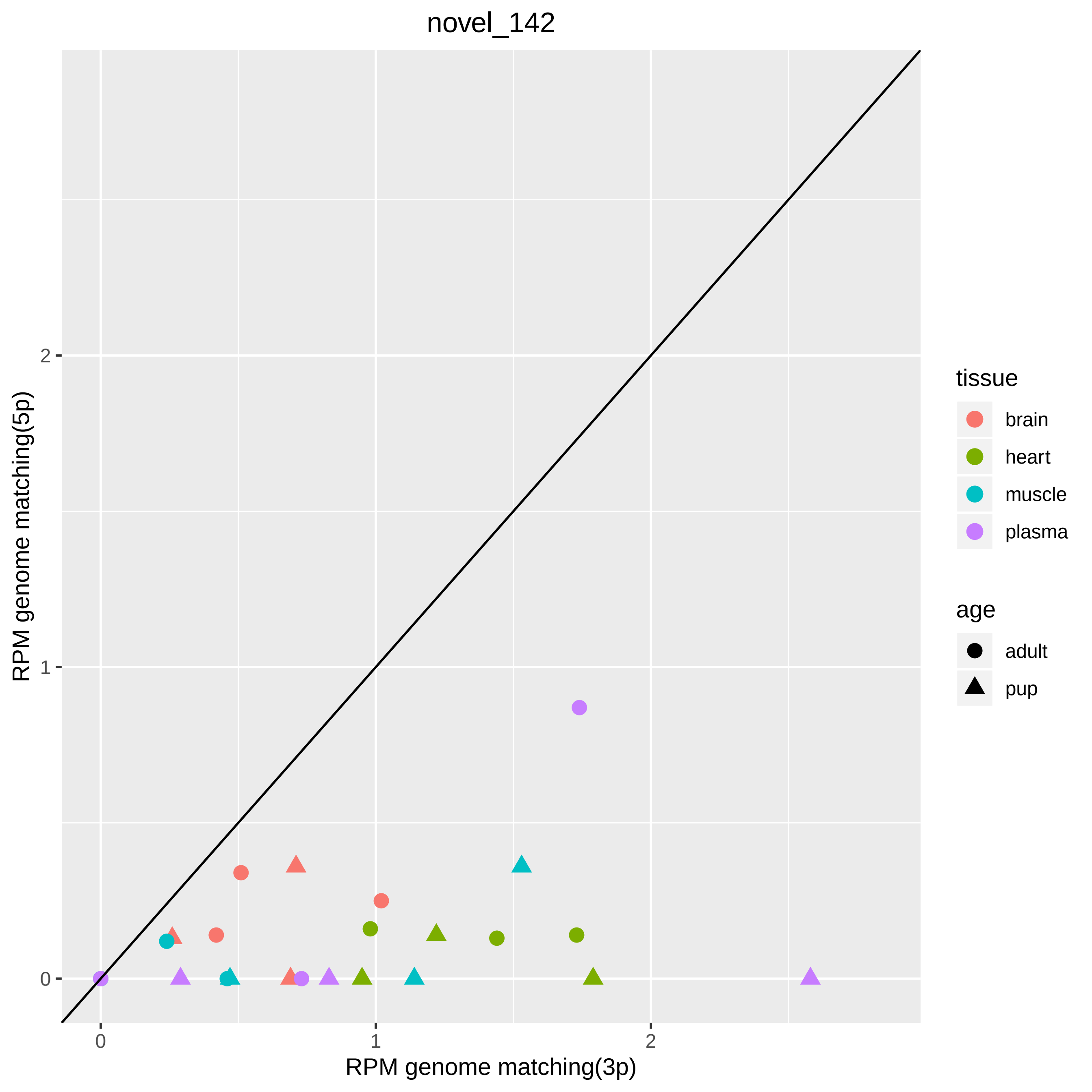
novel_142
RPM genome matching(5p)
RPM genome matching(3p)
0
1
2
0
1
2
tissue
brain
heart
muscle
plasma
age
adult
pup

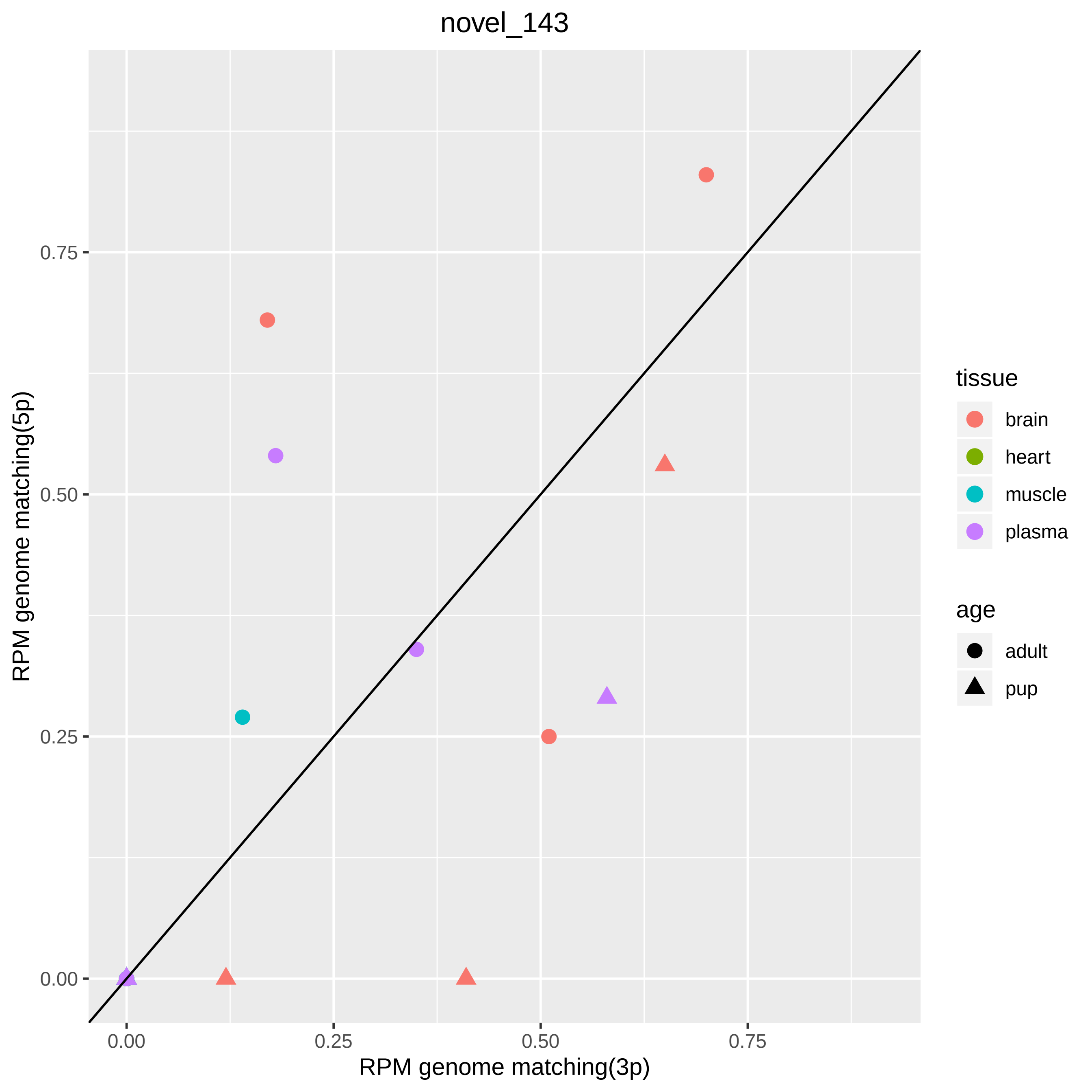

novel_143
RPM genome matching(5p)
RPM genome matching(3p)
0.00
0.25
0.50
0.75
tissue
brain
heart
muscle
plasma
age
adult
pup

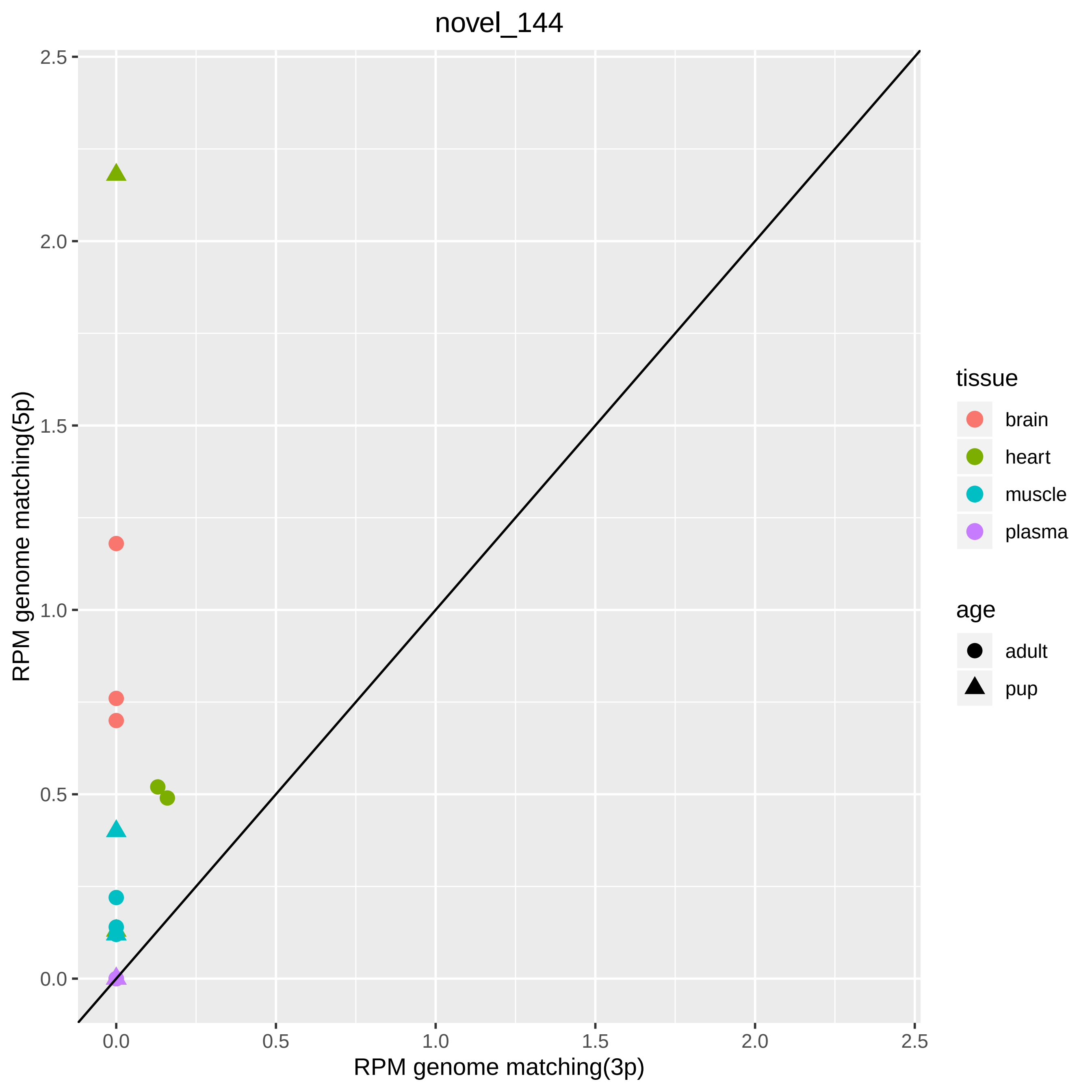

novel_144
RPM genome matching(5p)
RPM genome matching(3p)
0.0
0.5
1.0
1.5
2.0
2.5
tissue
brain
heart
muscle
plasma
age
adult
pup

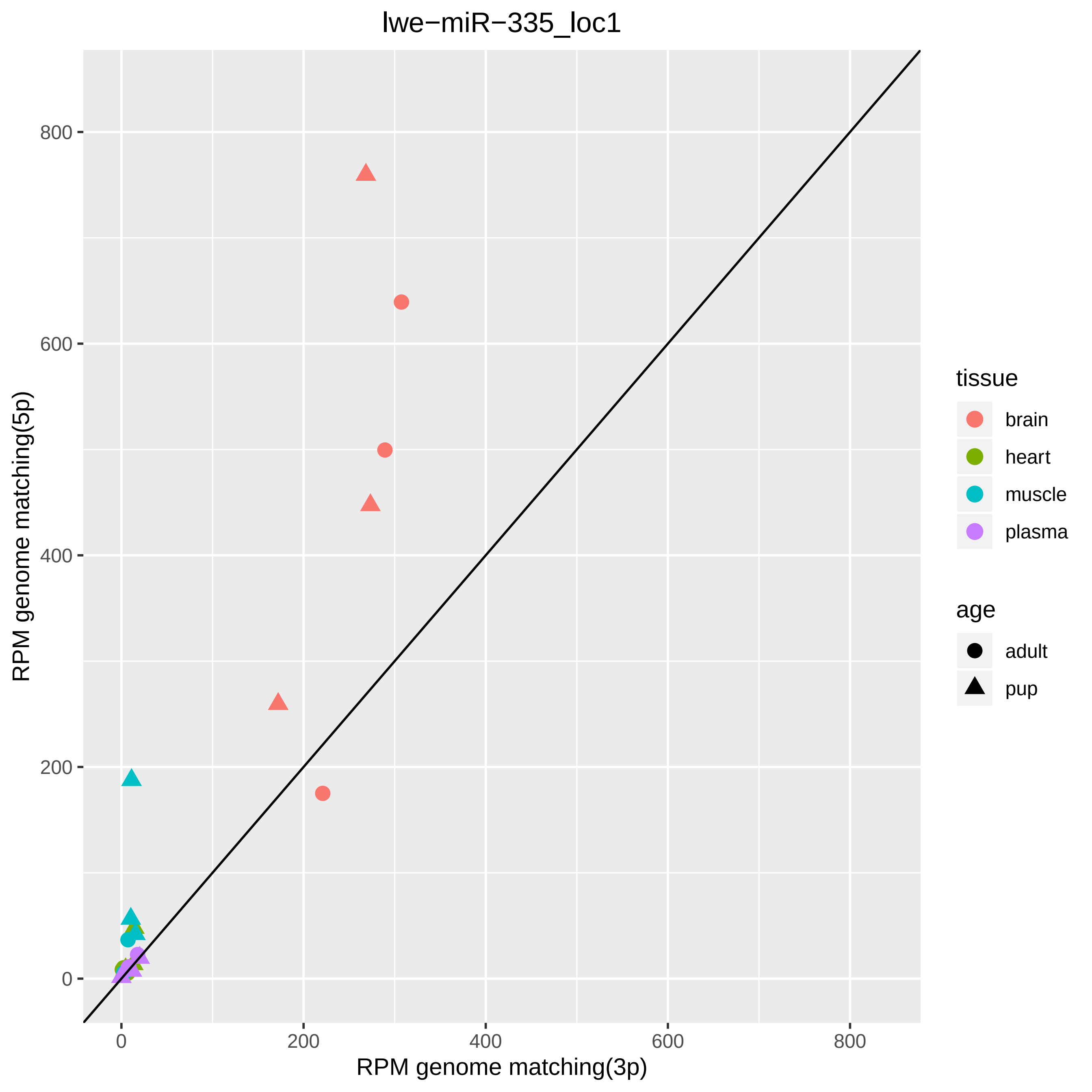

lwe−miR−335_loc1
RPM genome matching(5p)
RPM genome matching(3p)
0
200
400
600
800
tissue
brain
heart
muscle
plasma
age
adult
pup

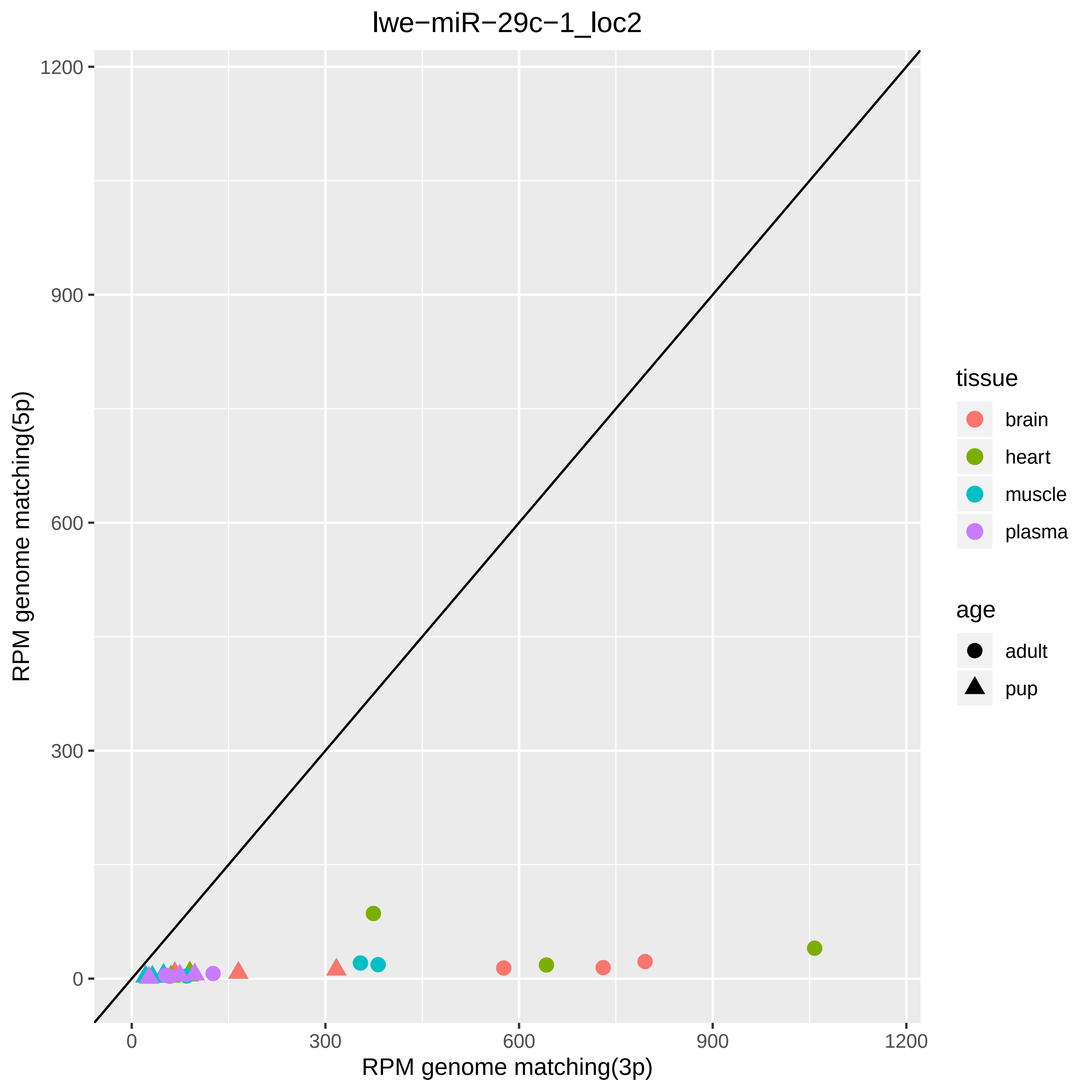

lwe−miR−29c−1_loc2
RPM genome matching(5p)
RPM genome matching(3p)
0
300
600
900
1200
0
300
600
900
1200
tissue
brain
heart
muscle
plasma
age
adult
pup

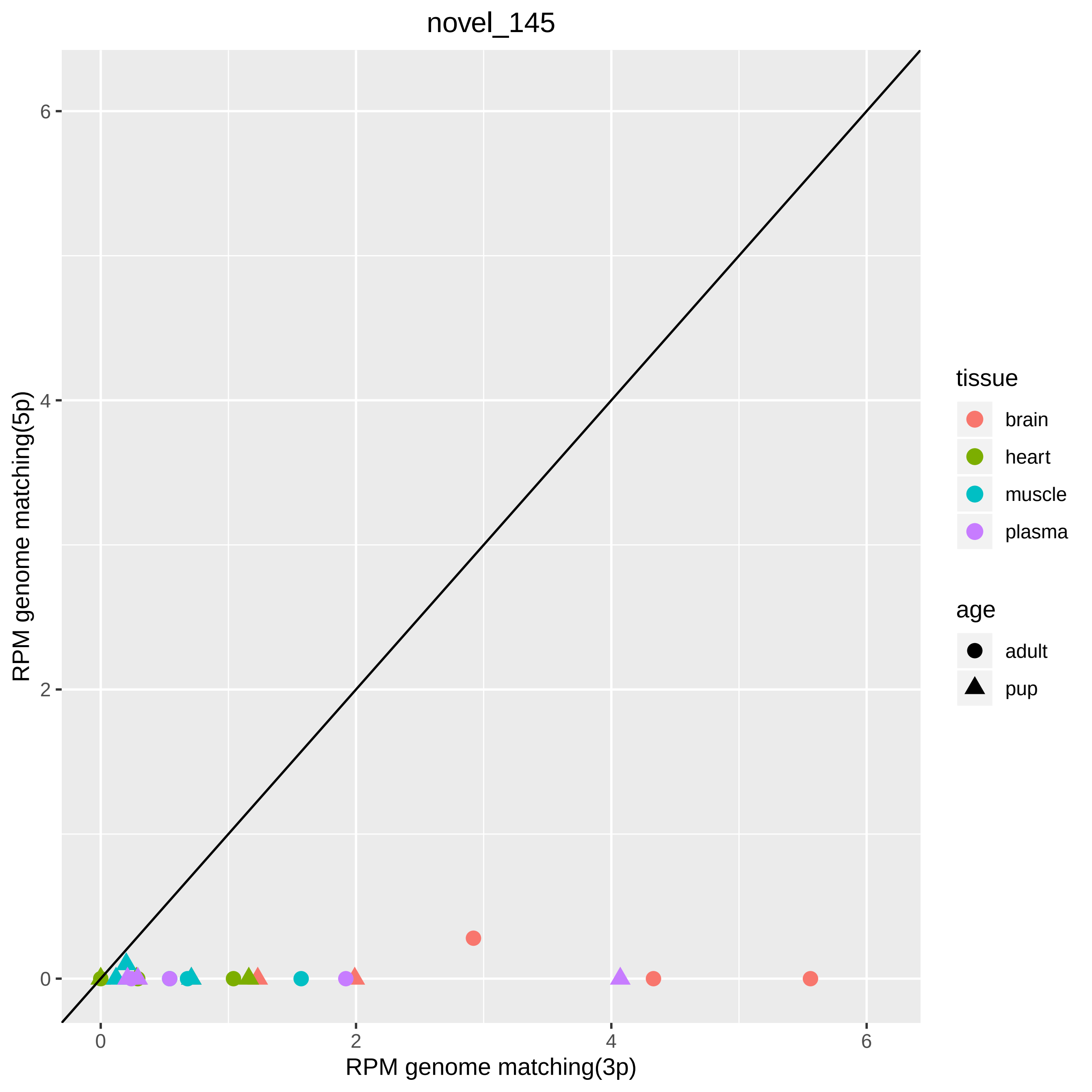

novel_145
RPM genome matching(5p)
RPM genome matching(3p)
0
2
4
6
0
2
4
6
tissue
brain
heart
muscle
plasma
age
adult
pup

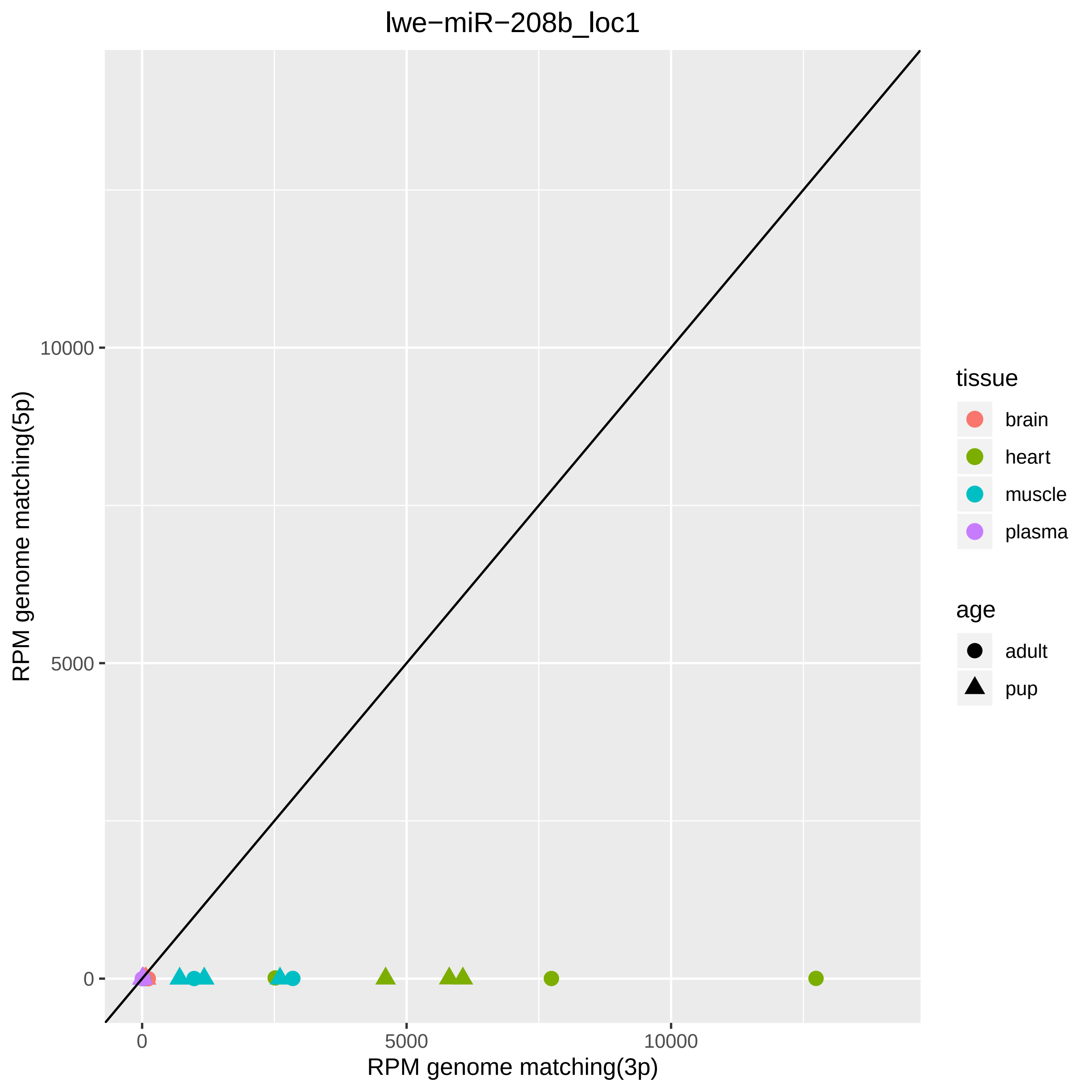

lwe−miR−208b_loc1
RPM genome matching(5p)
10000
5000
0
0
5000
10000
RPM genome matching(3p)
tissue
brain
heart
muscle
plasma
age
adult
pup

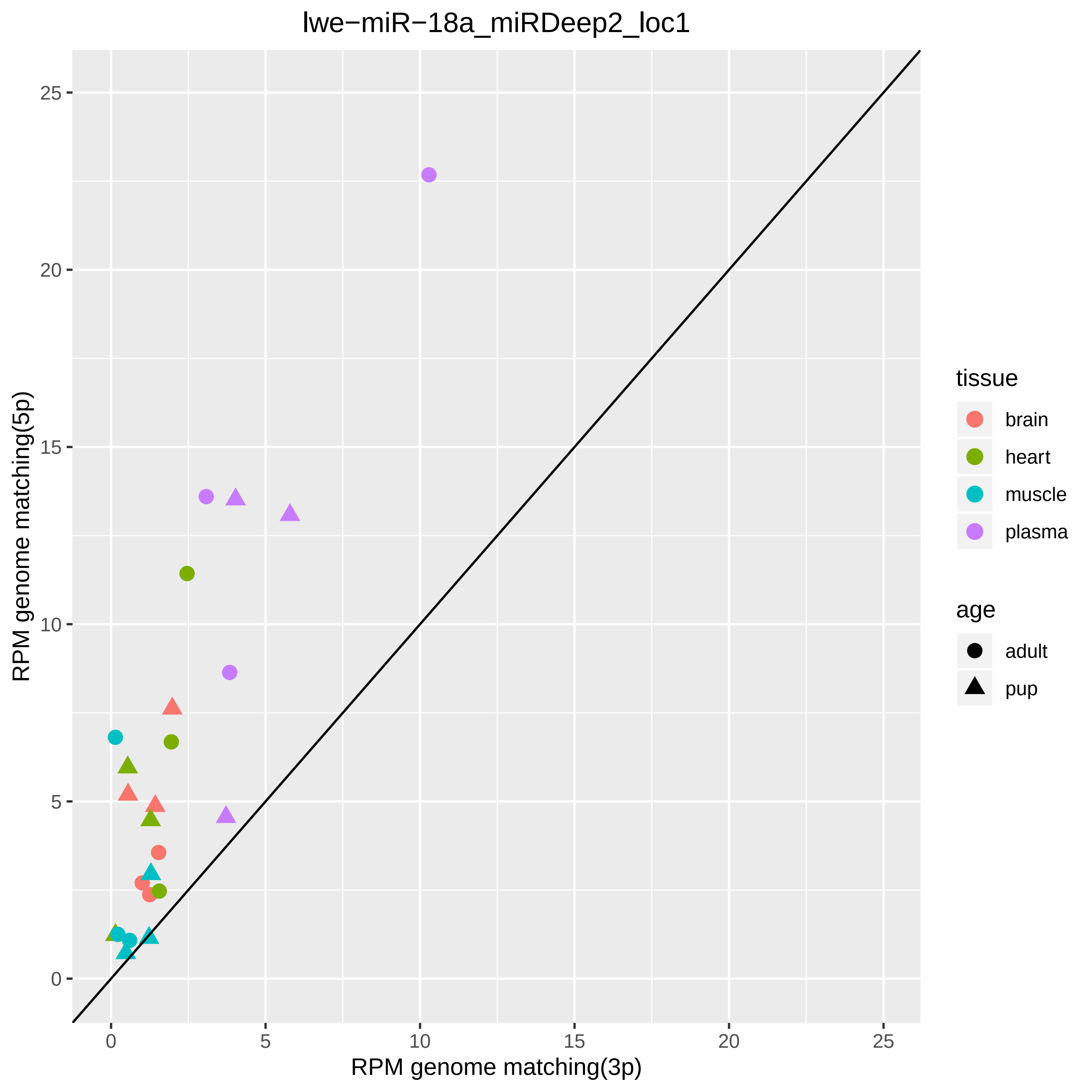
lwe−miR−18a_miRDeep2_loc1
RPM genome matching(5p)
RPM genome matching(3p)
0
5
10
15
20
25
tissue
brain
heart
muscle
plasma
age
adult
pup

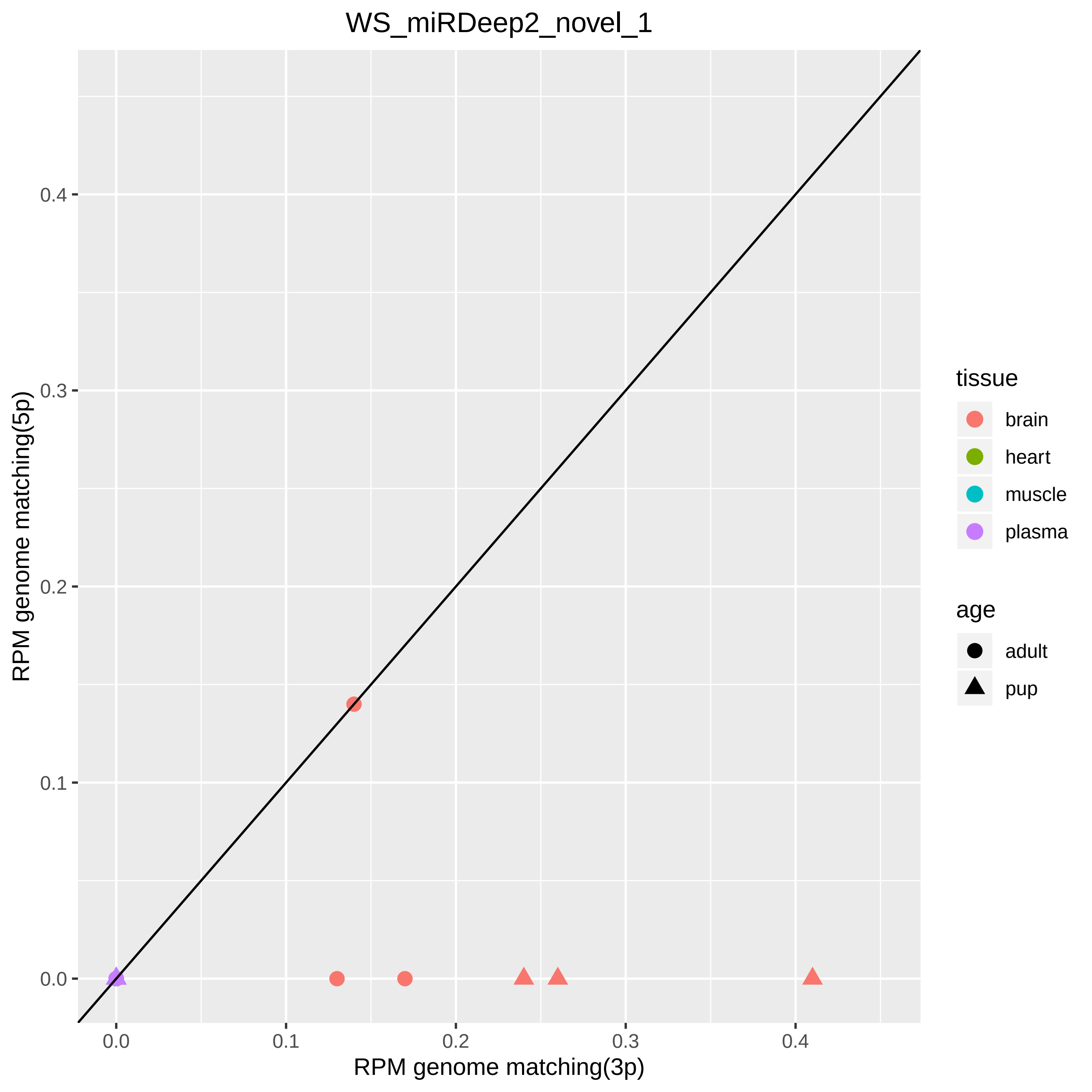
WS_miRDeep2_novel_1
RPM genome matching(5p)
RPM genome matching(3p)
0.0
0.1
0.2
0.3
0.4
tissue
brain
heart
muscle
plasma
age
adult
pup

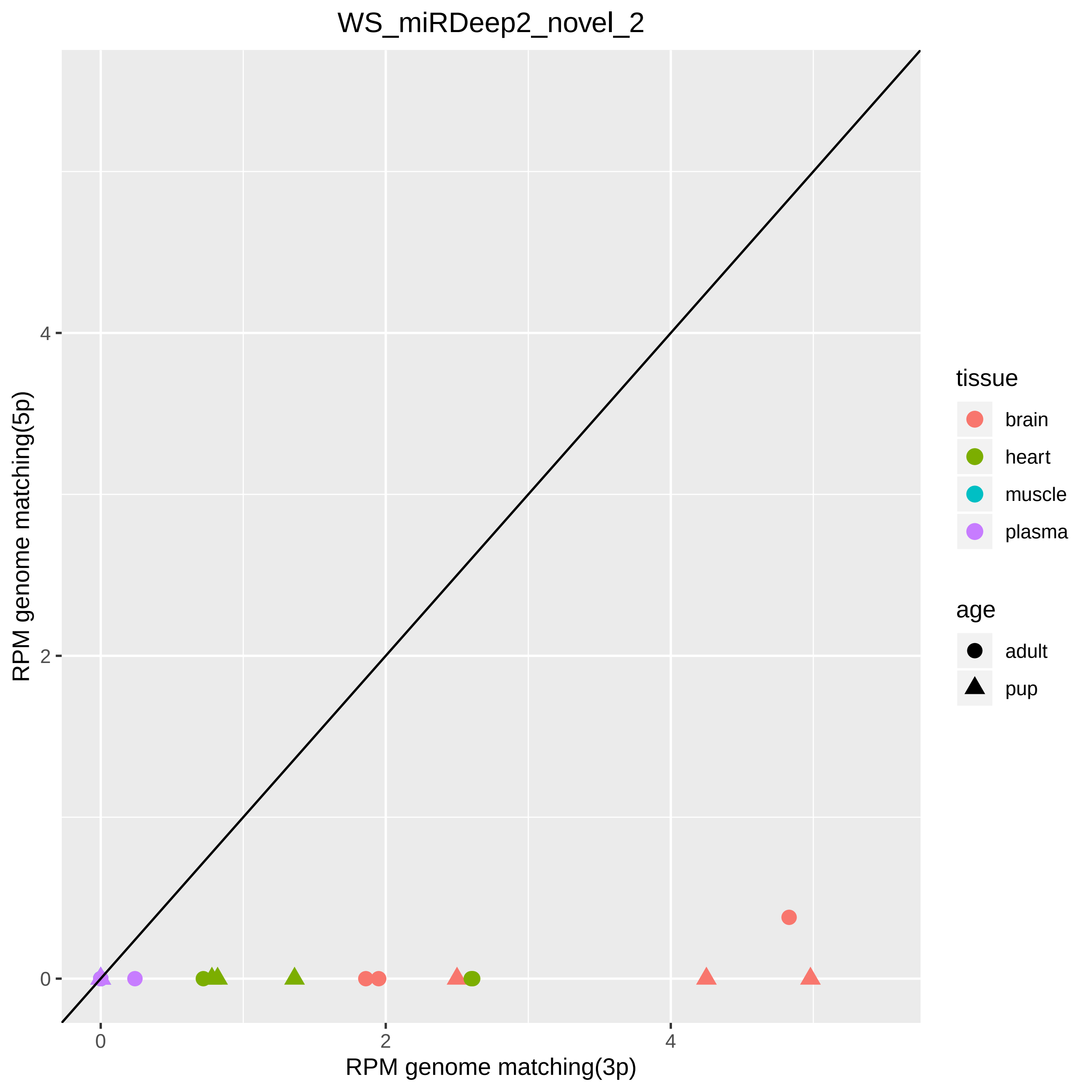
WS_miRDeep2_novel_2
RPM genome matching(5p)
RPM genome matching(3p)
0
2
4
tissue
brain
heart
muscle
plasma
age
adult
pup

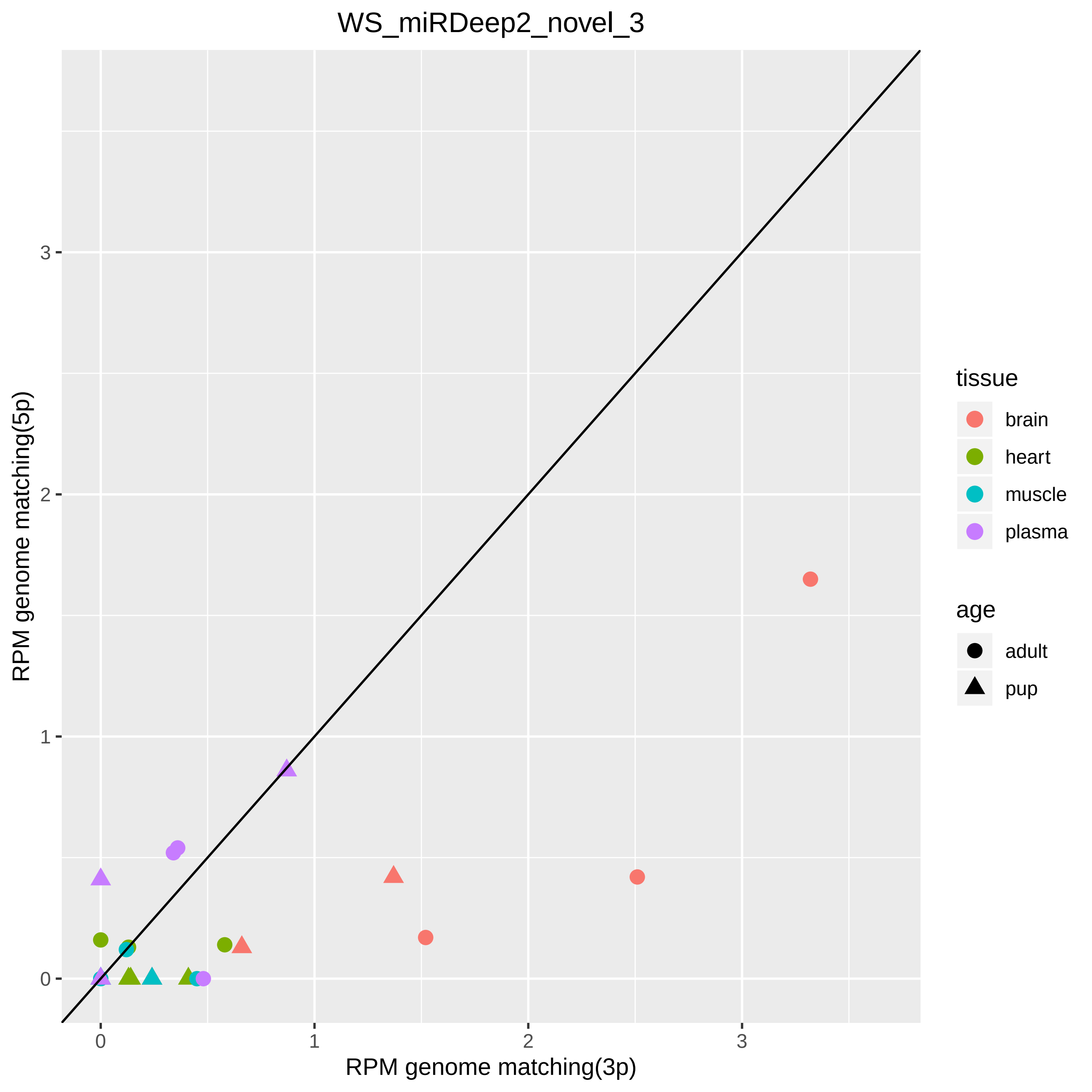
WS_miRDeep2_novel_3
RPM genome matching(5p)
RPM genome matching(3p)
0
1
2
3
tissue
brain
heart
muscle
plasma
age
adult
pup

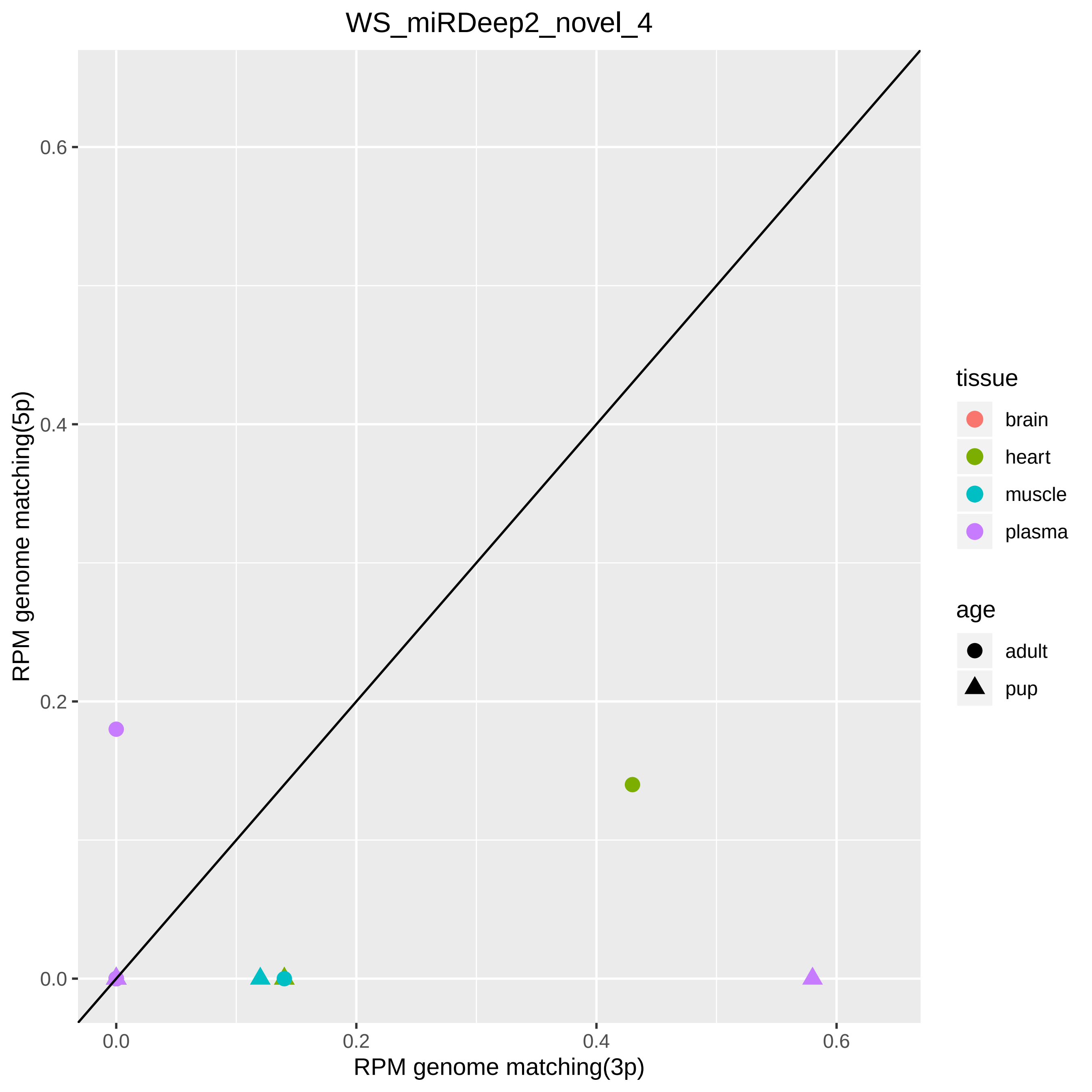

WS_miRDeep2_novel_4
RPM genome matching(5p)
RPM genome matching(3p)
0.0
0.2
0.4
0.6
tissue
brain
heart
muscle
plasma
age
adult
pup

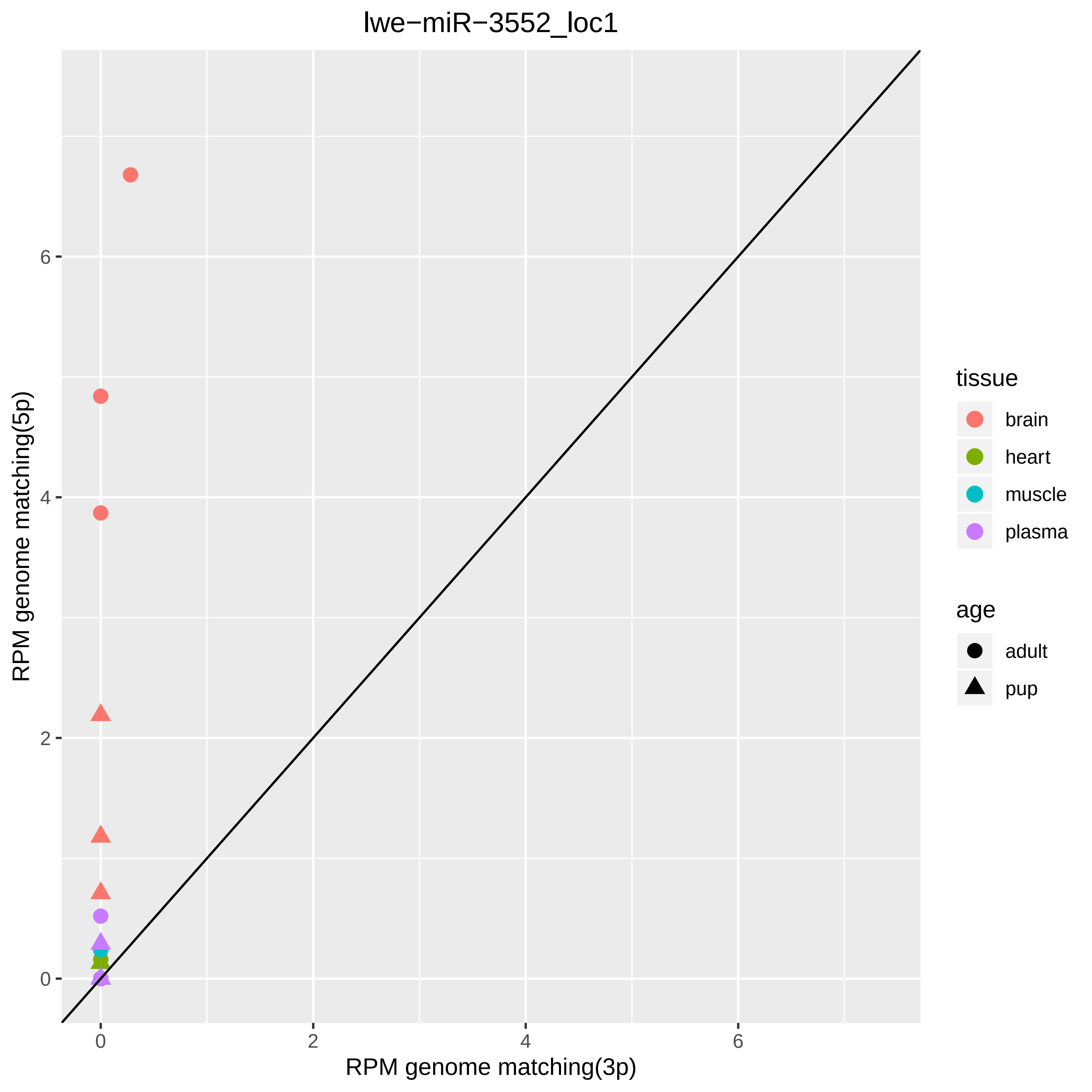
lwe−miR−3552_loc1
RPM genome matching(5p)
RPM genome matching(3p)
0
2
4
6
tissue
brain
heart
muscle
plasma
age
adult
pup

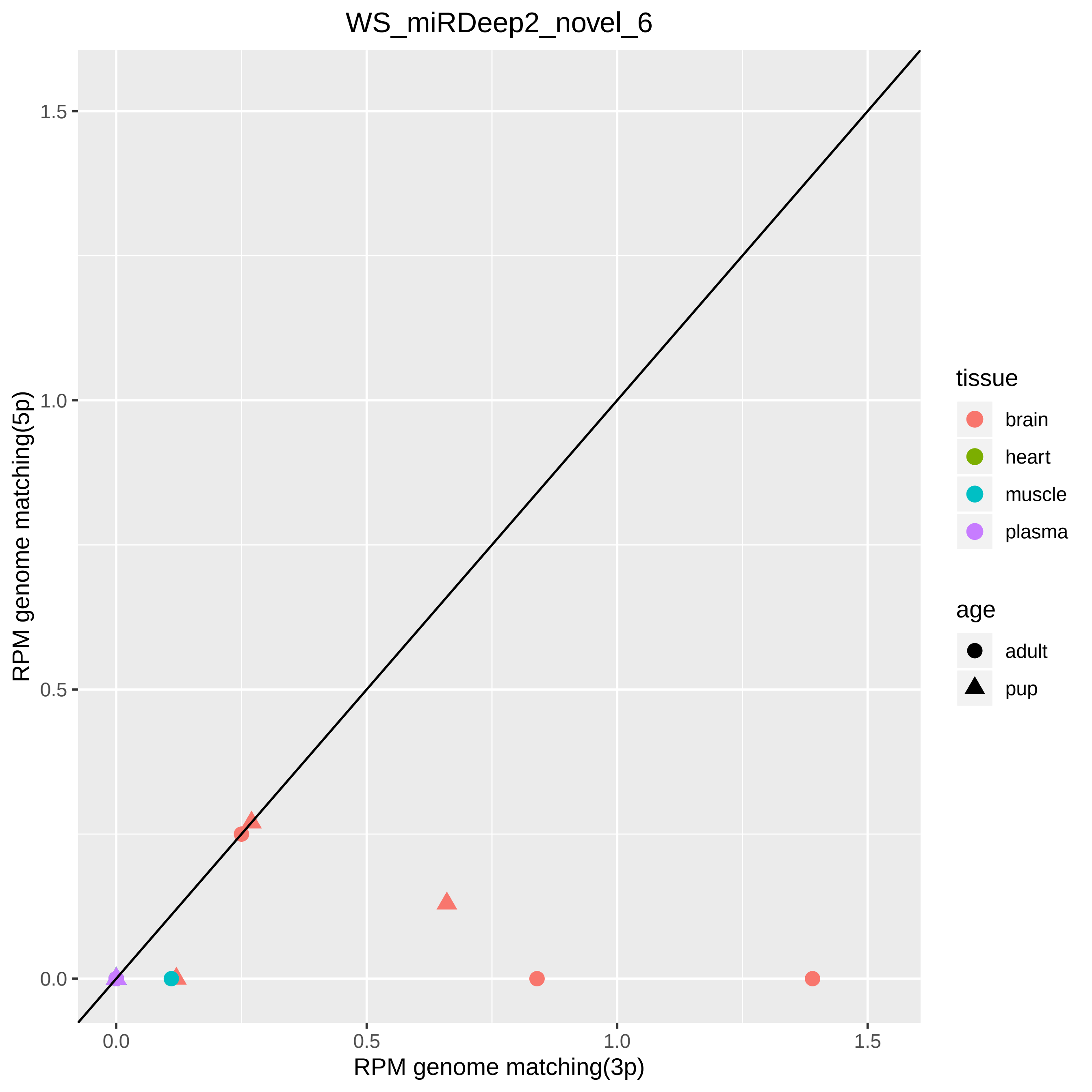

WS_miRDeep2_novel_6
RPM genome matching(5p)
RPM genome matching(3p)
0.0
0.5
1.0
1.5
tissue
brain
heart
muscle
plasma
age
adult
pup

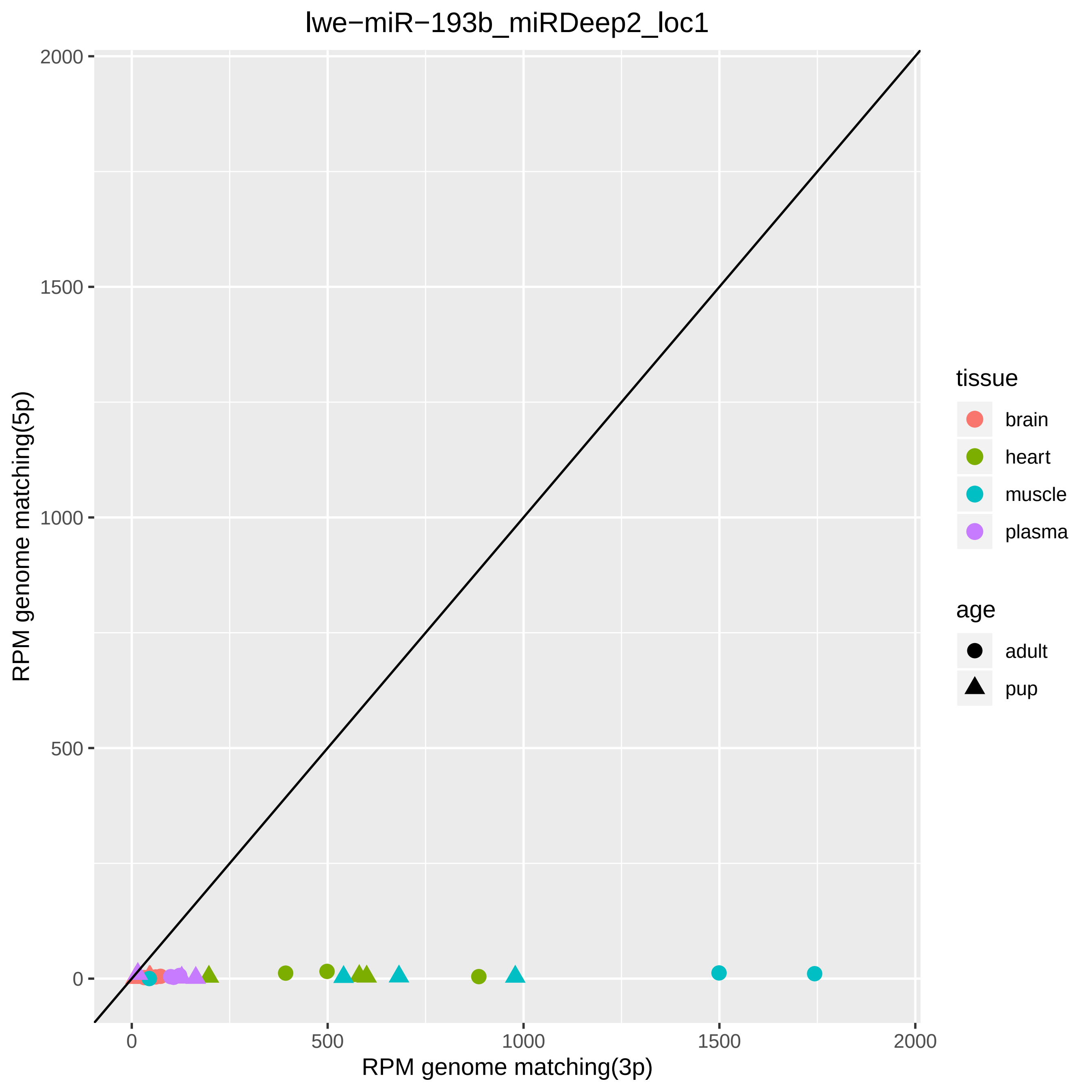

lwe−miR−193b_miRDeep2_loc1
2000
1500
1000
500
0
RPM genome matching(5p)
0
500
1000
1500
2000
RPM genome matching(3p)
tissue
brain
heart
muscle
plasma
age
adult
pup

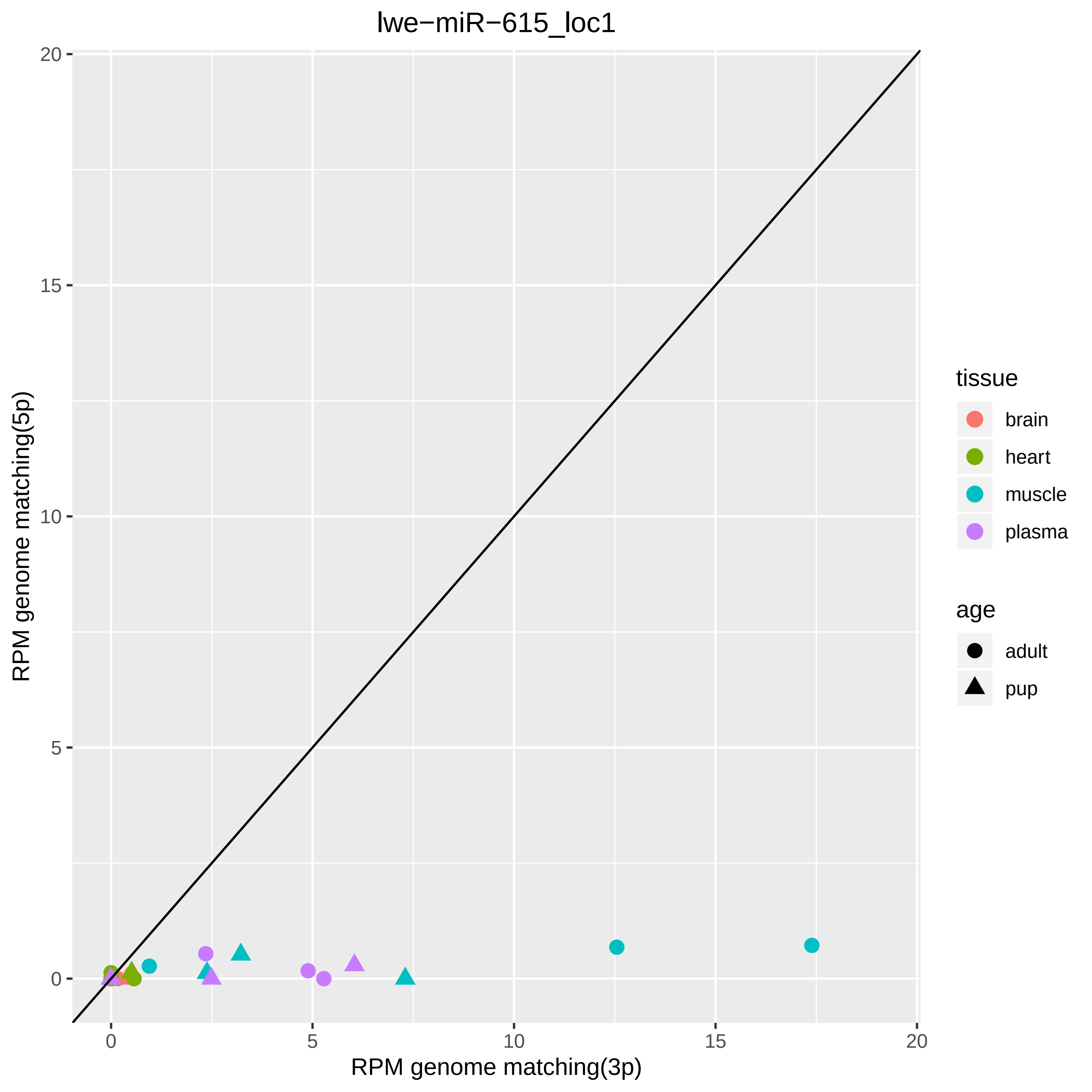

lwe−miR−615_loc1
RPM genome matching(5p)
RPM genome matching(3p)
0
5
10
15
20
tissue
brain
heart
muscle
plasma
age
adult
pup

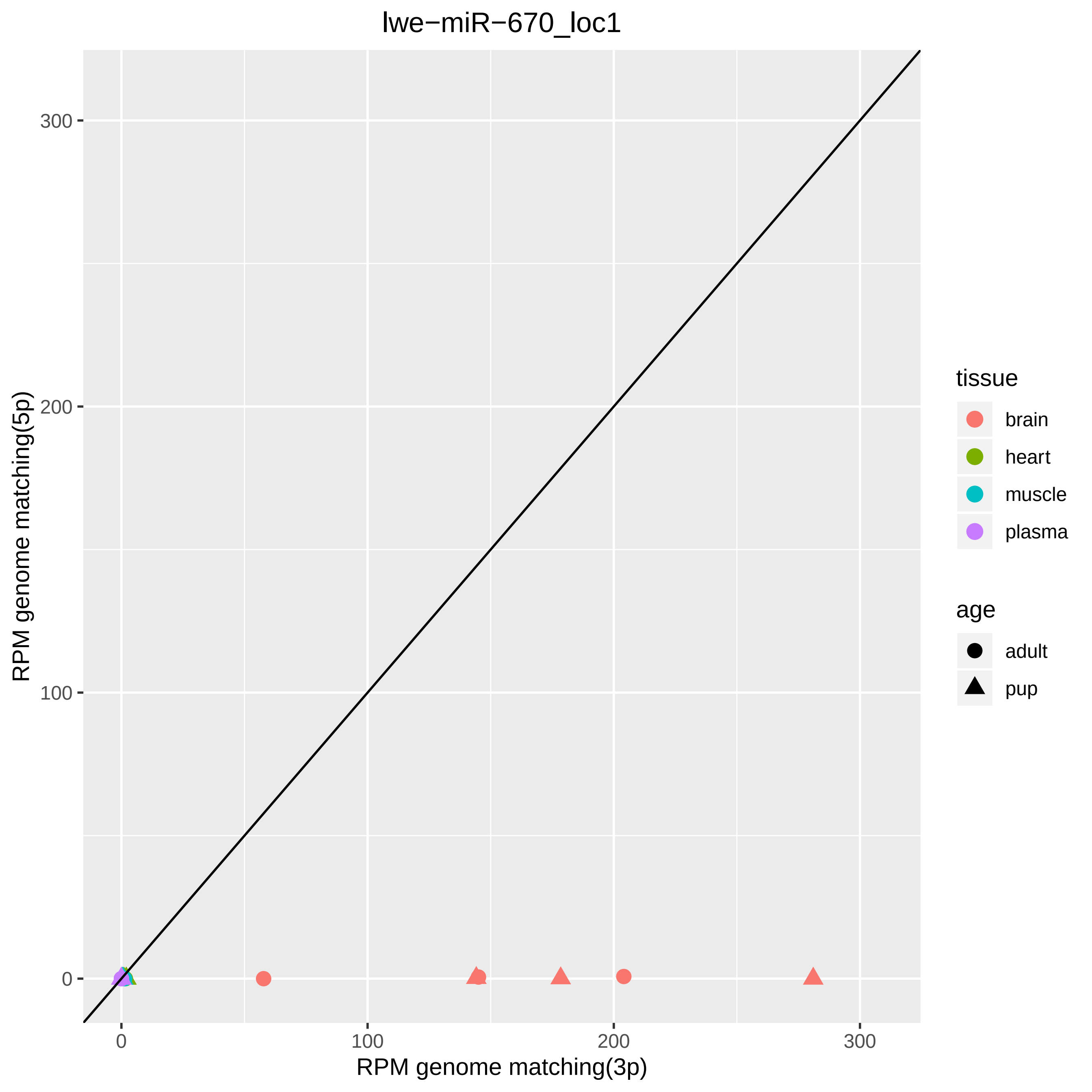
lwe−miR−670_loc1
300
200
100
0
0
100
200
300
RPM genome matching(5p)
RPM genome matching(3p)
tissue
brain
heart
muscle
plasma
age
adult
pup

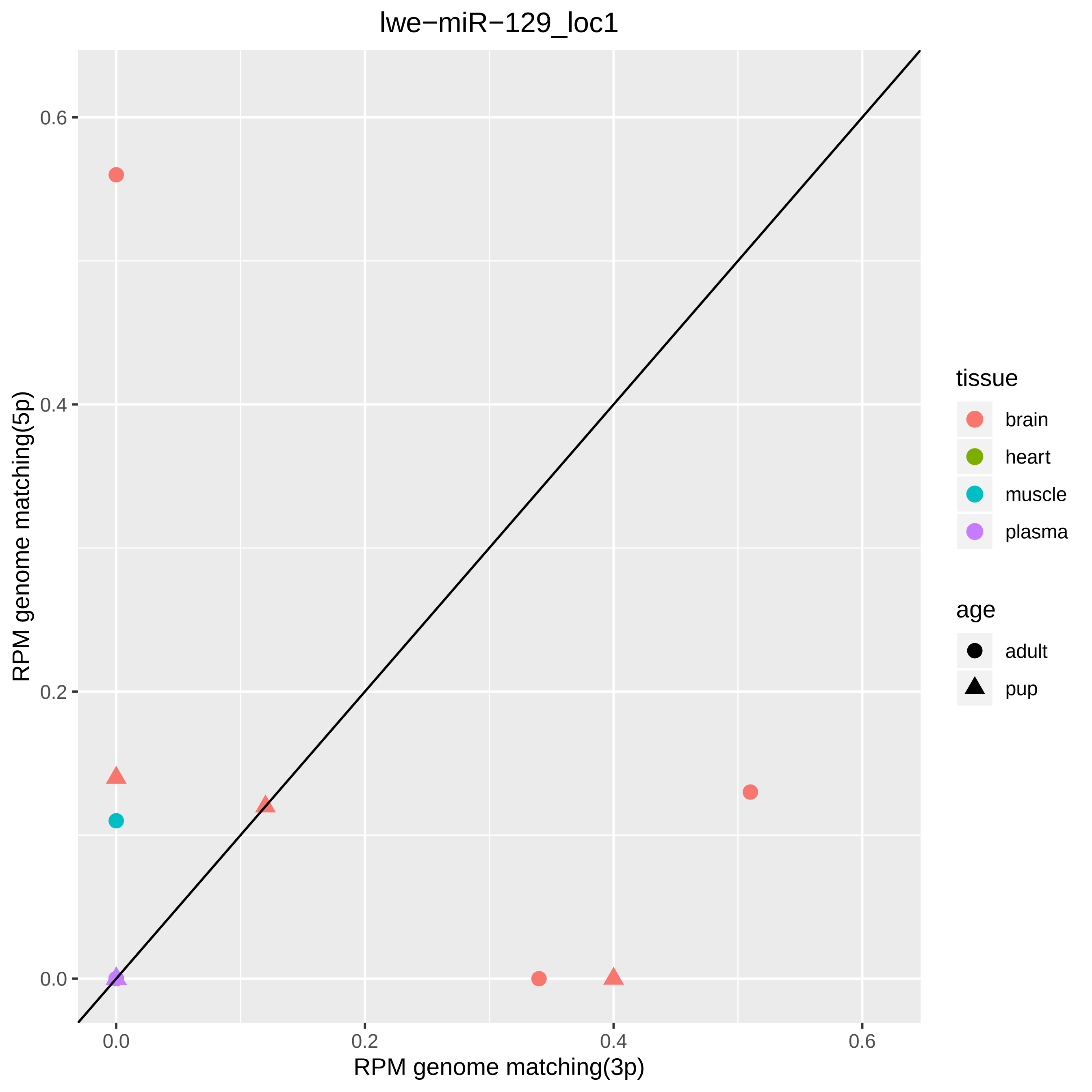

lwe−miR−129_loc1
RPM genome matching(5p)
RPM genome matching(3p)
0.0
0.2
0.4
0.6
0.0
0.2
0.4
0.6
tissue
brain
heart
muscle
plasma
age
adult
pup

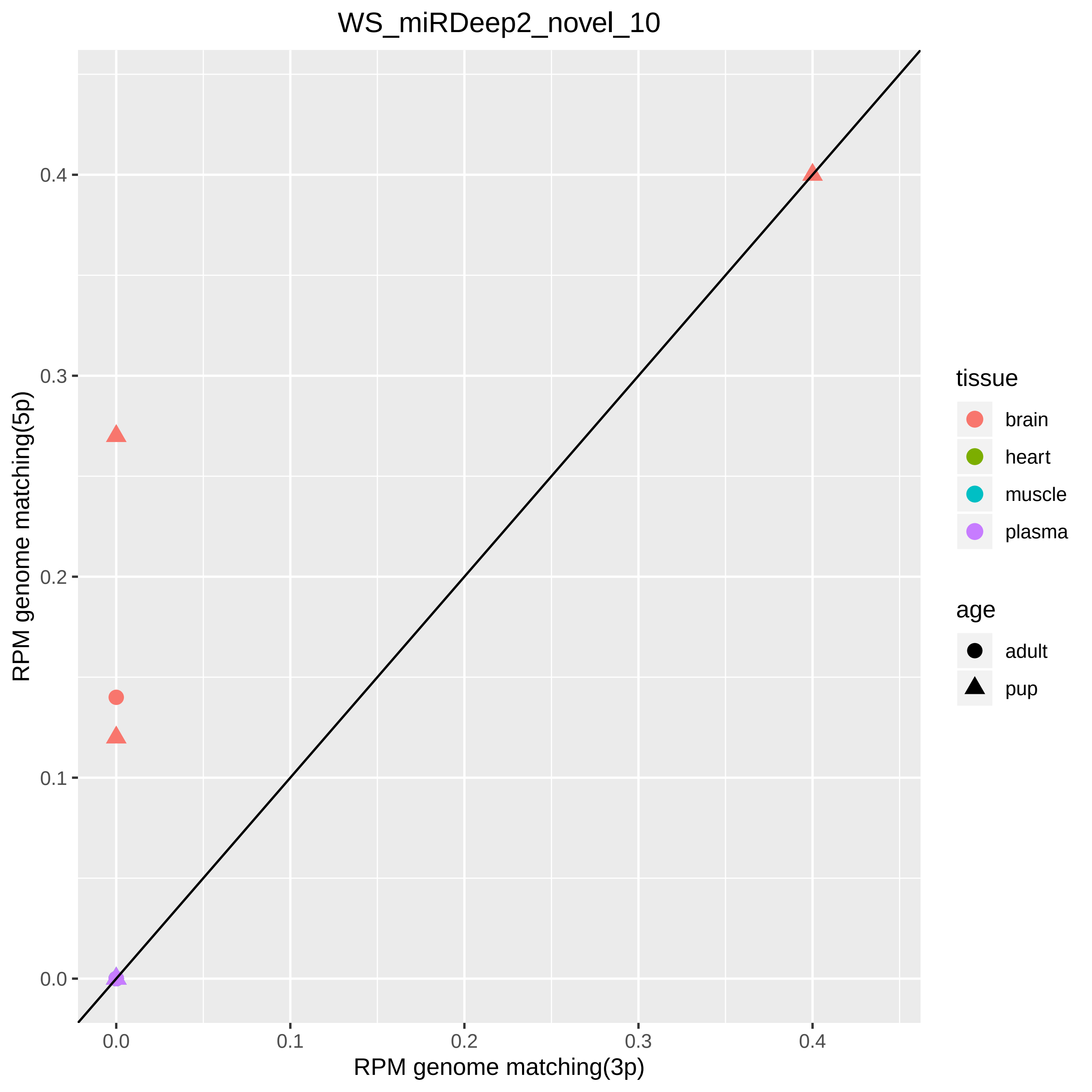

WS_miRDeep2_novel_10
RPM genome matching(5p)
RPM genome matching(3p)
0.0
0.1
0.2
0.3
0.4
tissue
brain
heart
muscle
plasma
age
adult
pup

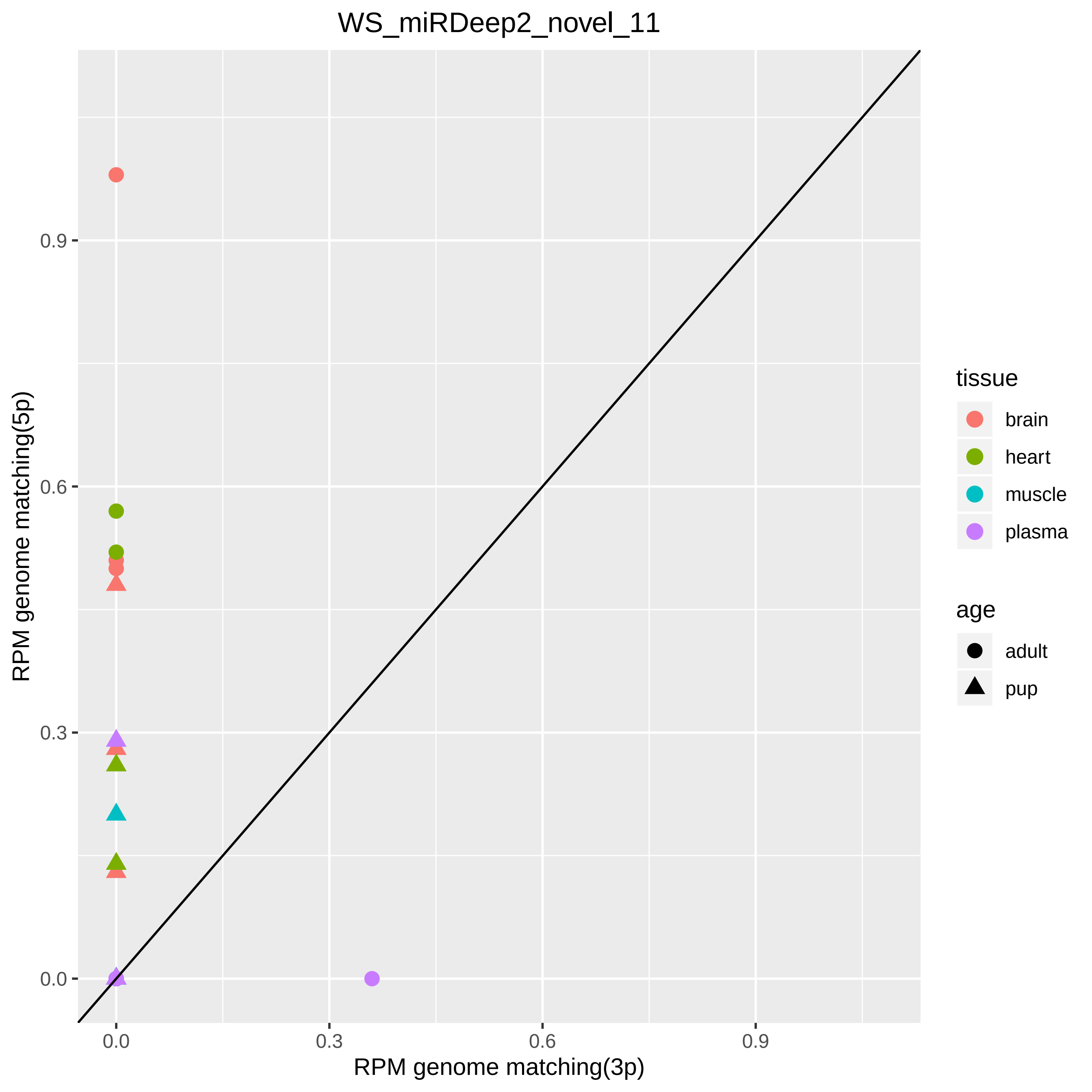

WS_miRDeep2_novel_11
RPM genome matching(5p)
RPM genome matching(3p)
0.0
0.3
0.6
0.9
0.0
0.3
0.6
0.9
tissue
brain
heart
muscle
plasma
age
adult
pup

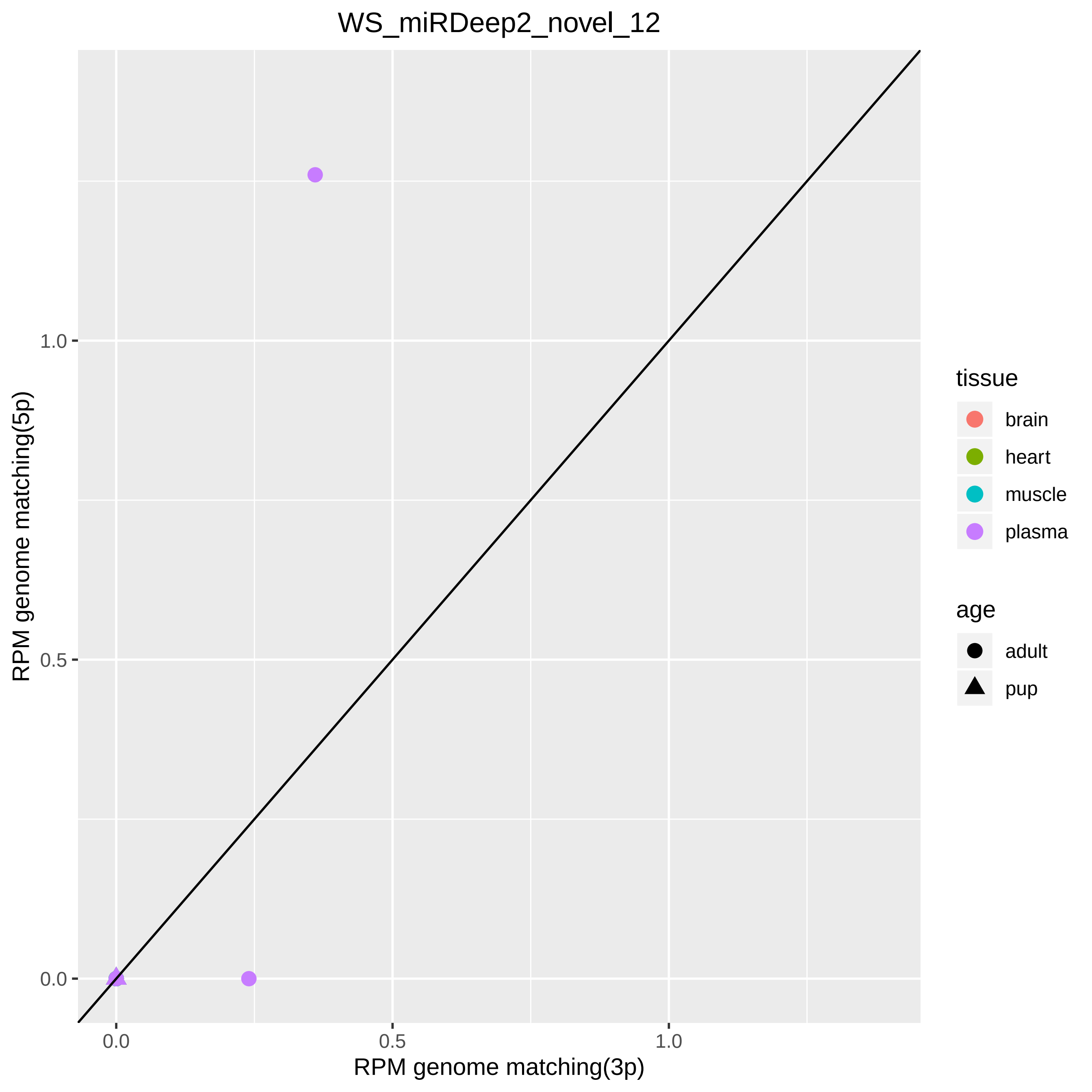

WS_miRDeep2_novel_12
RPM genome matching(5p)
RPM genome matching(3p)
1.0
0.5
0.0
0.0
0.5
1.0
tissue
brain
heart
muscle
plasma
age
adult
pup

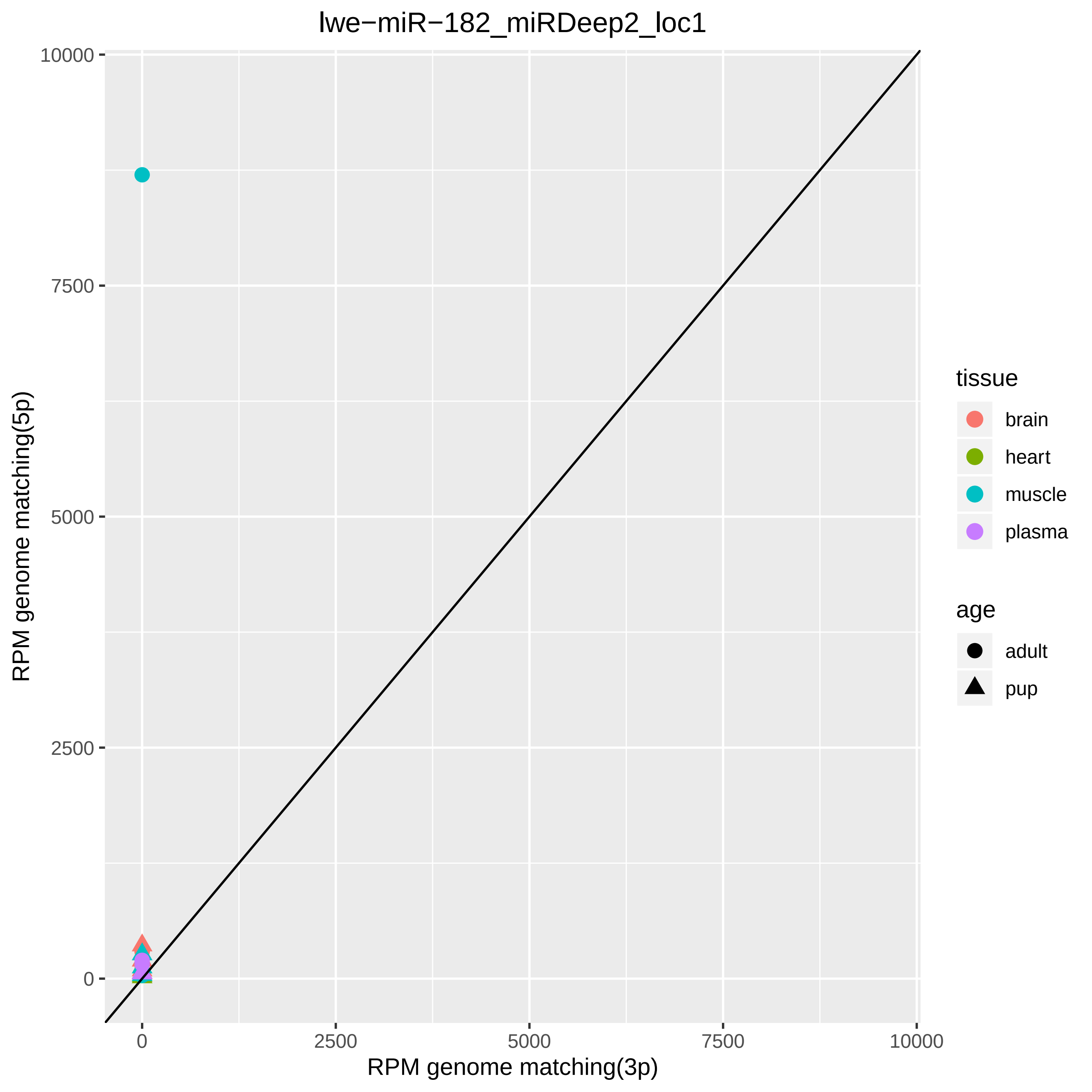
lwe−miR−182_miRDeep2_loc1
10000
7500
5000
2500
0
0
2500
5000
7500
10000
RPM genome matching(5p)
RPM genome matching(3p)
tissue
brain
heart
muscle
plasma
age
adult
pup

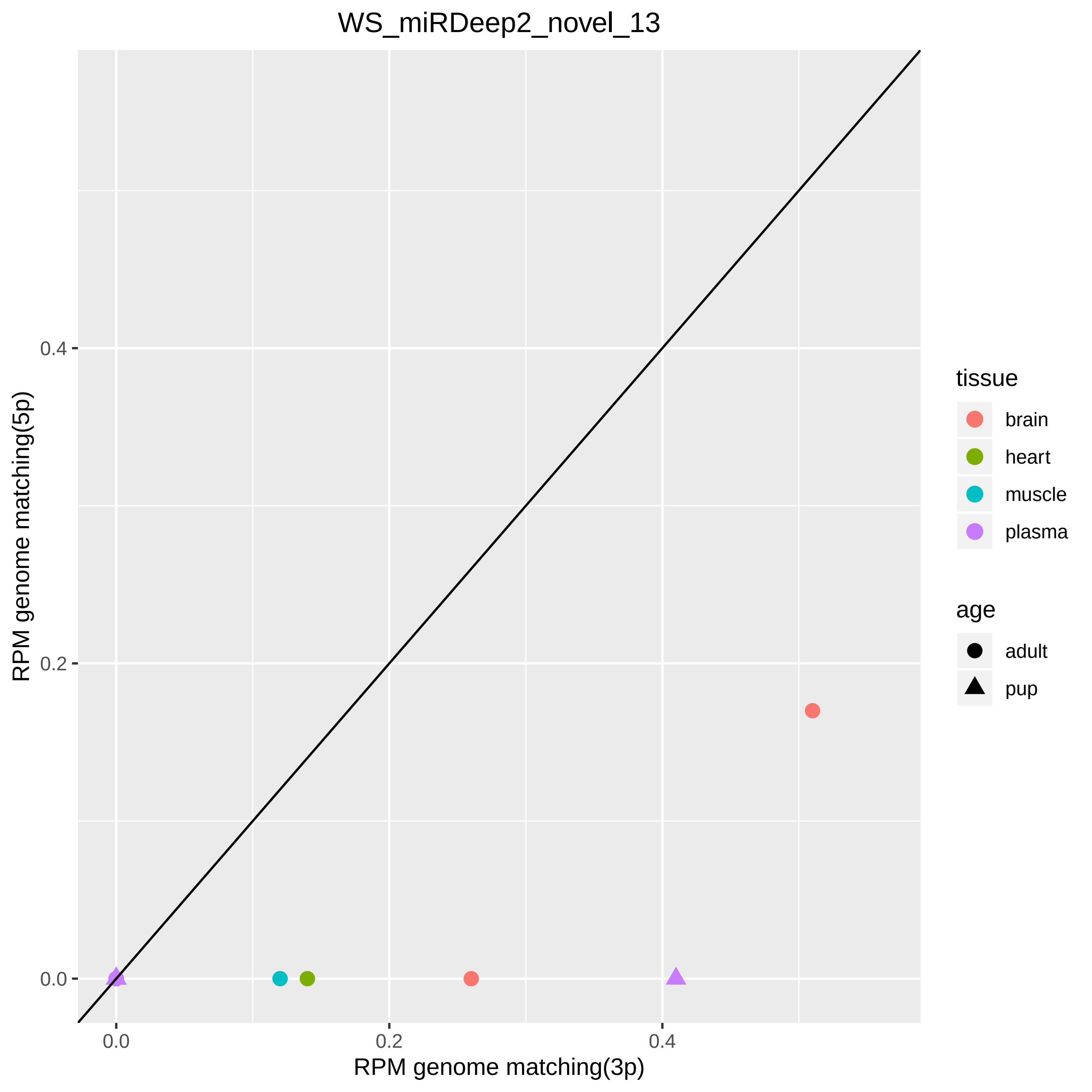

WS_miRDeep2_novel_13
RPM genome matching(5p)
RPM genome matching(3p)
0.0
0.2
0.4
0.0
0.2
0.4
tissue
brain
heart
muscle
plasma
age
adult
pup

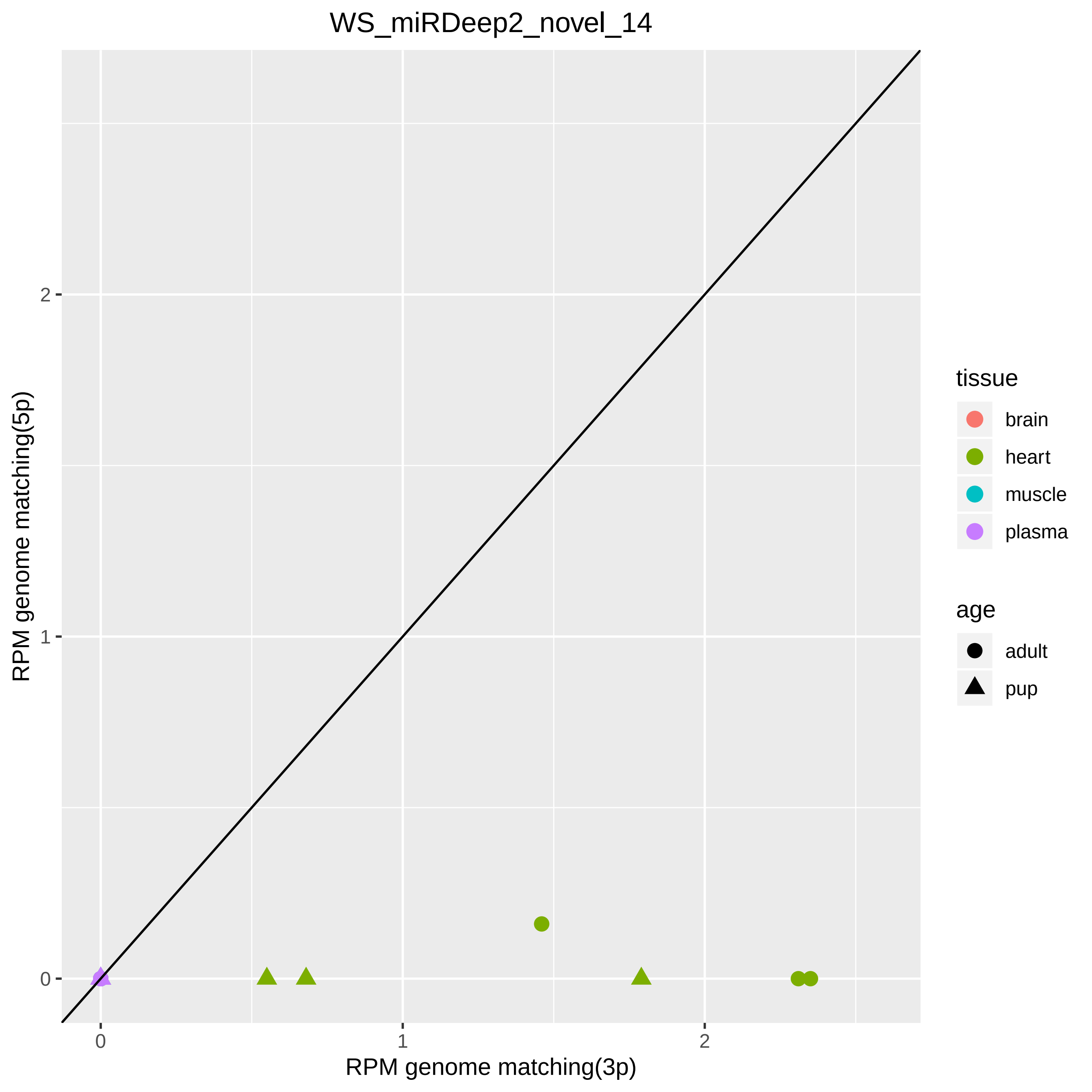

WS_miRDeep2_novel_14
RPM genome matching(5p)
RPM genome matching(3p)
0
1
2
0
1
2
tissue
brain
heart
muscle
plasma
age
adult
pup

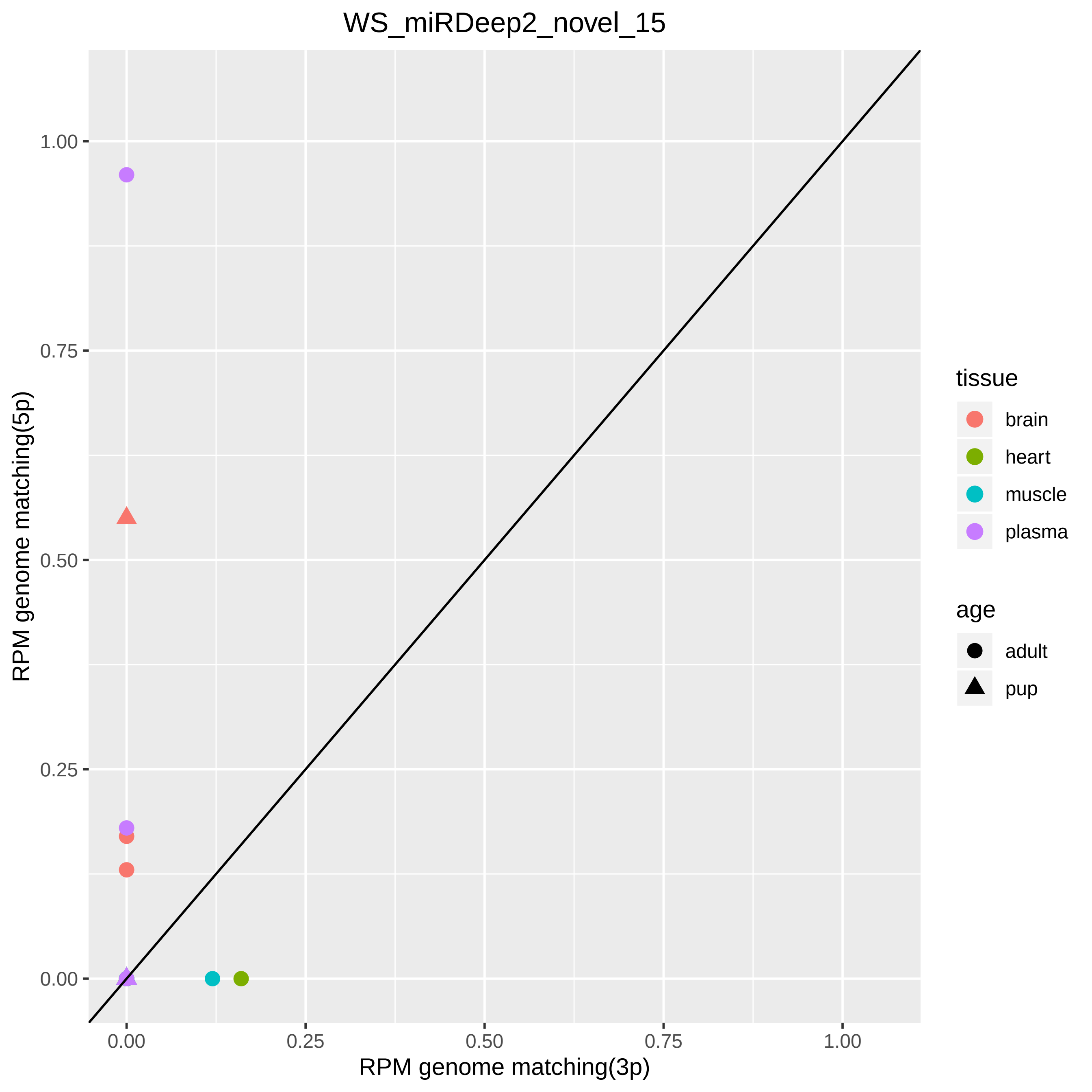
WS_miRDeep2_novel_15
RPM genome matching(5p)
RPM genome matching(3p)
1.00
0.75
0.50
0.25
0.00
0.00
0.25
0.50
0.75
1.00
tissue
brain
heart
muscle
plasma
age
adult
pup

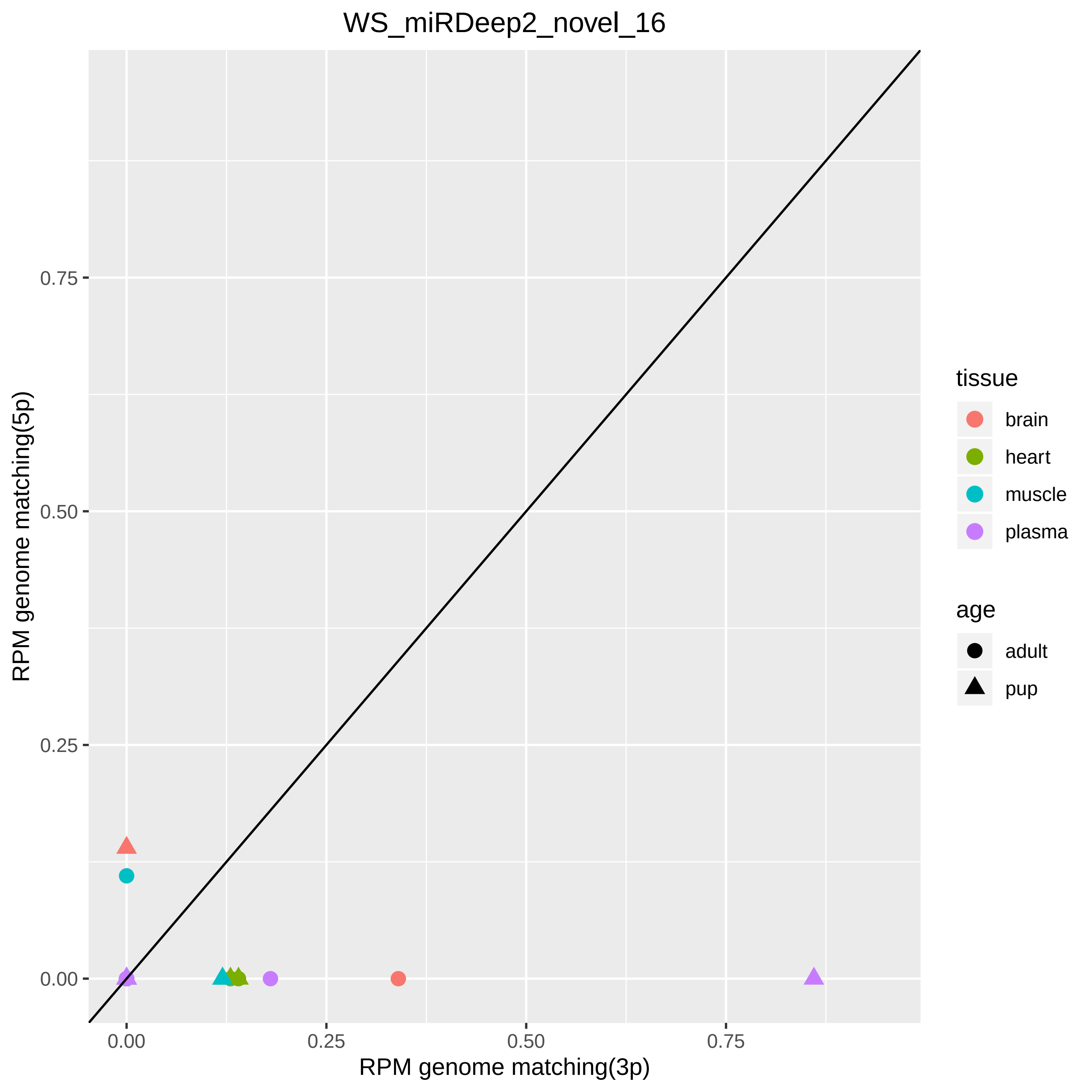

WS_miRDeep2_novel_16
RPM genome matching(5p)
RPM genome matching(3p)
0.00
0.25
0.50
0.75
tissue
brain
heart
muscle
plasma
age
adult
pup

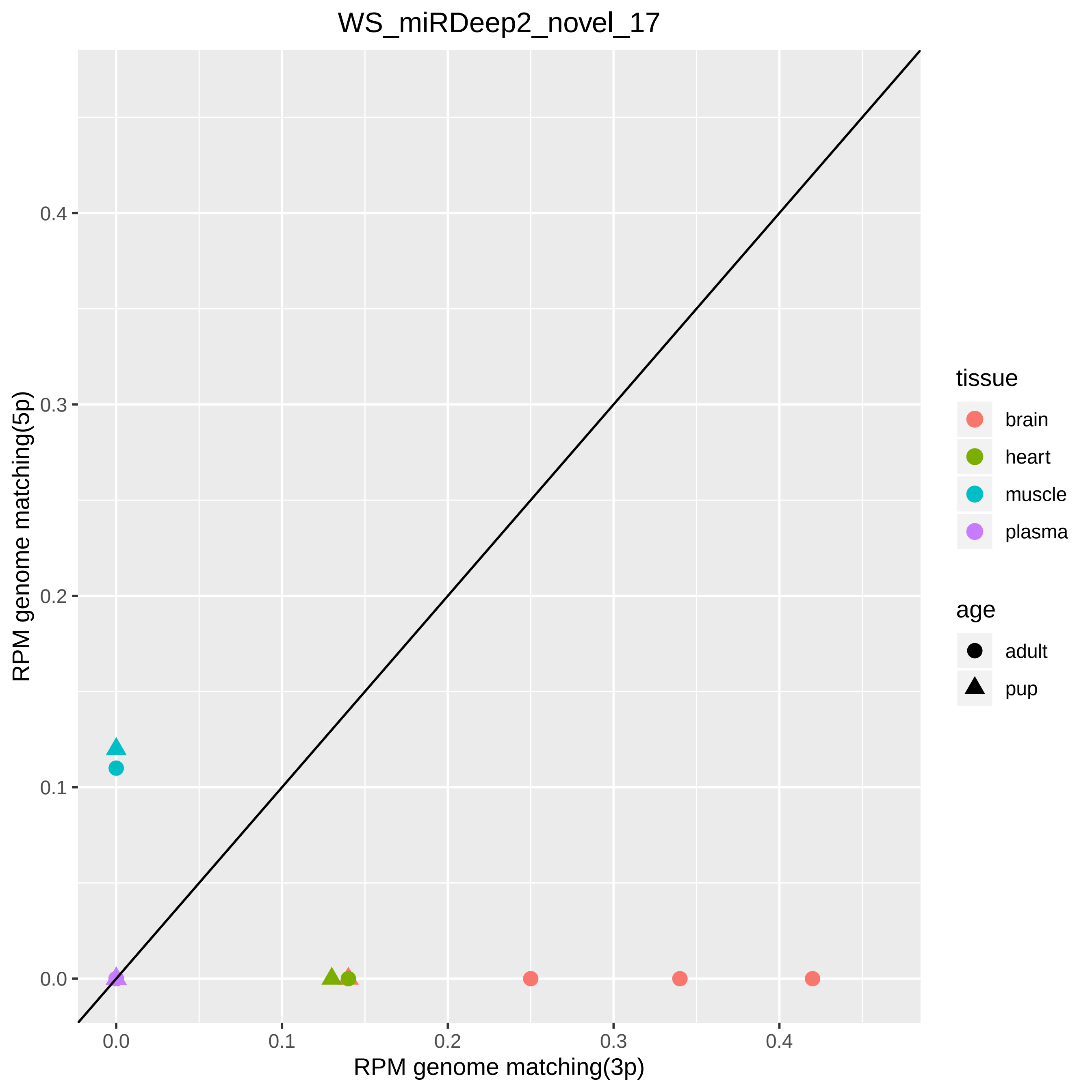
WS_miRDeep2_novel_17
RPM genome matching(5p)
RPM genome matching(3p)
0.0
0.1
0.2
0.3
0.4
tissue
brain
heart
muscle
plasma
age
adult
pup

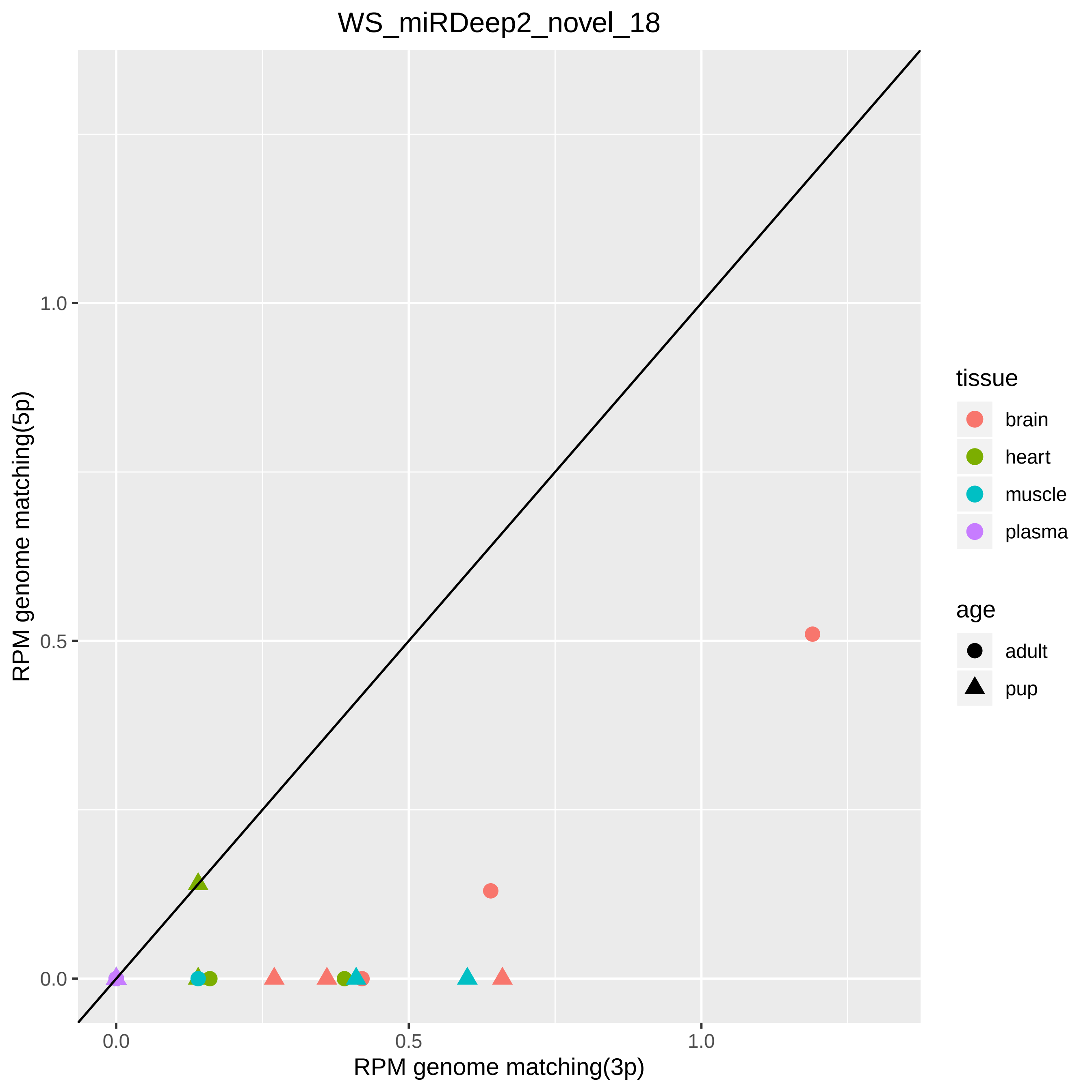

WS_miRDeep2_novel_18
RPM genome matching(5p)
RPM genome matching(3p)
0.0
0.5
1.0
tissue
brain
heart
muscle
plasma
age
adult
pup

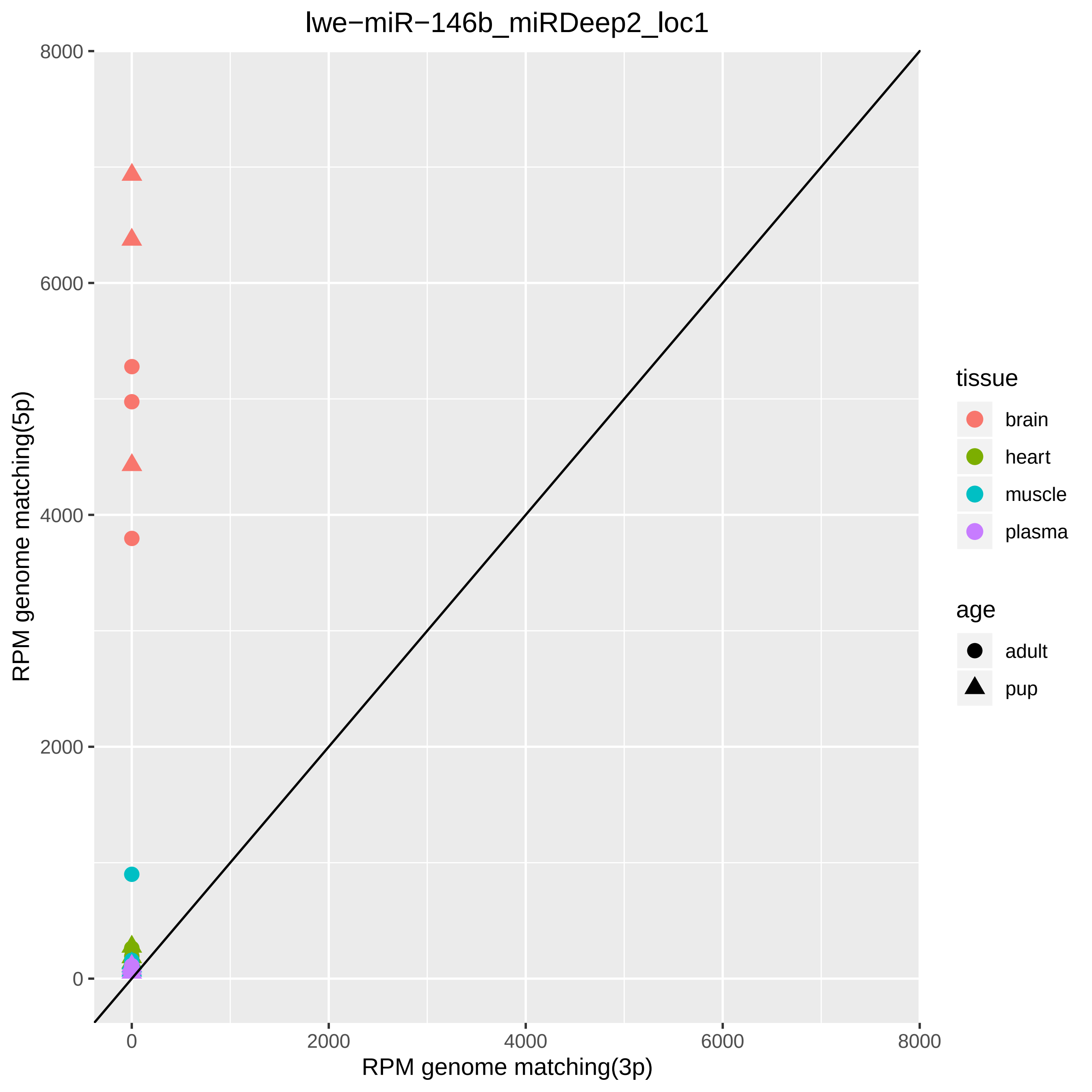
lwe−miR−146b_miRDeep2_loc1
8000
6000
4000
2000
0
RPM genome matching(5p)
0
2000
4000
6000
8000
RPM genome matching(3p)
tissue
brain
heart
muscle
plasma
age
adult
pup

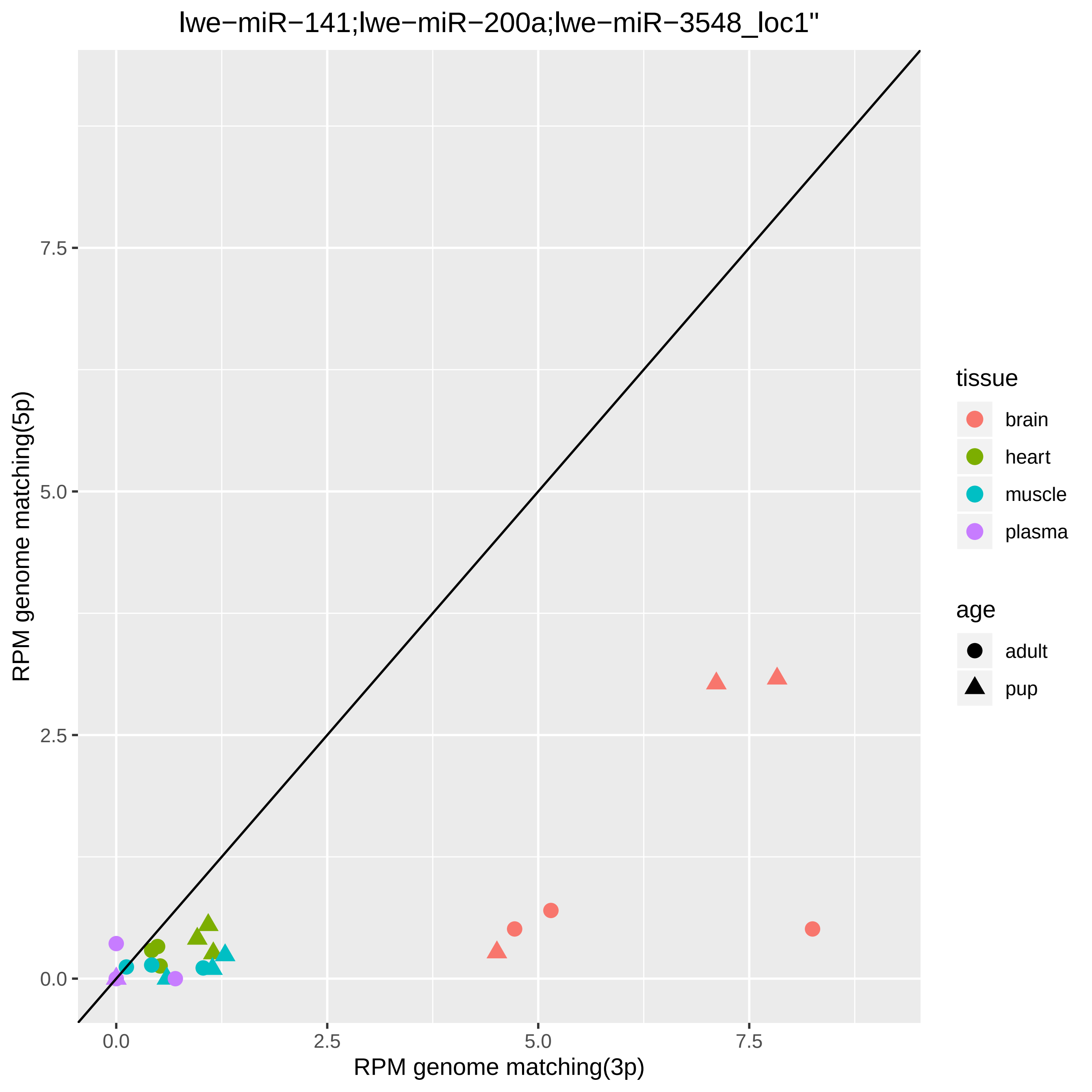

lwe−miR−141;lwe−miR−200a;lwe−miR−3548_loc1"
RPM genome matching(5p)
RPM genome matching(3p)
0.0
2.5
5.0
7.5
tissue
brain
heart
muscle
plasma
age
adult
pup

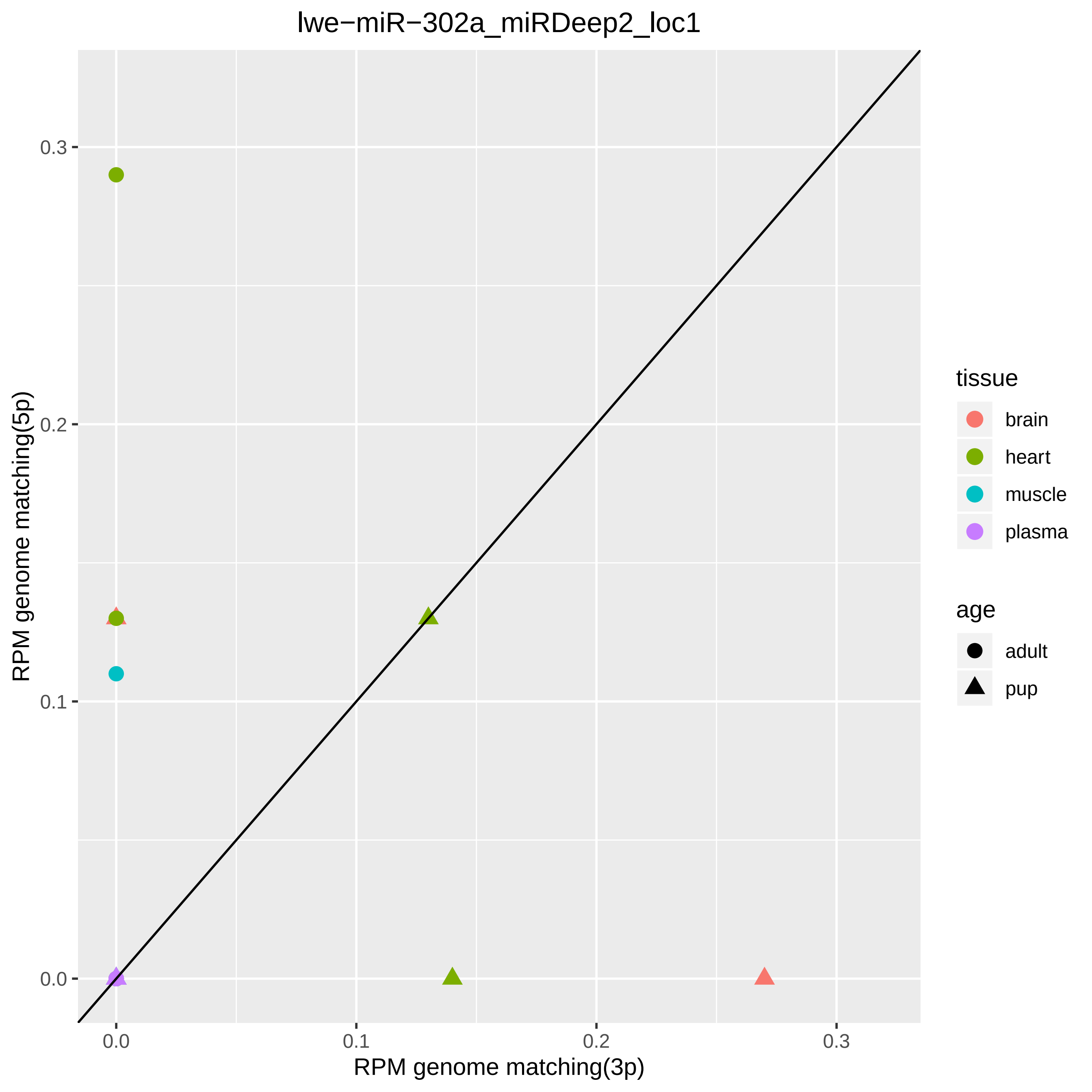
lwe−miR−302a_miRDeep2_loc1
RPM genome matching(5p)
0.3
0.2
0.1
0.0
0.0
0.1
0.2
0.3
RPM genome matching(3p)
tissue
brain
heart
muscle
plasma
age
adult
pup

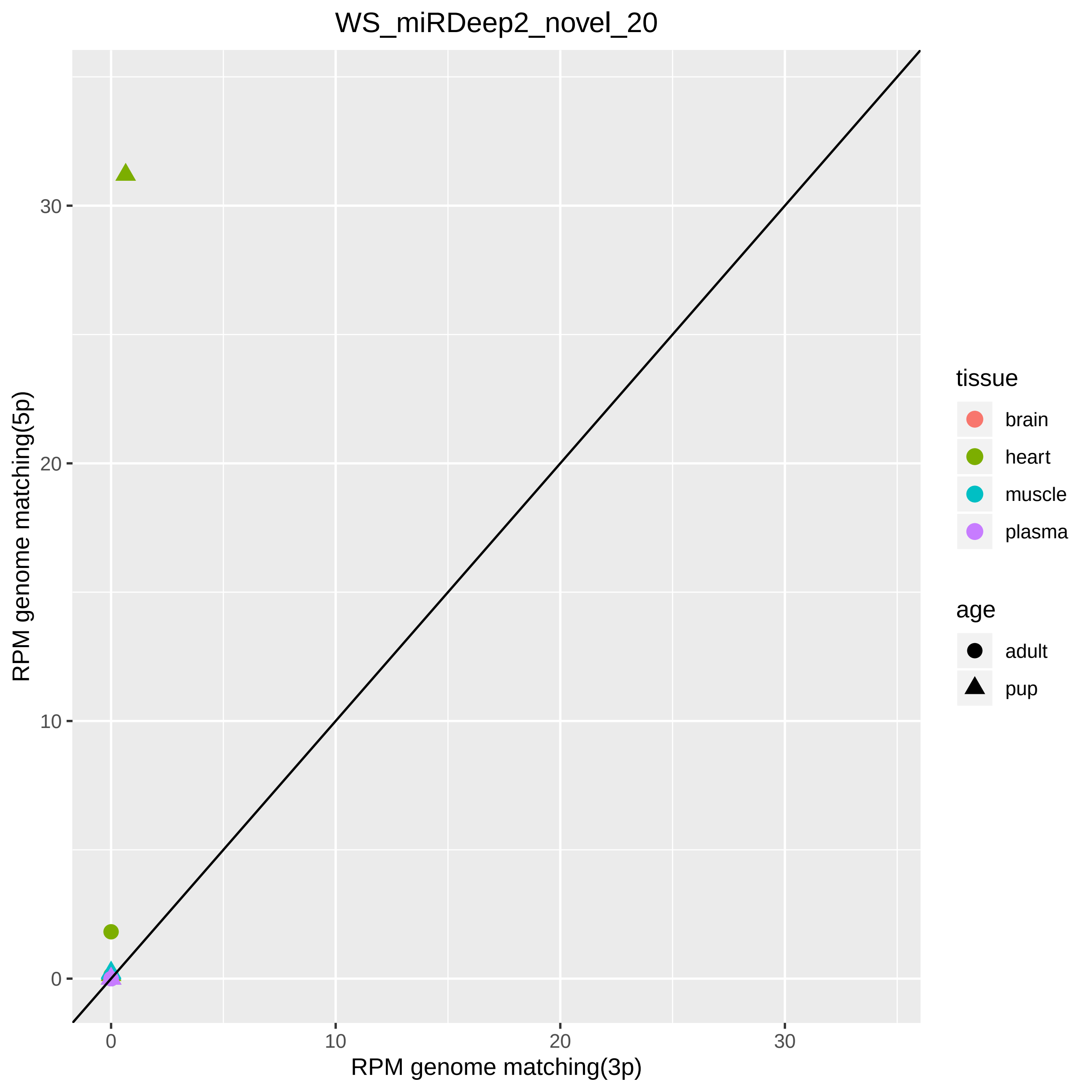
WS_miRDeep2_novel_20
RPM genome matching(5p)
RPM genome matching(3p)
0
10
20
30
0
10
20
30
tissue
brain
heart
muscle
plasma
age
adult
pup

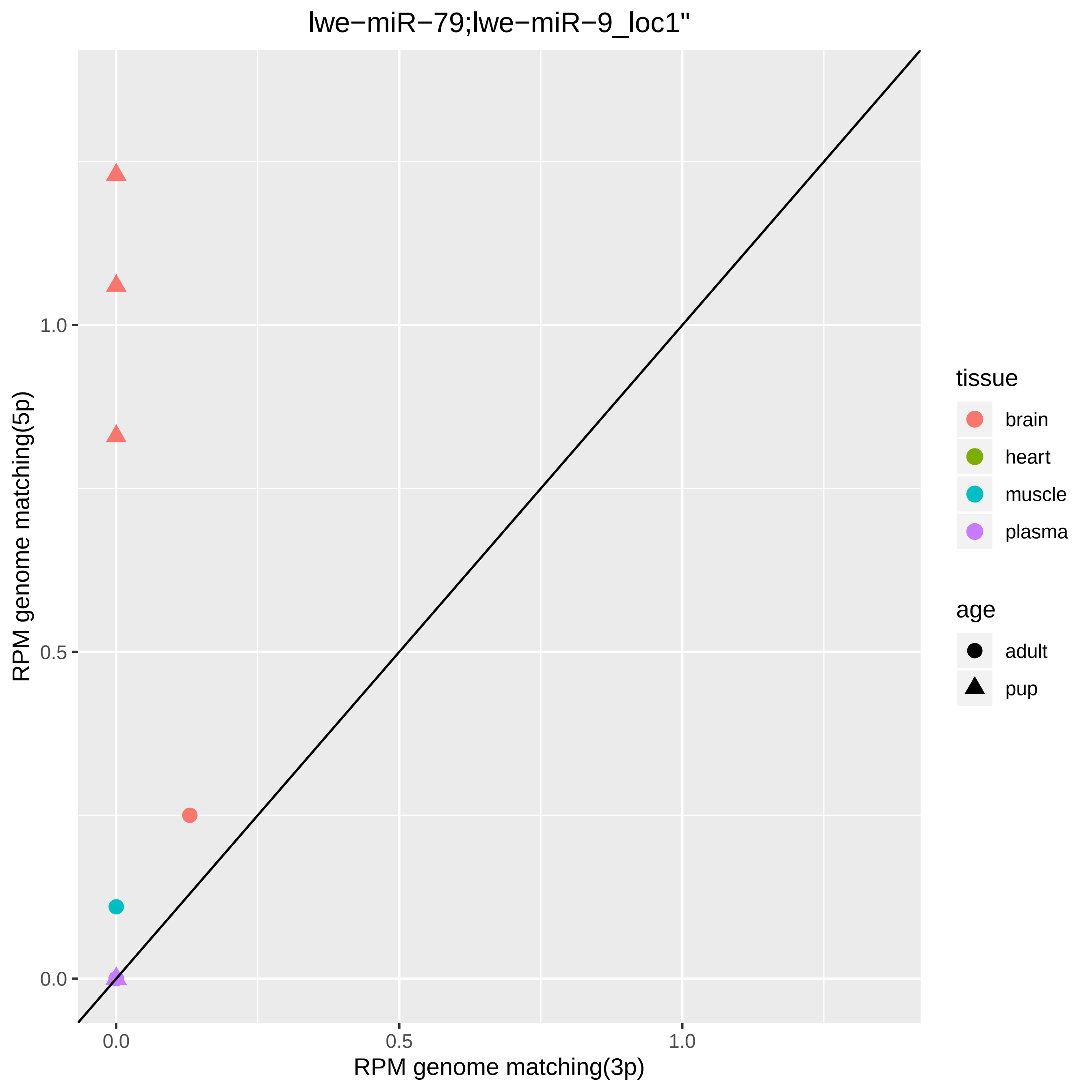
lwe−miR−79;lwe−miR−9_loc1"
RPM genome matching(5p)
RPM genome matching(3p)
1.0
0.5
0.0
0.0
0.5
1.0
tissue
brain
heart
muscle
plasma
age
adult
pup

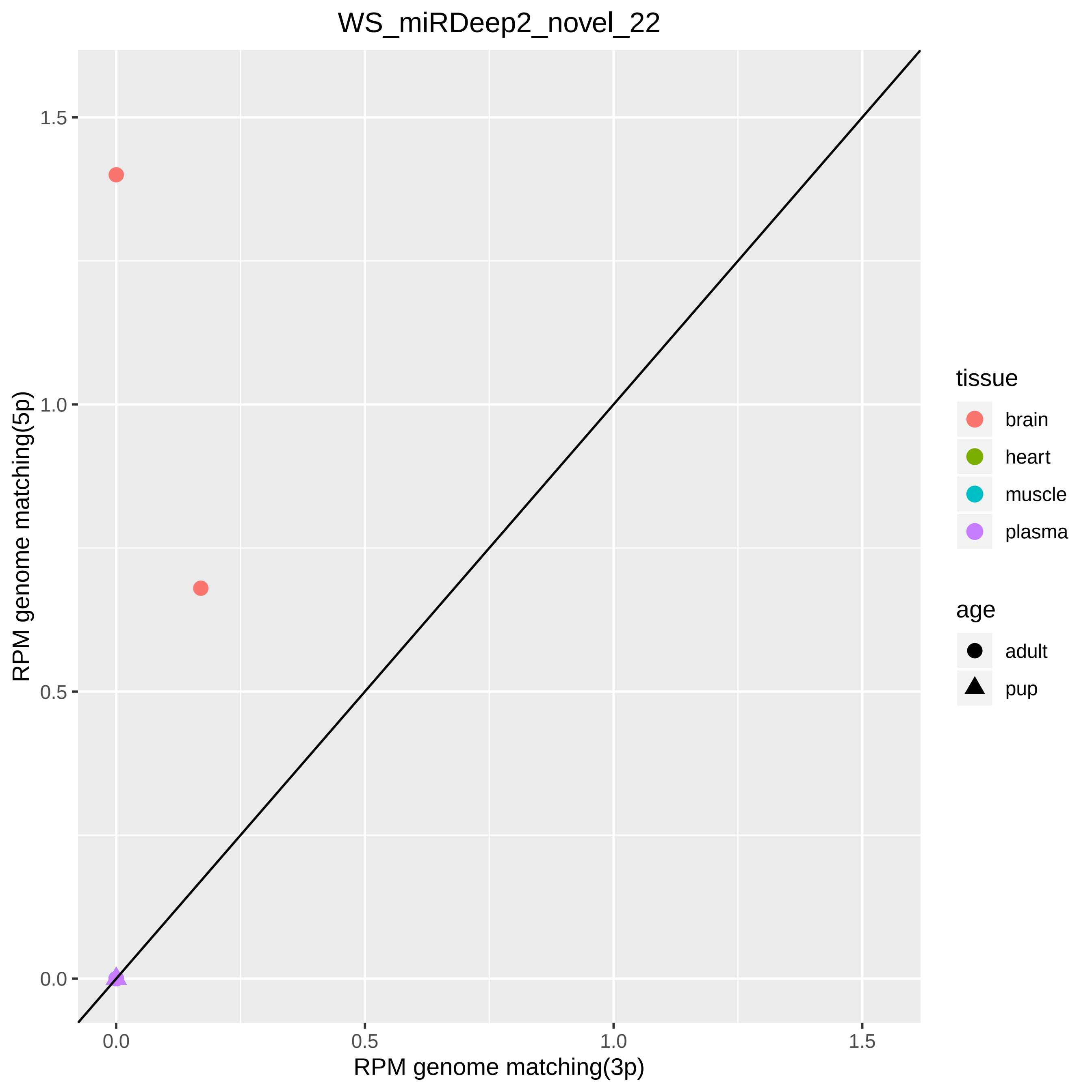

WS_miRDeep2_novel_22
RPM genome matching(5p)
RPM genome matching(3p)
0.0
0.5
1.0
1.5
tissue
brain
heart
muscle
plasma
age
adult
pup

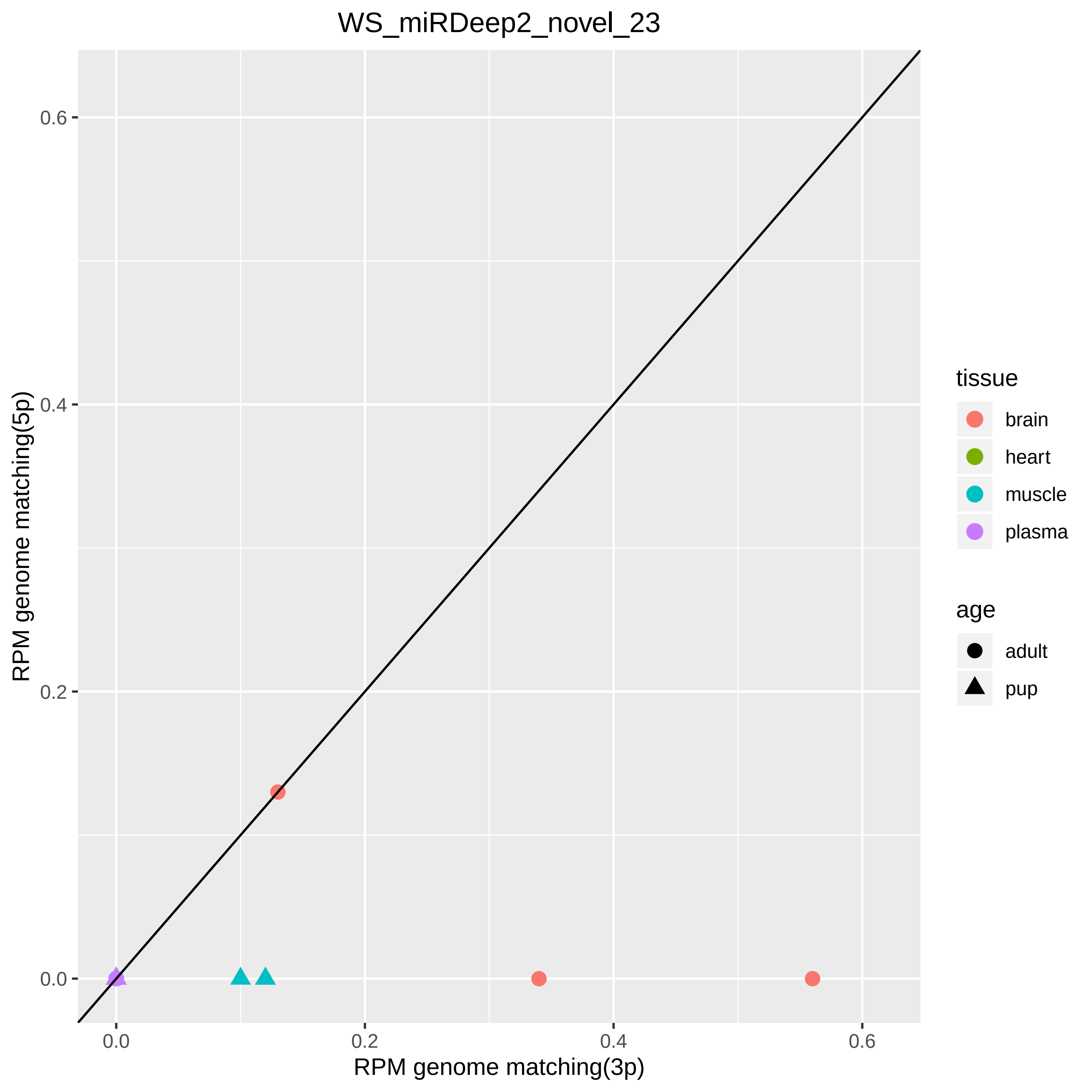
WS_miRDeep2_novel_23
RPM genome matching(5p)
RPM genome matching(3p)
0.0
0.2
0.4
0.6
0.0
0.2
0.4
0.6
tissue
brain
heart
muscle
plasma
age
adult
pup

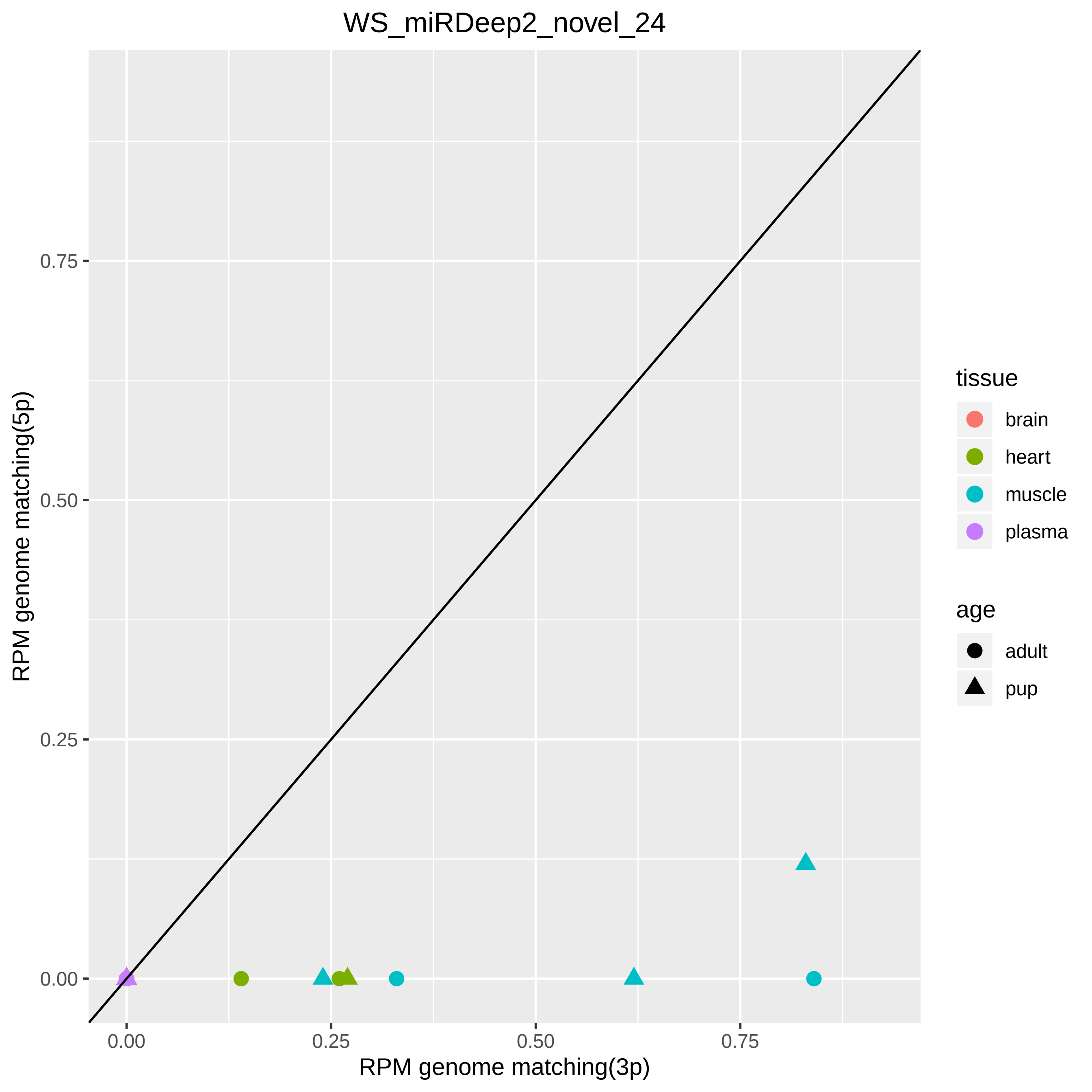

WS_miRDeep2_novel_24
RPM genome matching(5p)
RPM genome matching(3p)
0.00
0.25
0.50
0.75
tissue
brain
heart
muscle
plasma
age
adult
pup

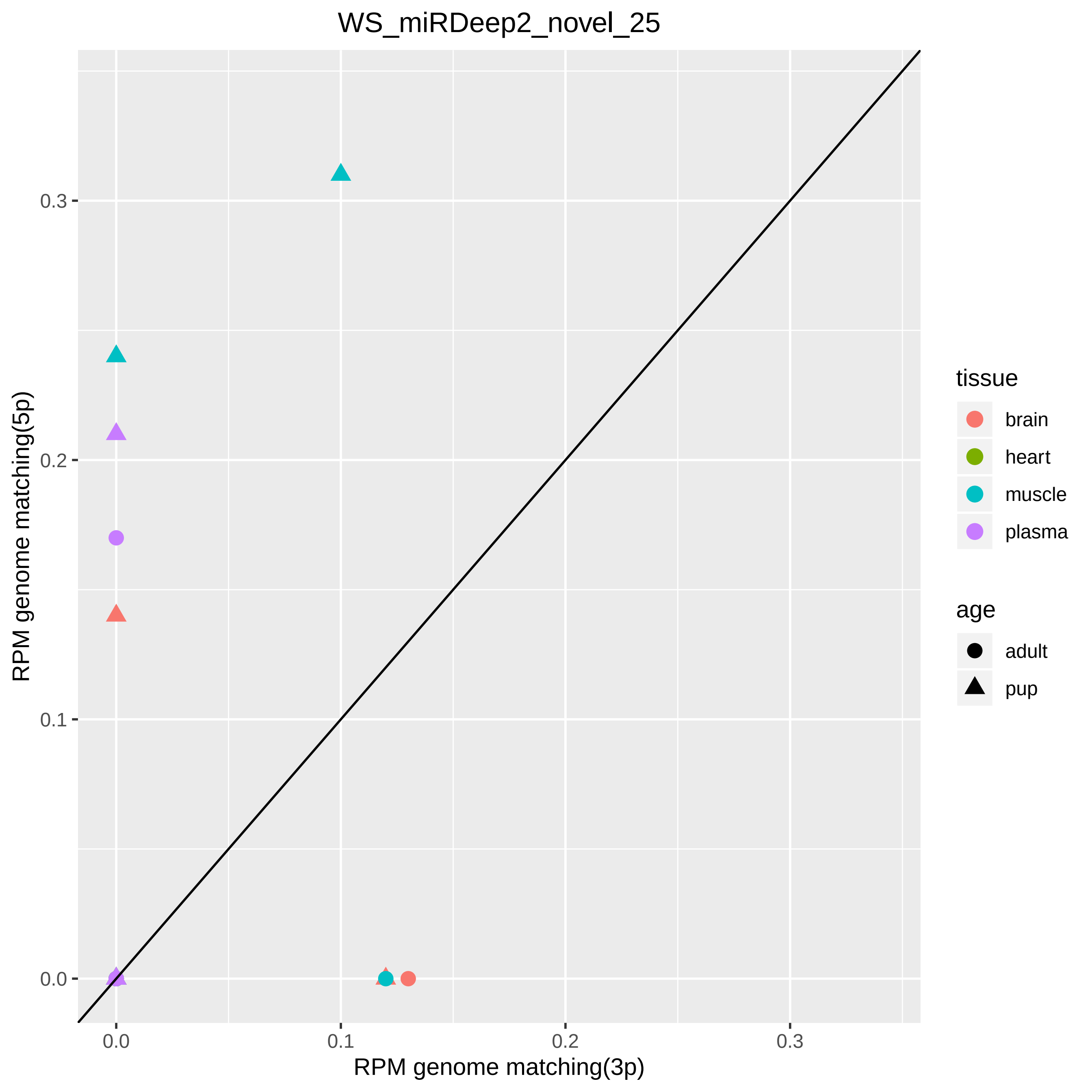
WS_miRDeep2_novel_25
RPM genome matching(5p)
RPM genome matching(3p)
0.0
0.1
0.2
0.3
tissue
brain
heart
muscle
plasma
age
adult
pup

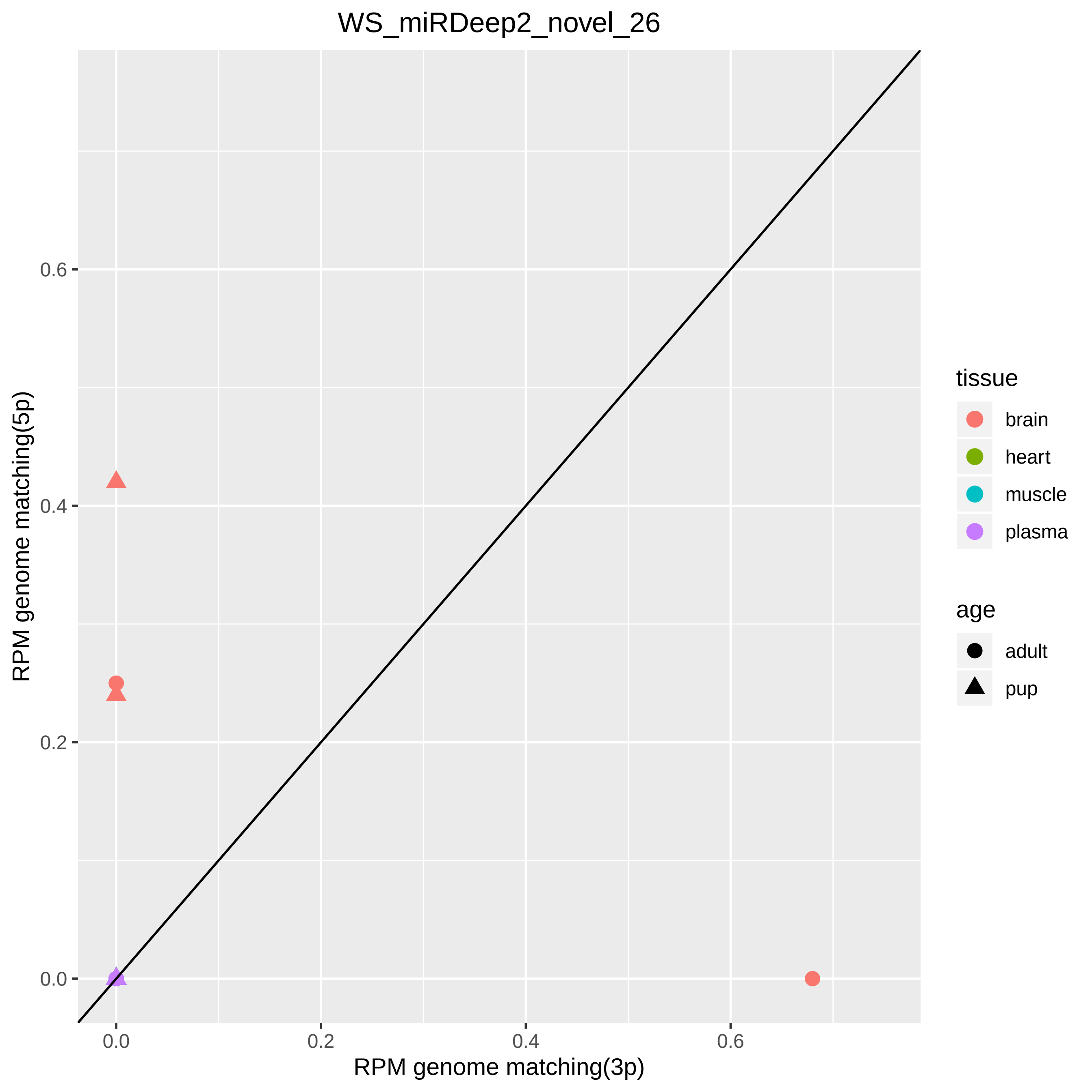

WS_miRDeep2_novel_26
RPM genome matching(5p)
RPM genome matching(3p)
0.0
0.2
0.4
0.6
tissue
brain
heart
muscle
plasma
age
adult
pup

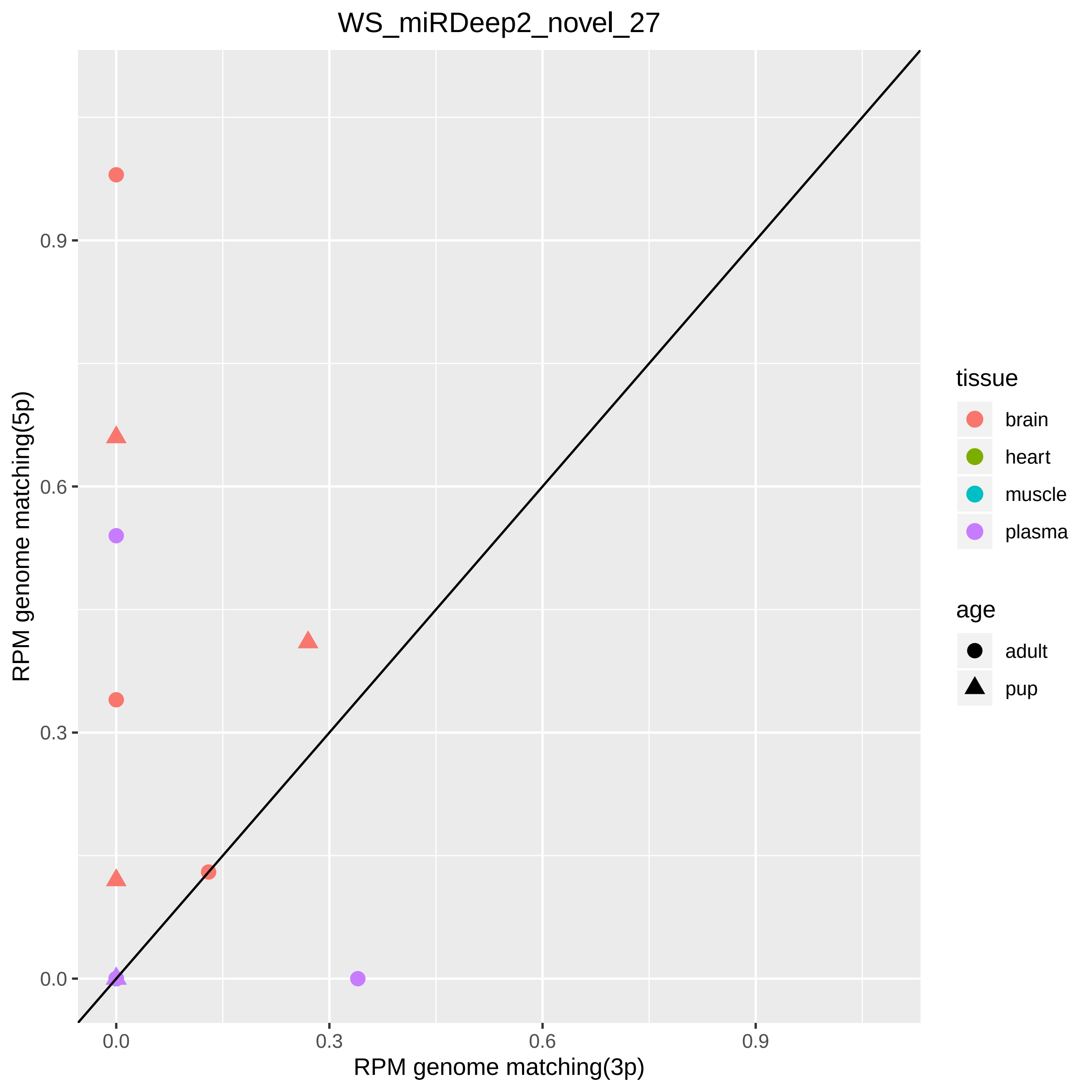

WS_miRDeep2_novel_27
RPM genome matching(5p)
RPM genome matching(3p)
0.0
0.3
0.6
0.9
tissue
brain
heart
muscle
plasma
age
adult
pup

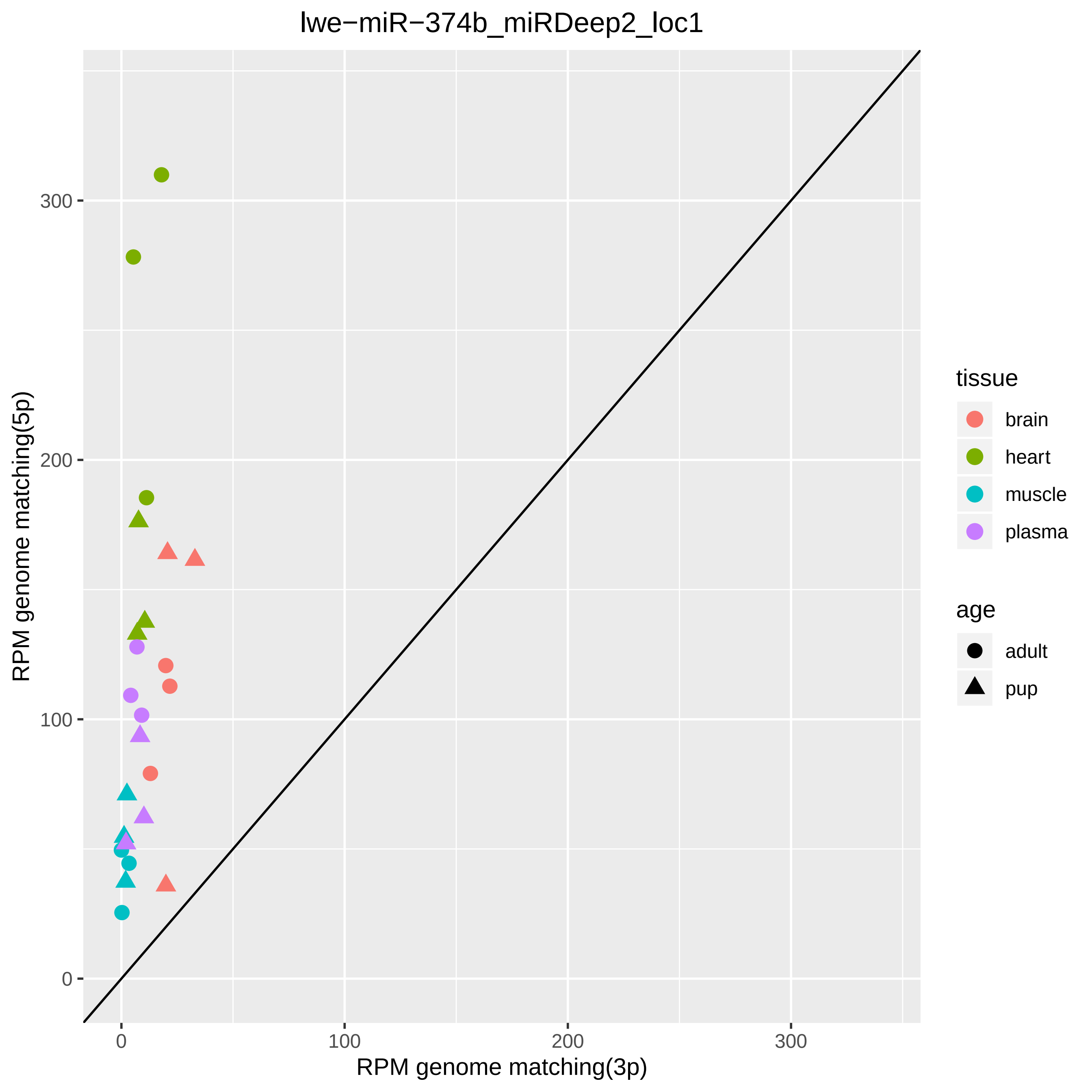
lwe−miR−374b_miRDeep2_loc1
RPM genome matching(5p)
RPM genome matching(3p)
0
100
200
300
0
100
200
300
tissue
brain
heart
muscle
plasma
age
adult
pup

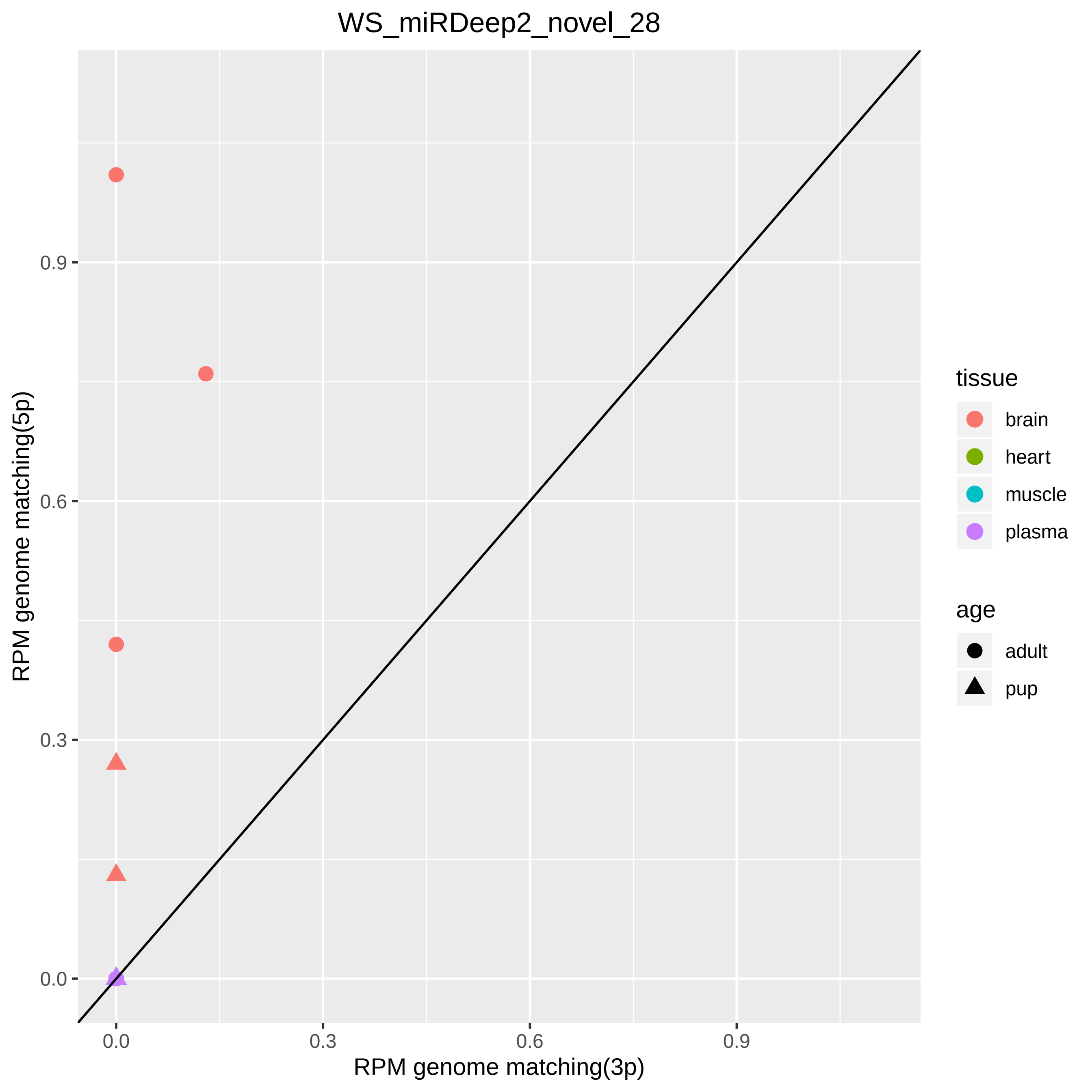

WS_miRDeep2_novel_28
RPM genome matching(5p)
RPM genome matching(3p)
0.0
0.3
0.6
0.9
tissue
brain
heart
muscle
plasma
age
adult
pup

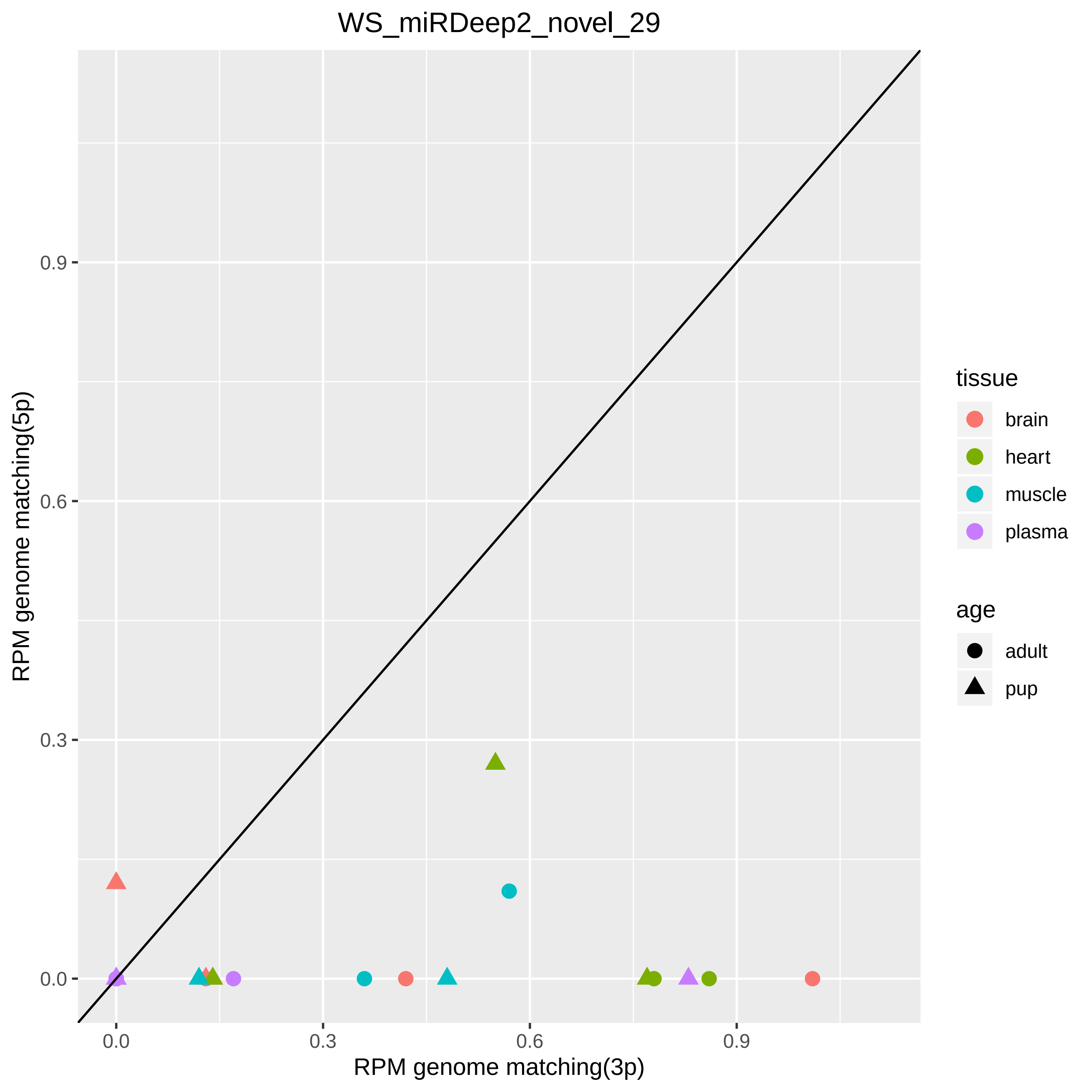

WS_miRDeep2_novel_29
RPM genome matching(5p)
RPM genome matching(3p)
0.0
0.3
0.6
0.9
tissue
brain
heart
muscle
plasma
age
adult
pup

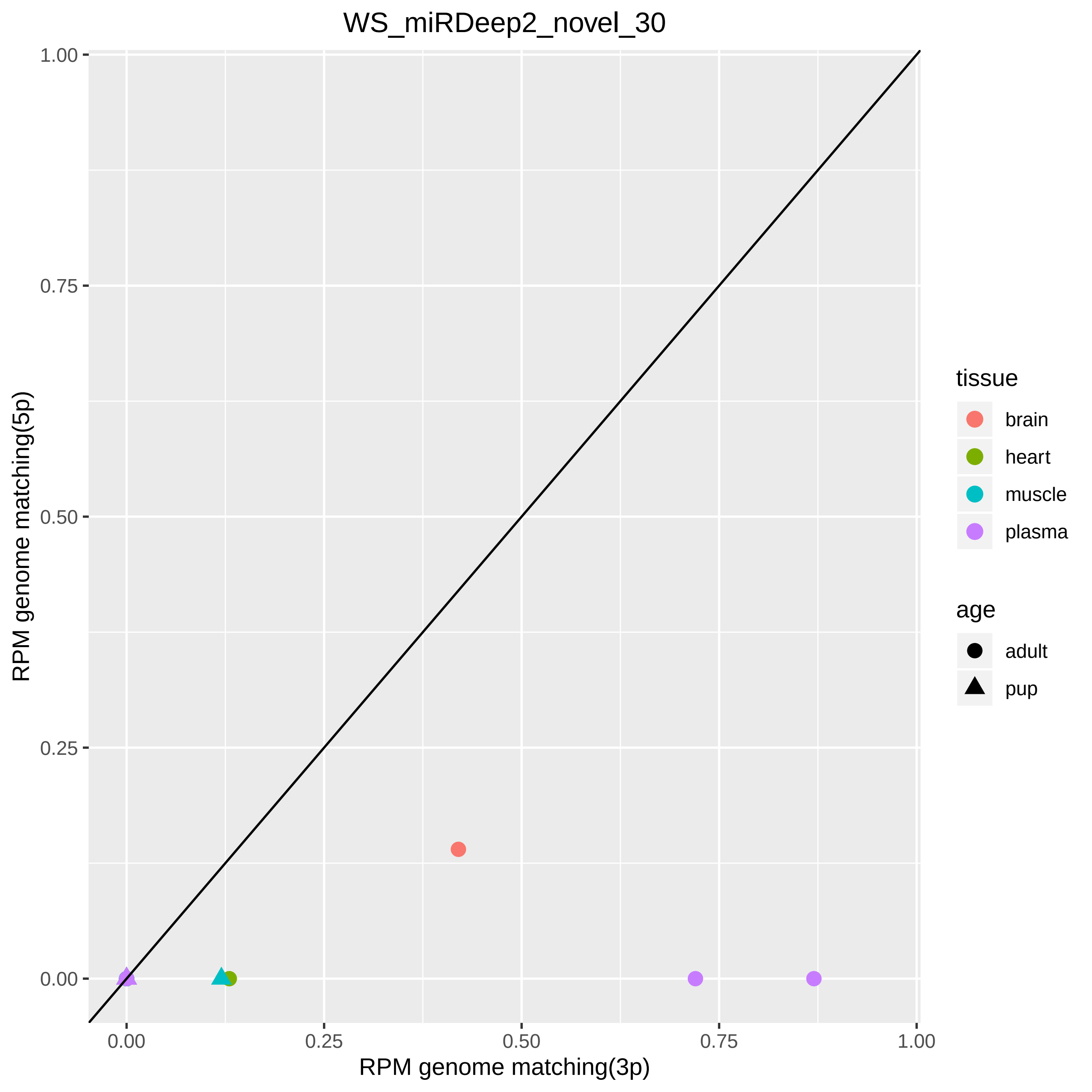
WS_miRDeep2_novel_30
RPM genome matching(5p)
RPM genome matching(3p)
0.00
0.25
0.50
0.75
1.00
tissue
brain
heart
muscle
plasma
age
adult
pup

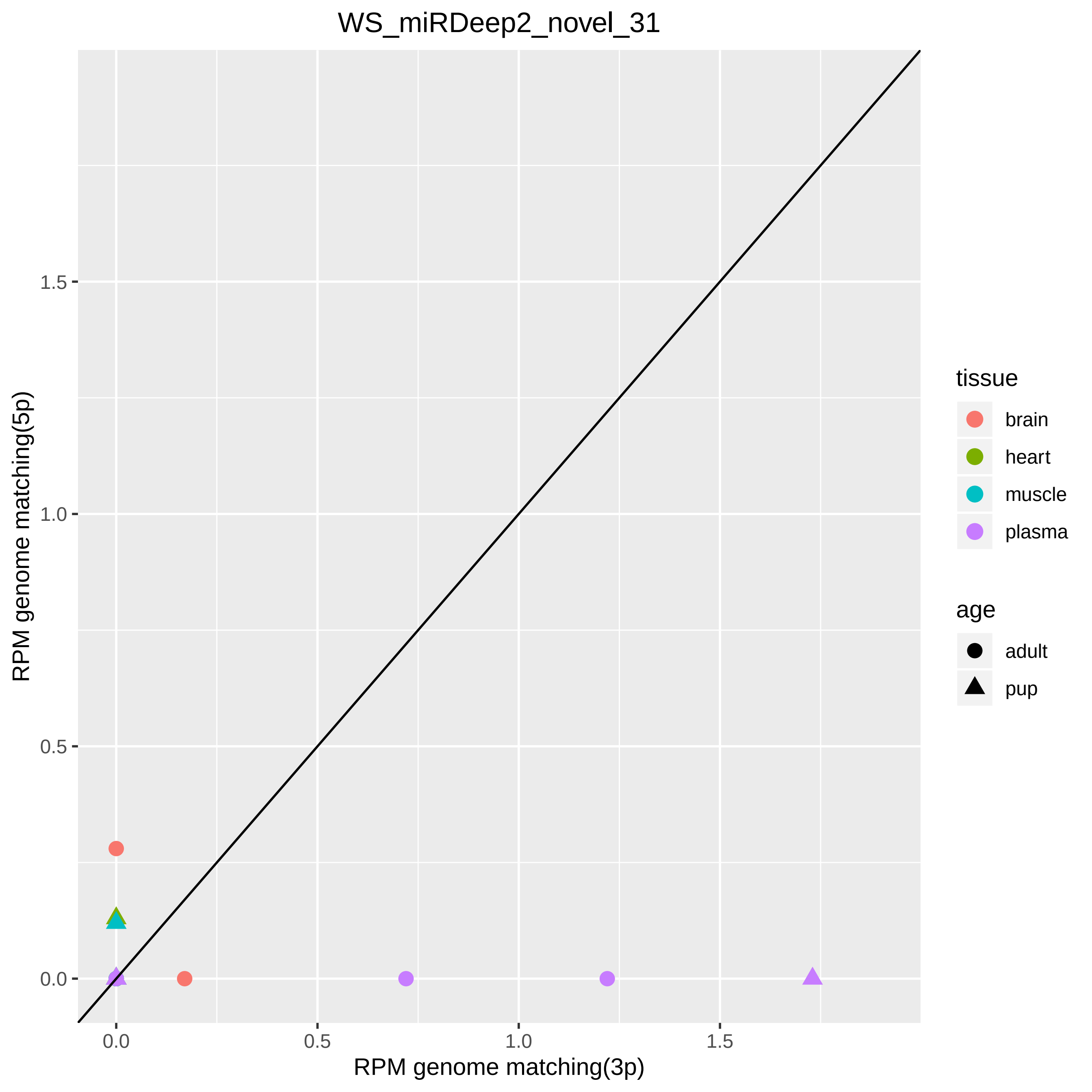

WS_miRDeep2_novel_31
RPM genome matching(5p)
RPM genome matching(3p)
0.0
0.5
1.0
1.5
tissue
brain
heart
muscle
plasma
age
adult
pup

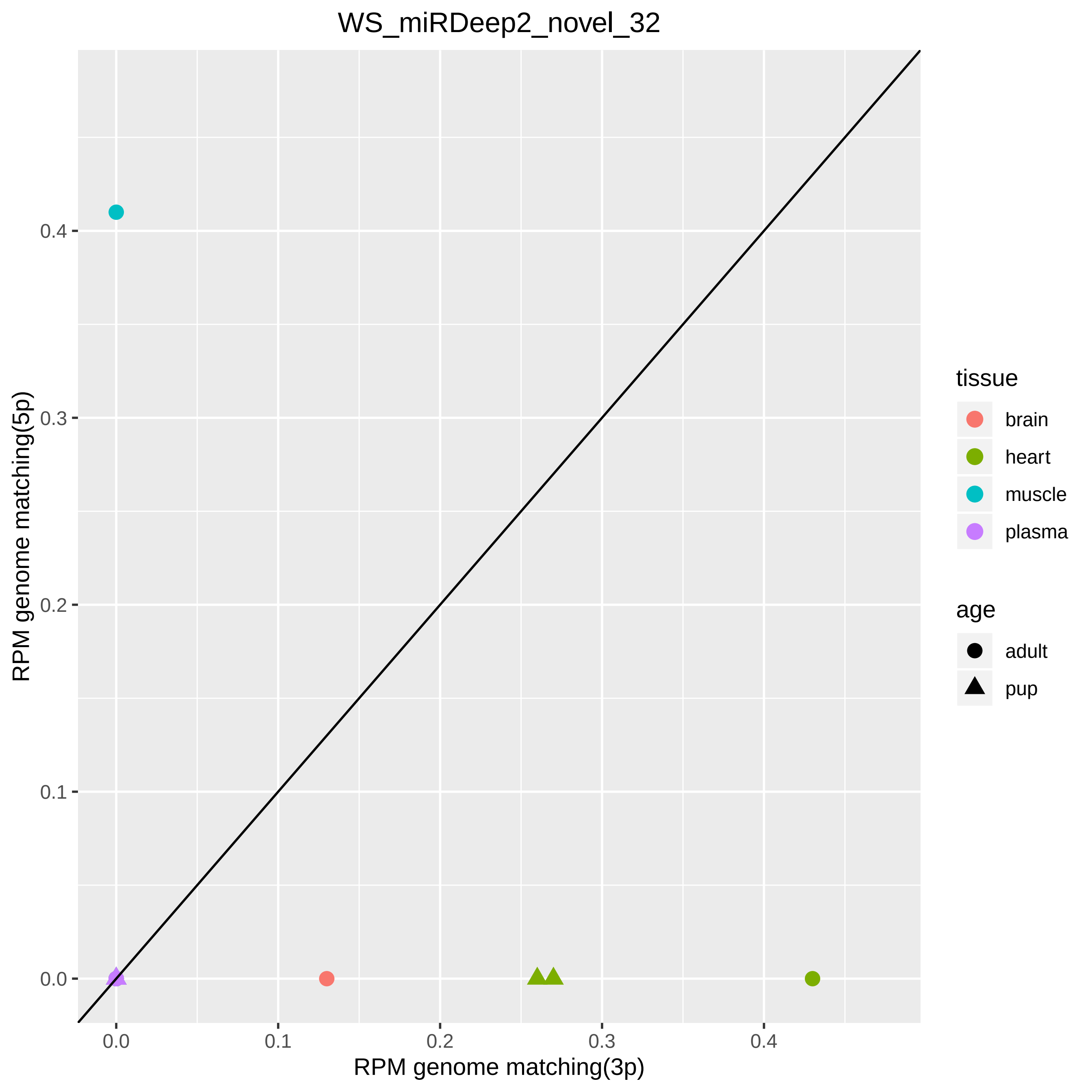

WS_miRDeep2_novel_32
RPM genome matching(5p)
RPM genome matching(3p)
0.0
0.1
0.2
0.3
0.4
tissue
brain
heart
muscle
plasma
age
adult
pup

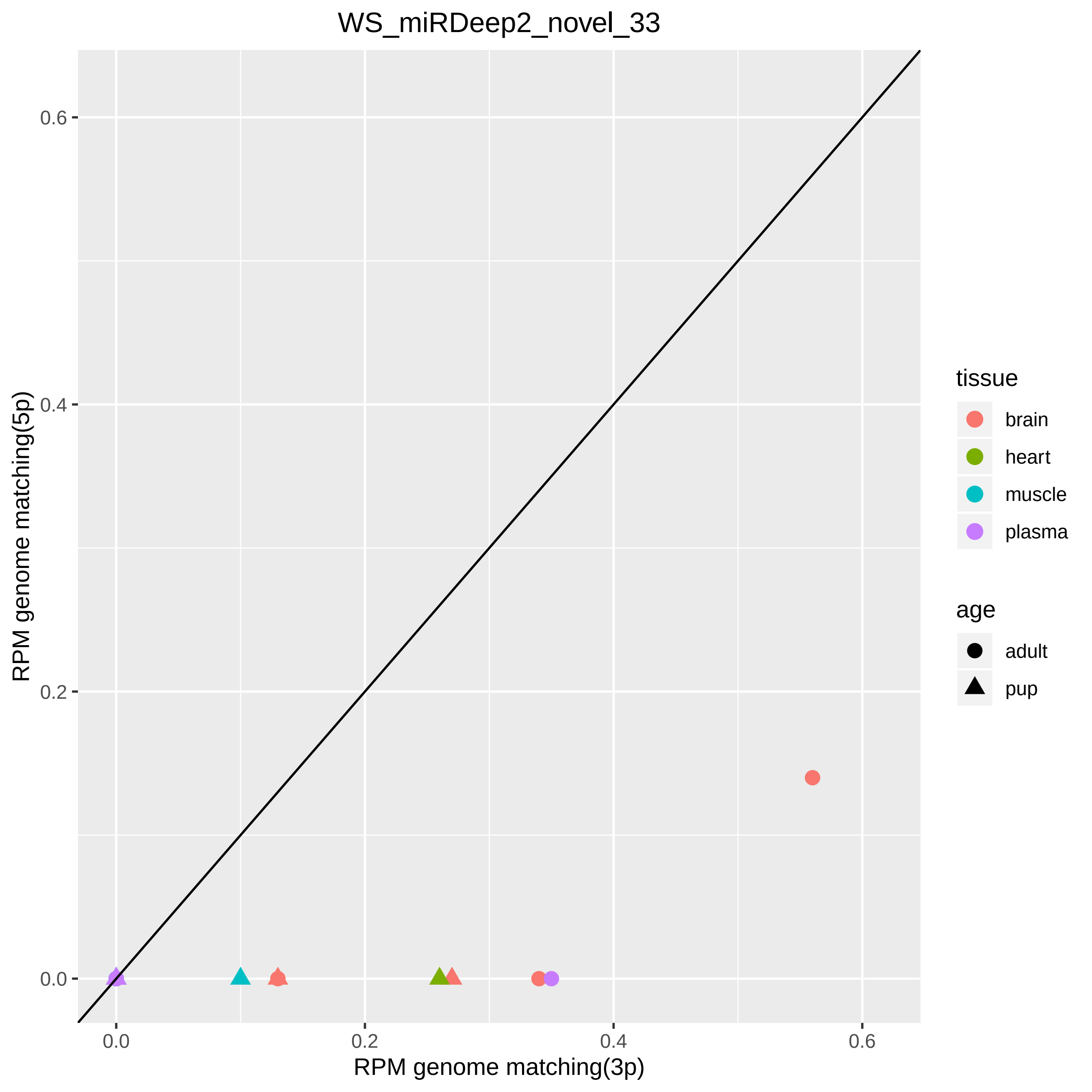
WS_miRDeep2_novel_33
RPM genome matching(5p)
RPM genome matching(3p)
0.0
0.2
0.4
0.6
tissue
brain
heart
muscle
plasma
age
adult
pup

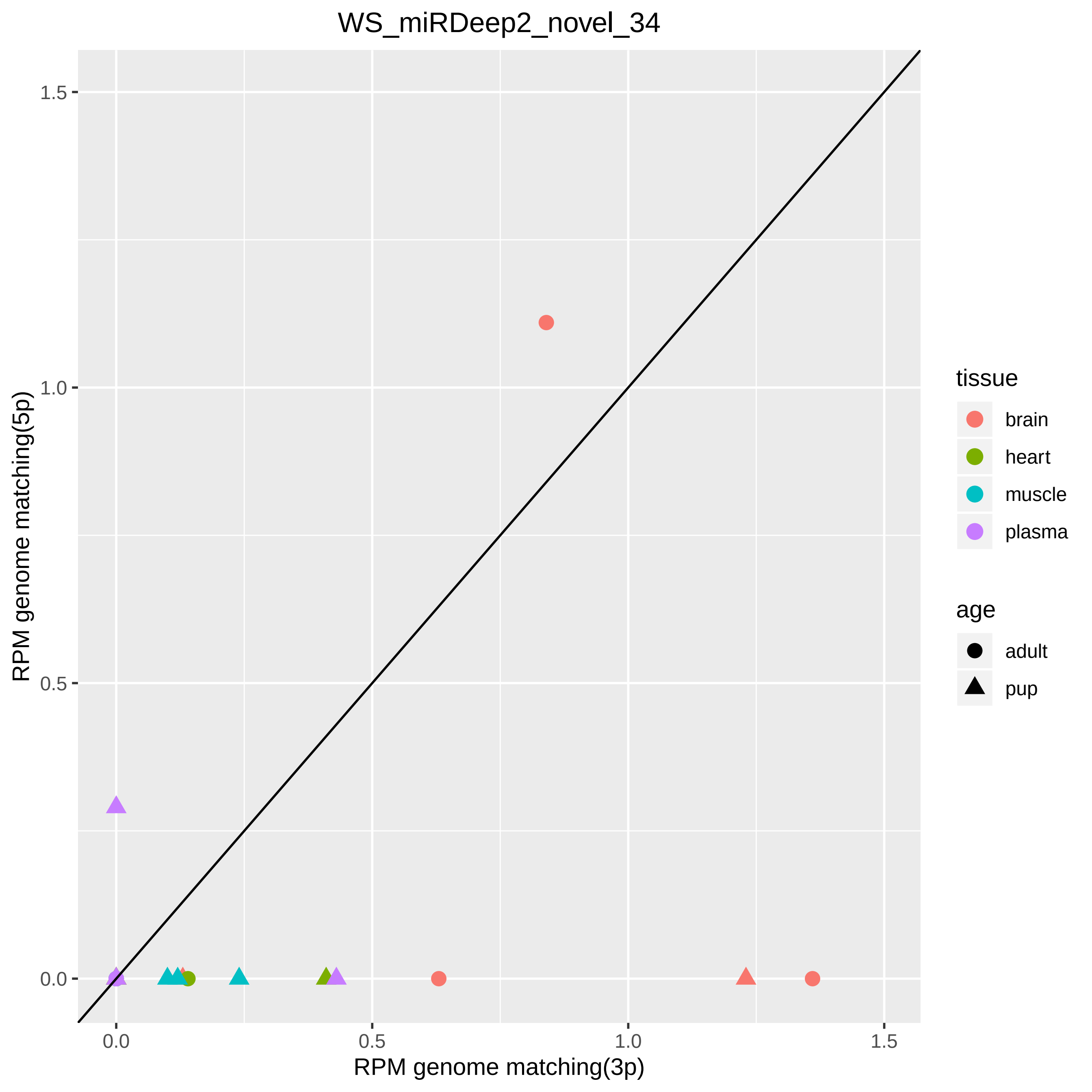

WS_miRDeep2_novel_34
RPM genome matching(5p)
RPM genome matching(3p)
0.0
0.5
1.0
1.5
tissue
brain
heart
muscle
plasma
age
adult
pup

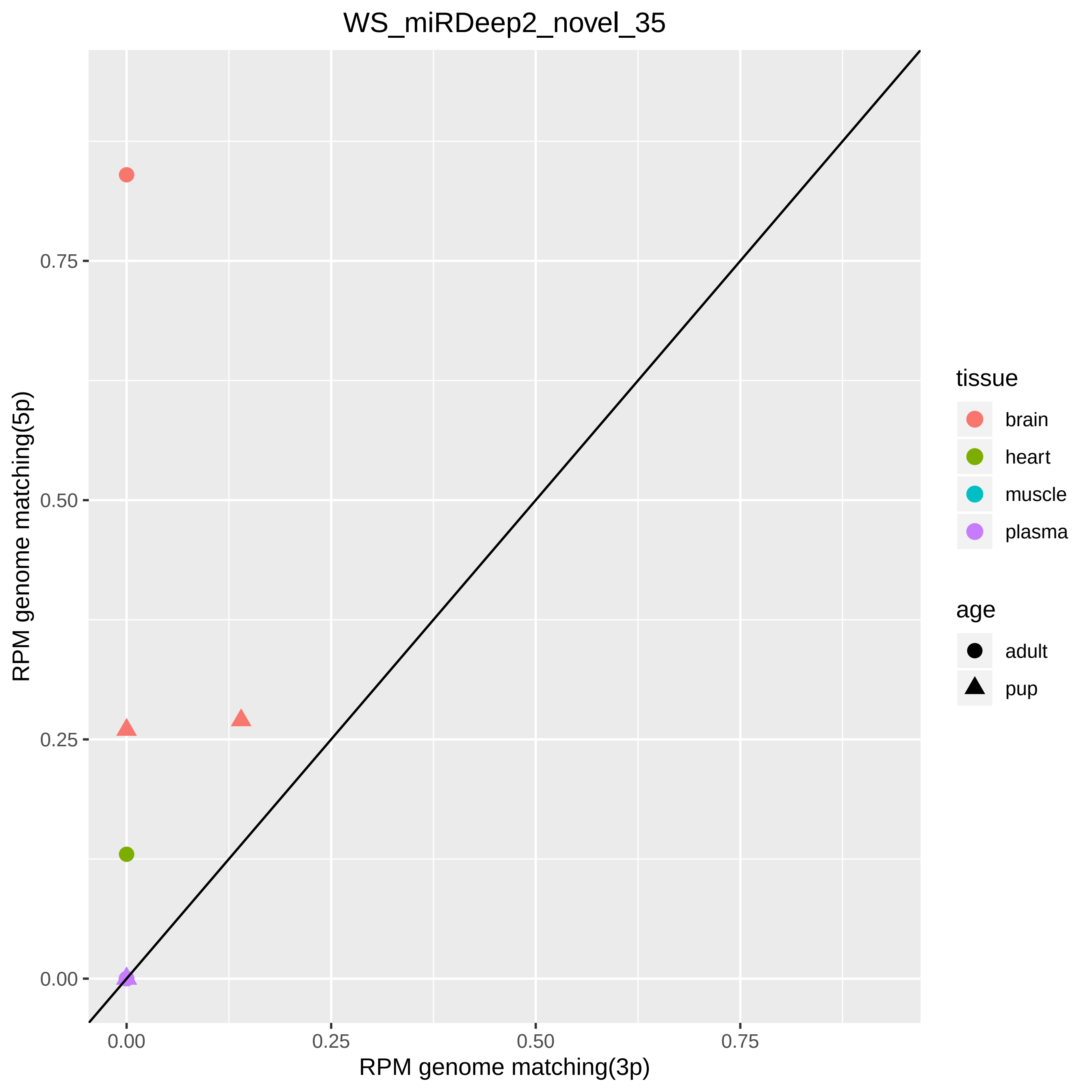

WS_miRDeep2_novel_35
RPM genome matching(5p)
RPM genome matching(3p)
0.00
0.25
0.50
0.75
tissue
brain
heart
muscle
plasma
age
adult
pup

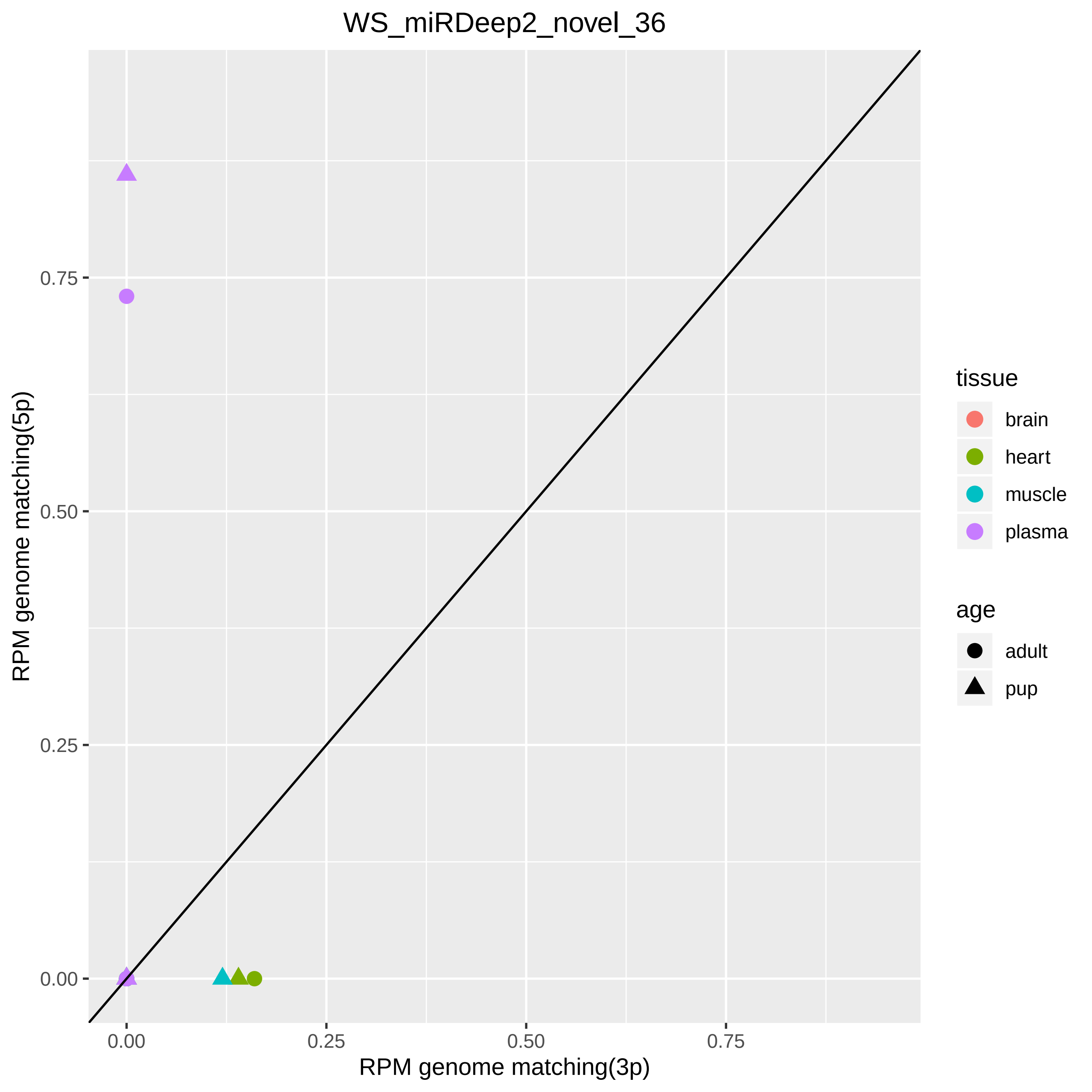
WS_miRDeep2_novel_36
RPM genome matching(5p)
RPM genome matching(3p)
0.00
0.25
0.50
0.75
tissue
brain
heart
muscle
plasma
age
adult
pup

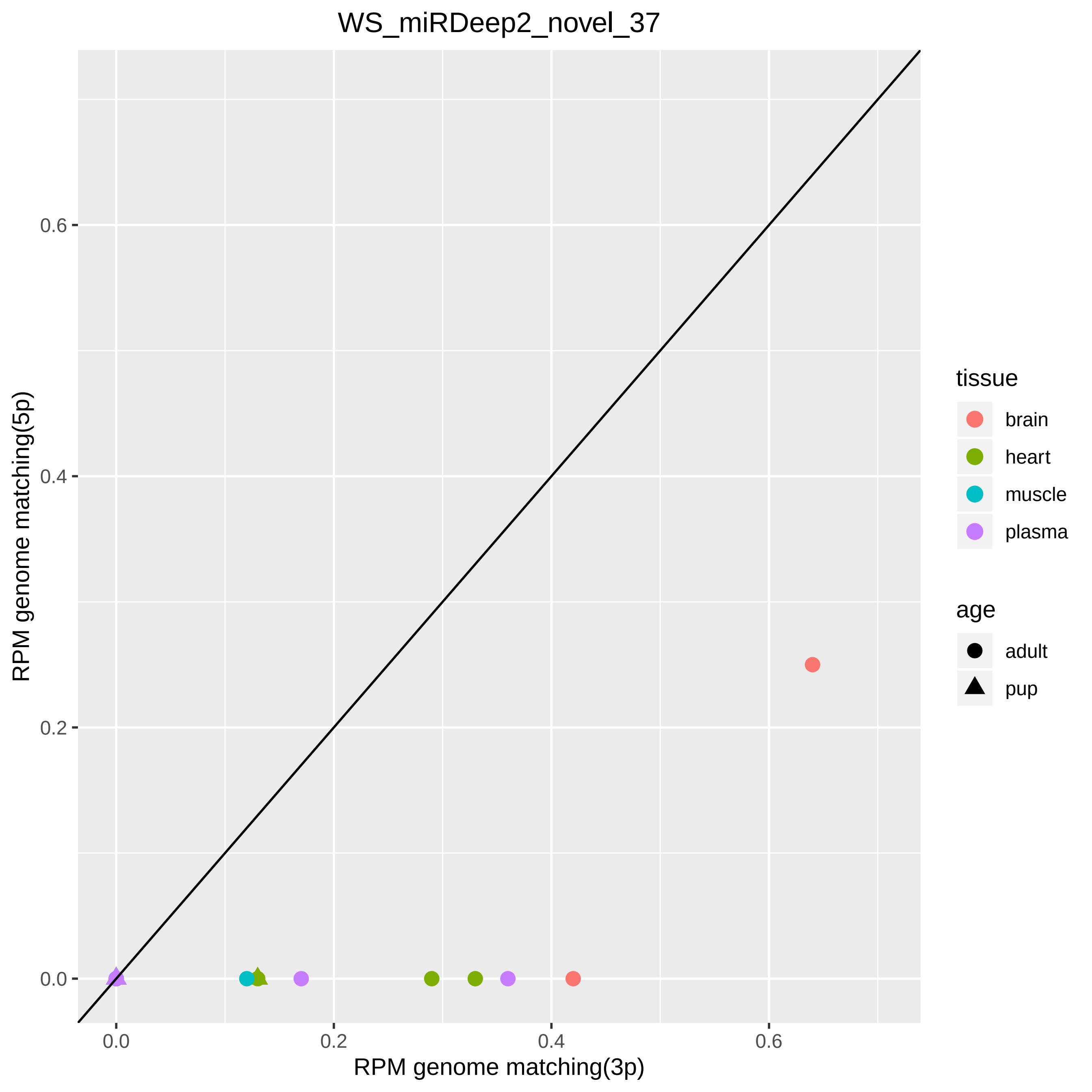
WS_miRDeep2_novel_37
RPM genome matching(5p)
RPM genome matching(3p)
0.0
0.2
0.4
0.6
tissue
brain
heart
muscle
plasma
age
adult
pup

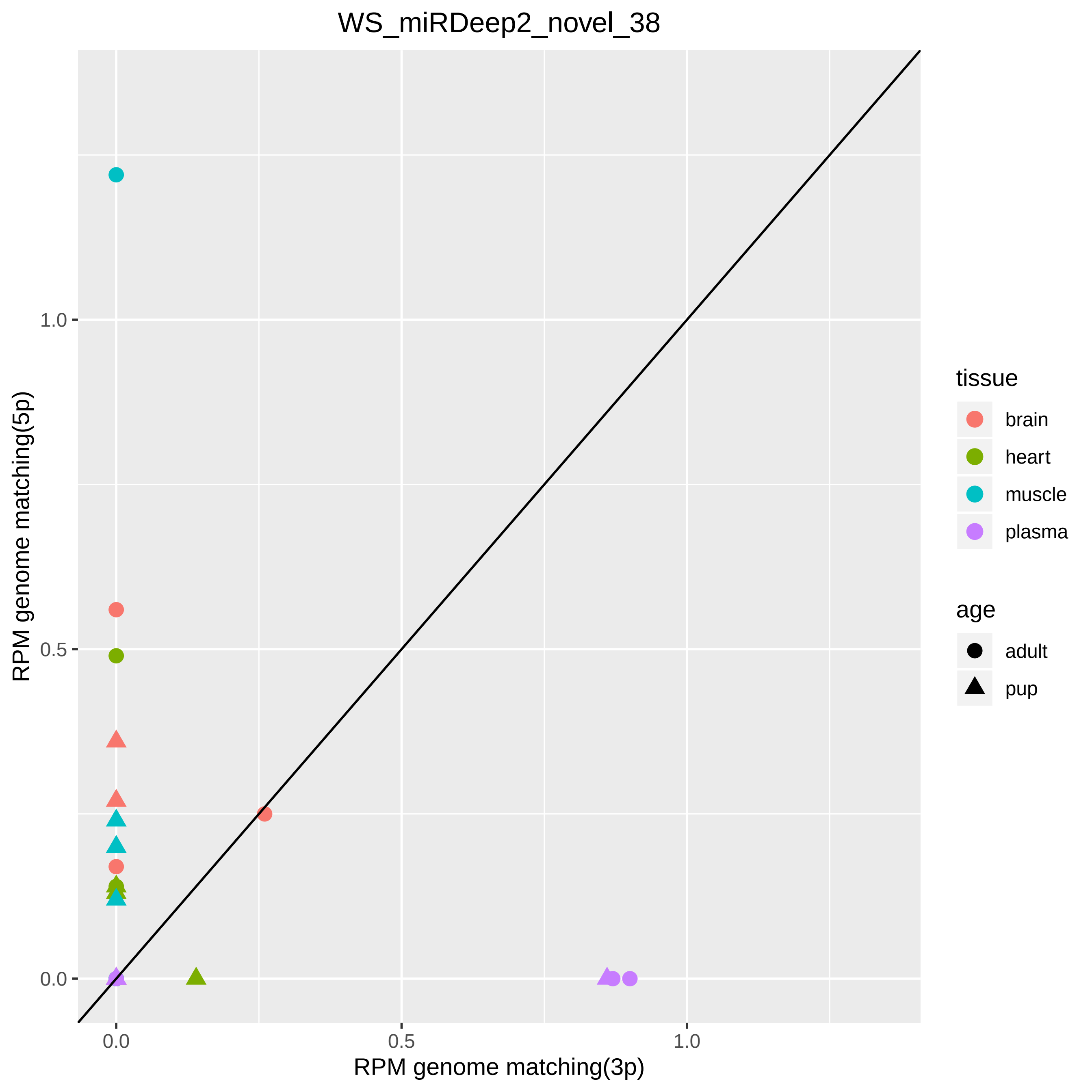

WS_miRDeep2_novel_38
RPM genome matching(5p)
RPM genome matching(3p)
1.0
0.5
0.0
0.0
0.5
1.0
tissue
brain
heart
muscle
plasma
age
adult
pup

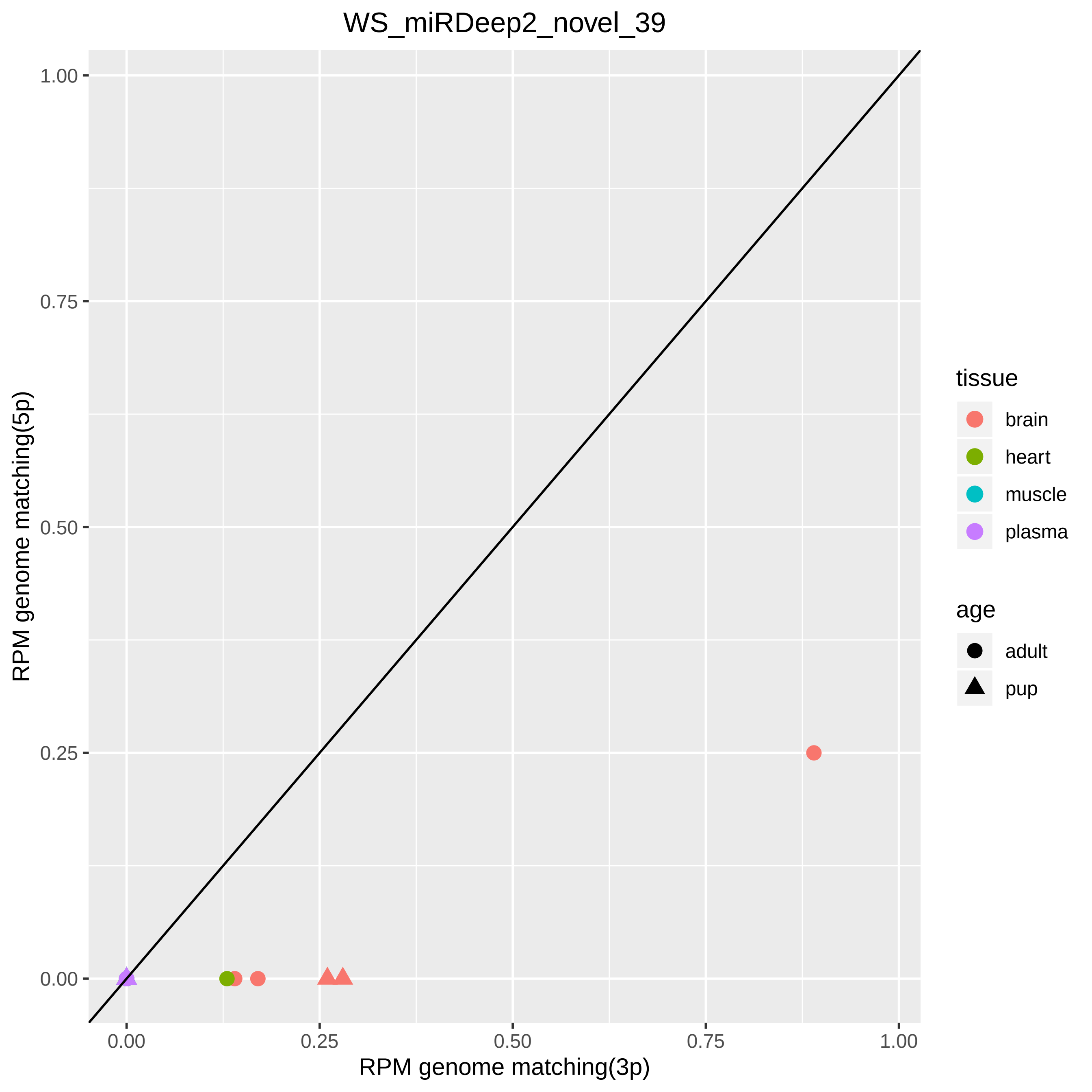

WS_miRDeep2_novel_39
RPM genome matching(5p)
RPM genome matching(3p)
1.00
0.75
0.50
0.25
0.00
0.00
0.25
0.50
0.75
1.00
tissue
brain
heart
muscle
plasma
age
adult
pup

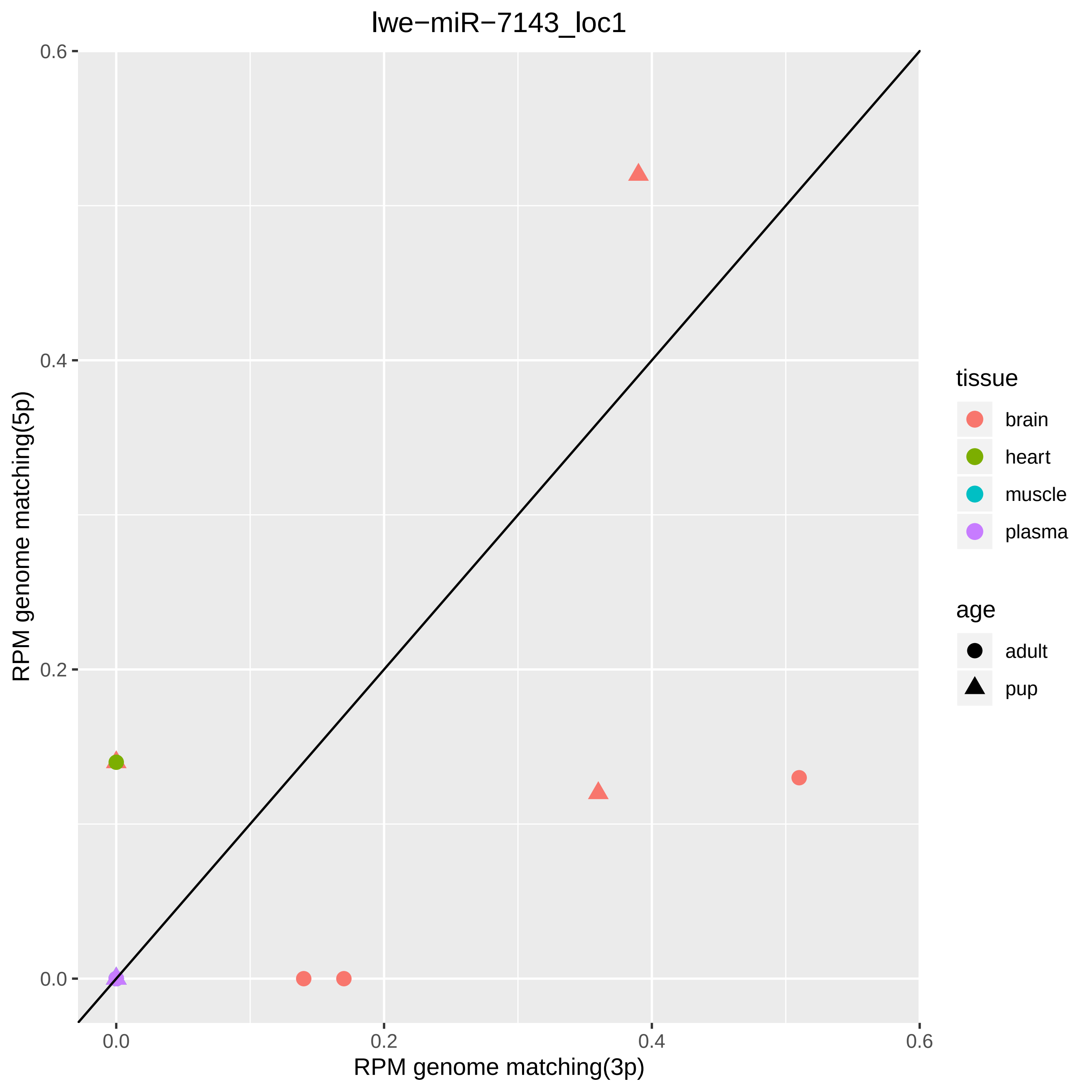

lwe−miR−7143_loc1
RPM genome matching(5p)
RPM genome matching(3p)
0.0
0.2
0.4
0.6
tissue
brain
heart
muscle
plasma
age
adult
pup

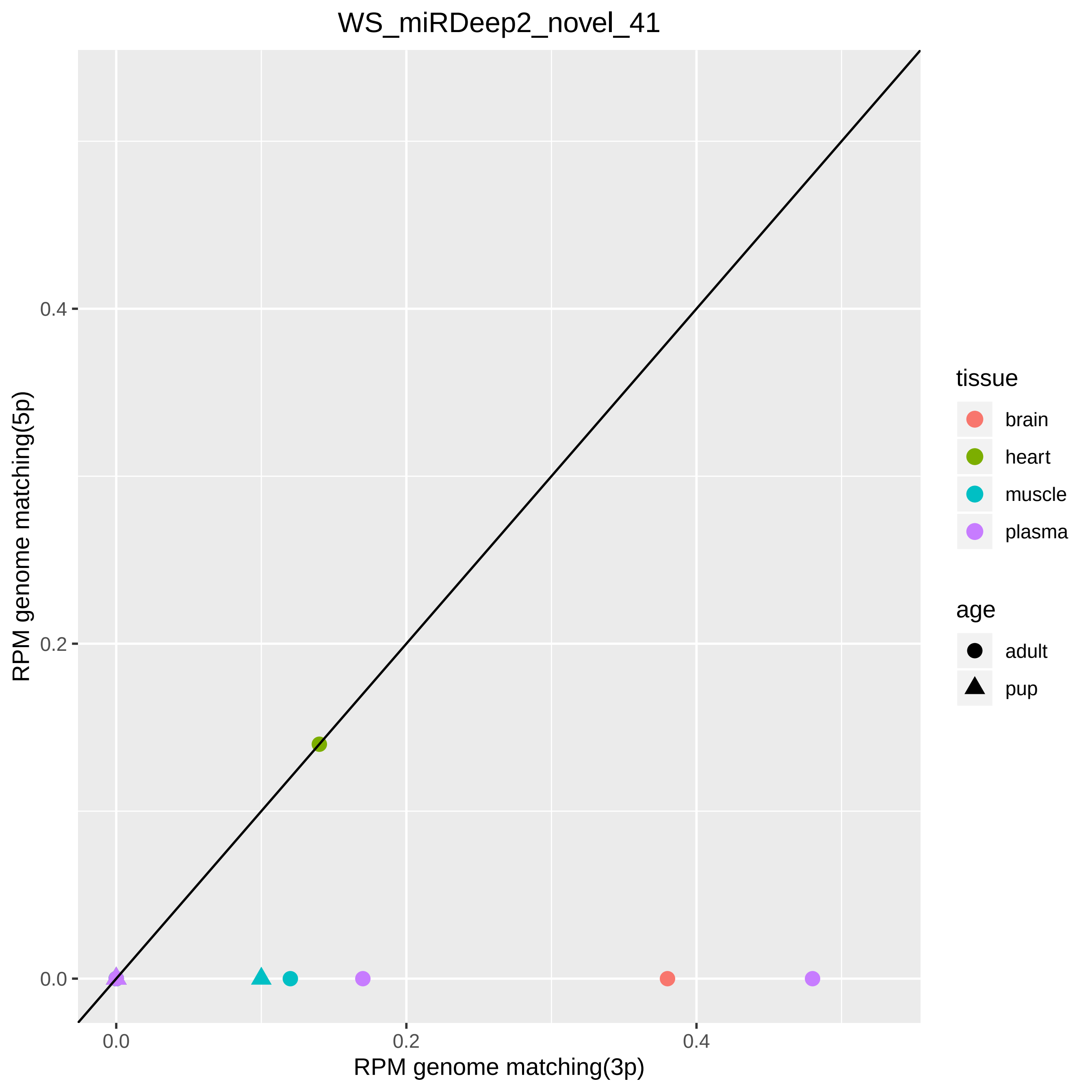

WS_miRDeep2_novel_41
RPM genome matching(5p)
RPM genome matching(3p)
0.0
0.2
0.4
0.0
0.2
0.4
tissue
brain
heart
muscle
plasma
age
adult
pup

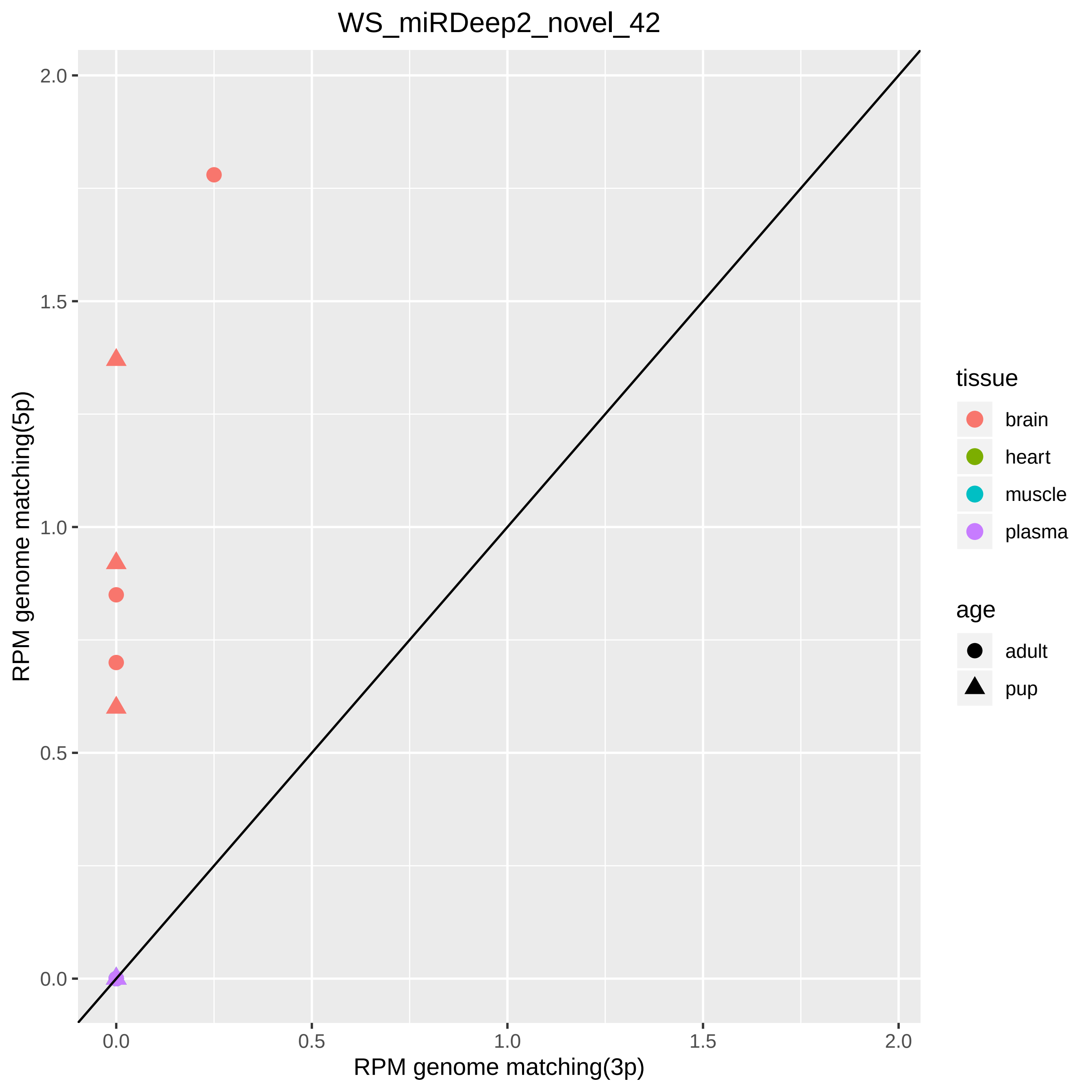

WS_miRDeep2_novel_42
RPM genome matching(5p)
RPM genome matching(3p)
0.0
0.5
1.0
1.5
2.0
tissue
brain
heart
muscle
plasma
age
adult
pup

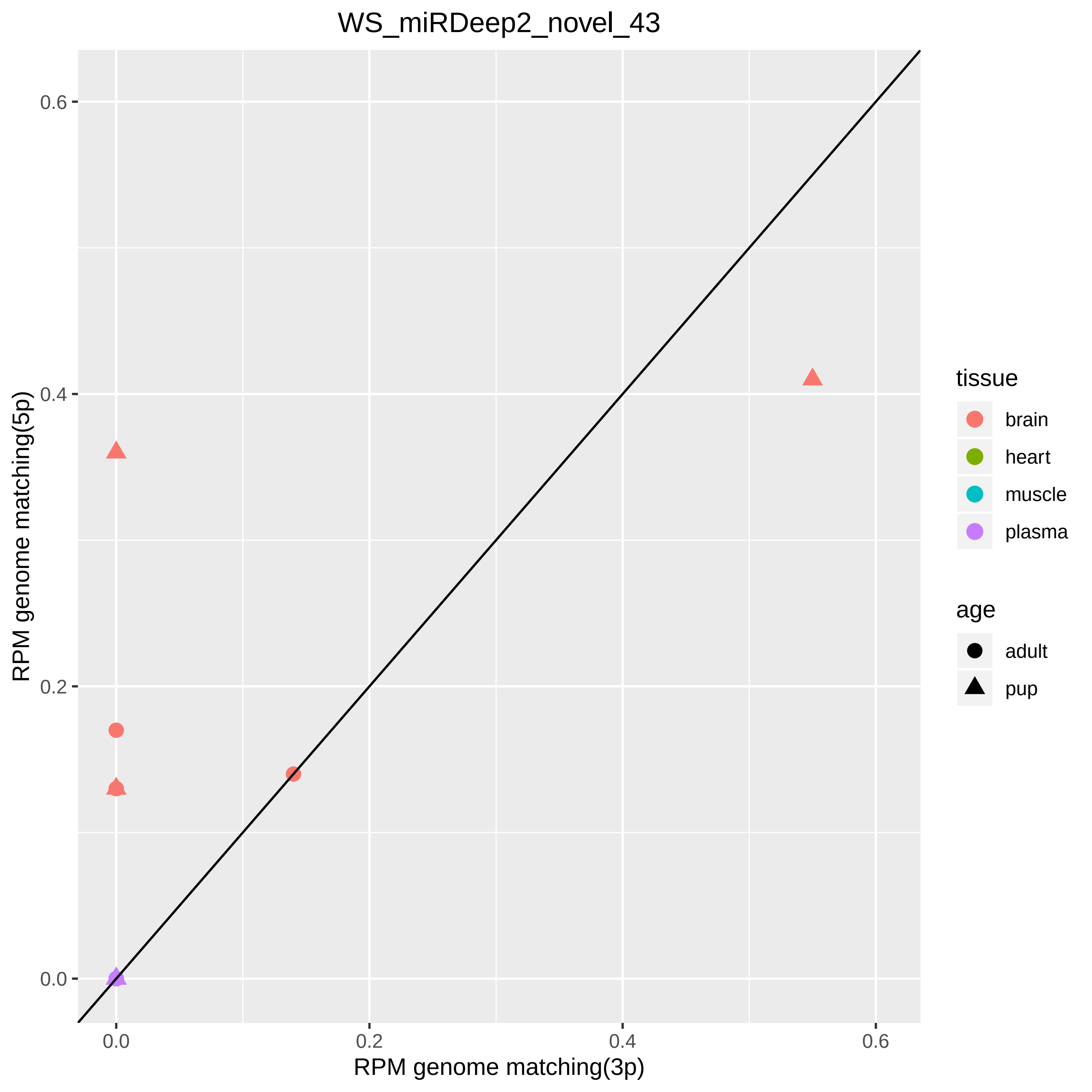

WS_miRDeep2_novel_43
RPM genome matching(5p)
RPM genome matching(3p)
0.0
0.2
0.4
0.6
tissue
brain
heart
muscle
plasma
age
adult
pup

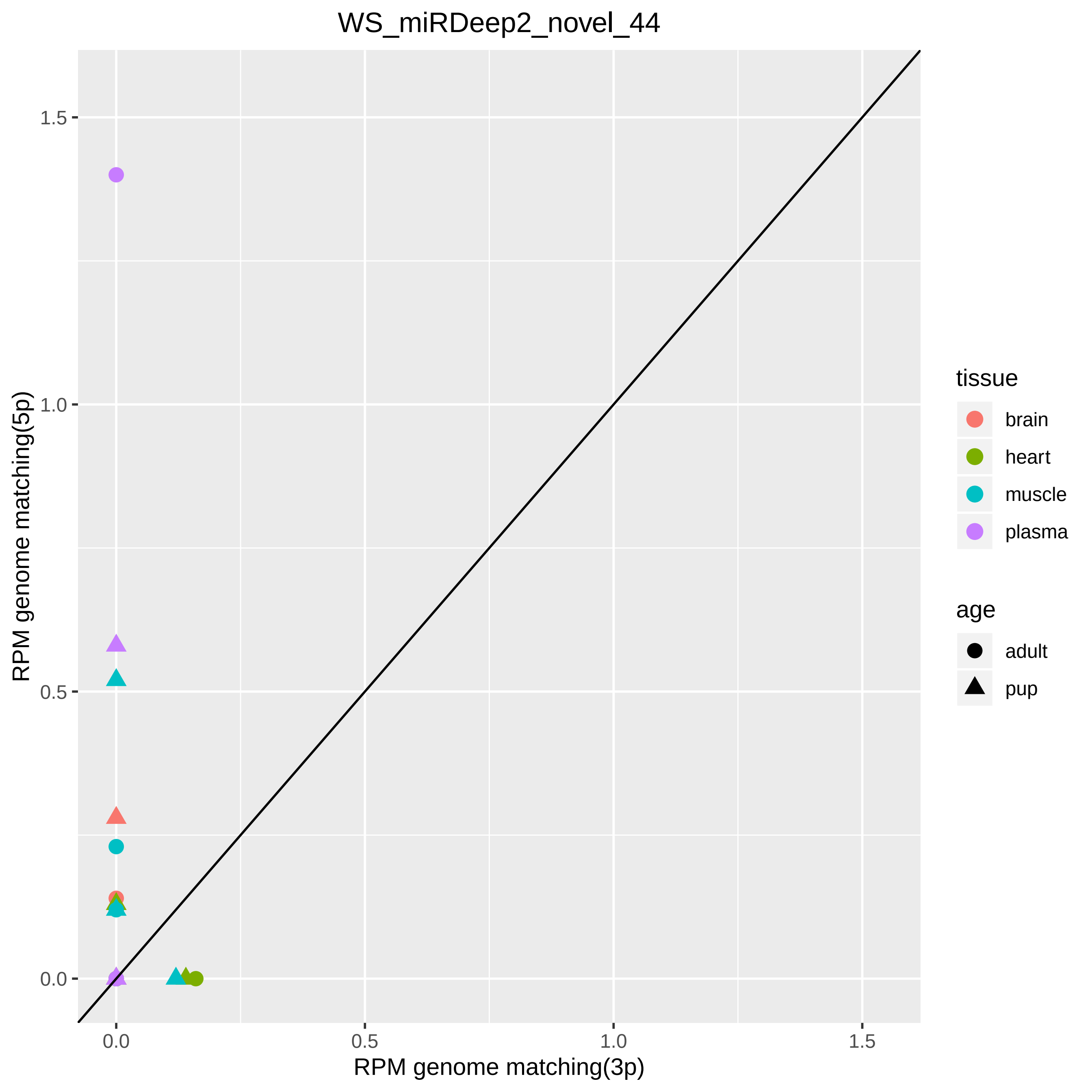
WS_miRDeep2_novel_44
RPM genome matching(5p)
RPM genome matching(3p)
0.0
0.5
1.0
1.5
0.0
0.5
1.0
1.5
tissue
brain
heart
muscle
plasma
age
adult
pup

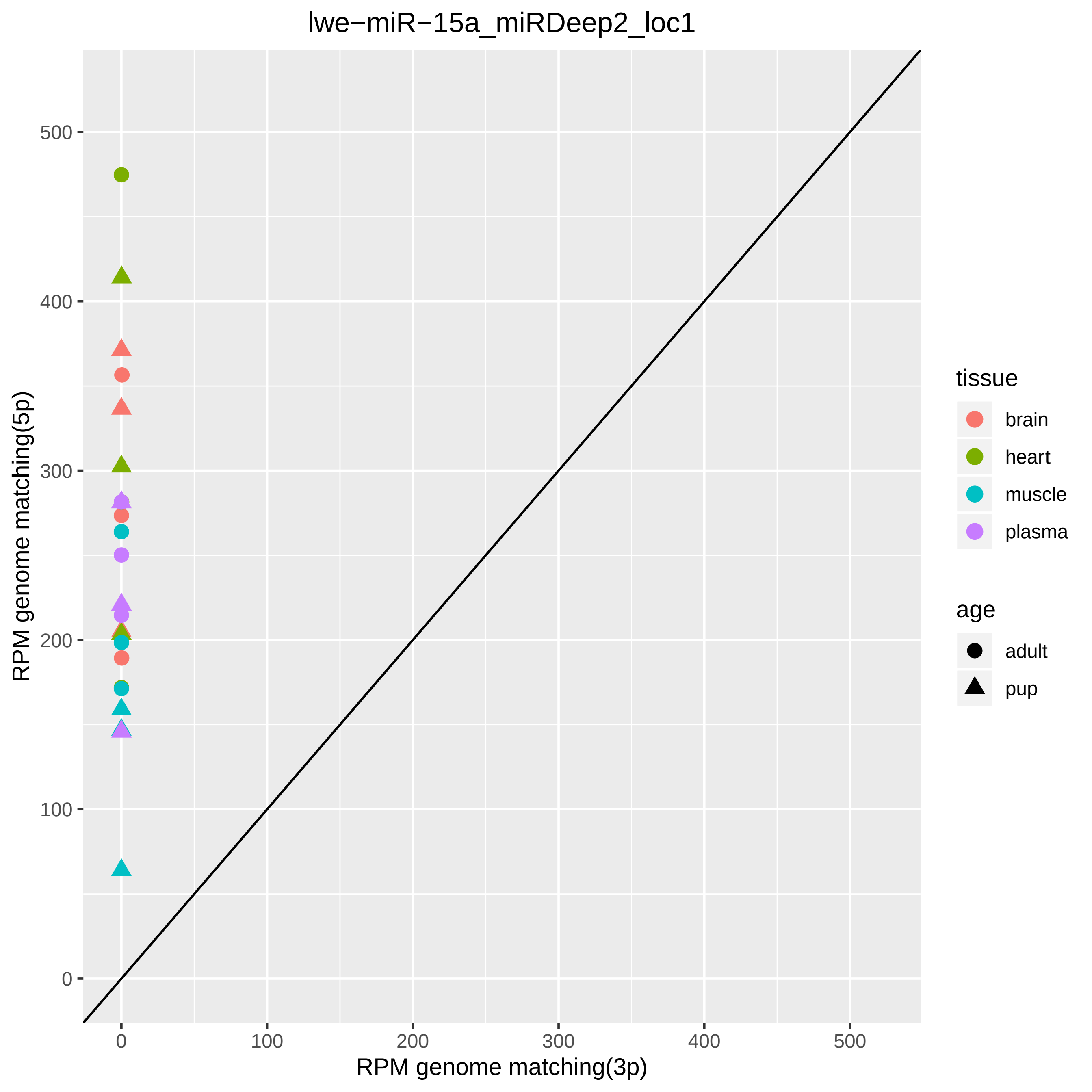
lwe−miR−15a_miRDeep2_loc1
RPM genome matching(5p)
RPM genome matching(3p)
0
100
200
300
400
500
tissue
brain
heart
muscle
plasma
age
adult
pup

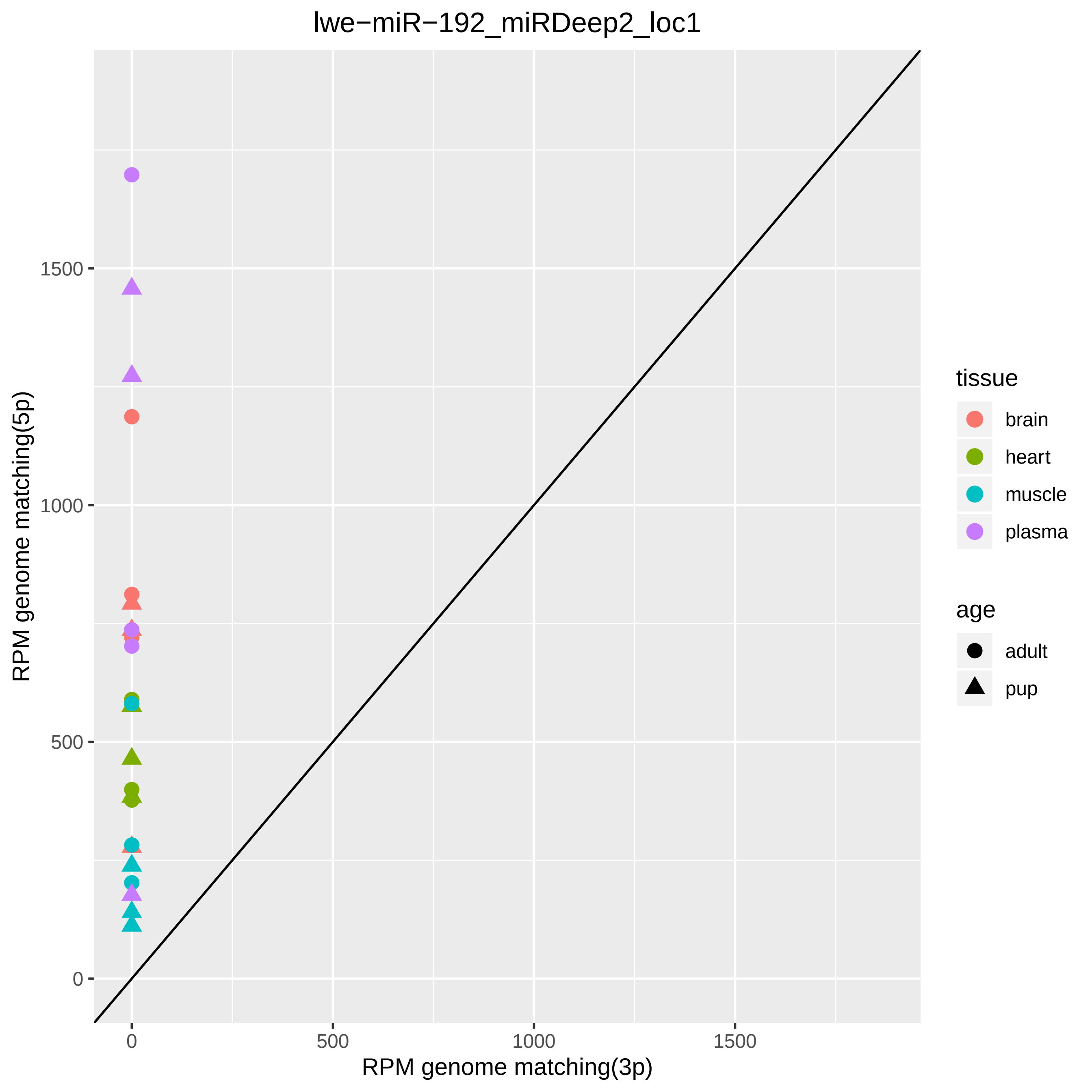
lwe−miR−192_miRDeep2_loc1
RPM genome matching(5p)
RPM genome matching(3p)
0
500
1000
1500
0
500
1000
1500
tissue
brain
heart
muscle
plasma
age
adult
pup

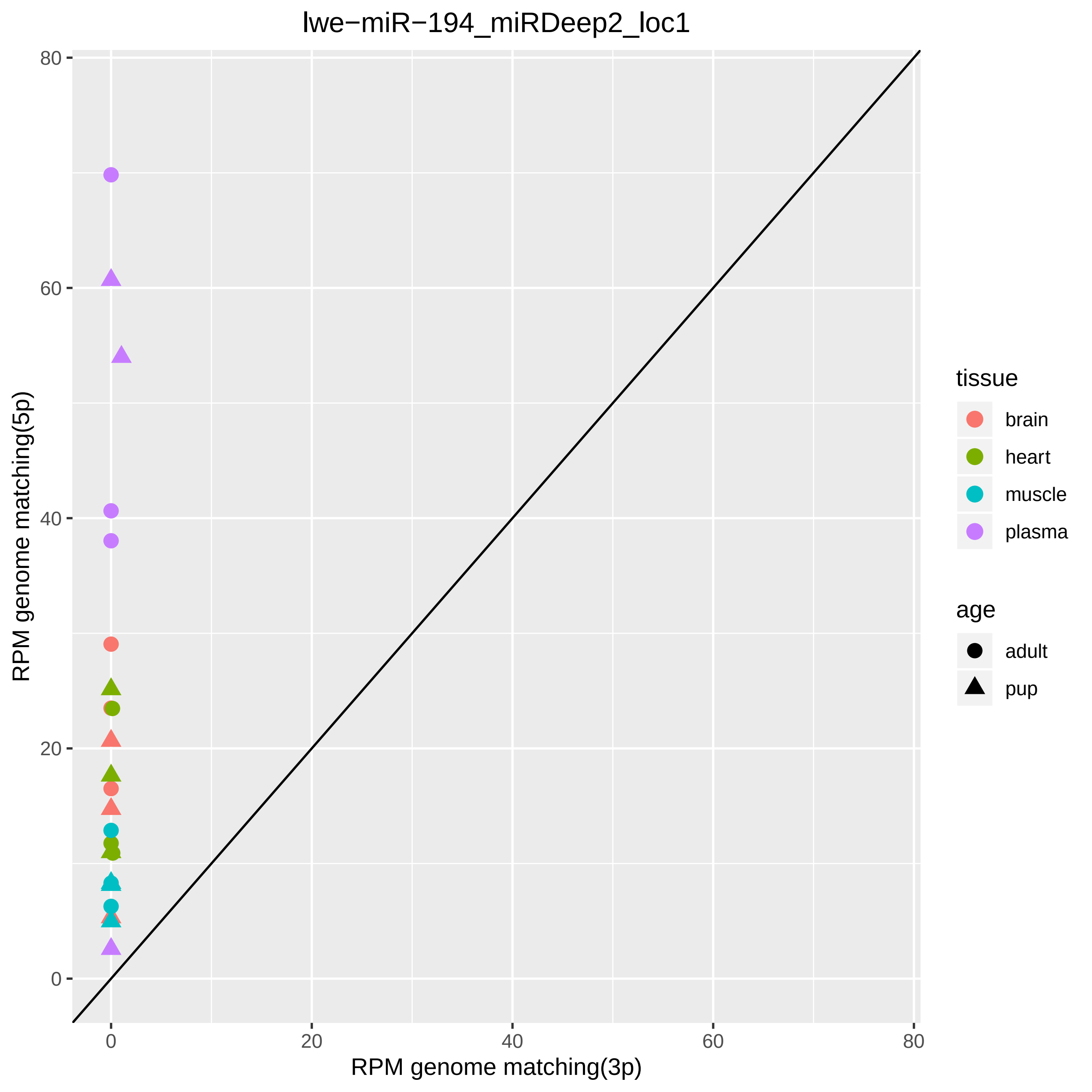

lwe−miR−194_miRDeep2_loc1
RPM genome matching(5p)
RPM genome matching(3p)
0
20
40
60
80
0
20
40
60
80
tissue
brain
heart
muscle
plasma
age
adult
pup

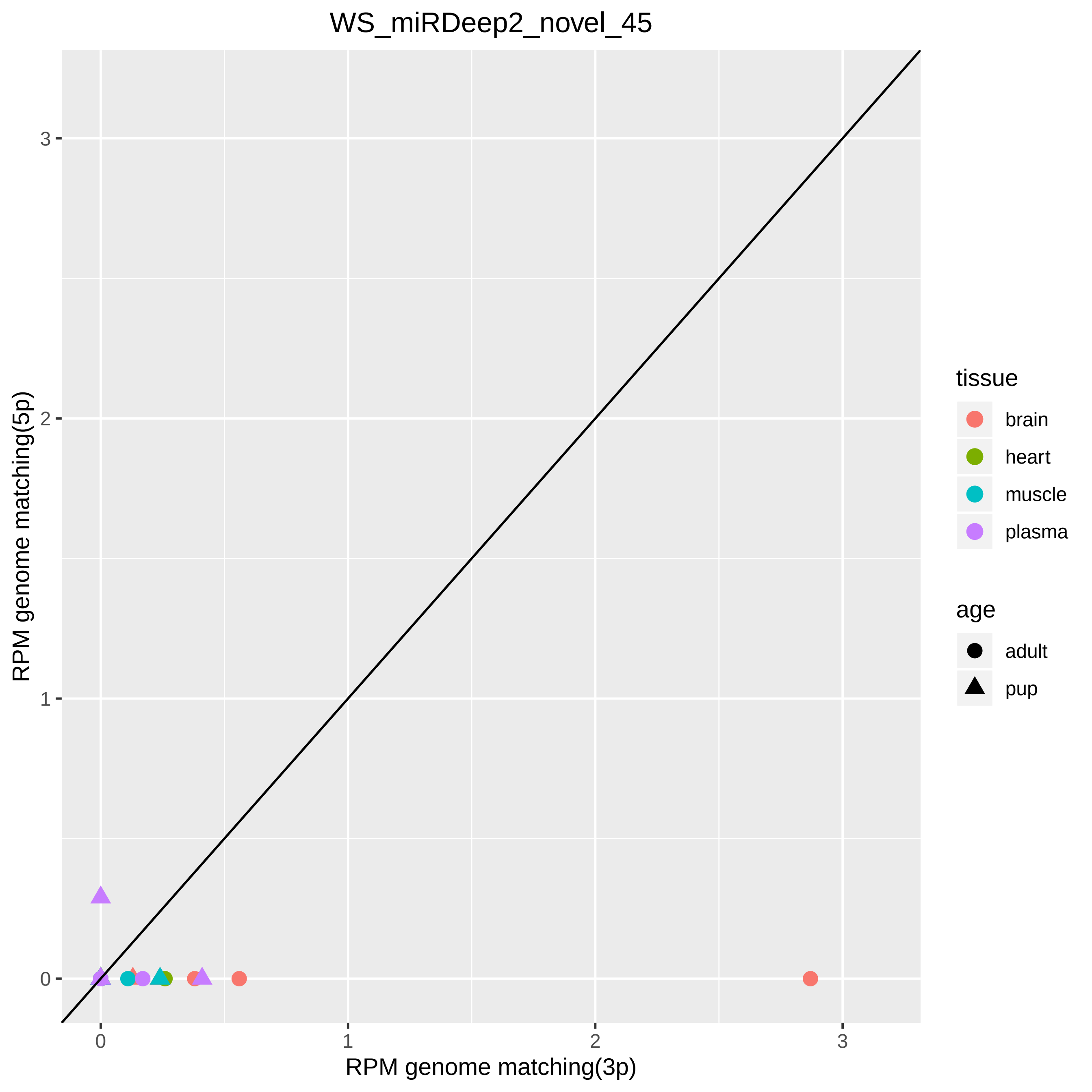

WS_miRDeep2_novel_45
RPM genome matching(5p)
RPM genome matching(3p)
0
1
2
3
0
1
2
3
tissue
brain
heart
muscle
plasma
age
adult
pup

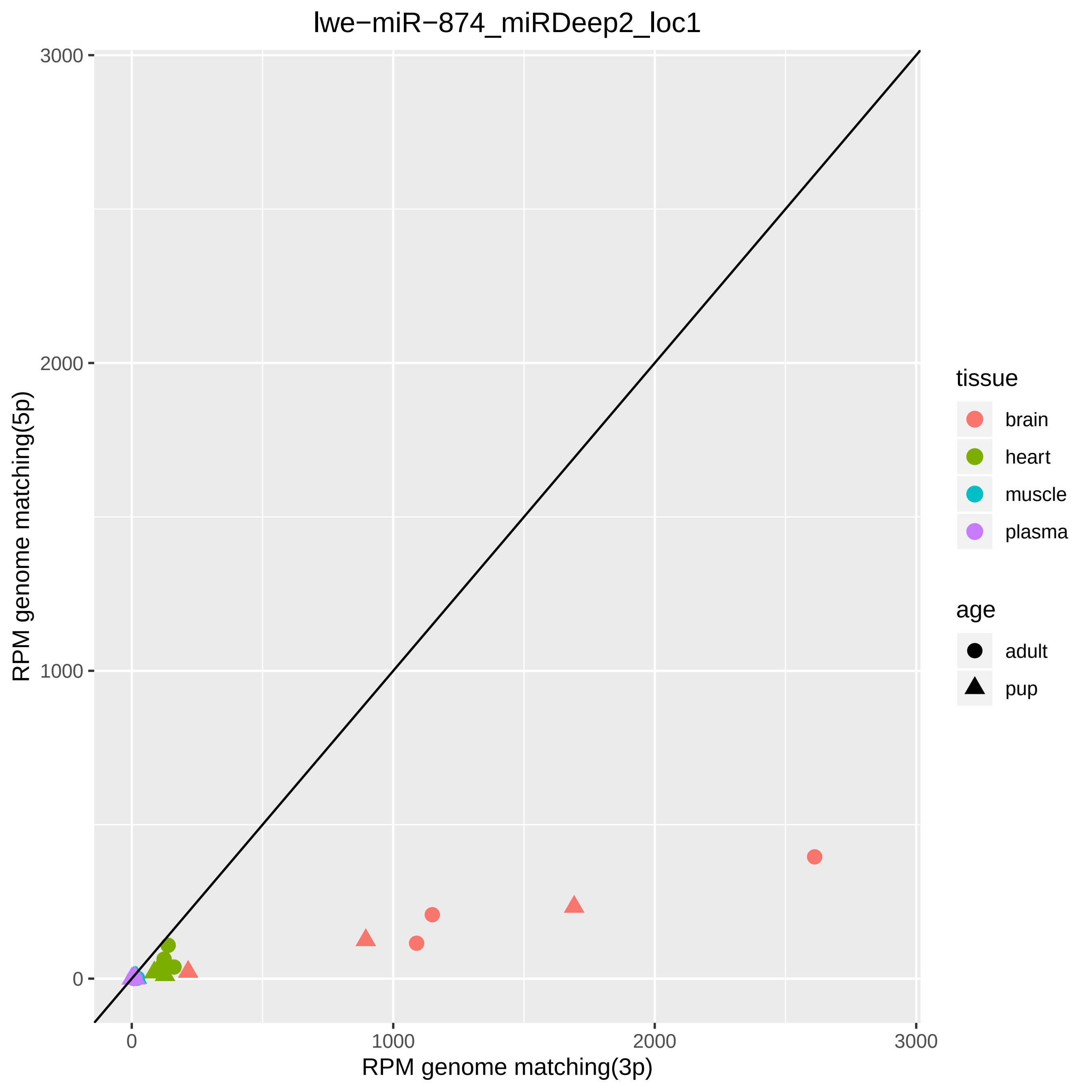
lwe−miR−874_miRDeep2_loc1
3000
2000
1000
0
RPM genome matching(5p)
0
1000
2000
3000
RPM genome matching(3p)
tissue
brain
heart
muscle
plasma
age
adult
pup

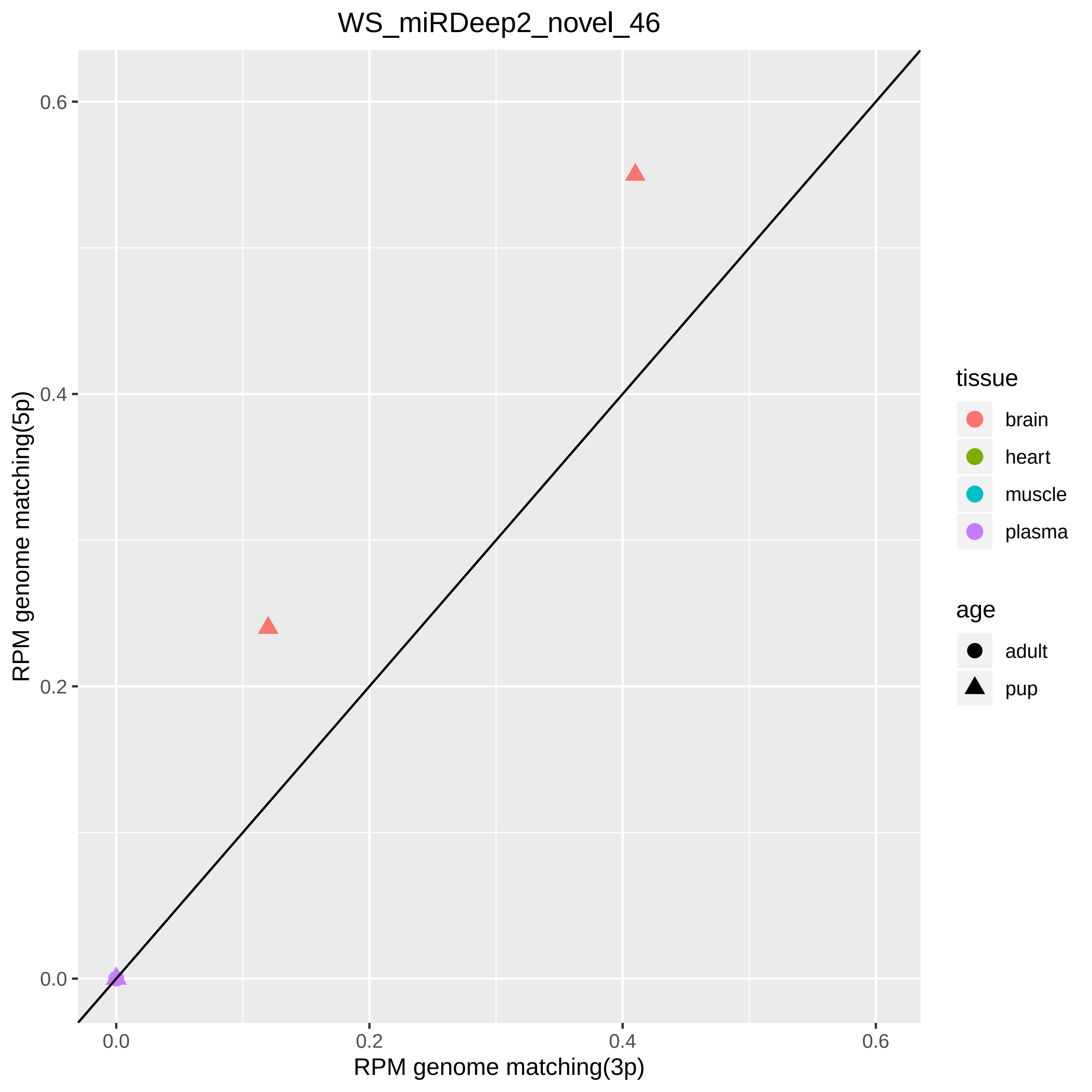
WS_miRDeep2_novel_46
RPM genome matching(5p)
RPM genome matching(3p)
0.0
0.2
0.4
0.6
tissue
brain
heart
muscle
plasma
age
adult
pup

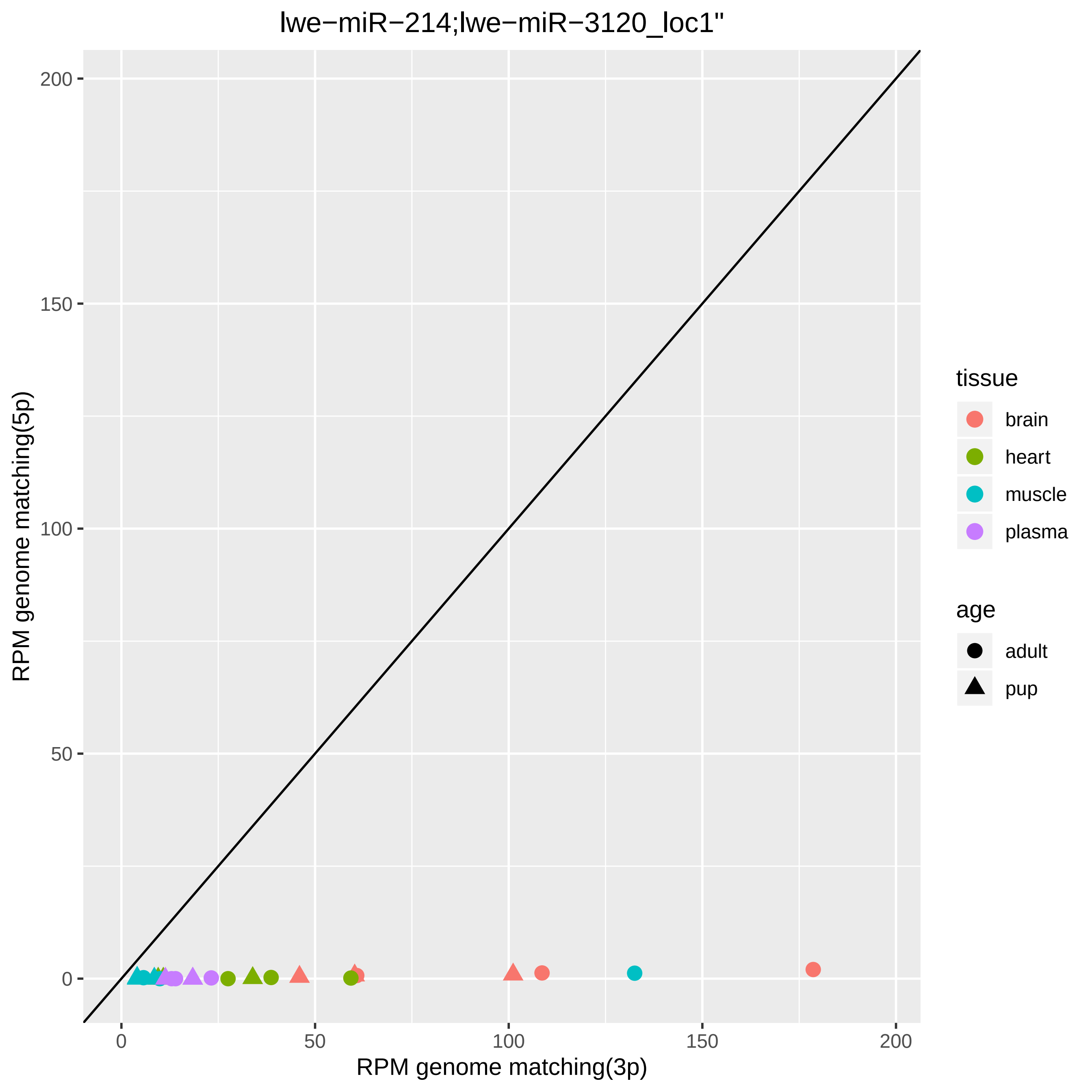

lwe−miR−214;lwe−miR−3120_loc1"
RPM genome matching(5p)
RPM genome matching(3p)
0
50
100
150
200
tissue
brain
heart
muscle
plasma
age
adult
pup

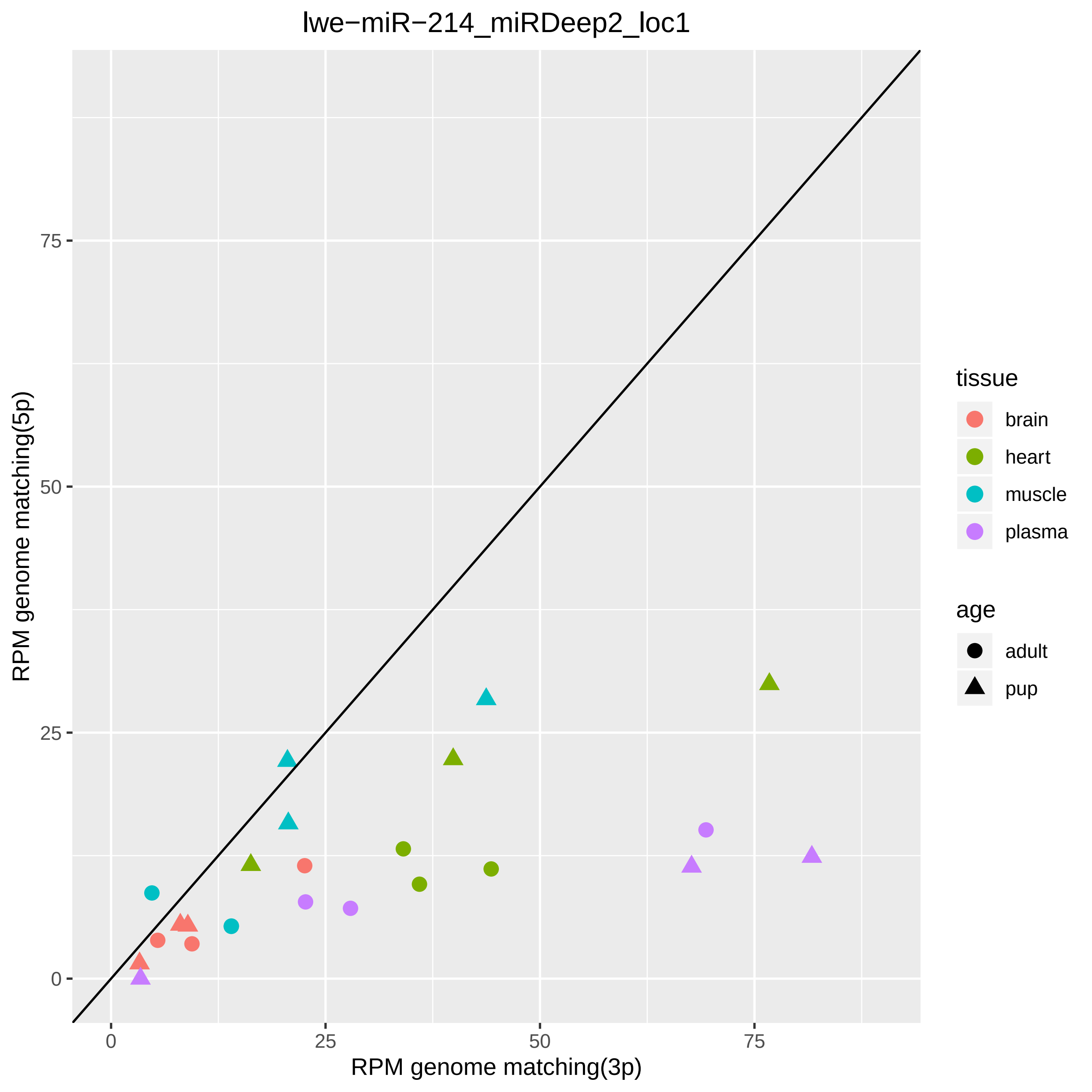
lwe−miR−214_miRDeep2_loc1
RPM genome matching(5p)
RPM genome matching(3p)
0
25
50
75
tissue
brain
heart
muscle
plasma
age
adult
pup

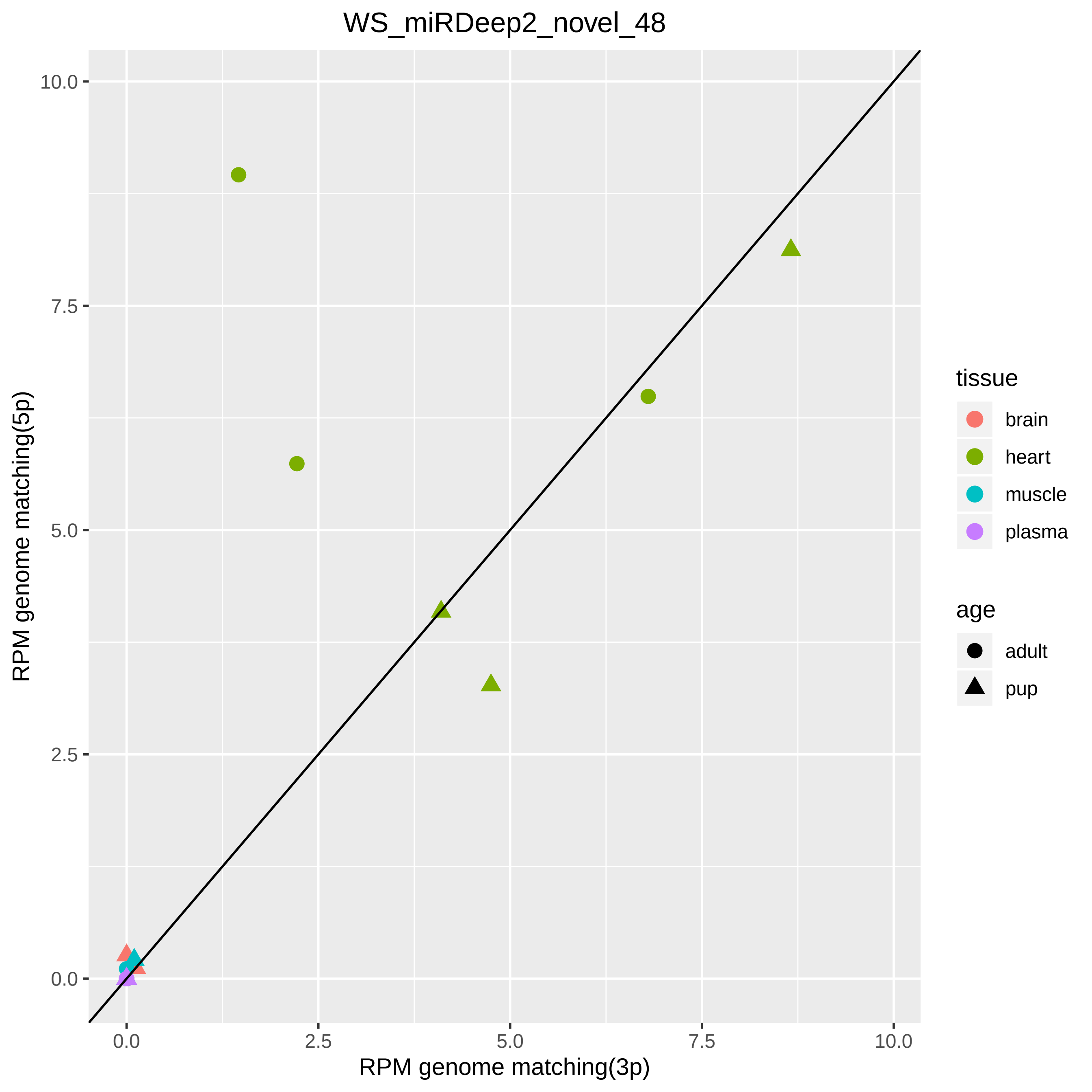

WS_miRDeep2_novel_48
RPM genome matching(5p)
RPM genome matching(3p)
10.0
7.5
5.0
2.5
0.0
0.0
2.5
5.0
7.5
10.0
tissue
brain
heart
muscle
plasma
age
adult
pup

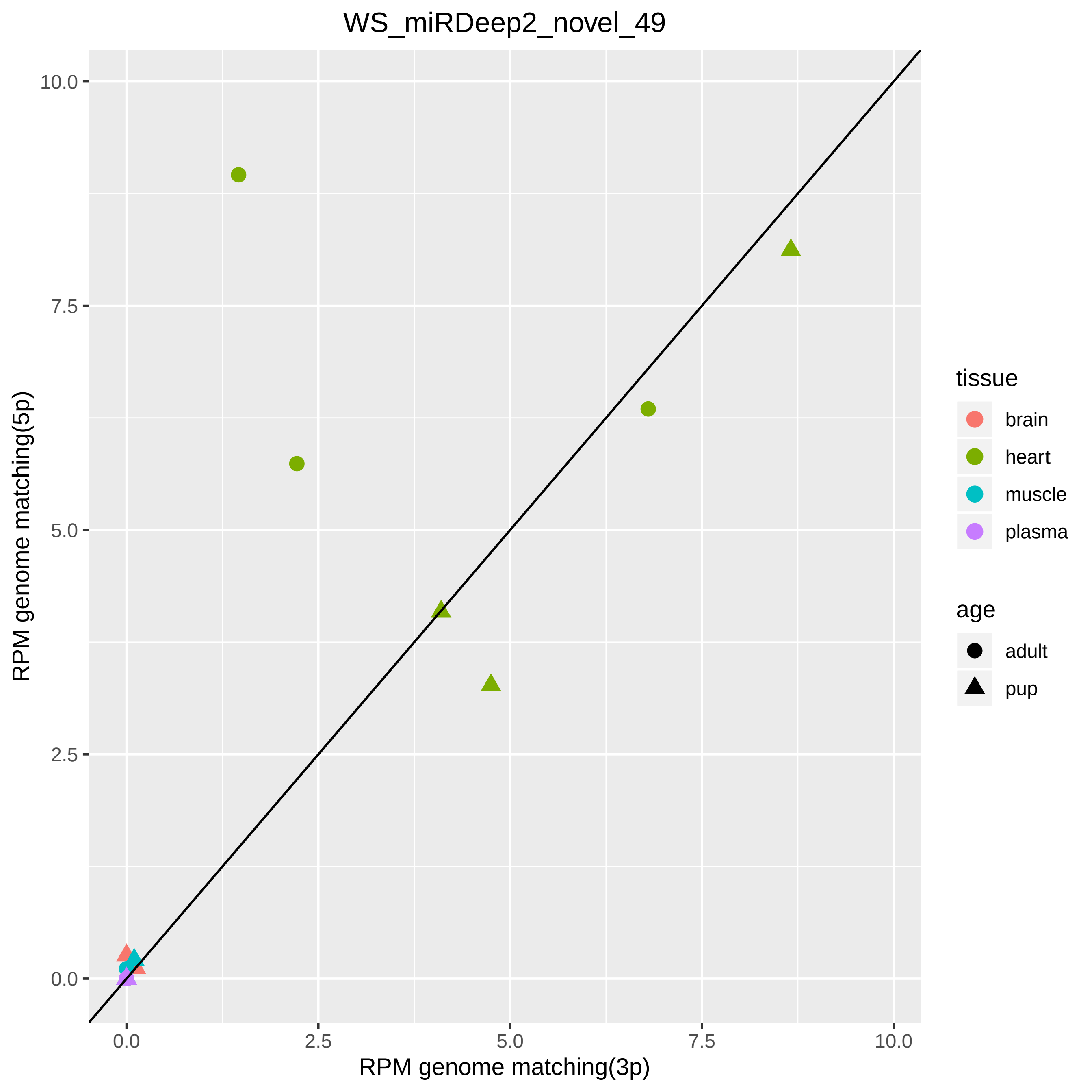

WS_miRDeep2_novel_49
RPM genome matching(5p)
RPM genome matching(3p)
0.0
2.5
5.0
7.5
10.0
tissue
brain
heart
muscle
plasma
age
adult
pup

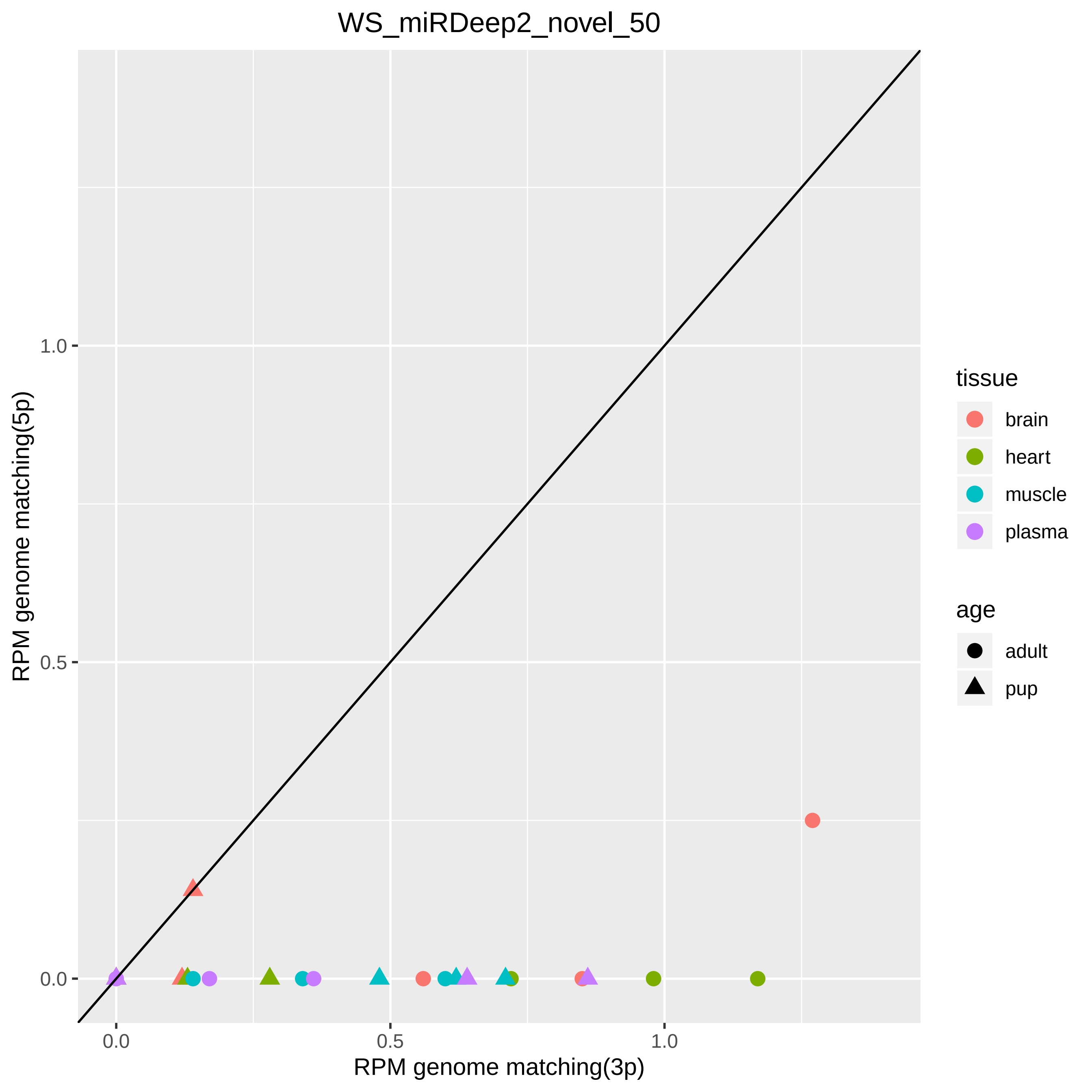
WS_miRDeep2_novel_50
RPM genome matching(5p)
RPM genome matching(3p)
0.0
0.5
1.0
tissue
brain
heart
muscle
plasma
age
adult
pup

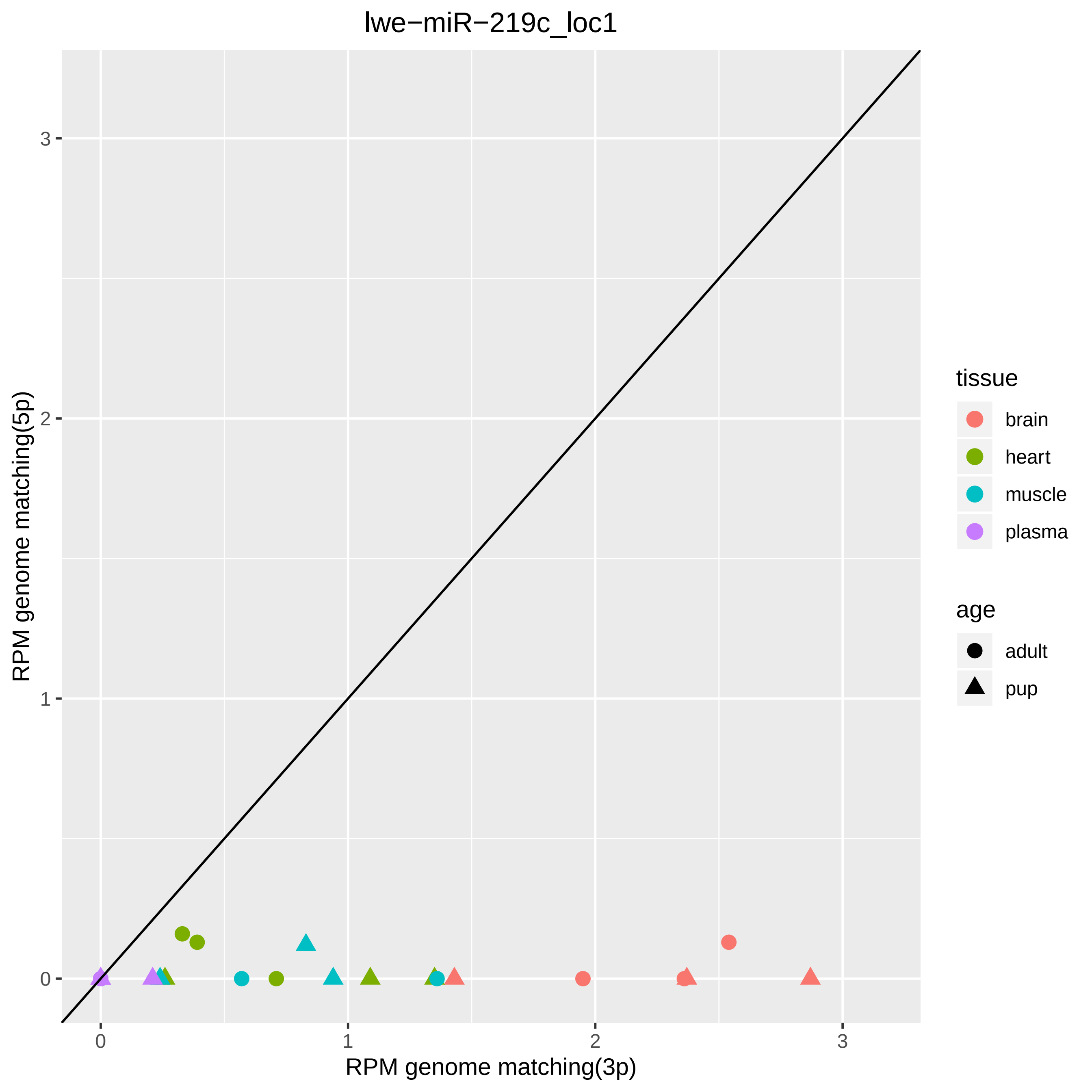

lwe−miR−219c_loc1
RPM genome matching(5p)
RPM genome matching(3p)
0
1
2
3
0
1
2
3
tissue
brain
heart
muscle
plasma
age
adult
pup

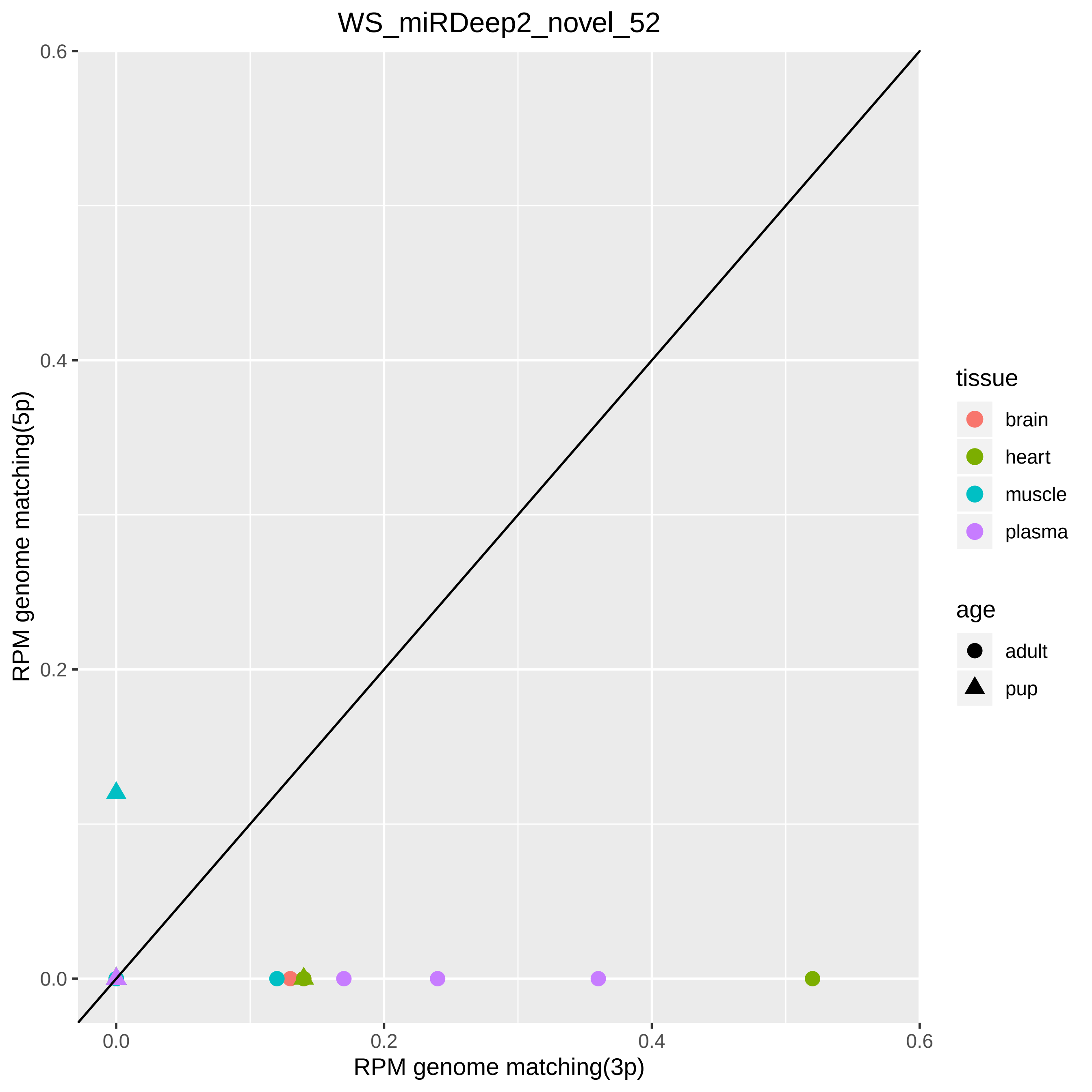

WS_miRDeep2_novel_52
RPM genome matching(5p)
RPM genome matching(3p)
0.0
0.2
0.4
0.6
tissue
brain
heart
muscle
plasma
age
adult
pup

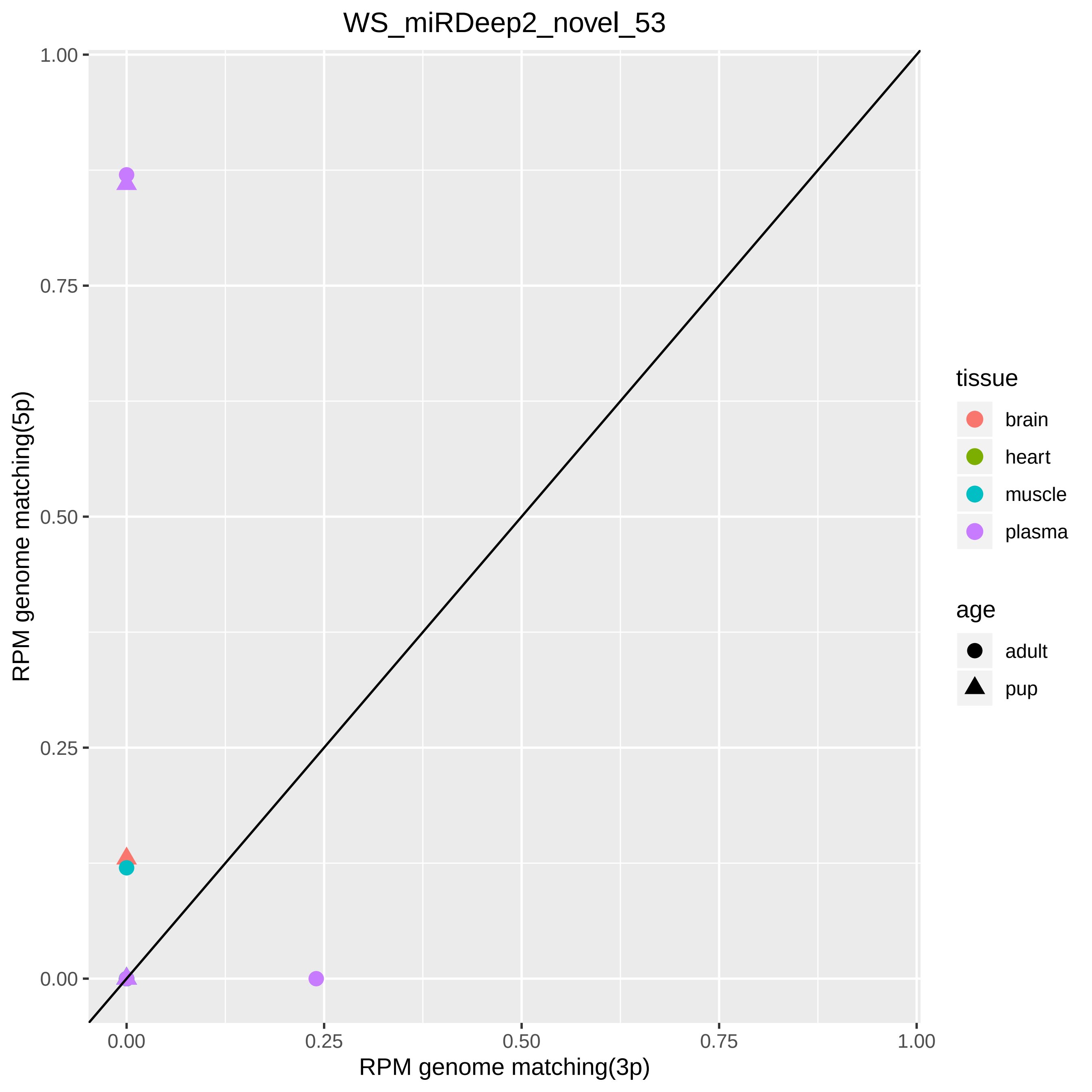

WS_miRDeep2_novel_53
RPM genome matching(5p)
RPM genome matching(3p)
0.00
0.25
0.50
0.75
1.00
tissue
brain
heart
muscle
plasma
age
adult
pup

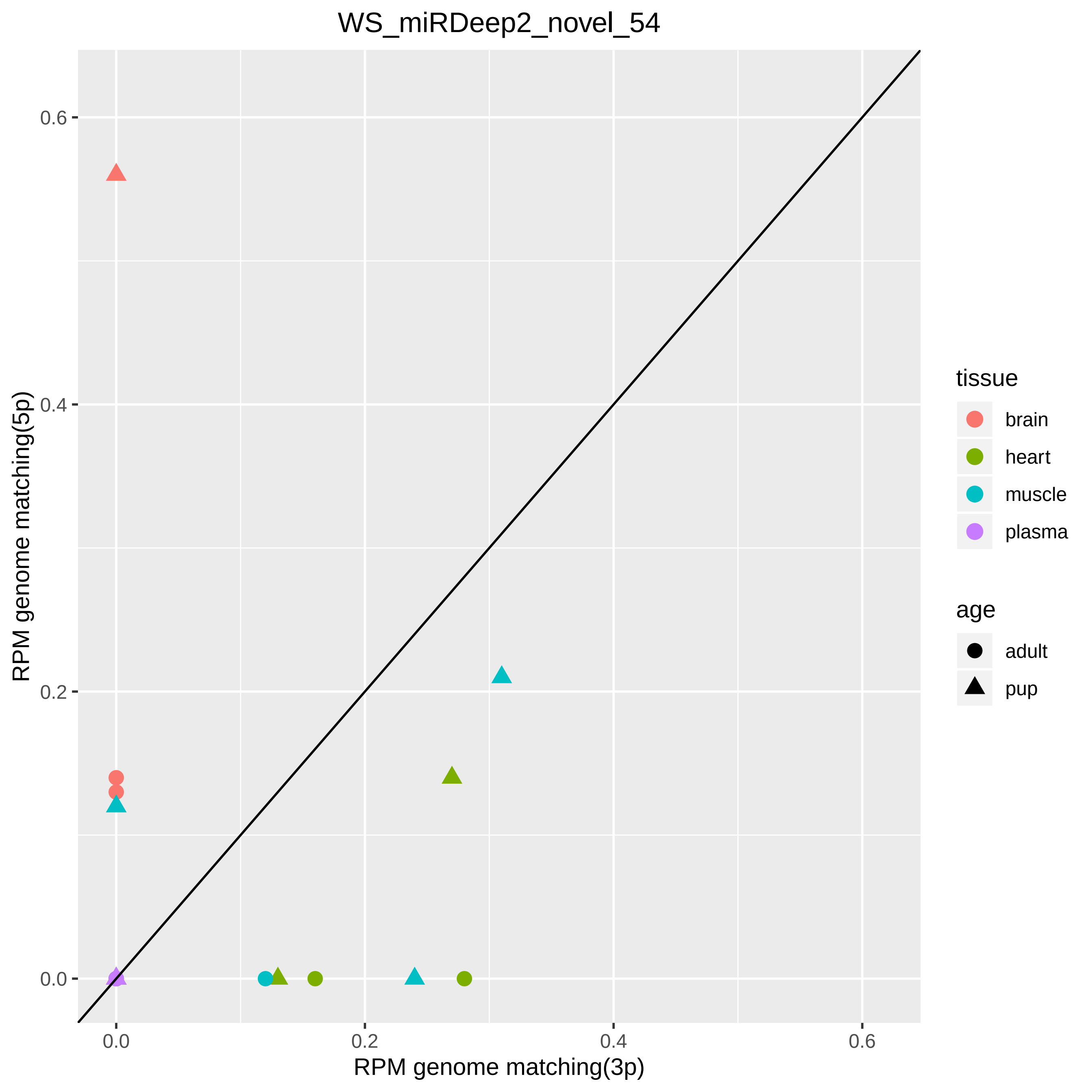

WS_miRDeep2_novel_54
RPM genome matching(5p)
RPM genome matching(3p)
0.0
0.2
0.4
0.6
tissue
brain
heart
muscle
plasma
age
adult
pup

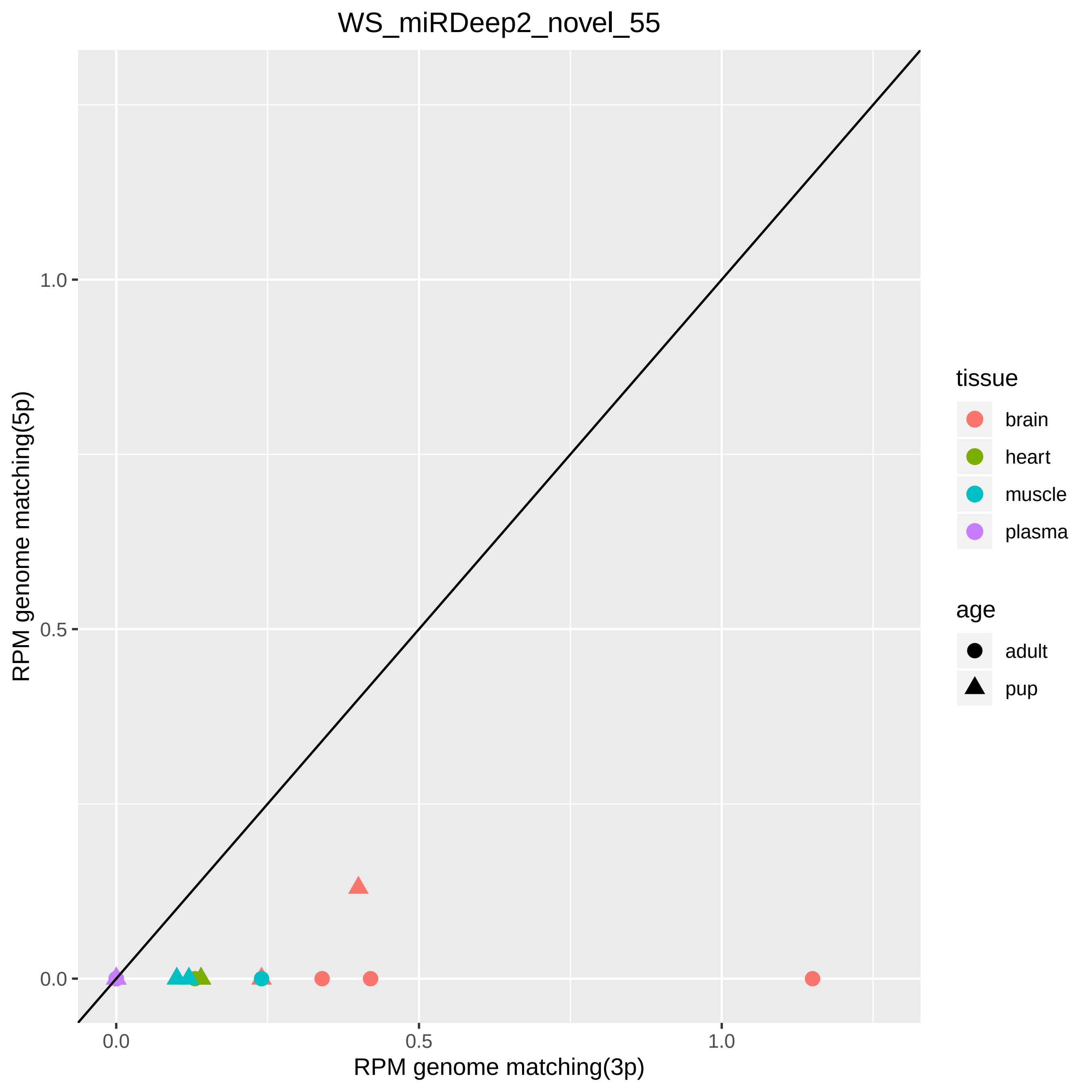

WS_miRDeep2_novel_55
RPM genome matching(5p)
RPM genome matching(3p)
0.0
0.5
1.0
tissue
brain
heart
muscle
plasma
age
adult
pup

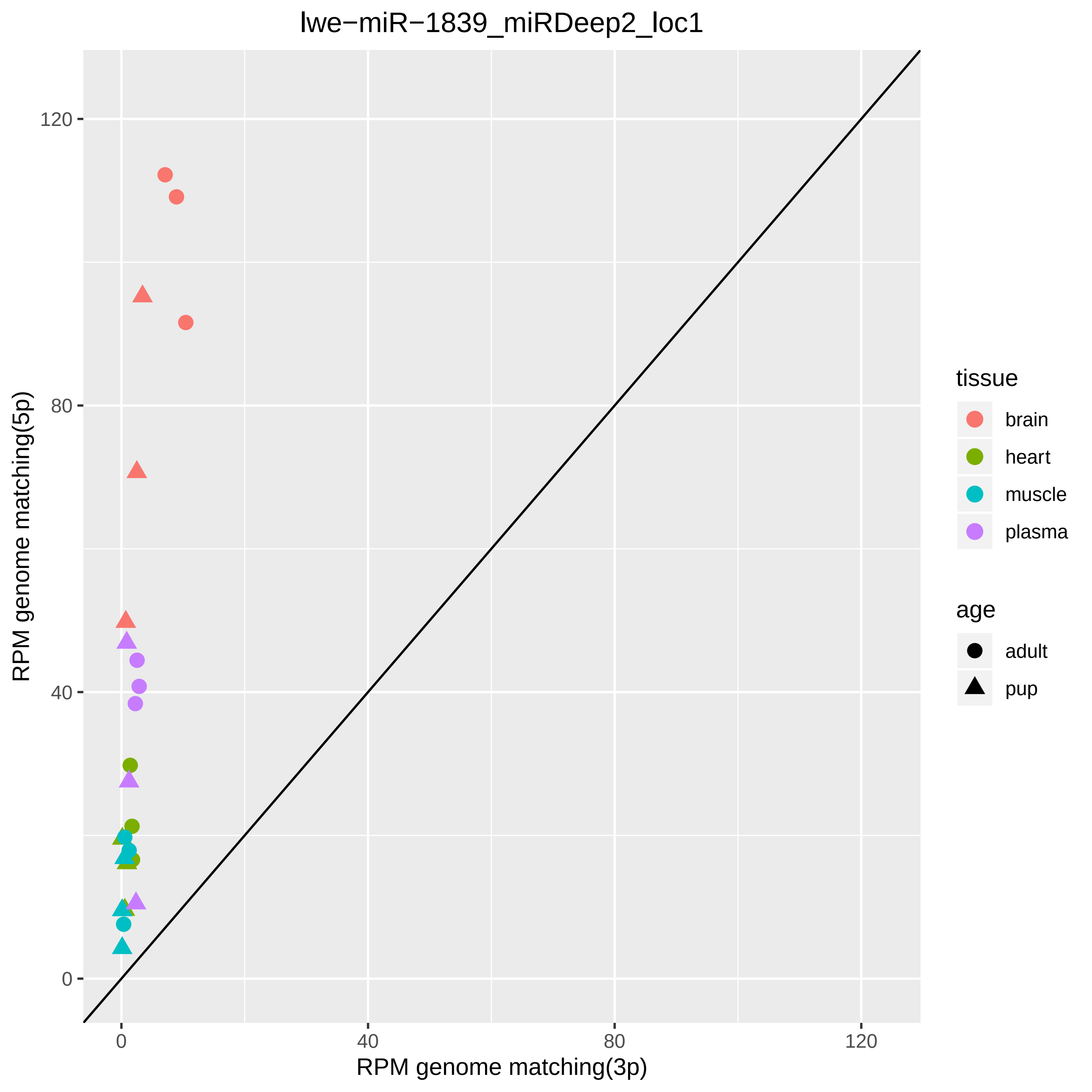
lwe−miR−1839_miRDeep2_loc1
RPM genome matching(5p)
RPM genome matching(3p)
0
40
80
120
0
40
80
120
tissue
brain
heart
muscle
plasma
age
adult
pup

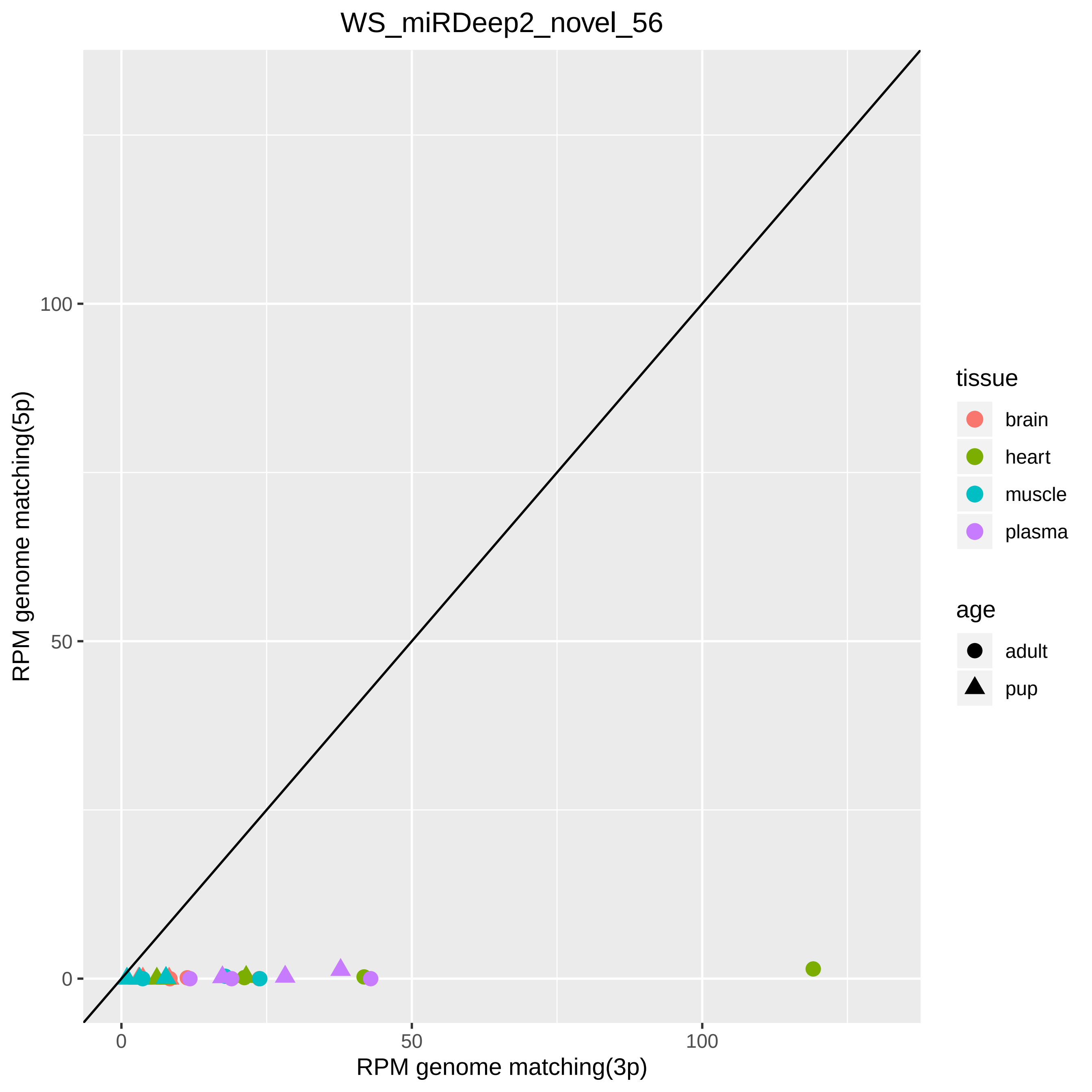
WS_miRDeep2_novel_56
RPM genome matching(5p)
RPM genome matching(3p)
0
50
100
0
50
100
tissue
brain
heart
muscle
plasma
age
adult
pup

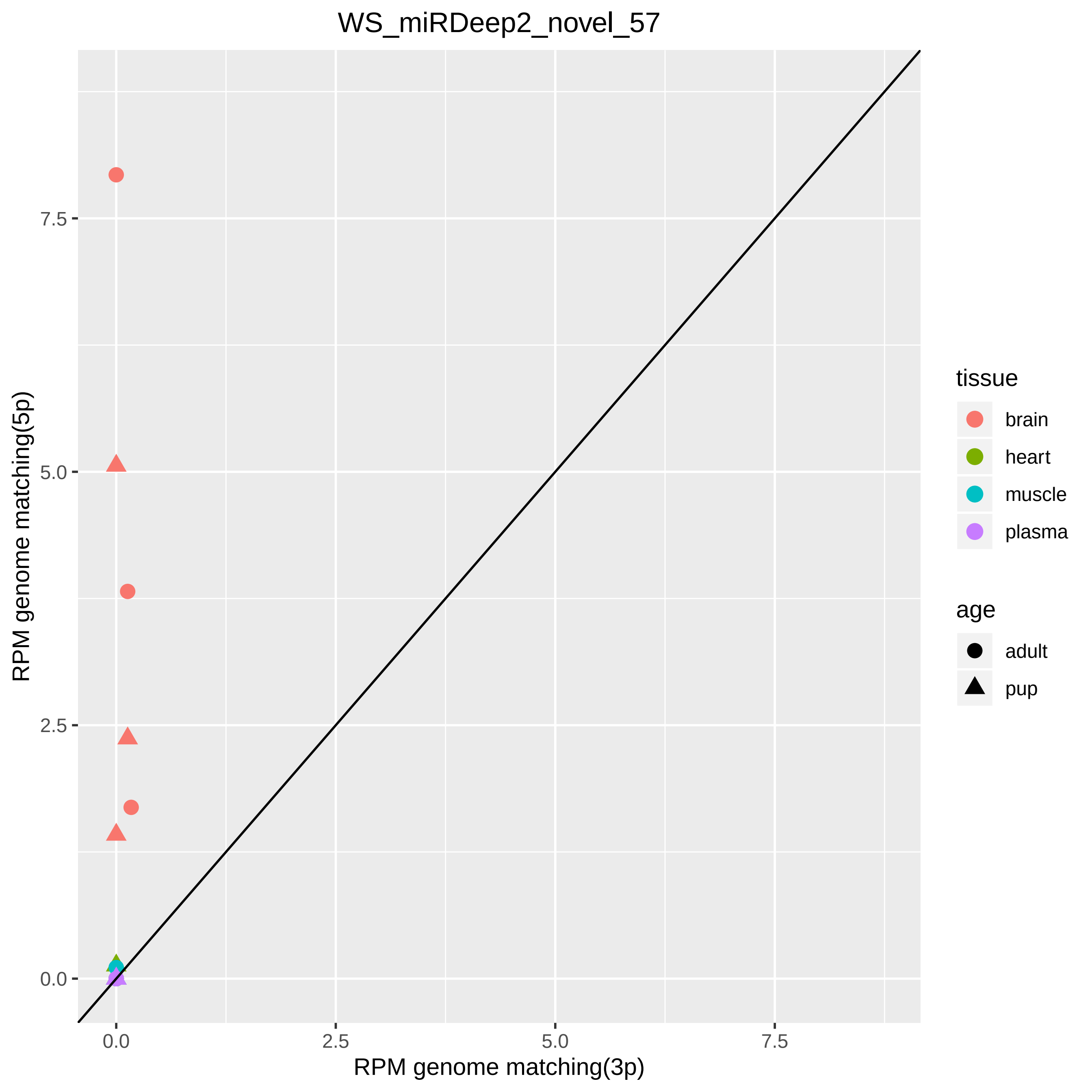

WS_miRDeep2_novel_57
RPM genome matching(5p)
RPM genome matching(3p)
0.0
2.5
5.0
7.5
tissue
brain
heart
muscle
plasma
age
adult
pup

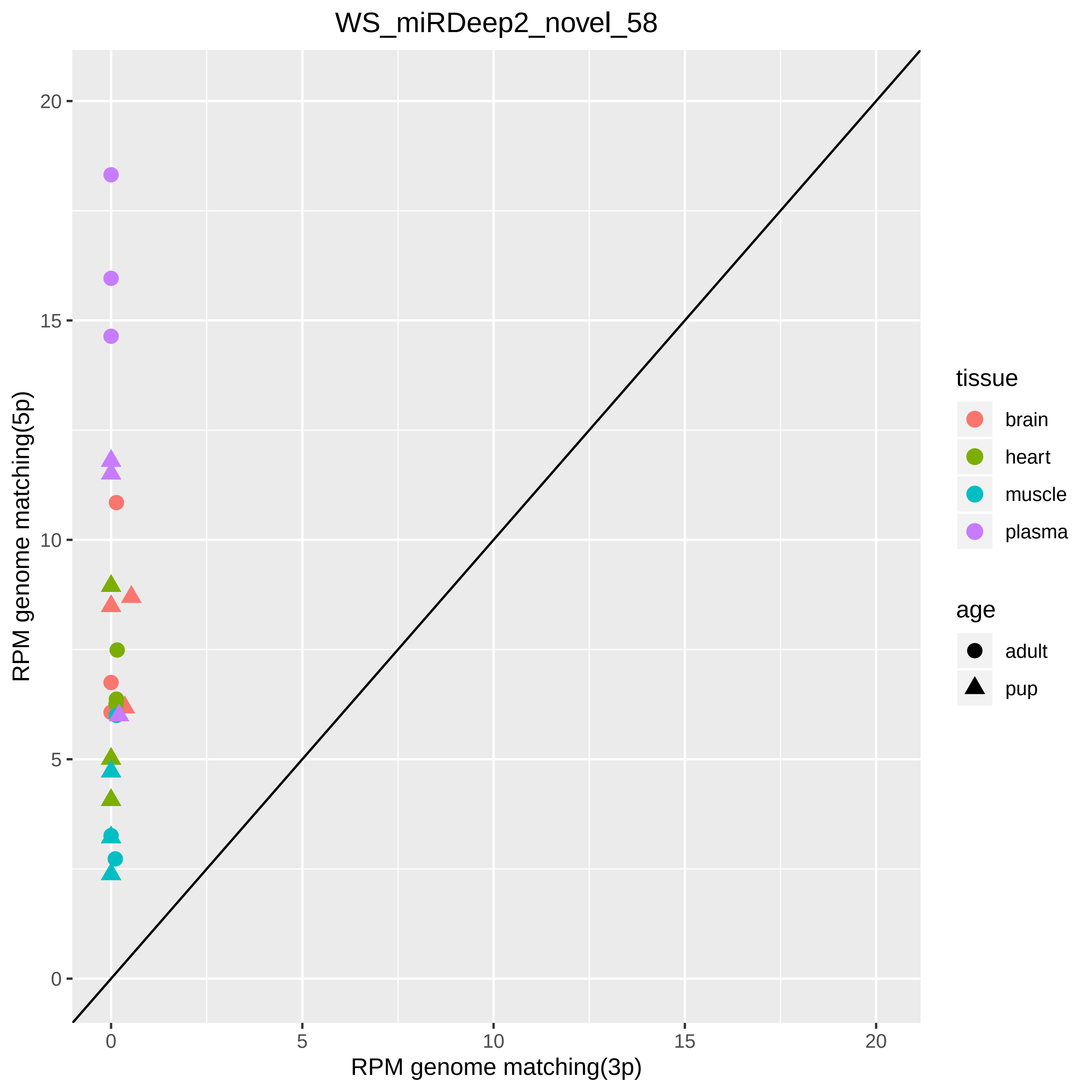

WS_miRDeep2_novel_58
RPM genome matching(5p)
RPM genome matching(3p)
0
5
10
15
20
tissue
brain
heart
muscle
plasma
age
adult
pup

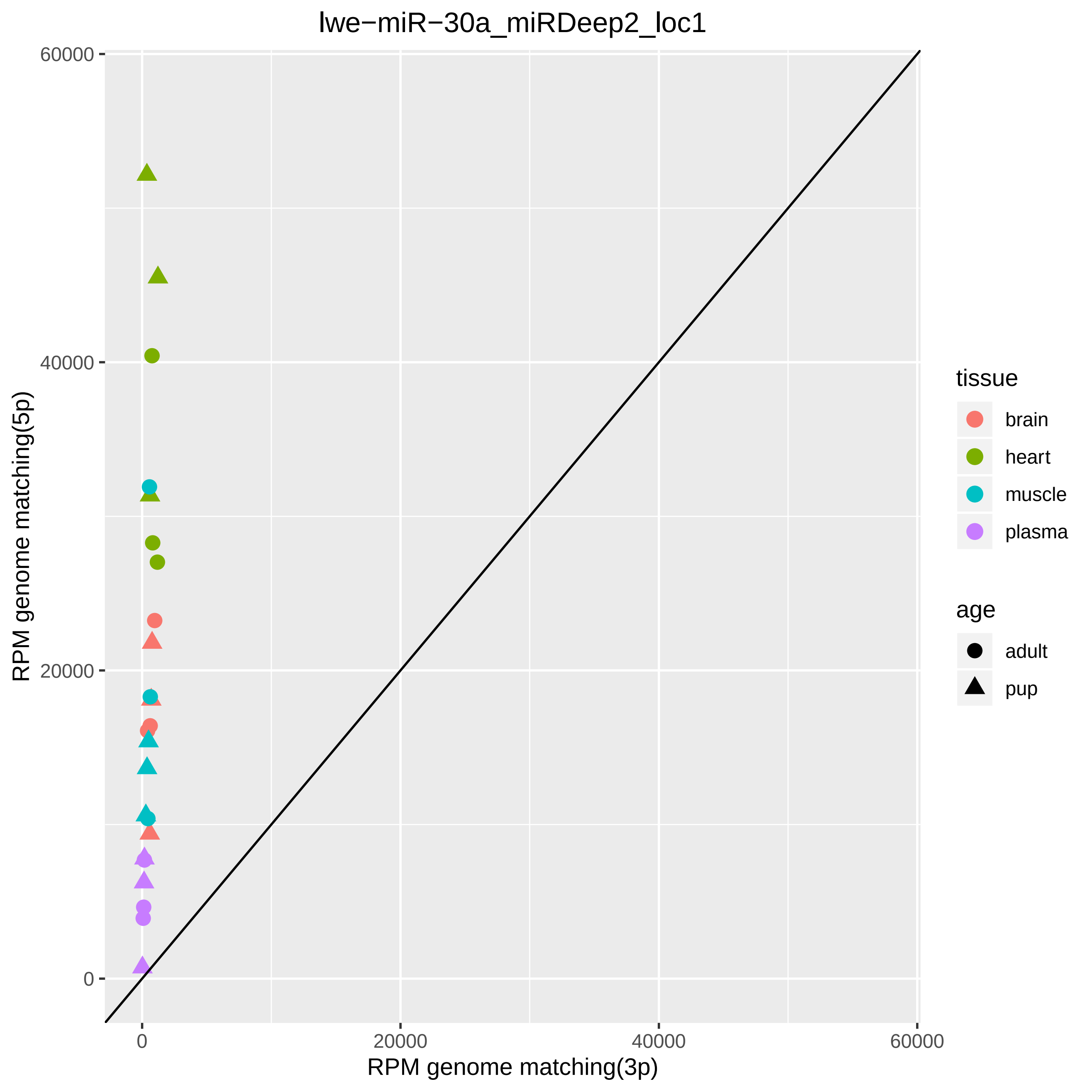

lwe−miR−30a_miRDeep2_loc1
RPM genome matching(5p)
RPM genome matching(3p)
0
20000
40000
60000
0
20000
40000
60000
tissue
brain
heart
muscle
plasma
age
adult
pup

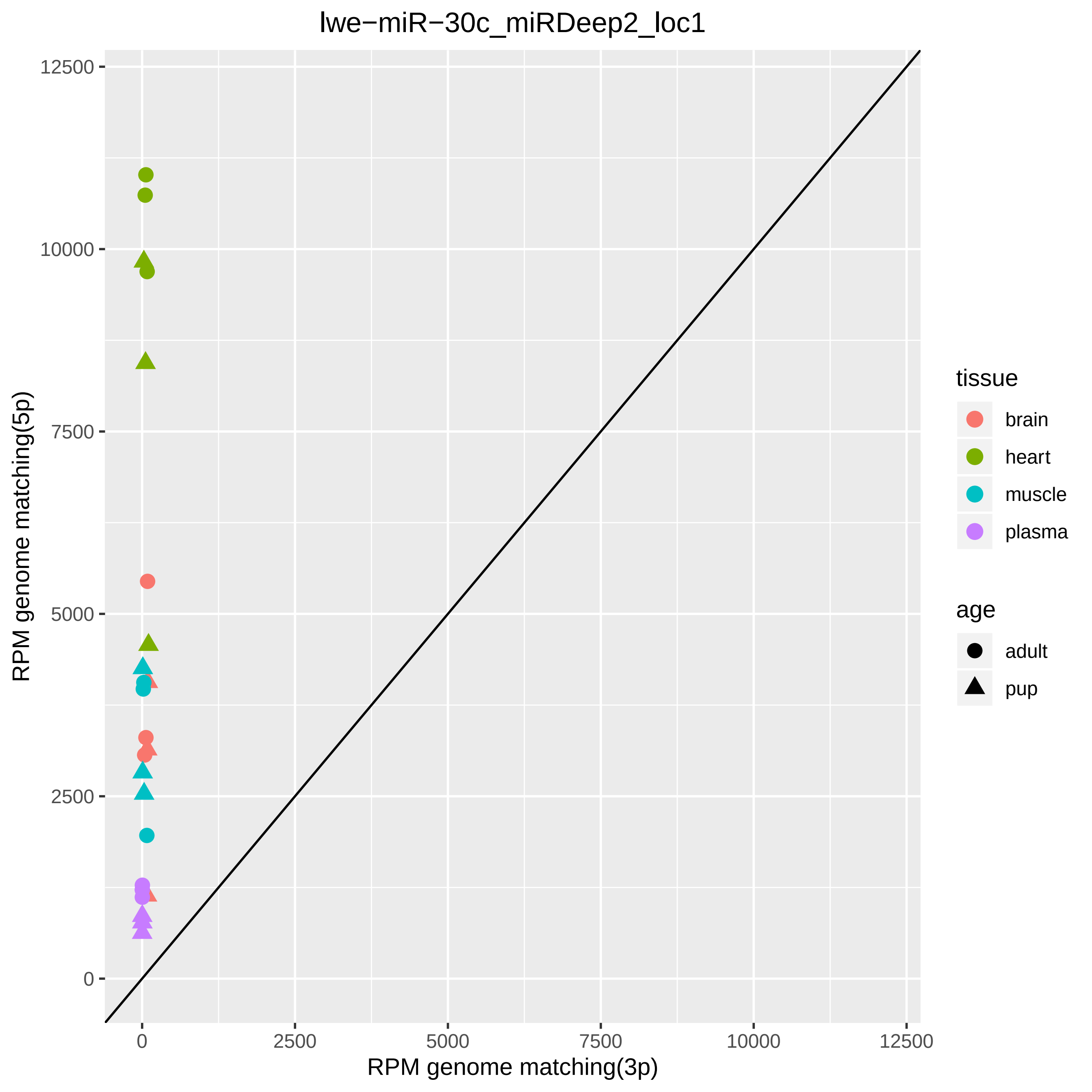

lwe−miR−30c_miRDeep2_loc1
RPM genome matching(5p)
12500
10000
7500
5000
2500
0
RPM genome matching(3p)
0
2500
5000
7500
10000
12500
tissue
brain
heart
muscle
plasma
age
adult
pup

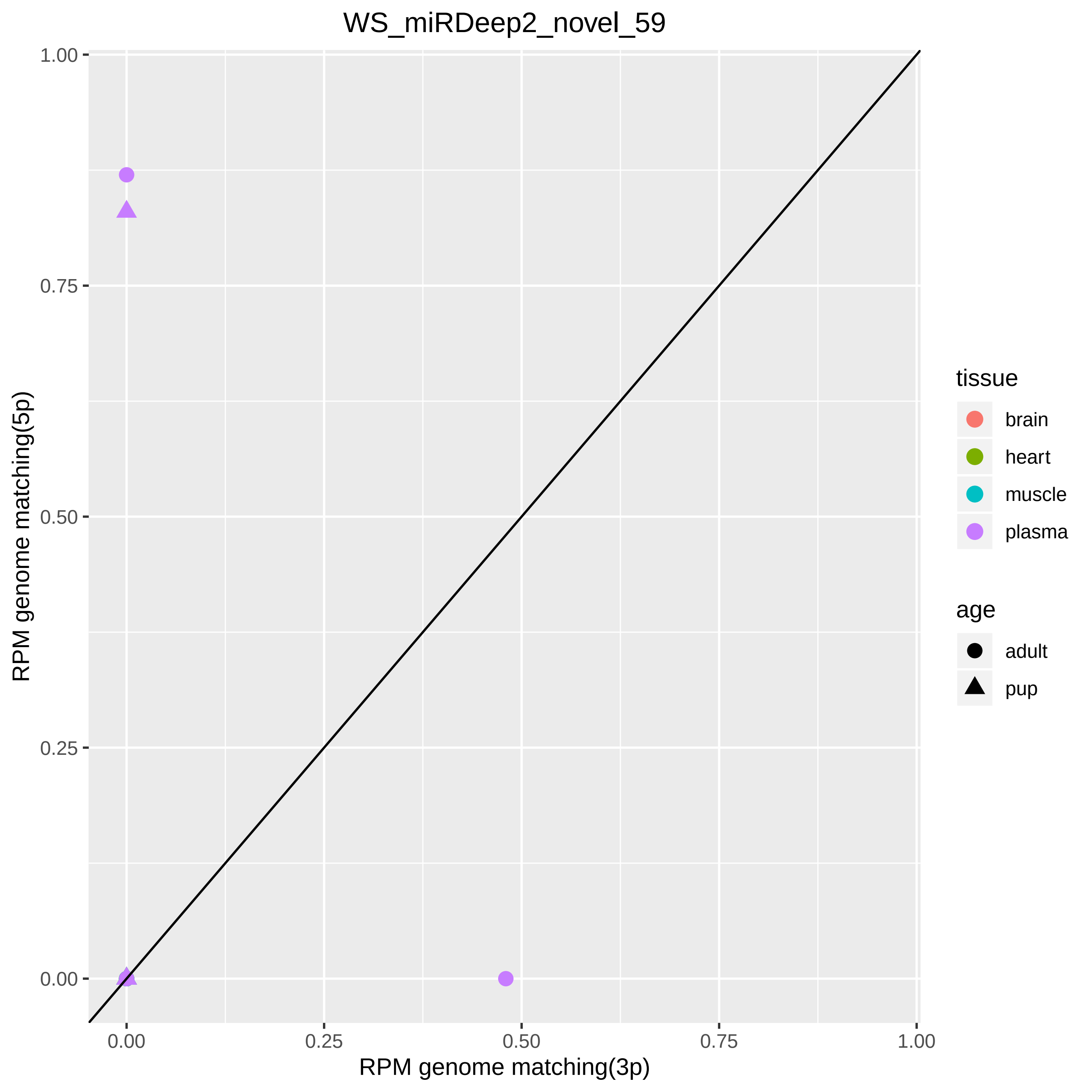
WS_miRDeep2_novel_59
RPM genome matching(5p)
RPM genome matching(3p)
0.00
0.25
0.50
0.75
1.00
tissue
brain
heart
muscle
plasma
age
adult
pup

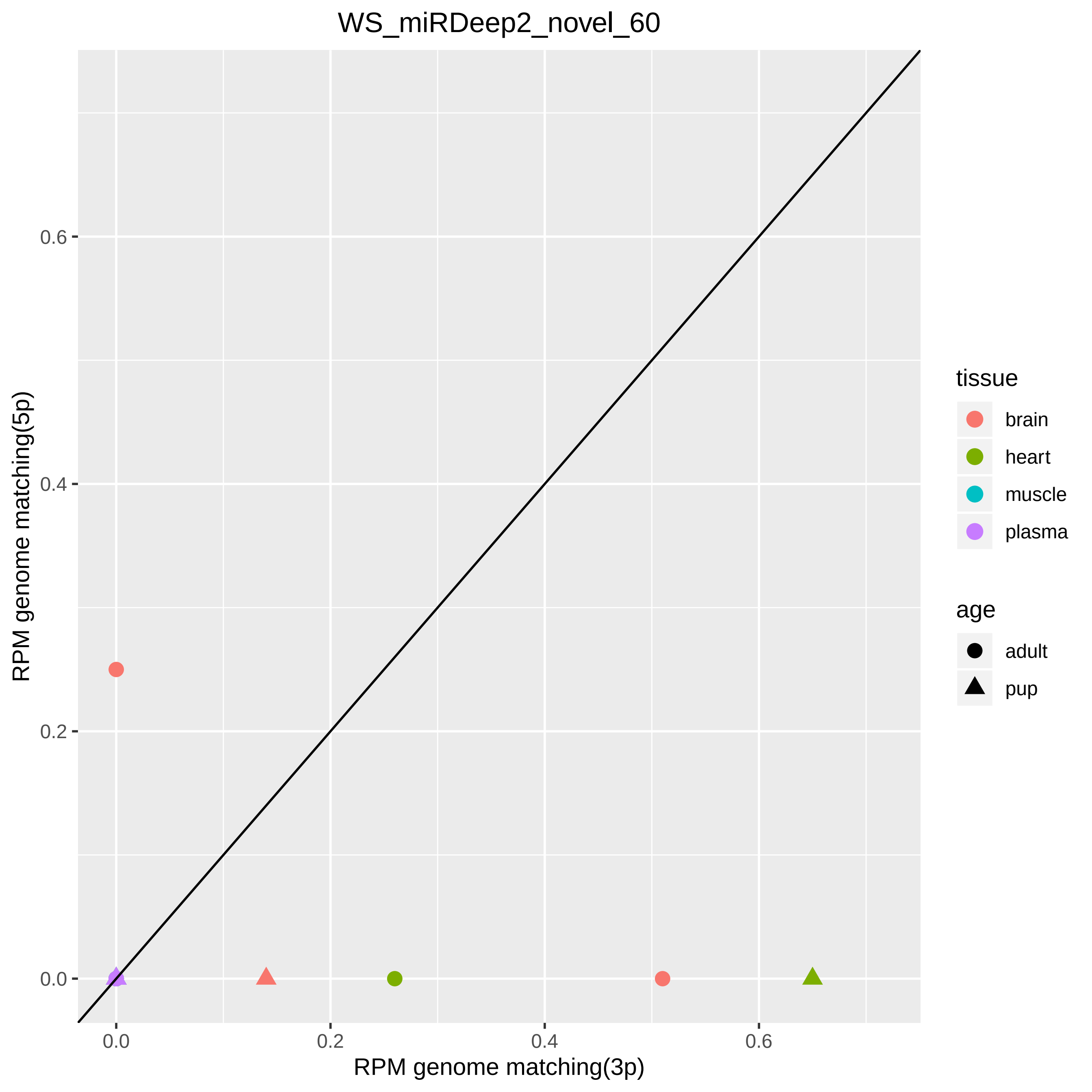

WS_miRDeep2_novel_60
RPM genome matching(5p)
RPM genome matching(3p)
0.0
0.2
0.4
0.6
tissue
brain
heart
muscle
plasma
age
adult
pup

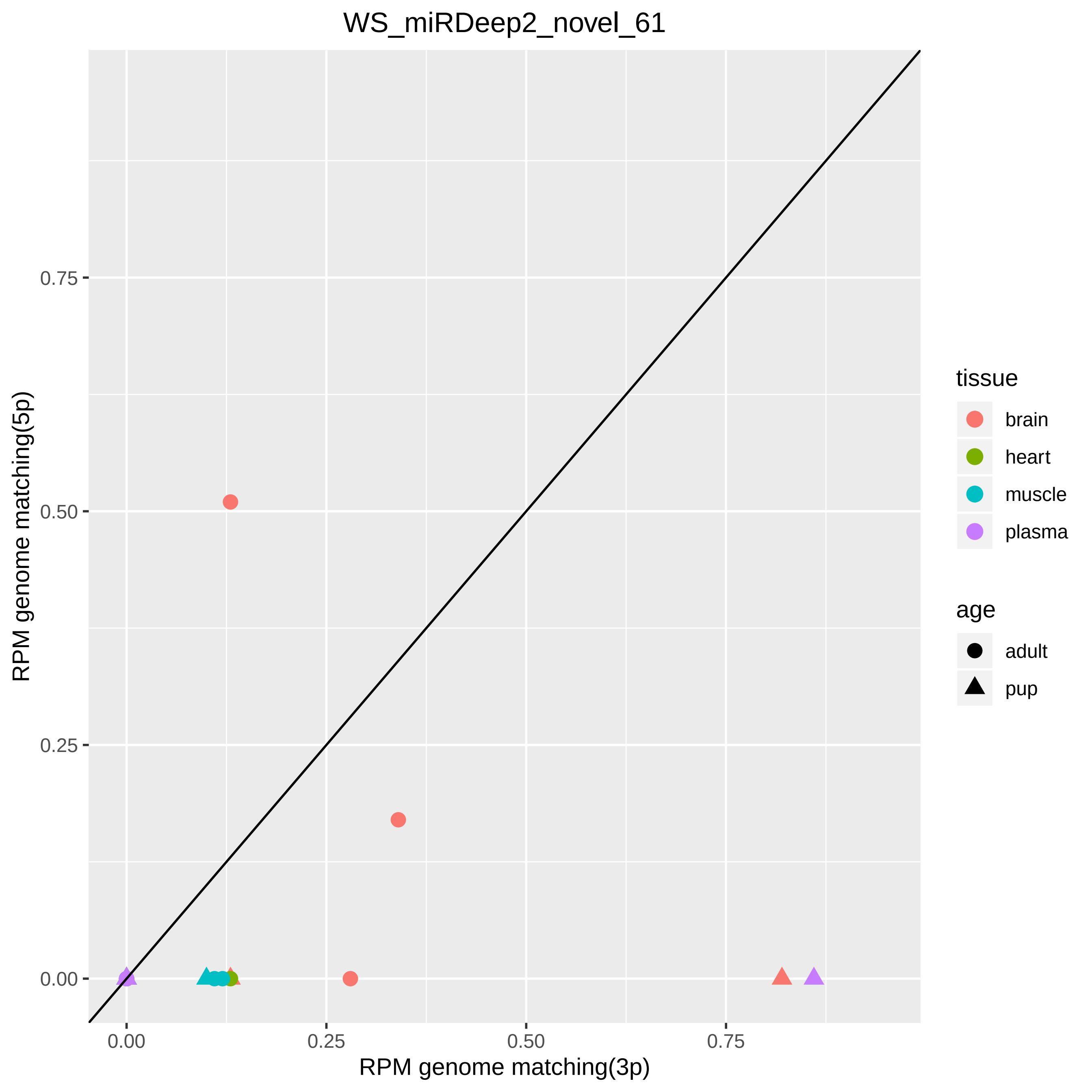

WS_miRDeep2_novel_61
RPM genome matching(5p)
RPM genome matching(3p)
0.00
0.25
0.50
0.75
tissue
brain
heart
muscle
plasma
age
adult
pup

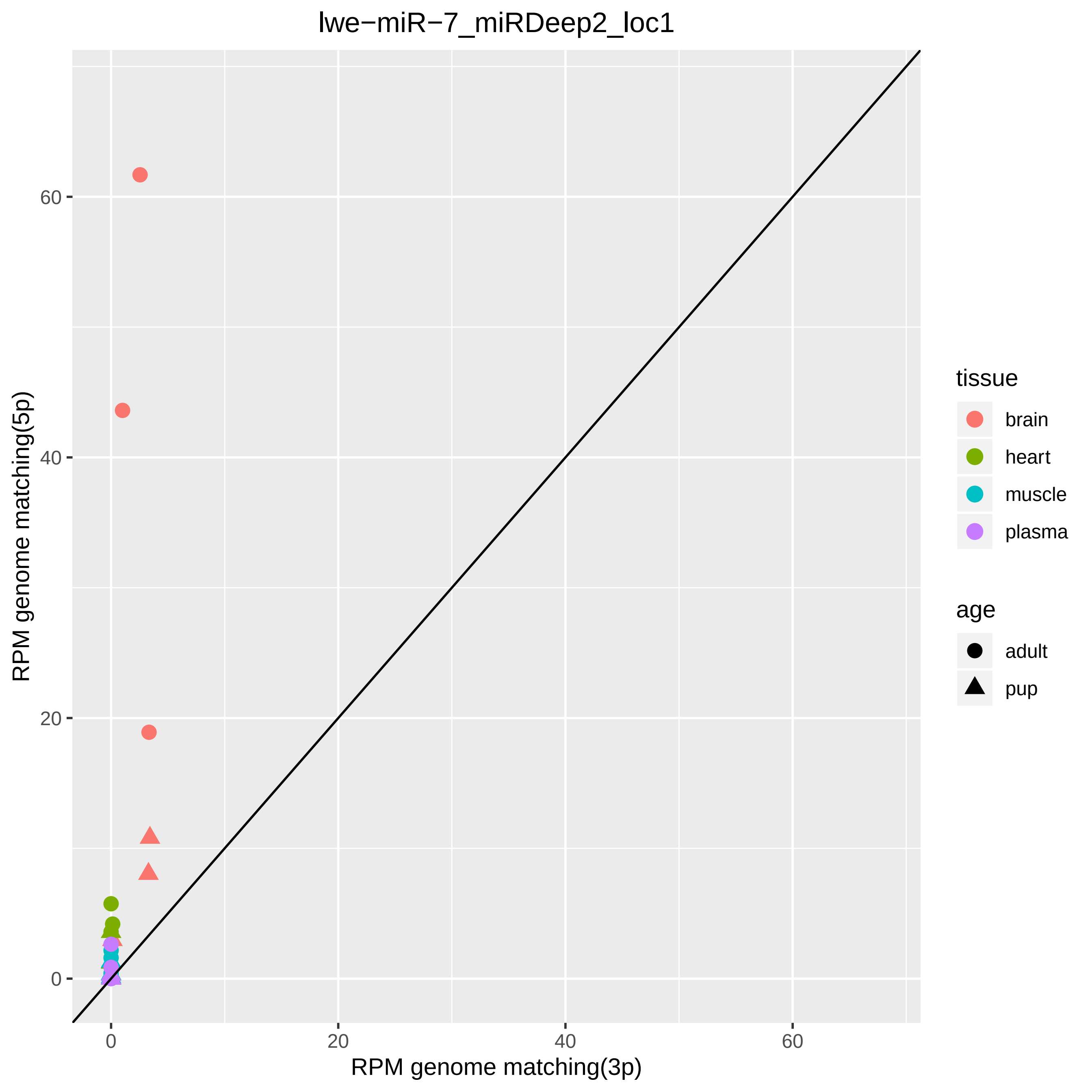

lwe−miR−7_miRDeep2_loc1
RPM genome matching(5p)
60
40
20
0
RPM genome matching(3p)
0
20
40
60
tissue
brain
heart
muscle
plasma
age
adult
pup

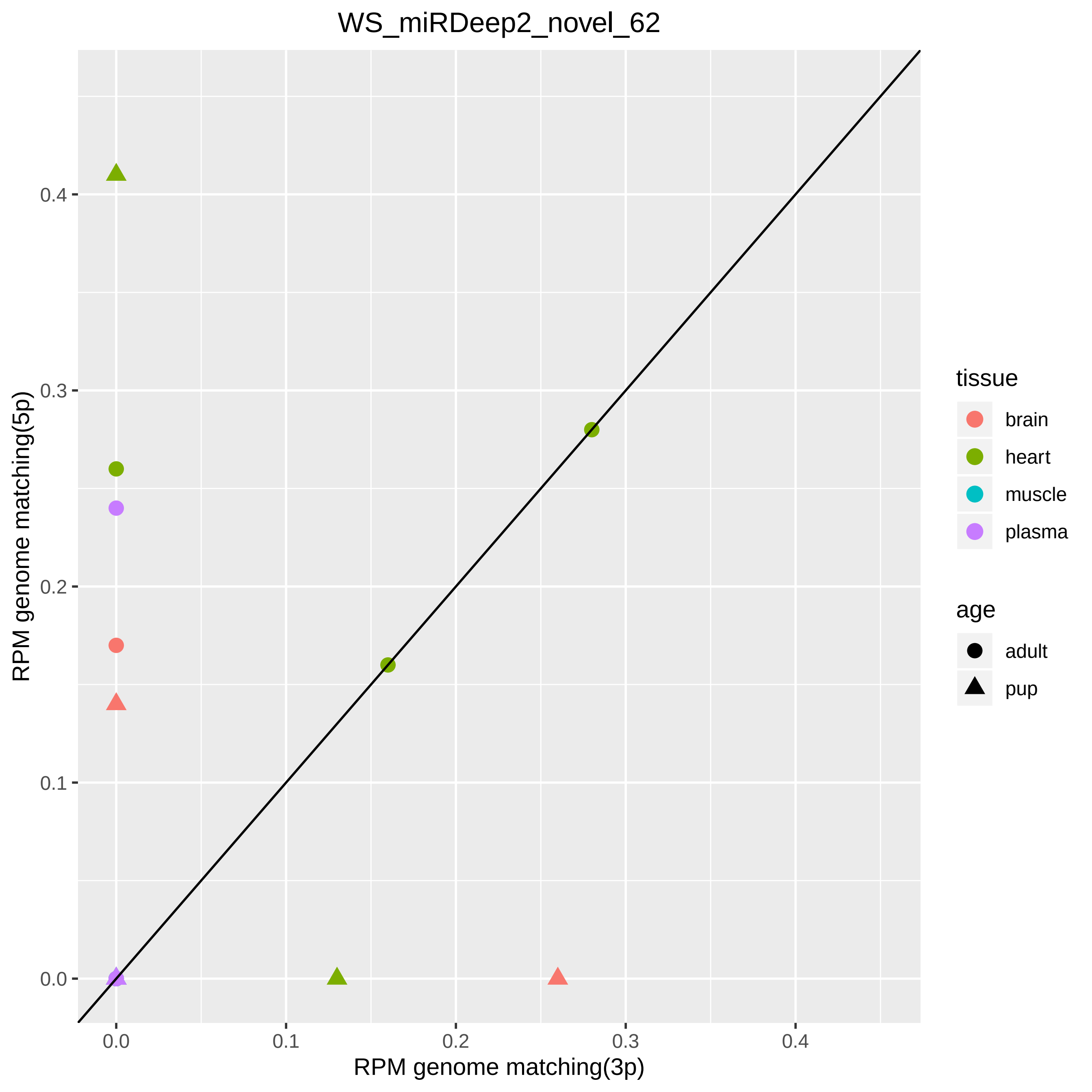

WS_miRDeep2_novel_62
RPM genome matching(5p)
RPM genome matching(3p)
0.0
0.1
0.2
0.3
0.4
tissue
brain
heart
muscle
plasma
age
adult
pup

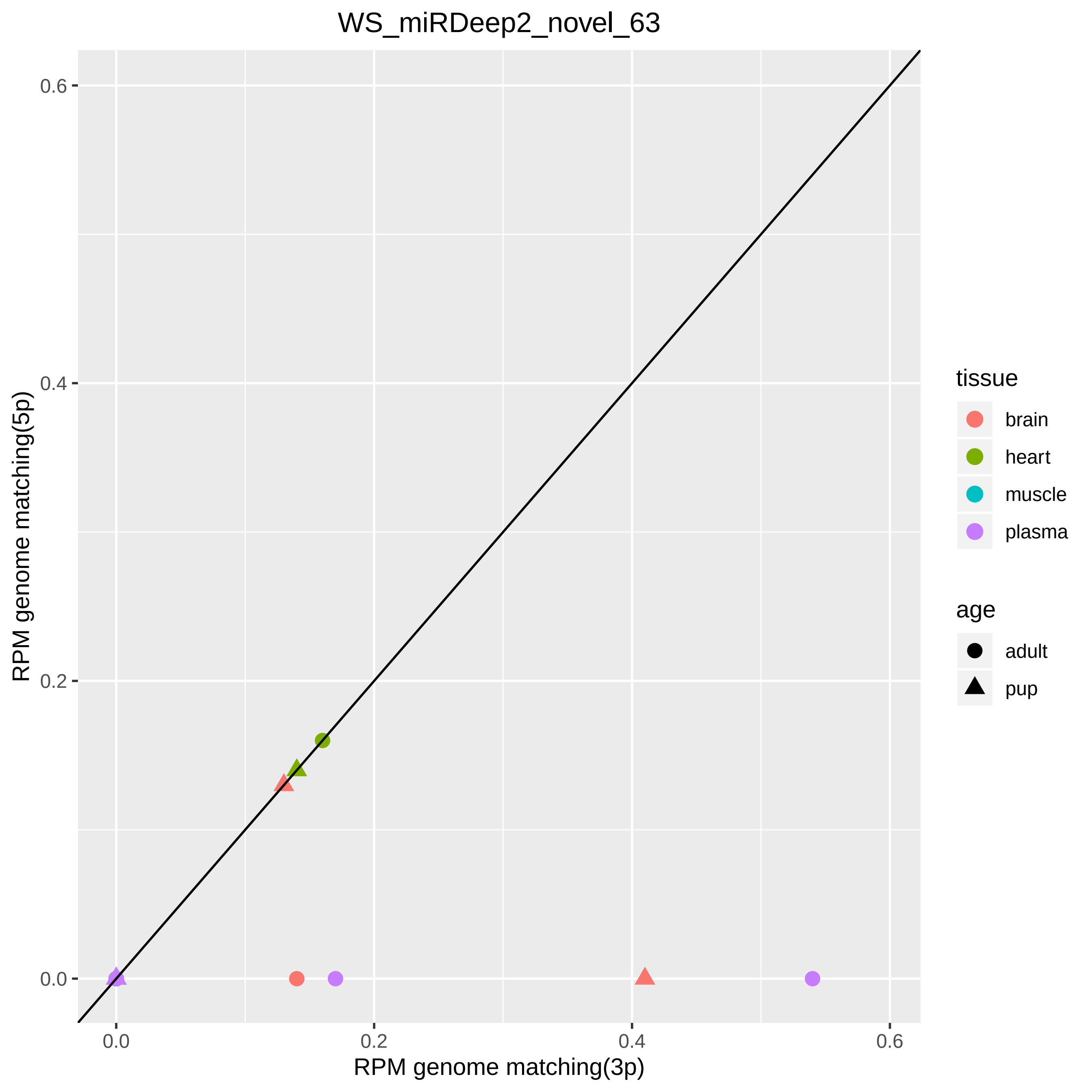
WS_miRDeep2_novel_63
RPM genome matching(5p)
RPM genome matching(3p)
0.0
0.2
0.4
0.6
0.0
0.2
0.4
0.6
tissue
brain
heart
muscle
plasma
age
adult
pup

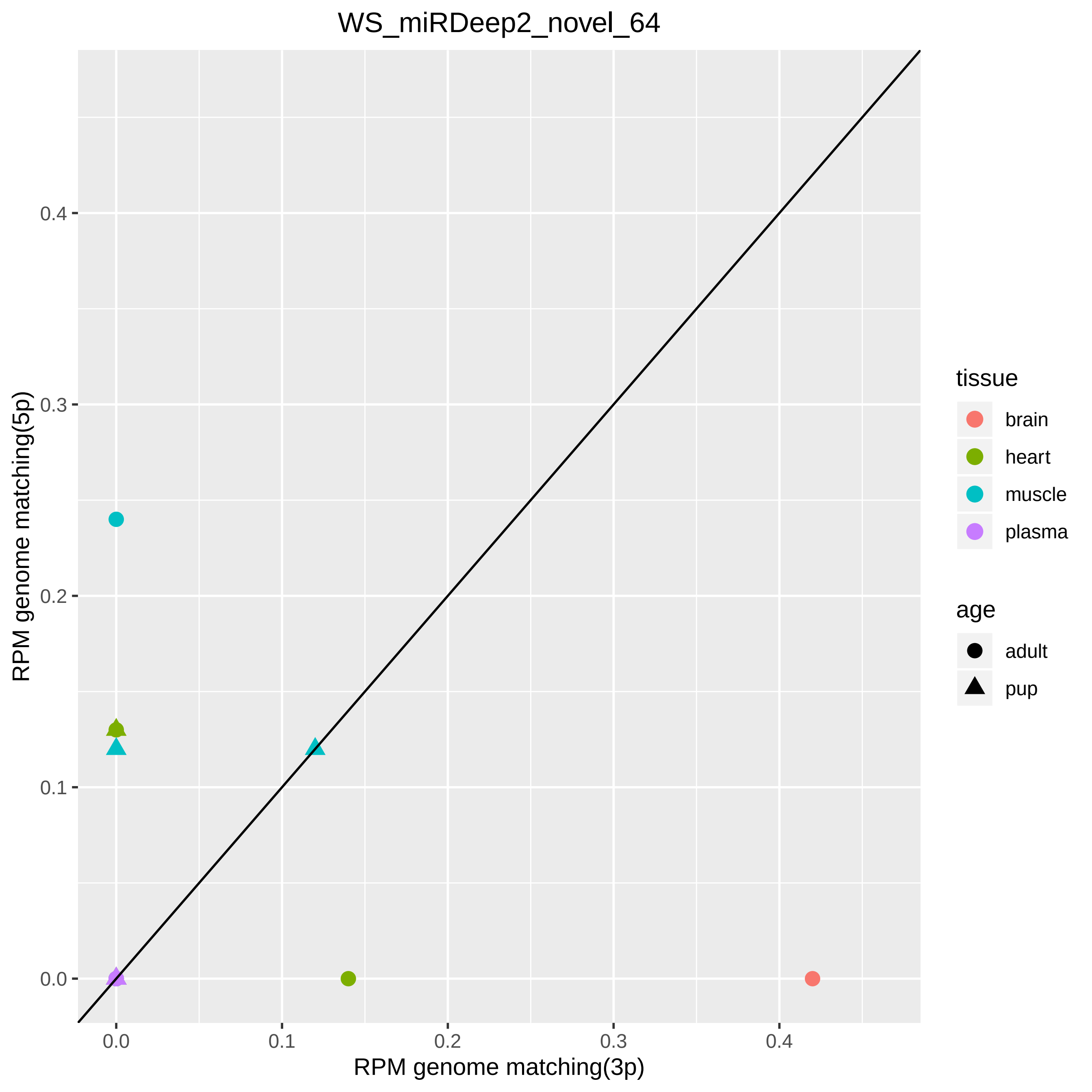

WS_miRDeep2_novel_64
RPM genome matching(5p)
RPM genome matching(3p)
0.0
0.1
0.2
0.3
0.4
tissue
brain
heart
muscle
plasma
age
adult
pup

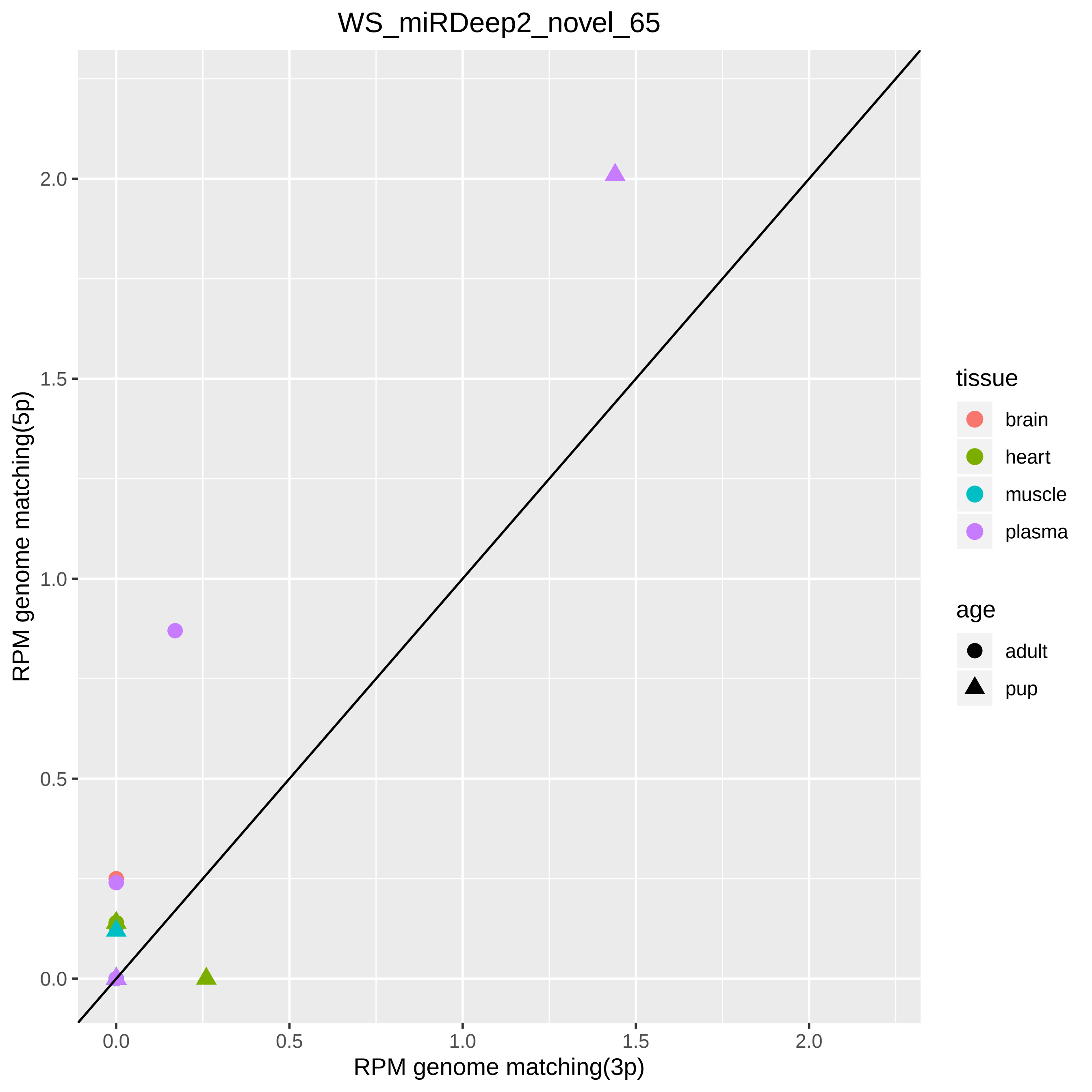
WS_miRDeep2_novel_65
RPM genome matching(5p)
RPM genome matching(3p)
0.0
0.5
1.0
1.5
2.0
tissue
brain
heart
muscle
plasma
age
adult
pup

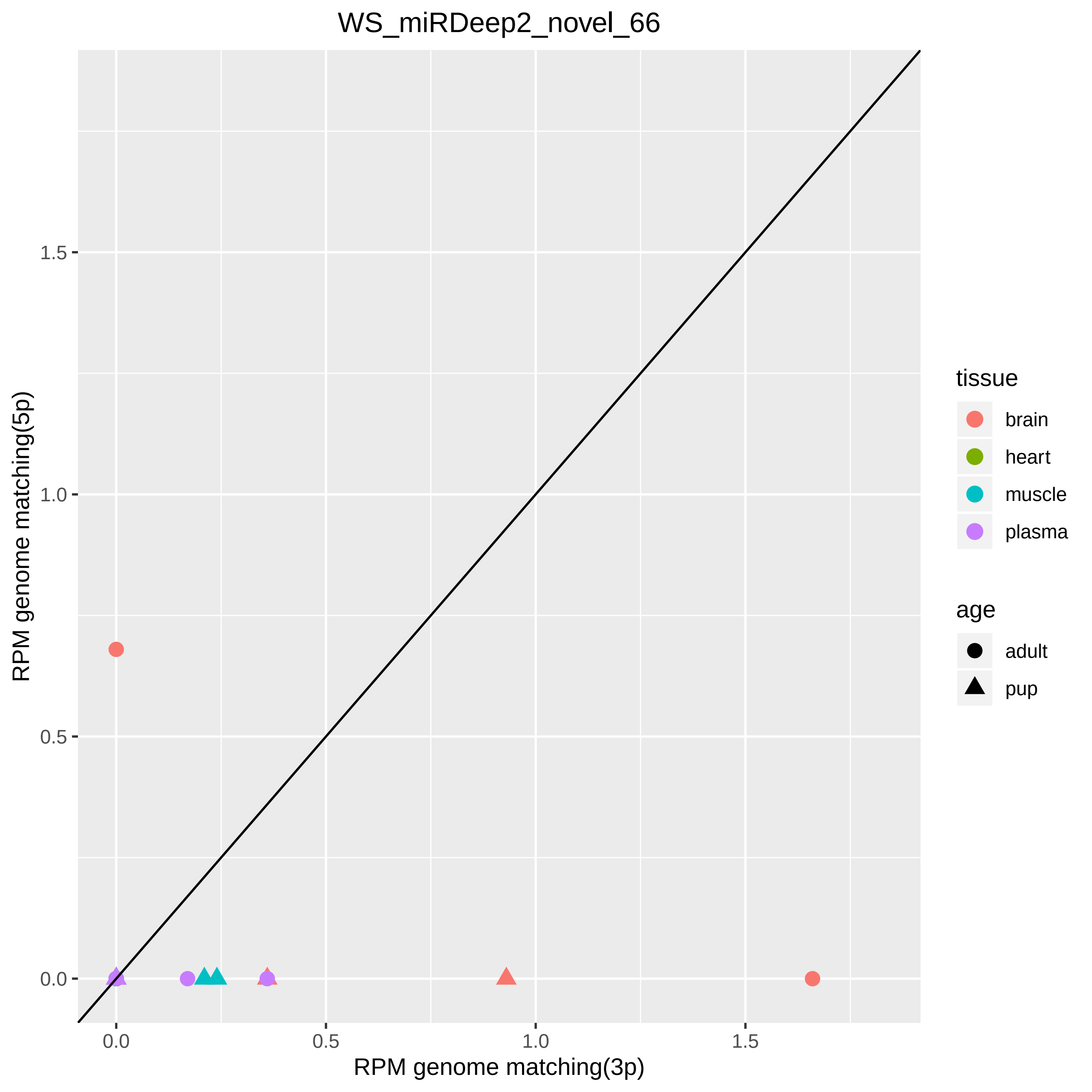
WS_miRDeep2_novel_66
RPM genome matching(5p)
RPM genome matching(3p)
0.0
0.5
1.0
1.5
tissue
brain
heart
muscle
plasma
age
adult
pup

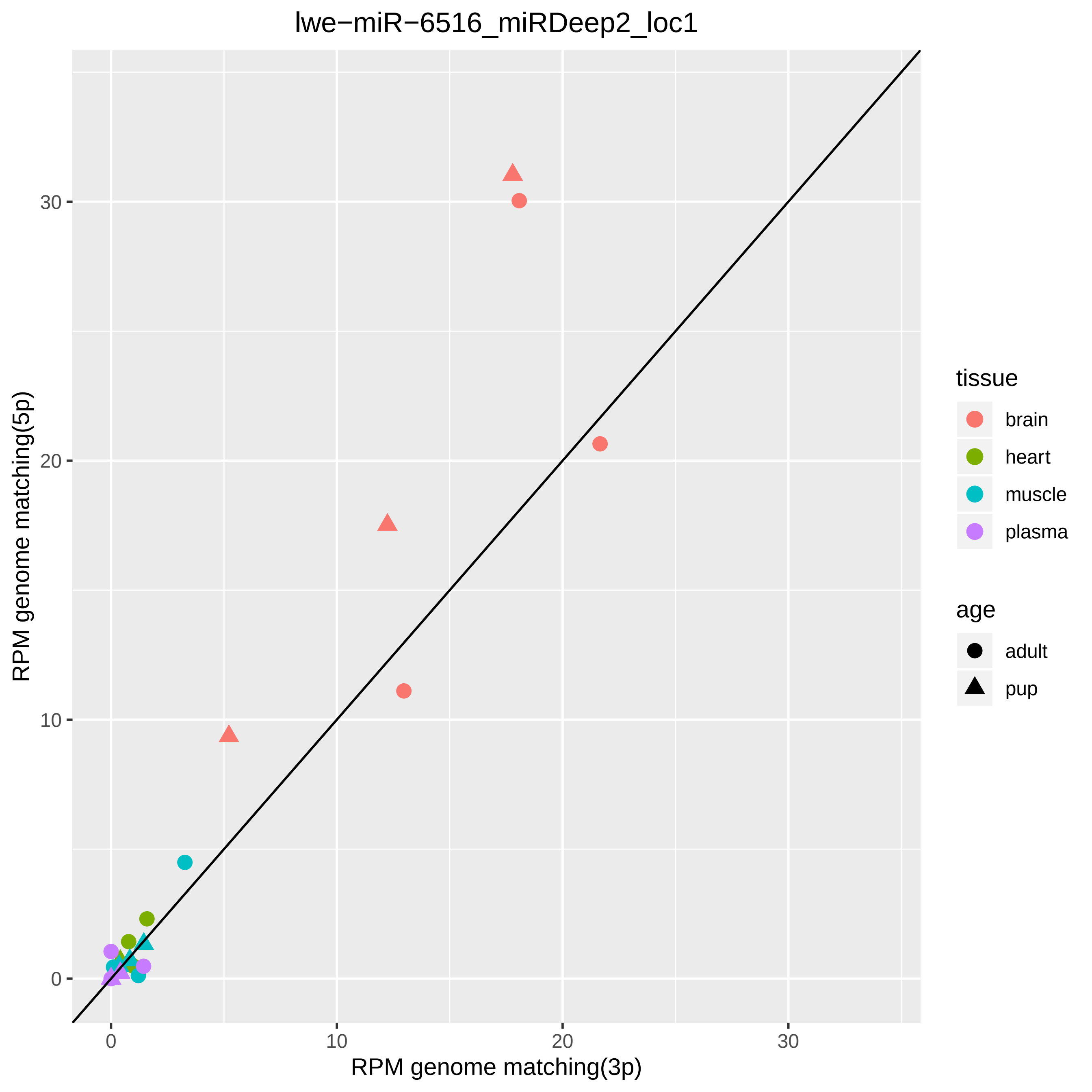
lwe−miR−6516_miRDeep2_loc1
RPM genome matching(5p)
RPM genome matching(3p)
0
10
20
30
tissue
brain
heart
muscle
plasma
age
adult
pup

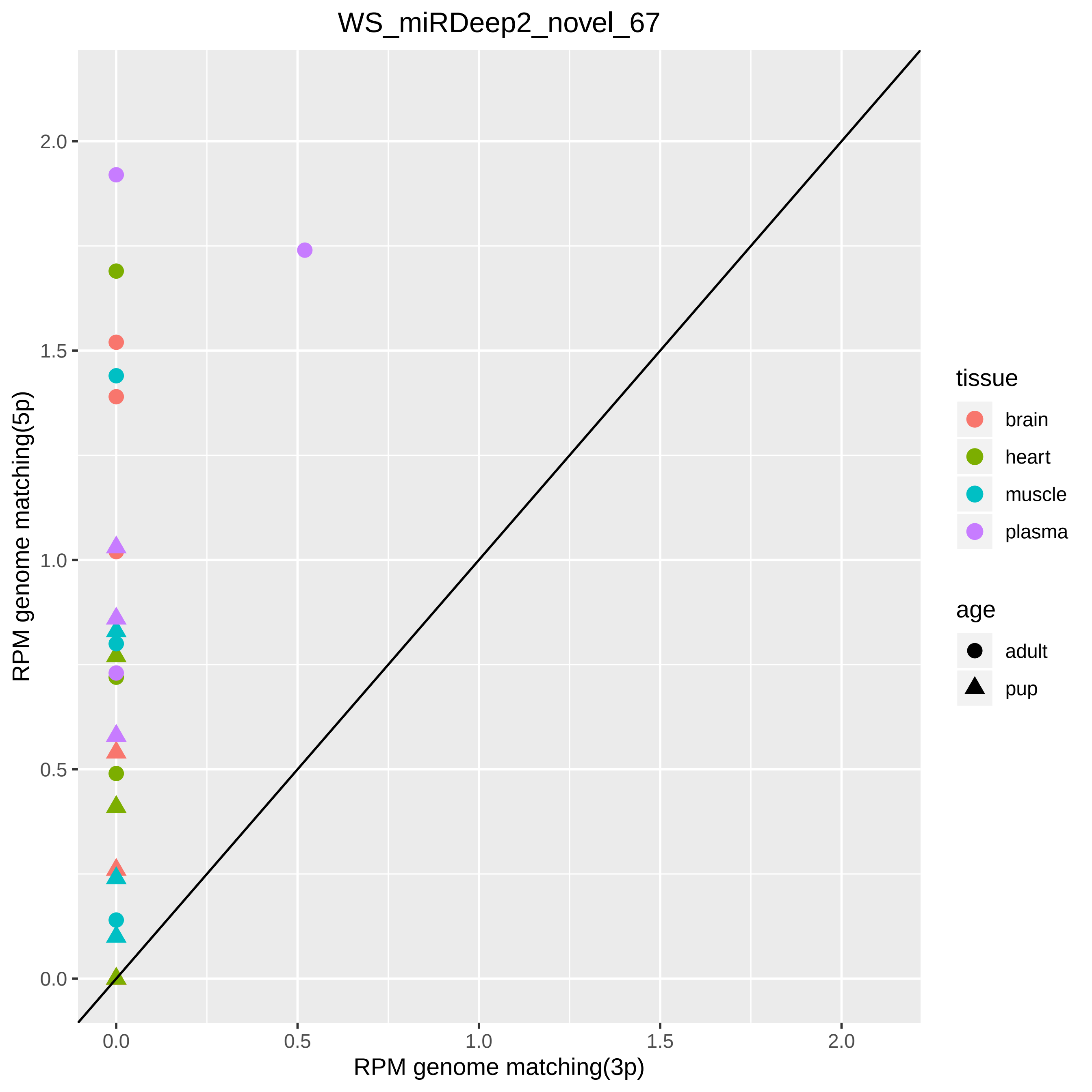

WS_miRDeep2_novel_67
RPM genome matching(5p)
RPM genome matching(3p)
0.0
0.5
1.0
1.5
2.0
tissue
brain
heart
muscle
plasma
age
adult
pup

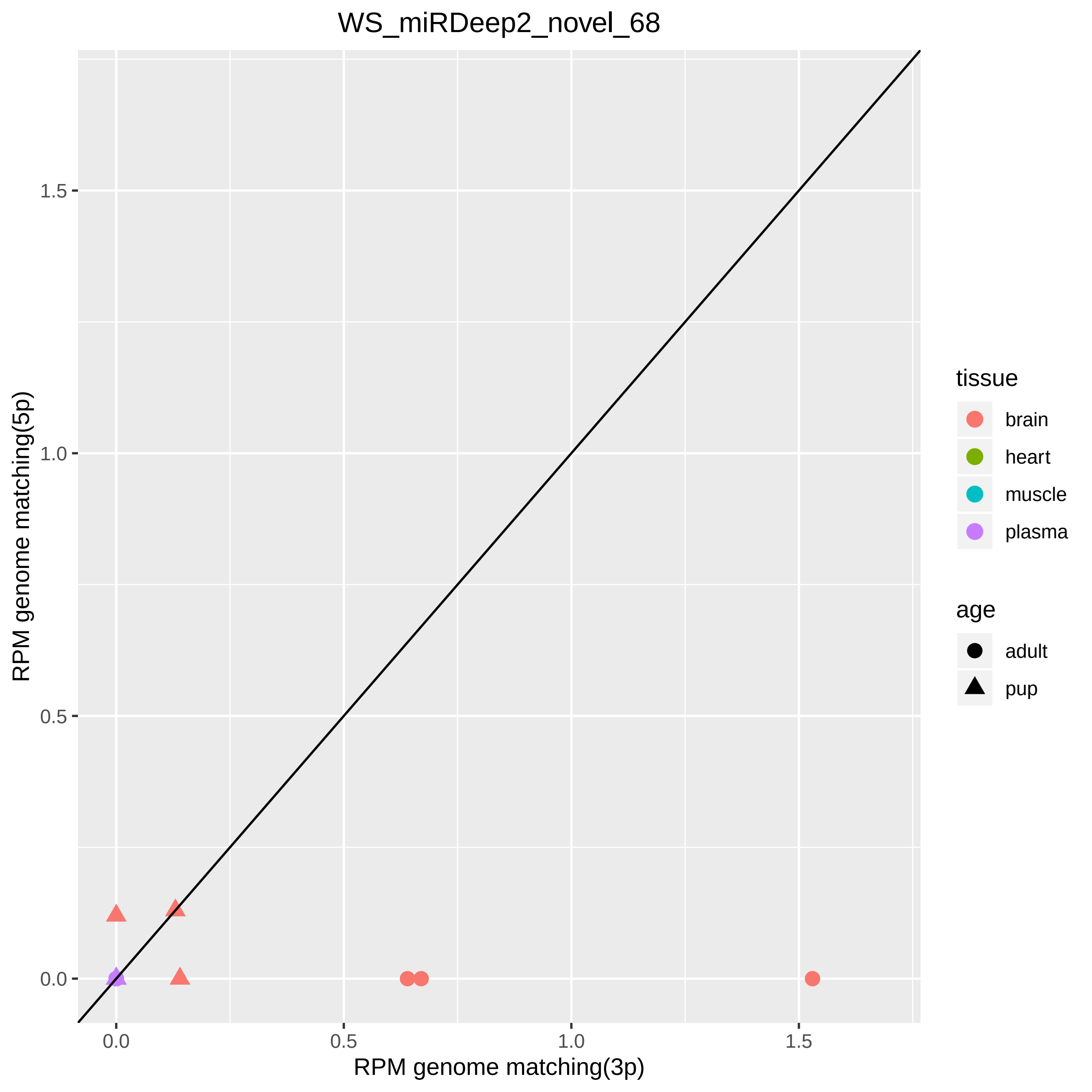

WS_miRDeep2_novel_68
RPM genome matching(5p)
RPM genome matching(3p)
0.0
0.5
1.0
1.5
tissue
brain
heart
muscle
plasma
age
adult
pup

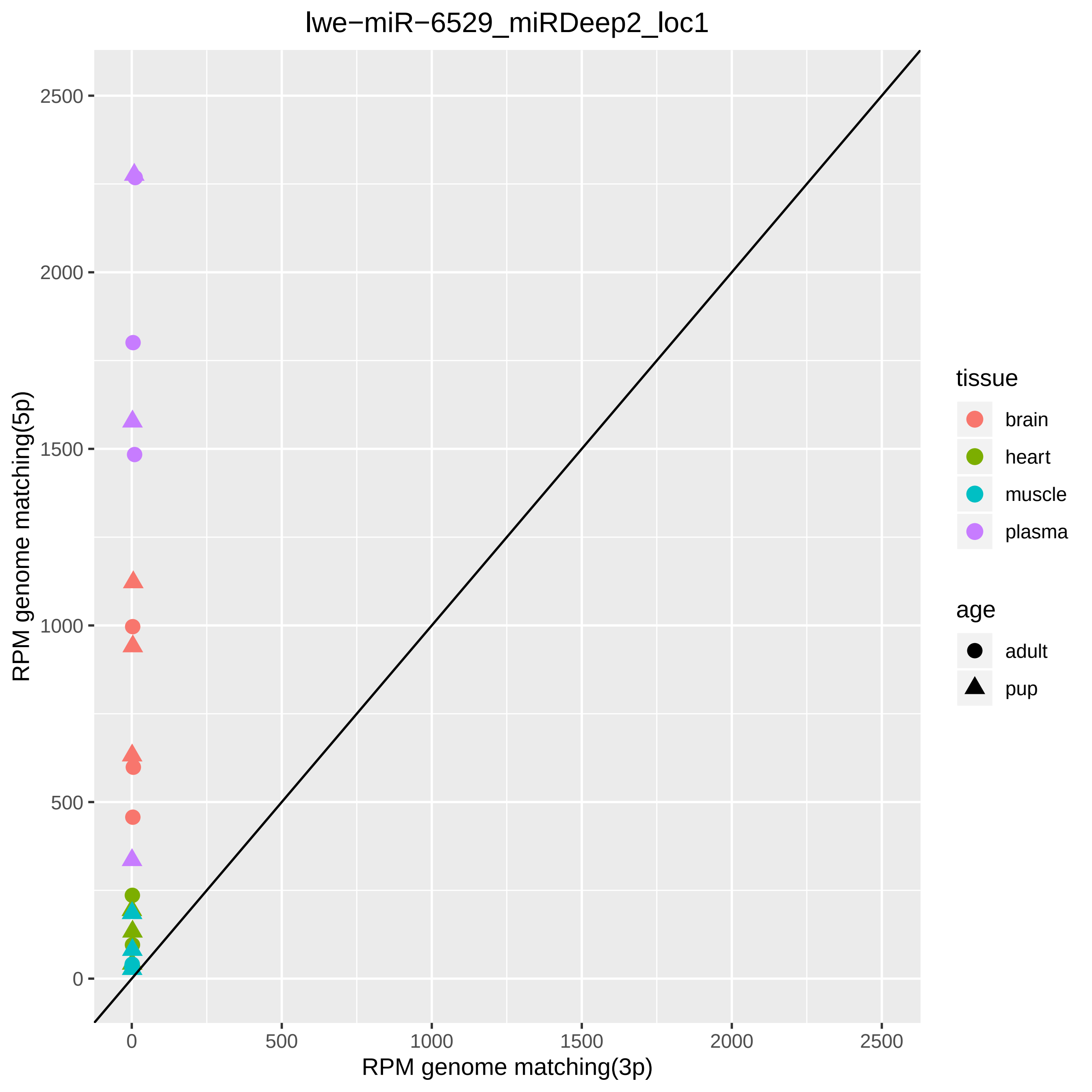
lwe−miR−6529_miRDeep2_loc1
RPM genome matching(5p)
RPM genome matching(3p)
2500
2000
1500
1000
500
0
0
500
1000
1500
2000
2500
tissue
brain
heart
muscle
plasma
age
adult
pup

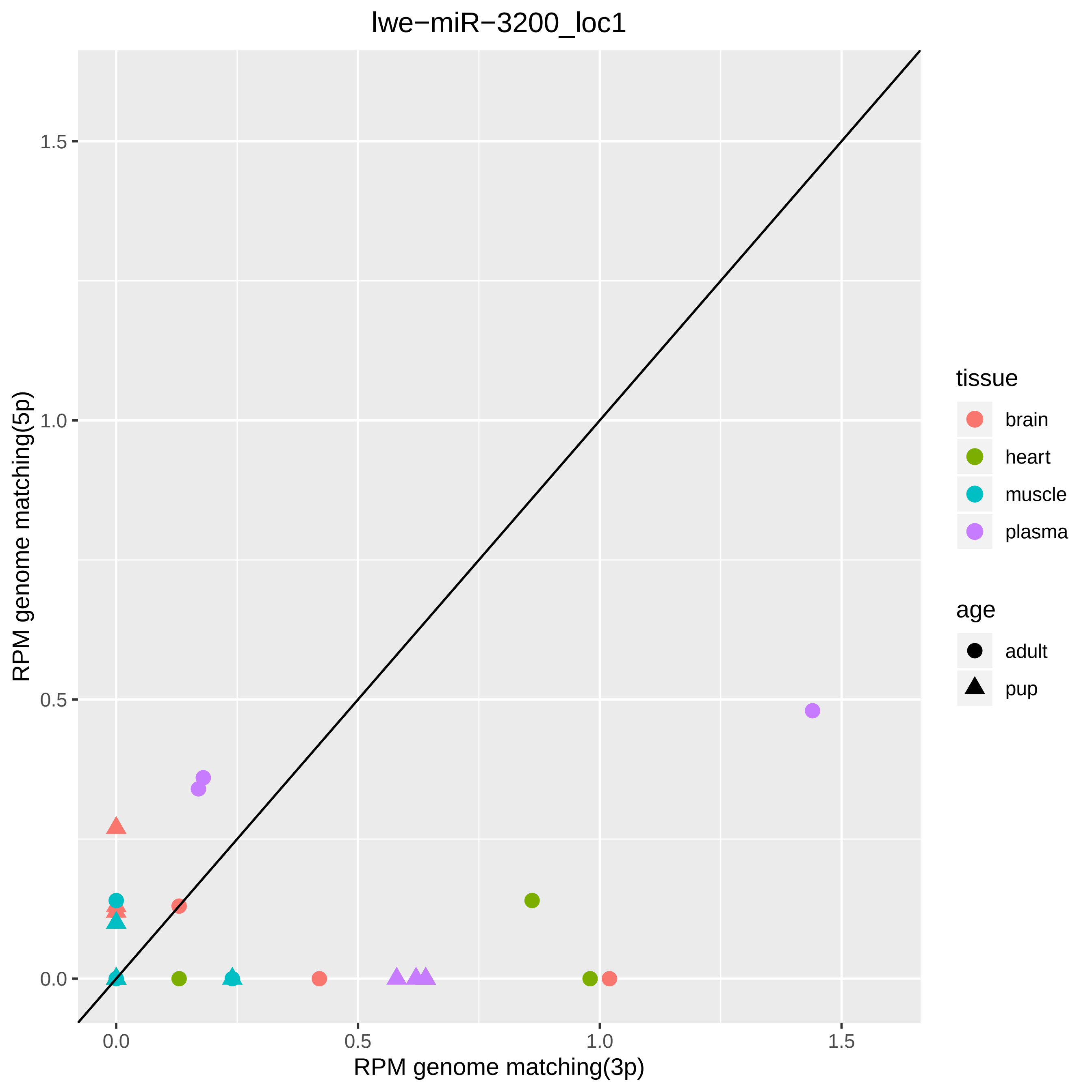
lwe−miR−3200_loc1
RPM genome matching(5p)
RPM genome matching(3p)
0.0
0.5
1.0
1.5
tissue
brain
heart
muscle
plasma
age
adult
pup

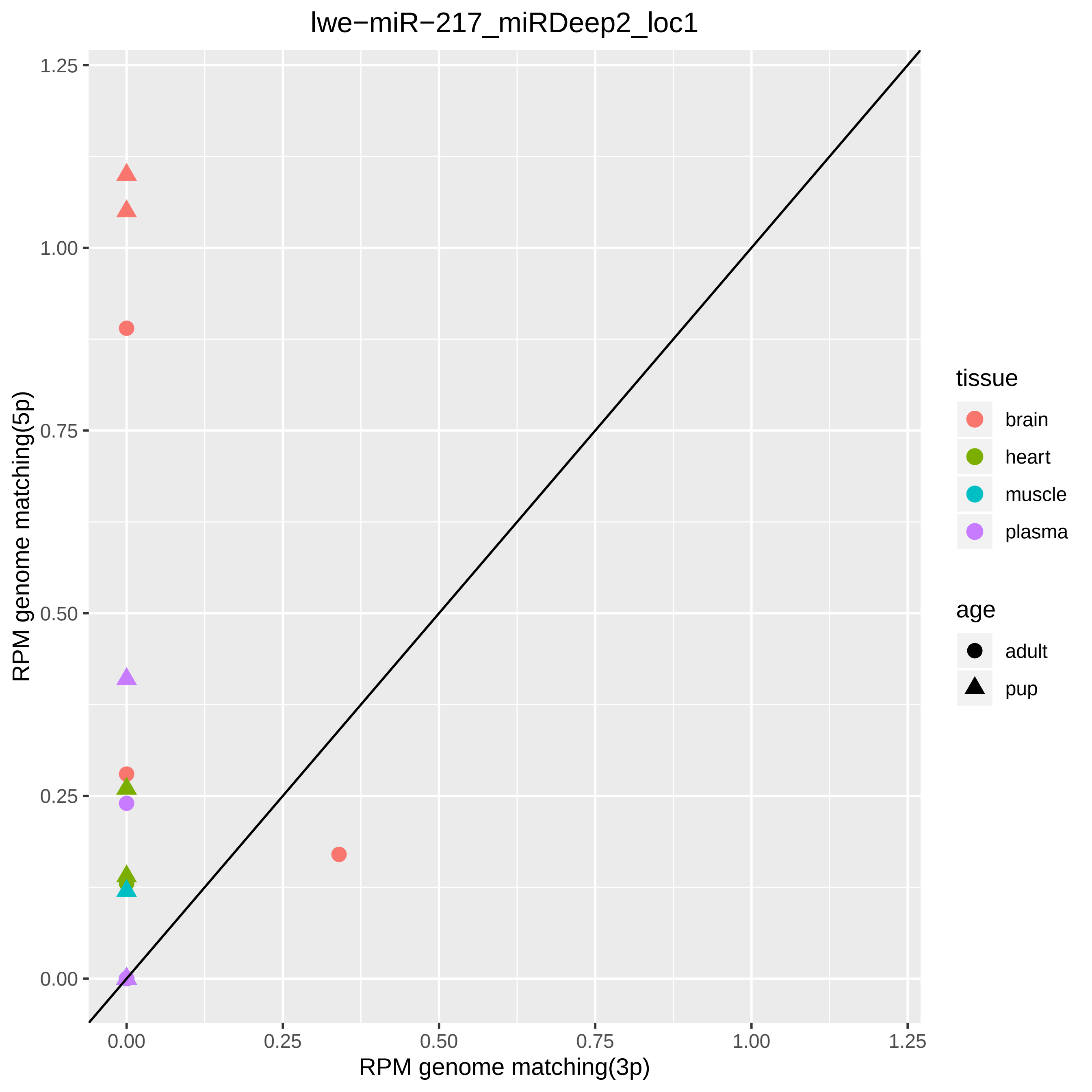

lwe−miR−217_miRDeep2_loc1
RPM genome matching(5p)
RPM genome matching(3p)
1.25
1.00
0.75
0.50
0.25
0.00
0.00
0.25
0.50
0.75
1.00
1.25
tissue
brain
heart
muscle
plasma
age
adult
pup

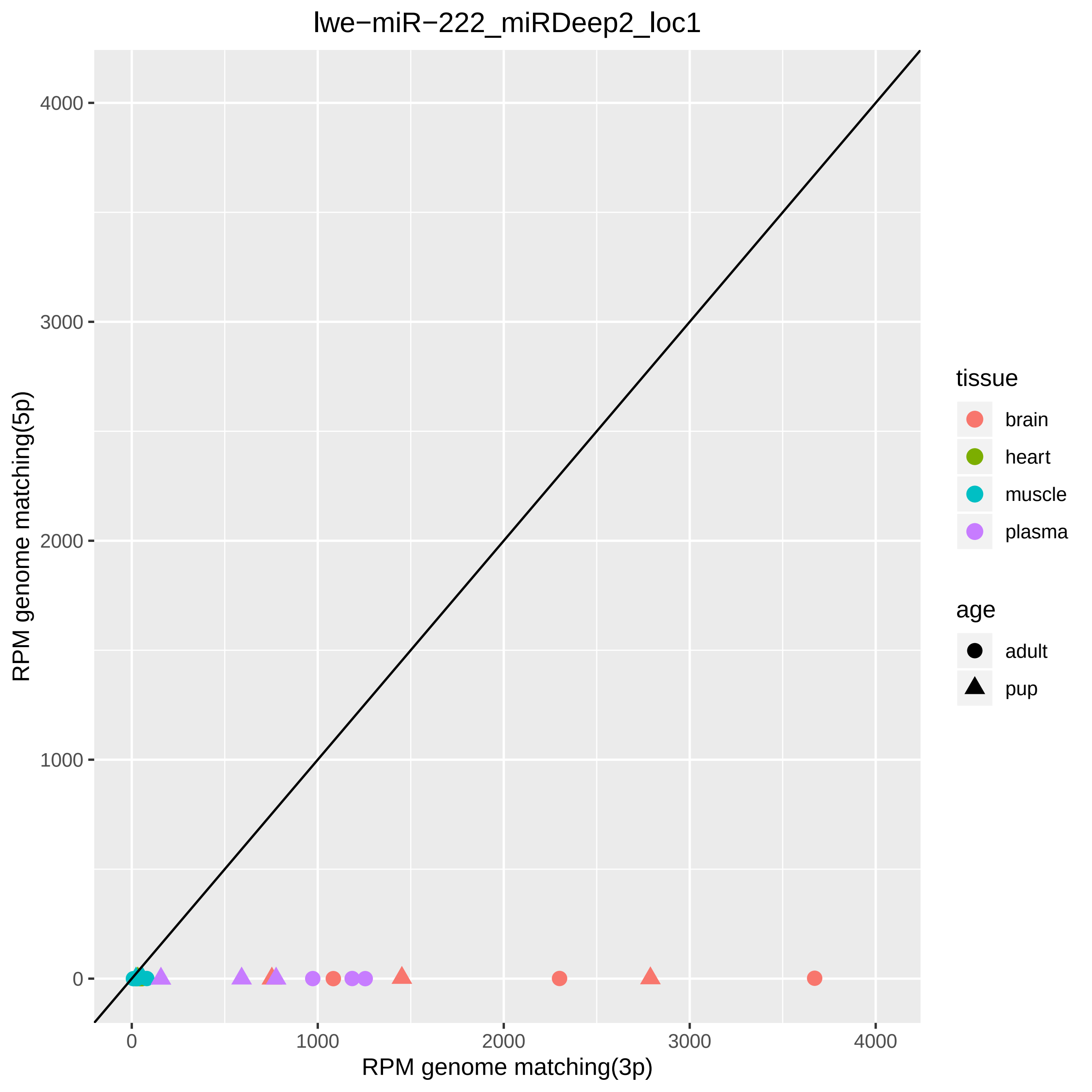

lwe−miR−222_miRDeep2_loc1
RPM genome matching(5p)
RPM genome matching(3p)
0
1000
2000
3000
4000
tissue
brain
heart
muscle
plasma
age
adult
pup

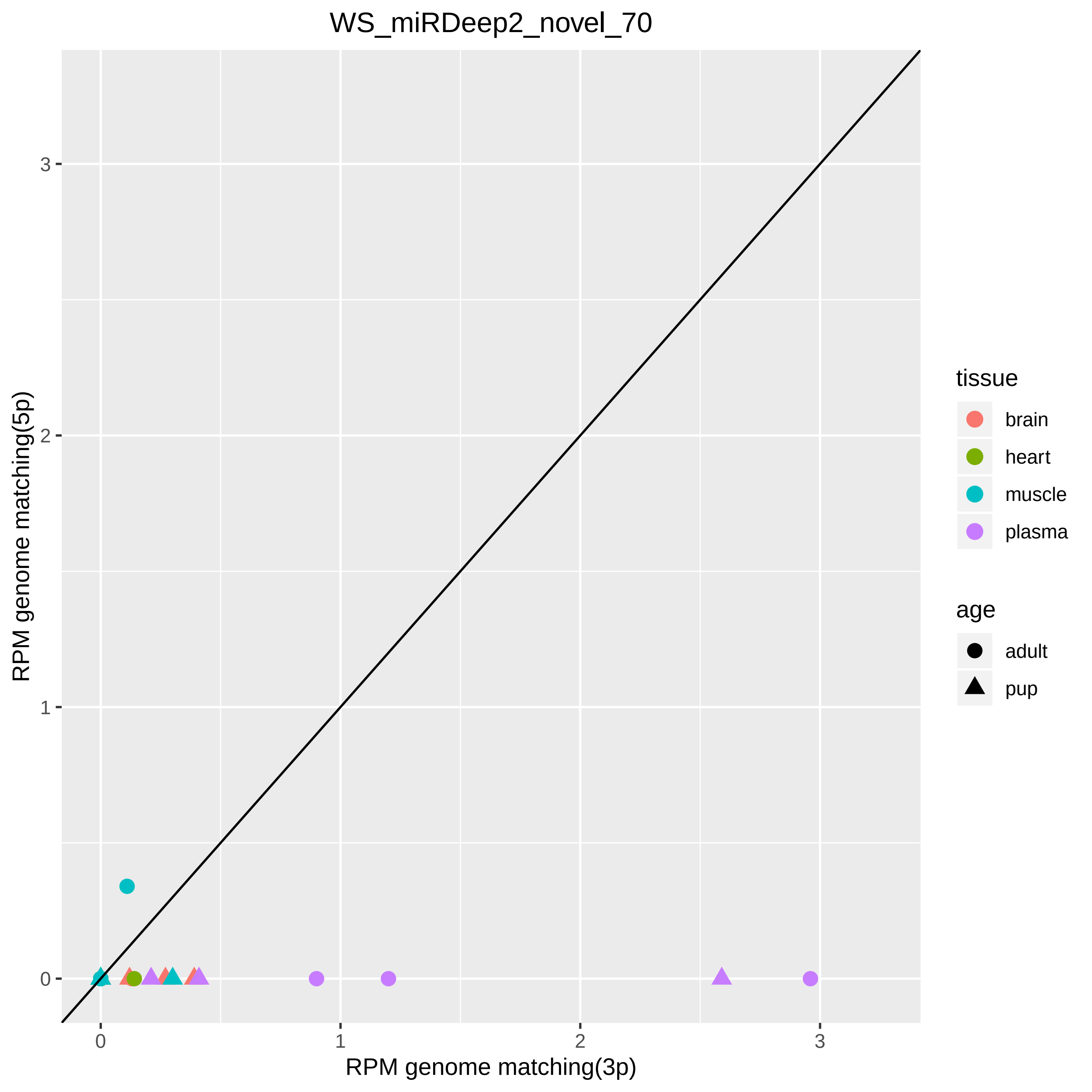

WS_miRDeep2_novel_70
RPM genome matching(5p)
RPM genome matching(3p)
0
1
2
3
0
1
2
3
tissue
brain
heart
muscle
plasma
age
adult
pup

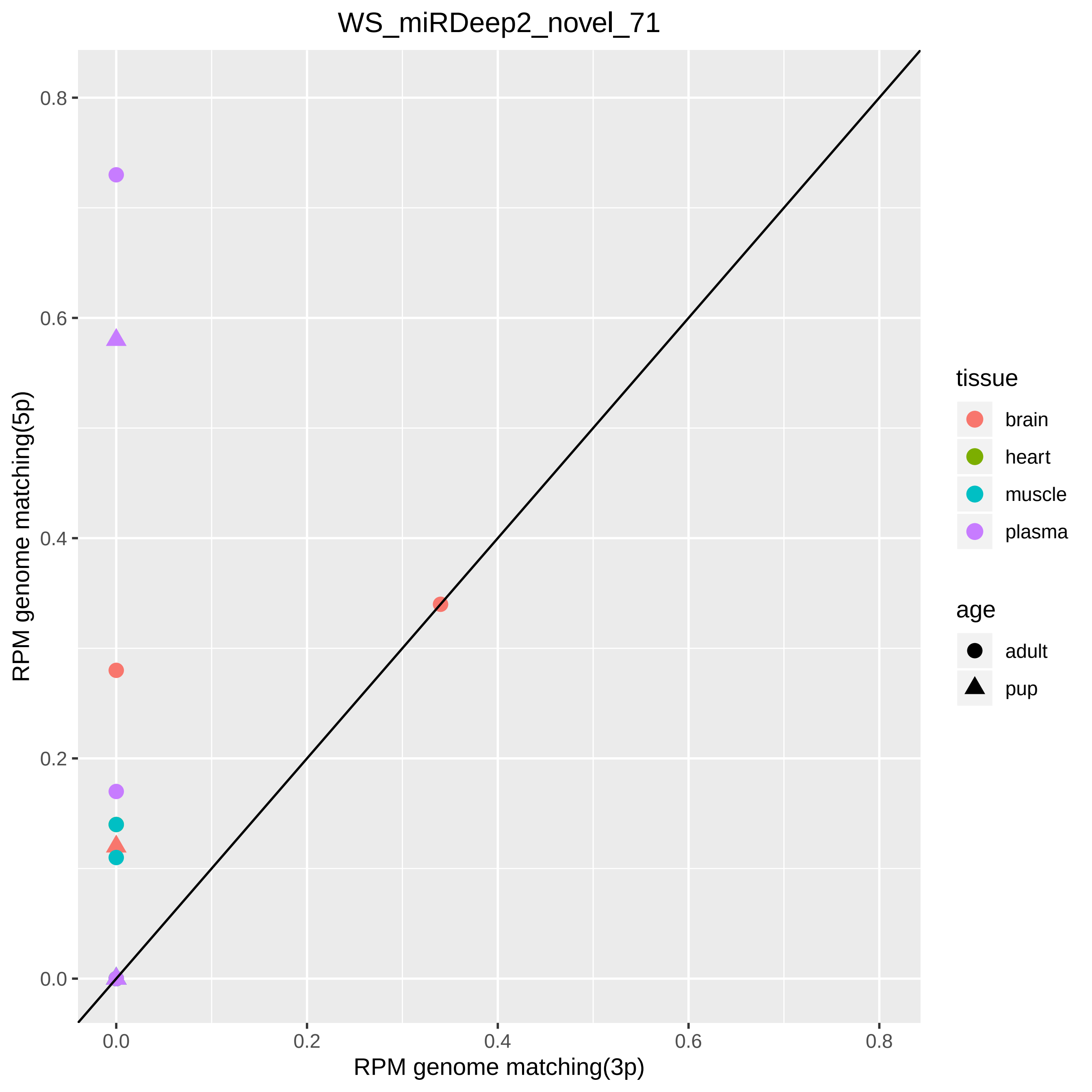

WS_miRDeep2_novel_71
RPM genome matching(5p)
RPM genome matching(3p)
0.0
0.2
0.4
0.6
0.8
0.0
0.2
0.4
0.6
0.8
tissue
brain
heart
muscle
plasma
age
adult
pup

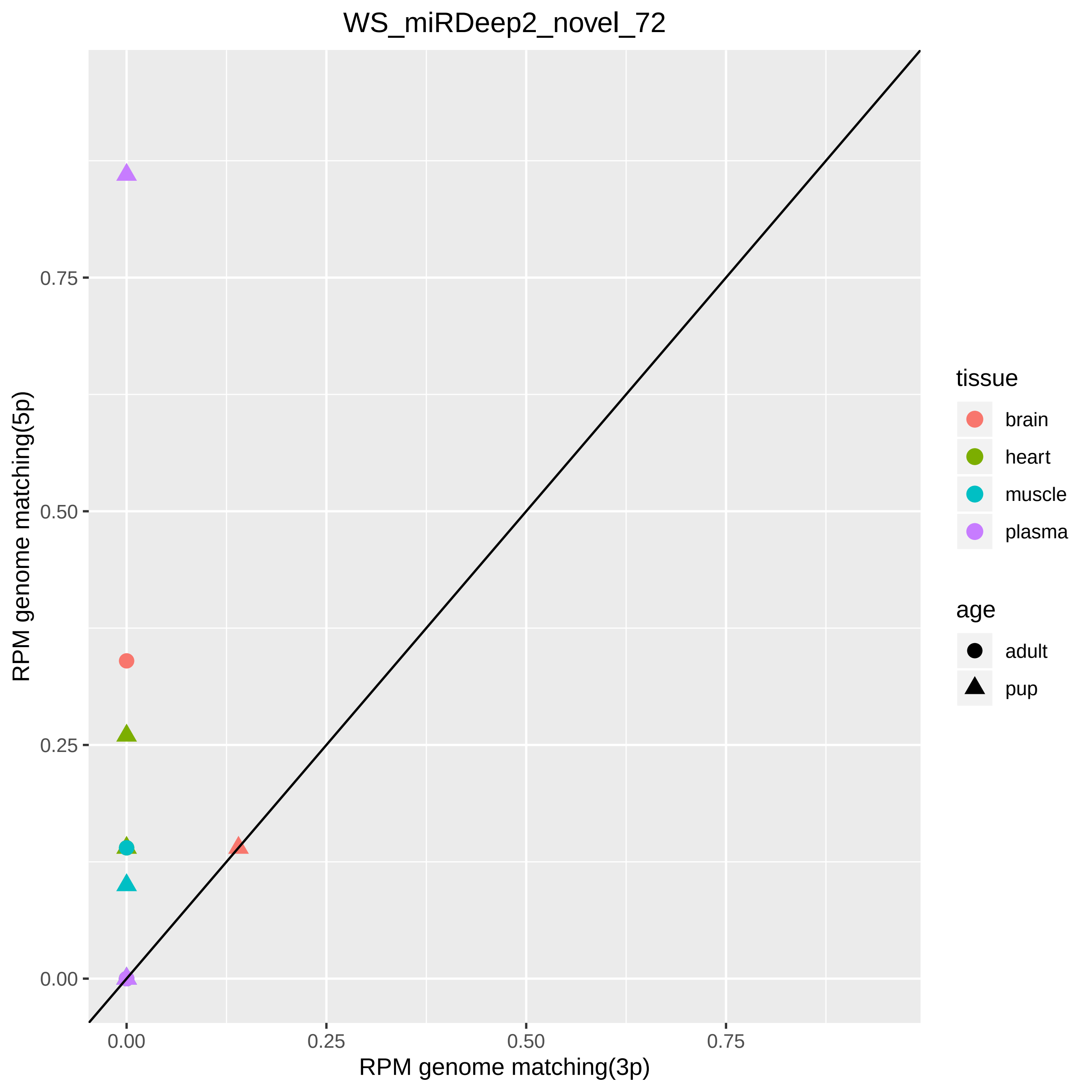

WS_miRDeep2_novel_72
RPM genome matching(5p)
RPM genome matching(3p)
0.00
0.25
0.50
0.75
tissue
brain
heart
muscle
plasma
age
adult
pup

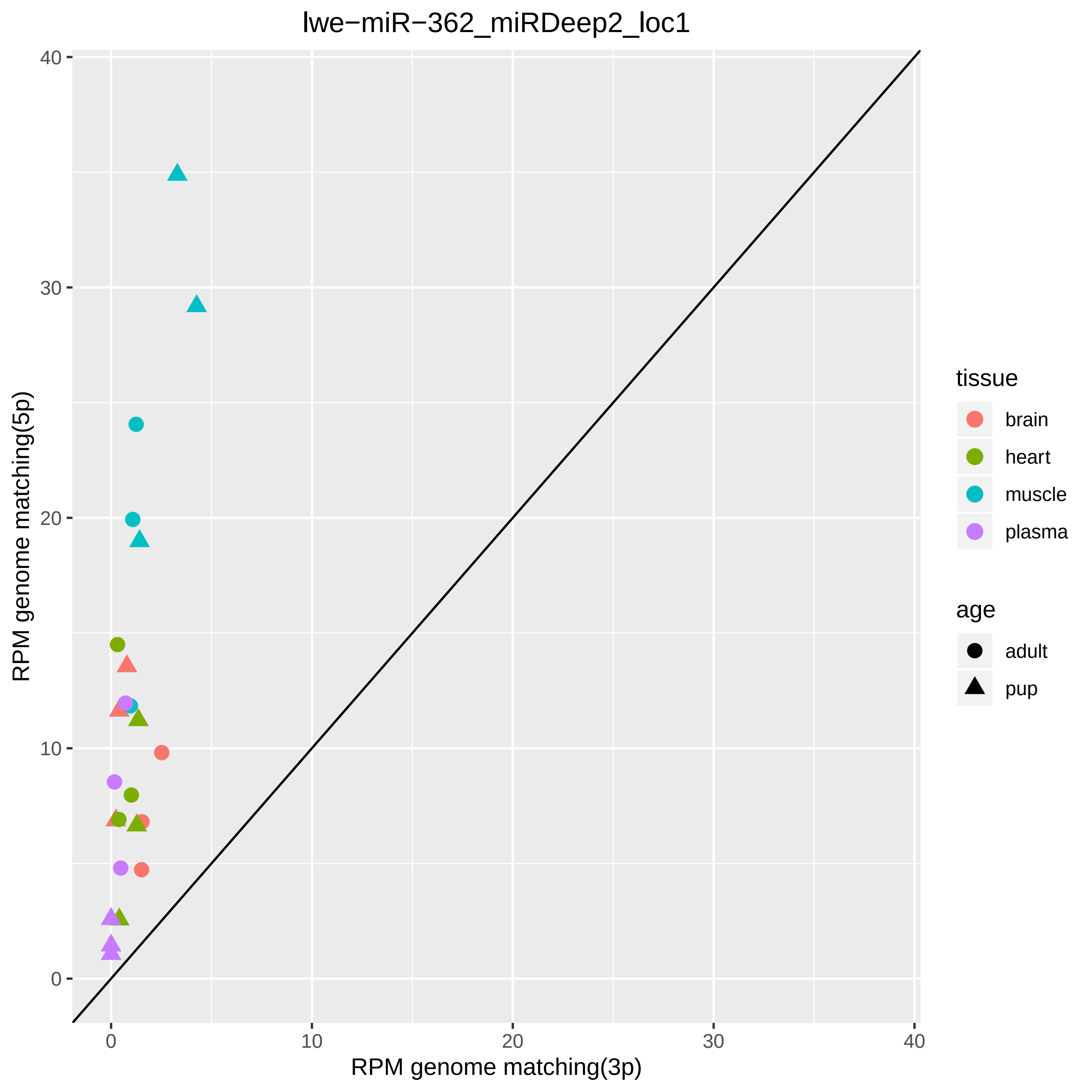

lwe−miR−362_miRDeep2_loc1
RPM genome matching(5p)
RPM genome matching(3p)
0
10
20
30
40
0
10
20
30
40
tissue
brain
heart
muscle
plasma
age
adult
pup

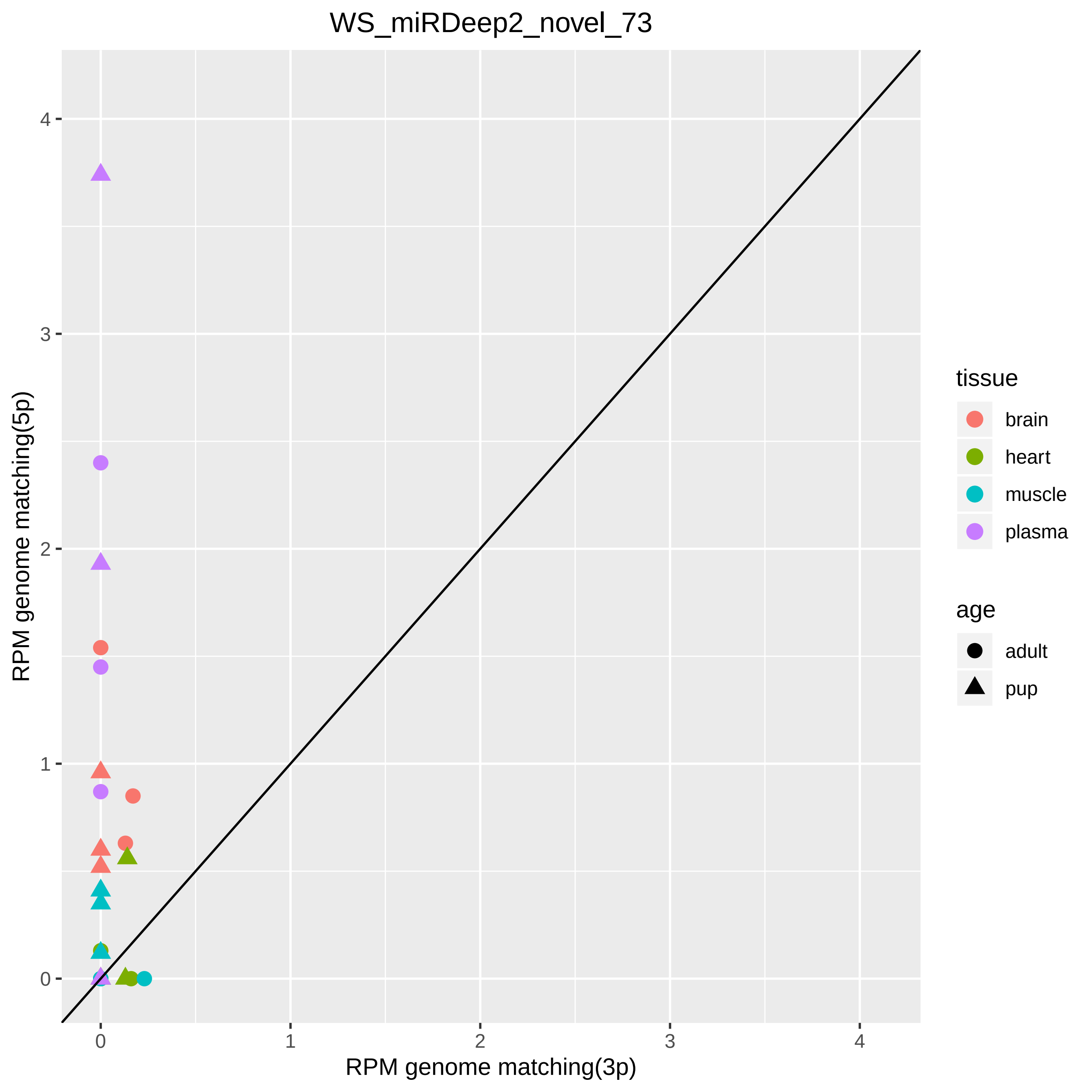
WS_miRDeep2_novel_73
RPM genome matching(5p)
RPM genome matching(3p)
0
1
2
3
4
tissue
brain
heart
muscle
plasma
age
adult
pup

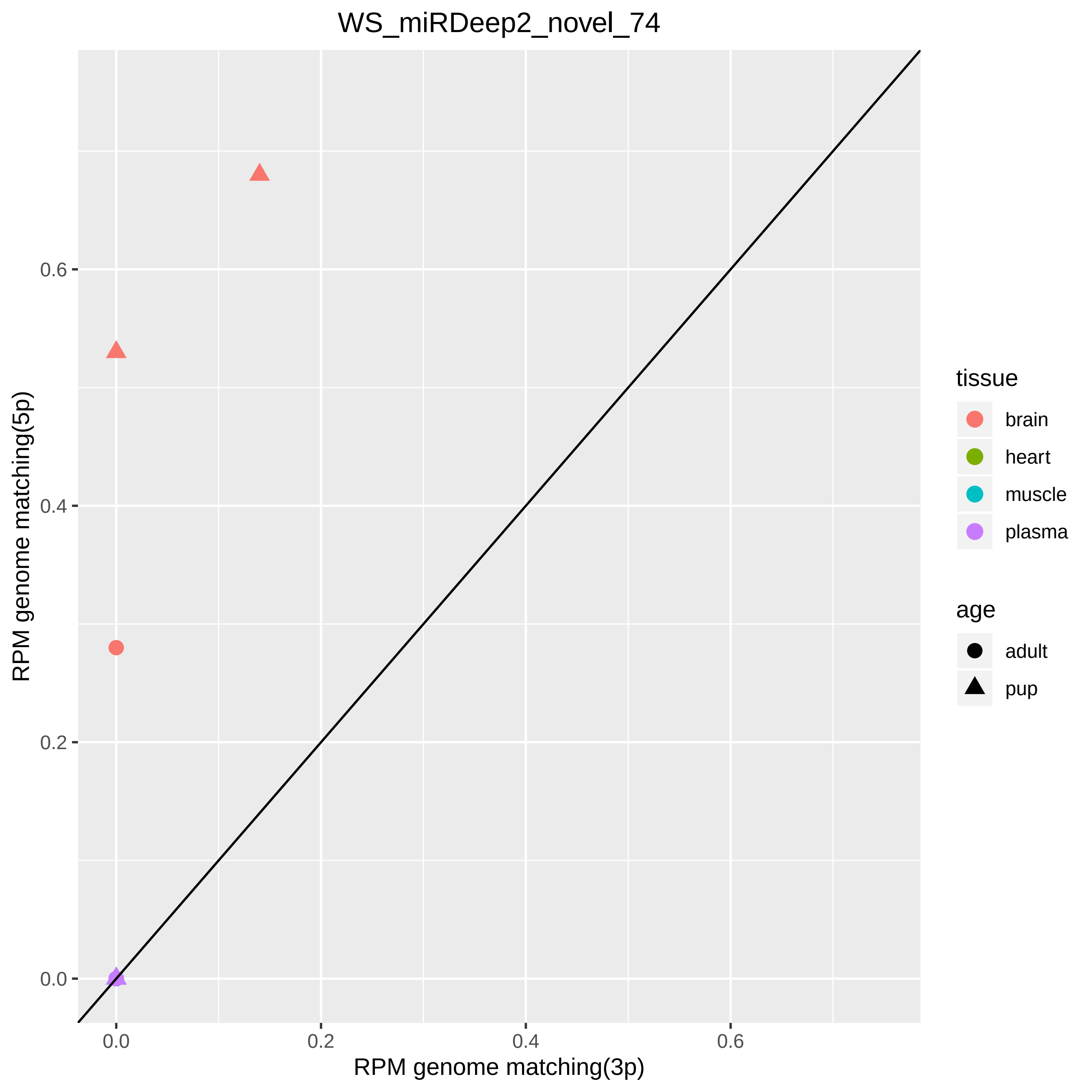

WS_miRDeep2_novel_74
RPM genome matching(5p)
RPM genome matching(3p)
0.0
0.2
0.4
0.6
tissue
brain
heart
muscle
plasma
age
adult
pup

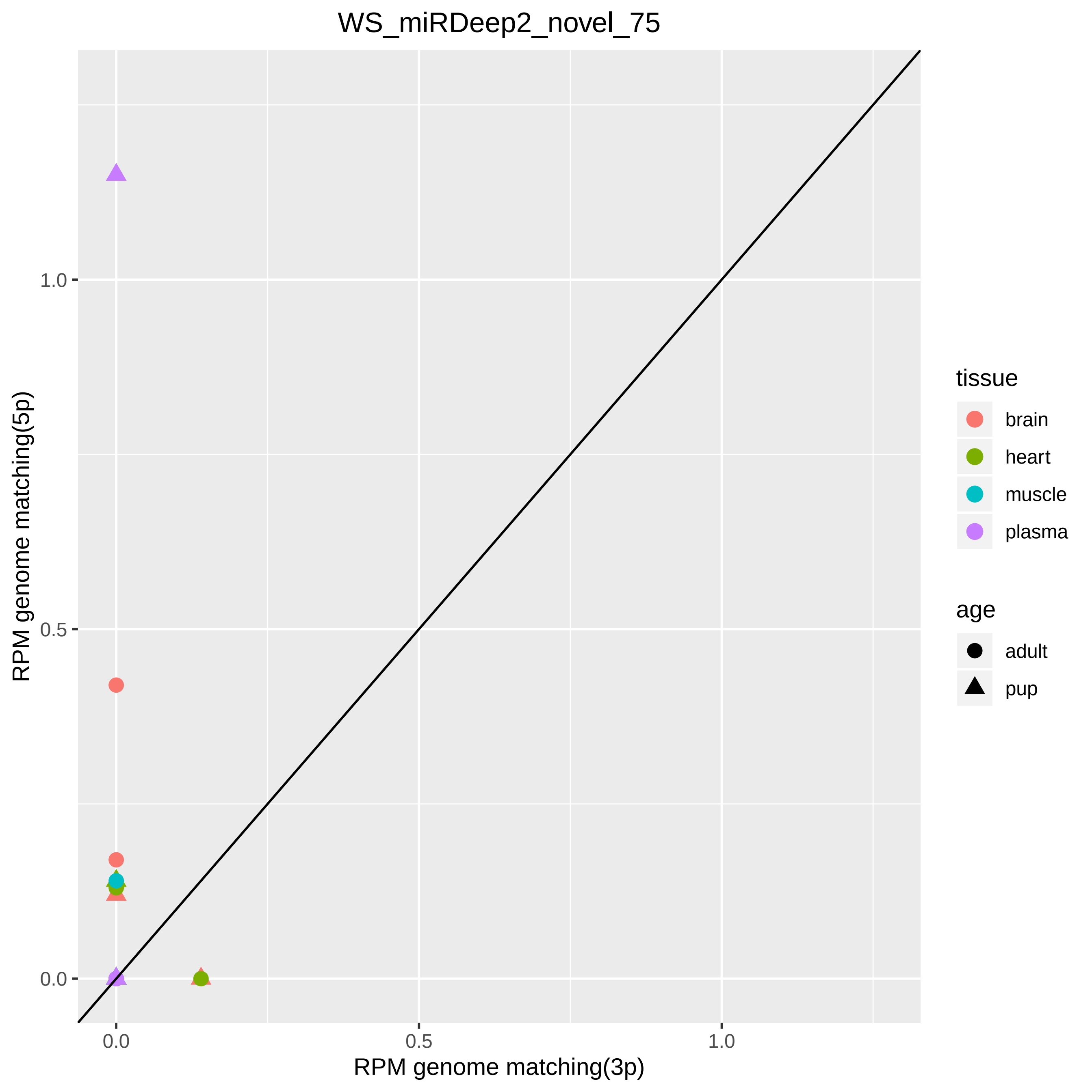
WS_miRDeep2_novel_75
RPM genome matching(5p)
RPM genome matching(3p)
1.0
0.5
0.0
0.0
0.5
1.0
tissue
brain
heart
muscle
plasma
age
adult
pup

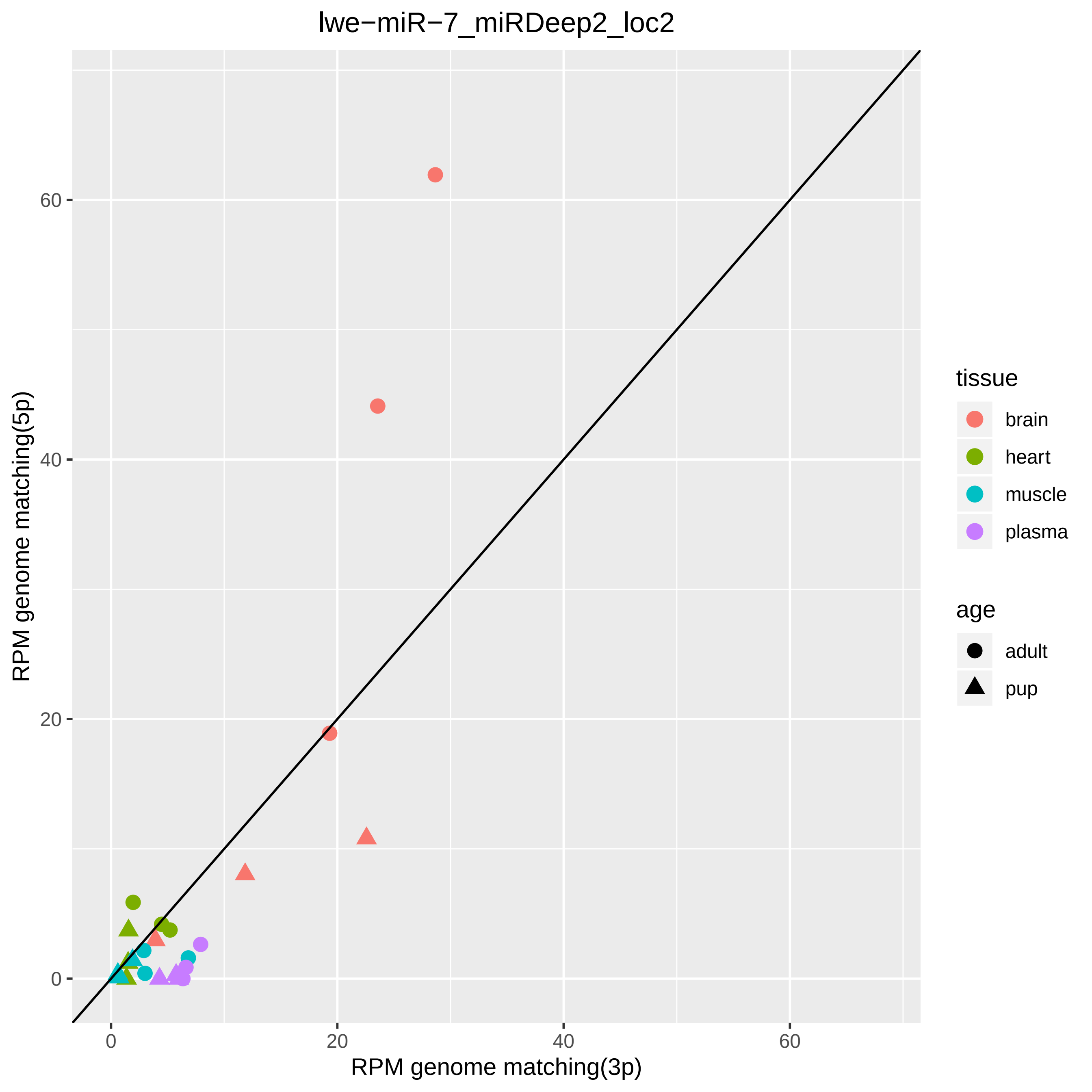

lwe−miR−7_miRDeep2_loc2
RPM genome matching(5p)
RPM genome matching(3p)
0
20
40
60
0
20
40
60
tissue
brain
heart
muscle
plasma
age
adult
pup

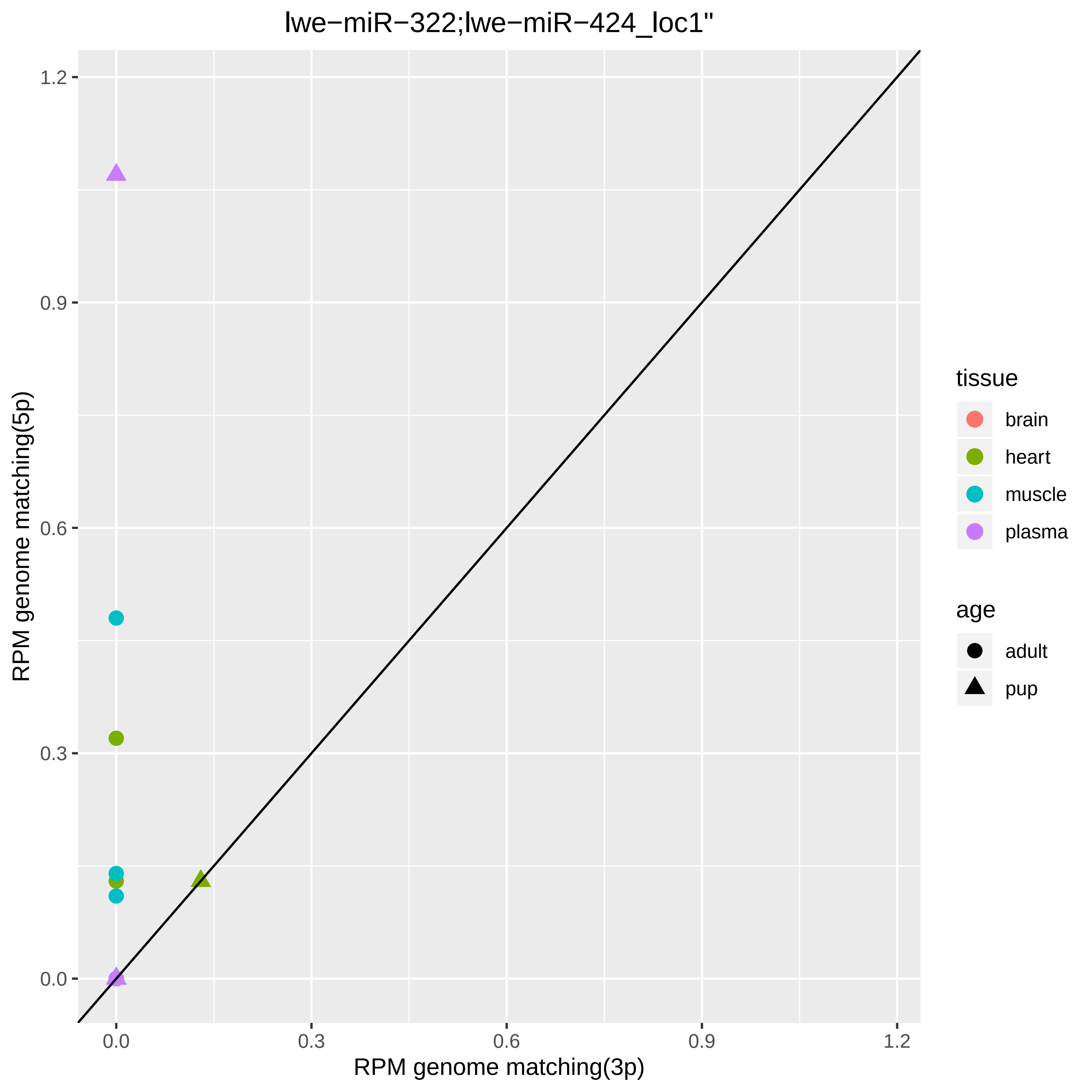

lwe−miR−322;lwe−miR−424_loc1"
RPM genome matching(5p)
RPM genome matching(3p)
1.2
0.9
0.6
0.3
0.0
0.0
0.3
0.6
0.9
1.2
tissue
brain
heart
muscle
plasma
age
adult
pup

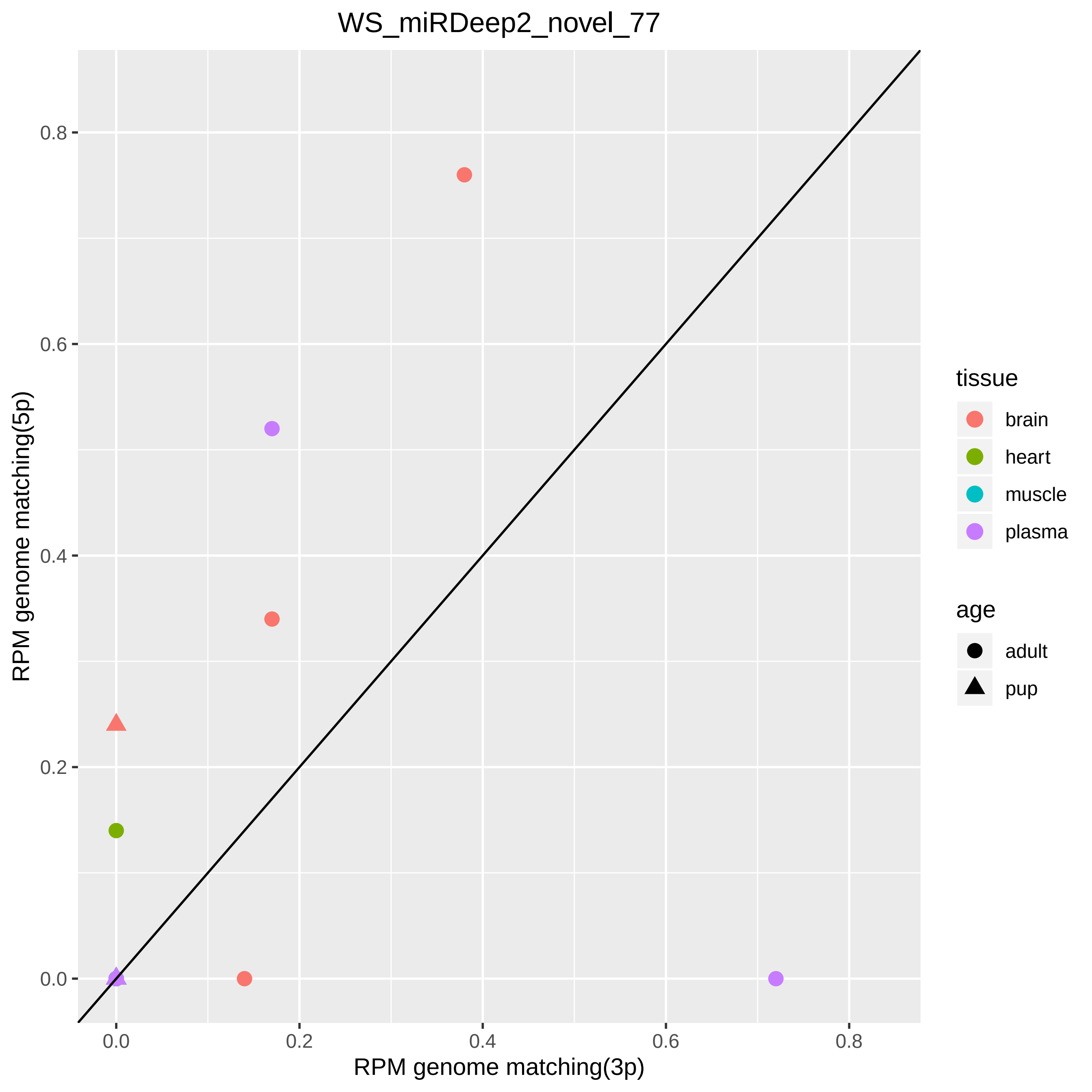

WS_miRDeep2_novel_77
RPM genome matching(5p)
RPM genome matching(3p)
0.0
0.2
0.4
0.6
0.8
tissue
brain
heart
muscle
plasma
age
adult
pup

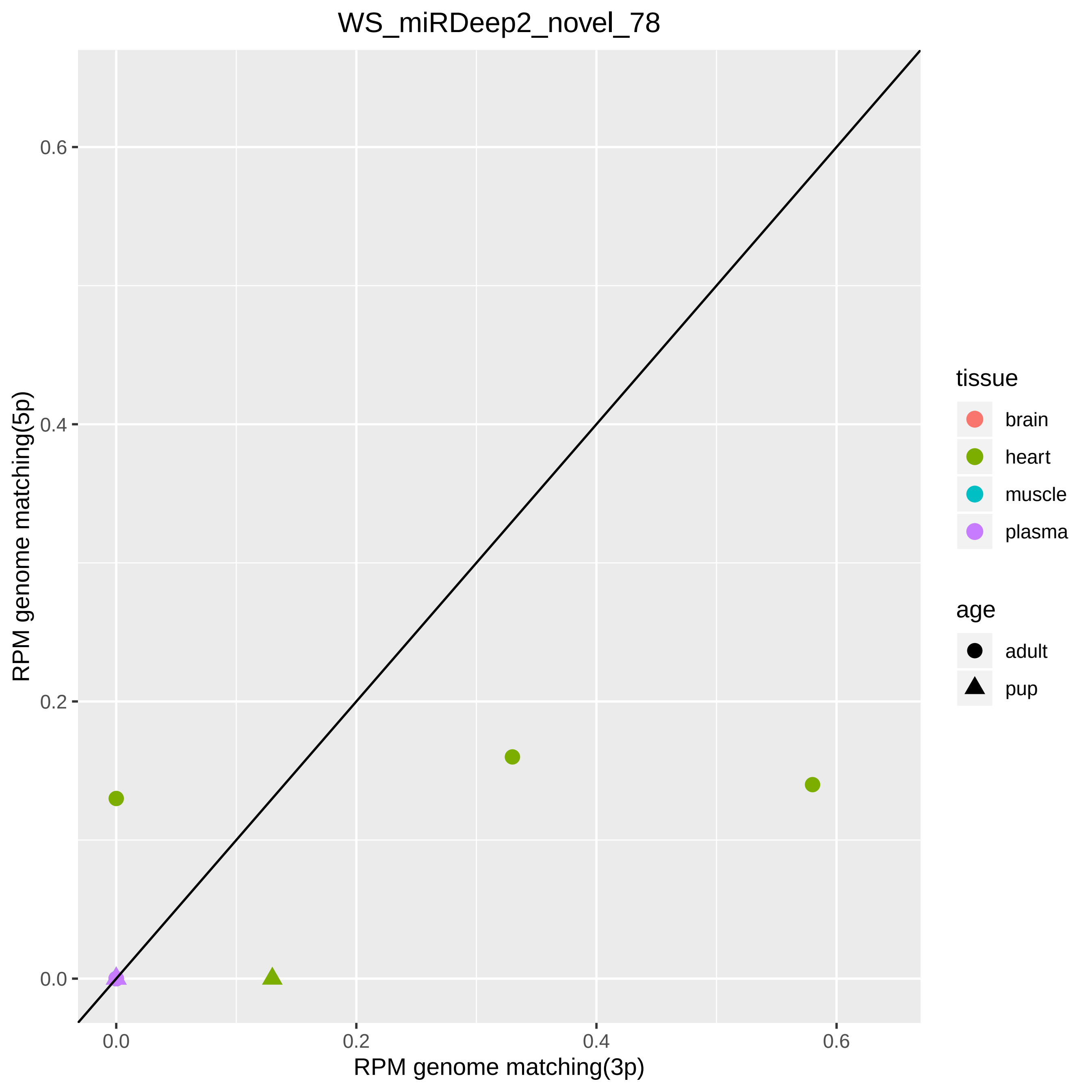
WS_miRDeep2_novel_78
RPM genome matching(5p)
RPM genome matching(3p)
0.0
0.2
0.4
0.6
0.0
0.2
0.4
0.6
tissue
brain
heart
muscle
plasma
age
adult
pup

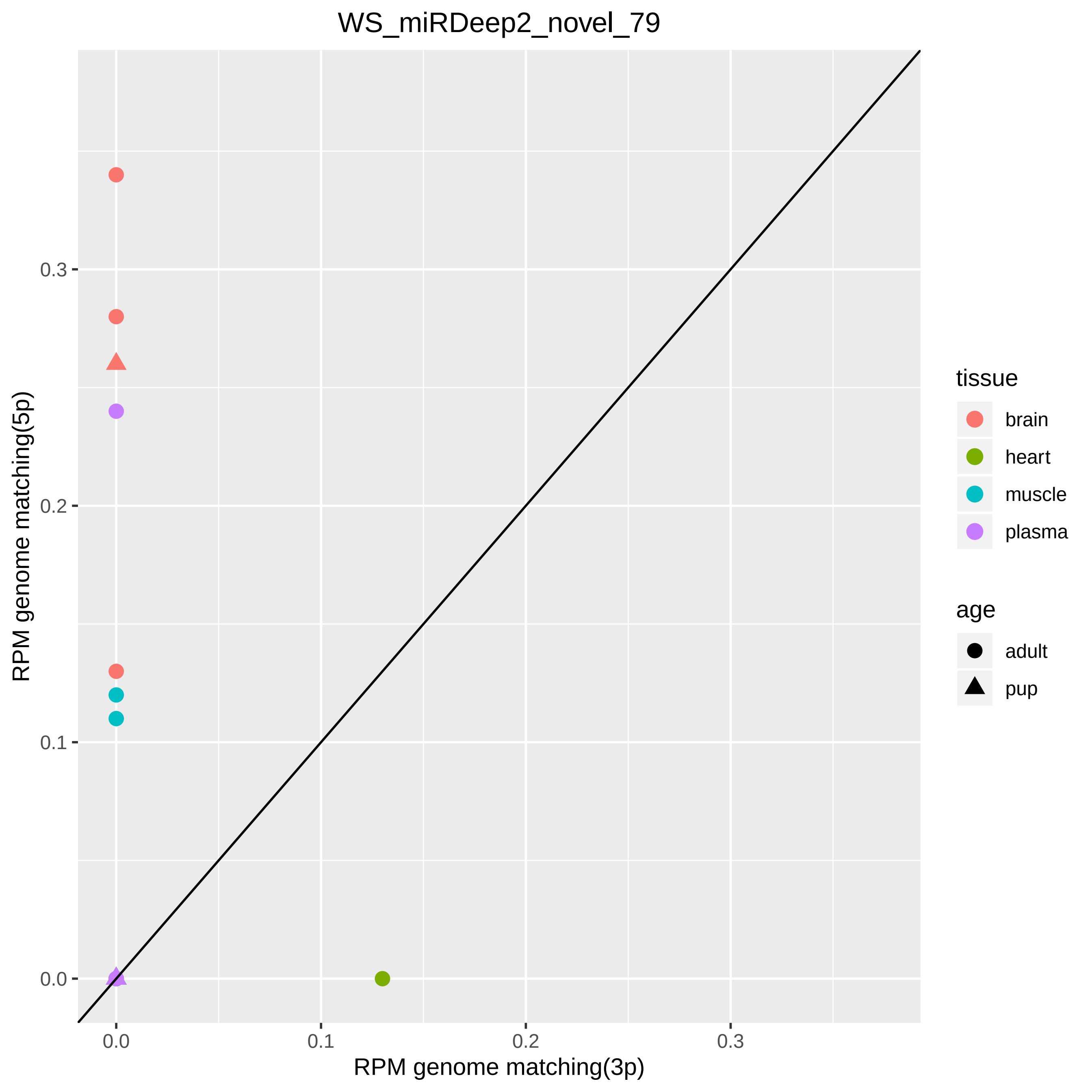
WS_miRDeep2_novel_79
RPM genome matching(5p)
RPM genome matching(3p)
0.0
0.1
0.2
0.3
tissue
brain
heart
muscle
plasma
age
adult
pup

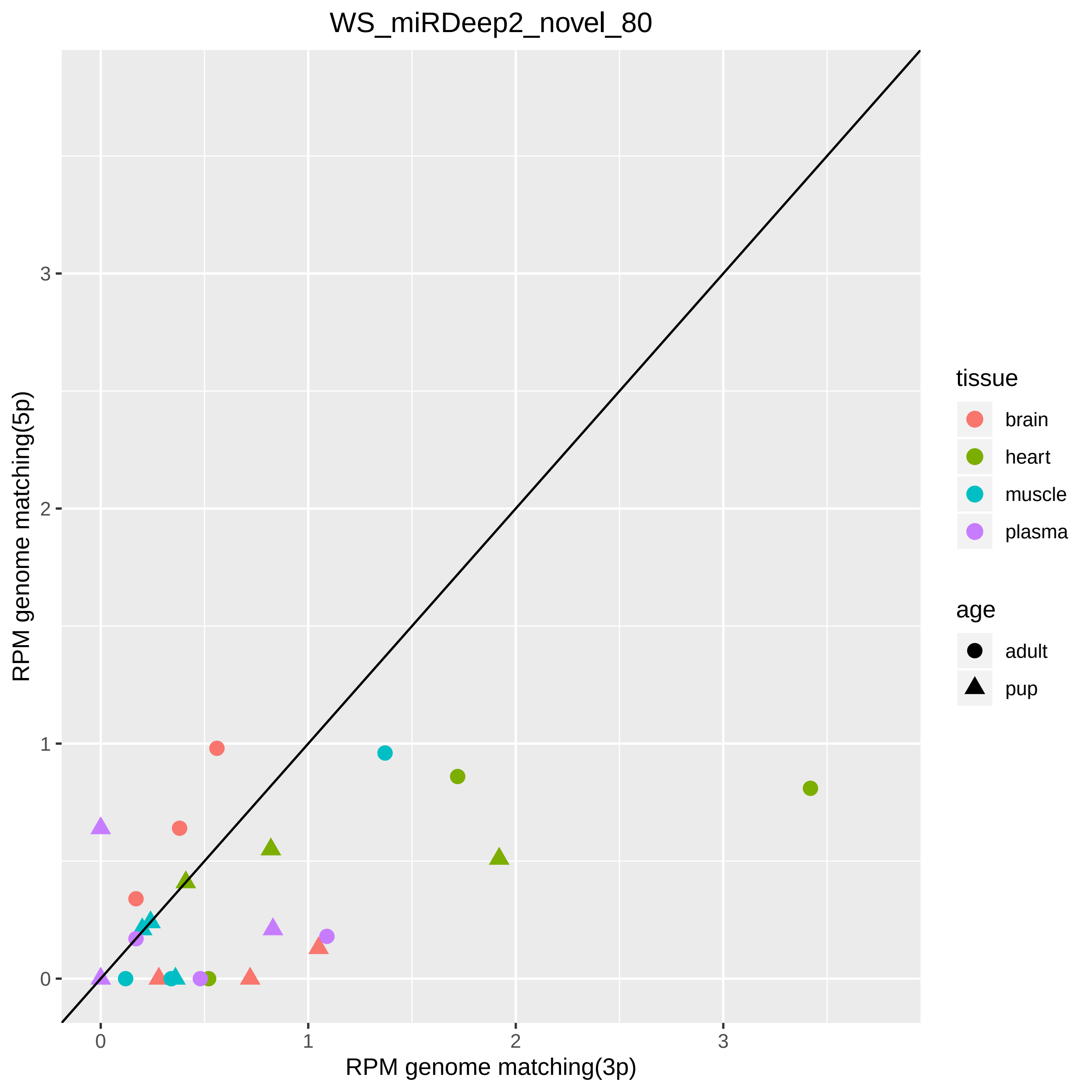
WS_miRDeep2_novel_80
RPM genome matching(5p)
RPM genome matching(3p)
0
1
2
3
0
1
2
3
tissue
brain
heart
muscle
plasma
age
adult
pup

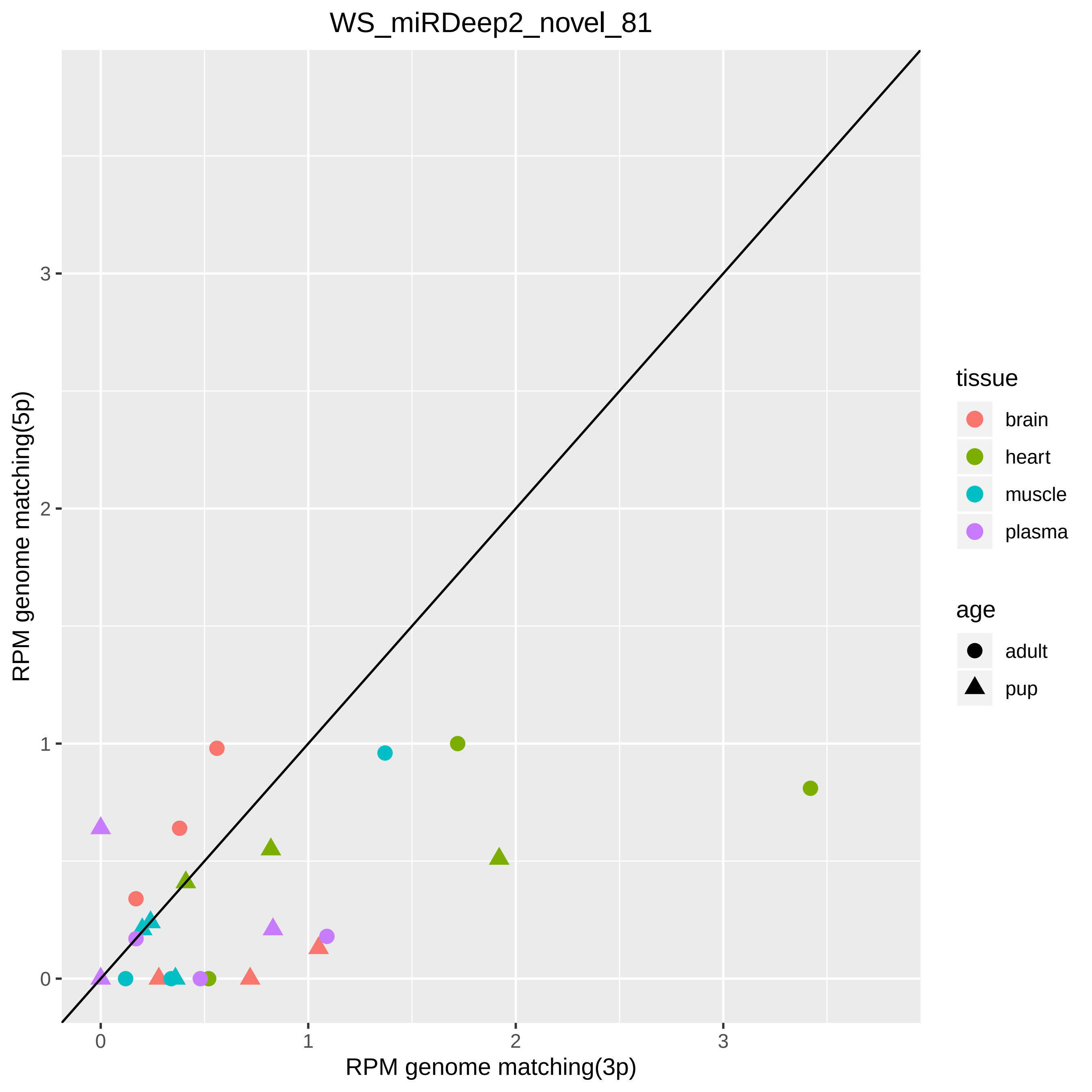

WS_miRDeep2_novel_81
RPM genome matching(5p)
RPM genome matching(3p)
0
1
2
3
tissue
brain
heart
muscle
plasma
age
adult
pup

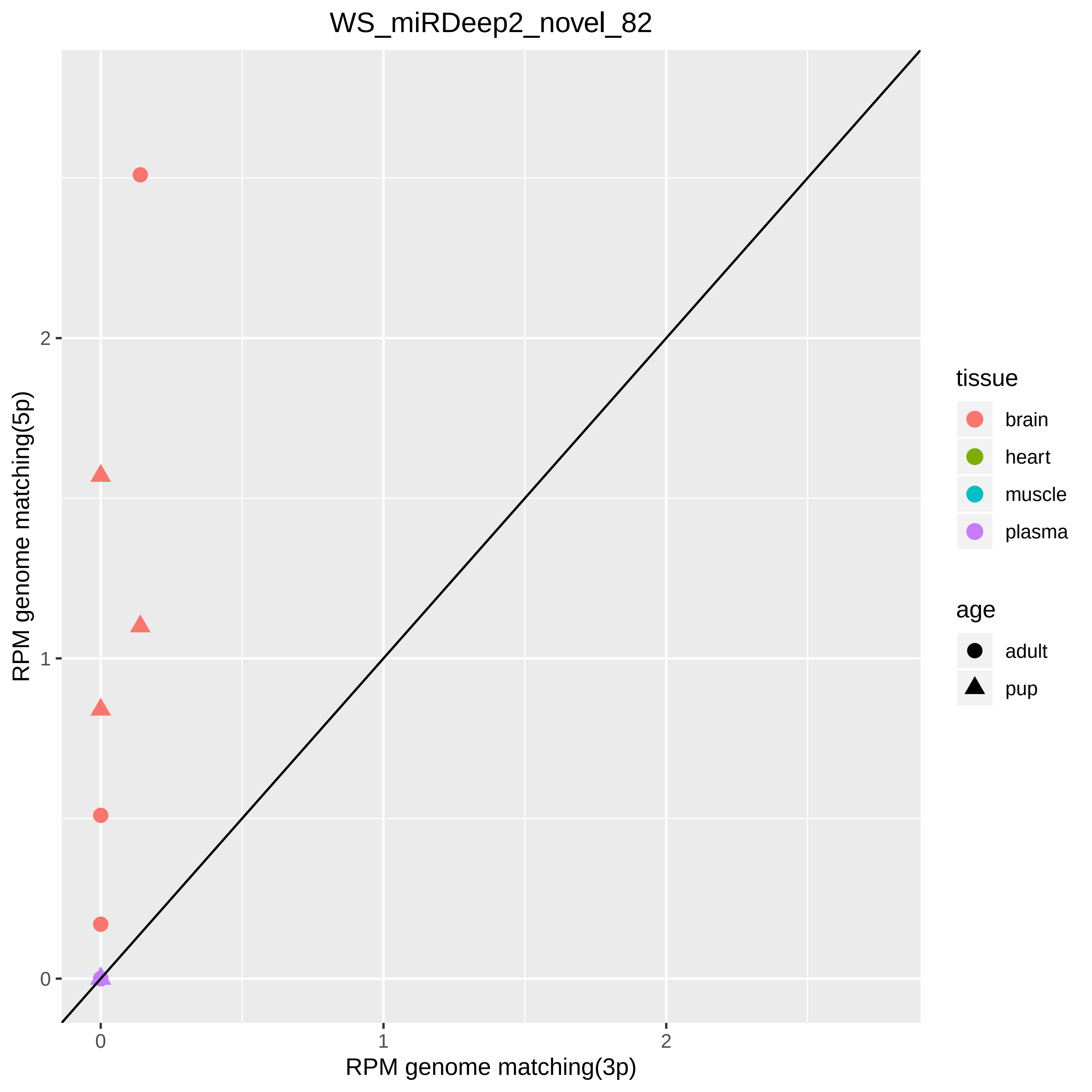
WS_miRDeep2_novel_82
RPM genome matching(5p)
RPM genome matching(3p)
0
1
2
0
1
2
tissue
brain
heart
muscle
plasma
age
adult
pup

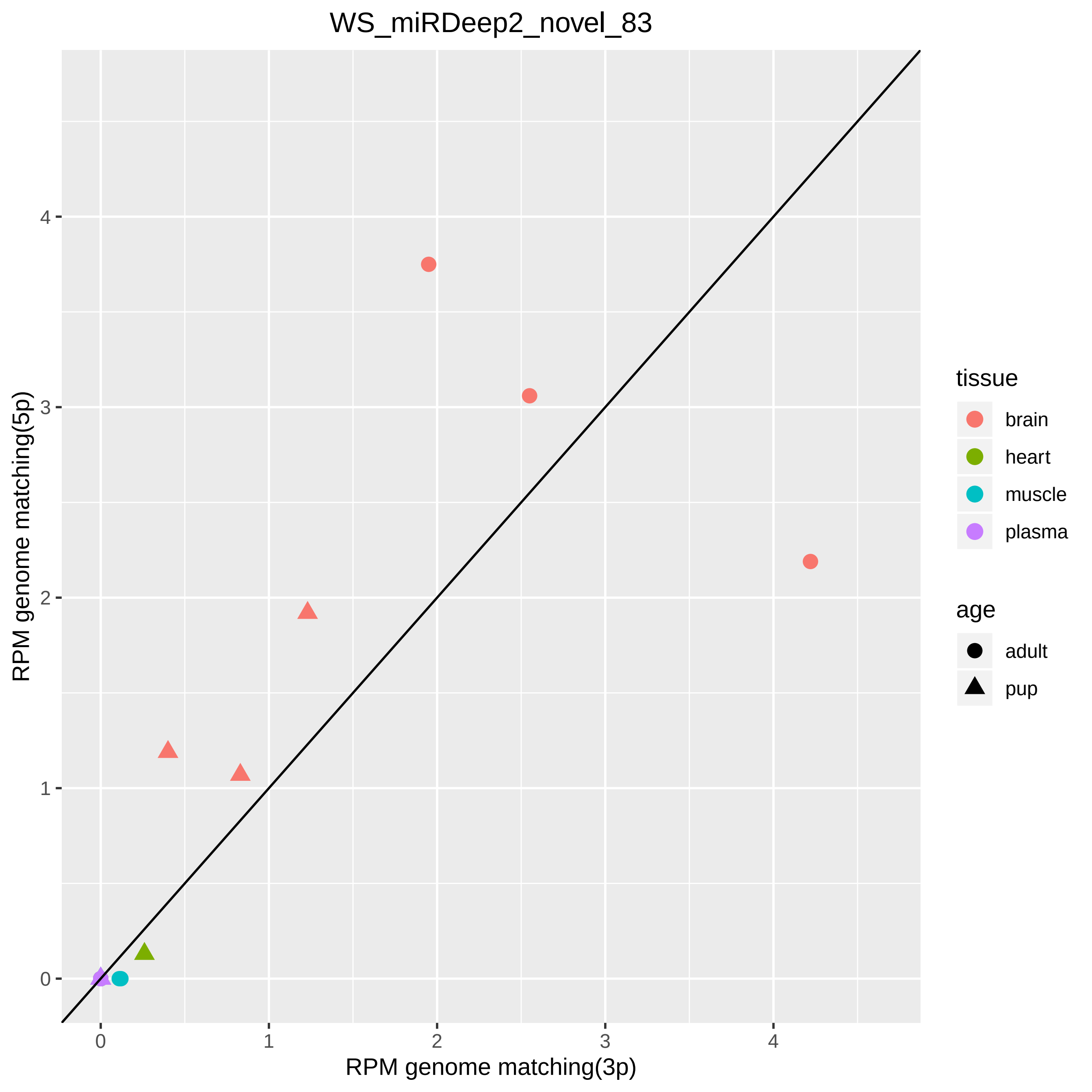
WS_miRDeep2_novel_83
RPM genome matching(5p)
RPM genome matching(3p)
0
1
2
3
4
tissue
brain
heart
muscle
plasma
age
adult
pup

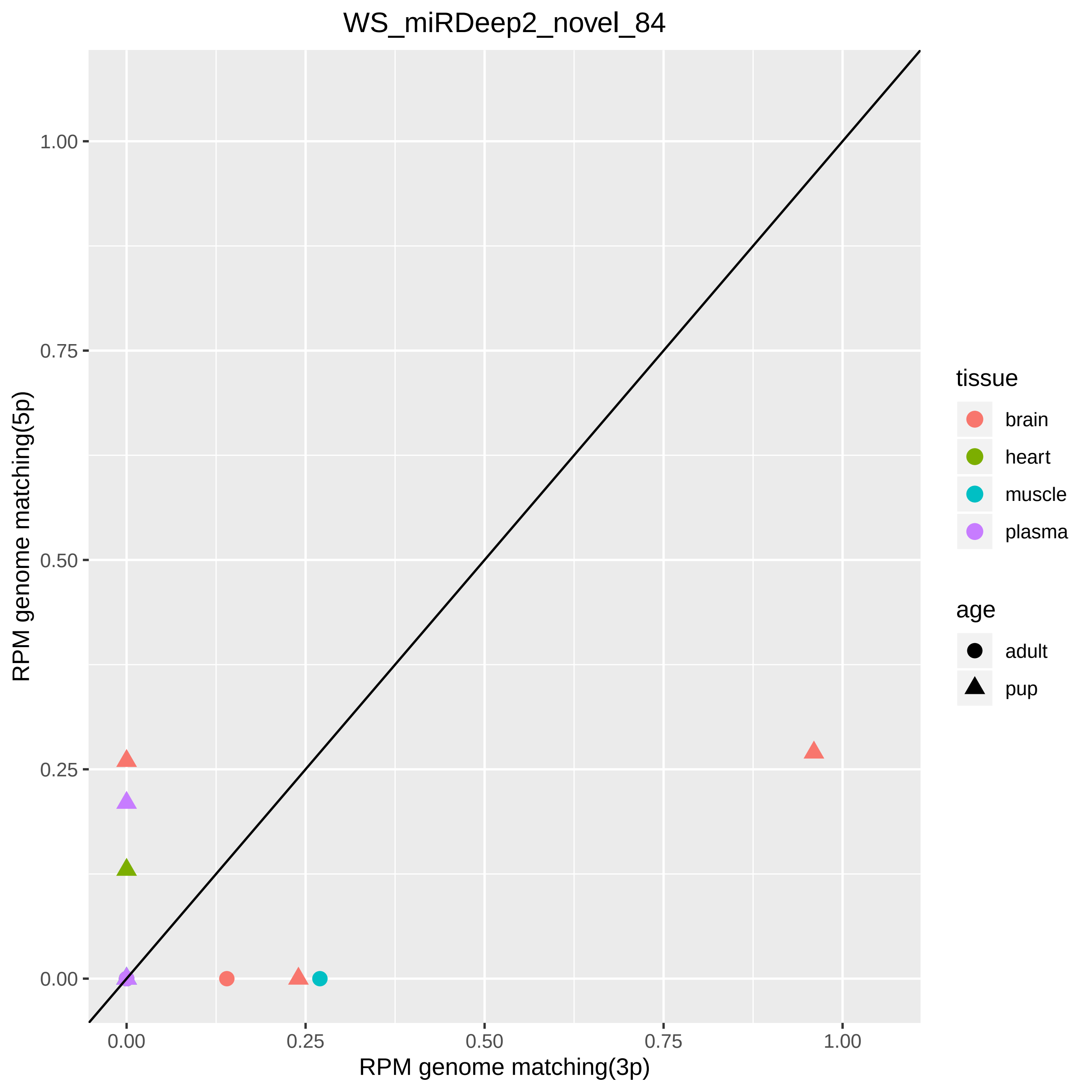

WS_miRDeep2_novel_84
RPM genome matching(5p)
RPM genome matching(3p)
0.00
0.25
0.50
0.75
1.00
tissue
brain
heart
muscle
plasma
age
adult
pup

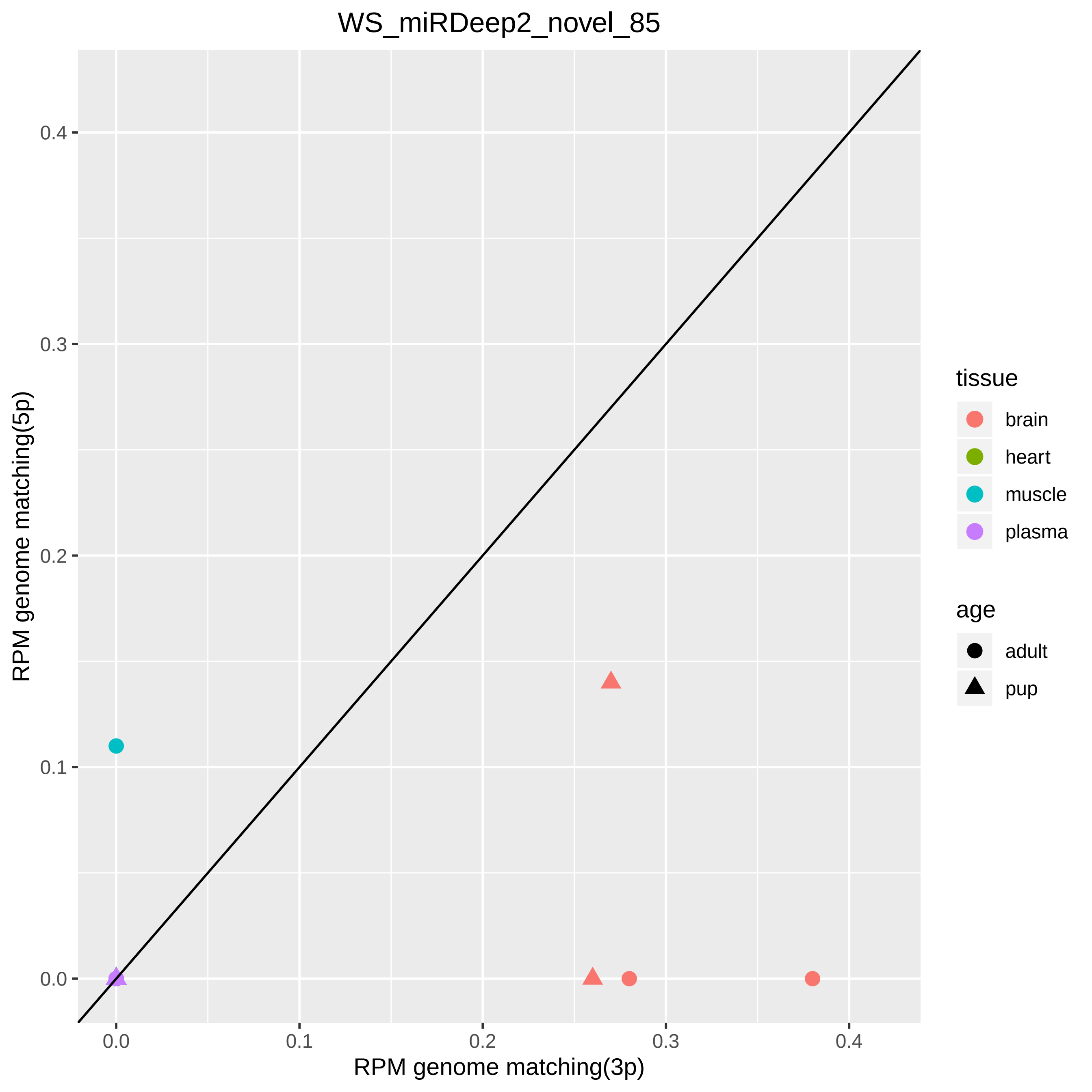

WS_miRDeep2_novel_85
RPM genome matching(5p)
RPM genome matching(3p)
0.0
0.1
0.2
0.3
0.4
tissue
brain
heart
muscle
plasma
age
adult
pup

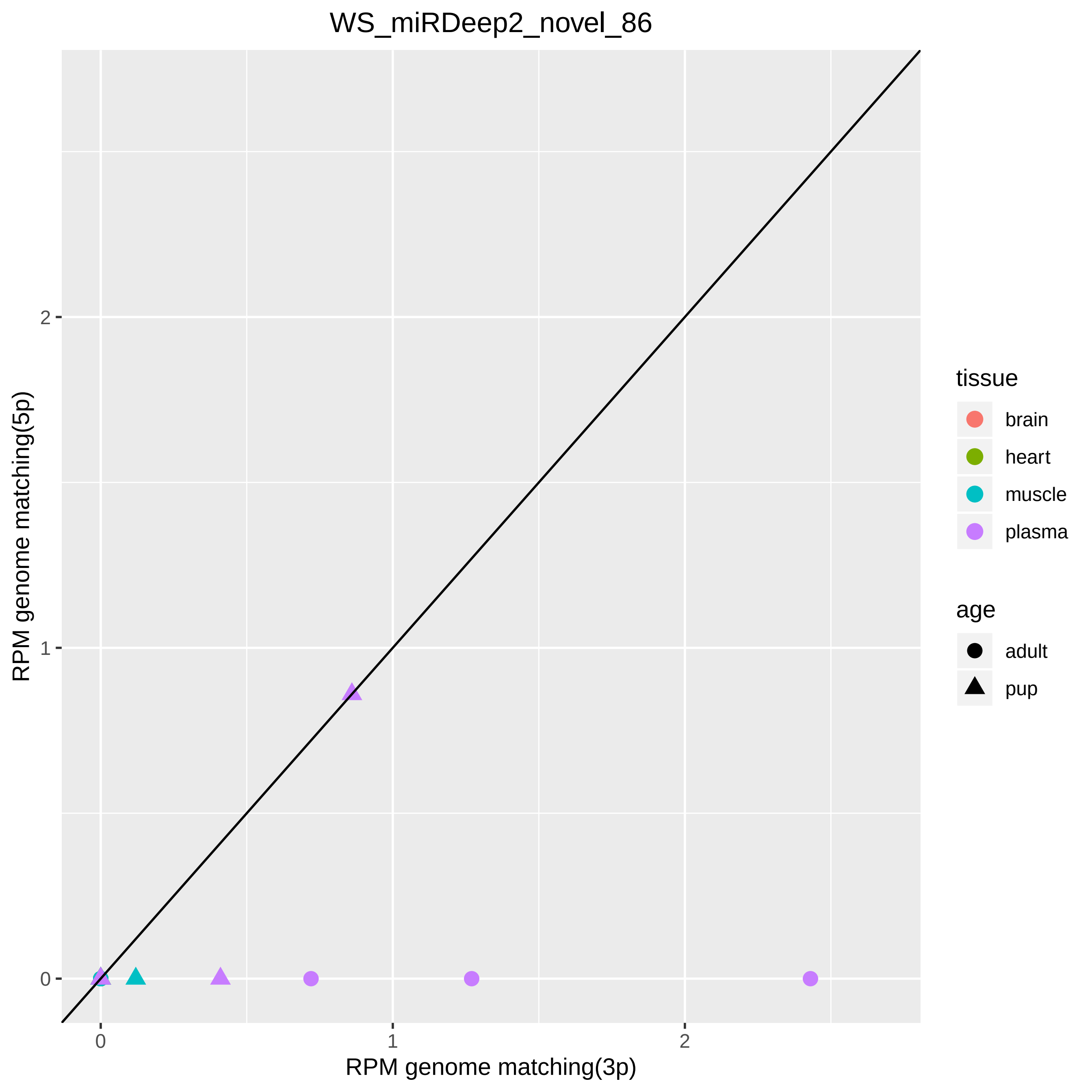

WS_miRDeep2_novel_86
RPM genome matching(5p)
RPM genome matching(3p)
0
1
2
0
1
2
tissue
brain
heart
muscle
plasma
age
adult
pup

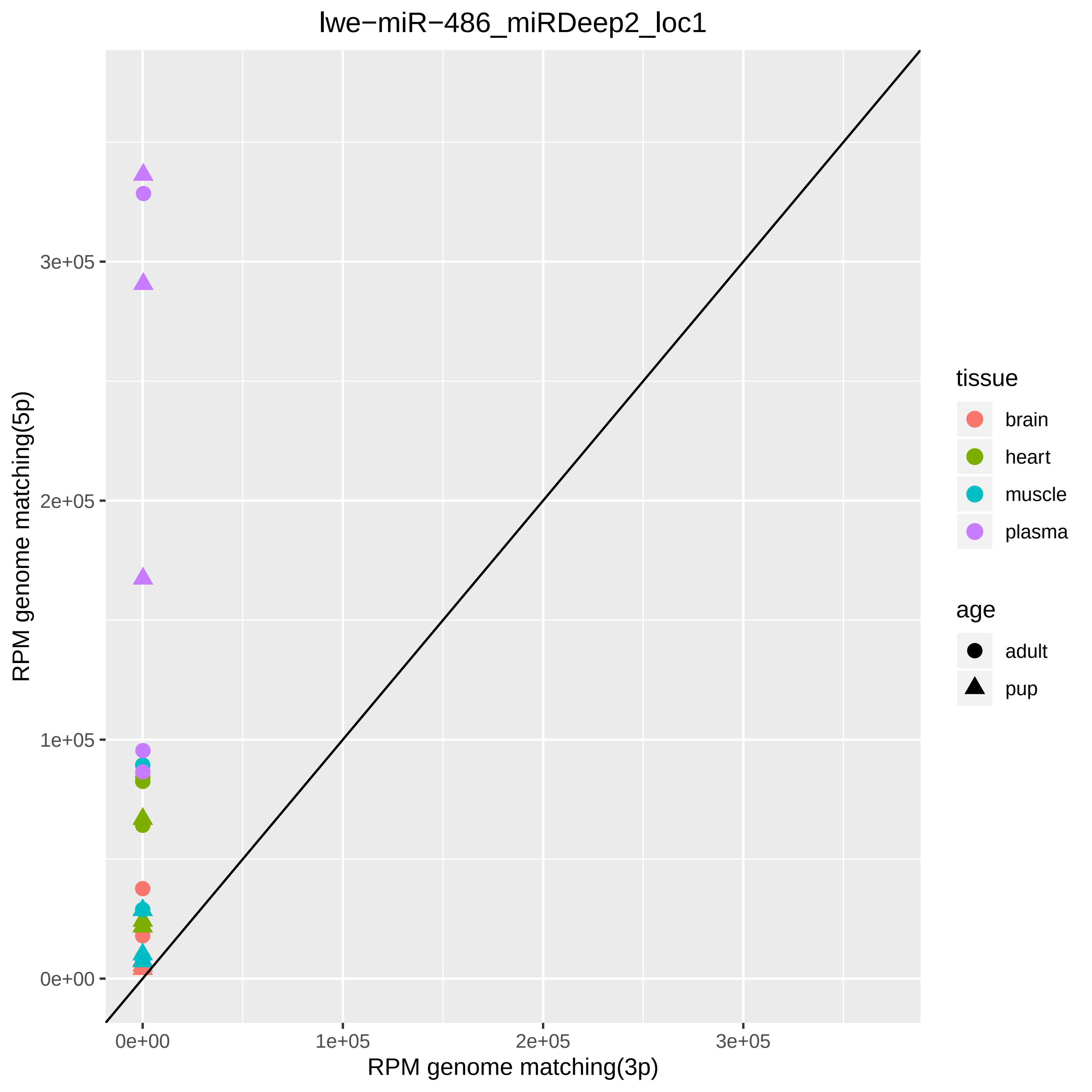
lwe−miR−486_miRDeep2_loc1
RPM genome matching(5p)
RPM genome matching(3p)
3e+05
2e+05
1e+05
0e+00
0e+00
1e+05
2e+05
3e+05
tissue
brain
heart
muscle
plasma
age
adult
pup

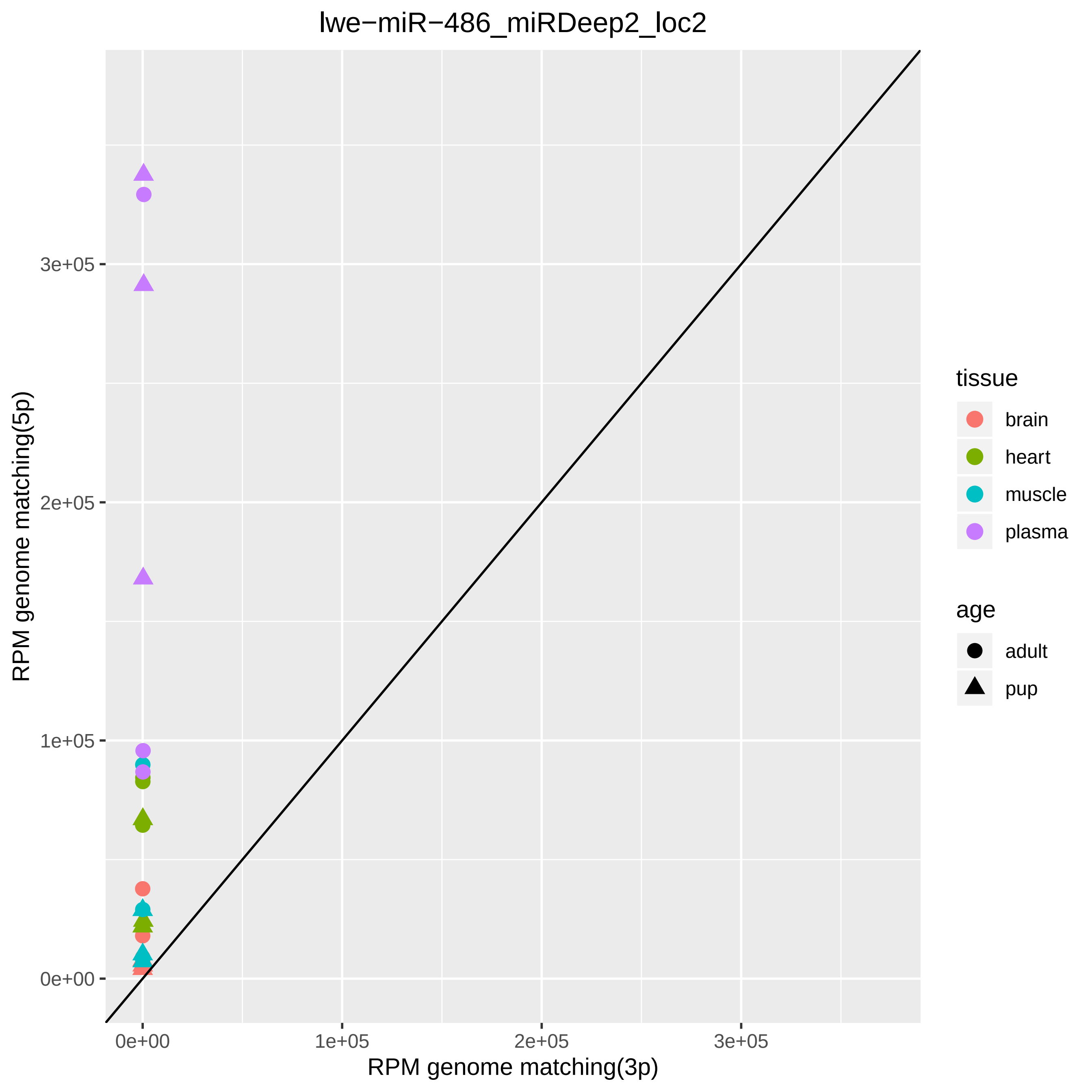
lwe−miR−486_miRDeep2_loc2
RPM genome matching(5p)
3e+05
2e+05
1e+05
0e+00
0e+00
1e+05
2e+05
3e+05
RPM genome matching(3p)
tissue
brain
heart
muscle
plasma
age
adult
pup

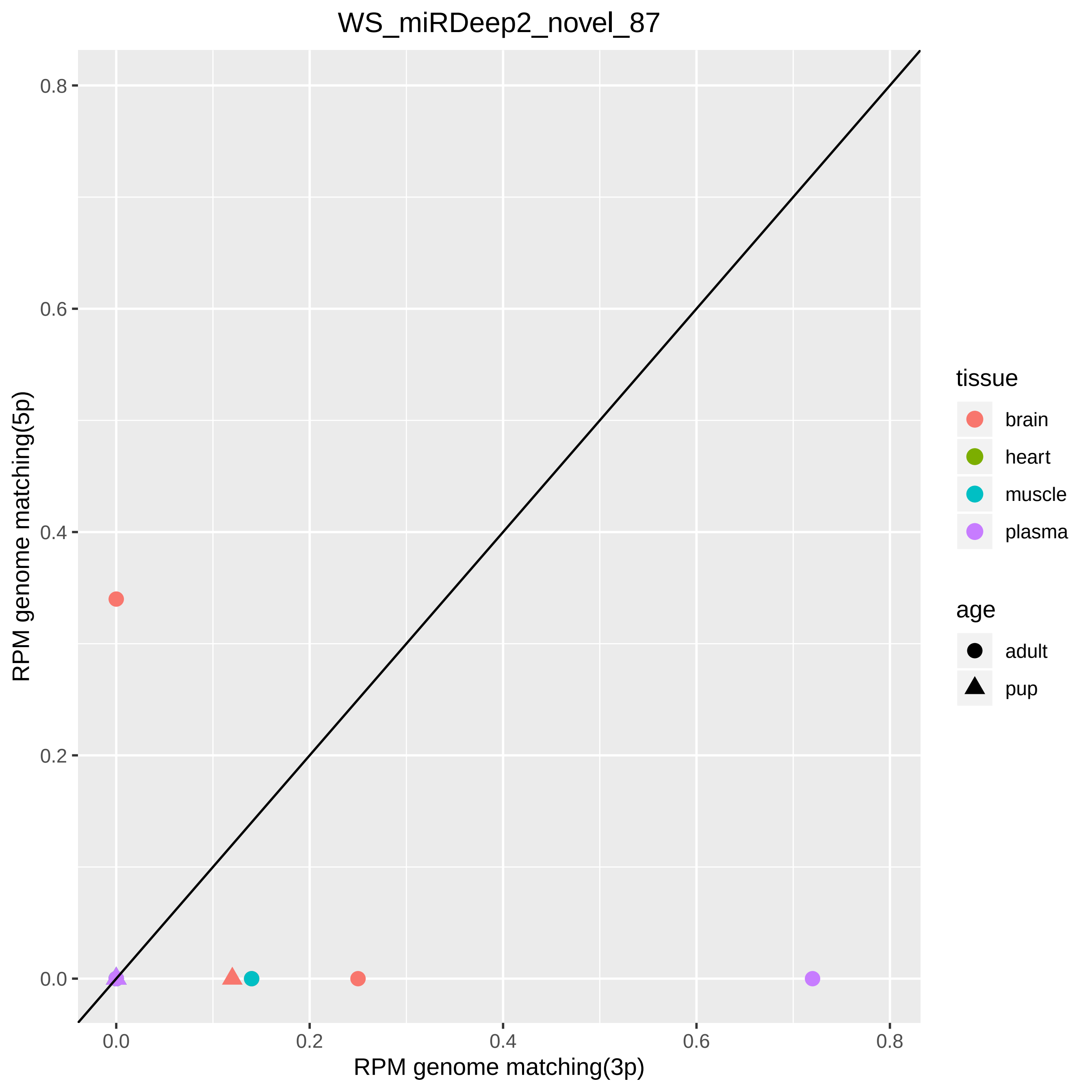
WS_miRDeep2_novel_87
RPM genome matching(5p)
RPM genome matching(3p)
0.0
0.2
0.4
0.6
0.8
tissue
brain
heart
muscle
plasma
age
adult
pup

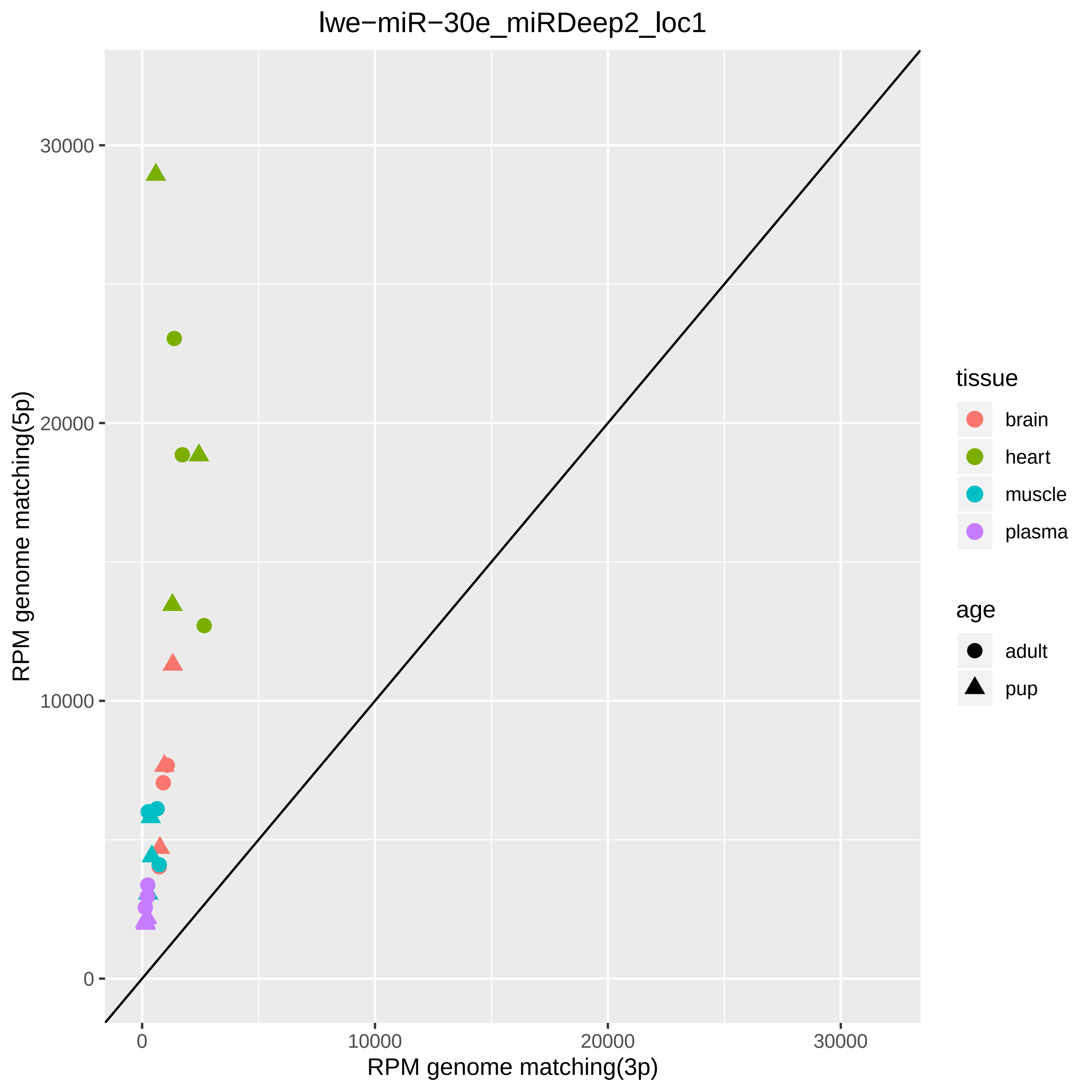
lwe−miR−30e_miRDeep2_loc1
RPM genome matching(5p)
RPM genome matching(3p)
0
10000
20000
30000
tissue
brain
heart
muscle
plasma
age
adult
pup

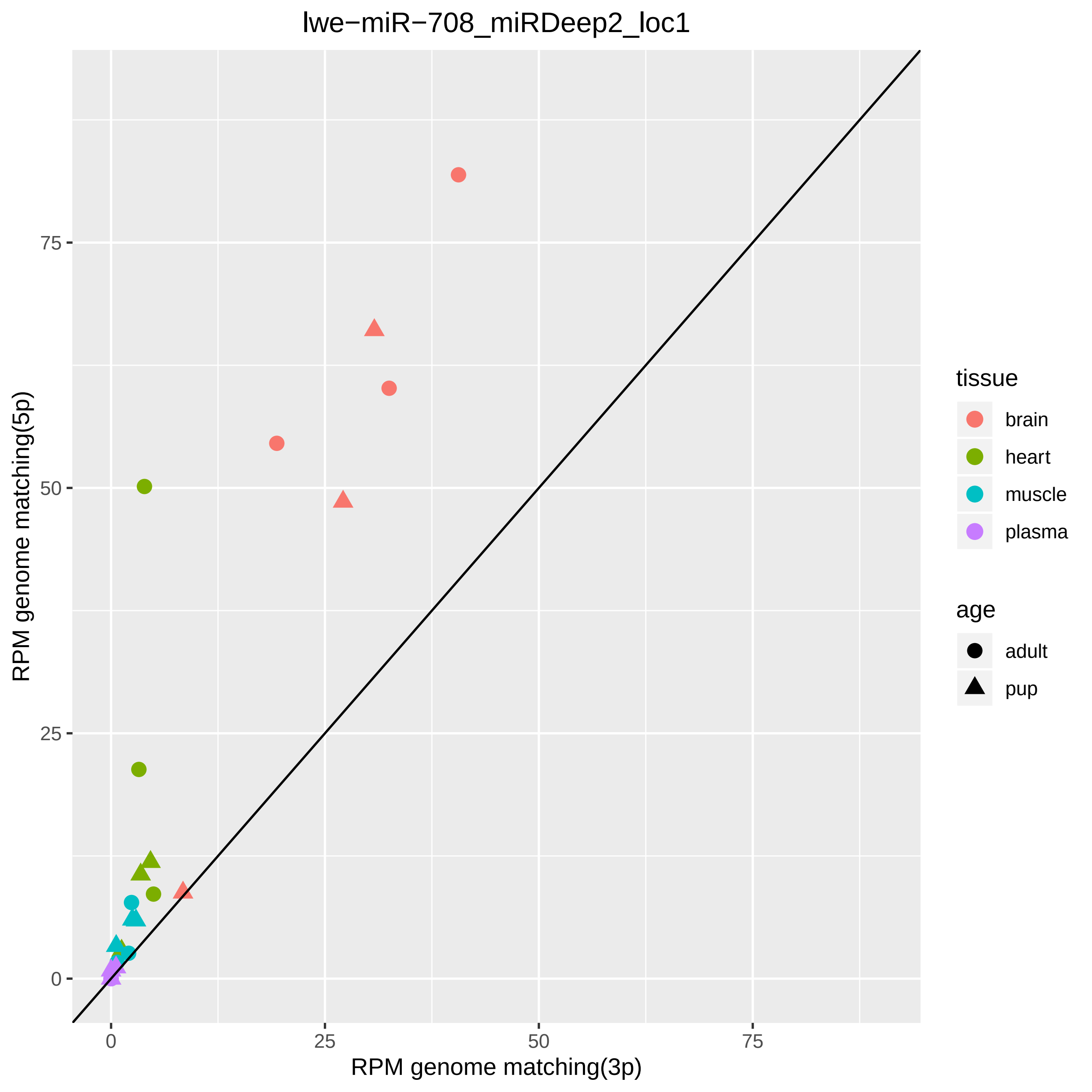
lwe−miR−708_miRDeep2_loc1
RPM genome matching(5p)
RPM genome matching(3p)
0
25
50
75
0
25
50
75
tissue
brain
heart
muscle
plasma
age
adult
pup

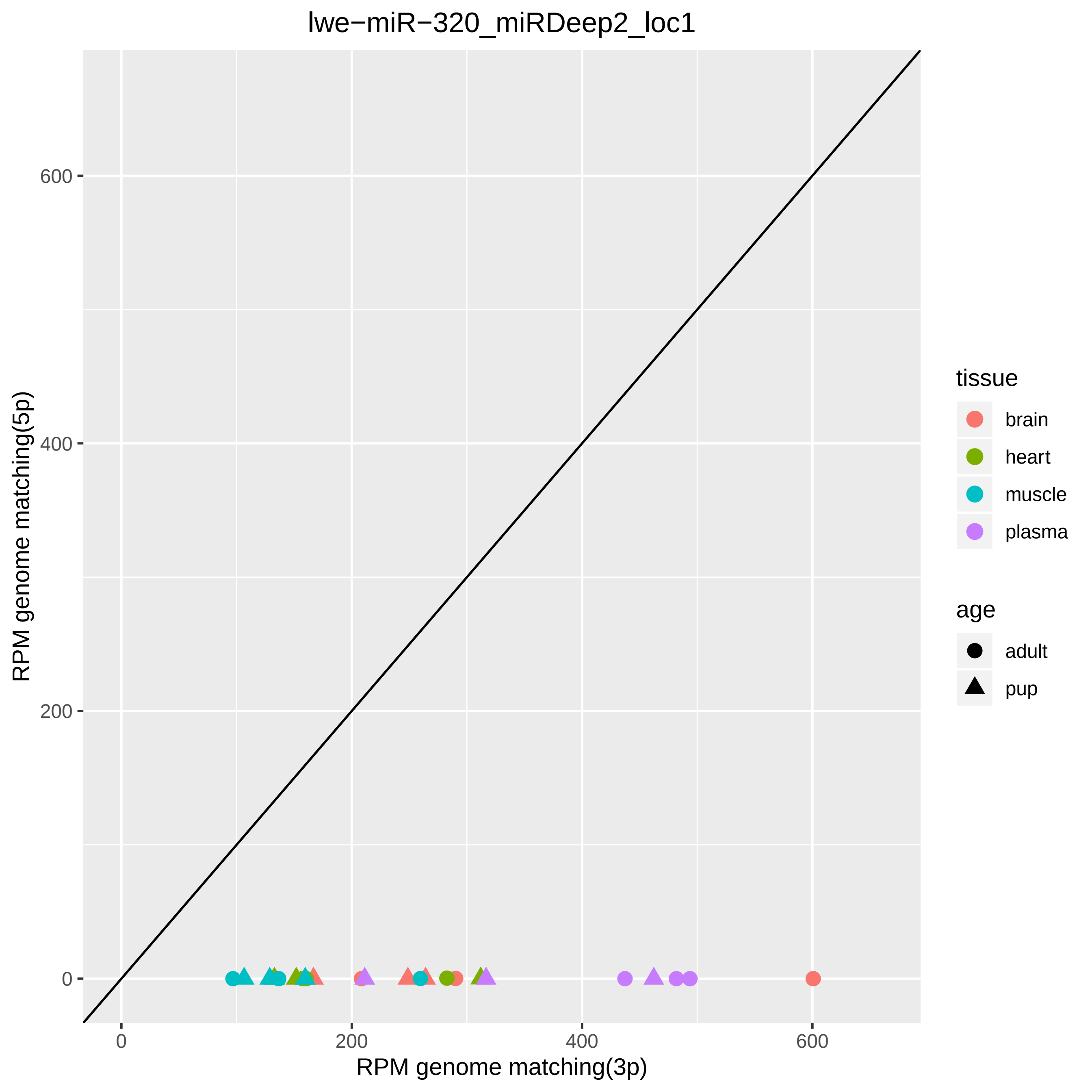

lwe−miR−320_miRDeep2_loc1
RPM genome matching(5p)
RPM genome matching(3p)
0
200
400
600
tissue
brain
heart
muscle
plasma
age
adult
pup

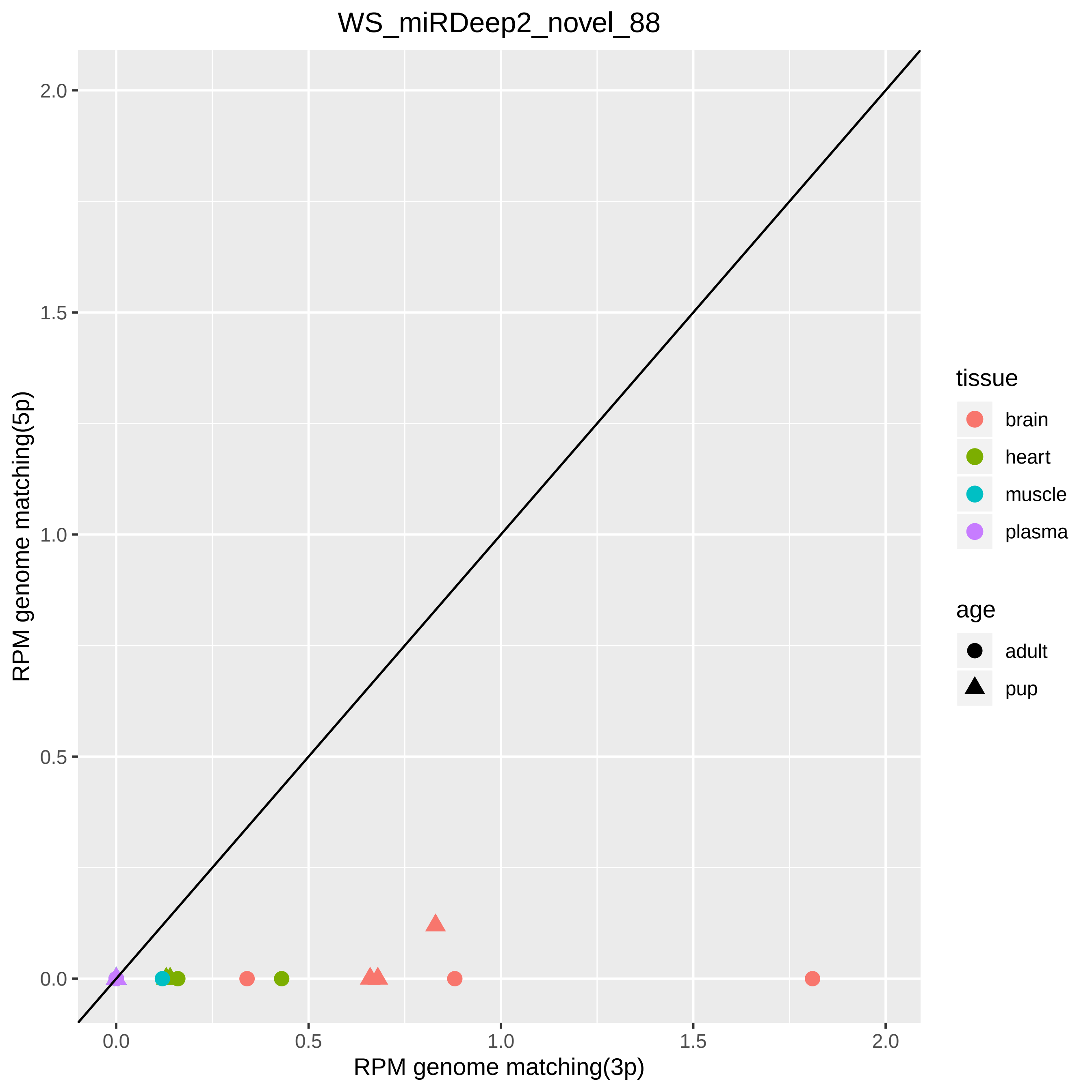

WS_miRDeep2_novel_88
RPM genome matching(5p)
RPM genome matching(3p)
0.0
0.5
1.0
1.5
2.0
tissue
brain
heart
muscle
plasma
age
adult
pup

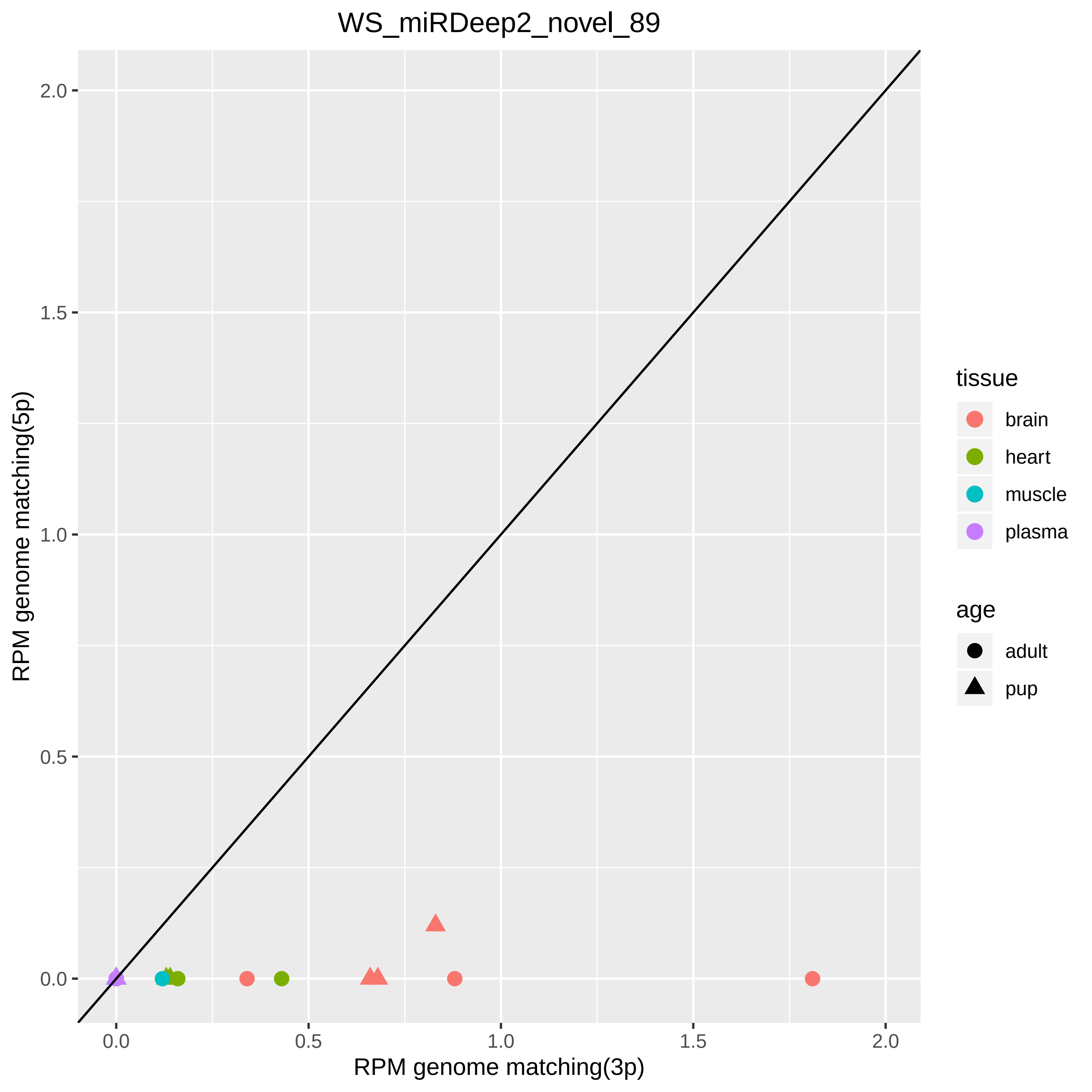

WS_miRDeep2_novel_89
RPM genome matching(5p)
RPM genome matching(3p)
0.0
0.5
1.0
1.5
2.0
tissue
brain
heart
muscle
plasma
age
adult
pup

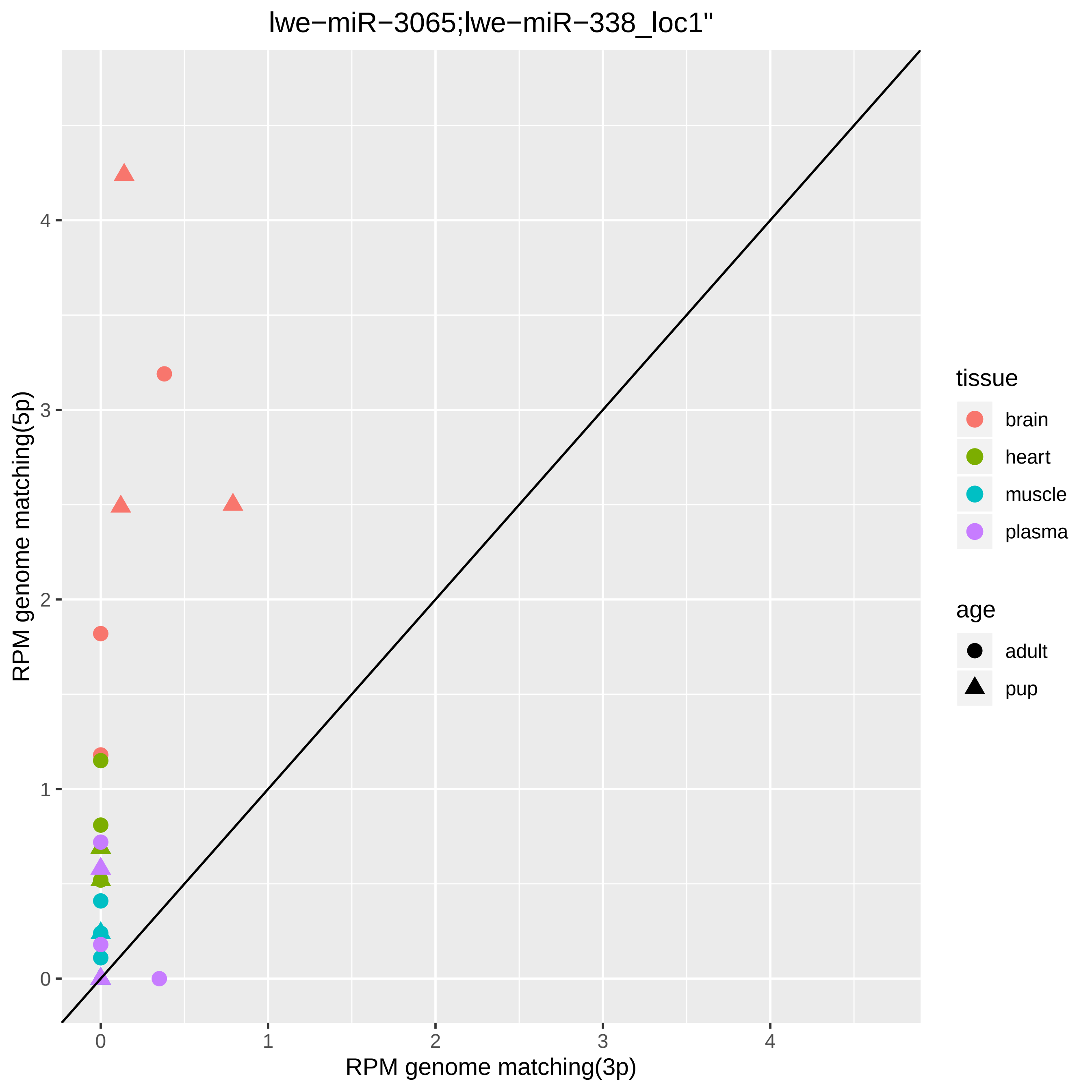

lwe−miR−3065;lwe−miR−338_loc1"
RPM genome matching(5p)
RPM genome matching(3p)
0
1
2
3
4
0
1
2
3
4
tissue
brain
heart
muscle
plasma
age
adult
pup

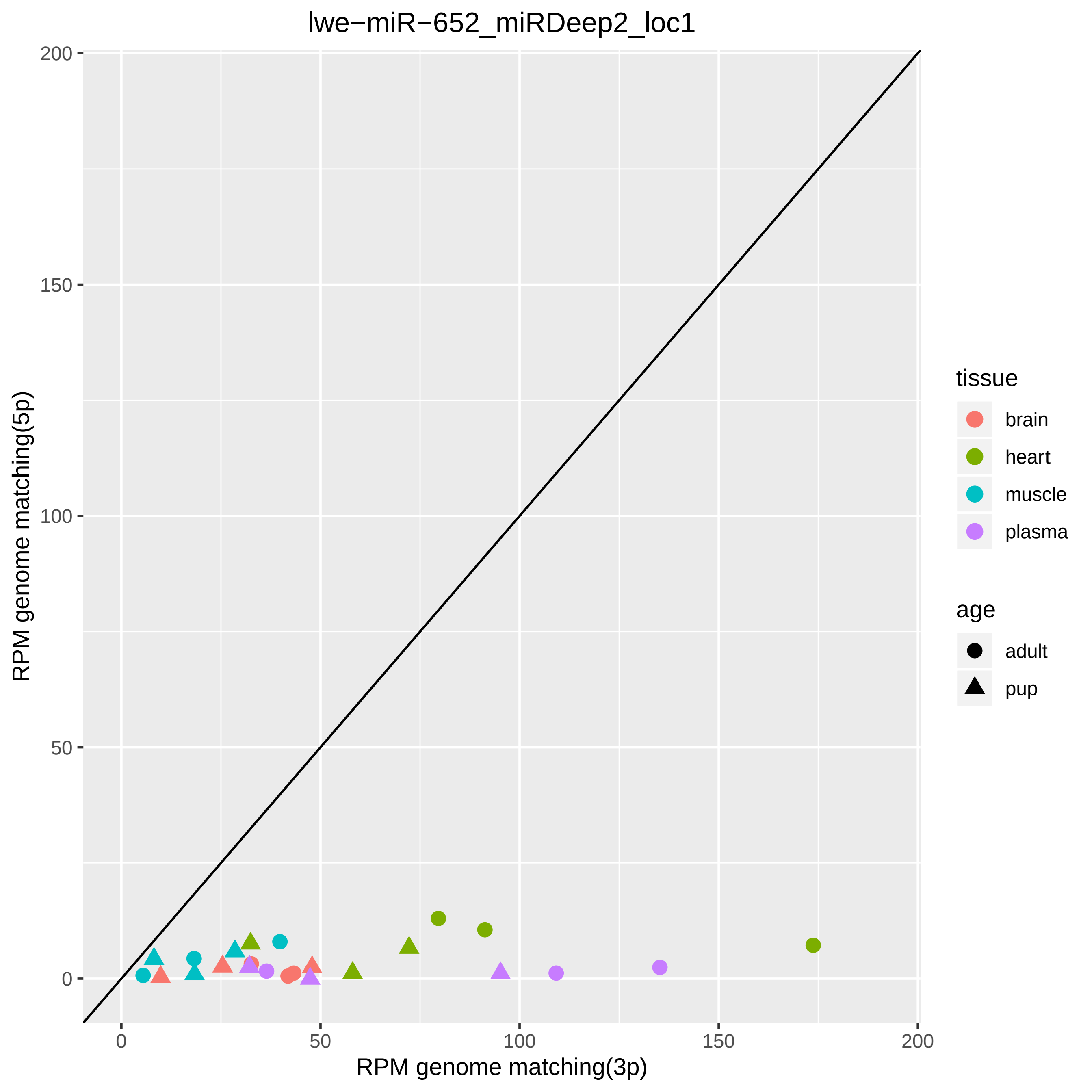
lwe−miR−652_miRDeep2_loc1
RPM genome matching(5p)
RPM genome matching(3p)
0
50
100
150
200
0
50
100
150
200
tissue
brain
heart
muscle
plasma
age
adult
pup

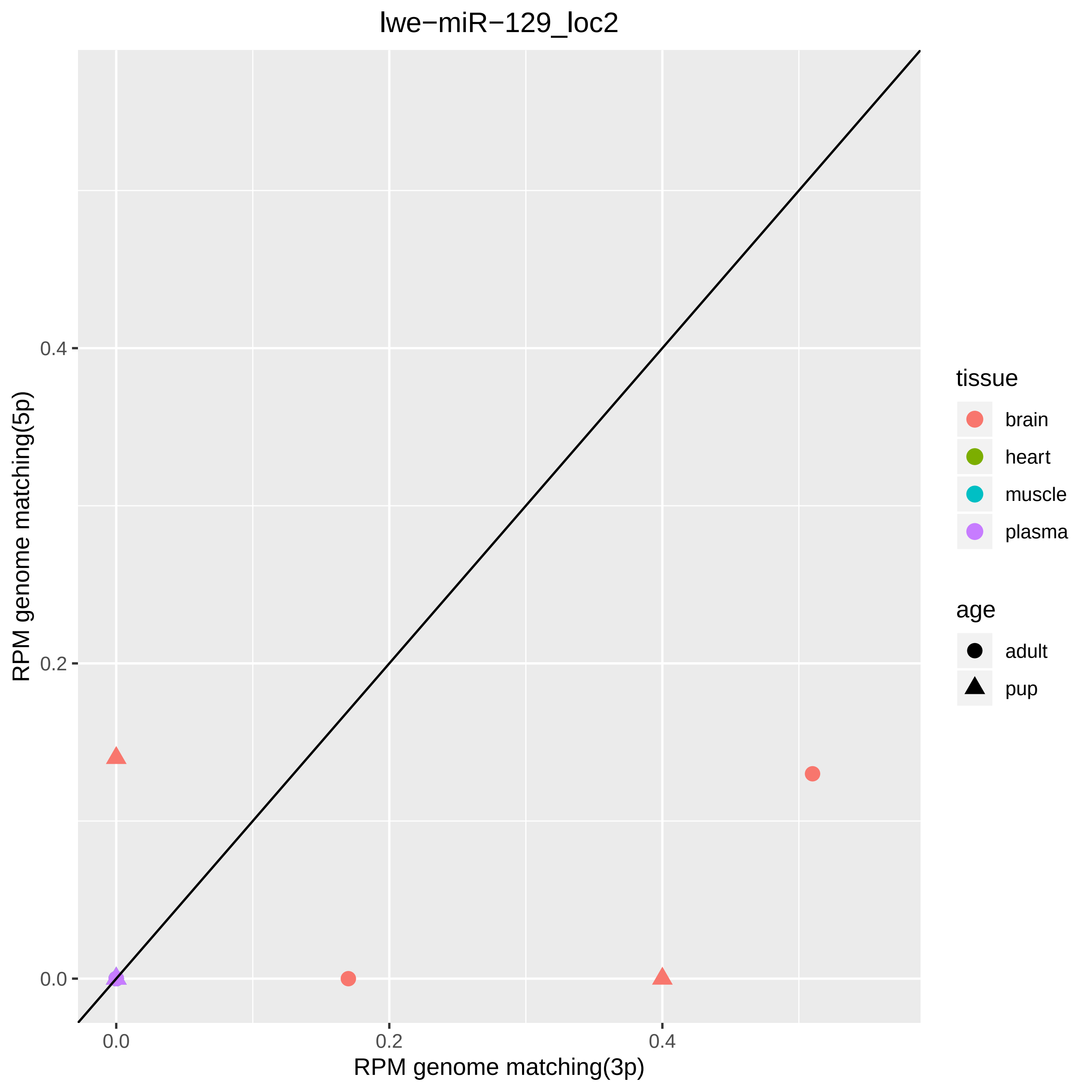
lwe−miR−129_loc2
RPM genome matching(5p)
RPM genome matching(3p)
0.0
0.2
0.4
0.0
0.2
0.4
tissue
brain
heart
muscle
plasma
age
adult
pup

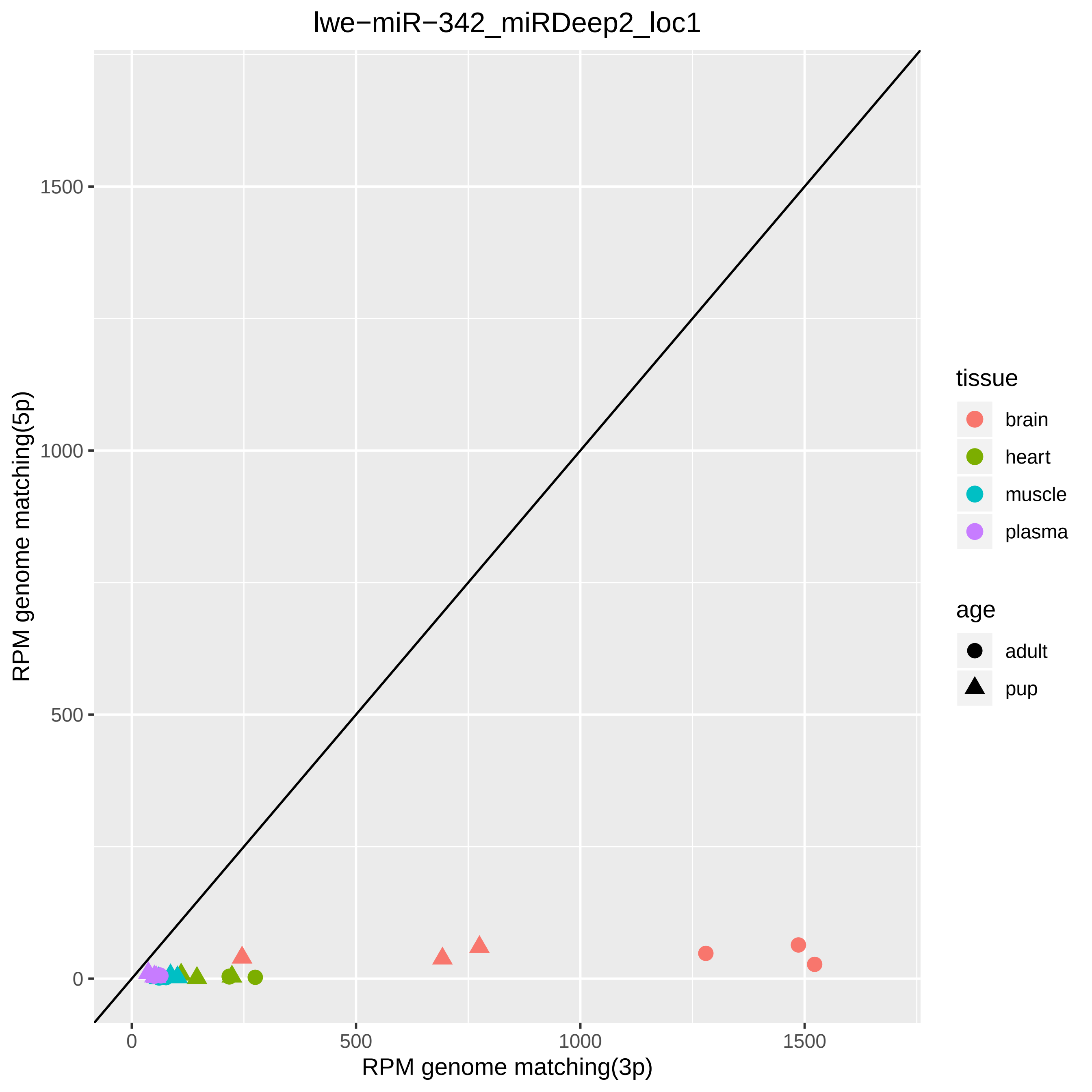
lwe−miR−342_miRDeep2_loc1
RPM genome matching(5p)
RPM genome matching(3p)
0
500
1000
1500
0
500
1000
1500
tissue
brain
heart
muscle
plasma
age
adult
pup

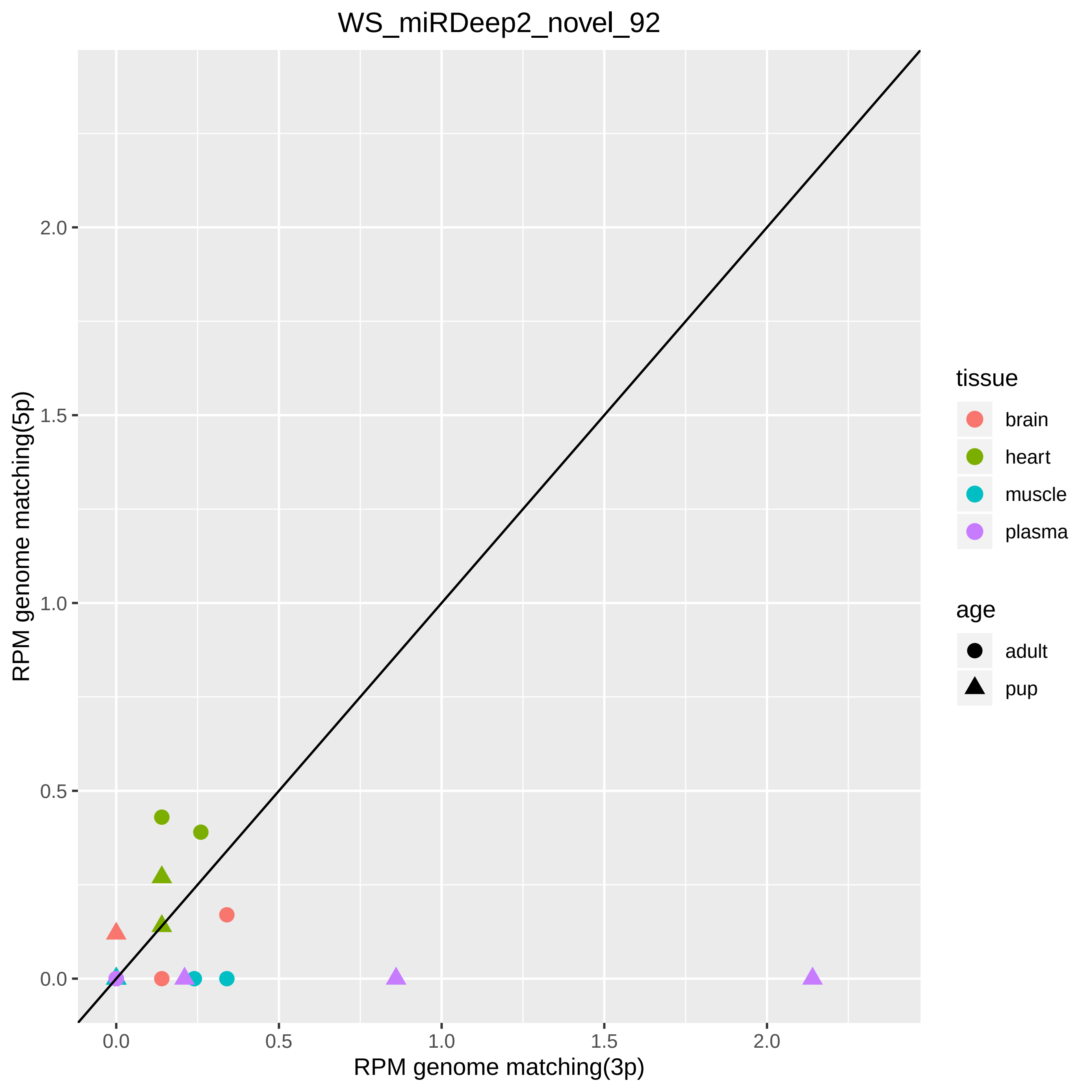

WS_miRDeep2_novel_92
RPM genome matching(5p)
RPM genome matching(3p)
0.0
0.5
1.0
1.5
2.0
tissue
brain
heart
muscle
plasma
age
adult
pup

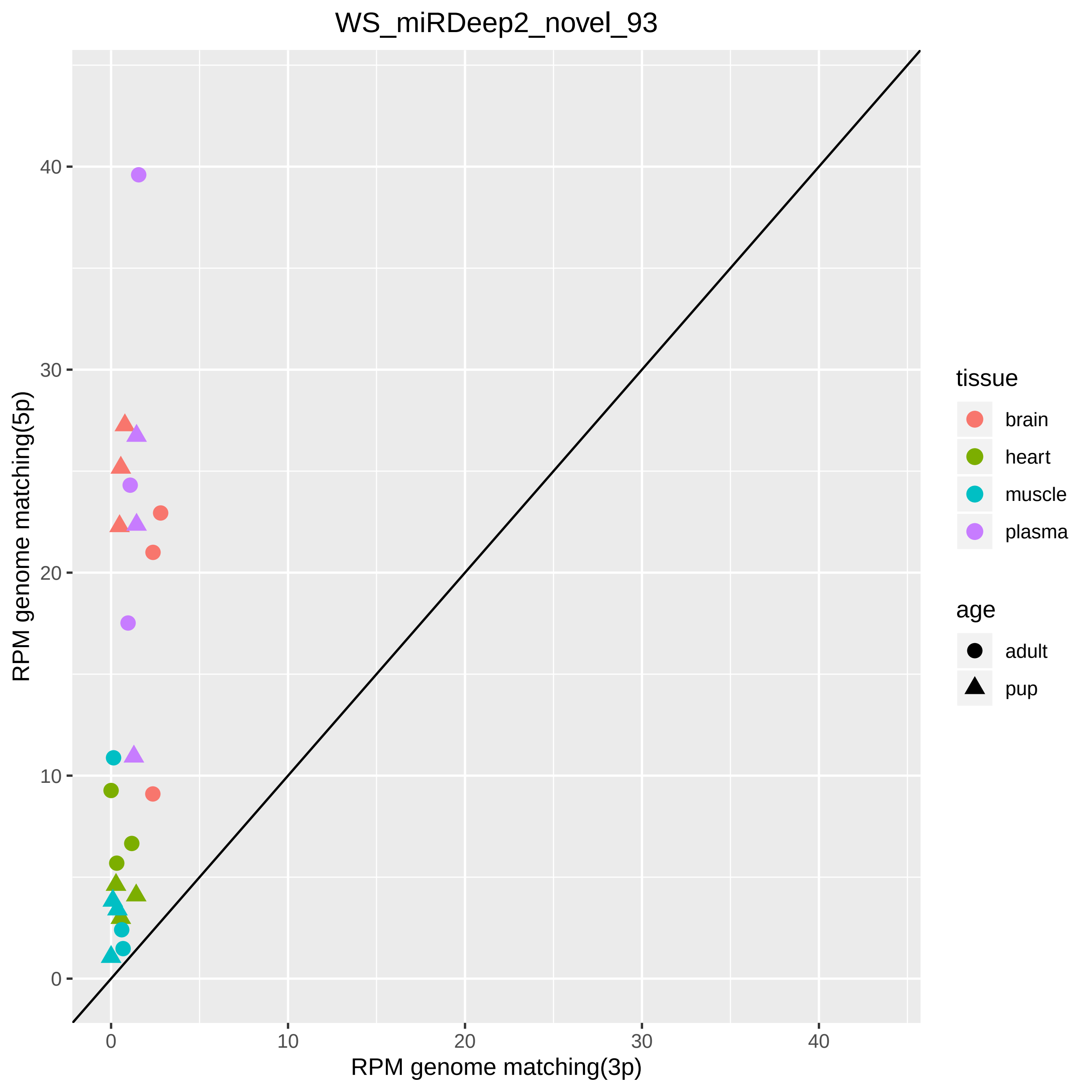

WS_miRDeep2_novel_93
RPM genome matching(5p)
RPM genome matching(3p)
0
10
20
30
40
0
10
20
30
40
tissue
brain
heart
muscle
plasma
age
adult
pup

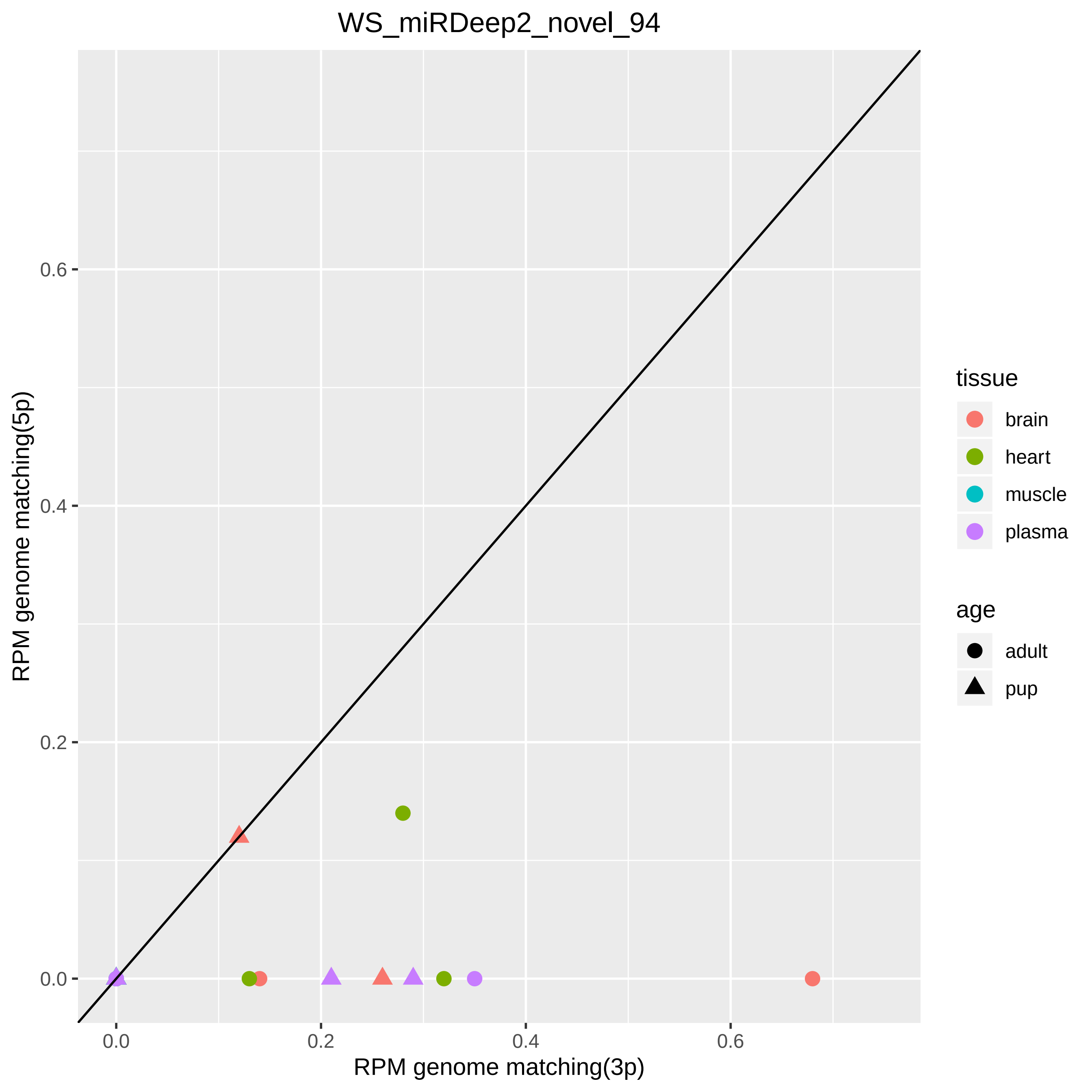

WS_miRDeep2_novel_94
RPM genome matching(5p)
RPM genome matching(3p)
0.0
0.2
0.4
0.6
tissue
brain
heart
muscle
plasma
age
adult
pup

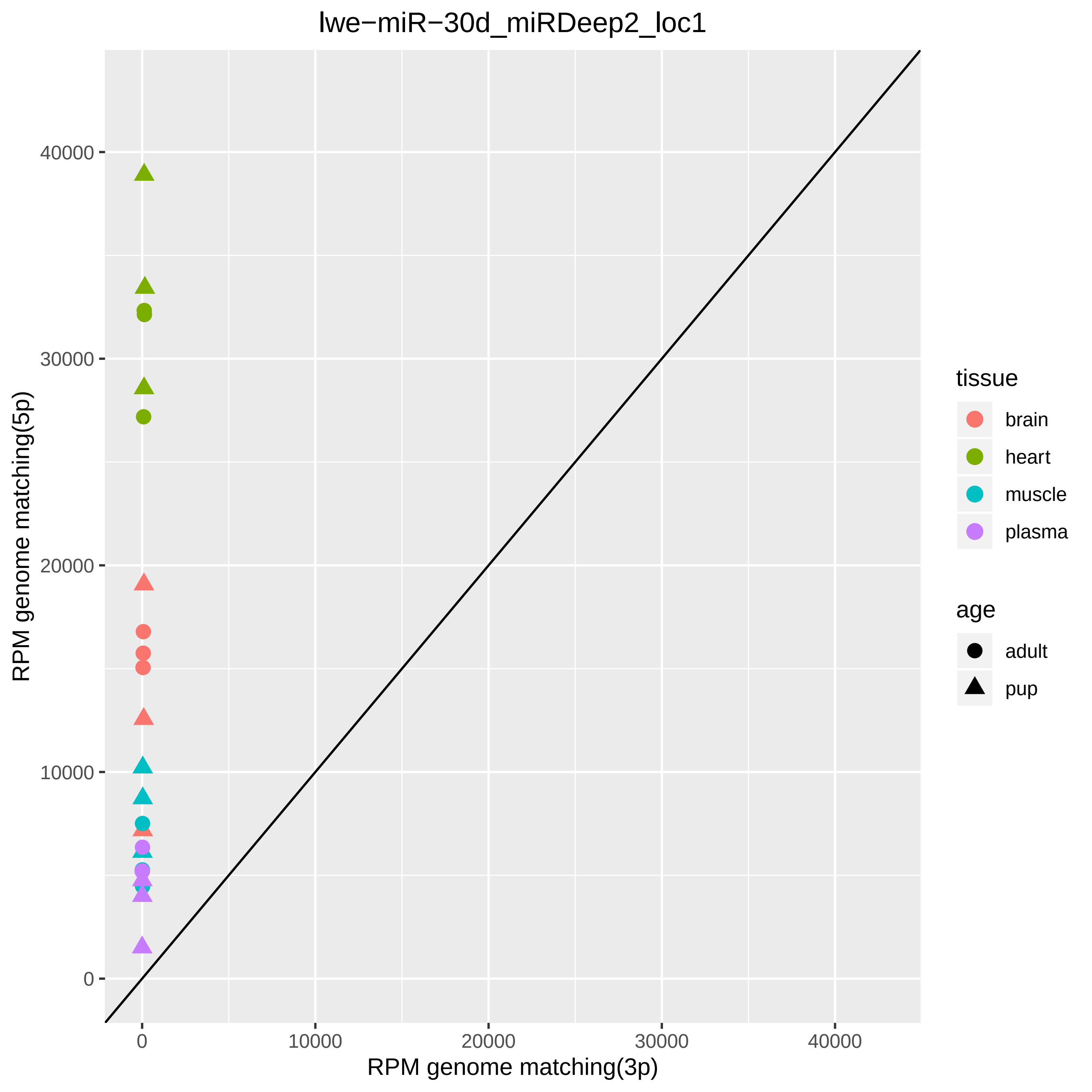
lwe−miR−30d_miRDeep2_loc1
RPM genome matching(5p)
RPM genome matching(3p)
0
10000
20000
30000
40000
tissue
brain
heart
muscle
plasma
age
adult
pup

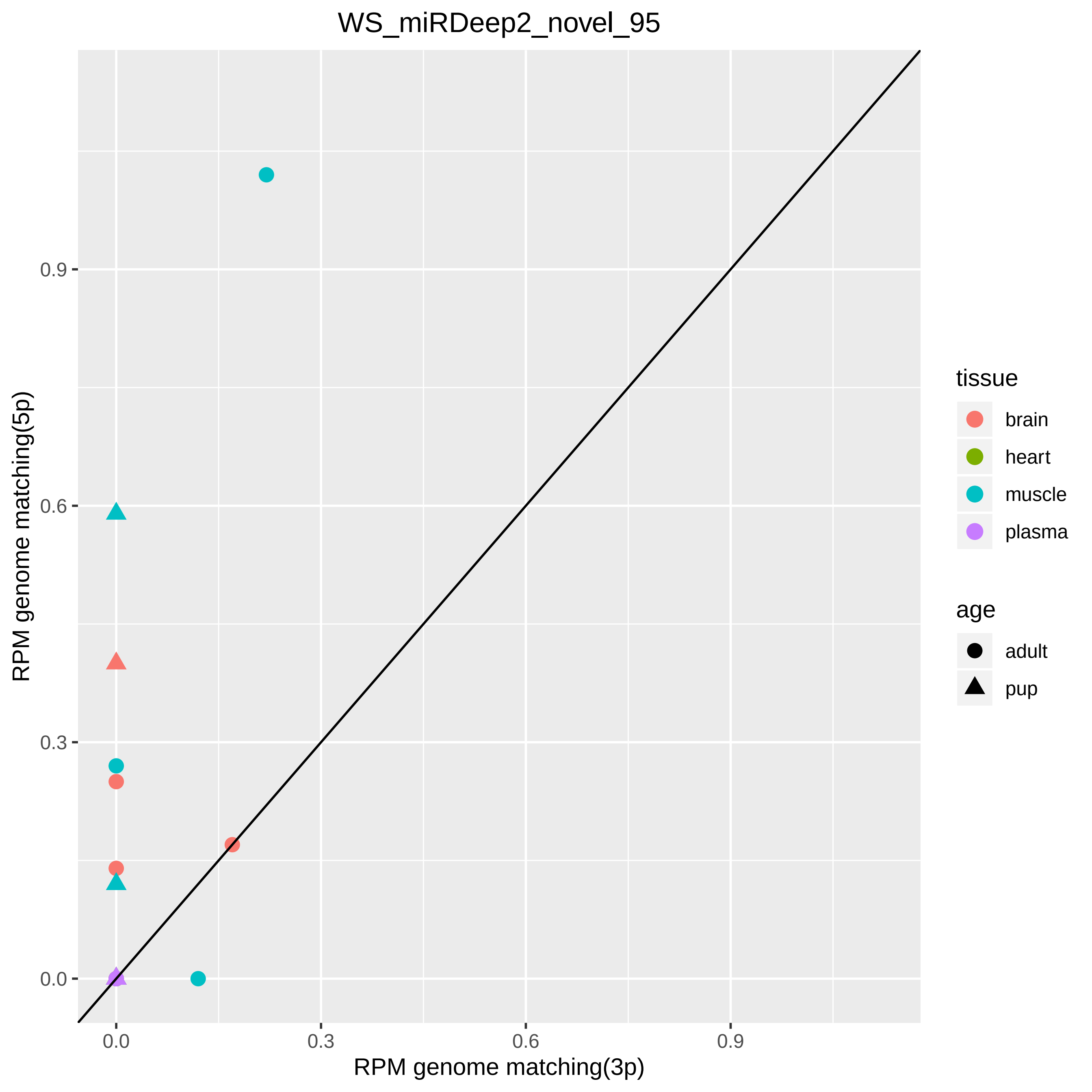

WS_miRDeep2_novel_95
RPM genome matching(5p)
RPM genome matching(3p)
0.0
0.3
0.6
0.9
0.0
0.3
0.6
0.9
tissue
brain
heart
muscle
plasma
age
adult
pup

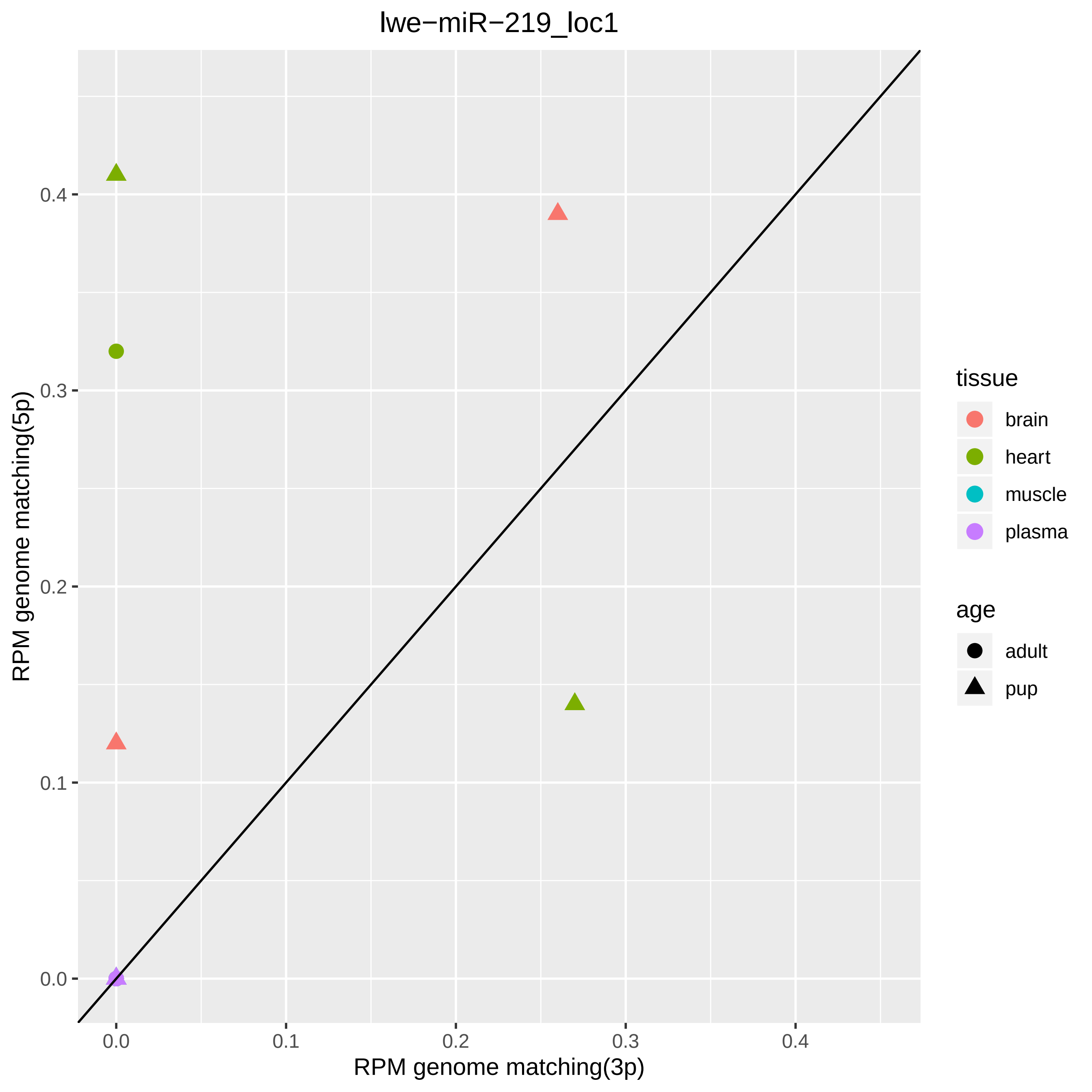
lwe−miR−219_loc1
RPM genome matching(5p)
RPM genome matching(3p)
0.0
0.1
0.2
0.3
0.4
tissue
brain
heart
muscle
plasma
age
adult
pup

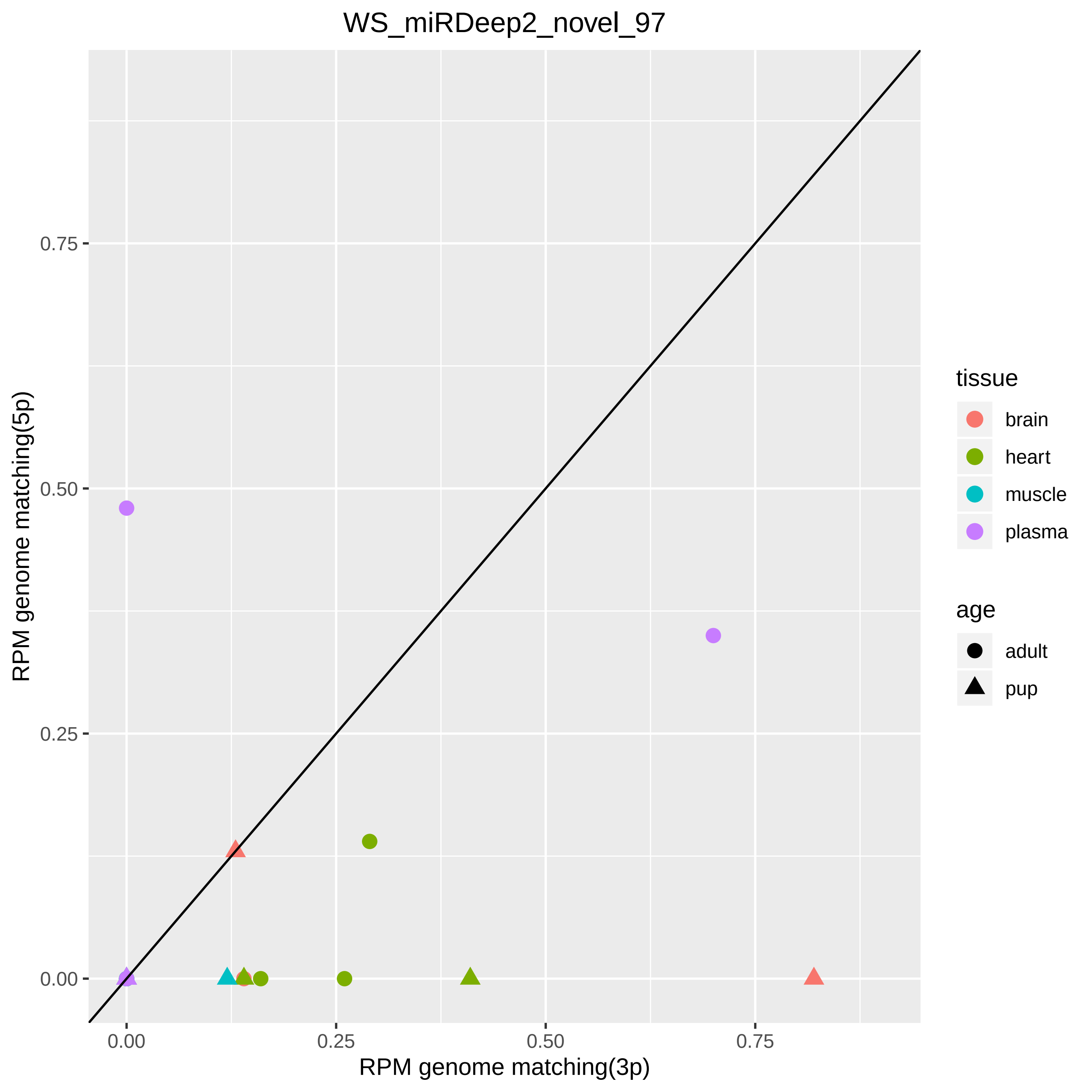
WS_miRDeep2_novel_97
RPM genome matching(5p)
RPM genome matching(3p)
0.00
0.25
0.50
0.75
tissue
brain
heart
muscle
plasma
age
adult
pup

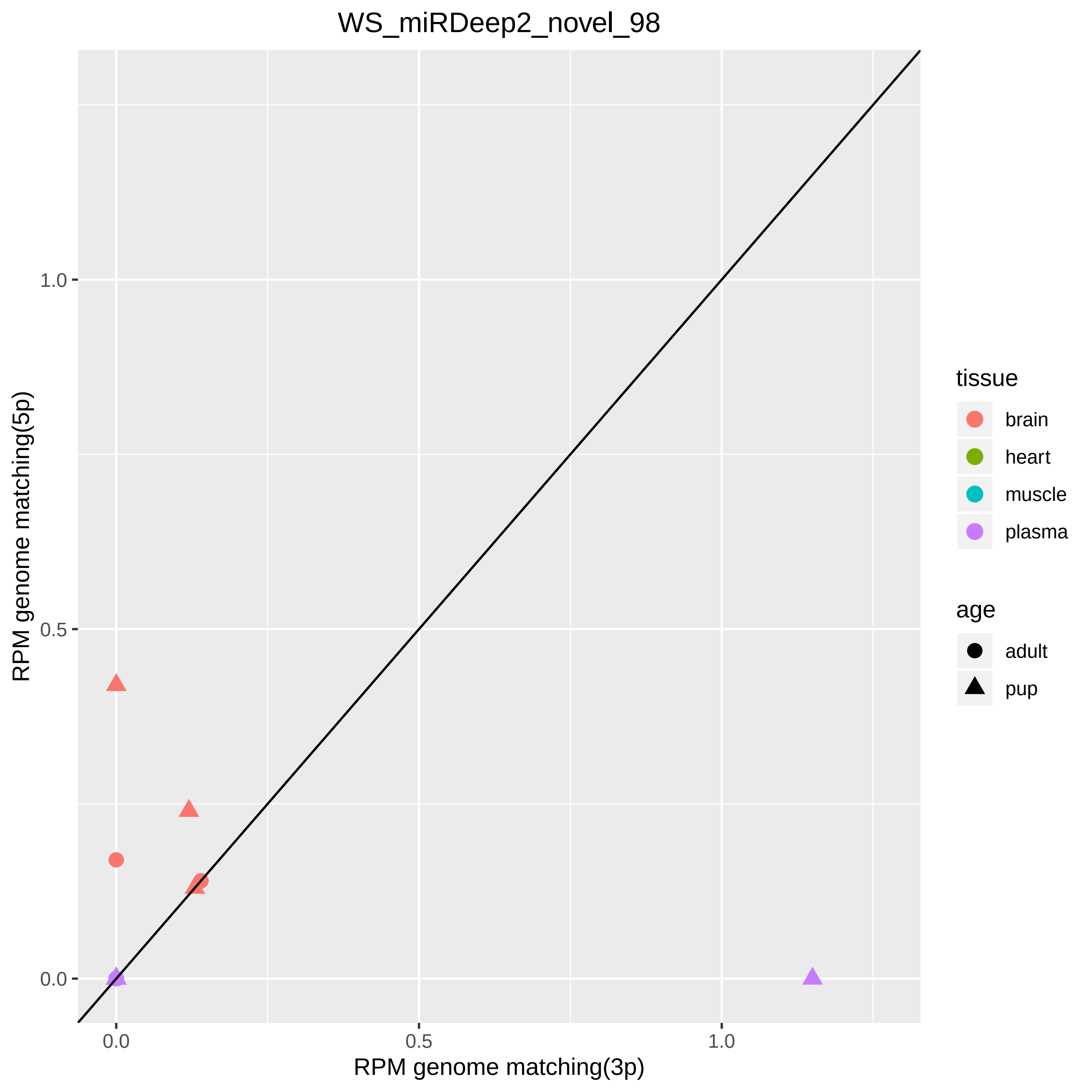

WS_miRDeep2_novel_98
RPM genome matching(5p)
1.0
0.5
0.0
0.0
0.5
1.0
RPM genome matching(3p)
tissue
brain
heart
muscle
plasma
age
adult
pup

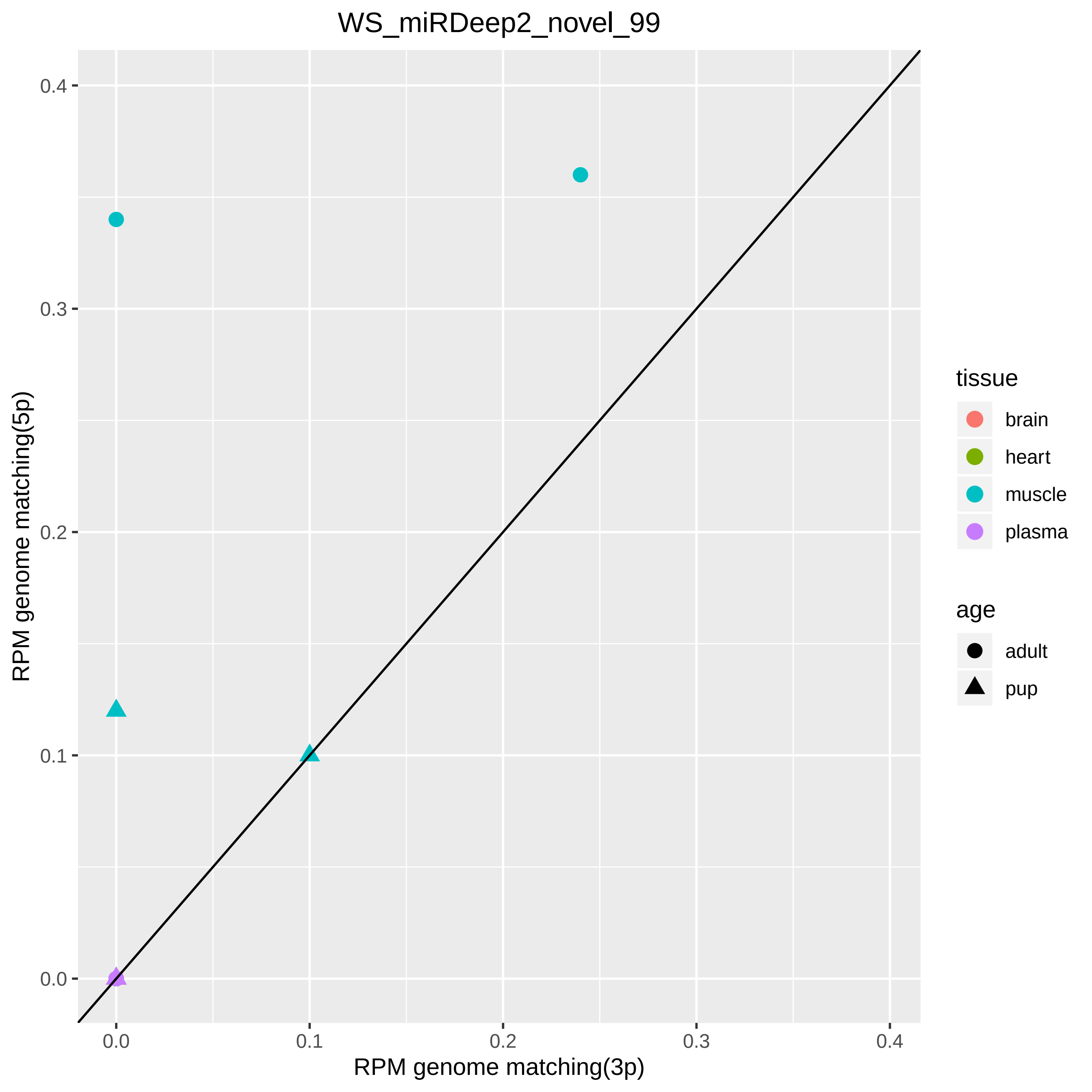

WS_miRDeep2_novel_99
RPM genome matching(5p)
RPM genome matching(3p)
0.0
0.1
0.2
0.3
0.4
tissue
brain
heart
muscle
plasma
age
adult
pup

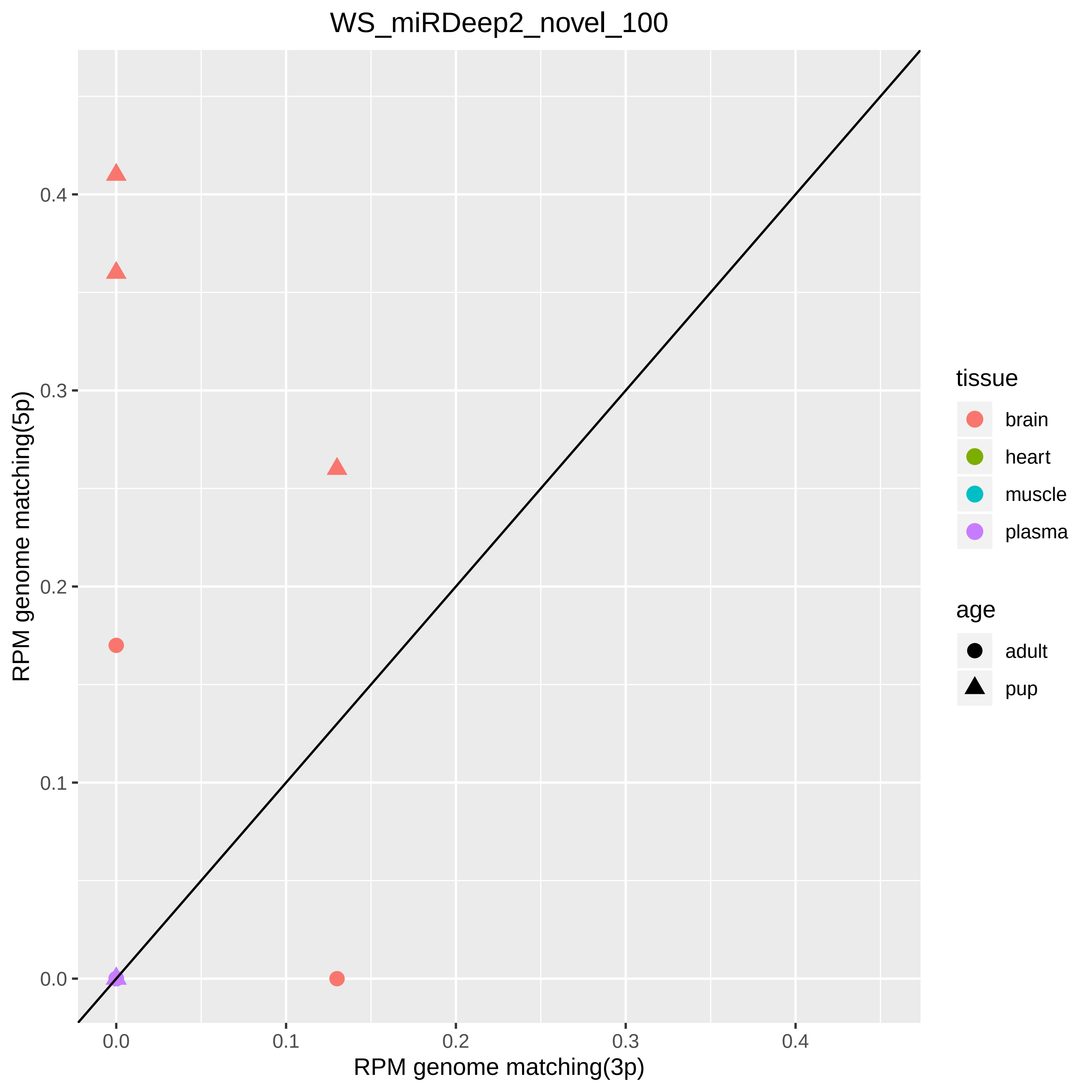
WS_miRDeep2_novel_100
RPM genome matching(5p)
RPM genome matching(3p)
0.0
0.1
0.2
0.3
0.4
tissue
brain
heart
muscle
plasma
age
adult
pup

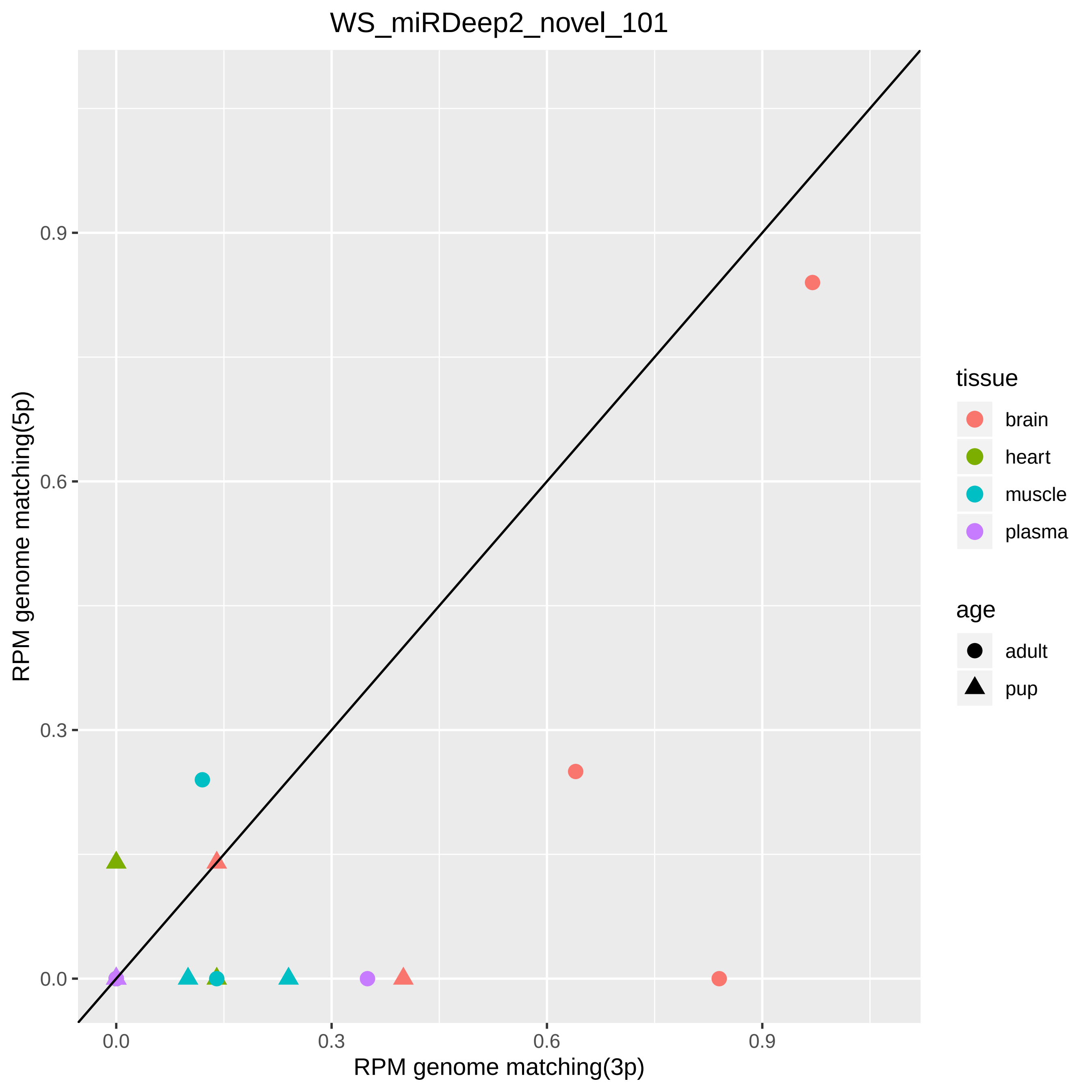

WS_miRDeep2_novel_101
RPM genome matching(5p)
RPM genome matching(3p)
0.0
0.3
0.6
0.9
tissue
brain
heart
muscle
plasma
age
adult
pup

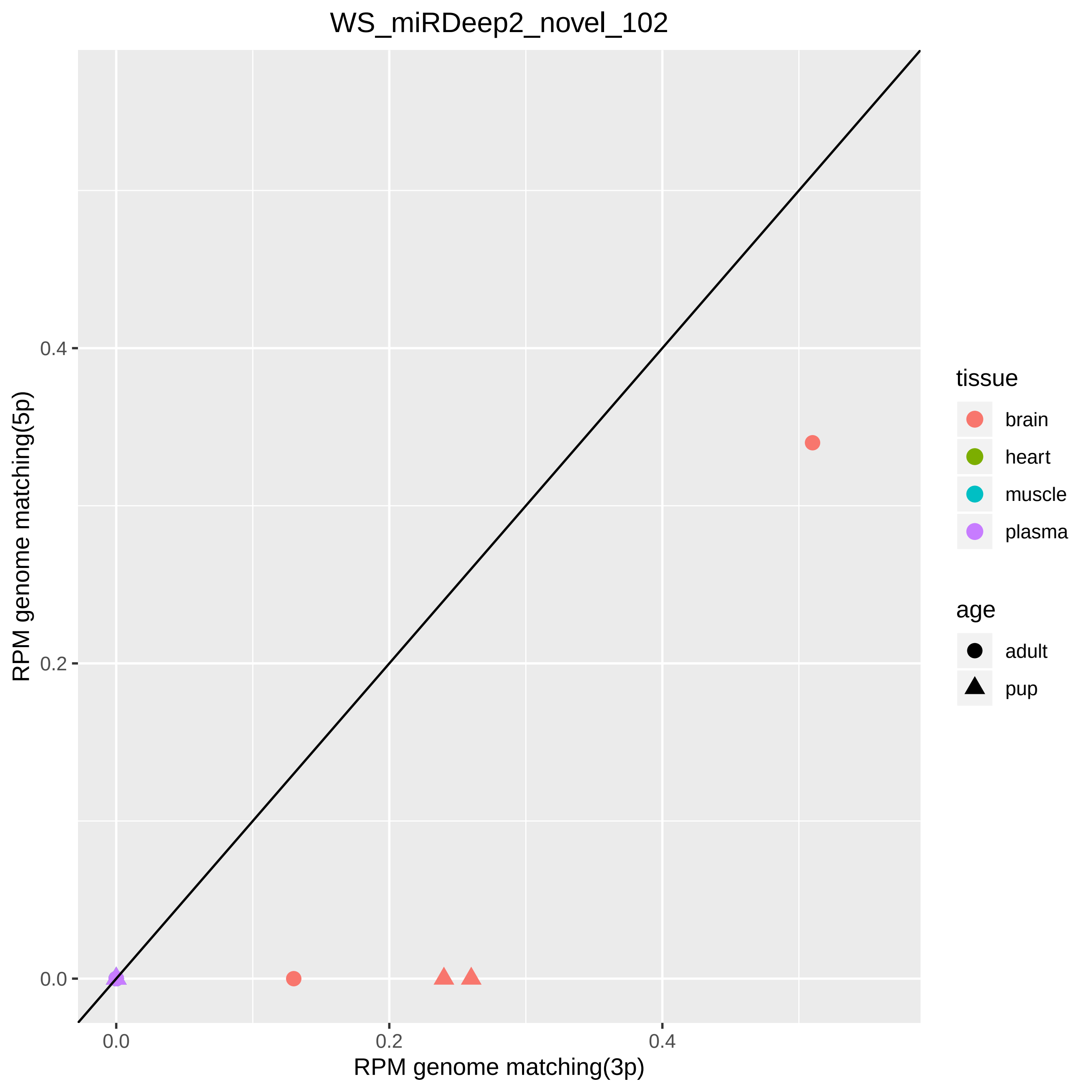

WS_miRDeep2_novel_102
RPM genome matching(5p)
RPM genome matching(3p)
0.0
0.2
0.4
0.0
0.2
0.4
tissue
brain
heart
muscle
plasma
age
adult
pup

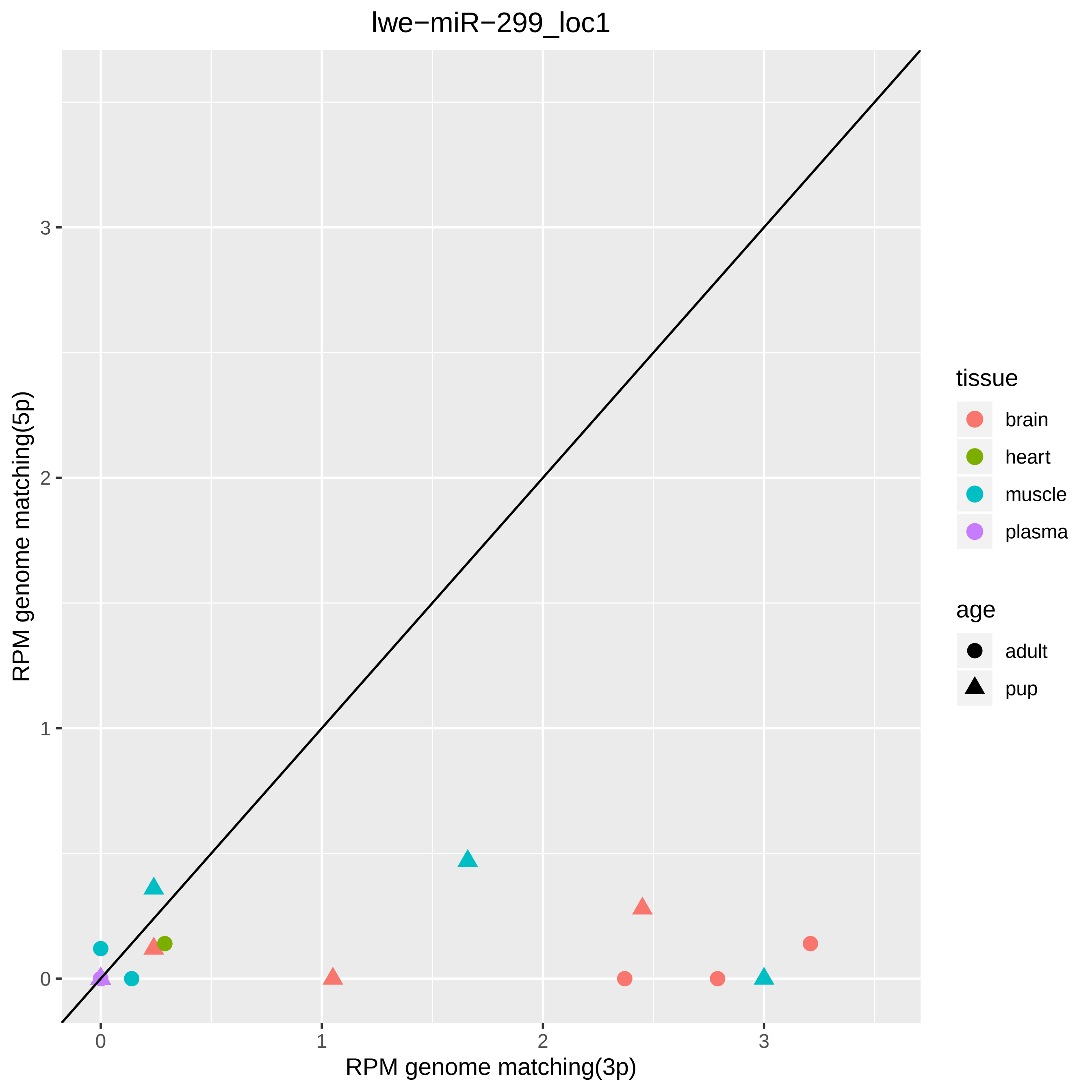
lwe−miR−299_loc1
RPM genome matching(5p)
RPM genome matching(3p)
0
1
2
3
0
1
2
3
tissue
brain
heart
muscle
plasma
age
adult
pup

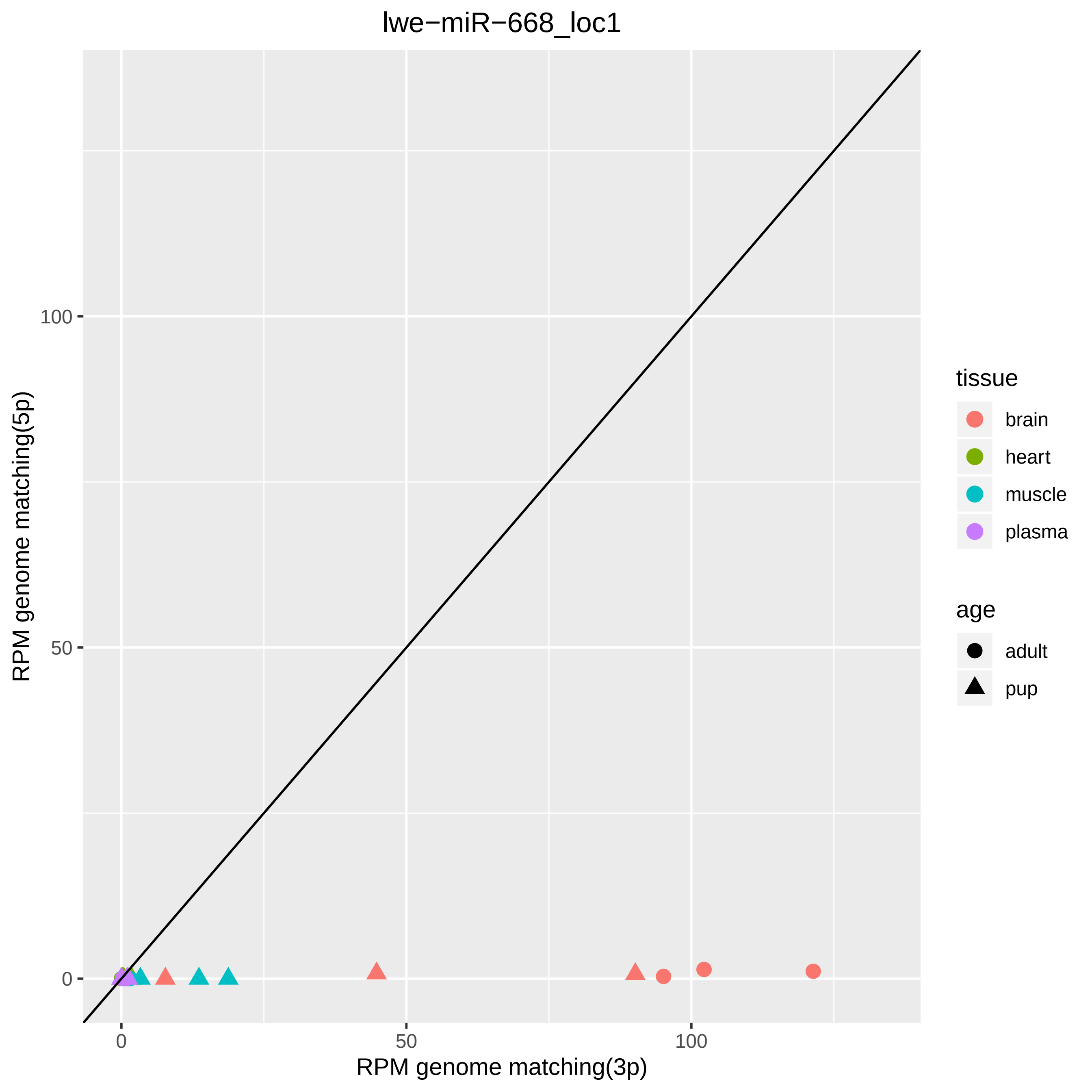
lwe−miR−668_loc1
RPM genome matching(5p)
RPM genome matching(3p)
0
50
100
0
50
100
tissue
brain
heart
muscle
plasma
age
adult
pup

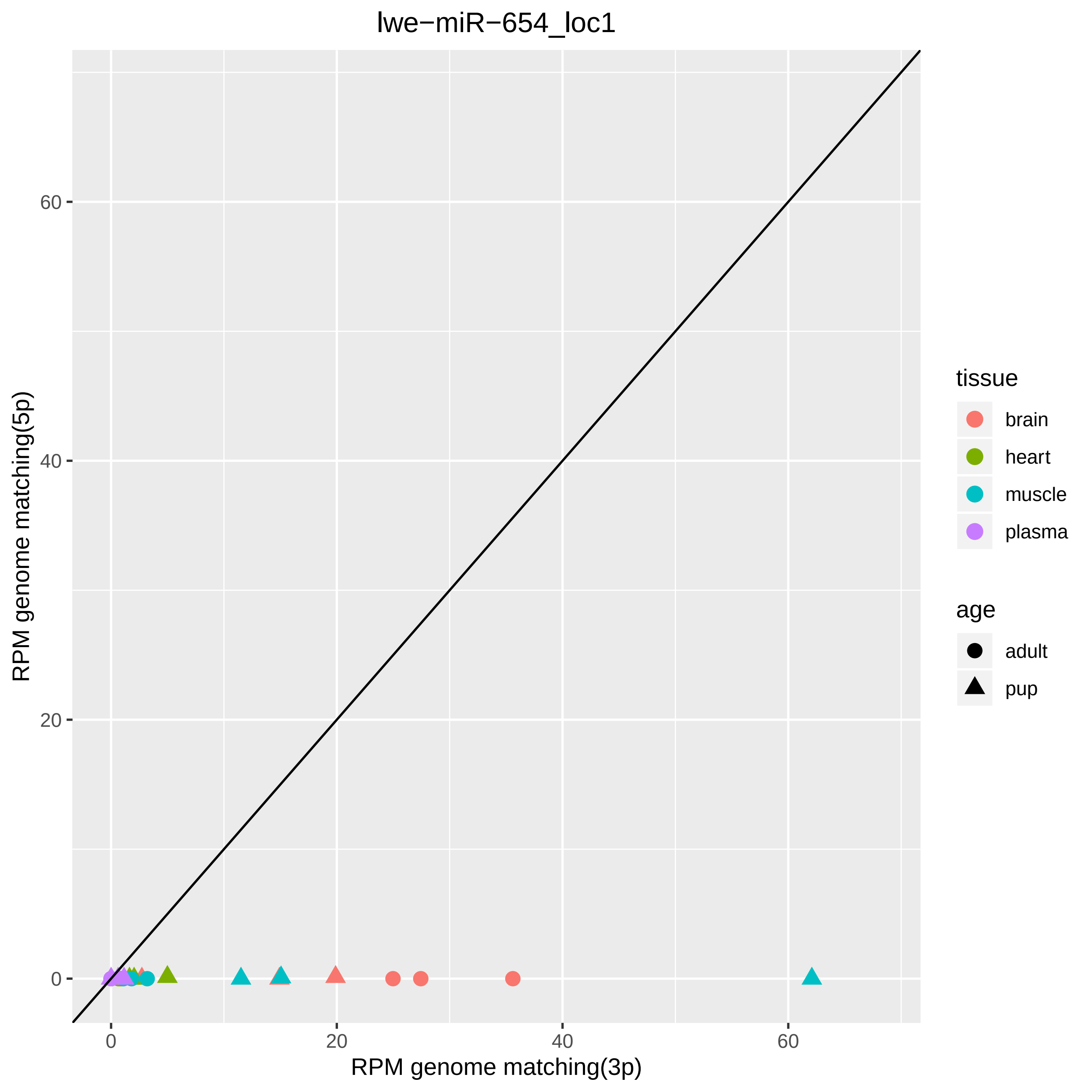

lwe−miR−654_loc1
RPM genome matching(5p)
RPM genome matching(3p)
0
20
40
60
tissue
brain
heart
muscle
plasma
age
adult
pup

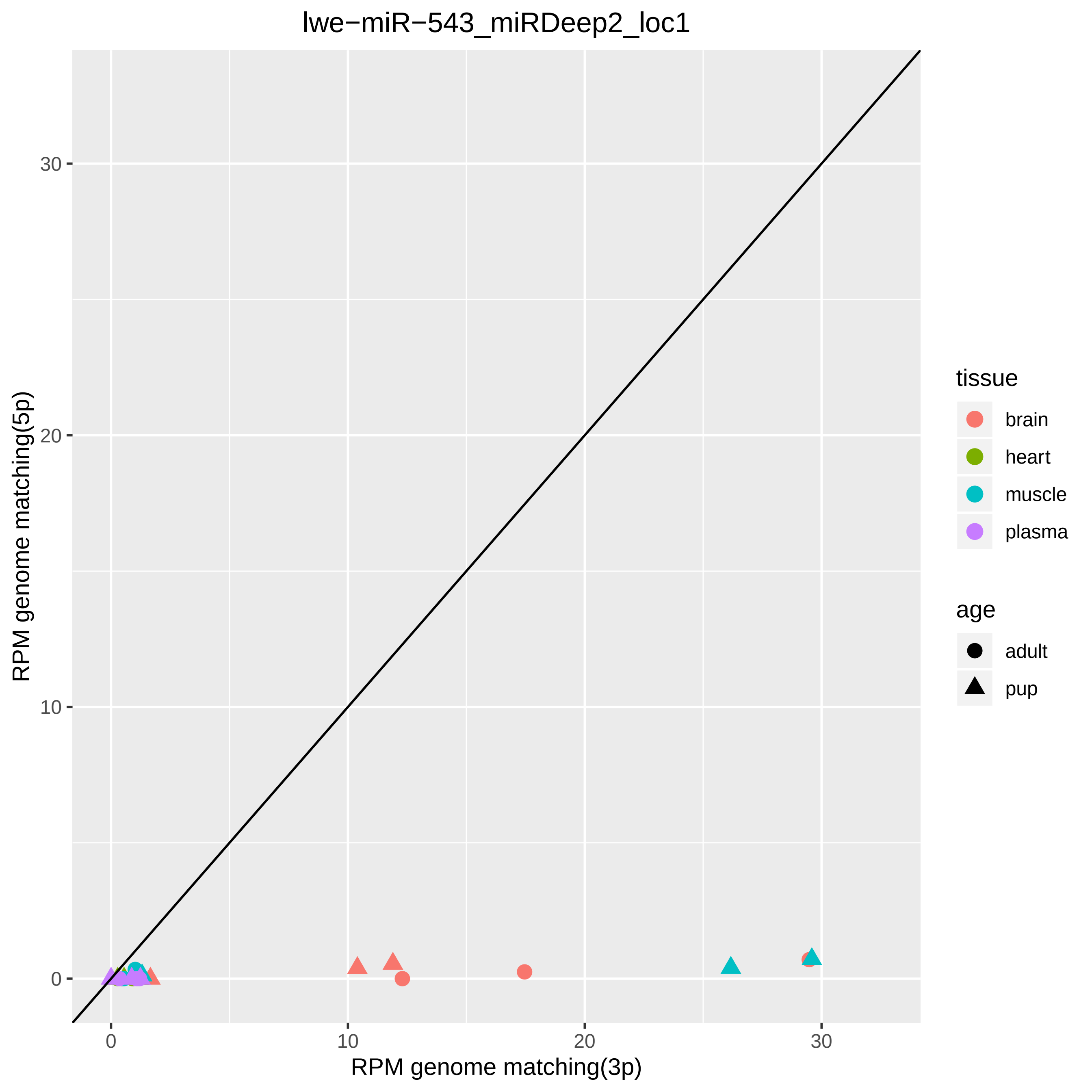
lwe−miR−543_miRDeep2_loc1
RPM genome matching(5p)
RPM genome matching(3p)
0
10
20
30
0
10
20
30
tissue
brain
heart
muscle
plasma
age
adult
pup

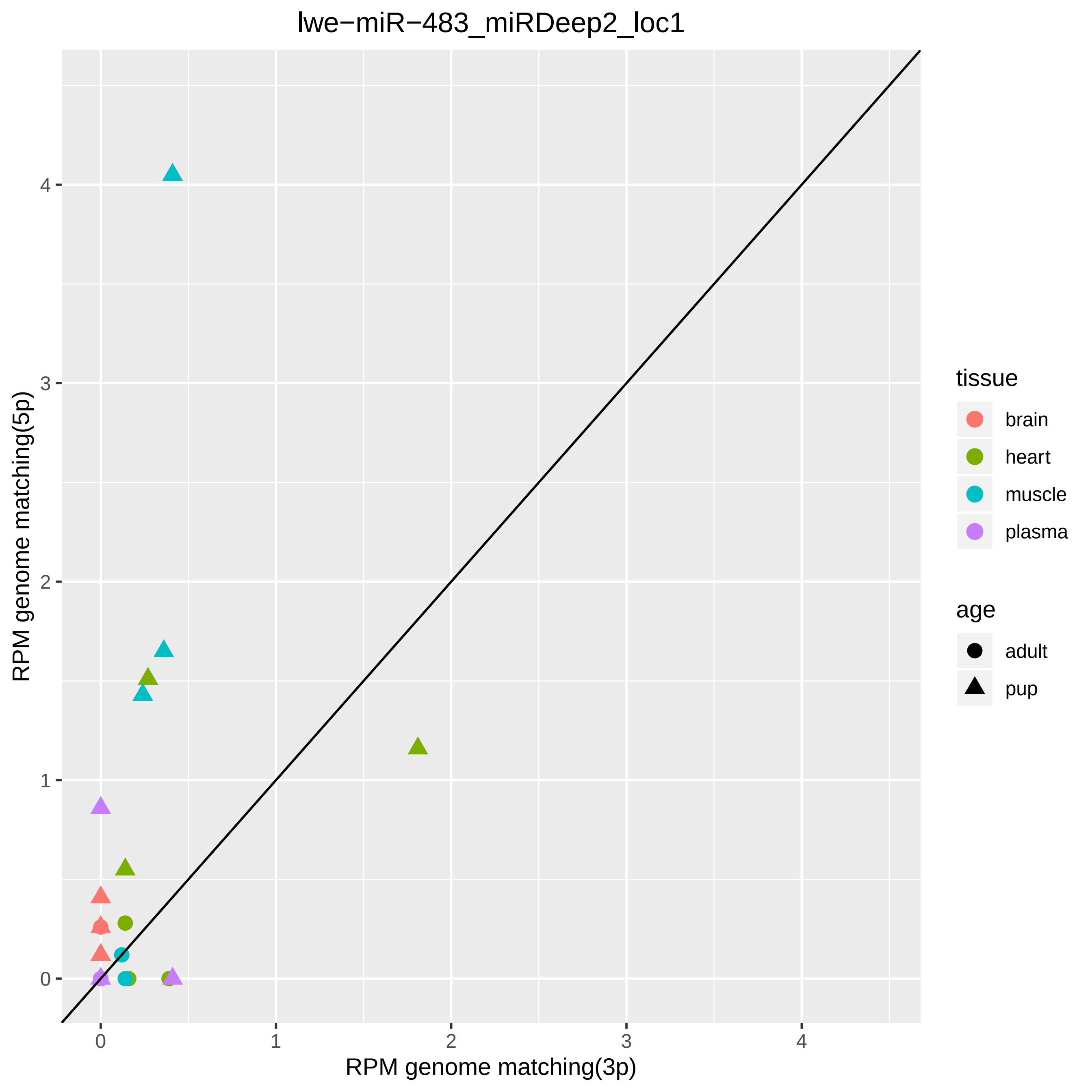

lwe−miR−483_miRDeep2_loc1
RPM genome matching(5p)
RPM genome matching(3p)
0
1
2
3
4
tissue
brain
heart
muscle
plasma
age
adult
pup

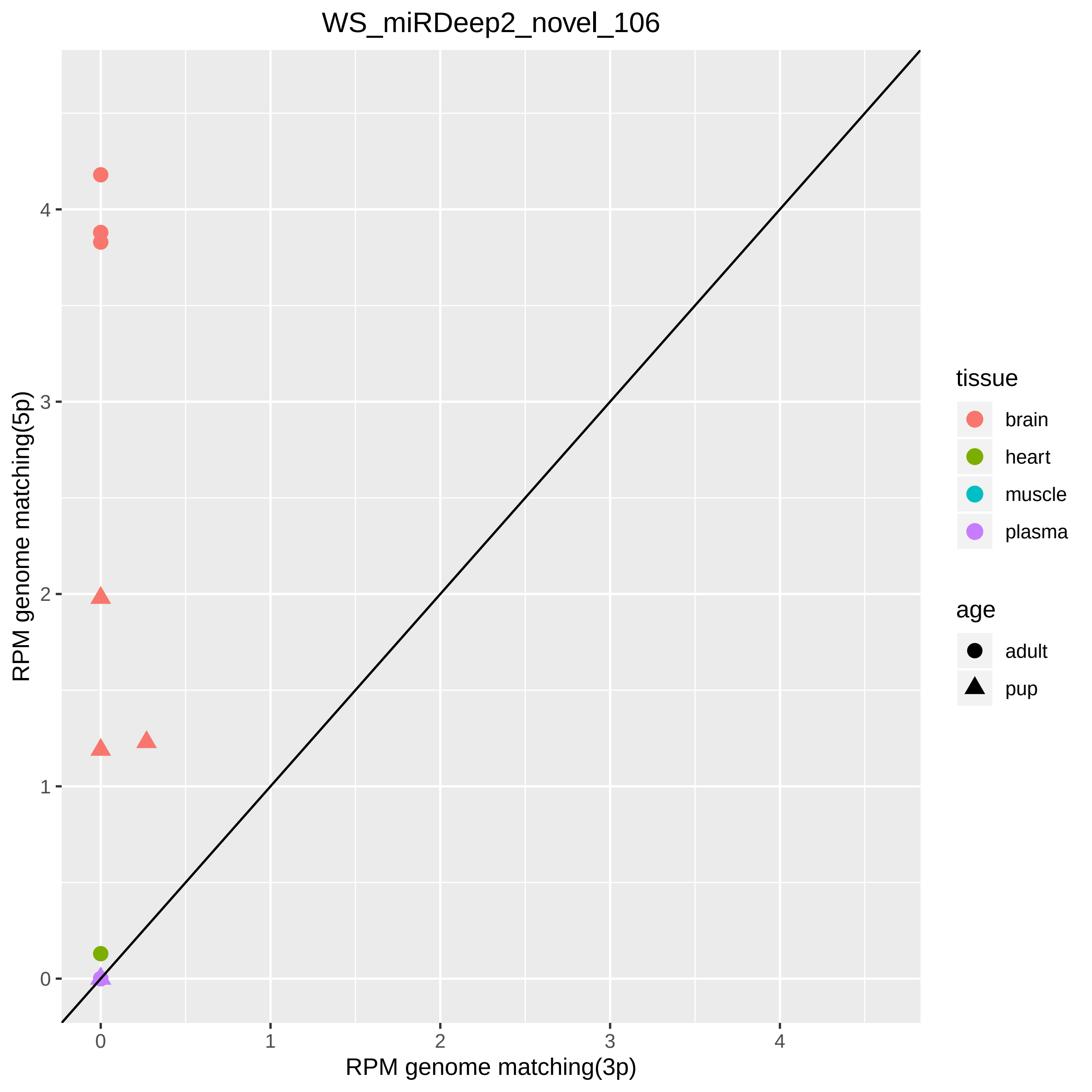

WS_miRDeep2_novel_106
RPM genome matching(5p)
RPM genome matching(3p)
0
1
2
3
4
tissue
brain
heart
muscle
plasma
age
adult
pup

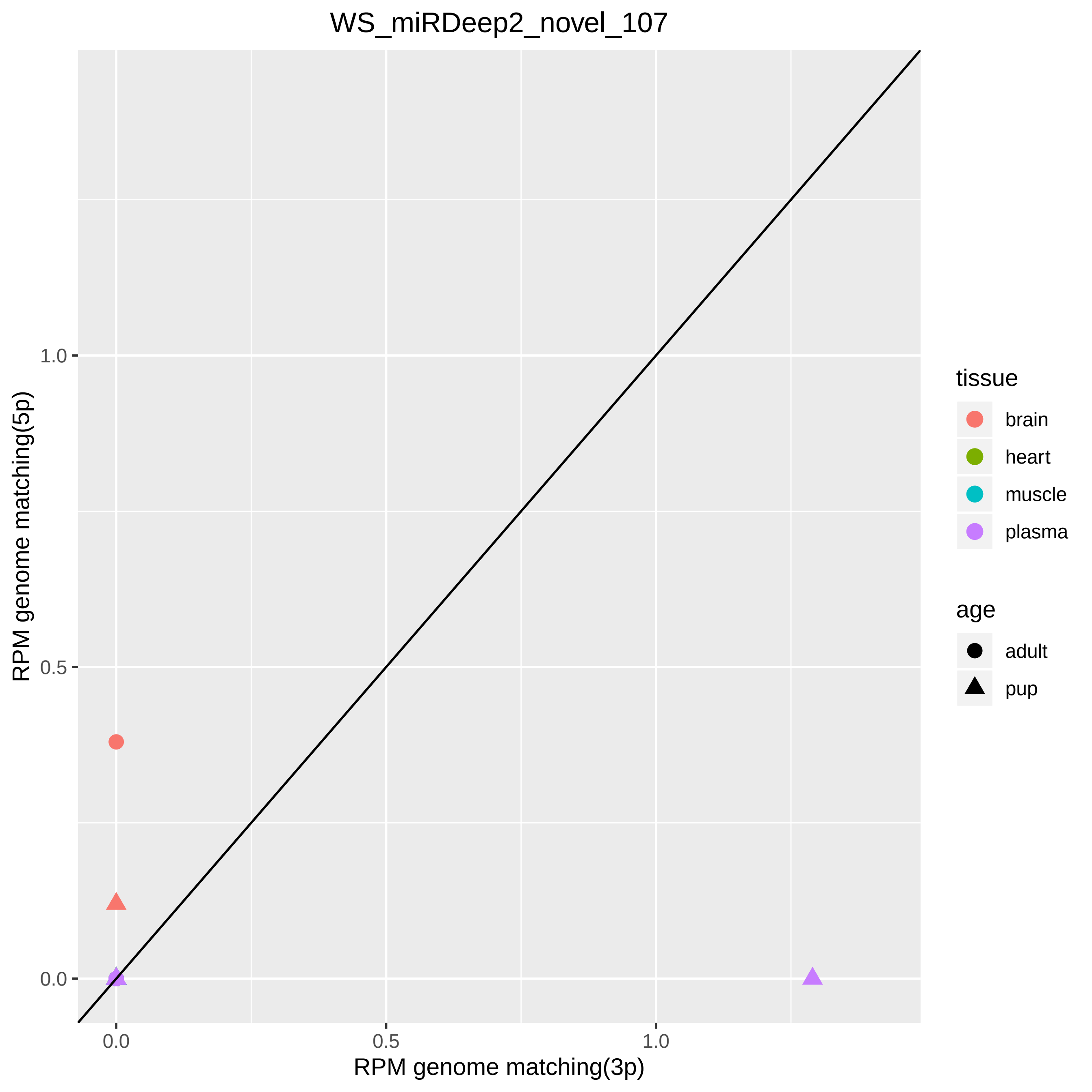
WS_miRDeep2_novel_107
RPM genome matching(5p)
RPM genome matching(3p)
0.0
0.5
1.0
tissue
brain
heart
muscle
plasma
age
adult
pup

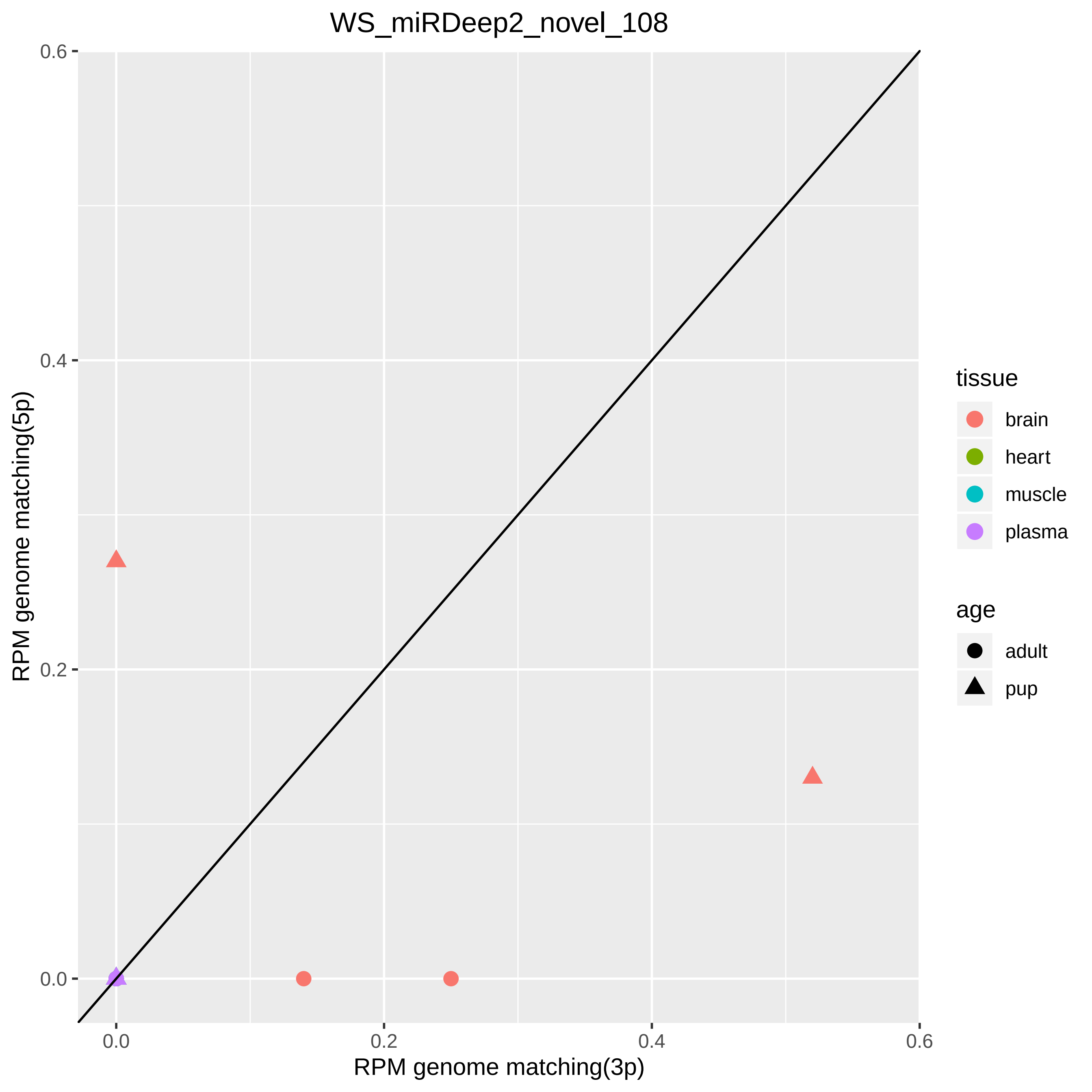

WS_miRDeep2_novel_108
RPM genome matching(5p)
RPM genome matching(3p)
0.0
0.2
0.4
0.6
tissue
brain
heart
muscle
plasma
age
adult
pup

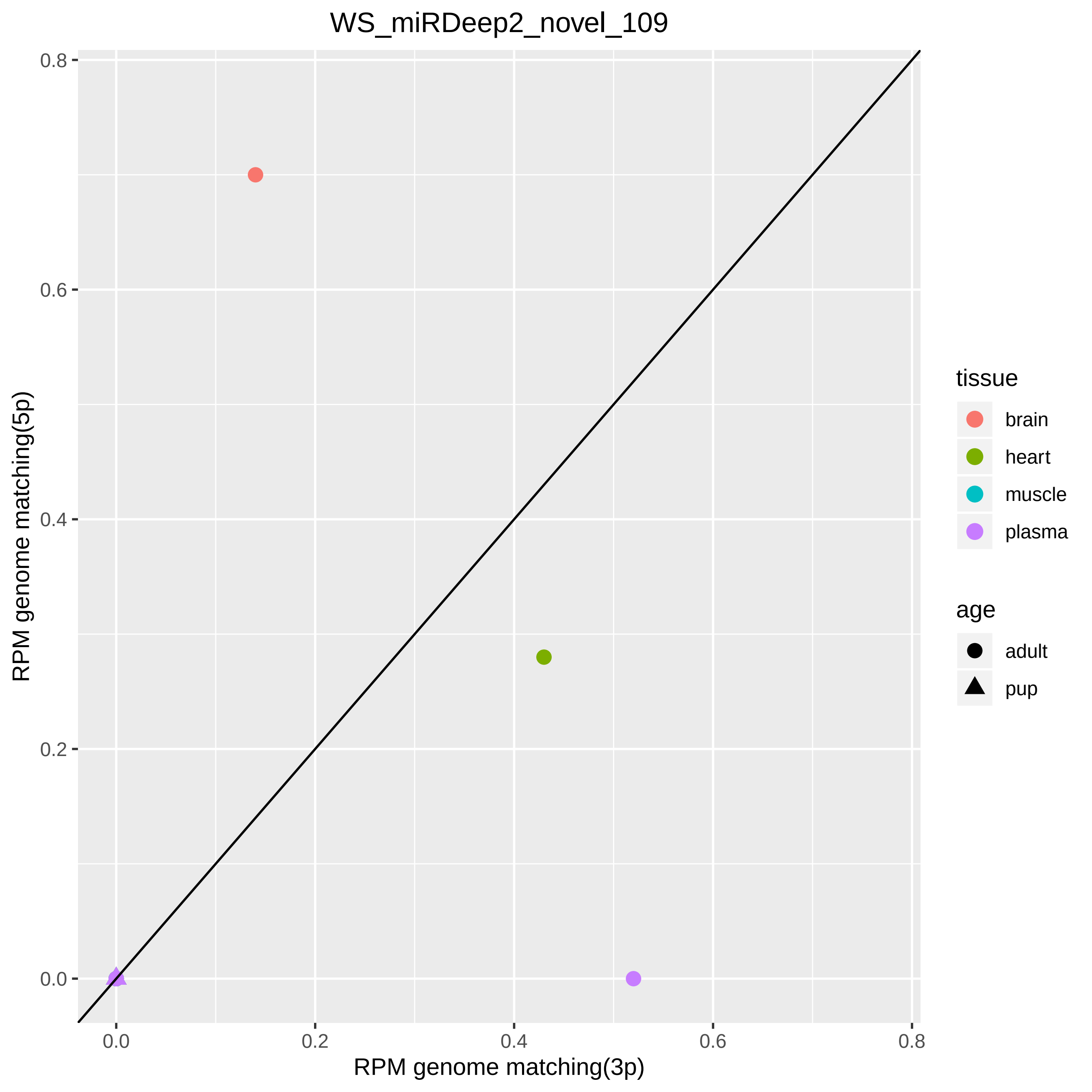
WS_miRDeep2_novel_109
RPM genome matching(5p)
RPM genome matching(3p)
0.0
0.2
0.4
0.6
0.8
tissue
brain
heart
muscle
plasma
age
adult
pup

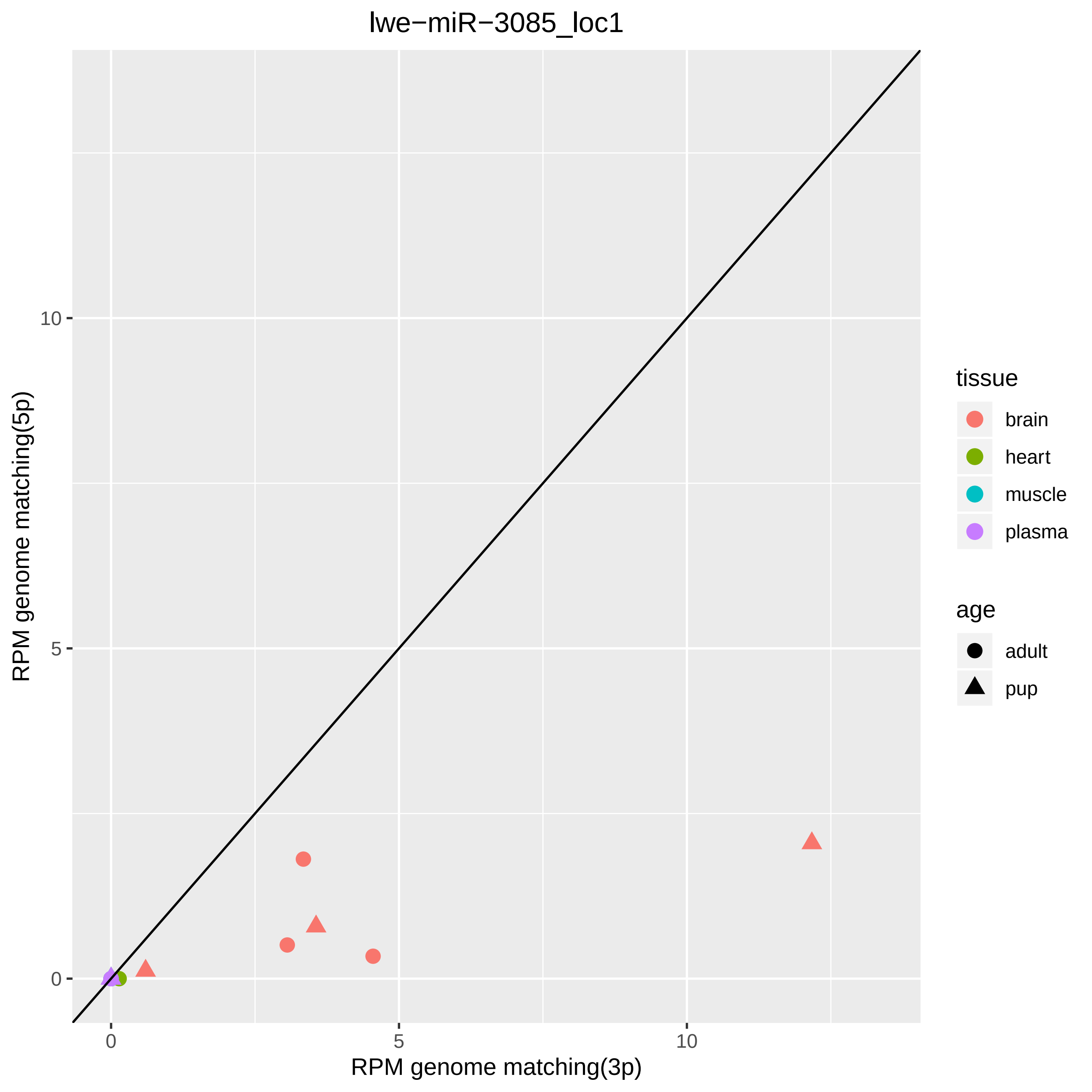

lwe−miR−3085_loc1
RPM genome matching(5p)
RPM genome matching(3p)
0
5
10
tissue
brain
heart
muscle
plasma
age
adult
pup

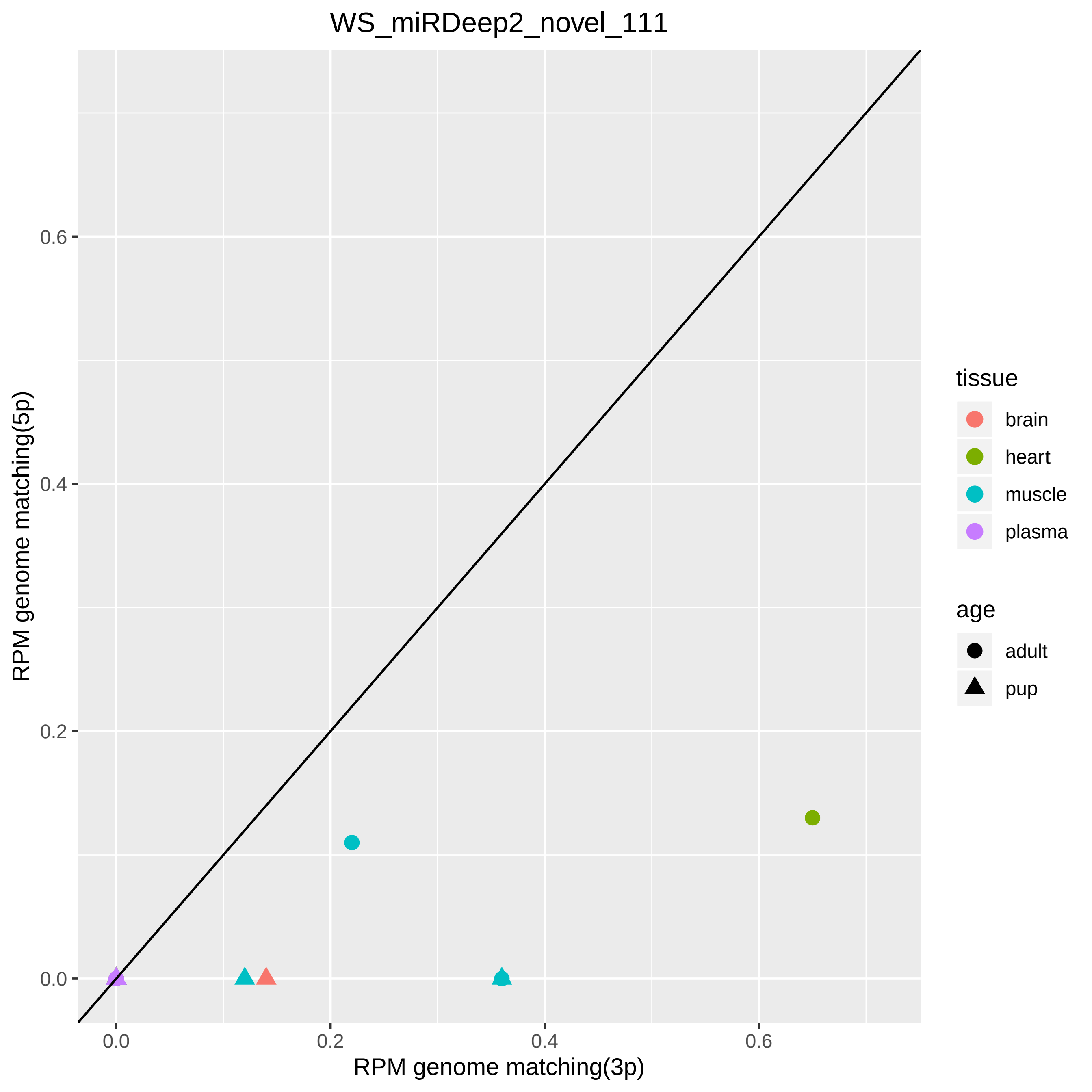

WS_miRDeep2_novel_111
RPM genome matching(5p)
RPM genome matching(3p)
0.0
0.2
0.4
0.6
tissue
brain
heart
muscle
plasma
age
adult
pup

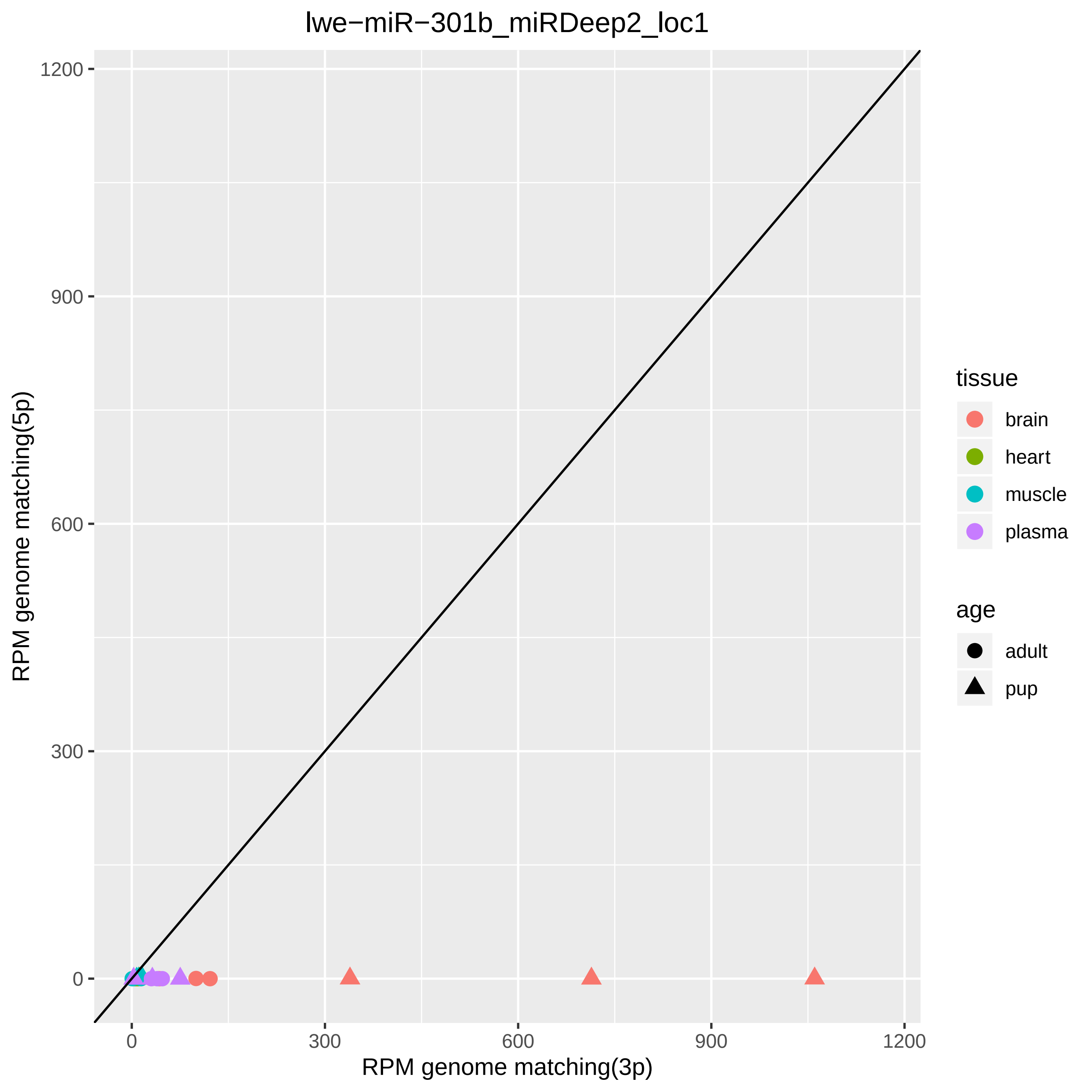
lwe−miR−301b_miRDeep2_loc1
RPM genome matching(5p)
RPM genome matching(3p)
0
300
600
900
1200
tissue
brain
heart
muscle
plasma
age
adult
pup

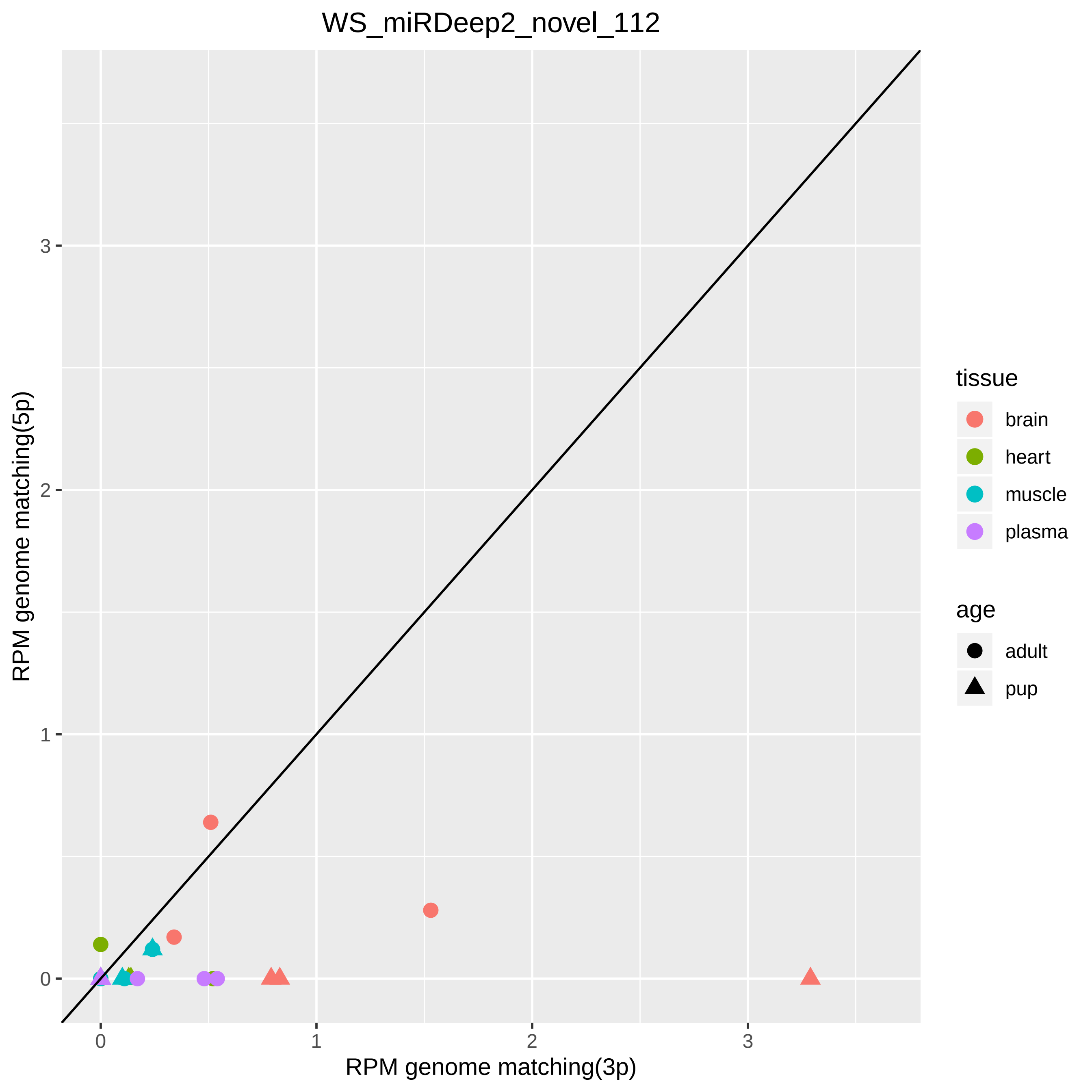

WS_miRDeep2_novel_112
RPM genome matching(5p)
RPM genome matching(3p)
0
1
2
3
0
1
2
3
tissue
brain
heart
muscle
plasma
age
adult
pup

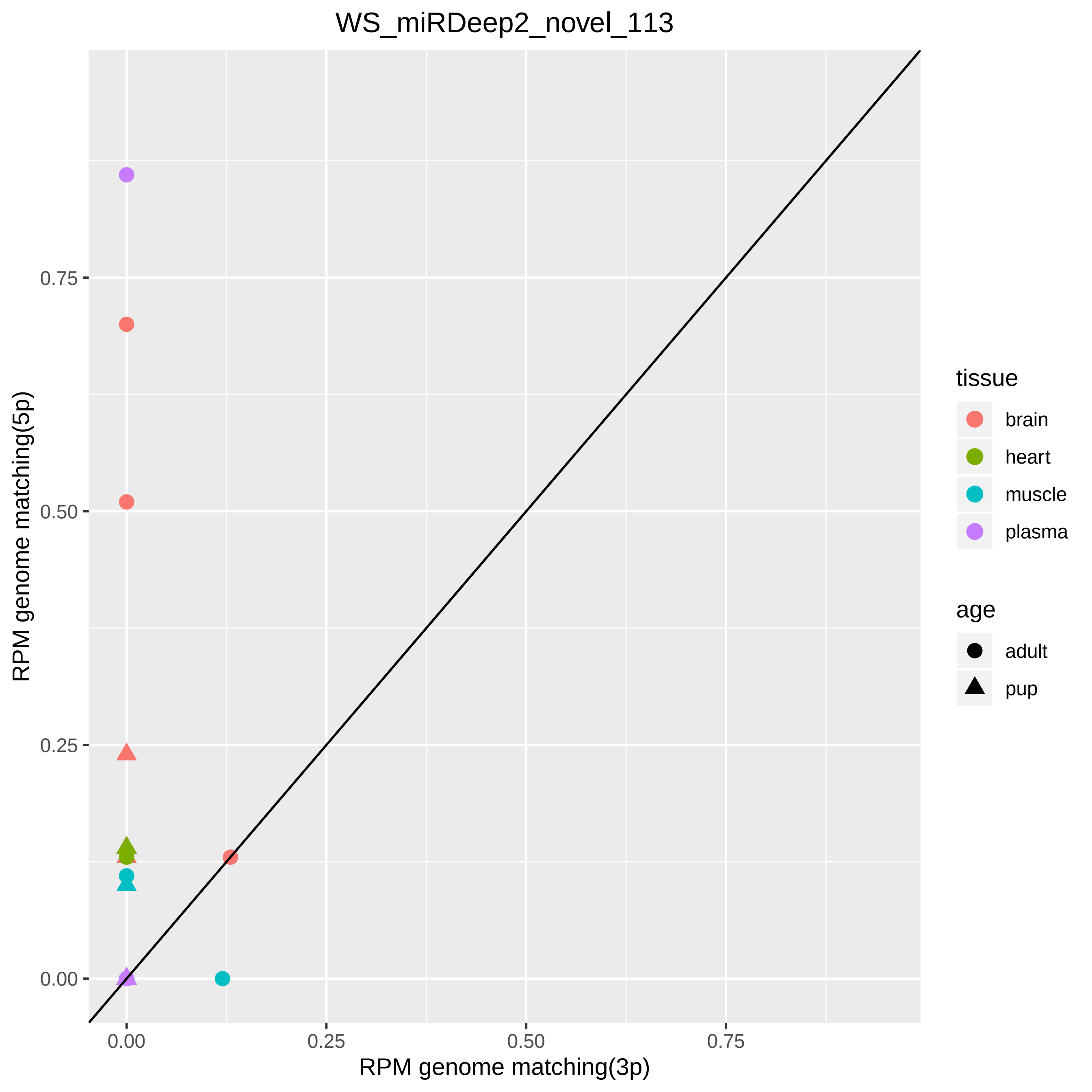

WS_miRDeep2_novel_113
RPM genome matching(5p)
RPM genome matching(3p)
0.00
0.25
0.50
0.75
0.00
0.25
0.50
0.75
tissue
brain
heart
muscle
plasma
age
adult
pup

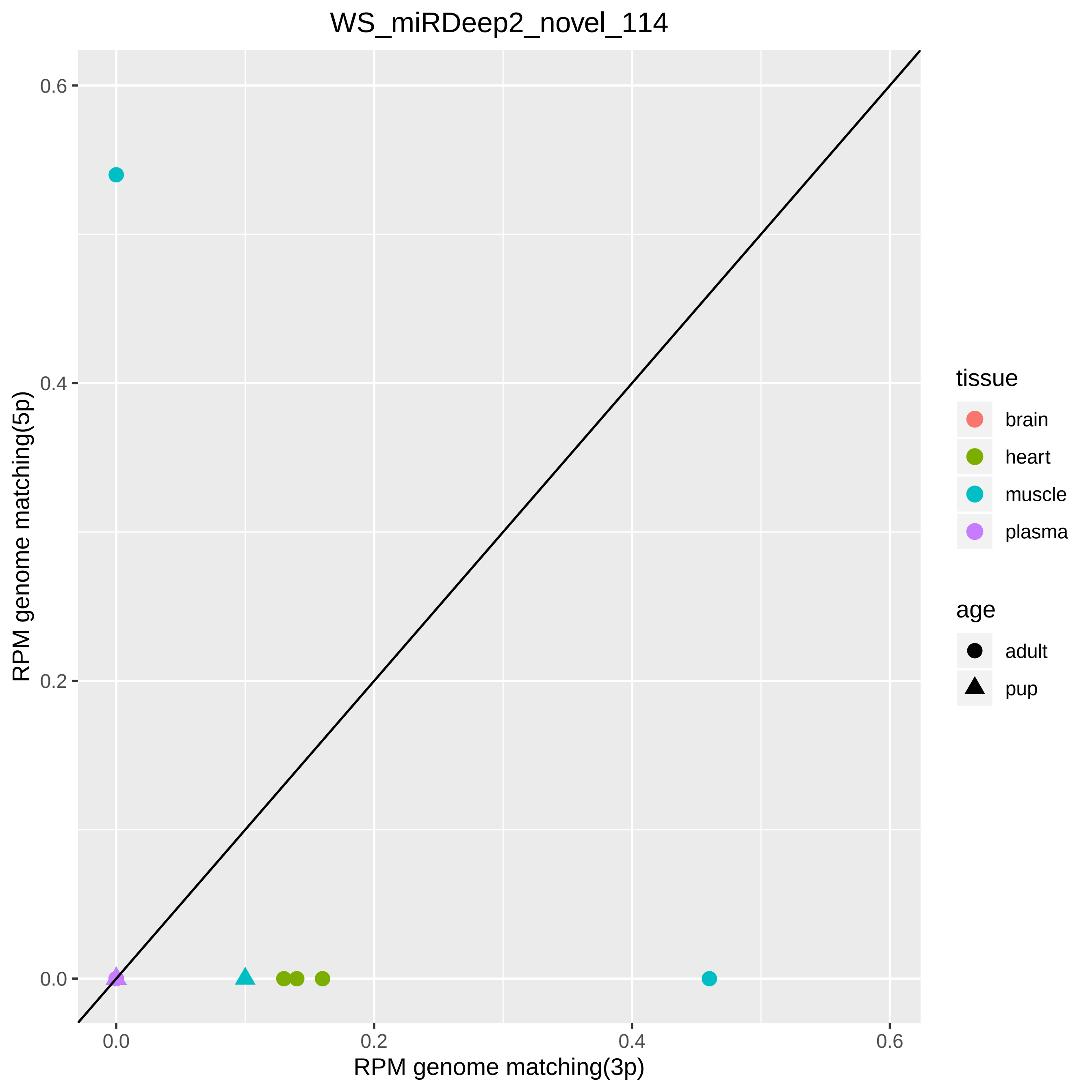
WS_miRDeep2_novel_114
RPM genome matching(5p)
RPM genome matching(3p)
0.0
0.2
0.4
0.6
0.0
0.2
0.4
0.6
tissue
brain
heart
muscle
plasma
age
adult
pup

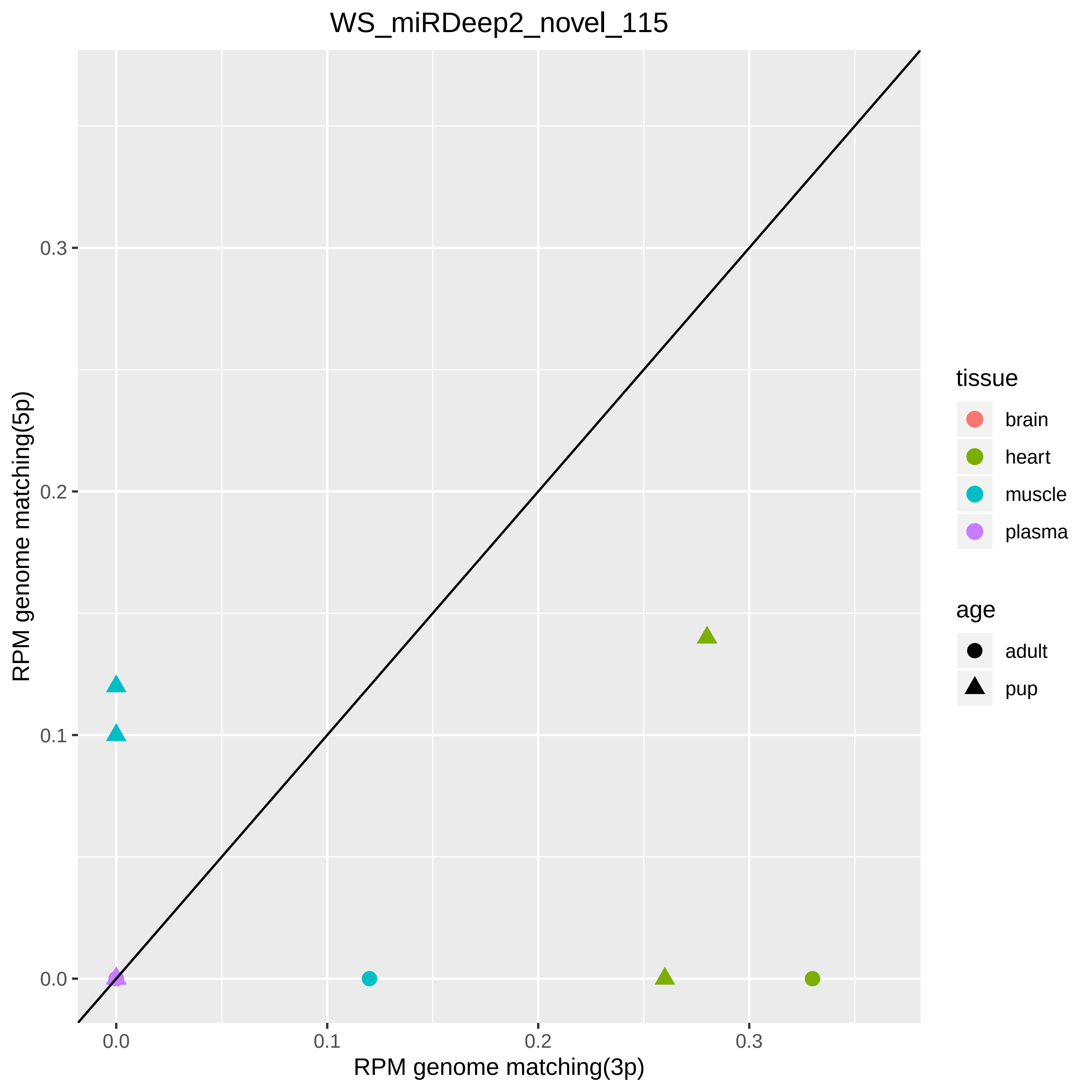

WS_miRDeep2_novel_115
RPM genome matching(5p)
RPM genome matching(3p)
0.0
0.1
0.2
0.3
tissue
brain
heart
muscle
plasma
age
adult
pup

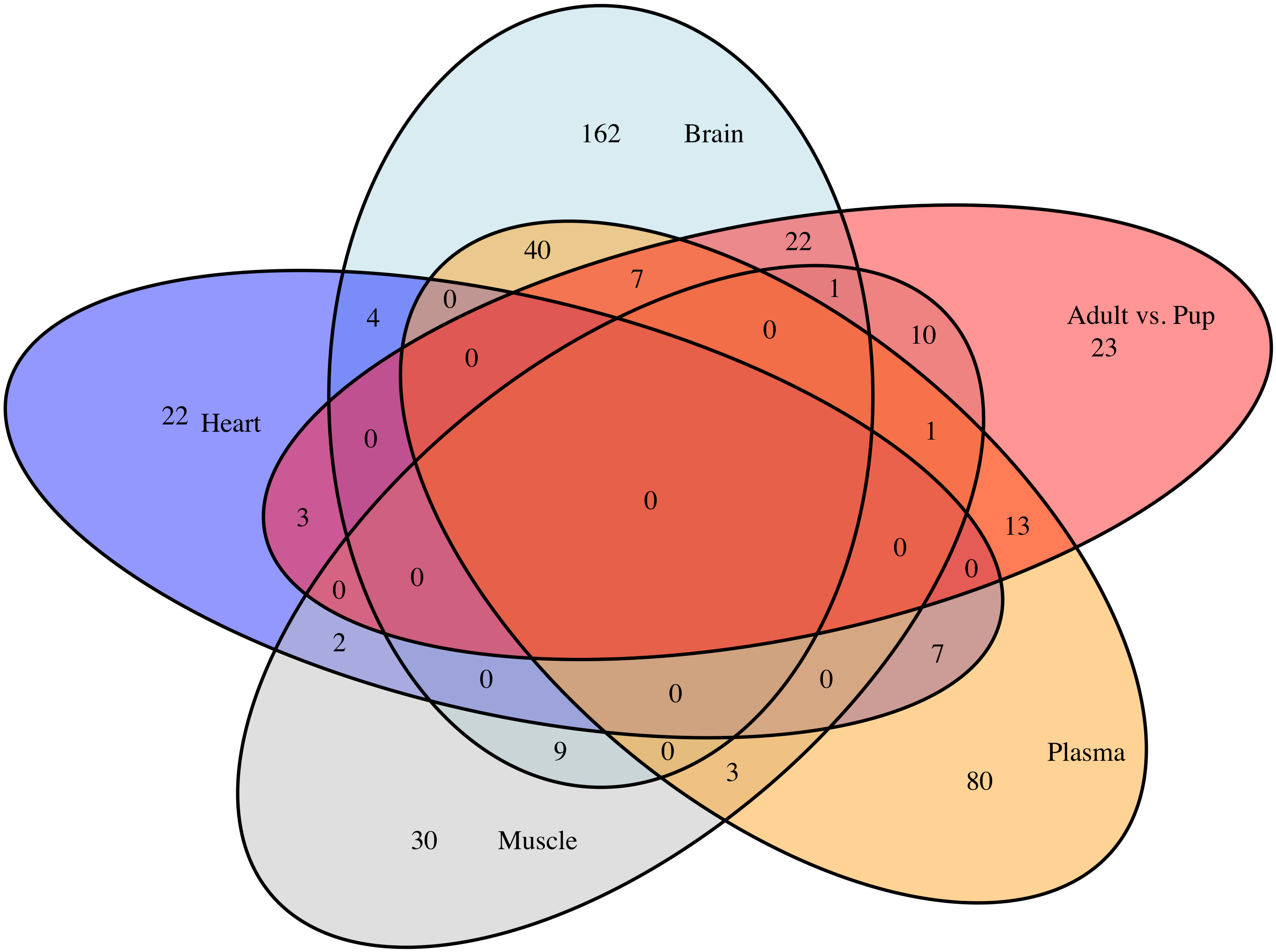


Supplementary Figure S7. Venn diagrams showing the overlap between sets of DE miRNAs for each comparison: “Brain”, “Heart”, “Muscle”, “Plasma” and “Adult vs. Pup”.

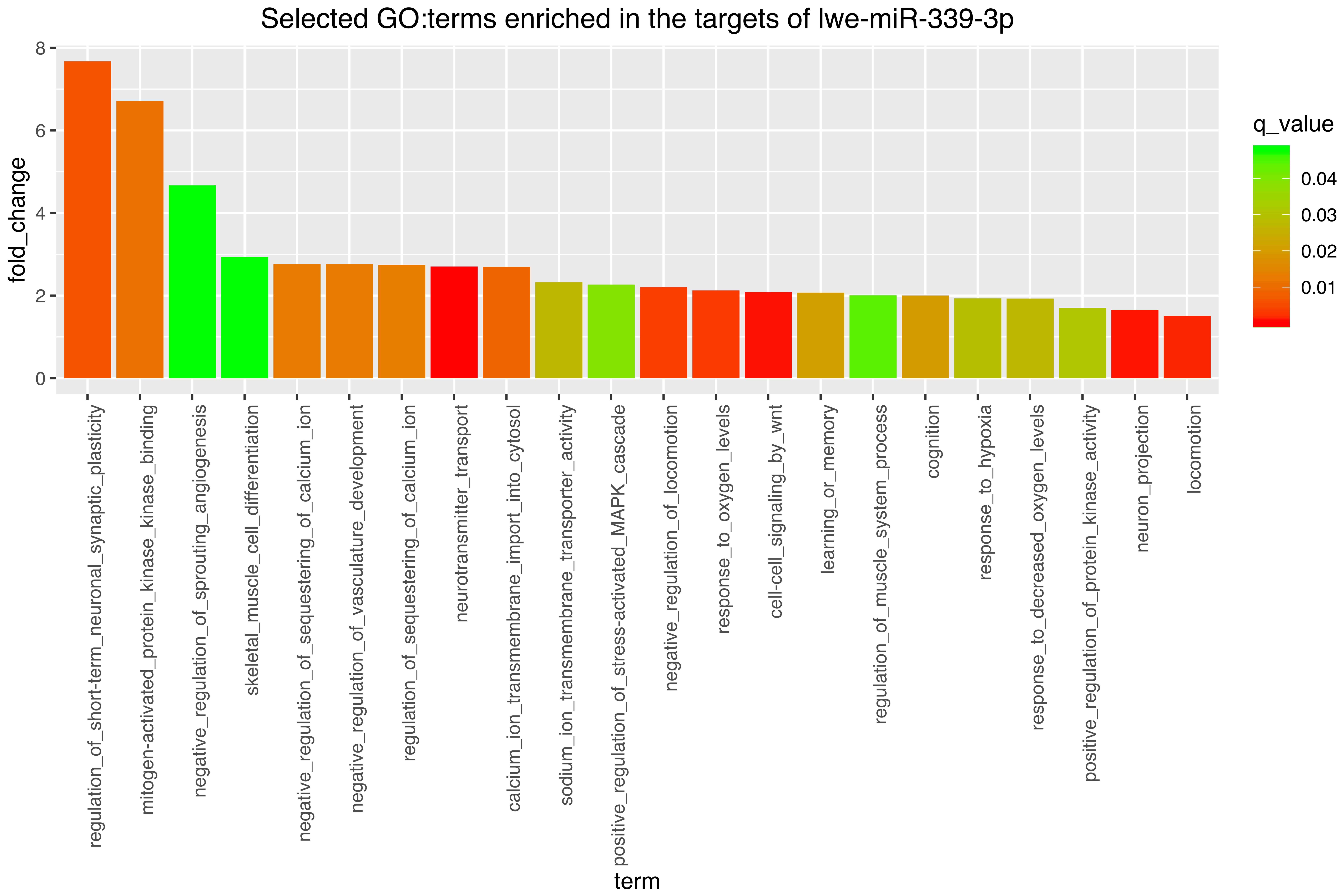


Supplementary Figure S8. Selected GO accessions enriched in the targets of miR-339-3p. The color gradient reflects the calculated q-value; different GO accessions are listed along the x-axis, while the fold change enrichment is shown on the y-axis.

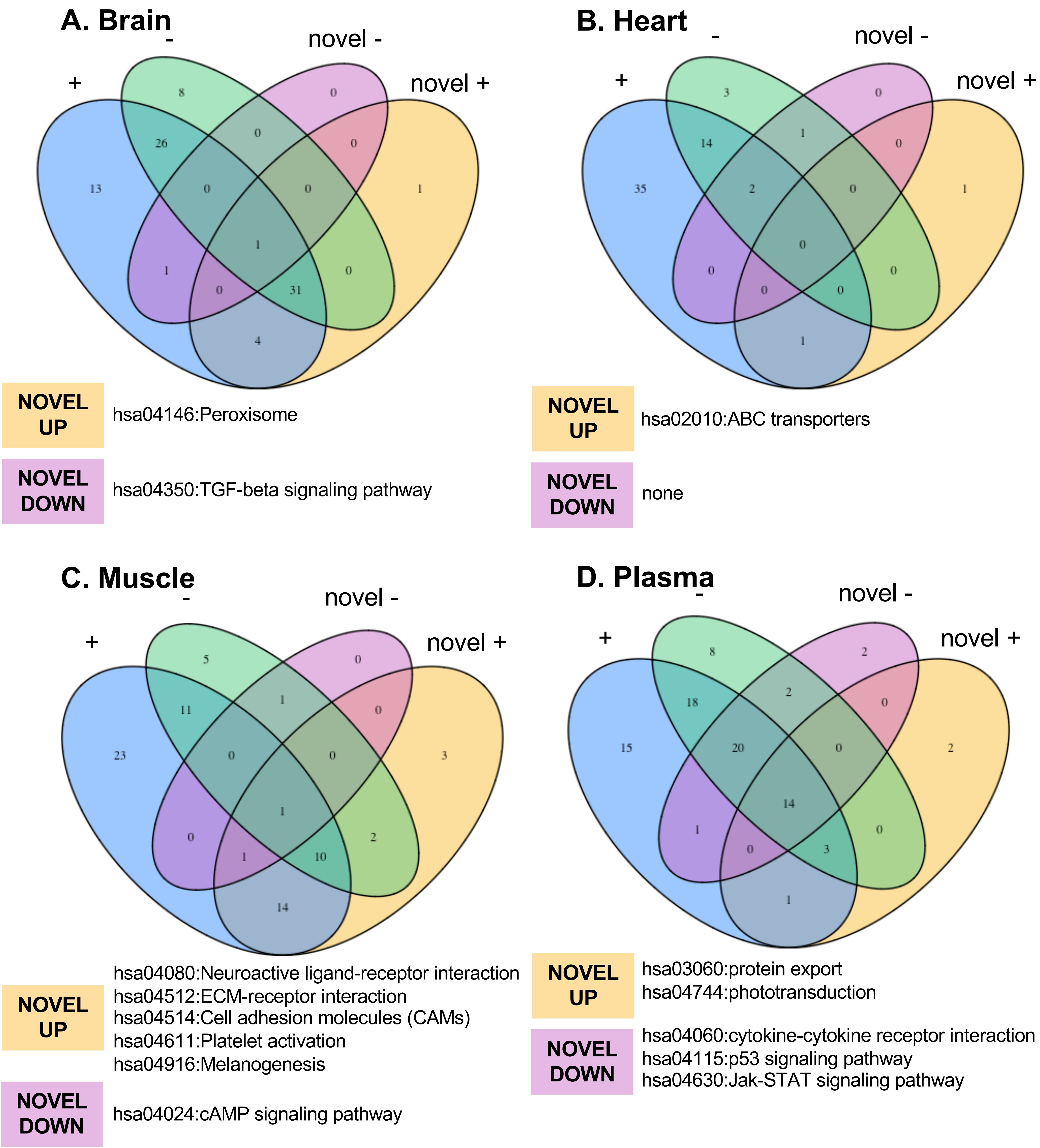


Supplementary Figure S9. Venn diagrams demonstrate overlap between enriched pathways of mRNA targets predicted to vary across 4 sample types in Weddell seals. Enriched KEGG pathways were identified based on miRNAs significantly up (+) or down (-) regulated in each tissue, compared to all others. They were then categorized as pathways related to previously annotated, miRBase miRNAs versus novel miRNAs identified in the Weddell seal (novel +, novel -). For each direction of regulation, the identities of enriched pathways targeted by novel miRNAs that were not detected from annotated miRNAs with the same direction of differential expression are listed for each tissue.
